# Science should be machine-readable

**DOI:** 10.64898/2026.01.30.702911

**Authors:** A. Sina Booeshaghi, Laura Luebbert, Lior Pachter

## Abstract

We develop a machine-automated approach for extracting results from papers, which we assess via a comprehensive review of the entire eLife corpus. Our method facilitates a direct comparison of machine and peer review, and sheds light on key challenges that must be overcome in order to facilitate AI-assisted science. In particular, the results point the way towards a machine-readable framework for disseminating scientific information. We therefore argue that publication systems should optimize separately for the dissemination of data and results versus the conveying of novel ideas, and the former should be machine-readable.

## Introduction

Science is primarily communicated through stories. Journal articles and preprints present scientific results as figures, tables, and prose organized for human readers. This format has served the scientific enterprise well. However, the number of scientific papers is growing rapidly (1). The resulting volume makes it increasingly difficult for scientists to systematically search, evaluate, and synthesize existing work, while effective tools for navigating the scientific literature remain limited.

The core challenge lies in how scientific results are presented. Results are typically reported in narrative form, expressed using domain-specific and often hard-to-parse language, and linked through citations that are coarse and at times incomplete. Data is presented in figures that may not be machine-readable, and raw data is not always available or annotated in a way that makes it usable. As a result, answering basic questions—^1^such as which data and studies support a given claim or how methods differ between studies—remains labor-intensive, even when the relevant information exists.

Machine-readable systems offer a way to address this challenge. When scientific content is structured, it becomes possible to index, compare, and analyze effectively (2). Some progress has been made as journals transition from PDF manuscripts to XML-based representations, stored in indexed file systems such as GitHub. The Journal Article Tag Suite (JATS) XML format provides a machine-readable encoding of journal articles that enables automated rendering, version control, and large-scale corpus management (3). While these formats and systems improve how papers are stored and distributed, they do not make scientific results explicit. As a consequence, synthesizing results across the literature remains difficult, even when the underlying data and code are available.

Recent advances in large language models highlight the importance of this limitation. LLMs are general-purpose parsers of natural language. Any scientific artifact that can be represented as text—papers, code, methods, and reviews—can be ingested, transformed, and reorganized by an LLM. This makes them well-suited to structuring scientific information (4). Their ability to generate, execute, and evaluate code also allows them to verify aspects of structured scientific outputs, providing a mechanism for consistency checks and validation. These capabilities make LLMs candidates for changing how scientific results are organized, evaluated, and disseminated.

LLMs can partially address the representation problem by extracting claims and results from existing papers and converting them into structured, machine-readable formats. However, they may be imperfect and limited by how well authors have organized the information in their articles. This raises the question of whether structure should be made explicit at publication.

Human-readable narratives remain essential. They support interpretation, creativity, and the communication of ideas. But they need not be the sole or primary representation of scientific output. Machine-readable repositories of results, code, and evidence could coexist with narrative papers, providing a substrate on top of which human-facing views are constructed. In this model, the narrative can become a highly curated and specific view of validated objects, rather than a single, fixed narrative.

In this work, we demonstrate the utility of machine-readable science by undertaking a journal-scale evaluation of peer review. Peer review is a core mechanism by which scientific claims are assessed, yet it is traditionally opaque, qualitative, and difficult to analyze at scale. By representing manuscripts and reviews in a shared, machine-readable framework of claims and results, we are able to compare reviews produced by large language models and by human reviewers across an entire journal corpus.

This comparison is not an argument for replacing human reviewers. Rather, it serves as a concrete demonstration of how machine-readable representations make previously inaccessible analyses accessible. The same framework that enables peer review comparison also supports other tasks, such as tracking claims across papers, comparing methods, auditing experiments, and synthesizing results across the literature. In this sense, the peer review analysis we present is one example of a broader class of evaluations that follow naturally from making scientific results explicit and machine-readable.

Taken together, these observations argue for scientific publication systems that should distinguish between the dissemination of results and the communication of ideas, and optimize them separately. Results—claims, evidence, and evaluations— should be published in explicit, machine-readable form, while narrative text serves as an interpretive layer for human readers. We treat peer review as a concrete case study of this principle: by making results and evaluations machine-readable, it becomes possible to analyze how scientific claims are assessed at scale.

## Machine-Readable Evaluation of Peer Review

Peer review is a relatively recent innovation in the scientific enterprise. First established in 1752, peer review became a systematic, institutionalized process only in the 20th century (5, 6).

The etymology of “peer” is derived from the Latin “par,” meaning an equal or matched companion, and the rationale for peer review was that peer experts are best informed to critique manuscripts and improve the research of their colleagues through the publication process (7). While the peer review process has been, and continues to be, widely critiqued (8), the process is not without merit. A famous example of the benefit of peer review is Nick Katz’s questioning of Andrew Wiles during the review of Wiles’ proof of Fermat’s Last Theorem, leading to the identification of a gap later fixed by Wiles and Taylor (9). Similarly, Howard Percy Robertson’s review of Albert Einstein’s 1936 paper rejecting gravitational waves—later revised and published—identified errors in the non-existence proof, which relied on singularities in the Einstein-Rosen metric (10). Robertson showed, in a 10-page referee report submitted to The Physical Review—a report that Einstein ignored—that recasting the metric in cylindrical coordinates removed these singularities, illustrating the central role of peer review: identifying mistakes before publication.

Peer review has contributed to improving research publications (11), although the small number of reviewers can result in mistakes (12), helps in dissemination of results (13), although the proliferation of preprints has partly displaced that purpose (14), and provides confidence in published results (15), although there is evidence that the public may be losing faith in the scientific enterprise (16).

Automated AI approaches to reviewing and analyzing manuscripts have been proposed in the past (17), but interest has recently increased (18) as a result of major advances in natural language processing (NLP) through the use of large language models (LLMs). In theory, LLMs can parse large amounts of complex technical writing, reason over empirical evidence, and generate structured critiques. Numerous private and public efforts have been announced that seek to leverage LLMs for these purposes, e.g., [Edison Scientific’s Kosmos](https://edisonscientific.com/articles/announcing-kosmos), refine.ink, [qedscience](https://www.qedscience.com/), [Elicit](https://elicit.com/), [Perplexity](https://www.perplexity.ai/), [Semantic Scholar](https://www.semanticscholar.org/), [ResearchAgent](https://github.com/JinheonBaek/ResearchAgent), and many more. However, the accuracy of AI-driven reviews is a topic of debate (19-23), and there have been several high-profile failures of machine reviews (22, 24). The question of what role artificial intelligence will play in advancing science, specifically regarding the evaluation and publication of research, remains open.

To address this question, we introduce a machine-readable, systematic comparison between LLM-based and human-based reviews to benchmark current model performance and understand how AI systems might augment, complement, or replace traditional review processes. We embarked on a large-scale assessment of AI review grounded in the open-access eLife corpus (Supplementary References), which, due to its publication model (25), is an exceptional resource providing not only the full text of papers under a non-restrictive CC-BY license but also invaluable peer reviews of those papers, Fig. 1 (green).

**Fig. 1.**
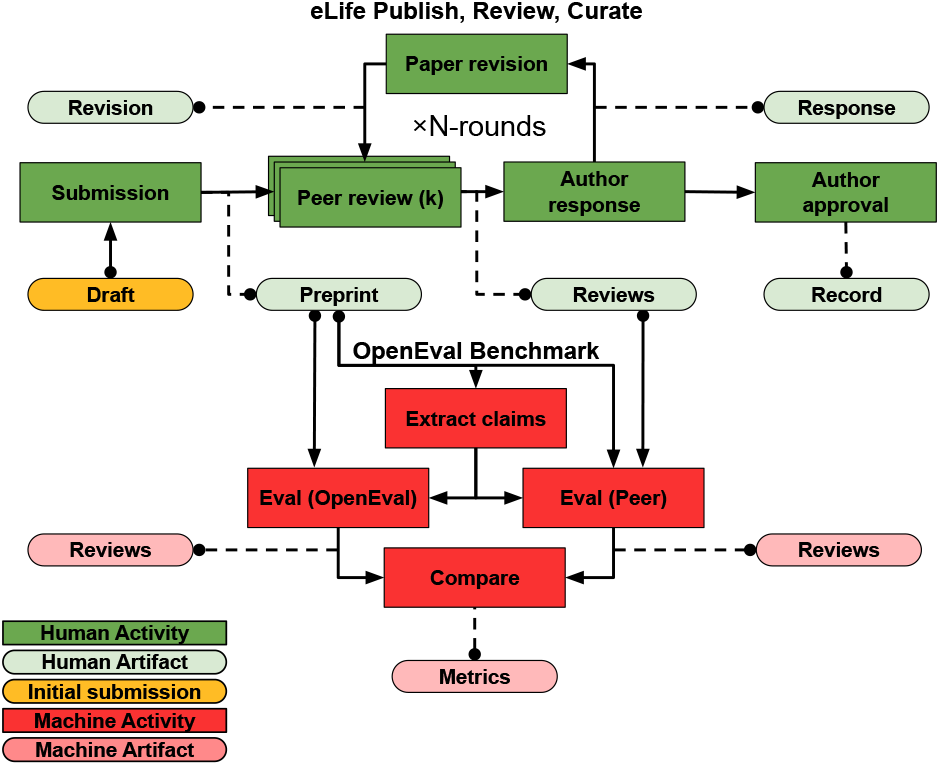
OpenEval Benchmark of the OpenEval Dataset. The eLife Publish, Review, Curate (PRC) model is represented as a block diagram. Scientists submit a draft either directly to eLife or to a preprint server. eLife decides to send the preprint for peer review, where it gets evaluated by *k* reviewers (*k* is typically three). Reviewers send their reviews to the authors, who respond and decide to either make their preprint (and reviews and responses) part of the record, or choose to respond and revise their paper for another round of peer review. This process can occur indefinitely. The OpenEval Benchmark takes, from eLife, the preprint and reviews of the preprint to construct the claims, results, and evaluations for comparison and analysis.

Our approach, which we term OpenEval, is open-source and constitutes a framework for evaluating and comparing the performance of machine versus human peer reviewers in assessing scientific results. OpenEval implements a structured, five-stage review: (1) extraction of atomic, factual, explicit and implicit claims from a manuscript, (2) grouping of those claims into research results, (3) evaluation of those research results by an LLM, (4) evaluation of the research results made by peer review commentary, and (5) comparison of the OpenEval and peer evaluations, Fig. 1 (red). Each stage outputs machine-readable, structured JSON data, enabling transparent, reproducible, and fine-grained analyses of agreement and divergence across reviewer types.

By aligning machine and peer reasoning on matched results based on the same set of possible claims, OpenEval provides a principled framework to compare LLM to peer reviews. Perhaps most importantly, the OpenEval results showcase how the framework wherein claims are extracted, subsets of them form results, and subsets of results are evaluated against each other, may be useful more generally for direct machine-readable dissemination of results in the future.

### OpenEval Dataset and Benchmark

Scientific papers are increasingly available online through two main channels. Preprint servers like arXiv, bioRxiv, and medRxiv host hundreds of thousands of papers ([arXiv](https://arxiv.org/stats/monthly_submissions): 2,935,336; [bioRxiv and medRxiv](https://openrxiv.org/2025-year-in-review/): >389,000, as of January 16, 2026) with minimal barriers to publication and no peer review requirement. Openaccess journals like Cell, Nature, and Science publish peer-reviewed papers, but charge authors fees (for example, [up to $12,690 per paper at Nature](https://www.nature.com/nature/for-authors/publishing-options)), and most of their papers remain behind paywalls.

Even when papers are freely available, three problems remain in benchmarking machine review. First, peer reviews are rarely published alongside articles. Nature only began requiring authors to release peer reviews in June 2025 (26) after offering this as an optional feature only five years prior. Second, even when peer reviews are available, they can be assessed only alongside the published articles, and not with respect to the submissions on which they were based. Third, restrictive licenses like [CC BY-NC-ND](https://www.nature.com/nature-portfolio/editorial-policies/self-archiving-and-license-to-publish) prohibit derivative works, posing a legal barrier to the use of LLMs in assessing manuscripts or their peer reviews.

The recently adopted publication system at eLife addresses all three of these issues. The eLife journal, which was [founded in 2012](https://elifesciences.org/inside-elife/12dd10ad/welcome-to-elife), adopted the permissive [CC BY license](https://elifesciences.org/terms) from the start and gave authors the option to publish peer reviews along with their papers (25). In 2023, eLife changed its publication model (27) to review selected preprints, along with their peer reviews and editorial assessments^1^ (Fig. 1). This new system, called the Publish, Review, Curate (PRC) publishing model, treats manuscripts as [“an open ‘conversation’… without the threat of rejection”](https://elifesciences.org/inside-elife/036d99a7/publishing-with-elife-an-open-conversation-without-the-threat-of-rejection). This process, shown in Fig. 1, starts with a `draft`. The submission passes through *N* rounds of peer review, during which the `preprint` receives `reviews` from *k* reviewers, with the authors given an opportunity to `respond` and make `revisions`. Once satisfied, the authors can decide to approve the state of the `preprint` and declare a version of `record` (subject to Editorial approval). Throughout this process, the journal publishes updates to `draft`, now called a `reviewed preprint`, in a machine-readable format (JATS XML) and makes them available via a GitHub repository (28).

JATS (Journal Article Tag Suite) is an XML format developed by the National Information Standard Organization that provides a standard way to annotate scientific articles (3). Like HTML, it uses tags to mark structural elements— title, abstract, author list, sections, figures—giving publishers a consistent vocabulary across journals and preprint servers. JATS files contain more information than language models need. The format includes metadata, formatting instructions, and cross-references that are useful for rendering articles but are currently unnecessary for text analysis with LLMs. These metadata inflate file size and consume context window space. We therefore built a command-line tool to convert JATS files to plain markdown. The tool, called `jats` (Methods), parses the eLife XML and outputs the article text as a markdown file, with optional export of peer reviews and author responses. This conversion reduces file size by about 65%, making the eLife corpus practical for use with language models.

We selected these eLife papers (post-2023, reviewed and disseminated using this new system) for our evaluation dataset, which we call the OpenEval Dataset (Fig. 2). The combination of permissive licensing, published submissions, peer reviews, and machine-readable format makes eLife uniquely suited for comparing LLM evaluations to human reviewers. Our dataset includes 2,487 papers with their associated peer reviews. We then selected the remaining 13,600 papers for OpenEval-only analysis.

**Fig. 2.**
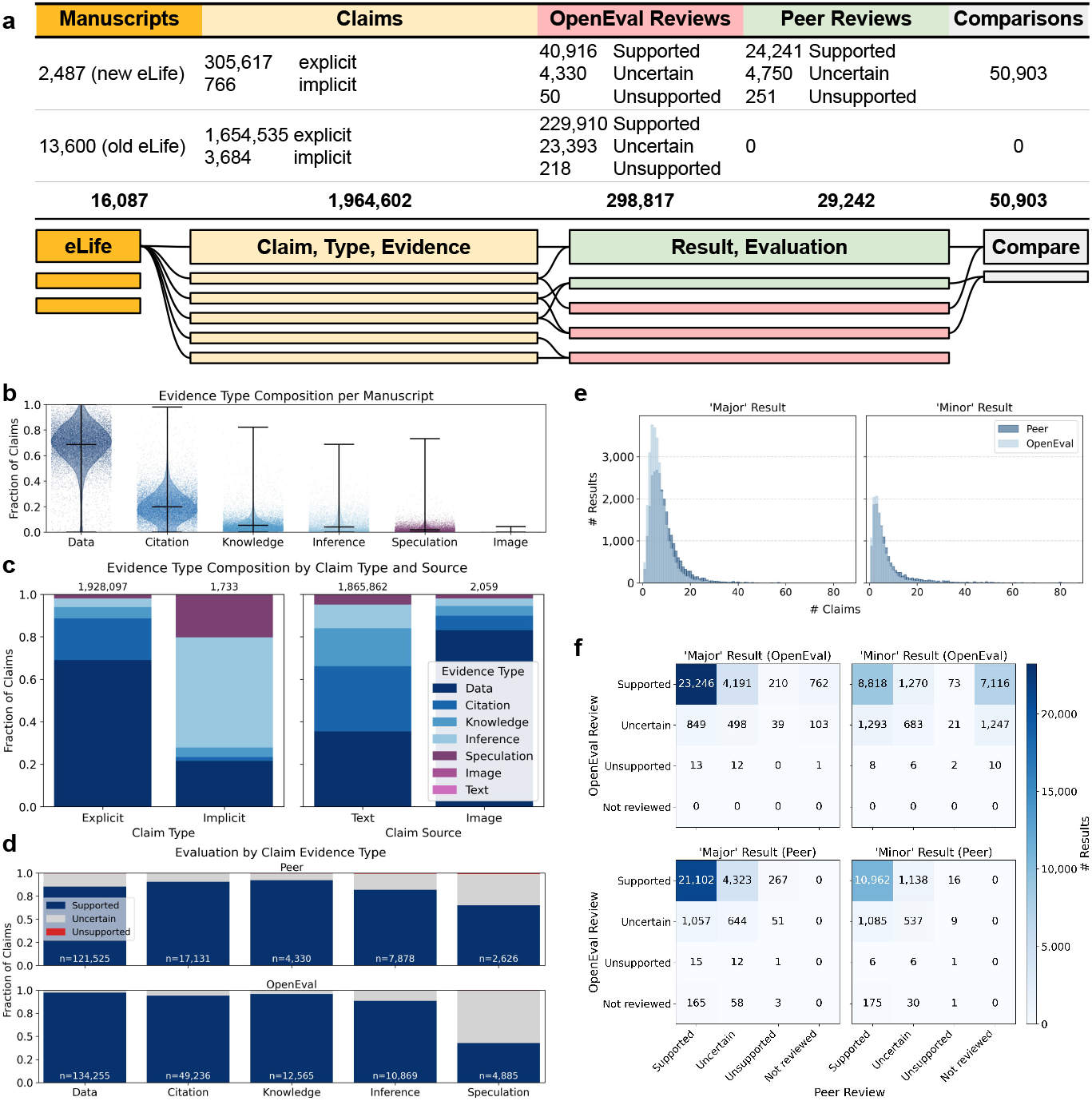
Claims, results, and evaluations of the OpenEval Dataset. **(a)** The OpenEval Dataset consists of claims, results, and evaluations made by OpenEval and peer reviewers on 2,487 eLife manuscripts. The OpenEval Dataset further includes a post-publication review of 13,600 eLife manuscripts performed with OpenEval. The OpenEval Benchmark consists of a comparison between the peer and OpenEval evaluations (including claims and results) from the OpenEval Dataset. **(b)** The fraction of claims, classified by their type, per manuscript. **(c)** The distribution of evidence types for explicit and implicit claims (left). The distribution of evidence types for text vs. image-based claims (right). This figure includes only claims associated with a single evidence type; the full breakdown is shown in Fig. S1e. **(d)** Fraction of claims per evidence type that were assigned to a supported, uncertain, or unsupported result by peer (top) and OpenEval (bottom). **(e)** The distribution of the number of claims per result for OpenEval and peers, for major and minor results. **f)** The number of supported, uncertain, unsupported, and not reviewed results evaluated by OpenEval and peer reviews, split by major or minor types by either OpenEval or peer review.

To compare machine review to peer review, we developed the OpenEval Benchmark, Fig. 1 (red). The benchmark consists of claim extraction, result collation and evaluation, followed by comparison. The benchmark starts with a reviewed `preprint` and associated peer `reviews`. LLMs extract claims linked to the `preprint`, evaluate groups of claims as results, and output these evaluations in a machine-readable structure for comparison. We generate machine `reviews` with Claude Sonnet 4.5 (in the OpenEval framework) and structure peer `reviews` (from the peer review file) using the same framework. We anticipate that the OpenEval Dataset and Benchmark will be useful for training AI-based peer review systems.

### Claim extraction

The task of claim extraction from scientific texts is a long-standing NLP problem (29). Techniques include rule-based and pattern-matching approaches, supervised sequence-labeling models (e.g., CRFs and BiLSTM-CRFs), neural span-extraction and relation-classification frameworks. In recent years, the efficacy of transformers and LLMs for claim extraction has been investigated, with promising results (30). LLMs have the ability to parse large amounts of text and to accurately extract precise scientific results.

To extract claims from the literature using current state-of-the-art LLMs, we developed a prompt guided by repeated experiments with respect to published peer review in the eLife corpus (see Methods). OpenEval extracted 1,964,602 claims in total from the eLife corpus (16,087 manuscripts), consisting of 1,960,309 explicit and 4,450 implicit claims. To prevent hallucinations, we ensured that extracted claims were traceable. Claims consisted of literal excerpts from the original manuscripts that are mapped back to the text itself (https://openevalproject.com).

We packaged claim extraction into a Python command-line tool, `cllm` (short for claim extraction with LLMs, Methods). `cllm` takes as input the markdown version of an eLife manuscript and generates a JSON file with extracted claims. The CLI allows users to set an API key for Anthropic’s Claude and set the specific LLM model to run claim extraction and verification.

Using `cllm`, OpenEval extracted an average of ∼112 claims per paper. The number of claims was not correlated with the length of the paper (*R*^2^ = 0.039; Fig. S1a). OpenEval associated each claim with a claim type, evidence type, and source type based on a predefined set of categories (see Methods). Across manuscripts, the most prominent evidence type was `data`, which accounted for approximately ∼75% of all evidence types, followed by `citations` (∼20%) and explicit `knowledge` claims (∼10%), with `inference` and `speculation` comprising smaller fractions. This distribution was consistent across manuscripts (Fig. 2b), indicating a strong reliance on empirical and referenced support in scientific writing.

We next examined how evidence composition varied by claim type and source. Explicit claims were overwhelmingly supported by `data` and `citations`, whereas implicit claims showed a shift toward `inference` and `speculation`. As expected, image-based claims were predominantly grounded in `data` (Fig. 2c).

### Result collation and evaluation

Extracted claims are literal statements from a manuscript, each representing one factual assertion. A paper contains many such claims, often closely related. We used an LLM to group claims into “results,” where each result is a set of related claims that together support a single finding.

Results form a subset of the power set of claims; let *C* be the set of claims. A result *r* is a subset of *C* (*r* ⊆ *C*), and the same claim may appear in multiple results—although in practice this was infrequent (less than one percent of claims). The set of all results *R* is therefore a subset of the power set of *C*, i.e., *R* ⊆ 2^*C*^. Through this grouping, a result is associated with claims, each of which is annotated with an evidence type. Each result is classified as major or minor and is given an evaluation (supported, unsupported, or uncertain). These attributes are both assigned by the evaluating system, either by peer reviewers or OpenEval.

We generated result-claim sets for both peer reviewers and OpenEval using `cllm`. The 1,964,602 claims were grouped into 298,817 results, and the LLM provided reasoning for each grouping decision. OpenEval evaluated 18.2 results per paper; peer reviewers evaluated 11.7 results per paper. Peer reviewer results contained more claims on average (8.5 claims per result) than OpenEval results (6.5 claims per result) (Fig. 2e). Overall, OpenEval identified 29,924 major and 20,547 minor results. Of OpenEval’s major results, 28,409 (95.0%) were supported, 1,489 (5.0%) were uncertain, and 26 (0.1%) were unsupported. Of its minor results, 17,277 (84.1%) were supported, 3,244 (15.8%) were uncertain, and 26 (0.1%) were unsupported.

Peer reviewers identified 27,698 major and 13,966 minor results. Peer reviewers marked a smaller fraction of major results as supported compared to OpenEval: 22,339 (80.7%) supported, 5,037 (18.2%) uncertain, and 322 (1.2%) unsupported. Their minor results showed 12,228 (87.5%) supported, 1,711 (12.3%) uncertain, and 27 (0.2%) unsupported (Fig. 2f).

The evidence type underlying each claim was predictive of the evaluation outcome of its associated results. Results with claims backed by `data` or `citations` were most often marked as supported by both systems (Fig. 2d). Results with claims based on `inference` or `speculation` showed higher rates of uncertainty or non-support. OpenEval supported a smaller fraction of `speculation`-based results than peer reviewers did (Fig. 2d).

To highlight the utility of machine review for presenting reviewed content, we built https://openevalproject.com, where extracted claims and results for all manuscripts can be explored interactively.

### OpenEval and peer review comparison

We compared OpenEval’s automated assessments to human peer reviews for eLife manuscripts where both were available, and examined how often the two approaches agreed, whether agreement changed over time, and whether agreements depended on the use of related claims.

OpenEval and human reviewers agreed on most research results (Fig. 3a). Of the 41,232 results evaluated by both, 33,247 (81%) received matching assessments. In 7,681 cases (19%), the two evaluations partially agreed (one expressed uncertainty while the other gave a definitive judgment). Most partial agreements reflected uncertainty from the human reviewer (5,497 cases). Outright disagreement was rare, occurring in only 304 cases (0.7%). Results where OpenEval and peer reviewers agreed showed consistently higher overlap in their underlying claims than results with partial agreement or disagreement (Fig. 3b). OpenEval and peer evaluations also agreed in in 32,320 (78.4%) of cases when classifying results as major or minor (Fig. S1b).

**Fig. 3.**
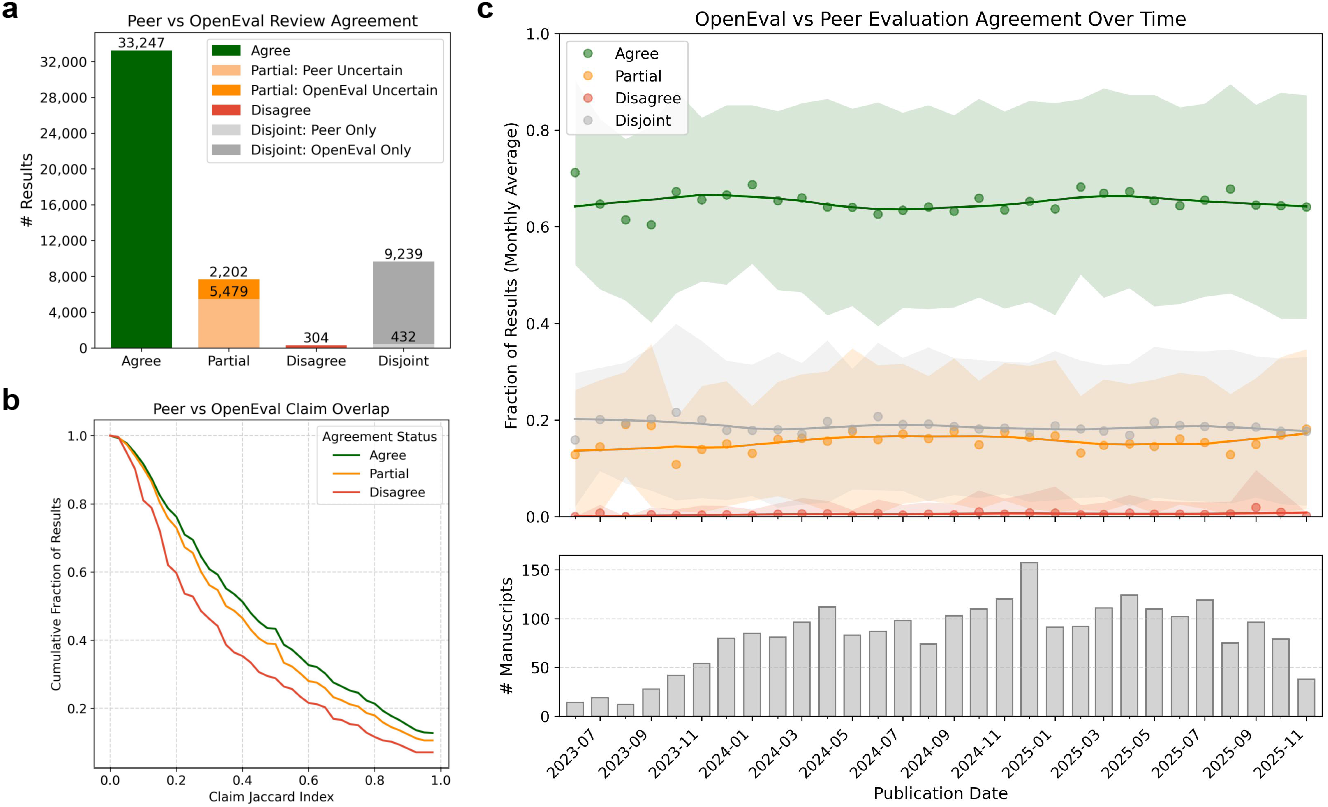
Comparison of machine and peer review. **(a)** The number of results where OpenEval and peer reviewers agreed, partially agreed, or disagreed in their evaluation of the result. Results that were not evaluated by peer reviewers but were evaluated by OpenEval (disjoint) are in dark gray, the converse are shown in light gray. **(b)** The Jaccard index on the set of claims per result between peer reviewers and OpenEval for comparisons where they both agree (green), partially agree (orange), and disagree (red). **(c)** Fraction of results where OpenEval and peer reviewers agreed, partially agreed, or disagreed in their evaluation, over time. Results are presented as monthly averages since the start of eLife’s PRC model, with the number of manuscripts given in the lower bar plot.

The two evaluations differed substantially in how many research results they assessed (Fig. 2f). In total, OpenEval found results that peer reviewers did not address (866 major and 8,373 minor), some of which were evaluated as “Uncertain” (103 major and 1,247 minor) or “Unsupported” (1 major and 10 minor) by OpenEval. The reverse was less common—peer reviewers rarely evaluated results that OpenEval missed (226 major and 206 minor). This difference is likely due to OpenEval’s usage of an LLM with a large context size, giving it the ability to systematically examine almost every result in a manuscript. In our comparison, OpenEval assessed 92.8% of research claims in a manuscript on average, while human reviewers were more selective, assessing only 68.1% of claims on average, leaving many statements unevaluated.

Evaluation and agreement patterns remained stable over time (Fig. 3c, Fig. S1c). The proportion of matching assessments did not change meaningfully across publication dates, nor did rates of partial agreement or disagreement. They did, however, exhibit high variability (Fig. 3c), likely due to variability in peer judgements (Fig. S1c). This stability in agreement rates over time, albeit with high variability, suggests that our findings are not artifacts of changing editorial practices or model behavior.

### OpenEval review of the entire eLife corpus

Having established that machine review has utility in assessing eLife publications using the benchmark dataset, we reviewed the remaining 13,600 manuscripts, made available under the old publishing model, with OpenEval.

Claim extraction yielded 1,654,535 claims (121.7 claims per manuscript on average). Results collation grouped these into 253,521 results (18.6 per manuscript). Of these results, OpenEval marked 229,910 (90.7%) as supported, 23,393 (9.2%) as uncertain, and 218 (0.09%) as unsupported. The low unsupported rate is expected for published work that has been peer-reviewed. 39.77% of uncertain and 39.51% of unsupported results primarily involved claims based on `inference` or `speculation`, compared to only 6.08% of supported results, consistent with patterns observed in the benchmark dataset (Fig. 2d).

### Connecting results across manuscripts

Machine readability makes it possible to connect papers at the level of individual results, rather than entire manuscripts. To explore this, we embedded all 298,817 results extracted by OpenEval using a text embedding model. Each result was mapped to a numerical vector. We then constructed a *k*-Nearest-Neighbor graph over these embeddings and identified pairs of results from different papers that were related under a cosine similarity metric (see Methods).

This analysis revealed several distinct classes of related result pairs. These included trivial pairs, where shared vocabulary led to superficial similarity; explicitly cited pairs, where two papers reported the same or closely related results and the more recent one cited the other; implicitly cited pairs, where a paper cited relevant prior work but not for the specific result identified by OpenEval; and uncited pairs, where related results appeared in papers that did not reference each other at all (Table S1).

Trivial pairs primarily reflected methodological overlap. For example, two studies describing RNA-seq experiments clustered together due to shared terminology around differential expression, “up- and down-regulation”, and “enrichment analysis”, even when the biological systems differed (31, 32). These connections appear to arise solely from the use of a shared vocabulary instead of from a deeper conceptual link. In analogy with LLM hallucinations, these LLM-induced false positive connections are paranormal citations, i.e., paracites.

Explicitly cited pairs often corresponded to closely related findings reported across papers from the same research group or line of work. In one such pair, two studies described oscillatory neural circuits in crustaceans, reporting consistent properties of pyloric and gastric mill rhythms and overlapping neuron populations (including identifying the number of Anterior Burster and Pyloric Dilator neurons involved in the circuit) (33, 34). In these cases, OpenEval recovered relationships that were already present in the citation graph.

Implicitly cited pairs revealed more subtle connections. In these cases, two papers cited are connected by a citation, but not for the specific result identified by OpenEval. For example, two studies characterized concentration-dependent oligomerization behavior of related centrosomal proteins, identifying similar domain-driven assembly mechanisms (35, 36). While one paper referenced the other, the specific connection between the results was not explicitly acknowledged in the context of the citation. It was recovered only through result-level comparison with OpenEval.

Finally, uncited pairs can reveal links between related studies with no direct citation link. In one case, two papers independently examined the mechanisms of timing-dependent long-term depression (tLTD) in different brain circuits. One showed that tLTD exists either with or without NMDA receptors, depending on the synaptic connection (37). The other showed that NMDA receptors are required and that they mediate tLTD through non-ionotropic signaling (38). Although the studies do not cite each other—they worked on different circuits—OpenEval links them by their specific results. This example highlights how AI agents could identify connected and related results leading to the novel synthesis of ideas if scientific results were machine-readable. Of course, determining the significance and utility of connections is non-trivial and requires domain expertise, in this case neuroscience expertise.

## Call to Action

Machine-readable science is enabling. The eLife journal was the first to make its manuscript corpus machine-readable by adopting the JATS format. Representing each paper as a single structured XML file made it straightforward to store the entire corpus on GitHub, including peer reviews. This, in turn, enabled automatic rendering of HTML views, version differences across manuscript revisions, and lower publication costs, since each paper could be tracked as a versioned file rather than a rendered document with scattered metadata.

We have taken the eLife approach one step further by representing papers as structured claims and results, which allowed us to compare machine and human peer review. The same representation supports other tasks that we do not perform here, but that follow naturally from machine readability. These include citation tracking, method comparison, literature search, and fact-checking. At the same time, our study highlights the limits of post hoc restructuring. Retrofitting machine readability onto existing papers using LLMs is costly, resource-intensive, and imperfect. More importantly, grouping claims into results required an additional inferential step by the model, which may not align with how the authors or reviewers understood the work.

The clear implication is that machine-readable science should be generated at the point of publication, not after the fact. Science should be disseminated in a structured and machine-readable form, along with traditional human-readable narratives. We do not propose a single, universal framework for organizing all of science. The diversity of scientific practice likely demands multiple, domain-specific solutions. This leads us to a clear call to action for the scientific community: (1) Develop shared nomenclature standards. (2) Build novel open-publication, reproducible, and transparent infrastructure. (3) Train models on structured scientific outputs.

Pursuing these goals will require a departure from current publication practices, which treat the scientific narrative as the primary research artifact. The machine learning community is well-positioned to lead this change, as it has the cultural influence to normalize machine-readable dissemination of scientific results, beginning in well-defined domains with existing standards and evaluation mechanisms.

The mathlib project (39) is a concrete example of this approach. Work in the mathematics community has shown how formal structure, through the Lean proof assistant, enables verification, reuse, and cumulative progress. Lean is thus complementing, not replacing, mathematics publications, which are essential for communicating narratives of ideas.

## The Utility of Machine-Readable Science

Scientific papers are narratives. They transmit ideas from one scientist to another by weaving together data, observations, hypotheses, inferences, and speculation in a form optimized for human reading. In this sense, a paper functions as an API (application programming interface) for human understanding. But it is a narrow one: rich in meaning for people, but opaque to machines, and distilled to only one fixed narrative.

A shift to machine-readable representations alongside narratives optimized for humans enables science to be indexed. Claims, results, methods, and citations can be extracted and organized into a structured graph that reflects how scientific ideas relate to one another across papers. Claims can be linked to results, results to citations, and citations to papers, across thousands of manuscripts. As this index fills in, navigation becomes easier for both humans and automated agents to discover relevant work, generate precise (and not hallucinated) citations, and to identify gaps in the citation record. Importantly, this graph need not be limited to explicit citations between papers. In fact, as we’ve shown here, there are likely many instances of connections between studies that do not cite each other but address related scientific questions that can be made in an automated way. Such an index could serve as the foundation on which machine-readable science delivers its benefits: it makes science easier to produce, evaluate, and consume.

### Producing Science

An indexed scientific record lowers the cost of engaging with prior work. Claims, results, and methods can be searched and compared directly. A researcher can quickly identify related findings, check whether an idea has been tested before, and generate new hypotheses by inspecting patterns across many papers, rather than relying on memory or manual reading alone. Literature review, a core scientific activity, becomes faster, systematic, and reproducible when the record itself is machine-readable.

### Evaluating Science

The same index helps scientists evaluate the growing volume of human- and machine-generated science. Claims and results can be grounded explicitly in the data that support them, as well as be put into the context of the scientific record. In this way, evaluations can help determine the utility of new work at a scale not possible by any one human.

### Consuming Science

Scientific narratives are essential. They are the interface through which humans reason, learn, communicate, and create. Narratives should not be replaced, but rather complemented. Our work shows that language models can help build out an index post hoc, enabling summaries, comparisons, and explanations that update as the underlying science evolves. Consumption of scientific knowledge becomes more flexible, reflecting the current state of the record rather than a static snapshot.

### AI-assisted Science

Finally, AI-assisted science depends on machine-readable scientific records. To *produce* hypotheses, explanations, or analyses, an agent must first *consume* and then *evaluate* existing knowledge. Machine-readable science enables this by transforming the literature into structured findings that can be composed at inference time. When scientific outputs are grounded in an explicit index rather than implicit text alone, AI systems can navigate the record in a way that is both scalable and faithful to the underlying evidence.

## Alternative Views

Our position on machine-readable science builds on a long history of efforts to improve reproducibility in scientific research. Like earlier advocates of reproducible science, our goal is pragmatic: to make it easier for scientists to build on prior work and to reduce friction in verification, comparison, and reuse. Efforts to formalize reproducibility have often been met with valid objections. These include increased burden on researchers, loss of creativity, and the difficulty of enforcing standards across fields. Similar concerns are likely to arise in response to proposals for machine-readable science.

**Objection: Efforts to make science machine-readable increase the burden on researchers**. While there is an upfront cost in organizing data and results in a machine-readable format, the extra work will yield numerous dividends.

**Objection: Imposing machine-readability risks rigidity and loss of scientific creativity**. While this is a serious and potentially valid concern, we are not proposing to make all aspects of publications machine-readable, and believe that there is an important role for communication of ideas and concepts, an aspect of science that may be difficult or impossible to formalize.

**Objection: Scientific claims and results are inseparable from the narrative that describes them**. Enumerating the types of claims and results that may appear in publications from all scientific fields is beyond the scope of this paper, however we believe that for many areas, most claims and results could be structured for machine readability.

**Objection: Writing a scientific narrative, after synthesizing results from an experiment, is crucial to the development of those ideas**. We agree with this and we are advocating for narratives to be written by scientists, rather than outsourcing writing to LLMs. Our paper is about machine readability of claims and results.

**Objection: Prior literature, by virtue of not being machine-readable, is inaccessible under the proposed system**. As we have demonstrated, LLMs can be used to make science machine-readable post hoc.

**Objection: Machine-readable science complicates evaluation and credit**. We do not claim to solve this challenge. However, we note that existing metrics already struggle to accurately reflect scientific contributions.

**Objection: Automation shifts authority from scientists to machines**. In our framework, LLMs function as tools for parsing and organizing scientific content, not as arbiters of truth. Human judgment remains central at every stage: in designing experiments, interpreting results, evaluating claims, and formulating narratives that transmit ideas.

## Supporting information

Supplementary References

## Acknowledgments

We thank Jonah Cool and Anthropic for providing funding that facilitated the deployment of OpenEval on the eLife corpus. We thank eLife for making its content accessible and usable via GitHub, as well as the authors who have submitted their work and the reviewers and editors who evaluated it. We also thank the Howard Hughes Medical Institute for supporting ASB through the Hanna H. Gray Fellows program. Work at the Eric and Wendy Schmidt Center is supported by the generosity of Eric and Wendy Schmidt.

## Impact Statement

This work highlights how representing scientific claims, evidence, and evaluations in machine-readable form can enable new forms of large-scale analysis of the scientific literature. Such representations make it easier to compare results across papers, track how claims are supported or revised over time, and identify relationships that are difficult to detect otherwise. While not a substitute for human judgment or scientific creativity, machine-readable science has significant potential to improve verification, synthesis, and reuse of existing results across many areas of research.

## Author Contributions

ASB conceived the idea of structuring scientific literature into a machine-readable form for AI-based peer review. ASB and LL jointly conceived the comparative study between AI-generated and human peer reviews using this structure. ASB developed the OpenEval framework with input from LL and LP. LL performed the analyses with input from ASB and LP. All authors contributed to writing the manuscript.

## Data and Code Availability

The OpenEval Dataset and Benchmark can be viewed interactively at https://openevalproject.com.

The `cllm` tool can be found here: https://github.com/OpenEvalProject/cllm.

The `jats` tool can be found here: https://github.com/OpenEvalProject/jats.

The OpenEval Dataset, consisting of the manuscripts, extracted claims, and results from 16,087 eLife papers—including peer reviews and comparisons, where available—has been organized to facilitate benchmarking of AI tools, and can be found here: https://github.com/OpenEvalProject/evals.

## Methods

### OpenEval Dataset and Benchmark

We generated the OpenEval Dataset from the public `elife-article-xml` GitHub repository (28). We cloned the repository and treated each XML file as one manuscript version (for a given manuscript ID). We restricted the analysis to `articles`.

For the OpenEval Benchmark (machine vs. peer comparison), we further restricted to manuscripts in the Publish, Review, Curate (PRC) era where peer review sub-articles were available and could be reliably associated with a specific reviewed preprint version. For the OpenEval-only corpus analysis, we included all remaining eligible eLife research manuscripts, including the pre-PRC era, where peer reviews were not consistently available.

A single eLife article may have multiple XML versions (e.g., `v1, v2`) corresponding to multiple rounds of revision. In the benchmark dataset, we selected versions depending on the publishing model used for the manuscript. For PRC manuscripts, we selected the earliest version of the manuscript. For non-PRC manuscripts, we selected the most recent version.

### Journal Article Tag Suite (JATS)

eLife articles are stored in the JATS file format. JATS is an XML standard for representing the structure of scholarly articles with additional metadata.

We developed a Python library and command-line tool called `jats`. The main function of `jats` is to convert eLife XML files to plain text. The tool parses JATS XML and outputs three markdown files: the manuscript body, peer reviews, and author responses. JATS files contain structural markup, cross-references, and metadata that inflate file size without adding content for language model analysis. Our conversion strips this overhead while preserving the text, figures, tables, and mathematical notation. The `jats` tool extracts the following article components:

- Title
- Authors
- Affiliations
- Abstract
- References
- Figures
- Tables
- Body sections
- Sub-articles containing peer reviews

For mathematical content, `jats` converts MathML to LaTeX notation. For citations, it preserves reference text and links to DOIs. The output follows standard markdown syntax, with HTML tables.

We classified sub-articles by their JATS article-type attribute. Decision letters, editor reports, and referee reports became peer review files. Author replies, and comments became response files. The tool extracts reviewer metadata when available, including names, affiliations, and ORCID identifiers, though many reviews are anonymous.

### Claim extraction

We developed a command-line tool `cllm` (claim extraction with Anthropic’s Claude Sonnet 4.5, claude-sonnet-4-5-20250929) to extract atomic factual claims from the eLife manuscripts. `cllm` tool takes the JATS converted markdown files as input and sends them to an LLM with a structured prompt. It expects a structured JSON array of claims. Each claim contains six fields:

1. The claim text states a single factual assertion.
2. The claim type is either explicit (stated directly) or implicit (logically entailed).
3. The source field contains the exact excerpt from the manuscript, preserving inline citations.
4. The source type indicates whether the claim derives from text, images, or both.
5. The evidence field explains how the claim is supported.
6. The evidence type classifies support as data, citation, knowledge, inference, or speculation.

We required claims to be atomic: one discrete assertion per claim. We required sources to be verbatim excerpts that could be located in the original text. This traceability constraint reduces hallucination; any claim whose source cannot be found in the manuscript is discarded. This step removed fewer than 1% of claims in practice.

### Result collation

Claims describe individual facts, and results describe scientific findings. A result is a grouping of related claims that together support one conclusion. We used Anthropic’s Claude Sonnet 4.5 to group claims into results and evaluate whether each result is supported by the evidence. Each result has three attributes:

1. The evaluation type is supported, unsupported, or uncertain.
2. The result type is major (core finding) or minor (supporting detail).
3. The evaluation field contains a brief explanation of the assessment.

We generated results with the `cllm` tool in two ways: by LLM evaluation and by peer review interpretation.

#### LLM Evaluation

For LLM evaluation, `cllm` sends the manuscript and extracted claims to Anthropic’s Claude Sonnet 4.5. The prompt instructs the model to act as a critical scientific reviewer. The prompt specifies evaluation criteria: statistical soundness, experimental rigor, computational integrity, logical consistency, and accuracy of prior work claims.

The model groups related claims and assigns each group an evaluation. It provides reasoning that references specific claims by their identifiers (C1, C2, etc.). It classifies each result as major or minor based on scientific significance.

#### Peer review interpretation

For peer review evaluation, `cllm` sends the manuscript, claims, and peer review text to Anthropic’s Claude Sonnet 4.5. The prompt instructs the model to identify which claims the reviewers addressed and how.

Anthropic’s Claude Sonnet 4.5 groups claims according to how they were discussed by the reviewers. It assigns evaluations based on reviewer statements: affirmation yields “supported”, explicit critique yields “unsupported”, expressed doubt, or requests for additional experiments yield “uncertain”. The model quotes or paraphrases reviewer feedback in its reasoning. This design separates what the reviewers said from the independent assessment. The LLM interprets peer commentary; it does not substitute its own judgment for theirs.

### Result comparisons

To compare OpenEval evaluations with human peer review, we aligned results within each manuscript using a two-step procedure. First, we computed the Jaccard index between all OpenEval and peer review results based on their associated claim identifiers. For an OpenEval result and a peer result, the Jaccard index is defined as the size of the intersection of their claim sets divided by the size of their union. This provided a simple measure of claim-level overlap and was used to identify candidate result pairings. Second, we passed the candidate pairings to an LLM for comparison. For each candidate pair, the model was given the two results, their evaluation labels, and the claims associated with each result. The model was instructed to determine whether the two results addressed the same underlying scientific finding and, if so, to briefly describe how the OpenEval and peer evaluations aligned or differed. If a result appeared in only one system, the model recorded it as evaluated by OpenEval only or by peer review only.

The output of this step was a linking of OpenEval results to peer results, along with a short explanation of agreement, partial agreement, or disagreement. These concordance metrics were used to compute agreement statistics and to analyze differences in claim coverage.

### The cllm workflow

We processed the eLife corpus using a batch script that coordinates the workflow. The script queries a SQLite database for manuscripts with status QUEUED, processes them in parallel (up to 10 concurrent jobs), and updates their status to PROCESSED or FAILED. The workflow has five stages per manuscript:

1. Claim extraction from manuscript markdown
2. LLM evaluation of claims (OpenEval)
3. Peer review evaluation of claims (if reviews exist)
4. Comparison of evaluations (if peer reviews exist)
5. Export to database format

Each stage writes output to a JSON file. The final stage consolidates all outputs into a single JSON file with normalized tables for database import.

We tracked manuscript versions separately. A paper may have multiple versions (v1, v2, etc.) corresponding to revision cycles. Each version has its own XML file, markdown conversion, and evaluation outputs.

We classified papers by publication model. Papers published under eLife’s Publish, Review, Curate model (post-2023) have peer reviews available. Papers published under the traditional model have manuscript text only. We processed both, but generated comparisons only for papers with peer reviews.

### Result embeddings

To enable similarity analysis across claims and results, we generated vector embeddings for all extracted claims and all OpenEval results. For each manuscript version, we embedded the text of each claim and each result independently using a fixed text embedding model (OpenAI `text-embedding-3-large`). Claims were embedded using their claim text, and results were embedded using the short natural-language result summaries produced by OpenEval.

We loaded precomputed embeddings for all results and normalized them to unit length, so that the inner product corresponds to cosine similarity. To identify related results at scale, we constructed a *k*-Nearest-Neighbor (kNN) index over all result embeddings using FAISS with an HNSW index and inner-product similarity. For each result, we retrieved its top 50 nearest neighbors, excluding the result itself. Similarity scores and neighbor indices were stored to disk to allow downstream analysis without re-running the index.

We first examined how often nearest neighbors came from the same manuscript versus different manuscripts. While many high-similarity neighbors originated from the same paper, a substantial fraction linked results across different papers, motivating cross-manuscript analysis.

To summarize similarity at the manuscript level, we aggregated result-level similarities into a directed paper-paper graph. For each pair of papers, we summed the similarity scores of all result pairs connecting them, excluding within-paper links. To prevent papers with many results from dominating the graph, outgoing edge weights were normalized by the total outgoing similarity from each paper.

To inspect individual cross-paper relationships, we identified, for a given result, the most similar result originating from a different manuscript and examined the paired result texts directly.

### OpenEval resource usage

To understand the resource consumption of LLMs for claim extraction and machine review, we measured the token usage, time, and cost of running OpenEval on the entire eLife corpus. Extracting claims from the entire eLife corpus cost a total of $6,844.77 ($0.43 per manuscript) and evaluation with OpenEval cost $5,287.94 ($0.33 per manuscript), with a total single-thread processing time of 1,252.9 h (280.38 s per manuscript) and 422.7 h (94.58 s per manuscript), respectively (Fig. S1d).

## Supplementary Information

**Fig. S1.**
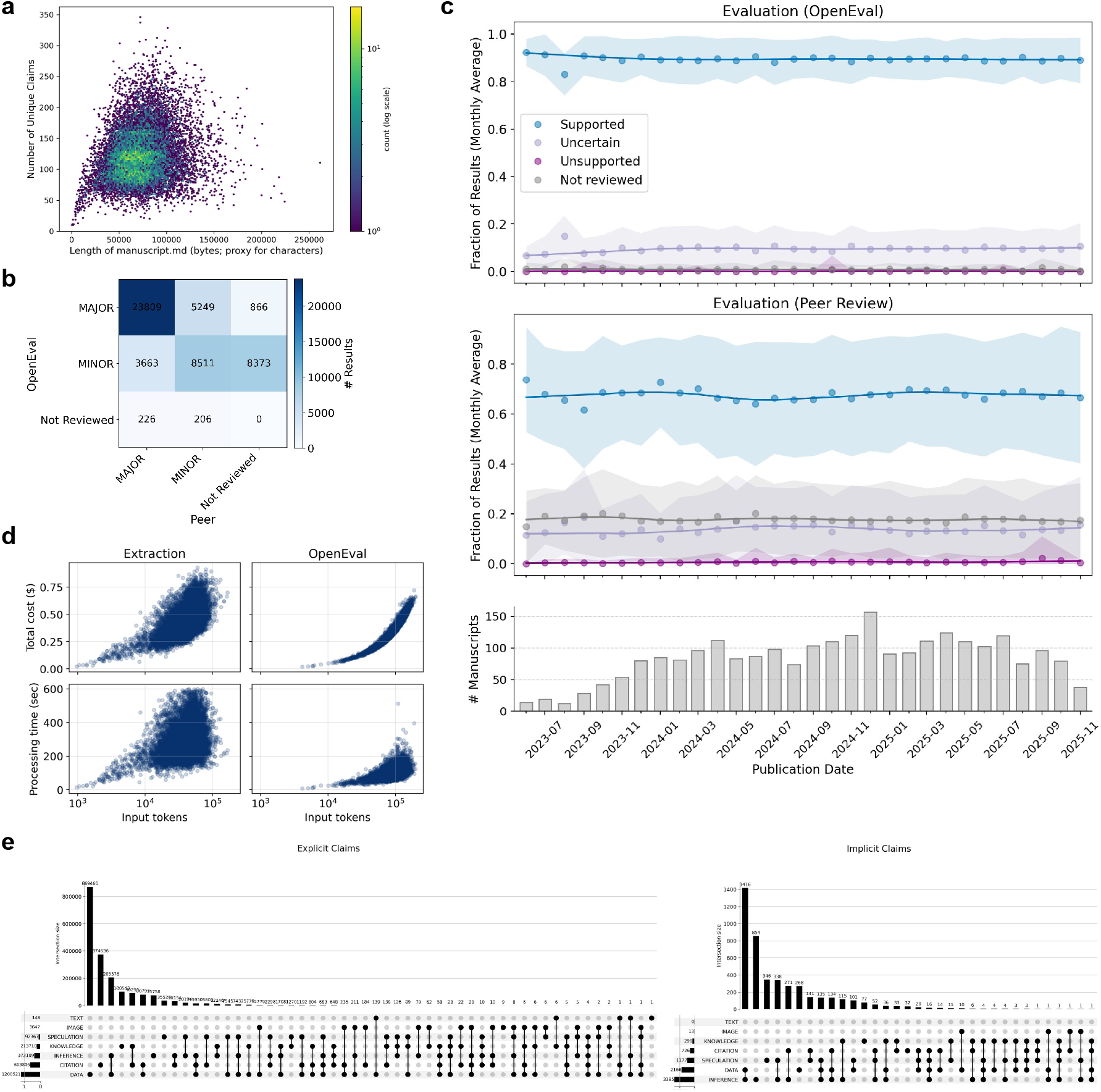
**(a)** The number of claims per manuscript as a function of manuscript length. **(b)** Agreement between peer and OpenEval categorization of results as major and minor. **(c)** Fraction of results evaluated by OpenEval or peer reviewers as supported, uncertain, or unsupported over time. Results are shown as monthly averages since the start of eLife’s PRC model, with the number of manuscripts indicated in the lower bar plot. **(d)** Processing time and cost per input token for claim extraction and OpenEval evaluation of each manuscript in the entire eLife corpus. **(e)** The number of explicit (left) and implicit (right) claims, grouped by evidence type

**Table S1.**
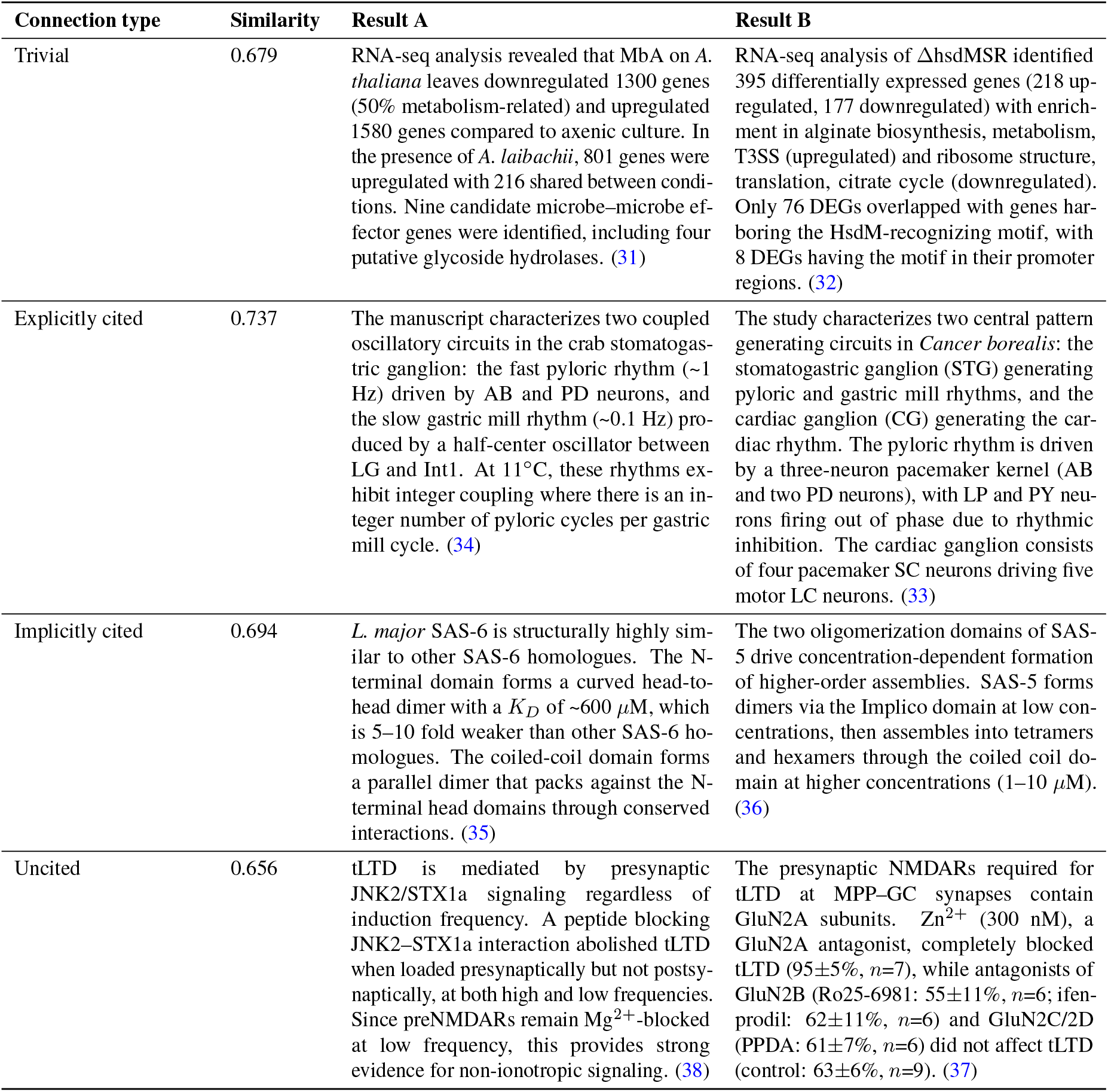
Examples of result-level connections across manuscripts. Representative pairs of related results identified via embedding-based similarity, illustrating different types of connections between papers. Results are inferred by OpenEval from the reviewed papers. The similarity refers to the cosine similarity on the embedded result text (see Methods).

## Footnotes

1 The em dashes reflect our stylistic preference and should not be interpreted as evidence of AI-assisted text generation.

1 Note that while most papers in this new publishing era have peer reviews by default, eLife does not explicitly mandate that papers get published in that way. Authors still have the option of publishing on eLife under the old system.

