## Supplementary References for "Science should be machine-readable"

### The eLife Corpus

The papers below were analyzed by OpenEval. They are distributed by eLife under the CC-BY license.

[2] Ron Ammar, Dax Torti, Kyle Tsui, Marinella Gebbia, Tanja Durbic, Gary D Bader, Guri Giaever, and Corey Nislow. Chromatin is an ancient innovation conserved between Archaea and Eukarya. *eLife*, 1, 2012.

[3] Preetha Anand, Silvia Cermelli, Zhihuan Li, Adam Kassan, Marta Bosch, Robilyn Sigua, Lan Huang, Andre J Ouellette, Albert Pol, Michael A Welte, and Steven P Gross. A novel role for lipid droplets in the organismal antibacterial response. *eLife*, 1, 2012.

[4] Tsutomu Aoki, Ali Sarkeshik, John Yates, and Paul Schedl. Elba, a novel developmentally regulated chromatin boundary factor is a hetero-tripartite DNA binding complex. *eLife*, 1, 2012.

[5] Yael Artzy-Randrup, Mary M Rorick, Karen Day, Donald Chen, Andrew P Dobson, and Mercedes Pascual. Population structuring of multi-copy, antigen-encoding genes in Plasmodium falciparum. *eLife*, 1, 2012.

[6] B Tom Burnley, Pavel V Afonine, Paul D Adams, and Piet Gros. Modelling dynamics in protein crystal structures by ensemble refinement. *eLife*, 1, 2012.

[7] Claudio Ciferri, Gabriel C Lander, Alessio Maiolica, Franz Herzog, Ruedi Aebersold, and Eva Nogales. Molecular architecture of human polycomb repressive complex 2. *eLife*, 1, 2012.

[8] Bray Denard, Ching Lee, and Jin Ye. Doxorubicin blocks proliferation of cancer cells through proteolytic activation of CREB3L1. *eLife*, 1, 2012.

[9] Jiajie Diao, Patricia Grob, Daniel J Cipriano, Minjoung Kyoung, Yunxiang Zhang, Sachi Shah, Amie Nguyen, Mark Padolina, Ankita Srivastava, Marija Vrljic, Ankita Shah, Eva Nogales, Steven Chu, and Axel T Brunger. Synaptic proteins promote calcium-triggered fast transition from point contact to full fusion. *eLife*, 1, 2012.

[10] Anca M Farcas, Neil P Blackledge, Ian Sudbery, Hannah K Long, Joanna F McGouran, Nathan R Rose, Sheena Lee, David Sims, Andrea Cerase, Thomas W Sheahan, Haruhiko Koseki, Neil Brockdorff, Chris P Ponting, Benedikt M Kessler, and Robert J Klose. KDM2B links the Polycomb Repressive Complex 1 (PRC1) to recognition of CpG islands. *eLife*, 1, 2012.

[11] Brian J Ferguson, Daniel S Mansur, Nicholas E Peters, Hongwei Ren, and Geoffrey L Smith. DNA-PK is a DNA sensor for IRF-3-dependent innate immunity. *eLife*, 1, 2012.

[12] Tim Grotjohann, Ilaria Testa, Matthias Reuss, Tanja Brakemann, Christian Eggeling, Stefan W Hell, and Stefan Jakobs. rsEGFP2 enables fast RESOLFT nanoscopy of living cells. *eLife*, 1, 2012.

[13] Ryan J Hum, Prabhat Jha, Anita M McGahan, and Yu-Ling Cheng. Global divergence in critical income for adult and childhood survival: analyses of mortality using Michaelis–Menten. *eLife*, 1, 2012.

[14] Philipp Kimmig, Marcy Diaz, Jiashun Zheng, Christopher C Williams, Alexander Lang, Tomas Aragón, Hao Li, and Peter Walter. The unfolded protein response in fission yeast modulates stability of select mRNAs to maintain protein homeostasis. *eLife*, 1, 2012.

[15] Chunguang Kong, Weizhong Zeng, Sheng Ye, Liping Chen, David Bryant Sauer, Yeeling Lam, Mehabaw Getahun Derebe, and Youxing Jiang. Distinct gating mechanisms revealed by the structures of a multi-ligand gated K+ channel. *eLife*, 1, 2012.

[16] Xiao-Dong Li and Zhijian J Chen. Sequence specific detection of bacterial 23S ribosomal RNA by TLR13. *eLife*, 1, 2012.

[17] Glenn A Maston, Lihua Julie Zhu, Lynn Chamberlain, Ling Lin, Minggang Fang, and Michael R Green. Non-canonical TAF complexes regulate active promoters in human embryonic stem cells. *eLife*, 1, 2012.

[18] Jerome S Menet, Joseph Rodriguez, Katharine C Abruzzi, and Michael Rosbash. Nascent-Seq reveals novel features of mouse circadian transcriptional regulation. *eLife*, 1, 2012.

[19] Matthew P Miller, ElÇin ünal, Gloria A Brar, and Angelika Amon. Meiosis I chromosome segregation is established through regulation of microtubule–kinetochore interactions. *eLife*, 1, 2012.

[20] Syed M Moin and Sinisa Urban. Membrane immersion allows rhomboid proteases to achieve specificity by reading transmembrane segment dynamics. *eLife*, 1, 2012.

[21] Paolo Pretto, Jean-Pierre Bresciani, Gregor Rainer, and Heinrich H Bülthoff. Foggy perception slows us down. *eLife*, 1, 2012.

[22] Kaushik Ragunathan, Cheng Liu, and Taekjip Ha. RecA filament sliding on DNA facilitates homology search. *eLife*, 1, 2012.

[23] Stephen W Santoro and Catherine Dulac. The activity-dependent histone variant H2BE modulates the life span of olfactory neurons. *eLife*, 1, 2012.

[24] Meredith C Schuman, Kathleen Barthel, and Ian T Baldwin. Herbivory-induced volatiles function as defenses increasing fitness of the native plant Nicotiana attenuata in nature. *eLife*, 1, 2012.

[25] Hao Wu, John Williams, and Jeremy Nathans. Morphologic diversity of cutaneous sensory afferents revealed by genetically directed sparse labeling. *eLife*, 1, 2012.

[26] Huan Yan, Guocai Zhong, Guangwei Xu, Wenhui He, Zhiyi Jing, Zhenchao Gao, Yi Huang, Yonghe Qi, Bo Peng, Haimin Wang, Liran Fu, Mei Song, Pan Chen, Wenqing Gao, Bijie Ren, Yinyan Sun, Tao Cai, Xiaofeng Feng, Jianhua Sui, and Wenhui Li. Sodium taurocholate cotransporting polypeptide is a functional receptor for human hepatitis B and D virus. *eLife*, 1, 2012.

[27] Yuan Zhang, Yang Xie, Eric D Berglund, Katie Colbert Coate, Tian Teng He, Takeshi Katafuchi, Guanghua Xiao, Matthew J Potthoff, Wei Wei, Yihong Wan, Ruth T Yu, Ronald M Evans, Steven A Kliewer, and David J Mangelsdorf. The starvation hormone, fibroblast growth factor-21, extends lifespan in mice. *eLife*, 1, 2012.

[28] Sadeem Ahmad, Satya Brata Routh, Venu Kamarthapu, Jisha Chalissery, Sowndarya Muthukumar, Tanweer Hussain, Shobha P Kruparani, Mandar V Deshmukh, and Rajan Sankaranarayanan. Mechanism of chiral proofreading during translation of the genetic code. *eLife*, 2, 2013.

[29] Benjamin Allen, Jeff Gore, and Martin A Nowak. Spatial dilemmas of diffusible public goods. *eLife*, 2, 2013.

[30] Silke Allmann, Anna Späthe, Sonja Bisch-Knaden, Mario Kallenbach, Andreas Reinecke, Silke Sachse, Ian T Baldwin, and Bill S Hansson. Feeding-induced rearrangement of green leaf volatiles reduces moth oviposition. *eLife*, 2, 2013.

[31] Kasper R Andersen, Evgeny Onischenko, Jeffrey H Tang, Pravin Kumar, James Z Chen, Alexander Ulrich, Jan T Liphardt, Karsten Weis, and Thomas U Schwartz. Scaffold nucleoporins Nup188 and Nup192 share structural and functional properties with nuclear transport receptors. *eLife*, 2, 2013.

[32] Keigo Araki, Keiko Kawauchi, Hiroaki Hirata, Mie Yamamoto, and Yoichi Taya. Cytoplasmic translocation of the retinoblastoma protein disrupts sarcomeric organization. *eLife*, 2, 2013.

[33] Anton Arkhipov, Yibing Shan, Eric T Kim, Ron O Dror, and David E Shaw. Her2 activation mechanism reflects evolutionary preservation of asymmetric ectodomain dimers in the human EGFR family. *eLife*, 2, 2013.

[34] Moritz Armbruster, Mirko Messa, Shawn M Ferguson, Pietro De Camilli, and Timothy A Ryan. Dynamin phosphorylation controls optimization of endocytosis for brief action potential bursts. *eLife*, 2, 2013.

[35] Alyson Ashe, Tony Bélicard, Jérémie Le Pen, Peter Sarkies, Lise Frézal, Nicolas J Lehrbach, Marie-Anne Félix, and Eric A Miska. A deletion polymorphism in the Caenorhabditis elegans RIG-I homolog disables viral RNA dicing and antiviral immunity. *eLife*, 2, 2013.

[36] Xiao-chen Bai, Israel S Fernandez, Greg McMullan, and Sjors HW Scheres. Ribosome structures to near-atomic resolution from thirty thousand cryo-EM particles. *eLife*, 2, 2013.

[37] Minna D Balbas, Michael J Evans, David J Hosfield, John Wongvipat, Vivek K Arora, Philip A Watson, Yu Chen, Geoffrey L Greene, Yang Shen, and Charles L Sawyers. Overcoming mutation-based resistance to antiandrogens with rational drug design. *eLife*, 2, 2013.

[38] Anirban Banerjee, Alice Lee, Ernest Campbell, and Roderick MacKinnon. Structure of a pore-blocking toxin in complex with a eukaryotic voltage-dependent K+ channel. *eLife*, 2, 2013.

[39] István Bartha, Jonathan M Carlson, Chanson J Brumme, Paul J McLaren, Zabrina L Brumme, Mina John, David W Haas, Javier Martinez-Picado, Judith Dalmau, Cecilio López-Galíndez, Concepción Casado, Andri Rauch, Huldrych F Günthard, Enos Bernasconi, Pietro Vernazza, Thomas Klimkait, Sabine Yerly, Stephen J O'Brien, Jennifer Listgarten, Nico Pfeifer, Christoph Lippert, Nicolo Fusi, Zoltán Kutalik, Todd M Allen, Viktor Müller, P Richard Harrigan, David Heckerman, Amalio Telenti, and Jacques Fellay. A genome-to-genome analysis of associations between human genetic variation, HIV-1 sequence diversity, and viral control. *eLife*, 2, 2013.

[40] Franziska Bleichert, Maxim Balasov, Igor Chesnokov, Eva Nogales, Michael R Botchan, and James M Berger. A Meier-Gorlin syndrome mutation in a conserved C-terminal helix of Orc6 impedes origin recognition complex formation. *eLife*, 2, 2013.

[41] Hannah G Blitzblau and Andreas Hochwagen. ATR/Mec1 prevents lethal meiotic recombination initiation on partially replicated chromosomes in budding yeast. *eLife*, 2, 2013.

[42] David Bolduc, Meghdad Rahdar, Becky Tu-Sekine, Sindhu Carmen Sivakumaren, Daniel Raben, L Mario Amzel, Peter Devreotes, Sandra B Gabelli, and Philip Cole. Phosphorylation-mediated PTEN conformational closure and deactivation revealed with protein semisynthesis. *eLife*, 2, 2013.

[43] Henry R Bourne. A fair deal for PhD students and postdocs. *eLife*, 2, 2013.

[44] Henry R Bourne. A recipe for mediocrity and disaster, in five axioms. *eLife*, 2, 2013.

[45] Henry R Bourne. The writing on the wall. *eLife*, 2, 2013.

[46] Sean Michael Boyle, Shane McInally, and Anandasankar Ray. Expanding the olfactory code by in silico decoding of odor-receptor chemical space. *eLife*, 2, 2013.

[47] Ivana Bozic, Johannes G Reiter, Benjamin Allen, Tibor Antal, Krishnendu Chatterjee, Preya Shah, Yo Sup Moon, Amin Yaqubie, Nicole Kelly, Dung T Le, Evan J Lipson, Paul B Chapman, Luis A Diaz, Bert Vogelstein, and Martin A Nowak. Evolutionary dynamics of cancer in response to targeted combination therapy. *eLife*, 2, 2013.

[48] Amy Brand. Faculty appointments and the record of scholarship. *eLife*, 2, 2013.

[49] Nicolas Brandes, Heather Tienson, Antje Lindemann, Victor Vitvitsky, Dana Reichmann, Ruma Banerjee, and Ursula Jakob. Time line of redox events in aging postmitotic cells. *eLife*, 2, 2013.

[50] Katherine Noelani Chang, Shan Zhong, Matthew T Weirauch, Gary Hon, Mattia Pelizzola, Hai Li, Shao-shan Carol Huang, Robert J Schmitz, Mark A Urich, Dwight Kuo, Joseph R Nery, Hong Qiao, Ally Yang, Abdullah Jamali, Huaming Chen, Trey Ideker, Bing Ren, Ziv Bar-Joseph, Timothy R Hughes, and Joseph R Ecker. Temporal transcriptional response to ethylene gas drives growth hormone cross-regulation in Arabidopsis. *eLife*, 2, 2013.

[51] Alex Chattwood, Koki Nagayama, Parvin Bolourani, Lauren Harkin, Marzieh Kamjoo, Gerald Weeks, and Christopher RL Thompson. Developmental lineage priming in Dictyostelium by heterogeneous Ras activation. *eLife*, 2, 2013.

[52] Dipayan Chaudhuri, Yasemin Sancak, Vamsi K Mootha, and David E Clapham. MCU encodes the pore conducting mitochondrial calcium currents. *eLife*, 2, 2013.

[53] Kai Chen, Jeff Johnston, Wanqing Shao, Samuel Meier, Cynthia Staber, and Julia Zeitlinger. A global change in RNA polymerase II pausing during the Drosophila midblastula transition. *eLife*, 2, 2013.

[54] Poshen B Chen, Jui-Hung Hung, Taylor L Hickman, Andrew H Coles, James F Carey, Zhiping Weng, Feixia Chu, and Thomas G Fazzio. Hdac6 regulates Tip60-p400 function in stem cells. *eLife*, 2, 2013.

[55] Xiao-Wei Chen, He Wang, Kanika Bajaj, Pengcheng Zhang, Zhuo-Xian Meng, Danjun Ma, Yongsheng Bai, Hui-Hui Liu, Elizabeth Adams, Andrea Baines, Genggeng Yu, Maureen A Sartor, Bin Zhang, Zhengping Yi, Jiandie Lin, Stephen G Young, Randy Schekman, and David Ginsburg. SEC24A deficiency lowers plasma cholesterol through reduced PCSK9 secretion. *eLife*, 2, 2013.

[56] Jingheng Cheng and Daoyun Ji. Rigid firing sequences undermine spatial memory codes in a neurodegenerative mouse model. *eLife*, 2, 2013.

[57] Smita Cherry, Eugene Jennifer Jin, Mehmet Neset özel, Zhiyuan Lu, Egemen Agi, Dong Wang, Wei-Hung Jung, Daniel Epstein, Ian A Meinertzhagen, Chih-Chiang Chan, and P Robin Hiesinger. Charcot-Marie-Tooth 2B mutations in rab7 cause dosage-dependent neurodegeneration due to partial loss of function. *eLife*, 2, 2013.

[58] Hong Y Choi, Yun Liu, Christian Tennert, Yoshie Sugiura, Andromachi Karakatsani, Stephan Kröger, Eric B Johnson, Robert E Hammer, Weichun Lin, and Joachim Herz. APP interacts with LRP4 and agrin to coordinate the development of the neuromuscular junction in mice. *eLife*, 2, 2013.

[59] Thomas S Churcher, Teun Bousema, Martin Walker, Chris Drakeley, Petra Schneider, André Lin Ouédraogo, and María-Gloria Basáñez. Predicting mosquito infection from Plasmodium falciparum gametocyte density and estimating the reservoir of infection. *eLife*, 2, 2013.

[60] Matthias Corrotte, Patricia E Almeida, Christina Tam, Thiago Castro-Gomes, Maria Cecilia Fernandes, Bryan A Millis, Mauro Cortez, Heather Miller, Wenxia Song, Timothy K Maugel, and Norma W Andrews. Caveolae internalization repairs wounded cells and muscle fibers. *eLife*, 2, 2013.

[61] Matthew A Cottee, Nadine Muschalik, Yao Liang Wong, Christopher M Johnson, Steven Johnson, Antonina Andreeva, Karen Oegema, Susan M Lea, Jordan W Raff, and Mark van Breugel. Crystal structures of the CPAP/STIL complex reveal its role in centriole assembly and human microcephaly. *eLife*, 2, 2013.

[62] Christoph D Dahl and Ikuma Adachi. Conceptual metaphorical mapping in chimpanzees (Pan troglodytes). *eLife*, 2, 2013.

[63] Stephen R Daley, Kristen M Coakley, Daniel Y Hu, Katrina L Randall, Craig N Jenne, Andre Limnander, Darienne R Myers, Noelle K Polakos, Anselm Enders, Carla Roots, Bhavani Balakishnan, Lisa A Miosge, Geoff Sjollema, Edward M Bertram, Matthew A Field, Yunli Shao, T Daniel Andrews, Belinda Whittle, S Whitney Barnes, John R Walker, Jason G Cyster, Christopher C Goodnow, and Jeroen P Roose. Rasgrp1 mutation increases naïve T-cell CD44 expression and drives mTOR-dependent accumulation of Helios+ T cells and autoantibodies. *eLife*, 2, 2013.

[64] Manchuta Dangkulwanich, Toyotaka Ishibashi, Shixin Liu, Maria L Kireeva, Lucyna Lubkowska, Mikhail Kashlev, and Carlos J Bustamante. Complete dissection of transcription elongation reveals slow translocation of RNA polymerase II in a linear ratchet mechanism. *eLife*, 2, 2013.

[65] Roberto De Michele, Cindy Ast, Dominique Loqué, Cheng-Hsun Ho, Susana LA Andrade, Viviane Lanquar, Guido Grossmann, Sören Gehne, Michael U Kumke, and Wolf B Frommer. Fluorescent sensors reporting the activity of ammonium transceptors in live cells. *eLife*, 2, 2013.

[66] Nikki Dellas, Suzanne T Thomas, Gerard Manning, and Joseph P Noel. Discovery of a metabolic alternative to the classical mevalonate pathway. *eLife*, 2, 2013.

[67] Wei Deng, Mark Mayford, and Fred H Gage. Selection of distinct populations of dentate granule cells in response to inputs as a mechanism for pattern separation in mice. *eLife*, 2, 2013.

[68] Sara C Di Rienzi, Itai Sharon, Kelly C Wrighton, Omry Koren, Laura A Hug, Brian C Thomas, Julia K Goodrich, Jordana T Bell, Timothy D Spector, Jillian F Banfield, and Ruth E Ley. The human gut and groundwater harbor non-photosynthetic bacteria belonging to a new candidate phylum sibling to Cyanobacteria. *eLife*, 2, 2013.

[69] Jiajie Diao, Jacqueline Burré, Sandro Vivona, Daniel J Cipriano, Manu Sharma, Minjoung Kyoung, Thomas C Südhof, and Axel T Brunger. Native $\alpha$-synuclein induces clustering of synaptic-vesicle mimics via binding to phospholipids and synaptobrevin-2/VAMP2. *eLife*, 2, 2013.

[70] Ana I Domingos, Aylesse Sordillo, Marcelo O Dietrich, Zhong-Wu Liu, Luis A Tellez, Jake Vaynshteyn, Jozelia G Ferreira, Mats I Ekstrand, Tamas L Horvath, Ivan E de Araujo, and Jeffrey M Friedman. Hypothalamic melanin concentrating hormone neurons communicate the nutrient value of sugar. *eLife*, 2, 2013.

[71] George Dragoi and Susumu Tonegawa. Development of schemas revealed by prior experience and NMDA receptor knock-out. *eLife*, 2, 2013.

[72] Adrien Ducret, Betty Fleuchot, Ptissam Bergam, and Tâm Mignot. Direct live imaging of cell–cell protein transfer by transient outer membrane fusion in Myxococcus xanthus. *eLife*, 2, 2013.

[73] Joshua G Dunn, Catherine K Foo, Nicolette G Belletier, Elizabeth R Gavis, and Jonathan S Weissman. Ribosome profiling reveals pervasive and regulated stop codon readthrough in Drosophila melanogaster. *eLife*, 2, 2013.

[74] Caitlin Engelhard, Sarah Sarsfield, Janna Merte, Qiang Wang, Peng Li, Hideyuki Beppu, Alex L Kolodkin, Henry M Sucov, and David D Ginty. MEGF8 is a modifier of BMP signaling in trigeminal sensory neurons. *eLife*, 2, 2013.

[75] Taesun Eom, Chaolin Zhang, Huidong Wang, Kenneth Lay, John Fak, Jeffrey L Noebels, and Robert B Darnell. NOVA-dependent regulation of cryptic NMD exons controls synaptic protein levels after seizure. *eLife*, 2, 2013.

[76] Ombretta Foresti, Annamaria Ruggiano, Hans K Hannibal-Bach, Christer S Ejsing, and Pedro Carvalho. Sterol homeostasis requires regulated degradation of squalene monooxygenase by the ubiquitin ligase Doa10/Teb4. *eLife*, 2, 2013.

[77] Luis M Franco, Kristine L Bucasas, Janet M Wells, Diane Niño, Xueqing Wang, Gladys E Zapata, Nancy Arden, Alexander Renwick, Peng Yu, John M Quarles, Molly S Bray, Robert B Couch, John W Belmont, and Chad A Shaw. Integrative genomic analysis of the human immune response to influenza vaccination. *eLife*, 2, 2013.

[78] Jason P Gallivan, D Adam McLean, Kenneth F Valyear, and Jody C Culham. Decoding the neural mechanisms of human tool use. *eLife*, 2, 2013.

[79] Michael A Gaviño, Danielle Wenemoser, Irving E Wang, and Peter W Reddien. Tissue absence initiates regeneration through Follistatin-mediated inhibition of Activin signaling. *eLife*, 2, 2013.

[80] Liang Ge, David Melville, Min Zhang, and Randy Schekman. The ER–Golgi intermediate compartment is a key membrane source for the LC3 lipidation step of autophagosome biogenesis. *eLife*, 2, 2013.

[81] Lizhi Ian Gong, Marc A Suchard, and Jesse D Bloom. Stability-mediated epistasis constrains the evolution of an influenza protein. *eLife*, 2, 2013.

[82] Dan FM Goodman, Victor Benichoux, and Romain Brette. Decoding neural responses to temporal cues for sound localization. *eLife*, 2, 2013.

[83] Mingyu Gu, Qiang Liu, Shigeki Watanabe, Lin Sun, Gunther Hollopeter, Barth D Grant, and Erik M Jorgensen. AP2 hemicomplexes contribute independently to synaptic vesicle endocytosis. *eLife*, 2, 2013.

[84] Yusong Guo, Giulia Zanetti, and Randy Schekman. A novel GTP-binding protein–adaptor protein complex responsible for export of Vangl2 from the trans Golgi network. *eLife*, 2, 2013.

[85] Maria Gutierrez-Arcelus, Tuuli Lappalainen, Stephen B Montgomery, Alfonso Buil, Halit Ongen, Alisa Yurovsky, Julien Bryois, Thomas Giger, Luciana Romano, Alexandra Planchon, Emilie Falconnet, Deborah Bielser, Maryline Gagnebin, Ismael Padioleau, Christelle Borel, Audrey Letourneau, Periklis Makrythanasis, Michel Guipponi, Corinne Gehrig, Stylianos E Antonarakis, and Emmanouil T Dermitzakis. Passive and active DNA methylation and the interplay with genetic variation in gene regulation. *eLife*, 2, 2013.

[86] Svasti Haricharan, Jie Dong, Sarah Hein, Jay P Reddy, Zhijun Du, Michael Toneff, Kimberly Holloway, Susan G Hilsenbeck, Shixia Huang, Rachel Atkinson, Wendy Woodward, Sonali Jindal, Virginia F Borges, Carolina Gutierrez, Hong Zhang, Pepper J Schedin, C Kent Osborne, David J Tweardy, and Yi Li. Mechanism and preclinical prevention of increased breast cancer risk caused by pregnancy. *eLife*, 2, 2013.

[87] Mark Hedglin, Senthil K Perumal, Zhenxin Hu, and Stephen Benkovic. Stepwise assembly of the human replicative polymerase holoenzyme. *eLife*, 2, 2013.

[88] Amy Heidersbach, Chris Saxby, Karen Carver-Moore, Yu Huang, Yen-Sin Ang, Pieter J de Jong, Kathryn N Ivey, and Deepak Srivastava. microRNA-1 regulates sarcomere formation and suppresses smooth muscle gene expression in the mammalian heart. *eLife*, 2, 2013.

[89] Stefan Heinrichs, Lillian F Conover, Carlos E Bueso-Ramos, Outi Kilpivaara, Kristen Stevenson, Donna Neuberg, Mignon L Loh, Wen-Shu Wu, Scott J Rodig, Guillermo Garcia-Manero, Hagop M Kantarjian, and A Thomas Look. MYBL2 is a sub-haploinsufficient tumor suppressor gene in myeloid malignancy. *eLife*, 2, 2013.

[90] Luke A Helgeson and Brad J Nolen. Mechanism of synergistic activation of Arp2/3 complex by cortactin and N-WASP. *eLife*, 2, 2013.

[91] Victor J Hernandez, Jian Weng, Peter Ly, Shanica Pompey, Hongyun Dong, Lopa Mishra, Margaret Schwarz, Richard GW Anderson, and Peter Michaely. Cavin-3 dictates the balance between ERK and Akt signaling. *eLife*, 2, 2013.

[92] Donald W Hilgemann, Michael Fine, Maurine E Linder, Benjamin C Jennings, and Mei-Jung Lin. Massive endocytosis triggered by surface membrane palmitoylation under mitochondrial control in BHK fibroblasts. *eLife*, 2, 2013.

[93] Samuel Andrew Hires, Lorenz Pammer, Karel Svoboda, and David Golomb. Tapered whiskers are required for active tactile sensation. *eLife*, 2, 2013.

[94] Polly Yingshan Hsu, Upendra K Devisetty, and Stacey L Harmer. Accurate timekeeping is controlled by a cycling activator in Arabidopsis. *eLife*, 2, 2013.

[95] Shaohui Hu, Jun Wan, Yijing Su, Qifeng Song, Yaxue Zeng, Ha Nam Nguyen, Jaehoon Shin, Eric Cox, Hee Sool Rho, Crystal Woodard, Shuli Xia, Shuang Liu, Huibin Lyu, Guo-Li Ming, Herschel Wade, Hongjun Song, Jiang Qian, and Heng Zhu. DNA methylation presents distinct binding sites for human transcription factors. *eLife*, 2, 2013.

[96] Zhitao Hu, Xia-Jing Tong, and Joshua M Kaplan. UNC-13L, UNC-13S, and Tomosyn form a protein code for fast and slow neurotransmitter release in Caenorhabditis elegans. *eLife*, 2, 2013.

[97] Zhong L Hua, Philip M Smallwood, and Jeremy Nathans. Frizzled3 controls axonal development in distinct populations of cranial and spinal motor neurons. *eLife*, 2, 2013.

[98] Cheng-Chiu Huang, Ken Sugino, Yasuyuki Shima, Caiying Guo, Suxia Bai, Brett D Mensh, Sacha B Nelson, and Adam W Hantman. Convergence of pontine and proprioceptive streams onto multimodal cerebellar granule cells. *eLife*, 2, 2013.

[99] Garth R Ilsley, Jasmin Fisher, Rolf Apweiler, Angela H DePace, and Nicholas M Luscombe. Cellular resolution models for even skipped regulation in the entire Drosophila embryo. *eLife*, 2, 2013.

[100] Jeffrey S Iwig, Yvonne Vercoulen, Rahul Das, Tiago Barros, Andre Limnander, Yan Che, Jeffrey G Pelton, David E Wemmer, Jeroen P Roose, and John Kuriyan. Structural analysis of autoinhibition in the Ras-specific exchange factor RasGRP1. *eLife*, 2, 2013.

[101] Lin Jiang, Cong Liu, David Leibly, Meytal Landau, Minglei Zhao, Michael P Hughes, and David S Eisenberg. Structure-based discovery of fiber-binding compounds that reduce the cytotoxicity of amyloid beta. *eLife*, 2, 2013.

[102] Yunyun Jin, Jinjin Xu, Meng-Xin Yin, Yi Lu, Lianxin Hu, Peixue Li, Peng Zhang, Zengqiang Yuan, Margaret S Ho, Hongbin Ji, Yun Zhao, and Lei Zhang. Brahma is essential for Drosophila intestinal stem cell proliferation and regulated by Hippo signaling. *eLife*, 2, 2013.

[103] Martin Jinek, Alexandra East, Aaron Cheng, Steven Lin, Enbo Ma, and Jennifer Doudna. RNA-programmed genome editing in human cells. *eLife*, 2, 2013.

[104] Bindu Joseph, Jason A Corwin, Baohua Li, Suzi Atwell, and Daniel J Kliebenstein. Cytoplasmic genetic variation and extensive cytonuclear interactions influence natural variation in the metabolome. *eLife*, 2, 2013.

[105] Hari B Kamadurai, Yu Qiu, Alan Deng, Joseph S Harrison, Chris MacDonald, Marcelo Actis, Patrick Rodrigues, Darcie J Miller, Judith Souphron, Steven M Lewis, Igor Kurinov, Naoaki Fujii, Michal Hammel, Robert Piper, Brian Kuhlman, and Brenda A Schulman. Mechanism of ubiquitin ligation and lysine prioritization by a HECT E3. *eLife*, 2, 2013.

[106] Jason Karch, Jennifer Q Kwong, Adam R Burr, Michelle A Sargent, John W Elrod, Pablo M Peixoto, Sonia Martinez-Caballero, Hanna Osinska, Emily H-Y Cheng, Jeffrey Robbins, Kathleen W Kinnally, and Jeffery D Molkentin. Bax and Bak function as the outer membrane component of the mitochondrial permeability pore in regulating necrotic cell death in mice. *eLife*, 2, 2013.

[107] Marc A Kerenyi, Zhen Shao, Yu-Jung Hsu, Guoji Guo, Sidinh Luc, Kassandra O'Brien, Yuko Fujiwara, Cong Peng, Minh Nguyen, and Stuart H Orkin. Histone demethylase Lsd1 represses hematopoietic stem and progenitor cell signatures during blood cell maturation. *eLife*, 2, 2013.

[108] Elizabeth D Kirby, Sandra E Muroy, Wayne G Sun, David Covarrubias, Megan J Leong, Laurel A Barchas, and Daniela Kaufer. Acute stress enhances adult rat hippocampal neurogenesis and activation of newborn neurons via secreted astrocytic FGF2. *eLife*, 2, 2013.

[109] John H Koschwanez, Kevin R Foster, and Andrew W Murray. Improved use of a public good selects for the evolution of undifferentiated multicellularity. *eLife*, 2, 2013.

[110] William S Kruesi, Leighton J Core, Colin T Waters, John T Lis, and Barbara J Meyer. Condensin controls recruitment of RNA polymerase II to achieve nematode X-chromosome dosage compensation. *eLife*, 2, 2013.

[111] Mikhail Kudryashev, Marco Stenta, Stefan Schmelz, Marlise Amstutz, Ulrich Wiesand, Daniel Castaño-Díez, Matteo T Degiacomi, Stefan Münnich, Christopher KE Bleck, Julia Kowal, Andreas Diepold, Dirk W Heinz, Matteo Dal Peraro, Guy R Cornelis, and Henning Stahlberg. In situ structural analysis of the Yersinia enterocolitica injectisome. *eLife*, 2, 2013.

[112] Bridget R Kulasekara, Cassandra Kamischke, Hemantha D Kulasekara, Matthias Christen, Paul A Wiggins, and Samuel I Miller. c-di-GMP heterogeneity is generated by the chemotaxis machinery to regulate flagellar motility. *eLife*, 2, 2013.

[113] Pooja Kumari, Patrick C Gilligan, Shimin Lim, Long Duc Tran, Sylke Winkler, Robin Philp, and Karuna Sampath. An essential role for maternal control of Nodal signaling. *eLife*, 2, 2013.

[114] Marion Langen, Marta Koch, Jiekun Yan, Natalie De Geest, Maria-Luise Erfurth, Barret D Pfeiffer, Dietmar Schmucker, Yves Moreau, and Bassem A Hassan. Mutual inhibition among postmitotic neurons regulates robustness of brain wiring in Drosophila. *eLife*, 2, 2013.

[115] Gregory Lavieu, Hong Zheng, and James E Rothman. Stapled Golgi cisternae remain in place as cargo passes through the stack. *eLife*, 2, 2013.

[116] Bettina L Lee, Joanne E Moon, Jeffrey H Shu, Lin Yuan, Zachary R Newman, Randy Schekman, and Gregory M Barton. UNC93B1 mediates differential trafficking of endosomal TLRs. *eLife*, 2, 2013.

[117] Ji-Eun Lee, Chaochen Wang, Shiliyang Xu, Young-Wook Cho, Lifeng Wang, Xuesong Feng, Anne Baldridge, Vittorio Sartorelli, Lenan Zhuang, Weiqun Peng, and Kai Ge. H3K4 mono- and dimethyltransferase MLL4 is required for enhancer activation during cell differentiation. *eLife*, 2, 2013.

[118] Jonghyeob Lee, Takuya Sugiyama, Yinghua Liu, Jing Wang, Xueying Gu, Ji Lei, James F Markmann, Satsuki Miyazaki, Jun-ichi Miyazaki, Gregory L Szot, Rita Bottino, and Seung K Kim. Expansion and conversion of human pancreatic ductal cells into insulin-secreting endocrine cells. *eLife*, 2, 2013.

[119] Pengpeng Li, Kevin M Collins, Michael R Koelle, and Kang Shen. LIN-12/Notch signaling instructs postsynaptic muscle arm development by regulating UNC-40/DCC and MADD-2 in Caenorhabditis elegans. *eLife*, 2, 2013.

[120] Jingtao Lilue, Urs Benedikt Müller, Tobias Steinfeldt, and Jonathan C Howard. Reciprocal virulence and resistance polymorphism in the relationship between Toxoplasma gondii and the house mouse. *eLife*, 2, 2013.

[121] Mei-Jung Lin, Michael Fine, Jui-Yun Lu, Sandra L Hofmann, Gary Frazier, and Donald W Hilgemann. Massive palmitoylation-dependent endocytosis during reoxygenation of anoxic cardiac muscle. *eLife*, 2, 2013.

[122] Siqi Liu, Jueqi Chen, Xin Cai, Jiaxi Wu, Xiang Chen, You-Tong Wu, Lijun Sun, and Zhijian J Chen. MAVS recruits multiple ubiquitin E3 ligases to activate antiviral signaling cascades. *eLife*, 2, 2013.

[123] Hannah K Long, David Sims, Andreas Heger, Neil P Blackledge, Claudia Kutter, Megan L Wright, Frank Grützner, Duncan T Odom, Roger Patient, Chris P Ponting, and Robert J Klose. Epigenetic conservation at gene regulatory elements revealed by non-methylated DNA profiling in seven vertebrates. *eLife*, 2, 2013.

[124] Rosa Lozano-Durán, Alberto P Macho, Freddy Boutrot, Cécile Segonzac, Imre E Somssich, and Cyril Zipfel. The transcriptional regulator BZR1 mediates trade-off between plant innate immunity and growth. *eLife*, 2, 2013.

[125] Ling Luo, Hironori Uehara, Xiaohui Zhang, Subrata K Das, Thomas Olsen, Derick Holt, Jacquelyn M Simonis, Kyle Jackman, Nirbhai Singh, Tadashi R Miya, Wei Huang, Faisal Ahmed, Ana Bastos-Carvalho, Yun Zheng Le, Christina Mamalis, Vince A Chiodo, William W Hauswirth, Judit Baffi, Pedro M Lacal, Angela Orecchia, Napoleone Ferrara, Guangping Gao, Kim Young-hee, Yingbin Fu, Leah Owen, Romulo Albuquerque, Wolfgang Baehr, Kirk Thomas, Dean Y Li, Kakarla V Chalam, Masabumi Shibuya, Salvatore Grisanti, David J Wilson, Jayakrishna Ambati, and Balamurali K Ambati. Photoreceptor avascular privilege is shielded by soluble VEGF receptor-1. *eLife*, 2, 2013.

[126] Tamar R Makin, Alona O Cramer, Jan Scholz, Avital Hahamy, David Henderson Slater, Irene Tracey, and Heidi Johansen-Berg. Deprivation-related and use-dependent plasticity go hand in hand. *eLife*, 2, 2013.

[127] Mattias Malaguti, Paul A Nistor, Guillaume Blin, Amy Pegg, Xinzhi Zhou, and Sally Lowell. Bone morphogenic protein signalling suppresses differentiation of pluripotent cells by maintaining expression of E-Cadherin. *eLife*, 2, 2013.

[128] Nadja Mannowetz, Natasha M Naidoo, Seung-A Sara Choo, James F Smith, and Polina V Lishko. Slo1 is the principal potassium channel of human spermatozoa. *eLife*, 2, 2013.

[129] Alexis Marianes and Allan C Spradling. Physiological and stem cell compartmentalization within the Drosophila midgut. *eLife*, 2, 2013.

[130] Alexandre Martinière, Aurélie Bak, Jean-Luc Macia, Nicole Lautredou, Daniel Gargani, Juliette Doumayrou, Elisa Garzo, Aranzazu Moreno, Alberto Fereres, Stéphane Blanc, and Martin Drucker. A virus responds instantly to the presence of the vector on the host and forms transmission morphs. *eLife*, 2, 2013.

[131] Meghan M McLean, Yong Chang, Gautam Dhar, John K Heiss, and Reid C Johnson. Multiple interfaces between a serine recombinase and an enhancer control site-specific DNA inversion. *eLife*, 2, 2013.

[132] Louise Meigh, Sophie A Greenhalgh, Thomas L Rodgers, Martin J Cann, David I Roper, and Nicholas Dale. CO2 directly modulates connexin 26 by formation of carbamate bridges between subunits. *eLife*, 2, 2013.

[133] César S Mendes, Imre Bartos, Turgay Akay, Szabolcs Márka, and Richard S Mann. Quantification of gait parameters in freely walking wild type and sensory deprived Drosophila melanogaster. *eLife*, 2, 2013.

[134] Jessica L Metcalf, Laura Wegener Parfrey, Antonio Gonzalez, Christian L Lauber, Dan Knights, Gail Ackermann, Gregory C Humphrey, Matthew J Gebert, Will Van Treuren, Donna Berg-Lyons, Kyle Keepers, Yan Guo, James Bullard, Noah Fierer, David O Carter, and Rob Knight. A microbial clock provides an accurate estimate of the postmortem interval in a mouse model system. *eLife*, 2, 2013.

[135] Yong Miao, Cathrine Miner, Lei Zhang, Phyllis I Hanson, Adish Dani, and Monika Vig. An essential and NSF independent role for $\alpha$-SNAP in store-operated calcium entry. *eLife*, 2, 2013.

[136] Deryck J Mills, Stella Vitt, Mike Strauss, Seigo Shima, and Janet Vonck. De novo modeling of the F420-reducing [NiFe]-hydrogenase from a methanogenic archaeon by cryo-electron microscopy. *eLife*, 2, 2013.

[137] Sandra Mitrovic, Cristina Nogueira, Gerard Cantero-Recasens, Kerstin Kiefer, José M Fernández-Fernández, Jean-FranÇois Popoff, Laetitia Casano, Frederic A Bard, Raul Gomez, Miguel A Valverde, and Vivek Malhotra. TRPM5-mediated calcium uptake regulates mucin secretion from human colon goblet cells. *eLife*, 2, 2013.

[138] Babak Momeni, Kristen A Brileya, Matthew W Fields, and Wenying Shou. Strong inter-population cooperation leads to partner intermixing in microbial communities. *eLife*, 2, 2013.

[139] Babak Momeni, Adam James Waite, and Wenying Shou. Spatial self-organization favors heterotypic cooperation over cheating. *eLife*, 2, 2013.

[140] Andrew Morin, Ben Eisenbraun, Jason Key, Paul C Sanschagrin, Michael A Timony, Michelle Ottaviano, and Piotr Sliz. Collaboration gets the most out of software. *eLife*, 2, 2013.

[141] Jonathan Mowers, Maeran Uhm, Shannon M Reilly, Joshua Simon, Dara Leto, Shian-Huey Chiang, Louise Chang, and Alan R Saltiel. Inflammation produces catecholamine resistance in obesity via activation of PDE3B by the protein kinases IKK$\epsilon$ and TBK1. *eLife*, 2, 2013.

[142] Andrew Murley, Laura L Lackner, Christof Osman, Matthew West, Gia K Voeltz, Peter Walter, and Jodi Nunnari. ER-associated mitochondrial division links the distribution of mitochondria and mitochondrial DNA in yeast. *eLife*, 2, 2013.

[143] Sigrid Nachtergaele, Daniel M Whalen, Laurel K Mydock, Zhonghua Zhao, Tomas Malinauskas, Kathiresan Krishnan, Philip W Ingham, Douglas F Covey, Christian Siebold, and Rajat Rohatgi. Structure and function of the Smoothened extracellular domain in vertebrate Hedgehog signaling. *eLife*, 2, 2013.

[144] Stanislav Nagy, Charles Wright, Nora Tramm, Nicholas Labello, Stanislav Burov, and David Biron. A longitudinal study of Caenorhabditis elegans larvae reveals a novel locomotion switch, regulated by G$\alpha$s signaling. *eLife*, 2, 2013.

[145] Jinsuke Nishino, Sunjung Kim, Yuan Zhu, Hao Zhu, and Sean J Morrison. A network of heterochronic genes including Imp1 regulates temporal changes in stem cell properties. *eLife*, 2, 2013.

[146] Mari Nishino, Jonathan W Choy, Nathan N Gushwa, Juan A Oses-Prieto, Kyriacos Koupparis, Alma L Burlingame, Adam R Renslo, James H McKerrow, and Jack Taunton. Hypothemycin, a fungal natural product, identifies therapeutic targets in Trypanosoma brucei. *eLife*, 2, 2013.

[147] Robert Noble, Zóe Christodoulou, Sue Kyes, Robert Pinches, Chris I Newbold, and Mario Recker. The antigenic switching network of Plasmodium falciparum and its implications for the immunoepidemiology of malaria. *eLife*, 2, 2013.

[148] Katarzyna Nowak, Gerhard Seisenbacher, Ernst Hafen, and Hugo Stocker. Nutrient restriction enhances the proliferative potential of cells lacking the tumor suppressor PTEN in mitotic tissues. *eLife*, 2, 2013.

[149] Geoff P O'Donoghue, Rafal M Pielak, Alexander A Smoligovets, Jenny J Lin, and Jay T Groves. Direct single molecule measurement of TCR triggering by agonist pMHC in living primary T cells. *eLife*, 2, 2013.

[150] Virginie Olive, Erich Sabio, Margaux J Bennett, Caitlin S De Jong, Anne Biton, James C McGann, Samantha K Greaney, Nicole M Sodir, Alicia Y Zhou, Asha Balakrishnan, Mona Foth, Micah A Luftig, Andrei Goga, Terence P Speed, Zhenyu Xuan, Gerard I Evan, Ying Wan, Alex C Minella, and Lin He. A component of the mir-17-92 polycistronic oncomir promotes oncogene-dependent apoptosis. *eLife*, 2, 2013.

[151] Calla M Olson, Marissa R Donovan, Michael J Spellberg, and Michael T Marr. The insulin receptor cellular IRES confers resistance to eIF4A inhibition. *eLife*, 2, 2013.

[152] Dana E Orange, Nathalie E Blachere, John Fak, Salina Parveen, Mayu O Frank, Margo Herre, Suyan Tian, Sebastien Monette, and Robert B Darnell. Dendritic cells loaded with FK506 kill T cells in an antigen-specific manner and prevent autoimmunity in vivo. *eLife*, 2, 2013.

[153] Jesper Pallesen, Yaser Hashem, Gürkan Korkmaz, Ravi Kiran Koripella, Chenhui Huang, Måns Ehrenberg, Suparna Sanyal, and Joachim Frank. Cryo-EM visualization of the ribosome in termination complex with apo-RF3 and RF1. *eLife*, 2, 2013.

[154] Natalya N Pavlova, Christian Pallasch, Andrew EH Elia, Christian J Braun, Thomas F Westbrook, Michael Hemann, and Stephen J Elledge. A role for PVRL4-driven cell–cell interactions in tumorigenesis. *eLife*, 2, 2013.

[155] Verena Pawlak, David S Greenberg, Henning Sprekeler, Wulfram Gerstner, and Jason ND Kerr. Changing the responses of cortical neurons from sub- to suprathreshold using single spikes in vivo. *eLife*, 2, 2013.

[156] Hugh Pelham. Building for the future. *eLife*, 2, 2013.

[157] Patrina A Pellett, Felix Dietrich, Jörg Bewersdorf, James E Rothman, and Grégory Lavieu. Inter-Golgi transport mediated by COPI-containing vesicles carrying small cargoes. *eLife*, 2, 2013.

[158] Rajika L Perera, Rubben Torella, Sebastian Klinge, Mairi L Kilkenny, Joseph D Maman, and Luca Pellegrini. Mechanism for priming DNA synthesis by yeast DNA Polymerase $\alpha$. *eLife*, 2, 2013.

[159] Ainhoa Pérez-Garijo, Yaron Fuchs, and Hermann Steller. Apoptotic cells can induce non-autonomous apoptosis through the TNF pathway. *eLife*, 2, 2013.

[160] Stephan A Pless, Jason D Galpin, Ana P Niciforovic, Harley T Kurata, and Christopher A Ahern. Hydrogen bonds as molecular timers for slow inactivation in voltage-gated potassium channels. *eLife*, 2, 2013.

[161] Ivana Primorac, John R Weir, Elena Chiroli, Fridolin Gross, Ingrid Hoffmann, Suzan van Gerwen, Andrea Ciliberto, and Andrea Musacchio. Bub3 reads phosphorylated MELT repeats to promote spindle assembly checkpoint signaling. *eLife*, 2, 2013.

[162] Zhi Qi, Robert A Pugh, Maria Spies, and Yann R Chemla. Sequence-dependent base pair stepping dynamics in XPD helicase unwinding. *eLife*, 2, 2013.

[163] Nicole A Rapicavoli, Kun Qu, Jiajing Zhang, Megan Mikhail, Remi-Martin Laberge, and Howard Y Chang. A mammalian pseudogene lncRNA at the interface of inflammation and anti-inflammatory therapeutics. *eLife*, 2, 2013.

[164] Ahmed Rattani, Magda Wolna, Mickael Ploquin, Wolfgang Helmhart, Seamus Morrone, Bernd Mayer, Jonathan Godwin, Wenqing Xu, Olaf Stemmann, Alberto Pendas, and Kim Nasmyth. Sgol2 provides a regulatory platform that coordinates essential cell cycle processes during meiosis I in oocytes. *eLife*, 2, 2013.

[165] Ricardo Antonio Rosselló, Chun-Chun Chen, Rui Dai, Jason T Howard, Ute Hochgeschwender, and Erich D Jarvis. Mammalian genes induce partially reprogrammed pluripotent stem cells in non-mammalian vertebrate and invertebrate species. *eLife*, 2, 2013.

[166] Tal Rousso, Annette M Shewan, Keith E Mostov, Eyal D Schejter, and Ben-Zion Shilo. Apical targeting of the formin Diaphanous in Drosophila tubular epithelia. *eLife*, 2, 2013.

[167] Yashar Sadian, Christos Gatsogiannis, Csilla Patasi, Oliver Hofnagel, Roger S Goody, Marian FarkaŠovský, and Stefan Raunser. The role of Cdc42 and Gic1 in the regulation of septin filament formation and dissociation. *eLife*, 2, 2013.

[168] Miriam Samstein, Heidi A Schreiber, Ingrid M Leiner, Bože SuŠac, Michael S Glickman, and Eric G Pamer. Essential yet limited role for CCR2+ inflammatory monocytes during Mycobacterium tuberculosis-specific T cell priming. *eLife*, 2, 2013.

[169] Andrea Sánchez-Vallet, Raspudin Saleem-Batcha, Anja Kombrink, Guido Hansen, Dirk-Jan Valkenburg, Bart PHJ Thomma, and Jeroen R Mesters. Fungal effector Ecp6 outcompetes host immune receptor for chitin binding through intrachain LysM dimerization. *eLife*, 2, 2013.

[170] Martin Sauvageau, Loyal A Goff, Simona Lodato, Boyan Bonev, Abigail F Groff, Chiara Gerhardinger, Diana B Sanchez-Gomez, Ezgi Hacisuleyman, Eric Li, Matthew Spence, Stephen C Liapis, William Mallard, Michael Morse, Mavis R Swerdel, Michael F D'Ecclessis, Jennifer C Moore, Venus Lai, Guochun Gong, George D Yancopoulos, David Frendewey, Manolis Kellis, Ronald P Hart, David M Valenzuela, Paola Arlotta, and John L Rinn. Multiple knockout mouse models reveal lincRNAs are required for life and brain development. *eLife*, 2, 2013.

[171] Ankur Saxena, Brian N Peng, and Marianne E Bronner. Sox10-dependent neural crest origin of olfactory microvillous neurons in zebrafish. *eLife*, 2, 2013.

[172] Jose U Scher, Andrew Sczesnak, Randy S Longman, Nicola Segata, Carles Ubeda, Craig Bielski, Tim Rostron, Vincenzo Cerundolo, Eric G Pamer, Steven B Abramson, Curtis Huttenhower, and Dan R Littman. Expansion of intestinal Prevotella copri correlates with enhanced susceptibility to arthritis. *eLife*, 2, 2013.

[173] Joshua T Schiffer, David Swan, Ramzi Al Sallaq, Amalia Magaret, Christine Johnston, Karen E Mark, Stacy Selke, Negusse Ocbamichael, Steve Kuntz, Jia Zhu, Barry Robinson, Meei-Li Huang, Keith R Jerome, Anna Wald, and Lawrence Corey. Rapid localized spread and immunologic containment define Herpes simplex virus-2 reactivation in the human genital tract. *eLife*, 2, 2013.

[174] Ursula Schulze-Gahmen, Heather Upton, Andrew Birnberg, Katherine Bao, Seemay Chou, Nevan J Krogan, Qiang Zhou, and Tom Alber. The AFF4 scaffold binds human P-TEFb adjacent to HIV Tat. *eLife*, 2, 2013.

[175] Gregory W Schwartz and Fred Rieke. Controlling gain one photon at a time. *eLife*, 2, 2013.

[176] Antonio Scialdone, Sam T Mugford, Doreen Feike, Alastair Skeffington, Philippa Borrill, Alexander Graf, Alison M Smith, and Martin Howard. Arabidopsis plants perform arithmetic division to prevent starvation at night. *eLife*, 2, 2013.

[177] Arnau Sebé-Pedrós, Manuel Irimia, Javier del Campo, Helena Parra-Acero, Carsten Russ, Chad Nusbaum, Benjamin J Blencowe, and Iñaki Ruiz-Trillo. Regulated aggregative multicellularity in a close unicellular relative of metazoa. *eLife*, 2, 2013.

[178] Sarah H Shahmoradian, Jesus G Galaz-Montoya, Michael F Schmid, Yao Cong, Boxue Ma, Christoph Spiess, Judith Frydman, Steven J Ludtke, and Wah Chiu. TRiC's tricks inhibit huntingtin aggregation. *eLife*, 2, 2013.

[179] Chandan Shee, Ben D Cox, Franklin Gu, Elizabeth M Luengas, Mohan C Joshi, Li-Ya Chiu, David Magnan, Jennifer A Halliday, Ryan L Frisch, Janet L Gibson, Ralf Bernd Nehring, Huong G Do, Marcos Hernandez, Lei Li, Christophe Herman, PJ Hastings, David Bates, Reuben S Harris, Kyle M Miller, and Susan M Rosenberg. Engineered proteins detect spontaneous DNA breakage in human and bacterial cells. *eLife*, 2, 2013.

[180] Yair Shemesh, Yehezkel Sztainberg, Oren Forkosh, Tamar Shlapobersky, Alon Chen, and Elad Schneidman. High-order social interactions in groups of mice. *eLife*, 2, 2013.

[181] Dan Shi, Brent L Nannenga, Matthew G Iadanza, and Tamir Gonen. Three-dimensional electron crystallography of protein microcrystals. *eLife*, 2, 2013.

[182] Sheng Min Shih, Benjamin D Engel, Fatih Kocabas, Thomas Bilyard, Arne Gennerich, Wallace F Marshall, and Ahmet Yildiz. Intraflagellar transport drives flagellar surface motility. *eLife*, 2, 2013.

[183] Kazuhiro Shimomura, Vivek Kumar, Nobuya Koike, Tae-Kyung Kim, Jason Chong, Ethan D Buhr, Andrew R Whiteley, Sharon S Low, Chiaki Omura, Deborah Fenner, Joseph R Owens, Marc Richards, Seung-Hee Yoo, Hee-Kyung Hong, Martha H Vitaterna, Joseph Bass, Mathew T Pletcher, Tim Wiltshire, John Hogenesch, Phillip L Lowrey, and Joseph S Takahashi. Usf1, a suppressor of the circadian Clock mutant, reveals the nature of the DNA-binding of the CLOCK:BMAL1 complex in mice. *eLife*, 2, 2013.

[184] Carmela Sidrauski, Diego Acosta-Alvear, Arkady Khoutorsky, Punitha Vedantham, Brian R Hearn, Han Li, Karine Gamache, Ciara M Gallagher, Kenny K-H Ang, Chris Wilson, Voytek Okreglak, Avi Ashkenazi, Byron Hann, Karim Nader, Michelle R Arkin, Adam R Renslo, Nahum Sonenberg, and Peter Walter. Pharmacological brake-release of mRNA translation enhances cognitive memory. *eLife*, 2, 2013.

[185] Benjamin A Smith, Shae B Padrick, Lynda K Doolittle, Karen Daugherty-Clarke, Ivan R Corrêa, Ming-Qun Xu, Bruce L Goode, Michael K Rosen, and Jeff Gelles. Three-color single molecule imaging shows WASP detachment from Arp2/3 complex triggers actin filament branch formation. *eLife*, 2, 2013.

[186] F Donelson Smith, Steve L Reichow, Jessica L Esseltine, Dan Shi, Lorene K Langeberg, John D Scott, and Tamir Gonen. Intrinsic disorder within an AKAP-protein kinase A complex guides local substrate phosphorylation. *eLife*, 2, 2013.

[187] Bo-mi Song, Serge Faumont, Shawn Lockery, and Leon Avery. Recognition of familiar food activates feeding via an endocrine serotonin signal in Caenorhabditis elegans. *eLife*, 2, 2013.

[188] Se Jin Song, Christian Lauber, Elizabeth K Costello, Catherine A Lozupone, Gregory Humphrey, Donna Berg-Lyons, J Gregory Caporaso, Dan Knights, Jose C Clemente, Sara Nakielny, Jeffrey I Gordon, Noah Fierer, and Rob Knight. Cohabiting family members share microbiota with one another and with their dogs. *eLife*, 2, 2013.

[189] Ashley L St John, Abhay PS Rathore, Bhuvanakantham Raghavan, Mah-Lee Ng, and Soman N Abraham. Contributions of mast cells and vasoactive products, leukotrienes and chymase, to dengue virus-induced vascular leakage. *eLife*, 2, 2013.

[190] Randy B Stockbridge, Janice L Robertson, Ludmila Kolmakova-Partensky, and Christopher Miller. A family of fluoride-specific ion channels with dual-topology architecture. *eLife*, 2, 2013.

[191] Anja S Strauss, Sven Peters, Wilhelm Boland, and Antje Burse. ABC transporter functions as a pacemaker for sequestration of plant glucosides in leaf beetles. *eLife*, 2, 2013.

[192] Hume Stroud, Bo Ding, Stacey A Simon, Suhua Feng, Maria Bellizzi, Matteo Pellegrini, Guo-Liang Wang, Blake C Meyers, and Steven E Jacobsen. Plants regenerated from tissue culture contain stable epigenome changes in rice. *eLife*, 2, 2013.

[193] Arvind R Subramaniam, Aaron DeLoughery, Niels Bradshaw, Yun Chen, Erin O'Shea, Richard Losick, and Yunrong Chai. A serine sensor for multicellularity in a bacterium. *eLife*, 2, 2013.

[194] Hani Suleiman, Lei Zhang, Robyn Roth, John E Heuser, Jeffrey H Miner, Andrey S Shaw, and Adish Dani. Nanoscale protein architecture of the kidney glomerular basement membrane. *eLife*, 2, 2013.

[195] Jianjun Sun and Allan C Spradling. Ovulation in Drosophila is controlled by secretory cells of the female reproductive tract. *eLife*, 2, 2013.

[196] Liou Y Sun, Adam Spong, William R Swindell, Yimin Fang, Cristal Hill, Joshua A Huber, Jacob D Boehm, Reyhan Westbrook, Roberto Salvatori, and Andrzej Bartke. Growth hormone-releasing hormone disruption extends lifespan and regulates response to caloric restriction in mice. *eLife*, 2, 2013.

[197] Yaping Sun, Zhiqiang Dong, Taihao Jin, Kean-Hooi Ang, Miller Huang, Kelly M Haston, Jisong Peng, Tao P Zhong, Steven Finkbeiner, William A Weiss, Michelle R Arkin, Lily Y Jan, and Su Guo. Imaging-based chemical screening reveals activity-dependent neural differentiation of pluripotent stem cells. *eLife*, 2, 2013.

[198] Susumu Takahashi. Hierarchical organization of context in the hippocampal episodic code. *eLife*, 2, 2013.

[199] Marvin E Tanenbaum, Ronald D Vale, and Richard J McKenney. Cytoplasmic dynein crosslinks and slides anti-parallel microtubules using its two motor domains. *eLife*, 2, 2013.

[200] Li Tao, Qi Xie, Yue-He Ding, Shang-Tong Li, Shengyi Peng, Yan-Ping Zhang, Dan Tan, Zengqiang Yuan, and Meng-Qiu Dong. CAMKII and Calcineurin regulate the lifespan of Caenorhabditis elegans through the FOXO transcription factor DAF-16. *eLife*, 2, 2013.

[201] Benjamin JM Taylor, Serena Nik-Zainal, Yee Ling Wu, Lucy A Stebbings, Keiran Raine, Peter J Campbell, Cristina Rada, Michael R Stratton, and Michael S Neuberger. DNA deaminases induce break-associated mutation showers with implication of APOBEC3B and 3A in breast cancer kataegis. *eLife*, 2, 2013.

[202] Vitor H Teixeira, Parthiban Nadarajan, Trevor A Graham, Christodoulos P Pipinikas, James M Brown, Mary Falzon, Emma Nye, Richard Poulsom, David Lawrence, Nicholas A Wright, Stuart McDonald, Adam Giangreco, Benjamin D Simons, and Sam M Janes. Stochastic homeostasis in human airway epithelium is achieved by neutral competition of basal cell progenitors. *eLife*, 2, 2013.

[203] Sundeep Teki, Maria Chait, Sukhbinder Kumar, Shihab Shamma, and Timothy D Griffiths. Segregation of complex acoustic scenes based on temporal coherence. *eLife*, 2, 2013.

[204] Dawn A Thompson, Sushmita Roy, Michelle Chan, Mark P Styczynsky, Jenna Pfiffner, Courtney French, Amanda Socha, Anne Thielke, Sara Napolitano, Paul Muller, Manolis Kellis, Jay H Konieczka, Ilan Wapinski, and Aviv Regev. Evolutionary principles of modular gene regulation in yeasts. *eLife*, 2, 2013.

[205] Sharon E Torigoe, Ashok Patel, Mai T Khuong, Gregory D Bowman, and James T Kadonaga. ATP-dependent chromatin assembly is functionally distinct from chromatin remodeling. *eLife*, 2, 2013.

[206] Budd A Tucker, Robert F Mullins, Luan M Streb, Kristin Anfinson, Mari E Eyestone, Emily Kaalberg, Megan J Riker, Arlene V Drack, Terry A Braun, and Edwin M Stone. Patient-specific iPSC-derived photoreceptor precursor cells as a means to investigate retinitis pigmentosa. *eLife*, 2, 2013.

[207] Seralynne D Vann. Dismantling the Papez circuit for memory in rats. *eLife*, 2, 2013.

[208] Rati Verma, Robert S Oania, Natalie J Kolawa, and Raymond J Deshaies. Cdc48/p97 promotes degradation of aberrant nascent polypeptides bound to the ribosome. *eLife*, 2, 2013.

[209] S Nahuel Villegas, Michaela Rothová, Martin E Barrios-Llerena, Maria Pulina, Anna-Katerina Hadjantonakis, Thierry Le Bihan, Sophie Astrof, and Joshua M Brickman. PI3K/Akt1 signalling specifies foregut precursors by generating regionalized extra-cellular matrix. *eLife*, 2, 2013.

[210] Harina Vin, Sandra S Ojeda, Grace Ching, Marco L Leung, Vida Chitsazzadeh, David W Dwyer, Charles H Adelmann, Monica Restrepo, Kristen N Richards, Larissa R Stewart, Lili Du, Scarlett B Ferguson, Deepavali Chakravarti, Karin Ehrenreiter, Manuela Baccarini, Rosamaria Ruggieri, Jonathan L Curry, Kevin B Kim, Ana M Ciurea, Madeleine Duvic, Victor G Prieto, Stephen E Ullrich, Kevin N Dalby, Elsa R Flores, and Kenneth Y Tsai. BRAF inhibitors suppress apoptosis through off-target inhibition of JNK signaling. *eLife*, 2, 2013.

[211] Sven K Vogel, Zdenek Petrasek, Fabian Heinemann, and Petra Schwille. Myosin motors fragment and compact membrane-bound actin filaments. *eLife*, 2, 2013.

[212] Ayelet Voskoboynik, Norma F Neff, Debashis Sahoo, Aaron M Newman, Dmitry Pushkarev, Winston Koh, Benedetto Passarelli, H Christina Fan, Gary L Mantalas, Karla J Palmeri, Katherine J Ishizuka, Carmela Gissi, Francesca Griggio, Rachel Ben-Shlomo, Daniel M Corey, Lolita Penland, Richard A White, Irving L Weissman, and Stephen R Quake. The genome sequence of the colonial chordate, Botryllus schlosseri. *eLife*, 2, 2013.

[213] Lamia Wahba, Steven K Gore, and Douglas Koshland. The homologous recombination machinery modulates the formation of RNA–DNA hybrids and associated chromosome instability. *eLife*, 2, 2013.

[214] Bo Wang, James J Collins, and Phillip A Newmark. Functional genomic characterization of neoblast-like stem cells in larval Schistosoma mansoni. *eLife*, 2, 2013.

[215] Yuxiao Wang, Heath G Pascoe, Chad A Brautigam, Huawei He, and Xuewu Zhang. Structural basis for activation and non-canonical catalysis of the Rap GTPase activating protein domain of plexin. *eLife*, 2, 2013.

[216] Shigeki Watanabe, Qiang Liu, M Wayne Davis, Gunther Hollopeter, Nikita Thomas, Nels B Jorgensen, and Erik M Jorgensen. Ultrafast endocytosis at Caenorhabditis elegans neuromuscular junctions. *eLife*, 2, 2013.

[217] Hua Wen, Jeffrey M Hubbard, Benjamin Rakela, Michael W Linhoff, Gail Mandel, and Paul Brehm. Synchronous and asynchronous modes of synaptic transmission utilize different calcium sources. *eLife*, 2, 2013.

[218] Ruifen Weng, Jacqueline SR Chin, Joanne Y Yew, Natascha Bushati, and Stephen M Cohen. miR-124 controls male reproductive success in Drosophila. *eLife*, 2, 2013.

[219] Tobias Wiestler and Jörn Diedrichsen. Skill learning strengthens cortical representations of motor sequences. *eLife*, 2, 2013.

[220] Arun P Wiita, Etay Ziv, Paul J Wiita, Anatoly Urisman, Olivier Julien, Alma L Burlingame, Jonathan S Weissman, and James A Wells. Global cellular response to chemotherapy-induced apoptosis. *eLife*, 2, 2013.

[221] Erik Wijnker, Geo Velikkakam James, Jia Ding, Frank Becker, Jonas R Klasen, Vimal Rawat, Beth A Rowan, Daniël F de Jong, C Bastiaan de Snoo, Luis Zapata, Bruno Huettel, Hans de Jong, Stephan Ossowski, Detlef Weigel, Maarten Koornneef, Joost JB Keurentjes, and Korbinian Schneeberger. The genomic landscape of meiotic crossovers and gene conversions in Arabidopsis thaliana. *eLife*, 2, 2013.

[222] Jeremy D Wilbur and Rebecca Heald. Mitotic spindle scaling during Xenopus development by kif2a and importin $\alpha$. *eLife*, 2, 2013.

[223] Ben P Williams, Iain G Johnston, Sarah Covshoff, and Julian M Hibberd. Phenotypic landscape inference reveals multiple evolutionary paths to C4 photosynthesis. *eLife*, 2, 2013.

[224] Shawniqua T Williams, Mary M Conte, Andrew M Goldfine, Quentin Noirhomme, Olivia Gosseries, Marie Thonnard, Bradley Beattie, Jennifer Hersh, Douglas I Katz, Jonathan D Victor, Steven Laureys, and Nicholas D Schiff. Common resting brain dynamics indicate a possible mechanism underlying zolpidem response in severe brain injury. *eLife*, 2, 2013.

[225] Xiu Ming Wong, Susan Younger, Christian J Peters, Yuh Nung Jan, and Lily Y Jan. Subdued, a TMEM16 family Ca2+−activated Cl− channel in Drosophila melanogaster with an unexpected role in host defense. *eLife*, 2, 2013.

[226] Jing Yan, Dan L Chao, Shiori Toba, Kotaro Koyasako, Takuo Yasunaga, Shinji Hirotsune, and Kang Shen. Kinesin-1 regulates dendrite microtubule polarity in Caenorhabditis elegans. *eLife*, 2, 2013.

[227] Li Yang, Mingli Xu, Yeonjong Koo, Jia He, and R Scott Poethig. Sugar promotes vegetative phase change in Arabidopsis thaliana by repressing the expression of MIR156A and MIR156C. *eLife*, 2, 2013.

[228] Yan Yang and Stephen G Lisberger. Interaction of plasticity and circuit organization during the acquisition of cerebellum-dependent motor learning. *eLife*, 2, 2013.

[229] Fan Ye, David K Breslow, Elena F Koslover, Andrew J Spakowitz, W James Nelson, and Maxence V Nachury. Single molecule imaging reveals a major role for diffusion in the exploration of ciliary space by signaling receptors. *eLife*, 2, 2013.

[230] Tangsheng Yi and Jason G Cyster. EBI2-mediated bridging channel positioning supports splenic dendritic cell homeostasis and particulate antigen capture. *eLife*, 2, 2013.

[231] Avihu H Yona, Zohar Bloom-Ackermann, Idan Frumkin, Victor Hanson-Smith, Yoav Charpak-Amikam, Qinghua Feng, Jef D Boeke, Orna Dahan, and Yitzhak Pilpel. tRNA genes rapidly change in evolution to meet novel translational demands. *eLife*, 2, 2013.

[232] Ihab Younis, Kimberly Dittmar, Wei Wang, Shawn W Foley, Michael G Berg, Karen Y Hu, Zhi Wei, Lili Wan, and Gideon Dreyfuss. Minor introns are embedded molecular switches regulated by highly unstable U6atac snRNA. *eLife*, 2, 2013.

[233] Sha Yu, Li Cao, Chuan-Miao Zhou, Tian-Qi Zhang, Heng Lian, Yue Sun, Jianqiang Wu, Jirong Huang, Guodong Wang, and Jia-Wei Wang. Sugar is an endogenous cue for juvenile-to-adult phase transition in plants. *eLife*, 2, 2013.

[234] Tian Yu, Linda C Meiners, Katrin Danielsen, Monica TY Wong, Timothy Bowler, Danny Reinberg, Peter J Scambler, Conny MA van Ravenswaaij-Arts, and M Albert Basson. Deregulated FGF and homeotic gene expression underlies cerebellar vermis hypoplasia in CHARGE syndrome. *eLife*, 2, 2013.

[235] Wei-Ming Yu, Jessica M Appler, Ye-Hyun Kim, Allison M Nishitani, Jeffrey R Holt, and Lisa V Goodrich. A Gata3–Mafb transcriptional network directs post-synaptic differentiation in synapses specialized for hearing. *eLife*, 2, 2013.

[236] Giulia Zanetti, Simone Prinz, Sebastian Daum, Annette Meister, Randy Schekman, Kirsten Bacia, and John AG Briggs. The structure of the COPII transport-vesicle coat assembled on membranes. *eLife*, 2, 2013.

[237] Xing Zhang, Huatao Guo, Lei Jin, Elizabeth Czornyj, Asher Hodes, Wong H Hui, Angela W Nieh, Jeff F Miller, and Z Hong Zhou. A new topology of the HK97-like fold revealed in Bordetella bacteriophage by cryoEM at 3.5 Å resolution. *eLife*, 2, 2013.

[238] Jimmy L Zhao, Dinesh S Rao, Ryan M O'Connell, Yvette Garcia-Flores, and David Baltimore. MicroRNA-146a acts as a guardian of the quality and longevity of hematopoietic stem cells in mice. *eLife*, 2, 2013.

[239] Yingying Zhao, Jason A MacGurn, Max Liu, and Scott Emr. The ART-Rsp5 ubiquitin ligase network comprises a plasma membrane quality control system that protects yeast cells from proteotoxic stress. *eLife*, 2, 2013.

[240] Haiying Zhou, Tommy Kaplan, Yan Li, Ivan Grubisic, Zhengjian Zhang, P Jeremy Wang, Michael B Eisen, and Robert Tjian. Dual functions of TAF7L in adipocyte differentiation. *eLife*, 2, 2013.

[241] Keming Zhou, Tamara M Stawicki, Alexandr Goncharov, and Yishi Jin. Position of UNC-13 in the active zone regulates synaptic vesicle release probability and release kinetics. *eLife*, 2, 2013.

[242] Huanhu Zhu, Huali Shen, Aileen K Sewell, Marina Kniazeva, and Min Han. A novel sphingolipid-TORC1 pathway critically promotes postembryonic development in Caenorhabditis elegans. *eLife*, 2, 2013.

[243] Nixon M Abraham, Roberto Vincis, Samuel Lagier, Ivan Rodriguez, and Alan Carleton. Long term functional plasticity of sensory inputs mediated by olfactory learning. *eLife*, 3, 2014.

[244] Enas Abu Shah and Kinneret Keren. Symmetry breaking in reconstituted actin cortices. *eLife*, 3, 2014.

[245] Carolyn E Adler, Chris W Seidel, Sean A McKinney, and Alejandro Sánchez Alvarado. Selective amputation of the pharynx identifies a FoxA-dependent regeneration program in planaria. *eLife*, 3, 2014.

[246] Oluwaseun Akeju, Marco L Loggia, Ciprian Catana, Kara J Pavone, Rafael Vazquez, James Rhee, Violeta Contreras Ramirez, Daniel B Chonde, David Izquierdo-Garcia, Grae Arabasz, Shirley Hsu, Kathleen Habeeb, Jacob M Hooker, Vitaly Napadow, Emery N Brown, and Patrick L Purdon. Disruption of thalamic functional connectivity is a neural correlate of dexmedetomidine-induced unconsciousness. *eLife*, 3, 2014.

[247] Luca Albergante, J Julian Blow, and Timothy J Newman. Buffered Qualitative Stability explains the robustness and evolvability of transcriptional networks. *eLife*, 3, 2014.

[248] Matteo Allegretti, Deryck J Mills, Greg McMullan, Werner Kühlbrandt, and Janet Vonck. Atomic model of the F420-reducing [NiFe] hydrogenase by electron cryo-microscopy using a direct electron detector. *eLife*, 3, 2014.

[249] Mary Ann Allen, Zdenek Andrysik, Veronica L Dengler, Hestia S Mellert, Anna Guarnieri, Justin A Freeman, Kelly D Sullivan, Matthew D Galbraith, Xin Luo, W Lee Kraus, Robin D Dowell, and Joaquin M Espinosa. Global analysis of p53-regulated transcription identifies its direct targets and unexpected regulatory mechanisms. *eLife*, 3, 2014.

[250] Daniil Alpern, Diana Langer, Benoit Ballester, Stephanie Le Gras, Christophe Romier, Gabrielle Mengus, and Irwin Davidson. TAF4, a subunit of transcription factor II D, directs promoter occupancy of nuclear receptor HNF4A during post-natal hepatocyte differentiation. *eLife*, 3, 2014.

[251] Till F M Andlauer, Sabrina Scholz-Kornehl, Rui Tian, Marieluise Kirchner, Husam A Babikir, Harald Depner, Bernhard Loll, Christine Quentin, Varun K Gupta, Matthew G Holt, Shubham Dipt, Michael Cressy, Markus C Wahl, André Fiala, Matthias Selbach, Martin Schwärzel, and Stephan J Sigrist. Drep-2 is a novel synaptic protein important for learning and memory. *eLife*, 3, 2014.

[252] Muhammad Usman Anwer, Eleni Boikoglou, Eva Herrero, Marc Hallstein, Amanda Melaragno Davis, Geo Velikkakam James, Ferenc Nagy, and Seth Jon Davis. Natural variation reveals that intracellular distribution of ELF3 protein is associated with function in the circadian clock. *eLife*, 3, 2014.

[253] Itaru Arai and Peter Jonas. Nanodomain coupling explains Ca2+ independence of transmitter release time course at a fast central synapse. *eLife*, 3, 2014.

[254] Silvia Ardissone, Coralie Fumeaux, Matthieu Bergé, Audrey Beaussart, Laurence Théraulaz, Sunish Kumar Radhakrishnan, Yves F Dufrêne, and Patrick H Viollier. Cell cycle constraints on capsulation and bacteriophage susceptibility. *eLife*, 3, 2014.

[255] Armando Arias, Lucy Thorne, and Ian Goodfellow. Favipiravir elicits antiviral mutagenesis during virus replication in vivo. *eLife*, 3, 2014.

[256] Yoshinori Aso, Daisuke Hattori, Yang Yu, Rebecca M Johnston, Nirmala A Iyer, Teri-TB Ngo, Heather Dionne, LF Abbott, Richard Axel, Hiromu Tanimoto, and Gerald M Rubin. The neuronal architecture of the mushroom body provides a logic for associative learning. *eLife*, 3, 2014.

[257] Yoshinori Aso, Divya Sitaraman, Toshiharu Ichinose, Karla R Kaun, Katrin Vogt, Ghislain Belliart-Guérin, Pierre-Yves PlaÇais, Alice A Robie, Nobuhiro Yamagata, Christopher Schnaitmann, William J Rowell, Rebecca M Johnston, Teri-T B Ngo, Nan Chen, Wyatt Korff, Michael N Nitabach, Ulrike Heberlein, Thomas Preat, Kristin M Branson, Hiromu Tanimoto, and Gerald M Rubin. Mushroom body output neurons encode valence and guide memory-based action selection in Drosophila. *eLife*, 3, 2014.

[258] Julie L Aspden, Ying Chen Eyre-Walker, Rose J Phillips, Unum Amin, Muhammad Ali S Mumtaz, Michele Brocard, and Juan-Pablo Couso. Extensive translation of small Open Reading Frames revealed by Poly-Ribo-Seq. *eLife*, 3, 2014.

[259] Sabrina Asteriti, Claudia Gargini, and Lorenzo Cangiano. Mouse rods signal through gap junctions with cones. *eLife*, 3, 2014.

[260] Joseph Atherton, Irene Farabella, I-Mei Yu, Steven S Rosenfeld, Anne Houdusse, Maya Topf, and Carolyn A Moores. Conserved mechanisms of microtubule-stimulated ADP release, ATP binding, and force generation in transport kinesins. *eLife*, 3, 2014.

[261] Magda Luciana Atilano, Pedro Matos Pereira, Filipa Vaz, Maria João Catalão, Patricia Reed, Inês Ramos Grilo, Rita GonÇalves Sobral, Petros Ligoxygakis, Mariana Gomes Pinho, and Sérgio Raposo Filipe. Bacterial autolysins trim cell surface peptidoglycan to prevent detection by the Drosophila innate immune system. *eLife*, 3, 2014.

[262] Andrei S Avanesov, Siming Ma, Kerry A Pierce, Sun Hee Yim, Byung Cheon Lee, Clary B Clish, and Vadim N Gladyshev. Age- and diet-associated metabolome remodeling characterizes the aging process driven by damage accumulation. *eLife*, 3, 2014.

[263] Pelin Ayaz, Sarah Munyoki, Elisabeth A Geyer, Felipe-Andrés Piedra, Emily S Vu, Raquel Bromberg, Zbyszek Otwinowski, Nick V Grishin, Chad A Brautigam, and Luke M Rice. A tethered delivery mechanism explains the catalytic action of a microtubule polymerase. *eLife*, 3, 2014.

[264] Yun Bai, Akshay Tambe, Kaihong Zhou, and Jennifer A Doudna. RNA-guided assembly of Rev-RRE nuclear export complexes. *eLife*, 3, 2014.

[265] Adam P Baker, Matthew J Brookes, Iead A Rezek, Stephen M Smith, Timothy Behrens, Penny J Probert Smith, and Mark Woolrich. Fast transient networks in spontaneous human brain activity. *eLife*, 3, 2014.

[266] Benoit Ballester, Alejandra Medina-Rivera, Dominic Schmidt, Mar Gonzàlez-Porta, Matthew Carlucci, Xiaoting Chen, Kyle Chessman, Andre J Faure, Alister PW Funnell, Angela Goncalves, Claudia Kutter, Margus Lukk, Suraj Menon, William M McLaren, Klara Stefflova, Stephen Watt, Matthew T Weirauch, Merlin Crossley, John C Marioni, Duncan T Odom, Paul Flicek, and Michael D Wilson. Multi-species, multi-transcription factor binding highlights conserved control of tissue-specific biological pathways. *eLife*, 3, 2014.

[267] Sudeep Banjade and Michael K Rosen. Phase transitions of multivalent proteins can promote clustering of membrane receptors. *eLife*, 3, 2014.

[268] Xiucong Bao, Yi Wang, Xin Li, Xiao-Meng Li, Zheng Liu, Tangpo Yang, Chi Fat Wong, Jiangwen Zhang, Quan Hao, and Xiang David Li. Identification of 'erasers' for lysine crotonylated histone marks using a chemical proteomics approach. *eLife*, 3, 2014.

[269] Antonio P Baptista, Ramon Roozendaal, Rogier M Reijmers, Jasper J Koning, Wendy W Unger, Mascha Greuter, Eelco D Keuning, Rosalie Molenaar, Gera Goverse, Marlous M S Sneeboer, Joke M M den Haan, Marianne Boes, and Reina E Mebius. Lymph node stromal cells constrain immunity via MHC class II self-antigen presentation. *eLife*, 3, 2014.

[270] Rachael M Barry, Anne-Florence Bitbol, Alexander Lorestani, Emeric J Charles, Chris H Habrian, Jesse M Hansen, Hsin-Jung Li, Enoch P Baldwin, Ned S Wingreen, Justin M Kollman, and Zemer Gitai. Large-scale filament formation inhibits the activity of CTP synthetase. *eLife*, 3, 2014.

[271] Robert L Barry, Seth A Smith, Adrienne N Dula, and John C Gore. Resting state functional connectivity in the human spinal cord. *eLife*, 3, 2014.

[272] Federica Basilico, Stefano Maffini, John R Weir, Daniel Prumbaum, Ana M Rojas, Tomasz Zimniak, Anna De Antoni, Sadasivam Jeganathan, Beate Voss, Suzan van Gerwen, Veronica Krenn, Lucia Massimiliano, Alfonso Valencia, Ingrid R Vetter, Franz Herzog, Stefan Raunser, Sebastiano Pasqualato, and Andrea Musacchio. The pseudo GTPase CENP-M drives human kinetochore assembly. *eLife*, 3, 2014.

[273] Sulochanadevi Baskaran, Lars-Anders Carlson, Goran Stjepanovic, Lindsey N Young, Do Jin Kim, Patricia Grob, Robin E Stanley, Eva Nogales, and James H Hurley. Architecture and dynamics of the autophagic phosphatidylinositol 3-kinase complex. *eLife*, 3, 2014.

[274] Andrew R Bassett, Asifa Akhtar, Denise P Barlow, Adrian P Bird, Neil Brockdorff, Denis Duboule, Anne Ephrussi, Anne C Ferguson-Smith, Thomas R Gingeras, Wilfried Haerty, Douglas R Higgs, Eric A Miska, and Chris P Ponting. Considerations when investigating lncRNA function in vivo. *eLife*, 3, 2014.

[275] Christopher Batram, Nicola G Jones, Christian J Janzen, Sebastian M Markert, and Markus Engstler. Expression site attenuation mechanistically links antigenic variation and development in Trypanosoma brucei. *eLife*, 3, 2014.

[276] Michel Becuwe and Sébastien Léon. Integrated control of transporter endocytosis and recycling by the arrestin-related protein Rod1 and the ubiquitin ligase Rsp5. *eLife*, 3, 2014.

[277] Trevor Bedford, Marc A Suchard, Philippe Lemey, Gytis Dudas, Victoria Gregory, Alan J Hay, John W McCauley, Colin A Russell, Derek J Smith, and Andrew Rambaut. Integrating influenza antigenic dynamics with molecular evolution. *eLife*, 3, 2014.

[278] Philip Bejon, Thomas N Williams, Christopher Nyundo, Simon I Hay, David Benz, Peter W Gething, Mark Otiende, Judy Peshu, Mahfudh Bashraheil, Bryan Greenhouse, Teun Bousema, Evasius Bauni, Kevin Marsh, David L Smith, and Steffen Borrmann. A micro-epidemiological analysis of febrile malaria in Coastal Kenya showing hotspots within hotspots. *eLife*, 3, 2014.

[279] Nicholas W Bellono, Iliana E Escobar, Ariel J Lefkovith, Michael S Marks, and Elena Oancea. An intracellular anion channel critical for pigmentation. *eLife*, 3, 2014.

[280] Inbal Ben-Ami Bartal, David A Rodgers, Maria Sol Bernardez Sarria, Jean Decety, and Peggy Mason. Pro-social behavior in rats is modulated by social experience. *eLife*, 3, 2014.

[281] Joseph F Bergan, Yoram Ben-Shaul, and Catherine Dulac. Sex-specific processing of social cues in the medial amygdala. *eLife*, 3, 2014.

[282] Galina V Beznoussenko, Seetharaman Parashuraman, Riccardo Rizzo, Roman Polishchuk, Oliviano Martella, Daniele Di Giandomenico, Aurora Fusella, Alexander Spaar, Michele Sallese, Maria Grazia Capestrano, Margit Pavelka, Matthijn R Vos, Yuri GM Rikers, Volkhard Helms, Alexandre A Mironov, and Alberto Luini. Transport of soluble proteins through the Golgi occurs by diffusion via continuities across cisternae. *eLife*, 3, 2014.

[283] Vassilis Bitsikas, Ivan R Corrêa, and Benjamin J Nichols. Clathrin-independent pathways do not contribute significantly to endocytic flux. *eLife*, 3, 2014.

[284] Barak Blum, Adam N Roose, Ornella Barrandon, René Maehr, Anthony C Arvanites, Lance S Davidow, Jeffrey C Davis, Quinn P Peterson, Lee L Rubin, and Douglas A Melton. Reversal of $\beta$ cell de-differentiation by a small molecule inhibitor of the TGF$\beta$ pathway. *eLife*, 3, 2014.

[285] David Blum and Samuel LaBarge. Registered report: Tumour micro-environment elicits innate resistance to RAF inhibitors through HGF secretion. *eLife*, 3, 2014.

[286] Ian D Blum, Lei Zhu, Luc Moquin, Maia V Kokoeva, Alain Gratton, Bruno Giros, and Kai-Florian Storch. A highly tunable dopaminergic oscillator generates ultradian rhythms of behavioral arousal. *eLife*, 3, 2014.

[287] Dani L Bodor, João F Mata, Mikhail Sergeev, Ana Filipa David, Kevan J Salimian, Tanya Panchenko, Don W Cleveland, Ben E Black, Jagesh V Shah, and Lars ET Jansen. The quantitative architecture of centromeric chromatin. *eLife*, 3, 2014.

[288] Roberto Bonasio, Emilio Lecona, Varun Narendra, Philipp Voigt, Fabio Parisi, Yuval Kluger, and Danny Reinberg. Interactions with RNA direct the Polycomb group protein SCML2 to chromatin where it represses target genes. *eLife*, 3, 2014.

[289] Daniel G Booth, Masatoshi Takagi, Luis Sanchez-Pulido, Elizabeth Petfalski, Giulia Vargiu, Kumiko Samejima, Naoko Imamoto, Chris P Ponting, David Tollervey, William C Earnshaw, and Paola Vagnarelli. Ki-67 is a PP1-interacting protein that organises the mitotic chromosome periphery. *eLife*, 3, 2014.

[290] David S Booth, Yifan Cheng, and Alan D Frankel. The export receptor Crm1 forms a dimer to promote nuclear export of HIV RNA. *eLife*, 3, 2014.

[291] Justin A Bosch, Taryn M Sumabat, Yassi Hafezi, Brett J Pellock, Kevin D Gandhi, and Iswar K Hariharan. The Drosophila F-box protein Fbxl7 binds to the protocadherin Fat and regulates Dachs localization and Hippo signaling. *eLife*, 3, 2014.

[292] Jeffrey I Boucher, Joseph R Jacobowitz, Brian C Beckett, Scott Classen, and Douglas L Theobald. An atomic-resolution view of neofunctionalization in the evolution of apicomplexan lactate dehydrogenases. *eLife*, 3, 2014.

[293] Sarah K Bowman, Aimee M Deaton, Heber Domingues, Peggy I Wang, Ruslan I Sadreyev, Robert E Kingston, and Welcome Bender. H3K27 modifications define segmental regulatory domains in the Drosophila bithorax complex. *eLife*, 3, 2014.

[294] Marko Brankatschk, Sebastian Dunst, Linda Nemetschke, and Suzanne Eaton. Delivery of circulating lipoproteins to specific neurons in the Drosophila brain regulates systemic insulin signaling. *eLife*, 3, 2014.

[295] Christoph Brenker, Yu Zhou, Astrid Müller, Fabio Andres Echeverry, Christian Trötschel, Ansgar Poetsch, Xiao-Ming Xia, Wolfgang Bönigk, Christopher J Lingle, U Benjamin Kaupp, and Timo Strünker. The Ca2+-activated K+ current of human sperm is mediated by Slo3. *eLife*, 3, 2014.

[296] Caitlin A Brennan, Jason R Hunt, Natacha Kremer, Benjamin C Krasity, Michael A Apicella, Margaret J McFall-Ngai, and Edward G Ruby. A model symbiosis reveals a role for sheathed-flagellum rotation in the release of immunogenic lipopolysaccharide. *eLife*, 3, 2014.

[297] Ariane Briegel, Mark S Ladinsky, Catherine Oikonomou, Christopher W Jones, Michael J Harris, Daniel J Fowler, Yi-Wei Chang, Lynmarie K Thompson, Judith P Armitage, and Grant J Jensen. Structure of bacterial cytoplasmic chemoreceptor arrays and implications for chemotactic signaling. *eLife*, 3, 2014.

[298] Andrew Anand Brown, Alfonso Buil, Ana Viñuela, Tuuli Lappalainen, Hou-Feng Zheng, J Brent Richards, Kerrin S Small, Timothy D Spector, Emmanouil T Dermitzakis, and Richard Durbin. Genetic interactions affecting human gene expression identified by variance association mapping. *eLife*, 3, 2014.

[299] Jennifer Brown, Wei-Xing Pan, and Joshua Tate Dudman. The inhibitory microcircuit of the substantia nigra provides feedback gain control of the basal ganglia output. *eLife*, 3, 2014.

[300] Douglas R Brumley, Kirsty Y Wan, Marco Polin, and Raymond E Goldstein. Flagellar synchronization through direct hydrodynamic interactions. *eLife*, 3, 2014.

[301] János Brunner, Máté Neubrandt, Susan Van-Weert, Tibor Andrási, Felix B Kleine Borgmann, Sebastian Jessberger, and János Szabadics. Adult-born granule cells mature through two functionally distinct states. *eLife*, 3, 2014.

[302] Jean-Marie Buerstedde, Noel Lowndes, and David G Schatz. Induction of homologous recombination between sequence repeats by the activation induced cytidine deaminase (AID) protein. *eLife*, 3, 2014.

[303] Tammy J Bullwinkle, Noah M Reynolds, Medha Raina, Adil Moghal, Eleftheria Matsa, Andrei Rajkovic, Huseyin Kayadibi, Farbod Fazlollahi, Christopher Ryan, Nathaniel Howitz, Kym F Faull, Beth A Lazazzera, and Michael Ibba. Oxidation of cellular amino acid pools leads to cytotoxic mistranslation of the genetic code. *eLife*, 3, 2014.

[304] Megan L Burger, Kenneth K Leung, Margaux J Bennett, and Astar Winoto. T cell-specific inhibition of multiple apoptotic pathways blocks negative selection and causes autoimmunity. *eLife*, 3, 2014.

[305] Rose T Byrne, Audrey J Klingele, Eric L Cabot, Wendy S Schackwitz, Jeffrey A Martin, Joel Martin, Zhong Wang, Elizabeth A Wood, Christa Pennacchio, Len A Pennacchio, Nicole T Perna, John R Battista, and Michael M Cox. Evolution of extreme resistance to ionizing radiation via genetic adaptation of DNA repair. *eLife*, 3, 2014.

[306] Joseph JE Caesar, Hayley Lavender, Philip N Ward, Rachel M Exley, Jack Eaton, Emily Chittock, Talat H Malik, Elena Goiecoechea De Jorge, Matthew C Pickering, Christoph M Tang, and Susan M Lea. Competition between antagonistic complement factors for a single protein on N. meningitidis rules disease susceptibility. *eLife*, 3, 2014.

[307] Yujia Cai, Rasmus O Bak, and Jacob Giehm Mikkelsen. Targeted genome editing by lentiviral protein transduction of zinc-finger and TAL-effector nucleases. *eLife*, 3, 2014.

[308] Benjamin Cain, Jessica M Budke, Kelsey J Wood, Neal T Sweeney, and Benjamin Schwessinger. How postdocs benefit from building a union. *eLife*, 3, 2014.

[309] Adam J Calhoun, Sreekanth H Chalasani, and Tatyana O Sharpee. Maximally informative foraging by Caenorhabditis elegans. *eLife*, 3, 2014.

[310] Sinan Can, Mark A Dewitt, and Ahmet Yildiz. Bidirectional helical motility of cytoplasmic dynein around microtubules. *eLife*, 3, 2014.

[311] Yangrong Cao, Yan Liang, Kiwamu Tanaka, Cuong T Nguyen, Robert P Jedrzejczak, Andrzej Joachimiak, and Gary Stacey. The kinase LYK5 is a major chitin receptor in Arabidopsis and forms a chitin-induced complex with related kinase CERK1. *eLife*, 3, 2014.

[312] Florence Caro, Vida Ahyong, Miguel Betegon, and Joseph L DeRisi. Genome-wide regulatory dynamics of translation in the Plasmodium falciparum asexual blood stages. *eLife*, 3, 2014.

[313] Marta Carroni, Eva Kummer, Yuki Oguchi, Petra Wendler, Daniel K Clare, Irmgard Sinning, Jürgen Kopp, Axel Mogk, Bernd Bukau, and Helen R Saibil. Head-to-tail interactions of the coiled-coil domains regulate ClpB activity and cooperation with Hsp70 in protein disaggregation. *eLife*, 3, 2014.

[314] Sílvia Carvalho, Alexandra C Vítor, Sreerama C Sridhara, Filipa B Martins, Ana C Raposo, Joana MP Desterro, João Ferreira, and Sérgio F de Almeida. SETD2 is required for DNA double-strand break repair and activation of the p53-mediated checkpoint. *eLife*, 3, 2014.

[315] Anil G Cashikar, Soomin Shim, Robyn Roth, Michael R Maldazys, John E Heuser, and Phyllis I Hanson. Structure of cellular ESCRT-III spirals and their relationship to HIV budding. *eLife*, 3, 2014.

[316] Angela Castanieto, Michael J Johnston, and Todd G Nystul. EGFR signaling promotes self-renewal through the establishment of cell polarity in Drosophila follicle stem cells. *eLife*, 3, 2014.

[317] Vincenzo Cavalieri and Giovanni Spinelli. Early asymmetric cues triggering the dorsal/ventral gene regulatory network of the sea urchin embryo. *eLife*, 3, 2014.

[318] Fanny Cazettes, Brian J Fischer, and Jose L Pena. Spatial cue reliability drives frequency tuning in the barn Owl's midbrain. *eLife*, 3, 2014.

[319] Vladislava Chalei, Stephen N Sansom, Lesheng Kong, Sheena Lee, Juan F Montiel, Keith W Vance, and Chris P Ponting. The long non-coding RNA Dali is an epigenetic regulator of neural differentiation. *eLife*, 3, 2014.

[320] Elik Chapnik, Natalia Rivkin, Alexander Mildner, Gilad Beck, Ronit Pasvolsky, Eyal Metzl-Raz, Yehudit Birger, Gail Amir, Itay Tirosh, Ziv Porat, Liron L Israel, Emmanuel Lellouche, Shulamit Michaeli, Jean-Paul M Lellouche, Shai Izraeli, Steffen Jung, and Eran Hornstein. miR-142 orchestrates a network of actin cytoskeleton regulators during megakaryopoiesis. *eLife*, 3, 2014.

[321] Debashree Chatterjee, Richard B Cooley, Chelsea D Boyd, Ryan A Mehl, George A O'Toole, and Holger Sondermann. Mechanistic insight into the conserved allosteric regulation of periplasmic proteolysis by the signaling molecule cyclic-di-GMP. *eLife*, 3, 2014.

[322] Rishidev Chaudhuri, Alberto Bernacchia, and Xiao-Jing Wang. A diversity of localized timescales in network activity. *eLife*, 3, 2014.

[323] Tomasz Chelmicki, Friederike Dündar, Matthew James Turley, Tasneem Khanam, Tugce Aktas, Fidel Ramírez, Anne-Valerie Gendrel, Patrick Rudolf Wright, Pavankumar Videm, Rolf Backofen, Edith Heard, Thomas Manke, and Asifa Akhtar. MOF-associated complexes ensure stem cell identity and Xist repression. *eLife*, 3, 2014.

[324] Guo-Yun Chen, Nicholas K Brown, Wei Wu, Zahra Khedri, Hai Yu, Xi Chen, Diantha van de Vlekkert, Alessandra D'Azzo, Pan Zheng, and Yang Liu. Broad and direct interaction between TLR and Siglec families of pattern recognition receptors and its regulation by Neu1. *eLife*, 3, 2014.

[325] Jia Chen, Brendan F Miller, and Anthony V Furano. Repair of naturally occurring mismatches can induce mutations in flanking DNA. *eLife*, 3, 2014.

[326] Jun Chen, Bo Wang, Michael S Engel, Torsten Wappler, Edmund A Jarzembowski, Haichun Zhang, Xiaoli Wang, Xiaoting Zheng, and Jes Rust. Extreme adaptations for aquatic ectoparasitism in a Jurassic fly larva. *eLife*, 3, 2014.

[327] Lin Chen, Yibin Xu, Julie Healer, Jenny K Thompson, Brian J Smith, Michael C Lawrence, and Alan F Cowman. Crystal structure of PfRh5, an essential P. falciparum ligand for invasion of human erythrocytes. *eLife*, 3, 2014.

[328] Shanping Chen, Diancai Cai, Kaycey Pearce, Philip Y-W Sun, Adam C Roberts, and David L Glanzman. Reinstatement of long-term memory following erasure of its behavioral and synaptic expression in Aplysia. *eLife*, 3, 2014.

[329] Guo Cheng, Ming Zhong, Riki Kawaguchi, Miki Kassai, Muayyad Al-Ubaidi, Jun Deng, Mariam Ter-Stepanian, and Hui Sun. Identification of PLXDC1 and PLXDC2 as the transmembrane receptors for the multifunctional factor PEDF. *eLife*, 3, 2014.

[330] Yinwei Cheng, Wenjiao Zhu, Yuxiao Chen, Shinsaku Ito, Tadao Asami, and Xuelu Wang. Brassinosteroids control root epidermal cell fate via direct regulation of a MYB-bHLH-WD40 complex by GSK3-like kinases. *eLife*, 3, 2014.

[331] Joanne Chia, Keit Min Tham, David James Gill, Emilie Anne Bard-Chapeau, and Frederic A Bard. ERK8 is a negative regulator of O-GalNAc glycosylation and cell migration. *eLife*, 3, 2014.

[332] Shiho Chiba, Hiroaki Ikushima, Hiroshi Ueki, Hideyuki Yanai, Yoshitaka Kimura, Sho Hangai, Junko Nishio, Hideo Negishi, Tomohiko Tamura, Shinobu Saijo, Yoichiro Iwakura, and Tadatsugu Taniguchi. Recognition of tumor cells by Dectin-1 orchestrates innate immune cells for anti-tumor responses. *eLife*, 3, 2014.

[333] Jacqueline SR Chin, Shane R Ellis, Huong T Pham, Stephen J Blanksby, Kenji Mori, Qi Ling Koh, William J Etges, and Joanne Y Yew. Sex-specific triacylglycerides are widely conserved in Drosophila and mediate mating behavior. *eLife*, 3, 2014.

[334] Isaac M Chiu, Lee B Barrett, Erika K Williams, David E Strochlic, Seungkyu Lee, Andy D Weyer, Shan Lou, Gregory S Bryman, David P Roberson, Nader Ghasemlou, Cara Piccoli, Ezgi Ahat, Victor Wang, Enrique J Cobos, Cheryl L Stucky, Qiufu Ma, Stephen D Liberles, and Clifford J Woolf. Transcriptional profiling at whole population and single cell levels reveals somatosensory neuron molecular diversity. *eLife*, 3, 2014.

[335] Young Eun Choi, Yunfeng Pan, Eunmi Park, Panagiotis Konstantinopoulos, Subhajyoti De, Alan D'Andrea, and Dipanjan Chowdhury. MicroRNAs down-regulate homologous recombination in the G1 phase of cycling cells to maintain genomic stability. *eLife*, 3, 2014.

[336] Denise Chroscinski, Darryl Sampey, and Alex Hewitt. Registered report: Melanoma genome sequencing reveals frequent PREX2 mutations. *eLife*, 3, 2014.

[337] Mei-I Chung, Taejoon Kwon, Fan Tu, Eric R Brooks, Rakhi Gupta, Matthew Meyer, Julie C Baker, Edward M Marcotte, and John B Wallingford. Coordinated genomic control of ciliogenesis and cell movement by RFX2. *eLife*, 3, 2014.

[338] Jessica K Cinkornpumin, Dona R Wisidagama, Veronika Rapoport, James L Go, Christoph Dieterich, Xiaoyue Wang, Ralf J Sommer, and Ray L Hong. A host beetle pheromone regulates development and behavior in the nematode Pristionchus pacificus. *eLife*, 3, 2014.

[339] Sara Cipolat, Esther Hoste, Ken Natsuga, Sven R Quist, and Fiona M Watt. Epidermal barrier defects link atopic dermatitis with altered skin cancer susceptibility. *eLife*, 3, 2014.

[340] Lori Clay, Fabrice Caudron, Annina Denoth-Lippuner, Barbara Boettcher, Stéphanie Buvelot Frei, Erik Lee Snapp, and Yves Barral. A sphingolipid-dependent diffusion barrier confines ER stress to the yeast mother cell. *eLife*, 3, 2014.

[341] Carina C Clingman, Laura M Deveau, Samantha A Hay, Ryan M Genga, Shivender MD Shandilya, Francesca Massi, and Sean P Ryder. Allosteric inhibition of a stem cell RNA-binding protein by an intermediary metabolite. *eLife*, 3, 2014.

[342] Paul T Conduit, Jennifer H Richens, Alan Wainman, James Holder, Catarina C Vicente, Metta B Pratt, Carly I Dix, Zsofia A Novak, Ian M Dobbie, Lothar Schermelleh, and Jordan W Raff. A molecular mechanism of mitotic centrosome assembly in Drosophila. *eLife*, 3, 2014.

[343] Alessandro Costa, Ludovic Renault, Paolo Swuec, Tatjana Petojevic, James J Pesavento, Ivar Ilves, Kirsty MacLellan-Gibson, Roland A Fleck, Michael R Botchan, and James M Berger. DNA binding polarity, dimerization, and ATPase ring remodeling in the CMG helicase of the eukaryotic replisome. *eLife*, 3, 2014.

[344] Antoine Coulon, Matthew L Ferguson, Valeria de Turris, Murali Palangat, Carson C Chow, and Daniel R Larson. Kinetic competition during the transcription cycle results in stochastic RNA processing. *eLife*, 3, 2014.

[345] Cristina Cruz and Jonathan Houseley. Endogenous RNA interference is driven by copy number. *eLife*, 3, 2014.

[346] Akash Das, Michael S Brown, Donald D Anderson, Joseph L Goldstein, and Arun Radhakrishnan. Three pools of plasma membrane cholesterol and their relation to cholesterol homeostasis. *eLife*, 3, 2014.

[347] Paolo De Los Rios and Alessandro Barducci. Hsp70 chaperones are non-equilibrium machines that achieve ultra-affinity by energy consumption. *eLife*, 3, 2014.

[348] Paul G DeCaen, Yuka Takahashi, Terry A Krulwich, Masahiro Ito, and David E Clapham. Ionic selectivity and thermal adaptations within the voltage-gated sodium channel family of alkaliphilic Bacillus. *eLife*, 3, 2014.

[349] Safia Deddouche, Delphine Goubau, Jan Rehwinkel, Probir Chakravarty, Sharmin Begum, Pierre V Maillard, Annabel Borg, Nik Matthews, Qian Feng, Frank J M van Kuppeveld, and Caetano Reis e Sousa. Identification of an LGP2-associated MDA5 agonist in picornavirus-infected cells. *eLife*, 3, 2014.

[350] David Degen, Yu Feng, Yu Zhang, Katherine Y Ebright, Yon W Ebright, Matthew Gigliotti, Hanif Vahedian-Movahed, Sukhendu Mandal, Meliza Talaue, Nancy Connell, Eddy Arnold, William Fenical, and Richard H Ebright. Transcription inhibition by the depsipeptide antibiotic salinamide A. *eLife*, 3, 2014.

[351] Annina Denoth-Lippuner, Marek Konrad Krzyzanowski, Catherine Stober, and Yves Barral. Role of SAGA in the asymmetric segregation of DNA circles during yeast ageing. *eLife*, 3, 2014.

[352] Noah Dephoure, Sunyoung Hwang, Ciara O'Sullivan, Stacie E Dodgson, Steven P Gygi, Angelika Amon, and Eduardo M Torres. Quantitative proteomic analysis reveals posttranslational responses to aneuploidy in yeast. *eLife*, 3, 2014.

[353] Mehabaw G Derebe, Clare M Zlatkov, Sureka Gattu, Kelly A Ruhn, Shipra Vaishnava, Gretchen E Diehl, John B MacMillan, Noelle S Williams, and Lora V Hooper. Serum amyloid A is a retinol binding protein that transports retinol during bacterial infection. *eLife*, 3, 2014.

[354] Nicolas Descostes, Martin Heidemann, Lionel Spinelli, Roland Schüller, Muhammad Ahmad Maqbool, Romain Fenouil, Frederic Koch, Charlène Innocenti, Marta Gut, Ivo Gut, Dirk Eick, and Jean-Christophe Andrau. Tyrosine phosphorylation of RNA polymerase II CTD is associated with antisense promoter transcription and active enhancers in mammalian cells. *eLife*, 3, 2014.

[355] W Patrick Devine, Joshua D Wythe, Matthew George, Kazuko Koshiba-Takeuchi, and Benoit G Bruneau. Early patterning and specification of cardiac progenitors in gastrulating mesoderm. *eLife*, 3, 2014.

[356] Vincenzo Di Cerbo, Fabio Mohn, Daniel P Ryan, Emilie Montellier, Salim Kacem, Philipp Tropberger, Eleni Kallis, Monika Holzner, Leslie Hoerner, Angelika Feldmann, Florian Martin Richter, Andrew J Bannister, Gerhard Mittler, Jens Michaelis, Saadi Khochbin, Robert Feil, Dirk Schuebeler, Tom Owen-Hughes, Sylvain Daujat, and Robert Schneider. Acetylation of histone H3 at lysine 64 regulates nucleosome dynamics and facilitates transcription. *eLife*, 3, 2014.

[357] Daniela Dimastrogiovanni, Kathrin S Fröhlich, Katarzyna J Bandyra, Heather A Bruce, Susann Hohensee, Jörg Vogel, and Ben F Luisi. Recognition of the small regulatory RNA RydC by the bacterial Hfq protein. *eLife*, 3, 2014.

[358] Scott J Dixon, Darpan N Patel, Matthew Welsch, Rachid Skouta, Eric D Lee, Miki Hayano, Ajit G Thomas, Caroline E Gleason, Nicholas P Tatonetti, Barbara S Slusher, and Brent R Stockwell. Pharmacological inhibition of cystine–glutamate exchange induces endoplasmic reticulum stress and ferroptosis. *eLife*, 3, 2014.

[359] David W Dowdy, Jason R Andrews, Peter J Dodd, and Robert H Gilman. A user-friendly, open-source tool to project impact and cost of diagnostic tests for tuberculosis. *eLife*, 3, 2014.

[360] Hauke Drechsler, Toni McHugh, Martin R Singleton, Nicholas J Carter, and Andrew D McAinsh. The Kinesin-12 Kif15 is a processive track-switching tetramer. *eLife*, 3, 2014.

[361] Ines A Drinnenberg, Dakota deYoung, Steven Henikoff, and Harmit Singh Malik. Recurrent loss of CenH3 is associated with independent transitions to holocentricity in insects. *eLife*, 3, 2014.

[362] Jan Drugowitsch, Gregory C DeAngelis, Eliana M Klier, Dora E Angelaki, and Alexandre Pouget. Optimal multisensory decision-making in a reaction-time task. *eLife*, 3, 2014.

[363] Ngoc-Hien Du, Alaaddin Bulak Arpat, Mara De Matos, and David Gatfield. MicroRNAs shape circadian hepatic gene expression on a transcriptome-wide scale. *eLife*, 3, 2014.

[364] Yuntao Duan, Shih-Hsiu Wang, Juan Song, Yevgeniya Mironova, Guo-li Ming, Alex L Kolodkin, and Roman J Giger. Semaphorin 5A inhibits synaptogenesis in early postnatal- and adult-born hippocampal dentate granule cells. *eLife*, 3, 2014.

[365] Martial A Dufour, Adele Woodhouse, Julien Amendola, and Jean-Marc Goaillard. Non-linear developmental trajectory of electrical phenotype in rat substantia nigra pars compacta dopaminergic neurons. *eLife*, 3, 2014.

[366] Edward H Egelman. Ambiguities in helical reconstruction. *eLife*, 3, 2014.

[367] Thomas Eicher, Markus A Seeger, Claudio Anselmi, Wenchang Zhou, Lorenz Brandstätter, FranÇois Verrey, Kay Diederichs, José D Faraldo-Gómez, and Klaas M Pos. Coupling of remote alternating-access transport mechanisms for protons and substrates in the multidrug efflux pump AcrB. *eLife*, 3, 2014.

[368] Marcella L Erb, James A Kraemer, Joanna K C Coker, Vorrapon Chaikeeratisak, Poochit Nonejuie, David A Agard, and Joe Pogliano. A bacteriophage tubulin harnesses dynamic instability to center DNA in infected cells. *eLife*, 3, 2014.

[369] Aysen L Erdem, Malgorzata Jaszczur, Jeffrey G Bertram, Roger Woodgate, Michael M Cox, and Myron F Goodman. DNA polymerase V activity is autoregulated by a novel intrinsic DNA-dependent ATPase. *eLife*, 3, 2014.

[370] Timothy M Errington, Elizabeth Iorns, William Gunn, Fraser Elisabeth Tan, Joelle Lomax, and Brian A Nosek. An open investigation of the reproducibility of cancer biology research. *eLife*, 3, 2014.

[371] Kevin M Esvelt, Andrea L Smidler, Flaminia Catteruccia, and George M Church. Concerning RNA-guided gene drives for the alteration of wild populations. *eLife*, 3, 2014.

[372] Pietro Fazzari, An Snellinx, Victor Sabanov, Tariq Ahmed, Lutgarde Serneels, Annette Gartner, S Ali M Shariati, Detlef Balschun, and Bart De Strooper. Cell autonomous regulation of hippocampal circuitry via Aph1b-$\gamma$-secretase/neuregulin 1 signalling. *eLife*, 3, 2014.

[373] Quentin Ferry, Julia Steinberg, Caleb Webber, David R FitzPatrick, Chris P Ponting, Andrew Zisserman, and Christoffer Nellåker. Diagnostically relevant facial gestalt information from ordinary photos. *eLife*, 3, 2014.

[374] Diasynou Fioravante, YunXiang Chu, Arthur PH de Jong, Michael Leitges, Pascal S Kaeser, and Wade G Regehr. RETRACTED: Protein kinase C is a calcium sensor for presynaptic short-term plasticity. *eLife*, 3, 2014.

[375] Nir Fluman, Sivan Navon, Eitan Bibi, and Yitzhak Pilpel. mRNA-programmed translation pauses in the targeting of E. coli membrane proteins. *eLife*, 3, 2014.

[376] Yick W Fong, Jaclyn J Ho, Carla Inouye, and Robert Tjian. The dyskerin ribonucleoprotein complex as an OCT4/SOX2 coactivator in embryonic stem cells. *eLife*, 3, 2014.

[377] Matteo Forloni, Shaillay Kumar Dogra, Yuying Dong, Darryl Conte, Jianhong Ou, Lihua Julie Zhu, April Deng, Meera Mahalingam, Michael R Green, and Narendra Wajapeyee. miR-146a promotes the initiation and progression of melanoma by activating Notch signaling. *eLife*, 3, 2014.

[378] Nicholas W Frankel, William Pontius, Yann S Dufour, Junjiajia Long, Luis Hernandez-Nunez, and Thierry Emonet. Adaptability of non-genetic diversity in bacterial chemotaxis. *eLife*, 3, 2014.

[379] Clifford A Froelich, Sukhyun Kang, Leslie B Epling, Stephen P Bell, and Eric J Enemark. A conserved MCM single-stranded DNA binding element is essential for replication initiation. *eLife*, 3, 2014.

[380] Wenxian Fu, Julia Farache, Susan M Clardy, Kimie Hattori, Palwinder Mander, Kevin Lee, Inmaculada Rioja, Ralph Weissleder, Rab K Prinjha, Christophe Benoist, and Diane Mathis. Epigenetic modulation of type-1 diabetes via a dual effect on pancreatic macrophages and $\beta$ cells. *eLife*, 3, 2014.

[381] Hotaka Fukushima, Yue Zhang, Georgia Archbold, Rie Ishikawa, Karim Nader, and Satoshi Kida. Enhancement of fear memory by retrieval through reconsolidation. *eLife*, 3, 2014.

[382] Milos Galic, Feng-Chiao Tsai, Sean R Collins, Maja Matis, Samuel Bandara, and Tobias Meyer. Dynamic recruitment of the curvature-sensitive protein ArhGAP44 to nanoscale membrane deformations limits exploratory filopodia initiation in neurons. *eLife*, 3, 2014.

[383] Christian M Gallardo, Martin Darvas, Mia Oviatt, Chris H Chang, Mateusz Michalik, Timothy F Huddy, Emily E Meyer, Scott A Shuster, Antonio Aguayo, Elizabeth M Hill, Karun Kiani, Jonathan Ikpeazu, Johan S Martinez, Mari Purpura, Andrea N Smit, Danica F Patton, Ralph E Mistlberger, Richard D Palmiter, and Andrew D Steele. Dopamine receptor 1 neurons in the dorsal striatum regulate food anticipatory circadian activity rhythms in mice. *eLife*, 3, 2014.

[384] Yann Gambin, Nicholas Ariotti, Kerrie-Ann McMahon, Michele Bastiani, Emma Sierecki, Oleksiy Kovtun, Mark E Polinkovsky, Astrid Magenau, WooRam Jung, Satomi Okano, Yong Zhou, Natalya Leneva, Sergey Mureev, Wayne Johnston, Katharina Gaus, John F Hancock, Brett M Collins, Kirill Alexandrov, and Robert G Parton. Single-molecule analysis reveals self assembly and nanoscale segregation of two distinct cavin subcomplexes on caveolae. *eLife*, 3, 2014.

[385] Don B Gammon, Sophie Duraffour, Daniel K Rozelle, Heidi Hehnly, Rita Sharma, Michael E Sparks, Cara C West, Ying Chen, James J Moresco, Graciela Andrei, John H Connor, Darryl Conte, Dawn E Gundersen-Rindal, William L Marshall, John R Yates, Neal Silverman, and Craig C Mello. A single vertebrate DNA virus protein disarms invertebrate immunity to RNA virus infection. *eLife*, 3, 2014.

[386] Justin Gardin, Rukhsana Yeasmin, Alisa Yurovsky, Ying Cai, Steve Skiena, and Bruce Futcher. Measurement of average decoding rates of the 61 sense codons in vivo. *eLife*, 3, 2014.

[387] Liora Garion, Uri Dubin, Yoav Rubin, Mohamed Khateb, Yitzhak Schiller, Rony Azouz, and Jackie Schiller. Texture coarseness responsive neurons and their mapping in layer 2–3 of the rat barrel cortex in vivo. *eLife*, 3, 2014.

[388] Jonathan Garst-Orozco, Baktash Babadi, and Bence P ölveczky. A neural circuit mechanism for regulating vocal variability during song learning in zebra finches. *eLife*, 3, 2014.

[389] Liang Ge, Min Zhang, and Randy Schekman. Phosphatidylinositol 3-kinase and COPII generate LC3 lipidation vesicles from the ER-Golgi intermediate compartment. *eLife*, 3, 2014.

[390] Moritz Gegg, Anika Böttcher, Ingo Burtscher, Stefan Hasenoeder, Claude Van Campenhout, Michaela Aichler, Axel Walch, Seth G N Grant, and Heiko Lickert. Flattop regulates basal body docking and positioning in mono- and multiciliated cells. *eLife*, 3, 2014.

[391] Maria V Gelfand, Nellwyn Hagan, Aleksandra Tata, Won-Jong Oh, Baptiste Lacoste, Kyu-Tae Kang, Justyna Kopycinska, Joyce Bischoff, Jia-Huai Wang, and Chenghua Gu. Neuropilin-1 functions as a VEGFR2 co-receptor to guide developmental angiogenesis independent of ligand binding. *eLife*, 3, 2014.

[392] özgür GenÇ, Olexiy Kochubey, Ruud F Toonen, Matthijs Verhage, and Ralf Schneggenburger. Munc18-1 is a dynamically regulated PKC target during short-term enhancement of transmitter release. *eLife*, 3, 2014.

[393] Adrien Georges, David L'Hôte, Anne Laure Todeschini, Aurélie Auguste, Bérangère Legois, Alain Zider, and Reiner A Veitia. The transcription factor FOXL2 mobilizes estrogen signaling to maintain the identity of ovarian granulosa cells. *eLife*, 3, 2014.

[394] Sebastian A Giusti, Annette M Vogl, Marisa M Brockmann, Claudia A Vercelli, Martin L Rein, Dietrich Trümbach, Wolfgang Wurst, Demian Cazalla, Valentin Stein, Jan M Deussing, and Damian Refojo. MicroRNA-9 controls dendritic development by targeting REST. *eLife*, 3, 2014.

[395] Judith R Glynn, Ndoliwe Kayuni, Levie Gondwe, Alison J Price, and Amelia C Crampin. Earlier menarche is associated with a higher prevalence of Herpes simplex type-2 (HSV-2) in young women in rural Malawi. *eLife*, 3, 2014.

[396] Andrea M Gomez, Robert C Froemke, and Steven J Burden. Synaptic plasticity and cognitive function are disrupted in the absence of Lrp4. *eLife*, 3, 2014.

[397] Marcin Grabowicz, Dorothee Andres, Matthew D Lebar, Goran Malojčić, Daniel Kahne, and Thomas J Silhavy. A mutant Escherichia coli that attaches peptidoglycan to lipopolysaccharide and displays cell wall on its surface. *eLife*, 3, 2014.

[398] Arnulf BA Graf and Richard A Andersen. Inferring eye position from populations of lateral intraparietal neurons. *eLife*, 3, 2014.

[399] Edward Greenfield and Erin Griner. Registered report: Widespread potential for growth factor-driven resistance to anticancer kinase inhibitors. *eLife*, 3, 2014.

[400] William N Grimes, Mrinalini Hoon, Kevin L Briggman, Rachel O Wong, and Fred Rieke. Cross-synaptic synchrony and transmission of signal and noise across the mouse retina. *eLife*, 3, 2014.

[401] Yong Gu, Dora E Angelaki, and Gregory C DeAngelis. Contribution of correlated noise and selective decoding to choice probability measurements in extrastriate visual cortex. *eLife*, 3, 2014.

[402] Carine Gubelmann, Petra C Schwalie, Sunil K Raghav, Eva Röder, Tenagne Delessa, Elke Kiehlmann, Sebastian M Waszak, Andrea Corsinotti, Gilles Udin, Wiebke Holcombe, Gottfried Rudofsky, Didier Trono, Christian Wolfrum, and Bart Deplancke. Identification of the transcription factor ZEB1 as a central component of the adipogenic gene regulatory network. *eLife*, 3, 2014.

[403] Leonardo Gastón Guilgur, Pedro Prudêncio, Daniel Sobral, Denisa Liszekova, André Rosa, and Rui GonÇalo Martinho. Requirement for highly efficient pre-mRNA splicing during Drosophila early embryonic development. *eLife*, 3, 2014.

[404] Rajesh D Gunage, Heinrich Reichert, and K VijayRaghavan. Identification of a new stem cell population that generates Drosophila flight muscles. *eLife*, 3, 2014.

[405] Ufuk Günesdogan, Herbert Jäckle, and Alf Herzig. Histone supply regulates S phase timing and cell cycle progression. *eLife*, 3, 2014.

[406] Fang Guo, Isadora Cerullo, Xiao Chen, and Michael Rosbash. PDF neuron firing phase-shifts key circadian activity neurons in Drosophila. *eLife*, 3, 2014.

[407] Tingqing Guo, Patricia Marmol, Annalena Moliner, Marie Björnholm, Chao Zhang, Kevan M Shokat, and Carlos F Ibanez. Adipocyte ALK7 links nutrient overload to catecholamine resistance in obesity. *eLife*, 3, 2014.

[408] Xiaoyan Guo and L René García. SIR-2.1 integrates metabolic homeostasis with the reproductive neuromuscular excitability in early aging male Caenorhabditis elegans. *eLife*, 3, 2014.

[409] Sachiko Haga-Yamanaka, Limei Ma, Jie He, Qiang Qiu, Luke D Lavis, Loren L Looger, and C Ron Yu. Integrated action of pheromone signals in promoting courtship behavior in male mice. *eLife*, 3, 2014.

[410] Parthiv Haldipur, Gwendolyn S Gillies, Olivia K Janson, Victor V Chizhikov, Divakar S Mithal, Richard J Miller, and Kathleen J Millen. Foxc1 dependent mesenchymal signalling drives embryonic cerebellar growth. *eLife*, 3, 2014.

[411] Kosuke Hamaguchi, Katherine A Tschida, Inho Yoon, Bruce R Donald, and Richard Mooney. Auditory synapses to song premotor neurons are gated off during vocalization in zebra finches. *eLife*, 3, 2014.

[412] Brenda Y Han, Shuang Wu, Chuan-Sheng Foo, Robert M Horton, Craig N Jenne, Susan R Watson, Belinda Whittle, Chris C Goodnow, and Jason G Cyster. Zinc finger protein Zfp335 is required for the formation of the naïve T cell compartment. *eLife*, 3, 2014.

[413] Timothy Hanks, Roozbeh Kiani, and Michael N Shadlen. A neural mechanism of speed-accuracy tradeoff in macaque area LIP. *eLife*, 3, 2014.

[414] Max Emanuel Harner, Ann-Katrin Unger, Toshiaki Izawa, Dirk M Walther, Cagakan özbalci, Stefan Geimer, Fulvio Reggiori, Britta Brügger, Matthias Mann, Benedikt Westermann, and Walter Neupert. Aim24 and MICOS modulate respiratory function, tafazzin-related cardiolipin modification and mitochondrial architecture. *eLife*, 3, 2014.

[415] Megumi Hatori, Shubhroz Gill, Ludovic S Mure, Martyn Goulding, Dennis D M O'Leary, and Satchidananda Panda. Lhx1 maintains synchrony among circadian oscillator neurons of the SCN. *eLife*, 3, 2014.

[416] Laura Dean Heckman, Maria H Chahrour, and Huda Y Zoghbi. Rett-causing mutations reveal two domains critical for MeCP2 function and for toxicity in MECP2 duplication syndrome mice. *eLife*, 3, 2014.

[417] Kiersten A Henderson, Adam L Hughes, and Daniel E Gottschling. Mother-daughter asymmetry of pH underlies aging and rejuvenation in yeast. *eLife*, 3, 2014.

[418] Steven Henikoff, Srinivas Ramachandran, Kristina Krassovsky, Terri D Bryson, Christine A Codomo, Kristin Brogaard, Jonathan Widom, Ji-Ping Wang, and Jorja G Henikoff. The budding yeast Centromere DNA Element II wraps a stable Cse4 hemisome in either orientation in vivo. *eLife*, 3, 2014.

[419] Alexander M Herman, Longwen Huang, Dona K Murphey, Isabella Garcia, and Benjamin R Arenkiel. Cell type-specific and time-dependent light exposure contribute to silencing in neurons expressing Channelrhodopsin-2. *eLife*, 3, 2014.

[420] Jeremy K Herren, Juan C Paredes, Fanny Schüpfer, Karim Arafah, Philippe Bulet, and Bruno Lemaitre. Insect endosymbiont proliferation is limited by lipid availability. *eLife*, 3, 2014.

[421] Francisco J Herrera, Teppei Yamaguchi, Henk Roelink, and Robert Tjian. Core promoter factor TAF9B regulates neuronal gene expression. *eLife*, 3, 2014.

[422] Salvador C Herrera and Ginés Morata. Transgressions of compartment boundaries and cell reprogramming during regeneration in Drosophila. *eLife*, 3, 2014.

[423] Jennifer Hirst, Alexander Schlacht, John P Norcott, David Traynor, Gareth Bloomfield, Robin Antrobus, Robert R Kay, Joel B Dacks, and Margaret S Robinson. Characterization of TSET, an ancient and widespread membrane trafficking complex. *eLife*, 3, 2014.

[424] Richard K Hite, Joel A Butterwick, and Roderick MacKinnon. Phosphatidic acid modulation of Kv channel voltage sensor function. *eLife*, 3, 2014.

[425] Cheng-Hsun Ho and Wolf B Frommer. Fluorescent sensors for activity and regulation of the nitrate transceptor CHL1/NRT1.1 and oligopeptide transporters. *eLife*, 3, 2014.

[426] Jan-Erik Hoffmann, Yessica Fermin, Ruth LO Stricker, Katja Ickstadt, and Eli Zamir. Symmetric exchange of multi-protein building blocks between stationary focal adhesions and the cytosol. *eLife*, 3, 2014.

[427] Gunther Hollopeter, Jeffrey J Lange, Ying Zhang, Thien N Vu, Mingyu Gu, Michael Ailion, Eric J Lambie, Brian D Slaughter, Jay R Unruh, Laurence Florens, and Erik M Jorgensen. The membrane-associated proteins FCHo and SGIP are allosteric activators of the AP2 clathrin adaptor complex. *eLife*, 3, 2014.

[428] Alf Honigmann, Sina Sadeghi, Jan Keller, Stefan W Hell, Christian Eggeling, and Richard Vink. A lipid bound actin meshwork organizes liquid phase separation in model membranes. *eLife*, 3, 2014.

[429] Johanna L Höög, Sylvain Lacomble, Eileen T O'Toole, Andreas Hoenger, J Richard McIntosh, and Keith Gull. Modes of flagellar assembly in Chlamydomonas reinhardtii and Trypanosoma brucei. *eLife*, 3, 2014.

[430] Thomas A Hopf, Charlotta P I Schärfe, João P G L M Rodrigues, Anna G Green, Oliver Kohlbacher, Chris Sander, Alexandre M J J Bonvin, and Debora S Marks. Sequence co-evolution gives 3D contacts and structures of protein complexes. *eLife*, 3, 2014.

[431] Gene Hopping, Jackson Kellock, Ravi Pratap Barnwal, Peter Law, James Bryers, Gabriele Varani, Byron Caughey, and Valerie Daggett. Designed $\alpha$-sheet peptides inhibit amyloid formation by targeting toxic oligomers. *eLife*, 3, 2014.

[432] Shauna L Houlihan and Yuanyi Feng. The scaffold protein Nde1 safeguards the brain genome during S phase of early neural progenitor differentiation. *eLife*, 3, 2014.

[433] Thomas House. Epidemiological dynamics of Ebola outbreaks. *eLife*, 3, 2014.

[434] Conor J Howard, Victor Hanson-Smith, Kristopher J Kennedy, Chad J Miller, Hua Jane Lou, Alexander D Johnson, Benjamin E Turk, and Liam J Holt. Ancestral resurrection reveals evolutionary mechanisms of kinase plasticity. *eLife*, 3, 2014.

[435] Jing-Ping Hsin, Wencheng Li, Mainul Hoque, Bin Tian, and James L Manley. RNAP II CTD tyrosine 1 performs diverse functions in vertebrate cells. *eLife*, 3, 2014.

[436] Bruno Hudry, Morgane Thomas-Chollier, Yael Volovik, Marilyne Duffraisse, Amélie Dard, Dale Frank, Ulrich Technau, and Samir Merabet. Molecular insights into the origin of the Hox-TALE patterning system. *eLife*, 3, 2014.

[437] Diego Huet, Thierry Blisnick, Sylvie Perrot, and Philippe Bastin. The GTPase IFT27 is involved in both anterograde and retrograde intraflagellar transport. *eLife*, 3, 2014.

[438] William W Hwang, Ryan D Salinas, Jason J Siu, Kevin W Kelley, Ryan N Delgado, Mercedes F Paredes, Arturo Alvarez-Buylla, Michael C Oldham, and Daniel A Lim. Distinct and separable roles for EZH2 in neurogenic astroglia. *eLife*, 3, 2014.

[439] Uwe Irion, Hans Georg Frohnhöfer, Jana Krauss, Tuğba Çolak Champollion, Hans-Martin Maischein, Silke Geiger-Rudolph, Christian Weiler, and Christiane Nüsslein-Volhard. Gap junctions composed of connexins 41.8 and 39.4 are essential for colour pattern formation in zebrafish. *eLife*, 3, 2014.

[440] Joan Isern, Andrés García-García, Ana M Martín, Lorena Arranz, Daniel Martín-Pérez, Carlos Torroja, Fátima Sánchez-Cabo, and Simón Méndez-Ferrer. The neural crest is a source of mesenchymal stem cells with specialized hematopoietic stem cell niche function. *eLife*, 3, 2014.

[441] Hiroaki Ishikawa, Takahiro Ide, Toshiki Yagi, Xue Jiang, Masafumi Hirono, Hiroyuki Sasaki, Haruaki Yanagisawa, Kimberly A Wemmer, Didier YR Stainier, Hongmin Qin, Ritsu Kamiya, and Wallace F Marshall. TTC26/DYF13 is an intraflagellar transport protein required for transport of motility-related proteins into flagella. *eLife*, 3, 2014.

[442] Taichi Isobe, Shigeo Hisamori, Daniel J Hogan, Maider Zabala, David G Hendrickson, Piero Dalerba, Shang Cai, Ferenc Scheeren, Angera H Kuo, Shaheen S Sikandar, Jessica S Lam, Dalong Qian, Frederick M Dirbas, George Somlo, Kaiqin Lao, Patrick O Brown, Michael F Clarke, and Yohei Shimono. miR-142 regulates the tumorigenicity of human breast cancer stem cells through the canonical WNT signaling pathway. *eLife*, 3, 2014.

[443] Motoko Iwashita. Phasic activation of ventral tegmental neurons increases response and pattern similarity in prefrontal cortex neurons. *eLife*, 3, 2014.

[444] Ignacio Izeddin, Vincent Récamier, Lana Bosanac, Ibrahim I Cissé, Lydia Boudarene, Claire Dugast-Darzacq, Florence Proux, Olivier Bénichou, Raphaël Voituriez, Olivier Bensaude, Maxime Dahan, and Xavier Darzacq. Single-molecule tracking in live cells reveals distinct target-search strategies of transcription factors in the nucleus. *eLife*, 3, 2014.

[445] Mariko Izumo, Martina Pejchal, Andrew C Schook, Ryan P Lange, Jacqueline A Walisser, Takashi R Sato, Xiaozhong Wang, Christopher A Bradfield, and Joseph S Takahashi. Differential effects of light and feeding on circadian organization of peripheral clocks in a forebrain Bmal1 mutant. *eLife*, 3, 2014.

[446] Chris G C Jacobs, Herman P Spaink, and Maurijn van der Zee. The extraembryonic serosa is a frontier epithelium providing the insect egg with a full-range innate immune response. *eLife*, 3, 2014.

[447] Maciej M Jankowski, Md Nurul Islam, Nicholas F Wright, Seralynne D Vann, Jonathan T Erichsen, John P Aggleton, and Shane M O'Mara. Nucleus reuniens of the thalamus contains head direction cells. *eLife*, 3, 2014.

[448] Bhargavi Jayaraman, David C Crosby, Christina Homer, Isabel Ribeiro, David Mavor, and Alan D Frankel. RNA-directed remodeling of the HIV-1 protein Rev orchestrates assembly of the Rev–Rev response element complex. *eLife*, 3, 2014.

[449] Misty R Jenkins, Jane C Stinchcombe, Byron B Au-Yeung, Yukako Asano, Alex T Ritter, Arthur Weiss, and Gillian M Griffiths. Distinct structural and catalytic roles for Zap70 in formation of the immunological synapse in CTL. *eLife*, 3, 2014.

[450] Xiaodan Ji, Huasong Lu, Qiang Zhou, and Kunxin Luo. LARP7 suppresses P-TEFb activity to inhibit breast cancer progression and metastasis. *eLife*, 3, 2014.

[451] Xiaofei Jia, Erin Weber, Andrey Tokarev, Mary Lewinski, Maryan Rizk, Marissa Suarez, John Guatelli, and Yong Xiong. Structural basis of HIV-1 Vpu-mediated BST2 antagonism via hijacking of the clathrin adaptor protein complex 1. *eLife*, 3, 2014.

[452] Zachary Lee Johnson, Jun-Ho Lee, Kiyoun Lee, Minhee Lee, Do-Yeon Kwon, Jiyong Hong, and Seok-Yong Lee. Structural basis of nucleoside and nucleoside drug selectivity by concentrative nucleoside transporters. *eLife*, 3, 2014.

[453] Alexander M Jones, Jonas ÅH Danielson, Shruti N ManojKumar, Viviane Lanquar, Guido Grossmann, and Wolf B Frommer. Abscisic acid dynamics in roots detected with genetically encoded FRET sensors. *eLife*, 3, 2014.

[454] Iris Jonkers, Hojoong Kwak, and John T Lis. Genome-wide dynamics of Pol II elongation and its interplay with promoter proximal pausing, chromatin, and exons. *eLife*, 3, 2014.

[455] Young Seok Ju, Ludmil B Alexandrov, Moritz Gerstung, Inigo Martincorena, Serena Nik-Zainal, Manasa Ramakrishna, Helen R Davies, Elli Papaemmanuil, Gunes Gundem, Adam Shlien, Niccolo Bolli, Sam Behjati, Patrick S Tarpey, Jyoti Nangalia, Charles E Massie, Adam P Butler, Jon W Teague, George S Vassiliou, Anthony R Green, Ming-Qing Du, Ashwin Unnikrishnan, John E Pimanda, Bin Tean Teh, Nikhil Munshi, Mel Greaves, Paresh Vyas, Adel K El-Naggar, Tom Santarius, V Peter Collins, Richard Grundy, Jack A Taylor, D Neil Hayes, David Malkin, Christopher S Foster, Anne Y Warren, Hayley C Whitaker, Daniel Brewer, Rosalind Eeles, Colin Cooper, David Neal, Tapio Visakorpi, William B Isaacs, G Steven Bova, Adrienne M Flanagan, P Andrew Futreal, Andy G Lynch, Patrick F Chinnery, Ultan McDermott, Michael R Stratton, and Peter J Campbell. Origins and functional consequences of somatic mitochondrial DNA mutations in human cancer. *eLife*, 3, 2014.

[456] Megumi Kaneko and Michael P Stryker. Sensory experience during locomotion promotes recovery of function in adult visual cortex. *eLife*, 3, 2014.

[457] Vasily Kantsler, Jörn Dunkel, Martyn Blayney, and Raymond E Goldstein. Rheotaxis facilitates upstream navigation of mammalian sperm cells. *eLife*, 3, 2014.

[458] Mihoko Kato, Tsui-Fen Chou, Collin Z Yu, John DeModena, and Paul W Sternberg. LINKIN, a new transmembrane protein necessary for cell adhesion. *eLife*, 3, 2014.

[459] Yarden Katz, Feifei Li, Nicole J Lambert, Ethan S Sokol, Wai-Leong Tam, Albert W Cheng, Edoardo M Airoldi, Christopher J Lengner, Piyush B Gupta, Zhengquan Yu, Rudolf Jaenisch, and Christopher B Burge. Musashi proteins are post-transcriptional regulators of the epithelial-luminal cell state. *eLife*, 3, 2014.

[460] Satdip Kaur, Andrew B Fielding, Gisela Gassner, Nicholas J Carter, and Stephen J Royle. An unmet actin requirement explains the mitotic inhibition of clathrin-mediated endocytosis. *eLife*, 3, 2014.

[461] Jennifer M Kavran, Jacqueline M McCabe, Patrick O Byrne, Mary Katherine Connacher, Zhihong Wang, Alexander Ramek, Sarvenaz Sarabipour, Yibing Shan, David E Shaw, Kalina Hristova, Philip A Cole, and Daniel J Leahy. How IGF-1 activates its receptor. *eLife*, 3, 2014.

[462] Tomokazu Kawashima, Daisuke Maruyama, Murat Shagirov, Jing Li, Yuki Hamamura, Ramesh Yelagandula, Yusuke Toyama, and Frédéric Berger. Dynamic F-actin movement is essential for fertilization in Arabidopsis thaliana. *eLife*, 3, 2014.

[463] Roberto A Keller, Christian Peeters, and Patrícia Beldade. Evolution of thorax architecture in ant castes highlights trade-off between flight and ground behaviors. *eLife*, 3, 2014.

[464] Mehdi Keramati and Boris Gutkin. Homeostatic reinforcement learning for integrating reward collection and physiological stability. *eLife*, 3, 2014.

[465] Edward Kim, Yuan Wang, Sun-Jung Kim, Miriam Bornhorst, Emmanuelle S Jecrois, Todd E Anthony, Chenran Wang, Yi E Li, Jun-Lin Guan, Geoffrey G Murphy, and Yuan Zhu. Transient inhibition of the ERK pathway prevents cerebellar developmental defects and improves long-term motor functions in murine models of neurofibromatosis type 1. *eLife*, 3, 2014.

[466] Il Hwan Kim, Hong Wang, Scott H Soderling, and Ryohei Yasuda. Loss of Cdc42 leads to defects in synaptic plasticity and remote memory recall. *eLife*, 3, 2014.

[467] Namsuk Kim, Kwang Wook Min, Kyung Hwa Kang, Eun Jung Lee, Hyoung-Tai Kim, Kyunghwan Moon, Jiheon Choi, Dai Le, Sang-Hee Lee, and Jin Woo Kim. Regulation of retinal axon growth by secreted Vax1 homeodomain protein. *eLife*, 3, 2014.

[468] Rhea R Kimpo, Jacob M Rinaldi, Christina K Kim, Hannah L Payne, and Jennifer L Raymond. Gating of neural error signals during motor learning. *eLife*, 3, 2014.

[469] Courtney L Klaips, Megan L Hochstrasser, Christine R Langlois, and Tricia R Serio. Spatial quality control bypasses cell-based limitations on proteostasis to promote prion curing. *eLife*, 3, 2014.

[470] Kathrin Kläsener, Palash C Maity, Elias Hobeika, Jianying Yang, and Michael Reth. B cell activation involves nanoscale receptor reorganizations and inside-out signaling by Syk. *eLife*, 3, 2014.

[471] Zachary A Knight, Sarah F Schmidt, Kivanc Birsoy, Keith Tan, and Jeffrey M Friedman. A critical role for mTORC1 in erythropoiesis and anemia. *eLife*, 3, 2014.

[472] Chwee Tat Koe, Song Li, Fabrizio Rossi, Jack Jing Lin Wong, Yan Wang, Zhizhuo Zhang, Keng Chen, Sherry Shiying Aw, Helena E Richardson, Paul Robson, Wing-Kin Sung, Fengwei Yu, Cayetano Gonzalez, and Hongyan Wang. The Brm-HDAC3-Erm repressor complex suppresses dedifferentiation in Drosophila type II neuroblast lineages. *eLife*, 3, 2014.

[473] Hideyuki Komori, Qi Xiao, Derek H Janssens, Yali Dou, and Cheng-Yu Lee. Trithorax maintains the functional heterogeneity of neural stem cells through the transcription factor Buttonhead. *eLife*, 3, 2014.

[474] Katja Kornysheva and Jörn Diedrichsen. Human premotor areas parse sequences into their spatial and temporal features. *eLife*, 3, 2014.

[475] Takashi Koyama, Marisa A Rodrigues, Alekos Athanasiadis, Alexander W Shingleton, and Christen K Mirth. Nutritional control of body size through FoxO-Ultraspiracle mediated ecdysone biosynthesis. *eLife*, 3, 2014.

[476] Dima Kozakov, Keyong Li, David R Hall, Dmitri Beglov, Jiefu Zheng, Pirooz Vakili, Ora Schueler-Furman, Ioannis Ch Paschalidis, G Marius Clore, and Sandor Vajda. Encounter complexes and dimensionality reduction in protein–protein association. *eLife*, 3, 2014.

[477] Michal Krawczyk and Beverly M Emerson. p50-associated COX-2 extragenic RNA (PACER) activates COX-2 gene expression by occluding repressive NF-$\kappa$B complexes. *eLife*, 3, 2014.

[478] Arnaud R Krebs, Sophie Dessus-Babus, Lukas Burger, and Dirk Schübeler. High-throughput engineering of a mammalian genome reveals building principles of methylation states at CG rich regions. *eLife*, 3, 2014.

[479] Pavan Kumar P, Uchenna Emechebe, Richard Smith, Sarah Franklin, Barry Moore, Mark Yandell, Stephen L Lessnick, and Anne M Moon. Coordinated control of senescence by lncRNA and a novel T-box3 co-repressor complex. *eLife*, 3, 2014.

[480] Snehlata Kumari, Younes Redouane, Jaime Lopez-Mosqueda, Ryoko Shiraishi, Malgorzata Romanowska, Stefan Lutzmayer, Jan Kuiper, Conception Martinez, Ivan Dikic, Manolis Pasparakis, and Fumiyo Ikeda. Sharpin prevents skin inflammation by inhibiting TNFR1-induced keratinocyte apoptosis. *eLife*, 3, 2014.

[481] Sen-Lin Lai and Chris Q Doe. Transient nuclear Prospero induces neural progenitor quiescence. *eLife*, 3, 2014.

[482] Ying Lai, Jiajie Diao, Daniel J Cipriano, Yunxiang Zhang, Richard A Pfuetzner, Mark S Padolina, and Axel T Brunger. Complexin inhibits spontaneous release and synchronizes Ca2+-triggered synaptic vesicle fusion by distinct mechanisms. *eLife*, 3, 2014.

[483] Harriet Lane-Serff, Paula MacGregor, Edward D Lowe, Mark Carrington, and Matthew K Higgins. Structural basis for ligand and innate immunity factor uptake by the trypanosome haptoglobin-haemoglobin receptor. *eLife*, 3, 2014.

[484] Lars Langemeyer, Ricardo Nunes Bastos, Yiying Cai, Aymelt Itzen, Karin M Reinisch, and Francis A Barr. Diversity and plasticity in Rab GTPase nucleotide release mechanism has consequences for Rab activation and inactivation. *eLife*, 3, 2014.

[485] Liana F Lareau, Dustin H Hite, Gregory J Hogan, and Patrick O Brown. Distinct stages of the translation elongation cycle revealed by sequencing ribosome-protected mRNA fragments. *eLife*, 3, 2014.

[486] Sofía Lavista-Llanos, AleŠ SvatoŠ, Marco Kai, Thomas Riemensperger, Serge Birman, Marcus C Stensmyr, and Bill S Hansson. Dopamine drives Drosophila sechellia adaptation to its toxic host. *eLife*, 3, 2014.

[487] Brigitte LeBoeuf, Paola Correa, Changhoon Jee, and L René García. Caenorhabditis elegans male sensory-motor neurons and dopaminergic support cells couple ejaculation and post-ejaculatory behaviors. *eLife*, 3, 2014.

[488] Lauren LeBon, Tom V Lee, David Sprinzak, Hamed Jafar-Nejad, and Michael B Elowitz. Fringe proteins modulate Notch-ligand cis and trans interactions to specify signaling states. *eLife*, 3, 2014.

[489] Jeremy Leitz and Ege T Kavalali. Fast retrieval and autonomous regulation of single spontaneously recycling synaptic vesicles. *eLife*, 3, 2014.

[490] Félix Leroy, Boris Lamotte d'Incamps, Rebecca D Imhoff-Manuel, and Daniel Zytnicki. Early intrinsic hyperexcitability does not contribute to motoneuron degeneration in amyotrophic lateral sclerosis. *eLife*, 3, 2014.

[491] Daine R Lesniak, Kara L Marshall, Scott A Wellnitz, Blair A Jenkins, Yoshichika Baba, Matthew N Rasband, Gregory J Gerling, and Ellen A Lumpkin. Computation identifies structural features that govern neuronal firing properties in slowly adapting touch receptors. *eLife*, 3, 2014.

[492] Carl Leung, Natalya V Dudkina, Natalya Lukoyanova, Adrian W Hodel, Irene Farabella, Arun P Pandurangan, Nasrin Jahan, Mafalda Pires Damaso, Dino Osmanović, Cyril F Reboul, Michelle A Dunstone, Peter W Andrew, Rana Lonnen, Maya Topf, Helen R Saibil, and Bart W Hoogenboom. Stepwise visualization of membrane pore formation by suilysin, a bacterial cholesterol-dependent cytolysin. *eLife*, 3, 2014.

[493] Tera C Levin, Allison J Greaney, Laura Wetzel, and Nicole King. The rosetteless gene controls development in the choanoflagellate S. rosetta. *eLife*, 3, 2014.

[494] Erin D Lew, Jennifer Oh, Patrick G Burrola, Irit Lax, Anna Zagórska, Paqui G Través, Joseph Schlessinger, and Greg Lemke. Differential TAM receptor–ligand–phospholipid interactions delimit differential TAM bioactivities. *eLife*, 3, 2014.

[495] Cedric Leyrat, Max Renner, Karl Harlos, Juha T Huiskonen, and Jonathan M Grimes. Drastic changes in conformational dynamics of the antiterminator M2-1 regulate transcription efficiency in Pneumovirinae. *eLife*, 3, 2014.

[496] Lishi Li and David D Ginty. The structure and organization of lanceolate mechanosensory complexes at mouse hair follicles. *eLife*, 3, 2014.

[497] Qin Li, Sika Zheng, Areum Han, Chia-Ho Lin, Peter Stoilov, Xiang-Dong Fu, and Douglas L Black. The splicing regulator PTBP2 controls a program of embryonic splicing required for neuronal maturation. *eLife*, 3, 2014.

[498] Weida Li, Mio Nakanishi, Adrian Zumsteg, Matthew Shear, Christopher Wright, Douglas A Melton, and Qiao Zhou. In vivo reprogramming of pancreatic acinar cells to three islet endocrine subtypes. *eLife*, 3, 2014.

[499] Xiao-Yong Li, Melissa M Harrison, Jacqueline E Villalta, Tommy Kaplan, and Michael B Eisen. Establishment of regions of genomic activity during the Drosophila maternal to zygotic transition. *eLife*, 3, 2014.

[500] Hoong Chuin Lim, Ivan Vladimirovich Surovtsev, Bruno Gabriel Beltran, Fang Huang, Jörg Bewersdorf, and Christine Jacobs-Wagner. Evidence for a DNA-relay mechanism in ParABS-mediated chromosome segregation. *eLife*, 3, 2014.

[501] Shu-Ting Lin, Luoying Zhang, Xiaoyan Lin, Linda Chen Zhang, Valentina Elizabeth Garcia, Chen-Wei Tsai, Louis Ptáček, and Ying-Hui Fu. Nuclear envelope protein MAN1 regulates clock through BMAL1. *eLife*, 3, 2014.

[502] Steven Lin, Brett T Staahl, Ravi K Alla, and Jennifer A Doudna. Enhanced homology-directed human genome engineering by controlled timing of CRISPR/Cas9 delivery. *eLife*, 3, 2014.

[503] Tien-chen Lin, Annett Neuner, Yvonne T Schlosser, Annette ND Scharf, Lisa Weber, and Elmar Schiebel. Cell-cycle dependent phosphorylation of yeast pericentrin regulates $\gamma$-TuSC-mediated microtubule nucleation. *eLife*, 3, 2014.

[504] Michelle A Linterman, Alice E Denton, Devina P Divekar, Ilona Zvetkova, Leanne Kane, Cristina Ferreira, Marc Veldhoen, Simon Clare, Gordon Dougan, Marion Espéli, and Kenneth GC Smith. CD28 expression is required after T cell priming for helper T cell responses and protective immunity to infection. *eLife*, 3, 2014.

[505] Huisheng Liu, Hua Bai, Enfu Hui, Lu Yang, Chantell S Evans, Zhao Wang, Sung E Kwon, and Edwin R Chapman. Synaptotagmin 7 functions as a Ca2+-sensor for synaptic vesicle replenishment. *eLife*, 3, 2014.

[506] Tingting Liu, Ariel Kaplan, Lisa Alexander, Shannon Yan, Jin-Der Wen, Laura Lancaster, Charles E Wickersham, Kurt Fredrick, Harry Noller, Ignacio Tinoco, and Carlos J Bustamante. Direct measurement of the mechanical work during translocation by the ribosome. *eLife*, 3, 2014.

[507] Xiaokun Liu, Heini M Grabherr, Roland Willmann, Dagmar Kolb, Frédéric Brunner, Ute Bertsche, Daniel Kühner, Mirita Franz-Wachtel, Bushra Amin, Georg Felix, Marc Ongena, Thorsten Nürnberger, and Andrea A Gust. Host-induced bacterial cell wall decomposition mediates pattern-triggered immunity in Arabidopsis. *eLife*, 3, 2014.

[508] Yanfen Liu, Nia Soetandyo, Jin-gu Lee, Liping Liu, Yue Xu, William M Clemons, and Yihong Ye. USP13 antagonizes gp78 to maintain functionality of a chaperone in ER-associated degradation. *eLife*, 3, 2014.

[509] Zhe Liu, Wesley R Legant, Bi-Chang Chen, Li Li, Jonathan B Grimm, Luke D Lavis, Eric Betzig, and Robert Tjian. 3D imaging of Sox2 enhancer clusters in embryonic stem cells. *eLife*, 3, 2014.

[510] Wan-Lin Lo, Benjamin D Solomon, David L Donermeyer, Chyi-Song Hsieh, and Paul M Allen. T cell immunodominance is dictated by the positively selecting self-peptide. *eLife*, 3, 2014.

[511] Braden T Lobingier, Daniel P Nickerson, Sheng-Ying Lo, and Alexey J Merz. SM proteins Sly1 and Vps33 co-assemble with Sec17 and SNARE complexes to oppose SNARE disassembly by Sec18. *eLife*, 3, 2014.

[512] Irakli Loladze. Hidden shift of the ionome of plants exposed to elevated CO2 depletes minerals at the base of human nutrition. *eLife*, 3, 2014.

[513] David R Lorenz, Lauren F Meyer, Patrick J R Grady, Michelle M Meyer, and Hugh P Cam. Heterochromatin assembly and transcriptome repression by Set1 in coordination with a class II histone deacetylase. *eLife*, 3, 2014.

[514] Tony Ly, Yasmeen Ahmad, Adam Shlien, Dominique Soroka, Allie Mills, Michael J Emanuele, Michael R Stratton, and Angus I Lamond. A proteomic chronology of gene expression through the cell cycle in human myeloid leukemia cells. *eLife*, 3, 2014.

[515] Benjamin R Macadangdang, Amit Oberai, Tanya Spektor, Oscar A Campos, Fang Sheng, Michael F Carey, Maria Vogelauer, and Siavash K Kurdistani. Evolution of histone 2A for chromatin compaction in eukaryotes. *eLife*, 3, 2014.

[516] Andrew L MacKinnon, Ville O Paavilainen, Ajay Sharma, Ramanujan S Hegde, and Jack Taunton. An allosteric Sec61 inhibitor traps nascent transmembrane helices at the lateral gate. *eLife*, 3, 2014.

[517] Anna L Mallam, David J Sidote, and Alan M Lambowitz. Molecular insights into RNA and DNA helicase evolution from the determinants of specificity for a DEAD-box RNA helicase. *eLife*, 3, 2014.

[518] Julia Mallmann, David Heckmann, Andrea Bräutigam, Martin J Lercher, Andreas PM Weber, Peter Westhoff, and Udo Gowik. The role of photorespiration during the evolution of C4 photosynthesis in the genus Flaveria. *eLife*, 3, 2014.

[519] Jennifer L Mamrosh, Jae Man Lee, Martin Wagner, Peter J Stambrook, Richard J Whitby, Richard N Sifers, San-Pin Wu, Ming-Jer Tsai, Francesco J DeMayo, and David D Moore. Nuclear receptor LRH-1/NR5A2 is required and targetable for liver endoplasmic reticulum stress resolution. *eLife*, 3, 2014.

[520] Yael Mandelblat-Cerf, Liora Las, Natalia Denisenko, and Michale S Fee. A role for descending auditory cortical projections in songbird vocal learning. *eLife*, 3, 2014.

[521] Niall M Mangan and Michael P Brenner. Systems analysis of the CO2 concentrating mechanism in cyanobacteria. *eLife*, 3, 2014.

[522] Martial Marbouty, Axel Cournac, Jean-FranÇois Flot, Hervé Marie-Nelly, Julien Mozziconacci, and Romain Koszul. Metagenomic chromosome conformation capture (meta3C) unveils the diversity of chromosome organization in microorganisms. *eLife*, 3, 2014.

[523] Jana Dorfman Marcette, Jessica Jie Chen, and Michael L Nonet. The Caenorhabditis elegans microtubule minus-end binding homolog PTRN-1 stabilizes synapses and neurites. *eLife*, 3, 2014.

[524] Karolina Marciniak, Artin Atabaki, Peter W Dicke, and Peter Thier. Disparate substrates for head gaze following and face perception in the monkey superior temporal sulcus. *eLife*, 3, 2014.

[525] Magdalena M Maslon, Sara R Heras, Nicolas Bellora, Eduardo Eyras, and Javier F Cáceres. The translational landscape of the splicing factor SRSF1 and its role in mitosis. *eLife*, 3, 2014.

[526] Joshua D Mast, Consuelo M De Moraes, Hans T Alborn, Luke D Lavis, and David L Stern. Evolved differences in larval social behavior mediated by novel pheromones. *eLife*, 3, 2014.

[527] Maja Matis, David A Russler-Germain, Qie Hu, Claire J Tomlin, and Jeffrey D Axelrod. Microtubules provide directional information for core PCP function. *eLife*, 3, 2014.

[528] Juliana L Matos, On Sun Lau, Charles Hachez, Alfredo Cruz-Ramírez, Ben Scheres, and Dominique C Bergmann. Irreversible fate commitment in the Arabidopsis stomatal lineage requires a FAMA and RETINOBLASTOMA-RELATED module. *eLife*, 3, 2014.

[529] Yuval Mazor, Daniel Nataf, Hila Toporik, and Nathan Nelson. Crystal structures of virus-like photosystem I complexes from the mesophilic cyanobacterium Synechocystis PCC 6803. *eLife*, 3, 2014.

[530] Shawn McClelland, Gary P Brennan, Celine Dubé, Seeta Rajpara, Shruti Iyer, Cristina Richichi, Christophe Bernard, and Tallie Z Baram. The transcription factor NRSF contributes to epileptogenesis by selective repression of a subset of target genes. *eLife*, 3, 2014.

[531] Michael A McDannald, Guillem R Esber, Meredyth A Wegener, Heather M Wied, Tzu-Lan Liu, Thomas A Stalnaker, Joshua L Jones, Jason Trageser, and Geoffrey Schoenbaum. Orbitofrontal neurons acquire responses to 'valueless' Pavlovian cues during unblocking. *eLife*, 3, 2014.

[532] James C McGann, Jon A Oyer, Saurabh Garg, Huilan Yao, Jun Liu, Xin Feng, Lujian Liao, John R Yates, and Gail Mandel. Polycomb- and REST-associated histone deacetylases are independent pathways toward a mature neuronal phenotype. *eLife*, 3, 2014.

[533] Rhoanne C McPherson, Joanne E Konkel, Catriona T Prendergast, John P Thomson, Raffaele Ottaviano, Melanie D Leech, Oliver Kay, Stephanie E J Zandee, Claire H Sweenie, David C Wraith, Richard R Meehan, Amanda J Drake, and Stephen M Anderton. Epigenetic modification of the PD-1 (Pdcd1) promoter in effector CD4+ T cells tolerized by peptide immunotherapy. *eLife*, 3, 2014.

[534] Patrick J Mears, Santosh Koirala, Chris V Rao, Ido Golding, and Yann R Chemla. Escherichia coli swimming is robust against variations in flagellar number. *eLife*, 3, 2014.

[535] Sohum Mehta, Nwe-Nwe Aye-Han, Ambhighainath Ganesan, Laurel Oldach, Kirill Gorshkov, and Jin Zhang. Calmodulin-controlled spatial decoding of oscillatory Ca2+ signals by calcineurin. *eLife*, 3, 2014.

[536] Louise Meigh, Naveed Hussain, Daniel K Mulkey, and Nicholas Dale. Connexin26 hemichannels with a mutation that causes KID syndrome in humans lack sensitivity to CO2. *eLife*, 3, 2014.

[537] Romain Mercier, Yoshikazu Kawai, and Jeff Errington. General principles for the formation and proliferation of a wall-free (L-form) state in bacteria. *eLife*, 3, 2014.

[538] Mirko Messa, Rubén Fernández-Busnadiego, Elizabeth Wen Sun, Hong Chen, Heather Czapla, Kristie Wrasman, Yumei Wu, Genevieve Ko, Theodora Ross, Beverly Wendland, and Pietro De Camilli. Epsin deficiency impairs endocytosis by stalling the actin-dependent invagination of endocytic clathrin-coated pits. *eLife*, 3, 2014.

[539] Jason A Metcalf, Lisa J Funkhouser-Jones, Kristen Brileya, Anna-Louise Reysenbach, and Seth R Bordenstein. Antibacterial gene transfer across the tree of life. *eLife*, 3, 2014.

[540] John K Mich, Robert AJ Signer, Daisuke Nakada, André Pineda, Rebecca J Burgess, Tou Yia Vue, Jane E Johnson, and Sean J Morrison. Prospective identification of functionally distinct stem cells and neurosphere-initiating cells in adult mouse forebrain. *eLife*, 3, 2014.

[541] Joshua K Michener, Aline A Camargo Neves, Stéphane Vuilleumier, FranÇoise Bringel, and Christopher J Marx. Effective use of a horizontally-transferred pathway for dichloromethane catabolism requires post–transfer refinement. *eLife*, 3, 2014.

[542] Oliver H Miller, Lingling Yang, Chih-Chieh Wang, Elizabeth A Hargroder, Yihui Zhang, Eric Delpire, and Benjamin J Hall. GluN2B-containing NMDA receptors regulate depression-like behavior and are critical for the rapid antidepressant actions of ketamine. *eLife*, 3, 2014.

[543] Smriti Mishra, Brandy Verhalen, Richard A Stein, Po-Chao Wen, Emad Tajkhorshid, and Hassane S Mchaourab. Conformational dynamics of the nucleotide binding domains and the power stroke of a heterodimeric ABC transporter. *eLife*, 3, 2014.

[544] Christine Missbach, Hany KM Dweck, Heiko Vogel, Andreas Vilcinskas, Marcus C Stensmyr, Bill S Hansson, and Ewald Grosse-Wilde. Evolution of insect olfactory receptors. *eLife*, 3, 2014.

[545] Mehrab N Modi, Ashesh K Dhawale, and Upinder S Bhalla. CA1 cell activity sequences emerge after reorganization of network correlation structure during associative learning. *eLife*, 3, 2014.

[546] Bama Charan Mondal, Jiwon Shim, Cory J Evans, and Utpal Banerjee. Pvr expression regulators in equilibrium signal control and maintenance of Drosophila blood progenitors. *eLife*, 3, 2014.

[547] Felipe Mora-Bermúdez, Fumio Matsuzaki, and Wieland B Huttner. Specific polar subpopulations of astral microtubules control spindle orientation and symmetric neural stem cell division. *eLife*, 3, 2014.

[548] Meghan Morgan-Smith, Yaohong Wu, Xiaoqin Zhu, Julia Pringle, and William D Snider. GSK-3 signaling in developing cortical neurons is essential for radial migration and dendritic orientation. *eLife*, 3, 2014.

[549] Sean J Morrison. Time to do something about reproducibility. *eLife*, 3, 2014.

[550] Timothy J Mosca and Liqun Luo. Synaptic organization of the Drosophila antennal lobe and its regulation by the Teneurins. *eLife*, 3, 2014.

[551] Alexander Muir, Subramaniam Ramachandran, FranÇoise M Roelants, Garrett Timmons, and Jeremy Thorner. TORC2-dependent protein kinase Ypk1 phosphorylates ceramide synthase to stimulate synthesis of complex sphingolipids. *eLife*, 3, 2014.

[552] Shankar Mukherji and Erin K O'Shea. Mechanisms of organelle biogenesis govern stochastic fluctuations in organelle abundance. *eLife*, 3, 2014.

[553] Alexandra Naba, Karl R Clauser, John M Lamar, Steven A Carr, and Richard O Hynes. Extracellular matrix signatures of human mammary carcinoma identify novel metastasis promoters. *eLife*, 3, 2014.

[554] Sundar Ram Naganathan, Sebastian Fürthauer, Masatoshi Nishikawa, Frank Jülicher, and Stephan W Grill. Active torque generation by the actomyosin cell cortex drives left–right symmetry breaking. *eLife*, 3, 2014.

[555] Stanislav Nagy, Nora Tramm, Jarred Sanders, Shachar Iwanir, Ian A Shirley, Erel Levine, and David Biron. Homeostasis in C. elegans sleep is characterized by two behaviorally and genetically distinct mechanisms. *eLife*, 3, 2014.

[556] Farzaneh Najafi, Andrea Giovannucci, Samuel S-H Wang, and Javier F Medina. Coding of stimulus strength via analog calcium signals in Purkinje cell dendrites of awake mice. *eLife*, 3, 2014.

[557] Ramiro Nández, Daniel M Balkin, Mirko Messa, Liang Liang, Summer Paradise, Heather Czapla, Marco Y Hein, James S Duncan, Matthias Mann, and Pietro De Camilli. A role of OCRL in clathrin-coated pit dynamics and uncoating revealed by studies of Lowe syndrome cells. *eLife*, 3, 2014.

[558] Shannon N Nangle, Clark Rosensweig, Nobuya Koike, Hajime Tei, Joseph S Takahashi, Carla B Green, and Ning Zheng. Molecular assembly of the period-cryptochrome circadian transcriptional repressor complex. *eLife*, 3, 2014.

[559] Brent L Nannenga, Dan Shi, Johan Hattne, Francis E Reyes, and Tamir Gonen. Structure of catalase determined by MicroED. *eLife*, 3, 2014.

[560] Anil M Narasimha, Manuel Kaulich, Gary S Shapiro, Yoon J Choi, Piotr Sicinski, and Steven F Dowdy. Cyclin D activates the Rb tumor suppressor by mono-phosphorylation. *eLife*, 3, 2014.

[561] Elin Násström, Nga Tran Vu Thieu, Sabina Dongol, Abhilasha Karkey, Phat Voong Vinh, Tuyen Ha Thanh, Anders Johansson, Amit Arjyal, Guy Thwaites, Christiane Dolecek, Buddha Basnyat, Stephen Baker, and Henrik Antti. Salmonella Typhi and Salmonella Paratyphi A elaborate distinct systemic metabolite signatures during enteric fever. *eLife*, 3, 2014.

[562] Mei Rosa Ng, Achim Besser, Joan S Brugge, and Gaudenz Danuser. Mapping the dynamics of force transduction at cell–cell junctions of epithelial clusters. *eLife*, 3, 2014.

[563] David P Nguyen and Shih-Chieh Lin. A frontal cortex event-related potential driven by the basal forebrain. *eLife*, 3, 2014.

[564] Tania Nguyen, Harry Fischl, FranÇoise S Howe, Ronja Woloszczuk, Ana Serra Barros, Zhenyu Xu, David Brown, Struan C Murray, Simon Haenni, James M Halstead, Leigh O'Connor, Gergana Shipkovenska, Lars M Steinmetz, and Jane Mellor. Transcription mediated insulation and interference direct gene cluster expression switches. *eLife*, 3, 2014.

[565] Simone A Nish, Dominik Schenten, F Thomas Wunderlich, Scott D Pope, Yan Gao, Namiko Hoshi, Shuang Yu, Xiting Yan, Heung Kyu Lee, Lesley Pasman, Igor Brodsky, Brian Yordy, Hongyu Zhao, Jens Brüning, and Ruslan Medzhitov. T cell-intrinsic role of IL-6 signaling in primary and memory responses. *eLife*, 3, 2014.

[566] Cristina Nogueira, Patrik Erlmann, Julien Villeneuve, António JM Santos, Emma Martínez-Alonso, José ángel Martínez-Menárguez, and Vivek Malhotra. SLY1 and Syntaxin 18 specify a distinct pathway for procollagen VII export from the endoplasmic reticulum. *eLife*, 3, 2014.

[567] Daan Noordermeer, Marion Leleu, Patrick Schorderet, Elisabeth Joye, Fabienne Chabaud, and Denis Duboule. Temporal dynamics and developmental memory of 3D chromatin architecture at Hox gene loci. *eLife*, 3, 2014.

[568] Thomas R Noriega, Jin Chen, Peter Walter, and Joseph D Puglisi. Real-time observation of signal recognition particle binding to actively translating ribosomes. *eLife*, 3, 2014.

[569] Jerome C Nwachukwu, Sathish Srinivasan, Nelson E Bruno, Alexander A Parent, Travis S Hughes, Julie A Pollock, Olsi Gjyshi, Valerie Cavett, Jason Nowak, Ruben D Garcia-Ordonez, René Houtman, Patrick R Griffin, Douglas J Kojetin, John A Katzenellenbogen, Michael D Conkright, and Kendall W Nettles. Resveratrol modulates the inflammatory response via an estrogen receptor-signal integration network. *eLife*, 3, 2014.

[570] Kamila Oglęcka, Padmini Rangamani, Bo Liedberg, Rachel S Kraut, and Atul N Parikh. Oscillatory phase separation in giant lipid vesicles induced by transmembrane osmotic differentials. *eLife*, 3, 2014.

[571] Eunkyoo Oh, Jia-Ying Zhu, Ming-Yi Bai, Rafael Augusto Arenhart, Yu Sun, and Zhi-Yong Wang. Cell elongation is regulated through a central circuit of interacting transcription factors in the Arabidopsis hypocotyl. *eLife*, 3, 2014.

[572] William H Olds and Tian Xu. Regulation of food intake by mechanosensory ion channels in enteric neurons. *eLife*, 3, 2014.

[573] Llwyd David Orton and Adrian Rees. Intercollicular commissural connections refine the representation of sound frequency and level in the auditory midbrain. *eLife*, 3, 2014.

[574] Olga Ostrovskaya, Keqiang Xie, Ikuo Masuho, Ana Fajardo-Serrano, Rafael Lujan, Kevin Wickman, and Kirill A Martemyanov. RGS7/G$\beta$5/R7BP complex regulates synaptic plasticity and memory by modulating hippocampal GABABR-GIRK signaling. *eLife*, 3, 2014.

[575] Sergey Ovchinnikov, Hetunandan Kamisetty, and David Baker. Robust and accurate prediction of residue–residue interactions across protein interfaces using evolutionary information. *eLife*, 3, 2014.

[576] William Hunt Palmer, Dongyu Jia, and Wu-Min Deng. Cis-interactions between Notch and its ligands block ligand-independent Notch activity. *eLife*, 3, 2014.

[577] Nicola Palmieri, Carolin Kosiol, and Christian Schlötterer. The life cycle of Drosophila orphan genes. *eLife*, 3, 2014.

[578] Kally Z Pan, Timothy E Saunders, Ignacio Flor-Parra, Martin Howard, and Fred Chang. Cortical regulation of cell size by a sizer cdr2p. *eLife*, 3, 2014.

[579] Romeo Papazyan, Ekaterina Voronina, Jessica R Chapman, Teresa R Luperchio, Tonya M Gilbert, Elizabeth Meier, Samuel G Mackintosh, Jeffrey Shabanowitz, Alan J Tackett, Karen L Reddy, Robert S Coyne, Donald F Hunt, Yifan Liu, and Sean D Taverna. Methylation of histone H3K23 blocks DNA damage in pericentric heterochromatin during meiosis. *eLife*, 3, 2014.

[580] Joanne L Parker, Joseph A Mindell, and Simon Newstead. Thermodynamic evidence for a dual transport mechanism in a POT peptide transporter. *eLife*, 3, 2014.

[581] James R Partridge, Laura A Lavery, Daniel Elnatan, Nariman Naber, Roger Cooke, and David A Agard. A novel N-terminal extension in mitochondrial TRAP1 serves as a thermal regulator of chaperone activity. *eLife*, 3, 2014.

[582] Cristina Paulino and Werner Kühlbrandt. pH- and sodium-induced changes in a sodium/proton antiporter. *eLife*, 3, 2014.

[583] Cristina Paulino, David Wöhlert, Ekaterina Kapotova, özkan Yildiz, and Werner Kühlbrandt. Structure and transport mechanism of the sodium/proton antiporter MjNhaP1. *eLife*, 3, 2014.

[584] Christopher J Peck and C Daniel Salzman. Amygdala neural activity reflects spatial attention towards stimuli promising reward or threatening punishment. *eLife*, 3, 2014.

[585] Ke Peng, Walter Muranyi, Bärbel Glass, Vibor Laketa, Stephen R Yant, Luong Tsai, Tomas Cihlar, Barbara Müller, and Hans-Georg Kräusslich. Quantitative microscopy of functional HIV post-entry complexes reveals association of replication with the viral capsid. *eLife*, 3, 2014.

[586] Ivana Petrovska, Elisabeth Nüske, Matthias C Munder, Gayathrie Kulasegaran, Liliana Malinovska, Sonja Kroschwald, Doris Richter, Karim Fahmy, Kimberley Gibson, Jean-Marc Verbavatz, and Simon Alberti. Filament formation by metabolic enzymes is a specific adaptation to an advanced state of cellular starvation. *eLife*, 3, 2014.

[587] Alexandre Pfister, Marie Barberon, Julien Alassimone, Lothar Kalmbach, Yuree Lee, Joop EM Vermeer, Misako Yamazaki, Guowei Li, Christophe Maurel, Junpei Takano, Takehiro Kamiya, David E Salt, Daniele Roppolo, and Niko Geldner. A receptor-like kinase mutant with absent endodermal diffusion barrier displays selective nutrient homeostasis defects. *eLife*, 3, 2014.

[588] Hyewon Phee, Byron B Au-Yeung, Olga Pryshchep, Kyle Leonard O'Hagan, Stephanie Grace Fairbairn, Maria Radu, Rachelle Kosoff, Marianne Mollenauer, Debra Cheng, Jonathan Chernoff, and Arthur Weiss. Pak2 is required for actin cytoskeleton remodeling, TCR signaling, and normal thymocyte development and maturation. *eLife*, 3, 2014.

[589] Daniela Pignatta, Robert M Erdmann, Elias Scheer, Colette L Picard, George W Bell, and Mary Gehring. Natural epigenetic polymorphisms lead to intraspecific variation in Arabidopsis gene imprinting. *eLife*, 3, 2014.

[590] David M Pigott, Samir Bhatt, Nick Golding, Kirsten A Duda, Katherine E Battle, Oliver J Brady, Jane P Messina, Yves Balard, Patrick Bastien, Francine Pratlong, John S Brownstein, Clark C Freifeld, Sumiko R Mekaru, Peter W Gething, Dylan B George, Monica F Myers, Richard Reithinger, and Simon I Hay. Global distribution maps of the leishmaniases. *eLife*, 3, 2014.

[591] David M Pigott, Nick Golding, Adrian Mylne, Zhi Huang, Andrew J Henry, Daniel J Weiss, Oliver J Brady, Moritz UG Kraemer, David L Smith, Catherine L Moyes, Samir Bhatt, Peter W Gething, Peter W Horby, Isaac I Bogoch, John S Brownstein, Sumiko R Mekaru, Andrew J Tatem, Kamran Khan, and Simon I Hay. Mapping the zoonotic niche of Ebola virus disease in Africa. *eLife*, 3, 2014.

[592] Ivan KH Poon, Amy A Baxter, Fung T Lay, Grant D Mills, Christopher G Adda, Jennifer AE Payne, Thanh Kha Phan, Gemma F Ryan, Julie A White, Prem K Veneer, Nicole L van der Weerden, Marilyn A Anderson, Marc Kvansakul, and Mark D Hulett. Phosphoinositide-mediated oligomerization of a defensin induces cell lysis. *eLife*, 3, 2014.

[593] Jeremy L Praissman, David H Live, Shuo Wang, Annapoorani Ramiah, Zoeisha S Chinoy, Geert-Jan Boons, Kelley W Moremen, and Lance Wells. B4GAT1 is the priming enzyme for the LARGE-dependent functional glycosylation of $\alpha$-dystroglycan. *eLife*, 3, 2014.

[594] Daniel J Puleston, Hanlin Zhang, Timothy J Powell, Elina Lipina, Stuart Sims, Isabel Panse, Alexander S Watson, Vincenzo Cerundolo, Alain RM Townsend, Paul Klenerman, and Anna Katharina Simon. Autophagy is a critical regulator of memory CD8+ T cell formation. *eLife*, 3, 2014.

[595] Vladimir Purvanov, Manuel Holst, Jameel Khan, Christian Baarlink, and Robert Grosse. G-protein-coupled receptor signaling and polarized actin dynamics drive cell-in-cell invasion. *eLife*, 3, 2014.

[596] Yun Qing, Feng Yin, Wei Wang, Yonggang Zheng, Pengfei Guo, Frederick Schozer, Hua Deng, and Duojia Pan. The Hippo effector Yorkie activates transcription by interacting with a histone methyltransferase complex through Ncoa6. *eLife*, 3, 2014.

[597] Shu Quan, Lili Wang, Evgeniy V Petrotchenko, Karl AT Makepeace, Scott Horowitz, Jianyi Yang, Yang Zhang, Christoph H Borchers, and James CA Bardwell. Super Spy variants implicate flexibility in chaperone action. *eLife*, 3, 2014.

[598] Delphine Quénet and Yamini Dalal. A long non-coding RNA is required for targeting centromeric protein A to the human centromere. *eLife*, 3, 2014.

[599] Senthil K Radhakrishnan, Willem den Besten, and Raymond J Deshaies. p97-dependent retrotranslocation and proteolytic processing govern formation of active Nrf1 upon proteasome inhibition. *eLife*, 3, 2014.

[600] Michael B Radke, Manuel H Taft, Britta Stapel, Denise Hilfiker-Kleiner, Matthias Preller, and Dietmar J Manstein. Small molecule-mediated refolding and activation of myosin motor function. *eLife*, 3, 2014.

[601] Daniel T Radoff, Yongming Dong, David Snead, Jihong Bai, David Eliezer, and Jeremy S Dittman. The accessory helix of complexin functions by stabilizing central helix secondary structure. *eLife*, 3, 2014.

[602] Nadine Randel, Albina Asadulina, Luis A Bezares-Calderón, Csaba Verasztó, Elizabeth A Williams, Markus Conzelmann, Réza Shahidi, and Gáspár Jékely. Neuronal connectome of a sensory-motor circuit for visual navigation. *eLife*, 3, 2014.

[603] Rajnish P Rao, Falk Mielke, Evgeny Bobrov, and Michael Brecht. Vocalization–whisking coordination and multisensory integration of social signals in rat auditory cortex. *eLife*, 3, 2014.

[604] Stéphanie Ratté, Yi Zhu, Kwan Yeop Lee, and Steven A Prescott. Criticality and degeneracy in injury-induced changes in primary afferent excitability and the implications for neuropathic pain. *eLife*, 3, 2014.

[605] Sarina Ravens, Marjorie Fournier, Tao Ye, Matthieu Stierle, Doulaye Dembele, Virginie Chavant, and Làszlò Tora. Mof-associated complexes have overlapping and unique roles in regulating pluripotency in embryonic stem cells and during differentiation. *eLife*, 3, 2014.

[606] Xuefeng Ren, Sang Yoon Park, Juan S Bonifacino, and James H Hurley. How HIV-1 Nef hijacks the AP-2 clathrin adaptor to downregulate CD4. *eLife*, 3, 2014.

[607] Claire E Richardson, Kerri A Spilker, Juan G Cueva, John Perrino, Miriam B Goodman, and Kang Shen. PTRN-1, a microtubule minus end-binding CAMSAP homolog, promotes microtubule function in Caenorhabditis elegans neurons. *eLife*, 3, 2014.

[608] Sandra R Richardson, Iñigo Narvaiza, Randy A Planegger, Matthew D Weitzman, and John V Moran. APOBEC3A deaminates transiently exposed single-strand DNA during LINE-1 retrotransposition. *eLife*, 3, 2014.

[609] Sandra Richter, Marika Kientz, Sabine Brumm, Mads Eggert Nielsen, Misoon Park, Richard Gavidia, Cornelia Krause, Ute Voss, Hauke Beckmann, Ulrike Mayer, York-Dieter Stierhof, and Gerd Jürgens. Delivery of endocytosed proteins to the cell–division plane requires change of pathway from recycling to secretion. *eLife*, 3, 2014.

[610] James A Rickard, Holly Anderton, Nima Etemadi, Ueli Nachbur, Maurice Darding, Nieves Peltzer, Najoua Lalaoui, Kate E Lawlor, Hannah Vanyai, Cathrine Hall, Aleks Bankovacki, Lahiru Gangoda, Wendy Wei-Lynn Wong, Jason Corbin, Chunzi Huang, Edward S Mocarski, James M Murphy, Warren S Alexander, Anne K Voss, David L Vaux, William J Kaiser, Henning Walczak, and John Silke. TNFR1-dependent cell death drives inflammation in Sharpin-deficient mice. *eLife*, 3, 2014.

[611] Martina Katharina Ried, Meritxell Antolín-Llovera, and Martin Parniske. Spontaneous symbiotic reprogramming of plant roots triggered by receptor-like kinases. *eLife*, 3, 2014.

[612] W Christopher Risher, Sagar Patel, Il Hwan Kim, Akiyoshi Uezu, Srishti Bhagat, Daniel K Wilton, Louis-Jan Pilaz, Jonnathan Singh Alvarado, Osman Y Calhan, Debra L Silver, Beth Stevens, Nicole Calakos, Scott H Soderling, and Cagla Eroglu. Astrocytes refine cortical connectivity at dendritic spines. *eLife*, 3, 2014.

[613] Anthony J Roberts, Brian S Goodman, and Samara L Reck-Peterson. Reconstitution of dynein transport to the microtubule plus end by kinesin. *eLife*, 3, 2014.

[614] Frederic A Roemschied, Monika JB Eberhard, Jan-Hendrik Schleimer, Bernhard Ronacher, and Susanne Schreiber. Cell-intrinsic mechanisms of temperature compensation in a grasshopper sensory receptor neuron. *eLife*, 3, 2014.

[615] Alexandre B Roland, Ana Ricobaraza, Damien Carrel, Benjamin M Jordan, Felix Rico, Anne Simon, Marie Humbert-Claude, Jeremy Ferrier, Maureen H McFadden, Simon Scheuring, and Zsolt Lenkei. Cannabinoid-induced actomyosin contractility shapes neuronal morphology and growth. *eLife*, 3, 2014.

[616] Giselle Roman-Hernandez, Janine H Peterson, and Harris D Bernstein. Reconstitution of bacterial autotransporter assembly using purified components. *eLife*, 3, 2014.

[617] Miriam I Rosenberg, Ava E Brent, FranÇois Payre, and Claude Desplan. Dual mode of embryonic development is highlighted by expression and function of Nasonia pair-rule genes. *eLife*, 3, 2014.

[618] Simon Roux, Alyse K Hawley, Monica Torres Beltran, Melanie Scofield, Patrick Schwientek, Ramunas Stepanauskas, Tanja Woyke, Steven J Hallam, and Matthew B Sullivan. Ecology and evolution of viruses infecting uncultivated SUP05 bacteria as revealed by single-cell- and meta-genomics. *eLife*, 3, 2014.

[619] Jorge Ruiz-Orera, Xavier Messeguer, Juan Antonio Subirana, and M Mar Alba. Long non-coding RNAs as a source of new peptides. *eLife*, 3, 2014.

[620] Colin A Russell, Peter M Kasson, Ruben O Donis, Steven Riley, John Dunbar, Andrew Rambaut, Jason Asher, Stephen Burke, C Todd Davis, Rebecca J Garten, Sandrasegaram Gnanakaran, Simon I

Hay, Sander Herfst, Nicola S Lewis, James O Lloyd-Smith, Catherine A Macken, Sebastian Maurer-Stroh, Elizabeth Neuhaus, Colin R Parrish, Kim M Pepin, Samuel S Shepard, David L Smith, David L Suarez, Susan C Trock, Marc-Alain Widdowson, Dylan B George, Marc Lipsitch, and Jesse D Bloom. Improving pandemic influenza risk assessment. *eLife*, 3, 2014.

[621] Owen W Ryan, Jeffrey M Skerker, Matthew J Maurer, Xin Li, Jordan C Tsai, Snigdha Poddar, Michael E Lee, Will DeLoache, John E Dueber, Adam P Arkin, and Jamie HD Cate. Selection of chromosomal DNA libraries using a multiplex CRISPR system. *eLife*, 3, 2014.

[622] Pedro Saavedra, Jean-Paul Vincent, Isabel M Palacios, Peter A Lawrence, and José Casal. Plasticity of both planar cell polarity and cell identity during the development of Drosophila. *eLife*, 3, 2014.

[623] Akira Sakurai, Arianna N Tamvacakis, and Paul S Katz. Hidden synaptic differences in a neural circuit underlie differential behavioral susceptibility to a neural injury. *eLife*, 3, 2014.

[624] Arun Sampathkumar, Pawel Krupinski, Raymond Wightman, Pascale Milani, Alexandre Berquand, Arezki Boudaoud, Olivier Hamant, Henrik Jönsson, and Elliot M Meyerowitz. Subcellular and supracellular mechanical stress prescribes cytoskeleton behavior in Arabidopsis cotyledon pavement cells. *eLife*, 3, 2014.

[625] Hector Sandoval, Chi-Kuang Yao, Kuchuan Chen, Manish Jaiswal, Taraka Donti, Yong Qi Lin, Vafa Bayat, Bo Xiong, Ke Zhang, Gabriela David, Wu-Lin Charng, Shinya Yamamoto, Lita Duraine, Brett H Graham, and Hugo J Bellen. Mitochondrial fusion but not fission regulates larval growth and synaptic development through steroid hormone production. *eLife*, 3, 2014.

[626] Martial Sankar, Kaisa Nieminen, Laura Ragni, Ioannis Xenarios, and Christian S Hardtke. Automated quantitative histology reveals vascular morphodynamics during Arabidopsis hypocotyl secondary growth. *eLife*, 3, 2014.

[627] Dipali G Sashital, Candacia A Greeman, Dmitry Lyumkis, Clinton S Potter, Bridget Carragher, and James R Williamson. A combined quantitative mass spectrometry and electron microscopy analysis of ribosomal 30S subunit assembly in E. coli. *eLife*, 3, 2014.

[628] Sjors HW Scheres. Beam-induced motion correction for sub-megadalton cryo-EM particles. *eLife*, 3, 2014.

[629] Jens C Schmidt, Andrew B Dalby, and Thomas R Cech. Identification of human TERT elements necessary for telomerase recruitment to telomeres. *eLife*, 3, 2014.

[630] Jessica E Scholey, Stanley Nithianantham, Jonathan M Scholey, and Jawdat Al-Bassam. Structural basis for the assembly of the mitotic motor Kinesin-5 into bipolar tetramers. *eLife*, 3, 2014.

[631] Courtney M Schroeder, Jonathan ML Ostrem, Nicholas T Hertz, and Ronald D Vale. A Ras-like domain in the light intermediate chain bridges the dynein motor to a cargo-binding region. *eLife*, 3, 2014.

[632] Fabian Schueren, Thomas Lingner, Rosemol George, Julia Hofhuis, Corinna Dickel, Jutta Gärtner, and Sven Thoms. Peroxisomal lactate dehydrogenase is generated by translational readthrough in mammals. *eLife*, 3, 2014.

[633] Ursula Schulze-Gahmen, Huasong Lu, Qiang Zhou, and Tom Alber. AFF4 binding to Tat-P-TEFb indirectly stimulates TAR recognition of super elongation complexes at the HIV promoter. *eLife*, 3, 2014.

[634] Molly Schumer, Rongfeng Cui, Daniel L Powell, Rebecca Dresner, Gil G Rosenthal, and Peter Andolfatto. High-resolution mapping reveals hundreds of genetic incompatibilities in hybridizing fish species. *eLife*, 3, 2014.

[635] Sabina Schütz, Ute Fischer, Martin Altvater, Purnima Nerurkar, Cohue Peña, Michaela Gerber, Yiming Chang, Stefanie Caesar, Olga T Schubert, Gabriel Schlenstedt, and Vikram G Panse. A RanGTP-independent mechanism allows ribosomal protein nuclear import for ribosome assembly. *eLife*, 3, 2014.

[636] Kimberley D Seed, Minmin Yen, B Jesse Shapiro, Isabelle J Hilaire, Richelle C Charles, Jessica E Teng, Louise C Ivers, Jacques Boncy, Jason B Harris, and Andrew Camilli. Evolutionary consequences of intra-patient phage predation on microbial populations. *eLife*, 3, 2014.

[637] Andrew M Seeds, Primoz Ravbar, Phuong Chung, Stefanie Hampel, Frank M Midgley, Brett D Mensh, and Julie H Simpson. A suppression hierarchy among competing motor programs drives sequential grooming in Drosophila. *eLife*, 3, 2014.

[638] Julia L Semmelhack, Joseph C Donovan, Tod R Thiele, Enrico Kuehn, Eva Laurell, and Herwig Baier. A dedicated visual pathway for prey detection in larval zebrafish. *eLife*, 3, 2014.

[639] Sonia Sen, Deshou Cao, Ramveer Choudhary, Silvia Biagini, Jing W Wang, Heinrich Reichert, and K VijayRaghavan. Genetic transformation of structural and functional circuitry rewires the Drosophila brain. *eLife*, 3, 2014.

[640] Ryan W Serra, Minggang Fang, Sung Mi Park, Lloyd Hutchinson, and Michael R Green. A KRAS-directed transcriptional silencing pathway that mediates the CpG island methylator phenotype. *eLife*, 3, 2014.

[641] Zhiguo Shang, Kaifeng Zhou, Chen Xu, Roseann Csencsits, Jared C Cochran, and Charles V Sindelar. High-resolution structures of kinesin on microtubules provide a basis for nucleotide-gated force-generation. *eLife*, 3, 2014.

[642] Ehud Shapiro. Correcting the bias against interdisciplinary research. *eLife*, 3, 2014.

[643] Lincoln T Shenje, Peter Andersen, Hideki Uosaki, Laviel Fernandez, Peter P Rainer, Gun-sik Cho, Dong-ik Lee, Weimin Zhong, Richard P Harvey, David A Kass, and Chulan Kwon. Precardiac deletion of Numb and Numblike reveals renewal of cardiac progenitors. *eLife*, 3, 2014.

[644] Alexander A Shestov, Xiaojing Liu, Zheng Ser, Ahmad A Cluntun, Yin P Hung, Lei Huang, Dongsung Kim, Anne Le, Gary Yellen, John G Albeck, and Jason W Locasale. Quantitative determinants of aerobic glycolysis identify flux through the enzyme GAPDH as a limiting step. *eLife*, 3, 2014.

[645] Mi Shi, Zhifeng Yue, Alexandre Kuryatov, Jon M Lindstrom, and Amita Sehgal. Identification of Redeye, a new sleep-regulating protein whose expression is modulated by sleep amount. *eLife*, 3, 2014.

[646] Yi Shi, Xiaoling Xu, Qian Zhang, Guangsen Fu, Zhongying Mo, George S Wang, Shuji Kishi, and Xiang-Lei Yang. tRNA synthetase counteracts c-Myc to develop functional vasculature. *eLife*, 3, 2014.

[647] Joshua H Siegle and Matthew A Wilson. Enhancement of encoding and retrieval functions through theta phase-specific manipulation of hippocampus. *eLife*, 3, 2014.

[648] Peter J Skene, Aaron E Hernandez, Mark Groudine, and Steven Henikoff. The nucleosomal barrier to promoter escape by RNA polymerase II is overcome by the chromatin remodeler Chd1. *eLife*, 3, 2014.

[649] Dorothy K Sojka, Beatrice Plougastel-Douglas, Liping Yang, Melissa A Pak-Wittel, Maxim N Artyomov, Yulia Ivanova, Chao Zhong, Julie M Chase, Paul B Rothman, Jenny Yu, Joan K Riley, Jinfang Zhu, Zhigang Tian, and Wayne M Yokoyama. Tissue-resident natural killer (NK) cells are cell lineages distinct from thymic and conventional splenic NK cells. *eLife*, 3, 2014.

[650] Brian A Sosa, F Esra Demircioglu, James Z Chen, Jessica Ingram, Hidde L Ploegh, and Thomas U Schwartz. How lamina-associated polypeptide 1 (LAP1) activates Torsin. *eLife*, 3, 2014.

[651] Harish Chandra Soundararajan and Simon L Bullock. The influence of dynein processivity control, MAPs, and microtubule ends on directional movement of a localising mRNA. *eLife*, 3, 2014.

[652] Charlotte J Stagg, Velicia Bachtiar, Ugwechi Amadi, Christel A Gudberg, Andrei S Ilie, Cassandra Sampaio-Baptista, Jacinta O'Shea, Mark Woolrich, Stephen M Smith, Nicola Filippini, Jamie Near, and Heidi Johansen-Berg. Local GABA concentration is related to network-level resting functional connectivity. *eLife*, 3, 2014.

[653] Jennifer L Stamos, Matthew Ling-Hon Chu, Michael D Enos, Niket Shah, and William I Weis. Structural basis of GSK-3 inhibition by N-terminal phosphorylation and by the Wnt receptor LRP6. *eLife*, 3, 2014.

[654] Edward Stanek, Steven Cheng, Jun Takatoh, Bao-Xia Han, and Fan Wang. Monosynaptic premotor circuit tracing reveals neural substrates for oro-motor coordination. *eLife*, 3, 2014.

[655] Florian A Steiner and Steven Henikoff. Holocentromeres are dispersed point centromeres localized at transcription factor hotspots. *eLife*, 3, 2014.

[656] Andrew M Stern, Arturo Casadevall, R Grant Steen, and Ferric C Fang. Financial costs and personal consequences of research misconduct resulting in retracted publications. *eLife*, 3, 2014.

[657] Jonathan M Stokes, Joseph H Davis, Chand S Mangat, James R Williamson, and Eric D Brown. Discovery of a small molecule that inhibits bacterial ribosome biogenesis. *eLife*, 3, 2014.

[658] Alberto Stolfi, Elijah K Lowe, Claudia Racioppi, Filomena Ristoratore, C Titus Brown, Billie J Swalla, and Lionel Christiaen. Divergent mechanisms regulate conserved cardiopharyngeal development and gene expression in distantly related ascidians. *eLife*, 3, 2014.

[659] Margaret Stratton, Il-Hyung Lee, Moitrayee Bhattacharyya, Sune M Christensen, Luke H Chao, Howard Schulman, Jay T Groves, and John Kuriyan. Activation-triggered subunit exchange between CaMKII holoenzymes facilitates the spread of kinase activity. *eLife*, 3, 2014.

[660] Antonia Strutz, Jan Soelter, Amelie Baschwitz, Abu Farhan, Veit Grabe, Jürgen Rybak, Markus Knaden, Michael Schmuker, Bill S Hansson, and Silke Sachse. Decoding odor quality and intensity in the Drosophila brain. *eLife*, 3, 2014.

[661] Arul Subramanian and Thomas F Schilling. Thrombospondin-4 controls matrix assembly during development and repair of myotendinous junctions. *eLife*, 3, 2014.

[662] Qian Sun, Kalyan V Srinivas, Alaba Sotayo, and Steven A Siegelbaum. Dendritic Na+ spikes enable cortical input to drive action potential output from hippocampal CA2 pyramidal neurons. *eLife*, 3, 2014.

[663] Syuan-Jyun Sun, Dustin R Rubenstein, Bo-Fei Chen, Shih-Fan Chan, Jian-Nan Liu, Mark Liu, Wenbe Hwang, Ping-Shih Yang, and Sheng-Feng Shen. Climate-mediated cooperation promotes niche expansion in burying beetles. *eLife*, 3, 2014.

[664] Jan Sundquist, Kristina Sundquist, and Jianguang Ji. Autism and attention-deficit/hyperactivity disorder among individuals with a family history of alcohol use disorders. *eLife*, 3, 2014.

[665] Catherine M Sweeney-Reed, Tino Zaehle, Juergen Voges, Friedhelm C Schmitt, Lars Buentjen, Klaus Kopitzki, Christine Esslinger, Hermann Hinrichs, Hans-Jochen Heinze, Robert T Knight, and Alan Richardson-Klavehn. Corticothalamic phase synchrony and cross-frequency coupling predict human memory formation. *eLife*, 3, 2014.

[666] Johanna Liinamaria Syrjänen, Luca Pellegrini, and Owen Richard Davies. A molecular model for the role of SYCP3 in meiotic chromosome organisation. *eLife*, 3, 2014.

[667] Piotr Szwedziak, Qing Wang, Tanmay A M Bharat, Matthew Tsim, and Jan Löwe. Architecture of the ring formed by the tubulin homologue FtsZ in bacterial cell division. *eLife*, 3, 2014.

[668] Shigeyuki Tanaka, Thomas Brefort, Nina Neidig, Armin Djamei, Jörg Kahnt, Wilfred Vermerris, Stefanie Koenig, Kirstin Feussner, Ivo Feussner, and Regine Kahmann. A secreted Ustilago maydis effector promotes virulence by targeting anthocyanin biosynthesis in maize. *eLife*, 3, 2014.

[669] Benjamin JM Taylor, Yee Ling Wu, and Cristina Rada. Active RNAP pre-initiation sites are highly mutated by cytidine deaminases in yeast, with AID targeting small RNA genes. *eLife*, 3, 2014.

[670] Aiguo Tian and Jin Jiang. Intestinal epithelium-derived BMP controls stem cell self-renewal in Drosophila adult midgut. *eLife*, 3, 2014.

[671] Jason Tien, Christian J Peters, Xiu Ming Wong, Tong Cheng, Yuh Nung Jan, Lily Yeh Jan, and Huanghe Yang. A comprehensive search for calcium binding sites critical for TMEM16A calcium-activated chloride channel activity. *eLife*, 3, 2014.

[672] Karen E Tkach, Debashis Barik, Guillaume Voisinne, Nicole Malandro, Matthew M Hathorn, Jesse W Cotari, Robert Vogel, Taha Merghoub, Jedd Wolchok, Oleg Krichevsky, and Grégoire Altan-Bonnet. T cells translate individual, quantal activation into collective, analog cytokine responses via time-integrated feedbacks. *eLife*, 3, 2014.

[673] Angela Torlopp, Mohsin A F Khan, Nidia M M Oliveira, Ingrid Lekk, Luz Mayela Soto-Jiménez, Alona Sosinsky, and Claudio D Stern. The transcription factor Pitx2 positions the embryonic axis and regulates twinning. *eLife*, 3, 2014.

[674] Katerina Toropova, Sirui Zou, Anthony J Roberts, William B Redwine, Brian S Goodman, Samara L Reck-Peterson, and Andres E Leschziner. Lis1 regulates dynein by sterically blocking its mechanochemical cycle. *eLife*, 3, 2014.

[675] Thorsten Trimbuch, Junjie Xu, David Flaherty, Diana R Tomchick, Josep Rizo, and Christian Rosenmund. Re-examining how complexin inhibits neurotransmitter release. *eLife*, 3, 2014.

[676] Nicolas X Tritsch, Won-Jong Oh, Chenghua Gu, and Bernardo L Sabatini. Midbrain dopamine neurons sustain inhibitory transmission using plasma membrane uptake of GABA, not synthesis. *eLife*, 3, 2014.

[677] Satoshi Tsunoda, Edward Avezov, Alisa Zyryanova, Tasuku Konno, Leonardo Mendes-Silva, Eduardo Pinho Melo, Heather P Harding, and David Ron. Intact protein folding in the glutathione-depleted endoplasmic reticulum implicates alternative protein thiol reductants. *eLife*, 3, 2014.

[678] Srinivasan Tupal, Wei-Hsiang Huang, Maria Cristina D Picardo, Guang-Yi Ling, Christopher A Del Negro, Huda Y Zoghbi, and Paul A Gray. Atoh1-dependent rhombic lip neurons are required for temporal delay between independent respiratory oscillators in embryonic mice. *eLife*, 3, 2014.

[679] Leslie M Turner and Bettina Harr. Genome-wide mapping in a house mouse hybrid zone reveals hybrid sterility loci and Dobzhansky-Muller interactions. *eLife*, 3, 2014.

[680] Perunthottathu K Umasankar, Li Ma, James R Thieman, Anupma Jha, Balraj Doray, Simon C Watkins, and Linton M Traub. A clathrin coat assembly role for the muniscin protein central linker revealed by TALEN-mediated gene editing. *eLife*, 3, 2014.

[681] Federico Vaggi, Tommaso Schiavinotto, Jonathan LD Lawson, Anatole Chessel, James Dodgson, Marco Geymonat, Masamitsu Sato, Rafael Edgardo Carazo Salas, and Attila Csikász-Nagy. A network approach to mixing delegates at meetings. *eLife*, 3, 2014.

[682] Eelco van Anken, David Pincus, Scott Coyle, Tomás Aragón, Christof Osman, Federica Lari, Silvia Gómez Puerta, Alexei V Korennykh, and Peter Walter. Specificity in endoplasmic reticulum-stress signaling in yeast entails a step-wise engagement of HAC1 mRNA to clusters of the stress sensor Ire1. *eLife*, 3, 2014.

[683] Maaike PA van Bragt, Xin Hu, Ying Xie, and Zhe Li. RUNX1, a transcription factor mutated in breast cancer, controls the fate of ER-positive mammary luminal cells. *eLife*, 3, 2014.

[684] Mark van Breugel, Rainer Wilcken, Stephen H McLaughlin, Trevor J Rutherford, and Christopher M Johnson. Structure of the SAS-6 cartwheel hub from Leishmania major. *eLife*, 3, 2014.

[685] Fusinita van den Ent, Thierry Izoré, Tanmay AM Bharat, Christopher M Johnson, and Jan Löwe. Bacterial actin MreB forms antiparallel double filaments. *eLife*, 3, 2014.

[686] Ben C L van Schaijk, Ivo H J Ploemen, Takeshi Annoura, Martijn W Vos, Lander Foquet, Geert-Jan van Gemert, Severine Chevalley-Maurel, Marga van de Vegte-Bolmer, Mohammed Sajid, Jean-Francois Franetich, Audrey Lorthiois, Geert Leroux-Roels, Philip Meuleman, Cornelius C Hermsen, Dominique Mazier, Stephen L Hoffman, Chris J Janse, Shahid M Khan, and Robert W Sauerwein. A genetically attenuated malaria vaccine candidate based on P. falciparum b9/slarp gene-deficient sporozoites. *eLife*, 3, 2014.

[687] Hannes Vanhaeren, Nathalie Gonzalez, Frederik Coppens, Liesbeth De Milde, Twiggy Van Daele, Mattias Vermeersch, Nubia B Eloy, Veronique Storme, and Dirk Inzé. Combining growth-promoting genes leads to positive epistasis in Arabidopsis thaliana. *eLife*, 3, 2014.

[688] Sakari Vanharanta, Christina B Marney, Weiping Shu, Manuel Valiente, Yilong Zou, Aldo Mele, Robert B Darnell, and Joan Massagué. Loss of the multifunctional RNA-binding protein RBM47 as a source of selectable metastatic traits in breast cancer. *eLife*, 3, 2014.

[689] Csaba Varga, Mikko Oijala, Jonathan Lish, Gergely G Szabo, Marianne Bezaire, Ivan Marchionni, Peyman Golshani, and Ivan Soltesz. Functional fission of parvalbumin interneuron classes during fast network events. *eLife*, 3, 2014.

[690] Maria Vera, Bibhusita Pani, Lowri A Griffiths, Christian Muchardt, Catherine M Abbott, Robert H Singer, and Evgeny Nudler. The translation elongation factor eEF1A1 couples transcription to translation during heat shock response. *eLife*, 3, 2014.

[691] Grégory Verdon, SeCheol Oh, Ryan N Serio, and Olga Boudker. Coupled ion binding and structural transitions along the transport cycle of glutamate transporters. *eLife*, 3, 2014.

[692] Kitty F Verzijlbergen, Olga O Nerusheva, David Kelly, Alastair Kerr, Dean Clift, Flavia de Lima Alves, Juri Rappsilber, and Adele L Marston. Shugoshin biases chromosomes for biorientation through condensin recruitment to the pericentromere. *eLife*, 3, 2014.

[693] Igor Vivanco, Zhi C Chen, Barbara Tanos, Barbara Oldrini, Wan-Ying Hsieh, Nicolas Yannuzzi, Carl Campos, and Ingo K Mellinghoff. A kinase-independent function of AKT promotes cancer cell survival. *eLife*, 3, 2014.

[694] Katrin Vogt, Christopher Schnaitmann, Kristina V Dylla, Stephan Knapek, Yoshinori Aso, Gerald M Rubin, and Hiromu Tanimoto. Shared mushroom body circuits underlie visual and olfactory memories in Drosophila. *eLife*, 3, 2014.

[695] Octavian Voiculescu, Lawrence Bodenstein, I-Jun Lau, and Claudio D Stern. Local cell interactions and self-amplifying individual cell ingression drive amniote gastrulation. *eLife*, 3, 2014.

[696] Rainer Waadt, Kenichi Hitomi, Noriyuki Nishimura, Chiharu Hitomi, Stephen R Adams, Elizabeth D Getzoff, and Julian I Schroeder. FRET-based reporters for the direct visualization of abscisic acid concentration changes and distribution in Arabidopsis. *eLife*, 3, 2014.

[697] Setsuko Wakao, Brian L Chin, Heidi K Ledford, Rachel M Dent, David Casero, Matteo Pellegrini, Sabeeha S Merchant, and Krishna K Niyogi. Phosphoprotein SAK1 is a regulator of acclimation to singlet oxygen in Chlamydomonas reinhardtii. *eLife*, 3, 2014.

[698] Guoqiang Wan, Maria E Gómez-Casati, Angelica R Gigliello, M Charles Liberman, and Gabriel Corfas. Neurotrophin-3 regulates ribbon synapse density in the cochlea and induces synapse regeneration after acoustic trauma. *eLife*, 3, 2014.

[699] Chao Wang, Zhiyi Wei, Keyu Chen, Fei Ye, Cong Yu, Vann Bennett, and Mingjie Zhang. Structural basis of diverse membrane target recognitions by ankyrins. *eLife*, 3, 2014.

[700] Jennifer T Wang, Jarrett Smith, Bi-Chang Chen, Helen Schmidt, Dominique Rasoloson, Alexandre Paix, Bramwell G Lambrus, Deepika Calidas, Eric Betzig, and Geraldine Seydoux. Regulation of RNA granule dynamics by phosphorylation of serine-rich, intrinsically disordered proteins in C. elegans. *eLife*, 3, 2014.

[701] Jie Wang, Kristeen A Pareja, Chris A Kaiser, and Carolyn S Sevier. Redox signaling via the molecular chaperone BiP protects cells against endoplasmic reticulum-derived oxidative stress. *eLife*, 3, 2014.

[702] Jiguang Wang, Hossein Khiabanian, Davide Rossi, Giulia Fabbri, Valter Gattei, Francesco Forconi, Luca Laurenti, Roberto Marasca, Giovanni Del Poeta, Robin Foà, Laura Pasqualucci, Gianluca Gaidano, and Raul Rabadan. Tumor evolutionary directed graphs and the history of chronic lymphocytic leukemia. *eLife*, 3, 2014.

[703] Weiping Wang, Tao Wu, and Marc W Kirschner. The master cell cycle regulator APC-Cdc20 regulates ciliary length and disassembly of the primary cilium. *eLife*, 3, 2014.

[704] Weiwei Wang, Matthew R Whorton, and Roderick MacKinnon. Quantitative analysis of mammalian GIRK2 channel regulation by G proteins, the signaling lipid PIP2 and Na+ in a reconstituted system. *eLife*, 3, 2014.

[705] Xueying Wang, John A Hayes, Ann L Revill, Hanbing Song, Andrew Kottick, Nikolas C Vann, M Drew LaMar, Maria Cristina D Picardo, Victoria T Akins, Gregory D Funk, and Christopher A Del Negro. Laser ablation of Dbx1 neurons in the pre-Bötzinger complex stops inspiratory rhythm and impairs output in neonatal mice. *eLife*, 3, 2014.

[706] Yong Wang, Yanxin Liu, Hannah A DeBerg, Takeshi Nomura, Melinda Tonks Hoffman, Paul R Rohde, Klaus Schulten, Boris Martinac, and Paul R Selvin. Single molecule FRET reveals pore size and opening mechanism of a mechano-sensitive ion channel. *eLife*, 3, 2014.

[707] Yubao Wang, Young-Mi Lee, Lukas Baitsch, Alan Huang, Yi Xiang, Haoxuan Tong, Ana Lako, Thanh Von, Christine Choi, Elgene Lim, Junxia Min, Li Li, Frank Stegmeier, Robert Schlegel, Michael J Eck, Nathanael S Gray, Timothy J Mitchison, and Jean J Zhao. MELK is an oncogenic kinase essential for mitotic progression in basal-like breast cancer cells. *eLife*, 3, 2014.

[708] Jonathan J Ward, Hélio Roque, Claude Antony, and FranÇois Nédélec. Mechanical design principles of a mitotic spindle. *eLife*, 3, 2014.

[709] Alexander A Warkentin, Michael S Lopez, Elisabeth A Lasater, Kimberly Lin, Bai-Liang He, Anskar YH Leung, Catherine C Smith, Neil P Shah, and Kevan M Shokat. Overcoming myelosuppression due to synthetic lethal toxicity for FLT3-targeted acute myeloid leukemia therapy. *eLife*, 3, 2014.

[710] Benjamin P Weaver, Rebecca Zabinsky, Yi M Weaver, Eui Seung Lee, Ding Xue, and Min Han. CED-3 caspase acts with miRNAs to regulate non-apoptotic gene expression dynamics for robust development in C. elegans. *eLife*, 3, 2014.

[711] William W Wheaton, Samuel E Weinberg, Robert B Hamanaka, Saul Soberanes, Lucas B Sullivan, Elena Anso, Andrea Glasauer, Eric Dufour, Gokhan M Mutlu, GR Scott Budigner, and Navdeep S Chandel. Metformin inhibits mitochondrial complex I of cancer cells to reduce tumorigenesis. *eLife*, 3, 2014.

[712] Michael T White, Stephan Karl, Katherine E Battle, Simon I Hay, Ivo Mueller, and Azra C Ghani. Modelling the contribution of the hypnozoite reservoir to Plasmodium vivax transmission. *eLife*, 3, 2014.

[713] Stephan Wickles, Abhishek Singharoy, Jessica Andreani, Stefan Seemayer, Lukas Bischoff, Otto Berninghausen, Johannes Soeding, Klaus Schulten, Eli O van der Sluis, and Roland Beckmann. A structural model of the active ribosome-bound membrane protein insertase YidC. *eLife*, 3, 2014.

[714] Gregg A Wildenberg and Andrew W Murray. Evolving a 24-hr oscillator in budding yeast. *eLife*, 3, 2014.

[715] Florian Wilfling, Abdou Rachid Thiam, Maria-Jesus Olarte, Jing Wang, Rainer Beck, Travis J Gould, Edward S Allgeyer, Frederic Pincet, Jörg Bewersdorf, Robert V Farese, and Tobias C Walther. Arf1/COPI machinery acts directly on lipid droplets and enables their connection to the ER for protein targeting. *eLife*, 3, 2014.

[716] Tobias Willer, Kei-ichiro Inamori, David Venzke, Corinne Harvey, Greg Morgensen, Yuji Hara, Daniel Beltrán Valero de Bernabé, Liping Yu, Kevin M Wright, and Kevin P Campbell. The glucuronyltransferase B4GAT1 is required for initiation of LARGE-mediated $\alpha$-dystroglycan functional glycosylation. *eLife*, 3, 2014.

[717] Byron C Williams, Joshua J Filter, Kristina A Blake-Hodek, Brian E Wadzinski, Nicholas J Fuda, David Shalloway, and Michael L Goldberg. Greatwall-phosphorylated Endosulfine is both an inhibitor and a substrate of PP2A-B55 heterotrimers. *eLife*, 3, 2014.

[718] James P Williams, Jacob Hurst, Wolfgang Stöhr, Nicola Robinson, Helen Brown, Martin Fisher, Sabine Kinloch, David Cooper, Mauro Schechter, Giuseppe Tambussi, Sarah Fidler, Mary Carrington, Abdel Babiker, Jonathan Weber, Kersten K Koelsch, Anthony D Kelleher, Rodney E Phillips, and John Frater. HIV-1 DNA predicts disease progression and post-treatment virological control. *eLife*, 3, 2014.

[719] Jan Wisniewski, Bassam Hajj, Jiji Chen, Gaku Mizuguchi, Hua Xiao, Debbie Wei, Maxime Dahan, and Carl Wu. Imaging the fate of histone Cse4 reveals de novo replacement in S phase and subsequent stable residence at centromeres. *eLife*, 3, 2014.

[720] David Wöhlert, Werner Kühlbrandt, and özkan Yildiz. Structure and substrate ion binding in the sodium/proton antiporter PaNhaP. *eLife*, 3, 2014.

[721] Wilson Wong, Xiao-chen Bai, Alan Brown, Israel S Fernandez, Eric Hanssen, Melanie Condron, Yan Hong Tan, Jake Baum, and Sjors HW Scheres. Cryo-EM structure of the Plasmodium falciparum 80S ribosome bound to the anti-protozoan drug emetine. *eLife*, 3, 2014.

[722] Hao Wu, John Williams, and Jeremy Nathans. Complete morphologies of basal forebrain cholinergic neurons in the mouse. *eLife*, 3, 2014.

[723] Shu-Zon Wu and Magdalena Bezanilla. Myosin VIII associates with microtubule ends and together with actin plays a role in guiding plant cell division. *eLife*, 3, 2014.

[724] Yumei Wu, Eileen T O'Toole, Martine Girard, Brigitte Ritter, Mirko Messa, Xinran Liu, Peter S McPherson, Shawn M Ferguson, and Pietro De Camilli. A dynamin 1-, dynamin 3- and clathrin-independent pathway of synaptic vesicle recycling mediated by bulk endocytosis. *eLife*, 3, 2014.

[725] Nicky Wybouw, Wannes Dermauw, Luc Tirry, Christian Stevens, Miodrag Grbić, René Feyereisen, and Thomas Van Leeuwen. A gene horizontally transferred from bacteria protects arthropods from host plant cyanide poisoning. *eLife*, 3, 2014.

[726] Guiran Xiao, Zhihui Wan, Qiangwang Fan, Xiaona Tang, and Bing Zhou. The metal transporter ZIP13 supplies iron into the secretory pathway in Drosophila melanogaster. *eLife*, 3, 2014.

[727] Yonggang Xie, Xiaosu Li, Xian Zhang, Shaolin Mei, Hongyu Li, Andreacarola Urso, and Sijun Zhu. The Drosophila Sp8 transcription factor Buttonhead prevents premature differentiation of intermediate neural progenitors. *eLife*, 3, 2014.

[728] Xiufang Xiong, Yongchao Zhao, Fei Tang, Dongping Wei, Daffyd Thomas, Xiang Wang, Yang Liu, Pan Zheng, and Yi Sun. Ribosomal protein S27-like is a physiological regulator of p53 that suppresses genomic instability and tumorigenesis. *eLife*, 3, 2014.

[729] Hui Xu, Xiaojing He, Hui Zheng, Lily J Huang, Fajian Hou, Zhiheng Yu, Michael Jason de la Cruz, Brian Borkowski, Xuewu Zhang, Zhijian J Chen, and Qiu-Xing Jiang. Structural basis for the prion-like MAVS filaments in antiviral innate immunity. *eLife*, 3, 2014.

[730] Hiroshi Yamaguchi, Toshihiko Maruyama, Yoshihiro Urade, and Shigekazu Nagata. Immunosuppression via adenosine receptor activation by adenosine monophosphate released from apoptotic cells. *eLife*, 3, 2014.

[731] Koji Yamano, Adam I Fogel, Chunxin Wang, Alexander M van der Bliek, and Richard J Youle. Mitochondrial Rab GAPs govern autophagosome biogenesis during mitophagy. *eLife*, 3, 2014.

[732] Naoki Yamawaki, Katharine Borges, Benjamin A Suter, Kenneth D Harris, and Gordon M G Shepherd. A genuine layer 4 in motor cortex with prototypical synaptic circuit connectivity. *eLife*, 3, 2014.

[733] Haidun Yan, Juan L Pablo, Chaojian Wang, and Geoffrey S Pitt. FGF14 modulates resurgent sodium current in mouse cerebellar Purkinje neurons. *eLife*, 3, 2014.

[734] Fan Yang and Jie Zheng. High temperature sensitivity is intrinsic to voltage-gated potassium channels. *eLife*, 3, 2014.

[735] Yul W Yang, Ryan A Flynn, Yong Chen, Kun Qu, Bingbing Wan, Kevin C Wang, Ming Lei, and Howard Y Chang. Essential role of lncRNA binding for WDR5 maintenance of active chromatin and embryonic stem cell pluripotency. *eLife*, 3, 2014.

[736] Zhiping Yang, Yuh-Chin T Huang, Henry Koziel, Rini de Crom, Hartmut Ruetten, Paulus Wohlfart, Reimar W Thomsen, Johnny A Kahlert, Henrik Toft Sørensen, Szczepan Jozefowski, Amy Colby, and Lester Kobzik. Female resistance to pneumonia identifies lung macrophage nitric oxide synthase-3 as a therapeutic target. *eLife*, 3, 2014.

[737] Marianna Yanike and Vincent P Ferrera. Interpretive monitoring in the caudate nucleus. *eLife*, 3, 2014.

[738] Qing Yao, Qiuhe Lu, Xiaobo Wan, Feng Song, Yue Xu, Mo Hu, Alla Zamyatina, Xiaoyun Liu, Niu Huang, Ping Zhu, and Feng Shao. A structural mechanism for bacterial autotransporter glycosylation by a dodecameric heptosyltransferase family. *eLife*, 3, 2014.

[739] Risheng Ye, William L Holland, Ruth Gordillo, Miao Wang, Qiong A Wang, Mengle Shao, Thomas S Morley, Rana K Gupta, Andreas Stahl, and Philipp E Scherer. Adiponectin is essential for lipid homeostasis and survival under insulin deficiency and promotes $\beta$-cell regeneration. *eLife*, 3, 2014.

[740] Keren Yizhak, Edoardo Gaude, Sylvia Le Dévédec, Yedael Y Waldman, Gideon Y Stein, Bob van de Water, Christian Frezza, and Eytan Ruppin. Phenotype-based cell-specific metabolic modeling reveals metabolic liabilities of cancer. *eLife*, 3, 2014.

[741] Xiaofei Yu, Yuhao Wang, Mi Deng, Yun Li, Kelly A Ruhn, Cheng Cheng Zhang, and Lora V Hooper. The basic leucine zipper transcription factor NFIL3 directs the development of a common innate lymphoid cell precursor. *eLife*, 3, 2014.

[742] Andy H Yuan, Sean J Garrity, Entela Nako, and Ann Hochschild. Prion propagation can occur in a prokaryote and requires the ClpB chaperone. *eLife*, 3, 2014.

[743] Shen Yue, Liu-Ya Tang, Ying Tang, Yi Tang, Qiu-Hong Shen, Jie Ding, Yan Chen, Zengdi Zhang, Ting-Ting Yu, Ying E Zhang, and Steven Y Cheng. Requirement of Smurf-mediated endocytosis of Patched1 in sonic hedgehog signal reception. *eLife*, 3, 2014.

[744] Jina Yun, Rajat Puri, Huan Yang, Michael A Lizzio, Chunlai Wu, Zu-Hang Sheng, and Ming Guo. MUL1 acts in parallel to the PINK1/parkin pathway in regulating mitofusin and compensates for loss of PINK1/parkin. *eLife*, 3, 2014.

[745] Margarita Zacharogianni, Angelica Aguilera-Gomez, Tineke Veenendaal, Jan Smout, and Catherine Rabouille. A stress assembly that confers cell viability by preserving ERES components during amino-acid starvation. *eLife*, 3, 2014.

[746] Sarah E Zanders, Michael T Eickbush, Jonathan S Yu, Ji-Won Kang, Kyle R Fowler, Gerald R Smith, and Harmit Singh Malik. Genome rearrangements and pervasive meiotic drive cause hybrid infertility in fission yeast. *eLife*, 3, 2014.

[747] Mark A Zaydman, Marina A Kasimova, Kelli McFarland, Zachary Beller, Panpan Hou, Holly E Kinser, Hongwu Liang, Guohui Zhang, Jingyi Shi, Mounir Tarek, and Jianmin Cui. Domain–domain interactions determine the gating, permeation, pharmacology, and subunit modulation of the IKs ion channel. *eLife*, 3, 2014.

[748] Xun Zeng, Christina Meyer, Jun Huang, Evan W Newell, Brian A Kidd, Yu-Ling Wei, and Yueh-hsiu Chien. Gamma delta T cells recognize haptens and mount a hapten-specific response. *eLife*, 3, 2014.

[749] Jianbo Zhang, Tao Zuo, Dafang Wang, and Thomas Peterson. Transposition-mediated DNA re-replication in maize. *eLife*, 3, 2014.

[750] Qi Zhang, Xiang Zhou, RuiZhi Wu, Amber Mosley, Shelya X Zeng, Zhen Xing, and Hua Lu. The role of IMP dehydrogenase 2 in Inauhzin-induced ribosomal stress. *eLife*, 3, 2014.

[751] Wei Zhang, Zhiqiang Yan, Bingxue Li, Lily Yeh Jan, and Yuh Nung Jan. Identification of motor neurons and a mechanosensitive sensory neuron in the defecation circuitry of Drosophila larvae. *eLife*, 3, 2014.

[752] Zhe Zhang, Tianlong Zhang, Shanshan Wang, Zhou Gong, Chun Tang, Jiangye Chen, and Jianping Ding. Molecular mechanism for Rabex-5 GEF activation by Rabaptin-5. *eLife*, 3, 2014.

[753] Zhengjian Zhang, Andrey Revyakin, Jonathan B Grimm, Luke D Lavis, and Robert Tjian. Single-molecule tracking of the transcription cycle by sub-second RNA detection. *eLife*, 3, 2014.

[754] Hui Zhao, Bernard Thienpont, Betül Tuba Yesilyurt, Matthieu Moisse, Joke Reumers, Lieve Coenegrachts, Xavier Sagaert, Stefanie Schrauwen, Dominiek Smeets, Gert Matthijs, Stein Aerts, Jan Cools, Alex Metcalf, Amanda Spurdle, Frederic Amant, and Diether Lambrechts. Mismatch repair deficiency endows tumors with a unique mutation signature and sensitivity to DNA double-strand breaks. *eLife*, 3, 2014.

[755] Suwen Zhao, Ayano Sakai, Xinshuai Zhang, Matthew W Vetting, Ritesh Kumar, Brandan Hillerich, Brian San Francisco, Jose Solbiati, Adam Steves, Shoshana Brown, Eyal Akiva, Alan Barber, Ronald D Seidel, Patricia C Babbitt, Steven C Almo, John A Gerlt, and Matthew P Jacobson. Prediction and characterization of enzymatic activities guided by sequence similarity and genome neighborhood networks. *eLife*, 3, 2014.

[756] Ya Zhao, Jimin Lin, Beiying Xu, Sida Hu, Xue Zhang, and Ligang Wu. MicroRNA-mediated repression of nonsense mRNAs. *eLife*, 3, 2014.

[757] Xingguo Zheng, Vera Valakh, Aaron DiAntonio, and Yehuda Ben-Shahar. Natural antisense transcripts regulate the neuronal stress response and excitability. *eLife*, 3, 2014.

[758] Guisheng Zhong, Jiang He, Ruobo Zhou, Damaris Lorenzo, Hazen P Babcock, Vann Bennett, and Xiaowei Zhuang. Developmental mechanism of the periodic membrane skeleton in axons. *eLife*, 3, 2014.

[759] Haibo Zhou, Zhanmin Lin, Kai Voges, Chiheng Ju, Zhenyu Gao, Laurens WJ Bosman, Tom JH Ruigrok, Freek E Hoebeek, Chris I De Zeeuw, and Martijn Schonewille. Cerebellar modules operate at different frequencies. *eLife*, 3, 2014.

[760] Haiying Zhou, Bo Wan, Ivan Grubisic, Tommy Kaplan, and Robert Tjian. TAF7L modulates brown adipose tissue formation. *eLife*, 3, 2014.

[761] Ruobo Zhou, Jichuan Zhang, Matthew L Bochman, Virginia A Zakian, and Taekjip Ha. Periodic DNA patrolling underlies diverse functions of Pif1 on R-loops and G-rich DNA. *eLife*, 3, 2014.

[762] Qingqing Zhu, Lu Song, Guangdun Peng, Na Sun, Jun Chen, Ting Zhang, Nengyin Sheng, Wei Tang, Cheng Qian, Yunbo Qiao, Ke Tang, Jing-Dong Jackie Han, Jinsong Li, and Naihe Jing. The transcription factor Pou3f1 promotes neural fate commitment via activation of neural lineage genes and inhibition of external signaling pathways. *eLife*, 3, 2014.

[763] Michael Zick, Christopher Stroupe, Amy Orr, Deborah Douville, and William T Wickner. Membranes linked by trans-SNARE complexes require lipids prone to non-bilayer structure for progression to fusion. *eLife*, 3, 2014.

[764] Michael Zick and William T Wickner. A distinct tethering step is vital for vacuole membrane fusion. *eLife*, 3, 2014.

[765] Katja Zieske and Petra Schwille. Reconstitution of self-organizing protein gradients as spatial cues in cell-free systems. *eLife*, 3, 2014.

[766] Adelajda Zorba, Vanessa Buosi, Steffen Kutter, Nadja Kern, Francesco Pontiggia, Young-Jin Cho, and Dorothee Kern. Molecular mechanism of Aurora A kinase autophosphorylation and its allosteric activation by TPX2. *eLife*, 3, 2014.

[767] Sylvain Zorman, Aleksander A Rebane, Lu Ma, Guangcan Yang, Matthew A Molski, Jeff Coleman, Frederic Pincet, James E Rothman, and Yongli Zhang. Common intermediates and kinetics, but different energetics, in the assembly of SNARE proteins. *eLife*, 3, 2014.

[768] Melina Zourelidou, Birgit Absmanner, Benjamin Weller, Inês CR Barbosa, Björn C Willige, Astrid Fastner, Verena Streit, Sarah A Port, Jean Colcombet, Sergio de la Fuente van Bentem, Heribert Hirt, Bernhard Kuster, Waltraud X Schulze, Ulrich Z Hammes, and Claus Schwechheimer. Auxin efflux by PIN-FORMED proteins is activated by two different protein kinases, D6 PROTEIN KINASE and PINOID. *eLife*, 3, 2014.

[769] Marlen Zschätzsch, Carlos Oliva, Marion Langen, Natalie De Geest, Mehmet Neset özel, W Ryan Williamson, William C Lemon, Alessia Soldano, Sebastian Munck, P Robin Hiesinger, Natalia Sanchez-Soriano, and Bassem A Hassan. Regulation of branching dynamics by axon-intrinsic asymmetries in Tyrosine Kinase Receptor signaling. *eLife*, 3, 2014.

[770] Diego Acosta-Alvear, Min Y Cho, Thomas Wild, Tonia J Buchholz, Alana G Lerner, Olga Simakova, Jamie Hahn, Neha Korde, Ola Landgren, Irina Maric, Chunaram Choudhary, Peter Walter, Jonathan S Weissman, and Martin Kampmann. Paradoxical resistance of multiple myeloma to proteasome inhibitors by decreased levels of 19S proteasomal subunits. *eLife*, 4, 2015.

[771] Chinnakkaruppan Adaikkan and Kobi Rosenblum. A molecular mechanism underlying gustatory memory trace for an association in the insular cortex. *eLife*, 4, 2015.

[772] Vikram Agarwal, George W Bell, Jin-Wu Nam, and David P Bartel. Predicting effective microRNA target sites in mammalian mRNAs. *eLife*, 4, 2015.

[773] Sobhika Agarwala, Sandra Duquesne, Kun Liu, Anton Boehm, Lin Grimm, Sandra Link, Sabine König, Stefan Eimer, Olaf Ronneberger, and Virginie Lecaudey. Amotl2a interacts with the Hippo effector Yap1 and the Wnt/$\beta$-catenin effector Lef1 to control tissue size in zebrafish. *eLife*, 4, 2015.

[774] Tarjani Agrawal and Gaiti Hasan. Maturation of a central brain flight circuit in Drosophila requires Fz2/Ca2+ signaling. *eLife*, 4, 2015.

[775] Sara Ajina, Franco Pestilli, Ariel Rokem, Christopher Kennard, and Holly Bridge. Human blindsight is mediated by an intact geniculo-extrastriate pathway. *eLife*, 4, 2015.

[776] Andrei R Akhmetzhanov and Michael E Hochberg. Dynamics of preventive vs post-diagnostic cancer control using low-impact measures. *eLife*, 4, 2015.

[777] Koichiro Akiyama, Shinya Mizuno, Yohei Hizukuri, Hiroyuki Mori, Terukazu Nogi, and Yoshinori Akiyama. Roles of the membrane-reentrant $\beta$-hairpin-like loop of RseP protease in selective substrate cleavage. *eLife*, 4, 2015.

[778] Cécile Alanio, Francesco Nicoli, Philippe Sultanik, Tobias Flecken, Brieuc Perot, Darragh Duffy, Elisabetta Bianchi, Annick Lim, Emmanuel Clave, Marit M van Buuren, Aurélie Schnuriger, Kerstin Johnsson, Jeremy Boussier, Antoine Garbarg-Chenon, Laurence Bousquet, Estelle Mottez, Ton N Schumacher, Antoine Toubert, Victor Appay, Farhad Heshmati, Robert Thimme, Stanislas Pol, Vincent Mallet, and Matthew L Albert. Bystander hyperactivation of preimmune CD8+ T cells in chronic HCV patients. *eLife*, 4, 2015.

[779] FranÇois Alberge, Leon Espinosa, Farida Seduk, Léa Sylvi, René Toci, Anne Walburger, and Axel Magalon. Dynamic subcellular localization of a respiratory complex controls bacterial respiration. *eLife*, 4, 2015.

[780] Jane Alfred and Ian T Baldwin. New opportunities at the wild frontier. *eLife*, 4, 2015.

[781] Shahar Alon, Sandra C Garrett, Erez Y Levanon, Sara Olson, Brenton R Graveley, Joshua J C Rosenthal, and Eli Eisenberg. The majority of transcripts in the squid nervous system are extensively recoded by A-to-I RNA editing. *eLife*, 4, 2015.

[782] Meghan O Altman, Jack R Bennink, Jonathan W Yewdell, and Brantley R Herrin. Lamprey VLRB response to influenza virus supports universal rules of immunogenicity and antigenicity. *eLife*, 4, 2015.

[783] Hermann C Altmeppen, Johannes Prox, Susanne Krasemann, Berta Puig, Katharina Kruszewski, Frank Dohler, Christian Bernreuther, Ana Hoxha, Luise Linsenmeier, Beata Sikorska, Pawel P Liberski, Udo Bartsch, Paul Saftig, and Markus Glatzel. The sheddase ADAM10 is a potent modulator of prion disease. *eLife*, 4, 2015.

[784] Vikram Alva, Johannes Söding, and Andrei N Lupas. A vocabulary of ancient peptides at the origin of folded proteins. *eLife*, 4, 2015.

[785] Abhijit A Ambegaonkar and Kenneth D Irvine. Coordination of planar cell polarity pathways through Spiny-legs. *eLife*, 4, 2015.

[786] Dave W Anderson, Alesia N McKeown, and Joseph W Thornton. Intermolecular epistasis shaped the function and evolution of an ancient transcription factor and its DNA binding sites. *eLife*, 4, 2015.

[787] Johan OL Andreasson, Bojan Milic, Geng-Yuan Chen, Nicholas R Guydosh, William O Hancock, and Steven M Block. Examining kinesin processivity within a general gating framework. *eLife*, 4, 2015.

[788] Joanna Andrecka, Jaime Ortega Arroyo, Yasuharu Takagi, Gabrielle de Wit, Adam Fineberg, Lachlan MacKinnon, Gavin Young, James R Sellers, and Philipp Kukura. Structural dynamics of myosin 5 during processive motion revealed by interferometric scattering microscopy. *eLife*, 4, 2015.

[789] Samuel Andrew Hires, Diego A Gutnisky, Jianing Yu, Daniel H O'Connor, and Karel Svoboda. Low-noise encoding of active touch by layer 4 in the somatosensory cortex. *eLife*, 4, 2015.

[790] Carmen Andrikou, Chih-Yu Pai, Yi-Hsien Su, and Maria Ina Arnone. Logics and properties of a genetic regulatory program that drives embryonic muscle development in an echinoderm. *eLife*, 4, 2015.

[791] Lea Ankri, Zoé Husson, Katarzyna Pietrajtis, Rémi Proville, Clément Léna, Yosef Yarom, Stéphane Dieudonné, and Marylka Yoe Uusisaari. A novel inhibitory nucleo-cortical circuit controls cerebellar Golgi cell activity. *eLife*, 4, 2015.

[792] Rajaguru Aradhya, Monika Zmojdzian, Jean Philippe Da Ponte, and Krzysztof Jagla. Muscle niche-driven Insulin-Notch-Myc cascade reactivates dormant Adult Muscle Precursors in Drosophila. *eLife*, 4, 2015.

[793] Michael J Arcaro, Christopher J Honey, Ryan EB Mruczek, Sabine Kastner, and Uri Hasson. Widespread correlation patterns of fMRI signal across visual cortex reflect eccentricity organization. *eLife*, 4, 2015.

[794] Yasunobu Arima, Daisuke Kamimura, Toru Atsumi, Masaya Harada, Tadafumi Kawamoto, Naoki Nishikawa, Andrea Stofkova, Takuto Ohki, Kotaro Higuchi, Yuji Morimoto, Peter Wieghofer, Yuka Okada, Yuki Mori, Saburo Sakoda, Shizuya Saika, Yoshichika Yoshioka, Issei Komuro, Toshihide Yamashita, Toshio Hirano, Marco Prinz, and Masaaki Murakami. A pain-mediated neural signal induces relapse in murine autoimmune encephalomyelitis, a multiple sclerosis model. *eLife*, 4, 2015.

[795] Christian Arquint, Anna-Maria Gabryjonczyk, Stefan Imseng, Raphael Böhm, Evelyn Sauer, Sebastian Hiller, Erich A Nigg, and Timm Maier. STIL binding to Polo-box 3 of PLK4 regulates centriole duplication. *eLife*, 4, 2015.

[796] Aaron Arvey, Joris van der Veeken, George Plitas, Stephen S Rich, Patrick Concannon, and Alexander Y Rudensky. Genetic and epigenetic variation in the lineage specification of regulatory T cells. *eLife*, 4, 2015.

[797] David George Ashbrook, Beatrice Gini, and Reinmar Hager. Genetic variation in offspring indirectly influences the quality of maternal behaviour in mice. *eLife*, 4, 2015.

[798] Cindy Ast, Roberto De Michele, Michael U Kumke, and Wolf B Frommer. Single-fluorophore membrane transport activity sensors with dual-emission read-out. *eLife*, 4, 2015.

[799] Sabrina Asteriti, Sten Grillner, and Lorenzo Cangiano. A Cambrian origin for vertebrate rods. *eLife*, 4, 2015.

[800] Thomas O Auer, Tong Xiao, Valerie Bercier, Christoph Gebhardt, Karine Duroure, Jean-Paul Concordet, Claire Wyart, Maximiliano Suster, Koichi Kawakami, Joachim Wittbrodt, Herwig Baier, and Filippo Del Bene. Deletion of a kinesin I motor unmasks a mechanism of homeostatic branching control by neurotrophin-3. *eLife*, 4, 2015.

[801] Stephen D Auger, Peter Zeidman, and Eleanor A Maguire. A central role for the retrosplenial cortex in de novo environmental learning. *eLife*, 4, 2015.

[802] Kevin Axelrod, Alvaro Sanchez, and Jeff Gore. Phenotypic states become increasingly sensitive to perturbations near a bifurcation in a synthetic gene network. *eLife*, 4, 2015.

[803] Anthony W Azevedo, Thuy Doan, Hormoz Moaven, Iza Sokal, Faiza Baameur, Sergey A Vishnivetskiy, Kristoff T Homan, John JG Tesmer, Vsevolod V Gurevich, Jeannie Chen, and Fred Rieke. C-terminal threonines and serines play distinct roles in the desensitization of rhodopsin, a G protein-coupled receptor. *eLife*, 4, 2015.

[804] Nicolas Aznar, Krishna K Midde, Ying Dunkel, Inmaculada Lopez-Sanchez, Yelena Pavlova, Arthur Marivin, Jorge Barbazán, Fiona Murray, Ulrich Nitsche, Klaus-Peter Janssen, Karl Willert, Ajay Goel, Miguel Abal, Mikel Garcia-Marcos, and Pradipta Ghosh. Daple is a novel non-receptor GEF required for trimeric G protein activation in Wnt signaling. *eLife*, 4, 2015.

[805] Velicia Bachtiar, Jamie Near, Heidi Johansen-Berg, and Charlotte J Stagg. Modulation of GABA and resting state functional connectivity by transcranial direct current stimulation. *eLife*, 4, 2015.

[806] Brian Bae, James Chen, Elizabeth Davis, Katherine Leon, Seth A Darst, and Elizabeth A Campbell. CarD uses a minor groove wedge mechanism to stabilize the RNA polymerase open promoter complex. *eLife*, 4, 2015.

[807] Brian Bae, Andrey Feklistov, Agnieszka Lass-Napiorkowska, Robert Landick, and Seth A Darst. Structure of a bacterial RNA polymerase holoenzyme open promoter complex. *eLife*, 4, 2015.

[808] Nicolas Baeyens, Stefania Nicoli, Brian G Coon, Tyler D Ross, Koen Van den Dries, Jinah Han, Holly M Lauridsen, Cecile O Mejean, Anne Eichmann, Jean-Leon Thomas, Jay D Humphrey, and Martin A Schwartz. Vascular remodeling is governed by a VEGFR3-dependent fluid shear stress set point. *eLife*, 4, 2015.

[809] Manon Baëza, Séverine Viala, Marjorie Heim, Amélie Dard, Bruno Hudry, Marilyne Duffraisse, Ana Rogulja-Ortmann, Christine Brun, and Samir Merabet. Inhibitory activities of short linear motifs underlie Hox interactome specificity in vivo. *eLife*, 4, 2015.

[810] Xiao-chen Bai, Eeson Rajendra, Guanghui Yang, Yigong Shi, and Sjors HW Scheres. Sampling the conformational space of the catalytic subunit of human $\gamma$-secretase. *eLife*, 4, 2015.

[811] Fernanda Bajanca, Vinicio Gonzalez-Perez, Sean J Gillespie, Cyriaque Beley, Luis Garcia, Eric Theveneau, Richard P Sear, and Simon M Hughes. In vivo dynamics of skeletal muscle Dystrophin in zebrafish embryos revealed by improved FRAP analysis. *eLife*, 4, 2015.

[812] Christa A Baker, Kevin R Huck, and Bruce A Carlson. Peripheral sensory coding through oscillatory synchrony in weakly electric fish. *eLife*, 4, 2015.

[813] Maxim Y Balakirev, James E Mullally, Adrien Favier, Nicole Assard, Eric Sulpice, David F Lindsey, Anastasia V Rulina, Xavier Gidrol, and Keith D Wilkinson. Wss1 metalloprotease partners with Cdc48/Doa1 in processing genotoxic SUMO conjugates. *eLife*, 4, 2015.

[814] Rupa Banerjee, Christina Gladkova, Koyeli Mapa, Gregor Witte, and Dejana Mokranjac. Protein translocation channel of mitochondrial inner membrane and matrix-exposed import motor communicate via two-domain coupling protein. *eLife*, 4, 2015.

[815] Balaji Banoth, Budhaditya Chatterjee, Bharath Vijayaragavan, MVR Prasad, Payel Roy, and Soumen Basak. Stimulus-selective crosstalk via the NF-$\kappa$B signaling system reinforces innate immune response to alleviate gut infection. *eLife*, 4, 2015.

[816] Pinglei Bao, Christopher J Purington, and Bosco S Tjan. Using an achiasmic human visual system to quantify the relationship between the fMRI BOLD signal and neural response. *eLife*, 4, 2015.

[817] Pierre Barbier de Reuille, Anne-Lise Routier-Kierzkowska, Daniel Kierzkowski, George W Bassel, Thierry Schüpbach, Gerardo Tauriello, Namrata Bajpai, Sören Strauss, Alain Weber, Annamaria Kiss, Agata Burian, Hugo Hofhuis, Aleksandra Sapala, Marcin Lipowczan, Maria B Heimlicher, Sarah Robinson, Emmanuelle M Bayer, Konrad Basler, Petros Koumoutsakos, Adrienne HK Roeder, Tinri Aegerter-Wilmsen, Naomi Nakayama, Miltos Tsiantis, Angela Hay, Dorota Kwiatkowska, Ioannis Xenarios, Cris Kuhlemeier, and Richard S Smith. MorphoGraphX: A platform for quantifying morphogenesis in 4D. *eLife*, 4, 2015.

[818] David Barneda, Joan Planas-Iglesias, Maria L Gaspar, Dariush Mohammadyani, Sunil Prasannan, Dirk Dormann, Gil-Soo Han, Stephen A Jesch, George M Carman, Valerian Kagan, Malcolm G Parker, Nicholas T Ktistakis, Judith Klein-Seetharaman, Ann M Dixon, Susan A Henry, and Mark Christian. The brown adipocyte protein CIDEA promotes lipid droplet fusion via a phosphatidic acid-binding amphipathic helix. *eLife*, 4, 2015.

[819] Steven P Barrett, Peter L Wang, and Julia Salzman. Circular RNA biogenesis can proceed through an exon-containing lariat precursor. *eLife*, 4, 2015.

[820] Thomas M Bartol, Cailey Bromer, Justin Kinney, Michael A Chirillo, Jennifer N Bourne, Kristen M Harris, and Terrence J Sejnowski. Nanoconnectomic upper bound on the variability of synaptic plasticity. *eLife*, 4, 2015.

[821] Simon Baumann, Olivier Joly, Adrian Rees, Christopher I Petkov, Li Sun, Alexander Thiele, and Timothy D Griffiths. The topography of frequency and time representation in primate auditory cortices. *eLife*, 4, 2015.

[822] Kyle M Baumbauer, Jennifer J DeBerry, Peter C Adelman, Richard H Miller, Junichi Hachisuka, Kuan Hsien Lee, Sarah E Ross, H Richard Koerber, Brian M Davis, and Kathryn M Albers. Keratinocytes can modulate and directly initiate nociceptive responses. *eLife*, 4, 2015.

[823] Martin Baumdick, Yannick Brüggemann, Malte Schmick, Georgia Xouri, Ola Sabet, Lloyd Davis, Jason W Chin, and Philippe IH Bastiaens. EGF-dependent re-routing of vesicular recycling switches spontaneous phosphorylation suppression to EGFR signaling. *eLife*, 4, 2015.

[824] Thomas W Bebee, Juw Won Park, Katherine I Sheridan, Claude C Warzecha, Benjamin W Cieply, Alex M Rohacek, Yi Xing, and Russ P Carstens. The splicing regulators Esrp1 and Esrp2 direct an epithelial splicing program essential for mammalian development. *eLife*, 4, 2015.

[825] Nicole L Bedford and Hopi E Hoekstra. Peromyscus mice as a model for studying natural variation. *eLife*, 4, 2015.

[826] Brittany J Belin, Terri Lee, and R Dyche Mullins. DNA damage induces nuclear actin filament assembly by Formin-2 and Spire-1/2 that promotes efficient DNA repair. *eLife*, 4, 2015.

[827] Jason C Bell, Bian Liu, and Stephen C Kowalczykowski. Imaging and energetics of single SSB-ssDNA molecules reveal intramolecular condensation and insight into RecOR function. *eLife*, 4, 2015.

[828] Timothy Bellay, Andreas Klaus, Saurav Seshadri, and Dietmar Plenz. Irregular spiking of pyramidal neurons organizes as scale-invariant neuronal avalanches in the awake state. *eLife*, 4, 2015.

[829] Victor Benichoux, Bertrand Fontaine, Tom P Franken, Shotaro Karino, Philip X Joris, and Romain Brette. Neural tuning matches frequency-dependent time differences between the ears. *eLife*, 4, 2015.

[830] Robert A Benson, Megan KL MacLeod, Benjamin G Hale, Agapitos Patakas, Paul Garside, and James M Brewer. Antigen presentation kinetics control T cell/dendritic cell interactions and follicular helper T cell generation in vivo. *eLife*, 4, 2015.

[831] Lee R Berger, John Hawks, Darryl J de Ruiter, Steven E Churchill, Peter Schmid, Lucas K Delezene, Tracy L Kivell, Heather M Garvin, Scott A Williams, Jeremy M DeSilva, Matthew M Skinner, Charles M Musiba, Noel Cameron, Trenton W Holliday, William Harcourt-Smith, Rebecca R Ackermann, Markus Bastir, Barry Bogin, Debra Bolter, Juliet Brophy, Zachary D Cofran, Kimberly A Congdon, Andrew S Deane, Mana Dembo, Michelle Drapeau, Marina C Elliott, Elen M Feuerriegel, Daniel Garcia-Martinez, David J Green, Alia Gurtov, Joel D Irish, Ashley Kruger, Myra F Laird, Damiano Marchi, Marc R Meyer, Shahed Nalla, Enquye W Negash, Caley M Orr, Davorka Radovcic, Lauren Schroeder, Jill E Scott, Zachary Throckmorton, Matthew W Tocheri, Caroline VanSickle, Christopher S Walker, Pianpian Wei, and Bernhard Zipfel. Homo naledi, a new species of the genus Homo from the Dinaledi Chamber, South Africa. *eLife*, 4, 2015.

[832] Scott Berry, Matthew Hartley, Tjelvar S G Olsson, Caroline Dean, and Martin Howard. Local chromatin environment of a Polycomb target gene instructs its own epigenetic inheritance. *eLife*, 4, 2015.

[833] Samir Bhatt, Daniel J Weiss, Bonnie Mappin, Ursula Dalrymple, Ewan Cameron, Donal Bisanzio, David L Smith, Catherine L Moyes, Andrew J Tatem, Michael Lynch, Cristin A Fergus, Joshua Yukich, Adam Bennett, Thomas P Eisele, Jan Kolaczinski, Richard E Cibulskis, Simon I Hay, and Peter W Gething. Coverage and system efficiencies of insecticide-treated nets in Africa from 2000 to 2017. *eLife*, 4, 2015.

[834] Chunjing Bian, Qiang Chen, and Xiaochun Yu. The zinc finger proteins ZNF644 and WIZ regulate the G9a/GLP complex for gene repression. *eLife*, 4, 2015.

[835] Beyenech Binotti, Nathan J Pavlos, Dietmar Riedel, Dirk Wenzel, Gerd Vorbrüggen, Amanda M Schalk, Karin Kühnel, Janina Boyken, Christian Erck, Henrik Martens, John JE Chua, and Reinhard Jahn. The GTPase Rab26 links synaptic vesicles to the autophagy pathway. *eLife*, 4, 2015.

[836] Steven M Blaauboer, Samira Mansouri, Heidi R Tucker, Hatti L Wang, Vincent D Gabrielle, and Lei Jin. The mucosal adjuvant cyclic di-GMP enhances antigen uptake and selectively activates pinocytosis-efficient cells in vivo. *eLife*, 4, 2015.

[837] Todd Blevins, Ram Podicheti, Vibhor Mishra, Michelle Marasco, Jing Wang, Doug Rusch, Haixu Tang, and Craig S Pikaard. Identification of Pol IV and RDR2-dependent precursors of 24 nt siRNAs guiding de novo DNA methylation in Arabidopsis. *eLife*, 4, 2015.

[838] Fabian Blombach, Enrico Salvadori, Thomas Fouqueau, Jun Yan, Julia Reimann, Carol Sheppard, Katherine L Smollett, Sonja V Albers, Christopher WM Kay, Konstantinos Thalassinos, and Finn Werner. Archaeal TFE$\alpha/\beta$ is a hybrid of TFIIE and the RNA polymerase III subcomplex hRPC62/39. *eLife*, 4, 2015.

[839] Gareth Bloomfield, David Traynor, Sophia P Sander, Douwe M Veltman, Justin A Pachebat, and Robert R Kay. Neurofibromin controls macropinocytosis and phagocytosis in Dictyostelium. *eLife*, 4, 2015.

[840] Zachary D Blount. The unexhausted potential of E. coli. *eLife*, 4, 2015.

[841] David Blum, Haiping Hao, and Michael McCarthy. Registered report: Transcriptional amplification in tumor cells with elevated c-Myc. *eLife*, 4, 2015.

[842] Batiste Boëda and Sandrine Etienne-Manneville. Spectrin binding motifs regulate Scribble cortical dynamics and polarity function. *eLife*, 4, 2015.

[843] Nicholas A Bokulich, Jordyn Bergsveinson, Barry Ziola, and David A Mills. Mapping microbial ecosystems and spoilage-gene flow in breweries highlights patterns of contamination and resistance. *eLife*, 4, 2015.

[844] Jacques P Bothma, Hernan G Garcia, Samuel Ng, Michael W Perry, Thomas Gregor, and Michael Levine. Enhancer additivity and non-additivity are determined by enhancer strength in the Drosophila embryo. *eLife*, 4, 2015.

[845] Mouloud Bouhadfane, Attila Kaszás, Balázs Rózsa, Ronald M Harris-Warrick, Laurent Vinay, and Frédéric Brocard. Sensitization of neonatal rat lumbar motoneuron by the inflammatory pain mediator bradykinin. *eLife*, 4, 2015.

[846] Romaric Bouveret, Ashley J Waardenberg, Nicole Schonrock, Mirana Ramialison, Tram Doan, Danielle de Jong, Antoine Bondue, Gurpreet Kaur, Stephanie Mohamed, Hananeh Fonoudi, Chiannmun Chen, Merridee A Wouters, Shoumo Bhattacharya, Nicolas Plachta, Sally L Dunwoodie, Gavin Chapman, Cédric Blanpain, and Richard P Harvey. NKX2-5 mutations causative for congenital heart disease retain functionality and are directed to hundreds of targets. *eLife*, 4, 2015.

[847] Gunther Boysen, Christopher E Barbieri, Davide Prandi, Mirjam Blattner, Sung-Suk Chae, Arun Dahija, Srilakshmi Nataraj, Dennis Huang, Clarisse Marotz, Limei Xu, Julie Huang, Paola Lecca,

Sagar Chhangawala, Deli Liu, Pengbo Zhou, Andrea Sboner, Johann S de Bono, Francesca Demichelis, Yariv Houvras, and Mark A Rubin. SPOP mutation leads to genomic instability in prostate cancer. *eLife*, 4, 2015.

[848] Brian Bradshaw, Kerry Thompson, and Uri Frank. Distinct mechanisms underlie oral vs aboral regeneration in the cnidarian Hydractinia echinata. *eLife*, 4, 2015.

[849] Niels Bradshaw and Richard Losick. Asymmetric division triggers cell-specific gene expression through coupled capture and stabilization of a phosphatase. *eLife*, 4, 2015.

[850] Olivier Britz, Jingming Zhang, Katja S Grossmann, Jason Dyck, Jun C Kim, Susan Dymecki, Simon Gosgnach, and Martyn Goulding. A genetically defined asymmetry underlies the inhibitory control of flexor–extensor locomotor movements. *eLife*, 4, 2015.

[851] Boris Brumshtein, Shannon R Esswein, Lukasz Salwinski, Martin L Phillips, Alan T Ly, Duilio Cascio, Michael R Sawaya, and David S Eisenberg. Inhibition by small-molecule ligands of formation of amyloid fibrils of an immunoglobulin light chain variable domain. *eLife*, 4, 2015.

[852] Martina Brunati, Simone Perucca, Ling Han, Angela Cattaneo, Francesco Consolato, Annapaola Andolfo, Céline Schaeffer, Eric Olinger, Jianhao Peng, Sara Santambrogio, Romain Perrier, Shuo Li, Marcel Bokhove, Angela Bachi, Edith Hummler, Olivier Devuyst, Qingyu Wu, Luca Jovine, and Luca Rampoldi. The serine protease hepsin mediates urinary secretion and polymerisation of Zona Pellucida domain protein uromodulin. *eLife*, 4, 2015.

[853] Zuzanna Brzosko, Wolfram Schultz, and Ole Paulsen. Retroactive modulation of spike timing-dependent plasticity by dopamine. *eLife*, 4, 2015.

[854] Michael G Buhnerkempe, Katelyn Gostic, Miran Park, Prianna Ahsan, Jessica A Belser, and James O Lloyd-Smith. Mapping influenza transmission in the ferret model to transmission in humans. *eLife*, 4, 2015.

[855] Brandon D Bunker, Tittu T Nellimoottil, Ryan M Boileau, Anne K Classen, and David Bilder. The transcriptional response to tumorigenic polarity loss in Drosophila. *eLife*, 4, 2015.

[856] Alondra Schweizer Burguete, Sandra Almeida, Fen-Biao Gao, Robert Kalb, Michael R Akins, and Nancy M Bonini. GGGGCC microsatellite RNA is neuritically localized, induces branching defects, and perturbs transport granule function. *eLife*, 4, 2015.

[857] Shannon Burns, Jennifer S Avena, Jay R Unruh, Zulin Yu, Sarah E Smith, Brian D Slaughter, Mark Winey, and Sue L Jaspersen. Structured illumination with particle averaging reveals novel roles for yeast centrosome components during duplication. *eLife*, 4, 2015.

[858] Nigel J Burroughs, Edward F Harry, and Andrew D McAinsh. Super-resolution kinetochore tracking reveals the mechanisms of human sister kinetochore directional switching. *eLife*, 4, 2015.

[859] Diane M Bushman, Gwendolyn E Kaeser, Benjamin Siddoway, Jurgen W Westra, Richard R Rivera, Stevens K Rehen, Yun C Yung, and Jerold Chun. Genomic mosaicism with increased amyloid precursor protein (APP) gene copy number in single neurons from sporadic Alzheimer's disease brains. *eLife*, 4, 2015.

[860] Agata Butryn, Jan M Schuller, Gabriele Stoehr, Petra Runge-Wollmann, Friedrich Förster, David T Auble, and Karl-Peter Hopfner. Structural basis for recognition and remodeling of the TBP:DNA:NC2 complex by Mot1. *eLife*, 4, 2015.

[861] Anna Caballe, Dawn M Wenzel, Monica Agromayor, Steven L Alam, Jack J Skalicky, Magdalena Kloc, Jeremy G Carlton, Leticia Labrador, Wesley I Sundquist, and Juan Martin-Serrano. ULK3 regulates cytokinetic abscission by phosphorylating ESCRT-III proteins. *eLife*, 4, 2015.

[862] Pauline Cabochette, Guillermo Vega-Lopez, Juliette Bitard, Karine Parain, Romain Chemouny, Christel Masson, Caroline Borday, Marie Hedderich, Kristine A Henningfeld, Morgane Locker, Odile Bronchain, and Muriel Perron. YAP controls retinal stem cell DNA replication timing and genomic stability. *eLife*, 4, 2015.

[863] Stuart M Cahalan, Viktor Lukacs, Sanjeev S Ranade, Shu Chien, Michael Bandell, and Ardem Patapoutian. Piezo1 links mechanical forces to red blood cell volume. *eLife*, 4, 2015.

[864] Nathaniel Calloway, Géraldine Gouzer, Mingyu Xue, and Timothy A Ryan. The active-zone protein Munc13 controls the use-dependence of presynaptic voltage-gated calcium channels. *eLife*, 4, 2015.

[865] Kate Campbell, Jakob Vowinckel, Michael Mülleder, Silke Malmsheimer, Nicola Lawrence, Enrica Calvani, Leonor Miller-Fleming, Mohammad T Alam, Stefan Christen, Markus A Keller, and Markus Ralser. Self-establishing communities enable cooperative metabolite exchange in a eukaryote. *eLife*, 4, 2015.

[866] Melody G Campbell, David Veesler, Anchi Cheng, Clinton S Potter, and Bridget Carragher. 2.8 Å resolution reconstruction of the Thermoplasma acidophilum 20S proteasome using cryo-electron microscopy. *eLife*, 4, 2015.

[867] Muqing Cao, Jue Ning, Carmen I Hernandez-Lara, Olivier Belzile, Qian Wang, Susan K Dutcher, Yanjie Liu, and William J Snell. Uni-directional ciliary membrane protein trafficking by a cytoplasmic retrograde IFT motor and ciliary ectosome shedding. *eLife*, 4, 2015.

[868] Jordan D Carelli, Steven G Sethofer, Geoffrey A Smith, Howard R Miller, Jillian L Simard, William C Merrick, Rishi K Jain, Nathan T Ross, and Jack Taunton. Ternatin and improved synthetic variants kill cancer cells by targeting the elongation factor-1A ternary complex. *eLife*, 4, 2015.

[869] Lucas B Carey. RNA polymerase errors cause splicing defects and can be regulated by differential expression of RNA polymerase subunits. *eLife*, 4, 2015.

[870] Etienne Caron, Lucia Espona, Daniel J Kowalewski, Heiko Schuster, Nicola Ternette, Adán Alpízar, Ralf B Schittenhelm, Sri H Ramarathinam, Cecilia S Lindestam Arlehamn, Ching Chiek Koh, Ludovic C Gillet, Armin Rabsteyn, Pedro Navarro, Sangtae Kim, Henry Lam, Theo Sturm, Miguel Marcilla, Alessandro Sette, David S Campbell, Eric W Deutsch, Robert L Moritz, Anthony W Purcell, Hans-Georg Rammensee, Stefan Stevanovic, and Ruedi Aebersold. An open-source computational and data resource to analyze digital maps of immunopeptidomes. *eLife*, 4, 2015.

[871] Marta Carrara, Filippo Prischi, Piotr R Nowak, Megan C Kopp, and Maruf MU Ali. Noncanonical binding of BiP ATPase domain to Ire1 and Perk is dissociated by unfolded protein CH1 to initiate ER stress signaling. *eLife*, 4, 2015.

[872] Anne-Ruxandra Carvunis, Tina Wang, Dylan Skola, Alice Yu, Jonathan Chen, Jason F Kreisberg, and Trey Ideker. Evidence for a common evolutionary rate in metazoan transcriptional networks. *eLife*, 4, 2015.

[873] C Keith Cassidy, Benjamin A Himes, Frances J Alvarez, Jun Ma, Gongpu Zhao, Juan R Perilla, Klaus Schulten, and Peijun Zhang. CryoEM and computer simulations reveal a novel kinase conformational switch in bacterial chemotaxis signaling. *eLife*, 4, 2015.

[874] Claudia Castillo-González, Xiuying Liu, Changjun Huang, Changjiang Zhao, Zeyang Ma, Tao Hu, Feng Sun, Yijun Zhou, Xueping Zhou, Xiu-Jie Wang, and Xiuren Zhang. Geminivirus-encoded TrAP suppressor inhibits the histone methyltransferase SUVH4/KYP to counter host defense. *eLife*, 4, 2015.

[875] Gamze Çamdere, Vincent Guacci, Jeremiah Stricklin, and Douglas Koshland. The ATPases of cohesin interface with regulators to modulate cohesin-mediated DNA tethering. *eLife*, 4, 2015.

[876] Diana J Cha, Jeffrey L Franklin, Yongchao Dou, Qi Liu, James N Higginbotham, Michelle Demory Beckler, Alissa M Weaver, Kasey Vickers, Nirpesh Prasad, Shawn Levy, Bing Zhang, Robert J Coffey, and James G Patton. KRAS-dependent sorting of miRNA to exosomes. *eLife*, 4, 2015.

[877] Angus Chadwick, Mark CW van Rossum, and Matthew F Nolan. Independent theta phase coding accounts for CA1 population sequences and enables flexible remapping. *eLife*, 4, 2015.

[878] Joseph E Chambers, Lucy E Dalton, Hanna J Clarke, Elke Malzer, Caia S Dominicus, Vruti Patel, Greg Moorhead, David Ron, and Stefan J Marciniak. Actin dynamics tune the integrated stress response by regulating eukaryotic initiation factor 2$\alpha$ dephosphorylation. *eLife*, 4, 2015.

[879] FuJung Chang, Alberto Riera, Cecile Evrin, Jingchuan Sun, Huilin Li, Christian Speck, and Michael Weinreich. Cdc6 ATPase activity disengages Cdc6 from the pre-replicative complex to promote DNA replication. *eLife*, 4, 2015.

[880] Tao-Hsin Chang, Fu-Lien Hsieh, Matthias Zebisch, Karl Harlos, Jonathan Elegheert, and E Yvonne Jones. Structure and functional properties of Norrin mimic Wnt for signalling with Frizzled4, Lrp5/6, and proteoglycan. *eLife*, 4, 2015.

[881] Ying-Ju Chang, Scott Pownall, Thomas E Jensen, Samar Mouaaz, Warren Foltz, Lily Zhou, Nicole Liadis, Minna Woo, Zhenyue Hao, Previn Dutt, Philip J Bilan, Amira Klip, Tak Mak, and Vuk Stambolic. The Rho-guanine nucleotide exchange factor PDZ-RhoGEF governs susceptibility to diet-induced obesity and type 2 diabetes. *eLife*, 4, 2015.

[882] Zenas C Chao, Yasuo Nagasaka, and Naotaka Fujii. Cortical network architecture for context processing in primate brain. *eLife*, 4, 2015.

[883] Paul E Chappell, El Kahina Meziane, Michael Harrison, Łukasz Magiera, Clemens Hermann, Laura Mears, Antoni G Wrobel, Charlotte Durant, Lise Lotte Nielsen, Søren Buus, Nicola Ternette, William Mwangi, Colin Butter, Venugopal Nair, Trudy Ahyee, Richard Duggleby, Alejandro Madrigal, Pietro Roversi, Susan M Lea, and Jim Kaufman. Expression levels of MHC class I molecules are inversely correlated with promiscuity of peptide binding. *eLife*, 4, 2015.

[884] Zachary Charlop-Powers, Jeremy G Owen, Boojala Vijay B Reddy, Melinda A Ternei, Denise O Guimarães, Ulysses A de Frias, Monica T Pupo, Prudy Seepe, Zhiyang Feng, and Sean F Brady. Global biogeographic sampling of bacterial secondary metabolism. *eLife*, 4, 2015.

[885] Christina Chatzi, Eric Schnell, and Gary L Westbrook. Localized hypoxia within the subgranular zone determines the early survival of newborn hippocampal granule cells. *eLife*, 4, 2015.

[886] Lizhen Chen, Marian Chuang, Thijs Koorman, Mike Boxem, Yishi Jin, and Andrew D Chisholm. Axon injury triggers EFA-6 mediated destabilization of axonal microtubules via TACC and doublecortin like kinase. *eLife*, 4, 2015.

[887] Ming Chen, Yanfang Zhao, Hualan Yang, Wenjie Luan, Jiaojiao Song, Dongyang Cui, Yi Dong, Bin Lai, Lan Ma, and Ping Zheng. Morphine disinhibits glutamatergic input to VTA dopamine neurons and promotes dopamine neuron excitation. *eLife*, 4, 2015.

[888] Ruming Chen, Cláudia Rato, Yahui Yan, Ana Crespillo-Casado, Hanna J Clarke, Heather P Harding, Stefan J Marciniak, Randy J Read, and David Ron. G-actin provides substrate-specificity to eukaryotic initiation factor 2$\alpha$ holophosphatases. *eLife*, 4, 2015.

[889] Wei-Ta Chen, Nancy D Ebelt, Travis H Stracker, Blerta Xhemalce, Carla L Van Den Berg, and Kyle M Miller. ATM regulation of IL-8 links oxidative stress to cancer cell migration and invasion. *eLife*, 4, 2015.

[890] Xiaomo Chen and Veit Stuphorn. Sequential selection of economic good and action in medial frontal cortex of macaques during value-based decisions. *eLife*, 4, 2015.

[891] Yu-Chen Chen, Xiaowei Li, Lijie Liu, Jian Wang, Chun-Qiang Lu, Ming Yang, Yun Jiao, Feng-Chao Zang, Kelly Radziwon, Guang-Di Chen, Wei Sun, Vijaya Prakash Krishnan Muthaiah, Richard Salvi, and Gao-Jun Teng. Tinnitus and hyperacusis involve hyperactivity and enhanced connectivity in auditory-limbic-arousal-cerebellar network. *eLife*, 4, 2015.

[892] Clarence Yu Cheng, Fang-Chieh Chou, Wipapat Kladwang, Siqi Tian, Pablo Cordero, and Rhiju Das. Consistent global structures of complex RNA states through multidimensional chemical mapping. *eLife*, 4, 2015.

[893] Tammy MK Cheng, Sebastian Heeger, Raphaël AG Chaleil, Nik Matthews, Aengus Stewart, Jon Wright, Carmay Lim, Paul A Bates, and Frank Uhlmann. A simple biophysical model emulates budding yeast chromosome condensation. *eLife*, 4, 2015.

[894] Yunsheng Cheng, Jiaming Wang, Yu Wang, and Mei Ding. Synaptotagmin 1 directs repetitive release by coupling vesicle exocytosis to the Rab3 cycle. *eLife*, 4, 2015.

[895] Mi Cheong Cheong, Alexander B Artyukhin, Young-Jai You, and Leon Avery. An opioid-like system regulating feeding behavior in C. elegans. *eLife*, 4, 2015.

[896] Lydia WT Cheung, Katarzyna W Walkiewicz, Tabot MD Besong, Huifang Guo, David H Hawke, Stefan T Arold, and Gordon B Mills. Regulation of the PI3K pathway through a p85$\alpha$ monomer–homodimer equilibrium. *eLife*, 4, 2015.

[897] Pak-yan Patricia Cheung, Charles Limouse, Hideo Mabuchi, and Suzanne R Pfeffer. Protein flexibility is required for vesicle tethering at the Golgi. *eLife*, 4, 2015.

[898] Kunitoshi Chiba, Joshua Z Johnson, Jacob M Vogan, Tina Wagner, John M Boyle, and Dirk Hockemeyer. Cancer-associated TERT promoter mutations abrogate telomerase silencing. *eLife*, 4, 2015.

[899] Youngshik Choe, Trung Huynh, and Samuel J Pleasure. Epithelial cells supply Sonic Hedgehog to the perinatal dentate gyrus via transport by platelets. *eLife*, 4, 2015.

[900] Hongyoon Choi, Yoori Choi, Kyu Wan Kim, Hyejin Kang, Do Won Hwang, E Edmund Kim, June-Key Chung, and Dong Soo Lee. Maturation of metabolic connectivity of the adolescent rat brain. *eLife*, 4, 2015.

[901] Alexandre Chojnowski, Peh Fern Ong, Esther SM Wong, John SY Lim, Rafidah A Mutalif, Raju Navasankari, Bamaprasad Dutta, Henry Yang, Yi Y Liow, Siu K Sze, Thomas Boudier, Graham D Wright, Alan Colman, Brian Burke, Colin L Stewart, and Oliver Dreesen. Progerin reduces LAP2$\alpha$-telomere association in Hutchinson-Gilford progeria. *eLife*, 4, 2015.

[902] Jean-Paul Chretien, Steven Riley, and Dylan B George. Mathematical modeling of the West Africa Ebola epidemic. *eLife*, 4, 2015.

[903] Ryan Patrick Christensen, Alexandra Bokinsky, Anthony Santella, Yicong Wu, Javier Marquina-Solis, Min Guo, Ismar Kovacevic, Abhishek Kumar, Peter W Winter, Nicole Tashakkori, Evan McCreedy, Huafeng Liu, Matthew McAuliffe, William Mohler, Daniel A Colón-Ramos, Zhirong Bao, and Hari Shroff. Untwisting the Caenorhabditis elegans embryo. *eLife*, 4, 2015.

[904] Denise Chroscinski, Nimet Maherali, and Erin Griner. Registered report: The CD47-signal regulated protein alpha (SIRPa) interaction is a therapeutic target for human solid tumors. *eLife*, 4, 2015.

[905] Denise Chroscinski, Darryl Sampey, and Nimet Maherali. Registered report: Tumour vascularization via endothelial differentiation of glioblastoma stem-like cells. *eLife*, 4, 2015.

[906] Michael A Cianfrocco and Andres E Leschziner. Low cost, high performance processing of single particle cryo-electron microscopy data in the cloud. *eLife*, 4, 2015.

[907] Christopher M Ciarleglio, Arseny S Khakhalin, Angelia F Wang, Alexander C Constantino, Sarah P Yip, and Carlos D Aizenman. Multivariate analysis of electrophysiological diversity of Xenopus visual neurons during development and plasticity. *eLife*, 4, 2015.

[908] Nela Cicmil, Bruce G Cumming, Andrew J Parker, and Kristine Krug. Reward modulates the effect of visual cortical microstimulation on perceptual decisions. *eLife*, 4, 2015.

[909] Elizabeth L Clare and Marc W Holderied. Acoustic shadows help gleaning bats find prey, but may be defeated by prey acoustic camouflage on rough surfaces. *eLife*, 4, 2015.

[910] Thomas M Coate, Nathalie A Spita, Kaidi D Zhang, Kevin T Isgrig, and Matthew W Kelley. Neuropilin-2/Semaphorin-3F-mediated repulsion promotes inner hair cell innervation by spiral ganglion neurons. *eLife*, 4, 2015.

[911] Michael J Coffey, Brad E Sleebs, Alessandro D Uboldi, Alexandra Garnham, Magdalena Franco, Nicole D Marino, Michael W Panas, David JP Ferguson, Marta Enciso, Matthew T O'Neill, Sash Lopaticki, Rebecca J Stewart, Grant Dewson, Gordon K Smyth, Brian J Smith, Seth L Masters, John C Boothroyd, Justin A Boddey, and Christopher J Tonkin. An aspartyl protease defines a novel pathway for export of Toxoplasma proteins into the host cell. *eLife*, 4, 2015.

[912] Daniel M Cohen, Kyoung-Jae Won, Nha Nguyen, Mitchell A Lazar, Christopher S Chen, and David J Steger. ATF4 licenses C/EBP$\beta$ activity in human mesenchymal stem cells primed for adipogenesis. *eLife*, 4, 2015.

[913] Jeremiah Y Cohen, Mackenzie W Amoroso, and Naoshige Uchida. Serotonergic neurons signal reward and punishment on multiple timescales. *eLife*, 4, 2015.

[914] Caroline Colijn and Ted Cohen. How competition governs whether moderate or aggressive treatment minimizes antibiotic resistance. *eLife*, 4, 2015.

[915] Mayeul Collot, Christian D Wilms, Asma Bentkhayet, Païkan Marcaggi, Kiri Couchman, Serge Charpak, Stéphane Dieudonné, Michael Häusser, Anne Feltz, and Jean-Maurice Mallet. CaRuby-Nano: a novel high affinity calcium probe for dual color imaging. *eLife*, 4, 2015.

[916] Paul T Conduit, Alan Wainman, Zsofia A Novak, Timothy T Weil, and Jordan W Raff. Re-examining the role of Drosophila Sas-4 in centrosome assembly using two-colour-3D-SIM FRAP. *eLife*, 4, 2015.

[917] Thomas R Connor, Clare R Barker, Kate S Baker, FranÇois-Xavier Weill, Kaisar Ali Talukder, Anthony M Smith, Stephen Baker, Malika Gouali, Duy Pham Thanh, Ishrat Jahan Azmi, Wanderley Dias da Silveira, Torsten Semmler, Lothar H Wieler, Claire Jenkins, Alejandro Cravioto, Shah M Faruque, Julian Parkhill, Dong Wook Kim, Karen H Keddy, and Nicholas R Thomson. Species-wide whole genome sequencing reveals historical global spread and recent local persistence in Shigella flexneri. *eLife*, 4, 2015.

[918] Román A Corfas and Leslie B Vosshall. The cation channel TRPA1 tunes mosquito thermotaxis to host temperatures. *eLife*, 4, 2015.

[919] Laura Cornelissen, Seong-Eun Kim, Patrick L Purdon, Emery N Brown, and Charles B Berde. Age-dependent electroencephalogram (EEG) patterns during sevoflurane general anesthesia in infants. *eLife*, 4, 2015.

[920] Daniel B Cortes, Karen L McNally, Paul E Mains, and Francis J McNally. The asymmetry of female meiosis reduces the frequency of inheritance of unpaired chromosomes. *eLife*, 4, 2015.

[921] Rui Ponte Costa, Robert C Froemke, P Jesper Sjöström, and Mark CW van Rossum. Unified pre- and postsynaptic long-term plasticity enables reliable and flexible learning. *eLife*, 4, 2015.

[922] Matthew A Cottee, Nadine Muschalik, Steven Johnson, Joanna Leveson, Jordan W Raff, and Susan M Lea. The homo-oligomerisation of both Sas-6 and Ana2 is required for efficient centriole assembly in flies. *eLife*, 4, 2015.

[923] Yoan Coudert, Wojtek Palubicki, Karin Ljung, Ondrej Novak, Ottoline Leyser, and C Jill Harrison. Three ancient hormonal cues co-ordinate shoot branching in a moss. *eLife*, 4, 2015.

[924] Alvaro H Crevenna, Marcelino Arciniega, Aurélie Dupont, Naoko Mizuno, Kaja Kowalska, Oliver F Lange, Roland Wedlich-Söldner, and Don C Lamb. Side-binding proteins modulate actin filament dynamics. *eLife*, 4, 2015.

[925] Ben M Crittenden, Daniel J Mitchell, and John Duncan. Recruitment of the default mode network during a demanding act of executive control. *eLife*, 4, 2015.

[926] Jixin Cui, Junyu Xiao, Vincent S Tagliabracci, Jianzhong Wen, Meghdad Rahdar, and Jack E Dixon. A secretory kinase complex regulates extracellular protein phosphorylation. *eLife*, 4, 2015.

[927] Daniel M Czajkowsky, Jielin Sun, Yi Shen, and Zhifeng Shao. Single molecule compression reveals intra-protein forces drive cytotoxin pore formation. *eLife*, 4, 2015.

[928] Zoulfia Darieva, Aaron Webber, Stacey Warwood, and Andrew D Sharrocks. Protein kinase C coordinates histone H3 phosphorylation and acetylation. *eLife*, 4, 2015.

[929] Alex de Mendoza, Hiroshi Suga, Jon Permanyer, Manuel Irimia, and Iñaki Ruiz-Trillo. Complex transcriptional regulation and independent evolution of fungal-like traits in a relative of animals. *eLife*, 4, 2015.

[930] Alfonso De Simone, Francesco A Aprile, Anne Dhulesia, Christopher M Dobson, and Michele Vendruscolo. Structure of a low-population intermediate state in the release of an enzyme product. *eLife*, 4, 2015.

[931] Urko del Castillo, Michael Winding, Wen Lu, and Vladimir I Gelfand. Interplay between kinesin-1 and cortical dynein during axonal outgrowth and microtubule organization in Drosophila neurons. *eLife*, 4, 2015.

[932] Rosa Della Monica, Roberta Visconti, Nando Cervone, Angela Flavia Serpico, and Domenico Grieco. Fcp1 phosphatase controls Greatwall kinase to promote PP2A-B55 activation and mitotic progression. *eLife*, 4, 2015.

[933] Lauren M Dembeck, Katalin Böröczky, Wen Huang, Coby Schal, Robert R H Anholt, and Trudy F C Mackay. Genetic architecture of natural variation in cuticular hydrocarbon composition in Drosophila melanogaster. *eLife*, 4, 2015.

[934] Nicolas Denans, Tadahiro Iimura, and Olivier Pourquié. Hox genes control vertebrate body elongation by collinear Wnt repression. *eLife*, 4, 2015.

[935] Hua Deng, Wei Wang, Jianzhong Yu, Yonggang Zheng, Yun Qing, and Duojia Pan. Spectrin regulates Hippo signaling by modulating cortical actomyosin activity. *eLife*, 4, 2015.

[936] Neha R Deshpande, Heather L Parrish, and Michael S Kuhns. Self-recognition drives the preferential accumulation of promiscuous CD4+ T-cells in aged mice. *eLife*, 4, 2015.

[937] Amandine Di-Cicco, Valérie Petit, Aurélie Chiche, Laura Bresson, Mathilde Romagnoli, Véronique Orian-Rousseau, Maria dM Vivanco, Daniel Medina, Marisa M Faraldo, Marina A Glukhova, and Marie-Ange Deugnier. Paracrine Met signaling triggers epithelial–mesenchymal transition in mammary luminal progenitors, affecting their fate. *eLife*, 4, 2015.

[938] João D Dias, Tiago Rito, Elena Torlai Triglia, Alexander Kukalev, Carmelo Ferrai, Mita Chotalia, Emily Brookes, Hiroshi Kimura, and Ana Pombo. Methylation of RNA polymerase II non-consensus Lysine residues marks early transcription in mammalian cells. *eLife*, 4, 2015.

[939] Wieneke Dijk, Markus Heine, Laurent Vergnes, Mariëtte R Boon, Gert Schaart, Matthijs KC Hesselink, Karen Reue, Wouter D van Marken Lichtenbelt, Gunilla Olivecrona, Patrick CN Rensen, Joerg Heeren, and Sander Kersten. ANGPTL4 mediates shuttling of lipid fuel to brown adipose tissue during sustained cold exposure. *eLife*, 4, 2015.

[940] Paul HGM Dirks, Lee R Berger, Eric M Roberts, Jan D Kramers, John Hawks, Patrick S Randolph-Quinney, Marina Elliott, Charles M Musiba, Steven E Churchill, Darryl J de Ruiter, Peter Schmid, Lucinda R Backwell, Georgy A Belyanin, Pedro Boshoff, K Lindsay Hunter, Elen M Feuerriegel, Alia Gurtov, James du G Harrison, Rick Hunter, Ashley Kruger, Hannah Morris, Tebogo V Makhubela, Becca Peixotto, and Steven Tucker. Geological and taphonomic context for the new hominin species Homo naledi from the Dinaledi Chamber, South Africa. *eLife*, 4, 2015.

[941] Rose E Dixon, Claudia M Moreno, Can Yuan, Ximena Opitz-Araya, Marc D Binder, Manuel F Navedo, and Luis F Santana. Graded Ca2+/calmodulin-dependent coupling of voltage-gated CaV1.2 channels. *eLife*, 4, 2015.

[942] Anne E Dodson and Jasper Rine. Heritable capture of heterochromatin dynamics in Saccharomyces cerevisiae. *eLife*, 4, 2015.

[943] Julia F Doerner, Markus Delling, and David E Clapham. Ion channels and calcium signaling in motile cilia. *eLife*, 4, 2015.

[944] Shi-Hui Dong, Weixin Tang, Tiit Lukk, Yi Yu, Satish K Nair, and Wilfred A van der Donk. The enterococcal cytolysin synthetase has an unanticipated lipid kinase fold. *eLife*, 4, 2015.

[945] Yongming Dong, Yueyang Gou, Yi Li, Yan Liu, and Jihong Bai. Synaptojanin cooperates in vivo with endophilin through an unexpected mechanism. *eLife*, 4, 2015.

[946] Jefferson J Doyle, Alexander J Doyle, Nicole K Wilson, Jennifer P Habashi, Djahida Bedja, Ryan E Whitworth, Mark E Lindsay, Florian Schoenhoff, Loretha Myers, Nick Huso, Suha Bachir, Oliver Squires, Benjamin Rusholme, Hamid Ehsan, David Huso, Craig J Thomas, Mark J Caulfield, Jennifer E Van Eyk, Daniel P Judge, and Harry C Dietz. A deleterious gene-by-environment interaction imposed by calcium channel blockers in Marfan syndrome. *eLife*, 4, 2015.

[947] Jan Drugowitsch, Gregory C DeAngelis, Dora E Angelaki, and Alexandre Pouget. Tuning the speed-accuracy trade-off to maximize reward rate in multisensory decision-making. *eLife*, 4, 2015.

[948] Manu J Dubin, Pei Zhang, Dazhe Meng, Marie-Stanislas Remigereau, Edward J Osborne, Francesco Paolo Casale, Philipp Drewe, André Kahles, Geraldine Jean, Bjarni Vilhjálmsson, Joanna Jagoda, Selen Irez, Viktor Voronin, Qiang Song, Quan Long, Gunnar Rätsch, Oliver Stegle, Richard M Clark, and Magnus Nordborg. DNA methylation in Arabidopsis has a genetic basis and shows evidence of local adaptation. *eLife*, 4, 2015.

[949] Julien Dubrulle, Benjamin M Jordan, Laila Akhmetova, Jeffrey A Farrell, Seok-Hyung Kim, Lilianna Solnica-Krezel, and Alexander F Schier. Response to Nodal morphogen gradient is determined by the kinetics of target gene induction. *eLife*, 4, 2015.

[950] Van K Duesterberg, Irena T Fischer-Hwang, Christian F Perez, Daniel W Hogan, and Steven M Block. Observation of long-range tertiary interactions during ligand binding by the TPP riboswitch aptamer. *eLife*, 4, 2015.

[951] Susan Duncan, Svante Holm, Julia Questa, Judith Irwin, Alastair Grant, and Caroline Dean. Seasonal shift in timing of vernalization as an adaptation to extreme winter. *eLife*, 4, 2015.

[952] Kyle Dunovan, Brighid Lynch, Tara Molesworth, and Timothy Verstynen. Competing basal ganglia pathways determine the difference between stopping and deciding not to go. *eLife*, 4, 2015.

[953] Briana R Dye, David R Hill, Michael AH Ferguson, Yu-Hwai Tsai, Melinda S Nagy, Rachel Dyal, James M Wells, Christopher N Mayhew, Roy Nattiv, Ophir D Klein, Eric S White, Gail H Deutsch, and Jason R Spence. In vitro generation of human pluripotent stem cell derived lung organoids. *eLife*, 4, 2015.

[954] Kate Eaton and Wanwan Yang. Registered report: Intestinal inflammation targets cancer-inducing activity of the microbiota. *eLife*, 4, 2015.

[955] Marwa El Zein, Valentin Wyart, and Julie Grèzes. Anxiety dissociates the adaptive functions of sensory and motor response enhancements to social threats. *eLife*, 4, 2015.

[956] Christopher E Ellison and Doris Bachtrog. Non-allelic gene conversion enables rapid evolutionary change at multiple regulatory sites encoded by transposable elements. *eLife*, 4, 2015.

[957] Benjamin D Engel, Miroslava Schaffer, Luis Kuhn Cuellar, Elizabeth Villa, Jürgen M Plitzko, and Wolfgang Baumeister. Native architecture of the Chlamydomonas chloroplast revealed by in situ cryo-electron tomography. *eLife*, 4, 2015.

[958] Eugeni V Entchev, Dhaval S Patel, Mei Zhan, Andrew J Steele, Hang Lu, and QueeLim Ch'ng. A gene-expression-based neural code for food abundance that modulates lifespan. *eLife*, 4, 2015.

[959] Dae Seok Eom, Emily J Bain, Larissa B Patterson, Megan E Grout, and David M Parichy. Long-distance communication by specialized cellular projections during pigment pattern development and evolution. *eLife*, 4, 2015.

[960] Jeffrey C Erlich, Bingni W Brunton, Chunyu A Duan, Timothy D Hanks, and Carlos D Brody. Distinct effects of prefrontal and parietal cortex inactivations on an accumulation of evidence task in the rat. *eLife*, 4, 2015.

[961] Nima Etemadi, Michael Chopin, Holly Anderton, Maria C Tanzer, James A Rickard, Waruni Abeysekera, Cathrine Hall, Sukhdeep K Spall, Bing Wang, Yuquan Xiong, Timothy Hla, Stuart M Pitson, Claudine S Bonder, Wendy Wei-Lynn Wong, Matthias Ernst, Gordon K Smyth, David L Vaux, Stephen L Nutt, Ueli Nachbur, and John Silke. TRAF2 regulates TNF and NF-$\kappa$B signalling to suppress apoptosis and skin inflammation independently of Sphingosine kinase 1. *eLife*, 4, 2015.

[962] Raphaël Etournay, Marko Popović, Matthias Merkel, Amitabha Nandi, Corinna Blasse, Benoît Aigouy, Holger Brandl, Gene Myers, Guillaume Salbreux, Frank Jülicher, and Suzanne Eaton. Interplay of cell dynamics and epithelial tension during morphogenesis of the Drosophila pupal wing. *eLife*, 4, 2015.

[963] Brad Evans and Erin Griner. Registered report: Oncometabolite 2-hydroxyglutarate is a competitive inhibitor of $\alpha$-ketoglutarate-dependent dioxygenases. *eLife*, 4, 2015.

[964] James Evans, Anthony Essex, Hong Xin, Nurith Amitai, Lindsey Brinton, and Erin Griner. Registered report: Wnt activity defines colon cancer stem cells and is regulated by the microenvironment. *eLife*, 4, 2015.

[965] Timothy A Evans, Celine Santiago, Elise Arbeille, and Greg J Bashaw. Robo2 acts in trans to inhibit Slit-Robo1 repulsion in pre-crossing commissural axons. *eLife*, 4, 2015.

[966] Andrey Fadeev, Jana Krauss, Hans Georg Frohnhöfer, Uwe Irion, and Christiane Nüsslein-Volhard. Tight Junction Protein 1a regulates pigment cell organisation during zebrafish colour patterning. *eLife*, 4, 2015.

[967] Qing Fang, Artur A Indzhykulian, Mirna Mustapha, Gavin P Riordan, David F Dolan, Thomas B Friedman, Inna A Belyantseva, Gregory I Frolenkov, Sally A Camper, and Jonathan E Bird. The 133-kDa N-terminal domain enables myosin 15 to maintain mechanotransducing stereocilia and is essential for hearing. *eLife*, 4, 2015.

[968] Tomas Fanutza, Dolores Del Prete, Michael J Ford, Pablo E Castillo, and Luciano D'Adamio. APP and APLP2 interact with the synaptic release machinery and facilitate transmitter release at hippocampal synapses. *eLife*, 4, 2015.

[969] Margherita Farina, Rhea van de Bospoort, Enqi He, Claudia M Persoon, Jan RT van Weering, Jurjen H Broeke, Matthijs Verhage, and Ruud F Toonen. CAPS-1 promotes fusion competence of stationary dense-core vesicles in presynaptic terminals of mammalian neurons. *eLife*, 4, 2015.

[970] Paul Farrow, Konstantin Khodosevich, Yechiam Sapir, Anton Schulmann, Muhammad Aslam, Yael Stern-Bach, Hannah Monyer, and Jakob von Engelhardt. Auxiliary subunits of the CKAMP family differentially modulate AMPA receptor properties. *eLife*, 4, 2015.

[971] Robin Fears and Volker ter Meulen. What next for gain-of-function research in Europe? *eLife*, 4, 2015.

[972] Sylvia Fechner, Luis Alvarez, Wolfgang Bönigk, Astrid Müller, Thomas K Berger, Rene Pascal, Christian Trötschel, Ansgar Poetsch, Gabriel Stölting, Kellee R Siegfried, Elisabeth Kremmer, Reinhard Seifert, and U Benjamin Kaupp. A K+-selective CNG channel orchestrates Ca2+ signalling in zebrafish sperm. *eLife*, 4, 2015.

[973] Justin Feng, Benjamin W Jester, Christine E Tinberg, Daniel J Mandell, Mauricio S Antunes, Raj Chari, Kevin J Morey, Xavier Rios, June I Medford, George M Church, Stanley Fields, and David Baker. A general strategy to construct small molecule biosensors in eukaryotes. *eLife*, 4, 2015.

[974] Qing Feng, Lauren Snider, Sujatha Jagannathan, Rabi Tawil, Silvère M van der Maarel, Stephen J Tapscott, and Robert K Bradley. A feedback loop between nonsense-mediated decay and the retrogene DUX4 in facioscapulohumeral muscular dystrophy. *eLife*, 4, 2015.

[975] Rafael Fernandez-Leiro, Julian Conrad, Sjors HW Scheres, and Meindert H Lamers. cryo-EM structures of the E. coli replicative DNA polymerase reveal its dynamic interactions with the DNA sliding clamp, exonuclease and $\tau$. *eLife*, 4, 2015.

[976] Marc Fiedler, Michael Graeb, Juliusz Mieszczanek, Trevor J Rutherford, Christopher M Johnson, and Mariann Bienz. An ancient Pygo-dependent Wnt enhanceosome integrated by Chip/LDB-SSDP. *eLife*, 4, 2015.

[977] Steven Fiering, Lay-Hong Ang, Judith Lacoste, Tim D Smith, and Erin Griner. Registered report: Biomechanical remodeling of the microenvironment by stromal caveolin-1 favors tumor invasion and metastasis. *eLife*, 4, 2015.

[978] Bernd Fischer, Thomas Sandmann, Thomas Horn, Maximilian Billmann, Varun Chaudhary, Wolfgang Huber, and Michael Boutros. A map of directional genetic interactions in a metazoan cell. *eLife*, 4, 2015.

[979] Ute Fischer, Nico Schäuble, Sabina Schütz, Martin Altvater, Yiming Chang, Marius Boulos Faza, and Vikram Govind Panse. A non-canonical mechanism for Crm1-export cargo complex assembly. *eLife*, 4, 2015.

[980] James E Fitzgerald and Damon A Clark. Nonlinear circuits for naturalistic visual motion estimation. *eLife*, 4, 2015.

[981] Michael S Fleming, Anna Vysochan, Sónia Paixão, Jingwen Niu, Rüdiger Klein, Joseph M Savitt, and Wenqin Luo. Cis and trans RET signaling control the survival and central projection growth of rapidly adapting mechanoreceptors. *eLife*, 4, 2015.

[982] David Flores-Benitez and Elisabeth Knust. Crumbs is an essential regulator of cytoskeletal dynamics and cell-cell adhesion during dorsal closure in Drosophila. *eLife*, 4, 2015.

[983] Christopher B Ford, Jason M Funt, Darren Abbey, Luca Issi, Candace Guiducci, Diego A Martinez, Toni Delorey, Bi yu Li, Theodore C White, Christina Cuomo, Reeta P Rao, Judith Berman, Dawn A Thompson, and Aviv Regev. The evolution of drug resistance in clinical isolates of Candida albicans. *eLife*, 4, 2015.

[984] Peter J Foster, Sebastian Fürthauer, Michael J Shelley, and Daniel J Needleman. Active contraction of microtubule networks. *eLife*, 4, 2015.

[985] Silke Frahm, Beatriz Antolin-Fontes, Andreas Görlich, Johannes-Friedrich Zander, Gudrun Ahnert-Hilger, and Ines Ibañez-Tallon. An essential role of acetylcholine-glutamate synergy at habenular synapses in nicotine dependence. *eLife*, 4, 2015.

[986] Nicholas T Franklin and Michael J Frank. A cholinergic feedback circuit to regulate striatal population uncertainty and optimize reinforcement learning. *eLife*, 4, 2015.

[987] Eleonora Franzoni, Sam A Booker, Srinivas Parthasarathy, Frederick Rehfeld, Sabine Grosser, Swathi Srivatsa, Heiko R Fuchs, Victor Tarabykin, Imre Vida, and F Gregory Wulczyn. miR-128 regulates neuronal migration, outgrowth and intrinsic excitability via the intellectual disability gene Phf6. *eLife*, 4, 2015.

[988] Tanya S Freedman, Ying X Tan, Katarzyna M Skrzypczynska, Boryana N Manz, Frances V Sjaastad, Helen S Goodridge, Clifford A Lowell, and Arthur Weiss. LynA regulates an inflammation-sensitive signaling checkpoint in macrophages. *eLife*, 4, 2015.

[989] Jeremy Freeman, Greg D Field, Peter H Li, Martin Greschner, Deborah E Gunning, Keith Mathieson, Alexander Sher, Alan M Litke, Liam Paninski, Eero P Simoncelli, and EJ Chichilnisky. Mapping nonlinear receptive field structure in primate retina at single cone resolution. *eLife*, 4, 2015.

[990] Lise Frézal and Marie-Anne Félix. C. elegans outside the Petri dish. *eLife*, 4, 2015.

[991] Jonathan R Friedman, Arnaud Mourier, Justin Yamada, J Michael McCaffery, and Jodi Nunnari. MICOS coordinates with respiratory complexes and lipids to establish mitochondrial inner membrane architecture. *eLife*, 4, 2015.

[992] Florian Fröhlich, Constance Petit, Nora Kory, Romain Christiano, Hans-Kristian Hannibal-Bach, Morven Graham, Xinran Liu, Christer S Ejsing, Robert V Farese, and Tobias C Walther. The GARP complex is required for cellular sphingolipid homeostasis. *eLife*, 4, 2015.

[993] Idan Frumin, Ofer Perl, Yaara Endevelt-Shapira, Ami Eisen, Neetai Eshel, Iris Heller, Maya Shemesh, Aharon Ravia, Lee Sela, Anat Arzi, and Noam Sobel. A social chemosignaling function for human handshaking. *eLife*, 4, 2015.

[994] Yu Fu, Megumi Kaneko, Yunshuo Tang, Arturo Alvarez-Buylla, and Michael P Stryker. A cortical disinhibitory circuit for enhancing adult plasticity. *eLife*, 4, 2015.

[995] Ho Yee Joyce Fung, Szu-Chin Fu, Chad A Brautigam, and Yuh Min Chook. Structural determinants of nuclear export signal orientation in binding to exportin CRM1. *eLife*, 4, 2015.

[996] Juan José Fung, Alan Kosaka, Xiaochuan Shan, Gwenn Danet-Desnoyers, Michael Gormally, and Kate Owen. Registered report: Inhibition of BET recruitment to chromatin as an effective treatment for MLL-fusion leukemia. *eLife*, 4, 2015.

[997] Irene Gallego Romero, Bryan J Pavlovic, Irene Hernando-Herraez, Xiang Zhou, Michelle C Ward, Nicholas E Banovich, Courtney L Kagan, Jonathan E Burnett, Constance H Huang, Amy Mitrano, Claudia I Chavarria, Inbar Friedrich Ben-Nun, Yingchun Li, Karen Sabatini, Trevor R Leonardo, Mana Parast, Tomas Marques-Bonet, Louise C Laurent, Jeanne F Loring, and Yoav Gilad. A panel of induced pluripotent stem cells from chimpanzees: a resource for comparative functional genomics. *eLife*, 4, 2015.

[998] Elad Ganmor, Ronen Segev, and Elad Schneidman. A thesaurus for a neural population code. *eLife*, 4, 2015.

[999] Stephanie C Gantz, Brooks G Robinson, David C Buck, James R Bunzow, Rachael L Neve, John T Williams, and Kim A Neve. Distinct regulation of dopamine D2S and D2L autoreceptor signaling by calcium. *eLife*, 4, 2015.

[1000] Chenxi Gao, Guangming Chen, Shih-Fan Kuan, Dennis Han Zhang, David D Schlaepfer, and Jing Hu. FAK/PYK2 promotes the Wnt/$\beta$-catenin pathway and intestinal tumorigenesis by phosphorylating GSK3$\beta$. *eLife*, 4, 2015.

[1001] Jie Gao, Shannon M Buckley, Luisa Cimmino, Maria Guillamot, Alexandros Strikoudis, Yong Cang, Stephen P Goff, and Iannis Aifantis. The CUL4-DDB1 ubiquitin ligase complex controls adult and embryonic stem cell differentiation and homeostasis. *eLife*, 4, 2015.

[1002] Xing-Huang Gao, Dawid Krokowski, Bo-Jhih Guan, Ilya Bederman, Mithu Majumder, Marc Parisien, Luda Diatchenko, Omer Kabil, Belinda Willard, Ruma Banerjee, Benlian Wang, Gurkan Bebek, Charles R. Evans, Paul L. Fox, Stanton L. Gerson, Charles L. Hoppel, Ming Liu, Peter Arvan, and Maria Hatzoglou. Quantitative H2S-mediated protein sulfhydration reveals metabolic reprogramming during the integrated stress response. *eLife*, 4, 2015.

[1003] Thomas Garcia, Guilhem Doulcier, and Silvia De Monte. The evolution of adhesiveness as a social adaptation. *eLife*, 4, 2015.

[1004] Ivana Gasic, Purnima Nerurkar, and Patrick Meraldi. Centrosome age regulates kinetochore–microtubule stability and biases chromosome mis-segregation. *eLife*, 4, 2015.

[1005] Alberto T Gatta, Louise H Wong, Yves Y Sere, Diana M Calderón-Noreña, Shamshad Cockcroft, Anant K Menon, and Tim P Levine. A new family of StART domain proteins at membrane contact sites has a role in ER-PM sterol transport. *eLife*, 4, 2015.

[1006] Michael A Gaviño, Kevin J Ford, Santiago Archila, and Graeme W Davis. Homeostatic synaptic depression is achieved through a regulated decrease in presynaptic calcium channel abundance. *eLife*, 4, 2015.

[1007] Xuecai Ge, Ljiljana Milenkovic, Kaye Suyama, Tom Hartl, Teresa Purzner, Amy Winans, Tobias Meyer, and Matthew P Scott. Phosphodiesterase 4D acts downstream of Neuropilin to control Hedgehog signal transduction and the growth of medulloblastoma. *eLife*, 4, 2015.

[1008] Matthew Gemberling, Ravi Karra, Amy L Dickson, and Kenneth D Poss. Nrg1 is an injury-induced cardiomyocyte mitogen for the endogenous heart regeneration program in zebrafish. *eLife*, 4, 2015.

[1009] Vincenzo A Gennarino, Callison E Alcott, Chun-An Chen, Arindam Chaudhury, Madelyn A Gillentine, Jill A Rosenfeld, Sumit Parikh, James W Wheless, Elizabeth R Roeder, Dafne DG Horovitz, Erin K Roney, Janice L Smith, Sau W Cheung, Wei Li, Joel R Neilson, Christian P Schaaf, and Huda Y Zoghbi. NUDT21-spanning CNVs lead to neuropsychiatric disease and altered MeCP2 abundance via alternative polyadenylation. *eLife*, 4, 2015.

[1010] Joju George, Cary Soares, Audrey Montersino, Jean-Claude Beique, and Gareth M Thomas. Palmitoylation of LIM Kinase-1 ensures spine-specific actin polymerization and morphological plasticity. *eLife*, 4, 2015.

[1011] Roxana E Georgescu, Grant D Schauer, Nina Y Yao, Lance D Langston, Olga Yurieva, Dan Zhang, Jeff Finkelstein, and Mike E O'Donnell. Reconstitution of a eukaryotic replisome reveals suppression mechanisms that define leading/lagging strand operation. *eLife*, 4, 2015.

[1012] Ruben Gepner, Mirna Mihovilovic Skanata, Natalie M Bernat, Margarita Kaplow, and Marc Gershow. Computations underlying Drosophila photo-taxis, odor-taxis, and multi-sensory integration. *eLife*, 4, 2015.

[1013] Anna C Geraghty, Sandra E Muroy, Sheng Zhao, George E Bentley, Lance J Kriegsfeld, and Daniela Kaufer. Knockdown of hypothalamic RFRP3 prevents chronic stress-induced infertility and embryo resorption. *eLife*, 4, 2015.

[1014] Richard C Gerkin and Jason B Castro. The number of olfactory stimuli that humans can discriminate is still unknown. *eLife*, 4, 2015.

[1015] Elisabeth A Geyer, Alexander Burns, Beth A Lalonde, Xuecheng Ye, Felipe-Andres Piedra, Tim C Huffaker, and Luke M Rice. A mutation uncouples the tubulin conformational and GTPase cycles, revealing allosteric control of microtubule dynamics. *eLife*, 4, 2015.

[1016] Lisa F Gill, Wolfgang Goymann, Andries Ter Maat, and Manfred Gahr. Patterns of call communication between group-housed zebra finches change during the breeding cycle. *eLife*, 4, 2015.

[1017] Sumanprava Giri, Vasudha Aggarwal, Julien Pontis, Zhen Shen, Arindam Chakraborty, Abid Khan, Craig Mizzen, Kannanganattu V Prasanth, Slimane Ait-Si-Ali, Taekjip Ha, and Supriya G Prasanth. The preRC protein ORCA organizes heterochromatin by assembling histone H3 lysine 9 methyltransferases on chromatin. *eLife*, 4, 2015.

[1018] Christopher M Glaze, Joseph W Kable, and Joshua I Gold. Normative evidence accumulation in unpredictable environments. *eLife*, 4, 2015.

[1019] Sezgi Goksan, Caroline Hartley, Faith Emery, Naomi Cockrill, Ravi Poorun, Fiona Moultrie, Richard Rogers, Jon Campbell, Michael Sanders, Eleri Adams, Stuart Clare, Mark Jenkinson, Irene Tracey, and Rebeccah Slater. fMRI reveals neural activity overlap between adult and infant pain. *eLife*, 4, 2015.

[1020] Vicki AM Gold, Ralf Salzer, Beate Averhoff, and Werner Kühlbrandt. Structure of a type IV pilus machinery in the open and closed state. *eLife*, 4, 2015.

[1021] Seth R Goldman, Nikhil U Nair, Christopher D Wells, Bryce E Nickels, and Ann Hochschild. The primary $\sigma$ factor in Escherichia coli can access the transcription elongation complex from solution in vivo. *eLife*, 4, 2015.

[1022] Matthew D Golub, Byron M Yu, and Steven M Chase. Internal models for interpreting neural population activity during sensorimotor control. *eLife*, 4, 2015.

[1023] Stephen N Gomperts, Fabian Kloosterman, and Matthew A Wilson. VTA neurons coordinate with the hippocampal reactivation of spatial experience. *eLife*, 4, 2015.

[1024] Katelyn M Gostic, Adam J Kucharski, and James O Lloyd-Smith. Effectiveness of traveller screening for emerging pathogens is shaped by epidemiology and natural history of infection. *eLife*, 4, 2015.

[1025] Katja Gotthardt, Mandy Lokaj, Carolin Koerner, Nathalie Falk, Andreas Gießl, and Alfred Wittinghofer. A G-protein activation cascade from Arl13B to Arl3 and implications for ciliary targeting of lipidated proteins. *eLife*, 4, 2015.

[1026] Thiago S Gouvêa, Tiago Monteiro, Asma Motiwala, Sofia Soares, Christian Machens, and Joseph J Paton. Striatal dynamics explain duration judgments. *eLife*, 4, 2015.

[1027] Benjamin G Gowen, Bryan Chim, Caleb D Marceau, Trever T Greene, Patrick Burr, Jeanmarie R Gonzalez, Charles R Hesser, Peter A Dietzen, Teal Russell, Alexandre Iannello, Laurent Coscoy, Charles L Sentman, Jan E Carette, Stefan A Muljo, and David H Raulet. A forward genetic screen reveals novel independent regulators of ULBP1, an activating ligand for natural killer cells. *eLife*, 4, 2015.

[1028] Katharine R Grabek, Cecilia Diniz Behn, Gregory S Barsh, Jay R Hesselberth, and Sandra L Martin. Enhanced stability and polyadenylation of select mRNAs support rapid thermogenesis in the brown fat of a hibernator. *eLife*, 4, 2015.

[1029] Timothy Grant and Nikolaus Grigorieff. Measuring the optimal exposure for single particle cryo-EM using a 2.6 Å reconstruction of rotavirus VP6. *eLife*, 4, 2015.

[1030] Andrew N Gray, Alexander JF Egan, Inge L van't Veer, Jolanda Verheul, Alexandre Colavin, Alexandra Koumoutsi, Jacob Biboy, A F Maarten Altelaar, Mirjam J Damen, Kerwyn Casey Huang, Jean-Pierre Simorre, Eefjan Breukink, Tanneke den Blaauwen, Athanasios Typas, Carol A Gross, and Waldemar Vollmer. Coordination of peptidoglycan synthesis and outer membrane constriction during Escherichia coli cell division. *eLife*, 4, 2015.

[1031] Bree K Grillo-Hill, Changhoon Choi, Maite Jimenez-Vidal, and Diane L Barber. Increased H+ efflux is sufficient to induce dysplasia and necessary for viability with oncogene expression. *eLife*, 4, 2015.

[1032] William N Grimes, Logan R Graves, Mathew T Summers, and Fred Rieke. A simple retinal mechanism contributes to perceptual interactions between rod- and cone-mediated responses in primates. *eLife*, 4, 2015.

[1033] Derek Groen and Ben Calderhead. Science hackathons for developing interdisciplinary research and collaborations. *eLife*, 4, 2015.

[1034] Flora S Groothuizen, Ines Winkler, Michele Cristóvão, Alexander Fish, Herrie HK Winterwerp, Annet Reumer, Andreas D Marx, Nicolaas Hermans, Robert A Nicholls, Garib N Murshudov, Joyce HG Lebbink, Peter Friedhoff, and Titia K Sixma. MutS/MutL crystal structure reveals that the MutS sliding clamp loads MutL onto DNA. *eLife*, 4, 2015.

[1035] Ana R Grosso, Ana P Leite, Sílvia Carvalho, Mafalda R Matos, Filipa B Martins, Alexandra C Vítor, Joana MP Desterro, Maria Carmo-Fonseca, and Sérgio F de Almeida. Pervasive transcription read-through promotes aberrant expression of oncogenes and RNA chimeras in renal carcinoma. *eLife*, 4, 2015.

[1036] Tina M Gruene, Katelyn Flick, Alexis Stefano, Stephen D Shea, and Rebecca M Shansky. Sexually divergent expression of active and passive conditioned fear responses in rats. *eLife*, 4, 2015.

[1037] Yapeng Gu, Jill Shea, Gloria Slattum, Matthew A Firpo, Margaret Alexander, Sean J Mulvihill, Vita M Golubovskaya, and Jody Rosenblatt. Defective apical extrusion signaling contributes to aggressive tumor hallmarks. *eLife*, 4, 2015.

[1038] Jikui Guan, Ganesh Umapathy, Yasuo Yamazaki, Georg Wolfstetter, Patricia Mendoza, Kathrin Pfeifer, Ateequrrahman Mohammed, Fredrik Hugosson, Hongbing Zhang, Amy W Hsu, Robert Halenbeck, Bengt Hallberg, and Ruth H Palmer. FAM150A and FAM150B are activating ligands for anaplastic lymphoma kinase. *eLife*, 4, 2015.

[1039] Virginia Guarani, Elizabeth M McNeill, Joao A Paulo, Edward L Huttlin, Florian Fröhlich, Steven P Gygi, David Van Vactor, and J Wade Harper. QIL1 is a novel mitochondrial protein required for MICOS complex stability and cristae morphology. *eLife*, 4, 2015.

[1040] Jian-Zhong Guo, Austin R Graves, Wendy W Guo, Jihong Zheng, Allen Lee, Juan Rodríguez-González, Nuo Li, John J Macklin, James W Phillips, Brett D Mensh, Kristin Branson, and Adam W Hantman. Cortex commands the performance of skilled movement. *eLife*, 4, 2015.

[1041] Kanchan Gupta, Maryam Zamanian, Chanhyung Bae, Mirela Milescu, Dmitriy Krepkiy, Drew C Tilley, Jon T Sack, Vladimir Yarov-Yarovoy, Jae Il Kim, and Kenton J Swartz. Tarantula toxins use common surfaces for interacting with Kv and ASIC ion channels. *eLife*, 4, 2015.

[1042] Kevin C Hadley, Rishi Rakhit, Hongbo Guo, Yulong Sun, James EN Jonkman, Joanne McLaurin, Lili-Naz Hazrati, Andrew Emili, and Avijit Chakrabartty. Determining composition of micron-scale protein deposits in neurodegenerative disease by spatially targeted optical microproteomics. *eLife*, 4, 2015.

[1043] Daniel Christopher Haggerty and Daoyun Ji. Activities of visual cortical and hippocampal neurons co-fluctuate in freely moving rats during spatial behavior. *eLife*, 4, 2015.

[1044] Avital Hahamy, Stamatios N Sotiropoulos, David Henderson Slater, Rafael Malach, Heidi Johansen-Berg, and Tamar R Makin. Normalisation of brain connectivity through compensatory behaviour, despite congenital hand absence. *eLife*, 4, 2015.

[1045] Sarah Hake and Jeffrey Ross-Ibarra. Genetic, evolutionary and plant breeding insights from the domestication of maize. *eLife*, 4, 2015.

[1046] Rosalind Hale, Amy L Brittle, Katherine H Fisher, Nicholas A M Monk, and David Strutt. Cellular interpretation of the long-range gradient of Four-jointed activity in the Drosophila wing. *eLife*, 4, 2015.

[1047] L Shahul Hameed, Daniel A Berg, Laure Belnoue, Lasse D Jensen, Yihai Cao, and András Simon. Environmental changes in oxygen tension reveal ROS-dependent neurogenesis and regeneration in the adult newt brain. *eLife*, 4, 2015.

[1048] Stefanie Hampel, Romain Franconville, Julie H Simpson, and Andrew M Seeds. A neural command circuit for grooming movement control. *eLife*, 4, 2015.

[1049] L Naomi Handly, Anna Pilko, and Roy Wollman. Paracrine communication maximizes cellular response fidelity in wound signaling. *eLife*, 4, 2015.

[1050] Anders S Hansen and Erin K O'Shea. Limits on information transduction through amplitude and frequency regulation of transcription factor activity. *eLife*, 4, 2015.

[1051] Jody M Hansen, Daniela R Chavez, and Gillian M Stanfield. COMP-1 promotes competitive advantage of nematode sperm. *eLife*, 4, 2015.

[1052] Scott D Hansen and R Dyche Mullins. Lamellipodin promotes actin assembly by clustering Ena/VASP proteins and tethering them to actin filaments. *eLife*, 4, 2015.

[1053] Rizal F Hariadi, Ruth F Sommese, and Sivaraj Sivaramakrishnan. Tuning myosin-driven sorting on cellular actin networks. *eLife*, 4, 2015.

[1054] Robin M Harris, Barret D Pfeiffer, Gerald M Rubin, and James W Truman. Neuron hemilineages provide the functional ground plan for the Drosophila ventral nervous system. *eLife*, 4, 2015.

[1055] Tomomi Hashidate-Yoshida, Takeshi Harayama, Daisuke Hishikawa, Ryo Morimoto, Fumie Hamano, Suzumi M Tokuoka, Miki Eto, Miwa Tamura-Nakano, Rieko Yanobu-Takanashi, Yoshiko Mukumoto, Hiroshi Kiyonari, Tadashi Okamura, Yoshihiro Kita, Hideo Shindou, and Takao Shimizu. Fatty acid remodeling by LPCAT3 enriches arachidonate in phospholipid membranes and regulates triglyceride transport. *eLife*, 4, 2015.

[1056] Evan P Hass and David C Zappulla. The Ku subunit of telomerase binds Sir4 to recruit telomerase to lengthen telomeres in S. cerevisiae. *eLife*, 4, 2015.

[1057] Melissa A Hausburg, Jason D Doles, Sandra L Clement, Adam B Cadwallader, Monica N Hall, Perry J Blackshear, Jens Lykke-Andersen, and Bradley B Olwin. Post-transcriptional regulation of satellite cell quiescence by TTP-mediated mRNA decay. *eLife*, 4, 2015.

[1058] Paula R Haynes, Bethany L Christmann, and Leslie C Griffith. A single pair of neurons links sleep to memory consolidation in Drosophila melanogaster. *eLife*, 4, 2015.

[1059] Lian He, Yuanwei Zhang, Guolin Ma, Peng Tan, Zhanjun Li, Shengbing Zang, Xiang Wu, Ji Jing, Shaohai Fang, Lijuan Zhou, Youjun Wang, Yun Huang, Patrick G Hogan, Gang Han, and Yubin Zhou. Near-infrared photoactivatable control of Ca2+ signaling and optogenetic immunomodulation. *eLife*, 4, 2015.

[1060] Stephan Heermann, Lucas Schütz, Steffen Lemke, Kerstin Krieglstein, and Joachim Wittbrodt. Eye morphogenesis driven by epithelial flow into the optic cup facilitated by modulation of bone morphogenetic protein. *eLife*, 4, 2015.

[1061] Ramanath Narayana Hegde, Seetharaman Parashuraman, Francesco Iorio, Fabiana Ciciriello, Fabrizio Capuani, Annamaria Carissimo, Diego Carrella, Vincenzo Belcastro, Advait Subramanian, Laura Bounti, Maria Persico, Graeme Carlile, Luis Galietta, David Y Thomas, Diego Di Bernardo, and Alberto Luini. Unravelling druggable signalling networks that control F508del-CFTR proteostasis. *eLife*, 4, 2015.

[1062] Heidi Hehnly, David Canton, Paula Bucko, Lorene K Langeberg, Leah Ogier, Irwin Gelman, L Fernando Santana, Linda Wordeman, and John D Scott. A mitotic kinase scaffold depleted in testicular seminomas impacts spindle orientation in germ line stem cells. *eLife*, 4, 2015.

[1063] Christian SM Helker, Annika Schuermann, Cathrin Pollmann, Serene C Chng, Friedemann Kiefer, Bruno Reversade, and Wiebke Herzog. The hormonal peptide Elabela guides angioblasts to the midline during vasculogenesis. *eLife*, 4, 2015.

[1064] Valerie C Henderson, Nadine Demko, Amanda Hakala, Nathalie MacKinnon, Carole A Federico, Dean Fergusson, and Jonathan Kimmelman. A meta-analysis of threats to valid clinical inference in preclinical research of sunitinib. *eLife*, 4, 2015.

[1065] Fredrick E Henry, Ken Sugino, Adam Tozer, Tiago Branco, and Scott M Sternson. Cell type-specific transcriptomics of hypothalamic energy-sensing neuron responses to weight-loss. *eLife*, 4, 2015.

[1066] Anton G Henssen, Elizabeth Henaff, Eileen Jiang, Amy R Eisenberg, Julianne R Carson, Camila M Villasante, Mondira Ray, Eric Still, Melissa Burns, Jorge Gandara, Cedric Feschotte, Christopher E Mason, and Alex Kentsis. Genomic DNA transposition induced by human PGBD5. *eLife*, 4, 2015.

[1067] Clemens Hermann, Andy van Hateren, Nico Trautwein, Andreas Neerincx, Patrick J Duriez, Stefan Stevanović, John Trowsdale, Janet E Deane, Tim Elliott, and Louise H Boyle. TAPBPR alters MHC class I peptide presentation by functioning as a peptide exchange catalyst. *eLife*, 4, 2015.

[1068] Robert G Hesse, Gayle K Kouklis, Nadav Ahituv, and Jason H Pomerantz. The human ARF tumor suppressor senses blastema activity and suppresses epimorphic tissue regeneration. *eLife*, 4, 2015.

[1069] Simone Hettmer, Anna C Schinzel, Daria Tchessalova, Michaela Schneider, Christina L Parker, Roderick T Bronson, Nigel GJ Richards, William C Hahn, and Amy J Wagers. Functional genomic screening reveals asparagine dependence as a metabolic vulnerability in sarcoma. *eLife*, 4, 2015.

[1070] Anneke Hibbel, Aliona Bogdanova, Mohammed Mahamdeh, Anita Jannasch, Marko Storch, Erik Schäffer, Dimitris Liakopoulos, and Jonathon Howard. Kinesin Kip2 enhances microtubule growth in vitro through length-dependent feedback on polymerization and catastrophe. *eLife*, 4, 2015.

[1071] Kevin G Hicks, Scott P Delbecq, Enea Sancho-Vaello, Marie-Pierre Blanc, Katja K Dove, Lynne R Prost, Margaret E Daley, Kornelius Zeth, Rachel E Klevit, and Samuel I Miller. Acidic pH and divalent cation sensing by PhoQ are dispensable for systemic salmonellae virulence. *eLife*, 4, 2015.

[1072] Stephen M Hinshaw, Vasso Makrantoni, Alastair Kerr, Adèle L Marston, and Stephen C Harrison. Structural evidence for Scc4-dependent localization of cohesin loading. *eLife*, 4, 2015.

[1073] Susumu Hirabayashi and Ross L Cagan. Salt-inducible kinases mediate nutrient-sensing to link dietary sugar and tumorigenesis in Drosophila. *eLife*, 4, 2015.

[1074] Toru Hiratsuka, Yoshihisa Fujita, Honda Naoki, Kazuhiro Aoki, Yuji Kamioka, and Michiyuki Matsuda. Intercellular propagation of extracellular signal-regulated kinase activation revealed by in vivo imaging of mouse skin. *eLife*, 4, 2015.

[1075] Natsuko Hitora-Imamura, Yuki Miura, Chie Teshirogi, Yuji Ikegaya, Norio Matsuki, and Hiroshi Nomura. Prefrontal dopamine regulates fear reinstatement through the downregulation of extinction circuits. *eLife*, 4, 2015.

[1076] Florian Hladik, Adam Burgener, Lamar Ballweber, Raphael Gottardo, Lucia Vojtech, Slim Fourati, James Y Dai, Mark J Cameron, Johanna Strobl, Sean M Hughes, Craig Hoesley, Philip Andrew, Sherri Johnson, Jeanna Piper, David R Friend, T Blake Ball, Ross D Cranston, Kenneth H Mayer, M Juliana McElrath, and Ian McGowan. Mucosal effects of tenofovir 1% gel. *eLife*, 4, 2015.

[1077] Diana M Ho, SK Pallavi, and Spyros Artavanis-Tsakonas. The Notch-mediated hyperplasia circuitry in Drosophila reveals a Src-JNK signaling axis. *eLife*, 4, 2015.

[1078] Carl J Hodgetts, Mark Postans, Jonathan P Shine, Derek K Jones, Andrew D Lawrence, and Kim S Graham. Dissociable roles of the inferior longitudinal fasciculus and fornix in face and place perception. *eLife*, 4, 2015.

[1079] Doris Höglinger, Per Haberkant, Auxiliadora Aguilera-Romero, Howard Riezman, Forbes D Porter, Frances M Platt, Antony Galione, and Carsten Schultz. Intracellular sphingosine releases calcium from lysosomes. *eLife*, 4, 2015.

[1080] Maria-Graciela Hollm-Delgado, Frédéric B Piel, Daniel J Weiss, Rosalind E Howes, Elizabeth A Stuart, Simon I Hay, and Robert E Black. Vitamin A supplements, routine immunization, and the subsequent risk of Plasmodium infection among children under 5 years in sub-Saharan Africa. *eLife*, 4, 2015.

[1081] Eric D Hoopfer, Yonil Jung, Hidehiko K Inagaki, Gerald M Rubin, and David J Anderson. P1 interneurons promote a persistent internal state that enhances inter-male aggression in Drosophila. *eLife*, 4, 2015.

[1082] Christine S Hopp, Kevin Chiou, Daniel RT Ragheb, Ahmed M Salman, Shahid M Khan, Andrea J Liu, and Photini Sinnis. Longitudinal analysis of Plasmodium sporozoite motility in the dermis reveals component of blood vessel recognition. *eLife*, 4, 2015.

[1083] James Hose, Chris Mun Yong, Maria Sardi, Zhishi Wang, Michael A Newton, and Audrey P Gasch. Dosage compensation can buffer copy-number variation in wild yeast. *eLife*, 4, 2015.

[1084] Loren E Hough, Kaushik Dutta, Samuel Sparks, Deniz B Temel, Alia Kamal, Jaclyn Tetenbaum-Novatt, Michael P Rout, and David Cowburn. The molecular mechanism of nuclear transport revealed by atomic-scale measurements. *eLife*, 4, 2015.

[1085] Ted M Hsu, Joel D Hahn, Vaibhav R Konanur, Emily E Noble, Andrea N Suarez, Jessica Thai, Emily M Nakamoto, and Scott E Kanoski. Hippocampus ghrelin signaling mediates appetite through lateral hypothalamic orexin pathways. *eLife*, 4, 2015.

[1086] Hai Huang and Thomas B Kornberg. Myoblast cytonemes mediate Wg signaling from the wing imaginal disc and Delta-Notch signaling to the air sac primordium. *eLife*, 4, 2015.

[1087] Hung-Hsiang Huang, Antti Hassinen, Subha Sundaram, Andrej-Nikolai Spiess, Sakari Kellokumpu, and Pamela Stanley. GnT1IP-L specifically inhibits MGAT1 in the Golgi via its luminal domain. *eLife*, 4, 2015.

[1088] Siyi Huang, Kevin J O'Donovan, Eric E Turner, Jian Zhong, and David D Ginty. Extrinsic and intrinsic signals converge on the Runx1/CBF$\beta$ transcription factor for nonpeptidergic nociceptor maturation. *eLife*, 4, 2015.

[1089] Jessica R Hughes, Vladimir A Levchenko, Stephen J Blanksby, Todd W Mitchell, Alan Williams, and Roger JW Truscott. No turnover in lens lipids for the entire human lifespan. *eLife*, 4, 2015.

[1090] James E Hughes Hallett, Xiangxia Luo, and Andrew P Capaldi. Snf1/AMPK promotes the formation of Kog1/Raptor-bodies to increase the activation threshold of TORC1 in budding yeast. *eLife*, 4, 2015.

[1091] Sung-Ho Huh, Mark E Warchol, and David M Ornitz. Cochlear progenitor number is controlled through mesenchymal FGF receptor signaling. *eLife*, 4, 2015.

[1092] Simon T Hui, Brian W Parks, Elin Org, Frode Norheim, Nam Che, Calvin Pan, Lawrence W Castellani, Sarada Charugundla, Darwin L Dirks, Nikolaos Psychogios, Isaac Neuhaus, Robert E Gerszten, Todd Kirchgessner, Peter S Gargalovic, and Aldons J Lusis. The genetic architecture of NAFLD among inbred strains of mice. *eLife*, 4, 2015.

[1093] Laurence T Hunt, Timothy EJ Behrens, Takayuki Hosokawa, Jonathan D Wallis, and Steven W Kennerley. Capturing the temporal evolution of choice across prefrontal cortex. *eLife*, 4, 2015.

[1094] Alexandre Hyafil, Lorenzo Fontolan, Claire Kabdebon, Boris Gutkin, and Anne-Lise Giraud. Speech encoding by coupled cortical theta and gamma oscillations. *eLife*, 4, 2015.

[1095] Toshiharu Ichinose, Yoshinori Aso, Nobuhiro Yamagata, Ayako Abe, Gerald M Rubin, and Hiromu Tanimoto. Reward signal in a recurrent circuit drives appetitive long-term memory formation. *eLife*, 4, 2015.

[1096] Kinga Igloi, Giulia Gaggioni, Virginie Sterpenich, and Sophie Schwartz. A nap to recap or how reward regulates hippocampal-prefrontal memory networks during daytime sleep in humans. *eLife*, 4, 2015.

[1097] Seol Hee Im, Kendra Takle, Juyeon Jo, Daniel T Babcock, Zhiguo Ma, Yang Xiang, and Michael J Galko. Tachykinin acts upstream of autocrine Hedgehog signaling during nociceptive sensitization in Drosophila. *eLife*, 4, 2015.

[1098] Takashi Imai, Hidekazu Ishida, Kazutomo Suzue, Tomoyo Taniguchi, Hiroko Okada, Chikako Shimokawa, and Hajime Hisaeda. Cytotoxic activities of CD8+ T cells collaborate with macrophages to protect against blood-stage murine malaria. *eLife*, 4, 2015.

[1099] Mayu Inaba, Zsolt G Venkei, and Yukiko M Yamashita. The polarity protein Baz forms a platform for the centrosome orientation during asymmetric stem cell division in the Drosophila male germline. *eLife*, 4, 2015.

[1100] Francesca Incardona, M Mehdi Doroudchi, Nawfal Ismail, Alberto Carreno, Erin Griner, and Minyoung Anna Lim. Registered report: Interactions between cancer stem cells and their niche govern metastatic colonization. *eLife*, 4, 2015.

[1101] Taro Ishikawa, Misa Shimuta, and Michael Häusser. Multimodal sensory integration in single cerebellar granule cells in vivo. *eLife*, 4, 2015.

[1102] Ayaka Ito, Cynthia Hong, Xin Rong, Xuewei Zhu, Elizabeth J Tarling, Per Niklas Hedde, Enrico Gratton, John Parks, and Peter Tontonoz. LXRs link metabolism to inflammation through Abca1-dependent regulation of membrane composition and TLR signaling. *eLife*, 4, 2015.

[1103] Shingo Iwami, Junko S Takeuchi, Shinji Nakaoka, Fabrizio Mammano, FranÇois Clavel, Hisashi Inaba, Tomoko Kobayashi, Naoko Misawa, Kazuyuki Aihara, Yoshio Koyanagi, and Kei Sato. Cell-to-cell infection by HIV contributes over half of virus infection. *eLife*, 4, 2015.

[1104] Etai Jacob, Ron Unger, and Amnon Horovitz. Codon-level information improves predictions of inter-residue contacts in proteins by correlated mutation analysis. *eLife*, 4, 2015.

[1105] Pablo D Jadzinsky and Stephen A Baccus. Synchronized amplification of local information transmission by peripheral retinal input. *eLife*, 4, 2015.

[1106] Vera Jansen, Luis Alvarez, Melanie Balbach, Timo Strünker, Peter Hegemann, U Benjamin Kaupp, and Dagmar Wachten. Controlling fertilization and cAMP signaling in sperm by optogenetics. *eLife*, 4, 2015.

[1107] Georges E Janssens, Anne C Meinema, Javier González, Justina C Wolters, Alexander Schmidt, Victor Guryev, Rainer Bischoff, Ernst C Wit, Liesbeth M Veenhoff, and Matthias Heinemann. Protein biogenesis machinery is a driver of replicative aging in yeast. *eLife*, 4, 2015.

[1108] Sébastien Jeay, Swann Gaulis, Stéphane Ferretti, Hans Bitter, Moriko Ito, Thérèse Valat, Masato Murakami, Stephan Ruetz, Daniel A Guthy, Caroline Rynn, Michael R Jensen, Marion Wiesmann, Joerg Kallen, Pascal Furet, FranÇois Gessier, Philipp Holzer, Keiichi Masuya, Jens Würthner, Ensar Halilovic, Francesco Hofmann, William R Sellers, and Diana Graus Porta. A distinct p53 target gene set predicts for response to the selective p53–HDM2 inhibitor NVP-CGM097. *eLife*, 4, 2015.

[1109] Wei-ke Ji, Anna L Hatch, Ronald A Merrill, Stefan Strack, and Henry N Higgs. Actin filaments target the oligomeric maturation of the dynamin GTPase Drp1 to mitochondrial fission sites. *eLife*, 4, 2015.

[1110] Zhe Ji, Ruisheng Song, Aviv Regev, and Kevin Struhl. Many lncRNAs, 5′UTRs, and pseudogenes are translated and some are likely to express functional proteins. *eLife*, 4, 2015.

[1111] Chimari Jiko, Karen M Davies, Kyoko Shinzawa-Itoh, Kazutoshi Tani, Shintaro Maeda, Deryck J Mills, Tomitake Tsukihara, Yoshinori Fujiyoshi, Werner Kühlbrandt, and Christoph Gerle. Bovine F1Fo ATP synthase monomers bend the lipid bilayer in 2D membrane crystals. *eLife*, 4, 2015.

[1112] Yui Jin and Lois S Weisman. The vacuole/lysosome is required for cell-cycle progression. *eLife*, 4, 2015.

[1113] Markus A Jobst, Lukas F Milles, Constantin Schoeler, Wolfgang Ott, Daniel B Fried, Edward A Bayer, Hermann E Gaub, and Michael A Nash. Resolving dual binding conformations of cellulosome cohesin-dockerin complexes using single-molecule force spectroscopy. *eLife*, 4, 2015.

[1114] Vanessa M Johnen, Franz-Xaver Neubert, Ethan R Buch, Lennart Verhagen, Jill X O'Reilly, Rogier B Mars, and Matthew F S Rushworth. Causal manipulation of functional connectivity in a specific neural pathway during behaviour and at rest. *eLife*, 4, 2015.

[1115] Alyssa E Johnson, Huidy Shu, Anna G Hauswirth, Amy Tong, and Graeme W Davis. VCP-dependent muscle degeneration is linked to defects in a dynamic tubular lysosomal network in vivo. *eLife*, 4, 2015.

[1116] Stuart L Johnson. Membrane properties specialize mammalian inner hair cells for frequency or intensity encoding. *eLife*, 4, 2015.

[1117] Iain G Johnston, Joerg P Burgstaller, Vitezslav Havlicek, Thomas Kolbe, Thomas Rülicke, Gottfried Brem, Jo Poulton, and Nick S Jones. Stochastic modelling, Bayesian inference, and new in vivo measurements elucidate the debated mtDNA bottleneck mechanism. *eLife*, 4, 2015.

[1118] Jamie Johnston and Leon Lagnado. General features of the retinal connectome determine the computation of motion anticipation. *eLife*, 4, 2015.

[1119] Eric Jonas and Konrad Kording. Automatic discovery of cell types and microcircuitry from neural connectomics. *eLife*, 4, 2015.

[1120] Graham L Jones, Marta E Soden, Cerise R Knakal, Heather Lee, Amanda S Chung, Elliott B Merriam, and Larry S Zweifel. A genetic link between discriminative fear coding by the lateral amygdala, dopamine, and fear generalization. *eLife*, 4, 2015.

[1121] Seung-Hye Jung, Catherine Hueston, and Vikas Bhandawat. Odor-identity dependent motor programs underlie behavioral responses to odors. *eLife*, 4, 2015.

[1122] Balint Z Kacsoh, Julianna Bozler, Mani Ramaswami, and Giovanni Bosco. Social communication of predator-induced changes in Drosophila behavior and germ line physiology. *eLife*, 4, 2015.

[1123] Benjamin R Kallman, Heesoo Kim, and Kristin Scott. Excitation and inhibition onto central courtship neurons biases Drosophila mate choice. *eLife*, 4, 2015.

[1124] Nicholas S Kalson, Yinhui Lu, Susan H Taylor, Tobias Starborg, David F Holmes, and Karl E Kadler. A structure-based extracellular matrix expansion mechanism of fibrous tissue growth. *eLife*, 4, 2015.

[1125] Irawati Kandela, James Chou, and Kartoa Chow. Registered report: Coadministration of a tumor-penetrating peptide enhances the efficacy of cancer drugs. *eLife*, 4, 2015.

[1126] Irawati Kandela, Hyun Yong Jin, and Katherine Owen. Registered report: BET bromodomain inhibition as a therapeutic strategy to target c-Myc. *eLife*, 4, 2015.

[1127] Irawati Kandela and Ioannis Zervantonakis. Registered report: Discovery and preclinical validation of drug indications using compendia of public gene expression data. *eLife*, 4, 2015.

[1128] Woo Kyu Kang, Yeong Hyeock Kim, Hyun Ah Kang, Ki-Sun Kwon, and Jeong-Yoon Kim. Sir2 phosphorylation through cAMP-PKA and CK2 signaling inhibits the lifespan extension activity of Sir2 in yeast. *eLife*, 4, 2015.

[1129] Paulina Kaniewska, Shahar Alon, Sarit Karako-Lampert, Ove Hoegh-Guldberg, and Oren Levy. Signaling cascades and the importance of moonlight in coral broadcast mass spawning. *eLife*, 4, 2015.

[1130] Rebeccah J Katzenberger, Stanislava Chtarbanova, Stacey A Rimkus, Julie A Fischer, Gulpreet Kaur, Jocelyn M Seppala, Laura C Swanson, Jocelyn E Zajac, Barry Ganetzky, and David A Wassarman. Death following traumatic brain injury in Drosophila is associated with intestinal barrier dysfunction. *eLife*, 4, 2015.

[1131] Matthew T Kaufman, Mark M Churchland, Stephen I Ryu, and Krishna V Shenoy. Vacillation, indecision and hesitation in moment-by-moment decoding of monkey motor cortex. *eLife*, 4, 2015.

[1132] Matthew S Kayser, Benjamin Mainwaring, Zhifeng Yue, and Amita Sehgal. Sleep deprivation suppresses aggression in Drosophila. *eLife*, 4, 2015.

[1133] Daniel A Keedy, Lillian R Kenner, Matthew Warkentin, Rahel A Woldeyes, Jesse B Hopkins, Michael C Thompson, Aaron S Brewster, Andrew H Van Benschoten, Elizabeth L Baxter, Monarin Uervirojnangkoorn, Scott E McPhillips, Jinhu Song, Roberto Alonso-Mori, James M Holton, William I Weis, Axel T Brunger, S Michael Soltis, Henrik Lemke, Ana Gonzalez, Nicholas K Sauter, Aina E Cohen, Henry van den Bedem, Robert E Thorne, and James S Fraser. Mapping the conformational landscape of a dynamic enzyme by multitemperature and XFEL crystallography. *eLife*, 4, 2015.

[1134] Rachel Kerwin, Julie Feusier, Jason Corwin, Matthew Rubin, Catherine Lin, Alise Muok, Brandon Larson, Baohua Li, Bindu Joseph, Marta Francisco, Daniel Copeland, Cynthia Weinig, and Daniel J Kliebenstein. Natural genetic variation in Arabidopsis thaliana defense metabolism genes modulates field fitness. *eLife*, 4, 2015.

[1135] Israr Khan, John Kerwin, Kate Owen, and Erin Griner. Registered report: A coding-independent function of gene and pseudogene mRNAs regulates tumour biology. *eLife*, 4, 2015.

[1136] Bernard Khor, John D Gagnon, Gautam Goel, Marly I Roche, Kara L Conway, Khoa Tran, Leslie N Aldrich, Thomas B Sundberg, Alison M Paterson, Scott Mordecai, David Dombkowski, Melanie Schirmer, Pauline H Tan, Atul K Bhan, Rahul Roychoudhuri, Nicholas P Restifo, John J O'Shea, Benjamin D Medoff, Alykhan F Shamji, Stuart L Schreiber, Arlene H Sharpe, Stanley Y Shaw, and Ramnik J Xavier. The kinase DYRK1A reciprocally regulates the differentiation of Th17 and regulatory T cells. *eLife*, 4, 2015.

[1137] Arkady Khoutorsky, Robert P Bonin, Robert E Sorge, Christos G Gkogkas, Sophie Anne Pawlowski, Seyed Mehdi Jafarnejad, Mark H Pitcher, Tommy Alain, Jimena Perez-Sanchez, Eric W Salter, Loren Martin, Alfredo Ribeiro-da Silva, Yves De Koninck, Fernando Cervero, Jeffrey S Mogil, and Nahum Sonenberg. Translational control of nociception via 4E-binding protein 1. *eLife*, 4, 2015.

[1138] Philipp Khuc Trong, Hélène Doerflinger, Jörn Dunkel, Daniel St Johnston, and Raymond E Goldstein. Cortical microtubule nucleation can organise the cytoskeleton of Drosophila oocytes to define the anteroposterior axis. *eLife*, 4, 2015.

[1139] Rebecca M Kilner, Giuseppe Boncoraglio, Jonathan M Henshaw, Benjamin JM Jarrett, Ornela De Gasperin, Alfredo Attisano, and Hanna Kokko. Parental effects alter the adaptive value of an adult behavioural trait. *eLife*, 4, 2015.

[1140] Anmo J Kim, Aurel A Lazar, and Yevgeniy B Slutskiy. Projection neurons in Drosophila antennal lobes signal the acceleration of odor concentrations. *eLife*, 4, 2015.

[1141] Sangmok Kim and Kelsey C Martin. Neuron-wide RNA transport combines with netrin-mediated local translation to spatially regulate the synaptic proteome. *eLife*, 4, 2015.

[1142] Tae-Jin Kim, Chirlmin Joo, Jihye Seong, Reza Vafabakhsh, Elliot L Botvinick, Michael W Berns, Amy E Palmer, Ning Wang, Taekjip Ha, Eric Jakobsson, Jie Sun, and Yingxiao Wang. Distinct mechanisms regulating mechanical force-induced Ca2+ signals at the plasma membrane and the ER in human MSCs. *eLife*, 4, 2015.

[1143] Tahnbee Kim, Florentina Soto, and Daniel Kerschensteiner. An excitatory amacrine cell detects object motion and provides feature-selective input to ganglion cells in the mouse retina. *eLife*, 4, 2015.

[1144] Yujin Kim, Ching-Lung Hsu, Mark S Cembrowski, Brett D Mensh, and Nelson Spruston. Dendritic sodium spikes are required for long-term potentiation at distal synapses on hippocampal pyramidal neurons. *eLife*, 4, 2015.

[1145] Judith Kimble, William M Bement, Qiang Chang, Benjamin L Cox, Norman R Drinkwater, Richard L Gourse, Aaron A Hoskins, Anna Huttenlocher, Pamela K Kreeger, Paul F Lambert, Marsha R Mailick, Shigeki Miyamoto, Richard L Moss, Kate M O'Connor-Giles, Avtar Roopra, Krishanu Saha, and Hannah S Seidel. Strategies from UW-Madison for rescuing biomedical research in the US. *eLife*, 4, 2015.

[1146] Indigo Chris King, James Gleixner, Lindsey Doyle, Alexandre Kuzin, John F Hunt, Rong Xiao, Gaetano T Montelione, Barry L Stoddard, Frank DiMaio, and David Baker. Precise assembly of complex beta sheet topologies from de novo designed building blocks. *eLife*, 4, 2015.

[1147] Koray Kırlı, Samir Karaca, Heinz Jürgen Dehne, Matthias Samwer, Kuan Ting Pan, Christof Lenz, Henning Urlaub, and Dirk Görlich. A deep proteomics perspective on CRM1-mediated nuclear export and nucleocytoplasmic partitioning. *eLife*, 4, 2015.

[1148] Alexander D Kloth, Aleksandra Badura, Amy Li, Adriana Cherskov, Sara G Connolly, Andrea Giovannucci, M Ali Bangash, Giorgio Grasselli, Olga Peñagarikano, Claire Piochon, Peter T Tsai, Daniel H Geschwind, Christian Hansel, Mustafa Sahin, Toru Takumi, Paul F Worley, and Samuel S-H Wang. Cerebellar associative sensory learning defects in five mouse autism models. *eLife*, 4, 2015.

[1149] Kang I Ko, Cory M Root, Scott A Lindsay, Orel A Zaninovich, Andrew K Shepherd, Steven A Wasserman, Susy M Kim, and Jing W Wang. Starvation promotes concerted modulation of appetitive olfactory behavior via parallel neuromodulatory circuits. *eLife*, 4, 2015.

[1150] Andrew Kodani, Timothy W Yu, Jeffrey R Johnson, Divya Jayaraman, Tasha L Johnson, Lihadh Al-Gazali, Lāszló Sztriha, Jennifer N Partlow, Hanjun Kim, Alexis L Krup, Alexander Dammermann, Nevan J Krogan, Christopher A Walsh, and Jeremy F Reiter. Centriolar satellites assemble centrosomal microcephaly proteins to recruit CDK2 and promote centriole duplication. *eLife*, 4, 2015.

[1151] Katia Koelle and David A Rasmussen. The effects of a deleterious mutation load on patterns of influenza A/H3N2's antigenic evolution in humans. *eLife*, 4, 2015.

[1152] Kurtulus Kok, Ahmet Ay, Li M Li, and David N Arnosti. Genome-wide errant targeting by Hairy. *eLife*, 4, 2015.

[1153] Agnieszka Konopacka, Mingkwan Greenwood, Su-Yi Loh, Julian Paton, and David Murphy. RNA binding protein Caprin-2 is a pivotal regulator of the central osmotic defense response. *eLife*, 4, 2015.

[1154] Anil Korkut, Weiqing Wang, Emek Demir, Bülent Arman Aksoy, Xiaohong Jing, Evan J Molinelli, özgün Babur, Debra L Bemis, Selcuk Onur Sumer, David B Solit, Christine A Pratilas, and Chris Sander. Perturbation biology nominates upstream–downstream drug combinations in RAF inhibitor resistant melanoma cells. *eLife*, 4, 2015.

[1155] Natalya Korogod, Carl CH Petersen, and Graham W Knott. Ultrastructural analysis of adult mouse neocortex comparing aldehyde perfusion with cryo fixation. *eLife*, 4, 2015.

[1156] Dimitar Kostadinov and Joshua R Sanes. Protocadherin-dependent dendritic self-avoidance regulates neural connectivity and circuit function. *eLife*, 4, 2015.

[1157] Matthew V Kotlajich, Daniel R Hron, Beth A Boudreau, Zhiqiang Sun, Yuri L Lyubchenko, and Robert Landick. Bridged filaments of histone-like nucleoid structuring protein pause RNA polymerase and aid termination in bacteria. *eLife*, 4, 2015.

[1158] Kristin S Koutmou, Anthony P Schuller, Julie L Brunelle, Aditya Radhakrishnan, Sergej Djuranovic, and Rachel Green. Ribosomes slide on lysine-encoding homopolymeric A stretches. *eLife*, 4, 2015.

[1159] Yevgenia Kozorovitskiy, Rui Peixoto, Wengang Wang, Arpiar Saunders, and Bernardo L Sabatini. Neuromodulation of excitatory synaptogenesis in striatal development. *eLife*, 4, 2015.

[1160] Moritz UG Kraemer, Marianne E Sinka, Kirsten A Duda, Adrian QN Mylne, Freya M Shearer, Christopher M Barker, Chester G Moore, Roberta G Carvalho, Giovanini E Coelho, Wim Van Bortel, Guy Hendrickx, Francis Schaffner, Iqbal RF Elyazar, Hwa-Jen Teng, Oliver J Brady, Jane P Messina, David M Pigott, Thomas W Scott, David L Smith, GR William Wint, Nick Golding, and Simon I Hay. The global distribution of the arbovirus vectors Aedes aegypti and Ae. albopictus. *eLife*, 4, 2015.

[1161] Nathan M Krah, Jean-Paul De La O, Galvin H Swift, Chinh Q Hoang, Spencer G Willet, Fong Chen Pan, Gabriela M Cash, Mary P Bronner, Christopher VE Wright, Raymond J MacDonald, and L Charles Murtaugh. The acinar differentiation determinant PTF1A inhibits initiation of pancreatic ductal adenocarcinoma. *eLife*, 4, 2015.

[1162] Ute Krämer. Planting molecular functions in an ecological context with Arabidopsis thaliana. *eLife*, 4, 2015.

[1163] Shyam S Krishnakumar, Feng Li, Jeff Coleman, Curtis M Schauder, Daniel Kümmel, Frederic Pincet, James E Rothman, and Karin M Reinisch. Re-visiting the trans insertion model for complexin clamping. *eLife*, 4, 2015.

[1164] Sonja Kroschwald, Shovamayee Maharana, Daniel Mateju, Liliana Malinovska, Elisabeth Nüske, Ina Poser, Doris Richter, and Simon Alberti. Promiscuous interactions and protein disaggregases determine the material state of stress-inducible RNP granules. *eLife*, 4, 2015.

[1165] Tom Kruitwagen, Annina Denoth-Lippuner, Bryan J Wilkins, Heinz Neumann, and Yves Barral. Axial contraction and short-range compaction of chromatin synergistically promote mitotic chromosome condensation. *eLife*, 4, 2015.

[1166] Yoshiyuki Kubota, Satoru Kondo, Masaki Nomura, Sayuri Hatada, Noboru Yamaguchi, Alsayed A Mohamed, Fuyuki Karube, Joachim Lübke, and Yasuo Kawaguchi. Functional effects of distinct innervation styles of pyramidal cells by fast spiking cortical interneurons. *eLife*, 4, 2015.

[1167] Michael J Kuiper, Craig J Morton, Sneha E Abraham, and Angus Gray-Weale. The biological function of an insect antifreeze protein simulated by molecular dynamics. *eLife*, 4, 2015.

[1168] Nathan Kumar, Jenna Richter, Josh Cutts, Kevin T Bush, Cleber Trujillo, Sanjay K Nigam, Terry Gaasterland, David Brafman, and Karl Willert. Generation of an expandable intermediate mesoderm restricted progenitor cell line from human pluripotent stem cells. *eLife*, 4, 2015.

[1169] Sudha Kumari, David Depoil, Roberta Martinelli, Edward Judokusumo, Guillaume Carmona, Frank B Gertler, Lance C Kam, Christopher V Carman, Janis K Burkhardt, Darrell J Irvine, and Michael L Dustin. Actin foci facilitate activation of the phospholipase C-$\gamma$ in primary T lymphocytes via the WASP pathway. *eLife*, 4, 2015.

[1170] Prabhat S Kunwar, Moriel Zelikowsky, Ryan Remedios, Haijiang Cai, Melis Yilmaz, Markus Meister, and David J Anderson. Ventromedial hypothalamic neurons control a defensive emotion state. *eLife*, 4, 2015.

[1171] Zeb Kurth-Nelson, Gareth Barnes, Dino Sejdinovic, Ray Dolan, and Peter Dayan. Temporal structure in associative retrieval. *eLife*, 4, 2015.

[1172] Colin T Kyle, Jared D Stokes, Jennifer S Lieberman, Abdul S Hassan, and Arne D Ekstrom. Successful retrieval of competing spatial environments in humans involves hippocampal pattern separation mechanisms. *eLife*, 4, 2015.

[1173] Liedewij Laan, John H Koschwanez, and Andrew W Murray. Evolutionary adaptation after crippling cell polarization follows reproducible trajectories. *eLife*, 4, 2015.

[1174] Cathleen M Lake, Rachel J Nielsen, Fengli Guo, Jay R Unruh, Brian D Slaughter, and R Scott Hawley. Vilya, a component of the recombination nodule, is required for meiotic double-strand break formation in Drosophila. *eLife*, 4, 2015.

[1175] Dominic Landgraf, Anthony H Tsang, Alexei Leliavski, Christiane E Koch, Johanna L Barclay, Daniel J Drucker, and Henrik Oster. Oxyntomodulin regulates resetting of the liver circadian clock by food. *eLife*, 4, 2015.

[1176] Benoît Landrein, Annamaria Kiss, Massimiliano Sassi, Aurélie Chauvet, Pradeep Das, Millan Cortizo, Patrick Laufs, Seiji Takeda, Mitsuhiro Aida, Jan Traas, Teva Vernoux, Arezki Boudaoud, and Olivier Hamant. Mechanical stress contributes to the expression of the STM homeobox gene in Arabidopsis shoot meristems. *eLife*, 4, 2015.

[1177] Kelly M LaRue, Jan Clemens, Gordon J Berman, and Mala Murthy. Acoustic duetting in Drosophila virilis relies on the integration of auditory and tactile signals. *eLife*, 4, 2015.

[1178] Cristian A Lasagna-Reeves, Maxime WC Rousseaux, Marcos J Guerrero-Munoz, Luis Vilanova-Velez, Jeehye Park, Lauren See, Paymaan Jafar-Nejad, Ronald Richman, Harry T Orr, Rakez Kayed, and Huda Y Zoghbi. Ataxin-1 oligomers induce local spread of pathology and decreasing them by passive immunization slows Spinocerebellar ataxia type 1 phenotypes. *eLife*, 4, 2015.

[1179] Cristian A Lasagna-Reeves, Maxime WC Rousseaux, Marcos J Guerrero-Muñoz, Jeehye Park, Paymaan Jafar-Nejad, Ronald Richman, Nan Lu, Urmi Sengupta, Alexandra Litvinchuk, Harry T Orr, Rakez Kayed, and Huda Y Zoghbi. A native interactor scaffolds and stabilizes toxic ATAXIN-1 oligomers in SCA1. *eLife*, 4, 2015.

[1180] Charlotte Lässig, Sarah Matheisl, Konstantin MJ Sparrer, Carina C de Oliveira Mann, Manuela Moldt, Jenish R Patel, Marion Goldeck, Gunther Hartmann, Adolfo García-Sastre, Veit Hornung, Karl-Klaus Conzelmann, Roland Beckmann, and Karl-Peter Hopfner. ATP hydrolysis by the viral RNA sensor RIG-I prevents unintentional recognition of self-RNA. *eLife*, 4, 2015.

[1181] Patrick Laurent, Zoltan Soltesz, Geoffrey M Nelson, Changchun Chen, Fausto Arellano-Carbajal, Emmanuel Levy, and Mario de Bono. Decoding a neural circuit controlling global animal state in C. elegans. *eLife*, 4, 2015.

[1182] Patrick Laurette, Thomas Strub, Dana Koludrovic, Céline Keime, Stéphanie Le Gras, Hannah Seberg, Eric Van Otterloo, Hana Imrichova, Robert Siddaway, Stein Aerts, Robert A Cornell, Gabrielle Mengus, and Irwin Davidson. Transcription factor MITF and remodeller BRG1 define chromatin organisation at regulatory elements in melanoma cells. *eLife*, 4, 2015.

[1183] Alison Leaf and Mark Von Zastrow. Dopamine receptors reveal an essential role of IFT-B, KIF17, and Rab23 in delivering specific receptors to primary cilia. *eLife*, 4, 2015.

[1184] Ho-Jin Lee, De-Li Shi, and Jie J Zheng. Conformational change of Dishevelled plays a key regulatory role in the Wnt signaling pathways. *eLife*, 4, 2015.

[1185] Peter L Lee, Maikke B Ohlson, and Suzanne R Pfeffer. The Rab6-regulated KIF1C kinesin motor domain contributes to Golgi organization. *eLife*, 4, 2015.

[1186] Shun-Hsiao Lee, Lissa Nicola Princz, Maren Felizitas Klügel, Bianca Habermann, Boris Pfander, and Christian Biertümpfel. Human Holliday junction resolvase GEN1 uses a chromodomain for efficient DNA recognition and cleavage. *eLife*, 4, 2015.

[1187] Yool Lee, A Reum Jang, Lauren J Francey, Amita Sehgal, and John B Hogenesch. KPNB1 mediates PER/CRY nuclear translocation and circadian clock function. *eLife*, 4, 2015.

[1188] Yu-Ling Lee, Yuan-Ching Li, Chia-Hsin Su, Chun-Hui Chiao, I-Hsuan Lin, and Ming-Ta Hsu. MAF1 represses CDKN1A through a Pol III-dependent mechanism. *eLife*, 4, 2015.

[1189] Sarah G Leinwand, Claire J Yang, Daphne Bazopoulou, Nikos Chronis, Jagan Srinivasan, and Sreekanth H Chalasani. Circuit mechanisms encoding odors and driving aging-associated behavioral declines in Caenorhabditis elegans. *eLife*, 4, 2015.

[1190] Alexandre B Leitão and élio Sucena. Drosophila sessile hemocyte clusters are true hematopoietic tissues that regulate larval blood cell differentiation. *eLife*, 4, 2015.

[1191] Michele LeRoux, Robin L Kirkpatrick, Elena I Montauti, Bao Q Tran, S Brook Peterson, Brittany N Harding, John C Whitney, Alistair B Russell, Beth Traxler, Young Ah Goo, David R Goodlett, Paul A Wiggins, and Joseph D Mougous. Kin cell lysis is a danger signal that activates antibacterial pathways of Pseudomonas aeruginosa. *eLife*, 4, 2015.

[1192] Mor Levi-Ferber, Hai Gian, Reut Dudkevich, and Sivan Henis-Korenblit. Transdifferentiation mediated tumor suppression by the endoplasmic reticulum stress sensor IRE-1 in C. elegans. *eLife*, 4, 2015.

[1193] Mia T Levine, Helen M Vander Wende, and Harmit S Malik. Mitotic fidelity requires transgenerational action of a testis-restricted HP1. *eLife*, 4, 2015.

[1194] Amanda H Lewis and Jörg Grandl. Mechanical sensitivity of Piezo1 ion channels can be tuned by cellular membrane tension. *eLife*, 4, 2015.

[1195] Laura D Lewis, Jakob Voigts, Francisco J Flores, L Ian Schmitt, Matthew A Wilson, Michael M Halassa, and Emery N Brown. Thalamic reticular nucleus induces fast and local modulation of arousal state. *eLife*, 4, 2015.

[1196] Chao Li, Fang-Ling Yeh, Alice Y Cheung, Qiaohong Duan, Daniel Kita, Ming-Che Liu, Jacob Maman, Emily J Luu, Brendan W Wu, Laura Gates, Methun Jalal, Amy Kwong, Hunter Carpenter, and Hen-Ming Wu. Glycosylphosphatidylinositol-anchored proteins as chaperones and co-receptors for FERONIA receptor kinase signaling in Arabidopsis. *eLife*, 4, 2015.

[1197] Ci-Xiu Li, Mang Shi, Jun-Hua Tian, Xian-Dan Lin, Yan-Jun Kang, Liang-Jun Chen, Xin-Cheng Qin, Jianguo Xu, Edward C Holmes, and Yong-Zhen Zhang. Unprecedented genomic diversity of RNA viruses in arthropods reveals the ancestry of negative-sense RNA viruses. *eLife*, 4, 2015.

[1198] Jia Li and Matthew Lam. Registered report: the microRNA miR-34a inhibits prostate cancer stem cells and metastasis by directly repressing CD44. *eLife*, 4, 2015.

[1199] Li Li, Silvia Sanchez Martinez, Wenxin Hu, Zhe Liu, and Robert Tjian. A specific E3 ligase/deubiquitinase pair modulates TBP protein levels during muscle differentiation. *eLife*, 4, 2015.

[1200] Ming Li, Arjan Hada, Payel Sen, Lola Olufemi, Michael A Hall, Benjamin Y Smith, Scott Forth, Jeffrey N McKnight, Ashok Patel, Gregory D Bowman, Blaine Bartholomew, and Michelle D Wang. Dynamic regulation of transcription factors by nucleosome remodeling. *eLife*, 4, 2015.

[1201] Qian Li, Yaw Tachie-Baffour, Zhikai Liu, Maude W Baldwin, Andrew C Kruse, and Stephen D Liberles. Non-classical amine recognition evolved in a large clade of olfactory receptors. *eLife*, 4, 2015.

[1202] Ran Li, Jin Zhang, Jiancai Li, Guoxin Zhou, Qi Wang, Wenbo Bian, Matthias Erb, and Yonggen Lou. Prioritizing plant defence over growth through WRKY regulation facilitates infestation by non-target herbivores. *eLife*, 4, 2015.

[1203] Shuang Li, Bopanna I Kalappa, and Thanos Tzounopoulos. Noise-induced plasticity of KCNQ2/3 and HCN channels underlies vulnerability and resilience to tinnitus. *eLife*, 4, 2015.

[1204] Xiaoxiao Li, Wei Wei, HoangDinh Huynh, Hao Zuo, Xueqian Wang, and Yihong Wan. Nur77 prevents excessive osteoclastogenesis by inducing ubiquitin ligase Cbl-b to mediate NFATc1 self-limitation. *eLife*, 4, 2015.

[1205] Xin Li, Vivian Yaci Yu, Yuping Lin, Kulika Chomvong, Raíssa Estrela, Annsea Park, Julie M Liang, Elizabeth A Znameroski, Joanna Feehan, Soo Rin Kim, Yong-Su Jin, N Louise Glass, and Jamie HD Cate. Expanding xylose metabolism in yeast for plant cell wall conversion to biofuels. *eLife*, 4, 2015.

[1206] Xue Liang, Kyle Bittinger, Xuanwen Li, Darrell R Abernethy, Frederic D Bushman, and Garret A FitzGerald. Bidirectional interactions between indomethacin and the murine intestinal microbiota. *eLife*, 4, 2015.

[1207] Juliane Liepe, Hermann-Georg Holzhütter, Elena Bellavista, Peter M Kloetzel, Michael PH Stumpf, and Michele Mishto. Quantitative time-resolved analysis reveals intricate, differential regulation of standard- and immuno-proteasomes. *eLife*, 4, 2015.

[1208] Yoong Wearn Lim, Lionel A Sanz, Xiaoqin Xu, Stella R Hartono, and Frédéric Chédin. Genome-wide DNA hypomethylation and RNA:DNA hybrid accumulation in Aicardi–Goutières syndrome. *eLife*, 4, 2015.

[1209] Chun-Chieh Lin, Katharine A Prokop-Prigge, George Preti, and Christopher J Potter. Food odors trigger Drosophila males to deposit a pheromone that guides aggregation and female oviposition decisions. *eLife*, 4, 2015.

[1210] David Tse Shen Lin and Elizabeth Conibear. ABHD17 proteins are novel protein depalmitoylases that regulate N-Ras palmitate turnover and subcellular localization. *eLife*, 4, 2015.

[1211] Michelle I Lin, Emily N Price, Sonja Boatman, Elliott J Hagedorn, Eirini Trompouki, Sruthi Satishchandran, Charles W Carspecken, Audrey Uong, Anthony DiBiase, Song Yang, Matthew C Canver, Ann Dahlberg, Zhigang Lu, Cheng Cheng Zhang, Stuart H Orkin, Irwin D Bernstein, Jon C Aster, Richard M White, and Leonard I Zon. Angiopoietin-like proteins stimulate HSPC development through interaction with notch receptor signaling. *eLife*, 4, 2015.

[1212] Zhi Lin, Jason Y Tann, Eddy TH Goh, Claire Kelly, Kim Buay Lim, Jian Fang Gao, and Carlos F Ibanez. Structural basis of death domain signaling in the p75 neurotrophin receptor. *eLife*, 4, 2015.

[1213] Anthony J Linares, Chia-Ho Lin, Andrey Damianov, Katrina L Adams, Bennett G Novitch, and Douglas L Black. The splicing regulator PTBP1 controls the activity of the transcription factor Pbx1 during neuronal differentiation. *eLife*, 4, 2015.

[1214] Peter A Lind, Andrew D Farr, and Paul B Rainey. Experimental evolution reveals hidden diversity in evolutionary pathways. *eLife*, 4, 2015.

[1215] Nils O Lindström, Melanie L Lawrence, Sally F Burn, Jeanette A Johansson, Elvira RM Bakker, Rachel A Ridgway, C-Hong Chang, Michele J Karolak, Leif Oxburgh, Denis J Headon, Owen J Sansom, Ron Smits, Jamie A Davies, and Peter Hohenstein. Integrated $\beta$-catenin, BMP, PTEN, and Notch signalling patterns the nephron. *eLife*, 4, 2015.

[1216] Gianni Liti. The fascinating and secret wild life of the budding yeast S. cerevisiae. *eLife*, 4, 2015.

[1217] Albert Liu, Neeraj Jain, Ajai Vyas, and Lee Wei Lim. Ventromedial prefrontal cortex stimulation enhances memory and hippocampal neurogenesis in the middle-aged rats. *eLife*, 4, 2015.

[1218] Jia Liu, Hyun Joo Lee, Andrew J Weitz, Zhongnan Fang, Peter Lin, ManKin Choy, Robert Fisher, Vadim Pinskiy, Alexander Tolpygo, Partha Mitra, Nicholas Schiff, and Jin Hyung Lee. Frequency-selective control of cortical and subcortical networks by central thalamus. *eLife*, 4, 2015.

[1219] Ninning Liu, Gheorghe Chistol, and Carlos Bustamante. Two-subunit DNA escort mechanism and inactive subunit bypass in an ultra-fast ring ATPase. *eLife*, 4, 2015.

[1220] Shouan Liu, Barbara Kracher, Jörg Ziegler, Rainer P Birkenbihl, and Imre E Somssich. Negative regulation of ABA signaling by WRKY33 is critical for Arabidopsis immunity towards Botrytis cinerea 2100. *eLife*, 4, 2015.

[1221] Sunbin Liu, Sina Mozaffari-Jovin, Jan Wollenhaupt, Karine F Santos, Matthias Theuser, Stanislaw Dunin-Horkawicz, Patrizia Fabrizio, Janusz M Bujnicki, Reinhard Lührmann, and Markus C Wahl. A composite double-/single-stranded RNA-binding region in protein Prp3 supports tri-snRNP stability and splicing. *eLife*, 4, 2015.

[1222] Wenxuan Liu, Lan Wei-LaPierre, Alanna Klose, Robert T Dirksen, and Joe V Chakkalakal. Inducible depletion of adult skeletal muscle stem cells impairs the regeneration of neuromuscular junctions. *eLife*, 4, 2015.

[1223] Xili Liu, Xin Wang, Xiaojing Yang, Sen Liu, Lingli Jiang, Yimiao Qu, Lufeng Hu, Qi Ouyang, and Chao Tang. Reliable cell cycle commitment in budding yeast is ensured by signal integration. *eLife*, 4, 2015.

[1224] Yi Bessie Liu, Ambika Tewari, Johnny Salameh, Elena Arystarkhova, Thomas G Hampton, Allison Brashear, Laurie J Ozelius, Kamran Khodakhah, and Kathleen J Sweadner. A dystonia-like movement disorder with brain and spinal neuronal defects is caused by mutation of the mouse laminin $\beta$1 subunit, Lamb1. *eLife*, 4, 2015.

[1225] Zhu Liu, Zhou Gong, Wen-Xue Jiang, Ju Yang, Wen-Kai Zhu, Da-Chuan Guo, Wei-Ping Zhang, Mai-Li Liu, and Chun Tang. Lys63-linked ubiquitin chain adopts multiple conformational states for specific target recognition. *eLife*, 4, 2015.

[1226] Christian Löfke, Kai Dünser, David Scheuring, and Jürgen Kleine-Vehn. Auxin regulates SNARE-dependent vacuolar morphology restricting cell size. *eLife*, 4, 2015.

[1227] Jeremy S Logue, Alexander X Cartagena-Rivera, Michelle A Baird, Michael W Davidson, Richard S Chadwick, and Clare M Waterman. Erk regulation of actin capping and bundling by Eps8 promotes cortex tension and leader bleb-based migration. *eLife*, 4, 2015.

[1228] Dawn H Loh, Shekib A Jami, Richard E Flores, Danny Truong, Cristina A Ghiani, Thomas J O'Dell, and Christopher S Colwell. Misaligned feeding impairs memories. *eLife*, 4, 2015.

[1229] Alex John London and Jonathan Kimmelman. Why clinical translation cannot succeed without failure. *eLife*, 4, 2015.

[1230] Nina Lopatina, Michael A McDannald, Clay V Styer, Brian F Sadacca, Joseph F Cheer, and Geoffrey Schoenbaum. Lateral orbitofrontal neurons acquire responses to upshifted, downshifted, or blocked cues during unblocking. *eLife*, 4, 2015.

[1231] Stilianos Louca and Michael Doebeli. Calibration and analysis of genome-based models for microbial ecology. *eLife*, 4, 2015.

[1232] Alan R Lowe, Jeffrey H Tang, Jaime Yassif, Michael Graf, William YC Huang, Jay T Groves, Karsten Weis, and Jan T Liphardt. Importin-$\beta$ modulates the permeability of the nuclear pore complex in a Ran-dependent manner. *eLife*, 4, 2015.

[1233] Feiran Lu, Qiren Liang, Lina Abi-Mosleh, Akash Das, Jef K De Brabander, Joseph L Goldstein, and Michael S Brown. Identification of NPC1 as the target of U18666A, an inhibitor of lysosomal cholesterol export and Ebola infection. *eLife*, 4, 2015.

[1234] Huasong Lu, Yuhua Xue, Guoying K Yu, Carolina Arias, Julie Lin, Susan Fong, Michel Faure, Ben Weisburd, Xiaodan Ji, Alexandre Mercier, James Sutton, Kunxin Luo, Zhenhai Gao, and Qiang Zhou. Compensatory induction of MYC expression by sustained CDK9 inhibition via a BRD4-dependent mechanism. *eLife*, 4, 2015.

[1235] Edina Lump, Laura M Castellano, Christoph Meier, Janine Seeliger, Nelli Erwin, Benjamin Sperlich, Christina M Stürzel, Shariq Usmani, Rebecca M Hammond, Jens von Einem, Gisa Gerold, Florian Kreppel, Kenny Bravo-Rodriguez, Thomas Pietschmann, Veronica M Holmes, David Palesch, Onofrio Zirafi, Drew Weissman, Andrea Sowislok, Burkhard Wettig, Christian Heid, Frank Kirchhoff, Tanja Weil, Frank-Gerrit Klärner, Thomas Schrader, Gal Bitan, Elsa Sanchez-Garcia, Roland Winter, James Shorter, and Jan Münch. A molecular tweezer antagonizes seminal amyloids and HIV infection. *eLife*, 4, 2015.

[1236] Tony Ly, Aki Endo, and Angus I Lamond. Proteomic analysis of the response to cell cycle arrests in human myeloid leukemia cells. *eLife*, 4, 2015.

[1237] Hansong Ma and Patrick H O'Farrell. Selections that isolate recombinant mitochondrial genomes in animals. *eLife*, 4, 2015.

[1238] Lu Ma, Aleksander A Rebane, Guangcan Yang, Zhiqun Xi, Yuhao Kang, Ying Gao, and Yongli Zhang. Munc18-1-regulated stage-wise SNARE assembly underlying synaptic exocytosis. *eLife*, 4, 2015.

[1239] Anne Maass, David Berron, Laura A Libby, Charan Ranganath, and Emrah Düzel. Functional subregions of the human entorhinal cortex. *eLife*, 4, 2015.

[1240] Donna A MacDuff, Tiffany A Reese, Jacqueline M Kimmey, Leslie A Weiss, Christina Song, Xin Zhang, Amal Kambal, Erning Duan, Javier A Carrero, Bertrand Boisson, Emmanuel Laplantine, Alain Israel, Capucine Picard, Marco Colonna, Brian T Edelson, L David Sibley, Christina L Stallings, Jean-Laurent Casanova, Kazuhiro Iwai, and Herbert W Virgin. Phenotypic complementation of genetic immunodeficiency by chronic herpesvirus infection. *eLife*, 4, 2015.

[1241] Ana S Machado, Dana M Darmohray, João Fayad, Hugo G Marques, and Megan R Carey. A quantitative framework for whole-body coordination reveals specific deficits in freely walking ataxic mice. *eLife*, 4, 2015.

[1242] Lauran Madden, Mark Juhas, William E Kraus, George A Truskey, and Nenad Bursac. Bioengineered human myobundles mimic clinical responses of skeletal muscle to drugs. *eLife*, 4, 2015.

[1243] Ross K Maddox, Huriye Atilgan, Jennifer K Bizley, and Adrian KC Lee. Auditory selective attention is enhanced by a task-irrelevant temporally coherent visual stimulus in human listeners. *eLife*, 4, 2015.

[1244] Andrea Mair, Lorenzo Pedrotti, Bernhard Wurzinger, Dorothea Anrather, Andrea Simeunovic, Christoph Weiste, Concetta Valerio, Katrin Dietrich, Tobias Kirchler, Thomas Nägele, Jesús Vicente Carbajosa, Johannes Hanson, Elena Baena-González, Christina Chaban, Wolfram Weckwerth, Wolfgang Dröge-Laser, and Markus Teige. SnRK1-triggered switch of bZIP63 dimerization mediates the low-energy response in plants. *eLife*, 4, 2015.

[1245] Florian Mair, Stefanie Joller, Romy Hoeppli, Lucas Onder, Matthias Hahn, Burkhard Ludewig, Ari Waisman, and Burkhard Becher. The NF$\kappa$B-inducing kinase is essential for the developmental programming of skin-resident and IL-17-producing $\gamma\delta$ T cells. *eLife*, 4, 2015.

[1246] Siu-Shan Mak, Cantas Alev, Hiroki Nagai, Anna Wrabel, Yoko Matsuoka, Akira Honda, Guojun Sheng, and Raj K Ladher. Characterization of the finch embryo supports evolutionary conservation of the naive stage of development in amniotes. *eLife*, 4, 2015.

[1247] Christophe Malabat, Frank Feuerbach, Laurence Ma, Cosmin Saveanu, and Alain Jacquier. Quality control of transcription start site selection by nonsense-mediated-mRNA decay. *eLife*, 4, 2015.

[1248] Eva Malatinkova, Ward De Spiegelaere, Pawel Bonczkowski, Maja Kiselinova, Karen Vervisch, Wim Trypsteen, Margaret Johnson, Chris Verhofstede, Danny de Looze, Charles Murray, Sabine Kinloch-de Loes, and Linos Vandekerckhove. Impact of a decade of successful antiretroviral therapy initiated at HIV-1 seroconversion on blood and rectal reservoirs. *eLife*, 4, 2015.

[1249] Kwun Nok M Man, Cordelia Imig, Alexander M Walter, Paulo S Pinheiro, David R Stevens, Jens Rettig, Jakob B Sørensen, Benjamin H Cooper, Nils Brose, and Sonja M Wojcik. Identification of a Munc13-sensitive step in chromaffin cell large dense-core vesicle exocytosis. *eLife*, 4, 2015.

[1250] Joseph D Mancias, Laura Pontano Vaites, Sahar Nissim, Douglas E Biancur, Andrew J Kim, Xiaoxu Wang, Yu Liu, Wolfram Goessling, Alec C Kimmelman, and J Wade Harper. Ferritinophagy via NCOA4 is required for erythropoiesis and is regulated by iron dependent HERC2-mediated proteolysis. *eLife*, 4, 2015.

[1251] Yael Mandelblat-Cerf, Rohan N Ramesh, Christian R Burgess, Paola Patella, Zongfang Yang, Bradford B Lowell, and Mark L Andermann. Arcuate hypothalamic AgRP and putative POMC neurons show opposite changes in spiking across multiple timescales. *eLife*, 4, 2015.

[1252] Nicola Manfrini, Michela Clerici, Maxime Wery, Chiara Vittoria Colombo, Marc Descrimes, Antonin Morillon, Fabrizio d'Adda di Fagagna, and Maria Pia Longhese. Resection is responsible for loss of transcription around a double-strand break in Saccharomyces cerevisiae. *eLife*, 4, 2015.

[1253] Niall M Mangan and Michael P Brenner. Correction: Systems analysis of the CO2 concentrating mechanism in cyanobacteria. *eLife*, 4, 2015.

[1254] Uri Manor, Sadie Bartholomew, Gonen Golani, Eric Christenson, Michael Kozlov, Henry Higgs, James Spudich, and Jennifer Lippincott-Schwartz. A mitochondria-anchored isoform of the actin-nucleating spire protein regulates mitochondrial division. *eLife*, 4, 2015.

[1255] Virginie Mansuy-Aubert, Laurent Gautron, Syann Lee, Angie L Bookout, Christine M Kusminski, Kai Sun, Yuan Zhang, Philipp E Scherer, David J Mangelsdorf, and Joel K Elmquist. Loss of the liver X receptor LXR$\alpha/\beta$ in peripheral sensory neurons modifies energy expenditure. *eLife*, 4, 2015.

[1256] Boryana N Manz, Ying Xim Tan, Adam H Courtney, Florentine Rutaganira, Ed Palmer, Kevan M Shokat, and Arthur Weiss. Small molecule inhibition of Csk alters affinity recognition by T cells. *eLife*, 4, 2015.

[1257] A Maqbool, H Saitoh, M Franceschetti, CEM Stevenson, A Uemura, H Kanzaki, S Kamoun, R Terauchi, and MJ Banfield. Structural basis of pathogen recognition by an integrated HMA domain in a plant NLR immune receptor. *eLife*, 4, 2015.

[1258] Therese Ann Markow. The secret lives of Drosophila flies. *eLife*, 4, 2015.

[1259] Laura Marroqui, Miguel Lopes, Reinaldo S dos Santos, Fabio A Grieco, Merja Roivainen, Sarah J Richardson, Noel G Morgan, Anne Op de beeck, and Decio L Eizirik. Differential cell autonomous responses determine the outcome of coxsackievirus infections in murine pancreatic $\alpha$ and $\beta$ cells. *eLife*, 4, 2015.

[1260] Heather D Marshall, John P Ray, Brian J Laidlaw, Nianzhi Zhang, Dipika Gawande, Matthew M Staron, Joe Craft, and Susan M Kaech. The transforming growth factor beta signaling pathway is critical for the formation of CD4 T follicular helper cells and isotype-switched antibody responses in the lung mucosa. *eLife*, 4, 2015.

[1261] Megan L Martik and David R McClay. Deployment of a retinal determination gene network drives directed cell migration in the sea urchin embryo. *eLife*, 4, 2015.

[1262] E Anne Martin, Shruti Muralidhar, Zhirong Wang, Diégo Cordero Cervantes, Raunak Basu, Matthew R Taylor, Jennifer Hunter, Tyler Cutforth, Scott A Wilke, Anirvan Ghosh, and Megan E Williams. The intellectual disability gene Kirrel3 regulates target-specific mossy fiber synapse development in the hippocampus. *eLife*, 4, 2015.

[1263] Pedro L Martinez-Espinosa, Jianping Wu, Chengtao Yang, Vivian Gonzalez-Perez, Huifang Zhou, Hongwu Liang, Xiao-Ming Xia, and Christopher J Lingle. Knockout of Slo2.2 enhances itch, abolishes KNa current, and increases action potential firing frequency in DRG neurons. *eLife*, 4, 2015.

[1264] Nicholas GD Masca, Elizabeth MA Hensor, Victoria R Cornelius, Francesca M Buffa, Helen M Marriott, James M Eales, Michael P Messenger, Amy E Anderson, Chris Boot, Catey Bunce, Robert D Goldin, Jessica Harris, Rod F Hinchliffe, Hiba Junaid, Shaun Kingston, Carmen Martin-Ruiz, Christopher P Nelson, Janet Peacock, Paul T Seed, Bethany Shinkins, Karl J Staples, Jamie Toombs, Adam KA Wright, and M Dawn Teare. RIPOSTE: a framework for improving the design and analysis of laboratory-based research. *eLife*, 4, 2015.

[1265] Philippe Materne, Jayamani Anandhakumar, Valerie Migeot, Ignacio Soriano, Carlo Yague-Sanz, Elena Hidalgo, Carole Mignion, Luis Quintales, Francisco Antequera, and Damien Hermand. Promoter nucleosome dynamics regulated by signalling through the CTD code. *eLife*, 4, 2015.

[1266] Alexander Mathis, Martin B Stemmler, and Andreas VM Herz. Probable nature of higher-dimensional symmetries underlying mammalian grid-cell activity patterns. *eLife*, 4, 2015.

[1267] Ryo Matsuda, Chie Hosono, Christos Samakovlis, and Kaoru Saigo. Multipotent versus differentiated cell fate selection in the developing Drosophila airways. *eLife*, 4, 2015.

[1268] Yuval Mazor, Anna Borovikova, and Nathan Nelson. The structure of plant photosystem I super-complex at 2.8 Å resolution. *eLife*, 4, 2015.

[1269] Ryan M McCormack, Lesley R de Armas, Motoaki Shiratsuchi, Desiree G Fiorentino, Melissa L Olsson, Mathias G Lichtenheld, Alejo Morales, Kirill Lyapichev, Louis E Gonzalez, Natasa Strbo, Neelima Sukumar, Olivera Stojadinovic, Gregory V Plano, George P Munson, Marjana Tomic-Canic, Robert S Kirsner, David G Russell, and Eckhard R Podack. Perforin-2 is essential for intracellular defense of parenchymal cells and phagocytes against pathogenic bacteria. *eLife*, 4, 2015.

[1270] Ryan M McCormack, Kirill Lyapichev, Melissa L Olsson, Eckhard R Podack, and George P Munson. Enteric pathogens deploy cell cycle inhibiting factors to block the bactericidal activity of Perforin-2. *eLife*, 4, 2015.

[1271] Aaron P McGrath, Elise L Laming, G Patricia Casas Garcia, Marc Kvansakul, J Mitchell Guss, Jill Trewhella, Benoit Calmes, Paul V Bernhardt, Graeme R Hanson, Ulrike Kappler, and Megan J Maher. Structural basis of interprotein electron transfer in bacterial sulfite oxidation. *eLife*, 4, 2015.

[1272] Markus Meister. On the dimensionality of odor space. *eLife*, 4, 2015.

[1273] Aaron S Mendez, Jennifer Alfaro, Marisol A Morales-Soto, Arvin C Dar, Emma McCullagh, Katja Gotthardt, Han Li, Diego Acosta-Alvear, Carmela Sidrauski, Alexei V Korennykh, Sebastian Bernales, Kevan M Shokat, and Peter Walter. Endoplasmic reticulum stress-independent activation of unfolded protein response kinases by a small molecule ATP-mimic. *eLife*, 4, 2015.

[1274] William Menegas, Joseph F Bergan, Sachie K Ogawa, Yoh Isogai, Kannan Umadevi Venkataraju, Pavel Osten, Naoshige Uchida, and Mitsuko Watabe-Uchida. Dopamine neurons projecting to the posterior striatum form an anatomically distinct subclass. *eLife*, 4, 2015.

[1275] Gaelle Mercenne, Steven L Alam, Jun Arii, Matthew S Lalonde, and Wesley I Sundquist. Angiomotin functions in HIV-1 assembly and budding. *eLife*, 4, 2015.

[1276] Dana Meyen, Katsiaryna Tarbashevich, Torsten U Banisch, Carolina Wittwer, Michal Reichman-Fried, Benoît Maugis, Cecilia Grimaldi, Esther-Maria Messerschmidt, and Erez Raz. Dynamic filopodia are required for chemokine-dependent intracellular polarization during guided cell migration in vivo. *eLife*, 4, 2015.

[1277] Christine Mieck, Maxim I Molodtsov, Katarzyna Drzewicka, Babet van der Vaart, Gabriele Litos, Gerald Schmauss, Alipasha Vaziri, and Stefan Westermann. Non-catalytic motor domains enable processive movement and functional diversification of the kinesin-14 Kar3. *eLife*, 4, 2015.

[1278] Alexander S Mikheyev and Timothy A Linksvayer. Genes associated with ant social behavior show distinct transcriptional and evolutionary patterns. *eLife*, 4, 2015.

[1279] Shilpi Minocha, Delphine Valloton, Isabelle Brunet, Anne Eichmann, Jean-Pierre Hornung, and Cecile Lebrand. NG2 glia are required for vessel network formation during embryonic development. *eLife*, 4, 2015.

[1280] Anish Mitra, Abraham Z Snyder, Enzo Tagliazucchi, Helmut Laufs, and Marcus E Raichle. Propagated infra-slow intrinsic brain activity reorganizes across wake and slow wave sleep. *eLife*, 4, 2015.

[1281] Matthew A Mitsche, Jeffrey G McDonald, Helen H Hobbs, and Jonathan C Cohen. Flux analysis of cholesterol biosynthesis in vivo reveals multiple tissue and cell-type specific pathways. *eLife*, 4, 2015.

[1282] Yusuke Miyazaki, Ling-chun Chen, Bernard W Chu, Tomek Swigut, and Thomas J Wandless. Distinct transcriptional responses elicited by unfolded nuclear or cytoplasmic protein in mammalian cells. *eLife*, 4, 2015.

[1283] Binyam Mogessie, Daniel Roth, Zainab Rahil, and Anne Straube. A novel isoform of MAP4 organises the paraxial microtubule array required for muscle cell differentiation. *eLife*, 4, 2015.

[1284] Jorrit S Montijn, Pieter M Goltstein, and Cyriel MA Pennartz. Mouse V1 population correlates of visual detection rely on heterogeneity within neuronal response patterns. *eLife*, 4, 2015.

[1285] Hernán Morales-Navarrete, Fabián Segovia-Miranda, Piotr Klukowski, Kirstin Meyer, Hidenori Nonaka, Giovanni Marsico, Mikhail Chernykh, Alexander Kalaidzidis, Marino Zerial, and Yannis Kalaidzidis. A versatile pipeline for the multi-scale digital reconstruction and quantitative analysis of 3D tissue architecture. *eLife*, 4, 2015.

[1286] Marco Morselli, William A Pastor, Barbara Montanini, Kevin Nee, Roberto Ferrari, Kai Fu, Giancarlo Bonora, Liudmilla Rubbi, Amander T Clark, Simone Ottonello, Steven E Jacobsen, and Matteo Pellegrini. In vivo targeting of de novo DNA methylation by histone modifications in yeast and mouse. *eLife*, 4, 2015.

[1287] Yangling Mu, Chunmei Zhao, Nicolas Toni, Jun Yao, and Fred H Gage. Distinct roles of NMDA receptors at different stages of granule cell development in the adult brain. *eLife*, 4, 2015.

[1288] Alexander Muir, FranÇoise M Roelants, Garrett Timmons, Kristin L Leskoske, and Jeremy Thorner. Down-regulation of TORC2-Ypk1 signaling promotes MAPK-independent survival under hyperosmotic stress. *eLife*, 4, 2015.

[1289] Martin Müller, Oliver Schmidt, Mihaela Angelova, Klaus Faserl, Sabine Weys, Leopold Kremser, Thaddäus Pfaffenwimmer, Thomas Dalik, Claudine Kraft, Zlatko Trajanoski, Herbert Lindner, and David Teis. The coordinated action of the MVB pathway and autophagy ensures cell survival during starvation. *eLife*, 4, 2015.

[1290] Peter R Murphy, Ian H Robertson, Siobhán Harty, and Redmond G O'Connell. Neural evidence accumulation persists after choice to inform metacognitive judgments. *eLife*, 4, 2015.

[1291] Elisabeth A Murray, Emily J Moylan, Kadharbatcha S Saleem, Benjamin M Basile, and Janita Turchi. Specialized areas for value updating and goal selection in the primate orbitofrontal cortex. *eLife*, 4, 2015.

[1292] Christina C Muscat, Keila M Torre-Santiago, Michael V Tran, James A Powers, and Sarah M Wignall. Kinetochore-independent chromosome segregation driven by lateral microtubule bundles. *eLife*, 4, 2015.

[1293] Nicholas E Myers, Gustavo Rohenkohl, Valentin Wyart, Mark W Woolrich, Anna C Nobre, and Mark G Stokes. Testing sensory evidence against mnemonic templates. *eLife*, 4, 2015.

[1294] Nataliya Nady, Ankit Gupta, Ziyang Ma, Tomek Swigut, Akiko Koide, Shohei Koide, and Joanna Wysocka. ETO family protein Mtgr1 mediates Prdm14 functions in stem cell maintenance and primordial germ cell formation. *eLife*, 4, 2015.

[1295] Sonal Nagarkar-Jaiswal, Steven Z DeLuca, Pei-Tseng Lee, Wen-Wen Lin, Hongling Pan, Zhongyuan Zuo, Jiangxing Lv, Allan C Spradling, and Hugo J Bellen. A genetic toolkit for tagging intronic MiMIC containing genes. *eLife*, 4, 2015.

[1296] Wiebke Nahrendorf, Philip J Spence, Irene Tumwine, Prisca Lévy, William Jarra, Robert W Sauerwein, and Jean Langhorne. Blood-stage immunity to Plasmodium chabaudi malaria following chemoprophylaxis and sporozoite immunization. *eLife*, 4, 2015.

[1297] Divya Nandakumar, Manjula Pandey, and Smita S Patel. Cooperative base pair melting by helicase and polymerase positioned one nucleotide from each other. *eLife*, 4, 2015.

[1298] Kamesh Narasimhan, Samuel A Lambert, Ally WH Yang, Jeremy Riddell, Sanie Mnaimneh, Hong Zheng, Mihai Albu, Hamed S Najafabadi, John S Reece-Hoyes, Juan I Fuxman Bass, Albertha JM Walhout, Matthew T Weirauch, and Timothy R Hughes. Mapping and analysis of Caenorhabditis elegans transcription factor sequence specificities. *eLife*, 4, 2015.

[1299] Ryan G Natan, John J Briguglio, Laetitia Mwilambwe-Tshilobo, Sara I Jones, Mark Aizenberg, Ethan M Goldberg, and Maria Neimark Geffen. Complementary control of sensory adaptation by two types of cortical interneurons. *eLife*, 4, 2015.

[1300] Tobias Navarro Schröder, Koen V Haak, Nestor I Zaragoza Jimenez, Christian F Beckmann, and Christian F Doeller. Functional topography of the human entorhinal cortex. *eLife*, 4, 2015.

[1301] Scott J Neal, Asuka Takeishi, Michael P O'Donnell, JiSoo Park, Myeongjin Hong, Rebecca A Butcher, Kyuhyung Kim, and Piali Sengupta. Feeding state-dependent regulation of developmental plasticity via CaMKI and neuroendocrine signaling. *eLife*, 4, 2015.

[1302] Joshua P Neunuebel, Adam L Taylor, Ben J Arthur, and SE Roian Egnor. Female mice ultrasonically interact with males during courtship displays. *eLife*, 4, 2015.

[1303] Aitana Neves, Coralie Busso, and Pierre Gönczy. Cellular hallmarks reveal restricted aerobic metabolism at thermal limits. *eLife*, 4, 2015.

[1304] Jonathan P Newman, Ming-fai Fong, Daniel C Millard, Clarissa J Whitmire, Garrett B Stanley, and Steve M Potter. Optogenetic feedback control of neural activity. *eLife*, 4, 2015.

[1305] Erin Newman-Smith, Matthew J Kourakis, Wendy Reeves, Michael Veeman, and William C Smith. Reciprocal and dynamic polarization of planar cell polarity core components and myosin. *eLife*, 4, 2015.

[1306] Kien Xuan Ngo, Noriyuki Kodera, Eisaku Katayama, Toshio Ando, and Taro QP Uyeda. Cofilin-induced unidirectional cooperative conformational changes in actin filaments revealed by high-speed atomic force microscopy. *eLife*, 4, 2015.

[1307] Michael O Nguyen, Manisha Jalan, Carl A Morrow, Fekret Osman, and Matthew C Whitby. Recombination occurs within minutes of replication blockage by RTS1 producing restarted forks that are prone to collapse. *eLife*, 4, 2015.

[1308] Mai Nguyen-Chi, Béryl Laplace-Builhe, Jana Travnickova, Patricia Luz-Crawford, Gautier Tejedor, Quang Tien Phan, Isabelle Duroux-Richard, Jean-Pierre Levraud, Karima Kissa, Georges Lutfalla, Christian Jorgensen, and Farida Djouad. Identification of polarized macrophage subsets in zebrafish. *eLife*, 4, 2015.

[1309] Joshua M Nicholson, Joana C Macedo, Aaron J Mattingly, Darawalee Wangsa, Jordi Camps, Vera Lima, Ana M Gomes, Sofia Dória, Thomas Ried, Elsa Logarinho, and Daniela Cimini. Chromosome mis-segregation and cytokinesis failure in trisomic human cells. *eLife*, 4, 2015.

[1310] Henrike Niederholtmeyer, Zachary Z Sun, Yutaka Hori, Enoch Yeung, Amanda Verpoorte, Richard M Murray, and Sebastian J Maerkl. Rapid cell-free forward engineering of novel genetic ring oscillators. *eLife*, 4, 2015.

[1311] Stanley Nithianantham, Sinh Le, Elbert Seto, Weitao Jia, Julie Leary, Kevin D Corbett, Jeffrey K Moore, and Jawdat Al-Bassam. Tubulin cofactors and Arl2 are cage-like chaperones that regulate the soluble $\alpha\beta$-tubulin pool for microtubule dynamics. *eLife*, 4, 2015.

[1312] Kazuhiro R Nitta, Arttu Jolma, Yimeng Yin, Ekaterina Morgunova, Teemu Kivioja, Junaid Akhtar, Korneel Hens, Jarkko Toivonen, Bart Deplancke, Eileen E M Furlong, and Jussi Taipale. Conservation of transcription factor binding specificities across 600 million years of bilateria evolution. *eLife*, 4, 2015.

[1313] Faye M Nixon, Cristina Gutiérrez-Caballero, Fiona E Hood, Daniel G Booth, Ian A Prior, and Stephen J Royle. The mesh is a network of microtubule connectors that stabilizes individual kinetochore fibers of the mitotic spindle. *eLife*, 4, 2015.

[1314] Natasha M O'Brown, Brian R Summers, Felicity C Jones, Shannon D Brady, and David M Kingsley. A recurrent regulatory change underlying altered expression and Wnt response of the stickleback armor plates gene EDA. *eLife*, 4, 2015.

[1315] Benson Ogunjimi, Lander Willem, Philippe Beutels, and Niel Hens. Integrating between-host transmission and within-host immunity to analyze the impact of varicella vaccination on zoster. *eLife*, 4, 2015.

[1316] H Freyja ólafsdóttir, Caswell Barry, Aman B Saleem, Demis Hassabis, and Hugo J Spiers. Hippocampal place cells construct reward related sequences through unexplored space. *eLife*, 4, 2015.

[1317] Enno R Oldewurtel, Nadzeya Kouzel, Lena Dewenter, Katja Henseler, and Berenike Maier. Differential interaction forces govern bacterial sorting in early biofilms. *eLife*, 4, 2015.

[1318] Jason W Olejarz, Benjamin Allen, Carl Veller, and Martin A Nowak. The evolution of non-reproductive workers in insect colonies with haplodiploid genetics. *eLife*, 4, 2015.

[1319] Koh Onimaru, Shigehiro Kuraku, Wataru Takagi, Susumu Hyodo, James Sharpe, and Mikiko Tanaka. A shift in anterior–posterior positional information underlies the fin-to-limb evolution. *eLife*, 4, 2015.

[1320] David Orduz, Paloma P Maldonado, Maddalena Balia, Mateo Vélez-Fort, Vincent de Sars, Yuchio Yanagawa, Valentina Emiliani, and Maria Cecilia Angulo. Interneurons and oligodendrocyte progenitors form a structured synaptic network in the developing neocortex. *eLife*, 4, 2015.

[1321] Yael Oren, Aharon Nachshon, Amit Frishberg, Roni Wilentzik, and Irit Gat-Viks. Linking traits based on their shared molecular mechanisms. *eLife*, 4, 2015.

[1322] Andrew R Osborn, Khaled H Almabruk, Garrett Holzwarth, Shumpei Asamizu, Jane LaDu, Kelsey M Kean, P Andrew Karplus, Robert L Tanguay, Alan T Bakalinsky, and Taifo Mahmud. De novo synthesis of a sunscreen compound in vertebrates. *eLife*, 4, 2015.

[1323] Radim Osicka, Adriana Osickova, Shakir Hasan, Ladislav Bumba, Jiri Cerny, and Peter Sebo. Bordetella adenylate cyclase toxin is a unique ligand of the integrin complement receptor 3. *eLife*, 4, 2015.

[1324] Sergey Ovchinnikov, Lisa Kinch, Hahnbeom Park, Yuxing Liao, Jimin Pei, David E Kim, Hetunandan Kamisetty, Nick V Grishin, and David Baker. Large-scale determination of previously unsolved protein structures using evolutionary information. *eLife*, 4, 2015.

[1325] Katharina Overlack, Ivana Primorac, Mathijs Vleugel, Veronica Krenn, Stefano Maffini, Ingrid Hoffmann, Geert J P L Kops, and Andrea Musacchio. A molecular basis for the differential roles of Bub1 and BubR1 in the spindle assembly checkpoint. *eLife*, 4, 2015.

[1326] Mehmet Neset özel, Marion Langen, Bassem A Hassan, and P Robin Hiesinger. Filopodial dynamics and growth cone stabilization in Drosophila visual circuit development. *eLife*, 4, 2015.

[1327] Annalise B Paaby, Amelia G White, David D Riccardi, Kristin C Gunsalus, Fabio Piano, and Matthew V Rockman. Wild worm embryogenesis harbors ubiquitous polygenic modifier variation. *eLife*, 4, 2015.

[1328] Marta Pallotto, Paul V Watkins, Boma Fubara, Joshua H Singer, and Kevin L Briggman. Extracellular space preservation aids the connectomic analysis of neural circuits. *eLife*, 4, 2015.

[1329] Chethan Pandarinath, Vikash Gilja, Christine H Blabe, Paul Nuyujukian, Anish A Sarma, Brittany L Sorice, Emad N Eskandar, Leigh R Hochberg, Jaimie M Henderson, and Krishna V Shenoy. Neural population dynamics in human motor cortex during movements in people with ALS. *eLife*, 4, 2015.

[1330] Effrosyni Papanikou, Kasey J Day, Jotham Austin, and Benjamin S Glick. COPI selectively drives maturation of the early Golgi. *eLife*, 4, 2015.

[1331] Stephanie J Papp, Anne-Laure Huber, Sabine D Jordan, Anna Kriebs, Madelena Nguyen, James J Moresco, John R Yates, and Katja A Lamia. DNA damage shifts circadian clock time via Hausp-dependent Cry1 stabilization. *eLife*, 4, 2015.

[1332] Samuel S Pappas, Katherine Darr, Sandra M Holley, Carlos Cepeda, Omar S Mabrouk, Jenny-Marie T Wong, Tessa M LeWitt, Reema Paudel, Henry Houlden, Robert T Kennedy, Michael S Levine, and William T Dauer. Forebrain deletion of the dystonia protein torsinA causes dystonic-like movements and loss of striatal cholinergic neurons. *eLife*, 4, 2015.

[1333] María Belén Pardi, Mora Belén Ogando, Alejandro F Schinder, and Antonia Marin-Burgin. Differential inhibition onto developing and mature granule cells generates high-frequency filters with variable gain. *eLife*, 4, 2015.

[1334] David M Parichy. Advancing biology through a deeper understanding of zebrafish ecology and evolution. *eLife*, 4, 2015.

[1335] Chung Park, James Arthos, Claudia Cicala, and John H Kehrl. The HIV-1 envelope protein gp120 is captured and displayed for B cell recognition by SIGN-R1+ lymph node macrophages. *eLife*, 4, 2015.

[1336] Timothy J Parnell, Alisha Schlichter, Boris G Wilson, and Bradley R Cairns. The chromatin remodelers RSC and ISW1 display functional and chromatin-based promoter antagonism. *eLife*, 4, 2015.

[1337] Viktoria Désirée Paul, Ulrich Mühlenhoff, Martin Stümpfig, Jan Seebacher, Karl G Kugler, Christian Renicke, Christof Taxis, Anne-Claude Gavin, Antonio J Pierik, and Roland Lill. The deca-GX3 proteins Yae1-Lto1 function as adaptors recruiting the ABC protein Rli1 for iron-sulfur cluster insertion. *eLife*, 4, 2015.

[1338] Jillian N Pearring, William J Spencer, Eric C Lieu, and Vadim Y Arshavsky. Guanylate cyclase 1 relies on rhodopsin for intracellular stability and ciliary trafficking. *eLife*, 4, 2015.

[1339] Nicolás Peláez, Arnau Gavalda-Miralles, Bao Wang, Heliodoro Tejedor Navarro, Herman Gudjonson, Ilaria Rebay, Aaron R Dinner, Aggelos K Katsaggelos, Luís AN Amaral, and Richard W Carthew. Dynamics and heterogeneity of a fate determinant during transition towards cell differentiation. *eLife*, 4, 2015.

[1340] William G Pembroke, Arran Babbs, Kay E Davies, Chris P Ponting, and Peter L Oliver. Temporal transcriptomics suggest that twin-peaking genes reset the clock. *eLife*, 4, 2015.

[1341] Laura Pereira, Paschalis Kratsios, Esther Serrano-Saiz, Hila Sheftel, Avi E Mayo, David H Hall, John G White, Brigitte LeBoeuf, L Rene Garcia, Uri Alon, and Oliver Hobert. A cellular and regulatory map of the cholinergic nervous system of C. elegans. *eLife*, 4, 2015.

[1342] Julio D Perez, Nimrod D Rubinstein, Daniel E Fernandez, Stephen W Santoro, Leigh A Needleman, Olivia Ho-Shing, John J Choi, Mariela Zirlinger, Shau-Kwaun Chen, Jun S Liu, and Catherine Dulac. Quantitative and functional interrogation of parent-of-origin allelic expression biases in the brain. *eLife*, 4, 2015.

[1343] Christopher Petro, Pablo A González, Natalia Cheshenko, Thomas Jandl, Nazanin Khajoueinejad, Angèle Bénard, Mayami Sengupta, Betsy C Herold, and William R Jacobs. Herpes simplex type 2 virus deleted in glycoprotein D protects against vaginal, skin and neural disease. *eLife*, 4, 2015.

[1344] Megan Phifer-Rixey and Michael W Nachman. Insights into mammalian biology from the wild house mouse Mus musculus. *eLife*, 4, 2015.

[1345] Andrea Picco, Markus Mund, Jonas Ries, FranÇois Nédélec, and Marko Kaksonen. Visualizing the functional architecture of the endocytic machinery. *eLife*, 4, 2015.

[1346] Justyna Pielecka-Fortuna, Evgenia Kalogeraki, Michal G Fortuna, and Siegrid Löwel. Optimal level activity of matrix metalloproteinases is critical for adult visual plasticity in the healthy and stroke-affected brain. *eLife*, 4, 2015.

[1347] Francisco J Piña and Maho Niwa. The ER Stress Surveillance (ERSU) pathway regulates daughter cell ER protein aggregate inheritance. *eLife*, 4, 2015.

[1348] André Plagens, Michael Daume, Julia Wiegel, and Lennart Randau. Circularization restores signal recognition particle RNA functionality in Thermoproteus. *eLife*, 4, 2015.

[1349] Tino Pleiner, Mark Bates, Sergei Trakhanov, Chung-Tien Lee, Jan Erik Schliep, Hema Chug, Marc Böhning, Holger Stark, Henning Urlaub, and Dirk Görlich. Nanobodies: site-specific labeling for super-resolution imaging, rapid epitope-mapping and native protein complex isolation. *eLife*, 4, 2015.

[1350] Anne Plessis, Christoph Hafemeister, Olivia Wilkins, Zennia Jean Gonzaga, Rachel Sarah Meyer, Inês Pires, Christian Müller, Endang M Septiningsih, Richard Bonneau, and Michael Purugganan. Multiple abiotic stimuli are integrated in the regulation of rice gene expression under field conditions. *eLife*, 4, 2015.

[1351] Rachel Plumb, Zai-Rong Zhang, Suhila Appathurai, and Malaiyalam Mariappan. A functional link between the co-translational protein translocation pathway and the UPR. *eLife*, 4, 2015.

[1352] Thomas Pohlmann, Sebastian Baumann, Carl Haag, Mario Albrecht, and Michael Feldbrügge. A FYVE zinc finger domain protein specifically links mRNA transport to endosome trafficking. *eLife*, 4, 2015.

[1353] Sebastian Poliak, Daniel Morales, Louis-Philippe Croteau, Dayana Krawchuk, Elena Palmesino, Susan Morton, Jean-FranÇois Cloutier, Frederic Charron, Matthew B Dalva, Susan L Ackerman, Tzu-Jen Kao, and Artur Kania. Synergistic integration of Netrin and ephrin axon guidance signals by spinal motor neurons. *eLife*, 4, 2015.

[1354] Welkin H Pope, Charles A Bowman, Daniel A Russell, Deborah Jacobs-Sera, David J Asai, Steven G Cresawn, William R Jacobs, Roger W Hendrix, Jeffrey G Lawrence, and Graham F Hatfull. Whole genome comparison of a large collection of mycobacteriophages reveals a continuum of phage genetic diversity. *eLife*, 4, 2015.

[1355] Silvia Portugal, Christopher M Tipton, Haewon Sohn, Younoussou Kone, Jing Wang, Shanping Li, Jeff Skinner, Kimmo Virtaneva, Daniel E Sturdevant, Stephen F Porcella, Ogobara K Doumbo, Safiatou Doumbo, Kassoum Kayentao, Aissata Ongoiba, Boubacar Traore, Inaki Sanz, Susan K Pierce, and Peter D Crompton. Malaria-associated atypical memory B cells exhibit markedly reduced B cell receptor signaling and effector function. *eLife*, 4, 2015.

[1356] Kate Powell, Alexandre Mathy, Ian Duguid, and Michael Häusser. Synaptic representation of locomotion in single cerebellar granule cells. *eLife*, 4, 2015.

[1357] Steffen Preissler, Joseph E Chambers, Ana Crespillo-Casado, Edward Avezov, Elena Miranda, Juan Perez, Linda M Hendershot, Heather P Harding, and David Ron. Physiological modulation of BiP activity by trans-protomer engagement of the interdomain linker. *eLife*, 4, 2015.

[1358] Steffen Preissler, Cláudia Rato, Ruming Chen, Robin Antrobus, Shujing Ding, Ian M Fearnley, and David Ron. AMPylation matches BiP activity to client protein load in the endoplasmic reticulum. *eLife*, 4, 2015.

[1359] Mira I Pronobis, Nasser M Rusan, and Mark Peifer. A novel GSK3-regulated APC:Axin interaction regulates Wnt signaling by driving a catalytic cycle of efficient $\beta$catenin destruction. *eLife*, 4, 2015.

[1360] Erik M Quandt, Jimmy Gollihar, Zachary D Blount, Andrew D Ellington, George Georgiou, and Jeffrey E Barrick. Fine-tuning citrate synthase flux potentiates and refines metabolic innovation in the Lenski evolution experiment. *eLife*, 4, 2015.

[1361] Neil C Rabinowitz, Robbe L Goris, Marlene Cohen, and Eero P Simoncelli. Attention stabilizes the shared gain of V4 populations. *eLife*, 4, 2015.

[1362] Lilliana Radoshevich, Francis Impens, David Ribet, Juan J Quereda, To Nam Tham, Marie-Anne Nahori, Hélène Bierne, Olivier Dussurget, Javier Pizarro-Cerdá, Klaus-Peter Knobeloch, and Pascale Cossart. ISG15 counteracts Listeria monocytogenes infection. *eLife*, 4, 2015.

[1363] Ponni Rajagopal, Eric Tse, Andrew J Borst, Scott P Delbecq, Lei Shi, Daniel R Southworth, and Rachel E Klevit. A conserved histidine modulates HSPB5 structure to trigger chaperone activity in response to stress-related acidosis. *eLife*, 4, 2015.

[1364] Ghanasyam Rallapalli, Diane GO Saunders, Kentaro Yoshida, Anne Edwards, Carlos A Lugo, Steve Collin, Bernardo Clavijo, Manuel Corpas, David Swarbreck, Matthew Clark, J Allan Downie, Sophien Kamoun, and Dan MacLean. Lessons from Fraxinus, a crowd-sourced citizen science game in genomics. *eLife*, 4, 2015.

[1365] Indira M Raman. Triaging Shakespeare. *eLife*, 4, 2015.

[1366] Roybel R Ramiscal, Ian A Parish, Robert S Lee-Young, Jeffrey J Babon, Julianna Blagih, Alvin Pratama, Jaime Martin, Naomi Hawley, Jean Y Cappello, Pablo F Nieto, Julia I Ellyard, Nadia J Kershaw, Rebecca A Sweet, Christopher C Goodnow, Russell G Jones, Mark A Febbraio, Carola G Vinuesa, and Vicki Athanasopoulos. Attenuation of AMPK signaling by ROQUIN promotes T follicular helper cell formation. *eLife*, 4, 2015.

[1367] Nadine Randel, Réza Shahidi, Csaba Verasztó, Luis A Bezares-Calderón, Steffen Schmidt, and Gáspár Jékely. Inter-individual stereotypy of the Platynereis larval visual connectome. *eLife*, 4, 2015.

[1368] Sunitha Rangaraju, Gregory M Solis, Ryan C Thompson, Rafael L Gomez-Amaro, Leo Kurian, Sandra E Encalada, Alexander B Niculescu, Daniel R Salomon, and Michael Petrascheck. Suppression of transcriptional drift extends C. elegans lifespan by postponing the onset of mortality. *eLife*, 4, 2015.

[1369] Anand Ranjan, Feng Wang, Gaku Mizuguchi, Debbie Wei, Yingzi Huang, and Carl Wu. H2A histone-fold and DNA elements in nucleosome activate SWR1-mediated H2A.Z replacement in budding yeast. *eLife*, 4, 2015.

[1370] Prashanth R Rao, Li Lin, Hai Huang, Arjun Guha, Sougata Roy, and Thomas B Kornberg. Developmental compartments in the larval trachea of Drosophila. *eLife*, 4, 2015.

[1371] Samrrah Raouf, Claire Weston, and Nora Yucel. Registered report: Senescence surveillance of pre-malignant hepatocytes limits liver cancer development. *eLife*, 4, 2015.

[1372] Benjamin Rauwel, Suk Min Jang, Marco Cassano, Adamandia Kapopoulou, Isabelle Barde, and Didier Trono. Release of human cytomegalovirus from latency by a KAP1/TRIM28 phosphorylation switch. *eLife*, 4, 2015.

[1373] Tali Raveh-Sadka, Brian C Thomas, Andrea Singh, Brian Firek, Brandon Brooks, Cindy J Castelle, Itai Sharon, Robyn Baker, Misty Good, Michael J Morowitz, and Jillian F Banfield. Gut bacteria are rarely shared by co-hospitalized premature infants, regardless of necrotizing enterocolitis development. *eLife*, 4, 2015.

[1374] David C Rawling, Megan E Fitzgerald, and Anna Marie Pyle. Establishing the role of ATP for the function of the RIG-I innate immune sensor. *eLife*, 4, 2015.

[1375] Jonathan E Reeder, Youn-Tae Kwak, Ryan P McNamara, Christian V Forst, and Iván D'Orso. HIV Tat controls RNA Polymerase II and the epigenetic landscape to transcriptionally reprogram target immune cells. *eLife*, 4, 2015.

[1376] Austin L Reese and Ege T Kavalali. Spontaneous neurotransmission signals through store-driven Ca2+ transients to maintain synaptic homeostasis. *eLife*, 4, 2015.

[1377] Tobias Reiff, Jake Jacobson, Paola Cognigni, Zeus Antonello, Esther Ballesta, Kah Junn Tan, Joanne Y Yew, Maria Dominguez, and Irene Miguel-Aliaga. Endocrine remodelling of the adult intestine sustains reproduction in Drosophila. *eLife*, 4, 2015.

[1378] Robert C Reiner, Arnaud Le Menach, Simon Kunene, Nyasatu Ntshalintshali, Michelle S Hsiang, T Alex Perkins, Bryan Greenhouse, Andrew J Tatem, Justin M Cohen, and David L Smith. Mapping residual transmission for malaria elimination. *eLife*, 4, 2015.

[1379] Rubén Rellán-álvarez, Guillaume Lobet, Heike Lindner, Pierre-Luc Pradier, Jose Sebastian, Muh-Ching Yee, Yu Geng, Charlotte Trontin, Therese LaRue, Amanda Schrager-Lavelle, Cara H Haney, Rita Nieu, Julin Maloof, John P Vogel, and José R Dinneny. GLO-Roots: an imaging platform enabling multidimensional characterization of soil-grown root systems. *eLife*, 4, 2015.

[1380] Ann L Revill, Nikolas C Vann, Victoria T Akins, Andrew Kottick, Paul A Gray, Christopher A Del Negro, and Gregory D Funk. Dbx1 precursor cells are a source of inspiratory XII premotoneurons. *eLife*, 4, 2015.

[1381] Kent Riemondy, Xiao-jing Wang, Enrique C Torchia, Dennis R Roop, and Rui Yi. MicroRNA-203 represses selection and expansion of oncogenic Hras transformed tumor initiating cells. *eLife*, 4, 2015.

[1382] Todd R Riley, Allan Lazarovici, Richard S Mann, and Harmen J Bussemaker. Building accurate sequence-to-affinity models from high-throughput in vitro protein-DNA binding data using FeatureREDUCE. *eLife*, 4, 2015.

[1383] Maureen Ritchey, Maria E Montchal, Andrew P Yonelinas, and Charan Ranganath. Delay-dependent contributions of medial temporal lobe regions to episodic memory retrieval. *eLife*, 4, 2015.

[1384] Philip J Robinson, Michael J Trnka, Riccardo Pellarin, Charles H Greenberg, David A Bushnell, Ralph Davis, Alma L Burlingame, Andrej Sali, and Roger D Kornberg. Molecular architecture of the yeast Mediator complex. *eLife*, 4, 2015.

[1385] Mathieu P Rodero, Lucie Poupel, Pierre-Louis Loyher, Pauline Hamon, Fabrice Licata, Charlotte Pessel, David A Hume, Christophe Combadière, and Alexandre Boissonnas. Immune surveillance of the lung by migrating tissue monocytes. *eLife*, 4, 2015.

[1386] Aida Rodrigo Albors, Akira Tazaki, Fabian Rost, Sergej Nowoshilow, Osvaldo Chara, and Elly M Tanaka. Planar cell polarity-mediated induction of neural stem cell expansion during axolotl spinal cord regeneration. *eLife*, 4, 2015.

[1387] Kacper B Rogala, Nicola J Dynes, Georgios N Hatzopoulos, Jun Yan, Sheng Kai Pong, Carol V Robinson, Charlotte M Deane, Pierre Gönczy, and Ioannis Vakonakis. The Caenorhabditis elegans protein SAS-5 forms large oligomeric assemblies critical for centriole formation. *eLife*, 4, 2015.

[1388] Clare Rollie, Stefanie Schneider, Anna Sophie Brinkmann, Edward L Bolt, and Malcolm F White. Intrinsic sequence specificity of the Cas1 integrase directs new spacer acquisition. *eLife*, 4, 2015.

[1389] Xin Rong, Bo Wang, Merlow M Dunham, Per Niklas Hedde, Jinny S Wong, Enrico Gratton, Stephen G Young, David A Ford, and Peter Tontonoz. Lpcat3-dependent production of arachidonoyl phospholipids is a key determinant of triglyceride secretion. *eLife*, 4, 2015.

[1390] Susanne Roosing, Matan Hofree, Sehyun Kim, Eric Scott, Brett Copeland, Marta Romani, Jennifer L Silhavy, Rasim O Rosti, Jana Schroth, Tommaso Mazza, Elide Miccinilli, Maha S Zaki, Kathryn J Swoboda, Joanne Milisa-Drautz, William B Dobyns, Mohamed A Mikati, Faruk İncecik, Matloob Azam, Renato Borgatti, Romina Romaniello, Rose-Mary Boustany, Carol L Clericuzio, Stefano D'Arrigo, Petter Strømme, Eugen Boltshauser, Franco Stanzial, Marisol Mirabelli-Badenier, Isabella Moroni, Enrico Bertini, Francesco Emma, Maja Steinlin, Friedhelm Hildebrandt, Colin A Johnson, Michael Freilinger, Keith K Vaux, Stacey B Gabriel, Pedro Aza-Blanc, Susanne Heynen-Genel, Trey Ideker, Brian D Dynlacht, Ji Eun Lee, Enza Maria Valente, Joon Kim, and Joseph G Gleeson. Functional genome-wide siRNA screen identifies KIAA0586 as mutated in Joubert syndrome. *eLife*, 4, 2015.

[1391] Juliana M Rosa, Rémi Bos, Georgeann S Sack, Cécile Fortuny, Amit Agarwal, Dwight E Bergles, John G Flannery, and Marla B Feller. Neuron-glia signaling in developing retina mediated by neurotransmitter spillover. *eLife*, 4, 2015.

[1392] Alicia Rosello, Mathias Mossoko, Stefan Flasche, Albert Jan Van Hoek, Placide Mbala, Anton Camacho, Sebastian Funk, Adam Kucharski, Benoit Kebela Ilunga, W John Edmunds, Peter Piot, Marc Baguelin, and Jean-Jacques Muyembe Tamfum. Ebola virus disease in the Democratic Republic of the Congo, 1976-2014. *eLife*, 4, 2015.

[1393] S Michael Rothenberg, Kyle Concannon, Sarah Cullen, Gaylor Boulay, Alexa B Turke, Anthony C Faber, Elizabeth L Lockerman, Miguel N Rivera, Jeffrey A Engelman, Shyamala Maheswaran, and Daniel A Haber. Inhibition of mutant EGFR in lung cancer cells triggers SOX2-FOXO6-dependent survival pathways. *eLife*, 4, 2015.

[1394] Simon Roux, Steven J Hallam, Tanja Woyke, and Matthew B Sullivan. Viral dark matter and virus–host interactions resolved from publicly available microbial genomes. *eLife*, 4, 2015.

[1395] Miguel Rovira, Pedro Saavedra, José Casal, and Peter A Lawrence. Regions within a single epidermal cell of Drosophila can be planar polarised independently. *eLife*, 4, 2015.

[1396] Achira Roy, Jonathan Skibo, Franck Kalume, Jing Ni, Sherri Rankin, Yiling Lu, William B Dobyns, Gordon B Mills, Jean J Zhao, Suzanne J Baker, and Kathleen J Millen. Mouse models of human PIK3CA-related brain overgrowth have acutely treatable epilepsy. *eLife*, 4, 2015.

[1397] Christian K Roy, Sara Olson, Brenton R Graveley, Phillip D Zamore, and Melissa J Moore. Assessing long-distance RNA sequence connectivity via RNA-templated DNA–DNA ligation. *eLife*, 4, 2015.

[1398] Alon Rubin, Nitzan Geva, Liron Sheintuch, and Yaniv Ziv. Hippocampal ensemble dynamics timestamp events in long-term memory. *eLife*, 4, 2015.

[1399] Marina Rudan, Dominique Schneider, Tobias Warnecke, and Anita Krisko. RNA chaperones buffer deleterious mutations in E. coli. *eLife*, 4, 2015.

[1400] Marisa D Ruehle, Haibo Zhang, Ryan M Sheridan, Somdeb Mitra, Yuanwei Chen, Ruben L Gonzalez, Barry S Cooperman, and Jeffrey S Kieft. A dynamic RNA loop in an IRES affects multiple steps of elongation factor-mediated translation initiation. *eLife*, 4, 2015.

[1401] Pierre-Louis Ruffault, Fabien D'Autréaux, John A Hayes, Marc Nomaksteinsky, Sandra Autran, Tomoyuki Fujiyama, Mikio Hoshino, Martin Hägglund, Ole Kiehn, Jean-FranÇois Brunet, Gilles Fortin, and Christo Goridis. The retrotrapezoid nucleus neurons expressing Atoh1 and Phox2b are essential for the respiratory response to CO2. *eLife*, 4, 2015.

[1402] Joanna P Rys, Christopher C DuFort, David A Monteiro, Michelle A Baird, Juan A Oses-Prieto, Shreya Chand, Alma L Burlingame, Michael W Davidson, and Tamara N Alliston. Discrete spatial organization of TGF$\beta$ receptors couples receptor multimerization and signaling to cellular tension. *eLife*, 4, 2015.

[1403] Hannes P Saal, Michael A Harvey, and Sliman J Bensmaia. Rate and timing of cortical responses driven by separate sensory channels. *eLife*, 4, 2015.

[1404] Juha Saarikangas and Yves Barral. Protein aggregates are associated with replicative aging without compromising protein quality control. *eLife*, 4, 2015.

[1405] Lena Sachs, Yen-Ta Chen, Axel Drechsler, Jeremy A Lynch, Kristen A Panfilio, Michael Lässig, Johannes Berg, and Siegfried Roth. Dynamic BMP signaling polarized by Toll patterns the dorsoventral axis in a hemimetabolous insect. *eLife*, 4, 2015.

[1406] Anusmita Sahoo, Shruti Khare, Sivasankar Devanarayanan, Pankaj C. Jain, and Raghavan Varadarajan. Residue proximity information and protein model discrimination using saturation-suppressor mutagenesis. *eLife*, 4, 2015.

[1407] Monica Sala-Rabanal, Zeynep Yurtsever, Colin G Nichols, and Tom J Brett. Secreted CLCA1 modulates TMEM16A to activate Ca2+-dependent chloride currents in human cells. *eLife*, 4, 2015.

[1408] Moti Salti, Simo Monto, Lucie Charles, Jean-Remi King, Lauri Parkkonen, and Stanislas Dehaene. Distinct cortical codes and temporal dynamics for conscious and unconscious percepts. *eLife*, 4, 2015.

[1409] António JM Santos, Ishier Raote, Margherita Scarpa, Nathalie Brouwers, and Vivek Malhotra. TANGO1 recruits ERGIC membranes to the endoplasmic reticulum for procollagen export. *eLife*, 4, 2015.

[1410] Fernando J Santos, Rodrigo F Oliveira, Xin Jin, and Rui M Costa. Corticostriatal dynamics encode the refinement of specific behavioral variability during skill learning. *eLife*, 4, 2015.

[1411] Ignacio Sarria, Johan Pahlberg, Yan Cao, Alexander V Kolesnikov, Vladimir J Kefalov, Alapakkam P Sampath, and Kirill A Martemyanov. Sensitivity and kinetics of signal transmission at the first visual synapse differentially impact visually-guided behavior. *eLife*, 4, 2015.

[1412] Takunori Satoh, Aya Ohba, Ziguang Liu, Tsuyoshi Inagaki, and Akiko K Satoh. dPob/EMC is essential for biosynthesis of rhodopsin and other multi-pass membrane proteins in Drosophila photoreceptors. *eLife*, 4, 2015.

[1413] Jonas-Frederic Sauer, Michael Strüber, and Marlene Bartos. Impaired fast-spiking interneuron function in a genetic mouse model of depression. *eLife*, 4, 2015.

[1414] Arpiar Saunders, Adam J Granger, and Bernardo L Sabatini. Corelease of acetylcholine and GABA from cholinergic forebrain neurons. *eLife*, 4, 2015.

[1415] Lizbeth Sayavedra, Manuel Kleiner, Ruby Ponnudurai, Silke Wetzel, Eric Pelletier, Valerie Barbe, Nori Satoh, Eiichi Shoguchi, Dennis Fink, Corinna Breusing, Thorsten BH Reusch, Philip Rosenstiel, Markus B Schilhabel, Dörte Becher, Thomas Schweder, Stephanie Markert, Nicole Dubilier, and Jillian M Petersen. Abundant toxin-related genes in the genomes of beneficial symbionts from deep-sea hydrothermal vent mussels. *eLife*, 4, 2015.

[1416] Nicole T Schirle, Jessica Sheu-Gruttadauria, Stanley D Chandradoss, Chirlmin Joo, and Ian J MacRae. Water-mediated recognition of t1-adenosine anchors Argonaute2 to microRNA targets. *eLife*, 4, 2015.

[1417] Michael Schleyer, Daisuke Miura, Teiichi Tanimura, and Bertram Gerber. Learning the specific quality of taste reinforcement in larval Drosophila. *eLife*, 4, 2015.

[1418] Hermann Broder Schmidt and Dirk Görlich. Nup98 FG domains from diverse species spontaneously phase-separate into particles with nuclear pore-like permselectivity. *eLife*, 4, 2015.

[1419] Ute I Scholl, Gabriel Stölting, Carol Nelson-Williams, Alfred A Vichot, Murim Choi, Erin Loring, Manju L Prasad, Gerald Goh, Tobias Carling, C Christofer Juhlin, Ivo Quack, Lars C Rump, Anne Thiel, Marc Lande, Britney G Frazier, Majid Rasoulpour, David L Bowlin, Christine B Sethna, Howard Trachtman, Christoph Fahlke, and Richard P Lifton. Recurrent gain of function mutation in calcium channel CACNA1H causes early-onset hypertension with primary aldosteronism. *eLife*, 4, 2015.

[1420] Sebastiaan Schotten, Marieke Meijer, Alexander Matthias Walter, Vincent Huson, Lauren Mamer, Lawrence Kalogreades, Mirelle ter Veer, Marvin Ruiter, Nils Brose, Christian Rosenmund, Jakob Balslev Sørensen, Matthijs Verhage, and Lennart Niels Cornelisse. Additive effects on the energy barrier for synaptic vesicle fusion cause supralinear effects on the vesicle fusion rate. *eLife*, 4, 2015.

[1421] Dietmar Schreiner, Jovan Simicevic, Erik Ahrné, Alexander Schmidt, and Peter Scheiffele. Quantitative isoform-profiling of highly diversified recognition molecules. *eLife*, 4, 2015.

[1422] Aljoscha Schulze, Alex Gomez-Marin, Vani G Rajendran, Gus Lott, Marco Musy, Parvez Ahammad, Ajinkya Deogade, James Sharpe, Julia Riedl, David Jarriault, Eric T Trautman, Christopher Werner, Madhusudhan Venkadesan, Shaul Druckmann, Vivek Jayaraman, and Matthieu Louis. Dynamical feature extraction at the sensory periphery guides chemotaxis. *eLife*, 4, 2015.

[1423] Marc M Schumacher, Rania Elsabrouty, Joachim Seemann, Youngah Jo, and Russell A DeBose-Boyd. The prenyltransferase UBIAD1 is the target of geranylgeraniol in degradation of HMG CoA reductase. *eLife*, 4, 2015.

[1424] Meredith C Schuman, Silke Allmann, and Ian T Baldwin. Plant defense phenotypes determine the consequences of volatile emission for individuals and neighbors. *eLife*, 4, 2015.

[1425] Flavio Schwarz, Oliver MT Pearce, Xiaoxia Wang, Annie N Samraj, Heinz Läubli, Javier O Garcia, Hongqiao Lin, Xiaoming Fu, Andrea Garcia-Bingman, Patrick Secrest, Casey E Romanoski, Charles Heyser, Christopher K Glass, Stanley L Hazen, Nissi Varki, Ajit Varki, and Pascal Gagneux. Siglec receptors impact mammalian lifespan by modulating oxidative stress. *eLife*, 4, 2015.

[1426] Benjamin B Scott, Christine M Constantinople, Jeffrey C Erlich, David W Tank, and Carlos D Brody. Sources of noise during accumulation of evidence in unrestrained and voluntarily head-restrained rats. *eLife*, 4, 2015.

[1427] David Secco, Chuang Wang, Huixia Shou, Matthew D Schultz, Serge Chiarenza, Laurent Nussaume, Joseph R Ecker, James Whelan, and Ryan Lister. Stress induced gene expression drives transient DNA methylation changes at adjacent repetitive elements. *eLife*, 4, 2015.

[1428] Paolo S Segre, Roslyn Dakin, Victor B Zordan, Michael H Dickinson, Andrew D Straw, and Douglas L Altshuler. Burst muscle performance predicts the speed, acceleration, and turning performance of Anna's hummingbirds. *eLife*, 4, 2015.

[1429] Ivonne M Sehring, Pierre Recho, Elsa Denker, Matthew Kourakis, Birthe Mathiesen, Edouard Hannezo, Bo Dong, and Di Jiang. Assembly and positioning of actomyosin rings by contractility and planar cell polarity. *eLife*, 4, 2015.

[1430] Hannah S Seidel and Judith Kimble. Cell-cycle quiescence maintains Caenorhabditis elegans germline stem cells independent of GLP-1/Notch. *eLife*, 4, 2015.

[1431] Ashok Sekhar, Jessica AO Rumfeldt, Helen R Broom, Colleen M Doyle, Guillaume Bouvignies, Elizabeth M Meiering, and Lewis E Kay. Thermal fluctuations of immature SOD1 lead to separate folding and misfolding pathways. *eLife*, 4, 2015.

[1432] Mohini Sengupta and Vatsala Thirumalai. AMPA receptor mediated synaptic excitation drives state-dependent bursting in Purkinje neurons of zebrafish larvae. *eLife*, 4, 2015.

[1433] Eric N Senning and Sharona E Gordon. Activity and Ca2+ regulate the mobility of TRPV1 channels in the plasma membrane of sensory neurons. *eLife*, 4, 2015.

[1434] Suhagi Shah, Gijsbert M Grotenbreg, Amariliz Rivera, and George S Yap. An extrafollicular pathway for the generation of effector CD8+ T cells driven by the proinflammatory cytokine, IL-12. *eLife*, 4, 2015.

[1435] Réza Shahidi, Elizabeth A Williams, Markus Conzelmann, Albina Asadulina, Csaba Verasztó, Sanja Jasek, Luis A Bezares-Calderón, and Gáspár Jékely. A serial multiplex immunogold labeling method for identifying peptidergic neurons in connectomes. *eLife*, 4, 2015.

[1436] Shruti Shankar, Jia Yi Chua, Kah Junn Tan, Meredith EK Calvert, Ruifen Weng, Wan Chin Ng, Kenji Mori, and Joanne Y Yew. The neuropeptide tachykinin is essential for pheromone detection in a gustatory neural circuit. *eLife*, 4, 2015.

[1437] Nengyin Sheng, Yun S Shi, Richa Madan Lomash, Katherine W Roche, and Roger A Nicoll. Neto auxiliary proteins control both the trafficking and biophysical properties of the kainate receptor GluK1. *eLife*, 4, 2015.

[1438] Hirohito Shimizu, Johann Schredelseker, Jie Huang, Kui Lu, Shamim Naghdi, Fei Lu, Sarah Franklin, Hannah DG Fiji, Kevin Wang, Huanqi Zhu, Cheng Tian, Billy Lin, Haruko Nakano, Amy Ehrlich, Junichi Nakai, Adam Z Stieg, James K Gimzewski, Atsushi Nakano, Joshua I Goldhaber, Thomas M Vondriska, György Hajnóczky, Ohyun Kwon, and Jau-Nian Chen. Mitochondrial Ca2+ uptake by the voltage-dependent anion channel 2 regulates cardiac rhythmicity. *eLife*, 4, 2015.

[1439] Jae Yen Shin, Javier Lopez-Garrido, Sang-Hyuk Lee, Cesar Diaz-Celis, Tinya Fleming, Carlos Bustamante, and Kit Pogliano. Visualization and functional dissection of coaxial paired SpoIIIE channels across the sporulation septum. *eLife*, 4, 2015.

[1440] Wenying Shou. Acknowledging selection at sub-organismal levels resolves controversy on pro-cooperation mechanisms. *eLife*, 4, 2015.

[1441] Prerana Shrestha, Awni Mousa, and Nathaniel Heintz. Layer 2/3 pyramidal cells in the medial prefrontal cortex moderate stress induced depressive behaviors. *eLife*, 4, 2015.

[1442] Carmela Sidrauski, Anna M McGeachy, Nicholas T Ingolia, and Peter Walter. The small molecule ISRIB reverses the effects of eIF2$\alpha$ phosphorylation on translation and stress granule assembly. *eLife*, 4, 2015.

[1443] Carmela Sidrauski, Jordan C Tsai, Martin Kampmann, Brian R Hearn, Punitha Vedantham, Priyadarshini Jaishankar, Masaaki Sokabe, Aaron S Mendez, Billy W Newton, Edward L Tang, Erik Verschueren, Jeffrey R Johnson, Nevan J Krogan, Christopher S Fraser, Jonathan S Weissman, Adam R Renslo, and Peter Walter. Pharmacological dimerization and activation of the exchange factor eIF2B antagonizes the integrated stress response. *eLife*, 4, 2015.

[1444] Matthias Siebert, Mathias A Böhme, Jan H Driller, Husam Babikir, Malou M Mampell, Ulises Rey, Niraja Ramesh, Tanja Matkovic, Nicole Holton, Suneel Reddy-Alla, Fabian Göttfert, Dirk Kamin, Christine Quentin, Susan Klinedinst, Till FM Andlauer, Stefan W Hell, Catherine A Collins, Markus C Wahl, Bernhard Loll, and Stephan J Sigrist. A high affinity RIM-binding protein/Aplip1 interaction prevents the formation of ectopic axonal active zones. *eLife*, 4, 2015.

[1445] Paul A Sigala, Jan R Crowley, Jeffrey P Henderson, and Daniel E Goldberg. Deconvoluting heme biosynthesis to target blood-stage malaria parasites. *eLife*, 4, 2015.

[1446] Chanpreet Singh, Grigorios Oikonomou, and David A Prober. Norepinephrine is required to promote wakefulness and for hypocretin-induced arousal in zebrafish. *eLife*, 4, 2015.

[1447] Peter J Skene and Steven Henikoff. A simple method for generating high-resolution maps of genome-wide protein binding. *eLife*, 4, 2015.

[1448] Edden Slomowitz, Boaz Styr, Irena Vertkin, Hila Milshtein-Parush, Israel Nelken, Michael Slutsky, and Inna Slutsky. Interplay between population firing stability and single neuron dynamics in hippocampal networks. *eLife*, 4, 2015.

[1449] Peter Soba, Chun Han, Yi Zheng, Daniel Perea, Irene Miguel-Aliaga, Lily Yeh Jan, and Yuh Nung Jan. The Ret receptor regulates sensory neuron dendrite growth and integrin mediated adhesion. *eLife*, 4, 2015.

[1450] Nicholas James Sofroniew, Yurii A Vlasov, Samuel Andrew Hires, Jeremy Freeman, and Karel Svoboda. Neural coding in barrel cortex during whisker-guided locomotion. *eLife*, 4, 2015.

[1451] Lukas Solanka, Mark CW van Rossum, and Matthew F Nolan. Noise promotes independent control of gamma oscillations and grid firing within recurrent attractor networks. *eLife*, 4, 2015.

[1452] Maja Šolman, Alessio Ligabue, Olga BlaževitŠ, Alok Jaiswal, Yong Zhou, Hong Liang, Benoit Lectez, Kari Kopra, Camilo Guzmán, Harri Härmä, John F Hancock, Tero Aittokallio, and Daniel Abankwa. Specific cancer-associated mutations in the switch III region of Ras increase tumorigenicity by nanocluster augmentation. *eLife*, 4, 2015.

[1453] Dmitriy Sonkin. Expression signature based on TP53 target genes doesn't predict response to TP53-MDM2 inhibitor in wild type TP53 tumors. *eLife*, 4, 2015.

[1454] Konstantinos Sousounis, Feng Qi, Manisha C Yadav, José Luis Millán, Fubito Toyama, Chikafumi Chiba, Yukiko Eguchi, Goro Eguchi, and Panagiotis A Tsonis. A robust transcriptional program in newts undergoing multiple events of lens regeneration throughout their lifespan. *eLife*, 4, 2015.

[1455] Thomas Spatzal, Kathryn A Perez, James B Howard, and Douglas C Rees. Catalysis-dependent selenium incorporation and migration in the nitrogenase active site iron-molybdenum cofactor. *eLife*, 4, 2015.

[1456] Noah Spies, Cheryl L Smith, Jesse M Rodriguez, Julie C Baker, Serafim Batzoglou, and Arend Sidow. Constraint and divergence of global gene expression in the mammalian embryo. *eLife*, 4, 2015.

[1457] Vicki Springthorpe and Steven Penfield. Flowering time and seed dormancy control use external coincidence to generate life history strategy. *eLife*, 4, 2015.

[1458] Michael J Steinbaugh, Sri Devi Narasimhan, Stacey Robida-Stubbs, Lorenza E Moronetti Mazzeo, Jonathan M Dreyfuss, John M Hourihan, Prashant Raghavan, Theresa N Operaña, Reza Esmaillie, and T Keith Blackwell. Lipid-mediated regulation of SKN-1/Nrf in response to germ cell absence. *eLife*, 4, 2015.

[1459] Federico Stella and Alessandro Treves. The self-organization of grid cells in 3D. *eLife*, 4, 2015.

[1460] Drew Stenesen, Andrew T Moehlman, and Helmut Krämer. The carcinine transporter CarT is required in Drosophila photoreceptor neurons to sustain histamine recycling. *eLife*, 4, 2015.

[1461] Louise A Stephen, Hasan Tawamie, Gemma M Davis, Lars Tebbe, Peter Nürnberg, Gudrun Nürnberg, Holger Thiele, Michaela Thoenes, Eugen Boltshauser, Steffen Uebe, Oliver Rompel, André Reis, Arif B Ekici, Lynn McTeir, Amy M Fraser, Emma A Hall, Pleasantine Mill, Nicolas Daudet, Courtney Cross, Uwe Wolfrum, Rami Abou Jamra, Megan G Davey, and Hanno J Bolz. TALPID3 controls centrosome and cell polarity and the human ortholog KIAA0586 is mutated in Joubert syndrome (JBTS23). *eLife*, 4, 2015.

[1462] Gabriella R Sterne, Jung Hwan Kim, and Bing Ye. Dysregulated Dscam levels act through Abelson tyrosine kinase to enlarge presynaptic arbors. *eLife*, 4, 2015.

[1463] Sarah A Stratmann, Seamus R Morrone, Antoine M van Oijen, and Jungsan Sohn. The innate immune sensor IFI16 recognizes foreign DNA in the nucleus by scanning along the duplex. *eLife*, 4, 2015.

[1464] Tian Su, Tanya Bondar, Xu Zhou, Cuiling Zhang, Hang He, and Ruslan Medzhitov. Two-signal requirement for growth-promoting function of Yap in hepatocytes. *eLife*, 4, 2015.

[1465] Kenji Sugiyama, Atsushi Kawaguchi, Mitsuru Okuwaki, and Kyosuke Nagata. pp32 and APRIL are host cell-derived regulators of influenza virus RNA synthesis from cRNA. *eLife*, 4, 2015.

[1466] Sukrit Suksombat, Rustem Khafizov, Alexander G Kozlov, Timothy M Lohman, and Yann R Chemla. Structural dynamics of E. coli single-stranded DNA binding protein reveal DNA wrapping and unwrapping pathways. *eLife*, 4, 2015.

[1467] Fanny Sunden, Ariana Peck, Julia Salzman, Susanne Ressl, and Daniel Herschlag. Extensive site-directed mutagenesis reveals interconnected functional units in the alkaline phosphatase active site. *eLife*, 4, 2015.

[1468] Adhira Sunkara, Gregory C DeAngelis, and Dora E Angelaki. Role of visual and non-visual cues in constructing a rotation-invariant representation of heading in parietal cortex. *eLife*, 4, 2015.

[1469] Vladislav Susoy, Erik J Ragsdale, Natsumi Kanzaki, and Ralf J Sommer. Rapid diversification associated with a macroevolutionary pulse of developmental plasticity. *eLife*, 4, 2015.

[1470] Miyuki Suzawa, Diego A Miranda, Karmela A Ramos, Kenny K-H Ang, Emily J Faivre, Christopher G Wilson, Laura Caboni, Michelle R Arkin, Yeong-Sang Kim, Robert J Fletterick, Aaron Diaz, John S Schneekloth, and Holly A Ingraham. A gene-expression screen identifies a non-toxic sumoylation inhibitor that mimics SUMO-less human LRH-1 in liver. *eLife*, 4, 2015.

[1471] Catherine M Sweeney-Reed, Tino Zaehle, Jürgen Voges, Friedhelm C Schmitt, Lars Buentjen, Klaus Kopitzki, Hermann Hinrichs, Hans-Jochen Heinze, Michael D Rugg, Robert T Knight, and Alan Richardson-Klavehn. Thalamic theta phase alignment predicts human memory formation and anterior thalamic cross-frequency coupling. *eLife*, 4, 2015.

[1472] Ruhma Syeda, Jie Xu, Adrienne E Dubin, Bertrand Coste, Jayanti Mathur, Truc Huynh, Jason Matzen, Jianmin Lao, David C Tully, Ingo H Engels, H Michael Petrassi, Andrew M Schumacher, Mauricio Montal, Michael Bandell, and Ardem Patapoutian. Chemical activation of the mechanotransduction channel Piezo1. *eLife*, 4, 2015.

[1473] Ayesa Syenina, Cyril J Jagaraj, Siti AB Aman, Aishwarya Sridharan, and Ashley L St John. Dengue vascular leakage is augmented by mast cell degranulation mediated by immunoglobulin Fc$\gamma$ receptors. *eLife*, 4, 2015.

[1474] Susumu Takahashi. Episodic-like memory trace in awake replay of hippocampal place cell activity sequences. *eLife*, 4, 2015.

[1475] Akira Takano, Takuya Kajita, Makoto Mochizuki, Toshiya Endo, and Tohru Yoshihisa. Cytosolic Hsp70 and co-chaperones constitute a novel system for tRNA import into the nucleus. *eLife*, 4, 2015.

[1476] Sandeep K Talapatra, Bethany Harker, and Julie PI Welburn. The C-terminal region of the motor protein MCAK controls its structure and activity through a conformational switch. *eLife*, 4, 2015.

[1477] Chia Huei Tan, Ivana Gasic, Sabina P Huber-Reggi, Damian Dudka, Marin Barisic, Helder Maiato, and Patrick Meraldi. The equatorial position of the metaphase plate ensures symmetric cell divisions. *eLife*, 4, 2015.

[1478] Ek Han Tan, Isabelle M Henry, Maruthachalam Ravi, Keith R Bradnam, Terezie Mandakova, Mohan PA Marimuthu, Ian Korf, Martin A Lysak, Luca Comai, and Simon WL Chan. Catastrophic chromosomal restructuring during genome elimination in plants. *eLife*, 4, 2015.

[1479] Marvin E Tanenbaum, Noam Stern-Ginossar, Jonathan S Weissman, and Ronald D Vale. Regulation of mRNA translation during mitosis. *eLife*, 4, 2015.

[1480] Shaogeng Tang, W Mike Henne, Peter P Borbat, Nicholas J Buchkovich, Jack H Freed, Yuxin Mao, J Christopher Fromme, and Scott D Emr. Structural basis for activation, assembly and membrane binding of ESCRT-III Snf7 filaments. *eLife*, 4, 2015.

[1481] Susan H Taylor, Ching-Yan Chloé Yeung, Nicholas S Kalson, Yinhui Lu, Paola Zigrino, Tobias Starborg, Stacey Warwood, David F Holmes, Elizabeth G Canty-Laird, Cornelia Mauch, and Karl E Kadler. Matrix metalloproteinase 14 is required for fibrous tissue expansion. *eLife*, 4, 2015.

[1482] Alex Tendler and Shlomo Wagner. Different types of theta rhythmicity are induced by social and fearful stimuli in a network associated with social memory. *eLife*, 4, 2015.

[1483] Palaniraja Thandapani, Jingwen Song, Valentina Gandin, Yutian Cai, Samuel G Rouleau, Jean-Michel Garant, Francois-Michel Boisvert, Zhenbao Yu, Jean-Pierre Perreault, Ivan Topisirovic, and Stéphane Richard. Aven recognition of RNA G-quadruplexes regulates translation of the mixed lineage leukemia protooncogenes. *eLife*, 4, 2015.

[1484] Hanh Thi-Kim Vu, Jochen C Rink, Sean A McKinney, Melainia McClain, Naharajan Lakshmanaperumal, Richard Alexander, and Alejandro Sánchez Alvarado. Stem cells and fluid flow drive cyst formation in an invertebrate excretory organ. *eLife*, 4, 2015.

[1485] Karen J Thompson, Cynthia M Harley, Grant M Barthel, Mark A Sanders, and Karen A Mesce. Plasmon resonance and the imaging of metal-impregnated neurons with the laser scanning confocal microscope. *eLife*, 4, 2015.

[1486] Juan Tian, Libo Han, Zhidi Feng, Guangda Wang, Weiwei Liu, Yinping Ma, Yanjun Yu, and Zhaosheng Kong. Orchestration of microtubules and the actin cytoskeleton in trichome cell shape determination by a plant-unique kinesin. *eLife*, 4, 2015.

[1487] Tiffany W Todd, Hiroshi Kokubu, Helen C Miranda, Constanza J Cortes, Albert R La Spada, and Janghoo Lim. Nemo-like kinase is a novel regulator of spinal and bulbar muscular atrophy. *eLife*, 4, 2015.

[1488] Sari Tojkander, Gergana Gateva, Amjad Husain, Ramaswamy Krishnan, and Pekka Lappalainen. Generation of contractile actomyosin bundles depends on mechanosensitive actin filament assembly and disassembly. *eLife*, 4, 2015.

[1489] Xia-Jing Tong, Zhitao Hu, Yu Liu, Dorian Anderson, and Joshua M Kaplan. A network of autism linked genes stabilizes two pools of synaptic GABAA receptors. *eLife*, 4, 2015.

[1490] Jeremy S Treger, Michael F Priest, and Francisco Bezanilla. Single-molecule fluorimetry and gating currents inspire an improved optical voltage indicator. *eLife*, 4, 2015.

[1491] Peter Tsvetkov, Marc L Mendillo, Jinghui Zhao, Jan E Carette, Parker H Merrill, Domagoj Cikes, Malini Varadarajan, Ferdy R van Diemen, Josef M Penninger, Alfred L Goldberg, Thijn R Brummelkamp, Sandro Santagata, and Susan Lindquist. Compromising the 19S proteasome complex protects cells from reduced flux through the proteasome. *eLife*, 4, 2015.

[1492] Kimberly C Tu, Li-Chun Cheng, Hanh TK Vu, Jeffrey J Lange, Sean A McKinney, Chris W Seidel, and Alejandro Sánchez Alvarado. Egr-5 is a post-mitotic regulator of planarian epidermal differentiation. *eLife*, 4, 2015.

[1493] Shun Tu, Shu-Juan Guo, Chien-Sheng Chen, Cheng-Xi Liu, He-Wei Jiang, Feng Ge, Jiao-Yu Deng, Yi-Ming Zhou, Daniel M Czajkowsky, Yang Li, Bang-Ruo Qi, Young-Hoon Ahn, Philip A Cole, Heng Zhu, and Sheng-Ce Tao. YcgC represents a new protein deacetylase family in prokaryotes. *eLife*, 4, 2015.

[1494] Jenny Tung, Luis B Barreiro, Michael B Burns, Jean-Christophe Grenier, Josh Lynch, Laura E Grieneisen, Jeanne Altmann, Susan C Alberts, Ran Blekhman, and Elizabeth A Archie. Social networks predict gut microbiome composition in wild baboons. *eLife*, 4, 2015.

[1495] Jenny Tung, Xiang Zhou, Susan C Alberts, Matthew Stephens, and Yoav Gilad. The genetic architecture of gene expression levels in wild baboons. *eLife*, 4, 2015.

[1496] Monarin Uervirojnangkoorn, Oliver B Zeldin, Artem Y Lyubimov, Johan Hattne, Aaron S Brewster, Nicholas K Sauter, Axel T Brunger, and William I Weis. Enabling X-ray free electron laser crystallography for challenging biological systems from a limited number of crystals. *eLife*, 4, 2015.

[1497] Balázs B Ujfalussy, Judit K Makara, Tiago Branco, and Máté Lengyel. Dendritic nonlinearities are tuned for efficient spike-based computations in cortical circuits. *eLife*, 4, 2015.

[1498] Arun M Unni, William W Lockwood, Kreshnik Zejnullahu, Shih-Queen Lee-Lin, and Harold Varmus. Evidence that synthetic lethality underlies the mutual exclusivity of oncogenic KRAS and EGFR mutations in lung adenocarcinoma. *eLife*, 4, 2015.

[1499] Reid C Van Lehn, Bin Zhang, and Thomas F Miller. Regulation of multispanning membrane protein topology via post-translational annealing. *eLife*, 4, 2015.

[1500] Anna Vangone and Alexandre MJJ Bonvin. Contacts-based prediction of binding affinity in protein–protein complexes. *eLife*, 4, 2015.

[1501] Benoit Vanhollebeke, Oliver A Stone, Naguissa Bostaille, Chris Cho, Yulian Zhou, Emilie Maquet, Anne Gauquier, Pauline Cabochette, Shigetomo Fukuhara, Naoki Mochizuki, Jeremy Nathans, and Didier YR Stainier. Tip cell-specific requirement for an atypical Gpr124- and Reck-dependent Wnt/$\beta$-catenin pathway during brain angiogenesis. *eLife*, 4, 2015.

[1502] Harish N Vasudevan, Pierre Mazot, Fenglei He, and Philippe Soriano. Receptor tyrosine kinases modulate distinct transcriptional programs by differential usage of intracellular pathways. *eLife*, 4, 2015.

[1503] Jorge Vera, Lydia Lartigue, Suzanne Vigneron, Gilles Gadea, Veronique Gire, Maguy Del Rio, Isabelle Soubeyran, Frederic Chibon, Thierry Lorca, and Anna Castro. Greatwall promotes cell transformation by hyperactivating AKT in human malignancies. *eLife*, 4, 2015.

[1504] Pushpa Verma and Stephen M Cohen. miR-965 controls cell proliferation and migration during tissue morphogenesis in the Drosophila abdomen. *eLife*, 4, 2015.

[1505] Andrés Vidal-Gadea, Kristi Ward, Celia Beron, Navid Ghorashian, Sertan Gokce, Joshua Russell, Nicholas Truong, Adhishri Parikh, Otilia Gadea, Adela Ben-Yakar, and Jonathan Pierce-Shimomura. Magnetosensitive neurons mediate geomagnetic orientation in Caenorhabditis elegans. *eLife*, 4, 2015.

[1506] Dhanasekaran Vijaykrishna, Edward C Holmes, Udayan Joseph, Mathieu Fourment, Yvonne CF Su, Rebecca Halpin, Raphael TC Lee, Yi-Mo Deng, Vithiagaran Gunalan, Xudong Lin, Timothy B Stockwell, Nadia B Fedorova, Bin Zhou, Natalie Spirason, Denise Kühnert, Veronika BoŠková, Tanja Stadler, Anna-Maria Costa, Dominic E Dwyer, Q Sue Huang, Lance C Jennings, William Rawlinson, Sheena G Sullivan, Aeron C Hurt, Sebastian Maurer-Stroh, David E Wentworth, Gavin JD Smith, and Ian G Barr. The contrasting phylodynamics of human influenza B viruses. *eLife*, 4, 2015.

[1507] Roberto Villaseñor, Hidenori Nonaka, Perla Del Conte-Zerial, Yannis Kalaidzidis, and Marino Zerial. Regulation of EGFR signal transduction by analogue-to-digital conversion in endosomes. *eLife*, 4, 2015.

[1508] Philippe FY Vincent, Yohan Bouleau, Christine Petit, and Didier Dulon. A synaptic F-actin network controls otoferlin-dependent exocytosis in auditory inner hair cells. *eLife*, 4, 2015.

[1509] Nadine Vincenten, Lisa-Marie Kuhl, Isabel Lam, Ashwini Oke, Alastair RW Kerr, Andreas Hochwagen, Jennifer Fung, Scott Keeney, Gerben Vader, and Adèle L Marston. The kinetochore prevents centromere-proximal crossover recombination during meiosis. *eLife*, 4, 2015.

[1510] Yaron Vinik, Hadas Shatz-Azoulay, Alessia Vivanti, Navit Hever, Yifat Levy, Rotem Karmona, Vlad Brumfeld, Saja Baraghithy, Malka Attar-Lamdar, Sigalit Boura-Halfon, Itai Bab, and Yehiel Zick. The mammalian lectin galectin-8 induces RANKL expression, osteoclastogenesis, and bone mass reduction in mice. *eLife*, 4, 2015.

[1511] Jasper J Visser, Yolanda Cheng, Steven C Perry, Andrew Benjamin Chastain, Bayan Parsa, Shatha S Masri, Thomas A Ray, Jeremy N Kay, and Woj M Wojtowicz. An extracellular biochemical screen reveals that FLRTs and Unc5s mediate neuronal subtype recognition in the retina. *eLife*, 4, 2015.

[1512] Jyothsna Visweswaraiah, Yvette Pittman, Thomas E Dever, and Alan G Hinnebusch. The $\beta$-hairpin of 40S exit channel protein Rps5/uS7 promotes efficient and accurate translation initiation in vivo. *eLife*, 4, 2015.

[1513] Katharina S Volz, Andrew H Jacobs, Heidi I Chen, Aruna Poduri, Andrew S McKay, Daniel P Riordan, Natalie Kofler, Jan Kitajewski, Irving Weissman, and Kristy Red-Horse. Pericytes are progenitors for coronary artery smooth muscle. *eLife*, 4, 2015.

[1514] Rebecca M Voorhees and Ramanujan S Hegde. Structures of the scanning and engaged states of the mammalian SRP-ribosome complex. *eLife*, 4, 2015.

[1515] Miriam Walden, John M Edwards, Aleksandra M Dziewulska, Rene Bergmann, Gerhard Saalbach, Su-Yin Kan, Ona K Miller, Miriam Weckener, Rosemary J Jackson, Sally L Shirran, Catherine H Botting, Gordon J Florence, Manfred Rohde, Mark J Banfield, and Ulrich Schwarz-Linek. An internal thioester in a pathogen surface protein mediates covalent host binding. *eLife*, 4, 2015.

[1516] Zhengpeng Wan, Xiangjun Chen, Haodong Chen, Qinghua Ji, Yingjia Chen, Jing Wang, Yiyun Cao, Fei Wang, Jizhong Lou, Zhuo Tang, and Wanli Liu. The activation of IgM- or isotype-switched IgG- and IgE-BCR exhibits distinct mechanical force sensitivity and threshold. *eLife*, 4, 2015.

[1517] Amberlyn M Wands, Akiko Fujita, Janet E McCombs, Jakob Cervin, Benjamin Dedic, Andrea C Rodriguez, Nicole Nischan, Michelle R Bond, Marcel Mettlen, David C Trudgian, Andrew Lemoff, Marianne Quiding-Järbrink, Bengt Gustavsson, Catharina Steentoft, Henrik Clausen, Hamid Mirzaei, Susann Teneberg, Ulf Yrlid, and Jennifer J Kohler. Fucosylation and protein glycosylation create functional receptors for cholera toxin. *eLife*, 4, 2015.

[1518] Bo Wang, Fangyuan Xia, Torsten Wappler, Ewa Simon, Haichun Zhang, Edmund A Jarzembowski, and Jacek Szwedo. Brood care in a 100-million-year-old scale insect. *eLife*, 4, 2015.

[1519] Eric Wang, Shinpei Kawaoka, Jae-Seok Roe, Junwei Shi, Anja F Hohmann, Yali Xu, Anand S Bhagwat, Yutaka Suzuki, Justin B Kinney, and Christopher R Vakoc. The transcriptional cofactor TRIM33 prevents apoptosis in B lymphoblastic leukemia by deactivating a single enhancer. *eLife*, 4, 2015.

[1520] Guangliang Wang, Surendra K Rajpurohit, Fabien Delaspre, Steven L Walker, David T White, Alexis Ceasrine, Rejji Kuruvilla, Ruo-jing Li, Joong S Shim, Jun O Liu, Michael J Parsons, and Jeff S Mumm. First quantitative high-throughput screen in zebrafish identifies novel pathways for increasing pancreatic $\beta$-cell mass. *eLife*, 4, 2015.

[1521] Gui-Zhen Wang, Xin Cheng, Bo Zhou, Zhe-Sheng Wen, Yun-Chao Huang, Hao-Bin Chen, Gao-Feng Li, Zhi-Liang Huang, Yong-Chun Zhou, Lin Feng, Ming-Ming Wei, Li-Wei Qu, Yi Cao, and Guang-Biao Zhou. The chemokine CXCL13 in lung cancers associated with environmental polycyclic aromatic hydrocarbons pollution. *eLife*, 4, 2015.

[1522] Jia Wang, Wen Yao, Dan Zhu, Weibo Xie, and Qifa Zhang. Genetic basis of sRNA quantitative variation analyzed using an experimental population derived from an elite rice hybrid. *eLife*, 4, 2015.

[1523] Jiyong Wang, Bharat D Reddy, and Songtao Jia. Rapid epigenetic adaptation to uncontrolled heterochromatin spreading. *eLife*, 4, 2015.

[1524] Qi Wang, Erik M Vogan, Laura M Nocka, Connor E Rosen, Julie A Zorn, Stephen C Harrison, and John Kuriyan. Autoinhibition of Bruton's tyrosine kinase (Btk) and activation by soluble inositol hexakisphosphate. *eLife*, 4, 2015.

[1525] Shaohe Wang, Di Wu, Sophie Quintin, Rebecca A Green, Dhanya K Cheerambathur, Stacy D Ochoa, Arshad Desai, and Karen Oegema. NOCA-1 functions with $\gamma$-tubulin and in parallel to Patronin to assemble non-centrosomal microtubule arrays in C. elegans. *eLife*, 4, 2015.

[1526] Su Wang, Yuan Gao, Xiaoqing Song, Xing Ma, Xiujuan Zhu, Ying Mao, Zhihao Yang, Jianquan Ni, Hua Li, Kathryn E Malanowski, Perera Anoja, Jungeun Park, Jeff Haug, and Ting Xie. Wnt signaling-mediated redox regulation maintains the germ line stem cell differentiation niche. *eLife*, 4, 2015.

[1527] Won-Jing Wang, Devrim Acehan, Chien-Han Kao, Wann-Neng Jane, Kunihiro Uryu, and Meng-Fu Bryan Tsou. De novo centriole formation in human cells is error-prone and does not require SAS-6 self-assembly. *eLife*, 4, 2015.

[1528] Ye Wang and Valentin Dragoi. Rapid learning in visual cortical networks. *eLife*, 4, 2015.

[1529] Yi-Zhi Wang, Jing-Jing Wang, Yu Huang, Fan Liu, Wei-Zheng Zeng, Ying Li, Zhi-Gang Xiong, Michael X Zhu, and Tian-Le Xu. Tissue acidosis induces neuronal necroptosis via ASIC1a channel independent of its ionic conduction. *eLife*, 4, 2015.

[1530] Masashi Watanabe, Hidehisa Takahashi, Yasushi Saeki, Takashi Ozaki, Shihori Itoh, Masanobu Suzuki, Wataru Mizushima, Keiji Tanaka, and Shigetsugu Hatakeyama. The E3 ubiquitin ligase TRIM23 regulates adipocyte differentiation via stabilization of the adipogenic activator PPAR$\gamma$. *eLife*, 4, 2015.

[1531] Andrew J Watrous, Lorena Deuker, Juergen Fell, and Nikolai Axmacher. Phase-amplitude coupling supports phase coding in human ECoG. *eLife*, 4, 2015.

[1532] Christopher R Weber, Guo Hua Liang, Yitang Wang, Sudipto Das, Le Shen, Alan S L Yu, Deborah J Nelson, and Jerrold R Turner. Claudin-2-dependent paracellular channels are dynamically gated. *eLife*, 4, 2015.

[1533] Xing Wei, Audrey S Howell, Xintong Dong, Caitlin A Taylor, Roshni C Cooper, Jianqi Zhang, Wei Zou, David R Sherwood, and Kang Shen. The unfolded protein response is required for dendrite morphogenesis. *eLife*, 4, 2015.

[1534] Yongjie Wei, Zhenyi An, Zhongju Zou, Rhea Sumpter, Minfei Su, Xiao Zang, Sangita Sinha, Matthias Gaestel, and Beth Levine. The stress-responsive kinases MAPKAPK2/MAPKAPK3 activate starvation-induced autophagy through Beclin 1 phosphorylation. *eLife*, 4, 2015.

[1535] Clemens L Weiß, Michael Dannemann, Kay Prüfer, and Hernán A Burbano. Contesting the presence of wheat in the British Isles 8,000 years ago by assessing ancient DNA authenticity from low-coverage data. *eLife*, 4, 2015.

[1536] Suzanne PM Welten, Anke Redeker, Kees LMC Franken, Jennifer D Oduro, Ferry Ossendorp, Luka čičin Šain, Cornelis JM Melief, Peter Aichele, and Ramon Arens. The viral context instructs the redundancy of costimulatory pathways in driving CD8+ T cell expansion. *eLife*, 4, 2015.

[1537] Benjamin Werner, Fabian Beier, Sebastian Hummel, Stefan Balabanov, Lisa Lassay, Thorsten Orlikowsky, David Dingli, Tim H Brümmendorf, and Arne Traulsen. Reconstructing the in vivo dynamics of hematopoietic stem cells from telomere length distributions. *eLife*, 4, 2015.

[1538] Diek W Wheeler, Charise M White, Christopher L Rees, Alexander O Komendantov, David J Hamilton, and Giorgio A Ascoli. Hippocampome.org: a knowledge base of neuron types in the rodent hippocampus. *eLife*, 4, 2015.

[1539] Glen Wheeler, Takahiro Ishikawa, Varissa Pornsaksit, and Nicholas Smirnoff. Evolution of alternative biosynthetic pathways for vitamin C following plastid acquisition in photosynthetic eukaryotes. *eLife*, 4, 2015.

[1540] Guy Wiedermann, Robert Alexander Bone, Joana Clara Silva, Mia Bjorklund, Philip J Murray, and J Kim Dale. A balance of positive and negative regulators determines the pace of the segmentation clock. *eLife*, 4, 2015.

[1541] Larissa Wilhelm, Frank Bürmann, Anita Minnen, Ho-Chul Shin, Christopher P Toseland, Byung-Ha Oh, and Stephan Gruber. SMC condensin entraps chromosomal DNA by an ATP hydrolysis dependent loading mechanism in Bacillus subtilis. *eLife*, 4, 2015.

[1542] Amy L Williams, Giulio Genovese, Thomas Dyer, Nicolas Altemose, Katherine Truax, Goo Jun, Nick Patterson, Simon R Myers, Joanne E Curran, Ravi Duggirala, John Blangero, David Reich, and Molly Przeworski. Non-crossover gene conversions show strong GC bias and unexpected clustering in humans. *eLife*, 4, 2015.

[1543] Rory P Wilson, Iwan W Griffiths, Michael GL Mills, Chris Carbone, John W Wilson, and David M Scantlebury. Mass enhances speed but diminishes turn capacity in terrestrial pursuit predators. *eLife*, 4, 2015.

[1544] Jennifer Winstanley, Annick Sawala, Clair Baldock, and Hilary L Ashe. Synthetic enzyme-substrate tethering obviates the Tolloid-ECM interaction during Drosophila BMP gradient formation. *eLife*, 4, 2015.

[1545] Wipawee Winuthayanon, Miranda L Bernhardt, Elizabeth Padilla-Banks, Page H Myers, Matthew L Edin, Fred B Lih, Sylvia C Hewitt, Kenneth S Korach, and Carmen J Williams. Oviductal estrogen receptor $\alpha$ signaling prevents protease-mediated embryo death. *eLife*, 4, 2015.

[1546] Danuta M Wisniewska, John M Ratcliffe, Kristian Beedholm, Christian B Christensen, Mark Johnson, Jens C Koblitz, Magnus Wahlberg, and Peter T Madsen. Range-dependent flexibility in the acoustic field of view of echolocating porpoises (Phocoena phocoena). *eLife*, 4, 2015.

[1547] David Wöhlert, Maria J Grötzinger, Werner Kühlbrandt, and özkan Yildiz. Mechanism of Na+-dependent citrate transport from the structure of an asymmetrical CitS dimer. *eLife*, 4, 2015.

[1548] Luise Wolf, Olin K Silander, and Erik van Nimwegen. Expression noise facilitates the evolution of gene regulation. *eLife*, 4, 2015.

[1549] Philip Wolff, Hua Jiang, Guifeng Wang, Juan Santos-González, and Claudia Köhler. Paternally expressed imprinted genes establish postzygotic hybridization barriers in Arabidopsis thaliana. *eLife*, 4, 2015.

[1550] Jasmine C Wong, Kelley M Weinfurtner, Maria del pilar Alzamora, Scott C Kogan, Michael R Burgess, Yan Zhang, Joy Nakitandwe, Jing Ma, Jinjun Cheng, Shann-Ching Chen, Theodore T Ho, Johanna Flach, Damien Reynaud, Emmanuelle Passegué, James R Downing, and Kevin Shannon. Functional evidence implicating chromosome 7q22 haploinsufficiency in myelodysplastic syndrome pathogenesis. *eLife*, 4, 2015.

[1551] Yong H Woo, Hifzur Ansari, Thomas D Otto, Christen M Klinger, Martin Kolisko, Jan Michálek, Alka Saxena, Dhanasekaran Shanmugam, Annageldi Tayyrov, Alaguraj Veluchamy, Shahjahan Ali, Axel Bernal, Javier del Campo, Jaromír Cihlář, Pavel Flegontov, Sebastian G Gornik, Eva HajduŠková, AleŠ Horák, Jan JanouŠkovec, Nicholas J Katris, Fred D Mast, Diego Miranda-Saavedra, Tobias Mourier, Raeece Naeem, Mridul Nair, Aswini K Panigrahi, Neil D Rawlings, Eriko Padron-Regalado, Abhinay Ramaprasad, Nadira Samad, AleŠ Tomčala, Jon Wilkes, Daniel E Neafsey, Christian Doerig, Chris Bowler, Patrick J Keeling, David S Roos, Joel B Dacks, Thomas J Templeton, Ross F Waller, Julius LukeŠ, Miroslav Oborník, and Arnab Pain. Chromerid genomes reveal the evolutionary path from photosynthetic algae to obligate intracellular parasites. *eLife*, 4, 2015.

[1552] Karl R Wotton, Eva Jiménez-Guri, Anton Crombach, Hilde Janssens, Anna Alcaine-Colet, Steffen Lemke, Urs Schmidt-Ott, and Johannes Jaeger. Quantitative system drift compensates for altered maternal inputs to the gap gene network of the scuttle fly Megaselia abdita. *eLife*, 4, 2015.

[1553] Chia-Hung Wu, Maja Bialecka-Fornal, and Dianne K Newman. Methylation at the C-2 position of hopanoids increases rigidity in native bacterial membranes. *eLife*, 4, 2015.

[1554] Haitao Wu, Arnab Barik, Yisheng Lu, Chengyong Shen, Andrew Bowman, Lei Li, Anupama Sathyamurthy, Thiri W Lin, Wen-Cheng Xiong, and Lin Mei. Slit2 as a $\beta$-catenin/Ctnnb1-dependent retrograde signal for presynaptic differentiation. *eLife*, 4, 2015.

[1555] Jianguo Wu, Zhirui Yang, Yu Wang, Lijia Zheng, Ruiqiang Ye, Yinghua Ji, Shanshan Zhao, Shaoyi Ji, Ruofei Liu, Le Xu, Hong Zheng, Yijun Zhou, Xin Zhang, Xiaofeng Cao, Lianhui Xie, Zujian Wu, Yijun Qi, and Yi Li. Viral-inducible Argonaute18 confers broad-spectrum virus resistance in rice by sequestering a host microRNA. *eLife*, 4, 2015.

[1556] Linwei Wu, Liem H Nguyen, Kejin Zhou, T Yvanka de Soysa, Lin Li, Jason B Miller, Jianmin Tian, Joseph Locker, Shuyuan Zhang, Gen Shinoda, Marc T Seligson, Lauren R Zeitels, Asha Acharya, Sam C Wang, Joshua T Mendell, Xiaoshun He, Jinsuke Nishino, Sean J Morrison, Daniel J Siegwart, George Q Daley, Ng Shyh-Chang, and Hao Zhu. Precise let-7 expression levels balance organ regeneration against tumor suppression. *eLife*, 4, 2015.

[1557] Robert Alexander Wu, Yavuz S Dagdas, S Tunc Yilmaz, Ahmet Yildiz, and Kathleen Collins. Single-molecule imaging of telomerase reverse transcriptase in human telomerase holoenzyme and minimal RNP complexes. *eLife*, 4, 2015.

[1558] Bettina Wurzer, Gabriele Zaffagnini, Dorotea Fracchiolla, Eleonora Turco, Christine Abert, Julia Romanov, and Sascha Martens. Oligomerization of p62 allows for selection of ubiquitinated cargo and isolation membrane during selective autophagy. *eLife*, 4, 2015.

[1559] Keqiang Xie, Ikuo Masuho, Chien-Cheng Shih, Yan Cao, Keita Sasaki, Chun Wan J Lai, Pyung-Lim Han, Hiroshi Ueda, Carmen W Dessauer, Michelle E Ehrlich, Baoji Xu, Barry M Willardson, and Kirill A Martemyanov. Stable G protein-effector complexes in striatal neurons: mechanism of assembly and role in neurotransmitter signaling. *eLife*, 4, 2015.

[1560] Daichao Xu, Bing Shan, Byung-Hoon Lee, Kezhou Zhu, Tao Zhang, Huawang Sun, Min Liu, Linyu Shi, Wei Liang, Lihui Qian, Juan Xiao, Lili Wang, Lifeng Pan, Daniel Finley, and Junying Yuan. Phosphorylation and activation of ubiquitin-specific protease-14 by Akt regulates the ubiquitin-proteasome system. *eLife*, 4, 2015.

[1561] Hui Xu, Xiaojing He, Hui Zheng, Lily J Huang, Fajian Hou, Zhiheng Yu, Michael Jason de la Cruz, Brian Borkowski, Xuewu Zhang, Zhijian J Chen, and Qiu-Xing Jiang. Correction: Structural basis for the prion-like MAVS filaments in antiviral innate immunity. *eLife*, 4, 2015.

[1562] Ming Xu, Allyson K Palmer, Husheng Ding, Megan M Weivoda, Tamar Pirtskhalava, Thomas A White, Anna Sepe, Kurt O Johnson, Michael B Stout, Nino Giorgadze, Michael D Jensen, Nathan K LeBrasseur, Tamar Tchkonia, and James L Kirkland. Targeting senescent cells enhances adipogenesis and metabolic function in old age. *eLife*, 4, 2015.

[1563] Zijian Xu, Wenjie Wang, Kaiju Jiang, Zhou Yu, Huanwei Huang, Fengchao Wang, Bin Zhou, and Ting Chen. Embryonic attenuated Wnt/$\beta$-catenin signaling defines niche location and long-term stem cell fate in hair follicle. *eLife*, 4, 2015.

[1564] Guy Yachdav, Tatyana Goldberg, Sebastian Wilzbach, David Dao, Iris Shih, Saket Choudhary, Steve Crouch, Max Franz, Alexander García, Leyla J García, Björn A Grüning, Devasena Inupakutika, Ian Sillitoe, Anil S Thanki, Bruno Vieira, José M Villaveces, Maria V Schneider, Suzanna Lewis, Steve Pettifer, Burkhard Rost, and Manuel Corpas. Anatomy of BioJS, an open source community for the life sciences. *eLife*, 4, 2015.

[1565] Yfat Yahalom-Ronen, Dana Rajchman, Rachel Sarig, Benjamin Geiger, and Eldad Tzahor. Reduced matrix rigidity promotes neonatal cardiomyocyte dedifferentiation, proliferation and clonal expansion. *eLife*, 4, 2015.

[1566] Bo Yang, Mingliang Qu, Rengang Wang, Jon E Chatterton, Xiao-Bo Liu, Bing Zhu, Sonoko Narisawa, Jose Luis Millan, Nobuki Nakanishi, Kathryn Swoboda, Stuart A Lipton, and Dongxian Zhang. The critical role of membralin in postnatal motor neuron survival and disease. *eLife*, 4, 2015.

[1567] Hanseul Yang, Daniel Schramek, Rene C Adam, Brice E Keyes, Ping Wang, Deyou Zheng, and Elaine Fuchs. ETS family transcriptional regulators drive chromatin dynamics and malignancy in squamous cell carcinomas. *eLife*, 4, 2015.

[1568] Rui Yang, Dario Lirussi, Tina M Thornton, Dawn M Jelley-Gibbs, Sean A Diehl, Laure K Case, Muniswamy Madesh, Douglas J Taatjes, Cory Teuscher, Laura Haynes, and Mercedes Rincón. Mitochondrial Ca2+ and membrane potential, an alternative pathway for Interleukin 6 to regulate CD4 cell effector function. *eLife*, 4, 2015.

[1569] Yang Yang, Chunlin Wang, Qunying Yang, Aaron B Kantor, Hiutung Chu, Eliver EB Ghosn, Guang Qin, Sarkis K Mazmanian, Jian Han, and Leonore A Herzenberg. Distinct mechanisms define murine B cell lineage immunoglobulin heavy chain (IgH) repertoires. *eLife*, 4, 2015.

[1570] Yunlei Yang, Peter Lee, and Scott M Sternson. Cell type-specific pharmacology of NMDA receptors using masked MK801. *eLife*, 4, 2015.

[1571] Yuwei Yao, Xiaotian Cui, Ismael Al-Ramahi, Xiaoli Sun, Bo Li, Jiapeng Hou, Marian Difiglia, James Palacino, Zhi-Ying Wu, Lixiang Ma, Juan Botas, and Boxun Lu. A striatal-enriched intronic GPCR modulates huntingtin levels and toxicity. *eLife*, 4, 2015.

[1572] Laura Yelin-Bekerman, Idan Elbaz, Alex Diber, Dvir Dahary, Liron Gibbs-Bar, Shahar Alon, Tali Lerer-Goldshtein, and Lior Appelbaum. Hypocretin neuron-specific transcriptome profiling identifies the sleep modulator Kcnh4a. *eLife*, 4, 2015.

[1573] Yasuhiro Yokota, Hiroyuki Nakajima, Yuki Wakayama, Akira Muto, Koichi Kawakami, Shigetomo Fukuhara, and Naoki Mochizuki. Endothelial Ca2+ oscillations reflect VEGFR signaling-regulated angiogenic capacity in vivo. *eLife*, 4, 2015.

[1574] Ran You, Wen Lu, Ming Shan, Jacob M Berlin, Errol LG Samuel, Daniela C Marcano, Zhengzong Sun, William KA Sikkema, Xiaoyi Yuan, Lizhen Song, Amanda Y Hendrix, James M Tour, David B Corry, and Farrah Kheradmand. Nanoparticulate carbon black in cigarette smoke induces DNA cleavage and Th17-mediated emphysema. *eLife*, 4, 2015.

[1575] Kuai Yu, Jarred M Whitlock, Kyleen Lee, Eric A Ortlund, Yuan Yuan Cui, and H Criss Hartzell. Identification of a lipid scrambling domain in ANO6/TMEM16F. *eLife*, 4, 2015.

[1576] Xuekui Yu, Jiansen Jiang, Jingchen Sun, and Z Hong Zhou. A putative ATPase mediates RNA transcription and capping in a dsRNA virus. *eLife*, 4, 2015.

[1577] Yunguo Yu, Anita M Schmid, and Jonathan D Victor. Visual processing of informative multipoint correlations arises primarily in V2. *eLife*, 4, 2015.

[1578] Xuetian Yue, Yuhan Zhao, Juan Liu, Cen Zhang, Haiyang Yu, Jiabei Wang, Tongsen Zheng, Lianxin Liu, Jun Li, Zhaohui Feng, and Wenwei Hu. BAG2 promotes tumorigenesis through enhancing mutant p53 protein levels and function. *eLife*, 4, 2015.

[1579] Maximina H Yun, Hongorzul Davaapil, and Jeremy P Brockes. Recurrent turnover of senescent cells during regeneration of a complex structure. *eLife*, 4, 2015.

[1580] Amanda D Yzaguirre, Arun Padmanabhan, Eric D de Groh, Kurt A Engleka, Jun Li, Nancy A Speck, and Jonathan A Epstein. Loss of neurofibromin Ras-GAP activity enhances the formation of cardiac blood islands in murine embryos. *eLife*, 4, 2015.

[1581] Yiran Zang, Ming Wan, Min Liu, Hongmei Ke, Shuangchun Ma, Lu-Ping Liu, Jian-Quan Ni, and José Carlos Pastor-Pareja. Plasma membrane overgrowth causes fibrotic collagen accumulation and immune activation in Drosophila adipocytes. *eLife*, 4, 2015.

[1582] Fabio Zanini, Johanna Brodin, Lina Thebo, Christa Lanz, Göran Bratt, Jan Albert, and Richard A Neher. Population genomics of intrapatient HIV-1 evolution. *eLife*, 4, 2015.

[1583] David C Zebrowski, Silvia Vergarajauregui, Chi-Chung Wu, Tanja Piatkowski, Robert Becker, Marina Leone, Sofia Hirth, Filomena Ricciardi, Nathalie Falk, Andreas Giessl, Steffen Just, Thomas Braun, Gilbert Weidinger, and Felix B Engel. Developmental alterations in centrosome integrity contribute to the post-mitotic state of mammalian cardiomyocytes. *eLife*, 4, 2015.

[1584] Yashar Zeighami, Miguel Ulla, Yasser Iturria-Medina, Mahsa Dadar, Yu Zhang, Kevin Michel-Herve Larcher, Vladimir Fonov, Alan C Evans, D Louis Collins, and Alain Dagher. Network structure of brain atrophy in de novo Parkinson's disease. *eLife*, 4, 2015.

[1585] Andreas Zembrzycki, Adam M Stocker, Axel Leingärtner, Setsuko Sahara, Shen-Ju Chou, Valery Kalatsky, Scott R May, Michael P Stryker, and Dennis DM O'Leary. Genetic mechanisms control the linear scaling between related cortical primary and higher order sensory areas. *eLife*, 4, 2015.

[1586] Donglei Zhang and Michael Glotzer. The RhoGAP activity of CYK-4/MgcRacGAP functions non-canonically by promoting RhoA activation during cytokinesis. *eLife*, 4, 2015.

[1587] Jun Zhang, Xijiang Pan, Kaige Yan, Shan Sun, Ning Gao, and Sen-Fang Sui. Mechanisms of ribosome stalling by SecM at multiple elongation steps. *eLife*, 4, 2015.

[1588] Li Zhang, Ngoc-Tung Tran, Hairui Su, Rui Wang, Yuheng Lu, Haiping Tang, Sayura Aoyagi, Ailan Guo, Alireza Khodadadi-Jamayran, Dewang Zhou, Kun Qian, Todd Hricik, Jocelyn Côté, Xiaosi Han, Wenping Zhou, Suparna Laha, Omar Abdel-Wahab, Ross L Levine, Glen Raffel, Yanyan Liu, Dongquan Chen, Haitao Li, Tim Townes, Hengbin Wang, Haiteng Deng, Y George Zheng, Christina Leslie, Minkui Luo, and Xinyang Zhao. Cross-talk between PRMT1-mediated methylation and ubiquitylation on RBM15 controls RNA splicing. *eLife*, 4, 2015.

[1589] Min Zhang, Samuel J Kenny, Liang Ge, Ke Xu, and Randy Schekman. Translocation of interleukin-$1\beta$ into a vesicle intermediate in autophagy-mediated secretion. *eLife*, 4, 2015.

[1590] Tao Zhang, Kangyun Dong, Wei Liang, Daichao Xu, Hongguang Xia, Jiefei Geng, Ayaz Najafov, Min Liu, Yanxia Li, Xiaoran Han, Juan Xiao, Zhenzhen Jin, Ting Peng, Yang Gao, Yu Cai, Chunting Qi, Qing Zhang, Anyang Sun, Marta Lipinski, Hong Zhu, Yue Xiong, Pier Paolo Pandolfi, He Li, Qiang Yu, and Junying Yuan. G-protein-coupled receptors regulate autophagy by ZBTB16-mediated ubiquitination and proteasomal degradation of Atg14L. *eLife*, 4, 2015.

[1591] Zhenghua Zhang, Caihua Li, Fei Wu, Ruixiao Ma, Jing Luan, Feng Yang, Weida Liu, Li Wang, Shoumin Zhang, Yan Liu, Jun Gu, Wenlian Hua, Min Fan, Hua Peng, Xuemei Meng, Ningjing Song, Xinling Bi, Chaoying Gu, Zhen Zhang, Qiong Huang, Lianjun Chen, Leihong Xiang, Jinhua Xu, Zhizhong Zheng, and Zhengwen Jiang. Genomic variations of the mevalonate pathway in porokeratosis. *eLife*, 4, 2015.

[1592] Zhengjian Zhang, Zarko Boskovic, Mahmud M Hussain, Wenxin Hu, Carla Inouye, Han-Je Kim, A Katherine Abole, Mary K Doud, Timothy A Lewis, Angela N Koehler, Stuart L Schreiber, and Robert Tjian. Chemical perturbation of an intrinsically disordered region of TFIID distinguishes two modes of transcription initiation. *eLife*, 4, 2015.

[1593] Jie Zhao, Sarah Siddiqui, Shaobin Shang, Yao Bian, Sreya Bagchi, Ying He, and Chyung-Ru Wang. Mycolic acid-specific T cells protect against Mycobacterium tuberculosis infection in a humanized transgenic mouse model. *eLife*, 4, 2015.

[1594] Jing Zhao, Quan Lin, Kevin J Kim, Faranak D Dardashti, Jennifer Kim, Fei He, and Yi Sun. Ngn1 inhibits astrogliogenesis through induction of miR-9 during neuronal fate specification. *eLife*, 4, 2015.

[1595] Ning Zheng, Okunola Jeyifous, Charlotte Munro, Johanna M Montgomery, and William N Green. Synaptic activity regulates AMPA receptor trafficking through different recycling pathways. *eLife*, 4, 2015.

[1596] Xunhai Zheng, Lalith Perera, Geoffrey A Mueller, Eugene F DeRose, and Robert E London. Asymmetric conformational maturation of HIV-1 reverse transcriptase. *eLife*, 4, 2015.

[1597] Anna Zhou, Alexis Rohou, Daniel G Schep, John V Bason, Martin G Montgomery, John E Walker, Nikolaus Grigorieff, and John L Rubinstein. Structure and conformational states of the bovine mitochondrial ATP synthase by cryo-EM. *eLife*, 4, 2015.

[1598] Bo O Zhou, Lei Ding, and Sean J Morrison. Hematopoietic stem and progenitor cells regulate the regeneration of their niche by secreting Angiopoietin-1. *eLife*, 4, 2015.

[1599] Chuan Zhou, Romain Franconville, Alexander G Vaughan, Carmen C Robinett, Vivek Jayaraman, and Bruce S Baker. Central neural circuitry mediating courtship song perception in male Drosophila. *eLife*, 4, 2015.

[1600] Chun Zhou, Sergei Pourmal, and Nikola P Pavletich. Dna2 nuclease-helicase structure, mechanism and regulation by Rpa. *eLife*, 4, 2015.

[1601] Shu Jun Zhu, Stephanie E Hallows, Ko W Currie, ChangJiang Xu, and Bret J Pearson. A mex3 homolog is required for differentiation during planarian stem cell lineage development. *eLife*, 4, 2015.

[1602] Agata P Zielinska, Zuzana Holubcova, Martyn Blayney, Kay Elder, and Melina Schuh. Sister kinetochore splitting and precocious disintegration of bivalents could explain the maternal age effect. *eLife*, 4, 2015.

[1603] Piotr A Ziolkowski, Luke E Berchowitz, Christophe Lambing, Nataliya E Yelina, Xiaohui Zhao, Krystyna A Kelly, Kyuha Choi, Liliana Ziolkowska, Viviana June, Eugenio Sanchez-Moran, Chris Franklin, Gregory P Copenhaver, and Ian R Henderson. Juxtaposition of heterozygous and homozygous regions causes reciprocal crossover remodelling via interference during Arabidopsis meiosis. *eLife*, 4, 2015.

[1604] Jun Zou, Diana Tran, Mai Baalbaki, Ling Fung Tang, Annie Poon, Angelo Pelonero, Erron W Titus, Christiana Yuan, Chenxu Shi, Shruthi Patchava, Elizabeth Halper, Jasmine Garg, Irina Movsesyan, Chaoying Yin, Roland Wu, Lisa D Wilsbacher, Jiandong Liu, Ronald L Hager, Shaun R Coughlin, Martin Jinek, Clive R Pullinger, John P Kane, Daniel O Hart, Pui-Yan Kwok, and Rahul C Deo. An internal promoter underlies the difference in disease severity between N- and C-terminal truncation mutations of Titin in zebrafish. *eLife*, 4, 2015.

[1605] Jan J Zylicz, Sabine Dietmann, Ufuk Günesdogan, Jamie A Hackett, Delphine Cougot, Caroline Lee, and M Azim Surani. Chromatin dynamics and the role of G9a in gene regulation and enhancer silencing during early mouse development. *eLife*, 4, 2015.

[1606] Andisheh Abedini, Annette Plesner, Ping Cao, Zachary Ridgway, Jinghua Zhang, Ling-Hsien Tu, Chris T Middleton, Brian Chao, Daniel J Sartori, Fanling Meng, Hui Wang, Amy G Wong, Martin T Zanni, C Bruce Verchere, Daniel P Raleigh, and Ann Marie Schmidt. Time-resolved studies define the nature of toxic IAPP intermediates, providing insight for anti-amyloidosis therapeutics. *eLife*, 5, 2016.

[1607] Priyanka D Abeyrathne, Cha San Koh, Timothy Grant, Nikolaus Grigorieff, and Andrei A Korostelev. Ensemble cryo-EM uncovers inchworm-like translocation of a viral IRES through the ribosome. *eLife*, 5, 2016.

[1608] Katie Abley, Susanna Sauret-Güeto, Athanasius FM Marée, and Enrico Coen. Formation of polarity convergences underlying shoot outgrowths. *eLife*, 5, 2016.

[1609] Rhys M Adams, Thierry Mora, Aleksandra M Walczak, and Justin B Kinney. Measuring the sequence-affinity landscape of antibodies with massively parallel titration curves. *eLife*, 5, 2016.

[1610] Sonia Afroz, Julie Parato, Hui Shen, and Sheryl Sue Smith. Synaptic pruning in the female hippocampus is triggered at puberty by extrasynaptic GABAA receptors on dendritic spines. *eLife*, 5, 2016.

[1611] Angelica Aguilera-Gomez, Marinke M van Oorschot, Tineke Veenendaal, and Catherine Rabouille. In vivo vizualisation of mono-ADP-ribosylation by dPARP16 upon amino-acid starvation. *eLife*, 5, 2016.

[1612] Ehud Ahissar and Eldad Assa. Perception as a closed-loop convergence process. *eLife*, 5, 2016.

[1613] Moloud Ahmadi and Richard Roy. AMPK acts as a molecular trigger to coordinate glutamatergic signals and adaptive behaviours during acute starvation. *eLife*, 5, 2016.

[1614] Colin Echeverría Aitken, Petra Beznosková, Vladislava Vlčkova, Wen-Ling Chiu, Fujun Zhou, LeoŠ Shivaya ValáŠek, Alan G Hinnebusch, and Jon R Lorsch. Eukaryotic translation initiation factor 3 plays distinct roles at the mRNA entry and exit channels of the ribosomal preinitiation complex. *eLife*, 5, 2016.

[1615] Zohreh AkhavanAghdam, Joydeb Sinha, Omar P Tabbaa, and Nan Hao. Dynamic control of gene regulatory logic by seemingly redundant transcription factors. *eLife*, 5, 2016.

[1616] Hessameddin Akhlaghpour, Joost Wiskerke, Jung Yoon Choi, Joshua P Taliaferro, Jennifer Au, and Ilana B Witten. Dissociated sequential activity and stimulus encoding in the dorsomedial striatum during spatial working memory. *eLife*, 5, 2016.

[1617] Orkun Akin and S Lawrence Zipursky. Frazzled promotes growth cone attachment at the source of a Netrin gradient in the Drosophila visual system. *eLife*, 5, 2016.

[1618] Felicity Alcock, Phillip J Stansfeld, Hajra Basit, Johann Habersetzer, Matthew AB Baker, Tracy Palmer, Mark I Wallace, and Ben C Berks. Assembling the Tat protein translocase. *eLife*, 5, 2016.

[1619] Zehra Ali-Murthy and Thomas B Kornberg. Bicoid gradient formation and function in the Drosophila pre-syncytial blastoderm. *eLife*, 5, 2016.

[1620] Micah Allen, Darya Frank, D Samuel Schwarzkopf, Francesca Fardo, Joel S Winston, Tobias U Hauser, and Geraint Rees. Unexpected arousal modulates the influence of sensory noise on confidence. *eLife*, 5, 2016.

[1621] William John Allen, Robin Adam Corey, Peter Oatley, Richard Barry Sessions, Steve A Baldwin, Sheena E Radford, Roman Tuma, and Ian Collinson. Two-way communication between SecY and SecA suggests a Brownian ratchet mechanism for protein translocation. *eLife*, 5, 2016.

[1622] Karmel A Allison, Eniko Sajti, Jana G Collier, David Gosselin, Ty Dale Troutman, Erica L Stone, Stephen M Hedrick, and Christopher K Glass. Affinity and dose of TCR engagement yield proportional enhancer and gene activity in CD4+ T cells. *eLife*, 5, 2016.

[1623] John Paul Alvarez, Chihiro Furumizu, Idan Efroni, Yuval Eshed, and John L Bowman. Active suppression of a leaf meristem orchestrates determinate leaf growth. *eLife*, 5, 2016.

[1624] Frederike Alwes, Camille Enjolras, and Michalis Averof. Live imaging reveals the progenitors and cell dynamics of limb regeneration. *eLife*, 5, 2016.

[1625] Ryoji Amamoto, Violeta Gisselle Lopez Huerta, Emi Takahashi, Guangping Dai, Aaron K Grant, Zhanyan Fu, and Paola Arlotta. Adult axolotls can regenerate original neuronal diversity in response to brain injury. *eLife*, 5, 2016.

[1626] Jeremy D Amon and Douglas Koshland. RNase H enables efficient repair of R-loop induced DNA damage. *eLife*, 5, 2016.

[1627] Katia Ancelin, Laurène Syx, Maud Borensztein, Noémie Ranisavljevic, Ivaylo Vassilev, Luis Briseño-Roa, Tao Liu, Eric Metzger, Nicolas Servant, Emmanuel Barillot, Chong-Jian Chen, Roland Schüle, and Edith Heard. Maternal LSD1/KDM1A is an essential regulator of chromatin and transcription landscapes during zygotic genome activation. *eLife*, 5, 2016.

[1628] Douglas P Anderson, Dustin S Whitney, Victor Hanson-Smith, Arielle Woznica, William Campodonico-Burnett, Brian F Volkman, Nicole King, Joseph W Thornton, and Kenneth E Prehoda. Correction: Evolution of an ancient protein function involved in organized multicellularity in animals. *eLife*, 5, 2016.

[1629] Douglas P Anderson, Dustin S Whitney, Victor Hanson-Smith, Arielle Woznica, William Campodonico-Burnett, Brian F Volkman, Nicole King, Joseph W Thornton, and Kenneth E Prehoda. Evolution of an ancient protein function involved in organized multicellularity in animals. *eLife*, 5, 2016.

[1630] John M Ankers, Raheela Awais, Nicholas A Jones, James Boyd, Sheila Ryan, Antony D Adamson, Claire V Harper, Lloyd Bridge, David G Spiller, Dean A Jackson, Pawel Paszek, Violaine Sée, and Michael RH White. Dynamic NF-$\kappa$B and E2F interactions control the priority and timing of inflammatory signalling and cell proliferation. *eLife*, 5, 2016.

[1631] Giovanni Anobile, Roberto Arrighi, Irene Togoli, and David Charles Burr. A shared numerical representation for action and perception. *eLife*, 5, 2016.

[1632] Luisa CS Antunes, Daniel Poppleton, Andreas Klingl, Alexis Criscuolo, Bruno Dupuy, Céline Brochier-Armanet, Christophe Beloin, and Simonetta Gribaldo. Phylogenomic analysis supports the ancestral presence of LPS-outer membranes in the Firmicutes. *eLife*, 5, 2016.

[1633] Evan G Antzoulatos and Earl K Miller. Synchronous beta rhythms of frontoparietal networks support only behaviorally relevant representations. *eLife*, 5, 2016.

[1634] David Arkadir, Angela Radulescu, Deborah Raymond, Naomi Lubarr, Susan B Bressman, Pietro Mazzoni, and Yael Niv. DYT1 dystonia increases risk taking in humans. *eLife*, 5, 2016.

[1635] Christopher P Arnold, M Shane Merryman, Aleishia Harris-Arnold, Sean A McKinney, Chris W Seidel, Sydney Loethen, Kylie N Proctor, Longhua Guo, and Alejandro Sánchez Alvarado. Pathogenic shifts in endogenous microbiota impede tissue regeneration via distinct activation of TAK1/MKK/p38. *eLife*, 5, 2016.

[1636] Amjad Askary, Joanna Smeeton, Sandeep Paul, Simone Schindler, Ingo Braasch, Nicholas A Ellis, John Postlethwait, Craig T Miller, and J Gage Crump. Ancient origin of lubricated joints in bony vertebrates. *eLife*, 5, 2016.

[1637] Yoshinori Aso and Gerald M Rubin. Dopaminergic neurons write and update memories with cell-type-specific rules. *eLife*, 5, 2016.

[1638] Jeremy F Atherton, Eileen L McIver, Matthew RM Mullen, David L Wokosin, D James Surmeier, and Mark D Bevan. Early dysfunction and progressive degeneration of the subthalamic nucleus in mouse models of Huntington's disease. *eLife*, 5, 2016.

[1639] Lauren Y Atlas, Bradley B Doll, Jian Li, Nathaniel D Daw, and Elizabeth A Phelps. Instructed knowledge shapes feedback-driven aversive learning in striatum and orbitofrontal cortex, but not the amygdala. *eLife*, 5, 2016.

[1640] Jan Attig, Igor Ruiz de los Mozos, Nejc Haberman, Zhen Wang, Warren Emmett, Kathi Zarnack, Julian König, and Jernej Ule. Splicing repression allows the gradual emergence of new Alu-exons in primate evolution. *eLife*, 5, 2016.

[1641] Nicolas Aznar, Arjun Patel, Cristina C Rohena, Ying Dunkel, Linda P Joosen, Vanessa Taupin, Irina Kufareva, Marilyn G Farquhar, and Pradipta Ghosh. AMP-activated protein kinase fortifies epithelial tight junctions during energetic stress via its effector GIV/Girdin. *eLife*, 5, 2016.

[1642] Marije Baas, Alix BesanÇon, Tania Goncalves, Fabrice Valette, Hideo Yagita, Birgit Sawitzki, Hans-Dieter Volk, Emmanuelle Waeckel-Enée, Benedita Rocha, Lucienne Chatenoud, and Sylvaine You. TGF$\beta$-dependent expression of PD-1 and PD-L1 controls CD8+ T cell anergy in transplant tolerance. *eLife*, 5, 2016.

[1643] Bartholomew J Bacak, Taegyo Kim, Jeffrey C Smith, Jonathan E Rubin, and Ilya A Rybak. Mixed-mode oscillations and population bursting in the pre-Bötzinger complex. *eLife*, 5, 2016.

[1644] Chanhyung Bae, Claudio Anselmi, Jeet Kalia, Andres Jara-Oseguera, Charles D Schwieters, Dmitriy Krepkiy, Chul Won Lee, Eun-Hee Kim, Jae Il Kim, José D Faraldo-Gómez, and Kenton J Swartz. Structural insights into the mechanism of activation of the TRPV1 channel by a membrane-bound tarantula toxin. *eLife*, 5, 2016.

[1645] Parikshit Bagchi, Takamasa Inoue, and Billy Tsai. EMC1-dependent stabilization drives membrane penetration of a partially destabilized non-enveloped virus. *eLife*, 5, 2016.

[1646] Christopher A Baker, Yishai M Elyada, Andres Parra, and M McLean Bolton. Cellular resolution circuit mapping with temporal-focused excitation of soma-targeted channelrhodopsin. *eLife*, 5, 2016.

[1647] Xiaojun Bao, Louise LG Raguet, Sydni M Cole, James D Howard, and Jay A Gottfried. The role of piriform associative connections in odor categorization. *eLife*, 5, 2016.

[1648] Xiaoyan Robert Bao, Shao-En Ong, Olga Goldberger, Jun Peng, Rohit Sharma, Dawn A Thompson, Scott B Vafai, Andrew G Cox, Eizo Marutani, Fumito Ichinose, Wolfram Goessling, Aviv Regev, Steven A Carr, Clary B Clish, and Vamsi K Mootha. Mitochondrial dysfunction remodels one-carbon metabolism in human cells. *eLife*, 5, 2016.

[1649] Anna P Baron, Conrad von Schubert, Fabien Cubizolles, Gerhard Siemeister, Marion Hitchcock, Anne Mengel, Jens Schröder, Amaury Fernández-Montalván, Franz von Nussbaum, Dominik Mumberg, and Erich A Nigg. Probing the catalytic functions of Bub1 kinase using the small molecule inhibitors BAY-320 and BAY-524. *eLife*, 5, 2016.

[1650] Olga Barreiro, Danay Cibrian, Cristina Clemente, David Alvarez, Vanessa Moreno, íñigo Valiente, Antonio Bernad, Dietmar Vestweber, Alicia G Arroyo, Pilar Martín, Ulrich H von Andrian, and Francisco Sánchez Madrid. Pivotal role for skin transendothelial radio-resistant anti-inflammatory macrophages in tissue repair. *eLife*, 5, 2016.

[1651] David GT Barrett, Sophie Denève, and Christian K Machens. Optimal compensation for neuron loss. *eLife*, 5, 2016.

[1652] Francisco J Barrionuevo, Alicia Hurtado, Gwang-Jin Kim, Francisca M Real, Mohammed Bakkali, Janel L Kopp, Maike Sander, Gerd Scherer, Miguel Burgos, and Rafael Jiménez. Sox9 and Sox8 protect the adult testis from male-to-female genetic reprogramming and complete degeneration. *eLife*, 5, 2016.

[1653] William E Barry and Carl S Thummel. The Drosophila HNF4 nuclear receptor promotes glucose-stimulated insulin secretion and mitochondrial function in adults. *eLife*, 5, 2016.

[1654] Martin L Basch, Rogers M Brown, Hsin-I Jen, Fatih Semerci, Frederic Depreux, Renée K Edlund, Hongyuan Zhang, Christine R Norton, Thomas Gridley, Susan E Cole, Angelika Doetzlhofer, Mirjana Maletic-Savatic, Neil Segil, and Andrew K Groves. Fine-tuning of Notch signaling sets the boundary of the organ of Corti and establishes sensory cell fates. *eLife*, 5, 2016.

[1655] Erin A Bassett, Nicholas Tokarew, Ema A Allemano, Chantal Mazerolle, Katy Morin, Alan J Mears, Brian McNeill, Randy Ringuette, Charles Campbell, Sheila Smiley, Neno T Pokrajac, Adrian M Dubuc, Vijay Ramaswamy, Paul A Northcott, Marc Remke, Philippe P Monnier, David Potter, Kim Paes, Laura L Kirkpatrick, Kenneth J Coker, Dennis S Rice, Carol Perez-Iratxeta, Michael D Taylor, and Valerie A Wallace. Norrin/Frizzled4 signalling in the preneoplastic niche blocks medulloblastoma initiation. *eLife*, 5, 2016.

[1656] Aakash Basu, Samuel Lagier, Maria Vologodskaia, Brian A Fabella, and AJ Hudspeth. Direct mechanical stimulation of tip links in hair cells through DNA tethers. *eLife*, 5, 2016.

[1657] Gervasio Batista, Jennifer Leigh Johnson, Elena Dominguez, Mauro Costa-Mattioli, and Jose L Pena. Translational control of auditory imprinting and structural plasticity by eIF2$\alpha$. *eLife*, 5, 2016.

[1658] Aleksandra Bebel, Ezgi Karaca, Banushree Kumar, W Marshall Stark, and Orsolya Barabas. Structural snapshots of Xer recombination reveal activation by synaptic complex remodeling and DNA bending. *eLife*, 5, 2016.

[1659] Christian Behrens, Timm Schubert, Silke Haverkamp, Thomas Euler, and Philipp Berens. Connectivity map of bipolar cells and photoreceptors in the mouse retina. *eLife*, 5, 2016.

[1660] Reza Behrouzi, Chenning Lu, Mark A Currie, Gloria Jih, Nahid Iglesias, and Danesh Moazed. Heterochromatin assembly by interrupted Sir3 bridges across neighboring nucleosomes. *eLife*, 5, 2016.

[1661] Jacob LS Bellmund, Lorena Deuker, Tobias Navarro Schröder, and Christian F Doeller. Grid-cell representations in mental simulation. *eLife*, 5, 2016.

[1662] Stephanie Beltz, Jens Bassler, and Joachim E Schultz. Regulation by the quorum sensor from Vibrio indicates a receptor function for the membrane anchors of adenylate cyclases. *eLife*, 5, 2016.

[1663] Blair W Benham-Pyle, Joo Yong Sim, Kevin C Hart, Beth L Pruitt, and William James Nelson. Increasing $\beta$-catenin/Wnt3A activity levels drive mechanical strain-induced cell cycle progression through mitosis. *eLife*, 5, 2016.

[1664] Nuria Benito, Gonzalo Martín-Vázquez, Julia Makarova, Valeri A Makarov, and Oscar Herreras. The right hippocampus leads the bilateral integration of gamma-parsed lateralized information. *eLife*, 5, 2016.

[1665] Matthew E Berck, Avinash Khandelwal, Lindsey Claus, Luis Hernandez-Nunez, Guangwei Si, Christopher J Tabone, Feng Li, James W Truman, Rick D Fetter, Matthieu Louis, Aravinthan DT Samuel, and Albert Cardona. The wiring diagram of a glomerular olfactory system. *eLife*, 5, 2016.

[1666] Adi Bercovich-Kinori, Julie Tai, Idit Anna Gelbart, Alina Shitrit, Shani Ben-Moshe, Yaron Drori, Shalev Itzkovitz, Michal Mandelboim, and Noam Stern-Ginossar. A systematic view on influenza induced host shutoff. *eLife*, 5, 2016.

[1667] Matthieu Bergé, Sébastien Campagne, Johann Mignolet, Seamus Holden, Laurence Théraulaz, Suliana Manley, Frédéric H-T Allain, and Patrick H Viollier. Modularity and determinants of a (bi-)polarization control system from free-living and obligate intracellular bacteria. *eLife*, 5, 2016.

[1668] Stephanie Berger, Erik Procko, Daciana Margineantu, Erinna F Lee, Betty W Shen, Alex Zelter, Daniel-Adriano Silva, Kusum Chawla, Marco J Herold, Jean-Marc Garnier, Richard Johnson, Michael J MacCoss, Guillaume Lessene, Trisha N Davis, Patrick S Stayton, Barry L Stoddard, W Douglas Fairlie, David M Hockenbery, and David Baker. Computationally designed high specificity inhibitors delineate the roles of BCL2 family proteins in cancer. *eLife*, 5, 2016.

[1669] Shai Berlin, Stephanie Szobota, Andreas Reiner, Elizabeth C Carroll, Michael A Kienzler, Alice Guyon, Tong Xiao, Dirk Trauner, and Ehud Y Isacoff. A family of photoswitchable NMDA receptors. *eLife*, 5, 2016.

[1670] Lisa K Berry, Guðjón ólafsson, Elena Ledesma-Fernández, and Peter H Thorpe. Synthetic protein interactions reveal a functional map of the cell. *eLife*, 5, 2016.

[1671] Michael T Bethune, Marvin H Gee, Mario Bunse, Mark S Lee, Eric H Gschweng, Meghana S Pagadala, Jing Zhou, Donghui Cheng, James R Heath, Donald B Kohn, Michael S Kuhns, Wolfgang Uckert, and David Baltimore. Domain-swapped T cell receptors improve the safety of TCR gene therapy. *eLife*, 5, 2016.

[1672] Marianne J Bezaire, Ivan Raikov, Kelly Burk, Dhrumil Vyas, and Ivan Soltesz. Interneuronal mechanisms of hippocampal theta oscillations in a full-scale model of the rodent CA1 circuit. *eLife*, 5, 2016.

[1673] Ajay Bhargava, Madan Anant, and Hildegard Mack. Registered report: Kinase-dead BRAF and oncogenic RAS cooperate to drive tumor progression through CRAF. *eLife*, 5, 2016.

[1674] Ajay Bhargava, Steven Pelech, Ben Woodard, John Kerwin, and Nimet Maherali. Registered report: RAF inhibitors prime wild-type RAF to activate the MAPK pathway and enhance growth. *eLife*, 5, 2016.

[1675] Debashish Bhattacharya, Shobhit Agrawal, Manuel Aranda, Sebastian Baumgarten, Mahdi Belcaid, Jeana L Drake, Douglas Erwin, Sylvian Foret, Ruth D Gates, David F Gruber, Bishoy Kamel, Michael P Lesser, Oren Levy, Yi Jin Liew, Matthew MacManes, Tali Mass, Monica Medina, Shaadi Mehr, Eli Meyer, Dana C Price, Hollie M Putnam, Huan Qiu, Chuya Shinzato, Eiichi Shoguchi, Alexander J Stokes, Sylvie Tambutté, Dan Tchernov, Christian R Voolstra, Nicole Wagner, Charles W Walker, Andreas PM Weber, Virginia Weis, Ehud Zelzion, Didier Zoccola, and Paul G Falkowski. Comparative genomics explains the evolutionary success of reef-forming corals. *eLife*, 5, 2016.

[1676] Moitrayee Bhattacharyya, Margaret M Stratton, Catherine C Going, Ethan D McSpadden, Yongjian Huang, Anna C Susa, Anna Elleman, Yumeng Melody Cao, Nishant Pappireddi, Pawel Burkhardt, Christine L Gee, Tiago Barros, Howard Schulman, Evan R Williams, and John Kuriyan. Molecular mechanism of activation-triggered subunit exchange in Ca2+/calmodulin-dependent protein kinase II. *eLife*, 5, 2016.

[1677] Sanchari Bhattacharyya, Shimon Bershtein, Jin Yan, Tijda Argun, Amy I Gilson, Sunia A Trauger, and Eugene I Shakhnovich. Transient protein-protein interactions perturb E. coli metabolome and cause gene dosage toxicity. *eLife*, 5, 2016.

[1678] Pengpeng Bi, Feng Yue, Yusuke Sato, Sara Wirbisky, Weiyi Liu, Tizhong Shan, Yefei Wen, Daoguo Zhou, Jennifer Freeman, and Shihuan Kuang. Stage-specific effects of Notch activation during skeletal myogenesis. *eLife*, 5, 2016.

[1679] Alessandro Bitto, Takashi K Ito, Victor V Pineda, Nicolas J LeTexier, Heather Z Huang, Elissa Sutlief, Herman Tung, Nicholas Vizzini, Belle Chen, Kaleb Smith, Daniel Meza, Masanao Yajima, Richard P Beyer, Kathleen F Kerr, Daniel J Davis, Catherine H Gillespie, Jessica M Snyder, Piper M Treuting, and Matt Kaeberlein. Transient rapamycin treatment can increase lifespan and healthspan in middle-aged mice. *eLife*, 5, 2016.

[1680] E Rebecka Björnfors and Abdeljabbar El Manira. Functional diversity of excitatory commissural interneurons in adult zebrafish. *eLife*, 5, 2016.

[1681] Sophia M Blake, Stefan H Stricker, Hanna Halavach, Anna R Poetsch, George Cresswell, Gavin Kelly, Nnennaya Kanu, Silvia Marino, Nicholas M Luscombe, Steven M Pollard, and Axel Behrens. Inactivation of the ATMIN/ATM pathway protects against glioblastoma formation. *eLife*, 5, 2016.

[1682] FranÇois Blanquart, Mary Kate Grabowski, Joshua Herbeck, Fred Nalugoda, David Serwadda, Michael A Eller, Merlin L Robb, Ronald Gray, Godfrey Kigozi, Oliver Laeyendecker, Katrina A Lythgoe, Gertrude Nakigozi, Thomas C Quinn, Steven J Reynolds, Maria J Wawer, and Christophe Fraser. A transmission-virulence evolutionary trade-off explains attenuation of HIV-1 in Uganda. *eLife*, 5, 2016.

[1683] Kim R Blasdell, Veasna Duong, Marc Eloit, Fabrice Chretien, Sowath Ly, Vibol Hul, Vincent Deubel, Serge Morand, and Philippe Buchy. Evidence of human infection by a new mammarenavirus endemic to Southeastern Asia. *eLife*, 5, 2016.

[1684] Shelby A Blythe and Eric F Wieschaus. Establishment and maintenance of heritable chromatin structure during early Drosophila embryogenesis. *eLife*, 5, 2016.

[1685] Gudrun Böhmdorfer, Shriya Sethuraman, M Jordan Rowley, Michal Krzyszton, M Hafiz Rothi, Lilia Bouzit, and Andrzej T Wierzbicki. Long non-coding RNA produced by RNA polymerase V determines boundaries of heterochromatin. *eLife*, 5, 2016.

[1686] David M Bolduc, Daniel R Montagna, Matthew C Seghers, Michael S Wolfe, and Dennis J Selkoe. The amyloid-beta forming tripeptide cleavage mechanism of $\gamma$-secretase. *eLife*, 5, 2016.

[1687] James J Bonaiuto, Archy de Berker, and Sven Bestmann. Response repetition biases in human perceptual decisions are explained by activity decay in competitive attractor models. *eLife*, 5, 2016.

[1688] Benjamin Bonneau, Hideaki Ando, Katsuhiro Kawaai, Matsumi Hirose, Hiromi Takahashi-Iwanaga, and Katsuhiko Mikoshiba. IRBIT controls apoptosis by interacting with the Bcl-2 homolog, Bcl2l10, and by promoting ER-mitochondria contact. *eLife*, 5, 2016.

[1689] Kirsten I Bos, Alexander Herbig, Jason Sahl, Nicholas Waglechner, Mathieu Fourment, Stephen A Forrest, Jennifer Klunk, Verena J Schuenemann, Debi Poinar, Melanie Kuch, G Brian Golding, Olivier Dutour, Paul Keim, David M Wagner, Edward C Holmes, Johannes Krause, and Hendrik N Poinar. Eighteenth century Yersinia pestis genomes reveal the long-term persistence of an historical plague focus. *eLife*, 5, 2016.

[1690] Salvatore Botta, Elena Marrocco, Nicola de Prisco, Fabiola Curion, Mario Renda, Martina Sofia, Mariangela Lupo, Annamaria Carissimo, Maria Laura Bacci, Carlo Gesualdo, Settimio Rossi, Francesca Simonelli, and Enrico Maria Surace. Rhodopsin targeted transcriptional silencing by DNA-binding. *eLife*, 5, 2016.

[1691] Benjamin P Bouchet, Rosemarie E Gough, York-Christoph Ammon, Dieudonnée van de Willige, Harm Post, Guillaume Jacquemet, AF Maarten Altelaar, Albert JR Heck, Benjamin T Goult, and Anna Akhmanova. Talin-KANK1 interaction controls the recruitment of cortical microtubule stabilizing complexes to focal adhesions. *eLife*, 5, 2016.

[1692] Fabrice Bouilloux, Jérôme Thireau, Stéphanie Ventéo, Charlotte Farah, Sarah Karam, Yves Dauvilliers, Jean Valmier, Neal G Copeland, Nancy A Jenkins, Sylvain Richard, and Frédéric Marmigère. Loss of the transcription factor Meis1 prevents sympathetic neurons target-field innervation and increases susceptibility to sudden cardiac death. *eLife*, 5, 2016.

[1693] Cara C Boutte, Christina E Baer, Kadamba Papavinasasundaram, Weiru Liu, Michael R Chase, Xavier Meniche, Sarah M Fortune, Christopher M Sassetti, Thomas R Ioerger, and Eric J Rubin. A cytoplasmic peptidoglycan amidase homologue controls mycobacterial cell wall synthesis. *eLife*, 5, 2016.

[1694] Tobias Brandt, Laetitia Cavellini, Werner Kühlbrandt, and Mickaël M Cohen. A mitofusin-dependent docking ring complex triggers mitochondrial fusion in vitro. *eLife*, 5, 2016.

[1695] Michael O Breckwoldt, Julia Bode, Felix T Kurz, Angelika Hoffmann, Katharina Ochs, Martina Ott, Katrin Deumelandt, Thomas Krüwel, Daniel Schwarz, Manuel Fischer, Xavier Helluy, David Milford, Klara Kirschbaum, Gergely Solecki, Sara Chiblak, Amir Abdollahi, Frank Winkler, Wolfgang Wick, Michael Platten, Sabine Heiland, Martin Bendszus, and Björn Tews. Correlated magnetic resonance imaging and ultramicroscopy (MR-UM) is a tool kit to assess the dynamics of glioma angiogenesis. *eLife*, 5, 2016.

[1696] Johanna Brodin, Fabio Zanini, Lina Thebo, Christa Lanz, Göran Bratt, Richard A Neher, and Jan Albert. Establishment and stability of the latent HIV-1 DNA reservoir. *eLife*, 5, 2016.

[1697] Thibaut Brunet, Antje HL Fischer, Patrick RH Steinmetz, Antonella Lauri, Paola Bertucci, and Detlev Arendt. The evolutionary origin of bilaterian smooth and striated myocytes. *eLife*, 5, 2016.

[1698] Krzysztof Brzezinka, Simone Altmann, Hjördis Czesnick, Philippe Nicolas, Michal Gorka, Eileen Benke, Tina Kabelitz, Felix Jähne, Alexander Graf, Christian Kappel, and Isabel Bäurle. Arabidopsis FORGETTER1 mediates stress-induced chromatin memory through nucleosome remodeling. *eLife*, 5, 2016.

[1699] Tuan V Bui, Nicolas Stifani, Turgay Akay, and Robert M Brownstone. Spinal microcircuits comprising dI3 interneurons are necessary for motor functional recovery following spinal cord transection. *eLife*, 5, 2016.

[1700] George BJ Busby, Gavin Band, Quang Si Le, Muminatou Jallow, Edith Bougama, Valentina D Mangano, Lucas N Amenga-Etego, Anthony Enimil, Tobias Apinjoh, Carolyne M Ndila, Alphaxard Manjurano, Vysaul Nyirongo, Ogobara Doumba, Kirk A Rockett, Dominic P Kwiatkowski, and Chris CA Spencer. Admixture into and within sub-Saharan Africa. *eLife*, 5, 2016.

[1701] Nicholas E Bush, Christopher L Schroeder, Jennifer A Hobbs, Anne ET Yang, Lucie A Huet, Sara A Solla, and Mitra JZ Hartmann. Decoupling kinematics and mechanics reveals coding properties of trigeminal ganglion neurons in the rat vibrissal system. *eLife*, 5, 2016.

[1702] Alexandra B Byrne, Rebecca D McWhirter, Yuichi Sekine, Stephen M Strittmatter, David M Miller, and Marc Hammarlund. Inhibiting poly(ADP-ribosylation) improves axon regeneration. *eLife*, 5, 2016.

[1703] Yujia Cai, Anders Laustsen, Yan Zhou, Chenglong Sun, Mads Valdemar Anderson, Shengting Li, Niels Uldbjerg, Yonglun Luo, Martin R Jakobsen, and Jacob Giehm Mikkelsen. Targeted, homology-driven gene insertion in stem cells by ZFN-loaded 'all-in-one' lentiviral vectors. *eLife*, 5, 2016.

[1704] Alessia Calcagnì, Lotte kors, Eric Verschuren, Rossella De Cegli, Nicolina Zampelli, Edoardo Nusco, Stefano Confalonieri, Giovanni Bertalot, Salvatore Pece, Carmine Settembre, Gabriel G Malouf, Jaklien C Leemans, Emile de Heer, Marco Salvatore, Dorien JM Peters, Pier Paolo Di Fiore, and Andrea Ballabio. Modelling TFE renal cell carcinoma in mice reveals a critical role of WNT signaling. *eLife*, 5, 2016.

[1705] Margarita Calvo, Natalie Richards, Annina B Schmid, Alejandro Barroso, Lan Zhu, Dinka Ivulic, Ning Zhu, Philipp Anwandter, Manzoor A Bhat, Felipe A Court, Stephen B McMahon, and David LH Bennett. Altered potassium channel distribution and composition in myelinated axons suppresses hyperexcitability following injury. *eLife*, 5, 2016.

[1706] Dario Campagner, Mathew Hywel Evans, Michael Ross Bale, Andrew Erskine, and Rasmus Strange Petersen. Prediction of primary somatosensory neuron activity during active tactile exploration. *eLife*, 5, 2016.

[1707] Bo Cao, Kebing Wang, Jun-Ming Liao, Xiang Zhou, Peng Liao, Shelya X Zeng, Meifang He, Lianzhou Chen, Yulong He, Wen Li, and Hua Lu. Inactivation of oncogenic cAMP-specific phosphodiesterase 4D by miR-139-5p in response to p53 activation. *eLife*, 5, 2016.

[1708] Paul Capewell, Christelle Cren-Travaillé, Francesco Marchesi, Pamela Johnston, Caroline Clucas, Robert A Benson, Taylor-Anne Gorman, Estefania Calvo-Alvarez, Aline Crouzols, Grégory Jouvion, Vincent Jamonneau, William Weir, M Lynn Stevenson, Kerry O'Neill, Anneli Cooper, Nonoraymond Kuispond Swar, Bruno Bucheton, Dieudonné Mumba Ngoyi, Paul Garside, Brice Rotureau, and Annette MacLeod. The skin is a significant but overlooked anatomical reservoir for vector-borne African trypanosomes. *eLife*, 5, 2016.

[1709] Carmen Capone, Fabrice Dabertrand, Celine Baron-Menguy, Athena Chalaris, Lamia Ghezali, Valérie Domenga-Denier, Stefanie Schmidt, Clément Huneau, Stefan Rose-John, Mark T Nelson, and Anne Joutel. Mechanistic insights into a TIMP3-sensitive pathway constitutively engaged in the regulation of cerebral hemodynamics. *eLife*, 5, 2016.

[1710] Lars-Anders Carlson, Yun Bai, Sarah C Keane, Jennifer A Doudna, and James H Hurley. Reconstitution of selective HIV-1 RNA packaging in vitro by membrane-bound Gag assemblies. *eLife*, 5, 2016.

[1711] Michelle A Carmell, Gregoriy A Dokshin, Helen Skaletsky, Yueh-Chiang Hu, Josien C van Wolfswinkel, Kyomi J Igarashi, Daniel W Bellott, Michael Nefedov, Peter W Reddien, George C Enders, Vladimir N Uversky, Craig C Mello, and David C Page. A widely employed germ cell marker is an ancient disordered protein with reproductive functions in diverse eukaryotes. *eLife*, 5, 2016.

[1712] Rita Carmona, Ana Cañete, Elena Cano, Laura Ariza, Anabel Rojas, and Ramon Muñoz-Chápuli. Conditional deletion of WT1 in the septum transversum mesenchyme causes congenital diaphragmatic hernia in mice. *eLife*, 5, 2016.

[1713] Bernadette Carroll, Dorothea Maetzel, Oliver DK Maddocks, Gisela Otten, Matthew Ratcliff, Graham R Smith, Elaine A Dunlop, João F Passos, Owen R Davies, Rudolf Jaenisch, Andrew R Tee, Sovan Sarkar, and Viktor I Korolchuk. Control of TSC2-Rheb signaling axis by arginine regulates mTORC1 activity. *eLife*, 5, 2016.

[1714] Alecia J Carter, Miquel Torrents Ticó, and Guy Cowlishaw. Sequential phenotypic constraints on social information use in wild baboons. *eLife*, 5, 2016.

[1715] Yaron Caspi and Cees Dekker. Mapping out Min protein patterns in fully confined fluidic chambers. *eLife*, 5, 2016.

[1716] Sean E Cavanagh, Joni D Wallis, Steven W Kennerley, and Laurence T Hunt. Autocorrelation structure at rest predicts value correlates of single neurons during reward-guided choice. *eLife*, 5, 2016.

[1717] Mark S Cembrowski, Lihua Wang, Ken Sugino, Brenda C Shields, and Nelson Spruston. Hipposeq: a comprehensive RNA-seq database of gene expression in hippocampal principal neurons. *eLife*, 5, 2016.

[1718] Rahul Chadda, Venkatramanan Krishnamani, Kacey Mersch, Jason Wong, Marley Brimberry, Ankita Chadda, Ludmila Kolmakova-Partensky, Larry J Friedman, Jeff Gelles, and Janice L Robertson. The dimerization equilibrium of a ClC Cl−/H+ antiporter in lipid bilayers. *eLife*, 5, 2016.

[1719] Angus Chadwick, Mark CW van Rossum, and Matthew F Nolan. Flexible theta sequence compression mediated via phase precessing interneurons. *eLife*, 5, 2016.

[1720] Subhojit Chakraborty, Nils Kolling, Mark E Walton, and Anna S Mitchell. Critical role for the mediodorsal thalamus in permitting rapid reward-guided updating in stochastic reward environments. *eLife*, 5, 2016.

[1721] Matthew Chalk, Boris Gutkin, and Sophie Denève. Neural oscillations as a signature of efficient coding in the presence of synaptic delays. *eLife*, 5, 2016.

[1722] Anna L Chaly, Dollada Srisai, Ellen E Gardner, and Julien A Sebag. The Melanocortin Receptor Accessory Protein 2 promotes food intake through inhibition of the Prokineticin Receptor-1. *eLife*, 5, 2016.

[1723] Clara Champagne, David Georges Salthouse, Richard Paul, Van-Mai Cao-Lormeau, Benjamin Roche, and Bernard Cazelles. Structure in the variability of the basic reproductive number (R0) for Zika epidemics in the Pacific islands. *eLife*, 5, 2016.

[1724] Ming Chao, Hao Wu, Kai Jin, Bin Li, Jianjun Wu, Guangqiang Zhang, Gong Yang, and Xun Hu. A nonrandomized cohort and a randomized study of local control of large hepatocarcinoma by targeting intratumoral lactic acidosis. *eLife*, 5, 2016.

[1725] Karthikeyani Chellappa, Poonamjot Deol, Jane R Evans, Linh M Vuong, Gang Chen, Nadege BrianÇon, Eugene Bolotin, Christian Lytle, Meera G Nair, and Frances M Sladek. Opposing roles of nuclear receptor HNF4$\alpha$ isoforms in colitis and colitis-associated colon cancer. *eLife*, 5, 2016.

[1726] Cuie Chen, Mayu Inaba, Zsolt G Venkei, and Yukiko M Yamashita. Klp10A, a stem cell centrosome-enriched kinesin, balances asymmetries in Drosophila male germline stem cell division. *eLife*, 5, 2016.

[1727] Jerry L Chen, Fabian F Voigt, Mitra Javadzadeh, Roland Krueppel, and Fritjof Helmchen. Long-range population dynamics of anatomically defined neocortical networks. *eLife*, 5, 2016.

[1728] Jun Chen, Na Xu, Huanwei Huang, Tao Cai, and Rongwen Xi. A feedback amplification loop between stem cells and their progeny promotes tissue regeneration and tumorigenesis. *eLife*, 5, 2016.

[1729] Kuchuan Chen, Tammy Szu-Yu Ho, Guang Lin, Kai Li Tan, Matthew N Rasband, and Hugo J Bellen. Loss of Frataxin activates the iron/sphingolipid/PDK1/Mef2 pathway in mammals. *eLife*, 5, 2016.

[1730] Kuchuan Chen, Guang Lin, Nele A Haelterman, Tammy Szu-Yu Ho, Tongchao Li, Zhihong Li, Lita Duraine, Brett H Graham, Manish Jaiswal, Shinya Yamamoto, Matthew N Rasband, and Hugo J Bellen. Loss of Frataxin induces iron toxicity, sphingolipid synthesis, and Pdk1/Mef2 activation, leading to neurodegeneration. *eLife*, 5, 2016.

[1731] Lizhen Chen, Zhijie Liu, Bing Zhou, Chaoliang Wei, Yu Zhou, Michael G Rosenfeld, Xiang-Dong Fu, Andrew D Chisholm, and Yishi Jin. CELF RNA binding proteins promote axon regeneration in C. elegans and mammals through alternative splicing of Syntaxins. *eLife*, 5, 2016.

[1732] Naiyan Chen, Hiroki Sugihara, Jinah Kim, Zhanyan Fu, Boaz Barak, Mriganka Sur, Guoping Feng, and Weiping Han. Direct modulation of GFAP-expressing glia in the arcuate nucleus bi-directionally regulates feeding. *eLife*, 5, 2016.

[1733] Qiang Chen, Zhe Pei, David Koren, and Wei Wei. Stimulus-dependent recruitment of lateral inhibition underlies retinal direction selectivity. *eLife*, 5, 2016.

[1734] Susu Chen, George J Augustine, and Paul Chadderton. The cerebellum linearly encodes whisker position during voluntary movement. *eLife*, 5, 2016.

[1735] Xiangsong Chen, Li Lu, Kevin S Mayer, Mark Scalf, Shuiming Qian, Aaron Lomax, Lloyd M Smith, and Xuehua Zhong. POWERDRESS interacts with HISTONE DEACETYLASE 9 to promote aging in Arabidopsis. *eLife*, 5, 2016.

[1736] Xiao Chen, Reazur Rahman, Fang Guo, and Michael Rosbash. Genome-wide identification of neuronal activity-regulated genes in Drosophila. *eLife*, 5, 2016.

[1737] Yiming Chen, Yen-Chu Lin, Christopher A Zimmerman, Rachel A Essner, and Zachary A Knight. Hunger neurons drive feeding through a sustained, positive reinforcement signal. *eLife*, 5, 2016.

[1738] Kaiying Cheng, Hong Xu, Xuanyi Chen, Liangyan Wang, Bing Tian, Ye Zhao, and Yuejin Hua. Structural basis for DNA 5´-end resection by RecJ. *eLife*, 5, 2016.

[1739] Tat Cheung Cheng, Chuan Hong, Ildikó V Akey, Shujun Yuan, and Christopher W Akey. A near atomic structure of the active human apoptosome. *eLife*, 5, 2016.

[1740] Wing Yin Cheng, HoangDinh Huynh, Peiwen Chen, Samuel Peña-Llopis, and Yihong Wan. Macrophage PPAR$\gamma$ inhibits Gpr132 to mediate the anti-tumor effects of rosiglitazone. *eLife*, 5, 2016.

[1741] Connie Cheung, Liberty S Hamilton, Keith Johnson, and Edward F Chang. The auditory representation of speech sounds in human motor cortex. *eLife*, 5, 2016.

[1742] Marc Chevalier, Natalia Toporikova, John Simmers, and Muriel Thoby-Brisson. Development of pacemaker properties and rhythmogenic mechanisms in the mouse embryonic respiratory network. *eLife*, 5, 2016.

[1743] Christopher Chidley, Sunia A Trauger, KıvanÇ Birsoy, and Erin K O'Shea. The anticancer natural product ophiobolin A induces cytotoxicity by covalent modification of phosphatidylethanolamine. *eLife*, 5, 2016.

[1744] Christine E Cho, Chantal Brueggemann, Noelle D L'Etoile, and Cornelia I Bargmann. Parallel encoding of sensory history and behavioral preference during Caenorhabditis elegans olfactory learning. *eLife*, 5, 2016.

[1745] Won-Ki Cho, Namrata Jayanth, Brian P English, Takuma Inoue, J Owen Andrews, William Conway, Jonathan B Grimm, Jan-Hendrik Spille, Luke D Lavis, Timothée Lionnet, and Ibrahim I Cisse. RNA Polymerase II cluster dynamics predict mRNA output in living cells. *eLife*, 5, 2016.

[1746] Ucheor B Choi, Minglei Zhao, Yunxiang Zhang, Ying Lai, and Axel T Brunger. Complexin induces a conformational change at the membrane-proximal C-terminal end of the SNARE complex. *eLife*, 5, 2016.

[1747] Subhendu Roy Choudhury, Anand K Singh, Tina McLeod, Marco Blanchette, Boyun Jang, Paul Badenhorst, Aditi Kanhere, and Saverio Brogna. Exon junction complex proteins bind nascent transcripts independently of pre-mRNA splicing in Drosophila melanogaster. *eLife*, 5, 2016.

[1748] Chih-Wen Chu and Sergei Y Sokol. Wnt proteins can direct planar cell polarity in vertebrate ectoderm. *eLife*, 5, 2016.

[1749] Man Chu, Taotao Li, Bin Shen, Xudong Cao, Haoyu Zhong, Luqing Zhang, Fei Zhou, Wenjuan Ma, Haijuan Jiang, Pancheng Xie, Zhengzheng Liu, Ningzheng Dong, Ying Xu, Yun Zhao, Guoqiang Xu, Peirong Lu, Jincai Luo, Qingyu Wu, Kari Alitalo, Gou Young Koh, Ralf H Adams, and Yulong He. Angiopoietin receptor Tie2 is required for vein specification and maintenance via regulating COUP-TFII. *eLife*, 5, 2016.

[1750] Marian Chuang, Tiffany I Hsiao, Amy Tong, Suhong Xu, and Andrew D Chisholm. DAPK interacts with Patronin and the microtubule cytoskeleton in epidermal development and wound repair. *eLife*, 5, 2016.

[1751] Vladimir Chubanov, Silvia Ferioli, Annika Wisnowsky, David G Simmons, Christin Leitzinger, Claudia Einer, Wenke Jonas, Yuriy Shymkiv, Harald Bartsch, Attila Braun, Banu Akdogan, Lorenz Mittermeier, Ludmila Sytik, Friedrich Torben, Vindi Jurinovic, Emiel PC van der Vorst, Christian Weber, önder A Yildirim, Karl Sotlar, Annette Schürmann, Susanna Zierler, Hans Zischka, Alexey G Ryazanov, and Thomas Gudermann. Epithelial magnesium transport by TRPM6 is essential for prenatal development and adult survival. *eLife*, 5, 2016.

[1752] Ho-Ryun Chung, Chao Xu, Alisa Fuchs, Andreas Mund, Martin Lange, Hannah Staege, Tobias Schubert, Chuanbing Bian, Ilona Dunkel, Anton Eberharter, Catherine Regnard, Henrike Klinker, David Meierhofer, Luca Cozzuto, Andreas Winterpacht, Luciano Di Croce, Jinrong Min, Hans Will, and Sarah Kinkley. PHF13 is a molecular reader and transcriptional co-regulator of H3K4me2/3. *eLife*, 5, 2016.

[1753] Onur Cizmecioglu, Jing Ni, Shaozhen Xie, Jean J Zhao, and Thomas M Roberts. Rac1-mediated membrane raft localization of PI3K/p110$\beta$ is required for its activation by GPCRs or PTEN loss. *eLife*, 5, 2016.

[1754] Erik Clark and Michael Akam. Odd-paired controls frequency doubling in Drosophila segmentation by altering the pair-rule gene regulatory network. *eLife*, 5, 2016.

[1755] Natalie M Clark, Elizabeth Hinde, Cara M Winter, Adam P Fisher, Giuseppe Crosti, Ikram Blilou, Enrico Gratton, Philip N Benfey, and Rosangela Sozzani. Tracking transcription factor mobility and interaction in Arabidopsis roots with fluorescence correlation spectroscopy. *eLife*, 5, 2016.

[1756] Paul A Clarke, Maria-Jesus Ortiz-Ruiz, Robert TePoele, Olajumoke Adeniji-Popoola, Gary Box, Will Court, Stephanie Czasch, Samer El Bawab, Christina Esdar, Ken Ewan, Sharon Gowan, Alexis De Haven Brandon, Phillip Hewitt, Stephen M Hobbs, Wolfgang Kaufmann, Aurélie Mallinger, Florence Raynaud, Toby Roe, Felix Rohdich, Kai Schiemann, Stephanie Simon, Richard Schneider, Melanie Valenti, Stefan Weigt, Julian Blagg, Andree Blaukat, Trevor C Dale, Suzanne A Eccles, Stefan Hecht, Klaus Urbahns, Paul Workman, and Dirk Wienke. Assessing the mechanism and therapeutic potential of modulators of the human Mediator complex-associated protein kinases. *eLife*, 5, 2016.

[1757] Shaun L Cloherty, Nicholas J Hughes, Markus A Hietanen, Partha S Bhagavatula, Geoffrey J Goodhill, and Michael R Ibbotson. Sensory experience modifies feature map relationships in visual cortex. *eLife*, 5, 2016.

[1758] Luca Cocchi, Martin V Sale, Leonardo L Gollo, Peter T Bell, Vinh T Nguyen, Andrew Zalesky, Michael Breakspear, and Jason B Mattingley. A hierarchy of timescales explains distinct effects of local inhibition of primary visual cortex and frontal eye fields. *eLife*, 5, 2016.

[1759] James J Collins, George R Wendt, Harini Iyer, and Phillip A Newmark. Stem cell progeny contribute to the schistosome host-parasite interface. *eLife*, 5, 2016.

[1760] Kevin M Collins, Addys Bode, Robert W Fernandez, Jessica E Tanis, Jacob C Brewer, Matthew S Creamer, and Michael R Koelle. Activity of the C. elegans egg-laying behavior circuit is controlled by competing activation and feedback inhibition. *eLife*, 5, 2016.

[1761] Andrew R Conery, Richard C Centore, Adrianne Neiss, Patricia J Keller, Shivangi Joshi, Kerry L Spillane, Peter Sandy, Charlie Hatton, Eneida Pardo, Laura Zawadzke, Archana Bommi-Reddy, Karen E Gascoigne, Barbara M Bryant, Jennifer A Mertz, and Robert J Sims. Bromodomain inhibition of the transcriptional coactivators CBP/EP300 as a therapeutic strategy to target the IRF4 network in multiple myeloma. *eLife*, 5, 2016.

[1762] Erin G Conlon, Lei Lu, Aarti Sharma, Takashi Yamazaki, Timothy Tang, Neil A Shneider, and James L Manley. The C9ORF72 GGGGCC expansion forms RNA G-quadruplex inclusions and sequesters hnRNP H to disrupt splicing in ALS brains. *eLife*, 5, 2016.

[1763] Richard B Cooley, John P O'Donnell, and Holger Sondermann. Coincidence detection and bi-directional transmembrane signaling control a bacterial second messenger receptor. *eLife*, 5, 2016.

[1764] Deani L Cooper and Susan T Lovett. Recombinational branch migration by the RadA/Sms paralog of RecA in Escherichia coli. *eLife*, 5, 2016.

[1765] Verónica María Corrales-Carvajal, Aldo A Faisal, and Carlos Ribeiro. Internal states drive nutrient homeostasis by modulating exploration-exploitation trade-off. *eLife*, 5, 2016.

[1766] Susana S Correia, Anna G McGrath, Allison Lee, Ann M Graybiel, and Ki A Goosens. Amygdala-ventral striatum circuit activation decreases long-term fear. *eLife*, 5, 2016.

[1767] Adam M Corrigan, Edward Tunnacliffe, Danielle Cannon, and Jonathan R Chubb. A continuum model of transcriptional bursting. *eLife*, 5, 2016.

[1768] Anthony J Covarrubias, Halil Ibrahim Aksoylar, Jiujiu Yu, Nathaniel W Snyder, Andrew J Worth, Shankar S Iyer, Jiawei Wang, Issam Ben-Sahra, Vanessa Byles, Tiffany Polynne-Stapornkul, Erika C Espinosa, Dudley Lamming, Brendan D Manning, Yijing Zhang, Ian A Blair, and Tiffany Horng. Akt-mTORC1 signaling regulates Acly to integrate metabolic input to control of macrophage activation. *eLife*, 5, 2016.

[1769] Scott M Coyle and Wendell A Lim. Mapping the functional versatility and fragility of Ras GTPase signaling circuits through in vitro network reconstitution. *eLife*, 5, 2016.

[1770] Dina Cramer, Luis Serrano, and Martin H Schaefer. A network of epigenetic modifiers and DNA repair genes controls tissue-specific copy number alteration preference. *eLife*, 5, 2016.

[1771] C Jeremy Craven. Evaluation of predictions of the stochastic model of organelle production based on exact distributions. *eLife*, 5, 2016.

[1772] Oliver Crawley, Consuelo Barroso, Sarah Testori, Nuria Ferrandiz, Nicola Silva, Maikel Castellano-Pozo, Angel Luis Jaso-Tamame, and Enrique Martinez-Perez. Cohesin-interacting protein WAPL-1 regulates meiotic chromosome structure and cohesion by antagonizing specific cohesin complexes. *eLife*, 5, 2016.

[1773] Frank Criscione, Yumin Qi, and Zhijian Tu. GUY1 confers complete female lethality and is a strong candidate for a male-determining factor in Anopheles stephensi. *eLife*, 5, 2016.

[1774] Yihui Cui, Ilya Prokin, Hao Xu, Bruno Delord, Stephane Genet, Laurent Venance, and Hugues Berry. Endocannabinoid dynamics gate spike-timing dependent depression and potentiation. *eLife*, 5, 2016.

[1775] Yuwei Cui, Yanbin V Wang, Silvia J H Park, Jonathan B Demb, and Daniel A Butts. Divisive suppression explains high-precision firing and contrast adaptation in retinal ganglion cells. *eLife*, 5, 2016.

[1776] Ko W Currie, Alyssa M Molinaro, and Bret J Pearson. Neuronal sources of hedgehog modulate neurogenesis in the adult planarian brain. *eLife*, 5, 2016.

[1777] Amy J Curwin, Nathalie Brouwers, Manuel Alonso Y Adell, David Teis, Gabriele Turacchio, Seetharaman Parashuraman, Paolo Ronchi, and Vivek Malhotra. ESCRT-III drives the final stages of CUPS maturation for unconventional protein secretion. *eLife*, 5, 2016.

[1778] Yasin F Dagdas, Khaoula Belhaj, Abbas Maqbool, Angela Chaparro-Garcia, Pooja Pandey, Benjamin Petre, Nadra Tabassum, Neftaly Cruz-Mireles, Richard K Hughes, Jan Sklenar, Joe Win, Frank Menke, Kim Findlay, Mark J Banfield, Sophien Kamoun, and Tolga O Bozkurt. An effector of the Irish potato famine pathogen antagonizes a host autophagy cargo receptor. *eLife*, 5, 2016.

[1779] Giuseppe D'Agostino, David J Lyons, Claudia Cristiano, Luke K Burke, Joseph C Madara, John N Campbell, Ana Paula Garcia, Benjamin B Land, Bradford B Lowell, Ralph J Dileone, and Lora K Heisler. Appetite controlled by a cholecystokinin nucleus of the solitary tract to hypothalamus neurocircuit. *eLife*, 5, 2016.

[1780] Chiraj K Dalal, Ignacio A Zuleta, Kaitlin F Mitchell, David R Andes, Hana El-Samad, and Alexander D Johnson. Transcriptional rewiring over evolutionary timescales changes quantitative and qualitative properties of gene expression. *eLife*, 5, 2016.

[1781] Corey M Dambacher, Evan J Worden, Mark A Herzik, Andreas Martin, and Gabriel C Lander. Atomic structure of the 26S proteasome lid reveals the mechanism of deubiquitinase inhibition. *eLife*, 5, 2016.

[1782] Nicolas Damond, Fabrizio Thorel, Julie S Moyers, Maureen J Charron, Patricia M Vuguin, Alvin C Powers, and Pedro L Herrera. Blockade of glucagon signaling prevents or reverses diabetes onset only if residual $\beta$-cells persist. *eLife*, 5, 2016.

[1783] Hod Dana, Boaz Mohar, Yi Sun, Sujatha Narayan, Andrew Gordus, Jeremy P Hasseman, Getahun Tsegaye, Graham T Holt, Amy Hu, Deepika Walpita, Ronak Patel, John J Macklin, Cornelia I Bargmann, Misha B Ahrens, Eric R Schreiter, Vivek Jayaraman, Loren L Looger, Karel Svoboda, and Douglas S Kim. Sensitive red protein calcium indicators for imaging neural activity. *eLife*, 5, 2016.

[1784] Radostin Danev and Wolfgang Baumeister. Cryo-EM single particle analysis with the Volta phase plate. *eLife*, 5, 2016.

[1785] Benjamin Dann, Jonathan A Michaels, Stefan Schaffelhofer, and Hansjörg Scherberger. Uniting functional network topology and oscillations in the fronto-parietal single unit network of behaving primates. *eLife*, 5, 2016.

[1786] Samir Das, John Gilchrist, Frank Bosmans, and Filip Van Petegem. Binary architecture of the Nav1.2-$\beta$2 signaling complex. *eLife*, 5, 2016.

[1787] Matthew D Daugherty, Aaron M Schaller, Adam P Geballe, and Harmit S Malik. Evolution-guided functional analyses reveal diverse antiviral specificities encoded by IFIT1 genes in mammals. *eLife*, 5, 2016.

[1788] Geoffry A Davis, Atsuko Kanazawa, Mark Aurel Schöttler, Kaori Kohzuma, John E Froehlich, A William Rutherford, Mio Satoh-Cruz, Deepika Minhas, Stefanie Tietz, Amit Dhingra, and David M Kramer. Limitations to photosynthesis by proton motive force-induced photosystem II photodamage. *eLife*, 5, 2016.

[1789] Saralin Davis, Juan Wang, Ming Zhu, Kyle Stahmer, Ramya Lakshminarayan, Majid Ghassemian, Yu Jiang, Elizabeth A Miller, and Susan Ferro-Novick. Sec24 phosphorylation regulates autophagosome abundance during nutrient deprivation. *eLife*, 5, 2016.

[1790] Bart-Jan de Kreuk, Alexandre R Gingras, James DR Knight, Jian J Liu, Anne-Claude Gingras, and Mark H Ginsberg. Heart of glass anchors Rasip1 at endothelial cell-cell junctions to support vascular integrity. *eLife*, 5, 2016.

[1791] Aimee M Deaton, Mariluz Gómez-Rodríguez, Jakub Mieczkowski, Michael Y Tolstorukov, Sharmistha Kundu, Ruslan I Sadreyev, Lars ET Jansen, and Robert E Kingston. Enhancer regions show high histone H3.3 turnover that changes during differentiation. *eLife*, 5, 2016.

[1792] Paul G DeCaen, Xiaowen Liu, Sunday Abiria, and David E Clapham. Atypical calcium regulation of the PKD2-L1 polycystin ion channel. *eLife*, 5, 2016.

[1793] Marc Deffains, Liliya Iskhakova, Shiran Katabi, Suzanne N Haber, Zvi Israel, and Hagai Bergman. Subthalamic, not striatal, activity correlates with basal ganglia downstream activity in normal and parkinsonian monkeys. *eLife*, 5, 2016.

[1794] Rowena DeJesus, Francesca Moretti, Gregory McAllister, Zuncai Wang, Phil Bergman, Shanming Liu, Elizabeth Frias, John Alford, John S Reece-Hoyes, Alicia Lindeman, Jennifer Kelliher, Carsten Russ, Judith Knehr, Walter Carbone, Martin Beibel, Guglielmo Roma, Aylwin Ng, John A Tallarico, Jeffery A Porter, Ramnik J Xavier, Craig Mickanin, Leon O Murphy, Gregory R Hoffman, and Beat Nyfeler. Functional CRISPR screening identifies the ufmylation pathway as a regulator of SQSTM1/p62. *eLife*, 5, 2016.

[1795] Brian M Dekleva, Pavan Ramkumar, Paul A Wanda, Konrad P Kording, and Lee E Miller. Uncertainty leads to persistent effects on reach representations in dorsal premotor cortex. *eLife*, 5, 2016.

[1796] Francesco della Gatta, Francesca Garbarini, Guglielmo Puglisi, Antonella Leonetti, Annamaria Berti, and Paola Borroni. Decreased motor cortex excitability mirrors own hand disembodiment during the rubber hand illusion. *eLife*, 5, 2016.

[1797] Rebecca Delventhal and John R Carlson. Bitter taste receptors confer diverse functions to neurons. *eLife*, 5, 2016.

[1798] Beatrice Demarchi, Shaun Hall, Teresa Roncal-Herrero, Colin L Freeman, Jos Woolley, Molly K Crisp, Julie Wilson, Anna Fotakis, Roman Fischer, Benedikt M Kessler, Rosa Rakownikow Jersie-Christensen, Jesper V Olsen, James Haile, Jessica Thomas, Curtis W Marean, John Parkington, Samantha Presslee, Julia Lee-Thorp, Peter Ditchfield, Jacqueline F Hamilton, Martyn W Ward, Chunting Michelle Wang, Marvin D Shaw, Terry Harrison, Manuel Domínguez-Rodrigo, Ross DE MacPhee, Amandus Kwekason, Michaela Ecker, Liora Kolska Horwitz, Michael Chazan, Roland Kröger, Jane Thomas-Oates, John H Harding, Enrico Cappellini, Kirsty Penkman, and Matthew J Collins. Protein sequences bound to mineral surfaces persist into deep time. *eLife*, 5, 2016.

[1799] F Esra Demircioglu, Brian A Sosa, Jessica Ingram, Hidde L Ploegh, and Thomas U Schwartz. Structures of TorsinA and its disease-mutant complexed with an activator reveal the molecular basis for primary dystonia. *eLife*, 5, 2016.

[1800] Stuti K Desai, Ricksen S Winardhi, Saravanan Periasamy, Michal M Dykas, Yan Jie, and Linda J Kenney. The horizontally-acquired response regulator SsrB drives a Salmonella lifestyle switch by relieving biofilm silencing. *eLife*, 5, 2016.

[1801] Lorena Deuker, Jacob LS Bellmund, Tobias Navarro Schröder, and Christian F Doeller. An event map of memory space in the hippocampus. *eLife*, 5, 2016.

[1802] Rebecca Devlin, Catarina A Marques, Daniel Paape, Marko Prorocic, Andrea C Zurita-Leal, Samantha J Campbell, Craig Lapsley, Nicholas Dickens, and Richard McCulloch. Mapping replication dynamics in Trypanosoma brucei reveals a link with telomere transcription and antigenic variation. *eLife*, 5, 2016.

[1803] Nidhi Sharma Dey, Parvathy Ramesh, Mayank Chugh, Sudip Mandal, and Lolitika Mandal. Dpp dependent Hematopoietic stem cells give rise to Hh dependent blood progenitors in larval lymph gland of Drosophila. *eLife*, 5, 2016.

[1804] Carole Deyts, Mary Clutter, Stacy Herrera, Natalia Jovanovic, Anna Goddi, and Angèle T Parent. Loss of presenilin function is associated with a selective gain of APP function. *eLife*, 5, 2016.

[1805] Madhurima Dhara, Antonio Yarzagaray, Mazen Makke, Barbara Schindeldecker, Yvonne Schwarz, Ahmed Shaaban, Satyan Sharma, Rainer A Böckmann, Manfred Lindau, Ralf Mohrmann, and Dieter Bruns. v-SNARE transmembrane domains function as catalysts for vesicle fusion. *eLife*, 5, 2016.

[1806] Madhur S Dhingra, Jean Artois, Timothy P Robinson, Catherine Linard, Celia Chaiban, Ioannis Xenarios, Robin Engler, Robin Liechti, Dmitri Kuznetsov, Xiangming Xiao, Sophie Von Dobschuetz, Filip Claes, Scott H Newman, Gwenaëlle Dauphin, and Marius Gilbert. Global mapping of highly pathogenic avian influenza H5N1 and H5Nx clade 2.3.4.4 viruses with spatial cross-validation. *eLife*, 5, 2016.

[1807] Rachael Di Santo, Soufiane Aboulhouda, and David E Weinberg. The fail-safe mechanism of post-transcriptional silencing of unspliced HAC1 mRNA. *eLife*, 5, 2016.

[1808] Maria Diamantaki, Markus Frey, Philipp Berens, Patricia Preston-Ferrer, and Andrea Burgalossi. Sparse activity of identified dentate granule cells during spatial exploration. *eLife*, 5, 2016.

[1809] William E Diehl, Nirali Patel, Kate Halm, and Welkin E Johnson. Tracking interspecies transmission and long-term evolution of an ancient retrovirus using the genomes of modern mammals. *eLife*, 5, 2016.

[1810] Margarita C Dinamarca, Francesca Guzzetti, Anna Karpova, Dmitry Lim, Nico Mitro, Stefano Musardo, Manuela Mellone, Elena Marcello, Jennifer Stanic, Tanmoy Samaddar, Adeline Burguière, Antonio Caldarelli, Armando A Genazzani, Julie Perroy, Laurent Fagni, Pier Luigi Canonico, Michael R Kreutz, Fabrizio Gardoni, and Monica Di Luca. Ring finger protein 10 is a novel synaptonuclear messenger encoding activation of NMDA receptors in hippocampus. *eLife*, 5, 2016.

[1811] Johnny Phong Do, Min Xu, Seung-Hee Lee, Wei-Cheng Chang, Siyu Zhang, Shinjae Chung, Tyler J Yung, Jiang Lan Fan, Kazunari Miyamichi, Liqun Luo, and Yang Dan. Cell type-specific long-range connections of basal forebrain circuit. *eLife*, 5, 2016.

[1812] Daniel Dominguez, Yi-Hsuan Tsai, Robert Weatheritt, Yang Wang, Benjamin J Blencowe, and Zefeng Wang. An extensive program of periodic alternative splicing linked to cell cycle progression. *eLife*, 5, 2016.

[1813] Eric T Domyan, Zev Kronenberg, Carlos R Infante, Anna I Vickrey, Sydney A Stringham, Rebecca Bruders, Michael W Guernsey, Sungdae Park, Jason Payne, Robert B Beckstead, Gabrielle Kardon, Douglas B Menke, Mark Yandell, and Michael D Shapiro. Molecular shifts in limb identity underlie development of feathered feet in two domestic avian species. *eLife*, 5, 2016.

[1814] Xintong Dong, Hui Chiu, Yeonhee Jenny Park, Wei Zou, Yan Zou, Engin özkan, Chieh Chang, and Kang Shen. Precise regulation of the guidance receptor DMA-1 by KPC-1/Furin instructs dendritic branching decisions. *eLife*, 5, 2016.

[1815] Yedidyah Dordek, Daniel Soudry, Ron Meir, and Dori Derdikman. Extracting grid cell characteristics from place cell inputs using non-negative principal component analysis. *eLife*, 5, 2016.

[1816] James R Dotimas, Austin W Lee, Angela B Schmider, Shannon H Carroll, Anu Shah, Julide Bilen, Kayla R Elliott, Ronald B Myers, Roy J Soberman, Jun Yoshioka, and Richard T Lee. Diabetes regulates fructose absorption through thioredoxin-interacting protein. *eLife*, 5, 2016.

[1817] Joanna M Dragich, Takaaki Kuwajima, Megumi Hirose-Ikeda, Michael S Yoon, Evelien Eenjes, Joan R Bosco, Leora M Fox, Alf H Lystad, Tinmarla F Oo, Olga Yarygina, Tomohiro Mita, Satoshi Waguri, Yoshinobu Ichimura, Masaaki Komatsu, Anne Simonsen, Robert E Burke, Carol A Mason, and Ai Yamamoto. Autophagy linked FYVE (Alfy/WDFY3) is required for establishing neuronal connectivity in the mammalian brain. *eLife*, 5, 2016.

[1818] Rinaldo David D'Souza, Andrew Max Meier, Pawan Bista, Quanxin Wang, and Andreas Burkhalter. Recruitment of inhibition and excitation across mouse visual cortex depends on the hierarchy of interconnecting areas. *eLife*, 5, 2016.

[1819] Eun Jo Du, Tae Jung Ahn, Xianlan Wen, Dae-Won Seo, Duk L Na, Jae Young Kwon, Myunghwan Choi, Hyung-Wook Kim, Hana Cho, and KyeongJin Kang. Nucleophile sensitivity of Drosophila TRPA1 underlies light-induced feeding deterrence. *eLife*, 5, 2016.

[1820] Wen Du, Maoge Zhou, Wei Zhao, Dongwan Cheng, Lifen Wang, Jingze Lu, Eli Song, Wei Feng, Yanhong Xue, Pingyong Xu, and Tao Xu. HID-1 is required for homotypic fusion of immature secretory granules during maturation. *eLife*, 5, 2016.

[1821] Laurence Dubois, Jean-Louis Frendo, Hélène Chanut-Delalande, Michèle Crozatier, and Alain Vincent. Genetic dissection of the Transcription Factor code controlling serial specification of muscle identities in Drosophila. *eLife*, 5, 2016.

[1822] Christian Duellberg, Nicholas I Cade, David Holmes, and Thomas Surrey. The size of the EB cap determines instantaneous microtubule stability. *eLife*, 5, 2016.

[1823] Timothy W Dunn, Yu Mu, Sujatha Narayan, Owen Randlett, Eva A Naumann, Chao-Tsung Yang, Alexander F Schier, Jeremy Freeman, Florian Engert, and Misha B Ahrens. Brain-wide mapping of neural activity controlling zebrafish exploratory locomotion. *eLife*, 5, 2016.

[1824] Agustina D'Urso, Yoh-hei Takahashi, Bin Xiong, Jessica Marone, Robert Coukos, Carlo Randise-Hinchliff, Ji-Ping Wang, Ali Shilatifard, and Jason H Brickner. Set1/COMPASS and Mediator are repurposed to promote epigenetic transcriptional memory. *eLife*, 5, 2016.

[1825] Hany KM Dweck, Shimaa AM Ebrahim, Mohammed A Khallaf, Christopher Koenig, Abu Farhan, Regina Stieber, Jerrit Weißflog, AleŠ SvatoŠ, Ewald Grosse-Wilde, Markus Knaden, and Bill S Hansson. Olfactory channels associated with the Drosophila maxillary palp mediate short- and long-range attraction. *eLife*, 5, 2016.

[1826] Chiara M Eandi, Hugo Charles Messance, Sébastien Augustin, Elisa Dominguez, Sophie Lavalette, Valérie Forster, Shulong Justin Hu, Lourdes Siquieros, Cheryl Mae Craft, José-Alain Sahel, Ramin Tadayoni, Michel Paques, Xavier Guillonneau, and Florian Sennlaub. Subretinal mononuclear phagocytes induce cone segment loss via IL-1$\beta$. *eLife*, 5, 2016.

[1827] Michael N Economo, Nathan G Clack, Luke D Lavis, Charles R Gerfen, Karel Svoboda, Eugene W Myers, and Jayaram Chandrashekar. A platform for brain-wide imaging and reconstruction of individual neurons. *eLife*, 5, 2016.

[1828] James R Edgar, Paul T Manna, Shinichi Nishimura, George Banting, and Margaret S Robinson. Tetherin is an exosomal tether. *eLife*, 5, 2016.

[1829] Dawn Z Eichenfield, Ty Dale Troutman, Verena M Link, Michael T Lam, Han Cho, David Gosselin, Nathanael J Spann, Hanna P Lesch, Jenhan Tao, Jun Muto, Richard L Gallo, Ronald M Evans, and Christopher K Glass. Tissue damage drives co-localization of NF-$\kappa$B, Smad3, and Nrf2 to direct Rev-erb sensitive wound repair in mouse macrophages. *eLife*, 5, 2016.

[1830] Stephen W Eichhorn, Alexander O Subtelny, Iva Kronja, Jamie C Kwasnieski, Terry L Orr-Weaver, and David P Bartel. mRNA poly(A)-tail changes specified by deadenylation broadly reshape translation in Drosophila oocytes and early embryos. *eLife*, 5, 2016.

[1831] Raya Eilam, Rina Aharoni, Ruth Arnon, and Rafael Malach. Astrocyte morphology is confined by cortical functional boundaries in mammals ranging from mice to human. *eLife*, 5, 2016.

[1832] Assaf Elazar, Jonathan Weinstein, Ido Biran, Yearit Fridman, Eitan Bibi, and Sarel Jacob Fleishman. Mutational scanning reveals the determinants of protein insertion and association energetics in the plasma membrane. *eLife*, 5, 2016.

[1833] Vegard Eldholm, Adrien Rieux, Johana Monteserin, Julia Montana Lopez, Domingo Palmero, Beatriz Lopez, Viviana Ritacco, Xavier Didelot, and Francois Balloux. Impact of HIV co-infection on the evolution and transmission of multidrug-resistant tuberculosis. *eLife*, 5, 2016.

[1834] Ellen N Elliott, Karyn L Sheaffer, and Klaus H Kaestner. The 'de novo' DNA methyltransferase Dnmt3b compensates the Dnmt1-deficient intestinal epithelium. *eLife*, 5, 2016.

[1835] Uchenna Emechebe, Pavan Kumar P, Julian M Rozenberg, Bryn Moore, Ashley Firment, Tooraj Mirshahi, and Anne M Moon. T-box3 is a ciliary protein and regulates stability of the Gli3 transcription factor to control digit number. *eLife*, 5, 2016.

[1836] David Enard, Le Cai, Carina Gwennap, and Dmitri A Petrov. Viruses are a dominant driver of protein adaptation in mammals. *eLife*, 5, 2016.

[1837] Elif Engin, Kiersten S Smith, Yudong Gao, David Nagy, Rachel A Foster, Evgeny Tsvetkov, Ruth Keist, Florence Crestani, Jean-Marc Fritschy, Vadim Y Bolshakov, Mihaly Hajos, Scott A Heldt, and Uwe Rudolph. Modulation of anxiety and fear via distinct intrahippocampal circuits. *eLife*, 5, 2016.

[1838] Renske Erion, Anna N King, Gang Wu, John B Hogenesch, and Amita Sehgal. Neural clocks and Neuropeptide F/Y regulate circadian gene expression in a peripheral metabolic tissue. *eLife*, 5, 2016.

[1839] Cihan Erkut, Vamshidhar R Gade, Sunil Laxman, and Teymuras V Kurzchalia. The glyoxylate shunt is essential for desiccation tolerance in C. elegans and budding yeast. *eLife*, 5, 2016.

[1840] Christina Ernst, Jeremy Pike, Sarah J Aitken, Hannah K Long, Nils Eling, Lovorka Stojic, Michelle C Ward, Frances Connor, Timothy F Rayner, Margus Lukk, Robert J Klose, Claudia Kutter, and Duncan T Odom. Successful transmission and transcriptional deployment of a human chromosome via mouse male meiosis. *eLife*, 5, 2016.

[1841] Joana Esteves de Lima, Marie-Ange Bonnin, Carmen Birchmeier, and Delphine Duprez. Muscle contraction is required to maintain the pool of muscle progenitors via YAP and NOTCH during fetal myogenesis. *eLife*, 5, 2016.

[1842] Raphaël Etournay, Matthias Merkel, Marko Popović, Holger Brandl, Natalie A Dye, Benoît Aigouy, Guillaume Salbreux, Suzanne Eaton, and Frank Jülicher. TissueMiner: A multiscale analysis toolkit to quantify how cellular processes create tissue dynamics. *eLife*, 5, 2016.

[1843] Gilad D Evrony, Eunjung Lee, Peter J Park, and Christopher A Walsh. Resolving rates of mutation in the brain using single-neuron genomics. *eLife*, 5, 2016.

[1844] Michelle C Faits, Chunmeng Zhang, Florentina Soto, and Daniel Kerschensteiner. Dendritic mitochondria reach stable positions during circuit development. *eLife*, 5, 2016.

[1845] MalariaGEN Plasmodium falciparum Community Project. Genomic epidemiology of artemisinin resistant malaria. *eLife*, 5:e08714, mar 2016.

[1846] Jill Elaine Falk, Dai Tsuchiya, Jolien Verdaasdonk, Soni Lacefield, Kerry Bloom, and Angelika Amon. Spatial signals link exit from mitosis to spindle position. *eLife*, 5, 2016.

[1847] Ferric C Fang, Anthony Bowen, and Arturo Casadevall. NIH peer review percentile scores are poorly predictive of grant productivity. *eLife*, 5, 2016.

[1848] Qianglin Fang, Ping Chen, Mingzhu Wang, Junnan Fang, Na Yang, Guohong Li, and Rui-Ming Xu. Human cytomegalovirus IE1 protein alters the higher-order chromatin structure by targeting the acidic patch of the nucleosome. *eLife*, 5, 2016.

[1849] Sean W Fanning, Christopher G Mayne, Venkatasubramanian Dharmarajan, Kathryn E Carlson, Teresa A Martin, Scott J Novick, Weiyi Toy, Bradley Green, Srinivas Panchamukhi, Benita S Katzenellenbogen, Emad Tajkhorshid, Patrick R Griffin, Yang Shen, Sarat Chandarlapaty, John A Katzenellenbogen, and Geoffrey L Greene. Estrogen receptor alpha somatic mutations Y537S and D538G confer breast cancer endocrine resistance by stabilizing the activating function-2 binding conformation. *eLife*, 5, 2016.

[1850] Jordan S Farrell, Ismael Gaxiola-Valdez, Marshal D Wolff, Laurence S David, Haruna I Dika, Bryce L Geeraert, X Rachel Wang, Shaily Singh, Simon C Spanswick, Jeff F Dunn, Michael C Antle, Paolo Federico, and G Campbell Teskey. Postictal behavioural impairments are due to a severe prolonged hypoperfusion/hypoxia event that is COX-2 dependent. *eLife*, 5, 2016.

[1851] Olivia K Faull, Mark Jenkinson, Martyn Ezra, and Kyle TS Pattinson. Conditioned respiratory threat in the subdivisions of the human periaqueductal gray. *eLife*, 5, 2016.

[1852] Karen Featherstone, Kirsty Hey, Hiroshi Momiji, Anne V McNamara, Amanda L Patist, Joanna Woodburn, David G Spiller, Helen C Christian, Alan S McNeilly, John J Mullins, Bärbel F Finkenstädt, David A Rand, Michael RH White, and Julian RE Davis. Spatially coordinated dynamic gene transcription in living pituitary tissue. *eLife*, 5, 2016.

[1853] Alison F Feder, Soo-Yon Rhee, Susan P Holmes, Robert W Shafer, Dmitri A Petrov, and Pleuni S Pennings. More effective drugs lead to harder selective sweeps in the evolution of drug resistance in HIV-1. *eLife*, 5, 2016.

[1854] Daniel L Felch, Arseny S Khakhalin, and Carlos D Aizenman. Multisensory integration in the developing tectum is constrained by the balance of excitation and inhibition. *eLife*, 5, 2016.

[1855] MatyáŠ Fendrych, Jeffrey Leung, and Jiří Friml. TIR1/AFB-Aux/IAA auxin perception mediates rapid cell wall acidification and growth of Arabidopsis hypocotyls. *eLife*, 5, 2016.

[1856] Delphine Fessart, Charlotte Domblides, Tony Avril, Leif A Eriksson, Hugues Begueret, Raphael Pineau, Camille Malrieux, Nathalie Dugot-Senant, Carlo Lucchesi, Eric Chevet, and Frederic Delom. Secretion of protein disulphide isomerase AGR2 confers tumorigenic properties. *eLife*, 5, 2016.

[1857] Oliver Fiehn, Megan Reed Showalter, and Christine E Schaner-Tooley. Registered report: The common feature of leukemia-associated IDH1 and IDH2 mutations is a neomorphic enzyme activity converting alpha-ketoglutarate to 2-hydroxyglutarate. *eLife*, 5, 2016.

[1858] Duarte D Figueiredo, Rita A Batista, Pawel J Roszak, Lars Hennig, and Claudia Köhler. Auxin production in the endosperm drives seed coat development in Arabidopsis. *eLife*, 5, 2016.

[1859] Stephen N Floor and Jennifer A Doudna. Tunable protein synthesis by transcript isoforms in human cells. *eLife*, 5, 2016.

[1860] Oscar Flórez-Vargas, Andy Brass, George Karystianis, Michael Bramhall, Robert Stevens, Sheena Cruickshank, and Goran Nenadic. Bias in the reporting of sex and age in biomedical research on mouse models. *eLife*, 5, 2016.

[1861] Chii Shyang Fong, Gregory Mazo, Tuhin Das, Joshua Goodman, Minhee Kim, Brian P O'Rourke, Denisse Izquierdo, and Meng-Fu Bryan Tsou. 53BP1 and USP28 mediate p53-dependent cell cycle arrest in response to centrosome loss and prolonged mitosis. *eLife*, 5, 2016.

[1862] Ehud Fonio, Yael Heyman, Lucas Boczkowski, Aviram Gelblum, Adrian Kosowski, Amos Korman, and Ofer Feinerman. A locally-blazed ant trail achieves efficient collective navigation despite limited information. *eLife*, 5, 2016.

[1863] David Forsberg, Zachi Horn, Evangelia Tserga, Erik Smedler, Gilad Silberberg, Yuri Shvarev, Kai Kaila, Per Uhlén, and Eric Herlenius. CO2-evoked release of PGE2 modulates sighs and inspiration as demonstrated in brainstem organotypic culture. *eLife*, 5, 2016.

[1864] Jean-Baptiste Fourmann, Olexandr Dybkov, Dmitry E Agafonov, Marcel J Tauchert, Henning Urlaub, Ralf Ficner, Patrizia Fabrizio, and Reinhard Lührmann. The target of the DEAH-box NTP triphosphatase Prp43 in Saccharomyces cerevisiae spliceosomes is the U2 snRNP-intron interaction. *eLife*, 5, 2016.

[1865] Dorotea Fracchiolla, Justyna Sawa-Makarska, Bettina Zens, Anita de Ruiter, Gabriele Zaffagnini, Andrea Brezovich, Julia Romanov, Kathrin Runggatscher, Claudine Kraft, Bojan Zagrovic, and Sascha Martens. Mechanism of cargo-directed Atg8 conjugation during selective autophagy. *eLife*, 5, 2016.

[1866] Claudio A Franco, Martin L Jones, Miguel O Bernabeu, Anne-Clemence Vion, Pedro Barbacena, Jieqing Fan, Thomas Mathivet, Catarina G Fonseca, Anan Ragab, Terry P Yamaguchi, Peter V Coveney, Richard A Lang, and Holger Gerhardt. Non-canonical Wnt signalling modulates the endothelial shear stress flow sensor in vascular remodelling. *eLife*, 5, 2016.

[1867] Kai Fu, Xin Sun, Eric M Wier, Andrea Hodgson, Ryan P Hobbs, and Fengyi Wan. Sam68/KHDRBS1-dependent NF-$\kappa$B activation confers radioprotection to the colon epithelium in $\gamma$-irradiated mice. *eLife*, 5, 2016.

[1868] Kai Fu, Xin Sun, Eric M Wier, Andrea Hodgson, Yue Liu, Cynthia L Sears, and Fengyi Wan. Sam68/KHDRBS1 is critical for colon tumorigenesis by regulating genotoxic stress-induced NF-$\kappa$B activation. *eLife*, 5, 2016.

[1869] Lin Fu, Hailong Wu, Steven Y Cheng, Daming Gao, Lei Zhang, and Yun Zhao. Set7 mediated Gli3 methylation plays a positive role in the activation of Sonic Hedgehog pathway in mammals. *eLife*, 5, 2016.

[1870] Paulina Fuentes, Fei Zhou, Alexander Erban, Daniel Karcher, Joachim Kopka, and Ralph Bock. A new synthetic biology approach allows transfer of an entire metabolic pathway from a medicinal plant to a biomass crop. *eLife*, 5, 2016.

[1871] Franziska Fuhrmann, Timo Lischke, Fridolin Gross, Tobias Scheel, Laura Bauer, Khalid Wasim Kalim, Andreas Radbruch, Hanspeter Herzel, Andreas Hutloff, and Ria Baumgrass. Adequate immune response ensured by binary IL-2 and graded CD25 expression in a murine transfer model. *eLife*, 5, 2016.

[1872] Teppei Fujikawa, Carlos M Castorena, Mackenzie Pearson, Christine M Kusminski, Newaz Ahmed, Pavan K Battiprolu, Ki Woo Kim, Syann Lee, Joseph A Hill, Philipp E Scherer, William L Holland, and Joel K Elmquist. SF-1 expression in the hypothalamus is required for beneficial metabolic effects of exercise. *eLife*, 5, 2016.

[1873] Matthew J Fuxjager, Franz Goller, Annika Dirkse, Gloria D Sanin, and Sarah Garcia. Select forelimb muscles have evolved superfast contractile speed to support acrobatic social displays. *eLife*, 5, 2016.

[1874] Gilles Gadea, Nikola Arsic, Kenneth Fernandes, Alexandra Diot, Sébastien M Joruiz, Samer Abdallah, Valerie Meuray, Stéphanie Vinot, Christelle Anguille, Judit Remenyi, Marie P Khoury, Philip R Quinlan, Colin A Purdie, Lee B Jordan, Frances V Fuller-Pace, Marion de Toledo, Maïlys Cren, Alastair M Thompson, Jean-Christophe Bourdon, and Pierre Roux. TP53 drives invasion through expression of its $\delta$133p53$\beta$ variant. *eLife*, 5, 2016.

[1875] Dahai Gai, Damian Wang, Shu-Xing Li, and Xiaojiang S Chen. RETRACTED: The structure of SV40 large T hexameric helicase in complex with AT-rich origin DNA. *eLife*, 5, 2016.

[1876] Ciara M Gallagher, Carolina Garri, Erica L Cain, Kenny Kean-Hooi Ang, Christopher G Wilson, Steven Chen, Brian R Hearn, Priyadarshini Jaishankar, Andres Aranda-Diaz, Michelle R Arkin, Adam R Renslo, and Peter Walter. Ceapins are a new class of unfolded protein response inhibitors, selectively targeting the ATF6$\alpha$ branch. *eLife*, 5, 2016.

[1877] Ciara M Gallagher and Peter Walter. Ceapins inhibit ATF6$\alpha$ signaling by selectively preventing transport of ATF6$\alpha$ to the Golgi apparatus during ER stress. *eLife*, 5, 2016.

[1878] Joyonna Carrie Gamble-George, Rita Baldi, Lindsay Halladay, Adrina Kocharian, Nolan Hartley, Carolyn Grace Silva, Holly Roberts, Andre Haymer, Lawrence J Marnett, Andrew Holmes, and Sachin Patel. Cyclooxygenase-2 inhibition reduces stress-induced affective pathology. *eLife*, 5, 2016.

[1879] Liang Gao, Gülce Sila Gülcüler, Lieke Golbach, Helena Block, Alexander Zarbock, and Ana Martin-Villalba. Endothelial cell-derived CD95 ligand serves as a chemokine in induction of neutrophil slow rolling and adhesion. *eLife*, 5, 2016.

[1880] Zhaofeng Gao, Andrea A Putnam, Heath A Bowers, Ulf-Peter Guenther, Xuan Ye, Audrey Kindsfather, Angela K Hilliker, and Eckhard Jankowsky. Coupling between the DEAD-box RNA helicases Ded1p and eIF4A. *eLife*, 5, 2016.

[1881] Farid N Garas, Rahul S Shah, Eszter Kormann, Natalie M Doig, Federica Vinciati, Kouichi C Nakamura, Matthijs C Dorst, Yoland Smith, Peter J Magill, and Andrew Sharott. Secretagogin expression delineates functionally-specialized populations of striatal parvalbumin-containing interneurons. *eLife*, 5, 2016.

[1882] David M Garcia, David Dietrich, Jon Clardy, and Daniel F Jarosz. A common bacterial metabolite elicits prion-based bypass of glucose repression. *eLife*, 5, 2016.

[1883] Andrew M Garrett, Abigail LD Tadenev, Yuna T Hammond, Peter G Fuerst, and Robert W Burgess. Replacing the PDZ-interacting C-termini of DSCAM and DSCAML1 with epitope tags causes different phenotypic severity in different cell populations. *eLife*, 5, 2016.

[1884] Abigail G Garrity, Wuyang Wang, Crystal MD Collier, Sara A Levey, Qiong Gao, and Haoxing Xu. The endoplasmic reticulum, not the pH gradient, drives calcium refilling of lysosomes. *eLife*, 5, 2016.

[1885] José Ignacio Garzón, Lei Deng, Diana Murray, Sagi Shapira, Donald Petrey, and Barry Honig. A computational interactome and functional annotation for the human proteome. *eLife*, 5, 2016.

[1886] Audrey P Gasch, James Hose, Michael A Newton, Maria Sardi, Mun Yong, and Zhishi Wang. Further support for aneuploidy tolerance in wild yeast and effects of dosage compensation on gene copy-number evolution. *eLife*, 5, 2016.

[1887] Jeremie Gaveau, Bastien Berret, Dora E Angelaki, and Charalambos Papaxanthis. Direction-dependent arm kinematics reveal optimal integration of gravity cues. *eLife*, 5, 2016.

[1888] Zhiyun Ge, Bao Lin Quek, Karen L Beemon, and J Robert Hogg. Polypyrimidine tract binding protein 1 protects mRNAs from recognition by the nonsense-mediated mRNA decay pathway. *eLife*, 5, 2016.

[1889] Lena Gehre, Olivier Gorgette, Stéphanie Perrinet, Marie-Christine Prevost, Mathieu Ducatez, Amanda M Giebel, David E Nelson, Steven G Ball, and Agathe Subtil. Sequestration of host metabolism by an intracellular pathogen. *eLife*, 5, 2016.

[1890] Marie Gendrel, Emily G Atlas, and Oliver Hobert. A cellular and regulatory map of the GABAergic nervous system of C. elegans. *eLife*, 5, 2016.

[1891] Yong Geng, Lidia Mosyak, Igor Kurinov, Hao Zuo, Emmanuel Sturchler, Tat Cheung Cheng, Prakash Subramanyam, Alice P Brown, Sarah C Brennan, Hee-chang Mun, Martin Bush, Yan Chen, Trang X Nguyen, Baohua Cao, Donald D Chang, Matthias Quick, Arthur D Conigrave, Henry M Colecraft, Patricia McDonald, and Qing R Fan. Structural mechanism of ligand activation in human calcium-sensing receptor. *eLife*, 5, 2016.

[1892] Matthew A Geramita, Shawn D Burton, and Nathan N Urban. Distinct lateral inhibitory circuits drive parallel processing of sensory information in the mammalian olfactory bulb. *eLife*, 5, 2016.

[1893] Dave T Gerrard, Andrew A Berry, Rachel E Jennings, Karen Piper Hanley, Nicoletta Bobola, and Neil A Hanley. An integrative transcriptomic atlas of organogenesis in human embryos. *eLife*, 5, 2016.

[1894] Kenneth D Gibbs, Jacob Basson, Imam M Xierali, and David A Broniatowski. Decoupling of the minority PhD talent pool and assistant professor hiring in medical school basic science departments in the US. *eLife*, 5, 2016.

[1895] Claire M Gillan, Michal Kosinski, Robert Whelan, Elizabeth A Phelps, and Nathaniel D Daw. Characterizing a psychiatric symptom dimension related to deficits in goal-directed control. *eLife*, 5, 2016.

[1896] Michael J Goard, Gerald N Pho, Jonathan Woodson, and Mriganka Sur. Distinct roles of visual, parietal, and frontal motor cortices in memory-guided sensorimotor decisions. *eLife*, 5, 2016.

[1897] Onur GökÇe, Tobias Bonhoeffer, and Volker Scheuss. Clusters of synaptic inputs on dendrites of layer 5 pyramidal cells in mouse visual cortex. *eLife*, 5, 2016.

[1898] Tal Golan, Ido Davidesco, Meir Meshulam, David M Groppe, Pierre Mégevand, Erin M Yeagle, Matthew S Goldfinger, Michal Harel, Lucia Melloni, Charles E Schroeder, Leon Y Deouell, Ashesh D Mehta, and Rafael Malach. Human intracranial recordings link suppressed transients rather than 'filling-in' to perceptual continuity across blinks. *eLife*, 5, 2016.

[1899] Marcel P Goldschen-Ohm, Vadim A Klenchin, David S White, John B Cowgill, Qiang Cui, Randall H Goldsmith, and Baron Chanda. Structure and dynamics underlying elementary ligand binding events in human pacemaking channels. *eLife*, 5, 2016.

[1900] Anton Goloborodko, Maxim V Imakaev, John F Marko, and Leonid Mirny. Compaction and segregation of sister chromatids via active loop extrusion. *eLife*, 5, 2016.

[1901] Javier A Gomez and D Thomas Rutkowski. Experimental reconstitution of chronic ER stress in the liver reveals feedback suppression of BiP mRNA expression. *eLife*, 5, 2016.

[1902] Kerry M Goodman, Masahito Yamagata, Xiangshu Jin, Seetha Mannepalli, Phinikoula S Katsamba, Göran Ahlsén, Alina P Sergeeva, Barry Honig, Joshua R Sanes, and Lawrence Shapiro. Molecular basis of sidekick-mediated cell-cell adhesion and specificity. *eLife*, 5, 2016.

[1903] Kerry Marie Goodman, Rotem Rubinstein, Chan Aye Thu, Seetha Mannepalli, Fabiana Bahna, Göran Ahlsén, Chelsea Rittenhouse, Tom Maniatis, Barry Honig, and Lawrence Shapiro. $\gamma$-Protocadherin structural diversity and functional implications. *eLife*, 5, 2016.

[1904] Aniket V Gore, Brett Athans, James R Iben, Kristin Johnson, Valya Russanova, Daniel Castranova, Van N Pham, Matthew G Butler, Lisa Williams-Simons, James T Nichols, Erica Bresciani, Bejamin Feldman, Charles B Kimmel, Paul P Liu, and Brant M Weinstein. Epigenetic regulation of hematopoiesis by DNA methylation. *eLife*, 5, 2016.

[1905] Fabian Grabenhorst, Istvan Hernadi, and Wolfram Schultz. Primate amygdala neurons evaluate the progress of self-defined economic choice sequences. *eLife*, 5, 2016.

[1906] Joshua S Greene, May Dobosiewicz, Rebecca A Butcher, Patrick T McGrath, and Cornelia I Bargmann. Regulatory changes in two chemoreceptor genes contribute to a Caenorhabditis elegans QTL for foraging behavior. *eLife*, 5, 2016.

[1907] Trever T Greene, Maria Tokuyama, Giselle M Knudsen, Michele Kunz, James Lin, Alexander L Greninger, Victor R DeFilippis, Joseph L DeRisi, David H Raulet, and Laurent Coscoy. A Herpesviral induction of RAE-1 NKG2D ligand expression occurs through release of HDAC mediated repression. *eLife*, 5, 2016.

[1908] Edward JD Greenwood, Nicholas J Matheson, Kim Wals, Dick JH van den Boomen, Robin Antrobus, James C Williamson, and Paul J Lehner. Temporal proteomic analysis of HIV infection reveals remodelling of the host phosphoproteome by lentiviral Vif variants. *eLife*, 5, 2016.

[1909] Joaquim Grego-Bessa, Joshua Bloomekatz, Pau Castel, Tatiana Omelchenko, José Baselga, and Kathryn V Anderson. The tumor suppressor PTEN and the PDK1 kinase regulate formation of the columnar neural epithelium. *eLife*, 5, 2016.

[1910] Roddy M Grieves, Emma R Wood, and Paul A Dudchenko. Place cells on a maze encode routes rather than destinations. *eLife*, 5, 2016.

[1911] Benjamin Groves, Arjun Khakhar, Cory M Nadel, Richard G Gardner, and Georg Seelig. Rewiring MAP kinases in Saccharomyces cerevisiae to regulate novel targets through ubiquitination. *eLife*, 5, 2016.

[1912] Thibaud Gruber, Klaus Zuberbühler, and Christof Neumann. Travel fosters tool use in wild chimpanzees. *eLife*, 5, 2016.

[1913] Magda Grudniewska, Stijn Mouton, Daniil Simanov, Frank Beltman, Margriet Grelling, Katrien de Mulder, Wibowo Arindrarto, Philipp M. Weissert, Stefan van der Elst, and Eugene Berezikov. Transcriptional signatures of somatic neoblasts and germline cells in Macrostomum lignano. *eLife*, 5, 2016.

[1914] Matthias Gruhn, Philipp Rosenbaum, Till Bockemühl, and Ansgar Büschges. Body side-specific control of motor activity during turning in a walking animal. *eLife*, 5, 2016.

[1915] Yuliya Gryaznova, Ayse Koca Caydasi, Gabriele Malengo, Victor Sourjik, and Gislene Pereira. A FRET-based study reveals site-specific regulation of spindle position checkpoint proteins at yeast centrosomes. *eLife*, 5, 2016.

[1916] Michaela Gschweitl, Anna Ulbricht, Christopher A Barnes, Radoslav I Enchev, Ingrid Stoffel-Studer, Nathalie Meyer-Schaller, Jatta Huotari, Yohei Yamauchi, Urs F Greber, Ari Helenius, and Matthias Peter. A SPOPL/Cullin-3 ubiquitin ligase complex regulates endocytic trafficking by targeting EPS15 at endosomes. *eLife*, 5, 2016.

[1917] Yu Gu and Jianhua Cang. Binocular matching of thalamocortical and intracortical circuits in the mouse visual cortex. *eLife*, 5, 2016.

[1918] Virginia Guarani, Claude Jardel, Dominique Chrétien, Anne Lombès, Paule Bénit, Clémence Labasse, Emmanuelle Lacène, Agnès Bourillon, Apolline Imbard, Jean-FranÇois Benoist, Imen Dorboz, Mylène Gilleron, Eric S Goetzman, Pauline Gaignard, Abdelhamid Slama, Monique Elmaleh-Bergès, Norma B Romero, Pierre Rustin, Hélène Ogier de Baulny, Joao A Paulo, J Wade Harper, and Manuel Schiff. QIL1 mutation causes MICOS disassembly and early onset fatal mitochondrial encephalopathy with liver disease. *eLife*, 5, 2016.

[1919] Isabel Guerreiro, Sandra Gitto, Ana Novoa, Julien Codourey, Thi Hanh Nguyen Huynh, Federico Gonzalez, Michel C Milinkovitch, Moises Mallo, and Denis Duboule. Reorganisation of Hoxd regulatory landscapes during the evolution of a snake-like body plan. *eLife*, 5, 2016.

[1920] Matthias Guggenmos, Gregor Wilbertz, Martin N Hebart, and Philipp Sterzer. Mesolimbic confidence signals guide perceptual learning in the absence of external feedback. *eLife*, 5, 2016.

[1921] Jordina Guillén-Boixet, Víctor Buzon, Xavier Salvatella, and Raúl Méndez. CPEB4 is regulated during cell cycle by ERK2/Cdk1-mediated phosphorylation and its assembly into liquid-like droplets. *eLife*, 5, 2016.

[1922] Jingjing Guo and Huan-Xiang Zhou. Allosteric activation of SENP1 by SUMO1 $\beta$-grasp domain involves a dock-and-coalesce mechanism. *eLife*, 5, 2016.

[1923] Xinzheng Guo, William D Snider, and Bo Chen. GSK3$\beta$ regulates AKT-induced central nervous system axon regeneration via an eIF2B$\epsilon$-dependent, mTORC1-independent pathway. *eLife*, 5, 2016.

[1924] Romi Gupta, Matteo Forloni, Malik Bisserier, Shaillay Kumar Dogra, Qiaohong Yang, and Narendra Wajapeyee. Interferon alpha-inducible protein 6 regulates NRASQ61K-induced melanomagenesis and growth. *eLife*, 5, 2016.

[1925] Juergen Haag, Alexander Arenz, Etienne Serbe, Fabrizio Gabbiani, and Alexander Borst. Complementary mechanisms create direction selectivity in the fly. *eLife*, 5, 2016.

[1926] Chloé Habermacher, Adeline Martz, Nicolas Calimet, Damien Lemoine, Laurie Peverini, Alexandre Specht, Marco Cecchini, and Thomas Grutter. Photo-switchable tweezers illuminate pore-opening motions of an ATP-gated P2X ion channel. *eLife*, 5, 2016.

[1927] Junichi Hachisuka, Kyle M Baumbauer, Yu Omori, Lindsey M Snyder, H Richard Koerber, and Sarah E Ross. Semi-intact ex vivo approach to investigate spinal somatosensory circuits. *eLife*, 5, 2016.

[1928] Travis A Hage, Yujie Sun, and Zayd M Khaliq. Electrical and Ca2+ signaling in dendritic spines of substantia nigra dopaminergic neurons. *eLife*, 5, 2016.

[1929] Sarah J Hainer, Kurtis N McCannell, Jun Yu, Ly-Sha Ee, Lihua J Zhu, Oliver J Rando, and Thomas G Fazzio. DNA methylation directs genomic localization of Mbd2 and Mbd3 in embryonic stem cells. *eLife*, 5, 2016.

[1930] Benjamin Hallier, Ronja Schiemann, Eva Cordes, Jessica Vitos-Faleato, Stefan Walter, Jürgen J Heinisch, Anders Malmendal, Achim Paululat, and Heiko Meyer. Drosophila neprilysins control insulin signaling and food intake via cleavage of regulatory peptides. *eLife*, 5, 2016.

[1931] Eileen P Hamilton, Aurélie Kapusta, Piroska E Huvos, Shelby L Bidwell, Nikhat Zafar, Haibao Tang, Michalis Hadjithomas, Vivek Krishnakumar, Jonathan H Badger, Elisabet V Caler, Carsten Russ, Qiandong Zeng, Lin Fan, Joshua Z Levin, Terrance Shea, Sarah K Young, Ryan Hegarty, Riza Daza, Sharvari Gujja, Jennifer R Wortman, Bruce W Birren, Chad Nusbaum, Jainy Thomas, Clayton M Carey, Ellen J Pritham, Cédric Feschotte, Tomoko Noto, Kazufumi Mochizuki, Romeo Papazyan, Sean D Taverna, Paul H Dear, Donna M Cassidy-Hanley, Jie Xiong, Wei Miao, Eduardo Orias, and Robert S Coyne. Structure of the germline genome of Tetrahymena thermophila and relationship to the massively rearranged somatic genome. *eLife*, 5, 2016.

[1932] Ali S Hamodi, Zhenyu Liu, and Kara G Pratt. An NMDA receptor-dependent mechanism for subcellular segregation of sensory inputs in the tadpole optic tectum. *eLife*, 5, 2016.

[1933] Erik C Hansen, Monica Ransom, Jay R Hesselberth, Nina N Hosmane, Adam A Capoferri, Katherine M Bruner, Ross A Pollack, Hao Zhang, Michael Bradley Drummond, Janet M Siliciano, Robert Siliciano, and James T Stivers. Diverse fates of uracilated HIV-1 DNA during infection of myeloid lineage cells. *eLife*, 5, 2016.

[1934] Cyril Hanus, Helene Geptin, Georgi Tushev, Sakshi Garg, Beatriz Alvarez-Castelao, Sivakumar Sambandan, Lisa Kochen, Anne-Sophie Hafner, Julian D Langer, and Erin M Schuman. Unconventional secretory processing diversifies neuronal ion channel properties. *eLife*, 5, 2016.

[1935] Bryan T Harada, William L Hwang, Sebastian Deindl, Nilanjana Chatterjee, Blaine Bartholomew, and Xiaowei Zhuang. Stepwise nucleosome translocation by RSC remodeling complexes. *eLife*, 5, 2016.

[1936] Kawssar Harb, Elia Magrinelli, Céline S Nicolas, Nikita Lukianets, Laura Frangeul, Mariel Pietri, Tao Sun, Guillaume Sandoz, Franck Grammont, Denis Jabaudon, Michèle Studer, and Christian Alfano. Area-specific development of distinct projection neuron subclasses is regulated by postnatal epigenetic modifications. *eLife*, 5, 2016.

[1937] Max E Harner, Ann-Katrin Unger, Willie JC Geerts, Muriel Mari, Toshiaki Izawa, Maria Stenger, Stefan Geimer, Fulvio Reggiori, Benedikt Westermann, and Walter Neupert. An evidence based hypothesis on the existence of two pathways of mitochondrial crista formation. *eLife*, 5, 2016.

[1938] Adam J Harrington, Aram Raissi, Kacey Rajkovich, Stefano Berto, Jaswinder Kumar, Gemma Molinaro, Jonathan Raduazzo, Yuhong Guo, Kris Loerwald, Genevieve Konopka, Kimberly M Huber, and Christopher W Cowan. MEF2C regulates cortical inhibitory and excitatory synapses and behaviors relevant to neurodevelopmental disorders. *eLife*, 5, 2016.

[1939] Robin E Harris, Linda Setiawan, Josh Saul, and Iswar K Hariharan. Localized epigenetic silencing of a damage-activated WNT enhancer limits regeneration in mature Drosophila imaginal discs. *eLife*, 5, 2016.

[1940] Joseph S Harrison, Evan M Cornett, Dennis Goldfarb, Paul A DaRosa, Zimeng M Li, Feng Yan, Bradley M Dickson, Angela H Guo, Daniel V Cantu, Lilia Kaustov, Peter J Brown, Cheryl H Arrowsmith, Dorothy A Erie, Michael B Major, Rachel E Klevit, Krzysztof Krajewski, Brian Kuhlman, Brian D Strahl, and Scott B Rothbart. Hemi-methylated DNA regulates DNA methylation inheritance through allosteric activation of H3 ubiquitylation by UHRF1. *eLife*, 5, 2016.

[1941] Bianca Hartmann, Timothy Wai, Hao Hu, Thomas MacVicar, Luciana Musante, Björn Fischer-Zirnsak, Werner Stenzel, Ralph Gräf, Lambert van den Heuvel, Hans-Hilger Ropers, Thomas F Wienker, Christoph Hübner, Thomas Langer, and Angela M Kaindl. Homozygous YME1L1 mutation causes mitochondriopathy with optic atrophy and mitochondrial network fragmentation. *eLife*, 5, 2016.

[1942] Marcus D Hartmann, Claudia T Mendler, Jens Bassler, Ioanna Karamichali, Oswin Ridderbusch, Andrei N Lupas, and Birte Hernandez Alvarez. $\alpha/\beta$ coiled coils. *eLife*, 5, 2016.

[1943] Catherine A Hartzell, Katarzyna I Jankowska, Janis K Burkhardt, and Richard S Lewis. Calcium influx through CRAC channels controls actin organization and dynamics at the immune synapse. *eLife*, 5, 2016.

[1944] Yehudit Hasin-Brumshtein, Arshad H Khan, Farhad Hormozdiari, Calvin Pan, Brian W Parks, Vladislav A Petyuk, Paul D Piehowski, Anneke Brümmer, Matteo Pellegrini, Xinshu Xiao, Eleazar Eskin, Richard D Smith, Aldons J Lusis, and Desmond J Smith. Hypothalamic transcriptomes of 99 mouse strains reveal trans eQTL hotspots, splicing QTLs and novel non-coding genes. *eLife*, 5, 2016.

[1945] Kirsten Hattermann, Henrike Gebhardt, Sebastian Krossa, Andreas Ludwig, Ralph Lucius, Janka Held-Feindt, and Rolf Mentlein. Transmembrane chemokines act as receptors in a novel mechanism termed inverse signaling. *eLife*, 5, 2016.

[1946] Julia Hatzold, Filippo Beleggia, Hannah Herzig, Janine Altmüller, Peter Nürnberg, Wilhelm Bloch, Bernd Wollnik, and Matthias Hammerschmidt. Tumor suppression in basal keratinocytes via dual non-cell-autonomous functions of a Na,K-ATPase beta subunit. *eLife*, 5, 2016.

[1947] Robbert Havekes, Alan J Park, Jennifer C Tudor, Vincent G Luczak, Rolf T Hansen, Sarah L Ferri, Vibeke M Bruinenberg, Shane G Poplawski, Jonathan P Day, Sara J Aton, Kasia Radwańska, Peter Meerlo, Miles D Houslay, George S Baillie, and Ted Abel. Sleep deprivation causes memory deficits by negatively impacting neuronal connectivity in hippocampal area CA1. *eLife*, 5, 2016.

[1948] Babette Haven, Elysia Heilig, Cristine Donham, Michael Settles, Nicole Vasilevsky, and Katherine Owen. Registered report: A chromatin-mediated reversible drug-tolerant state in cancer cell subpopulations. *eLife*, 5, 2016.

[1949] Alexander Haverkamp, Felipe Yon, Ian W Keesey, Christine Mißbach, Christopher Koenig, Bill S Hansson, Ian T Baldwin, Markus Knaden, and Danny Kessler. Hawkmoths evaluate scenting flowers with the tip of their proboscis. *eLife*, 5, 2016.

[1950] Shinichiro Hayashi, Ichiro Manabe, Yumi Suzuki, Frédéric Relaix, and Yumiko Oishi. Klf5 regulates muscle differentiation by directly targeting muscle-specific genes in cooperation with MyoD in mice. *eLife*, 5, 2016.

[1951] Didi He, Sam Hughes, Sally Vanden-Hehir, Atanas Georgiev, Kirsten Altenbach, Emma Tarrant, C Logan Mackay, Kevin J Waldron, David J Clarke, and Jon Marles-Wright. Structural characterization of encapsulated ferritin provides insight into iron storage in bacterial nanocompartments. *eLife*, 5, 2016.

[1952] Shuning He, Marc R Mansour, Mark W Zimmerman, Dong Hyuk Ki, Hillary M Layden, Koshi Akahane, Evisa Gjini, Eric D de Groh, Antonio R Perez-Atayde, Shizhen Zhu, Jonathan A Epstein, and A Thomas Look. Synergy between loss of NF1 and overexpression of MYCN in neuroblastoma is mediated by the GAP-related domain. *eLife*, 5, 2016.

[1953] Mark Hedglin, Binod Pandey, and Stephen J Benkovic. Characterization of human translesion DNA synthesis across a UV-induced DNA lesion. *eLife*, 5, 2016.

[1954] Merja Heinäniemi, Tapio Vuorenmaa, Susanna Teppo, Minna U Kaikkonen, Maria Bouvy-Liivrand, Juha Mehtonen, Henri Niskanen, Vasilios Zachariadis, Saara Laukkanen, Thomas Liuksiala, Kaisa Teittinen, and Olli Lohi. Transcription-coupled genetic instability marks acute lymphoblastic leukemia structural variation hotspots. *eLife*, 5, 2016.

[1955] Udo Heintz and Ilme Schlichting. Blue light-induced LOV domain dimerization enhances the affinity of Aureochrome 1a for its target DNA sequence. *eLife*, 5, 2016.

[1956] Richard G Held, Changliang Liu, and Pascal S Kaeser. ELKS controls the pool of readily releasable vesicles at excitatory synapses through its N-terminal coiled-coil domains. *eLife*, 5, 2016.

[1957] Sarah R Hengel, Eva Malacaria, Laura Folly da Silva Constantino, Fletcher E Bain, Andrea Diaz, Brandon G Koch, Liping Yu, Meng Wu, Pietro Pichierri, M Ashley Spies, and Maria Spies. Small-molecule inhibitors identify the RAD52-ssDNA interaction as critical for recovery from replication stress and for survival of BRCA2 deficient cells. *eLife*, 5, 2016.

[1958] Sid Henriksen, Rich Pang, and Mark Wronkiewicz. A simple generative model of the mouse mesoscale connectome. *eLife*, 5, 2016.

[1959] Nikolai Hentze, Laura Le Breton, Jan Wiesner, Georg Kempf, and Matthias P Mayer. Molecular mechanism of thermosensory function of human heat shock transcription factor Hsf1. *eLife*, 5, 2016.

[1960] Su-Jin Heo, Tristan P Driscoll, Stephen D Thorpe, Nandan L Nerurkar, Brendon M Baker, Michael T Yang, Christopher S Chen, David A Lee, and Robert L Mauck. Differentiation alters stem cell nuclear architecture, mechanics, and mechano-sensitivity. *eLife*, 5, 2016.

[1961] Lars Hering, Jamal-Eddine Bouameur, Julian Reichelt, Thomas M Magin, and Georg Mayer. Novel origin of lamin-derived cytoplasmic intermediate filaments in tardigrades. *eLife*, 5, 2016.

[1962] Alexander Heuck, Sonja Schitter-Sollner, Marcin Józef Suskiewicz, Robert Kurzbauer, Juliane Kley, Alexander Schleiffer, Pascaline Rombaut, Franz Herzog, and Tim Clausen. Structural basis for the disaggregase activity and regulation of Hsp104. *eLife*, 5, 2016.

[1963] Maarten Hilbrant, Thorsten Horn, Stefan Koelzer, and Kristen A Panfilio. The beetle amnion and serosa functionally interact as apposed epithelia. *eLife*, 5, 2016.

[1964] Emily R Hildebrandt, Michael Cheng, Peng Zhao, June H Kim, Lance Wells, and Walter K Schmidt. A shunt pathway limits the CaaX processing of Hsp40 Ydj1p and regulates Ydj1p-dependent phenotypes. *eLife*, 5, 2016.

[1965] Jennifer Hampton Hill, Eric A Franzosa, Curtis Huttenhower, and Karen Guillemin. A conserved bacterial protein induces pancreatic beta cell expansion during zebrafish development. *eLife*, 5, 2016.

[1966] Patrick Hillenbrand, Kerstin C Maier, Patrick Cramer, and Ulrich Gerland. Inference of gene regulation functions from dynamic transcriptome data. *eLife*, 5, 2016.

[1967] Arisa Hirano, Guangsen Shi, Christopher R Jones, Anna Lipzen, Len A Pennacchio, Ying Xu, William C Hallows, Thomas McMahon, Maya Yamazaki, Louis J Ptáček, and Ying-Hui Fu. A Cryptochrome 2 mutation yields advanced sleep phase in humans. *eLife*, 5, 2016.

[1968] Sungho Hong, Mario Negrello, Marc Junker, Aleksandra Smilgin, Peter Thier, and Erik De Schutter. Multiplexed coding by cerebellar Purkinje neurons. *eLife*, 5, 2016.

[1969] Iris V Hood and James M Berger. Viral hijacking of a replicative helicase loader and its implications for helicase loading control and phage replication. *eLife*, 5, 2016.

[1970] Max A Horlbeck, Luke A Gilbert, Jacqueline E Villalta, Britt Adamson, Ryan A Pak, Yuwen Chen, Alexander P Fields, Chong Yon Park, Jacob E Corn, Martin Kampmann, and Jonathan S Weissman. Compact and highly active next-generation libraries for CRISPR-mediated gene repression and activation. *eLife*, 5, 2016.

[1971] Max A Horlbeck, Lea B Witkowsky, Benjamin Guglielmi, Joseph M Replogle, Luke A Gilbert, Jacqueline E Villalta, Sharon E Torigoe, Robert Tjian, and Jonathan S Weissman. Nucleosomes impede Cas9 access to DNA in vivo and in vitro. *eLife*, 5, 2016.

[1972] Felix Horns, Christopher Vollmers, Derek Croote, Sally F Mackey, Gary E Swan, Cornelia L Dekker, Mark M Davis, and Stephen R Quake. Lineage tracing of human B cells reveals the in vivo landscape of human antibody class switching. *eLife*, 5, 2016.

[1973] Aaron A Hoskins, Margaret L Rodgers, Larry J Friedman, Jeff Gelles, and Melissa J Moore. Single molecule analysis reveals reversible and irreversible steps during spliceosome activation. *eLife*, 5, 2016.

[1974] Shauna L Houlihan, Alison A Lanctot, Yan Guo, and Yuanyi Feng. Upregulation of neurovascular communication through filamin abrogation promotes ectopic periventricular neurogenesis. *eLife*, 5, 2016.

[1975] Gerta Hoxhaj, Edward Caddye, Ayaz Najafov, Vanessa P Houde, Catherine Johnson, Kumara Dissanayake, Rachel Toth, David G Campbell, Alan R Prescott, and Carol MacKintosh. The E3 ubiquitin ligase ZNRF2 is a substrate of mTORC1 and regulates its activation by amino acids. *eLife*, 5, 2016.

[1976] Xichan Hu, Jinqiang Liu, Hyun-IK Jun, Jin-Kwang Kim, and Feng Qiao. Multi-step coordination of telomerase recruitment in fission yeast through two coupled telomere-telomerase interfaces. *eLife*, 5, 2016.

[1977] Hai Huang and Thomas B Kornberg. Cells must express components of the planar cell polarity system and extracellular matrix to support cytonemes. *eLife*, 5, 2016.

[1978] He Huang, Chan Yul Yoo, Rebecca Bindbeutel, Jessica Goldsworthy, Allison Tielking, Sophie Alvarez, Michael J Naldrett, Bradley S Evans, Meng Chen, and Dmitri A Nusinow. PCH1 integrates circadian and light-signaling pathways to control photoperiod-responsive growth in Arabidopsis. *eLife*, 5, 2016.

[1979] Junqi Huang, Ting Gang Chew, Ying Gu, Saravanan Palani, Anton Kamnev, Douglas S Martin, Nicholas J Carter, Robert Anthony Cross, Snezhana Oliferenko, and Mohan K Balasubramanian. Curvature-induced expulsion of actomyosin bundles during cytokinetic ring contraction. *eLife*, 5, 2016.

[1980] Wei Huang, Andon N Placzek, Gonzalo Viana Di Prisco, Sanjeev Khatiwada, Carmela Sidrauski, KreŠimir Krnjević, Peter Walter, John A Dani, and Mauro Costa-Mattioli. Translational control by eIF2$\alpha$ phosphorylation regulates vulnerability to the synaptic and behavioral effects of cocaine. *eLife*, 5, 2016.

[1981] Wen Huang, Richard F Lyman, Rachel A Lyman, Mary Anna Carbone, Susan T Harbison, Michael M Magwire, and Trudy FC Mackay. Spontaneous mutations and the origin and maintenance of quantitative genetic variation. *eLife*, 5, 2016.

[1982] Ying Huang, Artur Matysiak, Peter Heil, Reinhard König, and Michael Brosch. Persistent neural activity in auditory cortex is related to auditory working memory in humans and nonhuman primates. *eLife*, 5, 2016.

[1983] Yongjian Huang, Shashank Bharill, Deepti Karandur, Sean M Peterson, Morgan Marita, Xiaojun Shi, Megan J Kaliszewski, Adam W Smith, Ehud Y Isacoff, and John Kuriyan. Molecular basis for multimerization in the activation of the epidermal growth factor receptor. *eLife*, 5, 2016.

[1984] Robert TR Huckstepp, Lauren E Henderson, Kathryn P Cardoza, and Jack L Feldman. Interactions between respiratory oscillators in adult rats. *eLife*, 5, 2016.

[1985] Clare Hudson, Cathy Sirour, and Hitoyoshi Yasuo. Co-expression of Foxa.a, Foxd and Fgf9/16/20 defines a transient mesendoderm regulatory state in ascidian embryos. *eLife*, 5, 2016.

[1986] Adam L Hughes, Casey E Hughes, Kiersten A Henderson, Nina Yazvenko, and Daniel E Gottschling. Selective sorting and destruction of mitochondrial membrane proteins in aged yeast. *eLife*, 5, 2016.

[1987] Jean-Emmanuel Hugonnet, Dominique Mengin-Lecreulx, Alejandro Monton, Tanneke den Blaauwen, Etienne Carbonnelle, Carole Veckerlé, V. Brun, Yves, Michael van Nieuwenhze, Christiane Bouchier, Kuyek Tu, Louis B Rice, and Michel Arthur. Factors essential for L,D-transpeptidase-mediated peptidoglycan cross-linking and $\beta$-lactam resistance in Escherichia coli. *eLife*, 5, 2016.

[1988] Christine J Huh, Bo Zhang, Matheus B Victor, Sonika Dahiya, Luis FZ Batista, Steve Horvath, and Andrew S Yoo. Maintenance of age in human neurons generated by microRNA-based neuronal conversion of fibroblasts. *eLife*, 5, 2016.

[1989] Pim J Huis in 't Veld, Sadasivam Jeganathan, Arsen Petrovic, Priyanka Singh, Juliane John, Veronica Krenn, Florian Weissmann, Tanja Bange, and Andrea Musacchio. Molecular basis of outer kinetochore assembly on CENP-T. *eLife*, 5, 2016.

[1990] Ingrid Hums, Julia Riedl, Fanny Mende, Saul Kato, Harris S Kaplan, Richard Latham, Michael Sonntag, Lisa Traunmüller, and Manuel Zimmer. Regulation of two motor patterns enables the gradual adjustment of locomotion strategy in Caenorhabditis elegans. *eLife*, 5, 2016.

[1991] Barbara J Hunnicutt, Bart C Jongbloets, William T Birdsong, Katrina J Gertz, Haining Zhong, and Tianyi Mao. A comprehensive excitatory input map of the striatum reveals novel functional organization. *eLife*, 5, 2016.

[1992] Cuong Nguyen Huu, Christian Kappel, Barbara Keller, Adrien Sicard, Yumiko Takebayashi, Holger Breuninger, Michael D Nowak, Isabel Bäurle, Axel Himmelbach, Michael Burkart, Thomas Ebbing-Lohaus, Hitoshi Sakakibara, Lothar Altschmied, Elena Conti, and Michael Lenhard. Presence versus absence of CYP734A50 underlies the style-length dimorphism in primroses. *eLife*, 5, 2016.

[1993] Kiyohito Iigaya. Adaptive learning and decision-making under uncertainty by metaplastic synapses guided by a surprise detection system. *eLife*, 5, 2016.

[1994] Robert S Illingworth, Jurriaan J Hölzenspies, Fabian V Roske, Wendy A Bickmore, and Joshua M Brickman. Polycomb enables primitive endoderm lineage priming in embryonic stem cells. *eLife*, 5, 2016.

[1995] Hideo Imamura, Tim Downing, Frederik Van den Broeck, Mandy J Sanders, Suman Rijal, Shyam Sundar, An Mannaert, Manu Vanaerschot, Maya Berg, Géraldine De Muylder, Franck Dumetz, Bart Cuypers, Ilse Maes, Malgorzata Domagalska, Saskia Decuypere, Keshav Rai, Surendra Uranw, Narayan Raj Bhattarai, Basudha Khanal, Vijay Kumar Prajapati, Smriti Sharma, Olivia Stark, Gabriele Schönian, Harry P De Koning, Luca Settimo, Benoit Vanhollebeke, Syamal Roy, Bart Ostyn, Marleen Boelaert, Louis Maes, Matthew Berriman, Jean-Claude Dujardin, and James A Cotton. Evolutionary genomics of epidemic visceral leishmaniasis in the Indian subcontinent. *eLife*, 5, 2016.

[1996] Iordan Iordanov, Csaba Mihályi, Balázs Tóth, and László Csanády. The proposed channel-enzyme transient receptor potential melastatin 2 does not possess ADP ribose hydrolase activity. *eLife*, 5, 2016.

[1997] Asif J Iqbal, Tessa J Barrett, Lewis Taylor, Eileen McNeill, Arun Manmadhan, Carlota Recio, Alfredo Carmineri, Maximillian H Brodermann, Gemma E White, Dianne Cooper, Joseph A DiDonato, Maryam Zamanian-Daryoush, Stanley L Hazen, Keith M Channon, David R Greaves, and Edward A Fisher. Acute exposure to apolipoprotein A1 inhibits macrophage chemotaxis in vitro and monocyte recruitment in vivo. *eLife*, 5, 2016.

[1998] R Stefan Isaac, Fuguo Jiang, Jennifer A Doudna, Wendell A Lim, Geeta J Narlikar, and Ricardo Almeida. Nucleosome breathing and remodeling constrain CRISPR-Cas9 function. *eLife*, 5, 2016.

[1999] Luke Isbel, Lexie Prokopuk, Haoyu Wu, Lucia Daxinger, Harald Oey, Alex Spurling, Adam J Lawther, Matthew W Hale, and Emma Whitelaw. Wiz binds active promoters and CTCF-binding sites and is required for normal behaviour in the mouse. *eLife*, 5, 2016.

[2000] Keisuke Ishihara, Kirill S Korolev, and Timothy J Mitchison. Physical basis of large microtubule aster growth. *eLife*, 5, 2016.

[2001] Rie Ishikawa, Hotaka Fukushima, Paul W Frankland, and Satoshi Kida. Hippocampal neurogenesis enhancers promote forgetting of remote fear memory after hippocampal reactivation by retrieval. *eLife*, 5, 2016.

[2002] Ryuta Ishimura, Gabor Nagy, Ivan Dotu, Jeffrey H Chuang, and Susan L Ackerman. Activation of GCN2 kinase by ribosome stalling links translation elongation with translation initiation. *eLife*, 5, 2016.

[2003] Daniel N Itzhak, Stefka Tyanova, Jürgen Cox, and Georg HH Borner. Global, quantitative and dynamic mapping of protein subcellular localization. *eLife*, 5, 2016.

[2004] Thierry Izoré, Danguole Kureisaite-Ciziene, Stephen H McLaughlin, and Jan Löwe. Crenactin forms actin-like double helical filaments regulated by arcadin-2. *eLife*, 5, 2016.

[2005] Zane A Jaafar, Akihiro Oguro, Yoshikazu Nakamura, and Jeffrey S Kieft. Translation initiation by the hepatitis C virus IRES requires eIF1A and ribosomal complex remodeling. *eLife*, 5, 2016.

[2006] Joshua A Jadwin, Dongmyung Oh, Timothy G Curran, Mari Ogiue-Ikeda, Lin Jia, Forest M White, Kazuya Machida, Ji Yu, and Bruce J Mayer. Time-resolved multimodal analysis of Src Homology 2 (SH2) domain binding in signaling by receptor tyrosine kinases. *eLife*, 5, 2016.

[2007] Clemens Jaitner, Chethan Reddy, Andreas Abentung, Nigel Whittle, Dietmar Rieder, Andrea Delekate, Martin Korte, Gaurav Jain, Andre Fischer, Farahnaz Sananbenesi, Isabella Cera, Nicolas Singewald, Georg Dechant, and Galina Apostolova. Satb2 determines miRNA expression and long-term memory in the adult central nervous system. *eLife*, 5, 2016.

[2008] Annelies Janssens, Maarten Gees, Balazs Istvan Toth, Debapriya Ghosh, Marie Mulier, Rudi Vennekens, Joris Vriens, Karel Talavera, and Thomas Voets. Definition of two agonist types at the mammalian cold-activated channel TRPM8. *eLife*, 5, 2016.

[2009] Andres Jara-Oseguera, Chanhyung Bae, and Kenton J Swartz. An external sodium ion binding site controls allosteric gating in TRPV1 channels. *eLife*, 5, 2016.

[2010] Aline Jelenkovic, Yoon-Mi Hur, Reijo Sund, Yoshie Yokoyama, Sisira H Siribaddana, Matthew Hotopf, Athula Sumathipala, Fruhling Rijsdijk, Qihua Tan, Dongfeng Zhang, Zengchang Pang, Sari Aaltonen, Kauko Heikkilä, Sevgi Y öncel, Fazil Aliev, Esther Rebato, Adam D Tarnoki, David L Tarnoki, Kaare Christensen, Axel Skytthe, Kirsten O Kyvik, Judy L Silberg, Lindon J Eaves, Hermine H Maes, Tessa L Cutler, John L Hopper, Juan R Ordoñana, Juan F Sánchez-Romera, Lucia Colodro-Conde, Wendy Cozen, Amie E Hwang, Thomas M Mack, Joohon Sung, Yun-Mi Song, Sarah Yang, Kayoung Lee, Carol E Franz, William S Kremen, Michael J Lyons, Andreas Busjahn, Tracy L Nelson, Keith E Whitfield, Christian Kandler, Kerry L Jang, Margaret Gatz, David A Butler, Maria A Stazi, Corrado Fagnani, Cristina D'Ippolito, Glen E Duncan, Dedra Buchwald, Catherine A Derom, Robert F Vlietinck, Ruth JF Loos, Nicholas G Martin, Sarah E Medland, Grant W Montgomery, Hoe-Uk Jeong, Gary E Swan, Ruth Krasnow, Patrik KE Magnusson, Nancy L Pedersen, Anna K Dahl-Aslan, Tom A McAdams, Thalia C Eley, Alice M Gregory, Per Tynelius, Laura A Baker, Catherine Tuvblad, Gombojav Bayasgalan, Danshiitsoodol Narandalai, Paul Lichtenstein, Timothy D Spector, Massimo Mangino, Genevieve Lachance, Meike Bartels, Toos CEM van Beijsterveldt, Gonneke Willemsen, S Alexandra Burt, Kelly L Klump, Jennifer R Harris, Ingunn Brandt,

Thomas Sevenius Nilsen, Robert F Krueger, Matt McGue, Shandell Pahlen, Robin P Corley, Jacob v B Hjelmborg, Jack H Goldberg, Yoshinori Iwatani, Mikio Watanabe, Chika Honda, Fujio Inui, Finn Rasmussen, Brooke M Huibregtse, Dorret I Boomsma, Thorkild I A Sørensen, Jaakko Kaprio, and Karri Silventoinen. Genetic and environmental influences on adult human height across birth cohorts from 1886 to 1994. *eLife*, 5, 2016.

[2011] Danique Jeurissen, Matthew W Self, and Pieter R Roelfsema. Serial grouping of 2D-image regions with object-based attention in humans. *eLife*, 5, 2016.

[2012] John R Jimah, Nichole D Salinas, Monica Sala-Rabanal, Nathaniel G Jones, L David Sibley, Colin G Nichols, Paul H Schlesinger, and Niraj H Tolia. Malaria parasite CelTOS targets the inner leaflet of cell membranes for pore-dependent disruption. *eLife*, 5, 2016.

[2013] Maximilian Joesch, David Mankus, Masahito Yamagata, Ali Shahbazi, Richard Schalek, Adi Suissa-Peleg, Markus Meister, Jeff W Lichtman, Walter J Scheirer, and Joshua R Sanes. Reconstruction of genetically identified neurons imaged by serial-section electron microscopy. *eLife*, 5, 2016.

[2014] Brian W Jones, Jennifer Deem, Thomas J Younts, Michael Weisenhaus, Christina A Sanford, Margaret C Slack, Jenesa Chin, Daniela Nachmanson, Alex McKennon, Pablo E Castillo, and G Stanley McKnight. Targeted deletion of AKAP7 in dentate granule cells impairs spatial discrimination. *eLife*, 5, 2016.

[2015] James D Joseph, Beatrice Darimont, Wei Zhou, Alfonso Arrazate, Amy Young, Ellen Ingalla, Kimberly Walter, Robert A Blake, Jim Nonomiya, Zhengyu Guan, Lorna Kategaya, Steven P Govek, Andiliy G Lai, Mehmet Kahraman, Dan Brigham, John Sensintaffar, Nhin Lu, Gang Shao, Jing Qian, Kate Grillot, Michael Moon, Rene Prudente, Eric Bischoff, Kyoung-Jin Lee, Celine Bonnefous, Karensa L Douglas, Jackaline D Julien, Johnny Y Nagasawa, Anna Aparicio, Josh Kaufman, Benjamin Haley, Jennifer M Giltnane, Ingrid E Wertz, Mark R Lackner, Michelle A Nannini, Deepak Sampath, Luis Schwarz, Henry Charles Manning, Mohammed Noor Tantawy, Carlos L Arteaga, Richard A Heyman, Peter J Rix, Lori Friedman, Nicholas D Smith, Ciara Metcalfe, and Jeffrey H Hager. The selective estrogen receptor downregulator GDC-0810 is efficacious in diverse models of ER+ breast cancer. *eLife*, 5, 2016.

[2016] Ryan P Joyner, Jeffrey H Tang, Jonne Helenius, Elisa Dultz, Christiane Brune, Liam J Holt, Sebastien Huet, Daniel J Müller, and Karsten Weis. A glucose-starvation response regulates the diffusion of macromolecules. *eLife*, 5, 2016.

[2017] Lining Ju, Yunfeng Chen, Lingzhou Xue, Xiaoping Du, and Cheng Zhu. Cooperative unfolding of distinctive mechanoreceptor domains transduces force into signals. *eLife*, 5, 2016.

[2018] Xiang-Chun Ju, Qiong-Qiong Hou, Ai-Li Sheng, Kong-Yan Wu, Yang Zhou, Ying Jin, Tieqiao Wen, Zhengang Yang, Xiaoqun Wang, and Zhen-Ge Luo. The hominoid-specific gene TBC1D3 promotes generation of basal neural progenitors and induces cortical folding in mice. *eLife*, 5, 2016.

[2019] Daniela Justa-Schuch, Maria Silva-Garcia, Esther Pilla, Michael Engelke, Markus Kilisch, Christof Lenz, Ulrike Möller, Fumihiko Nakamura, Henning Urlaub, and Ruth Geiss-Friedlander. DPP9 is a novel component of the N-end rule pathway targeting the tyrosine kinase Syk. *eLife*, 5, 2016.

[2020] Agnieszka M Kabat, Oliver J Harrison, Thomas Riffelmacher, Amin E Moghaddam, Claire F Pearson, Adam Laing, Lucie Abeler-Dörner, Simon P Forman, Richard K Grencis, Quentin Sattentau, Anna Katharina Simon, Johanna Pott, and Kevin J Maloy. The autophagy gene Atg16l1 differentially regulates Treg and TH2 cells to control intestinal inflammation. *eLife*, 5, 2016.

[2021] Ivan Kadurin, Laurent Ferron, Simon W Rothwell, James O Meyer, Leon R Douglas, Claudia S Bauer, Beatrice Lana, Wojciech Margas, Orpheas Alexopoulos, Manuela Nieto-Rostro, Wendy S Pratt, and Annette C Dolphin. Proteolytic maturation of $\alpha 2\delta$ represents a checkpoint for activation and neuronal trafficking of latent calcium channels. *eLife*, 5, 2016.

[2022] Pinhas Kafri, Sarah E Hasenson, Itamar Kanter, Jonathan Sheinberger, Noa Kinor, Sharon Yunger, and Yaron Shav-Tal. Quantifying $\beta$-catenin subcellular dynamics and cyclin D1 mRNA transcription during Wnt signaling in single living cells. *eLife*, 5, 2016.

[2023] Thorsten Kahnt and Philippe N Tobler. Dopamine regulates stimulus generalization in the human hippocampus. *eLife*, 5, 2016.

[2024] Tae-In Kam, Hyejin Park, Youngdae Gwon, Sungmin Song, Seo-Hyun Kim, Seo Won Moon, Dong-Gyu Jo, and Yong-Keun Jung. Fc$\gamma$RIIb-SHIP2 axis links A$\beta$ to tau pathology by disrupting phosphoinositide metabolism in Alzheimer's disease model. *eLife*, 5, 2016.

[2025] Kenji Kamimoto, Kota Kaneko, Cindy Yuet-Yin Kok, Hajime Okada, Atsushi Miyajima, and Tohru Itoh. Heterogeneity and stochastic growth regulation of biliary epithelial cells dictate dynamic epithelial tissue remodeling. *eLife*, 5, 2016.

[2026] Yilin Kang, Michael James Baker, Michael Liem, Jade Louber, Matthew McKenzie, Ishara Atukorala, Ching-Seng Ang, Shivakumar Keerthikumar, Suresh Mathivanan, and Diana Stojanovski. Tim29 is a novel subunit of the human TIM22 translocase and is involved in complex assembly and stability. *eLife*, 5, 2016.

[2027] Satomi Kanno, Jean-FranÇois Arrighi, Serge Chiarenza, Vincent Bayle, Richard Berthomé, Benjamin Péret, Hélène Javot, Etienne Delannoy, Elena Marin, Tomoko M Nakanishi, Marie-Christine Thibaud, and Laurent Nussaume. A novel role for the root cap in phosphate uptake and homeostasis. *eLife*, 5, 2016.

[2028] Ravi Kant, Baisen Zeng, Celestine J Thomas, Brian Bothner, and Stephen R Sprang. Ric-8A, a G protein chaperone with nucleotide exchange activity induces long-range secondary structure changes in G$\alpha$. *eLife*, 5, 2016.

[2029] Damian Kao, Alvina G Lai, Evangelia Stamataki, Silvana Rosic, Nikolaos Konstantinides, Erin Jarvis, Alessia Di Donfrancesco, Natalia Pouchkina-Stancheva, Marie Sémon, Marco Grillo, Heather Bruce, Suyash Kumar, Igor Siwanowicz, Andy Le, Andrew Lemire, Michael B Eisen, Cassandra Extavour, William E Browne, Carsten Wolff, Michalis Averof, Nipam H Patel, Peter Sarkies, Anastasios Pavlopoulos, and Aziz Aboobaker. The genome of the crustacean Parhyale hawaiensis, a model for animal development, regeneration, immunity and lignocellulose digestion. *eLife*, 5, 2016.

[2030] Akira Karasawa and Toshimitsu Kawate. Structural basis for subtype-specific inhibition of the P2X7 receptor. *eLife*, 5, 2016.

[2031] Andreas A Kardamakis, Juan Pérez-Fernández, and Sten Grillner. Spatiotemporal interplay between multisensory excitation and recruited inhibition in the lamprey optic tectum. *eLife*, 5, 2016.

[2032] Amit Karmon and Yitzhak Pilpel. Biological causal links on physiological and evolutionary time scales. *eLife*, 5, 2016.

[2033] Claudia Kathe, Thomas Haynes Hutson, Stephen Brendan McMahon, and Lawrence David Falcon Moon. Intramuscular Neurotrophin-3 normalizes low threshold spinal reflexes, reduces spasms and improves mobility after bilateral corticospinal tract injury in rats. *eLife*, 5, 2016.

[2034] Shachi Katira, Kranthi K Mandadapu, Suriyanarayanan Vaikuntanathan, Berend Smit, and David Chandler. Pre-transition effects mediate forces of assembly between transmembrane proteins. *eLife*, 5, 2016.

[2035] Zachary B Katz, Brian P English, Timothée Lionnet, Young J Yoon, Nilah Monnier, Ben Ovryn, Mark Bathe, and Robert H Singer. Mapping translation 'hot-spots' in live cells by tracking single molecules of mRNA and ribosomes. *eLife*, 5, 2016.

[2036] Yoshitaka Kawasoe, Toshiki Tsurimoto, Takuro Nakagawa, Hisao Masukata, and Tatsuro S Takahashi. MutS$\alpha$ maintains the mismatch repair capability by inhibiting PCNA unloading. *eLife*, 5, 2016.

[2037] Peter Keating, Onayomi Rosenior-Patten, Johannes C Dahmen, Olivia Bell, and Andrew J King. Behavioral training promotes multiple adaptive processes following acute hearing loss. *eLife*, 5, 2016.

[2038] William Thomas Keenan, Alan C Rupp, Rachel A Ross, Preethi Somasundaram, Suja Hiriyanna, Zhijian Wu, Tudor C Badea, Phyllis R Robinson, Bradford B Lowell, and Samer S Hattar. A visual circuit uses complementary mechanisms to support transient and sustained pupil constriction. *eLife*, 5, 2016.

[2039] Christian Keine, Rudolf Rübsamen, and Bernhard Englitz. Inhibition in the auditory brainstem enhances signal representation and regulates gain in complex acoustic environments. *eLife*, 5, 2016.

[2040] Tatiana Kent, Pedro A Mateos-Gomez, Agnel Sfeir, and Richard T Pomerantz. Polymerase $\theta$ is a robust terminal transferase that oscillates between three different mechanisms during end-joining. *eLife*, 5, 2016.

[2041] Envel Kerdaffrec, Danièle L Filiault, Arthur Korte, Eriko Sasaki, Viktoria Nizhynska, ümit Seren, and Magnus Nordborg. Multiple alleles at a single locus control seed dormancy in Swedish Arabidopsis. *eLife*, 5, 2016.

[2042] Katalin Kerti-Szigeti and Zoltan Nusser. Similar GABAA receptor subunit composition in somatic and axon initial segment synapses of hippocampal pyramidal cells. *eLife*, 5, 2016.

[2043] Amin Khalifeh-Soltani, Arnold Ha, Michael J Podolsky, Donald A McCarthy, William McKleroy, Saeedeh Azary, Stephen Sakuma, Kevin M Tharp, Nanyan Wu, Yasuyuki Yokosaki, Daniel Hart, Andreas Stahl, and Kamran Atabai. $\alpha 8\beta 1$ integrin regulates nutrient absorption through an Mfge8-PTEN dependent mechanism. *eLife*, 5, 2016.

[2044] Chandra M Khantwal, Sherwin J Abraham, Wei Han, Tao Jiang, Tanmay S Chavan, Ricky C Cheng, Shelley M Elvington, Corey W Liu, Irimpan I Mathews, Richard A Stein, Hassane S Mchaourab, Emad Tajkhorshid, and Merritt Maduke. Revealing an outward-facing open conformational state in a CLC Cl–/H+ exchange transporter. *eLife*, 5, 2016.

[2045] Christina Kiel, Hannah Benisty, Veronica Lloréns-Rico, and Luis Serrano. The yin–yang of kinase activation and unfolding explains the peculiarity of Val600 in the activation segment of BRAF. *eLife*, 5, 2016.

[2046] Chintan K Kikani, Xiaoying Wu, Litty Paul, Hana Sabic, Zuolian Shen, Arvind Shakya, Alexandra Keefe, Claudio Villanueva, Gabrielle Kardon, Barbara Graves, Dean Tantin, and Jared Rutter. Pask integrates hormonal signaling with histone modification via Wdr5 phosphorylation to drive myogenesis. *eLife*, 5, 2016.

[2047] Sanne Kikkert, James Kolasinski, Saad Jbabdi, Irene Tracey, Christian F Beckmann, Heidi Johansen-Berg, and Tamar R Makin. Revealing the neural fingerprints of a missing hand. *eLife*, 5, 2016.

[2048] Gukhan Kim, Rafael Luján, Jochen Schwenk, Melissa H Kelley, Carolina Aguado, Masahiko Watanabe, Bernd Fakler, James Maylie, and John P Adelman. Membrane palmitoylated protein 2 is a synaptic scaffold protein required for synaptic SK2-containing channel function. *eLife*, 5, 2016.

[2049] Hyun-Ji Kim, Myong-Ho Jeong, Kyung-Ran Kim, Chang-Yun Jung, Seul-Yi Lee, Hanna Kim, Jewoo Koh, Tuan Anh Vuong, Seungmoon Jung, Hyunwoo Yang, Su-Kyung Park, Dahee Choi, Sung Hun Kim, KyeongJin Kang, Jong-Woo Sohn, Joo Min Park, Daejong Jeon, Seung-Hoi Koo, Won-Kyung Ho, Jong-Sun Kang, Seong-Tae Kim, and Hana Cho. Protein arginine methylation facilitates KCNQ channel-PIP2 interaction leading to seizure suppression. *eLife*, 5, 2016.

[2050] Myungjin Kim, Erin Sandford, Damian Gatica, Yu Qiu, Xu Liu, Yumei Zheng, Brenda A Schulman, Jishu Xu, Ian Semple, Seung-Hyun Ro, Boyoung Kim, R Nehir Mavioglu, Aslıhan Tolun, Andras Jipa, Szabolcs Takats, Manuela Karpati, Jun Z Li, Zuhal Yapici, Gabor Juhasz, Jun Hee Lee, Daniel J Klionsky, and Margit Burmeister. Mutation in ATG5 reduces autophagy and leads to ataxia with developmental delay. *eLife*, 5, 2016.

[2051] Dari Kimanius, Björn O Forsberg, Sjors HW Scheres, and Erik Lindahl. Accelerated cryo-EM structure determination with parallelisation using GPUs in RELION-2. *eLife*, 5, 2016.

[2052] Maxime J Kinet, Jennifer A Malin, Mary C Abraham, Elyse S Blum, Melanie R Silverman, Yun Lu, and Shai Shaham. HSF-1 activates the ubiquitin proteasome system to promote non-apoptotic developmental cell death in C. elegans. *eLife*, 5, 2016.

[2053] Julius B Kirkegaard, Ambre Bouillant, Alan O Marron, Kyriacos C Leptos, and Raymond E Goldstein. Aerotaxis in the closest relatives of animals. *eLife*, 5, 2016.

[2054] Zachary A Knecht, Ana F Silbering, Lina Ni, Mason Klein, Gonzalo Budelli, Rati Bell, Liliane Abuin, Anggie J Ferrer, Aravinthan DT Samuel, Richard Benton, and Paul A Garrity. Distinct combinations of variant ionotropic glutamate receptors mediate thermosensation and hygrosensation in Drosophila. *eLife*, 5, 2016.

[2055] Michael Knoblauch, Jan Knoblauch, Daniel L Mullendore, Jessica A Savage, Benjamin A Babst, Sierra D Beecher, Adam C Dodgen, Kaare H Jensen, and N Michele Holbrook. Testing the Münch hypothesis of long distance phloem transport in plants. *eLife*, 5, 2016.

[2056] Hyo Rim Ko, Il-Sun Kwon, Inwoo Hwang, Eun-Ju Jin, Joo-Ho Shin, Angela M Brennan-Minnella, Raymond Swanson, Sung-Woo Cho, Kyung-Hoon Lee, and Jee-Yin Ahn. Akt1-Inhibitor of DNA binding2 is essential for growth cone formation and axon growth and promotes central nervous system axon regeneration. *eLife*, 5, 2016.

[2057] Dmitry Kobak, Wieland Brendel, Christos Constantinidis, Claudia E Feierstein, Adam Kepecs, Zachary F Mainen, Xue-Lian Qi, Ranulfo Romo, Naoshige Uchida, and Christian K Machens. Demixed principal component analysis of neural population data. *eLife*, 5, 2016.

[2058] Daniel L Kober, Jennifer M Alexander-Brett, Celeste M Karch, Carlos Cruchaga, Marco Colonna, Michael J Holtzman, and Thomas J Brett. Neurodegenerative disease mutations in TREM2 reveal a functional surface and distinct loss-of-function mechanisms. *eLife*, 5, 2016.

[2059] Mahdokht Kohansal-Nodehi, John JE Chua, Henning Urlaub, Reinhard Jahn, and Dominika Czernik. Analysis of protein phosphorylation in nerve terminal reveals extensive changes in active zone proteins upon exocytosis. *eLife*, 5, 2016.

[2060] Shun Koizumi, Taro Irie, Shoshiro Hirayama, Yasuyuki Sakurai, Hideki Yashiroda, Isao Naguro, Hidenori Ichijo, Jun Hamazaki, and Shigeo Murata. The aspartyl protease DDI2 activates Nrf1 to compensate for proteasome dysfunction. *eLife*, 5, 2016.

[2061] James Kolasinski, Tamar R Makin, John P Logan, Saad Jbabdi, Stuart Clare, Charlotte J Stagg, and Heidi Johansen-Berg. Perceptually relevant remapping of human somatotopy in 24 hours. *eLife*, 5, 2016.

[2062] Anna Konovalova, Angela M Mitchell, and Thomas J Silhavy. A lipoprotein/$\beta$-barrel complex monitors lipopolysaccharide integrity transducing information across the outer membrane. *eLife*, 5, 2016.

[2063] Ippei Kotera, Nhat Anh Tran, Donald Fu, Jimmy HJ Kim, Jarlath Byrne Rodgers, and William S Ryu. Pan-neuronal screening in Caenorhabditis elegans reveals asymmetric dynamics of AWC neurons is critical for thermal avoidance behavior. *eLife*, 5, 2016.

[2064] Elisabeth Kravets, Daniel Degrandi, Qijun Ma, Thomas-Otavio Peulen, Verena Klümpers, Suren Felekyan, Ralf Kühnemuth, Stefanie Weidtkamp-Peters, Claus AM Seidel, and Klaus Pfeffer. Guanylate binding proteins directly attack Toxoplasma gondii via supramolecular complexes. *eLife*, 5, 2016.

[2065] Anjali Krishnan, Choong-Wan Woo, Luke J Chang, Luka Ruzic, Xiaosi Gu, Marina López-Solà, Philip L Jackson, Jesús Pujol, Jin Fan, and Tor D Wager. Somatic and vicarious pain are represented by dissociable multivariate brain patterns. *eLife*, 5, 2016.

[2066] Giri P Krishnan, Sylvain Chauvette, Isaac Shamie, Sara Soltani, Igor Timofeev, Sydney S Cash, Eric Halgren, and Maxim Bazhenov. Cellular and neurochemical basis of sleep stages in the thalamocortical network. *eLife*, 5, 2016.

[2067] Oliver A Krupke, Ivona Zysk, Dan O Mellott, and Robert D Burke. Eph and Ephrin function in dispersal and epithelial insertion of pigmented immunocytes in sea urchin embryos. *eLife*, 5, 2016.

[2068] Benjamin Krusche, Cristina Ottone, Melanie P Clements, Ewan R Johnstone, Katrin Goetsch, Huang Lieven, Silvia G Mota, Poonam Singh, Sanjay Khadayate, Azhaar Ashraf, Timothy Davies, Steven M Pollard, Vincenzo De Paola, Federico Roncaroli, Jorge Martinez-Torrecuadrada, Paul Bertone, and Simona Parrinello. EphrinB2 drives perivascular invasion and proliferation of glioblastoma stem-like cells. *eLife*, 5, 2016.

[2069] Amy W Ku, Jason B Muhitch, Colin A Powers, Michael Diehl, Minhyung Kim, Daniel T Fisher, Anand P Sharda, Virginia K Clements, Kieran O'Loughlin, Hans Minderman, Michelle N Messmer, Jing Ma, Joseph J Skitzki, Douglas A Steeber, Bruce Walcheck, Suzanne Ostrand-Rosenberg, Scott I Abrams, and Sharon S Evans. Tumor-induced MDSC act via remote control to inhibit L-selectin-dependent adaptive immunity in lymph nodes. *eLife*, 5, 2016.

[2070] Meihua Christina Kuang, Paul D Hutchins, Jason D Russell, Joshua J Coon, and Chris Todd Hittinger. Ongoing resolution of duplicate gene functions shapes the diversification of a metabolic network. *eLife*, 5, 2016.

[2071] Daniel Kueh, William H Barnett, Gennady S Cymbalyuk, and Ronald L Calabrese. Na+/K+ pump interacts with the h-current to control bursting activity in central pattern generator neurons of leeches. *eLife*, 5, 2016.

[2072] Peer-Hendrik Kuhn, Alessio Vittorio Colombo, Benjamin Schusser, Daniela Dreymueller, Sebastian Wetzel, Ute Schepers, Julia Herber, Andreas Ludwig, Elisabeth Kremmer, Dirk Montag, Ulrike Müller, Michaela Schweizer, Paul Saftig, Stefan Bräse, and Stefan F Lichtenthaler. Systematic substrate identification indicates a central role for the metalloprotease ADAM10 in axon targeting and synapse function. *eLife*, 5, 2016.

[2073] Abhishek Kulkarni, Deniz Ertekin, Chi-Hon Lee, and Thomas Hummel. Birth order dependent growth cone segregation determines synaptic layer identity in the Drosophila visual system. *eLife*, 5, 2016.

[2074] Yosuke Kumamoto, Toshiro Hirai, Patrick W Wong, Daniel H Kaplan, and Akiko Iwasaki. CD301b+ dendritic cells suppress T follicular helper cells and antibody responses to protein antigens. *eLife*, 5, 2016.

[2075] Ganesan Senthil Kumar, Ezgi Gokhan, Sofie De Munter, Mathieu Bollen, Paola Vagnarelli, Wolfgang Peti, and Rebecca Page. The Ki-67 and RepoMan mitotic phosphatases assemble via an identical, yet novel mechanism. *eLife*, 5, 2016.

[2076] Sandra Kümper, Faraz K Mardakheh, Afshan McCarthy, Maggie Yeo, Gordon W Stamp, Angela Paul, Jonathan Worboys, Amine Sadok, Claus Jørgensen, Sabrina Guichard, and Christopher J Marshall. Rho-associated kinase (ROCK) function is essential for cell cycle progression, senescence and tumorigenesis. *eLife*, 5, 2016.

[2077] Esther Kur, Jiha Kim, Aleksandra Tata, Cesar H Comin, Kyle I Harrington, Luciano da F Costa, Katie Bentley, and Chenghua Gu. Temporal modulation of collective cell behavior controls vascular network topology. *eLife*, 5, 2016.

[2078] Toshihide Kurihara, Peter D Westenskow, Marin L Gantner, Yoshihiko Usui, Andrew Schultz, Stephen Bravo, Edith Aguilar, Carli Wittgrove, Mollie SH Friedlander, Liliana P Paris, Emily Chew, Gary Siuzdak, and Martin Friedlander. Hypoxia-induced metabolic stress in retinal pigment epithelial cells is sufficient to induce photoreceptor degeneration. *eLife*, 5, 2016.

[2079] Chun Wai Kwan, Jackie Gavin-Smyth, Edwin L Ferguson, and Urs Schmidt-Ott. Functional evolution of a morphogenetic gradient. *eLife*, 5, 2016.

[2080] Agnieszka Kłosowska, Tomasz Chamera, and Krzysztof Liberek. Adenosine diphosphate restricts the protein remodeling activity of the Hsp104 chaperone to Hsp70 assisted disaggregation. *eLife*, 5, 2016.

[2081] Piergiorgio La Rosa, Pamela Bielli, Claudia Compagnucci, Eleonora Cesari, Elisabetta Volpe, Stefano Farioli Vecchioli, and Claudio Sette. Sam68 promotes self-renewal and glycolytic metabolism in mouse neural progenitor cells by modulating Aldh1a3 pre-mRNA 3′-end processing. *eLife*, 5, 2016.

[2082] Armin Lak, William R Stauffer, and Wolfram Schultz. Dopamine neurons learn relative chosen value from probabilistic rewards. *eLife*, 5, 2016.

[2083] Samira Lakhal-Littleton, Magda Wolna, Yu Jin Chung, Helen C Christian, Lisa C Heather, Marcella Brescia, Vicky Ball, Rebeca Diaz, Ana Santos, Daniel Biggs, Kieran Clarke, Benjamin Davies, and Peter A Robbins. An essential cell-autonomous role for hepcidin in cardiac iron homeostasis. *eLife*, 5, 2016.

[2084] Manoj M Lalu, Katrina J Sullivan, Shirley HJ Mei, David Moher, Alexander Straus, Dean A Fergusson, Duncan J Stewart, Mazen Jazi, Malcolm MacLeod, Brent Winston, John Marshall, Brian Hutton, Keith R Walley, and Lauralyn McIntyre. Evaluating mesenchymal stem cell therapy for sepsis with preclinical meta-analyses prior to initiating a first-in-human trial. *eLife*, 5, 2016.

[2085] Eva Lana-Elola, Sheona Watson-Scales, Amy Slender, Dorota Gibbins, Alexandrine Martineau, Charlotte Douglas, Timothy Mohun, Elizabeth MC Fisher, and Victor LJ Tybulewicz. Genetic dissection of Down syndrome-associated congenital heart defects using a new mouse mapping panel. *eLife*, 5, 2016.

[2086] Rachel Lander and Christian P Petersen. Wnt, Ptk7, and FGFRL expression gradients control trunk positional identity in planarian regeneration. *eLife*, 5, 2016.

[2087] Brian J Lane, Pranit Samarth, Joseph L Ransdell, Satish S Nair, and David J Schulz. Synergistic plasticity of intrinsic conductance and electrical coupling restores synchrony in an intact motor network. *eLife*, 5, 2016.

[2088] Harriet Lane-Serff, Paula MacGregor, Lori Peacock, Olivia JS Macleod, Christopher Kay, Wendy Gibson, Matthew K Higgins, and Mark Carrington. Evolutionary diversification of the trypanosome haptoglobin-haemoglobin receptor from an ancestral haemoglobin receptor. *eLife*, 5, 2016.

[2089] Damien Lanfray, Alexandre Caron, Marie-Claude Roy, Mathieu Laplante, Fabrice Morin, Jérôme Leprince, Marie-Christine Tonon, and Denis Richard. Involvement of the Acyl-CoA binding domain containing 7 in the control of food intake and energy expenditure in mice. *eLife*, 5, 2016.

[2090] Nikki K Larter, Jennifer S Sun, and John R Carlson. Organization and function of Drosophila odorant binding proteins. *eLife*, 5, 2016.

[2091] Nicholas B Last, Ludmila Kolmakova-Partensky, Tania Shane, and Christopher Miller. Mechanistic signs of double-barreled structure in a fluoride ion channel. *eLife*, 5, 2016.

[2092] Meirav Lavy, Michael J Prigge, Sibo Tao, Stephanie Shain, April Kuo, Kerstin Kirchsteiger, and Mark Estelle. Constitutive auxin response in Physcomitrella reveals complex interactions between Aux/IAA and ARF proteins. *eLife*, 5, 2016.

[2093] Gurunathan Laxmikanthan, Chen Xu, Axel F Brilot, David Warren, Lindsay Steele, Nicole Seah, Wenjun Tong, Nikolaus Grigorieff, Arthur Landy, and Gregory D Van Duyne. Structure of a Holliday junction complex reveals mechanisms governing a highly regulated DNA transaction. *eLife*, 5, 2016.

[2094] Andres M Lebensohn, Ramin Dubey, Leif R Neitzel, Ofelia Tacchelly-Benites, Eungi Yang, Caleb D Marceau, Eric M Davis, Bhaven B Patel, Zahra Bahrami-Nejad, Kyle J Travaglini, Yashi Ahmed, Ethan Lee, Jan E Carette, and Rajat Rohatgi. Comparative genetic screens in human cells reveal new regulatory mechanisms in WNT signaling. *eLife*, 5, 2016.

[2095] ChangHwan Lee, Erika B Sorensen, Tina R Lynch, and Judith Kimble. C. elegans GLP-1/Notch activates transcription in a probability gradient across the germline stem cell pool. *eLife*, 5, 2016.

[2096] GaYoung Lee and Thomas L Schwarz. Filamin, a synaptic organizer in Drosophila, determines glutamate receptor composition and membrane growth. *eLife*, 5, 2016.

[2097] Wei-Hua Lee, Hitoshi Higuchi, Sakae Ikeda, Erica L Macke, Tetsuya Takimoto, Bikash R Pattnaik, Che Liu, Li-Fang Chu, Sandra M Siepka, Kathleen J Krentz, C Dustin Rubinstein, Robert F Kalejta, James A Thomson, Robert F Mullins, Joseph S Takahashi, Lawrence H Pinto, and Akihiro Ikeda. Mouse Tmem135 mutation reveals a mechanism involving mitochondrial dynamics that leads to age-dependent retinal pathologies. *eLife*, 5, 2016.

[2098] Janelle C Leggere, Yuhki Saito, Robert B Darnell, Marc Tessier-Lavigne, Harald J Junge, and Zhe Chen. NOVA regulates Dcc alternative splicing during neuronal migration and axon guidance in the spinal cord. *eLife*, 5, 2016.

[2099] Nicolas J Lehrbach and Gary Ruvkun. Proteasome dysfunction triggers activation of SKN-1A/Nrf1 by the aspartic protease DDI-1. *eLife*, 5, 2016.

[2100] Lilia Leisle, Rahul Chadda, John D Lueck, Daniel T Infield, Jason D Galpin, Venkatramanan Krishnamani, Janice L Robertson, and Christopher A Ahern. Cellular encoding of Cy dyes for single-molecule imaging. *eLife*, 5, 2016.

[2101] Andrea Leo, Giacomo Handjaras, Matteo Bianchi, Hamal Marino, Marco Gabiccini, Andrea Guidi, Enzo Pasquale Scilingo, Pietro Pietrini, Antonio Bicchi, Marco Santello, and Emiliano Ricciardi. A synergy-based hand control is encoded in human motor cortical areas. *eLife*, 5, 2016.

[2102] Jake Lesnik, Travis Antes, Jeewon Kim, Erin Griner, and Luisa Pedro. Registered report: Melanoma exosomes educate bone marrow progenitor cells toward a pro-metastatic phenotype through MET. *eLife*, 5, 2016.

[2103] Robert F Levendosky, Anton Sabantsev, Sebastian Deindl, and Gregory D Bowman. The Chd1 chromatin remodeler shifts hexasomes unidirectionally. *eLife*, 5, 2016.

[2104] Roi Levy, David Levitan, and Abraham J Susswein. New learning while consolidating memory during sleep is actively blocked by a protein synthesis dependent process. *eLife*, 5, 2016.

[2105] Nicola S Lewis, Colin A Russell, Pinky Langat, Tavis K Anderson, Kathryn Berger, Filip Bielejec, David F Burke, Gytis Dudas, Judith M Fonville, Ron AM Fouchier, Paul Kellam, Bjorn F Koel, Philippe Lemey, Tung Nguyen, Bundit Nuansrichy, JS Malik Peiris, Takehiko Saito, Gaelle Simon, Eugene Skepner, Nobuhiro Takemae, Richard J Webby, Kristien Van Reeth, Sharon M Brookes, Lars Larsen, Simon J Watson, Ian H Brown, and Amy L Vincent. The global antigenic diversity of swine influenza A viruses. *eLife*, 5, 2016.

[2106] Jian Li and Suzanne R Pfeffer. Lysosomal membrane glycoproteins bind cholesterol and contribute to lysosomal cholesterol export. *eLife*, 5, 2016.

[2107] Li Li, Hui Liu, Peng Dong, Dong Li, Wesley R Legant, Jonathan B Grimm, Luke D Lavis, Eric Betzig, Robert Tjian, and Zhe Liu. Real-time imaging of Huntingtin aggregates diverting target search and gene transcription. *eLife*, 5, 2016.

[2108] Peng Li, Giomar Rivera-Cancel, Lisa N Kinch, Dor Salomon, Diana R Tomchick, Nick V Grishin, and Kim Orth. Bile salt receptor complex activates a pathogenic type III secretion system. *eLife*, 5, 2016.

[2109] Ruo-Jing Li, Jing Xu, Chenglai Fu, Jing Zhang, Yujun George Zheng, Hao Jia, and Jun O Liu. Regulation of mTORC1 by lysosomal calcium and calmodulin. *eLife*, 5, 2016.

[2110] Shengben Li, Brandon Le, Xuan Ma, Shaofang Li, Chenjiang You, Yu Yu, Bailong Zhang, Lin Liu, Lei Gao, Ting Shi, Yonghui Zhao, Beixin Mo, Xiaofeng Cao, and Xuemei Chen. Biogenesis of phased siRNAs on membrane-bound polysomes in Arabidopsis. *eLife*, 5, 2016.

[2111] Tongchao Li, Nikolaos Giagtzoglou, Daniel F Eberl, Sonal Nagarkar Jaiswal, Tiantian Cai, Dorothea Godt, Andrew K Groves, and Hugo J Bellen. The E3 ligase Ubr3 regulates Usher syndrome and MYH9 disorder proteins in the auditory organs of Drosophila and mammals. *eLife*, 5, 2016.

[2112] Xuan Li and Jessica K Tyler. Nucleosome disassembly during human non-homologous end joining followed by concerted HIRA- and CAF-1-dependent reassembly. *eLife*, 5, 2016.

[2113] Yen-Li Li, Viswanathan Chandrasekaran, Stephen D Carter, Cora L Woodward, Devin E Christensen, Kelly A Dryden, Owen Pornillos, Mark Yeager, Barbie K Ganser-Pornillos, Grant J Jensen, and Wesley I Sundquist. Primate TRIM5 proteins form hexagonal nets on HIV-1 capsids. *eLife*, 5, 2016.

[2114] Yubing Li, Dianyi Liu, Cristina López-Paz, Bradley JSC Olson, and James G Umen. A new class of cyclin dependent kinase in Chlamydomonas is required for coupling cell size to cell division. *eLife*, 5, 2016.

[2115] Xiangxiu Liang, Pingtao Ding, Kehui Lian, Jinlong Wang, Miaomiao Ma, Lin Li, Lei Li, Meng Li, Xiaojuan Zhang, She Chen, Yuelin Zhang, and Jian-Min Zhou. Arabidopsis heterotrimeric G proteins regulate immunity by directly coupling to the FLS2 receptor. *eLife*, 5, 2016.

[2116] Sara I Liin, Johan E Larsson, Rene Barro-Soria, Bo Hjorth Bentzen, and H Peter Larsson. Fatty acid analogue N-arachidonoyl taurine restores function of IKs channels with diverse long QT mutations. *eLife*, 5, 2016.

[2117] Cherry Lim, Emi Takahashi, Maliwan Hongsuwan, Vanaporn Wuthiekanun, Visanu Thamlikitkul, Soawapak Hinjoy, Nicholas PJ Day, Sharon J Peacock, and Direk Limmathurotsakul. Epidemiology and burden of multidrug-resistant bacterial infection in a developing country. *eLife*, 5, 2016.

[2118] Yoonseob Lim, Ryan Lagoy, Barbara G Shinn-Cunningham, and Timothy J Gardner. Transformation of temporal sequences in the zebra finch auditory system. *eLife*, 5, 2016.

[2119] Tsung-Chieh Lin, Chia-Yi Su, Pei-Yu Wu, Tsung-Ching Lai, Wen-An Pan, Yi-Hua Jan, Yu-Chang Chang, Chi-Tai Yeh, Chi-Long Chen, Luo-Ping Ger, Hong-Tai Chang, Chih-Jen Yang, Ming-Shyan Huang, Yu-Peng Liu, Yuan-Feng Lin, John Y-J Shyy, Ming-Daw Tsai, and Michael Hsiao. The nucleolar protein NIFK promotes cancer progression via CK1$\alpha$/$\beta$-catenin in metastasis and Ki-67-dependent cell proliferation. *eLife*, 5, 2016.

[2120] Mario J Lintz and Gidon Felsen. Basal ganglia output reflects internally-specified movements. *eLife*, 5, 2016.

[2121] Chan-Wang Lio, Jiayuan Zhang, Edahí González-Avalos, Patrick G Hogan, Xing Chang, and Anjana Rao. Tet2 and Tet3 cooperate with B-lineage transcription factors to regulate DNA modification and chromatin accessibility. *eLife*, 5, 2016.

[2122] Marc Lipsitch, Wendy Barclay, Rahul Raman, Charles J Russell, Jessica A Belser, Sarah Cobey, Peter M Kasson, James O Lloyd-Smith, Sebastian Maurer-Stroh, Steven Riley, Catherine AA Beauchemin, Trevor Bedford, Thomas C Friedrich, Andreas Handel, Sander Herfst, Pablo R Murcia, Benjamin Roche, Claus O Wilke, and Colin A Russell. Viral factors in influenza pandemic risk assessment. *eLife*, 5, 2016.

[2123] Haiyang Liu, Jianchao Li, Manmeet H Raval, Ningning Yao, Xiaoying Deng, Qing Lu, Si Nie, Wei Feng, Jun Wan, Christopher M Yengo, Wei Liu, and Mingjie Zhang. Myosin III-mediated cross-linking and stimulation of actin bundling activity of Espin. *eLife*, 5, 2016.

[2124] Libin Liu and Paul F Pilch. PTRF/Cavin-1 promotes efficient ribosomal RNA transcription in response to metabolic challenges. *eLife*, 5, 2016.

[2125] Liu D Liu, Ralf M Haefner, and Christopher C Pack. A neural basis for the spatial suppression of visual motion perception. *eLife*, 5, 2016.

[2126] Min Liu, Yajuan Li, Aiguo Liu, Ruifeng Li, Ying Su, Juan Du, Cheng Li, and Alan Jian Zhu. The exon junction complex regulates the splicing of cell polarity gene dlg1 to control Wingless signaling in development. *eLife*, 5, 2016.

[2127] Qingqing Liu, Xing Yang, Jingsong Tian, Zhongbao Gao, Meng Wang, Yan Li, and Aike Guo. Gap junction networks in mushroom bodies participate in visual learning and memory in Drosophila. *eLife*, 5, 2016.

[2128] Tie Liu, Adam D Longhurst, Franklin Talavera-Rauh, Samuel A Hokin, and M Kathryn Barton. The Arabidopsis transcription factor ABIG1 relays ABA signaled growth inhibition and drought induced senescence. *eLife*, 5, 2016.

[2129] Wallace H Liu, Sarah C Roemer, Yeyun Zhou, Zih-Jie Shen, Briana K Dennehey, Jeremy L Balsbaugh, Jennifer C Liddle, Travis Nemkov, Natalie G Ahn, Kirk C Hansen, Jessica K Tyler, and Mair EA Churchill. The Cac1 subunit of histone chaperone CAF-1 organizes CAF-1-H3/H4 architecture and tetramerizes histones. *eLife*, 5, 2016.

[2130] Xiaoxia Liu, Alpay Burak Seven, Marcial Camacho, Victoria Esser, Junjie Xu, Thorsten Trimbuch, Bradley Quade, Lijing Su, Cong Ma, Christian Rosenmund, and Josep Rizo. Functional synergy between the Munc13 C-terminal C1 and C2 domains. *eLife*, 5, 2016.

[2131] Zhong-Yu Liu, Xiao-Feng Li, Tao Jiang, Yong-Qiang Deng, Qing Ye, Hui Zhao, Jiu-Yang Yu, and Cheng-Feng Qin. Viral RNA switch mediates the dynamic control of flavivirus replicase recruitment by genome cyclization. *eLife*, 5, 2016.

[2132] Melissa A Lopes Pinheiro, Alwin Kamermans, Juan J Garcia-Vallejo, Bert van het Hof, Laura Wierts, Tom O'Toole, Daniël Boeve, Marleen Verstege, Susanne MA van der Pol, Yvette van Kooyk, Helga E de Vries, and Wendy WJ Unger. Internalization and presentation of myelin antigens by the brain endothelium guides antigen-specific T cell migration. *eLife*, 5, 2016.

[2133] Jaime Lopez-Mosqueda, Karthik Maddi, Stefan Prgomet, Sissy Kalayil, Ivana Marinovic-Terzic, Janos Terzic, and Ivan Dikic. SPRTN is a mammalian DNA-binding metalloprotease that resolves DNA-protein crosslinks. *eLife*, 5, 2016.

[2134] Alizée Lopez-Persem, Philippe Domenech, and Mathias Pessiglione. How prior preferences determine decision-making frames and biases in the human brain. *eLife*, 5, 2016.

[2135] David S Lorberbaum, Andrea I Ramos, Kevin A Peterson, Brandon S Carpenter, David S Parker, Sandip De, Lauren E Hillers, Victoria M Blake, Yuichi Nishi, Matthew R McFarlane, Ason CY Chiang, Judith A Kassis, Benjamin L Allen, Andrew P McMahon, and Scott Barolo. An ancient yet flexible cis-regulatory architecture allows localized Hedgehog tuning by patched/Ptch1. *eLife*, 5, 2016.

[2136] Francesca Lorenzin, Uwe Benary, Apoorva Baluapuri, Susanne Walz, Lisa Anna Jung, Björn von Eyss, Caroline Kisker, Jana Wolf, Martin Eilers, and Elmar Wolf. Different promoter affinities account for specificity in MYC-dependent gene regulation. *eLife*, 5, 2016.

[2137] Olga Lositsky, Janice Chen, Daniel Toker, Christopher J Honey, Michael Shvartsman, Jordan L Poppenk, Uri Hasson, and Kenneth A Norman. Neural pattern change during encoding of a narrative predicts retrospective duration estimates. *eLife*, 5, 2016.

[2138] Anna B Loveland, Eugene Bah, Rohini Madireddy, Ying Zhang, Axel F Brilot, Nikolaus Grigorieff, and Andrei A Korostelev. Ribosome•RelA structures reveal the mechanism of stringent response activation. *eLife*, 5, 2016.

[2139] Rachel Lowe, Caio AS Coelho, Christovam Barcellos, Marilia Sá Carvalho, Rafael De Castro Catão, Giovanini E Coelho, Walter Massa Ramalho, Trevor C Bailey, David B Stephenson, and Xavier Rodó. Evaluating probabilistic dengue risk forecasts from a prototype early warning system for Brazil. *eLife*, 5, 2016.

[2140] Fang-Min Lu, Christine Deisl, and Donald W Hilgemann. Profound regulation of Na/K pump activity by transient elevations of cytoplasmic calcium in murine cardiac myocytes. *eLife*, 5, 2016.

[2141] Zhenwei Lu, Sijo Mathew, Jiang Chen, Arina Hadziselimovic, Riya Palamuttam, Billy G Hudson, Reinhard Fässler, Ambra Pozzi, Charles R Sanders, and Roy Zent. Implications of the differing roles of the $\beta$1 and $\beta$3 transmembrane and cytoplasmic domains for integrin function. *eLife*, 5, 2016.

[2142] Giovanni Luchetti, Ria Sircar, Jennifer H Kong, Sigrid Nachtergaele, Andreas Sagner, Eamon FX Byrne, Douglas F Covey, Christian Siebold, and Rajat Rohatgi. Cholesterol activates the G-protein coupled receptor Smoothened to promote Hedgehog signaling. *eLife*, 5, 2016.

[2143] John D Lueck, Adam L Mackey, Daniel T Infield, Jason D Galpin, Jing Li, Benoît Roux, and Christopher A Ahern. Atomic mutagenesis in ion channels with engineered stoichiometry. *eLife*, 5, 2016.

[2144] Andrew Lutas, Carolina Lahmann, Magali Soumillon, and Gary Yellen. The leak channel NALCN controls tonic firing and glycolytic sensitivity of substantia nigra pars reticulata neurons. *eLife*, 5, 2016.

[2145] Penelope Anne Lynch and Mike Boots. Using evolution to generate sustainable malaria control with spatial repellents. *eLife*, 5, 2016.

[2146] Declan G Lyons, Alexandre Parpaleix, Morgane Roche, and Serge Charpak. Mapping oxygen concentration in the awake mouse brain. *eLife*, 5, 2016.

[2147] Artem Y Lyubimov, Monarin Uervirojnangkoorn, Oliver B Zeldin, Qiangjun Zhou, Minglei Zhao, Aaron S Brewster, Tara Michels-Clark, James M Holton, Nicholas K Sauter, William I Weis, and Axel T Brunger. Advances in X-ray free electron laser (XFEL) diffraction data processing applied to the crystal structure of the synaptotagmin-1 / SNARE complex. *eLife*, 5, 2016.

[2148] Péter Lőrincz, Zsolt Lakatos, ágnes Varga, Tamás Maruzs, Zsófia Simon-Vecsei, Zsuzsanna Darula, Péter Benkő, Gábor Csordás, Mónika Lippai, István Andó, Krisztina Hegedűs, Katalin F Medzihradszky, Szabolcs Takáts, and Gábor Juhász. MiniCORVET is a Vps8-containing early endosomal tether in Drosophila. *eLife*, 5, 2016.

[2149] Siming Ma, Akhil Upneja, Andrzej Galecki, Yi-Miau Tsai, Charles F Burant, Sasha Raskind, Quanwei Zhang, Zhengdong D Zhang, Andrei Seluanov, Vera Gorbunova, Clary B Clish, Richard A Miller, and Vadim N Gladyshev. Cell culture-based profiling across mammals reveals DNA repair and metabolism as determinants of species longevity. *eLife*, 5, 2016.

[2150] Paul M Macdonald, Matt Kanke, and Andrew Kenny. Community effects in regulation of translation. *eLife*, 5, 2016.

[2151] Ricardo AR Machado, Mark McClure, Maxime R Hervé, Ian T Baldwin, and Matthias Erb. Benefits of jasmonate-dependent defenses against vertebrate herbivores in nature. *eLife*, 5, 2016.

[2152] Maud Magdeleine, Romain Gautier, Pierre Gounon, Hélène Barelli, Stefano Vanni, and Bruno Antonny. A filter at the entrance of the Golgi that selects vesicles according to size and bulk lipid composition. *eLife*, 5, 2016.

[2153] Rebekah Elizabeth Mahoney, Jorge Azpurua, and Benjamin A Eaton. Insulin signaling controls neurotransmission via the 4eBP-dependent modification of the exocytotic machinery. *eLife*, 5, 2016.

[2154] Lara Maldanis, Murilo Carvalho, Mariana Ramos Almeida, Francisco Idalécio Freitas, José Artur Ferreira Gomes de Andrade, Rafael Silva Nunes, Carlos Eduardo Rochitte, Ronei Jesus Poppi, Raul Oliveira Freitas, Fábio Rodrigues, Sandra Siljeström, Frederico Alves Lima, Douglas Galante, Ismar S Carvalho, Carlos Alberto Perez, Marcelo Rodrigues de Carvalho, Jefferson Bettini, Vincent Fernandez, and José Xavier-Neto. Heart fossilization is possible and informs the evolution of cardiac outflow tract in vertebrates. *eLife*, 5, 2016.

[2155] Barbora Malecova, Alessandra Dall'Agnese, Luca Madaro, Sole Gatto, Paula Coutinho Toto, Sonia Albini, Tammy Ryan, Làszlò Tora, and Pier Lorenzo Puri. TBP/TFIID-dependent activation of MyoD target genes in skeletal muscle cells. *eLife*, 5, 2016.

[2156] Somya Mani and Mukund Thattai. Stacking the odds for Golgi cisternal maturation. *eLife*, 5, 2016.

[2157] Maria Manifava, Matthew Smith, Sergio Rotondo, Simon Walker, Izabella Niewczas, Roberto Zoncu, Jonathan Clark, and Nicholas T Ktistakis. Dynamics of mTORC1 activation in response to amino acids. *eLife*, 5, 2016.

[2158] Moutusi Manna, Miia Niemelä, Joona Tynkkynen, Matti Javanainen, Waldemar Kulig, Daniel J Müller, Tomasz Rog, and Ilpo Vattulainen. Mechanism of allosteric regulation of $\beta$2-adrenergic receptor by cholesterol. *eLife*, 5, 2016.

[2159] Sabrina F Mansilla, Agustina P Bertolin, Valérie Bergoglio, Marie-Jeanne Pillaire, Marina A González Besteiro, Carlos Luzzani, Santiago G Miriuka, Christophe Cazaux, Jean-Sébastien Hoffmann, and Vanesa Gottifredi. Cyclin Kinase-independent role of p21CDKN1A in the promotion of nascent DNA elongation in unstressed cells. *eLife*, 5, 2016.

[2160] Nimi Marcel and Apurva Sarin. Notch1 regulated autophagy controls survival and suppressor activity of activated murine T-regulatory cells. *eLife*, 5, 2016.

[2161] Luciano Marcon, Xavier Diego, James Sharpe, and Patrick Müller. High-throughput mathematical analysis identifies Turing networks for patterning with equally diffusing signals. *eLife*, 5, 2016.

[2162] Alvaro Martinez Barrio, Sangeet Lamichhaney, Guangyi Fan, Nima Rafati, Mats Pettersson, He Zhang, Jacques Dainat, Diana Ekman, Marc Höppner, Patric Jern, Marcel Martin, Björn Nystedt, Xin Liu, Wenbin Chen, Xinming Liang, Chengcheng Shi, Yuanyuan Fu, Kailong Ma, Xiao Zhan, Chungang Feng, Ulla Gustafson, Carl-Johan Rubin, Markus Sällman Almén, Martina Blass, Michele Casini, Arild Folkvord, Linda Laikre, Nils Ryman, Simon Ming-Yuen Lee, Xun Xu, and Leif Andersson. The genetic basis for ecological adaptation of the Atlantic herring revealed by genome sequencing. *eLife*, 5, 2016.

[2163] Fidelis T Masao, Elgidius B Ichumbaki, Marco Cherin, Angelo Barili, Giovanni Boschian, Dawid A Iurino, Sofia Menconero, Jacopo Moggi-Cecchi, and Giorgio Manzi. New footprints from Laetoli (Tanzania) provide evidence for marked body size variation in early hominins. *eLife*, 5, 2016.

[2164] Philippe Materne, Enrique Vázquez, Mar Sánchez, Carlo Yague-Sanz, Jayamani Anandhakumar, Valerie Migeot, Francisco Antequera, and Damien Hermand. Histone H2B ubiquitylation represses gametogenesis by opposing RSC-dependent chromatin remodeling at the ste11 master regulator locus. *eLife*, 5, 2016.

[2165] Rebecca S Mathew, Antonis Tatarakis, Andrii Rudenko, Erin M Johnson-Venkatesh, Yawei J Yang, Elisabeth A Murphy, Travis P Todd, Scott T Schepers, Nertila Siuti, Anthony J Martorell, William A Falls, Sayamwong E Hammack, Christopher A Walsh, Li-Huei Tsai, Hisashi Umemori, Mark E Bouton, and Danesh Moazed. A microRNA negative feedback loop downregulates vesicle transport and inhibits fear memory. *eLife*, 5, 2016.

[2166] Hideyuki Matsumoto, Ju Tian, Naoshige Uchida, and Mitsuko Watabe-Uchida. Midbrain dopamine neurons signal aversion in a reward-context-dependent manner. *eLife*, 5, 2016.

[2167] Madlen Matz-Soja, Christiane Rennert, Kristin Schönefeld, Susanne Aleithe, Jan Boettger, Wolfgang Schmidt-Heck, Thomas S Weiss, Amalya Hovhannisyan, Sebastian Zellmer, Nora Klöting, Angela Schulz, Jürgen Kratzsch, Reinhardt Guthke, and Rolf Gebhardt. Hedgehog signaling is a potent regulator of liver lipid metabolism and reveals a GLI-code associated with steatosis. *eLife*, 5, 2016.

[2168] David Mavor, Kyle Barlow, Samuel Thompson, Benjamin A Barad, Alain R Bonny, Clinton L Cario, Garrett Gaskins, Zairan Liu, Laura Deming, Seth D Axen, Elena Caceres, Weilin Chen, Adolfo Cuesta, Rachel E Gate, Evan M Green, Kaitlin R Hulce, Weiyue Ji, Lillian R Kenner, Bruk Mensa, Leanna S Morinishi, Steven M Moss, Marco Mravic, Ryan K Muir, Stefan Niekamp, Chimno I Nnadi, Eugene Palovcak, Erin M Poss, Tyler D Ross, Eugenia C Salcedo, Stephanie K See, Meena Subramaniam, Allison W Wong, Jennifer Li, Kurt S Thorn, Shane ó Conchúir, Benjamin P Roscoe, Eric D Chow, Joseph L DeRisi, Tanja Kortemme, Daniel N Bolon, and James S Fraser. Determination of ubiquitin fitness landscapes under different chemical stresses in a classroom setting. *eLife*, 5, 2016.

[2169] Kathleen L McCann, Takamasa Teramoto, Jun Zhang, Traci M Tanaka Hall, and Susan J Baserga. The molecular basis for ANE syndrome revealed by the large ribosomal subunit processome interactome. *eLife*, 5, 2016.

[2170] Shelly TH McClatchey, Zheng Wang, Lara M Linden, Eric L Hastie, Lin Wang, Wanqing Shen, Alan Chen, Qiuyi Chi, and David R Sherwood. Boundary cells restrict dystroglycan trafficking to control basement membrane sliding during tissue remodeling. *eLife*, 5, 2016.

[2171] Gail McConnell, Johanna Trägårdh, Rumelo Amor, John Dempster, Es Reid, and William Bradshaw Amos. A novel optical microscope for imaging large embryos and tissue volumes with sub-cellular resolution throughout. *eLife*, 5, 2016.

[2172] Kristopher McEown, Yohko Takata, Yoan Cherasse, Nanae Nagata, Kosuke Aritake, and Michael Lazarus. Chemogenetic inhibition of the medial prefrontal cortex reverses the effects of REM sleep loss on sucrose consumption. *eLife*, 5, 2016.

[2173] John P McGinnis, Huoqing Jiang, Moutaz Ali Agha, Consuelo Perez Sanchez, Jeff Lange, Zulin Yu, Frederic Marion-Poll, and Kausik Si. Immediate perception of a reward is distinct from the reward's long-term salience. *eLife*, 5, 2016.

[2174] Skye C McIver, Koichi R Katsumura, Elsa Davids, Peng Liu, Yoon-A Kang, David Yang, and Emery H Bresnick. Exosome complex orchestrates developmental signaling to balance proliferation and differentiation during erythropoiesis. *eLife*, 5, 2016.

[2175] Alison C McKelvey, Travis B Lear, Sarah R Dunn, John Evankovich, James D Londino, Joseph S Bednash, Yingze Zhang, Bryan J McVerry, Yuan Liu, and Bill B Chen. RING finger E3 ligase PPP1R11 regulates TLR2 signaling and innate immunity. *eLife*, 5, 2016.

[2176] Erin C McKiernan, Philip E Bourne, C Titus Brown, Stuart Buck, Amye Kenall, Jennifer Lin, Damon McDougall, Brian A Nosek, Karthik Ram, Courtney K Soderberg, Jeffrey R Spies, Kaitlin Thaney, Andrew Updegrove, Kara H Woo, and Tal Yarkoni. How open science helps researchers succeed. *eLife*, 5, 2016.

[2177] Curtis McMurtrey, Thomas Trolle, Tiffany Sansom, Soumya G Remesh, Thomas Kaever, Wilfried Bardet, Kenneth Jackson, Rima McLeod, Alessandro Sette, Morten Nielsen, Dirk M Zajonc, Ira J Blader, Bjoern Peters, and William Hildebrand. Toxoplasma gondii peptide ligands open the gate of the HLA class I binding groove. *eLife*, 5, 2016.

[2178] Darpan Medhi, Alastair SH Goldman, and Michael Lichten. Local chromosome context is a major determinant of crossover pathway biochemistry during budding yeast meiosis. *eLife*, 5, 2016.

[2179] Edgar M Medina, Jonathan J Turner, Raluca Gordân, Jan M Skotheim, and Nicolas E Buchler. Punctuated evolution and transitional hybrid network in an ancestral cell cycle of fungi. *eLife*, 5, 2016.

[2180] Feng Mei, Klaus Lehmann-Horn, Yun-An A Shen, Kelsey A Rankin, Karin J Stebbins, Daniel S Lorrain, Kara Pekarek, Sharon A Sagan, Lan Xiao, Cory Teuscher, H-Christian von Büdingen, Jürgen Wess, J Josh Lawrence, Ari J Green, Stephen PJ Fancy, Scott S Zamvil, and Jonah R Chan. Accelerated remyelination during inflammatory demyelination prevents axonal loss and improves functional recovery. *eLife*, 5, 2016.

[2181] Markus Meister. Physical limits to magnetogenetics. *eLife*, 5, 2016.

[2182] Andrew Melber, Un Na, Ajay Vashisht, Benjamin D Weiler, Roland Lill, James A Wohlschlegel, and Dennis R Winge. Role of Nfu1 and Bol3 in iron-sulfur cluster transfer to mitochondrial clients. *eLife*, 5, 2016.

[2183] Ignacio Mena, Martha I Nelson, Francisco Quezada-Monroy, Jayeeta Dutta, Refugio Cortes-Fernández, J Horacio Lara-Puente, Felipa Castro-Peralta, Luis F Cunha, Nídia S Trovão, Bernardo Lozano-Dubernard, Andrew Rambaut, Harm van Bakel, and Adolfo García-Sastre. Origins of the 2009 H1N1 influenza pandemic in swine in Mexico. *eLife*, 5, 2016.

[2184] Wilson Mena, Sören Diegelmann, Christian Wegener, and John Ewer. Stereotyped responses of Drosophila peptidergic neuronal ensemble depend on downstream neuromodulators. *eLife*, 5, 2016.

[2185] Philipe RF MendonÇa, Mariana Vargas-Caballero, Ferenc Erdélyi, Gábor Szabó, Ole Paulsen, and Hugh PC Robinson. Stochastic and deterministic dynamics of intrinsically irregular firing in cortical inhibitory interneurons. *eLife*, 5, 2016.

[2186] Lingfeng Meng, Albert Zhang, Yishi Jin, and Dong Yan. Regulation of neuronal axon specification by glia-neuron gap junctions in C. elegans. *eLife*, 5, 2016.

[2187] Xiangling Meng, Wei Wang, Hui Lu, Ling-jie He, Wu Chen, Eugene S Chao, Marta L Fiorotto, Bin Tang, Jose A Herrera, Michelle L Seymour, Jeffrey L Neul, Fred A Pereira, Jianrong Tang, Mingshan Xue, and Huda Y Zoghbi. Manipulations of MeCP2 in glutamatergic neurons highlight their contributions to Rett and other neurological disorders. *eLife*, 5, 2016.

[2188] Audrey A Mercer, Kristin J Palarz, Nino Tabatadze, Catherine S Woolley, and Indira M Raman. Sex differences in cerebellar synaptic transmission and sex-specific responses to autism-linked Gabrb3 mutations in mice. *eLife*, 5, 2016.

[2189] Jane P Messina, Moritz UG Kraemer, Oliver J Brady, David M Pigott, Freya M Shearer, Daniel J Weiss, Nick Golding, Corrine W Ruktanonchai, Peter W Gething, Emily Cohn, John S Brownstein, Kamran Khan, Andrew J Tatem, Thomas Jaenisch, Christopher JL Murray, Fatima Marinho, Thomas W Scott, and Simon I Hay. Mapping global environmental suitability for Zika virus. *eLife*, 5, 2016.

[2190] Michael G Metzen, Volker Hofmann, and Maurice J Chacron. Neural correlations enable invariant coding and perception of natural stimuli in weakly electric fish. *eLife*, 5, 2016.

[2191] Florent Meyniel, Guy M Goodwin, JF William Deakin, Corinna Klinge, Christine MacFadyen, Holly Milligan, Emma Mullings, Mathias Pessiglione, and Raphaël Gaillard. A specific role for serotonin in overcoming effort cost. *eLife*, 5, 2016.

[2192] Linqing Miao, Liu Yang, Haoliang Huang, Feisi Liang, Chen Ling, and Yang Hu. mTORC1 is necessary but mTORC2 and GSK3$\beta$ are inhibitory for AKT3-induced axon regeneration in the central nervous system. *eLife*, 5, 2016.

[2193] Kristina D Micheva, Dylan Wolman, Brett D Mensh, Elizabeth Pax, JoAnn Buchanan, Stephen J Smith, and Davi D Bock. A large fraction of neocortical myelin ensheathes axons of local inhibitory neurons. *eLife*, 5, 2016.

[2194] Johann Mignolet, Seamus Holden, Matthieu Bergé, Gaël Panis, Ezgi Eroglu, Laurence Théraulaz, Suliana Manley, and Patrick H Viollier. Functional dichotomy and distinct nanoscale assemblies of a cell cycle-controlled bipolar zinc-finger regulator. *eLife*, 5, 2016.

[2195] Csaba Mihályi, Beáta Töröcsik, and László Csanády. Obligate coupling of CFTR pore opening to tight nucleotide-binding domain dimerization. *eLife*, 5, 2016.

[2196] Emiko Mihara, Hidenori Hirai, Hideki Yamamoto, Keiko Tamura-Kawakami, Mami Matano, Akira Kikuchi, Toshiro Sato, and Junichi Takagi. Active and water-soluble form of lipidated Wnt protein is maintained by a serum glycoprotein afamin/$\alpha$-albumin. *eLife*, 5, 2016.

[2197] Christopher J Millard, Niranjan Varma, Almutasem Saleh, Kyle Morris, Peter J Watson, Andrew R Bottrill, Louise Fairall, Corinne J Smith, and John WR Schwabe. The structure of the core NuRD repression complex provides insights into its interaction with chromatin. *eLife*, 5, 2016.

[2198] Tyne W Miller-Fleming, Sarah C Petersen, Laura Manning, Cristina Matthewman, Megan Gornet, Allison Beers, Sayaka Hori, Shohei Mitani, Laura Bianchi, Janet Richmond, and David M Miller. The DEG/ENaC cation channel protein UNC-8 drives activity-dependent synapse removal in remodeling GABAergic neurons. *eLife*, 5, 2016.

[2199] Loïs S Miraucourt, Jennifer Tsui, Delphine Gobert, Jean-FranÇois Desjardins, Anne Schohl, Mari Sild, Perry Spratt, Annie Castonguay, Yves De Koninck, Nicholas Marsh-Armstrong, Paul W Wiseman, and Edward S Ruthazer. Endocannabinoid signaling enhances visual responses through modulation of intracellular chloride levels in retinal ganglion cells. *eLife*, 5, 2016.

[2200] Yevgeniya A Mironova, Guy M Lenk, Jing-Ping Lin, Seung Joon Lee, Jeffery L Twiss, Ilaria Vaccari, Alessandra Bolino, Leif A Havton, Sang H Min, Charles S Abrams, Peter Shrager, Miriam H Meisler, and Roman J Giger. PI(3,5)P2 biosynthesis regulates oligodendrocyte differentiation by intrinsic and extrinsic mechanisms. *eLife*, 5, 2016.

[2201] Diana M Mitrea, Jaclyn A Cika, Clifford S Guy, David Ban, Priya R Banerjee, Christopher B Stanley, Amanda Nourse, Ashok A Deniz, and Richard W Kriwacki. Nucleophosmin integrates within the nucleolus via multi-modal interactions with proteins displaying R-rich linear motifs and rRNA. *eLife*, 5, 2016.

[2202] Haik Mkhikian, Christie-Lynn Mortales, Raymond W Zhou, Khachik Khachikyan, Gang Wu, Stuart M Haslam, Patil Kavarian, Anne Dell, and Michael Demetriou. Golgi self-correction generates bioequivalent glycans to preserve cellular homeostasis. *eLife*, 5, 2016.

[2203] Alisa Mo, Chongyuan Luo, Fred P Davis, Eran A Mukamel, Gilbert L Henry, Joseph R Nery, Mark A Urich, Serge Picard, Ryan Lister, Sean R Eddy, Michael A Beer, Joseph R Ecker, and Jeremy Nathans. Epigenomic landscapes of retinal rods and cones. *eLife*, 5, 2016.

[2204] Jeffrey R Moffitt, Shristi Pandey, Alistair N Boettiger, Siyuan Wang, and Xiaowei Zhuang. Spatial organization shapes the turnover of a bacterial transcriptome. *eLife*, 5, 2016.

[2205] Gábor Molnár, Márton Rózsa, Judith Baka, Noémi Holderith, Pál Barzó, Zoltan Nusser, and Gábor Tamás. Human pyramidal to interneuron synapses are mediated by multi-vesicular release and multiple docked vesicles. *eLife*, 5, 2016.

[2206] Felipe Mora-Bermúdez, Farhath Badsha, Sabina Kanton, J Gray Camp, Benjamin Vernot, Kathrin Köhler, Birger Voigt, Keisuke Okita, Tomislav Maricic, Zhisong He, Robert Lachmann, Svante Pääbo, Barbara Treutlein, and Wieland B Huttner. Differences and similarities between human and chimpanzee neural progenitors during cerebral cortex development. *eLife*, 5, 2016.

[2207] Deborah Morena, Nicola Maestro, Francesca Bersani, Paolo Emanuele Forni, Marcello Francesco Lingua, Valentina Foglizzo, Petar Šćepanović, Silvia Miretti, Alessandro Morotti, Jack F Shern, Javed Khan, Ugo Ala, Paolo Provero, Valentina Sala, Tiziana Crepaldi, Patrizia Gasparini, Michela Casanova, Andrea Ferrari, Gabriella Sozzi, Roberto Chiarle, Carola Ponzetto, and Riccardo Taulli. Hepatocyte Growth Factor-mediated satellite cells niche perturbation promotes development of distinct sarcoma subtypes. *eLife*, 5, 2016.

[2208] Claudia M Moreno, Rose E Dixon, Sendoa Tajada, Can Yuan, Ximena Opitz-Araya, Marc D Binder, and Luis F Santana. Ca2+ entry into neurons is facilitated by cooperative gating of clustered CaV1.3 channels. *eLife*, 5, 2016.

[2209] Clive P Morgan, Jocelyn F Krey, M'hamed Grati, Bo Zhao, Shannon Fallen, Abhiraami Kannan-Sundhari, Xue Zhong Liu, Dongseok Choi, Ulrich Müller, and Peter G Barr-Gillespie. PDZD7-MYO7A complex identified in enriched stereocilia membranes. *eLife*, 5, 2016.

[2210] Shane J Morley, Yanmei Qi, Loredana Iovino, Laura Andolfi, Da Guo, Nereo Kalebic, Laura Castaldi, Christian Tischer, Carla Portulano, Giulia Bolasco, Kalyanee Shirlekar, Claudia M Fusco, Antonino Asaro, Federica Fermani, Mayya Sundukova, Ulf Matti, Luc Reymond, Adele De Ninno, Luca Businaro, Kai Johnsson, Marco Lazzarino, Jonas Ries, Yannick Schwab, Jing Hu, and Paul A Heppenstall. Acetylated tubulin is essential for touch sensation in mice. *eLife*, 5, 2016.

[2211] Elizabeth R Morris, Heather Grey, Grant McKenzie, Anita C Jones, and Julia M Richardson. A bend, flip and trap mechanism for transposon integration. *eLife*, 5, 2016.

[2212] Ruzbeh Mosadeghi, Kurt M Reichermeier, Martin Winkler, Anne Schreiber, Justin M Reitsma, Yaru Zhang, Florian Stengel, Junyue Cao, Minsoo Kim, Michael J Sweredoski, Sonja Hess, Alexander Leitner, Ruedi Aebersold, Matthias Peter, Raymond J Deshaies, and Radoslav I Enchev. Structural and kinetic analysis of the COP9-Signalosome activation and the cullin-RING ubiquitin ligase deneddylation cycle. *eLife*, 5, 2016.

[2213] Xiang Mou and Daoyun Ji. Social observation enhances cross-environment activation of hippocampal place cell patterns. *eLife*, 5, 2016.

[2214] Zineb Mounir, Joshua M Korn, Thomas Westerling, Fallon Lin, Christina A Kirby, Markus Schirle, Gregg McAllister, Greg Hoffman, Nadire Ramadan, Anke Hartung, Yan Feng, David Randal Kipp, Christopher Quinn, Michelle Fodor, Jason Baird, Marie Schoumacher, Ronald Meyer, James Deeds, Gilles Buchwalter, Travis Stams, Nicholas Keen, William R Sellers, Myles Brown, and Raymond A Pagliarini. ERG signaling in prostate cancer is driven through PRMT5-dependent methylation of the Androgen Receptor. *eLife*, 5, 2016.

[2215] André Mourão, Sophie Bonnal, Komal Soni, Lisa Warner, Rémy Bordonné, Juan Valcárcel, and Michael Sattler. Structural basis for the recognition of spliceosomal SmN/B/B' proteins by the RBM5 OCRE domain in splicing regulation. *eLife*, 5, 2016.

[2216] Sandrine Moutel, Nicolas Bery, Virginie Bernard, Laura Keller, Emilie Lemesre, Ario de Marco, Laetitia Ligat, Jean-Christophe Rain, Gilles Favre, Aurélien Olichon, and Franck Perez. NaLi-H1: A universal synthetic library of humanized nanobodies providing highly functional antibodies and intrabodies. *eLife*, 5, 2016.

[2217] Louise A Moyle, Eric Blanc, Oihane Jaka, Johanna Prueller, Christopher RS Banerji, Francesco Saverio Tedesco, Stephen DR Harridge, Robert D Knight, and Peter S Zammit. Ret function in muscle stem cells points to tyrosine kinase inhibitor therapy for facioscapulohumeral muscular dystrophy. *eLife*, 5, 2016.

[2218] Shatanik Mukherjee, Vera Jansen, Jan F Jikeli, Hussein Hamzeh, Luis Alvarez, Marco Dombrowski, Melanie Balbach, Timo Strünker, Reinhard Seifert, U Benjamin Kaupp, and Dagmar Wachten. A novel biosensor to study cAMP dynamics in cilia and flagella. *eLife*, 5, 2016.

[2219] Lyle Muller, Giovanni Piantoni, Dominik Koller, Sydney S Cash, Eric Halgren, and Terrence J Sejnowski. Rotating waves during human sleep spindles organize global patterns of activity that repeat precisely through the night. *eLife*, 5, 2016.

[2220] Yasunobu Murata and Matthew T Colonnese. An excitatory cortical feedback loop gates retinal wave transmission in rodent thalamus. *eLife*, 5, 2016.

[2221] Keith R Murphy, Sonali A Deshpande, Maria E Yurgel, James P Quinn, Jennifer L Weissbach, Alex C Keene, Ken Dawson-Scully, Robert Huber, Seth M Tomchik, and William W Ja. Postprandial sleep mechanics in Drosophila. *eLife*, 5, 2016.

[2222] Sean C Murphy, Lucy M Palmer, Thomas Nyffeler, René M Müri, and Matthew E Larkum. Transcranial magnetic stimulation (TMS) inhibits cortical dendrites. *eLife*, 5, 2016.

[2223] Christopher I Murray, Maartje Westhoff, Jodene Eldstrom, Emely Thompson, Robert Emes, and David Fedida. Unnatural amino acid photo-crosslinking of the IKs channel complex demonstrates a KCNE1:KCNQ1 stoichiometry of up to 4:4. *eLife*, 5, 2016.

[2224] Jason Murray, Christos G Savva, Byung-Sik Shin, Thomas E Dever, V Ramakrishnan, and Israel S Fernández. Structural characterization of ribosome recruitment and translocation by type IV IRES. *eLife*, 5, 2016.

[2225] Samuel A Myers, Sailaja Peddada, Nilanjana Chatterjee, Tara Friedrich, Kiichrio Tomoda, Gregor Krings, Sean Thomas, Jason Maynard, Michael Broeker, Matthew Thomson, Katherine Pollard, Shinya Yamanaka, Alma L Burlingame, and Barbara Panning. SOX2 O-GlcNAcylation alters its protein-protein interactions and genomic occupancy to modulate gene expression in pluripotent cells. *eLife*, 5, 2016.

[2226] Simon Mysling, Kristian Kølby Kristensen, Mikael Larsson, Anne P Beigneux, Henrik Gårdsvoll, Loren G Fong, André Bensadouen, Thomas JD Jørgensen, Stephen G Young, and Michael Ploug. The acidic domain of the endothelial membrane protein GPIHBP1 stabilizes lipoprotein lipase activity by preventing unfolding of its catalytic domain. *eLife*, 5, 2016.

[2227] Simon Mysling, Kristian Kølby Kristensen, Mikael Larsson, Oleg Kovrov, André Bensadouen, Thomas JD Jørgensen, Gunilla Olivecrona, Stephen G Young, and Michael Ploug. The angiopoietin-like protein ANGPTL4 catalyzes unfolding of the hydrolase domain in lipoprotein lipase and the endothelial membrane protein GPIHBP1 counteracts this unfolding. *eLife*, 5, 2016.

[2228] Saravanapriah Nadarajan, Firaz Mohideen, Yonatan B Tzur, Nuria Ferrandiz, Oliver Crawley, Alex Montoya, Peter Faull, Ambrosius P Snijders, Pedro R Cutillas, Ashwini Jambhekar, Michael D Blower, Enrique Martinez-Perez, J Wade Harper, and Monica P Colaiacovo. The MAP kinase pathway coordinates crossover designation with disassembly of synaptonemal complex proteins during meiosis. *eLife*, 5, 2016.

[2229] Karine Narbonne-Reveau, Elodie Lanet, Caroline Dillard, Sophie Foppolo, Ching-Huan Chen, Hugues Parrinello, Stéphanie Rialle, Nicholas S Sokol, and Cédric Maurange. Neural stem cell-encoded temporal patterning delineates an early window of malignant susceptibility in Drosophila. *eLife*, 5, 2016.

[2230] Christopher A Natale, Elizabeth K Duperret, Junqian Zhang, Rochelle Sadeghi, Ankit Dahal, Kevin Tyler O'Brien, Rosa Cookson, Jeffrey D Winkler, and Todd W Ridky. Sex steroids regulate skin pigmentation through nonclassical membrane-bound receptors. *eLife*, 5, 2016.

[2231] Jeremy Nathans and Peter Sterling. How scientists can reduce their carbon footprint. *eLife*, 5, 2016.

[2232] NCD Risk Factor Collaboration (NCD-RisC). A century of trends in adult human height. *eLife*, 5:e13410, jul 2016.

[2233] Tamilla Nechiporuk, James McGann, Karin Mullendorff, Jenny Hsieh, Wolfgang Wurst, Thomas Floss, and Gail Mandel. The REST remodeling complex protects genomic integrity during embryonic neurogenesis. *eLife*, 5, 2016.

[2234] Derek Evan Nee and Mark D'Esposito. The hierarchical organization of the lateral prefrontal cortex. *eLife*, 5, 2016.

[2235] Takefumi Negishi, Naoyuki Miyazaki, Kazuyoshi Murata, Hitoyoshi Yasuo, and Naoto Ueno. Physical association between a novel plasma-membrane structure and centrosome orients cell division. *eLife*, 5, 2016.

[2236] Jonathan O Nelson, Kristin A Moore, Alex Chapin, Julie Hollien, and Mark M Metzstein. Degradation of Gadd45 mRNA by nonsense-mediated decay is essential for viability. *eLife*, 5, 2016.

[2237] Rafik Neme and Diethard Tautz. Fast turnover of genome transcription across evolutionary time exposes entire non-coding DNA to de novo gene emergence. *eLife*, 5, 2016.

[2238] Huyen Nguyen, Peter Dayan, Zac Pujic, Justin Cooper-White, and Geoffrey J Goodhill. RETRACTED: A mathematical model explains saturating axon guidance responses to molecular gradients. *eLife*, 5, 2016.

[2239] Quynh-Anh Nguyen, Meryl E Horn, and Roger A Nicoll. Distinct roles for extracellular and intracellular domains in neuroligin function at inhibitory synapses. *eLife*, 5, 2016.

[2240] Thi-Minh Nguyen, Dietmar Schreiner, Le Xiao, Lisa Traunmüller, Caroline Bornmann, and Peter Scheiffele. An alternative splicing switch shapes neurexin repertoires in principal neurons versus interneurons in the mouse hippocampus. *eLife*, 5, 2016.

[2241] Kun-Ming Ni, Xiao-Jun Hou, Ci-Hang Yang, Ping Dong, Yue Li, Ying Zhang, Ping Jiang, Darwin K Berg, Shumin Duan, and Xiao-Ming Li. Selectively driving cholinergic fibers optically in the thalamic reticular nucleus promotes sleep. *eLife*, 5, 2016.

[2242] Lina Ni, Mason Klein, Kathryn V Svec, Gonzalo Budelli, Elaine C Chang, Anggie J Ferrer, Richard Benton, Aravinthan DT Samuel, and Paul A Garrity. The Ionotropic Receptors IR21a and IR25a mediate cool sensing in Drosophila. *eLife*, 5, 2016.

[2243] Thomas K Ni and Charlotte Kuperwasser. Premature polyadenylation of MAGI3 produces a dominantly-acting oncogene in human breast cancer. *eLife*, 5, 2016.

[2244] María Nicolás-Pérez, Franz Kuchling, Joaquín Letelier, Rocío Polvillo, Jochen Wittbrodt, and Juan R Martínez-Morales. Analysis of cellular behavior and cytoskeletal dynamics reveal a constriction mechanism driving optic cup morphogenesis. *eLife*, 5, 2016.

[2245] John M Nicoludis, Bennett E Vogt, Anna G Green, Charlotta PI Schärfe, Debora S Marks, and Rachelle Gaudet. Antiparallel protocadherin homodimers use distinct affinity- and specificity-mediating regions in cadherin repeats 1-4. *eLife*, 5, 2016.

[2246] Keisuke Nimura, Masamichi Yamamoto, Makiko Takeichi, Kotaro Saga, Katsuyoshi Takaoka, Norihiko Kawamura, Hirohisa Nitta, Hiromichi Nagano, Saki Ishino, Tatsuya Tanaka, Robert J Schwartz, Hiroyuki Aburatani, and Yasufumi Kaneda. Regulation of alternative polyadenylation by Nkx2-5 and Xrn2 during mouse heart development. *eLife*, 5, 2016.

[2247] Silas Boye Nissen, Tali Magidson, Kevin Gross, and Carl T Bergstrom. Publication bias and the canonization of false facts. *eLife*, 5, 2016.

[2248] Marie-Julie Nokin, Florence Durieux, Paul Peixoto, Barbara Chiavarina, Olivier Peulen, Arnaud Blomme, Andrei Turtoi, Brunella Costanza, Nicolas Smargiasso, Dominique Baiwir, Jean L Scheijen, Casper G Schalkwijk, Justine Leenders, Pascal De Tullio, Elettra Bianchi, Marc Thiry, Koji Uchida, David A Spiegel, James R Cochrane, Craig A Hutton, Edwin De Pauw, Philippe Delvenne, Dominique Belpomme, Vincent Castronovo, and Akeila Bellahcène. Methylglyoxal, a glycolysis side-product, induces Hsp90 glycation and YAP-mediated tumor growth and metastasis. *eLife*, 5, 2016.

[2249] Mark Norman, Robin Vuilleumier, Alexander Springhorn, Jennifer Gawlik, and George Pyrowolakis. Pentagone internalises glypicans to fine-tune multiple signalling pathways. *eLife*, 5, 2016.

[2250] Noelle R Noyes, Xiang Yang, Lyndsey M Linke, Roberta J Magnuson, Adam Dettenwanger, Shaun Cook, Ifigenia Geornaras, Dale E Woerner, Sheryl P Gow, Tim A McAllister, Hua Yang, Jaime Ruiz, Kenneth L Jones, Christina A Boucher, Paul S Morley, and Keith E Belk. Resistome diversity in cattle and the environment decreases during beef production. *eLife*, 5, 2016.

[2251] Daichi Nozaki, Atsushi Yokoi, Takahiro Kimura, Masaya Hirashima, and Jean-Jacques Orban de Xivry. Tagging motor memories with transcranial direct current stimulation allows later artificially-controlled retrieval. *eLife*, 5, 2016.

[2252] Illyce Nuñez, Marbelys Rodriguez Pino, David J Wiley, Maitreyi E Das, Chuan Chen, Tetsuya Goshima, Kazunori Kume, Dai Hirata, Takashi Toda, and Fulvia Verde. Spatial control of translation repression and polarized growth by conserved NDR kinase Orb6 and RNA-binding protein Sts5. *eLife*, 5, 2016.

[2253] Daniel Oberfeld and Felicitas Klöckner-Nowotny. Individual differences in selective attention predict speech identification at a cocktail party. *eLife*, 5, 2016.

[2254] Calogero Maria Oddo, Stanisa Raspopovic, Fiorenzo Artoni, Alberto Mazzoni, Giacomo Spigler, Francesco Petrini, Federica Giambattistelli, Fabrizio Vecchio, Francesca Miraglia, Loredana Zollo, Giovanni Di Pino, Domenico Camboni, Maria Chiara Carrozza, Eugenio Guglielmelli, Paolo Maria Rossini, Ugo Faraguna, and Silvestro Micera. Intraneural stimulation elicits discrimination of textural features by artificial fingertip in intact and amputee humans. *eLife*, 5, 2016.

[2255] Yasuhiro Ogawa, Yoko Irukayama-Tomobe, Nobuyuki Murakoshi, Maiko Kiyama, Yui Ishikawa, Naoto Hosokawa, Hiromu Tominaga, Shuntaro Uchida, Saki Kimura, Mika Kanuka, Miho Morita, Michito Hamada, Satoru Takahashi, Yu Hayashi, and Masashi Yanagisawa. Peripherally administered orexin improves survival of mice with endotoxin shock. *eLife*, 5, 2016.

[2256] Yoko Ogawa, Satoru Morikawa, Hideyuki Okano, Yo Mabuchi, Sadafumi Suzuki, Tomonori Yaguchi, Yukio Sato, Shin Mukai, Saori Yaguchi, Takaaki Inaba, Shinichiro Okamoto, Yutaka Kawakami, Kazuo Tsubota, Yumi Matsuzaki, and Shigeto Shimmura. MHC-compatible bone marrow stromal/stem cells trigger fibrosis by activating host T cells in a scleroderma mouse model. *eLife*, 5, 2016.

[2257] Koji Oishi, Nao Nakagawa, Kashiko Tachikawa, Shinji Sasaki, Michihiko Aramaki, Shinji Hirano, Nobuhiko Yamamoto, Yumiko Yoshimura, and Kazunori Nakajima. Identity of neocortical layer 4 neurons is specified through correct positioning into the cortex. *eLife*, 5, 2016.

[2258] Nikola Ojkic, Javier López-Garrido, Kit Pogliano, and Robert G Endres. Cell-wall remodeling drives engulfment during Bacillus subtilis sporulation. *eLife*, 5, 2016.

[2259] Masahiro Oka, Sonoko Mura, Kohji Yamada, Percival Sangel, Saki Hirata, Kazumitsu Maehara, Koichi Kawakami, Taro Tachibana, Yasuyuki Ohkawa, Hiroshi Kimura, and Yoshihiro Yoneda. Chromatin-prebound Crm1 recruits Nup98-HoxA9 fusion to induce aberrant expression of Hox cluster genes. *eLife*, 5, 2016.

[2260] Yuji Okamoto, Noa Lipstein, Yunfeng Hua, Kun-Han Lin, Nils Brose, Takeshi Sakaba, and Mitsuharu Midorikawa. Distinct modes of endocytotic presynaptic membrane and protein uptake at the calyx of Held terminal of rats and mice. *eLife*, 5, 2016.

[2261] Michael L Oldham, Nikolaus Grigorieff, and Jue Chen. Structure of the transporter associated with antigen processing trapped by herpes simplex virus. *eLife*, 5, 2016.

[2262] Saleh Omairi, Antonios Matsakas, Hans Degens, Oliver Kretz, Kenth-Arne Hansson, Andreas Våvang Solbrå, Jo C Bruusgaard, Barbara Joch, Roberta Sartori, Natasa Giallourou, Robert Mitchell, Henry Collins-Hooper, Keith Foster, Arja Pasternack, Olli Ritvos, Marco Sandri, Vihang Narkar, Jonathan R Swann, Tobias B Huber, and Ketan Patel. Enhanced exercise and regenerative capacity in a mouse model that violates size constraints of oxidative muscle fibres. *eLife*, 5, 2016.

[2263] Mohsen Omrani, Chantelle D Murnaghan, J Andrew Pruszynski, and Stephen H Scott. Distributed task-specific processing of somatosensory feedback for voluntary motor control. *eLife*, 5, 2016.

[2264] Shotaro Otsuka, Khanh Huy Bui, Martin Schorb, M Julius Hossain, Antonio Z Politi, Birgit Koch, Mikhail Eltsov, Martin Beck, and Jan Ellenberg. Nuclear pore assembly proceeds by an inside-out extrusion of the nuclear envelope. *eLife*, 5, 2016.

[2265] Gilles Ouanounou, Gérard Baux, and Thierry Bal. A novel synaptic plasticity rule explains homeostasis of neuromuscular transmission. *eLife*, 5, 2016.

[2266] Arzu öztürk Çolak, Bernard Moussian, Sofia J Araújo, and Jordi Casanova. A feedback mechanism converts individual cell features into a supracellular ECM structure in Drosophila trachea. *eLife*, 5, 2016.

[2267] Yamuna D Paila, Lynn GL Richardson, Hitoshi Inoue, Elizabeth S Parks, James McMahon, Kentaro Inoue, and Danny J Schnell. Multi-functional roles for the polypeptide transport associated domains of Toc75 in chloroplast protein import. *eLife*, 5, 2016.

[2268] Janelle MP Pakan, Scott C Lowe, Evelyn Dylda, Sander W Keemink, Stephen P Currie, Christopher A Coutts, and Nathalie L Rochefort. Behavioral-state modulation of inhibition is context-dependent and cell type specific in mouse visual cortex. *eLife*, 5, 2016.

[2269] Richard Panayiotou, Francesc Miralles, Rafal Pawlowski, Jessica Diring, Helen R Flynn, Mark Skehel, and Richard Treisman. Phosphorylation acts positively and negatively to regulate MRTF-A subcellular localisation and activity. *eLife*, 5, 2016.

[2270] Alessandro Papale, Ilaria Maria Morella, Marzia Tina Indrigo, Rick Eugene Bernardi, Livia Marrone, Francesca Marchisella, Andrea Brancale, Rainer Spanagel, Riccardo Brambilla, and Stefania Fasano. Impairment of cocaine-mediated behaviours in mice by clinically relevant Ras-ERK inhibitors. *eLife*, 5, 2016.

[2271] Nicole D Paris, Andrew Soroka, Alanna Klose, Wenxuan Liu, and Joe V Chakkalakal. Smad4 restricts differentiation to promote expansion of satellite cell derived progenitors during skeletal muscle regeneration. *eLife*, 5, 2016.

[2272] Eun Chan Park and Christopher Rongo. The p38 MAP kinase pathway modulates the hypoxia response and glutamate receptor trafficking in aging neurons. *eLife*, 5, 2016.

[2273] Hyojin Park, Christoph Kayser, Gregor Thut, and Joachim Gross. Lip movements entrain the observers' low-frequency brain oscillations to facilitate speech intelligibility. *eLife*, 5, 2016.

[2274] Sunmee Park, Akhil Bandi, Christian R Lee, and David J Margolis. Peripheral optogenetic stimulation induces whisker movement and sensory perception in head-fixed mice. *eLife*, 5, 2016.

[2275] Maulik R Patel, Ganesh K Miriyala, Aimee J Littleton, Heiko Yang, Kien Trinh, Janet M Young, Scott R Kennedy, Yukiko M Yamashita, Leo J Pallanck, and Harmit S Malik. A mitochondrial DNA hypomorph of cytochrome oxidase specifically impairs male fertility in Drosophila melanogaster. *eLife*, 5, 2016.

[2276] Stephen Patterson, Susan Wyllie, Suzanne Norval, Laste Stojanovski, Frederick RC Simeons, Jennifer L Auer, Maria Osuna-Cabello, Kevin D Read, and Alan H Fairlamb. The anti-tubercular drug delamanid as a potential oral treatment for visceral leishmaniasis. *eLife*, 5, 2016.

[2277] Julia Patzig, Michelle S Erwig, Stefan Tenzer, Kathrin Kusch, Payam Dibaj, Wiebke Möbius, Sandra Goebbels, Nicole Schaeren-Wiemers, Klaus-Armin Nave, and Hauke B Werner. Septin/anillin filaments scaffold central nervous system myelin to accelerate nerve conduction. *eLife*, 5, 2016.

[2278] Hania J Pavlou, Andrew C Lin, Megan C Neville, Tetsuya Nojima, Fengqiu Diao, Brian E Chen, Benjamin H White, and Stephen F Goodwin. Neural circuitry coordinating male copulation. *eLife*, 5, 2016.

[2279] Claire Pearson, Emily E Thornton, Brent McKenzie, Anna-Lena Schaupp, Nicky Huskens, Thibault Griseri, Nathaniel West, Sim Tung, Benedict P Seddon, Holm H Uhlig, and Fiona Powrie. ILC3 GM-CSF production and mobilisation orchestrate acute intestinal inflammation. *eLife*, 5, 2016.

[2280] Joel Pearson, Rocco Chiou, Sebastian Rogers, Marcus Wicken, Stewart Heitmann, and Bard Ermentrout. Sensory dynamics of visual hallucinations in the normal population. *eLife*, 5, 2016.

[2281] Victoria E Pedanou, Stéphane Gobeil, Sébastien Tabariès, Tessa M Simone, Lihua Julie Zhu, Peter M Siegel, and Michael R Green. The histone H3K9 demethylase KDM3A promotes anoikis by transcriptionally activating pro-apoptotic genes BNIP3 and BNIP3L. *eLife*, 5, 2016.

[2282] Cengiz Pehlevan, Paolo Paoletti, and L Mahadevan. Integrative neuromechanics of crawling in D. melanogaster larvae. *eLife*, 5, 2016.

[2283] Valentina Peleh, Emmanuelle Cordat, and Johannes M Herrmann. Mia40 is a trans-site receptor that drives protein import into the mitochondrial intermembrane space by hydrophobic substrate binding. *eLife*, 5, 2016.

[2284] Gertrudis Perea, Ricardo Gómez, Sara Mederos, Ana Covelo, Jesús J Ballesteros, Laura Schlosser, Alicia Hernández-Vivanco, Mario Martín-Fernández, Ruth Quintana, Abdelrahman Rayan, Adolfo Díez, Marco Fuenzalida, Amit Agarwal, Dwight E Bergles, Bernhard Bettler, Denise Manahan-Vaughan, Eduardo D Martín, Frank Kirchhoff, and Alfonso Araque. Activity-dependent switch of GABAergic inhibition into glutamatergic excitation in astrocyte-neuron networks. *eLife*, 5, 2016.

[2285] Elliot A Perens, Zayra V Garavito-Aguilar, Gina P Guio-Vega, Karen T Peña, Yocheved L Schindler, and Deborah Yelon. Hand2 inhibits kidney specification while promoting vein formation within the posterior mesoderm. *eLife*, 5, 2016.

[2286] Mireia Perez-Camps, Jing Tian, Serene C Chng, Kai Pin Sem, Thankiah Sudhaharan, Cathleen Teh, Malte Wachsmuth, Vladimir Korzh, Sohail Ahmed, and Bruno Reversade. Quantitative imaging reveals real-time Pou5f3–Nanog complexes driving dorsoventral mesendoderm patterning in zebrafish. *eLife*, 5, 2016.

[2287] José Antonio Pérez-Escobar, Olga Kornienko, Patrick Latuske, Laura Kohler, and Kevin Allen. Visual landmarks sharpen grid cell metric and confer context specificity to neurons of the medial entorhinal cortex. *eLife*, 5, 2016.

[2288] ángel Pérez-Lara, Anusa Thapa, Sarah B Nyenhuis, David A Nyenhuis, Partho Halder, Michael Tietzel, Kai Tittmann, David S Cafiso, and Reinhard Jahn. PtdInsP2 and PtdSer cooperate to trap synaptotagmin-1 to the plasma membrane in the presence of calcium. *eLife*, 5, 2016.

[2289] María Inés Pérez Millán, Michelle L Brinkmeier, Amanda H Mortensen, and Sally A Camper. PROP1 triggers epithelial-mesenchymal transition-like process in pituitary stem cells. *eLife*, 5, 2016.

[2290] Jason D Perlmutter, Farzaneh Mohajerani, and Michael F Hagan. Many-molecule encapsulation by an icosahedral shell. *eLife*, 5, 2016.

[2291] Jan Perner, Roman Sobotka, Radek Sima, Jitka Konvickova, Daniel Sojka, Pedro Lagerblad de Oliveira, Ondrej Hajdusek, and Petr Kopacek. Acquisition of exogenous haem is essential for tick reproduction. *eLife*, 5, 2016.

[2292] Elsa Perrody, Laurence Abrami, Michal Feldman, Beatrice Kunz, Sylvie Urbé, and F Gisou van der Goot. Ubiquitin-dependent folding of the Wnt signaling coreceptor LRP6. *eLife*, 5, 2016.

[2293] Peter C Petersen and Rune W Berg. Lognormal firing rate distribution reveals prominent fluctuation–driven regime in spinal motor networks. *eLife*, 5, 2016.

[2294] Cohue Peña, Sabina Schütz, Ute Fischer, Yiming Chang, and Vikram G Panse. Prefabrication of a ribosomal protein subcomplex essential for eukaryotic ribosome formation. *eLife*, 5, 2016.

[2295] Anne Pfeiffer, Denis Janocha, Yihan Dong, Anna Medzihradszky, Stefanie Schöne, Gabor Daum, Takuya Suzaki, Joachim Forner, Tobias Langenecker, Eugen Rempel, Markus Schmid, Markus Wirtz, Rüdiger Hell, and Jan U Lohmann. Integration of light and metabolic signals for stem cell activation at the shoot apical meristem. *eLife*, 5, 2016.

[2296] Barbara Pfister, Antoni Sánchez-Ferrer, Ana Diaz, Kuanjen Lu, Caroline Otto, Mirko Holler, Farooque Razvi Shaik, Florence Meier, Raffaele Mezzenga, and Samuel C Zeeman. Recreating the synthesis of starch granules in yeast. *eLife*, 5, 2016.

[2297] Duy Pham Thanh, Abhilasha Karkey, Sabina Dongol, Nhan Ho Thi, Corinne N Thompson, Maia A Rabaa, Amit Arjyal, Kathryn E Holt, Vanessa Wong, Nga Tran Vu Thieu, Phat Voong Vinh, Tuyen Ha Thanh, Ashish Pradhan, Saroj Kumar Shrestha, Damoder Gajurel, Derek Pickard, Christopher M Parry, Gordon Dougan, Marcel Wolbers, Christiane Dolecek, Guy E Thwaites, Buddha Basnyat, and Stephen Baker. A novel ciprofloxacin-resistant subclade of H58 Salmonella Typhi is associated with fluoroquinolone treatment failure. *eLife*, 5, 2016.

[2298] Mitch Phelps, Chris Coss, Hongyan Wang, and Matthew Cook. Registered report: Coding-independent regulation of the tumor suppressor PTEN by competing endogenous mRNAs. *eLife*, 5, 2016.

[2299] Claude Philippe, Dulce B Vargas-Landin, Aurélien J Doucet, Dominic van Essen, Jorge Vera-Otarola, Monika Kuciak, Antoine Corbin, Pilvi Nigumann, and Gaël Cristofari. Activation of individual L1 retrotransposon instances is restricted to cell-type dependent permissive loci. *eLife*, 5, 2016.

[2300] Elizabeth AK Phillips and Andrea R Hasenstaub. Asymmetric effects of activating and inactivating cortical interneurons. *eLife*, 5, 2016.

[2301] Nick E Phillips, Cerys S Manning, Tom Pettini, Veronica Biga, Elli Marinopoulou, Peter Stanley, James Boyd, James Bagnall, Pawel Paszek, David G Spiller, Michael RH White, Marc Goodfellow, Tobias Galla, Magnus Rattray, and Nancy Papalopulu. Stochasticity in the miR-9/Hes1 oscillatory network can account for clonal heterogeneity in the timing of differentiation. *eLife*, 5, 2016.

[2302] Cosimo Pinto, Kristina Kasaciunaite, Ralf Seidel, and Petr Cejka. Human DNA2 possesses a cryptic DNA unwinding activity that functionally integrates with BLM or WRN helicases. *eLife*, 5, 2016.

[2303] Camille Piponiot, Plinio Sist, Lucas Mazzei, Marielos Peña-Claros, Francis E Putz, Ervan Rutishauser, Alexander Shenkin, Nataly Ascarrunz, Celso P de Azevedo, Christopher Baraloto, Mabiane FranÇa, Marcelino Guedes, Eurídice N Honorio Coronado, Marcus VN d'Oliveira, Ademir R Ruschel, Kátia E da Silva, Eleneide Doff Sotta, Cintia R de Souza, Edson Vidal, Thales AP West, and Bruno Hérault. Carbon recovery dynamics following disturbance by selective logging in Amazonian forests. *eLife*, 5, 2016.

[2304] Urszula Piskurewicz, Mayumi Iwasaki, Daichi Susaki, Christian Megies, Tetsu Kinoshita, and Luis Lopez-Molina. Dormancy-specific imprinting underlies maternal inheritance of seed dormancy in Arabidopsis thaliana. *eLife*, 5, 2016.

[2305] Andon N Placzek, David L Molfese, Sanjeev Khatiwada, Gonzalo Viana Di Prisco, Wei Huang, Carmela Sidrauski, KreŠimir Krnjević, Christopher L Amos, Russell Ray, John A Dani, Peter Walter, Ramiro Salas, and Mauro Costa-Mattioli. Translational control of nicotine-evoked synaptic potentiation in mice and neuronal responses in human smokers by eIF2$\alpha$. *eLife*, 5, 2016.

[2306] Andon N Placzek, Gonzalo Viana Di Prisco, Sanjeev Khatiwada, Martina Sgritta, Wei Huang, KreŠimir Krnjević, Randal J Kaufman, John A Dani, Peter Walter, and Mauro Costa-Mattioli. eIF2$\alpha$-mediated translational control regulates the persistence of cocaine-induced LTP in midbrain dopamine neurons. *eLife*, 5, 2016.

[2307] Leigh D Plant, Jeremy D Marks, and Steve AN Goldstein. SUMOylation of NaV1.2 channels mediates the early response to acute hypoxia in central neurons. *eLife*, 5, 2016.

[2308] Lars Plate, Christina B Cooley, John J Chen, Ryan J Paxman, Ciara M Gallagher, Franck Madoux, Joseph C Genereux, Wesley Dobbs, Dan Garza, Timothy P Spicer, Louis Scampavia, Steven J Brown, Hugh Rosen, Evan T Powers, Peter Walter, Peter Hodder, R Luke Wiseman, and Jeffery W Kelly. Small molecule proteostasis regulators that reprogram the ER to reduce extracellular protein aggregation. *eLife*, 5, 2016.

[2309] Shaina N Porter, Andrew S Cluster, Wei Yang, Kelsey A Busken, Riddhi M Patel, Jiyeon Ryoo, and Jeffrey A Magee. Fetal and neonatal hematopoietic progenitors are functionally and transcriptionally resistant to Flt3-ITD mutations. *eLife*, 5, 2016.

[2310] Sandra Postel, Daniel Deredge, Daniel A Bonsor, Xiong Yu, Kay Diederichs, Saskia Helmsing, Aviv Vromen, Assaf Friedler, Michael Hust, Edward H Egelman, Dorothy Beckett, Patrick L Wintrode, and Eric J Sundberg. Bacterial flagellar capping proteins adopt diverse oligomeric states. *eLife*, 5, 2016.

[2311] Shahram Jevin Poureetezadi, Christina N Cheng, Joseph M Chambers, Bridgette E Drummond, and Rebecca A Wingert. Prostaglandin signaling regulates nephron segment patterning of renal progenitors during zebrafish kidney development. *eLife*, 5, 2016.

[2312] Ehsan Pourkarimi, James M Bellush, and Iestyn Whitehouse. Spatiotemporal coupling and decoupling of gene transcription with DNA replication origins during embryogenesis in C. elegans. *eLife*, 5, 2016.

[2313] Jeremy L Praissman, Tobias Willer, M Osman Sheikh, Ants Toi, David Chitayat, Yung-Yao Lin, Hane Lee, Stephanie H Stalnaker, Shuo Wang, Pradeep Kumar Prabhakar, Stanley F Nelson, Derek L Stemple, Steven A Moore, Kelley W Moremen, Kevin P Campbell, and Lance Wells. The functional O-mannose glycan on $\alpha$-dystroglycan contains a phospho-ribitol primed for matriglycan addition. *eLife*, 5, 2016.

[2314] Patricia Preston-Ferrer, Stefano Coletta, Markus Frey, and Andrea Burgalossi. Anatomical organization of presubicular head-direction circuits. *eLife*, 5, 2016.

[2315] Noam Prywes, J Craig Blain, Francesca Del Frate, and Jack W Szostak. Nonenzymatic copying of RNA templates containing all four letters is catalyzed by activated oligonucleotides. *eLife*, 5, 2016.

[2316] Alicja Puścian, Szymon Łęski, Grzegorz Kasprowicz, Maciej Winiarski, Joanna Borowska, Tomasz Nikolaev, Paweł M Boguszewski, Hans-Peter Lipp, and Ewelina Knapska. Eco-HAB as a fully automated and ecologically relevant assessment of social impairments in mouse models of autism. *eLife*, 5, 2016.

[2317] Olena Pylypenko, Tobias Welz, Janine Tittel, Martin Kollmar, Florian Chardon, Gilles Malherbe, Sabine Weiss, Carina Ida Luise Michel, Annette Samol-Wolf, Andreas Till Grasskamp, Alistair Hume, Bruno Goud, Bruno Baron, Patrick England, Margaret A Titus, Petra Schwille, Thomas Weidemann, Anne Houdusse, and Eugen Kerkhoff. Coordinated recruitment of Spir actin nucleators and myosin V motors to Rab11 vesicle membranes. *eLife*, 5, 2016.

[2318] Shuhong Qi, Hui Li, Lisen Lu, Zhongyang Qi, Lei Liu, Lu Chen, Guanxin Shen, Ling Fu, Qingming Luo, and Zhihong Zhang. Long-term intravital imaging of the multicolor-coded tumor microenvironment during combination immunotherapy. *eLife*, 5, 2016.

[2319] Yi-xiang Qi, Jia Huang, Meng-qi Li, Ya-su Wu, Ren-ying Xia, and Gong-yin Ye. Serotonin modulates insect hemocyte phagocytosis via two different serotonin receptors. *eLife*, 5, 2016.

[2320] Jian Qiu, Casey C Nestor, Chunguang Zhang, Stephanie L Padilla, Richard D Palmiter, Martin J Kelly, and Oline K Rønnekleiv. High-frequency stimulation-induced peptide release synchronizes arcuate kisspeptin neurons and excites GnRH neurons. *eLife*, 5, 2016.

[2321] Jing Qiu, Jamie McQueen, Bilada Bilican, Owen Dando, Dario Magnani, Karolina Punovuori, Bhuvaneish T Selvaraj, Matthew Livesey, Ghazal Haghi, Samuel Heron, Karen Burr, Rickie Patani, Rinku Rajan, Olivia Sheppard, Peter C Kind, T Ian Simpson, Victor LJ Tybulewicz, David JA Wyllie, Elizabeth MC Fisher, Sally Lowell, Siddharthan Chandran, and Giles E Hardingham. Evidence for evolutionary divergence of activity-dependent gene expression in developing neurons. *eLife*, 5, 2016.

[2322] Fangfei Qu, Damaris N Lorenzo, Samantha J King, Rebecca Brooks, James E Bear, and Vann Bennett. Ankyrin-B is a PI3P effector that promotes polarized $\alpha5\beta1$-integrin recycling via recruiting RabGAP1L to early endosomes. *eLife*, 5, 2016.

[2323] Zhipeng Qu, Hai Zhang, Moli Huang, Guangsen Shi, Zhiwei Liu, Pancheng Xie, Hui Li, Wei Wang, Guoqiang Xu, Yang Zhang, Ling Yang, Guocun Huang, Joseph S Takahashi, Weiping J Zhang, and Ying Xu. Loss of ZBTB20 impairs circadian output and leads to unimodal behavioral rhythms. *eLife*, 5, 2016.

[2324] Leandro Quadrana, Amanda Bortolini Silveira, George F Mayhew, Chantal LeBlanc, Robert A Martienssen, Jeffrey A Jeddeloh, and Vincent Colot. The Arabidopsis thaliana mobilome and its impact at the species level. *eLife*, 5, 2016.

[2325] Josipa Raguz, Ines Jeric, Theodora Niault, Joanna Daniela Nowacka, Sanya Eduarda Kuzet, Christian Rupp, Irmgard Fischer, Silvia Biggi, Tiziana Borsello, and Manuela Baccarini. Epidermal RAF prevents allergic skin disease. *eLife*, 5, 2016.

[2326] Amrita Rai, Anastasia Oprisko, Jeremy Campos, Yangxue Fu, Timon Friese, Aymelt Itzen, Roger S Goody, Emerich Mihai Gazdag, and Matthias P Müller. bMERB domains are bivalent Rab8 family effectors evolved by gene duplication. *eLife*, 5, 2016.

[2327] Anil Raj, Sidney H Wang, Heejung Shim, Arbel Harpak, Yang I Li, Brett Engelmann, Matthew Stephens, Yoav Gilad, and Jonathan K Pritchard. Thousands of novel translated open reading frames in humans inferred by ribosome footprint profiling. *eLife*, 5, 2016.

[2328] Prithvi Raj, Ekta Rai, Ran Song, Shaheen Khan, Benjamin E Wakeland, Kasthuribai Viswanathan, Carlos Arana, Chaoying Liang, Bo Zhang, Igor Dozmorov, Ferdicia Carr-Johnson, Mitja Mitrovic, Graham B Wiley, Jennifer A Kelly, Bernard R Lauwerys, Nancy J Olsen, Chris Cotsapas, Christine K Garcia, Carol A Wise, John B Harley, Swapan K Nath, Judith A James, Chaim O Jacob, Betty P Tsao, Chandrashekhar Pasare, David R Karp, Quan Zhen Li, Patrick M Gaffney, and Edward K Wakeland. Regulatory polymorphisms modulate the expression of HLA class II molecules and promote autoimmunity. *eLife*, 5, 2016.

[2329] Kasmir Ramo, Koichi Sugamura, Siobhan Craige, John F Keaney, and Roger J Davis. Suppression of ischemia in arterial occlusive disease by JNK-promoted native collateral artery development. *eLife*, 5, 2016.

[2330] Gianpaolo Rando, Chek Kun Tan, Nourhène Khaled, Alexandra Montagner, Nicolas Leuenberger, Justine Bertrand-Michel, Eeswari Paramalingam, Hervé Guillou, and Walter Wahli. Glucocorticoid receptor-PPAR$\alpha$ axis in fetal mouse liver prepares neonates for milk lipid catabolism. *eLife*, 5, 2016.

[2331] Lara M Rangel, Jon W Rueckemann, Pamela D Riviere, Katherine R Keefe, Blake S Porter, Ian S Heimbuch, Carl H Budlong, and Howard Eichenbaum. Rhythmic coordination of hippocampal neurons during associative memory processing. *eLife*, 5, 2016.

[2332] Meera Rao, Voytek Okreglak, Un Seng Chio, Hyunju Cho, Peter Walter, and Shu-ou Shan. Multiple selection filters ensure accurate tail-anchored membrane protein targeting. *eLife*, 5, 2016.

[2333] Saikat Ray and Michael Brecht. Structural development and dorsoventral maturation of the medial entorhinal cortex. *eLife*, 5, 2016.

[2334] Austin L Reese and Ege T Kavalali. Single synapse evaluation of the postsynaptic NMDA receptors targeted by evoked and spontaneous neurotransmission. *eLife*, 5, 2016.

[2335] Max Renner, Mattia Bertinelli, Cédric Leyrat, Guido C Paesen, Laura Freitas Saraiva de Oliveira, Juha T Huiskonen, and Jonathan M Grimes. Nucleocapsid assembly in pneumoviruses is regulated by conformational switching of the N protein. *eLife*, 5, 2016.

[2336] John Repass, Nimet Maherali, and Kate Owen. Registered report: Fusobacterium nucleatum infection is prevalent in human colorectal carcinoma. *eLife*, 5, 2016.

[2337] Shanna L Resendez, Piper C Keyes, Jeremy J Day, Caely Hambro, Curtis J Austin, Francis K Maina, Lori N Eidson, Kirsten A Porter-Stransky, Natalie Nevárez, J William McLean, Morgan A Kuhnmuench, Anne Z Murphy, Tiffany A Mathews, and Brandon J Aragona. Dopamine and opioid systems interact within the nucleus accumbens to maintain monogamous pair bonds. *eLife*, 5, 2016.

[2338] Anne-Cecile Reymann, Fabio Staniscia, Anna Erzberger, Guillaume Salbreux, and Stephan W Grill. Cortical flow aligns actin filaments to form a furrow. *eLife*, 5, 2016.

[2339] Ed Reznik, Martin L Miller, Yasin şenbabaoğlu, Nadeem Riaz, Judy Sarungbam, Satish K Tickoo, Hikmat A Al-Ahmadie, William Lee, Venkatraman E Seshan, A Ari Hakimi, and Chris Sander. Mitochondrial DNA copy number variation across human cancers. *eLife*, 5, 2016.

[2340] Brian C Richardson, Steve L Halaby, Margaret A Gustafson, and J Christopher Fromme. The Sec7 N-terminal regulatory domains facilitate membrane-proximal activation of the Arf1 GTPase. *eLife*, 5, 2016.

[2341] Adam D Richarson, David A Scott, Olga Zagnitko, Pedro Aza-Blanc, Chih-Cheng Chang, and David A Russler-Germain. Registered report: IDH mutation impairs histone demethylation and results in a block to cell differentiation. *eLife*, 5, 2016.

[2342] Franziska R Richter, Rose A Cooper, Paul M Bays, and Jon S Simons. Distinct neural mechanisms underlie the success, precision, and vividness of episodic memory. *eLife*, 5, 2016.

[2343] Nick Riddiford and Gerhard Schlosser. Dissecting the pre-placodal transcriptome to reveal presumptive direct targets of Six1 and Eya1 in cranial placodes. *eLife*, 5, 2016.

[2344] Francesco Rigoli, Karl J Friston, Cristina Martinelli, Mirjana Selaković, Sukhwinder S Shergill, and Raymond J Dolan. A Bayesian model of context-sensitive value attribution. *eLife*, 5, 2016.

[2345] Pablo Ripollés, Josep Marco-Pallarés, Helena Alicart, Claus Tempelmann, Antoni Rodríguez-Fornells, and Toemme Noesselt. Intrinsic monitoring of learning success facilitates memory encoding via the activation of the SN/VTA-Hippocampal loop. *eLife*, 5, 2016.

[2346] Jennifer Ro, Gloria Pak, Paige A Malec, Yang Lyu, David B Allison, Robert T Kennedy, and Scott D Pletcher. Serotonin signaling mediates protein valuation and aging. *eLife*, 5, 2016.

[2347] Seung-Hyun Ro, Xiang Xue, Sadeesh K Ramakrishnan, Chun-Seok Cho, Sim Namkoong, Insook Jang, Ian A Semple, Allison Ho, Hwan-Woo Park, Yatrik M Shah, and Jun Hee Lee. Tumor suppressive role of sestrin2 during colitis and colon carcinogenesis. *eLife*, 5, 2016.

[2348] Brock Roberts, Catalina Casillas, Astrid C Alfaro, Carina Jägers, and Henk Roelink. Patched1 and Patched2 inhibit Smoothened non-cell autonomously. *eLife*, 5, 2016.

[2349] William M Roberts, Steven B Augustine, Kristy J Lawton, Theodore H Lindsay, Tod R Thiele, Eduardo J Izquierdo, Serge Faumont, Rebecca A Lindsay, Matthew Cale Britton, Navin Pokala, Cornelia I Bargmann, and Shawn R Lockery. A stochastic neuronal model predicts random search behaviors at multiple spatial scales in C. elegans. *eLife*, 5, 2016.

[2350] Rachel H Roberts-Galbraith, John L Brubacher, and Phillip A Newmark. A functional genomics screen in planarians reveals regulators of whole-brain regeneration. *eLife*, 5, 2016.

[2351] Crystal Rock, Hector Zurita, Charles Wilson, and Alfonso junior Apicella. An inhibitory corticostriatal pathway. *eLife*, 5, 2016.

[2352] Rebecca Rojansky, Moon-Yong Cha, and David C Chan. Elimination of paternal mitochondria in mouse embryos occurs through autophagic degradation dependent on PARKIN and MUL1. *eLife*, 5, 2016.

[2353] Benjamin Roland, Rebecca Jordan, Dara L Sosulski, Assunta Diodato, Izumi Fukunaga, Ian Wickersham, Kevin M Franks, Andreas T Schaefer, and Alexander Fleischmann. Massive normalization of olfactory bulb output in mice with a 'monoclonal nose'. *eLife*, 5, 2016.

[2354] Ana Rolo, Dawn Savery, Sarah Escuin, Sandra C de Castro, Hannah EJ Armer, Peter MG Munro, Matteo A Molè, Nicholas DE Greene, and Andrew J Copp. Regulation of cell protrusions by small GTPases during fusion of the neural folds. *eLife*, 5, 2016.

[2355] Michael Rosbash. Five suggestions for substantial NIH reforms. *eLife*, 5, 2016.

[2356] Nathan R Rose, Hamish W King, Neil P Blackledge, Nadezda A Fursova, Katherine JI Ember, Roman Fischer, Benedikt M Kessler, and Robert J Klose. RYBP stimulates PRC1 to shape chromatin-based communication between Polycomb repressive complexes. *eLife*, 5, 2016.

[2357] Fabian Rost, Aida Rodrigo Albors, Vladimir Mazurov, Lutz Brusch, Andreas Deutsch, Elly M Tanaka, and Osvaldo Chara. Accelerated cell divisions drive the outgrowth of the regenerating spinal cord in axolotls. *eLife*, 5, 2016.

[2358] Jason Seth Rothman, Laszlo Kocsis, Etienne Herzog, Zoltan Nusser, and Robin Angus Silver. Physical determinants of vesicle mobility and supply at a central synapse. *eLife*, 5, 2016.

[2359] Maxime WC Rousseaux, Maria de Haro, Cristian A Lasagna-Reeves, Antonia De Maio, Jeehye Park, Paymaan Jafar-Nejad, Ismael Al-Ramahi, Ajay Sharma, Lauren See, Nan Lu, Luis Vilanova-Velez, Tiemo J Klisch, Thomas F Westbrook, Juan C Troncoso, Juan Botas, and Huda Y Zoghbi. TRIM28 regulates the nuclear accumulation and toxicity of both alpha-synuclein and tau. *eLife*, 5, 2016.

[2360] Sébastien Roux, Frédéric Matonti, Florent Dupont, Louis Hoffart, Sylvain Takerkart, Serge Picaud, Pascale Pham, and Frédéric Chavane. Probing the functional impact of sub-retinal prosthesis. *eLife*, 5, 2016.

[2361] Taruni Roy Khurana and Sanjay P Sane. Airflow and optic flow mediate antennal positioning in flying honeybees. *eLife*, 5, 2016.

[2362] Huan Rui, Pablo Artigas, and Benoît Roux. The selectivity of the Na+/K+-pump is controlled by binding site protonation and self-correcting occlusion. *eLife*, 5, 2016.

[2363] Kerrianne Ryan, Zhiyuan Lu, and Ian A Meinertzhagen. The CNS connectome of a tadpole larva of Ciona intestinalis (L.) highlights sidedness in the brain of a chordate sibling. *eLife*, 5, 2016.

[2364] Brian F Sadacca, Joshua L Jones, and Geoffrey Schoenbaum. Midbrain dopamine neurons compute inferred and cached value prediction errors in a common framework. *eLife*, 5, 2016.

[2365] Yuhki Saito, Soledad Miranda-Rottmann, Matteo Ruggiu, Christopher Y Park, John J Fak, Ru Zhong, Jeremy S Duncan, Brian A Fabella, Harald J Junge, Zhe Chen, Roberto Araya, Bernd Fritzsch, A J Hudspeth, and Robert B Darnell. NOVA2-mediated RNA regulation is required for axonal pathfinding during development. *eLife*, 5, 2016.

[2366] Charanya Sampathkumar, Yuan-Ju Wu, Mayur Vadhvani, Thorsten Trimbuch, Britta Eickholt, and Christian Rosenmund. Loss of MeCP2 disrupts cell autonomous and autocrine BDNF signaling in mouse glutamatergic neurons. *eLife*, 5, 2016.

[2367] Raul Y Sanchez David, Chantal Combredet, Odile Sismeiro, Marie-Agnès Dillies, Bernd Jagla, Jean-Yves Coppée, Marie Mura, Mathilde Guerbois Galla, Philippe Despres, Frédéric Tangy, and Anastassia V Komarova. Comparative analysis of viral RNA signatures on different RIG-I-like receptors. *eLife*, 5, 2016.

[2368] Laura E Sanman, Yu Qian, Nicholas A Eisele, Tessie M Ng, Wouter A van der Linden, Denise M Monack, Eranthie Weerapana, and Matthew Bogyo. Disruption of glycolytic flux is a signal for inflammasome signaling and pyroptotic cell death. *eLife*, 5, 2016.

[2369] Julia Santiago, Benjamin Brandt, Mari Wildhagen, Ulrich Hohmann, Ludwig A Hothorn, Melinka A Butenko, and Michael Hothorn. Mechanistic insight into a peptide hormone signaling complex mediating floral organ abscission. *eLife*, 5, 2016.

[2370] Sarvenaz Sarabipour, Kurt Ballmer-Hofer, and Kalina Hristova. VEGFR-2 conformational switch in response to ligand binding. *eLife*, 5, 2016.

[2371] Derya Sargin, David K Oliver, and Evelyn K Lambe. Chronic social isolation reduces 5-HT neuronal activity via upregulated SK3 calcium-activated potassium channels. *eLife*, 5, 2016.

[2372] Mihail Sarov, Christiane Barz, Helena Jambor, Marco Y Hein, Christopher Schmied, Dana Suchold, Bettina Stender, Stephan Janosch, Vinay Vikas KJ, RT Krishnan, Aishwarya Krishnamoorthy, Irene RS Ferreira, Radoslaw K Ejsmont, Katja Finkl, Susanne Hasse, Philipp Kämpfer, Nicole Plewka, Elisabeth Vinis, Siegfried Schloissnig, Elisabeth Knust, Volker Hartenstein, Matthias Mann, Mani Ramaswami, K VijayRaghavan, Pavel Tomancak, and Frank Schnorrer. A genome-wide resource for the analysis of protein localisation in Drosophila. *eLife*, 5, 2016.

[2373] Pablo Sartori, Veikko F Geyer, Andre Scholich, Frank Jülicher, and Jonathon Howard. Dynamic curvature regulation accounts for the symmetric and asymmetric beats of Chlamydomonas flagella. *eLife*, 5, 2016.

[2374] Yo Sasaki, Takashi Nakagawa, Xianrong Mao, Aaron DiAntonio, and Jeffrey Milbrandt. NMNAT1 inhibits axon degeneration via blockade of SARM1-mediated NAD+ depletion. *eLife*, 5, 2016.

[2375] Satoru M Sato and Catherine S Woolley. Acute inhibition of neurosteroid estrogen synthesis suppresses status epilepticus in an animal model. *eLife*, 5, 2016.

[2376] Rupinder Sayal, Jacqueline M Dresch, Irina Pushel, Benjamin R Taylor, and David N Arnosti. Quantitative perturbation-based analysis of gene expression predicts enhancer activity in early Drosophila embryo. *eLife*, 5, 2016.

[2377] Dorothy P Schafer, Christopher T Heller, Georgia Gunner, Molly Heller, Christopher Gordon, Timothy Hammond, Yochai Wolf, Steffen Jung, and Beth Stevens. Microglia contribute to circuit defects in Mecp2 null mice independent of microglia-specific loss of Mecp2 expression. *eLife*, 5, 2016.

[2378] Louise Scharf, Anthony P West, Stuart A Sievers, Courtney Chen, Siduo Jiang, Han Gao, Matthew D Gray, Andrew T McGuire, Johannes F Scheid, Michel C Nussenzweig, Leonidas Stamatatos, and Pamela J Bjorkman. Structural basis for germline antibody recognition of HIV-1 immunogens. *eLife*, 5, 2016.

[2379] Claudia Scheckel, Elodie Drapeau, Maria A Frias, Christopher Y Park, John Fak, Ilana Zucker-Scharff, Yan Kou, Vahram Haroutunian, Avi Ma'ayan, Joseph D Buxbaum, and Robert B Darnell. Regulatory consequences of neuronal ELAV-like protein binding to coding and non-coding RNAs in human brain. *eLife*, 5, 2016.

[2380] Robson Scheffer-Teixeira and Adriano BL Tort. On cross-frequency phase-phase coupling between theta and gamma oscillations in the hippocampus. *eLife*, 5, 2016.

[2381] Sven Schenk, Christian Krauditsch, Peter Frühauf, Christopher Gerner, and Florian Raible. Discovery of methylfarnesoate as the annelid brain hormone reveals an ancient role of sesquiterpenoids in reproduction. *eLife*, 5, 2016.

[2382] Philipp Schlegel, Michael J Texada, Anton Miroschnikow, Andreas Schoofs, Sebastian Hückesfeld, Marc Peters, Casey M Schneider-Mizell, Haluk Lacin, Feng Li, Richard D Fetter, James W Truman, Albert Cardona, and Michael J Pankratz. Synaptic transmission parallels neuromodulation in a central food-intake circuit. *eLife*, 5, 2016.

[2383] Edward T Schmid, Iris K Pang, Eugenio A Carrera Silva, Lidia Bosurgi, Jonathan J Miner, Michael S Diamond, Akiko Iwasaki, and Carla V Rothlin. AXL receptor tyrosine kinase is required for T cell priming and antiviral immunity. *eLife*, 5, 2016.

[2384] Dominik Schmiedel, Julie Tai, Rachel Yamin, Orit Berhani, Yoav Bauman, and Ofer Mandelboim. The RNA binding protein IMP3 facilitates tumor immune escape by downregulating the stress-induced ligands ULPB2 and MICB. *eLife*, 5, 2016.

[2385] Casey M Schneider-Mizell, Stephan Gerhard, Mark Longair, Tom Kazimiers, Feng Li, Maarten F Zwart, Andrew Champion, Frank M Midgley, Richard D Fetter, Stephan Saalfeld, and Albert Cardona. Quantitative neuroanatomy for connectomics in Drosophila. *eLife*, 5, 2016.

[2386] Sebastian Schnorrenberg, Tim Grotjohann, Gerd Vorbrüggen, Alf Herzig, Stefan W Hell, and Stefan Jakobs. In vivo super-resolution RESOLFT microscopy of Drosophila melanogaster. *eLife*, 5, 2016.

[2387] Nils Schuergers, Tchern Lenn, Ronald Kampmann, Markus V Meissner, Tiago Esteves, Maja Temerinac-Ott, Jan G Korvink, Alan R Lowe, Conrad W Mullineaux, and Annegret Wilde. Cyanobacteria use micro-optics to sense light direction. *eLife*, 5, 2016.

[2388] Anne K Schuetz and Lewis E Kay. A Dynamic molecular basis for malfunction in disease mutants of p97/VCP. *eLife*, 5, 2016.

[2389] Ursula Schulze-Gahmen, Ignacia Echeverria, Goran Stjepanovic, Yun Bai, Huasong Lu, Dina Schneidman-Duhovny, Jennifer A Doudna, Qiang Zhou, Andrej Sali, and James H Hurley. Insights into HIV-1 proviral transcription from integrative structure and dynamics of the Tat:AFF4:P-TEFb:TAR complex. *eLife*, 5, 2016.

[2390] Judith Schütte, Huange Wang, Stella Antoniou, Andrew Jarratt, Nicola K Wilson, Joey Riepsaame, Fernando J Calero-Nieto, Victoria Moignard, Silvia Basilico, Sarah J Kinston, Rebecca L Hannah, Mun Chiang Chan, Sylvia T Nürnberg, Willem H Ouwehand, Nicola Bonzanni, Marella FTR de Bruijn, and Berthold Göttgens. An experimentally validated network of nine haematopoietic transcription factors reveals mechanisms of cell state stability. *eLife*, 5, 2016.

[2391] Achim H Schwermann, Tomy dos Santos Rolo, Michael S Caterino, Günter Bechly, Heiko Schmied, Tilo Baumbach, and Thomas van de Kamp. Preservation of three-dimensional anatomy in phosphatized fossil arthropods enriches evolutionary inference. *eLife*, 5, 2016.

[2392] Caspar M Schwiedrzik, Benjamin Bernstein, and Lucia Melloni. Motion along the mental number line reveals shared representations for numerosity and space. *eLife*, 5, 2016.

[2393] M Lucila Scimone, Lauren E Cote, Travis Rogers, and Peter W Reddien. Two FGFRL-Wnt circuits organize the planarian anteroposterior axis. *eLife*, 5, 2016.

[2394] William Sedley, Phillip E Gander, Sukhbinder Kumar, Christopher K Kovach, Hiroyuki Oya, Hiroto Kawasaki, Matthew A Howard, and Timothy D Griffiths. Neural signatures of perceptual inference. *eLife*, 5, 2016.

[2395] Stefanie Seelk, Irene Adrian-Kalchhauser, Balázs Hargitai, Martina Hajduskova, Silvia Gutnik, Baris Tursun, and Rafal Ciosk. Increasing Notch signaling antagonizes PRC2-mediated silencing to promote reprograming of germ cells into neurons. *eLife*, 5, 2016.

[2396] Elizabeth M Sefton, Bhart-Anjan S Bhullar, Zahra Mohaddes, and James Hanken. Evolution of the head-trunk interface in tetrapod vertebrates. *eLife*, 5, 2016.

[2397] Einat Segev, Thomas P Wyche, Ki Hyun Kim, Jörn Petersen, Claire Ellebrandt, Hera Vlamakis, Natasha Barteneva, Joseph N Paulson, Liraz Chai, Jon Clardy, and Roberto Kolter. Dynamic metabolic exchange governs a marine algal-bacterial interaction. *eLife*, 5, 2016.

[2398] Eyal Seidemann, Yuzhi Chen, Yoon Bai, Spencer C Chen, Preeti Mehta, Bridget L Kajs, Wilson S Geisler, and Boris V Zemelman. Calcium imaging with genetically encoded indicators in behaving primates. *eLife*, 5, 2016.

[2399] Lindsey Seldin, Andrew Muroyama, and Terry Lechler. NuMA-microtubule interactions are critical for spindle orientation and the morphogenesis of diverse epidermal structures. *eLife*, 5, 2016.

[2400] Melih Sener, Johan Strumpfer, Abhishek Singharoy, C Neil Hunter, and Klaus Schulten. Overall energy conversion efficiency of a photosynthetic vesicle. *eLife*, 5, 2016.

[2401] Laura Senst, Dinara Baimoukhametova, Toni-Lee Sterley, and Jaideep Singh Bains. Sexually dimorphic neuronal responses to social isolation. *eLife*, 5, 2016.

[2402] Irene Serrano, Pierre Buscaill, Corinne Audran, Cécile Pouzet, Alain Jauneau, and Susana Rivas. A non canonical subtilase attenuates the transcriptional activation of defence responses in Arabidopsis thaliana. *eLife*, 5, 2016.

[2403] Neel H Shah, Qi Wang, Qingrong Yan, Deepti Karandur, Theresa A Kadlecek, Ian R Fallahee, William P Russ, Rama Ranganathan, Arthur Weiss, and John Kuriyan. An electrostatic selection mechanism controls sequential kinase signaling downstream of the T cell receptor. *eLife*, 5, 2016.

[2404] Chun-Min Shan, Jiyong Wang, Ke Xu, Huijie Chen, Jia-Xing Yue, Stuart Andrews, James J Moresco, John R Yates, Peter L Nagy, Liang Tong, and Songtao Jia. A histone H3K9M mutation traps histone methyltransferase Clr4 to prevent heterochromatin spreading. *eLife*, 5, 2016.

[2405] Kevin Q Shan, Evgueniy V Lubenov, Maria Papadopoulou, and Athanassios G Siapas. Spatial tuning and brain state account for dorsal hippocampal CA1 activity in a non-spatial learning task. *eLife*, 5, 2016.

[2406] Vidhu Sharma, Lisa Young, Anne B Allison, and Kate Owen. Registered report: Diverse somatic mutation patterns and pathway alterations in human cancers. *eLife*, 5, 2016.

[2407] Vidhu Sharma, Lisa Young, Miguel Cavadas, and Kate Owen. Registered report: COT drives resistance to RAF inhibition through MAP kinase pathway reactivation. *eLife*, 5, 2016.

[2408] Koning Shen, Barbara Calamini, Jonathan A Fauerbach, Boxue Ma, Sarah H Shahmoradian, Ivana L Serrano Lachapel, Wah Chiu, Donald C Lo, and Judith Frydman. Control of the structural landscape and neuronal proteotoxicity of mutant Huntingtin by domains flanking the polyQ tract. *eLife*, 5, 2016.

[2409] Yang Shen, Marilia Barros, Tarek Vennemann, D Travis Gallagher, Yizhou Yin, Sara B Linden, Ryan D Heselpoth, Dennis J Spencer, David M Donovan, John Moult, Vincent A Fischetti, Frank Heinrich, Mathias Lösche, and Daniel C Nelson. A bacteriophage endolysin that eliminates intracellular streptococci. *eLife*, 5, 2016.

[2410] Etsuko Shibata, Manjari Kiran, Yoshiyuki Shibata, Samarendra Singh, Shashi Kiran, and Anindya Dutta. Two subunits of human ORC are dispensable for DNA replication and proliferation. *eLife*, 5, 2016.

[2411] Yasuyuki Shima, Ken Sugino, Chris Martin Hempel, Masami Shima, Praveen Taneja, James B Bullis, Sonam Mehta, Carlos Lois, and Sacha B Nelson. A Mammalian enhancer trap resource for discovering and manipulating neuronal cell types. *eLife*, 5, 2016.

[2412] Jaeyoung Shin, Shuichi Watanabe, Soraya Hoelper, Marcus Krüger, Sawa Kostin, Jochen Pöling, Thomas Kubin, and Thomas Braun. BRAF activates PAX3 to control muscle precursor cell migration during forelimb muscle development. *eLife*, 5, 2016.

[2413] Jihoon Shin, Tae Wan Kim, Hyunsoo Kim, Hye Ji Kim, Min Young Suh, Sangho Lee, Han-Teo Lee, Sojung Kwak, Sang-Eun Lee, Jong-Hyuk Lee, Hyonchol Jang, Eun-Jung Cho, and Hong-Duk Youn. Aurkb/PP1-mediated resetting of Oct4 during the cell cycle determines the identity of embryonic stem cells. *eLife*, 5, 2016.

[2414] Kotaro Shirakawa, Lan Wang, Na Man, Jasna Maksimoska, Alexander W Sorum, Hyung W Lim, Intelly S Lee, Tadahiro Shimazu, John C Newman, Sebastian Schröder, Melanie Ott, Ronen Marmorstein, Jordan Meier, Stephen Nimer, and Eric Verdin. Salicylate, diflunisal and their metabolites inhibit CBP/p300 and exhibit anticancer activity. *eLife*, 5, 2016.

[2415] Nitin H Shirole, Debjani Pal, Edward R Kastenhuber, Serif Senturk, Joseph Boroda, Paola Pisterzi, Madison Miller, Gustavo Munoz, Marko Anderluh, Marc Ladanyi, Scott W Lowe, and Raffaella Sordella. TP53 exon-6 truncating mutations produce separation of function isoforms with protumorigenic functions. *eLife*, 5, 2016.

[2416] Rabindra V Shivnaraine, Brendan Kelly, Krishana S Sankar, Dar'ya S Redka, Yi Rang Han, Fei Huang, Gwendolynne Elmslie, Daniel Pinto, Yuchong Li, Jonathan V Rocheleau, Claudiu C Gradinaru, John Ellis, and James W Wells. Allosteric modulation in monomers and oligomers of a G protein-coupled receptor. *eLife*, 5, 2016.

[2417] Matthew J Shurtleff, Morayma M Temoche-Diaz, Kate V Karfilis, Sayaka Ri, and Randy Schekman. Y-box protein 1 is required to sort microRNAs into exosomes in cells and in a cell-free reaction. *eLife*, 5, 2016.

[2418] Felix Siebenhühner, Sheng H Wang, J Matias Palva, and Satu Palva. Cross-frequency synchronization connects networks of fast and slow oscillations during visual working memory maintenance. *eLife*, 5, 2016.

[2419] Daniel Sieiro, Anne C Rios, Claire E Hirst, and Christophe Marcelle. Cytoplasmic NOTCH and membrane-derived $\beta$-catenin link cell fate choice to epithelial-mesenchymal transition during myogenesis. *eLife*, 5, 2016.

[2420] Jennie R Sims, Maria C Ow, Mailyn A Nishiguchi, Kyuhyung Kim, Piali Sengupta, and Sarah E Hall. Developmental programming modulates olfactory behavior in C. elegans via endogenous RNAi pathways. *eLife*, 5, 2016.

[2421] Shalini Singh, Danielle Howell, Niraj Trivedi, Ketty Kessler, Taren Ong, Pedro Rosmaninho, Alexandre ASF Raposo, Giles Robinson, Martine F Roussel, Diogo S Castro, and David J Solecki. Zeb1 controls neuron differentiation and germinal zone exit by a mesenchymal-epithelial-like transition. *eLife*, 5, 2016.

[2422] Abhishek Singharoy, Ivan Teo, Ryan McGreevy, John E Stone, Jianhua Zhao, and Klaus Schulten. Molecular dynamics-based refinement and validation for sub-5 Å cryo-electron microscopy maps. *eLife*, 5, 2016.

[2423] Katarzyna Siuda-Krzywicka, Łukasz Bola, Małgorzata Paplińska, Ewa Sumera, Katarzyna Jednoróg, Artur Marchewka, Magdalena W Śliwińska, Amir Amedi, and Marcin Szwed. Massive cortical reorganization in sighted Braille readers. *eLife*, 5, 2016.

[2424] Sushama Sivakumar, Paweł Ł Janczyk, Qianhui Qu, Chad A Brautigam, P Todd Stukenberg, Hongtao Yu, and Gary J Gorbsky. The human SKA complex drives the metaphase-anaphase cell cycle transition by recruiting protein phosphatase 1 to kinetochores. *eLife*, 5, 2016.

[2425] Samuel O Skinner, Heng Xu, Sonal Nagarkar-Jaiswal, Pablo R Freire, Thomas P Zwaka, and Ido Golding. Single-cell analysis of transcription kinetics across the cell cycle. *eLife*, 5, 2016.

[2426] Sama F Sleiman, Jeffrey Henry, Rami Al-Haddad, Lauretta El Hayek, Edwina Abou Haidar, Thomas Stringer, Devyani Ulja, Saravanan S Karuppagounder, Edward B Holson, Rajiv R Ratan, Ipe Ninan, and Moses V Chao. Exercise promotes the expression of brain derived neurotrophic factor (BDNF) through the action of the ketone body $\beta$-hydroxybutyrate. *eLife*, 5, 2016.

[2427] Chris A Smith, Andrew D McAinsh, and Nigel J Burroughs. Human kinetochores are swivel joints that mediate microtubule attachments. *eLife*, 5, 2016.

[2428] Heather L Smith, Jennifer N Bourne, Guan Cao, Michael A Chirillo, Linnaea E Ostroff, Deborah J Watson, and Kristen M Harris. Mitochondrial support of persistent presynaptic vesicle mobilization with age-dependent synaptic growth after LTP. *eLife*, 5, 2016.

[2429] Jarrett Smith, Deepika Calidas, Helen Schmidt, Tu Lu, Dominique Rasoloson, and Geraldine Seydoux. Spatial patterning of P granules by RNA-induced phase separation of the intrinsically-disordered protein MEG-3. *eLife*, 5, 2016.

[2430] Kristen M Smith-Edwards, Jennifer J DeBerry, Jami L Saloman, Brian M Davis, and C Jeffery Woodbury. Profound alteration in cutaneous primary afferent activity produced by inflammatory mediators. *eLife*, 5, 2016.

[2431] Michal Sobecki, Karim Mrouj, Alain Camasses, Nikolaos Parisis, Emilien Nicolas, David Llères, FranÇois Gerbe, Susana Prieto, Liliana Krasinska, Alexandre David, Manuel Eguren, Marie-Christine Birling, Serge Urbach, Sonia Hem, Jérôme Déjardin, Marcos Malumbres, Philippe Jay, Vjekoslav Dulic, Denis LJ Lafontaine, Robert Feil, and Daniel Fisher. The cell proliferation antigen Ki-67 organises heterochromatin. *eLife*, 5, 2016.

[2432] Meghna Sobti, Callum Smits, Andrew SW Wong, Robert Ishmukhametov, Daniela Stock, Sara Sandin, and Alastair G Stewart. Cryo-EM structures of the autoinhibited E. coli ATP synthase in three rotational states. *eLife*, 5, 2016.

[2433] Nicholas James Sofroniew, Daniel Flickinger, Jonathan King, and Karel Svoboda. A large field of view two-photon mesoscope with subcellular resolution for in vivo imaging. *eLife*, 5, 2016.

[2434] Ediz Sohoglu and Maria Chait. Detecting and representing predictable structure during auditory scene analysis. *eLife*, 5, 2016.

[2435] Jordi Solana, Manuel Irimia, Salah Ayoub, Marta Rodriguez Orejuela, Vera Zywitza, Marvin Jens, Javier Tapial, Debashish Ray, Quaid Morris, Timothy R Hughes, Benjamin J Blencowe, and Nikolaus Rajewsky. Conserved functional antagonism of CELF and MBNL proteins controls stem cell-specific alternative splicing in planarians. *eLife*, 5, 2016.

[2436] Alessia Soldano, Yeranddy A Alpizar, Brett Boonen, Luis Franco, Alejandro López-Requena, Guangda Liu, Natalia Mora, Emre Yaksi, Thomas Voets, Rudi Vennekens, Bassem A Hassan, and Karel Talavera. Gustatory-mediated avoidance of bacterial lipopolysaccharides via TRPA1 activation in Drosophila. *eLife*, 5, 2016.

[2437] Elliott D SoRelle, Orly Liba, Jos L Campbell, Roopa Dalal, Cristina L Zavaleta, and Adam de la Zerda. A hyperspectral method to assay the microphysiological fates of nanomaterials in histological samples. *eLife*, 5, 2016.

[2438] Julian Sosnik, Likun Zheng, Christopher V Rackauckas, Michelle Digman, Enrico Gratton, Qing Nie, and Thomas F Schilling. Noise modulation in retinoic acid signaling sharpens segmental boundaries of gene expression in the embryonic zebrafish hindbrain. *eLife*, 5, 2016.

[2439] Joana S Sousa, Deryck J Mills, Janet Vonck, and Werner Kühlbrandt. Functional asymmetry and electron flow in the bovine respirasome. *eLife*, 5, 2016.

[2440] Sheryl Southard, Ju-Ryoung Kim, SiewHui Low, Richard W Tsika, and Christoph Lepper. Myofiber-specific TEAD1 overexpression drives satellite cell hyperplasia and counters pathological effects of dystrophin deficiency. *eLife*, 5, 2016.

[2441] Naren Srinivasan, Oliver Gordon, Susan Ahrens, Anna Franz, Safia Deddouche, Probir Chakravarty, David Phillips, Ali A Yunus, Michael K Rosen, Rita S Valente, Luis Teixeira, Barry Thompson, Marc S Dionne, Will Wood, and Caetano Reis e Sousa. Actin is an evolutionarily-conserved damage-associated molecular pattern that signals tissue injury in Drosophila melanogaster. *eLife*, 5, 2016.

[2442] Shekhar Srivastava, Saswati Panda, Zhai Li, Stephen R Fuhs, Tony Hunter, Dennis J Thiele, Stevan R Hubbard, and Edward Y Skolnik. Histidine phosphorylation relieves copper inhibition in the mammalian potassium channel KCa3.1. *eLife*, 5, 2016.

[2443] Bernhard P Staresina, Sebastian Michelmann, Mathilde Bonnefond, Ole Jensen, Nikolai Axmacher, and Juergen Fell. Hippocampal pattern completion is linked to gamma power increases and alpha power decreases during recollection. *eLife*, 5, 2016.

[2444] Martin Steger, Francesca Tonelli, Genta Ito, Paul Davies, Matthias Trost, Melanie Vetter, Stefanie Wachter, Esben Lorentzen, Graham Duddy, Stephen Wilson, Marco AS Baptista, Brian K Fiske, Matthew J Fell, John A Morrow, Alastair D Reith, Dario R Alessi, and Matthias Mann. Phosphoproteomics reveals that Parkinson's disease kinase LRRK2 regulates a subset of Rab GTPases. *eLife*, 5, 2016.

[2445] Claire E Stelly, Matthew B Pomrenze, Jason B Cook, and Hitoshi Morikawa. Repeated social defeat stress enhances glutamatergic synaptic plasticity in the VTA and cocaine place conditioning. *eLife*, 5, 2016.

[2446] Caroline Townsend Stork, Michael Bocek, Madzia P Crossley, Julie Sollier, Lionel A Sanz, Frédéric Chédin, Tomek Swigut, and Karlene A Cimprich. Co-transcriptional R-loops are the main cause of estrogen-induced DNA damage. *eLife*, 5, 2016.

[2447] Benjamin M Stormo and Donald T Fox. Distinct responses to reduplicated chromosomes require distinct Mad2 responses. *eLife*, 5, 2016.

[2448] Andrea Strakova, Máire Ní Leathlobhair, Guo-Dong Wang, Ting-Ting Yin, Ilona Airikkala-Otter, Janice L Allen, Karen M Allum, Leontine Bansse-Issa, Jocelyn L Bisson, Artemio Castillo Domracheva, Karina F de Castro, Anne M Corrigan, Hugh R Cran, Jane T Crawford, Stephen M Cutter, Laura Delgadillo Keenan, Edward M Donelan, Ibikunle A Faramade, Erika Flores Reynoso, Eleni Fotopoulou, Skye N Fruean, Fanny Gallardo-Arrieta, Olga Glebova, Rodrigo F Häfelin Manrique, Joaquim JGP Henriques, Natalia Ignatenko, Debbie Koenig, Marta Lanza-Perea, Remo Lobetti, Adriana M Lopez Quintana, Thibault Losfelt, Gabriele Marino, Inigo Martincorena, Simón Martínez Castañeda, Mayra F Martínez-López, Michael Meyer, Berna Nakanwagi, Andrigo B De Nardi, Winifred Neunzig, Sally J Nixon, Marsden M Onsare, Antonio Ortega-Pacheco, Maria C Peleteiro, Ruth J Pye, John F Reece, Jose Rojas Gutierrez, Haleema Sadia, Sheila K Schmeling, Olga Shamanova, Richard K Ssuna, Audrey E Steenland-Smit, Alla Svitich, Ismail Thoya Ngoka, Bogdan A Vițălaru, Anna P de Vos, Johan P de Vos, Oliver Walkinton, David C Wedge, Alvaro S Wehrle-Martinez, Mirjam G van der Wel, Sophie AE Widdowson, and Elizabeth P Murchison. Mitochondrial genetic diversity, selection and recombination in a canine transmissible cancer. *eLife*, 5, 2016.

[2449] Johannes Stratmann, Hugo Gabilondo, Jonathan Benito-Sipos, and Stefan Thor. Neuronal cell fate diversification controlled by sub-temporal action of Kruppel. *eLife*, 5, 2016.

[2450] Ingeborg Streng-Ouwehand, Nataschja I Ho, Manja Litjens, Hakan Kalay, Martine Annemarie Boks, Lenneke AM Cornelissen, Satwinder Kaur Singh, Eirikur Saeland, Juan J Garcia-Vallejo, Ferry A Ossendorp, Wendy WJ Unger, and Yvette van Kooyk. Glycan modification of antigen alters its intracellular routing in dendritic cells, promoting priming of T cells. *eLife*, 5, 2016.

[2451] Carsen Stringer, Marius Pachitariu, Nicholas A Steinmetz, Michael Okun, Peter Bartho, Kenneth D Harris, Maneesh Sahani, and Nicholas A Lesica. Inhibitory control of correlated intrinsic variability in cortical networks. *eLife*, 5, 2016.

[2452] Tim Stuart, Steven R Eichten, Jonathan Cahn, Yuliya V Karpievitch, Justin O Borevitz, and Ryan Lister. Population scale mapping of transposable element diversity reveals links to gene regulation and epigenomic variation. *eLife*, 5, 2016.

[2453] Bradford M Stubenhaus, John P Dustin, Emily R Neverett, Megan S Beaudry, Leanna E Nadeau, Ethan Burk-McCoy, Xinwen He, Bret J Pearson, and Jason Pellettieri. Light-induced depigmentation in planarians models the pathophysiology of acute porphyrias. *eLife*, 5, 2016.

[2454] Saranya Subramani, Harmonie Perdreau-Dahl, and Jens Preben Morth. The magnesium transporter A is activated by cardiolipin and is highly sensitive to free magnesium in vitro. *eLife*, 5, 2016.

[2455] Jørgen Sugar and Menno P Witter. Postnatal development of retrosplenial projections to the parahippocampal region of the rat. *eLife*, 5, 2016.

[2456] Michael Sulak, Lindsey Fong, Katelyn Mika, Sravanthi Chigurupati, Lisa Yon, Nigel P Mongan, Richard D Emes, and Vincent J Lynch. TP53 copy number expansion is associated with the evolution of increased body size and an enhanced DNA damage response in elephants. *eLife*, 5, 2016.

[2457] Kelly D Sullivan, Hannah C Lewis, Amanda A Hill, Ahwan Pandey, Leisa P Jackson, Joseph M Cabral, Keith P Smith, L Alexander Liggett, Eliana B Gomez, Matthew D Galbraith, James DeGregori, and Joaquín M Espinosa. Trisomy 21 consistently activates the interferon response. *eLife*, 5, 2016.

[2458] Wenjing Sun, Elizabeth A Matthews, Vicky Nicolas, Susanne Schoch, and Dirk Dietrich. NG2 glial cells integrate synaptic input in global and dendritic calcium signals. *eLife*, 5, 2016.

[2459] Yishan Sun, Sergiu P Paşca, Thomas Portmann, Carleton Goold, Kathleen A Worringer, Wendy Guan, Karen C Chan, Hui Gai, Daniel Vogt, Ying-Jiun J Chen, Rong Mao, Karrie Chan, John LR Rubenstein, Daniel V Madison, Joachim Hallmayer, Wendy M Froehlich-Santino, Jonathan A Bernstein, and Ricardo E Dolmetsch. A deleterious Nav1.1 mutation selectively impairs telencephalic inhibitory neurons derived from Dravet Syndrome patients. *eLife*, 5, 2016.

[2460] Min-Kyung Sung, Tanya R Porras-Yakushi, Justin M Reitsma, Ferdinand M Huber, Michael J Sweredoski, André Hoelz, Sonja Hess, and Raymond J Deshaies. A conserved quality-control pathway that mediates degradation of unassembled ribosomal proteins. *eLife*, 5, 2016.

[2461] Nobuyoshi Suto, Amanda Laque, Genna L De Ness, Grant E Wagner, Debbie Watry, Tony Kerr, Eisuke Koya, Mark R Mayford, Bruce T Hope, and Friedbert Weiss. Distinct memory engrams in the infralimbic cortex of rats control opposing environmental actions on a learned behavior. *eLife*, 5, 2016.

[2462] Joel M Swenson, Serafin U Colmenares, Amy R Strom, Sylvain V Costes, and Gary H Karpen. The composition and organization of Drosophila heterochromatin are heterogeneous and dynamic. *eLife*, 5, 2016.

[2463] Durafshan Sakeena Syed, Swetha B.M. Gowda, O Venkateswara Reddy, Heinrich Reichert, and K VijayRaghavan. Glial and neuronal Semaphorin signaling instruct the development of a functional myotopic map for Drosophila walking. *eLife*, 5, 2016.

[2464] Andreas T Sørensen, Yonatan A Cooper, Michael V Baratta, Feng-Ju Weng, Yuxiang Zhang, Kartik Ramamoorthi, Robin Fropf, Emily LaVerriere, Jian Xue, Andrew Young, Colleen Schneider, Casper René Gøtzsche, Martin Hemberg, Jerry CP Yin, Steven F Maier, and Yingxi Lin. A robust activity marking system for exploring active neuronal ensembles. *eLife*, 5, 2016.

[2465] Erica Tagliatti, Manuela Fadda, Antonio Falace, Fabio Benfenati, and Anna Fassio. Arf6 regulates the cycling and the readily releasable pool of synaptic vesicles at hippocampal synapse. *eLife*, 5, 2016.

[2466] Seika Takayanagi-Kiya, Keming Zhou, and Yishi Jin. Release-dependent feedback inhibition by a presynaptically localized ligand-gated anion channel. *eLife*, 5, 2016.

[2467] Akira Takeda, Daichi Kobayashi, Keita Aoi, Naoko Sasaki, Yuki Sugiura, Hidemitsu Igarashi, Kazuo Tohya, Asuka Inoue, Erina Hata, Noriyuki Akahoshi, Haruko Hayasaka, Junichi Kikuta, Elke Scandella, Burkhard Ludewig, Satoshi Ishii, Junken Aoki, Makoto Suematsu, Masaru Ishii, Kiyoshi Takeda, Sirpa Jalkanen, Masayuki Miyasaka, and Eiji Umemoto. Fibroblastic reticular cell-derived lysophosphatidic acid regulates confined intranodal T-cell motility. *eLife*, 5, 2016.

[2468] Dan Tan, Qiang Li, Mei-Jun Zhang, Chao Liu, Chengying Ma, Pan Zhang, Yue-He Ding, Sheng-Bo Fan, Li Tao, Bing Yang, Xiangke Li, Shoucai Ma, Junjie Liu, Boya Feng, Xiaohui Liu, Hong-Wei Wang, Si-Min He, Ning Gao, Keqiong Ye, Meng-Qiu Dong, and Xiaoguang Lei. Trifunctional cross-linker for mapping protein-protein interaction networks and comparing protein conformational states. *eLife*, 5, 2016.

[2469] Huiling Tan, Alek Pogosyan, Keyoumars Ashkan, Alexander L Green, Tipu Aziz, Thomas Foltynie, Patricia Limousin, Ludvic Zrinzo, Marwan Hariz, and Peter Brown. Decoding gripping force based on local field potentials recorded from subthalamic nucleus in humans. *eLife*, 5, 2016.

[2470] Hanlin Tang, Hsiang-Yu Yu, Chien-Chen Chou, Nathan E Crone, Joseph R Madsen, William S Anderson, and Gabriel Kreiman. Cascade of neural processing orchestrates cognitive control in human frontal cortex. *eLife*, 5, 2016.

[2471] Jonathan CY Tang, Eugene Drokhlyansky, Behzad Etemad, Stephanie Rudolph, Binggege Guo, Sui Wang, Emily G Ellis, Jonathan Z Li, and Constance L Cepko. Detection and manipulation of live antigen-expressing cells using conditionally stable nanobodies. *eLife*, 5, 2016.

[2472] Shaogeng Tang, Nicholas J Buchkovich, W Mike Henne, Sudeep Banjade, Yun Jung Kim, and Scott D Emr. ESCRT-III activation by parallel action of ESCRT-I/II and ESCRT-0/Bro1 during MVB biogenesis. *eLife*, 5, 2016.

[2473] Trinh To Tat, Patricia A Maroney, Sangpen Chamnongpol, Jeff Coller, and Timothy W Nilsen. Cotranslational microRNA mediated messenger RNA destabilization. *eLife*, 5, 2016.

[2474] Anjali Teckchandani and Jonathan A Cooper. The ubiquitin-proteasome system regulates focal adhesions at the leading edge of migrating cells. *eLife*, 5, 2016.

[2475] Hanno Teeling, Bernhard M Fuchs, Christin M Bennke, Karen Krüger, Meghan Chafee, Lennart Kappelmann, Greta Reintjes, Jost Waldmann, Christian Quast, Frank Oliver Glöckner, Judith Lucas, Antje Wichels, Gunnar Gerdts, Karen H Wiltshire, and Rudolf I Amann. Recurring patterns in bacterioplankton dynamics during coastal spring algae blooms. *eLife*, 5, 2016.

[2476] Kai Wen Teng, Yuji Ishitsuka, Pin Ren, Yeoan Youn, Xiang Deng, Pinghua Ge, Sang Hak Lee, Andrew S Belmont, and Paul R Selvin. Labeling proteins inside living cells using external fluorophores for microscopy. *eLife*, 5, 2016.

[2477] Shin-Ichiro Terada, Daisuke Matsubara, Koun Onodera, Masanori Matsuzaki, Tadashi Uemura, and Tadao Usui. Neuronal processing of noxious thermal stimuli mediated by dendritic Ca2+ influx in Drosophila somatosensory neurons. *eLife*, 5, 2016.

[2478] Akiko Terauchi, Erin M Johnson-Venkatesh, Brenna Bullock, Maria K Lehtinen, and Hisashi Umemori. Retrograde fibroblast growth factor 22 (FGF22) signaling regulates insulin-like growth factor 2 (IGF2) expression for activity-dependent synapse stabilization in the mammalian brain. *eLife*, 5, 2016.

[2479] Robert J Tetley, Guy B Blanchard, Alexander G Fletcher, Richard J Adams, and Bénédicte Sanson. Unipolar distributions of junctional Myosin II identify cell stripe boundaries that drive cell intercalation throughout Drosophila axis extension. *eLife*, 5, 2016.

[2480] Sheila S Teves, Luye An, Anders S Hansen, Liangqi Xie, Xavier Darzacq, and Robert Tjian. A dynamic mode of mitotic bookmarking by transcription factors. *eLife*, 5, 2016.

[2481] Rajan Thakur, Aniruddha Panda, Elise Coessens, Nikita Raj, Shweta Yadav, Sruthi Balakrishnan, Qifeng Zhang, Plamen Georgiev, Bishal Basak, Renu Pasricha, Michael JO Wakelam, Nicholas T Ktistakis, and Padinjat Raghu. Phospholipase D activity couples plasma membrane endocytosis with retromer dependent recycling. *eLife*, 5, 2016.

[2482] Marina Theodosiou, Moritz Widmaier, Ralph T Böttcher, Emanuel Rognoni, Maik Veelders, Mitasha Bharadwaj, Armin Lambacher, Katharina Austen, Daniel J Müller, Roy Zent, and Reinhard Fässler. Kindlin-2 cooperates with talin to activate integrins and induces cell spreading by directly binding paxillin. *eLife*, 5, 2016.

[2483] John A Thompson, Jamie D Costabile, and Gidon Felsen. Mesencephalic representations of recent experience influence decision making. *eLife*, 5, 2016.

[2484] Mary K Thompson, Maria F Rojas-Duran, Paritosh Gangaramani, and Wendy V Gilbert. The ribosomal protein Asc1/RACK1 is required for efficient translation of short mRNAs. *eLife*, 5, 2016.

[2485] Shuhua Thong, Bilge Ercan, Federico Torta, Zhen Yang Fong, Hui Yi Alvina Wong, Markus R Wenk, and Shu-Sin Chng. Defining key roles for auxiliary proteins in an ABC transporter that maintains bacterial outer membrane lipid asymmetry. *eLife*, 5, 2016.

[2486] Mikhail Tikhonov. Community-level cohesion without cooperation. *eLife*, 5, 2016.

[2487] Andrew T Timberlake, Jungmin Choi, Samir Zaidi, Qiongshi Lu, Carol Nelson-Williams, Eric D Brooks, Kaya Bilguvar, Irina Tikhonova, Shrikant Mane, Jenny F Yang, Rajendra Sawh-Martinez, Sarah Persing, Elizabeth G Zellner, Erin Loring, Carolyn Chuang, Amy Galm, Peter W Hashim, Derek M Steinbacher, Michael L DiLuna, Charles C Duncan, Kevin A Pelphrey, Hongyu Zhao, John A Persing, and Richard P Lifton. Two locus inheritance of non-syndromic midline craniosynostosis via rare SMAD6 and common BMP2 alleles. *eLife*, 5, 2016.

[2488] Alexandre Tiriac and Mark S Blumberg. Gating of reafference in the external cuneate nucleus during self-generated movements in wake but not sleep. *eLife*, 5, 2016.

[2489] Nagaraja Tirumuru, Boxuan Simen Zhao, Wuxun Lu, Zhike Lu, Chuan He, and Li Wu. N6-methyladenosine of HIV-1 RNA regulates viral infection and HIV-1 Gag protein expression. *eLife*, 5, 2016.

[2490] Marketa Tomkova, Michael McClellan, Skirmantas Kriaucionis, and Benjamin Schuster-Boeckler. 5-hydroxymethylcytosine marks regions with reduced mutation frequency in human DNA. *eLife*, 5, 2016.

[2491] Matthew B Toomey, Olle Lind, Rikard Frederiksen, Robert W Curley, Ken M Riedl, David Wilby, Steven J Schwartz, Christopher C Witt, Earl H Harrison, Nicholas W Roberts, Misha Vorobyev, Kevin J McGraw, M Carter Cornwall, Almut Kelber, and Joseph C Corbo. Complementary shifts in photoreceptor spectral tuning unlock the full adaptive potential of ultraviolet vision in birds. *eLife*, 5, 2016.

[2492] Eduardo M Torres, Michael Springer, and Angelika Amon. No current evidence for widespread dosage compensation in S. cerevisiae. *eLife*, 5, 2016.

[2493] Manon Torres, Denis Becquet, Marie-Pierre Blanchard, Séverine Guillen, Bénédicte Boyer, Mathias Moreno, Jean-Louis Franc, and Anne-Marie FranÇois-Bellan. Circadian RNA expression elicited by 3′-UTR IRAlu-paraspeckle associated elements. *eLife*, 5, 2016.

[2494] Junko Y Toshima, Eri Furuya, Makoto Nagano, Chisa Kanno, Yuta Sakamoto, Masashi Ebihara, Daria Elisabeth Siekhaus, and Jiro Toshima. Yeast Eps15-like endocytic protein Pan1p regulates the interaction between endocytic vesicles, endosomes and the actin cytoskeleton. *eLife*, 5, 2016.

[2495] Kouki K Touhara, Weiwei Wang, and Roderick MacKinnon. The GIRK1 subunit potentiates G protein activation of cardiac GIRK1/4 hetero-tetramers. *eLife*, 5, 2016.

[2496] Felipe Trajtenberg, Juan A Imelio, Matías R Machado, Nicole Larrieux, Marcelo A Marti, Gonzalo Obal, Ariel E Mechaly, and Alejandro Buschiazzo. Regulation of signaling directionality revealed by 3D snapshots of a kinase:regulator complex in action. *eLife*, 5, 2016.

[2497] Michael Tramantano, Lu Sun, Christy Au, Daniel Labuz, Zhimin Liu, Mindy Chou, Chen Shen, and Ed Luk. Constitutive turnover of histone H2A.Z at yeast promoters requires the preinitiation complex. *eLife*, 5, 2016.

[2498] Vítor Trovisco, Katsiaryna Belaya, Dmitry Nashchekin, Uwe Irion, George Sirinakis, Richard Butler, Jack J Lee, Elizabeth R Gavis, and Daniel St Johnston. bicoid mRNA localises to the Drosophila oocyte anterior by random Dynein-mediated transport and anchoring. *eLife*, 5, 2016.

[2499] Ming-Feng Tsai, Charles B Phillips, Matthew Ranaghan, Chen-Wei Tsai, Yujiao Wu, Carole Williams, and Christopher Miller. Dual functions of a small regulatory subunit in the mitochondrial calcium uniporter complex. *eLife*, 5, 2016.

[2500] Pei-Ling Tsai, Chenguang Zhao, Elizabeth Turner, and Christian Schlieker. The Lamin B receptor is essential for cholesterol synthesis and perturbed by disease-causing mutations. *eLife*, 5, 2016.

[2501] Panayiotis Tsokas, Changchi Hsieh, Yudong Yao, Edith Lesburguères, Emma Jane Claire Wallace, Andrew Tcherepanov, Desingarao Jothianandan, Benjamin Rush Hartley, Ling Pan, Bruno Rivard, Robert V Farese, Mini P Sajan, Peter John Bergold, Alejandro Iván Hernández, James E Cottrell, Harel Z Shouval, André Antonio Fenton, and Todd Charlton Sacktor. Compensation for PKM$\zeta$in long-term potentiation and spatial long-term memory in mutant mice. *eLife*, 5, 2016.

[2502] Michal Turek, Judith Besseling, Jan-Philipp Spies, Sabine König, and Henrik Bringmann. Sleep-active neuron specification and sleep induction require FLP-11 neuropeptides to systemically induce sleep. *eLife*, 5, 2016.

[2503] Matti Turtola and Georgiy A Belogurov. NusG inhibits RNA polymerase backtracking by stabilizing the minimal transcription bubble. *eLife*, 5, 2016.

[2504] Richard CV Tyser, Antonio MA Miranda, Chiann-mun Chen, Sean M Davidson, Shankar Srinivas, and Paul R Riley. Calcium handling precedes cardiac differentiation to initiate the first heartbeat. *eLife*, 5, 2016.

[2505] Benedetta Ubezio, Raquel Agudo Blanco, Ilse Geudens, Fabio Stanchi, Thomas Mathivet, Martin L Jones, Anan Ragab, Katie Bentley, and Holger Gerhardt. Synchronization of endothelial Dll4-Notch dynamics switch blood vessels from branching to expansion. *eLife*, 5, 2016.

[2506] Hana Uhlirova, Kıvılcım KılıÇ, Peifang Tian, Martin Thunemann, Michèle Desjardins, Payam A Saisan, Sava Sakadžić, Torbjørn V Ness, Celine Mateo, Qun Cheng, Kimberly L Weldy, Florence Razoux, Matthieu Vandenberghe, Jonathan A Cremonesi, Christopher GL Ferri, Krystal Nizar, Vishnu B Sridhar, Tyler C Steed, Maxim Abashin, Yeshaiahu Fainman, Eliezer Masliah, Srdjan Djurovic, Ole A Andreassen, Gabriel A Silva, David A Boas, David Kleinfeld, Richard B Buxton, Gaute T Einevoll, Anders M Dale, and Anna Devor. Cell type specificity of neurovascular coupling in cerebral cortex. *eLife*, 5, 2016.

[2507] Sinan Uğur Umu, Anthony M Poole, Renwick CJ Dobson, and Paul P Gardner. Avoidance of stochastic RNA interactions can be harnessed to control protein expression levels in bacteria and archaea. *eLife*, 5, 2016.

[2508] Silvia Urbansky, Paula González Avalos, Maike Wosch, and Steffen Lemke. Folded gastrulation and T48 drive the evolution of coordinated mesoderm internalization in flies. *eLife*, 5, 2016.

[2509] Kerstin Ure, Hui Lu, Wei Wang, Aya Ito-Ishida, Zhenyu Wu, Ling-jie He, Yehezkel Sztainberg, Wu Chen, Jianrong Tang, and Huda Y Zoghbi. Restoration of Mecp2 expression in GABAergic neurons is sufficient to rescue multiple disease features in a mouse model of Rett syndrome. *eLife*, 5, 2016.

[2510] Marta A Uzarska, Veronica Nasta, Benjamin D Weiler, Farah Spantgar, Simone Ciofi-Baffoni, Maria Rosaria Saviello, Leonardo Gonnelli, Ulrich Mühlenhoff, Lucia Banci, and Roland Lill. Mitochondrial Bol1 and Bol3 function as assembly factors for specific iron-sulfur proteins. *eLife*, 5, 2016.

[2511] Thirumalini Vaithianathan, Diane Henry, Wendy Akmentin, and Gary Matthews. Nanoscale dynamics of synaptic vesicle trafficking and fusion at the presynaptic active zone. *eLife*, 5, 2016.

[2512] Ronald D Vale and Anthony A Hyman. Priority of discovery in the life sciences. *eLife*, 5, 2016.

[2513] Antoine M Valera, Francesca Binda, Sophie A Pawlowski, Jean-Luc Dupont, Jean-FranÇois Casella, Jeffrey D Rothstein, Bernard Poulain, and Philippe Isope. Stereotyped spatial patterns of functional synaptic connectivity in the cerebellar cortex. *eLife*, 5, 2016.

[2514] Ronald van den Berg, Kavitha Anandalingam, Ariel Zylberberg, Roozbeh Kiani, Michael N Shadlen, and Daniel M Wolpert. A common mechanism underlies changes of mind about decisions and confidence. *eLife*, 5, 2016.

[2515] An Van den Bossche, Steven W Hardwick, Pieter-Jan Ceyssens, Hanne Hendrix, Marleen Voet, Tom Dendooven, Katarzyna J Bandyra, Marc De Maeyer, Abram Aertsen, Jean-Paul Noben, Ben F Luisi, and Rob Lavigne. Structural elucidation of a novel mechanism for the bacteriophage-based inhibition of the RNA degradosome. *eLife*, 5, 2016.

[2516] Jonathan G Van Vranken, Mi-Young Jeong, Peng Wei, Yu-Chan Chen, Steven P Gygi, Dennis R Winge, and Jared Rutter. The mitochondrial acyl carrier protein (ACP) coordinates mitochondrial fatty acid synthesis with iron sulfur cluster biogenesis. *eLife*, 5, 2016.

[2517] John P Vanden Heuvel and Jessica Bullenkamp. Registered report: Systematic identification of genomic markers of drug sensitivity in cancer cells. *eLife*, 5, 2016.

[2518] Dieter Vanderelst, Jan Steckel, Andre Boen, Herbert Peremans, and Marc W Holderied. Place recognition using batlike sonar. *eLife*, 5, 2016.

[2519] Davy Vanhoutte, Tobias G Schips, Jennifer Q Kwong, Jennifer Davis, Andoria Tjondrokoesoemo, Matthew J Brody, Michelle A Sargent, Onur Kanisicak, Hong Yi, Quan Q Gao, Joseph E Rabinowitz, Talila Volk, Elizabeth M McNally, and Jeffery D Molkentin. Thrombospondin expression in myofibers stabilizes muscle membranes. *eLife*, 5, 2016.

[2520] Jorge Vaquero-Garcia, Alejandro Barrera, Matthew R Gazzara, Juan González-Vallinas, Nicholas F Lahens, John B Hogenesch, Kristen W Lynch, and Yoseph Barash. A new view of transcriptome complexity and regulation through the lens of local splicing variations. *eLife*, 5, 2016.

[2521] Claudia G Vasquez, Sarah M Heissler, Neil Billington, James R Sellers, and Adam C Martin. Drosophila non-muscle myosin II motor activity determines the rate of tissue folding. *eLife*, 5, 2016.

[2522] Alejandro Vélez and Bruce A Carlson. Detection of transient synchrony across oscillating receptors by the central electrosensory system of mormyrid fish. *eLife*, 5, 2016.

[2523] Douwe M Veltman, Thomas D Williams, Gareth Bloomfield, Bi-Chang Chen, Eric Betzig, Robert H Insall, and Robert R Kay. A plasma membrane template for macropinocytic cups. *eLife*, 5, 2016.

[2524] Bram-Ernst Verhoef and John HR Maunsell. Attention operates uniformly throughout the classical receptive field and the surround. *eLife*, 5, 2016.

[2525] Carla S Verissimo, René M Overmeer, Bas Ponsioen, Jarno Drost, Sander Mertens, Ingrid Verlaan-Klink, Bastiaan van Gerwen, Marieke van der Ven, Marc van de Wetering, David A Egan, René Bernards, Hans Clevers, Johannes L Bos, and Hugo J Snippert. Targeting mutant RAS in patient-derived colorectal cancer organoids by combinatorial drug screening. *eLife*, 5, 2016.

[2526] Santiago Vernia, Yvonne JK Edwards, Myoung Sook Han, Julie Cavanagh-Kyros, Tamera Barrett, Jason K Kim, and Roger J Davis. An alternative splicing program promotes adipose tissue thermogenesis. *eLife*, 5, 2016.

[2527] Santiago Vernia, Caroline Morel, Joseph C Madara, Julie Cavanagh-Kyros, Tamera Barrett, Kathryn Chase, Norman J Kennedy, Dae Young Jung, Jason K Kim, Neil Aronin, Richard A Flavell, Bradford B Lowell, and Roger J Davis. Excitatory transmission onto AgRP neurons is regulated by cJun NH2-terminal kinase 3 in response to metabolic stress. *eLife*, 5, 2016.

[2528] Andreu Viader, Daisuke Ogasawara, Christopher M Joslyn, Manuel Sanchez-Alavez, Simone Mori, William Nguyen, Bruno Conti, and Benjamin F Cravatt. A chemical proteomic atlas of brain serine hydrolases identifies cell type-specific pathways regulating neuroinflammation. *eLife*, 5, 2016.

[2529] Ravi Vijayvargia, Raquel Epand, Alexander Leitner, Tae-Yang Jung, Baehyun Shin, Roy Jung, Alejandro Lloret, Randy Singh Atwal, Hyeongseok Lee, Jong-Min Lee, Ruedi Aebersold, Hans Hebert, Ji-Joon Song, and Ihn Sik Seong. Huntingtin's spherical solenoid structure enables polyglutamine tract-dependent modulation of its structure and function. *eLife*, 5, 2016.

[2530] Claudia Villani, Giuseppina Sacchetti, Renzo Bagnati, Alice Passoni, Federica Fusco, Mirjana Carli, and Roberto William Invernizzi. Lovastatin fails to improve motor performance and survival in methyl-CpG-binding protein2-null mice. *eLife*, 5, 2016.

[2531] Alejandro Villarino, Arian Laurence, Gertraud W Robinson, Michael Bonelli, Barbara Dema, Behdad Afzali, Han-Yu Shih, Hong-Wei Sun, Stephen R Brooks, Lothar Hennighausen, Yuka Kanno, and John J O'Shea. Signal transducer and activator of transcription 5 (STAT5) paralog dose governs T cell effector and regulatory functions. *eLife*, 5, 2016.

[2532] Roberto Vincis and Alfredo Fontanini. Associative learning changes cross-modal representations in the gustatory cortex. *eLife*, 5, 2016.

[2533] Hanneke Vlaming, Thom M Molenaar, Tibor van Welsem, Deepani W Poramba-Liyanage, Desiree E Smith, Arno Velds, Liesbeth Hoekman, Tessy Korthout, Sjoerd Hendriks, AF Maarten Altelaar, and Fred van Leeuwen. Direct screening for chromatin status on DNA barcodes in yeast delineates the regulome of H3K79 methylation by Dot1. *eLife*, 5, 2016.

[2534] Andre Voelzmann, Pilar Okenve-Ramos, Yue Qu, Monika Chojnowska-Monga, Manuela del Caño-Espinel, Andreas Prokop, and Natalia Sanchez-Soriano. Tau and spectraplakins promote synapse formation and maintenance through Jun kinase and neuronal trafficking. *eLife*, 5, 2016.

[2535] Jacob M Vogan, Xiaozhu Zhang, Daniel T Youmans, Samuel G Regalado, Joshua Z Johnson, Dirk Hockemeyer, and Kathleen Collins. Minimized human telomerase maintains telomeres and resolves endogenous roles of H/ACA proteins, TCAB1, and Cajal bodies. *eLife*, 5, 2016.

[2536] Katrin Vogt, Yoshinori Aso, Toshihide Hige, Stephan Knapek, Toshiharu Ichinose, Anja B Friedrich, Glenn C Turner, Gerald M Rubin, and Hiromu Tanimoto. Direct neural pathways convey distinct visual information to Drosophila mushroom bodies. *eLife*, 5, 2016.

[2537] Oleg A Volkov, Lisa Kinch, Carson Ariagno, Xiaoyi Deng, Shihua Zhong, Nick Grishin, Diana R Tomchick, Zhe Chen, and Margaret A Phillips. Relief of autoinhibition by conformational switch explains enzyme activation by a catalytically dead paralog. *eLife*, 5, 2016.

[2538] Yannick von Grabowiecki, Paula Abreu, Orphee Blanchard, Lavinia Palamiuc, Samir Benosman, Sophie Mériaux, Véronique Devignot, Isabelle Gross, Georg Mellitzer, José L Gonzalez de Aguilar, and Christian Gaiddon. Transcriptional activator TAp63 is upregulated in muscular atrophy during ALS and induces the pro-atrophic ubiquitin ligase Trim63. *eLife*, 5, 2016.

[2539] Seychelle M Vos, David Pöllmann, Livia Caizzi, Katharina B Hofmann, Pascaline Rombaut, Tomasz Zimniak, Franz Herzog, and Patrick Cramer. Architecture and RNA binding of the human negative elongation factor. *eLife*, 5, 2016.

[2540] Andrei Vovk, Chad Gu, Michael G Opferman, Larisa E Kapinos, Roderick YH Lim, Rob D Coalson, David Jasnow, and Anton Zilman. Simple biophysics underpins collective conformations of the intrinsically disordered proteins of the Nuclear Pore Complex. *eLife*, 5, 2016.

[2541] Nicholas P Vyleta, Carolina Borges-Merjane, and Peter Jonas. Plasticity-dependent, full detonation at hippocampal mossy fiber–CA3 pyramidal neuron synapses. *eLife*, 5, 2016.

[2542] Jonathan M Wagner, Marcin D Roganowicz, Katarzyna Skorupka, Steven L Alam, Devin Christensen, Ginna Doss, Yueping Wan, Gabriel A Frank, Barbie K Ganser-Pornillos, Wesley I Sundquist, and Owen Pornillos. Mechanism of B-box 2 domain-mediated higher-order assembly of the retroviral restriction factor TRIM5$\alpha$. *eLife*, 5, 2016.

[2543] Jonathan M Wagner, Kaneil K Zadrozny, Jakub Chrustowicz, Michael D Purdy, Mark Yeager, Barbie K Ganser-Pornillos, and Owen Pornillos. Crystal structure of an HIV assembly and maturation switch. *eLife*, 5, 2016.

[2544] Peter Walentek, Ian K Quigley, Dingyuan I Sun, Umeet K Sajjan, Christopher Kintner, and Richard M Harland. Ciliary transcription factors and miRNAs precisely regulate Cp110 levels required for ciliary adhesions and ciliogenesis. *eLife*, 5, 2016.

[2545] Pauline Wales, Christian E Schuberth, Roland Aufschnaiter, Johannes Fels, Ireth García-Aguilar, Annette Janning, Christopher P Dlugos, Marco Schäfer-Herte, Christoph Klingner, Mike Wälte, Julian Kuhlmann, Ekaterina Menis, Laura Hockaday Kang, Kerstin C Maier, Wenya Hou, Antonella Russo, Henry N Higgs, Hermann Pavenstädt, Thomas Vogl, Johannes Roth, Britta Qualmann, Michael M Kessels, Dietmar E Martin, Bela Mulder, and Roland Wedlich-Söldner. Calcium-mediated actin reset (CaAR) mediates acute cell adaptations. *eLife*, 5, 2016.

[2546] Mannu K Walia, Patricia MW Ho, Scott Taylor, Alvin JM Ng, Ankita Gupte, Alistair M Chalk, Andrew CW Zannettino, T John Martin, and Carl R Walkley. Activation of PTHrP-cAMP-CREB1 signaling following p53 loss is essential for osteosarcoma initiation and maintenance. *eLife*, 5, 2016.

[2547] Ward G Walkup, Tara L Mastro, Leslie T Schenker, Jost Vielmetter, Rebecca Hu, Ariella Iancu, Meera Reghunathan, Barry Dylan Bannon, and Mary B Kennedy. A model for regulation by SynGAP-$\alpha$1 of binding of synaptic proteins to PDZ-domain 'Slots' in the postsynaptic density. *eLife*, 5, 2016.

[2548] Marius Walter, Aurélie Teissandier, Raquel Pérez-Palacios, and Déborah Bourc'his. An epigenetic switch ensures transposon repression upon dynamic loss of DNA methylation in embryonic stem cells. *eLife*, 5, 2016.

[2549] Cheng-Wei Wang, Arunima Purkayastha, Kevin T Jones, Shivani K Thaker, and Utpal Banerjee. In vivo genetic dissection of tumor growth and the Warburg effect. *eLife*, 5, 2016.

[2550] Doudou Wang, Ryo Nitta, Manatsu Morikawa, Hiroaki Yajima, Shigeyuki Inoue, Hideki Shigematsu, Masahide Kikkawa, and Nobutaka Hirokawa. Motility and microtubule depolymerization mechanisms of the Kinesin-8 motor, KIF19A. *eLife*, 5, 2016.

[2551] Feng Wang, JongDae Shin, Jeremy M Shea, Jun Yu, Ana BoŠković, Meg Byron, Xiaochun Zhu, Alex K Shalek, Aviv Regev, Jeanne B Lawrence, Eduardo M Torres, Lihua J Zhu, Oliver J Rando, and Ingolf Bach. Regulation of X-linked gene expression during early mouse development by Rlim. *eLife*, 5, 2016.

[2552] Gordon X Wang, Stephen J Smith, and Philippe Mourrain. Sub-synaptic, multiplexed analysis of proteins reveals Fragile X related protein 2 is mislocalized in Fmr1 KO synapses. *eLife*, 5, 2016.

[2553] Huajin Wang, Michel Becuwe, Benjamin E Housden, Chandramohan Chitraju, Ashley J Porras, Morven M Graham, Xinran N Liu, Abdou Rachid Thiam, David B Savage, Anil K Agarwal, Abhimanyu Garg, Maria-Jesus Olarte, Qingqing Lin, Florian Fröhlich, Hans Kristian Hannibal-Bach, Srigokul Upadhyayula, Norbert Perrimon, Tomas Kirchhausen, Christer S Ejsing, Tobias C Walther, and Robert V Farese. Seipin is required for converting nascent to mature lipid droplets. *eLife*, 5, 2016.

[2554] Irving E Wang, Sylvain W Lapan, M Lucila Scimone, Thomas R Clandinin, and Peter W Reddien. Hedgehog signaling regulates gene expression in planarian glia. *eLife*, 5, 2016.

[2555] Kang Wang, Pedro Mateos-Aparicio, Christoph Hönigsperger, Vijeta Raghuram, Wendy W Wu, Margreet C Ridder, Pankaj Sah, Jim Maylie, Johan F Storm, and John P Adelman. IK1 channels do not contribute to the slow afterhyperpolarization in pyramidal neurons. *eLife*, 5, 2016.

[2556] Lei Wang, Kwanwoo Lee, Ryan Malonis, Irma Sanchez, and Brian D Dynlacht. Tethering of an E3 ligase by PCM1 regulates the abundance of centrosomal KIAA0586/Talpid3 and promotes ciliogenesis. *eLife*, 5, 2016.

[2557] Lihua Wang, Pengcheng Bu, Yiwei Ai, Tara Srinivasan, Huanhuan Joyce Chen, Kun Xiang, Steven M Lipkin, and Xiling Shen. A long non-coding RNA targets microRNA miR-34a to regulate colon cancer stem cell asymmetric division. *eLife*, 5, 2016.

[2558] Ray Yu-Ruei Wang, Yifan Song, Benjamin A Barad, Yifan Cheng, James S Fraser, and Frank DiMaio. Automated structure refinement of macromolecular assemblies from cryo-EM maps using Rosetta. *eLife*, 5, 2016.

[2559] Shen Wang, Yun Li, and Cong Ma. Synaptotagmin-1 C2B domain interacts simultaneously with SNAREs and membranes to promote membrane fusion. *eLife*, 5, 2016.

[2560] Songyu Wang, Hanna Tukachinsky, Fabian B Romano, and Tom A Rapoport. Cooperation of the ER-shaping proteins atlastin, lunapark, and reticulons to generate a tubular membrane network. *eLife*, 5, 2016.

[2561] Xinchen Wang, Nathan R Tucker, Gizem Rizki, Robert Mills, Peter HL Krijger, Elzo de Wit, Vidya Subramanian, Eric Bartell, Xinh-Xinh Nguyen, Jiangchuan Ye, Jordan Leyton-Mange, Elena V Dolmatova, Pim van der Harst, Wouter de Laat, Patrick T Ellinor, Christopher Newton-Cheh, David J Milan, Manolis Kellis, and Laurie A Boyer. Discovery and validation of sub-threshold genome-wide association study loci using epigenomic signatures. *eLife*, 5, 2016.

[2562] Yin Wang, Xi Wang, Thorsten Wohland, and Karuna Sampath. Extracellular interactions and ligand degradation shape the nodal morphogen gradient. *eLife*, 5, 2016.

[2563] Yingxue Wang, Zachary Roth, and Eva Pastalkova. Synchronized excitability in a network enables generation of internal neuronal sequences. *eLife*, 5, 2016.

[2564] Yong Wang, Elena Papaleo, and Kresten Lindorff-Larsen. Mapping transiently formed and sparsely populated conformations on a complex energy landscape. *eLife*, 5, 2016.

[2565] Yuxuan Wang, Karin Sundfeldt, Constantina Mateoiu, Ie-Ming Shih, Robert J Kurman, Joy Schaefer, Natalie Silliman, Isaac Kinde, Simeon Springer, Michael Foote, Björg Kristjansdottir, Nathan James, Kenneth W Kinzler, Nickolas Papadopoulos, Luis A Diaz, and Bert Vogelstein. Diagnostic potential of tumor DNA from ovarian cyst fluid. *eLife*, 5, 2016.

[2566] Jadiel A Wasson, Ashley K Simon, Dexter A Myrick, Gernot Wolf, Shawn Driscoll, Samuel L Pfaff, Todd S Macfarlan, and David J Katz. Maternally provided LSD1/KDM1A enables the maternal-to-zygotic transition and prevents defects that manifest postnatally. *eLife*, 5, 2016.

[2567] Emma Watson, Viridiana Olin-Sandoval, Michael J Hoy, Chi-Hua Li, Timo Louisse, Victoria Yao, Akihiro Mori, Amy D Holdorf, Olga G Troyanskaya, Markus Ralser, and Albertha JM Walhout. Metabolic network rewiring of propionate flux compensates vitamin B12 deficiency in C. elegans. *eLife*, 5, 2016.

[2568] Robbyn L Weaver, Jazeel F Limzerwala, Ryan M Naylor, Karthik B Jeganathan, Darren J Baker, and Jan M van Deursen. BubR1 alterations that reinforce mitotic surveillance act against aneuploidy and cancer. *eLife*, 5, 2016.

[2569] Alexis B Webb, Iván M Lengyel, David J Jörg, Guillaume Valentin, Frank Jülicher, Luis G Morelli, and Andrew C Oates. Persistence, period and precision of autonomous cellular oscillators from the zebrafish segmentation clock. *eLife*, 5, 2016.

[2570] Janine Weber, Han Bao, Christoph Hartlmüller, Zhiqin Wang, Almut Windhager, Robert Janowski, Tobias Madl, Peng Jin, and Dierk Niessing. Structural basis of nucleic-acid recognition and double-strand unwinding by the essential neuronal protein Pur-alpha. *eLife*, 5, 2016.

[2571] Chase A Weidmann, Chen Qiu, René M Arvola, Tzu-Fang Lou, Jordan Killingsworth, Zachary T Campbell, Traci M Tanaka Hall, and Aaron C Goldstrohm. Drosophila Nanos acts as a molecular clamp that modulates the RNA-binding and repression activities of Pumilio. *eLife*, 5, 2016.

[2572] Luise Westernberg, Claire Conche, Yina Hsing Huang, Stephanie Rigaud, Yisong Deng, Sabine Siegemund, Sayak Mukherjee, Lyn'Al Nosaka, Jayajit Das, and Karsten Sauer. Non-canonical antagonism of PI3K by the kinase Itpkb delays thymocyte $\beta$-selection and renders it Notch-dependent. *eLife*, 5, 2016.

[2573] Bayly S Wheeler, Erika Anderson, Christian Frøkjær-Jensen, Qian Bian, Erik Jorgensen, and Barbara J Meyer. Chromosome-wide mechanisms to decouple gene expression from gene dose during sex-chromosome evolution. *eLife*, 5, 2016.

[2574] Joshua R Wheeler, Tyler Matheny, Saumya Jain, Robert Abrisch, and Roy Parker. Distinct stages in stress granule assembly and disassembly. *eLife*, 5, 2016.

[2575] Anjar Wibowo, Claude Becker, Gianpiero Marconi, Julius Durr, Jonathan Price, Jorg Hagmann, Ranjith Papareddy, Hadi Putra, Jorge Kageyama, Jorg Becker, Detlef Weigel, and Jose Gutierrez-Marcos. Hyperosmotic stress memory in Arabidopsis is mediated by distinct epigenetically labile sites in the genome and is restricted in the male germline by DNA glycosylase activity. *eLife*, 5, 2016.

[2576] Candice L Wike, Hillary K Graves, Reva Hawkins, Matthew D Gibson, Michelle B Ferdinand, Tao Zhang, Zhihong Chen, Damien F Hudson, Jennifer J Ottesen, Michael G Poirier, Jill Schumacher, and Jessica K Tyler. Aurora-A mediated histone H3 phosphorylation of threonine 118 controls condensin I and cohesin occupancy in mitosis. *eLife*, 5, 2016.

[2577] Martin Wilkinson, Yuriy Chaban, and Dale B Wigley. Mechanism for nuclease regulation in RecBCD. *eLife*, 5, 2016.

[2578] Martin Wilkinson, Luca A Troman, Wan AK Wan Nur Ismah, Yuriy Chaban, Matthew B Avison, Mark S Dillingham, and Dale B Wigley. Structural basis for the inhibition of RecBCD by Gam and its synergistic antibacterial effect with quinolones. *eLife*, 5, 2016.

[2579] Alex H Williams, Cian O'Donnell, Terrence J Sejnowski, and Timothy O'Leary. Dendritic trafficking faces physiologically critical speed-precision tradeoffs. *eLife*, 5, 2016.

[2580] Helen Rankin Willsey, Xiaoyan Zheng, José Carlos Pastor-Pareja, A Jeremy Willsey, Philip A Beachy, and Tian Xu. Localized JNK signaling regulates organ size during development. *eLife*, 5, 2016.

[2581] Amy M Winans, Sean R Collins, and Tobias Meyer. Waves of actin and microtubule polymerization drive microtubule-based transport and neurite growth before single axon formation. *eLife*, 5, 2016.

[2582] Somsakul Pop Wongpalee, Ajay Vashisht, Shalini Sharma, Darryl Chui, James A Wohlschlegel, and Douglas L Black. Large-scale remodeling of a repressed exon ribonucleoprotein to an exon definition complex active for splicing. *eLife*, 5, 2016.

[2583] C David Wood, Hildegonda Veenstra, Sarika Khasnis, Andrea Gunnell, Helen M Webb, Claire Shannon-Lowe, Simon Andrews, Cameron S Osborne, and Michelle J West. MYC activation and BCL2L11 silencing by a tumour virus through the large-scale reconfiguration of enhancer-promoter hubs. *eLife*, 5, 2016.

[2584] Dalei Wu, Xiaoyu Su, Nalini Potluri, Youngchang Kim, and Fraydoon Rastinejad. NPAS1-ARNT and NPAS3-ARNT crystal structures implicate the bHLH-PAS family as multi-ligand binding transcription factors. *eLife*, 5, 2016.

[2585] Jianguo Wu, Andrei I Ivanov, Paul B Fisher, and Zheng Fu. Polo-like kinase 1 induces epithelial-to-mesenchymal transition and promotes epithelial cell motility by activating CRAF/ERK signaling. *eLife*, 5, 2016.

[2586] Ming Wu, Aljoscha Nern, W Ryan Williamson, Mai M Morimoto, Michael B Reiser, Gwyneth M Card, and Gerald M Rubin. Visual projection neurons in the Drosophila lobula link feature detection to distinct behavioral programs. *eLife*, 5, 2016.

[2587] Nicholas C Wu, Lei Dai, C Anders Olson, James O Lloyd-Smith, and Ren Sun. Adaptation in protein fitness landscapes is facilitated by indirect paths. *eLife*, 5, 2016.

[2588] Filip J Wymeersch, Yali Huang, Guillaume Blin, Noemí Cambray, Ron Wilkie, Frederick CK Wong, and Valerie Wilson. Position-dependent plasticity of distinct progenitor types in the primitive streak. *eLife*, 5, 2016.

[2589] Antoine Wystrach, Konstantinos Lagogiannis, and Barbara Webb. Continuous lateral oscillations as a core mechanism for taxis in Drosophila larvae. *eLife*, 5, 2016.

[2590] Shuting Xia, Zikai Zhou, Celeste Leung, Yuehua Zhu, Xingxiu Pan, Junxia Qi, Maria Morena, Matthew N Hill, Wei Xie, and Zhengping Jia. p21-activated kinase 1 restricts tonic endocannabinoid signaling in the hippocampus. *eLife*, 5, 2016.

[2591] Ying-Zi Xiong, Jun-Yun Zhang, and Cong Yu. Bottom-up and top-down influences at untrained conditions determine perceptual learning specificity and transfer. *eLife*, 5, 2016.

[2592] Kenta Yamamoto, Jiguang Wang, Lisa Sprinzen, Jun Xu, Christopher J Haddock, Chen Li, Brian J Lee, Denis G Loredan, Wenxia Jiang, Alessandro Vindigni, Dong Wang, Raul Rabadan, and Shan Zha. Kinase-dead ATM protein is highly oncogenic and can be preferentially targeted by Topo-isomerase I inhibitors. *eLife*, 5, 2016.

[2593] Satoshi Yamashita, Haruhiko Yamaguchi, Toshiyuki Waki, Yuichi Aoki, Makie Mizuno, Fumihiro Yanbe, Tomoki Ishii, Ayuta Funaki, Yuzuru Tozawa, Yukino Miyagi-Inoue, Kazuhisa Fushihara, Toru Nakayama, and Seiji Takahashi. Identification and reconstitution of the rubber biosynthetic machinery on rubber particles from Hevea brasiliensis. *eLife*, 5, 2016.

[2594] Takayuki Yamashita and Carl CH Petersen. Target-specific membrane potential dynamics of neocortical projection neurons during goal-directed behavior. *eLife*, 5, 2016.

[2595] Jialong Yang, Xingguang Lin, Yun Pan, Jinli Wang, Pengcheng Chen, Hongxiang Huang, Hai-Hui Xue, Jimin Gao, and Xiao-Ping Zhong. Critical roles of mTOR Complex 1 and 2 for T follicular helper cell differentiation and germinal center responses. *eLife*, 5, 2016.

[2596] Junhua Yang, Hongbin Yang, Yali Liu, Xia Li, Liming Qin, Huifang Lou, Shumin Duan, and Hao Wang. Astrocytes contribute to synapse elimination via type 2 inositol 1,4,5-trisphosphate receptor-dependent release of ATP. *eLife*, 5, 2016.

[2597] Junsheng Yang, Xinxin Hao, Xiuling Cao, Beidong Liu, and Thomas Nyström. Spatial sequestration and detoxification of Huntingtin by the ribosome quality control complex. *eLife*, 5, 2016.

[2598] Scott Cheng-Hsin Yang, Máté Lengyel, and Daniel M Wolpert. Active sensing in the categorization of visual patterns. *eLife*, 5, 2016.

[2599] Zhiyong Yang, Marcus J Robinson, Xiangjun Chen, Geoffrey A Smith, Jack Taunton, Wanli Liu, and Christopher D C Allen. Regulation of B cell fate by chronic activity of the IgE B cell receptor. *eLife*, 5, 2016.

[2600] Tao Yao, Madhura Ketkar, Stefan Treue, and B Suresh Krishna. Visual attention is available at a task-relevant location rapidly after a saccade. *eLife*, 5, 2016.

[2601] Shi Yun Yeo, Yoko Itahana, Alvin Kunyao Guo, Rachel Han, Kozue Iwamoto, Hung Thanh Nguyen, Yi Bao, Kai Kleiber, Ya Jun Wu, Boon Huat Bay, Mathijs Voorhoeve, and Koji Itahana. Transglutaminase 2 contributes to a TP53-induced autophagy program to prevent oncogenic transformation. *eLife*, 5, 2016.

[2602] Xiao Yi and Antony M Dean. Phenotypic plasticity as an adaptation to a functional trade-off. *eLife*, 5, 2016.

[2603] Mariko Yokouchi, Toru Atsugi, Mark van Logtestijn, Reiko J Tanaka, Mayumi Kajimura, Makoto Suematsu, Mikio Furuse, Masayuki Amagai, and Akiharu Kubo. Epidermal cell turnover across tight junctions based on Kelvin's tetrakaidecahedron cell shape. *eLife*, 5, 2016.

[2604] Ryo Yonashiro, Erich B Tahara, Mario H Bengtson, Maria Khokhrina, Holger Lorenz, Kai-Chun Chen, Yu Kigoshi-Tansho, Jeffrey N Savas, John R Yates, Steve A Kay, Elizabeth A Craig, Axel Mogk, Bernd Bukau, and Claudio AP Joazeiro. The Rqc2/Tae2 subunit of the ribosome-associated quality control (RQC) complex marks ribosome-stalled nascent polypeptide chains for aggregation. *eLife*, 5, 2016.

[2605] Isseki Yu, Takaharu Mori, Tadashi Ando, Ryuhei Harada, Jaewoon Jung, Yuji Sugita, and Michael Feig. Biomolecular interactions modulate macromolecular structure and dynamics in atomistic model of a bacterial cytoplasm. *eLife*, 5, 2016.

[2606] Jessica C Yu and Rodrigo Fernandez-Gonzalez. Local mechanical forces promote polarized junctional assembly and axis elongation in Drosophila. *eLife*, 5, 2016.

[2607] Yue Yu, Rui Huang, Jie Ye, Vivian Zhang, Chao Wu, Guo Cheng, Junling Jia, and Liming Wang. Regulation of starvation-induced hyperactivity by insulin and glucagon signaling in adult Drosophila. *eLife*, 5, 2016.

[2608] Zhou Yu, Taoyong Chen, Xuelian Li, Mingjin Yang, Songqing Tang, Xuhui Zhu, Yan Gu, Xiaoping Su, Meng Xia, Weihua Li, Xuemin Zhang, Qingqing Wang, Xuetao Cao, and Jianli Wang. Lys29-linkage of ASK1 by Skp1−Cullin 1−Fbxo21 ubiquitin ligase complex is required for antiviral innate response. *eLife*, 5, 2016.

[2609] Meng Yuan, Yinggen Ke, Renyan Huang, Ling Ma, Zeyu Yang, Zhaohui Chu, Jinghua Xiao, Xianghua Li, and Shiping Wang. A host basal transcription factor is a key component for infection of rice by TALE-carrying bacteria. *eLife*, 5, 2016.

[2610] Rui Yue, Bo Shen, and Sean J Morrison. Clec11a/osteolectin is an osteogenic growth factor that promotes the maintenance of the adult skeleton. *eLife*, 5, 2016.

[2611] Sophie Zaaijer and Yaniv Erlich. Using mobile sequencers in an academic classroom. *eLife*, 5, 2016.

[2612] Raphael Zahn, Dino Osmanović, Severin Ehret, Carolina Araya Callis, Steffen Frey, Murray Stewart, Changjiang You, Dirk Görlich, Bart W Hoogenboom, and Ralf P Richter. A physical model describing the interaction of nuclear transport receptors with FG nucleoporin domain assemblies. *eLife*, 5, 2016.

[2613] Samuel Zambrano, Ilario De Toma, Arianna Piffer, Marco E Bianchi, and Alessandra Agresti. NF-$\kappa$B oscillations translate into functionally related patterns of gene expression. *eLife*, 5, 2016.

[2614] Valeria Zampini, Jian K Liu, Marco A Diana, Paloma P Maldonado, Nicolas Brunel, and Stéphane Dieudonné. Mechanisms and functional roles of glutamatergic synapse diversity in a cerebellar circuit. *eLife*, 5, 2016.

[2615] Maria N Zanetti, Oscar D Bello, Jing Wang, Jeff Coleman, Yiying Cai, Charles V Sindelar, James E Rothman, and Shyam S Krishnakumar. Ring-like oligomers of Synaptotagmins and related C2 domain proteins. *eLife*, 5, 2016.

[2616] Juan Pablo Zanin, Elizabeth Abercrombie, and Wilma J Friedman. Proneurotrophin-3 promotes cell cycle withdrawal of developing cerebellar granule cell progenitors via the p75 neurotrophin receptor. *eLife*, 5, 2016.

[2617] Shota Zarnadze, Peter Bäuerle, Julio Santos-Torres, Claudia Böhm, Dietmar Schmitz, Jörg RP Geiger, Tamar Dugladze, and Tengis Gloveli. Cell-specific synaptic plasticity induced by network oscillations. *eLife*, 5, 2016.

[2618] Anatoly V Zaytsev, Dario Segura-Peña, Maxim Godzi, Abram Calderon, Edward R Ballister, Rumen Stamatov, Alyssa M Mayo, Laura Peterson, Ben E Black, Fazly I Ataullakhanov, Michael A Lampson, and Ekaterina L Grishchuk. Bistability of a coupled Aurora B kinase-phosphatase system in cell division. *eLife*, 5, 2016.

[2619] Sabrina Zechel, Yasushi Nakagawa, and Carlos F Ibáñez. Thalamo-cortical axons regulate the radial dispersion of neocortical GABAergic interneurons. *eLife*, 5, 2016.

[2620] Cen Zhang, Juan Liu, Yuhan Zhao, Xuetian Yue, Yu Zhu, Xiaolong Wang, Hao Wu, Felix Blanco, Shaohua Li, Gyan Bhanot, Bruce G Haffty, Wenwei Hu, and Zhaohui Feng. Glutaminase 2 is a novel negative regulator of small GTPase Rac1 and mediates p53 function in suppressing metastasis. *eLife*, 5, 2016.

[2621] Feng Zhang, Sonya M Hanson, Andres Jara-Oseguera, Dmitriy Krepkiy, Chanhyung Bae, Larry V Pearce, Peter M Blumberg, Simon Newstead, and Kenton J Swartz. Engineering vanilloid-sensitivity into the rat TRPV2 channel. *eLife*, 5, 2016.

[2622] Haibo Zhang, Martin Y Ng, Yuanwei Chen, and Barry S Cooperman. Kinetics of initiating polypeptide elongation in an IRES-dependent system. *eLife*, 5, 2016.

[2623] Hongkang Zhang, Elaine Reichert, and Adam E Cohen. Optical electrophysiology for probing function and pharmacology of voltage-gated ion channels. *eLife*, 5, 2016.

[2624] Peng Zhang, Yan Bai, Ling Lu, Yun Li, and Cunming Duan. An oxygen-insensitive Hif-3$\alpha$ isoform inhibits Wnt signaling by destabilizing the nuclear $\beta$-catenin complex. *eLife*, 5, 2016.

[2625] Wei Zhang, Paul Jong Kim, Zhongcan Chen, Hidayat Lokman, Lifeng Qiu, Ke Zhang, Steven George Rozen, Eng King Tan, Hyunsoo Shawn Je, and Li Zeng. MiRNA-128 regulates the proliferation and neurogenesis of neural precursors by targeting PCM1 in the developing cortex. *eLife*, 5, 2016.

[2626] Weilin Zhang, He Ren, Chunling Xu, Chongzhuo Zhu, Hao Wu, Dong Liu, Jun Wang, Lei Liu, Wei Li, Qi Ma, Lei Du, Ming Zheng, Chuanmao Zhang, Junling Liu, and Quan Chen. Hypoxic mitophagy regulates mitochondrial quality and platelet activation and determines severity of I/R heart injury. *eLife*, 5, 2016.

[2627] Xiaonan Zhang and Quentin Gaudry. Functional integration of a serotonergic neuron in the Drosophila antennal lobe. *eLife*, 5, 2016.

[2628] Yang Zhang, Theodore L Roth, Elizabeth E Gray, Hsin Chen, Lauren B Rodda, Yin Liang, Patrick Ventura, Saul Villeda, Paul R Crocker, and Jason G Cyster. Migratory and adhesive cues controlling innate-like lymphocyte surveillance of the pathogen-exposed surface of the lymph node. *eLife*, 5, 2016.

[2629] Yifei Zhang, Xing Wang, Hitoshi Matakatsu, Richard Fehon, and Seth S Blair. The novel SH3 domain protein Dlish/CG10933 mediates fat signaling in Drosophila by binding and regulating Dachs. *eLife*, 5, 2016.

[2630] Bo Zhao, Zizhen Wu, and Ulrich Müller. Murine Fam65b forms ring-like structures at the base of stereocilia critical for mechanosensory hair cell function. *eLife*, 5, 2016.

[2631] Hongyun Zhao, Lifeng Yang, Joelle Baddour, Abhinav Achreja, Vincent Bernard, Tyler Moss, Juan C Marini, Thavisha Tudawe, Elena G Seviour, F Anthony San Lucas, Hector Alvarez, Sonal Gupta, Sourindra N Maiti, Laurence Cooper, Donna Peehl, Prahlad T Ram, Anirban Maitra, and Deepak Nagrath. Tumor microenvironment derived exosomes pleiotropically modulate cancer cell metabolism. *eLife*, 5, 2016.

[2632] Huaying Zhao, Yan Fu, Carla Glasser, Eric J Andrade Alba, Mark L Mayer, George Patterson, and Peter Schuck. Monochromatic multicomponent fluorescence sedimentation velocity for the study of high-affinity protein interactions. *eLife*, 5, 2016.

[2633] Juan Zhao and Rikard Blunck. The isolated voltage sensing domain of the Shaker potassium channel forms a voltage-gated cation channel. *eLife*, 5, 2016.

[2634] Chao Yu Zhen, Roubina Tatavosian, Thao Ngoc Huynh, Huy Nguyen Duc, Raibatak Das, Marko Kokotovic, Jonathan B Grimm, Luke D Lavis, Jun Lee, Frances J Mejia, Yang Li, Tingting Yao, and Xiaojun Ren. Live-cell single-molecule tracking reveals co-recognition of H3K27me3 and DNA targets polycomb Cbx7-PRC1 to chromatin. *eLife*, 5, 2016.

[2635] Xinde Zheng, Leah Boyer, Mingji Jin, Yongsung Kim, Weiwei Fan, Cedric Bardy, Travis Berggren, Ronald M Evans, Fred H Gage, and Tony Hunter. Alleviation of neuronal energy deficiency by mTOR inhibition as a treatment for mitochondria-related neurodegeneration. *eLife*, 5, 2016.

[2636] Xinde Zheng, Leah Boyer, Mingji Jin, Jerome Mertens, Yongsung Kim, Li Ma, Li Ma, Michael Hamm, Fred H Gage, and Tony Hunter. Metabolic reprogramming during neuronal differentiation from aerobic glycolysis to neuronal oxidative phosphorylation. *eLife*, 5, 2016.

[2637] Xu Zheng, Joanna Krakowiak, Nikit Patel, Ali Beyzavi, Jideofor Ezike, Ahmad S Khalil, and David Pincus. Dynamic control of Hsf1 during heat shock by a chaperone switch and phosphorylation. *eLife*, 5, 2016.

[2638] Guocai Zhong, Haimin Wang, Charles C Bailey, Guangping Gao, and Michael Farzan. Rational design of aptazyme riboswitches for efficient control of gene expression in mammalian cells. *eLife*, 5, 2016.

[2639] Miou Zhou, Stuart Greenhill, Shan Huang, Tawnie K Silva, Yoshitake Sano, Shumin Wu, Ying Cai, Yoshiko Nagaoka, Megha Sehgal, Denise J Cai, Yong-Seok Lee, Kevin Fox, and Alcino J Silva. CCR5 is a suppressor for cortical plasticity and hippocampal learning and memory. *eLife*, 5, 2016.

[2640] Wenwen Zhou, Qiuping He, Chunxia Zhang, Xin He, Zongbin Cui, Feng Liu, and Wei Li. BLOS2 negatively regulates Notch signaling during neural and hematopoietic stem and progenitor cell development. *eLife*, 5, 2016.

[2641] Wenwu Zhou, Thomas Brockmöller, Zhihao Ling, Ashton Omdahl, Ian T Baldwin, and Shuqing Xu. Evolution of herbivore-induced early defense signaling was shaped by genome-wide duplications in Nicotiana. *eLife*, 5, 2016.

[2642] Xiang Zhou, Qian Hao, Peng Liao, Shiwen Luo, Minhong Zhang, Guohui Hu, Hongbing Liu, Yiwei Zhang, Bo Cao, Melody Baddoo, Erik K Flemington, Shelya X Zeng, and Hua Lu. Nerve growth factor receptor negates the tumor suppressor p53 as a feedback regulator. *eLife*, 5, 2016.

[2643] Xu Zhou, Alexander W Blocker, Edoardo M Airoldi, and Erin K O'Shea. A computational approach to map nucleosome positions and alternative chromatin states with base pair resolution. *eLife*, 5, 2016.

[2644] Yang Zhou, Yining Liu, Haidong Lu, Si Wu, and Mingsha Zhang. Neuronal representation of saccadic error in macaque posterior parietal cortex (PPC). *eLife*, 5, 2016.

[2645] Hongbo Zhu, Edgardo Sepulveda, Marcus D Hartmann, Manjunatha Kogenaru, Astrid Ursinus, Eva Sulz, Reinhard Albrecht, Murray Coles, Jörg Martin, and Andrei N Lupas. Origin of a folded repeat protein from an intrinsically disordered ancestor. *eLife*, 5, 2016.

[2646] Qinyu Zhu, David Venzke, Ameya S Walimbe, Mary E Anderson, Qiuyu Fu, Lisa N Kinch, Wei Wang, Xing Chen, Nick V Grishin, Niu Huang, Liping Yu, Jack E Dixon, Kevin P Campbell, and Junyu Xiao. Structure of protein O-mannose kinase reveals a unique active site architecture. *eLife*, 5, 2016.

[2647] Jia-Min Zhuo, Hua-an Tseng, Mitul Desai, Mark E Bucklin, Ali I Mohammed, Nick TM Robinson, Edward S Boyden, Lara M Rangel, Alan P Jasanoff, Howard J Gritton, and Xue Han. Young adult born neurons enhance hippocampal dependent performance via influences on bilateral networks. *eLife*, 5, 2016.

[2648] Christina M Zimanyi, Percival Yang-Ting Chen, Gyunghoon Kang, Michael A Funk, and Catherine L Drennan. Molecular basis for allosteric specificity regulation in class Ia ribonucleotide reductase from Escherichia coli. *eLife*, 5, 2016.

[2649] Wei Zou, Ao Shen, Xintong Dong, Madina Tugizova, Yang K Xiang, and Kang Shen. A multi-protein receptor-ligand complex underlies combinatorial dendrite guidance choices in C. elegans. *eLife*, 5, 2016.

[2650] Ariel Zylberberg, Christopher R Fetsch, and Michael N Shadlen. The influence of evidence volatility on choice, reaction time and confidence in a perceptual decision. *eLife*, 5, 2016.

[2651] Paweł Śledź and Martin Jinek. Structural insights into the molecular mechanism of the m6A writer complex. *eLife*, 5, 2016.

[2652] John Aach, Jeantine Lunshof, Eswar Iyer, and George M Church. Addressing the ethical issues raised by synthetic human entities with embryo-like features. *eLife*, 6, 2017.

[2653] Laurence Abrami, Tiziano Dallavilla, Patrick A Sandoz, Mustafa Demir, Béatrice Kunz, Georgios Savoglidis, Vassily Hatzimanikatis, and F Gisou van der Goot. Identification and dynamics of the human ZDHHC16-ZDHHC6 palmitoylation cascade. *eLife*, 6, 2017.

[2654] Martina Absinta, Seung-Kwon Ha, Govind Nair, Pascal Sati, Nicholas J Luciano, Maryknoll Palisoc, Antoine Louveau, Kareem A Zaghloul, Stefania Pittaluga, Jonathan Kipnis, and Daniel S Reich. Human and nonhuman primate meninges harbor lymphatic vessels that can be visualized noninvasively by MRI. *eLife*, 6, 2017.

[2655] Jennifer M Achiro, John Shen, and Sarah W Bottjer. Neural activity in cortico-basal ganglia circuits of juvenile songbirds encodes performance during goal-directed learning. *eLife*, 6, 2017.

[2656] Manuel Alonso Y Adell, Simona M Migliano, Srigokul Upadhyayula, Yury S Bykov, Simon Sprenger, Mehrshad Pakdel, Georg F Vogel, Gloria Jih, Wesley Skillern, Reza Behrouzi, Markus Babst, Oliver Schmidt, Michael W Hess, John AG Briggs, Tomas Kirchhausen, and David Teis. Recruitment dynamics of ESCRT-III and Vps4 to endosomes and implications for reverse membrane budding. *eLife*, 6, 2017.

[2657] Elena W Adlaf, Ryan J Vaden, Anastasia J Niver, Allison F Manuel, Vincent C Onyilo, Matheus T Araujo, Cristina V Dieni, Hai T Vo, Gwendalyn D King, Jacques I Wadiche, and Linda Overstreet-Wadiche. Adult-born neurons modify excitatory synaptic transmission to existing neurons. *eLife*, 6, 2017.

[2658] Ji-Eun Ahn, Yan Chen, and Hubert Amrein. Molecular basis of fatty acid taste in Drosophila. *eLife*, 6, 2017.

[2659] Shanshan Ai, Yong Peng, Chen Li, Fei Gu, Xianhong Yu, Yanzhu Yue, Qing Ma, Jinghai Chen, Zhiqiang Lin, Pingzhu Zhou, Huafeng Xie, Terence W Prendiville, Wen Zheng, Yuli Liu, Stuart H Orkin, Da-Zhi Wang, Jia Yu, William T Pu, and Aibin He. EED orchestration of heart maturation through interaction with HDACs is H3K27me3-independent. *eLife*, 6, 2017.

[2660] Fraser Aird, Irawati Kandela, and Christine Mantis. Replication Study: BET bromodomain inhibition as a therapeutic strategy to target c-Myc. *eLife*, 6, 2017.

[2661] Takahiko Akematsu, Yasuhiro Fukuda, Jyoti Garg, Jeffrey S Fillingham, Ronald E Pearlman, and Josef Loidl. Post-meiotic DNA double-strand breaks occur in Tetrahymena, and require Topoisomerase II and Spo11. *eLife*, 6, 2017.

[2662] Ismael Al-Ramahi, Sai Srinivas Panapakkam Giridharan, Yu-Chi Chen, Samarjit Patnaik, Nathaniel Safren, Junya Hasegawa, Maria de Haro, Amanda K Wagner Gee, Steven A Titus, Hyunkyung Jeong, Jonathan Clarke, Dimitri Krainc, Wei Zheng, Robin F Irvine, Sami Barmada, Marc Ferrer, Noel Southall, Lois S Weisman, Juan Botas, and Juan Jose Marugan. Inhibition of PIP4K$\gamma$ ameliorates the pathological effects of mutant huntingtin protein. *eLife*, 6, 2017.

[2663] Lorenzo Alamo, James S Ware, Antonio Pinto, Richard E Gillilan, Jonathan G Seidman, Christine E Seidman, and Raúl Padrón. Effects of myosin variants on interacting-heads motif explain distinct hypertrophic and dilated cardiomyopathy phenotypes. *eLife*, 6, 2017.

[2664] Sarah Elizabeth Albritton, Anna-Lena Kranz, Lara Heermans Winterkorn, Lena Annika Street, and Sevinc Ercan. Cooperation between a hierarchical set of recruitment sites targets the X chromosome for dosage compensation. *eLife*, 6, 2017.

[2665] Felicity Alcock, Merel PM Damen, Jesper Levring, and Ben C Berks. In vivo experiments do not support the charge zipper model for Tat translocase assembly. *eLife*, 6, 2017.

[2666] Sarah E Ali-Khan, Liam W Harris, and E Richard Gold. Motivating participation in open science by examining researcher incentives. *eLife*, 6, 2017.

[2667] Amin Allalou, Yuelong Wu, Mostafa Ghannad-Rezaie, Peter M Eimon, and Mehmet Fatih Yanik. Automated deep-phenotyping of the vertebrate brain. *eLife*, 6, 2017.

[2668] Nicolas Altemose, Nudrat Noor, Emmanuelle Bitoun, Afidalina Tumian, Michael Imbeault, J Ross Chapman, A Radu Aricescu, and Simon R Myers. A map of human PRDM9 binding provides evidence for novel behaviors of PRDM9 and other zinc-finger proteins in meiosis. *eLife*, 6, 2017.

[2669] Alejandra Alvarado, Andreas Kjær, Wen Yang, Petra Mann, Ariane Briegel, Matthew K Waldor, and Simon Ringgaard. Coupling chemosensory array formation and localization. *eLife*, 6, 2017.

[2670] Katherine Amberg-Johnson, Sanjay B Hari, Suresh M Ganesan, Hernan A Lorenzi, Robert T Sauer, Jacquin C Niles, and Ellen Yeh. Small molecule inhibition of apicomplexan FtsH1 disrupts plastid biogenesis in human pathogens. *eLife*, 6, 2017.

[2671] Ariel Amir. Is cell size a spandrel? *eLife*, 6, 2017.

[2672] Veronique Amor, Chuansheng Zhang, Anna Vainshtein, Ao Zhang, Daniel R Zollinger, Yael Eshed-Eisenbach, Peter J Brophy, Matthew N Rasband, and Elior Peles. The paranodal cytoskeleton clusters Na+ channels at nodes of Ranvier. *eLife*, 6, 2017.

[2673] Bobae An, Jihye Kim, Kyungjoon Park, Sukwon Lee, Sukwoon Song, and Sukwoo Choi. Amount of fear extinction changes its underlying mechanisms. *eLife*, 6, 2017.

[2674] Sojin An, Jungmin Yoon, Hanseong Kim, Ji-Joon Song, and Uhn-soo Cho. Structure-based nuclear import mechanism of histones H3 and H4 mediated by Kap123. *eLife*, 6, 2017.

[2675] Yu A An, Kai Sun, Nolwenn Joffin, Fang Zhang, Yingfeng Deng, Olivier Donzé, Christine M Kusminski, and Philipp E Scherer. Angiopoietin-2 in white adipose tissue improves metabolic homeostasis through enhanced angiogenesis. *eLife*, 6, 2017.

[2676] Rachel W Anantha, Srilatha Simhadri, Tzeh Keong Foo, Susanna Miao, Jingmei Liu, Zhiyuan Shen, Shridar Ganesan, and Bing Xia. Functional and mutational landscapes of BRCA1 for homology-directed repair and therapy resistance. *eLife*, 6, 2017.

[2677] Daniel Andergassen, Christoph P Dotter, Daniel Wenzel, Verena Sigl, Philipp C Bammer, Markus Muckenhuber, Daniela Mayer, Tomasz M Kulinski, Hans-Christian Theussl, Josef M Penninger, Christoph Bock, Denise P Barlow, Florian M Pauler, and Quanah J Hudson. Mapping the mouse Allelome reveals tissue-specific regulation of allelic expression. *eLife*, 6, 2017.

[2678] Caitlin Anderson, India Reiss, Cyrus Zhou, Annie Cho, Haziq Siddiqi, Benjamin Mormann, Cameron M Avelis, Peter Deford, Alan Bergland, Elijah Roberts, James Taylor, Daniel Vasiliauskas, and Robert J Johnston. Natural variation in stochastic photoreceptor specification and color preference in Drosophila. *eLife*, 6, 2017.

[2679] Charles T Anderson, Manoj Kumar, Shanshan Xiong, and Thanos Tzounopoulos. Cell-specific gain modulation by synaptically released zinc in cortical circuits of audition. *eLife*, 6, 2017.

[2680] Madeline G Andrews, Lorenzo M del Castillo, Eliana Ochoa-Bolton, Ken Yamauchi, Jan Smogorzewski, and Samantha J Butler. BMPs direct sensory interneuron identity in the developing spinal cord using signal-specific not morphogenic activities. *eLife*, 6, 2017.

[2681] Viviana Anelli, Jacques A Villefranc, Sagar Chhangawala, Raul Martinez-McFaline, Eleonora Riva, Anvy Nguyen, Akanksha Verma, Rohan Bareja, Zhengming Chen, Theresa Scognamiglio, Olivier Elemento, and Yariv Houvras. Oncogenic BRAF disrupts thyroid morphogenesis and function via twist expression. *eLife*, 6, 2017.

[2682] Yesseinia I Angleró-Rodríguez, Octavio AC Talyuli, Benjamin J Blumberg, Seokyoung Kang, Celia Demby, Alicia Shields, Jenny Carlson, Natapong Jupatanakul, and George Dimopoulos. An Aedes aegypti-associated fungus increases susceptibility to dengue virus by modulating gut trypsin activity. *eLife*, 6, 2017.

[2683] Manabu Aoki, Hironori Hayashi, Kalapala Venkateswara Rao, Debananda Das, Nobuyo Higashi-Kuwata, Haydar Bulut, Hiromi Aoki-Ogata, Yuki Takamatsu, Ravikiran S Yedidi, David A Davis, Shin-ichiro Hattori, Noriko Nishida, Kazuya Hasegawa, Nobutoki Takamune, Prasanth R Nyalapatla, Heather L Osswald, Hirofumi Jono, Hideyuki Saito, Robert Yarchoan, Shogo Misumi, Arun K Ghosh, and Hiroaki Mitsuya. A novel central nervous system-penetrating protease inhibitor overcomes human immunodeficiency virus 1 resistance with unprecedented aM to pM potency. *eLife*, 6, 2017.

[2684] I Lorena Arancibia-Cárcamo, Marc C Ford, Lee Cossell, Kinji Ishida, Koujiro Tohyama, and David Attwell. Node of Ranvier length as a potential regulator of myelinated axon conduction speed. *eLife*, 6, 2017.

[2685] Lindsey Araujo, Phillip Khim, Haik Mkhikian, Christie-Lynn Mortales, and Michael Demetriou. Glycolysis and glutaminolysis cooperatively control T cell function by limiting metabolite supply to N-glycosylation. *eLife*, 6, 2017.

[2686] Michael J Arcaro and Margaret S Livingstone. A hierarchical, retinotopic proto-organization of the primate visual system at birth. *eLife*, 6, 2017.

[2687] Tomoki Arichi, Kimberley Whitehead, Giovanni Barone, Ronit Pressler, Francesco Padormo, A David Edwards, and Lorenzo Fabrizi. Localization of spontaneous bursting neuronal activity in the preterm human brain with simultaneous EEG-fMRI. *eLife*, 6, 2017.

[2688] Jaisy Arikkatt, Md Ashik Ullah, Kirsty Renfree Short, Vivan Zhang, Wan Jun Gan, Zhixuan Loh, Rhiannon B Werder, Jennifer Simpson, Peter D Sly, Stuart B Mazzone, Kirsten M Spann, Manuel AR Ferreira, John W Upham, Maria B Sukkar, and Simon Phipps. RAGE deficiency predisposes mice to virus-induced paucigranulocytic asthma. *eLife*, 6, 2017.

[2689] Yasunobu Arima, Takuto Ohki, Naoki Nishikawa, Kotaro Higuchi, Mitsutoshi Ota, Yuki Tanaka, Junko Nio-Kobayashi, Mohamed Elfeky, Ryota Sakai, Yuki Mori, Tadafumi Kawamoto, Andrea Stofkova, Yukihiro Sakashita, Yuji Morimoto, Masaki Kuwatani, Toshihiko Iwanaga, Yoshichika Yoshioka, Naoya Sakamoto, Akihiko Yoshimura, Mitsuyoshi Takiguchi, Saburo Sakoda, Marco Prinz, Daisuke Kamimura, and Masaaki Murakami. Brain micro-inflammation at specific vessels dysregulates organ-homeostasis via the activation of a new neural circuit. *eLife*, 6, 2017.

[2690] Ana Paula Arruda, Benedicte Mengel Pers, Günes Parlakgul, Ekin Güney, Ted Goh, Erika Cagampan, Grace Yankun Lee, Renata L Goncalves, and Gökhan S Hotamisligil. Defective STIM-mediated store operated Ca2+ entry in hepatocytes leads to metabolic dysfunction in obesity. *eLife*, 6, 2017.

[2691] Filippo Artoni, Rebecca E Kreipke, Ondina Palmeira, Connor Dixon, Zachary Goldberg, and Hannele Ruohola-Baker. Loss of foxo rescues stem cell aging in Drosophila germ line. *eLife*, 6, 2017.

[2692] Joseph Atherton, I-Mei Yu, Alexander Cook, Joseph M Muretta, Agnel Joseph, Jennifer Major, Yannick Sourigues, Jeffrey Clause, Maya Topf, Steven S Rosenfeld, Anne Houdusse, and Carolyn A Moores. The divergent mitotic kinesin MKLP2 exhibits atypical structure and mechanochemistry. *eLife*, 6, 2017.

[2693] Jegath C Athilingam, Roy Ben-Shalom, Caroline M Keeshen, Vikaas S Sohal, and Kevin J Bender. Serotonin enhances excitability and gamma frequency temporal integration in mouse prefrontal fast-spiking interneurons. *eLife*, 6, 2017.

[2694] Ishara F Azmi, Shinya Watanabe, Michael F Maloney, Sukhyun Kang, Jason A Belsky, David M MacAlpine, Craig L Peterson, and Stephen P Bell. Nucleosomes influence multiple steps during replication initiation. *eLife*, 6, 2017.

[2695] Thomas Badet, Remi Peyraud, Malick Mbengue, Olivier Navaud, Mark Derbyshire, Richard P Oliver, Adelin Barbacci, and Sylvain Raffaele. Codon optimization underpins generalist parasitism in fungi. *eLife*, 6, 2017.

[2696] Doreen Badheka, Yevgen Yudin, Istvan Borbiro, Cassandra M Hartle, Aysenur Yazici, Tooraj Mirshahi, and Tibor Rohacs. Inhibition of Transient Receptor Potential Melastatin 3 ion channels by G-protein $\beta\gamma$ subunits. *eLife*, 6, 2017.

[2697] Sung Jun Bae, Lisheng Ni, Adam Osinski, Diana R Tomchick, Chad A Brautigam, and Xuelian Luo. SAV1 promotes Hippo kinase activation through antagonizing the PP2A phosphatase STRIPAK. *eLife*, 6, 2017.

[2698] Alexis S Bailey, Pedro J Batista, Rebecca S Gold, Y Grace Chen, Dirk G de Rooij, Howard Y Chang, and Margaret T Fuller. The conserved RNA helicase YTHDC2 regulates the transition from proliferation to differentiation in the germline. *eLife*, 6, 2017.

[2699] Rasmus O Bak, Daniel P Dever, Andreas Reinisch, David Cruz Hernandez, Ravindra Majeti, and Matthew H Porteus. Multiplexed genetic engineering of human hematopoietic stem and progenitor cells using CRISPR/Cas9 and AAV6. *eLife*, 6, 2017.

[2700] Zachary Baker, Molly Schumer, Yuki Haba, Lisa Bashkirova, Chris Holland, Gil G Rosenthal, and Molly Przeworski. Repeated losses of PRDM9-directed recombination despite the conservation of PRDM9 across vertebrates. *eLife*, 6, 2017.

[2701] Anush Bakunts, Andrea Orsi, Milena Vitale, Angela Cattaneo, Federica Lari, Laura Tadè, Roberto Sitia, Andrea Raimondi, Angela Bachi, and Eelco van Anken. Ratiometric sensing of BiP-client versus BiP levels by the unfolded protein response determines its signaling amplitude. *eLife*, 6, 2017.

[2702] Nicholas L Balderston, Elizabeth Hale, Abigail Hsiung, Salvatore Torrisi, Tom Holroyd, Frederick W Carver, Richard Coppola, Monique Ernst, and Christian Grillon. Threat of shock increases excitability and connectivity of the intraparietal sulcus. *eLife*, 6, 2017.

[2703] Sandro Baldi, Alessio Bolognesi, Anne Cornelis Meinema, and Yves Barral. Heat stress promotes longevity in budding yeast by relaxing the confinement of age-promoting factors in the mother cell. *eLife*, 6, 2017.

[2704] Michael R Bale, Malamati Bitzidou, Anna Pitas, Leonie S Brebner, Lina Khazim, Stavros T Anagnou, Caitlin D Stevenson, and Miguel Maravall. Learning and recognition of tactile temporal sequences by mice and humans. *eLife*, 6, 2017.

[2705] Pradeep Bandaru, Neel H Shah, Moitrayee Bhattacharyya, John P Barton, Yasushi Kondo, Joshua C Cofsky, Christine L Gee, Arup K Chakraborty, Tanja Kortemme, Rama Ranganathan, and John Kuriyan. Deconstruction of the Ras switching cycle through saturation mutagenesis. *eLife*, 6, 2017.

[2706] Jennifer J Banerjee, Birgit L Aerne, Maxine V Holder, Simon Hauri, Matthias Gstaiger, and Nicolas Tapon. Meru couples planar cell polarity with apical-basal polarity during asymmetric cell division. *eLife*, 6, 2017.

[2707] Susanne CS Bantele, Pedro Ferreira, Dalia Gritenaite, Dominik Boos, and Boris Pfander. Targeting of the Fun30 nucleosome remodeller by the Dpb11 scaffold facilitates cell cycle-regulated DNA end resection. *eLife*, 6, 2017.

[2708] Lara Barrio and Marco Milán. Boundary Dpp promotes growth of medial and lateral regions of the Drosophila wing. *eLife*, 6, 2017.

[2709] Kevin C Barry, Nicholas T Ingolia, and Russell E Vance. Global analysis of gene expression reveals mRNA superinduction is required for the inducible immune response to a bacterial pathogen. *eLife*, 6, 2017.

[2710] Michael J Bartlett, Tammy E Steeves, Neil J Gemmell, and Patrice C Rosengrave. Sperm competition risk drives rapid ejaculate adjustments mediated by seminal fluid. *eLife*, 6, 2017.

[2711] Sandip Basak, Nicolaus Schmandt, Yvonne Gicheru, and Sudha Chakrapani. Crystal structure and dynamics of a lipid-induced potential desensitized-state of a pentameric ligand-gated channel. *eLife*, 6, 2017.

[2712] Sofia Battaglia, Michael Lidschreiber, Carlo Baejen, Phillipp Torkler, Seychelle M Vos, and Patrick Cramer. RNA-dependent chromatin association of transcription elongation factors and Pol II CTD kinases. *eLife*, 6, 2017.

[2713] Philippe J Batut and Thomas R Gingeras. Conserved noncoding transcription and core promoter regulatory code in early Drosophila development. *eLife*, 6, 2017.

[2714] Matthias Bauer, Johannes Knebel, Matthias Lechner, Peter Pickl, and Erwin Frey. Ecological feedback in quorum-sensing microbial populations can induce heterogeneous production of autoinducers. *eLife*, 6, 2017.

[2715] Fabian Baumann, Magnus Sebastian Bauer, Martin Rees, Alexander Alexandrovich, Mathias Gautel, Diana Angela Pippig, and Hermann Eduard Gaub. Increasing evidence of mechanical force as a functional regulator in smooth muscle myosin light chain kinase. *eLife*, 6, 2017.

[2716] Janina Baumbach, Andal Murthy, Mark A McClintock, Carly I Dix, Ruta Zalyte, Ha Thi Hoang, and Simon L Bullock. Lissencephaly-1 is a context-dependent regulator of the human dynein complex. *eLife*, 6, 2017.

[2717] Thomas R Beattie, Nitin Kapadia, Emilien Nicolas, Stephan Uphoff, Adam JM Wollman, Mark C Leake, and Rodrigo Reyes-Lamothe. Frequent exchange of the DNA polymerase during bacterial chromosome replication. *eLife*, 6, 2017.

[2718] Esteban J Beckwith, Quentin Geissmann, Alice S French, and Giorgio F Gilestro. Regulation of sleep homeostasis by sexual arousal. *eLife*, 6, 2017.

[2719] Muheeb Beg, Nazish Abdullah, Fathima Shazna Thowfeik, Nasser K Altorki, and Timothy E McGraw. Distinct Akt phosphorylation states are required for insulin regulated Glut4 and Glut1-mediated glucose uptake. *eLife*, 6, 2017.

[2720] Marcel Beining, Lucas Alberto Mongiat, Stephan Wolfgang Schwarzacher, Hermann Cuntz, and Peter Jedlicka. T2N as a new tool for robust electrophysiological modeling demonstrated for mature and adult-born dentate granule cells. *eLife*, 6, 2017.

[2721] Gillian Morven Belbin, Jacqueline Odgis, Elena P Sorokin, Muh-Ching Yee, Sumita Kohli, Benjamin S Glicksberg, Christopher R Gignoux, Genevieve L Wojcik, Tielman Van Vleck, Janina M Jeff, Michael Linderman, Claudia Schurmann, Douglas Ruderfer, Xiaoqiang Cai, Amanda Merkelson, Anne E Justice, Kristin L Young, Misa Graff, Kari E North, Ulrike Peters, Regina James, Lucia Hindorff, Ruth Kornreich, Lisa Edelmann, Omri Gottesman, Eli EA Stahl, Judy H Cho, Ruth JF Loos, Erwin P Bottinger, Girish N Nadkarni, Noura S Abul-Husn, and Eimear E Kenny. Genetic identification of a common collagen disease in Puerto Ricans via identity-by-descent mapping in a health system. *eLife*, 6, 2017.

[2722] Carmelo Bellardita, Vittorio Caggiano, Roberto Leiras, Vanessa Caldeira, Andrea Fuchs, Julien Bouvier, Peter Löw, and Ole Kiehn. Spatiotemporal correlation of spinal network dynamics underlying spasms in chronic spinalized mice. *eLife*, 6, 2017.

[2723] Stephanie F Bellmaine, Dmitry A Ovchinnikov, David T Manallack, Claire E Cuddy, Andrew G Elefanty, Edouard G Stanley, Ernst J Wolvetang, Spencer J Williams, and Martin Pera. Inhibition of DYRK1A disrupts neural lineage specificationin human pluripotent stem cells. *eLife*, 6, 2017.

[2724] Maria Benito-Jardón, Sarah Klapproth, Irene Gimeno-LLuch, Tobias Petzold, Mitasha Bharadwaj, Daniel J Müller, Gabriele Zuchtriegel, Christoph A Reichel, and Mercedes Costell. The fibronectin synergy site re-enforces cell adhesion and mediates a crosstalk between integrin classes. *eLife*, 6, 2017.

[2725] Joshua B Benoit, Aurélien Vigneron, Nichole A Broderick, Yineng Wu, Jennifer S Sun, John R Carlson, Serap Aksoy, and Brian L Weiss. Symbiont-induced odorant binding proteins mediate insect host hematopoiesis. *eLife*, 6, 2017.

[2726] Lee R Berger, John Hawks, Paul HGM Dirks, Marina Elliott, and Eric M Roberts. Homo naledi and Pleistocene hominin evolution in subequatorial Africa. *eLife*, 6, 2017.

[2727] Gustavo S Betini, Andrew G McAdam, Cortland K Griswold, and D Ryan Norris. A fitness trade-off between seasons causes multigenerational cycles in phenotype and population size. *eLife*, 6, 2017.

[2728] Irene Beusch, Pierre Barraud, Ahmed Moursy, Antoine Cléry, and Frédéric Hai-Trieu Allain. Tandem hnRNP A1 RNA recognition motifs act in concert to repress the splicing of survival motor neuron exon 7. *eLife*, 6, 2017.

[2729] Upinder Singh Bhalla. Synaptic input sequence discrimination on behavioral timescales mediated by reaction-diffusion chemistry in dendrites. *eLife*, 6, 2017.

[2730] Matthew J Bick, Per J Greisen, Kevin J Morey, Mauricio S Antunes, David La, Banumathi Sankaran, Luc Reymond, Kai Johnsson, June I Medford, and David Baker. Computational design of environmental sensors for the potent opioid fentanyl. *eLife*, 6, 2017.

[2731] Brent W Bisgrove, Yi-Chu Su, and H Joseph Yost. Maternal Gdf3 is an obligatory cofactor in Nodal signaling for embryonic axis formation in zebrafish. *eLife*, 6, 2017.

[2732] Paul-Yannick Bitome-Essono, Benjamin Ollomo, Céline Arnathau, Patrick Durand, Nancy Diamella Mokoudoum, Lauriane Yacka-Mouele, Alain-Prince Okouga, Larson Boundenga, Bertrand Mve-Ondo, Judicaël Obame-Nkoghe, Philippe Mbehang-Nguema, Flobert Njiokou, Boris Makanga, Rémi Wattier, Diego Ayala, Francisco J Ayala, Francois Renaud, Virginie Rougeron, Francois Bretagnolle, Franck Prugnolle, and Christophe Paupy. Tracking zoonotic pathogens using blood-sucking flies as 'flying syringes'. *eLife*, 6, 2017.

[2733] Daniel Blanco-Melo, Robert J Gifford, and Paul D Bieniasz. Co-option of an endogenous retrovirus envelope for host defense in hominid ancestors. *eLife*, 6, 2017.

[2734] Oriane Blanquie, Jenq-Wei Yang, Werner Kilb, Salim Sharopov, Anne Sinning, and Heiko J Luhmann. Electrical activity controls area-specific expression of neuronal apoptosis in the mouse developing cerebral cortex. *eLife*, 6, 2017.

[2735] Dvir Blivis, Gal Haspel, Philip Z Mannes, Michael J O'Donovan, and Michael J Iadarola. Identification of a novel spinal nociceptive-motor gate control for A$\delta$ pain stimuli in rats. *eLife*, 6, 2017.

[2736] Bernard Bloem, Rafiq Huda, Mriganka Sur, and Ann M Graybiel. Two-photon imaging in mice shows striosomes and matrix have overlapping but differential reinforcement-related responses. *eLife*, 6, 2017.

[2737] Kevin A Bolding and Kevin M Franks. Complementary codes for odor identity and intensity in olfactory cortex. *eLife*, 6, 2017.

[2738] Tobias Boothe, Lennart Hilbert, Michael Heide, Lea Berninger, Wieland B Huttner, Vasily Zaburdaev, Nadine L Vastenhouw, Eugene W Myers, David N Drechsel, and Jochen C Rink. A tunable refractive index matching medium for live imaging cells, tissues and model organisms. *eLife*, 6, 2017.

[2739] Martin Borch Jensen, Yanyan Qi, Rebeccah Riley, Liya Rabkina, and Heinrich Jasper. PGAM5 promotes lasting FoxO activation after developmental mitochondrial stress and extends lifespan in Drosophila. *eLife*, 6, 2017.

[2740] Alexander Borodavka, Eric C Dykeman, Waldemar Schrimpf, and Don C Lamb. Protein-mediated RNA folding governs sequence-specific interactions between rotavirus genome segments. *eLife*, 6, 2017.

[2741] Pablo Sanchez Bosch, Ruta Ziukaite, Cyrille Alexandre, Konrad Basler, and Jean-Paul Vincent. Dpp controls growth and patterning in Drosophila wing precursors through distinct modes of action. *eLife*, 6, 2017.

[2742] Yves Boubenec, Jennifer Lawlor, Urszula Górska, Shihab Shamma, and Bernhard Englitz. Detecting changes in dynamic and complex acoustic environments. *eLife*, 6, 2017.

[2743] Aaron B Bowen, Ashley M Bourke, Brian G Hiester, Cyril Hanus, and Matthew J Kennedy. Golgi-independent secretory trafficking through recycling endosomes in neuronal dendrites and spines. *eLife*, 6, 2017.

[2744] Janine Bowring, Maan M Neamah, Jorge Donderis, Ignacio Mir-Sanchis, Christian Alite, J Rafael Ciges-Tomas, Elisa Maiques, Iltyar Medmedov, Alberto Marina, and José R Penadés. Pirating conserved phage mechanisms promotes promiscuous staphylococcal pathogenicity island transfer. *eLife*, 6, 2017.

[2745] Niels Bradshaw, Vladimir M Levdikov, Christina M Zimanyi, Rachelle Gaudet, Anthony J Wilkinson, and Richard Losick. A widespread family of serine/threonine protein phosphatases shares a common regulatory switch with proteasomal proteases. *eLife*, 6, 2017.

[2746] Tobias Brandt, Arnaud Mourier, Luke S Tain, Linda Partridge, Nils-Göran Larsson, and Werner Kühlbrandt. Changes of mitochondrial ultrastructure and function during ageing in mice and Drosophila. *eLife*, 6, 2017.

[2747] Colin S Brent, John A Byers, and Anat Levi-Zada. An insect anti-antiaphrodisiac. *eLife*, 6, 2017.

[2748] James Alexander Briggs, Victor C Li, Seungkyu Lee, Clifford J Woolf, Allon Klein, and Marc W Kirschner. Mouse embryonic stem cells can differentiate via multiple paths to the same state. *eLife*, 6, 2017.

[2749] Paul J Brighton, Yojiro Maruyama, Katherine Fishwick, Pavle Vrljicak, Shreeya Tewary, Risa Fujihara, Joanne Muter, Emma S Lucas, Taihei Yamada, Laura Woods, Raffaella Lucciola, Yie Hou Lee, Satoru Takeda, Sascha Ott, Myriam Hemberger, Siobhan Quenby, and Jan Joris Brosens. Clearance of senescent decidual cells by uterine natural killer cells in cycling human endometrium. *eLife*, 6, 2017.

[2750] Amy E Brinegar, Zheng Xia, James Anthony Loehr, Wei Li, George Gerald Rodney, and Thomas A Cooper. Extensive alternative splicing transitions during postnatal skeletal muscle development are required for calcium handling functions. *eLife*, 6, 2017.

[2751] Graham J Britton, Rachel Ambler, Danielle J Clark, Elaine V Hill, Helen M Tunbridge, Kerrie E McNally, Bronwen R Burton, Philomena Butterweck, Catherine Sabatos-Peyton, Lea A Hampton-O'Neil, Paul Verkade, Christoph Wülfing, and David Cameron Wraith. PKC$\theta$ links proximal T cell and Notch signaling through localized regulation of the actin cytoskeleton. *eLife*, 6, 2017.

[2752] Angela M Bruno, William N Frost, and Mark D Humphries. A spiral attractor network drives rhythmic locomotion. *eLife*, 6, 2017.

[2753] Zuzanna Brzosko, Sara Zannone, Wolfram Schultz, Claudia Clopath, and Ole Paulsen. Sequential neuromodulation of Hebbian plasticity offers mechanism for effective reward-based navigation. *eLife*, 6, 2017.

[2754] Christoph A Bücherl, Iris K Jarsch, Christian Schudoma, Cécile Segonzac, Malick Mbengue, Silke Robatzek, Daniel MacLean, Thomas Ott, and Cyril Zipfel. Plant immune and growth receptors share common signalling components but localise to distinct plasma membrane nanodomains. *eLife*, 6, 2017.

[2755] Katherine M Buckley, Eric Chun Hei Ho, Taku Hibino, Catherine S Schrankel, Nicholas W Schuh, Guizhi Wang, and Jonathan P Rast. IL17 factors are early regulators in the gut epithelium during inflammatory response to Vibrio in the sea urchin larva. *eLife*, 6, 2017.

[2756] Luke K Burke, Tamana Darwish, Althea R Cavanaugh, Sam Virtue, Emma Roth, Joanna Morro, Shun-Mei Liu, Jing Xia, Jeffrey W Dalley, Keith Burling, Streamson Chua, Toni Vidal-Puig, Gary J Schwartz, and Clémence Blouet. mTORC1 in AGRP neurons integrates exteroceptive and interoceptive food-related cues in the modulation of adaptive energy expenditure in mice. *eLife*, 6, 2017.

[2757] David H Burkhardt, Silvi Rouskin, Yan Zhang, Gene-Wei Li, Jonathan S Weissman, and Carol A Gross. Operon mRNAs are organized into ORF-centric structures that predict translation efficiency. *eLife*, 6, 2017.

[2758] John A Burns, Huanjia Zhang, Elizabeth Hill, Eunsoo Kim, and Ryan Kerney. Transcriptome analysis illuminates the nature of the intracellular interaction in a vertebrate-algal symbiosis. *eLife*, 6, 2017.

[2759] Yury S Bykov, Miroslava Schaffer, Svetlana O Dodonova, Sahradha Albert, Jürgen M Plitzko, Wolfgang Baumeister, Benjamin D Engel, and John AG Briggs. The structure of the COPI coat determined within the cell. *eLife*, 6, 2017.

[2760] Monica Pia Caggiano, Xiulian Yu, Neha Bhatia, André Larsson, Hasthi Ram, Carolyn K Ohno, Pia Sappl, Elliot M Meyerowitz, Henrik Jönsson, and Marcus G Heisler. Cell type boundaries organize plant development. *eLife*, 6, 2017.

[2761] Vladimir Camarena, David W Sant, Tyler C Huff, Sushmita Mustafi, Ryan K Muir, Allegra T Aron, Christopher J Chang, Adam R Renslo, Paula V Monje, and Gaofeng Wang. cAMP signaling regulates DNA hydroxymethylation by augmenting the intracellular labile ferrous iron pool. *eLife*, 6, 2017.

[2762] Felix Campelo, Josse van Galen, Gabriele Turacchio, Seetharaman Parashuraman, Michael M Kozlov, María F García-Parajo, and Vivek Malhotra. Sphingomyelin metabolism controls the shape and function of the Golgi cisternae. *eLife*, 6, 2017.

[2763] Sebastian Canovas, Elena Ivanova, Raquel Romar, Soledad García-Martínez, Cristina Soriano-Úbeda, Francisco A García-Vázquez, Heba Saadeh, Simon Andrews, Gavin Kelsey, and Pilar Coy. DNA methylation and gene expression changes derived from assisted reproductive technologies can be decreased by reproductive fluids. *eLife*, 6, 2017.

[2764] Limian Cao, Pengfei Zhang, Jinming Li, and Mian Wu. LAST, a c-Myc-inducible long noncoding RNA, cooperates with CNBP to promote CCND1 mRNA stability in human cells. *eLife*, 6, 2017.

[2765] Juliana S Capitanio, Ben Montpetit, and Richard W Wozniak. Human Nup98 regulates the localization and activity of DExH/D-box helicase DHX9. *eLife*, 6, 2017.

[2766] Paulina Carriba and Alun M Davies. CD40 is a major regulator of dendrite growth from developing excitatory and inhibitory neurons. *eLife*, 6, 2017.

[2767] Marta Carroni, Kamila B Franke, Michael Maurer, Jasmin Jäger, Ingo Hantke, Felix Gloge, Daniela Linder, Sebastian Gremer, Kürşad Turgay, Bernd Bukau, and Axel Mogk. Regulatory coiled-coil domains promote head-to-head assemblies of AAA+ chaperones essential for tunable activity control. *eLife*, 6, 2017.

[2768] Ian M Cartwright, Xinjian Liu, Min Zhou, Fang Li, and Chuan-Yuan Li. Essential roles of Caspase-3 in facilitating Myc-induced genetic instability and carcinogenesis. *eLife*, 6, 2017.

[2769] Elisabeth Cassier, Nathalie Gallay, Thomas Bourquard, Sylvie Claeysen, Joël Bockaert, Pascale Crépieux, Anne Poupon, Eric Reiter, Philippe Marin, and Franck Vandermoere. Phosphorylation of $\beta$-arrestin2 at Thr383 by MEK underlies $\beta$-arrestin-dependent activation of Erk1/2 by GPCRs. *eLife*, 6, 2017.

[2770] Laura Cato, Antje Neeb, Adam Sharp, Victor Buzón, Scott B Ficarro, Linxiao Yang, Claudia Muhle-Goll, Nane C Kuznik, Ruth Riisnaes, Daniel Nava Rodrigues, Olivier Armant, Victor Gourain, Guillaume Adelmant, Emmanuel A Ntim, Thomas Westerling, David Dolling, Pasquale Rescigno, Ines Figueiredo, Friedrich Fauser, Jennifer Wu, Jaice T Rottenberg, Liubov Shatkina, Claudia Ester, Burkhard Luy, Holger Puchta, Jakob Troppmair, Nicole Jung, Stefan Bräse, Uwe Strähle, Jarrod A Marto, Gerd Ulrich Nienhaus, Bissan Al-Lazikani, Xavier Salvatella, Johann S de Bono, Andrew CB Cato, and Myles Brown. Development of Bag-1L as a therapeutic target in androgen receptor-dependent prostate cancer. *eLife*, 6, 2017.

[2771] Magali Cazade, Isabelle Bidaud, Philippe Lory, and Jean Chemin. Activity-dependent regulation of T-type calcium channels by submembrane calcium ions. *eLife*, 6, 2017.

[2772] Cemre Celen, Jen-Chieh Chuang, Xin Luo, Nadine Nijem, Angela K Walker, Fei Chen, Shuyuan Zhang, Andrew S Chung, Liem H Nguyen, Ibrahim Nassour, Albert Budhipramono, Xuxu Sun, Levinus A Bok, Meriel McEntagart, Evelien F Gevers, Shari G Birnbaum, Amelia J Eisch, Craig M Powell, Woo-Ping Ge, Gijs WE Santen, Maria Chahrour, and Hao Zhu. Arid1b haploinsufficient mice reveal neuropsychiatric phenotypes and reversible causes of growth impairment. *eLife*, 6, 2017.

[2773] Barbara Celona, John von Dollen, Sarat C Vatsavayai, Risa Kashima, Jeffrey R Johnson, Amy A Tang, Akiko Hata, Bruce L Miller, Eric J Huang, Nevan J Krogan, William W Seeley, and Brian L Black. Suppression of C9orf72 RNA repeat-induced neurotoxicity by the ALS-associated RNA-binding protein Zfp106. *eLife*, 6, 2017.

[2774] Isaac Cervantes-Sandoval, Anna Phan, Molee Chakraborty, and Ronald L Davis. Reciprocal synapses between mushroom body and dopamine neurons form a positive feedback loop required for learning. *eLife*, 6, 2017.

[2775] Rima S Chakrabarti, Sally A Ingham, Julia Kozlitina, Austin Gay, Jonathan C Cohen, Arun Radhakrishnan, and Helen H Hobbs. Variability of cholesterol accessibility in human red blood cells measured using a bacterial cholesterol-binding toxin. *eLife*, 6, 2017.

[2776] Kasturi Chakraborty, KaHo Leung, and Yamuna Krishnan. High lumenal chloride in the lysosome is critical for lysosome function. *eLife*, 6, 2017.

[2777] Simon Chamberland, Helen H Yang, Michael M Pan, Stephen W Evans, Sihui Guan, Mariya Chavarha, Ying Yang, Charleen Salesse, Haodi Wu, Joseph C Wu, Thomas R Clandinin, Katalin Toth, Michael Z Lin, and FranÇois St-Pierre. Fast two-photon imaging of subcellular voltage dynamics in neuronal tissue with genetically encoded indicators. *eLife*, 6, 2017.

[2778] Eunice HoYee Chan, Pruthvi Chavadimane Shivakumar, Raphaël Clément, Edith Laugier, and Pierre-FranÇois Lenne. Patterned cortical tension mediated by N-cadherin controls cell geometric order in the Drosophila eye. *eLife*, 6, 2017.

[2779] Vijayendran Chandran, Kun Gao, Vivek Swarup, Revital Versano, Hongmei Dong, Maria C Jordan, and Daniel H Geschwind. Inducible and reversible phenotypes in a novel mouse model of Friedreich's Ataxia. *eLife*, 6, 2017.

[2780] Chia-Wei Chang, Julia R Wilkerson, Carly F Hale, Jay R Gibson, and Kimberly M Huber. Distinct stages of synapse elimination are induced by burst firing of CA1 neurons and differentially require MEF2A/D. *eLife*, 6, 2017.

[2781] Ting-Ya Chang, Chen Chen, Min Lee, Ya-Chu Chang, Chi-Huan Lu, Shao-Tzu Lu, De-Yao Wang, Aijun Wang, Chin-Lin Guo, and Pei-Lin Cheng. Paxillin facilitates timely neurite initiation on soft-substrate environments by interacting with the endocytic machinery. *eLife*, 6, 2017.

[2782] Camille Charoy, Sarah Dinvaut, Yohan Chaix, Laurette Morlé, Isabelle Sanyas, Muriel Bozon, Karine Kindbeiter, Bénédicte Durand, Jennifer M Skidmore, Lies De Groef, Motoaki Seki, Lieve Moons, Christiana Ruhrberg, James F Martin, Donna M Martin, Julien Falk, and Valerie Castellani. Genetic specification of left–right asymmetry in the diaphragm muscles and their motor innervation. *eLife*, 6, 2017.

[2783] Dhananjay Chaturvedi, Heinrich Reichert, Rajesh D Gunage, and K VijayRaghavan. Identification and functional characterization of muscle satellite cells in Drosophila. *eLife*, 6, 2017.

[2784] Ritu Chaudhary, Berkley Gryder, Wendy S Woods, Murugan Subramanian, Matthew F Jones, Xiao Ling Li, Lisa M Jenkins, Svetlana A Shabalina, Min Mo, Mary Dasso, Yuan Yang, Lalage M Wakefield, Yuelin Zhu, Susan M Frier, Branden S Moriarity, Kannanganattu V Prasanth, Pablo Perez-Pinera, and Ashish Lal. Prosurvival long noncoding RNA PINCR regulates a subset of p53 targets in human colorectal cancer cells by binding to Matrin 3. *eLife*, 6, 2017.

[2785] Baoyu Chen, Hui-Ting Chou, Chad A Brautigam, Wenmin Xing, Sheng Yang, Lisa Henry, Lynda K Doolittle, Thomas Walz, and Michael K Rosen. Rac1 GTPase activates the WAVE regulatory complex through two distinct binding sites. *eLife*, 6, 2017.

[2786] Huaibin Chen, William M Marsiglia, Min-Kyu Cho, Zhifeng Huang, Jingjing Deng, Steven P Blais, Weiming Gai, Shibani Bhattacharya, Thomas A Neubert, Nathaniel J Traaseth, and Moosa Mohammadi. Elucidation of a four-site allosteric network in fibroblast growth factor receptor tyrosine kinases. *eLife*, 6, 2017.

[2787] Jie Chen, Yuhui Ou, Yi Li, Shumei Hu, Li-Wa Shao, and Ying Liu. Metformin extends C. elegans lifespan through lysosomal pathway. *eLife*, 6, 2017.

[2788] Jingxun Chen, Amy Tresenrider, Minghao Chia, David T McSwiggen, Gianpiero Spedale, Victoria Jorgensen, Hanna Liao, Folkert Jacobus van Werven, and ElÇin ünal. Kinetochore inactivation by expression of a repressive mRNA. *eLife*, 6, 2017.

[2789] Keyu Chen, Jianchao Li, Chao Wang, Zhiyi Wei, and Mingjie Zhang. Autoinhibition of ankyrin-B/G membrane target bindings by intrinsically disordered segments from the tail regions. *eLife*, 6, 2017.

[2790] Lin Chen, Yibin Xu, Wilson Wong, Jennifer K Thompson, Julie Healer, Ethan D Goddard-Borger, Michael C Lawrence, and Alan F Cowman. Structural basis for inhibition of erythrocyte invasion by antibodies to Plasmodium falciparum protein CyRPA. *eLife*, 6, 2017.

[2791] Meiting Chen, Ziyi Zhao, Jin Yang, Kai Peng, Matthew AB Baker, Fan Bai, and Chien-Jung Lo. Length-dependent flagellar growth of Vibrio alginolyticus revealed by real time fluorescent imaging. *eLife*, 6, 2017.

[2792] Ming Chen, Jinzhao Wang, Jinxiang Jiang, Xingzhi Zheng, Nicholas J Justice, Kun Wang, Xiangqian Ran, Yi Li, Qingwei Huo, Jiajia Zhang, Hongmei Li, Nannan Lu, Ying Wang, Hui Zheng, Cheng Long, and Li Yang. APP modulates KCC2 expression and function in hippocampal GABAergic inhibition. *eLife*, 6, 2017.

[2793] Mary Hongying Cheng, Delany Torres-Salazar, Aneysis D Gonzalez-Suarez, Susan G Amara, and Ivet Bahar. Substrate transport and anion permeation proceed through distinct pathways in glutamate transporters. *eLife*, 6, 2017.

[2794] Kylie S Chew, Jordan M Renna, David S McNeill, Diego C Fernandez, William T Keenan, Michael B Thomsen, Jennifer L Ecker, Gideon S Loevinsohn, Cassandra VanDunk, Daniel C Vicarel, Adele Tufford, Shijun Weng, Paul A Gray, Michel Cayouette, Erik D Herzog, Haiqing Zhao, David M Berson, and Samer Hattar. A subset of ipRGCs regulates both maturation of the circadian clock and segregation of retinogeniculate projections in mice. *eLife*, 6, 2017.

[2795] Kim Hou Chia, Tomoyuki Fukuda, Fajar Sofyantoro, Takato Matsuda, Takamitsu Amai, and Kazuhiro Shiozaki. Ragulator and GATOR1 complexes promote fission yeast growth by attenuating TOR complex 1 through Rag GTPases. *eLife*, 6, 2017.

[2796] Minghao Chia, Amy Tresenrider, Jingxun Chen, Gianpiero Spedale, Victoria Jorgensen, ElÇin ünal, and Folkert Jacobus van Werven. Transcription of a 5′ extended mRNA isoform directs dynamic chromatin changes and interference of a downstream promoter. *eLife*, 6, 2017.

[2797] David M Chiasson, Kristina Haage, Katharina Sollweck, Andreas Brachmann, Petra Dietrich, and Martin Parniske. A quantitative hypermorphic CNGC allele confers ectopic calcium flux and impairs cellular development. *eLife*, 6, 2017.

[2798] Alexander Chien, Sheng Min Shih, Raqual Bower, Douglas Tritschler, Mary E Porter, and Ahmet Yildiz. Dynamics of the IFT machinery at the ciliary tip. *eLife*, 6, 2017.

[2799] Yashodhan Chinchore, Tedi Begaj, David Wu, Eugene Drokhlyansky, and Constance L Cepko. Glycolytic reliance promotes anabolism in photoreceptors. *eLife*, 6, 2017.

[2800] Krishna Chinthalapudi, Sarah M Heissler, Matthias Preller, James R Sellers, and Dietmar J Manstein. Mechanistic insights into the active site and allosteric communication pathways in human nonmuscle myosin-2C. *eLife*, 6, 2017.

[2801] Jungnam Cho and Jerzy Paszkowski. Regulation of rice root development by a retrotransposon acting as a microRNA sponge. *eLife*, 6, 2017.

[2802] Sandip Chorghade, Joseph Seimetz, Russell Emmons, Jing Yang, Stefan M Bresson, Michael De Lisio, Gianni Parise, Nicholas K Conrad, and Auinash Kalsotra. Poly(A) tail length regulates PABPC1 expression to tune translation in the heart. *eLife*, 6, 2017.

[2803] Jenna R Christensen, Glen M Hocky, Kaitlin E Homa, Alisha N Morganthaler, Sarah E Hitchcock-DeGregori, Gregory A Voth, and David R Kovar. Competition between Tropomyosin, Fimbrin, and ADF/Cofilin drives their sorting to distinct actin filament networks. *eLife*, 6, 2017.

[2804] Jacky Chung, Johannes G Wittig, Alireza Ghamari, Manami Maeda, Tamara A Dailey, Hector Bergonia, Martin D Kafina, Emma E Coughlin, Catherine E Minogue, Alexander S Hebert, Liangtao Li, Jerry Kaplan, Harvey F Lodish, Daniel E Bauer, Stuart H Orkin, Alan B Cantor, Takahiro Maeda, John D Phillips, Joshua J Coon, David J Pagliarini, Harry A Dailey, and Barry H Paw. Erythropoietin signaling regulates heme biosynthesis. *eLife*, 6, 2017.

[2805] Jean-Ju Chung, Kiyoshi Miki, Doory Kim, Sang-Hee Shim, Huanan F Shi, Jae Yeon Hwang, Xinjiang Cai, Yusuf Iseri, Xiaowei Zhuang, and David E Clapham. CatSper$\zeta$regulates the structural continuity of sperm Ca2+ signaling domains and is required for normal fertility. *eLife*, 6, 2017.

[2806] SeYeon Chung, Sangjoon Kim, and Deborah J Andrew. Uncoupling apical constriction from tissue invagination. *eLife*, 6, 2017.

[2807] Angela J Churchill, Giselle Dominguez Gutiérrez, Ruth A Singer, David S Lorberbaum, Kevin A Fischer, and Lori Sussel. Genetic evidence that Nkx2.2 acts primarily downstream of Neurog3 in pancreatic endocrine lineage development. *eLife*, 6, 2017.

[2808] Matthew A Churgin, Sang-Kyu Jung, Chih-Chieh Yu, Xiangmei Chen, David M Raizen, and Christopher Fang-Yen. Longitudinal imaging of Caenorhabditis elegans in a microfabricated device reveals variation in behavioral decline during aging. *eLife*, 6, 2017.

[2809] Lindsay D Clark, Igor Dikiy, Karen Chapman, Karin EJ Rödström, James Aramini, Michael V LeVine, George Khelashvili, Søren GF Rasmussen, Kevin H Gardner, and Daniel M Rosenbaum. Ligand modulation of sidechain dynamics in a wild-type human GPCR. *eLife*, 6, 2017.

[2810] Geraldine M Clarke, Kirk Rockett, Katja Kivinen, Christina Hubbart, Anna E Jeffreys, Kate Rowlands, Muminatou Jallow, David J Conway, Kalifa A Bojang, Margaret Pinder, Stanley Usen, Fatoumatta Sisay-Joof, Giorgio Sirugo, Ousmane Toure, Mahamadou A Thera, Salimata Konate, Sibiry Sissoko, Amadou Niangaly, Belco Poudiougou, Valentina D Mangano, Edith C Bougouma, Sodiomon B Sirima, David Modiano, Lucas N Amenga-Etego, Anita Ghansah, Kwadwo A Koram, Michael D Wilson, Anthony Enimil, Jennifer Evans, Olukemi K Amodu, Subulade Olaniyan, Tobias Apinjoh, Regina Mugri, Andre Ndi, Carolyne M Ndila, Sophie Uyoga, Alexander Macharia, Norbert Peshu, Thomas N Williams, Alphaxard Manjurano, Nuno Sepúlveda, Taane G Clark, Eleanor Riley, Chris Drakeley, Hugh Reyburn, Vysaul Nyirongo, David Kachala, Malcolm Molyneux, Sarah J Dunstan, Nguyen Hoan Phu, Nguyen Ngoc Quyen, Cao Quang Thai, Tran Tinh Hien, Laurens Manning, Moses Laman, Peter Siba, Harin Karunajeewa, Steve Allen, Angela Allen, Timothy ME Davis, Pascal Michon, Ivo Mueller, Síle F Molloy, Susana Campino, Angeliki Kerasidou, Victoria J Cornelius, Lee Hart, Shivang S Shah, Gavin Band, Chris CA Spencer, Tsiri Agbenyega, Eric Achidi, Ogobara K Doumbo, Jeremy Farrar, Kevin Marsh, Terrie Taylor, and Dominic P Kwiatkowski. Characterisation of the opposing effects of G6PD deficiency on cerebral malaria and severe malarial anaemia. *eLife*, 6, 2017.

[2811] Sara J Clasen, Wei Shao, He Gu, and Peter J Espenshade. Prolyl dihydroxylation of unassembled uS12/Rps23 regulates fungal hypoxic adaptation. *eLife*, 6, 2017.

[2812] Clémence Claussin, David Porubský, Diana CJ Spierings, Nancy Halsema, Stefan Rentas, Victor Guryev, Peter M Lansdorp, and Michael Chang. Genome-wide mapping of sister chromatid exchange events in single yeast cells using Strand-seq. *eLife*, 6, 2017.

[2813] Marcello Clerici, Marco Faini, Ruedi Aebersold, and Martin Jinek. Structural insights into the assembly and polyA signal recognition mechanism of the human CPSF complex. *eLife*, 6, 2017.

[2814] Davide M Coda, Tessa Gaarenstroom, Philip East, Harshil Patel, Daniel S J Miller, Anna Lobley, Nik Matthews, Aengus Stewart, and Caroline S Hill. Distinct modes of SMAD2 chromatin binding and remodeling shape the transcriptional response to NODAL/Activin signaling. *eLife*, 6, 2017.

[2815] Jeremy D Cohen, Mark Bolstad, and Albert K Lee. Experience-dependent shaping of hippocampal CA1 intracellular activity in novel and familiar environments. *eLife*, 6, 2017.

[2816] Bárbara Coimbra, Carina Soares-Cunha, Sónia Borges, Nivaldo AP Vasconcelos, Nuno Sousa, and Ana João Rodrigues. Impairments in laterodorsal tegmentum to VTA projections underlie glucocorticoid-triggered reward deficits. *eLife*, 6, 2017.

[2817] Giles L Colclough, Stephen M Smith, Thomas E Nichols, Anderson M Winkler, Stamatios N Sotiropoulos, Matthew F Glasser, David C Van Essen, and Mark W Woolrich. The heritability of multi-modal connectivity in human brain activity. *eLife*, 6, 2017.

[2818] Kate E Coleman, Miklós Békés, Jessica R Chapman, Sarah B Crist, Mathew JK Jones, Beatrix M Ueberheide, and Tony T Huang. SENP8 limits aberrant neddylation of NEDD8 pathway components to promote cullin-RING ubiquitin ligase function. *eLife*, 6, 2017.

[2819] Sebastien Colin, Luis Pedro Coelho, Shinichi Sunagawa, Chris Bowler, Eric Karsenti, Peer Bork, Rainer Pepperkok, and Colomban de Vargas. Quantitative 3D-imaging for cell biology and ecology of environmental microbial eukaryotes. *eLife*, 6, 2017.

[2820] Sven Collette, Wolfgang M Pauli, Peter Bossaerts, and John O'Doherty. Neural computations underlying inverse reinforcement learning in the human brain. *eLife*, 6, 2017.

[2821] Mauricio Comas-Garcia, Siddhartha AK Datta, Laura Baker, Rajat Varma, Prabhakar R Gudla, and Alan Rein. Dissection of specific binding of HIV-1 Gag to the 'packaging signal' in viral RNA. *eLife*, 6, 2017.

[2822] Heather S Comstra, Jacob McArthy, Samantha Rudin-Rush, Cortnie Hartwig, Avanti Gokhale, Stephanie A Zlatic, Jessica B Blackburn, Erica Werner, Michael Petris, Priya D'Souza, Parinya Panuwet, Dana Boyd Barr, Vladimir Lupashin, Alysia Vrailas-Mortimer, and Victor Faundez. The interactome of the copper transporter ATP7A belongs to a network of neurodevelopmental and neurodegeneration factors. *eLife*, 6, 2017.

[2823] Lin Cong, Zeguan Wang, Yuming Chai, Wei Hang, Chunfeng Shang, Wenbin Yang, Lu Bai, Jiulin Du, Kai Wang, and Quan Wen. Rapid whole brain imaging of neural activity in freely behaving larval zebrafish (Danio rerio). *eLife*, 6, 2017.

[2824] Marisa Connell, Helen Chen, Jihong Jiang, Chia-Wei Kuan, Abbas Fotovati, Tony LH Chu, Zhengcheng He, Tess C Lengyell, Huaibiao Li, Torsten Kroll, Amanda M Li, Daniel Goldowitz, Lucien Frappart, Aspasia Ploubidou, Millan S Patel, Linda M Pilarski, Elizabeth M Simpson, Philipp F Lange, Douglas W Allan, and Christopher A Maxwell. HMMR acts in the PLK1-dependent spindle positioning pathway and supports neural development. *eLife*, 6, 2017.

[2825] Anneli Cooper, Hamidou Ilboudo, V Pius Alibu, Sophie Ravel, John Enyaru, William Weir, Harry Noyes, Paul Capewell, Mamadou Camara, Jacqueline Milet, Vincent Jamonneau, Oumou Camara, Enock Matovu, Bruno Bucheton, and Annette MacLeod. APOL1 renal risk variants have contrasting resistance and susceptibility associations with African trypanosomiasis. *eLife*, 6, 2017.

[2826] Robert M Cooper, Lev Tsimring, and Jeff Hasty. Inter-species population dynamics enhance microbial horizontal gene transfer and spread of antibiotic resistance. *eLife*, 6, 2017.

[2827] Patrícia A Correia, Eran Lottem, Dhruba Banerjee, Ana S Machado, Megan R Carey, and Zachary F Mainen. Transient inhibition and long-term facilitation of locomotion by phasic optogenetic activation of serotonin neurons. *eLife*, 6, 2017.

[2828] Andrea Corsinotti, Frederick CK Wong, Tülin Tatar, Iwona Szczerbinska, Florian Halbritter, Douglas Colby, Sabine Gogolok, Raphaël Pantier, Kirsten Liggat, Elham S Mirfazeli, Elisa Hall-Ponsele, Nicholas P Mullin, Valerie Wilson, and Ian Chambers. Distinct SoxB1 networks are required for naïve and primed pluripotency. *eLife*, 6, 2017.

[2829] Francis Corson and Eric D Siggia. Gene-free methodology for cell fate dynamics during development. *eLife*, 6, 2017.

[2830] Natalie A Counihan, Scott A Chisholm, Hayley E Bullen, Anubhav Srivastava, Paul R Sanders, Thorey K Jonsdottir, Greta E Weiss, Sreejoyee Ghosh, Brendan S Crabb, Darren J Creek, Paul R Gilson, and Tania F de Koning-Ward. Plasmodium falciparum parasites deploy RhopH2 into the host erythrocyte to obtain nutrients, grow and replicate. *eLife*, 6, 2017.

[2831] Andrew S Cowburn, David Macias, Charlotte Summers, Edwin R Chilvers, and Randall S Johnson. Cardiovascular adaptation to hypoxia and the role of peripheral resistance. *eLife*, 6, 2017.

[2832] Ana Crespillo-Casado, Joseph E Chambers, Peter M Fischer, Stefan J Marciniak, and David Ron. PPP1R15A-mediated dephosphorylation of eIF2$\alpha$ is unaffected by Sephin1 or Guanabenz. *eLife*, 6, 2017.

[2833] Justin Crest, Alba Diz-Muñoz, Dong-Yuan Chen, Daniel A Fletcher, and David Bilder. Organ sculpting by patterned extracellular matrix stiffness. *eLife*, 6, 2017.

[2834] Frédéric Crevecoeur and Konrad P Kording. Saccadic suppression as a perceptual consequence of efficient sensorimotor estimation. *eLife*, 6, 2017.

[2835] Kristina Crona, Alex Gavryushkin, Devin Greene, and Niko Beerenwinkel. Inferring genetic interactions from comparative fitness data. *eLife*, 6, 2017.

[2836] Dorottya Cserpán, Domokos Meszéna, Lucia Wittner, Kinga Tóth, István Ulbert, Zoltán Somogyvári, and Daniel K Wójcik. Revealing the distribution of transmembrane currents along the dendritic tree of a neuron from extracellular recordings. *eLife*, 6, 2017.

[2837] Luis G Cuello, D Marien Cortes, and Eduardo Perozo. The gating cycle of a K+ channel at atomic resolution. *eLife*, 6, 2017.

[2838] Tie-Zhong Cui, Tabitha A Peterson, and Christopher G Burd. A CDC25 family protein phosphatase gates cargo recognition by the Vps26 retromer subunit. *eLife*, 6, 2017.

[2839] Christopher L Cunningham, Zizhen Wu, Aria Jafari, Bo Zhao, Kat Schrode, Sarah Harkins-Perry, Amanda Lauer, and Ulrich Müller. The murine catecholamine methyltransferase mTOMT is essential for mechanotransduction by cochlear hair cells. *eLife*, 6, 2017.

[2840] Gucan Dai and William N Zagotta. Molecular mechanism of voltage-dependent potentiation of KCNH potassium channels. *eLife*, 6, 2017.

[2841] Ning Dai, Fei Ji, Jason Wright, Liliana Minichiello, Ruslan Sadreyev, and Joseph Avruch. IGF2 mRNA binding protein-2 is a tumor promoter that drives cancer proliferation through its client mRNAs IGF2 and HMGA1. *eLife*, 6, 2017.

[2842] Ursula Dalrymple, Ewan Cameron, Samir Bhatt, Daniel J Weiss, Sunetra Gupta, and Peter W Gething. Quantifying the contribution of Plasmodium falciparum malaria to febrile illness amongst African children. *eLife*, 6, 2017.

[2843] Lola Danet, Jérémie Pariente, Pierre Eustache, Nicolas Raposo, Igor Sibon, Jean-FranÇois Albucher, Fabrice Bonneville, Patrice Péran, and Emmanuel J Barbeau. Medial thalamic stroke and its impact on familiarity and recollection. *eLife*, 6, 2017.

[2844] Radostin Danev, Dimitry Tegunov, and Wolfgang Baumeister. Using the Volta phase plate with defocus for cryo-EM single particle analysis. *eLife*, 6, 2017.

[2845] James A Daniel, Benjamin H Cooper, Jorma J Palvimo, Fu-Ping Zhang, Nils Brose, and Marilyn Tirard. Analysis of SUMO1-conjugation at synapses. *eLife*, 6, 2017.

[2846] Simon M Danner, Natalia A Shevtsova, Alain Frigon, and Ilya A Rybak. Computational modeling of spinal circuits controlling limb coordination and gaits in quadrupeds. *eLife*, 6, 2017.

[2847] Benoit Daubech, Philippe Remigi, Ginaini Doin de Moura, Marta Marchetti, Cécile Pouzet, Marie-Christine Auriac, Chaitanya S Gokhale, Catherine Masson-Boivin, and Delphine Capela. Spatio-temporal control of mutualism in legumes helps spread symbiotic nitrogen fixation. *eLife*, 6, 2017.

[2848] Janine M Daum, özkan Keles, Sjoerd JB Holwerda, Hubertus Kohler, Filippo M Rijli, Michael Stadler, and Botond Roska. The formation of the light-sensing compartment of cone photoreceptors coincides with a transcriptional switch. *eLife*, 6, 2017.

[2849] Kashyap Dave, Inderpreet Sur, Jian Yan, Jilin Zhang, Eevi Kaasinen, Fan Zhong, Leander Blaas, Xiaoze Li, Shabnam Kharazi, Charlotte Gustafsson, Ayla De Paepe, Robert Månsson, and Jussi Taipale. Mice deficient of Myc super-enhancer region reveal differential control mechanism between normal and pathological growth. *eLife*, 6, 2017.

[2850] Erin L Davies, Kai Lei, Christopher W Seidel, Amanda E Kroesen, Sean A McKinney, Longhua Guo, Sofia MC Robb, Eric J Ross, Kirsten Gotting, and Alejandro Sánchez Alvarado. Embryonic origin of adult stem cells required for tissue homeostasis and regeneration. *eLife*, 6, 2017.

[2851] Valentina De Col, Philippe Fuchs, Thomas Nietzel, Marlene Elsässer, Chia Pao Voon, Alessia Candeo, Ingo Seeliger, Mark D Fricker, Christopher Grefen, Ian Max Møller, Andrea Bassi, Boon Leong Lim, Marco Zancani, Andreas J Meyer, Alex Costa, Stephan Wagner, and Markus Schwarzländer. ATP sensing in living plant cells reveals tissue gradients and stress dynamics of energy physiology. *eLife*, 6, 2017.

[2852] Jan Willem de Gee, Olympia Colizoli, Niels A Kloosterman, Tomas Knapen, Sander Nieuwenhuis, and Tobias H Donner. Dynamic modulation of decision biases by brainstem arousal systems. *eLife*, 6, 2017.

[2853] Lylah D Deady, Wei Li, and Jianjun Sun. The zinc-finger transcription factor Hindsight regulates ovulation competency of Drosophila follicles. *eLife*, 6, 2017.

[2854] Altaira D Dearborn, Erin A Wall, James L Kizziah, Laura Klenow, Laura K Parker, Keith A Manning, Michael S Spilman, John M Spear, Gail E Christie, and Terje Dokland. Competing scaffolding proteins determine capsid size during mobilization of Staphylococcus aureus pathogenicity islands. *eLife*, 6, 2017.

[2855] Sarah N Deffit, Brian A Yee, Aidan C Manning, Suba Rajendren, Pranathi Vadlamani, Emily C Wheeler, Alain Domissy, Michael C Washburn, Gene W Yeo, and Heather A Hundley. The C. elegans neural editome reveals an ADAR target mRNA required for proper chemotaxis. *eLife*, 6, 2017.

[2856] M Joaquina Delás, Leah R Sabin, Egor Dolzhenko, Simon RV Knott, Ester Munera Maravilla, Benjamin T Jackson, Sophia A Wild, Tatjana Kovacevic, Eva Maria Stork, Meng Zhou, Nicolas Erard, Emily Lee, David R Kelley, Mareike Roth, Inês AM Barbosa, Johannes Zuber, John L Rinn, Andrew D Smith, and Gregory J Hannon. lncRNA requirements for mouse acute myeloid leukemia and normal differentiation. *eLife*, 6, 2017.

[2857] Sandeep Dembla, Marc Behrendt, Florian Mohr, Christian Goecke, Julia Sondermann, Franziska M Schneider, Marlene Schmidt, Julia Stab, Raissa Enzeroth, Michael G Leitner, Paulina Nuñez-Badinez, Jochen Schwenk, Bernd Nürnberg, Alejandro Cohen, Stephan E Philipp, Wolfgang Greffrath, Moritz Bünemann, Dominik Oliver, Eleonora Zakharian, Manuela Schmidt, and Johannes Oberwinkler. Anti-nociceptive action of peripheral mu-opioid receptors by G-beta-gamma protein-mediated inhibition of TRPM3 channels. *eLife*, 6, 2017.

[2858] Keren Demishtein-Zohary, Umut Günsel, Milit Marom, Rupa Banerjee, Walter Neupert, Abdussalam Azem, and Dejana Mokranjac. Role of Tim17 in coupling the import motor to the translocation channel of the mitochondrial presequence translocase. *eLife*, 6, 2017.

[2859] Gabriel Demo, Aviram Rasouly, Nikita Vasilyev, Vladimir Svetlov, Anna B Loveland, Ruben Diaz-Avalos, Nikolaus Grigorieff, Evgeny Nudler, and Andrei A Korostelev. Structure of RNA polymerase bound to ribosomal 30S subunit. *eLife*, 6, 2017.

[2860] Gabriel Demo, Egor Svidritskiy, Rohini Madireddy, Ruben Diaz-Avalos, Timothy Grant, Nikolaus Grigorieff, Duncan Sousa, and Andrei A Korostelev. Mechanism of ribosome rescue by ArfA and RF2. *eLife*, 6, 2017.

[2861] Niels Denkert, Alexander Benjamin Schendzielorz, Mariam Barbot, Lennart Versemann, Frank Richter, Peter Rehling, and Michael Meinecke. Cation selectivity of the presequence translocase channel Tim23 is crucial for efficient protein import. *eLife*, 6, 2017.

[2862] An-Sofie Desmet, Carla Cirillo, Jan Tack, Wim Vandenberghe, and Pieter Vanden Berghe. Live calcium and mitochondrial imaging in the enteric nervous system of Parkinson patients and controls. *eLife*, 6, 2017.

[2863] Alison M Devault, Tatum D Mortimer, Andrew Kitchen, Henrike Kiesewetter, Jacob M Enk, G Brian Golding, John Southon, Melanie Kuch, Ana T Duggan, William Aylward, Shea N Gardner, Jonathan E Allen, Andrew M King, Gerard Wright, Makoto Kuroda, Kengo Kato, Derek EG Briggs, Gino Fornaciari, Edward C Holmes, Hendrik N Poinar, and Caitlin S Pepperell. A molecular portrait of maternal sepsis from Byzantine Troy. *eLife*, 6, 2017.

[2864] Ashesh K Dhawale, Rajesh Poddar, Steffen BE Wolff, Valentin A Normand, Evi Kopelowitz, and Bence P ölveczky. Automated long-term recording and analysis of neural activity in behaving animals. *eLife*, 6, 2017.

[2865] Giovanni Diana, Dhaval S Patel, Eugeni V Entchev, Mei Zhan, Hang Lu, and QueeLim Ch'ng. Genetic control of encoding strategy in a food-sensing neural circuit. *eLife*, 6, 2017.

[2866] Feici Diao, Amicia D Elliott, Fengqiu Diao, Sarav Shah, and Benjamin H White. Neuromodulatory connectivity defines the structure of a behavioral neural network. *eLife*, 6, 2017.

[2867] Luis Miguel Díaz-Santín, Natasha Lukoyanova, Emir Aciyan, and Alan CM Cheung. Cryo-EM structure of the SAGA and NuA4 coactivator subunit Tra1 at 3.7 angstrom resolution. *eLife*, 6, 2017.

[2868] Shanaz Diessler, Corinne Kostic, Yvan Arsenijevic, Aki Kawasaki, and Paul Franken. Rai1 frees mice from the repression of active wake behaviors by light. *eLife*, 6, 2017.

[2869] Edoardo D'Imprima, Ralf Salzer, Ramachandra M Bhaskara, Ricardo Sánchez, Ilona Rose, Lennart Kirchner, Gerhard Hummer, Werner Kühlbrandt, Janet Vonck, and Beate Averhoff. Cryo-EM structure of the bifunctional secretin complex of Thermus thermophilus. *eLife*, 6, 2017.

[2870] Paul HGM Dirks, Eric M Roberts, Hannah Hilbert-Wolf, Jan D Kramers, John Hawks, Anthony Dosseto, Mathieu Duval, Marina Elliott, Mary Evans, Rainer Grün, John Hellstrom, Andy IR Herries, Renaud Joannes-Boyau, Tebogo V Makhubela, Christa J Placzek, Jessie Robbins, Carl Spandler, Jelle Wiersma, Jon Woodhead, and Lee R Berger. The age of Homo naledi and associated sediments in the Rising Star Cave, South Africa. *eLife*, 6, 2017.

[2871] Marina Diskowski, Ahmad Reza Mehdipour, Dorith Wunnicke, Deryck J Mills, Vedrana Mikusevic, Natalie Bärland, Jan Hoffmann, Nina Morgner, Heinz-Jürgen Steinhoff, Gerhard Hummer, Janet Vonck, and Inga Hänelt. Helical jackknives control the gates of the double-pore K+ uptake system KtrAB. *eLife*, 6, 2017.

[2872] Svetlana O Dodonova, Patrick Aderhold, Juergen Kopp, Iva Ganeva, Simone Röhling, Wim J H Hagen, Irmgard Sinning, Felix Wieland, and John A G Briggs. 9Å structure of the COPI coat reveals that the Arf1 GTPase occupies two contrasting molecular environments. *eLife*, 6, 2017.

[2873] Michael Doebeli, Yaroslav Ispolatov, and Burt Simon. Towards a mechanistic foundation of evolutionary theory. *eLife*, 6, 2017.

[2874] Sunil Kumar Dogga, Budhaditya Mukherjee, Damien Jacot, Tobias Kockmann, Luca Molino, Pierre-Mehdi Hammoudi, Ruben C Hartkoorn, Adrian B Hehl, and Dominique Soldati-Favre. A druggable secretory protein maturase of Toxoplasma essential for invasion and egress. *eLife*, 6, 2017.

[2875] Lan-Feng Dong, Jaromira Kovarova, Martina Bajzikova, Ayenachew Bezawork-Geleta, David Svec, Berwini Endaya, Karishma Sachaphibulkij, Ana R Coelho, Natasa Sebkova, Anna Ruzickova, An S Tan, Katarina Kluckova, Kristyna Judasova, Katerina Zamecnikova, Zuzana Rychtarcikova, Vinod Gopalan, Ladislav Andera, Margarita Sobol, Bing Yan, Bijay Pattnaik, Naveen Bhatraju, Jaroslav Truksa, Pavel Stopka, Pavel Hozak, Alfred K Lam, Radislav Sedlacek, Paulo J Oliveira, Mikael Kubista, Anurag Agrawal, Katerina Dvorakova-Hortova, Jakub Rohlena, Michael V Berridge, and Jiri Neuzil. Horizontal transfer of whole mitochondria restores tumorigenic potential in mitochondrial DNA-deficient cancer cells. *eLife*, 6, 2017.

[2876] Tobias X Dong, Shivashankar Othy, Milton L Greenberg, Amit Jairaman, Chijioke Akunwafo, Sabrina Leverrier, Ying Yu, Ian Parker, Joseph L Dynes, and Michael D Cahalan. Intermittent Ca2+ signals mediated by Orai1 regulate basal T cell motility. *eLife*, 6, 2017.

[2877] Tobias X Dong, Shivashankar Othy, Amit Jairaman, Jonathan Skupsky, Angel Zavala, Ian Parker, Joseph L Dynes, and Michael D Cahalan. T-cell calcium dynamics visualized in a ratiometric tdTomato-GCaMP6f transgenic reporter mouse. *eLife*, 6, 2017.

[2878] Richard G Dorrell, Gillian Gile, Giselle McCallum, Raphaël Méheust, Eric P Bapteste, Christen M Klinger, Loraine Brillet-Guéguen, Katalina D Freeman, Daniel J Richter, and Chris Bowler. Chimeric origins of ochrophytes and haptophytes revealed through an ancient plastid proteome. *eLife*, 6, 2017.

[2879] Frédéric Doussau, Hartmut Schmidt, Kevin Dorgans, Antoine M Valera, Bernard Poulain, and Philippe Isope. Frequency-dependent mobilization of heterogeneous pools of synaptic vesicles shapes presynaptic plasticity. *eLife*, 6, 2017.

[2880] Catherine M Drerup, Amy L Herbert, Kelly R Monk, and Alex V Nechiporuk. Regulation of mitochondria-dynactin interaction and mitochondrial retrograde transport in axons. *eLife*, 6, 2017.

[2881] Jonathan W Driver, Elisabeth A Geyer, Megan E Bailey, Luke M Rice, and Charles L Asbury. Direct measurement of conformational strain energy in protofilaments curling outward from disassembling microtubule tips. *eLife*, 6, 2017.

[2882] Jianyang Du, Margaret P Price, Rebecca J Taugher, Daniel Grigsby, Jamison J Ash, Austin C Stark, Md Zubayer Hossain Saad, Kritika Singh, Juthika Mandal, John A Wemmie, and Michael J Welsh. Transient acidosis while retrieving a fear-related memory enhances its lability. *eLife*, 6, 2017.

[2883] Guillaume P Dugué, Matthieu Tihy, Boris Gourévitch, and Clément Léna. Cerebellar re-encoding of self-generated head movements. *eLife*, 6, 2017.

[2884] Jason G Dumelie and Samie R Jaffrey. Defining the location of promoter-associated R-loops at near-nucleotide resolution using bisDRIP-seq. *eLife*, 6, 2017.

[2885] Ana Maria G Dumitru, Scott F Rusin, Amber E M Clark, Arminja N Kettenbach, and Duane A Compton. Cyclin A/Cdk1 modulates Plk1 activity in prometaphase to regulate kinetochore-microtubule attachment stability. *eLife*, 6, 2017.

[2886] Rémi Dumollard, Nicolas Minc, Gregory Salez, Sameh Ben Aicha, Faisal Bekkouche, Céline Hebras, Lydia Besnardeau, and Alex McDougall. The invariant cleavage pattern displayed by ascidian embryos depends on spindle positioning along the cell's longest axis in the apical plane and relies on asynchronous cell divisions. *eLife*, 6, 2017.

[2887] David Duneau, Jean-Baptiste Ferdy, Jonathan Revah, Hannah Kondolf, Gerardo A Ortiz, Brian P Lazzaro, and Nicolas Buchon. Stochastic variation in the initial phase of bacterial infection predicts the probability of survival in D. melanogaster. *eLife*, 6, 2017.

[2888] Sylvain Durand, Frédérique Braun, Anne-Catherine Helfer, Pascale Romby, and Ciarán Condon. sRNA-mediated activation of gene expression by inhibition of $5'$-$3'$ exonucleolytic mRNA degradation. *eLife*, 6, 2017.

[2889] Joanne Durgan, Yun-Yu Tseng, Jens C Hamann, Marie-Charlotte Domart, Lucy Collinson, Alan Hall, Michael Overholtzer, and Oliver Florey. Mitosis can drive cell cannibalism through entosis. *eLife*, 6, 2017.

[2890] Yann Duroc, Rajeev Kumar, Lepakshi Ranjha, Céline Adam, Raphaël Guérois, Khan Md Muntaz, Marie-Claude Marsolier-Kergoat, Florent Dingli, Raphaëlle Laureau, Damarys Loew, Bertrand Llorente, Jean-Baptiste Charbonnier, Petr Cejka, and Valérie Borde. Concerted action of the MutL$\beta$ heterodimer and Mer3 helicase regulates the global extent of meiotic gene conversion. *eLife*, 6, 2017.

[2891] Sylvia Dyballa, Thierry Savy, Philipp Germann, Karol Mikula, Mariana Remesikova, Róbert Špir, Andrea Zecca, Nadine Peyriéras, and Cristina Pujades. Distribution of neurosensory progenitor pools during inner ear morphogenesis unveiled by cell lineage reconstruction. *eLife*, 6, 2017.

[2892] Brittany M Edens, Jianhua Yan, Nimrod Miller, Han-Xiang Deng, Teepu Siddique, and Yongchao C Ma. A novel ALS-associated variant in UBQLN4 regulates motor axon morphogenesis. *eLife*, 6, 2017.

[2893] Benedikt V Ehinger, Katja Häusser, José P Ossandón, and Peter König. Humans treat unreliable filled-in percepts as more real than veridical ones. *eLife*, 6, 2017.

[2894] J Christopher Ehlen, Allison J Brager, Julie Baggs, Lennisha Pinckney, Cloe L Gray, Jason P DeBruyne, Karyn A Esser, Joseph S Takahashi, and Ketema N Paul. Bmal1 function in skeletal muscle regulates sleep. *eLife*, 6, 2017.

[2895] Kevin M Elias, Wojciech Fendler, Konrad Stawiski, Stephen J Fiascone, Allison F Vitonis, Ross S Berkowitz, Gyorgy Frendl, Panagiotis Konstantinopoulos, Christopher P Crum, Magdalena Kedzierska, Daniel W Cramer, and Dipanjan Chowdhury. Diagnostic potential for a serum miRNA neural network for detection of ovarian cancer. *eLife*, 6, 2017.

[2896] Daniel Elnatan, Miguel Betegon, Yanxin Liu, Theresa Ramelot, Michael A Kennedy, and David A Agard. Symmetry broken and rebroken during the ATP hydrolysis cycle of the mitochondrial Hsp90 TRAP1. *eLife*, 6, 2017.

[2897] Cherilyn A Elwell, Nadine Czudnochowski, John von Dollen, Jeffrey R Johnson, Rachel Nakagawa, Kathleen Mirrashidi, Nevan J Krogan, Joanne N Engel, and Oren S Rosenberg. Chlamydia interfere with an interaction between the mannose-6-phosphate receptor and sorting nexins to counteract host restriction. *eLife*, 6, 2017.

[2898] Elaine Emmerson, Alison J May, Sara Nathan, Noel Cruz-Pacheco, Carlos O Lizama, Lenka Maliskova, Ann C Zovein, Yin Shen, Marcus O Muench, and Sarah M Knox. SOX2 regulates acinar cell development in the salivary gland. *eLife*, 6, 2017.

[2899] Mitsuhiro Endoh, Takaho A Endo, Jun Shinga, Katsuhiko Hayashi, Anca Farcas, Kit-Wan Ma, Shinsuke Ito, Jafar Sharif, Tamie Endoh, Naoko Onaga, Manabu Nakayama, Tomoyuki Ishikura, Osamu Masui, Benedikt M Kessler, Toshio Suda, Osamu Ohara, Akihiko Okuda, Robert Klose, and Haruhiko Koseki. PCGF6-PRC1 suppresses premature differentiation of mouse embryonic stem cells by regulating germ cell-related genes. *eLife*, 6, 2017.

[2900] Timothy Erickson, Clive P Morgan, Jennifer Olt, Katherine Hardy, Elisabeth Busch-Nentwich, Reo Maeda, Rachel Clemens, Jocelyn F Krey, Alex Nechiporuk, Peter G Barr-Gillespie, Walter Marcotti, and Teresa Nicolson. Integration of Tmc1/2 into the mechanotransduction complex in zebrafish hair cells is regulated by Transmembrane O-methyltransferase (Tomt). *eLife*, 6, 2017.

[2901] Simon Erlendsson, Kamil Gotfryd, Flemming Hofmann Larsen, Jonas Sigurd Mortensen, Michel-Andreas Geiger, Barth-Jan van Rossum, Hartmut Oschkinat, Ulrik Gether, Kaare Teilum, and Claus J Loland. Direct assessment of substrate binding to the Neurotransmitter:Sodium Symporter LeuT by solid state NMR. *eLife*, 6, 2017.

[2902] Michelle V Evans, Tad A Dallas, Barbara A Han, Courtney C Murdock, and John M Drake. Data-driven identification of potential Zika virus vectors. *eLife*, 6, 2017.

[2903] Collin Yvès Ewald, John M Hourihan, Monet S Bland, Carolin Obieglo, Iskra Katic, Lorenza E Moronetti Mazzeo, Joy Alcedo, T Keith Blackwell, and Nancy E Hynes. NADPH oxidase-mediated redox signaling promotes oxidative stress resistance and longevity through memo-1 in C. elegans. *eLife*, 6, 2017.

[2904] Melanie Falgairolle, Joshua G Puhl, Avinash Pujala, Wenfang Liu, and Michael J O'Donovan. Motoneurons regulate the central pattern generator during drug-induced locomotor-like activity in the neonatal mouse. *eLife*, 6, 2017.

[2905] Fushun Fan, Kohichi Matsunaga, Hao Wang, Ray Ishizaki, Eri Kobayashi, Hiroshi Kiyonari, Yoshiko Mukumoto, Katsuhide Okunishi, and Tetsuro Izumi. Exophilin-8 assembles secretory granules for exocytosis in the actin cortex via interaction with RIM-BP2 and myosin-VIIa. *eLife*, 6, 2017.

[2906] Yichao Fan, Jiao Yue, Mengtao Xiao, Han Han-Zhang, Yao Vickie Wang, Chun Ma, Zhilin Deng, Yingxiang Li, Yanyan Yu, Xinghao Wang, Shen Niu, Youjia Hua, Zhiping Weng, Peter Atadja, En Li, and Bin Xiang. FXR1 regulates transcription and is required for growth of human cancer cells with TP53/FXR2 homozygous deletion. *eLife*, 6, 2017.

[2907] Hanwei Fang, Natacha Klages, Bastien Baechler, Evelyn Hillner, Lu Yu, Mercedes Pardo, Jyoti Choudhary, and Mathieu Brochet. Multiple short windows of calcium-dependent protein kinase 4 activity coordinate distinct cell cycle events during Plasmodium gametogenesis. *eLife*, 6, 2017.

[2908] Olivia K Faull and Kyle TS Pattinson. The cortical connectivity of the periaqueductal gray and the conditioned response to the threat of breathlessness. *eLife*, 6, 2017.

[2909] Paola Favuzza, Elena Guffart, Marco Tamborrini, Bianca Scherer, Anita M Dreyer, Arne C Rufer, Johannes Erny, Joerg Hoernschemeyer, Ralf Thoma, Georg Schmid, Bernard Gsell, Araceli Lamelas, Joerg Benz, Catherine Joseph, Hugues Matile, Gerd Pluschke, and Markus G Rudolph. Structure of the malaria vaccine candidate antigen CyRPA and its complex with a parasite invasion inhibitory antibody. *eLife*, 6, 2017.

[2910] Robin Fears and Volker ter Meulen. How should the applications of genome editing be assessed and regulated? *eLife*, 6, 2017.

[2911] Chungang Feng, Mats Pettersson, Sangeet Lamichhaney, Carl-Johan Rubin, Nima Rafati, Michele Casini, Arild Folkvord, and Leif Andersson. Moderate nucleotide diversity in the Atlantic herring is associated with a low mutation rate. *eLife*, 6, 2017.

[2912] Shiran Ferber, Galia Tiram, Ana Sousa-Herves, Anat Eldar-Boock, Adva Krivitsky, Anna Scomparin, Eilam Yeini, Paula Ofek, Dikla Ben-Shushan, Laura Isabel Vossen, Kai Licha, Rachel Grossman, Zvi Ram, Jack Henkin, Eytan Ruppin, Noam Auslander, Rainer Haag, Marcelo Calderón, and Ronit Satchi-Fainaro. Co-targeting the tumor endothelium and P-selectin-expressing glioblastoma cells leads to a remarkable therapeutic outcome. *eLife*, 6, 2017.

[2913] Joana S Ferreira, Thomas Papouin, Laurent Ladépêche, Andrea Yao, Valentin C Langlais, Delphine Bouchet, Jérôme Dulong, Jean-Pierre Mothet, Silvia Sacchi, Loredano Pollegioni, Pierre Paoletti, Stéphane Henri Richard Oliet, and Laurent Groc. Co-agonists differentially tune GluN2B-NMDA receptor trafficking at hippocampal synapses. *eLife*, 6, 2017.

[2914] Rita R Ferreira, Andrej Vilfan, Frank Jülicher, Willy Supatto, and Julien Vermot. Physical limits of flow sensing in the left-right organizer. *eLife*, 6, 2017.

[2915] Michael A Ferrin and Arvind R Subramaniam. Kinetic modeling predicts a stimulatory role for ribosome collisions at elongation stall sites in bacteria. *eLife*, 6, 2017.

[2916] Giulia Fianco, Maria Patrizia Mongiardi, Andrea Levi, Teresa De Luca, Marianna Desideri, Daniela Trisciuoglio, Donatella Del Bufalo, Irene Cinà, Anna Di Benedetto, Marcella Mottolese, Antonietta Gentile, Diego Centonze, Fabrizio Ferrè, and Daniela Barilà. Caspase-8 contributes to angiogenesis and chemotherapy resistance in glioblastoma. *eLife*, 6, 2017.

[2917] Aaron L Fidler, Carl E Darris, Sergei V Chetyrkin, Vadim K Pedchenko, Sergei P Boudko, Kyle L Brown, W Gray Jerome, Julie K Hudson, Antonis Rokas, and Billy G Hudson. Collagen IV and basement membrane at the evolutionary dawn of metazoan tissues. *eLife*, 6, 2017.

[2918] Felix Fiederling, Markus Weschenfelder, Martin Fritz, Anne von Philipsborn, Martin Bastmeyer, and Franco Weth. Ephrin-A/EphA specific co-adaptation as a novel mechanism in topographic axon guidance. *eLife*, 6, 2017.

[2919] Ceri A Fielding, Michael P Weekes, Luis V Nobre, Eva Ruckova, Gavin S Wilkie, Joao A Paulo, Chiwen Chang, Nicolás M Suárez, James A Davies, Robin Antrobus, Richard J Stanton, Rebecca J Aicheler, Hester Nichols, Borek Vojtesek, John Trowsdale, Andrew J Davison, Steven P Gygi, Peter Tomasec, Paul J Lehner, and Gavin W G Wilkinson. Control of immune ligands by members of a cytomegalovirus gene expansion suppresses natural killer cell activation. *eLife*, 6, 2017.

[2920] Kathryn H Fife, Navarre A Gutierrez-Reed, Vivien Zell, Julie Bailly, Christina M Lewis, Adam R Aron, and Thomas S Hnasko. Causal role for the subthalamic nucleus in interrupting behavior. *eLife*, 6, 2017.

[2921] Gianluca Figlia, Camilla Norrmén, Jorge A Pereira, Daniel Gerber, and Ueli Suter. Dual function of the PI3K-Akt-mTORC1 axis in myelination of the peripheral nervous system. *eLife*, 6, 2017.

[2922] Joshua J Filter, Byron C Williams, Masumi Eto, David Shalloway, and Michael L Goldberg. Unfair competition governs the interaction of pCPI-17 with myosin phosphatase (PP1-MYPT1). *eLife*, 6, 2017.

[2923] Nicolas Fiorini, David J Lipman, and Zhiyong Lu. Towards PubMed 2.0. *eLife*, 6, 2017.

[2924] Annette Fischer, Kelly S Harrison, Yesid Ramirez, Daniela Auer, Suvagata Roy Chowdhury, Bhupesh K Prusty, Florian Sauer, Zoe Dimond, Caroline Kisker, P Scott Hefty, and Thomas Rudel. Chlamydia trachomatis-containing vacuole serves as deubiquitination platform to stabilize Mcl-1 and to interfere with host defense. *eLife*, 6, 2017.

[2925] Caleb N Fischer, Eric P Trautman, Jason M Crawford, Eric V Stabb, Jo Handelsman, and Nichole A Broderick. Metabolite exchange between microbiome members produces compounds that influence Drosophila behavior. *eLife*, 6, 2017.

[2926] Petra Fischer, Alek Pogosyan, Damian M Herz, Binith Cheeran, Alexander L Green, James Fitzgerald, Tipu Z Aziz, Jonathan Hyam, Simon Little, Thomas Foltynie, Patricia Limousin, Ludvic Zrinzo, Peter Brown, and Huiling Tan. Subthalamic nucleus gamma activity increases not only during movement but also during movement inhibition. *eLife*, 6, 2017.

[2927] Gemma LM Fisher, César L Pastrana, Victoria A Higman, Alan Koh, James A Taylor, Annika Butterer, Timothy Craggs, Frank Sobott, Heath Murray, Matthew P Crump, Fernando Moreno-Herrero, and Mark S Dillingham. The structural basis for dynamic DNA binding and bridging interactions which condense the bacterial centromere. *eLife*, 6, 2017.

[2928] Yvette E Fisher, Helen H Yang, Jesse Isaacman-Beck, Marjorie Xie, Daryl M Gohl, and Thomas R Clandinin. FlpStop, a tool for conditional gene control in Drosophila. *eLife*, 6, 2017.

[2929] Gabriel A Fitzgerald, Christopher Mulligan, and Joseph A Mindell. A general method for determining secondary active transporter substrate stoichiometry. *eLife*, 6, 2017.

[2930] Grant Parker Flowers, Lucas D Sanor, and Craig M Crews. Lineage tracing of genome-edited alleles reveals high fidelity axolotl limb regeneration. *eLife*, 6, 2017.

[2931] Jennifer L Fogel, Daniel L Lakeland, In Kyoung Mah, and Francesca V Mariani. A minimally sufficient model for rib proximal-distal patterning based on genetic analysis and agent-based simulations. *eLife*, 6, 2017.

[2932] Floris Foijer, Lee A Albacker, Bjorn Bakker, Diana C Spierings, Ying Yue, Stephanie Z Xie, Stephanie Davis, Annegret Lutum-Jehle, Darin Takemoto, Brian Hare, Brinley Furey, Roderick T Bronson, Peter M Lansdorp, Allan Bradley, and Peter K Sorger. Deletion of the MAD2L1 spindle assembly checkpoint gene is tolerated in mouse models of acute T-cell lymphoma and hepatocellular carcinoma. *eLife*, 6, 2017.

[2933] Jiunn CN Fong, Andrew Rogers, Alicia K Michael, Nicole C Parsley, William-Cole Cornell, Yu-Cheng Lin, Praveen K Singh, Raimo Hartmann, Knut Drescher, Evgeny Vinogradov, Lars EP Dietrich, Carrie L Partch, and Fitnat H Yildiz. Structural dynamics of RbmA governs plasticity of Vibrio cholerae biofilms. *eLife*, 6, 2017.

[2934] Joan Font, Marc López-Cano, Serena Notartomaso, Pamela Scarselli, Paola Di Pietro, Roger Bresolí-Obach, Giuseppe Battaglia, Fanny Malhaire, Xavier Rovira, Juanlo Catena, Jesús Giraldo, Jean-Philippe Pin, Víctor Fernández-Dueñas, Cyril Goudet, Santi Nonell, Ferdinando Nicoletti, Amadeu Llebaria, and Francisco Ciruela. Optical control of pain in vivo with a photoactive mGlu5 receptor negative allosteric modulator. *eLife*, 6, 2017.

[2935] Pietro Fontana, Juan José Bonfiglio, Luca Palazzo, Edward Bartlett, Ivan Matic, and Ivan Ahel. Serine ADP-ribosylation reversal by the hydrolase ARH3. *eLife*, 6, 2017.

[2936] David Forsberg, Thomas Ringstedt, and Eric Herlenius. Astrocytes release prostaglandin E2 to modify respiratory network activity. *eLife*, 6, 2017.

[2937] Antonio Elia Forte, Octave Etard, and Tobias Reichenbach. The human auditory brainstem response to running speech reveals a subcortical mechanism for selective attention. *eLife*, 6, 2017.

[2938] Baptiste Fouquet, Patrycja Pawlikowska, Sandrine Caburet, Celine Guigon, Marika Mäkinen, Laura Tanner, Marja Hietala, Kaja Urbanska, Laura Bellutti, Bérangère Legois, Bettina Bessieres, Alain Gougeon, Alexandra Benachi, Gabriel Livera, Filippo Rosselli, Reiner A Veitia, and Micheline Misrahi. A homozygous FANCM mutation underlies a familial case of non-syndromic primary ovarian insufficiency. *eLife*, 6, 2017.

[2939] David T Fraebel, Harry Mickalide, Diane Schnitkey, Jason Merritt, Thomas E Kuhlman, and Seppe Kuehn. Environment determines evolutionary trajectory in a constrained phenotypic space. *eLife*, 6, 2017.

[2940] Camila Tenorio FranÇa, Michael T White, Wen-Qiang He, Jessica B Hostetler, Jessica Brewster, Gabriel Frato, Indu Malhotra, Jakub Gruszczyk, Christele Huon, Enmoore Lin, Benson Kiniboro, Anjali Yadava, Peter Siba, Mary R Galinski, Julie Healer, Chetan Chitnis, Alan F Cowman, Eizo Takashima, Takafumi Tsuboi, Wai-Hong Tham, Rick M Fairhurst, Julian C Rayner, Christopher L King, and Ivo Mueller. Identification of highly-protective combinations of Plasmodium vivax recombinant proteins for vaccine development. *eLife*, 6, 2017.

[2941] Janusz Franco-Barraza, Ralph Francescone, Tiffany Luong, Neelima Shah, Raj Madhani, Gil Cukierman, Essel Dulaimi, Karthik Devarajan, Brian L Egleston, Emmanuelle Nicolas, R Katherine Alpaugh, Ruchi Malik, Robert G Uzzo, John P Hoffman, Erica A Golemis, and Edna Cukierman. Matrix-regulated integrin $\alpha v\beta 5$ maintains $\alpha 5\beta 1$-dependent desmoplastic traits prognostic of neoplastic recurrence. *eLife*, 6, 2017.

[2942] Paula Freire-Pritchett, Stefan Schoenfelder, Csilla Várnai, Steven W Wingett, Jonathan Cairns, Amanda J Collier, Raquel García-Vílchez, Mayra Furlan-Magaril, Cameron S Osborne, Peter Fraser,

Peter J Rugg-Gunn, and Mikhail Spivakov. Global reorganisation of cis-regulatory units upon lineage commitment of human embryonic stem cells. *eLife*, 6, 2017.

[2943] Rachel Fremont, Ambika Tewari, Chantal Angueyra, and Kamran Khodakhah. A role for cerebellum in the hereditary dystonia DYT1. *eLife*, 6, 2017.

[2944] Erez Freud, Jody C Culham, David C Plaut, and Marlene Behrmann. The large-scale organization of shape processing in the ventral and dorsal pathways. *eLife*, 6, 2017.

[2945] Lillian K Fritz-Laylin, Megan Riel-Mehan, Bi-Chang Chen, Samuel J Lord, Thomas D Goddard, Thomas E Ferrin, Susan M Nicholson-Dykstra, Henry Higgs, Graham T Johnson, Eric Betzig, and R Dyche Mullins. Actin-based protrusions of migrating neutrophils are intrinsically lamellar and facilitate direction changes. *eLife*, 6, 2017.

[2946] Jelke Jan Fros, Isabelle Dietrich, Kinda Alshaikhahmed, Tim Casper Passchier, David John Evans, and Peter Simmonds. CpG and UpA dinucleotides in both coding and non-coding regions of echovirus 7 inhibit replication initiation post-entry. *eLife*, 6, 2017.

[2947] Sean Froudist-Walsh, Michael AP Bloomfield, Mattia Veronese, Jasmin Kroll, Vyacheslav R Karolis, Sameer Jauhar, Ilaria Bonoldi, Philip K McGuire, Shitij Kapur, Robin M Murray, Chiara Nosarti, and Oliver Howes. The effect of perinatal brain injury on dopaminergic function and hippocampal volume in adult life. *eLife*, 6, 2017.

[2948] Guo Fu, Li-Chun Tu, Anton Zilman, and Siegfried M Musser. Investigating molecular crowding within nuclear pores using polarization-PALM. *eLife*, 6, 2017.

[2949] Naonobu Fujita, Wilson Huang, Tzu-han Lin, Jean-Francois Groulx, Steve Jean, Jen Nguyen, Yoshihiko Kuchitsu, Ikuko Koyama-Honda, Noboru Mizushima, Mitsunori Fukuda, and Amy A Kiger. Genetic screen in Drosophila muscle identifies autophagy-mediated T-tubule remodeling and a Rab2 role in autophagy. *eLife*, 6, 2017.

[2950] James R Fuller and Phoebe A Rice. Target DNA bending by the Mu transpososome promotes careful transposition and prevents its reversal. *eLife*, 6, 2017.

[2951] Ho Yee Joyce Fung, Szu-Chin Fu, and Yuh Min Chook. Nuclear export receptor CRM1 recognizes diverse conformations in nuclear export signals. *eLife*, 6, 2017.

[2952] Leon A Furchtgott, Samuel Melton, Vilas Menon, and Sharad Ramanathan. Discovering sparse transcription factor codes for cell states and state transitions during development. *eLife*, 6, 2017.

[2953] Amber M Fyall, Yasmine El-Shamayleh, Hannah Choi, Eric Shea-Brown, and Anitha Pasupathy. Dynamic representation of partially occluded objects in primate prefrontal and visual cortex. *eLife*, 6, 2017.

[2954] Christophe Gaillochet, Thomas Stiehl, Christian Wenzl, Juan-José Ripoll, Lindsay J Bailey-Steinitz, Lanxin Li, Anne Pfeiffer, Andrej Miotk, Jana P Hakenjos, Joachim Forner, Martin F Yanofsky, Anna Marciniak-Czochra, and Jan U Lohmann. Control of plant cell fate transitions by transcriptional and hormonal signals. *eLife*, 6, 2017.

[2955] Federico Gaiti, Katia Jindrich, Selene L Fernandez-Valverde, Kathrein E Roper, Bernard M Degnan, and MiloŠ Tanurdžić. Landscape of histone modifications in a sponge reveals the origin of animal cis-regulatory complexity. *eLife*, 6, 2017.

[2956] Joshua M Galanter, Christopher R Gignoux, Sam S Oh, Dara Torgerson, Maria Pino-Yanes, Neeta Thakur, Celeste Eng, Donglei Hu, Scott Huntsman, Harold J Farber, Pedro C Avila, Emerita Brigino-Buenaventura, Michael A LeNoir, Kelly Meade, Denise Serebrisky, William Rodríguez-Cintrón, Rajesh Kumar, Jose R Rodríguez-Santana, Max A Seibold, Luisa N Borrell, Esteban G Burchard, and Noah Zaitlen. Differential methylation between ethnic sub-groups reflects the effect of genetic ancestry and environmental exposures. *eLife*, 6, 2017.

[2957] Marina Venero Galanternik, Daniel Castranova, Aniket V Gore, Nathan H Blewett, Hyun Min Jung, Amber N Stratman, Martha R Kirby, James Iben, Mayumi F Miller, Koichi Kawakami, Richard J Maraia, and Brant M Weinstein. A novel perivascular cell population in the zebrafish brain. *eLife*, 6, 2017.

[2958] Marco Galardini, Alexandra Koumoutsi, Lucia Herrera-Dominguez, Juan Antonio Cordero Varela, Anja Telzerow, Omar Wagih, Morgane Wartel, Olivier Clermont, Erick Denamur, Athanasios Typas, and Pedro Beltrao. Phenotype inference in an Escherichia coli strain panel. *eLife*, 6, 2017.

[2959] Antonio Galeone, Seung Yeop Han, Chengcheng Huang, Akira Hosomi, Tadashi Suzuki, and Hamed Jafar-Nejad. Tissue-specific regulation of BMP signaling by Drosophila N-glycanase 1. *eLife*, 6, 2017.

[2960] Valentina Galli, Rafael Sebastian, Sandrine Moutel, Jason Ecard, Franck Perez, and Aurélien Roux. Uncoupling of dynamin polymerization and GTPase activity revealed by the conformation-specific nanobody dynab. *eLife*, 6, 2017.

[2961] Daniel J Galtieri, Chad M Estep, David L Wokosin, Stephen Traynelis, and D James Surmeier. Pedunculopontine glutamatergic neurons control spike patterning in substantia nigra dopaminergic neurons. *eLife*, 6, 2017.

[2962] Serge Gangloff, Guillaume Achaz, Stefania Francesconi, Adrien Villain, Samia Miled, Claire Denis, and Benoit Arcangioli. Quiescence unveils a novel mutational force in fission yeast. *eLife*, 6, 2017.

[2963] Jie Gao, Mate Marosi, Jinkuk Choi, Jennifer M Achiro, Sangmok Kim, Sandy Li, Klara Otis, Kelsey C Martin, Carlos Portera-Cailliau, and Peter Tontonoz. The E3 ubiquitin ligase IDOL regulates synaptic ApoER2 levels and is important for plasticity and learning. *eLife*, 6, 2017.

[2964] Yuan Gao, Carl Budlong, Emily Durlacher, and Ian G Davison. Neural mechanisms of social learning in the female mouse. *eLife*, 6, 2017.

[2965] Leonor García-Bayona, Monica S Guo, and Michael T Laub. Contact-dependent killing by Caulobacter crescentus via cell surface-associated, glycine zipper proteins. *eLife*, 6, 2017.

[2966] Juan-Carlos García-Betancur, Angel Goñi-Moreno, Thomas Horger, Melanie Schott, Malvika Sharan, Julian Eikmeier, Barbara Wohlmuth, Alma Zernecke, Knut Ohlsen, Christina Kuttler, and Daniel Lopez. Cell differentiation defines acute and chronic infection cell types in Staphylococcus aureus. *eLife*, 6, 2017.

[2967] Luis Carlos Garcia del Molino, Guangyu Robert Yang, Jorge F Mejias, and Xiao-Jing Wang. Paradoxical response reversal of top-down modulation in cortical circuits with three interneuron types. *eLife*, 6, 2017.

[2968] Stefano Garofalo, Alessandra Porzia, Fabrizio Mainiero, Silvia Di Angelantonio, Barbara Cortese, Bernadette Basilico, Francesca Pagani, Giorgio Cignitti, Giuseppina Chece, Roberta Maggio, Marie-Eve Tremblay, Julie Savage, Kanchan Bisht, Vincenzo Esposito, Giovanni Bernardini, Thomas Seyfried, Jakub Mieczkowski, Karolina Stepniak, Bozena Kaminska, Angela Santoni, and Cristina Limatola. Environmental stimuli shape microglial plasticity in glioma. *eLife*, 6, 2017.

[2969] Mona M Garvert, Raymond J Dolan, and Timothy EJ Behrens. A map of abstract relational knowledge in the human hippocampal–entorhinal cortex. *eLife*, 6, 2017.

[2970] Ludmila Gaspar, Cedric Howald, Konstantin Popadin, Bert Maier, Daniel Mauvoisin, Ermanno Moriggi, Maria Gutierrez-Arcelus, Emilie Falconnet, Christelle Borel, Dieter Kunz, Achim Kramer, Frederic Gachon, Emmanouil T Dermitzakis, Stylianos E Antonarakis, and Steven A Brown. The genomic landscape of human cellular circadian variation points to a novel role for the signalosome. *eLife*, 6, 2017.

[2971] özgür GenÇ, Dion K Dickman, Wenpei Ma, Amy Tong, Richard D Fetter, and Graeme W Davis. MCTP is an ER-resident calcium sensor that stabilizes synaptic transmission and homeostatic plasticity. *eLife*, 6, 2017.

[2972] Stephan Gerhard, Ingrid Andrade, Richard D Fetter, Albert Cardona, and Casey M Schneider-Mizell. Conserved neural circuit structure across Drosophila larval development revealed by comparative connectomics. *eLife*, 6, 2017.

[2973] Samuel J Gershman, Marie-H Monfils, Kenneth A Norman, and Yael Niv. The computational nature of memory modification. *eLife*, 6, 2017.

[2974] Stephan Geuter, Sabrina Boll, Falk Eippert, and Christian Büchel. Functional dissociation of stimulus intensity encoding and predictive coding of pain in the insula. *eLife*, 6, 2017.

[2975] Erinn S Gideons, Pei-Yi Lin, Melissa Mahgoub, Ege T Kavalali, and Lisa M Monteggia. Chronic lithium treatment elicits its antimanic effects via BDNF-TrkB dependent synaptic downscaling. *eLife*, 6, 2017.

[2976] Aditya Gilra and Wulfram Gerstner. Predicting non-linear dynamics by stable local learning in a recurrent spiking neural network. *eLife*, 6, 2017.

[2977] Donato Giovannelli, Stefan M Sievert, Michael Hügler, Stephanie Markert, Dörte Becher, Thomas Schweder, and Costantino Vetriani. Insight into the evolution of microbial metabolism from the deep-branching bacterium, Thermovibrio ammonificans. *eLife*, 6, 2017.

[2978] Ksenia Gnedeva, Adrian Jacobo, Joshua D Salvi, Aleksandra A Petelski, and A J Hudspeth. Elastic force restricts growth of the murine utricle. *eLife*, 6, 2017.

[2979] Qingnian Goh and Douglas P Millay. Requirement of myomaker-mediated stem cell fusion for skeletal muscle hypertrophy. *eLife*, 6, 2017.

[2980] Tal Golan, Ido Davidesco, Meir Meshulam, David M Groppe, Pierre Mégevand, Erin M Yeagle, Matthew S Goldfinger, Michal Harel, Lucia Melloni, Charles E Schroeder, Leon Y Deouell, Ashesh D Mehta, and Rafael Malach. Increasing suppression of saccade-related transients along the human visual hierarchy. *eLife*, 6, 2017.

[2981] Ognjen Golub, Brett Wee, Rhonda A Newman, Nicole M Paterson, and Kenneth E Prehoda. Activation of Discs large by aPKC aligns the mitotic spindle to the polarity axis during asymmetric cell division. *eLife*, 6, 2017.

[2982] James E Gomez, Benjamin B Kaufmann-Malaga, Carl N Wivagg, Peter B Kim, Melanie R Silvis, Nikolai Renedo, Thomas R Ioerger, Rushdy Ahmad, Jonathan Livny, Skye Fishbein, James C Sacchettini, Steven A Carr, and Deborah T Hung. Ribosomal mutations promote the evolution of antibiotic resistance in a multidrug environment. *eLife*, 6, 2017.

[2983] Bo Gong, Weimin Shen, Wanghua Xiao, Yaping Meng, Anming Meng, and Shunji Jia. The Sec14-like phosphatidylinositol transfer proteins Sec14l3/SEC14L2 act as GTPase proteins to mediate Wnt/Ca2+ signaling. *eLife*, 6, 2017.

[2984] Laura González-Méndez, Irene Seijo-Barandiarán, and Isabel Guerrero. Cytoneme-mediated cell-cell contacts for Hedgehog reception. *eLife*, 6, 2017.

[2985] Sara González-Rodríguez, Mohiuddin A Quadir, Shilpi Gupta, Karolina A Walker, Xuejiao Zhang, Viola Spahn, Dominika Labuz, Antonio Rodriguez-Gaztelumendi, Martin Schmelz, Jan Joseph, Maria K Parr, Halina Machelska, Rainer Haag, and Christoph Stein. Polyglycerol-opioid conjugate produces analgesia devoid of side effects. *eLife*, 6, 2017.

[2986] Noam Gordon, Roger Koenig-Robert, Naotsugu Tsuchiya, Jeroen JA van Boxtel, and Jakob Hohwy. Neural markers of predictive coding under perceptual uncertainty revealed with Hierarchical Frequency Tagging. *eLife*, 6, 2017.

[2987] Bryan B Gore, Samara M Miller, Yong Sang Jo, Madison A Baird, Mrinalini Hoon, Christina A Sanford, Avery Hunker, Weining Lu, Rachel O Wong, and Larry S Zweifel. Roundabout receptor 2 maintains inhibitory control of the adult midbrain. *eLife*, 6, 2017.

[2988] Srinivas Gorur-Shandilya, Mahmut Demir, Junjiajia Long, Damon A Clark, and Thierry Emonet. Olfactory receptor neurons use gain control and complementary kinetics to encode intermittent odorant stimuli. *eLife*, 6, 2017.

[2989] Graeme Gossel, Thea Hogan, Daniel Cownden, Benedict Seddon, and Andrew J Yates. Memory CD4 T cell subsets are kinetically heterogeneous and replenished from naive T cells at high levels. *eLife*, 6, 2017.

[2990] Youlian Goulev, Sandrine Morlot, Audrey Matifas, Bo Huang, Mikael Molin, Michel B Toledano, and Gilles Charvin. Nonlinear feedback drives homeostatic plasticity in H2O2 stress response. *eLife*, 6, 2017.

[2991] Anitha P Govind, Yolanda F Vallejo, Jacob R Stolz, Jing-Zhi Yan, Geoffrey T Swanson, and William N Green. Selective and regulated trapping of nicotinic receptor weak base ligands and relevance to smoking cessation. *eLife*, 6, 2017.

[2992] Alejandro A Granados, Matthew M Crane, Luis F Montano-Gutierrez, Reiko J Tanaka, Margaritis Voliotis, and Peter S Swain. Distributing tasks via multiple input pathways increases cellular survival in stress. *eLife*, 6, 2017.

[2993] James G Granneman, Vickie A Kimler, Huamei Zhang, Xiangqun Ye, Xixia Luo, John H Postlethwait, and Ryan Thummel. Lipid droplet biology and evolution illuminated by the characterization of a novel perilipin in teleost fish. *eLife*, 6, 2017.

[2994] Xavier Grau-Bové, Guifré Torruella, Stuart Donachie, Hiroshi Suga, Guy Leonard, Thomas A Richards, and Iñaki Ruiz-Trillo. Dynamics of genomic innovation in the unicellular ancestry of animals. *eLife*, 6, 2017.

[2995] Lucas T Gray, Zizhen Yao, Thuc Nghi Nguyen, Tae Kyung Kim, Hongkui Zeng, and Bosiljka Tasic. Layer-specific chromatin accessibility landscapes reveal regulatory networks in adult mouse visual cortex. *eLife*, 6, 2017.

[2996] Kathleen Greenham, Carmela Rosaria Guadagno, Malia A Gehan, Todd C Mockler, Cynthia Weinig, Brent E Ewers, and C Robertson McClung. Temporal network analysis identifies early physiological and transcriptomic indicators of mild drought in Brassica rapa. *eLife*, 6, 2017.

[2997] Saskia Gressel, Björn Schwalb, Tim Michael Decker, Weihua Qin, Heinrich Leonhardt, Dirk Eick, and Patrick Cramer. CDK9-dependent RNA polymerase II pausing controls transcription initiation. *eLife*, 6, 2017.

[2998] Adam Graham Grieve, Hongmei Xu, Ulrike Künzel, Paul Bambrough, Boris Sieber, and Matthew Freeman. Phosphorylation of iRhom2 at the plasma membrane controls mammalian TACE-dependent inflammatory and growth factor signalling. *eLife*, 6, 2017.

[2999] Joscha Griger, Robin Schneider, Ines Lahmann, Verena Schöwel, Charles Keller, Simone Spuler, Marc Nazare, and Carmen Birchmeier. Loss of Ptpn11 (Shp2) drives satellite cells into quiescence. *eLife*, 6, 2017.

[3000] Christopher M Groen, Cormac McGrath, Katherine A Campbell, Cecilia Götherström, Anthony J Windebank, and Natalia Landázuri. Promoting international collaboration and creativity in doctoral students. *eLife*, 6, 2017.

[3001] John P Grogan, Demitra Tsivos, Laura Smith, Brogan E Knight, Rafal Bogacz, Alan Whone, and Elizabeth J Coulthard. Effects of dopamine on reinforcement learning and consolidation in Parkinson's disease. *eLife*, 6, 2017.

[3002] Julien Gronnier, Jean-Marc Crowet, Birgit Habenstein, Mehmet Nail Nasir, Vincent Bayle, Eric Hosy, Matthieu Pierre Platre, Paul Gouguet, Sylvain Raffaele, Denis Martinez, Axelle Grelard, Antoine Loquet, FranÇoise Simon-Plas, Patricia Gerbeau-Pissot, Christophe Der, Emmanuelle M Bayer, Yvon Jaillais, Magali Deleu, Véronique Germain, Laurence Lins, and Sébastien Mongrand. Structural basis for plant plasma membrane protein dynamics and organization into functional nanodomains. *eLife*, 6, 2017.

[3003] Matthew Grove, Hyukmin Kim, Maryline Santerre, Alexander J Krupka, Seung Baek Han, Jinbin Zhai, Jennifer Y Cho, Raehee Park, Michele Harris, Seonhee Kim, Bassel E Sawaya, Shin H Kang, Mary F Barbe, Seo-Hee Cho, Michel A Lemay, and Young-Jin Son. YAP/TAZ initiate and maintain Schwann cell myelination. *eLife*, 6, 2017.

[3004] Paolo Grumati, Giulio Morozzi, Soraya Hölper, Muriel Mari, Marie-Lena IE Harwardt, Riqiang Yan, Stefan Müller, Fulvio Reggiori, Mike Heilemann, and Ivan Dikic. Full length RTN3 regulates turnover of tubular endoplasmic reticulum via selective autophagy. *eLife*, 6, 2017.

[3005] Benedikt Grünewald, Maren D Lange, Christian Werner, Aet O'Leary, Andreas Weishaupt, Sandy Popp, David A Pearce, Heinz Wiendl, Andreas Reif, Hans C Pape, Klaus V Toyka, Claudia Sommer, and Christian Geis. Defective synaptic transmission causes disease signs in a mouse model of juvenile neuronal ceroid lipofuscinosis. *eLife*, 6, 2017.

[3006] Bin Gu, Jean-Philippe Lambert, Katie Cockburn, Anne-Claude Gingras, and Janet Rossant. AIRE is a critical spindle-associated protein in embryonic stem cells. *eLife*, 6, 2017.

[3007] Wuqiang Guan, Jun-Wei Cao, Lin-Yun Liu, Zhi-Hao Zhao, Yinghui Fu, and Yong-Chun Yu. Eye opening differentially modulates inhibitory synaptic transmission in the developing visual cortex. *eLife*, 6, 2017.

[3008] Zhuo Guan, Maria Bykhovskaia, Ramon A Jorquera, Roger Bryan Sutton, Yulia Akbergenova, and J Troy Littleton. A synaptotagmin suppressor screen indicates SNARE binding controls the timing and Ca2+ cooperativity of vesicle fusion. *eLife*, 6, 2017.

[3009] Iva Gudernova, Silvie Foldynova-Trantirkova, Barbora El Ghannamova, Bohumil Fafilek, Miroslav Varecha, Lukas Balek, Eva Hruba, Lucie Jonatova, Iva Jelinkova, Michaela Kunova Bosakova, Lukas Trantirek, Jiri Mayer, and Pavel Krejci. One reporter for in-cell activity profiling of majority of protein kinase oncogenes. *eLife*, 6, 2017.

[3010] Michael W Guernsey, Edward B Chuong, Guillaume Cornelis, Marilyn B Renfree, and Julie C Baker. Molecular conservation of marsupial and eutherian placentation and lactation. *eLife*, 6, 2017.

[3011] Olivia Guest and Bradley C Love. What the success of brain imaging implies about the neural code. *eLife*, 6, 2017.

[3012] Vincent Guichard, Nelly Bonilla, Aurélie Durand, Alexandra Audemard-Verger, Thomas Guilbert, Bruno Martin, Bruno Lucas, and Cédric Auffray. Calcium-mediated shaping of naive CD4 T-cell phenotype and function. *eLife*, 6, 2017.

[3013] Laurent Guillaud, Dimitar Dimitrov, and Tomoyuki Takahashi. Presynaptic morphology and vesicular composition determine vesicle dynamics in mouse central synapses. *eLife*, 6, 2017.

[3014] Michael Guo, Zun Liu, Jessie Willen, Cameron P Shaw, Daniel Richard, Evelyn Jagoda, Andrew C Doxey, Joel Hirschhorn, and Terence D Capellini. Epigenetic profiling of growth plate chondrocytes sheds insight into regulatory genetic variation influencing height. *eLife*, 6, 2017.

[3015] Yusong R Guo and Roderick MacKinnon. Structure-based membrane dome mechanism for Piezo mechanosensitivity. *eLife*, 6, 2017.

[3016] Kapil Gupta, Aleksandra A Watson, Tiago Baptista, Elisabeth Scheer, Anna L Chambers, Christine Koehler, Juan Zou, Ima Obong-Ebong, Eaazhisai Kandiah, Arturo Temblador, Adam Round, Eric Forest, Petr Man, Christoph Bieniossek, Ernest D Laue, Edward A Lemke, Juri Rappsilber, Carol V Robinson, Didier Devys, Làszlò Tora, and Imre Berger. Architecture of TAF11/TAF13/TBP complex suggests novel regulation properties of general transcription factor TFIID. *eLife*, 6, 2017.

[3017] Pinar S Gurel, Laura Y Kim, Paul V Ruijgrok, Tosan Omabegho, Zev Bryant, and Gregory M Alushin. Cryo-EM structures reveal specialization at the myosin VI-actin interface and a mechanism of force sensitivity. *eLife*, 6, 2017.

[3018] Nicholas R Guydosh, Philipp Kimmig, Peter Walter, and Rachel Green. Regulated Ire1-dependent mRNA decay requires no-go mRNA degradation to maintain endoplasmic reticulum homeostasis in S. pombe. *eLife*, 6, 2017.

[3019] Juergen Haag, Abhishek Mishra, and Alexander Borst. A common directional tuning mechanism of Drosophila motion-sensing neurons in the ON and in the OFF pathway. *eLife*, 6, 2017.

[3020] Nobuhiro Hagura, Patrick Haggard, and Jörn Diedrichsen. Perceptual decisions are biased by the cost to act. *eLife*, 6, 2017.

[3021] Tang Hai, Chunwei Cao, Haitao Shang, Weiwei Guo, Yanshuang Mu, Shulin Yang, Ying Zhang, Qiantao Zheng, Tao Zhang, Xianlong Wang, Yu Liu, Qingran Kong, Kui Li, Dayu Wang, Meng Qi, Qianlong Hong, Rui Zhang, Xiupeng Wang, Qitao Jia, Xiao Wang, Guosong Qin, Yongshun Li, Ailing Luo, Weiwu Jin, Jing Yao, Jiaojiao Huang, Hongyong Zhang, Menghua Li, Xiangmo Xie, Xuejuan Zheng, Kenan Guo, Qinghua Wang, Shibin Zhang, Liang Li, Fei Xie, Yu Zhang, Xiaogang Weng, Zhi Yin, Kui Hu, Yimei Cong, Peng Zheng, Hailong Zou, Leilei Xin, Jihan Xia, Jinxue Ruan, Hegang Li, Weiming Zhao, Jing Yuan, Zizhan Liu, Weiwang Gu, Ming Li, Yong Wang, Hongmei Wang, Shiming Yang, Zhonghua Liu, Hong Wei, Jianguo Zhao, Qi Zhou, and Anming Meng. Pilot study of large-scale production of mutant pigs by ENU mutagenesis. *eLife*, 6, 2017.

[3022] Parthiv Haldipur, Derek Dang, Kimberly A Aldinger, Olivia K Janson, Fabien Guimiot, Homa Adle-Biasette, William B Dobyns, Joseph R Siebert, Rosa Russo, and Kathleen J Millen. Phenotypic outcomes in Mouse and Human Foxc1 dependent Dandy-Walker cerebellar malformation suggest shared mechanisms. *eLife*, 6, 2017.

[3023] Angela M Halstead, Chiraag D Kapadia, Jennifer Bolzenius, Clarence E Chu, Andrew Schriefer, Lukas D Wartman, Gregory R Bowman, and Vivek K Arora. Bladder-cancer-associated mutations in RXRA activate peroxisome proliferator-activated receptors to drive urothelial proliferation. *eLife*, 6, 2017.

[3024] Stefanie Hampel, Claire E McKellar, Julie H Simpson, and Andrew M Seeds. Simultaneous activation of parallel sensory pathways promotes a grooming sequence in Drosophila. *eLife*, 6, 2017.

[3025] Bicheng Han, Yongming Dong, Lin Zhang, Yan Liu, Ithai Rabinowitch, and Jihong Bai. Dopamine signaling tunes spatial pattern selectivity in C. elegans. *eLife*, 6, 2017.

[3026] Han Han, Nicole Monroe, Wesley I Sundquist, Peter S Shen, and Christopher P Hill. The AAA ATPase Vps4 binds ESCRT-III substrates through a repeating array of dipeptide-binding pockets. *eLife*, 6, 2017.

[3027] Yan Han, Chunli Yan, Thi Hoang Duong Nguyen, Ashleigh J Jackobel, Ivaylo Ivanov, Bruce A Knutson, and Yuan He. Structural mechanism of ATP-independent transcription initiation by RNA polymerase I. *eLife*, 6, 2017.

[3028] Yongchao Han, Liangyao Xiong, Ying Xu, Tian Tian, and Tao Wang. The $\beta$-alanine transporter BalaT is required for visual neurotransmission in Drosophila. *eLife*, 6, 2017.

[3029] Colleen E Hannon, Shelby A Blythe, and Eric F Wieschaus. Concentration dependent chromatin states induced by the bicoid morphogen gradient. *eLife*, 6, 2017.

[3030] Anders S Hansen, Iryna Pustova, Claudia Cattoglio, Robert Tjian, and Xavier Darzacq. CTCF and cohesin regulate chromatin loop stability with distinct dynamics. *eLife*, 6, 2017.

[3031] Yangguang Hao and Li Hua Jin. Dual role for Jumu in the control of hematopoietic progenitors in the Drosophila lymph gland. *eLife*, 6, 2017.

[3032] Masatoshi Hara, Boryana Petrova, and Terry L Orr-Weaver. Control of PNG kinase, a key regulator of mRNA translation, is coupled to meiosis completion at egg activation. *eLife*, 6, 2017.

[3033] Stefan Harmansa, Ilaria Alborelli, Dimitri Bieli, Emmanuel Caussinus, and Markus Affolter. A nanobody-based toolset to investigate the role of protein localization and dispersal in Drosophila. *eLife*, 6, 2017.

[3034] Thomas C Harmon, Uri Magaram, David L McLean, and Indira M Raman. Distinct responses of Purkinje neurons and roles of simple spikes during associative motor learning in larval zebrafish. *eLife*, 6, 2017.

[3035] Tyler S Harmon, Alex S Holehouse, Michael K Rosen, and Rohit V Pappu. Intrinsically disordered linkers determine the interplay between phase separation and gelation in multivalent proteins. *eLife*, 6, 2017.

[3036] Hala Harony-Nicolas, Maya Kay, Johann du Hoffmann, Matthew E Klein, Ozlem Bozdagi-Gunal, Mohammed Riad, Nikolaos P Daskalakis, Sankalp Sonar, Pablo E Castillo, Patrick R Hof, Matthew L Shapiro, Mark G Baxter, Shlomo Wagner, and Joseph D Buxbaum. Oxytocin improves behavioral and electrophysiological deficits in a novel Shank3-deficient rat. *eLife*, 6, 2017.

[3037] Michael JM Harrap, Sean A Rands, Natalie Hempel de Ibarra, and Heather M Whitney. The diversity of floral temperature patterns, and their use by pollinators. *eLife*, 6, 2017.

[3038] Kelley Harris and Jonathan K Pritchard. Rapid evolution of the human mutation spectrum. *eLife*, 6, 2017.

[3039] Gesa Hartwigsen, Danilo Bzdok, Maren Klein, Max Wawrzyniak, Anika Stockert, Katrin Wrede, Joseph Classen, and Dorothee Saur. Rapid short-term reorganization in the language network. *eLife*, 6, 2017.

[3040] Tomoya Hasegawa, Christopher J Hall, Philip S Crosier, Gembu Abe, Koichi Kawakami, Akira Kudo, and Atsushi Kawakami. Transient inflammatory response mediated by interleukin-1$\beta$ is required for proper regeneration in zebrafish fin fold. *eLife*, 6, 2017.

[3041] Elizabeth S Haswell. The sustainable professor. *eLife*, 6, 2017.

[3042] Tobias U Hauser, Micah Allen, Nina Purg, Michael Moutoussis, Geraint Rees, and Raymond J Dolan. Noradrenaline blockade specifically enhances metacognitive performance. *eLife*, 6, 2017.

[3043] Virginia E Hawkins, Ana C Takakura, Ashley Trinh, Milene R Malheiros-Lima, Colin M Cleary, Ian C Wenker, Todd Dubreuil, Elliot M Rodriguez, Mark T Nelson, Thiago S Moreira, and Daniel K Mulkey. Purinergic regulation of vascular tone in the retrotrapezoid nucleus is specialized to support the drive to breathe. *eLife*, 6, 2017.

[3044] Yakupjan Haxim, Asigul Ismayil, Qi Jia, Yan Wang, Xiyin Zheng, Tianyuan Chen, Lichao Qian, Na Liu, Yunjing Wang, Shaojie Han, Jiaxuan Cheng, Yijun Qi, Yiguo Hong, and Yule Liu. Autophagy functions as an antiviral mechanism against geminiviruses in plants. *eLife*, 6, 2017.

[3045] Naoto Hayasaka, Arisa Hirano, Yuka Miyoshi, Isao T Tokuda, Hikari Yoshitane, Junichiro Matsuda, and Yoshitaka Fukada. Salt-inducible kinase 3 regulates the mammalian circadian clock by destabilizing PER2 protein. *eLife*, 6, 2017.

[3046] Bin Z He, Xu Zhou, and Erin K O'Shea. Evolution of reduced co-activator dependence led to target expansion of a starvation response pathway. *eLife*, 6, 2017.

[3047] Li He, Helena Steinocher, Ashish Shelar, Emily B Cohen, Erin N Heim, Birthe B Kragelund, Gevorg Grigoryan, and Daniel DiMaio. Single methyl groups can act as toggle switches to specify transmembrane Protein-protein interactions. *eLife*, 6, 2017.

[3048] Florian T Heer, David J Posson, Wojciech Wojtas-Niziurski, Crina M Nimigean, and Simon Bernèche. Mechanism of activation at the selectivity filter of the KcsA K+ channel. *eLife*, 6, 2017.

[3049] Peter D Heintzman, Grant D Zazula, Ross DE MacPhee, Eric Scott, James A Cahill, Brianna K McHorse, Joshua D Kapp, Mathias Stiller, Matthew J Wooller, Ludovic Orlando, John Southon, Duane G Froese, and Beth Shapiro. A new genus of horse from Pleistocene North America. *eLife*, 6, 2017.

[3050] Sebastian Carsten Johannes Helle, Qian Feng, Mathias J Aebersold, Luca Hirt, Raphael R Grüter, Afshin Vahid, Andrea Sirianni, Serge Mostowy, Jess G Snedeker, Anđela Šarić, Timon Idema, Tomaso Zambelli, and Benoît Kornmann. Mechanical force induces mitochondrial fission. *eLife*, 6, 2017.

[3051] Markus Helmer, Manuel Schottdorf, Andreas Neef, and Demian Battaglia. Gender bias in scholarly peer review. *eLife*, 6, 2017.

[3052] Xin Heng, Qiuxia Guo, Alan W Leung, and James YH Li. Analogous mechanism regulating formation of neocortical basal radial glia and cerebellar Bergmann glia. *eLife*, 6, 2017.

[3053] Erik Henrich, Oliver Peetz, Christopher Hein, Aisha Laguerre, Beate Hoffmann, Jan Hoffmann, Volker Dötsch, Frank Bernhard, and Nina Morgner. Analyzing native membrane protein assembly in nanodiscs by combined non-covalent mass spectrometry and synthetic biology. *eLife*, 6, 2017.

[3054] Chelsea Hepler, Mengle Shao, Jonathan Y Xia, Alexandra L Ghaben, Mackenzie J Pearson, Lavanya Vishvanath, Ankit X Sharma, Thomas S Morley, William L Holland, and Rana K Gupta. Directing visceral white adipocyte precursors to a thermogenic adipocyte fate improves insulin sensitivity in obese mice. *eLife*, 6, 2017.

[3055] Damian M Herz, Huiling Tan, John-Stuart Brittain, Petra Fischer, Binith Cheeran, Alexander L Green, James FitzGerald, Tipu Z Aziz, Keyoumars Ashkan, Simon Little, Thomas Foltynie, Patricia Limousin, Ludvic Zrinzo, Rafal Bogacz, and Peter Brown. Distinct mechanisms mediate speed-accuracy adjustments in cortico-subthalamic networks. *eLife*, 6, 2017.

[3056] André Heuer, Emma Thomson, Christian Schmidt, Otto Berninghausen, Thomas Becker, Ed Hurt, and Roland Beckmann. Cryo-EM structure of a late pre-40S ribosomal subunit from Saccharomyces cerevisiae. *eLife*, 6, 2017.

[3057] Jonathan D Hibshman, Alexander E Doan, Brad T Moore, Rebecca EW Kaplan, Anthony Hung, Amy K Webster, Dhaval P Bhatt, Rojin Chitrakar, Matthew D Hirschey, and L Ryan Baugh. daf-16/FoxO promotes gluconeogenesis and trehalose synthesis during starvation to support survival. *eLife*, 6, 2017.

[3058] Zoë A Hilbert and Dennis H Kim. Sexually dimorphic control of gene expression in sensory neurons regulates decision-making behavior in C. elegans. *eLife*, 6, 2017.

[3059] David R Hill, Sha Huang, Melinda S Nagy, Veda K Yadagiri, Courtney Fields, Dishari Mukherjee, Brooke Bons, Priya H Dedhia, Alana M Chin, Yu-Hwai Tsai, Shrikar Thodla, Thomas M Schmidt, Seth Walk, Vincent B Young, and Jason R Spence. Bacterial colonization stimulates a complex physiological response in the immature human intestinal epithelium. *eLife*, 6, 2017.

[3060] Daniel Scott Himmelstein, Antoine Lizee, Christine Hessler, Leo Brueggeman, Sabrina L Chen, Dexter Hadley, Ari Green, Pouya Khankhanian, and Sergio E Baranzini. Systematic integration of biomedical knowledge prioritizes drugs for repurposing. *eLife*, 6, 2017.

[3061] Stefan Hirschberg, Yong Li, Andrew Randall, Eric J Kremer, and Anthony E Pickering. Functional dichotomy in spinal- vs prefrontal-projecting locus coeruleus modules splits descending noradrenergic analgesia from ascending aversion and anxiety in rats. *eLife*, 6, 2017.

[3062] Peter Hitchcock, Ambika Mathur, Jabbar Bennett, Patricia Cameron, Christine Chow, Philip Clifford, Robert Duvoisin, Andrew Feig, Kevin Finneran, Diane M Klotz, Richard McGee, Mary O'Riordan, Christine Pfund, Christopher Pickett, Nancy Schwartz, Nancy E Street, Elizabeth Watkins, Jonathan Wiest, and David Engelke. The future of graduate and postdoctoral training in the biosciences. *eLife*, 6, 2017.

[3063] Jeroen NA Hoffer, Janine Mariën, Jacintha Ellers, and Joris M Koene. Sexual selection gradients change over time in a simultaneous hermaphrodite. *eLife*, 6, 2017.

[3064] Natalie E Hofmann, Stephan Karl, Rahel Wampfler, Benson Kiniboro, Albina Teliki, Jonah Iga, Andreea Waltmann, Inoni Betuela, Ingrid Felger, Leanne J Robinson, and Ivo Mueller. The complex relationship of exposure to new Plasmodium infections and incidence of clinical malaria in Papua New Guinea. *eLife*, 6, 2017.

[3065] Nicholas T Hogan, Michael B Whalen, Lindsey K Stolze, Nizar K Hadeli, Michael T Lam, James R Springstead, Christopher K Glass, and Casey E Romanoski. Transcriptional networks specifying homeostatic and inflammatory programs of gene expression in human aortic endothelial cells. *eLife*, 6, 2017.

[3066] Annika Hohendahl, Nathaniel Talledge, Valentina Galli, Peter S Shen, Frédéric Humbert, Pietro De Camilli, Adam Frost, and Aurélien Roux. Structural inhibition of dynamin-mediated membrane fission by endophilin. *eLife*, 6, 2017.

[3067] Esteban Hoijman, L Fargas, Patrick Blader, and Berta Alsina. Pioneer neurog1 expressing cells ingress into the otic epithelium and instruct neuronal specification. *eLife*, 6, 2017.

[3068] Sebastian C Holst, Alexandra Sousek, Katharina Hefti, Sohrab Saberi-Moghadam, Alfred Buck, Simon M Ametamey, Milan Scheidegger, Paul Franken, Anke Henning, Erich Seifritz, Mehdi Tafti, and Hans-Peter Landolt. Cerebral mGluR5 availability contributes to elevated sleep need and behavioral adjustment after sleep deprivation. *eLife*, 6, 2017.

[3069] Guillaume Holzer, Marc Besson, Anne Lambert, Loïc FranÇois, Paul Barth, Benjamin Gillet, Sandrine Hughes, Gwenaël Piganeau, Francois Leulier, Laurent Viriot, David Lecchini, and Vincent Laudet. Fish larval recruitment to reefs is a thyroid hormone-mediated metamorphosis sensitive to the pesticide chlorpyrifos. *eLife*, 6, 2017.

[3070] Keiji Honda, Sung Huhn Kim, Michael C Kelly, Joseph C Burns, Laura Constance, Xiangming Li, Fei Zhou, Michael Hoa, Matthew W Kelley, Philine Wangemann, Robert J Morell, and Andrew J Griffith. Molecular architecture underlying fluid absorption by the developing inner ear. *eLife*, 6, 2017.

[3071] Sungki Hong, Mallory A Freeberg, Ting Han, Avani Kamath, Yao Yao, Tomoko Fukuda, Tsukasa Suzuki, John K Kim, and Ken Inoki. LARP1 functions as a molecular switch for mTORC1-mediated translation of an essential class of mRNAs. *eLife*, 6, 2017.

[3072] Stephen K Horrigan. Replication Study: The CD47-signal regulatory protein alpha (SIRPa) interaction is a therapeutic target for human solid tumors. *eLife*, 6, 2017.

[3073] Stephen K Horrigan, Pascal Courville, Darryl Sampey, Faren Zhou, and Steve Cai. Replication Study: Melanoma genome sequencing reveals frequent PREX2 mutations. *eLife*, 6, 2017.

[3074] Li Hou, Pengcheng Yang, Feng Jiang, Qing Liu, Xianhui Wang, and Le Kang. The neuropeptide F/nitric oxide pathway is essential for shaping locomotor plasticity underlying locust phase transition. *eLife*, 6, 2017.

[3075] Sébastien Houy, Alexander J Groffen, Iwona Ziomkiewicz, Matthijs Verhage, Paulo S Pinheiro, and Jakob Balslev Sørensen. Doc2B acts as a calcium sensor for vesicle priming requiring synaptotagmin-1, Munc13-2 and SNAREs. *eLife*, 6, 2017.

[3076] FranÇoise S Howe, Andrew Russell, Anna R Lamstaes, Afaf El-Sagheer, Anitha Nair, Tom Brown, and Jane Mellor. CRISPRi is not strand-specific at all loci and redefines the transcriptional landscape. *eLife*, 6, 2017.

[3077] Jen-Chun Hsiang, Keith P Johnson, Linda Madisen, Hongkui Zeng, and Daniel Kerschensteiner. Local processing in neurites of VGluT3-expressing amacrine cells differentially organizes visual information. *eLife*, 6, 2017.

[3078] Ching-Lin Hsieh, Christopher P Ptak, Andrew Tseng, Igor Massahiro de Souza Suguiura, Sean P McDonough, Tepyuda Sritrakul, Ting Li, Yi-Pin Lin, Richard E Gillilan, Robert E Oswald, and Yung-Fu Chang. Extended low-resolution structure of a Leptospira antigen offers high bactericidal antibody accessibility amenable to vaccine design. *eLife*, 6, 2017.

[3079] Fu-Lien Hsieh and Matthew K Higgins. The structure of a LAIR1-containing human antibody reveals a novel mechanism of antigen recognition. *eLife*, 6, 2017.

[3080] Yen-Ping Hsueh, Matthew R Gronquist, Erich M Schwarz, Ravi David Nath, Ching-Han Lee, Shalha Gharib, Frank C Schroeder, and Paul W Sternberg. Nematophagous fungus Arthrobotrys oligospora mimics olfactory cues of sex and food to lure its nematode prey. *eLife*, 6, 2017.

[3081] Wen Hu, Zhao-Di Jiang, Fang Suo, Jin-Xin Zheng, Wan-Zhong He, and Li-Lin Du. A large gene family in fission yeast encodes spore killers that subvert Mendel's law. *eLife*, 6, 2017.

[3082] Anqi Huang, Christopher Amourda, Shaobo Zhang, Nicholas S Tolwinski, and Timothy E Saunders. Decoding temporal interpretation of the morphogen Bicoid in the early Drosophila embryo. *eLife*, 6, 2017.

[3083] Hai-Tsang Huang, Hyuk-Soo Seo, Tinghu Zhang, Yubao Wang, Baishan Jiang, Qing Li, Dennis L Buckley, Behnam Nabet, Justin M Roberts, Joshiawa Paulk, Shiva Dastjerdi, Georg E Winter, Hilary McLauchlan, Jennifer Moran, James E Bradner, Michael J Eck, Sirano Dhe-Paganon, Jean J Zhao, and Nathanael S Gray. MELK is not necessary for the proliferation of basal-like breast cancer cells. *eLife*, 6, 2017.

[3084] Ting-hao Huang, Peter Niesman, Deepshika Arasu, Donghyung Lee, Aubrie L De La Cruz, Antuca Callejas, Elizabeth J Hong, and Carlos Lois. Tracing neuronal circuits in transgenic animals by transneuronal control of transcription (TRACT). *eLife*, 6, 2017.

[3085] Xin Huang, Sophie Balmer, Fan Yang, Miguel Fidalgo, Dan Li, Diana Guallar, Anna-Katerina Hadjantonakis, and Jianlong Wang. Zfp281 is essential for mouse epiblast maturation through transcriptional and epigenetic control of Nodal signaling. *eLife*, 6, 2017.

[3086] Yu Huang, Anli A Liu, Belen Lafon, Daniel Friedman, Michael Dayan, Xiuyuan Wang, Marom Bikson, Werner K Doyle, Orrin Devinsky, and Lucas C Parra. Measurements and models of electric fields in the in vivo human brain during transcranial electric stimulation. *eLife*, 6, 2017.

[3087] Yun Huang, Jong Kyoung Kim, Dang Vinh Do, Caroline Lee, Christopher A Penfold, Jan J Zylicz, John C Marioni, Jamie A Hackett, and M Azim Surani. Stella modulates transcriptional and endogenous retrovirus programs during maternal-to-zygotic transition. *eLife*, 6, 2017.

[3088] Elizabeth A Hubin, Allison Fay, Catherine Xu, James M Bean, Ruth M Saecker, Michael S Glickman, Seth A Darst, and Elizabeth A Campbell. Structure and function of the mycobacterial transcription initiation complex with the essential regulator RbpA. *eLife*, 6, 2017.

[3089] Christina L Hueschen, Samuel J Kenny, Ke Xu, and Sophie Dumont. NuMA recruits dynein activity to microtubule minus-ends at mitosis. *eLife*, 6, 2017.

[3090] Karthik Hullahalli, Marinelle Rodrigues, and Kelli L Palmer. Exploiting CRISPR-Cas to manipulate Enterococcus faecalis populations. *eLife*, 6, 2017.

[3091] Greta Hultqvist, Emma Åberg, Carlo Camilloni, Gustav N Sundell, Eva Andersson, Jakob Dogan, Celestine N Chi, Michele Vendruscolo, and Per Jemth. Emergence and evolution of an interaction between intrinsically disordered proteins. *eLife*, 6, 2017.

[3092] Victoria Hung, Stephanie S Lam, Namrata D Udeshi, Tanya Svinkina, Gaelen Guzman, Vamsi K Mootha, Steven A Carr, and Alice Y Ting. Proteomic mapping of cytosol-facing outer mitochondrial and ER membranes in living human cells by proximity biotinylation. *eLife*, 6, 2017.

[3093] Yin P Hung, Carolyn Teragawa, Nont Kosaisawe, Taryn E Gillies, Michael Pargett, Marta Minguet, Kevin Distor, Briana L Rocha-Gregg, Jonathan L Coloff, Mark A Keibler, Gregory Stephanopoulos, Gary Yellen, Joan S Brugge, and John G Albeck. Akt regulation of glycolysis mediates bioenergetic stability in epithelial cells. *eLife*, 6, 2017.

[3094] Matthew Hur, Charlotte A Gistelinck, Philippe Huber, Jane Lee, Marjorie H Thompson, Adrian T Monstad-Rios, Claire J Watson, Sarah K McMenamin, Andy Willaert, David M Parichy, Paul Coucke, and Ronald Y Kwon. MicroCT-based phenomics in the zebrafish skeleton reveals virtues of deep phenotyping in a distributed organ system. *eLife*, 6, 2017.

[3095] Wonmuk Hwang, Matthew J Lang, and Martin Karplus. Kinesin motility is driven by subdomain dynamics. *eLife*, 6, 2017.

[3096] Yu-Hsien Hwang Fu, William Y C Huang, Kuang Shen, Jay T Groves, Thomas Miller, and Shu-ou Shan. Two-step membrane binding by the bacterial SRP receptor enable efficient and accurate Co-translational protein targeting. *eLife*, 6, 2017.

[3097] Alexandre Hyafil and Rubén Moreno-Bote. Breaking down hierarchies of decision-making in primates. *eLife*, 6, 2017.

[3098] Michael J Iannacone, Isabel Beets, Lindsey E Lopes, Matthew A Churgin, Christopher Fang-Yen, Matthew D Nelson, Liliane Schoofs, and David M Raizen. The RFamide receptor DMSR-1 regulates stress-induced sleep in C. elegans. *eLife*, 6, 2017.

[3099] Ximena Ibarra-Soria, Thiago S Nakahara, Jingtao Lilue, Yue Jiang, Casey Trimmer, Mateus AA Souza, Paulo HM Netto, Kentaro Ikegami, Nicolle R Murphy, Mairi Kusma, Andrea Kirton, Luis R Saraiva, Thomas M Keane, Hiroaki Matsunami, Joel Mainland, Fabio Papes, and Darren W Logan. Variation in olfactory neuron repertoires is genetically controlled and environmentally modulated. *eLife*, 6, 2017.

[3100] Guilhem Ibos and David J Freedman. Sequential sensory and decision processing in posterior parietal cortex. *eLife*, 6, 2017.

[3101] Yuichi Ichikawa, Caitlin F Connelly, Alon Appleboim, Thomas CR Miller, Hadas Jacobi, Nebiyu A Abshiru, Hsin-Jung Chou, Yuanyuan Chen, Upasna Sharma, Yupeng Zheng, Paul M Thomas, Hsuiyi V Chen, Vineeta Bajaj, Christoph W Müller, Neil L Kelleher, Nir Friedman, Daniel NA Bolon, Oliver J Rando, and Paul D Kaufman. A synthetic biology approach to probing nucleosome symmetry. *eLife*, 6, 2017.

[3102] Ronit Ilouz, Varda Lev-Ram, Eric A Bushong, Travis L Stiles, Dinorah Friedmann-Morvinski, Christopher Douglas, Jeffrey L Goldberg, Mark H Ellisman, and Susan S Taylor. Isoform-specific subcellular localization and function of protein kinase A identified by mosaic imaging of mouse brain. *eLife*, 6, 2017.

[3103] Rodney Elwood Infante and Arun Radhakrishnan. Continuous transport of a small fraction of plasma membrane cholesterol to endoplasmic reticulum regulates total cellular cholesterol. *eLife*, 6, 2017.

[3104] Tokiro Ishikawa, Makoto Kashima, Atsushi J Nagano, Tomoko Ishikawa-Fujiwara, Yasuhiro Kamei, Takeshi Todo, and Kazutoshi Mori. Unfolded protein response transducer IRE1-mediated signaling independent of XBP1 mRNA splicing is not required for growth and development of medaka fish. *eLife*, 6, 2017.

[3105] Daisuke Ito, Marc A Schureck, and Sanjay A Desai. An essential dual-function complex mediates erythrocyte invasion and channel-mediated nutrient uptake in malaria parasites. *eLife*, 6, 2017.

[3106] Kenzo Ivanovitch, Susana Temiño, and Miguel Torres. Live imaging of heart tube development in mouse reveals alternating phases of cardiac differentiation and morphogenesis. *eLife*, 6, 2017.

[3107] Eiko Iwakoshi-Ukena, Kenshiro Shikano, Kunihiro Kondo, Shusuke Taniuchi, Megumi Furumitsu, Yuta Ochi, Tsutomu Sasaki, Shiki Okamoto, George E Bentley, Lance J Kriegsfeld, Yasuhiko Minokoshi, and Kazuyoshi Ukena. Neurosecretory protein GL stimulates food intake, de novo lipogenesis, and onset of obesity. *eLife*, 6, 2017.

[3108] Matthew DB Jackson, Hao Xu, Salva Duran-Nebreda, Petra Stamm, and George W Bassel. Topological analysis of multicellular complexity in the plant hypocotyl. *eLife*, 6, 2017.

[3109] Sagi Jaffe-Dax, Or Frenkel, and Merav Ahissar. Dyslexics' faster decay of implicit memory for sounds and words is manifested in their shorter neural adaptation. *eLife*, 6, 2017.

[3110] Uday K Jagadisan and Neeraj J Gandhi. Removal of inhibition uncovers latent movement potential during preparation. *eLife*, 6, 2017.

[3111] Arjen J Jakobi, Matthias Wilmanns, and Carsten Sachse. Model-based local density sharpening of cryo-EM maps. *eLife*, 6, 2017.

[3112] Ryan Janke, Anthony T Iavarone, and Jasper Rine. Oncometabolite D-2-Hydroxyglutarate enhances gene silencing through inhibition of specific H3K36 histone demethylases. *eLife*, 6, 2017.

[3113] Philipp Janz, Niels Schwaderlapp, Katharina Heining, Ute Häussler, Jan G Korvink, Dominik von Elverfeldt, Jürgen Hennig, Ulrich Egert, Pierre LeVan, and Carola A Haas. Early tissue damage and microstructural reorganization predict disease severity in experimental epilepsy. *eLife*, 6, 2017.

[3114] Arman Javadi, Ravi K Deevi, Emma Evergren, Elodie Blondel-Tepaz, George S Baillie, Mark GH Scott, and Frederick C Campbell. PTEN controls glandular morphogenesis through a juxtamembrane $\beta$-Arrestin1/ARHGAP21 scaffolding complex. *eLife*, 6, 2017.

[3115] Pauline Jeannot, Ada Nowosad, Renaud T Perchey, Caroline Callot, Evangeline Bennana, Takanori Katsube, Patrick Mayeux, FranÇois Guillonneau, Stéphane Manenti, and Arnaud Besson. p27Kip1 promotes invadopodia turnover and invasion through the regulation of the PAK1/Cortactin pathway. *eLife*, 6, 2017.

[3116] Martin D Jennings, Christopher J Kershaw, Tomas Adomavicius, and Graham D Pavitt. Fail-safe control of translation initiation by dissociation of eIF2$\alpha$ phosphorylated ternary complexes. *eLife*, 6, 2017.

[3117] Justin M Jenson, Jeremy A Ryan, Robert A Grant, Anthony Letai, and Amy E Keating. Epistatic mutations in PUMA BH3 drive an alternate binding mode to potently and selectively inhibit anti-apoptotic Bfl-1. *eLife*, 6, 2017.

[3118] Daniel Jercog, Alex Roxin, Peter Barthó, Artur Luczak, Albert Compte, and Jaime de la Rocha. UP-DOWN cortical dynamics reflect state transitions in a bistable network. *eLife*, 6, 2017.

[3119] Elizabeth R Jerison, Sergey Kryazhimskiy, James Kameron Mitchell, Joshua S Bloom, Leonid Kruglyak, and Michael M Desai. Genetic variation in adaptability and pleiotropy in budding yeast. *eLife*, 6, 2017.

[3120] Zhejian Ji, Haishan Gao, Luying Jia, Bing Li, and Hongtao Yu. A sequential multi-target Mps1 phosphorylation cascade promotes spindle checkpoint signaling. *eLife*, 6, 2017.

[3121] Tao Jiang, Katie Kindt, and Doris K Wu. Transcription factor Emx2 controls stereociliary bundle orientation of sensory hair cells. *eLife*, 6, 2017.

[3122] Tao Jiang, Kuai Yu, H Criss Hartzell, and Emad Tajkhorshid. Lipids and ions traverse the membrane by the same physical pathway in the nhTMEM16 scramblase. *eLife*, 6, 2017.

[3123] Yuwei Jiang, Daniel C Berry, and Jonathan M Graff. Distinct cellular and molecular mechanisms for $\beta$3 adrenergic receptor-induced beige adipocyte formation. *eLife*, 6, 2017.

[3124] Hui Jing, Xiaoyu Zhang, Stephanie A Wisner, Xiao Chen, Nicole A Spiegelman, Maurine E Linder, and Hening Lin. SIRT2 and lysine fatty acylation regulate the transforming activity of K-Ras4a. *eLife*, 6, 2017.

[3125] Jeanyoung Jo, Krista L Cortez, William Cole Cornell, Alexa Price-Whelan, and Lars EP Dietrich. An orphan cbb3-type cytochrome oxidase subunit supports Pseudomonas aeruginosa biofilm growth and virulence. *eLife*, 6, 2017.

[3126] Whitney L Johnson, William T Yewdell, Jason C Bell, Shannon M McNulty, Zachary Duda, Rachel J O'Neill, Beth A Sullivan, and Aaron F Straight. RNA-dependent stabilization of SUV39H1 at constitutive heterochromatin. *eLife*, 6, 2017.

[3127] Scott L Jones, Minh-Son To, and Greg J Stuart. Dendritic small conductance calcium-activated potassium channels activated by action potentials suppress EPSPs and gate spike-timing dependent synaptic plasticity. *eLife*, 6, 2017.

[3128] Stephanie E Jones, Louis Ho, Christiaan A Rees, Jane E Hill, Justin R Nodwell, and Marie A Elliot. Streptomyces exploration is triggered by fungal interactions and volatile signals. *eLife*, 6, 2017.

[3129] Ryan M Joseph, Jennifer S Sun, Edric Tam, and John R Carlson. A receptor and neuron that activate a circuit limiting sucrose consumption. *eLife*, 6, 2017.

[3130] Shai R Joseph, Máté Pálfy, Lennart Hilbert, Mukesh Kumar, Jens Karschau, Vasily Zaburdaev, Andrej Shevchenko, and Nadine L Vastenhouw. Competition between histone and transcription factor binding regulates the onset of transcription in zebrafish embryos. *eLife*, 6, 2017.

[3131] Ana Lilia Juárez-Vázquez, Janaka N Edirisinghe, Ernesto A Verduzco-Castro, Karolina Michalska, Chenggang Wu, Lianet Noda-García, Gyorgy Babnigg, Michael Endres, Sofía Medina-Ruíz, Julián Santoyo-Flores, Mauricio Carrillo-Tripp, Hung Ton-That, Andrzej Joachimiak, Christopher S Henry, and Francisco Barona-Gómez. Evolution of substrate specificity in a retained enzyme driven by gene loss. *eLife*, 6, 2017.

[3132] Jennifer Jung, Arnab Nayak, Véronique Schaeffer, Tatjana Starzetz, Achim K Kirsch, Stefan Müller, Ivan Dikic, Michel Mittelbronn, and Christian Behrends. Multiplex image-based autophagy RNAi screening identifies SMCR8 as ULK1 kinase activity and gene expression regulator. *eLife*, 6, 2017.

[3133] Victoria Juskaite, David S Corcoran, and Birgit Leitinger. Collagen induces activation of DDR1 through lateral dimer association and phosphorylation between dimers. *eLife*, 6, 2017.

[3134] Malene Juul, Johanna Bertl, Qianyun Guo, Morten Muhlig Nielsen, Michał Świtnicki, Henrik Hornshøj, Tobias Madsen, Asger Hobolth, and Jakob Skou Pedersen. Non-coding cancer driver candidates identified with a sample- and position-specific model of the somatic mutation rate. *eLife*, 6, 2017.

[3135] Mikko Juusola, An Dau, Zhuoyi Song, Narendra Solanki, Diana Rien, David Jaciuch, Sidhartha Anil Dongre, Florence Blanchard, Gonzalo G de Polavieja, Roger C Hardie, and Jouni Takalo. Microsaccadic sampling of moving image information provides Drosophila hyperacute vision. *eLife*, 6, 2017.

[3136] Morten Egevang Jørgensen, Deyang Xu, Christoph Crocoll, Heidi Asschenfeldt Ernst, David Ramírez, Mohammed Saddik Motawia, Carl Erik Olsen, Osman Mirza, Hussam Hassan Nour-Eldin, and Barbara Ann Halkier. Origin and evolution of transporter substrate specificity within the NPF family. *eLife*, 6, 2017.

[3137] Aashiq H Kachroo, Jon M Laurent, Azat Akhmetov, Madelyn Szilagyi-Jones, Claire D McWhite, Alice Zhao, and Edward M Marcotte. Systematic bacterialization of yeast genes identifies a near-universally swappable pathway. *eLife*, 6, 2017.

[3138] Pornchai Kaewsapsak, David Michael Shechner, William Mallard, John L Rinn, and Alice Y Ting. Live-cell mapping of organelle-associated RNAs via proximity biotinylation combined with protein-RNA crosslinking. *eLife*, 6, 2017.

[3139] Irawati Kandela and Fraser Aird. Replication Study: Discovery and preclinical validation of drug indications using compendia of public gene expression data. *eLife*, 6, 2017.

[3140] Jin Young Kang, Paul Dominic B Olinares, James Chen, Elizabeth A Campbell, Arkady Mustaev, Brian T Chait, Max E Gottesman, and Seth A Darst. Structural basis of transcription arrest by coliphage HK022 Nun in an Escherichia coli RNA polymerase elongation complex. *eLife*, 6, 2017.

[3141] Scott A Kanner, Travis Morgenstern, and Henry M Colecraft. Sculpting ion channel functional expression with engineered ubiquitin ligases. *eLife*, 6, 2017.

[3142] Mark A Kanow, Michelle M Giarmarco, Connor SR Jankowski, Kristine Tsantilas, Abbi L Engel, Jianhai Du, Jonathan D Linton, Christopher C Farnsworth, Stephanie R Sloat, Austin Rountree, Ian R Sweet, Ken J Lindsay, Edward D Parker, Susan E Brockerhoff, Martin Sadilek, Jennifer R Chao, and James B Hurley. Biochemical adaptations of the retina and retinal pigment epithelium support a metabolic ecosystem in the vertebrate eye. *eLife*, 6, 2017.

[3143] G Elif Karagöz, Diego Acosta-Alvear, Hieu T Nguyen, Crystal P Lee, Feixia Chu, and Peter Walter. An unfolded protein-induced conformational switch activates mammalian IRE1. *eLife*, 6, 2017.

[3144] Akira Karasawa, Kevin Michalski, Polina Mikhelzon, and Toshimitsu Kawate. The P2X7 receptor forms a dye-permeable pore independent of its intracellular domain but dependent on membrane lipid composition. *eLife*, 6, 2017.

[3145] Jason Karch, Tobias G Schips, Bryan D Maliken, Matthew J Brody, Michelle A Sargent, Onur Kanisicak, and Jeffery D Molkentin. Autophagic cell death is dependent on lysosomal membrane permeability through Bax and Bak. *eLife*, 6, 2017.

[3146] Matthias Karnahl, Misoon Park, Ulrike Mayer, Ulrike Hiller, and Gerd Jürgens. ER assembly of SNARE complexes mediating formation of partitioning membrane in Arabidopsis cytokinesis. *eLife*, 6, 2017.

[3147] Ioannis Kasioulis, Raman M Das, and Kate G Storey. Inter-dependent apical microtubule and actin dynamics orchestrate centrosome retention and neuronal delamination. *eLife*, 6, 2017.

[3148] Peter Kasson, Frank DiMaio, Xiong Yu, Soizick Lucas-Staat, Mart Krupovic, Stefan Schouten, David Prangishvili, and Edward H Egelman. Model for a novel membrane envelope in a filamentous hyperthermophilic virus. *eLife*, 6, 2017.

[3149] Marketa Kaucka, Tomas Zikmund, Marketa Tesarova, Daniel Gyllborg, Andreas Hellander, Josef Jaros, Jozef Kaiser, Julian Petersen, Bara Szarowska, Phillip T Newton, Vyacheslav Dyachuk, Lei Li, Hong Qian, Anne-Sofie Johansson, Yuji Mishina, Joshua D Currie, Elly M Tanaka, Alek Erickson, Andrew Dudley, Hjalmar Brismar, Paul Southam, Enrico Coen, Min Chen, Lee S Weinstein, Ales Hampl, Ernest Arenas, Andrei S Chagin, Kaj Fried, and Igor Adameyko. Oriented clonal cell dynamics enables accurate growth and shaping of vertebrate cartilage. *eLife*, 6, 2017.

[3150] Kendrick N Kay and Jason D Yeatman. Bottom-up and top-down computations in word- and face-selective cortex. *eLife*, 6, 2017.

[3151] Lukasz Kedzierski, Michelle D Tate, Alan C Hsu, Tatiana B Kolesnik, Edmond M Linossi, Laura Dagley, Zhaoguang Dong, Sarah Freeman, Giuseppe Infusini, Malcolm R Starkey, Nicola L Bird, Simon M Chatfield, Jeffrey J Babon, Nicholas Huntington, Gabrielle Belz, Andrew Webb, Peter AB Wark, Nicos A Nicola, Jianqing Xu, Katherine Kedzierska, Philip M Hansbro, and Sandra E Nicholson. Suppressor of cytokine signaling (SOCS)5 ameliorates influenza infection via inhibition of EGFR signaling. *eLife*, 6, 2017.

[3152] Jutta Keicher, Nina Jaspert, Katrin Weckermann, Claudia Möller, Christian Throm, Aaron Kintzi, and Claudia Oecking. Arabidopsis 14-3-3 epsilon members contribute to polarity of PIN auxin carrier and auxin transport-related development. *eLife*, 6, 2017.

[3153] Christian Keine, Rudolf Rübsamen, and Bernhard Englitz. Signal integration at spherical bushy cells enhances representation of temporal structure but limits its range. *eLife*, 6, 2017.

[3154] Andreas J Keller, Rachael Houlton, Björn M Kampa, Nicholas A Lesica, Thomas D Mrsic-Flogel, Georg B Keller, and Fritjof Helmchen. Stimulus relevance modulates contrast adaptation in visual cortex. *eLife*, 6, 2017.

[3155] Timothy W R Kelso, Devin K Porter, Maria Luisa Amaral, Maxim N Shokhirev, Christopher Benner, and Diana C Hargreaves. Chromatin accessibility underlies synthetic lethality of SWI/SNF subunits in ARID1A-mutant cancers. *eLife*, 6, 2017.

[3156] Elaine F Kenny, Alf Herzig, Renate Krüger, Aaron Muth, Santanu Mondal, Paul R Thompson, Volker Brinkmann, Horst von Bernuth, and Arturo Zychlinsky. Diverse stimuli engage different neutrophil extracellular trap pathways. *eLife*, 6, 2017.

[3157] David M Kern, Julie K Monda, Kuan-Chung Su, Elizabeth M Wilson-Kubalek, and Iain M Cheeseman. Astrin-SKAP complex reconstitution reveals its kinetochore interaction with microtubule-bound Ndc80. *eLife*, 6, 2017.

[3158] Andreas Keymer, Priya Pimprikar, Vera Wewer, Claudia Huber, Mathias Brands, Simone L Bucerius, Pierre-Marc Delaux, Verena Klingl, Edda von Röpenack-Lahaye, Trevor L Wang, Wolfgang Eisenreich, Peter Dörmann, Martin Parniske, and Caroline Gutjahr. Lipid transfer from plants to arbuscular mycorrhiza fungi. *eLife*, 6, 2017.

[3159] Rohan J Khadilkar, Wayne Vogl, Katharine Goodwin, and Guy Tanentzapf. Modulation of occluding junctions alters the hematopoietic niche to trigger immune activation. *eLife*, 6, 2017.

[3160] Preeya Khanna and Jose M Carmena. Beta band oscillations in motor cortex reflect neural population signals that delay movement onset. *eLife*, 6, 2017.

[3161] Mohamed Khateb, Jackie Schiller, and Yitzhak Schiller. Feedforward motor information enhances somatosensory responses and sharpens angular tuning of rat S1 barrel cortex neurons. *eLife*, 6, 2017.

[3162] Vikram Khedgikar, Genevieve Abbruzzese, Ketan Mathavan, Hannah Szydlo, Helene Cousin, and Dominique Alfandari. Dual control of pcdh8l/PCNS expression and function in Xenopus laevis neural crest cells by adam13/33 via the transcription factors tfap2$\alpha$ and arid3a. *eLife*, 6, 2017.

[3163] Jian Ming Khor and Charles A Ettensohn. Functional divergence of paralogous transcription factors supported the evolution of biomineralization in echinoderms. *eLife*, 6, 2017.

[3164] Pierre Khoueiry, Charles Girardot, Lucia Ciglar, Pei-Chen Peng, E Hilary Gustafson, Saurabh Sinha, and Eileen EM Furlong. Uncoupling evolutionary changes in DNA sequence, transcription factor occupancy and enhancer activity. *eLife*, 6, 2017.

[3165] Collin Kieffer, Mark S Ladinsky, Allen Ninh, Rachel P Galimidi, and Pamela J Bjorkman. Longitudinal imaging of HIV-1 spread in humanized mice with parallel 3D immunofluorescence and electron tomography. *eLife*, 6, 2017.

[3166] Byunghyuk Kim and Scott W Emmons. Multiple conserved cell adhesion protein interactions mediate neural wiring of a sensory circuit in C. elegans. *eLife*, 6, 2017.

[3167] Daeyeon Kim, Mar Alvarez, Laura M Lechuga, and Matthieu Louis. Species-specific modulation of food-search behavior by respiration and chemosensation in Drosophila larvae. *eLife*, 6, 2017.

[3168] Heesoo Kim, Colleen Kirkhart, and Kristin Scott. Long-range projection neurons in the taste circuit of Drosophila. *eLife*, 6, 2017.

[3169] Jae ook Kim, Alex Zelter, Neil T Umbreit, Athena Bollozos, Michael Riffle, Richard Johnson, Michael J MacCoss, Charles L Asbury, and Trisha N Davis. The Ndc80 complex bridges two Dam1 complex rings. *eLife*, 6, 2017.

[3170] Min Woo Kim, Wenjing Wang, Mateo I Sanchez, Robert Coukos, Mark von Zastrow, and Alice Y Ting. Time-gated detection of protein-protein interactions with transcriptional readout. *eLife*, 6, 2017.

[3171] Seungsoo Kim, Ivan Liachko, Donna G Brickner, Kate Cook, William S Noble, Jason H Brickner, Jay Shendure, and Maitreya J Dunham. The dynamic three-dimensional organization of the diploid yeast genome. *eLife*, 6, 2017.

[3172] Yeha Kim, Soyeon Lim, Taejeong Ha, You-Hyang Song, Young-In Sohn, Dae-Jin Park, Sun-Sook Paik, Joo-ri Kim-Kaneyama, Mi-Ryoung Song, Amanda Leung, Edward M Levine, In-Beom Kim, Yong Sook Goo, Seung-Hee Lee, Kyung Hwa Kang, and Jin Woo Kim. The LIM protein complex establishes a retinal circuitry of visual adaptation by regulating Pax6 $\alpha$-enhancer activity. *eLife*, 6, 2017.

[3173] David Kimelman, Natalie L Smith, Jason Kuan Han Lai, and Didier YR Stainier. Regulation of posterior body and epidermal morphogenesis in zebrafish by localized Yap1 and Wwtr1. *eLife*, 6, 2017.

[3174] Makoto Kimura, Yuriko Morinaka, Kenichiro Imai, Shingo Kose, Paul Horton, and Naoko Imamoto. Extensive cargo identification reveals distinct biological roles of the 12 importin pathways. *eLife*, 6, 2017.

[3175] Hamish W King and Robert J Klose. The pioneer factor OCT4 requires the chromatin remodeller BRG1 to support gene regulatory element function in mouse embryonic stem cells. *eLife*, 6, 2017.

[3176] Stuart RF King. Consultative review is worth the wait. *eLife*, 6, 2017.

[3177] Stuart RF King, Emma Pewsey, and Sarah Shailes. An inside guide to eLife digests. *eLife*, 6, 2017.

[3178] Julianne Kitevski-LeBlanc, Amélie Fradet-Turcotte, Predrag Kukic, Marcus D Wilson, Guillem Portella, Tairan Yuwen, Stephanie Panier, Shili Duan, Marella D Canny, Hugo van Ingen, Cheryl H Arrowsmith, John L Rubinstein, Michele Vendruscolo, Daniel Durocher, and Lewis E Kay. The RNF168 paralog RNF169 defines a new class of ubiquitylated histone reader involved in the response to DNA damage. *eLife*, 6, 2017.

[3179] Matthew P Klassen, Christian J Peters, Shiwei Zhou, Hannah H Williams, Lily Yeh Jan, and Yuh Nung Jan. Age-dependent diastolic heart failure in an in vivo Drosophila model. *eLife*, 6, 2017.

[3180] Mason Klein, Sergei V Krivov, Anggie J Ferrer, Linjiao Luo, Aravinthan DT Samuel, and Martin Karplus. Exploratory search during directed navigation in C. elegans and Drosophila larva. *eLife*, 6, 2017.

[3181] Sandra Kleinecke, Sarah Richert, Livia de Hoz, Britta Brügger, Theresa Kungl, Ebrahim Asadollahi, Susanne Quintes, Judith Blanz, Rhona McGonigal, Kobra Naseri, Michael W Sereda, Timo Sachsenheimer, Christian Lüchtenborg, Wiebke Möbius, Hugh Willison, Myriam Baes, Klaus-Armin Nave, and Celia Michèle Kassmann. Peroxisomal dysfunctions cause lysosomal storage and axonal Kv1 channel redistribution in peripheral neuropathy. *eLife*, 6, 2017.

[3182] Michael Kleyman, Emre Sefer, Teodora Nicola, Celia Espinoza, Divya Chhabra, James S Hagood, Naftali Kaminski, Namasivayam Ambalavanan, and Ziv Bar-Joseph. Selecting the most appropriate time points to profile in high-throughput studies. *eLife*, 6, 2017.

[3183] Björn Udo Klink, Eldar Zent, Puneet Juneja, Anne Kuhlee, Stefan Raunser, and Alfred Wittinghofer. A recombinant BBSome core complex and how it interacts with ciliary cargo. *eLife*, 6, 2017.

[3184] Viktoria Klippenstein, Christian Hoppmann, Shixin Ye, Lei Wang, and Pierre Paoletti. Optocontrol of glutamate receptor activity by single side-chain photoisomerization. *eLife*, 6, 2017.

[3185] Tiemo J Klisch, Anna Vainshtein, Akash J Patel, and Huda Y Zoghbi. Jak2-mediated phosphorylation of Atoh1 is critical for medulloblastoma growth. *eLife*, 6, 2017.

[3186] Peter J Klover, Rajesh L Thangapazham, Jiro Kato, Ji-an Wang, Stasia A Anderson, Victoria Hoffmann, Wendy K Steagall, Shaowei Li, Elizabeth McCart, Neera Nathan, Joshua D Bernstock, Matthew D Wilkerson, Clifton L Dalgard, Joel Moss, and Thomas N Darling. Tsc2 disruption in mesenchymal progenitors results in tumors with vascular anomalies overexpressing Lgals3. *eLife*, 6, 2017.

[3187] Dennis Klug and Friedrich Frischknecht. Motility precedes egress of malaria parasites from oocysts. *eLife*, 6, 2017.

[3188] Niklas Klusch, Bonnie J Murphy, Deryck J Mills, özkan Yildiz, and Werner Kühlbrandt. Structural basis of proton translocation and force generation in mitochondrial ATP synthase. *eLife*, 6, 2017.

[3189] Steven Knafo, Kevin Fidelin, Andrew Prendergast, Po-En Brian Tseng, Alexandre Parrin, Charles Dickey, Urs Lucas Böhm, Sophie Nunes Figueiredo, Olivier Thouvenin, Hugues Pascal-Moussellard, and Claire Wyart. Mechanosensory neurons control the timing of spinal microcircuit selection during locomotion. *eLife*, 6, 2017.

[3190] Zachary A Knecht, Ana F Silbering, Joyner Cruz, Ludi Yang, Vincent Croset, Richard Benton, and Paul A Garrity. Ionotropic Receptor-dependent moist and dry cells control hygrosensation in Drosophila. *eLife*, 6, 2017.

[3191] Masashi Kondo, Kenta Kobayashi, Masamichi Ohkura, Junichi Nakai, and Masanori Matsuzaki. Two-photon calcium imaging of the medial prefrontal cortex and hippocampus without cortical invasion. *eLife*, 6, 2017.

[3192] Yannick Kopp, Wei-Han Lang, Tobias B Schuster, Adrián Martínez-Limón, Harald F Hofbauer, Robert Ernst, Giulia Calloni, and R Martin Vabulas. CHIP as a membrane-shuttling proteostasis sensor. *eLife*, 6, 2017.

[3193] Jörgen Kornfeld, Sam E Benezra, Rajeevan T Narayanan, Fabian Svara, Robert Egger, Marcel Oberlaender, Winfried Denk, and Michael A Long. EM connectomics reveals axonal target variation in a sequence-generating network. *eLife*, 6, 2017.

[3194] Hiroaki Koseki, Matteo Donegá, Brian YH Lam, Veselina Petrova, Susan van Erp, Giles SH Yeo, Jessica CF Kwok, Charles ffrench Constant, Richard Eva, and James W Fawcett. Selective rab11 transport and the intrinsic regenerative ability of CNS axons. *eLife*, 6, 2017.

[3195] Eleni Kougioumtzidou, Takahiro Shimizu, Nicola B Hamilton, Koujiro Tohyama, Rolf Sprengel, Hannah Monyer, David Attwell, and William D Richardson. Signalling through AMPA receptors on oligodendrocyte precursors promotes myelination by enhancing oligodendrocyte survival. *eLife*, 6, 2017.

[3196] Onur Ozan Koyluoglu, Yoni Pertzov, Sanjay Manohar, Masud Husain, and Ila R Fiete. Fundamental bound on the persistence and capacity of short-term memory stored as graded persistent activity. *eLife*, 6, 2017.

[3197] Agnė Kozlovskaja-Gumbrienė, Ren Yi, Richard Alexander, Andy Aman, Ryan Jiskra, Danielle Nagelberg, Holger Knaut, Melainia McClain, and Tatjana Piotrowski. Proliferation-independent regulation of organ size by Fgf/Notch signaling. *eLife*, 6, 2017.

[3198] Elsa B Krall, Belinda Wang, Diana M Munoz, Nina Ilic, Srivatsan Raghavan, Matthew J Niederst, Kristine Yu, David A Ruddy, Andrew J Aguirre, Jong Wook Kim, Amanda J Redig, Justin F Gainor, Juliet A Williams, John M Asara, John G Doench, Pasi A Janne, Alice T Shaw, Robert E McDonald III, Jeffrey A Engelman, Frank Stegmeier, Michael R Schlabach, and William C Hahn. KEAP1 loss modulates sensitivity to kinase targeted therapy in lung cancer. *eLife*, 6, 2017.

[3199] Paschalis Kratsios, Sze Yen Kerk, Catarina Catela, Joseph Liang, Berta Vidal, Emily A Bayer, Weidong Feng, Estanisla Daniel De La Cruz, Laura Croci, G Giacomo Consalez, Kota Mizumoto, and Oliver Hobert. An intersectional gene regulatory strategy defines subclass diversity of C. elegans motor neurons. *eLife*, 6, 2017.

[3200] Michael Krieg, Jan Stühmer, Juan G Cueva, Richard Fetter, Kerri Spilker, Daniel Cremers, Kang Shen, Alexander R Dunn, and Miriam B Goodman. Genetic defects in $\beta$-spectrin and tau sensitize C. elegans axons to movement-induced damage via torque-tension coupling. *eLife*, 6, 2017.

[3201] Pascal Krotee, Jose A Rodriguez, Michael R Sawaya, Duilio Cascio, Francis E Reyes, Dan Shi, Johan Hattne, Brent L Nannenga, Marie E Oskarsson, Stephan Philipp, Sarah Griner, Lin Jiang, Charles G Glabe, Gunilla T Westermark, Tamir Gonen, and David S Eisenberg. Atomic structures of fibrillar segments of hIAPP suggest tightly mated $\beta$-sheets are important for cytotoxicity. *eLife*, 6, 2017.

[3202] Amy T Ku, Timothy M Shaver, Ajay S Rao, Jeffrey M Howard, Christine N Rodriguez, Qi Miao, Gloria Garcia, Diep Le, Diane Yang, Malgorzata Borowiak, Daniel N Cohen, Vida Chitsazzadeh, Abdul H Diwan, Kenneth Y Tsai, and Hoang Nguyen. TCF7L1 promotes skin tumorigenesis independently of $\beta$-catenin through induction of LCN2. *eLife*, 6, 2017.

[3203] Zheng Kuang, Sudarshan Pinglay, Hongkai Ji, and Jef D Boeke. Msn2/4 regulate expression of glycolytic enzymes and control transition from quiescence to growth. *eLife*, 6, 2017.

[3204] Bridget M Kuehn. Rooting out bias. *eLife*, 6, 2017.

[3205] Bridget M Kuehn. The value of a healthy relationship. *eLife*, 6, 2017.

[3206] Inge Kühl, Maria Miranda, Ilian Atanassov, Irina Kuznetsova, Yvonne Hinze, Arnaud Mourier, Aleksandra Filipovska, and Nils-Göran Larsson. Transcriptomic and proteomic landscape of mitochondrial dysfunction reveals secondary coenzyme Q deficiency in mammals. *eLife*, 6, 2017.

[3207] Atul Kumar, Jevgenia Tamjar, Andrew D Waddell, Helen I Woodroof, Olawale G Raimi, Andrew M Shaw, Mark Peggie, Miratul MK Muqit, and Daan MF van Aalten. Structure of PINK1 and mechanisms of Parkinson's disease-associated mutations. *eLife*, 6, 2017.

[3208] Dhivya Kumar, Daniela Strenkert, Ramila S Patel-King, Michael T Leonard, Sabeeha S Merchant, Richard E Mains, Stephen M King, and Betty A Eipper. A bioactive peptide amidating enzyme is required for ciliogenesis. *eLife*, 6, 2017.

[3209] C Leopold Kurz, Bernard Charroux, Delphine Chaduli, Annelise Viallat-Lieutaud, and Julien Royet. Peptidoglycan sensing by octopaminergic neurons modulates Drosophila oviposition. *eLife*, 6, 2017.

[3210] Antonija Kuzmanic, Ludovico Sutto, Giorgio Saladino, Angel R Nebreda, Francesco L Gervasio, and Modesto Orozco. Changes in the free-energy landscape of p38$\alpha$ MAP kinase through its canonical activation and binding events as studied by enhanced molecular dynamics simulations. *eLife*, 6, 2017.

[3211] Mato Lagator, Tiago Paixão, Nicholas H Barton, Jonathan P Bollback, and Călin C Guet. On the mechanistic nature of epistasis in a canonical cis-regulatory element. *eLife*, 6, 2017.

[3212] Mato Lagator, Srdjan Sarikas, Hande Acar, Jonathan P Bollback, and Călin C Guet. Regulatory network structure determines patterns of intermolecular epistasis. *eLife*, 6, 2017.

[3213] Roni M Lahr, Bruno D Fonseca, Gabrielle E Ciotti, Hiba A Al-Ashtal, Jian-Jun Jia, Marius R Niklaus, Sarah P Blagden, Tommy Alain, and Andrea J Berman. La-related protein 1 (LARP1) binds the mRNA cap, blocking eIF4F assembly on TOP mRNAs. *eLife*, 6, 2017.

[3214] Garet P Lahvis. Unbridle biomedical research from the laboratory cage. *eLife*, 6, 2017.

[3215] Shih-Lei Lai, Rubén Marín-Juez, Pedro Luís Moura, Carsten Kuenne, Jason Kuan Han Lai, Ayele Taddese Tsedeke, Stefan Guenther, Mario Looso, and Didier YR Stainier. Reciprocal analyses in zebrafish and medaka reveal that harnessing the immune response promotes cardiac regeneration. *eLife*, 6, 2017.

[3216] Emma E Laing, Carla S Möller-Levet, Norman Poh, Nayantara Santhi, Simon N Archer, and Derk-Jan Dijk. Blood transcriptome based biomarkers for human circadian phase. *eLife*, 6, 2017.

[3217] Lance Langston and Mike O'Donnell. Action of CMG with strand-specific DNA blocks supports an internal unwinding mode for the eukaryotic replicative helicase. *eLife*, 6, 2017.

[3218] Lance D Langston, Ryan Mayle, Grant D Schauer, Olga Yurieva, Daniel Zhang, Nina Y Yao, Roxana E Georgescu, and Mike E O'Donnell. Mcm10 promotes rapid isomerization of CMG-DNA for replisome bypass of lagging strand DNA blocks. *eLife*, 6, 2017.

[3219] Pierre Lapaquette, Sabrina Fritah, Nouara Lhocine, Alexandra Andrieux, Giulia Nigro, Joëlle Mounier, Philippe Sansonetti, and Anne Dejean. Shigella entry unveils a calcium/calpain-dependent mechanism for inhibiting sumoylation. *eLife*, 6, 2017.

[3220] Ivan Larderet, Pauline MJ Fritsch, Nanae Gendre, G Larisa Neagu-Maier, Richard D Fetter, Casey M Schneider-Mizell, James W Truman, Marta Zlatic, Albert Cardona, and Simon G Sprecher. Organization of the Drosophila larval visual circuit. *eLife*, 6, 2017.

[3221] Martin Larhammar, Sarah Huntwork-Rodriguez, Zhiyu Jiang, Hilda Solanoy, Arundhati Sengupta Ghosh, Bei Wang, Joshua S Kaminker, Kevin Huang, Jeffrey Eastham-Anderson, Michael Siu, Zora Modrusan, Madeline M Farley, Marc Tessier-Lavigne, Joseph W Lewcock, and Trent A Watkins. Dual leucine zipper kinase-dependent PERK activation contributes to neuronal degeneration following insult. *eLife*, 6, 2017.

[3222] Joshua Donald Larson and Aaron A Hoskins. Dynamics and consequences of spliceosome E complex formation. *eLife*, 6, 2017.

[3223] Nicholas B Last, Senmiao Sun, Minh C Pham, and Christopher Miller. Molecular determinants of permeation in a fluoride-specific ion channel. *eLife*, 6, 2017.

[3224] Jean Laurens, Sheng Liu, Xiong-Jie Yu, Raymond Chan, David Dickman, Gregory C DeAngelis, and Dora E Angelaki. Transformation of spatiotemporal dynamics in the macaque vestibular system from otolith afferents to cortex. *eLife*, 6, 2017.

[3225] My-Tra Le, Wojciech K Kasprzak, Taejin Kim, Feng Gao, Megan YL Young, Xuefeng Yuan, Bruce A Shapiro, Joonil Seog, and Anne E Simon. Folding behavior of a T-shaped, ribosome-binding translation enhancer implicated in a wide-spread conformational switch. *eLife*, 6, 2017.

[3226] Johanne Le Coq, Marta Camacho-Artacho, José Vicente Velázquez, Clara M Santiveri, Luis Heredia Gallego, Ramón Campos-Olivas, Nicole Dölker, and Daniel Lietha. Structural basis for interdomain communication in SHIP2 providing high phosphatase activity. *eLife*, 6, 2017.

[3227] Jean-FranÇois Le Garrec, Jorge N Domínguez, Audrey Desgrange, Kenzo D Ivanovitch, Etienne Raphaël, J Andrew Bangham, Miguel Torres, Enrico Coen, Timothy J Mohun, and Sigolène M Meilhac. A predictive model of asymmetric morphogenesis from 3D reconstructions of mouse heart looping dynamics. *eLife*, 6, 2017.

[3228] Stephanie L Leal, Susan M Landau, Rachel K Bell, and William J Jagust. Hippocampal activation is associated with longitudinal amyloid accumulation and cognitive decline. *eLife*, 6, 2017.

[3229] Salvatore Lecca, Frank Julius Meye, Massimo Trusel, Anna Tchenio, Julia Harris, Martin Karl Schwarz, Denis Burdakov, Francois Georges, and Manuel Mameli. Aversive stimuli drive hypothalamus-to-habenula excitation to promote escape behavior. *eLife*, 6, 2017.

[3230] Chih-Yung Sean Lee, Tu Lu, and Geraldine Seydoux. Nanos promotes epigenetic reprograming of the germline by down-regulation of the THAP transcription factor LIN-15B. *eLife*, 6, 2017.

[3231] Daniel A Lee, Andrey Andreev, Thai V Truong, Audrey Chen, Andrew J Hill, Grigorios Oikonomou, Uyen Pham, Young K Hong, Steven Tran, Laura Glass, Viveca Sapin, Jae Engle, Scott E Fraser, and David A Prober. Genetic and neuronal regulation of sleep by neuropeptide VF. *eLife*, 6, 2017.

[3232] Jennifer K Lee, Peter T Hallock, and Steven J Burden. Abelson tyrosine-protein kinase 2 regulates myoblast proliferation and controls muscle fiber length. *eLife*, 6, 2017.

[3233] Kenneth Pak Kin Lee, Jue Chen, and Roderick MacKinnon. Molecular structure of human KATP in complex with ATP and ADP. *eLife*, 6, 2017.

[3234] Kunwoo Lee, Vanessa A Mackley, Anirudh Rao, Anthony T Chong, Mark A Dewitt, Jacob E Corn, and Niren Murthy. Synthetically modified guide RNA and donor DNA are a versatile platform for CRISPR-Cas9 engineering. *eLife*, 6, 2017.

[3235] Norman Lee and Andrew C Mason. How spatial release from masking may fail to function in a highly directional auditory system. *eLife*, 6, 2017.

[3236] Pedro Lee, Rupali Gund, Abhik Dutta, Neha Pincha, Isha Rana, Subhasri Ghosh, Deborah Witherden, Eve Kandyba, Amanda MacLeod, Krzysztof Kobielak, Wendy L Havran, and Colin Jamora. Stimulation of hair follicle stem cell proliferation through an IL-1 dependent activation of $\gamma\delta$T-cells. *eLife*, 6, 2017.

[3237] Sora Lee, Jessica M Tumolo, Aaron C Ehlinger, Kristin K Jernigan, Susan J Qualls-Histed, Pi-Chiang Hsu, W Hayes McDonald, Walter J Chazin, and Jason A MacGurn. Ubiquitin turnover and endocytic trafficking in yeast are regulated by Ser57 phosphorylation of ubiquitin. *eLife*, 6, 2017.

[3238] Yuh Chwen G Lee and Gary H Karpen. Pervasive epigenetic effects of Drosophila euchromatic transposable elements impact their evolution. *eLife*, 6, 2017.

[3239] John A Lees, Nicholas J Croucher, David Goldblatt, FranÇois Nosten, Julian Parkhill, Claudia Turner, Paul Turner, and Stephen D Bentley. Genome-wide identification of lineage and locus specific variation associated with pneumococcal carriage duration. *eLife*, 6, 2017.

[3240] Jérémie Lefebvre, Axel Hutt, and Flavio Frohlich. Stochastic resonance mediates the state-dependent effect of periodic stimulation on cortical alpha oscillations. *eLife*, 6, 2017.

[3241] Marco Lepore, Artem Kalinichenko, Salvatore Calogero, Pavanish Kumar, Bhairav Paleja, Mathias Schmaler, Vipin Narang, Francesca Zolezzi, Michael Poidinger, Lucia Mori, and Gennaro De Libero. Functionally diverse human T cells recognize non-microbial antigens presented by MR1. *eLife*, 6, 2017.

[3242] Josien Levenga, Helen Wong, Ryan A Milstead, Bailey N Keller, Lauren E LaPlante, and Charles A Hoeffer. AKT isoforms have distinct hippocampal expression and roles in synaptic plasticity. *eLife*, 6, 2017.

[3243] Jacob S Lewis, Lisanne M Spenkelink, Slobodan Jergic, Elizabeth A Wood, Enrico Monachino, Nicholas P Horan, Karl E Duderstadt, Michael M Cox, Andrew Robinson, Nicholas E Dixon, and Antoine M van Oijen. Single-molecule visualization of fast polymerase turnover in the bacterial replisome. *eLife*, 6, 2017.

[3244] Eugene Leypunskiy, Jenny Lin, Haneul Yoo, UnJin Lee, Aaron R Dinner, and Michael J Rust. The cyanobacterial circadian clock follows midday in vivo and in vitro. *eLife*, 6, 2017.

[3245] Bingshuo Li, Juha P Virtanen, Axel Oeltermann, Cornelius Schwarz, Martin A Giese, Ulf Ziemann, and Alia Benali. Lifting the veil on the dynamics of neuronal activities evoked by transcranial magnetic stimulation. *eLife*, 6, 2017.

[3246] Dan Li, Wenhui He, Ximing Liu, Sanduo Zheng, Yonghe Qi, Huiyu Li, Fengfeng Mao, Juan Liu, Yinyan Sun, Lijing Pan, Kaixin Du, Keqiong Ye, Wenhui Li, and Jianhua Sui. A potent human neutralizing antibody Fc-dependently reduces established HBV infections. *eLife*, 6, 2017.

[3247] Dianrong Li, Lingjun Meng, Tao Xu, Yaning Su, Xiao Liu, Zhiyuan Zhang, and Xiaodong Wang. RIPK1-RIPK3-MLKL-dependent necrosis promotes the aging of mouse male reproductive system. *eLife*, 6, 2017.

[3248] Dongming Li, Ana Marie S Palanca, So Youn Won, Lei Gao, Ying Feng, Ajay A Vashisht, Li Liu, Yuanyuan Zhao, Xigang Liu, Xiuyun Wu, Shaofang Li, Brandon Le, Yun Ju Kim, Guodong Yang, Shengben Li, Jinyuan Liu, James A Wohlschlegel, Hongwei Guo, Beixin Mo, Xuemei Chen, and Julie A Law. The MBD7 complex promotes expression of methylated transgenes without significantly altering their methylation status. *eLife*, 6, 2017.

[3249] Hanqing Li, Ash Watson, Agnieszka Olechwier, Michael Anaya, Siamak K Sorooshyari, Dermott P Harnett, Hyung-Kook (Peter) Lee, Jost Vielmetter, Mario A Fares, K Christopher Garcia, Engin özkan, Juan-Pablo Labrador, and Kai Zinn. Deconstruction of the beaten Path-Sidestep interaction network provides insights into neuromuscular system development. *eLife*, 6, 2017.

[3250] Jianneng Li, Mohammad Alyamani, Ao Zhang, Kai-Hsiung Chang, Michael Berk, Zhenfei Li, Ziqi Zhu, Marianne Petro, Cristina Magi-Galluzzi, Mary-Ellen Taplin, Jorge A Garcia, Kevin Courtney, Eric A Klein, and Nima Sharifi. Aberrant corticosteroid metabolism in tumor cells enables GR takeover in enzalutamide resistant prostate cancer. *eLife*, 6, 2017.

[3251] Jiaxing Li, Yao V Zhang, Elham Asghari Adib, Doychin T Stanchev, Xin Xiong, Susan Klinedinst, Pushpanjali Soppina, Thomas Robert Jahn, Richard I Hume, Tobias M Rasse, and Catherine A Collins. Restraint of presynaptic protein levels by Wnd/DLK signaling mediates synaptic defects associated with the kinesin-3 motor Unc-104. *eLife*, 6, 2017.

[3252] Jing Li, Jordan T White, Harry Saavedra, James O Wrabl, Hesam N Motlagh, Kaixian Liu, James Sowers, Trina A Schroer, E Brad Thompson, and Vincent J Hilser. Genetically tunable frustration controls allostery in an intrinsically disordered transcription factor. *eLife*, 6, 2017.

[3253] Liyi Li, Sébastien Sultan, Stefanie Heigele, Charlotte Schmidt-Salzmann, Nicolas Toni, and Josef Bischofberger. Silent synapses generate sparse and orthogonal action potential firing in adult-born hippocampal granule cells. *eLife*, 6, 2017.

[3254] Meng Amy Li, Paulo P Amaral, Priscilla Cheung, Jan H Bergmann, Masaki Kinoshita, Tüzer Kalkan, Meryem Ralser, Sam Robson, Ferdinand von Meyenn, Maike Paramor, Fengtang Yang, Caifu Chen, Jennifer Nichols, David L Spector, Tony Kouzarides, Lin He, and Austin Smith. A lncRNA fine tunes the dynamics of a cell state transition involving Lin28, let-7 and de novo DNA methylation. *eLife*, 6, 2017.

[3255] Xiaolei Li, Zhiqiang Wu, Xiaojing An, Qian Mei, Miaomiao Bai, Leena Hanski, Xiang Li, Tero Ahola, and Weidong Han. Blockade of the LRP16-PKR-NF-$\kappa$B signaling axis sensitizes colorectal carcinoma cells to DNA-damaging cytotoxic therapy. *eLife*, 6, 2017.

[3256] Yao Li, Yu Zhang, Yu Xu, Alec Kittredge, Nancy Ward, Shoudeng Chen, Stephen H Tsang, and Tingting Yang. Patient-specific mutations impair BESTROPHIN1's essential role in mediating Ca2+-dependent Cl- currents in human RPE. *eLife*, 6, 2017.

[3257] Yize Li, Shuvojit Banerjee, Stephen A Goldstein, Beihua Dong, Christina Gaughan, Sneha Rath, Jesse Donovan, Alexei Korennykh, Robert H Silverman, and Susan R Weiss. Ribonuclease L mediates the cell-lethal phenotype of double-stranded RNA editing enzyme ADAR1 deficiency in a human cell line. *eLife*, 6, 2017.

[3258] Yuwei Li, Ang Li, Jason Junge, and Marianne Bronner. Planar cell polarity signaling coordinates oriented cell division and cell rearrangement in clonally expanding growth plate cartilage. *eLife*, 6, 2017.

[3259] Ziwei Liang, Karen E Brown, Thomas Carroll, Benjamin Taylor, Isabel Ferreirós Vidal, Brian Hendrich, David Rueda, Amanda G Fisher, and Matthias Merkenschlager. A high-resolution map of transcriptional repression. *eLife*, 6, 2017.

[3260] Peng Liao, Weichao Wang, Yu Li, Rui Wang, Jiali Jin, Weijuan Pang, Yunfei Chen, Mingyue Shen, Xinbo Wang, Dongyang Jiang, Jinjiang Pang, Mingyao Liu, Xia Lin, Xin-Hua Feng, Ping Wang, and Xin Ge. Palmitoylated SCP1 is targeted to the plasma membrane and negatively regulates angiogenesis. *eLife*, 6, 2017.

[3261] Ann Lin, Christopher J Giuliano, Nicole M Sayles, and Jason M Sheltzer. CRISPR/Cas9 mutagenesis invalidates a putative cancer dependency targeted in on-going clinical trials. *eLife*, 6, 2017.

[3262] Audrie Lin, Benjamin F Arnold, Andrew N Mertens, Jue Lin, Jade Benjamin-Chung, Shahjahan Ali, Alan E Hubbard, Christine P Stewart, Abul K Shoab, Md Ziaur Rahman, Md Saheen Hossen, Palash Mutsuddi, Syeda L Famida, Salma Akther, Mahbubur Rahman, Leanne Unicomb, Firdaus S Dhabhar, Lia C H Fernald, John M Colford, and Stephen P Luby. Effects of water, sanitation, handwashing, and nutritional interventions on telomere length among children in a cluster-randomized controlled trial in rural Bangladesh. *eLife*, 6, 2017.

[3263] Chuwen Lin, Erica Yao, Kuan Zhang, Xuan Jiang, Stacey Croll, Katherine Thompson-Peer, and Pao-Tien Chuang. YAP is essential for mechanical force production and epithelial cell proliferation during lung branching morphogenesis. *eLife*, 6, 2017.

[3264] Jing-Ping Lin, Yevgeniya A Mironova, Peter Shrager, and Roman J Giger. LRP1 regulates peroxisome biogenesis and cholesterol homeostasis in oligodendrocytes and is required for proper CNS myelin development and repair. *eLife*, 6, 2017.

[3265] Julieta E Lischinsky, Katie Sokolowski, Peijun Li, Shigeyuki Esumi, Yasmin Kamal, Meredith Goodrich, Livio Oboti, Timothy R Hammond, Meera Krishnamoorthy, Daniel Feldman, Molly Huntsman, Judy Liu, and Joshua G Corbin. Embryonic transcription factor expression in mice predicts medial amygdala neuronal identity and sex-specific responses to innate behavioral cues. *eLife*, 6, 2017.

[3266] Michael J Litt, G Donald Okoye, Daniel Lark, Isin Cakir, Christy Moore, Mary C Barber, James Atkinson, Josh Fessel, Javid Moslehi, and Roger D Cone. Loss of the melanocortin-4 receptor in mice causes dilated cardiomyopathy. *eLife*, 6, 2017.

[3267] Daifei Liu, Xueqi Liu, Zhiguo Shang, and Charles V Sindelar. Structural basis of cooperativity in kinesin revealed by 3D reconstruction of a two-head-bound state on microtubules. *eLife*, 6, 2017.

[3268] Jie Liu, Christopher Ede, William D Wright, Steven K Gore, Shirin S Jenkins, Bret D Freudenthal, M Todd Washington, Xavier Veaute, and Wolf-Dietrich Heyer. Srs2 promotes synthesis-dependent strand annealing by disrupting DNA polymerase $\delta$-extending D-loops. *eLife*, 6, 2017.

[3269] Junke Liu, Zongyong Zhang, David Moreno-Delgado, James AR Dalton, Xavier Rovira, Ana Trapero, Cyril Goudet, Amadeu Llebaria, Jesús Giraldo, Qilin Yuan, Philippe Rondard, Siluo Huang, Jianfeng Liu, and Jean-Philippe Pin. Allosteric control of an asymmetric transduction in a G protein-coupled receptor heterodimer. *eLife*, 6, 2017.

[3270] Nan Liu, Yaxiong Yang, Lin Ge, Min Liu, Henry M Colecraft, and Xiaodong Liu. Cooperative and acute inhibition by multiple C-terminal motifs of L-type Ca2+ channels. *eLife*, 6, 2017.

[3271] Song Liu, Sangeeta Kumari, Qiang Hu, Dhirodatta Senapati, Varadha Balaji Venkadakrishnan, Dan Wang, Adam D DePriest, Simon E Schlanger, Salma Ben-Salem, Malyn May Valenzuela, Belinda Willard, Shaila Mudambi, Wendy M Swetzig, Gokul M Das, Mojgan Shourideh, Shahriah Koochekpour, Sara Moscovita Falzarano, Cristina Magi-Galluzzi, Neelu Yadav, Xiwei Chen, Changshi Lao, Jianmin Wang, Jean-Noel Billaud, and Hannelore V Heemers. A comprehensive analysis of coregulator recruitment, androgen receptor function and gene expression in prostate cancer. *eLife*, 6, 2017.

[3272] Wei Liu, Jing Li, Weinan Zheng, Yingli Shang, Zhendong Zhao, Shanshan Wang, Yuhai Bi, Shuang Zhang, Chongfeng Xu, Ziyuan Duan, Lianfeng Zhang, Yue L Wang, Zhengfan Jiang, Wenjun Liu, and Lei Sun. Cyclophilin A-regulated ubiquitination is critical for RIG-I-mediated antiviral immune responses. *eLife*, 6, 2017.

[3273] Wenxuan Liu, Alanna Klose, Sophie Forman, Nicole D Paris, Lan Wei-LaPierre, Mariela Cortés-Lopéz, Aidi Tan, Morgan Flaherty, Pedro Miura, Robert T Dirksen, and Joe V Chakkalakal. Loss of adult skeletal muscle stem cells drives age-related neuromuscular junction degeneration. *eLife*, 6, 2017.

[3274] Yi Liu, Mike Visetsouk, Michelle Mynlieff, Hongmin Qin, Karl F Lechtreck, and Pinfen Yang. H+- and Na+- elicited rapid changes of the microtubule cytoskeleton in the biflagellated green alga Chlamydomonas. *eLife*, 6, 2017.

[3275] Peter M LoCoco, April L Risinger, Hudson R Smith, Teresa S Chavera, Kelly A Berg, and William P Clarke. Pharmacological augmentation of nicotinamide phosphoribosyltransferase (NAMPT) protects against paclitaxel-induced peripheral neuropathy. *eLife*, 6, 2017.

[3276] Yicheng Long, Ben Bolanos, Lihu Gong, Wei Liu, Karen J Goodrich, Xin Yang, Siming Chen, Anne R Gooding, Karen A Maegley, Ketan S Gajiwala, Alexei Brooun, Thomas R Cech, and Xin Liu. Conserved RNA-binding specificity of polycomb repressive complex 2 is achieved by dispersed amino acid patches in EZH2. *eLife*, 6, 2017.

[3277] Marta Losa, Victor Latorre, Munazah Andrabi, Franck Ladam, Charles Sagerström, Ana Novoa, Peyman Zarrineh, Laure Bridoux, Neil A Hanley, Moises Mallo, and Nicoletta Bobola. A tissue-specific, Gata6-driven transcriptional program instructs remodeling of the mature arterial tree. *eLife*, 6, 2017.

[3278] Isabelle Louradour, Anurag Sharma, Ismael Morin-Poulard, Manon Letourneau, Alain Vincent, Michèle Crozatier, and Nathalie Vanzo. Reactive oxygen species-dependent Toll/NF-$\kappa$B activation in the Drosophila hematopoietic niche confers resistance to wasp parasitism. *eLife*, 6, 2017.

[3279] José LourenÇo, Maricelia Maia de Lima, Nuno Rodrigues Faria, Andrew Walker, Moritz UG Kraemer, Christian Julian Villabona-Arenas, Ben Lambert, Erenilde Marques de Cerqueira, Oliver G Pybus, Luiz CJ Alcantara, and Mario Recker. Epidemiological and ecological determinants of Zika virus transmission in an urban setting. *eLife*, 6, 2017.

[3280] Eric Lowet, Mark J Roberts, Alina Peter, Bart Gips, and Peter De Weerd. A quantitative theory of gamma synchronization in macaque V1. *eLife*, 6, 2017.

[3281] Olga Loza, Idse Heemskerk, Nadav Gordon-Bar, Liat Amir-Zilberstein, Yunmin Jung, and David Sprinzak. A synthetic planar cell polarity system reveals localized feedback on Fat4-Ds1 complexes. *eLife*, 6, 2017.

[3282] Kevin L Lu and Yukiko M Yamashita. Germ cell connectivity enhances cell death in response to DNA damage in the Drosophila testis. *eLife*, 6, 2017.

[3283] Matthias Lübbert, R Oliver Goral, Rachel Satterfield, Travis Putzke, Arn MJM van den Maagdenberg, Naomi Kamasawa, and Samuel M Young. A novel region in the CaV2.1 $\alpha1$ subunit C-terminus regulates fast synaptic vesicle fusion and vesicle docking at the mammalian presynaptic active zone. *eLife*, 6, 2017.

[3284] Elodie Luche, Virginie Robert, Vincent Cuminetti, Celine Pomié, Quentin Sastourné-Arrey, Aurélie Waget, Emmanuelle Arnaud, Audrey Varin, Elodie Labit, Patrick Laharrague, Remy Burcelin, Louis Casteilla, and Beatrice Cousin. Corrupted adipose tissue endogenous myelopoiesis initiates diet-induced metabolic disease. *eLife*, 6, 2017.

[3285] Pauline Luczynski, Monica Tramullas, Maria Viola, Fergus Shanahan, Gerard Clarke, Siobhain O'Mahony, Timothy G Dinan, and John F Cryan. Microbiota regulates visceral pain in the mouse. *eLife*, 6, 2017.

[3286] Johanna Ludwigsen, Sabrina Pfennig, Ashish K Singh, Christina Schindler, Nadine Harrer, Ignasi Forné, Martin Zacharias, and Felix Mueller-Planitz. Concerted regulation of ISWI by an autoinhibitory domain and the H4 N-terminal tail. *eLife*, 6, 2017.

[3287] Martijn S Luijsterburg, Dimitris Typas, Marie-Christine Caron, Wouter W Wiegant, Diana van den Heuvel, Rick A Boonen, Anthony M Couturier, Leon H Mullenders, Jean-Yves Masson, and Haico van Attikum. A PALB2-interacting domain in RNF168 couples homologous recombination to DNA break-induced chromatin ubiquitylation. *eLife*, 6, 2017.

[3288] Lydia V Luncz, Amanda Tan, Michael Haslam, Lars Kulik, Tomos Proffitt, Suchinda Malaivijitnond, and Michael Gumert. Resource depletion through primate stone technology. *eLife*, 6, 2017.

[3289] Ulrich Lutz, Thomas Nussbaumer, Manuel Spannagl, Julia Diener, Klaus FX Mayer, and Claus Schwechheimer. Natural haplotypes of FLM non-coding sequences fine-tune flowering time in ambient spring temperatures in Arabidopsis. *eLife*, 6, 2017.

[3290] Tal Luzzatto-Knaan, Neha Garg, Mingxun Wang, Evgenia Glukhov, Yao Peng, Gail Ackermann, Amnon Amir, Brendan M Duggan, Sergey Ryazanov, Lena Gerwick, Rob Knight, Theodore Alexandrov, Nuno Bandeira, William H Gerwick, and Pieter C Dorrestein. Digitizing mass spectrometry data to explore the chemical diversity and distribution of marine cyanobacteria and algae. *eLife*, 6, 2017.

[3291] Tony Ly, Arlene Whigham, Rosemary Clarke, Alejandro J Brenes-Murillo, Brett Estes, Diana Madhessian, Emma Lundberg, Patricia Wadsworth, and Angus I Lamond. Proteomic analysis of cell cycle progression in asynchronous cultures, including mitotic subphases, using PRIMMUS. *eLife*, 6, 2017.

[3292] Timothy Lynagh, Emelie Flood, Céline Boiteux, Matthias Wulf, Vitaly V Komnatnyy, Janne M Colding, Toby W Allen, and Stephan A Pless. A selectivity filter at the intracellular end of the acid-sensing ion channel pore. *eLife*, 6, 2017.

[3293] Michael Lynch and Georgi K Marinov. Membranes, energetics, and evolution across the prokaryote-eukaryote divide. *eLife*, 6, 2017.

[3294] David B Lyons and Daniel Zilberman. DDM1 and Lsh remodelers allow methylation of DNA wrapped in nucleosomes. *eLife*, 6, 2017.

[3295] Lu Ma, Yiying Cai, Yanghui Li, Junyi Jiao, Zhenyong Wu, Ben O'Shaughnessy, Pietro De Camilli, Erdem Karatekin, and Yongli Zhang. Single-molecule force spectroscopy of protein-membrane interactions. *eLife*, 6, 2017.

[3296] Wenfu Ma, Elena Goldberg, and Jonathan Goldberg. ER retention is imposed by COPII protein sorting and attenuated by 4-phenylbutyrate. *eLife*, 6, 2017.

[3297] Daniel R Machado, Dinis JS Afonso, Alexandra R Kenny, Arzu öztürk Çolak, Emilia H Moscato, Benjamin Mainwaring, Matthew Kayser, and Kyunghee Koh. Identification of octopaminergic neurons that modulate sleep suppression by male sex drive. *eLife*, 6, 2017.

[3298] Mateusz Maciejewski, Eugen Lounkine, Steven Whitebread, Pierre Farmer, William DuMouchel, Brian K Shoichet, and Laszlo Urban. Reverse translation of adverse event reports paves the way for de-risking preclinical off-targets. *eLife*, 6, 2017.

[3299] Marie MacLennan, Marta García-Cañadas, Judith Reichmann, Elena Khazina, Gabriele Wagner, Christopher J Playfoot, Carmen Salvador-Palomeque, Abigail R Mann, Paula Peressini, Laura Sanchez, Karen Dobie, David Read, Chao-Chun Hung, Ragnhild Eskeland, Richard R Meehan, Oliver Weichenrieder, Jose Luis García-Pérez, and Ian R Adams. Mobilization of LINE-1 retrotransposons is restricted by Tex19.1 in mouse embryonic stem cells. *eLife*, 6, 2017.

[3300] Asaf Madi, Asaf Poran, Eric Shifrut, Shlomit Reich-Zeliger, Erez Greenstein, Irena Zaretsky, Tomer Arnon, Francois Van Laethem, Alfred Singer, Jinghua Lu, Peter D Sun, Irun R Cohen, and Nir Friedman. T cell receptor repertoires of mice and humans are clustered in similarity networks around conserved public CDR3 sequences. *eLife*, 6, 2017.

[3301] Hiromi Maekawa, Annett Neuner, Diana Rüthnick, Elmar Schiebel, Gislene Pereira, and Yoshinobu Kaneko. Polo-like kinase Cdc5 regulates Spc72 recruitment to spindle pole body in the methylotrophic yeast Ogataea polymorpha. *eLife*, 6, 2017.

[3302] Masato Maesako, Jana Horlacher, Katarzyna M Zoltowska, Ksenia V Kastanenka, Eleanna Kara, Sarah Svirsky, Laura J Keller, Xuejing Li, Bradley T Hyman, Brian J Bacskai, and Oksana Berezovska. Pathogenic PS1 phosphorylation at Ser367. *eLife*, 6, 2017.

[3303] Lukas Franz Mager, Viktor Hendrik Koelzer, Regula Stuber, Lester Thoo, Irene Keller, Ivonne Koeck, Maya Langenegger, Cedric Simillion, Simona P Pfister, Martin Faderl, Vera Genitsch, Irina Tcymbarevich, Pascal Juillerat, Xiaohong Li, Yu Xia, Eva Karamitopoulou, Ruth Lyck, Inti Zlobec, Siegfried Hapfelmeier, Rémy Bruggmann, Kathy D McCoy, Andrew J Macpherson, Christoph Müller, Bruce Beutler, and Philippe Krebs. The ESRP1-GPR137 axis contributes to intestinal pathogenesis. *eLife*, 6, 2017.

[3304] Christina FS Mages, Axel Wintsche, Stephan H Bernhart, and Gerd A Müller. The DREAM complex through its subunit Lin37 cooperates with Rb to initiate quiescence. *eLife*, 6, 2017.

[3305] Vivek Mahadevan, C Sahara Khademullah, Zahra Dargaei, Jonah Chevrier, Pavel Uvarov, Julian Kwan, Richard D Bagshaw, Tony Pawson, Andrew Emili, Yves De Koninck, Victor Anggono, Matti Airaksinen, and Melanie A Woodin. Native KCC2 interactome reveals PACSIN1 as a critical regulator of synaptic inhibition. *eLife*, 6, 2017.

[3306] Deeqa Mahamed, Mikael Boulle, Yashica Ganga, Chanelle Mc Arthur, Steven Skroch, Lance Oom, Oana Catinas, Kelly Pillay, Myshnee Naicker, Sanisha Rampersad, Colisile Mathonsi, Jessica Hunter, Emily B Wong, Moosa Suleman, Gopalkrishna Sreejit, Alexander S Pym, Gila Lustig, and Alex Sigal. Intracellular growth of Mycobacterium tuberculosis after macrophage cell death leads to serial killing of host cells. *eLife*, 6, 2017.

[3307] Lara Maister, Teresa Tang, and Manos Tsakiris. Neurobehavioral evidence of interoceptive sensitivity in early infancy. *eLife*, 6, 2017.

[3308] Frederikke Gro Malinovsky, Marie-Louise F Thomsen, Sebastian J Nintemann, Lea Møller Jagd, Baptiste Bourgine, Meike Burow, and Daniel J Kliebenstein. An evolutionarily young defense metabolite influences the root growth of plants via the ancient TOR signaling pathway. *eLife*, 6, 2017.

[3309] Duccio Malinverni, Alfredo Jost Lopez, Paolo De Los Rios, Gerhard Hummer, and Alessandro Barducci. Modeling Hsp70/Hsp40 interaction by multi-scale molecular simulations and coevolutionary sequence analysis. *eLife*, 6, 2017.

[3310] Sarah M Mangiameli, Christopher N Merrikh, Paul A Wiggins, and Houra Merrikh. Transcription leads to pervasive replisome instability in bacteria. *eLife*, 6, 2017.

[3311] Zoe F Mann, Héctor Gálvez, David Pedreno, Ziqi Chen, Elena Chrysostomou, Magdalena Żak, Miso Kang, Elachumee Canden, and Nicolas Daudet. Shaping of inner ear sensory organs through antagonistic interactions between Notch signalling and Lmx1a. *eLife*, 6, 2017.

[3312] Anitha Manohar, Andre L Curtis, Stephen A Zderic, and Rita J Valentino. Brainstem network dynamics underlying the encoding of bladder information. *eLife*, 6, 2017.

[3313] Anitha Manohar, Guglielmo Foffani, Patrick D Ganzer, John R Bethea, and Karen A Moxon. Cortex-dependent recovery of unassisted hindlimb locomotion after complete spinal cord injury in adult rats. *eLife*, 6, 2017.

[3314] Christine Mantis, Irawati Kandela, and Fraser Aird. Replication Study: Coadministration of a tumor-penetrating peptide enhances the efficacy of cancer drugs. *eLife*, 6, 2017.

[3315] Giulia Manzoni, Carine Marinach, Selma TopÇu, Sylvie Briquet, Morgane Grand, Matthieu Tolle, Marion Gransagne, Julien Lescar, Chiara Andolina, Jean-FranÇois Franetich, Mirjam B Zeisel, Thierry Huby, Eric Rubinstein, Georges Snounou, Dominique Mazier, FranÇois Nosten, Thomas F Baumert, and Olivier Silvie. Plasmodium P36 determines host cell receptor usage during sporozoite invasion. *eLife*, 6, 2017.

[3316] Antoine MarÇais, Marie Marotel, Sophie Degouve, Alice Koenig, Sébastien Fauteux-Daniel, Annabelle Drouillard, Heinrich Schlums, Sébastien Viel, Laurie Besson, Omran Allatif, Mathieu Bléry, Eric Vivier, Yenan Bryceson, Olivier Thaunat, and Thierry Walzer. High mTOR activity is a hallmark of reactive natural killer cells and amplifies early signaling through activating receptors. *eLife*, 6, 2017.

[3317] Ricardo Marchante, David M Beal, Nadejda Koloteva-Levine, Tracey J Purton, Mick F Tuite, and Wei-Feng Xue. The physical dimensions of amyloid aggregates control their infective potential as prion particles. *eLife*, 6, 2017.

[3318] Daniel S Marigold and Trevor Drew. Posterior parietal cortex estimates the relationship between object and body location during locomotion. *eLife*, 6, 2017.

[3319] Welber Marinovic, Eugene Poh, Aymar de Rugy, and Timothy J Carroll. Action history influences subsequent movement via two distinct processes. *eLife*, 6, 2017.

[3320] Philippa Marrack, Sai Harsha Krovi, Daniel Silberman, Janice White, Eleanor Kushnir, Maki Nakayama, James Crooks, Thomas Danhorn, Sonia Leach, Randy Anselment, James Scott-Browne, Laurent Gapin, and John Kappler. The somatically generated portion of T cell receptor CDR3$\alpha$ contributes to the MHC allele specificity of the T cell receptor. *eLife*, 6, 2017.

[3321] James S Martenson, Tokiwa Yamasaki, Nashid H Chaudhury, David Albrecht, and Susumu Tomita. Assembly rules for GABAA receptor complexes in the brain. *eLife*, 6, 2017.

[3322] Gregory M Martin, Balamurugan Kandasamy, Frank DiMaio, Craig Yoshioka, and Show-Ling Shyng. Anti-diabetic drug binding site in a mammalian KATP channel revealed by Cryo-EM. *eLife*, 6, 2017.

[3323] Gregory M Martin, Craig Yoshioka, Emily A Rex, Jonathan F Fay, Qing Xie, Matthew R Whorton, James Z Chen, and Show-Ling Shyng. Cryo-EM structure of the ATP-sensitive potassium channel illuminates mechanisms of assembly and gating. *eLife*, 6, 2017.

[3324] Pilar Martin-Marcos, Fujun Zhou, Charm Karunasiri, Fan Zhang, Jinsheng Dong, Jagpreet Nanda, Shardul D Kulkarni, Neelam Dabas Sen, Mercedes Tamame, Michael Zeschnigk, Jon R Lorsch, and Alan G Hinnebusch. eIF1A residues implicated in cancer stabilize translation preinitiation complexes and favor suboptimal initiation sites in yeast. *eLife*, 6, 2017.

[3325] Olga Martzoukou, Sotiris Amillis, Amalia Zervakou, Savvas Christoforidis, and George Diallinas. The AP-2 complex has a specialized clathrin-independent role in apical endocytosis and polar growth in fungi. *eLife*, 6, 2017.

[3326] Ameya A Mashruwala, Adriana van de Guchte, and Jeffrey M Boyd. Impaired respiration elicits SrrAB-dependent programmed cell lysis and biofilm formation in Staphylococcus aureus. *eLife*, 6, 2017.

[3327] Miriam Matamales, Zala Skrbis, Matthew R Bailey, Peter D Balsam, Bernard W Balleine, Jürgen Götz, and Jesus Bertran-Gonzalez. A corticostriatal deficit promotes temporal distortion of automatic action in ageing. *eLife*, 6, 2017.

[3328] Valéry Matarazzo, Laura Caccialupi, Fabienne Schaller, Yuri Shvarev, Nazim Kourdougli, Alessandra Bertoni, Clément Menuet, Nicolas Voituron, Evan Deneris, Patricia Gaspar, Laurent Bezin, Pascale Durbec, Gérard Hilaire, and FranÇoise Muscatelli. Necdin shapes serotonergic development and SERT activity modulating breathing in a mouse model for Prader-Willi syndrome. *eLife*, 6, 2017.

[3329] Sara Matias, Eran Lottem, Guillaume P Dugué, and Zachary F Mainen. Activity patterns of serotonin neurons underlying cognitive flexibility. *eLife*, 6, 2017.

[3330] Jacob Peter Matson, Raluca Dumitru, Philip Coryell, Ryan M Baxley, Weili Chen, Kirk Twaroski, Beau R Webber, Jakub Tolar, Anja-Katrin Bielinsky, Jeremy E Purvis, and Jeanette Gowen Cook. Rapid DNA replication origin licensing protects stem cell pluripotency. *eLife*, 6, 2017.

[3331] Shinya Matsuda and Markus Affolter. Dpp from the anterior stripe of cells is crucial for the growth of the Drosophila wing disc. *eLife*, 6, 2017.

[3332] Naoyuki Matsumoto, Yohei Shinmyo, Yoshie Ichikawa, and Hiroshi Kawasaki. Gyrification of the cerebral cortex requires FGF signaling in the mammalian brain. *eLife*, 6, 2017.

[3333] Sandy Mattijssen, Aneeshkumar G Arimbasseri, James R Iben, Sergei Gaidamakov, Joowon Lee, Markus Hafner, and Richard J Maraia. LARP4 mRNA codon-tRNA match contributes to LARP4 activity for ribosomal protein mRNA poly(A) tail length protection. *eLife*, 6, 2017.

[3334] Francesca Mattiroli, Yajie Gu, Tejas Yadav, Jeremy L Balsbaugh, Michael R Harris, Eileen S Findlay, Yang Liu, Catherine A Radebaugh, Laurie A Stargell, Natalie G Ahn, Iestyn Whitehouse, and Karolin Luger. DNA-mediated association of two histone-bound complexes of yeast Chromatin Assembly Factor-1 (CAF-1) drives tetrasome assembly in the wake of DNA replication. *eLife*, 6, 2017.

[3335] Jordan G McCall, Edward R Siuda, Dionnet L Bhatti, Lamley A Lawson, Zoe A McElligott, Garret D Stuber, and Michael R Bruchas. Locus coeruleus to basolateral amygdala noradrenergic projections promote anxiety-like behavior. *eLife*, 6, 2017.

[3336] Heather A McCauley, Véronique Chevrier, Daniel Birnbaum, and Géraldine Guasch. De-repression of the RAC activator ELMO1 in cancer stem cells drives progression of TGF$\beta$-deficient squamous cell carcinoma from transition zones. *eLife*, 6, 2017.

[3337] Kevin McCluskey, Kyria Boundy-Mills, Greg Dye, Erin Ehmke, Gregg F Gunnell, Hippokratis Kiaris, Maxi Polihronakis Richmond, Anne D Yoder, Daniel R Zeigler, Sarah Zehr, and Erich Grotewold. The challenges faced by living stock collections in the USA. *eLife*, 6, 2017.

[3338] Thomas McColgan, Ji Liu, Paula Tuulia Kuokkanen, Catherine Emily Carr, Hermann Wagner, and Richard Kempter. Dipolar extracellular potentials generated by axonal projections. *eLife*, 6, 2017.

[3339] Nathan A McDonald, Abigail L Lind, Sarah E Smith, Rong Li, and Kathleen L Gould. Nanoscale architecture of the Schizosaccharomyces pombe contractile ring. *eLife*, 6, 2017.

[3340] Alicia N McMurchy, Przemyslaw Stempor, Tessa Gaarenstroom, Brian Wysolmerski, Yan Dong, Darya Aussianikava, Alex Appert, Ni Huang, Paulina Kolasinska-Zwierz, Alexandra Sapetschnig, Eric A Miska, and Julie Ahringer. A team of heterochromatin factors collaborates with small RNA pathways to combat repetitive elements and germline stress. *eLife*, 6, 2017.

[3341] Jamie McQueen, Tomás J Ryan, Sean McKay, Katie Marwick, Paul Baxter, Sarah M Carpanini, Thomas M Wishart, Thomas H Gillingwater, Jean C Manson, David J A Wyllie, Seth G N Grant, Barry W McColl, Noboru H Komiyama, and Giles E Hardingham. Pro-death NMDA receptor signaling is promoted by the GluN2B C-terminus independently of Dapk1. *eLife*, 6, 2017.

[3342] Andrew F Mead, Nerea Osinalde, Niels Ørtenblad, Joachim Nielsen, Jonathan Brewer, Michiel Vellema, Iris Adam, Constance Scharff, Yafeng Song, Ulrik Frandsen, Blagoy Blagoev, Irina Kratchmarova, and Coen PH Elemans. Fundamental constraints in synchronous muscle limit superfast motor control in vertebrates. *eLife*, 6, 2017.

[3343] Netra Pal Meena and Alan R Kimmel. Chemotactic network responses to live bacteria show independence of phagocytosis from chemoreceptor sensing. *eLife*, 6, 2017.

[3344] Pratap Meera, Stefan Pulst, and Thomas Otis. A positive feedback loop linking enhanced mGluR function and basal calcium in spinocerebellar ataxia type 2. *eLife*, 6, 2017.

[3345] Michael P Meers, Telmo Henriques, Christopher A Lavender, Daniel J McKay, Brian D Strahl, Robert J Duronio, Karen Adelman, and A Gregory Matera. Histone gene replacement reveals a post-transcriptional role for H3K36 in maintaining metazoan transcriptome fidelity. *eLife*, 6, 2017.

[3346] Karen Mendelson, Suveg Pandey, Yu Hisano, Frank Carellini, Bhaskar C Das, Timothy Hla, and Todd Evans. The ceramide synthase 2b gene mediates genomic sensing and regulation of sphingosine levels during zebrafish embryogenesis. *eLife*, 6, 2017.

[3347] William Menegas, Benedicte M Babayan, Naoshige Uchida, and Mitsuko Watabe-Uchida. Opposite initialization to novel cues in dopamine signaling in ventral and posterior striatum in mice. *eLife*, 6, 2017.

[3348] Anaïs Menny, Solène N Lefebvre, Philipp AM Schmidpeter, Emmanuelle Drège, Zaineb Fourati, Marc Delarue, Stuart J Edelstein, Crina M Nimigean, Delphine Joseph, and Pierre-Jean Corringer. Identification of a pre-active conformation of a pentameric channel receptor. *eLife*, 6, 2017.

[3349] Elisa Merklinger, Jan-Gero Schloetel, Pascal Weber, Helena Batoulis, Sarah Holz, Nora Karnowski, Jérôme Finke, and Thorsten Lang. The packing density of a supramolecular membrane protein cluster is controlled by cytoplasmic interactions. *eLife*, 6, 2017.

[3350] Yannick Mesrouze, Fedir Bokhovchuk, Marco Meyerhofer, Patrizia Fontana, Catherine Zimmermann, Typhaine Martin, Clara Delaunay, Dirk Erdmann, Tobias Schmelzle, and Patrick Chène. Dissection of the interaction between the intrinsically disordered YAP protein and the transcription factor TEAD. *eLife*, 6, 2017.

[3351] Michael G Metzen and Maurice J Chacron. Stimulus background influences phase invariant coding by correlated neural activity. *eLife*, 6, 2017.

[3352] Eyal Metzl-Raz, Moshe Kafri, Gilad Yaakov, Ilya Soifer, Yonat Gurvich, and Naama Barkai. Principles of cellular resource allocation revealed by condition-dependent proteome profiling. *eLife*, 6, 2017.

[3353] Heather M Meyer, José Teles, Pau Formosa-Jordan, Yassin Refahi, Rita San-Bento, Gwyneth Ingram, Henrik Jönsson, James C W Locke, and Adrienne H K Roeder. Fluctuations of the transcription factor ATML1 generate the pattern of giant cells in the Arabidopsis sepal. *eLife*, 6, 2017.

[3354] Matthias Meyer, Eleftheria Palkopoulou, Sina Baleka, Mathias Stiller, Kirsty E H Penkman, Kurt W Alt, Yasuko Ishida, Dietrich Mania, Swapan Mallick, Tom Meijer, Harald Meller, Sarah Nagel, Birgit Nickel, Sven Ostritz, Nadin Rohland, Karol Schauer, Tim Schüler, Alfred L Roca, David Reich, Beth Shapiro, and Michael Hofreiter. Palaeogenomes of Eurasian straight-tusked elephants challenge the current view of elephant evolution. *eLife*, 6, 2017.

[3355] Natasha Mhatre, Robert Malkin, Rittik Deb, Rohini Balakrishnan, and Daniel Robert. Tree crickets optimize the acoustics of baffles to exaggerate their mate-attraction signal. *eLife*, 6, 2017.

[3356] Yong Miao, Jaya Bhushan, Adish Dani, and Monika Vig. Na+ influx via Orai1 inhibits intracellular ATP-induced mTORC2 signaling to disrupt CD4 T cell gene expression and differentiation. *eLife*, 6, 2017.

[3357] Nicolas Michalski, Juan D Goutman, Sarah Marie Auclair, Jacques Boutet de Monvel, Margot Tertrais, Alice Emptoz, Alexandre Parrin, Sylvie Nouaille, Marc Guillon, Martin Sachse, Danica Ciric, Amel Bahloul, Jean-Pierre Hardelin, Roger Bryan Sutton, Paul Avan, Shyam S Krishnakumar, James E Rothman, Didier Dulon, Saaid Safieddine, and Christine Petit. Otoferlin acts as a Ca2+ sensor for vesicle fusion and vesicle pool replenishment at auditory hair cell ribbon synapses. *eLife*, 6, 2017.

[3358] Agnès H Michel, Riko Hatakeyama, Philipp Kimmig, Meret Arter, Matthias Peter, Joao Matos, Claudio De Virgilio, and Benoît Kornmann. Functional mapping of yeast genomes by saturated transposition. *eLife*, 6, 2017.

[3359] Thomas Miconi. Biologically plausible learning in recurrent neural networks reproduces neural dynamics observed during cognitive tasks. *eLife*, 6, 2017.

[3360] Anna L Miles, Stephen P Burr, Guinevere L Grice, and James A Nathan. The vacuolar-ATPase complex and assembly factors, TMEM199 and CCDC115, control HIF1$\alpha$ prolyl hydroxylation by regulating cellular iron levels. *eLife*, 6, 2017.

[3361] Adam C Miller, Alex C Whitebirch, Arish N Shah, Kurt C Marsden, Michael Granato, John O'Brien, and Cecilia B Moens. A genetic basis for molecular asymmetry at vertebrate electrical synapses. *eLife*, 6, 2017.

[3362] Jeremy A Miller, Angela Guillozet-Bongaarts, Laura E Gibbons, Nadia Postupna, Anne Renz, Allison E Beller, Susan M Sunkin, Lydia Ng, Shannon E Rose, Kimberly A Smith, Aaron Szafer, Chris Barber, Darren Bertagnolli, Kristopher Bickley, Krissy Brouner, Shiella Caldejon, Mike Chapin, Mindy L Chua, Natalie M Coleman, Eiron Cudaback, Christine Cuhaciyan, Rachel A Dalley, Nick Dee, Tsega Desta, Tim A Dolbeare, Nadezhda I Dotson, Michael Fisher, Nathalie Gaudreault, Garrett Gee, Terri L Gilbert, Jeff Goldy, Fiona Griffin, Caroline Habel, Zeb Haradon, Nika Hejazinia, Leanne L Hellstern, Steve Horvath, Kim Howard, Robert Howard, Justin Johal, Nikolas L Jorstad, Samuel R Josephsen, Chihchau L Kuan, Florence Lai, Eric Lee, Felix Lee, Tracy Lemon, Xianwu Li, Desiree A Marshall, Jose Melchor, Shubhabrata Mukherjee, Julie Nyhus, Julie Pendergraft, Lydia Potekhina, Elizabeth Y Rha, Samantha Rice, David Rosen, Abharika Sapru, Aimee Schantz, Elaine Shen, Emily Sherfield, Shu Shi, Andy J Sodt, Nivretta Thatra, Michael Tieu, Angela M Wilson, Thomas J Montine, Eric B Larson, Amy Bernard, Paul K Crane, Richard G Ellenbogen, C Dirk Keene, and Ed Lein. Neuropathological and transcriptomic characteristics of the aged brain. *eLife*, 6, 2017.

[3363] Laura Milligan, Camille Sayou, Alex Tuck, Tatsiana Auchynnikava, Jane EA Reid, Ross Alexander, Flavia de Lima Alves, Robin Allshire, Christos Spanos, Juri Rappsilber, Jean D Beggs, Grzegorz Kudla, and David Tollervey. RNA polymerase II stalling at pre-mRNA splice sites is enforced by ubiquitination of the catalytic subunit. *eLife*, 6, 2017.

[3364] Susana Mingote, Nao Chuhma, Abigail Kalmbach, Gretchen M Thomsen, Yvonne Wang, Andra Mihali, Caroline Sferrazza, Ilana Zucker-Scharff, Anna-Claire Siena, Martha G Welch, José Lizardi-Ortiz, David Sulzer, Holly Moore, Inna Gaisler-Salomon, and Stephen Rayport. Dopamine neuron dependent behaviors mediated by glutamate cotransmission. *eLife*, 6, 2017.

[3365] Ajay Mishra, Bénédicte Oulès, Angela Oliveira Pisco, Tony Ly, Kifayathullah Liakath-Ali, Gernot Walko, Priyalakshmi Viswanathan, Matthieu Tihy, Jagdeesh Nijjher, Sara-Jane Dunn, Angus I Lamond, and Fiona M Watt. A protein phosphatase network controls the temporal and spatial dynamics of differentiation commitment in human epidermis. *eLife*, 6, 2017.

[3366] Saurabh Mishra, Prashant Shukla, Ashima Bhaskar, Kushi Anand, Priyanka Baloni, Rajiv Kumar Jha, Abhilash Mohan, Raju S Rajmani, Valakunja Nagaraja, Nagasuma Chandra, and Amit Singh. Efficacy of $\beta$-lactam/$\beta$-lactamase inhibitor combination is linked to WhiB4-mediated changes in redox physiology of Mycobacterium tuberculosis. *eLife*, 6, 2017.

[3367] Melinda S Modrell, Mike Lyne, Adrian R Carr, Harold H Zakon, David Buckley, Alexander S Campbell, Marcus C Davis, Gos Micklem, and Clare VH Baker. Insights into electrosensory organ development, physiology and evolution from a lateral line-enriched transcriptome. *eLife*, 6, 2017.

[3368] Gaurav D Moghe, Bryan J Leong, Steven M Hurney, A Daniel Jones, and Robert L Last. Evolutionary routes to biochemical innovation revealed by integrative analysis of a plant-defense related specialized metabolic pathway. *eLife*, 6, 2017.

[3369] Susana Moleirinho, Sany Hoxha, Vinay Mandati, Graziella Curtale, Scott Troutman, Ursula Ehmer, and Joseph L Kissil. Regulation of localization and function of the transcriptional co-activator YAP by angiomotin. *eLife*, 6, 2017.

[3370] Babak Momeni, Li Xie, and Wenying Shou. Lotka-Volterra pairwise modeling fails to capture diverse pairwise microbial interactions. *eLife*, 6, 2017.

[3371] Bishakha Mona, Ana Uruena, Rahul K Kollipara, Zhenzhong Ma, Mark D Borromeo, Joshua C Chang, and Jane E Johnson. Repression by PRDM13 is critical for generating precision in neuronal identity. *eLife*, 6, 2017.

[3372] Kevin Monahan, Ira Schieren, Jonah Cheung, Alice Mumbey-Wafula, Edwin S Monuki, and Stavros Lomvardas. Cooperative interactions enable singular olfactory receptor expression in mouse olfactory neurons. *eLife*, 6, 2017.

[3373] Arindam Mondal, Anthony R Dawson, Gregory K Potts, Elyse C Freiberger, Steven F Baker, Lindsey A Moser, Kristen A Bernard, Joshua J Coon, and Andrew Mehle. Influenza virus recruits host protein kinase C to control assembly and activity of its replication machinery. *eLife*, 6, 2017.

[3374] Nicole Monroe, Han Han, Peter S Shen, Wesley I Sundquist, and Christopher P Hill. Structural basis of protein translocation by the Vps4-Vta1 AAA ATPase. *eLife*, 6, 2017.

[3375] Tessa G Montague and Alexander F Schier. Vg1-Nodal heterodimers are the endogenous inducers of mesendoderm. *eLife*, 6, 2017.

[3376] Stefania Monterisi, Miguel J Lobo, Craig Livie, John C Castle, Michael Weinberger, George Baillie, Nicoletta C Surdo, Nshunge Musheshe, Alessandra Stangherlin, Eyal Gottlieb, Rory Maizels, Mario Bortolozzi, Massimo Micaroni, and Manuela Zaccolo. PDE2A2 regulates mitochondria morphology and apoptotic cell death via local modulation of cAMP/PKA signalling. *eLife*, 6, 2017.

[3377] David Florentino Montez, Finnegan J Calabro, and Beatriz Luna. The expression of established cognitive brain states stabilizes with working memory development. *eLife*, 6, 2017.

[3378] Ignacio Moraga, Jamie B Spangler, Juan L Mendoza, Milica Gakovic, Tom S Wehrman, Peter Krutzik, and K Christopher Garcia. Synthekines are surrogate cytokine and growth factor agonists that compel signaling through non-natural receptor dimers. *eLife*, 6, 2017.

[3379] Abigail J Morales, Javier A Carrero, Putzer J Hung, Anthony T Tubbs, Jared M Andrews, Brian T Edelson, Boris Calderon, Cynthia L Innes, Richard S Paules, Jacqueline E Payton, and Barry P Sleckman. A type I IFN-dependent DNA damage response regulates the genetic program and inflammasome activation in macrophages. *eLife*, 6, 2017.

[3380] David Moreau, Ian J Kirk, and Karen E Waldie. High-intensity training enhances executive function in children in a randomized, placebo-controlled trial. *eLife*, 6, 2017.

[3381] David Moreno Delgado, Thor C Møller, Jeanne Ster, Jesús Giraldo, Damien Maurel, Xavier Rovira, Pauline Scholler, Jurriaan M Zwier, Julie Perroy, Thierry Durroux, Eric Trinquet, Laurent Prezeau, Philippe Rondard, and Jean-Philippe Pin. Pharmacological evidence for a metabotropic glutamate receptor heterodimer in neuronal cells. *eLife*, 6, 2017.

[3382] Julie Morin-Rivat, Adeline Fayolle, Charly Favier, Laurent Bremond, Sylvie Gourlet-Fleury, Nicolas Bayol, Philippe Lejeune, Hans Beeckman, and Jean-Louis Doucet. Present-day central African forest is a legacy of the 19th century human history. *eLife*, 6, 2017.

[3383] Jason A Morrison, Rebecca McLennan, Lauren A Wolfe, Madelaine M Gogol, Samuel Meier, Mary C McKinney, Jessica M Teddy, Laura Holmes, Craig L Semerad, Andrew C Box, Hua Li, Kathryn E Hall, Anoja G Perera, and Paul M Kulesa. Single-cell transcriptome analysis of avian neural crest migration reveals signatures of invasion and molecular transitions. *eLife*, 6, 2017.

[3384] Mark D Morrissey, Nathan Insel, and Kaori Takehara-Nishiuchi. Generalizable knowledge outweighs incidental details in prefrontal ensemble code over time. *eLife*, 6, 2017.

[3385] Carl A Morrow, Michael O Nguyen, Andrew Fower, Io Nam Wong, Fekret Osman, Claire Bryer, and Matthew C Whitby. Inter-Fork Strand Annealing causes genomic deletions during the termination of DNA replication. *eLife*, 6, 2017.

[3386] Timothy J Mosca, David J Luginbuhl, Irving E Wang, and Liqun Luo. Presynaptic LRP4 promotes synapse number and function of excitatory CNS neurons. *eLife*, 6, 2017.

[3387] Margarida Moura, Mariana Osswald, Nelson LeÇa, João Barbosa, António J Pereira, Helder Maiato, Claudio E Sunkel, and Carlos Conde. Protein Phosphatase 1 inactivates Mps1 to ensure efficient Spindle Assembly Checkpoint silencing. *eLife*, 6, 2017.

[3388] Frauke Mücksch, Vibor Laketa, Barbara Müller, Carsten Schultz, and Hans-Georg Kräusslich. Synchronized HIV assembly by tunable PIP2 changes reveals PIP2 requirement for stable Gag anchoring. *eLife*, 6, 2017.

[3389] Alexander Muir, Laura V Danai, Dan Y Gui, Chiara Y Waingarten, Caroline A Lewis, and Matthew G Vander Heiden. Environmental cystine drives glutamine anaplerosis and sensitizes cancer cells to glutaminase inhibition. *eLife*, 6, 2017.

[3390] Haripriya Mukundarajan, Felix Jan Hein Hol, Erica Araceli Castillo, Cooper Newby, and Manu Prakash. Using mobile phones as acoustic sensors for high-throughput mosquito surveillance. *eLife*, 6, 2017.

[3391] Jean M Mulcahy Levy, Shadi Zahedi, Andrea M Griesinger, Andrew Morin, Kurtis D Davies, Dara L Aisner, BK Kleinschmidt-DeMasters, Brent E Fitzwalter, Megan L Goodall, Jacqueline Thorburn, Vladimir Amani, Andrew M Donson, Diane K Birks, David M Mirsky, Todd C Hankinson, Michael H Handler, Adam L Green, Rajeev Vibhakar, Nicholas K Foreman, and Andrew Thorburn. Autophagy inhibition overcomes multiple mechanisms of resistance to BRAF inhibition in brain tumors. *eLife*, 6, 2017.

[3392] Wahid A Mulla, Chris W Seidel, Jin Zhu, Hung-Ji Tsai, Sarah E Smith, Pushpendra Singh, William D Bradford, Scott McCroskey, Anjali R Nelliat, Juliana Conkright, Allison Peak, Kathryn E Malanowski, Anoja G Perera, and Rong Li. Aneuploidy as a cause of impaired chromatin silencing and mating-type specification in budding yeast. *eLife*, 6, 2017.

[3393] Rahul Munshi, Shahnaz M Qadri, Qian Zhang, Idoia Castellanos Rubio, Pablo del Pino, and Arnd Pralle. Magnetothermal genetic deep brain stimulation of motor behaviors in awake, freely moving mice. *eLife*, 6, 2017.

[3394] Sachiko Murase, Crystal L Lantz, and Elizabeth M Quinlan. Light reintroduction after dark exposure reactivates plasticity in adults via perisynaptic activation of MMP-9. *eLife*, 6, 2017.

[3395] Andrew Muroyama and Terry Lechler. A transgenic toolkit for visualizing and perturbing microtubules reveals unexpected functions in the epidermis. *eLife*, 6, 2017.

[3396] James M Murray and G Sean Escola. Learning multiple variable-speed sequences in striatum via cortical tutoring. *eLife*, 6, 2017.

[3397] Veronica Musante, Lu Li, Jean Kanyo, Tukiet T Lam, Christopher M Colangelo, Shuk Kei Cheng, A Harrison Brody, Paul Greengard, Nicolas Le Novère, and Angus C Nairn. Reciprocal regulation of ARPP-16 by PKA and MAST3 kinases provides a cAMP-regulated switch in protein phosphatase 2A inhibition. *eLife*, 6, 2017.

[3398] Saravanapriah Nadarajan, Talley J Lambert, Elisabeth Altendorfer, Jinmin Gao, Michael D Blower, Jennifer C Waters, and Monica P Colaiácovo. Polo-like kinase-dependent phosphorylation of the synaptonemal complex protein SYP-4 regulates double-strand break formation through a negative feedback loop. *eLife*, 6, 2017.

[3399] Carey D Nadell, Deirdre Ricaurte, Jing Yan, Knut Drescher, and Bonnie L Bassler. Flow environment and matrix structure interact to determine spatial competition in Pseudomonas aeruginosa biofilms. *eLife*, 6, 2017.

[3400] Sonal Nagarkar-Jaiswal, Sathiya N Manivannan, Zhongyuan Zuo, and Hugo J Bellen. A cell cycle-independent, conditional gene inactivation strategy for differentially tagging wild-type and mutant cells. *eLife*, 6, 2017.

[3401] Nicole Ann Najor, Gillian Nicole Fitz, Jennifer Leigh Koetsier, Lisa Marie Godsel, Lauren Veronica Albrecht, Robert Harmon, and Kathleen Janee Green. Epidermal Growth Factor Receptor neddylation is regulated by a desmosomal-COP9 (Constitutive Photomorphogenesis 9) signalosome complex. *eLife*, 6, 2017.

[3402] Paul A Nakamura, Andy A Shimchuk, Shibing Tang, Zhizhi Wang, Kole DeGolier, Sheng Ding, and Thomas A Reh. Small molecule Photoregulin3 prevents retinal degeneration in the RhoP23H mouse model of retinitis pigmentosa. *eLife*, 6, 2017.

[3403] Haruko Nakano, Itsunari Minami, Daniel Braas, Herman Pappoe, Xiuju Wu, Addelynn Sagadevan, Laurent Vergnes, Kai Fu, Marco Morselli, Christopher Dunham, Xueqin Ding, Adam Z Stieg, James K Gimzewski, Matteo Pellegrini, Peter M Clark, Karen Reue, Aldons J Lusis, Bernard Ribalet, Siavash K Kurdistani, Heather Christofk, Norio Nakatsuji, and Atsushi Nakano. Glucose inhibits cardiac muscle maturation through nucleotide biosynthesis. *eLife*, 6, 2017.

[3404] Kei Nakayama, Rie Ohashi, Yo Shinoda, Maya Yamazaki, Manabu Abe, Akihiro Fujikawa, Shuji Shigenobu, Akira Futatsugi, Masaharu Noda, Katsuhiko Mikoshiba, Teiichi Furuichi, Kenji Sakimura, and Nobuyuki Shiina. RNG105/caprin1, an RNA granule protein for dendritic mRNA localization, is essential for long-term memory formation. *eLife*, 6, 2017.

[3405] Nilay Nandi, Lauren K Tyra, Drew Stenesen, and Helmut Krämer. Stress-induced Cdk5 activity enhances cytoprotective basal autophagy in Drosophila melanogaster by phosphorylating acinus at serine437. *eLife*, 6, 2017.

[3406] Drew M Nassal, Xiaoping Wan, Haiyan Liu, Danielle Maleski, Angelina Ramirez-Navarro, Christine S Moravec, Eckhard Ficker, Kenneth R Laurita, and Isabelle Deschênes. KChIP2 is a core transcriptional regulator of cardiac excitability. *eLife*, 6, 2017.

[3407] Elin Näsström, Christopher M Parry, Nga Tran Vu Thieu, Rapeephan R Maude, Hanna K de Jong, Masako Fukushima, Olena Rzhepishevska, Florian Marks, Ursula Panzner, Justin Im, Hyonjin Jeon, Seeun Park, Zabeen Chaudhury, Aniruddha Ghose, Rasheda Samad, Tan Trinh Van, Anders Johansson, Arjen M Dondorp, Guy E Thwaites, Abul Faiz, Henrik Antti, and Stephen Baker. Reproducible diagnostic metabolites in plasma from typhoid fever patients in Asia and Africa. *eLife*, 6, 2017.

[3408] Dipti D Nayak, Nilkamal Mahanta, Douglas A Mitchell, and William W Metcalf. Post-translational thioamidation of methyl-coenzyme M reductase, a key enzyme in methanogenic and methanotrophic Archaea. *eLife*, 6, 2017.

[3409] Evgeniya V Nazarova, Christine R Montague, Thuy La, Kaley M Wilburn, Neelima Sukumar, Wonsik Lee, Shannon Caldwell, David G Russell, and Brian C VanderVen. Rv3723/LucA coordinates fatty acid and cholesterol uptake in Mycobacterium tuberculosis. *eLife*, 6, 2017.

[3410] Inna V Nechipurenko, Cristina Berciu, Piali Sengupta, and Daniela Nicastro. Centriolar remodeling underlies basal body maturation during ciliogenesis in Caenorhabditis elegans. *eLife*, 6, 2017.

[3411] Derek Evan Nee and Mark D'Esposito. Causal evidence for lateral prefrontal cortex dynamics supporting cognitive control. *eLife*, 6, 2017.

[3412] Andreas Neerincx, Clemens Hermann, Robin Antrobus, Andy van Hateren, Huan Cao, Nico Trautwein, Stefan Stevanović, Tim Elliott, Janet E Deane, and Louise H Boyle. TAPBPR bridges UDP-glucose:glycoprotein glucosyltransferase 1 onto MHC class I to provide quality control in the antigen presentation pathway. *eLife*, 6, 2017.

[3413] David R Nelson, Basel Khraiwesh, Weiqi Fu, Saleh Alseekh, Ashish Jaiswal, Amphun Chaiboonchoe, Khaled M Hazzouri, Matthew J O'Connor, Glenn L Butterfoss, Nizar Drou, Jillian D Rowe, Jamil Harb, Alisdair R Fernie, Kristin C Gunsalus, and Kourosh Salehi-Ashtiani. The genome and phenome of the green alga Chloroidium sp. UTEX 3007 reveal adaptive traits for desert acclimatization. *eLife*, 6, 2017.

[3414] Jutta Nesper, Isabelle Hug, Setsu Kato, Chee-Seng Hee, Judith Maria Habazettl, Pablo Manfredi, Stephan Grzesiek, Tilman Schirmer, Thierry Emonet, and Urs Jenal. Cyclic di-GMP differentially tunes a bacterial flagellar motor through a novel class of CheY-like regulators. *eLife*, 6, 2017.

[3415] Emilie-Fleur Neubauer, Angela Z Poole, Philipp Neubauer, Olivier Detournay, Kenneth Tan, Simon K Davy, and Virginia M Weis. A diverse host thrombospondin-type-1 repeat protein repertoire promotes symbiont colonization during establishment of cnidarian-dinoflagellate symbiosis. *eLife*, 6, 2017.

[3416] TD Barbara Nguyen-Vu, Grace Q Zhao, Subhaneil Lahiri, Rhea R Kimpo, Hanmi Lee, Surya Ganguli, Carla J Shatz, and Jennifer L Raymond. A saturation hypothesis to explain both enhanced and impaired learning with enhanced plasticity. *eLife*, 6, 2017.

[3417] Yanxiang Ni, Bo Cao, Tszshan Ma, Gang Niu, Yingdong Huo, Jiandong Huang, Danni Chen, Yi Liu, Bin Yu, Michael Q Zhang, and Hanben Niu. Super-resolution imaging of a 2.5 kb non-repetitive DNA in situ in the nuclear genome using molecular beacon probes. *eLife*, 6, 2017.

[3418] Angela K Nietz, Jada H Vaden, Luke T Coddington, Linda Overstreet-Wadiche, and Jacques I Wadiche. Non-synaptic signaling from cerebellar climbing fibers modulates Golgi cell activity. *eLife*, 6, 2017.

[3419] Ross Nieuwburg, Dmitry Nashchekin, Maximilian Jakobs, Andrew P Carter, Philipp Khuc Trong, Raymond E Goldstein, and Daniel St Johnston. Localised dynactin protects growing microtubules to deliver oskar mRNA to the posterior cortex of the Drosophila oocyte. *eLife*, 6, 2017.

[3420] Marko Z Nikolić, Oriol Caritg, Quitz Jeng, Jo-Anne Johnson, Dawei Sun, Kate J Howell, Jane L Brady, Usua Laresgoiti, George Allen, Richard Butler, Matthias Zilbauer, Adam Giangreco, and Emma L Rawlins. Human embryonic lung epithelial tips are multipotent progenitors that can be expanded in vitro as long-term self-renewing organoids. *eLife*, 6, 2017.

[3421] Nadinath B Nillegoda, Antonia Stank, Duccio Malinverni, Niels Alberts, Anna Szlachcic, Alessandro Barducci, Paolo De Los Rios, Rebecca C Wade, and Bernd Bukau. Evolution of an intricate J-protein network driving protein disaggregation in eukaryotes. *eLife*, 6, 2017.

[3422] Simon Nimpf, Erich Pascal Malkemper, Mattias Lauwers, Lyubov Ushakova, Gregory Nordmann, Andrea Wenninger-Weinzierl, Thomas R Burkard, Sonja Jacob, Thomas Heuser, Guenter P Resch, and David A Keays. Subcellular analysis of pigeon hair cells implicates vesicular trafficking in cuticulosome formation and maintenance. *eLife*, 6, 2017.

[3423] Masatoshi Nishikawa, Sundar Ram Naganathan, Frank Jülicher, and Stephan W Grill. Controlling contractile instabilities in the actomyosin cortex. *eLife*, 6, 2017.

[3424] Yang Niu, Zhonghua Dai, Wenxue Liu, Cheng Zhang, Yanrui Yang, Zhenzhen Guo, Xiaoyu Li, Chenchang Xu, Xiahe Huang, Yingchun Wang, Yun S Shi, and Jia-Jia Liu. Ablation of SNX6 leads to defects in synaptic function of CA1 pyramidal neurons and spatial memory. *eLife*, 6, 2017.

[3425] Isabel Nocedal, Eugenio Mancera, and Alexander D Johnson. Gene regulatory network plasticity predates a switch in function of a conserved transcription regulator. *eLife*, 6, 2017.

[3426] Kentaro Noma, Alexandr Goncharov, Mark H Ellisman, and Yishi Jin. Microtubule-dependent ribosome localization in C. elegans neurons. *eLife*, 6, 2017.

[3427] Kaori Noridomi, Go Watanabe, Melissa N Hansen, Gye Won Han, and Lin Chen. Structural insights into the molecular mechanisms of myasthenia gravis and their therapeutic implications. *eLife*, 6, 2017.

[3428] Davide Normanno, Aurélie Négrel, Abinadabe J de Melo, Stéphane Betzi, Katheryn Meek, and Mauro Modesti. Mutational phospho-mimicry reveals a regulatory role for the XRCC4 and XLF C-terminal tails in modulating DNA bridging during classical non-homologous end joining. *eLife*, 6, 2017.

[3429] Adam D Norris, Xicotencatl Gracida, and John A Calarco. CRISPR-mediated genetic interaction profiling identifies RNA binding proteins controlling metazoan fitness. *eLife*, 6, 2017.

[3430] Megan L Norris, Andrea Pauli, James A Gagnon, Nathan D Lord, Katherine W Rogers, Christian Mosimann, Leonard I Zon, and Alexander F Schier. Toddler signaling regulates mesodermal cell migration downstream of Nodal signaling. *eLife*, 6, 2017.

[3431] Brian A Nosek and Timothy M Errington. Making sense of replications. *eLife*, 6, 2017.

[3432] Nicole L Nuckolls, María Angélica Bravo Núñez, Michael T Eickbush, Janet M Young, Jeffrey J Lange, Jonathan S Yu, Gerald R Smith, Sue L Jaspersen, Harmit S Malik, and Sarah E Zanders. wtf genes are prolific dual poison-antidote meiotic drivers. *eLife*, 6, 2017.

[3433] Patrick J Nygren, Sohum Mehta, Devin K Schweppe, Lorene K Langeberg, Jennifer L Whiting, Chad R Weisbrod, James E Bruce, Jin Zhang, David Veesler, and John D Scott. Intrinsic disorder within AKAP79 fine-tunes anchored phosphatase activity toward substrates and drug sensitivity. *eLife*, 6, 2017.

[3434] Sebastian Ocklenburg, Judith Schmitz, Zahra Moinfar, Dirk Moser, Rena Klose, Stephanie Lor, Georg Kunz, Martin Tegenthoff, Pedro Faustmann, Clyde Francks, Jörg T Epplen, Robert Kumsta, and Onur Güntürkün. Epigenetic regulation of lateralized fetal spinal gene expression underlies hemispheric asymmetries. *eLife*, 6, 2017.

[3435] Devin Lee O'Connor, Samuel Elton, Fabrizio Ticchiarelli, Mon Mandy Hsia, John P Vogel, and Ottoline Leyser. Cross-species functional diversity within the PIN auxin efflux protein family. *eLife*, 6, 2017.

[3436] Cian O'Donnell, J Tiago GonÇalves, Carlos Portera-Cailliau, and Terrence J Sejnowski. Beyond excitation/inhibition imbalance in multidimensional models of neural circuit changes in brain disorders. *eLife*, 6, 2017.

[3437] Fergus M O'Farrell, Svetlana Mastitskaya, Matthew Hammond-Haley, Felipe Freitas, Wen Rui Wah, and David Attwell. Capillary pericytes mediate coronary no-reflow after myocardial ischaemia. *eLife*, 6, 2017.

[3438] Kelly H Oh, James J Haney, Xiaohong Wang, Chiou-Fen Chuang, Janet E Richmond, and Hongkyun Kim. ERG-28 controls BK channel trafficking in the ER to regulate synaptic function and alcohol response in C. elegans. *eLife*, 6, 2017.

[3439] Justin K O'Hare, Haofang Li, Namsoo Kim, Erin Gaidis, Kristen Ade, Jeff Beck, Henry Yin, and Nicole Calakos. Striatal fast-spiking interneurons selectively modulate circuit output and are required for habitual behavior. *eLife*, 6, 2017.

[3440] Patrick J O'Hern, Inês do Carmo G. GonÇalves, Johanna Brecht, Eduardo Javier López Soto, Jonah Simon, Natalie Chapkis, Diane Lipscombe, Min Jeong Kye, and Anne C Hart. Decreased microRNA levels lead to deleterious increases in neuronal M2 muscarinic receptors in Spinal Muscular Atrophy models. *eLife*, 6, 2017.

[3441] Hayao Ohno, Morikatsu Yoshida, Takahiro Sato, Johji Kato, Mikiya Miyazato, Masayasu Kojima, Takanori Ida, and Yuichi Iino. Luqin-like RYamide peptides regulate food-evoked responses in C. elegans. *eLife*, 6, 2017.

[3442] Hironobu Okuno, Francois Renault Mihara, Shigeki Ohta, Kimiko Fukuda, Kenji Kurosawa, Wado Akamatsu, Tsukasa Sanosaka, Jun Kohyama, Kanehiro Hayashi, Kazunori Nakajima, Takao Takahashi, Joanna Wysocka, Kenjiro Kosaki, and Hideyuki Okano. CHARGE syndrome modeling using patient-iPSCs reveals defective migration of neural crest cells harboring CHD7 mutations. *eLife*, 6, 2017.

[3443] Ana-Carolina Oliveira, João Francisco Gomes-Neto, Carlos-Henrique Dantas Barbosa, Alessandra Granato, Bernardo S Reis, Bruno Maia Santos, Rita Fucs, Fábio B Canto, Helder I Nakaya, Alberto Nóbrega, and Maria Bellio. Crucial role for T cell-intrinsic IL-18R-MyD88 signaling in cognate immune response to intracellular parasite infection. *eLife*, 6, 2017.

[3444] William Olson, Ishmail Abdus-Saboor, Lian Cui, Justin Burdge, Tobias Raabe, Minghong Ma, and Wenqin Luo. Sparse genetic tracing reveals regionally specific functional organization of mammalian nociceptors. *eLife*, 6, 2017.

[3445] M Teague O'Mara, Martin Wikelski, Christian C Voigt, Andries Ter Maat, Henry S Pollock, Gary Burness, Lanna M Desantis, and Dina KN Dechmann. Cyclic bouts of extreme bradycardia counteract the high metabolism of frugivorous bats. *eLife*, 6, 2017.

[3446] Keisuke Onishi and Yimin Zou. Sonic Hedgehog switches on Wnt/planar cell polarity signaling in commissural axon growth cones by reducing levels of Shisa2. *eLife*, 6, 2017.

[3447] Koun Onodera, Shumpei Baba, Akira Murakami, Tadashi Uemura, and Tadao Usui. Small conductance Ca2+-activated K+ channels induce the firing pause periods during the activation of Drosophila nociceptive neurons. *eLife*, 6, 2017.

[3448] Jacinta O'Shea, Patrice Revol, Helena Cousijn, Jamie Near, Pierre Petitet, Sophie Jacquin-Courtois, Heidi Johansen-Berg, Gilles Rode, and Yves Rossetti. Induced sensorimotor cortex plasticity remediates chronic treatment-resistant visual neglect. *eLife*, 6, 2017.

[3449] Beatriz A Osuna, Conor J Howard, Subheksha KC, Adam Frost, and David E Weinberg. In vitro analysis of RQC activities provides insights into the mechanism and function of CAT tailing. *eLife*, 6, 2017.

[3450] Adriane G Otopalik, Marie L Goeritz, Alexander C Sutton, Ted Brookings, Cosmo Guerini, and Eve Marder. Sloppy morphological tuning in identified neurons of the crustacean stomatogastric ganglion. *eLife*, 6, 2017.

[3451] Adriane G Otopalik, Alexander C Sutton, Matthew Banghart, and Eve Marder. When complex neuronal structures may not matter. *eLife*, 6, 2017.

[3452] Jeroen Overman, Frank Fontaine, Mehdi Moustaqil, Deepak Mittal, Emma Sierecki, Natalia Sacilotto, Johannes Zuegg, Avril AB Robertson, Kelly Holmes, Angela A Salim, Sreeman Mamidyala, Mark S Butler, Ashley S Robinson, Emmanuelle Lesieur, Wayne Johnston, Kirill Alexandrov, Brian L Black, Benjamin M Hogan, Sarah De Val, Robert J Capon, Jason S Carroll, Timothy L Bailey, Peter Koopman, Ralf Jauch, Matthew A Cooper, Yann Gambin, and Mathias Francois. Pharmacological targeting of the transcription factor SOX18 delays breast cancer in mice. *eLife*, 6, 2017.

[3453] B Duygu özpolat, Mette Handberg-Thorsager, Michel Vervoort, and Guillaume Balavoine. Cell lineage and cell cycling analyses of the 4d micromere using live imaging in the marine annelid Platynereis dumerilii. *eLife*, 6, 2017.

[3454] Nicolas A Pabon and Carlos J Camacho. Probing protein flexibility reveals a mechanism for selective promiscuity. *eLife*, 6, 2017.

[3455] Zahid Padamsey, Rudi Tong, and Nigel Emptage. Glutamate is required for depression but not potentiation of long-term presynaptic function. *eLife*, 6, 2017.

[3456] Lars Paeger, Ismene Karakasilioti, Janine Altmüller, Peter Frommolt, Jens Brüning, and Peter Kloppenburg. Antagonistic modulation of NPY/AgRP and POMC neurons in the arcuate nucleus by noradrenalin. *eLife*, 6, 2017.

[3457] Lars Paeger, Andreas Pippow, Simon Hess, Moritz Paehler, Andreas C Klein, Andreas Husch, Christophe Pouzat, Jens C Brüning, and Peter Kloppenburg. Energy imbalance alters Ca2+ handling and excitability of POMC neurons. *eLife*, 6, 2017.

[3458] Athma A Pai, Telmo Henriques, Kayla McCue, Adam Burkholder, Karen Adelman, and Christopher B Burge. The kinetics of pre-mRNA splicing in the Drosophila genome and the influence of gene architecture. *eLife*, 6, 2017.

[3459] Madhukar Pai and Jennifer Furin. Tuberculosis innovations mean little if they cannot save lives. *eLife*, 6, 2017.

[3460] Debjani Pal, Anja Pertot, Nitin H Shirole, Zhan Yao, Naishitha Anaparthy, Tyler Garvin, Hilary Cox, Kenneth Chang, Fred Rollins, Jude Kendall, Leyla Edwards, Vijay A Singh, Gary C Stone, Michael C Schatz, James Hicks, Gregory J Hannon, and Raffaella Sordella. TGF-$\beta$ reduces DNA ds-break repair mechanisms to heighten genetic diversity and adaptability of CD44+/CD24$-$ cancer cells. *eLife*, 6, 2017.

[3461] Luisa F Pallares, Ronan Ledevin, Sophie Pantalacci, Leslie M Turner, Eirikur Steingrimsson, and Sabrina Renaud. Genomic regions controlling shape variation in the first upper molar of the house mouse. *eLife*, 6, 2017.

[3462] Dongqing Pan, Kerstin Klare, Arsen Petrovic, Annika Take, Kai Walstein, Priyanka Singh, Arnaud Rondelet, Alexander W Bird, and Andrea Musacchio. CDK-regulated dimerization of M18BP1 on a Mis18 hexamer is necessary for CENP-A loading. *eLife*, 6, 2017.

[3463] Chethan Pandarinath, Paul Nuyujukian, Christine H Blabe, Brittany L Sorice, Jad Saab, Francis R Willett, Leigh R Hochberg, Krishna V Shenoy, and Jaimie M Henderson. High performance communication by people with paralysis using an intracortical brain-computer interface. *eLife*, 6, 2017.

[3464] M Sabrina Pankey, Randi L Foxall, Ian M Ster, Lauren A Perry, Brian M Schuster, Rachel A Donner, Matthew Coyle, Vaughn S Cooper, and Cheryl A Whistler. Host-selected mutations converging on a global regulator drive an adaptive leap towards symbiosis in bacteria. *eLife*, 6, 2017.

[3465] Brenton R Paolella, William J Gibson, Laura M Urbanski, John A Alberta, Travis I Zack, Pratiti Bandopadhayay, Caitlin A Nichols, Pankaj K Agarwalla, Meredith S Brown, Rebecca Lamothe, Yong Yu, Peter S Choi, Esther A Obeng, Dirk Heckl, Guo Wei, Belinda Wang, Aviad Tsherniak, Francisca Vazquez, Barbara A Weir, David E Root, Glenn S Cowley, Sara J Buhrlage, Charles D Stiles, Benjamin L Ebert, William C Hahn, Robin Reed, and Rameen Beroukhim. Copy-number and gene dependency analysis reveals partial copy loss of wild-type SF3B1 as a novel cancer vulnerability. *eLife*, 6, 2017.

[3466] Alan Jung Park, Robbert Havekes, Xiuping Fu, Rolf Hansen, Jennifer C Tudor, Lucia Peixoto, Zhi Li, Yen-Ching Wu, Shane G Poplawski, Jay M Baraban, and Ted Abel. Learning induces the translin/trax RNase complex to express activin receptors for persistent memory. *eLife*, 6, 2017.

[3467] Eun-Hye Park, Jae-Rong Ahn, and Inah Lee. Interactions between stimulus and response types are more strongly represented in the entorhinal cortex than in its upstream regions in rats. *eLife*, 6, 2017.

[3468] Hyun Bong Park, Corey E Perez, Karl W Barber, Jesse Rinehart, and Jason M Crawford. Genome mining unearths a hybrid nonribosomal peptide synthetase-like-pteridine synthase biosynthetic gene cluster. *eLife*, 6, 2017.

[3469] Junchol Park and Bita Moghaddam. Risk of punishment influences discrete and coordinated encoding of reward-guided actions by prefrontal cortex and VTA neurons. *eLife*, 6, 2017.

[3470] Raghavendran Partha, Bharesh K Chauhan, Zelia Ferreira, Joseph D Robinson, Kira Lathrop, Ken K Nischal, Maria Chikina, and Nathan L Clark. Subterranean mammals show convergent regression in ocular genes and enhancers, along with adaptation to tunneling. *eLife*, 6, 2017.

[3471] Benjamin Pasquereau and Robert S Turner. A selective role for ventromedial subthalamic nucleus in inhibitory control. *eLife*, 6, 2017.

[3472] Tulsi Patel and Oliver Hobert. Coordinated control of terminal differentiation and restriction of cellular plasticity. *eLife*, 6, 2017.

[3473] Irini Pateraki, Johan Andersen-Ranberg, Niels Bjerg Jensen, Sileshi Gizachew Wubshet, Allison Maree Heskes, Victor Forman, Björn Hallström, Britta Hamberger, Mohammed Saddik Motawia, Carl Erik Olsen, Dan Staerk, Jørgen Hansen, Birger Lindberg Møller, and Björn Hamberger. Total biosynthesis of the cyclic AMP booster forskolin from Coleus forskohlii. *eLife*, 6, 2017.

[3474] Ardan Patwardhan, Robert Brandt, Sarah J Butcher, Lucy Collinson, David Gault, Kay Grünewald, Corey Hecksel, Juha T Huiskonen, Andrii Iudin, Martin L Jones, Paul K Korir, Abraham J Koster, Ingvar Lagerstedt, Catherine L Lawson, David Mastronarde, Matthew McCormick, Helen Parkinson, Peter B Rosenthal, Stephan Saalfeld, Helen R Saibil, Sirarat Sarntivijai, Irene Solanes Valero, Sriram Subramaniam, Jason R Swedlow, Ilinca Tudose, Martyn Winn, and Gerard J Kleywegt. Building bridges between cellular and molecular structural biology. *eLife*, 6, 2017.

[3475] Blessy Paul, Hyun Sung Kim, Markus C Kerr, Wilhelmina M Huston, Rohan D Teasdale, and Brett M Collins. Structural basis for the hijacking of endosomal sorting nexin proteins by Chlamydia trachomatis. *eLife*, 6, 2017.

[3476] Anja Paulick, Vladimir Jakovljevic, SiMing Zhang, Michael Erickstad, Alex Groisman, Yigal Meir, William S Ryu, Ned S Wingreen, and Victor Sourjik. Mechanism of bidirectional thermotaxis in Escherichia coli. *eLife*, 6, 2017.

[3477] Cristina Paulino, Yvonne Neldner, Andy KM Lam, Valeria Kalienkova, Janine Denise Brunner, Stephan Schenck, and Raimund Dutzler. Structural basis for anion conduction in the calcium-activated chloride channel TMEM16A. *eLife*, 6, 2017.

[3478] Matthew D Pauly, Megan C Procario, and Adam S Lauring. A novel twelve class fluctuation test reveals higher than expected mutation rates for influenza A viruses. *eLife*, 6, 2017.

[3479] Komal Ishwar Pawar, Katta Suma, Ayshwarya Seenivasan, Santosh Kumar Kuncha, Satya Brata Routh, Shobha P Kruparani, and Rajan Sankaranarayanan. Role of D-aminoacyl-tRNA deacylase beyond chiral proofreading as a cellular defense against glycine mischarging by AlaRS. *eLife*, 6, 2017.

[3480] Sumit Pawar, Rosemarie Ungricht, Peter Tiefenboeck, Jean-Christophe Leroux, and Ulrike Kutay. Efficient protein targeting to the inner nuclear membrane requires Atlastin-dependent maintenance of ER topology. *eLife*, 6, 2017.

[3481] Andrea Pawellek, Ursula Ryder, Triin Tammsalu, Lewis J King, Helmi Kreinin, Tony Ly, Ronald T Hay, Richard C Hartley, and Angus I Lamond. Characterisation of the biflavonoid hinokiflavone as a pre-mRNA splicing modulator that inhibits SENP. *eLife*, 6, 2017.

[3482] Hannah L Payne and Jennifer L Raymond. Magnetic eye tracking in mice. *eLife*, 6, 2017.

[3483] Kaycey Pearce, Diancai Cai, Adam C Roberts, and David L Glanzman. Role of protein synthesis and DNA methylation in the consolidation and maintenance of long-term memory in Aplysia. *eLife*, 6, 2017.

[3484] Sarah F Pearce, Joanna Rorbach, Lindsey Van Haute, Aaron R D'Souza, Pedro Rebelo-Guiomar, Christopher A Powell, Ian Brierley, Andrew E Firth, and Michal Minczuk. Maturation of selected human mitochondrial tRNAs requires deadenylation. *eLife*, 6, 2017.

[3485] Matthias Pechmann, Matthew A Benton, Nathan J Kenny, Nico Posnien, and Siegfried Roth. A novel role for Ets4 in axis specification and cell migration in the spider Parasteatoda tepidariorum. *eLife*, 6, 2017.

[3486] Stano Pekár, Lenka Petráková, Matthew W Bulbert, Martin J Whiting, and Marie E Herberstein. The golden mimicry complex uses a wide spectrum of defence to deter a community of predators. *eLife*, 6, 2017.

[3487] Thaher Pelaseyed, Raghuvir Viswanatha, Cécile Sauvanet, Joshua J Filter, Michael L Goldberg, and Anthony Bretscher. Ezrin activation by LOK phosphorylation involves a PIP2-dependent wedge mechanism. *eLife*, 6, 2017.

[3488] Jose L Pelliccia, Granton A Jindal, and Rebecca D Burdine. Gdf3 is required for robust Nodal signaling during germ layer formation and left-right patterning. *eLife*, 6, 2017.

[3489] Silvia Peluso, Adam Douglas, Alison Hill, Carlo De Angelis, Benjamin L Moore, Graeme Grimes, Giulia Petrovich, Abdelkader Essafi, and Robert E Hill. Fibroblast growth factors (FGFs) prime the limb specific Shh enhancer for chromatin changes that balance histone acetylation mediated by E26 transformation-specific (ETS) factors. *eLife*, 6, 2017.

[3490] Satyakrishna Pentakota, Keda Zhou, Charlotte Smith, Stefano Maffini, Arsen Petrovic, Garry P Morgan, John R Weir, Ingrid R Vetter, Andrea Musacchio, and Karolin Luger. Decoding the centromeric nucleosome through CENP-N. *eLife*, 6, 2017.

[3491] Ernesto Pérez, Jillian L Lindblad, and Andreas Bergmann. Tumor-promoting function of apoptotic caspases by an amplification loop involving ROS, macrophages and JNK in Drosophila. *eLife*, 6, 2017.

[3492] Javier Periz, Jamie Whitelaw, Clare Harding, Simon Gras, Mario Igor Del Rosario Minina, Fernanda Latorre-Barragan, Leandro Lemgruber, Madita Alice Reimer, Robert Insall, Aoife Heaslip, and Markus Meissner. Toxoplasma gondii F-actin forms an extensive filamentous network required for material exchange and parasite maturation. *eLife*, 6, 2017.

[3493] Natalia Petrenko, Yi Jin, Koon Ho Wong, and Kevin Struhl. Evidence that Mediator is essential for Pol II transcription, but is not a required component of the preinitiation complex in vivo. *eLife*, 6, 2017.

[3494] Angela M Phillips, Luna O Gonzalez, Emmanuel E Nekongo, Anna I Ponomarenko, Sean M McHugh, Vincent L Butty, Stuart S Levine, Yu-Shan Lin, Leonid A Mirny, and Matthew D Shoulders. Host proteostasis modulates influenza evolution. *eLife*, 6, 2017.

[3495] Aurèle Piazza, Xiaojie Cui, Michael Adrian, Frédéric Samazan, Brahim Heddi, Anh-Tuan Phan, and Alain G Nicolas. Non-Canonical G-quadruplexes cause the hCEB1 minisatellite instability in Saccharomyces cerevisiae. *eLife*, 6, 2017.

[3496] Joseph Robert Piechura, Kapil Amarnath, and Erin K O'Shea. Natural changes in light interact with circadian regulation at promoters to control gene expression in cyanobacteria. *eLife*, 6, 2017.

[3497] Maryna Pilkiw, Nathan Insel, Younghua Cui, Caitlin Finney, Mark D Morrissey, and Kaori Takehara-Nishiuchi. Phasic and tonic neuron ensemble codes for stimulus-environment conjunctions in the lateral entorhinal cortex. *eLife*, 6, 2017.

[3498] Itziar Pinilla-Macua, Alexandre Grassart, Umamaheswar Duvvuri, Simon C Watkins, and Alexander Sorkin. EGF receptor signaling, phosphorylation, ubiquitylation and endocytosis in tumors in vivo. *eLife*, 6, 2017.

[3499] Ewa Piskadlo, Alexandra Tavares, and Raquel A Oliveira. Metaphase chromosome structure is dynamically maintained by condensin I-directed DNA (de)catenation. *eLife*, 6, 2017.

[3500] Pontus Plavén-Sigray, Granville James Matheson, Björn Christian Schiffler, and William Hedley Thompson. The readability of scientific texts is decreasing over time. *eLife*, 6, 2017.

[3501] William F Podlaski, Alexander Seeholzer, Lukas N Groschner, Gero Miesenböck, Rajnish Ranjan, and Tim P Vogels. Mapping the function of neuronal ion channels in model and experiment. *eLife*, 6, 2017.

[3502] Abigail M Polter, Kelsey Barcomb, Rudy W Chen, Paige M Dingess, Nicholas M Graziane, Travis E Brown, and Julie A Kauer. Constitutive activation of kappa opioid receptors at ventral tegmental area inhibitory synapses following acute stress. *eLife*, 6, 2017.

[3503] Autumn P Pomreinke, Gary H Soh, Katherine W Rogers, Jennifer K Bergmann, Alexander J Bläßle, and Patrick Müller. Dynamics of BMP signaling and distribution during zebrafish dorsal-ventral patterning. *eLife*, 6, 2017.

[3504] Wannarat Pornsiriwong, Gonzalo M Estavillo, Kai Xun Chan, Estee E Tee, Diep Ganguly, Peter A Crisp, Su Yin Phua, Chenchen Zhao, Jiaen Qiu, Jiyoung Park, Miing Tiem Yong, Nazia Nisar, Arun Kumar Yadav, Benjamin Schwessinger, John Rathjen, Christopher I Cazzonelli, Philippa B Wilson, Matthew Gilliham, Zhong-Hua Chen, and Barry J Pogson. A chloroplast retrograde signal, 3′-phosphoadenosine 5′-phosphate, acts as a secondary messenger in abscisic acid signaling in stomatal closure and germination. *eLife*, 6, 2017.

[3505] Eszter Posfai, Sophie Petropoulos, Flavia Regina Oliveira de Barros, John Paul Schell, Igor Jurisica, Rickard Sandberg, Fredrik Lanner, and Janet Rossant. Position- and Hippo signaling-dependent plasticity during lineage segregation in the early mouse embryo. *eLife*, 6, 2017.

[3506] Sebastian Pott. Simultaneous measurement of chromatin accessibility, DNA methylation, and nucleosome phasing in single cells. *eLife*, 6, 2017.

[3507] Paula A Pousinha, Xavier Mouska, Elisabeth F Raymond, Carole Gwizdek, Gihen Dhib, Gwenola Poupon, Laure-Emmanuelle Zaragosi, Camilla Giudici, Ingrid Bethus, Emilie Pacary, Michael Willem, and Hélène Marie. Physiological and pathophysiological control of synaptic GluN2B-NMDA receptors by the C-terminal domain of amyloid precursor protein. *eLife*, 6, 2017.

[3508] Fanny Pouyet, Dominique Mouchiroud, Laurent Duret, and Marie Sémon. Recombination, meiotic expression and human codon usage. *eLife*, 6, 2017.

[3509] Steffen Preissler, Lukas Rohland, Yahui Yan, Ruming Chen, Randy J Read, and David Ron. AMPylation targets the rate-limiting step of BiP's ATPase cycle for its functional inactivation. *eLife*, 6, 2017.

[3510] Alexander M Price, Joanne Dai, Quentin Bazot, Luv Patel, Pavel A Nikitin, Reza Djavadian, Peter S Winter, Cristina A Salinas, Ashley Perkins Barry, Kris C Wood, Eric C Johannsen, Anthony Letai, Martin J Allday, and Micah A Luftig. Epstein-Barr virus ensures B cell survival by uniquely modulating apoptosis at early and late times after infection. *eLife*, 6, 2017.

[3511] Eric A Prinslow, Chad A Brautigam, and Josep Rizo. Reconciling isothermal titration calorimetry analyses of interactions between complexin and truncated SNARE complexes. *eLife*, 6, 2017.

[3512] Ashutosh Pudasaini, Jae Sung Shim, Young Hun Song, Hua Shi, Takatoshi Kiba, David E Somers, Takato Imaizumi, and Brian D Zoltowski. Kinetics of the LOV domain of ZEITLUPE determine its circadian function in Arabidopsis. *eLife*, 6, 2017.

[3513] Erdem Pulcu and Michael Browning. Affective bias as a rational response to the statistics of rewards and punishments. *eLife*, 6, 2017.

[3514] Maria D Purice, Arpita Ray, Eva Jolanda Münzel, Bernard J Pope, Daniel J Park, Sean D Speese, and Mary A Logan. A novel Drosophila injury model reveals severed axons are cleared through a Draper/MMP-1 signaling cascade. *eLife*, 6, 2017.

[3515] Anna M Puszynska and Erin K O'Shea. Switching of metabolic programs in response to light availability is an essential function of the cyanobacterial circadian output pathway. *eLife*, 6, 2017.

[3516] William Putzbach, Quan Q Gao, Monal Patel, Stijn van Dongen, Ashley Haluck-Kangas, Aishe A Sarshad, Elizabeth T Bartom, Kwang-Youn A Kim, Denise M Scholtens, Markus Hafner, Jonathan C Zhao, Andrea E Murmann, and Marcus E Peter. Many si/shRNAs can kill cancer cells by targeting multiple survival genes through an off-target mechanism. *eLife*, 6, 2017.

[3517] Daniela Puzzo, Roberto Piacentini, Mauro Fá, Walter Gulisano, Domenica D Li Puma, Agnes Staniszewski, Hong Zhang, Maria Rosaria Tropea, Sara Cocco, Agostino Palmeri, Paul Fraser, Luciano D'Adamio, Claudio Grassi, and Ottavio Arancio. LTP and memory impairment caused by extracellular A$\beta$ and Tau oligomers is APP-dependent. *eLife*, 6, 2017.

[3518] Edward Pym, Nikhil Sasidharan, Katherine L Thompson-Peer, David J Simon, Anthony Anselmo, Ruslan Sadreyev, Qi Hall, Stephen Nurrish, and Joshua M Kaplan. Shank is a dose-dependent regulator of Cav1 calcium current and CREB target expression. *eLife*, 6, 2017.

[3519] Bin Qi, Marina Kniazeva, and Min Han. A vitamin-B2-sensing mechanism that regulates gut protease activity to impact animal's food behavior and growth. *eLife*, 6, 2017.

[3520] Xingyun Qi, Soon-Ki Han, Jonathan H Dang, Jacqueline M Garrick, Masaki Ito, Alex K Hofstetter, and Keiko U Torii. Autocrine regulation of stomatal differentiation potential by EPF1 and ERECTA-LIKE1 ligand-receptor signaling. *eLife*, 6, 2017.

[3521] Yongjun Qian, Yue Cao, Bowen Deng, Guang Yang, Jiayun Li, Rui Xu, Dandan zhang, Juan Huang, and Yi Rao. Sleep homeostasis regulated by 5HT2b receptor in a small subset of neurons in the dorsal fan-shaped body of drosophila. *eLife*, 6, 2017.

[3522] C Alisha Quandt, Denis Beaudet, Daniele Corsaro, Julia Walochnik, Rolf Michel, Nicolas Corradi, and Timothy Y James. The genome of an intranuclear parasite, Paramicrosporidium saccamoebae, reveals alternative adaptations to obligate intracellular parasitism. *eLife*, 6, 2017.

[3523] Jaclyn E Quin, Ioana Bujila, Mariama Chérif, Guillaume S Sanou, Ying Qu, Manijeh Vafa Homann, Anna Rolicka, Sodiomon B Sirima, Mary A O'Connell, Andreas Lennartsson, Marita Troye-Blomberg, Issa Nebie, and Ann-Kristin östlund Farrants. Major transcriptional changes observed in the Fulani, an ethnic group less susceptible to malaria. *eLife*, 6, 2017.

[3524] Maia A Rabaa, Yves Girerd-Chambaz, Kien Duong Thi Hue, Trung Vu Tuan, Bridget Wills, Matthew Bonaparte, Diane van der Vliet, Edith Langevin, Margarita Cortes, Betzana Zambrano, Corinne Dunod, Anh Wartel-Tram, Nicholas Jackson, and Cameron P Simmons. Genetic epidemiology of dengue viruses in phase III trials of the CYD tetravalent dengue vaccine and implications for efficacy. *eLife*, 6, 2017.

[3525] Julien Racle, Kaat de Jonge, Petra Baumgaertner, Daniel E Speiser, and David Gfeller. Simultaneous enumeration of cancer and immune cell types from bulk tumor gene expression data. *eLife*, 6, 2017.

[3526] Mohammed Mostafizur Rahman, Sonal Kedia, Giselle Fernandes, and Sumantra Chattarji. Activation of the same mGluR5 receptors in the amygdala causes divergent effects on specific versus indiscriminate fear. *eLife*, 6, 2017.

[3527] Jean-Baptiste Raina, Peta L Clode, Soshan Cheong, Jeremy Bougoure, Matt R Kilburn, Anthony Reeder, Sylvain Forêt, Michael Stat, Victor Beltran, Peter Thomas-Hall, Dianne Tapiolas, Cherie M Motti, Bill Gong, Mathieu Pernice, Christopher E Marjo, Justin R Seymour, Bette L Willis, and David G Bourne. Subcellular tracking reveals the location of dimethylsulfoniopropionate in microalgae and visualises its uptake by marine bacteria. *eLife*, 6, 2017.

[3528] Ganesh V Raj, Gangadhara Reddy Sareddy, Shihong Ma, Tae-Kyung Lee, Suryavathi Viswanadhapalli, Rui Li, Xihui Liu, Shino Murakami, Chien-Cheng Chen, Wan-Ru Lee, Monica Mann, Samaya Rajeshwari Krishnan, Bikash Manandhar, Vijay K Gonugunta, Douglas Strand, Rajeshwar Rao Tekmal, Jung-Mo Ahn, and Ratna K Vadlamudi. Estrogen receptor coregulator binding modulators (ERXs) effectively target estrogen receptor positive human breast cancers. *eLife*, 6, 2017.

[3529] Andres Ramirez-Martinez, Bercin Kutluk Cenik, Svetlana Bezprozvannaya, Beibei Chen, Rhonda Bassel-Duby, Ning Liu, and Eric N Olson. KLHL41 stabilizes skeletal muscle sarcomeres by nonproteolytic ubiquitination. *eLife*, 6, 2017.

[3530] Cyrille Ramond, Nicolas Glaser, Claire Berthault, Jacqueline Ameri, Jeannette Schlichting Kirkegaard, Mattias Hansson, Christian Honoré, Henrik Semb, and Raphaël Scharfmann. Reconstructing human pancreatic differentiation by mapping specific cell populations during development. *eLife*, 6, 2017.

[3531] Michal Ramot, Sara Kimmich, Javier Gonzalez-Castillo, Vinai Roopchansingh, Haroon Popal, Emily White, Stephen J Gotts, and Alex Martin. Direct modulation of aberrant brain network connectivity through real-time NeuroFeedback. *eLife*, 6, 2017.

[3532] Péter Rapali, Romain Mitteau, Craig Braun, Aurèlie Massoni-Laporte, Caner ünlü, Laure Bataille, Floriane Saint Arramon, Steven P Gygi, and Derek McCusker. Scaffold-mediated gating of Cdc42 signalling flux. *eLife*, 6, 2017.

[3533] Fahad Rashid, Paul D Harris, Manal S Zaher, Mohamed A Sobhy, Luay I Joudeh, Chunli Yan, Hubert Piwonski, Susan E Tsutakawa, Ivaylo Ivanov, John A Tainer, Satoshi Habuchi, and Samir M Hamdan. Single-molecule FRET unveils induced-fit mechanism for substrate selectivity in flap endonuclease 1. *eLife*, 6, 2017.

[3534] Robert G Rasmussen, Andrew Schwartz, and Steven M Chase. Dynamic range adaptation in primary motor cortical populations. *eLife*, 6, 2017.

[3535] Sarah Rauscher and Régis Pomès. The liquid structure of elastin. *eLife*, 6, 2017.

[3536] Alexandra B Rebocho, Paul Southam, J Richard Kennaway, J Andrew Bangham, and Enrico Coen. Generation of shape complexity through tissue conflict resolution. *eLife*, 6, 2017.

[3537] Naresh S Redhu, Vasudevan Bakthavatchalu, Evan A Conaway, Dror S Shouval, Amy Tsou, Jeremy A Goettel, Amlan Biswas, Chuanwu Wang, Michael Field, Werner Muller, Andre Bleich, Ning Li, Georg K Gerber, Lynn Bry, James G Fox, Scott B Snapper, and Bruce H Horwitz. Macrophage dysfunction initiates colitis during weaning of infant mice lacking the interleukin-10 receptor. *eLife*, 6, 2017.

[3538] William B Redwine, Morgan E DeSantis, Ian Hollyer, Zaw Min Htet, Phuoc Tien Tran, Selene K Swanson, Laurence Florens, Michael P Washburn, and Samara L Reck-Peterson. The human cytoplasmic dynein interactome reveals novel activators of motility. *eLife*, 6, 2017.

[3539] Aviv Regev, Sarah A Teichmann, Eric S Lander, Ido Amit, Christophe Benoist, Ewan Birney, Bernd Bodenmiller, Peter Campbell, Piero Carninci, Menna Clatworthy, Hans Clevers, Bart Deplancke, Ian Dunham, James Eberwine, Roland Eils, Wolfgang Enard, Andrew Farmer, Lars Fugger, Berthold Göttgens, Nir Hacohen, Muzlifah Haniffa, Martin Hemberg, Seung Kim, Paul Klenerman, Arnold Kriegstein, Ed Lein, Sten Linnarsson, Emma Lundberg, Joakim Lundeberg, Partha Majumder, John C Marioni, Miriam Merad, Musa Mhlanga, Martijn Nawijn, Mihai Netea, Garry Nolan, Dana Pe'er, Anthony Phillipakis, Chris P Ponting, Stephen Quake, Wolf Reik, Orit Rozenblatt-Rosen, Joshua Sanes, Rahul Satija, Ton N Schumacher, Alex Shalek, Ehud Shapiro, Padmanee Sharma, Jay W Shin, Oliver Stegle, Michael Stratton, Michael J T Stubbington, Fabian J Theis, Matthias Uhlen, Alexander van Oudenaarden, Allon Wagner, Fiona Watt, Jonathan Weissman, Barbara Wold, Ramnik Xavier, and Nir Yosef. The Human Cell Atlas. *eLife*, 6, 2017.

[3540] Sylvie Remaud, Fernando C Ortiz, Marine Perret-Jeanneret, Marie-Stéphane Aigrot, Jean-David Gothié, Csaba Fekete, Zsuzsanna Kvárta-Papp, Balázs Gereben, Dominique Langui, Catherine Lubetzki, Maria Cecilia Angulo, Bernard Zalc, and Barbara Demeneix. Transient hypothyroidism favors oligodendrocyte generation providing functional remyelination in the adult mouse brain. *eLife*, 6, 2017.

[3541] Yi Ren, Philip Schmiege, and Günter Blobel. Structural and biochemical analyses of the DEAD-box ATPase Sub2 in association with THO or Yra1. *eLife*, 6, 2017.

[3542] Thibaud T Renault, Anthony O Abraham, Tobias Bergmiller, Guillaume Paradis, Simon Rainville, Emmanuelle Charpentier, Călin C Guet, Yuhai Tu, Keiichi Namba, James P Keener, Tohru Minamino, and Marc Erhardt. Bacterial flagella grow through an injection-diffusion mechanism. *eLife*, 6, 2017.

[3543] Nicolas Renier, Chloé Dominici, Reha S Erzurumlu, Claudius F Kratochwil, Filippo M Rijli, Patricia Gaspar, and Alain Chédotal. A mutant with bilateral whisker to barrel inputs unveils somatosensory mapping rules in the cerebral cortex. *eLife*, 6, 2017.

[3544] Maria C Renner, Eva HH Albers, Nicolas Gutierrez-Castellanos, Niels R Reinders, Aile N van Huijstee, Hui Xiong, Tessa R Lodder, and Helmut W Kessels. Synaptic plasticity through activation of GluA3-containing AMPA-receptors. *eLife*, 6, 2017.

[3545] Ronen Reshef, Elena Kudryavitskaya, Haran Shani-Narkiss, Batya Isaacson, Neta Rimmerman, Adi Mizrahi, and Raz Yirmiya. The role of microglia and their CX3CR1 signaling in adult neurogenesis in the olfactory bulb. *eLife*, 6, 2017.

[3546] Jennifer Resnik and Daniel B Polley. Fast-spiking GABA circuit dynamics in the auditory cortex predict recovery of sensory processing following peripheral nerve damage. *eLife*, 6, 2017.

[3547] Kévin Rey, Romain Amiot, FranÇois Fourel, Fernando Abdala, Frédéric Fluteau, Nour-Eddine Jalil, Jun Liu, Bruce S Rubidge, Roger MH Smith, J Sébastien Steyer, Pia A Viglietti, Xu Wang, and Christophe Lécuyer. Oxygen isotopes suggest elevated thermometabolism within multiple Permo-Triassic therapsid clades. *eLife*, 6, 2017.

[3548] Ed Reznik, Qingguo Wang, Konnor La, Nikolaus Schultz, and Chris Sander. Mitochondrial respiratory gene expression is suppressed in many cancers. *eLife*, 6, 2017.

[3549] James Rhodes, Davide Mazza, Kim Nasmyth, and Stephan Uphoff. Scc2/Nipbl hops between chromosomal cohesin rings after loading. *eLife*, 6, 2017.

[3550] Elizabeth Rickenbacher, Rosemarie E Perry, Regina M Sullivan, and Marta A Moita. Freezing suppression by oxytocin in central amygdala allows alternate defensive behaviours and mother-pup interactions. *eLife*, 6, 2017.

[3551] J Peter Rickgauer, Nikolaus Grigorieff, and Winfried Denk. Single-protein detection in crowded molecular environments in cryo-EM images. *eLife*, 6, 2017.

[3552] Michelle M Riehle, Tullu Bukhari, Awa Gneme, Wamdaogo M Guelbeogo, Boubacar Coulibaly, Abdrahamane Fofana, Adrien Pain, Emmanuel Bischoff, Francois Renaud, Abdoul H Beavogui, Sekou F Traore, N'Fale Sagnon, and Kenneth D Vernick. The Anopheles gambiae 2La chromosome inversion is associated with susceptibility to Plasmodium falciparum in Africa. *eLife*, 6, 2017.

[3553] Jacopo Rigosa, Alessandro Lucantonio, Giovanni Noselli, Arash Fassihi, Erik Zorzin, Fabrizio Manzino, Francesca Pulecchi, and Mathew E Diamond. Dye-enhanced visualization of rat whiskers for behavioral studies. *eLife*, 6, 2017.

[3554] Lorenzo Rinaldi, Alexandra Avgustinova, Mercè Martín, Debayan Datta, Guiomar Solanas, Neus Prats, and Salvador Aznar Benitah. Loss of Dnmt3a and Dnmt3b does not affect epidermal homeostasis but promotes squamous transformation through PPAR-$\gamma$. *eLife*, 6, 2017.

[3555] Silvia Ripamonti, Mateusz C Ambrozkiewicz, Francesca Guzzi, Marta Gravati, Gerardo Biella, Ingo Bormuth, Matthieu Hammer, Liam P Tuffy, Albrecht Sigler, Hiroshi Kawabe, Katsuhiko Nishimori, Mauro Toselli, Nils Brose, Marco Parenti, and JeongSeop Rhee. Transient oxytocin signaling primes the development and function of excitatory hippocampal neurons. *eLife*, 6, 2017.

[3556] Zev A Ripstein, Rui Huang, Rafal Augustyniak, Lewis E Kay, and John L Rubinstein. Structure of a AAA+ unfoldase in the process of unfolding substrate. *eLife*, 6, 2017.

[3557] Irene Riva, Clarissa Eibl, Rudolf Volkmer, Anna L Carbone, and Andrew JR Plested. Control of AMPA receptor activity by the extracellular loops of auxiliary proteins. *eLife*, 6, 2017.

[3558] Nadia R Roan, Nathallie Sandi-Monroy, Nargis Kohgadai, Shariq M Usmani, Katherine G Hamil, Jason Neidleman, Mauricio Montano, Ludger Ständker, Annika Röcker, Marielle Cavrois, Jared Rosen, Kara Marson, James F Smith, Christopher D Pilcher, Friedrich Gagsteiger, Olena Sakk, Michael O'Rand, Polina V Lishko, Frank Kirchhoff, Jan Münch, and Warner C Greene. Semen amyloids participate in spermatozoa selection and clearance. *eLife*, 6, 2017.

[3559] Christelle AM Robert, Xi Zhang, Ricardo AR Machado, Stefanie Schirmer, Martina Lori, Pierre Mateo, Matthias Erb, and Jonathan Gershenzon. Sequestration and activation of plant toxins protect the western corn rootworm from enemies at multiple trophic levels. *eLife*, 6, 2017.

[3560] Jerome Robert, Emily B Button, Brian Yuen, Megan Gilmour, Kevin Kang, Arvin Bahrabadi, Sophie Stukas, Wenchen Zhao, Iva Kulic, and Cheryl L Wellington. Clearance of beta-amyloid is facilitated by apolipoprotein E and circulating high-density lipoproteins in bioengineered human vessels. *eLife*, 6, 2017.

[3561] Brooks G Robinson, Alec F Condon, Daniela Radl, Emiliana Borrelli, John T Williams, and Kim A Neve. Cocaine-induced adaptation of dopamine D2S, but not D2L autoreceptors. *eLife*, 6, 2017.

[3562] Nuno Rocha, David A Bulger, Andrea Frontini, Hannah Titheradge, Sigrid Bjerge Gribsholt, Rachel Knox, Matthew Page, Julie Harris, Felicity Payne, Claire Adams, Alison Sleigh, John Crawford, Anette Prior Gjesing, Jette Bork-Jensen, Oluf Pedersen, Inês Barroso, Torben Hansen, Helen Cox, Mary Reilly, Alex Rossor, Rebecca J Brown, Simeon I Taylor, Duncan McHale, Martin Armstrong, Elif A Oral, Vladimir Saudek, Stephen O'Rahilly, Eamonn R Maher, Bjørn Richelsen, David B Savage, and Robert K Semple. Human biallelic MFN2 mutations induce mitochondrial dysfunction, upper body adipose hyperplasia, and suppression of leptin expression. *eLife*, 6, 2017.

[3563] Peter Rodgers. Decisions, decisions. *eLife*, 6, 2017.

[3564] Diana C Rodriguez Camargo, Kyle J Korshavn, Alexander Jussupow, Kolio Raltchev, David Goricanec, Markus Fleisch, Riddhiman Sarkar, Kai Xue, Michaela Aichler, Gabriele Mettenleiter, Axel Karl Walch, Carlo Camilloni, Franz Hagn, Bernd Reif, and Ayyalusamy Ramamoorthy. Stabilization and structural analysis of a membrane-associated hIAPP aggregation intermediate. *eLife*, 6, 2017.

[3565] José A Rodríguez-Martínez, Aaron W Reinke, Devesh Bhimsaria, Amy E Keating, and Aseem Z Ansari. Combinatorial bZIP dimers display complex DNA-binding specificity landscapes. *eLife*, 6, 2017.

[3566] Daniela Roellig, Johanna Tan-Cabugao, Sevan Esaian, and Marianne E Bronner. Dynamic transcriptional signature and cell fate analysis reveals plasticity of individual neural plate border cells. *eLife*, 6, 2017.

[3567] Ofer Rog, Simone Köhler, and Abby F Dernburg. The synaptonemal complex has liquid crystalline properties and spatially regulates meiotic recombination factors. *eLife*, 6, 2017.

[3568] Katherine W Rogers, Nathan D Lord, James A Gagnon, Andrea Pauli, Steven Zimmerman, Deniz C Aksel, Deepak Reyon, Shengdar Q Tsai, J Keith Joung, and Alexander F Schier. Nodal patterning without Lefty inhibitory feedback is functional but fragile. *eLife*, 6, 2017.

[3569] Mohammad Hossein Rohban, Shantanu Singh, Xiaoyun Wu, Julia B Berthet, Mark-Anthony Bray, Yashaswi Shrestha, Xaralabos Varelas, Jesse S Boehm, and Anne E Carpenter. Systematic morphological profiling of human gene and allele function via Cell Painting. *eLife*, 6, 2017.

[3570] Sophie Rohlf, Boukje Habets, Marco von Frieling, and Brigitte Röder. Infants are superior in implicit crossmodal learning and use other learning mechanisms than adults. *eLife*, 6, 2017.

[3571] Benjamin Roland, Thomas Deneux, Kevin M Franks, Brice Bathellier, and Alexander Fleischmann. Odor identity coding by distributed ensembles of neurons in the mouse olfactory cortex. *eLife*, 6, 2017.

[3572] Shunxing Rong, Víctor A Cortés, Shirya Rashid, Norma N Anderson, Jeffrey G McDonald, Guosheng Liang, Young-Ah Moon, Robert E Hammer, and Jay D Horton. Expression of SREBP-1c Requires SREBP-2-mediated Generation of a Sterol Ligand for LXR in Livers of Mice. *eLife*, 6, 2017.

[3573] Irine Ronin, Naama Katsowich, Ilan Rosenshine, and Nathalie Q Balaban. A long-term epigenetic memory switch controls bacterial virulence bimodality. *eLife*, 6, 2017.

[3574] Alberto Roselló-Díez, Daniel Stephen, and Alexandra L Joyner. Altered paracrine signaling from the injured knee joint impairs postnatal long bone growth. *eLife*, 6, 2017.

[3575] Gideon Rosenthal, Michal Tanzer, Erez Simony, Uri Hasson, Marlene Behrmann, and Galia Avidan. Altered topology of neural circuits in congenital prosopagnosia. *eLife*, 6, 2017.

[3576] Rina Rosenzweig, Ashok Sekhar, Jayashree Nagesh, and Lewis E Kay. Promiscuous binding by Hsp70 results in conformational heterogeneity and fuzzy chaperone-substrate ensembles. *eLife*, 6, 2017.

[3577] Timothy J Ross-Elliott, Kaare H Jensen, Katrine S Haaning, Brittney M Wager, Jan Knoblauch, Alexander H Howell, Daniel L Mullendore, Alexander G Monteith, Danae Paultre, Dawei Yan, Sofia Otero, Matthieu Bourdon, Ross Sager, Jung-Youn Lee, Ykä Helariutta, Michael Knoblauch, and Karl J Oparka. Phloem unloading in Arabidopsis roots is convective and regulated by the phloem-pole pericycle. *eLife*, 6, 2017.

[3578] Ian A Roundtree, Guan-Zheng Luo, Zijie Zhang, Xiao Wang, Tao Zhou, Yiquang Cui, Jiahao Sha, Xingxu Huang, Laura Guerrero, Phil Xie, Emily He, Bin Shen, and Chuan He. YTHDC1 mediates nuclear export of N6-methyladenosine methylated mRNAs. *eLife*, 6, 2017.

[3579] Eleonora Russo and Daniel Durstewitz. Cell assemblies at multiple time scales with arbitrary lag constellations. *eLife*, 6, 2017.

[3580] Srivatsun Sadagopan, Wilbert Zarco, and Winrich A Freiwald. A causal relationship between face-patch activity and face-detection behavior. *eLife*, 6, 2017.

[3581] Michael Saitakis, Stéphanie Dogniaux, Christel Goudot, Nathalie Bufi, Sophie Asnacios, Mathieu Maurin, Clotilde Randriamampita, Atef Asnacios, and Claire Hivroz. Different TCR-induced T lymphocyte responses are potentiated by stiffness with variable sensitivity. *eLife*, 6, 2017.

[3582] Naoya Sakamoto, Ying Feng, Carmine Stolfi, Yuki Kurosu, Maranne Green, Jeffry Lin, Megan E Green, Kazuhiro Sentani, Wataru Yasui, Martin McMahon, Karin M Hardiman, Jason R Spence, Nobukatsu Horita, Joel K Greenson, Rork Kuick, Kathleen R Cho, and Eric R Fearon. BRAFV600E cooperates with CDX2 inactivation to promote serrated colorectal tumorigenesis. *eLife*, 6, 2017.

[3583] Sondos Samandi, Annie V Roy, Vivian Delcourt, Jean-FranÇois Lucier, Jules Gagnon, Maxime C Beaudoin, Benoît Vanderperre, Marc-André Breton, Julie Motard, Jean-FranÇois Jacques, Mylène Brunelle, Isabelle Gagnon-Arsenault, Isabelle Fournier, Aida Ouangraoua, Darel J Hunting, Alan A Cohen, Christian R Landry, Michelle S Scott, and Xavier Roucou. Deep transcriptome annotation enables the discovery and functional characterization of cryptic small proteins. *eLife*, 6, 2017.

[3584] Biswajit Samanta and Gerald F Joyce. A reverse transcriptase ribozyme. *eLife*, 6, 2017.

[3585] Bodo Sander, Wenshan Xu, Martin Eilers, Nikita Popov, and Sonja Lorenz. A conformational switch regulates the ubiquitin ligase HUWE1. *eLife*, 6, 2017.

[3586] Imelda T Sandoval, Richard Glenn C Delacruz, Braden N Miller, Shauna Hill, Kristofor A Olson, Ana E Gabriel, Kevin Boyd, Christeena Satterfield, Holly Van Remmen, Jared Rutter, and David A Jones. A metabolic switch controls intestinal differentiation downstream of Adenomatous polyposis coli (APC). *eLife*, 6, 2017.

[3587] Andrew Santiago-Frangos, Jeliazko R Jeliazkov, Jeffrey J Gray, and Sarah A Woodson. Acidic C-terminal domains autoregulate the RNA chaperone Hfq. *eLife*, 6, 2017.

[3588] Joseph M Santin, Mauricio Vallejo, and Lynn K Hartzler. Synaptic up-scaling preserves motor circuit output after chronic, natural inactivity. *eLife*, 6, 2017.

[3589] Jorge M Santos, Saskia Egarter, Vanessa Zuzarte-Luís, Hirdesh Kumar, Catherine A Moreau, Jessica Kehrer, Andreia Pinto, Mário da Costa, Blandine Franke-Fayard, Chris J Janse, Friedrich Frischknecht, and Gunnar R Mair. Malaria parasite LIMP protein regulates sporozoite gliding motility and infectivity in mosquito and mammalian hosts. *eLife*, 6, 2017.

[3590] Paul Victor Sauer, Jennifer Timm, Danni Liu, David Sitbon, Elisabetta Boeri-Erba, Christophe Velours, Norbert Mücke, Jörg Langowski, FranÇoise Ochsenbein, Geneviève Almouzni, and Daniel Panne. Insights into the molecular architecture and histone H3-H4 deposition mechanism of yeast Chromatin assembly factor 1. *eLife*, 6, 2017.

[3591] Francesco Savelli, JD Luck, and James J Knierim. Framing of grid cells within and beyond navigation boundaries. *eLife*, 6, 2017.

[3592] Elise Savier, Stephen J Eglen, Amélie Bathélémy, Martine Perraut, Frank W Pfrieger, Greg Lemke, and Michael Reber. A molecular mechanism for the topographic alignment of convergent neural maps. *eLife*, 6, 2017.

[3593] Elizabeth A Savory, Skylar L Fuller, Alexandra J Weisberg, William J Thomas, Michael I Gordon, Danielle M Stevens, Allison L Creason, Michael S Belcher, Maryna Serdani, Michele S Wiseman, Niklaus J Grünwald, Melodie L Putnam, and Jeff H Chang. Evolutionary transitions between beneficial and phytopathogenic Rhodococcus challenge disease management. *eLife*, 6, 2017.

[3594] Shogo Sawaguchi, Shweta Varshney, Mitsutaka Ogawa, Yuta Sakaidani, Hirokazu Yagi, Kyosuke Takeshita, Toyoaki Murohara, Koichi Kato, Subha Sundaram, Pamela Stanley, and Tetsuya Okajima. O-GlcNAc on NOTCH1 EGF repeats regulates ligand-induced Notch signaling and vascular development in mammals. *eLife*, 6, 2017.

[3595] Volkan I Sayin, Sarah E LeBoeuf, Simranjit X Singh, Shawn M Davidson, Douglas Biancur, Betul S Guzelhan, Samantha W Alvarez, Warren L Wu, Triantafyllia R Karakousi, Anastasia Maria Zavitsanou, Julian Ubriaco, Alexander Muir, Dimitris Karagiannis, Patrick J Morris, Craig J Thomas, Richard Possemato, Matthew G Vander Heiden, and Thales Papagiannakopoulos. Activation of the NRF2 antioxidant program generates an imbalance in central carbon metabolism in cancer. *eLife*, 6, 2017.

[3596] Pierluigi Scerbo, Leslie Marchal, and Laurent Kodjabachian. Lineage commitment of embryonic cells involves MEK1-dependent clearance of pluripotency regulator Ventx2. *eLife*, 6, 2017.

[3597] Michael D Schaller, Gary McDowell, André Porter, Dorothy Shippen, Katherine L Friedman, Matthew S Gentry, Tricia R Serio, and Wesley I Sundquist. What's in a name? *eLife*, 6, 2017.

[3598] Paul E Schavemaker, Wojciech M Śmigiel, and Bert Poolman. Ribosome surface properties may impose limits on the nature of the cytoplasmic proteome. *eLife*, 6, 2017.

[3599] Gisela Schimmack, Kenji Schorpp, Kerstin Kutzner, Torben Gehring, Jara Kerstin Brenke, Kamyar Hadian, and Daniel Krappmann. YOD1/TRAF6 association balances p62-dependent IL-1 signaling to NF-$\kappa$B. *eLife*, 6, 2017.

[3600] Leanne J M Schmitz, Remco V Klaassen, Marta Ruiperez-Alonso, Azra Elia Zamri, Jasper Stroeder, Priyanka Rao-Ruiz, Johannes C Lodder, Rolinka J van der Loo, Huib D Mansvelder, August B Smit, and Sabine Spijker. The AMPA receptor-associated protein Shisa7 regulates hippocampal synaptic function and contextual memory. *eLife*, 6, 2017.

[3601] Christina Schoenherr, Adam Byron, Emma Sandilands, Ketevan Paliashvili, George S Baillie, Amaya Garcia-Munoz, Cristina Valacca, Francesco Cecconi, Bryan Serrels, and Margaret C Frame. Ambra1 spatially regulates Src activity and Src/FAK-mediated cancer cell invasion via trafficking networks. *eLife*, 6, 2017.

[3602] Jacqueline Scholl, Nils Kolling, Natalie Nelissen, Charlotte J Stagg, Catherine J Harmer, and Matthew FS Rushworth. Excitation and inhibition in anterior cingulate predict use of past experiences. *eLife*, 6, 2017.

[3603] Nicole Scholz, Chonglin Guan, Matthias Nieberler, Alexander Grotemeyer, Isabella Maiellaro, Shiqiang Gao, Sebastian Beck, Matthias Pawlak, Markus Sauer, Esther Asan, Sven Rothemund, Jana Winkler, Simone Prömel, Georg Nagel, Tobias Langenhan, and Robert J Kittel. Mechano-dependent signaling by Latrophilin/CIRL quenches cAMP in proprioceptive neurons. *eLife*, 6, 2017.

[3604] Willemijn D Schot, Eli Brenner, and Jeroen BJ Smeets. Unusual prism adaptation reveals how grasping is controlled. *eLife*, 6, 2017.

[3605] Sarah Schuster, Timothy Krüger, Ines Subota, Sina Thusek, Brice Rotureau, Andreas Beilhack, and Markus Engstler. Developmental adaptations of trypanosome motility to the tsetse fly host environments unravel a multifaceted in vivo microswimmer system. *eLife*, 6, 2017.

[3606] Miriam Schwalm, Florian Schmid, Lydia Wachsmuth, Hendrik Backhaus, Andrea Kronfeld, Felipe Aedo Jury, Pierre-Hugues Prouvot, Consuelo Fois, Franziska Albers, Timo van Alst, Cornelius Faber, and Albrecht Stroh. Cortex-wide BOLD fMRI activity reflects locally-recorded slow oscillation-associated calcium waves. *eLife*, 6, 2017.

[3607] Matthew L Schwartz, Daniel P Nickerson, Braden T Lobingier, Rachael L Plemel, Mengtong Duan, Cortney G Angers, Michael Zick, and Alexey J Merz. Sec17 ($\alpha$-SNAP) and an SM-tethering complex regulate the outcome of SNARE zippering in vitro and in vivo. *eLife*, 6, 2017.

[3608] Juliane Schwarz and Henrik Bringmann. Analysis of the NK2 homeobox gene ceh-24 reveals sublateral motor neuron control of left-right turning during sleep. *eLife*, 6, 2017.

[3609] Olivia Schwarz, Ali Asgar Bohra, Xinyu Liu, Heinrich Reichert, Krishnaswamy VijayRaghavan, and Jan Pielage. Motor control of Drosophila feeding behavior. *eLife*, 6, 2017.

[3610] Matthew A Sdano, James M Fulcher, Sowmiya Palani, Mahesh B Chandrasekharan, Timothy J Parnell, Frank G Whitby, Tim Formosa, and Christopher P Hill. A novel SH2 recognition mechanism recruits Spt6 to the doubly phosphorylated RNA polymerase II linker at sites of transcription. *eLife*, 6, 2017.

[3611] Eric Seemann, Minxuan Sun, Sarah Krueger, Jessica Tröger, Wenya Hou, Natja Haag, Susann Schüler, Martin Westermann, Christian A Huebner, Bernd Romeike, Michael M Kessels, and Britta Qualmann. Deciphering caveolar functions by syndapin III KO-mediated impairment of caveolar invagination. *eLife*, 6, 2017.

[3612] Kerstin Seidel, Pauline Marangoni, Cynthia Tang, Bahar Houshmand, Wen Du, Richard L Maas, Steven Murray, Michael C Oldham, and Ophir D Klein. Resolving stem and progenitor cells in the adult mouse incisor through gene co-expression analysis. *eLife*, 6, 2017.

[3613] Lisa Seidel, Barbara Zarzycka, Saheem A Zaidi, Vsevolod Katritch, and Irene Coin. Structural insight into the activation of a class B G-protein-coupled receptor by peptide hormones in live human cells. *eLife*, 6, 2017.

[3614] Vladislav Sekulić and Frances K Skinner. Computational models of O-LM cells are recruited by low or high theta frequency inputs depending on h-channel distributions. *eLife*, 6, 2017.

[3615] Ali Seleit, Isabel Krämer, Bea F Riebesehl, Elizabeth M Ambrosio, Julian S Stolper, Colin Q Lischik, Nicolas Dross, and Lazaro Centanin. Neural stem cells induce the formation of their physical niche during organogenesis. *eLife*, 6, 2017.

[3616] Fatih Semerci, William Tin-Shing Choi, Aleksandar Bajic, Aarohi Thakkar, Juan Manuel Encinas, Frederic Depreux, Neil Segil, Andrew K Groves, and Mirjana Maletic-Savatic. Lunatic fringe-mediated Notch signaling regulates adult hippocampal neural stem cell maintenance. *eLife*, 6, 2017.

[3617] Arnold Y Seo, Pick-Wei Lau, Daniel Feliciano, Prabuddha Sengupta, Mark A Le Gros, Bertrand Cinquin, Carolyn A Larabell, and Jennifer Lippincott-Schwartz. AMPK and vacuole-associated Atg14p orchestrate $\mu$-lipophagy for energy production and long-term survival under glucose starvation. *eLife*, 6, 2017.

[3618] Sachin Sethi and Jing W Wang. A versatile genetic tool for post-translational control of gene expression in Drosophila melanogaster. *eLife*, 6, 2017.

[3619] Sema K Sgaier, Maria Eletskaya, Elisabeth Engl, Owen Mugurungi, Bushimbwa Tambatamba, Gertrude Ncube, Sinokuthemba Xaba, Alice Nanga, Svetlana Gogolina, Patrick Odawo, Sehlulekile Gumede-Moyo, and Steve Kretschmer. A case study for a psychographic-behavioral segmentation approach for targeted demand generation in voluntary medical male circumcision. *eLife*, 6, 2017.

[3620] Neel Shah, Ping Wang, John Wongvipat, Wouter R Karthaus, Wassim Abida, Joshua Armenia, Shira Rockowitz, Yotam Drier, Bradley E Bernstein, Henry W Long, Matthew L Freedman, Vivek K Arora, Deyou Zheng, and Charles L Sawyers. Regulation of the glucocorticoid receptor via a BET-dependent enhancer drives antiandrogen resistance in prostate cancer. *eLife*, 6, 2017.

[3621] Samina Shaheen, Zhengpeng Wan, Zongyu Li, Alicia Chau, Xinxin Li, Shaosen Zhang, Yang Liu, Junyang Yi, Yingyue Zeng, Jing Wang, Xiangjun Chen, Liling Xu, Wei Chen, Fei Wang, Yun Lu, Wenjie Zheng, Yan Shi, Xiaolin Sun, Zhanguo Li, Chunyang Xiong, and Wanli Liu. Substrate stiffness governs the initiation of B cell activation by the concerted signaling of PKC$\beta$ and focal adhesion kinase. *eLife*, 6, 2017.

[3622] Sarah Shailes. Something for everyone. *eLife*, 6, 2017.

[3623] Sarah Shailes. To fund or not to fund? *eLife*, 6, 2017.

[3624] Xiaochuan Shan, Juan Jose Fung, Alan Kosaka, and Gwenn Danet-Desnoyers. Replication Study: Inhibition of BET recruitment to chromatin as an effective treatment for MLL-fusion leukaemia. *eLife*, 6, 2017.

[3625] Guijun Shang, Chad A Brautigam, Rui Chen, Defen Lu, Jesús Torres-Vázquez, and Xuewu Zhang. Structure analyses reveal a regulated oligomerization mechanism of the PlexinD1/GIPC/myosin VI complex. *eLife*, 6, 2017.

[3626] Yongliang Shang, Fuxi Zhu, Lina Wang, Ying-Chun Ouyang, Ming-Zhe Dong, Chao Liu, Haichao Zhao, Xiuhong Cui, Dongyuan Ma, Zhiguo Zhang, Xiaoyu Yang, Yueshuai Guo, Feng Liu, Li Yuan, Fei Gao, Xuejiang Guo, Qing-Yuan Sun, Yunxia Cao, and Wei Li. Essential role for SUN5 in anchoring sperm head to the tail. *eLife*, 6, 2017.

[3627] Ruchira Sharma, Yoshiro Ishimaru, Ian Davison, Kentaro Ikegami, Ming-Shan Chien, Helena You, Quiyi Chi, Momoka Kubota, Masafumi Yohda, Michael Ehlers, and Hiroaki Matsunami. Olfactory receptor accessory proteins play crucial roles in receptor function and gene choice. *eLife*, 6, 2017.

[3628] Melissa J Sharpe, Hannah M Batchelor, and Geoffrey Schoenbaum. Preconditioned cues have no value. *eLife*, 6, 2017.

[3629] Emma S Sherling, Ellen Knuepfer, Joseph A Brzostowski, Louis H Miller, Michael J Blackman, and Christiaan van Ooij. The Plasmodium falciparum rhoptry protein RhopH3 plays essential roles in host cell invasion and nutrient uptake. *eLife*, 6, 2017.

[3630] Shu-Hsien Sheu, Juan Carlos Tapia, Shlomo Tsuriel, and Jeff W Lichtman. Similar synapse elimination motifs at successive relays in the same efferent pathway during development in mice. *eLife*, 6, 2017.

[3631] Cheng Shi, Alexi M Runnels, and Coleen T Murphy. Mating and male pheromone kill Caenorhabditis males through distinct mechanisms. *eLife*, 6, 2017.

[3632] Yu Shi and Fanxin Long. Hedgehog signaling via Gli2 prevents obesity induced by high-fat diet in adult mice. *eLife*, 6, 2017.

[3633] Hirohito Shimizu, Adam D Langenbacher, Jie Huang, Kevin Wang, Georg Otto, Robert Geisler, Yibin Wang, and Jau-Nian Chen. The Calcineurin-FoxO-MuRF1 signaling pathway regulates myofibril integrity in cardiomyocytes. *eLife*, 6, 2017.

[3634] Donghyuk Shin, Wooju Na, Ji-Hyung Lee, Gyuhee Kim, Jiseok Baek, Seok Hee Park, Cheol Yong Choi, and Sangho Lee. Site-specific monoubiquitination downregulates Rab5 by disrupting effector binding and guanine nucleotide conversion. *eLife*, 6, 2017.

[3635] Hyeyoung Shin, Robert Law, Shawn Tsutsui, Christopher I Moore, and Stephanie R Jones. The rate of transient beta frequency events predicts behavior across tasks and species. *eLife*, 6, 2017.

[3636] Atsuko Shirai, Takayuki Kawaguchi, Hideaki Shimojo, Daisuke Muramatsu, Mayumi Ishida-Yonetani, Yoshifumi Nishimura, Hiroshi Kimura, Jun-ichi Nakayama, and Yoichi Shinkai. Impact of nucleic acid and methylated H3K9 binding activities of Suv39h1 on its heterochromatin assembly. *eLife*, 6, 2017.

[3637] Megan Reed Showalter, Jason Hatakeyama, Tomas Cajka, Kacey VanderVorst, Kermit L Carraway, and Oliver Fiehn. Replication Study: The common feature of leukemia-associated IDH1 and IDH2 mutations is a neomorphic enzyme activity converting alpha-ketoglutarate to 2-hydroxyglutarate. *eLife*, 6, 2017.

[3638] Nataly Shtraizent, Charles DeRossi, Shikha Nayar, Ravi Sachidanandam, Liora S Katz, Adam Prince, Anna P Koh, Adam Vincek, Yoav Hadas, Yujin Hoshida, Donald K Scott, Efrat Eliyahu, Hudson H Freeze, Kirsten C Sadler, and Jaime Chu. MPI depletion enhances O-GlcNAcylation of p53 and suppresses the Warburg effect. *eLife*, 6, 2017.

[3639] Leia C Shuhaibar, Jerid W Robinson, Giulia Vigone, Ninna P Shuhaibar, Jeremy R Egbert, Valentina Baena, Tracy F Uliasz, Deborah Kaback, Siu-Pok Yee, Robert Feil, Melanie C Fisher, Caroline N Dealy, Lincoln R Potter, and Laurinda A Jaffe. Dephosphorylation of the NPR2 guanylyl cyclase contributes to inhibition of bone growth by fibroblast growth factor. *eLife*, 6, 2017.

[3640] Anzhalika Sidarovich, Cindy L Will, Maria M Anokhina, Javier Ceballos, Sonja Sievers, Dmitry E Agafonov, Timur Samatov, Penghui Bao, Berthold Kastner, Henning Urlaub, Herbert Waldmann, and Reinhard Lührmann. Identification of a small molecule inhibitor that stalls splicing at an early step of spliceosome activation. *eLife*, 6, 2017.

[3641] Jaydeep Sidhaye and Caren Norden. Concerted action of neuroepithelial basal shrinkage and active epithelial migration ensures efficient optic cup morphogenesis. *eLife*, 6, 2017.

[3642] Kevin D Siegenthaler, Kristeen A Pareja, Jie Wang, and Carolyn S Sevier. An unexpected role for the yeast nucleotide exchange factor Sil1 as a reductant acting on the molecular chaperone BiP. *eLife*, 6, 2017.

[3643] Jennifer Simkin, Thomas R Gawriluk, John C Gensel, and Ashley W Seifert. Macrophages are necessary for epimorphic regeneration in African spiny mice. *eLife*, 6, 2017.

[3644] Fabrizio Simonetti, Tito Candelli, Sebastien Leon, Domenico Libri, and Mathieu Rougemaille. Ubiquitination-dependent control of sexual differentiation in fission yeast. *eLife*, 6, 2017.

[3645] Fabien Sindikubwabo, Shuai Ding, Tahir Hussain, Philippe Ortet, Mohamed Barakat, Sebastian Baumgarten, Dominique Cannella, Andrés Palencia, Alexandre Bougdour, Lucid Belmudes, Yohann Couté, Isabelle Tardieux, Cyrille Y Botté, Artur Scherf, and Mohamed-ali Hakimi. Modifications at K31 on the lateral surface of histone H4 contribute to genome structure and expression in apicomplexan parasites. *eLife*, 6, 2017.

[3646] Esti Singer, Yardena BH Silas, Sigal Ben-Yehuda, and Ophry Pines. Bacterial fumarase and L-malic acid are evolutionary ancient components of the DNA damage response. *eLife*, 6, 2017.

[3647] Conor W Sipe and Sarah E Siegrist. Eyeless uncouples mushroom body neuroblast proliferation from dietary amino acids in Drosophila. *eLife*, 6, 2017.

[3648] Xavier Sisquella, Thomas Nebl, Jennifer K Thompson, Lachlan Whitehead, Brian M Malpede, Nichole D Salinas, Kelly Rogers, Niraj H Tolia, Andrea Fleig, Joseph O'Neill, Wai-Hong Tham, F David Horgen, and Alan F Cowman. Plasmodium falciparum ligand binding to erythrocytes induce alterations in deformability essential for invasion. *eLife*, 6, 2017.

[3649] Ewa Sitarska, Junjie Xu, Seungmee Park, Xiaoxia Liu, Bradley Quade, Karolina Stepien, Kyoko Sugita, Chad A Brautigam, Shuzo Sugita, and Josep Rizo. Autoinhibition of Munc18-1 modulates synaptobrevin binding and helps to enable Munc13-dependent regulation of membrane fusion. *eLife*, 6, 2017.

[3650] Nathan G Skene, Marcia Roy, and Seth GN Grant. A genomic lifespan program that reorganises the young adult brain is targeted in schizophrenia. *eLife*, 6, 2017.

[3651] Peter J Skene and Steven Henikoff. An efficient targeted nuclease strategy for high-resolution mapping of DNA binding sites. *eLife*, 6, 2017.

[3652] Dylan S Small, Terrie E Taylor, Douglas G Postels, Nicholas AV Beare, Jing Cheng, Ian JC MacCormick, and Karl B Seydel. Evidence from a natural experiment that malaria parasitemia is pathogenic in retinopathy-negative cerebral malaria. *eLife*, 6, 2017.

[3653] Daniel Šmít, Coralie Fouquet, Frédéric Pincet, Martin Zapotocky, and Alain Trembleau. Axon tension regulates fasciculation/defasciculation through the control of axon shaft zippering. *eLife*, 6, 2017.

[3654] Alex J Smith, Xiaoming Yao, James A Dix, Byung-Ju Jin, and Alan S Verkman. Test of the 'glymphatic' hypothesis demonstrates diffusive and aquaporin-4-independent solute transport in rodent brain parenchyma. *eLife*, 6, 2017.

[3655] Patrick Smith, David Willemsen, Miriam Popkes, Franziska Metge, Edson Gandiwa, Martin Reichard, and Dario Riccardo Valenzano. Regulation of life span by the gut microbiota in the short-lived African turquoise killifish. *eLife*, 6, 2017.

[3656] H Francis Song, Guangyu R Yang, and Xiao-Jing Wang. Reward-based training of recurrent neural networks for cognitive and value-based tasks. *eLife*, 6, 2017.

[3657] Hongki Song, Amy Orr, Mengtong Duan, Alexey J Merz, and William Wickner. Sec17/Sec18 act twice, enhancing membrane fusion and then disassembling cis-SNARE complexes. *eLife*, 6, 2017.

[3658] Lina Song and Adam D Linstedt. Inhibitor of ppGalNAc-T3-mediated O-glycosylation blocks cancer cell invasiveness and lowers FGF23 levels. *eLife*, 6, 2017.

[3659] Ben Sorum, Beáta Töröcsik, and László Csanády. Asymmetry of movements in CFTR's two ATP sites during pore opening serves their distinct functions. *eLife*, 6, 2017.

[3660] Naoyuki Sotta, Susan Duncan, Mayuki Tanaka, Takafumi Sato, Athanasius FM Marée, Toru Fujiwara, and Verônica A Grieneisen. Rapid transporter regulation prevents substrate flow traffic jams in boron transport. *eLife*, 6, 2017.

[3661] Cristina Spalletti, Claudia Alia, Stefano Lai, Alessandro Panarese, Sara Conti, Silvestro Micera, and Matteo Caleo. Combining robotic training and inactivation of the healthy hemisphere restores pre-stroke motor patterns in mice. *eLife*, 6, 2017.

[3662] Felix Spira, Sara Cuylen-Haering, Shalin Mehta, Matthias Samwer, Anne Reversat, Amitabh Verma, Rudolf Oldenbourg, Michael Sixt, and Daniel W Gerlich. Cytokinesis in vertebrate cells initiates by contraction of an equatorial actomyosin network composed of randomly oriented filaments. *eLife*, 6, 2017.

[3663] Eric C Spivey, Stephen K Jones, James R Rybarski, Fatema A Saifuddin, and Ilya J Finkelstein. An aging-independent replicative lifespan in a symmetrically dividing eukaryote. *eLife*, 6, 2017.

[3664] Michael R Stadler, Jenna E Haines, and Michael B Eisen. Convergence of topological domain boundaries, insulators, and polytene interbands revealed by high-resolution mapping of chromatin contacts in the early Drosophila melanogaster embryo. *eLife*, 6, 2017.

[3665] Benedikt Staffler, Manuel Berning, Kevin M Boergens, Anjali Gour, Patrick van der Smagt, and Moritz Helmstaedter. SynEM, automated synapse detection for connectomics. *eLife*, 6, 2017.

[3666] Martin Steger, Federico Diez, Herschel S Dhekne, Pawel Lis, Raja S Nirujogi, Ozge Karayel, Francesca Tonelli, Terina N Martinez, Esben Lorentzen, Suzanne R Pfeffer, Dario R Alessi, and Matthias Mann. Systematic proteomic analysis of LRRK2-mediated Rab GTPase phosphorylation establishes a connection to ciliogenesis. *eLife*, 6, 2017.

[3667] Ximena Steinberg, Marina A Kasimova, Deny Cabezas-Bratesco, Jason D Galpin, Ernesto Ladron-de Guevara, Federica Villa, Vincenzo Carnevale, Leon Islas, Christopher A Ahern, and Sebastian E Brauchi. Conformational dynamics in TRPV1 channels reported by an encoded coumarin amino acid. *eLife*, 6, 2017.

[3668] Hannah Steinert, Florian Sochor, Anna Wacker, Janina Buck, Christina Helmling, Fabian Hiller, Sara Keyhani, Jonas Noeske, Steffen Grimm, Martin M Rudolph, Heiko Keller, Rachel Anne Mooney, Robert Landick, Beatrix Suess, Boris Fürtig, Jens Wöhnert, and Harald Schwalbe. Pausing guides RNA folding to populate transiently stable RNA structures for riboswitch-based transcription regulation. *eLife*, 6, 2017.

[3669] Jonathan B Steinman, Cristina C Santarossa, Rand M Miller, Lola S Yu, Anna S Serpinskaya, Hideki Furukawa, Sachie Morimoto, Yuta Tanaka, Mitsuyoshi Nishitani, Moriteru Asano, Ruta Zalyte, Alison E Ondrus, Alex G Johnson, Fan Ye, Maxence V Nachury, Yoshiyuki Fukase, Kazuyoshi Aso, Michael A Foley, Vladimir I Gelfand, James K Chen, Andrew P Carter, and Tarun M Kapoor. Chemical structure-guided design of dynapyrazoles, cell-permeable dynein inhibitors with a unique mode of action. *eLife*, 6, 2017.

[3670] Magdalena Steinrueck and Călin C Guet. Complex chromosomal neighborhood effects determine the adaptive potential of a gene under selection. *eLife*, 6, 2017.

[3671] Julia P Steringer, Sascha Lange, Sabína čujová, Radek Šachl, Chetan Poojari, Fabio Lolicato, Oliver Beutel, Hans-Michael Müller, Sebastian Unger, ünal Coskun, Alf Honigmann, Ilpo Vattulainen, Martin Hof, Christian Freund, and Walter Nickel. Key steps in unconventional secretion of fibroblast growth factor 2 reconstituted with purified components. *eLife*, 6, 2017.

[3672] Matthew J Sternfeld, Christopher A Hinckley, Niall J Moore, Matthew T Pankratz, Kathryn L Hilde, Shawn P Driscoll, Marito Hayashi, Neal D Amin, Dario Bonanomi, Wesley D Gifford, Kamal Sharma, Martyn Goulding, and Samuel L Pfaff. Speed and segmentation control mechanisms characterized in rhythmically-active circuits created from spinal neurons produced from genetically-tagged embryonic stem cells. *eLife*, 6, 2017.

[3673] Francesca Stillitano, Jens Hansen, Chi-Wing Kong, Ioannis Karakikes, Christian Funck-Brentano, Lin Geng, Stuart Scott, Stephan Reynier, Ma Wu, Yannick Valogne, Carole Desseaux, Joe-Elie Salem, Dorota Jeziorowska, Noël Zahr, Ronald Li, Ravi Iyengar, Roger J Hajjar, and Jean-Sébastien Hulot. Modeling susceptibility to drug-induced long QT with a panel of subject-specific induced pluripotent stem cells. *eLife*, 6, 2017.

[3674] Alexandra Stolyarova and Alicia Izquierdo. Complementary contributions of basolateral amygdala and orbitofrontal cortex to value learning under uncertainty. *eLife*, 6, 2017.

[3675] Matthew B Stone, Sarah A Shelby, Marcos F Núñez, Kathleen Wisser, and Sarah L Veatch. Protein sorting by lipid phase-like domains supports emergent signaling function in B lymphocyte plasma membranes. *eLife*, 6, 2017.

[3676] Ariana Strandburg-Peshkin, Damien R Farine, Margaret C Crofoot, and Iain D Couzin. Habitat and social factors shape individual decisions and emergent group structure during baboon collective movement. *eLife*, 6, 2017.

[3677] Jonathan Strecker, Sonia Stinus, Mariana Pliego Caballero, Rachel K Szilard, Michael Chang, and Daniel Durocher. A sharp Pif1-dependent threshold separates DNA double-strand breaks from critically short telomeres. *eLife*, 6, 2017.

[3678] Ting Su, Jingdong Cheng, Daniel Sohmen, Rickard Hedman, Otto Berninghausen, Gunnar von Heijne, Daniel N Wilson, and Roland Beckmann. The force-sensing peptide VemP employs extreme compaction and secondary structure formation to induce ribosomal stalling. *eLife*, 6, 2017.

[3679] Kotaro Sugimoto, Subhra P Hui, Delicia Z Sheng, and Kazu Kikuchi. Dissection of zebrafish shha function using site-specific targeting with a Cre-dependent genetic switch. *eLife*, 6, 2017.

[3680] Kenji Sugioka, Danielle R Hamill, Joshua B Lowry, Marie E McNeely, Molly Enrick, Alyssa C Richter, Lauren E Kiebler, James R Priess, and Bruce Bowerman. Centriolar SAS-7 acts upstream of SPD-2 to regulate centriole assembly and pericentriolar material formation. *eLife*, 6, 2017.

[3681] Liou Y Sun, Yimin Fang, Amit Patki, Jacob JE Koopman, David B Allison, Cristal M Hill, Michal M Masternak, Justin Darcy, Jian Wang, Samuel McFadden, and Andrzej Bartke. Longevity is impacted by growth hormone action during early postnatal period. *eLife*, 6, 2017.

[3682] Qi Sun, Xing Zhu, Jia Qi, Weidong An, Pengfei Lan, Dan Tan, Rongchang Chen, Bing Wang, Sanduo Zheng, Cheng Zhang, Xining Chen, Wei Zhang, Jing Chen, Meng-Qiu Dong, and Keqiong Ye. Molecular architecture of the 90S small subunit pre-ribosome. *eLife*, 6, 2017.

[3683] Yidi Sun, Nicole T Leong, Tommy Jiang, Astou Tangara, Xavier Darzacq, and David G Drubin. Switch-like Arp2/3 activation upon WASP and WIP recruitment to an apparent threshold level by multivalent linker proteins in vivo. *eLife*, 6, 2017.

[3684] Arunkumar Sundaram, Rachel Plumb, Suhila Appathurai, and Malaiyalam Mariappan. The Sec61 translocon limits IRE1$\alpha$ signaling during the unfolded protein response. *eLife*, 6, 2017.

[3685] Ramasubramanian Sundaramoorthy, Amanda L Hughes, Vijender Singh, Nicola Wiechens, Daniel P Ryan, Hassane El-Mkami, Maxim Petoukhov, Dmitri I Svergun, Barbara Treutlein, Salina Quack, Monika Fischer, Jens Michaelis, Bettina Böttcher, David G Norman, and Tom Owen-Hughes. Structural reorganization of the chromatin remodeling enzyme Chd1 upon engagement with nucleosomes. *eLife*, 6, 2017.

[3686] Michael W Susman, Edith P Karuna, Ryan C Kunz, Taranjit S Gujral, Andrea V Cantú, Shannon S Choi, Brigette Y Jong, Kyoko Okada, Michael K Scales, Jennie Hum, Linda S Hu, Marc W Kirschner, Ryuichi Nishinakamura, Soichiro Yamada, Diana J Laird, Li-En Jao, Steven P Gygi, Michael E Greenberg, and Hsin-Yi Henry Ho. Kinesin superfamily protein Kif26b links Wnt5a-Ror signaling to the control of cell and tissue behaviors in vertebrates. *eLife*, 6, 2017.

[3687] Shigetomo Suyama, Alexandra Ralevski, Zhong-Wu Liu, Marcelo O Dietrich, Toshihiko Yada, Stephanie E Simonds, Michael A Cowley, Xiao-Bing Gao, Sabrina Diano, and Tamas L Horvath. Plasticity of calcium-permeable AMPA glutamate receptors in Pro-opiomelanocortin neurons. *eLife*, 6, 2017.

[3688] Jennifer C Swart, Monja I Froböse, Jennifer L Cook, Dirk EM Geurts, Michael J Frank, Roshan Cools, and Hanneke EM den Ouden. Catecholaminergic challenge uncovers distinct Pavlovian and instrumental mechanisms of motivated (in)action. *eLife*, 6, 2017.

[3689] Toon Swings, Bram Van den Bergh, Sander Wuyts, Eline Oeyen, Karin Voordeckers, Kevin J Verstrepen, Maarten Fauvart, Natalie Verstraeten, and Jan Michiels. Adaptive tuning of mutation rates allows fast response to lethal stress in Escherichia coli. *eLife*, 6, 2017.

[3690] Mubarak Hussain Syed, Brandon Mark, and Chris Q Doe. Steroid hormone induction of temporal gene expression in Drosophila brain neuroblasts generates neuronal and glial diversity. *eLife*, 6, 2017.

[3691] Johanna L Syrjänen, Iddo Heller, Andrea Candelli, Owen R Davies, Erwin J G Peterman, Gijs J L Wuite, and Luca Pellegrini. Single-molecule observation of DNA compaction by meiotic protein SYCP3. *eLife*, 6, 2017.

[3692] Aleksander D Szymaniak, Rongjuan Mi, Shannon E McCarthy, Adam C Gower, Taylor L Reynolds, Michael Mingueneau, Maria Kukuruzinska, and Xaralabos Varelas. The Hippo pathway effector YAP is an essential regulator of ductal progenitor patterning in the mouse submandibular gland. *eLife*, 6, 2017.

[3693] Elin Sørhus, John P Incardona, Tomasz Furmanek, Giles W Goetz, Nathaniel L Scholz, Sonnich Meier, Rolf B Edvardsen, and Sissel Jentoft. Novel adverse outcome pathways revealed by chemical genetics in a developing marine fish. *eLife*, 6, 2017.

[3694] Jacqueline M Tabler, Maggie M Rigney, Gordon J Berman, Swetha Gopalakrishnan, Eglantine Heude, Hadeel Adel Al-lami, Basil Z Yannakoudakis, Rebecca D Fitch, Christopher Carter, Steven Vokes, Karen J Liu, Shahragim Tajbakhsh, SE Roian Egnor, and John B Wallingford. Cilia-mediated Hedgehog signaling controls form and function in the mammalian larynx. *eLife*, 6, 2017.

[3695] Sina Tafazoli, Houman Safaai, Gioia De Franceschi, Federica Bianca Rosselli, Walter Vanzella, Margherita Riggi, Federica Buffolo, Stefano Panzeri, and Davide Zoccolan. Emergence of transformation-tolerant representations of visual objects in rat lateral extrastriate cortex. *eLife*, 6, 2017.

[3696] Thibaud Taillefumier, Anna Posfai, Yigal Meir, and Ned S Wingreen. Microbial consortia at steady supply. *eLife*, 6, 2017.

[3697] Satohiro Tajima, Kowa Koida, Chihiro I Tajima, Hideyuki Suzuki, Kazuyuki Aihara, and Hidehiko Komatsu. Task-dependent recurrent dynamics in visual cortex. *eLife*, 6, 2017.

[3698] Norihiro Takakuwa, Rikako Kato, Peter Redgrave, and Tadashi Isa. Emergence of visually-evoked reward expectation signals in dopamine neurons via the superior colliculus in V1 lesioned monkeys. *eLife*, 6, 2017.

[3699] Shin-ya Takemura, Yoshinori Aso, Toshihide Hige, Allan Wong, Zhiyuan Lu, C Shan Xu, Patricia K Rivlin, Harald Hess, Ting Zhao, Toufiq Parag, Stuart Berg, Gary Huang, William Katz, Donald J Olbris, Stephen Plaza, Lowell Umayam, Roxanne Aniceto, Lei-Ann Chang, Shirley Lauchie, Omotara Ogundeyi, Christopher Ordish, Aya Shinomiya, Christopher Sigmund, Satoko Takemura, Julie Tran, Glenn C Turner, Gerald M Rubin, and Louis K Scheffer. A connectome of a learning and memory center in the adult Drosophila brain. *eLife*, 6, 2017.

[3700] Shin-ya Takemura, Aljoscha Nern, Dmitri B Chklovskii, Louis K Scheffer, Gerald M Rubin, and Ian A Meinertzhagen. The comprehensive connectome of a neural substrate for 'ON' motion detection in Drosophila. *eLife*, 6, 2017.

[3701] Nicolas Talarek, Elisabeth Gueydon, and Etienne Schwob. Homeostatic control of START through negative feedback between Cln3-Cdk1 and Rim15/Greatwall kinase in budding yeast. *eLife*, 6, 2017.

[3702] Ana Tamarkin-Ben-Harush, Jean-Jacques Vasseur, FranÇoise Debart, Igor Ulitsky, and Rivka Dikstein. Cap-proximal nucleotides via differential eIF4E binding and alternative promoter usage mediate translational response to energy stress. *eLife*, 6, 2017.

[3703] Prasad Tammineni, Xuan Ye, Tuancheng Feng, Daniyal Aikal, and Qian Cai. Impaired retrograde transport of axonal autophagosomes contributes to autophagic stress in Alzheimer's disease neurons. *eLife*, 6, 2017.

[3704] Zhi Xuan Tan and Kang Hao Cheong. Nomadic-colonial life strategies enable paradoxical survival and growth despite habitat destruction. *eLife*, 6, 2017.

[3705] Fei Tang, Peng Zhang, Peiying Ye, Christopher A Lazarski, Qi Wu, Ingrid L Bergin, Timothy P Bender, Michael N Hall, Ya Cui, Liguo Zhang, Taijiao Jiang, Yang Liu, and Pan Zheng. A population of innate myelolymphoblastoid effector cell expanded by inactivation of mTOR complex 1 in mice. *eLife*, 6, 2017.

[3706] Xin Tang, Sanne Roessingh, Sean E Hayley, Michelle L Chu, Nobuaki K Tanaka, Werner Wolfgang, Seongho Song, Ralf Stanewsky, and Fumika N Hamada. The role of PDF neurons in setting the preferred temperature before dawn in Drosophila. *eLife*, 6, 2017.

[3707] Yuki Tanimoto, Akiko Yamazoe-Umemoto, Kosuke Fujita, Yuya Kawazoe, Yosuke Miyanishi, Shuhei J Yamazaki, Xianfeng Fei, Karl Emanuel Busch, Keiko Gengyo-Ando, Junichi Nakai, Yuichi Iino, Yuishi Iwasaki, Koichi Hashimoto, and Koutarou D Kimura. Calcium dynamics regulating the timing of decision-making in C. elegans. *eLife*, 6, 2017.

[3708] Hisashi Tatebe, Shinichi Murayama, Toshiya Yonekura, Tomoyuki Hatano, David Richter, Tomomi Furuya, Saori Kataoka, Kyoko Furuita, Chojiro Kojima, and Kazuhiro Shiozaki. Substrate specificity of TOR complex 2 is determined by a ubiquitin-fold domain of the Sin1 subunit. *eLife*, 6, 2017.

[3709] Marcel J Tauchert, Jean-Baptiste Fourmann, Reinhard Lührmann, and Ralf Ficner. Structural insights into the mechanism of the DEAH-box RNA helicase Prp43. *eLife*, 6, 2017.

[3710] Barbara Tavares, Raquel Jacinto, Pedro Sampaio, Sara Pestana, Andreia Pinto, Andreia Vaz, Mónica Roxo-Rosa, Rui Gardner, Telma Lopes, Britta Schilling, Ian Henry, Leonor Saúde, and Susana Santos Lopes. Notch/Her12 signalling modulates, motile/immotile cilia ratio downstream of Foxj1a in zebrafish left-right organizer. *eLife*, 6, 2017.

[3711] Tiberiu Teşileanu, Bence ölveczky, and Vijay Balasubramanian. Rules and mechanisms for efficient two-stage learning in neural circuits. *eLife*, 6, 2017.

[3712] Michiel M ten Brinke, Shane A Heiney, Xiaolu Wang, Martina Proietti-Onori, Henk-Jan Boele, Jacob Bakermans, Javier F Medina, Zhenyu Gao, and Chris I De Zeeuw. Dynamic modulation of activity in cerebellar nuclei neurons during pavlovian eyeblink conditioning in mice. *eLife*, 6, 2017.

[3713] Inês M Tenente, Madeline N Hayes, Myron S Ignatius, Karin McCarthy, Marielle Yohe, Sivasish Sindiri, Berkley Gryder, Mariana L Oliveira, Ashwin Ramakrishnan, Qin Tang, Eleanor Y Chen, G Petur Nielsen, Javed Khan, and David M Langenau. Myogenic regulatory transcription factors regulate growth in rhabdomyosarcoma. *eLife*, 6, 2017.

[3714] Yayoi Teramoto, Daniel Y Takahashi, Philip Holmes, and Asif A Ghazanfar. Vocal development in a Waddington landscape. *eLife*, 6, 2017.

[3715] Liku B Tezera, Magdalena K Bielecka, Andrew Chancellor, Michaela T Reichmann, Basim Al Shammari, Patience Brace, Alex Batty, Annie Tocheva, Sanjay Jogai, Ben G Marshall, Marc Tebruegge, Suwan N Jayasinghe, Salah Mansour, and Paul T Elkington. Dissection of the host-pathogen interaction in human tuberculosis using a bioengineered 3-dimensional model. *eLife*, 6, 2017.

[3716] J Terrig Thomas, D Eric Dollins, Kristin R Andrykovich, Tehyen Chu, Brian G Stultz, Deborah A Hursh, and Malcolm Moos. SMOC can act as both an antagonist and an expander of BMP signaling. *eLife*, 6, 2017.

[3717] Thornton W Thompson, Alexander Byungsuk Kim, P Jonathan Li, Jiaxi Wang, Benjamin T Jackson, Kristen Ting Hui Huang, Lily Zhang, and David H Raulet. Endothelial cells express NKG2D ligands and desensitize antitumor NK responses. *eLife*, 6, 2017.

[3718] Qinghai Tian, Lars Kaestner, Laura Schröder, Jia Guo, and Peter Lipp. An adaptation of astronomical image processing enables characterization and functional 3D mapping of individual sites of excitation-contraction coupling in rat cardiac muscle. *eLife*, 6, 2017.

[3719] Yuhua Tian, Xianghui Ma, Cong Lv, Xiaole Sheng, Xiang Li, Ran Zhao, Yongli Song, Thomas Andl, Maksim V Plikus, Jinyue Sun, Fazheng Ren, Jianwei Shuai, Christopher J Lengner, Wei Cui, and Zhengquan Yu. Stress responsive miR-31 is a major modulator of mouse intestinal stem cells during regeneration and tumorigenesis. *eLife*, 6, 2017.

[3720] Christian Tiede, Robert Bedford, Sophie J Heseltine, Gina Smith, Imeshi Wijetunga, Rebecca Ross, Danah AlQallaf, Ashley PE Roberts, Alexander Balls, Alistair Curd, Ruth E Hughes, Heather Martin, Sarah R Needham, Laura C Zanetti-Domingues, Yashar Sadigh, Thomas P Peacock, Anna A Tang, Naomi Gibson, Hannah Kyle, Geoffrey W Platt, Nicola Ingram, Thomas Taylor, Louise P Coletta, Iain Manfield, Margaret Knowles, Sandra Bell, Filomena Esteves, Azhar Maqbool, Raj K Prasad, Mark Drinkhill, Robin S Bon, Vikesh Patel, Sarah A Goodchild, Marisa Martin-Fernandez, Ray J Owens, Joanne E Nettleship, Michael E Webb, Michael Harrison, Jonathan D Lippiat, Sreenivasan Ponnambalam, Michelle Peckham, Alastair Smith, Paul Ko Ferrigno, Matt Johnson, Michael J McPherson, and Darren Charles Tomlinson. Affimer proteins are versatile and renewable affinity reagents. *eLife*, 6, 2017.

[3721] M Hadi Timachi, Cedric AJ Hutter, Michael Hohl, Tufa Assafa, Simon Böhm, Anshumali Mittal, Markus A Seeger, and Enrica Bordignon. Exploring conformational equilibria of a heterodimeric ABC transporter. *eLife*, 6, 2017.

[3722] See-Yeun Ting, Nicholas L Yan, Brenda A Schilke, and Elizabeth A Craig. Dual interaction of scaffold protein Tim44 of mitochondrial import motor with channel-forming translocase subunit Tim23. *eLife*, 6, 2017.

[3723] Ali S Titiz, Michael R H Hill, Emily A Mankin, Zahra M Aghajan, Dawn Eliashiv, Natalia Tchemodanov, Uri Maoz, John Stern, Michelle E Tran, Peter Schuette, Eric Behnke, Nanthia A Suthana, and Itzhak Fried. Theta-burst microstimulation in the human entorhinal area improves memory specificity. *eLife*, 6, 2017.

[3724] William F Tobin, Rachel I Wilson, and Wei-Chung Allen Lee. Wiring variations that enable and constrain neural computation in a sensory microcircuit. *eLife*, 6, 2017.

[3725] Ante Tocilj, Kin Fan On, Zuanning Yuan, Jingchuan Sun, Elad Elkayam, Huilin Li, Bruce Stillman, and Leemor Joshua-Tor. Structure of the active form of human origin recognition complex and its ATPase motor module. *eLife*, 6, 2017.

[3726] Kirill Tokarev, Julia Hyland Bruno, Iva Ljubičić, Paresh J Kothari, Santosh A Helekar, Ofer Tchernichovski, and Henning U Voss. Sexual dimorphism in striatal dopaminergic responses promotes monogamy in social songbirds. *eLife*, 6, 2017.

[3727] Romana Tomasoni, Raffaella Morini, Jose P Lopez-Atalaya, Irene Corradini, Alice Canzi, Marco Rasile, Cristina Mantovani, Davide Pozzi, Cecilia Garlanda, Alberto Mantovani, Elisabetta Menna, Angel Barco, and Michela Matteoli. Lack of IL-1R8 in neurons causes hyperactivation of IL-1 receptor pathway and induces MECP2-dependent synaptic defects. *eLife*, 6, 2017.

[3728] Alice Tomassini, Luca Ambrogioni, W Pieter Medendorp, and Eric Maris. Theta oscillations locked to intended actions rhythmically modulate perception. *eLife*, 6, 2017.

[3729] Yusuke Tomina and Daniel A Wagenaar. A double-sided microscope to realize whole-ganglion imaging of membrane potential in the medicinal leech. *eLife*, 6, 2017.

[3730] Fiona J Tooke, Marion Babot, Govind Chandra, Grant Buchanan, and Tracy Palmer. A unifying mechanism for the biogenesis of membrane proteins co-operatively integrated by the Sec and Tat pathways. *eLife*, 6, 2017.

[3731] Eva Torreira, Jaime Alegrio Louro, Irene Pazos, Noelia González-Polo, David Gil-Carton, Ana Garcia Duran, Sébastien Tosi, Oriol Gallego, Olga Calvo, and Carlos Fernández-Tornero. The dynamic assembly of distinct RNA polymerase I complexes modulates rDNA transcription. *eLife*, 6, 2017.

[3732] Alan Tran, Angelina Tang, Colleen T O'Loughlin, Anthony Balistreri, Eric Chang, Doris Coto Villa, Joy Li, Aruna Varshney, Vanessa Jimenez, Jacqueline Pyle, Bryan Tsujimoto, Christopher Wellbrook, Christopher Vargas, Alex Duong, Nebat Ali, Sarah Y Matthews, Samantha Levinson, Sarah Woldemariam, Sami Khuri, Martina Bremer, Daryl K Eggers, Noelle L'Etoile, Laura C Miller Conrad, and Miri K VanHoven. C. elegans avoids toxin-producing Streptomyces using a seven transmembrane domain chemosensory receptor. *eLife*, 6, 2017.

[3733] Christoph D Treiber and Scott Waddell. Resolving the prevalence of somatic transposition in Drosophila. *eLife*, 6, 2017.

[3734] Andrea Trevisiol, Aiman S Saab, Ulrike Winkler, Grit Marx, Hiromi Imamura, Wiebke Möbius, Kathrin Kusch, Klaus-Armin Nave, and Johannes Hirrlinger. Monitoring ATP dynamics in electrically active white matter tracts. *eLife*, 6, 2017.

[3735] Darinka Trübutschek, Sébastien Marti, Andrés Ojeda, Jean-Rémi King, Yuanyuan Mi, Misha Tsodyks, and Stanislas Dehaene. A theory of working memory without consciousness or sustained activity. *eLife*, 6, 2017.

[3736] Torrey LS Truszkowski, Oscar A Carrillo, Julia Bleier, Carolina M Ramirez-Vizcarrondo, Daniel L Felch, Molly McQuillan, Christopher P Truszkowski, Arseny S Khakhalin, and Carlos D Aizenman. A cellular mechanism for inverse effectiveness in multisensory integration. *eLife*, 6, 2017.

[3737] Albert Tsai, Anand K Muthusamy, Mariana RP Alves, Luke D Lavis, Robert H Singer, David L Stern, and Justin Crocker. Nuclear microenvironments modulate transcription from low-affinity enhancers. *eLife*, 6, 2017.

[3738] Teja Turk, Nadine Bachmann, Claus Kadelka, Jürg Böni, Sabine Yerly, Vincent Aubert, Thomas Klimkait, Manuel Battegay, Enos Bernasconi, Alexandra Calmy, Matthias Cavassini, Hansjakob Furrer, Matthias Hoffmann, Huldrych F Günthard, Roger D Kouyos, V Aubert, V Aubert, M Battegay, E Bernasconi, J Böni, DL Braun, HC Bucher, A Calmy, M Cavassini, A Ciuffi, G Dollenmaier, M Egger, L Elzi, J Fehr, J Fellay, H Furrer, CA Fux, HF Günthard, D Haerry, B Hasse, HH Hirsch, M Hoffmann, I Hösli, C Kahlert, L Kaiser, O Keiser, T Klimkait, RD Kouyos, H Kovari, B Ledergerber, G Martinetti, B Martinez de Tejada, C Marzolini, KJ Metzner, N Müller, D Nicca, G Pantaleo, P Paioni, A Rauch, C Rudin, AU Scherrer, P Schmid, R Speck, M Stöckle, P Tarr, A Trkola, P Vernazza, G Wandeler, R Weber, and S Yerly. Assessing the danger of self-sustained HIV epidemics in heterosexuals by population based phylogenetic cluster analysis. *eLife*, 6, 2017.

[3739] Daniel Turner-Evans, Stephanie Wegener, Hervé Rouault, Romain Franconville, Tanya Wolff, Johannes D Seelig, Shaul Druckmann, and Vivek Jayaraman. Angular velocity integration in a fly heading circuit. *eLife*, 6, 2017.

[3740] Akira Uchida, Divakaran Murugesapillai, Markus Kastner, Yao Wang, Maria F Lodeiro, Shaan Prabhakar, Guinevere V Oliver, Jamie J Arnold, L James Maher, Mark C Williams, and Craig E Cameron. Unexpected sequences and structures of mtDNA required for efficient transcription from the first heavy-strand promoter. *eLife*, 6, 2017.

[3741] Kohei Ueno, Ema Suzuki, Shintaro Naganos, Kyoko Ofusa, Junjiro Horiuchi, and Minoru Saitoe. Coincident postsynaptic activity gates presynaptic dopamine release to induce plasticity in Drosophila mushroom bodies. *eLife*, 6, 2017.

[3742] Giovanni Stefano Ugolini, Andrea Pavesi, Marco Rasponi, Gianfranco Beniamino Fiore, Roger Kamm, and Monica Soncini. Human cardiac fibroblasts adaptive responses to controlled combined mechanical strain and oxygen changes in vitro. *eLife*, 6, 2017.

[3743] Rachel T Uren, Martin O'Hely, Sweta Iyer, Ray Bartolo, Melissa X Shi, Jason M Brouwer, Amber E Alsop, Grant Dewson, and Ruth M Kluck. Disordered clusters of Bak dimers rupture mitochondria during apoptosis. *eLife*, 6, 2017.

[3744] James R Usherwood and Zoe T Self Davies. Work minimization accounts for footfall phasing in slow quadrupedal gaits. *eLife*, 6, 2017.

[3745] Shahabeddin Vahdat, Stuart Fogel, Habib Benali, and Julien Doyon. Network-wide reorganization of procedural memory during NREM sleep revealed by fMRI. *eLife*, 6, 2017.

[3746] Markus P Vallaster, Shweta Kukreja, Xin Y Bing, Jennifer Ngolab, Rubing Zhao-Shea, Paul D Gardner, Andrew R Tapper, and Oliver J Rando. Paternal nicotine exposure alters hepatic xenobiotic metabolism in offspring. *eLife*, 6, 2017.

[3747] Björn van der Hoort, Maria Reingardt, and H Henrik Ehrsson. Body ownership promotes visual awareness. *eLife*, 6, 2017.

[3748] Petra van der Lelij, Simone Lieb, Julian Jude, Gordana Wutz, Catarina P Santos, Katrina Falkenberg, Andreas Schlattl, Jozef Ban, Raphaela Schwentner, Thomas Hoffmann, Heinrich Kovar, Francisco X Real, Todd Waldman, Mark A Pearson, Norbert Kraut, Jan-Michael Peters, Johannes Zuber, and Mark Petronczki. Synthetic lethality between the cohesin subunits STAG1 and STAG2 in diverse cancer contexts. *eLife*, 6, 2017.

[3749] Ramon A van der Valk, Jocelyne Vreede, Liang Qin, Geri F Moolenaar, Andreas Hofmann, Nora Goosen, and Remus T Dame. Mechanism of environmentally driven conformational changes that modulate H-NS DNA-bridging activity. *eLife*, 6, 2017.

[3750] Katharina van Pee, Alexander Neuhaus, Edoardo D'Imprima, Deryck J Mills, Werner Kühlbrandt, and özkan Yildiz. CryoEM structures of membrane pore and prepore complex reveal cytolytic mechanism of Pneumolysin. *eLife*, 6, 2017.

[3751] Wilhelmina E van Riel, Ankit Rai, Sarah Bianchi, Eugene A Katrukha, Qingyang Liu, Albert JR Heck, Casper C Hoogenraad, Michel O Steinmetz, Lukas C Kapitein, and Anna Akhmanova. Kinesin-4 KIF21B is a potent microtubule pausing factor. *eLife*, 6, 2017.

[3752] Laurens M van Tienen, Juliusz Mieszczanek, Marc Fiedler, Trevor J Rutherford, and Mariann Bienz. Constitutive scaffolding of multiple Wnt enhanceosome components by Legless/BCL9. *eLife*, 6, 2017.

[3753] Michiel van Veen, Elisa Matas-Rico, Koen van de Wetering, Daniela Leyton-Puig, Katarzyna M Kedziora, Valentina De Lorenzi, Yvette Stijf-Bultsma, Bram van den Broek, Kees Jalink, Nicolai Sidenius, Anastassis Perrakis, and Wouter H Moolenaar. Negative regulation of urokinase receptor activity by a GPI-specific phospholipase C in breast cancer cells. *eLife*, 6, 2017.

[3754] Sebastian A Vasquez-Lopez, Yves Weissenberger, Michael Lohse, Peter Keating, Andrew J King, and Johannes C Dahmen. Thalamic input to auditory cortex is locally heterogeneous but globally tonotopic. *eLife*, 6, 2017.

[3755] Christopher N Vassallo, Pengbo Cao, Austin Conklin, Hayley Finkelstein, Christopher S Hayes, and Daniel Wall. Infectious polymorphic toxins delivered by outer membrane exchange discriminate kin in myxobacteria. *eLife*, 6, 2017.

[3756] Pavan Vedula, Satoshi Kurosaka, Nicolae Adrian Leu, Yuri I Wolf, Svetlana A Shabalina, Junling Wang, Stephanie Sterling, Dawei W Dong, and Anna Kashina. Diverse functions of homologous actin isoforms are defined by their nucleotide, rather than their amino acid sequence. *eLife*, 6, 2017.

[3757] Oscar Velazquez Camacho, Carmen Galan, Kalina Swist-Rosowska, Reagan Ching, Michael Gamalinda, Fethullah Karabiber, Inti De La Rosa-Velazquez, Bettina Engist, Birgit Koschorz, Nicholas Shukeir, Megumi Onishi-Seebacher, Suzanne van de Nobelen, and Thomas Jenuwein. Major satellite repeat RNA stabilize heterochromatin retention of Suv39h enzymes by RNA-nucleosome association and RNA:DNA hybrid formation. *eLife*, 6, 2017.

[3758] A Catalina Vélez-Ortega, Mary J Freeman, Artur A Indzhykulian, Jonathan M Grossheim, and Gregory I Frolenkov. Mechanotransduction current is essential for stability of the transducing stereocilia in mammalian auditory hair cells. *eLife*, 6, 2017.

[3759] Donovan Ventimiglia and Cornelia I Bargmann. Diverse modes of synaptic signaling, regulation, and plasticity distinguish two classes of C. elegans glutamatergic neurons. *eLife*, 6, 2017.

[3760] Csaba Verasztó, Nobuo Ueda, Luis A Bezares-Calderón, Aurora Panzera, Elizabeth A Williams, Réza Shahidi, and Gáspár Jékely. Ciliomotor circuitry underlying whole-body coordination of ciliary activity in the Platynereis larva. *eLife*, 6, 2017.

[3761] Daniëlle Rianne José Verboogen, Natalia González Mancha, Martin ter Beest, and Geert van den Bogaart. Fluorescence Lifetime Imaging Microscopy reveals rerouting of SNARE trafficking driving dendritic cell activation. *eLife*, 6, 2017.

[3762] Yvonne Vercoulen, Yasushi Kondo, Jeffrey S Iwig, Axel B Janssen, Katharine A White, Mojtaba Amini, Diane L Barber, John Kuriyan, and Jeroen P Roose. A Histidine pH sensor regulates activation of the Ras-specific guanine nucleotide exchange factor RasGRP1. *eLife*, 6, 2017.

[3763] Judit M Veres, Gergő A Nagy, and Norbert Hájos. Perisomatic GABAergic synapses of basket cells effectively control principal neuron activity in amygdala networks. *eLife*, 6, 2017.

[3764] Sujith Vijayan, Kyle Q Lepage, Nancy J Kopell, and Sydney S Cash. Frontal beta-theta network during REM sleep. *eLife*, 6, 2017.

[3765] Frans Vinberg, Tian Wang, Alicia De Maria, Haiqing Zhao, Steven Bassnett, Jeannie Chen, and Vladimir J Kefalov. The Na+/Ca2+, K+ exchanger NCKX4 is required for efficient cone-mediated vision. *eLife*, 6, 2017.

[3766] Ines R Violante, Lucia M Li, David W Carmichael, Romy Lorenz, Robert Leech, Adam Hampshire, John C Rothwell, and David J Sharp. Externally induced frontoparietal synchronization modulates network dynamics and enhances working memory performance. *eLife*, 6, 2017.

[3767] Jyothsna Visweswaraiah and Alan G Hinnebusch. Interface between 40S exit channel protein uS7/Rps5 and eIF2$\alpha$ modulates start codon recognition in vivo. *eLife*, 6, 2017.

[3768] Oscar Vivas, Claudia M Moreno, Luis F Santana, and Bertil Hille. Proximal clustering between BK and CaV1.3 channels promotes functional coupling and BK channel activation at low voltage. *eLife*, 6, 2017.

[3769] Oliver Vöcking, Ioannis Kourtesis, Sharat Chandra Tumu, and Harald Hausen. Co-expression of xenopsin and rhabdomeric opsin in photoreceptors bearing microvilli and cilia. *eLife*, 6, 2017.

[3770] Sven Kenjiro Vogel, Ferdinand Greiss, Alena Khmelinskaia, and Petra Schwille. Control of lipid domain organization by a biomimetic contractile actomyosin cortex. *eLife*, 6, 2017.

[3771] Mattia Volta, Dayne A Beccano-Kelly, Sarah A Paschall, Stefano Cataldi, Sarah E MacIsaac, Naila Kuhlmann, Chelsie A Kadgien, Igor Tatarnikov, Jesse Fox, Jaskaran Khinda, Emma Mitchell, Sabrina Bergeron, Heather Melrose, Matthew J Farrer, and Austen J Milnerwood. Initial elevations in glutamate and dopamine neurotransmission decline with age, as does exploratory behavior, in LRRK2 G2019S knock-in mice. *eLife*, 6, 2017.

[3772] Daniel von Wangenheim, Robert Hauschild, MatyáŠ Fendrych, Vanessa Barone, Eva Benková, and Jiří Friml. Live tracking of moving samples in confocal microscopy for vertically grown roots. *eLife*, 6, 2017.

[3773] Annemarie Voorberg-van der Wel, Guglielmo Roma, Devendra Kumar Gupta, Sven Schuierer, Florian Nigsch, Walter Carbone, Anne-Marie Zeeman, Boon Heng Lee, Sam O Hofman, Bart W Faber, Judith Knehr, Erica Pasini, Bernd Kinzel, Pablo Bifani, Ghislain M C Bonamy, Tewis Bouwmeester, Clemens H M Kocken, and Thierry Tidiane Diagana. A comparative transcriptomic analysis of replicating and dormant liver stages of the relapsing malaria parasite Plasmodium cynomolgi. *eLife*, 6, 2017.

[3774] Thanh Thi Kim Vuong-Brender, Martine Ben Amar, Julien Pontabry, and Michel Labouesse. The interplay of stiffness and force anisotropies drives embryo elongation. *eLife*, 6, 2017.

[3775] Takeshi Wakida, Masae Ikura, Kenji Kuriya, Shinji Ito, Yoshiharu Shiroiwa, Toshiyuki Habu, Takuo Kawamoto, Katsuzumi Okumura, Tsuyoshi Ikura, and Kanji Furuya. The CDK-PLK1 axis targets the DNA damage checkpoint sensor protein RAD9 to promote cell proliferation and tolerance to genotoxic stress. *eLife*, 6, 2017.

[3776] Lauren J Walker, Daniel W Summers, Yo Sasaki, EJ Brace, Jeffrey Milbrandt, and Aaron DiAntonio. MAPK signaling promotes axonal degeneration by speeding the turnover of the axonal maintenance factor NMNAT2. *eLife*, 6, 2017.

[3777] Alexander M Walter, Rainer Müller, Bassam Tawfik, Keimpe DB Wierda, Paulo S Pinheiro, André Nadler, Anthony W McCarthy, Iwona Ziomkiewicz, Martin Kruse, Gregor Reither, Jens Rettig, Martin Lehmann, Volker Haucke, Bertil Hille, Carsten Schultz, and Jakob Balslev Sørensen. Phosphatidylinositol 4,5-bisphosphate optical uncaging potentiates exocytosis. *eLife*, 6, 2017.

[3778] Jun Wan, Yijing Su, Qifeng Song, Brian Tung, Olutobi Oyinlade, Sheng Liu, Mingyao Ying, Guo-li Ming, Hongjun Song, Jiang Qian, Heng Zhu, and Shuli Xia. Methylated cis-regulatory elements mediate KLF4-dependent gene transactivation and cell migration. *eLife*, 6, 2017.

[3779] Chenggang Wang, Mingqi Zhou, Xudong Zhang, Jin Yao, Yanping Zhang, and Zhonglin Mou. A lectin receptor kinase as a potential sensor for extracellular nicotinamide adenine dinucleotide in Arabidopsis thaliana. *eLife*, 6, 2017.

[3780] Chien-Hsiang Wang, Yi-Chun Huang, Pei-Yi Chen, Ying-Ju Cheng, Hsiu-Hua Kao, Haiwei Pi, and Cheng-Ting Chien. USP5/Leon deubiquitinase confines postsynaptic growth by maintaining ubiquitin homeostasis through Ubiquilin. *eLife*, 6, 2017.

[3781] Hansen Wang, Lisa D Muiznieks, Punam Ghosh, Declan Williams, Michael Solarski, Andrew Fang, Alejandro Ruiz-Riquelme, Régis Pomès, Joel C Watts, Avi Chakrabartty, Holger Wille, Simon Sharpe, and Gerold Schmitt-Ulms. Somatostatin binds to the human amyloid $\beta$ peptide and favors the formation of distinct oligomers. *eLife*, 6, 2017.

[3782] Haoqing Wang, Harry B Gristick, Louise Scharf, Anthony P West, Rachel P Galimidi, Michael S Seaman, Natalia T Freund, Michel C Nussenzweig, and Pamela J Bjorkman. Asymmetric recognition of HIV-1 Envelope trimer by V1V2 loop-targeting antibodies. *eLife*, 6, 2017.

[3783] Hui Qin Wang, Ensar Halilovic, Xiaoyan Li, Jinsheng Liang, Yichen Cao, Daniel P Rakiec, David A Ruddy, Sebastien Jeay, Jens U Wuerthner, Noelito Timple, Shailaja Kasibhatla, Nanxin Li, Juliet A Williams, William R Sellers, Alan Huang, and Fang Li. Combined ALK and MDM2 inhibition increases antitumor activity and overcomes resistance in human ALK mutant neuroblastoma cell lines and xenograft models. *eLife*, 6, 2017.

[3784] Jennifer T Wang, Dong Kong, Christian R Hoerner, Jadranka Loncarek, and Tim Stearns. Centriole triplet microtubules are required for stable centriole formation and inheritance in human cells. *eLife*, 6, 2017.

[3785] Jing Wang, Feng Li, Oscar D Bello, Charles Vaughn Sindelar, Frédéric Pincet, Shyam S Krishnakumar, and James E Rothman. Circular oligomerization is an intrinsic property of synaptotagmin. *eLife*, 6, 2017.

[3786] Miranda Wang, Michael Ly, Andrew Lugowski, John D Laver, Howard D Lipshitz, Craig A Smibert, and Olivia S Rissland. ME31B globally represses maternal mRNAs by two distinct mechanisms during the Drosophila maternal-to-zygotic transition. *eLife*, 6, 2017.

[3787] Mu-Yun Wang and Hideaki Takeuchi. Individual recognition and the 'face inversion effect' in medaka fish (Oryzias latipes). *eLife*, 6, 2017.

[3788] Qixuan Wang, Ji Won Oh, Hye-Lim Lee, Anukriti Dhar, Tao Peng, Raul Ramos, Christian Fernando Guerrero-Juarez, Xiaojie Wang, Ran Zhao, Xiaoling Cao, Jonathan Le, Melisa A Fuentes, Shelby C Jocoy, Antoni R Rossi, Brian Vu, Kim Pham, Xiaoyang Wang, Nanda Maya Mali, Jung Min Park, June-Hyug Choi, Hyunsu Lee, Julien M D Legrand, Eve Kandyba, Jung Chul Kim, Moonkyu Kim, John Foley, Zhengquan Yu, Krzysztof Kobielak, Bogi Andersen, Kiarash Khosrotehrani, Qing Nie, and Maksim V Plikus. A multi-scale model for hair follicles reveals heterogeneous domains driving rapid spatiotemporal hair growth patterning. *eLife*, 6, 2017.

[3789] Wei Wang, Wan Zhao, Jing Li, Lan Luo, Le Kang, and Feng Cui. The c-Jun N-terminal kinase pathway of a vector insect is activated by virus capsid protein and promotes viral replication. *eLife*, 6, 2017.

[3790] Xianhua Wang, Xing Zhang, Di Wu, Zhanglong Huang, Tingting Hou, Chongshu Jian, Peng Yu, Fujian Lu, Rufeng Zhang, Tao Sun, Jinghang Li, Wenfeng Qi, Yanru Wang, Feng Gao, and Heping Cheng. Mitochondrial flashes regulate ATP homeostasis in the heart. *eLife*, 6, 2017.

[3791] Xinbo Wang, Shanshan Li, Haicheng Wang, Wenqing Shui, and Junjie Hu. Quantitative proteomics reveal proteins enriched in tubular endoplasmic reticulum of Saccharomyces cerevisiae. *eLife*, 6, 2017.

[3792] Yi Wang, Xu Zhang, Huihui Huang, Yin Xia, YiFei Yao, Arthur Fuk-Tat Mak, Patrick Shu-Hang Yung, Kai-Ming Chan, Li Wang, Chenglin Zhang, Yu Huang, and Kingston King-Lun Mak. Osteocalcin expressing cells from tendon sheaths in mice contribute to tendon repair by activating Hedgehog signaling. *eLife*, 6, 2017.

[3793] Zhao Wang, Guizhen Fan, Corey F Hryc, James N Blaza, Irina I Serysheva, Michael F Schmid, Wah Chiu, Ben F Luisi, and Dijun Du. An allosteric transport mechanism for the AcrAB-TolC multidrug efflux pump. *eLife*, 6, 2017.

[3794] Zhaowei Wang, Xiaoling Xia, Xueli Yang, Xueyi Zhang, Yongxiang Liu, Di Wu, Yuan Fang, Yujie Liu, Jiuyue Xu, Yang Qiu, and Xi Zhou. A picorna-like virus suppresses the N-end rule pathway to inhibit apoptosis. *eLife*, 6, 2017.

[3795] Elizabeth V Wasmuth, John C Zinder, Dimitrios Zattas, Mom Das, and Christopher D Lima. Structure and reconstitution of yeast Mpp6-nuclear exosome complexes reveals that Mpp6 stimulates RNA decay and recruits the Mtr4 helicase. *eLife*, 6, 2017.

[3796] Mika Watanabe, Ken Natsuga, Wataru Nishie, Yasuaki Kobayashi, Giacomo Donati, Shotaro Suzuki, Yu Fujimura, Tadasuke Tsukiyama, Hideyuki Ujiie, Satoru Shinkuma, Hideki Nakamura, Masamoto Murakami, Michitaka Ozaki, Masaharu Nagayama, Fiona M Watt, and Hiroshi Shimizu. Type XVII collagen coordinates proliferation in the interfollicular epidermis. *eLife*, 6, 2017.

[3797] Jake F Watson, Hinze Ho, and Ingo H Greger. Synaptic transmission and plasticity require AMPA receptor anchoring via its N-terminal domain. *eLife*, 6, 2017.

[3798] James Watson, Walter RJ Taylor, Didier Menard, Sim Kheng, and Nicholas J White. Modelling primaquine-induced haemolysis in G6PD deficiency. *eLife*, 6, 2017.

[3799] Michael Weber, Nico Scherf, Alexander M Meyer, Daniela Panáková, Peter Kohl, and Jan Huisken. Cell-accurate optical mapping across the entire developing heart. *eLife*, 6, 2017.

[3800] Pascal Weber, Helena Batoulis, Kerstin M Rink, Stefan Dahlhoff, Kerstin Pinkwart, Thomas H Söllner, and Thorsten Lang. Electrostatic anchoring precedes stable membrane attachment of SNAP25/SNAP23 to the plasma membrane. *eLife*, 6, 2017.

[3801] Andrew Weems and Michael McMurray. The step-wise pathway of septin hetero-octamer assembly in budding yeast. *eLife*, 6, 2017.

[3802] Simon Weinberger, Matthew P Topping, Jiekun Yan, Annelies Claeys, Natalie De Geest, Duru Ozbay, Talah Hassan, Xiaoli He, Joerg T Albert, Bassem A Hassan, and Ariane Ramaekers. Evolutionary changes in transcription factor coding sequence quantitatively alter sensory organ development and function. *eLife*, 6, 2017.

[3803] Nicholas R Weir, Roarke A Kamber, James S Martenson, and Vladimir Denic. The AAA protein Msp1 mediates clearance of excess tail-anchored proteins from the peroxisomal membrane. *eLife*, 6, 2017.

[3804] Alexandria C Wells, Keith A Daniels, Constance C Angelou, Eric Fagerberg, Amy S Burnside, Michele Markstein, Dominique Alfandari, Raymond M Welsh, Elena L Pobezinskaya, and Leonid A Pobezinsky. Modulation of let-7 miRNAs controls the differentiation of effector CD8 T cells. *eLife*, 6, 2017.

[3805] Brent S Wells, Daniela Pistillo, Erin Barnhart, and Claude Desplan. Parallel Activin and BMP signaling coordinates R7/R8 photoreceptor subtype pairing in the stochastic Drosophila retina. *eLife*, 6, 2017.

[3806] Jung-Kun Wen, Yi-Ting Wang, Chih-Chiang Chan, Cheng-Wen Hsieh, Hsiao-Man Liao, Chin-Chun Hung, and Guang-Chao Chen. Atg9 antagonizes TOR signaling to regulate intestinal cell growth and epithelial homeostasis in Drosophila. *eLife*, 6, 2017.

[3807] Max Werth, Kai M Schmidt-Ott, Thomas Leete, Andong Qiu, Christian Hinze, Melanie Viltard, Neal Paragas, Carrie J Shawber, Wenqiang Yu, Peter Lee, Xia Chen, Abby Sarkar, Weiyi Mu, Alexander Rittenberg, Chyuan-Sheng Lin, Jan Kitajewski, Qais Al-Awqati, and Jonathan Barasch. Transcription factor TFCP2L1 patterns cells in the mouse kidney collecting ducts. *eLife*, 6, 2017.

[3808] Richard J White, John E Collins, Ian M Sealy, Neha Wali, Christopher M Dooley, Zsofia Digby, Derek L Stemple, Daniel N Murphy, Konstantinos Billis, Thibaut Hourlier, Anja Füllgrabe, Matthew P Davis, Anton J Enright, and Elisabeth M Busch-Nentwich. A high-resolution mRNA expression time course of embryonic development in zebrafish. *eLife*, 6, 2017.

[3809] Alexandra M Whiteley, Miguel A Prado, Ivan Peng, Alexander R Abbas, Benjamin Haley, Joao A Paulo, Mike Reichelt, Anand Katakam, Meredith Sagolla, Zora Modrusan, Dong Yun Lee, Merone Roose-Girma, Donald S Kirkpatrick, Brent S McKenzie, Steven P Gygi, Daniel Finley, and Eric J Brown. Ubiquilin1 promotes antigen-receptor mediated proliferation by eliminating mislocalized mitochondrial proteins. *eLife*, 6, 2017.

[3810] Thomas J Whitford, Bradley N Jack, Daniel Pearson, Oren Griffiths, David Luque, Anthony WF Harris, Kevin M Spencer, and Mike E Le Pelley. Neurophysiological evidence of efference copies to inner speech. *eLife*, 6, 2017.

[3811] John C Whitney, S Brook Peterson, Jungyun Kim, Manuel Pazos, Adrian J Verster, Matthew C Radey, Hemantha D Kulasekara, Mary Q Ching, Nathan P Bullen, Diane Bryant, Young Ah Goo, Michael G Surette, Elhanan Borenstein, Waldemar Vollmer, and Joseph D Mougous. A broadly distributed toxin family mediates contact-dependent antagonism between gram-positive bacteria. *eLife*, 6, 2017.

[3812] Steven D Wiederman, Joseph M Fabian, James R Dunbier, and David C O'Carroll. A predictive focus of gain modulation encodes target trajectories in insect vision. *eLife*, 6, 2017.

[3813] Lukasz Wieteska, Saeid Shahidi, and Anastasia Zhuravleva. Allosteric fine-tuning of the conformational equilibrium poises the chaperone BiP for post-translational regulation. *eLife*, 6, 2017.

[3814] Geert-Jan Will, Robb B Rutledge, Michael Moutoussis, and Raymond J Dolan. Neural and computational processes underlying dynamic changes in self-esteem. *eLife*, 6, 2017.

[3815] Elizabeth A Williams, Csaba Verasztó, Sanja Jasek, Markus Conzelmann, Réza Shahidi, Philipp Bauknecht, Olivier Mirabeau, and Gáspár Jékely. Synaptic and peptidergic connectome of a neurosecretory center in the annelid brain. *eLife*, 6, 2017.

[3816] Jenna L Wingfield, Ilaria Mengoni, Heather Bomberger, Yu-Yang Jiang, Jonathon D Walsh, Jason M Brown, Tyler Picariello, Deborah A Cochran, Bing Zhu, Junmin Pan, Jonathan Eggenschwiler, Jacek Gaertig, George B Witman, Peter Kner, and Karl Lechtreck. IFT trains in different stages of assembly queue at the ciliary base for consecutive release into the cilium. *eLife*, 6, 2017.

[3817] Michael B Winter, Florencia La Greca, Shirin Arastu-Kapur, Francesco Caiazza, Peter Cimermancic, Tonia J Buchholz, Janet L Anderl, Matthew Ravalin, Markus F Bohn, Andrej Sali, Anthony J O'Donoghue, and Charles S Craik. Immunoproteasome functions explained by divergence in cleavage specificity and regulation. *eLife*, 6, 2017.

[3818] Bonnie C Wintle, Christian R Boehm, Catherine Rhodes, Jennifer C Molloy, Piers Millett, Laura Adam, Rainer Breitling, Rob Carlson, Rocco Casagrande, Malcolm Dando, Robert Doubleday, Eric Drexler, Brett Edwards, Tom Ellis, Nicholas G Evans, Richard Hammond, Jim Haseloff, Linda Kahl, Todd Kuiken, Benjamin R Lichman, Colette A Matthewman, Johnathan A Napier, Seán S óhéigeartaigh, Nicola J Patron, Edward Perello, Philip Shapira, Joyce Tait, Eriko Takano, and

William J Sutherland. A transatlantic perspective on 20 emerging issues in biological engineering. *eLife*, 6, 2017.

[3819] Einav Wircer, Janna Blechman, Nataliya Borodovsky, Michael Tsoory, Ana Rita Nunes, Rui F Oliveira, and Gil Levkowitz. Homeodomain protein Otp affects developmental neuropeptide switching in oxytocin neurons associated with a long-term effect on social behavior. *eLife*, 6, 2017.

[3820] Kristen Witte, Devin Strickland, and Michael Glotzer. Cell cycle entry triggers a switch between two modes of Cdc42 activation during yeast polarization. *eLife*, 6, 2017.

[3821] Sharon Grayer Wolf, Yael Mutsafi, Tali Dadosh, Tal Ilani, Zipora Lansky, Ben Horowitz, Sarah Rubin, Michael Elbaum, and Deborah Fass. 3D visualization of mitochondrial solid-phase calcium stores in whole cells. *eLife*, 6, 2017.

[3822] Gabriella Hannah Wolff, Hanne Halkinrud Thoen, Justin Marshall, Marcel E Sayre, and Nicholas James Strausfeld. An insect-like mushroom body in a crustacean brain. *eLife*, 6, 2017.

[3823] Jonci Nikolai Wolff, Neil J Gemmell, Daniel M Tompkins, and Damian K Dowling. Introduction of a male-harming mitochondrial haplotype via 'Trojan Females' achieves population suppression in fruit flies. *eLife*, 6, 2017.

[3824] Adam JM Wollman, Sviatlana Shashkova, Erik G Hedlund, Rosmarie Friemann, Stefan Hohmann, and Mark C Leake. Transcription factor clusters regulate genes in eukaryotic cells. *eLife*, 6, 2017.

[3825] Aaron L Wong, Jeff Goldsmith, Alexander D Forrence, Adrian M Haith, and John W Krakauer. Reaction times can reflect habits rather than computations. *eLife*, 6, 2017.

[3826] Wing Tak Wong, Gianfranco Matrone, XiaoYu Tian, Simion Alin Tomoiaga, Kin Fai Au, Shu Meng, Sayumi Yamazoe, Daniel Sieveking, Kaifu Chen, David M Burns, James K Chen, Helen M Blau, and John P Cooke. Discovery of novel determinants of endothelial lineage using chimeric heterokaryons. *eLife*, 6, 2017.

[3827] Elliot C Woods, FuiBoon Kai, J Matthew Barnes, Kayvon Pedram, Michael W Pickup, Michael J Hollander, Valerie M Weaver, and Carolyn R Bertozzi. A bulky glycocalyx fosters metastasis formation by promoting G1 cell cycle progression. *eLife*, 6, 2017.

[3828] Bei Wu, Alex J McDonald, Kathleen Markham, Celeste B Rich, Kyle P McHugh, Jörg Tatzelt, David W Colby, Glenn L Millhauser, and David A Harris. The N-terminus of the prion protein is a toxic effector regulated by the C-terminus. *eLife*, 6, 2017.

[3829] David Wu, Ru-Ting Huang, Robert B Hamanaka, Matt Krause, Myung-Jin Oh, Cheng-Hsiang Kuo, Recep Nigdelioglu, Angelo Y Meliton, Leah Witt, Guohao Dai, Mete Civelek, Nanduri R Prabhakar, Yun Fang, and Gökhan M Mutlu. HIF-1$\alpha$ is required for disturbed flow-induced metabolic reprogramming in human and porcine vascular endothelium. *eLife*, 6, 2017.

[3830] Fuqing Wu, Ri-Qi Su, Ying-Cheng Lai, and Xiao Wang. Engineering of a synthetic quadrastable gene network to approach Waddington landscape and cell fate determination. *eLife*, 6, 2017.

[3831] Zhenyong Wu, Oscar D Bello, Sathish Thiyagarajan, Sarah Marie Auclair, Wensi Vennekate, Shyam S Krishnakumar, Ben O'Shaughnessy, and Erdem Karatekin. Dilation of fusion pores by crowding of SNARE proteins. *eLife*, 6, 2017.

[3832] Zhongshou Wu, Shuai Huang, Xiaobo Zhang, Di Wu, Shitou Xia, and Xin Li. Regulation of plant immune receptor accumulation through translational repression by a glycine-tyrosine-phenylalanine (GYF) domain protein. *eLife*, 6, 2017.

[3833] Frances Xia, Blake A Richards, Matthew M Tran, Sheena A Josselyn, Kaori Takehara-Nishiuchi, and Paul W Frankland. Parvalbumin-positive interneurons mediate neocortical-hippocampal interactions that are necessary for memory consolidation. *eLife*, 6, 2017.

[3834] Xunde Xian, Yinyuan Ding, Marco Dieckmann, Li Zhou, Florian Plattner, Mingxia Liu, John S Parks, Robert E Hammer, Philippe Boucher, Shirling Tsai, and Joachim Herz. LRP1 integrates murine macrophage cholesterol homeostasis and inflammatory responses in atherosclerosis. *eLife*, 6, 2017.

[3835] Dongsheng Xiao, Matthieu P Vanni, Catalin C Mitelut, Allen W Chan, Jeffrey M LeDue, Yicheng Xie, Andrew CN Chen, Nicholas V Swindale, and Timothy H Murphy. Mapping cortical mesoscopic networks of single spiking cortical or sub-cortical neurons. *eLife*, 6, 2017.

[3836] Mei-Fang Xiao, Desheng Xu, Michael T Craig, Kenneth A Pelkey, Chun-Che Chien, Yang Shi, Juhong Zhang, Susan Resnick, Olga Pletnikova, David Salmon, James Brewer, Steven Edland, Jerzy Wegiel, Benjamin Tycko, Alena Savonenko, Roger H Reeves, Juan C Troncoso, Chris J McBain, Douglas Galasko, and Paul F Worley. NPTX2 and cognitive dysfunction in Alzheimer's Disease. *eLife*, 6, 2017.

[3837] Zhi-Jie Xiao, Jing Liu, Si-Qi Wang, Yun Zhu, Xu-Yuan Gao, Vicky Pui-Chi Tin, Jing Qin, Jun-Wen Wang, and Maria Pik Wong. NFATc2 enhances tumor-initiating phenotypes through the NFATc2/SOX2/ALDH axis in lung adenocarcinoma. *eLife*, 6, 2017.

[3838] Xiaojun Xie, Masashi Tabuchi, Matthew P Brown, Sarah P Mitchell, Mark N Wu, and Alex L Kolodkin. The laminar organization of the Drosophila ellipsoid body is semaphorin-dependent and prevents the formation of ectopic synaptic connections. *eLife*, 6, 2017.

[3839] Guangyan Xiong, Sajedah M Hindi, Aman K Mann, Yann S Gallot, Kyle R Bohnert, Douglas R Cavener, Scott R Whittemore, and Ashok Kumar. The PERK arm of the unfolded protein response regulates satellite cell-mediated skeletal muscle regeneration. *eLife*, 6, 2017.

[3840] C Shan Xu, Kenneth J Hayworth, Zhiyuan Lu, Patricia Grob, Ahmed M Hassan, José G García-Cerdán, Krishna K Niyogi, Eva Nogales, Richard J Weinberg, and Harald F Hess. Enhanced FIB-SEM systems for large-volume 3D imaging. *eLife*, 6, 2017.

[3841] Chiwei Xu, Junjie Luo, Li He, Craig Montell, and Norbert Perrimon. Oxidative stress induces stem cell proliferation via TRPA1/RyR-mediated Ca2+ signaling in the Drosophila midgut. *eLife*, 6, 2017.

[3842] Jin Xu, Xinxin Peng, Yuxin Chen, Yuezheng Zhang, Qin Ma, Liang Liang, Ava C Carter, Xuemei Lu, and Chung-I Wu. Free-living human cells reconfigure their chromosomes in the evolution back to uni-cellularity. *eLife*, 6, 2017.

[3843] Junjie Xu, Marcial Camacho, Yibin Xu, Victoria Esser, Xiaoxia Liu, Thorsten Trimbuch, Yun-Zu Pan, Cong Ma, Diana R Tomchick, Christian Rosenmund, and Josep Rizo. Mechanistic insights into neurotransmitter release and presynaptic plasticity from the crystal structure of Munc13-1 C1C2BMUN. *eLife*, 6, 2017.

[3844] Peng Xu, Hannah M Hankins, Chris MacDonald, Samuel J Erlinger, Meredith N Frazier, Nicholas S Diab, Robert C Piper, Lauren P Jackson, Jason A MacGurn, and Todd R Graham. COPI mediates recycling of an exocytic SNARE by recognition of a ubiquitin sorting signal. *eLife*, 6, 2017.

[3845] Xiang-Ru Shannon Xu, Valentino Matteo Gantz, Natalia Siomava, and Ethan Bier. CRISPR/Cas9 and active genetics-based trans-species replacement of the endogenous Drosophila kni-L2 CRM reveals unexpected complexity. *eLife*, 6, 2017.

[3846] Yixi Xu, Shaokai Ning, Zheng Wei, Ran Xu, Xinlin Xu, Mengtan Xing, Rong Guo, and Dongyi Xu. 53BP1 and BRCA1 control pathway choice for stalled replication restart. *eLife*, 6, 2017.

[3847] Zhao Xuan, Laura Manning, Jessica Nelson, Janet E Richmond, Daniel A Colón-Ramos, Kang Shen, and Peri T Kurshan. Clarinet (CLA-1), a novel active zone protein required for synaptic vesicle clustering and release. *eLife*, 6, 2017.

[3848] Katherine S Xue, Terry Stevens-Ayers, Angela P Campbell, Janet A Englund, Steven A Pergam, Michael Boeckh, and Jesse D Bloom. Parallel evolution of influenza across multiple spatiotemporal scales. *eLife*, 6, 2017.

[3849] Rajesh K Yadav, Carolyn M Jablonowski, Alfonso G Fernandez, Brandon R Lowe, Ryan A Henry, David Finkelstein, Kevin J Barnum, Alison L Pidoux, Yin-Ming Kuo, Jie Huang, Matthew J O'Connell, Andrew J Andrews, Arzu Onar-Thomas, Robin C Allshire, and Janet F Partridge. Histone H3G34R mutation causes replication stress, homologous recombination defects and genomic instability in S. pombe. *eLife*, 6, 2017.

[3850] Belgin YalÇın, Lu Zhao, Martin Stofanko, Niamh C O'Sullivan, Zi Han Kang, Annika Roost, Matthew R Thomas, Sophie Zaessinger, Olivier Blard, Alex L Patto, Anood Sohail, Valentina Baena, Mark Terasaki, and Cahir J O'Kane. Modeling of axonal endoplasmic reticulum network by spastic paraplegia proteins. *eLife*, 6, 2017.

[3851] Qing Yan, Benjamin Philmus, Jeff H Chang, and Joyce E Loper. Novel mechanism of metabolic co-regulation coordinates the biosynthesis of secondary metabolites in Pseudomonas protegens. *eLife*, 6, 2017.

[3852] Aimin Yang, Supansa Pantoom, and Yao-Wen Wu. Elucidation of the anti-autophagy mechanism of the Legionella effector RavZ using semisynthetic LC3 proteins. *eLife*, 6, 2017.

[3853] Ke Yang, Ling-Qiao Huang, Chao Ning, and Chen-Zhu Wang. Two single-point mutations shift the ligand selectivity of a pheromone receptor between two closely related moth species. *eLife*, 6, 2017.

[3854] Lihua Yang and Yong Gu. Distinct spatial coordinate of visual and vestibular heading signals in macaque FEFsem and MSTd. *eLife*, 6, 2017.

[3855] Xinan H Yang, Rangarajan D Nadadur, Catharina RE Hilvering, Valerio Bianchi, Michael Werner, Stefan R Mazurek, Margaret Gadek, Kaitlyn M Shen, Joseph Aaron Goldman, Leonid Tyan, Jenna Bekeny, Johnathon M Hall, Nutishia Lee, Carlos Perez-Cervantes, Ozanna Burnicka-Turek, Kenneth D Poss, Christopher R Weber, Wouter de Laat, Alexander J Ruthenburg, and Ivan P Moskowitz. Transcription-factor-dependent enhancer transcription defines a gene regulatory network for cardiac rhythm. *eLife*, 6, 2017.

[3856] Yan Yang, Qian Wang, Shu-Rong Wang, Yi Wang, and Qian Xiao. Representation of time interval entrained by periodic stimuli in the visual thalamus of pigeons. *eLife*, 6, 2017.

[3857] Shenqin Yao, Joseph Bergan, Anne Lanjuin, and Catherine Dulac. Oxytocin signaling in the medial amygdala is required for sex discrimination of social cues. *eLife*, 6, 2017.

[3858] Kelvin Yeung, Ann Boija, Edvin Karlsson, Per-Henrik Holmqvist, Yonit Tsatskis, Ilaria Nisoli, Damian Yap, Alireza Lorzadeh, Michelle Moksa, Martin Hirst, Samuel Aparicio, Manolis Fanto, Per Stenberg, Mattias Mannervik, and Helen McNeill. Atrophin controls developmental signaling pathways via interactions with Trithorax-like. *eLife*, 6, 2017.

[3859] Seung-Woo Yoo and Inah Lee. Functional double dissociation within the entorhinal cortex for visual scene-dependent choice behavior. *eLife*, 6, 2017.

[3860] Dongjoo You, Emma Nilsson, Danielle E Tenen, Anna Lyubetskaya, James C Lo, Rencong Jiang, Jasmine Deng, Brian A Dawes, Allan Vaag, Charlotte Ling, Evan D Rosen, and Sona Kang. Dnmt3a is an epigenetic mediator of adipose insulin resistance. *eLife*, 6, 2017.

[3861] Bernadette C Young, Chieh-Hsi Wu, N Claire Gordon, Kevin Cole, James R Price, Elian Liu, Anna E Sheppard, Sanuki Perera, Jane Charlesworth, Tanya Golubchik, Zamin Iqbal, Rory Bowden, Ruth C Massey, John Paul, Derrick W Crook, Timothy E Peto, A Sarah Walker, Martin J Llewelyn, David H Wyllie, and Daniel J Wilson. Severe infections emerge from commensal bacteria by adaptive evolution. *eLife*, 6, 2017.

[3862] Paul Yourik, Colin Echeverría Aitken, Fujun Zhou, Neha Gupta, Alan G Hinnebusch, and Jon R Lorsch. Yeast eIF4A enhances recruitment of mRNAs regardless of their structural complexity. *eLife*, 6, 2017.

[3863] Feiqiao Brian Yu, Paul C Blainey, Frederik Schulz, Tanja Woyke, Mark A Horowitz, and Stephen R Quake. Microfluidic-based mini-metagenomics enables discovery of novel microbial lineages from complex environmental samples. *eLife*, 6, 2017.

[3864] Jai Y Yu, Kenneth Kay, Daniel F Liu, Irene Grossrubatscher, Adrianna Loback, Marielena Sosa, Jason E Chung, Mattias P Karlsson, Margaret C Larkin, and Loren M Frank. Distinct hippocampal-cortical memory representations for experiences associated with movement versus immobility. *eLife*, 6, 2017.

[3865] Libing Yu, Jared T Winkelman, Chirangini Pukhrambam, Terence R Strick, Bryce E Nickels, and Richard H Ebright. The mechanism of variability in transcription start site selection. *eLife*, 6, 2017.

[3866] Xiao-Wen Yu, Daniel M Curlik, M Matthew Oh, Jerry CP Yin, and John F Disterhoft. CREB overexpression in dorsal CA1 ameliorates long-term memory deficits in aged rats. *eLife*, 6, 2017.

[3867] Mei Yuan, Thomas Meyer, Christoph Benkowitz, Shakuntala Savanthrapadian, Laura Ansel-Bollepalli, Angelica Foggetti, Peer Wulff, Pepe Alcami, Claudio Elgueta, and Marlene Bartos. Somatostatin-positive interneurons in the dentate gyrus of mice provide local- and long-range septal synaptic inhibition. *eLife*, 6, 2017.

[3868] Xiang-Shan Yuan, Lu Wang, Hui Dong, Wei-Min Qu, Su-Rong Yang, Yoan Cherasse, Michael Lazarus, Serge N Schiffmann, Alban de Kerchove d'Exaerde, Rui-Xi Li, and Zhi-Li Huang. Striatal adenosine A2A receptor neurons control active-period sleep via parvalbumin neurons in external globus pallidus. *eLife*, 6, 2017.

[3869] Xiaomei Yuan, Bi-Huei Yang, Yi Dong, Asami Yamamura, and Wenxian Fu. CRIg, a tissue-resident macrophage specific immune checkpoint molecule, promotes immunological tolerance in NOD mice, via a dual role in effector and regulatory T cells. *eLife*, 6, 2017.

[3870] ZY Yuan, F Jiao, XR Shi, Jordi Sardans, Fernando T Maestre, Manuel Delgado-Baquerizo, Peter B Reich, and Josep Peñuelas. Experimental and observational studies find contrasting responses of soil nutrients to climate change. *eLife*, 6, 2017.

[3871] Wendy Wing Sze Yue, Rikard Frederiksen, Xiaozhi Ren, Dong-Gen Luo, Takahiro Yamashita, Yoshinori Shichida, M Carter Cornwall, and King-Wai Yau. Spontaneous activation of visual pigments in relation to openness/closedness of chromophore-binding pocket. *eLife*, 6, 2017.

[3872] Sophie Zaaijer, Assaf Gordon, Daniel Speyer, Robert Piccone, Simon Cornelis Groen, and Yaniv Erlich. Rapid re-identification of human samples using portable DNA sequencing. *eLife*, 6, 2017.

[3873] Hiba Ahmad Zahreddine, Biljana Culjkovic-Kraljacic, Audrey Emond, Filippa Pettersson, Ronald Midura, Mark Lauer, Sonia Del Rincon, Valbona Cali, Sarit Assouline, Wilson H Miller, Vincent Hascall, and Katherine LB Borden. The eukaryotic translation initiation factor eIF4E harnesses hyaluronan production to drive its malignant activity. *eLife*, 6, 2017.

[3874] Kara E Zang, Elver Ho, and Niels Ringstad. Inhibitory peptidergic modulation of C. elegans serotonin neurons is gated by T-type calcium channels. *eLife*, 6, 2017.

[3875] Assaf Zaritsky, Uri Obolski, Zhuo Gan, Carlos R Reis, Zuzana Kadlecova, Yi Du, Sandra L Schmid, and Gaudenz Danuser. Decoupling global biases and local interactions between cell biological variables. *eLife*, 6, 2017.

[3876] Baltazar A Zavala, Anthony I Jang, and Kareem A Zaghloul. Human subthalamic nucleus activity during non-motor decision making. *eLife*, 6, 2017.

[3877] Longhui Zeng, Li Zhang, Pengran Wang, and Guoyu Meng. Structural basis of host recognition and biofilm formation by Salmonella Saf pili. *eLife*, 6, 2017.

[3878] Bo Zhang, Mattias Holmlund, Severine Lorrain, Mikael Norberg, László Bakó, Christian Fankhauser, and Ove Nilsson. BLADE-ON-PETIOLE proteins act in an E3 ubiquitin ligase complex to regulate PHYTOCHROME INTERACTING FACTOR 4 abundance. *eLife*, 6, 2017.

[3879] Haoran Zhang, Li Wang, Elaine Yee Man Wong, Sze Lan Tsang, Pin-Xian Xu, Urban Lendahl, and Mai Har Sham. An Eya1-Notch axis specifies bipotential epibranchial differentiation in mammalian craniofacial morphogenesis. *eLife*, 6, 2017.

[3880] Lei Zhang, Yubin Hou, Zhiyuan Li, Xinmiao Ji, Ze Wang, Huizhen Wang, Xiaofei Tian, Fazhi Yu, Zhenye Yang, Li Pi, Timothy J Mitchison, Qingyou Lu, and Xin Zhang. 27 T ultra-high static magnetic field changes orientation and morphology of mitotic spindles in human cells. *eLife*, 6, 2017.

[3881] Li Zhang, Prashant Rajbhandari, Christina Priest, Jaspreet Sandhu, Xiaohui Wu, Ryan Temel, Antonio Castrillo, Thomas Q de Aguiar Vallim, Tamer Sallam, and Peter Tontonoz. Inhibition of cholesterol biosynthesis through RNF145-dependent ubiquitination of SCAP. *eLife*, 6, 2017.

[3882] Miao Zhang, Martin Bommer, Ruchira Chatterjee, Rana Hussein, Junko Yano, Holger Dau, Jan Kern, Holger Dobbek, and Athina Zouni. Structural insights into the light-driven auto-assembly process of the water-oxidizing Mn4CaO5-cluster in photosystem II. *eLife*, 6, 2017.

[3883] Qiaosheng Zhang, Toby Manders, Ai Phuong Tong, Runtao Yang, Arpan Garg, Erik Martinez, Haocheng Zhou, Jahrane Dale, Abhinav Goyal, Louise Urien, Guang Yang, Zhe Chen, and Jing Wang. Chronic pain induces generalized enhancement of aversion. *eLife*, 6, 2017.

[3884] Rui Zhang, Johanna Roostalu, Thomas Surrey, and Eva Nogales. Structural insight into TPX2-stimulated microtubule assembly. *eLife*, 6, 2017.

[3885] Ting Zhang, Prashant Mishra, Bruce A Hay, David Chan, and Ming Guo. Valosin-containing protein (VCP/p97) inhibitors relieve Mitofusin-dependent mitochondrial defects due to VCP disease mutants. *eLife*, 6, 2017.

[3886] Ting-ting Zhang, David G Gonzalez, Christine M Cote, Steven M Kerfoot, Shaoli Deng, Yuqing Cheng, Masaki Magari, and Ann M Haberman. Germinal center B cell development has distinctly regulated stages completed by disengagement from T cell help. *eLife*, 6, 2017.

[3887] Wei Zhang, Clarence Hue Lok Yeung, Liwen Wu, and Karen Wing Yee Yuen. E3 ubiquitin ligase Bre1 couples sister chromatid cohesion establishment to DNA replication in Saccharomyces cerevisiae. *eLife*, 6, 2017.

[3888] Xiaodong Zhang, Alicia M Saarinen, Taro Hitosugi, Zhenghe Wang, Liguo Wang, Thai H Ho, and Jun Liu. Inhibition of intracellular lipolysis promotes human cancer cell adaptation to hypoxia. *eLife*, 6, 2017.

[3889] Xiaoyu Zhang, Nicole A Spiegelman, Ornella D Nelson, Hui Jing, and Hening Lin. SIRT6 regulates Ras-related protein R-Ras2 by lysine defatty-acylation. *eLife*, 6, 2017.

[3890] Xiulin Zhang, Birgit T Priest, Inna Belfer, and Michael S Gold. Voltage-gated Na+ currents in human dorsal root ganglion neurons. *eLife*, 6, 2017.

[3891] Xu Zhang, Xiaoli Chen, Qiuying Liu, Shaojie Zhang, and Wenqian Hu. Translation repression via modulation of the cytoplasmic poly(A)-binding protein in the inflammatory response. *eLife*, 6, 2017.

[3892] Xuemeng Zhang, Thomas Kampourakis, Ziqian Yan, Ivanka Sevrieva, Malcolm Irving, and Yin-Biao Sun. Distinct contributions of the thin and thick filaments to length-dependent activation in heart muscle. *eLife*, 6, 2017.

[3893] Yaqing Zhang, Wei Yan, Esha Mathew, Kevin T Kane, Arthur Brannon, Maeva Adoumie, Alekya Vinta, Howard C Crawford, and Marina Pasca di Magliano. Epithelial-Myeloid cell crosstalk regulates acinar cell plasticity and pancreatic remodeling in mice. *eLife*, 6, 2017.

[3894] Yu Zhang, Kevin Michel-Herve Larcher, Bratislav Misic, and Alain Dagher. Anatomical and functional organization of the human substantia nigra and its connections. *eLife*, 6, 2017.

[3895] Zhijian Zhang, Qing Liu, Pengjie Wen, Jiaozhen Zhang, Xiaoping Rao, Ziming Zhou, Hongruo Zhang, Xiaobin He, Juan Li, Zheng Zhou, Xiaoran Xu, Xueyi Zhang, Rui Luo, Guanghui Lv, Haohong Li, Pei Cao, Liping Wang, and Fuqiang Xu. Activation of the dopaminergic pathway from VTA to the medial olfactory tubercle generates odor-preference and reward. *eLife*, 6, 2017.

[3896] Zhonghui Zhang, Fuqu Hu, Min Woo Sung, Chang Shu, Claudia Castillo-González, Hisashi Koiwa, Guiliang Tang, Martin Dickman, Pingwei Li, and Xiuren Zhang. RISC-interacting clearing $3'$- $5'$ exoribonucleases (RICEs) degrade uridylated cleavage fragments to maintain functional RISC in Arabidopsis thaliana. *eLife*, 6, 2017.

[3897] Gengjing Zhao, Emma S Gleave, and Meindert Hugo Lamers. Single-molecule studies contrast ordered DNA replication with stochastic translesion synthesis. *eLife*, 6, 2017.

[3898] Huaying Zhao, Suvendu Lomash, Sagar Chittori, Carla Glasser, Mark L Mayer, and Peter Schuck. Preferential assembly of heteromeric kainate and AMPA receptor amino terminal domains. *eLife*, 6, 2017.

[3899] Shanshan Zhao, Wei Hong, Jianguo Wu, Yu Wang, Shaoyi Ji, Shuyi Zhu, Chunhong Wei, Jinsong Zhang, and Yi Li. A viral protein promotes host SAMS1 activity and ethylene production for the benefit of virus infection. *eLife*, 6, 2017.

[3900] Zhaoqing Zheng, Ganesh Ambigapathy, and Joyce Keifer. MeCP2 regulates Tet1-catalyzed demethylation, CTCF binding, and learning-dependent alternative splicing of the BDNF gene in Turtle. *eLife*, 6, 2017.

[3901] Yongwang Zhong, Jiou Wang, Mark J Henderson, Peixin Yang, Brian M Hagen, Teepu Siddique, Bruce E Vogel, Han-Xiang Deng, and Shengyun Fang. Nuclear export of misfolded SOD1 mediated by a normally buried NES-like sequence reduces proteotoxicity in the nucleus. *eLife*, 6, 2017.

[3902] Hao Zhou, Katarzyna Bulek, Xiao Li, Tomasz Herjan, Minjia Yu, Wen Qian, Han Wang, Gao Zhou, Xing Chen, Hui Yang, Lingzi Hong, Junjie Zhao, Luke Qin, Koichi Fukuda, Annette Flotho, Ji Gao, Ashok Dongre, Julie A Carman, Zizhen Kang, Bing Su, Timothy S Kern, Jonathan D Smith, Thomas A Hamilton, Frauke Melchior, Paul L Fox, and Xiaoxia Li. IRAK2 directs stimulus-dependent nuclear export of inflammatory mRNAs. *eLife*, 6, 2017.

[3903] Pingzhu Zhou, Fei Gu, Lina Zhang, Brynn N Akerberg, Qing Ma, Kai Li, Aibin He, Zhiqiang Lin, Sean M Stevens, Bin Zhou, and William T Pu. Mapping cell type-specific transcriptional enhancers using high affinity, lineage-specific Ep300 bioChIP-seq. *eLife*, 6, 2017.

[3904] Wenchang Zhou, Fabrizio Marinelli, Corrine Nief, and José D Faraldo-Gómez. Atomistic simulations indicate the c-subunit ring of the F1Fo ATP synthase is not the mitochondrial permeability transition pore. *eLife*, 6, 2017.

[3905] Zhen Zhou, Yu-Ting Liu, Li Ma, Ting Gong, Ya-Nan Hu, Hong-Tao Li, Chen Cai, Ling-Li Zhang, Gang Wei, and Jin-Qiu Zhou. Independent manipulation of histone H3 modifications in individual nucleosomes reveals the contributions of sister histones to transcription. *eLife*, 6, 2017.

[3906] Jun-yi Zhu, Yulong Fu, Margaret Nettleton, Adam Richman, and Zhe Han. High throughput in vivo functional validation of candidate congenital heart disease genes in Drosophila. *eLife*, 6, 2017.

[3907] Lu Zhu, Jeff R Jorgensen, Ming Li, Ya-Shan Chuang, and Scott D Emr. ESCRTs function directly on the lysosome membrane to downregulate ubiquitinated lysosomal membrane proteins. *eLife*, 6, 2017.

[3908] Wenyi Zhu, Saritha Krishna, Cristina Garcia, Chia-Ching John Lin, Bartley D Mitchell, Kenneth L Scott, Carrie A Mohila, Chad J Creighton, Seung-Hee Yoo, Hyun Kyoung Lee, and Benjamin Deneen. Daam2 driven degradation of VHL promotes gliomagenesis. *eLife*, 6, 2017.

[3909] Jun Zhuang, Lydia Ng, Derric Williams, Matthew Valley, Yang Li, Marina Garrett, and Jack Waters. An extended retinotopic map of mouse cortex. *eLife*, 6, 2017.

[3910] Joseph Zinski, Ye Bu, Xu Wang, Wei Dou, David Umulis, and Mary C Mullins. Systems biology derived source-sink mechanism of BMP gradient formation. *eLife*, 6, 2017.

[3911] Stefano Zucca, Giulia D'Urso, Valentina Pasquale, Dania Vecchia, Giuseppe Pica, Serena Bovetti, Claudio Moretti, Stefano Varani, Manuel Molano-Mazón, Michela Chiappalone, Stefano Panzeri, and Tommaso Fellin. An inhibitory gate for state transition in cortex. *eLife*, 6, 2017.

[3912] Yaara Zwang, Oliver Jonas, Casandra Chen, Mikael L Rinne, John G Doench, Federica Piccioni, Li Tan, Hai-Tsang Huang, Jinhua Wang, Young Jin Ham, Joyce O'Connell, Patrick Bhola, Mihir Doshi, Matthew Whitman, Michael Cima, Anthony Letai, David E Root, Robert S Langer, Nathanael Gray, and William C Hahn. Synergistic interactions with PI3K inhibition that induce apoptosis. *eLife*, 6, 2017.

[3913] Peter-Paul Zwetsloot, Mira Van Der Naald, Emily S Sena, David W Howells, Joanna IntHout, Joris AH De Groot, Steven AJ Chamuleau, Malcolm R MacLeod, and Kimberley E Wever. Standardized mean differences cause funnel plot distortion in publication bias assessments. *eLife*, 6, 2017.

[3914] Ariel Zylberberg, Jeannette AM Lorteije, Brian G Ouellette, Chris I De Zeeuw, Mariano Sigman, and Pieter Roelfsema. Serial, parallel and hierarchical decision making in primates. *eLife*, 6, 2017.

[3915] Daniel Zúñiga, Yann Gager, Hanna Kokko, Adam Michael Fudickar, Andreas Schmidt, Beat Naef-Daenzer, Martin Wikelski, and Jesko Partecke. Migration confers winter survival benefits in a partially migratory songbird. *eLife*, 6, 2017.

[3916] Ethan V Abel, Masashi Goto, Brian Magnuson, Saji Abraham, Nikita Ramanathan, Emily Hotaling, Anthony A Alaniz, Chandan Kumar-Sinha, Michele L Dziubinski, Sumithra Urs, Lidong Wang, Jiaqi Shi, Meghna Waghray, Mats Ljungman, Howard C Crawford, and Diane M Simeone. HNF1A is a novel oncogene that regulates human pancreatic cancer stem cell properties. *eLife*, 7, 2018.

[3917] Diego Acosta-Alvear, G Elif Karagöz, Florian Fröhlich, Han Li, Tobias C Walther, and Peter Walter. The unfolded protein response and endoplasmic reticulum protein targeting machineries converge on the stress sensor IRE1. *eLife*, 7, 2018.

[3918] Sarah Adio, Heena Sharma, Tamara Senyushkina, Prajwal Karki, Cristina Maracci, Ingo Wohlgemuth, Wolf Holtkamp, Frank Peske, and Marina V Rodnina. Dynamics of ribosomes and release factors during translation termination in E. coli. *eLife*, 7, 2018.

[3919] Anamika Agrawal, Gulcin Pekkurnaz, and Elena F Koslover. Spatial control of neuronal metabolism through glucose-mediated mitochondrial transport regulation. *eLife*, 7, 2018.

[3920] Raphaël Aguillon, Julie Batut, Arul Subramanian, Romain Madelaine, Pascale Dufourcq, Thomas F Schilling, and Patrick Blader. Cell-type heterogeneity in the early zebrafish olfactory epithelium is generated from progenitors within preplacodal ectoderm. *eLife*, 7, 2018.

[3921] Joachim Ahlbeck, Lingzhen Song, Mattia Chini, Sebastian H Bitzenhofer, and Ileana L Hanganu-Opatz. Glutamatergic drive along the septo-temporal axis of hippocampus boosts prelimbic oscillations in the neonatal mouse. *eLife*, 7, 2018.

[3922] Saumel Ahmadi, Sunny Xia, Yu-Sheng Wu, Michelle Di Paola, Randolph Kissoon, Catherine Luk, Fan Lin, Kai Du, Johanna Rommens, and Christine E Bear. SLC6A14, an amino acid transporter, modifies the primary CF defect in fluid secretion. *eLife*, 7, 2018.

[3923] Sizhi Ai, Yunlu Yin, Yu Chen, Cong Wang, Yan Sun, Xiangdong Tang, Lin Lu, Lusha Zhu, and Jie Shi. Promoting subjective preferences in simple economic choices during nap. *eLife*, 7, 2018.

[3924] Abhishek Aich, Cong Wang, Arpita Chowdhury, Christin Ronsör, David Pacheu-Grau, Ricarda Richter-Dennerlein, Sven Dennerlein, and Peter Rehling. COX16 promotes COX2 metallation and assembly during respiratory complex IV biogenesis. *eLife*, 7, 2018.

[3925] Yulia Akbergenova, Karen L Cunningham, Yao V Zhang, Shirley Weiss, and J Troy Littleton. Characterization of developmental and molecular factors underlying release heterogeneity at Drosophila synapses. *eLife*, 7, 2018.

[3926] Takuya Akiyama, Sırma D User, and Matthew C Gibson. Somatic clones heterozygous for recessive disease alleles of BMPR1A exhibit unexpected phenotypes in Drosophila. *eLife*, 7, 2018.

[3927] Ream Al-Hasani, Jenny-Marie T Wong, Omar S Mabrouk, Jordan G McCall, Gavin P Schmitz, Kirsten A Porter-Stransky, Brandon J Aragona, Robert T Kennedy, and Michael R Bruchas. In vivo detection of optically-evoked opioid peptide release. *eLife*, 7, 2018.

[3928] Luis Alarcon-Martinez, Sinem Yilmaz-Ozcan, Muge Yemisci, Jesse Schallek, Kıvılcım KılıÇ, Alp Can, Adriana Di Polo, and Turgay Dalkara. Capillary pericytes express $\alpha$-smooth muscle actin, which requires prevention of filamentous-actin depolymerization for detection. *eLife*, 7, 2018.

[3929] Frank Wolfgang Albert, Joshua S Bloom, Jake Siegel, Laura Day, and Leonid Kruglyak. Genetics of trans-regulatory variation in gene expression. *eLife*, 7, 2018.

[3930] Fabien Alcaraz, Virginie Fresno, Alain R Marchand, Eric J Kremer, Etienne Coutureau, and Mathieu Wolff. Thalamocortical and corticothalamic pathways differentially contribute to goal-directed behaviors in the rat. *eLife*, 7, 2018.

[3931] Hervé Alégot, Pierre Pouchin, Olivier Bardot, and Vincent Mirouse. Jak-Stat pathway induces Drosophila follicle elongation by a gradient of apical contractility. *eLife*, 7, 2018.

[3932] Cristiano Alessandro, Benjamin A Rellinger, Filipe Oliveira Barroso, and Matthew C Tresch. Adaptation after vastus lateralis denervation in rats demonstrates neural regulation of joint stresses and strains. *eLife*, 7, 2018.

[3933] Georgia M Alexander, Logan Y Brown, Shannon Farris, Daniel Lustberg, Caroline Pantazis, Bernd Gloss, Nicholas W Plummer, Patricia Jensen, and Serena M Dudek. CA2 neuronal activity controls hippocampal low gamma and ripple oscillations. *eLife*, 7, 2018.

[3934] Alexandre Alié, Lucie Devos, Jorge Torres-Paz, Lise Prunier, Fanny Boulet, Maryline Blin, Yannick Elipot, and Sylvie Retaux. Developmental evolution of the forebrain in cavefish, from natural variations in neuropeptides to behavior. *eLife*, 7, 2018.

[3935] Tom Alisch, James D Crall, Albert B Kao, Dave Zucker, and Benjamin L de Bivort. MAPLE (modular automated platform for large-scale experiments), a robot for integrated organism-handling and phenotyping. *eLife*, 7, 2018.

[3936] Sonia Alonso-Martin, Frédéric Auradé, Despoina Mademtzoglou, Anne Rochat, Peter S Zammit, and Frédéric Relaix. SOXF factors regulate murine satellite cell self-renewal and function through inhibition of $\beta$-catenin activity. *eLife*, 7, 2018.

[3937] Drew M Altschul, William D Hopkins, Elizabeth S Herrelko, Miho Inoue-Murayama, Tetsuro Matsuzawa, James E King, Stephen R Ross, and Alexander Weiss. Personality links with lifespan in chimpanzees. *eLife*, 7, 2018.

[3938] Efrén álvarez Salvado, Angela M Licata, Erin G Connor, Margaret K McHugh, Benjamin MN King, Nicholas Stavropoulos, Jonathan D Victor, John P Crimaldi, and Katherine I Nagel. Elementary sensory-motor transformations underlying olfactory navigation in walking fruit-flies. *eLife*, 7, 2018.

[3939] Daiane S Alves, Justin M Westerfield, Xiaojun Shi, Vanessa P Nguyen, Katherine M Stefanski, Kristen R Booth, Soyeon Kim, Jennifer Morrell-Falvey, Bing-Cheng Wang, Steven M Abel, Adam W Smith, and Francisco N Barrera. A novel pH-dependent membrane peptide that binds to EphA2 and inhibits cell migration. *eLife*, 7, 2018.

[3940] Andrew J Aman, Alexis N Fulbright, and David M Parichy. Wnt/$\beta$-catenin regulates an ancient signaling network during zebrafish scale development. *eLife*, 7, 2018.

[3941] Adithya N Ananth, Ankur Mishra, Steffen Frey, Arvind Dwarkasing, Roderick Versloot, Erik van der Giessen, Dirk Görlich, Patrick Onck, and Cees Dekker. Spatial structure of disordered proteins dictates conductance and selectivity in nuclear pore complex mimics. *eLife*, 7, 2018.

[3942] Sandra Breum Andersen, Melanie Ghoul, Rasmus L Marvig, Zhuo-Bin Lee, Søren Molin, Helle Krogh Johansen, and Ashleigh S Griffin. Privatisation rescues function following loss of cooperation. *eLife*, 7, 2018.

[3943] Dmitry E Andreev, Maxim Arnold, Stephen J Kiniry, Gary Loughran, Audrey M Michel, Dmitrii Rachinskii, and Pavel V Baranov. TASEP modelling provides a parsimonious explanation for the ability of a single uORF to derepress translation during the integrated stress response. *eLife*, 7, 2018.

[3944] Jean-Michel Arbona, Arach Goldar, Olivier Hyrien, Alain Arneodo, and Benjamin Audit. The eukaryotic bell-shaped temporal rate of DNA replication origin firing emanates from a balance between origin activation and passivation. *eLife*, 7, 2018.

[3945] Michelle Armenta Salas, Luke Bashford, Spencer Kellis, Matiar Jafari, HyeongChan Jo, Daniel Kramer, Kathleen Shanfield, Kelsie Pejsa, Brian Lee, Charles Y Liu, and Richard A Andersen. Proprioceptive and cutaneous sensations in humans elicited by intracortical microstimulation. *eLife*, 7, 2018.

[3946] Isabelle Arnoux, Michael Willam, Nadine Griesche, Jennifer Krummeich, Hirofumi Watari, Nina Offermann, Stephanie Weber, Partha Narayan Dey, Changwei Chen, Olivia Monteiro, Sven Buettner, Katharina Meyer, Daniele Bano, Konstantin Radyushkin, Rosamund Langston, Jeremy J Lambert, Erich Wanker, Axel Methner, Sybille Krauss, Susann Schweiger, and Albrecht Stroh. Metformin reverses early cortical network dysfunction and behavior changes in Huntington's disease. *eLife*, 7, 2018.

[3947] Gunjan Arora, Geoffrey T Hart, Javier Manzella-Lapeira, Justin YA Doritchamou, David L Narum, L Michael Thomas, Joseph Brzostowski, Sumati Rajagopalan, Ogobara K Doumbo, Boubacar Traore, Louis H Miller, Susan K Pierce, Patrick E Duffy, Peter D Crompton, Sanjay A Desai, and Eric O Long. NK cells inhibit Plasmodium falciparum growth in red blood cells via antibody-dependent cellular cytotoxicity. *eLife*, 7, 2018.

[3948] Joshua A Arribere and Andrew Z Fire. Nonsense mRNA suppression via nonstop decay. *eLife*, 7, 2018.

[3949] Gregory Artiushin, Shirley L Zhang, Hervé Tricoire, and Amita Sehgal. Endocytosis at the Drosophila blood–brain barrier as a function for sleep. *eLife*, 7, 2018.

[3950] Seksiri Arttamangkul, Daniel A Heinz, James R Bunzow, Xianqiang Song, and John T Williams. Cellular tolerance at the $\mu$-opioid receptor is phosphorylation dependent. *eLife*, 7, 2018.

[3951] Zahra Assur Sanghai, Qun Liu, Oliver B Clarke, Meagan Belcher-Dufrisne, Pattama Wiriyasermkul, M Hunter Giese, Edgar Leal-Pinto, Brian Kloss, Shantelle Tabuso, James Love, Marco Punta, Surajit Banerjee, Kanagalaghatta R Rajashankar, Burkhard Rost, Diomedes Logothetis, Matthias Quick, Wayne A Hendrickson, and Filippo Mancia. Structure-based analysis of CysZ-mediated cellular uptake of sulfate. *eLife*, 7, 2018.

[3952] James Attwater, Aditya Raguram, Alexey S Morgunov, Edoardo Gianni, and Philipp Holliger. Ribozyme-catalysed RNA synthesis using triplet building blocks. *eLife*, 7, 2018.

[3953] Gabriel N Aughey, Alicia Estacio Gomez, Jamie Thomson, Hang Yin, and Tony D Southall. CATaDa reveals global remodelling of chromatin accessibility during stem cell differentiation in vivo. *eLife*, 7, 2018.

[3954] Prachee Avasthi, Alice Soragni, and Joshua N Bembenek. Journal clubs in the time of preprints. *eLife*, 7, 2018.

[3955] Inna Averbukh, Sen-Lin Lai, Chris Q Doe, and Naama Barkai. A repressor-decay timer for robust temporal patterning in embryonic Drosophila neuroblast lineages. *eLife*, 7, 2018.

[3956] Fikret Aydin, Naomi Courtemanche, Thomas D Pollard, and Gregory A Voth. Gating mechanisms during actin filament elongation by formins. *eLife*, 7, 2018.

[3957] Sayyed M Azimi, Steven D Sheridan, Mostafa Ghannad-Rezaie, Peter M Eimon, and Mehmet Fatih Yanik. Combinatorial programming of human neuronal progenitors using magnetically-guided stoichiometric mRNA delivery. *eLife*, 7, 2018.

[3958] Kentarou Baba, Wataru Yoshida, Michinori Toriyama, Tadayuki Shimada, Colleen F Manning, Michiko Saito, Kenji Kohno, James S Trimmer, Rikiya Watanabe, and Naoyuki Inagaki. Gradient-reading and mechano-effector machinery for netrin-1-induced axon guidance. *eLife*, 7, 2018.

[3959] Aleksandra Badura, Jessica L Verpeut, Julia W Metzger, Talmo D Pereira, Thomas J Pisano, Ben Deverett, Dariya E Bakshinskaya, and Samuel S-H Wang. Normal cognitive and social development require posterior cerebellar activity. *eLife*, 7, 2018.

[3960] Thomas D Baird, Ken Chih-Chien Cheng, Yu-Chi Chen, Eugen Buehler, Scott E Martin, James Inglese, and J Robert Hogg. ICE1 promotes the link between splicing and nonsense-mediated mRNA decay. *eLife*, 7, 2018.

[3961] Diego Balboa, Jonna Saarimäki-Vire, Daniel Borshagovski, Mantas Survila, Päivi Lindholm, Emilia Galli, Solja Eurola, Jarkko Ustinov, Heli Grym, Hanna Huopio, Juha Partanen, Kirmo Wartiovaara, and Timo Otonkoski. Insulin mutations impair beta-cell development in a patient-derived iPSC model of neonatal diabetes. *eLife*, 7, 2018.

[3962] Angela Ballesteros, Cristina Fenollar-Ferrer, and Kenton Jon Swartz. Structural relationship between the putative hair cell mechanotransduction channel TMC1 and TMEM16 proteins. *eLife*, 7, 2018.

[3963] Hugo Balleza-Tapia, Sophie Crux, Yuniesky Andrade-Talavera, Pablo Dolz-Gaiton, Daniela Papadia, Gefei Chen, Jan Johansson, and André Fisahn. TrpV1 receptor activation rescues neuronal function and network gamma oscillations from A$\beta$-induced impairment in mouse hippocampus in vitro. *eLife*, 7, 2018.

[3964] Emily T Baltz, Ege A Yalcinbas, Rafael Renteria, and Christina M Gremel. Orbital frontal cortex updates state-induced value change for decision-making. *eLife*, 7, 2018.

[3965] Xun X Bao, Christos Spanos, Tomoko Kojidani, Eric M Lynch, Juri Rappsilber, Yasushi Hiraoka, Tokuko Haraguchi, and Kenneth E Sawin. Exportin Crm1 is repurposed as a docking protein to generate microtubule organizing centers at the nuclear pore. *eLife*, 7, 2018.

[3966] Anissa Bara, Antonia Manduca, Axel Bernabeu, Milene Borsoi, Michela Serviado, Olivier Lassalle, Michelle Murphy, Jim Wager-Miller, Ken Mackie, Anne-Laure Pelissier-Alicot, Viviana Trezza, and Olivier J Manzoni. Sex-dependent effects of in utero cannabinoid exposure on cortical function. *eLife*, 7, 2018.

[3967] Jelena Baranovic and Andrew JR Plested. Auxiliary subunits keep AMPA receptors compact during activation and desensitization. *eLife*, 7, 2018.

[3968] Catherine Baranowski, Michael A Welsh, Lok-To Sham, Haig A Eskandarian, Hoong Chuin Lim, Karen J Kieser, Jeffrey C Wagner, John D McKinney, Georg E Fantner, Thomas R Ioerger, Suzanne Walker, Thomas G Bernhardt, Eric J Rubin, and E Hesper Rego. Maturing Mycobacterium smegmatis peptidoglycan requires non-canonical crosslinks to maintain shape. *eLife*, 7, 2018.

[3969] Jad Barchini, Xuefeng Shi, Hui Chen, and Jianhua Cang. Bidirectional encoding of motion contrast in the mouse superior colliculus. *eLife*, 7, 2018.

[3970] Valentina Barrera, Ian James Callum MacCormick, Gabriela Czanner, Paul Stephenson Hiscott, Valerie Ann White, Alister Gordon Craig, Nicholas Alexander Venton Beare, Lucy Hazel Culshaw, Yalin Zheng, Simon Charles Biddolph, Danny Arnold Milner, Steve Kamiza, Malcolm Edward Molyneux, Terrie Ellen Taylor, and Simon Peter Harding. Neurovascular sequestration in paediatric P. falciparum malaria is visible clinically in the retina. *eLife*, 7, 2018.

[3971] Emilie Battivelli, Matthew S Dahabieh, Mohamed Abdel-Mohsen, J Peter Svensson, Israel Tojal Da Silva, Lillian B Cohn, Andrea Gramatica, Steven Deeks, Warner C Greene, Satish K Pillai, and Eric Verdin. Distinct chromatin functional states correlate with HIV latency reactivation in infected primary CD4+ T cells. *eLife*, 7, 2018.

[3972] N Sumru Bayin, Alexandre Wojcinski, Aurelien Mourton, Hiromitsu Saito, Noboru Suzuki, and Alexandra L Joyner. Age-dependent dormant resident progenitors are stimulated by injury to regenerate Purkinje neurons. *eLife*, 7, 2018.

[3973] Wael Bazzi, Pierre B Cattenoz, Claude Delaporte, Vasanthi Dasari, Rosy Sakr, Yoshihiro Yuasa, and Angela Giangrande. Embryonic hematopoiesis modulates the inflammatory response and larval hematopoiesis in Drosophila. *eLife*, 7, 2018.

[3974] Gwendolyn M Beacham, Edward A Partlow, Jeffrey J Lange, and Gunther Hollopeter. NECAPs are negative regulators of the AP2 clathrin adaptor complex. *eLife*, 7, 2018.

[3975] Robin Beaven and Barry Denholm. Release and spread of Wingless is required to pattern the proximo-distal axis of Drosophila renal tubules. *eLife*, 7, 2018.

[3976] Lars Becker, Michael E Schnee, Mamiko Niwa, Willy Sun, Stephan Maxeiner, Sara Talaei, Bechara Kachar, Mark A Rutherford, and Anthony J Ricci. The presynaptic ribbon maintains vesicle populations at the hair cell afferent fiber synapse. *eLife*, 7, 2018.

[3977] Jason C Bell, David Jukam, Nicole A Teran, Viviana I Risca, Owen K Smith, Whitney L Johnson, Jan M Skotheim, William James Greenleaf, and Aaron F Straight. Chromatin-associated RNA sequencing (ChAR-seq) maps genome-wide RNA-to-DNA contacts. *eLife*, 7, 2018.

[3978] Bente Benedict, Tanja van Harn, Marleen Dekker, Simone Hermsen, Asli Kucukosmanoglu, Wietske Pieters, Elly Delzenne-Goette, Josephine C Dorsman, Eva Petermann, Floris Foijer, and Hein te Riele. Loss of p53 suppresses replication-stress-induced DNA breakage in G1/S checkpoint deficient cells. *eLife*, 7, 2018.

[3979] Noah C Benson and Jonathan Winawer. Bayesian analysis of retinotopic maps. *eLife*, 7, 2018.

[3980] Isabelle Bergiers, Tallulah Andrews, özge Vargel Bölükbaşı, Andreas Buness, Ewa Janosz, Natalia Lopez-Anguita, Kerstin Ganter, Kinga Kosim, Cemre Celen, Gülce Itır PerÇin, Paul Collier, Bianka Baying, Vladimir Benes, Martin Hemberg, and Christophe Lancrin. Single-cell transcriptomics reveals a new dynamical function of transcription factors during embryonic hematopoiesis. *eLife*, 7, 2018.

[3981] Giulia Bertolin, Anne-Laure Bulteau, Marie-Clotilde Alves-Guerra, Agnes Burel, Marie-Thérèse Lavault, Olivia Gavard, Stephanie Le Bras, Jean-Philippe Gagné, Guy G Poirier, Roland Le Borgne, Claude Prigent, and Marc Tramier. Aurora kinase A localises to mitochondria to control organelle dynamics and energy production. *eLife*, 7, 2018.

[3982] Nicole L Bertschi, Annemarie Voorberg-van der Wel, Anne-Marie Zeeman, Sven Schuierer, Florian Nigsch, Walter Carbone, Judith Knehr, Devendra K Gupta, Sam O Hofman, Nicole van der Werff, Ivonne Nieuwenhuis, Els Klooster, Bart W Faber, Erika L Flannery, Sebastian A Mikolajczak, Vorada Chuenchob, Binesh Shrestha, Martin Beibel, Tewis Bouwmeester, Niwat Kangwanrangsan, Jetsumon Sattabongkot, Thierry T Diagana, Clemens HM Kocken, and Guglielmo Roma. Transcriptomic analysis reveals reduced transcriptional activity in the malaria parasite Plasmodium cynomolgi during progression into dormancy. *eLife*, 7, 2018.

[3983] Nicolas Bery, Abimael Cruz-Migoni, Carole JR Bataille, Camilo E Quevedo, Hanna Tulmin, Ami Miller, Angela Russell, Simon EV Phillips, Stephen B Carr, and Terence H Rabbitts. BRET-based RAS biosensors that show a novel small molecule is an inhibitor of RAS-effector protein-protein interactions. *eLife*, 7, 2018.

[3984] Marina Besprozvannaya, Eamonn Dickson, Hao Li, Kenneth S Ginburg, Donald M Bers, Johan Auwerx, and Jodi Nunnari. GRAM domain proteins specialize functionally distinct ER-PM contact sites in human cells. *eLife*, 7, 2018.

[3985] Ewelina Betleja, Rashmi Nanjundappa, Tao Cheng, and Moe R Mahjoub. A novel Cep120-dependent mechanism inhibits centriole maturation in quiescent cells. *eLife*, 7, 2018.

[3986] Marie Bettonville, Stefania d'Aria, Kathleen Weatherly, Paolo E Porporato, Jinyu Zhang, Sabrina Bousbata, Pierre Sonveaux, and Michel Y Braun. Long-term antigen exposure irreversibly modifies metabolic requirements for T cell function. *eLife*, 7, 2018.

[3987] Luis A Bezares-Calderón, Jürgen Berger, Sanja Jasek, Csaba Verasztó, Sara Mendes, Martin Gühmann, Rodrigo Almeda, Réza Shahidi, and Gáspár Jékely. Neural circuitry of a polycystin-mediated hydrodynamic startle response for predator avoidance. *eLife*, 7, 2018.

[3988] Debadrita Bhattacharya, Megan Rothstein, Ana Paula Azambuja, and Marcos Simoes-Costa. Control of neural crest multipotency by Wnt signaling and the Lin28/let-7 axis. *eLife*, 7, 2018.

[3989] Andrej Bicanski and Neil Burgess. A neural-level model of spatial memory and imagery. *eLife*, 7, 2018.

[3990] Vincent T Bicocca, Tereza Ormsby, Keyur K Adhvaryu, Shinji Honda, and Eric U Selker. ASH1-catalyzed H3K36 methylation drives gene repression and marks H3K27me2/3-competent chromatin. *eLife*, 7, 2018.

[3991] Janine Diane Bijsterbosch, Mark W Woolrich, Matthew F Glasser, Emma C Robinson, Christian F Beckmann, David C Van Essen, Samuel J Harrison, and Stephen M Smith. The relationship between spatial configuration and functional connectivity of brain regions. *eLife*, 7, 2018.

[3992] Célian Bimbard, Charlie Demene, Constantin Girard, Susanne Radtke-Schuller, Shihab Shamma, Mickael Tanter, and Yves Boubenec. Multi-scale mapping along the auditory hierarchy using high-resolution functional UltraSound in the awake ferret. *eLife*, 7, 2018.

[3993] Paola Binda, Jan W Kurzawski, Claudia Lunghi, Laura Biagi, Michela Tosetti, and Maria Concetta Morrone. Response to short-term deprivation of the human adult visual cortex measured with 7T BOLD. *eLife*, 7, 2018.

[3994] Jeremy G Bird, Urmimala Basu, David Kuster, Aparna Ramachandran, Ewa Grudzien-Nogalska, Atif Towheed, Douglas C Wallace, Megerditch Kiledjian, Dmitry Temiakov, Smita S Patel, Richard H Ebright, and Bryce E Nickels. Highly efficient $5'$ capping of mitochondrial RNA with NAD+ and NADH by yeast and human mitochondrial RNA polymerase. *eLife*, 7, 2018.

[3995] Julien Bischerour, Simran Bhullar, Cyril Denby Wilkes, Vinciane Régnier, Nathalie Mathy, Emeline Dubois, Aditi Singh, Estienne Swart, Olivier Arnaiz, Linda Sperling, Mariusz Nowacki, and Mireille Bétermier. Six domesticated PiggyBac transposases together carry out programmed DNA elimination in Paramecium. *eLife*, 7, 2018.

[3996] Johannes Bischof, Marilyne Duffraisse, Edy Furger, Leiore Ajuria, Guillaume Giraud, Solene Vanderperre, Rachel Paul, Mikael Björklund, Damien Ahr, Alexis W Ahmed, Lionel Spinelli, Christine Brun, Konrad Basler, and Samir Merabet. Generation of a versatile BiFC ORFeome library for analyzing protein–protein interactions in live Drosophila. *eLife*, 7, 2018.

[3997] Adam J Bisogni, Shila Ghazanfar, Eric O Williams, Heather M Marsh, Jean YH Yang, and David M Lin. Tuning of delta-protocadherin adhesion through combinatorial diversity. *eLife*, 7, 2018.

[3998] Leonora S Bittleston, Charles J Wolock, Bakhtiar E Yahya, Xin Yue Chan, Kok Gan Chan, Naomi E Pierce, and Anne Pringle. Convergence between the microcosms of Southeast Asian and North American pitcher plants. *eLife*, 7, 2018.

[3999] Landry Blanc, Isaac B Daudelin, Brendan K Podell, Pei-Yu Chen, Matthew Zimmerman, Amanda J Martinot, Rada M Savic, Brendan Prideaux, and Véronique Dartois. High-resolution mapping of fluoroquinolones in TB rabbit lesions reveals specific distribution in immune cell types. *eLife*, 7, 2018.

[4000] Marta Bogacz and R Luise Krauth-Siegel. Tryparedoxin peroxidase-deficiency commits trypanosomes to ferroptosis-type cell death. *eLife*, 7, 2018.

[4001] James J Bonaiuto, Sofie S Meyer, Simon Little, Holly Rossiter, Martina F Callaghan, Frederic Dick, Gareth R Barnes, and Sven Bestmann. Lamina-specific cortical dynamics in human visual and sensorimotor cortices. *eLife*, 7, 2018.

[4002] Frédéric Bonnet, Angie Molina, Mélanie Roussat, Manon Azais, Sophie Bel-Vialar, Jacques Gautrais, Fabienne Pituello, and Eric Agius. Neurogenic decisions require a cell cycle independent function of the CDC25B phosphatase. *eLife*, 7, 2018.

[4003] Alena Boos, Jutta Distler, Heike Rudolf, Martin Klingler, and Ezzat El-Sherif. A re-inducible gap gene cascade patterns the anterior–posterior axis of insects in a threshold-free fashion. *eLife*, 7, 2018.

[4004] Andrew J Borst, Connor E Weidle, Matthew D Gray, Brandon Frenz, Joost Snijder, M Gordon Joyce, Ivelin S Georgiev, Guillaume BE Stewart-Jones, Peter D Kwong, Andrew T McGuire, Frank DiMaio, Leonidas Stamatatos, Marie Pancera, and David Veesler. Germline VRC01 antibody recognition of a modified clade C HIV-1 envelope trimer and a glycosylated HIV-1 gp120 core. *eLife*, 7, 2018.

[4005] Firas Bou Daher, Yuanjie Chen, Behruz Bozorg, Jack Clough, Henrik Jönsson, and Siobhan A Braybrook. Anisotropic growth is achieved through the additive mechanical effect of material anisotropy and elastic asymmetry. *eLife*, 7, 2018.

[4006] Hadi Boukhatmi and Sarah Bray. A population of adult satellite-like cells in Drosophila is maintained through a switch in RNA-isoforms. *eLife*, 7, 2018.

[4007] Guy Bouvier, Johnatan Aljadeff, Claudia Clopath, Célian Bimbard, Jonas Ranft, Antonin Blot, Jean-Pierre Nadal, Nicolas Brunel, Vincent Hakim, and Boris Barbour. Cerebellar learning using perturbations. *eLife*, 7, 2018.

[4008] Hande Boyaci, James Chen, Mirjana Lilic, Margaret Palka, Rachel Anne Mooney, Robert Landick, Seth A Darst, and Elizabeth A Campbell. Fidaxomicin jams Mycobacterium tuberculosis RNA polymerase motions needed for initiation via RbpA contacts. *eLife*, 7, 2018.

[4009] John Bradley, Will Stone, Dari F Da, Isabelle Morlais, Alassane Dicko, Anna Cohuet, Wamdaogo M Guelbeogo, Almahamoudou Mahamar, Sandrine Nsango, Harouna M Soumaré, Halimatou Diawara, Kjerstin Lanke, Wouter Graumans, Rianne Siebelink-Stoter, Marga van de Vegte-Bolmer, Ingrid Chen, Alfred Tiono, Bronner Pamplona GonÇalves, Roland Gosling, Robert W Sauerwein, Chris Drakeley, Thomas S Churcher, and Teun Bousema. Predicting the likelihood and intensity of mosquito infection from sex specific Plasmodium falciparum gametocyte density. *eLife*, 7, 2018.

[4010] Owen A Brady, Eutteum Jeong, José A Martina, Mehdi Pirooznia, Ilker Tunc, and Rosa Puertollano. The transcription factors TFE3 and TFEB amplify p53 dependent transcriptional programs in response to DNA damage. *eLife*, 7, 2018.

[4011] Philipp Brand, Hugh M Robertson, Wei Lin, Ratnasri Pothula, William E Klingeman, Juan Luis Jurat-Fuentes, and Brian R Johnson. The origin of the odorant receptor gene family in insects. *eLife*, 7, 2018.

[4012] Andreas M Brandmaier, Elisabeth Wenger, Nils C Bodammer, Simone Kühn, Naftali Raz, and Ulman Lindenberger. Assessing reliability in neuroimaging research through intra-class effect decomposition (ICED). *eLife*, 7, 2018.

[4013] Joerg E Braun, Larry J Friedman, Jeff Gelles, and Melissa J Moore. Synergistic assembly of human pre-spliceosomes across introns and exons. *eLife*, 7, 2018.

[4014] Gerard L Brien, David Remillard, Junwei Shi, Matthew L Hemming, Jonathon Chabon, Kieran Wynne, Eugène T Dillon, Gerard Cagney, Guido Van Mierlo, Marijke P Baltissen, Michiel Vermeulen, Jun Qi, Stefan Fröhling, Nathanael S Gray, James E Bradner, Christopher R Vakoc, and Scott A Armstrong. Targeted degradation of BRD9 reverses oncogenic gene expression in synovial sarcoma. *eLife*, 7, 2018.

[4015] Edward J Brignole, Kuang-Lei Tsai, Johnathan Chittuluru, Haoran Li, Yimon Aye, Pawel A Penczek, JoAnne Stubbe, Catherine L Drennan, and Francisco Asturias. 3.3-Å resolution cryo-EM structure of human ribonucleotide reductase with substrate and allosteric regulators bound. *eLife*, 7, 2018.

[4016] Alan Brown, Matthew R Baird, Matthew CJ Yip, Jason Murray, and Sichen Shao. Structures of translationally inactive mammalian ribosomes. *eLife*, 7, 2018.

[4017] Vanessa M Brown, Lusha Zhu, John M Wang, B Christopher Frueh, Brooks King-Casas, and Pearl H Chiu. Associability-modulated loss learning is increased in posttraumatic stress disorder. *eLife*, 7, 2018.

[4018] Christoph Brütting, Cristina Maria Crava, Martin Schäfer, Meredith C Schuman, Stefan Meldau, Nora Adam, and Ian T Baldwin. Cytokinin transfer by a free-living mirid to Nicotiana attenuata recapitulates a strategy of endophytic insects. *eLife*, 7, 2018.

[4019] Ambjorn Brynnel, Yaeren Hernandez, Balazs Kiss, Johan Lindqvist, Maya Adler, Justin Kolb, Robbert van der Pijl, Jochen Gohlke, Joshua Strom, John Smith, Coen Ottenheijm, and Henk L Granzier. Downsizing the molecular spring of the giant protein titin reveals that skeletal muscle titin determines passive stiffness and drives longitudinal hypertrophy. *eLife*, 7, 2018.

[4020] Christian Buchel, Stephan Miedl, and Christian Sprenger. Hedonic processing in humans is mediated by an opioidergic mechanism in a mesocorticolimbic system. *eLife*, 7, 2018.

[4021] Simon Bulley, Carlos Fernández-Peña, Raquibul Hasan, M Dennis Leo, Padmapriya Muralidharan, Charles E Mackay, Kirk W Evanson, Luiz Moreira-Junior, Alejandro Mata-Daboin, Sarah K Burris, Qian Wang, Korah P Kuruvilla, and Jonathan H Jaggar. Arterial smooth muscle cell PKD2 (TRPP1) channels regulate systemic blood pressure. *eLife*, 7, 2018.

[4022] Philippe Bun, Serge Dmitrieff, Julio M Belmonte, FranÇois J Nédélec, and Péter Lénárt. A disassembly-driven mechanism explains F-actin-mediated chromosome transport in starfish oocytes. *eLife*, 7, 2018.

[4023] Marianne Burbage, Francesca Gasparrini, Shweta Aggarwal, Mauro Gaya, Johan Arnold, Usha Nair, Michael Way, Andreas Bruckbauer, and Facundo D Batista. Tuning of in vivo cognate B-T cell interactions by Intersectin 2 is required for effective anti-viral B cell immunity. *eLife*, 7, 2018.

[4024] Anita Burgos, Ken Honjo, Tomoko Ohyama, Cheng Sam Qian, Grace Ji-eun Shin, Daryl M Gohl, Marion Silies, W Daniel Tracey, Marta Zlatic, Albert Cardona, and Wesley B Grueber. Nociceptive interneurons control modular motor pathways to promote escape behavior in Drosophila. *eLife*, 7, 2018.

[4025] Zachary Daniel Burkett, Nancy F Day, Todd Haswell Kimball, Caitlin M Aamodt, Jonathan B Heston, Austin T Hilliard, Xinshu Xiao, and Stephanie A White. FoxP2 isoforms delineate spatiotemporal transcriptional networks for vocal learning in the zebra finch. *eLife*, 7, 2018.

[4026] Silja Barbara Burkhard and Jeroen Bakkers. Spatially resolved RNA-sequencing of the embryonic heart identifies a role for Wnt/$\beta$-catenin signaling in autonomic control of heart rate. *eLife*, 7, 2018.

[4027] Nikolay Burnaevskiy, Shengying Chen, Miguel Mailig, Anthony Reynolds, Shruti Karanth, Alexander Mendenhall, Marc Van Gilst, and Matt Kaeberlein. Reactivation of RNA metabolism underlies somatic restoration after adult reproductive diapause in C. elegans. *eLife*, 7, 2018.

[4028] Bronwen R Burton, Richard K Tennant, John Love, Richard W Titball, David C Wraith, and Harry N White. Variant proteins stimulate more IgM+ GC B-cells revealing a mechanism of cross-reactive recognition by antibody memory. *eLife*, 7, 2018.

[4029] Verena Burtscher, Matej Hotka, Yang Li, Michael Freissmuth, and Walter Sandtner. A label-free approach to detect ligand binding to cell surface proteins in real time. *eLife*, 7, 2018.

[4030] Mitchell T Butler and John B Wallingford. Spatial and temporal analysis of PCP protein dynamics during neural tube closure. *eLife*, 7, 2018.

[4031] Agata Butryn, Stephan Woike, Savera J Shetty, David T Auble, and Karl-Peter Hopfner. Crystal structure of the full Swi2/Snf2 remodeler Mot1 in the resting state. *eLife*, 7, 2018.

[4032] Joseph M Cabral, Hyung Suk Oh, and David M Knipe. ATRX promotes maintenance of herpes simplex virus heterochromatin during chromatin stress. *eLife*, 7, 2018.

[4033] Jaime Cadena-Valencia, Otto García-Garibay, Hugo Merchant, Mehrdad Jazayeri, and Victor de Lafuente. Entrainment and maintenance of an internal metronome in supplementary motor area. *eLife*, 7, 2018.

[4034] Alison Caldwell, Rachel Coleby, Cesar Tovar, Maximilian R Stammnitz, Young Mi Kwon, Rachel S Owen, Marios Tringides, Elizabeth P Murchison, Karsten Skjødt, Gareth J Thomas, Jim Kaufman, Tim Elliott, Gregory M Woods, and Hannah VT Siddle. The newly-arisen Devil facial tumour disease 2 (DFT2) reveals a mechanism for the emergence of a contagious cancer. *eLife*, 7, 2018.

[4035] Sara Calhoun, Magdalena Korczynska, Daniel J Wichelecki, Brian San Francisco, Suwen Zhao, Dmitry A Rodionov, Matthew W Vetting, Nawar F Al-Obaidi, Henry Lin, Matthew J O'Meara, David A Scott, John H Morris, Daniel Russel, Steven C Almo, Andrei L Osterman, John A Gerlt, Matthew P Jacobson, Brian K Shoichet, and Andrej Sali. Prediction of enzymatic pathways by integrative pathway mapping. *eLife*, 7, 2018.

[4036] Soledad A Camolotto, Shrivatsav Pattabiraman, Timothy L Mosbruger, Alex Jones, Veronika K Belova, Grace Orstad, Mitchell Streiff, Lydia Salmond, Chris Stubben, Klaus H Kaestner, and Eric L Snyder. FoxA1 and FoxA2 drive gastric differentiation and suppress squamous identity in NKX2-1-negative lung cancer. *eLife*, 7, 2018.

[4037] Amy E Campbell, Sean C Shadle, Sujatha Jagannathan, Jong-Won Lim, Rebecca Resnick, Rabi Tawil, Silvère M van der Maarel, and Stephen J Tapscott. NuRD and CAF-1-mediated silencing of the D4Z4 array is modulated by DUX4-induced MBD3L proteins. *eLife*, 7, 2018.

[4038] Sharon M Campbell, Johanna A Knipper, Dominik Ruckerl, Conor M Finlay, Nicola Logan, Carlos M Minutti, Matthias Mack, Stephen J Jenkins, Matthew D Taylor, and Judith E Allen. Myeloid cell recruitment versus local proliferation differentiates susceptibility from resistance to filarial infection. *eLife*, 7, 2018.

[4039] Joseph J Campo, Timothy Q Le, Jozelyn V Pablo, Christopher Hung, Andy A Teng, Hervé Tettelin, Andrea Tate, William P Hanage, Mark R Alderson, Xiaowu Liang, Richard Malley, Marc Lipsitch, and Nicholas J Croucher. Panproteome-wide analysis of antibody responses to whole cell pneumococcal vaccination. *eLife*, 7, 2018.

[4040] Jessica Cande, Shigehiro Namiki, Jirui Qiu, Wyatt Korff, Gwyneth M Card, Joshua W Shaevitz, David L Stern, and Gordon J Berman. Optogenetic dissection of descending behavioral control in Drosophila. *eLife*, 7, 2018.

[4041] Tito Candelli, Julien Gros, and Domenico Libri. Pervasive transcription fine-tunes replication origin activity. *eLife*, 7, 2018.

[4042] Gerard Cantero-Recasens, Cristian M Butnaru, Miguel A Valverde, José R Naranjo, Nathalie Brouwers, and Vivek Malhotra. KChIP3 coupled to Ca2+ oscillations exerts a tonic brake on baseline mucin release in the colon. *eLife*, 7, 2018.

[4043] Sarah Cantor, Wei Zhang, Nicolas Delestrée, Leonor Remédio, George Z Mentis, and Steven J Burden. Preserving neuromuscular synapses in ALS by stimulating MuSK with a therapeutic agonist antibody. *eLife*, 7, 2018.

[4044] Luyan Cao, Mikael Kerleau, Emiko L. Suzuki, Hugo Wioland, Sandy Jouet, Berengere Guichard, Martin Lenz, Guillaume Romet-Lemonne, and Antoine Jegou. Modulation of formin processivity by profilin and mechanical tension. *eLife*, 7, 2018.

[4045] Shu-Xia Cao, Ying Zhang, Xing-Yue Hu, Bin Hong, Peng Sun, Hai-Yang He, Hong-Yan Geng, Ai-Min Bao, Shu-Min Duan, Jian-Ming Yang, Tian-Ming Gao, Hong Lian, and Xiao-Ming Li. ErbB4 deletion in noradrenergic neurons in the locus coeruleus induces mania-like behavior via elevated catecholamines. *eLife*, 7, 2018.

[4046] Xinyun Cao, Tobias Koch, Lydia Steffens, Julia Finkensieper, Renate Zigann, John E Cronan, and Christiane Dahl. Lipoate-binding proteins and specific lipoate-protein ligases in microbial sulfur oxidation reveal an atpyical role for an old cofactor. *eLife*, 7, 2018.

[4047] Daiana A Capdevila, Fidel Huerta, Katherine A Edmonds, My Tra Le, Hongwei Wu, and David P Giedroc. Tuning site-specific dynamics to drive allosteric activation in a pneumococcal zinc uptake regulator. *eLife*, 7, 2018.

[4048] Kevin Caref and Saleem M Nicola. Endogenous opioids in the nucleus accumbens promote approach to high-fat food in the absence of caloric need. *eLife*, 7, 2018.

[4049] Alexandre Caron, Heather M Dungan Lemko, Carlos M Castorena, Teppei Fujikawa, Syann Lee, Caleb C Lord, Newaz Ahmed, Charlotte E Lee, William L Holland, Chen Liu, and Joel K Elmquist. POMC neurons expressing leptin receptors coordinate metabolic responses to fasting via suppression of leptin levels. *eLife*, 7, 2018.

[4050] Francesca Caroti, Everardo González Avalos, Viola Noeske, Paula González Avalos, Dimitri Kromm, Maike Wosch, Lucas Schütz, Lars Hufnagel, and Steffen Lemke. Decoupling from yolk sac is required for extraembryonic tissue spreading in the scuttle fly Megaselia abdita. *eLife*, 7, 2018.

[4051] Arnaldo Carreira-Rosario, Aref Arzan Zarin, Matthew Q Clark, Laurina Manning, Richard D Fetter, Albert Cardona, and Chris Q Doe. MDN brain descending neurons coordinately activate backward and inhibit forward locomotion. *eLife*, 7, 2018.

[4052] Isaac Casanovas-Vilar, Joan Garcia-Porta, Josep Fortuny, óscar Sanisidro, Jérôme Prieto, Marina Querejeta, Sergio Llácer, Josep M Robles, Federico Bernardini, and David M Alba. Oldest skeleton of a fossil flying squirrel casts new light on the phylogeny of the group. *eLife*, 7, 2018.

[4053] Gianni M Castiglione and Belinda SW Chang. Functional trade-offs and environmental variation shaped ancient trajectories in the evolution of dim-light vision. *eLife*, 7, 2018.

[4054] Giovanna Catavitello, Yury Ivanenko, and Francesco Lacquaniti. A kinematic synergy for terrestrial locomotion shared by mammals and birds. *eLife*, 7, 2018.

[4055] Alexis M Ceasrine, Eugene E Lin, David N Lumelsky, Radhika Iyer, and Rejji Kuruvilla. Adrb2 controls glucose homeostasis by developmental regulation of pancreatic islet vasculature. *eLife*, 7, 2018.

[4056] Mark S Cembrowski, Lihua Wang, Andrew L Lemire, Monique Copeland, Salvatore F DiLisio, Jody Clements, and Nelson Spruston. The subiculum is a patchwork of discrete subregions. *eLife*, 7, 2018.

[4057] Bram Cerulus, Abbas Jariani, Gemma Perez-Samper, Lieselotte Vermeersch, Julian MJ Pietsch, Matthew M Crane, Aaron M New, Brigida Gallone, Miguel Roncoroni, Maria C Dzialo, Sander K Govers, Jhana O Hendrickx, Eva Galle, Maarten Coomans, Pieter Berden, Sara Verbandt, Peter S Swain, and Kevin J Verstrepen. Transition between fermentation and respiration determines history-dependent behavior in fluctuating carbon sources. *eLife*, 7, 2018.

[4058] Hicham Chahtane, Thanise Nogueira Füller, Pierre-Marie Allard, Laurence Marcourt, Emerson Ferreira Queiroz, Venkatasalam Shanmugabalaji, Jacques Falquet, Jean-Luc Wolfender, and Luis Lopez-Molina. The plant pathogen Pseudomonas aeruginosa triggers a DELLA-dependent seed germination arrest in Arabidopsis. *eLife*, 7, 2018.

[4059] Andrea Chai, Ana M Mateus, Fazal Oozeer, and Rita Sousa-Nunes. Spatiotemporally controlled genetic perturbation for efficient large-scale studies of cell non-autonomous effects. *eLife*, 7, 2018.

[4060] Joseph M Chambers, Shahram Jevin Poureetezadi, Amanda Addiego, Manuela Lahne, and Rebecca A Wingert. ppargc1a controls nephron segmentation during zebrafish embryonic kidney ontogeny. *eLife*, 7, 2018.

[4061] John D Chan, Timothy A Day, and Jonathan S Marchant. Coalescing beneficial host and deleterious antiparasitic actions as an antischistosomal strategy. *eLife*, 7, 2018.

[4062] Leon Y Chan, Christopher F Mugler, Stephanie Heinrich, Pascal Vallotton, and Karsten Weis. Non-invasive measurement of mRNA decay reveals translation initiation as the major determinant of mRNA stability. *eLife*, 7, 2018.

[4063] Natali L Chanaday and Ege T Kavalali. Optical detection of three modes of endocytosis at hippocampal synapses. *eLife*, 7, 2018.

[4064] Roger Chang, Jacob Eriksen, and Robert H Edwards. The dual role of chloride in synaptic vesicle glutamate transport. *eLife*, 7, 2018.

[4065] Luke H Chao, Jaebong Jang, Adam Johnson, Anthony Nguyen, Nathanael S Gray, Priscilla L Yang, and Stephen C Harrison. How small-molecule inhibitors of dengue-virus infection interfere with viral membrane fusion. *eLife*, 7, 2018.

[4066] Richard H Chapple, Tianyuan Hu, Yu-Jung Tseng, Lu Liu, Ayumi Kitano, Victor Luu, Kevin A Hoegenauer, Takao Iwawaki, Qing Li, and Daisuke Nakada. ER$\alpha$ promotes murine hematopoietic regeneration through the Ire1$\alpha$-mediated unfolded protein response. *eLife*, 7, 2018.

[4067] Jillian Chase, Andrew Catalano, Alex J Noble, Edward T Eng, Paul DB Olinares, Kelly Molloy, Danaya Pakotiprapha, Martin Samuels, Brian Chait, Amedee des Georges, and David Jeruzalmi. Mechanisms of opening and closing of the bacterial replicative helicase. *eLife*, 7, 2018.

[4068] Ishanu Chattopadhyay, Emre Kiciman, Joshua W Elliott, Jeffrey L Shaman, and Andrey Rzhetsky. Conjunction of factors triggering waves of seasonal influenza. *eLife*, 7, 2018.

[4069] Kevin F Chau, Morgan L Shannon, Ryann M Fame, Erin Fonseca, Hillary Mullan, Matthew B Johnson, Anoop K Sendamarai, Mark W Springel, Benoit Laurent, and Maria K Lehtinen. Down-regulation of ribosome biogenesis during early forebrain development. *eLife*, 7, 2018.

[4070] Swarnava Chaudhuri, Liping Li, Matthew Zimmerman, Yuemeng Chen, Yu-Xiang Chen, Melody N Toosky, Michelle Gardner, Miaomiao Pan, Yang-Yang Li, Qingwen Kawaji, Jun-Hao Zhu, Hong-Wei Su, Amanda J Martinot, Eric J Rubin, Veronique Anne Dartois, and Babak Javid. Kasugamycin potentiates rifampicin and limits emergence of resistance in Mycobacterium tuberculosis by specifically decreasing mycobacterial mistranslation. *eLife*, 7, 2018.

[4071] Cuie Chen, Ryan Cummings, Aghapi Mordovanakis, Alan J Hunt, Michael Mayer, David Sept, and Yukiko M Yamashita. Cytokine receptor-Eb1 interaction couples cell polarity and fate during asymmetric cell division. *eLife*, 7, 2018.

[4072] Eric CH Chen, Stephanie Mathieu, Anne Hoffrichter, Kinga Sedzielewska-Toro, Max Peart, Adrian Pelin, Steve Ndikumana, Jeanne Ropars, Steven Dreissig, Jorg Fuchs, Andreas Brachmann, and Nicolas Corradi. Single nucleus sequencing reveals evidence of inter-nucleus recombination in arbuscular mycorrhizal fungi. *eLife*, 7, 2018.

[4073] Guifen Chen, John Andrew King, Yi Lu, Francesca Cacucci, and Neil Burgess. Spatial cell firing during virtual navigation of open arenas by head-restrained mice. *eLife*, 7, 2018.

[4074] Wenyang Chen, Sridhar Mandali, Stephen P Hancock, Pramod Kumar, Michael Collazo, Duilio Cascio, and Reid C Johnson. Multiple serine transposase dimers assemble the transposon-end synaptic complex during IS607-family transposition. *eLife*, 7, 2018.

[4075] Xi Chen, Akihiro CE Shibata, Ardalan Hendi, Mizuki Kurashina, Ethan Fortes, Nicholas L Weilinger, Brian A MacVicar, Hideji Murakoshi, and Kota Mizumoto. Rap2 and TNIK control Plexin-dependent tiled synaptic innervation in C. elegans. *eLife*, 7, 2018.

[4076] Yu-Chen Chen, Pei-Heng Jiang, Hsuan-Ming Chen, Chang-Han Chen, Yi-Ting Wang, Yu-Ju Chen, Chia-Jung Yu, and Shu-Chun Teng. Glucose intake hampers PKA-regulated HSP90 chaperone activity. *eLife*, 7, 2018.

[4077] Chun-Chun Cheng, Ko Tsutsui, Toru Taguchi, Noriko Sanzen, Asako Nakagawa, Kisa Kakiguchi, Shigenobu Yonemura, Chiharu Tanegashima, Sean D Keeley, Hiroshi Kiyonari, Yasuhide Furuta, Yasuko Tomono, Fiona M Watt, and Hironobu Fujiwara. Hair follicle epidermal stem cells define a niche for tactile sensation. *eLife*, 7, 2018.

[4078] Kaiying Cheng and Dale B Wigley. DNA translocation mechanism of an XPD family helicase. *eLife*, 7, 2018.

[4079] Leonard Cheung, Paul Le Tissier, Sam GJ Goldsmith, Mathias Treier, Robin Lovell-Badge, and Karine Rizzoti. NOTCH activity differentially affects alternative cell fate acquisition and maintenance. *eLife*, 7, 2018.

[4080] Kelda Chia, Julie Mazzolini, Marina Mione, and Dirk Sieger. Tumor initiating cells induce Cxcr4-mediated infiltration of pro-tumoral macrophages into the brain. *eLife*, 7, 2018.

[4081] Poh Hui Chia, Franklin Lei Zhong, Shinsuke Niwa, Carine Bonnard, Kagistia Hana Utami, Ruizhu Zeng, Hane Lee, Ascia Eskin, Stanley F Nelson, William H Xie, Samah Al-Tawalbeh, Mohammad El-Khateeb, Mohammad Shboul, Mahmoud A Pouladi, Mohammed Al-Raqad, and Bruno Reversade. A homozygous loss-of-function CAMK2A mutation causes growth delay, frequent seizures and severe intellectual disability. *eLife*, 7, 2018.

[4082] Nicholas Chim, Lynnette N Jackson, Anh M Trinh, and John C Chaput. Crystal structures of DNA polymerase I capture novel intermediates in the DNA synthesis pathway. *eLife*, 7, 2018.

[4083] Ucheor B Choi, Minglei Zhao, K Ian White, Richard A Pfuetzner, Luis Esquivies, Qiangjun Zhou, and Axel T Brunger. NSF-mediated disassembly of on- and off-pathway SNARE complexes and inhibition by complexin. *eLife*, 7, 2018.

[4084] Lasse Christiansen, MA Urbin, Gordon S Mitchell, and Monica A Perez. Acute intermittent hypoxia enhances corticospinal synaptic plasticity in humans. *eLife*, 7, 2018.

[4085] Chee Wai Chua, Nusrat J Epsi, Eva Y Leung, Shouhong Xuan, Ming Lei, Bo I Li, Sarah K Bergren, Hanina Hibshoosh, Antonina Mitrofanova, and Michael M Shen. Differential requirements of androgen receptor in luminal progenitors during prostate regeneration and tumor initiation. *eLife*, 7, 2018.

[4086] Nao Chuhma, Susana Mingote, Leora Yetnikoff, Abigail Kalmbach, Thong Ma, Samira Ztaou, Anna-Claire Sienna, Sophia Tepler, Jean-Francois Poulin, Mark Ansorge, Rajeshwar Awatramani, Un Jung Kang, and Stephen Rayport. Dopamine neuron glutamate cotransmission evokes a delayed excitation in lateral dorsal striatal cholinergic interneurons. *eLife*, 7, 2018.

[4087] Ilya Chuykin, Olga Ossipova, and Sergei Y Sokol. Par3 interacts with Prickle3 to generate apical PCP complexes in the vertebrate neural plate. *eLife*, 7, 2018.

[4088] Davide Ciliberti, Frédéric Michon, and Fabian Kloosterman. Real-time classification of experience-related ensemble spiking patterns for closed-loop applications. *eLife*, 7, 2018.

[4089] Erica Cirri, Sébastien Brier, Reda Assal, Juan Carlos Canul-Tec, Julia Chamot-Rooke, and Nicolas Reyes. Consensus designs and thermal stability determinants of a human glutamate transporter. *eLife*, 7, 2018.

[4090] Sarah Clark, Janette B Myers, Ashleigh King, Radovan Fiala, Jiri Novacek, Grant Pearce, Jörg Heierhorst, Steve L Reichow, and Elisar J Barbar. Multivalency regulates activity in an intrinsically disordered transcription factor. *eLife*, 7, 2018.

[4091] Jeroen Claus, Gargi Patel, Flavia Autore, Audrey Colomba, Gregory Weitsman, Tanya N Soliman, Selene Roberts, Laura C Zanetti-Domingues, Michael Hirsch, Francesca Collu, Roger George, Elena Ortiz-Zapater, Paul R Barber, Boris Vojnovic, Yosef Yarden, Marisa L Martin-Fernandez, Angus Cameron, Franca Fraternali, Tony Ng, and Peter J Parker. Inhibitor-induced HER2-HER3 heterodimerisation promotes proliferation through a novel dimer interface. *eLife*, 7, 2018.

[4092] Alice M Clement, Benedict King, Sam Giles, Brian Choo, Per E Ahlberg, Gavin C Young, and John A Long. Neurocranial anatomy of an enigmatic Early Devonian fish sheds light on early osteichthyan evolution. *eLife*, 7, 2018.

[4093] Jessica Coates, Bo Ryoung Park, Dai Le, Emrah şimşek, Waqas Chaudhry, and Minsu Kim. Antibiotic-induced population fluctuations and stochastic clearance of bacteria. *eLife*, 7, 2018.

[4094] Erez Cohen, Scott R Allen, Jessica K Sawyer, and Donald T Fox. Fizzy-Related dictates A cell cycle switch during organ repair and tissue growth responses in the Drosophila hindgut. *eLife*, 7, 2018.

[4095] Michelle M Collins, Hans-Martin Maischein, Pascale Dufourcq, Marine Charpentier, Patrick Blader, and Didier YR Stainier. Pitx2c orchestrates embryonic axis extension via mesendodermal cell migration. *eLife*, 7, 2018.

[4096] Tamica N Collins, Yingyu Mao, Hongge Li, Michael Bouaziz, Angela Hong, Gen-Sheng Feng, Fen Wang, Lawrence A Quilliam, Lin Chen, Taeju Park, Tom Curran, and Xin Zhang. Crk proteins transduce FGF signaling to promote lens fiber cell elongation. *eLife*, 7, 2018.

[4097] Yaara Y Columbus-Shenkar, Maria Y Sachkova, Jason Macrander, Arie Fridrich, Vengamanaidu Modepalli, Adam M Reitzel, Kartik Sunagar, and Yehu Moran. Dynamics of venom composition across a complex life cycle. *eLife*, 7, 2018.

[4098] Mauricio Comas-Garcia, Tomas Kroupa, Siddhartha AK Datta, Demetria P Harvin, Wei-Shau Hu, and Alan Rein. Efficient support of virus-like particle assembly by the HIV-1 packaging signal. *eLife*, 7, 2018.

[4099] Joan Comenge, Jack Sharkey, Oihane Fragueiro, Bettina Wilm, Mathias Brust, Patricia Murray, Raphael Levy, and Antonius Plagge. Multimodal cell tracking from systemic administration to tumour growth by combining gold nanorods and reporter genes. *eLife*, 7, 2018.

[4100] Barbara Commisso, Lingjun Ding, Karl Varadi, Martin Gorges, David Bayer, Tobias M Boeckers, Albert C Ludolph, Jan Kassubek, Oliver J Müller, and Francesco Roselli. Stage-dependent remodeling of projections to motor cortex in ALS mouse model revealed by a new variant retrograde-AAV9. *eLife*, 7, 2018.

[4101] Andrew James Kerr Conlan, Martin Vordermeier, Mart CM de Jong, and James LN Wood. The intractable challenge of evaluating cattle vaccination as a control for bovine Tuberculosis. *eLife*, 7, 2018.

[4102] Erin G Conlon, Delphine Fagegaltier, Phaedra Agius, Julia Davis-Porada, James Gregory, Isabel Hubbard, Kristy Kang, Duyang Kim, Hemali Phatnani, Hemali Phatnani, Justin Kwan, Dhruv Sareen, James R Broach, Zachary Simmons, Ximena Arcila-Londono, Edward B Lee, Vivianna M Van Deerlin, Neil A Shneider, Ernest Fraenkel, Lyle W Ostrow, Frank Baas, Noah Zaitlen, James D Berry, Andrea Malaspina, Pietro Fratta, Gregory A Cox, Leslie M Thompson, Steve Finkbeiner, Efthimios Dardiotis, Timothy M Miller, Siddharthan Chandran, Suvankar Pal, Eran Hornstein, Daniel J MacGowan, Terry Heiman-Patterson, Molly G Hammell, Nikolaos A Patsopoulos, Joshua Dubnau, Avindra Nath, Hemali Phatnani, Neil A Shneider, and James L Manley. Unexpected similarities between C9ORF72 and sporadic forms of ALS/FTD suggest a common disease mechanism. *eLife*, 7, 2018.

[4103] William D Constance, Amrita Mukherjee, Yvette E Fisher, Sinziana Pop, Eric Blanc, Yusuke Toyama, and Darren W Williams. Neurexin and Neuroligin-based adhesion complexes drive axonal arborisation growth independent of synaptic activity. *eLife*, 7, 2018.

[4104] Samuel T Coradetti, Dominic Pinel, Gina M Geiselman, Masakazu Ito, Stephen J Mondo, Morgann C Reilly, Ya-Fang Cheng, Stefan Bauer, Igor V Grigoriev, John M Gladden, Blake A Simmons, Rachel B Brem, Adam P Arkin, and Jeffrey M Skerker. Functional genomics of lipid metabolism in the oleaginous yeast Rhodosporidium toruloides. *eLife*, 7, 2018.

[4105] Tom Cornelissen, Sven Vilain, Katlijn Vints, Natalia Gounko, Patrik Verstreken, and Wim Vandenberghe. Deficiency of parkin and PINK1 impairs age-dependent mitophagy in Drosophila. *eLife*, 7, 2018.

[4106] Julie Corre, Ruud van Zessen, Michaël Loureiro, Tommaso Patriarchi, Lin Tian, Vincent Pascoli, and Christian Lüscher. Dopamine neurons projecting to medial shell of the nucleus accumbens drive heroin reinforcement. *eLife*, 7, 2018.

[4107] Kevin C Courtney, Karen YY Fung, Frederick R Maxfield, Gregory D Fairn, and Xiaohui Zha. Comment on 'Orthogonal lipid sensors identify transbilayer asymmetry of plasma membrane cholesterol'. *eLife*, 7, 2018.

[4108] Ana Covelo and Alfonso Araque. Neuronal activity determines distinct gliotransmitter release from a single astrocyte. *eLife*, 7, 2018.

[4109] Sophie Croizier, Soyoung Park, Julien Maillard, and Sebastien G Bouret. Central Dicer-miR-103/107 controls developmental switch of POMC progenitors into NPY neurons and impacts glucose homeostasis. *eLife*, 7, 2018.

[4110] Cristina Cruz, Monica Della Rosa, Christel Krueger, Qian Gao, Dorottya Horkai, Michelle King, Lucy Field, and Jonathan Houseley. Tri-methylation of histone H3 lysine 4 facilitates gene expression in ageing cells. *eLife*, 7, 2018.

[4111] Bridgette M Cumming, Kelvin W Addicott, John H Adamson, and Adrie JC Steyn. Mycobacterium tuberculosis induces decelerated bioenergetic metabolism in human macrophages. *eLife*, 7, 2018.

[4112] Yasin F Dagdas, Pooja Pandey, Yasin Tumtas, Nattapong Sanguankiattichai, Khaoula Belhaj, Cian Duggan, Alexandre Y Leary, Maria E Segretin, Mauricio P Contreras, Zachary Savage, Virendrasinh S Khandare, Sophien Kamoun, and Tolga O Bozkurt. Host autophagy machinery is diverted to the pathogen interface to mediate focal defense responses against the Irish potato famine pathogen. *eLife*, 7, 2018.

[4113] Ajinkya Dahake, Anna L Stöckl, James J Foster, Sanjay P Sane, and Almut Kelber. The roles of vision and antennal mechanoreception in hawkmoth flight control. *eLife*, 7, 2018.

[4114] Agnik Dasgupta, Matthias Merkel, Madeline J Clark, Andrew E Jacob, Jonathan Edward Dawson, M Lisa Manning, and Jeffrey D Amack. Cell volume changes contribute to epithelial morphogenesis in zebrafish Kupffer's vesicle. *eLife*, 7, 2018.

[4115] Matthew J Davidson, David Alais, Jeroen JA van Boxtel, and Naotsugu Tsuchiya. Attention periodically samples competing stimuli during binocular rivalry. *eLife*, 7, 2018.

[4116] Tim Davies, Han X Kim, Natalia Romano Spica, Benjamin J Lesea-Pringle, Julien Dumont, Mimi Shirasu-Hiza, and Julie C Canman. Cell-intrinsic and -extrinsic mechanisms promote cell-type-specific cytokinetic diversity. *eLife*, 7, 2018.

[4117] Antonio Davila, Ling Liu, Karthikeyani Chellappa, Philip Redpath, Eiko Nakamaru-Ogiso, Lauren M Paolella, Zhigang Zhang, Marie E Migaud, Joshua D Rabinowitz, and Joseph A Baur. Nicotinamide adenine dinucleotide is transported into mammalian mitochondria. *eLife*, 7, 2018.

[4118] Gregory M Davis, Shikui Tu, Joshua WT Anderson, Rhys N Colson, Menachem J Gunzburg, Michelle A Francisco, Debashish Ray, Sean P Shrubsole, Julia A Sobotka, Uri Seroussi, Robert X Lao, Tuhin Maity, Monica Z Wu, Katherine McJunkin, Quaid D Morris, Timothy R Hughes, Jacqueline A Wilce, Julie M Claycomb, Zhiping Weng, and Peter R Boag. The TRIM-NHL protein NHL-2 is a co-factor in the nuclear and somatic RNAi pathways in C. elegans. *eLife*, 7, 2018.

[4119] Matthew Day, Mathieu Rappas, Katie Ptasinska, Dominik Boos, Antony W Oliver, and Laurence H Pearl. BRCT domains of the DNA damage checkpoint proteins TOPBP1/Rad4 display distinct specificities for phosphopeptide ligands. *eLife*, 7, 2018.

[4120] Benjamin J De Corte, Rebecca R Della Valle, and Matthew S Matell. Recalibrating timing behavior via expected covariance between temporal cues. *eLife*, 7, 2018.

[4121] Janaina de Freitas Nascimento, Steven Kelly, Jack Sunter, and Mark Carrington. Codon choice directs constitutive mRNA levels in trypanosomes. *eLife*, 7, 2018.

[4122] Andrea Cristina de Lima-Pardini, Daniel Boari Coelho, Carolina Pinto Souza, Carolina Oliveira Souza, Maria Gabriela dos Santos Ghilardi, Tiago Garcia, Mariana Voos, Matija Milosevic, Clement Hamani, Luis Augusto Teixeira, and Erich Talamoni Fonoff. Effects of spinal cord stimulation on postural control in Parkinson's disease patients with freezing of gait. *eLife*, 7, 2018.

[4123] Franziska Decker, David Oriola, Benjamin Dalton, and Jan Brugués. Autocatalytic microtubule nucleation determines the size and mass of Xenopus laevis egg extract spindles. *eLife*, 7, 2018.

[4124] Christoph M Deeg, Cheryl-Emiliane T Chow, and Curtis A Suttle. The kinetoplastid-infecting Bodo saltans virus (BsV), a window into the most abundant giant viruses in the sea. *eLife*, 7, 2018.

[4125] Adriana Degrossoli, Alexandra Müller, Kaibo Xie, Jannis F Schneider, Verian Bader, Konstanze F Winklhofer, Andreas J Meyer, and Lars I Leichert. Neutrophil-generated HOCl leads to non-specific thiol oxidation in phagocytized bacteria. *eLife*, 7, 2018.

[4126] Josefina Inés del Mármol, Kouki K Touhara, Gist Croft, and Roderick MacKinnon. Piezo1 forms a slowly-inactivating mechanosensory channel in mouse embryonic stem cells. *eLife*, 7, 2018.

[4127] Daniel J Denman, Jennifer A Luviano, Douglas R Ollerenshaw, Sissy Cross, Derric Williams, Michael A Buice, Shawn R Olsen, and R Clay Reid. Mouse color and wavelength-specific luminance contrast sensitivity are non-uniform across visual space. *eLife*, 7, 2018.

[4128] Ben Deverett, Sue Ann Koay, Marlies Oostland, and Samuel S-H Wang. Cerebellar involvement in an evidence-accumulation decision-making task. *eLife*, 7, 2018.

[4129] Arun Devotta, Chang-Soo Hong, and Jean-Pierre Saint-Jeannet. Dkk2 promotes neural crest specification by activating Wnt/$\beta$-catenin signaling in a GSK3$\beta$ independent manner. *eLife*, 7, 2018.

[4130] Pooran Singh Dewari, Benjamin Southgate, Katrina Mccarten, German Monogarov, Eoghan O'Duibhir, Niall Quinn, Ashley Tyrer, Marie-Christin Leitner, Colin Plumb, Maria Kalantzaki, Carla Blin, Rebecca Finch, Raul Bardini Bressan, Gillian Morrison, Ashley M Jacobi, Mark A Behlke, Alex von Kriegsheim, Simon Tomlinson, Jeroen Krijgsveld, and Steven M Pollard. An efficient and scalable pipeline for epitope tagging in mammalian stem cells using Cas9 ribonucleoprotein. *eLife*, 7, 2018.

[4131] Richard Burkett Dewell and Fabrizio Gabbiani. Biophysics of object segmentation in a collision-detecting neuron. *eLife*, 7, 2018.

[4132] William S DeWitt, Anajane Smith, Gary Schoch, John A Hansen, Frederick A Matsen, and Philip Bradley. Human T cell receptor occurrence patterns encode immune history, genetic background, and receptor specificity. *eLife*, 7, 2018.

[4133] Herschel S Dhekne, Izumi Yanatori, Rachel C Gomez, Francesca Tonelli, Federico Diez, Birgitt Schüle, Martin Steger, Dario R Alessi, and Suzanne R Pfeffer. A pathway for Parkinson's Disease LRRK2 kinase to block primary cilia and Sonic hedgehog signaling in the brain. *eLife*, 7, 2018.

[4134] Kevin D'hoe, Stefan Vet, Karoline Faust, Frédéric Moens, Gwen Falony, Didier Gonze, Verónica Lloréns-Rico, Lendert Gelens, Jan Danckaert, Luc De Vuyst, and Jeroen Raes. Integrated culturing, modeling and transcriptomics uncovers complex interactions and emergent behavior in a three-species synthetic gut community. *eLife*, 7, 2018.

[4135] Sara C Di Rienzi, Juliet Jacobson, Elizabeth A Kennedy, Mary E Bell, Qiaojuan Shi, Jillian L Waters, Peter Lawrence, J Thomas Brenna, Robert A Britton, Jens Walter, and Ruth E Ley. Resilience of small intestinal beneficial bacteria to the toxicity of soybean oil fatty acids. *eLife*, 7, 2018.

[4136] Min Diao, Sulin Ren, Qiannan Wang, Lichao Qian, Jiangfeng Shen, Yule Liu, and Shanjin Huang. Arabidopsis formin 2 regulates cell-to-cell trafficking by capping and stabilizing actin filaments at plasmodesmata. *eLife*, 7, 2018.

[4137] Claire Dickson, Adam J Fletcher, Marina Vaysburd, Ji-Chun Yang, Donna L Mallery, Jingwei Zeng, Christopher M Johnson, Stephen H McLaughlin, Mark Skehel, Sarah Maslen, James Cruickshank, Nicolas Huguenin-Dezot, Jason W Chin, David Neuhaus, and Leo C James. Intracellular antibody signalling is regulated by phosphorylation of the Fc receptor TRIM21. *eLife*, 7, 2018.

[4138] Maria M Diehl, Christian Bravo-Rivera, Jose Rodriguez-Romaguera, Pablo A Pagan-Rivera, Anthony Burgos-Robles, Ciorana Roman-Ortiz, and Gregory J Quirk. Active avoidance requires inhibitory signaling in the rodent prelimbic prefrontal cortex. *eLife*, 7, 2018.

[4139] Nadine Dijkstra, Pim Mostert, Floris P de Lange, Sander Bosch, and Marcel AJ van Gerven. Differential temporal dynamics during visual imagery and perception. *eLife*, 7, 2018.

[4140] Emilia Dimitrova, Takashi Kondo, Angelika Feldmann, Manabu Nakayama, Yoko Koseki, Rebecca Konietzny, Benedikt M Kessler, Haruhiko Koseki, and Robert J Klose. FBXL19 recruits CDK-Mediator to CpG islands of developmental genes priming them for activation during lineage commitment. *eLife*, 7, 2018.

[4141] Chen Ding and Marc Hammarlund. Aberrant information transfer interferes with functional axon regeneration. *eLife*, 7, 2018.

[4142] Siyuan Ding, Shu Zhu, Lili Ren, Ningguo Feng, Yanhua Song, Xiaomei Ge, Bin Li, Richard A Flavell, and Harry B Greenberg. Rotavirus VP3 targets MAVS for degradation to inhibit type III interferon expression in intestinal epithelial cells. *eLife*, 7, 2018.

[4143] Cindy Dirscherl, Zeynep Hein, Venkat Raman Ramnarayan, Catherine Jacob-Dolan, and Sebastian Springer. A two-hybrid antibody micropattern assay reveals specific in cis interactions of MHC I heavy chains at the cell surface. *eLife*, 7, 2018.

[4144] Guillaume Diss and Ben Lehner. The genetic landscape of a physical interaction. *eLife*, 7, 2018.

[4145] Garron T Dodd, Natalie J Michael, Robert S Lee-Young, Salvatore P Mangiafico, Jack T Pryor, Astrid C Munder, Stephanie E Simonds, Jens Claus Brüning, Zhong-Yin Zhang, Michael A Cowley, Sofianos Andrikopoulos, Tamas L Horvath, David Spanswick, and Tony Tiganis. Insulin regulates POMC neuronal plasticity to control glucose metabolism. *eLife*, 7, 2018.

[4146] Stefan Donat, Marta LourenÇo, Alessio Paolini, Cécile Otten, Marc Renz, and Salim Abdelilah-Seyfried. Heg1 and Ccm1/2 proteins control endocardial mechanosensitivity during zebrafish valvulogenesis. *eLife*, 7, 2018.

[4147] Qiping Dong, Qing Liu, Ronghui Li, Anxin Wang, Qian Bu, Kuan Hong Wang, and Qiang Chang. Mechanism and consequence of abnormal calcium homeostasis in Rett syndrome astrocytes. *eLife*, 7, 2018.

[4148] Katherine A Donovan, Jian An, Radosław P Nowak, Jingting C Yuan, Emma C Fink, Bethany C Berry, Benjamin L Ebert, and Eric S Fischer. Thalidomide promotes degradation of SALL4, a transcription factor implicated in Duane Radial Ray syndrome. *eLife*, 7, 2018.

[4149] James C Dooley and Mark S Blumberg. Developmental 'awakening' of primary motor cortex to the sensory consequences of movement. *eLife*, 7, 2018.

[4150] Daniel B Dorman, Joanna Jędrzejewska-Szmek, and Kim T Blackwell. Inhibition enhances spatially-specific calcium encoding of synaptic input patterns in a biologically constrained model. *eLife*, 7, 2018.

[4151] Aline R Dörrbaum, Lisa Kochen, Julian D Langer, and Erin M Schuman. Local and global influences on protein turnover in neurons and glia. *eLife*, 7, 2018.

[4152] Eliot Dow, Adrian Jacobo, Sajjad Hossain, Kimberly Siletti, and A J Hudspeth. Connectomics of the zebrafish's lateral-line neuromast reveals wiring and miswiring in a simple microcircuit. *eLife*, 7, 2018.

[4153] Florian Drawitsch, Ali Karimi, Kevin M Boergens, and Moritz Helmstaedter. FluoEM, virtual labeling of axons in three-dimensional electron microscopy data for long-range connectomics. *eLife*, 7, 2018.

[4154] Terri M Driessen, Paul J Lee, and Janghoo Lim. Molecular pathway analysis towards understanding tissue vulnerability in spinocerebellar ataxia type 1. *eLife*, 7, 2018.

[4155] Lijuan Du, Alex Sohr, Ge Yan, and Sougata Roy. Feedback regulation of cytoneme-mediated transport shapes a tissue-specific FGF morphogen gradient. *eLife*, 7, 2018.

[4156] Dorine Jeanne Mariëtte du Mee, Maxim Ivanov, Joseph Paul Parker, Stephen Buratowski, and Sebastian Marquardt. Efficient termination of nuclear lncRNA transcription promotes mitochondrial genome maintenance. *eLife*, 7, 2018.

[4157] Gytis Dudas, Luiz Max Carvalho, Andrew Rambaut, and Trevor Bedford. MERS-CoV spillover at the camel-human interface. *eLife*, 7, 2018.

[4158] Camille Dumon, Diabe Diabira, Ilona Chudotvorova, Francesca Bader, Semra Sahin, Jinwei Zhang, Christophe Porcher, Gary Wayman, Igor Medina, and Jean-Luc Gaiarsa. The adipocyte hormone leptin sets the emergence of hippocampal inhibition in mice. *eLife*, 7, 2018.

[4159] Romain Durand-de Cuttoli, Sarah Mondoloni, Fabio Marti, Damien Lemoine, Claire Nguyen, Jérémie Naudé, Thibaut d'Izarny Gargas, Stéphanie Pons, Uwe Maskos, Dirk Trauner, Richard H Kramer, Philippe Faure, and Alexandre Mourot. Manipulating midbrain dopamine neurons and reward-related behaviors with light-controllable nicotinic acetylcholine receptors. *eLife*, 7, 2018.

[4160] Kira M Düsterwald, Christopher B Currin, Richard J Burman, Colin J Akerman, Alan R Kay, and Joseph V Raimondo. Biophysical models reveal the relative importance of transporter proteins and impermeant anions in chloride homeostasis. *eLife*, 7, 2018.

[4161] Fabien Duveau, Andrea Hodgins-Davis, Brian PH Metzger, Bing Yang, Stephen Tryban, Elizabeth A Walker, Tricia Lybrook, and Patricia J Wittkopp. Fitness effects of altering gene expression noise in Saccharomyces cerevisiae. *eLife*, 7, 2018.

[4162] Jason J Early, Katy LH Marshall-Phelps, Jill M Williamson, Matthew Swire, Hari Kamadurai, Marc Muskavitch, and David A Lyons. An automated high-resolution in vivo screen in zebrafish to identify chemical regulators of myelination. *eLife*, 7, 2018.

[4163] Kathryn Eaton, Ali Pirani, Evan S Snitkin, Elizabeth Iorns, Elizabeth Iorns, Rachel Tsui, Alexandria Denis, Nicole Perfito, Timothy M Errington, Elizabeth Iorns, Rachel Tsui, Alexandria Denis, Nicole Perfito, and Timothy M Errington. Replication Study: Intestinal inflammation targets cancer-inducing activity of the microbiota. *eLife*, 7, 2018.

[4164] Hani Ebrahimi, Hirohisa Masuda, Devanshi Jain, and Julia Promisel Cooper. Distinct 'safe zones' at the nuclear envelope ensure robust replication of heterochromatic chromosome regions. *eLife*, 7, 2018.

[4165] Frances Edwards, Gilliane Maton, Nelly Gareil, Julie C Canman, and Julien Dumont. BUB-1 promotes amphitelic chromosome biorientation via multiple activities at the kinetochore. *eLife*, 7, 2018.

[4166] Yuichi Eguchi, Koji Makanae, Tomohisa Hasunuma, Yuko Ishibashi, Keiji Kito, and Hisao Moriya. Estimating the protein burden limit of yeast cells by measuring the expression limits of glycolytic proteins. *eLife*, 7, 2018.

[4167] Nico Eisenhauer, Jes Hines, Forest Isbell, Fons van der Plas, Sarah E Hobbie, Clare E Kazanski, Anika Lehmann, Mengyun Liu, Alfred Lochner, Matthias C Rillig, Anja Vogel, Kally Worm, and Peter B Reich. Plant diversity maintains multiple soil functions in future environments. *eLife*, 7, 2018.

[4168] Stephan A Eisler, Filipa Curado, Gisela Link, Sarah Schulz, Melanie Noack, Maren Steinke, Monilola A Olayioye, and Angelika Hausser. A Rho signaling network links microtubules to PKD controlled carrier transport to focal adhesions. *eLife*, 7, 2018.

[4169] Krystel El Hage, Florent Hédin, Prashant K Gupta, Markus Meuwly, and Martin Karplus. Valid molecular dynamics simulations of human hemoglobin require a surprisingly large box size. *eLife*, 7, 2018.

[4170] Harri Elamaa, Minna Kihlström, Emmi Kapiainen, Mika Kaakinen, Ilkka Miinalainen, Symantas Ragauskas, Marc Cerrada-Gimenez, Satu Mering, Marjut Nätynki, and Lauri Eklund. Angiopoietin-4-dependent venous maturation and fluid drainage in the peripheral retina. *eLife*, 7, 2018.

[4171] Lotem Elber-Dorozko and Yonatan Loewenstein. Striatal action-value neurons reconsidered. *eLife*, 7, 2018.

[4172] Mark AG Eldridge, Narihisa Matsumoto, John H Wittig, Evan C Masseau, Richard C Saunders, and Barry J Richmond. Perceptual processing in the ventral visual stream requires area TE but not rhinal cortex. *eLife*, 7, 2018.

[4173] Lorriane Eley, Ahlam MS Alqahtani, Donal MacGrogan, Rachel V Richardson, Lindsay Murphy, Alejandro Salguero-Jimenez, Marcos Sintes Rodriguez San Pedro, Shindi Tiurma, Lauren McCutcheon, Adam Gilmore, José Luis de La Pompa, Bill Chaudhry, and Deborah J Henderson. A novel source of arterial valve cells linked to bicuspid aortic valve without raphe in mice. *eLife*, 7, 2018.

[4174] Brian T Emmer, Geoffrey G Hesketh, Emilee Kotnik, Vi T Tang, Paul J Lascuna, Jie Xiang, Anne-Claude Gingras, Xiao-Wei Chen, and David Ginsburg. The cargo receptor SURF4 promotes the efficient cellular secretion of PCSK9. *eLife*, 7, 2018.

[4175] Sonya Entova, Jean-Marc Billod, Jean-Marie Swiecicki, Sonsoles Martín-Santamaría, and Barbara Imperiali. Insights into the key determinants of membrane protein topology enable the identification of new monotopic folds. *eLife*, 7, 2018.

[4176] Kali Esancy, Logan Condon, Jing Feng, Corinna Kimball, Andrew Curtright, and Ajay Dhaka. A zebrafish and mouse model for selective pruritus via direct activation of TRPA1. *eLife*, 7, 2018.

[4177] Prahathees J Eswara, Robert S Brzozowski, Marissa G Viola, Gianni Graham, Catherine Spanoudis, Catherine Trebino, Jyoti Jha, Joseph I Aubee, Karl M Thompson, Jodi L Camberg, and Kumaran S Ramamurthi. An essential Staphylococcus aureus cell division protein directly regulates FtsZ dynamics. *eLife*, 7, 2018.

[4178] Alex Evilevitch. The mobility of packaged phage genome controls ejection dynamics. *eLife*, 7, 2018.

[4179] Giulia Faini, Andrea Aguirre, Silvia Landi, Didi Lamers, Tommaso Pizzorusso, Gian Michele Ratto, Charlotte Deleuze, and Alberto Bacci. Perineuronal nets control visual input via thalamic recruitment of cortical PV interneurons. *eLife*, 7, 2018.

[4180] Timothy R Fallon, Sarah E Lower, Ching-Ho Chang, Manabu Bessho-Uehara, Gavin J Martin, Adam J Bewick, Megan Behringer, Humberto J Debat, Isaac Wong, John C Day, Anton Suvorov, Christian J Silva, Kathrin F Stanger-Hall, David W Hall, Robert J Schmitz, David R Nelson, Sara M Lewis, Shuji Shigenobu, Seth M Bybee, Amanda M Larracuente, Yuichi Oba, and Jing-Ke Weng. Firefly genomes illuminate parallel origins of bioluminescence in beetles. *eLife*, 7, 2018.

[4181] Chen Fan, Wooyoung Choi, Weinan Sun, Juan Du, and Wei Lü. Structure of the human lipid-gated cation channel TRPC3. *eLife*, 7, 2018.

[4182] Li Fan, Yichao Lu, Xiulian Shen, Hong Shao, Lun Suo, and Qiang Wu. Alpha protocadherins and Pyk2 kinase regulate cortical neuron migration and cytoskeletal dynamics via Rac1 GTPase and WAVE complex in mice. *eLife*, 7, 2018.

[4183] Yunshu Fan, Joshua I Gold, and Long Ding. Ongoing, rational calibration of reward-driven perceptual biases. *eLife*, 7, 2018.

[4184] Dong Fang, Haiyun Gan, Liang Cheng, Jeong-Heon Lee, Hui Zhou, Jann N Sarkaria, David J Daniels, and Zhiguo Zhang. H3.3K27M mutant proteins reprogram epigenome by sequestering the PRC2 complex to poised enhancers. *eLife*, 7, 2018.

[4185] Ziling Fang, Bo Cao, Jun-Ming Liao, Jun Deng, Kevin D Plummer, Peng Liao, Tao Liu, Wensheng Zhang, Kun Zhang, Li Li, David Margolin, Shelya X Zeng, Jianping Xiong, and Hua Lu. SPIN1 promotes tumorigenesis by blocking the uL18 (universal large ribosomal subunit protein 18)-MDM2-p53 pathway in human cancer. *eLife*, 7, 2018.

[4186] Sean W Fanning, Rinath Jeselsohn, Venkatasubramanian Dharmarajan, Christopher G Mayne, Mostafa Karimi, Gilles Buchwalter, René Houtman, Weiyi Toy, Colin E Fowler, Ross Han, Muriel Lainé, Kathryn E Carlson, Teresa A Martin, Jason Nowak, Jerome C Nwachukwu, David J Hosfield,

Sarat Chandarlapaty, Emad Tajkhorshid, Kendall W Nettles, Patrick R Griffin, Yang Shen, John A Katzenellenbogen, Myles Brown, and Geoffrey L Greene. The SERM/SERD bazedoxifene disrupts ESR1 helix 12 to overcome acquired hormone resistance in breast cancer cells. *eLife*, 7, 2018.

[4187] Farnoush Farahpour, Mohammadkarim Saeedghalati, Verena S Brauer, and Daniel Hoffmann. Trade-off shapes diversity in eco-evolutionary dynamics. *eLife*, 7, 2018.

[4188] Jamshid Faraji, Mitra Karimi, Nabiollah Soltanpour, Alireza Moharrerie, Zahra Rouhzadeh, Hamid lotfi, S Abedin Hosseini, S Yaghoob Jafari, Shabnam Roudaki, Reza Moeeini, and Gerlinde AS Metz. Oxytocin-mediated social enrichment promotes longer telomeres and novelty seeking. *eLife*, 7, 2018.

[4189] Zoltán Farkas, Dorottya Kalapis, Zoltán Bódi, Béla Szamecz, Andreea Daraba, Karola Almási, Károly Kovács, Gábor Boross, Ferenc Pál, Péter Horváth, Tamás Balassa, Csaba Molnár, Aladár Pettkó-Szandtner, éva Klement, Edit Rutkai, Attila Szvetnik, Balázs Papp, and Csaba Pál. Hsp70-associated chaperones have a critical role in buffering protein production costs. *eLife*, 7, 2018.

[4190] Ali Farshchian, Alessandra Sciutti, Assaf Pressman, Ilana Nisky, and Ferdinando A Mussa-Ivaldi. Energy exchanges at contact events guide sensorimotor integration. *eLife*, 7, 2018.

[4191] Zohreh Farsi, Sindhuja Gowrisankaran, Matija Krunic, Burkhard Rammner, Andrew Woehler, Eileen M Lafer, Carsten Mim, Reinhard Jahn, and Ira Milosevic. Clathrin coat controls synaptic vesicle acidification by blocking vacuolar ATPase activity. *eLife*, 7, 2018.

[4192] Aitak Farzi, Jackie Lau, Chi Kin Ip, Yue Qi, Yan-Chuan Shi, Lei Zhang, Ramon Tasan, Günther Sperk, and Herbert Herzog. Arcuate nucleus and lateral hypothalamic CART neurons in the mouse brain exert opposing effects on energy expenditure. *eLife*, 7, 2018.

[4193] Jean-Denis Faure and Johnathan A Napier. Europe's first and last field trial of gene-edited plants? *eLife*, 7, 2018.

[4194] Tyler B Faust, Yang Li, Curtis W Bacon, Gwendolyn M Jang, Amit Weiss, Bhargavi Jayaraman, Billy W Newton, Nevan J Krogan, Iván D'Orso, and Alan D Frankel. The HIV-1 Tat protein recruits a ubiquitin ligase to reorganize the 7SK snRNP for transcriptional activation. *eLife*, 7, 2018.

[4195] Daniel J Fazakerley, Rima Chaudhuri, Pengyi Yang, Ghassan J Maghzal, Kristen C Thomas, James R Krycer, Sean J Humphrey, Benjamin L Parker, Kelsey H Fisher-Wellman, Christopher C Meoli, Nolan J Hoffman, Ciana Diskin, James G Burchfield, Mark J Cowley, Warren Kaplan, Zora Modrusan, Ganesh Kolumam, Jean YH Yang, Daniel L Chen, Dorit Samocha-Bonet, Jerry R Greenfield, Kyle L Hoehn, Roland Stocker, and David E James. Mitochondrial CoQ deficiency is a common driver of mitochondrial oxidants and insulin resistance. *eLife*, 7, 2018.

[4196] Juan Feng, Xianchi Dong, Jennifer Pinello, Jun Zhang, Chafen Lu, Roxana E Iacob, John R Engen, William J Snell, and Timothy A Springer. Fusion surface structure, function, and dynamics of gamete fusogen HAP2. *eLife*, 7, 2018.

[4197] Suihan Feng, Takeshi Harayama, Sylvie Montessuit, Fabrice PA David, Nicolas Winssinger, Jean-Claude Martinou, and Howard Riezman. Mitochondria-specific photoactivation to monitor local sphingosine metabolism and function. *eLife*, 7, 2018.

[4198] Xi Feng, Sharon Liu, David Chen, Susanna Rosi, and Nalin Gupta. Rescue of cognitive function following fractionated brain irradiation in a novel preclinical glioma model. *eLife*, 7, 2018.

[4199] Aidan M Fenix, Abigail C Neininger, Nilay Taneja, Karren Hyde, Mike R Visetsouk, Ryan J Garde, Baohong Liu, Benjamin R Nixon, Annabelle E Manalo, Jason R Becker, Scott W Crawley, David M Bader, Matthew J Tyska, Qi Liu, Jennifer H Gutzman, and Dylan T Burnette. Muscle-specific stress fibers give rise to sarcomeres in cardiomyocytes. *eLife*, 7, 2018.

[4200] Chrysanthi Fergani, Silvia Leon, Stephanie L Padilla, Anne MJ Verstegen, Richard D Palmiter, and Victor M Navarro. NKB signaling in the posterodorsal medial amygdala stimulates gonadotropin release in a kisspeptin-independent manner in female mice. *eLife*, 7, 2018.

[4201] Anne C Ferguson-Smith and Deborah Bourc'his. The discovery and importance of genomic imprinting. *eLife*, 7, 2018.

[4202] Laura MJ Fernandez, Gil Vantomme, Alejandro Osorio-Forero, Romain Cardis, Elidie Béard, and Anita Lüthi. Thalamic reticular control of local sleep in mouse sensory cortex. *eLife*, 7, 2018.

[4203] Ryan D Ferrao, Heidi JA Wallweber, and Patrick J Lupardus. Receptor-mediated dimerization of JAK2 FERM domains is required for JAK2 activation. *eLife*, 7, 2018.

[4204] Andrea Ferrario, Robert Merrison-Hort, Stephen R Soffe, and Roman Borisyuk. Structural and functional properties of a probabilistic model of neuronal connectivity in a simple locomotor network. *eLife*, 7, 2018.

[4205] Laurent Ferron, Ivan Kadurin, and Annette C Dolphin. Proteolytic maturation of $\alpha 2\delta$ controls the probability of synaptic vesicular release. *eLife*, 7, 2018.

[4206] Tomas Fessl, Daniel Watkins, Peter Oatley, William John Allen, Robin Adam Corey, Jim Horne, Steve A Baldwin, Sheena E Radford, Ian Collinson, and Roman Tuma. Dynamic action of the Sec machinery during initiation, protein translocation and termination. *eLife*, 7, 2018.

[4207] Christopher R Fetsch, Naomi N Odean, Danique Jeurissen, Yasmine El-Shamayleh, Gregory D Horwitz, and Michael N Shadlen. Focal optogenetic suppression in macaque area MT biases direction discrimination and decision confidence, but only transiently. *eLife*, 7, 2018.

[4208] Lars-Eric Fielmich, Ruben Schmidt, Daniel J Dickinson, Bob Goldstein, Anna Akhmanova, and Sander van den Heuvel. Optogenetic dissection of mitotic spindle positioning in vivo. *eLife*, 7, 2018.

[4209] Danièle L Filiault, Evangeline S Ballerini, Terezie Mandáková, GökÇe Aköz, Nathan J Derieg, Jeremy Schmutz, Jerry Jenkins, Jane Grimwood, Shengqiang Shu, Richard D Hayes, Uffe Hellsten, Kerrie Barry, Juying Yan, Sirma Mihaltcheva, Miroslava Karafiátová, Viktoria Nizhynska, Elena M Kramer, Martin A Lysak, Scott A Hodges, and Magnus Nordborg. The Aquilegia genome provides insight into adaptive radiation and reveals an extraordinarily polymorphic chromosome with a unique history. *eLife*, 7, 2018.

[4210] Daniel J Firl, Soren E Degn, Timothy Padera, and Michael C Carroll. Capturing change in clonal composition amongst single mouse germinal centers. *eLife*, 7, 2018.

[4211] Juliane Fischer, Sebastian Y Müller, Tina Netzker, Nils Jäger, Agnieszka Gacek-Matthews, Kirstin Scherlach, Maria C Stroe, María García-Altares, Francesco Pezzini, Hanno Schoeler, Michael Reichelt, Jonathan Gershenzon, Mario KC Krespach, Ekaterina Shelest, Volker Schroeckh, Vito Valiante, Thorsten Heinzel, Christian Hertweck, Joseph Strauss, and Axel A Brakhage. Chromatin mapping identifies BasR, a key regulator of bacteria-triggered production of fungal secondary metabolites. *eLife*, 7, 2018.

[4212] Marta Florio, Michael Heide, Anneline Pinson, Holger Brandl, Mareike Albert, Sylke Winkler, Pauline Wimberger, Wieland B Huttner, and Michael Hiller. Evolution and cell-type specificity of human-specific genes preferentially expressed in progenitors of fetal neocortex. *eLife*, 7, 2018.

[4213] Esmeralda Fonseca, Victor de Lafuente, Sidney A Simon, and Ranier Gutierrez. Sucrose intensity coding and decision-making in rat gustatory cortices. *eLife*, 7, 2018.

[4214] Talitha L Forcier, Andalus Ayaz, Manraj S Gill, Daniel Jones, Rob Phillips, and Justin B Kinney. Measuring cis-regulatory energetics in living cells using allelic manifolds. *eLife*, 7, 2018.

[4215] Anthony D Fouad, Shelly Teng, Julian R Mark, Alice Liu, Pilar Alvarez-Illera, Hongfei Ji, Angelica Du, Priya D Bhirgoo, Eli Cornblath, Sihui Asuka Guan, and Christopher Fang-Yen. Distributed rhythm generators underlie Caenorhabditis elegans forward locomotion. *eLife*, 7, 2018.

[4216] Juan Jose Fraire-Zamora, Johannes Jaeger, and Jérôme Solon. Two consecutive microtubule-based epithelial seaming events mediate dorsal closure in the scuttle fly Megaselia abdita. *eLife*, 7, 2018.

[4217] Romain Franconville, Celia Beron, and Vivek Jayaraman. Building a functional connectome of the Drosophila central complex. *eLife*, 7, 2018.

[4218] Polytimi Frangou, Marta Correia, and Zoe Kourtzi. GABA, not BOLD, reveals dissociable learning-dependent plasticity mechanisms in the human brain. *eLife*, 7, 2018.

[4219] Tom P Franken, Philip X Joris, and Philip H Smith. Principal cells of the brainstem's interaural sound level detector are temporal differentiators rather than integrators. *eLife*, 7, 2018.

[4220] Meghan Whitney Franklin, Sergey Nepomnyachyi, Ryan Feehan, Nir Ben-Tal, Rachel Kolodny, and Joanna SG Slusky. Evolutionary pathways of repeat protein topology in bacterial outer membrane proteins. *eLife*, 7, 2018.

[4221] Lydia Freddolino, Jamie Yang, Amir Momen-Roknabadi, and Saeed Tavazoie. Stochastic tuning of gene expression enables cellular adaptation in the absence of pre-existing regulatory circuitry. *eLife*, 7, 2018.

[4222] Jacob W Freimer, TJ Hu, and Robert Blelloch. Decoupling the impact of microRNAs on translational repression versus RNA degradation in embryonic stem cells. *eLife*, 7, 2018.

[4223] Thomas JR Frith, Ilaria Granata, Matthew Wind, Erin Stout, Oliver Thompson, Katrin Neumann, Dylan Stavish, Paul R Heath, Daniel Ortmann, James OS Hackland, Konstantinos Anastassiadis, Mina Gouti, James Briscoe, Valerie Wilson, Stuart L Johnson, Marysia Placzek, Mario R Guarracino, Peter W Andrews, and Anestis Tsakiridis. Human axial progenitors generate trunk neural crest cells in vitro. *eLife*, 7, 2018.

[4224] Sean Froudist-Walsh, Philip GF Browning, James J Young, Kathy L Murphy, Rogier B Mars, Lazar Fleysher, and Paula L Croxson. Macro-connectomics and microstructure predict dynamic plasticity patterns in the non-human primate brain. *eLife*, 7, 2018.

[4225] Tristan Frum, Tayler M Murphy, and Amy Ralston. HIPPO signaling resolves embryonic cell fate conflicts during establishment of pluripotency in vivo. *eLife*, 7, 2018.

[4226] Qiang Fu, Jose Bienvenido Diez, Mike Pole, Manuel García ávila, Zhong-Jian Liu, Hang Chu, Yemao Hou, Pengfei Yin, Guo-Qiang Zhang, Kaihe Du, and Xin Wang. An unexpected noncarpellate epigynous flower from the Jurassic of China. *eLife*, 7, 2018.

[4227] Daniel R Fuentes, Tomek Swigut, and Joanna Wysocka. Systematic perturbation of retroviral LTRs reveals widespread long-range effects on human gene regulation. *eLife*, 7, 2018.

[4228] Hajime Fukui, Takahiro Miyazaki, Renee Wei-Yan Chow, Hiroyuki Ishikawa, Hiroyuki Nakajima, Julien Vermot, and Naoki Mochizuki. Hippo signaling determines the number of venous pole cells that originate from the anterior lateral plate mesoderm in zebrafish. *eLife*, 7, 2018.

[4229] Saori Furuta, Gang Ren, Jian-Hua Mao, and Mina J Bissell. Laminin signals initiate the reciprocal loop that informs breast-specific gene expression and homeostasis by activating NO, p53 and microRNAs. *eLife*, 7, 2018.

[4230] Michael A Gaffield, Matthew J M Rowan, Samantha B Amat, Hirokazu Hirai, and Jason M Christie. Inhibition gates supralinear Ca2+ signaling in Purkinje cell dendrites during practiced movements. *eLife*, 7, 2018.

[4231] Sandra Galic, Kim Loh, Lisa Murray-Segal, Gregory R Steinberg, Zane B Andrews, and Bruce E Kemp. AMPK signaling to acetyl-CoA carboxylase is required for fasting- and cold-induced appetite but not thermogenesis. *eLife*, 7, 2018.

[4232] Elisa Galliano, Eleonora Franzoni, Marine Breton, Annisa N Chand, Darren J Byrne, Venkatesh N Murthy, and Matthew S Grubb. Embryonic and postnatal neurogenesis produce functionally distinct subclasses of dopaminergic neuron. *eLife*, 7, 2018.

[4233] Selene Gallo, Riccardo Paracampo, Laura Müller-Pinzler, Mario Carlo Severo, Laila Blömer, Carolina Fernandes-Henriques, Anna Henschel, Balint Kalista Lammes, Tatjana Maskaljunas, Judith Suttrup, Alessio Avenanti, Christian Keysers, and Valeria Gazzola. The causal role of the somatosensory cortex in prosocial behaviour. *eLife*, 7, 2018.

[4234] Nathan Gamarra, Stephanie L Johnson, Michael J Trnka, Alma L Burlingame, and Geeta J Narlikar. The nucleosomal acidic patch relieves auto-inhibition by the ISWI remodeler SNF2h. *eLife*, 7, 2018.

[4235] Patrick D Ganzer, Michael J Darrow, Eric C Meyers, Bleyda R Solorzano, Andrea D Ruiz, Nicole M Robertson, Katherine S Adcock, Justin T James, Han S Jeong, April M Becker, Mark P Goldberg, David T Pruitt, Seth A Hays, Michael P Kilgard, and Robert L Rennaker. Closed-loop neuromodulation restores network connectivity and motor control after spinal cord injury. *eLife*, 7, 2018.

[4236] Jieqiong Gao, Lars Langemeyer, Daniel Kümmel, Fulvio Reggiori, and Christian Ungermann. Molecular mechanism to target the endosomal Mon1-Ccz1 GEF complex to the pre-autophagosomal structure. *eLife*, 7, 2018.

[4237] Shangbang Gao, Sihui Asuka Guan, Anthony D Fouad, Jun Meng, Taizo Kawano, Yung-Chi Huang, Yi Li, Salvador Alcaire, Wesley Hung, Yangning Lu, Yingchuan Billy Qi, Yishi Jin, Mark Alkema, Christopher Fang-Yen, and Mei Zhen. Excitatory motor neurons are local oscillators for backward locomotion. *eLife*, 7, 2018.

[4238] Maria Daniela Garcia-Castillo, Daniel J-F Chinnapen, Yvonne M te Welscher, Rodrigo J Gonzalez, Samir Softic, Michele Pacheco, Randall J Mrsny, C Ronald Kahn, Ulrich H von Andrian, Jesper Lau, Bradley L Pentelute, and Wayne I Lencer. Mucosal absorption of therapeutic peptides by harnessing the endogenous sorting of glycosphingolipids. *eLife*, 7, 2018.

[4239] Alam García-Heredia, Amol Arunrao Pohane, Emily S Melzer, Caleb R Carr, Taylor J Fiolek, Sarah R Rundell, Hoong Chuin Lim, Jeffrey C Wagner, Yasu S Morita, Benjamin M Swarts, and M Sloan Siegrist. Peptidoglycan precursor synthesis along the sidewall of pole-growing mycobacteria. *eLife*, 7, 2018.

[4240] Javier García-Nafría, Yang Lee, Xiaochen Bai, Byron Carpenter, and Christopher G Tate. Cryo-EM structure of the adenosine A2A receptor coupled to an engineered heterotrimeric G protein. *eLife*, 7, 2018.

[4241] Matthew PH Gardner, Jessica C Conroy, Clay V Styer, Timothy Huynh, Leslie R Whitaker, and Geoffrey Schoenbaum. Medial orbitofrontal inactivation does not affect economic choice. *eLife*, 7, 2018.

[4242] Cecilia Garmendia-Torres, Olivier Tassy, Audrey Matifas, Nacho Molina, and Gilles Charvin. Multiple inputs ensure yeast cell size homeostasis during cell cycle progression. *eLife*, 7, 2018.

[4243] Jingpeng Ge, Johannes Elferich, April Goehring, Huaying Zhao, Peter Schuck, and Eric Gouaux. Structure of mouse protocadherin 15 of the stereocilia tip link in complex with LHFPL5. *eLife*, 7, 2018.

[4244] Lauren Geary and Carole LaBonne. FGF mediated MAPK and PI3K/Akt Signals make distinct contributions to pluripotency and the establishment of Neural Crest. *eLife*, 7, 2018.

[4245] Ron S Gejman, Aaron Y Chang, Heather F Jones, Krysta DiKun, Abraham Ari Hakimi, Andrea Schietinger, and David A Scheinberg. Rejection of immunogenic tumor clones is limited by clonal fraction. *eLife*, 7, 2018.

[4246] Jie Geng, John D Altman, Sujatha Krishnakumar, and Malini Raghavan. Empty conformers of HLA-B preferentially bind CD8 and regulate CD8+ T cell function. *eLife*, 7, 2018.

[4247] Ruben Gepner, Jason Wolk, Digvijay Shivaji Wadekar, Sophie Dvali, and Marc Gershow. Variance adaptation in navigational decision making. *eLife*, 7, 2018.

[4248] Piotr Gerlach, Jan M Schuller, Fabien Bonneau, Jérôme Basquin, Peter Reichelt, Sebastian Falk, and Elena Conti. Distinct and evolutionary conserved structural features of the human nuclear exosome complex. *eLife*, 7, 2018.

[4249] Federico Germani, Daniel Hain, Denise Sternlicht, Eduardo Moreno, and Konrad Basler. The Toll pathway inhibits tissue growth and regulates cell fitness in an infection-dependent manner. *eLife*, 7, 2018.

[4250] Elisabeth A Geyer, Matthew P Miller, Chad A Brautigam, Sue Biggins, and Luke M Rice. Design principles of a microtubule polymerase. *eLife*, 7, 2018.

[4251] Sabina Gherman and Marios G. Philiastides. Human VMPFC encodes early signatures of confidence in perceptual decisions. *eLife*, 7, 2018.

[4252] Shrestha Ghosh, Sheung Kin Wong, Zhixin Jiang, Baohua Liu, Yi Wang, Quan Hao, Vera Gorbunova, Xinguang Liu, and Zhongjun Zhou. Haploinsufficiency of Trp53 dramatically extends the lifespan of Sirt6-deficient mice. *eLife*, 7, 2018.

[4253] Stefanie Giera, Rong Luo, Yanqin Ying, Sarah D Ackerman, Sung-Jin Jeong, Hannah M Stoveken, Christopher J Folts, Christina A Welsh, Gregory G Tall, Beth Stevens, Kelly R Monk, and Xianhua Piao. Microglial transglutaminase-2 drives myelination and myelin repair via GPR56/ADGRG1 in oligodendrocyte precursor cells. *eLife*, 7, 2018.

[4254] Sarah Gilbertson, Joel D Federspiel, Ella Hartenian, Ileana M Cristea, and Britt Glaunsinger. Changes in mRNA abundance drive shuttling of RNA binding proteins, linking cytoplasmic RNA degradation to transcription. *eLife*, 7, 2018.

[4255] Miriam Bracha Ginzberg, Nancy Chang, Heather D'Souza, Nish Patel, Ran Kafri, and Marc W Kirschner. Cell size sensing in animal cells coordinates anabolic growth rates and cell cycle progression to maintain cell size uniformity. *eLife*, 7, 2018.

[4256] Nomeda Girnius, Yvonne JK Edwards, David S Garlick, and Roger J Davis. The cJUN NH2-terminal kinase (JNK) signaling pathway promotes genome stability and prevents tumor initiation. *eLife*, 7, 2018.

[4257] Christopher J Giuliano, Ann Lin, Joan C Smith, Ann C Palladino, and Jason M Sheltzer. MELK expression correlates with tumor mitotic activity but is not required for cancer growth. *eLife*, 7, 2018.

[4258] Adrian G Glover, Helena Wiklund, Chong Chen, and Thomas G Dahlgren. Managing a sustainable deep-sea 'blue economy' requires knowledge of what actually lives there. *eLife*, 7, 2018.

[4259] Sebastian Gluth, Mikhail S Spektor, and Jörg Rieskamp. Value-based attentional capture affects multi-alternative decision making. *eLife*, 7, 2018.

[4260] Jimmie M Gmaz, James E Carmichael, and Matthijs AA van der Meer. Persistent coding of outcome-predictive cue features in the rat nucleus accumbens. *eLife*, 7, 2018.

[4261] Sezgi Goksan, Luke Baxter, Fiona Moultrie, Eugene Duff, Gareth Hathway, Caroline Hartley, Irene Tracey, and Rebeccah Slater. The influence of the descending pain modulatory system on infant pain-related brain activity. *eLife*, 7, 2018.

[4262] Raymond E Goldstein. Are theoretical results 'Results'? *eLife*, 7, 2018.

[4263] Pieter M Goltstein, Guido T Meijer, and Cyriel MA Pennartz. Conditioning sharpens the spatial representation of rewarded stimuli in mouse primary visual cortex. *eLife*, 7, 2018.

[4264] Carla GonÇalves, Jennifer H Wisecaver, Jacek Kominek, Madalena Salema Oom, Maria José Leandro, Xing-Xing Shen, Dana A Opulente, Xiaofan Zhou, David Peris, Cletus P Kurtzman, Chris Todd Hittinger, Antonis Rokas, and Paula GonÇalves. Evidence for loss and reacquisition of alcoholic fermentation in a fructophilic yeast lineage. *eLife*, 7, 2018.

[4265] Estibaliz González-Fernández, Hey-Kyeong Jeong, Masahiro Fukaya, Hyukmin Kim, Rabia R Khawaja, Isha N Srivastava, Ari Waisman, Young-Jin Son, and Shin H Kang. PTEN negatively regulates the cell lineage progression from NG2+ glial progenitor to oligodendrocyte via mTOR-independent signaling. *eLife*, 7, 2018.

[4266] Carlos González-García, Matthew W Flounders, Raymond Chang, Alexis T Baria, and Biyu J He. Content-specific activity in frontoparietal and default-mode networks during prior-guided visual perception. *eLife*, 7, 2018.

[4267] Oliver Gordon, Conor M Henry, Naren Srinivasan, Susan Ahrens, Anna Franz, Safia Deddouche, Probir Chakravarty, David Phillips, Roger George, Svend Kjaer, David Frith, Ambrosius P Snijders, Rita S Valente, Carolina J Simoes da Silva, Luis Teixeira, Barry Thompson, Marc S Dionne, Will Wood, and Caetano Reis e Sousa. $\alpha$-actinin accounts for the bioactivity of actin preparations in inducing STAT target genes in Drosophila melanogaster. *eLife*, 7, 2018.

[4268] Sharona E Gordon, Mika Munari, and William N Zagotta. Visualizing conformational dynamics of proteins in solution and at the cell membrane. *eLife*, 7, 2018.

[4269] Natalia A Goriounova, Djai B Heyer, René Wilbers, Matthijs B Verhoog, Michele Giugliano, Christophe Verbist, Joshua Obermayer, Amber Kerkhofs, Harriët Smeding, Maaike Verberne, Sander Idema, Johannes C Baayen, Anton W Pieneman, Christiaan PJ de Kock, Martin Klein, and Huibert D Mansvelder. Large and fast human pyramidal neurons associate with intelligence. *eLife*, 7, 2018.

[4270] So Goto, Akishi Onishi, Kazuyo Misaki, Shigenobu Yonemura, Sunao Sugita, Hiromi Ito, Yoko Ohigashi, Masatsugu Ema, Hirokazu Sakaguchi, Kohji Nishida, and Masayo Takahashi. Neural retina-specific Aldh1a1 controls dorsal choroidal vascular development via Sox9 expression in retinal pigment epithelial cells. *eLife*, 7, 2018.

[4271] Vishwa Goudar and Dean V Buonomano. Encoding sensory and motor patterns as time-invariant trajectories in recurrent neural networks. *eLife*, 7, 2018.

[4272] Elizabeth A Gould, Nicolas Busquet, Douglas Shepherd, Robert M Dietz, Paco S Herson, Fabio M Simoes de Souza, Anan Li, Nicholas M George, Diego Restrepo, and Wendy B Macklin. Mild myelin disruption elicits early alteration in behavior and proliferation in the subventricular zone. *eLife*, 7, 2018.

[4273] Peter D Gould, Mirela Domijan, Mark Greenwood, Isao T Tokuda, Hannah Rees, Laszlo Kozma-Bognar, Anthony JW Hall, and James CW Locke. Coordination of robust single cell rhythms in the Arabidopsis circadian clock via spatial waves of gene expression. *eLife*, 7, 2018.

[4274] Geoffrey Gourinchas, Udo Heintz, and Andreas Winkler. Asymmetric activation mechanism of a homodimeric red light-regulated photoreceptor. *eLife*, 7, 2018.

[4275] Arvina Grahl, Selim Onat, and Christian Büchel. The periaqueductal gray and Bayesian integration in placebo analgesia. *eLife*, 7, 2018.

[4276] Timothy Grant, Alexis Rohou, and Nikolaus Grigorieff. cisTEM, user-friendly software for single-particle image processing. *eLife*, 7, 2018.

[4277] Hannah J Green, Annabel GM Griffiths, Jari Ylänne, and Nicholas H Brown. Novel functions for integrin-associated proteins revealed by analysis of myofibril attachment in Drosophila. *eLife*, 7, 2018.

[4278] Garrett A Greenan, Bettina Keszthelyi, Ronald D Vale, and David A Agard. Insights into centriole geometry revealed by cryotomography of doublet and triplet centrioles. *eLife*, 7, 2018.

[4279] R A Greenstein, Stephen K Jones, Eric C Spivey, James R Rybarski, Ilya J Finkelstein, and Bassem Al-Sady. Noncoding RNA-nucleated heterochromatin spreading is intrinsically labile and requires accessory elements for epigenetic stability. *eLife*, 7, 2018.

[4280] Efrat Esther Greenwald, Lior Baltiansky, and Ofer Feinerman. Individual crop loads provide local control for collective food intake in ant colonies. *eLife*, 7, 2018.

[4281] Sona Gregorova, Vaclav Gergelits, Irena Chvatalova, Tanmoy Bhattacharyya, Barbora Valiskova, Vladana Fotopulosova, Petr Jansa, Diana Wiatrowska, and Jiri Forejt. Modulation of Prdm9-controlled meiotic chromosome asynapsis overrides hybrid sterility in mice. *eLife*, 7, 2018.

[4282] Tineke Grent-'t Jong, Joachim Gross, Jozien Goense, Michael Wibral, Ruchika Gajwani, Andrew I Gumley, Stephen M Lawrie, Matthias Schwannauer, Frauke Schultze-Lutter, Tobias Navarro Schröder, Dagmar Koethe, F Markus Leweke, Wolf Singer, and Peter J Uhlhaas. Resting-state gamma-band power alterations in schizophrenia reveal E/I-balance abnormalities across illness-stages. *eLife*, 7, 2018.

[4283] William N Grimes, Jacob Baudin, Anthony W Azevedo, and Fred Rieke. Range, routing and kinetics of rod signaling in primate retina. *eLife*, 7, 2018.

[4284] Iris IA Groen, Michelle R Greene, Christopher Baldassano, Li Fei-Fei, Diane M Beck, and Chris I Baker. Distinct contributions of functional and deep neural network features to representational similarity of scenes in human brain and behavior. *eLife*, 7, 2018.

[4285] Fenghui Guan, Jiayu Yu, Jie Yu, Yang Liu, Ying Li, Xin-Hua Feng, Kerwyn Casey Huang, Zengyi Chang, and Sheng Ye. Lateral interactions between protofilaments of the bacterial tubulin homolog FtsZ are essential for cell division. *eLife*, 7, 2018.

[4286] Ye Guan, Zhengda Li, Shiyuan Wang, Patrick M Barnes, Xuwen Liu, Haotian Xu, Minjun Jin, Allen P Liu, and Qiong Yang. A robust and tunable mitotic oscillator in artificial cells. *eLife*, 7, 2018.

[4287] Wamdaogo M Guelbéogo, Bronner Pamplona GonÇalves, Lynn Grignard, John Bradley, Samuel S Serme, Joel Hellewell, Kjerstin Lanke, Soumanaba Zongo, Nuno Sepúlveda, Issiaka Soulama, Dimitri W Wangrawa, Laith Yakob, N'Falé Sagnon, Teun Bousema, and Chris Drakeley. Variation in natural exposure to anopheles mosquitoes and its effects on malaria transmission. *eLife*, 7, 2018.

[4288] Xavier Guell, Jeremy D Schmahmann, John DE Gabrieli, and Satrajit S Ghosh. Functional gradients of the cerebellum. *eLife*, 7, 2018.

[4289] Antoine Guérin, Gaspard Kerner, Nico Marr, Janet G Markle, Florence Fenollar, Natalie Wong, Sabri Boughorbel, Danielle T Avery, Cindy S Ma, Salim Bougarn, Matthieu Bouaziz, Vivien Béziat, Erika Della Mina, Carmen Oleaga-Quintas, Tomi Lazarov, Lisa Worley, Tina Nguyen, Etienne Patin, Caroline Deswarte, Rubén Martinez-Barricarte, Soraya Boucherit, Xavier Ayral, Sophie Edouard, Stéphanie Boisson-Dupuis, Vimel Rattina, Benedetta Bigio, Guillaume Vogt, Frédéric Geissmann, Lluis Quintana-Murci, Damien Chaussabel, Stuart G Tangye, Didier Raoult, Laurent Abel, Jacinta Bustamante, and Jean-Laurent Casanova. IRF4 haploinsufficiency in a family with Whipple's disease. *eLife*, 7, 2018.

[4290] Almary Guerra, Raoul FV Germano, Oliver Stone, Rima Arnaout, Stefan Guenther, Suchit Ahuja, Verónica Uribe, Benoit Vanhollebeke, Didier YR Stainier, and Sven Reischauer. Distinct myocardial lineages break atrial symmetry during cardiogenesis in zebrafish. *eLife*, 7, 2018.

[4291] Ricardo Guerrero-Ferreira, Nicholas MI Taylor, Daniel Mona, Philippe Ringler, Matthias E Lauer, Roland Riek, Markus Britschgi, and Henning Stahlberg. Cryo-EM structure of alpha-synuclein fibrils. *eLife*, 7, 2018.

[4292] Judith Gunzelmann, Diana Rüthnick, Tien-chen Lin, Wanlu Zhang, Annett Neuner, Ursula Jäkle, and Elmar Schiebel. The microtubule polymerase Stu2 promotes oligomerization of the $\gamma$-TuSC for cytoplasmic microtubule nucleation. *eLife*, 7, 2018.

[4293] Neha Gupta, Jon R Lorsch, and Alan G Hinnebusch. Yeast Ded1 promotes 48S translation pre-initiation complex assembly in an mRNA-specific and eIF4F-dependent manner. *eLife*, 7, 2018.

[4294] Jose Arturo Gutierrez-Triana, Tinatini Tavhelidse, Thomas Thumberger, Isabelle Thomas, Beate Wittbrodt, Tanja Kellner, Kerim Anlas, Erika Tsingos, and Joachim Wittbrodt. Efficient single-copy HDR by 5′ modified long dsDNA donors. *eLife*, 7, 2018.

[4295] Ngoc T Ha and Kimberly J Dougherty. Spinal Shox2 interneuron interconnectivity related to function and development. *eLife*, 7, 2018.

[4296] Kalina T Haas, Benjamin Compans, Mathieu Letellier, Thomas M Bartol, Dolors Grillo-Bosch, Terrence J Sejnowski, Matthieu Sainlos, Daniel Choquet, Olivier Thoumine, and Eric Hosy. Pre-post synaptic alignment through neuroligin-1 tunes synaptic transmission efficiency. *eLife*, 7, 2018.

[4297] Jodi A Hadden, Juan R Perilla, Christopher John Schlicksup, Balasubramanian Venkatakrishnan, Adam Zlotnick, and Klaus Schulten. All-atom molecular dynamics of the HBV capsid reveals insights into biological function and cryo-EM resolution limits. *eLife*, 7, 2018.

[4298] Hugh K Haddox, Adam S Dingens, Sarah K Hilton, Julie Overbaugh, and Jesse D Bloom. Mapping mutational effects along the evolutionary landscape of HIV envelope. *eLife*, 7, 2018.

[4299] Tatsuya Haga and Tomoki Fukai. Recurrent network model for learning goal-directed sequences through reverse replay. *eLife*, 7, 2018.

[4300] Richard Hakim, Kiarash Shamardani, and Hillel Adesnik. A neural circuit for gamma-band coherence across the retinotopic map in mouse visual cortex. *eLife*, 7, 2018.

[4301] Jessica A Haley, David Hampton, and Eve Marder. Two central pattern generators from the crab, Cancer borealis, respond robustly and differentially to extreme extracellular pH. *eLife*, 7, 2018.

[4302] Zachary Jonas Hall and Vincent Tropepe. Movement maintains forebrain neurogenesis via peripheral neural feedback in larval zebrafish. *eLife*, 7, 2018.

[4303] Arda Halu, Jian-Guo Wang, Hiroshi Iwata, Alexander Mojcher, Ana Luisa Abib, Sasha A Singh, Masanori Aikawa, and Amitabh Sharma. Context-enriched interactome powered by proteomics helps the identification of novel regulators of macrophage activation. *eLife*, 7, 2018.

[4304] Mayuko Hamada, Katja Schröder, Jay Bathia, Ulrich Kürn, Sebastian Fraune, Mariia Khalturina, Konstantin Khalturin, Chuya Shinzato, Nori Satoh, and Thomas CG Bosch. Metabolic co-dependence drives the evolutionarily ancient Hydra–Chlorella symbiosis. *eLife*, 7, 2018.

[4305] Shuting Han, Ekaterina Taralova, Christophe Dupre, and Rafael Yuste. Comprehensive machine learning analysis of Hydra behavior reveals a stable basal behavioral repertoire. *eLife*, 7, 2018.

[4306] Yang Han, Monika Eipel, Julia Franzen, Vadim Sakk, Bertien Dethmers-Ausema, Laura Yndriago, Ander Izeta, Gerald de Haan, Hartmut Geiger, and Wolfgang Wagner. Epigenetic age-predictor for mice based on three CpG sites. *eLife*, 7, 2018.

[4307] Matthew Hancock, Anne D Hafstad, Adam A Nabeebaccus, Norman Catibog, Angela Logan, Ioannis Smyrnias, Synne S Hansen, Johanna Lanner, Katrin Schröder, Michael P Murphy, Ajay M Shah, and Min Zhang. Myocardial NADPH oxidase-4 regulates the physiological response to acute exercise. *eLife*, 7, 2018.

[4308] Matthew Robert Hannaford, Anne Ramat, Nicolas Loyer, and Jens Januschke. aPKC-mediated displacement and actomyosin-mediated retention polarize Miranda in Drosophila neuroblasts. *eLife*, 7, 2018.

[4309] Anders S Hansen, Maxime Woringer, Jonathan B Grimm, Luke D Lavis, Robert Tjian, and Xavier Darzacq. Robust model-based analysis of single-particle tracking experiments with Spot-On. *eLife*, 7, 2018.

[4310] Yingsong Hao, Wenxing Yang, Jing Ren, Qi Hall, Yun Zhang, and Joshua M Kaplan. Thioredoxin shapes the C. elegans sensory response to Pseudomonas produced nitric oxide. *eLife*, 7, 2018.

[4311] Masatoshi Hara, Sebastian Lourido, Boryana Petrova, Hua Jane Lou, Jessica R Von Stetina, Helena Kashevsky, Benjamin E Turk, and Terry L Orr-Weaver. Identification of PNG kinase substrates uncovers interactions with the translational repressor TRAL in the oocyte-to-embryo transition. *eLife*, 7, 2018.

[4312] Osama F Harraz, Thomas A Longden, David Hill-Eubanks, and Mark T Nelson. PIP2 depletion promotes TRPV4 channel activity in mouse brain capillary endothelial cells. *eLife*, 7, 2018.

[4313] Ian F Harrison, Bernard Siow, Aisha B Akilo, Phoebe G Evans, Ozama Ismail, Yolanda Ohene, Payam Nahavandi, David L Thomas, Mark F Lythgoe, and Jack A Wells. Non-invasive imaging of CSF-mediated brain clearance pathways via assessment of perivascular fluid movement with diffusion tensor MRI. *eLife*, 7, 2018.

[4314] Andrea L Hartzell, Kelly M Martyniuk, G Stefano Brigidi, Daniel A Heinz, Nathalie A Djaja, Anja Payne, and Brenda L Bloodgood. NPAS4 recruits CCK basket cell synapses and enhances cannabinoid-sensitive inhibition in the mouse hippocampus. *eLife*, 7, 2018.

[4315] Christoph A Haselwandter and Roderick MacKinnon. Piezo's membrane footprint and its contribution to mechanosensitivity. *eLife*, 7, 2018.

[4316] Christopher K Hauser, Dantong Zhu, Terrence R Stanford, and Emilio Salinas. Motor selection dynamics in FEF explain the reaction time variance of saccades to single targets. *eLife*, 7, 2018.

[4317] Anna G Hauswirth, Kevin J Ford, Tingting Wang, Richard D Fetter, Amy Tong, and Graeme W Davis. A postsynaptic PI3K-cII dependent signaling controller for presynaptic homeostatic plasticity. *eLife*, 7, 2018.

[4318] Michael H Hayes, Elizabeth H Peuchen, Norman J Dovichi, and Daniel L Weeks. Dual roles for ATP in the regulation of phase separated protein aggregates in Xenopus oocyte nucleoli. *eLife*, 7, 2018.

[4319] Joel Haywood, Jason W Schmidberger, Amy M James, Samuel G Nonis, Kirill V Sukhoverkov, Mikael Elias, Charles S Bond, and Joshua S Mylne. Structural basis of ribosomal peptide macrocyclization in plants. *eLife*, 7, 2018.

[4320] Feng He, Alper Celik, Chan Wu, and Allan Jacobson. General decapping activators target different subsets of inefficiently translated mRNAs. *eLife*, 7, 2018.

[4321] Huan He, Marie-Pierre Brenier-Pinchart, Laurence Braun, Alexandra Kraut, Bastien Touquet, Yohann Couté, Isabelle Tardieux, Mohamed-Ali Hakimi, and Alexandre Bougdour. Characterization of a Toxoplasma effector uncovers an alternative GSK3/$\beta$-catenin-regulatory pathway of inflammation. *eLife*, 7, 2018.

[4322] Sicong He, Jiahao Chen, Yunyun Jiang, Yi Wu, Lu Zhu, Wan Jin, Changlong Zhao, Tao Yu, Tienan Wang, Shuting Wu, Xi Lin, Jianan Y Qu, Zilong Wen, Wenqing Zhang, and Jin Xu. Adult zebrafish Langerhans cells arise from hematopoietic stem/progenitor cells. *eLife*, 7, 2018.

[4323] Wenxuan He, David Kemp, and Tianying Ren. Timing of the reticular lamina and basilar membrane vibration in living gerbil cochleae. *eLife*, 7, 2018.

[4324] Michael D Healy, Manuela K Hospenthal, Ryan J Hall, Mintu Chandra, Molly Chilton, Vikas Tillu, Kai-En Chen, Dion J Celligoi, Fiona J McDonald, Peter J Cullen, J Shaun Lott, Brett M Collins, and Rajesh Ghai. Structural insights into the architecture and membrane interactions of the conserved COMMD proteins. *eLife*, 7, 2018.

[4325] Martin N Hebart, Brett B Bankson, Assaf Harel, Chris I Baker, and Radoslaw M Cichy. The representational dynamics of task and object processing in humans. *eLife*, 7, 2018.

[4326] Florian Heigwer, Christian Scheeder, Thilo Miersch, Barbara Schmitt, Claudia Blass, Mischan Vali Pour Jamnani, and Michael Boutros. Time-resolved mapping of genetic interactions to model rewiring of signaling pathways. *eLife*, 7, 2018.

[4327] Gibran Hemani, Jie Zheng, Benjamin Elsworth, Kaitlin H Wade, Valeriia Haberland, Denis Baird, Charles Laurin, Stephen Burgess, Jack Bowden, Ryan Langdon, Vanessa Y Tan, James Yarmolinsky, Hashem A Shihab, Nicholas J Timpson, David M Evans, Caroline Relton, Richard M Martin, George Davey Smith, Tom R Gaunt, and Philip C Haycock. The MR-Base platform supports systematic causal inference across the human phenome. *eLife*, 7, 2018.

[4328] David G Hendrickson, Ilya Soifer, Bernd J Wranik, Griffin Kim, Michael Robles, Patrick A Gibney, and R Scott McIsaac. A new experimental platform facilitates assessment of the transcriptional and chromatin landscapes of aging yeast. *eLife*, 7, 2018.

[4329] Jay A Hennig, Matthew D Golub, Peter J Lund, Patrick T Sadtler, Emily R Oby, Kristin M Quick, Stephen I Ryu, Elizabeth C Tyler-Kabara, Aaron P Batista, Byron M Yu, and Steven M Chase. Constraints on neural redundancy. *eLife*, 7, 2018.

[4330] Chelsea Hepler, Bo Shan, Qianbin Zhang, Gervaise H Henry, Mengle Shao, Lavanya Vishvanath, Alexandra L Ghaben, Angela B Mobley, Douglas Strand, Gary C Hon, and Rana K Gupta. Identification of functionally distinct fibro-inflammatory and adipogenic stromal subpopulations in visceral adipose tissue of adult mice. *eLife*, 7, 2018.

[4331] Dante J Heredia, Cheng-Yuan Feng, Grant W Hennig, Robert B Renden, and Thomas W Gould. Activity-induced Ca2+ signaling in perisynaptic Schwann cells of the early postnatal mouse is mediated by P2Y1 receptors and regulates muscle fatigue. *eLife*, 7, 2018.

[4332] Salvador C Herrera and Erika A Bach. JNK signaling triggers spermatogonial dedifferentiation during chronic stress to maintain the germline stem cell pool in the Drosophila testis. *eLife*, 7, 2018.

[4333] Haley Hieronymus, Rajmohan Murali, Amy Tin, Kamlesh Yadav, Wassim Abida, Henrik Moller, Daniel Berney, Howard Scher, Brett Carver, Peter Scardino, Nikolaus Schultz, Barry Taylor, Andrew Vickers, Jack Cuzick, and Charles L Sawyers. Tumor copy number alteration burden is a pan-cancer prognostic factor associated with recurrence and death. *eLife*, 7, 2018.

[4334] Zoë A Hilbert and Dennis H Kim. PDF-1 neuropeptide signaling regulates sexually dimorphic gene expression in shared sensory neurons of C. elegans. *eLife*, 7, 2018.

[4335] Eric M Hill and Christian P Petersen. Positional information specifies the site of organ regeneration and not tissue maintenance in planarians. *eLife*, 7, 2018.

[4336] Rose Z Hill, Benjamin U Hoffman, Takeshi Morita, Stephanie M Campos, Ellen A Lumpkin, Rachel B Brem, and Diana M Bautista. The signaling lipid sphingosine 1-phosphate regulates mechanical pain. *eLife*, 7, 2018.

[4337] Daniel S Himmelstein, Ariel Rodriguez Romero, Jacob G Levernier, Thomas Anthony Munro, Stephen Reid McLaughlin, Bastian Greshake Tzovaras, and Casey S Greene. Sci-Hub provides access to nearly all scholarly literature. *eLife*, 7, 2018.

[4338] Clémence Hocquet, Xavier Robellet, Laurent Modolo, Xi-Ming Sun, Claire Burny, Sara Cuylen-Haering, Esther Toselli, Sandra Clauder-Münster, Lars Steinmetz, Christian H Haering, Samuel Marguerat, and Pascal Bernard. Condensin controls cellular RNA levels through the accurate segregation of chromosomes instead of directly regulating transcription. *eLife*, 7, 2018.

[4339] Karl P Hodel, Richard de Borja, Erin E Henninger, Brittany B Campbell, Nathan Ungerleider, Nicholas Light, Tong Wu, Kimberly G LeCompte, A Yasemin Goksenin, Bruce A Bunnell, Uri Tabori, Adam Shlien, and Zachary F Pursell. Explosive mutation accumulation triggered by heterozygous human Pol $\epsilon$ proofreading-deficiency is driven by suppression of mismatch repair. *eLife*, 7, 2018.

[4340] Jackson A Hoffman, Kevin W Trotter, James M Ward, and Trevor K Archer. BRG1 governs glucocorticoid receptor interactions with chromatin and pioneer factors across the genome. *eLife*, 7, 2018.

[4341] Paul Hoffman, Ekaterina Loginova, and Asatta Russell. Poor coherence in older people's speech is explained by impaired semantic and executive processes. *eLife*, 7, 2018.

[4342] Lukas Hofmann, Dorothea Hose, Anne Grießhammer, Robert Blum, Frank Döring, Sulayman Dib-Hajj, Stephen Waxman, Claudia Sommer, Erhard Wischmeyer, and Nurcan üÇeyler. Characterization of small fiber pathology in a mouse model of Fabry disease. *eLife*, 7, 2018.

[4343] Andrew N Holding, Amy E Cullen, and Florian Markowetz. Genome-wide Estrogen Receptor-$\alpha$ activation is sustained, not cyclical. *eLife*, 7, 2018.

[4344] Devon Hominick, Asitha Silva, Noor Khurana, Ying Liu, Paul C Dechow, Jian Q Feng, Bronislaw Pytowski, Joseph M Rutkowski, Kari Alitalo, and Michael T Dellinger. VEGF-C promotes the development of lymphatics in bone and bone loss. *eLife*, 7, 2018.

[4345] Noriko Homma, Ruyun Zhou, Muhammad Imran Naseer, Adeel G Chaudhary, Mohammed H Al-Qahtani, and Nobutaka Hirokawa. KIF2A regulates the development of dentate granule cells and postnatal hippocampal wiring. *eLife*, 7, 2018.

[4346] Jungeui Hong, Nathan Brandt, Farah Abdul-Rahman, Ally Yang, Tim Hughes, and David Gresham. An incoherent feedforward loop facilitates adaptive tuning of gene expression. *eLife*, 7, 2018.

[4347] Xuehui Hong, He Huang, Xingfeng Qiu, Zhijie Ding, Xing Feng, Yuekun Zhu, Huiqin Zhuo, Jingjing Hou, Jiabao Zhao, Wangyu Cai, Ruihua Sha, Xinya Hong, Yongxiang Li, Hongjiang Song, and Zhiyong Zhang. RETRACTED: Targeting posttranslational modifications of RIOK1 inhibits the progression of colorectal and gastric cancers. *eLife*, 7, 2018.

[4348] Suvi Honkanen, Anna Thamm, Mario A Arteaga-Vazquez, and Liam Dolan. Negative regulation of conserved RSL class I bHLH transcription factors evolved independently among land plants. *eLife*, 7, 2018.

[4349] Richard M Hooy and Jungsan Sohn. The allosteric activation of cGAS underpins its dynamic signaling landscape. *eLife*, 7, 2018.

[4350] C Matthew Hope, Jemma L Webber, Sherzod A Tokamov, and Ilaria Rebay. Tuned polymerization of the transcription factor Yan limits off-DNA sequestration to confer context-specific repression. *eLife*, 7, 2018.

[4351] Jane Anne Horne, Carlie Langille, Sari McLin, Meagan Wiederman, Zhiyuan Lu, C Shan Xu, Stephen M Plaza, Louis K Scheffer, Harald F Hess, and Ian A Meinertzhagen. A resource for the Drosophila antennal lobe provided by the connectome of glomerulus VA1v. *eLife*, 7, 2018.

[4352] Xiaowei Hou, Shana R Burstein, and Stephen Barstow Long. Structures reveal opening of the store-operated calcium channel Orai. *eLife*, 7, 2018.

[4353] Peter Hraber, Bette Korber, Kshitij Wagh, David Montefiori, and Mario Roederer. A single, continuous metric to define tiered serum neutralization potency against HIV. *eLife*, 7, 2018.

[4354] Joel Hrit, Leeanne Goodrich, Cheng Li, Bang-An Wang, Ji Nie, Xiaolong Cui, Elizabeth Allene Martin, Eric Simental, Jenna Fernandez, Monica Yun Liu, Joseph R Nery, Rosa Castanon, Rahul M Kohli, Natalia Tretyakova, Chuan He, Joseph R Ecker, Mary Goll, and Barbara Panning. OGT binds a conserved C-terminal domain of TET1 to regulate TET1 activity and function in development. *eLife*, 7, 2018.

[4355] FoSheng Hsu, Stephanie Spannl, Charles Ferguson, Anthony A Hyman, Robert G Parton, and Marino Zerial. Rab5 and Alsin regulate stress-activated cytoprotective signaling on mitochondria. *eLife*, 7, 2018.

[4356] Marian Y Hu, Jia-Jiun Yan, Inga Petersen, Nina Himmerkus, Markus Bleich, and Meike Stumpp. A SLC4 family bicarbonate transporter is critical for intracellular pH regulation and biomineralization in sea urchin embryos. *eLife*, 7, 2018.

[4357] Zheng Hu, Bo Xia, Spike DL Postnikoff, Zih-Jie Shen, Alin S Tomoiaga, Troy A Harkness, Ja Hwan Seol, Wei Li, Kaifu Chen, and Jessica K Tyler. Ssd1 and Gcn2 suppress global translation efficiency in replicatively aged yeast while their activation extends lifespan. *eLife*, 7, 2018.

[4358] Chengjie G Huang, Michael G Metzen, and Maurice J Chacron. Feedback optimizes neural coding and perception of natural stimuli. *eLife*, 7, 2018.

[4359] Qiaoli Huang, Jianrong Jia, Qiming Han, and Huan Luo. Fast-backward replay of sequentially memorized items in humans. *eLife*, 7, 2018.

[4360] Xu Huang, Qian Zhang, Yupei Jiang, Chuanwei Yang, Qianyue Wang, and Lin Li. Shade-induced nuclear localization of PIF7 is regulated by phosphorylation and 14-3-3 proteins in Arabidopsis. *eLife*, 7, 2018.

[4361] Yan Huang, Sheng Huang, Coralie Di Scala, Qifu Wang, Hans H Wandall, Jacques Fantini, and Yong Q Zhang. The glycosphingolipid MacCer promotes synaptic bouton formation in Drosophila by interacting with Wnt. *eLife*, 7, 2018.

[4362] Ziqiang Huang, Houda G Khaled, Moritz Kirschmann, Sharon MH Gobes, and Richard HR Hahnloser. Excitatory and inhibitory synapse reorganization immediately after critical sensory experience in a vocal learner. *eLife*, 7, 2018.

[4363] Diego Huet, Esther Rajendran, Giel G van Dooren, and Sebastian Lourido. Identification of cryptic subunits from an apicomplexan ATP synthase. *eLife*, 7, 2018.

[4364] Ryan L Huizar, Chanjae Lee, Alexander A Boulgakov, Amjad Horani, Fan Tu, Edward M Marcotte, Steven L Brody, and John B Wallingford. A liquid-like organelle at the root of motile ciliopathy. *eLife*, 7, 2018.

[4365] Raymond E Hulse, Zongli Li, Rick K Huang, Jin Zhang, and David E Clapham. Cryo-EM structure of the polycystin 2-l1 ion channel. *eLife*, 7, 2018.

[4366] Ashiq Hussain, Atefeh Pooryasin, Mo Zhang, Laura F Loschek, Marco La Fortezza, Anja B Friedrich, Catherine-Marie Blais, Habibe K üÇpunar, Vicente A Yépez, Martin Lehmann, Nicolas Gompel, Julien Gagneur, Stephan J Sigrist, and Ilona C Grunwald Kadow. Inhibition of oxidative stress in cholinergic projection neurons fully rescues aging-associated olfactory circuit degeneration in Drosophila. *eLife*, 7, 2018.

[4367] Saman Hussain, Carl N Wivagg, Piotr Szwedziak, Felix Wong, Kaitlin Schaefer, Thierry Izoré, Lars D Renner, Matthew J Holmes, Yingjie Sun, Alexandre W Bisson-Filho, Suzanne Walker, Ariel Amir, Jan Löwe, and Ethan C Garner. MreB filaments align along greatest principal membrane curvature to orient cell wall synthesis. *eLife*, 7, 2018.

[4368] Young Sun Hwang, Minseok Seo, Sang Kyung Kim, Sohyun Bang, Heebal Kim, and Jae Yong Han. Zygotic gene activation in the chicken occurs in two waves, the first involving only maternally derived genes. *eLife*, 7, 2018.

[4369] Yuichi Ichikawa, Noriko Saitoh, and Paul D Kaufman. An asymmetric centromeric nucleosome. *eLife*, 7, 2018.

[4370] Maxim Igaev and Helmut Grubmüller. Microtubule assembly governed by tubulin allosteric gain in flexibility and lattice induced fit. *eLife*, 7, 2018.

[4371] Myron S Ignatius, Madeline N Hayes, Finola E Moore, Qin Tang, Sara P Garcia, Patrick R Blackburn, Kunal Baxi, Long Wang, Alexander Jin, Ashwin Ramakrishnan, Sophia Reeder, Yidong Chen, Gunnlaugur Petur Nielsen, Eleanor Y Chen, Robert P Hasserjian, Franck Tirode, Stephen C Ekker, and David M Langenau. tp53 deficiency causes a wide tumor spectrum and increases embryonal rhabdomyosarcoma metastasis in zebrafish. *eLife*, 7, 2018.

[4372] Tsuyoshi Ikegami, Gowrishankar Ganesh, Tatsuya Takeuchi, and Hiroki Nakamoto. Prediction error induced motor contagions in human behaviors. *eLife*, 7, 2018.

[4373] F Tudor Ilca, Andreas Neerincx, Clemens Hermann, Ana Marcu, Stefan Stevanović, Janet E Deane, and Louise H Boyle. TAPBPR mediates peptide dissociation from MHC class I using a leucine lever. *eLife*, 7, 2018.

[4374] Mikiko Inaki, Ryo Hatori, Naotaka Nakazawa, Takashi Okumura, Tomoki Ishibashi, Junichi Kikuta, Masaru Ishii, Kenji Matsuno, and Hisao Honda. Chiral cell sliding drives left-right asymmetric organ twisting. *eLife*, 7, 2018.

[4375] Takuma Inoshita and Tomoo Hirano. Occurrence of long-term depression in the cerebellar flocculus during adaptation of optokinetic response. *eLife*, 7, 2018.

[4376] Ran Inoue, Kareem Abdou, Ayumi Hayashi-Tanaka, Shin-ichi Muramatsu, Kaori Mino, Kaoru Inokuchi, and Hisashi Mori. Glucocorticoid receptor-mediated amygdalar metaplasticity underlies adaptive modulation of fear memory by stress. *eLife*, 7, 2018.

[4377] Elias B Issa, Charles F Cadieu, and James J DiCarlo. Neural dynamics at successive stages of the ventral visual stream are consistent with hierarchical error signals. *eLife*, 7, 2018.

[4378] Emi Ito, Kazuo Ebine, Seung-won Choi, Sakura Ichinose, Tomohiro Uemura, Akihiko Nakano, and Takashi Ueda. Integration of two RAB5 groups during endosomal transport in plants. *eLife*, 7, 2018.

[4379] Roland Ivanyi-Nagy, Syed Moiz Ahmed, Sabrina Peter, Priya Dharshana Ramani, Peh Fern Ong, Oliver Dreesen, and Peter Dröge. The RNA interactome of human telomerase RNA reveals a coding-independent role for a histone mRNA in telomere homeostasis. *eLife*, 7, 2018.

[4380] Laurelle Jackson, Jessica Hunter, Sandile Cele, Isabella Markham Ferreira, Andrew C Young, Farina Karim, Rajhmun Madansein, Kaylesh J Dullabh, Chih-Yuan Chen, Noel J Buckels, Yashica Ganga, Khadija Khan, Mikael Boulle, Gila Lustig, Richard A Neher, and Alex Sigal. Incomplete inhibition of HIV infection results in more HIV infected lymph node cells by reducing cell death. *eLife*, 7, 2018.

[4381] Alexandria H Jaeger, Molly Stanley, Zachary F Weiss, Pierre-Yves Musso, Rachel CW Chan, Han Zhang, Damian Feldman-Kiss, and Michael D Gordon. A complex peripheral code for salt taste in Drosophila. *eLife*, 7, 2018.

[4382] Seyed Mehdi Jafarnejad, Clément Chapat, Edna Matta-Camacho, Idit Anna Gelbart, Geoffrey G Hesketh, Meztli Arguello, Aitor Garzia, Sung-Hoon Kim, Jan Attig, Maayan Shapiro, Masahiro Morita, Arkady Khoutorsky, Tommy Alain, G Gkogkas, Christos, Noam Stern-Ginossar, Thomas Tuschl, Anne-Claude Gingras, Thomas F Duchaine, and Nahum Sonenberg. Translational control of ERK signaling through miRNA/4EHP-directed silencing. *eLife*, 7, 2018.

[4383] Sagi Jaffe-Dax, Eva Kimel, and Merav Ahissar. Shorter cortical adaptation in dyslexia is broadly distributed in the superior temporal lobe and includes the primary auditory cortex. *eLife*, 7, 2018.

[4384] Madhav Jagannathan, Ryan Cummings, and Yukiko M Yamashita. A conserved function for pericentromeric satellite DNA. *eLife*, 7, 2018.

[4385] James B Jaggard, Bethany A Stahl, Evan Lloyd, David A Prober, Erik R Duboue, and Alex C Keene. Hypocretin underlies the evolution of sleep loss in the Mexican cavefish. *eLife*, 7, 2018.

[4386] Devanshi Jain, M Rhyan Puno, Cem Meydan, Nathalie Lailler, Christopher E Mason, Christopher D Lima, Kathryn V Anderson, and Scott Keeney. ketu mutant mice uncover an essential meiotic function for the ancient RNA helicase YTHDC2. *eLife*, 7, 2018.

[4387] Kirsty Jamieson, Kevin J McNaught, Tereza Ormsby, Neena A Leggett, Shinji Honda, and Eric U Selker. Telomere repeats induce domains of H3K27 methylation in Neurospora. *eLife*, 7, 2018.

[4388] Jürgen Jänes, Yan Dong, Michael Schoof, Jacques Serizay, Alex Appert, Chiara Cerrato, Carson Woodbury, Ron Chen, Carolina Gemma, Ni Huang, Djem Kissiov, Przemyslaw Stempor, Annette Steward, Eva Zeiser, Sascha Sauer, and Julie Ahringer. Chromatin accessibility dynamics across C. elegans development and ageing. *eLife*, 7, 2018.

[4389] Sharan Janjuha, Sumeet Pal Singh, Anastasia Tsakmaki, S Neda Mousavy Gharavy, Priyanka Murawala, Judith Konantz, Sarah Birke, David J Hodson, Guy A Rutter, Gavin A Bewick, and Nikolay Ninov. Age-related islet inflammation marks the proliferative decline of pancreatic beta-cells in zebrafish. *eLife*, 7, 2018.

[4390] Byoung Il Je, Fang Xu, Qingyu Wu, Lei Liu, Robert Meeley, Joseph P Gallagher, Leo Corcilius, Richard J Payne, Madelaine E Bartlett, and David Jackson. The CLAVATA receptor FASCIATED EAR2 responds to distinct CLE peptides by signaling through two downstream effectors. *eLife*, 7, 2018.

[4391] Laura Jeacock, Joana Faria, and David Horn. Codon usage bias controls mRNA and protein abundance in trypanosomatids. *eLife*, 7, 2018.

[4392] Philippe Jean, David Lopez de la Morena, Susann Michanski, Lina María Jaime Tobón, Rituparna Chakrabarti, Maria Magdalena Picher, Jakob Neef, SangYong Jung, Mehmet Gültas, Stephan Maxeiner, Andreas Neef, Carolin Wichmann, Nicola Strenzke, Chad Grabner, and Tobias Moser. The synaptic ribbon is critical for sound encoding at high rates and with temporal precision. *eLife*, 7, 2018.

[4393] Matthew Jemielita, Ned S Wingreen, and Bonnie L Bassler. Quorum sensing controls Vibrio cholerae multicellular aggregate formation. *eLife*, 7, 2018.

[4394] Anders B Jensen, Thorkild IA Sørensen, Oluf Pedersen, Tine Jess, Søren Brunak, and Kristine H Allin. Increase in clinically recorded type 2 diabetes after colectomy. *eLife*, 7, 2018.

[4395] Bjarke Jensen, Bastiaan J Boukens, Dane A Crossley, Justin Conner, Rajiv A Mohan, Karel van Duijvenboden, Alex V Postma, Christopher R Gloschat, Ruth M Elsey, David Sedmera, Igor R Efimov, and Vincent M Christoffels. Specialized impulse conduction pathway in the alligator heart. *eLife*, 7, 2018.

[4396] Yeon-Tae Jeong, Daniele Simoneschi, Sarah Keegan, David Melville, Natalia S Adler, Anita Saraf, Laurence Florens, Michael P Washburn, Claudio N Cavasotto, David Fenyö, Ana Maria Cuervo, Mario Rossi, and Michele Pagano. The ULK1-FBXW5-SEC23B nexus controls autophagy. *eLife*, 7, 2018.

[4397] SaŠa Jereb, Hun-Way Hwang, Eric Van Otterloo, Eve-Ellen Govek, John J Fak, Yuan Yuan, Mary E Hatten, and Robert B Darnell. Differential 3′ processing of specific transcripts expands regulatory and protein diversity across neuronal cell types. *eLife*, 7, 2018.

[4398] Young Rae Ji, Sunita Warrier, Tao Jiang, Doris K Wu, and Katie S Kindt. Directional selectivity of afferent neurons in zebrafish neuromasts is regulated by Emx2 in presynaptic hair cells. *eLife*, 7, 2018.

[4399] Jiefeng Jiang, Anthony D Wagner, and Tobias Egner. Integrated externally and internally generated task predictions jointly guide cognitive control in prefrontal cortex. *eLife*, 7, 2018.

[4400] Wei Jiang, Yuehua Wei, Yong Long, Arthur Owen, Bingying Wang, Xuebing Wu, Shuo Luo, Yongjun Dang, and Dengke K Ma. A genetic program mediates cold-warming response and promotes stress-induced phenoptosis in C. elegans. *eLife*, 7, 2018.

[4401] Junyi Jiao, Mengze He, Sarah A Port, Richard W Baker, Yonggang Xu, Hong Qu, Yujian Xiong, Yukun Wang, Huaizhou Jin, Travis J Eisemann, Frederick M Hughson, and Yongli Zhang. Munc18-1 catalyzes neuronal SNARE assembly by templating SNARE association. *eLife*, 7, 2018.

[4402] Manuel Jiménez-Zaragoza, Marina PL Yubero, Esther Martín-Forero, Jose R Castón, David Reguera, Daniel Luque, Pedro J de Pablo, and Javier M Rodríguez. Biophysical properties of single rotavirus particles account for the functions of protein shells in a multilayered virus. *eLife*, 7, 2018.

[4403] Daniel S Johnson, Marina Bleck, and Sanford M Simon. Timing of ESCRT-III protein recruitment and membrane scission during HIV-1 assembly. *eLife*, 7, 2018.

[4404] Matthew Richard Johnson, Roxan Amanda Stephenson, Sina Ghaemmaghami, and Michael Andreas Welte. Developmentally regulated H2Av buffering via dynamic sequestration to lipid droplets in Drosophila embryos. *eLife*, 7, 2018.

[4405] Adam B Johnston, Denise M Hilton, Patrick McConnell, Britney Johnson, Meghan T Harris, Avital Simone, Gaya K Amarasinghe, John A Cooper, and Bruce L Goode. A novel mode of capping protein-regulation by twinfilin. *eLife*, 7, 2018.

[4406] Sebastian Jojoa-Cruz, Kei Saotome, Swetha E Murthy, Che Chun Alex Tsui, Mark SP Sansom, Ardem Patapoutian, and Andrew B Ward. Cryo-EM structure of the mechanically activated ion channel OSCA1.2. *eLife*, 7, 2018.

[4407] Mark G Jones, Orestis G Andriotis, James JW Roberts, Kerry Lunn, Victoria J Tear, Lucy Cao, Kjetil Ask, David E Smart, Alessandra Bonfanti, Peter Johnson, Aiman Alzetani, Franco Conforti, Regan Doherty, Chester Y Lai, Benjamin Johnson, Konstantinos N Bourdakos, Sophie V Fletcher, Ben G Marshall, Sanjay Jogai, Christopher J Brereton, Serena J Chee, Christian H Ottensmeier, Patricia Sime, Jack Gauldie, Martin Kolb, Sumeet Mahajan, Aurelie Fabre, Atul Bhaskar, Wolfgang Jarolimek, Luca Richeldi, Katherine MA O'Reilly, Phillip D Monk, Philipp J Thurner, and Donna E Davies. Nanoscale dysregulation of collagen structure-function disrupts mechano-homeostasis and mediates pulmonary fibrosis. *eLife*, 7, 2018.

[4408] Sheena A Josselyn. The past, present and future of light-gated ion channels and optogenetics. *eLife*, 7, 2018.

[4409] Karina Kaczmarek-Hajek, Jiong Zhang, Robin Kopp, Antje Grosche, Björn Rissiek, Anika Saul, Santina Bruzzone, Tobias Engel, Tina Jooss, Anna Krautloher, Stefanie Schuster, Tim Magnus, Christine Stadelmann, Swetlana Sirko, Friedrich Koch-Nolte, Volker Eulenburg, and Annette Nicke. Re-evaluation of neuronal P2X7 expression using novel mouse models and a P2X7-specific nanobody. *eLife*, 7, 2018.

[4410] Preetish Kadur Lakshminarasimha Murthy, Tara Srinivasan, Matthew S Bochter, Rui Xi, Anastasia Kristine Varanko, Kuei-Ling Tung, Fatih Semerci, Keli Xu, Mirjana Maletic-Savatic, Susan E Cole, and Xiling Shen. Radical and lunatic fringes modulate notch ligands to support mammalian intestinal homeostasis. *eLife*, 7, 2018.

[4411] Nereo Kalebic, Carlotta Gilardi, Mareike Albert, Takashi Namba, Katherine R Long, Milos Kostic, Barbara Langen, and Wieland B Huttner. Human-specific ARHGAP11B induces hallmarks of neocortical expansion in developing ferret neocortex. *eLife*, 7, 2018.

[4412] Povilas Karvelis, Aaron R Seitz, Stephen M Lawrie, and Peggy Seriès. Autistic traits, but not schizotypy, predict increased weighting of sensory information in Bayesian visual integration. *eLife*, 7, 2018.

[4413] Herbert G Kasler, Intelly S Lee, Hyung W Lim, and Eric Verdin. Histone Deacetylase 7 mediates tissue-specific autoimmunity via control of innate effector function in invariant Natural Killer T Cells. *eLife*, 7, 2018.

[4414] Jakob Nikolas Kather, Meggy Suarez-Carmona, Pornpimol Charoentong, Cleo-Aron Weis, Daniela Hirsch, Peter Bankhead, Marcel Horning, Dyke Ferber, Ivan Kel, Esther Herpel, Sarah Schott, Inka Zörnig, Jochen Utikal, Alexander Marx, Timo Gaiser, Herrmann Brenner, Jenny Chang-Claude, Michael Hoffmeister, Dirk Jäger, and Niels Halama. Topography of cancer-associated immune cells in human solid tumors. *eLife*, 7, 2018.

[4415] Junya Kato, Supratim Dey, Jose E Soto, Carmen Butan, Mason C Wilkinson, Roberto N De Guzman, and Jorge E Galan. A protein secreted by the Salmonella type III secretion system controls needle filament assembly. *eLife*, 7, 2018.

[4416] Marketa Kaucka, Julian Petersen, Marketa Tesarova, Bara Szarowska, Maria Eleni Kastriti, Meng Xie, Anna Kicheva, Karl Annusver, Maria Kasper, Orsolya Symmons, Leslie Pan, Francois Spitz, Jozef Kaiser, Maria Hovorakova, Tomas Zikmund, Kazunori Sunadome, Michael P Matise, Hui Wang, Ulrika Marklund, Hind Abdo, Patrik Ernfors, Pascal Maire, Maud Wurmser, Andrei S Chagin, Kaj Fried, and Igor Adameyko. Signals from the brain and olfactory epithelium control shaping of the mammalian nasal capsule cartilage. *eLife*, 7, 2018.

[4417] Akira Kawanabe, Masaki Hashimoto, Manami Nishizawa, Kazuhisa Nishizawa, Hirotaka Narita, Tomoko Yonezawa, Yuka Jinno, Souhei Sakata, Atsushi Nakagawa, and Yasushi Okamura. The hydrophobic nature of a novel membrane interface regulates the enzyme activity of a voltage-sensing phosphatase. *eLife*, 7, 2018.

[4418] Kerim B Kaylan, Ian C Berg, Matthew J Biehl, Aidan Brougham-Cook, Ishita Jain, Sameed M Jamil, Lauren H Sargeant, Nicholas J Cornell, Lori T Raetzman, and Gregory H Underhill. Spatial patterning of liver progenitor cell differentiation mediated by cellular contractility and Notch signaling. *eLife*, 7, 2018.

[4419] Sara E Kee, Xiang Mou, Huda Y Zoghbi, and Daoyun Ji. Impaired spatial memory codes in a mouse model of Rett syndrome. *eLife*, 7, 2018.

[4420] Daniel A Keedy, Zachary B Hill, Justin T Biel, Emily Kang, T Justin Rettenmaier, José Brandão Neto, Nicholas M Pearce, Frank von Delft, James A Wells, and James S Fraser. An expanded allosteric network in PTP1B by multitemperature crystallography, fragment screening, and covalent tethering. *eLife*, 7, 2018.

[4421] Ruedeerat Keerativittayayut, Ryuta Aoki, Mitra Taghizadeh Sarabi, Koji Jimura, and Kiyoshi Nakahara. Large-scale network integration in the human brain tracks temporal fluctuations in memory encoding performance. *eLife*, 7, 2018.

[4422] Jennifer M Kefauver, Kei Saotome, Adrienne E Dubin, Jesper Pallesen, Christopher A Cottrell, Stuart M Cahalan, Zhaozhu Qiu, Gunhee Hong, Christopher S Crowley, Tess Whitwam, Wen-Hsin Lee, Andrew B Ward, and Ardem Patapoutian. Structure of the human volume regulated anion channel. *eLife*, 7, 2018.

[4423] Alexandra T Keinath, Russell A Epstein, and Vijay Balasubramanian. Environmental deformations dynamically shift the grid cell spatial metric. *eLife*, 7, 2018.

[4424] Genevieve C Kendall, Sarah Watson, Lin Xu, Collette A LaVigne, Whitney Murchison, Dinesh Rakheja, Stephen X Skapek, Franck Tirode, Olivier Delattre, and James F Amatruda. PAX3-FOXO1 transgenic zebrafish models identify HES3 as a mediator of rhabdomyosarcoma tumorigenesis. *eLife*, 7, 2018.

[4425] Arjun Khakhar, Alexander R Leydon, Andrew C Lemmex, Eric Klavins, and Jennifer L Nemhauser. Synthetic hormone-responsive transcription factors can monitor and re-program plant development. *eLife*, 7, 2018.

[4426] Renat N Khaliullin, Rebecca A Green, Linda Z Shi, J Sebastian Gomez-Cavazos, Michael W Berns, Arshad Desai, and Karen Oegema. A positive-feedback-based mechanism for constriction rate acceleration during cytokinesis in Caenorhabditis elegans. *eLife*, 7, 2018.

[4427] Elena Khazina and Oliver Weichenrieder. Human LINE-1 retrotransposition requires a metastable coiled coil and a positively charged N-terminus in L1ORF1p. *eLife*, 7, 2018.

[4428] Andrei Khilkevich, Juan Zambrano, Molly-Marie Richards, and Michael Dean Mauk. Cerebellar implementation of movement sequences through feedback. *eLife*, 7, 2018.

[4429] Atsushi Kikumoto and Ulrich Mayr. Decoding hierarchical control of sequential behavior in oscillatory EEG activity. *eLife*, 7, 2018.

[4430] Jeewon Kim, Amirali Afshari, Ranjita Sengupta, Vittorio Sebastiano, Archana Gupta, Young H Kim, Elizabeth Iorns, Elizabeth Iorns, Rachel Tsui, Alexandria Denis, Nicole Perfito, Timothy M Errington, Elizabeth Iorns, Rachel Tsui, Alexandria Denis, Nicole Perfito, and Timothy M Errington. Replication Study: Melanoma exosomes educate bone marrow progenitor cells toward a pro-metastatic phenotype through MET. *eLife*, 7, 2018.

[4431] Kyung Won Kim, Ngang Heok Tang, Christopher A Piggott, Matthew G Andrusiak, Seungmee Park, Ming Zhu, Naina Kurup, Salvatore J Cherra, Zilu Wu, Andrew D Chisholm, and Yishi Jin. Expanded genetic screening in Caenorhabditis elegans identifies new regulators and an inhibitory role for NAD+ in axon regeneration. *eLife*, 7, 2018.

[4432] Seha Kim and Johannes Burge. The lawful imprecision of human surface tilt estimation in natural scenes. *eLife*, 7, 2018.

[4433] Skylar Xantus Kim, Gamze Çamdere, Xuchen Hu, Douglas Koshland, and Hugo Tapia. Synergy between the small intrinsically disordered protein Hsp12 and trehalose sustain viability after severe desiccation. *eLife*, 7, 2018.

[4434] Sooyun Kim, Yoonsub Kim, Suk-Ho Lee, and Won-Kyung Ho. Dendritic spikes in hippocampal granule cells are necessary for long-term potentiation at the perforant path synapse. *eLife*, 7, 2018.

[4435] Sung Hyun Kim, Mahipal Ganji, Eugene Kim, Jaco van der Torre, Elio Abbondanzieri, and Cees Dekker. DNA sequence encodes the position of DNA supercoils. *eLife*, 7, 2018.

[4436] Mushtak Kisko, Nadia Bouain, Alaeddine Safi, Anna Medici, Robert C Akkers, David Secco, Gilles Fouret, Gabriel Krouk, Mark GM Aarts, Wolfgang Busch, and Hatem Rouached. LPCAT1 controls phosphate homeostasis in a zinc-dependent manner. *eLife*, 7, 2018.

[4437] Kathryn E Kistler, Tatjana Trcek, Thomas R Hurd, Ruoyu Chen, Feng-Xia Liang, Joseph Sall, Masato Kato, and Ruth Lehmann. Phase transitioned nuclear Oskar promotes cell division of Drosophila primordial germ cells. *eLife*, 7, 2018.

[4438] Amrutha Kizhedathu, Archit V Bagul, and Arjun Guha. Negative regulation of G2-M by ATR (mei-41)/Chk1(Grapes) facilitates tracheoblast growth and tracheal hypertrophy in Drosophila. *eLife*, 7, 2018.

[4439] Jonas Klinkhammer, Daniel Schnepf, Liang Ye, Marilena Schwaderlapp, Hans Henrik Gad, Rune Hartmann, Dominique Garcin, Tanel Mahlakõiv, and Peter Staeheli. IFN-$\lambda$ prevents influenza virus spread from the upper airways to the lungs and limits virus transmission. *eLife*, 7, 2018.

[4440] Jason R Klug, Max D Engelhardt, Cara N Cadman, Hao Li, Jared B Smith, Sarah Ayala, Elora W Williams, Hilary Hoffman, and Xin Jin. Differential inputs to striatal cholinergic and parvalbumin interneurons imply functional distinctions. *eLife*, 7, 2018.

[4441] Andrew John Knappenberger, Caroline Wetherington Reiss, and Scott A Strobel. Structures of two aptamers with differing ligand specificity reveal ruggedness in the functional landscape of RNA. *eLife*, 7, 2018.

[4442] Gavin T Knight, Brady F Lundin, Nisha Iyer, Lydia MT Ashton, William A Sethares, Rebecca M Willett, and Randolph Scott Ashton. Engineering induction of singular neural rosette emergence within hPSC-derived tissues. *eLife*, 7, 2018.

[4443] Elisabeth R Knoll, Z Iris Zhu, Debasish Sarkar, David Landsman, and Randall H Morse. Role of the pre-initiation complex in Mediator recruitment and dynamics. *eLife*, 7, 2018.

[4444] David A Knowles, Courtney K Burrows, John D Blischak, Kristen M Patterson, Daniel J Serie, Nadine Norton, Carole Ober, Jonathan K Pritchard, and Yoav Gilad. Determining the genetic basis of anthracycline-cardiotoxicity by molecular response QTL mapping in induced cardiomyocytes. *eLife*, 7, 2018.

[4445] Suhasa B Kodandaramaiah, Francisco J Flores, Gregory L Holst, Annabelle C Singer, Xue Han, Emery N Brown, Edward S Boyden, and Craig R Forest. Multi-neuron intracellular recording in vivo via interacting autopatching robots. *eLife*, 7, 2018.

[4446] Chwee Tat Koe, Ye Sing Tan, Max Lönnfors, Seong Kwon Hur, Christine Siok Lan Low, Yingjie Zhang, Pakorn Kanchanawong, Vytas A Bankaitis, and Hongyan Wang. Vibrator and PI4KIII$\alpha$ govern neuroblast polarity by anchoring non-muscle myosin II. *eLife*, 7, 2018.

[4447] Fotini Kokou, Goor Sasson, Tali Nitzan, Adi Doron-Faigenboim, Sheenan Harpaz, Avner Cnaani, and Itzhak Mizrahi. Host genetic selection for cold tolerance shapes microbiome composition and modulates its response to temperature. *eLife*, 7, 2018.

[4448] Krishna Kalyan Kolluri, Constantine Alifrangis, Neelam Kumar, Yuki Ishii, Stacey Price, Magali Michaut, Steven Williams, Syd Barthorpe, Howard Lightfoot, Sara Busacca, Annabel Sharkey, Zhenqiang Yuan, Elizabeth K Sage, Sabarinath Vallath, John Le Quesne, David A Tice, Doraid Alrifai, Sylvia von Karstedt, Antonella Montinaro, Naomi Guppy, David A Waller, Apostolos Nakas, Robert Good, Alan Holmes, Henning Walczak, Dean A Fennell, Mathew Garnett, Francesco Iorio, Lodewyk Wessels, Ultan McDermott, and Samuel M Janes. Loss of functional BAP1 augments sensitivity to TRAIL in cancer cells. *eLife*, 7, 2018.

[4449] Terje R Kolstad, Jonas van den Brink, Niall MacQuaide, Per Kristian Lunde, Michael Frisk, Jan Magnus Aronsen, Einar S Norden, Alessandro Cataliotti, Ivar Sjaastad, Ole M Sejersted, Andrew G Edwards, Glenn Terje Lines, and William E Louch. Ryanodine receptor dispersion disrupts Ca2+ release in failing cardiac myocytes. *eLife*, 7, 2018.

[4450] Megan C Kopp, Piotr R Nowak, Natacha Larburu, Christopher J Adams, and Maruf MU Ali. In vitro FRET analysis of IRE1 and BiP association and dissociation upon endoplasmic reticulum stress. *eLife*, 7, 2018.

[4451] Nitzan Koppel, Jordan E Bisanz, Maria-Eirini Pandelia, Peter J Turnbaugh, and Emily P Balskus. Discovery and characterization of a prevalent human gut bacterial enzyme sufficient for the inactivation of a family of plant toxins. *eLife*, 7, 2018.

[4452] Julia C Körholz, Sara Zocher, Anna N Grzyb, Benjamin Morisse, Alexandra Poetzsch, Fanny Ehret, Christopher Schmied, and Gerd Kempermann. Selective increases in inter-individual variability in response to environmental enrichment in female mice. *eLife*, 7, 2018.

[4453] Olga Kornienko, Patrick Latuske, Mathis Bassler, Laura Kohler, and Kevin Allen. Non-rhythmic head-direction cells in the parahippocampal region are not constrained by attractor network dynamics. *eLife*, 7, 2018.

[4454] Evangelia-Georgia Kostaki, Timokratis Karamitros, Garyfallia Stefanou, Ioannis Mamais, Konstantinos Angelis, Angelos Hatzakis, Anna Kramvis, and Dimitrios Paraskevis. Unravelling the history of hepatitis B virus genotypes A and D infection using a full-genome phylogenetic and phylogeographic approach. *eLife*, 7, 2018.

[4455] Ninad B Kothari, Melville J Wohlgemuth, and Cynthia F Moss. Dynamic representation of 3D auditory space in the midbrain of the free-flying echolocating bat. *eLife*, 7, 2018.

[4456] Joanna Krakowiak, Xu Zheng, Nikit Patel, Zoë A Feder, Jayamani Anandhakumar, Kendra Valerius, David S Gross, Ahmad S Khalil, and David Pincus. Hsf1 and Hsp70 constitute a two-component feedback loop that regulates the yeast heat shock response. *eLife*, 7, 2018.

[4457] Ben Krause-Kyora, Julian Susat, Felix M Key, Denise Kühnert, Esther Bosse, Alexander Immel, Christoph Rinne, Sabin-Christin Kornell, Diego Yepes, Sören Franzenburg, Henrike O Heyne, Thomas Meier, Sandra Lösch, Harald Meller, Susanne Friederich, Nicole Nicklisch, Kurt W Alt, Stefan Schreiber, Andreas Tholey, Alexander Herbig, Almut Nebel, and Johannes Krause. Neolithic and medieval virus genomes reveal complex evolution of hepatitis B. *eLife*, 7, 2018.

[4458] Magdalena Kremer, Nora Kuhlmann, Marius Lechner, Linda Baldus, and Michael Lammers. Comment on 'YcgC represents a new protein deacetylase family in prokaryotes'. *eLife*, 7, 2018.

[4459] William A Kronert, Kaylyn M Bell, Meera C Viswanathan, Girish C Melkani, Adriana S Trujillo, Alice Huang, Anju Melkani, Anthony Cammarato, Douglas M Swank, and Sanford I Bernstein. Prolonged cross-bridge binding triggers muscle dysfunction in a Drosophila model of myosin-based hypertrophic cardiomyopathy. *eLife*, 7, 2018.

[4460] Michael Krumin, Julie J Lee, Kenneth D Harris, and Matteo Carandini. Decision and navigation in mouse parietal cortex. *eLife*, 7, 2018.

[4461] Adam J Kucharski, Mike Kama, Conall H Watson, Maite Aubry, Sebastian Funk, Alasdair D Henderson, Oliver J Brady, Jessica Vanhomwegen, Jean-Claude Manuguerra, Colleen L Lau, W John Edmunds, John Aaskov, Eric James Nilles, Van-Mai Cao-Lormeau, Stéphane Hué, and Martin L Hibberd. Using paired serology and surveillance data to quantify dengue transmission and control during a large outbreak in Fiji. *eLife*, 7, 2018.

[4462] Renuka Kudva, Pengfei Tian, Fátima Pardo-Avila, Marta Carroni, Robert B Best, Harris D Bernstein, and Gunnar von Heijne. The shape of the bacterial ribosome exit tunnel affects cotranslational protein folding. *eLife*, 7, 2018.

[4463] Bridget M Kuehn. Communities creating change. *eLife*, 7, 2018.

[4464] Bridget M Kuehn. Cultural shifts. *eLife*, 7, 2018.

[4465] Bridget M Kuehn. Looking for the best of both worlds. *eLife*, 7, 2018.

[4466] Anika Küken, Frederik Sommer, Liliya Yaneva-Roder, Luke CM Mackinder, Melanie Höhne, Stefan Geimer, Martin C Jonikas, Michael Schroda, Mark Stitt, Zoran Nikoloski, and Tabea Mettler-Altmann. Effects of microcompartmentation on flux distribution and metabolic pools in Chlamydomonas reinhardtii chloroplasts. *eLife*, 7, 2018.

[4467] Amit Kumar, Oded Schiff, Edi Barkai, Bartlett W Mel, Alon Poleg-Polsky, and Jackie Schiller. NMDA spikes mediate amplification of inputs in the rat piriform cortex. *eLife*, 7, 2018.

[4468] Amutha Sampath Kumar, Eunsook Park, Alexander Nedo, Ali Alqarni, Li Ren, Kyle Hoban, Shannon Modla, John H McDonald, Chandra Kambhamettu, Savithramma P Dinesh-Kumar, and Jeffrey Lewis Caplan. Stromule extension along microtubules coordinated with actin-mediated anchoring guides perinuclear chloroplast movement during innate immunity. *eLife*, 7, 2018.

[4469] Santosh Kumar Kuncha, Katta Suma, Komal Ishwar Pawar, Jotin Gogoi, Satya Brata Routh, Sambhavi Pottabathini, Shobha P Kruparani, and Rajan Sankaranarayanan. A discriminator code–based DTD surveillance ensures faithful glycine delivery for protein biosynthesis in bacteria. *eLife*, 7, 2018.

[4470] Jun Kunimatsu, Tomoki W Suzuki, Shogo Ohmae, and Masaki Tanaka. Different contributions of preparatory activity in the basal ganglia and cerebellum for self-timing. *eLife*, 7, 2018.

[4471] Ulrike Künzel, Adam Graham Grieve, Yao Meng, Boris Sieber, Sally A Cowley, and Matthew Freeman. FRMD8 promotes inflammatory and growth factor signalling by stabilising the iRhom/ADAM17 sheddase complex. *eLife*, 7, 2018.

[4472] Sebastian Kwiatkowski, Agnieszka K Seliga, Didier Vertommen, Marianna Terreri, Takao Ishikawa, Iwona Grabowska, Marcel Tiebe, Aurelio A Teleman, Adam K Jagielski, Maria Veiga-da Cunha, and Jakub Drozak. SETD3 protein is the actin-specific histidine N-methyltransferase. *eLife*, 7, 2018.

[4473] Andrew S LaCroix, Andrew D Lynch, Matthew E Berginski, and Brenton D Hoffman. Tunable molecular tension sensors reveal extension-based control of vinculin loading. *eLife*, 7, 2018.

[4474] Franck Ladam, William Stanney, Ian J Donaldson, Ozge Yildiz, Nicoletta Bobola, and Charles G Sagerström. TALE factors use two distinct functional modes to control an essential zebrafish gene expression program. *eLife*, 7, 2018.

[4475] Arnaud L Lalive, Anthony D Lien, Thomas K Roseberry, Christopher H Donahue, and Anatol C Kreitzer. Motor thalamus supports striatum-driven reinforcement. *eLife*, 7, 2018.

[4476] Andy KM Lam and Raimund Dutzler. Calcium-dependent electrostatic control of anion access to the pore of the calcium-activated chloride channel TMEM16A. *eLife*, 7, 2018.

[4477] Francois M Lambert, Laura Cardoit, Elric Courty, Marion Bougerol, Muriel Thoby-Brisson, John Simmers, Hervé Tostivint, and Didier Le Ray. Functional limb muscle innervation prior to cholinergic transmitter specification during early metamorphosis in Xenopus. *eLife*, 7, 2018.

[4478] Katherine Young Bezold Lamm, Maddison L Johnson, Julie Baker Phillips, Michael B Muntifering, Jeanne M James, Helen N Jones, Raymond W Redline, Antonis Rokas, and Louis J Muglia. Inverted formin 2 regulates intracellular trafficking, placentation, and pregnancy outcome. *eLife*, 7, 2018.

[4479] Fabienne Lampert, Diana Stafa, Algera Goga, Martin Varis Soste, Samuel Gilberto, Natacha Olieric, Paola Picotti, Markus Stoffel, and Matthias Peter. The multi-subunit GID/CTLH E3 ubiquitin ligase promotes cell proliferation and targets the transcription factor Hbp1 for degradation. *eLife*, 7, 2018.

[4480] Mylene Lancino, Sara Majello, Sebastien Herbert, Fabrice De Chaumont, Jean-Yves Tinevez, Jean-Christophe Olivo-Marin, Philippe Herbomel, and Anne Schmidt. Anisotropic organization of circumferential actomyosin characterizes hematopoietic stem cells emergence in the zebrafish. *eLife*, 7, 2018.

[4481] Lukas Landler, Simon Nimpf, Tobias Hochstoeger, Gregory C Nordmann, Artemis Papadaki-Anastasopoulou, and David A Keays. Comment on "Magnetosensitive neurons mediate geomagnetic orientation in Caenorhabditis elegans". *eLife*, 7, 2018.

[4482] Lisa Landskron, Victoria Steinmann, Francois Bonnay, Thomas R Burkard, Jonas Steinmann, Ilka Reichardt, Heike Harzer, Anne-Sophie Laurenson, Heinrich Reichert, and Jürgen A Knoblich. The asymmetrically segregating lncRNA cherub is required for transforming stem cells into malignant cells. *eLife*, 7, 2018.

[4483] Brian J Lane, Daniel R Kick, David K Wilson, Satish S Nair, and David J Schulz. Dopamine maintains network synchrony via direct modulation of gap junctions in the crustacean cardiac ganglion. *eLife*, 7, 2018.

[4484] Jackie Lang, Adrienne Barber, and Sue Biggins. An assay for de novo kinetochore assembly reveals a key role for the CENP-T pathway in budding yeast. *eLife*, 7, 2018.

[4485] Damien Laporte, Laëtitia Gouleme, Laure Jimenez, Ines Khemiri, and Isabelle Sagot. Mitochondria reorganization upon proliferation arrest predicts individual yeast cell fate. *eLife*, 7, 2018.

[4486] Antonio H Lara, Gamaleldin F Elsayed, Andrew J Zimnik, John P Cunningham, and Mark M Churchland. Conservation of preparatory neural events in monkey motor cortex regardless of how movement is initiated. *eLife*, 7, 2018.

[4487] Valerie A Larson, Yevgeniya Mironova, Kimberly G Vanderpool, Ari Waisman, John E Rash, Amit Agarwal, and Dwight E Bergles. Oligodendrocytes control potassium accumulation in white matter and seizure susceptibility. *eLife*, 7, 2018.

[4488] Johan E Larsson, H Peter Larsson, and Sara I Liin. KCNE1 tunes the sensitivity of KV7.1 to polyunsaturated fatty acids by moving turret residues close to the binding site. *eLife*, 7, 2018.

[4489] Charlotte Lässig, Katja Lammens, Jacob Lucián Gorenflos López, Sebastian Michalski, Olga Fettscher, and Karl-Peter Hopfner. Unified mechanisms for self-RNA recognition by RIG-I Singleton-Merten syndrome variants. *eLife*, 7, 2018.

[4490] Sarah Louise Latty, Jiro Sakai, Lee Hopkins, Brett Verstak, Teresa Paramo, Nils A Berglund, Eugenia Cammarota, Pietro Cicuta, Nicholas J Gay, Peter J Bond, David Klenerman, and Clare E Bryant. Activation of Toll-like receptors nucleates assembly of the MyDDosome signaling hub. *eLife*, 7, 2018.

[4491] Christopher E Laumer, Harald Gruber-Vodicka, Michael G Hadfield, Vicki B Pearse, Ana Riesgo, John C Marioni, and Gonzalo Giribet. Support for a clade of Placozoa and Cnidaria in genes with minimal compositional bias. *eLife*, 7, 2018.

[4492] Kenneth Lay, Shaopeng Yuan, Shiri Gur-Cohen, Yuxuan Miao, Tianxiao Han, Shruti Naik, H Amalia Pasolli, Samantha B Larsen, and Elaine Fuchs. Stem cells repurpose proliferation to contain a breach in their niche barrier. *eLife*, 7, 2018.

[4493] Amanda J Lea, Christopher M Vockley, Rachel A Johnston, Christina A Del Carpio, Luis B Barreiro, Timothy E Reddy, and Jenny Tung. Genome-wide quantification of the effects of DNA methylation on human gene regulation. *eLife*, 7, 2018.

[4494] Andres M Lebensohn and Rajat Rohatgi. R-spondins can potentiate WNT signaling without LGRs. *eLife*, 7, 2018.

[4495] Danny LeBert, Jayne M Squirrell, Chrissy Freisinger, Julie Rindy, Netta Golenberg, Grace Frecentese, Angela Gibson, Kevin W Eliceiri, and Anna Huttenlocher. Damage-induced reactive oxygen species regulate vimentin and dynamic collagen-based projections to mediate wound repair. *eLife*, 7, 2018.

[4496] Lucy LeBlanc, Bum-Kyu Lee, Andy C Yu, Mijeong Kim, Aparna V Kambhampati, Shannon M Dupont, Davide Seruggia, Byoung U Ryu, Stuart H Orkin, and Jonghwan Kim. Yap1 safeguards mouse embryonic stem cells from excessive apoptosis during differentiation. *eLife*, 7, 2018.

[4497] Chang Hoon Lee, Hongwei H Zhang, Satya P Singh, Lily Koo, Juraj Kabat, Hsinyi Tsang, Tej Pratap Singh, and Joshua M Farber. C/EBP$\delta$ drives interactions between human MAIT cells and endothelial cells that are important for extravasation. *eLife*, 7, 2018.

[4498] Christine C Lee, Bing Li, Hongtao Yu, and Michael J Matunis. Sumoylation promotes optimal APC/C activation and timely anaphase. *eLife*, 7, 2018.

[4499] Kian-Yong Lee, Rebecca A Green, Edgar Gutierrez, J Sebastian Gomez-Cavazos, Irina Kolotuev, Shaohe Wang, Arshad Desai, Alex Groisman, and Karen Oegema. CYK-4 functions independently of its centralspindlin partner ZEN-4 to cellularize oocytes in germline syncytia. *eLife*, 7, 2018.

[4500] Pei-Tseng Lee, Guang Lin, Wen-Wen Lin, Fengqiu Diao, Benjamin H White, and Hugo J Bellen. A kinase-dependent feedforward loop affects CREBB stability and long term memory formation. *eLife*, 7, 2018.

[4501] Pei-Tseng Lee, Jonathan Zirin, Oguz Kanca, Wen-Wen Lin, Karen L Schulze, David Li-Kroeger, Rong Tao, Colby Devereaux, Yanhui Hu, Verena Chung, Ying Fang, Yuchun He, Hongling Pan, Ming Ge, Zhongyuan Zuo, Benjamin E Housden, Stephanie E Mohr, Shinya Yamamoto, Robert W Levis, Allan C Spradling, Norbert Perrimon, and Hugo J Bellen. A gene-specific T2A-GAL4 library for Drosophila. *eLife*, 7, 2018.

[4502] Bryan A Leland, Angela C Chen, Amy Y Zhao, Robert C Wharton, and Megan C King. Rev7 and 53BP1/Crb2 prevent RecQ helicase-dependent hyper-resection of DNA double-strand breaks. *eLife*, 7, 2018.

[4503] Katerina Leonova, Alfiya Safina, Elimelech Nesher, Poorva Sandlesh, Rachel Pratt, Catherine Burkhart, Brittany Lipchick, Ilya Gitlin, Costakis Frangou, Igor Koman, Jianmin Wang, Kirill Kirsanov, Marianna G Yakubovskaya, Andrei V Gudkov, and Katerina Gurova. TRAIN (Transcription of Repeats Activates INterferon) in response to chromatin destabilization induced by small molecules in mammalian cells. *eLife*, 7, 2018.

[4504] L Michelle Lewis, Meredith C Edwards, Zachary R Meyers, C Conover Talbot, Haiping Hao, David Blum, Elizabeth Iorns, Elizabeth Iorns, Rachel Tsui, Alexandria Denis, Nicole Perfito, and Timothy M Errington. Replication Study: Transcriptional amplification in tumor cells with elevated c-Myc. *eLife*, 7, 2018.

[4505] Laura D Lewis, Giovanni Piantoni, Robert A Peterfreund, Emad N Eskandar, Priscilla Grace Harrell, Oluwaseun Akeju, Linda S Aglio, Sydney S Cash, Emery N Brown, Eran A Mukamel, and Patrick L Purdon. A transient cortical state with sleep-like sensory responses precedes emergence from general anesthesia in humans. *eLife*, 7, 2018.

[4506] Aurore L'honoré, Pierre-Henri Commère, Elisa Negroni, Giorgia Pallafacchina, Bertrand Friguet, Jacques Drouin, Margaret Buckingham, and Didier Montarras. The role of Pitx2 and Pitx3 in muscle stem cells gives new insights into P38$\alpha$ MAP kinase and redox regulation of muscle regeneration. *eLife*, 7, 2018.

[4507] Bo Li, Chouin Wong, Shihong Max Gao, Rulan Zhang, Rongbo Sun, Yulong Li, and Yan Song. The retromer complex safeguards against neural progenitor-derived tumorigenesis by regulating Notch receptor trafficking. *eLife*, 7, 2018.

[4508] Ji Li, Peter S Choi, Christine L Chaffer, Katherine Labella, Justin H Hwang, Andrew O Giacomelli, Jong Wook Kim, Nina Ilic, John G Doench, Seav Huong Ly, Chao Dai, Kimberly Hagel, Andrew L Hong, Ole Gjoerup, Shom Goel, Jennifer Y Ge, David E Root, Jean J Zhao, Angela N Brooks, Robert A Weinberg, and William C Hahn. An alternative splicing switch in FLNB promotes the mesenchymal cell state in human breast cancer. *eLife*, 7, 2018.

[4509] Jiancai Li, Meredith C Schuman, Rayko Halitschke, Xiang Li, Han Guo, Veit Grabe, Austin Hammer, and Ian T Baldwin. The decoration of specialized metabolites influences stylar development. *eLife*, 7, 2018.

[4510] Jiefu Li, Ricardo Guajardo, Chuanyun Xu, Bing Wu, Hongjie Li, Tongchao Li, David J Luginbuhl, Xiaojun Xie, and Liqun Luo. Stepwise wiring of the Drosophila olfactory map requires specific Plexin B levels. *eLife*, 7, 2018.

[4511] Rongbo Li, Ksenija Bernau, Nathan Sandbo, Jing Gu, Sebastian Preissl, and Xin Sun. Pdgfra marks a cellular lineage with distinct contributions to myofibroblasts in lung maturation and injury response. *eLife*, 7, 2018.

[4512] Wankun L Li, Monica W Chu, An Wu, Yusuke Suzuki, Itaru Imayoshi, and Takaki Komiyama. Adult-born neurons facilitate olfactory bulb pattern separation during task engagement. *eLife*, 7, 2018.

[4513] Weihan Li, Voytek Okreglak, Jirka Peschek, Philipp Kimmig, Meghan Zubradt, Jonathan S Weissman, and Peter Walter. Engineering ER-stress dependent non-conventional mRNA splicing. *eLife*, 7, 2018.

[4514] Xiaodong Li, Hiroshi Ishimoto, and Azusa Kamikouchi. Auditory experience controls the maturation of song discrimination and sexual response in Drosophila. *eLife*, 7, 2018.

[4515] Xiling Li, Pragya Goel, Catherine Chen, Varun Angajala, Xun Chen, and Dion K Dickman. Synapse-specific and compartmentalized expression of presynaptic homeostatic potentiation. *eLife*, 7, 2018.

[4516] Yan Li, Kyle W Muir, Matthew W Bowler, Jutta Metz, Christian H Haering, and Daniel Panne. Structural basis for Scc3-dependent cohesin recruitment to chromatin. *eLife*, 7, 2018.

[4517] David Li-Kroeger, Oguz Kanca, Pei-Tseng Lee, Sierra Cowan, Michael T Lee, Manish Jaiswal, Jose Luis Salazar, Yuchun He, Zhongyuan Zuo, and Hugo J Bellen. An expanded toolkit for gene tagging based on MiMIC and scarless CRISPR tagging in Drosophila. *eLife*, 7, 2018.

[4518] Gaojian Lian, JN Rashida Gnanaprakasam, Tingting Wang, Ruohan Wu, Xuyong Chen, Lingling Liu, Yuqing Shen, Mao Yang, Jun Yang, Ying Chen, Vasilis Vasiliou, Teresa A Cassel, Douglas R Green, Yusen Liu, Teresa WM Fan, and Ruoning Wang. Glutathione de novo synthesis but not recycling process coordinates with glutamine catabolism to control redox homeostasis and directs murine T cell differentiation. *eLife*, 7, 2018.

[4519] Lili Liao, Zongzhi Z Liu, Lauren Langbein, Weijia Cai, Eun-Ah Cho, Jie Na, Xiaohua Niu, Wei Jiang, Zhijiu Zhong, Wesley L Cai, Geetha Jagannathan, Essel Dulaimi, Joseph R Testa, Robert G Uzzo, Yuxin Wang, George R Stark, Jianxin Sun, Stephen Peiper, Yaomin Xu, Qin Yan, and Haifeng Yang. Multiple tumor suppressors regulate a HIF-dependent negative feedback loop via ISGF3 in human clear cell renal cancer. *eLife*, 7, 2018.

[4520] Ashley RG Libby, David A Joy, Po-Lin So, Mohammad A Mandegar, Jonathon M Muncie, Federico N Mendoza-Camacho, Valerie M Weaver, Bruce R Conklin, and Todd C McDevitt. Spatiotemporal mosaic self-patterning of pluripotent stem cells using CRISPR interference. *eLife*, 7, 2018.

[4521] Nina T Lichtenberg, Brian Lee, Vadim Kashtelyan, Bharadwaja S Chappa, Henok T Girma, Elizabeth A Green, Shir Kantor, Dave A Lagowala, Matthew A Myers, Danielle Potemri, Meredith G Pecukonis, Robel T Tesfay, Michael S Walters, Adam C Zhao, R James R Blair, Joseph F Cheer, and Matthew R Roesch. Rat behavior and dopamine release are modulated by conspecific distress. *eLife*, 7, 2018.

[4522] Helen Louise Lightfoot, Timo Hagen, Antoine Cléry, Frédéric Hai-Trieu Allain, and Jonathan Hall. Control of the polyamine biosynthesis pathway by G2-quadruplexes. *eLife*, 7, 2018.

[4523] Shion An Lim, Eric Richard Bolin, and Susan Marqusee. Tracing a protein's folding pathway over evolutionary time using ancestral sequence reconstruction and hydrogen exchange. *eLife*, 7, 2018.

[4524] Jia-Ren Lin, Benjamin Izar, Shu Wang, Clarence Yapp, Shaolin Mei, Parin M Shah, Sandro Santagata, and Peter K Sorger. Highly multiplexed immunofluorescence imaging of human tissues and tumors using t-CyCIF and conventional optical microscopes. *eLife*, 7, 2018.

[4525] Grace W Lindsay and Kenneth D Miller. How biological attention mechanisms improve task performance in a large-scale visual system model. *eLife*, 7, 2018.

[4526] Marcela Lipovsek and Richard JT Wingate. Conserved and divergent development of brainstem vestibular and auditory nuclei. *eLife*, 7, 2018.

[4527] Han-Hsuan Liu, Daniel B McClatchy, Lucio Schiapparelli, Wanhua Shen, John R Yates, and Hollis T Cline. Role of the visual experience-dependent nascent proteome in neuronal plasticity. *eLife*, 7, 2018.

[4528] Linxia Liu, Guang-Jun He, Lei Chen, Jiao Zheng, Yingying Chen, Lan Shen, Xiuyun Tian, Erwei Li, Ence Yang, Guojian Liao, and Linqi Wang. Genetic basis for coordination of meiosis and sexual structure maturation in Cryptococcus neoformans. *eLife*, 7, 2018.

[4529] Mochi Liu, Anuj K Sharma, Joshua W Shaevitz, and Andrew M Leifer. Temporal processing and context dependency in Caenorhabditis elegans response to mechanosensation. *eLife*, 7, 2018.

[4530] Qinwen Liu, Pinar Onal, Rhea R Datta, Julia M Rogers, Urs Schmidt-Ott, Martha L Bulyk, Stephen Small, and Joseph W Thornton. Ancient mechanisms for the evolution of the bicoid homeodomain's function in fly development. *eLife*, 7, 2018.

[4531] Rui Liu, Xiangyong Yuan, Kepu Chen, Yi Jiang, and Wen Zhou. Perception of social interaction compresses subjective duration in an oxytocin-dependent manner. *eLife*, 7, 2018.

[4532] Shixuan Liu, Miriam Bracha Ginzberg, Nish Patel, Marc Hild, Bosco Leung, Zhengda Li, Yen-Chi Chen, Nancy Chang, Yuan Wang, Ceryl Tan, Shulamit Diena, William Trimble, Larry Wasserman, Jeremy L Jenkins, Marc W Kirschner, and Ran Kafri. Size uniformity of animal cells is actively maintained by a p38 MAPK-dependent regulation of G1-length. *eLife*, 7, 2018.

[4533] Xiao-Man Liu, Akinori Yamasaki, Xiao-Min Du, Valerie C Coffman, Yoshinori Ohsumi, Hitoshi Nakatogawa, Jian-Qiu Wu, Nobuo N Noda, and Li-Lin Du. Lipidation-independent vacuolar functions of Atg8 rely on its noncanonical interaction with a vacuole membrane protein. *eLife*, 7, 2018.

[4534] Xiaowen Liu, Thuy Vien, Jingjing Duan, Shu-Hsien Sheu, Paul G DeCaen, and David E Clapham. Polycystin-2 is an essential ion channel subunit in the primary cilium of the renal collecting duct epithelium. *eLife*, 7, 2018.

[4535] Yang Liu, Noelle RB Stiles, and Markus Meister. Augmented reality powers a cognitive assistant for the blind. *eLife*, 7, 2018.

[4536] Yubing Liu, Wei Zou, Peiguo Yang, Li Wang, Yan Ma, Hong Zhang, and Xiaochen Wang. Autophagy-dependent ribosomal RNA degradation is essential for maintaining nucleotide homeostasis during C. elegans development. *eLife*, 7, 2018.

[4537] Yuchen Liu, Jianfa Li, Zhicong Chen, Weiren Huang, and Zhiming Cai. Synthesizing artificial devices that redirect cellular information at will. *eLife*, 7, 2018.

[4538] Kathryn E Livingston, Jacob P Mahoney, Aashish Manglik, Roger K Sunahara, and John R Traynor. Measuring ligand efficacy at the mu-opioid receptor using a conformational biosensor. *eLife*, 7, 2018.

[4539] Geulah Livshits, Direna Alonso-Curbelo, John P Morris, Richard Koche, Michael Saborowski, John Erby Wilkinson, and Scott W Lowe. Arid1a restrains Kras-dependent changes in acinar cell identity. *eLife*, 7, 2018.

[4540] Laura Lleras Forero, Rachna Narayanan, Leonie FA Huitema, Maaike VanBergen, Alexander Apschner, Josi Peterson-Maduro, Ive Logister, Guillaume Valentin, Luis G Morelli, Andrew C Oates, and Stefan Schulte-Merker. Segmentation of the zebrafish axial skeleton relies on notochord sheath cells and not on the segmentation clock. *eLife*, 7, 2018.

[4541] Carla Lloret-Fernández, Miren Maicas, Carlos Mora-Martínez, Alejandro Artacho, ángela Jimeno-Martín, Laura Chirivella, Peter Weinberg, and Nuria Flames. A transcription factor collective defines the HSN serotonergic neuron regulatory landscape. *eLife*, 7, 2018.

[4542] Pang-Kuo Lo, Yuan Yao, Ji Shin Lee, Yongshu Zhang, Weiliang Huang, Maureen A Kane, and Qun Zhou. LIPG signaling promotes tumor initiation and metastasis of human basal-like triple-negative breast cancer. *eLife*, 7, 2018.

[4543] James Anthony Loehr, Shang Wang, Tanya R Cully, Rituraj Pal, Irina V Larina, Kirill V Larin, and George G Rodney. NADPH oxidase mediates microtubule alterations and diaphragm dysfunction in dystrophic mice. *eLife*, 7, 2018.

[4544] Roxanne Lofredi, Wolf-Julian Neumann, Antje Bock, Andreas Horn, Julius Huebl, Sandy Siegert, Gerd-Helge Schneider, Joachim K Krauss, and Andrea A Kühn. Dopamine-dependent scaling of subthalamic gamma bursts with movement velocity in patients with Parkinson's disease. *eLife*, 7, 2018.

[4545] Juan Carlos Lopez-Baez, Daniel J Simpson, Laura LLeras Forero, Zhiqiang Zeng, Hannah Brunsdon, Angela Salzano, Alessandro Brombin, Cameron Wyatt, Witold Rybski, Leonie F A Huitema, Rodney M Dale, Koichi Kawakami, Christoph Englert, Tamir Chandra, Stefan Schulte-Merker, Nicholas D Hastie, and E Elizabeth Patton. Wilms Tumor 1b defines a wound-specific sheath cell subpopulation associated with notochord repair. *eLife*, 7, 2018.

[4546] Albert Lu, Paulina Wawro, David W Morgens, Fernando Portela, Michael C Bassik, and Suzanne R Pfeffer. Genome-wide interrogation of extracellular vesicle biology using barcoded miRNAs. *eLife*, 7, 2018.

[4547] Alex X Lu, Yolanda T Chong, Ian Shen Hsu, Bob Strome, Louis-Francois Handfield, Oren Kraus, Brenda J Andrews, and Alan M Moses. Integrating images from multiple microscopy screens reveals diverse patterns of change in the subcellular localization of proteins. *eLife*, 7, 2018.

[4548] Gang Lu, Stephanie Weng, Mary Matyskiela, Xinde Zheng, Wei Fang, Scott Wood, Christine Surka, Reina Mizukoshi, Chin-Chun Lu, Derek Mendy, In Sock Jang, Kai Wang, Mathieu Marella, Suzana Couto, Brian Cathers, James Carmichael, Philip Chamberlain, and Mark Rolfe. UBE2G1 governs the destruction of cereblon neomorphic substrates. *eLife*, 7, 2018.

[4549] Kevin L Lu, Jonathan O Nelson, George J Watase, Natalie Warsinger-Pepe, and Yukiko M Yamashita. Transgenerational dynamics of rDNA copy number in Drosophila male germline stem cells. *eLife*, 7, 2018.

[4550] Victoria A Lund, Katarzyna Wacnik, Robert D Turner, Bryony E Cotterell, Christa G Walther, Samuel J Fenn, Fabian Grein, Adam JM Wollman, Mark C Leake, Nicolas Olivier, Ashley Cadby, Stéphane Mesnage, Simon Jones, and Simon J Foster. Molecular coordination of Staphylococcus aureus cell division. *eLife*, 7, 2018.

[4551] Phi Luong, Matija Hedl, Jie Yan, Tao Zuo, Tian-Min Fu, Xiaomo Jiang, Jay R Thiagarajah, Steen H Hansen, Cammie F Lesser, Hao Wu, Clara Abraham, and Wayne I Lencer. INAVA-ARNO complexes bridge mucosal barrier function with inflammatory signaling. *eLife*, 7, 2018.

[4552] Katharina Lust and Joachim Wittbrodt. Activating the regenerative potential of Müller glia cells in a regeneration-deficient retina. *eLife*, 7, 2018.

[4553] Long Luu and Alan A Stocker. Post-decision biases reveal a self-consistency principle in perceptual inference. *eLife*, 7, 2018.

[4554] Michael Lynch. Phylogenetic divergence of cell biological features. *eLife*, 7, 2018.

[4555] Emilie Ma, Pauline Dupaigne, Laurent Maloisel, Raphaël Guerois, Eric Le Cam, and Eric Coïc. Rad52-Rad51 association is essential to protect Rad51 filaments against Srs2, but facultative for filament formation. *eLife*, 7, 2018.

[4556] Natalie Jing Ma, Colin F Hemez, Karl W Barber, Jesse Rinehart, and Farren J Isaacs. Organisms with alternative genetic codes resolve unassigned codons via mistranslation and ribosomal rescue. *eLife*, 7, 2018.

[4557] Wen Ma, Kevin D Whitley, Yann R Chemla, Zaida Luthey-Schulten, and Klaus Schulten. Free-energy simulations reveal molecular mechanism for functional switch of a DNA helicase. *eLife*, 7, 2018.

[4558] Xiaochu Ma, Maolin Lu, Jason Gorman, Daniel S Terry, Xinyu Hong, Zhou Zhou, Hong Zhao, Roger B Altman, James Arthos, Scott C Blanchard, Peter D Kwong, James B Munro, and Walther Mothes. HIV-1 Env trimer opens through an asymmetric intermediate in which individual protomers adopt distinct conformations. *eLife*, 7, 2018.

[4559] Zaijun Ma, Hui Wang, Yuping Cai, Han Wang, Kongyan Niu, Xiaofen Wu, Huanhuan Ma, Yun Yang, Wenhua Tong, Feng Liu, Zhandong Liu, Yaoyang Zhang, Rui Liu, Zheng-Jiang Zhu, and Nan Liu. Epigenetic drift of H3K27me3 in aging links glycolysis to healthy longevity in Drosophila. *eLife*, 7, 2018.

[4560] Zhiwei Ma and Nanyin Zhang. Temporal transitions of spontaneous brain activity. *eLife*, 7, 2018.

[4561] Kourosh Maboudi, Etienne Ackermann, Laurel Watkins de Jong, Brad E Pfeiffer, David Foster, Kamran Diba, and Caleb Kemere. Uncovering temporal structure in hippocampal output patterns. *eLife*, 7, 2018.

[4562] Joshua S MacCready, Pusparanee Hakim, Eric J Young, Longhua Hu, Jian Liu, Katherine W Osteryoung, Anthony G Vecchiarelli, and Daniel C Ducat. Protein gradients on the nucleoid position the carbon-fixing organelles of cyanobacteria. *eLife*, 7, 2018.

[4563] David Macias, Andrew S Cowburn, Hortensia Torres-Torrelo, Patricia Ortega-Sáenz, José López-Barneo, and Randall S Johnson. HIF-2$\alpha$ is essential for carotid body development and function. *eLife*, 7, 2018.

[4564] Despoina Mademtzoglou, Yoko Asakura, Matthew J Borok, Sonia Alonso-Martin, Philippos Mourikis, Yusaku Kodaka, Amrudha Mohan, Atsushi Asakura, and Frederic Relaix. Cellular localization of the cell cycle inhibitor Cdkn1c controls growth arrest of adult skeletal muscle stem cells. *eLife*, 7, 2018.

[4565] Britta Mahlert, Hanno Gerritsmann, Gabrielle Stalder, Thomas Ruf, Alexandre Zahariev, Stéphane Blanc, and Sylvain Giroud. Implications of being born late in the active season for growth, fattening, torpor use, winter survival and fecundity. *eLife*, 7, 2018.

[4566] Gita Mahmoudabadi and Rob Phillips. A comprehensive and quantitative exploration of thousands of viral genomes. *eLife*, 7, 2018.

[4567] Nguyen TH Mai, Nicholas Dobbs, Nguyen Hoan Phu, Romain A Colas, Le TP Thao, Nguyen TT Thuong, Ho DT Nghia, Nguyen HH Hanh, Nguyen T Hang, A Dorothee Heemskerk, Jeremy N Day, Lucy Ly, Do DA Thu, Laura Merson, Evelyne Kestelyn, Marcel Wolbers, Ronald Geskus, David Summers, Nguyen VV Chau, Jesmond Dalli, and Guy E Thwaites. A randomised double blind placebo controlled phase 2 trial of adjunctive aspirin for tuberculous meningitis in HIV-uninfected adults. *eLife*, 7, 2018.

[4568] Homa Majd, Martin S King, Shane M Palmer, Anthony C Smith, Liam DH Elbourne, Ian T Paulsen, David Sharples, Peter JF Henderson, and Edmund RS Kunji. Screening of candidate substrates and coupling ions of transporters by thermostability shift assays. *eLife*, 7, 2018.

[4569] Kinjal Majumder, Juexin Wang, Maria Boftsi, Matthew S Fuller, Jordan E Rede, Trupti Joshi, and David J Pintel. Parvovirus minute virus of mice interacts with sites of cellular DNA damage to establish and amplify its lytic infection. *eLife*, 7, 2018.

[4570] Rupamanjari Majumder, Iolanda Feola, Alexander S Teplenin, Antoine AF de Vries, Alexander V Panfilov, and Daniel A Pijnappels. Optogenetics enables real-time spatiotemporal control over spiral wave dynamics in an excitable cardiac system. *eLife*, 7, 2018.

[4571] Catherine A Makarewich, Amir Z Munir, Gabriele G Schiattarella, Svetlana Bezprozvannaya, Olga N Raguimova, Ellen E Cho, Alexander H Vidal, Seth L Robia, Rhonda Bassel-Duby, and Eric N Olson. The DWORF micropeptide enhances contractility and prevents heart failure in a mouse model of dilated cardiomyopathy. *eLife*, 7, 2018.

[4572] Nodar Makharashvili, Sucheta Arora, Yizhi Yin, Qiong Fu, Xuemei Wen, Ji-Hoon Lee, Chung-Hsuan Kao, Justin WC Leung, Kyle M Miller, and Tanya T Paull. Sae2/CtIP prevents R-loop accumulation in eukaryotic cells. *eLife*, 7, 2018.

[4573] Saori Maki-Yonekura, Rei Matsuoka, Yoshiki Yamashita, Hirofumi Shimizu, Maiko Tanaka, Fumie Iwabuki, and Koji Yonekura. Hexameric and pentameric complexes of the ExbBD energizer in the Ton system. *eLife*, 7, 2018.

[4574] Mazen Makke, Maria Mantero Martinez, Surya Gaya, Yvonne Schwarz, Walentina Frisch, Lina Silva-Bermudez, Martin Jung, Ralf Mohrmann, Madhurima Dhara, and Dieter Bruns. A mechanism for exocytotic arrest by the Complexin C-terminus. *eLife*, 7, 2018.

[4575] Divyanshu Malhotra, Jimann Shin, Lilianna Solnica-Krezel, and Erez Raz. Spatio-temporal regulation of concurrent developmental processes by generic signaling downstream of chemokine receptors. *eLife*, 7, 2018.

[4576] Girish R Mali, Patricia L Yeyati, Seiya Mizuno, Daniel O Dodd, Peter A Tennant, Margaret A Keighren, Petra zur Lage, Amelia Shoemark, Amaya Garcia-Munoz, Atsuko Shimada, Hiroyuki Takeda, Frank Edlich, Satoru Takahashi, Alex von Kreigsheim, Andrew P Jarman, and Pleasantine Mill. ZMYND10 functions in a chaperone relay during axonemal dynein assembly. *eLife*, 7, 2018.

[4577] Ahmed M Malik, Roberto A Miguez, Xingli Li, Ye-Shih Ho, Eva L Feldman, and Sami J Barmada. Matrin 3-dependent neurotoxicity is modified by nucleic acid binding and nucleocytoplasmic localization. *eLife*, 7, 2018.

[4578] Donna L Mallery, Chantal L Márquez, William A McEwan, Claire F Dickson, David A Jacques, Madhanagopal Anandapadamanaban, Katsiaryna Bichel, Gregory J Towers, Adolfo Saiardi, Till Böcking, and Leo C James. IP6 is an HIV pocket factor that prevents capsid collapse and promotes DNA synthesis. *eLife*, 7, 2018.

[4579] Anna Malolepszy, Simon Kelly, Kasper Kildegaard Sørensen, Euan Kevin James, Christina Kalisch, Zoltan Bozsoki, Michael Panting, Stig U Andersen, Shusei Sato, Ke Tao, Dorthe Bødker Jensen, Maria Vinther, Noor de Jong, Lene Heegaard Madsen, Yosuke Umehara, Kira Gysel, Mette U Berentsen, Mickael Blaise, Knud Jørgen Jensen, Mikkel B Thygesen, Niels Sandal, Kasper Røjkjær Andersen, and Simona Radutoiu. A plant chitinase controls cortical infection thread progression and nitrogen-fixing symbiosis. *eLife*, 7, 2018.

[4580] Marta Maltese, Jennifer Stanic, Annalisa Tassone, Giuseppe Sciamanna, Giulia Ponterio, Valentina Vanni, Giuseppina Martella, Paola Imbriani, Paola Bonsi, Nicola Biagio Mercuri, Fabrizio Gardoni, and Antonio Pisani. Early structural and functional plasticity alterations in a susceptibility period of DYT1 dystonia mouse striatum. *eLife*, 7, 2018.

[4581] Nathalie Mandairon, Nicola Kuczewski, Florence Kermen, Jérémy Forest, Maellie Midroit, Marion Richard, Marc Thevenet, Joelle Sacquet, Christiane Linster, and Anne Didier. Opposite regulation of inhibition by adult-born granule cells during implicit versus explicit olfactory learning. *eLife*, 7, 2018.

[4582] Dipak Manna, Christian Stephan Lentz, Gretchen Marie Ehrenkaufer, Susmitha Suresh, Amrita Bhat, and Upinder Singh. An NAD+-dependent novel transcription factor controls stage conversion in Entamoeba. *eLife*, 7, 2018.

[4583] Marco M Manni, Marion L Tiberti, Sophie Pagnotta, Hélène Barelli, Romain Gautier, and Bruno Antonny. Acyl chain asymmetry and polyunsaturation of brain phospholipids facilitate membrane vesiculation without leakage. *eLife*, 7, 2018.

[4584] Kelly A Manthei, Shyh-Ming Yang, Bolormaa Baljinnyam, Louise Chang, Alisa Glukhova, Wenmin Yuan, Lita A Freeman, David J Maloney, Anna Schwendeman, Alan T Remaley, Ajit Jadhav, and John JG Tesmer. Molecular basis for activation of lecithin:cholesterol acyltransferase by a compound that increases HDL cholesterol. *eLife*, 7, 2018.

[4585] Chantal L Márquez, Derrick Lau, James Walsh, Vaibhav Shah, Conall McGuinness, Andrew Wong, Anupriya Aggarwal, Michael W Parker, David A Jacques, Stuart Turville, and Till Böcking. Kinetics of HIV-1 capsid uncoating revealed by single-molecule analysis. *eLife*, 7, 2018.

[4586] Rogier B Mars, Stamatios N Sotiropoulos, Richard E Passingham, Jerome Sallet, Lennart Verhagen, Alexandre A Khrapitchev, Nicola Sibson, and Saad Jbabdi. Whole brain comparative anatomy using connectivity blueprints. *eLife*, 7, 2018.

[4587] Richard S Marshall and Richard D Vierstra. RETRACTED: Proteasome storage granules protect proteasomes from autophagic degradation upon carbon starvation. *eLife*, 7, 2018.

[4588] Chris B Martin, Danielle Douglas, Rachel N Newsome, Louisa LY Man, and Morgan D Barense. Integrative and distinctive coding of visual and conceptual object features in the ventral visual stream. *eLife*, 7, 2018.

[4589] Judy Lisette Martin, Erin Nicole Sanders, Paola Moreno-Roman, Leslie Ann Jaramillo Koyama, Shruthi Balachandra, XinXin Du, and Lucy Erin O'Brien. Long-term live imaging of the Drosophila adult midgut reveals real-time dynamics of division, differentiation and loss. *eLife*, 7, 2018.

[4590] Maud Martin, Alexandra Veloso, Jingchao Wu, Eugene A Katrukha, and Anna Akhmanova. Control of endothelial cell polarity and sprouting angiogenesis by non-centrosomal microtubules. *eLife*, 7, 2018.

[4591] éric Martineau, Adriana Di Polo, Christine Vande Velde, and Richard Robitaille. Dynamic neuromuscular remodeling precedes motor-unit loss in a mouse model of ALS. *eLife*, 7, 2018.

[4592] Inés Martínez, Maria X Maldonado-Gomez, João Carlos Gomes-Neto, Hatem Kittana, Hua Ding, Robert Schmaltz, Payal Joglekar, Roberto Jiménez Cardona, Nathan L Marsteller, Steven W Kembel, Andrew K Benson, Daniel A Peterson, Amanda E Ramer-Tait, and Jens Walter. Experimental evaluation of the importance of colonization history in early-life gut microbiota assembly. *eLife*, 7, 2018.

[4593] Neus Martínez-Abadías, Roger Mateu Estivill, Jaume Sastre Tomas, Susan Motch Perrine, Melissa Yoon, Alexandre Robert-Moreno, Jim Swoger, Lucia Russo, Kazuhiko Kawasaki, Joan Richtsmeier, and James Sharpe. Quantification of gene expression patterns to reveal the origins of abnormal morphogenesis. *eLife*, 7, 2018.

[4594] Juan Ramón Martínez-FranÇois, María Carmen Fernández-Agüera, Nidhi Nathwani, Carolina Lahmann, Veronica L Burnham, Nika N Danial, and Gary Yellen. BAD and KATP channels regulate neuron excitability and epileptiform activity. *eLife*, 7, 2018.

[4595] María de Lourdes Martínez-Silva, Rebecca D Imhoff-Manuel, Aarti Sharma, CJ Heckman, Neil A Shneider, Francesco Roselli, Daniel Zytnicki, and Marin Manuel. Hypoexcitability precedes denervation in the large fast-contracting motor units in two unrelated mouse models of ALS. *eLife*, 7, 2018.

[4596] Alexander J Martinko, Charles Truillet, Olivier Julien, Juan E Diaz, Max A Horlbeck, Gordon Whiteley, Josip Blonder, Jonathan S Weissman, Sourav Bandyopadhyay, Michael J Evans, and James A Wells. Targeting RAS-driven human cancer cells with antibodies to upregulated and essential cell-surface proteins. *eLife*, 7, 2018.

[4597] Tatsuro Maruyama, Shunsuke Imai, Tsukasa Kusakizako, Motoyuki Hattori, Ryuichiro Ishitani, Osamu Nureki, Koichi Ito, Andrès D Maturana, Ichio Shimada, and Masanori Osawa. Functional roles of Mg2+ binding sites in ion-dependent gating of a Mg2+ channel, MgtE, revealed by solution NMR. *eLife*, 7, 2018.

[4598] Dario Maschi, Michael W Gramlich, and Vitaly A Klyachko. Myosin V functions as a vesicle tether at the plasma membrane to control neurotransmitter release in central synapses. *eLife*, 7, 2018.

[4599] Ali Mashhoori, Saeedeh Hashemnia, Bruce L McNaughton, David R Euston, and Aaron J Gruber. Rat anterior cingulate cortex recalls features of remote reward locations after disfavoured reinforcements. *eLife*, 7, 2018.

[4600] Yasuhiro Matsunaga, Tsutomu Yamane, Tohru Terada, Kei Moritsugu, Hiroshi Fujisaki, Satoshi Murakami, Mitsunori Ikeguchi, and Akinori Kidera. Energetics and conformational pathways of functional rotation in the multidrug transporter AcrB. *eLife*, 7, 2018.

[4601] Benjamin Mattes, Yonglong Dang, Gediminas Greicius, Lilian Tamara Kaufmann, Benedikt Prunsche, Jakob Rosenbauer, Johannes Stegmaier, Ralf Mikut, Suat özbek, Gerd Ulrich Nienhaus, Alexander Schug, David M Virshup, and Steffen Scholpp. Wnt/PCP controls spreading of Wnt/$\beta$-catenin signals by cytonemes in vertebrates. *eLife*, 7, 2018.

[4602] Bryan J Matthews and David J Waxman. Computational prediction of CTCF/cohesin-based intra-TAD loops that insulate chromatin contacts and gene expression in mouse liver. *eLife*, 7, 2018.

[4603] Doreen Matthies, Chanhyung Bae, Gilman ES Toombes, Tara Fox, Alberto Bartesaghi, Sriram Subramaniam, and Kenton Jon Swartz. Single-particle cryo-EM structure of a voltage-activated potassium channel in lipid nanodiscs. *eLife*, 7, 2018.

[4604] Michelle E Maxson, Xenia Naj, Teresa R O'Meara, Jonathan D Plumb, Leah E Cowen, and Sergio Grinstein. Integrin-based diffusion barrier separates membrane domains enabling the formation of microbiostatic frustrated phagosomes. *eLife*, 7, 2018.

[4605] Douglas Maya Miles, Xenia Peñate, Trinidad Sanmartín Olmo, Frederic Jourquin, Maria Cruz Muñoz Centeno, Manuel Mendoza, Marie-Noelle Simon, Sebastian Chavez, and Vincent Geli. High levels of histones promote whole-genome-duplications and trigger a Swe1WEE1-dependent phosphorylation of Cdc28CDK1. *eLife*, 7, 2018.

[4606] Kristen R Maynard, John W Hobbs, BaDoi N Phan, Amolika Gupta, Sumita Rajpurohit, Courtney Williams, Anandita Rajpurohit, Joo Heon Shin, Andrew E Jaffe, and Keri Martinowich. BDNF-TrkB signaling in oxytocin neurons contributes to maternal behavior. *eLife*, 7, 2018.

[4607] Mark A McClintock, Carly I Dix, Christopher M Johnson, Stephen H McLaughlin, Rory J Maizels, Ha Thi Hoang, and Simon L Bullock. RNA-directed activation of cytoplasmic dynein-1 in reconstituted transport RNPs. *eLife*, 7, 2018.

[4608] Sean C McConnell, Erica L Westerman, Joseph F Pierre, Erin J Heckler, and Nancy B Schwartz. United States National Postdoc Survey results and the interaction of gender, career choice and mentor impact. *eLife*, 7, 2018.

[4609] John T McCrone, Robert J Woods, Emily T Martin, Ryan E Malosh, Arnold S Monto, and Adam S Lauring. Stochastic processes constrain the within and between host evolution of influenza virus. *eLife*, 7, 2018.

[4610] Kara L McKinley, Nico Stuurman, Loic A Royer, Christoph Schartner, David Castillo-Azofeifa, Markus Delling, Ophir D Klein, and Ronald D Vale. Cellular aspect ratio and cell division mechanics underlie the patterning of cell progeny in diverse mammalian epithelia. *eLife*, 7, 2018.

[4611] Gian-Luca McLelland, Thomas Goiran, Wei Yi, Geneviève Dorval, Carol X Chen, Nadine D Lauinger, Andrea I Krahn, Sepideh Valimehr, Aleksandar Rakovic, Isabelle Rouiller, Thomas M Durcan, Jean-FranÇois Trempe, and Edward A Fon. Mfn2 ubiquitination by PINK1/parkin gates the p97-dependent release of ER from mitochondria to drive mitophagy. *eLife*, 7, 2018.

[4612] Daniel Medina-Cano, Ekin Ucuncu, Lam Son Nguyen, Michael Nicouleau, Joanna Lipecka, Jean-Charles Bizot, Christian Thiel, FranÇois Foulquier, Nathalie Lefort, Catherine Faivre-Sarrailh, Laurence Colleaux, Ida Chiara Guerrera, and Vincent Cantagrel. High N-glycan multiplicity is critical for neuronal adhesion and sensitizes the developing cerebellum to N-glycosylation defect. *eLife*, 7, 2018.

[4613] Margarita V Meer, Dmitriy I Podolskiy, Alexander Tyshkovskiy, and Vadim N Gladyshev. A whole lifespan mouse multi-tissue DNA methylation clock. *eLife*, 7, 2018.

[4614] Colin D Meiklejohn, Emily L Landeen, Kathleen E Gordon, Thomas Rzatkiewicz, Sarah B Kingan, Anthony J Geneva, Jeffrey P Vedanayagam, Christina A Muirhead, Daniel Garrigan, David L Stern, and Daven C Presgraves. Gene flow mediates the role of sex chromosome meiotic drive during complex speciation. *eLife*, 7, 2018.

[4615] Anat Melamed, Hiroko Yaguchi, Michi Miura, Aviva Witkover, Tomas W Fitzgerald, Ewan Birney, and Charles RM Bangham. The human leukemia virus HTLV-1 alters the structure and transcription of host chromatin in cis. *eLife*, 7, 2018.

[4616] Luca Melli, Neil Billington, Sara A Sun, Jonathan E Bird, Attila Nagy, Thomas B Friedman, Yasuharu Takagi, and James R Sellers. Bipolar filaments of human nonmuscle myosin 2-A and 2-B have distinct motile and mechanical properties. *eLife*, 7, 2018.

[4617] Richard G Melvin, Nicole Lamichane, Essi Havula, Krista Kokki, Charles Soeder, Corbin D Jones, and Ville Hietakangas. Natural variation in sugar tolerance associates with changes in signaling and mitochondrial ribosome biogenesis. *eLife*, 7, 2018.

[4618] Edvin Memet, Feodor Hilitski, Margaret A Morris, Walter J Schwenger, Zvonimir Dogic, and L Mahadevan. Microtubules soften due to cross-sectional flattening. *eLife*, 7, 2018.

[4619] Fábio K Mendes, Jesualdo A Fuentes-González, Joshua G Schraiber, and Matthew W Hahn. A multispecies coalescent model for quantitative traits. *eLife*, 7, 2018.

[4620] Sam A Menzies, Norbert Volkmar, Dick JH van den Boomen, Richard T Timms, Anna S Dickson, James A Nathan, and Paul J Lehner. The sterol-responsive RNF145 E3 ubiquitin ligase mediates the degradation of HMG-CoA reductase together with gp78 and Hrd1. *eLife*, 7, 2018.

[4621] Matthias Merker, Maxime Barbier, Helen Cox, Jean-Philippe Rasigade, Silke Feuerriegel, Thomas Andreas Kohl, Roland Diel, Sonia Borrell, Sebastien Gagneux, Vladyslav Nikolayevskyy, Sönke Andres, Ulrich Nübel, Philip Supply, Thierry Wirth, and Stefan Niemann. Compensatory evolution drives multidrug-resistant tuberculosis in Central Asia. *eLife*, 7, 2018.

[4622] Jessica E Messier, Hongmei Chen, Zhao-Lin Cai, and Mingshan Xue. Targeting light-gated chloride channels to neuronal somatodendritic domain reduces their excitatory effect in the axon. *eLife*, 7, 2018.

[4623] Humberto Mestre, Lauren M Hablitz, Anna LR Xavier, Weixi Feng, Wenyan Zou, Tinglin Pu, Hiromu Monai, Giridhar Murlidharan, Ruth M Castellanos Rivera, Matthew J Simon, Martin M Pike, Virginia Plá, Ting Du, Benjamin T Kress, Xiaowen Wang, Benjamin A Plog, Alexander S Thrane, Iben Lundgaard, Yoichiro Abe, Masato Yasui, John H Thomas, Ming Xiao, Hajime Hirase, Aravind Asokan, Jeffrey J Iliff, and Maiken Nedergaard. Aquaporin-4-dependent glymphatic solute transport in the rodent brain. *eLife*, 7, 2018.

[4624] Jozsef Meszaros, Timothy Cheung, Maya M Erler, Un Jung Kang, Dalibor Sames, Christoph Kellendonk, and David Sulzer. Evoked transients of pH-sensitive fluorescent false neurotransmitter reveal dopamine hot spots in the globus pallidus. *eLife*, 7, 2018.

[4625] Travis Meyer and Nicole C Rust. Single-exposure visual memory judgments are reflected in inferotemporal cortex. *eLife*, 7, 2018.

[4626] Fernando Meza Gutierrez, Deniz Simsek, Arda Mizrak, Adam Deutschbauer, Hannes Braberg, Jeffrey Johnson, Jiewei Xu, Michael Shales, Michelle Nguyen, Raquel Tamse-Kuehn, Curt Palm, Lars M Steinmetz, Nevan J Krogan, and David P Toczyski. Genetic analysis reveals functions of atypical polyubiquitin chains. *eLife*, 7, 2018.

[4627] Nicholas Mikolajewicz, Elizabeth A Zimmermann, Bettina M Willie, and Svetlana V Komarova. Mechanically stimulated ATP release from murine bone cells is regulated by a balance of injury and repair. *eLife*, 7, 2018.

[4628] Meredith C Miles, Franz Goller, and Matthew J Fuxjager. Physiological constraint on acrobatic courtship behavior underlies rapid sympatric speciation in bearded manakins. *eLife*, 7, 2018.

[4629] Timothy F Miles, Katja Spiess, Kevin M Jude, Naotaka Tsutsumi, John S Burg, Jessica R Ingram, Deepa Waghray, Gertrud M Hjorto, Olav Larsen, Hidde L Ploegh, Mette M Rosenkilde, and K Christopher Garcia. Viral GPCR US28 can signal in response to chemokine agonists of nearly unlimited structural degeneracy. *eLife*, 7, 2018.

[4630] Rachel Millin, Tamar Kolodny, Anastasia V Flevaris, Alexander M Kale, Michael-Paul Schallmo, Jennifer Gerdts, Raphael A Bernier, and Scott Murray. Reduced auditory cortical adaptation in autism spectrum disorder. *eLife*, 7, 2018.

[4631] Daniel B Mills, Warren R Francis, Sergio Vargas, Morten Larsen, Coen PH Elemans, Donald E Canfield, and Gert Wörheide. The last common ancestor of animals lacked the HIF pathway and respired in low-oxygen environments. *eLife*, 7, 2018.

[4632] Gurdeep S Minhas, Daniel Bawdon, Reyme Herman, Michelle Rudden, Andrew P Stone, A Gordon James, Gavin H Thomas, and Simon Newstead. Structural basis of malodour precursor transport in the human axilla. *eLife*, 7, 2018.

[4633] Mustafa Mir, Michael R Stadler, Stephan A Ortiz, Colleen E Hannon, Melissa M Harrison, Xavier Darzacq, and Michael B Eisen. Dynamic multifactor hubs interact transiently with sites of active transcription in Drosophila embryos. *eLife*, 7, 2018.

[4634] Pablo Miranda, Miguel Holmgren, and Teresa Giraldez. Voltage-dependent dynamics of the BK channel cytosolic gating ring are coupled to the membrane-embedded voltage sensor. *eLife*, 7, 2018.

[4635] Hilda Mirbaha, Dailu Chen, Olga A Morazova, Kiersten M Ruff, Apurwa M Sharma, Xiaohua Liu, Mohammad Goodarzi, Rohit V Pappu, David W Colby, Hamid Mirzaei, Lukasz A Joachimiak, and Marc I Diamond. Inert and seed-competent tau monomers suggest structural origins of aggregation. *eLife*, 7, 2018.

[4636] Mitra L Miri, Martin Vinck, Rima Pant, and Jessica A Cardin. Altered hippocampal interneuron activity precedes ictal onset. *eLife*, 7, 2018.

[4637] Anton Miroschnikow, Philipp Schlegel, Andreas Schoofs, Sebastian Hueckesfeld, Feng Li, Casey M Schneider-Mizell, Richard D Fetter, James W Truman, Albert Cardona, and Michael J Pankratz. Convergence of monosynaptic and polysynaptic sensory paths onto common motor outputs in a Drosophila feeding connectome. *eLife*, 7, 2018.

[4638] Nathaniel J Miska, Leonidas MA Richter, Brian A Cary, Julijana Gjorgjieva, and Gina G Turrigiano. Sensory experience inversely regulates feedforward and feedback excitation-inhibition ratio in rodent visual cortex. *eLife*, 7, 2018.

[4639] Paolo Mita, Aleksandra Wudzinska, Xiaoji Sun, Joshua Andrade, Shruti Nayak, David J Kahler, Sana Badri, John LaCava, Beatrix Ueberheide, Chi Y Yun, David Fenyö, and Jef D Boeke. LINE-1 protein localization and functional dynamics during the cell cycle. *eLife*, 7, 2018.

[4640] Diana E Mitchell, Annie Kwan, Jerome Carriot, Maurice J Chacron, and Kathleen E Cullen. Neuronal variability and tuning are balanced to optimize naturalistic self-motion coding in primate vestibular pathways. *eLife*, 7, 2018.

[4641] Yasushi Miura, Satoshi Matsui, Naoko Miyata, Kenichi Harada, Yamato Kikkawa, Masaki Ohmuraya, Kimi Araki, Shinya Tsurusaki, Hitoshi Okochi, Nobuhito Goda, Atsushi Miyajima, and Minoru Tanaka. Differential expression of Lutheran/BCAM regulates biliary tissue remodeling in ductular reaction during liver regeneration. *eLife*, 7, 2018.

[4642] Andrew T Moehlman, Amanda K Casey, Kelly Servage, Kim Orth, and Helmut Krämer. Adaptation to constant light requires Fic-mediated AMPylation of BiP to protect against reversible photoreceptor degeneration. *eLife*, 7, 2018.

[4643] Francie Moehring, Ashley M Cowie, Anthony D Menzel, Andy D Weyer, Michael Grzybowski, Thiago Arzua, Aron M Geurts, Oleg Palygin, and Cheryl L Stucky. Keratinocytes mediate innocuous and noxious touch via ATP-P2X4 signaling. *eLife*, 7, 2018.

[4644] Chitra Mohan, Lisa M Kim, Nicole Hollar, Tailai Li, Eric Paulissen, Cheuk T Leung, and Ed Luk. VivosX, a disulfide crosslinking method to capture site-specific, protein-protein interactions in yeast and human cells. *eLife*, 7, 2018.

[4645] Yalda Mohsenzadeh, Sheng Qin, Radoslaw M Cichy, and Dimitrios Pantazis. Ultra-Rapid serial visual presentation reveals dynamics of feedforward and feedback processes in the ventral visual pathway. *eLife*, 7, 2018.

[4646] Lorna Moll, Noa Roitenberg, Michal Bejerano-Sagie, Hana Boocholez, Filipa Carvalhal Marques, Yuli Volovik, Tayir Elami, Atif Ahmed Siddiqui, Danielle Grushko, Adi Biram, Bar Lampert, Hana Achache, Tommer Ravid, Yonatan B Tzur, and Ehud Cohen. The insulin/IGF signaling cascade modulates SUMOylation to regulate aging and proteostasis in Caenorhabditis elegans. *eLife*, 7, 2018.

[4647] Ida Momennejad, A Ross Otto, Nathaniel D Daw, and Kenneth A Norman. Offline replay supports planning in human reinforcement learning. *eLife*, 7, 2018.

[4648] Aboozar Monavarfeshani, Gail Stanton, Jonathan Van Name, Kaiwen Su, William A Mills, Kenya Swilling, Alicia Kerr, Natalie A Huebschman, Jianmin Su, and Michael A Fox. LRRTM1 underlies synaptic convergence in visual thalamus. *eLife*, 7, 2018.

[4649] Marie Monniaux, Bjorn Pieper, Sarah M McKim, Anne-Lise Routier-Kierzkowska, Daniel Kierzkowski, Richard S Smith, and Angela Hay. The role of APETALA1 in petal number robustness. *eLife*, 7, 2018.

[4650] J Grey Monroe, Tyler Powell, Nicholas Price, Jack L Mullen, Anne Howard, Kyle Evans, John T Lovell, and John K McKay. Drought adaptation in Arabidopsis thaliana by extensive genetic loss-of-function. *eLife*, 7, 2018.

[4651] Lindsey E Montefiori, Debora R Sobreira, Noboru J Sakabe, Ivy Aneas, Amelia C Joslin, Grace T Hansen, Grazyna Bozek, Ivan P Moskowitz, Elizabeth M McNally, and Marcelo A Nóbrega. A promoter interaction map for cardiovascular disease genetics. *eLife*, 7, 2018.

[4652] Anna R Moore, Sarah E Richards, Katelyn Kenny, Leandro Royer, Urann Chan, Kelly Flavahan, Stephen D Van Hooser, and Suzanne Paradis. Rem2 stabilizes intrinsic excitability and spontaneous firing in visual circuits. *eLife*, 7, 2018.

[4653] Michael J Moore, Nathalie E Blachere, John J Fak, Christopher Y Park, Kirsty Sawicka, Salina Parveen, Ilana Zucker-Scharff, Bruno Moltedo, Alexander Y Rudensky, and Robert B Darnell. ZFP36 RNA-binding proteins restrain T cell activation and anti-viral immunity. *eLife*, 7, 2018.

[4654] Federico Andrea Moretti, Sarah Klapproth, Raphael Ruppert, Andreas Margraf, Jasmin Weber, Robert Pick, Christoph Scheiermann, Markus Sperandio, Reinhard Fässler, and Markus Moser. Differential requirement of kindlin-3 for T cell progenitor homing to the non-vascularized and vascularized thymus. *eLife*, 7, 2018.

[4655] Sophie M Morgani, Jakob J Metzger, Jennifer Nichols, Eric D Siggia, and Anna-Katerina Hadjantonakis. Micropattern differentiation of mouse pluripotent stem cells recapitulates embryo regionalized cell fate patterning. *eLife*, 7, 2018.

[4656] Ekaterina Morgunova, Yimeng Yin, Pratyush K Das, Arttu Jolma, Fangjie Zhu, Alexander Popov, You Xu, Lennart Nilsson, and Jussi Taipale. Two distinct DNA sequences recognized by transcription factors represent enthalpy and entropy optima. *eLife*, 7, 2018.

[4657] Juan Facundo Morici, Magdalena Miranda, Francisco Tomás Gallo, Belén Zanoni, Pedro Bekinschtein, and Noelia V Weisstaub. 5-HT2a receptor in mPFC influences context-guided reconsolidation of object memory in perirhinal cortex. *eLife*, 7, 2018.

[4658] Manon Morin, Emily C Pierce, and Rachel J Dutton. Changes in the genetic requirements for microbial interactions with increasing community complexity. *eLife*, 7, 2018.

[4659] Emma A Morrison, Samuel Bowerman, Kelli L Sylvers, Jeff Wereszczynski, and Catherine A Musselman. The conformation of the histone H3 tail inhibits association of the BPTF PHD finger with the nucleosome. *eLife*, 7, 2018.

[4660] Meghan A Morrissey, Adam P Williamson, Adriana M Steinbach, Edward W Roberts, Nadja Kern, Mark B Headley, and Ronald D Vale. Chimeric antigen receptors that trigger phagocytosis. *eLife*, 7, 2018.

[4661] Lucile Moynié, Ilaria Serra, Mariano A Scorciapino, Emilia Oueis, Malcolm GP Page, Matteo Ceccarelli, and James H Naismith. Preacinetobactin not acinetobactin is essential for iron uptake by the BauA transporter of the pathogen Acinetobacter baumannii. *eLife*, 7, 2018.

[4662] Didhiti Mukherjee, Greta Sokoloff, and Mark S Blumberg. Corollary discharge in precerebellar nuclei of sleeping infant rats. *eLife*, 7, 2018.

[4663] Diptendu Mukherjee, Bogna Marta Ignatowska-Jankowska, Eyal Itskovits, Ben Jerry Gonzales, Hagit Turm, Liz Izakson, Doron Haritan, Noa Bleistein, Chen Cohen, Ido Amit, Tal Shay, Brad Grueter, Alon Zaslaver, and Ami Citri. Salient experiences are represented by unique transcriptional signatures in the mouse brain. *eLife*, 7, 2018.

[4664] Sri Teja Mullapudi, Christian SM Helker, Giulia LM Boezio, Hans-Martin Maischein, Anna M Sokol, Stefan Guenther, Hiroki Matsuda, Stefan Kubicek, Johannes Graumann, Yu Hsuan Carol Yang, and Didier YR Stainier. Screening for insulin-independent pathways that modulate glucose homeostasis identifies androgen receptor antagonists. *eLife*, 7, 2018.

[4665] Christine Müller, Laura M Zidek, Tobias Ackermann, Tristan de Jong, Peng Liu, Verena Kliche, Mohamad Amr Zaini, Gertrud Kortman, Liesbeth Harkema, Dineke S Verbeek, Jan P Tuckermann, Julia von Maltzahn, Alain de Bruin, Victor Guryev, Zhao-Qi Wang, and Cornelis F Calkhoven. Reduced expression of C/EBP$\beta$-LIP extends health and lifespan in mice. *eLife*, 7, 2018.

[4666] Erinn M Muller, Erich Bartels, and Iliana B Baums. Bleaching causes loss of disease resistance within the threatened coral species Acropora cervicornis. *eLife*, 7, 2018.

[4667] Susanne Müller, Suzanne Ackloo, Cheryl H Arrowsmith, Marcus Bauser, Jeremy L Baryza, Julian Blagg, Jark Böttcher, Chas Bountra, Peter J Brown, Mark E Bunnage, Adrian J Carter, David Damerell, Volker Dötsch, David H Drewry, Aled M Edwards, James Edwards, Jon M Elkins, Christian Fischer, Stephen V Frye, Andreas Gollner, Charles E Grimshaw, Adriaan IJzerman, Thomas Hanke, Ingo V Hartung, Steve Hitchcock, Trevor Howe, Terry V Hughes, Stefan Laufer, Volkhart MJ Li, Spiros Liras, Brian D Marsden, Hisanori Matsui, John Mathias, Ronan C O'Hagan, Dafydd R Owen, Vineet Pande, Daniel Rauh, Saul H Rosenberg, Bryan L Roth, Natalie S Schneider, Cora Scholten, Kumar Singh Saikatendu, Anton Simeonov, Masayuki Takizawa, Chris Tse, Paul R Thompson, Daniel K Treiber, Amélia YI Viana, Carrow I Wells, Timothy M Willson, William J Zuercher, Stefan Knapp, and Anke Mueller-Fahrnow. Donated chemical probes for open science. *eLife*, 7, 2018.

[4668] Alexander Munden, Zhan Rong, Amanda Sun, Rama Gangula, Simon Mallal, and Jared T Nordman. Rif1 inhibits replication fork progression and controls DNA copy number in Drosophila. *eLife*, 7, 2018.

[4669] Nadine Mundt, Marc Spehr, and Polina V Lishko. TRPV4 is the temperature-sensitive ion channel of human sperm. *eLife*, 7, 2018.

[4670] Teona Munteanu, Katelyn J Noronha, Amanda C Leung, Simon Pan, Jasmine A Lucas, and Tiffany M Schmidt. Light-dependent pathways for dopaminergic amacrine cell development and function. *eLife*, 7, 2018.

[4671] Eiichi Murakami, Jeryl Cheng, Kira Gysel, Zoltan Bozsoki, Yasuyuki Kawaharada, Christian Toftegaard Hjuler, Kasper Kildegaard Sørensen, Ke Tao, Simon Kelly, Francesco Venice, Andrea Genre, Mikkel Boas Thygesen, Noor de Jong, Maria Vinther, Dorthe Bødker Jensen, Knud Jørgen Jensen, Michael Blaise, Lene Heegaard Madsen, Kasper Røjkjær Andersen, Jens Stougaard, and Simona Radutoiu. Epidermal LysM receptor ensures robust symbiotic signalling in Lotus japonicus. *eLife*, 7, 2018.

[4672] Patrick A Murphy, Vincent L Butty, Paul L Boutz, Shahinoor Begum, Amy L Kimble, Phillip A Sharp, Christopher B Burge, and Richard O Hynes. Alternative RNA splicing in the endothelium mediated in part by Rbfox2 regulates the arterial response to low flow. *eLife*, 7, 2018.

[4673] Swetha E Murthy, Adrienne E Dubin, Tess Whitwam, Sebastian Jojoa-Cruz, Stuart M Cahalan, Seyed Ali Reza Mousavi, Andrew B Ward, and Ardem Patapoutian. OSCA/TMEM63 are an evolutionarily conserved family of mechanically activated ion channels. *eLife*, 7, 2018.

[4674] Hannes Mutschler, Alexander I Taylor, Benjamin T Porebski, Alice Lightowlers, Gillian Houlihan, Mikhail Abramov, Piet Herdewijn, and Philipp Holliger. Random-sequence genetic oligomer pools display an innate potential for ligation and recombination. *eLife*, 7, 2018.

[4675] Sumanth K Mutte, Hirotaka Kato, Carl Rothfels, Michael Melkonian, Gane Ka-Shu Wong, and Dolf Weijers. Origin and evolution of the nuclear auxin response system. *eLife*, 7, 2018.

[4676] Martha M Muñoz, Y Hu, Philip S L Anderson, and SN Patek. Strong biomechanical relationships bias the tempo and mode of morphological evolution. *eLife*, 7, 2018.

[4677] Wiktor F Młynarski and Ann M Hermundstad. Adaptive coding for dynamic sensory inference. *eLife*, 7, 2018.

[4678] Ryo Nabeshima, Osamu Nishimura, Takako Maeda, Natsumi Shimizu, Takahiro Ide, Kenta Yashiro, Yasuo Sakai, Chikara Meno, Mitsutaka Kadota, Hidetaka Shiratori, Shigehiro Kuraku, and Hiroshi Hamada. Loss of Fam60a, a Sin3a subunit, results in embryonic lethality and is associated with aberrant methylation at a subset of gene promoters. *eLife*, 7, 2018.

[4679] Sundar Ram Naganathan, Sebastian Fürthauer, Josana Rodriguez, Bruno Thomas Fievet, Frank Jülicher, Julie Ahringer, Carlo Vittorio Cannistraci, and Stephan W Grill. Morphogenetic degeneracies in the actomyosin cortex. *eLife*, 7, 2018.

[4680] Monica Nagendran, Daniel P Riordan, Pehr B Harbury, and Tushar J Desai. Automated cell-type classification in intact tissues by single-cell molecular profiling. *eLife*, 7, 2018.

[4681] Takanori Nakane, Dari Kimanius, Erik Lindahl, and Sjors HW Scheres. Characterisation of molecular motions in cryo-EM single-particle data by multi-body refinement in RELION. *eLife*, 7, 2018.

[4682] Nagayasu Nakanishi and Mark Q Martindale. CRISPR knockouts reveal an endogenous role for ancient neuropeptides in regulating developmental timing in a sea anemone. *eLife*, 7, 2018.

[4683] Shigehiro Namiki, Michael H Dickinson, Allan M Wong, Wyatt Korff, and Gwyneth M Card. The functional organization of descending sensory-motor pathways in Drosophila. *eLife*, 7, 2018.

[4684] Pratibha Narayanan, Meike Hütte, Galina Kudryasheva, Francisco J Taberner, Stefan G Lechner, Florian Rehfeldt, David Gomez-Varela, and Manuela Schmidt. Myotubularin related protein-2 and its phospholipid substrate PIP2 control Piezo2-mediated mechanotransduction in peripheral sensory neurons. *eLife*, 7, 2018.

[4685] Amr Nassrallah, Martin Rougée, Clara Bourbousse, Stephanie Drevensek, Sandra Fonseca, Elisa Iniesto, Ouardia Ait-Mohamed, Anne-Flore Deton-Cabanillas, Gerald Zabulon, Ikhlak Ahmed, David Stroebel, Vanessa Masson, Berangere Lombard, Dominique Eeckhout, Kris Gevaert, Damarys Loew, Auguste Genovesio, Cecile Breyton, Geert De Jaeger, Chris Bowler, Vicente Rubio, and Fredy Barneche. DET1-mediated degradation of a SAGA-like deubiquitination module controls H2Bub homeostasis. *eLife*, 7, 2018.

[4686] Christopher A Natale, Jinyang Li, Junqian Zhang, Ankit Dahal, Tzvete Dentchev, Ben Z Stanger, and Todd W Ridky. Activation of G protein-coupled estrogen receptor signaling inhibits melanoma and improves response to immune checkpoint blockade. *eLife*, 7, 2018.

[4687] Richard W Naylor, Hao-Han G Chang, Sarah Qubisi, and Alan J Davidson. A novel mechanism of gland formation in zebrafish involving transdifferentiation of renal epithelial cells and live cell extrusion. *eLife*, 7, 2018.

[4688] Filipa Neto, Alexandra Klaus-Bergmann, Yu Ting Ong, Silvanus Alt, Anne-Clémence Vion, Anna Szymborska, Joana R Carvalho, Irene Hollfinger, Eireen Bartels-Klein, Claudio A Franco, Michael Potente, and Holger Gerhardt. YAP and TAZ regulate adherens junction dynamics and endothelial cell distribution during vascular development. *eLife*, 7, 2018.

[4689] Andrew F Neuwald, L Aravind, and Stephen F Altschul. Inferring joint sequence-structural determinants of protein functional specificity. *eLife*, 7, 2018.

[4690] Kenneth KH Ng, Mary A Yui, Arnav Mehta, Sharmayne Siu, Blythe Irwin, Shirley Pease, Satoshi Hirose, Michael B Elowitz, Ellen V Rothenberg, and Hao Yuan Kueh. A stochastic epigenetic switch controls the dynamics of T-cell lineage commitment. *eLife*, 7, 2018.

[4691] Elizabeth Nicholson, Dmitry A Kuzmin, Marco Leite, Thomas E Akam, and Dimitri Michael Kullmann. Analogue closed-loop optogenetic modulation of hippocampal pyramidal cells dissociates gamma frequency and amplitude. *eLife*, 7, 2018.

[4692] Mante S Nieuwland, Stephen Politzer-Ahles, Evelien Heyselaar, Katrien Segaert, Emily Darley, Nina Kazanina, Sarah Von Grebmer Zu Wolfsthurn, Federica Bartolozzi, Vita Kogan, Aine Ito, Diane Mézière, Dale J Barr, Guillaume A Rousselet, Heather J Ferguson, Simon Busch-Moreno, Xiao Fu, Jyrki Tuomainen, Eugenia Kulakova, E Matthew Husband, David I Donaldson, Zdenko Kohút, Shirley-Ann Rueschemeyer, and Falk Huettig. Large-scale replication study reveals a limit on probabilistic prediction in language comprehension. *eLife*, 7, 2018.

[4693] Mathieu Niquille, Greta Limoni, Foivos Markopoulos, Christelle Cadilhac, Julien Prados, Anthony Holtmaat, and Alexandre Dayer. Neurogliaform cortical interneurons derive from cells in the preoptic area. *eLife*, 7, 2018.

[4694] Silas Boye Nissen, Steven Rønhild, Ala Trusina, and Kim Sneppen. Theoretical tool bridging cell polarities with development of robust morphologies. *eLife*, 7, 2018.

[4695] Stanley Nithianantham, Brian D Cook, Madeleine Beans, Fei Guo, Fred Chang, and Jawdat Al-Bassam. Structural basis of tubulin recruitment and assembly by microtubule polymerases with tumor overexpressed gene (TOG) domain arrays. *eLife*, 7, 2018.

[4696] Jacqueline Niu, Ivy E Dick, Wanjun Yang, Moradeke A Bamgboye, David T Yue, Gordon Tomaselli, Takanari Inoue, and Manu Ben-Johny. Allosteric regulators selectively prevent Ca2+-feedback of CaV and NaV channels. *eLife*, 7, 2018.

[4697] Alex J Noble, Venkata P Dandey, Hui Wei, Julia Brasch, Jillian Chase, Priyamvada Acharya, Yong Zi Tan, Zhening Zhang, Laura Y Kim, Giovanna Scapin, Micah Rapp, Edward T Eng, William J Rice, Anchi Cheng, Carl J Negro, Lawrence Shapiro, Peter D Kwong, David Jeruzalmi, Amedee des Georges, Clinton S Potter, and Bridget Carragher. Routine single particle CryoEM sample and grid characterization by tomography. *eLife*, 7, 2018.

[4698] Charleston Noble, Ben Adlam, George M Church, Kevin M Esvelt, and Martin A Nowak. Current CRISPR gene drive systems are likely to be highly invasive in wild populations. *eLife*, 7, 2018.

[4699] Veronique Nogueira, Krushna C Patra, and Nissim Hay. Selective eradication of cancer displaying hyperactive Akt by exploiting the metabolic consequences of Akt activation. *eLife*, 7, 2018.

[4700] Sina Ibne Noor, Somayeh Jamali, Samantha Ames, Silke Langer, Joachim W Deitmer, and Holger M Becker. A surface proton antenna in carbonic anhydrase II supports lactate transport in cancer cells. *eLife*, 7, 2018.

[4701] Agnes Norbury, Trevor W Robbins, and Ben Seymour. Value generalization in human avoidance learning. *eLife*, 7, 2018.

[4702] Sigrid Noreng, Arpita Bharadwaj, Richard Posert, Craig Yoshioka, and Isabelle Baconguis. Structure of the human epithelial sodium channel by cryo-electron microscopy. *eLife*, 7, 2018.

[4703] Marco Novais-Cruz, Maria Alba Abad, Wilfred FJ van IJcken, Niels Galjart, A Arockia Jeyaprakash, Helder Maiato, and Cristina Ferrás. Mitotic progression, arrest, exit or death relies on centromere structural integrity, rather than de novo transcription. *eLife*, 7, 2018.

[4704] Mark Noviski, James L Mueller, Anne Satterthwaite, Lee Ann Garrett-Sinha, Frank Brombacher, and Julie Zikherman. IgM and IgD B cell receptors differentially respond to endogenous antigens and control B cell fate. *eLife*, 7, 2018.

[4705] Harry Nunns and Lea Goentoro. Signaling pathways as linear transmitters. *eLife*, 7, 2018.

[4706] Ignacio Obeso, Marius Moisa, Christian C Ruff, and Jean-Claude Dreher. A causal role for right temporo-parietal junction in signaling moral conflict. *eLife*, 7, 2018.

[4707] Sean R O'Bryan, Darrell A Worthy, Evan J Livesey, and Tyler Davis. Model-based fMRI reveals dissimilarity processes underlying base rate neglect. *eLife*, 7, 2018.

[4708] SeCheol Oh and Olga Boudker. Kinetic mechanism of coupled binding in sodium-aspartate symporter GltPh. *eLife*, 7, 2018.

[4709] Seunghee Oh, Tamaki Suganuma, Madelaine M Gogol, and Jerry L Workman. Histone H3 threonine 11 phosphorylation by Sch9 and CK2 regulates chronological lifespan by controlling the nutritional stress response. *eLife*, 7, 2018.

[4710] Molly OhAinle, Louisa Helms, Jolien Vermeire, Ferdinand Roesch, Daryl Humes, Ryan Basom, Jeffrey J Delrow, Julie Overbaugh, and Michael Emerman. A virus-packageable CRISPR screen identifies host factors mediating interferon inhibition of HIV. *eLife*, 7, 2018.

[4711] Ioanna Oikonomidi, Emma Burbridge, Miguel Cavadas, Graeme Sullivan, Blanka Collis, Heike Naegele, Danielle Clancy, Jana Brezinova, Tianyi Hu, Andrea Bileck, Christopher Gerner, Alfonso Bolado, Alex von Kriegsheim, Seamus J Martin, Florian Steinberg, Kvido Strisovsky, and Colin Adrain. iTAP, a novel iRhom interactor, controls TNF secretion by policing the stability of iRhom/TACE. *eLife*, 7, 2018.

[4712] Masako Okumura, Toyoaki Natsume, Masato T Kanemaki, and Tomomi Kiyomitsu. Dynein–Dynactin–NuMA clusters generate cortical spindle-pulling forces as a multi-arm ensemble. *eLife*, 7, 2018.

[4713] Scott W Olesen, Michael L Barnett, Derek R MacFadden, John S Brownstein, Sonia Hernández-Díaz, Marc Lipsitch, and Yonatan H Grad. The distribution of antibiotic use and its association with antibiotic resistance. *eLife*, 7, 2018.

[4714] Safia Omer, Samuel R Greenberg, and Wei-Lih Lee. Cortical dynein pulling mechanism is regulated by differentially targeted attachment molecule Num1. *eLife*, 7, 2018.

[4715] D Herbert Opi, Olivia Swann, Alexander Macharia, Sophie Uyoga, Gavin Band, Carolyne M Ndila, Ewen M Harrison, Mahamadou A Thera, Abdoulaye K Kone, Dapa A Diallo, Ogobara K Doumbo, Kirsten E Lyke, Christopher V Plowe, Joann M Moulds, Mohammed Shebbe, Neema Mturi, Norbert Peshu, Kathryn Maitland, Ahmed Raza, Dominic P Kwiatkowski, Kirk A Rockett, Thomas N Williams, and J Alexandra Rowe. Two complement receptor one alleles have opposing associations with cerebral malaria and interact with $\alpha$+thalassaemia. *eLife*, 7, 2018.

[4716] Jennifer M Ortega, özgür GenÇ, and Graeme W Davis. Molecular mechanisms that stabilize short term synaptic plasticity during presynaptic homeostatic plasticity. *eLife*, 7, 2018.

[4717] Matthew CW Oswald, Paul S Brooks, Maarten F Zwart, Amrita Mukherjee, Ryan JH West, Carlo NG Giachello, Khomgrit Morarach, Richard A Baines, Sean T Sweeney, and Matthias Landgraf. Reactive oxygen species regulate activity-dependent neuronal plasticity in Drosophila. *eLife*, 7, 2018.

[4718] Jeannine A Ott, Caitlin D Castro, Thaddeus C Deiss, Yuko Ohta, Martin F Flajnik, and Michael F Criscitiello. Somatic hypermutation of T cell receptor $\alpha$ chain contributes to selection in nurse shark thymus. *eLife*, 7, 2018.

[4719] Willem M Otte, Joeri K Tijdink, Paul L Weerheim, Herm J Lamberink, and Christiaan H Vinkers. Adequate statistical power in clinical trials is associated with the combination of a male first author and a female last author. *eLife*, 7, 2018.

[4720] Victor Ovchinnikov, Joy E Louveau, John P Barton, Martin Karplus, and Arup K Chakraborty. Role of framework mutations and antibody flexibility in the evolution of broadly neutralizing antibodies. *eLife*, 7, 2018.

[4721] Muge Ozker, Daniel Yoshor, and Michael S Beauchamp. Frontal cortex selects representations of the talker's mouth to aid in speech perception. *eLife*, 7, 2018.

[4722] Christian Louis Bonatto Paese, Anna Schoenauer, Daniel J Leite, Steven Russell, and Alistair P McGregor. A SoxB gene acts as an anterior gap gene and regulates posterior segment addition in a spider. *eLife*, 7, 2018.

[4723] May M Paing, Nichole D Salinas, Yvonne Adams, Anna Oksman, Anja TR Jensen, Daniel E Goldberg, and Niraj H Tolia. Shed EBA-175 mediates red blood cell clustering that enhances malaria parasite growth and enables immune evasion. *eLife*, 7, 2018.

[4724] Aurélie Pala and Carl CH Petersen. State-dependent cell-type-specific membrane potential dynamics and unitary synaptic inputs in awake mice. *eLife*, 7, 2018.

[4725] Luca Palazzo, Orsolya Leidecker, Evgeniia Prokhorova, Helen Dauben, Ivan Matic, and Ivan Ahel. Serine is the major residue for ADP-ribosylation upon DNA damage. *eLife*, 7, 2018.

[4726] Marios C Panayi and Simon Killcross. Functional heterogeneity within the rodent lateral orbitofrontal cortex dissociates outcome devaluation and reversal learning deficits. *eLife*, 7, 2018.

[4727] Ariel M Pani and Bob Goldstein. Direct visualization of a native Wnt in vivo reveals that a long-range Wnt gradient forms by extracellular dispersal. *eLife*, 7, 2018.

[4728] Martina Pannuzzo, Zachary A McDargh, and Markus Deserno. The role of scaffold reshaping and disassembly in dynamin driven membrane fission. *eLife*, 7, 2018.

[4729] Samuel S Pappas, Jay Li, Tessa M LeWitt, Jeong-Ki Kim, Umrao R Monani, and William T Dauer. A cell autonomous torsinA requirement for cholinergic neuron survival and motor control. *eLife*, 7, 2018.

[4730] Kristian Parey, Ulrich Brandt, Hao Xie, Deryck J Mills, Karin Siegmund, Janet Vonck, Werner Kühlbrandt, and Volker Zickermann. Cryo-EM structure of respiratory complex I at work. *eLife*, 7, 2018.

[4731] Donghyun Park, Maria Lara-Tejero, M Neal Waxham, Wenwei Li, Bo Hu, Jorge E Galán, and Jun Liu. Visualization of the type III secretion mediated Salmonella–host cell interface using cryo-electron tomography. *eLife*, 7, 2018.

[4732] Eunyong Park and Roderick MacKinnon. Structure of the CLC-1 chloride channel from Homo sapiens. *eLife*, 7, 2018.

[4733] Robbie M Parks, James E Bennett, Kyle J Foreman, Ralf Toumi, and Majid Ezzati. National and regional seasonal dynamics of all-cause and cause-specific mortality in the USA from 1980 to 2016. *eLife*, 7, 2018.

[4734] Arielle Waldstein Parsons, Tavis Forrester, Megan C Baker-Whatton, William J McShea, Christopher T Rota, Stephanie G Schuttler, Joshua J Millspaugh, and Roland Kays. Mammal communities are larger and more diverse in moderately developed areas. *eLife*, 7, 2018.

[4735] Nick R Parsons, M Dawn Teare, and Alice J Sitch. Unit of analysis issues in laboratory-based research. *eLife*, 7, 2018.

[4736] Freda Passam, Joyce Chiu, Lining Ju, Aster Pijning, Zeenat Jahan, Ronit Mor-Cohen, Adva Yeheskel, Katra KolŠek, Lena Thärichen, Camilo Aponte-Santamaría, Frauke Gräter, and Philip J Hogg. Mechano-redox control of integrin de-adhesion. *eLife*, 7, 2018.

[4737] Shaun R Patel, Todd M Herrington, Sameer A Sheth, Matthew Mian, Sarah K Bick, Jimmy C Yang, Alice W Flaherty, Michael J Frank, Alik S Widge, Darin Dougherty, and Emad N Eskandar. Intermittent subthalamic nucleus deep brain stimulation induces risk-aversive behavior in human subjects. *eLife*, 7, 2018.

[4738] Arnaud Pautrat, Marta Rolland, Margaux Barthelemy, Christelle Baunez, Valérie Sinniger, Brigitte Piallat, Marc Savasta, Paul G Overton, Olivier David, and Veronique Coizet. Revealing a novel nociceptive network that links the subthalamic nucleus to pain processing. *eLife*, 7, 2018.

[4739] Ryan Paxman, Lars Plate, Erik A Blackwood, Chris Glembotski, Evan T Powers, R Luke Wiseman, and Jeffery W Kelly. Pharmacologic ATF6 activating compounds are metabolically activated to selectively modify endoplasmic reticulum proteins. *eLife*, 7, 2018.

[4740] Pavel Payne, Lukas Geyrhofer, Nicholas H Barton, and Jonathan P Bollback. CRISPR-based herd immunity can limit phage epidemics in bacterial populations. *eLife*, 7, 2018.

[4741] Craig S Pearson, Caitlin P Mencio, Amanda C Barber, Keith R Martin, and Herbert M Geller. Identification of a critical sulfation in chondroitin that inhibits axonal regeneration. *eLife*, 7, 2018.

[4742] Thierry Pécot, Liu Zengzhen, Jérôme Boulanger, Jean Salamero, and Charles Kervrann. A quantitative approach for analyzing the spatio-temporal distribution of 3D intracellular events in fluorescence microscopy. *eLife*, 7, 2018.

[4743] Sen Pei, Flaviano Morone, Fredrik Liljeros, Hernán Makse, and Jeffrey L Shaman. Inference and control of the nosocomial transmission of methicillin-resistant Staphylococcus aureus. *eLife*, 7, 2018.

[4744] Pirita Pekkonen, Sanni Alve, Giuseppe Balistreri, Silvia Gramolelli, Olga Tatti-Bugaeva, Ilkka Paatero, Otso Niiranen, Krista Tuohinto, Nina Perälä, Adewale Taiwo, Nadezhda Zinovkina, Pauliina Repo, Katherine Icay, Johanna Ivaska, Pipsa Saharinen, Sampsa Hautaniemi, Kaisa Lehti, and Päivi M Ojala. Lymphatic endothelium stimulates melanoma metastasis and invasion via MMP14-dependent Notch3 and $\beta$1-integrin activation. *eLife*, 7, 2018.

[4745] Claudia Pellacani, Elisabetta Bucciarelli, Fioranna Renda, Daniel Hayward, Antonella Palena, Jack Chen, Silvia Bonaccorsi, James G Wakefield, Maurizio Gatti, and Maria Patrizia Somma. Splicing factors Sf3A2 and Prp31 have direct roles in mitotic chromosome segregation. *eLife*, 7, 2018.

[4746] Corinne L Pender and H Robert Horvitz. Hypoxia-inducible factor cell non-autonomously regulates C. elegans stress responses and behavior via a nuclear receptor. *eLife*, 7, 2018.

[4747] Jing Peng, Ivan J Santiago, Curie Ahn, Burak Gur, C Kimberly Tsui, Zhixiao Su, Chundi Xu, Aziz Karakhanyan, Marion Silies, and Matthew Y Pecot. Drosophila Fezf coordinates laminar-specific connectivity through cell-intrinsic and cell-extrinsic mechanisms. *eLife*, 7, 2018.

[4748] Xi Peng, Francesco Emiliani, Philip M Smallwood, Amir Rattner, Hong Lei, Mark F Sabbagh, and Jeremy Nathans. Affinity capture of polyribosomes followed by RNAseq (ACAPseq), a discovery platform for protein-protein interactions. *eLife*, 7, 2018.

[4749] Damián Pérez-Mazliah, Peter J Gardner, Edina Schweighoffer, Sarah McLaughlin, Caroline Hosking, Irene Tumwine, Randall S Davis, Alexandre J Potocnik, Victor LJ Tybulewicz, and Jean Langhorne. Plasmodium-specific atypical memory B cells are short-lived activated B cells. *eLife*, 7, 2018.

[4750] Stefano Pernigo, Magda S Chegkazi, Yan Y Yip, Conor Treacy, Giulia Glorani, Kjetil Hansen, Argyris Politis, Soi Bui, Mark P Dodding, and Roberto A Steiner. Structural basis for isoform-specific kinesin-1 recognition of Y-acidic cargo adaptors. *eLife*, 7, 2018.

[4751] Laurent Perrin, Ursula Loizides-Mangold, Stéphanie Chanon, Cédric Gobet, Nicolas Hulo, Laura Isenegger, Benjamin D Weger, Eugenia Migliavacca, Aline Charpagne, James A Betts, Jean-Philippe Walhin, Iain Templeman, Keith Stokes, Dylan Thompson, Kostas Tsintzas, Maud Robert, Cedric Howald, Howard Riezman, Jerome N Feige, Leonidas G Karagounis, Jonathan D Johnston, Emmanouil T Dermitzakis, Frédéric Gachon, Etienne Lefai, and Charna Dibner. Transcriptomic analyses reveal rhythmic and CLOCK-driven pathways in human skeletal muscle. *eLife*, 7, 2018.

[4752] Riley Perszyk, Brooke M Katzman, Hirofumi Kusumoto, Steven A Kell, Matthew P Epplin, Yesim A Tahirovic, Rhonda L Moore, David Menaldino, Pieter Burger, Dennis C Liotta, and Stephen F Traynelis. An NMDAR positive and negative allosteric modulator series share a binding site and are interconverted by methyl groups. *eLife*, 7, 2018.

[4753] Ilaria Pettinati, Pawel Grzechnik, Claudia Ribeiro de Almeida, Jurgen Brem, Michael A McDonough, Somdutta Dhir, Nick J Proudfoot, and Christopher J Schofield. Biosynthesis of histone messenger RNA employs a specific $3'$ end endonuclease. *eLife*, 7, 2018.

[4754] Emma Pewsey. Part of the research family. *eLife*, 7, 2018.

[4755] Thomas Pfeiffer, Stefanie Poll, Stephane Bancelin, Julie Angibaud, VVG Krishna Inavalli, Kevin Keppler, Manuel Mittag, Martin Fuhrmann, and U Valentin Nägerl. Chronic 2P-STED imaging reveals high turnover of dendritic spines in the hippocampus in vivo. *eLife*, 7, 2018.

[4756] Alison Philbrook, Shankar Ramachandran, Christopher M Lambert, Devyn Oliver, Jeremy Florman, Mark J Alkema, Michele Lemons, and Michael M Francis. Neurexin directs partner-specific synaptic connectivity in C. elegans. *eLife*, 7, 2018.

[4757] Fabian Philippart and Zayd M Khaliq. Gi/o protein-coupled receptors in dopamine neurons inhibit the sodium leak channel NALCN. *eLife*, 7, 2018.

[4758] David V Phizicky, Luke E Berchowitz, and Stephen P Bell. Multiple kinases inhibit origin licensing and helicase activation to ensure reductive cell division during meiosis. *eLife*, 7, 2018.

[4759] Swastik Phulera, Hongtao Zhu, Jie Yu, Derek P Claxton, Nate Yoder, Craig Yoshioka, and Eric Gouaux. Cryo-EM structure of the benzodiazepine-sensitive $\alpha 1\beta 1\gamma 2S$ tri-heteromeric GABAA receptor in complex with GABA. *eLife*, 7, 2018.

[4760] Marion A L Picard, Celine Cosseau, Sabrina Ferré, Thomas Quack, Christoph G Grevelding, Yohann Couté, and Beatriz Vicoso. Evolution of gene dosage on the Z-chromosome of schistosome parasites. *eLife*, 7, 2018.

[4761] Joseph Pickering, Constance A Rich, Holly Stainton, Cristina Aceituno, Kavitha Chinnaiya, Patricia Saiz-Lopez, Marian A Ros, and Matthew Towers. An intrinsic cell cycle timer terminates limb bud outgrowth. *eLife*, 7, 2018.

[4762] Sarah B Pickett, Eric D Thomas, Joy Y Sebe, Tor Linbo, Robert Esterberg, Dale W Hailey, and David W Raible. Cumulative mitochondrial activity correlates with ototoxin susceptibility in zebrafish mechanosensory hair cells. *eLife*, 7, 2018.

[4763] Ludivine Pidoux, Pascale Le Blanc, Carole Levenes, and Arthur Leblois. A subcortical circuit linking the cerebellum to the basal ganglia engaged in vocal learning. *eLife*, 7, 2018.

[4764] Paulo Wagner Pires and Scott Earley. Neuroprotective effects of TRPA1 channels in the cerebral endothelium following ischemic stroke. *eLife*, 7, 2018.

[4765] Vera P Pisareva, Andrey V Pisarev, and Israel S Fernández. Dual tRNA mimicry in the Cricket Paralysis Virus IRES uncovers an unexpected similarity with the Hepatitis C Virus IRES. *eLife*, 7, 2018.

[4766] Warintra Pitsawong, Vanessa Buosi, Renee Otten, Roman V Agafonov, Adelajda Zorba, Nadja Kern, Steffen Kutter, Gunther Kern, Ricardo AP Pádua, Xavier Meniche, and Dorothee Kern. Dynamics of human protein kinase Aurora A linked to drug selectivity. *eLife*, 7, 2018.

[4767] Weerapat Pittayakanchit, Zhiyue Lu, Justin Chew, Michael J Rust, and Arvind Murugan. Biophysical clocks face a trade-off between internal and external noise resistance. *eLife*, 7, 2018.

[4768] Andrew RG Plackett, Stephanie J Conway, Kristen D Hewett Hazelton, Ester H Rabbinowitsch, Jane A Langdale, and Verónica S Di Stilio. LEAFY maintains apical stem cell activity during shoot development in the fern Ceratopteris richardii. *eLife*, 7, 2018.

[4769] Eva Plesch, Cheng-Chang Chen, Elisabeth Butz, Anna Scotto Rosato, Einar K Krogsaeter, Hua Yinan, Karin Bartel, Marco Keller, Dina Robaa, Daniel Teupser, Lesca M Holdt, Angelika M Vollmar, Wolfgang Sippl, Rosa Puertollano, Diego Medina, Martin Biel, Christian Wahl-Schott, Franz Bracher, and Christian Grimm. Selective agonist of TRPML2 reveals direct role in chemokine release from innate immune cells. *eLife*, 7, 2018.

[4770] Mikhail V Pogorelyy, Anastasia A Minervina, Dmitriy M Chudakov, Ilgar Z Mamedov, Yuri B Lebedev, Thierry Mora, and Aleksandra M Walczak. Method for identification of condition-associated public antigen receptor sequences. *eLife*, 7, 2018.

[4771] Amy E Pohodich, Hari Yalamanchili, Ayush T Raman, Ying-Wooi Wan, Michael Gundry, Shuang Hao, Haijing Jin, Jianrong Tang, Zhandong Liu, and Huda Y Zoghbi. Forniceal deep brain stimulation induces gene expression and splicing changes that promote neurogenesis and plasticity. *eLife*, 7, 2018.

[4772] Adam J Pollak, Canzhao Liu, Aparna Gudlur, Joshua E Mayfield, Nancy D Dalton, Yusu Gu, Ju Chen, Joan Heller Brown, Patrick G Hogan, Sandra E Wiley, Kirk L Peterson, and Jack E Dixon. A secretory pathway kinase regulates sarcoplasmic reticulum Ca2+ homeostasis and protects against heart failure. *eLife*, 7, 2018.

[4773] Petr Popov, Yao Peng, Ling Shen, Raymond C Stevens, Vadim Cherezov, Zhi-Jie Liu, and Vsevolod Katritch. Computational design of thermostabilizing point mutations for G protein-coupled receptors. *eLife*, 7, 2018.

[4774] Dean A Pospisil, Anitha Pasupathy, and Wyeth Bair. 'Artiphysiology' reveals V4-like shape tuning in a deep network trained for image classification. *eLife*, 7, 2018.

[4775] Fanny Pouyet, Simon Aeschbacher, Alexandre Thiéry, and Laurent Excoffier. Background selection and biased gene conversion affect more than 95% of the human genome and bias demographic inferences. *eLife*, 7, 2018.

[4776] Juan Manuel Povedano, Paula Martinez, Rosa Serrano, águeda Tejera, Gonzalo Gómez-López, Maria Bobadilla, Juana Maria Flores, Fátima Bosch, and Maria A Blasco. Therapeutic effects of telomerase in mice with pulmonary fibrosis induced by damage to the lungs and short telomeres. *eLife*, 7, 2018.

[4777] Katrin H Preller, Joshua B Burt, Jie Lisa Ji, Charles H Schleifer, Brendan D Adkinson, Philipp Stämpfli, Erich Seifritz, Grega Repovs, John H Krystal, John D Murray, Franz X Vollenweider, and Alan Anticevic. Changes in global and thalamic brain connectivity in LSD-induced altered states of consciousness are attributable to the 5-HT2A receptor. *eLife*, 7, 2018.

[4778] J Andrew Pruszynski, J Randall Flanagan, and Roland S Johansson. Fast and accurate edge orientation processing during object manipulation. *eLife*, 7, 2018.

[4779] Christopher D Pull, Line V Ugelvig, Florian Wiesenhofer, Anna V Grasse, Simon Tragust, Thomas Schmitt, Mark JF Brown, and Sylvia Cremer. Destructive disinfection of infected brood prevents systemic disease spread in ant colonies. *eLife*, 7, 2018.

[4780] Will Putzbach, Ashley Haluck-Kangas, Quan Q Gao, Aishe A Sarshad, Elizabeth T Bartom, Austin Stults, Abdul S Qadir, Markus Hafner, and Marcus E Peter. CD95/Fas ligand mRNA is toxic to cells. *eLife*, 7, 2018.

[4781] Minxian Qian, Zuojun Liu, Linyuan Peng, Xiaolong Tang, Fanbiao Meng, Ying Ao, Mingyan Zhou, Ming Wang, Xinyue Cao, Baoming Qin, Zimei Wang, Zhongjun Zhou, Guangming Wang, Zhengliang Gao, Jun Xu, and Baohua Liu. Boosting ATM activity alleviates aging and extends lifespan in a mouse model of progeria. *eLife*, 7, 2018.

[4782] Bing Qiao, Chiyuan Li, Victoria W Allen, Mimi Shirasu-Hiza, and Sheyum Syed. Automated analysis of long-term grooming behavior in Drosophila using a k-nearest neighbors classifier. *eLife*, 7, 2018.

[4783] Jian Qiu, Heidi M Rivera, Martha A Bosch, Stephanie L Padilla, Todd L Stincic, Richard D Palmiter, Martin J Kelly, and Oline K Rønnekleiv. Estrogenic-dependent glutamatergic neurotransmission from kisspeptin neurons governs feeding circuits in females. *eLife*, 7, 2018.

[4784] Guosheng Qu, Carol Lyn Piazza, Dorie Smith, and Marlene Belfort. Group II intron inhibits conjugative relaxase expression in bacteria by mRNA targeting. *eLife*, 7, 2018.

[4785] Roberto Quaranta, Jakob Fell, Frank Rühle, Jyoti Rao, Ilaria Piccini, Marcos J Araúzo-Bravo, Arie O Verkerk, Monika Stoll, and Boris Greber. Revised roles of ISL1 in a hES cell-based model of human heart chamber specification. *eLife*, 7, 2018.

[4786] Gonzalo Quiroga Artigas, Pascal Lapébie, Lucas Leclère, Noriyo Takeda, Ryusaku Deguchi, Gáspár Jékely, Tsuyoshi Momose, and Evelyn Houliston. A gonad-expressed opsin mediates light-induced spawning in the jellyfish Clytia. *eLife*, 7, 2018.

[4787] Meytal Radzinski, Rosi Fassler, Ohad Yogev, William Breuer, Nadav Shai, Jenia Gutin, Sidra Ilyas, Yifat Geffen, Sabina Tsytkin-Kirschenzweig, Yaakov Nahmias, Tommer Ravid, Nir Friedman, Maya Schuldiner, and Dana Reichmann. Temporal profiling of redox-dependent heterogeneity in single cells. *eLife*, 7, 2018.

[4788] Atif Rahman, Ingileif Hallgrímsdóttir, Michael Eisen, and Lior Pachter. Association mapping from sequencing reads using k-mers. *eLife*, 7, 2018.

[4789] Mohammed Mostafizur Rahman, Ashutosh Shukla, and Sumantra Chattarji. Extinction recall of fear memories formed before stress is not affected despite higher theta activity in the amygdala. *eLife*, 7, 2018.

[4790] Srivarsha Rajshekar, Jun Yao, Paige K Arnold, Sara G Payne, Yinwen Zhang, Teresa V Bowman, Robert J Schmitz, John R Edwards, and Mary Goll. Pericentromeric hypomethylation elicits an interferon response in an animal model of ICF syndrome. *eLife*, 7, 2018.

[4791] Benjamin Rakela, Paul Brehm, and Gail Mandel. Astrocytic modulation of excitatory synaptic signaling in a mouse model of Rett syndrome. *eLife*, 7, 2018.

[4792] Anassuya Ramachandran, Pedro Vizán, Debipriya Das, Probir Chakravarty, Janis Vogt, Katherine W Rogers, Patrick Müller, Andrew P Hinck, Gopal P Sapkota, and Caroline S Hill. TGF-$\beta$ uses a novel mode of receptor activation to phosphorylate SMAD1/5 and induce epithelial-to-mesenchymal transition. *eLife*, 7, 2018.

[4793] David Ramos-Vicente, Jie Ji, Esther Gratacòs-Batlle, Gemma Gou, Rita Reig-Viader, Javier Luís, Demian Burguera, Enrique Navas-Perez, Jordi García-Fernández, Pablo Fuentes-Prior, Hector Escriva, Nerea Roher, David Soto, and Àlex Bayés. Metazoan evolution of glutamate receptors reveals unreported phylogenetic groups and divergent lineage-specific events. *eLife*, 7, 2018.

[4794] Ishier Raote, Maria Ortega-Bellido, António JM Santos, Ombretta Foresti, Chong Zhang, Maria F Garcia-Parajo, Felix Campelo, and Vivek Malhotra. TANGO1 builds a machine for collagen export by recruiting and spatially organizing COPII, tethers and membranes. *eLife*, 7, 2018.

[4795] Seyed Javad Rasouli, Mohamed El-Brolosy, Ayele Taddese Tsedeke, Anabela Bensimon-Brito, Parisa Ghanbari, Hans-Martin Maischein, Carsten Kuenne, and Didier Y Stainier. The flow responsive transcription factor Klf2 is required for myocardial wall integrity by modulating Fgf signaling. *eLife*, 7, 2018.

[4796] Madara Ratnadiwakara, Stuart K Archer, Craig I Dent, Igor Ruiz De Los Mozos, Traude H Beilharz, Anja S Knaupp, Christian M Nefzger, Jose M Polo, and Minna-Liisa Anko. SRSF3 promotes pluripotency through Nanog mRNA export and coordination of the pluripotency gene expression program. *eLife*, 7, 2018.

[4797] Matthieu Raveau, Denis Polygalov, Roman Boehringer, Kenji Amano, Kazuhiro Yamakawa, and Thomas J McHugh. Alterations of in vivo CA1 network activity in Dp(16)1Yey Down syndrome model mice. *eLife*, 7, 2018.

[4798] Thomas A Ray, Suva Roy, Christopher Kozlowski, Jingjing Wang, Jon Cafaro, Samuel W Hulbert, Christopher V Wright, Greg D Field, and Jeremy N Kay. Formation of retinal direction-selective circuitry initiated by starburst amacrine cell homotypic contact. *eLife*, 7, 2018.

[4799] Florian Razy-Krajka, Basile Gravez, Nicole Kaplan, Claudia Racioppi, Wei Wang, and Lionel Christiaen. An FGF-driven feed-forward circuit patterns the cardiopharyngeal mesoderm in space and time. *eLife*, 7, 2018.

[4800] Ignacio Rebollo, Anne-Dominique Devauchelle, Benoît Béranger, and Catherine Tallon-Baudry. Stomach-brain synchrony reveals a novel, delayed-connectivity resting-state network in humans. *eLife*, 7, 2018.

[4801] Gautam Reddy, Joseph D Zak, Massimo Vergassola, and Venkatesh N Murthy. Antagonism in olfactory receptor neurons and its implications for the perception of odor mixtures. *eLife*, 7, 2018.

[4802] Colin M Rees, Jun-Hai Yang, Marc Santolini, Aldons J Lusis, James N Weiss, and Alain Karma. The Ca2+ transient as a feedback sensor controlling cardiomyocyte ionic conductances in mouse populations. *eLife*, 7, 2018.

[4803] Aspen T Reese, Eugenia H Cho, Bruce Klitzman, Scott P Nichols, Natalie A Wisniewski, Max M Villa, Heather K Durand, Sharon Jiang, Firas S Midani, Sai N Nimmagadda, Thomas M O'Connell, Justin P Wright, Marc A Deshusses, and Lawrence A David. Antibiotic-induced changes in the microbiota disrupt redox dynamics in the gut. *eLife*, 7, 2018.

[4804] Patrick Reeson, Kevin Choi, and Craig E Brown. VEGF signaling regulates the fate of obstructed capillaries in mouse cortex. *eLife*, 7, 2018.

[4805] Adam J Reid, Arthur M Talman, Hayley M Bennett, Ana R Gomes, Mandy J Sanders, Christopher J R Illingworth, Oliver Billker, Matthew Berriman, and Mara KN Lawniczak. Single-cell RNA-seq reveals hidden transcriptional variation in malaria parasites. *eLife*, 7, 2018.

[4806] Justin Rendleman, Zhe Cheng, Shuvadeep Maity, Nicolai Kastelic, Mathias Munschauer, Kristina Allgoewer, Guoshou Teo, Yun Bin Matteo Zhang, Amy Lei, Brian Parker, Markus Landthaler, Lindsay Freeberg, Scott Kuersten, Hyungwon Choi, and Christine Vogel. New insights into the cellular temporal response to proteostatic stress. *eLife*, 7, 2018.

[4807] John Repass, Elizabeth Iorns, Elizabeth Iorns, Alexandria Denis, Stephen R Williams, Nicole Perfito, and Timothy M Errington. Replication Study: Fusobacterium nucleatum infection is prevalent in human colorectal carcinoma. *eLife*, 7, 2018.

[4808] Arbora Resulaj, Sarah Ruediger, Shawn R Olsen, and Massimo Scanziani. First spikes in visual cortex enable perceptual discrimination. *eLife*, 7, 2018.

[4809] Isaie J Reuling, Lisanne A van de Schans, Luc E Coffeng, Kjerstin Lanke, Lisette Meerstein-Kessel, Wouter Graumans, Geert-Jan van Gemert, Karina Teelen, Rianne Siebelink-Stoter, Marga van de Vegte-Bolmer, Quirijn de Mast, André J van der Ven, Karen Ivinson, Cornelus C Hermsen, Sake de Vlas, John Bradley, Katharine A Collins, Christian F Ockenhouse, James McCarthy, Robert W Sauerwein, and Teun Bousema. A randomized feasibility trial comparing four antimalarial drug regimens to induce Plasmodium falciparum gametocytemia in the controlled human malaria infection model. *eLife*, 7, 2018.

[4810] Jan Rheinberger, Xiaolong Gao, Philipp AM Schmidpeter, and Crina M Nimigean. Ligand discrimination and gating in cyclic nucleotide-gated ion channels from apo and partial agonist-bound cryo-EM structures. *eLife*, 7, 2018.

[4811] Yael Riahi, Tal Israeli, Roni Yeroslaviz, Shoshana Chimenez, Dana Avrahami, Miri Stolovich-Rain, Ido Alter, Marina Sebag, Nava Polin, Ernesto Bernal-Mizrachi, Yuval Dor, Erol Cerasi, and Gil Leibowitz. Inhibition of mTORC1 by ER stress impairs neonatal $\beta$-cell expansion and predisposes to diabetes in the Akita mouse. *eLife*, 7, 2018.

[4812] Benjamin L Ricca, Gautham Venugopalan, Saori Furuta, Kandice Tanner, Walter A Orellana, Clay D Reber, Douglas G Brownfield, Mina J Bissell, and Daniel A Fletcher. Transient external force induces phenotypic reversion of malignant epithelial structures via nitric oxide signaling. *eLife*, 7, 2018.

[4813] Jocelyn M Richard, Nakura Stout, Deanna Acs, and Patricia H Janak. Ventral pallidal encoding of reward-seeking behavior depends on the underlying associative structure. *eLife*, 7, 2018.

[4814] Daniel J Richter, Parinaz Fozouni, Michael B Eisen, and Nicole King. Gene family innovation, conservation and loss on the animal stem lineage. *eLife*, 7, 2018.

[4815] Erika A Riederer, Paul J Focke, Elka R Georgieva, Nurunisa Akyuz, Kimberly Matulef, Peter P Borbat, Jack H Freed, Scott C Blanchard, Olga Boudker, and Francis I Valiyaveetil. A facile approach for the in vitro assembly of multimeric membrane transport proteins. *eLife*, 7, 2018.

[4816] Kristina Riemer, Robert P Guralnick, and Ethan P White. No general relationship between mass and temperature in endothermic species. *eLife*, 7, 2018.

[4817] Pablo Ripollés, Laura Ferreri, Ernest Mas-Herrero, Helena Alicart, Alba Gómez-Andrés, Josep Marco-Pallares, Rosa Maria Antonijoan, Toemme Noesselt, Marta Valle, Jordi Riba, and Antoni Rodriguez-Fornells. Intrinsically regulated learning is modulated by synaptic dopamine signaling. *eLife*, 7, 2018.

[4818] Peter Rodgers. A question of time. *eLife*, 7, 2018.

[4819] Guillermo Rodrigo and Mario A Fares. Intrinsic adaptive value and early fate of gene duplication revealed by a bottom-up approach. *eLife*, 7, 2018.

[4820] Isabel Rodriguez-Barraquer, Emmanuel Arinaitwe, Prasanna Jagannathan, Moses R Kamya, Phillip J Rosenthal, John Rek, Grant Dorsey, Joaniter Nankabirwa, Sarah G Staedke, Maxwell Kilama, Chris Drakeley, Isaac Ssewanyana, David L Smith, and Bryan Greenhouse. Quantification of anti-parasite and anti-disease immunity to malaria as a function of age and exposure. *eLife*, 7, 2018.

[4821] Maxwell J Roeske, Eric M Camino, Sumant Grover, Mark Rebeiz, and Thomas Michael Williams. Cis-regulatory evolution integrated the Bric-à-brac transcription factors into a novel fruit fly gene regulatory network. *eLife*, 7, 2018.

[4822] Annah S Rolig, Emily Goers Sweeney, Lila E Kaye, Michael D DeSantis, Arden Perkins, Allison V Banse, M Kristina Hamilton, and Karen Guillemin. A bacterial immunomodulatory protein with lipocalin-like domains facilitates host–bacteria mutualism in larval zebrafish. *eLife*, 7, 2018.

[4823] Vincenzo Romano, Licia De Propris, Laurens WJ Bosman, Pascal Warnaar, Michiel M ten Brinke, Sander Lindeman, Chiheng Ju, Arthiha Velauthapillai, Jochen K Spanke, Emily Middendorp Guerra, Tycho M Hoogland, Mario Negrello, Egidio D'Angelo, and Chris I De Zeeuw. Potentiation of cerebellar Purkinje cells facilitates whisker reflex adaptation through increased simple spike activity. *eLife*, 7, 2018.

[4824] Gergely Rona, Domenico Roberti, Yandong Yin, Julia K Pagan, Harrison Homer, Elizabeth Sassani, Andras Zeke, Luca Busino, Eli Rothenberg, and Michele Pagano. PARP1-dependent recruitment of the FBXL10-RNF68-RNF2 ubiquitin ligase to sites of DNA damage controls H2A.Z loading. *eLife*, 7, 2018.

[4825] Jens-Christian Röper, Démosthène Mitrossilis, Guillaume Stirnemann, FranÇois Waharte, Isabel Brito, Maria-Elena Fernandez-Sanchez, Marc Baaden, Jean Salamero, and Emmanuel Farge. The major $\beta$-catenin/E-cadherin junctional binding site is a primary molecular mechano-transductor of differentiation in vivo. *eLife*, 7, 2018.

[4826] Adam Z Rosenthal, Yutao Qi, Sahand Hormoz, Jin Park, Sophia Hsin-Jung Li, and Michael B Elowitz. Metabolic interactions between dynamic bacterial subpopulations. *eLife*, 7, 2018.

[4827] Brian L Ross, Brian Tenner, Michele L Markwardt, Adam Zviman, Guoli Shi, Jaclyn P Kerr, Nicole E Snell, Jennifer J McFarland, Joseph R Mauban, Christopher W Ward, Megan A Rizzo, and Jin Zhang. Single-color, ratiometric biosensors for detecting signaling activities in live cells. *eLife*, 7, 2018.

[4828] Rachel A Ross, Silvia Leon, Joseph C Madara, Danielle Schafer, Chrysanthi Fergani, Caroline A Maguire, Anne MJ Verstegen, Emily Brengle, Dong Kong, Allan E Herbison, Ursula B Kaiser, Bradford B Lowell, and Victor M Navarro. PACAP neurons in the ventral premammillary nucleus regulate reproductive function in the female mouse. *eLife*, 7, 2018.

[4829] Zvi N Roth, David J Heeger, and Elisha P Merriam. Stimulus vignetting and orientation selectivity in human visual cortex. *eLife*, 7, 2018.

[4830] Christophe Rouillon, Januka S Athukoralage, Shirley Graham, Sabine Grüschow, and Malcolm F White. Control of cyclic oligoadenylate synthesis in a type III CRISPR system. *eLife*, 7, 2018.

[4831] Maxime WC Rousseaux, Jean-Pierre Revelli, Gabriel E Vázquez-Vélez, Ji-Yoen Kim, Evelyn Craigen, Kristyn Gonzales, Jaclyn Beckinghausen, and Huda Y Zoghbi. Depleting Trim28 in adult mice is well tolerated and reduces levels of $\alpha$-synuclein and tau. *eLife*, 7, 2018.

[4832] Richard H Row, Amy Pegg, Brian A Kinney, Gist H Farr, Lisa Maves, Sally Lowell, Valerie Wilson, and Benjamin L Martin. BMP and FGF signaling interact to pattern mesoderm by controlling basic helix-loop-helix transcription factor activity. *eLife*, 7, 2018.

[4833] David C Rowland, Horst A Obenhaus, Emilie R Skytøen, Qiangwei Zhang, Cliff G Kentros, Edvard I Moser, and May-Britt Moser. Functional properties of stellate cells in medial entorhinal cortex layer II. *eLife*, 7, 2018.

[4834] Sunetra Roy, Karl-Heinz Tomaszowski, Jessica W Luzwick, Soyoung Park, Jun Li, Maureen Murphy, and Katharina Schlacher. p53 orchestrates DNA replication restart homeostasis by suppressing mutagenic RAD52 and POL$\theta$ pathways. *eLife*, 7, 2018.

[4835] Inna Rozman Grinberg, Daniel Lundin, Mahmudul Hasan, Mikael Crona, Venkateswara Rao Jonna, Christoph Loderer, Margareta Sahlin, Natalia Markova, Ilya Borovok, Gustav Berggren, Anders Hofer, Derek T Logan, and Britt-Marie Sjöberg. Novel ATP-cone-driven allosteric regulation of ribonucleotide reductase via the radical-generating subunit. *eLife*, 7, 2018.

[4836] J Graham Ruby, Megan Smith, and Rochelle Buffenstein. Naked mole-rat mortality rates defy Gompertzian laws by not increasing with age. *eLife*, 7, 2018.

[4837] Marina Rudan, Peter Bou Dib, Marina Musa, Matea Kanunnikau, Sandra Sobočanec, David Rueda, Tobias Warnecke, and Anita KriŠko. Normal mitochondrial function in Saccharomyces cerevisiae has become dependent on inefficient splicing. *eLife*, 7, 2018.

[4838] Kathy Ruddy, Joshua Balsters, Dante Mantini, Quanying Liu, Pegah Kassraian-Fard, Nadja Enz, Ernest Mihelj, Bankim Subhash Chander, Surjo R Soekadar, and Nicole Wenderoth. Neural activity related to volitional regulation of cortical excitability. *eLife*, 7, 2018.

[4839] Martina Rudnicki, Ghoncheh Abdifarkosh, Emmanuel Nwadozi, Sofhia V Ramos, Armin Makki, Diane M Sepa-Kishi, Rolando B Ceddia, Christopher GR Perry, Emilie Roudier, and Tara L Haas. Endothelial-specific FoxO1 depletion prevents obesity-related disorders by increasing vascular metabolism and growth. *eLife*, 7, 2018.

[4840] Johannes Rudolph, Jyothi Mahadevan, Pamela Dyer, and Karolin Luger. Poly(ADP-ribose) polymerase 1 searches DNA via a 'monkey bar' mechanism. *eLife*, 7, 2018.

[4841] Emily F Ruff, Joseph M Muretta, Andrew R Thompson, Eric W Lake, Soreen Cyphers, Steven K Albanese, Sonya M Hanson, Julie M Behr, David D Thomas, John D Chodera, and Nicholas M Levinson. A dynamic mechanism for allosteric activation of Aurora kinase A by activation loop phosphorylation. *eLife*, 7, 2018.

[4842] Mario Ruiz, Rakesh Bodhicharla, Emma Svensk, Ranjan Devkota, Kiran Busayavalasa, Henrik Palmgren, Marcus Ståhlman, Jan Boren, and Marc Pilon. Membrane fluidity is regulated by the C. elegans transmembrane protein FLD-1 and its human homologs TLCD1/2. *eLife*, 7, 2018.

[4843] Alistair B Russell, Cole Trapnell, and Jesse D Bloom. Extreme heterogeneity of influenza virus infection in single cells. *eLife*, 7, 2018.

[4844] Mark F Sabbagh, Jacob S Heng, Chongyuan Luo, Rosa G Castanon, Joseph R Nery, Amir Rattner, Loyal A Goff, Joseph R Ecker, and Jeremy Nathans. Transcriptional and epigenomic landscapes of CNS and non-CNS vascular endothelial cells. *eLife*, 7, 2018.

[4845] Maria A Sacta, Bowranigan Tharmalingam, Maddalena Coppo, David A Rollins, Dinesh K Deochand, Bradley Benjamin, Li Yu, Bin Zhang, Xiaoyu Hu, Rong Li, Yurii Chinenov, and Inez Rogatsky. Gene-specific mechanisms direct glucocorticoid-receptor-driven repression of inflammatory response genes in macrophages. *eLife*, 7, 2018.

[4846] Brian F Sadacca, Heather M Wied, Nina Lopatina, Gurpreet K Saini, Daniel Nemirovsky, and Geoffrey Schoenbaum. Orbitofrontal neurons signal sensory associations underlying model-based inference in a sensory preconditioning task. *eLife*, 7, 2018.

[4847] Rochelle Shirin Sadeghi, Katarzyna Kulej, Rahul Singh Kathayat, Benjamin A Garcia, Bryan C Dickinson, Donita C Brady, and Eric S Witze. Wnt5a signaling induced phosphorylation increases APT1 activity and promotes melanoma metastatic behavior. *eLife*, 7, 2018.

[4848] Ehsan Sadeghipour, Miguel A Garcia, William James Nelson, and Beth L Pruitt. Shear-induced damped oscillations in an epithelium depend on actomyosin contraction and E-cadherin cell adhesion. *eLife*, 7, 2018.

[4849] Alexa Sadier, Kalina TJ Davies, Laurel R Yohe, Kun Yun, Paul Donat, Brandon P Hedrick, Elizabeth R Dumont, Liliana M Dávalos, Stephen J Rossiter, and Karen E Sears. Multifactorial processes underlie parallel opsin loss in neotropical bats. *eLife*, 7, 2018.

[4850] Tomohide Saio, Soichiro Kawagoe, Koichiro Ishimori, and Charalampos G Kalodimos. Oligomerization of a molecular chaperone modulates its activity. *eLife*, 7, 2018.

[4851] Richi Sakaguchi, Marcus N Leiwe, and Takeshi Imai. Bright multicolor labeling of neuronal circuits with fluorescent proteins and chemical tags. *eLife*, 7, 2018.

[4852] Elie Saliba, Minoas Evangelinos, Christos Gournas, Florent Corrillon, Isabelle Georis, and Bruno André. The yeast H+-ATPase Pma1 promotes Rag/Gtr-dependent TORC1 activation in response to H+-coupled nutrient uptake. *eLife*, 7, 2018.

[4853] Ximena I Salinas-Hernández, Pascal Vogel, Sebastian Betz, Raffael Kalisch, Torfi Sigurdsson, and Sevil Duvarci. Dopamine neurons drive fear extinction learning by signaling the omission of expected aversive outcomes. *eLife*, 7, 2018.

[4854] Miguel Salinas-Saavedra, Amber Q Rock, and Mark Q Martindale. Germ layer-specific regulation of cell polarity and adhesion gives insight into the evolution of mesoderm. *eLife*, 7, 2018.

[4855] Christi T Salisbury-Ruf, Clinton C Bertram, Aurelia Vergeade, Daniel S Lark, Qiong Shi, Marlene L Heberling, Niki L Fortune, G Donald Okoye, W Gray Jerome, Quinn S Wells, Josh Fessel, Javid Moslehi, Heidi Chen, L Jackson Roberts, Olivier Boutaud, Eric R Gamazon, and Sandra S Zinkel. Bid maintains mitochondrial cristae structure and function and protects against cardiac disease in an integrative genomics study. *eLife*, 7, 2018.

[4856] Olivier Sallin, Luc Reymond, Corentin Gondrand, Fabio Raith, Birgit Koch, and Kai Johnsson. Semisynthetic biosensors for mapping cellular concentrations of nicotinamide adenine dinucleotides. *eLife*, 7, 2018.

[4857] Yara E Sanchez-Corrales, Guy B Blanchard, and Katja Röper. Radially patterned cell behaviours during tube budding from an epithelium. *eLife*, 7, 2018.

[4858] Ana Sánchez-Moreno, Eduardo Guevara-Hernández, Ricardo Contreras-Cervera, Gisela Rangel-Yescas, Ernesto Ladrón-de Guevara, Tamara Rosenbaum, and León D Islas. Irreversible temperature gating in trpv1 sheds light on channel activation. *eLife*, 7, 2018.

[4859] Alfredo Sánchez-Tójar, Shinichi Nakagawa, Moisès Sánchez-Fortún, Dominic A Martin, Sukanya Ramani, Antje Girndt, Veronika Bókony, Bart Kempenaers, András Liker, David F Westneat, Terry Burke, and Julia Schroeder. Meta-analysis challenges a textbook example of status signalling and demonstrates publication bias. *eLife*, 7, 2018.

[4860] Fernando J Sánchez-Valdéz, Angel Padilla, Wei Wang, Dylan Orr, and Rick L Tarleton. Spontaneous dormancy protects Trypanosoma cruzi during extended drug exposure. *eLife*, 7, 2018.

[4861] Joana A Santos, Stephan Rempel, Sandra TM Mous, Cristiane T Pereira, Josy ter Beek, Jan-Willem de Gier, Albert Guskov, and Dirk J Slotboom. Functional and structural characterization of an ECF-type ABC transporter for vitamin B12. *eLife*, 7, 2018.

[4862] Aleksandra Sapala, Adam Runions, Anne-Lise Routier-Kierzkowska, Mainak Das Gupta, Lilan Hong, Hugo Hofhuis, Stéphane Verger, Gabriella Mosca, Chun-Biu Li, Angela Hay, Olivier Hamant, Adrienne HK Roeder, Miltos Tsiantis, Przemyslaw Prusinkiewicz, and Richard S Smith. Why plants make puzzle cells, and how their shape emerges. *eLife*, 7, 2018.

[4863] Andrea Saponaro, Francesca Cantini, Alessandro Porro, Annalisa Bucchi, Dario DiFrancesco, Vincenzo Maione, Chiara Donadoni, Bianca Introini, Pietro Mesirca, Matteo E Mangoni, Gerhard Thiel, Lucia Banci, Bina Santoro, and Anna Moroni. A synthetic peptide that prevents cAMP regulation in mammalian hyperpolarization-activated cyclic nucleotide-gated (HCN) channels. *eLife*, 7, 2018.

[4864] Panagiotis Sapountzis, Mariya Zhukova, Jonathan Z Shik, Morten Schiott, and Jacobus J Boomsma. Reconstructing the functions of endosymbiotic Mollicutes in fungus-growing ants. *eLife*, 7, 2018.

[4865] Shaarika Sarasija, Jocelyn T Laboy, Zahra Ashkavand, Jennifer Bonner, Yi Tang, and Kenneth R Norman. Presenilin mutations deregulate mitochondrial Ca2+ homeostasis and metabolic activity causing neurodegeneration in Caenorhabditis elegans. *eLife*, 7, 2018.

[4866] Mohsen Sarikhani, Sneha Mishra, Sangeeta Maity, Chaithanya Kotyada, Donald Wolfgeher, Mahesh P Gupta, Mahavir Singh, and Nagalingam R Sundaresan. SIRT2 deacetylase regulates the activity of GSK3 isoforms independent of inhibitory phosphorylation. *eLife*, 7, 2018.

[4867] Rashmi Sarnaik and Indira M Raman. Control of voluntary and optogenetically perturbed locomotion by spike rate and timing of neurons of the mouse cerebellar nuclei. *eLife*, 7, 2018.

[4868] Severin Sasso, Herwig Stibor, Maria Mittag, and Arthur R Grossman. From molecular manipulation of domesticated Chlamydomonas reinhardtii to survival in nature. *eLife*, 7, 2018.

[4869] Natasa Savic, Femke CAS Ringnalda, Helen Lindsay, Christian Berk, Katja Bargsten, Yizhou Li, Dario Neri, Mark D Robinson, Constance Ciaudo, Jonathan Hall, Martin Jinek, and Gerald Schwank. Covalent linkage of the DNA repair template to the CRISPR-Cas9 nuclease enhances homology-directed repair. *eLife*, 7, 2018.

[4870] Petter Säwen, Mohamed Eldeeb, Eva Erlandsson, Trine A Kristiansen, Cecilia Laterza, Zaal Kokaia, Göran Karlsson, Joan Yuan, Shamit Soneji, Pankaj K Mandal, Derrick J Rossi, and David Bryder. Murine HSCs contribute actively to native hematopoiesis but with reduced differentiation capacity upon aging. *eLife*, 7, 2018.

[4871] Matthew S Scarnati, Rahul Kataria, Mohana Biswas, and Kenneth G Paradiso. Active presynaptic ribosomes in the mammalian brain, and altered transmitter release after protein synthesis inhibition. *eLife*, 7, 2018.

[4872] Michael-Paul Schallmo, Alexander M Kale, Rachel Millin, Anastasia V Flevaris, Zoran Brkanac, Richard AE Edden, Raphael A Bernier, and Scott O Murray. Suppression and facilitation of human neural responses. *eLife*, 7, 2018.

[4873] Christoph T Schanzenbächer, Julian D Langer, and Erin M Schuman. Time- and polarity-dependent proteomic changes associated with homeostatic scaling at central synapses. *eLife*, 7, 2018.

[4874] Lucas Schirmer, Wiebke Möbius, Chao Zhao, Andrés Cruz-Herranz, Lucile Ben Haim, Christian Cordano, Lawrence R Shiow, Kevin W Kelley, Boguslawa Sadowski, Garrett Timmons, Anne-Katrin Pröbstel, Jackie N Wright, Jung Hyung Sin, Michael Devereux, Daniel E Morrison, Sandra M Chang, Khalida Sabeur, Ari J Green, Klaus-Armin Nave, Robin JM Franklin, and David H Rowitch. Oligodendrocyte-encoded Kir4.1 function is required for axonal integrity. *eLife*, 7, 2018.

[4875] Michael Schirner, Anthony Randal McIntosh, Viktor Jirsa, Gustavo Deco, and Petra Ritter. Inferring multi-scale neural mechanisms with brain network modelling. *eLife*, 7, 2018.

[4876] Katharina Schlereth, Dieter Weichenhan, Tobias Bauer, Tina Heumann, Evangelia Giannakouri, Daniel Lipka, Samira Jaeger, Matthias Schlesner, Patrick Aloy, Roland Eils, Christoph Plass, and Hellmut G Augustin. The transcriptomic and epigenetic map of vascular quiescence in the continuous lung endothelium. *eLife*, 7, 2018.

[4877] Christopher John Schlicksup, Joseph Che-Yen Wang, Samson Francis, Balasubramanian Venkatakrishnan, William W Turner, Michael VanNieuwenhze, and Adam Zlotnick. Hepatitis B virus core protein allosteric modulators can distort and disrupt intact capsids. *eLife*, 7, 2018.

[4878] Kerstin Schmidt, Qinyu Zhang, Alpaslan Tasdogan, Andreas Petzold, Andreas Dahl, Borros M Arneth, Robert Slany, Hans Jörg Fehling, Andrea Kranz, Adrian Francis Stewart, and Konstantinos Anastassiadis. The H3K4 methyltransferase Setd1b is essential for hematopoietic stem and progenitor cell homeostasis in mice. *eLife*, 7, 2018.

[4879] Evelyn Schubert, Ingrid R Vetter, Daniel Prumbaum, Pawel A Penczek, and Stefan Raunser. Membrane insertion of $\alpha$-xenorhabdolysin in near-atomic detail. *eLife*, 7, 2018.

[4880] Sabine Schürmann, Georg Steffes, Dominique Manikowski, Philipp Kastl, Ursula Malkus, Shyam Bandari, Stefanie Ohlig, Corinna Ortmann, Rocio Rebollido-Rios, Mandy Otto, Harald Nüsse, Daniel Hoffmann, Christian Klämbt, Milos Galic, Jürgen Klingauf, and Kay Grobe. Proteolytic processing of palmitoylated Hedgehog peptides specifies the 3-4 intervein region of the Drosophila wing. *eLife*, 7, 2018.

[4881] Benjamin Schwarz, Dominik Hollfelder, Katharina Scharf, Leonie Hartmann, and Ingolf Reim. Diversification of heart progenitor cells by EGF signaling and differential modulation of ETS protein activity. *eLife*, 7, 2018.

[4882] Cameron C Scott, Stefania Vossio, Jacques Rougemont, and Jean Gruenberg. TFAP2 transcription factors are regulators of lipid droplet biogenesis. *eLife*, 7, 2018.

[4883] Jermyn Z See, Craig A Atencio, Vikaas S Sohal, and Christoph E Schreiner. Coordinated neuronal ensembles in primary auditory cortical columns. *eLife*, 7, 2018.

[4884] Stephanie C Seeman, Luke Campagnola, Pasha A Davoudian, Alex Hoggarth, Travis A Hage, Alice Bosma-Moody, Christopher A Baker, Jung Hoon Lee, Stefan Mihalas, Corinne Teeter, Andrew L Ko, Jeffrey G Ojemann, Ryder P Gwinn, Daniel L Silbergeld, Charles Cobbs, John Phillips, Ed Lein, Gabe Murphy, Christof Koch, Hongkui Zeng, and Tim Jarsky. Sparse recurrent excitatory connectivity in the microcircuit of the adult mouse and human cortex. *eLife*, 7, 2018.

[4885] Mauricio Seguel, Felipe Montalva, Diego Perez-Venegas, Josefina Gutiérrez, Hector J Paves, Ananda Müller, Carola Valencia-Soto, Elizabeth Howerth, Victoria Mendiola, and Nicole Gottdenker. Immune-mediated hookworm clearance and survival of a marine mammal decrease with warmer ocean temperatures. *eLife*, 7, 2018.

[4886] Azadeh Seidi, Linden S Muellner-Wong, Esther Rajendran, Edwin T Tjhin, Laura F Dagley, Vincent YT Aw, Pierre Faou, Andrew I Webb, Christopher J Tonkin, and Giel G van Dooren. Elucidating the mitochondrial proteome of Toxoplasma gondii reveals the presence of a divergent cytochrome c oxidase. *eLife*, 7, 2018.

[4887] Ashok Sekhar, Algirdas Velyvis, Guy Zoltsman, Rina Rosenzweig, Guillaume Bouvignies, and Lewis E Kay. Conserved conformational selection mechanism of Hsp70 chaperone-substrate interactions. *eLife*, 7, 2018.

[4888] Ourania Semelidou, Summer F Acevedo, and Efthimios MC Skoulakis. Temporally specific engagement of distinct neuronal circuits regulating olfactory habituation in Drosophila. *eLife*, 7, 2018.

[4889] Mulugeta Semework, Sara C Steenrod, and Michael E Goldberg. A spatial memory signal shows that the parietal cortex has access to a craniotopic representation of space. *eLife*, 7, 2018.

[4890] Guadalupe Sepulveda, Mark Antkowiak, Ingrid Brust-Mascher, Karan Mahe, Tingyoung Ou, Noemi M Castro, Lana N Christensen, Lee Cheung, Xueer Jiang, Daniel Yoon, Bo Huang, and Li-En Jao. Co-translational protein targeting facilitates centrosomal recruitment of PCNT during centrosome maturation in vertebrates. *eLife*, 7, 2018.

[4891] Yasmina Serroukh, Chunyan Gu-Trantien, Baharak Hooshiar Kashani, Matthieu Defrance, Thien-Phong Vu Manh, Abdulkader Azouz, Aurélie Detavernier, Alice Hoyois, Jishnu Das, Martin Bizet, Emeline Pollet, Tressy Tabbuso, Emilie Calonne, Klaas van Gisbergen, Marc Dalod, FranÇois Fuks, Stanislas Goriely, and Arnaud Marchant. The transcription factors Runx3 and ThPOK cross-regulate acquisition of cytotoxic function by human Th1 lymphocytes. *eLife*, 7, 2018.

[4892] Omer Shafraz, Matthias Rübsam, Sara N Stahley, Amber L Caldara, Andrew P Kowalczyk, Carien M Niessen, and Sanjeevi Sivasankar. E-cadherin binds to desmoglein to facilitate desmosome assembly. *eLife*, 7, 2018.

[4893] Neel H Shah, Mark Löbel, Arthur Weiss, and John Kuriyan. Fine-tuning of substrate preferences of the Src-family kinase Lck revealed through a high-throughput specificity screen. *eLife*, 7, 2018.

[4894] Iryna Shakhmantsir, Soumyashant Nayak, Gregory R Grant, and Amita Sehgal. Spliceosome factors target timeless (tim) mRNA to control clock protein accumulation and circadian behavior in Drosophila. *eLife*, 7, 2018.

[4895] Daria N Shalaeva, Dmitry A Cherepanov, Michael Y Galperin, Andrey V Golovin, and Armen Y Mulkidjanian. Evolution of cation binding in the active sites of P-loop nucleoside triphosphatases in relation to the basic catalytic mechanism. *eLife*, 7, 2018.

[4896] Laura K Shanahan, Eva Gjorgieva, Ken A Paller, Thorsten Kahnt, and Jay A Gottfried. Odor-evoked category reactivation in human ventromedial prefrontal cortex during sleep promotes memory consolidation. *eLife*, 7, 2018.

[4897] Jinsai Shang, Richard Brust, Sarah A Mosure, Jared Bass, Paola Munoz-Tello, Hua Lin, Travis S Hughes, Miru Tang, Qingfeng Ge, Theodore M Kamenekca, and Douglas J Kojetin. Cooperative cobinding of synthetic and natural ligands to the nuclear receptor PPAR$\gamma$. *eLife*, 7, 2018.

[4898] Apurwa M Sharma, Talitha L Thomas, DaNae R Woodard, Omar M Kashmer, and Marc I Diamond. Tau monomer encodes strains. *eLife*, 7, 2018.

[4899] Chengyong Shen, Lei Li, Kai Zhao, Lei Bai, Ailian Wang, Xiaoqiu Shu, Yatao Xiao, Jianmin Zhang, Kejing Zhang, Tiankun Hui, Wenbing Chen, Bin Zhang, Wei Hsu, Wen-Cheng Xiong, and Lin Mei. Motoneuron Wnts regulate neuromuscular junction development. *eLife*, 7, 2018.

[4900] Madeline E Sherlock, Narasimhan Sudarsan, Shira Stav, and Ronald R Breaker. Tandem riboswitches form a natural Boolean logic gate to control purine metabolism in bacteria. *eLife*, 7, 2018.

[4901] Ellie Sherrard-Smith, Katarzyna A Sala, Michael Betancourt, Leanna M Upton, Fiona Angrisano, Merribeth J Morin, Azra C Ghani, Thomas S Churcher, and Andrew M Blagborough. Synergy in anti-malarial pre-erythrocytic and transmission-blocking antibodies is achieved by reducing parasite density. *eLife*, 7, 2018.

[4902] Zhimin Shi, Zoe Piccus, Xiaofang Zhang, Huidi Yang, Hannah Jarrell, Yan Ding, Zhaoqian Teng, Ofer Tchernichovski, and XiaoChing Li. miR-9 regulates basal ganglia-dependent developmental vocal learning and adult vocal performance in songbirds. *eLife*, 7, 2018.

[4903] Yuichi Shichino, Yoko Otsubo, Yoshitaka Kimori, Masayuki Yamamoto, and Akira Yamashita. YTH-RNA-binding protein prevents deleterious expression of meiotic proteins by tethering their mRNAs to nuclear foci. *eLife*, 7, 2018.

[4904] James M Shine, Matthew J Aburn, Michael Breakspear, and Russell A Poldrack. The modulation of neural gain facilitates a transition between functional segregation and integration in the brain. *eLife*, 7, 2018.

[4905] Marta Strecker Shocket, Sadie J Ryan, and Erin A Mordecai. Temperature explains broad patterns of Ross River virus transmission. *eLife*, 7, 2018.

[4906] David R Shook, Eric M Kasprowicz, Lance A Davidson, and Raymond Keller. Large, long range tensile forces drive convergence during Xenopus blastopore closure and body axis elongation. *eLife*, 7, 2018.

[4907] Kieran M Short, Alexander N Combes, Valerie Lisnyak, James G Lefevre, Lynelle K Jones, Melissa H Little, Nicholas A Hamilton, and Ian M Smyth. Branching morphogenesis in the developing kidney is not impacted by nephron formation or integration. *eLife*, 7, 2018.

[4908] Longfei Shu, Debra A Brock, Katherine S Geist, Jacob W Miller, David C Queller, Joan E Strassmann, and Susanne DiSalvo. Symbiont location, host fitness, and possible coadaptation in a symbiosis between social amoebae and bacteria. *eLife*, 7, 2018.

[4909] Matthew J Shurtleff, Daniel N Itzhak, Jeffrey A Hussmann, Nicole T Schirle Oakdale, Elizabeth A Costa, Martin Jonikas, Jimena Weibezahn, Katerina D Popova, Calvin H Jan, Pavel Sinitcyn, Shruthi S Vembar, Hilda Hernandez, Jürgen Cox, Alma L Burlingame, Jeffrey L Brodsky, Adam Frost, Georg HH Borner, and Jonathan S Weissman. The ER membrane protein complex interacts cotranslationally to enable biogenesis of multipass membrane proteins. *eLife*, 7, 2018.

[4910] Tânia Simões, Ramona Schuster, Fabian den Brave, and Mafalda Escobar-Henriques. Cdc48 regulates a deubiquitylase cascade critical for mitochondrial fusion. *eLife*, 7, 2018.

[4911] Yosef Singer, Yayoi Teramoto, Ben DB Willmore, Jan WH Schnupp, Andrew J King, and Nicol S Harper. Sensory cortex is optimized for prediction of future input. *eLife*, 7, 2018.

[4912] Dilawar Singh and Upinder Singh Bhalla. Subunit exchange enhances information retention by CaMKII in dendritic spines. *eLife*, 7, 2018.

[4913] Thomas E Sladewski, Neil Billington, M Yusuf Ali, Carol S Bookwalter, Hailong Lu, Elena B Krementsova, Trina A Schroer, and Kathleen M Trybus. Recruitment of two dyneins to an mRNA-dependent Bicaudal D transport complex. *eLife*, 7, 2018.

[4914] Jeroen B Smaers, Alan H Turner, Aida Gómez-Robles, and Chet C Sherwood. A cerebellar substrate for cognition evolved multiple times independently in mammals. *eLife*, 7, 2018.

[4915] Joan C Smith and Jason M Sheltzer. Systematic identification of mutations and copy number alterations associated with cancer patient prognosis. *eLife*, 7, 2018.

[4916] Katherine A Smith, Stephan Löser, Fumi Varyani, Yvonne Harcus, Henry J McSorley, Andrew NJ McKenzie, and Rick M Maizels. Concerted IL-25R and IL-4R$\alpha$ signaling drive innate type 2 effector immunity for optimal helminth expulsion. *eLife*, 7, 2018.

[4917] Mehdi Snoussi, John Paul Talledo, Nathan-Alexander Del Rosario, Salimeh Mohammadi, Bae-Yeun Ha, Andrej KoŠmrlj, and Sattar Taheri-Araghi. Heterogeneous absorption of antimicrobial peptide LL37 in Escherichia coli cells enhances population survivability. *eLife*, 7, 2018.

[4918] Katarzyna M Soczek, Tim Grant, Peter B Rosenthal, and Alfonso Mondragón. CryoEM structures of open dimers of gyrase A in complex with DNA illuminate mechanism of strand passage. *eLife*, 7, 2018.

[4919] Heun Soh, Suhyeorn Park, Kali Ryan, Kristen Springer, Atul Maheshwari, and Anastasios V Tzingounis. Deletion of KCNQ2/3 potassium channels from PV+ interneurons leads to homeostatic potentiation of excitatory transmission. *eLife*, 7, 2018.

[4920] Gregory M Solis, Rozina Kardakaris, Elizabeth R Valentine, Liron Bar-Peled, Alice L Chen, Megan M Blewett, Mark A McCormick, James R Williamson, Brian Kennedy, Benjamin F Cravatt, and Michael Petrascheck. Translation attenuation by minocycline enhances longevity and proteostasis in old post-stress-responsive organisms. *eLife*, 7, 2018.

[4921] Ming Song, Yi Yang, Jianghong He, Zhengyi Yang, Shan Yu, Qiuyou Xie, Xiaoyu Xia, Yuanyuan Dang, Qiang Zhang, Xinhuai Wu, Yue Cui, Bing Hou, Ronghao Yu, Ruxiang Xu, and Tianzi Jiang. Prognostication of chronic disorders of consciousness using brain functional networks and clinical characteristics. *eLife*, 7, 2018.

[4922] Tatiana Sorkina, Shiqi Ma, Mads Breum Larsen, Simon C Watkins, and Alexander Sorkin. Small molecule induced oligomerization, clustering and clathrin-independent endocytosis of the dopamine transporter. *eLife*, 7, 2018.

[4923] Trevor R Sorrells, Amanda N Johnson, Conor J Howard, Candace S Britton, Kyle R Fowler, Jordan T Feigerle, P Anthony Weil, and Alexander D Johnson. Intrinsic cooperativity potentiates parallel cis-regulatory evolution. *eLife*, 7, 2018.

[4924] Piotr Sosnowski, Linas Urnavicius, Andreas Boland, Robert Fagiewicz, Johan Busselez, Gabor Papai, and Helgo Schmidt. The CryoEM structure of the Saccharomyces cerevisiae ribosome maturation factor Rea1. *eLife*, 7, 2018.

[4925] Florentina Soto, Lei Zhao, and Daniel Kerschensteiner. Synapse maintenance and restoration in the retina by NGL2. *eLife*, 7, 2018.

[4926] Ruben Soto-Acosta, Xuping Xie, Chao Shan, Coleman K Baker, Pei-Yong Shi, Shannan L Rossi, Mariano A Garcia-Blanco, and Shelton Bradrick. Fragile X mental retardation protein is a Zika virus restriction factor that is antagonized by subgenomic flaviviral RNA. *eLife*, 7, 2018.

[4927] Valeria Souza, Alejandra Moreno-Letelier, Michael Travisano, Luis David Alcaraz, Gabriela Olmedo, and Luis Enrique Eguiarte. The lost world of Cuatro Ciénegas Basin, a relictual bacterial niche in a desert oasis. *eLife*, 7, 2018.

[4928] Caitlin N Spaulding, Henry Louis Schreiber, Weili Zheng, Karen W Dodson, Jennie E Hazen, Matt S Conover, Fengbin Wang, Pontus Svenmarker, Areli Luna-Rico, Olivera Francetic, Magnus Andersson, Scott Hultgren, and Edward H Egelman. Functional role of the type 1 pilus rod structure in mediating host-pathogen interactions. *eLife*, 7, 2018.

[4929] Pauline Spéder and Andrea H Brand. Systemic and local cues drive neural stem cell niche remodelling during neurogenesis in Drosophila. *eLife*, 7, 2018.

[4930] Holger Franz Sperdin, Ana Coito, Nada Kojovic, Tonia Anahi Rihs, Reem Kais Jan, Martina Franchini, Gijs Plomp, Serge Vulliemoz, Stephan Eliez, Christoph Martin Michel, and Marie Schaer. Early alterations of social brain networks in young children with autism. *eLife*, 7, 2018.

[4931] Corinna Speth, Emese Xochitl Szabo, Claudia Martinho, Silvio Collani, Sven zur Oven-Krockhaus, Sandra Richter, Irina Droste-Borel, Boris Macek, York-Dieter Stierhof, Markus Schmid, Chang Liu, and Sascha Laubinger. Arabidopsis RNA processing factor SERRATE regulates the transcription of intronless genes. *eLife*, 7, 2018.

[4932] Maria L Spletter, Christiane Barz, Assa Yeroslaviz, Xu Zhang, Sandra B Lemke, Adrien Bonnard, Erich Brunner, Giovanni Cardone, Konrad Basler, Bianca H Habermann, and Frank Schnorrer. A transcriptomics resource reveals a transcriptional transition during ordered sarcomere morphogenesis in flight muscle. *eLife*, 7, 2018.

[4933] Simeon U Springer, Chung-Hsin Chen, Maria Del Carmen Rodriguez Pena, Lu Li, Christopher Douville, Yuxuan Wang, Joshua David Cohen, Diana Taheri, Natalie Silliman, Joy Schaefer, Janine Ptak, Lisa Dobbyn, Maria Papoli, Isaac Kinde, Bahman Afsari, Aline C Tregnago, Stephania M Bezerra, Christopher VandenBussche, Kazutoshi Fujita, Dilek Ertoy, Isabela W Cunha, Lijia Yu, Trinity J Bivalacqua, Arthur P Grollman, Luis A Diaz, Rachel Karchin, Ludmila Danilova, Chao-Yuan Huang, Chia-Tung Shun, Robert J Turesky, Byeong Hwa Yun, Thomas A Rosenquist, Yeong-Shiau Pu, Ralph H Hruban, Cristian Tomasetti, Nickolas Papadopoulos, Ken W Kinzler, Bert Vogelstein, Kathleen G Dickman, and George J Netto. Non-invasive detection of urothelial cancer through the analysis of driver gene mutations and aneuploidy. *eLife*, 7, 2018.

[4934] Jordan W Squair, Seth Tigchelaar, Kyung-Mee Moon, Jie Liu, Wolfram Tetzlaff, Brian K Kwon, Andrei V Krassioukov, Christopher R West, Leonard J Foster, and Michael A Skinnider. Integrated systems analysis reveals conserved gene networks underlying response to spinal cord injury. *eLife*, 7, 2018.

[4935] Alex C Stabell, Nicholas R Meyerson, Rebekah C Gullberg, Alison R Gilchrist, Kristofor J Webb, William M Old, Rushika Perera, and Sara L Sawyer. Dengue viruses cleave STING in humans but not in nonhuman primates, their presumed natural reservoir. *eLife*, 7, 2018.

[4936] Stefanos Stagkourakis, Carolina Thörn Pérez, Arash Hellysaz, Rachida Ammari, and Christian Broberger. Network oscillation rules imposed by species-specific electrical coupling. *eLife*, 7, 2018.

[4937] Courtney W Stairs, Laura Eme, Sergio A Muñoz-Gómez, Alejandro Cohen, Graham Dellaire, Jennifer N Shepherd, James P Fawcett, and Andrew J Roger. Microbial eukaryotes have adapted to hypoxia by horizontal acquisitions of a gene involved in rhodoquinone biosynthesis. *eLife*, 7, 2018.

[4938] Kathrin Steck, Samuel J Walker, Pavel M Itskov, Célia Baltazar, José-Maria Moreira, and Carlos Ribeiro. Internal amino acid state modulates yeast taste neurons to support protein homeostasis in Drosophila. *eLife*, 7, 2018.

[4939] Richard R Stein, Takeshi Tanoue, Rose L Szabady, Shakti K Bhattarai, Bernat Olle, Jason M Norman, Wataru Suda, Kenshiro Oshima, Masahira Hattori, Georg K Gerber, Chris Sander, Kenya Honda, and Vanni Bucci. Computer-guided design of optimal microbial consortia for immune system modulation. *eLife*, 7, 2018.

[4940] Massimo Stella, Sanja Selakovic, Alberto Antonioni, and Cecilia S Andreazzi. Ecological multiplex interactions determine the role of species for parasite spread amplification. *eLife*, 7, 2018.

[4941] Peter Sterling. Predictive regulation and human design. *eLife*, 7, 2018.

[4942] Merav Stern, Kevin A Bolding, LF Abbott, and Kevin M Franks. A transformation from temporal to ensemble coding in a model of piriform cortex. *eLife*, 7, 2018.

[4943] Daniel P Stewart, Suresh Marada, William J Bodeen, Ashley Truong, Sadie Miki Sakurada, Tanushree Pandit, Shondra M Pruett-Miller, and Stacey K Ogden. Cleavage activates Dispatched for Sonic Hedgehog ligand release. *eLife*, 7, 2018.

[4944] Roman M Stilling, Gerard M Moloney, Feargal J Ryan, Alan E Hoban, Thomaz FS Bastiaanssen, Fergus Shanahan, Gerard Clarke, Marcus J Claesson, Timothy G Dinan, and John F Cryan. Social interaction-induced activation of RNA splicing in the amygdala of microbiome-deficient mice. *eLife*, 7, 2018.

[4945] Anastasiia Stratiievska, Sara Nelson, Eric N Senning, Jonathan D Lautz, Stephen EP Smith, and Sharona E Gordon. Reciprocal regulation among TRPV1 channels and phosphoinositide 3-kinase in response to nerve growth factor. *eLife*, 7, 2018.

[4946] Sebastian J Streichan, Matthew F Lefebvre, Nicholas Noll, Eric F Wieschaus, and Boris I Shraiman. Global morphogenetic flow is accurately predicted by the spatial distribution of myosin motors. *eLife*, 7, 2018.

[4947] Frederic Strobl, Anita Anderl, and Ernst HK Stelzer. A universal vector concept for a direct genotyping of transgenic organisms and a systematic creation of homozygous lines. *eLife*, 7, 2018.

[4948] Steven C Strutt, Rachel M Torrez, Emine Kaya, Oscar A Negrete, and Jennifer A Doudna. RNA-dependent RNA targeting by CRISPR-Cas9. *eLife*, 7, 2018.

[4949] Edward Stuttfeld, Christopher HS Aylett, Stefan Imseng, Daniel Boehringer, Alain Scaiola, Evelyn Sauer, Michael N Hall, Timm Maier, and Nenad Ban. Architecture of the human mTORC2 core complex. *eLife*, 7, 2018.

[4950] Arul Subramanian, Lauren Fallon Kanzaki, Jenna Lauren Galloway, and Thomas Friedrich Schilling. Mechanical force regulates tendon extracellular matrix organization and tenocyte morphogenesis through TGFbeta signaling. *eLife*, 7, 2018.

[4951] Jennifer S Sun, Nikki K Larter, J Sebastian Chahda, Douglas Rioux, Ankita Gumaste, and John R Carlson. Humidity response depends on the small soluble protein Obp59a in Drosophila. *eLife*, 7, 2018.

[4952] Jiandong Sun, Yan Liu, Yousheng Jia, Xiaoning Hao, Wei ju Lin, Jennifer Tran, Gary Lynch, Michel Baudry, and Xiaoning Bi. UBE3A-mediated p18/LAMTOR1 ubiquitination and degradation regulate mTORC1 activity and synaptic plasticity. *eLife*, 7, 2018.

[4953] Jiying Sun, Lin Shi, Aiko Kinomura, Atsuhiko Fukuto, Yasunori Horikoshi, Yukako Oma, Masahiko Harata, Masae Ikura, Tsuyoshi Ikura, Roland Kanaar, and Satoshi Tashiro. Distinct roles of ATM and ATR in the regulation of ARP8 phosphorylation to prevent chromosome translocations. *eLife*, 7, 2018.

[4954] Xianqiang Sun, Sukrit Singh, Kendall J Blumer, and Gregory R Bowman. Simulation of spontaneous G protein activation reveals a new intermediate driving GDP unbinding. *eLife*, 7, 2018.

[4955] Ramasubramanian Sundaramoorthy, Amanda L Hughes, Hassane El-Mkami, David G Norman, Helder Ferreira, and Tom Owen-Hughes. Structure of the chromatin remodelling enzyme Chd1 bound to a ubiquitinylated nucleosome. *eLife*, 7, 2018.

[4956] Megan Ann Supple, Jason G Bragg, Linda M Broadhurst, Adrienne B Nicotra, Margaret Byrne, Rose L Andrew, Abigail Widdup, Nicola C Aitken, and Justin O Borevitz. Landscape genomic prediction for restoration of a Eucalyptus foundation species under climate change. *eLife*, 7, 2018.

[4957] Laurie P Sutton, Cesare Orlandi, Chenghui Song, Won Chan Oh, Brian S Muntean, Keqiang Xie, Alice Filippini, Xiangyang Xie, Rachel Satterfield, Jazmine D W Yaeger, Kenneth J Renner, Samuel M Young, Baoji Xu, Hyungbae Kwon, and Kirill A Martemyanov. Orphan receptor GPR158 controls stress-induced depression. *eLife*, 7, 2018.

[4958] Aussie Suzuki, Sarah K Long, and Edward D Salmon. An optimized method for 3D fluorescence co-localization applied to human kinetochore protein architecture. *eLife*, 7, 2018.

[4959] Ian A Swinburne, Kishore R Mosaliganti, Srigokul Upadhyayula, Tsung-Li Liu, David G C Hildebrand, Tony Y C Tsai, Anzhi Chen, Ebaa Al-Obeidi, Anna K Fass, Samir Malhotra, Florian Engert, Jeff W Lichtman, Tomas Kirchhausen, Eric Betzig, and Sean G Megason. Lamellar projections in the endolymphatic sac act as a relief valve to regulate inner ear pressure. *eLife*, 7, 2018.

[4960] Andrew Michael Sydor, Etienne Coyaud, Cristina Rovelli, Estelle Laurent, Helen Liu, Brian Raught, and Vito Mennella. PPP1R35 is a novel centrosomal protein that regulates centriole length in concert with the microcephaly protein RTTN. *eLife*, 7, 2018.

[4961] Martin Szinte, Donatas Jonikaitis, Dragan Rangelov, and Heiner Deubel. Pre-saccadic remapping relies on dynamics of spatial attention. *eLife*, 7, 2018.

[4962] Milan Szuperak, Matthew A Churgin, Austin J Borja, David M Raizen, Christopher Fang-Yen, and Matthew S Kayser. A sleep state in Drosophila larvae required for neural stem cell proliferation. *eLife*, 7, 2018.

[4963] Tetsuya Takeda, Toshiya Kozai, Huiran Yang, Daiki Ishikuro, Kaho Seyama, Yusuke Kumagai, Tadashi Abe, Hiroshi Yamada, Takayuki Uchihashi, Toshio Ando, and Kohji Takei. Dynamic clustering of dynamin-amphiphysin helices regulates membrane constriction and fission coupled with GTP hydrolysis. *eLife*, 7, 2018.

[4964] Howard HF Tang, Shu Mei Teo, Danielle CM Belgrave, Michael D Evans, Daniel J Jackson, Marta Brozynska, Merci MH Kusel, Sebastian L Johnston, James E Gern, Robert F Lemanske, Angela Simpson, Adnan Custovic, Peter D Sly, Patrick G Holt, Kathryn E Holt, and Michael Inouye. Trajectories of childhood immune development and respiratory health relevant to asthma and allergy. *eLife*, 7, 2018.

[4965] Matthew F Tang, Cooper A Smout, Ehsan Arabzadeh, and Jason B Mattingley. Prediction error and repetition suppression have distinct effects on neural representations of visual information. *eLife*, 7, 2018.

[4966] Shiming Tang, Yimeng Zhang, Zhihao Li, Ming Li, Fang Liu, Hongfei Jiang, and Tai Sing Lee. Large-scale two-photon imaging revealed super-sparse population codes in the V1 superficial layer of awake monkeys. *eLife*, 7, 2018.

[4967] Frank Tanser, Daniel K Azongo, Alain Vandormael, Till Bärnighausen, and Christopher Appleton. Impact of the scale-up of piped water on urogenital schistosomiasis infection in rural South Africa. *eLife*, 7, 2018.

[4968] Heather J Tarbet, Lee Dolat, Timothy J Smith, Brett M Condon, E Timothy O'Brien, Raphael H Valdivia, and Michael Boyce. Site-specific glycosylation regulates the form and function of the intermediate filament cytoskeleton. *eLife*, 7, 2018.

[4969] Michael Taschner, Anna Lorentzen, André Mourão, Toby Collins, Grace M Freke, Dale Moulding, Jerome Basquin, Dagan Jenkins, and Esben Lorentzen. Crystal structure of intraflagellar transport protein 80 reveals a homo-dimer required for ciliogenesis. *eLife*, 7, 2018.

[4970] Ibrahim Tastekin, Avinash Khandelwal, David Tadres, Nico D Fessner, James W Truman, Marta Zlatic, Albert Cardona, and Matthieu Louis. Sensorimotor pathway controlling stopping behavior during chemotaxis in the Drosophila melanogaster larva. *eLife*, 7, 2018.

[4971] Martin S Taylor, Ilya Altukhov, Kelly R Molloy, Paolo Mita, Hua Jiang, Emily M Adney, Aleksandra Wudzinska, Sana Badri, Dmitry Ischenko, George Eng, Kathleen H Burns, David Fenyö, Brian T Chait, Dmitry Alexeev, Michael P Rout, Jef D Boeke, and John LaCava. Dissection of affinity captured LINE-1 macromolecular complexes. *eLife*, 7, 2018.

[4972] Ruth Rebecca Taylor, Anastasia Filia, Ursula Paredes, Yukako Asai, Jeffrey R Holt, Michael Lovett, and Andrew Forge. Regenerating hair cells in vestibular sensory epithelia from humans. *eLife*, 7, 2018.

[4973] Camilla S Teng, Man-chun Ting, D'Juan T Farmer, Mia Brockop, Robert E Maxson, and J Gage Crump. Altered bone growth dynamics prefigure craniosynostosis in a zebrafish model of Saethre-Chotzen syndrome. *eLife*, 7, 2018.

[4974] Erin E Terry, Xiping Zhang, Christy Hoffmann, Laura D Hughes, Scott A Lewis, Jiajia Li, Matthew J Wallace, Lance A Riley, Collin M Douglas, Miguel A Gutierrez-Monreal, Nicholas F Lahens, Ming C Gong, Francisco Andrade, Karyn A Esser, and Michael E Hughes. Transcriptional profiling reveals extraordinary diversity among skeletal muscle tissues. *eLife*, 7, 2018.

[4975] Fabian Tetzlaff, M Gordian Adam, Anja Feldner, Iris Moll, Amitai Menuchin, Juan Rodriguez-Vita, David Sprinzak, and Andreas Fischer. MPDZ promotes DLL4-induced Notch signaling during angiogenesis. *eLife*, 7, 2018.

[4976] Sheila S Teves, Luye An, Aarohi Bhargava-Shah, Liangqi Xie, Xavier Darzacq, and Robert Tjian. A stable mode of bookmarking by TBP recruits RNA polymerase II to mitotic chromosomes. *eLife*, 7, 2018.

[4977] Panagiota Theodoni, Bernat Rovira, Yingxue Wang, and Alex Roxin. Theta-modulation drives the emergence of connectivity patterns underlying replay in a network model of place cells. *eLife*, 7, 2018.

[4978] Abbey C Thompson, Terence D Capellini, Catherine A Guenther, Yingguang Frank Chan, Carlos R Infante, Douglas B Menke, and David M Kingsley. A novel enhancer near the Pitx1 gene influences development and evolution of pelvic appendages in vertebrates. *eLife*, 7, 2018.

[4979] Thornton W Thompson, Benjamin T Jackson, P Jonathan Li, Jiaxi Wang, Alexander Byungsuk Kim, Kristen Ting Hui Huang, Lily Zhang, and David H Raulet. Tumor-derived CSF-1 induces the NKG2D ligand RAE-1$\delta$ on tumor-infiltrating macrophages. *eLife*, 7, 2018.

[4980] Matthias Thurner, Martijn van de Bunt, Jason M Torres, Anubha Mahajan, Vibe Nylander, Amanda J Bennett, Kyle J Gaulton, Amy Barrett, Carla Burrows, Christopher G Bell, Robert Lowe, Stephan Beck, Vardhman K Rakyan, Anna L Gloyn, and Mark I McCarthy. Integration of human pancreatic islet genomic data refines regulatory mechanisms at Type 2 Diabetes susceptibility loci. *eLife*, 7, 2018.

[4981] Jing Tian, Jiaxi Wu, Xiang Chen, Tong Guo, Zhijian J Chen, Joseph L Goldstein, and Michael S Brown. BHLHE40, a third transcription factor required for insulin induction of SREBP-1c mRNA in rodent liver. *eLife*, 7, 2018.

[4982] Siqi Tian, Wipapat Kladwang, and Rhiju Das. Allosteric mechanism of the V. vulnificus adenine riboswitch resolved by four-dimensional chemical mapping. *eLife*, 7, 2018.

[4983] Hieng Chiong Tie, Alexander Ludwig, Sara Sandin, and Lei Lu. The spatial separation of processing and transport functions to the interior and periphery of the Golgi stack. *eLife*, 7, 2018.

[4984] Matthew Tien, Aretha Fiebig, and Sean Crosson. Gene network analysis identifies a central post-transcriptional regulator of cellular stress survival. *eLife*, 7, 2018.

[4985] Romain Tisserand, Christopher J Dakin, Machiel HF Van der Loos, Elizabeth A Croft, Timothy J Inglis, and Jean-Sébastien Blouin. Down regulation of vestibular balance stabilizing mechanisms to enable transition between motor states. *eLife*, 7, 2018.

[4986] Nicole Toepfner, Christoph Herold, Oliver Otto, Philipp Rosendahl, Angela Jacobi, Martin Kräter, Julia Stächele, Leonhard Menschner, Maik Herbig, Laura Ciuffreda, Lisa Ranford-Cartwright, Michal Grzybek, ünal Coskun, Elisabeth Reithuber, Geneviève Garriss, Peter Mellroth, Birgitta Henriques-Normark, Nicola Tregay, Meinolf Suttorp, Martin Bornhäuser, Edwin R Chilvers, Reinhard Berner, and Jochen Guck. Detection of human disease conditions by single-cell morpho-rheological phenotyping of blood. *eLife*, 7, 2018.

[4987] Oleg Tolstenkov, Petrus Van der Auwera, Wagner Steuer Costa, Olga Bazhanova, Tim M Gemeinhardt, Amelie CF Bergs, and Alexander Gottschalk. Functionally asymmetric motor neurons contribute to coordinating locomotion of Caenorhabditis elegans. *eLife*, 7, 2018.

[4988] Peter Tonzi, Yandong Yin, Chelsea Wei Ting Lee, Eli Rothenberg, and Tony T Huang. Translesion polymerase kappa-dependent DNA synthesis underlies replication fork recovery. *eLife*, 7, 2018.

[4989] Deniz Top, Jenna L O'Neil, Gregory E Merz, Kritika Dusad, Brian R Crane, and Michael W Young. CK1/Doubletime activity delays transcription activation in the circadian clock. *eLife*, 7, 2018.

[4990] Joana Torres, Remo Monti, Ariane L Moore, Makiko Seimiya, Yanrui Jiang, Niko Beerenwinkel, Christian Beisel, Jorge V Beira, and Renato Paro. A switch in transcription and cell fate governs the onset of an epigenetically-deregulated tumor in Drosophila. *eLife*, 7, 2018.

[4991] Kouki K Touhara and Roderick MacKinnon. Molecular basis of signaling specificity between GIRK channels and GPCRs. *eLife*, 7, 2018.

[4992] Jochen Triesch, Anh Duong Vo, and Anne-Sophie Hafner. Competition for synaptic building blocks shapes synaptic plasticity. *eLife*, 7, 2018.

[4993] Michael Nguyen Trinh, Michael S Brown, Joachim Seemann, Joseph L Goldstein, and Feiran Lu. Lysosomal cholesterol export reconstituted from fragments of Niemann-Pick C1. *eLife*, 7, 2018.

[4994] Michael Troup, Melvyn HW Yap, Chelsie Rohrscheib, Martyna J Grabowska, Deniz Ertekin, Roshini Randeniya, Benjamin Kottler, Aoife Larkin, Kelly Munro, Paul J Shaw, and Bruno van Swinderen. Acute control of the sleep switch in Drosophila reveals a role for gap junctions in regulating behavioral responsiveness. *eLife*, 7, 2018.

[4995] Feng-Ching Tsai, Aurelie Bertin, Hugo Bousquet, John Manzi, Yosuke Senju, Meng-Chen Tsai, Laura Picas, Stephanie Miserey-Lenkei, Pekka Lappalainen, Emmanuel Lemichez, Evelyne Coudrier, and Patricia Bassereau. Ezrin enrichment on curved membranes requires a specific conformation or interaction with a curvature-sensitive partner. *eLife*, 7, 2018.

[4996] Tin Ki Tsang, Eric A Bushong, Daniela Boassa, Junru Hu, Benedetto Romoli, Sebastien Phan, Davide Dulcis, Chih-Ying Su, and Mark H Ellisman. High-quality ultrastructural preservation using cryofixation for 3D electron microscopy of genetically labeled tissues. *eLife*, 7, 2018.

[4997] Chang-Hui Tsao, Chien-Chun Chen, Chen-Han Lin, Hao-Yu Yang, and Suewei Lin. Drosophila mushroom bodies integrate hunger and satiety signals to control innate food-seeking behavior. *eLife*, 7, 2018.

[4998] Marco Turi, David Charles Burr, and Paola Binda. Pupillometry reveals perceptual differences that are tightly linked to autistic traits in typical adults. *eLife*, 7, 2018.

[4999] Maxwell H Turner, Gregory W Schwartz, and Fred Rieke. Receptive field center-surround interactions mediate context-dependent spatial contrast encoding in the retina. *eLife*, 7, 2018.

[5000] Anita Tusche and Cendri A Hutcherson. Cognitive regulation alters social and dietary choice by changing attribute representations in domain-general and domain-specific brain circuits. *eLife*, 7, 2018.

[5001] Lena Tveriakhina, Karin Schuster-Gossler, Sanchez M Jarrett, Marie B Andrawes, Meike Rohrbach, Stephen C Blacklow, and Achim Gossler. The ectodomains determine ligand function in vivo and selectivity of DLL1 and DLL4 toward NOTCH1 and NOTCH2 in vitro. *eLife*, 7, 2018.

[5002] Stephen R Tymanskyj, Benjamin H Yang, Kristen J Verhey, and Le Ma. MAP7 regulates axon morphogenesis by recruiting kinesin-1 to microtubules and modulating organelle transport. *eLife*, 7, 2018.

[5003] Justyna Ulanska-Poutanen, Jakub Mieczkowski, Chao Zhao, Katarzyna Konarzewska, Beata Kaza, Hartmut BF Pohl, Lukasz Bugajski, Bozena Kaminska, Robin JM Franklin, and Malgorzata Zawadzka. Injury-induced perivascular niche supports alternative differentiation of adult rodent CNS progenitor cells. *eLife*, 7, 2018.

[5004] Niko Välimäki, Heli Kuisma, Annukka Pasanen, Oskari Heikinheimo, Jari Sjöberg, Ralf Bützow, Nanna Sarvilinna, Hanna-Riikka Heinonen, Jaana Tolvanen, Simona Bramante, Tomas Tanskanen, Juha Auvinen, Outi Uimari, Amjad Alkodsi, Rainer Lehtonen, Eevi Kaasinen, Kimmo Palin, and Lauri A Aaltonen. Genetic predisposition to uterine leiomyoma is determined by loci for genitourinary development and genome stability. *eLife*, 7, 2018.

[5005] Markus Johannes Van Ackeren, Francesca M Barbero, Stefania Mattioni, Roberto Bottini, and Olivier Collignon. Neuronal populations in the occipital cortex of the blind synchronize to the temporal dynamics of speech. *eLife*, 7, 2018.

[5006] Ronald van den Berg and Wei Ji Ma. A resource-rational theory of set size effects in human visual working memory. *eLife*, 7, 2018.

[5007] Meike E van der Heijden and Huda Y Zoghbi. Loss of Atoh1 from neurons regulating hypoxic and hypercapnic chemoresponses causes neonatal respiratory failure in mice. *eLife*, 7, 2018.

[5008] Daniel Marten van Es, Jan Theeuwes, and Tomas Knapen. Spatial sampling in human visual cortex is modulated by both spatial and feature-based attention. *eLife*, 7, 2018.

[5009] Anouk Mariette van Loon, Katya Olmos-Solis, Johannes Jacobus Fahrenfort, and Christian NL Olivers. Current and future goals are represented in opposite patterns in object-selective cortex. *eLife*, 7, 2018.

[5010] Matthijs van Veelen. Can Hamilton's rule be violated? *eLife*, 7, 2018.

[5011] John P Vanden Heuvel, Ewa Maddox, Samar W Maalouf, Elizabeth Iorns, Elizabeth Iorns, Rachel Tsui, Alexandria Denis, Nicole Perfito, and Timothy M Errington. Replication Study: Systematic identification of genomic markers of drug sensitivity in cancer cells. *eLife*, 7, 2018.

[5012] Timothy E Vanderleest, Celia M Smits, Yi Xie, Cayla E Jewett, J Todd Blankenship, and Dinah Loerke. Vertex sliding drives intercalation by radial coupling of adhesion and actomyosin networks during Drosophila germband extension. *eLife*, 7, 2018.

[5013] Mohamad-Javad Vaseghi, Kamel Chibani, Wilena Telman, Michael Florian Liebthal, Melanie Gerken, Helena Schnitzer, Sara Mareike Mueller, and Karl-Josef Dietz. The chloroplast 2-cysteine peroxiredoxin functions as thioredoxin oxidase in redox regulation of chloroplast metabolism. *eLife*, 7, 2018.

[5014] Rengasayee Veeraraghavan, Gregory S Hoeker, Anita Alvarez-Laviada, Daniel Hoagland, Xiaoping Wan, D Ryan King, Jose Sanchez-Alonso, Chunling Chen, Jane Jourdan, Lori L Isom, Isabelle Deschenes, James W Smyth, Julia Gorelik, Steven Poelzing, and Robert G Gourdie. The adhesion function of the sodium channel beta subunit ($\beta$1) contributes to cardiac action potential propagation. *eLife*, 7, 2018.

[5015] Csaba Verasztó, Martin Gühmann, Huiyong Jia, Vinoth Babu Veedin Rajan, Luis A Bezares-Calderón, Cristina Piñeiro-Lopez, Nadine Randel, Réza Shahidi, Nico K Michiels, Shozo Yokoyama, Kristin Tessmar-Raible, and Gáspár Jékely. Ciliary and rhabdomeric photoreceptor-cell circuits form a spectral depth gauge in marine zooplankton. *eLife*, 7, 2018.

[5016] Michael F Verderame, Victoria H Freedman, Lisa M Kozlowski, and Wayne T McCormack. Competency-based assessment for the training of PhD students and early-career scientists. *eLife*, 7, 2018.

[5017] Stéphane Verger, Yuchen Long, Arezki Boudaoud, and Olivier Hamant. A tension-adhesion feedback loop in plant epidermis. *eLife*, 7, 2018.

[5018] Robert McCoy Vernon, Paul Andrew Chong, Brian Tsang, Tae Hun Kim, Alaji Bah, Patrick Farber, Hong Lin, and Julie Deborah Forman-Kay. Pi-Pi contacts are an overlooked protein feature relevant to phase separation. *eLife*, 7, 2018.

[5019] Stephin J Vervoort, Olivier G de Jong, M Guy Roukens, Cynthia L Frederiks, Jeroen F Vermeulen, Ana Rita LourenÇo, Laura Bella, Ana Tufegdzic Vidakovic, José L Sandoval, Cathy Moelans, Miranda van Amersfoort, Margaret J Dallman, Alejandra Bruna, Carlos Caldas, Edward Nieuwenhuis, Elsken van der Wall, Patrick Derksen, Paul van Diest, Marianne C Verhaar, Eric W-F Lam, Michal Mokry, and Paul J Coffer. Global transcriptional analysis identifies a novel role for SOX4 in tumor-induced angiogenesis. *eLife*, 7, 2018.

[5020] Oriol Viader-Llargués, Valerio Lupperger, Laura Pola-Morell, Carsten Marr, and Hernán López-Schier. Live cell-lineage tracing and machine learning reveal patterns of organ regeneration. *eLife*, 7, 2018.

[5021] Anna I Vickrey, Rebecca Bruders, Zev Kronenberg, Emma Mackey, Ryan J Bohlender, Emily T Maclary, Raquel Maynez, Edward J Osborne, Kevin P Johnson, Chad D Huff, Mark Yandell, and Michael D Shapiro. Introgression of regulatory alleles and a missense coding mutation drive plumage pattern diversity in the rock pigeon. *eLife*, 7, 2018.

[5022] Andres Vidal-Gadea, Chance Bainbridge, Ben Clites, Bridgitte E Palacios, Layla Bakhtiari, Vernita Gordon, and Jonathan Pierce-Shimomura. Response to comment on "Magnetosensitive neurons mediate geomagnetic orientation in Caenorhabditis elegans". *eLife*, 7, 2018.

[5023] Deivanayagabarathy Vinayagam, Thomas Mager, Amir Apelbaum, Arne Bothe, Felipe Merino, Oliver Hofnagel, Christos Gatsogiannis, and Stefan Raunser. Electron cryo-microscopy structure of the canonical TRPC4 ion channel. *eLife*, 7, 2018.

[5024] Zoé Vincent-Mistiaen, Ahmed Elbediwy, Hannah Vanyai, Jennifer Cotton, Gordon Stamp, Emma Nye, Bradley Spencer-Dene, Gareth J Thomas, Junhao Mao, and Barry Thompson. YAP drives cutaneous squamous cell carcinoma formation and progression. *eLife*, 7, 2018.

[5025] Tim James Viney, Minas Salib, Abhilasha Joshi, Gunes Unal, Naomi Berry, and Peter Somogyi. Shared rhythmic subcortical GABAergic input to the entorhinal cortex and presubiculum. *eLife*, 7, 2018.

[5026] Raghuvir Viswanatha, Zhongchi Li, Yanhui Hu, and Norbert Perrimon. Pooled genome-wide CRISPR screening for basal and context-specific fitness gene essentiality in Drosophila cells. *eLife*, 7, 2018.

[5027] Daniela G Vitali, Sandro Käser, Antonia Kolb, Kai S Dimmer, Andre Schneider, and Doron Rapaport. Independent evolution of functionally exchangeable mitochondrial outer membrane import complexes. *eLife*, 7, 2018.

[5028] Eszter K Vladar, Miranda B Stratton, Maxwell L Saal, Glicella Salazar-De Simone, Xiangyuan Wang, Debra Wolgemuth, Tim Stearns, and Jeffrey D Axelrod. Cyclin-dependent kinase control of motile ciliogenesis. *eLife*, 7, 2018.

[5029] Vladimir A Volkov, Pim J Huis in 't Veld, Marileen Dogterom, and Andrea Musacchio. Multivalency of NDC80 in the outer kinetochore is essential to track shortening microtubules and generate forces. *eLife*, 7, 2018.

[5030] Laura Vuolo, Nicola L Stevenson, Kate J Heesom, and David J Stephens. Dynein-2 intermediate chains play crucial but distinct roles in primary cilia formation and function. *eLife*, 7, 2018.

[5031] Nickolai V Vysokov, John-Paul Silva, Vera G Lelianova, Jason Suckling, John Cassidy, Jennifer K Blackburn, Natalia Yankova, Mustafa BA Djamgoz, Serguei V Kozlov, Alexander G Tonevitsky, and Yuri A Ushkaryov. Proteolytically released Lasso/teneurin-2 induces axonal attraction by interacting with latrophilin-1 on axonal growth cones. *eLife*, 7, 2018.

[5032] Wayne P Wahls. The NIH must reduce disparities in funding to maximize its return on investments from taxpayers. *eLife*, 7, 2018.

[5033] Avner Wallach, Erik Harvey-Girard, James Jaeyoon Jun, André Longtin, and Len Maler. A time-stamp mechanism may provide temporal information necessary for egocentric to allocentric spatial transformations. *eLife*, 7, 2018.

[5034] Nian-Feng Wan, You-Ming Cai, Yan-Jun Shen, Xiang-Yun Ji, Xiang-Wen Wu, Xiang-Rong Zheng, Wei Cheng, Jun Li, Yao-Pei Jiang, Xin Chen, Jacob Weiner, Jie-Xian Jiang, Ming Nie, Rui-Ting Ju, Tao Yuan, Jian-Jun Tang, Wei-Dong Tian, Hao Zhang, and Bo Li. Increasing plant diversity with border crops reduces insecticide use and increases crop yield in urban agriculture. *eLife*, 7, 2018.

[5035] Bo Wang, Jayhun Lee, Pengyang Li, Amir Saberi, Huiying Yang, Chang Liu, Minglei Zhao, and Phillip A Newmark. Stem cell heterogeneity drives the parasitic life cycle of Schistosoma mansoni. *eLife*, 7, 2018.

[5036] Cheng Wang, Zhijia Tan, Ben Niu, Kwok Yeung Tsang, Andrew Tai, Wilson C W Chan, Rebecca L K Lo, Keith K H Leung, Nelson W F Dung, Nobuyuki Itoh, Michael Q Zhang, Danny Chan, and Kathryn Song Eng Cheah. Inhibiting the integrated stress response pathway prevents aberrant chondrocyte differentiation thereby alleviating chondrodysplasia. *eLife*, 7, 2018.

[5037] Congwei Wang, Fabian Schmich, Sumana Srivatsa, Julie Weidner, Niko Beerenwinkel, and Anne Spang. Context-dependent deposition and regulation of mRNAs in P-bodies. *eLife*, 7, 2018.

[5038] Haiguang Wang and Kristin A Hogquist. CCR7 defines a precursor for murine iNKT cells in thymus and periphery. *eLife*, 7, 2018.

[5039] Hao Wang, Shuo Li, Juejin Wang, Shenghan Chen, Xue-Long Sun, and Qingyu Wu. N-glycosylation in the protease domain of trypsin-like serine proteases mediates calnexin-assisted protein folding. *eLife*, 7, 2018.

[5040] Jiyu Wang, Jie Zhou, Qidi Yang, and Elizabeth J Grayhack. Multi-protein bridging factor 1(Mbf1), Rps3 and Asc1 prevent stalled ribosomes from frameshifting. *eLife*, 7, 2018.

[5041] Lihua Wang, Ergang Wang, Yi Wang, Robert Mines, Kun Xiang, Zhiguo Sun, Gaiting Zhou, Kai-Yuan Chen, Nikolai Rakhilin, Shanshan Chao, Gaoqi Ye, Zhenzhen Wu, Huiwen Yan, Hong Shen, Jeffrey Everitt, Pengcheng Bu, and Xiling Shen. miR-34a is a microRNA safeguard for Citrobacter-induced inflammatory colon oncogenesis. *eLife*, 7, 2018.

[5042] Lin Wang, Gina Kuperberg, and Ole Jensen. Specific lexico-semantic predictions are associated with unique spatial and temporal patterns of neural activity. *eLife*, 7, 2018.

[5043] Liu Wang, Barbora Valiskova, and Jiri Forejt. Cisplatin-induced DNA double-strand breaks promote meiotic chromosome synapsis in PRDM9-controlled mouse hybrid sterility. *eLife*, 7, 2018.

[5044] Ming Wang, Martin Schäfer, Dapeng Li, Rayko Halitschke, Chuanfu Dong, Erica McGale, Christian Paetz, Yuanyuan Song, Suhua Li, Junfu Dong, Sven Heiling, Karin Groten, Philipp Franken, Michael Bitterlich, Maria J Harrison, Uta Paszkowski, and Ian T Baldwin. Blumenols as shoot markers of root symbiosis with arbuscular mycorrhizal fungi. *eLife*, 7, 2018.

[5045] Qi Wang, Tae Hee Han, Peter Nguyen, Michal Jarnik, and Mihaela Serpe. Tenectin recruits integrin to stabilize bouton architecture and regulate vesicle release at the Drosophila neuromuscular junction. *eLife*, 7, 2018.

[5046] Qingqing Wang, Katharine C Abruzzi, Michael Rosbash, and Donald C Rio. Striking circadian neuron diversity and cycling of Drosophila alternative splicing. *eLife*, 7, 2018.

[5047] Wenke Wang, Amaresh Chaturbedi, Minghui Wang, Serim An, Satheeja Santhi Velayudhan, and Siu Sylvia Lee. SET-9 and SET-26 are H3K4me3 readers and play critical roles in germline development and longevity. *eLife*, 7, 2018.

[5048] Xiaoli Wang, Sytse J Piersma, Christopher A Nelson, Ya-Nan Dai, Ted Christensen, Eric Lazear, Liping Yang, Marjolein Sluijter, Thorbald van Hall, Ted H Hansen, Wayne M Yokoyama, and Daved H Fremont. A herpesvirus encoded Qa-1 mimic inhibits natural killer cell cytotoxicity through CD94/NKG2A receptor engagement. *eLife*, 7, 2018.

[5049] Xiaoming Wang, Stuart C White, Mairin Balisi, Jacob Biewer, Julia Sankey, Dennis Garber, and Z Jack Tseng. First bone-cracking dog coprolites provide new insight into bone consumption in Borophagus and their unique ecological niche. *eLife*, 7, 2018.

[5050] Ying Wang, Wei Wu, Xiangbing Wu, Yan Sun, Yi P Zhang, Ling-Xiao Deng, Melissa Jane Walker, Wenrui Qu, Chen Chen, Nai-Kui Liu, Qi Han, Heqiao Dai, Lisa BE Shields, Christopher B Shields, Dale R Sengelaub, Kathryn J Jones, George M Smith, and Xiao-Ming Xu. Remodeling of lumbar motor circuitry remote to a thoracic spinal cord injury promotes locomotor recovery. *eLife*, 7, 2018.

[5051] Yixuan Wang, Chengchen Zhao, Zhenzhen Hou, Yuanyuan Yang, Yan Bi, Hong Wang, Yong Zhang, and Shaorong Gao. Unique molecular events during reprogramming of human somatic cells to induced pluripotent stem cells (iPSCs) at naïve state. *eLife*, 7, 2018.

[5052] Yubao Wang, Young-Mi Li, Lukas Baitsch, Alan Huang, Yi Xiang, Haoxuan Tong, Ana Lako, Thanh Von, Christine Choi, Elgene Lim, Junxia Min, Li Li, Frank Stegmeier, Robert Schlegel, Michael J Eck, Nathanael S Gray, Timothy J Mitchison, and Jean J Zhao. Correction: MELK is an oncogenic kinase essential for mitotic progression in basal-like breast cancer cells. *eLife*, 7, 2018.

[5053] Yue Wang, Yuan Shang, Jianchao Li, Weidi Chen, Gang Li, Jun Wan, Wei Liu, and Mingjie Zhang. Specific Eph receptor-cytoplasmic effector signaling mediated by SAM–SAM domain interactions. *eLife*, 7, 2018.

[5054] Zheng Wang, Catherine Wu, Aaron Aslanian, John R Yates, and Tony Hunter. Defective RNA polymerase III is negatively regulated by the SUMO-Ubiquitin-Cdc48 pathway. *eLife*, 7, 2018.

[5055] Michelle C Ward, Siming Zhao, Kaixuan Luo, Bryan J Pavlovic, Mohammad M Karimi, Matthew Stephens, and Yoav Gilad. Silencing of transposable elements may not be a major driver of regulatory evolution in primate iPSCs. *eLife*, 7, 2018.

[5056] Sean C Warren, Max Nobis, Astrid Magenau, Yousuf H Mohammed, David Herrmann, Imogen Moran, Claire Vennin, James RW Conway, Pauline Mélénec, Thomas R Cox, Yingxiao Wang, Jennifer P Morton, Heidi CE Welch, Douglas Strathdee, Kurt I Anderson, Tri Giang Phan, Michael S Roberts, and Paul Timpson. Removing physiological motion from intravital and clinical functional imaging data. *eLife*, 7, 2018.

[5057] David Jon Wasilko, Qingqiu Huang, and Yuxin Mao. Insights into the ubiquitin transfer cascade catalyzed by the Legionella effector SidC. *eLife*, 7, 2018.

[5058] Andrew J Watrous, Jonathan Miller, Salman E Qasim, Itzhak Fried, and Joshua Jacobs. Phase-tuned neuronal firing encodes human contextual representations for navigational goals. *eLife*, 7, 2018.

[5059] Simon Nikolaus Weber and Henning Sprekeler. Learning place cells, grid cells and invariances with excitatory and inhibitory plasticity. *eLife*, 7, 2018.

[5060] Brian Webster, Scott W Werneke, Biljana Zafirova, Sébastien This, Séverin Coléon, Elodie Décembre, Helena Paidassi, Isabelle Bouvier, Pierre-Emmanuel Joubert, Darragh Duffy, Thierry Walzer, Matthew L Albert, and Marlène Dreux. Plasmacytoid dendritic cells control dengue and Chikungunya virus infections via IRF7-regulated interferon responses. *eLife*, 7, 2018.

[5061] Arne Weinhold, Elham Karimi Dorcheh, Ran Li, Natarajan Rameshkumar, and Ian T Baldwin. Antimicrobial peptide expression in a wild tobacco plant reveals the limits of host-microbe-manipulations in the field. *eLife*, 7, 2018.

[5062] Tracey L Weissgerber, Oscar Garcia-Valencia, Vesna D Garovic, Natasa M Milic, and Stacey J Winham. Why we need to report more than 'Data were Analyzed by t-tests or ANOVA'. *eLife*, 7, 2018.

[5063] Timothy J Wendorff and James M Berger. Topoisomerase VI senses and exploits both DNA crossings and bends to facilitate strand passage. *eLife*, 7, 2018.

[5064] George R Wendt, Julie NR Collins, Jimin Pei, Mark S Pearson, Hayley M Bennett, Alex Loukas, Matthew Berriman, Nick V Grishin, and James J Collins. Flatworm-specific transcriptional regulators promote the specification of tegumental progenitors in Schistosoma mansoni. *eLife*, 7, 2018.

[5065] Angela Wenning, Brian J Norris, Cengiz Günay, Daniel Kueh, and Ronald L Calabrese. Output variability across animals and levels in a motor system. *eLife*, 7, 2018.

[5066] Achim Werner, Regina Baur, Nia Teerikorpi, Deniz U Kaya, and Michael Rape. Multisite dependency of an E3 ligase controls monoubiquitylation-dependent cell fate decisions. *eLife*, 7, 2018.

[5067] Laura A Wetzel, Tera C Levin, Ryan E Hulett, Daniel Chan, Grant A King, Reef Aldayafleh, David S Booth, Monika Abedin Sigg, and Nicole King. Predicted glycosyltransferases promote development and prevent spurious cell clumping in the choanoflagellate S. rosetta. *eLife*, 7, 2018.

[5068] Aaron G Wexler, Whitman B Schofield, Patrick H Degnan, Ewa Folta-Stogniew, Natasha A Barry, and Andrew L Goodman. Human gut Bacteroides capture vitamin B12 via cell surface-exposed lipoproteins. *eLife*, 7, 2018.

[5069] K Ian White, Minglei Zhao, Ucheor B Choi, Richard A Pfuetzner, and Axel T Brunger. Structural principles of SNARE complex recognition by the AAA+ protein NSF. *eLife*, 7, 2018.

[5070] Ryan S Wible, Chidambaram Ramanathan, Carrie Hayes Sutter, Kristin M Olesen, Thomas W Kensler, Andrew C Liu, and Thomas R Sutter. NRF2 regulates core and stabilizing circadian clock loops, coupling redox and timekeeping in Mus musculus. *eLife*, 7, 2018.

[5071] Evan Wicker, Janita Turchi, Ludise Malkova, and Patrick A Forcelli. Mediodorsal thalamus is required for discrete phases of goal-directed behavior in macaques. *eLife*, 7, 2018.

[5072] Yves F Widmer, Cornelia Fritsch, Magali M Jungo, Silvia Almeida, Boris Egger, and Simon G Sprecher. Multiple neurons encode CrebB dependent appetitive long-term memory in the mushroom body circuit. *eLife*, 7, 2018.

[5073] J Simon Wiegert, Mauro Pulin, Christine Elizabeth Gee, and Thomas G Oertner. The fate of hippocampal synapses depends on the sequence of plasticity-inducing events. *eLife*, 7, 2018.

[5074] Sithara Wijeratne and Radhika Subramanian. Geometry of antiparallel microtubule bundles regulates relative sliding and stalling by PRC1 and Kif4A. *eLife*, 7, 2018.

[5075] Niklas Wilming, Peter König, Seth König, and Elizabeth A Buffalo. Entorhinal cortex receptive fields are modulated by spatial attention, even without movement. *eLife*, 7, 2018.

[5076] Jessica Winger, Ilana M Nodelman, Robert F Levendosky, and Gregory D Bowman. A twist defect mechanism for ATP-dependent translocation of nucleosomal DNA. *eLife*, 7, 2018.

[5077] Elizabeth C Wittenborn, Mériem Merrouch, Chie Ueda, Laura Fradale, Christophe Léger, Vincent Fourmond, Maria-Eirini Pandelia, Sébastien Dementin, and Catherine L Drennan. Redox-dependent rearrangements of the NiFeS cluster of carbon monoxide dehydrogenase. *eLife*, 7, 2018.

[5078] Josette J Wlaschin, Jacob M Gluski, Eileen Nguyen, Hanna Silberberg, James H Thompson, Alexander T Chesler, and Claire E Le Pichon. Dual leucine zipper kinase is required for mechanical allodynia and microgliosis after nerve injury. *eLife*, 7, 2018.

[5079] Carsten Wolff, Jean-Yves Tinevez, Tobias Pietzsch, Evangelia Stamataki, Benjamin Harich, Léo Guignard, Stephan Preibisch, Spencer Shorte, Philipp J Keller, Pavel Tomancak, and Anastasios Pavlopoulos. Multi-view light-sheet imaging and tracking with the MaMuT software reveals the cell lineage of a direct developing arthropod limb. *eLife*, 7, 2018.

[5080] Yao Liang Wong, Lauren LeBon, Rohinton Edalji, Hock Ben Lim, Chaohong Sun, and Carmela Sidrauski. The small molecule ISRIB rescues the stability and activity of Vanishing White Matter Disease eIF2B mutant complexes. *eLife*, 7, 2018.

[5081] Melanie I Worley, Larissa A Alexander, and Iswar K Hariharan. CtBP impedes JNK- and Upd/STAT-driven cell fate misspecifications in regenerating Drosophila imaginal discs. *eLife*, 7, 2018.

[5082] Mingxuan Wu, Dawn Hayward, Jay H Kalin, Yun Song, John WR Schwabe, and Philip A Cole. Lysine-14 acetylation of histone H3 in chromatin confers resistance to the deacetylase and demethylase activities of an epigenetic silencing complex. *eLife*, 7, 2018.

[5083] Xunde Xian, Theresa Pohlkamp, Murat S Durakoglugil, Connie H Wong, Jürgen K Beck, Courtney Lane-Donovan, Florian Plattner, and Joachim Herz. Reversal of ApoE4-induced recycling block as a novel prevention approach for Alzheimer's disease. *eLife*, 7, 2018.

[5084] Lei Xiao, Michael F Priest, and Yevgenia Kozorovitskiy. Oxytocin functions as a spatiotemporal filter for excitatory synaptic inputs to VTA dopamine neurons. *eLife*, 7, 2018.

[5085] Yang Xie, Chechang Nie, and Tianming Yang. Covert shift of attention modulates the value encoding in the orbitofrontal cortex. *eLife*, 7, 2018.

[5086] Guanglin Xing, Moyi Li, Yichen Sun, Menglong Rui, Yan Zhuang, Huihui Lv, Junhai Han, Zhengping Jia, and Wei Xie. Neurexin–Neuroligin 1 regulates synaptic morphology and functions via the WAVE regulatory complex in Drosophila neuromuscular junction. *eLife*, 7, 2018.

[5087] Yi Xiong, Adriana Souza Torsoni, Feihua Wu, Hong Shen, Yan Liu, Xiao Zhong, Mark J Canet, Yatrik M Shah, M Bishr Omary, Yong Liu, and Liangyou Rui. Hepatic NF-kB-inducing kinase (NIK) suppresses mouse liver regeneration in acute and chronic liver diseases. *eLife*, 7, 2018.

[5088] Ke Xu, Xiaodan Liu, Yuchun Wang, Chouin Wong, and Yan Song. Temporospatial induction of homeodomain gene cut dictates natural lineage reprogramming. *eLife*, 7, 2018.

[5089] Xiao Xu, Elitsa I Stoyanova, Agata E Lemiesz, Jie Xing, Deborah C Mash, and Nathaniel Heintz. Species and cell-type properties of classically defined human and rodent neurons and glia. *eLife*, 7, 2018.

[5090] Hiroshi Yamaguchi, Toshiyuki Oda, Masahide Kikkawa, and Hiroyuki Takeda. Systematic studies of all PIH proteins in zebrafish reveal their distinct roles in axonemal dynein assembly. *eLife*, 7, 2018.

[5091] Haruki Yamamoto, Mingxu Fang, Vladimira Dragnea, and Carl E Bauer. Differing isoforms of the cobalamin binding photoreceptor AerR oppositely regulate photosystem expression. *eLife*, 7, 2018.

[5092] Koji Yamano, Chunxin Wang, Shireen A Sarraf, Christian Münch, Reika Kikuchi, Nobuo N Noda, Yohei Hizukuri, Masato T Kanemaki, Wade Harper, Keiji Tanaka, Noriyuki Matsuda, and Richard J Youle. Endosomal Rab cycles regulate Parkin-mediated mitophagy. *eLife*, 7, 2018.

[5093] Masahiro Yamashita, Yujiro Yoshihara, Ryuichiro Hashimoto, Noriaki Yahata, Naho Ichikawa, Yuki Sakai, Takashi Yamada, Noriko Matsukawa, Go Okada, Saori C Tanaka, Kiyoto Kasai, Nobumasa Kato, Yasumasa Okamoto, Ben Seymour, Hidehiko Takahashi, Mitsuo Kawato, and Hiroshi Imamizu. A prediction model of working memory across health and psychiatric disease using whole-brain functional connectivity. *eLife*, 7, 2018.

[5094] Chao Yang, Shirng-Wern Tsaih, Angela Lemke, Michael J Flister, Monica S Thakar, and Subramaniam Malarkannan. mTORC1 and mTORC2 differentially promote natural killer cell development. *eLife*, 7, 2018.

[5095] Weijian Yang, Luis Carrillo-Reid, Yuki Bando, Darcy S Peterka, and Rafael Yuste. Simultaneous two-photon imaging and two-photon optogenetics of cortical circuits in three dimensions. *eLife*, 7, 2018.

[5096] Xi Yang, Felichi Mae Arines, Weichao Zhang, and Ming Li. Sorting of a multi-subunit ubiquitin ligase complex in the endolysosome system. *eLife*, 7, 2018.

[5097] Yang Yang, Dongwei Kang, Laura A Nguyen, Zachary B Smithline, Christophe Pannecouque, Peng Zhan, Xinyong Liu, and Thomas A Steitz. Structural basis for potent and broad inhibition of HIV-1 RT by thiophene[3,2-d]pyrimidine non-nucleoside inhibitors. *eLife*, 7, 2018.

[5098] Yu Hsuan Carol Yang, Koichi Kawakami, and Didier YR Stainier. A new mode of pancreatic islet innervation revealed by live imaging in zebrafish. *eLife*, 7, 2018.

[5099] Justin D Yao and Dan H Sanes. Developmental deprivation-induced perceptual and cortical processing deficits in awake-behaving animals. *eLife*, 7, 2018.

[5100] Lina Yao, Sarah Craven Seaton, Sula Ndousse-Fetter, Arijit A Adhikari, Nicholas DiBenedetto, Amir I Mina, Alexander S Banks, Lynn Bry, and A Sloan Devlin. A selective gut bacterial bile salt hydrolase alters host metabolism. *eLife*, 7, 2018.

[5101] Michael M Yartsev, Timothy D Hanks, Alice Misun Yoon, and Carlos D Brody. Causal contribution and dynamical encoding in the striatum during evidence accumulation. *eLife*, 7, 2018.

[5102] Brogan Yarzabek, Anita J Zaitouna, Eli Olson, Gayathri N Silva, Jie Geng, Aviva Geretz, Rasmi Thomas, Sujatha Krishnakumar, Daniel S Ramon, and Malini Raghavan. Variations in HLA-B cell surface expression, half-life and extracellular antigen receptivity. *eLife*, 7, 2018.

[5103] Azadeh Yazdan-Shahmorad, Daniel B Silversmith, Viktor Kharazia, and Philip N Sabes. Targeted cortical reorganization using optogenetics in non-human primates. *eLife*, 7, 2018.

[5104] Fei Ye, Yuxin Huang, Jianchao Li, Yuqian Ma, Chensu Xie, Zexu Liu, Xiaoying Deng, Jun Wan, Tian Xue, Wei Liu, and Mingjie Zhang. An unexpected INAD PDZ tandem-mediated plc$\beta$ binding in Drosophila photo receptors. *eLife*, 7, 2018.

[5105] Mengchen Ye, Kathryn M Lehigh, and David D Ginty. Multivesicular bodies mediate long-range retrograde NGF-TrkA signaling. *eLife*, 7, 2018.

[5106] Ya-Ping Yen, Wen-Fu Hsieh, Ya-Yin Tsai, Ya-Lin Lu, Ee Shan Liau, Ho-Chiang Hsu, Yen-Chung Chen, Ting-Chun Liu, Mien Chang, Joye Li, Shau-Ping Lin, Jui-Hung Hung, and Jun-An Chen. Dlk1-Dio3 locus-derived lncRNAs perpetuate postmitotic motor neuron cell fate and subtype identity. *eLife*, 7, 2018.

[5107] Pierre Yger, Giulia LB Spampinato, Elric Esposito, Baptiste Lefebvre, Stéphane Deny, Christophe Gardella, Marcel Stimberg, Florian Jetter, Guenther Zeck, Serge Picaud, Jens Duebel, and Olivier Marre. A spike sorting toolbox for up to thousands of electrodes validated with ground truth recordings in vitro and in vivo. *eLife*, 7, 2018.

[5108] Zhiqi Candice Yip and Maxwell G Heiman. Ordered arrangement of dendrites within a C. elegans sensory nerve bundle. *eLife*, 7, 2018.

[5109] Shigetoshi Yokoyama, Yan Cai, Miyuki Murata, Takeshi Tomita, Mitsuhiro Yoneda, Lei Xu, Aprile L Pilon, Raul E Cachau, and Shioko Kimura. A novel pathway of LPS uptake through syndecan-1 leading to pyroptotic cell death. *eLife*, 7, 2018.

[5110] Anna Yoney, Fred Etoc, Albert Ruzo, Thomas Carroll, Jakob J Metzger, Iain Martyn, Shu Li, Christoph Kirst, Eric D Siggia, and Ali H Brivanlou. WNT signaling memory is required for ACTIVIN to function as a morphogen in human gastruloids. *eLife*, 7, 2018.

[5111] Yosuke Yoneyama, Peter Lanzerstorfer, Hideaki Niwa, Takashi Umehara, Takashi Shibano, Shigeyuki Yokoyama, Kazuhiro Chida, Julian Weghuber, Fumihiko Hakuno, and Shin-Ichiro Takahashi. IRS-1 acts as an endocytic regulator of IGF-I receptor to facilitate sustained IGF signaling. *eLife*, 7, 2018.

[5112] Tae Yeon Yoo, Jeong-Mo Choi, William Conway, Che-Hang Yu, Rohit V Pappu, and Daniel J Needleman. Measuring NDC80 binding reveals the molecular basis of tension-dependent kinetochore-microtubule attachments. *eLife*, 7, 2018.

[5113] Harunori Yoshikawa, Mark Larance, Dylan J Harney, Ramasubramanian Sundaramoorthy, Tony Ly, Tom Owen-Hughes, and Angus I Lamond. Efficient analysis of mammalian polysomes in cells and tissues using Ribo Mega-SEC. *eLife*, 7, 2018.

[5114] Kazunori Yoshizawa, Yoshitaka Kamimura, Charles Lienhard, Rodrigo L Ferreira, and Alexander Blanke. A biological switching valve evolved in the female of a sex-role reversed cave insect to receive multiple seminal packages. *eLife*, 7, 2018.

[5115] Sangho Yu, Helia Cheng, Marie FranÇois, Emily Qualls-Creekmore, Clara Huesing, Yanlin He, Yanyan Jiang, Hong Gao, Yong Xu, Andrea Zsombok, Andrei V Derbenev, Eduardo A Nillni, David H Burk, Christopher D Morrison, Hans-Rudolf Berthoud, and Heike Münzberg. Preoptic leptin signaling modulates energy balance independent of body temperature regulation. *eLife*, 7, 2018.

[5116] Wenqi Yu, Dominique Missiakas, and Olaf Schneewind. Septal secretion of protein A in Staphylococcus aureus requires SecA and lipoteichoic acid synthesis. *eLife*, 7, 2018.

[5117] Zhou Yu, Lauren E Surface, Chong Yon Park, Max A Horlbeck, Gregory A Wyant, Monther Abu-Remaileh, Timothy R Peterson, David M Sabatini, Jonathan S Weissman, and Erin K O'Shea. Identification of a transporter complex responsible for the cytosolic entry of nitrogen-containing bisphosphonates. *eLife*, 7, 2018.

[5118] Fang Yuan, Xin Chen, Kai-Heng Fang, Yuanyuan Wang, Mingyan Lin, Shi-Bo Xu, Hai-Qin Huo, Min Xu, Lixiang Ma, Yuejun Chen, Shuijin He, and Yan Liu. Induction of human somatostatin and parvalbumin neurons by expressing a single transcription factor LIM homeobox 6. *eLife*, 7, 2018.

[5119] Giulia Zago, Irina Veith, Manish Kumar Singh, Laetitia Fuhrmann, Simon De Beco, Amanda Remorino, Saori Takaoka, Marjorie Palmeri, Frédérique Berger, Nathalie Brandon, Ahmed El Marjou, Anne Vincent-Salomon, Jacques Camonis, Mathieu Coppey, and Maria Carla Parrini. RalB directly triggers invasion downstream Ras by mobilizing the Wave complex. *eLife*, 7, 2018.

[5120] Christina Zakas, Jennifer M Deutscher, Alex D Kay, and Matthew V Rockman. Decoupled maternal and zygotic genetic effects shape the evolution of development. *eLife*, 7, 2018.

[5121] Fabio Zanini, Szu-Yuan Pu, Elena Bekerman, Shirit Einav, and Stephen R Quake. Single-cell transcriptional dynamics of flavivirus infection. *eLife*, 7, 2018.

[5122] Katarzyna Zawadzka, Pawel Zawadzki, Rachel Baker, Karthik V Rajasekar, Florence Wagner, David J Sherratt, and Lidia K Arciszewska. MukB ATPases are regulated independently by the N- and C-terminal domains of MukF kleisin. *eLife*, 7, 2018.

[5123] Mei Zeng, Nicholas P Kwiatkowski, Tinghu Zhang, Behnam Nabet, Mousheng Xu, Yanke Liang, Chunshan Quan, Jinhua Wang, Mingfeng Hao, Sangeetha Palakurthi, Shan Zhou, Qing Zeng, Paul T Kirschmeier, Khyati Meghani, Alan L Leggett, Jun Qi, Geoffrey I Shapiro, Joyce F Liu, Ursula A Matulonis, Charles Y Lin, Panagiotis A Konstantinopoulos, and Nathanael S Gray. Targeting MYC dependency in ovarian cancer through inhibition of CDK7 and CDK12/13. *eLife*, 7, 2018.

[5124] James P Zewe, Rachel C Wills, Sahana Sangappa, Brady D Goulden, and Gerald RV Hammond. SAC1 degrades its lipid substrate PtdIns4P in the endoplasmic reticulum to maintain a steep chemical gradient with donor membranes. *eLife*, 7, 2018.

[5125] Dao-Lai Zhang, Yu-Jing Sun, Ming-Liang Ma, Yi-jing Wang, Hui Lin, Rui-Rui Li, Zong-Lai Liang, Yuan Gao, Zhao Yang, Dong-Fang He, Amy Lin, Hui Mo, Yu-Jing Lu, Meng-Jing Li, Wei Kong, Ka Young Chung, Fan Yi, Jian-Yuan Li, Ying-Ying Qin, Jingxin Li, Alex R B Thomsen, Alem W Kahsai, Zi-Jiang Chen, Zhi-Gang Xu, Mingyao Liu, Dali Li, Xiao Yu, and Jin-Peng Sun. Gq activity- and $\beta$-arrestin-1 scaffolding-mediated ADGRG2/CFTR coupling are required for male fertility. *eLife*, 7, 2018.

[5126] Jiacheng Zhang, Bhaskar Chetnani, Eric D Cormack, Dulce Alonso, Wei Liu, Alfonso Mondragón, and Jingyi Fei. Specific structural elements of the T-box riboswitch drive the two-step binding of the tRNA ligand. *eLife*, 7, 2018.

[5127] Liangyu Zhang, Simone Köhler, Regina Rillo-Bohn, and Abby F Dernburg. A compartmentalized signaling network mediates crossover control in meiosis. *eLife*, 7, 2018.

[5128] Suyi Zhang, Hiroaki Mano, Michael Lee, Wako Yoshida, Mitsuo Kawato, Trevor W Robbins, and Ben Seymour. The control of tonic pain by active relief learning. *eLife*, 7, 2018.

[5129] Wen Zhang, Travis Walton, Li Li, and Jack W Szostak. Crystallographic observation of nonenzymatic RNA primer extension. *eLife*, 7, 2018.

[5130] Xianwei Zhang, Zicong Li, Huaqiang Yang, Dewu Liu, Gengyuan Cai, Guoling Li, Jianxin Mo, Dehua Wang, Cuili Zhong, Haoqiang Wang, Yue Sun, Junsong Shi, Enqin Zheng, Fanming Meng, Mao Zhang, Xiaoyan He, Rong Zhou, Jian Zhang, Miaorong Huang, Ran Zhang, Ning Li, Mingzhe Fan, Jinzeng Yang, and Zhenfang Wu. Novel transgenic pigs with enhanced growth and reduced environmental impact. *eLife*, 7, 2018.

[5131] Xue-Song Zhang, Jackie Li, Kimberly A Krautkramer, Michelle Badri, Thomas Battaglia, Timothy C Borbet, Hyunwook Koh, Sandy Ng, Rachel A Sibley, Yuanyuan Li, Wimal Pathmasiri, Shawn Jindal, Robin R Shields-Cutler, Ben Hillmann, Gabriel A Al-Ghalith, Victoria E Ruiz, Alexandra Livanos, Angélique B van 't Wout, Nabeetha Nagalingam, Arlin B Rogers, Susan Jenkins Sumner,

Dan Knights, John M Denu, Huilin Li, Kelly V Ruggles, Richard Bonneau, R Anthony Williamson, Marcus Rauch, and Martin J Blaser. Antibiotic-induced acceleration of type 1 diabetes alters maturation of innate intestinal immunity. *eLife*, 7, 2018.

[5132] Yingying Zhang, Jocelyn Charlton, Rahul Karnik, Isabel Beerman, Zachary D Smith, Hongcang Gu, Patrick Boyle, Xiaoli Mi, Kendell Clement, Ramona Pop, Andreas Gnirke, Derrick J Rossi, and Alexander Meissner. Targets and genomic constraints of ectopic Dnmt3b expression. *eLife*, 7, 2018.

[5133] Zhe Zhang, Balázs Tóth, Andras Szollosi, Jue Chen, and László Csanády. Structure of a TRPM2 channel in complex with Ca2+ explains unique gating regulation. *eLife*, 7, 2018.

[5134] Zhening Zhang, Wenguang G Liang, Lucas J Bailey, Yong Zi Tan, Hui Wei, Andrew Wang, Mara Farcasanu, Virgil A Woods, Lauren A McCord, David Lee, Weifeng Shang, Rebecca Deprez-Poulain, Benoit Deprez, David R Liu, Akiko Koide, Shohei Koide, Anthony A Kossiakoff, Sheng Li, Bridget Carragher, Clinton S Potter, and Wei-Jen Tang. Ensemble cryoEM elucidates the mechanism of insulin capture and degradation by human insulin degrading enzyme. *eLife*, 7, 2018.

[5135] Wei Zhao, Masayuki Oginuma, Rieko Ajima, Makoto Kiso, Akemi Okubo, and Yumiko Saga. Ripply2 recruits proteasome complex for Tbx6 degradation to define segment border during murine somitogenesis. *eLife*, 7, 2018.

[5136] Xiaoliang Zhao, Daniela Lenek, Ugur Dag, Barry J Dickson, and Krystyna Keleman. Persistent activity in a recurrent circuit underlies courtship memory in Drosophila. *eLife*, 7, 2018.

[5137] Yuehui Zhao, Lijiang Long, Wen Xu, Richard F Campbell, Edward E Large, Joshua S Greene, and Patrick T McGrath. Changes to social feeding behaviors are not sufficient for fitness gains of the Caenorhabditis elegans N2 reference strain. *eLife*, 7, 2018.

[5138] Yuhan Zhao, Lihua Wu, Xuetian Yue, Cen Zhang, Jianming Wang, Jun Li, Xiaohui Sun, Yiming Zhu, Zhaohui Feng, and Wenwei Hu. A polymorphism in the tumor suppressor p53 affects aging and longevity in mouse models. *eLife*, 7, 2018.

[5139] Ge Zheng, Mohammed Kanchwala, Chao Xing, and Hongtao Yu. MCM2–7-dependent cohesin loading during S phase promotes sister-chromatid cohesion. *eLife*, 7, 2018.

[5140] Peng Zhong, Casey R Vickstrom, Xiaojie Liu, Ying Hu, Laikang Yu, Han-Gang Yu, and Qing-song Liu. HCN2 channels in the ventral tegmental area regulate behavioral responses to chronic stress. *eLife*, 7, 2018.

[5141] Li Zhou, Mandy Yu Theng Lim, Prameet Kaur, Abil Saj, Diane Bortolamiol-Becet, Vikneswaran Gopal, Nicholas Tolwinski, Greg Tucker-Kellogg, and Katsutomo Okamura. Importance of miRNA stability and alternative primary miRNA isoforms in gene regulation during Drosophila development. *eLife*, 7, 2018.

[5142] Pengcheng Zhou, Shanna L Resendez, Jose Rodriguez-Romaguera, Jessica C Jimenez, Shay Q Neufeld, Andrea Giovannucci, Johannes Friedrich, Eftychios A Pnevmatikakis, Garret D Stuber, Rene Hen, Mazen A Kheirbek, Bernardo L Sabatini, Robert E Kass, and Liam Paninski. Efficient and accurate extraction of in vivo calcium signals from microendoscopic video data. *eLife*, 7, 2018.

[5143] Yu Zhou, Xiao-Ming Xia, and Christopher J Lingle. BK channel inhibition by strong extracellular acidification. *eLife*, 7, 2018.

[5144] Yue Zhou, Yuting Wang, Xinxing Zhang, Subhradeep Bhar, Rachel A Jones Lipinski, Jungsoo Han, Likui Feng, and Rebecca A Butcher. Biosynthetic tailoring of existing ascaroside pheromones alters their biological function in C. elegans. *eLife*, 7, 2018.

[5145] Zhipeng Zhou, Yunkun Dang, Mian Zhou, Haiyan Yuan, and Yi Liu. Codon usage biases co-evolve with transcription termination machinery to suppress premature cleavage and polyadenylation. *eLife*, 7, 2018.

[5146] Haichuan Zhu, Liuzhen Zhang, Yilin Wu, Bingjie Dong, Weilong Guo, Mei Wang, Lu Yang, Xiaoying Fan, Yuliang Tang, Ningshu Liu, Xiaoguang Lei, and Hong Wu. T-ALL leukemia stem cell 'stemness' is epigenetically controlled by the master regulator SPI1. *eLife*, 7, 2018.

[5147] Michal Ziemski, Ahmad Jomaa, Daniel Mayer, Sonja Rutz, Christoph Giese, Dmitry Veprintsev, and Eilika Weber-Ban. Cdc48-like protein of actinobacteria (Cpa) is a novel proteasome interactor in mycobacteria and related organisms. *eLife*, 7, 2018.

[5148] Iwan Zimmermann, Pascal Egloff, Cedric AJ Hutter, Fabian M Arnold, Peter Stohler, Nicolas Bocquet, Melanie N Hug, Sylwia Huber, Martin Siegrist, Lisa Hetemann, Jennifer Gera, Samira Gmür, Peter Spies, Daniel Gygax, Eric R Geertsma, Roger JP Dawson, and Markus A Seeger. Synthetic single domain antibodies for the conformational trapping of membrane proteins. *eLife*, 7, 2018.

[5149] Jasenko Zivanov, Takanori Nakane, Björn O Forsberg, Dari Kimanius, Wim JH Hagen, Erik Lindahl, and Sjors HW Scheres. New tools for automated high-resolution cryo-EM structure determination in RELION-3. *eLife*, 7, 2018.

[5150] Constantin M Zohner, Lidong Mo, and Susanne S Renner. Global warming reduces leaf-out and flowering synchrony among individuals. *eLife*, 7, 2018.

[5151] Philipp K Zuber, Irina Artsimovitch, Monali NandyMazumdar, Zhaokun Liu, Yuri Nedialkov, Kristian Schweimer, Paul Rösch, and Stefan H Knauer. The universally-conserved transcription factor RfaH is recruited to a hairpin structure of the non-template DNA strand. *eLife*, 7, 2018.

[5152] Stefano Zucca, Aya Zucca, Takashi Nakano, Sho Aoki, and Jeffery Wickens. Pauses in cholinergic interneuron firing exert an inhibitory control on striatal output in vivo. *eLife*, 7, 2018.

[5153] Jan J Zylicz, Maud Borensztein, Frederick CK Wong, Yun Huang, Caroline Lee, Sabine Dietmann, and M Azim Surani. G9a regulates temporal preimplantation developmental program and lineage segregation in blastocyst. *eLife*, 7, 2018.

[5154] Richard J Abdill and Ran Blekhman. Tracking the popularity and outcomes of all bioRxiv preprints. *eLife*, 8, 2019.

[5155] Merve Deniz Abdusselamoglu, Elif Eroglu, Thomas R Burkard, and Jürgen A Knoblich. The transcription factor odd-paired regulates temporal identity in transit-amplifying neural progenitors via an incoherent feed-forward loop. *eLife*, 8, 2019.

[5156] Amanda B Abildgaard, Amelie Stein, Sofie V Nielsen, Katrine Schultz-Knudsen, Elena Papaleo, Amruta Shrikhande, Eva R Hoffmann, Inge Bernstein, Anne-Marie Gerdes, Masanobu Takahashi, Chikashi Ishioka, Kresten Lindorff-Larsen, and Rasmus Hartmann-Petersen. Computational and cellular studies reveal structural destabilization and degradation of MLH1 variants in Lynch syndrome. *eLife*, 8, 2019.

[5157] Daniel Arthur Abrams, Aarthi Padmanabhan, Tianwen Chen, Paola Odriozola, Amanda E Baker, John Kochalka, Jennifer M Phillips, and Vinod Menon. Impaired voice processing in reward and salience circuits predicts social communication in children with autism. *eLife*, 8, 2019.

[5158] Enas Abu-Shah, Philippos Demetriou, Štefan Bálint, Viveka Mayya, Mikhail A Kutuzov, Omer Dushek, and Michael L Dustin. A tissue-like platform for studying engineered quiescent human T-cells' interactions with dendritic cells. *eLife*, 8, 2019.

[5159] Kevin Achberger, Christopher Probst, Jasmin Haderspeck, Sylvia Bolz, Julia Rogal, Johanna Chuchuy, Marina Nikolova, Virginia Cora, Lena Antkowiak, Wadood Haq, Nian Shen, Katja Schenke-Layland, Marius Ueffing, Stefan Liebau, and Peter Loskill. Merging organoid and organ-on-a-chip technology to generate complex multi-layer tissue models in a human retina-on-a-chip platform. *eLife*, 8, 2019.

[5160] Jessica E Ackerman, Anne EC Nichols, Valentina Studentsova, Katherine T Best, Emma Knapp, and Alayna E Loiselle. Cell non-autonomous functions of S100a4 drive fibrotic tendon healing. *eLife*, 8, 2019.

[5161] Frauke Ackermann, Kay Oliver Schink, Christine Bruns, Zsuzsanna Izsvák, F Kent Hamra, Christian Rosenmund, and Craig Curtis Garner. Critical role for Piccolo in synaptic vesicle retrieval. *eLife*, 8, 2019.

[5162] Sophie E Acton, Andrew JD Bell, Christopher P Toseland, and Alison Twelvetrees. A survey of new PIs in the UK. *eLife*, 8, 2019.

[5163] Hiroaki Adachi, Mauricio P Contreras, Adeline Harant, Chih-hang Wu, Lida Derevnina, Toshiyuki Sakai, Cian Duggan, Eleonora Moratto, Tolga O Bozkurt, Abbas Maqbool, Joe Win, and Sophien Kamoun. An N-terminal motif in NLR immune receptors is functionally conserved across distantly related plant species. *eLife*, 8, 2019.

[5164] Christina R Adams, Htet Htwe Htwe, Timothy Marsh, Aprilgate L Wang, Megan L Montoya, Lakshmipriya Subbaraj, Aaron D Tward, Nabeel Bardeesy, and Rushika M Perera. Transcriptional control of subtype switching ensures adaptation and growth of pancreatic cancer. *eLife*, 8, 2019.

[5165] Jenna N Adams, Anne Maass, Theresa M Harrison, Suzanne L Baker, and William J Jagust. Cortical tau deposition follows patterns of entorhinal functional connectivity in aging. *eLife*, 8, 2019.

[5166] Nicholas R Ader, Patrick C Hoffmann, Iva Ganeva, Alicia C Borgeaud, Chunxin Wang, Richard J Youle, and Wanda Kukulski. Molecular and topological reorganizations in mitochondrial architecture interplay during Bax-mediated steps of apoptosis. *eLife*, 8, 2019.

[5167] Evgenia Afanasieva, Indronil Chaudhuri, Jörg Martin, Eva Hertle, Astrid Ursinus, Vikram Alva, Marcus D Hartmann, and Andrei N Lupas. Structural diversity of oligomeric $\beta$-propellers with different numbers of identical blades. *eLife*, 8, 2019.

[5168] Olga Afonso, Colleen M Castellani, Liam P Cheeseman, Jorge G Ferreira, Bernardo Orr, Luisa T Ferreira, James J Chambers, Eurico Morais-de Sá, Thomas J Maresca, and Helder Maiato. Spatiotemporal control of mitotic exit during anaphase by an aurora B-Cdk1 crosstalk. *eLife*, 8, 2019.

[5169] Mohsen Afshar Bakooshli, Ethan S Lippmann, Ben Mulcahy, Nisha Iyer, Christine T Nguyen, Kayee Tung, Bryan A Stewart, Hubrecht van den Dorpel, Tobias Fuehrmann, Molly Shoichet, Anne Bigot, Elena Pegoraro, Henry Ahn, Howard Ginsberg, Mei Zhen, Randolph Scott Ashton, and Penney M Gilbert. A 3D culture model of innervated human skeletal muscle enables studies of the adult neuromuscular junction. *eLife*, 8, 2019.

[5170] Aleksandra Agapova, Agnese Serafini, Michael Petridis, Debbie M Hunt, Acely Garza-Garcia, Charles D Sohaskey, and Luiz Pedro Sório de Carvalho. Flexible nitrogen utilisation by the metabolic generalist pathogen Mycobacterium tuberculosis. *eLife*, 8, 2019.

[5171] Emil Dandanell Agerschou, Patrick Flagmeier, Theodora Saridaki, Céline Galvagnion, Daniel Komnig, Laetitia Heid, Vibha Prasad, Hamed Shaykhalishahi, Dieter Willbold, Christopher M Dobson, Aaron Voigt, Bjoern Falkenburger, Wolfgang Hoyer, and Alexander K Buell. An engineered monomer binding-protein for $\alpha$-synuclein efficiently inhibits the proliferation of amyloid fibrils. *eLife*, 8, 2019.

[5172] Shuchi Agrawal Singh, Mads Lerdrup, Ana-Luisa R Gomes, Harmen JG van de Werken, Jens Vilstrup Johansen, Robin Andersson, Albin Sandelin, Kristian Helin, and Klaus Hansen. PLZF targets developmental enhancers for activation during osteogenic differentiation of human mesenchymal stem cells. *eLife*, 8, 2019.

[5173] Zainab Ahdash, Euan Pyle, William John Allen, Robin A Corey, Ian Collinson, and Argyris Politis. HDX-MS reveals nucleotide-dependent, anti-correlated opening and closure of SecA and SecY channels of the bacterial translocon. *eLife*, 8, 2019.

[5174] Shivani Ahuja and Matthew R Whorton. Structural basis for mammalian nucleotide sugar transport. *eLife*, 8, 2019.

[5175] David O Ajasin, Vasudev R Rao, Xuhong Wu, Santhamani Ramasamy, Mario Pujato, Arthur P Ruiz, Andras Fiser, Anne R Bresnick, Ganjam V Kalpana, and Vinayaka R Prasad. CCL2 mobilizes ALIX to facilitate Gag-p6 mediated HIV-1 virion release. *eLife*, 8, 2019.

[5176] Kaur Alasoo, Julia Rodrigues, John Danesh, Daniel F Freitag, Dirk S Paul, and Daniel J Gaffney. Genetic effects on promoter usage are highly context-specific and contribute to complex traits. *eLife*, 8, 2019.

[5177] Mroj Alassaf, Emily C Daykin, Jaffna Mathiaparanam, and Marc A Wolman. Pregnancy-associated plasma protein-aa supports hair cell survival by regulating mitochondrial function. *eLife*, 8, 2019.

[5178] Benjamin Albert, Isabelle C Kos-Braun, Anthony K Henras, Christophe Dez, Maria Paula Rueda, Xu Zhang, Olivier Gadal, Martin Kos, and David Shore. A ribosome assembly stress response regulates transcription to maintain proteome homeostasis. *eLife*, 8, 2019.

[5179] Amal Alex, Valentina Piano, Soumitra Polley, Marchel Stuiver, Stephanie Voss, Giuseppe Ciossani, Katharina Overlack, Beate Voss, Sabine Wohlgemuth, Arsen Petrovic, Yaowen Wu, Philipp Selenko, Andrea Musacchio, and Stefano Maffini. Electroporated recombinant proteins as tools for in vivo functional complementation, imaging and chemical biology. *eLife*, 8, 2019.

[5180] Jeffrey M Alexander, Juan Guan, Bingkun Li, Lenka Maliskova, Michael Song, Yin Shen, Bo Huang, Stavros Lomvardas, and Orion D Weiner. Live-cell imaging reveals enhancer-dependent Sox2 transcription in the absence of enhancer proximity. *eLife*, 8, 2019.

[5181] Aminah T Ali, Lena Boehme, Guillermo Carbajosa, Vlad C Seitan, Kerrin S Small, and Alan Hodgkinson. Nuclear genetic regulation of the human mitochondrial transcriptome. *eLife*, 8, 2019.

[5182] Corey A H Allard, Hannah E Opalko, and James B Moseley. Stable Pom1 clusters form a glucose-modulated concentration gradient that regulates mitotic entry. *eLife*, 8, 2019.

[5183] Leandro M Alonso and Eve Marder. Visualization of currents in neural models with similar behavior and different conductance densities. *eLife*, 8, 2019.

[5184] Carolina Alvadia, Novandy K Lim, Vanessa Clerico Mosina, Gert T Oostergetel, Raimund Dutzler, and Cristina Paulino. Cryo-EM structures and functional characterization of the murine lipid scramblase TMEM16F. *eLife*, 8, 2019.

[5185] Ines Alvarez-Rodrigo, Thomas L Steinacker, Saroj Saurya, Paul T Conduit, Janina Baumbach, Zsofia A Novak, Mustafa G Aydogan, Alan Wainman, and Jordan W Raff. Evidence that a positive feedback loop drives centrosome maturation in fly embryos. *eLife*, 8, 2019.

[5186] Ryoji Amamoto, Mauricio D Garcia, Emma R West, Jiho Choi, Sylvain W Lapan, Elizabeth A Lane, Norbert Perrimon, and Constance L Cepko. Probe-Seq enables transcriptional profiling of specific cell types from heterogeneous tissue by RNA-based isolation. *eLife*, 8, 2019.

[5187] Olavo B Amaral, Kleber Neves, Ana P Wasilewska-Sampaio, and Clarissa FD Carneiro. The Brazilian Reproducibility Initiative. *eLife*, 8, 2019.

[5188] Katherine Cora Ames and Mark M Churchland. Motor cortex signals for each arm are mixed across hemispheres and neurons yet partitioned within the population response. *eLife*, 8, 2019.

[5189] Niko Amin-Wetzel, Lisa Neidhardt, Yahui Yan, Matthias P Mayer, and David Ron. Unstructured regions in IRE1$\alpha$ specify BiP-mediated destabilisation of the luminal domain dimer and repression of the UPR. *eLife*, 8, 2019.

[5190] Disi An, Ryosuke Fujiki, Dylan E Iannitelli, John W Smerdon, Shuvadeep Maity, Matthew F Rose, Alon Gelber, Elizabeth K Wanaselja, Ilona Yagudayeva, Joun Y Lee, Christine Vogel, Hynek Wichterle, Elizabeth C Engle, and Esteban Orlando Mazzoni. Stem cell-derived cranial and spinal motor neurons reveal proteostatic differences between ALS resistant and sensitive motor neurons. *eLife*, 8, 2019.

[5191] Savani Anbalagan, Janna Blechman, Michael Gliksberg, Ludmila Gordon, Ron Rotkopf, Tali Dadosh, Eyal Shimoni, and Gil Levkowitz. Robo2 regulates synaptic oxytocin content by affecting actin dynamics. *eLife*, 8, 2019.

[5192] Daniel Andergassen, Zachary D Smith, Jordan P Lewandowski, Chiara Gerhardinger, Alexander Meissner, and John L Rinn. In vivo Firre and Dxz4 deletion elucidates roles for autosomal gene regulation. *eLife*, 8, 2019.

[5193] Fredrik Andersen, Rani Lill Anjum, and Elena Rocca. Philosophical bias is the one bias that science cannot avoid. *eLife*, 8, 2019.

[5194] Jens Peter Andersen, Jesper Wiborg Schneider, Reshma Jagsi, and Mathias Wullum Nielsen. Gender variations in citation distributions in medicine are very small and due to self-citation and journal prestige. *eLife*, 8, 2019.

[5195] Brent W Anderson, Kuanqing Liu, Christine Wolak, Katarzyna Dubiel, Fukang She, Kenneth A Satyshur, James L Keck, and Jue D Wang. Evolution of (p)ppGpp-HPRT regulation through diversification of an allosteric oligomeric interaction. *eLife*, 8, 2019.

[5196] Gemma Anderson, John Dupré, and James G Wakefield. Drawing and the dynamic nature of living systems. *eLife*, 8, 2019.

[5197] Nicolas P Andrews, Justin X Boeckman, Colleen F Manning, Joe T Nguyen, Hannah Bechtold, Camelia Dumitras, Belvin Gong, Kimberly Nguyen, Deborah van der List, Karl D Murray, JoAnne Engebrecht, and James S Trimmer. A toolbox of IgG subclass-switched recombinant monoclonal antibodies for enhanced multiplex immunolabeling of brain. *eLife*, 8, 2019.

[5198] Cheen Euong Ang, Qing Ma, Orly L Wapinski, ShengHua Fan, Ryan A Flynn, Qian Yi Lee, Bradley Coe, Masahiro Onoguchi, Victor Hipolito Olmos, Brian T Do, Lynn Dukes-Rimsky, Jin Xu, Koji Tanabe, LiangJiang Wang, Ulrich Elling, Josef M Penninger, Yang Zhao, Kun Qu, Evan E Eichler, Anand Srivastava, Marius Wernig, and Howard Y Chang. The novel lncRNA lnc-NR2F1 is pro-neurogenic and mutated in human neurodevelopmental disorders. *eLife*, 8, 2019.

[5199] Francesca Angiolini, Elisa Belloni, Marco Giordano, Matteo Campioni, Federico Forneris, Maria Paola Paronetto, Michela Lupia, Chiara Brandas, Davide Pradella, Anna Di Matteo, Costanza Giampietro, Giovanna Jodice, Chiara Luise, Giovanni Bertalot, Stefano Freddi, Matteo Malinverno, Manuel Irimia, Jon D Moulton, James Summerton, Antonella Chiapparino, Carmen Ghilardi, Raffaella Giavazzi, Daniel Nyqvist, Davide Gabellini, Elisabetta Dejana, Ugo Cavallaro, and Claudia Ghigna. A novel L1CAM isoform with angiogenic activity generated by NOVA2-mediated alternative splicing. *eLife*, 8, 2019.

[5200] M Azim Ansari, Elihu Aranday-Cortes, Camilla LC Ip, Ana da Silva Filipe, Siu Hin Lau, Connor Bamford, David Bonsall, Amy Trebes, Paolo Piazza, Vattipally Sreenu, Vanessa M Cowton, J Ball, J Ball, E Barnes, G Burgess, G Cooke, J Dillon, G Foster, C Gore, N Guha, R Halford, C Holmes, E Hudson, S Hutchinson, W Irving, S Khakoo, P Klenerman, N Martin, T Mbisa, J McKeating, J McLauchlan, A Miners, A Murray, P Shaw, P Simmonds, S Smith, C Spencer, E Thomson, P Troke, P Vickerman, N Zitzmann, Emma Hudson, Rory Bowden, Arvind H Patel, Graham R Foster, William L Irving, Kosh Agarwal, Emma C Thomson, Peter Simmonds, Paul Klenerman, Chris Holmes, Eleanor Barnes, Chris CA Spencer, John McLauchlan, and Vincent Pedergnana. Interferon lambda 4 impacts the genetic diversity of hepatitis C virus. *eLife*, 8, 2019.

[5201] Paride Antinucci, Mónica Folgueira, and Isaac H Bianco. Pretectal neurons control hunting behaviour. *eLife*, 8, 2019.

[5202] Sho Aoki, Jared B Smith, Hao Li, Xunyi Yan, Masakazu Igarashi, Patrice Coulon, Jeffery R Wickens, Tom JH Ruigrok, and Xin Jin. An open cortico-basal ganglia loop allows limbic control over motor output via the nigrothalamic pathway. *eLife*, 8, 2019.

[5203] Miklos Argyelan, Leif Oltedal, Zhi-De Deng, Benjamin Wade, Marom Bikson, Andrea Joanlanne, Sohag Sanghani, Hauke Bartsch, Marta Cano, Anders M Dale, Udo Dannlowski, Annemiek Dols, Verena Enneking, Randall Espinoza, Ute Kessler, Katherine L Narr, Ketil J Oedegaard, Mardien L Oudega, Ronny Redlich, Max L Stek, Akihiro Takamiya, Louise Emsell, Filip Bouckaert, Pascal Sienaert, Jesus Pujol, Indira Tendolkar, Philip van Eijndhoven, Georgios Petrides, Anil K Malhotra, and Christopher Abbott. Electric field causes volumetric changes in the human brain. *eLife*, 8, 2019.

[5204] Georgios PD Argyropoulos, Clare Loane, Adriana Roca-Fernandez, Carmen Lage-Martinez, Oana Gurau, Sarosh R Irani, and Christopher R Butler. Network-wide abnormalities explain memory variability in hippocampal amnesia. *eLife*, 8, 2019.

[5205] Valentina Arkhipova, Gianluca Trinco, Thijs W Ettema, Sonja Jensen, Dirk J Slotboom, and Albert Guskov. Binding and transport of D-aspartate by the glutamate transporter homolog GltTk. *eLife*, 8, 2019.

[5206] Jean Paul Armache, Nathan Gamarra, Stephanie L Johnson, John D Leonard, Shenping Wu, Geeta J Narlikar, and Yifan Cheng. Cryo-EM structures of remodeler-nucleosome intermediates suggest allosteric control through the nucleosome. *eLife*, 8, 2019.

[5207] Catherine R Armbruster, Calvin K Lee, Jessica Parker-Gilham, Jaime de Anda, Aiguo Xia, Kun Zhao, Keiji Murakami, Boo Shan Tseng, Lucas R Hoffman, Fan Jin, Caroline S Harwood, Gerard CL Wong, and Matthew R Parsek. Heterogeneity in surface sensing suggests a division of labor in Pseudomonas aeruginosa populations. *eLife*, 8, 2019.

[5208] Lorena Armijo-Weingart, Andrea Ketschek, Rajiv Sainath, Almudena Pacheco, George M Smith, and Gianluca Gallo. Neurotrophins induce fission of mitochondria along embryonic sensory axons. *eLife*, 8, 2019.

[5209] Benjamin F Arnold, Diana L Martin, Jane Juma, Harran Mkocha, John B Ochieng, Gretchen M Cooley, Richard Omore, E Brook Goodhew, Jamae F Morris, Veronica Costantini, Jan Vinjé, Patrick J Lammie, and Jeffrey W Priest. Enteropathogen antibody dynamics and force of infection among children in low-resource settings. *eLife*, 8, 2019.

[5210] Torey R Arnold, Joseph H Shawky, Rachel E Stephenson, Kayla M Dinshaw, Tomohito Higashi, Farah Huq, Lance A Davidson, and Ann L Miller. Anillin regulates epithelial cell mechanics by structuring the medial-apical actomyosin network. *eLife*, 8, 2019.

[5211] Moises Arriaga and Edward B Han. Structured inhibitory activity dynamics in new virtual environments. *eLife*, 8, 2019.

[5212] Seksiri Arttamangkul, Andrew Plazek, Emily J Platt, Haihong Jin, Thomas F Murray, William T Birdsong, Kenner C Rice, David L Farrens, and John T Williams. Visualizing endogenous opioid receptors in living neurons using ligand-directed chemistry. *eLife*, 8, 2019.

[5213] Alok Arun, Susana M Coelho, Akira F Peters, Simon Bourdareau, Laurent Pérès, Delphine Scornet, Martina Strittmatter, Agnieszka P Lipinska, Haiqin Yao, Olivier Godfroy, Gabriel J Montecinos, Komlan Avia, Nicolas Macaisne, Christelle Troadec, Abdelhafid Bendahmane, and J Mark Cock. Convergent recruitment of TALE homeodomain life cycle regulators to direct sporophyte development in land plants and brown algae. *eLife*, 8, 2019.

[5214] Anoop Arunagiri, Leena Haataja, Anita Pottekat, Fawnnie Pamenan, Soohyun Kim, Lori M Zeltser, Adrienne W Paton, James C Paton, Billy Tsai, Pamela Itkin-Ansari, Randal J Kaufman, Ming Liu, and Peter Arvan. Proinsulin misfolding is an early event in the progression to type 2 diabetes. *eLife*, 8, 2019.

[5215] Patpicha Arunsan, Wannaporn Ittiprasert, Michael J Smout, Christina J Cochran, Victoria H Mann, Sujittra Chaiyadet, Shannon E Karinshak, Banchob Sripa, Neil David Young, Javier Sotillo, Alex Loukas, Paul J Brindley, and Thewarach Laha. Programmed knockout mutation of liver fluke granulin attenuates virulence of infection-induced hepatobiliary morbidity. *eLife*, 8, 2019.

[5216] James Ashley, Violet Sorrentino, Meike Lobb-Rabe, Sonal Nagarkar-Jaiswal, Liming Tan, Shuwa Xu, Qi Xiao, Kai Zinn, and Robert A Carrillo. Transsynaptic interactions between IgSF proteins DIP-$\alpha$ and Dpr10 are required for motor neuron targeting specificity. *eLife*, 8, 2019.

[5217] Yoshinori Aso, Robert P Ray, Xi Long, Daniel Bushey, Karol Cichewicz, Teri-TB Ngo, Brandi Sharp, Christina Christoforou, Amy Hu, Andrew L Lemire, Paul Tillberg, Jay Hirsh, Ashok Litwin-Kumar, and Gerald M Rubin. Nitric oxide acts as a cotransmitter in a subset of dopaminergic neurons to diversify memory dynamics. *eLife*, 8, 2019.

[5218] Salvatore Assenza, Alberto Stefano Sassi, Ruth Kellner, Benjamin Schuler, Paolo De Los Rios, and Alessandro Barducci. Efficient conversion of chemical energy into mechanical work by Hsp70 chaperones. *eLife*, 8, 2019.

[5219] Ruben M Atilho, Gayan Mirihana Arachchilage, Etienne B Greenlee, Kirsten M Knecht, and Ronald R Breaker. A bacterial riboswitch class for the thiamin precursor HMP-PP employs a terminator-embedded aptamer. *eLife*, 8, 2019.

[5220] Ayhan Atmanli, Dongjian Hu, Frederik Ernst Deiman, Annebel Marjolein van de Vrugt, FranÇois Cherbonneau, Lauren Deems Black, and Ibrahim John Domian. Multiplex live single-cell transcriptional analysis demarcates cellular functional heterogeneity. *eLife*, 8, 2019.

[5221] Jessica Ausborn, Natalia A Shevtsova, Vittorio Caggiano, Simon M Danner, and Ilya A Rybak. Computational modeling of brainstem circuits controlling locomotor frequency and gait. *eLife*, 8, 2019.

[5222] Emilie Auxerre-Plantié, Masayuki Nakamori, Yoan Renaud, Aline Huguet, Caroline Choquet, Cristiana Dondi, Lucile Miquerol, Masanori P Takahashi, Geneviève Gourdon, Guillaume Junion, Teresa Jagla, Monika Zmojdzian, and Krzysztof Jagla. Straightjacket/$\alpha 2\delta 3$ deregulation is associated with cardiac conduction defects in myotonic dystrophy type 1. *eLife*, 8, 2019.

[5223] Benjamin Auxier and Anna Bazzicalupo. Comment on 'Single nucleus sequencing reveals evidence of inter-nucleus recombination in arbuscular mycorrhizal fungi'. *eLife*, 8, 2019.

[5224] Maria Azkanaz, Aida Rodríguez López, Bauke de Boer, Wouter Huiting, Pierre-Olivier Angrand, Edo Vellenga, Harm H Kampinga, Steven Bergink, Joost HA Martens, Jan Jacob Schuringa, and Vincent van den Boom. Protein quality control in the nucleolus safeguards recovery of epigenetic regulators after heat shock. *eLife*, 8, 2019.

[5225] Maia Azubel, Stephen D Carter, Jennifer Weiszmann, Jun Zhang, Grant J Jensen, Yang Li, and Roger D Kornberg. FGF21 trafficking in intact human cells revealed by cryo-electron tomography with gold nanoparticles. *eLife*, 8, 2019.

[5226] Zeljana Babic, Amanda Capes-Davis, Maryann E Martone, Amos Bairoch, I Burak Ozyurt, Thomas H Gillespie, and Anita E Bandrowski. Incidences of problematic cell lines are lower in papers that use RRIDs to identify cell lines. *eLife*, 8, 2019.

[5227] Marc Bächinger, Rea Lehner, Felix Thomas, Samira Hanimann, Joshua Balsters, and Nicole Wenderoth. Human motor fatigability as evoked by repetitive movements results from a gradual breakdown of surround inhibition. *eLife*, 8, 2019.

[5228] Nathan A Baertsch and Jan-Marino Ramirez. Insights into the dynamic control of breathing revealed through cell-type-specific responses to substance P. *eLife*, 8, 2019.

[5229] Karen Baker, Irene A Gyamfi, Gregory I Mashanov, Justin E Molloy, Michael A Geeves, and Daniel P Mulvihill. TORC2-Gad8-dependent myosin phosphorylation modulates regulation by calcium. *eLife*, 8, 2019.

[5230] Lydia J Baker, Lindsay L Freed, Cole G Easson, Jose V Lopez, Danté Fenolio, Tracey T Sutton, Spencer V Nyholm, and Tory A Hendry. Diverse deep-sea anglerfishes share a genetically reduced luminous symbiont that is acquired from the environment. *eLife*, 8, 2019.

[5231] Akram Bakkour, Daniela J Palombo, Ariel Zylberberg, Yul HR Kang, Allison Reid, Mieke Verfaellie, Michael N Shadlen, and Daphna Shohamy. The hippocampus supports deliberation during value-based decisions. *eLife*, 8, 2019.

[5232] Timothy S Balmer and Laurence O Trussell. Selective targeting of unipolar brush cell subtypes by cerebellar mossy fibers. *eLife*, 8, 2019.

[5233] Sudeep Banjade, Shaogeng Tang, Yousuf H Shah, and Scott D Emr. Electrostatic lateral interactions drive ESCRT-III heteropolymer assembly. *eLife*, 8, 2019.

[5234] Belinda Baquero-Perez, Agne Antanaviciute, Ivaylo D Yonchev, Ian M Carr, Stuart A Wilson, and Adrian Whitehouse. The Tudor SND1 protein is an m6A RNA reader essential for replication of Kaposi's sarcoma-associated herpesvirus. *eLife*, 8, 2019.

[5235] Camilo Barbosa, Roderich Römhild, Philip Rosenstiel, and Hinrich Schulenburg. Evolutionary stability of collateral sensitivity to antibiotics in the model pathogen Pseudomonas aeruginosa. *eLife*, 8, 2019.

[5236] Paola Bardetti, Sónia Marisa Castanheira, Oliver Valerius, Gerhard H Braus, and José Pérez-Martín. Cytoplasmic retention and degradation of a mitotic inducer enable plant infection by a pathogenic fungus. *eLife*, 8, 2019.

[5237] Harihar Basnet, Lin Tian, Karuna Ganesh, Yun-Han Huang, Danilo G Macalinao, Edi Brogi, Lydia WS Finley, and Joan Massagué. Flura-seq identifies organ-specific metabolic adaptations during early metastatic colonization. *eLife*, 8, 2019.

[5238] Jens Bast, Kamil S Jaron, Donovan Schuseil, Denis Roze, and Tanja Schwander. Asexual reproduction reduces transposable element load in experimental yeast populations. *eLife*, 8, 2019.

[5239] Lucie Bastin-Héline, Arthur de Fouchier, Song Cao, Fotini Koutroumpa, Gabriela Caballero-Vidal, Stefania Robakiewicz, Christelle Monsempes, Marie-Christine FranÇois, Tatiana Ribeyre, Annick Maria, Thomas Chertemps, Anne de Cian, William B Walker, Guirong Wang, Emmanuelle Jacquin-Joly, and Nicolas Montagné. A novel lineage of candidate pheromone receptors for sex communication in moths. *eLife*, 8, 2019.

[5240] Rita A Batista, Jordi Moreno-Romero, Yichun Qiu, Joram van Boven, Juan Santos-González, Duarte D Figueiredo, and Claudia Köhler. The MADS-box transcription factor PHERES1 controls imprinting in the endosperm by binding to domesticated transposons. *eLife*, 8, 2019.

[5241] Rachel A Battaglia, Adriana S Beltran, Samed Delic, Raluca Dumitru, Jasmine A Robinson, Parijat Kabiraj, Laura E Herring, Victoria J Madden, Namritha Ravinder, Erik Willems, Rhonda A Newman, Roy A Quinlan, James E Goldman, Ming-Der Perng, Masaki Inagaki, and Natasha T Snider. Site-specific phosphorylation and caspase cleavage of GFAP are new markers of Alexander disease severity. *eLife*, 8, 2019.

[5242] Jacob Baudin, Juan M Angueyra, Raunak Sinha, and Fred Rieke. S-cone photoreceptors in the primate retina are functionally distinct from L and M cones. *eLife*, 8, 2019.

[5243] Samantha VH Bayer, Whitney R Grither, Audrey Brenot, Priscilla Y Hwang, Craig E Barcus, Melanie Ernst, Patrick Pence, Christopher Walter, Amit Pathak, and Gregory D Longmore. DDR2 controls breast tumor stiffness and metastasis by regulating integrin mediated mechanotransduction in CAFs. *eLife*, 8, 2019.

[5244] John Frederick Beckmann, Gagan Deep Sharma, Luis Mendez, Hongli Chen, and Mark Hochstrasser. The Wolbachia cytoplasmic incompatibility enzyme CidB targets nuclear import and protamine-histone exchange factors. *eLife*, 8, 2019.

[5245] David Alejandro Bejarano, Ke Peng, Vibor Laketa, Kathleen Börner, K Laurence Jost, Bojana Lucic, Bärbel Glass, Marina Lusic, Barbara Müller, and Hans-Georg Kräusslich. HIV-1 nuclear import in macrophages is regulated by CPSF6-capsid interactions at the nuclear pore complex. *eLife*, 8, 2019.

[5246] Tristan A Bell, Tania A Baker, and Robert T Sauer. Interactions between a subset of substrate side chains and AAA+ motor pore loops determine grip during protein unfolding. *eLife*, 8, 2019.

[5247] Jacob LS Bellmund, Lorena Deuker, and Christian F Doeller. Mapping sequence structure in the human lateral entorhinal cortex. *eLife*, 8, 2019.

[5248] Raphael M Bendriem, Shawn Singh, Alice Abdel Aleem, David A Antonetti, and M Elizabeth Ross. Tight junction protein occludin regulates progenitor Self-Renewal and survival in developing cortex. *eLife*, 8, 2019.

[5249] Fabienne Benz, Viraya Wichitnaowarat, Martin Lehmann, Raoul FV Germano, Diana Mihova, Jadranka Macas, Ralf H Adams, M Mark Taketo, Karl-Heinz Plate, Sylvaine Guérit, Benoit Vanhollebeke, and Stefan Liebner. Low wnt/$\beta$-catenin signaling determines leaky vessels in the subfornical organ and affects water homeostasis in mice. *eLife*, 8, 2019.

[5250] Flavio H Beraldo, Daniel Palmer, Sara Memar, David I Wasserman, Wai-Jane V Lee, Shuai Liang, Samantha D Creighton, Benjamin Kolisnyk, Matthew F Cowan, Justin Mels, Talal S Masood, Chris Fodor, Mohammed A Al-Onaizi, Robert Bartha, Tom Gee, Lisa M Saksida, Timothy J Bussey, Stephen S Strother, Vania F Prado, Boyer D Winters, and Marco AM Prado. MouseBytes, an open-access high-throughput pipeline and database for rodent touchscreen-based cognitive assessment. *eLife*, 8, 2019.

[5251] Jeremy J Berg, Arbel Harpak, Nasa Sinnott-Armstrong, Anja Moltke Joergensen, Hakhamanesh Mostafavi, Yair Field, Evan August Boyle, Xinjun Zhang, Fernando Racimo, Jonathan K Pritchard, and Graham Coop. Reduced signal for polygenic adaptation of height in UK Biobank. *eLife*, 8, 2019.

[5252] Kerryn Berndsen, Pawel Lis, Wondwossen M Yeshaw, Paulina S Wawro, Raja S Nirujogi, Melanie Wightman, Thomas Macartney, Mark Dorward, Axel Knebel, Francesca Tonelli, Suzanne R Pfeffer, and Dario R Alessi. PPM1H phosphatase counteracts LRRK2 signaling by selectively dephosphorylating Rab proteins. *eLife*, 8, 2019.

[5253] David B Bernstein, Floyd E Dewhirst, and Daniel Segrè. Metabolic network percolation quantifies biosynthetic capabilities across the human oral microbiome. *eLife*, 8, 2019.

[5254] Paul M Berube, Anna Rasmussen, Rogier Braakman, Ramunas Stepanauskas, and Sallie W Chisholm. Emergence of trait variability through the lens of nitrogen assimilation in Prochlorococcus. *eLife*, 8, 2019.

[5255] Tushar D Bhagat, Dagny Von Ahrens, Meelad Dawlaty, Yiyu Zou, Joelle Baddour, Abhinav Achreja, Hongyun Zhao, Lifeng Yang, Brijesh Patel, Changsoo Kwak, Gaurav S Choudhary, Shanisha Gordon-Mitchell, Srinivas Aluri, Sanchari Bhattacharyya, Srabani Sahu, Prafulla Bhagat, Yiting Yu, Matthias Bartenstein, Orsi Giricz, Masako Suzuki, Davendra Sohal, Sonal Gupta, Paola A Guerrero, Surinder Batra, Michael Goggins, Ulrich Steidl, John Greally, Beamon Agarwal, Kith Pradhan, Debabrata Banerjee, Deepak Nagrath, Anirban Maitra, and Amit Verma. Lactate-mediated epigenetic reprogramming regulates formation of human pancreatic cancer-associated fibroblasts. *eLife*, 8, 2019.

[5256] Vandita D Bhat, Kathleen L McCann, Yeming Wang, Dallas R Fonseca, Tarjani Shukla, Jacqueline C Alexander, Chen Qiu, Marv Wickens, Te-Wen Lo, Traci M Tanaka Hall, and Zachary T Campbell. Engineering a conserved RNA regulatory protein repurposes its biological function in vivo. *eLife*, 8, 2019.

[5257] Aanchal Bhatia, Sahil Moza, and Upinder Singh Bhalla. Precise excitation-inhibition balance controls gain and timing in the hippocampus. *eLife*, 8, 2019.

[5258] Neha Bhatia, Henrik Åhl, Henrik Jönsson, and Marcus G Heisler. Quantitative analysis of auxin sensing in leaf primordia argues against proposed role in regulating leaf dorsoventrality. *eLife*, 8, 2019.

[5259] Surabhi Bhutani, James D Howard, Rachel Reynolds, Phyllis C Zee, Jay Gottfried, and Thorsten Kahnt. Olfactory connectivity mediates sleep-dependent food choices in humans. *eLife*, 8, 2019.

[5260] Christopher Bianco and Ian Mohr. Ribosome biogenesis restricts innate immune responses to virus infection and DNA. *eLife*, 8, 2019.

[5261] John W Biddle, Maximilian Nguyen, and Jeremy Gunawardena. Negative reciprocity, not ordered assembly, underlies the interaction of Sox2 and Oct4 on DNA. *eLife*, 8, 2019.

[5262] Noa Bielopolski, Hoger Amin, Anthi A Apostolopoulou, Eyal Rozenfeld, Hadas Lerner, Wolf Huetteroth, Andrew C Lin, and Moshe Parnas. Inhibitory muscarinic acetylcholine receptors enhance aversive olfactory learning in adult Drosophila. *eLife*, 8, 2019.

[5263] Nicolas Bigot, Matthew Day, Robert A Baldock, Felicity Z Watts, Antony W Oliver, and Laurence H Pearl. Phosphorylation-mediated interactions with TOPBP1 couple 53BP1 and 9-1-1 to control the G1 DNA damage checkpoint. *eLife*, 8, 2019.

[5264] Janine Diane Bijsterbosch, Christian F Beckmann, Mark W Woolrich, Stephen M Smith, and Samuel J Harrison. The relationship between spatial configuration and functional connectivity of brain regions revisited. *eLife*, 8, 2019.

[5265] Loïc Binan, FranÇois Bélanger, Maxime Uriarte, Jean FranÇois Lemay, Jean Christophe Pelletier De Koninck, Joannie Roy, El Bachir Affar, Elliot Drobetsky, Hugo Wurtele, and Santiago Costantino. Opto-magnetic capture of individual cells based on visual phenotypes. *eLife*, 8, 2019.

[5266] William T Birdsong, Bart C Jongbloets, Kim A Engeln, Dong Wang, Grégory Scherrer, and Tianyi Mao. Synapse-specific opioid modulation of thalamo-cortico-striatal circuits. *eLife*, 8, 2019.

[5267] Ingvars Birznieks, Sarah McIntyre, Hanna Maria Nilsson, Saad S Nagi, Vaughan G Macefield, David A Mahns, and Richard M Vickery. Tactile sensory channels over-ruled by frequency decoding system that utilizes spike pattern regardless of receptor type. *eLife*, 8, 2019.

[5268] Avik Biswas, Allan Haldane, Eddy Arnold, and Ronald M Levy. Epistasis and entrenchment of drug resistance in HIV-1 subtype B. *eLife*, 8, 2019.

[5269] Mary-Ann Blätke and Andrea Bräutigam. Evolution of C4 photosynthesis predicted by constraint-based modelling. *eLife*, 8, 2019.

[5270] Isabelle Maria Blomfield, Brenda Rocamonde, Maria del Mar Masdeu, Eskeatnaf Mulugeta, Stefania Vaga, Debbie LC van den Berg, Emmanuelle Huillard, FranÇois Guillemot, and Noelia Urbán. Id4 promotes the elimination of the pro-activation factor Ascl1 to maintain quiescence of adult hippocampal stem cells. *eLife*, 8, 2019.

[5271] Joshua S Bloom, James Boocock, Sebastian Treusch, Meru J Sadhu, Laura Day, Holly Oates-Barker, and Leonid Kruglyak. Rare variants contribute disproportionately to quantitative trait variation in yeast. *eLife*, 8, 2019.

[5272] Michael AP Bloomfield, Robert A McCutcheon, Matthew Kempton, Tom P Freeman, and Oliver Howes. The effects of psychosocial stress on dopaminergic function and the acute stress response. *eLife*, 8, 2019.

[5273] Silvia C Bobeica, Shi-Hui Dong, Liujie Huo, Nuria Mazo, Martin I McLaughlin, Gonzalo Jiménez-Osés, Satish K Nair, and Wilfred A van der Donk. Insights into AMS/PCAT transporters from biochemical and structural characterization of a double Glycine motif protease. *eLife*, 8, 2019.

[5274] Irene Bolea, Alejandro Gella, Elisenda Sanz, Patricia Prada-Dacasa, Fabien Menardy, Angela M Bard, Pablo Machuca-Márquez, Abel Eraso-Pichot, Guillem Mòdol-Caballero, Xavier Navarro, Franck Kalume, and Albert Quintana. Defined neuronal populations drive fatal phenotype in a mouse model of Leigh syndrome. *eLife*, 8, 2019.

[5275] Florian Bolenz, Wouter Kool, Andrea MF Reiter, and Ben Eppinger. Metacontrol of decision-making strategies in human aging. *eLife*, 8, 2019.

[5276] Madison Bolger-Munro, Kate Choi, Joshua M Scurll, Libin Abraham, Rhys S Chappell, Duke Sheen, May Dang-Lawson, Xufeng Wu, John J Priatel, Daniel Coombs, John A Hammer, and Michael R Gold. Arp2/3 complex-driven spatial patterning of the BCR enhances immune synapse formation, BCR signaling and B cell activation. *eLife*, 8, 2019.

[5277] Andrew D Bolton, Martin Haesemeyer, Josua Jordi, Ulrich Schaechtle, Feras A Saad, Vikash K Mansinghka, Joshua B Tenenbaum, and Florian Engert. Elements of a stochastic 3D prediction engine in larval zebrafish prey capture. *eLife*, 8, 2019.

[5278] Valentina Bonnefil, Karen Dietz, Mario Amatruda, Maureen Wentling, Antonio V Aubry, Jeffrey L Dupree, Gary Temple, Hye-Jin Park, Nesha S Burghardt, Patrizia Casaccia, and Jia Liu. Region-specific myelin differences define behavioral consequences of chronic social defeat stress in mice. *eLife*, 8, 2019.

[5279] Alexander JR Booth, Zuojun Yue, John K Eykelenboom, Tom Stiff, GW Gant Luxton, Helfrid Hochegger, and Tomoyuki U Tanaka. Contractile acto-myosin network on nuclear envelope remnants positions human chromosomes for mitosis. *eLife*, 8, 2019.

[5280] Lauren N Booth, Travis J Maures, Robin W Yeo, Cindy Tantilert, and Anne Brunet. Self-sperm induce resistance to the detrimental effects of sexual encounters with males in hermaphroditic nematodes. *eLife*, 8, 2019.

[5281] Melissa A Borgen, Andrew C Giles, Dandan Wang, and Brock Grill. Synapse maintenance is impacted by ATAT-2 tubulin acetyltransferase activity and the RPM-1 signaling hub. *eLife*, 8, 2019.

[5282] Romain Bourboulou, Geoffrey Marti, FranÇois-Xavier Michon, Elissa El Feghaly, Morgane Nouguier, David Robbe, Julie Koenig, and Jerome Epsztein. Dynamic control of hippocampal spatial coding resolution by local visual cues. *eLife*, 8, 2019.

[5283] Kimberly Box, Bradley W Joyce, and Danelle Devenport. Epithelial geometry regulates spindle orientation and progenitor fate during formation of the mammalian epidermis. *eLife*, 8, 2019.

[5284] Julianna Bozler, Balint Z Kacsoh, and Giovanni Bosco. Transgenerational inheritance of ethanol preference is caused by maternal NPF repression. *eLife*, 8, 2019.

[5285] Aaron T Bozzi, Christina M Zimanyi, John M Nicoludis, Brandon K Lee, Casey H Zhang, and Rachelle Gaudet. Structures in multiple conformations reveal distinct transition metal and proton pathways in an Nramp transporter. *eLife*, 8, 2019.

[5286] Ryan Bracewell, Kamalakar Chatla, Matthew J Nalley, and Doris Bachtrog. Dynamic turnover of centromeres drives karyotype evolution in Drosophila. *eLife*, 8, 2019.

[5287] Alberto Brandariz-Nuñez, Ting Liu, Te Du, and Alex Evilevitch. Pressure-driven release of viral genome into a host nucleus is a mechanism leading to herpes infection. *eLife*, 8, 2019.

[5288] Meret Branscheidt, Panagiotis Kassavetis, Manuel Anaya, Davis Rogers, Han Debra Huang, Martin A Lindquist, and Pablo Celnik. Fatigue induces long-lasting detrimental changes in motor-skill learning. *eLife*, 8, 2019.

[5289] Andrea Brenna, Iwona Olejniczak, Rohit Chavan, Jürgen A Ripperger, Sonja Langmesser, Elisabetta Cameroni, Zehan Hu, Claudio De Virgilio, Jörn Dengjel, and Urs Albrecht. Cyclin-dependent kinase 5 (CDK5) regulates the circadian clock. *eLife*, 8, 2019.

[5290] Connor Brennan and Alexander Proekt. A quantitative model of conserved macroscopic dynamics predicts future motor commands. *eLife*, 8, 2019.

[5291] Marian Breuer, Emmy E Earnest, Chuck Merryman, Kim S Wise, Lijie Sun, Michaela R Lynott, Clyde A Hutchison, Hamilton O Smith, John D Lapek, David J Gonzalez, Valérie de Crécy-Lagard, Drago Haas, Andrew D Hanson, Piyush Labhsetwar, John I Glass, and Zaida Luthey-Schulten. Essential metabolism for a minimal cell. *eLife*, 8, 2019.

[5292] Helena Brezovjakova, Chris Tomlinson, Noor Mohd Naim, Pamela Swiatlowska, Jennifer C Erasmus, Stephan Huveneers, Julia Gorelik, Susann Bruche, and Vania MM Braga. Junction Mapper is a novel computer vision tool to decipher cell–cell contact phenotypes. *eLife*, 8, 2019.

[5293] Ben F Brian, Adrienne S Jolicoeur, Candace R Guerrero, Myra G Nunez, Zoi E Sychev, Siv A Hegre, Pål Sætrom, Nagy Habib, Justin M Drake, Kathryn L Schwertfeger, and Tanya S Freedman. Unique-region phosphorylation targets LynA for rapid degradation, tuning its expression and signaling in myeloid cells. *eLife*, 8, 2019.

[5294] Holly Bridge, Andrew H Bell, Matthew Ainsworth, Jerome Sallet, Elsie Premereur, Bashir Ahmed, Anna S Mitchell, Urs Schüffelgen, Mark Buckley, Benjamin C Tendler, Karla L Miller, Rogier B Mars, Andrew J Parker, and Kristine Krug. Preserved extrastriate visual network in a monkey with substantial, naturally occurring damage to primary visual cortex. *eLife*, 8, 2019.

[5295] Marisa M Brockmann, Marta Maglione, Claudia G Willmes, Alexander Stumpf, Boris A Bouazza, Laura M Velasquez, M Katharina Grauel, Prateep Beed, Martin Lehmann, Niclas Gimber, Jan Schmoranzer, Stephan J Sigrist, Christian Rosenmund, and Dietmar Schmitz. RIM-BP2 primes synaptic vesicles via recruitment of Munc13-1 at hippocampal mossy fiber synapses. *eLife*, 8, 2019.

[5296] Nichole A Broderick and Arturo Casadevall. Gender inequalities among authors who contributed equally. *eLife*, 8, 2019.

[5297] Stephen G Brohawn, Weiwei Wang, Annie Handler, Ernest B Campbell, Jürgen R Schwarz, and Roderick MacKinnon. The mechanosensitive ion channel TRAAK is localized to the mammalian node of Ranvier. *eLife*, 8, 2019.

[5298] D Garrett Brown, Raymond Soto, Soumya Yandamuri, Colleen Stone, Laura Dickey, Joao Carlos Gomes-Neto, Elissa D Pastuzyn, Rickesha Bell, Charisse Petersen, Kaitlin Buhrke, Robert S Fujinami, Ryan M O'Connell, W Zac Stephens, Jason D Shepherd, Thomas E Lane, and June L Round. The microbiota protects from viral-induced neurologic damage through microglia-intrinsic TLR signaling. *eLife*, 8, 2019.

[5299] Clarisse Brunet Avalos, G Larisa Maier, Rémy Bruggmann, and Simon G Sprecher. Single cell transcriptome atlas of the Drosophila larval brain. *eLife*, 8, 2019.

[5300] Jakub Witold Bubnicki, Marcin Churski, Krzysztof Schmidt, Tom A Diserens, and Dries PJ Kuijper. Linking spatial patterns of terrestrial herbivore community structure to trophic interactions. *eLife*, 8, 2019.

[5301] Abigail Buchwalter, Roberta Schulte, Hsiao Tsai, Juliana Capitanio, and Martin Hetzer. Selective clearance of the inner nuclear membrane protein emerin by vesicular transport during ER stress. *eLife*, 8, 2019.

[5302] Paula J Bucko, Chloe K Lombard, Lindsay Rathbun, Irvin Garcia, Akansha Bhat, Linda Wordeman, F Donelson Smith, Dustin J Maly, Heidi Hehnly, and John D Scott. Subcellular drug targeting illuminates local kinase action. *eLife*, 8, 2019.

[5303] Breane G Budaitis, Shashank Jariwala, Dana N Reinemann, Kristin I Schimert, Guido Scarabelli, Barry J Grant, David Sept, Matthew J Lang, and Kristen J Verhey. Neck linker docking is critical for Kinesin-1 force generation in cells but at a cost to motor speed and processivity. *eLife*, 8, 2019.

[5304] Julie G Burel, Mikhail Pomaznoy, Cecilia S Lindestam Arlehamn, Daniela Weiskopf, Ricardo da Silva Antunes, Yunmin Jung, Mariana Babor, Veronique Schulten, Gregory Seumois, Jason A Greenbaum, Sunil Premawansa, Gayani Premawansa, Ananda Wijewickrama, Dhammika Vidanagama, Bandu Gunasena, Rashmi Tippalagama, Aruna D deSilva, Robert H Gilman, Mayuko Saito, Randy Taplitz, Klaus Ley, Pandurangan Vijayanand, Alessandro Sette, and Bjoern Peters. Circulating T cell-monocyte complexes are markers of immune perturbations. *eLife*, 8, 2019.

[5305] C Joseph Burnett, Samuel C Funderburk, Jovana Navarrete, Alexander Sabol, Jing Liang-Guallpa, Theresa M Desrochers, and Michael J Krashes. Need-based prioritization of behavior. *eLife*, 8, 2019.

[5306] Niklas Byczkowicz, Abdelmoneim Eshra, Jacqueline Montanaro, Andrea Trevisiol, Johannes Hirrlinger, Maarten HP Kole, Ryuichi Shigemoto, and Stefan Hallermann. HCN channel-mediated neuromodulation can control action potential velocity and fidelity in central axons. *eLife*, 8, 2019.

[5307] Xiao-Chuan Cai, Tuo Zhang, Eui-jun Kim, Ming Jiang, Ke Wang, Junyi Wang, Shi Chen, Nawei Zhang, Hong Wu, Fengling Li, Carlo C dela Seña, Hong Zeng, Victor Vivcharuk, Xiang Niu, Weihong Zheng, Jonghan P Lee, Yuling Chen, Dalia Barsyte, Magda Szewczyk, Taraneh Hajian, Glorymar Ibáñez, Aiping Dong, Ludmila Dombrovski, Zhenyu Zhang, Haiteng Deng, Jinrong Min, Cheryl H Arrowsmith, Linas Mazutis, Lei Shi, Masoud Vedadi, Peter J Brown, Jenny Xiang, Li-Xuan Qin, Wei Xu, and Minkui Luo. A chemical probe of CARM1 alters epigenetic plasticity against breast cancer cell invasion. *eLife*, 8, 2019.

[5308] Axelle Calcus, Outi Tuomainen, Ana Campos, Stuart Rosen, and Lorna F Halliday. Functional brain alterations following mild-to-moderate sensorineural hearing loss in children. *eLife*, 8, 2019.

[5309] Rebecca A Callahan, Richard Roberts, Mohini Sengupta, Yukiko Kimura, Shin-ichi Higashijima, and Martha W Bagnall. Spinal V2b neurons reveal a role for ipsilateral inhibition in speed control. *eLife*, 8, 2019.

[5310] Ian Winsten Campbell, Xiaoxue Zhou, and Angelika Amon. The Mitotic Exit Network integrates temporal and spatial signals by distributing regulation across multiple components. *eLife*, 8, 2019.

[5311] Florent Campo-Paysaa, Jonathan DW Clarke, and Richard JT Wingate. Generation of the squamous epithelial roof of the 4th ventricle. *eLife*, 8, 2019.

[5312] Pengfei Cao, Sang-Jin Kim, Anqi Xing, Craig A Schenck, Lu Liu, Nan Jiang, Jie Wang, Robert L Last, and Federica Brandizzi. Homeostasis of branched-chain amino acids is critical for the activity of TOR signaling in Arabidopsis. *eLife*, 8, 2019.

[5313] Pengxing Cao, Katharine A Collins, Sophie Zaloumis, Thanaporn Wattanakul, Joel Tarning, Julie A Simpson, James McCarthy, and James M McCaw. Modeling the dynamics of Plasmodium falciparum gametocytes in humans during malaria infection. *eLife*, 8, 2019.

[5314] Christina M Caragine, Shannon C Haley, and Alexandra Zidovska. Nucleolar dynamics and interactions with nucleoplasm in living cells. *eLife*, 8, 2019.

[5315] Jeffrey N Carey, Erin L Mettert, Daniel R Fishman-Engel, Manuela Roggiani, Patricia J Kiley, and Mark Goulian. Phage integration alters the respiratory strategy of its host. *eLife*, 8, 2019.

[5316] Michael Luke Carlson, R Greg Stacey, John William Young, Irvinder Singh Wason, Zhiyu Zhao, David G Rattray, Nichollas Scott, Craig H Kerr, Mohan Babu, Leonard J Foster, and Franck Duong Van Hoa. Profiling the Escherichia coli membrane protein interactome captured in Peptidisc libraries. *eLife*, 8, 2019.

[5317] Jorge Carretero-Ortega, Zinal Chhangawala, Shane Hunt, Carlos Narvaez, Javier Menéndez-González, Carl M Gay, Tomasz Zygmunt, Xiaochun Li, and Jesús Torres-Vázquez. GIPC proteins negatively modulate Plexind1 signaling during vascular development. *eLife*, 8, 2019.

[5318] Luis Carrillo-Reid, Michelle Day, Zhong Xie, Alexandria E Melendez, Jyothisri Kondapalli, Joshua L Plotkin, David L Wokosin, Yu Chen, Geraldine J Kress, Michael Kaplitt, Ema Ilijic, Jaime N Guzman, C Savio Chan, and D James Surmeier. Mutant huntingtin enhances activation of dendritic Kv4 K+ channels in striatal spiny projection neurons. *eLife*, 8, 2019.

[5319] Joana R Carvalho, Isabela C Fortunato, Catarina G Fonseca, Anna Pezzarossa, Pedro Barbacena, Maria A Dominguez-Cejudo, Francisca F Vasconcelos, Nuno C Santos, Filomena A Carvalho, and Claudio A Franco. Non-canonical Wnt signaling regulates junctional mechanocoupling during angiogenic collective cell migration. *eLife*, 8, 2019.

[5320] Karine Casier, Valérie Delmarre, Nathalie Gueguen, Catherine Hermant, Elise Viodé, Chantal Vaury, Stéphane Ronsseray, Emilie Brasset, Laure Teysset, and Antoine Boivin. Environmentally-induced epigenetic conversion of a piRNA cluster. *eLife*, 8, 2019.

[5321] Antonino Casile, Jonathan D Victor, and Michele Rucci. Contrast sensitivity reveals an oculomotor strategy for temporally encoding space. *eLife*, 8, 2019.

[5322] Elisa Castaldi, Manuela Piazza, Stanislas Dehaene, Alexandre Vignaud, and Evelyn Eger. Attentional amplification of neural codes for number independent of other quantities along the dorsal visual stream. *eLife*, 8, 2019.

[5323] João PL Castro, Michelle N Yancoskie, Marta Marchini, Stefanie Belohlavy, Layla Hiramatsu, Marek Kučka, William H Beluch, Ronald Naumann, Isabella Skuplik, John Cobb, Nicholas H Barton, Campbell Rolian, and Yingguang Frank Chan. An integrative genomic analysis of the Longshanks selection experiment for longer limbs in mice. *eLife*, 8, 2019.

[5324] Claudia Cattoglio, Iryna Pustova, Nike Walther, Jaclyn J Ho, Merle Hantsche-Grininger, Carla J Inouye, M Julius Hossain, Gina M Dailey, Jan Ellenberg, Xavier Darzacq, Robert Tjian, and Anders S Hansen. Determining cellular CTCF and cohesin abundances to constrain 3D genome models. *eLife*, 8, 2019.

[5325] Jordi Cayuso, Qiling Xu, Megan Addison, and David G Wilkinson. Actomyosin regulation by Eph receptor signaling couples boundary cell formation to border sharpness. *eLife*, 8, 2019.

[5326] Adelaida M Celaya, Isabel Sánchez-Pérez, Jose M Bermúdez-Muñoz, Lourdes Rodríguez-de la Rosa, Laura Pintado-Berninches, Rosario Perona, Silvia Murillo-Cuesta, and Isabel Varela-Nieto. Deficit of mitogen-activated protein kinase phosphatase 1 (DUSP1) accelerates progressive hearing loss. *eLife*, 8, 2019.

[5327] Kanokwan Champasa, Caitlin Blank, Larry J Friedman, Jeff Gelles, and Stephen P Bell. A conserved Mcm4 motif is required for Mcm2-7 double-hexamer formation and origin DNA unwinding. *eLife*, 8, 2019.

[5328] Jackson Champer, Joan Chung, Yoo Lim Lee, Chen Liu, Emily Yang, Zhaoxin Wen, Andrew G Clark, and Philipp W Messer. Molecular safeguarding of CRISPR gene drive experiments. *eLife*, 8, 2019.

[5329] Hsiao-Han Chang, Amy Wesolowski, Ipsita Sinha, Christopher G Jacob, Ayesha Mahmud, Didar Uddin, Sazid Ibna Zaman, Md Amir Hossain, M Abul Faiz, Aniruddha Ghose, Abdullah Abu Sayeed, M Ridwanur Rahman, Akramul Islam, Mohammad Jahirul Karim, M Kamar Rezwan, Abul

Khair Mohammad Shamsuzzaman, Sanya Tahmina Jhora, M M Aktaruzzaman, Eleanor Drury, Sonia GonÇalves, Mihir Kekre, Mehul Dhorda, Ranitha Vongpromek, Olivo Miotto, Kenth Engø-Monsen, Dominic Kwiatkowski, Richard J Maude, and Caroline Buckee. Mapping imported malaria in Bangladesh using parasite genetic and human mobility data. *eLife*, 8, 2019.

[5330] Yunjie Chang, Ki Hwan Moon, Xiaowei Zhao, Steven J Norris, MD A Motaleb, and Jun Liu. Structural insights into flagellar stator–rotor interactions. *eLife*, 8, 2019.

[5331] Maxime Chantreau, Céline Poux, Marc F Lensink, Guillaume Brysbaert, Xavier Vekemans, and Vincent Castric. Asymmetrical diversification of the receptor-ligand interaction controlling self-incompatibility in Arabidopsis. *eLife*, 8, 2019.

[5332] Nimisha Chaturvedi, Evguenia S Svarovskaia, Hongmei Mo, Anu O Osinusi, Diana M Brainard, G Mani Subramanian, John G McHutchison, Stefan Zeuzem, and Jacques Fellay. Adaptation of hepatitis C virus to interferon lambda polymorphism across multiple viral genotypes. *eLife*, 8, 2019.

[5333] Christina Chatzi, Yingyu Zhang, Wiiliam D Hendricks, Yang Chen, Eric Schnell, Richard H Goodman, and Gary L Westbrook. Exercise-induced enhancement of synaptic function triggered by the inverse BAR protein, Mtss1L. *eLife*, 8, 2019.

[5334] Anwesha Chaudhury, Geoff D Miller, Daniel Eichner, and John M Higgins. Single-cell modeling of routine clinical blood tests reveals transient dynamics of human response to blood loss. *eLife*, 8, 2019.

[5335] Carolina Chavez, Grisel Cruz-Becerra, Jia Fei, George A Kassavetis, and James T Kadonaga. The tardigrade damage suppressor protein binds to nucleosomes and protects DNA from hydroxyl radicals. *eLife*, 8, 2019.

[5336] James Chen, Saumya Gopalkrishnan, Courtney Chiu, Albert Y Chen, Elizabeth A Campbell, Richard L Gourse, Wilma Ross, and Seth A Darst. E. coli TraR allosterically regulates transcription initiation by altering RNA polymerase conformation. *eLife*, 8, 2019.

[5337] Juyue Chen, Holly B Mandel, James E Fitzgerald, and Damon A Clark. Asymmetric ON-OFF processing of visual motion cancels variability induced by the structure of natural scenes. *eLife*, 8, 2019.

[5338] Ko-Fan Chen, Simon Lowe, Angélique Lamaze, Patrick Krätschmer, and James Jepson. Neurocalcin regulates nighttime sleep and arousal in Drosophila. *eLife*, 8, 2019.

[5339] Qingfeng Chen, Weizhong Zeng, Ji She, Xiao-chen Bai, and Youxing Jiang. Structural and functional characterization of an otopetrin family proton channel. *eLife*, 8, 2019.

[5340] Shi Chen, Rafal P Wiewiora, Fanwang Meng, Nicolas Babault, Anqi Ma, Wenyu Yu, Kun Qian, Hao Hu, Hua Zou, Junyi Wang, Shijie Fan, Gil Blum, Fabio Pittella-Silva, Kyle A Beauchamp, Wolfram Tempel, Hualiang Jiang, Kaixian Chen, Robert J Skene, Yujun George Zheng, Peter J Brown, Jian Jin, Cheng Luo, John D Chodera, and Minkui Luo. The dynamic conformational landscape of the protein methyltransferase SETD8. *eLife*, 8, 2019.

[5341] Tai-Heng Chen and Jun-An Chen. Multifaceted roles of microRNAs: From motor neuron generation in embryos to degeneration in spinal muscular atrophy. *eLife*, 8, 2019.

[5342] Xinlu Chen, Katrin Luck, Patrick Rabe, Christopher QD Dinh, Gad Shaulsky, David R Nelson, Jonathan Gershenzon, Jeroen S Dickschat, Tobias G Köllner, and Feng Chen. A terpene synthase-cytochrome P450 cluster in Dictyostelium discoideum produces a novel trisnorsesquiterpene. *eLife*, 8, 2019.

[5343] Xiuzhen Chen, Lukas A Widmer, Marcel M Stangier, Michel O Steinmetz, Jörg Stelling, and Yves Barral. Remote control of microtubule plus-end dynamics and function from the minus-end. *eLife*, 8, 2019.

[5344] Yiming Chen, Rachel A Essner, Seher Kosar, Oliver H Miller, Yen-Chu Lin, Sheyda Mesgarzadeh, and Zachary A Knight. Sustained NPY signaling enables AgRP neurons to drive feeding. *eLife*, 8, 2019.

[5345] Zhijie Chen, Ronen Gabizon, Aidan I Brown, Antony Lee, Aixin Song, César Díaz-Celis, Craig D Kaplan, Elena F Koslover, Tingting Yao, and Carlos Bustamante. High-resolution and high-accuracy topographic and transcriptional maps of the nucleosome barrier. *eLife*, 8, 2019.

[5346] Shouqiang Cheng, James Ashley, Justyna D Kurleto, Meike Lobb-Rabe, Yeonhee Jenny Park, Robert A Carrillo, and Engin özkan. Molecular basis of synaptic specificity by immunoglobulin superfamily receptors in Drosophila. *eLife*, 8, 2019.

[5347] Wei Cheng, Edmund T Rolls, Trevor W Robbins, Weikang Gong, Zhaowen Liu, Wujun Lv, Jingnan Du, Hongkai Wen, Liang Ma, Erin Burke Quinlan, Hugh Garavan, Eric Artiges, Dimitri Papadopoulos Orfanos, Michael N Smolka, Gunter Schumann, Keith Kendrick, and Jianfeng Feng. Decreased brain connectivity in smoking contrasts with increased connectivity in drinking. *eLife*, 8, 2019.

[5348] Ramesh Chennupati, Angela Wirth, Julie Favre, Rui Li, Rémy Bonnavion, Young-June Jin, Astrid Wietelmann, Frank Schweda, Nina Wettschureck, Daniel Henrion, and Stefan Offermanns. Myogenic vasoconstriction requires G12/G13 and LARG to maintain local and systemic vascular resistance. *eLife*, 8, 2019.

[5349] Patrick D Cherry, Sally E Peach, and Jay R Hesselberth. Multiple decay events target HAC1 mRNA during splicing to regulate the unfolded protein response. *eLife*, 8, 2019.

[5350] Kelda Chia, Marcus Keatinge, Julie Mazzolini, and Dirk Sieger. Brain tumours repurpose endogenous neuron to microglia signalling mechanisms to promote their own proliferation. *eLife*, 8, 2019.

[5351] Chris Cho, Yanshu Wang, Philip M Smallwood, John Williams, and Jeremy Nathans. Dlg1 activates beta-catenin signaling to regulate retinal angiogenesis and the blood-retina and blood-brain barriers. *eLife*, 8, 2019.

[5352] Chris Cho, Yanshu Wang, Philip M Smallwood, John Williams, and Jeremy Nathans. Molecular determinants in Frizzled, Reck, and Wnt7a for ligand-specific signaling in neurovascular development. *eLife*, 8, 2019.

[5353] Lucy Chou-Zheng and Asma Hatoum-Aslan. A type III-A CRISPR-Cas system employs degradosome nucleases to ensure robust immunity. *eLife*, 8, 2019.

[5354] Srikanta Chowdhury, Chi Jung Hung, Shuntaro Izawa, Ayumu Inutsuka, Meiko Kawamura, Takashi Kawashima, Haruhiko Bito, Itaru Imayoshi, Manabu Abe, Kenji Sakimura, and Akihiro Yamanaka. Dissociating orexin-dependent and -independent functions of orexin neurons using novel Orexin-Flp knock-in mice. *eLife*, 8, 2019.

[5355] Jenna R Christensen, Kaitlin E Homa, Alisha N Morganthaler, Rachel R Brown, Cristian Suarez, Alyssa J Harker, Meghan E O'Connell, and David R Kovar. Cooperation between tropomyosin and $\alpha$-actinin inhibits fimbrin association with actin filament networks in fission yeast. *eLife*, 8, 2019.

[5356] Zoé Christenson Wick, Madison R Tetzlaff, and Esther Krook-Magnuson. Novel long-range inhibitory nNOS-expressing hippocampal cells. *eLife*, 8, 2019.

[5357] Jeong Min Chung, Michael J Sheedlo, Anne M Campbell, Neha Sawhney, Arwen E Frick-Cheng, Dana Borden Lacy, Timothy L Cover, and Melanie D Ohi. Structure of the Helicobacter pylori Cag type IV secretion system. *eLife*, 8, 2019.

[5358] Dasan Mary Cibi, Masum M Mia, Shamini Guna Shekeran, Lim Sze Yun, Reddemma Sandireddy, Priyanka Gupta, Monalisa Hota, Lei Sun, Sujoy Ghosh, and Manvendra K Singh. Neural crest-specific deletion of Rbfox2 in mice leads to craniofacial abnormalities including cleft palate. *eLife*, 8, 2019.

[5359] Danielle J Clark, Laura E McMillan, Sin Lih Tan, Gaia Bellomo, Clementine Massoue, Harry Thompson, Lidiya Mykhaylechko, Dominic Alibhai, Xiongtao Ruan, Kentner L Singleton, Minna Du, Alan Hedges, Pamela L Schwartzberg, Paul Verkade, Robert F Murphy, and Christoph Wülfing. Transient protein accumulation at the center of the T cell antigen-presenting cell interface drives efficient IL-2 secretion. *eLife*, 8, 2019.

[5360] Benjamin A Clarke, Saurav Majumder, Hongling Zhu, Y Terry Lee, Mari Kono, Cuiling Li, Caroline Khanna, Hailey Blain, Ronit Schwartz, Vienna L Huso, Colleen Byrnes, Galina Tuymetova, Teresa M Dunn, Maria L Allende, and Richard L Proia. The Ormdl genes regulate the sphingolipid synthesis pathway to ensure proper myelination and neurologic function in mice. *eLife*, 8, 2019.

[5361] Veronica Cloud, Ada Thapa, Pedro Morales-Sosa, Tayla M Miller, Sara A Miller, Daniel Holsapple, Paige M Gerhart, Elaheh Momtahan, Jarrid L Jack, Edgardo Leiva, Sarah R Rapp, Lauren G Shelton, Richard A Pierce, Skylar Martin-Brown, Laurence Florens, Michael P Washburn, and Ryan D Mohan. Ataxin-7 and Non-stop coordinate SCAR protein levels, subcellular localization, and actin cytoskeleton organization. *eLife*, 8, 2019.

[5362] Amanda F Clouser, Hannah ER Baughman, Benjamin Basanta, Miklos Guttman, Abhinav Nath, and Rachel E Klevit. Interplay of disordered and ordered regions of a human small heat shock protein yields an ensemble of 'quasi-ordered' states. *eLife*, 8, 2019.

[5363] Matthew E Cockman, Kerstin Lippl, Ya-Min Tian, Hamish B Pegg, William D Figg, Martine I Abboud, Raphael Heilig, Roman Fischer, Johanna Myllyharju, Christopher J Schofield, and Peter J Ratcliffe. Lack of activity of recombinant HIF prolyl hydroxylases (PHDs) on reported non-HIF substrates. *eLife*, 8, 2019.

[5364] Ksenya Cohen Katsenelson, Joshua D Stender, Agnieszka T Kawashima, Gema Lordén, Satoshi Uchiyama, Victor Nizet, Christopher K Glass, and Alexandra C Newton. PHLPP1 counter-regulates STAT1-mediated inflammatory signaling. *eLife*, 8, 2019.

[5365] Glenda Comai, Eglantine Heude, Sebastian Mella, Sylvain Paisant, Francesca Pala, Mirialys Gallardo, Francina Langa, Gabrielle Kardon, Swetha Gopalakrishnan, and Shahragim Tajbakhsh. A distinct cardiopharyngeal mesoderm genetic hierarchy establishes antero-posterior patterning of esophagus striated muscle. *eLife*, 8, 2019.

[5366] Christine M Constantinople, Alex T Piet, Peter Bibawi, Athena Akrami, Charles Kopec, and Carlos D Brody. Lateral orbitofrontal cortex promotes trial-by-trial learning of risky, but not spatial, biases. *eLife*, 8, 2019.

[5367] Duccio Conti, Parveen Gul, Asifa Islam, José M Martín-Durán, Richard W Pickersgill, and Viji M Draviam. Kinetochores attached to microtubule-ends are stabilised by Astrin bound PP1 to ensure proper chromosome segregation. *eLife*, 8, 2019.

[5368] Eduardo J Contijoch, Graham J Britton, Chao Yang, Ilaria Mogno, Zhihua Li, Ruby Ng, Sean R Llewellyn, Sheela Hira, Crystal Johnson, Keren M Rabinowitz, Revital Barkan, Iris Dotan, Robert P Hirten, Shih-Chen Fu, Yuying Luo, Nancy Yang, Tramy Luong, Philippe R Labrias, Sergio Lira, Inga Peter, Ari Grinspan, Jose C Clemente, Roman Kosoy, Seunghee Kim-Schulze, Xiaochen Qin, Anabella Castillo, Amanda Hurley, Ashish Atreja, Jason Rogers, Farah Fasihuddin, Merjona Saliaj, Amy Nolan, Pamela Reyes-Mercedes, Carina Rodriguez, Sarah Aly, Kenneth Santa-Cruz, Lauren Peters, Mayte Suárez-Fariñas, Ruiqi Huang, Ke Hao, Jun Zhu, Bin Zhang, Bojan Losic, Haritz Irizar, Won-Min Song, Antonio Di Narzo, Wenhui Wang, Benjamin L Cohen, Christopher DiMaio, David Greenwald, Steven Itzkowitz, Aimee Lucas, James Marion, Elana Maser, Ryan Ungaro, Steven Naymagon, Joshua Novak, Brijen Shah, Thomas Ullman, Peter Rubin, James George, Peter Legnani, Shannon E Telesco, Joshua R Friedman, Carrie Brodmerkel, Scott Plevy, Judy H Cho, Jean-Frederic Colombel, Eric E Schadt, Carmen Argmann, Marla Dubinsky, Andrew Kasarskis, Bruce Sands, and Jeremiah J Faith. Gut microbiota density influences host physiology and is shaped by host and microbial factors. *eLife*, 8, 2019.

[5369] Jennifer L Cook, Jennifer C Swart, Monja I Froböse, Andreea O Diaconescu, Dirk EM Geurts, Hanneke EM den Ouden, and Roshan Cools. Catecholaminergic modulation of meta-learning. *eLife*, 8, 2019.

[5370] Aaron J Corcoran and Tyson L Hedrick. Compound-V formations in shorebird flocks. *eLife*, 8, 2019.

[5371] Robin A Corey, Zainab Ahdash, Anokhi Shah, Euan Pyle, William J Allen, Tomas Fessl, Janet E Lovett, Argyris Politis, and Ian Collinson. ATP-induced asymmetric pre-protein folding as a driver of protein translocation through the Sec machinery. *eLife*, 8, 2019.

[5372] Jonathan H Cornford, Marion S Mercier, Marco Leite, Vincent Magloire, Michael Häusser, and Dimitri M Kullmann. Dendritic NMDA receptors in parvalbumin neurons enable strong and stable neuronal assemblies. *eLife*, 8, 2019.

[5373] Andrea Cosolo, Janhvi Jaiswal, Gábor Csordás, Isabelle Grass, Mirka Uhlirova, and Anne-Kathrin Classen. JNK-dependent cell cycle stalling in G2 promotes survival and senescence-like phenotypes in tissue stress. *eLife*, 8, 2019.

[5374] Maïté Courel, Yves Clément, Clémentine Bossevain, Dominika Foretek, Olivia Vidal Cruchez, Zhou Yi, Marianne Bénard, Marie-Noëlle Benassy, Michel Kress, Caroline Vindry, Michèle Ernoult-Lange, Christophe Antoniewski, Antonin Morillon, Patrick Brest, Arnaud Hubstenberger, Hugues Roest Crollius, Nancy Standart, and Dominique Weil. GC content shapes mRNA storage and decay in human cells. *eLife*, 8, 2019.

[5375] Matthew M Crane, Adam E Russell, Brent J Schafer, Ben W Blue, Riley Whalen, Jared Almazan, Mung Gi Hong, Bao Nguyen, Joslyn E Goings, Kenneth L Chen, Ryan Kelly, and Matt Kaeberlein. DNA damage checkpoint activation impairs chromatin homeostasis and promotes mitotic catastrophe during aging. *eLife*, 8, 2019.

[5376] Thomas Crellen, Paul Turner, Sreymom Pol, Stephen Baker, To Nguyen Thi Nguyen, Nicole Stoesser, Nicholas PJ Day, Claudia Turner, and Ben S Cooper. Transmission dynamics and control of multidrug-resistant Klebsiella pneumoniae in neonates in a developing country. *eLife*, 8, 2019.

[5377] Thomas K Creson, Camilo Rojas, Ernie Hwaun, Thomas Vaissiere, Murat Kilinc, Andres Jimenez-Gomez, Jimmy Lloyd Holder, Jianrong Tang, Laura L Colgin, Courtney A Miller, and Gavin Rumbaugh. Re-expression of SynGAP protein in adulthood improves translatable measures of brain function and behavior. *eLife*, 8, 2019.

[5378] Tim A Crombie, Stefan Zdraljevic, Daniel E Cook, Robyn E Tanny, Shannon C Brady, Ye Wang, Kathryn S Evans, Steffen Hahnel, Daehan Lee, Briana C Rodriguez, Gaotian Zhang, Joost van der Zwagg, Karin Kiontke, and Erik C Andersen. Deep sampling of Hawaiian Caenorhabditis elegans reveals high genetic diversity and admixture with global populations. *eLife*, 8, 2019.

[5379] László Csanády and Beáta Töröcsik. Cystic fibrosis drug ivacaftor stimulates CFTR channels at picomolar concentrations. *eLife*, 8, 2019.

[5380] Andrea Cuentas-Condori, Ben Mulcahy, Siwei He, Sierra Palumbos, Mei Zhen, and David M Miller. C. elegans neurons have functional dendritic spines. *eLife*, 8, 2019.

[5381] Edward D Cui and Ben W Strowbridge. Selective attenuation of Ether-a-go-go related K+ currents by endogenous acetylcholine reduces spike-frequency adaptation and network correlation. *eLife*, 8, 2019.

[5382] Pierre Cunin, Rim Bouslama, Kellie R Machlus, Marta Martínez-Bonet, Pui Y Lee, Alexandra Wactor, Nathan Nelson-Maney, Allyn Morris, Li Guo, Andrew Weyrich, Martha Sola-Visner, Eric Boilard, Joseph E Italiano, and Peter A Nigrovic. Megakaryocyte emperipolesis mediates membrane transfer from intracytoplasmic neutrophils to platelets. *eLife*, 8, 2019.

[5383] Antonio Currais, Ling Huang, Joshua Goldberg, Michael Petrascheck, Gamze Ates, António Pinto-Duarte, Maxim N Shokhirev, David Schubert, and Pamela Maher. Elevating acetyl-CoA levels reduces aspects of brain aging. *eLife*, 8, 2019.

[5384] Ugur Dag, Zhengchang Lei, Jasmine Q Le, Allan Wong, Daniel Bushey, and Krystyna Keleman. Neuronal reactivation during post-learning sleep consolidates long-term memory in Drosophila. *eLife*, 8, 2019.

[5385] Wenli Dai, Brigitte Laforest, Leonid Tyan, Kaitlyn M Shen, Rangarajan D Nadadur, Francisco J Alvarado, Stefan R Mazurek, Sonja Lazarevic, Margaret Gadek, Yitang Wang, Ye Li, Hector H Valdivia, Le Shen, Michael T Broman, Ivan P Moskowitz, and Christopher R Weber. A calcium transport mechanism for atrial fibrillation in Tbx5-mutant mice. *eLife*, 8, 2019.

[5386] Vamsi K Daliparthi, Ryosuke O Tachibana, Brenton G Cooper, Richard HR Hahnloser, Satoshi Kojima, Samuel J Sober, and Todd F Roberts. Transitioning between preparatory and precisely sequenced neuronal activity in production of a skilled behavior. *eLife*, 8, 2019.

[5387] Philip Dammann, André Scherag, Nikolay Zak, Karol Szafranski, Susanne Holtze, Sabine Begall, Hynek Burda, Hans A Kestler, Thomas Hildebrandt, and Matthias Platzer. Comment on 'Naked mole-rat mortality rates defy Gompertzian laws by not increasing with age'. *eLife*, 8, 2019.

[5388] Christian Damsgaard, Henrik Lauridsen, Anette MD Funder, Jesper S Thomsen, Thomas Desvignes, Dane A Crossley, Peter R Møller, Do TT Huong, Nguyen T Phuong, H William Detrich, Annemarie Brüel, Horst Wilkens, Eric Warrant, Tobias Wang, Jens R Nyengaard, Michael Berenbrink, and Mark Bayley. Retinal oxygen supply shaped the functional evolution of the vertebrate eye. *eLife*, 8, 2019.

[5389] Valentin Danchev, Andrey Rzhetsky, and James A Evans. Centralized scientific communities are less likely to generate replicable results. *eLife*, 8, 2019.

[5390] Matteo D'Antonio, Joaquin Reyna, David Jakubosky, Margaret KR Donovan, Marc-Jan Bonder, Hiroko Matsui, Oliver Stegle, Naoki Nariai, Agnieszka D'Antonio-Chronowska, and Kelly A Frazer. Systematic genetic analysis of the MHC region reveals mechanistic underpinnings of HLA type associations with disease. *eLife*, 8, 2019.

[5391] Kristian Davidsen, Branden J Olson, William S DeWitt, Jean Feng, Elias Harkins, Philip Bradley, and Frederick A Matsen. Deep generative models for T cell receptor protein sequences. *eLife*, 8, 2019.

[5392] Neil Martin Davies, W David Hill, Emma L Anderson, Eleanor Sanderson, Ian J Deary, and George Davey Smith. Multivariable two-sample Mendelian randomization estimates of the effects of intelligence and education on health. *eLife*, 8, 2019.

[5393] Marijn de Boer, Giorgos Gouridis, Ruslan Vietrov, Stephanie L Begg, Gea K Schuurman-Wolters, Florence Husada, Nikolaos Eleftheriadis, Bert Poolman, Christopher A McDevitt, and Thorben Cordes. Conformational and dynamic plasticity in substrate-binding proteins underlies selective transport in ABC importers. *eLife*, 8, 2019.

[5394] Juan Carlos De la Concepcion, Marina Franceschetti, Dan MacLean, Ryohei Terauchi, Sophien Kamoun, and Mark J Banfield. Protein engineering expands the effector recognition profile of a rice NLR immune receptor. *eLife*, 8, 2019.

[5395] André Luiz de Oliveira, Andrew Calcino, and Andreas Wanninger. Ancient origins of arthropod moulting pathway components. *eLife*, 8, 2019.

[5396] Samuel Dean, Flavia Moreira-Leite, and Keith Gull. Basalin is an evolutionarily unconstrained protein revealed via a conserved role in flagellum basal plate function. *eLife*, 8, 2019.

[5397] Brian D DeAngelis, Jacob A Zavatone-Veth, and Damon A Clark. The manifold structure of limb coordination in walking Drosophila. *eLife*, 8, 2019.

[5398] Verena Dederer, Anton Khmelinskii, Anna Gesine Huhn, Voytek Okreglak, Michael Knop, and Marius K Lemberg. Cooperation of mitochondrial and ER factors in quality control of tail-anchored proteins. *eLife*, 8, 2019.

[5399] Samantha Del Borrello, Margot Lautens, Kathleen Dolan, June H Tan, Taylor Davie, Michael R Schertzberg, Mark A Spensley, Amy A Caudy, and Andrew G Fraser. Rhodoquinone biosynthesis in C. elegans requires precursors generated by the kynurenine pathway. *eLife*, 8, 2019.

[5400] Brian C Del Rosario, Andrea J Kriz, Amanda M Del Rosario, Anthony Anselmo, Christopher J Fry, Forest M White, Ruslan I Sadreyev, and Jeannie T Lee. Exploration of CTCF post-translation modifications uncovers Serine-224 phosphorylation by PLK1 at pericentric regions during the G2/M transition. *eLife*, 8, 2019.

[5401] Murilo Delgobo, Daniel AGB Mendes, Edgar Kozlova, Edroaldo Lummertz Rocha, Gabriela F Rodrigues-Luiz, Lucas Mascarin, Greicy Dias, Daniel O Patrício, Tim Dierckx, Maíra A Bicca, Gaëlle Bretton, Yonne Karoline Tenório de Menezes, Márick R Starick, Darcita Rovaris, Joanita Del Moral, Daniel S Mansur, Johan Van Weyenbergh, and André Báfica. An evolutionary recent IFN/IL-6/CEBP axis is linked to monocyte expansion and tuberculosis severity in humans. *eLife*, 8, 2019.

[5402] Tom O Delmont, Evan Kiefl, Ozsel Kilinc, Ozcan C Esen, Ismail Uysal, Michael S Rappé, Steven Giovannoni, and A Murat Eren. Single-amino acid variants reveal evolutionary processes that shape the biogeography of a global SAR11 subclade. *eLife*, 8, 2019.

[5403] Rebecca Delventhal, Reed M O'Connor, Meghan M Pantalia, Matthew Ulgherait, Han X Kim, Maylis K Basturk, Julie C Canman, and Mimi Shirasu-Hiza. Dissection of central clock function in Drosophila through cell-specific CRISPR-mediated clock gene disruption. *eLife*, 8, 2019.

[5404] Conor Dempsey, LF Abbott, and Nathaniel B Sawtell. Generalization of learned responses in the mormyrid electrosensory lobe. *eLife*, 8, 2019.

[5405] Bray Denard, Sungwon Han, JungYeon Kim, Elliott M Ross, and Jin Ye. Regulating G protein-coupled receptors by topological inversion. *eLife*, 8, 2019.

[5406] Eric Deneault, Muhammad Faheem, Sean H White, Deivid C Rodrigues, Song Sun, Wei Wei, Alina Piekna, Tadeo Thompson, Jennifer L Howe, Leon Chalil, Vickie Kwan, Susan Walker, Peter Pasceri, Frederick P Roth, Ryan KC Yuen, Karun K Singh, James Ellis, and Stephen W Scherer. CNTN5-/+or EHMT2-/+human iPSC-derived neurons from individuals with autism develop hyperactive neuronal networks. *eLife*, 8, 2019.

[5407] Thomas Deneux, Evan R Harrell, Alexandre Kempf, Sebastian Ceballo, Anton Filipchuk, and Brice Bathellier. Context-dependent signaling of coincident auditory and visual events in primary visual cortex. *eLife*, 8, 2019.

[5408] Qi Deng, Ping Li, Manju Che, Jiajia Liu, Soma Biswas, Gang Ma, Lin He, Zhanying Wei, Zhenlin Zhang, Yingzi Yang, Huijuan Liu, and Baojie Li. Activation of hedgehog signaling in mesenchymal stem cells induces cartilage and bone tumor formation via Wnt/$\beta$-Catenin. *eLife*, 8, 2019.

[5409] Yuqi Deng, Robert MG Reinhart, Inyong Choi, and Barbara G Shinn-Cunningham. Causal links between parietal alpha activity and spatial auditory attention. *eLife*, 8, 2019.

[5410] Kobe Desender, Annika Boldt, Tom Verguts, and Tobias H Donner. Confidence predicts speed-accuracy tradeoff for subsequent decisions. *eLife*, 8, 2019.

[5411] Jaigeeth Deveryshetty, Thibaut Peterlini, Mikhail Ryzhikov, Nadine Brahiti, Graham Dellaire, Jean-Yves Masson, and Sergey Korolev. Novel RNA and DNA strand exchange activity of the PALB2 DNA binding domain and its critical role for DNA repair in cells. *eLife*, 8, 2019.

[5412] Ciara A Devine, Christine Gaffney, Gerard M Loughnane, Simon P Kelly, and Redmond G O'Connell. The role of premature evidence accumulation in making difficult perceptual decisions under temporal uncertainty. *eLife*, 8, 2019.

[5413] Anita V Devineni, Bei Sun, Anna Zhukovskaya, and Richard Axel. Acetic acid activates distinct taste pathways in Drosophila to elicit opposing, state-dependent feeding responses. *eLife*, 8, 2019.

[5414] Riddhiman Dhar, Alsu M Missarova, Ben Lehner, and Lucas B Carey. Single cell functional genomics reveals the importance of mitochondria in cell-to-cell phenotypic variation. *eLife*, 8, 2019.

[5415] Elvira Diamantopoulou, Sarah Baxendale, Antonio de la Vega de León, Anzar Asad, Celia J Holdsworth, Leila Abbas, Valerie J Gillet, Giselle R Wiggin, and Tanya T Whitfield. Identification of compounds that rescue otic and myelination defects in the zebrafish adgrg6 (gpr126) mutant. *eLife*, 8, 2019.

[5416] Irene Díaz-López, René Toribio, Juan José Berlanga, and Iván Ventoso. An mRNA-binding channel in the ES6S region of the translation 48S-PIC promotes RNA unwinding and scanning. *eLife*, 8, 2019.

[5417] Carlos J Diaz Osterman, Duygu Ozmadenci, Elizabeth G Kleinschmidt, Kristin N Taylor, Allison M Barrie, Shulin Jiang, Lisa M Bean, Florian J Sulzmaier, Christine Jean, Isabelle Tancioni, Kristen Anderson, Sean Uryu, Edward A Cordasco, Jian Li, Xiao Lei Chen, Guo Fu, Marjaana Ojalill, Pekka Rappu, Jyrki Heino, Adam M Mark, Guorong Xu, Kathleen M Fisch, Vihren N Kolev, David T Weaver, Jonathan A Pachter, Balázs Győrffy, Michael T McHale, Denise C Connolly, Alfredo Molinolo, Dwayne G Stupack, and David D Schlaepfer. FAK activity sustains intrinsic and acquired ovarian cancer resistance to platinum chemotherapy. *eLife*, 8, 2019.

[5418] Penelope Dimas, Laura Montani, Jorge A Pereira, Daniel Moreno, Martin Trötzmüller, Joanne Gerber, Clay F Semenkovich, Harald C Köfeler, and Ueli Suter. CNS myelination and remyelination depend on fatty acid synthesis by oligodendrocytes. *eLife*, 8, 2019.

[5419] Edoardo D'Imprima, Davide Floris, Mirko Joppe, Ricardo Sánchez, Martin Grininger, and Werner Kühlbrandt. Protein denaturation at the air-water interface and how to prevent it. *eLife*, 8, 2019.

[5420] Keke Ding, Yifu Han, Taylor W Seid, Christopher Buser, Tomomi Karigo, Shishuo Zhang, Dion K Dickman, and David J Anderson. Imaging neuropeptide release at synapses with a genetically engineered reporter. *eLife*, 8, 2019.

[5421] Siyu Serena Ding, Linus J Schumacher, Avelino E Javer, Robert G Endres, and André EX Brown. Shared behavioral mechanisms underlie C. elegans aggregation and swarming. *eLife*, 8, 2019.

[5422] Yifu Ding, Daniel J Vanselow, Maksim A Yakovlev, Spencer R Katz, Alex Y Lin, Darin P Clark, Phillip Vargas, Xuying Xin, Jean E Copper, Victor A Canfield, Khai C Ang, Yuxin Wang, Xianghui Xiao, Francesco De Carlo, Damian B van Rossum, Patrick La Riviere, and Keith C Cheng. Computational 3D histological phenotyping of whole zebrafish by X-ray histotomography. *eLife*, 8, 2019.

[5423] Danielle DiTirro, Alison Philbrook, Kendrick Rubino, and Piali Sengupta. The Caenorhabditis elegans Tubby homolog dynamically modulates olfactory cilia membrane morphogenesis and phospholipid composition. *eLife*, 8, 2019.

[5424] Jonathon A Ditlev, Anthony R Vega, Darius Vasco Köster, Xiaolei Su, Tomomi Tani, Ashley M Lakoduk, Ronald D Vale, Satyajit Mayor, Khuloud Jaqaman, and Michael K Rosen. A composition-dependent molecular clutch between T cell signaling condensates and actin. *eLife*, 8, 2019.

[5425] Genevieve S Dobihal, Yannick R Brunet, Josué Flores-Kim, and David Z Rudner. Homeostatic control of cell wall hydrolysis by the WalRK two-component signaling pathway in Bacillus subtilis. *eLife*, 8, 2019.

[5426] May Dobosiewicz, Qiang Liu, and Cornelia I Bargmann. Reliability of an interneuron response depends on an integrated sensory state. *eLife*, 8, 2019.

[5427] Michael-John Dolan, Shahar Frechter, Alexander Shakeel Bates, Chuntao Dan, Paavo Huoviala, Ruairí JV Roberts, Philipp Schlegel, Serene Dhawan, Remy Tabano, Heather Dionne, Christina Christoforou, Kari Close, Ben Sutcliffe, Bianca Giuliani, Feng Li, Marta Costa, Gudrun Ihrke, Geoffrey Wilson Meissner, Davi D Bock, Yoshinori Aso, Gerald M Rubin, and Gregory SXE Jefferis. Neurogenetic dissection of the Drosophila lateral horn reveals major outputs, diverse behavioural functions, and interactions with the mushroom body. *eLife*, 8, 2019.

[5428] Katrin Domsch, Julie Carnesecchi, Vanessa Disela, Jana Friedrich, Nils Trost, Olga Ermakova, Maria Polychronidou, and Ingrid Lohmann. The Hox transcription factor Ubx stabilizes lineage commitment by suppressing cellular plasticity in Drosophila. *eLife*, 8, 2019.

[5429] Jie-Xian Dong, Yongam Lee, Michael Kirmiz, Stephanie Palacio, Camelia Dumitras, Claudia M Moreno, Richard Sando, L Fernando Santana, Thomas C Südhof, Belvin Gong, Karl D Murray, and James S Trimmer. A toolbox of nanobodies developed and validated for use as intrabodies and nanoscale immunolabels in mammalian brain neurons. *eLife*, 8, 2019.

[5430] Yibo Dong, Shichao Chen, Shifeng Cheng, Wenbin Zhou, Qing Ma, Zhiduan Chen, Cheng-Xin Fu, Xin Liu, Yun-peng Zhao, Pamela S Soltis, Gane Ka-Shu Wong, Douglas E Soltis, and Qiu-Yun(Jenny) Xiang. Natural selection and repeated patterns of molecular evolution following allopatric divergence. *eLife*, 8, 2019.

[5431] Benjamin T Donovan, Hengye Chen, Caroline Jipa, Lu Bai, and Michael G Poirier. Dissociation rate compensation mechanism for budding yeast pioneer transcription factors. *eLife*, 8, 2019.

[5432] Lauren J Donovan, William C Spencer, Meagan M Kitt, Brent A Eastman, Katherine J Lobur, Kexin Jiao, Jerry Silver, and Evan S Deneris. Lmx1b is required at multiple stages to build expansive serotonergic axon architectures. *eLife*, 8, 2019.

[5433] Karole N D'Orazio, Colin Chih-Chien Wu, Niladri Sinha, Raphael Loll-Krippleber, Grant W Brown, and Rachel Green. The endonuclease Cue2 cleaves mRNAs at stalled ribosomes during No Go Decay. *eLife*, 8, 2019.

[5434] Kevin Dorgans, Valérie Demais, Yannick Bailly, Bernard Poulain, Philippe Isope, and Frédéric Doussau. Short-term plasticity at cerebellar granule cell to molecular layer interneuron synapses expands information processing. *eLife*, 8, 2019.

[5435] éva Dóró, Sem H Jacobs, Ffion R Hammond, Henk Schipper, Remco PM Pieters, Mark Carrington, Geert F Wiegertjes, and Maria Forlenza. Visualizing trypanosomes in a vertebrate host reveals novel swimming behaviours, adaptations and attachment mechanisms. *eLife*, 8, 2019.

[5436] Michel Dosch, Joël Zindel, Fadi Jebbawi, Nicolas Melin, Daniel Sanchez-Taltavull, Deborah Stroka, Daniel Candinas, and Guido Beldi. Connexin-43-dependent ATP release mediates macrophage activation during sepsis. *eLife*, 8, 2019.

[5437] Nir Drayman, Parthiv Patel, Luke Vistain, and Savaş Tay. HSV-1 single-cell analysis reveals the activation of anti-viral and developmental programs in distinct sub-populations. *eLife*, 8, 2019.

[5438] Sarah E Dremel and Neal A DeLuca. Herpes simplex viral nucleoprotein creates a competitive transcriptional environment facilitating robust viral transcription and host shut off. *eLife*, 8, 2019.

[5439] Zhi-Yan Du, Krzysztof Zienkiewicz, Natalie Vande Pol, Nathaniel E Ostrom, Christoph Benning, and Gregory M Bonito. Algal-fungal symbiosis leads to photosynthetic mycelium. *eLife*, 8, 2019.

[5440] Veronika Dubinkina, Yulia Fridman, Parth Pratim Pandey, and Sergei Maslov. Multistability and regime shifts in microbial communities explained by competition for essential nutrients. *eLife*, 8, 2019.

[5441] Anne-Laure Duchemin, Hélène Vignes, and Julien Vermot. Mechanically activated piezo channels modulate outflow tract valve development through the Yap1 and Klf2-Notch signaling axis. *eLife*, 8, 2019.

[5442] Omaya Dudin, Andrej Ondracka, Xavier Grau-Bové, Arthur AB Haraldsen, Atsushi Toyoda, Hiroshi Suga, Jon Bråte, and Iñaki Ruiz-Trillo. A unicellular relative of animals generates a layer of polarized cells by actomyosin-dependent cellularization. *eLife*, 8, 2019.

[5443] Joseph G Duman, Shalaka Mulherkar, Yen-Kuei Tu, Kelly C Erikson, Christopher P Tzeng, Vasilis C Mavratsas, Tammy Szu-Yu Ho, and Kimberley F Tolias. The adhesion-GPCR BAI1 shapes dendritic arbors via Bcr-mediated RhoA activation causing late growth arrest. *eLife*, 8, 2019.

[5444] Anett Dunai, Réka Spohn, Zoltán Farkas, Viktória Lázár, ádám Györkei, Gábor Apjok, Gábor Boross, Balázs Szappanos, Gábor Grézal, Anikó Faragó, László Bodai, Balázs Papp, and Csaba Pál. Rapid decline of bacterial drug-resistance in an antibiotic-free environment through phenotypic reversion. *eLife*, 8, 2019.

[5445] Salvador Dura-Bernal, Benjamin A Suter, Padraig Gleeson, Matteo Cantarelli, Adrian Quintana, Facundo Rodriguez, David J Kedziora, George L Chadderdon, Cliff C Kerr, Samuel A Neymotin, Robert A McDougal, Michael Hines, Gordon MG Shepherd, and William W Lytton. NetPyNE, a tool for data-driven multiscale modeling of brain circuits. *eLife*, 8, 2019.

[5446] Natasha D Durham, Aditi Agrawal, Eric Waltari, Derek Croote, Fabio Zanini, Mallorie Fouch, Edgar Davidson, Olivia Smith, Esteban Carabajal, John E Pak, Benjamin J Doranz, Makeda Robinson, Ana M Sanz, Ludwig L Albornoz, Fernando Rosso, Shirit Einav, Stephen R Quake, Krista M McCutcheon, and Leslie Goo. Broadly neutralizing human antibodies against dengue virus identified by single B cell transcriptomics. *eLife*, 8, 2019.

[5447] Elizabeth ML Duxbury, Jonathan P Day, Davide Maria Vespasiani, Yannik Thüringer, Ignacio Tolosana, Sophia CL Smith, Lucia Tagliaferri, Altug Kamacioglu, Imogen Lindsley, Luca Love, Robert L Unckless, Francis M Jiggins, and Ben Longdon. Host-pathogen coevolution increases genetic variation in susceptibility to infection. *eLife*, 8, 2019.

[5448] David E Ehrlich and David Schoppik. A primal role for the vestibular sense in the development of coordinated locomotion. *eLife*, 8, 2019.

[5449] Ingrid Ehrmann, James H Crichton, Matthew R Gazzara, Katherine James, Yilei Liu, Sushma Nagaraja Grellscheid, Tomaž Curk, Dirk de Rooij, Jannetta S Steyn, Simon Cockell, Ian R Adams, Yoseph Barash, and David J Elliott. An ancient germ cell-specific RNA-binding protein protects the germline from cryptic splice site poisoning. *eLife*, 8, 2019.

[5450] Catherine A Eichel, Erick B Ríos-Pérez, Fang Liu, Margaret B Jameson, David K Jones, Jennifer J Knickelbine, and Gail A Robertson. A microtranslatome coordinately regulates sodium and potassium currents in the human heart. *eLife*, 8, 2019.

[5451] Krystel El Hage, Florent Hédin, Prashant K Gupta, Markus Meuwly, and Martin Karplus. Response to comment on 'Valid molecular dynamics simulations of human hemoglobin require a surprisingly large box size'. *eLife*, 8, 2019.

[5452] Mahmoud M Elguindy, Florian Kopp, Mohammad Goodarzi, Frederick Rehfeld, Anu Thomas, Tsung-Cheng Chang, and Joshua T Mendell. PUMILIO, but not RBMX, binding is required for regulation of genomic stability by noncoding RNA NORAD. *eLife*, 8, 2019.

[5453] Charles F Ericson, Fabian Eisenstein, João M Medeiros, Kyle E Malter, Giselle S Cavalcanti, Robert W Zeller, Dianne K Newman, Martin Pilhofer, and Nicholas J Shikuma. A contractile injection system stimulates tubeworm metamorphosis by translocating a proteinaceous effector. *eLife*, 8, 2019.

[5454] Simon Erlendsson, Thor Seneca Thorsen, Georges Vauquelin, Ina Ammendrup-Johnsen, Volker Wirth, Karen L Martinez, Kaare Teilum, Ulrik Gether, and Kenneth Lindegaard Madsen. Mechanisms of PDZ domain scaffold assembly illuminated by use of supported cell membrane sheets. *eLife*, 8, 2019.

[5455] Thomas Michael Ernst, Anna Evelina Brol, Marcel Gratz, Christoph Ritter, Ulrike Bingel, Marc Schlamann, Stefan Maderwald, Harald H Quick, Christian Josef Merz, and Dagmar Timmann. The cerebellum is involved in processing of predictions and prediction errors in a fear conditioning paradigm. *eLife*, 8, 2019.

[5456] Michelle S Erwig, Julia Patzig, Anna M Steyer, Payam Dibaj, Mareike Heilmann, Ingo Heilmann, Ramona B Jung, Kathrin Kusch, Wiebke Möbius, Olaf Jahn, Klaus-Armin Nave, and Hauke B Werner. Anillin facilitates septin assembly to prevent pathological outfoldings of central nervous system myelin. *eLife*, 8, 2019.

[5457] Anthony Essex, Javier Pineda, Grishma Acharya, Hong Xin, James Evans, Elizabeth Iorns, Elizabeth Iorns, Rachel Tsui, Alexandria Denis, Nicole Perfito, Timothy M Errington, Elizabeth Iorns, Rachel Tsui, Alexandria Denis, Nicole Perfito, and Timothy M Errington. Replication Study: Wnt activity defines colon cancer stem cells and is regulated by the microenvironment. *eLife*, 8, 2019.

[5458] Laura A Ewell, Kyle B Fischer, Christian Leibold, Stefan Leutgeb, and Jill K Leutgeb. The impact of pathological high-frequency oscillations on hippocampal network activity in rats with chronic epilepsy. *eLife*, 8, 2019.

[5459] Daphne Ezer and Kirstie Whitaker. Data science for the scientific life cycle. *eLife*, 8, 2019.

[5460] Martin H Fabry, Filippo Ciabrelli, Marzia Munafò, Evelyn L Eastwood, Emma Kneuss, Ilaria Falciatori, Federica A Falconio, Gregory J Hannon, and Benjamin Czech. piRNA-guided co-transcriptional silencing coopts nuclear export factors. *eLife*, 8, 2019.

[5461] Jocelyne Fadiga and Todd G Nystul. The follicle epithelium in the Drosophila ovary is maintained by a small number of stem cells. *eLife*, 8, 2019.

[5462] Thomas G Fai, Lishibanya Mohapatra, Prathitha Kar, Jane Kondev, and Ariel Amir. Length regulation of multiple flagella that self-assemble from a shared pool of components. *eLife*, 8, 2019.

[5463] Maria E Falzone, Jan Rheinberger, Byoung-Cheol Lee, Thasin Peyear, Linda Sasset, Ashleigh M Raczkowski, Edward T Eng, Annarita Di Lorenzo, Olaf S Andersen, Crina M Nimigean, and Alessio Accardi. Structural basis of Ca2+-dependent activation and lipid transport by a TMEM16 scramblase. *eLife*, 8, 2019.

[5464] Di Fan, Zoubida Chettouh, G Giacomo Consalez, and Jean-FranÇois Brunet. Taste bud formation depends on taste nerves. *eLife*, 8, 2019.

[5465] Li Fan, Ismar Kovacevic, Maxwell G Heiman, and Zhirong Bao. A multicellular rosette-mediated collective dendrite extension. *eLife*, 8, 2019.

[5466] Navid Farassat, Kauê Machado Costa, Strahinja Stojanovic, Stefan Albert, Lora Kovacheva, Josef Shin, Richard Egger, Mahalakshmi Somayaji, Sevil Duvarci, Gaby Schneider, and Jochen Roeper. In vivo functional diversity of midbrain dopamine neurons within identified axonal projections. *eLife*, 8, 2019.

[5467] Nona Farbehi, Ralph Patrick, Aude Dorison, Munira Xaymardan, Vaibhao Janbandhu, Katharina Wystub-Lis, Joshua WK Ho, Robert E Nordon, and Richard P Harvey. Single-cell expression profiling reveals dynamic flux of cardiac stromal, vascular and immune cells in health and injury. *eLife*, 8, 2019.

[5468] Chen Farhy, Santosh Hariharan, Jarkko Ylanko, Luis Orozco, Fu-Yue Zeng, Ian Pass, Fernando Ugarte, E Camilla Forsberg, Chun-Teng Huang, David W Andrews, and Alexey V Terskikh. Improving drug discovery using image-based multiparametric analysis of the epigenetic landscape. *eLife*, 8, 2019.

[5469] Daniel S Farrar, Shally Awasthi, Shaza A Fadel, Rajesh Kumar, Anju Sinha, Sze Hang Fu, Brian Wahl, Shaun K Morris, and Prabhat Jha. Seasonal variation and etiologic inferences of childhood pneumonia and diarrhea mortality in India. *eLife*, 8, 2019.

[5470] Michael Jan Fauth and Mark CW van Rossum. Self-organized reactivation maintains and reinforces memories despite synaptic turnover. *eLife*, 8, 2019.

[5471] Gareth W Fearnley, Katherine A Young, James R Edgar, Robin Antrobus, Iain M Hay, Wei-Ching Liang, Nadia Martinez-Martin, WeiYu Lin, Janet E Deane, and Hayley J Sharpe. The homophilic receptor PTPRK selectively dephosphorylates multiple junctional regulators to promote cell–cell adhesion. *eLife*, 8, 2019.

[5472] Ann Feke, Wei Liu, Jing Hong, Man-Wah Li, Chin-Mei Lee, Elton K Zhou, and Joshua M Gendron. Decoys provide a scalable platform for the identification of plant E3 ubiquitin ligases that regulate circadian function. *eLife*, 8, 2019.

[5473] Blake Ferguson, Herlina Y Handoko, Pamela Mukhopadhyay, Arash Chitsazan, Lois Balmer, Grant Morahan, and Graeme J Walker. Different genetic mechanisms mediate spontaneous versus UVR-induced malignant melanoma. *eLife*, 8, 2019.

[5474] Luke S Ferro, Sinan Can, Meghan A Turner, Mohamed M ElShenawy, and Ahmet Yildiz. Kinesin and dynein use distinct mechanisms to bypass obstacles. *eLife*, 8, 2019.

[5475] Mattia Ficarelli, Harry Wilson, Rui Pedro Galão, Michela Mazzon, Irati Antzin-Anduetza, Mark Marsh, Stuart JD Neil, and Chad M Swanson. KHNYN is essential for the zinc finger antiviral protein (ZAP) to restrict HIV-1 containing clustered CpG dinucleotides. *eLife*, 8, 2019.

[5476] Brandon D Fields, Son C Nguyen, Guy Nir, and Scott Kennedy. A multiplexed DNA FISH strategy for assessing genome architecture in Caenorhabditis elegans. *eLife*, 8, 2019.

[5477] Jonathan Fillatre, Jean-Daniel Fauny, Jasmine Alexandra Fels, Cheng Li, Mary Goll, Christine Thisse, and Bernard Thisse. TEADs, Yap, Taz, Vgll4s transcription factors control the establishment of Left-Right asymmetry in zebrafish. *eLife*, 8, 2019.

[5478] Andrew JP Fink, Richard Axel, and Carl E Schoonover. A virtual burrow assay for head–fixed mice measures habituation, discrimination, exploration and avoidance without training. *eLife*, 8, 2019.

[5479] Daniel Fisch, Artur Yakimovich, Barbara Clough, Joseph Wright, Monique Bunyan, Michael Howell, Jason Mercer, and Eva Frickel. Defining host–pathogen interactions employing an artificial intelligence workflow. *eLife*, 8, 2019.

[5480] Josef Fischböck-Halwachs, Sylvia Singh, Mia Potocnjak, Götz Hagemann, Victor Solis-Mezarino, Stephan Woike, Medini Ghodgaonkar-Steger, Florian Weissmann, Laura D Gallego, Julie Rojas, Jessica Andreani, Alwin Köhler, and Franz Herzog. The COMA complex interacts with Cse4 and positions Sli15/Ipl1 at the budding yeast inner kinetochore. *eLife*, 8, 2019.

[5481] Julia Fischer, James P Higham, Susan C Alberts, Louise Barrett, Jacinta C Beehner, Thore J Bergman, Alecia J Carter, Anthony Collins, Sarah Elton, Joël Fagot, Maria Joana Ferreira da Silva, Kurt Hammerschmidt, Peter Henzi, Clifford J Jolly, Sascha Knauf, Gisela H Kopp, Jeffrey Rogers, Christian Roos, Caroline Ross, Robert M Seyfarth, Joan Silk, Noah Snyder-Mackler, Veronika Staedele, Larissa Swedell, Michael L Wilson, and Dietmar Zinner. Insights into the evolution of social systems and species from baboon studies. *eLife*, 8, 2019.

[5482] Alexandra Fletcher-Jones, Keri L Hildick, Ashley J Evans, Yasuko Nakamura, Kevin A Wilkinson, and Jeremy M Henley. The C-terminal helix 9 motif in rat cannabinoid receptor type 1 regulates axonal trafficking and surface expression. *eLife*, 8, 2019.

[5483] Josué Flores-Kim, Genevieve S Dobihal, Andrew Fenton, David Z Rudner, and Thomas G Bernhardt. A switch in surface polymer biogenesis triggers growth-phase-dependent and antibiotic-induced bacteriolysis. *eLife*, 8, 2019.

[5484] Matthew W Flounders, Carlos González-García, Richard Hardstone, and Biyu J He. Neural dynamics of visual ambiguity resolution by perceptual prior. *eLife*, 8, 2019.

[5485] Lauren E Foley, Jinli Ling, Radhika Joshi, Naveh Evantal, Sebastian Kadener, and Patrick Emery. Drosophila PSI controls circadian period and the phase of circadian behavior under temperature cycle via tim splicing. *eLife*, 8, 2019.

[5486] Davide Folloni, Jerome Sallet, Alexandre A Khrapitchev, Nicola Sibson, Lennart Verhagen, and Rogier B Mars. Dichotomous organization of amygdala/temporal-prefrontal bundles in both humans and monkeys. *eLife*, 8, 2019.

[5487] Kimberly M Fornace, Neal Alexander, Tommy R Abidin, Paddy M Brock, Tock H Chua, Indra Vythilingam, Heather M Ferguson, Benny O Manin, Meng L Wong, Sui H Ng, Jon Cox, and Chris Drakeley. Local human movement patterns and land use impact exposure to zoonotic malaria in Malaysian Borneo. *eLife*, 8, 2019.

[5488] Kevin J Forsberg, Ishan V Bhatt, Danica T Schmidtke, Kamyab Javanmardi, Kaylee E Dillard, Barry L Stoddard, Ilya J Finkelstein, Brett K Kaiser, and Harmit S Malik. Functional metagenomics-guided discovery of potent Cas9 inhibitors in the human microbiome. *eLife*, 8, 2019.

[5489] Haleh Fotowat, Candice Lee, James Jaeyoon Jun, and Len Maler. Neural activity in a hippocampus-like region of the teleost pallium is associated with active sensing and navigation. *eLife*, 8, 2019.

[5490] Téo Fournier, Omar Abou Saada, Jing Hou, Jackson Peter, Elodie Caudal, and Joseph Schacherer. Extensive impact of low-frequency variants on the phenotypic landscape at population-scale. *eLife*, 8, 2019.

[5491] Mirko Francesconi, Bruno Di Stefano, Clara Berenguer, Luisa de Andrés-Aguayo, Marcos Plana-Carmona, Maria Mendez-Lago, Amy Guillaumet-Adkins, Gustavo Rodriguez-Esteban, Marta Gut, Ivo G Gut, Holger Heyn, Ben Lehner, and Thomas Graf. Single cell RNA-seq identifies the origins of heterogeneity in efficient cell transdifferentiation and reprogramming. *eLife*, 8, 2019.

[5492] Valerio Francioni, Zahid Padamsey, and Nathalie L Rochefort. High and asymmetric somato-dendritic coupling of V1 layer 5 neurons independent of visual stimulation and locomotion. *eLife*, 8, 2019.

[5493] Shahar Frechter, Alexander Shakeel Bates, Sina Tootoonian, Michael-John Dolan, James Manton, Arian Rokkum Jamasb, Johannes Kohl, Davi Bock, and Gregory Jefferis. Functional and anatomical specificity in a higher olfactory centre. *eLife*, 8, 2019.

[5494] Virginie Fresno, Shauna L Parkes, Angélique Faugère, Etienne Coutureau, and Mathieu Wolff. A thalamocortical circuit for updating action-outcome associations. *eLife*, 8, 2019.

[5495] Elias T Friman, Cédric Deluz, Antonio CA Meireles-Filho, Subashika Govindan, Vincent Gardeux, Bart Deplancke, and David M Suter. Dynamic regulation of chromatin accessibility by pluripotency transcription factors across the cell cycle. *eLife*, 8, 2019.

[5496] Darwin Y Fu and Jacob J Hughey. Releasing a preprint is associated with more attention and citations for the peer-reviewed article. *eLife*, 8, 2019.

[5497] Ryuichi Fukuda, Alla Aharonov, Yu Ting Ong, Oliver A Stone, Mohamed El-Brolosy, Hans-Martin Maischein, Michael Potente, Eldad Tzahor, and Didier YR Stainier. Metabolic modulation regulates cardiac wall morphogenesis in zebrafish. *eLife*, 8, 2019.

[5498] Sumiaki Fukuda, Akihiro Kaneshige, Takayuki Kaji, Yu-taro Noguchi, Yusei Takemoto, Lidan Zhang, Kazutake Tsujikawa, Hiroki Kokubo, Akiyoshi Uezumi, Kazumitsu Maehara, Akihito Harada, Yasuyuki Ohkawa, and So-ichiro Fukada. Sustained expression of HeyL is critical for the proliferation of muscle stem cells in overloaded muscle. *eLife*, 8, 2019.

[5499] Alexander D Fulford, Maxine V Holder, David Frith, Ambrosius P Snijders, Nicolas Tapon, and Paulo S Ribeiro. Casein kinase 1 family proteins promote Slimb-dependent Expanded degradation. *eLife*, 8, 2019.

[5500] Matteo Fumagalli, Stephane M Camus, Yoan Diekmann, Alice Burke, Marine D Camus, Paul J Norman, Agnel Joseph, Laurent Abi-Rached, Andrea Benazzo, Rita Rasteiro, Iain Mathieson, Maya Topf, Peter Parham, Mark G Thomas, and Frances M Brodsky. Genetic diversity of CHC22 clathrin impacts its function in glucose metabolism. *eLife*, 8, 2019.

[5501] Johanna Funk, Felipe Merino, Larisa Venkova, Lina Heydenreich, Jan Kierfeld, Pablo Vargas, Stefan Raunser, Matthieu Piel, and Peter Bieling. Profilin and formin constitute a pacemaker system for robust actin filament growth. *eLife*, 8, 2019.

[5502] Ryo Futahashi, Yumi Yamahama, Migaku Kawaguchi, Naoki Mori, Daisuke Ishii, Genta Okude, Yuji Hirai, Ryouka Kawahara-Miki, Kazutoshi Yoshitake, Shunsuke Yajima, Takahiko Hariyama, and Takema Fukatsu. Molecular basis of wax-based color change and UV reflection in dragonflies. *eLife*, 8, 2019.

[5503] Manuel Gálvez-Santisteban, Danni Chen, Ruilin Zhang, Ricardo Serrano, Cathleen Nguyen, Long Zhao, Laura Nerb, Evan M Masutani, Julien Vermot, Charles Geoffrey Burns, Caroline E Burns, Juan C del álamo, and Neil C Chi. Hemodynamic-mediated endocardial signaling controls in vivo myocardial reprogramming. *eLife*, 8, 2019.

[5504] Dana Gancz, Brian C Raftrey, Gal Perlmoter, Rubén Marín-Juez, Jonathan Semo, Ryota L Matsuoka, Ravi Karra, Hila Raviv, Noga Moshe, Yoseph Addadi, Ofra Golani, Kenneth D Poss, Kristy Red-Horse, Didier YR Stainier, and Karina Yaniv. Distinct origins and molecular mechanisms contribute to lymphatic formation during cardiac growth and regeneration. *eLife*, 8, 2019.

[5505] Kaiyu Gao, Yi Li, Shumei Hu, and Ying Liu. SUMO peptidase ULP-4 regulates mitochondrial UPR-mediated innate immunity and lifespan extension. *eLife*, 8, 2019.

[5506] Rongsui Gao, Wenhui Wei, Bachar H Hassan, Jun Li, Jiaoyu Deng, and Youjun Feng. A single regulator NrtR controls bacterial NAD+ homeostasis via its acetylation. *eLife*, 8, 2019.

[5507] Rui Gao, Anirban Chakraborty, Charlene Geater, Subrata Pradhan, Kara L Gordon, Jeffrey Snowden, Subo Yuan, Audrey S Dickey, Sanjeev Choudhary, Tetsuo Ashizawa, Lisa M Ellerby, Albert R La Spada, Leslie M Thompson, Tapas K Hazra, and Partha S Sarkar. Mutant huntingtin impairs PNKP and ATXN3, disrupting DNA repair and transcription. *eLife*, 8, 2019.

[5508] Vytautas Gapsys and Bert L de Groot. Comment on 'Valid molecular dynamics simulations of human hemoglobin require a surprisingly large box size'. *eLife*, 8, 2019.

[5509] Sara Garbarino, Marco Lorenzi, Neil P Oxtoby, Elisabeth J Vinke, Razvan V Marinescu, Arman Eshaghi, M Arfan Ikram, Wiro J Niessen, Olga Ciccarelli, Frederik Barkhof, Jonathan M Schott, Meike W Vernooij, and Daniel C Alexander. Differences in topological progression profile among neurodegenerative diseases from imaging data. *eLife*, 8, 2019.

[5510] Yasel Garcés Suárez, Jose L Martínez, David Torres Hernández, Haydee Olinca Hernández, Arianna Pérez-Delgado, Mayra Méndez, Christopher D Wood, Juan Manuel Rendon-Mancha, Daniela Silva-Ayala, Susana López, Adán Guerrero, and Carlos F Arias. Nanoscale organization of rotavirus replication machineries. *eLife*, 8, 2019.

[5511] Jacob M Garrigues, Brian V Tsu, Matthew D Daugherty, and Amy E Pasquinelli. Diversification of the Caenorhabditis heat shock response by Helitron transposable elements. *eLife*, 8, 2019.

[5512] Jenna M Gaska, Metodi Balev, Qiang Ding, Brigitte Heller, and Alexander Ploss. Differences across cyclophilin A orthologs contribute to the host range restriction of hepatitis C virus. *eLife*, 8, 2019.

[5513] Merse E Gáspár, Pierre-Olivier Polack, Peyman Golshani, Máté Lengyel, and Gergő Orbán. Representational untangling by the firing rate nonlinearity in V1 simple cells. *eLife*, 8, 2019.

[5514] Magdalena Gebala, Stephanie L Johnson, Geeta J Narlikar, and Dan Herschlag. Ion counting demonstrates a high electrostatic field generated by the nucleosome. *eLife*, 8, 2019.

[5515] Emma J Gehrke, Xiafei Zhang, Sheila M Pimentel-Elardo, Andrew R Johnson, Christiaan A Rees, Stephanie E Jones, Hindra, Sebastian S Gehrke, Sonya Turvey, Suzanne Boursalie, Jane E Hill, Erin E Carlson, Justin R Nodwell, and Marie A Elliot. Silencing cryptic specialized metabolism in Streptomyces by the nucleoid-associated protein Lsr2. *eLife*, 8, 2019.

[5516] Kathryn Geiger-Schuller, Jaba Mitra, Taekjip Ha, and Doug Barrick. Functional instability allows access to DNA in longer transcription Activator-Like effector (TALE) arrays. *eLife*, 8, 2019.

[5517] Sara Genovese, Raphaël Clément, Cassandra Gaultier, Florence Besse, Karine Narbonne-Reveau, Fabrice Daian, Sophie Foppolo, Nuno Miguel Luis, and Cédric Maurange. Coopted temporal patterning governs cellular hierarchy, heterogeneity and metabolism in Drosophila neuroblast tumors. *eLife*, 8, 2019.

[5518] Daniel Gerber, Monica Ghidinelli, Elisa Tinelli, Christian Somandin, Joanne Gerber, Jorge A Pereira, Andrea Ommer, Gianluca Figlia, Michaela Miehe, Lukas G Nägeli, Vanessa Suter, Valentina Tadini, Páris NM Sidiropoulos, Carsten Wessig, Klaus V Toyka, and Ueli Suter. Schwann cells, but not Oligodendrocytes, Depend Strictly on Dynamin 2 Function. *eLife*, 8, 2019.

[5519] Veneta Gerganova, Charlotte Floderer, Anna Archetti, Laetitia Michon, Lina Carlini, Thais Reichler, Suliana Manley, and Sophie G Martin. Multi-phosphorylation reaction and clustering tune Pom1 gradient mid-cell levels according to cell size. *eLife*, 8, 2019.

[5520] Swagata Ghatak, Nima Dolatabadi, Dorit Trudler, XiaoTong Zhang, Yin Wu, Madhav Mohata, Rajesh Ambasudhan, Maria Talantova, and Stuart A Lipton. Mechanisms of hyperexcitability in Alzheimer's disease hiPSC-derived neurons and cerebral organoids vs isogenic controls. *eLife*, 8, 2019.

[5521] Harshad Ghodke, Bishnu P Paudel, Jacob S Lewis, Slobodan Jergic, Kamya Gopal, Zachary J Romero, Elizabeth A Wood, Roger Woodgate, Michael M Cox, and Antoine M van Oijen. Spatial and temporal organization of RecA in the Escherichia coli DNA-damage response. *eLife*, 8, 2019.

[5522] Tobias W Giessen, Benjamin J Orlando, Andrew A Verdegaal, Melissa G Chambers, Jules Gardener, David C Bell, Gabriel Birrane, Maofu Liao, and Pamela A Silver. Large protein organelles form a new iron sequestration system with high storage capacity. *eLife*, 8, 2019.

[5523] Alison K Gillingham, Jessie Bertram, Farida Begum, and Sean Munro. In vivo identification of GTPase interactors by mitochondrial relocalization and proximity biotinylation. *eLife*, 8, 2019.

[5524] Dion Giovannone, Sandeep Paul, Simone Schindler, Claire Arata, D'Juan T Farmer, Punam Patel, Joanna Smeeton, and J Gage Crump. Programmed conversion of hypertrophic chondrocytes into osteoblasts and marrow adipocytes within zebrafish bones. *eLife*, 8, 2019.

[5525] Philipp Glock, Fridtjof Brauns, Jacob Halatek, Erwin Frey, and Petra Schwille. Design of biochemical pattern forming systems from minimal motifs. *eLife*, 8, 2019.

[5526] Sebastian Gluth and Nachshon Meiran. Leave-One-Trial-Out, LOTO, a general approach to link single-trial parameters of cognitive models to neural data. *eLife*, 8, 2019.

[5527] Kevin M Goff and Ethan M Goldberg. Vasoactive intestinal peptide-expressing interneurons are impaired in a mouse model of Dravet syndrome. *eLife*, 8, 2019.

[5528] Qingnian Goh, Taejeong Song, Michael J Petrany, Alyssa AW Cramer, Chengyi Sun, Sakthivel Sadayappan, Se-Jin Lee, and Douglas P Millay. Myonuclear accretion is a determinant of exercise-induced remodeling in skeletal muscle. *eLife*, 8, 2019.

[5529] David J Goldhamer and John B Lees-Shepard. Response to comment on 'Palovarotene reduces heterotopic ossification in juvenile FOP mice but exhibits pronounced skeletal toxicity'. *eLife*, 8, 2019.

[5530] Adriana E Golding, Ilaria Visco, Peter Bieling, and William M Bement. Extraction of active RhoGTPases by RhoGDI regulates spatiotemporal patterning of RhoGTPases. *eLife*, 8, 2019.

[5531] Rama Reddy Goluguri, Sreemantee Sen, and Jayant Udgaonkar. Microsecond sub-domain motions and the folding and misfolding of the mouse prion protein. *eLife*, 8, 2019.

[5532] Nicanor González-Morales, Yu Shu Xiao, Matthew Aaron Schilling, Océane Marescal, Kuo An Liao, and Frieder Schöck. Myofibril diameter is set by a finely tuned mechanism of protein oligomerization in Drosophila. *eLife*, 8, 2019.

[5533] Elena Gonzalo-Gil, Patrick B Rapuano, Uchenna Ikediobi, Rebecca Leibowitz, Sameet Mehta, Ayse K Coskun, J Zachary Porterfield, Teagan D Lampkin, Vincent C Marconi, David Rimland, Bruce D Walker, Steven Deeks, and Richard E Sutton. Transcriptional down-regulation of ccr5 in a subset of HIV+ controllers and their family members. *eLife*, 8, 2019.

[5534] Travis D Goode, Reed L Ressler, Gillian M Acca, Olivia W Miles, and Stephen Maren. Bed nucleus of the stria terminalis regulates fear to unpredictable threat signals. *eLife*, 8, 2019.

[5535] Carlos Gorbea, Tim Mosbruger, David A Nix, and Demián Cazalla. Viral miRNA adaptor differentially recruits miRNAs to target mRNAs through alternative base-pairing. *eLife*, 8, 2019.

[5536] Sergey Gorelick, Genevieve Buckley, Gediminas Gervinskas, Travis K Johnson, Ava Handley, Monica Pia Caggiano, James C Whisstock, Roger Pocock, and Alex de Marco. PIE-scope, integrated cryo-correlative light and FIB/SEM microscopy. *eLife*, 8, 2019.

[5537] Rachel A Gottschalk, Michael G Dorrington, Bhaskar Dutta, Kathleen S Krauss, Andrew J Martins, Stefan Uderhardt, Waipan Chan, John S Tsang, Parizad Torabi-Parizi, Iain DC Fraser, and Ronald N Germain. IFN-mediated negative feedback supports bacteria class-specific macrophage inflammatory responses. *eLife*, 8, 2019.

[5538] Mehdi Goudarzi, Kathryn Berg, Lindsey M Pieper, and Alexander F Schier. Individual long non-coding RNAs have no overt functions in zebrafish embryogenesis, viability and fertility. *eLife*, 8, 2019.

[5539] David Goyer, Marina A Silveira, Alexander P George, Nichole L Beebe, Ryan M Edelbrock, Peter T Malinski, Brett R Schofield, and Michael T Roberts. A novel class of inferior colliculus principal neurons labeled in vasoactive intestinal peptide-Cre mice. *eLife*, 8, 2019.

[5540] Fabian Grabenhorst, Ken-Ichiro Tsutsui, Shunsuke Kobayashi, and Wolfram Schultz. Primate prefrontal neurons signal economic risk derived from the statistics of recent reward experience. *eLife*, 8, 2019.

[5541] Matti Gralka and Oskar Hallatschek. Environmental heterogeneity can tip the population genetics of range expansions. *eLife*, 8, 2019.

[5542] Alec K Gramann, Arvind M Venkatesan, Melissa Guerin, and Craig J Ceol. Regulation of zebrafish melanocyte development by ligand-dependent BMP signaling. *eLife*, 8, 2019.

[5543] Justin J Grassmeyer, Asia L Cahill, Cassandra L Hays, Cody Barta, Rolen M Quadros, Channabasavaiah B Gurumurthy, and Wallace B Thoreson. Ca2+ sensor synaptotagmin-1 mediates exocytosis in mammalian photoreceptors. *eLife*, 8, 2019.

[5544] Basil J Greber, Daniel B Toso, Jie Fang, and Eva Nogales. The complete structure of the human TFIIH core complex. *eLife*, 8, 2019.

[5545] Maxim Greenberg, Aurélie Teissandier, Marius Walter, Daan Noordermeer, and Deborah Bourc'his. Dynamic enhancer partitioning instructs activation of a growth-related gene during exit from naïve pluripotency. *eLife*, 8, 2019.

[5546] Ethan J Greenblatt, Rebecca Obniski, Claire Mical, and Allan C Spradling. Prolonged ovarian storage of mature Drosophila oocytes dramatically increases meiotic spindle instability. *eLife*, 8, 2019.

[5547] Eric R Greene, Ellen A Goodall, Andres H de la Peña, Mary E Matyskiela, Gabriel C Lander, and Andreas Martin. Specific lid-base contacts in the 26s proteasome control the conformational switching required for substrate degradation. *eLife*, 8, 2019.

[5548] Benjamin James Griffiths, Stephen D Mayhew, Karen J Mullinger, João Jorge, Ian Charest, Maria Wimber, and Simon Hanslmayr. Alpha/beta power decreases track the fidelity of stimulus-specific information. *eLife*, 8, 2019.

[5549] Lin Grimm, Hiroyuki Nakajima, Smrita Chaudhury, Neil I Bower, Kazuhide S Okuda, Andrew G Cox, Natasha L Harvey, Katarzyna Koltowska, Naoki Mochizuki, and Benjamin M Hogan. Yap1 promotes sprouting and proliferation of lymphatic progenitors downstream of Vegfc in the zebrafish trunk. *eLife*, 8, 2019.

[5550] Maor Grinberg, Tomer Orevi, Shifra Steinberg, and Nadav Kashtan. Bacterial survival in microscopic surface wetness. *eLife*, 8, 2019.

[5551] Sarah L Griner, Paul Seidler, Jeannette Bowler, Kevin A Murray, Tianxiao Peter Yang, Shruti Sahay, Michael R Sawaya, Duilio Cascio, Jose A Rodriguez, Stephan Philipp, Justyna Sosna, Charles G Glabe, Tamir Gonen, and David S Eisenberg. Structure-based inhibitors of amyloid beta core suggest a common interface with tau. *eLife*, 8, 2019.

[5552] Mor Grinstein, Heather L Dingwall, Luke D O'Connor, Ken Zou, Terence Dante Capellini, and Jenna Lauren Galloway. A distinct transition from cell growth to physiological homeostasis in the tendon. *eLife*, 8, 2019.

[5553] Eyal Gruntman, Sandro Romani, and Michael B Reiser. The computation of directional selectivity in the Drosophila OFF motion pathway. *eLife*, 8, 2019.

[5554] Vincent Guacci, Fiona Chatterjee, Brett Robison, and Douglas E Koshland. Communication between distinct subunit interfaces of the cohesin complex promotes its topological entrapment of DNA. *eLife*, 8, 2019.

[5555] Ricardo Guajardo, David J Luginbuhl, Shuo Han, Liqun Luo, and Jiefu Li. Functional divergence of Plexin B structural motifs in distinct steps of Drosophila olfactory circuit assembly. *eLife*, 8, 2019.

[5556] Ricardo Guerrero-Ferreira, Nicholas MI Taylor, Ana-Andreea Arteni, Pratibha Kumari, Daniel Mona, Philippe Ringler, Markus Britschgi, Matthias E Lauer, Ali Makky, Joeri Verasdonck, Roland Riek, Ronald Melki, Beat H Meier, Anja Böckmann, Luc Bousset, and Henning Stahlberg. Two new polymorphic structures of human full-length alpha-synuclein fibrils solved by cryo-electron microscopy. *eLife*, 8, 2019.

[5557] Semih Günel, Helge Rhodin, Daniel Morales, João Campagnolo, Pavan Ramdya, and Pascal Fua. DeepFly3D, a deep learning-based approach for 3D limb and appendage tracking in tethered, adult Drosophila. *eLife*, 8, 2019.

[5558] Hui Guo, Toshiharu Suzuki, and John L Rubinstein. Structure of a bacterial ATP synthase. *eLife*, 8, 2019.

[5559] Pengfei Guo, Chang-Hyun Lee, Huiyan Lei, Yonggang Zheng, Katiuska Daniela Pulgar Prieto, and Duojia Pan. Nerfin-1 represses transcriptional output of Hippo signaling in cell competition. *eLife*, 8, 2019.

[5560] Qiong Guo, Shanhui Liao, Sebastian Kwiatkowski, Weronika Tomaka, Huijuan Yu, Gao Wu, Xiaoming Tu, Jinrong Min, Jakub Drozak, and Chao Xu. Structural insights into SETD3-mediated histidine methylation on $\beta$-actin. *eLife*, 8, 2019.

[5561] Wenna Guo, Liucun Zhu, Rui Zhu, Qihan Chen, Qiang Wang, and Jian-Qun Chen. A four-DNA methylation biomarker is a superior predictor of survival of patients with cutaneous melanoma. *eLife*, 8, 2019.

[5562] Devendra Kumar Gupta, Laurent Dembele, Annemarie Voorberg-van der Wel, Guglielmo Roma, Andy Yip, Vorada Chuenchob, Niwat Kangwanrangsan, Tomoko Ishino, Ashley M Vaughan, Stefan H Kappe, Erika L Flannery, Jetsumon Sattabongkot, Sebastian Mikolajczak, Pablo Bifani, Clemens HM Kocken, and Thierry Tidiane Diagana. The Plasmodium liver-specific protein 2 (LISP2) is an early marker of liver stage development. *eLife*, 8, 2019.

[5563] Kanchan Gupta, Gilman ES Toombes, and Kenton J Swartz. Exploring structural dynamics of a membrane protein by combining bioorthogonal chemistry and cysteine mutagenesis. *eLife*, 8, 2019.

[5564] Ritu Gupta, Adhish S Walvekar, Shun Liang, Zeenat Rashida, Premal Shah, and Sunil Laxman. A tRNA modification balances carbon and nitrogen metabolism by regulating phosphate homeostasis. *eLife*, 8, 2019.

[5565] Morgan L Gustison, Jeremy I Borjon, Daniel Y Takahashi, and Asif A Ghazanfar. Vocal and locomotor coordination develops in association with the autonomic nervous system. *eLife*, 8, 2019.

[5566] Leah Guthrie, Sarah Wolfson, and Libusha Kelly. The human gut chemical landscape predicts microbe-mediated biotransformation of foods and drugs. *eLife*, 8, 2019.

[5567] Gabrielle J Gutierrez and Sophie Denève. Population adaptation in efficient balanced networks. *eLife*, 8, 2019.

[5568] Vanessa A Gutzeit, Jordana Thibado, Daniel Starer Stor, Zhou Zhou, Scott C Blanchard, Olaf S Andersen, and Joshua Levitz. Conformational dynamics between transmembrane domains and allosteric modulation of a metabotropic glutamate receptor. *eLife*, 8, 2019.

[5569] Alenka Guček, Nikhil R Gandasi, Muhmmad Omar-Hmeadi, Marit Bakke, Stein O Døskeland, Anders Tengholm, and Sebastian Barg. Fusion pore regulation by cAMP/Epac2 controls cargo release during insulin exocytosis. *eLife*, 8, 2019.

[5570] Bat-Sheva Hadad and Sivan Schwartz. Perception in autism does not adhere to Weber's law. *eLife*, 8, 2019.

[5571] Zena Hadjivasiliou and Andrew Pomiankowski. Evolution of asymmetric gamete signaling and suppressed recombination at the mating type locus. *eLife*, 8, 2019.

[5572] Judith Hagenbuchner, Veronika Obsilova, Teresa Kaserer, Nora Kaiser, Bettina Rass, Katarina Psenakova, Vojtech Docekal, Miroslava Alblova, Klara Kohoutova, Daniela Schuster, Tatsiana Aneichyk, Jan Vesely, Petra Obexer, Tomas Obsil, and Michael J Ausserlechner. Modulating FOXO3 transcriptional activity by small, DBD-binding molecules. *eLife*, 8, 2019.

[5573] Nicole Hajicek, Nicholas C Keith, Edhriz Siraliev-Perez, Brenda RS Temple, Weigang Huang, Qisheng Zhang, T Kendall Harden, and John Sondek. Structural basis for the activation of PLC-$\gamma$ isozymes by phosphorylation and cancer-associated mutations. *eLife*, 8, 2019.

[5574] Briac Halbout, Andrew T Marshall, Ali Azimi, Mimi Liljeholm, Stephen V Mahler, Kate M Wassum, and Sean B Ostlund. Mesolimbic dopamine projections mediate cue-motivated reward seeking but not reward retrieval in rats. *eLife*, 8, 2019.

[5575] Matthew D Hall, Matthew TG Holden, Pramot Srisomang, Weera Mahavanakul, Vanaporn Wuthiekanun, Direk Limmathurotsakul, Kay Fountain, Julian Parkhill, Emma K Nickerson, Sharon J Peacock, and Christophe Fraser. Improved characterisation of MRSA transmission using within-host bacterial sequence diversity. *eLife*, 8, 2019.

[5576] Zachary F Hallberg, Chi Ho Chan, Todd A Wright, Philip J Kranzusch, Kevin W Doxzen, James J Park, Daniel R Bond, and Ming C Hammond. Structure and mechanism of a Hypr GGDEF enzyme that activates cGAMP signaling to control extracellular metal respiration. *eLife*, 8, 2019.

[5577] Machika Hamaguchi, Rieko Muramatsu, Harutoshi Fujimura, Hideki Mochizuki, Hirotoshi Kataoka, and Toshihide Yamashita. Circulating transforming growth factor-$\beta$1 facilitates remyelination in the adult central nervous system. *eLife*, 8, 2019.

[5578] Elizabeth Ann Hambleton, Victor Arnold Shivas Jones, Ira Maegele, David Kvaskoff, Timo Sachsenheimer, and Annika Guse. Sterol transfer by atypical cholesterol-binding NPC2 proteins in coral-algal symbiosis. *eLife*, 8, 2019.

[5579] Dandan Han, Lars Schomacher, Katrin M Schüle, Medhavi Mallick, Michael U Musheev, Emil Karaulanov, Laura Krebs, Annika von Seggern, and Christof Niehrs. NEIL1 and NEIL2 DNA glycosylases protect neural crest development against mitochondrial oxidative stress. *eLife*, 8, 2019.

[5580] Han Han, James M Fulcher, Venkata P Dandey, Janet H Iwasa, Wesley I Sundquist, Michael S Kay, Peter S Shen, and Christopher P Hill. Structure of Vps4 with circular peptides and implications for translocation of two polypeptide chains by AAA+ ATPases. *eLife*, 8, 2019.

[5581] Laura Hanson, Santhosh Sethuramanujam, Geoff deRosenroll, Varsha Jain, and Gautam B Awatramani. Retinal direction selectivity in the absence of asymmetric starburst amacrine cell responses. *eLife*, 8, 2019.

[5582] Mark Austin Hanson, Anna Dostálová, Camilla Ceroni, Mickael Poidevin, Shu Kondo, and Bruno Lemaitre. Synergy and remarkable specificity of antimicrobial peptides in vivo using a systematic knockout approach. *eLife*, 8, 2019.

[5583] Fransky Hantelys, Anne-Claire Godet, Florian David, Florence Tatin, Edith Renaud-Gabardos, FranÇoise Pujol, Leila H Diallo, Isabelle Ader, Laetitia Ligat, Anthony K Henras, Yasufumi Sato, Angelo Parini, Eric Lacazette, Barbara Garmy-Susini, and Anne-Catherine Prats. Vasohibin1, a new mouse cardiomyocyte IRES trans-acting factor that regulates translation in early hypoxia. *eLife*, 8, 2019.

[5584] Kumi Harada, Tetsuya Kotani, Hiromi Kirisako, Machiko Sakoh-Nakatogawa, Yu Oikawa, Yayoi Kimura, Hisashi Hirano, Hayashi Yamamoto, Yoshinori Ohsumi, and Hitoshi Nakatogawa. Two distinct mechanisms target the autophagy-related E3 complex to the pre-autophagosomal structure. *eLife*, 8, 2019.

[5585] Shogo Haraguchi, Masaki Kamata, Takuma Tokita, Kei-ichiro Tashiro, Miku Sato, Mitsuki Nozaki, Mayumi Okamoto-Katsuyama, Isao Shimizu, Guofeng Han, Vishwajit Sur Chowdhury, Xiao-Feng Lei, Takuro Miyazaki, Joo-ri Kim-Kaneyama, Tomoya Nakamachi, Kouhei Matsuda, Hirokazu Ohtaki, Toshinobu Tokumoto, Tetsuya Tachibana, Akira Miyazaki, and Kazuyoshi Tsutsui. Light-at-night exposure affects brain development through pineal allopregnanolone-dependent mechanisms. *eLife*, 8, 2019.

[5586] Eva Harde, LaShae Nicholson, Beatriz Furones Cuadrado, Diane Bissen, Sylvia Wigge, Severino Urban, Marta Segarra, Carmen Ruiz de Almodóvar, and Amparo Acker-Palmer. EphrinB2 regulates VEGFR2 during dendritogenesis and hippocampal circuitry development. *eLife*, 8, 2019.

[5587] Heather P Harding, Adriana Ordonez, Felicity Allen, Leopold Parts, Alison J Inglis, Roger L Williams, and David Ron. The ribosomal P-stalk couples amino acid starvation to GCN2 activation in mammalian cells. *eLife*, 8, 2019.

[5588] Holly Hardy, James GD Prendergast, Aara Patel, Sunit Dutta, Violeta Trejo-Reveles, Hannah Kroeger, Andrea R Yung, Lisa V Goodrich, Brian Brooks, Jane C Sowden, and Joe Rainger. Detailed analysis of chick optic fissure closure reveals Netrin-1 as an essential mediator of epithelial fusion. *eLife*, 8, 2019.

[5589] Jonathan Harnie, Adam Doelman, Emmanuelle de Vette, Johannie Audet, Etienne Desrochers, Nathaly Gaudreault, and Alain Frigon. The recovery of standing and locomotion after spinal cord injury does not require task-specific training. *eLife*, 8, 2019.

[5590] Clair Harris, Marissa Cloutier, Megan Trotter, Michael Hinten, Srimonta Gayen, Zhenhai Du, Wei Xie, and Sundeep Kalantry. Conversion of random X-inactivation to imprinted X-inactivation by maternal PRC2. *eLife*, 8, 2019.

[5591] Michael RM Harrison, Xidi Feng, Guqin Mo, Antonio Aguayo, Jessi Villafuerte, Tyler Yoshida, Caroline A Pearson, Stefan Schulte-Merker, and Ching-Ling Lien. Late developing cardiac lymphatic vasculature supports adult zebrafish heart function and regeneration. *eLife*, 8, 2019.

[5592] Samuel Frederick Mock Hart, Jose Mario Bello Pineda, Chi-Chun Chen, Robin Green, and Wenying Shou. Disentangling strictly self-serving mutations from win-win mutations in a mutualistic microbial community. *eLife*, 8, 2019.

[5593] Constantin Hays, Gérald Touak, Abdelouhab Bouaboud, Agnès Fouet, Julie Guignot, Claire Poyart, and Asmaa Tazi. Perinatal hormones favor CC17 group B Streptococcus intestinal translocation through M cells and hypervirulence in neonates. *eLife*, 8, 2019.

[5594] Amanda N Hayward, Eric J Aird, and Wendy R Gordon. A toolkit for studying cell surface shedding of diverse transmembrane receptors. *eLife*, 8, 2019.

[5595] Bicheng He, Marita Buescher, Max Stephen Farnworth, Frederic Strobl, Ernst HK Stelzer, Nikolaus DB Koniszewski, Dominik Muehlen, and Gregor Bucher. An ancestral apical brain region contributes to the central complex under the control of foxQ2 in the beetle Tribolium. *eLife*, 8, 2019.

[5596] Li He, Richard Binari, Jiuhong Huang, Julia Falo-Sanjuan, and Norbert Perrimon. In vivo study of gene expression with an enhanced dual-color fluorescent transcriptional timer. *eLife*, 8, 2019.

[5597] Shengbo He, Martin Vickers, Jingyi Zhang, and Xiaoqi Feng. Natural depletion of histone H1 in sex cells causes DNA demethylation, heterochromatin decondensation and transposon activation. *eLife*, 8, 2019.

[5598] Zhixiong He, Larry Young, Xin-Ming Ma, Qianqian Guo, Limin Wang, Yang Yang, Luo Luo, Wei Yuan, Laifu Li, Jing Zhang, Wenjuan Hou, Hui Qiao, Rui Jia, and Fadao Tai. Increased anxiety and decreased sociability induced by paternal deprivation involve the PVN-PrL OTergic pathway. *eLife*, 8, 2019.

[5599] Melissa Hebscher, Jed A Meltzer, and Asaf Gilboa. A causal role for the precuneus in network-wide theta and gamma oscillatory activity during complex memory retrieval. *eLife*, 8, 2019.

[5600] Idse Heemskerk, Kari Burt, Matthew Miller, Sapna Chhabra, M Cecilia Guerra, Lizhong Liu, and Aryeh Warmflash. Rapid changes in morphogen concentration control self-organized patterning in human embryonic stem cells. *eLife*, 8, 2019.

[5601] William Heffley and Court Hull. Classical conditioning drives learned reward prediction signals in climbing fibers across the lateral cerebellum. *eLife*, 8, 2019.

[5602] Ganapati V Hegde, Cecile de la Cruz, Jennifer M Giltnane, Lisa Crocker, Avinashnarayan Venkatanarayan, Gabriele Schaefer, Debra Dunlap, Joerg D Hoeck, Robert Piskol, Florian Gnad, Zora Modrusan, Frederic J de Sauvage, Christian W Siebel, and Erica L Jackson. NRG1 is a critical regulator of differentiation in TP63-driven squamous cell carcinoma. *eLife*, 8, 2019.

[5603] Robert N Helsley, Venkateshwari Varadharajan, Amanda L Brown, Anthony D Gromovsky, Rebecca C Schugar, Iyappan Ramachandiran, Kevin Fung, Mohammad Nasser Kabbany, Rakhee Banerjee, Chase K Neumann, Chelsea Finney, Preeti Pathak, Danny Orabi, Lucas J Osborn, William Massey, Renliang Zhang, Anagha Kadam, Brian E Sansbury, Calvin Pan, Jessica Sacks, Richard G Lee, Rosanne M Crooke, Mark J Graham, Madeleine E Lemieux, Valentin Gogonea, John P Kirwan, Daniela S Allende, Mete Civelek, Paul L Fox, Lawrence L Rudel, Aldons J Lusis, Matthew Spite, and J Mark Brown. Obesity-linked suppression of membrane-bound O-acyltransferase 7 (MBOAT7) drives non-alcoholic fatty liver disease. *eLife*, 8, 2019.

[5604] Ethan A Heming, Kevin P Cross, Tomohiko Takei, Douglas J Cook, and Stephen H Scott. Independent representations of ipsilateral and contralateral limbs in primary motor cortex. *eLife*, 8, 2019.

[5605] Nathan T Henderson, Sylvain J Le Marchand, Martin Hruska, Simon Hippenmeyer, Liqun Luo, and Matthew B Dalva. Ephrin-B3 controls excitatory synapse density through cell-cell competition for EphBs. *eLife*, 8, 2019.

[5606] Joseph Herdy, Simon Schafer, Yongsung Kim, Zoya Ansari, Dina Zangwill, Manching Ku, Apua Paquola, Hyungjun Lee, Jerome Mertens, and Fred H Gage. Chemical modulation of transcriptionally enriched signaling pathways to optimize the conversion of fibroblasts into neurons. *eLife*, 8, 2019.

[5607] Dora Hermes, Natalia Petridou, Kendrick N Kay, and Jonathan Winawer. An image-computable model for the stimulus selectivity of gamma oscillations. *eLife*, 8, 2019.

[5608] Caesar M Hernandez, Caitlin A Orsini, Chase C Labiste, Alexa-Rae Wheeler, Tyler W Ten Eyck, Matthew M Bruner, Todd J Sahagian, Scott W Harden, Charles J Frazier, Barry Setlow, and Jennifer L Bizon. Optogenetic dissection of basolateral amygdala contributions to intertemporal choice in young and aged rats. *eLife*, 8, 2019.

[5609] Diana Herrera-Perez, Alyson Haslam, Tyler Crain, Jennifer Gill, Catherine Livingston, Victoria Kaestner, Michael Hayes, Dan Morgan, Adam S Cifu, and Vinay Prasad. A comprehensive review of randomized clinical trials in three medical journals reveals 396 medical reversals. *eLife*, 8, 2019.

[5610] Sarah K Herring, Hee-Jung Moon, Punam Rawal, Anindit Chhibber, and Liqin Zhao. Brain clusterin protein isoforms and mitochondrial localization. *eLife*, 8, 2019.

[5611] Christina Hertel, Dmytro Fishman, Anna Lorenc, Annamari Ranki, Kai Krohn, Pärt Peterson, Kai Kisand, and Adrian Hayday. Response to comment on 'AIRE-deficient patients harbor unique high-affinity disease-ameliorating autoantibodies'. *eLife*, 8, 2019.

[5612] Mélina L Heuzé, Gautham Hari Narayana Sankara Narayana, Joseph D'Alessandro, Victor Cellerin, Tien Dang, David S Williams, Jan CM Van Hest, Philippe Marcq, René-Marc Mège, and Benoit Ladoux. Myosin II isoforms play distinct roles in adherens junction biogenesis. *eLife*, 8, 2019.

[5613] Leon Hilgers and Julia Schwarzer. The untapped potential of medaka and its wild relatives. *eLife*, 8, 2019.

[5614] Spencer Hill, Kurt Reichermeier, Daniel C Scott, Lorena Samentar, Jasmin Coulombe-Huntington, Luisa Izzi, Xiaojing Tang, Rebeca Ibarra, Thierry Bertomeu, Annie Moradian, Michael J Sweredoski, Nora Caberoy, Brenda A Schulman, Frank Sicheri, Mike Tyers, and Gary Kleiger. Robust cullin-RING ligase function is established by a multiplicity of poly-ubiquitylation pathways. *eLife*, 8, 2019.

[5615] Steven A Hill, Andrew S Blaeser, Austin A Coley, Yajun Xie, Katherine A Shepard, Corey C Harwell, Wen-Jun Gao, and A Denise R Garcia. Sonic hedgehog signaling in astrocytes mediates cell type-specific synaptic organization. *eLife*, 8, 2019.

[5616] Stephen M Hinshaw, Andrew N Dates, and Stephen C Harrison. The structure of the yeast Ctf3 complex. *eLife*, 8, 2019.

[5617] Stephen M Hinshaw and Stephen C Harrison. The structure of the Ctf19c/CCAN from budding yeast. *eLife*, 8, 2019.

[5618] Yoshinori Hirano, Yong-Guang Gao, Daniel J Stephenson, Ngoc T Vu, Lucy Malinina, Dhirendra K Simanshu, Charles E Chalfant, Dinshaw J Patel, and Rhoderick E Brown. Structural basis of phosphatidylcholine recognition by the C2–domain of cytosolic phospholipase A2$\alpha$. *eLife*, 8, 2019.

[5619] Antoine Hocher, Maria Rojec, Jacob B Swadling, Alexander Esin, and Tobias Warnecke. The DNA-binding protein HTa from Thermoplasma acidophilum is an archaeal histone analog. *eLife*, 8, 2019.

[5620] Austin D Hocker, Sarah A Beyeler, Alyssa N Gardner, Stephen M Johnson, Jyoti J Watters, and Adrianne G Huxtable. One bout of neonatal inflammation impairs adult respiratory motor plasticity in male and female rats. *eLife*, 8, 2019.

[5621] Carinna Hockham, Supachai Ekwattanakit, Samir Bhatt, Bridget S Penman, Sunetra Gupta, Vip Viprakasit, and Frédéric B Piel. Estimating the burden of $\alpha$-thalassaemia in Thailand using a comprehensive prevalence database for Southeast Asia. *eLife*, 8, 2019.

[5622] Brian A Hodge, Xiping Zhang, Miguel A Gutierrez-Monreal, Yi Cao, David W Hammers, Zizhen Yao, Christopher A Wolff, Ping Du, Denise Kemler, Andrew R Judge, and Karyn A Esser. MYOD1 functions as a clock amplifier as well as a critical co-factor for downstream circadian gene expression in muscle. *eLife*, 8, 2019.

[5623] Marieke MB Hoekstra, Yann Emmenegger, Jeffrey Hubbard, and Paul Franken. Cold-inducible RNA-binding protein (CIRBP) adjusts clock-gene expression and REM-sleep recovery following sleep deprivation. *eLife*, 8, 2019.

[5624] Andreas Hofmann, Jarno Mäkelä, David J Sherratt, Dieter Heermann, and Seán M Murray. Self-organised segregation of bacterial chromosomal origins. *eLife*, 8, 2019.

[5625] Thea Hogan, Maria Nowicka, Daniel Cownden, Claire F Pearson, Andrew J Yates, and Benedict Seddon. Differential impact of self and environmental antigens on the ontogeny and maintenance of CD4+ T cell memory. *eLife*, 8, 2019.

[5626] Mikael Holm, Chandra Sekhar Mandava, Måns Ehrenberg, and Suparna Sanyal. The mechanism of error induction by the antibiotic viomycin provides insight into the fidelity mechanism of translation. *eLife*, 8, 2019.

[5627] Leanne M Holt, Raymundo D Hernandez, Natasha L Pacheco, Beatriz Torres Ceja, Muhannah Hossain, and Michelle L Olsen. Astrocyte morphogenesis is dependent on BDNF signaling via astrocytic TrkB.T1. *eLife*, 8, 2019.

[5628] Johann Holzmann, Antonio Z Politi, Kota Nagasaka, Merle Hantsche-Grininger, Nike Walther, Birgit Koch, Johannes Fuchs, Gerhard Dürnberger, Wen Tang, Rene Ladurner, Roman R Stocsits, Georg A Busslinger, Béla Novák, Karl Mechtler, Iain Finley Davidson, Jan Ellenberg, and Jan-Michael Peters. Absolute quantification of cohesin, CTCF and their regulators in human cells. *eLife*, 8, 2019.

[5629] Andrew L Hong, Yuen-Yi Tseng, Jeremiah A Wala, Won-Jun Kim, Bryan D Kynnap, Mihir B Doshi, Guillaume Kugener, Gabriel J Sandoval, Thomas P Howard, Ji Li, Xiaoping Yang, Michelle Tillgren, Mahmhoud Ghandi, Abeer Sayeed, Rebecca Deasy, Abigail Ward, Brian McSteen, Katherine M Labella, Paula Keskula, Adam Tracy, Cora Connor, Catherine M Clinton, Alanna J Church, Brian D Crompton, Katherine A Janeway, Barbara Van Hare, David Sandak, Ole Gjoerup, Pratiti

Bandopadhayay, Paul A Clemons, Stuart L Schreiber, David E Root, Prafulla C Gokhale, Susan N Chi, Elizabeth A Mullen, Charles WM Roberts, Cigall Kadoch, Rameen Beroukhim, Keith L Ligon, Jesse S Boehm, and William C Hahn. Renal medullary carcinomas depend upon SMARCB1 loss and are sensitive to proteasome inhibition. *eLife*, 8, 2019.

[5630] Ray L Hong, Metta Riebesell, Daniel J Bumbarger, Steven J Cook, Heather R Carstensen, Tahmineh Sarpolaki, Luisa Cochella, Jessica Castrejon, Eduardo Moreno, Bogdan Sieriebriennikov, Oliver Hobert, and Ralf J Sommer. Evolution of neuronal anatomy and circuitry in two highly divergent nematode species. *eLife*, 8, 2019.

[5631] Hessel Honkoop, Dennis EM de Bakker, Alla Aharonov, Fabian Kruse, Avraham Shakked, Phong D Nguyen, Cecilia de Heus, Laurence Garric, Mauro J Muraro, Adam Shoffner, Federico Tessadori, Joshua Craiger Peterson, Wendy Noort, Alberto Bertozzi, Gilbert Weidinger, George Posthuma, Dominic Grün, Willem J van der Laarse, Judith Klumperman, Richard T Jaspers, Kenneth D Poss, Alexander van Oudenaarden, Eldad Tzahor, and Jeroen Bakkers. Single-cell analysis uncovers that metabolic reprogramming by ErbB2 signaling is essential for cardiomyocyte proliferation in the regenerating heart. *eLife*, 8, 2019.

[5632] Benjamin G Horst, Adam L Yokom, Daniel J Rosenberg, Kyle L Morris, Michal Hammel, James H Hurley, and Michael A Marletta. Allosteric activation of the nitric oxide receptor soluble guanylate cyclase mapped by cryo-electron microscopy. *eLife*, 8, 2019.

[5633] Myra Hosmillo, Jia Lu, Michael R McAllaster, James B Eaglesham, Xinjie Wang, Edward Emmott, Patricia Domingues, Yasmin Chaudhry, Tim J Fitzmaurice, Matthew KH Tung, Marc Dominik Panas, Gerald McInerney, Nicolas Locker, Craig B Wilen, and Ian G Goodfellow. Noroviruses subvert the core stress granule component G3BP1 to promote viral VPg-dependent translation. *eLife*, 8, 2019.

[5634] Panpan Hou, Jingyi Shi, Kelli McFarland White, Yuan Gao, and Jianmin Cui. ML277 specifically enhances the fully activated open state of KCNQ1 by modulating VSD-pore coupling. *eLife*, 8, 2019.

[5635] Nealia CM House, Erica J Polleys, Ishtiaque Quasem, Marjorie De la Rosa Mejia, Cailin E Joyce, Oliver Takacsi-Nagy, Jocelyn E Krebs, Stephen M Fuchs, and Catherine H Freudenreich. Distinct roles for S. cerevisiae H2A copies in recombination and repeat stability, with a role for H2A.1 threonine 126. *eLife*, 8, 2019.

[5636] Mark Howe, Imane Ridouh, Anna Letizia Allegra Mascaro, Alyssa Larios, Maite Azcorra, and Daniel A Dombeck. Coordination of rapid cholinergic and dopaminergic signaling in striatum during spontaneous movement. *eLife*, 8, 2019.

[5637] Sinisa Hrvatin, Christopher P Tzeng, M Aurel Nagy, Hume Stroud, Charalampia Koutsioumpa, Oren F Wilcox, Elena G Assad, Jonathan Green, Christopher D Harvey, Eric C Griffith, and Michael E Greenberg. A scalable platform for the development of cell-type-specific viral drivers. *eLife*, 8, 2019.

[5638] MengJie Hu, Keith E Schulze, Reena Ghildyal, Darren C Henstridge, Jacek L Kolanowski, Elizabeth J New, Yuning Hong, Alan C Hsu, Philip M Hansbro, Peter AB Wark, Marie A Bogoyevitch, and David A Jans. Respiratory syncytial virus co-opts host mitochondrial function to favour infectious virus production. *eLife*, 8, 2019.

[5639] Wei Hu, Yong Zhang, Xiaolin Sun, Tongtong Zhang, Liling Xu, Hengyi Xie, Zhanguo Li, Wanli Liu, Jizhong Lou, and Wei Chen. Fc$\gamma$RIIB-I232T polymorphic change allosterically suppresses ligand binding. *eLife*, 8, 2019.

[5640] Xuchen Hu, Ken Matsumoto, Rachel S Jung, Thomas A Weston, Patrick J Heizer, Cuiwen He, Norma P Sandoval, Christopher M Allan, Yiping Tu, Harry V Vinters, Linda M Liau, Rochelle M Ellison, Jazmin E Morales, Lynn J Baufeld, Nicholas A Bayley, Liqun He, Christer Betsholtz, Anne P Beigneux, David A Nathanson, Holger Gerhardt, Stephen G Young, Loren G Fong, and Haibo Jiang. GPIHBP1 expression in gliomas promotes utilization of lipoprotein-derived nutrients. *eLife*, 8, 2019.

[5641] Xuchen Hu, Thomas A Weston, Cuiwen He, Rachel S Jung, Patrick J Heizer, Brian D Young, Yiping Tu, Peter Tontonoz, James A Wohlschlegel, Haibo Jiang, Stephen G Young, and Loren G Fong. Release of cholesterol-rich particles from the macrophage plasma membrane during movement of filopodia and lamellipodia. *eLife*, 8, 2019.

[5642] Chaolie Huang, Sara Wagner-Valladolid, Amberley D Stephens, Raimund Jung, Chetan Poudel, Tessa Sinnige, Marie C Lechler, Nicole Schlörit, Meng Lu, Romain F Laine, Claire H Michel, Michele Vendruscolo, Clemens F Kaminski, Gabriele S Kaminski Schierle, and Della C David. Intrinsically aggregation-prone proteins form amyloid-like aggregates and contribute to tissue aging in Caenorhabditis elegans. *eLife*, 8, 2019.

[5643] Kee Wui Huang, Nicole E Ochandarena, Adrienne C Philson, Minsuk Hyun, Jaclyn E Birnbaum, Marcelo Cicconet, and Bernardo L Sabatini. Molecular and anatomical organization of the dorsal raphe nucleus. *eLife*, 8, 2019.

[5644] Yihe Huang, Becca Roth, Wei Lü, and Juan Du. Ligand recognition and gating mechanism through three ligand-binding sites of human TRPM2 channel. *eLife*, 8, 2019.

[5645] Ying Huang, Peter Heil, and Michael Brosch. Associations between sounds and actions in early auditory cortex of nonhuman primates. *eLife*, 8, 2019.

[5646] Yung-Chi Huang, Jennifer K Pirri, Diego Rayes, Shangbang Gao, Ben Mulcahy, Jeff Grant, Yasunori Saheki, Michael M Francis, Mei Zhen, and Mark J Alkema. Gain-of-function mutations in the UNC-2/CaV2$\alpha$ channel lead to excitation-dominant synaptic transmission in Caenorhabditis elegans. *eLife*, 8, 2019.

[5647] Taylor ET Hughes, John Smith Del Rosario, Abhijeet Kapoor, Aysenur Torun Yazici, Yevgen Yudin, Edwin C Fluck, Marta Filizola, Tibor Rohacs, and Vera Y Moiseenkova-Bell. Structure-based characterization of novel TRPV5 inhibitors. *eLife*, 8, 2019.

[5648] Pim J Huis in 't Veld, Vladimir A Volkov, Isabelle D Stender, Andrea Musacchio, and Marileen Dogterom. Molecular determinants of the Ska-Ndc80 interaction and their influence on microtubule tracking and force-coupling. *eLife*, 8, 2019.

[5649] Stevephen Hung, Alina Saiakhova, Zachary J Faber, Cynthia F Bartels, Devin Neu, Ian Bayles, Evelyn Ojo, Ellen S Hong, W Dean Pontius, Andrew R Morton, Ruifu Liu, Matthew F Kalady, David N Wald, Sanford Markowitz, and Peter C Scacheri. Mismatch repair-signature mutations activate gene enhancers across human colorectal cancer epigenomes. *eLife*, 8, 2019.

[5650] Mirjam Hunziker, Jonas Barandun, Olga Buzovetsky, Caitlin Steckler, Henrik Molina, and Sebastian Klinge. Conformational switches control early maturation of the eukaryotic small ribosomal subunit. *eLife*, 8, 2019.

[5651] Maxim Igaev, Carsten Kutzner, Lars V Bock, Andrea C Vaiana, and Helmut Grubmüller. Automated cryo-EM structure refinement using correlation-driven molecular dynamics. *eLife*, 8, 2019.

[5652] Antje Ihlefeld, Nima Alamatsaz, and Robert M Shapley. Population rate-coding predicts correctly that human sound localization depends on sound intensity. *eLife*, 8, 2019.

[5653] Ioannis Iliopoulos, Sophia Ananiadou, Antoine Danchin, John PA Ioannidis, Peter D Katsikis, Christos A Ouzounis, and Vasilis J Promponas. Hypothesis, analysis and synthesis, it's all Greek to me. *eLife*, 8, 2019.

[5654] Simon Imhof, Jiayan Zhang, Hui Wang, Khanh Huy Bui, Hoangkim Nguyen, Ivo Atanasov, Wong H Hui, Shun Kai Yang, Z Hong Zhou, and Kent L Hill. Cryo electron tomography with volta phase plate reveals novel structural foundations of the 96-nm axonemal repeat in the pathogen Trypanosoma brucei. *eLife*, 8, 2019.

[5655] Elliot Imler, Jin Sang Pyon, Selina Kindelay, Meaghan Torvund, Yong-quan Zhang, Sreeganga S Chandra, and Konrad E Zinsmaier. A Drosophila model of neuronal ceroid lipofuscinosis CLN4 reveals a hypermorphic gain of function mechanism. *eLife*, 8, 2019.

[5656] Ashley M Ingiosi, Hannah Schoch, Taylor Wintler, Kristan G Singletary, Dario Righelli, Leandro G Roser, Elizabeth Medina, Davide Risso, Marcos G Frank, and Lucia Peixoto. Shank3 modulates sleep and expression of circadian transcription factors. *eLife*, 8, 2019.

[5657] S Andrew Inkpen. Health, ecology and the microbiome. *eLife*, 8, 2019.

[5658] Michele N Insanally, Ioana Carcea, Rachel E Field, Chris C Rodgers, Brian DePasquale, Kanaka Rajan, Michael R DeWeese, Badr F Albanna, and Robert C Froemke. Spike-timing-dependent ensemble encoding by non-classically responsive cortical neurons. *eLife*, 8, 2019.

[5659] Iordan Iordanov, Balázs Tóth, Andras Szollosi, and László Csanády. Enzyme activity and selectivity filter stability of ancient TRPM2 channels were simultaneously lost in early vertebrates. *eLife*, 8, 2019.

[5660] Eric E Irons, Melissa M Lee-Sundlov, Yuqi Zhu, Sriram Neelamegham, Karin M Hoffmeister, and Joseph TY Lau. B cells suppress medullary granulopoiesis by an extracellular glycosylation-dependent mechanism. *eLife*, 8, 2019.

[5661] Eva S Istvan, Sudipta Das, Suyash Bhatnagar, Josh R Beck, Edward Owen, Manuel Llinas, Suresh M Ganesan, Jacquin C Niles, Elizabeth Winzeler, Akhil B Vaidya, and Daniel E Goldberg. Plasmodium Niemann-Pick type C1-related protein is a druggable target required for parasite membrane homeostasis. *eLife*, 8, 2019.

[5662] Daisuke Ito, Sihem Zitouni, Swadhin Chandra Jana, Paulo Duarte, Jaroslaw Surkont, Zita Carvalho-Santos, José B Pereira-Leal, Miguel Godinho Ferreira, and Mónica Bettencourt-Dias. Pericentrin-mediated SAS-6 recruitment promotes centriole assembly. *eLife*, 8, 2019.

[5663] Kie Itoh, Daisuke Murata, Takashi Kato, Tatsuya Yamada, Yoichi Araki, Atsushi Saito, Yoshihiro Adachi, Atsushi Igarashi, Shuo Li, Mikhail Pletnikov, Richard L Huganir, Shigeki Watanabe, Atsushi Kamiya, Miho Iijima, and Hiromi Sesaki. Brain-specific Drp1 regulates postsynaptic endocytosis and dendrite formation independently of mitochondrial division. *eLife*, 8, 2019.

[5664] Wannaporn Ittiprasert, Victoria H Mann, Shannon E Karinshak, Avril Coghlan, Gabriel Rinaldi, Geetha Sankaranarayanan, Apisit Chaidee, Toshihiko Tanno, Chutima Kumkhaek, Pannathee Prangtaworn, Margaret M Mentink-Kane, Christina J Cochran, Patrick Driguez, Nancy Holroyd, Alan Tracey, Rutchanee Rodpai, Bart Everts, Cornelis H Hokke, Karl F Hoffmann, Matthew Berriman, and Paul J Brindley. Programmed genome editing of the omega-1 ribonuclease of the blood fluke, Schistosoma mansoni. *eLife*, 8, 2019.

[5665] Mayumi Iwasaki, Lena Hyvärinen, Urszula Piskurewicz, and Luis Lopez-Molina. Non-canonical RNA-directed DNA methylation participates in maternal and environmental control of seed dormancy. *eLife*, 8, 2019.

[5666] Andrew Jaegle, Vahid Mehrpour, Yalda Mohsenzadeh, Travis Meyer, Aude Oliva, and Nicole Rust. Population response magnitude variation in inferotemporal cortex predicts image memorability. *eLife*, 8, 2019.

[5667] Daniyal J Jafree, Dale Moulding, Maria Kolatsi-Joannou, Nuria Perretta Tejedor, Karen L Price, Natalie J Milmoe, Claire L Walsh, Rosa Maria Correra, Paul JD Winyard, Peter C Harris, Christiana Ruhrberg, Simon Walker-Samuel, Paul R Riley, Adrian S Woolf, Peter J Scambler, and David A Long. Spatiotemporal dynamics and heterogeneity of renal lymphatics in mammalian development and cystic kidney disease. *eLife*, 8, 2019.

[5668] Madhav Jagannathan, Ryan Cummings, and Yukiko M Yamashita. The modular mechanism of chromocenter formation in Drosophila. *eLife*, 8, 2019.

[5669] Sujatha Jagannathan, Yuko Ogata, Philip R Gafken, Stephen J Tapscott, and Robert K Bradley. Quantitative proteomics reveals key roles for post-transcriptional gene regulation in the molecular pathology of facioscapulohumeral muscular dystrophy. *eLife*, 8, 2019.

[5670] Maximilian AH Jakobs, Andrea Dimitracopoulos, and Kristian Franze. KymoButler, a deep learning software for automated kymograph analysis. *eLife*, 8, 2019.

[5671] Manisha Jalan, Judith Oehler, Carl A Morrow, Fekret Osman, and Matthew C Whitby. Factors affecting template switch recombination associated with restarted DNA replication. *eLife*, 8, 2019.

[5672] Mohsen Jamali, Jerome Carriot, Maurice J Chacron, and Kathleen E Cullen. Coding strategies in the otolith system differ for translational head motion vs. static orientation relative to gravity. *eLife*, 8, 2019.

[5673] Thomas D James, Danielle J Zwiefelhofer, and C Andrew Frank. Maintenance of homeostatic plasticity at the Drosophila neuromuscular synapse requires continuous IP3-directed signaling. *eLife*, 8, 2019.

[5674] Mohammed Jamshad, Timothy J Knowles, Scott A White, Douglas G Ward, Fiyaz Mohammed, Kazi Fahmida Rahman, Max Wynne, Gareth W Hughes, Günter Kramer, Bernd Bukau, and Damon Huber. The C-terminal tail of the bacterial translocation ATPase SecA modulates its activity. *eLife*, 8, 2019.

[5675] Insook Jang, Anita Pottekat, Juthakorn Poothong, Jing Yong, Jacqueline Lagunas-Acosta, Adriana Charbono, Zhouji Chen, Donalyn L Scheuner, Ming Liu, Pamela Itkin-Ansari, Peter Arvan, and Randal J Kaufman. PDIA1/P4HB is required for efficient proinsulin maturation and ß cell health in response to diet induced obesity. *eLife*, 8, 2019.

[5676] Miae Jang, Elizabeth Gould, Jie Xu, Eun Jung Kim, and Jun Hee Kim. Oligodendrocytes regulate presynaptic properties and neurotransmission through BDNF signaling in the mouse brainstem. *eLife*, 8, 2019.

[5677] Jan JanouŠkovec, Gita G Paskerova, Tatiana S Miroliubova, Kirill V Mikhailov, Thomas Birley, Vladimir V Aleoshin, and Timur G Simdyanov. Apicomplexan-like parasites are polyphyletic and widely but selectively dependent on cryptic plastid organelles. *eLife*, 8, 2019.

[5678] Andrés Jara-Oseguera, Katherine E Huffer, and Kenton J Swartz. The ion selectivity filter is not an activation gate in TRPV1-3 channels. *eLife*, 8, 2019.

[5679] Nathalie Javidi-Sharifi, Jacqueline Martinez, Isabel English, Sunil K Joshi, Renata Scopim-Ribeiro, Shelton K Viola, David K Edwards, Anupriya Agarwal, Claudia Lopez, Danielle Jorgens, Jeffrey W Tyner, Brian J Druker, and Elie Traer. FGF2-FGFR1 signaling regulates release of Leukemia-Protective exosomes from bone marrow stromal cells. *eLife*, 8, 2019.

[5680] Míriam Javier-Torrent, Sergi Marco, Daniel Rocandio, Maria Pons-Vizcarra, Peter W Janes, Martin Lackmann, Joaquim Egea, and Carlos A Saura. Presenilin/$\gamma$-secretase-dependent EphA3 processing mediates axon elongation through non-muscle myosin IIA. *eLife*, 8, 2019.

[5681] Shobini Jayaraman, Marcus Fändrich, and Olga Gursky. Synergy between serum amyloid A and secretory phospholipase A2. *eLife*, 8, 2019.

[5682] Philip Jean-Richard-dit Bressel, Cassandra Ma, Laura A Bradfield, Simon Killcross, and Gavan P McNally. Punishment insensitivity emerges from impaired contingency detection, not aversion insensitivity or reward dominance. *eLife*, 8, 2019.

[5683] Hsin-I Jen, Matthew C Hill, Litao Tao, Kuanwei Sheng, Wenjian Cao, Hongyuan Zhang, Haoze V Yu, Juan Llamas, Chenghang Zong, James F Martin, Neil Segil, and Andrew K Groves. Transcriptomic and epigenetic regulation of hair cell regeneration in the mouse utricle and its potentiation by Atoh1. *eLife*, 8, 2019.

[5684] Blair A Jenkins, Natalia M Fontecilla, Catherine P Lu, Elaine Fuchs, and Ellen A Lumpkin. The cellular basis of mechanosensory Merkel-cell innervation during development. *eLife*, 8, 2019.

[5685] Jenna L Jewell, Vivian Fu, Audrey W Hong, Fa-Xing Yu, Delong Meng, Chase H Melick, Huanyu Wang, Wai-Ling Macrina Lam, Hai-Xin Yuan, Susan S Taylor, and Kun-Liang Guan. GPCR signaling inhibits mTORC1 via PKA phosphorylation of Raptor. *eLife*, 8, 2019.

[5686] Sawan Kumar Jha, Khushbu Rauniyar, Ewa Chronowska, Kenny Mattonet, Eunice Wairimu Maina, Hannu Koistinen, Ulf-Håkan Stenman, Kari Alitalo, and Michael Jeltsch. KLK3/PSA and cathepsin D activate VEGF-C and VEGF-D. *eLife*, 8, 2019.

[5687] Rui Jia and Juan S Bonifacino. Negative regulation of autophagy by UBA6-BIRC6–mediated ubiquitination of LC3. *eLife*, 8, 2019.

[5688] Zhihao Jia, Yaohui Nie, Feng Yue, Yifan Kong, Lijie Gu, Timothy P Gavin, Xiaoqi Liu, and Shihuan Kuang. A requirement of Polo-like kinase 1 in murine embryonic myogenesis and adult muscle regeneration. *eLife*, 8, 2019.

[5689] Hongbing Jiang, Christian Leung, Stephen Tahan, and David Wang. Entry by multiple picornaviruses is dependent on a pathway that includes TNK2, WASL, and NCK1. *eLife*, 8, 2019.

[5690] Nan Jiang, Jeffrey P Rasmussen, Joshua A Clanton, Marci F Rosenberg, Kory P Luedke, Mark R Cronan, Edward D Parker, Hyeon-Jin Kim, Joshua C Vaughan, Alvaro Sagasti, and Jay Z Parrish. A conserved morphogenetic mechanism for epidermal ensheathment of nociceptive sensory neurites. *eLife*, 8, 2019.

[5691] Rebeca Jimeno, Marta Lebrusant-Fernandez, Christian Margreitter, Beth Lucas, Natacha Veerapen, Gavin Kelly, Gurdyal S Besra, Franca Fraternali, Jo Spencer, Graham Anderson, and Patricia Barral. Tissue-specific shaping of the TCR repertoire and antigen specificity of iNKT cells. *eLife*, 8, 2019.

[5692] Junjun Jing, Jifan Feng, Jingyuan Li, Xia Han, Jinzhi He, Thach-Vu Ho, Jiahui Du, Xuedong Zhou, Mark Urata, and Yang Chai. Antagonistic interaction between Ezh2 and Arid1a coordinates root patterning and development via Cdkn2a in mouse molars. *eLife*, 8, 2019.

[5693] Youngah Jo, Jason S Hamilton, Seonghwan Hwang, Kristina Garland, Gennipher A Smith, Shan Su, Iris Fuentes, Sudha Neelam, Bonne M Thompson, Jeffrey G McDonald, and Russell A DeBose-Boyd. Schnyder corneal dystrophy-associated UBIAD1 inhibits ER-associated degradation of HMG CoA reductase in mice. *eLife*, 8, 2019.

[5694] Nathan R Johnson, Claude W dePamphilis, and Michael J Axtell. Compensatory sequence variation between trans-species small RNAs and their target sites. *eLife*, 8, 2019.

[5695] Verity Johnson, Harald J Junge, and Zhe Chen. Temporal regulation of axonal repulsion by alternative splicing of a conserved microexon in mammalian Robo1 and Robo2. *eLife*, 8, 2019.

[5696] David J Jörg, Elizabeth E Caygill, Anna E Hakes, Esteban G Contreras, Andrea H Brand, and Benjamin D Simons. The proneural wave in the Drosophila optic lobe is driven by an excitable reaction-diffusion mechanism. *eLife*, 8, 2019.

[5697] Aby Joseph, Andres Guevara-Torres, and Jesse Schallek. Imaging single-cell blood flow in the smallest to largest vessels in the living retina. *eLife*, 8, 2019.

[5698] Inokentijs Josts, Katharina Veith, and Henning Tidow. Ternary structure of the outer membrane transporter FoxA with resolved signalling domain provides insights into TonB-mediated siderophore uptake. *eLife*, 8, 2019.

[5699] Julie Jouette, Antoine Guichet, and Sandra B Claret. Dynein-mediated transport and membrane trafficking control PAR3 polarised distribution. *eLife*, 8, 2019.

[5700] Ashley L Juavinett, George Bekheet, and Anne K Churchland. Chronically implanted Neuropixels probes enable high-yield recordings in freely moving mice. *eLife*, 8, 2019.

[5701] Hyun Min Jung, Ciara T Hu, Alexandra M Fister, Andrew E Davis, Daniel Castranova, Van N Pham, Lisa M Price, and Brant M Weinstein. MicroRNA-mediated control of developmental lymphangiogenesis. *eLife*, 8, 2019.

[5702] Min Jung, Daniel Wells, Jannette Rusch, Suhaira Ahmad, Jonathan Marchini, Simon R Myers, and Donald F Conrad. Unified single-cell analysis of testis gene regulation and pathology in five mouse strains. *eLife*, 8, 2019.

[5703] Shaheen Kabir, Justin Cidado, Courtney Andersen, Cortni Dick, Pei-Chun Lin, Therese Mitros, Hong Ma, Seung Hyun Baik, Matthew A Belmonte, Lisa Drew, and Jacob E Corn. The CUL5 ubiquitin ligase complex mediates resistance to CDK9 and MCL1 inhibitors in lung cancer cells. *eLife*, 8, 2019.

[5704] Nirag Kadakia and Thierry Emonet. Front-end Weber-Fechner gain control enhances the fidelity of combinatorial odor coding. *eLife*, 8, 2019.

[5705] Daniel Kaiser, Jacopo Turini, and Radoslaw M Cichy. A neural mechanism for contextualizing fragmented inputs during naturalistic vision. *eLife*, 8, 2019.

[5706] Valeria Kalienkova, Vanessa Clerico Mosina, Laura Bryner, Gert T Oostergetel, Raimund Dutzler, and Cristina Paulino. Stepwise activation mechanism of the scramblase nhTMEM16 revealed by cryo-EM. *eLife*, 8, 2019.

[5707] Masashi Kameda, Shogo Ohmae, and Masaki Tanaka. Entrained neuronal activity to periodic visual stimuli in the primate striatum compared with the cerebellum. *eLife*, 8, 2019.

[5708] Cassandra Kamischke, Junping Fan, Julien Bergeron, Hemantha D Kulasekara, Zachary D Dalebroux, Anika Burrell, Justin M Kollman, and Samuel I Miller. The Acinetobacter baumannii Mla system and glycerophospholipid transport to the outer membrane. *eLife*, 8, 2019.

[5709] Lea Kampermann, Niklas Wilming, Arjen Alink, Christian Büchel, and Selim Onat. Fixation-pattern similarity analysis reveals adaptive changes in face-viewing strategies following aversive learning. *eLife*, 8, 2019.

[5710] Gary A Kane, Aaron M Bornstein, Amitai Shenhav, Robert C Wilson, Nathaniel D Daw, and Jonathan D Cohen. Rats exhibit similar biases in foraging and intertemporal choice tasks. *eLife*, 8, 2019.

[5711] Louis Kang and Vijay Balasubramanian. A geometric attractor mechanism for self-organization of entorhinal grid modules. *eLife*, 8, 2019.

[5712] Louis Kang and Michael R DeWeese. Replay as wavefronts and theta sequences as bump oscillations in a grid cell attractor network. *eLife*, 8, 2019.

[5713] Yilin Kang, Alexander J Anderson, Thomas Daniel Jackson, Catherine S Palmer, David P De Souza, Kenji M Fujihara, Tegan Stait, Ann E Frazier, Nicholas J Clemons, Deidreia Tull, David R Thorburn, Malcolm J McConville, Michael T Ryan, David A Stroud, and Diana Stojanovski. Function of hTim8a in complex IV assembly in neuronal cells provides insight into pathomechanism underlying Mohr-Tranebjærg syndrome. *eLife*, 8, 2019.

[5714] Yun Pyo Kang, Laura Torrente, Aimee Falzone, Cody M Elkins, Min Liu, John M Asara, Christian C Dibble, and Gina DeNicola. Correction: Cysteine dioxygenase 1 is a metabolic liability for non-small cell lung cancer. *eLife*, 8, 2019.

[5715] Yun Pyo Kang, Laura Torrente, Aimee Falzone, Cody M Elkins, Min Liu, John M Asara, Christian C Dibble, and Gina M DeNicola. Cysteine dioxygenase 1 is a metabolic liability for non-small cell lung cancer. *eLife*, 8, 2019.

[5716] Mohammed Kaplan, Debnath Ghosal, Poorna Subramanian, Catherine M Oikonomou, Andreas Kjaer, Sahand Pirbadian, Davi R Ortega, Ariane Briegel, Mohamed Y El-Naggar, and Grant J Jensen. The presence and absence of periplasmic rings in bacterial flagellar motors correlates with stator type. *eLife*, 8, 2019.

[5717] Theodoros K Karamanos, Matthew P Jackson, Antonio N Calabrese, Sophia C Goodchild, Emma E Cawood, Gary S Thompson, Arnout P Kalverda, Eric W Hewitt, and Sheena E Radford. Structural mapping of oligomeric intermediates in an amyloid assembly pathway. *eLife*, 8, 2019.

[5718] Patrick J Karas, John F Magnotti, Brian A Metzger, Lin L Zhu, Kristen B Smith, Daniel Yoshor, and Michael S Beauchamp. The visual speech head start improves perception and reduces superior temporal cortex responses to auditory speech. *eLife*, 8, 2019.

[5719] Vasileios Kargas, Pablo Castro-Hartmann, Norberto Escudero-Urquijo, Kyle Dent, Christine Hilcenko, Carolin Sailer, Gertrude Zisser, Maria J Marques-Carvalho, Simone Pellegrino, Leszek Wawiórka, Stefan MV Freund, Jane L Wagstaff, Antonina Andreeva, Alexandre Faille, Edwin Chen, Florian Stengel, Helmut Bergler, and Alan John Warren. Mechanism of completion of peptidyltransferase centre assembly in eukaryotes. *eLife*, 8, 2019.

[5720] Fuyuki Karube, Susumu Takahashi, Kenta Kobayashi, and Fumino Fujiyama. Motor cortex can directly drive the globus pallidus neurons in a projection neuron type-dependent manner in the rat. *eLife*, 8, 2019.

[5721] Marina A Kasimova, Debanjan Tewari, John B Cowgill, Willy Carrasquel Ursuleaz, Jenna L Lin, Lucie Delemotte, and Baron Chanda. Helix breaking transition in the S4 of HCN channel is critical for hyperpolarization-dependent gating. *eLife*, 8, 2019.

[5722] Ryan J Kast, Alexandra L Lanjewar, Colton D Smith, and Pat Levitt. FOXP2 exhibits projection neuron class specific expression, but is not required for multiple aspects of cortical histogenesis. *eLife*, 8, 2019.

[5723] Takeshi Katsuda, Juntaro Matsuzaki, Tomoko Yamaguchi, Yasuhiro Yamada, Marta Prieto-Vila, Kazunori Hosaka, Atsuko Takeuchi, Yoshimasa Saito, and Takahiro Ochiya. Generation of human hepatic progenitor cells with regenerative and metabolic capacities from primary hepatocytes. *eLife*, 8, 2019.

[5724] Diane Marie Keeling, Patricia Garza, Charisse Michelle Nartey, and Anne-Ruxandra Carvunis. The meanings of 'function' in biology and the problematic case of de novo gene emergence. *eLife*, 8, 2019.

[5725] Aaron Kerlin, Boaz Mohar, Daniel Flickinger, Bryan J MacLennan, Matthew B Dean, Courtney Davis, Nelson Spruston, and Karel Svoboda. Functional clustering of dendritic activity during decision-making. *eLife*, 8, 2019.

[5726] David M Kern, SeCheol Oh, Richard K Hite, and Stephen G Brohawn. Cryo-EM structures of the DCPIB-inhibited volume-regulated anion channel LRRC8A in lipid nanodiscs. *eLife*, 8, 2019.

[5727] Kanika Khanna, Javier Lopez-Garrido, Ziyi Zhao, Reika Watanabe, Yuan Yuan, Joseph Sugie, Kit Pogliano, and Elizabeth Villa. The molecular architecture of engulfment during Bacillus subtilis sporulation. *eLife*, 8, 2019.

[5728] Shahrzad Kharabian Masouleh, Simon B Eickhoff, Felix Hoffstaedter, and Sarah Genon. Empirical examination of the replicability of associations between brain structure and psychological variables. *eLife*, 8, 2019.

[5729] George Khelashvili, Neha Chauhan, Kalpana Pandey, David Eliezer, and Anant K Menon. Exchange of water for sterol underlies sterol egress from a StARkin domain. *eLife*, 8, 2019.

[5730] Yoonhee Ki and Chunghun Lim. Sleep-promoting effects of threonine link amino acid metabolism in Drosophila neuron to GABAergic control of sleep drive. *eLife*, 8, 2019.

[5731] David William Kikuchi, Anna Dornhaus, Vandana Gopeechund, and Thomas N Sherratt. Signal categorization by foraging animals depends on ecological diversity. *eLife*, 8, 2019.

[5732] Atsushi Kikumoto and Ulrich Mayr. Balancing model-based and memory-free action selection under competitive pressure. *eLife*, 8, 2019.

[5733] Tom Killelea, Adeline Palud, Farida Akcha, Mélanie Lemor, Stephane L'haridon, Anne Godfroy, and Ghislaine Henneke. The interplay at the replisome mitigates the impact of oxidative damage on the genetic integrity of hyperthermophilic Archaea. *eLife*, 8, 2019.

[5734] Konstantina Kilteni, Christian Houborg, and H Henrik Ehrsson. Rapid learning and unlearning of predicted sensory delays in self-generated touch. *eLife*, 8, 2019.

[5735] Byounghoon Kim, Shobha Channabasappa Kenchappa, Adhira Sunkara, Ting-Yu Chang, Lowell Thompson, Raymond Doudlah, and Ari Rosenberg. Real-time experimental control using network-based parallel processing. *eLife*, 8, 2019.

[5736] Byungchul Kim, Yen-Chih Wang, Charles W Hespen, Juliel Espinosa, Jeanne Salje, Kavita J Rangan, Deena A Oren, Jin Young Kang, Virginia A Pedicord, and Howard C Hang. Enterococcus faecium secreted antigen A generates muropeptides to enhance host immunity and limit bacterial pathogenesis. *eLife*, 8, 2019.

[5737] Christina K Kim, Kelvin F Cho, Min Woo Kim, and Alice Y Ting. Luciferase-LOV BRET enables versatile and specific transcriptional readout of cellular protein-protein interactions. *eLife*, 8, 2019.

[5738] Hyung-Seok Kim, Judith Neugebauer, Autumn McKnite, Anup Tilak, and Jan L Christian. BMP7 functions predominantly as a heterodimer with BMP2 or BMP4 during mammalian embryogenesis. *eLife*, 8, 2019.

[5739] Seungsoo Kim, Maitreya J Dunham, and Jay Shendure. A combination of transcription factors mediates inducible interchromosomal contacts. *eLife*, 8, 2019.

[5740] Soo-Kyoung Kim, Justin D Lormand, Cordelia A Weiss, Karin A Eger, Husan Turdiev, Asan Turdiev, Wade C Winkler, Holger Sondermann, and Vincent T Lee. A dedicated diribonuclease resolves a key bottleneck for the terminal step of RNA degradation. *eLife*, 8, 2019.

[5741] SungEun Kim, Yubin Kim, JungHo Kong, Eunjee Kim, Jae Hyeok Choi, Hyeong Dong Yuk, HyeSun Lee, Hwa-Ryeon Kim, Kyoung-Hwa Lee, Minyong Kang, Jae-Seok Roe, Kyung Chul Moon, Sanguk Kim, Ja Hyeon Ku, and Kunyoo Shin. Epigenetic regulation of mammalian Hedgehog signaling to the stroma determines the molecular subtype of bladder cancer. *eLife*, 8, 2019.

[5742] Grant A King, Jay S Goodman, Jennifer G Schick, Keerthana Chetlapalli, Danielle M Jorgens, Kent L McDonald, and ElÇin ünal. Meiotic cellular rejuvenation is coupled to nuclear remodeling in budding yeast. *eLife*, 8, 2019.

[5743] Maia Kinnebrew, Ellen J Iverson, Bhaven B Patel, Ganesh V Pusapati, Jennifer H Kong, Kristen A Johnson, Giovanni Luchetti, Kaitlyn M Eckert, Jeffrey G McDonald, Douglas F Covey, Christian Siebold, Arun Radhakrishnan, and Rajat Rohatgi. Cholesterol accessibility at the ciliary membrane controls hedgehog signaling. *eLife*, 8, 2019.

[5744] Charlotte Kirchhelle, Daniel Garcia-Gonzalez, Niloufer G Irani, Antoine Jérusalem, and Ian Moore. Two mechanisms regulate directional cell growth in Arabidopsis lateral roots. *eLife*, 8, 2019.

[5745] Isa Kristina Kirk, Christian Simon, Karina Banasik, Peter Christoffer Holm, Amalie Dahl Haue, Peter Bjødstrup Jensen, Lars Juhl Jensen, Cristina Leal Rodríguez, Mette Krogh Pedersen, Robert Eriksson, Henrik Ullits Andersen, Thomas Almdal, Jette Bork-Jensen, Niels Grarup, Knut Borch-Johnsen, Oluf Pedersen, Flemming Pociot, Torben Hansen, Regine Bergholdt, Peter Rossing, and Søren Brunak. Linking glycemic dysregulation in diabetes to symptoms, comorbidities, and genetics through EHR data mining. *eLife*, 8, 2019.

[5746] Shun Kishimoto, Jeffrey R Brender, Daniel R Crooks, Shingo Matsumoto, Tomohiro Seki, Nobu Oshima, Hellmut Merkle, Penghui Lin, Galen Reed, Albert P Chen, Jan Henrik Ardenkjaer-Larsen, Jeeva Munasinghe, Keita Saito, Kazutoshi Yamamoto, Peter L Choyke, James Mitchell, Andrew N Lane, Teresa WM Fan, W Marston Linehan, and Murali C Krishna. Imaging of glucose metabolism by 13C-MRI distinguishes pancreatic cancer subtypes in mice. *eLife*, 8, 2019.

[5747] Shunbun Kita, Shiro Fukuda, Norikazu Maeda, and Iichiro Shimomura. Native adiponectin in serum binds to mammalian cells expressing T-cadherin, but not AdipoRs or calreticulin. *eLife*, 8, 2019.

[5748] Keisuke Kitakaze, Shusuke Taniuchi, Eri Kawano, Yoshimasa Hamada, Masato Miyake, Miho Oyadomari, Hirotatsu Kojima, Hidetaka Kosako, Tomoko Kuribara, Suguru Yoshida, Takamitsu Hosoya, and Seiichi Oyadomari. Cell-based HTS identifies a chemical chaperone for preventing ER protein aggregation and proteotoxicity. *eLife*, 8, 2019.

[5749] Kerstin Klinkert, Nicolas Levernier, Peter Gross, Christian Gentili, Lukas von Tobel, Marie Pierron, Coralie Busso, Sarah Herrman, Stephan W Grill, Karsten Kruse, and Pierre Gönczy. Aurora A depletion reveals centrosome-independent polarization mechanism in Caenorhabditis elegans. *eLife*, 8, 2019.

[5750] Maja Kneissig, Kristina Keuper, Mirjam S de Pagter, Markus J van Roosmalen, Jana Martin, Hannah Otto, Verena Passerini, Aline Campos Sparr, Ivo Renkens, Fenna Kropveld, Anand Vasudevan, Jason M Sheltzer, Wigard P Kloosterman, and Zuzana Storchova. Micronuclei-based model system reveals functional consequences of chromothripsis in human cells. *eLife*, 8, 2019.

[5751] Laura D Knogler, Andreas M Kist, and Ruben Portugues. Motor context dominates output from purkinje cell functional regions during reflexive visuomotor behaviours. *eLife*, 8, 2019.

[5752] Gavin J Knott, Brady F Cress, Jun-Jie Liu, Brittney W Thornton, Rachel J Lew, Basem Al-Shayeb, Daniel J Rosenberg, Michal Hammel, Benjamin A Adler, Marco J Lobba, Michael Xu, Adam P Arkin, Christof Fellmann, and Jennifer A Doudna. Structural basis for AcrVA4 inhibition of specific CRISPR-Cas12a. *eLife*, 8, 2019.

[5753] Dmitry Kobak, Jose L Pardo-Vazquez, Mafalda Valente, Christian K Machens, and Alfonso Renart. State-dependent geometry of population activity in rat auditory cortex. *eLife*, 8, 2019.

[5754] Loes Koelewijn, Thomas M Lancaster, David Linden, Diana C Dima, Bethany C Routley, Lorenzo Magazzini, Kali Barawi, Lisa Brindley, Rachael Adams, Katherine E Tansey, Aline Bompas, Andrea Tales, Antony Bayer, and Krish Singh. Oscillatory hyperactivity and hyperconnectivity in young APOE-$\epsilon$4 carriers and hypoconnectivity in Alzheimer's disease. *eLife*, 8, 2019.

[5755] Daniel Koenig, Jörg Hagmann, Rachel Li, Felix Bemm, Tanja Slotte, Barbara Neuffer, Stephen I Wright, and Detlef Weigel. Long-term balancing selection drives evolution of immunity genes in Capsella. *eLife*, 8, 2019.

[5756] Shunshi Kohyama, Natsuhiko Yoshinaga, Miho Yanagisawa, Kei Fujiwara, and Nobuhide Doi. Cell-sized confinement controls generation and stability of a protein wave for spatiotemporal regulation in cells. *eLife*, 8, 2019.

[5757] Takuya Koike, Koshi Harada, Shu Horiuchi, and Daisuke Kitamura. The quantity of CD40 signaling determines the differentiation of B cells into functionally distinct memory cell subsets. *eLife*, 8, 2019.

[5758] Mina L Kojima, Dirk G de Rooij, and David C Page. Amplification of a broad transcriptional program by a common factor triggers the meiotic cell cycle in mice. *eLife*, 8, 2019.

[5759] Matthijs Kol, Ben Williams, Henry Toombs-Ruane, Henri G Franquelim, Sergei Korneev, Christian Schroeer, Petra Schwille, Dirk Trauner, Joost CM Holthuis, and James A Frank. Optical manipulation of sphingolipid biosynthesis using photoswitchable ceramides. *eLife*, 8, 2019.

[5760] Elif Kon, Elisa Calvo-Jiménez, Alexia Cossard, Youn Na, Jonathan A Cooper, and Yves Jossin. N-cadherin-regulated FGFR ubiquitination and degradation control mammalian neocortical projection neuron migration. *eLife*, 8, 2019.

[5761] Mariam M Konaté, Germán Plata, Jimin Park, Dinara R Usmanova, Harris Wang, and Dennis Vitkup. Molecular function limits divergent protein evolution on planetary timescales. *eLife*, 8, 2019.

[5762] Takefumi Kondo and Shigeo Hayashi. Two-step regulation of trachealess ensures tight coupling of cell fate with morphogenesis in the Drosophila trachea. *eLife*, 8, 2019.

[5763] Pratibha C Koneru, Ashwanth C Francis, Nanjie Deng, Stephanie V Rebensburg, Ashley C Hoyte, Jared Lindenberger, Daniel Adu-Ampratwum, Ross C Larue, Michael F Wempe, Alan N Engelman, Dmitry Lyumkis, James R Fuchs, Ronald M Levy, Gregory B Melikyan, and Mamuka Kvaratskhelia. HIV-1 integrase tetramers are the antiviral target of pyridine-based allosteric integrase inhibitors. *eLife*, 8, 2019.

[5764] Benjamin König, Yuchen Hao, Sophia Schwartz, Andrew JR Plested, and Tobias Stauber. A FRET sensor of C-terminal movement reveals VRAC activation by plasma membrane DAG signaling rather than ionic strength. *eLife*, 8, 2019.

[5765] Kalyn Kono, Shigeki Yoshiura, Ikumi Fujita, Yasushi Okada, Atsunori Shitamukai, Tatsuo Shibata, and Fumio Matsuzaki. Reconstruction of Par-dependent polarity in apolar cells reveals a dynamic process of cortical polarization. *eLife*, 8, 2019.

[5766] Florian Kopp, Mahmoud M Elguindy, Mehmet E Yalvac, He Zhang, Beibei Chen, Frank A Gillett, Sungyul Lee, Sushama Sivakumar, Hongtao Yu, Yang Xie, Prashant Mishra, Zarife Sahenk, and Joshua T Mendell. PUMILIO hyperactivity drives premature aging of Norad-deficient mice. *eLife*, 8, 2019.

[5767] Max Koppers, Roberta Cagnetta, Toshiaki Shigeoka, Lucia CS Wunderlich, Pedro Vallejo-Ramirez, Julie Qiaojin Lin, Sixian Zhao, Maximilian AH Jakobs, Asha Dwivedy, Michael S Minett, Anaïs Bellon, Clemens F Kaminski, William A Harris, John G Flanagan, and Christine E Holt. Receptor-specific interactome as a hub for rapid cue-induced selective translation in axons. *eLife*, 8, 2019.

[5768] Kevin P Koster, Walter Francesconi, Fulvia Berton, Sami Alahmadi, Roshan Srinivas, and Akira Yoshii. Developmental NMDA receptor dysregulation in the infantile neuronal ceroid lipofuscinosis mouse model. *eLife*, 8, 2019.

[5769] Dylan Kotliar, Adrian Veres, M Aurel Nagy, Shervin Tabrizi, Eran Hodis, Douglas A Melton, and Pardis C Sabeti. Identifying gene expression programs of cell-type identity and cellular activity with single-cell RNA-Seq. *eLife*, 8, 2019.

[5770] Matthew J Kourakis, Cezar Borba, Angela Zhang, Erin Newman-Smith, Priscilla Salas, B Manjunath, and William C Smith. Parallel visual circuitry in a basal chordate. *eLife*, 8, 2019.

[5771] Elena Kozgunova, Momoko Nishina, and Gohta Goshima. Kinetochore protein depletion underlies cytokinesis failure and somatic polyploidization in the moss Physcomitrella patens. *eLife*, 8, 2019.

[5772] Liesl A Krause and Susanna L Harris. Get online to support wellbeing of graduate students. *eLife*, 8, 2019.

[5773] Gabriela Krejčová, Adéla Danielová, Pavla Nedbalová, Michalina Kazek, LukáŠ Strych, Geetanjali Chawla, Jason M Tennessen, Jaroslava Lieskovská, Marek Jindra, TomáŠ Doležal, and Adam Bajgar. Drosophila macrophages switch to aerobic glycolysis to mount effective antibacterial defense. *eLife*, 8, 2019.

[5774] Lieselotte SM Kreuk, Meghan A Koch, Leianna C Slayden, Nicholas A Lind, Sophia Chu, Hannah P Savage, Aaron B Kantor, Nicole Baumgarth, and Gregory M Barton. B cell receptor and Toll-like receptor signaling coordinate to control distinct B-1 responses to both self and the microbiota. *eLife*, 8, 2019.

[5775] Lara Katharina Krüger, Jérémie-Luc Sanchez, Anne Paoletti, and Phong Thanh Tran. Kinesin-6 regulates cell-size-dependent spindle elongation velocity to keep mitosis duration constant in fission yeast. *eLife*, 8, 2019.

[5776] Bridget M Kuehn. Planning during pregnancy. *eLife*, 8, 2019.

[5777] Simone Kühn, Anna Mascharek, Tobias Banaschewski, Arun Bodke, Uli Bromberg, Christian Büchel, Erin Burke Quinlan, Sylvane Desrivieres, Herta Flor, Antoine Grigis, Hugh Garavan, Penny A Gowland, Andreas Heinz, Bernd Ittermann, Jean-Luc Martinot, Frauke Nees, Dimitri Papadopoulos Orfanos, Tomas Paus, Luise Poustka, Sabina Millenet, Juliane H Fröhner, Michael N Smolka, Henrik Walter, Robert Whelan, Gunter Schumann, Ulman Lindenberger, and Jürgen Gallinat. Predicting development of adolescent drinking behaviour from whole brain structure at 14 years of age. *eLife*, 8, 2019.

[5778] Yelena Kulik, Ryan Jones, Armen J Moughamian, Jenna Whippen, and Graeme W Davis. Dual separable feedback systems govern firing rate homeostasis. *eLife*, 8, 2019.

[5779] Yelena D Kulik, Deborah J Watson, Guan Cao, Masaaki Kuwajima, and Kristen M Harris. Structural plasticity of dendritic secretory compartments during LTP-induced synaptogenesis. *eLife*, 8, 2019.

[5780] Nathella Pavan Kumar, Kiyoshi F Fukutani, Basavaradhya S Shruthi, Thabata Alves, Paulo S Silveira-Mattos, Michael S Rocha, Kim West, Mohan Natarajan, Vijay Viswanathan, Subash Babu, Bruno B Andrade, and Hardy Kornfeld. Persistent inflammation during anti-tuberculosis treatment with diabetes comorbidity. *eLife*, 8, 2019.

[5781] Fu-Shan Kuo, Colin M Cleary, Joseph J LoTurco, Xinnian Chen, and Daniel K Mulkey. Disordered breathing in a mouse model of Dravet syndrome. *eLife*, 8, 2019.

[5782] Nadia S Kurd, Lydia K Lutes, Jaewon Yoon, Shiao Wei Chan, Ivan L Dzhagalov, Ashley R Hoover, and Ellen A Robey. A role for phagocytosis in inducing cell death during thymocyte negative selection. *eLife*, 8, 2019.

[5783] Yerbol Z Kurmangaliyev, Juyoun Yoo, Samuel A LoCascio, and S Lawrence Zipursky. Modular transcriptional programs separately define axon and dendrite connectivity. *eLife*, 8, 2019.

[5784] Shinae Kwak and Min Whan Jung. Distinct roles of striatal direct and indirect pathways in value-based decision making. *eLife*, 8, 2019.

[5785] Ursula Kwong-Brown, Martha L Tobias, Damian O Elias, Ian C Hall, Coen PH Elemans, and Darcy B Kelley. The return to water in ancestral Xenopus was accompanied by a novel mechanism for producing and shaping vocal signals. *eLife*, 8, 2019.

[5786] Samuel E Lacey, Shaoda He, Sjors HW Scheres, and Andrew P Carter. Cryo-EM of dynein microtubule-binding domains shows how an axonemal dynein distorts the microtubule. *eLife*, 8, 2019.

[5787] Haluk Lacin, Hui-Min Chen, Xi Long, Robert H Singer, Tzumin Lee, and James W Truman. Neurotransmitter identity is acquired in a lineage-restricted manner in the Drosophila CNS. *eLife*, 8, 2019.

[5788] Michael M Lacy, David Baddeley, and Julien Berro. Single-molecule turnover dynamics of actin and membrane coat proteins in clathrin-mediated endocytosis. *eLife*, 8, 2019.

[5789] Mark S Ladinsky, Wannisa Khamaikawin, Yujin Jung, Samantha Lin, Jennifer Lam, Dong Sung An, Pamela J Bjorkman, and Collin Kieffer. Mechanisms of virus dissemination in bone marrow of HIV-1–infected humanized BLT mice. *eLife*, 8, 2019.

[5790] Carl Laflamme, Paul M McKeever, Rahul Kumar, Julie Schwartz, Mahshad Kolahdouzan, Carol X Chen, Zhipeng You, Faiza Benaliouad, Opher Gileadi, Heidi M McBride, Thomas M Durcan, Aled M Edwards, Luke M Healy, Janice Robertson, and Peter S McPherson. Implementation of an antibody characterization procedure and application to the major ALS/FTD disease gene C9ORF72. *eLife*, 8, 2019.

[5791] Martina R Laidemitt, Larissa C Anderson, Helen J Wearing, Martin W Mutuku, Gerald M Mkoji, and Eric S Loker. Antagonism between parasites within snail hosts impacts the transmission of human schistosomiasis. *eLife*, 8, 2019.

[5792] Anna-Liisa Laine, Benoit Barrès, Elina Numminen, and Jukka P Siren. Variable opportunities for outcrossing result in hotspots of novel genetic variation in a pathogen metapopulation. *eLife*, 8, 2019.

[5793] Ismael Lamas-Toranzo, Noelia Fonseca Balvís, Ana Querejeta-Fernández, María José Izquierdo-Rico, Leopoldo González-Brusi, Pedro L Lorenzo, Pilar García-Rebollar, Manuel Avilés, and Pablo Bermejo-álvarez. ZP4 confers structural properties to the zona pellucida essential for embryo development. *eLife*, 8, 2019.

[5794] Nils Landegren, Lindsey B Rosen, Eva Freyhult, Daniel Eriksson, Tove Fall, Gustav Smith, Elise M N Ferre, Petter Brodin, Donald Sharon, Michael Snyder, Michail Lionakis, Mark Anderson, and Olle Kämpe. Comment on 'AIRE-deficient patients harbor unique high-affinity disease-ameliorating autoantibodies'. *eLife*, 8, 2019.

[5795] Lee F Langer, James M Ward, and Trevor K Archer. Tumor suppressor SMARCB1 suppresses super-enhancers to govern hESC lineage determination. *eLife*, 8, 2019.

[5796] Simon Langer, Christian Hammer, Kristina Hopfensperger, Lukas Klein, Dominik Hotter, Paul D De Jesus, Kristina M Herbert, Lars Pache, Nikaïa Smith, Johannes A van der Merwe, Sumit K Chanda, Jacques Fellay, Frank Kirchhoff, and Daniel Sauter. HIV-1 Vpu is a potent transcriptional suppressor of NF-$\kappa$B-elicited antiviral immune responses. *eLife*, 8, 2019.

[5797] Sasha A Langley, Karen H Miga, Gary H Karpen, and Charles H Langley. Haplotypes spanning centromeric regions reveal persistence of large blocks of archaic DNA. *eLife*, 8, 2019.

[5798] Lance D Langston and Michael E O'Donnell. An explanation for origin unwinding in eukaryotes. *eLife*, 8, 2019.

[5799] Jean-André Lapart, Marco Gottardo, Elisabeth Cortier, Jean-Luc Duteyrat, Céline Augière, Alain Mangé, Julie Jerber, Jérôme Solassol, Jay Gopalakrishnan, Joëlle Thomas, and Bénédicte Durand. Dzip1 and Fam92 form a ciliary transition zone complex with cell type specific roles in Drosophila. *eLife*, 8, 2019.

[5800] Lucie Laplane and Eric Solary. Towards a classification of stem cells. *eLife*, 8, 2019.

[5801] Azra Lari, Arvind Arul Nambi Rajan, Rima Sandhu, Taylor Reiter, Rachel Montpetit, Barry P Young, Chris JR Loewen, and Ben Montpetit. A nuclear role for the DEAD-box protein Dbp5 in tRNA export. *eLife*, 8, 2019.

[5802] Noga Larry, Merav Yarkoni, Adi Lixenberg, and Mati Joshua. Cerebellar climbing fibers encode expected reward size. *eLife*, 8, 2019.

[5803] Mads Breum Larsen, Mireia Perez Verdaguer, Brigitte F Schmidt, Marcel P Bruchez, Simon C Watkins, and Alexander Sorkin. Generation of endogenous pH-sensitive EGF receptor and its application in high-throughput screening for proteins involved in clathrin-mediated endocytosis. *eLife*, 8, 2019.

[5804] Kenneth W Latimer, Fred Rieke, and Jonathan W Pillow. Inferring synaptic inputs from spikes with a conductance-based neural encoding model. *eLife*, 8, 2019.

[5805] Johanna Lattner, Weihua Leng, Elisabeth Knust, Marko Brankatschk, and David Flores-Benitez. Crumbs organizes the transport machinery by regulating apical levels of PI(4,5)P2 in Drosophila. *eLife*, 8, 2019.

[5806] Janelle Lauer, Sandra Segeletz, Alice Cezanne, Giambattista Guaitoli, Francesco Raimondi, Marc Gentzel, Vikram Alva, Michael Habeck, Yannis Kalaidzidis, Marius Ueffing, Andrei N Lupas, Christian Johannes Gloeckner, and Marino Zerial. Auto-regulation of Rab5 GEF activity in Rabex5 by allosteric structural changes, catalytic core dynamics and ubiquitin binding. *eLife*, 8, 2019.

[5807] Melanie Laurin, Nicholas C Gomez, John Levorse, Ataman Sendoel, Megan Sribour, and Elaine Fuchs. An RNAi screen unravels the complexities of Rho GTPase networks in skin morphogenesis. *eLife*, 8, 2019.

[5808] Sydney Lavoie, Kara L Conway, Kara G Lassen, Humberto B Jijon, Hui Pan, Eunyoung Chun, Monia Michaud, Jessica K Lang, Carey Ann Gallini Comeau, Jonathan M Dreyfuss, Jonathan N Glickman, Hera Vlamakis, Ashwin Ananthakrishnan, Aleksander Kostic, Wendy S Garrett, and Ramnik J Xavier. The Crohn's disease polymorphism, ATG16L1 T300A, alters the gut microbiota and enhances the local Th1/Th17 response. *eLife*, 8, 2019.

[5809] Kynan T Lawlor, Luke Zappia, James Lefevre, Joo-Seop Park, Nicholas A Hamilton, Alicia Oshlack, Melissa H Little, and Alexander N Combes. Nephron progenitor commitment is a stochastic process influenced by cell migration. *eLife*, 8, 2019.

[5810] Samuel JD Lawrence, David G Norris, and Floris P de Lange. Dissociable laminar profiles of concurrent bottom-up and top-down modulation in the human visual cortex. *eLife*, 8, 2019.

[5811] Hannah Lawson, Edward Vuong, Renee M Miller, Karin Kiontke, David HA Fitch, and Douglas S Portman. The Makorin lep-2 and the lncRNA lep-5 regulate lin-28 to schedule sexual maturation of the C. elegans nervous system. *eLife*, 8, 2019.

[5812] Andrew K Lawton, Tyler Engstrom, Daniel Rohrbach, Masaaki Omura, Daniel H Turnbull, Jonathan Mamou, Teng Zhang, J M Schwarz, and Alexandra L Joyner. Cerebellar folding is initiated by mechanical constraints on a fluid-like layer without a cellular pre-pattern. *eLife*, 8, 2019.

[5813] Julia R Lazzari-Dean, Anneliese MM Gest, and Evan W Miller. Optical estimation of absolute membrane potential using fluorescence lifetime imaging. *eLife*, 8, 2019.

[5814] Amanda Lea, Meena Subramaniam, Arthur Ko, Terho Lehtimäki, Emma Raitoharju, Mika Kähönen, Ilkka Seppälä, Nina Mononen, Olli T Raitakari, Mika Ala-Korpela, Päivi Pajukanta, Noah Zaitlen, and Julien F Ayroles. Genetic and environmental perturbations lead to regulatory decoherence. *eLife*, 8, 2019.

[5815] Eleonore Lebeuf-Taylor, Nick McCloskey, Susan F Bailey, Aaron Hinz, and Rees Kassen. The distribution of fitness effects among synonymous mutations in a gene under directional selection. *eLife*, 8, 2019.

[5816] Adria C LeBoeuf, Patrice Waridel, Colin S Brent, Andre N Gonçalves, Laure Menin, Daniel Ortiz, Oksana Riba-Grognuz, Akiko Koto, Zamira G Soares, Eyal Privman, Eric A Miska, Richard Benton, and Laurent Keller. Oral transfer of chemical cues, growth proteins and hormones in social insects. *eLife*, 8, 2019.

[5817] Eugine Lee, John Wongvipat, Danielle Choi, Ping Wang, Young Sun Lee, Deyou Zheng, Philip A Watson, Anuradha Gopalan, and Charles L Sawyers. GREB1 amplifies androgen receptor output in human prostate cancer and contributes to antiandrogen resistance. *eLife*, 8, 2019.

[5818] Hayun Lee, Yukti Dhingra, and Dipali G Sashital. The Cas4-Cas1-Cas2 complex mediates precise prespacer processing during CRISPR adaptation. *eLife*, 8, 2019.

[5819] James Lee, David Tomasek, Thiago MA Santos, Mary D May, Ina Meuskens, and Daniel Kahne. Formation of a $\beta$-barrel membrane protein is catalyzed by the interior surface of the assembly machine protein BamA. *eLife*, 8, 2019.

[5820] Joo Lee, Caitlin A Taylor, Kristopher M Barnes, Ao Shen, Emerson V Stewart, Allison Chen, Yang K Xiang, Zhirong Bao, and Kang Shen. A Myt1 family transcription factor defines neuronal fate by repressing non-neuronal genes. *eLife*, 8, 2019.

[5821] Kwan Yeop Lee, Stéphanie Ratté, and Steven A Prescott. Excitatory neurons are more disinhibited than inhibitory neurons by chloride dysregulation in the spinal dorsal horn. *eLife*, 8, 2019.

[5822] Rachel S Lee, Marcelo G Mattar, Nathan F Parker, Ilana B Witten, and Nathaniel D Daw. Reward prediction error does not explain movement selectivity in DMS-projecting dopamine neurons. *eLife*, 8, 2019.

[5823] Sanghyeon Lee, Sunha Park, Hyojin Lee, Seulki Han, Jae-man Song, Dohyun Han, and Young Ho Suh. Nedd4 E3 ligase and beta-arrestins regulate ubiquitination, trafficking, and stability of the mGlu7 receptor. *eLife*, 8, 2019.

[5824] Seungkyu Lee, Sooyeon Jo, Sébastien Talbot, Han-Xiong Bear Zhang, Masakazu Kotoda, Nick A Andrews, Michelino Puopolo, Pin W Liu, Thomas Jacquemont, Maud Pascal, Laurel M Heckman, Aakanksha Jain, Jinbo Lee, Clifford J Woolf, and Bruce P Bean. Novel charged sodium and calcium channel inhibitor active against neurogenic inflammation. *eLife*, 8, 2019.

[5825] Teresa Wei-sy Lee, Heidi Shira David, Amanda Kathryn Engstrom, Brandon Scott Carpenter, and David John Katz. Repressive H3K9me2 protects lifespan against the transgenerational burden of COMPASS activity in C. elegans. *eLife*, 8, 2019.

[5826] Yerim Lee, Carey Phelps, Tao Huang, Barmak Mostofian, Lei Wu, Ying Zhang, Kai Tao, Young Hwan Chang, Philip JS Stork, Joe W Gray, Daniel M Zuckerman, and Xiaolin Nan. High-throughput, single-particle tracking reveals nested membrane domains that dictate KRasG12D diffusion and trafficking. *eLife*, 8, 2019.

[5827] Marco P Lehmann, He A Xu, Vasiliki Liakoni, Michael H Herzog, Wulfram Gerstner, and Kerstin Preuschoff. One-shot learning and behavioral eligibility traces in sequential decision making. *eLife*, 8, 2019.

[5828] Nicolas J Lehrbach and Gary Ruvkun. Endoplasmic reticulum-associated SKN-1A/Nrf1 mediates a cytoplasmic unfolded protein response and promotes longevity. *eLife*, 8, 2019.

[5829] Ingrid Lekk, Véronique Duboc, Ana Faro, Stephanos Nicolaou, Patrick Blader, and Stephen W Wilson. Sox1a mediates the ability of the parapineal to impart habenular left-right asymmetry. *eLife*, 8, 2019.

[5830] Amy M LeMessurier, Keven J Laboy-Juárez, Kathryn McClain, Shilin Chen, Theresa Nguyen, and Daniel E Feldman. Enrichment drives emergence of functional columns and improves sensory coding in the whisker map in L2/3 of mouse S1. *eLife*, 8, 2019.

[5831] Gábor Lengyel, Goda žalalytė, Alexandros Pantelides, James N Ingram, József Fiser, Máté Lengyel, and Daniel M Wolpert. Unimodal statistical learning produces multimodal object-like representations. *eLife*, 8, 2019.

[5832] Maureen Leninger, Ampon Sae Her, and Nathaniel J Traaseth. Inducing conformational preference of the membrane protein transporter EmrE through conservative mutations. *eLife*, 8, 2019.

[5833] Sabina Leonelli. The challenges of big data biology. *eLife*, 8, 2019.

[5834] Bluma J Lesch, Zuzana Tothova, Elizabeth A Morgan, Zhicong Liao, Roderick T Bronson, Benjamin L Ebert, and David C Page. Intergenerational epigenetic inheritance of cancer susceptibility in mammals. *eLife*, 8, 2019.

[5835] Itamar Lev, Hila Gingold, and Oded Rechavi. H3K9me3 is required for inheritance of small RNAs that target a unique subset of newly evolved genes. *eLife*, 8, 2019.

[5836] Robert F Levendosky and Gregory D Bowman. Asymmetry between the two acidic patches dictates the direction of nucleosome sliding by the ISWI chromatin remodeler. *eLife*, 8, 2019.

[5837] Baoqiang Li, Tatiana V Esipova, Ikbal Sencan, Kıvılcım KılıÇ, Buyin Fu, Michele Desjardins, Mohammad Moeini, Sreekanth Kura, Mohammad A Yaseen, Frederic Lesage, Leif Østergaard, Anna Devor, David A Boas, Sergei A Vinogradov, and Sava Sakadžić. More homogeneous capillary flow and oxygenation in deeper cortical layers correlate with increased oxygen extraction. *eLife*, 8, 2019.

[5838] Dayan J Li, Conor L McMann, and Peter W Reddien. Nuclear receptor NR4A is required for patterning at the ends of the planarian anterior-posterior axis. *eLife*, 8, 2019.

[5839] Hai Li, Chia-Ying Huang, Elena G Govorunova, Christopher T Schafer, Oleg A Sineshchekov, Meitian Wang, Lei Zheng, and John L Spudich. Crystal structure of a natural light-gated anion channelrhodopsin. *eLife*, 8, 2019.

[5840] Hao Li, Dominika Pullmann, Jennifer Y Cho, Maya Eid, and Thomas C Jhou. Generality and opponency of rostromedial tegmental (RMTg) roles in valence processing. *eLife*, 8, 2019.

[5841] Hao Li, Dominika Pullmann, and Thomas C Jhou. Valence-encoding in the lateral habenula arises from the entopeduncular region. *eLife*, 8, 2019.

[5842] Jie Li, Guijun Shang, Yu-Ju Chen, Chad A Brautigam, Jen Liou, Xuewu Zhang, and Xiao-chen Bai. Cryo-EM analyses reveal the common mechanism and diversification in the activation of RET by different ligands. *eLife*, 8, 2019.

[5843] Ke-Xin Li, Mu He, Wenlei Ye, Jeffrey Simms, Michael Gill, Xuaner Xiang, Yuh Nung Jan, and Lily Yeh Jan. TMEM16B regulates anxiety-related behavior and GABAergic neuronal signaling in the central lateral amygdala. *eLife*, 8, 2019.

[5844] Kefei Nina Li, Prachi Jain, Catherine Hua He, Flora Chae Eun, Sangjo Kang, and Tudorita Tumbar. Skin vasculature and hair follicle cross-talking associated with stem cell activation and tissue homeostasis. *eLife*, 8, 2019.

[5845] Mufeng Li, Yao Wang, Rahul Banerjee, Fabrizio Marinelli, Shai Silberberg, José D Faraldo-Gómez, Motoyuki Hattori, and Kenton Jon Swartz. Molecular mechanisms of human P2X3 receptor channel activation and modulation by divalent cation bound ATP. *eLife*, 8, 2019.

[5846] Nuo Li, Susu Chen, Zengcai V Guo, Han Chen, Yan Huo, Hidehiko K Inagaki, Guang Chen, Courtney Davis, David Hansel, Caiying Guo, and Karel Svoboda. Spatiotemporal constraints on optogenetic inactivation in cortical circuits. *eLife*, 8, 2019.

[5847] Sam Li, Jose-Jesus Fernandez, Wallace F Marshall, and David A Agard. Electron cryo-tomography provides insight into procentriole architecture and assembly mechanism. *eLife*, 8, 2019.

[5848] Weiwei Li, Haixia Wang, Xiaofei Xie, and Jian Li. Neural mediation of greed personality trait on economic risk-taking. *eLife*, 8, 2019.

[5849] Wenyuan Li, Yage Zhang, Bingzhou Han, Lianyan Li, Muhang Li, Xiaochan Lu, Cheng Chen, Mengjia Lu, Yujie Zhang, Xuefeng Jia, Zuoyan Zhu, Xiangjun Tong, and Bo Zhang. One-step efficient generation of dual-function conditional knockout and geno-tagging alleles in zebrafish. *eLife*, 8, 2019.

[5850] Xuehua Li, Li Han, Intawat Nookaew, Erin Mannen, Matthew J Silva, Maria Almeida, and Jinhu Xiong. Stimulation of Piezo1 by mechanical signals promotes bone anabolism. *eLife*, 8, 2019.

[5851] Yang Li, Kai-Xuan Li, Wei-Lin Hu, David M Ojcius, Jia-Qi Fang, Shi-Jun Li, Xu'ai Lin, and Jie Yan. RETRACTED: Endocytic recycling and vesicular transport systems mediate transcytosis of Leptospira interrogans across cell monolayer. *eLife*, 8, 2019.

[5852] Yu-Tung Li, Debashree Goswami, Melissa Follmer, Annette Artz, Mariana Pacheco-Blanco, and Dietmar Vestweber. Blood flow guides sequential support of neutrophil arrest and diapedesis by PILR-$\beta$1 and PILR-$\alpha$. *eLife*, 8, 2019.

[5853] Yuling Li, Jinhan Zhou, Sui Min, Yang Zhang, Yuqing Zhang, Qiao Zhou, Xiaofei Shen, Da Jia, Junhong Han, and Qingxiang Sun. Distinct RanBP1 nuclear export and cargo dissociation mechanisms between fungi and animals. *eLife*, 8, 2019.

[5854] Zhijie Li, Aidan CA Tomlinson, Alan HM Wong, Dongxia Zhou, Marc Desforges, Pierre J Talbot, Samir Benlekbir, John L Rubinstein, and James M Rini. The human coronavirus HCoV-229E S-protein structure and receptor binding. *eLife*, 8, 2019.

[5855] Zhiling Li, Marlene M Hao, Chris Van den Haute, Veerle Baekelandt, Werend Boesmans, and Pieter Vanden Berghe. Regional complexity in enteric neuron wiring reflects diversity of motility patterns in the mouse large intestine. *eLife*, 8, 2019.

[5856] Jianlin Liang, Nan Wang, Jing He, Jian Du, Yahui Guo, Lin Li, Wenbo Wu, Chencheng Yao, Zheng Li, and Kehkooi Kee. Induction of Sertoli-like cells from human fibroblasts by NR5A1 and GATA4. *eLife*, 8, 2019.

[5857] Rachel A Liberatore, Emily J Mastrocola, Elena Cassella, Fabian Schmidt, Jessie R Willen, Dennis Voronin, Trinity M Zang, Theodora Hatziioannou, and Paul D Bieniasz. Rhabdo-immunodeficiency virus, a murine model of acute HIV-1 infection. *eLife*, 8, 2019.

[5858] Simone Lieb, Silvia Blaha-Ostermann, Elisabeth Kamper, Janine Rippka, Cornelia Schwarz, Katharina Ehrenhöfer-Wölfer, Andreas Schlattl, Andreas Wernitznig, Jesse J Lipp, Kota Nagasaka, Petra van der Lelij, Gerd Bader, Minoru Koi, Ajay Goel, Ralph A Neumüller, Jan-Michael Peters, Norbert Kraut, Mark A Pearson, Mark Petronczki, and Simon Wöhrle. Werner syndrome helicase is a selective vulnerability of microsatellite instability-high tumor cells. *eLife*, 8, 2019.

[5859] Andrew Tze Fui Liew, Yong Hwee Foo, Yunfeng Gao, Parisa Zangoui, Moirangthem Kiran Singh, Ranjit Gulvady, and Linda J Kenney. Single cell, super-resolution imaging reveals an acid pH-dependent conformational switch in SsrB regulates SPI-2. *eLife*, 8, 2019.

[5860] Junghyun Lim, Hyunjoo Park, Jason Heisler, Timurs Maculins, Merone Roose-Girma, Min Xu, Brent Mckenzie, Menno van Lookeren Campagne, Kim Newton, and Aditya Murthy. Autophagy regulates inflammatory programmed cell death via turnover of RHIM-domain proteins. *eLife*, 8, 2019.

[5861] Sukbin Lim. Mechanisms underlying sharpening of visual response dynamics with familiarity. *eLife*, 8, 2019.

[5862] Dominique H Limoli, Elizabeth A Warren, Kaitlin D Yarrington, Niles P Donegan, Ambrose L Cheung, and George A O'Toole. Interspecies interactions induce exploratory motility in Pseudomonas aeruginosa. *eLife*, 8, 2019.

[5863] Cheng Yung Lin, Chia Lun Wu, Kok Zhi Lee, You Jei Chen, Po Hsiang Zhang, Chia Yu Chang, Horng Jyh Harn, Shinn Zong Lin, and Huai Jen Tsai. Extracellular Pgk1 enhances neurite outgrowth of motoneurons through Nogo66/NgR-independent targeting of NogoA. *eLife*, 8, 2019.

[5864] Meng Kuan Lin, Yeonsook Shin Takahashi, Bing-Xing Huo, Mitsutoshi Hanada, Jaimi Nagashima, Junichi Hata, Alexander S Tolpygo, Keerthi Ram, Brian C Lee, Michael I Miller, Marcello GP Rosa, Erika Sasaki, Atsushi Iriki, Hideyuki Okano, and Partha Mitra. A high-throughput neurohistological pipeline for brain-wide mesoscale connectivity mapping of the common marmoset. *eLife*, 8, 2019.

[5865] Zhijie Lin, Zhou Yang, Ruiling Xie, Zeyang Ji, Kunliang Guan, and Mingjie Zhang. Decoding WW domain tandem-mediated target recognitions in tissue growth and cell polarity. *eLife*, 8, 2019.

[5866] Daniel Linares, David Aguilar-Lleyda, and Joan López-Moliner. Decoupling sensory from decisional choice biases in perceptual decision making. *eLife*, 8, 2019.

[5867] Peter A Lind, Eric Libby, Jenny Herzog, and Paul B Rainey. Predicting mutational routes to new adaptive phenotypes. *eLife*, 8, 2019.

[5868] L Bailey Lindenmaier, Nicolas Parmentier, Caiying Guo, Fadel Tissir, and Kevin M Wright. Dystroglycan is a scaffold for extracellular axon guidance decisions. *eLife*, 8, 2019.

[5869] Shuo-Chien Ling, Somasish Ghosh Dastidar, Seiya Tokunaga, Wan Yun Ho, Kenneth Lim, Hristelina Ilieva, Philippe A Parone, Sheue-Houy Tyan, Tsemay M Tse, Jer-Cherng Chang, Oleksandr Platoshyn, Ngoc B Bui, Anh Bui, Anne Vetto, Shuying Sun, Melissa McAlonis-Downes, Joo Seok Han, Debbie Swing, Katannya Kapeli, Gene W Yeo, Lino Tessarollo, Martin Marsala, Christopher E Shaw, Greg Tucker-Kellogg, Albert R La Spada, Clotilde Lagier-Tourenne, Sandrine Da Cruz, and Don W Cleveland. Overriding FUS autoregulation in mice triggers gain-of-toxic dysfunctions in RNA metabolism and autophagy-lysosome axis. *eLife*, 8, 2019.

[5870] Maciej Lisicki, Marcos F Velho Rodrigues, Raymond E Goldstein, and Eric Lauga. Swimming eukaryotic microorganisms exhibit a universal speed distribution. *eLife*, 8, 2019.

[5871] Haowen Liu, Lei Li, Daniel Nedelcu, Qi Hall, Lijun Zhou, Wei Wang, Yi Yu, Joshua M Kaplan, and Zhitao Hu. Heterodimerization of UNC-13/RIM regulates synaptic vesicle release probability but not priming in C. elegans. *eLife*, 8, 2019.

[5872] Ling-Yu Liu, Xi Long, Ching-Po Yang, Rosa L Miyares, Ken Sugino, Robert H Singer, and Tzumin Lee. Mamo decodes hierarchical temporal gradients into terminal neuronal fate. *eLife*, 8, 2019.

[5873] Pan Liu, Seby Louis Edassery, Laith Ali, Benjamin R Thomson, Jeffrey N Savas, and Jing Jin. Long-lived metabolic enzymes in the crystalline lens identified by pulse-labeling of mice and mass spectrometry. *eLife*, 8, 2019.

[5874] Qian Liu, Elizabeth J Osterlund, Xiaoke Chi, Justin Pogmore, Brian Leber, and David William Andrews. Bim escapes displacement by BH3-mimetic anti-cancer drugs by double-bolt locking both Bcl-XL and Bcl-2. *eLife*, 8, 2019.

[5875] Shuang Liu, Shufeng Wang, Linzhi Zou, Jie Li, Chenmeng Song, Jiaofeng Chen, Qun Hu, Lian Liu, Pingbo Huang, and Wei Xiong. TMC1 is an essential component of a leak channel that modulates tonotopy and excitability of auditory hair cells in mice. *eLife*, 8, 2019.

[5876] Weiwei Liu, Anindya Ganguly, Jia Huang, Yijin Wang, Jinfei D Ni, Adishthi S Gurav, Morris A Aguilar, and Craig Montell. Neuropeptide F regulates courtship in Drosophila through a male-specific neuronal circuit. *eLife*, 8, 2019.

[5877] Yi Liu, Tobias Maierhofer, Katarzyna Rybak, Jan Sklenar, Andy Breakspear, Matthew G Johnston, Judith Fliegmann, Shouguang Huang, M Rob G Roelfsema, Georg Felix, Christine Faulkner, Frank LH Menke, Dietmar Geiger, Rainer Hedrich, and Silke Robatzek. Anion channel SLAH3 is a regulatory target of chitin receptor-associated kinase PBL27 in microbial stomatal closure. *eLife*, 8, 2019.

[5878] Alberto Llera, Thomas Wolfers, Peter Mulders, and Christian F Beckmann. Inter-individual differences in human brain structure and morphology link to variation in demographics and behavior. *eLife*, 8, 2019.

[5879] Alfredo Llorca, Gabriele Ciceri, Robert Beattie, Fong Kuan Wong, Giovanni Diana, Eleni Serafeimidou-Pouliou, Marian Fernández-Otero, Carmen Streicher, Sebastian J Arnold, Martin Meyer, Simon Hippenmeyer, Miguel Maravall, and Oscar Marin. A stochastic framework of neurogenesis underlies the assembly of neocortical cytoarchitecture. *eLife*, 8, 2019.

[5880] Emily J Lodge, Alice Santambrogio, John P Russell, Paraskevi Xekouki, Thomas S Jacques, Randy L Johnson, Selvam Thavaraj, Stefan R Bornstein, and Cynthia Lilian Andoniadou. Homeostatic and tumourigenic activity of SOX2+ pituitary stem cells is controlled by the LATS/YAP/TAZ cascade. *eLife*, 8, 2019.

[5881] Elsa Loissel. A question of support. *eLife*, 8, 2019.

[5882] Michael V Lombardo, Lisa Eyler, Adrienne Moore, Michael Datko, Cynthia Carter Barnes, Debra Cha, Eric Courchesne, and Karen Pierce. Default mode-visual network hypoconnectivity in an autism subtype with pronounced social visual engagement difficulties. *eLife*, 8, 2019.

[5883] Jason S Long, Alewo Idoko-Akoh, Bhakti Mistry, Daniel Goldhill, Ecco Staller, Jocelyn Schreyer, Craig Ross, Steve Goodbourn, Holly Shelton, Michael A Skinner, Helen Sang, Michael J McGrew, and Wendy Barclay. Species specific differences in use of ANP32 proteins by influenza A virus. *eLife*, 8, 2019.

[5884] Tina B Lonsdorf, Maren Klingelhöfer-Jens, Marta Andreatta, Tom Beckers, Anastasia Chalkia, Anna Gerlicher, Valerie L Jentsch, Shira Meir Drexler, Gaetan Mertens, Jan Richter, Rachel Sjouwerman, Julia Wendt, and Christian J Merz. Navigating the garden of forking paths for data exclusions in fear conditioning research. *eLife*, 8, 2019.

[5885] Tsui Han Loo, Xiaoqian Ye, Ruth Jinfen Chai, Mitsuteru Ito, Gisèle Bonne, Anne C Ferguson-Smith, and Colin L Stewart. The mammalian LINC complex component SUN1 regulates muscle regeneration by modulating drosha activity. *eLife*, 8, 2019.

[5886] Jorge López-Tello, Vicente Pérez-García, Jaspreet Khaira, Laura C Kusinski, Wendy N Cooper, Adam Andreani, Imogen Grant, Edurne Fernández de Liger, Brian YH Lam, Myriam Hemberger, Ionel Sandovici, Miguel Constancia, and Amanda N Sferruzzi-Perri. Fetal and trophoblast PI3K p110$\alpha$ have distinct roles in regulating resource supply to the growing fetus in mice. *eLife*, 8, 2019.

[5887] Yenisleidy Lorenzo-Ceballos, Willy Carrasquel-Ursulaez, Karen Castillo, Osvaldo Alvarez, and Ramon Latorre. Calcium-driven regulation of voltage-sensing domains in BK channels. *eLife*, 8, 2019.

[5888] Catrina M Loucks, Kwangjin Park, Denise S Walker, Andrea H McEwan, Tiffany A Timbers, Evan L Ardiel, Laura J Grundy, Chunmei Li, Jacque-Lynne Johnson, Julie Kennedy, Oliver E Blacque, William Schafer, Catharine H Rankin, and Michel R Leroux. EFHC1, implicated in juvenile myoclonic epilepsy, functions at the cilium and synapse to modulate dopamine signaling. *eLife*, 8, 2019.

[5889] Lingxi Lu, Qian Wang, Jingwei Sheng, Zhaowei Liu, Lang Qin, Liang Li, and Jia-Hong Gao. Neural tracking of speech mental imagery during rhythmic inner counting. *eLife*, 8, 2019.

[5890] Menghao Lu and Kota Mizumoto. Gradient-independent Wnt signaling instructs asymmetric neurite pruning in C. elegans. *eLife*, 8, 2019.

[5891] Robert Luck, Severino Urban, Andromachi Karakatsani, Eva Harde, Sivakumar Sambandan, LaShae Nicholson, Silke Haverkamp, Rebecca Mann, Ana Martin-Villalba, Erin Margaret Schuman, Amparo Acker-Palmer, and Carmen Ruiz de Almodóvar. VEGF/VEGFR2 signaling regulates hippocampal axon branching during development. *eLife*, 8, 2019.

[5892] Marshall Lukacs, Tia Roberts, Praneet Chatuverdi, and Rolf W Stottmann. Glycosylphosphatidylinositol biosynthesis and remodeling are required for neural tube closure, heart development, and cranial neural crest cell survival. *eLife*, 8, 2019.

[5893] Massimo Lumaca, Boris Kleber, Elvira Brattico, Peter Vuust, and Giosue Baggio. Functional connectivity in human auditory networks and the origins of variation in the transmission of musical systems. *eLife*, 8, 2019.

[5894] Lydia V Luncz, Mike Gill, Tomos Proffitt, Magdalena S Svensson, Lars Kulik, and Suchinda Malaivijitnond. Group-specific archaeological signatures of stone tool use in wild macaques. *eLife*, 8, 2019.

[5895] Chongyuan Luo, Qian Yi Lee, Orly Wapinski, Rosa Castanon, Joseph R Nery, Moritz Mall, Michael S Kareta, Sean M Cullen, Margaret A Goodell, Howard Y Chang, Marius Wernig, and Joseph R Ecker. Global DNA methylation remodeling during direct reprogramming of fibroblasts to neurons. *eLife*, 8, 2019.

[5896] Mark E Lush, Daniel C Diaz, Nina Koenecke, Sungmin Baek, Helena Boldt, Madeleine K St Peter, Tatiana Gaitan-Escudero, Andres Romero-Carvajal, Elisabeth M Busch-Nentwich, Anoja G Perera, Kathryn E Hall, Allison Peak, Jeffrey S Haug, and Tatjana Piotrowski. scRNA-Seq reveals distinct stem cell populations that drive hair cell regeneration after loss of Fgf and Notch signaling. *eLife*, 8, 2019.

[5897] Guillermo Luxán, Jonas Stewen, Noelia Díaz, Katsuhiro Kato, Sathish K Maney, Anusha Aravamudhan, Frank Berkenfeld, Nina Nagelmann, Hannes CA Drexler, Dagmar Zeuschner, Cornelius Faber, Hermann Schillers, Sven Hermann, John Wiseman, Juan M Vaquerizas, Mara E Pitulescu, and Ralf H Adams. Endothelial EphB4 maintains vascular integrity and transport function in adult heart. *eLife*, 8, 2019.

[5898] Fabrice Luyckx, Hamed Nili, Bernhard Spitzer, and Christopher Summerfield. Neural structure mapping in human probabilistic reward learning. *eLife*, 8, 2019.

[5899] Péter Lőrincz, Lili Anna Kenéz, Sarolta Tóth, Viktória Kiss, ágnes Varga, Tamás Csizmadia, Zsófia Simon-Vecsei, and Gábor Juhász. Vps8 overexpression inhibits HOPS-dependent trafficking routes by outcompeting Vps41/Lt. *eLife*, 8, 2019.

[5900] Wenxin Ma, Sean M Silverman, Lian Zhao, Rafael Villasmil, Maria M Campos, Juan Amaral, and Wai T Wong. Absence of TGF$\beta$ signaling in retinal microglia induces retinal degeneration and exacerbates choroidal neovascularization. *eLife*, 8, 2019.

[5901] Xiancai Ma, Tao Yang, Yuewen Luo, Liyang Wu, Yawen Jiang, Zheng Song, Ting Pan, Bingfeng Liu, Guangyan Liu, Jun Liu, Fei Yu, Zhangping He, Wanying Zhang, Jinyu Yang, Liting Liang, Yuanjun Guan, Xu Zhang, Linghua Li, Weiping Cai, Xiaoping Tang, Song Gao, Kai Deng, and Hui Zhang. TRIM28 promotes HIV-1 latency by SUMOylating CDK9 and inhibiting P-TEFb. *eLife*, 8, 2019.

[5902] Kate MacCord and Jane Maienschein. Understanding regeneration at different scales. *eLife*, 8, 2019.

[5903] Julia Machon, Jakob Krieger, Rebecca Meth, Magali Zbinden, Juliette Ravaux, Nicolas Montagné, Thomas Chertemps, and Steffen Harzsch. Neuroanatomy of a hydrothermal vent shrimp provides insights into the evolution of crustacean integrative brain centers. *eLife*, 8, 2019.

[5904] Emily L Mackevicius, Andrew H Bahle, Alex H Williams, Shijie Gu, Natalia I Denisenko, Mark S Goldman, and Michale S Fee. Unsupervised discovery of temporal sequences in high-dimensional datasets, with applications to neuroscience. *eLife*, 8, 2019.

[5905] Jamie A Macpherson, Alina Theisen, Laura Masino, Louise Fets, Paul C Driscoll, Vesela Encheva, Ambrosius P Snijders, Stephen R Martin, Jens Kleinjung, Perdita E Barran, Franca Fraternali, and Dimitrios Anastasiou. Functional cross-talk between allosteric effects of activating and inhibiting ligands underlies PKM2 regulation. *eLife*, 8, 2019.

[5906] Rajtarun Madangopal, Brendan J Tunstall, Lauren E Komer, Sophia J Weber, Jennifer K Hoots, Veronica A Lennon, Jennifer M Bossert, David H Epstein, Yavin Shaham, and Bruce T Hope. Discriminative stimuli are sufficient for incubation of cocaine craving. *eLife*, 8, 2019.

[5907] Julene Madariaga-Marcos, Cesar L Pastrana, Gemma LM Fisher, Mark Simon Dillingham, and Fernando Moreno-Herrero. ParB dynamics and the critical role of the CTD in DNA condensation unveiled by combined force-fluorescence measurements. *eLife*, 8, 2019.

[5908] Shintaro Maeda, Chinatsu Otomo, and Takanori Otomo. The autophagic membrane tether ATG2A transfers lipids between membranes. *eLife*, 8, 2019.

[5909] Jérémy Magescas, Jenny C Zonka, and Jessica L Feldman. A two-step mechanism for the inactivation of microtubule organizing center function at the centrosome. *eLife*, 8, 2019.

[5910] Maxime Maheu, Stanislas Dehaene, and Florent Meyniel. Brain signatures of a multiscale process of sequence learning in humans. *eLife*, 8, 2019.

[5911] Andrea Mair, Shou-Ling Xu, Tess C Branon, Alice Y Ting, and Dominique C Bergmann. Proximity labeling of protein complexes and cell-type-specific organellar proteomes in Arabidopsis enabled by TurboID. *eLife*, 8, 2019.

[5912] Peter Major, Kacper M Sendra, Paul Dean, Tom A Williams, Andrew K Watson, David T Thwaites, T Martin Embley, and Robert P Hirt. A new family of cell surface located purine transporters in Microsporidia and related fungal endoparasites. *eLife*, 8, 2019.

[5913] Tamar R Makin and Jean-Jacques Orban de Xivry. Ten common statistical mistakes to watch out for when writing or reviewing a manuscript. *eLife*, 8, 2019.

[5914] Samuel Andrew Malone, Georgios E Papadakis, Andrea Messina, Nour El Houda Mimouni, Sara Trova, Monica Imbernon, Cecile Allet, Irene Cimino, James Acierno, Daniele Cassatella, Cheng Xu, Richard Quinton, Gabor Szinnai, Pascal Pigny, Lur Alonso-Cotchico, Laura Masgrau, Jean-Didier Maréchal, Vincent Prevot, Nelly Pitteloud, and Paolo Giacobini. Defective AMH signaling disrupts GnRH neuron development and function and contributes to hypogonadotropic hypogonadism. *eLife*, 8, 2019.

[5915] Cristina Manatschal, Jonai Pujol-Giménez, Marion Poirier, Jean-Louis Reymond, Matthias A Hediger, and Raimund Dutzler. Mechanistic basis of the inhibition of SLC11/NRAMP-mediated metal ion transport by bis-isothiourea substituted compounds. *eLife*, 8, 2019.

[5916] Hetty E Manenschijn, Andrea Picco, Markus Mund, Anne-Sophie Rivier-Cordey, Jonas Ries, and Marko Kaksonen. Type-I myosins promote actin polymerization to drive membrane bending in endocytosis. *eLife*, 8, 2019.

[5917] Angelika Manhart, Téa Aleksandra Icheva, Christophe Guerin, Tobbias Klar, Rajaa Boujemaa-Paterski, Manuel Thery, Laurent Blanchoin, and Alex Mogilner. Quantitative regulation of the dynamic steady state of actin networks. *eLife*, 8, 2019.

[5918] Ernesto Manzo, Ileana Lorenzini, Dianne Barrameda, Abigail G O'Conner, Jordan M Barrows, Alexander Starr, Tina Kovalik, Benjamin E Rabichow, Erik M Lehmkuhl, Dakotah D Shreiner, Archi Joardar, Jean-Charles Liévens, Robert Bowser, Rita Sattler, and Daniela C Zarnescu. Glycolysis upregulation is neuroprotective as a compensatory mechanism in ALS. *eLife*, 8, 2019.

[5919] Uri Maoz, Gideon Yaffe, Christof Koch, and Liad Mudrik. Neural precursors of decisions that matter—an ERP study of deliberate and arbitrary choice. *eLife*, 8, 2019.

[5920] Aimee H Marceau, Caileen M Brison, Santrupti Nerli, Heather E Arsenault, Andrew C McShan, Eefei Chen, Hsiau-Wei Lee, Jennifer A Benanti, Nikolaos G Sgourakis, and Seth M Rubin. An order-to-disorder structural switch activates the FoxM1 transcription factor. *eLife*, 8, 2019.

[5921] Axelle Marchant, Angel F Cisneros, Alexandre K Dubé, Isabelle Gagnon-Arsenault, Diana Ascencio, Honey Jain, Simon Aubé, Chris Eberlein, Daniel Evans-Yamamoto, Nozomu Yachie, and Christian R Landry. The role of structural pleiotropy and regulatory evolution in the retention of heteromers of paralogs. *eLife*, 8, 2019.

[5922] Maria C Marchetto, Branka Hrvoj-Mihic, Bilal E Kerman, Diana X Yu, Krishna C Vadodaria, Sara B Linker, Iñigo Narvaiza, Renata Santos, Ahmet M Denli, Ana PD Mendes, Ruth Oefner, Jonathan

Cook, Lauren McHenry, Jaeson M Grasmick, Kelly Heard, Callie Fredlender, Lynne Randolph-Moore, Rijul Kshirsagar, Rea Xenitopoulos, Grace Chou, Nasun Hah, Alysson R Muotri, Krishnan Padmanabhan, Katerina Semendeferi, and Fred H Gage. Species-specific maturation profiles of human, chimpanzee and bonobo neural cells. *eLife*, 8, 2019.

[5923] Zoran S Marinkovic, Clément Vulin, Mislav Acman, Xiaohu Song, Jean-Marc Di Meglio, Ariel B Lindner, and Pascal Hersen. A microfluidic device for inferring metabolic landscapes in yeast monolayer colonies. *eLife*, 8, 2019.

[5924] Rosa María Marión, Juan J Montero, Isabel López de Silanes, Osvaldo Graña-Castro, Paula Martínez, Stefan Schoeftner, José Alejandro Palacios-Fábrega, and Maria A Blasco. TERRA regulate the transcriptional landscape of pluripotent cells through TRF1-dependent recruitment of PRC2. *eLife*, 8, 2019.

[5925] Andrew G Marshall, Manohar L Sharma, Kate Marley, Hakan Olausson, and Francis P McGlone. Spinal signalling of C-fiber mediated pleasant touch in humans. *eLife*, 8, 2019.

[5926] James AR Marshall, Ralf HJM Kurvers, Jens Krause, and Max Wolf. Quorums enable optimal pooling of independent judgements in biological systems. *eLife*, 8, 2019.

[5927] Carlotta Martelli and André Fiala. Slow presynaptic mechanisms that mediate adaptation in the olfactory pathway of Drosophila. *eLife*, 8, 2019.

[5928] Gregory M Martin, Min Woo Sung, Zhongying Yang, Laura M Innes, Balamurugan Kandasamy, Larry L David, Craig Yoshioka, and Show-Ling Shyng. Mechanism of pharmacochaperoning in a mammalian KATP channel revealed by cryo-EM. *eLife*, 8, 2019.

[5929] Ane Martin Anduaga, Naveh Evantal, Ines Lucia Patop, Osnat Bartok, Ron Weiss, and Sebastian Kadener. Thermosensitive alternative splicing senses and mediates temperature adaptation in Drosophila. *eLife*, 8, 2019.

[5930] Diana Martinez, Haroon Anwar, Amitabha Bose, Dirk M Bucher, and Farzan Nadim. Short-term synaptic dynamics control the activity phase of neurons in an oscillatory network. *eLife*, 8, 2019.

[5931] Jonathan Martinez-Fabregas, Stephan Wilmes, Luopin Wang, Maximillian Hafer, Elizabeth Pohler, Juliane Lokau, Christoph Garbers, Adeline Cozzani, Paul K Fyfe, Jacob Piehler, Majid Kazemian, Suman Mitra, and Ignacio Moraga. Kinetics of cytokine receptor trafficking determine signaling and functional selectivity. *eLife*, 8, 2019.

[5932] Valentina Marziano, Piero Poletti, Filippo Trentini, Alessia Melegaro, Marco Ajelli, and Stefano Merler. Parental vaccination to reduce measles immunity gaps in Italy. *eLife*, 8, 2019.

[5933] Matthew G Marzo, Jacqueline M Griswold, Kristina M Ruff, Rachel E Buchmeier, Colby P Fees, and Steven M Markus. Molecular basis for dyneinopathies reveals insight into dynein regulation and dysfunction. *eLife*, 8, 2019.

[5934] Sara Masachis, Nicolas J Tourasse, Claire Lays, Marion Faucher, Sandrine Chabas, Isabelle Iost, and Fabien Darfeuille. A genetic selection reveals functional metastable structures embedded in a toxin-encoding mRNA. *eLife*, 8, 2019.

[5935] David Massa López, Melanie Thelen, Felix Stahl, Christian Thiel, Arne Linhorst, Marc Sylvester, Irm Hermanns-Borgmeyer, Renate Lüllmann-Rauch, Winnie Eskild, Paul Saftig, and Markus Damme. The lysosomal transporter MFSD1 is essential for liver homeostasis and critically depends on its accessory subunit GLMP. *eLife*, 8, 2019.

[5936] Anna E Masser, Wenjing Kang, Joydeep Roy, Jayasankar Mohanakrishnan Kaimal, Jany Quintana-Cordero, Marc R Friedländer, and Claes Andréasson. Cytoplasmic protein misfolding titrates Hsp70 to activate nuclear Hsf1. *eLife*, 8, 2019.

[5937] Jonathan H Massey, Daayun Chung, Igor Siwanowicz, David L Stern, and Patricia J Wittkopp. The yellow gene influences Drosophila male mating success through sex comb melanization. *eLife*, 8, 2019.

[5938] Ambra Masuzzo, Gérard Manière, Annelise Viallat-Lieutaud, émilie Avazeri, Olivier Zugasti, Yaël Grosjean, C Léopold Kurz, and Julien Royet. Peptidoglycan-dependent NF-$\kappa$B activation in a small subset of brain octopaminergic neurons controls female oviposition. *eLife*, 8, 2019.

[5939] Mette Mathiesen Janiurek, Rana Soylu-Kucharz, Christina Christoffersen, Krzysztof Kucharz, and Martin Lauritzen. Apolipoprotein M-bound sphingosine-1-phosphate regulates blood–brain barrier paracellular permeability and transcytosis. *eLife*, 8, 2019.

[5940] Shin Matsubara, Akira Shiraishi, Tomohiro Osugi, Tsuyoshi Kawada, and Honoo Satake. The regulation of oocyte maturation and ovulation in the closest sister group of vertebrates. *eLife*, 8, 2019.

[5941] Benjamin J Matthews, Meg A Younger, and Leslie B Vosshall. The ion channel ppk301 controls freshwater egg-laying in the mosquito Aedes aegypti. *eLife*, 8, 2019.

[5942] Noémie Matthey, Sandrine Stutzmann, Candice Stoudmann, Nicolas Guex, Christian Iseli, and Melanie Blokesch. Neighbor predation linked to natural competence fosters the transfer of large genomic regions in Vibrio cholerae. *eLife*, 8, 2019.

[5943] Molly A Matty, Daphne R Knudsen, Eric M Walton, Rebecca W Beerman, Mark R Cronan, Charlie J Pyle, Rafael E Hernandez, and David M Tobin. Potentiation of P2RX7 as a host-directed strategy for control of mycobacterial infection. *eLife*, 8, 2019.

[5944] Joshua E Mayfield, Seema Irani, Edwin E Escobar, Zhao Zhang, Nathaniel T Burkholder, Michelle R Robinson, M Rachel Mehaffey, Sarah N Sipe, Wanjie Yang, Nicholas A Prescott, Karan R Kathuria, Zhijie Liu, Jennifer S Brodbelt, and Yan Zhang. Tyr1 phosphorylation promotes phosphorylation of Ser2 on the C-terminal domain of eukaryotic RNA polymerase II by P-TEFb. *eLife*, 8, 2019.

[5945] Leslie A McCauliff, Annette Langan, Ran Li, Olga Ilnytska, Debosreeta Bose, Miriam Waghalter, Kimberly Lai, Peter C Kahn, and Judith Storch. Intracellular cholesterol trafficking is dependent upon NPC2 interaction with lysobisphosphatidic acid. *eLife*, 8, 2019.

[5946] Gary S McDowell, John D Knutsen, June M Graham, Sarah K Oelker, and Rebeccah S Lijek. Co-reviewing and ghostwriting by early-career researchers in the peer review of manuscripts. *eLife*, 8, 2019.

[5947] Shane G McInally, Jane Kondev, and Scott C Dawson. Length-dependent disassembly maintains four different flagellar lengths in Giardia. *eLife*, 8, 2019.

[5948] Paul F McKay, Deniz Cizmeci, Yoann Aldon, Jeroen Maertzdorf, January Weiner, Stefan HE Kaufmann, David JM Lewis, Robert A van den Berg, Giuseppe Del Giudice, and Robin J Shattock. Identification of potential biomarkers of vaccine inflammation in mice. *eLife*, 8, 2019.

[5949] Erin C McKiernan, Lesley A Schimanski, Carol Muñoz Nieves, Lisa Matthias, Meredith T Niles, and Juan P Alperin. Use of the Journal Impact Factor in academic review, promotion, and tenure evaluations. *eLife*, 8, 2019.

[5950] Michael R McLaren, Amy D Willis, and Benjamin J Callahan. Consistent and correctable bias in metagenomic sequencing experiments. *eLife*, 8, 2019.

[5951] Mary McMahon, Adrian Contreras, Mikael Holm, Tamayo Uechi, Craig M Forester, Xiaming Pang, Cody Jackson, Meredith E Calvert, Bin Chen, David A Quigley, John M Luk, R Kate Kelley, John D Gordan, Ryan M Gill, Scott C Blanchard, and Davide Ruggero. A single H/ACA small nucleolar RNA mediates tumor suppression downstream of oncogenic RAS. *eLife*, 8, 2019.

[5952] David Trombley McSwiggen, Anders S Hansen, Sheila S Teves, Hervé Marie-Nelly, Yvonne Hao, Alec Basil Heckert, Kayla K Umemoto, Claire Dugast-Darzacq, Robert Tjian, and Xavier Darzacq. Evidence for DNA-mediated nuclear compartmentalization distinct from phase separation. *eLife*, 8, 2019.

[5953] Michael P Meers, Terri D Bryson, Jorja G Henikoff, and Steven Henikoff. Improved CUT&RUN chromatin profiling tools. *eLife*, 8, 2019.

[5954] Jessica U Meir, Julia M York, Bev A Chua, Wilhelmina Jardine, Lucy A Hawkes, and William K Milsom. Reduced metabolism supports hypoxic flight in the high-flying bar-headed goose (Anser indicus). *eLife*, 8, 2019.

[5955] James Meixiong, Chirag Vasavda, Dustin Green, Qin Zheng, Lijun Qi, Shawn G Kwatra, James P Hamilton, Solomon H Snyder, and Xinzhong Dong. Identification of a bilirubin receptor that may mediate a component of cholestatic itch. *eLife*, 8, 2019.

[5956] Sergio Menchero, Isabel Rollan, Antonio Lopez-Izquierdo, Maria Jose Andreu, Julio Sainz de Aja, Minjung Kang, Javier Adan, Rui Benedito, Teresa Rayon, Anna-Katerina Hadjantonakis, and Miguel Manzanares. Transitions in cell potency during early mouse development are driven by Notch. *eLife*, 8, 2019.

[5957] Guanghan Meng, Yajie Liang, Sarah Sarsfield, Wan-chen Jiang, Rongwen Lu, Joshua Tate Dudman, Yeka Aponte, and Na Ji. High-throughput synapse-resolving two-photon fluorescence microendoscopy for deep-brain volumetric imaging in vivo. *eLife*, 8, 2019.

[5958] Julia L Meng, Zarion D Marshall, Meike Lobb-Rabe, and Ellie S Heckscher. How prolonged expression of Hunchback, a temporal transcription factor, re-wires locomotor circuits. *eLife*, 8, 2019.

[5959] Xiangling Meng, Christopher M McGraw, Wei Wang, Junzhan Jing, Szu-Ying Yeh, Li Wang, Joanna Lopez, Amanda M Brown, Tao Lin, Wu Chen, Mingshan Xue, Roy V Sillitoe, Xiaolong Jiang, and Huda Y Zoghbi. Neurexophilin4 is a selectively expressed $\alpha$-neurexin ligand that modulates specific cerebellar synapses and motor functions. *eLife*, 8, 2019.

[5960] Kaushiki P Menon, Vivek Kulkarni, Shin-ya Takemura, Michael Anaya, and Kai Zinn. Interactions between Dpr11 and DIP-$\gamma$ control selection of amacrine neurons in Drosophila color vision circuits. *eLife*, 8, 2019.

[5961] David G Mets and Michael S Brainard. Learning is enhanced by tailoring instruction to individual genetic differences. *eLife*, 8, 2019.

[5962] Emily Mevers, Lin Su, Gleb Pishchany, Moshe Baruch, Jose Cornejo, Elissa Hobert, Eric Dimise, Caroline M Ajo-Franklin, and Jon Clardy. An elusive electron shuttle from a facultative anaerobe. *eLife*, 8, 2019.

[5963] Nancy L Meyer, Guiqing Hu, Omar Davulcu, Qing Xie, Alex J Noble, Craig Yoshioka, Drew S Gingerich, Andrew Trzynka, Larry David, Scott M Stagg, and Michael Stewart Chapman. Structure of the gene therapy vector, adeno-associated virus with its cell receptor, AAVR. *eLife*, 8, 2019.

[5964] Edwige Michel, Jean Gaudart, Samuel Beaulieu, Gregory Bulit, Martine Piarroux, Jacques Boncy, Patrick Dely, Renaud Piarroux, and Stanislas Rebaudet. Estimating effectiveness of case-area targeted response interventions against cholera in Haiti. *eLife*, 8, 2019.

[5965] Antonio Michelucci, Simona Boncompagni, Laura Pietrangelo, Maricela García-Castañeda, Takahiro Takano, Sundeep Malik, Robert T Dirksen, and Feliciano Protasi. Transverse tubule remodeling enhances Orai1-dependent Ca2+ entry in skeletal muscle. *eLife*, 8, 2019.

[5966] Teemu P Miettinen, Joon Ho Kang, Lucy F Yang, and Scott R Manalis. Mammalian cell growth dynamics in mitosis. *eLife*, 8, 2019.

[5967] Johann Mignolet, Guillaume Cerckel, Julien Damoczi, Laura Ledesma-Garcia, Andrea Sass, Tom Coenye, Sylvie Nessler, and Pascal Hols. Subtle selectivity in a pheromone sensor triumvirate desynchronizes competence and predation in a human gut commensal. *eLife*, 8, 2019.

[5968] Paul Glad Mihai, Michelle Moerel, Federico de Martino, Robert Trampel, Stefan Kiebel, and Katharina von Kriegstein. Modulation of tonotopic ventral medial geniculate body is behaviorally relevant for speech recognition. *eLife*, 8, 2019.

[5969] Snježana Mikuličić, Jérôme Finke, Fatima Boukhallouk, Elena Wüstenhagen, Dominik Sons, Yahya Homsi, Karina Reiss, Thorsten Lang, and Luise Florin. ADAM17-dependent signaling is required for oncogenic human papillomavirus entry platform assembly. *eLife*, 8, 2019.

[5970] Alexandria N Miller, George Vaisey, and Stephen B Long. Molecular mechanisms of gating in the calcium-activated chloride channel bestrophin. *eLife*, 8, 2019.

[5971] Brittany K Miller, Ryan Hughes, Lauren S Ligon, Nathan W Rigel, Seidu Malik, Brandon R Anjuwon-Foster, James C Sacchettini, and Miriam Braunstein. Mycobacterium tuberculosis SatS is a chaperone for the SecA2 protein export pathway. *eLife*, 8, 2019.

[5972] Ian F Miller, Robert A Barton, and Charles L Nunn. Quantitative uniqueness of human brain evolution revealed through phylogenetic comparative analysis. *eLife*, 8, 2019.

[5973] Steven W Miller, Artem Movsesyan, Sui Zhang, Rosa Fernández, and James W Posakony. Evolutionary emergence of Hairless as a novel component of the Notch signaling pathway. *eLife*, 8, 2019.

[5974] Wilbur Kyle Mills, Yuh Chwen G Lee, Antje M Kochendoerfer, Elaine M Dunleavy, and Gary H Karpen. RNA from a simple-tandem repeat is required for sperm maturation and male fertility in Drosophila melanogaster. *eLife*, 8, 2019.

[5975] Veronika Miskolci, Jayne Squirrell, Julie Rindy, William Vincent, John Demian Sauer, Angela Gibson, Kevin W Eliceiri, and Anna Huttenlocher. Distinct inflammatory and wound healing responses to complex caudal fin injuries of larval zebrafish. *eLife*, 8, 2019.

[5976] Avia Mizrachi, Shiri Graff van Creveld, Orr H Shapiro, Shilo Rosenwasser, and Assaf Vardi. Light-dependent single-cell heterogeneity in the chloroplast redox state regulates cell fate in a marine diatom. *eLife*, 8, 2019.

[5977] Andreas P Modlmeier, Ewan Colman, Ephraim M Hanks, Ryan Bringenberg, Shweta Bansal, and David P Hughes. Ant colonies maintain social homeostasis in the face of decreased density. *eLife*, 8, 2019.

[5978] Priyesh Mohanty, Rashmi Agrata, Batul Ismail Habibullah, Arun G S, and Ranabir Das. Deamidation disrupts native and transient contacts to weaken the interaction between UBC13 and RING-finger E3 ligases. *eLife*, 8, 2019.

[5979] Franziska Mohring, Melissa Natalie Hart, Thomas A Rawlinson, Ryan Henrici, James A Charleston, Ernest Diez Benavente, Avnish Patel, Joanna Hall, Neil Almond, Susana Campino, Taane G Clark, Colin J Sutherland, David A Baker, Simon J Draper, and Robert William Moon. Rapid and iterative genome editing in the malaria parasite Plasmodium knowlesi provides new tools for P. vivax research. *eLife*, 8, 2019.

[5980] Konstadinos Moissoglu, Kyota Yasuda, Tianhong Wang, George Chrisafis, and Stavroula Mili. Translational regulation of protrusion-localized RNAs involves silencing and clustering after transport. *eLife*, 8, 2019.

[5981] Hans Moldenhauer, Ignacio Díaz-Franulic, Horacio Poblete, and David Naranjo. Trans-toxin ion-sensitivity of charybdotoxin-blocked potassium-channels reveals unbinding transitional states. *eLife*, 8, 2019.

[5982] Sebastian Molina-Obando, Juan Felipe Vargas-Fique, Miriam Henning, Burak Gür, T Moritz Schladt, Junaid Akhtar, Thomas K Berger, and Marion Silies. ON selectivity in the Drosophila visual system is a multisynaptic process involving both glutamatergic and GABAergic inhibition. *eLife*, 8, 2019.

[5983] Sophie A Montandon, Anamarija Fofonjka, and Michel C Milinkovitch. Elastic instability during branchial ectoderm development causes folding of the Chlamydosaurus erectile frill. *eLife*, 8, 2019.

[5984] Scott WJ Mooney, Phillip J Marlow, and Barton L Anderson. The perception and misperception of optical defocus, shading, and shape. *eLife*, 8, 2019.

[5985] Hernán Morales-Navarrete, Hidenori Nonaka, André Scholich, Fabián Segovia-Miranda, Walter de Back, Kirstin Meyer, Roman L Bogorad, Victor Koteliansky, Lutz Brusch, Yannis Kalaidzidis, Frank Jülicher, Benjamin M Friedrich, and Marino Zerial. Liquid-crystal organization of liver tissue. *eLife*, 8, 2019.

[5986] Gideon J Mordecai, Kristina M Miller, Emiliano Di Cicco, Angela D Schulze, Karia H Kaukinen, Tobi J Ming, Shaorong Li, Amy Tabata, Amy Teffer, David A Patterson, Hugh W Ferguson, and Curtis A Suttle. Endangered wild salmon infected by newly discovered viruses. *eLife*, 8, 2019.

[5987] José-Maria Moreira, Pavel M Itskov, Dennis Goldschmidt, Celia Baltazar, Kathrin Steck, Ibrahim Tastekin, Samuel J Walker, and Carlos Ribeiro. optoPAD, a closed-loop optogenetics system to study the circuit basis of feeding behaviors. *eLife*, 8, 2019.

[5988] David F Moreno, Kirsten Jenkins, Sandrine Morlot, Gilles Charvin, Attila Csikasz-Nagy, and Martí Aldea. Proteostasis collapse, a hallmark of aging, hinders the chaperone-Start network and arrests cells in G1. *eLife*, 8, 2019.

[5989] Peter James Morgan, Romain Bourboulou, Caroline Filippi, Julie Koenig-Gambini, and Jérôme Epsztein. Kv1.1 contributes to a rapid homeostatic plasticity of intrinsic excitability in CA1 pyramidal neurons in vivo. *eLife*, 8, 2019.

[5990] David W Morgens, Charlene Chan, Andrew J Kane, Nicholas R Weir, Amy Li, Michael M Dubreuil, C Kimberly Tsui, Gaelen T Hess, Adam Lavertu, Kyuho Han, Nicole Polyakov, Jing Zhou, Emma L Handy, Philip Alabi, Amanda Dombroski, David Yao, Russ B Altman, Jason K Sello, Vladimir Denic, and Michael C Bassik. Retro-2 protects cells from ricin toxicity by inhibiting ASNA1-mediated ER targeting and insertion of tail-anchored proteins. *eLife*, 8, 2019.

[5991] Travis J Morgenstern, Jinseo Park, Qing R Fan, and Henry M Colecraft. A potent voltage-gated calcium channel inhibitor engineered from a nanobody targeted to auxiliary CaV$\beta$ subunits. *eLife*, 8, 2019.

[5992] Andrew B Morgenthaler, Wallis R Kinney, Christopher C Ebmeier, Corinne M Walsh, Daniel J Snyder, Vaughn S Cooper, William M Old, and Shelley D Copley. Mutations that improve efficiency of a weak-link enzyme are rare compared to adaptive mutations elsewhere in the genome. *eLife*, 8, 2019.

[5993] Hideaki Morishita, Yan G Zhao, Norito Tamura, Taki Nishimura, Yuki Kanda, Yuriko Sakamaki, Mitsuyo Okazaki, Dongfang Li, and Noboru Mizushima. A critical role of VMP1 in lipoprotein secretion. *eLife*, 8, 2019.

[5994] Nadege Morisot, Khanhky Phamluong, Yann Ehinger, Anthony L Berger, Jeffrey J Moffat, and Dorit Ron. mTORC1 in the orbitofrontal cortex promotes habitual alcohol seeking. *eLife*, 8, 2019.

[5995] Angel Morrow, Julie Underwood, Lindsey Seldin, Taylor Hinnant, and Terry Lechler. Regulated spindle orientation buffers tissue growth in the epidermis. *eLife*, 8, 2019.

[5996] Michael Morse, M Nabuan Naufer, Yuqing Feng, Linda Chelico, Ioulia Rouzina, and Mark C Williams. HIV restriction factor APOBEC3G binds in multiple steps and conformations to search and deaminate single-stranded DNA. *eLife*, 8, 2019.

[5997] Kishore R Mosaliganti, Ian A Swinburne, Chon U Chan, Nikolaus D Obholzer, Amelia A Green, Shreyas Tanksale, L Mahadevan, and Sean G Megason. Size control of the inner ear via hydraulic feedback. *eLife*, 8, 2019.

[5998] Noga Mosheiff and Yoram Burak. Velocity coupling of grid cell modules enables stable embedding of a low dimensional variable in a high dimensional neural attractor. *eLife*, 8, 2019.

[5999] Shannon N Mostyn, Katie A Wilson, Alexandra Schumann-Gillett, Zachary J Frangos, Susan Shimmon, Tristan Rawling, Renae M Ryan, Megan L O'Mara, and Robert J Vandenberg. Identification of an allosteric binding site on the human glycine transporter, GlyT2, for bioactive lipid analgesics. *eLife*, 8, 2019.

[6000] Christopher W Mount, Belgin YalÇın, Kennedy Cunliffe-Koehler, Shree Sundaresh, and Michelle Monje. Monosynaptic tracing maps brain-wide afferent oligodendrocyte precursor cell connectivity. *eLife*, 8, 2019.

[6001] Tyler Chistopher Moyer and Andrew Jon Holland. PLK4 promotes centriole duplication by phosphorylating STIL to link the procentriole cartwheel to the microtubule wall. *eLife*, 8, 2019.

[6002] Harriet Mpairwe, Milly Namutebi, Gyaviira Nkurunungi, Pius Tumwesige, Irene Nambuya, Mike Mukasa, Caroline Onen, Marble Nnaluwooza, Barbara Apule, Tonny Katongole, Gloria Oduru, Joseph Kahwa, Emily L Webb, Lawrence Lubyayi, Neil Pearce, and Alison M Elliott. Risk factors for asthma among schoolchildren who participated in a case-control study in urban Uganda. *eLife*, 8, 2019.

[6003] Elizabeth A Mueller, Alexander JF Egan, Eefjan Breukink, Waldemar Vollmer, and Petra Anne Levin. Plasticity of Escherichia coli cell wall metabolism promotes fitness and antibiotic resistance across environmental conditions. *eLife*, 8, 2019.

[6004] Alexander Mühleip, Sarah E McComas, and Alexey Amunts. Structure of a mitochondrial ATP synthase with bound native cardiolipin. *eLife*, 8, 2019.

[6005] Narendra Mukherjee, Joseph Wachutka, and Donald B Katz. Impact of precisely-timed inhibition of gustatory cortex on taste behavior depends on single-trial ensemble dynamics. *eLife*, 8, 2019.

[6006] Timothy H Muller, Rogier B Mars, Timothy E Behrens, and Jill X O'Reilly. Control of entropy in neural models of environmental state. *eLife*, 8, 2019.

[6007] Jennifer Munkley, Ling Li, S R Gokul Krishnan, Gerald Hysenaj, Emma Scott, Caroline Dalgliesh, Htoo Zarni Oo, Teresa Mendes Maia, Kathleen Cheung, Ingrid Ehrmann, Karen E Livermore, Hanna Zielinska, Oliver Thompson, Bridget Knight, Paul McCullagh, John McGrath, Malcolm Crundwell, Lorna W Harries, Mads Daugaard, Simon Cockell, Nuno L Barbosa-Morais, Sebastian Oltean, and David J Elliott. Androgen-regulated transcription of ESRP2 drives alternative splicing patterns in prostate cancer. *eLife*, 8, 2019.

[6008] David AD Munro, Yishay Wineberg, Julia Tarnick, Chris S Vink, Zhuan Li, Clare Pridans, Elaine Dzierzak, Tomer Kalisky, Peter Hohenstein, and Jamie A Davies. Macrophages restrict the nephrogenic field and promote endothelial connections during kidney development. *eLife*, 8, 2019.

[6009] Sachiko Murase, Dan Winkowski, Ji Liu, Patrick O Kanold, and Elizabeth M Quinlan. Homeostatic regulation of perisynaptic matrix metalloproteinase 9 (MMP9) activity in the amblyopic visual cortex. *eLife*, 8, 2019.

[6010] Daniel P Murphy, Andrew EO Hughes, Karen A Lawrence, Connie A Myers, and Joseph C Corbo. Cis-regulatory basis of sister cell type divergence in the vertebrate retina. *eLife*, 8, 2019.

[6011] Malia SQ Murphy, Steven Hawken, Wei Cheng, Lindsay A Wilson, Monica Lamoureux, Matthew Henderson, Jesmin Pervin, Azad Chowdhury, Courtney Gravett, Eve Lackritz, Beth K Potter, Mark

Walker, Julian Little, Anisur Rahman, Pranesh Chakraborty, and Kumanan Wilson. External validation of postnatal gestational age estimation using newborn metabolic profiles in Matlab, Bangladesh. *eLife*, 8, 2019.

[6012] Pierre-Yves Musso, Pierre Junca, Meghan Jelen, Damian Feldman-Kiss, Han Zhang, Rachel CW Chan, and Michael D Gordon. Closed-loop optogenetic activation of peripheral or central neurons modulates feeding in freely moving Drosophila. *eLife*, 8, 2019.

[6013] José Muñoz-Dorado, Aurelio Moraleda-Muñoz, Francisco Javier Marcos-Torres, Francisco Javier Contreras-Moreno, Ana Belen Martin-Cuadrado, Jared M Schrader, Penelope I Higgs, and Juana Pérez. Transcriptome dynamics of the Myxococcus xanthus multicellular developmental program. *eLife*, 8, 2019.

[6014] Sergio A Muñoz-Gómez, Sebastian Hess, Gertraud Burger, B Franz Lang, Edward Susko, Claudio H Slamovits, and Andrew J Roger. An updated phylogeny of the Alphaproteobacteria reveals that the parasitic Rickettsiales and Holosporales have independent origins. *eLife*, 8, 2019.

[6015] Adi Naamati, James C Williamson, Edward JD Greenwood, Sara Marelli, Paul J Lehner, and Nicholas J Matheson. Functional proteomic atlas of HIV infection in primary human CD4+ T cells. *eLife*, 8, 2019.

[6016] Jessica K Nadalin, Louis-Emmanuel Martinet, Ethan B Blackwood, Meng-Chen Lo, Alik S Widge, Sydney S Cash, Uri T Eden, and Mark A Kramer. A statistical framework to assess cross-frequency coupling while accounting for confounding analysis effects. *eLife*, 8, 2019.

[6017] Tomoki Naito, Bilge Ercan, Logesvaran Krshnan, Alexander Triebl, Dylan Hong Zheng Koh, Fan-Yan Wei, Kazuhito Tomizawa, Federico Tesio Torta, Markus R Wenk, and Yasunori Saheki. Movement of accessible plasma membrane cholesterol by the GRAMD1 lipid transfer protein complex. *eLife*, 8, 2019.

[6018] Alexander Naka, Julia Veit, Ben Shababo, Rebecca K Chance, Davide Risso, David Stafford, Benjamin Snyder, Andrew Egladyous, Desiree Chu, Savitha Sridharan, Daniel P Mossing, Liam Paninski, John Ngai, and Hillel Adesnik. Complementary networks of cortical somatostatin interneurons enforce layer specific control. *eLife*, 8, 2019.

[6019] Yoki Nakamura, Dilyan I Dryanovski, Yuriko Kimura, Shelley N Jackson, Amina S Woods, Yuko Yasui, Shang-Yi Tsai, Sachin Patel, Daniel P Covey, Tsung-Ping Su, and Carl R Lupica. Cocaine-induced endocannabinoid signaling mediated by sigma-1 receptors and extracellular vesicle secretion. *eLife*, 8, 2019.

[6020] Heejin Nam, Shin Jeon, Hyejin An, Jaeyoung Yoo, Hyo-Jong Lee, Soo-Kyung Lee, and Seunghee Lee. Critical roles of ARHGAP36 as a signal transduction mediator of Shh pathway in lateral motor columnar specification. *eLife*, 8, 2019.

[6021] Nagarajan Nandagopal, Leah A Santat, and Michael B Elowitz. Cis-activation in the Notch signaling pathway. *eLife*, 8, 2019.

[6022] Satish K Nandakumar, Sean K McFarland, Laura M Mateyka, Caleb A Lareau, Jacob C Ulirsch, Leif S Ludwig, Gaurav Agarwal, Jesse M Engreitz, Bartlomiej Przychodzen, Marie McConkey, Glenn S Cowley, John G Doench, Jaroslaw P Maciejewski, Benjamin L Ebert, David E Root, and Vijay G Sankaran. Gene-centric functional dissection of human genetic variation uncovers regulators of hematopoiesis. *eLife*, 8, 2019.

[6023] Anirvan Nandy, Jonathan J Nassi, Monika P Jadi, and John Reynolds. Optogenetically induced low-frequency correlations impair perception. *eLife*, 8, 2019.

[6024] Rashmi Nanjundappa, Dong Kong, Kyuhwan Shim, Tim Stearns, Steven L Brody, Jadranka Loncarek, and Moe R Mahjoub. Regulation of cilia abundance in multiciliated cells. *eLife*, 8, 2019.

[6025] Arjun Narayanan, Anatoli Meriin, J Owen Andrews, Jan-Hendrik Spille, Michael Y Sherman, and Ibrahim I Cisse. A first order phase transition mechanism underlies protein aggregation in mammalian cells. *eLife*, 8, 2019.

[6026] Craig A Nash, Wenhui Wei, Roshanak Irannejad, and Alan V Smrcka. Golgi localized $\beta$1-adrenergic receptors stimulate Golgi PI4P hydrolysis by PLC$\epsilon$ to regulate cardiac hypertrophy. *eLife*, 8, 2019.

[6027] Matthew R Nassar, Rasmus Bruckner, and Michael J Frank. Statistical context dictates the relationship between feedback-related EEG signals and learning. *eLife*, 8, 2019.

[6028] Evgeniya V Nazarova, Christine R Montague, Lu Huang, Thuy La, David Russell, and Brian C VanderVen. The genetic requirements of fatty acid import by Mycobacterium tuberculosis within macrophages. *eLife*, 8, 2019.

[6029] Camilla Nehammer, Patrick Ejlerskov, Sandeep Gopal, Ava Handley, Leelee Ng, Pedro Moreira, Huikyong Lee, Shohreh Issazadeh-Navikas, David C Rubinsztein, and Roger Pocock. Interferon-$\beta$-induced miR-1 alleviates toxic protein accumulation by controlling autophagy. *eLife*, 8, 2019.

[6030] Rebecca L Newcomer, Jason R Schrad, Eddie B Gilcrease, Sherwood R Casjens, Michael Feig, Carolyn M Teschke, Andrei T Alexandrescu, and Kristin N Parent. The phage L capsid decoration protein has a novel OB-fold and an unusual capsid binding strategy. *eLife*, 8, 2019.

[6031] Andrew Ying Hui Ng, Ziqing Li, Megan M Jones, Shuting Yang, Chunyi Li, Chuanyun Fu, Chengjian Tu, Merry Jo Oursler, Jun Qu, and Shuying Yang. Regulator of G protein signaling 12 enhances osteoclastogenesis by suppressing Nrf2-dependent antioxidant proteins to promote the generation of reactive oxygen species. *eLife*, 8, 2019.

[6032] Kevin W Ng, Jan Attig, George R Young, Eleonora Ottina, Spyros I Papamichos, Ioannis Kotsianidis, and George Kassiotis. Soluble PD-L1 generated by endogenous retroelement exaptation is a receptor antagonist. *eLife*, 8, 2019.

[6033] Ashley M Ngo, Matthew J Shurtleff, Katerina D Popova, Jessie Kulsuptrakul, Jonathan S Weissman, and Andreas S Puschnik. The ER membrane protein complex is required to ensure correct topology and stable expression of flavivirus polyproteins. *eLife*, 8, 2019.

[6034] Huy Quoc Nguyen, Jagoree Roy, Björn Harink, Nikhil P Damle, Naomi R Latorraca, Brian C Baxter, Kara Brower, Scott A Longwell, Tanja Kortemme, Kurt S Thorn, Martha S Cyert, and Polly Morrell Fordyce. Quantitative mapping of protein-peptide affinity landscapes using spectrally encoded beads. *eLife*, 8, 2019.

[6035] Minh Q Nguyen, Claire E Le Pichon, and Nicholas Ryba. Stereotyped transcriptomic transformation of somatosensory neurons in response to injury. *eLife*, 8, 2019.

[6036] Jinfei D Ni, Adishthi S Gurav, Weiwei Liu, Tyler H Ogunmowo, Hannah Hackbart, Ahmed Elsheikh, Andrew A Verdegaal, and Craig Montell. Differential regulation of the Drosophila sleep homeostat by circadian and arousal inputs. *eLife*, 8, 2019.

[6037] Isabel C Nimmo, Adrian C Barbrook, Imen Lassadi, Jit Ern Chen, Katrin Geisler, Alison G Smith, Manuel Aranda, Saul Purton, Ross F Waller, R Ellen R Nisbet, and Christopher J Howe. Genetic transformation of the dinoflagellate chloroplast. *eLife*, 8, 2019.

[6038] Luis V Nobre, Katie Nightingale, Benjamin J Ravenhill, Robin Antrobus, Lior Soday, Jenna Nichols, James A Davies, Sepehr Seirafian, Eddie CY Wang, Andrew J Davison, Gavin WG Wilkinson, Richard J Stanton, Edward L Huttlin, and Michael P Weekes. Human cytomegalovirus interactome analysis identifies degradation hubs, domain associations and viral protein functions. *eLife*, 8, 2019.

[6039] Hirofumi Noguchi, Jesse Garcia Castillo, Kinichi Nakashima, and Samuel J Pleasure. Suppressor of fused controls perinatal expansion and quiescence of future dentate adult neural stem cells. *eLife*, 8, 2019.

[6040] Jordan N Norwood, Qingguang Zhang, David Card, Amanda Craine, Timothy M Ryan, and Patrick J Drew. Anatomical basis and physiological role of cerebrospinal fluid transport through the murine cribriform plate. *eLife*, 8, 2019.

[6041] Matthew M Nour, Tarik Dahoun, Robert A McCutcheon, Rick A Adams, Matthew B Wall, and Oliver D Howes. Task-induced functional brain connectivity mediates the relationship between striatal D2/3 receptors and working memory. *eLife*, 8, 2019.

[6042] Melesse Nune, Michael T Morgan, Zaily Connell, Laura McCullough, Muhammad Jbara, Hao Sun, Ashraf Brik, Tim Formosa, and Cynthia Wolberger. FACT and Ubp10 collaborate to modulate H2B deubiquitination and nucleosome dynamics. *eLife*, 8, 2019.

[6043] Natasha M O'Brown, Sean G Megason, and Chenghua Gu. Suppression of transcytosis regulates zebrafish blood-brain barrier function. *eLife*, 8, 2019.

[6044] Bernadette O'Donovan, Adewale Adeluyi, Erin L Anderson, Robert D Cole, Jill R Turner, and Pavel I Ortinski. Altered gating of Kv1.4 in the nucleus accumbens suppresses motivation for reward. *eLife*, 8, 2019.

[6045] Hyung Suk Oh, Werner M Neuhausser, Pierce Eggan, Magdalena Angelova, Rory Kirchner, Kevin C Eggan, and David M Knipe. Herpesviral lytic gene functions render the viral genome susceptible to novel editing by CRISPR/Cas9. *eLife*, 8, 2019.

[6046] Nikola Ojkic, Diana Serbanescu, and Shiladitya Banerjee. Surface-to-volume scaling and aspect ratio preservation in rod-shaped bacteria. *eLife*, 8, 2019.

[6047] Masahiro Oka, Sonoko Mura, Mayumi Otani, Yoichi Miyamoto, Jumpei Nogami, Kazumitsu Maehara, Akihito Harada, Taro Tachibana, Yoshihiro Yoneda, and Yasuyuki Ohkawa. Chromatin-bound CRM1 recruits SET-Nup214 and NPM1c onto HOX clusters causing aberrant HOX expression in leukemia cells. *eLife*, 8, 2019.

[6048] Samora Okujeni and Ulrich Egert. Self-organization of modular network architecture by activity-dependent neuronal migration and outgrowth. *eLife*, 8, 2019.

[6049] Scott W Olesen, Marc Lipsitch, and Yonatan H Grad. Response to comment on 'The distribution of antibiotic use and its association with antibiotic resistance'. *eLife*, 8, 2019.

[6050] Chet Huan Oon and Kenneth E Prehoda. Asymmetric recruitment and actin-dependent cortical flows drive the neuroblast polarity cycle. *eLife*, 8, 2019.

[6051] Eduard Ort, Johannes Jacobus Fahrenfort, Tuomas ten Cate, Martin Eimer, and Christian NL Olivers. Humans can efficiently look for but not select multiple visual objects. *eLife*, 8, 2019.

[6052] Fabian E Ortega, Elena F Koslover, and Julie A Theriot. Listeria monocytogenes cell-to-cell spread in epithelia is heterogeneous and dominated by rare pioneer bacteria. *eLife*, 8, 2019.

[6053] Linnaea E Ostroff, Emanuela Santini, Robert Sears, Zachary Deane, Rahul N Kanadia, Joseph E LeDoux, Tenzin Lhakhang, Aristotelis Tsirigos, Adriana Heguy, and Eric Klann. Axon TRAP reveals learning-associated alterations in cortical axonal mRNAs in the lateral amygdala. *eLife*, 8, 2019.

[6054] Adriane G Otopalik, Jason Pipkin, and Eve Marder. Neuronal morphologies built for reliable physiology in a rhythmic motor circuit. *eLife*, 8, 2019.

[6055] Jeroen Overman, Frank Fontaine, Jill Wylie-Sears, Mehdi Moustaqil, Lan Huang, Marie Meurer, Ivy Kim Chiang, Emmanuelle Lesieur, Jatin Patel, Johannes Zuegg, Eddy Pasquier, Emma Sierecki, Yann Gambin, Mohamed Hamdan, Kiarash Khosrotehrani, Gregor Andelfinger, Joyce Bischoff, and Mathias Francois. R-propranolol is a small molecule inhibitor of the SOX18 transcription factor in a rare vascular syndrome and hemangioma. *eLife*, 8, 2019.

[6056] Allison M Owen, Samir P Patel, Jeffrey D Smith, Beverly K Balasuriya, Stephanie F Mori, Gregory S Hawk, Arnold J Stromberg, Naohide Kuriyama, Masao Kaneki, Alexander G Rabchevsky, Timothy A Butterfield, Karyn A Esser, Charlotte A Peterson, Marlene E Starr, and Hiroshi Saito. Chronic muscle weakness and mitochondrial dysfunction in the absence of sustained atrophy in a preclinical sepsis model. *eLife*, 8, 2019.

[6057] Nick Owens, Thaleia Papadopoulou, Nicola Festuccia, Alexandra Tachtsidi, Inma Gonzalez, Agnes Dubois, Sandrine Vandormael-Pournin, Elphège P Nora, Benoit G Bruneau, Michel Cohen-Tannoudji, and Pablo Navarro. CTCF confers local nucleosome resiliency after DNA replication and during mitosis. *eLife*, 8, 2019.

[6058] Maurizio Pacifici and Eileen M Shore. Comment on 'Palovarotene reduces heterotopic ossification in juvenile FOP mice but exhibits pronounced skeletal toxicity'. *eLife*, 8, 2019.

[6059] Abhilash Padavannil, Prithwijit Sarkar, Seung Joong Kim, Tolga Cagatay, Jenny Jiou, Chad A Brautigam, Diana R Tomchick, Andrej Sali, Sheena D'Arcy, and Yuh Min Chook. Importin-9 wraps around the H2A-H2B core to act as nuclear importer and histone chaperone. *eLife*, 8, 2019.

[6060] Pranesh Padmanabhan, Ramón Martínez-Mármol, Di Xia, Jürgen Götz, and Frédéric A Meunier. Frontotemporal dementia mutant Tau promotes aberrant Fyn nanoclustering in hippocampal dendritic spines. *eLife*, 8, 2019.

[6061] Petar Pajic, Pavlos Pavlidis, Kirsten Dean, Lubov Neznanova, Rose-Anne Romano, Danielle Garneau, Erin Daugherity, Anja Globig, Stefan Ruhl, and Omer Gokcumen. Independent amylase gene copy number bursts correlate with dietary preferences in mammals. *eLife*, 8, 2019.

[6062] Sander AL Palit, Daniel Vis, Suzan Stelloo, Cor Lieftink, Stefan Prekovic, Elise Bekers, Ingrid Hofland, Tonći ŠuŠtić, Liesanne Wolters, Roderick Beijersbergen, Andries M Bergman, Balázs Győrffy, Lodewyk FA Wessels, Wilbert Zwart, and Michiel S van der Heijden. TLE3 loss confers AR inhibitor resistance by facilitating GR-mediated human prostate cancer cell growth. *eLife*, 8, 2019.

[6063] Fabian Pallasdies, Sven Goedeke, Wilhelm Braun, and Raoul-Martin Memmesheimer. From single neurons to behavior in the jellyfish Aurelia aurita. *eLife*, 8, 2019.

[6064] Adam C Palmer, Christopher Chidley, and Peter K Sorger. A curative combination cancer therapy achieves high fractional cell killing through low cross-resistance and drug additivity. *eLife*, 8, 2019.

[6065] Marta Palomo-Guerrero, Rut Fadó, Maria Casas, Marta Pérez-Montero, Miguel Baena, Patrick O Helmer, José Luis Domínguez, Aina Roig, Dolors Serra, Heiko Hayen, Harald Stenmark, Camilla Raiborg, and Núria Casals. Sensing of nutrients by CPT1C regulates late endosome/lysosome anterograde transport and axon growth. *eLife*, 8, 2019.

[6066] Ting Pan, Zheng Song, Liyang Wu, Guangyan Liu, Xiancai Ma, Zhilin Peng, Mo Zhou, Liting Liang, Bingfeng Liu, Jun Liu, Junsong Zhang, Xuanhong Zhang, Ryan Huang, Jiacong Zhao, Yonghong Li, Xuemei Ling, Yuewen Luo, Xiaoping Tang, Weiping Cai, Kai Deng, Linghua Li, and Hui Zhang. USP49 potently stabilizes APOBEC3G protein by removing ubiquitin and inhibits HIV-1 replication. *eLife*, 8, 2019.

[6067] Georgia Panagiotakos, Christos Haveles, Arpana Arjun, Ralitsa Petrova, Anshul Rana, Thomas Portmann, Sergiu P Paşca, Theo D Palmer, and Ricardo E Dolmetsch. Aberrant calcium channel splicing drives defects in cortical differentiation in Timothy syndrome. *eLife*, 8, 2019.

[6068] Rich Pang and Adrienne L Fairhall. Fast and flexible sequence induction in spiking neural networks via rapid excitability changes. *eLife*, 8, 2019.

[6069] Ferenc Papp, Suvendu Lomash, Orsolya Szilagyi, Erika Babikow, Jaime Smith, Tsg-Hui Chang, Maria Isabel Bahamonde, Gilman Ewan Stephen Toombes, and Kenton Jon Swartz. TMEM266 is a functional voltage sensor regulated by extracellular Zn2+. *eLife*, 8, 2019.

[6070] Casey Paquola, Richard AI Bethlehem, Jakob Seidlitz, Konrad Wagstyl, Rafael Romero-Garcia, Kirstie J Whitaker, Reinder Vos de Wael, Guy B Williams, Petra E Vértes, Daniel S Margulies, Boris Bernhardt, and Edward T Bullmore. Shifts in myeloarchitecture characterise adolescent development of cortical gradients. *eLife*, 8, 2019.

[6071] Chung Park and John H Kehrl. An integrin/MFG-E8 shuttle loads HIV-1 viral-like particles onto follicular dendritic cells in mouse lymph node. *eLife*, 8, 2019.

[6072] Eugene Park, Swapneel Patel, Qiuling Wang, Prabhakar Andhey, Konstantin Zaitsev, Sophia Porter, Maxwell Hershey, Michael Bern, Beatrice Plougastel-Douglas, Patrick Collins, Marco Colonna, Kenneth M Murphy, Eugene Oltz, Maxim Artyomov, L David Sibley, and Wayne M Yokoyama. Toxoplasma gondii infection drives conversion of NK cells into ILC1-like cells. *eLife*, 8, 2019.

[6073] Hame Park and Christoph Kayser. Shared neural underpinnings of multisensory integration and trial-by-trial perceptual recalibration in humans. *eLife*, 8, 2019.

[6074] Jeehae Park, Javier Estrada, Gemma Johnson, Ben J Vincent, Chiara Ricci-Tam, Meghan DJ Bragdon, Yekaterina Shulgina, Anna Cha, Zeba Wunderlich, Jeremy Gunawardena, and Angela H DePace. Dissecting the sharp response of a canonical developmental enhancer reveals multiple sources of cooperativity. *eLife*, 8, 2019.

[6075] Jooyoung Park, Brinda Selvaraj, Andrew C McShan, Scott E Boyken, Kathy Y Wei, Gustav Oberdorfer, William DeGrado, Nikolaos G Sgourakis, Matthew J Cuneo, Dean AA Myles, and David Baker. De novo design of a homo-trimeric amantadine-binding protein. *eLife*, 8, 2019.

[6076] Myo-Hyeon Park, Ae kyung Kim, Sarala Manandhar, Su-Young Oh, Gun-Hyuk Jang, Li Kang, Dong-Won Lee, Do Young Hyeon, Sun-Hee Lee, Hye Eun Lee, Tae-Lin Huh, Sang Heon Suh, Daehee Hwang, Kyunghee Byun, Hae-Chul Park, and You Mie Lee. CCN1 interlinks integrin and hippo pathway to autoregulate tip cell activity. *eLife*, 8, 2019.

[6077] Daniel M Parker, Sai Thein Than Tun, Lisa J White, Ladda Kajeechiwa, May Myo Thwin, Jordi Landier, Victor Chaumeau, Vincent Corbel, Arjen M Dondorp, Lorenz von Seidlein, Nicholas J White, Richard J Maude, and FranÇois Nosten. Potential herd protection against Plasmodium falciparum infections conferred by mass antimalarial drug administrations. *eLife*, 8, 2019.

[6078] Margaret M Parker, Yuan Hao, Feng Guo, Betty Pham, Robert Chase, John Platig, Michael H Cho, Craig P Hersh, Victor J Thannickal, James Crapo, George Washko, Scott H Randell, Edwin K Silverman, Raúl San José Estépar, Xiaobo Zhou, and Peter J Castaldi. Identification of an emphysema-associated genetic variant near TGFB2 with regulatory effects in lung fibroblasts. *eLife*, 8, 2019.

[6079] Matthew W Parker, Maren Bell, Mustafa Mir, Jonchee A Kao, Xavier Darzacq, Michael R Botchan, and James M Berger. A new class of disordered elements controls DNA replication through initiator self-assembly. *eLife*, 8, 2019.

[6080] Edward A Partlow, Richard W Baker, Gwendolyn M Beacham, Joshua S Chappie, Andres E Leschziner, and Gunther Hollopeter. A structural mechanism for phosphorylation-dependent inactivation of the AP2 complex. *eLife*, 8, 2019.

[6081] Jean-Philippe Parvy, Yachuan Yu, Anna Dostalova, Shu Kondo, Alina Kurjan, Philippe Bulet, Bruno Lemaître, Marcos Vidal, and Julia B Cordero. The antimicrobial peptide defensin cooperates with tumour necrosis factor to drive tumour cell death in Drosophila. *eLife*, 8, 2019.

[6082] Avinash B Patel, Camille M Moore, Basil J Greber, Jie Luo, Stefan A Zukin, Jeff Ranish, and Eva Nogales. Architecture of the chromatin remodeler RSC and insights into its nucleosome engagement. *eLife*, 8, 2019.

[6083] Jay M Patel, Jessica Swanson, Kevin Ung, Alexander Herman, Elizabeth Hanson, Joshua Ortiz-Guzman, Jennifer Selever, Qingchun Tong, and Benjamin R Arenkiel. Sensory perception drives food avoidance through excitatory basal forebrain circuits. *eLife*, 8, 2019.

[6084] M Görkem Patir-Nebioglu, Zaida Andrés, Melanie Krebs, Fabian Fink, Katarzyna Drzewicka, Nicolas Stankovic-Valentin, Shoji Segami, Sebastian Schuck, Michael Büttner, Rüdiger Hell, Masayoshi Maeshima, Frauke Melchior, and Karin Schumacher. RETRACTED: Pyrophosphate modulates plant stress responses via SUMOylation. *eLife*, 8, 2019.

[6085] Hannah L Payne, Ranran L French, Christine C Guo, TD Barbara Nguyen-Vu, Tiina Manninen, and Jennifer L Raymond. Cerebellar Purkinje cells control eye movements with a rapid rate code that is invariant to spike irregularity. *eLife*, 8, 2019.

[6086] Matthew F Pech, Linda E Fong, Jacqueline E Villalta, Leanne JG Chan, Samir Kharbanda, Jonathon J O'Brien, Fiona E McAllister, Ari J Firestone, Calvin H Jan, and Jeffrey Settleman. Systematic identification of cancer cell vulnerabilities to natural killer cell-mediated immune surveillance. *eLife*, 8, 2019.

[6087] Michael Peer, Yorai Ron, Rotem Monsa, and Shahar Arzy. Processing of different spatial scales in the human brain. *eLife*, 8, 2019.

[6088] Xue Yuan Pei, Tom Dendooven, Elisabeth Sonnleitner, Shaoxia Chen, Udo Bläsi, and Ben F Luisi. Architectural principles for Hfq/Crc-mediated regulation of gene expression. *eLife*, 8, 2019.

[6089] Rini H Pek, Xiaojing Yuan, Nicole Rietzschel, Jianbing Zhang, Laurie Jackson, Eiji Nishibori, Ana Ribeiro, William Simmons, Jaya Jagadeesh, Hiroshi Sugimoto, Md Zahidul Alam, Lisa Garrett, Malay Haldar, Martina Ralle, John D Phillips, David M Bodine, and Iqbal Hamza. Hemozoin produced by mammals confers heme tolerance. *eLife*, 8, 2019.

[6090] Ma Teresa Pellicer Martinez, Jason C Crack, Melissa YY Stewart, Justin M Bradley, Dimitri A Svistunenko, Andrew WB Johnston, Myles R Cheesman, Jonathan D Todd, and Nick E Le Brun. Mechanisms of iron- and O2-sensing by the [4Fe-4S] cluster of the global iron regulator RirA. *eLife*, 8, 2019.

[6091] Jhan-Jie Peng, Shih-Han Lin, Yu-Tzu Liu, Hsin-Chieh Lin, Tsai-Ning Li, and Chi-Kuang Yao. A circuit-dependent ROS feedback loop mediates glutamate excitotoxicity to sculpt the Drosophila motor system. *eLife*, 8, 2019.

[6092] Yangfan Peng, Franz Xaver Mittermaier, Henrike Planert, Ulf Christoph Schneider, Henrik Alle, and Jörg Rolf Paul Geiger. High-throughput microcircuit analysis of individual human brains through next-generation multineuron patch-clamp. *eLife*, 8, 2019.

[6093] Alberto E Pereda. Science, art, society and Klimt's University of Vienna paintings. *eLife*, 8, 2019.

[6094] Joana B Pereira, Rik Ossenkoppele, Sebastian Palmqvist, Tor Olof Strandberg, Ruben Smith, Eric Westman, and Oskar Hansson. Amyloid and tau accumulate across distinct spatial networks and are differentially associated with brain connectivity. *eLife*, 8, 2019.

[6095] Laura Pereira, Florian Aeschimann, Chen Wang, Hannah Lawson, Esther Serrano-Saiz, Douglas S Portman, Helge Großhans, and Oliver Hobert. Timing mechanism of sexually dimorphic nervous system differentiation. *eLife*, 8, 2019.

[6096] Himasha M Perera and Michael A Trakselis. Amidst multiple binding orientations on fork DNA, Saccharolobus MCM helicase proceeds N-first for unwinding. *eLife*, 8, 2019.

[6097] Karina Perlaza, Hannah Toutkoushian, Morgane Boone, Mable Lam, Masakazu Iwai, Martin C Jonikas, Peter Walter, and Silvia Ramundo. The Mars1 kinase confers photoprotection through signaling in the chloroplast unfolded protein response. *eLife*, 8, 2019.

[6098] Maria Pernemalm, AnnSofi Sandberg, Yafeng Zhu, Jorrit Boekel, Davide Tamburro, Jochen M Schwenk, Albin Björk, Marie Wahren-Herlenius, Hanna Åmark, Claes-Göran östenson, Magnus Westgren, and Janne Lehtiö. In-depth human plasma proteome analysis captures tissue proteins and transfer of protein variants across the placenta. *eLife*, 8, 2019.

[6099] Jirka Peschek and Peter Walter. tRNA ligase structure reveals kinetic competition between non-conventional mRNA splicing and mRNA decay. *eLife*, 8, 2019.

[6100] Alina Peter, Cem Uran, Johanna Klon-Lipok, Rasmus Roese, Sylvia van Stijn, William Barnes, Jarrod R Dowdall, Wolf Singer, Pascal Fries, and Martin Vinck. Surface color and predictability determine contextual modulation of V1 firing and gamma oscillations. *eLife*, 8, 2019.

[6101] Brian G Peterson, Morgan L Glaser, Tom A Rapoport, and Ryan D Baldridge. Cycles of autoubiquitination and deubiquitination regulate the ERAD ubiquitin ligase Hrd1. *eLife*, 8, 2019.

[6102] Nikolai Petkau, Harun Budak, Xunlei Zhou, Henrik Oster, and Gregor Eichele. Acetylation of BMAL1 by TIP60 controls BRD4-P-TEFb recruitment to circadian promoters. *eLife*, 8, 2019.

[6103] Kasparas Petkevicius, Sam Virtue, Guillaume Bidault, Benjamin Jenkins, Cankut Çubuk, Cecilia Morgantini, Myriam Aouadi, Joaquin Dopazo, Mireille J Serlie, Albert Koulman, and Antonio Vidal-Puig. Accelerated phosphatidylcholine turnover in macrophages promotes adipose tissue inflammation in obesity. *eLife*, 8, 2019.

[6104] Natalia Petrenko, Yi Jin, Liguo Dong, Koon Ho Wong, and Kevin Struhl. Requirements for RNA polymerase II preinitiation complex formation in vivo. *eLife*, 8, 2019.

[6105] Anja Pflug, Florian Gompf, Muthuraman Muthuraman, Sergiu Groppa, and Christian Alexander Kell. Differential contributions of the two human cerebral hemispheres to action timing. *eLife*, 8, 2019.

[6106] Tung D Phan, Jessica A Wachter, Ethan A Solomon, and Michael J Kahana. Multivariate stochastic volatility modeling of neural data. *eLife*, 8, 2019.

[6107] Rachael Laura Philips, Jeong-Heon Lee, Krutika Gaonkar, Pritha Chanana, Ji Young Chung, Sinibaldo R Romero Arocha, Aaron Schwab, Tamas Ordog, and Virginia Smith Shapiro. HDAC3 restrains CD8-lineage genes to maintain a bi-potential state in CD4+CD8+ thymocytes for CD4-lineage commitment. *eLife*, 8, 2019.

[6108] Charles B Phillips, Chen-Wei Tsai, and Ming-Feng Tsai. The conserved aspartate ring of MCU mediates MICU1 binding and regulation in the mitochondrial calcium uniporter complex. *eLife*, 8, 2019.

[6109] Mary L Phillips, Holly Anne Robinson, and Lucas Pozzo-Miller. Ventral hippocampal projections to the medial prefrontal cortex regulate social memory. *eLife*, 8, 2019.

[6110] Ryan S Phillips, Tibin T John, Hidehiko Koizumi, Yaroslav I Molkov, and Jeffrey C Smith. Biophysical mechanisms in the mammalian respiratory oscillator re-examined with a new data-driven computational model. *eLife*, 8, 2019.

[6111] Francesco M Piccolo, Zhe Liu, Peng Dong, Ching-Lung Hsu, Elitsa I Stoyanova, Anjana Rao, Robert Tjian, and Nathaniel Heintz. MeCP2 nuclear dynamics in live neurons results from low and high affinity chromatin interactions. *eLife*, 8, 2019.

[6112] Joseph Pickering, Kavitha Chinnaiya, and Matthew Towers. An autoregulatory cell cycle timer integrates growth and specification in chick wing digit development. *eLife*, 8, 2019.

[6113] Mykola Pinkevych, Christine M Fennessey, Deborah Cromer, Carolyn Reid, Charles M Trubey, Jeffrey D Lifson, Brandon F Keele, and Miles P Davenport. Predictors of SIV recrudescence following antiretroviral treatment interruption. *eLife*, 8, 2019.

[6114] Basile Pinsard, Arnaud Boutin, Ella Gabitov, Ovidiu Lungu, Habib Benali, and Julien Doyon. Consolidation alters motor sequence-specific distributed representations. *eLife*, 8, 2019.

[6115] Benoît Pinson, Johanna Ceschin, Christelle Saint-Marc, and Bertrand Daignan-Fornier. Dual control of NAD+ synthesis by purine metabolites in yeast. *eLife*, 8, 2019.

[6116] Roger Pique-Regi, Roberto Romero, Adi L Tarca, Edward D Sendler, Yi Xu, Valeria Garcia-Flores, Yaozhu Leng, Francesca Luca, Sonia S Hassan, and Nardhy Gomez-Lopez. Single cell transcriptional signatures of the human placenta in term and preterm parturition. *eLife*, 8, 2019.

[6117] Matthew Piva, Kayla Velnoskey, Ruonan Jia, Amrita Nair, Ifat Levy, and Steve WC Chang. The dorsomedial prefrontal cortex computes task-invariant relative subjective value for self and other. *eLife*, 8, 2019.

[6118] Vanessa Plantier, Irene Sanchez-Brualla, Nejada Dingu, Cécile Brocard, Sylvie Liabeuf, Florian Gackière, and Frédéric Brocard. Calpain fosters the hyperexcitability of motoneurons after spinal cord injury and leads to spasticity. *eLife*, 8, 2019.

[6119] Jean-Claude Platel, Alexandra Angelova, Stephane Bugeon, Jenelle Wallace, Thibault Ganay, Ilona Chudotvorova, Jean-Christophe Deloulme, Christophe Béclin, Marie-Catherine Tiveron, Nathalie Coré, Venkatesh N Murthy, and Harold Cremer. Neuronal integration in the adult mouse olfactory bulb is a non-selective addition process. *eLife*, 8, 2019.

[6120] Laure-Anne Poissonnier, Sebastien Motsch, Jacques Gautrais, Camille Buhl, and Audrey Dussutour. Experimental investigation of ant traffic under crowded conditions. *eLife*, 8, 2019.

[6121] Andrey Poleshko, Cheryl L Smith, Son C Nguyen, Priya Sivaramakrishnan, Karen G Wong, John Isaac Murray, Melike Lakadamyali, Eric F Joyce, Rajan Jain, and Jonathan A Epstein. H3K9me2 orchestrates inheritance of spatial positioning of peripheral heterochromatin through mitosis. *eLife*, 8, 2019.

[6122] Denise M Poltavski, Pauline Colombier, Jianxin Hu, Alicia Duron, Brian L Black, and Takako Makita. Venous endothelin modulates responsiveness of cardiac sympathetic axons to arterial semaphorin. *eLife*, 8, 2019.

[6123] Alessandro Porro, Andrea Saponaro, Federica Gasparri, Daniel Bauer, Christine Gross, Matteo Pisoni, Gerardo Abbandonato, Kay Hamacher, Bina Santoro, Gerhard Thiel, and Anna Moroni. The HCN domain couples voltage gating and cAMP response in hyperpolarization-activated cyclic nucleotide-gated channels. *eLife*, 8, 2019.

[6124] Céline Portal, Panteleimos Rompolas, Peter Lwigale, and Carlo Iomini. Primary cilia deficiency in neural crest cells models anterior segment dysgenesis in mouse. *eLife*, 8, 2019.

[6125] Koen B Pouwels, Christopher C Butler, and Julie V Robotham. Comment on 'The distribution of antibiotic use and its association with antibiotic resistance'. *eLife*, 8, 2019.

[6126] Nicole Poweleit, Nadine Czudnochowski, Rachel Nakagawa, Donovan D Trinidad, Kenan C Murphy, Christopher M Sassetti, and Oren S Rosenberg. The structure of the endogenous ESX-3 secretion system. *eLife*, 8, 2019.

[6127] Maria Paz Prada, Arsalan U Syed, Olivia R Buonarati, Gopireddy R Reddy, Matthew A Nystoriak, Debapriya Ghosh, Sergi Simó, Daisuke Sato, Kent C Sasse, Sean M Ward, Luis F Santana, Yang K Xiang, Johannes W Hell, Madeline Nieves-Cintrón, and Manuel F Navedo. A Gs-coupled purinergic receptor boosts Ca2+ influx and vascular contractility during diabetic hyperglycemia. *eLife*, 8, 2019.

[6128] Thomas Pradeu. Immunology and individuality. *eLife*, 8, 2019.

[6129] Meenakshi Prajapati-DiNubila, Ana Benito-Gonzalez, Erin Jennifer Golden, Shuran Zhang, and Angelika Doetzlhofer. A counter gradient of Activin A and follistatin instructs the timing of hair cell differentiation in the murine cochlea. *eLife*, 8, 2019.

[6130] Thomas Cody Prang. The African ape-like foot of Ardipithecus ramidus and its implications for the origin of bipedalism. *eLife*, 8, 2019.

[6131] Asher Preska Steinberg, Sujit S Datta, Thomas Naragon, Justin C Rolando, Said R Bogatyrev, and Rustem F Ismagilov. High-molecular-weight polymers from dietary fiber drive aggregation of particulates in the murine small intestine. *eLife*, 8, 2019.

[6132] Eric A Prinslow, Karolina P Stepien, Yun-Zu Pan, Junjie Xu, and Josep Rizo. Multiple factors maintain assembled trans-SNARE complexes in the presence of NSF and $\alpha$SNAP. *eLife*, 8, 2019.

[6133] Thomas AA Prowse, Fatwa Adikusuma, Phillip Cassey, Paul Thomas, and Joshua V Ross. A Y-chromosome shredding gene drive for controlling pest vertebrate populations. *eLife*, 8, 2019.

[6134] Maria A Prusicki, Emma M Keizer, Rik P van Rosmalen, Shinichiro Komaki, Felix Seifert, Katja Müller, Erik Wijnker, Christian Fleck, and Arp Schnittger. Live cell imaging of meiosis in Arabidopsis thaliana. *eLife*, 8, 2019.

[6135] Avinash Pujala and Minoru Koyama. Chronology-based architecture of descending circuits that underlie the development of locomotor repertoire after birth. *eLife*, 8, 2019.

[6136] Michael Puljung, Natascia Vedovato, Samuel Usher, and Frances Ashcroft. Activation mechanism of ATP-sensitive K+ channels explored with real-time nucleotide binding. *eLife*, 8, 2019.

[6137] Mia C Pulos-Holmes, Daniel N Srole, Maria G Juarez, Amy S-Y Lee, David T McSwiggen, Nicholas T Ingolia, and Jamie H Cate. Repression of ferritin light chain translation by human eIF3. *eLife*, 8, 2019.

[6138] Ruth A Pumroy, Amrita Samanta, Yuhang Liu, Taylor ET Hughes, Siyuan Zhao, Yevgen Yudin, Tibor Rohacs, Seungil Han, and Vera Y Moiseenkova-Bell. Molecular mechanism of TRPV2 channel modulation by cannabidiol. *eLife*, 8, 2019.

[6139] Sruthi Purushothaman, Ahmed Elewa, and Ashley W Seifert. Fgf-signaling is compartmentalized within the mesenchyme and controls proliferation during salamander limb development. *eLife*, 8, 2019.

[6140] Yumin Qi, Yang Wu, Randy Saunders, Xiao-Guang Chen, Chunhong Mao, James Kite Biedler, and Zhijian Jake Tu. Guy1, a Y-linked embryonic signal, regulates dosage compensation in Anopheles stephensi by increasing X gene expression. *eLife*, 8, 2019.

[6141] Chen Qiu, Vandita D Bhat, Sanjana Rajeev, Chi Zhang, Alexa E Lasley, Robert N Wine, Zachary T Campbell, and Traci M Tanaka Hall. A crystal structure of a collaborative RNA regulatory complex reveals mechanisms to refine target specificity. *eLife*, 8, 2019.

[6142] Yue Qu, Ines Hahn, Meredith Lees, Jill Parkin, André Voelzmann, Karel Dorey, Alex Rathbone, Claire T Friel, Victoria J Allan, Pilar Okenve-Ramos, Natalia Sanchez-Soriano, and Andreas Prokop. Efa6 protects axons and regulates their growth and branching by inhibiting microtubule polymerisation at the cortex. *eLife*, 8, 2019.

[6143] Bradley Quade, Marcial Camacho, Xiaowei Zhao, Marta Orlando, Thorsten Trimbuch, Junjie Xu, Wei Li, Daniela Nicastro, Christian Rosenmund, and Josep Rizo. Membrane bridging by Munc13-1 is crucial for neurotransmitter release. *eLife*, 8, 2019.

[6144] Claudia Racioppi, Keira A Wiechecki, and Lionel Christiaen. Combinatorial chromatin dynamics foster accurate cardiopharyngeal fate choices. *eLife*, 8, 2019.

[6145] Nils Rademacher, Benno Kuropka, Stella-Amrei Kunde, Markus C Wahl, Christian Freund, and Sarah A Shoichet. Intramolecular domain dynamics regulate synaptic MAGUK protein interactions. *eLife*, 8, 2019.

[6146] Arunas L Radzvilavicius, Alexander J Stewart, and Joshua B Plotkin. Evolution of empathetic moral evaluation. *eLife*, 8, 2019.

[6147] Atiqur Rahman, Katherine M Henry, Kimberly D Herman, Alfred AR Thompson, Hannah M Isles, Claudia Tulotta, David Sammut, Julien JY Rougeot, Nika Khoshaein, Abigail E Reese, Kathryn Higgins, Caroline Tabor, Ian Sabroe, William J Zuercher, Caroline O Savage, Annemarie H Meijer, Moira KB Whyte, David H Dockrell, Stephen A Renshaw, and Lynne R Prince. Inhibition of ErbB kinase signalling promotes resolution of neutrophilic inflammation. *eLife*, 8, 2019.

[6148] Mathan K Raja, Julia Preobraschenski, Sergio Del Olmo-Cabrera, Rebeca Martinez-Turrillas, Reinhard Jahn, Isabel Perez-Otano, and John F Wesseling. Elevated synaptic vesicle release probability in synaptophysin/gyrin family quadruple knockouts. *eLife*, 8, 2019.

[6149] Malini Rajan, Cole P Anderson, Paul M Rindler, Steven Joshua Romney, Maria C Ferreira dos Santos, Jason Gertz, and Elizabeth A Leibold. NHR-14 loss of function couples intestinal iron uptake with innate immunity in C. elegans through PQM-1 signaling. *eLife*, 8, 2019.

[6150] Prashant Rajbhandari, Douglas Arneson, Sydney K Hart, In Sook Ahn, Graciel Diamante, Luis C Santos, Nima Zaghari, An-Chieh Feng, Brandon J Thomas, Laurent Vergnes, Stephen D Lee, Abha K Rajbhandari, Karen Reue, Stephen T Smale, Xia Yang, and Peter Tontonoz. Single cell analysis reveals immune cell–adipocyte crosstalk regulating the transcription of thermogenic adipocytes. *eLife*, 8, 2019.

[6151] Valeria Ramaglia, Salma Sheikh-Mohamed, Karen Legg, Calvin Park, Olga L Rojas, Stephanie Zandee, Fred Fu, Olga Ornatsky, Eric C Swanson, David Pitt, Alexandre Prat, Trevor D McKee, and Jennifer L Gommerman. Multiplexed imaging of immune cells in staged multiple sclerosis lesions by mass cytometry. *eLife*, 8, 2019.

[6152] Angela M Ramos-Lobo, Pryscila DS Teixeira, Isadora C Furigo, Helen M Melo, Natalia de M Lyra e Silva, Fernanda G De Felice, and Jose Donato. Long-term consequences of the absence of leptin signaling in early life. *eLife*, 8, 2019.

[6153] Martina Rangl, Nicolaus Schmandt, Eduardo Perozo, and Simon Scheuring. Real time dynamics of Gating-Related conformational changes in CorA. *eLife*, 8, 2019.

[6154] Olga Rapp and Ofer Yifrach. Evolutionary and functional insights into the mechanism underlying body-size-related adaptation of mammalian hemoglobin. *eLife*, 8, 2019.

[6155] David A Rasmussen and Tanja Stadler. Coupling adaptive molecular evolution to phylodynamics using fitness-dependent birth-death models. *eLife*, 8, 2019.

[6156] Arvind Ravichandran, özer Duman, Masoud Hoore, Guglielmo Saggiorato, Gerard A Vliegenthart, Thorsten Auth, and Gerhard Gompper. Chronology of motor-mediated microtubule streaming. *eLife*, 8, 2019.

[6157] Kate A Rawlinson, Francois Lapraz, Edward R Ballister, Mark Terasaki, Jessica Rodgers, Richard J McDowell, Johannes Girstmair, Katharine E Criswell, Miklos Boldogkoi, Fraser Simpson, David Goulding, Claire Cormie, Brian Hall, Robert J Lucas, and Maximilian J Telford. Extraocular, rod-like photoreceptors in a flatworm express xenopsin photopigment. *eLife*, 8, 2019.

[6158] Suparna Ray, Miriam I Rosenberg, Hélène Chanut-Delalande, Amélie Decaras, Barbara Schwertner, William Toubiana, Tzach Auman, Irene Schnellhammer, Matthias Teuscher, Philippe Valenti, Abderrahman Khila, Martin Klingler, and FranÇois Payre. The mlpt/Ubr3/Svb module comprises an ancient developmental switch for embryonic patterning. *eLife*, 8, 2019.

[6159] Yevgeniy Raynes and Daniel Weinreich. Selection on mutators is not frequency-dependent. *eLife*, 8, 2019.

[6160] Bharat Reddy, Navid Bavi, Allen Lu, Yeonwoo Park, and Eduardo Perozo. Molecular basis of force-from-lipids gating in the mechanosensitive channel MscS. *eLife*, 8, 2019.

[6161] Hemanth KN Reddy, Marta Carroni, Janos Hajdu, and Martin Svenda. Electron cryo-microscopy of bacteriophage PR772 reveals the elusive vertex complex and the capsid architecture. *eLife*, 8, 2019.

[6162] Martin Reichard and Matej Polačik. Nothobranchius furzeri, an 'instant' fish from an ephemeral habitat. *eLife*, 8, 2019.

[6163] Taylor A Reid, Courtney Coombes, Soumya Mukherjee, Rebecca R Goldblum, Kyle White, Sneha Parmar, Mark McClellan, Marija Zanic, Naomi Courtemanche, and Melissa K Gardner. Structural state recognition facilitates tip tracking of EB1 at growing microtubule ends. *eLife*, 8, 2019.

[6164] Katja Reinhard, Chen Li, Quan Do, Emily G Burke, Steven Heynderickx, and Karl Farrow. A projection specific logic to sampling visual inputs in mouse superior colliculus. *eLife*, 8, 2019.

[6165] Irina L Rempel, Matthew M Crane, David J Thaller, Ankur Mishra, Daniel PM Jansen, Georges Janssens, Petra Popken, Arman Akşit, Matt Kaeberlein, Erik van der Giessen, Anton Steen, Patrick R Onck, C Patrick Lusk, and Liesbeth M Veenhoff. Age-dependent deterioration of nuclear pore assembly in mitotic cells decreases transport dynamics. *eLife*, 8, 2019.

[6166] Jing Ren, Alina Isakova, Drew Friedmann, Jiawei Zeng, Sophie M Grutzner, Albert Pun, Grace Q Zhao, Sai Saroja Kolluru, Ruiyu Wang, Rui Lin, Pengcheng Li, Anan Li, Jennifer L Raymond, Qingming Luo, Minmin Luo, Stephen R Quake, and Liqun Luo. Single-cell transcriptomes and whole-brain projections of serotonin neurons in the mouse dorsal and median raphe nuclei. *eLife*, 8, 2019.

[6167] Tom G Richardson, Sean Harrison, Gibran Hemani, and George Davey Smith. An atlas of polygenic risk score associations to highlight putative causal relationships across the human phenome. *eLife*, 8, 2019.

[6168] Andreas S Richter, Claudia Banse, and Bernhard Grimm. The GluTR-binding protein is the heme-binding factor for feedback control of glutamyl-tRNA reductase. *eLife*, 8, 2019.

[6169] David Richter and Floris P de Lange. Statistical learning attenuates visual activity only for attended stimuli. *eLife*, 8, 2019.

[6170] Susanne Rinné, Aytug K Kiper, Kirsty S Vowinkel, David Ramírez, Marcus Schewe, Mauricio Bedoya, Diana Aser, Isabella Gensler, Michael F Netter, Phillip J Stansfeld, Thomas Baukrowitz, Wendy Gonzalez, and Niels Decher. The molecular basis for an allosteric inhibition of K+-flux gating in K2P channels. *eLife*, 8, 2019.

[6171] Martina Riva, Ioana Genescu, Chloé Habermacher, David Orduz, Fanny Ledonne, Filippo M Rijli, Guillermina López-Bendito, Eva Coppola, Sonia Garel, Maria Cecilia Angulo, and Alessandra Pierani. Activity-dependent death of transient Cajal-Retzius neurons is required for functional cortical wiring. *eLife*, 8, 2019.

[6172] Juan Rivera-Correa, Maria Sophia Mackroth, Thomas Jacobs, Julian Schulze zur Wiesch, Thierry Rolling, and Ana Rodriguez. Atypical memory B-cells are associated with Plasmodium falciparum anemia through anti-phosphatidylserine antibodies. *eLife*, 8, 2019.

[6173] Efraín E Rivera-Serrano, Olga González-López, Anshuman Das, and Stanley M Lemon. Cellular entry and uncoating of naked and quasi-enveloped human hepatoviruses. *eLife*, 8, 2019.

[6174] Allison W Roberts, Lauren M Popov, Gabriel Mitchell, Krystal L Ching, Daniel J Licht, Guillaume Golovkine, Gregory M Barton, and Jeffery S Cox. Cas9+ conditionally-immortalized macrophages as a tool for bacterial pathogenesis and beyond. *eLife*, 8, 2019.

[6175] Brooks G Robinson, Xintong Cai, Jiexin Wang, James R Bunzow, John T Williams, and Pascal S Kaeser. RIM is essential for stimulated but not spontaneous somatodendritic dopamine release in the midbrain. *eLife*, 8, 2019.

[6176] Cristina M Robinson, Kate T Snyder, and Nicole Creanza. Correlated evolution between repertoire size and song plasticity predicts that sexual selection on song promotes open-ended learning. *eLife*, 8, 2019.

[6177] J Elliott Robinson, Gerard M Coughlin, Acacia M Hori, Jounhong Ryan Cho, Elisha D Mackey, Zeynep Turan, Tommaso Patriarchi, Lin Tian, and Viviana Gradinaru. Optical dopamine monitoring with dLight1 reveals mesolimbic phenotypes in a mouse model of neurofibromatosis type 1. *eLife*, 8, 2019.

[6178] Morgane Roche, Emmanuelle Chaigneau, Ravi L Rungta, Davide Boido, Bruno Weber, and Serge Charpak. In vivo imaging with a water immersion objective affects brain temperature, blood flow and oxygenation. *eLife*, 8, 2019.

[6179] William RT Roderick, Diana D Chin, Mark R Cutkosky, and David Lentink. Birds land reliably on complex surfaces by adapting their foot-surface interactions upon contact. *eLife*, 8, 2019.

[6180] João V Rodrigues and Eugene I Shakhnovich. Adaptation to mutational inactivation of an essential gene converges to an accessible suboptimal fitness peak. *eLife*, 8, 2019.

[6181] Isabel Rodriguez-Barraquer, Henrik Salje, and Derek A Cummings. Opportunities for improved surveillance and control of dengue from age-specific case data. *eLife*, 8, 2019.

[6182] Kayla E Rohr, Harshida Pancholi, Shabi Haider, Christopher Karow, David Modert, Nicholas J Raddatz, and Jennifer Evans. Seasonal plasticity in GABAA signaling is necessary for restoring phase synchrony in the master circadian clock network. *eLife*, 8, 2019.

[6183] Maria Rojec, Antoine Hocher, Kathryn M Stevens, Matthias Merkenschlager, and Tobias Warnecke. Chromatinization of Escherichia coli with archaeal histones. *eLife*, 8, 2019.

[6184] Marco Romano, Paolo Citton, Isabella Salvador, Daniele Arobba, Ivano Rellini, Marco Firpo, Fabio Negrino, Marta Zunino, Elisabetta Starnini, and Marco Avanzini. A multidisciplinary approach to a unique palaeolithic human ichnological record from Italy (Bàsura Cave). *eLife*, 8, 2019.

[6185] Elena Rondeau, Nicolas Larmonier, Thomas Pradeu, and Andreas Bikfalvi. Characterizing causality in cancer. *eLife*, 8, 2019.

[6186] Sophie I Roper, Laabiah Wasim, Dessislava Malinova, Michael Way, Susan Cox, and Pavel Tolar. B cells extract antigens at Arp2/3-generated actin foci interspersed with linear filaments. *eLife*, 8, 2019.

[6187] Leah F Rosin, Olivia Crocker, Randi L Isenhart, Son C Nguyen, Zhuxuan Xu, and Eric F Joyce. Chromosome territory formation attenuates the translocation potential of cells. *eLife*, 8, 2019.

[6188] Achira Roy, Rory M Murphy, Mei Deng, James W MacDonald, Theo K Bammler, Kimberly A Aldinger, Ian A Glass, and Kathleen J Millen. PI3K-Yap activity drives cortical gyrification and hydrocephalus in mice. *eLife*, 8, 2019.

[6189] Arunava Roy, Anandita Ghosh, Binod Kumar, and Bala Chandran. IFI16, a nuclear innate immune DNA sensor, mediates epigenetic silencing of herpesvirus genomes by its association with H3K9 methyltransferases SUV39H1 and GLP. *eLife*, 8, 2019.

[6190] Derek J Royer, Jose Echegaray-Mendez, Liwen Lin, Grzegorz B Gmyrek, Rose Mathew, Daniel R Saban, Victor L Perez, and Daniel JJ Carr. Complement and CD4+ T cells drive context-specific corneal sensory neuropathy. *eLife*, 8, 2019.

[6191] Tania Rozario, Edward B Quinn, Jianbin Wang, Richard E Davis, and Phillip A Newmark. Region-specific regulation of stem cell-driven regeneration in tapeworms. *eLife*, 8, 2019.

[6192] Andrii Rozhok and James DeGregori. A generalized theory of age-dependent carcinogenesis. *eLife*, 8, 2019.

[6193] Gerald M Rubin and Erin K O'Shea. Looking back and looking forward at Janelia. *eLife*, 8, 2019.

[6194] J Graham Ruby, Megan Smith, and Rochelle Buffenstein. Response to comment on 'Naked mole-rat mortality rates defy Gompertzian laws by not increasing with age'. *eLife*, 8, 2019.

[6195] Hannes Ruge, Theo AJ Schäfer, Katharina Zwosta, Holger Mohr, and Uta Wolfensteller. Neural representation of newly instructed rule identities during early implementation trials. *eLife*, 8, 2019.

[6196] Mario Ruiz, Rakesh Bodhicharla, Marcus Ståhlman, Emma Svensk, Kiran Busayavalasa, Henrik Palmgren, Hanna Ruhanen, Jan Boren, and Marc Pilon. Evolutionarily conserved long-chain Acyl-CoA synthetases regulate membrane composition and fluidity. *eLife*, 8, 2019.

[6197] Alan C Rupp, Michelle Ren, Cara M Altimus, Diego C Fernandez, Melissa Richardson, Fred Turek, Samer Hattar, and Tiffany M Schmidt. Distinct ipRGC subpopulations mediate light's acute and circadian effects on body temperature and sleep. *eLife*, 8, 2019.

[6198] Hatice Duygu Saatcioglu, Motohiro Kano, Heiko Horn, Lihua Zhang, Wesley Samore, Nicholas Nagykery, Marie-Charlotte Meinsohn, Minsuk Hyun, Rana Suliman, Joy Poulo, Jennifer Hsu, Caitlin Sacha, Dan Wang, Guangping Gao, Kasper Lage, Esther Oliva, Mary E Morris Sabatini, Patricia K Donahoe, and David Pépin. Single-cell sequencing of neonatal uterus reveals an Misr2+ endometrial progenitor indispensable for fertility. *eLife*, 8, 2019.

[6199] Jason Saba, Xien Yu Chua, Tatiana V Mishanina, Dhananjaya Nayak, Tricia A Windgassen, Rachel Anne Mooney, and Robert Landick. The elemental mechanism of transcriptional pausing. *eLife*, 8, 2019.

[6200] Ruchika Sachdev, Maria Hondele, Miriam Linsenmeier, Pascal Vallotton, Christopher F Mugler, Paolo Arosio, and Karsten Weis. Pat1 promotes processing body assembly by enhancing the phase separation of the DEAD-box ATPase Dhh1 and RNA. *eLife*, 8, 2019.

[6201] Harleen Saini, Alicia A Bicknell, Sean R Eddy, and Melissa J Moore. Free circular introns with an unusual branchpoint in neuronal projections. *eLife*, 8, 2019.

[6202] Evan T Saitta, Renxing Liang, Maggie CY Lau, Caleb M Brown, Nicholas R Longrich, Thomas G Kaye, Ben J Novak, Steven L Salzberg, Mark A Norell, Geoffrey D Abbott, Marc R Dickinson, Jakob Vinther, Ian D Bull, Richard A Brooker, Peter Martin, Paul Donohoe, Timothy DJ Knowles, Kirsty EH Penkman, and Tullis Onstott. Cretaceous dinosaur bone contains recent organic material and provides an environment conducive to microbial communities. *eLife*, 8, 2019.

[6203] Jorune Sakalauskaite, Søren H Andersen, Paolo Biagi, Maria A Borrello, Théophile Cocquerez, André Carlo Colonese, Federica Dal Bello, Alberto Girod, Marion Heumüller, Hannah Koon, Giorgia Mandili, Claudio Medana, Kirsty EH Penkman, Laurent Plasseraud, Helmut Schlichtherle, Sheila Taylor, Caroline Tokarski, Jérôme Thomas, Julie Wilson, Frédéric Marin, and Beatrice Demarchi. 'Palaeoshellomics' reveals the use of freshwater mother-of-pearl in prehistory. *eLife*, 8, 2019.

[6204] Valerie S Salazar, Luciane P Capelo, Claudio Cantù, Dario Zimmerli, Nehal Gosalia, Steven Pregizer, Karen Cox, Satoshi Ohte, Marina Feigenson, Laura Gamer, Jeffry S Nyman, David J Carey, Aris Economides, Konrad Basler, and Vicki Rosen. Reactivation of a developmental Bmp2 signaling center is required for therapeutic control of the murine periosteal niche. *eLife*, 8, 2019.

[6205] Emily C Sales, Emily L Heckman, Timothy L Warren, and Chris Q Doe. Regulation of subcellular dendritic synapse specificity by axon guidance cues. *eLife*, 8, 2019.

[6206] David G Saliba, Pablo F Céspedes-Donoso, Štefan Bálint, Ewoud B Compeer, Kseniya Korobchevskaya, Salvatore Valvo, Viveka Mayya, Audun Kvalvaag, Yanchun Peng, Tao Dong, Maria-Laura Tognoli, Eric O'Neill, Sarah Bonham, Roman Fischer, Benedikt M Kessler, and Michael L Dustin. Composition and structure of synaptic ectosomes exporting antigen receptor linked to functional CD40 ligand from helper T cells. *eLife*, 8, 2019.

[6207] Emilio Salinas, Benjamin R Steinberg, Lauren A Sussman, Sophia M Fry, Christopher K Hauser, Denise D Anderson, and Terrence R Stanford. Voluntary and involuntary contributions to perceptually guided saccadic choices resolved with millisecond precision. *eLife*, 8, 2019.

[6208] Henrik Salje, Kishor Kumar Paul, Repon Paul, Isabel Rodriguez-Barraquer, Ziaur Rahman, Mohammad Shafiul Alam, Mahmadur Rahman, Hasan Mohammad Al-Amin, James Heffelfinger, and Emily Gurley. Nationally-representative serostudy of dengue in Bangladesh allows generalizable disease burden estimates. *eLife*, 8, 2019.

[6209] Lotta Salomies, Julia Eymann, Imran Khan, and Nicolas Di-Poï. The alternative regenerative strategy of bearded dragon unveils the key processes underlying vertebrate tooth renewal. *eLife*, 8, 2019.

[6210] Irepan Salvador-Martínez, Marco Grillo, Michalis Averof, and Maximilian J Telford. Is it possible to reconstruct an accurate cell lineage using CRISPR recorders? *eLife*, 8, 2019.

[6211] Luis Sánchez-Guardado and Carlos Lois. Lineage does not regulate the sensory synaptic input of projection neurons in the mouse olfactory bulb. *eLife*, 8, 2019.

[6212] Florian Sandhaeger, Constantin von Nicolai, Earl K Miller, and Markus Siegel. Monkey EEG links neuronal color and motion information across species and scales. *eLife*, 8, 2019.

[6213] Joshua R Sanes. Tell me a story. *eLife*, 8, 2019.

[6214] Dajun Sang, Sudarshan Pinglay, Rafal P Wiewiora, Myvizhi E Selvan, Hua Jane Lou, John D Chodera, Benjamin E Turk, Zeynep H Gümüş, and Liam J Holt. Ancestral reconstruction reveals mechanisms of ERK regulatory evolution. *eLife*, 8, 2019.

[6215] Ilaria Sani, Brent C McPherson, Heiko Stemmann, Franco Pestilli, and Winrich A Freiwald. Functionally defined white matter of the macaque monkey brain reveals a dorso-ventral attention network. *eLife*, 8, 2019.

[6216] Anita Santana-Sanchez, Daniel Solymosi, Henna Mustila, Luca Bersanini, Eva-Mari Aro, and Yagut Allahverdiyeva. Flavodiiron proteins 1–to-4 function in versatile combinations in O2 photoreduction in cyanobacteria. *eLife*, 8, 2019.

[6217] Alfonso Santos-Lopez, Christopher W Marshall, Michelle R Scribner, Daniel J Snyder, and Vaughn S Cooper. Evolutionary pathways to antibiotic resistance are dependent upon environmental structure and bacterial lifestyle. *eLife*, 8, 2019.

[6218] Alessandro Sanzeni, Samata Katta, Bryan Petzold, Beth L Pruitt, Miriam B Goodman, and Massimo Vergassola. Somatosensory neurons integrate the geometry of skin deformation and mechanotransduction channels to shape touch sensing. *eLife*, 8, 2019.

[6219] Hadas Sar Shalom, Ron Goldner, Yarden Golan-Vaishenker, and Avraham Yaron. Balance between BDNF and Semaphorins gates the innervation of the mammary gland. *eLife*, 8, 2019.

[6220] Thomas A Sasani, Brent S Pedersen, Ziyue Gao, Lisa Baird, Molly Przeworski, Lynn B Jorde, and Aaron R Quinlan. Large, three-generation human families reveal post-zygotic mosaicism and variability in germline mutation accumulation. *eLife*, 8, 2019.

[6221] Takashi R Sato, Takahide Itokazu, Hironobu Osaki, Makoto Ohtake, Tetsuya Yamamoto, Kazuhiro Sohya, Takakuni Maki, and Tatsuo K Sato. Interhemispherically dynamic representation of an eye movement-related activity in mouse frontal cortex. *eLife*, 8, 2019.

[6222] Miho Sato-Hashimoto, Tomomi Nozu, Riho Toriba, Ayano Horikoshi, Miho Akaike, Kyoko Kawamoto, Ayaka Hirose, Yuriko Hayashi, Hiromi Nagai, Wakana Shimizu, Ayaka Saiki, Tatsuya Ishikawa, Ruwaida Elhanbly, Takenori Kotani, Yoji Murata, Yasuyuki Saito, Masae Naruse, Koji Shibasaki, Per-Arne Oldenborg, Steffen Jung, Takashi Matozaki, Yugo Fukazawa, and Hiroshi Ohnishi. Microglial SIRP$\alpha$ regulates the emergence of CD11c+ microglia and demyelination damage in white matter. *eLife*, 8, 2019.

[6223] Julia Sauerwald, Wilko Backer, Till Matzat, Frank Schnorrer, and Stefan Luschnig. Matrix metalloproteinase 1 modulates invasive behavior of tracheal branches during entry into Drosophila flight muscles. *eLife*, 8, 2019.

[6224] Lauren M Saunders, Abhishek K Mishra, Andrew J Aman, Victor M Lewis, Matthew B Toomey, Jonathan S Packer, Xiaojie Qiu, Jose L McFaline-Figueroa, Joseph C Corbo, Cole Trapnell, and David M Parichy. Thyroid hormone regulates distinct paths to maturation in pigment cell lineages. *eLife*, 8, 2019.

[6225] Hannah P Savage, Kathrin Kläsener, Fauna L Smith, Zheng Luo, Michael Reth, and Nicole Baumgarth. TLR induces reorganization of the IgM-BCR complex regulating murine B-1 cell responses to infections. *eLife*, 8, 2019.

[6226] Kayle S Sawyer, Nasim Maleki, Trinity Urban, Ksenija Marinkovic, Steven Karson, Susan M Ruiz, Gordon J Harris, and Marlene Oscar-Berman. Alcoholism gender differences in brain responsivity to emotional stimuli. *eLife*, 8, 2019.

[6227] Daniel S Saxton and Jasper Rine. Epigenetic memory independent of symmetric histone inheritance. *eLife*, 8, 2019.

[6228] Bechara Saykali, Navrita Mathiah, Wallis Nahaboo, Marie-Lucie Racu, Latifa Hammou, Matthieu Defrance, and Isabelle Migeotte. Distinct mesoderm migration phenotypes in extra-embryonic and embryonic regions of the early mouse embryo. *eLife*, 8, 2019.

[6229] Thomas Scambler, Heledd H Jarosz-Griffiths, Samuel Lara-Reyna, Shelly Pathak, Chi Wong, Jonathan Holbrook, Fabio Martinon, Sinisa Savic, Daniel Peckham, and Michael F McDermott. ENaC-mediated sodium influx exacerbates NLRP3-dependent inflammation in cystic fibrosis. *eLife*, 8, 2019.

[6230] Emily Anne Scarborough, Trisha N Davis, and Charles L Asbury. Tight bending of the Ndc80 complex provides intrinsic regulation of its binding to microtubules. *eLife*, 8, 2019.

[6231] Jessie R Scheimann, Rachel D Moloney, Parinaz Mahbod, Rachel L Morano, Maureen Fitzgerald, Olivia Hoskins, Benjamin A Packard, Evelin M Cotella, Yueh-Chiang Hu, and James P Herman. Conditional deletion of glucocorticoid receptors in rat brain results in sex-specific deficits in fear and coping behaviors. *eLife*, 8, 2019.

[6232] Randy Schekman. Progress and promise. *eLife*, 8, 2019.

[6233] Randy Schekman and Ekemini AU Riley. Coordinating a new approach to basic research into Parkinson's disease. *eLife*, 8, 2019.

[6234] Sven Schenk, Stephanie C Bannister, Fritz J Sedlazeck, Dorothea Anrather, Bui Quang Minh, Andrea Bileck, Markus Hartl, Arndt von Haeseler, Christopher Gerner, Florian Raible, and Kristin Tessmar-Raible. Combined transcriptome and proteome profiling reveals specific molecular brain signatures for sex, maturation and circalunar clock phase. *eLife*, 8, 2019.

[6235] Torsten M Scheyer, John R Hutchinson, Olivier Strauss, Massimo Delfino, Jorge D Carrillo-Briceño, Rodolfo Sánchez, and Marcelo R Sánchez-Villagra. Giant extinct caiman breaks constraint on the axial skeleton of extant crocodylians. *eLife*, 8, 2019.

[6236] Matthias Schlichting, Madelen M Díaz, Jason Xin, and Michael Rosbash. Neuron-specific knockouts indicate the importance of network communication to Drosophila rhythmicity. *eLife*, 8, 2019.

[6237] Michael Schlierf, Ganggang Wang, Xiaojiang S Chen, and Taekjip Ha. Hexameric helicase G40P unwinds DNA in single base pair steps. *eLife*, 8, 2019.

[6238] Rolf M Schmidt, Julia P Schessner, Georg HH Borner, and Sebastian Schuck. The proteasome biogenesis regulator Rpn4 cooperates with the unfolded protein response to promote ER stress resistance. *eLife*, 8, 2019.

[6239] Thomas SB Schmidt, Matthew R Hayward, Luis P Coelho, Simone S Li, Paul I Costea, Anita Y Voigt, Jakob Wirbel, Oleksandr M Maistrenko, Renato JC Alves, Emma Bergsten, Carine de Beaufort, Iradj Sobhani, Anna Heintz-Buschart, Shinichi Sunagawa, Georg Zeller, Paul Wilmes, and Peer Bork. Extensive transmission of microbes along the gastrointestinal tract. *eLife*, 8, 2019.

[6240] Pia Schröder, Timo Torsten Schmidt, and Felix Blankenburg. Neural basis of somatosensory target detection independent of uncertainty, relevance, and reports. *eLife*, 8, 2019.

[6241] Johannes Schultz, Tom Willems, Maria Gädeke, Ghada Chakkour, Alexander Franke, Bernd Weber, and Rene Hurlemann. A human subcortical network underlying social avoidance revealed by risky economic choices. *eLife*, 8, 2019.

[6242] Angela Schulz, Nicola Victoria Müller, Nina Anne van de Lest, Andreas Eisenreich, Martina Schmidbauer, Andrei Barysenka, Bettina Purfürst, Anje Sporbert, Theodor Lorenzen, Alexander M Meyer, Laura Herlan, Anika Witten, Frank Rühle, Weibin Zhou, Emile de Heer, Marion Scharpfenecker, Daniela Panáková, Monika Stoll, and Reinhold Kreutz. Analysis of the genomic architecture of a complex trait locus in hypertensive rat models links Tmem63c to kidney damage. *eLife*, 8, 2019.

[6243] Philipp Schwartenbeck, Johannes Passecker, Tobias U Hauser, Thomas HB FitzGerald, Martin Kronbichler, and Karl J Friston. Computational mechanisms of curiosity and goal-directed exploration. *eLife*, 8, 2019.

[6244] Niklas Schwarz, Betül Uysal, Marc Welzer, Jacqueline C Bahr, Nikolas Layer, Heidi Löffler, Kornelijus Stanaitis, Harshad PA, Yvonne G Weber, Ulrike BS Hedrich, Jürgen B Honegger, Angelos Skodras, Albert J Becker, Thomas V Wuttke, and Henner Koch. Long-term adult human brain slice cultures as a model system to study human CNS circuitry and disease. *eLife*, 8, 2019.

[6245] Sarah Schwettmann, Joshua B Tenenbaum, and Nancy Kanwisher. Invariant representations of mass in the human brain. *eLife*, 8, 2019.

[6246] Madineh Sedigh-Sarvestani, Larry A Palmer, and Diego Contreras. Thalamocortical synapses in the cat visual system in vivo are weak and unreliable. *eLife*, 8, 2019.

[6247] Nidhi Seethapathi and Manoj Srinivasan. Step-to-step variations in human running reveal how humans run without falling. *eLife*, 8, 2019.

[6248] Sridhar Selvaraj, Ricardo Mondragon-Gonzalez, Bin Xu, Alessandro Magli, Hyunkee Kim, Jeanne Lainé, James Kiley, Holly Mckee, Fabrizio Rinaldi, Joy Aho, Nacira Tabti, Wei Shen, and Rita CR Perlingeiro. Screening identifies small molecules that enhance the maturation of human pluripotent stem cell-derived myotubes. *eLife*, 8, 2019.

[6249] Sonia Q Sen, Sachin Chanchani, Tony D Southall, and Chris Q Doe. Neuroblast-specific open chromatin allows the temporal transcription factor, Hunchback, to bind neuroblast-specific loci. *eLife*, 8, 2019.

[6250] Ishita Sengupta and Jayant Udgaonkar. Monitoring site-specific conformational changes in real-time reveals a misfolding mechanism of the prion protein. *eLife*, 8, 2019.

[6251] Yeonee Seol, Gábor M Harami, Mihály Kovács, and Keir C Neuman. Homology sensing via non-linear amplification of sequence-dependent pausing by RecQ helicase. *eLife*, 8, 2019.

[6252] B Semihcan Sermet, Pavel Truschow, Michael Feyerabend, Johannes M Mayrhofer, Tess B Oram, Ofer Yizhar, Jochen F Staiger, and Carl CH Petersen. Pathway-, layer- and cell-type-specific thalamic input to mouse barrel cortex. *eLife*, 8, 2019.

[6253] Kyle S Severson, Duo Xu, Hongdian Yang, and Daniel H O'Connor. Coding of whisker motion across the mouse face. *eLife*, 8, 2019.

[6254] Ahmed Shaaban, Madhurima Dhara, Walentina Frisch, Ali Harb, Ali H Shaib, Ute Becherer, Dieter Bruns, and Ralf Mohrmann. The SNAP-25 linker supports fusion intermediates by local lipid interactions. *eLife*, 8, 2019.

[6255] Vivekanandan Shanmuganathan, Nina Schiller, Anastasia Magoulopoulou, Jingdong Cheng, Katharina Braunger, Florian Cymer, Otto Berninghausen, Birgitta Beatrix, Kenji Kohno, Gunnar von Heijne, and Roland Beckmann. Structural and mutational analysis of the ribosome-arresting human XBP1u. *eLife*, 8, 2019.

[6256] Wanqing Shao, Sergio G-M Alcantara, and Julia Zeitlinger. Reporter-ChIP-nexus reveals strong contribution of the Drosophila initiator sequence to RNA polymerase pausing. *eLife*, 8, 2019.

[6257] Alexey Shapiguzov, Julia P Vainonen, Kerri Hunter, Helena Tossavainen, Arjun Tiwari, Sari Järvi, Maarit Hellman, Fayezeh Aarabi, Saleh Alseekh, Brecht Wybouw, Katrien Van Der Kelen, Lauri Nikkanen, Julia Krasensky-Wrzaczek, Nina Sipari, Markku Keinänen, Esa Tyystjärvi, Eevi Rintamäki, Bert De Rybel, Jarkko Salojärvi, Frank Van Breusegem, Alisdair R Fernie, Mikael Brosché, Perttu Permi, Eva-Mari Aro, Michael Wrzaczek, and Jaakko Kangasjärvi. Arabidopsis RCD1 coordinates chloroplast and mitochondrial functions through interaction with ANAC transcription factors. *eLife*, 8, 2019.

[6258] Arpit Sharma, Lalita Oonthonpan, Ryan D Sheldon, Adam J Rauckhorst, Zhiyong Zhu, Sean C Tompkins, Kevin Cho, Wojciech J Grzesik, Lawrence R Gray, Diego A Scerbo, Alvin D Pewa, Emily M Cushing, Michael C Dyle, James E Cox, Chris Adams, Brandon S Davies, Richard K Shields, Andrew W Norris, Gary Patti, Leonid V Zingman, and Eric B Taylor. Impaired skeletal muscle mitochondrial pyruvate uptake rewires glucose metabolism to drive whole-body leanness. *eLife*, 8, 2019.

[6259] Manju Sharma, Anuj Srivastava, Heather E Fairfield, David Bergstrom, William F Flynn, and Robert E Braun. Identification of EOMES-expressing spermatogonial stem cells and their regulation by PLZF. *eLife*, 8, 2019.

[6260] Aleksandra Shcherbakova, Matthias Preller, Manuel H Taft, Jordi Pujols, Salvador Ventura, Birgit Tiemann, Falk FR Buettner, and Hans Bakker. C-mannosylation supports folding and enhances stability of thrombospondin repeats. *eLife*, 8, 2019.

[6261] Ji She, Weizhong Zeng, Jiangtao Guo, Qingfeng Chen, Xiao-chen Bai, and Youxing Jiang. Structural mechanisms of phospholipid activation of the human TPC2 channel. *eLife*, 8, 2019.

[6262] Mee Rie Sheen, Jennifer L Fields, Brian Northan, Judith Lacoste, Lay-Hong Ang, Steven Fiering, Elizabeth Iorns, Elizabeth Iorns, Rachel Tsui, Alexandria Denis, Nicole Perfito, and Timothy M Errington. Replication Study: Biomechanical remodeling of the microenvironment by stromal caveolin-1 favors tumor invasion and metastasis. *eLife*, 8, 2019.

[6263] Anna Shemorry, Jonathan M Harnoss, Ofer Guttman, Scot A Marsters, László G Kőműves, David A Lawrence, and Avi Ashkenazi. Caspase-mediated cleavage of IRE1 controls apoptotic cell commitment during endoplasmic reticulum stress. *eLife*, 8, 2019.

[6264] Ao Shen, Dana Chen, Manpreet Kaur, Peter Bartels, Bing Xu, Qian Shi, Joseph M Martinez, Kwunnok Mimi Man, Madeline Nieves-Cintron, Johannes W Hell, Manuel F Navedo, Xi-Yong Yu, and Yang K Xiang. $\beta$-blockers augment L-type Ca2+ channel activity by targeting spatially restricted $\beta$2AR signaling in neurons. *eLife*, 8, 2019.

[6265] Bo Shen, Kristy Vardy, Payton Hughes, Alpaslan Tasdogan, Zhiyu Zhao, Rui Yue, Genevieve M Crane, and Sean J Morrison. Integrin alpha11 is an Osteolectin receptor and is required for the maintenance of adult skeletal bone mass. *eLife*, 8, 2019.

[6266] Zhixun Shen, Sichen Liu, Xinxin Li, Zhengpeng Wan, Youxiang Mao, Chunlai Chen, and Wanli Liu. Conformational change within the extracellular domain of B cell receptor in B cell activation upon antigen binding. *eLife*, 8, 2019.

[6267] Emily Shepard, Emma-Louise Cole, Andrew Neate, Emmanouil Lempidakis, and Andrew Ross. Wind prevents cliff-breeding birds from accessing nests through loss of flight control. *eLife*, 8, 2019.

[6268] Cheng Shi, Lauren N Booth, and Coleen T Murphy. Insulin-like peptides and the mTOR-TFEB pathway protect Caenorhabditis elegans hermaphrodites from mating-induced death. *eLife*, 8, 2019.

[6269] Kaixuan Shi, Xia Yin, Mei-Chun Cai, Ying Yan, Chenqiang Jia, Pengfei Ma, Shengzhe Zhang, Zhenfeng Zhang, Zhenyu Gu, Meiying Zhang, Wen Di, and Guanglei Zhuang. PAX8 regulon in human ovarian cancer links lineage dependency with epigenetic vulnerability to HDAC inhibitors. *eLife*, 8, 2019.

[6270] Daisuke Shimaoka, Nicholas A Steinmetz, Kenneth D Harris, and Matteo Carandini. The impact of bilateral ongoing activity on evoked responses in mouse cortex. *eLife*, 8, 2019.

[6271] Kazunori Shinomiya, Gary Huang, Zhiyuan Lu, Toufiq Parag, C Shan Xu, Roxanne Aniceto, Namra Ansari, Natasha Cheatham, Shirley Lauchie, Erika Neace, Omotara Ogundeyi, Christopher Ordish, David Peel, Aya Shinomiya, Claire Smith, Satoko Takemura, Iris Talebi, Patricia K Rivlin, Aljoscha Nern, Louis K Scheffer, Stephen M Plaza, and Ian A Meinertzhagen. Comparisons between the ON- and OFF-edge motion pathways in the Drosophila brain. *eLife*, 8, 2019.

[6272] Allison J Shultz and Timothy B Sackton. Immune genes are hotspots of shared positive selection across birds and mammals. *eLife*, 8, 2019.

[6273] Clara Sidor, Nerea Borreguero-Munoz, Georgina C Fletcher, Ahmed Elbediwy, Oriane Guillermin, and Barry J Thompson. Mask family proteins ANKHD1 and ANKRD17 regulate YAP nuclear import and stability. *eLife*, 8, 2019.

[6274] Luis F Sifuentes-Dominguez, Haiying Li, Ernesto Llano, Zhe Liu, Amika Singla, Ashish S Patel, Mahesh Kathania, Areen Khoury, Nicholas Norris, Jonathan J Rios, Petro Starokadomskyy, Jason Y Park, Purva Gopal, Qi Liu, Shuai Tan, Lillienne Chan, Theodora Ross, Steven Harrison, K Venuprasad, Linda A Baker, Da Jia, and Ezra Burstein. SCGN deficiency results in colitis susceptibility. *eLife*, 8, 2019.

[6275] Edward H Silson, Adam Steel, Alexis Kidder, Adrian W Gilmore, and Chris I Baker. Distinct subdivisions of human medial parietal cortex support recollection of people and places. *eLife*, 8, 2019.

[6276] M Catarina Silva, Fleur M Ferguson, Quan Cai, Katherine A Donovan, Ghata Nandi, Debasis Patnaik, Tinghu Zhang, Hai-Tsang Huang, Diane E Lucente, Bradford C Dickerson, Timothy J Mitchison, Eric S Fischer, Nathanael S Gray, and Stephen J Haggarty. Targeted degradation of aberrant tau in frontotemporal dementia patient-derived neuronal cell models. *eLife*, 8, 2019.

[6277] Cynthia B Silveira, Antoni Luque, Ty NF Roach, Helena Villela, Adam Barno, Kevin Green, Brandon Reyes, Esther Rubio-Portillo, Tram Le, Spencer Mead, Mark Hatay, Mark JA Vermeij, Yuichiro Takeshita, Andreas Haas, Barbara Bailey, and Forest Rohwer. Biophysical and physiological processes causing oxygen loss from coral reefs. *eLife*, 8, 2019.

[6278] Szandor Simmons, Naoko Sasaki, Eiji Umemoto, Yutaka Uchida, Shigetomo Fukuhara, Yusuke Kitazawa, Michiyo Okudaira, Asuka Inoue, Kazuo Tohya, Keita Aoi, Junken Aoki, Naoki Mochizuki, Kenjiro Matsuno, Kiyoshi Takeda, Masayuki Miyasaka, and Masaru Ishii. High-endothelial cell-derived S1P regulates dendritic cell localization and vascular integrity in the lymph node. *eLife*, 8, 2019.

[6279] David W Sims, Nicolas E Humphries, Nan Hu, Violeta Medan, and Jimena Berni. Optimal searching behaviour generated intrinsically by the central pattern generator for locomotion. *eLife*, 8, 2019.

[6280] Linda V Sinclair, Andrew JM Howden, Alejandro Brenes, Laura Spinelli, Jens L Hukelmann, Andrew N Macintyre, Xiaojing Liu, Sarah Thomson, Peter M Taylor, Jeffrey C Rathmell, Jason W Locasale, Angus I Lamond, and Doreen A Cantrell. Antigen receptor control of methionine metabolism in T cells. *eLife*, 8, 2019.

[6281] Anand K Singh, Subhendu Roy Choudhury, Sandip De, Jie Zhang, Stephen Kissane, Vibha Dwivedi, Preethi Ramanathan, Marija Petric, Luisa Orsini, Daniel Hebenstreit, and Saverio Brogna. The RNA helicase UPF1 associates with mRNAs co-transcriptionally and is required for the release of mRNAs from gene loci. *eLife*, 8, 2019.

[6282] Jogender Singh and Alejandro Aballay. Intestinal infection regulates behavior and learning via neuroendocrine signaling. *eLife*, 8, 2019.

[6283] Rohit Singh and Beom K Choi. Siglec1-expressing subcapsular sinus macrophages provide soil for melanoma lymph node metastasis. *eLife*, 8, 2019.

[6284] Anirban Sinha, René Lutter, Binbin Xu, Tamara Dekker, Barbara Dierdorp, Peter J Sterk, Urs Frey, and Edgar Delgado Eckert. Loss of adaptive capacity in asthmatic patients revealed by biomarker fluctuation dynamics after rhinovirus challenge. *eLife*, 8, 2019.

[6285] Tamara M Sirey, Kenny Roberts, Wilfried Haerty, Oscar Bedoya-Reina, Sebastian Rogatti-Granados, Jennifer Y Tan, Nick Li, Lisa C Heather, Roderick N Carter, Sarah Cooper, Andrew J Finch, Jimi Wills, Nicholas M Morton, Ana Claudia Marques, and Chris P Ponting. The long non-coding RNA Cerox1 is a post transcriptional regulator of mitochondrial complex I catalytic activity. *eLife*, 8, 2019.

[6286] Kevin R Sitek, Omer Faruk Gulban, Evan Calabrese, G Allan Johnson, Agustin Lage-Castellanos, Michelle Moerel, Satrajit S Ghosh, and Federico De Martino. Mapping the human subcortical auditory system using histology, postmortem MRI and in vivo MRI at 7T. *eLife*, 8, 2019.

[6287] Michael J Skelly, James J Furniss, Heather Grey, Ka-Wing Wong, and Steven H Spoel. Dynamic ubiquitination determines transcriptional activity of the plant immune coactivator NPR1. *eLife*, 8, 2019.

[6288] Alexander GJ Skorput, Stephanie M Lee, Pamela WL Yeh, and Hermes H Yeh. The NKCC1 antagonist bumetanide mitigates interneuronopathy associated with ethanol exposure in utero. *eLife*, 8, 2019.

[6289] Roman Sloutsky and Kristen M Naegle. ASPEN, a methodology for reconstructing protein evolution with improved accuracy using ensemble models. *eLife*, 8, 2019.

[6290] Kelly M Smith, Tyler J Browne, Olivia C Davis, A Coyle, Kieran A Boyle, Masahiko Watanabe, Sally A Dickinson, Jacqueline A Iredale, Mark A Gradwell, Phillip Jobling, Robert J Callister, Christopher V Dayas, David I Hughes, and Brett A Graham. Calretinin positive neurons form an excitatory amplifier network in the spinal cord dorsal horn. *eLife*, 8, 2019.

[6291] Stephen J Smith, Uygar Sümbül, Lucas T Graybuck, Forrest Collman, Sharmishtaa Seshamani, Rohan Gala, Olga Gliko, Leila Elabbady, Jeremy A Miller, Trygve E Bakken, Jean Rossier, Zizhen Yao, Ed Lein, Hongkui Zeng, Bosiljka Tasic, and Michael Hawrylycz. Single-cell transcriptomic evidence for dense intracortical neuropeptide networks. *eLife*, 8, 2019.

[6292] Mashaal Sohail, Robert M Maier, Andrea Ganna, Alex Bloemendal, Alicia R Martin, Michael C Turchin, Charleston WK Chiang, Joel Hirschhorn, Mark J Daly, Nick Patterson, Benjamin Neale, Iain Mathieson, David Reich, and Shamil R Sunyaev. Polygenic adaptation on height is overestimated due to uncorrected stratification in genome-wide association studies. *eLife*, 8, 2019.

[6293] Salma Sohrabi-Jahromi, Katharina B Hofmann, Andrea Boltendahl, Christian Roth, Saskia Gressel, Carlo Baejen, Johannes Soeding, and Patrick Cramer. Transcriptome maps of general eukaryotic RNA degradation factors. *eLife*, 8, 2019.

[6294] Meredith A Sommars, Krithika Ramachandran, Madhavi D Senagolage, Christopher R Futtner, Derrik M Germain, Amanda L Allred, Yasuhiro Omura, Ilya R Bederman, and Grant D Barish. Dynamic repression by BCL6 controls the genome-wide liver response to fasting and steatosis. *eLife*, 8, 2019.

[6295] Remi Sonneville, Rahul Bhowmick, Saskia Hoffmann, Niels Mailand, Ian D Hickson, and Karim Labib. TRAIP drives replisome disassembly and mitotic DNA repair synthesis at sites of incomplete DNA replication. *eLife*, 8, 2019.

[6296] Magda Spella, Ioannis Lilis, Mario AA Pepe, Yuanyuan Chen, Maria Armaka, Anne-Sophie Lamort, Dimitra E Zazara, Fani Roumelioti, Malamati Vreka, Nikolaos I Kanellakis, Darcy E Wagner, Anastasios D Giannou, Vasileios Armenis, Kristina AM Arendt, Laura V Klotz, Dimitrios Toumpanakis, Vassiliki Karavana, Spyros G Zakynthinos, Ioanna Giopanou, Antonia Marazioti, Vassilis Aidinis, Rocio Sotillo, and Georgios T Stathopoulos. Club cells form lung adenocarcinomas and maintain the alveoli of adult mice. *eLife*, 8, 2019.

[6297] Rosanne Spolski, Erin E West, Peng Li, Sharon Veenbergen, Sunny Yung, Majid Kazemian, Jangsuk Oh, Zu-Xi Yu, Alexandra F Freeman, Stephen M Holland, Philip M Murphy, and Warren J Leonard. IL-21/type I interferon interplay regulates neutrophil-dependent innate immune responses to Staphylococcus aureus. *eLife*, 8, 2019.

[6298] Megan E Spurgeon and Paul F Lambert. Sexual transmission of murine papillomavirus (MmuPV1) in Mus musculus. *eLife*, 8, 2019.

[6299] Madhusudhan Srinivasan, Naomi J Petela, Johanna C Scheinost, James Collier, Menelaos Voulgaris, Maurici B Roig, Frederic Beckouët, Bin Hu, and Kim A Nasmyth. Scc2 counteracts a Wapl-independent mechanism that releases cohesin from chromosomes during G1. *eLife*, 8, 2019.

[6300] Siddarth Srinivasan, C Nadir Kaplan, and L Mahadevan. A multiphase theory for spreading microbial swarms and films. *eLife*, 8, 2019.

[6301] Satyam P Srivastav, Reazur Rahman, Qicheng Ma, Jasmine Pierre, Saptaparni Bandyopadhyay, and Nelson C Lau. Har-P, a short P-element variant, weaponizes P-transposase to severely impair Drosophila development. *eLife*, 8, 2019.

[6302] Alexandra M Stafford, Cheryl Reed, Harue Baba, Nicole AR Walter, John RK Mootz, Robert W Williams, Kim A Neve, Lev M Fedorov, Aaron J Janowsky, and Tamara J Phillips. Taar1 gene variants have a causal role in methamphetamine intake and response and interact with Oprm1. *eLife*, 8, 2019.

[6303] Thomas A Stalnaker, James D Howard, Yuji K Takahashi, Samuel J Gershman, Thorsten Kahnt, and Geoffrey Schoenbaum. Dopamine neuron ensembles signal the content of sensory prediction errors. *eLife*, 8, 2019.

[6304] Justyna B Startek, Brett Boonen, Alejandro López-Requena, Ariel Talavera, Yeranddy A Alpizar, Debapriya Ghosh, Nele Van Ranst, Bernd Nilius, Thomas Voets, and Karel Talavera. Mouse TRPA1 function and membrane localization are modulated by direct interactions with cholesterol. *eLife*, 8, 2019.

[6305] Sergey D Stavisky, Francis R Willett, Guy H Wilson, Brian A Murphy, Paymon Rezaii, Donald T Avansino, William D Memberg, Jonathan P Miller, Robert F Kirsch, Leigh R Hochberg, A Bolu Ajiboye, Shaul Druckmann, Krishna V Shenoy, and Jaimie M Henderson. Neural ensemble dynamics in dorsal motor cortex during speech in people with paralysis. *eLife*, 8, 2019.

[6306] Jeffrey Stedehouder, Demi Brizee, Johan A Slotman, Maria Pascual-Garcia, Megan L Leyrer, Bibi LJ Bouwen, Clemens MF Dirven, Zhenyu Gao, David M Berson, Adriaan B Houtsmuller, and Steven A Kushner. Local axonal morphology guides the topography of interneuron myelination in mouse and human neocortex. *eLife*, 8, 2019.

[6307] Karoliina Stefanius, Kelly Servage, Marcela de Souza Santos, Hillery Fields Gray, Jason E Toombs, Suneeta Chimalapati, Min S Kim, Venkat S Malladi, Rolf Brekken, and Kim Orth. Human pancreatic cancer cell exosomes, but not human normal cell exosomes, act as an initiator in cell transformation. *eLife*, 8, 2019.

[6308] Jonathan Stieglitz, Benjamin C Trumble, Caleb E Finch, Dong Li, Matthew J Budoff, Hillard Kaplan, and Michael D Gurven. Computed tomography shows high fracture prevalence among physically active forager-horticulturalists with high fertility. *eLife*, 8, 2019.

[6309] Marcel Stimberg, Romain Brette, and Dan FM Goodman. Brian 2, an intuitive and efficient neural simulator. *eLife*, 8, 2019.

[6310] Arjen Stolk, Loek Brinkman, Mariska J Vansteensel, Erik Aarnoutse, Frans SS Leijten, Chris H Dijkerman, Robert T Knight, Floris P de Lange, and Ivan Toni. Electrocorticographic dissociation of alpha and beta rhythmic activity in the human sensorimotor system. *eLife*, 8, 2019.

[6311] Julian Stolper, Elizabeth Mayela Ambrosio, Diana-Patricia Danciu, Lorena Buono, David A Elliott, Kiyoshi Naruse, Juan R Martínez-Morales, Anna Marciniak-Czochra, and Lazaro Centanin. Stem cell topography splits growth and homeostatic functions in the fish gill. *eLife*, 8, 2019.

[6312] Kelly M Storek, Joyce Chan, Rajesh Vij, Nancy Chiang, Zhonghua Lin, Jack Bevers, Christopher M Koth, Jean-Michel Vernes, Y Gloria Meng, JianPing Yin, Heidi Wallweber, Olivier Dalmas, Stephanie Shriver, Christine Tam, Kellen Schneider, Dhaya Seshasayee, Gerald Nakamura, Peter A Smith, Jian Payandeh, James T Koerber, Laetitia Comps-Agrar, and Steven T Rutherford. Massive antibody discovery used to probe structure–function relationships of the essential outer membrane protein LptD. *eLife*, 8, 2019.

[6313] Federica Storti, Katrin Klee, Vyara Todorova, Regula Steiner, Alaa Othman, Saskia van der Velde-Visser, Marijana Samardzija, Isabelle Meneau, Maya Barben, Duygu Karademir, Valda Pauzuolyte, Sanford L Boye, Frank Blaser, Christoph Ullmer, Joshua L Dunaief, Thorsten Hornemann, Lucia Rohrer, Anneke den Hollander, Arnold von Eckardstein, Jürgen Fingerle, Cyrille Maugeais, and Christian Grimm. Impaired ABCA1/ABCG1-mediated lipid efflux in the mouse retinal pigment epithelium (RPE) leads to retinal degeneration. *eLife*, 8, 2019.

[6314] Johannes Felix Stortz, Mario Del Rosario, Mirko Singer, Jonathan M Wilkes, Markus Meissner, and Sujaan Das. Formin-2 drives polymerisation of actin filaments enabling segregation of apicoplasts and cytokinesis in Plasmodium falciparum. *eLife*, 8, 2019.

[6315] Cassandra L Stowe, Thomas A Burley, Helen Allan, Maria Vinci, Gabriela Kramer-Marek, Daniela M Ciobota, Gary N Parkinson, Tara L Southworth, Giulia Agliardi, Alastair Hotblack, Mark F Lythgoe, Bruce R Branchini, Tammy L Kalber, James C Anderson, and Martin A Pule. Near-infrared dual bioluminescence imaging in mouse models of cancer using infraluciferin. *eLife*, 8, 2019.

[6316] Erhard Strohm, Gudrun Herzner, Joachim Ruther, Martin Kaltenpoth, and Tobias Engl. Nitric oxide radicals are emitted by wasp eggs to kill mold fungi. *eLife*, 8, 2019.

[6317] Helen Strutt, Jessica Gamage, and David Strutt. Reciprocal action of Casein Kinase I$\epsilon$ on core planar polarity proteins regulates clustering and asymmetric localisation. *eLife*, 8, 2019.

[6318] Timothy S Strutzenberg, Ruben D Garcia-Ordonez, Scott J Novick, HaJeung Park, Mi Ra Chang, Christelle Doebellin, Yuanjun He, Rémi Patouret, Theodore M Kamenecka, and Patrick R Griffin. HDX-MS reveals structural determinants for ROR$\gamma$ hyperactivation by synthetic agonists. *eLife*, 8, 2019.

[6319] Sarah K Stumper, Harini Ravi, Larry J Friedman, Rachel Anne Mooney, Ivan R Corrêa, Anne Gershenson, Robert Landick, and Jeff Gelles. Delayed inhibition mechanism for secondary channel factor regulation of ribosomal RNA transcription. *eLife*, 8, 2019.

[6320] Jan Stundl, Anna Pospisilova, David Jandzik, Peter Fabian, Barbora Dobiasova, Martin Minarik, Brian D Metscher, Vladimir Soukup, and Robert Cerny. Bichir external gills arise via heterochronic shift that accelerates hyoid arch development. *eLife*, 8, 2019.

[6321] Maria R Stupnikov, Ying Yang, Munemasa Mori, Jining Lu, and Wellington V Cardoso. Jagged and Delta-like ligands control distinct events during airway progenitor cell differentiation. *eLife*, 8, 2019.

[6322] Javier A Suarez, James D Howard, Geoffrey Schoenbaum, and Thorsten Kahnt. Sensory prediction errors in the human midbrain signal identity violations independent of perceptual distance. *eLife*, 8, 2019.

[6323] Kaushikaram Subramanian, Martin Weigert, Oliver Borsch, Heike Petzold, Alfonso Garcia-Ulloa, Eugene W Myers, Marius Ader, Irina Solovei, and Moritz Kreysing. Rod nuclear architecture determines contrast transmission of the retina and behavioral sensitivity in mice. *eLife*, 8, 2019.

[6324] Ken Sugino, Erin Clark, Anton Schulmann, Yasuyuki Shima, Lihua Wang, David L Hunt, Bryan M Hooks, Dimitri Tränkner, Jayaram Chandrashekar, Serge Picard, Andrew L Lemire, Nelson Spruston, Adam W Hantman, and Sacha B Nelson. Mapping the transcriptional diversity of genetically and anatomically defined cell populations in the mouse brain. *eLife*, 8, 2019.

[6325] Luis F Sullivan, Timothy L Warren, and Chris Q Doe. Temporal identity establishes columnar neuron morphology, connectivity, and function in a Drosophila navigation circuit. *eLife*, 8, 2019.

[6326] Mark R Sullivan, Laura V Danai, Caroline A Lewis, Sze Ham Chan, Dan Y Gui, Tenzin Kunchok, Emily A Dennstedt, Matthew G Vander Heiden, and Alexander Muir. Quantification of microenvironmental metabolites in murine cancers reveals determinants of tumor nutrient availability. *eLife*, 8, 2019.

[6327] Alan Sulpizio, Marena E Minelli, Min Wan, Paul D Burrowes, Xiaochun Wu, Ethan J Sanford, Jung-Ho Shin, Byron C Williams, Michael L Goldberg, Marcus B Smolka, and Yuxin Mao. Protein polyglutamylation catalyzed by the bacterial calmodulin-dependent pseudokinase SidJ. *eLife*, 8, 2019.

[6328] Ruo Sun, Xingcong Jiang, Michael Reichelt, Jonathan Gershenzon, Sagar Subhash Pandit, and Daniel Giddings Vassão. Tritrophic metabolism of plant chemical defenses and its effects on herbivore and predator performance. *eLife*, 8, 2019.

[6329] Weijia Sun, Shaopeng Chi, Yuheng Li, Shukuan Ling, Yingjun Tan, Youjia Xu, Fan Jiang, Jianwei Li, Caizhi Liu, Guohui Zhong, Dengchao Cao, Xiaoyan Jin, Dingsheng Zhao, Xingcheng Gao, Zizhong Liu, Bailong Xiao, and Yingxian Li. The mechanosensitive Piezo1 channel is required for bone formation. *eLife*, 8, 2019.

[6330] Yidi Sun, Johannes Schöneberg, Xuyan Chen, Tommy Jiang, Charlotte Kaplan, Ke Xu, Thomas D Pollard, and David G Drubin. Direct comparison of clathrin-mediated endocytosis in budding and fission yeast reveals conserved and evolvable features. *eLife*, 8, 2019.

[6331] Ying Sun, Carlos Blanco-Centurion, Emmaline Bendell, Aurelio Vidal-Ortiz, Siwei Luo, and Meng Liu. Activity dynamics of amygdala GABAergic neurons during cataplexy of narcolepsy. *eLife*, 8, 2019.

[6332] Lalitta Suriya-Arunroj and Alexander Gail. Complementary encoding of priors in monkey frontoparietal network supports a dual process of decision-making. *eLife*, 8, 2019.

[6333] Damar Susilaradeya, Wei Xu, Thomas M Hall, Ferran Galán, Kai Alter, and Andrew Jackson. Extrinsic and intrinsic dynamics in movement intermittency. *eLife*, 8, 2019.

[6334] Nanoka Suzuki, Kodai Hirano, Hajime Ogino, and Haruki Ochi. Arid3a regulates nephric tubule regeneration via evolutionarily conserved regeneration signal-response enhancers. *eLife*, 8, 2019.

[6335] Egor Svidritskiy, Gabriel Demo, Anna B Loveland, Chen Xu, and Andrei A Korostelev. Extensive ribosome and RF2 rearrangements during translation termination. *eLife*, 8, 2019.

[6336] Nicole Swann. Are we there yet? *eLife*, 8, 2019.

[6337] Matthew Swire, Yuri Kotelevtsev, David J Webb, David A Lyons, and Charles ffrench Constant. Endothelin signalling mediates experience-dependent myelination in the CNS. *eLife*, 8, 2019.

[6338] Kathryn M Tabor, Gregory D Marquart, Christopher Hurt, Trevor S Smith, Alexandra K Geoca, Ashwin A Bhandiwad, Abhignya Subedi, Jennifer L Sinclair, Hannah M Rose, Nicholas F Polys, and Harold A Burgess. Brain-wide cellular resolution imaging of Cre transgenic zebrafish lines for functional circuit-mapping. *eLife*, 8, 2019.

[6339] Lucas Tafur, Yashar Sadian, Jonas Hanske, Rene Wetzel, Felix Weis, and Christoph W Müller. The cryo-EM structure of a 12-subunit variant of RNA polymerase I reveals dissociation of the A49-A34.5 heterodimer and rearrangement of subunit A12.2. *eLife*, 8, 2019.

[6340] Atsushi Takagi, Masaya Hirashima, Daichi Nozaki, and Etienne Burdet. Individuals physically interacting in a group rapidly coordinate their movement by estimating the collective goal. *eLife*, 8, 2019.

[6341] Naoki Takahashi, Nobuo Ogita, Tomonobu Takahashi, Shoji Taniguchi, Maho Tanaka, Motoaki Seki, and Masaaki Umeda. A regulatory module controlling stress-induced cell cycle arrest in Arabidopsis. *eLife*, 8, 2019.

[6342] Joshua S Talboom, Asta Håberg, Matthew D De Both, Marcus A Naymik, Isabelle Schrauwen, Candace R Lewis, Stacy F Bertinelli, Callie Hammersland, Mason A Fritz, Amanda J Myers, Meredith Hay, Carol A Barnes, Elizabeth Glisky, Lee Ryan, and Matthew J Huentelman. Family history of Alzheimer's disease alters cognition and is modified by medical and genetic factors. *eLife*, 8, 2019.

[6343] Jaya Talreja, Harvinder Talwar, Christian Bauerfeld, Lawrence I Grossman, Kezhong Zhang, Paul Tranchida, and Lobelia Samavati. HIF-1$\alpha$ regulates IL-1$\beta$ and IL-17 in sarcoidosis. *eLife*, 8, 2019.

[6344] Sanjeeta Tamang, Anastasiya Kishkevich, Carl A Morrow, Fekret Osman, Manisha Jalan, and Matthew C Whitby. The PCNA unloader Elg1 promotes recombination at collapsed replication forks in fission yeast. *eLife*, 8, 2019.

[6345] Leo TH Tang, Carlos A Diaz-Balzac, Maisha Rahman, Nelson J Ramirez-Suarez, Yehuda Salzberg, Maria I Lázaro-Peña, and Hannes E Bülow. TIAM-1/GEF can shape somatosensory dendrites independently of its GEF activity by regulating F-actin localization. *eLife*, 8, 2019.

[6346] Tianyu Tang, Timothy A Blenkinsop, and Eric J Lang. Complex spike synchrony dependent modulation of rat deep cerebellar nuclear activity. *eLife*, 8, 2019.

[6347] Wei Tang, Saad Jbabdi, Ziyi Zhu, Michiel Cottaar, Giorgia Grisot, Julia F Lehman, Anastasia Yendiki, and Suzanne N Haber. A connectional hub in the rostral anterior cingulate cortex links areas of emotion and cognitive control. *eLife*, 8, 2019.

[6348] Weiyi Tang, Megan L Martik, Yuwei Li, and Marianne E Bronner. Cardiac neural crest contributes to cardiomyocytes in amniotes and heart regeneration in zebrafish. *eLife*, 8, 2019.

[6349] Liangyu Tao, Siddhi Ozarkar, Jeffrey M Beck, and Vikas Bhandawat. Statistical structure of locomotion and its modulation by odors. *eLife*, 8, 2019.

[6350] Xiao Tao and Roderick MacKinnon. Cryo-EM structure of the KvAP channel reveals a non-domain-swapped voltage sensor topology. *eLife*, 8, 2019.

[6351] Xiao Tao and Roderick MacKinnon. Molecular structures of the human Slo1 K+ channel in complex with $\beta 4$. *eLife*, 8, 2019.

[6352] Yuyong Tao, Monika Mis, Levi Blazer, Mart Ustav, Zachary Steinhart, Rony Chidiac, Elli Kubarakos, Siobhan O'Brien, Xiaomei Wang, Nick Jarvik, Nish Patel, Jarrett Adams, Jason Moffat, Stephane Angers, and Sachdev S Sidhu. Tailored tetravalent antibodies potently and specifically activate Wnt/Frizzled pathways in cells, organoids and mice. *eLife*, 8, 2019.

[6353] Alexander J Tarashansky, Yuan Xue, Pengyang Li, Stephen R Quake, and Bo Wang. Self-assembling manifolds in single-cell RNA sequencing data. *eLife*, 8, 2019.

[6354] Oscar A Tarazona, Davys H Lopez, Leslie A Slota, and Martin J Cohn. Evolution of limb development in cephalopod mollusks. *eLife*, 8, 2019.

[6355] Michel Tassetto, Mark Kunitomi, Zachary J Whitfield, Patrick T Dolan, Irma Sánchez-Vargas, Miguel Garcia-Knight, Isabel Ribiero, Taotao Chen, Ken E Olson, and Raul Andino. Control of RNA viruses in mosquito cells through the acquisition of vDNA and endogenous viral elements. *eLife*, 8, 2019.

[6356] Andrew M Taverner, Lu Yang, Zachary J Barile, Becky Lin, Julie Peng, Ana P Pinharanda, Arya S Rao, Bartholomew P Roland, Aaron D Talsma, Daniel Wei, Georg Petschenka, Michael J Palladino, and Peter Andolfatto. Adaptive substitutions underlying cardiac glycoside insensitivity in insects exhibit epistasis in vivo. *eLife*, 8, 2019.

[6357] Gavin J Taylor, Pierre Tichit, Marie D Schmidt, Andrew J Bodey, Christoph Rau, and Emily Baird. Bumblebee visual allometry results in locally improved resolution and globally improved sensitivity. *eLife*, 8, 2019.

[6358] Tiberiu Teşileanu, Simona Cocco, Rémi Monasson, and Vijay Balasubramanian. Adaptation of olfactory receptor abundances for efficient coding. *eLife*, 8, 2019.

[6359] Ashley Tehranchi, Brian Hie, Michael Dacre, Irene Kaplow, Kade Pettie, Peter Combs, and Hunter B Fraser. Fine-mapping cis-regulatory variants in diverse human populations. *eLife*, 8, 2019.

[6360] Celso M Teixeira-Duarte, Fátima Fonseca, and João H Morais-Cabral. Activation of a nucleotide-dependent RCK domain requires binding of a cation cofactor to a conserved site. *eLife*, 8, 2019.

[6361] Morayma M Temoche-Diaz, Matthew J Shurtleff, Ryan M Nottingham, Jun Yao, Raj P Fadadu, Alan M Lambowitz, and Randy Schekman. Distinct mechanisms of microRNA sorting into cancer cell-derived extracellular vesicle subtypes. *eLife*, 8, 2019.

[6362] Camilla S Teng, Lionel Cavin, Robert E Maxson, Marcelo R Sánchez-Villagra, and J Gage Crump. Resolving homology in the face of shifting germ layer origins: Lessons from a major skull vault boundary. *eLife*, 8, 2019.

[6363] Emelyne Teo, Sudharshan Ravi, Diogo Barardo, Hyung-Seok Kim, Sheng Fong, Amaury Cazenave-Gassiot, Tsze Yin Tan, Jianhong Ching, Jean-Paul Kovalik, Markus R Wenk, Rudiyanto Gunawan, Philip K Moore, Barry Halliwell, Nicholas Tolwinski, and Jan Gruber. Metabolic stress is a primary pathogenic event in transgenic Caenorhabditis elegans expressing pan-neuronal human amyloid beta. *eLife*, 8, 2019.

[6364] Sofonias Tessema, Amy Wesolowski, Anna Chen, Maxwell Murphy, Jordan Wilheim, Anna-Rosa Mupiri, Nick W Ruktanonchai, Victor A Alegana, Andrew J Tatem, Munyaradzi Tambo, Bradley Didier, Justin M Cohen, Adam Bennett, Hugh JW Sturrock, Roland Gosling, Michelle S Hsiang, David L Smith, Davis R Mumbengegwi, Jennifer L Smith, and Bryan Greenhouse. Using parasite genetic and human mobility data to infer local and cross-border malaria connectivity in Southern Africa. *eLife*, 8, 2019.

[6365] David J Thaller, Matteo Allegretti, Sapan Borah, Paolo Ronchi, Martin Beck, and C Patrick Lusk. An ESCRT-LEM protein surveillance system is poised to directly monitor the nuclear envelope and nuclear transport system. *eLife*, 8, 2019.

[6366] Akanksha Thawani, Howard A Stone, Joshua W Shaevitz, and Sabine Petry. Spatiotemporal organization of branched microtubule networks. *eLife*, 8, 2019.

[6367] Eric D Thomas and David W Raible. Distinct progenitor populations mediate regeneration in the zebrafish lateral line. *eLife*, 8, 2019.

[6368] Jessica E Thomas, Gary R Carvalho, James Haile, Nicolas J Rawlence, Michael D Martin, Simon YW Ho, Arnór Þ Sigfússon, Vigfús A Jósefsson, Morten Frederiksen, Jannie F Linnebjerg, Jose A Samaniego Castruita, Jonas Niemann, Mikkel-Holger S Sinding, Marcela Sandoval-Velasco, André ER Soares, Robert Lacy, Christina Barilaro, Juila Best, Dirk Brandis, Chiara Cavallo, Mikelo Elorza, Kimball L Garrett, Maaike Groot, Friederike Johansson, Jan T Lifjeld, Göran Nilson, Dale Serjeanston, Paul Sweet, Errol Fuller, Anne Karin Hufthammer, Morten Meldgaard, Jon Fjeldså, Beth Shapiro, Michael Hofreiter, John R Stewart, M Thomas P Gilbert, and Michael Knapp. Demographic reconstruction from ancient DNA supports rapid extinction of the great auk. *eLife*, 8, 2019.

[6369] Albert Thommen, Steffen Werner, Olga Frank, Jenny Philipp, Oskar Knittelfelder, Yihui Quek, Karim Fahmy, Andrej Shevchenko, Benjamin M Friedrich, Frank Jülicher, and Jochen C Rink. Body size-dependent energy storage causes Kleiber's law scaling of the metabolic rate in planarians. *eLife*, 8, 2019.

[6370] Amelia J Thompson, Eva K Pillai, Ivan B Dimov, Sarah K Foster, Christine E Holt, and Kristian Franze. Rapid changes in tissue mechanics regulate cell behaviour in the developing embryonic brain. *eLife*, 8, 2019.

[6371] Morgan Thompson, Ryan Bixby, Robert Dalton, Alexa Vandenburg, John A Calarco, and Adam D Norris. Splicing in a single neuron is coordinately controlled by RNA binding proteins and transcription factors. *eLife*, 8, 2019.

[6372] Benjamin R Thomson, Isabel A Carota, Tomokazu Souma, Saily Soman, Dietmar Vestweber, and Susan E Quaggin. Targeting the vascular-specific phosphatase PTPRB protects against retinal ganglion cell loss in a pre-clinical model of glaucoma. *eLife*, 8, 2019.

[6373] Sushrut Thorat, Daria Proklova, and Marius V Peelen. The nature of the animacy organization in human ventral temporal cortex. *eLife*, 8, 2019.

[6374] Paul RHJ Timmers, Ninon Mounier, Kristi Lall, Krista Fischer, Zheng Ning, Xiao Feng, Andrew D Bretherick, David W Clark, M Agbessi, M Agbessi, H Ahsan, I Alves, A Andiappan, P Awadalla, A Battle, MJ Bonder, D Boomsma, M Christiansen, A Claringbould, P Deelen, J van Dongen, T Esko, M Favé, L Franke, T Frayling, SA Gharib, G Gibson, G Hemani, R Jansen, A Kalnapenkis, S Kasela, J Kettunen, Y Kim, H Kirsten, P Kovacs, K Krohn, J Kronberg-Guzman, V Kukushkina, Z Kutalik, M Kähönen, B Lee, T Lehtimäki, M Loeffler, U Marigorta, A Metspalu, J van Meurs, L Milani, M Müller-Nurasyid, M Nauck, M Nivard, B Penninx, M Perola, N Pervjakova, B Pierce, J Powell, H Prokisch, BM Psaty, O Raitakari, S Ring, S Ripatti, O Rotzschke, S Ruëger, A Saha, M Scholz, K Schramm, I Seppälä, M Stumvoll, P Sullivan, A Teumer, J Thiery, L Tong, A Tönjes, J Verlouw, PM Visscher, U Võsa, U Völker, H Yaghootkar, J Yang, B Zeng, F Zhang, M Agbessi, H Ahsan, I Alves, A Andiappan, P Awadalla, A Battle, MJ Bonder, D Boomsma, M Christiansen, A Claringbould, P Deelen, J van Dongen, T Esko, M Favé, L Franke, T Frayling, SA Gharib, G Gibson, G Hemani, R Jansen, A Kalnapenkis, S Kasela, J Kettunen, Y Kim, H Kirsten, P Kovacs, K Krohn, J Kronberg-Guzman, V Kukushkina, Z Kutalik, M Kähönen, B Lee, T Lehtimäki, M Loeffler, U Marigorta, A Metspalu, J van Meurs, L Milani, M Müller-Nurasyid, M Nauck, M Nivard, B Penninx, M Perola, N Pervjakova, B Pierce, J Powell, H Prokisch, BM Psaty, O Raitakari, S Ring, S Ripatti, O Rotzschke, S Ruëger, A Saha, M Scholz, K Schramm, I Seppälä, M Stumvoll, P Sullivan,

A Teumer, J Thiery, L Tong, A Tönjes, J Verlouw, PM Visscher, U Võsa, U Völker, H Yaghootkar, J Yang, B Zeng, F Zhang, Xia Shen, Tõnu Esko, Zoltán Kutalik, James F Wilson, and Peter K Joshi. Genomics of 1 million parent lifespans implicates novel pathways and common diseases and distinguishes survival chances. *eLife*, 8, 2019.

[6375] Doug K Tischer and Orion David Weiner. Light-based tuning of ligand half-life supports kinetic proofreading model of T cell signaling. *eLife*, 8, 2019.

[6376] Marco Tjioe, Saurabh Shukla, Rohit Vaidya, Alice Troitskaia, Carol S Bookwalter, Kathleen M Trybus, Yann R Chemla, and Paul R Selvin. Multiple kinesins induce tension for smooth cargo transport. *eLife*, 8, 2019.

[6377] Mélanie Tobin, Atitheb Chaiyasitdhi, Vincent Michel, Nicolas Michalski, and Pascal Martin. Stiffness and tension gradients of the hair cell's tip-link complex in the mammalian cochlea. *eLife*, 8, 2019.

[6378] Christopher D Todd, özgen Deniz, Darren Taylor, and Miguel R Branco. Functional evaluation of transposable elements as enhancers in mouse embryonic and trophoblast stem cells. *eLife*, 8, 2019.

[6379] Robert T Todd, Tyler D Wikoff, Anja Forche, and Anna Selmecki. Genome plasticity in Candida albicans is driven by long repeat sequences. *eLife*, 8, 2019.

[6380] Francesco Tomassoni-Ardori, Gianluca Fulgenzi, Jodi Becker, Colleen Barrick, Mary Ellen Palko, Skyler Kuhn, Vishal Koparde, Maggie Cam, Sudhirkumar Yanpallewar, Shalini Oberdoerffer, and Lino Tessarollo. Rbfox1 up-regulation impairs BDNF-dependent hippocampal LTP by dysregulating TrkB isoform expression levels. *eLife*, 8, 2019.

[6381] Jakub Tomek, Alfonso Bueno-Orovio, Elisa Passini, Xin Zhou, Ana Minchole, Oliver Britton, Chiara Bartolucci, Stefano Severi, Alvin Shrier, Laszlo Virag, Andras Varro, and Blanca Rodriguez. Development, calibration, and validation of a novel human ventricular myocyte model in health, disease, and drug block. *eLife*, 8, 2019.

[6382] Ailing Tong, John T Petroff, Fong-Fu Hsu, Philipp AM Schmidpeter, Crina M Nimigean, Liam Sharp, Grace Brannigan, and Wayland WL Cheng. Direct binding of phosphatidylglycerol at specific sites modulates desensitization of a ligand-gated ion channel. *eLife*, 8, 2019.

[6383] Sandra Elizabeth Torres, Ciara M Gallagher, Lars Plate, Meghna Gupta, Christina R Liem, Xiaoyan Guo, Ruilin Tian, Robert M Stroud, Martin Kampmann, Jonathan S Weissman, and Peter Walter. Ceapins block the unfolded protein response sensor ATF6$\alpha$ by inducing a neomorphic inter-organelle tether. *eLife*, 8, 2019.

[6384] Yamila N Torres Cleuren, Chee Kiang Ewe, Kyle C Chipman, Emily R Mears, Cricket G Wood, Coco Emma Alma Al-Alami, Melissa R Alcorn, Thomas L Turner, Pradeep M Joshi, Russell G Snell, and Joel H Rothman. Extensive intraspecies cryptic variation in an ancient embryonic gene regulatory network. *eLife*, 8, 2019.

[6385] Jorge Torres-Paz, Julien Leclercq, and Sylvie Rétaux. Maternally regulated gastrulation as a source of variation contributing to cavefish forebrain evolution. *eLife*, 8, 2019.

[6386] Nicolò Tosetti, Nicolas Dos Santos Pacheco, Dominique Soldati-Favre, and Damien Jacot. Three F-actin assembly centers regulate organelle inheritance, cell-cell communication and motility in Toxoplasma gondii. *eLife*, 8, 2019.

[6387] Shir Toubiana, Miriam Gagliardi, Mariarosaria Papa, Roberta Manco, Maty Tzukerman, Maria R Matarazzo, and Sara Selig. Persistent epigenetic memory impedes rescue of the telomeric phenotype in human ICF iPSCs following DNMT3B correction. *eLife*, 8, 2019.

[6388] Mai P Tran, Rio Tsutsumi, Joel M Erberich, Kevin D Chen, Michelle D Flores, and Kimberly L Cooper. Evolutionary loss of foot muscle during development with characteristics of atrophy and no evidence of cell death. *eLife*, 8, 2019.

[6389] Linton M Traub. A nanobody-based molecular toolkit provides new mechanistic insight into clathrin-coat initiation. *eLife*, 8, 2019.

[6390] Stanislao Igor Travisano, Vera Lucia Oliveira, Belén Prados, Joaquim Grego-Bessa, Rebeca Piñeiro-Sabarís, Vanesa Bou, Manuel J Gómez, Fátima Sánchez-Cabo, Donal MacGrogan, and José Luis de la Pompa. Coronary arterial development is regulated by a Dll4-Jag1-EphrinB2 signaling cascade. *eLife*, 8, 2019.

[6391] Anna S Trigos, Richard B Pearson, Anthony T Papenfuss, and David L Goode. Somatic mutations in early metazoan genes disrupt regulatory links between unicellular and multicellular genes in cancer. *eLife*, 8, 2019.

[6392] Albert Tsai, Mariana RP Alves, and Justin Crocker. Multi-enhancer transcriptional hubs confer phenotypic robustness. *eLife*, 8, 2019.

[6393] Ching-Ju Tsai, Jacopo Marino, Ricardo Adaixo, Filip Pamula, Jonas Muehle, Shoji Maeda, Tilman Flock, Nicholas MI Taylor, Inayatulla Mohammed, Hugues Matile, Roger JP Dawson, Xavier Deupi, Henning Stahlberg, and Gebhard Schertler. Cryo-EM structure of the rhodopsin-G$\alpha$i-$\beta\gamma$ complex reveals binding of the rhodopsin C-terminal tail to the g$\beta$ subunit. *eLife*, 8, 2019.

[6394] Erika Tsingos, Burkhard Höckendorf, Thomas Sütterlin, Stephan Kirchmaier, Niels Grabe, Lazaro Centanin, and Joachim Wittbrodt. Retinal stem cells modulate proliferative parameters to coordinate post-embryonic morphogenesis in the eye of fish. *eLife*, 8, 2019.

[6395] Joji Tsunada, Yale Cohen, and Joshua I Gold. Post-decision processing in primate prefrontal cortex influences subsequent choices on an auditory decision-making task. *eLife*, 8, 2019.

[6396] Shinichiro Tsutsumi, Naoki Hidaka, Yoshikazu Isomura, Masanori Matsuzaki, Kenji Sakimura, Masanobu Kano, and Kazuo Kitamura. Modular organization of cerebellar climbing fiber inputs during goal-directed behavior. *eLife*, 8, 2019.

[6397] Jérôme Tubiana, Simona Cocco, and Rémi Monasson. Learning protein constitutive motifs from sequence data. *eLife*, 8, 2019.

[6398] Raffaele Tucciarelli, Moritz Wurm, Elisa Baccolo, and Angelika Lingnau. The representational space of observed actions. *eLife*, 8, 2019.

[6399] Michal Tupec, AleŠ Buček, Václav JanouŠek, Heiko Vogel, Darina Prchalová, Jiří Kindl, Tereza Pavlíčková, Petra Wenzelová, Ullrich Jahn, Irena Valterová, and Iva Pichová. Expansion of the fatty acyl reductase gene family shaped pheromone communication in Hymenoptera. *eLife*, 8, 2019.

[6400] Rigney E Turnham, F Donelson Smith, Heidi L Kenerson, Mitchell H Omar, Martin Golkowski, Irvin Garcia, Renay Bauer, Ho-Tak Lau, Kevin M Sullivan, Lorene K Langeberg, Shao-En Ong, Kimberly J Riehle, Raymond S Yeung, and John D Scott. An acquired scaffolding function of the DNAJ-PKAc fusion contributes to oncogenic signaling in fibrolamellar carcinoma. *eLife*, 8, 2019.

[6401] Adam Tuttle, Catherine M Drerup, Molly Marra, Hillary McGraw, and Alex V Nechiporuk. Retrograde Ret signaling controls sensory pioneer axon outgrowth. *eLife*, 8, 2019.

[6402] Blake W Tye, Nicoletta Commins, Lillia V Ryazanova, Martin Wühr, Michael Springer, David Pincus, and L Stirling Churchman. Proteotoxicity from aberrant ribosome biogenesis compromises cell fitness. *eLife*, 8, 2019.

[6403] Ceereena Ubaida-Mohien, Alexey Lyashkov, Marta Gonzalez-Freire, Ravi Tharakan, Michelle Shardell, Ruin Moaddel, Richard D Semba, Chee W Chia, Myriam Gorospe, Ranjan Sen, and Luigi Ferrucci. Discovery proteomics in aging human skeletal muscle finds change in spliceosome, immunity, proteostasis and mitochondria. *eLife*, 8, 2019.

[6404] Emiko Uchikawa, Eunhee Choi, Guijun Shang, Hongtao Yu, and Xiao-chen Bai. Activation mechanism of the insulin receptor revealed by cryo-EM structure of the fully liganded receptor–ligand complex. *eLife*, 8, 2019.

[6405] SM Nashir Udden, Youn-Tae Kwak, Victoria Godfrey, Md Abdul Wadud Khan, Shahanshah Khan, Nicolas Loof, Lan Peng, Hao Zhu, and Hasan Zaki. NLRP12 suppresses hepatocellular carcinoma via downregulation of cJun N-terminal kinase activation in the hepatocyte. *eLife*, 8, 2019.

[6406] Akiyoshi Uezu, Erin Hisey, Yoshihiko Kobayashi, Yudong Gao, Tyler WA Bradshaw, Patrick Devlin, Ramona Rodriguiz, Purushothama Rao Tata, and Scott Soderling. Essential role for InSyn1 in dystroglycan complex integrity and cognitive behaviors in mice. *eLife*, 8, 2019.

[6407] Agnes Ulfig, Anton V Schulz, Alexandra Müller, Natalie Lupilov, and Lars I Leichert. N-chlorination mediates protective and immunomodulatory effects of oxidized human plasma proteins. *eLife*, 8, 2019.

[6408] Florian Ullrich, Sandy Blin, Katina Lazarow, Tony Daubitz, Jens Peter von Kries, and Thomas J Jentsch. Identification of TMEM206 proteins as pore of PAORAC/ASOR acid-sensitive chloride channels. *eLife*, 8, 2019.

[6409] Anne E Urai, Jan Willem de Gee, Konstantinos Tsetsos, and Tobias H Donner. Choice history biases subsequent evidence accumulation. *eLife*, 8, 2019.

[6410] Shinji Urata, Tadatsune Iida, Masamichi Yamamoto, Yu Mizushima, Chisato Fujimoto, Yu Matsumoto, Tatsuya Yamasoba, and Shigeo Okabe. Cellular cartography of the organ of Corti based on optical tissue clearing and machine learning. *eLife*, 8, 2019.

[6411] Jada H Vaden, Gokulakrishna Banumurthy, Eugeny S Gusarevich, Linda Overstreet-Wadiche, and Jacques I Wadiche. The readily-releasable pool dynamically regulates multivesicular release. *eLife*, 8, 2019.

[6412] Michael D Vahey and Daniel A Fletcher. Influenza A virus surface proteins are organized to help penetrate host mucus. *eLife*, 8, 2019.

[6413] Unnur A Valdimarsdóttir, Donghao Lu, Sigrún H Lund, Katja Fall, Fang Fang, Þórður Kristjánsson, Daníel Guðbjartsson, Agnar Helgason, and Kári Stefánsson. The mother's risk of premature death after child loss across two centuries. *eLife*, 8, 2019.

[6414] Giulia Vallardi, Lindsey A Allan, Lisa Crozier, and Adrian T Saurin. Division of labour between PP2A-B56 isoforms at the centromere and kinetochore. *eLife*, 8, 2019.

[6415] Katarina Valoskova, Julia Biebl, Marko Roblek, Shamsi Emtenani, Attila Gyoergy, Michaela Misova, Aparna Ratheesh, Patricia Reis-Rodrigues, Kateryna Shkarina, Ida Signe Bohse Larsen, Sergey Y Vakhrushev, Henrik Clausen, and Daria E Siekhaus. A conserved major facilitator superfamily member orchestrates a subset of O-glycosylation to aid macrophage tissue invasion. *eLife*, 8, 2019.

[6416] Bert van de Kooij, Pau Creixell, Anne van Vlimmeren, Brian A Joughin, Chad J Miller, Nasir Haider, Craig D Simpson, Rune Linding, Vuk Stambolic, Benjamin E Turk, and Michael B Yaffe. Comprehensive substrate specificity profiling of the human Nek kinome reveals unexpected signaling outputs. *eLife*, 8, 2019.

[6417] Jelle van den Ameele and Andrea H Brand. Neural stem cell temporal patterning and brain tumour growth rely on oxidative phosphorylation. *eLife*, 8, 2019.

[6418] Jochem van Kempen, Gerard M Loughnane, Daniel P Newman, Simon P Kelly, Alexander Thiele, Redmond G O'Connell, and Mark A Bellgrove. Behavioural and neural signatures of perceptual decision-making are modulated by pupil-linked arousal. *eLife*, 8, 2019.

[6419] Blaire Van Valkenburgh, Rolf O Peterson, Douglas W Smith, Daniel R Stahler, and John A Vucetich. Tooth fracture frequency in gray wolves reflects prey availability. *eLife*, 8, 2019.

[6420] J Edward van Veen, Michael Scherzer, Julia Boshuizen, Mollee Chu, Annie Liu, Allison Landman, Shon Green, Christy Trejo, and Martin McMahon. Mutationally-activated PI3′-kinase-$\alpha$ promotes de-differentiation of lung tumors initiated by the BRAFV600E oncoprotein kinase. *eLife*, 8, 2019.

[6421] Youna Vandaele, Nagaraj R Mahajan, David J Ottenheimer, Jocelyn M Richard, Shreesh P Mysore, and Patricia H Janak. Distinct recruitment of dorsomedial and dorsolateral striatum erodes with extended training. *eLife*, 8, 2019.

[6422] Sriram Varahan, Adhish Walvekar, Vaibhhav Sinha, Sandeep Krishna, and Sunil Laxman. Metabolic constraints drive self-organization of specialized cell groups. *eLife*, 8, 2019.

[6423] Grigor Varuzhanyan, Rebecca Rojansky, Michael J Sweredoski, Robert LJ Graham, Sonja Hess, Mark S Ladinsky, and David C Chan. Mitochondrial fusion is required for spermatogonial differentiation and meiosis. *eLife*, 8, 2019.

[6424] Ramakrishnan Vasudeva, Andreas Sutter, Kris Sales, Matthew E Dickinson, Alyson J Lumley, and Matthew JG Gage. Adaptive thermal plasticity enhances sperm and egg performance in a model insect. *eLife*, 8, 2019.

[6425] Ankita Ravi Vaswani, Beatrice Weykopf, Cathleen Hagemann, Hans-Ulrich Fried, Oliver Brüstle, and Sandra Blaess. Correct setup of the substantia nigra requires Reelin-mediated fast, laterally-directed migration of dopaminergic neurons. *eLife*, 8, 2019.

[6426] Anna Vavakou, Nigel P Cooper, and Marcel van der Heijden. The frequency limit of outer hair cell motility measured in vivo. *eLife*, 8, 2019.

[6427] Daniel čapek, Michael Smutny, Alexandra-Madelaine Tichy, Maurizio Morri, Harald Janovjak, and Carl-Philipp Heisenberg. Light-activated Frizzled7 reveals a permissive role of non-canonical wnt signaling in mesendoderm cell migration. *eLife*, 8, 2019.

[6428] Michiel Vellema, Mariana Diales Rocha, Sabrina Bascones, Sándor Zsebők, Jes Dreier, Stefan Leitner, Annemie Van der Linden, Jonathan Brewer, and Manfred Gahr. Accelerated redevelopment of vocal skills is preceded by lasting reorganization of the song motor circuitry. *eLife*, 8, 2019.

[6429] Lalanti Venkatasubramanian, Zhenhao Guo, Shuwa Xu, Liming Tan, Qi Xiao, Sonal Nagarkar-Jaiswal, and Richard S Mann. Stereotyped terminal axon branching of leg motor neurons mediated by IgSF proteins DIP-$\alpha$ and Dpr10. *eLife*, 8, 2019.

[6430] Frederick Verbruggen, Adam R Aron, Guido PH Band, Christian Beste, Patrick G Bissett, Adam T Brockett, Joshua W Brown, Samuel R Chamberlain, Christopher D Chambers, Hans Colonius, Lorenza S Colzato, Brian D Corneil, James P Coxon, Annie Dupuis, Dawn M Eagle, Hugh Garavan, Ian Greenhouse, Andrew Heathcote, René J Huster, Sara Jahfari, J Leon Kenemans, Inge Leunissen, Chiang-Shan R Li, Gordon D Logan, Dora Matzke, Sharon Morein-Zamir, Aditya Murthy, Martin Paré, Russell A Poldrack, K Richard Ridderinkhof, Trevor W Robbins, Matthew Roesch, Katya Rubia, Russell J Schachar, Jeffrey D Schall, Ann-Kathrin Stock, Nicole C Swann, Katharine N Thakkar, Maurits W van der Molen, Luc Vermeylen, Matthijs Vink, Jan R Wessel, Robert Whelan, Bram B Zandbelt, and C Nico Boehler. A consensus guide to capturing the ability to inhibit actions and impulsive behaviors in the stop-signal task. *eLife*, 8, 2019.

[6431] Berta Verd, Nicholas AM Monk, and Johannes Jaeger. Modularity, criticality, and evolvability of a developmental gene regulatory network. *eLife*, 8, 2019.

[6432] Lennart Verhagen, Cécile Gallea, Davide Folloni, Charlotte Constans, Daria EA Jensen, Harry Ahnine, Léa Roumazeilles, Mathieu Santin, Bashir Ahmed, Stéphane Lehericy, Miriam C Klein-Flügge, Kristine Krug, Rogier B Mars, Matthew FS Rushworth, Pierre Pouget, Jean-FranÇois Aubry, and Jerome Sallet. Offline impact of transcranial focused ultrasound on cortical activation in primates. *eLife*, 8, 2019.

[6433] Vikash Verma and Thomas J Maresca. Microtubule plus-ends act as physical signaling hubs to activate RhoA during cytokinesis. *eLife*, 8, 2019.

[6434] Carla Verna, Sree Janani Ravichandran, Megan G Sawchuk, Nguyen Manh Linh, and Enrico Scarpella. Coordination of tissue cell polarity by auxin transport and signaling. *eLife*, 8, 2019.

[6435] Cassondra L Vernier, Joshua J Krupp, Katelyn Marcus, Abraham Hefetz, Joel D Levine, and Yehuda Ben-Shahar. The cuticular hydrocarbon profiles of honey bee workers develop via a socially-modulated innate process. *eLife*, 8, 2019.

[6436] Jukka-Pekka Verta and Felicity C Jones. Predominance of cis-regulatory changes in parallel expression divergence of sticklebacks. *eLife*, 8, 2019.

[6437] Petra Vetter, Stephanie Badde, Elizabeth A Phelps, and Marisa Carrasco. Emotional faces guide the eyes in the absence of awareness. *eLife*, 8, 2019.

[6438] Nicholas C Vierra, Michael Kirmiz, Deborah van der List, L Fernando Santana, and James S Trimmer. Kv2.1 mediates spatial and functional coupling of L-type calcium channels and ryanodine receptors in mammalian neurons. *eLife*, 8, 2019.

[6439] Milena Vitale, Anush Bakunts, Andrea Orsi, Federica Lari, Laura Tadè, Alberto Danieli, Claudia Rato, Caterina Valetti, Roberto Sitia, Andrea Raimondi, John C Christianson, and Eelco van Anken. Inadequate BiP availability defines endoplasmic reticulum stress. *eLife*, 8, 2019.

[6440] Philip V'kovski, Markus Gerber, Jenna Kelly, Stephanie Pfaender, Nadine Ebert, Sophie Braga Lagache, Cedric Simillion, Jasmine Portmann, Hanspeter Stalder, Véronique Gaschen, Rémy Bruggmann, Michael H Stoffel, Manfred Heller, Ronald Dijkman, and Volker Thiel. Determination of host proteins composing the microenvironment of coronavirus replicase complexes by proximity-labeling. *eLife*, 8, 2019.

[6441] Aaron A Vogan, S Lorena Ament-Velásquez, Alexandra Granger-Farbos, Jesper Svedberg, Eric Bastiaans, Alfons JM Debets, Virginie Coustou, Hélène Yvanne, Corinne Clavé, Sven J Saupe, and Hanna Johannesson. Combinations of Spok genes create multiple meiotic drivers in Podospora. *eLife*, 8, 2019.

[6442] Joshua T Vogelstein, Eric W Bridgeford, Qing Wang, Carey E Priebe, Mauro Maggioni, and Cencheng Shen. Discovering and deciphering relationships across disparate data modalities. *eLife*, 8, 2019.

[6443] Timo Vögtle, Sumana Sharma, Jun Mori, Zoltan Nagy, Daniela Semeniak, Cyril Scandola, Mitchell J Geer, Christopher W Smith, Jordan Lane, Scott Pollack, Riitta Lassila, Annukka Jouppila, Alastair J Barr, Derek J Ogg, Tina D Howard, Helen J McMiken, Juli Warwicker, Catherine Geh, Rachel Rowlinson, W Mark Abbott, Anita Eckly, Harald Schulze, Gavin J Wright, Alexandra Mazharian, Klaus Fütterer, Sundaresan Rajesh, Michael R Douglas, and Yotis A Senis. Heparan sulfates are critical regulators of the inhibitory megakaryocyte-platelet receptor G6b-B. *eLife*, 8, 2019.

[6444] Hannah E Volkman, Stephanie Cambier, Elizabeth E Gray, and Daniel B Stetson. Tight nuclear tethering of cGAS is essential for preventing autoreactivity. *eLife*, 8, 2019.

[6445] James E Voss, Alicia Gonzalez-Martin, Raiees Andrabi, Roberta P Fuller, Ben Murrell, Laura E McCoy, Katelyn Porter, Deli Huang, Wenjuan Li, Devin Sok, Khoa Le, Bryan Briney, Morgan Chateau, Geoffrey Rogers, Lars Hangartner, Ann J Feeney, David Nemazee, Paula Cannon, and Dennis R Burton. Reprogramming the antigen specificity of B cells using genome-editing technologies. *eLife*, 8, 2019.

[6446] Leonhard Wachutka, Livia Caizzi, Julien Gagneur, and Patrick Cramer. Global donor and acceptor splicing site kinetics in human cells. *eLife*, 8, 2019.

[6447] Andrea Walens, Ashley V DiMarco, Ryan Lupo, Benjamin R Kroger, Jeffrey S Damrauer, and James V Alvarez. CCL5 promotes breast cancer recurrence through macrophage recruitment in residual tumors. *eLife*, 8, 2019.

[6448] Kerry MM Walker, Ray Gonzalez, Joe Z Kang, Josh H McDermott, and Andrew J King. Across-species differences in pitch perception are consistent with differences in cochlear filtering. *eLife*, 8, 2019.

[6449] Carolyn M Walsh, Rose Z Hill, Jamie Schwendinger-Schreck, Jacques Deguine, Emily C Brock, Natalie Kucirek, Ziad Rifi, Jessica Wei, Karsten Gronert, Rachel B Brem, Gregory M Barton, and Diana M Bautista. Neutrophils promote CXCR3-dependent itch in the development of atopic dermatitis. *eLife*, 8, 2019.

[6450] Justin D Walter, Marta Sawicka, and Raimund Dutzler. Cryo-EM structures and functional characterization of murine Slc26a9 reveal mechanism of uncoupled chloride transport. *eLife*, 8, 2019.

[6451] Chang-Zheng Wang, Jian Ma, Ye-Qian Xu, Shao-Na Jiang, Tian-Qi Chen, Zu-Liang Yuan, Xiao-Yi Mao, Shu-Qing Zhang, Lin-Yun Liu, Yinghui Fu, and Yong-Chun Yu. Early-generated interneurons regulate neuronal circuit formation during early postnatal development. *eLife*, 8, 2019.

[6452] Guiping Wang, David J Simon, Zhuhao Wu, Deanna M Belsky, Evan Heller, Melanie K O'Rourke, Nicholas T Hertz, Henrik Molina, Guisheng Zhong, Marc Tessier-Lavigne, and Xiaowei Zhuang. Structural plasticity of actin-spectrin membrane skeleton and functional role of actin and spectrin in axon degeneration. *eLife*, 8, 2019.

[6453] Luhong Wang, Charlotte Vanacker, Laura L Burger, Tammy Barnes, Yatrik M Shah, Martin G Myers, and Suzanne M Moenter. Genetic dissection of the different roles of hypothalamic kisspeptin neurons in regulating female reproduction. *eLife*, 8, 2019.

[6454] Qian Wang, Christin M Lepus, Harini Raghu, Laurent L Reber, Mindy M Tsai, Heidi H Wong, Ericka von Kaeppler, Nithya Lingampalli, Michelle S Bloom, Nick Hu, Eileen E Elliott, Francesca Oliviero, Leonardo Punzi, Nicholas J Giori, Stuart B Goodman, Constance R Chu, Jeremy Sokolove, Yoshihiro Fukuoka, Lawrence B Schwartz, Stephen J Galli, and William H Robinson. IgE-mediated mast cell activation promotes inflammation and cartilage destruction in osteoarthritis. *eLife*, 8, 2019.

[6455] Ruojun Wang, Kaj Kreutzfeldt, Helene Botella, Julien Vaubourgeix, Dirk Schnappinger, and Sabine Ehrt. Persistent Mycobacterium tuberculosis infection in mice requires PerM for successful cell division. *eLife*, 8, 2019.

[6456] Shixiang Wang, Zaoke He, Xuan Wang, Huimin Li, and Xue-Song Liu. Antigen presentation and tumor immunogenicity in cancer immunotherapy response prediction. *eLife*, 8, 2019.

[6457] Xingchao Wang, Qiong Wu, Laura Egan, Xiaosi Gu, Pinan Liu, Hong Gu, Yihong Yang, Jing Luo, Yanhong Wu, Zhixian Gao, and Jin Fan. Anterior insular cortex plays a critical role in interoceptive attention. *eLife*, 8, 2019.

[6458] Xiyue Wang, Xiaolin Chou, Bo Peng, Li Shen, Junxiang J Huang, Li I Zhang, and Huizhong W Tao. A cross-modality enhancement of defensive flight via parvalbumin neurons in zona incerta. *eLife*, 8, 2019.

[6459] Yan Wang, Menglong Rui, Quan Tang, Shufeng Bu, and Fengwei Yu. Patronin governs minus-end-out orientation of dendritic microtubules to promote dendrite pruning in Drosophila. *eLife*, 8, 2019.

[6460] Yanshu Wang, Mark F Sabbagh, Xiaowu Gu, Amir Rattner, John Williams, and Jeremy Nathans. Beta-catenin signaling regulates barrier-specific gene expression in circumventricular organ and ocular vasculatures. *eLife*, 8, 2019.

[6461] Yiyi Wang, Nicolas Bensaid, Pavan Tiruveedhula, Jianqiang Ma, Sowmya Ravikumar, and Austin Roorda. Human foveal cone photoreceptor topography and its dependence on eye length. *eLife*, 8, 2019.

[6462] Zong-Heng Wang, Yi Liu, Vijender Chaitankar, Mehdi Pirooznia, and Hong Xu. Electron transport chain biogenesis activated by a JNK-insulin-Myc relay primes mitochondrial inheritance in Drosophila. *eLife*, 8, 2019.

[6463] Michelle C Ward and Yoav Gilad. A generally conserved response to hypoxia in iPSC-derived cardiomyocytes from humans and chimpanzees. *eLife*, 8, 2019.

[6464] Kristen N Warren, Molly S Hermiller, Aneesha S Nilakantan, and Joel L Voss. Stimulating the hippocampal posterior-medial network enhances task-dependent connectivity and memory. *eLife*, 8, 2019.

[6465] Mya R Warren, Hui Sun, Yue Yan, Jonas Cremer, Bo Li, and Terence Hwa. Spatiotemporal establishment of dense bacterial colonies growing on hard agar. *eLife*, 8, 2019.

[6466] Leonhard Waschke, Sarah Tune, and Jonas Obleser. Local cortical desynchronization and pupil-linked arousal differentially shape brain states for optimal sensory performance. *eLife*, 8, 2019.

[6467] Samantha Washburn, Rachel Fremont, Maria Camila Moreno-Escobar, Chantal Angueyra, and Kamran Khodakhah. Acute cerebellar knockdown of Sgce reproduces salient features of myoclonus-dystonia (DYT11) in mice. *eLife*, 8, 2019.

[6468] Takamitsu Watanabe, Geraint Rees, and Naoki Masuda. Atypical intrinsic neural timescale in autism. *eLife*, 8, 2019.

[6469] James A Watson, Stije J Leopold, Julie A Simpson, Nicholas PJ Day, Arjen M Dondorp, and Nicholas J White. Collider bias and the apparent protective effect of glucose-6-phosphate dehydrogenase deficiency on cerebral malaria. *eLife*, 8, 2019.

[6470] Thomas Charles Watson, Pauline Obiang, Arturo Torres-Herraez, Aurélie Watilliaux, Patrice Coulon, Christelle Rochefort, and Laure Rondi-Reig. Anatomical and physiological foundations of cerebello-hippocampal interaction. *eLife*, 8, 2019.

[6471] Christoph Weber, Thomas Michaels, and L Mahadevan. Spatial control of irreversible protein aggregation. *eLife*, 8, 2019.

[6472] Michael W Webster, James AW Stowell, and Lori A Passmore. RNA-binding proteins distinguish between similar sequence motifs to promote targeted deadenylation by Ccr4-Not. *eLife*, 8, 2019.

[6473] Caroline Lei Wee, Erin Yue Song, Robert Evan Johnson, Deepak Ailani, Owen Randlett, Ji-Yoon Kim, Maxim Nikitchenko, Armin Bahl, Chao-Tsung Yang, Misha B Ahrens, Koichi Kawakami, Florian Engert, and Sam Kunes. A bidirectional network for appetite control in larval zebrafish. *eLife*, 8, 2019.

[6474] Martin Wegner, Valentina Diehl, Verena Bittl, Rahel de Bruyn, Svenja Wiechmann, Yves Matthess, Marie Hebel, Michael GB Hayes, Simone Schaubeck, Christopher Benner, Sven Heinz, Anja Bremm, Ivan Dikic, Andreas Ernst, and Manuel Kaulich. Circular synthesized CRISPR/Cas gRNAs for functional interrogations in the coding and noncoding genome. *eLife*, 8, 2019.

[6475] Fabian Wehnekamp, Gabriela Plucińska, Rachel Thong, Thomas Misgeld, and Don C Lamb. Nanoresolution real-time 3D orbital tracking for studying mitochondrial trafficking in vertebrate axons in vivo. *eLife*, 8, 2019.

[6476] Youheng Wei, Lucia Bettedi, Chun-Yuan Ting, Kuikwon Kim, Yingbiao Zhang, Jiadong Cai, and Mary A Lilly. The GATOR complex regulates an essential response to meiotic double-stranded breaks in Drosophila. *eLife*, 8, 2019.

[6477] Shirley Weiss, Jan E Melom, Kiel G Ormerod, Yao V Zhang, and J Troy Littleton. Glial Ca2+signaling links endocytosis to K+ buffering around neuronal somas to regulate excitability. *eLife*, 8, 2019.

[6478] Maaike Welling, Manuel Alexander Mohr, Aaron Ponti, Lluc Rullan Sabater, Andrea Boni, Yumiko K Kawamura, Prisca Liberali, Antoine HFM Peters, Pawel Pelczar, and Periklis Pantazis. Primed Track, high-fidelity lineage tracing in mouse pre-implantation embryos using primed conversion of photoconvertible proteins. *eLife*, 8, 2019.

[6479] Stephanie Wendt, Kim S Strunk, Jürgen Heinze, Andreas Roider, and Tomer J Czaczkes. Positive and negative incentive contrasts lead to relative value perception in ants. *eLife*, 8, 2019.

[6480] Jered M Wendte, Yinwen Zhang, Lexiang Ji, Xiuling Shi, Rashmi R Hazarika, Yadollah Shahryary, Frank Johannes, and Robert J Schmitz. Epimutations are associated with CHROMOMETHYLASE 3-induced de novo DNA methylation. *eLife*, 8, 2019.

[6481] Jan R Wessel, Darcy A Waller, and Jeremy DW Greenlee. Non-selective inhibition of inappropriate motor-tendencies during response-conflict by a fronto-subthalamic mechanism. *eLife*, 8, 2019.

[6482] Daan B Wesselink, Fiona MZ van den Heiligenberg, Naveed Ejaz, Harriet Dempsey-Jones, Lucilla Cardinali, Aurelie Tarall-Jozwiak, Jörn Diedrichsen, and Tamar R Makin. Obtaining and maintaining cortical hand representation as evidenced from acquired and congenital handlessness. *eLife*, 8, 2019.

[6483] Alan MV West, Scott C Rosenberg, Sarah N Ur, Madison K Lehmer, Qiaozhen Ye, Götz Hagemann, Iracema Caballero, Isabel Usón, Amy J MacQueen, Franz Herzog, and Kevin D Corbett. A conserved filamentous assembly underlies the structure of the meiotic chromosome axis. *eLife*, 8, 2019.

[6484] Jonathan R Whicher and Roderick MacKinnon. Regulation of Eag1 gating by its intracellular domains. *eLife*, 8, 2019.

[6485] Damien Beau Wilburn, Lisa M Tuttle, Rachel E Klevit, and Willie J Swanson. Indirect sexual selection drives rapid sperm protein evolution in abalone. *eLife*, 8, 2019.

[6486] Blake R Wilde, Zhizhou Ye, Tian-Yeh Lim, and Donald E Ayer. Cellular acidosis triggers human MondoA transcriptional activity by driving mitochondrial ATP production. *eLife*, 8, 2019.

[6487] Oliver J Wilkinson, Alejandro Martín-González, Haejoo Kang, Sarah J Northall, Dale B Wigley, Fernando Moreno-Herrero, and Mark Simon Dillingham. CtIP forms a tetrameric dumbbell-shaped particle which bridges complex DNA end structures for double-strand break repair. *eLife*, 8, 2019.

[6488] Amanda M Willard, Brian R Isett, Timothy C Whalen, Kevin J Mastro, Chris S Ki, Xiaobo Mao, and Aryn H Gittis. State transitions in the substantia nigra reticulata predict the onset of motor deficits in models of progressive dopamine depletion in mice. *eLife*, 8, 2019.

[6489] Ryan T Willett, N Sumru Bayin, Andrew S Lee, Anjana Krishnamurthy, Alexandre Wojcinski, Zhimin Lao, Daniel Stephen, Alberto Rosello-Diez, Katherine L Dauber-Decker, Grant D Orvis, Zhuhao Wu, Marc Tessier-Lavigne, and Alexandra L Joyner. Cerebellar nuclei excitatory neurons regulate developmental scaling of presynaptic Purkinje cell number and organ growth. *eLife*, 8, 2019.

[6490] Nathan H Williamson, Rea Ravin, Dan Benjamini, Hellmut Merkle, Melanie Falgairolle, Michael James O'Donovan, Dvir Blivis, Dave Ide, Teddy X Cai, Nima S Ghorashi, Ruiliang Bai, and Peter J Basser. Magnetic resonance measurements of cellular and sub-cellular membrane structures in live and fixed neural tissue. *eLife*, 8, 2019.

[6491] Ross S Williamson and Daniel B Polley. Parallel pathways for sound processing and functional connectivity among layer 5 and 6 auditory corticofugal neurons. *eLife*, 8, 2019.

[6492] Robert C Wilson and Anne GE Collins. Ten simple rules for the computational modeling of behavioral data. *eLife*, 8, 2019.

[6493] Evan Witt, Sigi Benjamin, Nicolas Svetec, and Li Zhao. Testis single-cell RNA-seq reveals the dynamics of de novo gene transcription and germline mutational bias in Drosophila. *eLife*, 8, 2019.

[6494] Jeroen Witteveldt, Lisanne I Knol, and Sara Macias. MicroRNA-deficient mouse embryonic stem cells acquire a functional interferon response. *eLife*, 8, 2019.

[6495] Guillaume Witz, Erik van Nimwegen, and Thomas Julou. Initiation of chromosome replication controls both division and replication cycles in E. coli through a double-adder mechanism. *eLife*, 8, 2019.

[6496] Dominik Witzigmann, Philipp Uhl, Sandro Sieber, Christina Kaufman, Tomaz Einfalt, Katrin Schöneweis, Philip Grossen, Jonas Buck, Yi Ni, Susanne H Schenk, Janine Hussner, Henriette E Meyer zu Schwabedissen, Gabriela Québatte, Walter Mier, Stephan Urban, and Jörg Huwyler. Optimization-by-design of hepatotropic lipid nanoparticles targeting the sodium-taurocholate co-transporting polypeptide. *eLife*, 8, 2019.

[6497] John W Wizeman, Qiuxia Guo, Elliott M Wilion, and James YH Li. Specification of diverse cell types during early neurogenesis of the mouse cerebellum. *eLife*, 8, 2019.

[6498] Aaron Benson Wong and J Gerard G Borst. Tonotopic and non-auditory organization of the mouse dorsal inferior colliculus revealed by two-photon imaging. *eLife*, 8, 2019.

[6499] Benjamin KC Wong, Shaza A Fadel, Shally Awasthi, Ajay Khera, Rajesh Kumar, Geetha Menon, and Prabhat Jha. The impact of measles immunization campaigns in India using a nationally representative sample of 27,000 child deaths. *eLife*, 8, 2019.

[6500] Felix Wong, Ethan C Garner, and Ariel Amir. Mechanics and dynamics of translocating MreB filaments on curved membranes. *eLife*, 8, 2019.

[6501] Francesca S Wong, R Fred Westbrook, and Nathan M Holmes. 'Online' integration of sensory and fear memories in the rat medial temporal lobe. *eLife*, 8, 2019.

[6502] Hiu-tung C Wong, Qiuxiang Zhang, Alisha J Beirl, Ronald S Petralia, Ya-Xian Wang, and Katie Kindt. Synaptic mitochondria regulate hair-cell synapse size and function. *eLife*, 8, 2019.

[6503] Io Nam Wong, Jacqueline PS Neo, Judith Oehler, Sophie Schafhauser, Fekret Osman, Stephen B Carr, and Matthew C Whitby. The Fml1-MHF complex suppresses inter-fork strand annealing in fission yeast. *eLife*, 8, 2019.

[6504] Kenneth Kin Lam Wong, Jenny Zhe Liao, and Esther M Verheyen. A positive feedback loop between Myc and aerobic glycolysis sustains tumor growth in a Drosophila tumor model. *eLife*, 8, 2019.

[6505] Yao Liang Wong, Lauren LeBon, Ana M Basso, Kathy L Kohlhaas, Arthur L Nikkel, Holly M Robb, Diana L Donnelly-Roberts, Janani Prakash, Andrew M Swensen, Nimrod D Rubinstein, Swathi Krishnan, Fiona E McAllister, Nicole V Haste, Jonathon J O'Brien, Margaret Roy, Andrea Ireland, Jennifer M Frost, Lei Shi, Stephan Riedmaier, Kathleen Martin, Michael J Dart, and Carmela Sidrauski. eIF2B activator prevents neurological defects caused by a chronic integrated stress response. *eLife*, 8, 2019.

[6506] Youngsik Woo, Soo Jeong Kim, Bo Kyoung Suh, Yongdo Kwak, Hyun-Jin Jung, Truong Thi My Nhung, Dong Jin Mun, Ji-Ho Hong, Su-Jin Noh, Seunghyun Kim, Ahryoung Lee, Seung Tae Baek, Minh Dang Nguyen, Youngshik Choe, and Sang Ki Park. Sequential phosphorylation of NDEL1 by the DYRK2-GSK3$\beta$ complex is critical for neuronal morphogenesis. *eLife*, 8, 2019.

[6507] Daniel Woods, Yinxin Dong, Frederic Bouche, Ryland Bednarek, Mark Rowe, Thomas Ream, and Richard Amasino. A florigen paralog is required for short-day vernalization in a pooid grass. *eLife*, 8, 2019.

[6508] Michael S Woody, Donald A Winkelmann, Marco Capitanio, E Michael Ostap, and Yale E Goldman. Single molecule mechanics resolves the earliest events in force generation by cardiac myosin. *eLife*, 8, 2019.

[6509] Oscar Woolnough, Kiefer James Forseth, Patrick Sarahan Rollo, and Nitin Tandon. Uncovering the functional anatomy of the human insula during speech. *eLife*, 8, 2019.

[6510] Kristina M Wright, Thomas C Jhou, Daniel Pimpinelli, and Michael A McDannald. Cue-inhibited ventrolateral periaqueductal gray neurons signal fear output and threat probability in male rats. *eLife*, 8, 2019.

[6511] Kristina M Wright and Michael A McDannald. Ventrolateral periaqueductal gray neurons prioritize threat probability over fear output. *eLife*, 8, 2019.

[6512] Bin Wu, FranÇois GC Blot, Aaron Benson Wong, Catarina Osório, Youri Adolfs, R Jeroen Pasterkamp, Jana Hartmann, Esther BE Becker, Henk-Jan Boele, Chris I De Zeeuw, and Martijn Schonewille. TRPC3 is a major contributor to functional heterogeneity of cerebellar Purkinje cells. *eLife*, 8, 2019.

[6513] Ling Wu, Ao Dong, Liting Dong, Shi-Qiang Wang, and Yulong Li. PARIS, an optogenetic method for functionally mapping gap junctions. *eLife*, 8, 2019.

[6514] Qiushuang Wu, Santiago Gerardo Medina, Gopal Kushawah, Michelle Lynn DeVore, Luciana A Castellano, Jacqelyn M Hand, Matthew Wright, and Ariel Alejandro Bazzini. Translation affects mRNA stability in a codon-dependent manner in human cells. *eLife*, 8, 2019.

[6515] Shun-Fan Wu, Ya-Long Ja, Yi-jie Zhang, and Chung-Hui Yang. Sweet neurons inhibit texture discrimination by signaling TMC-expressing mechanosensitive neurons in Drosophila. *eLife*, 8, 2019.

[6516] Yangyu Wu, Zhuyuan Chen, and Cecilia M Canessa. A valve-like mechanism controls desensitization of functional mammalian isoforms of acid-sensing ion channels. *eLife*, 8, 2019.

[6517] Shuke Xiao, Jennifer S Sun, and John R Carlson. Robust olfactory responses in the absence of odorant binding proteins. *eLife*, 8, 2019.

[6518] Chen Xie, Cemalettin Bekpen, Sven Künzel, Maryam Keshavarz, Rebecca Krebs-Wheaton, Neva Skrabar, Kristian Karsten Ullrich, and Diethard Tautz. A de novo evolved gene in the house mouse regulates female pregnancy cycles. *eLife*, 8, 2019.

[6519] Yi Xin, Allison Malick, Meiqin Hu, Chengdong Liu, Heya Batah, Haoxing Xu, and Cunming Duan. Cell-autonomous regulation of epithelial cell quiescence by calcium channel Trpv6. *eLife*, 8, 2019.

[6520] Guanglin Xing, Hongyang Jing, Lei Zhang, Yu Cao, Lei Li, Kai Zhao, Zhaoqi Dong, Wenbing Chen, Hongsheng Wang, Rangjuan Cao, Wen-Cheng Xiong, and Lin Mei. A mechanism in agrin signaling revealed by a prevalent Rapsyn mutation in congenital myasthenic syndrome. *eLife*, 8, 2019.

[6521] Ziyi Xiong, Gabriela Dankova, Laurence J Howe, Myoung Keun Lee, Pirro G Hysi, Markus A de Jong, Gu Zhu, Kaustubh Adhikari, Dan Li, Yi Li, Bo Pan, Eleanor Feingold, Mary L Marazita, John R Shaffer, Kerrie McAloney, Shu-Hua Xu, Li Jin, Sijia Wang, Femke MS de Vrij, Bas Lendemeijer, Stephen Richmond, Alexei Zhurov, Sarah Lewis, Gemma C Sharp, Lavinia Paternoster, Holly Thompson, Rolando Gonzalez-Jose, Maria Catira Bortolini, Samuel Canizales-Quinteros, Carla Gallo, Giovanni Poletti, Gabriel Bedoya, Francisco Rothhammer, André G Uitterlinden, M Arfan Ikram, Eppo Wolvius, Steven A Kushner, Tamar EC Nijsten, Robert-Jan TS Palstra, Stefan Boehringer, Sarah E Medland, Kun Tang, Andres Ruiz-Linares, Nicholas G Martin, Timothy D Spector, Evie Stergiakouli, Seth M Weinberg, Fan Liu, and Manfred Kayser. Novel genetic loci affecting facial shape variation in humans. *eLife*, 8, 2019.

[6522] Hui Xu, Shi-Jun Xu, Shu-Juan Xie, Yin Zhang, Jian-Hua Yang, Wei-Qi Zhang, Man-Ni Zheng, Hui Zhou, and Liang-Hu Qu. MicroRNA-122 supports robust innate immunity in hepatocytes by targeting the RTKs/STAT3 signaling pathway. *eLife*, 8, 2019.

[6523] Jiajia Xu, Yiyun Wang, Ching-Yun Hsu, Yongxing Gao, Carolyn Ann Meyers, Leslie Chang, Leititia Zhang, Kristen Broderick, Catherine Ding, Bruno Peault, Kenneth Witwer, and Aaron Watkins James. Human perivascular stem cell-derived extracellular vesicles mediate bone repair. *eLife*, 8, 2019.

[6524] Jianing Xu, Ed Reznik, Ho-Joon Lee, Gunes Gundem, Philip Jonsson, Judy Sarungbam, Anna Bialik, Francisco Sanchez-Vega, Chad J Creighton, Jake Hoekstra, Li Zhang, Peter Sajjakulnukit, Daniel Kremer, Zachary Tolstyka, Jozefina Casuscelli, Steve Stirdivant, Jie Tang, Nikolaus Schultz, Paul Jeng, Yiyu Dong, Wenjing Su, Emily H Cheng, Paul Russo, Jonathan A Coleman, Elli Papaemmanuil, Ying-Bei Chen, Victor E Reuter, Chris Sander, Scott R Kennedy, James J Hsieh, Costas A Lyssiotis, Satish K Tickoo, and A Ari Hakimi. Abnormal oxidative metabolism in a quiet genomic background underlies clear cell papillary renal cell carcinoma. *eLife*, 8, 2019.

[6525] Jin Xu, Kevin Nuno, Ulrike M Litzenburger, Yanyan Qi, M Ryan Corces, Ravindra Majeti, and Howard Y Chang. Single-cell lineage tracing by endogenous mutations enriched in transposase accessible mitochondrial DNA. *eLife*, 8, 2019.

[6526] Jingyue Xu, Han Liu, Yu Lan, Mike Adam, David E Clouthier, Steven Potter, and Rulang Jiang. Hedgehog signaling patterns the oral-aboral axis of the mandibular arch. *eLife*, 8, 2019.

[6527] Juncao Xu, Kaijie Cui, Liqiang Shen, Jing Shi, Lingting Li, Linlin You, Chengli Fang, Guoping Zhao, Yu Feng, Bei Yang, and Yu Zhang. Crl activates transcription by stabilizing active conformation of the master stress transcription initiation factor. *eLife*, 8, 2019.

[6528] Wen Xu, Lijiang Long, Yuehui Zhao, Lewis Stevens, Irene Felipe, Javier Munoz, Ronald E Ellis, and Patrick T McGrath. Evolution of Yin and Yang isoforms of a chromatin remodeling subunit precedes the creation of two genes. *eLife*, 8, 2019.

[6529] Xinhui Xu, Jinliang Gao, Wei Dai, Danyang Wang, Jian Wu, and Jinke Wang. Gene activation by a CRISPR-assisted trans enhancer. *eLife*, 8, 2019.

[6530] Koshika Yadava, Carlos Obed Medina, Heather Ishak, Irina Gurevich, Hedwich Kuipers, Elya Ali Shamskhou, Ievgen O Koliesnik, James J Moon, Casey Weaver, Kari Christine Nadeau, and Paul L Bollyky. Natural Tr1-like cells do not confer long-term tolerogenic memory. *eLife*, 8, 2019.

[6531] Oleksandr Yagensky, Mahdokht Kohansal-Nodehi, Saravanan Gunaseelan, Tamara Rabe, Saima Zafar, Inga Zerr, Wolfgang Härtig, Henning Urlaub, and John JE Chua. Increased expression of heme-binding protein 1 early in Alzheimer's disease is linked to neurotoxicity. *eLife*, 8, 2019.

[6532] Rachida Yakoubi, Astrid Rollenhagen, Marec von Lehe, Dorothea Miller, Bernd Walkenfort, Mike Hasenberg, Kurt Sätzler, and Joachim HR Lübke. Ultrastructural heterogeneity of layer 4 excitatory synaptic boutons in the adult human temporal lobe neocortex. *eLife*, 8, 2019.

[6533] Naoya Yamaguchi, Tugba Colak-Champollion, and Holger Knaut. zGrad is a nanobody-based degron system that inactivates proteins in zebrafish. *eLife*, 8, 2019.

[6534] Norihiro Yamaguchi, Ethan M Weinberg, Alexander Nguyen, Maria V Liberti, Hani Goodarzi, Yelena Y Janjigian, Philip B Paty, Leonard B Saltz, T Peter Kingham, Jia Min Loo, Elisa de Stanchina, and Sohail F Tavazoie. PCK1 and DHODH drive colorectal cancer liver metastatic colonization and hypoxic growth by promoting nucleotide synthesis. *eLife*, 8, 2019.

[6535] Kenta Yamamoto, Vikas Dubey, Katsumasa Irie, Hanayo Nakanishi, Himanshu Khandelia, Yoshinori Fujiyoshi, and Kazuhiro Abe. A single K+-binding site in the crystal structure of the gastric proton pump. *eLife*, 8, 2019.

[6536] King Faisal Yambire, Lorena Fernandez-Mosquera, Robert Steinfeld, Christiane Mühle, Elina Ikonen, Ira Milosevic, and Nuno Raimundo. Mitochondrial biogenesis is transcriptionally repressed in lysosomal lipid storage diseases. *eLife*, 8, 2019.

[6537] King Faisal Yambire, Christine Rostosky, Takashi Watanabe, David Pacheu-Grau, Sylvia Torres-Odio, Angela Sanchez-Guerrero, Ola Senderovich, Esther G Meyron-Holtz, Ira Milosevic, Jens Frahm, A Phillip West, and Nuno Raimundo. Impaired lysosomal acidification triggers iron deficiency and inflammation in vivo. *eLife*, 8, 2019.

[6538] Jing Yan, Chenyi Fei, Sheng Mao, Alexis Moreau, Ned S Wingreen, Andrej KoŠmrlj, Howard A Stone, and Bonnie L Bassler. Mechanical instability and interfacial energy drive biofilm morphogenesis. *eLife*, 8, 2019.

[6539] Le Yan, Richard A Neher, and Boris I Shraiman. Phylodynamic theory of persistence, extinction and speciation of rapidly adapting pathogens. *eLife*, 8, 2019.

[6540] Xuefei Yan, Beibei Tang, Biao Chen, Yongli Shan, Huajun Yang, Elizabeth Iorns, Elizabeth Iorns, Rachel Tsui, Alexandria Denis, Nicole Perfito, and Timothy M Errington. Replication Study: The microRNA miR-34a inhibits prostate cancer stem cells and metastasis by directly repressing CD44. *eLife*, 8, 2019.

[6541] Chi-Chun Yang, Hiroyuki Kato, Mayumi Shindo, and Hisao Masai. Cdc7 activates replication checkpoint by phosphorylating the Chk1-binding domain of Claspin in human cells. *eLife*, 8, 2019.

[6542] Dian Yang, Fangfei Qu, Hongchen Cai, Chen-Hua Chuang, Jing Shan Lim, Nadine Jahchan, Barbara M Grüner, Christin S Kuo, Christina Kong, Madeleine J Oudin, Monte M Winslow, and Julien Sage. Axon-like protrusions promote small cell lung cancer migration and metastasis. *eLife*, 8, 2019.

[6543] Fu Yang, Zhenghui Quan, Huanwei Huang, Minghui He, Xicheng Liu, Tao Cai, and Rongwen Xi. Ovaries absent links dLsd1 to HP1a for local H3K4 demethylation required for heterochromatic gene silencing. *eLife*, 8, 2019.

[6544] Meiling Yang, Yanli Wang, Qing Liu, Zhikang Liu, Feng Jiang, Huimin Wang, Xiaojiao Guo, Jianzhen Zhang, and Le Kang. A $\beta$-carotene-binding protein carrying a red pigment regulates body-color transition between green and black in locusts. *eLife*, 8, 2019.

[6545] Cong-Hui Yao, Rencheng Wang, Yahui Wang, Che-Pei Kung, Jason D Weber, and Gary J Patti. Mitochondrial fusion supports increased oxidative phosphorylation during cell proliferation. *eLife*, 8, 2019.

[6546] Mark Yarborough, Robert Nadon, and David G Karlin. Four erroneous beliefs thwarting more trustworthy research. *eLife*, 8, 2019.

[6547] Lihua Ye, Olaf Mueller, Jennifer Bagwell, Michel Bagnat, Rodger A Liddle, and John F Rawls. High fat diet induces microbiota-dependent silencing of enteroendocrine cells. *eLife*, 8, 2019.

[6548] Wenlei Ye, Tina W Han, Mu He, Yuh Nung Jan, and Lily Yeh Jan. Dynamic change of electrostatic field in TMEM16F permeation pathway shifts its ion selectivity. *eLife*, 8, 2019.

[6549] Wondwossen M Yeshaw, Marianne van der Zwaag, Francesco Pinto, Liza L Lahaye, Anita IE Faber, Rubén Gómez-Sánchez, Amalia M Dolga, Conor Poland, Anthony P Monaco, Sven CD van IJzendoorn, Nicola A Grzeschik, Antonio Velayos-Baeza, and Ody CM Sibon. Human VPS13A is associated with multiple organelles and influences mitochondrial morphology and lipid droplet motility. *eLife*, 8, 2019.

[6550] Marisa A Yonemitsu, Rachael M Giersch, Maria Polo-Prieto, Maurine Hammel, Alexis Simon, Florencia Cremonte, Fernando T Avilés, Nicolás Merino-Véliz, Erika AV Burioli, Annette F Muttray, James Sherry, Carol Reinisch, Susan A Baldwin, Stephen P Goff, Maryline Houssin, Gloria Arriagada, Nuria Vázquez, Nicolas Bierne, and Michael J Metzger. A single clonal lineage of transmissible cancer identified in two marine mussel species in South America and Europe. *eLife*, 8, 2019.

[6551] Jing Yong, Helmut Bischof, Sandra Burgstaller, Marina Siirin, Anne Murphy, Roland Malli, and Randal J Kaufman. Mitochondria supply ATP to the ER through a mechanism antagonized by cytosolic Ca2+. *eLife*, 8, 2019.

[6552] Yoseop Yoon, Jeff Klomp, Ines Martin-Martin, Frank Criscione, Eric Calvo, Jose Ribeiro, and Urs Schmidt-Ott. Embryo polarity in moth flies and mosquitoes relies on distinct old genes with localized transcript isoforms. *eLife*, 8, 2019.

[6553] Bernadette C Young, Sarah G Earle, Sona Soeng, Poda Sar, Varun Kumar, Songly Hor, Vuthy Sar, Rachel Bousfield, Nicholas D Sanderson, Leanne Barker, Nicole Stoesser, Katherine RW Emary, Christopher M Parry, Emma K Nickerson, Paul Turner, Rory Bowden, Derrick W Crook, David H Wyllie, Nicholas PJ Day, Daniel J Wilson, and Catrin E Moore. Panton–Valentine leucocidin is the key determinant of Staphylococcus aureus pyomyositis in a bacterial GWAS. *eLife*, 8, 2019.

[6554] Rodrigo M Young, Kenneth B Ewan, Veronica P Ferrer, Miguel L Allende, Jasminka Godovac-Zimmermann, Trevor C Dale, and Stephen W Wilson. Developmentally regulated Tcf7l2 splice variants mediate transcriptional repressor functions during eye formation. *eLife*, 8, 2019.

[6555] Rodrigo M Young, Thomas A Hawkins, Florencia Cavodeassi, Heather L Stickney, Quenten Schwarz, Lisa M Lawrence, Claudia Wierzbicki, Bowie YL Cheng, Jingyuan Luo, Elizabeth Mayela Ambrosio, Allison Klosner, Ian M Sealy, Jasmine Rowell, Chintan A Trivedi, Isaac H Bianco, Miguel L Allende, Elisabeth M Busch-Nentwich, Gaia Gestri, and Stephen W Wilson. Compensatory growth renders Tcf7l1a dispensable for eye formation despite its requirement in eye field specification. *eLife*, 8, 2019.

[6556] O Sascha Yousefi, Matthias Günther, Maximilian Hörner, Julia Chalupsky, Maximilian Wess, Simon M Brandl, Robert W Smith, Christian Fleck, Tim Kunkel, Matias D Zurbriggen, Thomas Höfer, Wilfried Weber, and Wolfgang WA Schamel. Optogenetic control shows that kinetic proofreading regulates the activity of the T cell receptor. *eLife*, 8, 2019.

[6557] Huasheng Yu, Tianjun Zhao, Simin Liu, Qinxue Wu, Omar Johnson, Zhaofa Wu, Zihao Zhuang, Yaocheng Shi, Luxin Peng, Renxi He, Yong Yang, Jianjun Sun, Xiaoqun Wang, Haifeng Xu, Zheng Zeng, Peng Zou, Xiaoguang Lei, Wenqin Luo, and Yulong Li. MRGPRX4 is a bile acid receptor for human cholestatic itch. *eLife*, 8, 2019.

[6558] Lu Yu, Valerie P O'Brien, Jonathan Livny, Denise Dorsey, Nirmalya Bandyopadhyay, Marco Colonna, Michael G Caparon, Elisha DO Roberson, Scott J Hultgren, and Thomas J Hannan. Mucosal infection rewires TNF$\alpha$ signaling dynamics to skew susceptibility to recurrence. *eLife*, 8, 2019.

[6559] Shuguang Yu and Jie He. Stochastic cell-cycle entry and cell-state-dependent fate outputs of injury-reactivated tectal radial glia in zebrafish. *eLife*, 8, 2019.

[6560] Xiaodi Yu, Olga Plotnikova, Paul D Bonin, Timothy A Subashi, Thomas J McLellan, Darren Dumlao, Ye Che, Yin Yao Dong, Elisabeth P Carpenter, Graham M West, Xiayang Qiu, Jeffrey S Culp, and Seungil Han. Cryo-EM structures of the human glutamine transporter SLC1A5 (ASCT2) in the outward-facing conformation. *eLife*, 8, 2019.

[6561] Zuanning Yuan, Roxana Georgescu, Ruda de Luna Almeida Santos, Daniel Zhang, Lin Bai, Nina Y Yao, Gongpu Zhao, Michael E O'Donnell, and Huilin Li. Ctf4 organizes sister replisomes and Pol $\alpha$ into a replication factory. *eLife*, 8, 2019.

[6562] Evanthia Zacharioudaki, Julia Falo Sanjuan, and Sarah Bray. Mi-2/NuRD complex protects stem cell progeny from mitogenic Notch signaling. *eLife*, 8, 2019.

[6563] Rani Zananiri, Omri Malik, Sergei Rudnizky, Vera Gaydar, Roman Kreiserman, Arnon Henn, and Ariel Kaplan. Synergy between RecBCD subunits is essential for efficient DNA unwinding. *eLife*, 8, 2019.

[6564] Mark Zander, Björn C Willige, Yupeng He, Thu A Nguyen, Amber E Langford, Ramlah Nehring, Elizabeth Howell, Robert McGrath, Anna Bartlett, Rosa Castanon, Joseph R Nery, Huaming Chen, Zhuzhu Zhang, Florian Jupe, Anna Stepanova, Robert J Schmitz, Mathew G Lewsey, Joanne Chory, and Joseph R Ecker. Epigenetic silencing of a multifunctional plant stress regulator. *eLife*, 8, 2019.

[6565] Aref Arzan Zarin, Brandon Mark, Albert Cardona, Ashok Litwin-Kumar, and Chris Q Doe. A multilayer circuit architecture for the generation of distinct locomotor behaviors in Drosophila. *eLife*, 8, 2019.

[6566] Taraneh Zarin, Bob Strome, Alex N Nguyen Ba, Simon Alberti, Julie D Forman-Kay, and Alan M Moses. Proteome-wide signatures of function in highly diverged intrinsically disordered regions. *eLife*, 8, 2019.

[6567] Eszter Zavodszky and Ramanujan S Hegde. Misfolded GPI-anchored proteins are escorted through the secretory pathway by ER-derived factors. *eLife*, 8, 2019.

[6568] TomáŠ Zavřel, Marjan Faizi, Cristina Loureiro, Gereon Poschmann, Kai Stühler, Maria Sinetova, Anna Zorina, Ralf Steuer, and Jan červený. Quantitative insights into the cyanobacterial cell economy. *eLife*, 8, 2019.

[6569] Stefan Zdraljevic, Bennett William Fox, Christine Strand, Oishika Panda, Francisco J Tenjo, Shannon C Brady, Tim A Crombie, John G Doench, Frank C Schroeder, and Erik C Andersen. Natural variation in C. elegans arsenic toxicity is explained by differences in branched chain amino acid metabolism. *eLife*, 8, 2019.

[6570] Hao Zeng, Johnny Castillo-Cabrera, Mika Manser, Bo Lu, Zinger Yang, Vaik Strande, Damien Begue, Raffaella Zamponi, Shumei Qiu, Frederic Sigoillot, Qiong Wang, Alicia Lindeman, John S Reece-Hoyes, Carsten Russ, Debora Bonenfant, Xiaomo Jiang, Youzhen Wang, and Feng Cong. Genome-wide CRISPR screening reveals genetic modifiers of mutant EGFR dependence in human NSCLC. *eLife*, 8, 2019.

[6571] Jingwei Zeng, Greg Slodkowicz, and Leo C James. Rare missense variants in the human cytosolic antibody receptor preserve antiviral function. *eLife*, 8, 2019.

[6572] Feng Zhang, Kenton Jon Swartz, and Andres Jara-Oseguera. Conserved allosteric pathways for activation of TRPV3 revealed through engineering vanilloid-sensitivity. *eLife*, 8, 2019.

[6573] Haiyang Zhang, Meghan V Petrie, Yiwei He, Jared M Peace, Irene E Chiolo, and Oscar M Aparicio. Dynamic relocalization of replication origins by Fkh1 requires execution of DDK function and Cdc45 loading at origins. *eLife*, 8, 2019.

[6574] Hejing Zhang, Jörg Gross, Carsten De Dreu, and Yina Ma. Oxytocin promotes coordinated out-group attack during intergroup conflict in humans. *eLife*, 8, 2019.

[6575] Hongsheng Zhang, Anupama Sathyamurthy, Fang Liu, Lei Li, Lei Zhang, Zhaoqi Dong, Wanpeng Cui, Xiangdong Sun, Kai Zhao, Hongsheng Wang, Hsin-Yi Henry Ho, Wen-Cheng Xiong, and Lin Mei. Agrin-Lrp4-Ror2 signaling regulates adult hippocampal neurogenesis in mice. *eLife*, 8, 2019.

[6576] Jie Zhang, Liang Li, Haoliang Huang, Fang Fang, Hannah C Webber, Pei Zhuang, Liang Liu, Roopa Dalal, Peter H Tang, Vinit B Mahajan, Yang Sun, Shaohua Li, Mingchang Zhang, Jeffrey L Goldberg, and Yang Hu. Silicone oil-induced ocular hypertension and glaucomatous neurodegeneration in mouse. *eLife*, 8, 2019.

[6577] Lu Zhang, John Lee, Christopher Rozell, and Annabelle C Singer. Sub-second dynamics of theta-gamma coupling in hippocampal CA1. *eLife*, 8, 2019.

[6578] Peipei Zhang, Baochang Fan, Peiguo Yang, Jamshid Temirov, James Messing, Hong Joo Kim, and J Paul Taylor. Chronic optogenetic induction of stress granules is cytotoxic and reveals the evolution of ALS-FTD pathology. *eLife*, 8, 2019.

[6579] Shuping Zhang, Alejandra Macias-Garcia, Jacob C Ulirsch, Jason Velazquez, Vincent L Butty, Stuart S Levine, Vijay G Sankaran, and Jane-Jane Chen. HRI coordinates translation necessary for protein homeostasis and mitochondrial function in erythropoiesis. *eLife*, 8, 2019.

[6580] Ting Zhang, Ling Zhu, Michele C Madigan, Wei Liu, Weiyong Shen, Svetlana Cherepanoff, Fanfan Zhou, Shaoxue Zeng, Jianhai Du, and Mark C Gillies. Human macular Müller cells rely more on serine biosynthesis to combat oxidative stress than those from the periphery. *eLife*, 8, 2019.

[6581] Wanying Zhang and Randy M Bruno. High-order thalamic inputs to primary somatosensory cortex are stronger and longer lasting than cortical inputs. *eLife*, 8, 2019.

[6582] Wei Zhang, Jason A Corwin, Daniel Harrison Copeland, Julie Feusier, Robert Eshbaugh, David E Cook, Suzi Atwell, and Daniel J Kliebenstein. Plant–necrotroph co-transcriptome networks illuminate a metabolic battlefield. *eLife*, 8, 2019.

[6583] Wen-Hao Zhang, He Wang, Aihua Chen, Yong Gu, Tai Sing Lee, KY Michael Wong, and Si Wu. Complementary congruent and opposite neurons achieve concurrent multisensory integration and segregation. *eLife*, 8, 2019.

[6584] Wenjuan Zhang, Benjamin Falcon, Alexey G Murzin, Juan Fan, R Anthony Crowther, Michel Goedert, and Sjors HW Scheres. Heparin-induced tau filaments are polymorphic and differ from those in Alzheimer's and Pick's diseases. *eLife*, 8, 2019.

[6585] Xi Zhang, Ricardo AR Machado, Cong Van Doan, Carla CM Arce, Lingfei Hu, and Christelle AM Robert. Entomopathogenic nematodes increase predation success by inducing cadaver volatiles that attract healthy herbivores. *eLife*, 8, 2019.

[6586] Xiao Min Zhang, Urielle FranÇois, Kätlin Silm, Maria Florencia Angelo, Maria Victoria Fernandez-Busch, Mona Maged, Christelle Martin, Véronique Bernard, Fabrice P Cordelières, Melissa Deshors, Stéphanie Pons, Uwe Maskos, Alexis Pierre Bemelmans, Sonja M Wojcik, Salah El Mestikawy, Yann Humeau, and Etienne Herzog. A proline-rich motif on VGLUT1 reduces synaptic vesicle super-pool and spontaneous release frequency. *eLife*, 8, 2019.

[6587] Xiaoli Zhang, Wei Chen, Ping Li, Raul Calvo, Noel Southall, Xin Hu, Melanie Bryant-Genevier, Xinghua Feng, Qi Geng, Chenlang Gao, Meimei Yang, Kaiyuan Tang, Marc Ferrer, Juan Jose Marugan, and Haoxing Xu. Agonist-specific voltage-dependent gating of lysosomal two-pore Na+ channels. *eLife*, 8, 2019.

[6588] Xiaonan Zhang, Kaylynn Coates, Andrew Dacks, Cengiz Günay, J Scott Lauritzen, Feng Li, Steven A Calle-Schuler, Davi Bock, and Quentin Gaudry. Local synaptic inputs support opposing, network-specific odor representations in a widely projecting modulatory neuron. *eLife*, 8, 2019.

[6589] Xiaoxing Zhang, Wenjun Yan, Wenliang Wang, Hongmei Fan, Ruiqing Hou, Yulei Chen, Zhaoqin Chen, Chaofan Ge, Shumin Duan, Albert Compte, and Chengyu T Li. Active information maintenance in working memory by a sensory cortex. *eLife*, 8, 2019.

[6590] Yue Zhang, Anne Lanjuin, Suvagata Roy Chowdhury, Meeta Mistry, Carlos G Silva-García, Heather J Weir, Chia-Lin Lee, Caroline C Escoubas, Emina Tabakovic, and William B Mair. Neuronal TORC1 modulates longevity via AMPK and cell nonautonomous regulation of mitochondrial dynamics in C. elegans. *eLife*, 8, 2019.

[6591] Jia Zhao, Weijian Zong, Yiwen Zhao, Dongzhou Gou, Shenghui Liang, Jiayu Shen, Yi Wu, Xuan Zheng, Runlong Wu, Xu Wang, Fuzeng Niu, Aimin Wang, Yunfeng Zhang, Jing-Wei Xiong, Liangyi Chen, and Yanmei Liu. In vivo imaging of $\beta$-cell function reveals glucose-mediated heterogeneity of $\beta$-cell functional development. *eLife*, 8, 2019.

[6592] Xiaocheng Zhao, Pavel Nedvetsky, Fabio Stanchi, Anne-Clemence Vion, Oliver Popp, Kerstin Zühlke, Gunnar Dittmar, Enno Klussmann, and Holger Gerhardt. Endothelial PKA activity regulates angiogenesis by limiting autophagy through phosphorylation of ATG16L1. *eLife*, 8, 2019.

[6593] Wang Zheng, Elena O Gracheva, and Sviatoslav N Bagriantsev. A hydrophobic gate in the inner pore helix is the major determinant of inactivation in mechanosensitive Piezo channels. *eLife*, 8, 2019.

[6594] Felix Y Zhou, Carlos Ruiz-Puig, Richard P Owen, Michael J White, Jens Rittscher, and Xin Lu. Motion sensing superpixels (MOSES) is a systematic computational framework to quantify and discover cellular motion phenotypes. *eLife*, 8, 2019.

[6595] Guangyu Zhou, Gregory Lane, Shiloh L Cooper, Thorsten Kahnt, and Christina Zelano. Characterizing functional pathways of the human olfactory system. *eLife*, 8, 2019.

[6596] Heng Zhou, Kevin R Neville, Nitsan Goldstein, Shushi Kabu, Naila Kausar, Rong Ye, Thuan Tinh Nguyen, Noah Gelwan, Bradley T Hyman, and Stephen N Gomperts. Cholinergic modulation of hippocampal calcium activity across the sleep-wake cycle. *eLife*, 8, 2019.

[6597] Lijun Zhou, Seohyun Chris Kim, Katherine H Ho, Derek K O'Flaherty, Constantin Giurgiu, Tom H Wright, and Jack W Szostak. Non-enzymatic primer extension with strand displacement. *eLife*, 8, 2019.

[6598] Qinbo Zhou, Bo Yu, Chastain Anderson, Zhan-Peng Huang, Jakub Hanus, Wensheng Zhang, Yu Han, Partha S Bhattacharjee, Sathish Srinivasan, Kun Zhang, Da-zhi Wang, and Shusheng Wang. LncEGFL7OS regulates human angiogenesis by interacting with MAX at the EGFL7/miR-126 locus. *eLife*, 8, 2019.

[6599] Qingtong Zhou, Dehua Yang, Meng Wu, Yu Guo, Wanjing Guo, Li Zhong, Xiaoqing Cai, Antao Dai, Wonjo Jang, Eugene I Shakhnovich, Zhi-Jie Liu, Raymond C Stevens, Nevin A Lambert, M Madan Babu, Ming-Wei Wang, and Suwen Zhao. Common activation mechanism of class A GPCRs. *eLife*, 8, 2019.

[6600] Wenchang Zhou, Giacomo Fiorin, Claudio Anselmi, Hossein Ali Karimi-Varzaneh, Horacio Poblete, Lucy R Forrest, and José D Faraldo-Gómez. Large-scale state-dependent membrane remodeling by a transporter protein. *eLife*, 8, 2019.

[6601] Ye Zhou, Can Cao, Lingli He, Xianping Wang, and Xuejun Cai Zhang. Crystal structure of dopamine receptor D4 bound to the subtype selective ligand, L745870. *eLife*, 8, 2019.

[6602] Yu-Lingzi Zhou, Adam Ślipiński, Dong Ren, and Joseph Parker. A Mesozoic clown beetle myrmecophile (Coleoptera: Histeridae). *eLife*, 8, 2019.

[6603] Jinsheng Zhu, Kelvin Lau, Robert Puschmann, Robert K Harmel, Youjun Zhang, Verena Pries, Philipp Gaugler, Larissa Broger, Amit K Dutta, Henning J Jessen, Gabriel Schaaf, Alisdair R Fernie, Ludwig A Hothorn, Dorothea Fiedler, and Michael Hothorn. Two bifunctional inositol pyrophosphate kinases/phosphatases control plant phosphate homeostasis. *eLife*, 8, 2019.

[6604] Sha Joe Zhu, Jason A Hendry, Jacob Almagro-Garcia, Richard D Pearson, Roberto Amato, Alistair Miles, Daniel J Weiss, Tim CD Lucas, Michele Nguyen, Peter W Gething, Dominic Kwiatkowski, and Gil McVean. The origins and relatedness structure of mixed infections vary with local prevalence of P. falciparum malaria. *eLife*, 8, 2019.

[6605] Ying Zhu, Mirko Scheibinger, Daniel Christian Ellwanger, Jocelyn F Krey, Dongseok Choi, Ryan T Kelly, Stefan Heller, and Peter G Barr-Gillespie. Single-cell proteomics reveals changes in expression during hair-cell development. *eLife*, 8, 2019.

[6606] Matthew N Zipple, Elizabeth A Archie, Jenny Tung, Jeanne Altmann, and Susan C Alberts. Intergenerational effects of early adversity on survival in wild baboons. *eLife*, 8, 2019.

[6607] Lejla Zubcevic, William F Borschel, Allen L Hsu, Mario J Borgnia, and Seok-Yong Lee. Regulatory switch at the cytoplasmic interface controls TRPV channel gating. *eLife*, 8, 2019.

[6608] Lejla Zubcevic, Allen L Hsu, Mario J Borgnia, and Seok-Yong Lee. Symmetry transitions during gating of the TRPV2 ion channel in lipid membranes. *eLife*, 8, 2019.

[6609] Esther Zumaquero, Sara L Stone, Christopher D Scharer, Scott A Jenks, Anoma Nellore, Betty Mousseau, Antonio Rosal-Vela, Davide Botta, John E Bradley, Wojciech Wojciechowski, Travis Ptacek, Maria I Danila, Jeffrey C Edberg, S Louis Bridges, Robert P Kimberly, W Winn Chatham, Trenton R Schoeb, Alexander F Rosenberg, Jeremy M Boss, Ignacio Sanz, and Frances E Lund. IFN$\gamma$ induces epigenetic programming of human T-bethi B cells and promotes TLR7/8 and IL-21 induced differentiation. *eLife*, 8, 2019.

[6610] Zhicheng Zuo, Ashwini Zolekar, Kesavan Babu, Victor JT Lin, Hamed S Hayatshahi, Rakhi Rajan, Yu-Chieh Wang, and Jin Liu. Structural and functional insights into the bona fide catalytic state of Streptococcus pyogenes Cas9 HNH nuclease domain. *eLife*, 8, 2019.

[6611] Perry Zurn and Danielle S Bassett. Seizing an opportunity. *eLife*, 8, 2019.

[6612] Hub Zwart. From primal scenes to synthetic cells. *eLife*, 8, 2019.

[6613] Katherine G Zyner, Darcie S Mulhearn, Santosh Adhikari, Sergio Martínez Cuesta, Marco Di Antonio, Nicolas Erard, Gregory J Hannon, David Tannahill, and Shankar Balasubramanian. Genetic interactions of G-quadruplexes in humans. *eLife*, 8, 2019.

[6614] Friso S Aalbers, Maximilian JLJ Fürst, Stefano Rovida, Milos Trajkovic, J Rubén Gómez Castellanos, Sebastian Bartsch, Andreas Vogel, Andrea Mattevi, and Marco W Fraaije. Approaching boiling point stability of an alcohol dehydrogenase through computationally-guided enzyme engineering. *eLife*, 9, 2020.

[6615] Syafiq Abd Wahab and Dirk Remus. Antagonistic control of DDK binding to licensed replication origins by Mcm2 and Rad53. *eLife*, 9, 2020.

[6616] Richard J Abdill, Elizabeth M Adamowicz, and Ran Blekhman. International authorship and collaboration across bioRxiv preprints. *eLife*, 9, 2020.

[6617] Nour J Abdulhay, Colin P McNally, Laura J Hsieh, Sivakanthan Kasinathan, Aidan Keith, Laurel S Estes, Mehran Karimzadeh, Jason G Underwood, Hani Goodarzi, Geeta J Narlikar, and Vijay Ramani. Massively multiplex single-molecule oligonucleosome footprinting. *eLife*, 9, 2020.

[6618] Sonia Accossato, Felix Kessler, and Venkatasalam Shanmugabalaji. SUMOylation contributes to proteostasis of the chloroplast protein import receptor TOC159 during early development. *eLife*, 9, 2020.

[6619] Titipat Achakulvisut, Tulakan Ruangrong, Isil Bilgin, Sofie Van Den Bossche, Brad Wyble, Dan FM Goodman, and Konrad P Kording. Improving on legacy conferences by moving online. *eLife*, 9, 2020.

[6620] Benjamin M Adams, Nathan P Canniff, Kevin P Guay, Ida Signe Bohse Larsen, and Daniel N Hebert. Quantitative glycoproteomics reveals cellular substrate selectivity of the ER protein quality control sensors UGGT1 and UGGT2. *eLife*, 9, 2020.

[6621] Nikolai M Adamski, Philippa Borrill, Jemima Brinton, Sophie A Harrington, Clémence Marchal, Alison R Bentley, William D Bovill, Luigi Cattivelli, James Cockram, Bruno Contreras-Moreira, Brett Ford, Sreya Ghosh, Wendy Harwood, Keywan Hassani-Pak, Sadiye Hayta, Lee T Hickey, Kostya Kanyuka, Julie King, Marco Maccaferrri, Guy Naamati, Curtis J Pozniak, Ricardo H Ramirez-Gonzalez, Carolina Sansaloni, Ben Trevaskis, Luzie U Wingen, Brande BH Wulff, and Cristobal Uauy. A roadmap for gene functional characterisation in crops with large genomes: Lessons from polyploid wheat. *eLife*, 9, 2020.

[6622] Naga Suresh Adapala, Gaurav Swarnkar, Manoj Arra, Jie Shen, Gabriel Mbalaviele, Ke Ke, and Yousef Abu-Amer. Inflammatory osteolysis is regulated by site-specific ISGylation of the scaffold protein NEMO. *eLife*, 9, 2020.

[6623] Rebecca C Adikes, Abraham Q Kohrman, Michael A Q Martinez, Nicholas J Palmisano, Jayson J Smith, Taylor N Medwig-Kinney, Mingwei Min, Maria D Sallee, Ononnah B Ahmed, Nuri Kim, Simeiyun Liu, Robert D Morabito, Nicholas Weeks, Qinyun Zhao, Wan Zhang, Jessica L Feldman, Michalis Barkoulas, Ariel M Pani, Sabrina L Spencer, Benjamin L Martin, and David Q Matus. Visualizing the metazoan proliferation-quiescence decision in vivo. *eLife*, 9, 2020.

[6624] Jeffrey R Adrion, Christopher B Cole, Noah Dukler, Jared G Galloway, Ariella L Gladstein, Graham Gower, Christopher C Kyriazis, Aaron P Ragsdale, Georgia Tsambos, Franz Baumdicker, Jedidiah Carlson, Reed A Cartwright, Arun Durvasula, Ilan Gronau, Bernard Y Kim, Patrick McKenzie, Philipp W Messer, Ekaterina Noskova, Diego Ortega-Del Vecchyo, Fernando Racimo, Travis J Struck, Simon Gravel, Ryan N Gutenkunst, Kirk E Lohmueller, Peter L Ralph, Daniel R Schrider, Adam Siepel, Jerome Kelleher, and Andrew D Kern. A community-maintained standard library of population genetic models. *eLife*, 9, 2020.

[6625] Felipe Aedo-Jury, Miriam Schwalm, Lara Hamzehpour, and Albrecht Stroh. Brain states govern the spatio-temporal dynamics of resting-state functional connectivity. *eLife*, 9, 2020.

[6626] Oshri Afanzar, Garrison K Buss, Tim Stearns, and James E Ferrell. The nucleus serves as the pacemaker for the cell cycle. *eLife*, 9, 2020.

[6627] Haggai Agmon and Yoram Burak. A theory of joint attractor dynamics in the hippocampus and the entorhinal cortex accounts for artificial remapping and grid cell field-to-field variability. *eLife*, 9, 2020.

[6628] Aakash Agrawal, KVS Hari, and SP Arun. A compositional neural code in high-level visual cortex can explain jumbled word reading. *eLife*, 9, 2020.

[6629] Sweta Agrawal, Evyn S Dickinson, Anne Sustar, Pralaksha Gurung, David Shepherd, James W Truman, and John C Tuthill. Central processing of leg proprioception in Drosophila. *eLife*, 9, 2020.

[6630] Noa Aharon-Hefetz, Idan Frumkin, Yoav Mayshar, Orna Dahan, Yitzhak Pilpel, and Roni Rak. Manipulation of the human tRNA pool reveals distinct tRNA sets that act in cellular proliferation or cell cycle arrest. *eLife*, 9, 2020.

[6631] Juraj Ahel, Anita Lehner, Antonia Vogel, Alexander Schleiffer, Anton Meinhart, David Haselbach, and Tim Clausen. Moyamoya disease factor RNF213 is a giant E3 ligase with a dynein-like core and a distinct ubiquitin-transfer mechanism. *eLife*, 9, 2020.

[6632] Shehryar Ahmad, Kara K Tsang, Kartik Sachar, Dennis Quentin, Tahmid M Tashin, Nathan P Bullen, Stefan Raunser, Andrew G McArthur, Gerd Prehna, and John C Whitney. Structural basis for effector transmembrane domain recognition by type VI secretion system chaperones. *eLife*, 9, 2020.

[6633] Shintaro Aibara, Vivek Singh, Angelika Modelska, and Alexey Amunts. Structural basis of mitochondrial translation. *eLife*, 9, 2020.

[6634] Timothy J Aikin, Amy F Peterson, Michael J Pokrass, Helen R Clark, and Sergi Regot. MAPK activity dynamics regulate non-cell autonomous effects of oncogene expression. *eLife*, 9, 2020.

[6635] Mattia Aime, Elisabete Augusto, Vladimir Kouskoff, Tiago Campelo, Christelle Martin, Yann Humeau, Nicolas Chenouard, and Frederic Gambino. The integration of Gaussian noise by long-range amygdala inputs in frontal circuit promotes fear learning in mice. *eLife*, 9, 2020.

[6636] Matthew Akamatsu, Ritvik Vasan, Daniel Serwas, Michael A Ferrin, Padmini Rangamani, and David G Drubin. Principles of self-organization and load adaptation by the actin cytoskeleton during clathrin-mediated endocytosis. *eLife*, 9, 2020.

[6637] Jyothi S Akella, Stephen P Carter, Ken Nguyen, Sofia Tsiropoulou, Ailis L Moran, Malan Silva, Fatima Rizvi, Breandan N Kennedy, David H Hall, Maureen M Barr, and Oliver E Blacque. Ciliary Rab28 and the BBSome negatively regulate extracellular vesicle shedding. *eLife*, 9, 2020.

[6638] Mazen Al Borno, Saurabh Vyas, Krishna V Shenoy, and Scott L Delp. High-fidelity musculoskeletal modeling reveals that motor planning variability contributes to the speed-accuracy tradeoff. *eLife*, 9, 2020.

[6639] Omar Al Rifai, Catherine Julien, Julie Lacombe, Denis Faubert, Erandi Lira-Navarrete, Yoshiki Narimatsu, Henrik Clausen, and Mathieu Ferron. The half-life of the bone-derived hormone osteocalcin is regulated through O-glycosylation in mice, but not in humans. *eLife*, 9, 2020.

[6640] Opeyemi O Alabi, M Felicia Davatolhagh, Mara Robinson, Michael P Fortunato, Luigim Vargas Cifuentes, Joseph W Kable, and Marc Vincent Fuccillo. Disruption of Nrxn1$\alpha$ within excitatory forebrain circuits drives value-based dysfunction. *eLife*, 9, 2020.

[6641] Andrea Alamia, Christopher Timmermann, David J Nutt, Rufin VanRullen, and Robin L Carhart-Harris. DMT alters cortical travelling waves. *eLife*, 9, 2020.

[6642] Cristina Alarcón-Vila, Alberto Baroja-Mazo, Carlos de Torre-Minguela, Carlos M Martínez, Juan J Martínez-García, Helios Martínez-Banaclocha, Carlos García-Palenciano, and Pablo Pelegrin. CD14 release induced by P2X7 receptor restricts inflammation and increases survival during sepsis. *eLife*, 9, 2020.

[6643] Catarina Albergaria, N Tatiana Silva, Dana M Darmohray, and Megan R Carey. Cannabinoids modulate associative cerebellar learning via alterations in behavioral state. *eLife*, 9, 2020.

[6644] Scott T Albert, Alkis M Hadjiosif, Jihoon Jang, Andrew J Zimnik, Demetris S Soteropoulos, Stuart N Baker, Mark M Churchland, John W Krakauer, and Reza Shadmehr. Postural control of arm and fingers through integration of movement commands. *eLife*, 9, 2020.

[6645] Callison E Alcott, Hari Krishna Yalamanchili, Ping Ji, Meike E van der Heijden, Alexander Saltzman, Nathan Elrod, Ai Lin, Mei Leng, Bhoomi Bhatt, Shuang Hao, Qi Wang, Afaf Saliba, Jianrong Tang, Anna Malovannaya, Eric J Wagner, Zhandong Liu, and Huda Y Zoghbi. Partial loss of CFIm25 causes learning deficits and aberrant neuronal alternative polyadenylation. *eLife*, 9, 2020.

[6646] Nicola Alessio, Tiziana Squillaro, Giovanni Di Bernardo, Giovanni Galano, Roberto De Rosa, Mariarosa AB Melone, Gianfranco Peluso, and Umberto Galderisi. Increase of circulating IGFBP-4 following genotoxic stress and its implication for senescence. *eLife*, 9, 2020.

[6647] Ryan K Alexander, Yae-Huei Liou, Nelson H Knudsen, Kyle A Starost, Chuanrui Xu, Alexander L Hyde, Sihao Liu, David Jacobi, Nan-Shih Liao, and Chih-Hao Lee. Bmal1 integrates mitochondrial metabolism and macrophage activation. *eLife*, 9, 2020.

[6648] Raymundo Alfaro-Aco, Akanksha Thawani, and Sabine Petry. Biochemical reconstitution of branching microtubule nucleation. *eLife*, 9, 2020.

[6649] Ali H Alghamdi, Jane C Munday, Gustavo Daniel Campagnaro, Dominik Gurvic, Fredrik Svensson, Chinyere E Okpara, Arvind Kumar, Juan Quintana, Maria Esther Martin Abril, Patrik Milić, Laura Watson, Daniel Paape, Luca Settimo, Anna Dimitriou, Joanna Wielinska, Graeme Smart, Laura F Anderson, Christopher M Woodley, Siu Pui Ying Kelly, Hasan MS Ibrahim, Fabian Hulpia, Mohammed I Al-Salabi, Anthonius A Eze, Teresa Sprenger, Ibrahim A Teka, Simon Gudin, Simone Weyand, Mark Field, Christophe Dardonville, Richard R Tidwell, Mark Carrington, Paul O'Neill, David W Boykin, Ulrich Zachariae, and Harry P De Koning. Positively selected modifications in the pore of TbAQP2 allow pentamidine to enter Trypanosoma brucei. *eLife*, 9, 2020.

[6650] Md Zulfikar Ali, Vinuselvi Parisutham, Sandeep Choubey, and Robert C Brewster. Inherent regulatory asymmetry emanating from network architecture in a prevalent autoregulatory motif. *eLife*, 9, 2020.

[6651] Tamer Ali and Phillip Grote. Beyond the RNA-dependent function of LncRNA genes. *eLife*, 9, 2020.

[6652] Ashna Alladin, Lucas Chaible, Lucia Garcia del Valle, Reither Sabine, Monika Loeschinger, Malte Wachsmuth, Jean-Karim Hériché, Christian Tischer, and Martin Jechlinger. Tracking cells in epithelial acini by light sheet microscopy reveals proximity effects in breast cancer initiation. *eLife*, 9, 2020.

[6653] Kathryn C Allaway, William Muñoz, Robin Tremblay, Mia Sherer, Jacob Herron, Bernardo Rudy, Robert Machold, and Gordon Fishell. Cellular birthdate predicts laminar and regional cholinergic projection topography in the forebrain. *eLife*, 9, 2020.

[6654] Aaron M Allen, Megan C Neville, Sebastian Birtles, Vincent Croset, Christoph Daniel Treiber, Scott Waddell, and Stephen F Goodwin. A single-cell transcriptomic atlas of the adult Drosophila ventral nerve cord. *eLife*, 9, 2020.

[6655] Robyn S Allen, Benjamin Tajer, Eileen M Shore, and Mary C Mullins. Fibrodysplasia ossificans progressiva mutant ACVR1 signals by multiple modalities in the developing zebrafish. *eLife*, 9, 2020.

[6656] Leandro M Alonso and Eve Marder. Temperature compensation in a small rhythmic circuit. *eLife*, 9, 2020.

[6657] Laura Alonso-Herranz, álvaro Sahún-Español, Ana Paredes, Pilar Gonzalo, Polyxeni Gkontra, Vanessa Núñez, Cristina Clemente, Marta Cedenilla, María Villalba-Orero, Javier Inserte, David García-Dorado, Alicia G Arroyo, and Mercedes Ricote. Macrophages promote endothelial-to-mesenchymal transition via MT1-MMP/TGF$\beta$1 after myocardial infarction. *eLife*, 9, 2020.

[6658] Ghada Alsaleh, Isabel Panse, Leo Swadling, Hanlin Zhang, Felix Clemens Richter, Alain Meyer, Janet Lord, Eleanor Barnes, Paul Klenerman, Christopher Green, and Anna Katharina Simon. Autophagy in T cells from aged donors is maintained by spermidine and correlates with function and vaccine responses. *eLife*, 9, 2020.

[6659] Beatriz Alvarez-Castelao, Susanne tom Dieck, Claudia M Fusco, Paul Donlin-Asp, Julio D Perez, and Erin M Schuman. The switch-like expression of heme-regulated kinase 1 mediates neuronal proteostasis following proteasome inhibition. *eLife*, 9, 2020.

[6660] Sara Alvira, Daniel W Watkins, Luca A Troman, William J Allen, James S Lorriman, Gianluca Degliesposti, Eli J Cohen, Morgan Beeby, Bertram Daum, Vicki AM Gold, J Mark Skehel, and Ian Collinson. Inter-membrane association of the Sec and BAM translocons for bacterial outer-membrane biogenesis. *eLife*, 9, 2020.

[6661] Hoger Amin, Anthi A Apostolopoulou, Raquel Suárez-Grimalt, Eleftheria Vrontou, and Andrew C Lin. Localized inhibition in the Drosophila mushroom body. *eLife*, 9, 2020.

[6662] Ana Rita Amândio, Lucille Lopez-Delisle, Christopher Chase Bolt, Bénédicte Mascrez, and Denis Duboule. A complex regulatory landscape involved in the development of mammalian external genitals. *eLife*, 9, 2020.

[6663] Jonathan Y An, Kristopher A Kerns, Andrew Ouellette, Laura Robinson, H Douglas Morris, Catherine Kaczorowski, So-Il Park, Title Mekvanich, Alex Kang, Jeffrey S McLean, Timothy C Cox, and Matt Kaeberlein. Rapamycin rejuvenates oral health in aging mice. *eLife*, 9, 2020.

[6664] Praveen Anand, Arjun Puranik, Murali Aravamudan, AJ Venkatakrishnan, and Venky Soundararajan. SARS-CoV-2 strategically mimics proteolytic activation of human ENaC. *eLife*, 9, 2020.

[6665] Jens Peter Andersen, Mathias Wullum Nielsen, Nicole L Simone, Resa E Lewiss, and Reshma Jagsi. COVID-19 medical papers have fewer women first authors than expected. *eLife*, 9, 2020.

[6666] Matthew J Anderson, Valentin Magidson, Ryoichiro Kageyama, and Mark Lewandoski. Fgf4 maintains Hes7 levels critical for normal somite segmentation clock function. *eLife*, 9, 2020.

[6667] Madeline G Andrews, Lakshmi Subramanian, and Arnold R Kriegstein. mTOR signaling regulates the morphology and migration of outer radial glia in developing human cortex. *eLife*, 9, 2020.

[6668] Gabriella Angiulli, Harveer Singh Dhupar, Hiroshi Suzuki, Irvinder Singh Wason, Franck Duong Van Hoa, and Thomas Walz. New approach for membrane protein reconstitution into peptidiscs and basis for their adaptability to different proteins. *eLife*, 9, 2020.

[6669] Neal Anthwal, Jane C Fenelon, Stephen D Johnston, Marilyn B Renfree, and Abigail S Tucker. Transient role of the middle ear as a lower jaw support across mammals. *eLife*, 9, 2020.

[6670] Paride Antinucci, Adna Dumitrescu, Charlotte Deleuze, Holly J Morley, Kristie Leung, Tom Hagley, Fumi Kubo, Herwig Baier, Isaac H Bianco, and Claire Wyart. A calibrated optogenetic toolbox of stable zebrafish opsin lines. *eLife*, 9, 2020.

[6671] Andrea Antonini, Andrea Sattin, Monica Moroni, Serena Bovetti, Claudio Moretti, Francesca Succol, Angelo Forli, Dania Vecchia, Vijayakumar P Rajamanickam, Andrea Bertoncini, Stefano Panzeri, Carlo Liberale, and Tommaso Fellin. Extended field-of-view ultrathin microendoscopes for high-resolution two-photon imaging with minimal invasiveness. *eLife*, 9, 2020.

[6672] Javier Antunez-Sanchez, Matthew Naish, Juan Sebastian Ramirez-Prado, Sho Ohno, Ying Huang, Alexander Dawson, Korawit Opassathian, Deborah Manza-Mianza, Federico Ariel, Cecile Raynaud, Anjar Wibowo, Josquin Daron, Minako Ueda, David Latrasse, R Keith Slotkin, Detlef Weigel, Moussa Benhamed, and Jose Gutierrez-Marcos. A new role for histone demethylases in the maintenance of plant genome integrity. *eLife*, 9, 2020.

[6673] Todd R Appleby and Michael B Manookin. Selectivity to approaching motion in retinal inputs to the dorsal visual pathway. *eLife*, 9, 2020.

[6674] Seiji Arai, Andreas Varkaris, Mannan Nouri, Sen Chen, Lisha Xie, and Steven P Balk. MARCH5 mediates NOXA-dependent MCL1 degradation driven by kinase inhibitors and integrated stress response activation. *eLife*, 9, 2020.

[6675] Yoichi Araki, Ingie Hong, Timothy R Gamache, Shaowen Ju, Leonardo Collado-Torres, Joo Heon Shin, and Richard L Huganir. SynGAP isoforms differentially regulate synaptic plasticity and dendritic development. *eLife*, 9, 2020.

[6676] Andrés Aranda-Díaz, Benjamin Obadia, Ren Dodge, Tani Thomsen, Zachary F Hallberg, Zehra Tüzün Güvener, William B Ludington, and Kerwyn Casey Huang. Bacterial interspecies interactions modulate pH-mediated antibiotic tolerance. *eLife*, 9, 2020.

[6677] Michael J Arcaro, Carlos Ponce, and Margaret Livingstone. The neurons that mistook a hat for a face. *eLife*, 9, 2020.

[6678] Robinson Arce-Molina, Francisca Cortés-Molina, Pamela Y Sandoval, Alex Galaz, Karin Alegría, Stefanie Schirmeier, L Felipe Barros, and Alejandro San Martín. A highly responsive pyruvate sensor reveals pathway-regulatory role of the mitochondrial pyruvate carrier MPC. *eLife*, 9, 2020.

[6679] Silvia Ardissone, Nicolas Kint, and Patrick H Viollier. Specificity in glycosylation of multiple flagellins by the modular and cell cycle regulated glycosyltransferase FlmG. *eLife*, 9, 2020.

[6680] Philip Arevalo, Huong Q McLean, Edward A Belongia, and Sarah Cobey. Earliest infections predict the age distribution of seasonal influenza A cases. *eLife*, 9, 2020.

[6681] Bilge Argunhan, Masayoshi Sakakura, Negar Afshar, Misato Kurihara, Kentaro Ito, Takahisa Maki, Shuji Kanamaru, Yasuto Murayama, Hideo Tsubouchi, Masayuki Takahashi, Hideo Takahashi, and Hiroshi Iwasaki. Cooperative interactions facilitate stimulation of Rad51 by the Swi5-Sfr1 auxiliary factor complex. *eLife*, 9, 2020.

[6682] Moritz Armbruster, Chris G Dulla, and Jeffrey S Diamond. Effects of fluorescent glutamate indicators on neurotransmitter diffusion and uptake. *eLife*, 9, 2020.

[6683] Laura L Arnold, Alessandra Cecchini, Danny A Stark, Jacqueline Ihnat, Rebecca N Craigg, Amory Carter, Sammy Zino, and DDW Cornelison. EphA7 promotes myogenic differentiation via cell-cell contact. *eLife*, 9, 2020.

[6684] Prateek Arora, Shivali Dongre, Renuka Raman, and Mahendra Sonawane. Stepwise polarisation of developing bilayered epidermis is mediated by aPKC and E-cadherin in zebrafish. *eLife*, 9, 2020.

[6685] Sebastian I Arriola Apelo, Amy Lin, Jacqueline A Brinkman, Emma Meyer, Mark Morrison, Jay L Tomasiewicz, Cassidy P Pumper, Emma L Baar, Nicole E Richardson, Mohammed Alotaibi, and Dudley W Lamming. Ovariectomy uncouples lifespan from metabolic health and reveals a sex-hormone-dependent role of hepatic mTORC2 in aging. *eLife*, 9, 2020.

[6686] Parisa Asghari, David RL Scriven, Myles Ng, Pankaj Panwar, Keng C Chou, Filip van Petegem, and Edwin DW Moore. Cardiac ryanodine receptor distribution is dynamic and changed by auxiliary proteins and post-translational modification. *eLife*, 9, 2020.

[6687] Sina Atashpaz, Sara Samadi Shams, Javier Martin Gonzalez, Endre Sebestyén, Negar Arghavanifard, Andrea Gnocchi, Eliene Albers, Simone Minardi, Giovanni Faga, Paolo Soffientini, Elisa Allievi, Valeria Cancila, Angela Bachi, óscar Fernández-Capetillo, Claudio Tripodo, Francesco Ferrari, Andrés Joaquin López-Contreras, and Vincenzo Costanzo. ATR expands embryonic stem cell fate potential in response to replication stress. *eLife*, 9, 2020.

[6688] Joseph Atherton, Jessica JA Hummel, Natacha Olieric, Julia Locke, Alejandro Peña, Steven S Rosenfeld, Michel O Steinmetz, Casper C Hoogenraad, and Carolyn A Moores. The mechanism of kinesin inhibition by kinesin-binding protein. *eLife*, 9, 2020.

[6689] Januka S Athukoralage, Shirley Graham, Christophe Rouillon, Sabine Grüschow, Clarissa M Czekster, and Malcolm F White. The dynamic interplay of host and viral enzymes in type III CRISPR-mediated cyclic nucleotide signalling. *eLife*, 9, 2020.

[6690] Januka S Athukoralage, Stuart McQuarrie, Sabine Grüschow, Shirley Graham, Tracey M Gloster, and Malcolm F White. Tetramerisation of the CRISPR ring nuclease Crn3/Csx3 facilitates cyclic oligoadenylate cleavage. *eLife*, 9, 2020.

[6691] Alisa Atkins, Michelle J Xu, Maggie Li, Nathaniel P Rogers, Marina V Pryzhkova, and Philip W Jordan. SMC5/6 is required for replication fork stability and faithful chromosome segregation during neurogenesis. *eLife*, 9, 2020.

[6692] CheukMan C Au, John B Furness, Kara Britt, Sofya Oshchepkova, Heta Ladumor, Kai Ying Soo, Brid Callaghan, Celine Gerard, Giorgio Inghirami, Vivek Mittal, Yufeng Wang, Xin Yun Huang, Jason A Spector, Eleni Andreopoulou, Paul Zumbo, Doron Betel, Lukas Dow, and Kristy A Brown. Three-dimensional growth of breast cancer cells potentiates the anti-tumor effects of unacylated ghrelin and AZP-531. *eLife*, 9, 2020.

[6693] Thomas G Aubier. Positive density dependence acting on mortality can help maintain species-rich communities. *eLife*, 9, 2020.

[6694] Arthur-Ervin Avramiea, Richard Hardstone, Jan-Matthis Lueckmann, Jan Bím, Huibert D Mansvelder, and Klaus Linkenkaer-Hansen. Pre-stimulus phase and amplitude regulation of phase-locked responses are maximized in the critical state. *eLife*, 9, 2020.

[6695] GökÇe B Ayan, Hye Jin Park, and Jenna Gallie. The birth of a bacterial tRNA gene by large-scale, tandem duplication events. *eLife*, 9, 2020.

[6696] Senem Aykul, Richard A Corpina, Erich J Goebel, Camille J Cunanan, Alexandra Dimitriou, Hyon Jong Kim, Qian Zhang, Ashique Rafique, Raymond Leidich, Xin Wang, Joyce McClain, Johanna Jimenez, Kalyan C Nannuru, Nyanza J Rothman, John B Lees-Shepard, Erik Martinez-Hackert, Andrew J Murphy, Thomas B Thompson, Aris N Economides, and Vincent Idone. Activin A forms a non-signaling complex with ACVR1 and type II Activin/BMP receptors via its finger 2 tip loop. *eLife*, 9, 2020.

[6697] Anthony W Azevedo, Evyn S Dickinson, Pralaksha Gurung, Lalanti Venkatasubramanian, Richard S Mann, and John C Tuthill. A size principle for recruitment of Drosophila leg motor neurons. *eLife*, 9, 2020.

[6698] Estefania P Azevedo, Bowen Tan, Lisa E Pomeranz, Violet Ivan, Robert Fetcho, Marc Schneeberger, Katherine R Doerig, Conor Liston, Jeffrey M Friedman, and Sarah A Stern. A limbic circuit selectively links active escape to food suppression. *eLife*, 9, 2020.

[6699] Zohre Azimi, Ruxandra Barzan, Katharina Spoida, Tatjana Surdin, Patric Wollenweber, Melanie D Mark, Stefan Herlitze, and Dirk Jancke. Separable gain control of ongoing and evoked activity in the visual cortex by serotonergic input. *eLife*, 9, 2020.

[6700] Afnan Azizi, Anne Herrmann, Yinan Wan, Salvador JRP Buse, Philipp J Keller, Raymond E Goldstein, and William A Harris. Nuclear crowding and nonlinear diffusion during interkinetic nuclear migration in the zebrafish retina. *eLife*, 9, 2020.

[6701] Chiara Azzi, Florian Aeschimann, Anca Neagu, and Helge Großhans. A branched heterochronic pathway directs juvenile-to-adult transition through two LIN-29 isoforms. *eLife*, 9, 2020.

[6702] Takashi Baba, Dai Miyazaki, Kodai Inata, Ryu Uotani, Hitomi Miyake, Shin-ichi Sasaki, Yumiko Shimizu, Yoshitsugu Inoue, and Kazuomi Nakamura. Role of IL-4 in bone marrow driven dysregulated angiogenesis and age-related macular degeneration. *eLife*, 9, 2020.

[6703] Travis A Babola, Calvin J Kersbergen, Han Chin Wang, and Dwight E Bergles. Purinergic signaling in cochlear supporting cells reduces hair cell excitability by increasing the extracellular space. *eLife*, 9, 2020.

[6704] Iris Bachmutsky, Xin Paul Wei, Eszter Kish, and Kevin Yackle. Opioids depress breathing through two small brainstem sites. *eLife*, 9, 2020.

[6705] Lindsey RF Backman, Yolanda Y Huang, Mary C Andorfer, Brian Gold, Ronald T Raines, Emily P Balskus, and Catherine L Drennan. Molecular basis for catabolism of the abundant metabolite trans-4-hydroxy-L-proline by a microbial glycyl radical enzyme. *eLife*, 9, 2020.

[6706] Stephanie Badde, Pia Ley, Siddhart S Rajendran, Idris Shareef, Ramesh Kekunnaya, and Brigitte Röder. Sensory experience during early sensitive periods shapes cross-modal temporal biases. *eLife*, 9, 2020.

[6707] Gaétan Bader, Ludovic Enkler, Yuhei Araiso, Marine Hemmerle, Krystyna Binko, Emilia Baranowska, Johan-Owen De Craene, Julie Ruer-Laventie, Jean Pieters, Déborah Tribouillard-Tanvier, Bruno Senger, Jean-Paul di Rago, Sylvie Friant, Roza Kucharczyk, and Hubert Dominique Becker. Assigning mitochondrial localization of dual localized proteins using a yeast Bi-Genomic Mitochondrial-Split-GFP. *eLife*, 9, 2020.

[6708] Sung Jun Bae, Lisheng Ni, and Xuelian Luo. STK25 suppresses Hippo signaling by regulating SAV1-STRIPAK antagonism. *eLife*, 9, 2020.

[6709] Pablo Baeza-Centurion, Belén Miñana, Juan Valcárcel, and Ben Lehner. Mutations primarily alter the inclusion of alternatively spliced exons. *eLife*, 9, 2020.

[6710] Jennifer Bagwell, James Norman, Kathryn Ellis, Brianna Peskin, James Hwang, Xiaoyan Ge, Stacy V Nguyen, Sarah K McMenamin, Didier YR Stainier, and Michel Bagnat. Notochord vacuoles absorb compressive bone growth during zebrafish spine formation. *eLife*, 9, 2020.

[6711] Lin Bai, Qinglong You, Bhawik K Jain, H Diessel Duan, Amanda Kovach, Todd R Graham, and Huilin Li. Transport mechanism of P4 ATPase phosphatidylcholine flippases. *eLife*, 9, 2020.

[6712] Xiaofei Bai, Jeff Bouffard, Avery Lord, Katherine Brugman, Paul W Sternberg, Erin J Cram, and Andy Golden. Caenorhabditis elegans PIEZO channel coordinates multiple reproductive tissues to govern ovulation. *eLife*, 9, 2020.

[6713] Yonghong Bai, Xinchao Yu, Hao Chen, Daniel Horne, Ryan White, Xiaosu Wu, Paul Lee, Yan Gu, Sudipa Ghimire-Rijal, Daniel C-H Lin, and Xin Huang. Structural basis for pharmacological modulation of the TRPC6 channel. *eLife*, 9, 2020.

[6714] Stacyann Bailey, Grazyna E Sroga, Betty Hoac, Orestis L Katsamenis, Zehai Wang, Nikolaos Bouropoulos, Marc D McKee, Esben S Sørensen, Philipp J Thurner, and Deepak Vashishth. The role of extracellular matrix phosphorylation on energy dissipation in bone. *eLife*, 9, 2020.

[6715] Konstantin I Bakhurin, Xiaoran Li, Alexander D Friedman, Nicholas A Lusk, Glenn DR Watson, Namsoo Kim, and Henry H Yin. Opponent regulation of action performance and timing by striatonigral and striatopallidal pathways. *eLife*, 9, 2020.

[6716] Rachael Bakker, Madhav Mani, and Richard W Carthew. The Wg and Dpp morphogens regulate gene expression by modulating the frequency of transcriptional bursts. *eLife*, 9, 2020.

[6717] Aurélia C Balestra, Mohammad Zeeshan, Edward Rea, Carla Pasquarello, Lorenzo Brusini, Tobias Mourier, Amit Kumar Subudhi, Natacha Klages, Patrizia Arboit, Rajan Pandey, Declan Brady, Sue Vaughan, Anthony A Holder, Arnab Pain, David JP Ferguson, Alexandre Hainard, Rita Tewari, and Mathieu Brochet. A divergent cyclin/cyclin-dependent kinase complex controls the atypical replication of a malaria parasite during gametogony and transmission. *eLife*, 9, 2020.

[6718] Jesus Javier Ballesteros, Jessica Blair Briscoe, and Yumiko Ishizawa. Neural signatures of $\alpha$2-Adrenergic agonist-induced unconsciousness and awakening by antagonist. *eLife*, 9, 2020.

[6719] Joshua Henk Balsters, Valerio Zerbi, Jerome Sallet, Nicole Wenderoth, and Rogier B Mars. Primate homologs of mouse cortico-striatal circuits. *eLife*, 9, 2020.

[6720] Connor J Balzer, Michael L James, Heidy Y Narvaez-Ortiz, Luke A Helgeson, Vladimir Sirotkin, and Brad J Nolen. Synergy between Wsp1 and Dip1 may initiate assembly of endocytic actin networks. *eLife*, 9, 2020.

[6721] Aditi Banerjee, Jinoh Lee, Paulina Nemcova, Changliang Liu, and Pascal S Kaeser. Synaptotagmin-1 is the Ca2+ sensor for fast striatal dopamine release. *eLife*, 9, 2020.

[6722] Dan Bang, Sara Ershadmanesh, Hamed Nili, and Stephen M Fleming. Private–public mappings in human prefrontal cortex. *eLife*, 9, 2020.

[6723] Edward J Banigan and Leonid A Mirny. The interplay between asymmetric and symmetric DNA loop extrusion. *eLife*, 9, 2020.

[6724] Edward J Banigan, Aafke A van den Berg, Hugo B Brandão, John F Marko, and Leonid A Mirny. Chromosome organization by one-sided and two-sided loop extrusion. *eLife*, 9, 2020.

[6725] Chen Bao, Sarah Loerch, Clarence Ling, Andrei A Korostelev, Nikolaus Grigorieff, and Dmitri N Ermolenko. mRNA stem-loops can pause the ribosome by hindering A-site tRNA binding. *eLife*, 9, 2020.

[6726] Jin Bao, Michael Graupner, Guadalupe Astorga, Thibault Collin, Abdelali Jalil, Dwi Wahyu Indriati, Jonathan Bradley, Ryuichi Shigemoto, and Isabel Llano. Synergism of type 1 metabotropic and ionotropic glutamate receptors in cerebellar molecular layer interneurons in vivo. *eLife*, 9, 2020.

[6727] Qiankun Bao, Bangying Zhang, Ya Suo, Chen Liu, Qian Yang, Kai Zhang, Ming Yuan, Meng Yuan, Yue Zhang, and Guangping Li. Intermittent hypoxia mediated by TSP1 dependent on STAT3 induces cardiac fibroblast activation and cardiac fibrosis. *eLife*, 9, 2020.

[6728] Yinon M Bar-On, Avi Flamholz, Rob Phillips, and Ron Milo. SARS-CoV-2 (COVID-19) by the numbers. *eLife*, 9, 2020.

[6729] Jeanette Baran-Gale, Michael D Morgan, Stefano Maio, Fatima Dhalla, Irene Calvo-Asensio, Mary E Deadman, Adam E Handel, Ashley Maynard, Steven Chen, Foad Green, Rene V Sit, Norma F Neff, Spyros Darmanis, Weilun Tan, Andy P May, John C Marioni, Chris P Ponting, and Georg A Holländer. Ageing compromises mouse thymus function and remodels epithelial cell differentiation. *eLife*, 9, 2020.

[6730] Kristopher M Barnes, Li Fan, Mark W Moyle, Christopher A Brittin, Yichi Xu, Daniel A Colón-Ramos, Anthony Santella, and Zhirong Bao. Cadherin preserves cohesion across involuting tissues during C. elegans neurulation. *eLife*, 9, 2020.

[6731] Richard Lee Barrett and Ellen Puré. Cancer-associated fibroblasts and their influence on tumor immunity and immunotherapy. *eLife*, 9, 2020.

[6732] Emilie Barruet, Steven M Garcia, Katharine Striedinger, Jake Wu, Solomon Lee, Lauren Byrnes, Alvin Wong, Sun Xuefeng, Stanley Tamaki, Andrew S Brack, and Jason H Pomerantz. Functionally heterogeneous human satellite cells identified by single cell RNA sequencing. *eLife*, 9, 2020.

[6733] Sandip Basak, Arvind Kumar, Steven Ramsey, Eric Gibbs, Abhijeet Kapoor, Marta Filizola, and Sudha Chakrapani. High-resolution structures of multiple 5-HT3AR-setron complexes reveal a novel mechanism of competitive inhibition. *eLife*, 9, 2020.

[6734] Timour Baslan, Jude Kendall, Konstantin Volyanskyy, Katherine McNamara, Hilary Cox, Sean D'Italia, Frank Ambrosio, Michael Riggs, Linda Rodgers, Anthony Leotta, Junyan Song, Yong Mao, Jie Wu, Ronak Shah, Rodrigo Gularte-Mérida, Kalyani Chadalavada, Gouri Nanjangud, Vinay Varadan, Assaf Gordon, Christina Curtis, Alex Krasnitz, Nevenka Dimitrova, Lyndsay Harris, Michael Wigler, and James Hicks. Novel insights into breast cancer copy number genetic heterogeneity revealed by single-cell genome sequencing. *eLife*, 9, 2020.

[6735] Nipun S Basrur, Maria Elena De Obaldia, Takeshi Morita, Margaret Herre, Ricarda K von Heynitz, Yael N Tsitohay, and Leslie B Vosshall. Fruitless mutant male mosquitoes gain attraction to human odor. *eLife*, 9, 2020.

[6736] Alexander Shakeel Bates, James D Manton, Sridhar R Jagannathan, Marta Costa, Philipp Schlegel, Torsten Rohlfing, and Gregory SXE Jefferis. The natverse, a versatile toolbox for combining and analysing neuroanatomical data. *eLife*, 9, 2020.

[6737] CJ Battey, Peter L Ralph, and Andrew D Kern. Predicting geographic location from genetic variation with deep neural networks. *eLife*, 9, 2020.

[6738] Nicolaas C Baudoin, Joshua M Nicholson, Kimberly Soto, Olga Martin, Jing Chen, and Daniela Cimini. Asymmetric clustering of centrosomes defines the early evolution of tetraploid cells. *eLife*, 9, 2020.

[6739] L Ryan Baugh and Troy Day. Nongenetic inheritance and multigenerational plasticity in the nematode C. elegans. *eLife*, 9, 2020.

[6740] Simranjot Bawa, David S Brooks, Kathryn E Neville, Marla Tipping, Md Abdul Sagar, Joseph A Kollhoff, Geetanjali Chawla, Brian V Geisbrecht, Jason M Tennessen, Kevin W Eliceiri, and Erika R Geisbrecht. Drosophila TRIM32 cooperates with glycolytic enzymes to promote cell growth. *eLife*, 9, 2020.

[6741] Matthew Baxter, Maria Voronkov, Toryn Poolman, Gina Galli, Christian Pinali, Laurence Goosey, Abigail Knight, Karolina Krakowiak, Robert Maidstone, Mudassar Iqbal, Min Zi, Sukhpal Prehar, Elizabeth J Cartwright, Julie Gibbs, Laura C Matthews, Antony D Adamson, Neil E Humphreys, Pedro Rebelo-Guiomar, Michal Minczuk, David A Bechtold, Andrew Loudon, and David Ray. Cardiac mitochondrial function depends on BUD23 mediated ribosome programming. *eLife*, 9, 2020.

[6742] Emily A Bayer, Rebecca C Stecky, Lauren Neal, Phinikoula S Katsamba, Goran Ahlsen, Vishnu Balaji, Thorsten Hoppe, Lawrence Shapiro, Meital Oren-Suissa, and Oliver Hobert. Ubiquitin-dependent regulation of a conserved DMRT protein controls sexually dimorphic synaptic connectivity and behavior. *eLife*, 9, 2020.

[6743] Oliver W Bayfield, Alasdair C Steven, and Alfred A Antson. Cryo-EM structure in situ reveals a molecular switch that safeguards virus against genome loss. *eLife*, 9, 2020.

[6744] Pierre-Louis Bazin, Anneke Alkemade, Martijn J Mulder, Amanda G Henry, and Birte U Forstmann. Multi-contrast anatomical subcortical structures parcellation. *eLife*, 9, 2020.

[6745] Léa Beaumelle, Frederik De Laender, and Nico Eisenhauer. Biodiversity mediates the effects of stressors but not nutrients on litter decomposition. *eLife*, 9, 2020.

[6746] Aleksandra Bebel, Melissa A Walsh, Ignacio Mir-Sanchis, and Phoebe A Rice. A novel DNA primase-helicase pair encoded by SCCmec elements. *eLife*, 9, 2020.

[6747] Ulrik Beierholm, Tim Rohe, Ambra Ferrari, Oliver Stegle, and Uta Noppeney. Using the past to estimate sensory uncertainty. *eLife*, 9, 2020.

[6748] Laura Bel Borja, Flavie Soubigou, Samuel J P Taylor, Conchita Fraguas Bringas, Jacqueline Budrewicz, Pablo Lara-Gonzalez, Christopher G Sorensen Turpin, Joshua N Bembenek, Dhanya K Cheerambathur, and Federico Pelisch. BUB-1 targets PP2A:B56 to regulate chromosome congression during meiosis I in C. elegans oocytes. *eLife*, 9, 2020.

[6749] Arnaud Belcour, Clémence Frioux, Méziane Aite, Anthony Bretaudeau, Falk Hildebrand, and Anne Siegel. Metage2Metabo, microbiota-scale metabolic complementarity for the identification of key species. *eLife*, 9, 2020.

[6750] Shilpak Bele, Shravan Babu Girada, Aramita Ray, Abhishek Gupta, Srinivas Oruganti, Phanithi Prakash Babu, Rahul SR Rayalla, Shashi Vardhan Kalivendi, Ahamed Ibrahim, Vishwajeet Puri, Venkateswar Adalla, Madhumohan R Katika, Richard DiMarchi, and Prasenjit Mitra. MS-275, a class 1 histone deacetylase inhibitor augments glucagon-like peptide-1 receptor agonism to improve glycemic control and reduce obesity in diet-induced obese mice. *eLife*, 9, 2020.

[6751] Daniel W Belsky, Avshalom Caspi, Louise Arseneault, Andrea Baccarelli, David L Corcoran, Xu Gao, Eiliss Hannon, Hona Lee Harrington, Line JH Rasmussen, Renate Houts, Kim Huffman, William E Kraus, Dayoon Kwon, Jonathan Mill, Carl F Pieper, Joseph A Prinz, Richie Poulton, Joel Schwartz, Karen Sugden, Pantel Vokonas, Benjamin S Williams, and Terrie E Moffitt. Quantification of the pace of biological aging in humans through a blood test, the DunedinPoAm DNA methylation algorithm. *eLife*, 9, 2020.

[6752] Francisca Benavente, Katja M Piltti, Mitra J Hooshmand, Aileen A Nava, Anita Lakatos, Brianna G Feld, Dana Creasman, Paul D Gershon, and Aileen Anderson. Novel C1q receptor-mediated signaling controls neural stem cell behavior and neurorepair. *eLife*, 9, 2020.

[6753] Lorena Benedetti, Jonathan S Marvin, Hanieh Falahati, Andres Guillén-Samander, Loren L Looger, and Pietro De Camilli. Optimized Vivid-derived Magnets photodimerizers for subcellular optogenetics in mammalian cells. *eLife*, 9, 2020.

[6754] Kateřina Benediktová, Jana Adámková, Jan Svoboda, Michael Scott Painter, Luděk BartoŠ, Petra Nováková, Lucie Vynikalová, Vlastimil Hart, John Phillips, and Hynek Burda. Magnetic alignment enhances homing efficiency of hunting dogs. *eLife*, 9, 2020.

[6755] Theresa Benezeder, Clemens Painsi, VijayKumar Patra, Saptaswa Dey, Martin Holcmann, Bernhard Lange-Asschenfeldt, Maria Sibilia, and Peter Wolf. Dithranol targets keratinocytes, their crosstalk with neutrophils and inhibits the IL-36 inflammatory loop in psoriasis. *eLife*, 9, 2020.

[6756] Tone Bengtsen, Viktor L Holm, Lisbeth Ravnkilde Kjølbye, Søren R Midtgaard, Nicolai Tidemand Johansen, Giulio Tesei, Sandro Bottaro, Birgit Schiøtt, Lise Arleth, and Kresten Lindorff-Larsen. Structure and dynamics of a nanodisc by integrating NMR, SAXS and SANS experiments with molecular dynamics simulations. *eLife*, 9, 2020.

[6757] Sabrina Bianca Bennstein, Sandra Weinhold, Angela Riccarda Manser, Nadine Scherenschlich, Angela Noll, Katharina Raba, Gesine Kögler, Lutz Walter, and Markus Uhrberg. Umbilical cord blood-derived ILC1-like cells constitute a novel precursor for mature KIR+NKG2A- NK cells. *eLife*, 9, 2020.

[6758] Klaas Bente, Sarah Mohammadinejad, Mohammad Avalin Charsooghi, Felix Bachmann, Agnese Codutti, Christopher T Lefèvre, Stefan Klumpp, and Damien Faivre. High-speed motility originates from cooperatively pushing and pulling flagella bundles in bilophotrichous bacteria. *eLife*, 9, 2020.

[6759] Richard Benton, Christophe Dessimoz, and David Moi. A putative origin of the insect chemosensory receptor superfamily in the last common eukaryotic ancestor. *eLife*, 9, 2020.

[6760] Yulia Bereshpolova, Xiaojuan Hei, Jose-Manuel Alonso, and Harvey A Swadlow. Three rules govern thalamocortical connectivity of fast-spike inhibitory interneurons in the visual cortex. *eLife*, 9, 2020.

[6761] Matthieu Bergé, Julian Pezzatti, Víctor González-Ruiz, Laurence Degeorges, Geneviève Mottet-Osman, Serge Rudaz, and Patrick H Viollier. Bacterial cell cycle control by citrate synthase independent of enzymatic activity. *eLife*, 9, 2020.

[6762] Michael Berger, Naubahar Shahryar Agha, and Alexander Gail. Wireless recording from unrestrained monkeys reveals motor goal encoding beyond immediate reach in frontoparietal cortex. *eLife*, 9, 2020.

[6763] Christopher Bergevin, Chandan Narayan, Joy Williams, Natasha Mhatre, Jennifer KE Steeves, Joshua GW Bernstein, and Brad Story. Overtone focusing in biphonic tuvan throat singing. *eLife*, 9, 2020.

[6764] Eva Berlot, Nicola J Popp, and Jörn Diedrichsen. A critical re-evaluation of fMRI signatures of motor sequence learning. *eLife*, 9, 2020.

[6765] Augusto Berrocal, Nicholas C Lammers, Hernan G Garcia, and Michael B Eisen. Kinetic sculpting of the seven stripes of the Drosophila even-skipped gene. *eLife*, 9, 2020.

[6766] Simone Bersini, Roberta Schulte, Ling Huang, Hannah Tsai, and Martin W Hetzer. Direct reprogramming of human smooth muscle and vascular endothelial cells reveals defects associated with aging and Hutchinson-Gilford progeria syndrome. *eLife*, 9, 2020.

[6767] Fabrice Besnard, Joao Picao-Osorio, Clément Dubois, and Marie-Anne Félix. A broad mutational target explains a fast rate of phenotypic evolution. *eLife*, 9, 2020.

[6768] Lisa R Beutler, Timothy V Corpuz, Jamie S Ahn, Seher Kosar, Weimin Song, Yiming Chen, and Zachary A Knight. Obesity causes selective and long-lasting desensitization of AgRP neurons to dietary fat. *eLife*, 9, 2020.

[6769] Amol K Bhandage, Gabriela C Olivera, Sachie Kanatani, Elizabeth Thompson, Karin Loré, Manuel Varas-Godoy, and Antonio Barragan. A motogenic GABAergic system of mononuclear phagocytes facilitates dissemination of coccidian parasites. *eLife*, 9, 2020.

[6770] Shivam Bhardwaj, Lim Si-Hui Jolander, Markus R Wenk, Jeffrey C Oliver, H Frederik Nijhout, and Antonia Monteiro. Origin of the mechanism of phenotypic plasticity in satyrid butterfly eyespots. *eLife*, 9, 2020.

[6771] Ashima Bhaskar, Santosh Kumar, Mehak Zahoor Khan, Amit Singh, Ved Prakash Dwivedi, and Vinay Kumar Nandicoori. Host sirtuin 2 as an immunotherapeutic target against tuberculosis. *eLife*, 9, 2020.

[6772] Pradyot Bhattacharya, Rada Ellegård, Mohammad Khalid, Cecilia Svanberg, Melissa Govender, Åsa V Keita, Johan D Söderholm, Pär Myrelid, Esaki M Shankar, Sofia Nyström, and Marie Larsson. Complement opsonization of HIV affects primary infection of human colorectal mucosa and subsequent activation of T cells. *eLife*, 9, 2020.

[6773] Moitrayee Bhattacharyya, Young Kwang Lee, Serena Muratcioglu, Baiyu Qiu, Priya Nyayapati, Howard Schulman, Jay T Groves, and John Kuriyan. Flexible linkers in CaMKII control the balance between activating and inhibitory autophosphorylation. *eLife*, 9, 2020.

[6774] Arjun Biddanda, Daniel P Rice, and John Novembre. A variant-centric perspective on geographic patterns of human allele frequency variation. *eLife*, 9, 2020.

[6775] Jessica E Biddinger, Roman M Lazarenko, Michael M Scott, and Richard Simerly. Leptin suppresses development of GLP-1 inputs to the paraventricular nucleus of the hypothalamus. *eLife*, 9, 2020.

[6776] Helena Bilandžija, Breanna Hollifield, Mireille Steck, Guanliang Meng, Mandy Ng, Andrew D Koch, Romana Gračan, Helena ćetković, Megan L Porter, Kenneth J Renner, and William Jeffery. Phenotypic plasticity as a mechanism of cave colonization and adaptation. *eLife*, 9, 2020.

[6777] Adrien Birot, Marta Tormos-Pérez, Sabine Vaur, Amélie Feytout, Julien Jaegy, Dácil Alonso Gil, Stéphanie Vazquez, Karl Ekwall, and Jean-Paul Javerzat. The CDK Pef1 and protein phosphatase 4 oppose each other for regulating cohesin binding to fission yeast chromosomes. *eLife*, 9, 2020.

[6778] Julia Bittmann, Rokas Grigaitis, Lorenzo Galanti, Silas Amarell, Florian Wilfling, Joao Matos, and Boris Pfander. An advanced cell cycle tag toolbox reveals principles underlying temporal control of structure-selective nucleases. *eLife*, 9, 2020.

[6779] Sebastian H Bitzenhofer, Jastyn A Pöpplau, and Ileana Hanganu-Opatz. Gamma activity accelerates during prefrontal development. *eLife*, 9, 2020.

[6780] Theresa E Bjorness, Ashwinikumar Kulkarni, Volodymyr Rybalchenko, Ayako Suzuki, Catherine Bridges, Adam J Harrington, Christopher W Cowan, Joseph S Takahashi, Genevieve Konopka, and Robert W Greene. An essential role for MEF2C in the cortical response to loss of sleep in mice. *eLife*, 9, 2020.

[6781] Jennifer M Blackwell, Alexandria MH Lesicko, Winnie Rao, Mariella De Biasi, and Maria N Geffen. Auditory cortex shapes sound responses in the inferior colliculus. *eLife*, 9, 2020.

[6782] Bastien Blain and Robb B Rutledge. Momentary subjective well-being depends on learning and not reward. *eLife*, 9, 2020.

[6783] Jorge Blanco, Jacob C Cooper, and Nicholas E Baker. Roles of C/EBP class bZip proteins in the growth and cell competition of Rp ('Minute') mutants in Drosophila. *eLife*, 9, 2020.

[6784] Adam R Blanden, Xin Yu, Alan J Blayney, Christopher Demas, Jeung-Hoi Ha, Yue Liu, Tracy Withers, Darren R Carpizo, and Stewart N Loh. Zinc shapes the folding landscape of p53 and establishes a pathway for reactivating structurally diverse cancer mutants. *eLife*, 9, 2020.

[6785] Cristina Blázquez, Andrea Ruiz-Calvo, Raquel Bajo-Grañeras, Jérôme M Baufreton, Eva Resel, Marjorie Varilh, Antonio C Pagano Zottola, Yamuna Mariani, Astrid Cannich, José A Rodríguez-Navarro, Giovanni Marsicano, Ismael Galve-Roperh, Luigi Bellocchio, and Manuel Guzmán. Inhibition of striatonigral autophagy as a link between cannabinoid intoxication and impairment of motor coordination. *eLife*, 9, 2020.

[6786] Solal Bloch, Hanako Hagio, Manon Thomas, Aurélie Heuzé, Jean-Michel Hermel, Elodie Lasserre, Ingrid Colin, Kimiko Saka, Pierre Affaticati, Arnim Jenett, Koichi Kawakami, Naoyuki Yamamoto, and Kei Yamamoto. Non-thalamic origin of zebrafish sensory nuclei implies convergent evolution of visual pathways in amniotes and teleosts. *eLife*, 9, 2020.

[6787] Leo Blondel, Tamsin EM Jones, and Cassandra G Extavour. Bacterial contribution to genesis of the novel germ line determinant oskar. *eLife*, 9, 2020.

[6788] Kyle P Blum, Kenneth S Campbell, Brian C Horslen, Paul Nardelli, Stephen N Housley, Timothy C Cope, and Lena H Ting. Diverse and complex muscle spindle afferent firing properties emerge from multiscale muscle mechanics. *eLife*, 9, 2020.

[6789] Daniel Blumenthal, Vidhi Chandra, Lyndsay Avery, and Janis K Burkhardt. Mouse T cell priming is enhanced by maturation-dependent stiffening of the dendritic cell cortex. *eLife*, 9, 2020.

[6790] Tatyana Bodrug, Elizabeth M Wilson-Kubalek, Stanley Nithianantham, Alex F Thompson, April Alfieri, Ignas Gaska, Jennifer Major, Garrett Debs, Sayaka Inagaki, Pedro Gutierrez, Larisa Gheber, Richard J McKenney, Charles Vaughn Sindelar, Ronald Milligan, Jason Stumpff, Steven S Rosenfeld, Scott T Forth, and Jawdat Al-Bassam. The kinesin-5 tail domain directly modulates the mechanochemical cycle of the motor domain for anti-parallel microtubule sliding. *eLife*, 9, 2020.

[6791] Giulia LM Boezio, Anabela Bensimon-Brito, Janett Piesker, Stefan Guenther, Christian SM Helker, and Didier YR Stainier. Endothelial TGF-$\beta$ signaling instructs smooth muscle cell development in the cardiac outflow tract. *eLife*, 9, 2020.

[6792] Rafal Bogacz. Dopamine role in learning and action inference. *eLife*, 9, 2020.

[6793] Katarzyna Bogucka, Malvika Pompaiah, Federico Marini, Harald Binder, Gregory Harms, Manuel Kaulich, Matthias Klein, Christian Michel, Markus P Radsak, Sebastian Rosigkeit, Peter Grimminger, Hansjörg Schild, and Krishnaraj Rajalingam. ERK3/MAPK6 controls IL-8 production and chemotaxis. *eLife*, 9, 2020.

[6794] Briana M Bohannon, Alicia de la Cruz, Xiaoan Wu, Jessica J Jowais, Marta E Perez, Derek M Dykxhoorn, Sara I Liin, and H Peter Larsson. Polyunsaturated fatty acid analogues differentially affect cardiac NaV, CaV, and KV channels through unique mechanisms. *eLife*, 9, 2020.

[6795] Claudia Böhm and Albert K Lee. Canonical goal-selective representations are absent from prefrontal cortex in a spatial working memory task requiring behavioral flexibility. *eLife*, 9, 2020.

[6796] Alfonso Bolado-Carrancio, Oleksii S Rukhlenko, Elena Nikonova, Mikhail A Tsyganov, Anne Wheeler, Amaya Garcia-Munoz, Walter Kolch, Alex von Kriegsheim, and Boris N Kholodenko. Periodic propagating waves coordinate RhoGTPase network dynamics at the leading and trailing edges during cell migration. *eLife*, 9, 2020.

[6797] Kevin A Bolding, Shivathmihai Nagappan, Bao-Xia Han, Fan Wang, and Kevin M Franks. Recurrent circuitry is required to stabilize piriform cortex odor representations across brain states. *eLife*, 9, 2020.

[6798] Mareike Bongers, Jordi Perez-Gil, Mark P Hodson, Lars Schrübbers, Tune Wulff, Morten OA Sommer, Lars K Nielsen, and Claudia E Vickers. Adaptation of hydroxymethylbutenyl diphosphate reductase enables volatile isoprenoid production. *eLife*, 9, 2020.

[6799] Sam A Booker, Harumi Harada, Claudio Elgueta, Julia Bank, Marlene Bartos, Akos Kulik, and Imre Vida. Presynaptic GABAB receptors functionally uncouple somatostatin interneurons from the active hippocampal network. *eLife*, 9, 2020.

[6800] Fivos Borbolis, John Rallis, George Kanatouris, Nikolitsa Kokla, Antonis Karamalegkos, Christina Vasileiou, Katerina M Vakaloglou, George Diallinas, Dimitrios J Stravopodis, Christos G Zervas, and Popi Syntichaki. mRNA decapping is an evolutionarily conserved modulator of neuroendocrine signaling that controls development and ageing. *eLife*, 9, 2020.

[6801] Charles L Bormann, Manoj Kumar Kanakasabapathy, Prudhvi Thirumalaraju, Raghav Gupta, Rohan Pooniwala, Hemanth Kandula, Eduardo Hariton, Irene Souter, Irene Dimitriadis, Leslie B Ramirez, Carol L Curchoe, Jason Swain, Lynn M Boehnlein, and Hadi Shafiee. Performance of a deep learning based neural network in the selection of human blastocysts for implantation. *eLife*, 9, 2020.

[6802] Benny Borremans, Amandine Gamble, KC Prager, Sarah K Helman, Abby M McClain, Caitlin Cox, Van Savage, and James O Lloyd-Smith. Quantifying antibody kinetics and RNA detection during early-phase SARS-CoV-2 infection by time since symptom onset. *eLife*, 9, 2020.

[6803] Francesca Maria Bosisio, Asier Antoranz, Yannick van Herck, Maddalena Maria Bolognesi, Lukas Marcelis, Clizia Chinello, Jasper Wouters, Fulvio Magni, Leonidas Alexopoulos, Marguerite Stas, Veerle Boecxstaens, Oliver Bechter, Giorgio Cattoretti, and Joost van den Oord. Functional heterogeneity of lymphocytic patterns in primary melanoma dissected through single-cell multiplexing. *eLife*, 9, 2020.

[6804] Flora Bouchacourt, Stefano Palminteri, Etienne Koechlin, and Srdjan Ostojic. Temporal chunking as a mechanism for unsupervised learning of task-sets. *eLife*, 9, 2020.

[6805] Christopher T Boughter, Marta T Borowska, Jenna J Guthmiller, Albert Bendelac, Patrick C Wilson, Benoit Roux, and Erin J Adams. Biochemical patterns of antibody polyreactivity revealed through a bioinformatics-based analysis of CDR loops. *eLife*, 9, 2020.

[6806] Rajaa Boujemaa-Paterski, Bruno Martins, Matthias Eibauer, Charlie T Beales, Benjamin Geiger, and Ohad Medalia. Talin-activated vinculin interacts with branched actin networks to initiate bundles. *eLife*, 9, 2020.

[6807] Alexandra Bova, Matt Gaidica, Amy Hurst, Yoshiko Iwai, Julia Hunter, and Daniel K Leventhal. Precisely timed dopamine signals establish distinct kinematic representations of skilled movements. *eLife*, 9, 2020.

[6808] Anne Ramsay Bowden, David A Morales-Juarez, Matylda Sczaniecka-Clift, Maria Martin Agudo, Natalia Lukashchuk, John Christopher Thomas, and Stephen P Jackson. Parallel CRISPR-Cas9 screens clarify impacts of p53 on screen performance. *eLife*, 9, 2020.

[6809] Anna J Bowen, Jane Y Chen, Y Waterlily Huang, Nathan A Baertsch, Sekun Park, and Richard D Palmiter. Dissociable control of unconditioned responses and associative fear learning by parabrachial CGRP neurons. *eLife*, 9, 2020.

[6810] Emily Bowie and Sarah C Goetz. TTBK2 and primary cilia are essential for the connectivity and survival of cerebellar Purkinje neurons. *eLife*, 9, 2020.

[6811] Caitlin R Bowman, Takako Iwashita, and Dagmar Zeithamova. Tracking prototype and exemplar representations in the brain across learning. *eLife*, 9, 2020.

[6812] Laura Bozal-Basterra, María Gonzalez-Santamarta, Veronica Muratore, Aitor Bermejo-Arteagabeitia, Carolina Da Fonseca, Orhi Barroso-Gomila, Mikel Azkargorta, Ibon Iloro, Olatz Pampliega, Ricardo Andrade, Natalia Martín-Martín, Tess C Branon, Alice Y Ting, Jose A Rodríguez, Arkaitz Carracedo, Felix Elortza, James D Sutherland, and Rosa Barrio. LUZP1, a novel regulator of primary cilia and the actin cytoskeleton, is a contributing factor in Townes-Brocks Syndrome. *eLife*, 9, 2020.

[6813] Nora Brackbill, Colleen Rhoades, Alexandra Kling, Nishal P Shah, Alexander Sher, Alan M Litke, and EJ Chichilnisky. Reconstruction of natural images from responses of primate retinal ganglion cells. *eLife*, 9, 2020.

[6814] Oliver Braganza, Daniel Mueller-Komorowska, Tony Kelly, and Heinz Beck. Quantitative properties of a feedback circuit predict frequency-dependent pattern separation. *eLife*, 9, 2020.

[6815] Marijke Brams, Cedric Govaerts, Kumiko Kambara, Kerry L Price, Radovan Spurny, Anant Gharpure, Els Pardon, Genevieve L Evans, Daniel Bertrand, Sarah CR Lummis, Ryan E Hibbs, Jan Steyaert, and Chris Ulens. Modulation of the Erwinia ligand-gated ion channel (ELIC) and the 5-HT3 receptor via a common vestibule site. *eLife*, 9, 2020.

[6816] Bob Bramson, Hanneke EM den Ouden, Ivan Toni, and Karin Roelofs. Improving emotional-action control by targeting long-range phase-amplitude neuronal coupling. *eLife*, 9, 2020.

[6817] Samuel PC Brand, Patrick Munywoki, David Walumbe, Matthew J Keeling, and David James Nokes. Reducing respiratory syncytial virus (RSV) hospitalization in a lower-income country by vaccinating mothers-to-be and their households. *eLife*, 9, 2020.

[6818] María Angélica Bravo Núñez, Ibrahim M Sabbarini, Lauren E Eide, Robert L Unckless, and Sarah E Zanders. Atypical meiosis can be adaptive in outcrossed Schizosaccharomyces pombe due to wtf meiotic drivers. *eLife*, 9, 2020.

[6819] Cedric Bressan, Alessandra Pecora, Dave Gagnon, Marina Snapyan, Simon Labrecque, Paul De Koninck, Martin Parent, and Armen Saghatelyan. The dynamic interplay between ATP/ADP levels and autophagy sustain neuronal migration in vivo. *eLife*, 9, 2020.

[6820] Jessica Briggs, Noam Teyssier, Joaniter I Nankabirwa, John Rek, Prasanna Jagannathan, Emmanuel Arinaitwe, Teun Bousema, Chris Drakeley, Margaret Murray, Emily Crawford, Nicholas Hathaway, Sarah G Staedke, David Smith, Phillip J Rosenthal, Moses Kamya, Grant Dorsey, Isabel Rodriguez-Barraquer, and Bryan Greenhouse. Sex-based differences in clearance of chronic Plasmodium falciparum infection. *eLife*, 9, 2020.

[6821] Christian Brion, Sheila M Lutz, and Frank Wolfgang Albert. Simultaneous quantification of mRNA and protein in single cells reveals post-transcriptional effects of genetic variation. *eLife*, 9, 2020.

[6822] Cara E Brook, Mike Boots, Kartik Chandran, Andrew P Dobson, Christian Drosten, Andrea L Graham, Bryan T Grenfell, Marcel A Müller, Melinda Ng, Lin-Fa Wang, and Anieke van Leeuwen. Accelerated viral dynamics in bat cell lines, with implications for zoonotic emergence. *eLife*, 9, 2020.

[6823] Eric R Brooks, Mohammed Tarek Islam, Kathryn V Anderson, and Jennifer A Zallen. Sonic hedgehog signaling directs patterned cell remodeling during cranial neural tube closure. *eLife*, 9, 2020.

[6824] William R Brothers, Steven Hebert, Claudia L Kleinman, and Marc R Fabian. A non-canonical role for the EDC4 decapping factor in regulating MARF1-mediated mRNA decay. *eLife*, 9, 2020.

[6825] Amanda M Brown, Joshua J White, Meike E van der Heijden, Joy Zhou, Tao Lin, and Roy V Sillitoe. Purkinje cell misfiring generates high-amplitude action tremors that are corrected by cerebellar deep brain stimulation. *eLife*, 9, 2020.

[6826] Stefan Brückner, Rajib Schubert, Timo Kraushaar, Raimo Hartmann, Daniel Hoffmann, Eric Jelli, Knut Drescher, Daniel J Müller, Lars Oliver Essen, and Hans-Ulrich Mösch. Kin discrimination in social yeast is mediated by cell surface receptors of the Flo11 adhesin family. *eLife*, 9, 2020.

[6827] Bart MH Bruininks, Paulo CT Souza, Helgi Ingolfsson, and Siewert J Marrink. A molecular view on the escape of lipoplexed DNA from the endosome. *eLife*, 9, 2020.

[6828] Giancarlo Noe Bruni and Joel M Kralj. Membrane voltage dysregulation driven by metabolic dysfunction underlies bactericidal activity of aminoglycosides. *eLife*, 9, 2020.

[6829] Andrä Brunner, Aldwin Suryo Rahmanto, Henrik Johansson, Marcela Franco, Johanna Viiliäinen, Mohiuddin Gazi, Oliver Frings, Erik Fredlund, Charles Spruck, Janne Lehtiö, Juha K Rantala, Lars-Gunnar Larsson, and Olle Sangfelt. PTEN and DNA-PK determine sensitivity and recovery in response to WEE1 inhibition in human breast cancer. *eLife*, 9, 2020.

[6830] Janine D Brunner, Roman P Jakob, Tobias Schulze, Yvonne Neldner, Anna Moroni, Gerhard Thiel, Timm Maier, and Stephan Schenck. Structural basis for ion selectivity in TMEM175 K+ channels. *eLife*, 9, 2020.

[6831] Julie Bruyère, Yah-Se Abada, Hélène Vitet, Gaëlle Fontaine, Jean-Christophe Deloulme, Aurélia Cès, Eric Denarier, Karin Pernet-Gallay, Annie Andrieux, Sandrine Humbert, Marie-Claude Potier, Benoît Delatour, and Frédéric Saudou. Presynaptic APP levels and synaptic homeostasis are regulated by Akt phosphorylation of huntingtin. *eLife*, 9, 2020.

[6832] Alessio P Buccino, Cole L Hurwitz, Samuel Garcia, Jeremy Magland, Joshua H Siegle, Roger Hurwitz, and Matthias H Hennig. SpikeInterface, a unified framework for spike sorting. *eLife*, 9, 2020.

[6833] Jonathan M Budzik, Danielle L Swaney, David Jimenez-Morales, Jeffrey R Johnson, Nicholas E Garelis, Teresa Repasy, Allison W Roberts, Lauren M Popov, Trevor J Parry, Dexter Pratt, Trey Ideker, Nevan J Krogan, and Jeffery S Cox. Dynamic post-translational modification profiling of Mycobacterium tuberculosis-infected primary macrophages. *eLife*, 9, 2020.

[6834] Carlos F Buen Abad Najar, Nir Yosef, and Liana F Lareau. Coverage-dependent bias creates the appearance of binary splicing in single cells. *eLife*, 9, 2020.

[6835] Ivan Buljan, Daniel Garcia-Costa, Francisco Grimaldo, Flaminio Squazzoni, and Ana MaruŠić. Large-scale language analysis of peer review reports. *eLife*, 9, 2020.

[6836] Kevin G Burfeind, Xinxia Zhu, Mason A Norgard, Peter R Levasseur, Christian Huisman, Abigail C Buenafe, Brennan Olson, Katherine A Michaelis, Eileen RS Torres, Sophia Jeng, Shannon McWeeney, Jacob Raber, and Daniel L Marks. Circulating myeloid cells invade the central nervous system to mediate cachexia during pancreatic cancer. *eLife*, 9, 2020.

[6837] Courtney A Burger, Jonathan Alevy, Anna K Casasent, Danye Jiang, Nicholas E Albrecht, Justine H Liang, Arlene A Hirano, Nicholas C Brecha, and Melanie A Samuel. LKB1 coordinates neurite remodeling to drive synapse layer emergence in the outer retina. *eLife*, 9, 2020.

[6838] Jeremy Carlos Burns, Bunny Cotleur, Dirk M Walther, Bekim Bajrami, Stephen J Rubino, Ru Wei, Nathalie Franchimont, Susan L Cotman, Richard M Ransohoff, and Michael Mingueneau. Differential accumulation of storage bodies with aging defines discrete subsets of microglia in the healthy brain. *eLife*, 9, 2020.

[6839] Leah Bury, Brittania Moodie, Jimmy Ly, Liliana S McKay, Karen HH Miga, and Iain M Cheeseman. Alpha-satellite RNA transcripts are repressed by centromere–nucleolus associations. *eLife*, 9, 2020.

[6840] Sean W Buskirk, Alecia B Rokes, and Gregory I Lang. Adaptive evolution of nontransitive fitness in yeast. *eLife*, 9, 2020.

[6841] Benjamin G Butt, Danielle J Owen, Cy M Jeffries, Lyudmila Ivanova, Chris H Hill, Jack W Houghton, Md Firoz Ahmed, Robin Antrobus, Dmitri I Svergun, John J Welch, Colin M Crump, and Stephen C Graham. Insights into herpesvirus assembly from the structure of the pUL7:pUL51 complex. *eLife*, 9, 2020.

[6842] Nicola Byrom. The challenges of lockdown for early-career researchers. *eLife*, 9, 2020.

[6843] David Cabañero, Angela Ramírez-López, Eva Drews, Anne Schmöle, David M Otte, Agnieszka Wawrzczak-Bargiela, Hector Huerga Encabo, Sami Kummer, Antonio Ferrer-Montiel, Ryszard Przewlocki, Andreas Zimmer, and Rafael Maldonado. Protective role of neuronal and lymphoid cannabinoid CB2 receptors in neuropathic pain. *eLife*, 9, 2020.

[6844] Cathryn R Cadwell, Federico Scala, Paul G Fahey, Dmitry Kobak, Shalaka Mulherkar, Fabian H Sinz, Stelios Papadopoulos, Zheng H Tan, Per Johnsson, Leonard Hartmanis, Shuang Li, Ronald J Cotton, Kimberley F Tolias, Rickard Sandberg, Philipp Berens, Xiaolong Jiang, and Andreas Savas Tolias. Cell type composition and circuit organization of clonally related excitatory neurons in the juvenile mouse neocortex. *eLife*, 9, 2020.

[6845] Lili X Cai, Katherine Pizano, Gregory W Gundersen, Cameron L Hayes, Weston T Fleming, Sebastian Holt, Julia M Cox, and Ilana B Witten. Distinct signals in medial and lateral VTA dopamine neurons modulate fear extinction at different times. *eLife*, 9, 2020.

[6846] CJ Cambier, Steven M Banik, Joseph A Buonomo, and Carolyn R Bertozzi. Spreading of a mycobacterial cell-surface lipid into host epithelial membranes promotes infectivity. *eLife*, 9, 2020.

[6847] Flavia Camponovo, Joseph J Campo, Timothy Q Le, Amit Oberai, Christopher Hung, Jozelyn V Pablo, Andy A Teng, Xiaowu Liang, B Kim Lee Sim, Said Jongo, Salim Abdulla, Marcel Tanner, Stephen L Hoffman, Claudia Daubenberger, and Melissa A Penny. Proteome-wide analysis of a malaria vaccine study reveals personalized humoral immune profiles in Tanzanian adults. *eLife*, 9, 2020.

[6848] Marta Canel, David Taggart, Andrew H Sims, David W Lonergan, Irene C Waizenegger, and Alan Serrels. T-cell co-stimulation in combination with targeting FAK drives enhanced anti-tumor immunity. *eLife*, 9, 2020.

[6849] Carla Cangalaya, Stoyan Stoyanov, Klaus-Dieter Fischer, and Alexander Dityatev. Light-induced engagement of microglia to focally remodel synapses in the adult brain. *eLife*, 9, 2020.

[6850] Yasmine Cantaut-Belarif, Adeline Orts Del'Immagine, Margot Penru, Guillaume Pézeron, Claire Wyart, and Pierre-Luc Bardet. Adrenergic activation modulates the signal from the Reissner fiber to cerebrospinal fluid-contacting neurons during development. *eLife*, 9, 2020.

[6851] Vincenza Caolo, Marjolaine Debant, Naima Endesh, T Simon Futers, Laeticia Lichtenstein, Fiona Bartoli, Gregory Parsonage, Elizabeth AV Jones, and David J Beech. Shear stress activates ADAM10 sheddase to regulate Notch1 via the Piezo1 force sensor in endothelial cells. *eLife*, 9, 2020.

[6852] Antonio Cappuccio, Shane T Jensen, Boris M Hartmann, Stuart C Sealfon, Vassili Soumelis, and Elena Zaslavsky. Deciphering the combinatorial landscape of immunity. *eLife*, 9, 2020.

[6853] Alexandre Carayon, Laetitia Bataillé, Gaëlle Lebreton, Laurence Dubois, Aurore Pelletier, Yannick Carrier, Antoine Wystrach, Alain Vincent, and Jean-Louis Frendo. Intrinsic control of muscle attachment sites matching. *eLife*, 9, 2020.

[6854] Sean M Carney, Andrew T Moreno, Sadie C Piatt, Metztli Cisneros-Aguirre, Felicia Wednesday Lopezcolorado, Jeremy M Stark, and Joseph J Loparo. XLF acts as a flexible connector during non-homologous end joining. *eLife*, 9, 2020.

[6855] Eli M Carrami, Sahand Sharifzadeh, Nina C Wietek, Mara Artibani, Salma El-Sahhar, Tatjana Sauka-Spengler, Christopher Yau, Volker Tresp, and Ahmed A Ahmed. A highly accurate platform for clone-specific mutation discovery enables the study of active mutational processes. *eLife*, 9, 2020.

[6856] Nicolas Carraro, Xavier Richard, Sandra Sulser, FranÇois Delavat, Christian Mazza, and Jan Roelof van der Meer. An analog to digital converter controls bistable transfer competence development of a widespread bacterial integrative and conjugative element. *eLife*, 9, 2020.

[6857] Rodrigo M Carrillo-Larco, C Joel Benites-Moya, Cecilia Anza-Ramirez, Leonardo Albitres-Flores, Diana Sánchez-Velazco, Niels Pacheco-Barrios, and Antonio Bernabe-Ortiz. A systematic review of population-based studies on lipid profiles in Latin America and the Caribbean. *eLife*, 9, 2020.

[6858] Brittany L Carroll, Tatsuro Nishikino, Wangbiao Guo, Shiwei Zhu, Seiji Kojima, Michio Homma, and Jun Liu. The flagellar motor of Vibrio alginolyticus undergoes major structural remodeling during rotational switching. *eLife*, 9, 2020.

[6859] Ava C Carter, Jin Xu, Meagan Y Nakamoto, Yuning Wei, Brian J Zarnegar, Quanming Shi, James P Broughton, Ryan C Ransom, Ankit Salhotra, Surya D Nagaraja, Rui Li, Diana R Dou, Kathryn E Yost, Seung-Woo Cho, Anil Mistry, Michael T Longaker, Paul A Khavari, Robert T Batey, Deborah S Wuttke, and Howard Y Chang. Spen links RNA-mediated endogenous retrovirus silencing and X chromosome inactivation. *eLife*, 9, 2020.

[6860] Paul Carter, Mathew Vithayathil, Siddhartha Kar, Rahul Potluri, Amy M Mason, Susanna C Larsson, and Stephen Burgess. Predicting the effect of statins on cancer risk using genetic variants from a Mendelian randomization study in the UK Biobank. *eLife*, 9, 2020.

[6861] Bruno Sylvain Carturan, Jason Pither, Jean-Philippe Maréchal, Corey JA Bradshaw, and Lael Parrott. Combining agent-based, trait-based and demographic approaches to model coral-community dynamics. *eLife*, 9, 2020.

[6862] Tanita Casci and Elizabeth Adams. Setting the right tone. *eLife*, 9, 2020.

[6863] Jason C Casler and Benjamin S Glick. A microscopy-based kinetic analysis of yeast vacuolar protein sorting. *eLife*, 9, 2020.

[6864] Victoria G Castiglioni, Helena R Pires, Rodrigo Rosas Bertolini, Amalia Riga, Jana Kerver, and Mike Boxem. Epidermal PAR-6 and PKC-3 are essential for larval development of C. elegans and organize non-centrosomal microtubules. *eLife*, 9, 2020.

[6865] Ryan Castro, Thomas Taetzsch, Sydney K Vaughan, Kerilyn Godbe, John Chappell, Robert E Settlage, and Gregorio Valdez. Specific labeling of synaptic schwann cells reveals unique cellular and molecular features. *eLife*, 9, 2020.

[6866] Sean Edward Cavanagh, Norman H Lam, John D Murray, Laurence Tudor Hunt, and Steven Wayne Kennerley. A circuit mechanism for decision-making biases and NMDA receptor hypofunction. *eLife*, 9, 2020.

[6867] Mathieu Cayla, Lindsay McDonald, Paula MacGregor, and Keith Matthews. An atypical DYRK kinase connects quorum-sensing with posttranscriptional gene regulation in Trypanosoma brucei. *eLife*, 9, 2020.

[6868] Christopher I Cazzonelli, Xin Hou, Yagiz Alagoz, John Rivers, Namraj Dhami, Jiwon Lee, Shashikanth Marri, and Barry J Pogson. A cis-carotene derived apocarotenoid regulates etioplast and chloroplast development. *eLife*, 9, 2020.

[6869] Jaroslaw Cendrowski, Marta Kaczmarek, Michał Mazur, Katarzyna Kuzmicz-Kowalska, Kamil Jastrzebski, Marta Brewinska-Olchowik, Agata Kominek, Katarzyna Piwocka, and Marta Miaczynska. Splicing variation of BMP2K balances abundance of COPII assemblies and autophagic degradation in erythroid cells. *eLife*, 9, 2020.

[6870] Nathan Cermak, Stephanie K Yu, Rebekah Clark, Yung-Chi Huang, Saba N Baskoylu, and Steven W Flavell. Whole-organism behavioral profiling reveals a role for dopamine in state-dependent motor program coupling in C. elegans. *eLife*, 9, 2020.

[6871] Alice Cezanne, Janelle Lauer, Anastasia Solomatina, Ivo F Sbalzarini, and Marino Zerial. A non-linear system patterns Rab5 GTPase on the membrane. *eLife*, 9, 2020.

[6872] Annie Wai Yeeng Chai, Pei San Yee, Stacey Price, Shi Mun Yee, Hui Mei Lee, Vivian KH Tiong, Emanuel GonÇalves, Fiona M Behan, Jessica Bateson, James Gilbert, Aik Choon Tan, Ultan McDermott, Mathew J Garnett, and Sok Ching Cheong. Genome-wide CRISPR screens of oral squamous cell carcinoma reveal fitness genes in the Hippo pathway. *eLife*, 9, 2020.

[6873] Leela Chakravarti Dilley, Milan Szuperak, Naihua N Gong, Charlette E Williams, Ricardo Linares Saldana, David S Garbe, Mubarak Hussain Syed, Rajan Jain, and Matthew S Kayser. Identification of a molecular basis for the juvenile sleep state. *eLife*, 9, 2020.

[6874] Karima Chakroun, David Mathar, Antonius Wiehler, Florian Ganzer, and Jan Peters. Dopaminergic modulation of the exploration/exploitation trade-off in human decision-making. *eLife*, 9, 2020.

[6875] Ka Man Carmen Chan, Sungmin Son, Eva M Schmid, and Daniel A Fletcher. A viral fusogen hijacks the actin cytoskeleton to drive cell-cell fusion. *eLife*, 9, 2020.

[6876] Zora Chui-Kuen Chan, Hiu-Lam Rachel Kwan, Yin Shun Wong, Zhixin Jiang, Zhongjun Zhou, Kin Wai Tam, Ying-Shing Chan, Chi Bun Chan, and Chi Wai Lee. Site-directed MT1-MMP trafficking and surface insertion regulate AChR clustering and remodeling at developing NMJs. *eLife*, 9, 2020.

[6877] Santosh Chandrasekaran, Ameya C Nanivadekar, Gina McKernan, Eric R Helm, Michael L Boninger, Jennifer L Collinger, Robert A Gaunt, and Lee E Fisher. Sensory restoration by epidural stimulation of the lateral spinal cord in upper-limb amputees. *eLife*, 9, 2020.

[6878] Ling-Shih Chang, Minseong Kim, Andrey Glinka, Carmen Reinhard, and Christof Niehrs. The tumor suppressor PTPRK promotes ZNRF3 internalization and is required for Wnt inhibition in the Spemann organizer. *eLife*, 9, 2020.

[6879] Ting-Yu Chang, Raymond Doudlah, Byounghoon Kim, Adhira Sunkara, Lowell W Thompson, Meghan E Lowe, and Ari Rosenberg. Functional links between sensory representations, choice activity, and sensorimotor associations in parietal cortex. *eLife*, 9, 2020.

[6880] Lynne Chantranupong, Jessica L Saulnier, Wengang Wang, Drew R Jones, Michael E Pacold, and Bernardo L Sabatini. Rapid purification and metabolomic profiling of synaptic vesicles from mammalian brain. *eLife*, 9, 2020.

[6881] Bolton KH Chau, Chun-Kit Law, Alizée Lopez-Persem, Miriam C Klein-Flügge, and Matthew FS Rushworth. Consistent patterns of distractor effects during decision making. *eLife*, 9, 2020.

[6882] Tanmay S Chavan, Ricky C Cheng, Tao Jiang, Irimpan I Mathews, Richard A Stein, Antoine Koehl, Hassane S Mchaourab, Emad Tajkhorshid, and Merritt Maduke. A CLC-ec1 mutant reveals global conformational change and suggests a unifying mechanism for the CLC Cl–/H+ transport cycle. *eLife*, 9, 2020.

[6883] Hui Che, Jie Li, You Li, Cheng Ma, Huan Liu, Jingyi Qin, Jianghui Dong, Zhen Zhang, Cory J Xian, Dengshun Miao, Liping Wang, and Yongxin Ren. p16 deficiency attenuates intervertebral disc degeneration by adjusting oxidative stress and nucleus pulposus cell cycle. *eLife*, 9, 2020.

[6884] Chen Chen, Todd D Murphey, and Malcolm A MacIver. Tuning movement for sensing in an uncertain world. *eLife*, 9, 2020.

[6885] Cheng-Yi Chen, Sean A McKinney, Lacey R Ellington, and Matthew C Gibson. Hedgehog signaling is required for endomesodermal patterning and germ cell development in the sea anemone Nematostella vectensis. *eLife*, 9, 2020.

[6886] Chuyu Chen, Giulia Soto, Vasin Dumrongprechachan, Nicholas Bannon, Shuo Kang, Yevgenia Kozorovitskiy, and Loukia Parisiadou. Pathway-specific dysregulation of striatal excitatory synapses by LRRK2 mutations. *eLife*, 9, 2020.

[6887] Cuie Chen and Yukiko M Yamashita. Alstrom syndrome gene is a stem-cell-specific regulator of centriole duplication in the Drosophila testis. *eLife*, 9, 2020.

[6888] Donglei Chen, Henning Blom, Sophie Sanchez, Paul Tafforeau, Tiiu Märss, and Per E Ahlberg. The developmental relationship between teeth and dermal odontodes in the most primitive bony fish Lophosteus. *eLife*, 9, 2020.

[6889] John Z Chen, Douglas M Fowler, and Nobuhiko Tokuriki. Comprehensive exploration of the translocation, stability and substrate recognition requirements in VIM-2 lactamase. *eLife*, 9, 2020.

[6890] Maorong Chen, Nathalia Amado, Jieqiong Tan, Alice Reis, Mengxu Ge, Jose Garcia Abreu, and Xi He. TMEM79/MATTRIN defines a pathway for Frizzled regulation and is required for Xenopus embryogenesis. *eLife*, 9, 2020.

[6891] Qiang Chen, Robert G Smith, Xiaolin Huang, and Wei Wei. Preserving inhibition with a disinhibitory microcircuit in the retina. *eLife*, 9, 2020.

[6892] Quanjing Chen, Adam Turnbull, Timothy M Baran, and Feng V Lin. Longitudinal stability of medial temporal lobe connectivity is associated with tau-related memory decline. *eLife*, 9, 2020.

[6893] Wu Chen, Zhao-Lin Cai, Eugene S Chao, Hongmei Chen, Colleen M Longley, Shuang Hao, Hsiao-Tuan Chao, Joo Hyun Kim, Jessica E Messier, Huda Y Zoghbi, Jianrong Tang, John W Swann, and Mingshan Xue. Stxbp1/Munc18-1 haploinsufficiency impairs inhibition and mediates key neurological features of STXBP1 encephalopathy. *eLife*, 9, 2020.

[6894] Chun-Chun Cheng, Joshua Wooten, Zane A Gibbs, Kathleen McGlynn, Prashant Mishra, and Angelique W Whitehurst. Sperm-specific COX6B2 enhances oxidative phosphorylation, proliferation, and survival in human lung adenocarcinoma. *eLife*, 9, 2020.

[6895] Ryan R Cheng, Vinicius G Contessoto, Erez Lieberman Aiden, Peter G Wolynes, Michele Di Pierro, and Jose N Onuchic. Exploring chromosomal structural heterogeneity across multiple cell lines. *eLife*, 9, 2020.

[6896] Yubo Cheng, Luke Saville, Babita Gollen, Christopher Isaac, Abel Belay, Jogender Mehla, Kush Patel, Nehal Thakor, Majid H Mohajerani, and Athanasios Zovoilis. Increased processing of SINE B2 ncRNAs unveils a novel type of transcriptome deregulation in amyloid beta neuropathology. *eLife*, 9, 2020.

[6897] Hosni Cherif, Daniel G Bisson, Matthew Mannarino, Oded Rabau, Jean A Ouellet, and Lisbet Haglund. Senotherapeutic drugs for human intervertebral disc degeneration and low back pain. *eLife*, 9, 2020.

[6898] Antoine Cherix, Thomas Larrieu, Jocelyn Grosse, João Rodrigues, Bruce McEwen, Carla Nasca, Rolf Gruetter, and Carmen Sandi. Metabolic signature in nucleus accumbens for anti-depressant-like effects of acetyl-L-carnitine. *eLife*, 9, 2020.

[6899] Xiaoke Chi, Dang Nguyen, James M Pemberton, Elizabeth J Osterlund, Qian Liu, Hetal Brahmbhatt, Zhi Zhang, Jialing Lin, Brian Leber, and David W Andrews. The carboxyl-terminal sequence of bim enables bax activation and killing of unprimed cells. *eLife*, 9, 2020.

[6900] Ying Ann Chiao, Huiliang Zhang, Mariya Sweetwyne, Jeremy Whitson, Ying Sonia Ting, Nathan Basisty, Lindsay K Pino, Ellen Quarles, Ngoc-Han Nguyen, Matthew D Campbell, Tong Zhang, Matthew J Gaffrey, Gennifer Merrihew, Lu Wang, Yongping Yue, Dongsheng Duan, Henk L Granzier, Hazel H Szeto, Wei-Jun Qian, David Marcinek, Michael J MacCoss, and Peter Rabinovitch. Late-life restoration of mitochondrial function reverses cardiac dysfunction in old mice. *eLife*, 9, 2020.

[6901] Antonella Chiapparino, Antonija Grbavac, Hendrik RA Jonker, Yvonne Hackmann, Sofia Mortensen, Ewa Zatorska, Andrea Schott, Gunter Stier, Krishna Saxena, Klemens Wild, Harald Schwalbe, Sabine Strahl, and Irmgard Sinning. Functional implications of MIR domains in protein O-mannosylation. *eLife*, 9, 2020.

[6902] Melissa A Chiasson, Nathan J Rollins, Jason J Stephany, Katherine A Sitko, Kenneth A Matreyek, Marta Verby, Song Sun, Frederick P Roth, Daniel DeSloover, Debora S Marks, Allan E Rettie, and Douglas M Fowler. Multiplexed measurement of variant abundance and activity reveals VKOR topology, active site and human variant impact. *eLife*, 9, 2020.

[6903] Suneeta Chimalapati, Marcela de Souza Santos, Alexander E Lafrance, Ann Ray, Wan-Ru Lee, Giomar Rivera-Cancel, GonÇalo Vale, Krzysztof Pawlowski, Matthew A Mitsche, Jeffrey G McDonald, Jen Liou, and Kim Orth. Vibrio deploys type 2 secreted lipase to esterify cholesterol with host fatty acids and mediate cell egress. *eLife*, 9, 2020.

[6904] Chue Vin Chin, Jisha Antony, Sarada Ketharnathan, Anastasia Labudina, Gregory Gimenez, Kate M Parsons, Jinshu He, Amee J George, Maria Michela Pallotta, Antonio Musio, Antony Braithwaite, Parry Guilford, Ross D Hannan, and Julia A Horsfield. Cohesin mutations are synthetic lethal with stimulation of WNT signaling. *eLife*, 9, 2020.

[6905] Praneeth Chitirala, Hsin-Fang Chang, Paloma Martzloff, Christiane Harenberg, Keerthana Ravichandran, Midhat H Abdulreda, Per-Olof Berggren, Elmar Krause, Claudia Schirra, Trese Leinders-Zufall, Fritz Benseler, Nils Brose, and Jens Rettig. Studying the biology of cytotoxic T lymphocytes in vivo with a fluorescent granzyme B-mTFP knock-in mouse. *eLife*, 9, 2020.

[6906] Ramesh Chittajallu, Kurt Auville, Vivek Mahadevan, Mandy Lai, Steven Hunt, Daniela Calvigioni, Kenneth A Pelkey, Kareem A Zaghloul, and Chris J McBain. Activity-dependent tuning of intrinsic excitability in mouse and human neurogliaform cells. *eLife*, 9, 2020.

[6907] Howard Choi, Deborah Simpson, Ding Wang, Mark Prescott, Andrew A Pitsillides, Jayesh Dudhia, Peter D Clegg, Peipei Ping, and Chavaunne T Thorpe. Heterogeneity of proteome dynamics between connective tissue phases of adult tendon. *eLife*, 9, 2020.

[6908] In Young Choi, Hotae Lim, Hyeon Jin Cho, Yohan Oh, Bin-Kuan Chou, Hao Bai, Linzhao Cheng, Yong Jun Kim, SangHwan Hyun, Hyesoo Kim, Joo Heon Shin, and Gabsang Lee. Transcriptional landscape of myogenesis from human pluripotent stem cells reveals a key role of TWIST1 in maintenance of skeletal muscle progenitors. *eLife*, 9, 2020.

[6909] Se Joon Choi, Thong C Ma, Yunmin Ding, Timothy Cheung, Neal Joshi, David Sulzer, Eugene V Mosharov, and Un Jung Kang. Alterations in the intrinsic properties of striatal cholinergic interneurons after dopamine lesion and chronic L-DOPA. *eLife*, 9, 2020.

[6910] Weng Man Chong, Won-Jing Wang, Chien-Hui Lo, Tzu-Yuan Chiu, Ting-Jui Chang, You-Pi Liu, Barbara Tanos, Gregory Mazo, Meng-Fu Bryan Tsou, Wann-Neng Jane, T Tony Yang, and Jung-Chi Liao. Super-resolution microscopy reveals coupling between mammalian centriole subdistal appendages and distal appendages. *eLife*, 9, 2020.

[6911] Xiao-lin Chou, Qi Fang, Linqing Yan, Wen Zhong, Bo Peng, Haifu Li, Jinxing Wei, Huizhong W Tao, and Li I Zhang. Contextual and cross-modality modulation of auditory cortical processing through pulvinar mediated suppression. *eLife*, 9, 2020.

[6912] Raeed H Chowdhury, Joshua I Glaser, and Lee E Miller. Area 2 of primary somatosensory cortex encodes kinematics of the whole arm. *eLife*, 9, 2020.

[6913] Nam Chu, Thibault Viennet, Hwan Bae, Antonieta Salguero, Andras Boeszoermenyi, Haribabu Arthanari, and Philip A Cole. The structural determinants of PH domain-mediated regulation of Akt revealed by segmental labeling. *eLife*, 9, 2020.

[6914] Xiakun Chu, Zucai Suo, and Jin Wang. Investigating the trade-off between folding and function in a multidomain Y-family DNA polymerase. *eLife*, 9, 2020.

[6915] Elin Claesson, Weixiao Yuan Wahlgren, Heikki Takala, Suraj Pandey, Leticia Castillon, Valentyna Kuznetsova, Léocadie Henry, Matthijs Panman, Melissa Carrillo, Joachim Kübel, Rahul Nanekar, Linnéa Isaksson, Amke Nimmrich, Andrea Cellini, Dmitry Morozov, Michał Maj, Moona Kurttila, Robert Bosman, Eriko Nango, Rie Tanaka, Tomoyuki Tanaka, Luo Fangjia, So Iwata, Shigeki Owada, Keith Moffat, Gerrit Groenhof, Emina A Stojković, Janne A Ihalainen, Marius Schmidt, and Sebastian Westenhoff. The primary structural photoresponse of phytochrome proteins captured by a femtosecond X-ray laser. *eLife*, 9, 2020.

[6916] Erin A Clark, Michael Rutlin, Lucia S Capano, Samuel Aviles, Jordan R Saadon, Praveen Taneja, Qiyu Zhang, James B Bullis, Timothy Lauer, Emma Myers, Anton Schulmann, Douglas Forrest, and Sacha B Nelson. Cortical ROR$\beta$ is required for layer 4 transcriptional identity and barrel integrity. *eLife*, 9, 2020.

[6917] Colin M Cleary, Thiago S Moreira, Ana C Takakura, Mark T Nelson, Thomas A Longden, and Daniel K Mulkey. Vascular control of the CO2/H+-dependent drive to breathe. *eLife*, 9, 2020.

[6918] Joshua C Cofsky, Deepti Karandur, Carolyn J Huang, Isaac P Witte, John Kuriyan, and Jennifer A Doudna. CRISPR-Cas12a exploits R-loop asymmetry to form double-strand breaks. *eLife*, 9, 2020.

[6919] Moran Cohen-Berkman, Reut Dudkevich, Shani Ben-Hamo, Alla Fishman, Yehuda Salzberg, Hiba Waldman Ben-Asher, Ayelet T Lamm, and Sivan Henis-Korenblit. Endogenous siRNAs promote proteostasis and longevity in germline-less Caenorhabditis elegans. *eLife*, 9, 2020.

[6920] Jonathan A Coleman, Vikas Navratna, Daniele Antermite, Dongxue Yang, James A Bull, and Eric Gouaux. Chemical and structural investigation of the paroxetine-human serotonin transporter complex. *eLife*, 9, 2020.

[6921] Enrico Sandro Colizzi, Renske MA Vroomans, and Roeland MH Merks. Evolution of multicellularity by collective integration of spatial information. *eLife*, 9, 2020.

[6922] James E Collier, Byung-Gil Lee, Maurici Brunet Roig, Stanislav Yatskevich, Naomi J Petela, Jean Metson, Menelaos Voulgaris, Andres Gonzalez Llamazares, Jan Löwe, and Kim A Nasmyth. Transport of DNA within cohesin involves clamping on top of engaged heads by Scc2 and entrapment within the ring by Scc3. *eLife*, 9, 2020.

[6923] Christine R Collins, Fiona Hackett, Steven A Howell, Ambrosius P Snijders, Matthew RG Russell, Lucy M Collinson, and Michael J Blackman. The malaria parasite sheddase SUB2 governs host red blood cell membrane sealing at invasion. *eLife*, 9, 2020.

[6924] Marco Colnaghi, Nick Lane, and Andrew Pomiankowski. Genome expansion in early eukaryotes drove the transition from lateral gene transfer to meiotic sex. *eLife*, 9, 2020.

[6925] Che C Colpitts, Sophie Ridewood, Bethany Schneiderman, Justin Warne, Keisuke Tabata, Caitlin F Ng, Ralf Bartenschlager, David L Selwood, and Greg J Towers. Hepatitis C virus exploits cyclophilin A to evade PKR. *eLife*, 9, 2020.

[6926] Alice Cont, Tamara Rossy, Zainebe Al-Mayyah, and Alexandre Persat. Biofilms deform soft surfaces and disrupt epithelia. *eLife*, 9, 2020.

[6927] Caroline Contat, Pierre-Benoit Ancey, Nadine Zangger, Silvia Sabatino, Justine Pascual, Stéphane Escrig, Louise Jensen, Christine Goepfert, Bernard Lanz, Mario Lepore, Rolf Gruetter, Anouk Rossier, Sabina Berezowska, Christina Neppl, Inti Zlobec, Stéphanie Clerc-Rosset, Graham William Knott, Jeffrey C Rathmell, E Dale Abel, Anders Meibom, and Etienne Meylan. Combined deletion of Glut1 and Glut3 impairs lung adenocarcinoma growth. *eLife*, 9, 2020.

[6928] Geoffrey MW Cook, Catia Sousa, Julia Schaeffer, Katherine Wiles, Prem Jareonsettasin, Asanish Kalyanasundaram, Eleanor Walder, Catharina Casper, Serena Patel, Pei Wei Chua, Gioia Riboni-Verri, Mansoor Raza, Nol Swaddiwudhipong, Andrew Hui, Ameer Abdullah, Saj Wajed, and Roger J Keynes. Regulation of nerve growth and patterning by cell surface protein disulphide isomerase. *eLife*, 9, 2020.

[6929] Nathalie Coré, Andrea Erni, Hanne M Hoffmann, Pamela L Mellon, Andrew J Saurin, Christophe Beclin, and Harold Cremer. Stem cell regionalization during olfactory bulb neurogenesis depends on regulatory interactions between Vax1 and Pax6. *eLife*, 9, 2020.

[6930] June K Corrigan, Deepti Ramachandran, Yuchen He, Colin J Palmer, Michael J Jurczak, Rui Chen, Bingshan Li, Randall H Friedline, Jason K Kim, Jon J Ramsey, Louise Lantier, Owen P McGuinness, and Alexander S Banks. A big-data approach to understanding metabolic rate and response to obesity in laboratory mice. *eLife*, 9, 2020.

[6931] Ana Rita Costa, Sara C Sousa, Rita Pinto-Costa, José C Mateus, Cátia DF Lopes, Ana Catarina Costa, David Rosa, Diana Machado, Luis Pajuelo, Xuewei Wang, Feng-quan Zhou, António J Pereira, Paula Sampaio, Boris Y Rubinstein, Inês Mendes Pinto, Marko Lampe, Paulo Aguiar, and Monica M Sousa. The membrane periodic skeleton is an actomyosin network that regulates axonal diameter and conduction. *eLife*, 9, 2020.

[6932] Nuno Costa-Borges, Katharina Spath, Irene Miguel-Escalada, Enric Mestres, Rosa Balmaseda, Anna Serafín, Maria Garcia-Jiménez, Ivette Vanrell, Jesús González, Klaus Rink, Dagan Wells, and Gloria Calderón. Maternal spindle transfer overcomes embryo developmental arrest caused by ooplasmic defects in mice. *eLife*, 9, 2020.

[6933] Lorenzo Costantino, Tsung-Han S Hsieh, Rebecca Lamothe, Xavier Darzacq, and Douglas Koshland. Cohesin residency determines chromatin loop patterns. *eLife*, 9, 2020.

[6934] Charles AR Cotton, Iria Bernhardsgrütter, Hai He, Simon Burgener, Luca Schulz, Nicole Paczia, Beau Dronsella, Alexander Erban, Stepan Toman, Marian Dempfle, Alberto De Maria, Joachim Kopka, Steffen N Lindner, Tobias J Erb, and Arren Bar-Even. Underground isoleucine biosynthesis pathways in E. coli. *eLife*, 9, 2020.

[6935] Nicolas Coudray, Georgia L Isom, Mark R MacRae, Mariyah N Saiduddin, Gira Bhabha, and Damian C Ekiert. Structure of bacterial phospholipid transporter MlaFEDB with substrate bound. *eLife*, 9, 2020.

[6936] Joshua F Coulcher, Agnès Roure, Rafath Chowdhury, Méryl Robert, Laury Lescat, Aurélie Bouin, Juliana Carvajal Cadavid, Hiroki Nishida, and Sébastien Darras. Conservation of peripheral nervous system formation mechanisms in divergent ascidian embryos. *eLife*, 9, 2020.

[6937] J Aaron Crapster, Paul G Rack, Zane J Hellmann, Austen D Le, Christopher M Adams, Ryan D Leib, Joshua E Elias, John Perrino, Barry Behr, Yanfeng Li, Jennifer Lin, Hong Zeng, and James K Chen. HIPK4 is essential for murine spermiogenesis. *eLife*, 9, 2020.

[6938] María Crespo, Barbara Gonzalez-Teran, Ivana Nikolic, Alfonso Mora, Cintia Folgueira, Elena Rodríguez, Luis Leiva-Vega, Aránzazu Pintor-Chocano, Macarena Fernández-Chacón, Irene Ruiz-Garrido, Beatriz Cicuéndez, Antonia Tomás-Loba, Noelia A-Gonzalez, Ainoa Caballero-Molano, Daniel Beiroa, Lourdes Hernández-Cosido, Jorge L Torres, Norman J Kennedy, Roger J Davis, Rui Benedito, Miguel Marcos, Ruben Nogueiras, Andrés Hidalgo, Nuria Matesanz, Magdalena Leiva, and Guadalupe Sabio. Neutrophil infiltration regulates clock-gene expression to organize daily hepatic metabolism. *eLife*, 9, 2020.

[6939] Mélanie Criqui, Aditi Qamra, Tsz Wai Chu, Monika Sharma, Julissa Tsao, Danielle A Henry, Dalia Barsyte-Lovejoy, Cheryl H Arrowsmith, Neil Winegarden, Mathieu Lupien, and Lea Harrington. Telomere dysfunction cooperates with epigenetic alterations to impair murine embryonic stem cell fate commitment. *eLife*, 9, 2020.

[6940] Katharine E Criswell and J Andrew Gillis. Resegmentation is an ancestral feature of the gnathostome vertebral skeleton. *eLife*, 9, 2020.

[6941] Kristina Crona. Rank orders and signed interactions in evolutionary biology. *eLife*, 9, 2020.

[6942] Richard B Crouse, Kristen Kim, Hannah M Batchelor, Eric M Girardi, Rufina Kamaletdinova, Justin Chan, Prithviraj Rajebhosale, Steven T Pittenger, Lorna W Role, David A Talmage, Miao Jing, Yulong Li, Xiao-Bing Gao, Yann S Mineur, and Marina R Picciotto. Acetylcholine is released in the basolateral amygdala in response to predictors of reward and enhances the learning of cue-reward contingency. *eLife*, 9, 2020.

[6943] Laura Crowley, Francesco Cambuli, Luis Aparicio, Maho Shibata, Brian D Robinson, Shouhong Xuan, Weiping Li, Hanina Hibshoosh, Massimo Loda, Raul Rabadan, and Michael M Shen. A single-cell atlas of the mouse and human prostate reveals heterogeneity and conservation of epithelial progenitors. *eLife*, 9, 2020.

[6944] Grisel Cruz-Becerra and James T Kadonaga. Enhancement of homology-directed repair with chromatin donor templates in cells. *eLife*, 9, 2020.

[6945] Gábor Csordás, Ferdinand Grawe, and Mirka Uhlirova. Eater cooperates with Multiplexin to drive the formation of hematopoietic compartments. *eLife*, 9, 2020.

[6946] Sean R Cuddy, Austin R Schinlever, Sara Dochnal, Philip V Seegren, Jon Suzich, Parijat Kundu, Taylor K Downs, Mina Farah, Bimal N Desai, Chris Boutell, and Anna R Cliffe. Neuronal hyperexcitability is a DLK-dependent trigger of herpes simplex virus reactivation that can be induced by IL-1. *eLife*, 9, 2020.

[6947] Xin Cui, Chao Ma, Varshini Vasudevaraja, Jonathan Serrano, Jie Tong, Yansong Peng, Michael Delorenzo, Guomiao Shen, Joshua Frenster, Renee-Tyler Tan Morales, Weiyi Qian, Aristotelis Tsirigos, Andrew S Chi, Rajan Jain, Sylvia C Kurz, Erik P Sulman, Dimitris G Placantonakis, Matija Snuderl, and Weiqiang Chen. Dissecting the immunosuppressive tumor microenvironments in Glioblastoma-on-a-Chip for optimized PD-1 immunotherapy. *eLife*, 9, 2020.

[6948] Yujun Cui, Chao Yang, Hongling Qiu, Hui Wang, Ruifu Yang, and Daniel Falush. The landscape of coadaptation in Vibrio parahaemolyticus. *eLife*, 9, 2020.

[6949] Zaixu Cui, Jennifer Stiso, Graham L Baum, Jason Z Kim, David R Roalf, Richard F Betzel, Shi Gu, Zhixin Lu, Cedric H Xia, Xiaosong He, Rastko Ciric, Desmond J Oathes, Tyler M Moore, Russell T Shinohara, Kosha Ruparel, Christos Davatzikos, Fabio Pasqualetti, Raquel E Gur, Ruben C Gur, Danielle S Bassett, and Theodore D Satterthwaite. Optimization of energy state transition trajectory supports the development of executive function during youth. *eLife*, 9, 2020.

[6950] Margaret M Cunniff, Eirene Markenscoff-Papadimitriou, Julia Ostrowski, John LR Rubenstein, and Vikaas Singh Sohal. Altered hippocampal-prefrontal communication during anxiety-related avoidance in mice deficient for the autism-associated gene Pogz. *eLife*, 9, 2020.

[6951] Kathleen M Cunningham, Kirstin Maulding, Kai Ruan, Mumine Senturk, Jonathan C Grima, Hyun Sung, Zhongyuan Zuo, Helen Song, Junli Gao, Sandeep Dubey, Jeffrey D Rothstein, Ke Zhang, Hugo J Bellen, and Thomas E Lloyd. TFEB/Mitf links impaired nuclear import to autophagolysosomal dysfunction in C9-ALS. *eLife*, 9, 2020.

[6952] Ryan R Cupo and James Shorter. Skd3 (human ClpB) is a potent mitochondrial protein disaggregase that is inactivated by 3-methylglutaconic aciduria-linked mutations. *eLife*, 9, 2020.

[6953] Timothy A Currier, Andrew MM Matheson, and Katherine I Nagel. Encoding and control of orientation to airflow by a set of Drosophila fan-shaped body neurons. *eLife*, 9, 2020.

[6954] Asher D Cutter and Joanna D Bundus. Speciation and the developmental alarm clock. *eLife*, 9, 2020.

[6955] Henry WP Dalgleish, Lloyd E Russell, Adam M Packer, Arnd Roth, Oliver M Gauld, Francesca Greenstreet, Emmett J Thompson, and Michael Häusser. How many neurons are sufficient for perception of cortical activity? *eLife*, 9, 2020.

[6956] Jeziel Dener Damasceno, Catarina A Marques, Dario Beraldi, Kathryn Crouch, Craig Lapsley, Ricardo Obonaga, Luiz RO Tosi, and Richard McCulloch. Genome duplication in Leishmania major relies on persistent subtelomeric DNA replication. *eLife*, 9, 2020.

[6957] Julia Damiano-Guercio, Laëtitia Kurzawa, Jan Mueller, Georgi Dimchev, Matthias Schaks, Maria Nemethova, Thomas Pokrant, Stefan Brühmann, Joern Linkner, Laurent Blanchoin, Michael Sixt, Klemens Rottner, and Jan Faix. Loss of Ena/VASP interferes with lamellipodium architecture, motility and integrin-dependent adhesion. *eLife*, 9, 2020.

[6958] James A D'Amour, Tyler Ekins, Stuti Ganatra, Xiaoqing Yuan, and Chris J McBain. Aberrant sorting of hippocampal complex pyramidal cells in type I lissencephaly alters topological innervation. *eLife*, 9, 2020.

[6959] Christian Damsgaard, Henrik Lauridsen, Till S Harter, Garfield T Kwan, Jesper S Thomsen, Anette MD Funder, Claudiu T Supuran, Martin Tresguerres, Philip GD Matthews, and Colin J Brauner. A novel acidification mechanism for greatly enhanced oxygen supply to the fish retina. *eLife*, 9, 2020.

[6960] Holger Dannenberg, Hallie Lazaro, Pranav Nambiar, Alec Hoyland, and Michael E Hasselmo. Effects of visual inputs on neural dynamics for coding of location and running speed in medial entorhinal cortex. *eLife*, 9, 2020.

[6961] Sven Dannhäuser, Thomas J Lux, Chun Hu, Mareike Selcho, Jeremy T-C Chen, Nadine Ehmann, Divya Sachidanandan, Sarah Stopp, Dennis Pauls, Matthias Pawlak, Tobias Langenhan, Peter Soba, Heike L Rittner, and Robert J Kittel. Antinociceptive modulation by the adhesion GPCR CIRL promotes mechanosensory signal discrimination. *eLife*, 9, 2020.

[6962] Heike Danzer, Joachim Glaesner, Anne Baerenwaldt, Carmen Reitinger, Anja Lux, Lukas Heger, Diana Dudziak, Thomas Harrer, André Gessner, and Falk Nimmerjahn. Human Fc$\gamma$-receptor IIb modulates pathogen-specific versus self-reactive antibody responses in lyme arthritis. *eLife*, 9, 2020.

[6963] Daniel Dar, Linda S Thomashow, David M Weller, and Dianne K Newman. Global landscape of phenazine biosynthesis and biodegradation reveals species-specific colonization patterns in agricultural soils and crop microbiomes. *eLife*, 9, 2020.

[6964] Timothy R Darlington and Stephen G Lisberger. Mechanisms that allow cortical preparatory activity without inappropriate movement. *eLife*, 9, 2020.

[6965] Jean-FranÇois Darrigrand, Mariana Valente, Glenda Comai, Pauline Martinez, Maxime Petit, Ryuichi Nishinakamura, Daniel S Osorio, Gilles Renault, Carmen Marchiol, Vanessa Ribes, and Bruno Cadot. Dullard-mediated Smad1/5/8 inhibition controls mouse cardiac neural crest cells condensation and outflow tract septation. *eLife*, 9, 2020.

[6966] Srijit Das, Felicia K Ooi, Johnny Cruz Corchado, Leah C Fuller, Joshua A Weiner, and Veena Prahlad. Serotonin signaling by maternal neurons upon stress ensures progeny survival. *eLife*, 9, 2020.

[6967] Suman G Das, Susana OL Direito, Bartlomiej Waclaw, Rosalind J Allen, and Joachim Krug. Predictable properties of fitness landscapes induced by adaptational tradeoffs. *eLife*, 9, 2020.

[6968] Ananya Dasgupta, Yu Jia Lim, Krishna Kumar, Nimmi Baby, Ka Lam Karen Pang, Amrita Benoy, Thomas Behnisch, and Sreedharan Sajikumar. Group III metabotropic glutamate receptors gate long-term potentiation and synaptic tagging/capture in rat hippocampal area CA2. *eLife*, 9, 2020.

[6969] Matthew J Davidson, Will Mithen, Hinze Hogendoorn, Jeroen JA van Boxtel, and Naotsugu Tsuchiya. The SSVEP tracks attention, not consciousness, during perceptual filling-in. *eLife*, 9, 2020.

[6970] Fred P Davis, Aljoscha Nern, Serge Picard, Michael B Reiser, Gerald M Rubin, Sean R Eddy, and Gilbert L Henry. A genetic, genomic, and computational resource for exploring neural circuit function. *eLife*, 9, 2020.

[6971] Stephanie M Davis, Harinder Singh, Cara M Weismann, Adriana Bankston, and Fátima Sancheznieto. Actionable recommendations from trainees to improve science training. *eLife*, 9, 2020.

[6972] Sejal Davla, Gregory Artiushin, Yongjun Li, Daryan Chitsaz, Sally Li, Amita Sehgal, and Donald J van Meyel. AANAT1 functions in astrocytes to regulate sleep homeostasis. *eLife*, 9, 2020.

[6973] Neal J Dawson, Luis Alza, Gabriele Nandal, Graham R Scott, and Kevin G McCracken. Convergent changes in muscle metabolism depend on duration of high-altitude ancestry across Andean waterfowl. *eLife*, 9, 2020.

[6974] Abhishek De, Yasmine El-Shamayleh, and Gregory D Horwitz. Fast and reversible neural inactivation in macaque cortex by optogenetic stimulation of GABAergic neurons. *eLife*, 9, 2020.

[6975] Jan Willem de Gee, Konstantinos Tsetsos, Lars Schwabe, Anne E Urai, David McCormick, Matthew J McGinley, and Tobias H Donner. Pupil-linked phasic arousal predicts a reduction of choice bias across species and decision domains. *eLife*, 9, 2020.

[6976] Peter C de Greef, Theres Oakes, Bram Gerritsen, Mazlina Ismail, James M Heather, Rutger Hermsen, Benjamin Chain, and Rob J de Boer. The naive T-cell receptor repertoire has an extremely broad distribution of clone sizes. *eLife*, 9, 2020.

[6977] Andres de Groot, Bastijn JG van den Boom, Romano M van Genderen, Joris Coppens, John van Veldhuijzen, Joop Bos, Hugo Hoedemaker, Mario Negrello, Ingo Willuhn, Chris I De Zeeuw, and Tycho M Hoogland. NINscope, a versatile miniscope for multi-region circuit investigations. *eLife*, 9, 2020.

[6978] Timothy J de Wet, Kristy R Winkler, Musa Mhlanga, Valerie Mizrahi, and Digby F Warner. Arrayed CRISPRi and quantitative imaging describe the morphotypic landscape of essential mycobacterial genes. *eLife*, 9, 2020.

[6979] Brian D DeAngelis, Jacob A Zavatone-Veth, and Damon A Clark. Correction: The manifold structure of limb coordination in walking Drosophila. *eLife*, 9, 2020.

[6980] Brian D DeAngelis, Jacob A Zavatone-Veth, Aneysis D Gonzalez-Suarez, and Damon A Clark. Spatiotemporally precise optogenetic activation of sensory neurons in freely walking Drosophila. *eLife*, 9, 2020.

[6981] Bisrat J Debebe, Lies Boelen, James C Lee, Eduard J Sanders, Eduard J Sanders, Omu Anzala, Anatoli Kamali, Pontiano Kaleebu, Etienne Karita, William Kilembe, Mubiana Inambao, Shabir Lakhi, Susan Allen, Eric Hunter, Vinodh A Edward, Pat E Fast, Matt A Price, Jill Gilmour, Jianming Tang, Chloe L Thio, Jacquie Astemborski, Gregory Kirk, Salim I Khakoo, Sharyne M Donfield, James J Goedert, and Becca Asquith. Identifying the immune interactions underlying HLA class I disease associations. *eLife*, 9, 2020.

[6982] Lukas Ded, Jae Yeon Hwang, Kiyoshi Miki, Huanan F Shi, and Jean-Ju Chung. 3D in situ imaging of the female reproductive tract reveals molecular signatures of fertilizing spermatozoa in mice. *eLife*, 9, 2020.

[6983] Tom D Deegan, Progya P Mukherjee, Ryo Fujisawa, Cristian Polo Rivera, and Karim Labib. CMG helicase disassembly is controlled by replication fork DNA, replisome components and a ubiquitin threshold. *eLife*, 9, 2020.

[6984] Jennifer D Deem, Chelsea L Faber, Christian Pedersen, Bao Anh Phan, Sarah A Larsen, Kayoko Ogimoto, Jarrell T Nelson, Vincent Damian, Megan A Tran, Richard D Palmiter, Karl J Kaiyala, Jarrad M Scarlett, Michael R Bruchas, Michael W Schwartz, and Gregory J Morton. Cold-induced hyperphagia requires AgRP neuron activation in mice. *eLife*, 9, 2020.

[6985] Kilian Dekoninck, Juliette Létoquart, Cédric Laguri, Pascal Demange, Robin Bevernaegie, Jean-Pierre Simorre, Olivia Dehu, Bogdan I Iorga, Benjamin Elias, Seung-Hyun Cho, and Jean-Francois Collet. Defining the function of OmpA in the Rcs stress response. *eLife*, 9, 2020.

[6986] Alexis Delabouglise, Nguyen Thi Le Thanh, Huynh Thi Ai Xuyen, Benjamin Nguyen-Van-Yen, Phung Ngoc Tuyet, Ha Minh Lam, and Maciej F Boni. Poultry farmer response to disease outbreaks in smallholder farming systems in southern Vietnam. *eLife*, 9, 2020.

[6987] Kira Delmore, Juan Carlos Illera, Javier Pérez-Tris, Gernot Segelbacher, Juan S Lugo Ramos, Gillian Durieux, Jun Ishigohoka, and Miriam Liedvogel. The evolutionary history and genomics of European blackcap migration. *eLife*, 9, 2020.

[6988] Steven Z DeLuca, Megha Ghildiyal, Liang-Yu Pang, and Allan C Spradling. Differentiating Drosophila female germ cells initiate Polycomb silencing by regulating PRC2-interacting proteins. *eLife*, 9, 2020.

[6989] Esin Demir, Y Ilker Yaman, Mustafa Basaran, and Askin Kocabas. Dynamics of pattern formation and emergence of swarming in Caenorhabditis elegans. *eLife*, 9, 2020.

[6990] Sunbin Deng, Buyan Pan, Leah Gottlieb, E James Petersson, and Ronen Marmorstein. Molecular basis for N-terminal alpha-synuclein acetylation by human NatB. *eLife*, 9, 2020.

[6991] Zhonghao Deng, Courtney Ng, Kazuki Inoue, Ziyu Chen, Yuhan Xia, Xiaoyu Hu, Matthew Greenblatt, Alessandra Pernis, and Baohong Zhao. Def6 regulates endogenous type-I interferon responses in osteoblasts and suppresses osteogenesis. *eLife*, 9, 2020.

[6992] Jonathan Desponds, Massimo Vergassola, and Aleksandra M Walczak. A mechanism for hunchback promoters to readout morphogenetic positional information in less than a minute. *eLife*, 9, 2020.

[6993] David Deutsch, Diego Pacheco, Lucas Encarnacion-Rivera, Talmo Pereira, Ramie Fathy, Jan Clemens, Cyrille Girardin, Adam Calhoun, Elise Ireland, Austin Burke, Sven Dorkenwald, Claire McKellar, Thomas Macrina, Ran Lu, Kisuk Lee, Nico Kemnitz, Dodam Ih, Manuel Castro, Akhilesh Halageri, Chris Jordan, William Silversmith, Jingpeng Wu, H Sebastian Seung, and Mala Murthy. The neural basis for a persistent internal state in Drosophila females. *eLife*, 9, 2020.

[6994] Danielle C DeVille, Rayus Kuplicki, Jennifer L Stewart, Robin L Aupperle, Robin L Aupperle, Jerzy Bodurka, Yoon-Hee Cha, Justin Feinstein, Jonathan B Savitz, Teresa A Victor, Martin P Paulus, and Sahib S Khalsa. Diminished responses to bodily threat and blunted interoception in suicide attempters. *eLife*, 9, 2020.

[6995] Renumathy Dhanasekaran, Virginie Baylot, Minsoon Kim, Sibu Kuruvilla, David I Bellovin, Nia Adeniji, Anand Rajan KD, Ian Lai, Meital Gabay, Ling Tong, Maya Krishnan, Jangho Park, Theodore Hu, Mustafa A Barbhuiya, Andrew J Gentles, Kasthuri Kannan, Phuoc T Tran, and Dean W Felsher. MYC and Twist1 cooperate to drive metastasis by eliciting crosstalk between cancer and innate immunity. *eLife*, 9, 2020.

[6996] Madhurima Dhara, Maria Mantero Martinez, Mazen Makke, Yvonne Schwarz, Ralf Mohrmann, and Dieter Bruns. Synergistic actions of v-SNARE transmembrane domains and membrane-curvature modifying lipids in neurotransmitter release. *eLife*, 9, 2020.

[6997] Giovanni M Di Liberto, Claire Pelofi, Roberta Bianco, Prachi Patel, Ashesh D Mehta, Jose L Herrero, Alain de Cheveigné, Shihab Shamma, and Nima Mesgarani. Cortical encoding of melodic expectations in human temporal cortex. *eLife*, 9, 2020.

[6998] André Dias, Anastasiia Lozovska, Filip J Wymeersch, Ana Nóvoa, Anahi Binagui-Casas, Daniel Sobral, Gabriel G Martins, Valerie Wilson, and Moises Mallo. A Tgfbr1/Snai1-dependent developmental module at the core of vertebrate axial elongation. *eLife*, 9, 2020.

[6999] Jennifer EL Diaz, Mehmet Eren Ahsen, Thomas Schaffter, Xintong Chen, Ronald B Realubit, Charles Karan, Andrea Califano, Bojan Losic, and Gustavo Stolovitzky. The transcriptomic response of cells to a drug combination is more than the sum of the responses to the monotherapies. *eLife*, 9, 2020.

[7000] Javier Díaz-Alonso, Wade Morishita, Salvatore Incontro, Jeffrey Simms, Julia Holtzman, Michael Gill, Lennart Mucke, Robert C Malenka, and Roger A Nicoll. Long-term potentiation is independent of the C-tail of the GluA1 AMPA receptor subunit. *eLife*, 9, 2020.

[7001] Maria M Diehl, Jorge M Iravedra-Garcia, Jonathan Morán-Sierra, Gabriel Rojas-Bowe, Fabiola N Gonzalez-Diaz, Viviana P Valentín-Valentín, and Gregory J Quirk. Divergent projections of the prelimbic cortex bidirectionally regulate active avoidance. *eLife*, 9, 2020.

[7002] Nadine Dijkstra, Luca Ambrogioni, Diego Vidaurre, and Marcel van Gerven. Neural dynamics of perceptual inference and its reversal during imagery. *eLife*, 9, 2020.

[7003] Jérôme Dockès, Russell A Poldrack, Romain Primet, Hande Gözükan, Tal Yarkoni, Fabian Suchanek, Bertrand Thirion, and Gaël Varoquaux. NeuroQuery, comprehensive meta-analysis of human brain mapping. *eLife*, 9, 2020.

[7004] Florelle Domart, Peter Cloetens, Stéphane Roudeau, Asuncion Carmona, Emeline Verdier, Daniel Choquet, and Richard Ortega. Correlating STED and synchrotron XRF nano-imaging unveils cosegregation of metals and cytoskeleton proteins in dendrites. *eLife*, 9, 2020.

[7005] Racquel Domingo-Gonzalez, Fabio Zanini, Xibing Che, Min Liu, Robert C Jones, Michael A Swift, Stephen R Quake, David N Cornfield, and Cristina M Alvira. Diverse homeostatic and immunomodulatory roles of immune cells in the developing mouse lung at single cell resolution. *eLife*, 9, 2020.

[7006] Ana Filipa Domingues, Rashmi Kulkarni, George Giotopoulos, Shikha Gupta, Laura Vinnenberg, Liliana Arede, Elena Foerner, Mitra Khalili, Rita Romano Adao, Ayona Johns, Shengjiang Tan, Keti Zeka, Brian J Huntly, Sudhakaran Prabakaran, and Cristina Pina. Loss of Kat2a enhances transcriptional noise and depletes acute myeloid leukemia stem-like cells. *eLife*, 9, 2020.

[7007] Nathaniel J Dominy, Salima Ikram, Gillian L Moritz, Patrick V Wheatley, John N Christensen, Jonathan W Chipman, and Paul L Koch. Mummified baboons reveal the far reach of early Egyptian mariners. *eLife*, 9, 2020.

[7008] Rafal Donczew, Linda Warfield, Derek Pacheco, Ariel Erijman, and Steven Hahn. Two roles for the yeast transcription coactivator SAGA and a set of genes redundantly regulated by TFIID and SAGA. *eLife*, 9, 2020.

[7009] Chen Dong, Mariana Beltcheva, Paul Gontarz, Bo Zhang, Pooja Popli, Laura A Fischer, Shafqat A Khan, Kyoung-mi Park, Eun-Ja Yoon, Xiaoyun Xing, Ramakrishna Kommagani, Ting Wang, Lilianna Solnica-Krezel, and Thorold W Theunissen. Derivation of trophoblast stem cells from naïve human pluripotent stem cells. *eLife*, 9, 2020.

[7010] Cheng Dong, Reiko Nakagawa, Kyohei Oyama, Yusuke Yamamoto, Weilian Zhang, Aiping Dong, Yanjun Li, Yuriko Yoshimura, Hiroyuki Kamiya, Jun-ichi Nakayama, Jun Ueda, and Jinrong Min. Structural basis for histone variant H3tK27me3 recognition by PHF1 and PHF19. *eLife*, 9, 2020.

[7011] Chuanfu Dong, Cameron J Weadick, Vincent Truffault, and Ralf J Sommer. Convergent evolution of small molecule pheromones in Pristionchus nematodes. *eLife*, 9, 2020.

[7012] Kirby M Donnelly, Olivia R DeLorenzo, Aprem DA Zaya, Gabrielle E Pisano, Wint M Thu, Liqun Luo, Ron R Kopito, and Margaret M Panning Pearce. Phagocytic glia are obligatory intermediates in transmission of mutant huntingtin aggregates across neuronal synapses. *eLife*, 9, 2020.

[7013] Andrew Dopheide, Andreas Makiola, Kate H Orwin, Robert J Holdaway, Jamie R Wood, and Ian A Dickie. Rarity is a more reliable indicator of land-use impacts on soil invertebrate communities than other diversity metrics. *eLife*, 9, 2020.

[7014] Aline Ricarda Dörrbaum, Beatriz Alvarez-Castelao, Belquis Nassim-Assir, Julian D Langer, and Erin M Schuman. Proteome dynamics during homeostatic scaling in cultured neurons. *eLife*, 9, 2020.

[7015] Steve Dos-Santos Carvalho, Maite M Moreau, Yeri Esther Hien, Mikael Garcia, Nathalie Aubailly, Deborah J Henderson, Vincent Studer, Nathalie Sans, Olivier Thoumine, and Mireille Montcouquiol. Vangl2 acts at the interface between actin and N-cadherin to modulate mammalian neuronal outgrowth. *eLife*, 9, 2020.

[7016] Guilhem Doulcier, Amaury Lambert, Silvia De Monte, and Paul B Rainey. Eco-evolutionary dynamics of nested Darwinian populations and the emergence of community-level heredity. *eLife*, 9, 2020.

[7017] Elizabeth B Draganova, Jiayan Zhang, Z Hong Zhou, and Ekaterina E Heldwein. Structural basis for capsid recruitment and coat formation during HSV-1 nuclear egress. *eLife*, 9, 2020.

[7018] Jefte M Drijvers, Arlene H Sharpe, and Marcia C Haigis. The effects of age and systemic metabolism on anti-tumor T cell responses. *eLife*, 9, 2020.

[7019] Adi Drori, Asaad Gammal, Shahar Azar, Liad Hinden, Rivka Hadar, Daniel Wesley, Alina Nemirovski, Gergő Szanda, Maayan Salton, Boaz Tirosh, and Joseph Tam. CB1R regulates soluble leptin receptor levels via CHOP, contributing to hepatic leptin resistance. *eLife*, 9, 2020.

[7020] Katarzyna Drożdżyk, Marta Sawicka, Maria-Isabel Bahamonde-Santos, Zaugg Jonas, Dawid Deneka, Christiane Albrecht, and Raimund Dutzler. Cryo-EM structures and functional properties of CALHM channels of the human placenta. *eLife*, 9, 2020.

[7021] Ryan S D'Souza, Jun Y Lim, Alper Turgut, Kelly Servage, Junmei Zhang, Kim Orth, Nisha G Sosale, Matthew J Lazzara, Jeremy Allegood, and James E Casanova. Calcium-stimulated disassembly of focal adhesions mediated by an ORP3/IQSec1 complex. *eLife*, 9, 2020.

[7022] Ramin Dubey, Peter van Kerkhof, Ingrid Jordens, Tomas Malinauskas, Ganesh V Pusapati, Joseph K McKenna, Dan Li, Jan E Carette, Mitchell Ho, Christian Siebold, Madelon Maurice, Andres M Lebensohn, and Rajat Rohatgi. R-spondins engage heparan sulfate proteoglycans to potentiate WNT signaling. *eLife*, 9, 2020.

[7023] Sushil Dubey, Nishita Bhembre, Shivani Bodas, Sukh Veer, Aurnab Ghose, Andrew Callan-Jones, and Pramod Pullarkat. The axonal actin-spectrin lattice acts as a tension buffering shock absorber. *eLife*, 9, 2020.

[7024] Hugo Ducuing, Thibault Gardette, Aurora Pignata, Karine Kindbeiter, Muriel Bozon, Olivier Thoumine, Céline Delloye-Bourgeois, Servane Tauszig-Delamasure, and Valerie Castellani. SlitC-PlexinA1 mediates iterative inhibition for orderly passage of spinal commissural axons through the floor plate. *eLife*, 9, 2020.

[7025] José M Duhart, Victoria Baccini, Yanan Zhang, Daniel R Machado, and Kyunghee Koh. Modulation of sleep-courtship balance by nutritional status in Drosophila. *eLife*, 9, 2020.

[7026] Florian Dunker, Adriana Trutzenberg, Jan S Rothenpieler, Sarah Kuhn, Reinhard Pröls, Tom Schreiber, Alain Tissier, Ariane Kemen, Eric Kemen, Ralph Hückelhoven, and Arne Weiberg. Oomycete small RNAs bind to the plant RNA-induced silencing complex for virulence. *eLife*, 9, 2020.

[7027] Timothy W Dunn and James E Fitzgerald. Correcting for physical distortions in visual stimuli improves reproducibility in zebrafish neuroscience. *eLife*, 9, 2020.

[7028] Leslie Duplaquet, Catherine Leroy, Audrey Vinchent, Sonia Paget, Jonathan Lefebvre, Fabien Vanden Abeele, Steve Lancel, Florence Giffard, Réjane Paumelle, Gabriel Bidaux, Laurent Heliot, Laurent Poulain, Alessandro Furlan, and David Tulasne. Control of cell death/survival balance by the MET dependence receptor. *eLife*, 9, 2020.

[7029] Maeva Dupont, Shanti Souriant, Luciana Balboa, Thien-Phong Vu Manh, Karine Pingris, Stella Rousset, Céline Cougoule, Yoann Rombouts, Renaud Poincloux, Myriam Ben Neji, Carolina Allers, Deepak Kaushal, Marcelo J Kuroda, Susana Benet, Javier Martinez-Picado, Nuria Izquierdo-Useros, Maria del Carmen Sasiain, Isabelle Maridonneau-Parini, Olivier Neyrolles, Christel Vérollet, and Geanncarlo Lugo-Villarino. Tuberculosis-associated IFN-I induces Siglec-1 on tunneling nanotubes and favors HIV-1 spread in macrophages. *eLife*, 9, 2020.

[7030] James B Eaglesham, Kacie L McCarty, and Philip J Kranzusch. Structures of diverse poxin cGAMP nucleases reveal a widespread role for cGAS-STING evasion in host–pathogen conflict. *eLife*, 9, 2020.

[7031] Alice Easton, Shenghan Gao, Scott P Lawton, Sasisekhar Bennuru, Asis Khan, Eric Dahlstrom, Rita G Oliveira, Stella Kepha, Stephen F Porcella, Joanne Webster, Roy Anderson, Michael E Grigg, Richard E Davis, Jianbin Wang, and Thomas B Nutman. Molecular evidence of hybridization between pig and human Ascaris indicates an interbred species complex infecting humans. *eLife*, 9, 2020.

[7032] Fanny E Eberhard, Sarah Cunze, Judith Kochmann, and Sven Klimpel. Modelling the climatic suitability of Chagas disease vectors on a global scale. *eLife*, 9, 2020.

[7033] Samantha Ebersoll, Marta Bogacz, Lina M Günter, Tobias P Dick, and R Luise Krauth-Siegel. A tryparedoxin-coupled biosensor reveals a mitochondrial trypanothione metabolism in trypanosomes. *eLife*, 9, 2020.

[7034] Elizabeth Eck, Jonathan Liu, Maryam Kazemzadeh-Atoufi, Sydney Ghoreishi, Shelby A Blythe, and Hernan G Garcia. Quantitative dissection of transcription in development yields evidence for transcription-factor-driven chromatin accessibility. *eLife*, 9, 2020.

[7035] Marina Eckermann, Jasper Frohn, Marius Reichardt, Markus Osterhoff, Michael Sprung, Fabian Westermeier, Alexandar Tzankov, Christopher Werlein, Mark Kühnel, Danny Jonigk, and Tim Salditt. 3D virtual pathohistology of lung tissue from Covid-19 patients based on phase contrast X-ray tomography. *eLife*, 9, 2020.

[7036] Antonia Franziska Eckert, Peng Gao, Janine Wesslowski, Xianxian Wang, Jasmijn Rath, Karin Nienhaus, Gary Davidson, and Gerd Ulrich Nienhaus. Measuring ligand-cell surface receptor affinities with axial line-scanning fluorescence correlation spectroscopy. *eLife*, 9, 2020.

[7037] Christopher Edelmaier, Adam R Lamson, Zachary R Gergely, Saad Ansari, Robert Blackwell, J Richard McIntosh, Matthew A Glaser, and Meredith D Betterton. Mechanisms of chromosome biorientation and bipolar spindle assembly analyzed by computational modeling. *eLife*, 9, 2020.

[7038] Tomás Edreira, Rubén Celador, Elvira Manjón, and Yolanda Sánchez. A novel checkpoint pathway controls actomyosin ring constriction trigger in fission yeast. *eLife*, 9, 2020.

[7039] Nicole Eichert, Emma C Robinson, Katherine L Bryant, Saad Jbabdi, Mark Jenkinson, Longchuan Li, Kristine Krug, Kate E Watkins, and Rogier B Mars. Cross-species cortical alignment identifies different types of anatomical reorganization in the primate temporal lobe. *eLife*, 9, 2020.

[7040] Catherine E Eichler, Anna C Hakes, Brooke Hull, and Elizabeth R Gavis. Compartmentalized oskar degradation in the germ plasm safeguards germline development. *eLife*, 9, 2020.

[7041] Travis J Eisemann, Frederick Allen, Kelly Lau, Gregory R Shimamura, Philip D Jeffrey, and Frederick M Hughson. The Sec1/Munc18 protein Vps45 holds the Qa-SNARE Tlg2 in an open conformation. *eLife*, 9, 2020.

[7042] Michael B Eisen, Anna Akhmanova, Timothy E Behrens, Diane M Harper, Detlef Weigel, and Mone Zaidi. Implementing a "publish, then review" model of publishing. *eLife*, 9, 2020.

[7043] Michael B Eisen, Anna Akhmanova, Timothy E Behrens, and Detlef Weigel. Publishing in the time of COVID-19. *eLife*, 9, 2020.

[7044] Tyler G Ekins, Vivek Mahadevan, Yajun Zhang, James A D'Amour, Gülcan Akgül, Timothy J Petros, and Chris J McBain. Emergence of non-canonical parvalbumin-containing interneurons in hippocampus of a murine model of type I lissencephaly. *eLife*, 9, 2020.

[7045] Lauretta El Hayek, Islam Oguz Tuncay, Nadine Nijem, Jamie Russell, Sara Ludwig, Kiran Kaur, Xiaohong Li, Priscilla Anderton, Miao Tang, Amanda Gerard, Anja Heinze, Pia Zacher, Hessa S Alsaif, Aboulfazl Rad, Kazem Hassanpour, Mohammad Reza Abbaszadegan, Camerun Washington, Barbara R DuPont, Raymond J Louie, Madeline Couse, Maha Faden, R Curtis Rogers, Rami Abou Jamra, Ellen R Elias, Reza Maroofian, Henry Houlden, Anna Lehman, Bruce Beutler, and Maria H Chahrour. KDM5A mutations identified in autism spectrum disorder using forward genetics. *eLife*, 9, 2020.

[7046] Eran Eldar, Gaëlle Lièvre, Peter Dayan, and Raymond J Dolan. The roles of online and offline replay in planning. *eLife*, 9, 2020.

[7047] Najia A Elkahlah, Jackson A Rogow, Maria Ahmed, and E Josephine Clowney. Presynaptic developmental plasticity allows robust sparse wiring of the Drosophila mushroom body. *eLife*, 9, 2020.

[7048] Jennifer L Elliott, Jenna E Eschbach, Pratibha C Koneru, Wen Li, Maritza Puray-Chavez, Dana Townsend, Dana Q Lawson, Alan N Engelman, Mamuka Kvaratskhelia, and Sebla B Kutluay. Integrase-RNA interactions underscore the critical role of integrase in HIV-1 virion morphogenesis. *eLife*, 9, 2020.

[7049] Kelsey H Elliott, Xiaoting Chen, Joseph Salomone, Praneet Chaturvedi, Preston A Schultz, Sai K Balchand, Jeffrey D Servetas, Aimée Zuniga, Rolf Zeller, Brian Gebelein, Matthew T Weirauch, Kevin A Peterson, and Samantha A Brugmann. Gli3 utilizes Hand2 to synergistically regulate tissue-specific transcriptional networks. *eLife*, 9, 2020.

[7050] Jon C Emery, Timothy W Russell, Yang Liu, Joel Hellewell, Carl AB Pearson, Katherine E Atkins, Katherine E Atkins, Petra Klepac, Akira Endo, Christopher I Jarvis, Nicholas G Davies, Eleanor M Rees, Sophie R Meakin, Alicia Rosello, Kevin van Zandvoort, James D Munday, W John Edmunds, Thibaut Jombart, Megan Auzenbergs, Emily S Nightingale, Mark Jit, Sam Abbott, David Simons, Nikos I Bosse, Quentin J Leclerc, Simon R Procter, C Julian Villabona-Arenas, Damien C Tully, Arminder K Deol, Fiona Yueqian Sun, Stéphane Hué, Anna M Foss, Kiesha Prem, Graham Medley, Amy Gimma, Rachel Lowe, Samuel Clifford, Matthew Quaife, Charlie Diamond, Hamish P Gibbs, Billy J Quilty, Kathleen OReilly, Gwenan M Knight, Rosalind M Eggo, Adam J Kucharski, Sebastian Funk, Stefan Flasche, and Rein MGJ Houben. The contribution of asymptomatic SARS-CoV-2 infections to transmission on the Diamond Princess cruise ship. *eLife*, 9, 2020.

[7051] Syed Usman Enam, Boris Zinshteyn, Daniel H Goldman, Madeline Cassani, Nathan M Livingston, Geraldine Seydoux, and Rachel Green. Puromycin reactivity does not accurately localize translation at the subcellular level. *eLife*, 9, 2020.

[7052] Pierre Enel, Joni D Wallis, and Erin L Rich. Stable and dynamic representations of value in the prefrontal cortex. *eLife*, 9, 2020.

[7053] Eric Engelbrecht, Michel V Levesque, Liqun He, Michael Vanlandewijck, Anja Nitzsche, Hira Niazi, Andrew Kuo, Sasha A Singh, Masanori Aikawa, Kristina Holton, Richard L Proia, Mari Kono, William T Pu, Eric Camerer, Christer Betsholtz, and Timothy Hla. Sphingosine 1-phosphate-regulated transcriptomes in heterogenous arterial and lymphatic endothelium of the aorta. *eLife*, 9, 2020.

[7054] Denis A Engemann, Oleh Kozynets, David Sabbagh, Guillaume Lemaître, Gael Varoquaux, Franziskus Liem, and Alexandre Gramfort. Combining magnetoencephalography with magnetic resonance imaging enhances learning of surrogate-biomarkers. *eLife*, 9, 2020.

[7055] Julia Erb, Lea-Maria Schmitt, and Jonas Obleser. Temporal selectivity declines in the aging human auditory cortex. *eLife*, 9, 2020.

[7056] Abril Angélica Escamilla-Ayala, Ragna Sannerud, Magali Mondin, Karin Poersch, Wendy Vermeire, Laura Paparelli, Caroline Berlage, Marcelle Koenig, Lucia Chavez-Gutierrez, Maximilian H Ulbrich, Sebastian Munck, Hideaki Mizuno, and Wim Annaert. Super-resolution microscopy reveals majorly mono- and dimeric presenilin1/$\gamma$-secretase at the cell surface. *eLife*, 9, 2020.

[7057] José Antonio Escudero, Aleksandra Nivina, Harry E Kemble, Céline Loot, Olivier Tenaillon, and Didier Mazel. Primary and promiscuous functions coexist during evolutionary innovation through whole protein domain acquisitions. *eLife*, 9, 2020.

[7058] Alejandra Escudero-Lara, Josep Argerich, David Cabañero, and Rafael Maldonado. Disease-modifying effects of natural $\delta$9-tetrahydrocannabinol in endometriosis-associated pain. *eLife*, 9, 2020.

[7059] Eben G Estell, Phuong T Le, Yosta Vegting, Hyeonwoo Kim, Christiane Wrann, Mary L Bouxsein, Kenichi Nagano, Roland Baron, Bruce M Spiegelman, and Clifford J Rosen. Irisin directly stimulates osteoclastogenesis and bone resorption in vitro and in vivo. *eLife*, 9, 2020.

[7060] Juan Estévez-Gallego, Fernando Josa-Prado, Siou Ku, Ruben M Buey, Francisco A Balaguer, Andrea E Prota, Daniel Lucena-Agell, Christina Kamma-Lorger, Toshiki Yagi, Hiroyuki Iwamoto, Laurence Duchesne, Isabel Barasoain, Michel O Steinmetz, Denis Chrétien, Shinji Kamimura, J Fernando Díaz, and Maria A Oliva. Structural model for differential cap maturation at growing microtubule ends. *eLife*, 9, 2020.

[7061] Chantell S Evans and Erika LF Holzbaur. Degradation of engulfed mitochondria is rate-limiting in Optineurin-mediated mitophagy in neurons. *eLife*, 9, 2020.

[7062] Daniel R Evans, Marissa P Griffith, Alexander J Sundermann, Kathleen A Shutt, Melissa I Saul, Mustapha M Mustapha, Jane W Marsh, Vaughn S Cooper, Lee H Harrison, and Daria Van Tyne. Systematic detection of horizontal gene transfer across genera among multidrug-resistant bacteria in a single hospital. *eLife*, 9, 2020.

[7063] Harrison Tudor Evans, Liviu-Gabriel Bodea, and Jürgen Götz. Cell-specific non-canonical amino acid labelling identifies changes in the de novo proteome during memory formation. *eLife*, 9, 2020.

[7064] David Exposito-Alonso, Catarina Osório, Clémence Bernard, Sandra Pascual-García, Isabel del Pino, Oscar Marín, and Beatriz Rico. Subcellular sorting of neuregulins controls the assembly of excitatory-inhibitory cortical circuits. *eLife*, 9, 2020.

[7065] Fanny Eyboulet, Célia Jeronimo, Jacques Côté, and FranÇois Robert. The deubiquitylase Ubp15 couples transcription to mRNA export. *eLife*, 9, 2020.

[7066] David W Eyre, Sheila F Lumley, Denise O'Donnell, Mark Campbell, Elizabeth Sims, Elaine Lawson, Fiona Warren, Tim James, Stuart Cox, Alison Howarth, George Doherty, Stephanie B Hatch, James Kavanagh, Kevin K Chau, Philip W Fowler, Jeremy Swann, Denis Volk, Fan Yang-Turner, Nicole Stoesser, Philippa C Matthews, Maria Dudareva, Timothy Davies, Robert H Shaw, Leon Peto, Louise O Downs, Alexander Vogt, Ali Amini, Bernadette C Young, Philip George Drennan, Alexander J Mentzer, Donal T Skelly, Fredrik Karpe, Matt J Neville, Monique Andersson, Andrew J Brent, Nicola Jones, Lucas Martins Ferreira, Thomas Christott, Brian D Marsden, Sarah Hoosdally, Richard Cornall, Derrick W Crook, David I Stuart, Gavin Screaton, Adam JR Watson, Adam JR Watson, Adan Taylor, Alan Chetwynd, Alexander Grassam-Rowe, Alexandra S Mighiu, Angus Livingstone, Annabel Killen, Caitlin Rigler, Callum Harries, Cameron East, Charlotte Lee, Chris JB Mason, Christian Holland, Connor Thompson, Conor Hennesey, Constantinos Savva, David S Kim, Edward WA Harris, Euan J McGivern, Evelyn Qian, Evie Rothwell, Francesca Back, Gabriella Kelly, Gareth Watson, Gregory Howgego, Hannah Chase, Hannah Danbury, Hannah Laurenson-Schafer, Harry L Ward, Holly Hendron, Imogen C Vorley, Isabel Tol, James Gunnell, Jocelyn LF Ward, Jonathan Drake, Joseph D Wilson, Joshua Morton, Julie Dequaire, Katherine O'Byrne, Kenzo Motohashi, Kirsty Harper, Krupa Ravi, Lancelot J Millar, Liam J Peck, Madeleine Oliver, Marcus Rex English, Mary Kumarendran, Matthew Wedlich, Olivia Ambler, Oscar T Deal, Owen Sweeney, Philip Cowie, Rebecca te Water Naudé, Rebecca Young, Rosie Freer, Samuel Scott, Samuel Sussmes, Sarah Peters, Saxon Pattenden, Seren Waite, Síle Ann Johnson, Stefan Kourdov, Stephanie Santos-Paulo, Stoyan Dimitrov, Sven Kerneis, Tariq Ahmed-Firani, Thomas B King, Thomas G Ritter, Thomas H Foord, Zoe De Toledo, Thomas Christie, Bernadett Gergely, David Axten, Emma-Jane Simons, Heather Nevard, Jane Philips, Justyna Szczurkowska, Kaisha Patel, Kyla Smit, Laura Warren, Lisa Morgan, Lucianne Smith, Maria Robles, Mary McKnight, Michael Luciw, Michelle Gates, Nellia Sande, Rachel Turford, Roshni Ray, Sonam Rughani, Tracey Mitchell, Trisha Bellinger, Vicki Wharton, Anita Justice, Gerald Jesuthasan, Susan Wareing, Nurul Huda Mohamad Fadzillah, Kathryn Cann, Richard Kirton, Claire Sutton, Claudia Salvagno, Gabriella DAmato, Gemma Pill, Lisa Butcher, Lydia Rylance-Knight, Merline Tabirao, Ruth Moroney, Sarah Wright, Timothy EA Peto, Bruno Holthof, Anne-Marie O'Donnell, Daniel Ebner, Christopher P

Conlon, Katie Jeffery, and Timothy M Walker. Differential occupational risks to healthcare workers from SARS-CoV-2 observed during a prospective observational study. *eLife*, 9, 2020.

[7067] Gunar Fabig, Robert Kiewisz, Norbert Lindow, James A Powers, Vanessa Cota, Luis J Quintanilla, Jan Brugués, Steffen Prohaska, Diana S Chu, and Thomas Müller-Reichert. Male meiotic spindle features that efficiently segregate paired and lagging chromosomes. *eLife*, 9, 2020.

[7068] Benjamin Jung Fair, Lauren E Blake, Abhishek Sarkar, Bryan J Pavlovic, Claudia Cuevas, and Yoav Gilad. Gene expression variability in human and chimpanzee populations share common determinants. *eLife*, 9, 2020.

[7069] Jiale Fan, Tingting Ji, Kai Wang, Jichang Huang, Mengqing Wang, Laura Manning, Xiaohua Dong, Yanjun Shi, Xumin Zhang, Zhiyong Shao, and Daniel A Colón-Ramos. A muscle-epidermis-glia signaling axis sustains synaptic specificity during allometric growth in Caenorhabditis elegans. *eLife*, 9, 2020.

[7070] Pengxiang Fan, Peipei Wang, Yann-Ru Lou, Bryan J Leong, Bethany M Moore, Craig A Schenck, Rachel Combs, Pengfei Cao, Federica Brandizzi, Shin-Han Shiu, and Robert L Last. Evolution of a plant gene cluster in Solanaceae and emergence of metabolic diversity. *eLife*, 9, 2020.

[7071] Xiaoxu Fan, Fan Wang, Hanyu Shao, Peng Zhang, and Sheng He. The bottom-up and top-down processing of faces in the human occipitotemporal cortex. *eLife*, 9, 2020.

[7072] Yunshu Fan, Joshua I Gold, and Long Ding. Frontal eye field and caudate neurons make different contributions to reward-biased perceptual decisions. *eLife*, 9, 2020.

[7073] Sean Fancher and Andrew Mugler. Diffusion vs. direct transport in the precision of morphogen readout. *eLife*, 9, 2020.

[7074] Hui Fang, Conrad C Labandeira, Yiming Ma, Bingyu Zheng, Dong Ren, Xinli Wei, Jiaxi Liu, and Yongjie Wang. Lichen mimesis in mid-Mesozoic lacewings. *eLife*, 9, 2020.

[7075] Kenneth T Farabaugh, Dawid Krokowski, Bo-Jhih Guan, Zhaofeng Gao, Xing-Huang Gao, Jing Wu, Raul Jobava, Greeshma Ray, Tristan J de Jesus, Massimiliano G Bianchi, Evelyn Chukwurah, Ovidio Bussolati, Michael Kilberg, David A Buchner, Ganes C Sen, Calvin Cotton, Christine McDonald, Michelle Longworth, Parameswaran Ramakrishnan, and Maria Hatzoglou. PACT-mediated PKR activation acts as a hyperosmotic stress intensity sensor weakening osmoadaptation and enhancing inflammation. *eLife*, 9, 2020.

[7076] Mor Farajian, Raz Leib, Hanna Kossowsky, Tomer Zaidenberg, Ferdinando A Mussa-Ivaldi, and Ilana Nisky. Stretching the skin immediately enhances perceived stiffness and gradually enhances the predictive control of grip force. *eLife*, 9, 2020.

[7077] Behnom Farboud, Catherine S Novak, Monique Nicoll, Alyssa Quiogue, and Barbara J Meyer. Dose-dependent action of the RNA binding protein FOX-1 to relay X-chromosome number and determine C. elegans sex. *eLife*, 9, 2020.

[7078] Reza Farhadifar, Che-Hang Yu, Gunar Fabig, Hai-Yin Wu, David B Stein, Matthew Rockman, Thomas Müller-Reichert, Michael J Shelley, and Daniel J Needleman. Stoichiometric interactions explain spindle dynamics and scaling across 100 million years of nematode evolution. *eLife*, 9, 2020.

[7079] Lucas Farnung, Moritz Ochmann, and Patrick Cramer. Nucleosome-CHD4 chromatin remodeler structure maps human disease mutations. *eLife*, 9, 2020.

[7080] Roman O Fedoryshchak, Magdalena Přechová, Abbey M Butler, Rebecca Lee, Nicola O'Reilly, Helen R Flynn, Ambrosius P Snijders, Noreen Eder, Sila Ultanir, Stephane Mouilleron, and Richard Treisman. Molecular basis for substrate specificity of the Phactr1/PP1 phosphatase holoenzyme. *eLife*, 9, 2020.

[7081] Xue Fei, Tristan A Bell, Sarah R Barkow, Tania A Baker, and Robert T Sauer. Structural basis of ClpXP recognition and unfolding of ssrA-tagged substrates. *eLife*, 9, 2020.

[7082] Xue Fei, Tristan A Bell, Simon Jenni, Benjamin M Stinson, Tania A Baker, Stephen C Harrison, and Robert T Sauer. Structures of the ATP-fueled ClpXP proteolytic machine bound to protein substrate. *eLife*, 9, 2020.

[7083] Nathalie Feiner, Illiam SC Jackson, Kirke L Munch, Reinder Radersma, and Tobias Uller. Plasticity and evolutionary convergence in the locomotor skeleton of Greater Antillean Anolis lizards. *eLife*, 9, 2020.

[7084] Viktor V Feketa, Yury A Nikolaev, Dana K Merriman, Sviatoslav N Bagriantsev, and Elena O Gracheva. CNGA3 acts as a cold sensor in hypothalamic neurons. *eLife*, 9, 2020.

[7085] Sebastian Felgner, Matthias Preusse, Ulrike Beutling, Stephanie Stahnke, Vinay Pawar, Manfred Rohde, Mark Brönstrup, Theresia Stradal, and Susanne Häussler. Host-induced spermidine production in motile Pseudomonas aeruginosa triggers phagocytic uptake. *eLife*, 9, 2020.

[7086] Natalia Felipe-Medina, Sandrine Caburet, Fernando Sánchez-Sáez, Yazmine B Condezo, Dirk G de Rooij, Laura Gómez-H, Rodrigo Garcia-Valiente, Anne Laure Todeschini, Paloma Duque, Manuel Adolfo Sánchez-Martin, Stavit A Shalev, Elena Llano, Reiner A Veitia, and Alberto M Pendás. A missense in HSF2BP causing primary ovarian insufficiency affects meiotic recombination by its novel interactor C19ORF57/BRME1. *eLife*, 9, 2020.

[7087] Sandra Fendl, Renee Marie Vieira, and Alexander Borst. Conditional protein tagging methods reveal highly specific subcellular distribution of ion channels in motion-sensing neurons. *eLife*, 9, 2020.

[7088] Emma J Fenech, Federica Lari, Philip D Charles, Roman Fischer, Marie Laétitia-Thézénas, Katrin Bagola, Adrienne W Paton, James C Paton, Mads Gyrd-Hansen, Benedikt M Kessler, and John C Christianson. Interaction mapping of endoplasmic reticulum ubiquitin ligases identifies modulators of innate immune signalling. *eLife*, 9, 2020.

[7089] Weidong Feng, Yinan Li, Pauline Dao, Jihad Aburas, Priota Islam, Benayahu Elbaz, Anna Kolarzyk, André EX Brown, and Paschalis Kratsios. A terminal selector prevents a Hox transcriptional switch to safeguard motor neuron identity throughout life. *eLife*, 9, 2020.

[7090] Xiaoxia Feng, Irene Altarelli, Karla Monzalvo, Guosheng Ding, Franck Ramus, Hua Shu, Stanislas Dehaene, Xiangzhi Meng, and Ghislaine Dehaene-Lambertz. A universal reading network and its modulation by writing system and reading ability in French and Chinese children. *eLife*, 9, 2020.

[7091] Ricardo A Fernandes, Chaoran Li, Gang Wang, Xinbo Yang, Christina S Savvides, Caleb R Glassman, Shen Dong, Eric Luxenberg, Leah V Sibener, Michael E Birnbaum, Christophe Benoist, Diane Mathis, and K Christopher Garcia. Discovery of surrogate agonists for visceral fat Treg cells that modulate metabolic indices in vivo. *eLife*, 9, 2020.

[7092] Sara Fernández-García, Sara Conde-Berriozabal, Esther García-García, Clara Gort-Paniello, David Bernal-Casas, Gerardo García-Díaz Barriga, Javier López-Gil, Emma Muñoz-Moreno, Guadalupe Soria, Leticia Campa, Francesc Artigas, Manuel José Rodríguez, Jordi Alberch, and Mercè Masana. M2 cortex-dorsolateral striatum stimulation reverses motor symptoms and synaptic deficits in Huntington's disease. *eLife*, 9, 2020.

[7093] André Ferreira Castro, Lothar Baltruschat, Tomke Stürner, Amirhoushang Bahrami, Peter Jedlicka, Gaia Tavosanis, and Hermann Cuntz. Achieving functional neuronal dendrite structure through sequential stochastic growth and retraction. *eLife*, 9, 2020.

[7094] Laura Ferrer-Font, Palak Mehta, Phoebe Harmos, Alfonso J Schmidt, Sally Chappell, Kylie M Price, Ian F Hermans, Franca Ronchese, Graham le Gros, and Johannes U Mayer. High-dimensional analysis of intestinal immune cells during helminth infection. *eLife*, 9, 2020.

[7095] Lindsay K Festa, Elena Irollo, Brian J Platt, Yuzen Tian, Stan Floresco, and Olimpia Meucci. CXCL12-induced rescue of cortical dendritic spines and cognitive flexibility. *eLife*, 9, 2020.

[7096] Reinhild Feuerstein, Aaron James Forde, Florens Lohrmann, Julia Kolter, Neftali Jose Ramirez, Jakob Zimmermann, Mercedes Gomez de Agüero, and Philipp Henneke. Resident macrophages acquire innate immune memory in staphylococcal skin infection. *eLife*, 9, 2020.

[7097] Alexandre LS Filipowicz, Christopher M Glaze, Joseph W Kable, and Joshua I Gold. Pupil diameter encodes the idiosyncratic, cognitive complexity of belief updating. *eLife*, 9, 2020.

[7098] Yaara Finkel, Dominik Schmiedel, Julie Tai-Schmiedel, Aharon Nachshon, Roni Winkler, Martina Dobesova, Michal Schwartz, Ofer Mandelboim, and Noam Stern-Ginossar. Comprehensive annotations of human herpesvirus 6A and 6B genomes reveal novel and conserved genomic features. *eLife*, 9, 2020.

[7099] Ksenia Finogenova, Jacques Bonnet, Simon Poepsel, Ingmar B Schäfer, Katja Finkl, Katharina Schmid, Claudia Litz, Mike Strauss, Christian Benda, and Jürg Müller. Structural basis for PRC2 decoding of active histone methylation marks H3K36me2/3. *eLife*, 9, 2020.

[7100] Lukas F Fischer, Raul Mojica Soto-Albors, Friederike Buck, and Mark T Harnett. Representation of visual landmarks in retrosplenial cortex. *eLife*, 9, 2020.

[7101] Petra Fischer, Witold J Lipski, Wolf-Julian Neumann, Robert S Turner, Pascal Fries, Peter Brown, and R Mark Richardson. Movement-related coupling of human subthalamic nucleus spikes to cortical gamma. *eLife*, 9, 2020.

[7102] Maximilian H Fitz-James, Pin Tong, Alison L Pidoux, Hakan Ozadam, Liyan Yang, Sharon A White, Job Dekker, and Robin C Allshire. Large domains of heterochromatin direct the formation of short mitotic chromosome loops. *eLife*, 9, 2020.

[7103] Avi I Flamholz, Eli Dugan, Cecilia Blikstad, Shmuel Gleizer, Roee Ben-Nissan, Shira Amram, Niv Antonovsky, Sumedha Ravishankar, Elad Noor, Arren Bar-Even, Ron Milo, and David F Savage. Functional reconstitution of a bacterial CO2 concentrating mechanism in Escherichia coli. *eLife*, 9, 2020.

[7104] Andrew I Flyak, Stormy E Ruiz, Jordan Salas, Semi Rho, Justin R Bailey, and Pamela J Bjorkman. An ultralong CDRH2 in HCV neutralizing antibody demonstrates structural plasticity of antibodies against E2 glycoprotein. *eLife*, 9, 2020.

[7105] Julia M Flynn, Ammeret Rossouw, Pamela Cote-Hammarlof, Inês Fragata, David Mavor, Carl Hollins, Claudia Bank, and Daniel NA Bolon. Comprehensive fitness maps of Hsp90 show widespread environmental dependence. *eLife*, 9, 2020.

[7106] István Fodor, Ahmed AA Hussein, Paul R Benjamin, Joris M Koene, and Zsolt Pirger. The unlimited potential of the great pond snail, Lymnaea stagnalis. *eLife*, 9, 2020.

[7107] Anton Fomenko, Kai-Hsiang Stanley Chen, Jean-FranÇois Nankoo, James Saravanamuttu, Yanqiu Wang, Mazen El-Baba, Xue Xia, Shakthi Sanjana Seerala, Kullervo Hynynen, Andres M Lozano, and Robert Chen. Systematic examination of low-intensity ultrasound parameters on human motor cortex excitability and behavior. *eLife*, 9, 2020.

[7108] Maria Fomicheva and Ian G Macara. Genome-wide CRISPR screen identifies noncanonical NF-$\kappa$B signaling as a regulator of density-dependent proliferation. *eLife*, 9, 2020.

[7109] Dominique Förster, Thomas O Helmbrecht, Duncan S Mearns, Linda Jordan, Nouwar Mokayes, and Herwig Baier. Retinotectal circuitry of larval zebrafish is adapted to detection and pursuit of prey. *eLife*, 9, 2020.

[7110] David J Forsthoefel, Nicholas I Cejda, Umair W Khan, and Phillip A Newmark. Cell-type diversity and regionalized gene expression in the planarian intestine. *eLife*, 9, 2020.

[7111] Neal P Fox, Matthew Leonard, Matthias J Sjerps, and Edward F Chang. Transformation of a temporal speech cue to a spatial neural code in human auditory cortex. *eLife*, 9, 2020.

[7112] Sebastià Franch-Expósito, Laia Bassaganyas, Maria Vila-Casadesús, Eva Hernández-Illán, Roger Esteban-Fabró, Marcos Díaz-Gay, Juan José Lozano, Antoni Castells, Josep Maria Llovet, Sergi Castellví-Bel, and Jordi Camps. CNApp, a tool for the quantification of copy number alterations and integrative analysis revealing clinical implications. *eLife*, 9, 2020.

[7113] Antonietta Franco, Xiawei Dang, Emily K Walton, Joshua N Ho, Barbara Zablocka, Cindy Ly, Timothy M Miller, Robert H Baloh, Michael E Shy, Andrew S Yoo, and Gerald W Dorn. Burst mitofusin activation reverses neuromuscular dysfunction in murine CMT2A. *eLife*, 9, 2020.

[7114] Jennifer L Fribourgh, Ashutosh Srivastava, Colby R Sandate, Alicia K Michael, Peter L Hsu, Christin Rakers, Leslee T Nguyen, Megan R Torgrimson, Gian Carlo G Parico, Sarvind Tripathi, Ning Zheng, Gabriel C Lander, Tsuyoshi Hirota, Florence Tama, and Carrie L Partch. Dynamics at the serine loop underlie differential affinity of cryptochromes for CLOCK:BMAL1 to control circadian timing. *eLife*, 9, 2020.

[7115] Matthias Fritsche, Eelke Spaak, and Floris P de Lange. A Bayesian and efficient observer model explains concurrent attractive and repulsive history biases in visual perception. *eLife*, 9, 2020.

[7116] Maike Frye, Simon Stritt, Henrik Ortsäter, Magda Hernandez Vasquez, Mika Kaakinen, Andres Vicente, John Wiseman, Lauri Eklund, Jorge L Martínez-Torrecuadrada, Dietmar Vestweber, and Taija Mäkinen. EphrinB2-EphB4 signalling provides Rho-mediated homeostatic control of lymphatic endothelial cell junction integrity. *eLife*, 9, 2020.

[7117] Hirofumi Fujita, Takashi Kodama, and Sascha du Lac. Modular output circuits of the fastigial nucleus for diverse motor and nonmotor functions of the cerebellar vermis. *eLife*, 9, 2020.

[7118] Tomoyuki Fukuda, Yuki Ebi, Tetsu Saigusa, Kentaro Furukawa, Shun-ichi Yamashita, Keiichi Inoue, Daiki Kobayashi, Yutaka Yoshida, and Tomotake Kanki. Atg43 tethers isolation membranes to mitochondria to promote starvation-induced mitophagy in fission yeast. *eLife*, 9, 2020.

[7119] Gregory G Fuller, Ting Han, Mallory A Freeberg, James J Moresco, Amirhossein Ghanbari Niaki, Nathan P Roach, John R Yates, Sua Myong, and John K Kim. RNA promotes phase separation of glycolysis enzymes into yeast G bodies in hypoxia. *eLife*, 9, 2020.

[7120] Marco Fumasoni and Andrew W Murray. The evolutionary plasticity of chromosome metabolism allows adaptation to constitutive DNA replication stress. *eLife*, 9, 2020.

[7121] Taro Furubayashi, Kensuke Ueda, Yohsuke Bansho, Daisuke Motooka, Shota Nakamura, Ryo Mizuuchi, and Norikazu Ichihashi. Emergence and diversification of a host-parasite RNA ecosystem through Darwinian evolution. *eLife*, 9, 2020.

[7122] Bjoern Gaertner, Sebastiaan van Heesch, Valentin Schneider-Lunitz, Jana Felicitas Schulz, Franziska Witte, Susanne Blachut, Steven Nguyen, Regina Wong, Ileana Matta, Norbert Hübner, and Maike Sander. A human ESC-based screen identifies a role for the translated lncRNA LINC00261 in pancreatic endocrine differentiation. *eLife*, 9, 2020.

[7123] Monica Gagliano, Vladyslav V Vyazovskiy, Alexander A Borbély, Martial Depczynski, and Ben Radford. Comment on 'Lack of evidence for associative learning in pea plants'. *eLife*, 9, 2020.

[7124] Christopher Gagne, Ondrej Zika, Peter Dayan, and Sonia J Bishop. Impaired adaptation of learning to contingency volatility in internalizing psychopathology. *eLife*, 9, 2020.

[7125] Mario U Gaimann, Maximilian Nguyen, Jonathan Desponds, and Andreas Mayer. Early life imprints the hierarchy of T cell clone sizes. *eLife*, 9, 2020.

[7126] Asaf Gal, Jonathan Saragosti, and Daniel JC Kronauer. anTraX, a software package for high-throughput video tracking of color-tagged insects. *eLife*, 9, 2020.

[7127] Jorge Luis Galeano Niño, Sophie V Pageon, Szun S Tay, Feyza Colakoglu, Daryan Kempe, Jack Hywood, Jessica K Mazalo, James Cremasco, Matt A Govendir, Laura F Dagley, Kenneth Hsu, Simone Rizzetto, Jerzy Zieba, Gregory Rice, Victoria Prior, Geraldine M O'Neill, Richard J Williams, David R Nisbet, Belinda Kramer, Andrew I Webb, Fabio Luciani, Mark N Read, and Maté Biro. Cytotoxic T cells swarm by homotypic chemokine signalling. *eLife*, 9, 2020.

[7128] Antonio Galeone, Joshua M Adams, Shinya Matsuda, Maximiliano F Presa, Ashutosh Pandey, Seung Yeop Han, Yuriko Tachida, Hiroto Hirayama, Thomas Vaccari, Tadashi Suzuki, Cathleen M Lutz, Markus Affolter, Aamir Zuberi, and Hamed Jafar-Nejad. Regulation of BMP4/Dpp retrotranslocation and signaling by deglycosylation. *eLife*, 9, 2020.

[7129] Alessandro R Galloni, Aeron Laffere, and Ede Rancz. Apical length governs computational diversity of layer 5 pyramidal neurons. *eLife*, 9, 2020.

[7130] Vahe Galstyan, Kabir Husain, Fangzhou Xiao, Arvind Murugan, and Rob Phillips. Proofreading through spatial gradients. *eLife*, 9, 2020.

[7131] Carlos S Galvan-Ampudia, Guillaume Cerutti, Jonathan Legrand, Géraldine Brunoud, Raquel Martin-Arevalillo, Romain Azais, Vincent Bayle, Steven Moussu, Christian Wenzl, Yvon Jaillais, Jan U Lohmann, Christophe Godin, and Teva Vernoux. Temporal integration of auxin information for the regulation of patterning. *eLife*, 9, 2020.

[7132] Melissa V Gammons, Miha Renko, Joshua E Flack, Juliusz Mieszczanek, and Mariann Bienz. Feedback control of Wnt signaling based on ultrastable histidine cluster co-aggregation between Naked/NKD and Axin. *eLife*, 9, 2020.

[7133] Jaba Gamrekelashvili, Tamar Kapanadze, Stefan Sablotny, Corina Ratiu, Khaled Dastagir, Matthias Lochner, Susanne Karbach, Philip Wenzel, Andre Sitnow, Susanne Fleig, Tim Sparwasser, Ulrich Kalinke, Bernhard Holzmann, Hermann Haller, and Florian P Limbourg. Notch and TLR signaling coordinate monocyte cell fate and inflammation. *eLife*, 9, 2020.

[7134] Massimo Ganassi, Sara Badodi, Kees Wanders, Peter S Zammit, and Simon M Hughes. Myogenin is an essential regulator of adult myofibre growth and muscle stem cell homeostasis. *eLife*, 9, 2020.

[7135] Shashank Gandhi, Erica J Hutchins, Krystyna Maruszko, Jong H Park, Matthew Thomson, and Marianne E Bronner. Bimodal function of chromatin remodeler Hmga1 in neural crest induction and Wnt-dependent emigration. *eLife*, 9, 2020.

[7136] Philippe Ganot, Eric Tambutté, Natacha Caminiti-Segonds, Gaëlle Toullec, Denis Allemand, and Sylvie Tambutté. Ubiquitous macropinocytosis in anthozoans. *eLife*, 9, 2020.

[7137] Stephanie C Gantz, Khaled Moussawi, and Holly S Hake. Delta glutamate receptor conductance drives excitation of mouse dorsal raphe neurons. *eLife*, 9, 2020.

[7138] Bo Gao, Sompob Saralamba, Yoel Lubell, Lisa J White, Arjen M Dondorp, and Ricardo Aguas. Determinants of MDA impact and designing MDAs towards malaria elimination. *eLife*, 9, 2020.

[7139] Richard Gao, Ruud L van den Brink, Thomas Pfeffer, and Bradley Voytek. Neuronal timescales are functionally dynamic and shaped by cortical microarchitecture. *eLife*, 9, 2020.

[7140] Vytautas Gapsys and Bert L de Groot. On the importance of statistics in molecular simulations for thermodynamics, kinetics and simulation box size. *eLife*, 9, 2020.

[7141] Judit García-González, Alistair J Brock, Matthew O Parker, Riva J Riley, David Joliffe, Ari Sudwarts, Muy-Teck Teh, Elisabeth M Busch-Nentwich, Derek L Stemple, Adrian R Martineau, Jaakko Kaprio, Teemu Palviainen, Valerie Kuan, Robert T Walton, and Caroline H Brennan. Identification of slit3 as a locus affecting nicotine preference in zebrafish and human smoking behaviour. *eLife*, 9, 2020.

[7142] Mikel Garcia-Marcos, Kshitij Parag-Sharma, Arthur Marivin, Marcin Maziarz, Alex Luebbers, and Lien T Nguyen. Optogenetic activation of heterotrimeric G-proteins by LOV2GIVe, a rationally engineered modular protein. *eLife*, 9, 2020.

[7143] Ankur Garg, Abdul Hannan, Qian Wang, Neoklis Makrides, Jian Zhong, Hongge Li, Sungtae Yoon, Yingyu Mao, and Xin Zhang. Etv transcription factors functionally diverge from their upstream FGF signaling in lens development. *eLife*, 9, 2020.

[7144] Marina Garrett, Sahar Manavi, Kate Roll, Douglas R Ollerenshaw, Peter A Groblewski, Nicholas D Ponvert, Justin T Kiggins, Linzy Casal, Kyla Mace, Ali Williford, Arielle Leon, Xiaoxuan Jia, Peter Ledochowitsch, Michael A Buice, Wayne Wakeman, Stefan Mihalas, and Shawn R Olsen. Experience shapes activity dynamics and stimulus coding of VIP inhibitory cells. *eLife*, 9, 2020.

[7145] Danielle M Garshott, Elayanambi Sundaramoorthy, Marilyn Leonard, and Eric J Bennett. Distinct regulatory ribosomal ubiquitylation events are reversible and hierarchically organized. *eLife*, 9, 2020.

[7146] Michael R Garvin, Christiane Alvarez, J Izaak Miller, Erica T Prates, Angelica M Walker, B Kirtley Amos, Alan E Mast, Amy Justice, Bruce Aronow, and Daniel Jacobson. A mechanistic model and therapeutic interventions for COVID-19 involving a RAS-mediated bradykinin storm. *eLife*, 9, 2020.

[7147] Remi Gau, Pierre-Louis Bazin, Robert Trampel, Robert Turner, and Uta Noppeney. Resolving multisensory and attentional influences across cortical depth in sensory cortices. *eLife*, 9, 2020.

[7148] Quentin Gaucher, Mariangela Panniello, Aleksandar Z Ivanov, Johannes C Dahmen, Andrew J King, and Kerry MM Walker. Complexity of frequency receptive fields predicts tonotopic variability across species. *eLife*, 9, 2020.

[7149] Marta Gawrys-Kopczynska, Marek Konop, Klaudia Maksymiuk, Katarzyna Kraszewska, Ladislav Derzsi, Krzysztof Sozanski, Robert Holyst, Marta Pilz, Emilia Samborowska, Leszek Dobrowolski, Kinga Jaworska, Izabella Mogilnicka, and Marcin Ufnal. TMAO, a seafood-derived molecule, produces diuresis and reduces mortality in heart failure rats. *eLife*, 9, 2020.

[7150] Katy AM Gaythorpe, Arran Hamlet, Laurence Cibrelus, Tini Garske, and Neil M Ferguson. The effect of climate change on yellow fever disease burden in Africa. *eLife*, 9, 2020.

[7151] Yifan Ge, Xiaojun Shi, Sivakumar Boopathy, Julie McDonald, Adam W Smith, and Luke H Chao. Two forms of Opa1 cooperate to complete fusion of the mitochondrial inner-membrane. *eLife*, 9, 2020.

[7152] Daniel A Gehrlach, Caroline Weiand, Thomas N Gaitanos, Eunjae Cho, Alexandra S Klein, Alexandru A Hennrich, Karl-Klaus Conzelmann, and Nadine Gogolla. A whole-brain connectivity map of mouse insular cortex. *eLife*, 9, 2020.

[7153] Tobias Geiger, Maria Lara-Tejero, Yong Xiong, and Jorge E Galán. Mechanisms of substrate recognition by a typhoid toxin secretion-associated muramidase. *eLife*, 9, 2020.

[7154] Joseph V Geisberg, Zarmik Moqtaderi, and Kevin Struhl. The transcriptional elongation rate regulates alternative polyadenylation in yeast. *eLife*, 9, 2020.

[7155] Aviram Gelblum, Ehud Fonio, Yoav Rodeh, Amos Korman, and Ofer Feinerman. Ant collective cognition allows for efficient navigation through disordered environments. *eLife*, 9, 2020.

[7156] özgür GenÇ, Joon-Yong An, Richard D Fetter, Yelena Kulik, Giulia Zunino, Stephan J Sanders, and Graeme W Davis. Homeostatic plasticity fails at the intersection of autism-gene mutations and a novel class of common genetic modifiers. *eLife*, 9, 2020.

[7157] Ajun Geng, Ting Wu, Cheguo Cai, Wenqian Song, Jiqiu Wang, Qing Cissy Yu, and Yi Arial Zeng. A novel function of R-spondin1 in regulating estrogen receptor expression independent of Wnt/$\beta$-catenin signaling. *eLife*, 9, 2020.

[7158] Ginto George, Satoshi Ninagawa, Hirokazu Yagi, Taiki Saito, Tokiro Ishikawa, Tetsushi Sakuma, Takashi Yamamoto, Koshi Imami, Yasushi Ishihama, Koichi Kato, Tetsuya Okada, and Kazutoshi Mori. EDEM2 stably disulfide-bonded to TXNDC11 catalyzes the first mannose trimming step in mammalian glycoprotein ERAD. *eLife*, 9, 2020.

[7159] Marco Geymonat, Qiuran Peng, Zhiang Guo, Zulin Yu, Jay R Unruh, Sue L Jaspersen, and Marisa Segal. Orderly assembly underpinning built-in asymmetry in the yeast centrosome duplication cycle requires cyclin-dependent kinase. *eLife*, 9, 2020.

[7160] Miruna Georgiana Ghinia Tegla, Diego F Buenaventura, Diana Y Kim, Cassandra Thakurdin, Kevin C Gonzalez, and Mark M Emerson. OTX2 represses sister cell fate choices in the developing retina to promote photoreceptor specification. *eLife*, 9, 2020.

[7161] Aniket Ghosh, Elise LV Malavasi, Diane L Sherman, and Peter J Brophy. Neurofascin and Kv7.3 are delivered to somatic and axon terminal surface membranes en route to the axon initial segment. *eLife*, 9, 2020.

[7162] Arpan C Ghosh, Sudhir Gopal Tattikota, Yifang Liu, Aram Comjean, Yanhui Hu, Victor Barrera, Shannan J Ho Sui, and Norbert Perrimon. Drosophila PDGF/VEGF signaling from muscles to hepatocyte-like cells protects against obesity. *eLife*, 9, 2020.

[7163] Tarini S Ghosh, Mrinmoy Das, Ian B Jeffery, and Paul W O'Toole. Adjusting for age improves identification of gut microbiome alterations in multiple diseases. *eLife*, 9, 2020.

[7164] Zane A Gibbs, Luis C Reza, Chun-Chun Cheng, Jill M Westcott, Kathleen McGlynn, and Angelique W Whitehurst. The testis protein ZNF165 is a SMAD3 cofactor that coordinates oncogenic TGF$\beta$ signaling in triple-negative breast cancer. *eLife*, 9, 2020.

[7165] Kimberley H Gibson, Felipe Trajtenberg, Elsio A Wunder, Megan R Brady, Fabiana San Martin, Ariel Mechaly, Zhiguo Shang, Jun Liu, Mathieu Picardeau, Albert Ko, Alejandro Buschiazzo, and Charles Vaughn Sindelar. An asymmetric sheath controls flagellar supercoiling and motility in the leptospira spirochete. *eLife*, 9, 2020.

[7166] Gabrielle E Giese, Melissa D Walker, Olga Ponomarova, Hefei Zhang, Xuhang Li, Gregory Minevich, and Albertha JM Walhout. Caenorhabditis elegans methionine/S-adenosylmethionine cycle activity is sensed and adjusted by a nuclear hormone receptor. *eLife*, 9, 2020.

[7167] Eduardo D Gigante, Megan R Taylor, Anna A Ivanova, Richard A Kahn, and Tamara Caspary. ARL13B regulates Sonic hedgehog signaling from outside primary cilia. *eLife*, 9, 2020.

[7168] Ariel Gilad, Ido Maor, and Adi Mizrahi. Learning-related population dynamics in the auditory thalamus. *eLife*, 9, 2020.

[7169] Joseph A Giovinazzo, Russell P Thomson, Nailya Khalizova, Patrick J Zager, Nirav Malani, Enrique Rodriguez-Boulan, Jayne Raper, and Ryan Schreiner. Apolipoprotein L-1 renal risk variants form active channels at the plasma membrane driving cytotoxicity. *eLife*, 9, 2020.

[7170] Alexandre Giraud-Gatineau, Juan Manuel Coya, Alexandra Maure, Anne Biton, Michael Thomson, Elliott M Bernard, Jade Marrec, Maximiliano G Gutierrez, Gérald Larrouy-Maumus, Roland Brosch, Brigitte Gicquel, and Ludovic Tailleux. The antibiotic bedaquiline activates host macrophage innate immune resistance to bacterial infection. *eLife*, 9, 2020.

[7171] Ritika Giri, Dimitrios K Papadopoulos, Diana M Posadas, Hemanth K Potluri, Pavel Tomancak, Madhav Mani, and Richard W Carthew. Ordered patterning of the sensory system is susceptible to stochastic features of gene expression. *eLife*, 9, 2020.

[7172] Bryan L Gitschlag, Ann T Tate, and Maulik R Patel. Nutrient status shapes selfish mitochondrial genome dynamics across different levels of selection. *eLife*, 9, 2020.

[7173] Lucas R Glover, Kerry M McFadden, Max Bjorni, Sawyer R Smith, Natalie G Rovero, Sarvar Oreizi-Esfahani, Takayuki Yoshida, Abagail F Postle, Mio Nonaka, Lindsay R Halladay, and Andrew Holmes. A prefrontal-bed nucleus of the stria terminalis circuit limits fear to uncertain threat. *eLife*, 9, 2020.

[7174] Keerthana Gnanapradeepan, Julia I-Ju Leu, Subhasree Basu, Thibaut Barnoud, Madeline Good, Joyce V Lee, William J Quinn, Che-Pei Kung, Rexford Ahima, Joseph A Baur, Kathryn E Wellen, Qin Liu, Zachary T Schug, Donna L George, and Maureen E Murphy. Increased mTOR activity and metabolic efficiency in mouse and human cells containing the African-centric tumor-predisposing p53 variant Pro47Ser. *eLife*, 9, 2020.

[7175] Raeann Goering, Laura I Hudish, Bryan B Guzman, Nisha Raj, Gary J Bassell, Holger A Russ, Daniel Dominguez, and J Matthew Taliaferro. FMRP promotes RNA localization to neuronal projections through interactions between its RGG domain and G-quadruplex RNA sequences. *eLife*, 9, 2020.

[7176] Sarah Goethals and Romain Brette. Theoretical relation between axon initial segment geometry and excitability. *eLife*, 9, 2020.

[7177] Stefan Golfier, Thomas Quail, Hiroshi Kimura, and Jan Brugués. Cohesin and condensin extrude DNA loops in a cell cycle-dependent manner. *eLife*, 9, 2020.

[7178] Elisa Gómez-Gil, Rebeca Martín-García, Jero Vicente-Soler, Alejandro Franco, Beatriz Vázquez-Marín, Francisco Prieto-Ruiz, Teresa Soto, Pilar Pérez, Marisa Madrid, and Jose Cansado. Stress-activated MAPK signaling controls fission yeast actomyosin ring integrity by modulating formin For3 levels. *eLife*, 9, 2020.

[7179] Pedro J GonÇalves, Jan-Matthis Lueckmann, Michael Deistler, Marcel Nonnenmacher, Kaan öcal, Giacomo Bassetto, Chaitanya Chintaluri, William F Podlaski, Sara A Haddad, Tim P Vogels, David S Greenberg, and Jakob H Macke. Training deep neural density estimators to identify mechanistic models of neural dynamics. *eLife*, 9, 2020.

[7180] Oscar C González, Yury Sokolov, Giri P Krishnan, Jean Erik Delanois, and Maxim Bazhenov. Can sleep protect memories from catastrophic forgetting? *eLife*, 9, 2020.

[7181] Christopher D Goodman, Taher Uddin, Natalie J Spillman, and Geoffrey I McFadden. A single point mutation in the Plasmodium falciparum FtsH1 metalloprotease confers actinonin resistance. *eLife*, 9, 2020.

[7182] Davis Goodnight and Jasper Rine. S-phase-independent silencing establishment in Saccharomyces cerevisiae. *eLife*, 9, 2020.

[7183] Joanne C Gordon, Natalie C Holt, Andrew Biewener, and Monica A Daley. Tuning of feedforward control enables stable muscle force-length dynamics after loss of autogenic proprioceptive feedback. *eLife*, 9, 2020.

[7184] Kacy L Gordon, Jay W Zussman, Xin Li, Camille Miller, and David R Sherwood. Stem cell niche exit in C. elegans via orientation and segregation of daughter cells by a cryptic cell outside the niche. *eLife*, 9, 2020.

[7185] Katelyn Gostic, Ana CR Gomez, Riley O Mummah, Adam J Kucharski, and James O Lloyd-Smith. Estimated effectiveness of symptom and risk screening to prevent the spread of COVID-19. *eLife*, 9, 2020.

[7186] Debanjan Goswami, De Chen, Yue Yang, Prabhakar R Gudla, John Columbus, Karen Worthy, Megan Rigby, Madeline Wheeler, Suman Mukhopadhyay, Katie Powell, William Burgan, Vanessa Wall, Dominic Esposito, Dhirendra K Simanshu, Felice C Lightstone, Dwight V Nissley, Frank McCormick, and Thomas Turbyville. Membrane interactions of the globular domain and the hypervariable region of KRAS4b define its unique diffusion behavior. *eLife*, 9, 2020.

[7187] Adam J Granger, Wengang Wang, Keiramarie Robertson, Mahmoud El-Rifai, Andrea F Zanello, Karina Bistrong, Arpiar Saunders, Brian W Chow, Vicente Nuñez, Miguel Turrero García, Corey C Harwell, Chenghua Gu, and Bernardo L Sabatini. Cortical ChAT+ neurons co-transmit acetylcholine and GABA in a target- and brain-region-specific manner. *eLife*, 9, 2020.

[7188] Andrea Graziadei, Frank Gabel, John Kirkpatrick, and Teresa Carlomagno. The guide sRNA sequence determines the activity level of box C/D RNPs. *eLife*, 9, 2020.

[7189] Jack P Green, Tessa Swanton, Lucy V Morris, Lina Y El-Sharkawy, James Cook, Shi Yu, James Beswick, Antony D Adamson, Neil E Humphreys, Richard Bryce, Sally Freeman, Catherine Lawrence, and David Brough. LRRC8A is essential for hypotonicity-, but not for DAMP-induced NLRP3 inflammasome activation. *eLife*, 9, 2020.

[7190] Kathleen Greenham, Ryan C Sartor, Stevan Zorich, Ping Lou, Todd C Mockler, and C Robertson McClung. Expansion of the circadian transcriptome in Brassica rapa and genome-wide diversification of paralog expression patterns. *eLife*, 9, 2020.

[7191] Michaela Gregorova, Daniel Morse, Tarcisio Brignoli, Joseph Steventon, Fergus Hamilton, Mahableshwar Albur, David Arnold, Matthew Thomas, Alice Halliday, Holly Baum, Christopher Rice, Matthew B Avison, Andrew D Davidson, Marianna Santopaolo, Elizabeth Oliver, Anu Goenka, Adam Finn, Linda Wooldridge, Borko Amulic, Rosemary J Boyton, Daniel M Altmann, David K Butler, Claire McMurray, Joanna Stockton, Sam Nicholls, Charles Cooper, Nicholas Loman, Michael J Cox, Laura Rivino, and Ruth C Massey. Post-acute COVID-19 associated with evidence of bystander T-cell activation and a recurring antibiotic-resistant bacterial pneumonia. *eLife*, 9, 2020.

[7192] Igor Gridchyn, Philipp Schoenenberger, Joseph O'Neill, and Jozsef Csicsvari. Optogenetic inhibition-mediated activity-dependent modification of CA1 pyramidal-interneuron connections during behavior. *eLife*, 9, 2020.

[7193] Percy Griffin, Patrick W Sheehan, Julie M Dimitry, Chun Guo, Michael F Kanan, Jiyeon Lee, Jinsong Zhang, and Erik S Musiek. REV-ERB$\alpha$ mediates complement expression and diurnal regulation of microglial synaptic phagocytosis. *eLife*, 9, 2020.

[7194] Linda Grigoraki, Xavier Grau-Bové, Henrietta Carrington Yates, Gareth J Lycett, and Hilary Ranson. Isolation and transcriptomic analysis of Anopheles gambiae oenocytes enables the delineation of hydrocarbon biosynthesis. *eLife*, 9, 2020.

[7195] John P Grogan, Timothy R Sandhu, Michele T Hu, and Sanjay G Manohar. Dopamine promotes instrumental motivation, but reduces reward-related vigour. *eLife*, 9, 2020.

[7196] Mor Gross-Vered, Sébastien Trzebanski, Anat Shemer, Biana Bernshtein, Caterina Curato, Gil Stelzer, Tomer-Meir Salame, Eyal David, Sigalit Boura-Halfon, Louise Chappell-Maor, Dena Leshkowitz, and Steffen Jung. Defining murine monocyte differentiation into colonic and ileal macrophages. *eLife*, 9, 2020.

[7197] Matthew Grove, Hyunkyoung Lee, Huaqing Zhao, and Young-Jin Son. Axon-dependent expression of YAP/TAZ mediates Schwann cell remyelination but not proliferation after nerve injury. *eLife*, 9, 2020.

[7198] Franz S Gruber, Zoe C Johnston, Christopher LR Barratt, and Paul D Andrews. A phenotypic screening platform utilising human spermatozoa identifies compounds with contraceptive activity. *eLife*, 9, 2020.

[7199] Stefanie Gryzik, Yen Hoang, Timo Lischke, Elodie Mohr, Melanie Venzke, Isabelle Kadner, Josephine Poetzsch, Detlef Groth, Andreas Radbruch, Andreas Hutloff, and Ria Baumgrass. Identification of a super-functional Tfh-like subpopulation in murine lupus by pattern perception. *eLife*, 9, 2020.

[7200] Bon-Mi Gu, Robert Schmidt, and Joshua D Berke. Globus pallidus dynamics reveal covert strategies for behavioral inhibition. *eLife*, 9, 2020.

[7201] Zhuo Guan, Monica C Quiñones-Frías, Yulia Akbergenova, and J Troy Littleton. Drosophila Synaptotagmin 7 negatively regulates synaptic vesicle release and replenishment in a dosage-dependent manner. *eLife*, 9, 2020.

[7202] Jeremy Guerin, Istvan Botos, Zijian Zhang, Karl Lundquist, James C Gumbart, and Susan K Buchanan. Structural insight into toxin secretion by contact-dependent growth inhibition transporters. *eLife*, 9, 2020.

[7203] Marion V Guerin, Veronica Finisguerra, Benoit J Van den Eynde, Nadege Bercovici, and Alain Trautmann. Preclinical murine tumor models: A structural and functional perspective. *eLife*, 9, 2020.

[7204] Emilie Guillon, Dipjyoti Das, Dörthe Jülich, Abdel-Rahman Hassan, Hannah Geller, and Scott Holley. Fibronectin is a smart adhesive that both influences and responds to the mechanics of early spinal column development. *eLife*, 9, 2020.

[7205] Krishnendu Guin, Yao Chen, Radha Mishra, Siti Rawaidah BM Muzaki, Bhagya C Thimmappa, Caoimhe E O'Brien, Geraldine Butler, Amartya Sanyal, and Kaustuv Sanyal. Spatial inter-centromeric interactions facilitated the emergence of evolutionary new centromeres. *eLife*, 9, 2020.

[7206] Suna Gulay, Neha Gupta, Jon R Lorsch, and Alan G Hinnebusch. Distinct interactions of eIF4A and eIF4E with RNA helicase Ded1 stimulate translation in vivo. *eLife*, 9, 2020.

[7207] Omer Faruk Gulban, Rainer Goebel, Michelle Moerel, Daniel Zachlod, Hartmut Mohlberg, Katrin Amunts, and Federico de Martino. Improving a probabilistic cytoarchitectonic atlas of auditory cortex using a novel method for inter-individual alignment. *eLife*, 9, 2020.

[7208] Diana Gumber, Myan Do, Neya Suresh Kumar, Pooja R Sonavane, Christina C N Wu, Luisjesus S Cruz, Stephanie Grainger, Dennis Carson, Terry Gaasterland, and Karl Willert. Selective activation of FZD7 promotes mesendodermal differentiation of human pluripotent stem cells. *eLife*, 9, 2020.

[7209] Justin T Gunesch, Amera L Dixon, Tasneem AM Ebrahim, Melissa M Berrien-Elliott, Swetha Tatineni, Tejas Kumar, Everardo Hegewisch-Solloa, Todd A Fehniger, and Emily M Mace. CD56 regulates human NK cell cytotoxicity through Pyk2. *eLife*, 9, 2020.

[7210] Eren Günseli and Mariam Aly. Preparation for upcoming attentional states in the hippocampus and medial prefrontal cortex. *eLife*, 9, 2020.

[7211] Syuan-Ming Guo, Li-Hao Yeh, Jenny Folkesson, Ivan E Ivanov, Anitha P Krishnan, Matthew G Keefe, Ezzat Hashemi, David Shin, Bryant B Chhun, Nathan H Cho, Manuel D Leonetti, May H Han, Tomasz J Nowakowski, and Shalin B Mehta. Revealing architectural order with quantitative label-free imaging and deep learning. *eLife*, 9, 2020.

[7212] Yajuan Guo, Catherine J Redmond, Krystynne A Leacock, Margarita V Brovkina, Suyun Ji, Vinod Jaskula-Ranga, and Pierre A Coulombe. Keratin 14-dependent disulfides regulate epidermal homeostasis and barrier function via 14-3-3$\sigma$ and YAP1. *eLife*, 9, 2020.

[7213] Avantika Gupta and Hugo Stocker. FoxO suppresses endoplasmic reticulum stress to inhibit growth of Tsc1-deficient tissues under nutrient restriction. *eLife*, 9, 2020.

[7214] E Mae Guthman, Joshua D Garcia, Ming Ma, Philip Chu, Serapio M Baca, Katharine R Smith, Diego Restrepo, and Molly M Huntsman. Cell-type-specific control of basolateral amygdala neuronal circuits via entorhinal cortex-driven feedforward inhibition. *eLife*, 9, 2020.

[7215] Ayse Güven, Nereo Kalebic, Katherine R Long, Marta Florio, Samir Vaid, Holger Brandl, Denise Stenzel, and Wieland B Huttner. Extracellular matrix-inducing Sox9 promotes both basal progenitor proliferation and gliogenesis in developing neocortex. *eLife*, 9, 2020.

[7216] Clotilde Guyon, Nada Jmari, Francine Padonou, Yen-Chin Li, Olga Ucar, Noriyuki Fujikado, Fanny Coulpier, Christophe Blanchet, David E Root, and Matthieu Giraud. Aire-dependent genes undergo Clp1-mediated 3′UTR shortening associated with higher transcript stability in the thymus. *eLife*, 9, 2020.

[7217] Chyi Yin Gwee, Kritika M Garg, Balaji Chattopadhyay, Keren R Sadanandan, Dewi M Prawiradilaga, Martin Irestedt, Fumin Lei, Luke M Bloch, Jessica GH Lee, Mohammad Irham, Tri Haryoko, Malcolm CK Soh, Kelvin S-H Peh, Karen MC Rowe, Teuku Reza Ferasyi, Shaoyuan Wu, Guinevere OU Wogan, Rauri CK Bowie, and Frank E Rheindt. Phylogenomics of white-eyes, a 'great speciator', reveals Indonesian archipelago as the center of lineage diversity. *eLife*, 9, 2020.

[7218] Laura Gwilliams and Jean-Remi King. Recurrent processes support a cascade of hierarchical decisions. *eLife*, 9, 2020.

[7219] Kotdaji Ha, Mai Nobuhara, Qinzhe Wang, Rebecca V Walker, Feng Qian, Christoph Schartner, Erhu Cao, and Markus Delling. The heteromeric PC-1/PC-2 polycystin complex is activated by the PC-1 N-terminus. *eLife*, 9, 2020.

[7220] Andrea Haag, Michael Walser, Adrian Henggeler, and Alex Hajnal. The CHORD protein CHP-1 regulates EGF receptor trafficking and signaling in C. elegans and in human cells. *eLife*, 9, 2020.

[7221] Verena Haage. A survey of travel behaviour among scientists in Germany and the potential for change. *eLife*, 9, 2020.

[7222] Amin Haghani, Mafalda Cacciottolo, Kevin R Doty, Carla D'Agostino, Max Thorwald, Nikoo Safi, Morgan E Levine, Constantinos Sioutas, Terrence C Town, Henry Jay Forman, Hongqiao Zhang, Todd E Morgan, and Caleb E Finch. Mouse brain transcriptome responses to inhaled nanoparticulate matter differed by sex and APOE in Nrf2-Nfkb interactions. *eLife*, 9, 2020.

[7223] Andreas Hahn, Michael Breakspear, Lucas Rischka, Wolfgang Wadsak, Godber M Godbersen, Verena Pichler, Paul Michenthaler, Thomas Vanicek, Marcus Hacker, Siegfried Kasper, Rupert Lanzenberger, and Luca Cocchi. Reconfiguration of functional brain networks and metabolic cost converge during task performance. *eLife*, 9, 2020.

[7224] Martin Hailstone, Dominic Waithe, Tamsin J Samuels, Lu Yang, Ita Costello, Yoav Arava, Elizabeth Robertson, Richard M Parton, and Ilan Davis. CytoCensus, mapping cell identity and division in tissues and organs using machine learning. *eLife*, 9, 2020.

[7225] Emma Hajaj, Galit Eisenberg, Shiri Klein, Shoshana Frankenburg, Sharon Merims, Inna Ben David, Thomas Eisenhaure, Sarah E Henrickson, Alexandra Chloé Villani, Nir Hacohen, Nathalie Abudi, Rinat Abramovich, Jonathan E Cohen, Tamar Peretz, Andre Veillette, and Michal Lotem. SLAMF6 deficiency augments tumor killing and skews toward an effector phenotype revealing it as a novel T cell checkpoint. *eLife*, 9, 2020.

[7226] Anna E Hakes and Andrea H Brand. Tailless/TLX reverts intermediate neural progenitors to stem cells driving tumourigenesis via repression of asense/ASCL1. *eLife*, 9, 2020.

[7227] Melissa S Haley, Stephen Bruno, Alfredo Fontanini, and Arianna Maffei. LTD at amygdalocortical synapses as a novel mechanism for hedonic learning. *eLife*, 9, 2020.

[7228] Kelsi R Hall, Katherine J Robins, Elsie M Williams, Michelle H Rich, Mark J Calcott, Janine N Copp, Rory F Little, Ralf Schwörer, Gary B Evans, Wayne M Patrick, and David F Ackerley. Intracellular complexities of acquiring a new enzymatic function revealed by mass-randomisation of active-site residues. *eLife*, 9, 2020.

[7229] Kelsey M Hallinen, Jason Karslake, and Kevin B Wood. Delayed antibiotic exposure induces population collapse in enterococcal communities with drug-resistant subpopulations. *eLife*, 9, 2020.

[7230] Julie Hamaide, Kristina Lukacova, Jasmien Orije, Georgios A Keliris, Marleen Verhoye, and Annemie Van der Linden. In vivo assessment of the neural substrate linked with vocal imitation accuracy. *eLife*, 9, 2020.

[7231] Daniel G Hamilton, Hannah Fraser, Rink Hoekstra, and Fiona Fidler. Journal policies and editors' opinions on peer review. *eLife*, 9, 2020.

[7232] Grace E Hamilton, Luke A Helgeson, Cameron L Noland, Charles L Asbury, Yoana N Dimitrova, and Trisha N Davis. Reconstitution reveals two paths of force transmission through the kinetochore. *eLife*, 9, 2020.

[7233] Ali S Hamodi, Aude Martinez Sabino, N Dalton Fitzgerald, Dionysia Moschou, and Michael C Crair. Transverse sinus injections drive robust whole-brain expression of transgenes. *eLife*, 9, 2020.

[7234] Stefanie Hampel, Katharina Eichler, Daichi Yamada, Davi D Bock, Azusa Kamikouchi, and Andrew M Seeds. Distinct subpopulations of mechanosensory chordotonal organ neurons elicit grooming of the fruit fly antennae. *eLife*, 9, 2020.

[7235] Fan Han, Minal Jamsandekar, Mats E Pettersson, Leyi Su, Angela P Fuentes-Pardo, Brian W Davis, Dorte Bekkevold, Florian Berg, Michele Casini, Geir Dahle, Edward D Farrell, Arild Folkvord, and Leif Andersson. Ecological adaptation in Atlantic herring is associated with large shifts in allele frequencies at hundreds of loci. *eLife*, 9, 2020.

[7236] Xiaochun Han, Michele J Gelfand, Bing Wu, Ting Zhang, Wenxin Li, Tianyu Gao, Chenyu Pang, Taoyu Wu, Yuqing Zhou, Shuai Zhou, Xinhuai Wu, and Shihui Han. A neurobiological association of revenge propensity during intergroup conflict. *eLife*, 9, 2020.

[7237] Naomi AK Hanemaaijer, Marko A Popovic, Xante Wilders, Sara Grasman, Oriol Pavón Arocas, and Maarten HP Kole. Ca2+ entry through NaV channels generates submillisecond axonal Ca2+ signaling. *eLife*, 9, 2020.

[7238] Jan N Hansen, Fabian Kaiser, Christina Klausen, Birthe Stüven, Raymond Chong, Wolfgang Bönigk, David U Mick, Andreas Möglich, Nathalie Jurisch-Yaksi, Florian I Schmidt, and Dagmar Wachten. Nanobody-directed targeting of optogenetic tools to study signaling in the primary cilium. *eLife*, 9, 2020.

[7239] Kristoffer B Hansen, Christian Staehr, Palle D Rohde, Casper Homilius, Sukhan Kim, Mette Nyegaard, Vladimir V Matchkov, and Ebbe Boedtkjer. PTPRG is an ischemia risk locus essential for HCO3—dependent regulation of endothelial function and tissue perfusion. *eLife*, 9, 2020.

[7240] Haley E Hanson, Noreen S Mathews, Mark E Hauber, and Lynn B Martin. The house sparrow in the service of basic and applied biology. *eLife*, 9, 2020.

[7241] Boran Hao, Shahabeddin Sotudian, Taiyao Wang, Tingting Xu, Yang Hu, Apostolos Gaitanidis, Kerry Breen, George C Velmahos, and Ioannis Ch Paschalidis. Early prediction of level-of-care requirements in patients with COVID-19. *eLife*, 9, 2020.

[7242] Qinyu Hao, Xinying Zong, Qinyu Sun, Yo-Chuen Lin, You Jin Song, Seyedsasan Hashemikhabir, Rosaline YC Hsu, Mohammad Kamran, Ritu Chaudhary, Vidisha Tripathi, Deepak Kumar Singh, Arindam Chakraborty, Xiao Ling Li, Yoon Jung Kim, Arturo V Orjalo, Maria Polycarpou-Schwarz, Branden S Moriarity, Lisa M Jenkins, Hans E Johansson, Yuelin J Zhu, Sven Diederichs, Anindya Bagchi, Tae Hoon Kim, Sarath C Janga, Ashish Lal, Supriya G Prasanth, and Kannanganattu V Prasanth. The S-phase-induced lncRNA SUNO1 promotes cell proliferation by controlling YAP1/Hippo signaling pathway. *eLife*, 9, 2020.

[7243] Xue Hao, Shimin Wang, Yi Lu, Wentao Yu, Pengyue Li, Dan Jiang, Tong Guo, Mengjie Li, Jinhui Li, Jinjin Xu, Wenqing Wu, Margaret S Ho, and Lei Zhang. Lola regulates Drosophila adult midgut homeostasis via non-canonical hippo signaling. *eLife*, 9, 2020.

[7244] Joseph L Harman, Andrea N Loes, Gus D Warren, Maureen C Heaphy, Kirsten J Lampi, and Michael J Harms. Evolution of multifunctionality through a pleiotropic substitution in the innate immune protein S100A9. *eLife*, 9, 2020.

[7245] Roy Harpaz and Elad Schneidman. Social interactions drive efficient foraging and income equality in groups of fish. *eLife*, 9, 2020.

[7246] Robin E Harris, Michael J Stinchfield, Spencer L Nystrom, Daniel J McKay, and Iswar K Hariharan. Damage-responsive, maturity-silenced enhancers regulate multiple genes that direct regeneration in Drosophila. *eLife*, 9, 2020.

[7247] Eric Hart and Alexander C Huk. Recurrent circuit dynamics underlie persistent activity in the macaque frontoparietal network. *eLife*, 9, 2020.

[7248] Evan E Hart, Melissa J Sharpe, Matthew PH Gardner, and Geoffrey Schoenbaum. Responding to preconditioned cues is devaluation sensitive and requires orbitofrontal cortex during cue-cue learning. *eLife*, 9, 2020.

[7249] Jonas Hartmann, Mie Wong, Elisa Gallo, and Darren Gilmour. An image-based data-driven analysis of cellular architecture in a developing tissue. *eLife*, 9, 2020.

[7250] Lucas A Hartsough, Mooncheol Park, Matthew V Kotlajich, John Tyler Lazar, Bing Han, Chih-Chun J Lin, Elena Musteata, Lauren Gambill, Meng C Wang, and Jeffrey J Tabor. Optogenetic control of gut bacterial metabolism to promote longevity. *eLife*, 9, 2020.

[7251] Gesa Hartwigsen, Anika Stockert, Louise Charpentier, Max Wawrzyniak, Julian Klingbeil, Katrin Wrede, Hellmuth Obrig, and Dorothee Saur. Short-term modulation of the lesioned language network. *eLife*, 9, 2020.

[7252] Md Rakibul Hasan, Maarit Takatalo, Hongqiang Ma, Ritva Rice, Tuija Mustonen, and David PC Rice. RAB23 coordinates early osteogenesis by repressing FGF10-pERK1/2 and GLI1. *eLife*, 9, 2020.

[7253] Kerstin Hasenpusch-Theil, Christine Laclef, Matt Colligan, Eamon Fitzgerald, Katherine Howe, Emily Carroll, Shaun R Abrams, Jeremy F Reiter, Sylvie Schneider-Maunoury, and Thomas Theil. A transient role of the ciliary gene Inpp5e in controlling direct versus indirect neurogenesis in cortical development. *eLife*, 9, 2020.

[7254] Saeedeh Hashemnia, David R Euston, and Aaron J Gruber. Amphetamine reduces reward encoding and stabilizes neural dynamics in rat anterior cingulate cortex. *eLife*, 9, 2020.

[7255] Amira Hassan, Pablo Araguas Rodriguez, Stefan K Heidmann, Emma L Walmsley, Gabriel N Aughey, and Tony D Southall. Condensin I subunit Cap-G is essential for proper gene expression during the maturation of post-mitotic neurons. *eLife*, 9, 2020.

[7256] Jordan F Hastings, Alvaro Gonzalez Rajal, Sharissa L Latham, Jeremy ZR Han, Rachael A McCloy, Yolande EI O'Donnell, Monica Phimmachanh, Alexander D Murphy, Adnan Nagrial, Dariush Daneshvar, Venessa Chin, D Neil Watkins, Andrew Burgess, and David R Croucher. Analysis of pulsed cisplatin signalling dynamics identifies effectors of resistance in lung adenocarcinoma. *eLife*, 9, 2020.

[7257] Anna Hatch and Stephen Curry. Changing how we evaluate research is difficult, but not impossible. *eLife*, 9, 2020.

[7258] Josué Haubrich, Matteo Bernabo, and Karim Nader. Noradrenergic projections from the locus coeruleus to the amygdala constrain fear memory reconsolidation. *eLife*, 9, 2020.

[7259] Vesa Havurinne and Esa Tyystjärvi. Photosynthetic sea slugs induce protective changes to the light reactions of the chloroplasts they steal from algae. *eLife*, 9, 2020.

[7260] James V Haxby, J Swaroop Guntupalli, Samuel A Nastase, and Ma Feilong. Hyperalignment: Modeling shared information encoded in idiosyncratic cortical topographies. *eLife*, 9, 2020.

[7261] Masayuki Hayakawa, Tetsuya Hiraiwa, Yuko Wada, Hidekazu Kuwayama, and Tatsuo Shibata. Polar pattern formation induced by contact following locomotion in a multicellular system. *eLife*, 9, 2020.

[7262] Luke Hayden, Katerina Lochovska, Marie Sémon, Sabrina Renaud, Marie-Laure Delignette-Muller, Maurine Vilcot, Renata Peterkova, Maria Hovorakova, and Sophie Pantalacci. Developmental variability channels mouse molar evolution. *eLife*, 9, 2020.

[7263] Lindsey R Hayes, Lauren Duan, Kelly Bowen, Petr Kalab, and Jeffrey D Rothstein. C9orf72 arginine-rich dipeptide repeat proteins disrupt karyopherin-mediated nuclear import. *eLife*, 9, 2020.

[7264] Mark David Hayes, Sophie Ward, Greg Crawford, Rocio Castro Seoane, William David Jackson, David Kipling, David Voehringer, Deborah Dunn-Walters, and Jessica Strid. Inflammation-induced IgE promotes epithelial hyperplasia and tumour growth. *eLife*, 9, 2020.

[7265] Michelle Hays, Janet M Young, Paula F Levan, and Harmit S Malik. A natural variant of the essential host gene MMS21 restricts the parasitic 2-micron plasmid in Saccharomyces cerevisiae. *eLife*, 9, 2020.

[7266] Stephanie G Hays and Kimberley D Seed. Dominant Vibrio cholerae phage exhibits lysis inhibition sensitive to disruption by a defensive phage satellite. *eLife*, 9, 2020.

[7267] Liu He, Robbelien Kooistra, Ravi Das, Ellen Oudejans, Eric van Leen, Johannes Ziegler, Sybren Portegies, Bart de Haan, Anna van Regteren Altena, Riccardo Stucchi, AF Maarten Altelaar, Stefan Wieser, Michael Krieg, Casper C Hoogenraad, and Martin Harterink. Cortical anchoring of the microtubule cytoskeleton is essential for neuron polarity. *eLife*, 9, 2020.

[7268] Mu He, Bing Wu, Wenlei Ye, Daniel D Le, Adriane W Sinclair, Valeria Padovano, Yuzhang Chen, Ke-Xin Li, Rene Sit, Michelle Tan, Michael J Caplan, Norma Neff, Yuh Nung Jan, Spyros Darmanis, and Lily Yeh Jan. Chloride channels regulate differentiation and barrier functions of the mammalian airway. *eLife*, 9, 2020.

[7269] Shenghong He, Abteen Mostofi, Emilie Syed, Flavie Torrecillos, Gerd Tinkhauser, Petra Fischer, Alek Pogosyan, Harutomo Hasegawa, Yuanqing Li, Keyoumars Ashkan, Erlick Pereira, Peter Brown, and Huiling Tan. Subthalamic beta-targeted neurofeedback speeds up movement initiation but increases tremor in Parkinsonian patients. *eLife*, 9, 2020.

[7270] Sicong He, Ye Tian, Shachuan Feng, Yi Wu, Xinwei Shen, Kani Chen, Yingzhu He, Qiqi Sun, Xuesong Li, Jin Xu, Zilong Wen, and Jianan Y Qu. In vivo single-cell lineage tracing in zebrafish using high-resolution infrared laser-mediated gene induction microscopy. *eLife*, 9, 2020.

[7271] Jade Hebras, Virginie Marty, Jean Personnaz, Pascale Mercier, Nicolai Krogh, Henrik Nielsen, Marion Aguirrebengoa, Hervé Seitz, Jean-Phillipe Pradere, Bruno P Guiard, and Jérôme Cavaille. Reassessment of the involvement of Snord115 in the serotonin 2c receptor pathway in a genetically relevant mouse model. *eLife*, 9, 2020.

[7272] Kristina Hedbacker, Yi-Hsueh Lu, Olof Dallner, Zhiying Li, Gulya Fayzikhodjaeva, KıvanÇ Birsoy, Chiayun Han, Chingwen Yang, and Jeffrey M Friedman. Limitation of adipose tissue by the number of embryonic progenitor cells. *eLife*, 9, 2020.

[7273] Matthew A Heinrich, Ricard Alert, Julienne M LaChance, Tom J Zajdel, Andrej KoŠmrlj, and Daniel J Cohen. Size-dependent patterns of cell proliferation and migration in freely-expanding epithelia. *eLife*, 9, 2020.

[7274] Clemens Heissenberger, Jarod A Rollins, Teresa L Krammer, Fabian Nagelreiter, Isabella Stocker, Ludivine Wacheul, Anton Shpylovyi, Koray Tav, Santina Snow, Johannes Grillari, Aric N Rogers, Denis L J Lafontaine, and Markus Schosserer. The ribosomal RNA m5C methyltransferase NSUN-1 modulates healthspan and oogenesis in Caenorhabditis elegans. *eLife*, 9, 2020.

[7275] Christian SM Helker, Jean Eberlein, Kerstin Wilhelm, Toshiya Sugino, Julian Malchow, Annika Schuermann, Stefan Baumeister, Hyouk-Bum Kwon, Hans-Martin Maischein, Michael Potente, Wiebke Herzog, and Didier YR Stainier. Apelin signaling drives vascular endothelial cells toward a pro-angiogenic state. *eLife*, 9, 2020.

[7276] Mathieu Hénault, Souhir Marsit, Guillaume Charron, and Christian R Landry. The effect of hybridization on transposable element accumulation in an undomesticated fungal species. *eLife*, 9, 2020.

[7277] Alasdair D Henderson, Maite Aubry, Mike Kama, Jessica Vanhomwegen, Anita Teissier, Teheipuaura Mariteragi-Helle, Tuterarii Paoaafaite, Yoann Teissier, Jean-Claude Manuguerra, John Edmunds, Jimmy Whitworth, Conall H Watson, Colleen L Lau, Van-Mai Cao-Lormeau, and Adam J Kucharski. Zika seroprevalence declines and neutralizing antibodies wane in adults following outbreaks in French Polynesia and Fiji. *eLife*, 9, 2020.

[7278] Joseph A Heng, Michael Woodford, and Rafael Polania. Efficient sampling and noisy decisions. *eLife*, 9, 2020.

[7279] Steven Henikoff, Jorja G Henikoff, Hatice S Kaya-Okur, and Kami Ahmad. Efficient chromatin accessibility mapping in situ by nucleosome-tethered tagmentation. *eLife*, 9, 2020.

[7280] Christopher A Henry, Mehrdad Jazayeri, Robert M Shapley, and Michael J Hawken. Distinct spatiotemporal mechanisms underlie extra-classical receptive field modulation in macaque V1 microcircuits. *eLife*, 9, 2020.

[7281] Julia U Henschke and Janelle MP Pakan. Disynaptic cerebrocerebellar pathways originating from multiple functionally distinct cortical areas. *eLife*, 9, 2020.

[7282] Nathaniel L Hepowit, Kevin N Pereira, Jessica M Tumolo, Walter J Chazin, and Jason A MacGurn. Identification of ubiquitin Ser57 kinases regulating the oxidative stress response in yeast. *eLife*, 9, 2020.

[7283] Jo Hepworth, Rea L Antoniou-Kourounioti, Kristina Berggren, Catja Selga, Eleri H Tudor, Bryony Yates, Deborah Cox, Barley Rose Collier Harris, Judith A Irwin, Martin Howard, Torbjörn Säll, Svante Holm, and Caroline Dean. Natural variation in autumn expression is the major adaptive determinant distinguishing Arabidopsis FLC haplotypes. *eLife*, 9, 2020.

[7284] Coralie Hérent, Séverine Diem, Gilles Fortin, and Julien Bouvier. Absent phasing of respiratory and locomotor rhythms in running mice. *eLife*, 9, 2020.

[7285] James P Herman, Fabrice Arcizet, and Richard J Krauzlis. Attention-related modulation of caudate neurons depends on superior colliculus activity. *eLife*, 9, 2020.

[7286] Caesar M Hernandez, Caitlin Orsini, Alexa-Rae Wheeler, Tyler W Ten Eyck, Sara M Betzhold, Chase C Labiste, Noelle G Wright, Barry Setlow, and Jennifer L Bizon. Testicular hormones mediate robust sex differences in impulsive choice in rats. *eLife*, 9, 2020.

[7287] J Jesús Hernández-Pérez, Keiland W Cooper, and Ehren L Newman. Medial entorhinal cortex activates in a traveling wave in the rat. *eLife*, 9, 2020.

[7288] Thurston Herricks, Magdalena Donczew, Fred D Mast, Tige Rustad, Robert Morrison, Timothy R Sterling, David R Sherman, and John D Aitchison. ODELAM, rapid sequence-independent detection of drug resistance in isolates of Mycobacterium tuberculosis. *eLife*, 9, 2020.

[7289] Sarah Hersman, David Allen, Mariko Hashimoto, Salvador Ignacio Brito, and Todd E Anthony. Stimulus salience determines defensive behaviors elicited by aversively conditioned serial compound auditory stimuli. *eLife*, 9, 2020.

[7290] Loreen Hertäg and Henning Sprekeler. Learning prediction error neurons in a canonical interneuron circuit. *eLife*, 9, 2020.

[7291] David J Herzfeld, Nathan J Hall, Marios Tringides, and Stephen G Lisberger. Principles of operation of a cerebellar learning circuit. *eLife*, 9, 2020.

[7292] Janis Karan Hesse and Doris Y Tsao. A new no-report paradigm reveals that face cells encode both consciously perceived and suppressed stimuli. *eLife*, 9, 2020.

[7293] Mark S Hibbins, Matthew JS Gibson, and Matthew W Hahn. Determining the probability of hemiplasy in the presence of incomplete lineage sorting and introgression. *eLife*, 9, 2020.

[7294] Alison B Hickman, Shweta Kailasan, Pavol Genzor, Astrid D Haase, and Fred Dyda. Casposase structure and the mechanistic link between DNA transposition and spacer acquisition by CRISPR-Cas. *eLife*, 9, 2020.

[7295] Allison L Hicks, Stephen M Kissler, Tatum D Mortimer, Kevin C Ma, George Taiaroa, Melinda Ashcroft, Deborah A Williamson, Marc Lipsitch, and Yonatan H Grad. Targeted surveillance strategies for efficient detection of novel antibiotic resistance variants. *eLife*, 9, 2020.

[7296] A Elizabeth Hildreth, Mitchell A Ellison, Alex M Francette, Julia M Seraly, Lauren M Lotka, and Karen M Arndt. The nucleosome DNA entry-exit site is important for transcription termination and prevention of pervasive transcription. *eLife*, 9, 2020.

[7297] Tom Hill and Robert L Unckless. Recurrent evolution of high virulence in isolated populations of a DNA virus. *eLife*, 9, 2020.

[7298] Sebastian Hillringhaus, Anil K Dasanna, Gerhard Gompper, and Dmitry A Fedosov. Stochastic bond dynamics facilitates alignment of malaria parasite at erythrocyte membrane upon invasion. *eLife*, 9, 2020.

[7299] Florian Hinte, Eelco van Anken, Boaz Tirosh, and Wolfram Brune. Repression of viral gene expression and replication by the unfolded protein response effector XBP1u. *eLife*, 9, 2020.

[7300] Ken-ichi Hirano, Akiko Suganami, Yutaka Tamura, Hideo Yagita, Sonoko Habu, Motoo Kitagawa, Takehito Sato, and Katsuto Hozumi. Delta-like 1 and Delta-like 4 differently require their extracellular domains for triggering Notch signaling in mice. *eLife*, 9, 2020.

[7301] Benjamin D Hobson, Linghao Kong, Erik W Hartwick, Ruben L Gonzalez, and Peter A Sims. Elongation inhibitors do not prevent the release of puromycylated nascent polypeptide chains from ribosomes. *eLife*, 9, 2020.

[7302] Andreas Hoehne, Maureen H McFadden, and David A DiGregorio. Feed-forward recruitment of electrical synapses enhances synchronous spiking in the mouse cerebellar cortex. *eLife*, 9, 2020.

[7303] Eric D Hoffer, Samuel Hong, S Sunita, Tatsuya Maehigashi, Ruben L Gonzalez, Paul C Whitford, and Christine M Dunham. Structural insights into mRNA reading frame regulation by tRNA modification and slippery codon–anticodon pairing. *eLife*, 9, 2020.

[7304] Sheila Hoffmann-Conaway, Marisa M Brockmann, Katharina Schneider, Anil Annamneedi, Kazi Atikur Rahman, Christine Bruns, Kathrin Textoris-Taube, Thorsten Trimbuch, Karl-Heinz Smalla, Christian Rosenmund, Eckart D Gundelfinger, Craig Curtis Garner, and Carolina Montenegro-Venegas. Parkin contributes to synaptic vesicle autophagy in Bassoon-deficient mice. *eLife*, 9, 2020.

[7305] Felix JH Hol, Louis Lambrechts, and Manu Prakash. BiteOscope, an open platform to study mosquito biting behavior. *eLife*, 9, 2020.

[7306] James Holder, Shabaz Mohammed, and Francis A Barr. Ordered dephosphorylation initiated by the selective proteolysis of cyclin B drives mitotic exit. *eLife*, 9, 2020.

[7307] Anne Kruse Hollensen, Henriette Sylvain Thomsen, Marta Lloret-Llinares, Andreas Bjerregaard Kamstrup, Jacob Malte Jensen, Majbritt Luckmann, Nanna Birkmose, Johan Palmfeldt, Torben Heick Jensen, Thomas B Hansen, and Christian Kroun Damgaard. circZNF827 nucleates a transcription inhibitory complex to balance neuronal differentiation. *eLife*, 9, 2020.

[7308] Jennifer A Honeycutt, Camila Demaestri, Shayna Peterzell, Marisa M Silveri, Xuezhu Cai, Praveen Kulkarni, Miles G Cunningham, Craig F Ferris, and Heather C Brenhouse. Altered corticolimbic connectivity reveals sex-specific adolescent outcomes in a rat model of early life adversity. *eLife*, 9, 2020.

[7309] Mingi Hong, Annabel Christ, Anna Christa, Thomas E Willnow, and Robert S Krauss. Cdon mutation and fetal alcohol converge on Nodal signaling in a mouse model of holoprosencephaly. *eLife*, 9, 2020.

[7310] Su Z Hong, Shiyong Huang, Daniel Severin, and Alfredo Kirkwood. Pull-push neuromodulation of cortical plasticity enables rapid bi-directional shifts in ocular dominance. *eLife*, 9, 2020.

[7311] Peter Jan Hooikaas, Hugo GJ Damstra, Oane J Gros, Wilhelmina E van Riel, Maud Martin, Yesper TH Smits, Jorg van Loosdregt, Lukas C Kapitein, Florian Berger, and Anna Akhmanova. Kinesin-4 KIF21B limits microtubule growth to allow rapid centrosome polarization in T cells. *eLife*, 9, 2020.

[7312] Lisa Horn, Thomas Bugnyar, Michael Griesser, Marietta Hengl, Ei-Ichi Izawa, Tim Oortwijn, Christiane Rössler, Clara Scheer, Martina Schiestl, Masaki Suyama, Alex H Taylor, Lisa-Claire Vanhooland, Auguste MP von Bayern, Yvonne Zürcher, and Jorg JM Massen. Sex-specific effects of cooperative breeding and colonial nesting on prosociality in corvids. *eLife*, 9, 2020.

[7313] Patricia M Horvath, Michelle K Piazza, Lisa M Monteggia, and Ege T Kavalali. Spontaneous and evoked neurotransmission are partially segregated at inhibitory synapses. *eLife*, 9, 2020.

[7314] James Hose, Leah E Escalante, Katie J Clowers, H Auguste Dutcher, DeElegant Robinson, Venera Bouriakov, Joshua J Coon, Evgenia Shishkova, and Audrey P Gasch. The genetic basis of aneuploidy tolerance in wild yeast. *eLife*, 9, 2020.

[7315] Xiaowei Hou, Ian R Outhwaite, Leanne Pedi, and Stephen Barstow Long. Cryo-EM structure of the calcium release-activated calcium channel Orai in an open conformation. *eLife*, 9, 2020.

[7316] Mona Hoyos, Michaela Huber, Konrad U Förstner, and Kai Papenfort. Gene autoregulation by $3'$ UTR-derived bacterial small RNAs. *eLife*, 9, 2020.

[7317] Jui-Yi Hsieh, Brittany N Ulrich, Fadi A Issa, Meng-chin A Lin, Brandon Brown, and Diane M Papazian. Infant and adult SCA13 mutations differentially affect Purkinje cell excitability, maturation, and viability in vivo. *eLife*, 9, 2020.

[7318] Sheng-Kai Hsu, Ana Marija JakŠić, Viola Nolte, Manolis Lirakis, Robert Kofler, Neda Barghi, Elisabetta Versace, and Christian Schlötterer. Rapid sex-specific adaptation to high temperature in Drosophila. *eLife*, 9, 2020.

[7319] Jia Ming Hu, Xue Mei Song, Qiannan Wang, and Anna Wang Roe. Curvature domains in V4 of macaque monkey. *eLife*, 9, 2020.

[7320] Kejia Hu, Mikael Eriksson, Yvonne Wengström, Kamila Czene, Per Hall, and Fang Fang. Sense of coherence and risk of breast cancer. *eLife*, 9, 2020.

[7321] Xiaoyun Hu, Shuangli Su, Qingsong Liu, Yaoyu Jiao, Yufa Peng, Yunhe Li, and Ted CJ Turlings. Caterpillar-induced rice volatiles provide enemy-free space for the offspring of the brown planthopper. *eLife*, 9, 2020.

[7322] Anqi Huang, Jean-FranÇois Rupprecht, and Timothy E Saunders. Embryonic geometry underlies phenotypic variation in decanalized conditions. *eLife*, 9, 2020.

[7323] Chun-Kai (Karl) Huang, Cameron Neylon, Richard Hosking, Lucy Montgomery, Katie S Wilson, Alkim Ozaygen, and Chloe Brookes-Kenworthy. Evaluating the impact of open access policies on research institutions. *eLife*, 9, 2020.

[7324] Nicholas Huang and Mounya Elhilali. Push-pull competition between bottom-up and top-down auditory attention to natural soundscapes. *eLife*, 9, 2020.

[7325] Rui Huang, Tingting Song, Haifeng Su, Zeliang Lai, Wusa Qin, Yinjun Tian, Xuan Dong, and Liming Wang. High-fat diet enhances starvation-induced hyperactivity via sensitizing hunger-sensing neurons in Drosophila. *eLife*, 9, 2020.

[7326] Tao Huang, Shenli Yuan, Lei Gao, Mengjing Li, Xiaochen Yu, Jianhong Zhan, Yingying Yin, Chao Liu, Chuanxin Zhang, Gang Lu, Wei Li, Jiang Liu, Zi-Jiang Chen, and Hongbin Liu. The histone modification reader ZCWPW1 links histone methylation to PRDM9-induced double-strand break repair. *eLife*, 9, 2020.

[7327] Yen-Lin Huang, Ching-Yeu Liang, Danilo Ritz, Ricardo Coelho, Dedy Septiadi, Manuela Estermann, Cécile Cumin, Natalie Rimmer, Andreas Schötzau, Mónica Núñez López, André Fedier, Martina Konantz, Tatjana Vlajnic, Diego Calabrese, Claudia Lengerke, Leonor David, Barbara Rothen-Rutishauser, Francis Jacob, and Viola Heinzelmann-Schwarz. Collagen-rich omentum is a premetastatic niche for integrin $\alpha$2-mediated peritoneal metastasis. *eLife*, 9, 2020.

[7328] Madlen Hubert, Elin Larsson, Naga Venkata Gayathri Vegesna, Maria Ahnlund, Annika I Johansson, Lindon WK Moodie, and Richard Lundmark. Lipid accumulation controls the balance between surface connection and scission of caveolae. *eLife*, 9, 2020.

[7329] John Huddleston, John R Barnes, Thomas Rowe, Xiyan Xu, Rebecca Kondor, David E Wentworth, Lynne Whittaker, Burcu Ermetal, Rodney Stuart Daniels, John W McCauley, Seiichiro Fujisaki, Kazuya Nakamura, Noriko Kishida, Shinji Watanabe, Hideki Hasegawa, Ian Barr, Kanta Subbarao, Pierre Barrat-Charlaix, Richard A Neher, and Trevor Bedford. Integrating genotypes and phenotypes improves long-term forecasts of seasonal influenza A/H3N2 evolution. *eLife*, 9, 2020.

[7330] Katherine E Huffer, Antoniya A Aleksandrova, Andrés Jara-Oseguera, Lucy R Forrest, and Kenton J Swartz. Global alignment and assessment of TRP channel transmembrane domain structures to explore functional mechanisms. *eLife*, 9, 2020.

[7331] Court Hull. Prediction signals in the cerebellum: Beyond supervised motor learning. *eLife*, 9, 2020.

[7332] Katina D Hulme, Limin Yan, Rebecca J Marshall, Conor J Bloxham, Kyle R Upton, Sumaira Z Hasnain, Helle Bielefeldt-Ohmann, Zhixuan Loh, Katharina Ronacher, Keng Yih Chew, Linda A Gallo, and Kirsty R Short. High glucose levels increase influenza-associated damage to the pulmonary epithelial-endothelial barrier. *eLife*, 9, 2020.

[7333] Ashley C Humphries, Sonali Narang, and Marek Mlodzik. Mutations associated with human neural tube defects display disrupted planar cell polarity in Drosophila. *eLife*, 9, 2020.

[7334] Chi Jung Hung, Daisuke Ono, Thomas S Kilduff, and Akihiro Yamanaka. Dual orexin and MCH neuron-ablated mice display severe sleep attacks and cataplexy. *eLife*, 9, 2020.

[7335] Harriet Hunter, Dana de Gracia Hahn, Amedine Duret, Yu Ri Im, Qinrong Cheah, Jiawen Dong, Madison Fairey, Clarissa Hjalmarsson, Alice Li, Hong Kai Lim, Lorcan McKeown, Claudia-Gabriela Mitrofan, Raunak Rao, Mrudula Utukuri, Ian A Rowe, and Jake P Mann. Weight loss, insulin resistance, and study design confound results in a meta-analysis of animal models of fatty liver. *eLife*, 9, 2020.

[7336] Vincent Huson, Marieke Meijer, Rien Dekker, Mirelle ter Veer, Marvin Ruiter, Jan RT van Weering, Matthijs Verhage, and Lennart Niels Cornelisse. Post-tetanic potentiation lowers the energy barrier for synaptic vesicle fusion independently of Synaptotagmin-1. *eLife*, 9, 2020.

[7337] Salik Hussain, Collin G Johnson, Joseph Sciurba, Xianglin Meng, Vandy P Stober, Caini Liu, Jaime M Cyphert-Daly, Katarzyna Bulek, Wen Qian, Alma Solis, Yosuke Sakamachi, Carol S Trempus, Jim J Aloor, Kym M Gowdy, W Michael Foster, John W Hollingsworth, Robert M Tighe, Xiaoxia Li, Michael B Fessler, and Stavros Garantziotis. TLR5 participates in the TLR4 receptor complex and promotes MyD88-dependent signaling in environmental lung injury. *eLife*, 9, 2020.

[7338] Nguyen PT Huynh, Catherine C Gloss, Jeremiah Lorentz, Ruhang Tang, Jonathan M Brunger, Audrey McAlinden, Bo Zhang, and Farshid Guilak. Long non-coding RNA GRASLND enhances chondrogenesis via suppression of the interferon type II signaling pathway. *eLife*, 9, 2020.

[7339] Lara S Hwa, Sofia Neira, Meghan E Flanigan, Christina M Stanhope, Melanie M Pina, Dipanwita Pati, Olivia J Hon, Waylin Yu, Emily Kokush, Rachel Calloway, Kristen Boyt, and Thomas L Kash. RETRACTED: Alcohol drinking alters stress response to predator odor via BNST kappa opioid receptor signaling in male mice. *eLife*, 9, 2020.

[7340] Jihee Hwang, Jae-Yeol Kim, Cheolhee Kim, Soojin Park, Sungmin Joo, Seong Keun Kim, and Nam Ki Lee. Single-molecule observation of ATP-independent SSB displacement by RecO in Deinococcus radiodurans. *eLife*, 9, 2020.

[7341] Noriko Ichino, MaKayla R Serres, Rhianna M Urban, Mark D Urban, Anthony J Treichel, Kyle J Schaefbauer, Lauren E Tallant, Gaurav K Varshney, Kimberly J Skuster, Melissa S McNulty, Camden L Daby, Ying Wang, Hsin-kai Liao, Suzan El-Rass, Yonghe Ding, Weibin Liu, Jennifer L Anderson, Mark D Wishman, Ankit Sabharwal, Lisa A Schimmenti, Sridhar Sivasubbu, Darius Balciunas, Matthias Hammerschmidt, Steven Arthur Farber, Xiao-Yan Wen, Xiaolei Xu, Maura McGrail, Jeffrey J Essner, Shawn M Burgess, Karl J Clark, and Stephen C Ekker. Building the vertebrate codex using the gene breaking protein trap library. *eLife*, 9, 2020.

[7342] Shintaro Ide, Yasuhito Yahara, Yoshihiko Kobayashi, Sarah A Strausser, Kana Ide, Anisha Watwe, Shengjie Xu-Vanpala, Jamie R Privratsky, Steven D Crowley, Mari L Shinohara, Benjamin A Alman, and Tomokazu Souma. Yolk-sac-derived macrophages progressively expand in the mouse kidney with age. *eLife*, 9, 2020.

[7343] Temitayo O Idowu, Valerie Etzrodt, Benjamin Seeliger, Patricia Bolanos-Palmieri, Kristina Thamm, Hermann Haller, and Sascha David. Identification of specific Tie2 cleavage sites and therapeutic modulation in experimental sepsis. *eLife*, 9, 2020.

[7344] Niv Ierushalmi, Maya Malik-Garbi, Angelika Manhart, Enas Abu Shah, Bruce L Goode, Alex Mogilner, and Kinneret Keren. Centering and symmetry breaking in confined contracting actomyosin networks. *eLife*, 9, 2020.

[7345] İbrahim Avşar Ilik, Michal Malszycki, Anna Katharina Lübke, Claudia Schade, David Meierhofer, and TuğÇe Aktaş. SON and SRRM2 are essential for nuclear speckle formation. *eLife*, 9, 2020.

[7346] Yukiko Imai, Mathilde Biot, Julie AJ Clément, Mariko Teragaki, Serge Urbach, Thomas Robert, Frédéric Baudat, Corinne Grey, and Bernard de Massy. PRDM9 activity depends on HELLS and promotes local 5-hydroxymethylcytosine enrichment. *eLife*, 9, 2020.

[7347] Hiromi Imamura, Shuichiro Sakamoto, Tomoki Yoshida, Yusuke Matsui, Silvia Penuela, Dale W Laird, Shin Mizukami, Kazuya Kikuchi, and Akira Kakizuka. Single-cell dynamics of pannexin-1-facilitated programmed ATP loss during apoptosis. *eLife*, 9, 2020.

[7348] Yasuko Inaba, Vasudha Chauhan, Aaron Paul van Loon, Lamia Saiyara Choudhury, and Alvaro Sagasti. Keratins and plakin family cytolinker proteins control the length of epithelial microridge protrusions. *eLife*, 9, 2020.

[7349] Samsoon Inayat, Surjeet Singh, Arashk Ghasroddashti, Qandeel, Pramuka Egodage, Ian Q Whishaw, and Majid H Mohajerani. A Matlab-based toolbox for characterizing behavior of rodents engaged in string-pulling. *eLife*, 9, 2020.

[7350] Aleksei Innokentev, Kentaro Furukawa, Tomoyuki Fukuda, Tetsu Saigusa, Keiichi Inoue, Shunichi Yamashita, and Tomotake Kanki. Association and dissociation between the mitochondrial Far complex and Atg32 regulate mitophagy. *eLife*, 9, 2020.

[7351] Ira Alexandra Iosub, Robert Willem van Nues, Stuart William McKellar, Karen Jule Nieken, Marta Marchioretto, Brandon Sy, Jai Justin Tree, Gabriella Viero, and Sander Granneman. Hfq CLASH uncovers sRNA-target interaction networks linked to nutrient availability adaptation. *eLife*, 9, 2020.

[7352] William T Ireland, Suzannah M Beeler, Emanuel Flores-Bautista, Nicholas S McCarty, Tom Röschinger, Nathan M Belliveau, Michael J Sweredoski, Annie Moradian, Justin B Kinney, and Rob Phillips. Deciphering the regulatory genome of Escherichia coli, one hundred promoters at a time. *eLife*, 9, 2020.

[7353] Kenichi Ishii, Margot Wohl, Andre DeSouza, and Kenta Asahina. Sex-determining genes distinctly regulate courtship capability and target preference via sexually dimorphic neurons. *eLife*, 9, 2020.

[7354] Hiroki Ito, Anna C Sales, Christopher H Fry, Anthony J Kanai, Marcus J Drake, and Anthony E Pickering. Probabilistic, spinally-gated control of bladder pressure and autonomous micturition by Barrington's nucleus CRH neurons. *eLife*, 9, 2020.

[7355] Sho Ito and Hiroaki Gomi. Visually-updated hand state estimates modulate the proprioceptive reflex independently of motor task requirements. *eLife*, 9, 2020.

[7356] Anna A Ivanova, Shashank Srikant, Yotaro Sueoka, Hope H Kean, Riva Dhamala, Una-May O'Reilly, Marina U Bers, and Evelina Fedorenko. Comprehension of computer code relies primarily on domain-general executive brain regions. *eLife*, 9, 2020.

[7357] Vasyl Ivashov, Johannes Zimmer, Sinead Schwabl, Jennifer Kahlhofer, Sabine Weys, Ronald Gstir, Thomas Jakschitz, Leopold Kremser, Günther K Bonn, Herbert Lindner, Lukas A Huber, Sebastien Leon, Oliver Schmidt, and David Teis. Complementary $\alpha$-arrestin-ubiquitin ligase complexes control nutrient transporter endocytosis in response to amino acids. *eLife*, 9, 2020.

[7358] Skyler L Jackman, Christopher H Chen, Heather L Offermann, Iain R Drew, Bailey M Harrison, Anna M Bowman, Katelyn M Flick, Isabella Flaquer, and Wade G Regehr. Cerebellar Purkinje cell activity modulates aggressive behavior. *eLife*, 9, 2020.

[7359] Christopher A Jackson, Dayanne M Castro, Giuseppe-Antonio Saldi, Richard Bonneau, and David Gresham. Gene regulatory network reconstruction using single-cell RNA sequencing of barcoded genotypes in diverse environments. *eLife*, 9, 2020.

[7360] Heidi IL Jacobs, Nikos Priovoulos, Benedikt A Poser, Linda HG Pagen, Dimo Ivanov, Frans RJ Verhey, and Kâmil Uludağ. Dynamic behavior of the locus coeruleus during arousal-related memory processing in a multi-modal 7T fMRI paradigm. *eLife*, 9, 2020.

[7361] Paul I Jaffe and Michael S Brainard. Acetylcholine acts on songbird premotor circuitry to invigorate vocal output. *eLife*, 9, 2020.

[7362] Anjali Jaiman and Mukund Thattai. Golgi compartments enable controlled biomolecular assembly using promiscuous enzymes. *eLife*, 9, 2020.

[7363] Payal Jain, Cecilia Ballare, Enrique Blanco, Pedro Vizan, and Luciano Di Croce. PHF19 mediated regulation of proliferation and invasiveness in prostate cancer cells. *eLife*, 9, 2020.

[7364] Varsha Jain, Benjamin L Murphy-Baum, Geoff deRosenroll, Santhosh Sethuramanujam, Mike Delsey, Kerry R Delaney, and Gautam Bhagwan Awatramani. The functional organization of excitation and inhibition in the dendrites of mouse direction-selective ganglion cells. *eLife*, 9, 2020.

[7365] Adam SB Jalal, Ngat T Tran, and Tung BK Le. ParB spreading on DNA requires cytidine triphosphate in vitro. *eLife*, 9, 2020.

[7366] Ankit Jambusaria, Zhigang Hong, Lianghui Zhang, Shubhi Srivastava, Arundhati Jana, Peter T Toth, Yang Dai, Asrar B Malik, and Jalees Rehman. Endothelial heterogeneity across distinct vascular beds during homeostasis and inflammation. *eLife*, 9, 2020.

[7367] Sebastian S James, Leah A Krubitzer, and Stuart P Wilson. Modelling the emergence of whisker barrels. *eLife*, 9, 2020.

[7368] Sumitash Jana, Ricci Hannah, Vignesh Muralidharan, and Adam R Aron. Temporal cascade of frontal, motor and muscle processes underlying human action-stopping. *eLife*, 9, 2020.

[7369] Matt J Jaremko, Kin Fan On, Dennis R Thomas, Bruce Stillman, and Leemor Joshua-Tor. The dynamic nature of the human origin recognition complex revealed through five cryoEM structures. *eLife*, 9, 2020.

[7370] Abbas Jariani, Lieselotte Vermeersch, Bram Cerulus, Gemma Perez-Samper, Karin Voordeckers, Thomas Van Brussel, Bernard Thienpont, Diether Lambrechts, and Kevin J Verstrepen. A new protocol for single-cell RNA-seq reveals stochastic gene expression during lag phase in budding yeast. *eLife*, 9, 2020.

[7371] Inga Jarmoskaite, Ishraq AlSadhan, Pavanapuresan P Vaidyanathan, and Daniel Herschlag. How to measure and evaluate binding affinities. *eLife*, 9, 2020.

[7372] Heledd H Jarosz-Griffiths, Thomas Scambler, Chi H Wong, Samuel Lara-Reyna, Jonathan Holbrook, Fabio Martinon, Sinisa Savic, Paul Whitaker, Christine Etherington, Giulia Spoletini, Ian Clifton, Anil Mehta, Michael F McDermott, and Daniel Peckham. Different CFTR modulator combinations downregulate inflammation differently in cystic fibrosis. *eLife*, 9, 2020.

[7373] Kyle Jasmin, Frederic Dick, Lauren Stewart, and Adam Taylor Tierney. Altered functional connectivity during speech perception in congenital amusia. *eLife*, 9, 2020.

[7374] Nicole L Jenkins, Simon A James, Agus Salim, Fransisca Sumardy, Terence P Speed, Marcus Conrad, Des R Richardson, Ashley I Bush, and Gawain McColl. Changes in ferrous iron and glutathione promote ferroptosis and frailty in aging Caenorhabditis elegans. *eLife*, 9, 2020.

[7375] Isaac J Jensen, Samantha N Jensen, Frances V Sjaastad, Katherine N Gibson-Corley, Thamothrampillai Dileepan, Thomas S Griffith, Ashutosh K Mangalam, and Vladimir P Badovinac. Sepsis impedes EAE disease development and diminishes autoantigen-specific naive CD4 T cells. *eLife*, 9, 2020.

[7376] Elizabeth R Jerison and Stephen R Quake. Heterogeneous T cell motility behaviors emerge from a coupling between speed and turning in vivo. *eLife*, 9, 2020.

[7377] Steven Jack Jerjian, Maneesh Sahani, and Alexander Kraskov. Movement initiation and grasp representation in premotor and primary motor cortex mirror neurons. *eLife*, 9, 2020.

[7378] Shan Jiang, Johan Bourghardt Fagman, Changyan Chen, Simon Alberti, and Beidong Liu. Protein phase separation and its role in tumorigenesis. *eLife*, 9, 2020.

[7379] Yihao Jiang, Vinay Idikuda, Sandipan Chowdhury, and Baron Chanda. Activation of the archaeal ion channel MthK is exquisitely regulated by temperature. *eLife*, 9, 2020.

[7380] Haojie Jin, Siying Wang, Esther A Zaal, Cun Wang, Haiqiu Wu, Astrid Bosma, Fleur Jochems, Nikita Isima, Guangzhi Jin, Cor Lieftink, Roderick Beijersbergen, Celia R Berkers, Wenxin Qin, and Rene Bernards. A powerful drug combination strategy targeting glutamine addiction for the treatment of human liver cancer. *eLife*, 9, 2020.

[7381] Peiqing Jin, Yuhan Lu, and Nai Ding. Low-frequency neural activity reflects rule-based chunking during speech listening. *eLife*, 9, 2020.

[7382] Youngah Jo, Steven S Kim, Kristina Garland, Iris Fuentes, Lisa M DiCarlo, Jessie L Ellis, Xueyan Fu, Sarah L Booth, Bret M Evers, and Russell A DeBose-Boyd. Enhanced ER-associated degradation of HMG CoA reductase causes embryonic lethality associated with Ubiad1 deficiency. *eLife*, 9, 2020.

[7383] Angelika Jochim, Lea Adolf, Darya Belikova, Nadine Anna Schilling, Inda Setyawati, Denny Chin, Severien Meyers, Peter Verhamme, David E Heinrichs, Dirk J Slotboom, and Simon Heilbronner. An ECF-type transporter scavenges heme to overcome iron-limitation in Staphylococcus lugdunensis. *eLife*, 9, 2020.

[7384] Courtney R Johnson, Marc G Steingesser, Andrew D Weems, Anum Khan, Amy Gladfelter, Aurélie Bertin, and Michael A McMurray. Guanidine hydrochloride reactivates an ancient septin hetero-oligomer assembly pathway in budding yeast. *eLife*, 9, 2020.

[7385] Matthew C Johnson and Justin M Kollman. Cryo-EM structures demonstrate human IMPDH2 filament assembly tunes allosteric regulation. *eLife*, 9, 2020.

[7386] Sean L Johnson, Bedri Ranxhi, Kozeta Libohova, Wei-Ling Tsou, and Sokol V Todi. Ubiquitin-interacting motifs of ataxin-3 regulate its polyglutamine toxicity through Hsc70-4-dependent aggregation. *eLife*, 9, 2020.

[7387] Calum HG Johnston, Anne-Lise Soulet, Matthieu Bergé, Marc Prudhomme, David De Lemos, and Patrice Polard. The alternative sigma factor $\sigma$X mediates competence shut-off at the cell pole in Streptococcus pneumoniae. *eLife*, 9, 2020.

[7388] Beryl M Jones, Vikyath D Rao, Tim Gernat, Tobias Jagla, Amy C Cash-Ahmed, Benjamin ER Rubin, Troy J Comi, Shounak Bhogale, Syed S Husain, Charles Blatti, Martin Middendorf, Saurabh Sinha, Sriram Chandrasekaran, and Gene E Robinson. Individual differences in honey bee behavior enabled by plasticity in brain gene regulatory networks. *eLife*, 9, 2020.

[7389] Eric M Jones, Nathan B Lubock, AJ Venkatakrishnan, Jeffrey Wang, Alex M Tseng, Joseph M Paggi, Naomi R Latorraca, Daniel Cancilla, Megan Satyadi, Jessica E Davis, M Madan Babu, Ron O Dror, and Sriram Kosuri. Structural and functional characterization of G protein–coupled receptors with deep mutational scanning. *eLife*, 9, 2020.

[7390] Jessica M Jones, William Foster, Colin R Twomey, Justin Burdge, Osama M Ahmed, Talmo D Pereira, Jessica A Wojick, Gregory Corder, Joshua B Plotkin, and Ishmail Abdus-Saboor. A machine-vision approach for automated pain measurement at millisecond timescales. *eLife*, 9, 2020.

[7391] Nick K Jones, Lucy Rivett, Dominic Sparkes, Sally Forrest, Sushmita Sridhar, Jamie Young, Joana Pereira-Dias, Claire Cormie, Harmeet Gill, Nicola Reynolds, Michelle Wantoch, Matthew Routledge, Ben Warne, Jack Levy, William David Córdova Jiménez, Fathima Nisha Begum Samad, Chris McNicholas, Mark Ferris, Jane Gray, Michael Gill, Stephen Baker, Stephen Baker, John Bradley, Gordon Dougan, Ian Goodfellow, Ravi Gupta, Paul J Lehner, Paul A Lyons, Nicholas J Matheson, Kenneth GC Smith, M Estee Torok, Mark Toshner, Michael P Weekes, Nicholas K Jones, Lucy Rivett, Matthew Routledge, Dominic Sparkes, Ben Warne, Claire Cormie, Sally Forrest, Harmeet Gill, Iain Kean, Joana Pereira-Dias, Nicola Reynolds, Sushmita Sridhar, Michelle Wantoch, Jamie Young, Sarah Caddy, Laura Caller, Theresa Feltwell, Grant Hall, William Hamilton, Myra Hosmillo, Charlotte Houldcroft, Aminu Jahun, Fahad Khokhar, Luke Meredith, Anna Yakovleva, Helen Butcher, Daniela Caputo, Debra Clapham-Riley, Helen Dolling, Anita Furlong, Barbara Graves, Emma Le Gresley, Nathalie Kingston, Sofia Papadia, Hannah Stark, Kathleen E Stirrups, Jennifer Webster, Joanna Calder, Julie Harris, Sarah Hewitt, Jane Kennet, Anne Meadows, Rebecca Rastall, Criona O Brien, Jo Price, Cherry Publico, Jane Rowlands, Valentina Ruffolo, Hugo Tordesillas, Michael Gill, Jane Gray, Greg Hannon, Karen Brookes, Laura Canna, Isabel Cruz, Katie Dempsey, Anne Elmer, Naidine Escoffery, Stewart Fuller, Heather Jones, Carla Ribeiro, Caroline Saunders, Angela Wright, Rutendo Nyagumbo, Anne Roberts, Ashlea Bucke, Simone Hargreaves, Danielle Johnson, Aileen Narcorda, Debbie Read, Christian Sparke, Lucy Worboys, Kirsty Lagadu, Lenette Mactavous, Tim Gould, Tim Raine, Ashley Shaw, Claire Mather, Nicola Ramenatte, Anne-Laure

Vallier, Mary Kasanicki, Penelope-Jane Eames, Chris McNicholas, Lisa Thake, Neil Bartholomew, Nick Brown, Martin Curran, Surendra Parmar, Hongyi Zhang, Ailsa Bowring, Mark Ferris, Geraldine Martell, Natalie Quinnell, Giles Wright, Jo Wright, Helen Murphy, Benjamin J Dunmore, Ekaterina Legchenko, Stefan Gräf, Christopher Huang, Josh Hodgson, Kelvin Hunter, Jennifer Martin, Federica Mescia, Ciara ODonnell, Linda Pointon, Joy Shih, Rachel Sutcliffe, Tobias Tilly, Zhen Tong, Carmen Treacy, Jennifer Wood, Laura Bergamaschi, Ariana Betancourt, Georgie Bowyer, Aloka De Sa, Maddie Epping, Andrew Hinch, Oisin Huhn, Isobel Jarvis, Daniel Lewis, Joe Marsden, Simon McCallum, Francescsa Nice, Ommar Omarjee, Marianne Perera, Nika Romashova, Mateusz Strezlecki, Natalia Savoinykh Yarkoni, Lori Turner, Barrie Bailey, Afzal Chaudhry, Rachel Doughton, Chris Workman, Caroline Trotter, William David, Cordova Jiménez, Jack Levy, Fatima NB Samad, Martin D Curran, Stewart Fuller, Afzal Chaudhry, Ashley Shaw, John R Bradley, Gregory J Hannon, Ian G Goodfellow, Gordon Dougan, Kenneth GC Smith, Paul J Lehner, Giles Wright, Nicholas J Matheson, Stephen Baker, and Michael P Weekes. Effective control of SARS-CoV-2 transmission between healthcare workers during a period of diminished community prevalence of COVID-19. *eLife*, 9, 2020.

[7392] Antony M Jose. The analysis of living systems can generate both knowledge and illusions. *eLife*, 9, 2020.

[7393] Aby Joseph, Colin J Chu, Guanping Feng, Kosha Dholakia, and Jesse Schallek. Label-free imaging of immune cell dynamics in the living retina using adaptive optics. *eLife*, 9, 2020.

[7394] Raji E Joseph, Neha Amatya, D Bruce Fulton, John R Engen, Thomas E Wales, and Amy Andreotti. Differential impact of BTK active site inhibitors on the conformational state of full-length BTK. *eLife*, 9, 2020.

[7395] Abidemi Junaid, Johannes Schoeman, Wei Yang, Wendy Stam, Alireza Mashaghi, Anton Jan van Zonneveld, and Thomas Hankemeier. Metabolic response of blood vessels to TNF$\alpha$. *eLife*, 9, 2020.

[7396] Da Hee Jung, Sung Min Ahn, Malk Eun Pak, Hong Ju Lee, Young Jin Jung, Ki Bong Kim, Yong-Il Shin, Hwa Kyoung Shin, and Byung Tae Choi. Therapeutic effects of anodal transcranial direct current stimulation in a rat model of ADHD. *eLife*, 9, 2020.

[7397] Szymon Juszkiewicz, Greg Slodkowicz, Zhewang Lin, Paula Freire-Pritchett, Sew-Yeu Peak-Chew, and Ramanujan S Hegde. Ribosome collisions trigger cis-acting feedback inhibition of translation initiation. *eLife*, 9, 2020.

[7398] Laurent Jutras-Dubé, Ezzat El-Sherif, and Paul FranÇois. Geometric models for robust encoding of dynamical information into embryonic patterns. *eLife*, 9, 2020.

[7399] Terese Sara Høj Jørgensen, Gunhild Tidemann Okholm, Kaare Christensen, Thorkild IA Sørensen, and Merere Osler. Body height in young adult men and risk of dementia later in adult life. *eLife*, 9, 2020.

[7400] Deepak A Kaji, Kristen L Howell, Zerina Balic, Dirk Hubmacher, and Alice H Huang. Tgf$\beta$ signaling is required for tenocyte recruitment and functional neonatal tendon regeneration. *eLife*, 9, 2020.

[7401] Anneke Dixie Kakebeen, Alexander Daniel Chitsazan, Madison Corinne Williams, Lauren M Saunders, and Andrea Elizabeth Wills. Chromatin accessibility dynamics and single cell RNA-Seq reveal new regulators of regeneration in neural progenitors. *eLife*, 9, 2020.

[7402] Ashley L Kalinski, Choya Yoon, Lucas D Huffman, Patrick C Duncker, Rafi Kohen, Ryan Passino, Hannah Hafner, Craig Johnson, Riki Kawaguchi, Kevin S Carbajal, Juan Sebastian Jara, Edmund Hollis, Daniel H Geschwind, Benjamin M Segal, and Roman J Giger. Analysis of the immune response to sciatic nerve injury identifies efferocytosis as a key mechanism of nerve debridement. *eLife*, 9, 2020.

[7403] Stephan Kamrad, María Rodríguez-López, Cristina Cotobal, Clara Correia-Melo, Markus Ralser, and Jürg Bähler. Pyphe, a python toolbox for assessing microbial growth and cell viability in high-throughput colony screens. *eLife*, 9, 2020.

[7404] Wei Kan, Michael D Enos, Elgin Korkmazhan, Stefan Muennich, Dong-Hua Chen, Melissa V Gammons, Mansi Vasishtha, Mariann Bienz, Alexander R Dunn, Georgios Skiniotis, and William I Weis. Limited dishevelled/Axin oligomerization determines efficiency of Wnt/$\beta$-catenin signal transduction. *eLife*, 9, 2020.

[7405] Gary A Kane, GonÇalo Lopes, Jonny L Saunders, Alexander Mathis, and Mackenzie W Mathis. Real-time, low-latency closed-loop feedback using markerless posture tracking. *eLife*, 9, 2020.

[7406] Joshua R Kane, Susan Fong, Jacob Shaul, Alexandra Frommlet, Andreas O Frank, Mark Knapp, Dirksen E Bussiere, Peter Kim, Elizabeth Ornelas, Carlos Cuellar, Anastasia Hyrina, Johanna R Abend, and Charles A Wartchow. A polyomavirus peptide binds to the capsid VP1 pore and has potent antiviral activity against BK and JC polyomaviruses. *eLife*, 9, 2020.

[7407] Husniye Kantarci, Yunzi Gou, and Bruce B Riley. The Warburg Effect and lactate signaling augment Fgf-MAPK to promote sensory-neural development in the otic vesicle. *eLife*, 9, 2020.

[7408] Chang-Hao Kao, Sangil Lee, Joshua I Gold, and Joseph W Kable. Neural encoding of task-dependent errors during adaptive learning. *eLife*, 9, 2020.

[7409] Iti Kapoor, Philip Olivares, and Satish K Nair. Biochemical basis for the regulation of biosynthesis of antiparasitics by bacterial hormones. *eLife*, 9, 2020.

[7410] Stoyo Karamihalev, Elena Brivio, Cornelia Flachskamm, Rainer Stoffel, Mathias V Schmidt, and Alon Chen. Social dominance mediates behavioral adaptation to chronic stress in a sex-specific manner. *eLife*, 9, 2020.

[7411] Deepti Karandur, Moitrayee Bhattacharyya, Zijie Xia, Young Kwang Lee, Serena Muratcioglu, Darren McAffee, Ethan D McSpadden, Baiyu Qiu, Jay T Groves, Evan R Williams, and John Kuriyan. Breakage of the oligomeric CaMKII hub by the regulatory segment of the kinase. *eLife*, 9, 2020.

[7412] Julia R Kardon, Jamie A Moroco, John R Engen, and Tania A Baker. Mitochondrial ClpX activates an essential biosynthetic enzyme through partial unfolding. *eLife*, 9, 2020.

[7413] Alexander Karge, Nicolle A Bonar, Scott Wood, and Christian P Petersen. tec-1 kinase negatively regulates regenerative neurogenesis in planarians. *eLife*, 9, 2020.

[7414] Dominic Kargl, Joanna Kaczanowska, Sophia Ulonska, Florian Groessl, Lukasz Piszczek, Jelena Lazovic, Katja Buehler, and Wulf Haubensak. The amygdala instructs insular feedback for affective learning. *eLife*, 9, 2020.

[7415] Ali Karimi, Jan Odenthal, Florian Drawitsch, Kevin M Boergens, and Moritz Helmstaedter. Cell-type specific innervation of cortical pyramidal cells at their apical dendrites. *eLife*, 9, 2020.

[7416] J Karimi Abadchi, Mojtaba Nazari-Ahangarkolaee, Sandra Gattas, Edgar Bermudez-Contreras, Artur Luczak, Bruce L McNaughton, and Majid H Mohajerani. Spatiotemporal patterns of neocortical activity around hippocampal sharp-wave ripples. *eLife*, 9, 2020.

[7417] Sofya A Kasatskaya, Kristin Ladell, Evgeniy S Egorov, Kelly L Miners, Alexey N Davydov, Maria Metsger, Dmitry B Staroverov, Elena K Matveyshina, Irina A Shagina, Ilgar Z Mamedov, Mark Izraelson, Pavel V Shelyakin, Olga V Britanova, David A Price, and Dmitriy M Chudakov. Functionally specialized human CD4+ T-cell subsets express physicochemically distinct TCRs. *eLife*, 9, 2020.

[7418] Magdalena Kasendra, Raymond Luc, Jianyi Yin, Dimitris V Manatakis, Gauri Kulkarni, Carolina Lucchesi, Josiah Sliz, Athanasia Apostolou, Laxmi Sunuwar, Jenifer Obrigewitch, Kyung-Jin Jang, Geraldine A Hamilton, Mark Donowitz, and Katia Karalis. Duodenum Intestine-Chip for preclinical drug assessment in a human relevant model. *eLife*, 9, 2020.

[7419] Bhavatharini Kasinathan, Serafin U Colmenares, Hannah McConnell, Janet M Young, Gary H Karpen, and Harmit S Malik. Innovation of heterochromatin functions drives rapid evolution of essential ZAD-ZNF genes in Drosophila. *eLife*, 9, 2020.

[7420] Daiki Kato, Tomonori Yaguchi, Takashi Iwata, Yuki Katoh, Kenji Morii, Kinya Tsubota, Yoshiaki Takise, Masaki Tamiya, Haruhiko Kamada, Hiroki Akiba, Kouhei Tsumoto, Satoshi Serada, Tetsuji Naka, Ryohei Nishimura, Takayuki Nakagawa, and Yutaka Kawakami. GPC1 specific CAR-T cells eradicate established solid tumor without adverse effects and synergize with anti-PD-1 Ab. *eLife*, 9, 2020.

[7421] Gurmeet Kaur, A Maxwell Burroughs, Lakshminarayan M Iyer, and L Aravind. Highly regulated, diversifying NTP-dependent biological conflict systems with implications for the emergence of multicellularity. *eLife*, 9, 2020.

[7422] Pamina Kazman, Marie-Theres Vielberg, María Daniela Pulido Cendales, Lioba Hunziger, Benedikt Weber, Ute Hegenbart, Martin Zacharias, Rolf Köhler, Stefan Schönland, Michael Groll, and Johannes Buchner. Fatal amyloid formation in a patient's antibody light chain is caused by a single point mutation. *eLife*, 9, 2020.

[7423] Yasuhiro Kazuki, Feng J Gao, Yicong Li, Anna J Moyer, Benjamin Devenney, Kei Hiramatsu, Sachiko Miyagawa-Tomita, Satoshi Abe, Kanako Kazuki, Naoyo Kajitani, Narumi Uno, Shoko Takehara, Masato Takiguchi, Miho Yamakawa, Atsushi Hasegawa, Ritsuko Shimizu, Satoko Matsukura, Naohiro Noda, Narumi Ogonuki, Kimiko Inoue, Shogo Matoba, Atsuo Ogura, Liliana D Florea, Alena Savonenko, Meifang Xiao, Dan Wu, Denise AS Batista, Junhua Yang, Zhaozhu Qiu, Nandini Singh, Joan T Richtsmeier, Takashi Takeuchi, Mitsuo Oshimura, and Roger H Reeves. A non-mosaic transchromosomic mouse model of Down syndrome carrying the long arm of human chromosome 21. *eLife*, 9, 2020.

[7424] Matthew D Keefe, Haille E Soderholm, Hung-Yu Shih, Tamara J Stevenson, Kathryn A Glaittli, D Miranda Bowles, Erika Scholl, Samuel Colby, Samer Merchant, Edward W Hsu, and Joshua L Bonkowsky. Vanishing white matter disease expression of truncated EIF2B5 activates induced stress response. *eLife*, 9, 2020.

[7425] Ian W Keesey, Veit Grabe, Markus Knaden, and Bill S Hansson. Divergent sensory investment mirrors potential speciation via niche partitioning across Drosophila. *eLife*, 9, 2020.

[7426] Anne Keitel, Joachim Gross, and Christoph Kayser. Shared and modality-specific brain regions that mediate auditory and visual word comprehension. *eLife*, 9, 2020.

[7427] Charlotte F Kelley, Thomas Litschel, Stephanie Schumacher, Dirk Dedden, Petra Schwille, and Naoko Mizuno. Phosphoinositides regulate force-independent interactions between talin, vinculin, and actin. *eLife*, 9, 2020.

[7428] Christina M Kelliher, Randy Lambreghts, Qijun Xiang, Christopher L Baker, Jennifer J Loros, and Jay C Dunlap. PRD-2 directly regulates casein kinase I and counteracts nonsense-mediated decay in the Neurospora circadian clock. *eLife*, 9, 2020.

[7429] Luke Kemp, Laura Adam, Christian R Boehm, Rainer Breitling, Rocco Casagrande, Malcolm Dando, Appolinaire Djikeng, Nicholas G Evans, Richard Hammond, Kelly Hills, Lauren A Holt, Todd Kuiken, Alemka Markotić, Piers Millett, Johnathan A Napier, Cassidy Nelson, Seán S óhéigeartaigh, Anne Osbourn, Megan J Palmer, Nicola J Patron, Edward Perello, Wibool Piyawattanametha, Vanessa Restrepo-Schild, Clarissa Rios-Rojas, Catherine Rhodes, Anna Roessing, Deborah Scott, Philip Shapira, Christopher Simuntala, Robert DJ Smith, Lalitha S Sundaram, Eriko Takano, Gwyn Uttmark, Bonnie C Wintle, Nadia B Zahra, and William J Sutherland. Bioengineering horizon scan 2020. *eLife*, 9, 2020.

[7430] Andrew S Kennard and Julie A Theriot. Osmolarity-independent electrical cues guide rapid response to injury in zebrafish epidermis. *eLife*, 9, 2020.

[7431] Rebecca Keogh, Johanna Bergmann, and Joel Pearson. Cortical excitability controls the strength of mental imagery. *eLife*, 9, 2020.

[7432] John Kerwin, Israr Khan, Elizabeth Iorns, Elizabeth Iorns, Rachel Tsui, Alexandria Denis, Nicole Perfito, and Timothy M Errington. Replication Study: A coding-independent function of gene and pseudogene mRNAs regulates tumour biology. *eLife*, 9, 2020.

[7433] Menoua Keshishian, Hassan Akbari, Bahar Khalighinejad, Jose L Herrero, Ashesh D Mehta, and Nima Mesgarani. Estimating and interpreting nonlinear receptive field of sensory neural responses with deep neural network models. *eLife*, 9, 2020.

[7434] Chun Kew, Wenming Huang, Julia Fischer, Raja Ganesan, Nirmal Robinson, and Adam Antebi. Evolutionarily conserved regulation of immunity by the splicing factor RNP-6/PUF60. *eLife*, 9, 2020.

[7435] Jeremiah Keyes, Ambhighainath Ganesan, Olivia Molinar-Inglis, Archer Hamidzadeh, Jinfan Zhang, Megan Ling, JoAnn Trejo, Andre Levchenko, and Jin Zhang. Signaling diversity enabled by Rap1-regulated plasma membrane ERK with distinct temporal dynamics. *eLife*, 9, 2020.

[7436] Arjun Khakhar, Colby G Starker, James C Chamness, Nayoung Lee, Sydney Stokke, Cecily Wang, Ryan Swanson, Furva Rizvi, Takato Imaizumi, and Daniel F Voytas. Building customizable auto-luminescent luciferase-based reporters in plants. *eLife*, 9, 2020.

[7437] Ahmad Abdelzaher Zaki Khalifa, Muneyoshi Ichikawa, Daniel Dai, Shintaroh Kubo, Corbin Steven Black, Katya Peri, Thomas S McAlear, Simon Veyron, Shun Kai Yang, Javier Vargas, Susanne Bechstedt, Jean-FranÇois Trempe, and Khanh Huy Bui. The inner junction complex of the cilia is an interaction hub that involves tubulin post-translational modifications. *eLife*, 9, 2020.

[7438] Aliaksandr Khaminets, Tal Ronnen-Oron, Maik Baldauf, Elke Meier, and Heinrich Jasper. Cohesin controls intestinal stem cell identity by maintaining association of Escargot with target promoters. *eLife*, 9, 2020.

[7439] Danish Khan, Dongju Lee, Gulcin Gulten, Anup Aggarwal, Joshua Wofford, Inna Krieger, Ashutosh Tripathi, John W Patrick, Debra M Eckert, Arthur Laganowsky, James Sacchettini, Paul Lindahl, and Vytas A Bankaitis. A Sec14-like phosphatidylinositol transfer protein paralog defines a novel class of heme-binding proteins. *eLife*, 9, 2020.

[7440] Hataf Khan, Rebecca P Sumner, Jane Rasaiyaah, Choon Ping Tan, Maria Teresa Rodriguez-Plata, Chris Van Tulleken, Douglas Fink, Lorena Zuliani-Alvarez, Lucy Thorne, David Stirling, Richard SB Milne, and Greg J Towers. HIV-1 Vpr antagonizes innate immune activation by targeting karyopherin-mediated NF-$\kappa$B/IRF3 nuclear transport. *eLife*, 9, 2020.

[7441] Mehak M Khan and Wade G Regehr. Loss of Doc2b does not influence transmission at Purkinje cell to deep nuclei synapses under physiological conditions. *eLife*, 9, 2020.

[7442] Shumayila Khan and James Gomes. Neuropathogenesis of SARS-CoV-2 infection. *eLife*, 9, 2020.

[7443] Khoosheh Khayati, Vrushank Bhatt, Zhixian Sherrie Hu, Sajid Fahumy, Xuefei Luo, and Jessie Yanxiang Guo. Autophagy compensates for Lkb1 loss to maintain adult mice homeostasis and survival. *eLife*, 9, 2020.

[7444] Kohki Kido, Satoshi Yamanaka, Shogo Nakano, Kou Motani, Souta Shinohara, Akira Nozawa, Hidetaka Kosako, Sohei Ito, and Tatsuya Sawasaki. AirID, a novel proximity biotinylation enzyme, for analysis of protein–protein interactions. *eLife*, 9, 2020.

[7445] Katrin Kierdorf, Fabian Hersperger, Jessica Sharrock, Crystal M Vincent, Pinar Ustaoglu, Jiawen Dou, Attila Gyoergy, Olaf Groß, Daria E Siekhaus, and Marc S Dionne. Muscle function and homeostasis require cytokine inhibition of AKT activity in Drosophila. *eLife*, 9, 2020.

[7446] Virapat Kieuvongngam, Paul Dominic B Olinares, Anthony Palillo, Michael L Oldham, Brian T Chait, and Jue Chen. Structural basis of substrate recognition by a polypeptide processing and secretion transporter. *eLife*, 9, 2020.

[7447] Chiho Kim, Xu-Dong Wang, and Yonghao Yu. PARP1 inhibitors trigger innate immunity via PARP1 trapping-induced DNA damage response. *eLife*, 9, 2020.

[7448] Eui Ho Kim, Matthew C Woodruff, Lilit Grigoryan, Barbara Maier, Song Hee Lee, Pratyusha Mandal, Mario Cortese, Muktha S Natrajan, Rajesh Ravindran, Huailiang Ma, Miriam Merad, Alexander D Gitlin, Edward S Mocarski, Joshy Jacob, and Bali Pulendran. Squalene emulsion-based vaccine adjuvants stimulate CD8 T cell, but not antibody responses, through a RIPK3-dependent pathway. *eLife*, 9, 2020.

[7449] Jong Wook Kim, Christian Berrios, Miju Kim, Amy E Schade, Guillaume Adelmant, Huwate Yeerna, Emily Damato, Amanda Balboni Iniguez, Laurence Florens, Michael P Washburn, Kim Stegmaier, Nathanael S Gray, Pablo Tamayo, Ole Gjoerup, Jarrod A Marto, James DeCaprio, and William C Hahn. STRIPAK directs PP2A activity toward MAP4K4 to promote oncogenic transformation of human cells. *eLife*, 9, 2020.

[7450] Joon T Kim, Stephanie M Terrell, Veronica L Li, Wei Wei, Curt R Fischer, and Jonathan Z Long. Cooperative enzymatic control of N-acyl amino acids by PM20D1 and FAAH. *eLife*, 9, 2020.

[7451] Kyungdeok Kim, Wangyong Shin, Muwon Kang, Suho Lee, Doyoun Kim, Ryeonghwa Kang, Yewon Jung, Yisul Cho, Esther Yang, Hyun Kim, Yong Chul Bae, and Eunjoon Kim. Presynaptic PTP$\sigma$ regulates postsynaptic NMDA receptor function through direct adhesion-independent mechanisms. *eLife*, 9, 2020.

[7452] Ok-Hee Kim, Carmen J Booth, Han Seok Choi, Jinwook Lee, Jinku Kang, June Hur, Woo Jin Jung, Yun-Shin Jung, Hyung Jin Choi, Hyeonjin Kim, Joong-Hyuck Auh, Jung-Wan Kim, Ji-Young Cha, Young Jae Lee, Cheol Soon Lee, Cheolsoo Choi, Yun Jae Jung, Jun-Young Yang, Seung-Soon Im, Dae Ho Lee, Sun Wook Cho, Young-Bum Kim, Kyong Soo Park, Young Joo Park, and Byung-Chul Oh. High-phytate/low-calcium diet is a risk factor for crystal nephropathies, renal phosphate wasting, and bone loss. *eLife*, 9, 2020.

[7453] Sora Kim, Kristin L Zuromski, Tristan A Bell, Robert T Sauer, and Tania A Baker. ClpAP proteolysis does not require rotation of the ClpA unfoldase relative to ClpP. *eLife*, 9, 2020.

[7454] Benedict King and Martin Rücklin. A Bayesian approach to dynamic homology of morphological characters and the ancestral phenotype of jawed vertebrates. *eLife*, 9, 2020.

[7455] Dana M King, Clarice Kit Yee Hong, James L Shepherdson, David M Granas, Brett B Maricque, and Barak A Cohen. Synthetic and genomic regulatory elements reveal aspects of cis-regulatory grammar in mouse embryonic stem cells. *eLife*, 9, 2020.

[7456] Amina A Kinkhabwala, Yi Gu, Dmitriy Aronov, and David W Tank. Visual cue-related activity of cells in the medial entorhinal cortex during navigation in virtual reality. *eLife*, 9, 2020.

[7457] Atsuko Kinoshita, Alice Vayssières, René Richter, Qing Sang, Adrian Roggen, Annabel D van Driel, Richard S Smith, and George Coupland. Regulation of shoot meristem shape by photoperiodic signaling and phytohormones during floral induction of Arabidopsis. *eLife*, 9, 2020.

[7458] Grant Kinsler, Kerry Geiler-Samerotte, and Dmitri A Petrov. Fitness variation across subtle environmental perturbations reveals local modularity and global pleiotropy of adaptation. *eLife*, 9, 2020.

[7459] Reiko Kintaka, Koji Makanae, Shotaro Namba, Hisaaki Kato, Keiji Kito, Shinsuke Ohnuki, Yoshikazu Ohya, Brenda J Andrews, Charles Boone, and Hisao Moriya. Genetic profiling of protein burden and nuclear export overload. *eLife*, 9, 2020.

[7460] Paul C Kirchberger and Howard Ochman. Resurrection of a global, metagenomically defined gokushovirus. *eLife*, 9, 2020.

[7461] Louise P Kirsch, Sahba Besharati, Christina Papadaki, Laura Crucianelli, Sara Bertagnoli, Nick Ward, Valentina Moro, Paul M Jenkinson, and Aikaterini Fotopoulou. Damage to the right insula disrupts the perception of affective touch. *eLife*, 9, 2020.

[7462] Jun-ichi Kishikawa, Atsuko Nakanishi, Aya Furuta, Takayuki Kato, Keiichi Namba, Masatada Tamakoshi, Kaoru Mitsuoka, and Ken Yokoyama. Mechanical inhibition of isolated Vo from V/A-ATPase for proton conductance. *eLife*, 9, 2020.

[7463] Nami Kitajima, Kenji Takikawa, Hiroshi Sekiya, Kaname Satoh, Daisuke Asanuma, Hirokazu Sakamoto, Shodai Takahashi, Kenjiro Hanaoka, Yasuteru Urano, Shigeyuki Namiki, Masamitsu Iino, and Kenzo Hirose. Real-time in vivo imaging of extracellular ATP in the brain with a hybrid-type fluorescent sensor. *eLife*, 9, 2020.

[7464] Amrutha Kizhedathu, Rose Sebastian Kunnappallil, Archit V Bagul, Puja Verma, and Arjun Guha. Multiple Wnts act synergistically to induce Chk1/Grapes expression and mediate G2 arrest in Drosophila tracheoblasts. *eLife*, 9, 2020.

[7465] PJ Klasse and John P Moore. Antibodies to SARS-CoV-2 and their potential for therapeutic passive immunization. *eLife*, 9, 2020.

[7466] Purnima Klingauf-Nerurkar, Ludovic C Gillet, Daniela Portugal-Calisto, Michaela Oborská-Oplová, Martin Jäger, Olga T Schubert, Agnese Pisano, Cohue Peña, Sanjana Rao, Martin Altvater, Yiming Chang, Ruedi Aebersold, and Vikram G Panse. The GTPase Nog1 co-ordinates the assembly, maturation and quality control of distant ribosomal functional centers. *eLife*, 9, 2020.

[7467] Björn Udo Klink, Christos Gatsogiannis, Oliver Hofnagel, Alfred Wittinghofer, and Stefan Raunser. Structure of the human BBSome core complex. *eLife*, 9, 2020.

[7468] Niels A Kloosterman, Julian Q Kosciessa, Ulman Lindenberger, Johannes Jacobus Fahrenfort, and Douglas D Garrett. Boosts in brain signal variability track liberal shifts in decision bias. *eLife*, 9, 2020.

[7469] Dennis Klug, Sarah Goellner, Jessica Kehrer, Julia Sattler, Léanne Strauss, Mirko Singer, Chafen Lu, Timothy A Springer, and Friedrich Frischknecht. Evolutionarily distant I domains can functionally replace the essential ligand-binding domain of Plasmodium TRAP. *eLife*, 9, 2020.

[7470] Elizabeth M Knapp, Wei Li, Vijender Singh, and Jianjun Sun. Nuclear receptor Ftz-f1 promotes follicle maturation and ovulation partly via bHLH/PAS transcription factor Sim. *eLife*, 9, 2020.

[7471] Athene Knüfer, Giovanni Diana, Gregory S Walsh, Jonathan DW Clarke, and Sarah Guthrie. Cadherins regulate nuclear topography and function of developing ocular motor circuitry. *eLife*, 9, 2020.

[7472] Janus RL Kobbersmed, Andreas T Grasskamp, Meida Jusyte, Mathias A Böhme, Susanne Ditlevsen, Jakob Balslev Sørensen, and Alexander M Walter. Rapid regulation of vesicle priming explains synaptic facilitation despite heterogeneous vesicle:Ca2+ channel distances. *eLife*, 9, 2020.

[7473] Robert Köchl, Lesley Vanes, Miriam Llorian Sopena, Probir Chakravarty, Harald Hartweger, Kathryn Fountain, Andrea White, Jennifer Cowan, Graham Anderson, and Victor LJ Tybulewicz. Critical role of WNK1 in MYC-dependent early mouse thymocyte development. *eLife*, 9, 2020.

[7474] Liis Kolberg, Nurlan Kerimov, Hedi Peterson, and Kaur Alasoo. Co-expression analysis reveals interpretable gene modules controlled by trans-acting genetic variants. *eLife*, 9, 2020.

[7475] Ljuvica R Kolich, Ya-Ting Chang, Nicolas Coudray, Sabrina I Giacometti, Mark R MacRae, Georgia L Isom, Evelyn M Teran, Gira Bhabha, and Damian C Ekiert. Structure of MlaFB uncovers novel mechanisms of ABC transporter regulation. *eLife*, 9, 2020.

[7476] Sebastian Korb, Sebastian J Götzendorfer, Claudia Massaccesi, Patrick Sezen, Irene Graf, Matthäus Willeit, Christoph Eisenegger, and Giorgia Silani. Dopaminergic and opioidergic regulation during anticipation and consumption of social and nonsocial rewards. *eLife*, 9, 2020.

[7477] Ilia Korobko, Hisham Mazal, Gilad Haran, and Amnon Horovitz. Measuring protein stability in the GroEL chaperonin cage reveals massive destabilization. *eLife*, 9, 2020.

[7478] Theodora Koromila, Fan Gao, Yasuno Iwasaki, Peng He, Lior Pachter, J Peter Gergen, and Angelike Stathopoulos. Odd-paired is a pioneer-like factor that coordinates with Zelda to control gene expression in embryos. *eLife*, 9, 2020.

[7479] Walter J Koroshetz, Shannon Behrman, Cynthia J Brame, Janet L Branchaw, Emery N Brown, Erin A Clark, David Dockterman, Jordan J Elm, Pamela L Gay, Katelyn M Green, Sherry Hsi, Michael G Kaplitt, Benedict J Kolber, Alex L Kolodkin, Diane Lipscombe, Malcolm R MacLeod, Caleb C McKinney, Marcus R Munafò, Barbara Oakley, Jeffrey T Olimpo, Nathalie Percie du Sert, Indira M Raman, Ceri Riley, Amy L Shelton, Stephen Miles Uzzo, Devon C Crawford, and Shai D Silberberg. Framework for advancing rigorous research. *eLife*, 9, 2020.

[7480] Isabelle C Kos-Braun, Björn Gerlach, and Claudia Pitzer. A survey of research quality in core facilities. *eLife*, 9, 2020.

[7481] Liang Kou, Lei Jiang, Stephan Hättenschwiler, Miaomiao Zhang, Shuli Niu, Xiaoli Fu, Xiaoqin Dai, Han Yan, Shenggong Li, and Huimin Wang. Diversity-decomposition relationships in forests worldwide. *eLife*, 9, 2020.

[7482] Marina Kovalenko, Serkan Erdin, Marissa A Andrew, Jason St Claire, Melissa Shaughnessey, Leroy Hubert, João Luís Neto, Alexei Stortchevoi, Daniel M Fass, Ricardo Mouro Pinto, Stephen J Haggarty, John H Wilson, Michael E Talkowski, and Vanessa C Wheeler. Histone deacetylase knockouts modify transcription, CAG instability and nuclear pathology in Huntington disease mice. *eLife*, 9, 2020.

[7483] Adrian Kovač, Csaba Miskey, Michael Menzel, Esther Grueso, Andreas Gogol-Döring, and Zoltán Ivics. RNA-guided retargeting of Sleeping Beauty transposition in human cells. *eLife*, 9, 2020.

[7484] Amanda Kowalczyk, Raghavendran Partha, Nathan L Clark, and Maria Chikina. Pan-mammalian analysis of molecular constraints underlying extended lifespan. *eLife*, 9, 2020.

[7485] Iryna Kozmikova and Zbynek Kozmik. Wnt/$\beta$-catenin signaling is an evolutionarily conserved determinant of chordate dorsal organizer. *eLife*, 9, 2020.

[7486] James E Kragel, Stephen VanHaerents, Jessica W Templer, Stephan Schuele, Joshua M Rosenow, Aneesha S Nilakantan, and Donna J Bridge. Hippocampal theta coordinates memory processing during visual exploration. *eLife*, 9, 2020.

[7487] Paul F Kramer, Emily L Twedell, Jung Hoon Shin, Renshu Zhang, and Zayd M Khaliq. Axonal mechanisms mediating $\gamma$-aminobutyric acid receptor type A (GABA-A) inhibition of striatal dopamine release. *eLife*, 9, 2020.

[7488] Robert Krautz, Dilan Khalili, and Ulrich Theopold. Tissue-autonomous immune response regulates stress signaling during hypertrophy. *eLife*, 9, 2020.

[7489] Alex J B Kreutzberger, Volker Kiessling, Catherine A Doyle, Noah Schenk, Clint M Upchurch, Margaret Elmer-Dixon, Amanda E Ward, Julia Preobraschenski, Syed S Hussein, Weronika Tomaka, Patrick Seelheim, Iman Kattan, Megan Harris, Binyong Liang, Anne K Kenworthy, Bimal N Desai, Norbert Leitinger, Arun Anantharam, J David Castle, and Lukas K Tamm. Distinct insulin granule subpopulations implicated in the secretory pathology of diabetes types 1 and 2. *eLife*, 9, 2020.

[7490] Ilja L Kruglikov, Manasi Shah, and Philipp E Scherer. Obesity and diabetes as comorbidities for COVID-19: Underlying mechanisms and the role of viral–bacterial interactions. *eLife*, 9, 2020.

[7491] Karen Krukowski, Amber Nolan, Elma S Frias, Morgane Boone, Gonzalo Ureta, Katherine Grue, Maria-Serena Paladini, Edward Elizarraras, Luz Delgado, Sebastian Bernales, Peter Walter, and Susanna Rosi. Small molecule cognitive enhancer reverses age-related memory decline in mice. *eLife*, 9, 2020.

[7492] Piotr Krzywkowski, Beatrice Penna, and Cornelius T Gross. Dynamic encoding of social threat and spatial context in the hypothalamus. *eLife*, 9, 2020.

[7493] Yi Kuang, Ohad Golan, Kristina Preusse, Brittany Cain, Collin J Christensen, Joseph Salomone, Ian Campbell, FearGod V Okwubido-Williams, Matthew R Hass, Zhenyu Yuan, Nathanel Eafergan, Kenneth H Moberg, Rhett A Kovall, Raphael Kopan, David Sprinzak, and Brian Gebelein. Enhancer architecture sensitizes cell specific responses to Notch gene dose via a bind and discard mechanism. *eLife*, 9, 2020.

[7494] Iga Kucharska, Elaine Thai, Ananya Srivastava, John L Rubinstein, Régis Pomès, and Jean-Philippe Julien. Structural ordering of the Plasmodium berghei circumsporozoite protein repeats by inhibitory antibody 3D11. *eLife*, 9, 2020.

[7495] Erika E Kuchen, Nils B Becker, Nina Claudino, and Thomas Höfer. Hidden long-range memories of growth and cycle speed correlate cell cycles in lineage trees. *eLife*, 9, 2020.

[7496] Georg Kuenze, Carlos G Vanoye, Reshma R Desai, Sneha Adusumilli, Kathryn R Brewer, Hope Woods, Eli F McDonald, Charles R Sanders, Alfred L George, and Jens Meiler. Allosteric mechanism for KCNE1 modulation of KCNQ1 potassium channel activation. *eLife*, 9, 2020.

[7497] Michael Kühl, Erik Trampe, Maria Mosshammer, Michael Johnson, Anthony WD Larkum, Niels-Ulrik Frigaard, and Klaus Koren. Substantial near-infrared radiation-driven photosynthesis of chlorophyll f-containing cyanobacteria in a natural habitat. *eLife*, 9, 2020.

[7498] André Kuhn, Sigurd Ramans Harborough, Heather M McLaughlin, Bhavani Natarajan, Inge Verstraeten, Jiří Friml, Stefan Kepinski, and Lars Østergaard. Direct ETTIN-auxin interaction controls chromatin states in gynoecium development. *eLife*, 9, 2020.

[7499] Laura E Kuil, Nynke Oosterhof, Giuliano Ferrero, Tereza MikuláŠová, Martina Hason, Jordy Dekker, Mireia Rovira, Herma C van der Linde, Paulina MH van Strien, Emma de Pater, Gerben Schaaf, Erik MJ Bindels, Valerie Wittamer, and Tjakko J van Ham. Zebrafish macrophage developmental arrest underlies depletion of microglia and reveals Csf1r-independent metaphocytes. *eLife*, 9, 2020.

[7500] Chaitanya A Kulkarni, Sergiy M Nadtochiy, Leslie Kennedy, Jimmy Zhang, Sophea Chhim, Hanan Alwaseem, Elizabeth Murphy, Dragony Fu, and Paul S Brookes. ALKBH7 mediates necrosis via rewiring of glyoxal metabolism. *eLife*, 9, 2020.

[7501] Alok Kumar, Kenji Chamoto, Partha S Chowdhury, and Tasuku Honjo. Tumors attenuating the mitochondrial activity in T cells escape from PD-1 blockade therapy. *eLife*, 9, 2020.

[7502] Ashutosh Kumar, Litao Xie, Chau My Ta, Antentor O Hinton, Susheel K Gunasekar, Rachel A Minerath, Karen Shen, Joshua M Maurer, Chad E Grueter, E Dale Abel, Gretchen Meyer, and Rajan Sah. SWELL1 regulates skeletal muscle cell size, intracellular signaling, adiposity and glucose metabolism. *eLife*, 9, 2020.

[7503] Tarun Kumar, Leo Blondel, and Cassandra G Extavour. Topology-driven protein-protein interaction network analysis detects genetic sub-networks regulating reproductive capacity. *eLife*, 9, 2020.

[7504] Sonya Kumar Bharathkar, Benjamin W Parker, Andrey G Malyutin, Nandan Haloi, Kathryn E Huey-Tubman, Emad Tajkhorshid, and Beth M Stadtmueller. The structures of secretory and dimeric immunoglobulin A. *eLife*, 9, 2020.

[7505] Santosh Kumar Kuncha, Vinitha Lakshmi Venkadasamy, Gurumoorthy Amudhan, Priyanka Dahate, Sankara Rao Kola, Sambhavi Pottabathini, Shobha P Kruparani, P Chandra Shekar, and Rajan Sankaranarayanan. Genomic innovation of ATD alleviates mistranslation associated with multicellularity in Animalia. *eLife*, 9, 2020.

[7506] Jun Kurushima, Nathalie Campo, Renske van Raaphorst, Guillaume Cerckel, Patrice Polard, and Jan-Willem Veening. Unbiased homeologous recombination during pneumococcal transformation allows for multiple chromosomal integration events. *eLife*, 9, 2020.

[7507] Masaru Kuwabara, Ningdong Kang, Timothy E Holy, and Camillo Padoa-Schioppa. Neural mechanisms of economic choices in mice. *eLife*, 9, 2020.

[7508] Akela Kuwahara, Ace E Lewis, Coohleen Coombes, Fang-Shiuan Leung, Michelle Percharde, and Jeffrey O Bush. Delineating the early transcriptional specification of the mammalian trachea and esophagus. *eLife*, 9, 2020.

[7509] Chiara La Rosa, Francesca Cavallo, Alessandra Pecora, Matteo Chincarini, Ugo Ala, Chris G Faulkes, Juan Nacher, Bruno Cozzi, Chet C Sherwood, Irmgard Amrein, and Luca Bonfanti. Phylogenetic variation in cortical layer II immature neuron reservoir of mammals. *eLife*, 9, 2020.

[7510] Loïc Labache, Bernard Mazoyer, Marc Joliot, Fabrice Crivello, Isabelle Hesling, and Nathalie Tzourio-Mazoyer. Typical and atypical language brain organization based on intrinsic connectivity and multitask functional asymmetries. *eLife*, 9, 2020.

[7511] Thibaut Laboute, Jorge Gandía, Lucie P Pellissier, Yannick Corde, Florian Rebeillard, Maria Gallo, Christophe Gauthier, Audrey Léauté, Jorge Diaz, Anne Poupon, Brigitte L Kieffer, Julie Le Merrer, and Jérôme AJ Becker. The orphan receptor GPR88 blunts the signaling of opioid receptors and multiple striatal GPCRs. *eLife*, 9, 2020.

[7512] Haluk Lacin, W Ryan Williamson, Gwyneth M Card, James B Skeath, and James W Truman. Unc-4 acts to promote neuronal identity and development of the take-off circuit in the Drosophila CNS. *eLife*, 9, 2020.

[7513] Mark S Ladinsky, Priyanthi NP Gnanapragasam, Zhi Yang, Anthony P West, Michael S Kay, and Pamela J Bjorkman. Electron tomography visualization of HIV-1 fusion with target cells using fusion inhibitors to trap the pre-hairpin intermediate. *eLife*, 9, 2020.

[7514] Juan Lafuente-Barquero, Maria Luisa García-Rubio, Marta San Martin-Alonso, Belén Gómez-González, and Andrés Aguilera. Harmful DNA:RNA hybrids are formed in cis and in a Rad51-independent manner. *eLife*, 9, 2020.

[7515] Vanessa Lage-Rupprecht, Li Zhou, Gaia Bianchini, S Sara Aghvami, Max Mueller, Balázs Rózsa, Marco Sassoè-Pognetto, and Veronica Egger. Presynaptic NMDARs cooperate with local spikes toward GABA release from the reciprocal olfactory bulb granule cell spine. *eLife*, 9, 2020.

[7516] Konstantinos Lagogiannis, Giovanni Diana, and Martin P Meyer. Learning steers the ontogeny of an efficient hunting sequence in zebrafish larvae. *eLife*, 9, 2020.

[7517] Mi Lai, Dana Al Rijjal, Hannes L Röst, Feihan F Dai, Erica P Gunderson, and Michael B Wheeler. Underlying dyslipidemia postpartum in women with a recent GDM pregnancy who develop type 2 diabetes. *eLife*, 9, 2020.

[7518] Armin Lak, Emily Hueske, Junya Hirokawa, Paul Masset, Torben Ott, Anne E Urai, Tobias H Donner, Matteo Carandini, Susumu Tonegawa, Naoshige Uchida, and Adam Kepecs. Reinforcement biases subsequent perceptual decisions when confidence is low, a widespread behavioral phenomenon. *eLife*, 9, 2020.

[7519] Mable Lam, Scot A Marsters, Avi Ashkenazi, and Peter Walter. Misfolded proteins bind and activate death receptor 5 to trigger apoptosis during unresolved endoplasmic reticulum stress. *eLife*, 9, 2020.

[7520] Assala Lamaa, Jonathan Humbert, Marion Aguirrebengoa, Xue Cheng, Estelle Nicolas, Jacques Côté, and Didier Trouche. Integrated analysis of H2A.Z isoforms function reveals a complex interplay in gene regulation. *eLife*, 9, 2020.

[7521] Ismael Lamas-Toranzo, Julieta G Hamze, Enrica Bianchi, Beatriz Fernández-Fuertes, Serafín Pérez-Cerezales, Ricardo Laguna-Barraza, Raúl Fernández-González, Pat Lonergan, Alfonso Gutiérrez-Adán, Gavin J Wright, María Jiménez-Movilla, and Pablo Bermejo-álvarez. TMEM95 is a sperm membrane protein essential for mammalian fertilization. *eLife*, 9, 2020.

[7522] W Marcus Lambert, Martin T Wells, Matthew F Cipriano, Jacob N Sneva, Juanita A Morris, and Linnie M Golightly. Career choices of underrepresented and female postdocs in the biomedical sciences. *eLife*, 9, 2020.

[7523] Shawn M Lamothe, Nazlee Sharmin, Grace Silver, Motoyasu Satou, Yubin Hao, Toru Tateno, Victoria A Baronas, and Harley T Kurata. Control of Slc7a5 sensitivity by the voltage-sensing domain of Kv1 channels. *eLife*, 9, 2020.

[7524] Lars Langemeyer, Ann-Christin Borchers, Eric Herrmann, Nadia Füllbrunn, Yaping Han, Angela Perz, Kathrin Auffarth, Daniel Kümmel, and Christian Ungermann. A conserved and regulated mechanism drives endosomal Rab transition. *eLife*, 9, 2020.

[7525] Lukas M Langer, Yair Gat, Fabien Bonneau, and Elena Conti. Structure of substrate-bound SMG1-8-9 kinase complex reveals molecular basis for phosphorylation specificity. *eLife*, 9, 2020.

[7526] Anthony B Lapsansky, Daniel Zatz, and Bret W Tobalske. Alcids 'fly' at efficient Strouhal numbers in both air and water but vary stroke velocity and angle. *eLife*, 9, 2020.

[7527] Hilal A Lashuel. What about faculty? *eLife*, 9, 2020.

[7528] Allison N Lau, Zhaoqi Li, Laura V Danai, Anna M Westermark, Alicia M Darnell, Raphael Ferreira, Vasilena Gocheva, Sharanya Sivanand, Evan C Lien, Kiera M Sapp, Jared R Mayers, Giulia Biffi, Christopher R Chin, Shawn M Davidson, David A Tuveson, Tyler Jacks, Nicholas J Matheson, Omer Yilmaz, and Matthew G Vander Heiden. Dissecting cell-type-specific metabolism in pancreatic ductal adenocarcinoma. *eLife*, 9, 2020.

[7529] Tatiana Lau, Samuel J Gershman, and Mina Cikara. Social structure learning in human anterior insula. *eLife*, 9, 2020.

[7530] Matthew D Lauver, Daniel J Goetschius, Colleen S Netherby-Winslow, Katelyn N Ayers, Ge Jin, Daniel G Haas, Elizabeth L Frost, Sung Hyun Cho, Carol M Bator, Stephanie M Bywaters, Neil D Christensen, Susan L Hafenstein, and Aron E Lukacher. Antibody escape by polyomavirus capsid mutation facilitates neurovirulence. *eLife*, 9, 2020.

[7531] Laura A Lavery, Kerstin Ure, Ying-Wooi Wan, Chongyuan Luo, Alexander J Trostle, Wei Wang, Haijing Jin, Joanna Lopez, Jacinta Lucero, Mark A Durham, Rosa Castanon, Joseph R Nery, Zhandong Liu, Margaret Goodell, Joseph R Ecker, M Margarita Behrens, and Huda Y Zoghbi. Losing Dnmt3a dependent methylation in inhibitory neurons impairs neural function by a mechanism impacting Rett syndrome. *eLife*, 9, 2020.

[7532] Belinda PP Lay, Audrey A Pitaru, Nathan Boulianne, Guillem R Esber, and Mihaela D Iordanova. Different methods of fear reduction are supported by distinct cortical substrates. *eLife*, 9, 2020.

[7533] André M Lazar, Roshanak Irannejad, Tanya A Baldwin, Aparna B Sundaram, J Silvio Gutkind, Asuka Inoue, Carmen W Dessauer, and Mark Von Zastrow. G protein-regulated endocytic trafficking of adenylyl cyclase type 9. *eLife*, 9, 2020.

[7534] Henry H Le, Chester JJ Wrobel, Sarah M Cohen, Jingfang Yu, Heenam Park, Maximilian J Helf, Brian J Curtis, Joseph C Kruempel, Pedro Reis Rodrigues, Patrick J Hu, Paul W Sternberg, and Frank C Schroeder. Modular metabolite assembly in Caenorhabditis elegans depends on carboxylesterases and formation of lysosome-related organelles. *eLife*, 9, 2020.

[7535] Chanjae Lee, Rachael M Cox, Ophelia Papoulas, Amjad Horani, Kevin Drew, Caitlin C Devitt, Steven L Brody, Edward M Marcotte, and John B Wallingford. Functional partitioning of a liquid-like organelle during assembly of axonemal dyneins. *eLife*, 9, 2020.

[7536] Chih-Yung S Lee, Andrea Putnam, Tu Lu, ShuaiXin He, John Paul T Ouyang, and Geraldine Seydoux. Recruitment of mRNAs to P granules by condensation with intrinsically-disordered proteins. *eLife*, 9, 2020.

[7537] Daniel A Lee, Grigorios Oikonomou, Tasha Cammidge, Andrey Andreev, Young Hong, Hannah Hurley, and David A Prober. Neuropeptide VF neurons promote sleep via the serotonergic raphe. *eLife*, 9, 2020.

[7538] Jihyung Lee, Junyan Zhang, Young-Jun Chung, Jun Hwan Kim, Chae Min Kook, José M González-Navajas, David S Herdman, Bernd Nürnberg, Paul A Insel, Maripat Corr, Ji-Hun Mo, Ailin Tao, Kei Yasuda, Ian R Rifkin, David H Broide, Roger Sciammas, Nicholas JG Webster, and Eyal Raz. Inhibition of IRF4 in dendritic cells by PRR-independent and -dependent signals inhibit Th2 and promote Th17 responses. *eLife*, 9, 2020.

[7539] Mi-Sun Lee, Jin Wan, and Daniel Goldman. Tgfb3 collaborates with PP2A and notch signaling pathways to inhibit retina regeneration. *eLife*, 9, 2020.

[7540] Moosung Lee, Young-Ho Lee, Jinyeop Song, Geon Kim, YoungJu Jo, HyunSeok Min, Chan Hyuk Kim, and YongKeun Park. Deep-learning-based three-dimensional label-free tracking and analysis of immunological synapses of CAR-T cells. *eLife*, 9, 2020.

[7541] Robyn S Lee, Jean-FranÇois Proulx, Fiona McIntosh, Marcel A Behr, and William P Hanage. Previously undetected super-spreading of Mycobacterium tuberculosis revealed by deep sequencing. *eLife*, 9, 2020.

[7542] Schuyler Lee, Haolin Liu, Ryan Hill, Chunjing Chen, Xia Hong, Fran Crawford, Molly Kingsley, Qianqian Zhang, Xinjian Liu, Zhongzhou Chen, Andreas Lengeling, Kathrin Maria Bernt, Philippa Marrack, John Kappler, Qiang Zhou, Chuan-Yuan Li, Yuhua Xue, Kirk Hansen, and Gongyi Zhang. JMJD6 cleaves MePCE to release positive transcription elongation factor b (P-TEFb) in higher eukaryotes. *eLife*, 9, 2020.

[7543] Wenn-Chyau Lee, Bruce Russell, Radoslaw Mikolaj Sobota, Khairunnisa Ghaffar, Shanshan W Howland, Zi Xin Wong, Alexander G Maier, Dominique Dorin-Semblat, Subhra Biswas, Benoit Gamain, Yee-Ling Lau, Benoit Malleret, Cindy Chu, FranÇois Nosten, and Laurent Renia. Plasmodium-infected erythrocytes induce secretion of IGFBP7 to form type II rosettes and escape phagocytosis. *eLife*, 9, 2020.

[7544] Ying-Jou Lee, Ching-Po Yang, Rosa L Miyares, Yu-Fen Huang, Yisheng He, Qingzhong Ren, Hui-Min Chen, Takashi Kawase, Masayoshi Ito, Hideo Otsuna, Ken Sugino, Yoshi Aso, Kei Ito, and Tzumin Lee. Conservation and divergence of related neuronal lineages in the Drosophila central brain. *eLife*, 9, 2020.

[7545] Nalin Leelatian, Justine Sinnaeve, Akshitkumar M Mistry, Sierra M Barone, Asa A Brockman, Kirsten E Diggins, Allison R Greenplate, Kyle D Weaver, Reid C Thompson, Lola B Chambless, Bret C Mobley, Rebecca A Ihrie, and Jonathan M Irish. Unsupervised machine learning reveals risk stratifying glioblastoma tumor cells. *eLife*, 9, 2020.

[7546] Yaara Lefler, Oren Amsalem, Nora Vrieler, Idan Segev, and Yosef Yarom. Using subthreshold events to characterize the functional architecture of the electrically coupled inferior olive network. *eLife*, 9, 2020.

[7547] Henri Leinonen, Nguyen C Pham, Taylor Boyd, Johanes Santoso, Krzysztof Palczewski, and Frans Vinberg. Homeostatic plasticity in the retina is associated with maintenance of night vision during retinal degenerative disease. *eLife*, 9, 2020.

[7548] Lilia Leisle, Yanyan Xu, Eva Fortea, Sangyun Lee, Jason D Galpin, Malvin Vien, Christopher A Ahern, Alessio Accardi, and Simon Bernèche. Divergent Cl- and H+ pathways underlie transport coupling and gating in CLC exchangers and channels. *eLife*, 9, 2020.

[7549] Alexandre B Leitão, Ramesh Arunkumar, Jonathan P Day, Emma M Geldman, Ismaël Morin-Poulard, Michèle Crozatier, and Francis M Jiggins. Constitutive activation of cellular immunity underlies the evolution of resistance to infection in Drosophila. *eLife*, 9, 2020.

[7550] Caroline Lekszas, Ombretta Foresti, Ishier Raote, Daniel Liedtke, Eva-Maria König, Indrajit Nanda, Barbara Vona, Peter De Coster, Rita Cauwels, Vivek Malhotra, and Thomas Haaf. Biallelic TANGO1 mutations cause a novel syndromal disease due to hampered cellular collagen secretion. *eLife*, 9, 2020.

[7551] Damien Lemoine, Sarah Mondoloni, Jérome Tange, Bertrand Lambolez, Philippe Faure, Antoine Taly, Ludovic Tricoire, and Alexandre Mourot. Probing the ionotropic activity of glutamate GluD2 receptor in HEK cells with genetically-engineered photopharmacology. *eLife*, 9, 2020.

[7552] Janna D Lendner, Randolph F Helfrich, Bryce A Mander, Luis Romundstad, Jack J Lin, Matthew P Walker, Pal G Larsson, and Robert T Knight. An electrophysiological marker of arousal level in humans. *eLife*, 9, 2020.

[7553] Martin Lenz. Reversal of contractility as a signature of self-organization in cytoskeletal bundles. *eLife*, 9, 2020.

[7554] Mathieu Letellier, Matthieu Lagardère, Béatrice Tessier, Harald Janovjak, and Olivier Thoumine. Optogenetic control of excitatory post-synaptic differentiation through neuroligin-1 tyrosine phosphorylation. *eLife*, 9, 2020.

[7555] Morgan Levine, Ross A McDevitt, Margarita Meer, Kathy Perdue, Andrea Di Francesco, Theresa Meade, Colin Farrell, Kyra Thrush, Meng Wang, Christopher Dunn, Matteo Pellegrini, Rafael de Cabo, and Luigi Ferrucci. A rat epigenetic clock recapitulates phenotypic aging and co-localizes with heterochromatin. *eLife*, 9, 2020.

[7556] David Levitan, Chenghao Liu, Tracy Yang, Yasuyuki Shima, Jian-You Lin, Joseph Wachutka, Yasmin Marrero, Ramin Ali Marandi Ghoddousi, Eduardo da Veiga Beltrame, Troy A Richter, Donald B Katz, and Sacha B Nelson. Deletion of Stk11 and Fos in mouse BLA projection neurons alters intrinsic excitability and impairs formation of long-term aversive memory. *eLife*, 9, 2020.

[7557] Rachel K Lex, Zhicheng Ji, Kristin N Falkenstein, Weiqiang Zhou, Joanna L Henry, Hongkai Ji, and Steven A Vokes. GLI transcriptional repression regulates tissue-specific enhancer activity in response to Hedgehog signaling. *eLife*, 9, 2020.

[7558] Bin-zhong Li, Christopher D Putnam, and Richard David Kolodner. Mechanisms underlying genome instability mediated by formation of foldback inversions in Saccharomyces cerevisiae. *eLife*, 9, 2020.

[7559] Binshuang Li, Ryan D Bickel, Benjamin J Parker, Omid Saleh Ziabari, Fangzhou Liu, Neetha Nanoth Vellichirammal, Jean-Christophe Simon, David L Stern, and Jennifer A Brisson. A large genomic insertion containing a duplicated follistatin gene is linked to the pea aphid male wing dimorphism. *eLife*, 9, 2020.

[7560] Chunmei Li, Carrie Barton, Katrin Henke, Jake Daane, Stephen Treaster, Joana Caetano-Lopes, Robyn L Tanguay, and Matthew P Harris. celsr1a is essential for tissue homeostasis and onset of aging phenotypes in the zebrafish. *eLife*, 9, 2020.

[7561] Dianrong Li, Youwei Ai, Jia Guo, Baijun Dong, Lin Li, Gaihong Cai, She Chen, Dan Xu, Fengchao Wang, and Xiaodong Wang. Casein kinase 1G2 suppresses necroptosis-promoted testis aging by inhibiting receptor-interacting kinase 3. *eLife*, 9, 2020.

[7562] Faxiang Li, Yang Li, Xuecheng Ye, Haishan Gao, Zhubing Shi, Xuelian Luo, Luke M Rice, and Hongtao Yu. Cryo-EM structure of VASH1-SVBP bound to microtubules. *eLife*, 9, 2020.

[7563] Guiyi Li, Manuel G Forero, Jill S Wentzell, Ilgim Durmus, Reinhard Wolf, Niki C Anthoney, Mieczyslaw Parker, Ruiying Jiang, Jacob Hasenauer, Nicholas James Strausfeld, Martin Heisenberg, and Alicia Hidalgo. A Toll-receptor map underlies structural brain plasticity. *eLife*, 9, 2020.

[7564] Ho Ling Li and Mark CW van Rossum. Energy efficient synaptic plasticity. *eLife*, 9, 2020.

[7565] Hongyu Li, Jacob R Hambrook, Emmanuel A Pila, Abdullah A Gharamah, Jing Fang, Xinzhong Wu, and Patrick Hanington. Coordination of humoral immune factors dictates compatibility between Schistosoma mansoni and Biomphalaria glabrata. *eLife*, 9, 2020.

[7566] Huirong Li, Lili Lian, Bo Liu, Yu Chen, Jinglei Yang, Shuhui Jian, Jiajia Zhou, Ying Xu, Xiaoyin Ma, Jia Qu, and Ling Hou. KIT ligand protects against both light-induced and genetic photoreceptor degeneration. *eLife*, 9, 2020.

[7567] Jay Li, Chun-Chi Liang, Samuel S Pappas, and William T Dauer. TorsinB overexpression prevents abnormal twisting in DYT1 dystonia mouse models. *eLife*, 9, 2020.

[7568] Jie Li, Rena Y Jiang, Kristin L Arendt, Yu-Tien Hsu, Sophia R Zhai, and Lu Chen. Defective memory engram reactivation underlies impaired fear memory recall in Fragile X syndrome. *eLife*, 9, 2020.

[7569] Ming Li, Zhongyu Xie, Jinteng Li, Jiajie Lin, Guan Zheng, Wenjie Liu, Su'an Tang, Shuizhong Cen, Guiwen Ye, Zhaofeng Li, Wenhui Yu, Peng Wang, Yanfeng Wu, and Huiyong Shen. GAS5 protects against osteoporosis by targeting UPF1/SMAD7 axis in osteoblast differentiation. *eLife*, 9, 2020.

[7570] Ming Li, Ting Yang, Nikolay P Kandul, Michelle Bui, Stephanie Gamez, Robyn Raban, Jared Bennett, Héctor M Sánchez C, Gregory C Lanzaro, Hanno Schmidt, Yoosook Lee, John M Marshall, and Omar S Akbari. Development of a confinable gene drive system in the human disease vector Aedes aegypti. *eLife*, 9, 2020.

[7571] Na Li, Stephanie Rignault-Clerc, Christelle Bielmann, Anne-Charlotte Bon-Mathier, Tamara Déglise, Alexia Carboni, Mégane Ducrest, and Nathalie Rosenblatt-Velin. Increasing heart vascularisation after myocardial infarction using brain natriuretic peptide stimulation of endothelial and WT1+ epicardial cells. *eLife*, 9, 2020.

[7572] Qiaochu Li, Daniel-Cosmin Marcu, Ottavia Palazzo, Frances Turner, Declan King, Tara L Spires-Jones, Melanie I Stefan, and Karl Emanuel Busch. High neural activity accelerates the decline of cognitive plasticity with age in Caenorhabditis elegans. *eLife*, 9, 2020.

[7573] Rui-Ting Li, Ling-Qiao Huang, Jun-Feng Dong, and Chen-Zhu Wang. A moth odorant receptor highly expressed in the ovipositor is involved in detecting host-plant volatiles. *eLife*, 9, 2020.

[7574] Shufen Li, Kirsty Y Wan, Wei Chen, Hui Tao, Xin Liang, and Junmin Pan. Functional exploration of heterotrimeric kinesin-II in IFT and ciliary length control in Chlamydomonas. *eLife*, 9, 2020.

[7575] Tsai-Ning Li, Yu-Jung Chen, Ting-Yi Lu, You-Tung Wang, Hsin-Chieh Lin, and Chi-Kuang Yao. A positive feedback loop between Flower and PI(4,5)P2 at periactive zones controls bulk endocytosis in Drosophila. *eLife*, 9, 2020.

[7576] Weiyi Li and Michael Lynch. Universally high transcript error rates in bacteria. *eLife*, 9, 2020.

[7577] Xia Li, Alexander Ploner, Yunzhang Wang, Patrik KE Magnusson, Chandra Reynolds, Deborah Finkel, Nancy L Pedersen, Juulia Jylhävä, and Sara Hägg. Longitudinal trajectories, correlations and mortality associations of nine biological ages across 20-years follow-up. *eLife*, 9, 2020.

[7578] Xiao Ling Li, Lőrinc Pongor, Wei Tang, Sudipto Das, Bruna R Muys, Matthew F Jones, Sarah B Lazar, Emily A Dangelmaier, Corrine CR Hartford, Ioannis Grammatikakis, Qinyu Hao, Qinyu Sun, Aaron Schetter, Jennifer L Martindale, BinWu Tang, Lisa M Jenkins, Ana I Robles, Robert L Walker, Stefan Ambs, Raj Chari, Svetlana A Shabalina, Myriam Gorospe, S Perwez Hussain, Curtis C Harris, Paul S Meltzer, Kannanganattu V Prasanth, Mirit I Aladjem, Thorkell Andresson, and Ashish Lal. A small protein encoded by a putative lncRNA regulates apoptosis and tumorigenicity in human colorectal cancer cells. *eLife*, 9, 2020.

[7579] Yadong Li, Hechen Bao, Yanjia Luo, Cherasse Yoan, Heather Anne Sullivan, Luis Quintanilla, Ian Wickersham, Michael Lazarus, Yen-Yu Ian Shih, and Juan Song. Supramammillary nucleus synchronizes with dentate gyrus to regulate spatial memory retrieval through glutamate release. *eLife*, 9, 2020.

[7580] Yan Li, Yongli Shan, Gokhul Krishna Kilaru, Stefano Berto, Guang-Zhong Wang, Kimberly H Cox, Seung-Hee Yoo, Shuzhang Yang, Genevieve Konopka, and Joseph S Takahashi. Epigenetic inheritance of circadian period in clonal cells. *eLife*, 9, 2020.

[7581] Yanjun Li, Katherine EM Tregillus, Qiongsha Luo, and Stephen A Engel. Visual mode switching learned through repeated adaptation to color. *eLife*, 9, 2020.

[7582] Yinan Li, Anthony Osuma, Edgar Correa, Munachiso A Okebalama, Pauline Dao, Olivia Gaylord, Jihad Aburas, Priota Islam, André EX Brown, and Paschalis Kratsios. Establishment and maintenance of motor neuron identity via temporal modularity in terminal selector function. *eLife*, 9, 2020.

[7583] Yingju Li, Amanda Dewar, Yeon Sun Kim, Sudhansu K Dey, and Xiaofei Sun. Pregnancy success in mice requires appropriate cannabinoid receptor signaling for primary decidua formation. *eLife*, 9, 2020.

[7584] Yong Li, Anthony L Hessel, Andreas Unger, David Ing, Jannik Recker, Franziska Koser, Johanna K Freundt, and Wolfgang A Linke. Graded titin cleavage progressively reduces tension and uncovers the source of A-band stability in contracting muscle. *eLife*, 9, 2020.

[7585] Yulan Li, Lixuan Li, Jintao Wu, Zhenggang Zhu, Xiang Feng, Liming Qin, Yuwei Zhu, Li Sun, Yijun Liu, Zilong Qiu, Shumin Duan, and Yan-Qin Yu. Activation of astrocytes in hippocampus decreases fear memory through adenosine A1 receptors. *eLife*, 9, 2020.

[7586] Xing Liang, Marcela Kokes, Richard D Fetter, Maria Danielle Sallee, Adrian W Moore, Jessica L Feldman, and Kang Shen. Growth cone-localized microtubule organizing center establishes microtubule orientation in dendrites. *eLife*, 9, 2020.

[7587] Tamar Licht, Esther Sasson, Batia Bell, Myriam Grunewald, Saran Kumar, Tirzah Kreisel, Ayal Ben-Zvi, and Eli Keshet. Hippocampal neural stem cells facilitate access from circulation via apical cytoplasmic processes. *eLife*, 9, 2020.

[7588] Ori J Lieberman, Micah D Frier, Avery F McGuirt, Christopher J Griffey, Elizabeth Rafikian, Mu Yang, Ai Yamamoto, Anders Borgkvist, Emanuela Santini, and David Sulzer. Cell-type-specific regulation of neuronal intrinsic excitability by macroautophagy. *eLife*, 9, 2020.

[7589] Tobias Lieblein, Rene Zangl, Janosch Martin, Jan Hoffmann, Marie J Hutchison, Tina Stark, Elke Stirnal, Thomas Schrader, Harald Schwalbe, and Nina Morgner. Structural rearrangement of amyloid-$\beta$ upon inhibitor binding suppresses formation of Alzheimer's disease related oligomers. *eLife*, 9, 2020.

[7590] Aaron R Lim, W Kimryn Rathmell, and Jeffrey C Rathmell. The tumor microenvironment as a metabolic barrier to effector T cells and immunotherapy. *eLife*, 9, 2020.

[7591] Chew Leng Lim, Yu Zuan Or, Zoe Ong, Hwa Hwa Chung, Hirohito Hayashi, Smeeta Shrestha, Shunsuke Chiba, Feng Lin, and Valerie Chun Ling Lin. Estrogen exacerbates mammary involution through neutrophil-dependent and -independent mechanism. *eLife*, 9, 2020.

[7592] Chu Hsien Lim, Prameet Kaur, Emelyne Teo, Vanessa Yuk Man Lam, Fangchen Zhu, Caroline Kibat, Jan Gruber, Ajay S Mathuru, and Nicholas S Tolwinski. Application of optogenetic Amyloid-$\beta$ distinguishes between metabolic and physical damages in neurodegeneration. *eLife*, 9, 2020.

[7593] Carmen Lin, Venus N Sherathiya, M Matthew Oh, and John F Disterhoft. Persistent firing in LEC III neurons is differentially modulated by learning and aging. *eLife*, 9, 2020.

[7594] Chih-Tung Lin, Daniela Lecca, Ling-Yu Yang, Weiming Luo, Michael T Scerba, David Tweedie, Pen-Sen Huang, Yoo-Jin Jung, Dong Seok Kim, Chih-Hao Yang, Barry J Hoffer, Jia-Yi Wang, and Nigel H Greig. 3,6'-dithiopomalidomide reduces neural loss, inflammation, behavioral deficits in brain injury and microglial activation. *eLife*, 9, 2020.

[7595] Feline W Lindhout, Robbelien Kooistra, Sybren Portegies, Lotte J Herstel, Riccardo Stucchi, Basten L Snoek, AF Maarten Altelaar, Harold D MacGillavry, Corette J Wierenga, and Casper C Hoogenraad. Quantitative mapping of transcriptome and proteome dynamics during polarization of human iPSC-derived neurons. *eLife*, 9, 2020.

[7596] Joachim Linssen, Anthony Ermens, Marvin Berrevoets, Michela Seghezzi, Giulia Previtali, Simone van der Sar-van der Brugge, Henk Russcher, Annelies Verbon, Judith Gillis, Jürgen Riedl, Eva de Jongh, Jarob Saker, Marion Münster, Imke CA Munnix, Anthonius Dofferhof, Volkher Scharnhorst, Heidi Ammerlaan, Kathleen Deiteren, Stephan JL Bakker, Lucas Joost Van Pelt, Yvette Kluiters-de Hingh, Mathie PG Leers, and Andre J van der Ven. A novel haemocytometric COVID-19 prognostic score developed and validated in an observational multicentre European hospital-based study. *eLife*, 9, 2020.

[7597] Jyun-you Liou, Elliot H Smith, Lisa M Bateman, Samuel L Bruce, Guy M McKhann, Robert R Goodman, Ronald G Emerson, Catherine A Schevon, and LF Abbott. A model for focal seizure onset, propagation, evolution, and progression. *eLife*, 9, 2020.

[7598] Michelle M Lissner, Katherine Cumnock, Nicole M Davis, José G Vilches-Moure, Priyanka Basak, Daniel J Navarrete, Jessica A Allen, and David Schneider. Metabolic profiling during malaria reveals the role of the aryl hydrocarbon receptor in regulating kidney injury. *eLife*, 9, 2020.

[7599] Thuy-Lan V Lite, Robert A Grant, Isabel Nocedal, Megan L Littlehale, Monica S Guo, and Michael T Laub. Uncovering the basis of protein-protein interaction specificity with a combinatorially complete library. *eLife*, 9, 2020.

[7600] Jamie C Little, Elisa Garcia-Garcia, Amanda Sul, and Daniel Kalderon. Drosophila hedgehog can act as a morphogen in the absence of regulated Ci processing. *eLife*, 9, 2020.

[7601] Nicole K Littlejohn, Nicolas Seban, Chung-Chih Liu, and Supriya Srinivasan. A feedback loop governs the relationship between lipid metabolism and longevity. *eLife*, 9, 2020.

[7602] Cheng-Hsin Liu, Ryan Seo, Tammy Szu-Yu Ho, Michael Stankewich, Peter J Mohler, Thomas J Hund, Jeffrey L Noebels, and Matthew N Rasband. $\beta$ spectrin-dependent and domain specific mechanisms for Na+ channel clustering. *eLife*, 9, 2020.

[7603] Cheng-Hsin Liu, Sharon R Stevens, Lindsay H Teliska, Michael Stankewich, Peter J Mohler, Thomas J Hund, and Matthew N Rasband. Nodal $\beta$ spectrins are required to maintain Na+ channel clustering and axon integrity. *eLife*, 9, 2020.

[7604] Chengdong Liu, Shuang Li, Pernille Rimmer Noer, Kasper Kjaer-Sorensen, Anna Karina Juhl, Allison Goldstein, Caihuan Ke, Claus Oxvig, and Cunming Duan. The metalloproteinase Papp-aa controls epithelial cell quiescence-proliferation transition. *eLife*, 9, 2020.

[7605] Huan Liu, Kaylia Duncan, Annika Helverson, Priyanka Kumari, Camille Mumm, Yao Xiao, Jenna Colavincenzo Carlson, Fabrice Darbellay, Axel Visel, Elizabeth Leslie, Patrick Breheny, Albert J Erives, and Robert A Cornell. Analysis of zebrafish periderm enhancers facilitates identification of a regulatory variant near human KRT8/18. *eLife*, 9, 2020.

[7606] Ran Liu, Joseph L Greenstein, James C Fackler, Melania M Bembea, and Raimond L Winslow. Spectral clustering of risk score trajectories stratifies sepsis patients by clinical outcome and interventions received. *eLife*, 9, 2020.

[7607] Xingchen Liu, Jordane Dimidschstein, Gordon Fishell, and Adam G Carter. Hippocampal inputs engage CCK+ interneurons to mediate endocannabinoid-modulated feed-forward inhibition in the prefrontal cortex. *eLife*, 9, 2020.

[7608] Yingqi Liu, Lingling Li, Xiaoqing Liu, Yibo Wang, Lingxia Liu, Lu Peng, Jiayuan Liu, Lian Zhang, Guannan Wang, Hongyuan Li, Dong-Xu Liu, Baiqu Huang, Jun Lu, and Yu Zhang. RETRACTED: Arginine methylation of SHANK2 by PRMT7 promotes human breast cancer metastasis through activating endosomal FAK signalling. *eLife*, 9, 2020.

[7609] Yun-Fei Liu, Judy Kim, Colin Wilson, and Marina Bedny. Computer code comprehension shares neural resources with formal logical inference in the fronto-parietal network. *eLife*, 9, 2020.

[7610] Zhimin Liu, Darach Miller, Fangfei Li, Xianan Liu, and Sasha F Levy. A large accessory protein interactome is rewired across environments. *eLife*, 9, 2020.

[7611] Marija Liutkute, Manisankar Maiti, Ekaterina Samatova, Jörg Enderlein, and Marina V Rodnina. Gradual compaction of the nascent peptide during cotranslational folding on the ribosome. *eLife*, 9, 2020.

[7612] Maria N Lo, Leah J Damon, Jian Wei Tay, Shang Jia, and Amy E Palmer. Single cell analysis reveals multiple requirements for zinc in the mammalian cell cycle. *eLife*, 9, 2020.

[7613] Savita Lochab, Yogendra Singh, Sagar Sengupta, and Vinay Kumar Nandicoori. Mycobacterium tuberculosis exploits host ATM kinase for survival advantage through SecA2 secretome. *eLife*, 9, 2020.

[7614] Jeffrey T Lock and Ian Parker. IP3 mediated global Ca2+ signals arise through two temporally and spatially distinct modes of Ca2+ release. *eLife*, 9, 2020.

[7615] Víctor J López-Madrona, Elena Pérez-Montoyo, Efrén álvarez Salvado, David Moratal, Oscar Herreras, Ernesto Pereda, Claudio R Mirasso, and Santiago Canals. Different theta frameworks coexist in the rat hippocampus and are coordinated during memory-guided and novelty tasks. *eLife*, 9, 2020.

[7616] Andres López-Perrote, Nele Hug, Ana González-Corpas, Carlos F Rodríguez, Marina Serna, Carmen García-Martín, Jasminka Boskovic, Rafael Fernandez-Leiro, Javier F Caceres, and Oscar Llorca. Regulation of RUVBL1-RUVBL2 AAA-ATPases by the nonsense-mediated mRNA decay factor DHX34, as evidenced by Cryo-EM. *eLife*, 9, 2020.

[7617] Eduardo Javier López Soto and Diane Lipscombe. Cell-specific exon methylation and CTCF binding in neurons regulate calcium ion channel splicing and function. *eLife*, 9, 2020.

[7618] Justin Losacco, Daniel Ramirez-Gordillo, Jesse Gilmer, and Diego Restrepo. Learning improves decoding of odor identity with phase-referenced oscillations in the olfactory bulb. *eLife*, 9, 2020.

[7619] Adam C Lu, Christine Kyuyoung Lee, Max Kleiman-Weiner, Brian Truong, Megan Wang, John R Huguenard, and Mark P Beenhakker. Nonlinearities between inhibition and T-type calcium channel activity bidirectionally regulate thalamic oscillations. *eLife*, 9, 2020.

[7620] Bo Lu, Liting Dong, Danyang Yi, Meiling Zhang, Chenxu Zhu, Xiaoyu Li, and Chengqi Yi. Transposase-assisted tagmentation of RNA/DNA hybrid duplexes. *eLife*, 9, 2020.

[7621] Kuan-Yi Lu, Charisse Flerida A Pasaje, Tamanna Srivastava, David R Loiselle, Jacquin C Niles, and Emily Derbyshire. Phosphatidylinositol 3-phosphate and Hsp70 protect Plasmodium falciparum from heat-induced cell death. *eLife*, 9, 2020.

[7622] Wen Lu, Margot Lakonishok, Rong Liu, Neil Billington, Ashley Rich, Michael Glotzer, James R Sellers, and Vladimir I Gelfand. Competition between kinesin-1 and myosin-V defines Drosophila posterior determination. *eLife*, 9, 2020.

[7623] Zhiwei Lu, Yuhua Xie, Huanwei Huang, Kaiju Jiang, Bin Zhou, Fengchao Wang, and Ting Chen. Hair follicle stem cells regulate retinoid metabolism to maintain the self-renewal niche for melanocyte stem cells. *eLife*, 9, 2020.

[7624] Haojiang Luan, Alexander Kuzin, Ward F Odenwald, and Benjamin H White. Cre-assisted fine-mapping of neural circuits using orthogonal split inteins. *eLife*, 9, 2020.

[7625] Jay Lubow, Maria C Virgilio, Madeline Merlino, David R Collins, Michael Mashiba, Brian G Peterson, Zana Lukic, Mark M Painter, Francisco Gomez-Rivera, Valeri Terry, Gretchen Zimmerman, and Kathleen L Collins. Mannose receptor is an HIV restriction factor counteracted by Vpr in macrophages. *eLife*, 9, 2020.

[7626] Naemi Luithle, Jelmi uit de Bos, Ruud Hovius, Daria Maslennikova, Renard TM Lewis, Rosemarie Ungricht, Beat Fierz, and Ulrike Kutay. Torsin ATPases influence chromatin interaction of the Torsin regulator LAP1. *eLife*, 9, 2020.

[7627] Casper K Lumby, Lei Zhao, Judith Breuer, and Christopher JR Illingworth. A large effective population size for established within-host influenza virus infection. *eLife*, 9, 2020.

[7628] Cheng Luo and Nai Ding. Cortical encoding of acoustic and linguistic rhythms in spoken narratives. *eLife*, 9, 2020.

[7629] Thomas Zhihao Luo, Adrian Gopnik Bondy, Diksha Gupta, Verity Alexander Elliott, Charles D Kopec, and Carlos D Brody. An approach for long-term, multi-probe Neuropixels recordings in unrestrained rats. *eLife*, 9, 2020.

[7630] Yicheng Luo, Elena Fefelova, Maria Ninova, Yung-Chia Ariel Chen, and Alexei A Aravin. Repression of interrupted and intact rDNA by the SUMO pathway in Drosophila melanogaster. *eLife*, 9, 2020.

[7631] Mirko Luoni, Serena Giannelli, Marzia Tina Indrigo, Antonio Niro, Luca Massimino, Angelo Iannielli, Laura Passeri, Fabio Russo, Giuseppe Morabito, Piera Calamita, Silvia Gregori, Benjamin Deverman, and Vania Broccoli. Whole brain delivery of an instability-prone Mecp2 transgene improves behavioral and molecular pathological defects in mouse models of Rett syndrome. *eLife*, 9, 2020.

[7632] Upendo Lupanga, Rachel Röhrich, Jana Askani, Stefan Hilmer, Christiane Kiefer, Melanie Krebs, Takehiko Kanazawa, Takashi Ueda, and Karin Schumacher. The Arabidopsis V-ATPase is localized to the TGN/EE via a seed plant-specific motif. *eLife*, 9, 2020.

[7633] Lu Lv, Peihao Chen, Longzhi Cao, Yamei Li, Zhi Zeng, Yue Cui, Qingcui Wu, Jiaojiao Li, Jian-Hua Wang, Meng-Qiu Dong, Xiangbing Qi, and Ting Han. Discovery of a molecular glue promoting CDK12-DDB1 interaction to trigger cyclin K degradation. *eLife*, 9, 2020.

[7634] Eugenia Lyashenko, Mario Niepel, Purushottam D Dixit, Sang Kyun Lim, Peter K Sorger, and Dennis Vitkup. Receptor-based mechanism of relative sensing and cell memory in mammalian signaling networks. *eLife*, 9, 2020.

[7635] Claire Y Ma, John C Marioni, Gillian M Griffiths, and Arianne C Richard. Stimulation strength controls the rate of initiation but not the molecular organisation of TCR-induced signalling. *eLife*, 9, 2020.

[7636] Ruihua Ma, Angelica S Martínez-Ramírez, Thomas L Borders, Fanding Gao, and Beatriz Sosa-Pineda. Metabolic and non-metabolic liver zonation is established non-synchronously and requires sinusoidal Wnts. *eLife*, 9, 2020.

[7637] Xiaojuan Ma, Yi Zhu, Jinxia Lu, Jingfei Xie, Chong Li, Woo Shik Shin, Jiali Qiang, Jiaqi Liu, Shuai Dou, Yi Xiao, Chuchu Wang, Chunyu Jia, Houfang Long, Juntao Yang, Yanshan Fang, Lin Jiang, Yaoyang Zhang, Shengnan Zhang, Rong Grace Zhai, Cong Liu, and Dan Li. Nicotinamide mononucleotide adenylyltransferase uses its NAD+ substrate-binding site to chaperone phosphorylated Tau. *eLife*, 9, 2020.

[7638] István Maák, Garyk Roelandt, and Patrizia d'Ettorre. A small number of workers with specific personality traits perform tool use in ants. *eLife*, 9, 2020.

[7639] Magali Macchi, Karine Magalon, Céline Zimmer, Elitsa Peeva, Bilal El Waly, Béatrice Brousse, Sarah Jaekel, Kay Grobe, Friedemann Kiefer, Anna Williams, Myriam Cayre, and Pascale Durbec. Mature oligodendrocytes bordering lesions limit demyelination and favor myelin repair via heparan sulfate production. *eLife*, 9, 2020.

[7640] Ana S Machado, Hugo G Marques, Diogo F Duarte, Dana M Darmohray, and Megan R Carey. Shared and specific signatures of locomotor ataxia in mutant mice. *eLife*, 9, 2020.

[7641] Charles E MacKay, M Dennis Leo, Carlos Fernández-Peña, Raquibul Hasan, Wen Yin, Alejandro Mata-Daboin, Simon Bulley, Jesse Gammons, Salvatore Mancarella, and Jonathan H Jaggar. Intravascular flow stimulates PKD2 (polycystin-2) channels in endothelial cells to reduce blood pressure. *eLife*, 9, 2020.

[7642] Isabelle Mackrous, Jérome Carriot, Kathleen E Cullen, and Maurice J Chacron. Neural variability determines coding strategies for natural self-motion in macaque monkeys. *eLife*, 9, 2020.

[7643] J Wesley Maddox, Kate L Randall, Ravi P Yadav, Brittany Williams, Jussara Hagen, Paul J Derr, Vasily Kerov, Luca Della Santina, Sheila A Baker, Nikolai Artemyev, Mrinalini Hoon, and Amy Lee. A dual role for Cav1.4 Ca2+ channels in the molecular and structural organization of the rod photoreceptor synapse. *eLife*, 9, 2020.

[7644] Maria-Bernadette Madel, Lidia Ibáñez, Thomas Ciucci, Julia Halper, Matthieu Rouleau, Antoine Boutin, Christophe Hue, Isabelle Duroux-Richard, Florence Apparailly, Henri-Jean Garchon, Abdelilah Wakkach, and Claudine Blin-Wakkach. Dissecting the phenotypic and functional heterogeneity of mouse inflammatory osteoclasts by the expression of Cx3cr1. *eLife*, 9, 2020.

[7645] Sukanya Madhwal, Mingyu Shin, Ankita Kapoor, Manisha Goyal, Manish K Joshi, Pirzada Mujeeb Ur Rehman, Kavan Gor, Jiwon Shim, and Tina Mukherjee. Metabolic control of cellular immune-competency by odors in Drosophila. *eLife*, 9, 2020.

[7646] Naïma Madi, Michiel Vos, Carmen Lia Murall, Pierre Legendre, and B Jesse Shapiro. Does diversity beget diversity in microbiomes? *eLife*, 9, 2020.

[7647] Lauren A Maggio, Ryan M Steinberg, Tiziano Piccardi, and John M Willinsky. Reader engagement with medical content on Wikipedia. *eLife*, 9, 2020.

[7648] Jeremy Magland, James J Jun, Elizabeth Lovero, Alexander J Morley, Cole Lincoln Hurwitz, Alessio Paolo Buccino, Samuel Garcia, and Alex H Barnett. SpikeForest, reproducible web-facing ground-truth validation of automated neural spike sorters. *eLife*, 9, 2020.

[7649] Jens P Magnusson, Margherita Zamboni, Giuseppe Santopolo, Jeff E Mold, Mauricio Barrientos-Somarribas, Carlos Talavera-Lopez, Björn Andersson, and Jonas Frisén. Activation of a neural stem cell transcriptional program in parenchymal astrocytes. *eLife*, 9, 2020.

[7650] Mohamed Mahgoub, Jacob Paiano, Melania Bruno, Wei Wu, Sarath Pathuri, Xing Zhang, Sherry Ralls, Xiaodong Cheng, André Nussenzweig, and Todd S Macfarlan. Dual histone methyl reader ZCWPW1 facilitates repair of meiotic double strand breaks in male mice. *eLife*, 9, 2020.

[7651] Keyvan Mahjoory, Jan-Mathijs Schoffelen, Anne Keitel, and Joachim Gross. The frequency gradient of human resting-state brain oscillations follows cortical hierarchies. *eLife*, 9, 2020.

[7652] Alexandre Mahrach, Guang Chen, Nuo Li, Carl van Vreeswijk, and David Hansel. Mechanisms underlying the response of mouse cortical networks to optogenetic manipulation. *eLife*, 9, 2020.

[7653] Femke Maij, Christian Seegelke, W Pieter Medendorp, and Tobias Heed. External location of touch is constructed post-hoc based on limb choice. *eLife*, 9, 2020.

[7654] Vayu Maini Rekdal, Paola Nol Bernadino, Michael U Luescher, Sina Kiamehr, Chip Le, Jordan E Bisanz, Peter J Turnbaugh, Elizabeth N Bess, and Emily P Balskus. A widely distributed metalloenzyme class enables gut microbial metabolism of host- and diet-derived catechols. *eLife*, 9, 2020.

[7655] Nairita Maitra, Chong He, Heidi M Blank, Mitsuhiro Tsuchiya, Birgit Schilling, Matt Kaeberlein, Rodolfo Aramayo, Brian K Kennedy, and Michael Polymenis. Translational control of one-carbon metabolism underpins ribosomal protein phenotypes in cell division and longevity. *eLife*, 9, 2020.

[7656] Rupamanjari Majumder, Tim De Coster, Nina Kudryashova, Arie O Verkerk, Ivan V Kazbanov, Balázs ördög, Niels Harlaar, Ronald Wilders, Antoine AF de Vries, Dirk L Ypey, Alexander V Panfilov, and Daniël A Pijnappels. Self-restoration of cardiac excitation rhythm by anti-arrhythmic ion channel gating. *eLife*, 9, 2020.

[7657] Cihan Makbul, Michael Nassal, and Bettina Böttcher. Slowly folding surface extension in the prototypic avian hepatitis B virus capsid governs stability. *eLife*, 9, 2020.

[7658] Tuomo Mäki-Marttunen, Nicolangelo Iannella, Andrew G Edwards, Gaute T Einevoll, and Kim T Blackwell. A unified computational model for cortical post-synaptic plasticity. *eLife*, 9, 2020.

[7659] Nathalia Luisa Sousa de Oliveira Malacco, Jessica Amanda Marques Souza, Flavia Rayssa Braga Martins, Milene Alvarenga Rachid, Janaina Aparecida Simplicio, Carlos Renato Tirapelli, Adriano de Paula Sabino, Celso Martins Queiroz-Junior, Grazielle Ribeiro Goes, Leda Quercia Vieira, Danielle Glória Souza, Vanessa Pinho, Mauro Martins Teixeira, and Frederico Marianetti Soriani. Chronic ethanol consumption compromises neutrophil function in acute pulmonary Aspergillus fumigatus infection. *eLife*, 9, 2020.

[7660] Gerardo Malagon, Takafumi Miki, Van Tran, Laura C Gomez, and Alain Marty. Incomplete vesicular docking limits synaptic strength under high release probability conditions. *eLife*, 9, 2020.

[7661] Maria Maldonado, Abhilash Padavannil, Long Zhou, Fei Guo, and James A Letts. Atomic structure of a mitochondrial complex I intermediate from vascular plants. *eLife*, 9, 2020.

[7662] Tatiana Malevich, Antimo Buonocore, and Ziad M Hafed. Rapid stimulus-driven modulation of slow ocular position drifts. *eLife*, 9, 2020.

[7663] Milene R Malheiros-Lima, Josiane N Silva, Felipe C Souza, Ana C Takakura, and Thiago S Moreira. C1 neurons are part of the circuitry that recruits active expiration in response to the activation of peripheral chemoreceptors. *eLife*, 9, 2020.

[7664] Jakob Malsy, Andrea C Alvarado, Joseph O Lamontagne, Karin Strittmatter, and Alexander G Marneros. Distinct effects of complement and of NLRP3- and non-NLRP3 inflammasomes for choroidal neovascularization. *eLife*, 9, 2020.

[7665] Kevin I Manage, Alicia K Rogers, Dylan C Wallis, Celja J Uebel, Dorian C Anderson, Dieu An H Nguyen, Katerina Arca, Kristen C Brown, Ricardo J Cordeiro Rodrigues, Bruno FM de Albuquerque, René F Ketting, Taiowa A Montgomery, and Carolyn Marie Phillips. A tudor domain protein, SIMR-1, promotes siRNA production at piRNA-targeted mRNAs in C. elegans. *eLife*, 9, 2020.

[7666] Walter R Mancia Leon, Julien Spatazza, Benjamin Rakela, Ankita Chatterjee, Viraj Pande, Tom Maniatis, Andrea R Hasenstaub, Michael P Stryker, and Arturo Alvarez-Buylla. Clustered gamma-protocadherins regulate cortical interneuron programmed cell death. *eLife*, 9, 2020.

[7667] Matteo Mancini, Agah Karakuzu, Julien Cohen-Adad, Mara Cercignani, Thomas E Nichols, and Nikola Stikov. An interactive meta-analysis of MRI biomarkers of myelin. *eLife*, 9, 2020.

[7668] Kyle Mangano, Tanja Florin, Xinhao Shao, Dorota Klepacki, Irina Chelysheva, Zoya Ignatova, Yu Gao, Alexander S Mankin, and Nora Vázquez-Laslop. Genome-wide effects of the antimicrobial peptide apidaecin on translation termination in bacteria. *eLife*, 9, 2020.

[7669] Joan Manils, Louise V Webb, Ashleigh Howes, Julia Janzen, Stefan Boeing, Anne M Bowcock, and Steven C Ley. CARD14E138A signalling in keratinocytes induces TNF-dependent skin and systemic inflammation. *eLife*, 9, 2020.

[7670] Shivani N Mann, Niran Hadad, Molly Nelson Holte, Alicia R Rothman, Roshini Sathiaseelan, Samim Ali Mondal, Martin-Paul Agbaga, Archana Unnikrishnan, Malayannan Subramaniam, John Hawse, Derek M Huffman, Willard M Freeman, and Michael B Stout. Health benefits attributed to 17$\alpha$-estradiol, a lifespan-extending compound, are mediated through estrogen receptor $\alpha$. *eLife*, 9, 2020.

[7671] Jorge Manuel, Natalia Färber, Darius A Gerlach, Karsten Heusser, Jens Jordan, Jens Tank, and Florian Beissner. Deciphering the neural signature of human cardiovascular regulation. *eLife*, 9, 2020.

[7672] Gabriela Manzano Nieves, Marilyn Bravo, Saba Baskoylu, and Kevin G Bath. Early life adversity decreases pre-adolescent fear expression by accelerating amygdala PV cell development. *eLife*, 9, 2020.

[7673] Yanbo Mao, Alexander Gabel, Thomas Nakel, Prisca Viehöver, Thomas Baum, Dawit Girma Tekleyohans, Dieu Vo, Ivo Grosse, and Rita Groß-Hardt. Selective egg cell polyspermy bypasses the triploid block. *eLife*, 9, 2020.

[7674] Zachary M March, Katelyn Sweeney, Hanna Kim, Xiaohui Yan, Laura M Castellano, Meredith E Jackrel, JiaBei Lin, Edward Chuang, Edward Gomes, Corey W Willicott, Karolina Michalska, Robert P Jedrzejczak, Andrzej Joachimiak, Kim A Caldwell, Guy A Caldwell, Ophir Shalem, and James Shorter. Therapeutic genetic variation revealed in diverse Hsp104 homologs. *eLife*, 9, 2020.

[7675] Julia M Marchingo, Linda V Sinclair, Andrew JM Howden, and Doreen A Cantrell. Quantitative analysis of how Myc controls T cell proteomes and metabolic pathways during T cell activation. *eLife*, 9, 2020.

[7676] Aleksandra Marconi, Amy Hancock-Ronemus, and J Andrew Gillis. Adult chondrogenesis and spontaneous cartilage repair in the skate, Leucoraja erinacea. *eLife*, 9, 2020.

[7677] Sara Marelli, James C Williamson, Anna V Protasio, Adi Naamati, Edward JD Greenwood, Janet E Deane, Paul J Lehner, and Nicholas J Matheson. Antagonism of PP2A is an independent and conserved function of HIV-1 Vif and causes cell cycle arrest. *eLife*, 9, 2020.

[7678] Kasey Markel. Lack of evidence for associative learning in pea plants. *eLife*, 9, 2020.

[7679] Kasey Markel. Response to comment on 'Lack of evidence for associative learning in pea plants'. *eLife*, 9, 2020.

[7680] Alexander L Markowitz and Radha Kalluri. Gradients in the biophysical properties of neonatal auditory neurons align with synaptic contact position and the intensity coding map of inner hair cells. *eLife*, 9, 2020.

[7681] Patricia Marmol, Favio Krapacher, and Carlos F Ibáñez. Control of brown adipose tissue adaptation to nutrient stress by the activin receptor ALK7. *eLife*, 9, 2020.

[7682] Ingo Marquardt, Peter De Weerd, Marian Schneider, Omer Faruk Gulban, Dimo Ivanov, Yawen Wang, and Kâmil Uludağ. Feedback contribution to surface motion perception in the human early visual cortex. *eLife*, 9, 2020.

[7683] Guillermo Marqués, Thomas Pengo, and Mark A Sanders. Imaging methods are vastly underreported in biomedical research. *eLife*, 9, 2020.

[7684] Joana G Marques, Berkley E Gryder, Blaz Pavlovic, Yeonjoo Chung, Quy A Ngo, Fabian Frommelt, Matthias Gstaiger, Young Song, Katharina Benischke, Dominik Laubscher, Marco Wachtel, Javed Khan, and Beat W Schäfer. NuRD subunit CHD4 regulates super-enhancer accessibility in rhabdomyosarcoma and represents a general tumor dependency. *eLife*, 9, 2020.

[7685] Bryan Marsh and Robert Blelloch. Single nuclei RNA-seq of mouse placental labyrinth development. *eLife*, 9, 2020.

[7686] Bryan A Martinez, Pedro Reis Rodrigues, Ricardo M Nuñez Medina, Prosenjit Mondal, Neale J Harrison, Museer A Lone, Amanda Webster, Aditi U Gurkar, Brock Grill, and Matthew S Gill. An alternatively spliced, non-signaling insulin receptor modulates insulin sensitivity via insulin peptide sequestration in C. elegans. *eLife*, 9, 2020.

[7687] Carlos Martinez-Ruiz, Rodrigo Pracana, Eckart Stolle, Carolina Ivon Paris, Richard A Nichols, and Yannick Wurm. Genomic architecture and evolutionary antagonism drive allelic expression bias in the social supergene of red fire ants. *eLife*, 9, 2020.

[7688] Daniel Martins, Anthony S Gabay, Mitul Mehta, and Yannis Paloyelis. Salivary and plasmatic oxytocin are not reliable trait markers of the physiology of the oxytocin system in humans. *eLife*, 9, 2020.

[7689] Dario Maschi and Vitaly A Klyachko. Spatiotemporal dynamics of multi-vesicular release is determined by heterogeneity of release sites within central synapses. *eLife*, 9, 2020.

[7690] Tara L Mastro, Anthony Preza, Shinjini Basu, Sumantra Chattarji, Sally M Till, Peter C Kind, and Mary B Kennedy. A sex difference in the response of the rodent postsynaptic density to synGAP haploinsufficiency. *eLife*, 9, 2020.

[7691] Akira Masuda, Chie Sano, Qi Zhang, Hiromichi Goto, Thomas J McHugh, Shigeyoshi Fujisawa, and Shigeyoshi Itohara. The hippocampus encodes delay and value information during delay-discounting decision making. *eLife*, 9, 2020.

[7692] Laura Matellán, Javier Manzano-López, and Fernando Monje-Casas. Polo-like kinase acts as a molecular timer that safeguards the asymmetric fate of spindle microtubule-organizing centers. *eLife*, 9, 2020.

[7693] Luana C Mathieu, Horace Cox, Angela M Early, Sachel Mok, Yassamine Lazrek, Jeanne-Celeste Paquet, Maria-Paz Ade, Naomi W Lucchi, Quacy Grant, Venkatachalam Udhayakumar, Jean SF Alexandre, Magalie Demar, Pascal Ringwald, Daniel E Neafsey, David A Fidock, and Lise Musset. Local emergence in Amazonia of Plasmodium falciparum k13 C580Y mutants associated with in vitro artemisinin resistance. *eLife*, 9, 2020.

[7694] Irina Matos, Amma Asare, John Levorse, Tamara Ouspenskaia, June de la Cruz-Racelis, Laura-Nadine Schuhmacher, and Elaine Fuchs. Progenitors oppositely polarize WNT activators and inhibitors to orchestrate tissue development. *eLife*, 9, 2020.

[7695] Naoyuki Matsumoto, Satoshi Tanaka, Toshihide Horiike, Yohei Shinmyo, and Hiroshi Kawasaki. A discrete subtype of neural progenitor crucial for cortical folding in the gyrencephalic mammalian brain. *eLife*, 9, 2020.

[7696] Sandy Mattijssen, James R Iben, Tianwei Li, Steven L Coon, and Richard J Maraia. Single molecule poly(A) tail-seq shows LARP4 opposes deadenylation throughout mRNA lifespan with most impact on short tails. *eLife*, 9, 2020.

[7697] Stefania Mattioni, Mohamed Rezk, Ceren Battal, Roberto Bottini, Karen E Cuculiza Mendoza, Nikolaas N Oosterhof, and Olivier Collignon. Categorical representation from sound and sight in the ventral occipito-temporal cortex of sighted and blind. *eLife*, 9, 2020.

[7698] William Mau, Michael E Hasselmo, and Denise J Cai. The brain in motion: How ensemble fluidity drives memory-updating and flexibility. *eLife*, 9, 2020.

[7699] Christina E May, Julia Rosander, Jennifer Gottfried, Evan Dennis, and Monica Dus. Dietary sugar inhibits satiation by decreasing the central processing of sweet taste. *eLife*, 9, 2020.

[7700] Omer Mazar and Yossi Yovel. A sensorimotor model shows why a spectral jamming avoidance response does not help bats deal with jamming. *eLife*, 9, 2020.

[7701] Matan Mazor, Karl J Friston, and Stephen M Fleming. Distinct neural contributions to metacognition for detecting, but not discriminating visual stimuli. *eLife*, 9, 2020.

[7702] Matthew P McCabe, Erin R Cullen, Caitlynn M Barrows, Amy N Shore, Katherine I Tooke, Kathryn A Laprade, James M Stafford, and Matthew C Weston. Genetic inactivation of mTORC1 or mTORC2 in neurons reveals distinct functions in glutamatergic synaptic transmission. *eLife*, 9, 2020.

[7703] Sarah McCallum, Yuuki Obata, Evangelia Fourli, Stefan Boeing, Christopher J Peddie, Qiling Xu, Stuart Horswell, Robert N Kelsh, Lucy Collinson, David Wilkinson, Carmen Pin, Vassilis Pachnis, and Tiffany A Heanue. Enteric glia as a source of neural progenitors in adult zebrafish. *eLife*, 9, 2020.

[7704] Erica McGale, Henrique Valim, Deepika Mittal, Jesús Morales Jimenez, Rayko Halitschke, Meredith C Schuman, and Ian T Baldwin. Determining the scale at which variation in a single gene changes population yields. *eLife*, 9, 2020.

[7705] Philip T McGilvray, S Andrei Anghel, Arunkumar Sundaram, Frank Zhong, Michael J Trnka, James R Fuller, Hong Hu, Alma L Burlingame, and Robert J Keenan. An ER translocon for multi-pass membrane protein biogenesis. *eLife*, 9, 2020.

[7706] Tracy L McGregor, Karen A Hunt, Elaine Yee, Dan Mason, Paul Nioi, Simina Ticau, Marissa Pelosi, Perry R Loken, Sarah Finer, Deborah A Lawlor, Eric B Fauman, Qin Qin Huang, Christopher J Griffiths, Daniel G MacArthur, Richard C Trembath, Devin Oglesbee, John C Lieske, David V Erbe, John Wright, and David A van Heel. Characterising a healthy adult with a rare HAO1 knockout to support a therapeutic strategy for primary hyperoxaluria. *eLife*, 9, 2020.

[7707] Claire E McKellar, Igor Siwanowicz, Barry J Dickson, and Julie H Simpson. Controlling motor neurons of every muscle for fly proboscis reaching. *eLife*, 9, 2020.

[7708] Scott McWilliams, Barbara Pierce, Andrea Wittenzellner, Lillie Langlois, Sophia Engel, John R Speakman, Olivia Fatica, Kristen DeMoranville, Wolfgang Goymann, Lisa Trost, Amadeusz Bryla, Maciej Dzialo, Edyta Sadowska, and Ulf Bauchinger. The energy savings-oxidative cost trade-off for migratory birds during endurance flight. *eLife*, 9, 2020.

[7709] William D Meador, Mrudang Mathur, Gabriella P Sugerman, Marcin Malinowski, Tomasz Jazwiec, Xinmei Wang, Carla MR Lacerda, Tomasz A Timek, and Manuel K Rausch. The tricuspid valve also maladapts as shown in sheep with biventricular heart failure. *eLife*, 9, 2020.

[7710] Edgar M Medina, Kristyn A Robinson, Kimberly Bellingham-Johnstun, Giuseppe Ianiri, Caroline Laplante, Lillian K Fritz-Laylin, and Nicolas E Buchler. Genetic transformation of Spizellomyces punctatus, a resource for studying chytrid biology and evolutionary cell biology. *eLife*, 9, 2020.

[7711] Stefan Meinke, Gesine Goldammer, A Ioana Weber, Victor Tarabykin, Alexander Neumann, Marco Preussner, and Florian Heyd. Srsf10 and the minor spliceosome control tissue-specific and dynamic SR protein expression. *eLife*, 9, 2020.

[7712] David Melamed and Daniel Kalderon. Opposing JAK-STAT and Wnt signaling gradients define a stem cell domain by regulating differentiation at two borders. *eLife*, 9, 2020.

[7713] Carlos I Mendoza and David Reguera. Shape selection and mis-assembly in viral capsid formation by elastic frustration. *eLife*, 9, 2020.

[7714] Juan L Mendoza, Suzanne Fischer, Marvin H Gee, Lilian H Lam, Simon Brackenridge, Fiona M Powrie, Michael Birnbaum, Andrew J McMichael, K Christopher Garcia, and Geraldine M Gillespie. Interrogating the recognition landscape of a conserved HIV-specific TCR reveals distinct bacterial peptide cross-reactivity. *eLife*, 9, 2020.

[7715] Louise Menendez, Talon Trecek, Suhasni Gopalakrishnan, Litao Tao, Alexander L Markowitz, Haoze V Yu, Xizi Wang, Juan Llamas, Chichou Huang, James Lee, Radha Kalluri, Justin Ichida, and Neil Segil. Generation of inner ear hair cells by direct lineage conversion of primary somatic cells. *eLife*, 9, 2020.

[7716] Fanbiao Meng, Minxian Qian, Bin Peng, Linyuan Peng, Xiaohui Wang, Kang Zheng, Zuojun Liu, Xiaolong Tang, Shuju Zhang, Shimin Sun, Xinyue Cao, Qiuxiang Pang, Bosheng Zhao, Wenbin Ma, Zhou Songyang, Bo Xu, Wei-Guo Zhu, Xingzhi Xu, and Baohua Liu. Synergy between SIRT1 and SIRT6 helps recognize DNA breaks and potentiates the DNA damage response and repair in humans and mice. *eLife*, 9, 2020.

[7717] Julia L Meng, Yupu Wang, Robert A Carrillo, and Ellie S Heckscher. Temporal transcription factors determine circuit membership by permanently altering motor neuron-to-muscle synaptic partnerships. *eLife*, 9, 2020.

[7718] Vinod Menon, Guillermo Gallardo, Mark A Pinsk, Van-Dang Nguyen, Jing-Rebecca Li, Weidong Cai, and Demian Wassermann. Microstructural organization of human insula is linked to its macrofunctional circuitry and predicts cognitive control. *eLife*, 9, 2020.

[7719] Clément Menuet, Angela A Connelly, Jaspreet K Bassi, Mariana R Melo, Sheng Le, Jessica Kamar, Natasha N Kumar, Stuart J McDougall, Simon McMullan, and Andrew M Allen. PreBötzinger complex neurons drive respiratory modulation of blood pressure and heart rate. *eLife*, 9, 2020.

[7720] Meredith B Metzger, Jessica L Scales, Mitchell F Dunklebarger, Jadranka Loncarek, and Allan M Weissman. A protein quality control pathway at the mitochondrial outer membrane. *eLife*, 9, 2020.

[7721] Renaud Mevel, Ivana Steiner, Susan Mason, Laura CA Galbraith, Rahima Patel, Muhammad ZH Fadlullah, Imran Ahmad, Hing Y Leung, Pedro Oliveira, Karen Blyth, Esther Baena, and Georges Lacaud. RUNX1 marks a luminal castration-resistant lineage established at the onset of prostate development. *eLife*, 9, 2020.

[7722] Valerie Michael, Jack Goffinet, John Pearson, Fan Wang, Katherine Tschida, and Richard Mooney. Circuit and synaptic organization of forebrain-to-midbrain pathways that promote and suppress vocalization. *eLife*, 9, 2020.

[7723] Thomas CT Michaels, Shuo Feng, Haiyi Liang, and L Mahadevan. Mechanics and kinetics of dynamic instability. *eLife*, 9, 2020.

[7724] Angie M Michaiel, Elliott TT Abe, and Cristopher M Niell. Dynamics of gaze control during prey capture in freely moving mice. *eLife*, 9, 2020.

[7725] FranÇois-Gaël Michalec, Itzhak Fouxon, Sami Souissi, and Markus Holzner. Efficient mate finding in planktonic copepods swimming in turbulence. *eLife*, 9, 2020.

[7726] Kevin Michalski, Johanna L Syrjanen, Erik Henze, Julia Kumpf, Hiro Furukawa, and Toshimitsu Kawate. The Cryo-EM structure of pannexin 1 reveals unique motifs for ion selection and inhibition. *eLife*, 9, 2020.

[7727] Eran Mick, Denis V Titov, Owen S Skinner, Rohit Sharma, Alexis A Jourdain, and Vamsi K Mootha. Distinct mitochondrial defects trigger the integrated stress response depending on the metabolic state of the cell. *eLife*, 9, 2020.

[7728] Laura E Mickelsen, William F Flynn, Kristen Springer, Lydia Wilson, Eric J Beltrami, Mohan Bolisetty, Paul Robson, and Alexander C Jackson. Cellular taxonomy and spatial organization of the murine ventral posterior hypothalamus. *eLife*, 9, 2020.

[7729] Federico Miguez-Cabello, Nuria Sánchez-Fernández, Natalia Yefimenko, Xavier Gasull, Esther Gratacòs-Batlle, and David Soto. AMPAR/TARP stoichiometry differentially modulates channel properties. *eLife*, 9, 2020.

[7730] Maria M Mikedis, Yuting Fan, Peter K Nicholls, Tsutomu Endo, Emily K Jackson, Sarah A Cobb, Dirk G de Rooij, and David C Page. DAZL mediates a broad translational program regulating expansion and differentiation of spermatogonial progenitors. *eLife*, 9, 2020.

[7731] Richard J Miles, Claire Kerridge, Laura Hilditch, Christopher Monit, David A Jacques, and Greg J Towers. MxB sensitivity of HIV-1 is determined by a highly variable and dynamic capsid surface. *eLife*, 9, 2020.

[7732] Hillary A Miller, Elizabeth S Dean, Scott D Pletcher, and Scott F Leiser. Cell non-autonomous regulation of health and longevity. *eLife*, 9, 2020.

[7733] Jeremy A Miller, Nathan W Gouwens, Bosiljka Tasic, Forrest Collman, Cindy TJ van Velthoven, Trygve E Bakken, Michael J Hawrylycz, Hongkui Zeng, Ed S Lein, and Amy Bernard. Common cell type nomenclature for the mammalian brain. *eLife*, 9, 2020.

[7734] Steve W Miller and James W Posakony. Disparate expression specificities coded by a shared Hox-C enhancer. *eLife*, 9, 2020.

[7735] Thomas D Miller, Trevor T-J Chong, Anne M Aimola Davies, Michael R Johnson, Sarosh R Irani, Masud Husain, Tammy WC Ng, Saiju Jacob, Paul Maddison, Christopher Kennard, Penny A Gowland, and Clive R Rosenthal. Human hippocampal CA3 damage disrupts both recent and remote episodic memories. *eLife*, 9, 2020.

[7736] Lakshmi E Miller-Vedam, Bastian Bräuning, Katerina D Popova, Nicole T Schirle Oakdale, Jessica L Bonnar, Jesuraj R Prabu, Elizabeth A Boydston, Natalia Sevillano, Matthew J Shurtleff, Robert M Stroud, Charles S Craik, Brenda A Schulman, Adam Frost, and Jonathan S Weissman. Structural and mechanistic basis of the EMC-dependent biogenesis of distinct transmembrane clients. *eLife*, 9, 2020.

[7737] Daniel J Millman, Gabriel Koch Ocker, Shiella Caldejon, India Kato, Josh D Larkin, Eric Kenji Lee, Jennifer Luviano, Chelsea Nayan, Thuyanh V Nguyen, Kat North, Sam Seid, Cassandra White, Jerome Lecoq, Clay Reid, Michael A Buice, and Saskia EJ de Vries. VIP interneurons in mouse primary visual cortex selectively enhance responses to weak but specific stimuli. *eLife*, 9, 2020.

[7738] Kirby L Mills and Nyeema C Harris. Humans disrupt access to prey for large African carnivores. *eLife*, 9, 2020.

[7739] Michael J Mina, C Jessica E Metcalf, Adrian B McDermott, Daniel C Douek, Jeremy Farrar, and Bryan T Grenfell. A Global lmmunological Observatory to meet a time of pandemics. *eLife*, 9, 2020.

[7740] Anastasia A Minervina, Mikhail V Pogorelyy, Ekaterina A Komech, Vadim K Karnaukhov, Petra Bacher, Elisa Rosati, Andre Franke, Dmitriy M Chudakov, Ilgar Z Mamedov, Yuri B Lebedev, Thierry Mora, and Aleksandra M Walczak. Primary and secondary anti-viral response captured by the dynamics and phenotype of individual T cell clones. *eLife*, 9, 2020.

[7741] Kyaw Thu Minn, Yuheng C Fu, Shenghua He, Sabine Dietmann, Steven C George, Mark A Anastasio, Samantha A Morris, and Lilianna Solnica-Krezel. High-resolution transcriptional and morphogenetic profiling of cells from micropatterned human ESC gastruloid cultures. *eLife*, 9, 2020.

[7742] Bogdan Andrei Mirauta, Daniel D Seaton, Dalila Bensaddek, Alejandro Brenes, Marc Jan Bonder, Helena Kilpinen, Chukwuma A Agu, Chukwuma A Agu, Alex Alderton, Petr Danecek, Rachel Denton, Richard Durbin, Daniel J Gaffney, Angela Goncalves, Reena Halai, Sarah Harper, Christopher M Kirton, Anja Kolb-Kokocinski, Andreas Leha, Shane A McCarthy, Yasin Memari, Minal Patel, Ewan Birney, Francesco Paolo Casale, Laura Clarke, Peter W Harrison, Helena Kilpinen, Ian Streeter, Davide Denovi, Oliver Stegle, Angus I Lamond, Ruta Meleckyte, Natalie Moens, Fiona M Watt, Willem H Ouwehand, Philip Beales, Oliver Stegle, and Angus I Lamond. Population-scale proteome variation in human induced pluripotent stem cells. *eLife*, 9, 2020.

[7743] Snigdha Misra and Mariana F Wolfner. Drosophila seminal sex peptide associates with rival as well as own sperm, providing SP function in polyandrous females. *eLife*, 9, 2020.

[7744] Patrick S Mitchell, Justin L Roncaioli, Elizabeth A Turcotte, Lisa Goers, Roberto A Chavez, Angus Y Lee, Cammie F Lesser, Isabella Rauch, and Russell E Vance. NAIP–NLRC4-deficient mice are susceptible to shigellosis. *eLife*, 9, 2020.

[7745] Briana E Mittleman, Sebastian Pott, Shane Warland, Tony Zeng, Zepeng Mu, Mayher Kaur, Yoav Gilad, and Yang Li. Alternative polyadenylation mediates genetic regulation of gene expression. *eLife*, 9, 2020.

[7746] Masatoshi Miyamoto, Takayuki Sassa, Megumi Sawai, and Akio Kihara. Lipid polarity gradient formed by $\omega$-hydroxy lipids in tear film prevents dry eye disease. *eLife*, 9, 2020.

[7747] Ryoji Miyazaki, Yoshinori Akiyama, and Hiroyuki Mori. Fine interaction profiling of VemP and mechanisms responsible for its translocation-coupled arrest-cancelation. *eLife*, 9, 2020.

[7748] Fredrick M Mobegi, Lex EX Leong, Fintan Thompson, Sean M Taylor, Linton R Harriss, Jocelyn M Choo, Steven L Taylor, Steve L Wesselingh, Robyn McDermott, Kerry L Ivey, and Geraint B Rogers. Intestinal microbiology shapes population health impacts of diet and lifestyle risk exposures in Torres Strait Islander communities. *eLife*, 9, 2020.

[7749] Samuel J Modlin, Derek Conkle-Gutierrez, Calvin Kim, Scott N Mitchell, Christopher Morrissey, Brian C Weinrick, William R Jacobs, Sarah M Ramirez-Busby, Sven E Hoffner, and Faramarz Valafar. Drivers and sites of diversity in the DNA adenine methylomes of 93 Mycobacterium tuberculosis complex clinical isolates. *eLife*, 9, 2020.

[7750] Klaudia Modlinska and Wojciech Pisula. The Norway rat, from an obnoxious pest to a laboratory pet. *eLife*, 9, 2020.

[7751] Polycarp Mogeni, Alain Vandormael, Diego Cuadros, Christopher Appleton, and Frank Tanser. Impact of community piped water coverage on re-infection with urogenital schistosomiasis in rural South Africa. *eLife*, 9, 2020.

[7752] Pedrum Mohammadi-Shemirani, Michael Chong, Marie Pigeyre, Robert W Morton, Hertzel C Gerstein, and Guillaume Paré. Effects of lifelong testosterone exposure on health and disease using Mendelian randomization. *eLife*, 9, 2020.

[7753] Laura Molina-García, Carla Lloret-Fernández, Steven J Cook, Byunghyuk Kim, Rachel C Bonnington, Michele Sammut, Jack M O'Shea, Sophie PR Gilbert, David J Elliott, David H Hall, Scott W Emmons, Arantza Barrios, and Richard J Poole. Direct glia-to-neuron transdifferentiation gives rise to a pair of male-specific neurons that ensure nimble male mating. *eLife*, 9, 2020.

[7754] Baisakhi Mondal, Hongjian Jin, Satish Kallappagoudar, Yurii Sedkov, Tanner Martinez, Monica F Sentmanat, Greg J Poet, Chunliang Li, Yiping Fan, Shondra M Pruett-Miller, and Hans-Martin Herz. The histone deacetylase complex MiDAC regulates a neurodevelopmental gene expression program to control neurite outgrowth. *eLife*, 9, 2020.

[7755] Hannah R Monday, Mathieu Bourdenx, Bryen A Jordan, and Pablo E Castillo. CB1-receptor-mediated inhibitory LTD triggers presynaptic remodeling via protein synthesis and ubiquitination. *eLife*, 9, 2020.

[7756] Viviana Monje-Galvan and Gregory A Voth. Binding mechanism of the matrix domain of HIV-1 gag on lipid membranes. *eLife*, 9, 2020.

[7757] Christian Montellese, Jasmin van den Heuvel, Caroline Ashiono, Kerstin Dörner, André Melnik, Stefanie Jonas, Ivo Zemp, Paola Picotti, Ludovic C Gillet, and Ulrike Kutay. USP16 counteracts mono-ubiquitination of RPS27a and promotes maturation of the 40S ribosomal subunit. *eLife*, 9, 2020.

[7758] Marta Montero-Crespo, Marta Dominguez-Alvaro, Patricia Rondon-Carrillo, Lidia Alonso-Nanclares, Javier DeFelipe, and Lidia Blazquez-Llorca. Three-dimensional synaptic organization of the human hippocampal CA1 field. *eLife*, 9, 2020.

[7759] Elisa Monzón-Casanova, Louise S Matheson, Kristina Tabbada, Kathi Zarnack, Christopher WJ Smith, and Martin Turner. Polypyrimidine tract-binding proteins are essential for B cell development. *eLife*, 9, 2020.

[7760] Hyang Mi Moon, Simon Hippenmeyer, Liqun Luo, and Anthony Wynshaw-Boris. LIS1 determines cleavage plane positioning by regulating actomyosin-mediated cell membrane contractility. *eLife*, 9, 2020.

[7761] Kyeong-Hye Moon, Ji-Hyun Ma, Hyehyun Min, Heiyeun Koo, HongKyung Kim, Hyuk Wan Ko, and Jinwoong Bok. Dysregulation of sonic hedgehog signaling causes hearing loss in ciliopathy mouse models. *eLife*, 9, 2020.

[7762] David Moreno-Delgado, Mar Puigdellívol, Estefanía Moreno, Mar Rodríguez-Ruiz, Joaquín Botta, Paola Gasperini, Anna Chiarlone, Lesley A Howell, Marco Scarselli, Vicent Casadó, Antoni Cortés, Sergi Ferré, Manuel Guzmán, Carmen Lluís, Jordi Alberch, Enric I Canela, Silvia Ginés, and Peter J McCormick. Modulation of dopamine D1 receptors via histamine H3 receptors is a novel therapeutic target for Huntington's disease. *eLife*, 9, 2020.

[7763] Mai M Morimoto, Aljoscha Nern, Arthur Zhao, Edward M Rogers, Allan M Wong, Mathew D Isaacson, Davi D Bock, Gerald M Rubin, and Michael B Reiser. Spatial readout of visual looming in the central brain of Drosophila. *eLife*, 9, 2020.

[7764] Valerie J Morley, Clare L Kinnear, Derek G Sim, Samantha N Olson, Lindsey M Jackson, Elsa Hansen, Grace A Usher, Scott A Showalter, Manjunath P Pai, Robert J Woods, and Andrew F Read. An adjunctive therapy administered with an antibiotic prevents enrichment of antibiotic-resistant clones of a colonizing opportunistic pathogen. *eLife*, 9, 2020.

[7765] Dylan H Morris, Velislava N Petrova, Fernando W Rossine, Edyth Parker, Bryan T Grenfell, Richard A Neher, Simon A Levin, and Colin A Russell. Asynchrony between virus diversity and antibody selection limits influenza virus evolution. *eLife*, 9, 2020.

[7766] Tim T Morris, Neil M Davies, and George Davey Smith. Can education be personalised using pupils' genetic data? *eLife*, 9, 2020.

[7767] Jeremiah K Morrow, Michael X Cohen, and Katalin M Gothard. Mesoscopic-scale functional networks in the primate amygdala. *eLife*, 9, 2020.

[7768] Jose F Moruno-Manchon, Pauline Lejault, Yaoxuan Wang, Brenna McCauley, Pedram Honarpisheh, Diego A Morales Scheihing, Shivani Singh, Weiwei Dang, Nayun Kim, Akihiko Urayama, Liang Zhu, David Monchaud, Louise D McCullough, and Andrey S Tsvetkov. Small-molecule G-quadruplex stabilizers reveal a novel pathway of autophagy regulation in neurons. *eLife*, 9, 2020.

[7769] James M Mossner, Renata Batista-Brito, Rima Pant, and Jessica A Cardin. Developmental loss of MeCP2 from VIP interneurons impairs cortical function and behavior. *eLife*, 9, 2020.

[7770] Hakhamanesh Mostafavi, Arbel Harpak, Ipsita Agarwal, Dalton Conley, Jonathan K Pritchard, and Molly Przeworski. Variable prediction accuracy of polygenic scores within an ancestry group. *eLife*, 9, 2020.

[7771] Dihia Moussaoui, James P Robblee, Daniel Auguin, Elena B Krementsova, Silvia Haase, Thomas CA Blake, Jake Baum, Julien Robert-Paganin, Kathleen M Trybus, and Anne Houdusse. Full-length Plasmodium falciparum myosin A and essential light chain PfELC structures provide new anti-malarial targets. *eLife*, 9, 2020.

[7772] Sara N Mouton, David J Thaller, Matthew M Crane, Irina L Rempel, Owen T Terpstra, Anton Steen, Matt Kaeberlein, C Patrick Lusk, Arnold J Boersma, and Liesbeth M Veenhoff. A physicochemical perspective of aging from single-cell analysis of pH, macromolecular and organellar crowding in yeast. *eLife*, 9, 2020.

[7773] Anusorn Mudla, Yanfei Jiang, Kei-ichiro Arimoto, Bingxian Xu, Adarsh Rajesh, Andy P Ryan, Wei Wang, Matthew D Daugherty, Dong-Er Zhang, and Nan Hao. Cell-cycle-gated feedback control mediates desensitization to interferon stimulation. *eLife*, 9, 2020.

[7774] Douaa Mugahid, Marian Kalocsay, Xili Liu, Jonathan Scott Gruver, Leonid Peshkin, and Marc W Kirschner. YAP regulates cell size and growth dynamics via non-cell autonomous mediators. *eLife*, 9, 2020.

[7775] Ananya Mukherjee, Randhir Singh, Sreeram Udayan, Sayan Biswas, Pothula Purushotham Reddy, Saumya Manmadhan, Geen George, Shilpa Kumar, Ranabir Das, Balaji M Rao, and Akash Gulyani. A Fyn biosensor reveals pulsatile, spatially localized kinase activity and signaling crosstalk in live mammalian cells. *eLife*, 9, 2020.

[7776] Arghya Mukherjee, Navdeep Bajwa, Norman H Lam, César Porrero, Francisco Clasca, and Michael M Halassa. Variation of connectivity across exemplar sensory and associative thalamocortical loops in the mouse. *eLife*, 9, 2020.

[7777] Shreyasi Mukherjee, Praneet Chaturvedi, Scott A Rankin, Margaret B Fish, Marcin Wlizla, Kitt D Paraiso, Melissa MacDonald, Xiaoting Chen, Matthew T Weirauch, Ira L Blitz, Ken WY Cho, and Aaron M Zorn. Sox17 and $\beta$-catenin co-occupy Wnt-responsive enhancers to govern the endoderm gene regulatory network. *eLife*, 9, 2020.

[7778] Marie Mulier, Nele Van Ranst, Nikky Corthout, Sebastian Munck, Pieter Vanden Berghe, Joris Vriens, Thomas Voets, and Lauri Moilanen. Upregulation of TRPM3 in nociceptors innervating inflamed tissue. *eLife*, 9, 2020.

[7779] Kenji Murata, Munehide Nakatsugawa, Muhammed A Rahman, Linh T Nguyen, Douglas G Millar, David T Mulder, Kenji Sugata, Hiroshi Saijo, Yukiko Matsunaga, Yuki Kagoya, Tingxi Guo, Mark Anczurowski, Chung-Hsi Wang, Brian D Burt, Dalam Ly, Kayoko Saso, Alexandra Easson, David P Goldstein, Michael Reedijk, Danny Ghazarian, Trevor J Pugh, Marcus O Butler, Tak W Mak, Pamela S Ohashi, and Naoto Hirano. Landscape mapping of shared antigenic epitopes and their cognate TCRs of tumor-infiltrating T lymphocytes in melanoma. *eLife*, 9, 2020.

[7780] Jonathan Muri, Helen Thut, Qian Feng, and Manfred Kopf. Thioredoxin-1 distinctly promotes NF-$\kappa$B target DNA binding and NLRP3 inflammasome activation independently of Txnip. *eLife*, 9, 2020.

[7781] Timothy H Murphy, Nicholas J Michelson, Jamie D Boyd, Tony Fong, Luis A Bolanos, David Bierbrauer, Teri Siu, Matilde Balbi, Federico Bolanos, Matthieu Vanni, and Jeff M LeDue. Automated task training and longitudinal monitoring of mouse mesoscale cortical circuits using home cages. *eLife*, 9, 2020.

[7782] Shreesh P Mysore and Ninad B Kothari. Mechanisms of competitive selection: A canonical neural circuit framework. *eLife*, 9, 2020.

[7783] Francisco M Nadal-Nicolás, Vincent P Kunze, John M Ball, Brian T Peng, Akshay Krishnan, Gaohui Zhou, Lijin Dong, and Wei Li. True S-cones are concentrated in the ventral mouse retina and wired for color detection in the upper visual field. *eLife*, 9, 2020.

[7784] Elina Nagaeva, Ivan Zubarev, Carolina Bengtsson Gonzales, Mikko Forss, Kasra Nikouei, Elena de Miguel, Lauri Elsilä, Anni-Maija Linden, Jens Hjerling-Leffler, George J Augustine, and Esa R Korpi. Heterogeneous somatostatin-expressing neuron population in mouse ventral tegmental area. *eLife*, 9, 2020.

[7785] Sumeda Nandadasa, Jason M Szafron, Vai Pathak, Sae-Il Murtada, Caroline M Kraft, Anna O'Donnell, Christian Norvik, Clare Hughes, Bruce Caterson, Miriam S Domowicz, Nancy B Schwartz, Karin Tran-Lundmark, Martina Veigl, David Sedwick, Elliot H Philipson, Jay D Humphrey, and Suneel S Apte. Vascular dimorphism ensured by regulated proteoglycan dynamics favors rapid umbilical artery closure at birth. *eLife*, 9, 2020.

[7786] Shyama Nandakumar, Olga Grushko, and Laura A Buttitta. Polyploidy in the adult Drosophila brain. *eLife*, 9, 2020.

[7787] Anirvan S Nandy, Jonathan J Nassi, Monika P Jadi, and John H Reynolds. Correction: Optogenetically induced low-frequency correlations impair perception. *eLife*, 9, 2020.

[7788] Madhumitha Narasimhan, Alexander Johnson, Roshan Prizak, Walter Anton Kaufmann, Shutang Tan, Barbara Casillas-Pérez, and Jiří Friml. Evolutionarily unique mechanistic framework of clathrin-mediated endocytosis in plants. *eLife*, 9, 2020.

[7789] Filipe Nascimento, Matthew James Broadhead, Efstathia Tetringa, Eirini Tsape, Laskaro Zagoraiou, and Gareth Brian Miles. Synaptic mechanisms underlying modulation of locomotor-related motoneuron output by premotor cholinergic interneurons. *eLife*, 9, 2020.

[7790] Zeyad D Nassar, Chui Yan Mah, Jonas Dehairs, Ingrid JG Burvenich, Swati Irani, Margaret M Centenera, Madison Helm, Raj K Shrestha, Max Moldovan, Anthony S Don, Jeff Holst, Andrew M Scott, Lisa G Horvath, David J Lynn, Luke A Selth, Andrew J Hoy, Johannes V Swinnen, and Lisa M Butler. Human DECR1 is an androgen-repressed survival factor that regulates PUFA oxidation to protect prostate tumor cells from ferroptosis. *eLife*, 9, 2020.

[7791] Marco A Navarro, Autoosa Salari, Jenna L Lin, Luke M Cowan, Nicholas J Penington, Mirela Milescu, and Lorin S Milescu. Sodium channels implement a molecular leaky integrator that detects action potentials and regulates neuronal firing. *eLife*, 9, 2020.

[7792] Jason Neidleman, Xiaoyu Luo, Julie Frouard, Guorui Xie, Feng Hsiao, Tongcui Ma, Vincent Morcilla, Ashley Lee, Sushama Telwatte, Reuben Thomas, Whitney Tamaki, Benjamin Wheeler, Rebecca Hoh, Ma Somsouk, Poonam Vohra, Jeffrey Milush, Katherine Sholtis James, Nancie M Archin, Peter W Hunt, Steven G Deeks, Steven A Yukl, Sarah Palmer, Warner C Greene, and Nadia R Roan. Phenotypic analysis of the unstimulated in vivo HIV CD4 T cell reservoir. *eLife*, 9, 2020.

[7793] Branden R Nelson, Rebecca D Hodge, Ray AM Daza, Prem Prakash Tripathi, Sebastian J Arnold, Kathleen J Millen, and Robert F Hevner. Intermediate progenitors support migration of neural stem cells into dentate gyrus outer neurogenic niches. *eLife*, 9, 2020.

[7794] Chase W Nelson, Zachary Ardern, Tony L Goldberg, Chen Meng, Chen-Hao Kuo, Christina Ludwig, Sergios-Orestis Kolokotronis, and Xinzhu Wei. Dynamically evolving novel overlapping gene as a factor in the SARS-CoV-2 pandemic. *eLife*, 9, 2020.

[7795] Elisa Nerli, Mauricio Rocha-Martins, and Caren Norden. Asymmetric neurogenic commitment of retinal progenitors involves Notch through the endocytic pathway. *eLife*, 9, 2020.

[7796] Ritam Neupane, Vera P Pisareva, Carlos F Rodriguez, Andrey V Pisarev, and Israel S Fernández. A complex IRES at the 5′-UTR of a viral mRNA assembles a functional 48S complex via an uAUG intermediate. *eLife*, 9, 2020.

[7797] Kleber Neves and Olavo B Amaral. Addressing selective reporting of experiments through predefined exclusion criteria. *eLife*, 9, 2020.

[7798] Samuel A Neymotin, Dylan S Daniels, Blake Caldwell, Robert A McDougal, Nicholas T Carnevale, Mainak Jas, Christopher I Moore, Michael L Hines, Matti Hämäläinen, and Stephanie R Jones. Human Neocortical Neurosolver (HNN), a new software tool for interpreting the cellular and network origin of human MEG/EEG data. *eLife*, 9, 2020.

[7799] Hong-Viet Ngo, Juergen Fell, and Bernhard Staresina. Sleep spindles mediate hippocampal-neocortical coupling during long-duration ripples. *eLife*, 9, 2020.

[7800] Tenzin Ngodup, Gabriel E Romero, and Laurence O Trussell. Identification of an inhibitory neuron subtype, the L-stellate cell of the cochlear nucleus. *eLife*, 9, 2020.

[7801] Giang T Nguyen, Lamyaa Shaban, Matthias Mack, Kenneth D Swanson, Stephen C Bunnell, David B Sykes, and Joan Mecsas. SKAP2 is required for defense against K. pneumoniae infection and neutrophil respiratory burst. *eLife*, 9, 2020.

[7802] John ME Nichols, Vlatka Antolović, Jacob D Reich, Sophie Brameyer, Peggy Paschke, and Jonathan R Chubb. Cell and molecular transitions during efficient dedifferentiation. *eLife*, 9, 2020.

[7803] Zuqin Nie, Chunhua Guo, Subhendu K Das, Carson C Chow, Eric Batchelor, S Stoney Simons, and David Levens. Dissecting transcriptional amplification by MYC. *eLife*, 9, 2020.

[7804] Rene Niehus, Esther van Kleef, Yin Mo, Agata Turlej-Rogacka, Christine Lammens, Yehuda Carmeli, Herman Goossens, Evelina Tacconelli, Biljana Carevic, Liliana Preotescu, Surbhi Malhotra-Kumar, and Ben S Cooper. Quantifying antibiotic impact on within-patient dynamics of extended-spectrum beta-lactamase resistance. *eLife*, 9, 2020.

[7805] Terren K Niethamer, Collin T Stabler, John P Leach, Jarod A Zepp, Michael P Morley, Apoorva Babu, Su Zhou, and Edward E Morrisey. Defining the role of pulmonary endothelial cell heterogeneity in the response to acute lung injury. *eLife*, 9, 2020.

[7806] Satoshi Ninagawa, Seiichiro Tada, Masaki Okumura, Kenta Inoguchi, Misaki Kinoshita, Shingo Kanemura, Koshi Imami, Hajime Umezawa, Tokiro Ishikawa, Robert B Mackin, Seiji Torii, Yasushi Ishihama, Kenji Inaba, Takayuki Anazawa, Takahiko Nagamine, and Kazutoshi Mori. Antipsychotic olanzapine-induced misfolding of proinsulin in the endoplasmic reticulum accounts for atypical development of diabetes. *eLife*, 9, 2020.

[7807] Long-Gang Niu, Ping Liu, Zhao-Wen Wang, and Bojun Chen. Slo2 potassium channel function depends on RNA editing-regulated expression of a SCYL1 protein. *eLife*, 9, 2020.

[7808] Meiqi Niu, Daniele Impieri, Lucija Rapan, Thomas Funck, Nicola Palomero-Gallagher, and Karl Zilles. Receptor-driven, multimodal mapping of cortical areas in the macaque monkey intraparietal sulcus. *eLife*, 9, 2020.

[7809] Yiming Niu, Xiao Tao, Kouki K Touhara, and Roderick MacKinnon. Cryo-EM analysis of PIP2 regulation in mammalian GIRK channels. *eLife*, 9, 2020.

[7810] Felix E Nolet, Alexandra Vandervelde, Arno Vanderbeke, Liliana Piñeros, Jeremy B Chang, and Lendert Gelens. Nuclei determine the spatial origin of mitotic waves. *eLife*, 9, 2020.

[7811] Pieter R Norden, Amélie Sabine, Ying Wang, Cansaran Saygili Demir, Ting Liu, Tatiana V Petrova, and Tsutomu Kume. Shear stimulation of FOXC1 and FOXC2 differentially regulates cytoskeletal activity during lymphatic valve maturation. *eLife*, 9, 2020.

[7812] Sigrid Noreng, Richard Posert, Arpita Bharadwaj, Alexandra Houser, and Isabelle Baconguis. Molecular principles of assembly, activation, and inhibition in epithelial sodium channel. *eLife*, 9, 2020.

[7813] Rosalind Norkett, Urko del Castillo, Wen Lu, and Vladimir I Gelfand. Ser/Thr kinase Trc controls neurite outgrowth in Drosophila by modulating microtubule-microtubule sliding. *eLife*, 9, 2020.

[7814] Sara M Nowinski, Ashley Solmonson, Scott F Rusin, J Alan Maschek, Claire L Bensard, Sarah Fogarty, Mi-Young Jeong, Sandra Lettlova, Jordan A Berg, Jeffrey T Morgan, Yeyun Ouyang, Bradley C Naylor, Joao A Paulo, Katsuhiko Funai, James E Cox, Steven P Gygi, Dennis R Winge, Ralph J DeBerardinis, and Jared Rutter. Mitochondrial fatty acid synthesis coordinates oxidative metabolism in mammalian mitochondria. *eLife*, 9, 2020.

[7815] Marlies P Noz, Siroon Bekkering, Laszlo Groh, Tim MJ Nielen, Evert JP Lamfers, Andreas Schlitzer, Saloua El Messaoudi, Niels van Royen, Erik HJPG Huys, Frank WMB Preijers, Esther MM Smeets, Erik HJG Aarntzen, Bowen Zhang, Yang Li, Manita EJ Bremmers, Walter JFM van der Velden, Harry Dolstra, Leo AB Joosten, Marc E Gomes, Mihai G Netea, and Niels P Riksen. Reprogramming of bone marrow myeloid progenitor cells in patients with severe coronary artery disease. *eLife*, 9, 2020.

[7816] Nicole L Nuckolls, Anthony C Mok, Jeffrey J Lange, Kexi Yi, Tejbir S Kandola, Andrew M Hunn, Scott McCroskey, Julia L Snyder, María Angélica Bravo Núñez, Melainia McClain, Sean A McKinney, Christopher Wood, Randal Halfmann, and Sarah E Zanders. The wtf4 meiotic driver utilizes controlled protein aggregation to generate selective cell death. *eLife*, 9, 2020.

[7817] John P O'Donnell, Ben P Phillips, Yuichi Yagita, Szymon Juszkiewicz, Armin Wagner, Duccio Malinverni, Robert J Keenan, Elizabeth A Miller, and Ramanujan S Hegde. The architecture of EMC reveals a path for membrane protein insertion. *eLife*, 9, 2020.

[7818] SeCheol Oh, Navid Paknejad, and Richard K Hite. Gating and selectivity mechanisms for the lysosomal K+ channel TMEM175. *eLife*, 9, 2020.

[7819] Seunghee Oh, Jaehyoun Lee, Selene K Swanson, Laurence Florens, Michael P Washburn, and Jerry L Workman. Yeast Nuak1 phosphorylates histone H3 threonine 11 in low glucose stress by the cooperation of AMPK and CK2 signaling. *eLife*, 9, 2020.

[7820] Yohei Ohashi, Shirley Tremel, Glenn Robert Masson, Lauren McGinney, Jerome Boulanger, Ksenia Rostislavleva, Christopher M Johnson, Izabella Niewczas, Jonathan Clark, and Roger L Williams. Membrane characteristics tune activities of endosomal and autophagic human VPS34 complexes. *eLife*, 9, 2020.

[7821] Machiko Ohbayashi. Inhibition of protein synthesis in M1 of monkeys disrupts performance of sequential movements guided by memory. *eLife*, 9, 2020.

[7822] Sho Ohta, Young Rae Ji, Daniel Martin, and Doris K Wu. Emx2 regulates hair cell rearrangement but not positional identity within neuromasts. *eLife*, 9, 2020.

[7823] Megan Okada, Ping Guo, Shai-anne Nalder, and Paul A Sigala. Doxycycline has distinct apicoplast-specific mechanisms of antimalarial activity. *eLife*, 9, 2020.

[7824] Benjamin W Okaty, Nikita Sturrock, Yasmin Escobedo Lozoya, YoonJeung Chang, Rebecca A Senft, Krissy A Lyon, Olga V Alekseyenko, and Susan M Dymecki. A single-cell transcriptomic and anatomic atlas of mouse dorsal raphe Pet1 neurons. *eLife*, 9, 2020.

[7825] Jordan G Okie, Amisha T Poret-Peterson, Zarraz MP Lee, Alexander Richter, Luis D Alcaraz, Luis E Eguiarte, Janet L Siefert, Valeria Souza, Chris L Dupont, and James J Elser. Genomic adaptations in information processing underpin trophic strategy in a whole-ecosystem nutrient enrichment experiment. *eLife*, 9, 2020.

[7826] Shinichiro Oku, Huijuan Feng, Steven Connor, Andrea Toledo, Peng Zhang, Yue Zhang, Olivier Thoumine, Chaolin Zhang, and Ann Marie Craig. Alternative splicing at neuroligin site A regulates glycan interaction and synaptogenic activity. *eLife*, 9, 2020.

[7827] Kanji Okumoto, Mahmoud El Shermely, Masanao Natsui, Hidetaka Kosako, Ryuichi Natsuyama, Toshihiro Marutani, and Yukio Fujiki. The peroxisome counteracts oxidative stresses by suppressing catalase import via Pex14 phosphorylation. *eLife*, 9, 2020.

[7828] Viktor János Oláh, David Lukacsovich, Jochen Winterer, Antónia Arszovszki, Andrea Lőrincz, Zoltan Nusser, Csaba Földy, and János Szabadics. Functional specification of CCK+ interneurons by alternative isoforms of Kv4.3 auxiliary subunits. *eLife*, 9, 2020.

[7829] Itsaso Olasagasti and Anne-Lise Giraud. Integrating prediction errors at two time scales permits rapid recalibration of speech sound categories. *eLife*, 9, 2020.

[7830] Claire S Oldfield, Irene Grossrubatscher, Mario Chávez, Adam Hoagland, Alex R Huth, Elizabeth C Carroll, Andrew Prendergast, Tony Qu, Jack L Gallant, Claire Wyart, and Ehud Y Isacoff. Experience, circuit dynamics, and forebrain recruitment in larval zebrafish prey capture. *eLife*, 9, 2020.

[7831] Timothy P O'Leary, Kaitlin E Sullivan, Lihua Wang, Jody Clements, Andrew L Lemire, and Mark S Cembrowski. Extensive and spatially variable within-cell-type heterogeneity across the basolateral amygdala. *eLife*, 9, 2020.

[7832] Erik Oliemuller, Richard Newman, Siu Man Tsang, Shane Foo, Gareth Muirhead, Farzana Noor, Syed Haider, Iskander Aurrekoetxea-Rodríguez, Maria dM Vivanco, and Beatrice A Howard. SOX11 promotes epithelial/mesenchymal hybrid state and alters tropism of invasive breast cancer cells. *eLife*, 9, 2020.

[7833] Cristina Olivieri, Yingjie Wang, Geoffrey C Li, Manu V S, Jonggul Kim, Benjamin R Stultz, Matthew Neibergall, Fernando Porcelli, Joseph M Muretta, David DT Thomas, Jiali Gao, Donald K Blumenthal, Susan S Taylor, and Gianluigi Veglia. Multi-state recognition pathway of the intrinsically disordered protein kinase inhibitor by protein kinase A. *eLife*, 9, 2020.

[7834] Lior Onn, Miguel Portillo, Stefan Ilic, Gal Cleitman, Daniel Stein, Shai Kaluski, Ido Shirat, Zeev Slobodnik, Monica Einav, Fabian Erdel, Barak Akabayov, and Debra Toiber. SIRT6 is a DNA double-strand break sensor. *eLife*, 9, 2020.

[7835] Nicola S Orefice, Owein Guillemot-Legris, Rosanna Capasso, Pauline Bottemanne, Philippe Hantraye, Michele Caraglia, Giuseppe Orefice, Mireille Alhouayek, and Giulio G Muccioli. miRNA profile is altered in a modified EAE mouse model of multiple sclerosis featuring cortical lesions. *eLife*, 9, 2020.

[7836] Krystal Ann Orlando, Amber K Douglas, Aierken Abudu, Yemin Wang, Basile Tessier-Cloutier, Weiping Su, Alec Peters, Larry S Sherman, Rayvon Moore, Vinh Nguyen, Gian Luca Negri, Shane Colborne, Gregg B Morin, Friedrich Kommoss, Jessica D Lang, William PD Hendricks, Elizabeth A Raupach, Patrick Pirrotte, David G Huntsman, Jeffrey M Trent, Joel S Parker, Jesse R Raab, and Bernard E Weissman. Re-expression of SMARCA4/BRG1 in small cell carcinoma of ovary, hypercalcemic type (SCCOHT) promotes an epithelial-like gene signature through an AP-1-dependent mechanism. *eLife*, 9, 2020.

[7837] Fabrizio Orsenigo, Lei Liu Conze, Suvi Jauhiainen, Monica Corada, Francesca Lazzaroni, Matteo Malinverno, Veronica Sundell, Sara Isabel Cunha, Johan Brännström, Maria Ascención Globisch, Claudio Maderna, Maria Grazia Lampugnani, Peetra Ulrica Magnusson, and Elisabetta Dejana. Mapping endothelial-cell diversity in cerebral cavernous malformations at single-cell resolution. *eLife*, 9, 2020.

[7838] Jennifer Orthmann-Murphy, Cody L Call, Gian C Molina-Castro, Yu Chen Hsieh, Matthew N Rasband, Peter A Calabresi, and Dwight E Bergles. Remyelination alters the pattern of myelin in the cerebral cortex. *eLife*, 9, 2020.

[7839] Lynda S Ostedgaard, Margaret P Price, Kristin M Whitworth, Mahmoud H Abou Alaiwa, Anthony J Fischer, Akshaya Warrier, Melissa Samuel, Lee D Spate, Patrick D Allen, Brieanna M Hilkin, Guillermo S Romano Ibarra, Miguel E Ortiz Bezara, Brian J Goodell, Steven E Mather, Linda S Powers, Mallory R Stroik, Nicholas D Gansemer, Camilla E Hippee, Keyan Zarei, J Adam Goeken, Thomas R Businga, Eric A Hoffman, David K Meyerholz, Randall S Prather, David A Stoltz, and Michael J Welsh. Lack of airway submucosal glands impairs respiratory host defenses. *eLife*, 9, 2020.

[7840] Minhui Ouyang, Qinmu Peng, Tina Jeon, Roy Heyne, Lina Chalak, and Hao Huang. Diffusion-MRI-based regional cortical microstructure at birth for predicting neurodevelopmental outcomes of 2-year-olds. *eLife*, 9, 2020.

[7841] Jin Rong Ow, Matias J Caldez, Gözde Zafer, Juat Chin Foo, Hong Yu Li, Soumita Ghosh, Heike Wollmann, Amaury Cazenave-Gassiot, Chee Bing Ong, Markus R Wenk, Weiping Han, Hyungwon Choi, and Philipp Kaldis. Remodeling of whole-body lipid metabolism and a diabetic-like phenotype caused by loss of CDK1 and hepatocyte division. *eLife*, 9, 2020.

[7842] Jennifer P Owen, Robert N Kelsh, and Christian A Yates. A quantitative modelling approach to zebrafish pigment pattern formation. *eLife*, 9, 2020.

[7843] Gizem özbaykal, Eva Wollrab, Francois Simon, Antoine Vigouroux, Baptiste Cordier, Andrey Aristov, Thibault Chaze, Mariette Matondo, and Sven van Teeffelen. The transpeptidase PBP2 governs initial localization and activity of the major cell-wall synthesis machinery in E. coli. *eLife*, 9, 2020.

[7844] Güliz Gürel özcan, Sumi Lim, Patricia LA Leighton, W Ted Allison, and Jason Rihel. Sleep is bi-directionally modified by amyloid beta oligomers. *eLife*, 9, 2020.

[7845] Giulia Paci, Tiantian Zheng, Joana Caria, Anton Zilman, and Edward A Lemke. Molecular determinants of large cargo transport into the nucleus. *eLife*, 9, 2020.

[7846] Jone Paesmans, Ella Martin, Babette Deckers, Marjolijn Berghmans, Ritika Sethi, Yannick Loeys, Els Pardon, Jan Steyaert, Patrik Verstreken, Christian Galicia, and Wim Versées. A structure of substrate-bound Synaptojanin1 provides new insights in its mechanism and the effect of disease mutations. *eLife*, 9, 2020.

[7847] Emily Ling-Lin Pai, Jin Chen, Siavash Fazel Darbandi, Frances S Cho, Jiapei Chen, Susan Lindtner, Julia S Chu, Jeanne T Paz, Daniel Vogt, Mercedes F Paredes, and John LR Rubenstein. Maf and Mafb control mouse pallial interneuron fate and maturation through neuropsychiatric disease gene regulation. *eLife*, 9, 2020.

[7848] Ananya Pal, Jia Yu Leung, Gareth Chin Khye Ang, Vinay Kumar Rao, Luca Pignata, Huey Jin Lim, Maxime Hebrard, Kenneth TE Chang, Victor KM Lee, Ernesto Guccione, and Reshma Taneja. EHMT2 epigenetically suppresses Wnt signaling and is a potential target in embryonal rhabdomyosarcoma. *eLife*, 9, 2020.

[7849] Hajnalka L Pálinkás, Angéla Békési, Gergely Róna, Lőrinc Pongor, Gábor Papp, Gergely Tihanyi, Eszter Holub, ádám Póti, Carolina Gemma, Simak Ali, Michael J Morten, Eli Rothenberg, Michele Pagano, Dávid Szűts, Balázs Győrffy, and Beáta G Vértessy. Genome-wide alterations of uracil distribution patterns in human DNA upon chemotherapeutic treatments. *eLife*, 9, 2020.

[7850] Suresh Pallikkuth, Sidhartha Chaudhury, Pinyi Lu, Li Pan, Erik Jongert, Ulrike Wille-Reece, and Savita Pahwa. A delayed fractionated dose RTS,S AS01 vaccine regimen mediates protection via improved T follicular helper and B cell responses. *eLife*, 9, 2020.

[7851] Li Pan, Madeleine E Lemieux, Tom Thomas, Julia M Rogers, Colin H Lipper, Winston Lee, Carl Johnson, Lynette M Sholl, Andrew P South, Jarrod A Marto, Guillaume O Adelmant, Stephen C Blacklow, and Jon C Aster. IER5, a DNA damage response gene, is required for Notch-mediated induction of squamous cell differentiation. *eLife*, 9, 2020.

[7852] Zhao-Qian Pan, Guang-Can Shao, Xiao-Man Liu, Quan Chen, Meng-Qiu Dong, and Li-Lin Du. Atg1 kinase in fission yeast is activated by Atg11-mediated dimerization and cis-autophosphorylation. *eLife*, 9, 2020.

[7853] Juanita Pang, Jennifer A Slyker, Sunando Roy, Josephine Bryant, Claire Atkinson, Juliana Cudini, Carey Farquhar, Paul Griffiths, James Kiarie, Sofia Morfopoulou, Alison C Roxby, Helena Tutil, Rachel Williams, Soren Gantt, Richard A Goldstein, and Judith Breuer. Mixed cytomegalovirus genotypes in HIV-positive mothers show compartmentalization and distinct patterns of transmission to infants. *eLife*, 9, 2020.

[7854] Vanja Panic, Stephanie Pearson, James Banks, Trevor S Tippetts, Jesse N Velasco-Silva, Sanghoon Lee, Judith Simcox, Gisela Geoghegan, Claire L Bensard, Tyler van Ry, Will L Holland, Scott A Summers, James Cox, Gregory S Ducker, Jared Rutter, and Claudio J Villanueva. Mitochondrial pyruvate carrier is required for optimal brown fat thermogenesis. *eLife*, 9, 2020.

[7855] Hendrik Pannen, Tim Rapp, and Thomas Klein. The ESCRT machinery regulates retromer-dependent transcytosis of septate junction components in Drosophila. *eLife*, 9, 2020.

[7856] Danai Papageorgiou and Damien Roger Farine. Group size and composition influence collective movement in a highly social terrestrial bird. *eLife*, 9, 2020.

[7857] Casey Paquola, Oualid Benkarim, Jordan DeKraker, Sara Larivière, Stefan Frässle, Jessica Royer, Shahin Tavakol, Sofie Valk, Andrea Bernasconi, Neda Bernasconi, Ali Khan, Alan C Evans, Adeel Razi, Jonathan Smallwood, and Boris C Bernhardt. Convergence of cortical types and functional motifs in the human mesiotemporal lobe. *eLife*, 9, 2020.

[7858] Cristina Parigini and Philip Greulich. Universality of clonal dynamics poses fundamental limits to identify stem cell self-renewal strategies. *eLife*, 9, 2020.

[7859] Annie Park, Tracy Tran, Elizabeth A Scheuermann, Dean P Smith, and Nigel S Atkinson. Alcohol potentiates a pheromone signal in flies. *eLife*, 9, 2020.

[7860] Hye Jin Park, Yuriy Pichugin, and Arne Traulsen. Why is cyclic dominance so rare? *eLife*, 9, 2020.

[7861] Jeong Hyang Park, Chang Geon Chung, Sung Soon Park, Davin Lee, Kyung Min Kim, Yeonjin Jeong, Eun Seon Kim, Jae Ho Cho, Yu-Mi Jeon, C-K James Shen, Hyung-Jun Kim, Daehee Hwang, and Sung Bae Lee. Cytosolic calcium regulates cytoplasmic accumulation of TDP-43 through Calpain-A and Importin $\alpha 3$. *eLife*, 9, 2020.

[7862] Silvia JH Park, Evan E Lieberman, Jiang-Bin Ke, Nao Rho, Padideh Ghorbani, Pouyan Rahmani, Na Young Jun, Hae-Lim Lee, In-Jung Kim, Kevin L Briggman, Jonathan B Demb, and Joshua H Singer. Connectomic analysis reveals an interneuron with an integral role in the retinal circuit for night vision. *eLife*, 9, 2020.

[7863] Matthew T Parker, Katarzyna Knop, Anna V Sherwood, Nicholas J Schurch, Katarzyna Mackinnon, Peter D Gould, Anthony JW Hall, Geoffrey J Barton, and Gordon G Simpson. Nanopore direct RNA sequencing maps the complexity of Arabidopsis mRNA processing and m6A modification. *eLife*, 9, 2020.

[7864] Andrey A Parkhitko, Divya Ramesh, Lin Wang, Dmitry Leshchiner, Elizabeth Filine, Richard Binari, Abby L Olsen, John M Asara, Valentin Cracan, Joshua D Rabinowitz, Axel Brockmann, and Norbert Perrimon. Downregulation of the tyrosine degradation pathway extends Drosophila lifespan. *eLife*, 9, 2020.

[7865] Aravindakshan Parthasarathy, Kenneth E Hancock, Kara Bennett, Victor DeGruttola, and Daniel B Polley. Bottom-up and top-down neural signatures of disordered multi-talker speech perception in adults with normal hearing. *eLife*, 9, 2020.

[7866] Enya Paschen, Claudio Elgueta, Katharina Heining, Diego M Vieira, Piret Kleis, Catarina Orcinha, Ute Häussler, Marlene Bartos, Ulrich Egert, Philipp Janz, and Carola A Haas. Hippocampal low-frequency stimulation prevents seizure generation in a mouse model of mesial temporal lobe epilepsy. *eLife*, 9, 2020.

[7867] Hugh Pastoll, Derek L Garden, Ioannis Papastathopoulos, Gülşen Sürmeli, and Matthew F Nolan. Inter- and intra-animal variation in the integrative properties of stellate cells in the medial entorhinal cortex. *eLife*, 9, 2020.

[7868] Nilkanth Patel, Xi Ping Huang, Jessica M Grandner, Linda C Johansson, Benjamin Stauch, John D McCorvy, Yongfeng Liu, Bryan Roth, and Vsevolod Katritch. Structure-based discovery of potent and selective melatonin receptor agonists. *eLife*, 9, 2020.

[7869] Yesha Patel, Heng Zhao, and John D Helmann. A regulatory pathway that selectively up-regulates elongasome function in the absence of class A PBPs. *eLife*, 9, 2020.

[7870] Trayambak Pathak, Maxime Gueguinou, Vonn Walter, Celine Delierneux, Martin T Johnson, Xuexin Zhang, Ping Xin, Ryan E Yoast, Scott M Emrich, Gregory S Yochum, Israel Sekler, Walter A Koltun, Donald L Gill, Nadine Hempel, and Mohamed Trebak. Dichotomous role of the human mitochondrial Na+/Ca2+/Li+ exchanger NCLX in colorectal cancer growth and metastasis. *eLife*, 9, 2020.

[7871] Sthitapranjya Pati, Kamal Saba, Sonali S Salvi, Praachi Tiwari, Pratik R Chaudhari, Vijaya Verma, Sourish Mukhopadhyay, Darshana Kapri, Shital Suryavanshi, James P Clement, Anant B Patel, and Vidita A Vaidya. Chronic postnatal chemogenetic activation of forebrain excitatory neurons evokes persistent changes in mood behavior. *eLife*, 9, 2020.

[7872] Bishnu P Paudel, Aaron Lavel Moye, Hala Abou Assi, Roberto El-Khoury, Scott B Cohen, Jessica K Holien, Monica L Birrento, Siritron Samosorn, Kamthorn Intharapichai, Christopher G Tomlinson, Marie-Paule Teulade-Fichou, Carlos González, Jennifer L Beck, Masad J Damha, Antoine M van Oijen, and Tracy M Bryan. A mechanism for the extension and unfolding of parallel telomeric G-quadruplexes by human telomerase at single-molecule resolution. *eLife*, 9, 2020.

[7873] Rodrigo Pavão, Elyse S Sussman, Brian J Fischer, and José L Peña. Natural ITD statistics predict human auditory spatial perception. *eLife*, 9, 2020.

[7874] Natalya N Pavlova, Bryan King, Rachel H Josselsohn, Sara Violante, Victoria L Macera, Santosha A Vardhana, Justin R Cross, and Craig B Thompson. Translation in amino-acid-poor environments is limited by tRNAGln charging. *eLife*, 9, 2020.

[7875] NataŠa Pavlović, Carlemi Calitz, Kess Thanapirom, Guiseppe Mazza, Krista Rombouts, Pär Gerwins, and Femke Heindryckx. Inhibiting IRE1$\alpha$-endonuclease activity decreases tumor burden in a mouse model for hepatocellular carcinoma. *eLife*, 9, 2020.

[7876] Slavica Pavlovic Djuranovic, Jessey Erath, Ryan J Andrews, Peter O Bayguinov, Joyce J Chung, Douglas L Chalker, James AJ Fitzpatrick, Walter N Moss, Pawel Szczesny, and Sergej Djuranovic. Plasmodium falciparum translational machinery condones polyadenosine repeats. *eLife*, 9, 2020.

[7877] Colin Pawlowski, Tyler Wagner, Arjun Puranik, Karthik Murugadoss, Liam Loscalzo, AJ Venkatakrishnan, Rajiv K Pruthi, Damon E Houghton, John C O'Horo, William G Morice, Amy W Williams, Gregory J Gores, John Halamka, Andrew D Badley, Elliot S Barnathan, Hideo Makimura, Najat Khan, and Venky Soundararajan. Inference from longitudinal laboratory tests characterizes temporal evolution of COVID-19-associated coagulopathy (CAC). *eLife*, 9, 2020.

[7878] James Peak, Billy Chieng, Genevra Hart, and Bernard W Balleine. Striatal direct and indirect pathway neurons differentially control the encoding and updating of goal-directed learning. *eLife*, 9, 2020.

[7879] Kenneth A Pelkey, Daniela Calvigioni, Calvin Fang, Geoffrey Vargish, Tyler Ekins, Kurt Auville, Jason C Wester, Mandy Lai, Connie Mackenzie-Gray Scott, Xiaoqing Yuan, Steven Hunt, Daniel Abebe, Qing Xu, Jordane Dimidschstein, Gordon Fishell, Ramesh Chittajallu, and Chris J McBain. Paradoxical network excitation by glutamate release from VGluT3+ GABAergic interneurons. *eLife*, 9, 2020.

[7880] James F Pelletier, Christine M Field, Sebastian Fürthauer, Matthew Sonnett, and Timothy J Mitchison. Co-movement of astral microtubules, organelles and F-actin by dynein and actomyosin forces in frog egg cytoplasm. *eLife*, 9, 2020.

[7881] Marie Pereira, Jeong-Hun Ko, John Logan, Hayley Protheroe, Kee-Beom Kim, Amelia Li Min Tan, Peter I Croucher, Kwon-Sik Park, Maxime Rotival, Enrico Petretto, JH Duncan Bassett, Graham R Williams, and Jacques Behmoaras. A trans-eQTL network regulates osteoclast multinucleation and bone mass. *eLife*, 9, 2020.

[7882] Margherita Perillo, Nathalie Oulhen, Stephany Foster, Maxwell Spurrell, Cristina Calestani, and Gary Wessel. Regulation of dynamic pigment cell states at single-cell resolution. *eLife*, 9, 2020.

[7883] Ofer Perl, Nahum Nahum, Katya Belelovsky, and Rafi Haddad. The contribution of temporal coding to odor coding and odor perception in humans. *eLife*, 9, 2020.

[7884] Nora G Peterson, Benjamin M Stormo, Kevin P Schoenfelder, Juliet S King, Rayson RS Lee, and Donald T Fox. Cytoplasmic sharing through apical membrane remodeling. *eLife*, 9, 2020.

[7885] Peristera-Ioanna Petropoulou, Ioanna Mosialou, Steven Shikhel, Lihong Hao, Konstantinos Panitsas, Brygida Bisikirska, Na Luo, Fabiana Bahna, Jongho Kim, Patrick Carberry, Francesca Zanderigo, Norman Simpson, Mihran Bakalian, Suham Kassir, Lawrence Shapiro, Mark D Underwood, Christina M May, Kiran Kumar Soligapuram Sai, Matthew J Jorgensen, Cyrille B Confavreux, Sue Shapses, Blandine Laferrère, Akiva Mintz, J John Mann, Mishaela Rubin, and Stavroula Kousteni. Lipocalin-2 is an anorexigenic signal in primates. *eLife*, 9, 2020.

[7886] Paul Pfeiffer, Alexei V Egorov, Franziska Lorenz, Jan-Hendrik Schleimer, Andreas Draguhn, and Susanne Schreiber. Clusters of cooperative ion channels enable a membrane-potential-based mechanism for short-term memory. *eLife*, 9, 2020.

[7887] Quan M Phan, Gracelyn M Fine, Lucia Salz, Gerardo G Herrera, Ben Wildman, Iwona M Driskell, and Ryan R Driskell. Lef1 expression in fibroblasts maintains developmental potential in adult skin to regenerate wounds. *eLife*, 9, 2020.

[7888] Ryan S Phillips, Ian Rosner, Aryn H Gittis, and Jonathan E Rubin. The effects of chloride dynamics on substantia nigra pars reticulata responses to pallidal and striatal inputs. *eLife*, 9, 2020.

[7889] Jonathan M Philpott, Rajesh Narasimamurthy, Clarisse G Ricci, Alfred M Freeberg, Sabrina R Hunt, Lauren E Yee, Rebecca S Pelofsky, Sarvind Tripathi, David M Virshup, and Carrie L Partch. Casein kinase 1 dynamics underlie substrate selectivity and the PER2 circadian phosphoswitch. *eLife*, 9, 2020.

[7890] Lokesh G Pimpale, Teije C Middelkoop, Alexander Mietke, and Stephan W Grill. Cell lineage-dependent chiral actomyosin flows drive cellular rearrangements in early Caenorhabditis elegans development. *eLife*, 9, 2020.

[7891] Damilola Pinheiro, Marie-Anne Mawhin, Maria Prendecki, and Kevin J Woollard. In-silico analysis of myeloid cells across the animal kingdom reveals neutrophil evolution by colony-stimulating factors. *eLife*, 9, 2020.

[7892] Roger Pique-Regi, Roberto Romero, Adi L Tarca, Francesca Luca, Yi Xu, Adnan Alazizi, Yaozhu Leng, Chaur-Dong Hsu, and Nardhy Gomez-Lopez. Does the human placenta express the canonical cell entry mediators for SARS-CoV-2? *eLife*, 9, 2020.

[7893] Ioannis Pisokas, Stanley Heinze, and Barbara Webb. The head direction circuit of two insect species. *eLife*, 9, 2020.

[7894] Anna Plana-Bonamaisó, Santiago López-Begines, David Fernández-Justel, Alexandra Junza, Ariadna Soler-Tapia, Jordi Andilla, Pablo Loza-Alvarez, Jose Luis Rosa, Esther Miralles, Isidre Casals, Oscar Yanes, Pedro de la Villa, Ruben M Buey, and Ana Méndez. Post-translational regulation of retinal IMPDH1 in vivo to adjust GTP synthesis to illumination conditions. *eLife*, 9, 2020.

[7895] Amanda M Pocratsky, Courtney T Shepard, Johnny R Morehouse, Darlene A Burke, Amberley S Riegler, Josiah T Hardin, Jason E Beare, Casey Hainline, Gregory JR States, Brandon L Brown, Scott R Whittemore, and David SK Magnuson. Long ascending propriospinal neurons provide flexible, context-specific control of interlimb coordination. *eLife*, 9, 2020.

[7896] Brandon Podyma, Dove-Anna Johnson, Laura Sipe, Thomas Parks Remcho, Katherine Battin, Yuxi Liu, Sung Ok Yoon, Christopher D Deppmann, and Ali Deniz Güler. The p75 neurotrophin receptor in AgRP neurons is necessary for homeostatic feeding and food anticipation. *eLife*, 9, 2020.

[7897] Amy R Poe, Yineng Xu, Christine Zhang, Joyce Lei, Kailyn Li, David Labib, and Chun Han. Low FoxO expression in Drosophila somatosensory neurons protects dendrite growth under nutrient restriction. *eLife*, 9, 2020.

[7898] Ana Rita Pombo Antunes, Isabelle Scheyltjens, Johnny Duerinck, Bart Neyns, Kiavash Movahedi, and Jo A Van Ginderachter. Understanding the glioblastoma immune microenvironment as basis for the development of new immunotherapeutic strategies. *eLife*, 9, 2020.

[7899] Adrian Ponce-Alvarez, Gabriela Mochol, Ainhoa Hermoso-Mendizabal, Jaime de la Rocha, and Gustavo Deco. Cortical state transitions and stimulus response evolve along stiff and sloppy parameter dimensions, respectively. *eLife*, 9, 2020.

[7900] Sinziana Pop, Chin-Lin Chen, Connor J Sproston, Shu Kondo, Pavan Ramdya, and Darren W Williams. Extensive and diverse patterns of cell death sculpt neural networks in insects. *eLife*, 9, 2020.

[7901] Bruno Popik, Felippe Espinelli Amorim, Olavo B Amaral, and Lucas De Oliveira Alvares. Shifting from fear to safety through deconditioning-update. *eLife*, 9, 2020.

[7902] Rosa Maria Porreca, Emilia Herrera-Moyano, Eleni Skourti, Pui Pik Law, Roser Gonzalez Franco, Alex Montoya, Peter Faull, Holger Kramer, and Jean-Baptiste Vannier. TRF1 averts chromatin remodelling, recombination and replication dependent-break induced replication at mouse telomeres. *eLife*, 9, 2020.

[7903] Fillip Port, Claudia Strein, Mona Stricker, Benedikt Rauscher, Florian Heigwer, Jun Zhou, Celine Beyersdörffer, Jana Frei, Amy Hess, Katharina Kern, Laura Lange, Nora Langner, Roberta Malamud, Bojana Pavlović, Kristin Rädecke, Lukas Schmitt, Lukas Voos, Erica Valentini, and Michael Boutros. A large-scale resource for tissue-specific CRISPR mutagenesis in Drosophila. *eLife*, 9, 2020.

[7904] Justin R Porter, Artur Meller, Maxwell I Zimmerman, Michael J Greenberg, and Gregory R Bowman. Conformational distributions of isolated myosin motor domains encode their mechanochemical properties. *eLife*, 9, 2020.

[7905] Harvie P Portugaliza, Shinya Miyazaki, Fiona JA Geurten, Christopher Pell, Anna Rosanas-Urgell, Chris J Janse, and Alfred Cortés. Artemisinin exposure at the ring or trophozoite stage impacts Plasmodium falciparum sexual conversion differently. *eLife*, 9, 2020.

[7906] Matthew J Powers, Brent W Simpson, and M Stephen Trent. The Mla pathway in Acinetobacter baumannii has no demonstrable role in anterograde lipid transport. *eLife*, 9, 2020.

[7907] Claudia Prahst, Parham Ashrafzadeh, Thomas Mead, Ana Figueiredo, Karen Chang, Douglas Richardson, Lakshmi Venkaraman, Mark Richards, Ana Martins Russo, Kyle Harrington, Marie Ouarné, Andreia Pena, Dong Feng Chen, Lena Claesson-Welsh, Kin-Sang Cho, Claudio A Franco, and Katie Bentley. Mouse retinal cell behaviour in space and time using light sheet fluorescence microscopy. *eLife*, 9, 2020.

[7908] Steffen Preissler, Claudia Rato, Yahui Yan, Luke A Perera, Aron Czako, and David Ron. Calcium depletion challenges endoplasmic reticulum proteostasis by destabilising BiP-substrate complexes. *eLife*, 9, 2020.

[7909] Michael J Prigge, Matthieu Platre, Nikita Kadakia, Yi Zhang, Kathleen Greenham, Whitnie Szutu, Bipin Kumar Pandey, Rahul Arvind Bhosale, Malcolm J Bennett, Wolfgang Busch, and Mark Estelle. Genetic analysis of the Arabidopsis TIR1/AFB auxin receptors reveals both overlapping and specialized functions. *eLife*, 9, 2020.

[7910] Kanella Prodromidou, Ioannis S Vlachos, Maria Gaitanou, Georgia Kouroupi, Artemis G Hatzigeorgiou, and Rebecca Matsas. MicroRNA-934 is a novel primate-specific small non-coding RNA with neurogenic function during early development. *eLife*, 9, 2020.

[7911] Myroslava Protsiv, Catherine Ley, Joanna Lankester, Trevor Hastie, and Julie Parsonnet. Decreasing human body temperature in the United States since the Industrial Revolution. *eLife*, 9, 2020.

[7912] Thomas Pühringer, Ulrich Hohmann, Laura Fin, Belén Pacheco-Fiallos, Ulla Schellhaas, Julius Brennecke, and Clemens Plaschka. Structure of the human core transcription-export complex reveals a hub for multivalent interactions. *eLife*, 9, 2020.

[7913] Joan Pulupa, Harriet Prior, Daniel S Johnson, and Sanford M Simon. Conformation of the nuclear pore in living cells is modulated by transport state. *eLife*, 9, 2020.

[7914] Xingyun Qi, Akira Yoshinari, Pengfei Bai, Michal Maes, Scott M Zeng, and Keiko U Torii. The manifold actions of signaling peptides on subcellular dynamics of a receptor specify stomatal cell fate. *eLife*, 9, 2020.

[7915] Zhenheng Qin, Lulu Bi, Xi-Miao Hou, Siqi Zhang, Xia Zhang, Ying Lu, Ming Li, Mauro Modesti, Xu-Guang Xi, and Bo Sun. Human RPA activates BLM's bidirectional DNA unwinding from a nick. *eLife*, 9, 2020.

[7916] Zhongwei Qu, Albert Zhang, and Dong Yan. Robo functions as an attractive cue for glial migration through SYG-1/Neph. *eLife*, 9, 2020.

[7917] Tran Minh Quan, Tran Thi Nhu Thao, Nguyen Manh Duy, Tran Minh Nhat, and Hannah Clapham. Estimates of the global burden of Japanese encephalitis and the impact of vaccination from 2000-2015. *eLife*, 9, 2020.

[7918] Daniel S Quintana. A synthetic dataset primer for the biobehavioural sciences to promote reproducibility and hypothesis generation. *eLife*, 9, 2020.

[7919] Kazi Rahman, Charles A Coomer, Saliha Majdoul, Selena Y Ding, Sergi Padilla-Parra, and Alex A Compton. Homology-guided identification of a conserved motif linking the antiviral functions of IFITM3 to its oligomeric state. *eLife*, 9, 2020.

[7920] Gulzhan Raiymbek, Sojin An, Nidhi Khurana, Saarang Gopinath, Ajay Larkin, Saikat Biswas, Raymond C Trievel, Uhn-soo Cho, and Kaushik Ragunathan. An H3K9 methylation-dependent protein interaction regulates the non-enzymatic functions of a putative histone demethylase. *eLife*, 9, 2020.

[7921] Sathish Ramakrishnan, Manindra Bera, Jeff Coleman, James E Rothman, and Shyam S Krishnakumar. Synergistic roles of Synaptotagmin-1 and complexin in calcium-regulated neuronal exocytosis. *eLife*, 9, 2020.

[7922] M Rosario Ramírez-Flores, Sergio Perez-Limon, Meng Li, Benjamín Barrales-Gamez, Doris Albinsky, Uta Paszkowski, Víctor Olalde-Portugal, and Ruairidh JH Sawers. The genetic architecture of host response reveals the importance of arbuscular mycorrhizae to maize cultivation. *eLife*, 9, 2020.

[7923] Héctor Vicente Ramírez-Gómez, Vilma Jimenez Sabinina, Martín Velázquez Pérez, Carmen Beltran, Jorge Carneiro, Christopher D Wood, Idan Tuval, Alberto Darszon, and Adán Guerrero. Sperm chemotaxis is driven by the slope of the chemoattractant concentration field. *eLife*, 9, 2020.

[7924] Srivastav Ranganathan and Eugene I Shakhnovich. Dynamic metastable long-living droplets formed by sticker-spacer proteins. *eLife*, 9, 2020.

[7925] Anand Ranjan, Vu Q Nguyen, Sheng Liu, Jan Wisniewski, Jee Min Kim, Xiaona Tang, Gaku Mizuguchi, Ejlal Elalaoui, Timothy J Nickels, Vivian Jou, Brian P English, Qinsi Zheng, Ed Luk, Luke D Lavis, Timothee Lionnet, and Carl Wu. Live-cell single particle imaging reveals the role of RNA polymerase II in histone H2A.Z eviction. *eLife*, 9, 2020.

[7926] Ishier Raote, Morgan Chabanon, Nikhil Walani, Marino Arroyo, Maria F Garcia-Parajo, Vivek Malhotra, and Felix Campelo. A physical mechanism of TANGO1-mediated bulky cargo export. *eLife*, 9, 2020.

[7927] Ishier Raote, Andreas M Ernst, Felix Campelo, James E Rothman, Frederic Pincet, and Vivek Malhotra. TANGO1 membrane helices create a lipid diffusion barrier at curved membranes. *eLife*, 9, 2020.

[7928] Amir Rattner, Chantelle E Terrillion, Claudia Jou, Tina Kleven, Shun Felix Hu, John Williams, Zhipeng Hou, Manisha Aggarwal, Susumu Mori, Gloria Shin, Loyal A Goff, Menno P Witter, Mikhail Pletnikov, André A Fenton, and Jeremy Nathans. Developmental, cellular, and behavioral phenotypes in a mouse model of congenital hypoplasia of the dentate gyrus. *eLife*, 9, 2020.

[7929] Subhasis Ray, Zane N Aldworth, and Mark A Stopfer. Feedback inhibition and its control in an insect olfactory circuit. *eLife*, 9, 2020.

[7930] Michelle S Reid, David M Kern, and Stephen Graf Brohawn. Cryo-EM structure of the potassium-chloride cotransporter KCC4 in lipid nanodiscs. *eLife*, 9, 2020.

[7931] Siqiang Ren, Wen Yao, Marc D Tambini, Tao Yin, Kelly A Norris, and Luciano D'Adamio. Microglia TREM2R47H Alzheimer-linked variant enhances excitatory transmission and reduces LTP via increased TNF-$\alpha$ levels. *eLife*, 9, 2020.

[7932] Kaushik Renganaath, Rockie Chong, Laura Day, Sriram Kosuri, Leonid Kruglyak, and Frank W Albert. Systematic identification of cis-regulatory variants that cause gene expression differences in a yeast cross. *eLife*, 9, 2020.

[7933] Henrik Renner, Martha Grabos, Katharina J Becker, Theresa E Kagermeier, Jie Wu, Mandy Otto, Stefan Peischard, Dagmar Zeuschner, Yaroslav TsyTsyura, Paul Disse, Jürgen Klingauf, Sebastian A Leidel, Guiscard Seebohm, Hans R Schöler, and Jan M Bruder. A fully automated high-throughput workflow for 3D-based chemical screening in human midbrain organoids. *eLife*, 9, 2020.

[7934] Nathan E Reticker-Flynn and Edgar G Engleman. Cancer systems immunology. *eLife*, 9, 2020.

[7935] Diogo Ribeiro, Ana Rita Nunes, Magda Teles, Savani Anbalagan, Janna Blechman, Gil Levkowitz, and Rui F Oliveira. Genetic variation in the social environment affects behavioral phenotypes of oxytocin receptor mutants in zebrafish. *eLife*, 9, 2020.

[7936] Biancamaria Ricci, Eric Tycksen, Hamza Celik, Jad I Belle, Francesca Fontana, Roberto Civitelli, and Roberta Faccio. Osterix-Cre marks distinct subsets of CD45- and CD45+ stromal populations in extra-skeletal tumors with pro-tumorigenic characteristics. *eLife*, 9, 2020.

[7937] Ashley Rich, Richard G Fehon, and Michael Glotzer. Rho1 activation recapitulates early gastrulation events in the ventral, but not dorsal, epithelium of Drosophila embryos. *eLife*, 9, 2020.

[7938] Nainoa Richardson, Isabelle Gillot, Elodie P Gregoire, Sameh A Youssef, Dirk de Rooij, Alain de Bruin, Marie-Cécile De Cian, and Marie-Christine Chaboissier. Sox8 and Sox9 act redundantly for ovarian-to-testicular fate reprogramming in the absence of R-spondin1 in mouse sex reversals. *eLife*, 9, 2020.

[7939] Delia Ricolo and Sofia J Araujo. Coordinated crosstalk between microtubules and actin by a spectraplakin regulates lumen formation and branching. *eLife*, 9, 2020.

[7940] Thomas Ridler, Jonathan Witton, Keith G Phillips, Andrew D Randall, and Jonathan T Brown. Impaired speed encoding and grid cell periodicity in a mouse model of tauopathy. *eLife*, 9, 2020.

[7941] Konstantin Riege, Helene Kretzmer, Arne Sahm, Simon S McDade, Steve Hoffmann, and Martin Fischer. Dissecting the DNA binding landscape and gene regulatory network of p63 and p53. *eLife*, 9, 2020.

[7942] Vera D Rinaldi, Elisa Donnard, Kyle Gellatly, Morten Rasmussen, Alper Kucukural, Onur Yukselen, Manuel Garber, Upasna Sharma, and Oliver J Rando. An atlas of cell types in the mouse epididymis and vas deferens. *eLife*, 9, 2020.

[7943] Rozenn Riou, Meriem Ladli, Sabine Gerbal-Chaloin, Pascale Bossard, Angélique Gougelet, Cécile Godard, Robin Loesch, Isabelle Lagoutte, Franck Lager, Julien Calderaro, Alexandre Dos Santos, Zhong Wang, Frédérique Verdier, and Sabine Colnot. ARID1A loss in adult hepatocytes activates $\beta$-catenin-mediated erythropoietin transcription. *eLife*, 9, 2020.

[7944] Zev A Ripstein, Siavash Vahidi, Walid A Houry, John L Rubinstein, and Lewis E Kay. A processive rotary mechanism couples substrate unfolding and proteolysis in the ClpXP degradation machinery. *eLife*, 9, 2020.

[7945] Elizabeth Rivers, Rajeev Rai, Jonas Lötscher, Michael Hollinshead, Gasper Markelj, James Thaventhiran, Austen Worth, Alessia Cavazza, Christoph Hess, Mona Bajaj-Elliott, and Adrian J Thrasher. Wiskott Aldrich syndrome protein regulates non-selective autophagy and mitochondrial homeostasis in human myeloid cells. *eLife*, 9, 2020.

[7946] Noemi Rives-Quinto, Hideyuki Komori, Cyrina M Ostgaard, Derek H Janssens, Shu Kondo, Qi Dai, Adrian W Moore, and Cheng-Yu Lee. Sequential activation of transcriptional repressors promotes progenitor commitment by silencing stem cell identity genes. *eLife*, 9, 2020.

[7947] Lucy Rivett, Sushmita Sridhar, Dominic Sparkes, Matthew Routledge, Nick K Jones, Sally Forrest, Jamie Young, Joana Pereira-Dias, William L Hamilton, Mark Ferris, M Estee Torok, Luke Meredith, Ravi Gupta, Ravi Gupta, Paul A Lyons, Mark Toshner, Ben Warne, Josefin Bartholdson Scott, Claire Cormie, Harmeet Gill, Iain Kean, Mailis Maes, Nicola Reynolds, Michelle Wantoch, Sarah Caddy, Laura Caller, Theresa Feltwell, Grant Hall, Myra Hosmillo, Charlotte Houldcroft, Aminu Jahun, Fahad Khokhar, Anna Yakovleva, Helen Butcher, Daniela Caputo, Debra Clapham-Riley, Helen Dolling, Anita Furlong, Barbara Graves, Emma Le Gresley, Nathalie Kingston, Sofia Papadia, Hannah Stark, Kathleen E Stirrups, Jennifer Webster, Joanna Calder, Julie Harris, Sarah Hewitt, Jane Kennet, Anne Meadows, Rebecca Rastall, Criona O Brien, Jo Price, Cherry Publico, Jane Rowlands, Valentina Ruffolo, Hugo Tordesillas, Karen Brookes, Laura Canna, Isabel Cruz, Katie Dempsey, Anne Elmer, Naidine Escoffery, Heather Jones, Carla Ribeiro, Caroline Saunders, Angela Wright, Rutendo Nyagumbo, Anne Roberts, Ashlea Bucke, Simone Hargreaves, Danielle Johnson, Aileen Narcorda, Debbie Read, Christian Sparke, Lucy Warboys, Kirsty Lagadu, Lenette Mactavous, Tim Gould, Tim Raine, Claire Mather, Nicola Ramenatte, Anne-Laure Vallier, Mary

Kasanicki, Penelope-Jane Eames, Chris McNicholas, Lisa Thake, Neil Bartholomew, Nick Brown, Surendra Parmar, Hongyi Zhang, Ailsa Bowring, Geraldine Martell, Natalie Quinnell, Jo Wright, Helen Murphy, Benjamin J Dunmore, Ekaterina Legchenko, Stefan Gräf, Christopher Huang, Josh Hodgson, Kelvin Hunter, Jennifer Martin, Federica Mescia, Ciara O'Donnell, Linda Pointon, Joy Shih, Rachel Sutcliffe, Tobias Tilly, Zhen Tong, Carmen Treacy, Jennifer Wood, Laura Bergamaschi, Ariana Betancourt, Georgie Bowyer, Aloka De Sa, Maddie Epping, Andrew Hinch, Oisin Huhn, Isobel Jarvis, Daniel Lewis, Joe Marsden, Simon McCallum, Francescsa Nice, Martin D Curran, Stewart Fuller, Afzal Chaudhry, Ashley Shaw, Richard J Samworth, John R Bradley, Gordon Dougan, Kenneth GC Smith, Paul J Lehner, Nicholas J Matheson, Giles Wright, Ian G Goodfellow, Stephen Baker, and Michael P Weekes. Screening of healthcare workers for SARS-CoV-2 highlights the role of asymptomatic carriage in COVID-19 transmission. *eLife*, 9, 2020.

[7948] Yvette Robbins, Sarah Greene, Jay Friedman, Paul E Clavijo, Carter Van Waes, Kellsye P Fabian, Michelle R Padget, Houssein Abdul Sater, John H Lee, Patrick Soon-Shiong, James Gulley, Jeffrey Schlom, James W Hodge, and Clint T Allen. Tumor control via targeting PD-L1 with chimeric antigen receptor modified NK cells. *eLife*, 9, 2020.

[7949] Nicolas Robinson-Garcia, Rodrigo Costas, Cassidy R Sugimoto, Vincent Larivière, and Gabriela F Nane. Task specialization across research careers. *eLife*, 9, 2020.

[7950] Pieter A Roelofs, Chai Yeen Goh, Boon Haow Chua, Matthew C Jarvis, Teneale A Stewart, Jennifer L McCann, Rebecca M McDougle, Michael A Carpenter, John WM Martens, Paul N Span, Dennis Kappei, and Reuben S Harris. Characterization of the mechanism by which the RB/E2F pathway controls expression of the cancer genomic DNA deaminase APOBEC3B. *eLife*, 9, 2020.

[7951] Friederike Roger, Cecilia Picazo, Wolfgang Reiter, Marouane Libiad, Chikako Asami, Sarah Hanzén, Chunxia Gao, Gilles Lagniel, Niek Welkenhuysen, Jean Labarre, Thomas Nyström, Morten Grøtli, Markus Hartl, Michel B Toledano, and Mikael Molin. Peroxiredoxin promotes longevity and H2O2-resistance in yeast through redox-modulation of protein kinase A. *eLife*, 9, 2020.

[7952] Katherine W Rogers, Mohammad ElGamacy, Benjamin M Jordan, and Patrick Müller. Optogenetic investigation of BMP target gene expression diversity. *eLife*, 9, 2020.

[7953] Connor Rogerson, Samuel Ogden, Edward Britton, Yeng Ang, and Andrew D Sharrocks. Repurposing of KLF5 activates a cell cycle signature during the progression from a precursor state to oesophageal adenocarcinoma. *eLife*, 9, 2020.

[7954] Seung-Eon Roh, Seung Ha Kim, Changhyeon Ryu, Chang-Eop Kim, Yong Gyu Kim, Paul F Worley, Sun Kwang Kim, and Sang Jeong Kim. Direct translation of climbing fiber burst-mediated sensory coding into post-synaptic Purkinje cell dendritic calcium. *eLife*, 9, 2020.

[7955] Antonis Rokas, Sam Mesiano, Ortal Tamam, Abigail LaBella, Ge Zhang, and Louis Muglia. Developing a theoretical evolutionary framework to solve the mystery of parturition initiation. *eLife*, 9, 2020.

[7956] Matthew L Rook, Abby Williamson, John D Lueck, Maria Musgaard, and David M Maclean. $\beta$11-12 linker isomerization governs acid-sensing ion channel desensitization and recovery. *eLife*, 9, 2020.

[7957] Christopher J Roome and Bernd Kuhn. Dendritic coincidence detection in Purkinje neurons of awake mice. *eLife*, 9, 2020.

[7958] Johanna Roostalu, Claire Thomas, Nicholas Ian Cade, Simone Kunzelmann, Ian A Taylor, and Thomas Surrey. The speed of GTP hydrolysis determines GTP cap size and controls microtubule stability. *eLife*, 9, 2020.

[7959] Stefania Rosano, Davide Corà, Sushant Parab, Serena Zaffuto, Claudio Isella, Roberta Porporato, Roxana Maria Hoza, Raffaele A Calogero, Chiara Riganti, Federico Bussolino, and Alessio Noghero. A regulatory microRNA network controls endothelial cell phenotypic switch during sprouting angiogenesis. *eLife*, 9, 2020.

[7960] Brittany Rosener, Serkan Sayin, Peter O Oluoch, Aurian P García González, Hirotada Mori, Albertha JM Walhout, and Amir Mitchell. Evolved bacterial resistance against fluoropyrimidines can lower chemotherapy impact in the Caenorhabditis elegans host. *eLife*, 9, 2020.

[7961] Anthony M Rossi and Claude Desplan. Extrinsic activin signaling cooperates with an intrinsic temporal program to increase mushroom body neuronal diversity. *eLife*, 9, 2020.

[7962] Gian Paolo Rossi, Viola Sanga, and Matthias Barton. Potential harmful effects of discontinuing ACE-inhibitors and ARBs in COVID-19 patients. *eLife*, 9, 2020.

[7963] Julien G Roth, Kristin L Muench, Aditya Asokan, Victoria M Mallett, Hui Gai, Yogendra Verma, Stephen Weber, Carol Charlton, Jonas L Fowler, Kyle M Loh, Ricardo E Dolmetsch, and Theo D Palmer. 16p11.2 microdeletion imparts transcriptional alterations in human iPSC-derived models of early neural development. *eLife*, 9, 2020.

[7964] Kelly A Rotstan, Michael M Abdelsayed, Luiz FM Passalacqua, Fabio Chizzolini, Kasireddy Sudarshan, A Richard Chamberlin, Jiří MíŠek, and Andrej Luptak. Regulation of mRNA translation by a photoriboswitch. *eLife*, 9, 2020.

[7965] Arani Roy, Jason J Osik, Benyamin Meschede-Krasa, Wesley T Alford, Daniel P Leman, and Stephen D Van Hooser. Synaptic and intrinsic mechanisms underlying development of cortical direction selectivity. *eLife*, 9, 2020.

[7966] Babhrubahan Roy, Simon JY Han, Adrienne Nicole Fontan, and Ajit P Joglekar. The copy-number and varied strengths of MELT motifs in Spc105 balance the strength and responsiveness of the spindle assembly checkpoint. *eLife*, 9, 2020.

[7967] Liron Rozenkrantz, Reut Weissgross, Tali Weiss, Inbal Ravreby, Idan Frumin, Sagit Shushan, Lior Gorodisky, Netta Reshef, Yael Holzman, Liron Pinchover, Yaara Endevelt-Shapira, Eva Mishor, Timna Soroka, Maya Finkel, Liav Tagania, Aharon Ravia, Ofer Perl, Edna Furman-Haran, Howard Carp, and Noam Sobel. Unexplained repeated pregnancy loss is associated with altered perceptual and brain responses to men's body-odor. *eLife*, 9, 2020.

[7968] Hang Ruan, Xiangyun Li, Xiang Xu, Brian J Leibowitz, Jingshan Tong, Lujia Chen, Luoquan Ao, Wei Xing, Jianhua Luo, Yanping Yu, Robert E Schoen, Nahum Sonenberg, Xinghua Lu, Lin Zhang, and Jian Yu. eIF4E S209 phosphorylation licenses myc- and stress-driven oncogenesis. *eLife*, 9, 2020.

[7969] Sarah Ruediger and Massimo Scanziani. Learning speed and detection sensitivity controlled by distinct cortico-fugal neurons in visual cortex. *eLife*, 9, 2020.

[7970] Rolando Ruiz-Vega, Chi-Fen Chen, Emaad Razzak, Priya Vasudeva, Tatiana B Krasieva, Jessica Shiu, Michael G Caldwell, Huaming Yan, John Lowengrub, Anand K Ganesan, and Arthur D Lander. Dynamics of nevus development implicate cell cooperation in the growth arrest of transformed melanocytes. *eLife*, 9, 2020.

[7971] Andrea Ruiz-Velasco, Min Zi, Susanne S Hille, Tayyiba Azam, Namrita Kaur, Juwei Jiang, Binh Nguyen, Karolina Sekeres, Pablo Binder, Lucy Collins, Fay Pu, Han Xiao, Kaomei Guan, Norbert Frey, Elizabeth J Cartwright, Oliver J Müller, Xin Wang, and Wei Liu. Targeting mir128-3p alleviates myocardial insulin resistance and prevents ischemia-induced heart failure. *eLife*, 9, 2020.

[7972] Michael E Rule, Adrianna R Loback, Dhruva V Raman, Laura N Driscoll, Christopher D Harvey, and Timothy O'Leary. Stable task information from an unstable neural population. *eLife*, 9, 2020.

[7973] Helene Rundqvist, Pedro VeliÇa, Laura Barbieri, Paulo A Gameiro, David Bargiela, Milos Gojkovic, Sara Mijwel, Stefan Markus Reitzner, David Wulliman, Emil Ahlstedt, Jernej Ule, Arne östman, and Randall S Johnson. Cytotoxic T-cells mediate exercise-induced reductions in tumor growth. *eLife*, 9, 2020.

[7974] Maria Ryazantseva, Jonas Englund, Alexandra Shintyapina, Johanna Huupponen, Vasilii Shteinikov, Asla Pitkänen, Juha M Partanen, and Sari E Lauri. Kainate receptors regulate development of glutamatergic synaptic circuitry in the rodent amygdala. *eLife*, 9, 2020.

[7975] Graham Rykiel, Claudia S López, Jessica L Riesterer, Ian Fries, Sanika Deosthali, Katherine Courchaine, Alina Maloyan, Kent Thornburg, and Sandra Rugonyi. Multiscale cardiac imaging spanning the whole heart and its internal cellular architecture in a small animal model. *eLife*, 9, 2020.

[7976] Mark F Sabbagh and Jeremy Nathans. A genome-wide view of the de-differentiation of central nervous system endothelial cells in culture. *eLife*, 9, 2020.

[7977] Nathanaëlle Saclier, Patrick Chardon, Florian Malard, Lara Konecny-Dupré, David Eme, Arnaud Bellec, Vincent Breton, Laurent Duret, Tristan Lefebure, and Christophe J Douady. Bedrock radioactivity influences the rate and spectrum of mutation. *eLife*, 9, 2020.

[7978] Katelyn E Sadler, Francie Moehring, and Cheryl L Stucky. Keratinocytes contribute to normal cold and heat sensation. *eLife*, 9, 2020.

[7979] Lina Sagert, Felix Hennig, Christoph Thomas, and Robert Tampé. A loop structure allows TAPBPR to exert its dual function as MHC I chaperone and peptide editor. *eLife*, 9, 2020.

[7980] Maria Luisa Saggio, Dakota Crisp, Jared M Scott, Philippa Karoly, Levin Kuhlmann, Mitsuyoshi Nakatani, Tomohiko Murai, Matthias Dümpelmann, Andreas Schulze-Bonhage, Akio Ikeda, Mark Cook, Stephen V Gliske, Jack Lin, Christophe Bernard, Viktor Jirsa, and William C Stacey. A taxonomy of seizure dynamotypes. *eLife*, 9, 2020.

[7981] Piyali Saha, Justin L Shumate, Jenna G Caldwell, Nadia Elghobashi-Meinhardt, Albert Lu, Lichao Zhang, Niclas E Olsson, Joshua E Elias, and Suzanne R Pfeffer. Inter-domain dynamics drive cholesterol transport by NPC1 and NPC1L1 proteins. *eLife*, 9, 2020.

[7982] Bastien Saint-Leandre, Courtney Christopher, and Mia T Levine. Adaptive evolution of an essential telomere protein restricts telomeric retrotransposons. *eLife*, 9, 2020.

[7983] Lily Goodyer Sait, Altin Sula, Mohammad-Reza Ghovanloo, David Hollingworth, Peter C Ruben, and BA Wallace. Cannabidiol interactions with voltage-gated sodium channels. *eLife*, 9, 2020.

[7984] Kazuki Saito, Rachel Green, and Allen R Buskirk. Ribosome recycling is not critical for translational coupling in Escherichia coli. *eLife*, 9, 2020.

[7985] Kazuki Saito, Rachel Green, and Allen R Buskirk. Translational initiation in E. coli occurs at the correct sites genome-wide in the absence of mRNA-rRNA base-pairing. *eLife*, 9, 2020.

[7986] Néstor Saiz, Laura Mora-Bitria, Shahadat Rahman, Hannah George, Jeremy P Herder, Jordi Garcia-Ojalvo, and Anna-Katerina Hadjantonakis. Growth-factor-mediated coupling between lineage size and cell fate choice underlies robustness of mammalian development. *eLife*, 9, 2020.

[7987] Miguel Salinas-Saavedra and Mark Q Martindale. Par protein localization during the early development of Mnemiopsis leidyi suggests different modes of epithelial organization in the metazoa. *eLife*, 9, 2020.

[7988] Timothy R Sampson, Collin Challis, Neha Jain, Anastasiya Moiseyenko, Mark S Ladinsky, Gauri G Shastri, Taren Thron, Brittany D Needham, Istvan Horvath, Justine W Debelius, Stefan Janssen, Rob Knight, Pernilla Wittung-Stafshede, Viviana Gradinaru, Matthew Chapman, and Sarkis K Mazmanian. A gut bacterial amyloid promotes $\alpha$-synuclein aggregation and motor impairment in mice. *eLife*, 9, 2020.

[7989] Tamsin J Samuels, Aino I Järvelin, David Ish-Horowicz, and Ilan Davis. Imp/IGF2BP levels modulate individual neural stem cell growth and division through myc mRNA stability. *eLife*, 9, 2020.

[7990] Liran Samuni, Franziska Wegdell, and Martin Surbeck. Behavioural diversity of bonobo prey preference as a potential cultural trait. *eLife*, 9, 2020.

[7991] Honi Sanders, Matthew A Wilson, and Samuel J Gershman. Hippocampal remapping as hidden state inference. *eLife*, 9, 2020.

[7992] Shaun S Sanders, Luiselys M Hernandez, Heun Soh, Santi Karnam, Randall S Walikonis, Anastasios V Tzingounis, and Gareth M Thomas. The palmitoyl acyltransferase ZDHHC14 controls Kv1-family potassium channel clustering at the axon initial segment. *eLife*, 9, 2020.

[7993] Eric M Sanford, Benjamin L Emert, Allison Coté, and Arjun Raj. Gene regulation gravitates toward either addition or multiplication when combining the effects of two signals. *eLife*, 9, 2020.

[7994] Smriti Sangwan, Shruti Sahay, Kevin A Murray, Sophie Morgan, Elizabeth L Guenther, Lin Jiang, Christopher K Williams, Harry V Vinters, Michel Goedert, and David S Eisenberg. Inhibition of synucleinopathic seeding by rationally designed inhibitors. *eLife*, 9, 2020.

[7995] Sundar Ram Sankaranarayanan, Giuseppe Ianiri, Marco A Coelho, Md Hashim Reza, Bhagya C Thimmappa, Promit Ganguly, Rakesh Netha Vadnala, Sheng Sun, Rahul Siddharthan, Christian Tellgren-Roth, Thomas L Dawson, Joseph Heitman, and Kaustuv Sanyal. Loss of centromere function drives karyotype evolution in closely related Malassezia species. *eLife*, 9, 2020.

[7996] Lucas D Sanor, Grant Parker Flowers, and Craig M Crews. Multiplex CRISPR/Cas screen in regenerating haploid limbs of chimeric Axolotls. *eLife*, 9, 2020.

[7997] Alessandro Sanzeni, Bradley Akitake, Hannah C Goldbach, Caitlin E Leedy, Nicolas Brunel, and Mark H Histed. Inhibition stabilization is a widespread property of cortical networks. *eLife*, 9, 2020.

[7998] Etai Sapoznik, Bo-Jui Chang, Jaewon Huh, Robert J Ju, Evgenia V Azarova, Theresa Pohlkamp, Erik S Welf, David Broadbent, Alexandre F Carisey, Samantha J Stehbens, Kyung-Min Lee, Arnaldo Marín, Ariella B Hanker, Jens C Schmidt, Carlos L Arteaga, Bin Yang, Yoshihiko Kobayashi, Purushothama Rao Tata, Rory Kruithoff, Konstantin Doubrovinski, Douglas P Shepherd, Alfred Millett-Sikking, Andrew G York, Kevin M Dean, and Reto P Fiolka. A versatile oblique plane microscope for large-scale and high-resolution imaging of subcellular dynamics. *eLife*, 9, 2020.

[7999] Sarvenaz Sarabipour. Virtual conferences raise standards for accessibility and interactions. *eLife*, 9, 2020.

[8000] Amijai Saragovi, Ifat Abramovich, Ibrahim Omar, Eliran Arbib, Ori Toker, Eyal Gottlieb, and Michael Berger. Systemic hypoxia inhibits T cell response by limiting mitobiogenesis via matrix substrate-level phosphorylation arrest. *eLife*, 9, 2020.

[8001] Jay F Sarthy, Michael P Meers, Derek H Janssens, Jorja G Henikoff, Heather Feldman, Patrick J Paddison, Christina M Lockwood, Nicholas A Vitanza, James M Olson, Kami Ahmad, and Steven Henikoff. Histone deposition pathways determine the chromatin landscapes of H3.1 and H3.3 K27M oncohistones. *eLife*, 9, 2020.

[8002] Yo Sasaki, Hiroki Kakita, Shunsuke Kubota, Abdoulaye Sene, Tae Jun Lee, Norimitsu Ban, Zhenyu Dong, Joseph B Lin, Sanford L Boye, Aaron DiAntonio, Shannon E Boye, Rajendra S Apte, and Jeffrey Milbrandt. SARM1 depletion rescues NMNAT1-dependent photoreceptor cell death and retinal degeneration. *eLife*, 9, 2020.

[8003] Sunetra Sase, Akshata A Almad, C Alexander Boecker, Pedro Guedes-Dias, Jian J Li, Asako Takanohashi, Akshilkumar Patel, Tara McCaffrey, Heta Patel, Divya Sirdeshpande, Julian Curiel, Judy Shih-Hwa Liu, Quasar Padiath, Erika LF Holzbaur, Steven S Scherer, and Adeline Vanderver. TUBB4A mutations result in both glial and neuronal degeneration in an H-ABC leukodystrophy mouse model. *eLife*, 9, 2020.

[8004] Tempei Sato, Kensuke Kataoka, Yoshiaki Ito, Shigetoshi Yokoyama, Masafumi Inui, Masaki Mori, Satoru Takahashi, Keiichi Akita, Shuji Takada, Hiroe Ueno-Kudoh, and Hiroshi Asahara. Lin28a/let-7 pathway modulates the Hox code via Polycomb regulation during axial patterning in vertebrates. *eLife*, 9, 2020.

[8005] David B Sauer, Noah Trebesch, Jennifer J Marden, Nicolette Cocco, Jinmei Song, Akiko Koide, Shohei Koide, Emad Tajkhorshid, and Da-Neng Wang. Structural basis for the reaction cycle of DASS dicarboxylate transporters. *eLife*, 9, 2020.

[8006] Elise Laura Savier, James Dunbar, Kyle Cheung, and Michael Reber. New insights on the modeling of the molecular mechanisms underlying neural maps alignment in the midbrain. *eLife*, 9, 2020.

[8007] Sansana Sawasdikosol and Steven Burakoff. A perspective on HPK1 as a novel immuno-oncology drug target. *eLife*, 9, 2020.

[8008] Alessandro Scacchetti, Tamas Schauer, Alexander Reim, Zivkos Apostolou, Aline Campos Sparr, Silke Krause, Patrick Heun, Michael Wierer, and Peter B Becker. Drosophila SWR1 and NuA4 complexes are defined by DOMINO isoforms. *eLife*, 9, 2020.

[8009] Kristin M Scaplen, Mustafa Talay, Kavin M Nunez, Sarah Salamon, Amanda G Waterman, Sydney Gang, Sophia L Song, Gilad Barnea, and Karla R Kaun. Circuits that encode and guide alcohol-associated preference. *eLife*, 9, 2020.

[8010] M Regina Scarpin, Samuel Leiboff, and Jacob O Brunkard. Parallel global profiling of plant TOR dynamics reveals a conserved role for LARP1 in translation. *eLife*, 9, 2020.

[8011] Matthew A Schaich, Samantha L Sanford, Griffin A Welfer, Samuel A Johnson, Thu H Khoang, Patricia L Opresko, and Bret D Freudenthal. Mechanisms of nucleotide selection by telomerase. *eLife*, 9, 2020.

[8012] Alexandra Schauer, Diana Pinheiro, Robert Hauschild, and Carl-Philipp Heisenberg. Zebrafish embryonic explants undergo genetically encoded self-assembly. *eLife*, 9, 2020.

[8013] Claudia Scheckel, Marigona Imeri, Petra Schwarz, and Adriano Aguzzi. Ribosomal profiling during prion disease uncovers progressive translational derangement in glia but not in neurons. *eLife*, 9, 2020.

[8014] Louis K Scheffer, C Shan Xu, Michal Januszewski, Zhiyuan Lu, Shin-ya Takemura, Kenneth J Hayworth, Gary B Huang, Kazunori Shinomiya, Jeremy Maitlin-Shepard, Stuart Berg, Jody Clements, Philip M Hubbard, William T Katz, Lowell Umayam, Ting Zhao, David Ackerman, Tim Blakely, John Bogovic, Tom Dolafi, Dagmar Kainmueller, Takashi Kawase, Khaled A Khairy, Laramie Leavitt, Peter H Li, Larry Lindsey, Nicole Neubarth, Donald J Olbris, Hideo Otsuna, Eric T Trautman, Masayoshi Ito, Alexander S Bates, Jens Goldammer, Tanya Wolff, Robert Svirskas, Philipp Schlegel, Erika Neace, Christopher J Knecht, Chelsea X Alvarado, Dennis A Bailey, Samantha Ballinger, Jolanta A Borycz, Brandon S Canino, Natasha Cheatham, Michael Cook, Marisa Dreher, Octave Duclos, Bryon Eubanks, Kelli Fairbanks, Samantha Finley, Nora Forknall, Audrey Francis, Gary Patrick Hopkins, Emily M Joyce, SungJin Kim, Nicole A Kirk, Julie Kovalyak, Shirley A Lauchie, Alanna Lohff, Charli Maldonado, Emily A Manley, Sari McLin, Caroline Mooney, Miatta Ndama, Omotara Ogundeyi, Nneoma Okeoma, Christopher Ordish, Nicholas Padilla, Christopher M Patrick, Tyler Paterson, Elliott E Phillips, Emily M Phillips, Neha Rampally, Caitlin Ribeiro, Madelaine K Robertson, Jon Thomson Rymer, Sean M Ryan, Megan Sammons, Anne K Scott, Ashley L Scott, Aya Shinomiya, Claire Smith, Kelsey Smith, Natalie L Smith, Margaret A Sobeski, Alia Suleiman, Jackie Swift, Satoko Takemura, Iris Talebi, Dorota Tarnogorska, Emily Tenshaw, Temour Tokhi, John J Walsh, Tansy Yang, Jane Anne Horne, Feng Li, Ruchi Parekh, Patricia K Rivlin, Vivek Jayaraman, Marta Costa, Gregory SXE Jefferis, Kei Ito, Stephan Saalfeld, Reed George, Ian A Meinertzhagen, Gerald M Rubin, Harald F Hess, Viren Jain, and Stephen M Plaza. A connectome and analysis of the adult Drosophila central brain. *eLife*, 9, 2020.

[8015] Jodie A Schiffer, Francesco A Servello, William R Heath, Francis Raj Gandhi Amrit, Stephanie V Stumbur, Matthias Eder, Olivier MF Martin, Sean B Johnsen, Julian A Stanley, Hannah Tam, Sarah J Brennan, Natalie G McGowan, Abigail L Vogelaar, Yuyan Xu, William T Serkin, Arjumand Ghazi, Nicholas Stroustrup, and Javier Apfeld. Caenorhabditis elegans processes sensory information to choose between freeloading and self-defense strategies. *eLife*, 9, 2020.

[8016] Christoph N Schlaffner, Konstantin Kahnert, Jan Muntel, Ruchi Chauhan, Bernhard Y Renard, Judith A Steen, and Hanno Steen. FLEXIQuant-LF to quantify protein modification extent in label-free proteomics data. *eLife*, 9, 2020.

[8017] Alisha Schlichter, Margaret M Kasten, Timothy J Parnell, and Bradley R Cairns. Specialization of the chromatin remodeler RSC to mobilize partially-unwrapped nucleosomes. *eLife*, 9, 2020.

[8018] Sonja Schmid and Thorsten Hugel. Controlling protein function by fine-tuning conformational flexibility. *eLife*, 9, 2020.

[8019] Claudia C Schmidt, Vedran Vasic, and Alexander Stein. Doa10 is a membrane protein retrotranslocase in ER-associated protein degradation. *eLife*, 9, 2020.

[8020] Isabelle Schmutz, Arjen R Mensenkamp, Kaori K Takai, Maaike Haadsma, Liesbeth Spruijt, Richarda M de Voer, Seunga Sara Choo, Franziska K Lorbeer, Emma J van Grinsven, Dirk Hockemeyer, Marjolijn CJ Jongmans, and Titia de Lange. TINF2 is a haploinsufficient tumor suppressor that limits telomere length. *eLife*, 9, 2020.

[8021] Sophie Schobesberger, Peter T Wright, Claire Poulet, Jose L Sanchez Alonso Mardones, Catherine Mansfield, Andreas Friebe, Sian E Harding, Jean-Luc Balligand, Viacheslav O Nikolaev, and Julia Gorelik. $\beta$3-Adrenoceptor redistribution impairs NO/cGMP/PDE2 signalling in failing cardiomyocytes. *eLife*, 9, 2020.

[8022] CM Schooling, JV Zhao, SL Au Yeung, and GM Leung. Investigating pleiotropic effects of statins on ischemic heart disease in the UK Biobank using Mendelian randomisation. *eLife*, 9, 2020.

[8023] Marina Schrecker, Julia Korobenko, and Richard K Hite. Cryo-EM structure of the lysosomal chloride-proton exchanger CLC-7 in complex with OSTM1. *eLife*, 9, 2020.

[8024] Catherine E Schretter, Yoshinori Aso, Alice A Robie, Marisa Dreher, Michael-John Dolan, Nan Chen, Masayoshi Ito, Tansy Yang, Ruchi Parekh, Kristin M Branson, and Gerald M Rubin. Cell types and neuronal circuitry underlying female aggression in Drosophila. *eLife*, 9, 2020.

[8025] Sandra K Schuller, Jan M Schuller, J Rajan Prabu, Marc Baumgärtner, Fabien Bonneau, Jérôme Basquin, and Elena Conti. Structural insights into the nucleic acid remodeling mechanisms of the yeast THO-Sub2 complex. *eLife*, 9, 2020.

[8026] Enrico Schulz, Anne Stankewitz, Anderson M Winkler, Stephanie Irving, Viktor Witkovský, and Irene Tracey. Ultra-high-field imaging reveals increased whole brain connectivity underpins cognitive strategies that attenuate pain. *eLife*, 9, 2020.

[8027] Lianna Schwartz-Orbach, Chenzhen Zhang, Simone Sidoli, Richa Amin, Diljeet Kaur, Anna Zhebrun, Julie Ni, and Sam G Gu. Caenorhabditis elegans nuclear RNAi factor SET-32 deposits the transgenerational histone modification, H3K23me3. *eLife*, 9, 2020.

[8028] Manon K Schweinfurth. The social life of Norway rats (Rattus norvegicus). *eLife*, 9, 2020.

[8029] Nicholas Sciascia, Wei Wu, Dali Zong, Yilun Sun, Nancy Wong, Sam John, Darawalee Wangsa, Thomas Ried, Samuel F Bunting, Yves Pommier, and André Nussenzweig. Suppressing proteasome mediated processing of topoisomerase II DNA-protein complexes preserves genome integrity. *eLife*, 9, 2020.

[8030] Brendon P Scicluna, Fabrice Uhel, Lonneke A van Vught, Maryse A Wiewel, Arie J Hoogendijk, Ingelore Baessman, Marek Franitza, Peter Nürnberg, Janneke Horn, Olaf L Cremer, Marc J Bonten, Marcus J Schultz, and Tom van der Poll. The leukocyte non-coding RNA landscape in critically ill patients with sepsis. *eLife*, 9, 2020.

[8031] Alessandra Sclip and Thomas C Südhof. LAR receptor phospho-tyrosine phosphatases regulate NMDA-receptor responses. *eLife*, 9, 2020.

[8032] Laura Sebastian Monasor, Stephan A Müller, Alessio Vittorio Colombo, Gaye Tanrioever, Jasmin König, Stefan Roth, Arthur Liesz, Anna Berghofer, Anke Piechotta, Matthias Prestel, Takashi Saito, Takaomi C Saido, Jochen Herms, Michael Willem, Christian Haass, Stefan F Lichtenthaler, and Sabina Tahirovic. Fibrillar A$\beta$ triggers microglial proteome alterations and dysfunction in Alzheimer mouse models. *eLife*, 9, 2020.

[8033] Dennis Segebarth, Matthias Griebel, Nikolai Stein, Cora R von Collenberg, Corinna Martin, Dominik Fiedler, Lucas B Comeras, Anupam Sah, Victoria Schoeffler, Teresa Lüffe, Alexander Dürr, Rohini Gupta, Manju Sasi, Christina Lillesaar, Maren D Lange, Ramon O Tasan, Nicolas Singewald, Hans-Christian Pape, Christoph M Flath, and Robert Blum. On the objectivity, reliability, and validity of deep learning enabled bioimage analyses. *eLife*, 9, 2020.

[8034] Paolo S Segre, Jean Potvin, David E Cade, John Calambokidis, Jacopo Di Clemente, Frank E Fish, Ari S Friedlaender, William T Gough, Shirel R Kahane-Rapport, Cláudia Oliveira, Susan E Parks, Gwenith S Penry, Malene Simon, Alison K Stimpert, David N Wiley, KC Bierlich, Peter T Madsen, and Jeremy A Goldbogen. Energetic and physical limitations on the breaching performance of large whales. *eLife*, 9, 2020.

[8035] Ki-Hyeon Seong, Taishi Matsumura, Yuko Shimada-Niwa, Ryusuke Niwa, and Siu Kang. The Drosophila Individual Activity Monitoring and Detection System (DIAMonDS). *eLife*, 9, 2020.

[8036] Pradyumna Sepulveda, Marius Usher, Ned Davies, Amy A Benson, Pietro Ortoleva, and Benedetto De Martino. Visual attention modulates the integration of goal-relevant evidence and not value. *eLife*, 9, 2020.

[8037] Lucas D Serdar, DaJuan L Whiteside, Sarah L Nock, David McGrath, and Kristian E Baker. Inhibition of post-termination ribosome recycling at premature termination codons in UPF1 ATPase mutants. *eLife*, 9, 2020.

[8038] Vahan Serobyan, Zacharias Kontarakis, Mohamed A El-Brolosy, Jordan M Welker, Oleg Tolstenkov, Amr M Saadeldein, Nicholas Retzer, Alexander Gottschalk, Ann M Wehman, and Didier YR Stainier. Transcriptional adaptation in Caenorhabditis elegans. *eLife*, 9, 2020.

[8039] Andrea Serra-Marques, Maud Martin, Eugene A Katrukha, Ilya Grigoriev, Cathelijn AE Peeters, Qingyang Liu, Peter Jan Hooikaas, Yao Yao, Veronika Solianova, Ihor Smal, Lotte B Pedersen, Erik Meijering, Lukas C Kapitein, and Anna Akhmanova. Concerted action of kinesins KIF5B and KIF13B promotes efficient secretory vesicle transport to microtubule plus ends. *eLife*, 9, 2020.

[8040] Olivier Serralbo, David Salgado, Nadège Véron, Caitlin Cooper, Marie-Julie Dejardin, Timothy Doran, Jérome Gros, and Christophe Marcelle. Transgenesis and web resources in quail. *eLife*, 9, 2020.

[8041] Inda Setyawati, Weronika K Stanek, Maria Majsnerowska, Lotteke J Y M Swier, Els Pardon, Jan Steyaert, Albert Guskov, and Dirk J Slotboom. In vitro reconstitution of dynamically interacting integral membrane subunits of energy-coupling factor transporters. *eLife*, 9, 2020.

[8042] Mark Shaaya, Jordan Fauser, Anastasia Zhurikhina, Jason E Conage-Pough, Vincent Huyot, Martin Brennan, Cameron T Flower, Jacob Matsche, Shahzeb Khan, Viswanathan Natarajan, Jalees Rehman, Pradeep Kota, Forest M White, Denis Tsygankov, and Andrei V Karginov. Light-regulated allosteric switch enables temporal and subcellular control of enzyme activity. *eLife*, 9, 2020.

[8043] Golia Shafiei, Ross D Markello, Reinder Vos de Wael, Boris C Bernhardt, Ben D Fulcher, and Bratislav Misic. Topographic gradients of intrinsic dynamics across neocortex. *eLife*, 9, 2020.

[8044] Nishal P Shah, Nora Brackbill, Colleen Rhoades, Alexandra Kling, Georges Goetz, Alan M Litke, Alexander Sher, Eero P Simoncelli, and EJ Chichilnisky. Inference of nonlinear receptive field subunits with spike-triggered clustering. *eLife*, 9, 2020.

[8045] Shanaya Shital Shah, Stella Hartono, Aurèle Piazza, Vanessa Som, William Wright, Frédéric Chédin, and Wolf-Dietrich Heyer. Rdh54/Tid1 inhibits Rad51-Rad54-mediated D-loop formation and limits D-loop length. *eLife*, 9, 2020.

[8046] Shanaya Shital Shah, Stella R Hartono, Frédéric Chédin, and Wolf-Dietrich Heyer. Bisulfite treatment and single-molecule real-time sequencing reveal D-loop length, position, and distribution. *eLife*, 9, 2020.

[8047] Or David Shahar and Erin Margaret Schuman. Large-scale cell-type-specific imaging of protein synthesis in a vertebrate brain. *eLife*, 9, 2020.

[8048] Julie Teresa Shapiro, Gilles Leboucher, Anne-Florence Myard-Dury, Pascale Girardo, Anatole Luzzati, Mélissa Mary, Jean-FranÇois Sauzon, Bénédicte Lafay, Olivier Dauwalder, Frédéric Laurent, Gerard Lina, Christian Chidiac, Sandrine Couray-Targe, FranÇois Vandenesch, Jean-Pierre Flandrois, and Jean-Philippe Rasigade. Metapopulation ecology links antibiotic resistance, consumption, and patient transfers in a network of hospital wards. *eLife*, 9, 2020.

[8049] Anamika Sharma and Gaiti Hasan. Modulation of flight and feeding behaviours requires presynaptic IP3Rs in dopaminergic neurons. *eLife*, 9, 2020.

[8050] Arun Sharma, Lauren K Wasson, Jon AL Willcox, Sarah U Morton, Joshua M Gorham, Daniel M DeLaughter, Meraj Neyazi, Manuel Schmid, Radhika Agarwal, Min Young Jang, Christopher N Toepfer, Tarsha Ward, Yuri Kim, Alexandre C Pereira, Steven R DePalma, Angela Tai, Seongwon Kim, David Conner, Daniel Bernstein, Bruce D Gelb, Wendy K Chung, Elizabeth Goldmuntz, George Porter, Martin Tristani-Firouzi, Deepak Srivastava, Jonathan G Seidman, and Christine E Seidman. GATA6 mutations in hiPSCs inform mechanisms for maldevelopment of the heart, pancreas, and diaphragm. *eLife*, 9, 2020.

[8051] Kriti Sharma, Márton Palatinszky, Georgi Nikolov, David Berry, and Elizabeth A Shank. Transparent soil microcosms for live-cell imaging and non-destructive stable isotope probing of soil microorganisms. *eLife*, 9, 2020.

[8052] Joseph Shaw, Rajendra Gosain, Monoj Mon Kalita, Toshana L Foster, Jayakanth Kankanala, D Ram Mahato, Sonia Abas, Barnabas J King, Claire Scott, Emma Brown, Matthew J Bentham, Laura Wetherill, Abigail Bloy, Adel Samson, Mark Harris, Jamel Mankouri, David J Rowlands, Andrew Macdonald, Alexander W Tarr, Wolfgang B Fischer, Richard Foster, and Stephen Griffin. Rationally derived inhibitors of hepatitis C virus (HCV) p7 channel activity reveal prospect for bimodal antiviral therapy. *eLife*, 9, 2020.

[8053] Rylan Shearn, Alison E Wright, Sylvain Mousset, Corinne Régis, Simon Penel, Jean-FranÇois Lemaitre, Guillaume Douay, Brigitte Crouau-Roy, Emilie Lecompte, and Gabriel AB Marais. Evolutionary stasis of the pseudoautosomal boundary in strepsirrhine primates. *eLife*, 9, 2020.

[8054] Birong Shen, Joseph H Chapman, Michael F Custance, Gianna M Tricola, Charles E Jones, and Anthony V Furano. Perturbation of base excision repair sensitizes breast cancer cells to APOBEC3 deaminase-mediated mutations. *eLife*, 9, 2020.

[8055] Hua Shen, Peiheng Gan, Kristy Wang, Ali Darehzereshki, Kai Wang, S Ram Kumar, Ching-Ling Lien, Michaela Patterson, Ge Tao, and Henry M Sucov. Mononuclear diploid cardiomyocytes support neonatal mouse heart regeneration in response to paracrine IGF2 signaling. *eLife*, 9, 2020.

[8056] Joel W Sher, Hoong Chuin Lim, and Thomas G Bernhardt. Global phenotypic profiling identifies a conserved actinobacterial cofactor for a bifunctional PBP-type cell wall synthase. *eLife*, 9, 2020.

[8057] Xiaoshan Shi, Chunmei Chang, Adam L Yokom, Liv E Jensen, and James H Hurley. The autophagy adaptor NDP52 and the FIP200 coiled-coil allosterically activate ULK1 complex membrane recruitment. *eLife*, 9, 2020.

[8058] Zhigang Shi, Nicole E Pelletier, Jennifer Wong, Baoxin Li, Andrei D Sdrulla, Christopher J Madden, Daniel L Marks, and Virginia L Brooks. Leptin increases sympathetic nerve activity via induction of its own receptor in the paraventricular nucleus. *eLife*, 9, 2020.

[8059] Takushi Shimomura, Yoshiki Yonekawa, Hitoshi Nagura, Michihiro Tateyama, Yoshinori Fujiyoshi, and Katsumasa Irie. A native prokaryotic voltage-dependent calcium channel with a novel selectivity filter sequence. *eLife*, 9, 2020.

[8060] Donghyuk Shin, Anshu Bhattacharya, Yi-Lin Cheng, Marta Campos Alonso, Ahmad Reza Mehdipour, Gerbrand J van der Heden van Noort, Huib Ovaa, Gerhard Hummer, and Ivan Dikic. Bacterial OTU deubiquitinases regulate substrate ubiquitination upon Legionella infection. *eLife*, 9, 2020.

[8061] Maggie M Shin, Catarina Catela, and Jeremy Dasen. Intrinsic control of neuronal diversity and synaptic specificity in a proprioceptive circuit. *eLife*, 9, 2020.

[8062] Sunhye Shin, Yiyu Pang, Jooman Park, Lifeng Liu, Brandon E Lukas, Seung Hyeon Kim, Ki-Wook Kim, Pingwen Xu, Daniel C Berry, and Yuwei Jiang. Dynamic control of adipose tissue development and adult tissue homeostasis by platelet-derived growth factor receptor alpha. *eLife*, 9, 2020.

[8063] Anastasia V Shindyapina, Aleksandr A Zenin, Andrei E Tarkhov, Didac Santesmasses, Peter O Fedichev, and Vadim N Gladyshev. Germline burden of rare damaging variants negatively affects human healthspan and lifespan. *eLife*, 9, 2020.

[8064] Mitsuru Shinohara, Takahisa Kanekiyo, Masaya Tachibana, Aishe Kurti, Motoko Shinohara, Yuan Fu, Jing Zhao, Xianlin Han, Patrick M Sullivan, G William Rebeck, John D Fryer, Michael G Heckman, and Guojun Bu. APOE2 is associated with longevity independent of Alzheimer's disease. *eLife*, 9, 2020.

[8065] Kazuki Shiotani, Yuta Tanisumi, Koshi Murata, Junya Hirokawa, Yoshio Sakurai, and Hiroyuki Manabe. Tuning of olfactory cortex ventral tenia tecta neurons to distinct task elements of goal-directed behavior. *eLife*, 9, 2020.

[8066] Gergana Shipkovenska, Alexander Durango, Marian Kalocsay, Steven P Gygi, and Danesh Moazed. A conserved RNA degradation complex required for spreading and epigenetic inheritance of heterochromatin. *eLife*, 9, 2020.

[8067] Miri Shnayder, Aharon Nachshon, Batsheva Rozman, Biana Bernshtein, Michael Lavi, Noam Fein, Emma Poole, Selmir Avdic, Emily Blyth, David Gottlieb, Allison Abendroth, Barry Slobedman, John Sinclair, Noam Stern-Ginossar, and Michal Schwartz. Single cell analysis reveals human cytomegalovirus drives latently infected cells towards an anergic-like monocyte state. *eLife*, 9, 2020.

[8068] Marta S Shocket, Anna B Verwillow, Mailo G Numazu, Hani Slamani, Jeremy M Cohen, Fadoua El Moustaid, Jason Rohr, Leah R Johnson, and Erin A Mordecai. Transmission of West Nile and five other temperate mosquito-borne viruses peaks at temperatures between 23°C and 26°C. *eLife*, 9, 2020.

[8069] Daniel Shriner, Amy R Bentley, Jie Zhou, Kenneth Ekoru, Ayo P Doumatey, Guanjie Chen, Adebowale Adeyemo, and Charles N Rotimi. Time-to-event modeling of hypertension reveals the nonexistence of true controls. *eLife*, 9, 2020.

[8070] Calla B Shubin and Carol W Greider. The role of Rif1 in telomere length regulation is separable from its role in origin firing. *eLife*, 9, 2020.

[8071] Sophie B Siems, Olaf Jahn, Maria A Eichel, Nirmal Kannaiyan, Lai Man N Wu, Diane L Sherman, Kathrin Kusch, Dörte Hesse, Ramona B Jung, Robert Fledrich, Michael W Sereda, Moritz J Rossner, Peter J Brophy, and Hauke B Werner. Proteome profile of peripheral myelin in healthy mice and in a neuropathy model. *eLife*, 9, 2020.

[8072] Sandeep K Singh, Miao Gui, Fujiet Koh, Matthew CJ Yip, and Alan Brown. Structure and activation mechanism of the BBSome membrane protein trafficking complex. *eLife*, 9, 2020.

[8073] Neel S Singhal, Meirong Bai, Evan M Lee, Shuo Luo, Kayleigh R Cook, and Dengke K Ma. Cytoprotection by a naturally occurring variant of ATP5G1 in Arctic ground squirrel neural progenitor cells. *eLife*, 9, 2020.

[8074] Niladri K Sinha, Alban Ordureau, Katharina Best, James A Saba, Boris Zinshteyn, Elayanambi Sundaramoorthy, Amit Fulzele, Danielle M Garshott, Timo Denk, Matthias Thoms, Joao A Paulo, J Wade Harper, Eric J Bennett, Roland Beckmann, and Rachel Green. EDF1 coordinates cellular responses to ribosome collisions. *eLife*, 9, 2020.

[8075] Chiara Sinigaglia, Sophie Peron, Jeanne Eichelbrenner, Sandra Chevalier, Julia Steger, Carine Barreau, Evelyn Houliston, and Lucas Leclère. Pattern regulation in a regenerating jellyfish. *eLife*, 9, 2020.

[8076] Kishor K Sivaraj, Backialakshmi Dharmalingam, Vishal Mohanakrishnan, Hyun-Woo Jeong, Katsuhiro Kato, Silke Schröder, Susanne Adams, Gou Young Koh, and Ralf H Adams. YAP1 and TAZ negatively control bone angiogenesis by limiting hypoxia-inducible factor signaling in endothelial cells. *eLife*, 9, 2020.

[8077] Andrew D Skol, Segun C Jung, Ana Marija Sokovic, Siquan Chen, Sarah Fazal, Olukayode Sosina, Poulami P Borkar, Amy Lin, Maria Sverdlov, Dingcai Cao, Anand Swaroop, Ionut Bebu, Barbara E Stranger, and Michael A Grassi. Integration of genomics and transcriptomics predicts diabetic retinopathy susceptibility genes. *eLife*, 9, 2020.

[8078] Victoria A Sleight and J Andrew Gillis. Embryonic origin and serial homology of gill arches and paired fins in the skate, Leucoraja erinacea. *eLife*, 9, 2020.

[8079] Ross O Smith, Takeshi Ninchoji, Emma Gordon, Helder André, Elisabetta Dejana, Dietmar Vestweber, Anders Kvanta, and Lena Claesson-Welsh. Vascular permeability in retinopathy is regulated by VEGFR2 Y949 signaling to VE-cadherin. *eLife*, 9, 2020.

[8080] Sarah Jacquelyn Smith, Lance A Davidson, and Mark Rebeiz. Evolutionary expansion of apical extracellular matrix is required for the elongation of cells in a novel structure. *eLife*, 9, 2020.

[8081] Stephen M Smith, Lloyd T Elliott, Fidel Alfaro-Almagro, Paul McCarthy, Thomas E Nichols, Gwenaëlle Douaud, and Karla L Miller. Brain aging comprises many modes of structural and functional change with distinct genetic and biophysical associations. *eLife*, 9, 2020.

[8082] Ediz Sohoglu and Matthew H Davis. Rapid computations of spectrotemporal prediction error support perception of degraded speech. *eLife*, 9, 2020.

[8083] Gabriel Sollberger, Robert Streeck, Falko Apel, Brian Edward Caffrey, Arthur I Skoultchi, and Arturo Zychlinsky. Linker histone H1.2 and H1.4 affect the neutrophil lineage determination. *eLife*, 9, 2020.

[8084] Isabella V Soluri, Lauren M Zumerling, Omar A Payan Parra, Eleanor G Clark, and Shelby A Blythe. Zygotic pioneer factor activity of Odd-paired/Zic is necessary for late function of the Drosophila segmentation network. *eLife*, 9, 2020.

[8085] Tim DD Somerville, Giulia Biffi, Juliane Daßler-Plenker, Stella K Hur, Xue-Yan He, Krysten E Vance, Koji Miyabayashi, Yali Xu, Diogo Maia-Silva, Olaf Klingbeil, Osama E Demerdash, Jonathan B Preall, Michael A Hollingsworth, Mikala Egeblad, David A Tuveson, and Christopher R Vakoc. Squamous trans-differentiation of pancreatic cancer cells promotes stromal inflammation. *eLife*, 9, 2020.

[8086] Hongki Song, Amy S Orr, Miriam Lee, Max E Harner, and William T Wickner. HOPS recognizes each SNARE, assembling ternary trans-complexes for rapid fusion upon engagement with the 4th SNARE. *eLife*, 9, 2020.

[8087] Huan Song, Henrik Larsson, Fang Fang, Catarina Almqvist, Nancy L Pedersen, Patrik KE Magnusson, and Unnur A Valdimarsdóttir. Risk of psychiatric disorders among the surviving twins after a co-twin loss. *eLife*, 9, 2020.

[8088] LouJin Song, Xian Chen, Terri A Swanson, Brianna LaViolette, Jincheng Pang, Teresa Cunio, Michael W Nagle, Shoh Asano, Katherine Hales, Arun Shipstone, Hanna Sobon, Sabra D Al-Harthy, Youngwook Ahn, Steven Kreuser, Andrew Robertson, Casey Ritenour, Frank Voigt, Magalie Boucher, Furong Sun, William C Sessa, and Rachel J Roth Flach. Lymphangiogenic therapy prevents cardiac dysfunction by ameliorating inflammation and hypertension. *eLife*, 9, 2020.

[8089] Konstantinos Sousounis, Donald M Bryant, Jose Martinez Fernandez, Samuel S Eddy, Stephanie L Tsai, Gregory C Gundberg, Jihee Han, Katharine Courtemanche, Michael Levin, and Jessica L Whited. Eya2 promotes cell cycle progression by regulating DNA damage response during vertebrate limb regeneration. *eLife*, 9, 2020.

[8090] Goffredina Spanò, Gloria Pizzamiglio, Cornelia McCormick, Ian A Clark, Sara De Felice, Thomas D Miller, Jamie O Edgin, Clive R Rosenthal, and Eleanor A Maguire. Dreaming with hippocampal damage. *eLife*, 9, 2020.

[8091] Melissa L Spear, Alex Diaz-Papkovich, Elad Ziv, Joseph M Yracheta, Simon Gravel, Dara G Torgerson, and Ryan D Hernandez. Recent shifts in the genomic ancestry of Mexican Americans may alter the genetic architecture of biomedical traits. *eLife*, 9, 2020.

[8092] Nicholas A Spidale, Nidhi Malhotra, Michela Frascoli, Katelyn Sylvia, Bing Miu, Coral Freeman, Brian D Stadinski, Eric Huseby, and Joonsoo Kang. Neonatal-derived IL-17 producing dermal $\gamma\delta$ T cells are required to prevent spontaneous atopic dermatitis. *eLife*, 9, 2020.

[8093] Michael Sporny, Julia Guez-Haddad, Tami Khazma, Avraham Yaron, Moshe Dessau, Yoel Shkolnisky, Carsten Mim, Michail N Isupov, Ran Zalk, Michael Hons, and Yarden Opatowsky. Structural basis for SARM1 inhibition and activation under energetic stress. *eLife*, 9, 2020.

[8094] Kate R Sprogis, Simone Videsen, and Peter T Madsen. Vessel noise levels drive behavioural responses of humpback whales with implications for whale-watching. *eLife*, 9, 2020.

[8095] Tiziana Squillaro, Gianfranco Peluso, Umberto Galderisi, and Giovanni Di Bernardo. Long noncoding RNAs in regulation of adipogenesis and adipose tissue function. *eLife*, 9, 2020.

[8096] Madhusudhan Srinivasan, Marco Fumasoni, Naomi J Petela, Andrew Murray, and Kim A Nasmyth. Cohesion is established during DNA replication utilising chromosome associated cohesin rings as well as those loaded de novo onto nascent DNAs. *eLife*, 9, 2020.

[8097] Elia Stahl, Théo Brillatz, Emerson Ferreira Queiroz, Laurence Marcourt, André Schmiesing, Olivier Hilfiker, Isabelle Riezman, Howard Riezman, Jean-Luc Wolfender, and Philippe Reymond. Phosphatidylcholines from Pieris brassicae eggs activate an immune response in Arabidopsis. *eLife*, 9, 2020.

[8098] Jessica J Stanisich, Dawid S Zyla, Pavel Afanasyev, Jingwei Xu, Anne Kipp, Eric Olinger, Olivier Devuyst, Martin Pilhofer, Daniel Boehringer, and Rudi Glockshuber. The cryo-EM structure of the human uromodulin filament core reveals a unique assembly mechanism. *eLife*, 9, 2020.

[8099] Todd A Starich, Xiaofei Bai, and David Greenstein. Gap junctions deliver malonyl-CoA from soma to germline to support embryogenesis in Caenorhabditis elegans. *eLife*, 9, 2020.

[8100] Marisa Stebegg, Alexandre Bignon, Danika Lea Hill, Alyssa Silva-Cayetano, Christel Krueger, Ine Vanderleyden, Silvia Innocentin, Louis Boon, Jiong Wang, Martin S Zand, James Dooley, Jonathan Clark, Adrian Liston, Edward Carr, and Michelle A Linterman. Rejuvenating conventional dendritic cells and T follicular helper cell formation after vaccination. *eLife*, 9, 2020.

[8101] Erich Stefan, Susanne Hofmann, and Robert Tampé. A single power stroke by ATP binding drives substrate translocation in a heterodimeric ABC transporter. *eLife*, 9, 2020.

[8102] Sonia Stefanovic, Brigitte Laforest, Jean-Pierre Desvignes, Fabienne Lescroart, Laurent Argiro, Corinne Maurel-Zaffran, David Salgado, Elise Plaindoux, Christopher De Bono, Kristijan Pazur, Magali Théveniau-Ruissy, Christophe Béroud, Michel Puceat, Anthony Gavalas, Robert G Kelly, and Stephane Zaffran. Hox-dependent coordination of mouse cardiac progenitor cell patterning and differentiation. *eLife*, 9, 2020.

[8103] Yannik Stegmann, Lea Ahrens, Paul Pauli, Andreas Keil, and Matthias J Wieser. Social aversive generalization learning sharpens the tuning of visuocortical neurons to facial identity cues. *eLife*, 9, 2020.

[8104] Emmanuelle Steib, Marine H Laporte, Davide Gambarotto, Natacha Olieric, Celine Zheng, Susanne Borgers, Vincent Olieric, Maeva Le Guennec, France Koll, Anne-Marie Tassin, Michel O Steinmetz, Paul Guichard, and Virginie Hamel. WDR90 is a centriolar microtubule wall protein important for centriole architecture integrity. *eLife*, 9, 2020.

[8105] Julia Steiner and Leonid Sazanov. Structure and mechanism of the Mrp complex, an ancient cation/proton antiporter. *eLife*, 9, 2020.

[8106] Janina Isabel Steinert, Shaukat Khan, Khudzie Mlambo, Fiona J Walsh, Emma Mafara, Charlotte Lejeune, Cebele Wong, Anita Hettema, Osondu Ogbuoji, Sebastian Vollmer, Jan-Walter De Neve, Sikhathele Mazibuko, Velephi Okello, Till Bärnighausen, and Pascal Geldsetzer. A stepped-wedge randomised trial on the impact of early ART initiation on HIV-patients' economic outcomes in Eswatini. *eLife*, 9, 2020.

[8107] Lukas S Stelzl, Despoina AI Mavridou, Emmanuel Saridakis, Diego Gonzalez, Andrew J Baldwin, Stuart J Ferguson, Mark SP Sansom, and Christina Redfield. Local frustration determines loop opening during the catalytic cycle of an oxidoreductase. *eLife*, 9, 2020.

[8108] Madlen Stephani, Lorenzo Picchianti, Alexander Gajic, Rebecca Beveridge, Emilio Skarwan, Victor Sanchez de Medina Hernandez, Azadeh Mohseni, Marion Clavel, Yonglun Zeng, Christin Naumann, Mateusz Matuszkiewicz, Eleonora Turco, Christian Loefke, Baiying Li, Gerhard Dürnberger, Michael Schutzbier, Hsiao Tieh Chen, Alibek Abdrakhmanov, Adriana Savova, Khong-Sam Chia, Armin Djamei, Irene Schaffner, Steffen Abel, Liwen Jiang, Karl Mechtler, Fumiyo Ikeda, Sascha Martens, Tim Clausen, and Yasin Dagdas. A cross-kingdom conserved ER-phagy receptor maintains endoplasmic reticulum homeostasis during stress. *eLife*, 9, 2020.

[8109] Zoe A Stephenson, Robert F Harvey, Kenneth R Pryde, Sarah Mistry, Rachel E Hardy, Riccardo Serreli, Injae Chung, Timothy EH Allen, Mark Stoneley, Marion MacFarlane, Peter M Fischer, Judy Hirst, Barrie Kellam, and Anne E Willis. Identification of a novel toxicophore in anti-cancer chemotherapeutics that targets mitochondrial respiratory complex I. *eLife*, 9, 2020.

[8110] Jonathan Stieglitz, Paul L Hooper, Benjamin C Trumble, Hillard Kaplan, and Michael D Gurven. Productivity loss associated with functional disability in a contemporary small-scale subsistence population. *eLife*, 9, 2020.

[8111] Gabriel M Stine, Ariel Zylberberg, Jochen Ditterich, and Michael N Shadlen. Differentiating between integration and non-integration strategies in perceptual decision making. *eLife*, 9, 2020.

[8112] Miriam Stoeber, Damien Jullié, Joy Li, Soumen Chakraborty, Susruta Majumdar, Nevin A Lambert, Aashish Manglik, and Mark von Zastrow. Agonist-selective recruitment of engineered protein probes and of GRK2 by opioid receptors in living cells. *eLife*, 9, 2020.

[8113] Thomas Stoeger and Luís A Nunes Amaral. COVID-19 research risks ignoring important host genes due to pre-established research patterns. *eLife*, 9, 2020.

[8114] David Q Stoye, Manuel Blesa, Gemma Sullivan, Paola Galdi, Gillian J Lamb, Gill S Black, Alan J Quigley, Michael J Thrippleton, Mark E Bastin, Rebecca M Reynolds, and James P Boardman. Maternal cortisol is associated with neonatal amygdala microstructure and connectivity in a sexually dimorphic manner. *eLife*, 9, 2020.

[8115] Isabelle Straub, Laurens Witter, Abdelmoneim Eshra, Miriam Hoidis, Niklas Byczkowicz, Sebastian Maas, Igor Delvendahl, Kevin Dorgans, Elise Savier, Ingo Bechmann, Martin Krueger, Philippe Isope, and Stefan Hallermann. Gradients in the mammalian cerebellar cortex enable Fourier-like transformation and improve storing capacity. *eLife*, 9, 2020.

[8116] Daniel J Strauss, Farah I Corona-Strauss, Andreas Schroeer, Philipp Flotho, Ronny Hannemann, and Steven A Hackley. Vestigial auriculomotor activity indicates the direction of auditory attention in humans. *eLife*, 9, 2020.

[8117] Ivana Strazic Geljic, Paola Kucan Brlic, Guillem Angulo, Ilija Brizic, Berislav Lisnic, Tina Jenus, Vanda Juranic Lisnic, Gian Pietro Pietri, Pablo Engel, Noa Kaynan, Jelena Zeleznjak, Peter Schu, Ofer Mandelboim, Astrid Krmpotic, Ana Angulo, Stipan Jonjic, and Tihana Lenac Rovis. Cytomegalovirus protein m154 perturbs the adaptor protein-1 compartment mediating broad-spectrum immune evasion. *eLife*, 9, 2020.

[8118] Anne M Stringer, Gabriele Baniulyte, Erica Lasek-Nesselquist, Kimberley D Seed, and Joseph T Wade. Transcription termination and antitermination of bacterial CRISPR arrays. *eLife*, 9, 2020.

[8119] Maria Cristina Stroe, Tina Netzker, Kirstin Scherlach, Thomas Krüger, Christian Hertweck, Vito Valiante, and Axel A Brakhage. Targeted induction of a silent fungal gene cluster encoding the bacteria-specific germination inhibitor fumigermin. *eLife*, 9, 2020.

[8120] Nils Stührwohldt, Stefan Scholl, Lisa Lang, Julia Katzenberger, Karin Schumacher, and Andreas Schaller. The biogenesis of CLEL peptides involves several processing events in consecutive compartments of the secretory pathway. *eLife*, 9, 2020.

[8121] Yusuke Sugasawa, Wayland WL Cheng, John R Bracamontes, Zi-Wei Chen, Lei Wang, Allison L Germann, Spencer R Pierce, Thomas C Senneff, Kathiresan Krishnan, David E Reichert, Douglas F Covey, Gustav Akk, and Alex S Evers. Site-specific effects of neurosteroids on GABAA receptor activation and desensitization. *eLife*, 9, 2020.

[8122] Demeng Sun, Sanling Liu, Siyu Li, Mengge Zhang, Fan Yang, Ming Wen, Pan Shi, Tao Wang, Man Pan, Shenghai Chang, Xing Zhang, Longhua Zhang, Changlin Tian, and Lei Liu. Structural insights into human acid-sensing ion channel 1a inhibition by snake toxin mambalgin1. *eLife*, 9, 2020.

[8123] Kaitlyn A Sun, Yan Li, Angelo Y Meliton, Parker S Woods, Lucas M Kimmig, Rengül Cetin-Atalay, Robert B Hamanaka, and Gökhan M Mutlu. Endogenous itaconate is not required for particulate matter-induced NRF2 expression or inflammatory response. *eLife*, 9, 2020.

[8124] Syuan-Jyun Sun and Rebecca M Kilner. Temperature stress induces mites to help their carrion beetle hosts by eliminating rival blowflies. *eLife*, 9, 2020.

[8125] Xiaoyan Sun, Alexandra Are, Karl Annusver, Unnikrishnan Sivan, Tina Jacob, Tim Dalessandri, Simon Joost, Anja Füllgrabe, Marco Gerling, and Maria Kasper. Coordinated hedgehog signaling induces new hair follicles in adult skin. *eLife*, 9, 2020.

[8126] Xuelong Sun, Shigang Yue, and Michael Mangan. A decentralised neural model explaining optimal integration of navigational strategies in insects. *eLife*, 9, 2020.

[8127] Yanan Sun, Meijiao Li, Dongfeng Zhao, Xin Li, Chonglin Yang, and Xiaochen Wang. Lysosome activity is modulated by multiple longevity pathways and is important for lifespan extension in C. elegans. *eLife*, 9, 2020.

[8128] Wilson Suraweera, David Warrell, Romulus Whitaker, Geetha Menon, Rashmi Rodrigues, Sze Hang Fu, Rehana Begum, Prabha Sati, Kapila Piyasena, Mehak Bhatia, Patrick Brown, and Prabhat Jha. Trends in snakebite deaths in India from 2000 to 2019 in a nationally representative mortality study. *eLife*, 9, 2020.

[8129] Aneesha K Suresh, James M Goodman, Elizaveta V Okorokova, Matthew Kaufman, Nicholas G Hatsopoulos, and Sliman J Bensmaia. Neural population dynamics in motor cortex are different for reach and grasp. *eLife*, 9, 2020.

[8130] Pooja Suresh, Alexandra F Long, and Sophie Dumont. Microneedle manipulation of the mammalian spindle reveals specialized, short-lived reinforcement near chromosomes. *eLife*, 9, 2020.

[8131] Katherine J Susa, Tom CM Seegar, Stephen C Blacklow, and Andrew C Kruse. A dynamic interaction between CD19 and the tetraspanin CD81 controls B cell co-receptor trafficking. *eLife*, 9, 2020.

[8132] Genjiro Suzuki, Sei Imura, Masato Hosokawa, Ryu Katsumata, Takashi Nonaka, Shin-Ichi Hisanaga, Yasushi Saeki, and Masato Hasegawa. $\alpha$-synuclein strains that cause distinct pathologies differentially inhibit proteasome. *eLife*, 9, 2020.

[8133] Yann Sweeney and Claudia Clopath. Population coupling predicts the plasticity of stimulus responses in cortical circuits. *eLife*, 9, 2020.

[8134] Marie E Sweet, Xihui Zhang, Hediye Erdjument-Bromage, Vikas Dubey, Himanshu Khandelia, Thomas A Neubert, Bjørn P Pedersen, and David L Stokes. Serine phosphorylation regulates the P-type potassium pump KdpFABC. *eLife*, 9, 2020.

[8135] Russell P Swift, Krithika Rajaram, Cyrianne Keutcha, Hans B Liu, Bobby Kwan, Amanda Dziedzic, Anne E Jedlicka, and Sean T Prigge. The NTP generating activity of pyruvate kinase II is critical for apicoplast maintenance in Plasmodium falciparum. *eLife*, 9, 2020.

[8136] Viktor Szegedi, Melinda Paizs, Judith Baka, Pál Barzó, Gábor Molnár, Gabor Tamas, and Karri Lamsa. Robust perisomatic GABAergic self-innervation inhibits basket cells in the human and mouse supragranular neocortex. *eLife*, 9, 2020.

[8137] Huy Q Ta, Natalia Dworak, Melissa L Ivey, Devin G Roller, and Daniel Gioeli. AR phosphorylation and CHK2 kinase activity regulates IR-stabilized AR–CHK2 interaction and prostate cancer survival. *eLife*, 9, 2020.

[8138] Alejandro Tabas, Glad Mihai, Stefan Kiebel, Robert Trampel, and Katharina von Kriegstein. Abstract rules drive adaptation in the subcortical sensory pathway. *eLife*, 9, 2020.

[8139] Clara Taffoni, Shizue Omi, Caroline Huber, Sébastien Mailfert, Mathieu Fallet, Jean-FranÇois Rupprecht, Jonathan J Ewbank, and Nathalie Pujol. Microtubule plus-end dynamics link wound repair to the innate immune response. *eLife*, 9, 2020.

[8140] Asuka Takeishi, Jihye Yeon, Nathan Harris, Wenxing Yang, and Piali Sengupta. Feeding state functionally reconfigures a sensory circuit to drive thermosensory behavioral plasticity. *eLife*, 9, 2020.

[8141] Dana M Talsness, Katie G Owings, Emily Coelho, Gaelle Mercenne, John M Pleinis, Raghavendran Partha, Kevin A Hope, Aamir R Zuberi, Nathan L Clark, Cathleen M Lutz, Aylin R Rodan, and Clement Y Chow. A Drosophila screen identifies NKCC1 as a modifier of NGLY1 deficiency. *eLife*, 9, 2020.

[8142] Vivian Tam, Peikai Chen, Anita Yee, Nestor Solis, Theo Klein, Mateusz Kudelko, Rakesh Sharma, Wilson CW Chan, Christopher M Overall, Lisbet Haglund, Pak C Sham, Kathryn Song Eng Cheah, and Danny Chan. DIPPER, a spatiotemporal proteomics atlas of human intervertebral discs for exploring ageing and degeneration dynamics. *eLife*, 9, 2020.

[8143] Hiromi Tamada, Jerome Blanc, Natalya Korogod, Carl CH Petersen, and Graham W Knott. Ultrastructural comparison of dendritic spine morphology preserved with cryo and chemical fixation. *eLife*, 9, 2020.

[8144] Marc D Tambini, Kelly A Norris, and Luciano D'Adamio. Opposite changes in APP processing and human A$\beta$ levels in rats carrying either a protective or a pathogenic APP mutation. *eLife*, 9, 2020.

[8145] Eshwar R Tammineni, Natalia Kraeva, Lourdes Figueroa, Carlo Manno, Carlos A Ibarra, Amira Klip, Sheila Riazi, and Eduardo Rios. Intracellular calcium leak lowers glucose storage in human muscle, promoting hyperglycemia and diabetes. *eLife*, 9, 2020.

[8146] Guak-Kim Tan, Brian A Pryce, Anna Stabio, John V Brigande, ChaoJie Wang, Zheng Xia, Sara F Tufa, Douglas R Keene, and Ronen Schweitzer. Tgf$\beta$ signaling is critical for maintenance of the tendon cell fate. *eLife*, 9, 2020.

[8147] Han L Tan, Richard H Roth, Austin R Graves, Robert H Cudmore, and Richard L Huganir. Lamina-specific AMPA receptor dynamics following visual deprivation in vivo. *eLife*, 9, 2020.

[8148] June H Tan, Margot Lautens, Laura Romanelli-Cedrez, Jianbin Wang, Michael R Schertzberg, Samantha R Reinl, Richard E Davis, Jennifer N Shepherd, Andrew G Fraser, and Gustavo Salinas. Alternative splicing of coq-2 controls the levels of rhodoquinone in animals. *eLife*, 9, 2020.

[8149] Kun Tan, Samantha H Jones, Blue B Lake, Jennifer N Chousal, Eleen Y Shum, Lingjuan Zhang, Song Chen, Abhishek Sohni, Shivam Pandya, Richard L Gallo, Kun Zhang, Heidi Cook-Andersen, and Miles F Wilkinson. The role of the NMD factor UPF3B in olfactory sensory neurons. *eLife*, 9, 2020.

[8150] Winnie Tan, Sylvie van Twest, Andrew Leis, Rohan Bythell-Douglas, Vincent J Murphy, Michael Sharp, Michael W Parker, Wayne Crismani, and Andrew J Deans. Monoubiquitination by the human Fanconi anemia core complex clamps FANCI:FANCD2 on DNA in filamentous arrays. *eLife*, 9, 2020.

[8151] Toshiko Tanaka, Nathan Basisty, Giovanna Fantoni, Julián Candia, Ann Z Moore, Angelique Biancotto, Birgit Schilling, Stefania Bandinelli, and Luigi Ferrucci. Plasma proteomic biomarker signature of age predicts health and life span. *eLife*, 9, 2020.

[8152] Cheng Tang, Roger Herikstad, Aishwarya Parthasarathy, Camilo Libedinsky, and Shih-Cheng Yen. Minimally dependent activity subspaces for working memory and motor preparation in the lateral prefrontal cortex. *eLife*, 9, 2020.

[8153] Rendong Tang, Qianling Song, Ying Li, Rui Zhang, Xingya Cai, and Haidong D Lu. Curvature-processing domains in primate V4. *eLife*, 9, 2020.

[8154] Rui Tang, Christopher W Murray, Ian L Linde, Nicholas J Kramer, Zhonglin Lyu, Min K Tsai, Leo C Chen, Hongchen Cai, Aaron D Gitler, Edgar Engleman, Wonjae Lee, and Monte M Winslow. A versatile system to record cell-cell interactions. *eLife*, 9, 2020.

[8155] Yizhe Tang, Juan Pablo Zuniga-Hertz, Cheng Han, Bin Yu, and Dongsheng Cai. Multifaceted secretion of htNSC-derived hypothalamic islets induces survival and antidiabetic effect via peripheral implantation in mice. *eLife*, 9, 2020.

[8156] Lili Tao, Andrew Lemoff, Guoxun Wang, Christina Zarek, Alexandria Lowe, Nan Yan, and Tiffany A Reese. Reactive oxygen species oxidize STING and suppress interferon production. *eLife*, 9, 2020.

[8157] Ammarah Tariq, Lucy Green, J Charles G Jeynes, Christian Soeller, and James G Wakefield. In vitro reconstitution of branching microtubule nucleation. *eLife*, 9, 2020.

[8158] Sudhir Gopal Tattikota, Bumsik Cho, Yifang Liu, Yanhui Hu, Victor Barrera, Michael J Steinbaugh, Sang-Ho Yoon, Aram Comjean, Fangge Li, Franz Dervis, Ruei-Jiun Hung, Jin-Wu Nam, Shannan Ho Sui, Jiwon Shim, and Norbert Perrimon. A single-cell survey of Drosophila blood. *eLife*, 9, 2020.

[8159] Rahil Taujale, Aarya Venkat, Liang-Chin Huang, Zhongliang Zhou, Wayland Yeung, Khaled M Rasheed, Sheng Li, Arthur S Edison, Kelley W Moremen, and Natarajan Kannan. Deep evolutionary analysis reveals the design principles of fold A glycosyltransferases. *eLife*, 9, 2020.

[8160] Andy Tay. Struggling to get started. *eLife*, 9, 2020.

[8161] Jennifer A Taylor, Benjamin P Bratton, Sophie R Sichel, Kris M Blair, Holly M Jacobs, Kristen E DeMeester, Erkin Kuru, Joe Gray, Jacob Biboy, Michael S VanNieuwenhze, Waldemar Vollmer, Catherine L Grimes, Joshua W Shaevitz, and Nina R Salama. Distinct cytoskeletal proteins define zones of enhanced cell wall synthesis in Helicobacter pylori. *eLife*, 9, 2020.

[8162] Keenan C Taylor, Po Wei Kang, Panpan Hou, Nien-Du Yang, Georg Kuenze, Jarrod A Smith, Jingyi Shi, Hui Huang, Kelli McFarland White, Dungeng Peng, Alfred L George, Jens Meiler, Robert L McFeeters, Jianmin Cui, and Charles R Sanders. Structure and physiological function of the human KCNQ1 channel voltage sensor intermediate state. *eLife*, 9, 2020.

[8163] Christine Tedijanto, Yonatan H Grad, and Marc Lipsitch. Potential impact of outpatient stewardship interventions on antibiotic exposures of common bacterial pathogens. *eLife*, 9, 2020.

[8164] Brian Tenner, Michael Getz, Brian Ross, Donya Ohadi, Christopher H Bohrer, Eric Greenwald, Sohum Mehta, Jie Xiao, Padmini Rangamani, and Jin Zhang. Spatially compartmentalized phase regulation of a Ca2+-cAMP-PKA oscillatory circuit. *eLife*, 9, 2020.

[8165] Jacob A Tennessen, Stephanie R Bollmann, Ekaterina Peremyslova, Brent A Kronmiller, Clint Sergi, Bulut Hamali, and Michael Scott Blouin. Clusters of polymorphic transmembrane genes control resistance to schistosomes in snail vectors. *eLife*, 9, 2020.

[8166] Jeannette L Tenthorey, Candice Young, Afeez Sodeinde, Michael Emerman, and Harmit S Malik. Mutational resilience of antiviral restriction favors primate TRIM5$\alpha$ in host-virus evolutionary arms races. *eLife*, 9, 2020.

[8167] Markus Terrey, Scott I Adamson, Alana L Gibson, Tianda Deng, Ryuta Ishimura, Jeffrey H Chuang, and Susan L Ackerman. GTPBP1 resolves paused ribosomes to maintain neuronal homeostasis. *eLife*, 9, 2020.

[8168] Tiberiu Tesileanu, Mary M Conte, John J Briguglio, Ann M Hermundstad, Jonathan D Victor, and Vijay Balasubramanian. Efficient coding of natural scene statistics predicts discrimination thresholds for grayscale textures. *eLife*, 9, 2020.

[8169] Liku B Tezera, Magdalena K Bielecka, Paul Ogongo, Naomi F Walker, Matthew Ellis, Diana J Garay-Baquero, Kristian Thomas, Michaela T Reichmann, David A Johnston, Katalin Andrea Wilkinson, Mohamed Ahmed, Sanjay Jogai, Suwan N Jayasinghe, Robert J Wilkinson, Salah Mansour, Gareth J Thomas, Christian H Ottensmeier, Alasdair Leslie, and Paul T Elkington. Anti-PD-1 immunotherapy leads to tuberculosis reactivation via dysregulation of TNF-$\alpha$. *eLife*, 9, 2020.

[8170] Vivek V Thacker, Neeraj Dhar, Kunal Sharma, Riccardo Barrile, Katia Karalis, and John D McKinney. A lung-on-chip model of early Mycobacterium tuberculosis infection reveals an essential role for alveolar epithelial cells in controlling bacterial growth. *eLife*, 9, 2020.

[8171] Akanksha Thawani, Michael J Rale, Nicolas Coudray, Gira Bhabha, Howard A Stone, Joshua W Shaevitz, and Sabine Petry. The transition state and regulation of $\gamma$-TuRC-mediated microtubule nucleation revealed by single molecule microscopy. *eLife*, 9, 2020.

[8172] Rachel C Thayer, Frances I Allen, and Nipam H Patel. Structural color in Junonia butterflies evolves by tuning scale lamina thickness. *eLife*, 9, 2020.

[8173] Patricia L Theard, Erin Sheffels, Nancy E Sealover, Amanda J Linke, David J Pratico, and Robert L Kortum. Marked synergy by vertical inhibition of EGFR signaling in NSCLC spheroids shows SOS1 is a therapeutic target in EGFR-mutated cancer. *eLife*, 9, 2020.

[8174] Jeanne L Theis, Georg Vogler, Maria A Missinato, Xing Li, Tanja Nielsen, Xin-Xin I Zeng, Almudena Martinez-Fernandez, Stanley M Walls, Anaïs Kervadec, James N Kezos, Katja Birker, Jared M Evans, Megan M O'Byrne, Zachary C Fogarty, André Terzic, Paul Grossfeld, Karen Ocorr, Timothy J Nelson, Timothy M Olson, Alexandre R Colas, and Rolf Bodmer. Patient-specific genomics and cross-species functional analysis implicate LRP2 in hypoplastic left heart syndrome. *eLife*, 9, 2020.

[8175] Christopher W Thomas, Mathilde CC Guillaumin, Laura E McKillop, Peter Achermann, and Vladyslav V Vyazovskiy. Global sleep homeostasis reflects temporally and spatially integrated local cortical neuronal activity. *eLife*, 9, 2020.

[8176] Erica N Thomas, Kyusik Q Kim, Emily P McHugh, Thomas Marcinkiewicz, and Hani S Zaher. Alkylative damage of mRNA leads to ribosome stalling and rescue by trans translation in bacteria. *eLife*, 9, 2020.

[8177] Connor J Thompson, Zhaoqian Su, Vinh H Vu, Yinghao Wu, Deborah E Leckband, and Daniel K Schwartz. Cadherin clusters stabilized by a combination of specific and nonspecific cis-interactions. *eLife*, 9, 2020.

[8178] Matthew G Thompson, Mark Dittmar, Michael J Mallory, Prasanna Bhat, Max B Ferretti, Beatriz MA Fontoura, Sara Cherry, and Kristen W Lynch. Viral-induced alternative splicing of host genes promotes influenza replication. *eLife*, 9, 2020.

[8179] Samuel Thompson, Yang Zhang, Christine Ingle, Kimberly A Reynolds, and Tanja Kortemme. Altered expression of a quality control protease in E. coli reshapes the in vivo mutational landscape of a model enzyme. *eLife*, 9, 2020.

[8180] Scott B Thompson, Adam M Sandor, Victor Lui, Jeffrey W Chung, Monique M Waldman, Robert A Long, Miriam L Estin, Jennifer L Matsuda, Rachel S Friedman, and Jordan Jacobelli. Formin-like 1 mediates effector T cell trafficking to inflammatory sites to enable T cell-mediated autoimmunity. *eLife*, 9, 2020.

[8181] William Hedley Thompson, Jessey Wright, and Patrick G Bissett. Open exploration. *eLife*, 9, 2020.

[8182] William Hedley Thompson, Jessey Wright, Patrick G Bissett, and Russell A Poldrack. Dataset decay and the problem of sequential analyses on open datasets. *eLife*, 9, 2020.

[8183] Johannes Thomsen, Magnus Berg Sletfjerding, Simon Bo Jensen, Stefano Stella, Bijoya Paul, Mette Galsgaard Malle, Guillermo Montoya, Troels Christian Petersen, and Nikos S Hatzakis. DeepFRET, a software for rapid and automated single-molecule FRET data classification using deep learning. *eLife*, 9, 2020.

[8184] Olivier Thouvenin, Ludovic Keiser, Yasmine Cantaut-Belarif, Martin Carbo-Tano, Frederik Verweij, Nathalie Jurisch-Yaksi, Pierre-Luc Bardet, Guillaume van Niel, Francois Gallaire, and Claire Wyart. Origin and role of the cerebrospinal fluid bidirectional flow in the central canal. *eLife*, 9, 2020.

[8185] Pascal N Timshel, Jonatan J Thompson, and Tune H Pers. Genetic mapping of etiologic brain cell types for obesity. *eLife*, 9, 2020.

[8186] Lauren C Tindale, Jessica E Stockdale, Michelle Coombe, Emma S Garlock, Wing Yin Venus Lau, Manu Saraswat, Louxin Zhang, Dongxuan Chen, Jacco Wallinga, and Caroline Colijn. Evidence for transmission of COVID-19 prior to symptom onset. *eLife*, 9, 2020.

[8187] Michael J Tisza, Diana V Pastrana, Nicole L Welch, Brittany Stewart, Alberto Peretti, Gabriel J Starrett, Yuk-Ying S Pang, Siddharth R Krishnamurthy, Patricia A Pesavento, David H McDermott,

Philip M Murphy, Jessica L Whited, Bess Miller, Jason Brenchley, Stephan P Rosshart, Barbara Rehermann, John Doorbar, Blake A Ta'ala, Olga Pletnikova, Juan C Troncoso, Susan M Resnick, Ben Bolduc, Matthew B Sullivan, Arvind Varsani, Anca M Segall, and Christopher B Buck. Discovery of several thousand highly diverse circular DNA viruses. *eLife*, 9, 2020.

[8188] Satish Kumar Tiwari, Ashish Ganeshlalji Toshniwal, Sudip Mandal, and Lolitika Mandal. Fatty acid $\beta$-oxidation is required for the differentiation of larval hematopoietic progenitors in Drosophila. *eLife*, 9, 2020.

[8189] Caroline Tizian, Annette Lahmann, Oliver Hölsken, Catalina Cosovanu, Michael Kofoed-Branzk, Frederik Heinrich, Mir-Farzin Mashreghi, Andrey Kruglov, Andreas Diefenbach, and Christian Neumann. c-Maf restrains T-bet-driven programming of CCR6-negative group 3 innate lymphoid cells. *eLife*, 9, 2020.

[8190] Victor Tobiasson and Alexey Amunts. Ciliate mitoribosome illuminates evolutionary steps of mitochondrial translation. *eLife*, 9, 2020.

[8191] Robert T Todd and Anna Selmecki. Expandable and reversible copy number amplification drives rapid adaptation to antifungal drugs. *eLife*, 9, 2020.

[8192] Yosuke Tona and Doris K Wu. Live imaging of hair bundle polarity acquisition demonstrates a critical timeline for transcription factor Emx2. *eLife*, 9, 2020.

[8193] Nicolò Tosetti, Nicolas Dos Santos Pacheco, Eloïse Bertiaux, Bohumil Maco, Lorène Bournonville, Virginie Hamel, Paul Guichard, and Dominique Soldati-Favre. Essential function of the alveolin network in the subpellicular microtubules and conoid assembly in Toxoplasma gondii. *eLife*, 9, 2020.

[8194] Simon Tragust, Claudia Herrmann, Jane Häfner, Ronja Braasch, Christina Tilgen, Maria Hoock, Margarita Artemis Milidakis, Roy Gross, and Heike Feldhaar. Formicine ants swallow their highly acidic poison for gut microbial selection and control. *eLife*, 9, 2020.

[8195] Stavros Trakoshis, Pablo Martínez-Cañada, Federico Rocchi, Carola Canella, Wonsang You, Bhismadev Chakrabarti, Amber NV Ruigrok, Edward T Bullmore, John Suckling, Marija Markicevic, Valerio Zerbi, Simon Baron-Cohen, Alessandro Gozzi, Meng-Chuan Lai, Stefano Panzeri, and Michael V Lombardo. Intrinsic excitation-inhibition imbalance affects medial prefrontal cortex differently in autistic men versus women. *eLife*, 9, 2020.

[8196] Alexandra N Trelle, Valerie A Carr, Scott A Guerin, Monica K Thieu, Manasi Jayakumar, Wanjia Guo, Ayesha Nadiadwala, Nicole K Corso, Madison P Hunt, Celia P Litovsky, Natalie J Tanner, Gayle K Deutsch, Jeffrey D Bernstein, Marc B Harrison, Anna M Khazenzon, Jiefeng Jiang, Sharon J Sha, Carolyn A Fredericks, Brian K Rutt, Elizabeth C Mormino, Geoffrey A Kerchner, and Anthony D Wagner. Hippocampal and cortical mechanisms at retrieval explain variability in episodic remembering in older adults. *eLife*, 9, 2020.

[8197] Mathieu Tremblay, Sophie Viala, Maxwell ER Shafer, Adda-Lee Graham-Paquin, Chloe Liu, and Maxime Bouchard. Regulation of stem/progenitor cell maintenance by BMP5 in prostate homeostasis and cancer initiation. *eLife*, 9, 2020.

[8198] Catherine G Triandafillou, Christopher D Katanski, Aaron R Dinner, and D Allan Drummond. Transient intracellular acidification regulates the core transcriptional heat shock response. *eLife*, 9, 2020.

[8199] Deepti Trivedi, Vinitha CM, Karishma Bisht, Vishnu Janardan, Awadhesh Pandit, Bishal Basak, Shwetha H, Navyashree Ramesh, and Padinjat Raghu. A genome engineering resource to uncover principles of cellular organization and tissue architecture by lipid signaling. *eLife*, 9, 2020.

[8200] Shubhanshi Trivedi, Daniel Labuz, Cole P Anderson, Claudia V Araujo, Antoinette Blair, Elizabeth A Middleton, Owen Jensen, Alexander Tran, Matthew A Mulvey, Robert A Campbell, J Scott Hale, Matthew T Rondina, and Daniel T Leung. Mucosal-associated invariant T (MAIT) cells mediate protective host responses in sepsis. *eLife*, 9, 2020.

[8201] Marie Trussart, Charis E Teh, Tania Tan, Lawrence Leong, Daniel HD Gray, and Terence P Speed. Removing unwanted variation with CytofRUV to integrate multiple CyTOF datasets. *eLife*, 9, 2020.

[8202] Hsiang-Yu Tsai, Dustin R Rubenstein, Bo-Fei Chen, Mark Liu, Shih-Fan Chan, De-Pei Chen, Syuan-Jyun Sun, Tzu-Neng Yuan, and Sheng-Feng Shen. Antagonistic effects of intraspecific cooperation and interspecific competition on thermal performance. *eLife*, 9, 2020.

[8203] Stephanie L Tsai, Clara Baselga-Garriga, and Douglas A Melton. Midkine is a dual regulator of wound epidermis development and inflammation during the initiation of limb regeneration. *eLife*, 9, 2020.

[8204] Anthony H Tsang, Christiane E Koch, Jana-Thabea Kiehn, Cosima X Schmidt, and Henrik Oster. An adipokine feedback regulating diurnal food intake rhythms in mice. *eLife*, 9, 2020.

[8205] Yiu Huen Tsang, Yumeng Wang, Kathleen Kong, Caitlin Grzeskowiak, Oksana Zagorodna, Turgut Dogruluk, Hengyu Lu, Nicole Villafane, Venkata Hemanjani Bhavana, Daniela Moreno, Sarah H Elsea, Han Liang, Gordon B Mills, and Kenneth L Scott. Differential expression of MAGEA6 toggles autophagy to promote pancreatic cancer progression. *eLife*, 9, 2020.

[8206] Tatsuhisa Tsuboi, Matheus P Viana, Fan Xu, Jingwen Yu, Raghav Chanchani, Ximena G Arceo, Evelina Tutucci, Joonhyuk Choi, Yang S Chen, Robert H Singer, Susanne M Rafelski, and Brian M Zid. Mitochondrial volume fraction and translation duration impact mitochondrial mRNA localization and protein synthesis. *eLife*, 9, 2020.

[8207] Taro Tsujimura, Osamu Takase, Masahiro Yoshikawa, Etsuko Sano, Matsuhiko Hayashi, Kazuto Hoshi, Tsuyoshi Takato, Atsushi Toyoda, Hideyuki Okano, and Keiichi Hishikawa. Controlling gene activation by enhancers through a drug-inducible topological insulator. *eLife*, 9, 2020.

[8208] Tomomi Tsunematsu, Amisha A Patel, Arno Onken, and Shuzo Sakata. State-dependent brainstem ensemble dynamics and their interactions with hippocampus across sleep states. *eLife*, 9, 2020.

[8209] Iku Tsutsui-Kimura, Hideyuki Matsumoto, Korleki Akiti, Melissa M Yamada, Naoshige Uchida, and Mitsuko Watabe-Uchida. Distinct temporal difference error signals in dopamine axons in three regions of the striatum in a decision-making task. *eLife*, 9, 2020.

[8210] Naotaka Tsutsumi, Somnath Mukherjee, Deepa Waghray, Claudia Y Janda, Kevin M Jude, Yi Miao, John S Burg, Nanda Gowtham Aduri, Anthony A Kossiakoff, Cornelius Gati, and K Christopher Garcia. Structure of human Frizzled5 by fiducial-assisted cryo-EM supports a heterodimeric mechanism of canonical Wnt signaling. *eLife*, 9, 2020.

[8211] Alzbeta Tuerkova, Ivo Kabelka, Tereza Králová, LukáŠ Sukeník, Šárka Pokorná, Martin Hof, and Robert Vácha. Effect of helical kink in antimicrobial peptides on membrane pore formation. *eLife*, 9, 2020.

[8212] Kirsteen M Tullett, Peck Szee Tan, Hae-Young Park, Ralf B Schittenhelm, Nicole Michael, Rong Li, Antonia N Policheni, Emily Gruber, Cheng Huang, Alex J Fulcher, Jillian C Danne, Peter E Czabotar, Linda M Wakim, Justine D Mintern, Georg Ramm, Kristen J Radford, Irina Caminschi, Meredith O'Keeffe, Jose A Villadangos, Mark D Wright, Marnie E Blewitt, William R Heath, Ken Shortman, Anthony W Purcell, Nicos A Nicola, Jian-Guo Zhang, and Mireille H Lahoud. RNF41 regulates the damage recognition receptor Clec9A and antigen cross-presentation in mouse dendritic cells. *eLife*, 9, 2020.

[8213] Hande Tunbak, Mireya Vazquez-Prada, Thomas Michael Ryan, Adam Raymond Kampff, and Elena Dreosti. Whole-brain mapping of socially isolated zebrafish reveals that lonely fish are not loners. *eLife*, 9, 2020.

[8214] Josef Turecek and Wade G Regehr. Cerebellar and vestibular nuclear synapses in the inferior olive have distinct release kinetics and neurotransmitters. *eLife*, 9, 2020.

[8215] Declan L Turner, Denis V Korneev, John G Purdy, Alex de Marco, and Rommel A Mathias. The host exosome pathway underpins biogenesis of the human cytomegalovirus virion. *eLife*, 9, 2020.

[8216] Kevin L Turner, Kyle W Gheres, Elizabeth A Proctor, and Patrick J Drew. Neurovascular coupling and bilateral connectivity during NREM and REM sleep. *eLife*, 9, 2020.

[8217] Motokazu Uchigashima, Kohtarou Konno, Emily Demchak, Amy Cheung, Takuya Watanabe, David G Keener, Manabu Abe, Timmy Le, Kenji Sakimura, Toshikuni Sasaoka, Takeshi Uemura, Yuka Imamura Kawasawa, Masahiko Watanabe, and Kensuke Futai. Specific Neuroligin3–$\alpha$Neurexin1 signaling regulates GABAergic synaptic function in mouse hippocampus. *eLife*, 9, 2020.

[8218] George Ueda, Aleksandar Antanasijevic, Jorge A Fallas, William Sheffler, Jeffrey Copps, Daniel Ellis, Geoffrey B Hutchinson, Adam Moyer, Anila Yasmeen, Yaroslav Tsybovsky, Young-Jun Park, Matthew J Bick, Banumathi Sankaran, Rebecca A Gillespie, Philip JM Brouwer, Peter H Zwart, David Veesler, Masaru Kanekiyo, Barney S Graham, Rogier W Sanders, John P Moore, Per Johan Klasse, Andrew B Ward, Neil P King, and David Baker. Tailored design of protein nanoparticle scaffolds for multivalent presentation of viral glycoprotein antigens. *eLife*, 9, 2020.

[8219] Rolf Ulrich and Jeff Miller. Questionable research practices may have little effect on replicability. *eLife*, 9, 2020.

[8220] Anthony D Umpierre, Lauren L Bystrom, Yanlu Ying, Yong U Liu, Gregory Worrell, and Long-Jun Wu. Microglial calcium signaling is attuned to neuronal activity in awake mice. *eLife*, 9, 2020.

[8221] Ambuj Upadhyay, Aidan J Peterson, Myung-Jun Kim, and Michael B O'Connor. Muscle-derived Myoglianin regulates Drosophila imaginal disc growth. *eLife*, 9, 2020.

[8222] Alessandro Urciuoli, Clément Zanolli, Amélie Beaudet, Jean Dumoncel, Frédéric Santos, Salvador Moyà-Solà, and David M Alba. The evolution of the vestibular apparatus in apes and humans. *eLife*, 9, 2020.

[8223] Samuel G Usher, Frances M Ashcroft, and Michael C Puljung. Nucleotide inhibition of the pancreatic ATP-sensitive K+ channel explored with patch-clamp fluorometry. *eLife*, 9, 2020.

[8224] Christopher E Vaaga, Spencer T Brown, and Indira M Raman. Cerebellar modulation of synaptic input to freezing-related neurons in the periaqueductal gray. *eLife*, 9, 2020.

[8225] Francesco Vacca, Caroline Chauché, Abhishek Jamwal, Elizabeth C Hinchy, Graham Heieis, Holly Webster, Adefunke Ogunkanbi, Zala Sekne, William F Gregory, Martin Wear, Georgia Perona-Wright, Matthew K Higgins, Josquin A Nys, E Suzanne Cohen, and Henry J McSorley. A helminth-derived suppressor of ST2 blocks allergic responses. *eLife*, 9, 2020.

[8226] Ryan J Vaden, Jose Carlos Gonzalez, Ming-Chi Tsai, Anastasia J Niver, Allison R Fusilier, Chelsea M Griffith, Richard H Kramer, Jacques I Wadiche, and Linda Overstreet-Wadiche. Parvalbumin interneurons provide spillover to newborn and mature dentate granule cells. *eLife*, 9, 2020.

[8227] Quentin Vagne, Jean-Patrick Vrel, and Pierre Sens. A minimal self-organisation model of the Golgi apparatus. *eLife*, 9, 2020.

[8228] Alicia N Vagnozzi, Kiran Garg, Carola Dewitz, Matthew T Moore, Jared M Cregg, Lucie Jeannotte, Niccolò Zampieri, Lynn T Landmesser, and Polyxeni Philippidou. Phrenic-specific transcriptional programs shape respiratory motor output. *eLife*, 9, 2020.

[8229] Horia Vais, Min Wang, Karthik Mallilankaraman, Riley Payne, Chris McKennan, Jeffrey T Lock, Lynn A Spruce, Carly Fiest, Matthew Yan-lok Chan, Ian Parker, Steven H Seeholzer, J Kevin Foskett, and Don-On Daniel Mak. ER-luminal [Ca2+] regulation of InsP3 receptor gating mediated by an ER-luminal peripheral Ca2+-binding protein. *eLife*, 9, 2020.

[8230] Nikolaos Vakirlis, Anne-Ruxandra Carvunis, and Aoife McLysaght. Synteny-based analyses indicate that sequence divergence is not the main source of orphan genes. *eLife*, 9, 2020.

[8231] Gabriele Valentini, Nobuaki Mizumoto, Stephen C Pratt, Theodore P Pavlic, and Sara I Walker. Revealing the structure of information flows discriminates similar animal social behaviors. *eLife*, 9, 2020.

[8232] Dan Valsky, Shai Heiman Grosberg, Zvi Israel, Thomas Boraud, Hagai Bergman, and Marc Deffains. What is the true discharge rate and pattern of the striatal projection neurons in Parkinson's disease and Dystonia? *eLife*, 9, 2020.

[8233] Sona Valuchova, Pavlina Mikulkova, Jana Pecinkova, Jana Klimova, Michal Krumnikl, Petr Bainar, Stefan Heckmann, Pavel Tomancak, and Karel Riha. Imaging plant germline differentiation within Arabidopsis flowers by light sheet microscopy. *eLife*, 9, 2020.

[8234] Frank L van de Veerdonk, Mihai G Netea, Marcel van Deuren, Jos WM van der Meer, Quirijn de Mast, Roger J Brüggemann, and Hans van der Hoeven. Kallikrein-kinin blockade in patients with COVID-19 to prevent acute respiratory distress syndrome. *eLife*, 9, 2020.

[8235] MGP van der Wijst, DH de Vries, HE Groot, G Trynka, CC Hon, MJ Bonder, O Stegle, MC Nawijn, Y Idaghdour, P van der Harst, CJ Ye, J Powell, FJ Theis, A Mahfouz, M Heinig, and L Franke. The single-cell eQTLGen consortium. *eLife*, 9, 2020.

[8236] Bram van Dijk, Paulien Hogeweg, Hilje M Doekes, and Nobuto Takeuchi. Slightly beneficial genes are retained by bacteria evolving DNA uptake despite selfish elements. *eLife*, 9, 2020.

[8237] Evelien Van Hoeymissen, Katharina Held, Ana Cristina Nogueira Freitas, Annelies Janssens, Thomas Voets, and Joris Vriens. Gain of channel function and modified gating properties in TRPM3 mutants causing intellectual disability and epilepsy. *eLife*, 9, 2020.

[8238] Dirk van Moorselaar, Eline Lampers, Elisa Cordesius, and Heleen A Slagter. Neural mechanisms underlying expectation-dependent inhibition of distracting information. *eLife*, 9, 2020.

[8239] Joury S van 't Klooster, Tan-Yun Cheng, Hendrik R Sikkema, Aike Jeucken, Branch Moody, and Bert Poolman. Periprotein lipidomes of Saccharomyces cerevisiae provide a flexible environment for conformational changes of membrane proteins. *eLife*, 9, 2020.

[8240] Jan Hendrik van Weerd, Rajiv A Mohan, Karel van Duijvenboden, Ingeborg B Hooijkaas, Vincent Wakker, Bastiaan J Boukens, Phil Barnett, and Vincent M Christoffels. Trait-associated noncoding variant regions affect TBX3 regulation and cardiac conduction. *eLife*, 9, 2020.

[8241] Joeri BG van Wijngaarden, Susanne S Babl, and Hiroshi T Ito. Entorhinal-retrosplenial circuits for allocentric-egocentric transformation of boundary coding. *eLife*, 9, 2020.

[8242] Hannes Vanhaeren, Ying Chen, Mattias Vermeersch, Liesbeth De Milde, Valerie De Vleeschhauwer, Annelore Natran, Geert Persiau, Dominique Eeckhout, Geert De Jaeger, Kris Gevaert, and Dirk Inzé. UBP12 and UBP13 negatively regulate the activity of the ubiquitin-dependent peptidases DA1, DAR1 and DAR2. *eLife*, 9, 2020.

[8243] Gilles Vannuscorps, Michael Andres, and Alfonso Caramazza. Efficient recognition of facial expressions does not require motor simulation. *eLife*, 9, 2020.

[8244] Calvin VanOpstall, Srikanth Perike, Hannah Brechka, Marc Gillard, Sophia Lamperis, Baizhen Zhu, Ryan Brown, Raj Bhanvadia, and Donald J Vander Griend. MEIS-mediated suppression of human prostate cancer growth and metastasis through HOXB13-dependent regulation of proteoglycans. *eLife*, 9, 2020.

[8245] Sriram Varahan, Vaibhhav Sinha, Adhish Walvekar, Sandeep Krishna, and Sunil Laxman. Resource plasticity-driven carbon-nitrogen budgeting enables specialization and division of labor in a clonal community. *eLife*, 9, 2020.

[8246] Carmen Varela and Matthew A Wilson. mPFC spindle cycles organize sparse thalamic activation and recently active CA1 cells during non-REM sleep. *eLife*, 9, 2020.

[8247] Yulia Vasianovich, Emmanuel Bajon, and Raymund J Wellinger. Telomerase biogenesis requires a novel Mex67 function and a cytoplasmic association with the Sm7 complex. *eLife*, 9, 2020.

[8248] Sara E Vazquez, Elise MN Ferré, David W Scheel, Sara Sunshine, Brenda Miao, Caleigh Mandel-Brehm, Zoe Quandt, Alice Y Chan, Mickie Cheng, Michael German, Michail Lionakis, Joseph L DeRisi, and Mark S Anderson. Identification of novel, clinically correlated autoantigens in the monogenic autoimmune syndrome APS1 by proteome-wide PhIP-Seq. *eLife*, 9, 2020.

[8249] Sapthaswaran Veerapathiran, Cathleen Teh, Shiwen Zhu, Indira Kartigayen, Vladimir Korzh, Paul T Matsudaira, and Thorsten Wohland. Wnt3 distribution in the zebrafish brain is determined by expression, diffusion and multiple molecular interactions. *eLife*, 9, 2020.

[8250] Griselda Velez-Aguilera, Sylvia Nkombo Nkoula, Batool Ossareh-Nazari, Jana Link, Dimitra Paouneskou, Lucie Van Hove, Nicolas Joly, Nicolas Tavernier, Jean-Marc Verbavatz, Verena Jantsch, and Lionel Pintard. PLK-1 promotes the merger of the parental genome into a single nucleus by triggering lamina disassembly. *eLife*, 9, 2020.

[8251] Valentina Vellani, Lianne P de Vries, Anne Gaule, and Tali Sharot. A selective effect of dopamine on information-seeking. *eLife*, 9, 2020.

[8252] AJ Venkatakrishnan, Arjun Puranik, Akash Anand, David Zemmour, Xiang Yao, Xiaoying Wu, Ramakrishna Chilaka, Dariusz K Murakowski, Kristopher Standish, Bharathwaj Raghunathan, Tyler Wagner, Enrique Garcia-Rivera, Hugo Solomon, Abhinav Garg, Rakesh Barve, Anuli Anyanwu-Ofili, Najat Khan, and Venky Soundararajan. Knowledge synthesis of 100 million biomedical documents augments the deep expression profiling of coronavirus receptors. *eLife*, 9, 2020.

[8253] Jelle Veraart, Daniel Nunes, Umesh Rudrapatna, Els Fieremans, Derek K Jones, Dmitry S Novikov, and Noam Shemesh. Noninvasive quantification of axon radii using diffusion MRI. *eLife*, 9, 2020.

[8254] Silvia Vergarajauregui, Robert Becker, Ulrike Steffen, Maria Sharkova, Tilman Esser, Jana Petzold, Florian Billing, Michael S Kapiloff, George Schett, Ingo Thievessen, and Felix B Engel. AKAP6 orchestrates the nuclear envelope microtubule-organizing center by linking golgi and nucleus via AKAP9. *eLife*, 9, 2020.

[8255] Jason D Vevea and Edwin R Chapman. Acute disruption of the synaptic vesicle membrane protein synaptotagmin 1 using knockoff in mouse hippocampal neurons. *eLife*, 9, 2020.

[8256] Sam Vickery, William D Hopkins, Chet C Sherwood, Steven J Schapiro, Robert D Latzman, Svenja Caspers, Christian Gaser, Simon B Eickhoff, Robert Dahnke, and Felix Hoffstaedter. Chimpanzee brain morphometry utilizing standardized MRI preprocessing and macroanatomical annotations. *eLife*, 9, 2020.

[8257] Valerie PI Vidal, Fariba Jian-Motamedi, Samah Rekima, Elodie P Gregoire, Emmanuelle Szenker-Ravi, Marc Leushacke, Bruno Reversade, Marie-Christine Chaboissier, and Andreas Schedl. R-spondin signalling is essential for the maintenance and differentiation of mouse nephron progenitors. *eLife*, 9, 2020.

[8258] Antoine Vigouroux, Baptiste Cordier, Andrey Aristov, Laura Alvarez, Gizem özbaykal, Thibault Chaze, Enno Rainer Oldewurtel, Mariette Matondo, Felipe Cava, David Bikard, and Sven van Teeffelen. Class-A penicillin binding proteins do not contribute to cell shape but repair cell-wall defects. *eLife*, 9, 2020.

[8259] Robin J Vigouroux, Quénol Cesar, Alain Chédotal, and Kim Tuyen Nguyen-Ba-Charvet. Revisiting the role of Dcc in visual system development with a novel eye clearing method. *eLife*, 9, 2020.

[8260] Aashutosh Vihani, Xiaoyang Serene Hu, Sivaji Gundala, Sachiko Koyama, Eric Block, and Hiroaki Matsunami. Semiochemical responsive olfactory sensory neurons are sexually dimorphic and plastic. *eLife*, 9, 2020.

[8261] Deivanayagabarathy Vinayagam, Dennis Quentin, Jing Yu-Strzelczyk, Oleg Sitsel, Felipe Merino, Markus Stabrin, Oliver Hofnagel, Maolin Yu, Mark W Ledeboer, Georg Nagel, Goran Malojcic, and Stefan Raunser. Structural basis of TRPC4 regulation by calmodulin and pharmacological agents. *eLife*, 9, 2020.

[8262] Todd A Vogel, Zachary M Savelson, A Ross Otto, and Mathieu Roy. Forced choices reveal a trade-off between cognitive effort and physical pain. *eLife*, 9, 2020.

[8263] Jakob Voigts, Christopher A Deister, and Christopher I Moore. Layer 6 ensembles can selectively regulate the behavioral impact and layer-specific representation of sensory deviants. *eLife*, 9, 2020.

[8264] Rashmi Voleti, Klaudia Jaczynska, and Josep Rizo. Ca2+-dependent release of synaptotagmin-1 from the SNARE complex on phosphatidylinositol 4,5-bisphosphate-containing membranes. *eLife*, 9, 2020.

[8265] Jonathan R Volpatti, Yukari Endo, Jessica Knox, Linda Groom, Stephanie Brennan, Ramil Noche, William J Zuercher, Peter Roy, Robert T Dirksen, and James J Dowling. Identification of drug modifiers for RYR1-related myopathy using a multi-species discovery pipeline. *eLife*, 9, 2020.

[8266] Corinne von Känel, Sergio A Muñoz-Gómez, Silke Oeljeklaus, Christoph Wenger, Bettina Warscheid, Jeremy G Wideman, Anke Harsman, and Andre Schneider. Homologue replacement in the import motor of the mitochondrial inner membrane of trypanosomes. *eLife*, 9, 2020.

[8267] Yuliya Voskobiynyk, Jonathan R Roth, J Nicholas Cochran, Travis Rush, Nancy VN Carullo, Jacob S Mesina, Mohammad Waqas, Rachael M Vollmer, Jeremy J Day, Lori L McMahon, and Erik D Roberson. Alzheimer's disease risk gene BIN1 induces Tau-dependent network hyperexcitability. *eLife*, 9, 2020.

[8268] Simon Vu, Vikrant Singh, Heike Wulff, Vladimir Yarov-Yarovoy, and Jie Zheng. New capsaicin analogs as molecular rulers to define the permissive conformation of the mouse TRPV1 ligand-binding pocket. *eLife*, 9, 2020.

[8269] Andra Waagmeester, Gregory Stupp, Sebastian Burgstaller-Muehlbacher, Benjamin M Good, Malachi Griffith, Obi L Griffith, Kristina Hanspers, Henning Hermjakob, Toby S Hudson, Kevin Hybiske, Sarah M Keating, Magnus Manske, Michael Mayers, Daniel Mietchen, Elvira Mitraka, Alexander R Pico, Timothy Putman, Anders Riutta, Nuria Queralt-Rosinach, Lynn M Schriml, Thomas Shafee, Denise Slenter, Ralf Stephan, Katherine Thornton, Ginger Tsueng, Roger Tu, Sabah Ul-Hasan, Egon Willighagen, Chunlei Wu, and Andrew I Su. Wikidata as a knowledge graph for the life sciences. *eLife*, 9, 2020.

[8270] Tyler Wagner, FNU Shweta, Karthik Murugadoss, Samir Awasthi, AJ Venkatakrishnan, Sairam Bade, Arjun Puranik, Martin Kang, Brian W Pickering, John C O'Horo, Philippe R Bauer, Raymund R Razonable, Paschalis Vergidis, Zelalem Temesgen, Stacey Rizza, Maryam Mahmood, Walter R Wilson, Douglas Challener, Praveen Anand, Matt Liebers, Zainab Doctor, Eli Silvert, Hugo Solomon, Akash Anand, Rakesh Barve, Gregory Gores, Amy W Williams, William G Morice, John Halamka, Andrew Badley, and Venky Soundararajan. Augmented curation of clinical notes from a massive EHR system reveals symptoms of impending COVID-19 diagnosis. *eLife*, 9, 2020.

[8271] Aboubacar Wague, Thomas T Joseph, Kellie A Woll, Weiming Bu, Kiran A Vaidya, Natarajan V Bhanu, Benjamin A Garcia, Crina M Nimigean, Roderic G Eckenhoff, and Paul M Riegelhaupt. Mechanistic insights into volatile anesthetic modulation of K2P channels. *eLife*, 9, 2020.

[8272] Ameya S Walimbe, Hidehiko Okuma, Soumya Joseph, Tiandi Yang, Takahiro Yonekawa, Jeffrey M Hord, David Venzke, Mary E Anderson, Silvia Torelli, Adnan Manzur, Megan Devereaux, Marco

Cuellar, Sally Prouty, Saul Ocampo Landa, Liping Yu, Junyu Xiao, Jack E Dixon, Francesco Muntoni, and Kevin P Campbell. POMK regulates dystroglycan function via LARGE1-mediated elongation of matriglycan. *eLife*, 9, 2020.

[8273] Denise S Walker and William R Schafer. Distinct roles for innexin gap junctions and hemichannels in mechanosensation. *eLife*, 9, 2020.

[8274] Emma C Walker, Kim Truong, Narelle E McGregor, Ingrid J Poulton, Tsuyoshi Isojima, Jonathan H Gooi, T John Martin, and Natalie A Sims. Cortical bone maturation in mice requires SOCS3 suppression of gp130/STAT3 signalling in osteocytes. *eLife*, 9, 2020.

[8275] Jenelle Wallace, Julia Lord, Lasse Dissing-Olesen, Beth Stevens, and Venkatesh N Murthy. Microglial depletion disrupts normal functional development of adult-born neurons in the olfactory bulb. *eLife*, 9, 2020.

[8276] Michael L Wallace, Kee Wui Huang, Daniel Hochbaum, Minsuk Hyun, Gianna Radeljic, and Bernardo L Sabatini. Anatomical and single-cell transcriptional profiling of the murine habenular complex. *eLife*, 9, 2020.

[8277] Jessica F Walls, Jeff J Subleski, Erika M Palmieri, Marieli Gonzalez-Cotto, Clair M Gardiner, Daniel W McVicar, and David K Finlay. Metabolic but not transcriptional regulation by PKM2 is important for natural killer cell responses. *eLife*, 9, 2020.

[8278] Morgan A Wambaugh, Steven T Denham, Magali Ayala, Brianna Brammer, Miekan A Stonhill, and Jessica CS Brown. Synergistic and antagonistic drug interactions in the treatment of systemic fungal infections. *eLife*, 9, 2020.

[8279] Sabrina Wamp, Zoe J Rutter, Jeanine Rismondo, Claire E Jennings, Lars Möller, Richard J Lewis, and Sven Halbedel. PrkA controls peptidoglycan biosynthesis through the essential phosphorylation of ReoM. *eLife*, 9, 2020.

[8280] Allen Wang, Joshua Chiou, Olivier B Poirion, Justin Buchanan, Michael J Valdez, Jamie M Verheyden, Xiaomeng Hou, Parul Kudtarkar, Sharvari Narendra, Jacklyn M Newsome, Minzhe Guo, Dina A Faddah, Kai Zhang, Randee E Young, Justinn Barr, Eniko Sajti, Ravi Misra, Heidie Huyck, Lisa Rogers, Cory Poole, Jeffery A Whitsett, Gloria Pryhuber, Yan Xu, Kyle J Gaulton, Sebastian Preissl, and Xin Sun. Single-cell multiomic profiling of human lungs reveals cell-type-specific and age-dynamic control of SARS-CoV2 host genes. *eLife*, 9, 2020.

[8281] Chenhui Wang and Allan C Spradling. An abundant quiescent stem cell population in Drosophila Malpighian tubules protects principal cells from kidney stones. *eLife*, 9, 2020.

[8282] Chongyuan Wang, Agata Jacewicz, Bryce D Delgado, Rozbeh Baradaran, and Stephen Barstow Long. Structures reveal gatekeeping of the mitochondrial Ca2+ uniporter by MICU1-MICU2. *eLife*, 9, 2020.

[8283] Dennis Wang, James Hensman, Ginte Kutkaite, Tzen S Toh, Ana Galhoz, Howard Lightfoot, Howard Lightfoot, Wanjuan Yang, Maryam Soleimani, Syd Barthorpe, Tatiana Mironenko, Alexandra Beck, Laura Richardson, Ermira Lleshi, James Hall, Charlotte Tolley, William Barendt, Jonathan R Dry, Julio Saez-Rodriguez, Mathew J Garnett, Michael P Menden, and Frank Dondelinger. A statistical framework for assessing pharmacological responses and biomarkers using uncertainty estimates. *eLife*, 9, 2020.

[8284] Hong Wang, Xueping Wan, Paul F Pilch, Leif W Ellisen, Susan K Fried, and Libin Liu. An AMPK-dependent, non-canonical p53 pathway plays a key role in adipocyte metabolic reprogramming. *eLife*, 9, 2020.

[8285] Hongyan Wang, Hanna S Radomska, Mitch A Phelps, Elizabeth Iorns, Elizabeth Iorns, Rachel Tsui, Alexandria Denis, Nicole Perfito, and Timothy M Errington. Replication Study: Coding-independent regulation of the tumor suppressor PTEN by competing endogenous mRNAs. *eLife*, 9, 2020.

[8286] Jing Wang, Eghbal Hosseini, Nicolas Meirhaeghe, Adam Akkad, and Mehrdad Jazayeri. Reinforcement regulates timing variability in thalamus. *eLife*, 9, 2020.

[8287] Lan Wang, Alexander Myasnikov, Xingjie Pan, and Peter Walter. Structure of the AAA protein Msp1 reveals mechanism of mislocalized membrane protein extraction. *eLife*, 9, 2020.

[8288] Li Wang, Wenya Guo, Xi Shen, Shel Yeo, Hui Long, Zhexuan Wang, Qifeng Lyu, Allan E Herbison, and Yanping Kuang. Different dendritic domains of the GnRH neuron underlie the pulse and surge modes of GnRH secretion in female mice. *eLife*, 9, 2020.

[8289] Ling Wang, Zachary Lee Johnson, Michael R Wasserman, Jesper Levring, Jue Chen, and Shixin Liu. Characterization of the kinetic cycle of an ABC transporter by single-molecule and cryo-EM analyses. *eLife*, 9, 2020.

[8290] Liping Wang, Yi Ding, Li He, Guiping Zhang, Jian-Kang Zhu, and Rosa Lozano-Duran. A virus-encoded protein suppresses methylation of the viral genome through its interaction with AGO4 in the Cajal body. *eLife*, 9, 2020.

[8291] Qin Wang, Huaxun Fan, Feng Li, Savanna S Skeeters, Vishnu V Krishnamurthy, Yuanquan Song, and Kai Zhang. Optical control of ERK and AKT signaling promotes axon regeneration and functional recovery of PNS and CNS in Drosophila. *eLife*, 9, 2020.

[8292] Shifan Wang, Huijuan Guo, Feng Ge, and Yucheng Sun. Apoptotic neurodegeneration in whitefly promotes the spread of TYLCV. *eLife*, 9, 2020.

[8293] Tian-Yi Wang, Jing Liu, and Haishan Yao. Control of adaptive action selection by secondary motor cortex during flexible visual categorization. *eLife*, 9, 2020.

[8294] Tianming Wang, Zheng Cao, Zhangfei Shen, Jingwen Yang, Xu Chen, Zhen Yang, Ke Xu, Xiaowei Xiang, Qiuhan Yu, Yimin Song, Weiwei Wang, Yanan Tian, Lina Sun, Libin Zhang, Su Guo, and Naiming Zhou. Existence and functions of a kisspeptin neuropeptide signaling system in a non-chordate deuterostome species. *eLife*, 9, 2020.

[8295] Tianyu Wang, Chunyan Wu, Dimitre G Ouzounov, Wenchao Gu, Fei Xia, Minsu Kim, Xusan Yang, Melissa R Warden, and Chris Xu. Quantitative analysis of 1300-nm three-photon calcium imaging in the mouse brain. *eLife*, 9, 2020.

[8296] Xiaobo Wang, Mary Ellenbecker, Benjamin Hickey, Nicholas J Day, Emily Osterli, Mikaya Terzo, and Ekaterina Voronina. Antagonistic control of Caenorhabditis elegans germline stem cell proliferation and differentiation by PUF proteins FBF-1 and FBF-2. *eLife*, 9, 2020.

[8297] Xiaoyu Wang and Olga Boudker. Large domain movements through the lipid bilayer mediate substrate release and inhibition of glutamate transporters. *eLife*, 9, 2020.

[8298] Xinru Wang, Dimitriya H Garvanska, Isha Nasa, Yumi Ueki, Gang Zhang, Arminja N Kettenbach, Wolfgang Peti, Jakob Nilsson, and Rebecca Page. A dynamic charge-charge interaction modulates PP2A:B56 substrate recruitment. *eLife*, 9, 2020.

[8299] Xinxin Wang, Zhiming Chen, Marcel Mettlen, Jungsik Noh, Sandra L Schmid, and Gaudenz Danuser. DASC, a sensitive classifier for measuring discrete early stages in clathrin-mediated endocytosis. *eLife*, 9, 2020.

[8300] Yan Wang, Yan Han, Ji She, Nam X Nguyen, Vamsi K Mootha, Xiao-chen Bai, and Youxing Jiang. Structural insights into the Ca2+-dependent gating of the human mitochondrial calcium uniporter. *eLife*, 9, 2020.

[8301] Ying Wang, Thomas S Chaffee, Rebecca S LaRue, Danielle N Huggins, Patrice M Witschen, Ayman M Ibrahim, Andrew C Nelson, Heather L Machado, and Kathryn L Schwertfeger. Tissue-resident macrophages promote extracellular matrix homeostasis in the mammary gland stroma of nulliparous mice. *eLife*, 9, 2020.

[8302] Yuan Wang, Xiaoqian Zhang, Qi Xin, Wesley Hung, Jeremy Florman, Jing Huo, Tianqi Xu, Yu Xie, Mark J Alkema, Mei Zhen, and Quan Wen. Flexible motor sequence generation during stereotyped escape responses. *eLife*, 9, 2020.

[8303] Zhipeng A Wang, Christopher J Millard, Chia-Liang Lin, Jennifer E Gurnett, Mingxuan Wu, Kwang-woon Lee, Louise Fairall, John WR Schwabe, and Philip A Cole. Diverse nucleosome Site-Selectivity among histone deacetylase complexes. *eLife*, 9, 2020.

[8304] Zijun Wang, Christopher O Barnes, Rajeev Gautam, Julio C Cetrulo Lorenzi, Christian T Mayer, Thiago Y Oliveira, Victor Ramos, Melissa Cipolla, Kristie M Gordon, Harry B Gristick, Anthony P West, Yoshiaki Nishimura, Henna Raina, Michael S Seaman, Anna Gazumyan, Malcolm Martin, Pamela J Bjorkman, Michel C Nussenzweig, and Amelia Escolano. A broadly neutralizing macaque monoclonal antibody against the HIV-1 V3-Glycan patch. *eLife*, 9, 2020.

[8305] Jamie R Wangen and Rachel Green. Stop codon context influences genome-wide stimulation of termination codon readthrough by aminoglycosides. *eLife*, 9, 2020.

[8306] Iromi Wanigasuriya, Quentin Gouil, Sarah A Kinkel, Andrés Tapia del Fierro, Tamara Beck, Ellise A Roper, Kelsey Breslin, Jessica Stringer, Karla Hutt, Heather J Lee, Andrew Keniry, Matthew E Ritchie, and Marnie E Blewitt. Smchd1 is a maternal effect gene required for genomic imprinting. *eLife*, 9, 2020.

[8307] Joanna A Warren, Shuntai Zhou, Yinyan Xu, Matthew J Moeser, Daniel R MacMillan, Olivia Council, Jennifer Kirchherr, Julia M Sung, Nadia R Roan, Adaora A Adimora, Sarah Joseph, JoAnn D Kuruc, Cynthia L Gay, David M Margolis, Nancie Archin, Zabrina L Brumme, Ronald Swanstrom, and Nilu Goonetilleke. The HIV-1 latent reservoir is largely sensitive to circulating T cells. *eLife*, 9, 2020.

[8308] Jack Waters. Sources of widefield fluorescence from the brain. *eLife*, 9, 2020.

[8309] James A Watson, Joel Tarning, Richard M Hoglund, Frederic J Baud, Bruno Megarbane, Jean-Luc Clemessy, and Nicholas J White. Concentration-dependent mortality of chloroquine in overdose. *eLife*, 9, 2020.

[8310] Zoe L Watson, Fred R Ward, Raphaël Méheust, Omer Ad, Alanna Schepartz, Jillian F Banfield, and Jamie HD Cate. Structure of the bacterial ribosome at 2 Å resolution. *eLife*, 9, 2020.

[8311] William J Watterson, Melikhan Tanyeri, Andrea R Watson, Candace M Cham, Yue Shan, Eugene B Chang, A Murat Eren, and Savaş Tay. Droplet-based high-throughput cultivation for accurate screening of antibiotic resistant gut microbes. *eLife*, 9, 2020.

[8312] Lotte P Watts, Toyoaki Natsume, Yuichiro Saito, Javier Garzon, Qianqian Dong, Lora Boteva, Nick Gilbert, Masato T Kanemaki, Shin-ichiro Hiraga, and Anne D Donaldson. The RIF1-long splice variant promotes G1 phase 53BP1 nuclear bodies to protect against replication stress. *eLife*, 9, 2020.

[8313] Rachel Waymack, Alvaro Fletcher, German Enciso, and Zeba Wunderlich. Shadow enhancers can suppress input transcription factor noise through distinct regulatory logic. *eLife*, 9, 2020.

[8314] Meltem Weger, Benjamin D Weger, Andrea Schink, Masanari Takamiya, Johannes Stegmaier, Cédric Gobet, Alice Parisi, Andrei Yu Kobitski, Jonas Mertes, Nils Krone, Uwe Strähle, Gerd Ulrich Nienhaus, Ralf Mikut, Frédéric Gachon, Philipp Gut, and Thomas Dickmeis. MondoA regulates gene expression in cholesterol biosynthesis-associated pathways required for zebrafish epiboly. *eLife*, 9, 2020.

[8315] Jessica Wei, Mary J Mattapallil, Reiko Horai, Yingyos Jittayasothorn, Arnav P Modi, H Nida Sen, Karsten Gronert, and Rachel R Caspi. A novel role for lipoxin A4 in driving a lymph node–eye axis that controls autoimmunity to the neuroretina. *eLife*, 9, 2020.

[8316] Chi G Weindel, Samantha L Bell, Krystal J Vail, Kelsi O West, Kristin L Patrick, and Robert O Watson. LRRK2 maintains mitochondrial homeostasis and regulates innate immune responses to Mycobacterium tuberculosis. *eLife*, 9, 2020.

[8317] Keiko Weir, Christophe Dupre, Lena van Giesen, Amy S-Y Lee, and Nicholas W Bellono. A molecular filter for the cnidarian stinging response. *eLife*, 9, 2020.

[8318] Tracey Weissgerber, Yaw Bediako, Charlotte M de Winde, Hedyeh Ebrahimi, Florencia Fernández-Chiappe, Vinodh Ilangovan, Devang Mehta, Carolina Paz Quezada, Julia L Riley, Shyam M Saladi, Sarvenaz Sarabipour, and Andy Tay. Mitigating the impact of conference and travel cancellations on researchers' futures. *eLife*, 9, 2020.

[8319] Nathan Weisz, Nadine Gabriele Kraft, and Gianpaolo Demarchi. Auditory cortical alpha/beta desynchronization prioritizes the representation of memory items during a retention period. *eLife*, 9, 2020.

[8320] Daniel Wells, Emmanuelle Bitoun, Daniela Moralli, Gang Zhang, Anjali Hinch, Julia Jankowska, Peter Donnelly, Catherine Green, and Simon R Myers. ZCWPW1 is recruited to recombination hotspots by PRDM9 and is essential for meiotic double strand break repair. *eLife*, 9, 2020.

[8321] Kristen L Wells, Corey N Miller, Andreas R Gschwind, Wu Wei, Jonah D Phipps, Mark S Anderson, and Lars M Steinmetz. Combined transient ablation and single-cell RNA-sequencing reveals the development of medullary thymic epithelial cells. *eLife*, 9, 2020.

[8322] Kenneth Wengler, Andrew T Goldberg, George Chahine, and Guillermo Horga. Distinct hierarchical alterations of intrinsic neural timescales account for different manifestations of psychosis. *eLife*, 9, 2020.

[8323] Jaime N Wertman, Nicole Melong, Matthew R Stoyek, Olivia Piccolo, Stewart Langley, Benno Orr, Shelby L Steele, Babak Razaghi, and Jason N Berman. The identification of dual protective agents against cisplatin-induced oto- and nephrotoxicity using the zebrafish model. *eLife*, 9, 2020.

[8324] Natalia Wesolowska, Ivan Avilov, Pedro Machado, Celina Geiss, Hiroshi Kondo, Masashi Mori, and Peter Lenart. Actin assembly ruptures the nuclear envelope by prying the lamina away from nuclear pores and nuclear membranes in starfish oocytes. *eLife*, 9, 2020.

[8325] Agata Wesolowska-Andersen, Grace Zhuo Yu, Vibe Nylander, Fernando Abaitua, Matthias Thurner, Jason M Torres, Anubha Mahajan, Anna L Gloyn, and Mark I McCarthy. Deep learning models predict regulatory variants in pancreatic islets and refine type 2 diabetes association signals. *eLife*, 9, 2020.

[8326] David S White, Marcel P Goldschen-Ohm, Randall H Goldsmith, and Baron Chanda. Top-down machine learning approach for high-throughput single-molecule analysis. *eLife*, 9, 2020.

[8327] Jordon D White, Tanzil M Arefin, Alexa Pugliese, Choong H Lee, Jeff Gassen, Jiangyang Zhang, and Arie Kaffman. Early life stress causes sex-specific changes in adult fronto-limbic connectivity that differentially drive learning. *eLife*, 9, 2020.

[8328] Kelly L Whiteford, Heather A Kreft, and Andrew J Oxenham. The role of cochlear place coding in the perception of frequency modulation. *eLife*, 9, 2020.

[8329] Wesley A Wierson, Jordan M Welker, Maira P Almeida, Carla M Mann, Dennis A Webster, Melanie E Torrie, Trevor J Weiss, Sekhar Kambakam, Macy K Vollbrecht, Merrina Lan, Kenna C McKeighan, Jacklyn Levey, Zhitao Ming, Alec Wehmeier, Christopher S Mikelson, Jeffrey A Haltom, Kristen M Kwan, Chi-Bin Chien, Darius Balciunas, Stephen C Ekker, Karl J Clark, Beau R Webber, Branden S Moriarity, Stacy L Solin, Daniel F Carlson, Drena L Dobbs, Maura McGrail, and Jeffrey Essner. Efficient targeted integration directed by short homology in zebrafish and mammalian cells. *eLife*, 9, 2020.

[8330] Wojciech Wietrzynski, Miroslava Schaffer, Dimitry Tegunov, Sahradha Albert, Atsuko Kanazawa, Jürgen M Plitzko, Wolfgang Baumeister, and Benjamin D Engel. Charting the native architecture of Chlamydomonas thylakoid membranes with single-molecule precision. *eLife*, 9, 2020.

[8331] David Willemsen, Rongfeng Cui, Martin Reichard, and Dario Riccardo Valenzano. Intra-species differences in population size shape life history and genome evolution. *eLife*, 9, 2020.

[8332] Allison Hillary Williams, Richard Wheeler, Ala-Eddine Deghmane, Ignacio Santecchia, Ryan E Schaub, Samia Hicham, Maryse Moya Nilges, Christian Malosse, Julia Chamot-Rooke, Ahmed Haouz, Joseph P Dillard, William P Robins, Muhamed-Kheir Taha, and Ivo Gomperts Boneca. Defective lytic transglycosylase disrupts cell morphogenesis by hindering cell wall de-O-acetylation in Neisseria meningitidis. *eLife*, 9, 2020.

[8333] Marc J Williams, Luis Zapata, Benjamin Werner, Chris P Barnes, Andrea Sottoriva, and Trevor A Graham. Measuring the distribution of fitness effects in somatic evolution by combining clonal dynamics with dN/dS ratios. *eLife*, 9, 2020.

[8334] Margot LK Williams and Lilianna Solnica-Krezel. Nodal and planar cell polarity signaling cooperate to regulate zebrafish convergence and extension gastrulation movements. *eLife*, 9, 2020.

[8335] Gordon Williamson, Giulia Tamburrino, Adriana Bizior, Mélanie Boeckstaens, Gaëtan Dias Mirandela, Marcus G Bage, Andrei Pisliakov, Callum M Ives, Eilidh Terras, Paul A Hoskisson, Anna Maria Marini, Ulrich Zachariae, and Arnaud Javelle. A two-lane mechanism for selective biological ammonium transport. *eLife*, 9, 2020.

[8336] Sandra Winters, William L Allen, and James P Higham. The structure of species discrimination signals across a primate radiation. *eLife*, 9, 2020.

[8337] Hannah S Wirtshafter and Matthew A Wilson. Differences in reward biased spatial representations in the lateral septum and hippocampus. *eLife*, 9, 2020.

[8338] Margot Wohl, Kenichi Ishii, and Kenta Asahina. Layered roles of fruitless isoforms in specification and function of male aggression-promoting neurons in Drosophila. *eLife*, 9, 2020.

[8339] Nicole C Woitowich, Annaliese Beery, and Teresa Woodruff. A 10-year follow-up study of sex inclusion in the biological sciences. *eLife*, 9, 2020.

[8340] Gernot Wolf, Alberto de Iaco, Ming-An Sun, Melania Bruno, Matthew Tinkham, Don Hoang, Apratim Mitra, Sherry Ralls, Didier Trono, and Todd S Macfarlan. KRAB-zinc finger protein gene expansion in response to active retrotransposons in the murine lineage. *eLife*, 9, 2020.

[8341] Adrian Wolny, Lorenzo Cerrone, Athul Vijayan, Rachele Tofanelli, Amaya Vilches Barro, Marion Louveaux, Christian Wenzl, Sören Strauss, David Wilson-Sánchez, Rena Lymbouridou, Susanne S Steigleder, Constantin Pape, Alberto Bailoni, Salva Duran-Nebreda, George W Bassel, Jan U Lohmann, Miltos Tsiantis, Fred A Hamprecht, Kay Schneitz, Alexis Maizel, and Anna Kreshuk. Accurate and versatile 3D segmentation of plant tissues at cellular resolution. *eLife*, 9, 2020.

[8342] Helen Wong, Josien Levenga, Lauren LaPlante, Bailey Keller, Andrew Cooper-Sansone, Curtis Borski, Ryan Milstead, Marissa Ehringer, and Charles Hoeffer. Isoform-specific roles for AKT in affective behavior, spatial memory, and extinction related to psychiatric disorders. *eLife*, 9, 2020.

[8343] Matthew WK Wong, Anbupalam Thalamuthu, Nady Braidy, Karen A Mather, Yue Liu, Liliana Ciobanu, Bernhardt T Baune, Nicola J Armstrong, John Kwok, Peter Schofield, Margaret J Wright, David Ames, Russell Pickford, Teresa Lee, Anne Poljak, and Perminder S Sachdev. Genetic and environmental determinants of variation in the plasma lipidome of older Australian twins. *eLife*, 9, 2020.

[8344] Evan J Worden, Xiangbin Zhang, and Cynthia Wolberger. Structural basis for COMPASS recognition of an H2B-ubiquitinated nucleosome. *eLife*, 9, 2020.

[8345] Amy A Worth, Rosemary Shoop, Katie Tye, Claire H Feetham, Giuseppe D'Agostino, Garron T Dodd, Frank Reimann, Fiona M Gribble, Emily C Beebe, James D Dunbar, Jesline T Alexander-Chacko, Dana K Sindelar, Tamer Coskun, Paul J Emmerson, and Simon M Luckman. The cytokine GDF15 signals through a population of brainstem cholecystokinin neurons to mediate anorectic signalling. *eLife*, 9, 2020.

[8346] Fengming Wu, Bowen Deng, Na Xiao, Tao Wang, Yining Li, Rencong Wang, Kai Shi, Dong-Gen Luo, Yi Rao, and Chuan Zhou. A neuropeptide regulates fighting behavior in Drosophila melanogaster. *eLife*, 9, 2020.

[8347] Jingjing Sherry Wu, Eunyoung Yi, Marco Manca, Hamad Javaid, Amanda M Lauer, and Elisabeth Glowatzki. Sound exposure dynamically induces dopamine synthesis in cholinergic LOC efferents for feedback to auditory nerve fibers. *eLife*, 9, 2020.

[8348] Yanying Wu, Yosuke Funato, Eleonora Meschi, Kristijan D Jovanoski, Hiroaki Miki, and Scott Waddell. Magnesium efflux from Drosophila Kenyon cells is critical for normal and diet-enhanced long-term memory. *eLife*, 9, 2020.

[8349] Zhi-Jing Wu, Jia-Cheng Liu, Xin Man, Xin Gu, Ting-Yi Li, Chen Cai, Ming-Hong He, Yangyang Shao, Ning Lu, Xiaoli Xue, Zhongjun Qin, and Jin-Qiu Zhou. Cdc13 is predominant over Stn1 and Ten1 in preventing chromosome end fusions. *eLife*, 9, 2020.

[8350] Linghe Xi, Thomas Carroll, Irina Matos, Ji-Dung Luo, Lisa Polak, H Amalia Pasolli, Samie R Jaffrey, and Elaine Fuchs. m6A RNA methylation impacts fate choices during skin morphogenesis. *eLife*, 9, 2020.

[8351] Kai Xie, Naixiang Wang, Xudong Lin, Zixun Wang, Xi Zhao, Peilin Fang, Haibing Yue, Junhwi Kim, Jing Luo, Shaoyang Cui, Feng Yan, and Peng Shi. Organic electrochemical transistor arrays for real-time mapping of evoked neurotransmitter release in vivo. *eLife*, 9, 2020.

[8352] Meng Xie, Pavel Gol'din, Anna Nele Herdina, Jordi Estefa, Ekaterina V Medvedeva, Lei Li, Phillip T Newton, Svetlana Kotova, Boris Shavkuta, Aditya Saxena, Lauren T Shumate, Brian D Metscher, Karl Großschmidt, Shigeki Nishimori, Anastasia Akovantseva, Anna P Usanova, Anastasiia D Kurenkova, Anoop Kumar, Irene Linares Arregui, Paul Tafforeau, Kaj Fried, Mattias Carlström, András Simon, Christian Gasser, Henry M Kronenberg, Murat Bastepe, Kimberly L Cooper, Peter Timashev, Sophie Sanchez, Igor Adameyko, Anders Eriksson, and Andrei S Chagin. Secondary ossification center induces and protects growth plate structure. *eLife*, 9, 2020.

[8353] Wenting Xie, Binxuan Jiao, Qing Bai, Vladimir A Ilin, Ming Sun, Charles E Burton, Dmytro Kolodieznyi, Michael J Calderon, Donna B Stolz, Patricia L Opresko, Claudette M St Croix, Simon Watkins, Bennett Van Houten, Marcel P Bruchez, and Edward A Burton. Chemoptogenetic ablation of neuronal mitochondria in vivo with spatiotemporal precision and controllable severity. *eLife*, 9, 2020.

[8354] Wenmin Xing, Denise Muhlrad, Roy Parker, and Michael K Rosen. A quantitative inventory of yeast P body proteins reveals principles of composition and specificity. *eLife*, 9, 2020.

[8355] Liyang Xiong, Yuansheng Cao, Robert Cooper, Wouter-Jan Rappel, Jeff Hasty, and Lev Tsimring. Flower-like patterns in multi-species bacterial colonies. *eLife*, 9, 2020.

[8356] Yan Xiong, M Julia Scerbo, Anett Seelig, Francesco Volta, Nils O'Brien, Andrea Dicker, Daniela Padula, Heiko Lickert, Jantje Mareike Gerdes, and Per-Olof Berggren. Islet vascularization is regulated by primary endothelial cilia via VEGF-A-dependent signaling. *eLife*, 9, 2020.

[8357] An Qi Xu, Rita R Barbosa, and Dinis Pedro Calado. Genetic timestamping of plasma cells in vivo reveals tissue-specific homeostatic population turnover. *eLife*, 9, 2020.

[8358] Jiajia Xu, Yiyun Wang, Ching-Yun Hsu, Stefano Negri, Robert J Tower, Yongxing Gao, Ye Tian, Takashi Sono, Carolyn A Meyers, Winters R Hardy, Leslie Chang, Shuaishuai Hu, Nusrat Kahn, Kristen Broderick, Bruno Péault, and Aaron W James. Lysosomal protein surface expression discriminates fat- from bone-forming human mesenchymal precursor cells. *eLife*, 9, 2020.

[8359] Kui Xu, Yanrong Zhou, Yulian Mu, Zhiguo Liu, Shaohua Hou, Yujian Xiong, Liurong Fang, Changli Ge, Yinghui Wei, Xiuling Zhang, Changjiang Xu, Jingjing Che, Ziyao Fan, Guangming Xiang, Jiankang Guo, Haitao Shang, Hua Li, Shaobo Xiao, Julang Li, and Kui Li. CD163 and pAPN double-knockout pigs are resistant to PRRSV and TGEV and exhibit decreased susceptibility to PDCoV while maintaining normal production performance. *eLife*, 9, 2020.

[8360] Lingna Xu, Xi Wang, Jia Zhou, Yunyi Qiu, Weina Shang, Jun-Ping Liu, Liquan Wang, and Chao Tong. Miga-mediated endoplasmic reticulum–mitochondria contact sites regulate neuronal homeostasis. *eLife*, 9, 2020.

[8361] Nannan Xu, Anne Oltmanns, Longsheng Zhao, Antoine Girot, Marzieh Karimi, Lara Hoepfner, Simon Kelterborn, Martin Scholz, Julia Beißel, Peter Hegemann, Oliver Bäumchen, Lu-Ning Liu, Kaiyao Huang, and Michael Hippler. Altered N-glycan composition impacts flagella-mediated adhesion in Chlamydomonas reinhardtii. *eLife*, 9, 2020.

[8362] Pengqi Xu, Volha U Chukhutsina, Wojciech J Nawrocki, Gert Schansker, Ludwik W Bielczynski, Yinghong Lu, Daniel Karcher, Ralph Bock, and Roberta Croce. Photosynthesis without $\beta$-carotene. *eLife*, 9, 2020.

[8363] Xiao-Ping Xu, Sabine Pokutta, Megan Torres, Mark F Swift, Dorit Hanein, Niels Volkmann, and William I Weis. Structural basis of $\alpha$E-catenin–F-actin catch bond behavior. *eLife*, 9, 2020.

[8364] Xiaomei Xu, Véronique Risoul, Deborah Byrne, Stéphanie Champ, Badreddine Douzi, and Amel Latifi. HetL, HetR and PatS form a reaction-diffusion system to control pattern formation in the cyanobacterium nostoc PCC 7120. *eLife*, 9, 2020.

[8365] Xin Xu, Huigang Shi, Xiaowen Gong, Pu Chen, Ying Gao, Xinzheng Zhang, and Song Xiang. Structural insights into sodium transport by the oxaloacetate decarboxylase sodium pump. *eLife*, 9, 2020.

[8366] Jing Xue, Tian Xie, Weizhong Zeng, Youxing Jiang, and Xiao-chen Bai. Cryo-EM structures of human ZnT8 in both outward- and inward-facing conformations. *eLife*, 9, 2020.

[8367] Yuan Xue, Terence C Theisen, Suchita Rastogi, Abel Ferrel, Stephen R Quake, and John C Boothroyd. A single-parasite transcriptional atlas of Toxoplasma Gondii reveals novel control of antigen expression. *eLife*, 9, 2020.

[8368] Junpei Yamashita, Akio Takeuchi, Kohei Hosono, Thomas Fleming, Yoshitaka Nagahama, and Kataaki Okubo. Male-predominant galanin mediates androgen-dependent aggressive chases in medaka. *eLife*, 9, 2020.

[8369] Chao Yang, Jason R Siebert, Robert Burns, Yongwei Zheng, Ao Mei, Benedetta Bonacci, Demin Wang, Raul A Urrutia, Matthew J Riese, Sridhar Rao, Karen-Sue Carlson, Monica S Thakar, and Subramaniam Malarkannan. Single-cell transcriptome reveals the novel role of T-bet in suppressing the immature NK gene signature. *eLife*, 9, 2020.

[8370] Hee Won Yang, Steven D Cappell, Ariel Jaimovich, Chad Liu, Mingyu Chung, Leighton H Daigh, Lindsey R Pack, Yilin Fan, Sergi Regot, Markus Covert, and Tobias Meyer. Stress-mediated exit to quiescence restricted by increasing persistence in CDK4/6 activation. *eLife*, 9, 2020.

[8371] Hua-Qian Yang, Marta Pérez-Hernández, Jose Sanchez-Alonso, Andriy Shevchuk, Julia Gorelik, Eli Rothenberg, Mario Delmar, and William A Coetzee. Ankyrin-G mediates targeting of both Na+ and KATP channels to the rat cardiac intercalated disc. *eLife*, 9, 2020.

[8372] Ke Yang, Xin-Lin Gong, Guo-Cheng Li, Ling-Qiao Huang, Chao Ning, and Chen-Zhu Wang. A gustatory receptor tuned to the steroid plant hormone brassinolide in Plutella xylostella (Lepidoptera: Plutellidae). *eLife*, 9, 2020.

[8373] Lei Yang, Michael Emerman, Harmit S Malik, and Richard N McLaughlin. Retrocopying expands the functional repertoire of APOBEC3 antiviral proteins in primates. *eLife*, 9, 2020.

[8374] Linlin Yang, Jessica A Jiménez, Alison M Earley, Victoria Hamlin, Victoria Kwon, Cameron T Dixon, and Celia E Shiau. Drainage of inflammatory macromolecules from the brain to periphery targets the liver for macrophage infiltration. *eLife*, 9, 2020.

[8375] Qi Yang, Thomas A Hughes, Anju Kelkar, Xinheng Yu, Kai Cheng, Sheldon Park, Wei-Chiao Huang, Jonathan F Lovell, and Sriram Neelamegham. Inhibition of SARS-CoV-2 viral entry upon blocking N- and O-glycan elaboration. *eLife*, 9, 2020.

[8376] Shuang Yang, Kriti Bahl, Hui-Ting Chou, Jonathan Woodsmith, Ulrich Stelzl, Thomas Walz, and Maxence V Nachury. Near-atomic structures of the BBSome reveal the basis for BBSome activation and binding to GPCR cargoes. *eLife*, 9, 2020.

[8377] Xiao Yang, Lian Zhao, Maria M Campos, Mones Abu-Asab, Davide Ortolan, Nathan Hotaling, Kapil Bharti, and Wai T Wong. CSF1R blockade induces macrophage ablation and results in mouse choroidal vascular atrophy and RPE disorganization. *eLife*, 9, 2020.

[8378] Yoonsun Yang, Joonyeol Lee, and Gunsoo Kim. Integration of locomotion and auditory signals in the mouse inferior colliculus. *eLife*, 9, 2020.

[8379] Zhe Yang, Hongcun Zhu, Kakit Kong, Xiaoxuan Wu, Jiayi Chen, Peiyao Li, Jialong Jiang, Jinchao Zhao, Bofei Cui, and Feng Liu. The dynamic transmission of positional information in stau- mutants during Drosophila embryogenesis. *eLife*, 9, 2020.

[8380] David Yanni, Shane Jacobeen, Pedro Márquez-Zacarías, Joshua S Weitz, William C Ratcliff, and Peter J Yunker. Topological constraints in early multicellularity favor reproductive division of labor. *eLife*, 9, 2020.

[8381] Jun Yao, Douglas C Wu, Ryan M Nottingham, and Alan M Lambowitz. Identification of protein-protected mRNA fragments and structured excised intron RNAs in human plasma by TGIRT-seq peak calling. *eLife*, 9, 2020.

[8382] Kenrick Yap, Alexander Drakew, Dinko Smilovic, Michael Rietsche, Mandy H Paul, Mario Vuksic, Domenico Del Turco, and Thomas Deller. The actin-modulating protein synaptopodin mediates long-term survival of dendritic spines. *eLife*, 9, 2020.

[8383] Mahdieh Yazdani, Guohui Zhang, Zhiguang Jia, Jingyi Shi, Jianmin Cui, and Jianhan Chen. Aromatic interactions with membrane modulate human BK channel activation. *eLife*, 9, 2020.

[8384] Luis Alfonso Yañez-Guerra, Xingxing Zhong, Ismail Moghul, Thomas Butts, Cleidiane G Zampronio, Alexandra M Jones, Olivier Mirabeau, and Maurice R Elphick. Echinoderms provide missing link in the evolution of PrRP/sNPF-type neuropeptide signalling. *eLife*, 9, 2020.

[8385] Fuzhou Ye, Ioly Kotta-Loizou, Milija Jovanovic, Xiaojiao Liu, David TF Dryden, Martin Buck, and Xiaodong Zhang. Structural basis of transcription inhibition by the DNA mimic protein Ocr of bacteriophage T7. *eLife*, 9, 2020.

[8386] Hui Ye, Shamsideen A Ojelade, David Li-Kroeger, Zhongyuan Zuo, Liping Wang, Yarong Li, Jessica YJ Gu, Ulrich Tepass, Avital Adah Rodal, Hugo J Bellen, and Joshua M Shulman. Retromer subunit, VPS29, regulates synaptic transmission and is required for endolysosomal function in the aging brain. *eLife*, 9, 2020.

[8387] Xinhai Ye, Shijiao Xiong, Ziwen Teng, Yi Yang, Jiale Wang, Kaili Yu, Huizi Wu, Yang Mei, Zhichao Yan, Sammy Cheng, Chuanlin Yin, Fang Wang, Hongwei Yao, Qi Fang, Qisheng Song, John H Werren, Gongyin Ye, and Fei Li. RETRACTED: Amino acid synthesis loss in parasitoid wasps and other hymenopterans. *eLife*, 9, 2020.

[8388] Zhian Ye, Zhongwu Su, Siyu Xie, Yuye Liu, Yongqiang Wang, Xi Xu, Yiqing Zheng, Meng Zhao, and Linjia Jiang. Yap-lin28a axis targets let7-Wnt pathway to restore progenitors for initiating regeneration. *eLife*, 9, 2020.

[8389] Zhifang Ye, Liang Shi, Anqi Li, Chuansheng Chen, and Gui Xue. Retrieval practice facilitates memory updating by enhancing and differentiating medial prefrontal cortex representations. *eLife*, 9, 2020.

[8390] Chia-An Yen, Dana L Ruter, Christian D Turner, Shanshan Pang, and Sean P Curran. Loss of flavin adenine dinucleotide (FAD) impairs sperm function and male reproductive advantage in C. elegans. *eLife*, 9, 2020.

[8391] Syn Kok Yeo, Xiaoting Zhu, Takako Okamoto, Mingang Hao, Cailian Wang, Peixin Lu, Long Jason Lu, and Jun-Lin Guan. Single-cell RNA-sequencing reveals distinct patterns of cell state heterogeneity in mouse models of breast cancer. *eLife*, 9, 2020.

[8392] Priscilla S-W Yeung, Christopher E Ing, Megumi Yamashita, Régis Pomès, and Murali Prakriya. A sulfur-aromatic gate latch is essential for opening of the Orai1 channel pore. *eLife*, 9, 2020.

[8393] Danielle Yi, Hai P Nguyen, Jennie Dinh, Jose A Viscarra, Ying Xie, Frances Lin, Madeleine Zhu, Jon M Dempersmier, Yuhui Wang, and Hei Sook Sul. Dot1l interacts with Zc3h10 to activate Ucp1 and other thermogenic genes. *eLife*, 9, 2020.

[8394] Rui Yin, Julia Siebert, Nico Eisenhauer, and Martin Schädler. Climate change and intensive land use reduce soil animal biomass via dissimilar pathways. *eLife*, 9, 2020.

[8395] Nate Yoder and Eric Gouaux. The His-Gly motif of acid-sensing ion channels resides in a reentrant 'loop' implicated in gating and ion selectivity. *eLife*, 9, 2020.

[8396] Saishu Yoshida, Katsuhiko Aoki, Ken Fujiwara, Takashi Nakakura, Akira Kawamura, Kohji Yamada, Masaya Ono, Satomi Yogosawa, and Kiyotsugu Yoshida. The novel ciliogenesis regulator DYRK2 governs Hedgehog signaling during mouse embryogenesis. *eLife*, 9, 2020.

[8397] Yuto Yoshinari, Tomotsune Ameku, Shu Kondo, Hiromu Tanimoto, Takayuki Kuraishi, Yuko Shimada-Niwa, and Ryusuke Niwa. Neuronal octopamine signaling regulates mating-induced germline stem cell increase in female Drosophila melanogaster. *eLife*, 9, 2020.

[8398] Chih-Chieh (Jay) Yu, Nicholas C Barry, Asmamaw T Wassie, Anubhav Sinha, Abhishek Bhattacharya, Shoh Asano, Chi Zhang, Fei Chen, Oliver Hobert, Miriam B Goodman, Gal Haspel, and Edward S Boyden. Expansion microscopy of C. elegans. *eLife*, 9, 2020.

[8399] Xiao-Wen Yu, Kiran Pandey, Aaron C Katzman, and Cristina M Alberini. A role for CIM6P/IGF2 receptor in memory consolidation and enhancement. *eLife*, 9, 2020.

[8400] Nora Yucel, Jessie Axsom, Yifan Yang, Li Li, Joshua H Rhoades, and Zoltan Arany. Cardiac endothelial cells maintain open chromatin and expression of cardiomyocyte myofibrillar genes. *eLife*, 9, 2020.

[8401] Tobias Zachs, Andreas Schertel, João Medeiros, Gregor L Weiss, Jannik Hugener, Joao Matos, and Martin Pilhofer. Fully automated, sequential focused ion beam milling for cryo-electron tomography. *eLife*, 9, 2020.

[8402] Susanne RK Zajitschek, Felix Zajitschek, Russell Bonduriansky, Robert C Brooks, Will Cornwell, Daniel S Falster, Malgorzata Lagisz, Jeremy Mason, Alistair M Senior, Daniel WA Noble, and Shinichi Nakagawa. Sexual dimorphism in trait variability and its eco-evolutionary and statistical implications. *eLife*, 9, 2020.

[8403] Masihuz Zaman and Maria Andreasen. Cross-talk between individual phenol-soluble modulins in Staphylococcus aureus biofilm enables rapid and efficient amyloid formation. *eLife*, 9, 2020.

[8404] Elisa Zamboni, Valentin G Kemper, Nuno Reis Goncalves, Ke Jia, Vasilis M Karlaftis, Samuel J Bell, Joseph Giorgio, Reuben Rideaux, Rainer Goebel, and Zoe Kourtzi. Fine-scale computations for adaptive processing in the human brain. *eLife*, 9, 2020.

[8405] Danae E Zamboulis, Chavaunne T Thorpe, Yalda Ashraf Kharaz, Helen L Birch, Hazel RC Screen, and Peter D Clegg. Postnatal mechanical loading drives adaptation of tissues primarily through modulation of the non-collagenous matrix. *eLife*, 9, 2020.

[8406] Natalia Zamorano Cuervo and Nathalie Grandvaux. ACE2: Evidence of role as entry receptor for SARS-CoV-2 and implications in comorbidities. *eLife*, 9, 2020.

[8407] Yunliang Zang, Sungho Hong, and Erik De Schutter. Firing rate-dependent phase responses of Purkinje cells support transient oscillations. *eLife*, 9, 2020.

[8408] Balyn W Zaro, Joseph J Noh, Victoria L Mascetti, Janos Demeter, Benson George, Monika Zukowska, Gunsagar S Gulati, Rahul Sinha, Ryan A Flynn, Allison Banuelos, Allison Zhang, Adam C Wilkinson, Peter Jackson, and Irving L Weissman. Proteomic analysis of young and old mouse hematopoietic stem cells and their progenitors reveals post-transcriptional regulation in stem cells. *eLife*, 9, 2020.

[8409] Michael Zavortink, Lauren N Rutt, Svetlana Dzitoyeva, Jesslyn C Henriksen, Chloe Barrington, Danielle Y Bilodeau, Miranda Wang, Xiao Xiao Lily Chen, and Olivia S Rissland. The E2 Marie Kondo and the CTLH E3 ligase clear deposited RNA binding proteins during the maternal-to-zygotic transition. *eLife*, 9, 2020.

[8410] Lihong Zhan, Li Fan, Lay Kodama, Peter Dongmin Sohn, Man Ying Wong, Gergey Alzaem Mousa, Yungui Zhou, Yaqiao Li, and Li Gan. A MAC2-positive progenitor-like microglial population is resistant to CSF1R inhibition in adult mouse brain. *eLife*, 9, 2020.

[8411] Ce Zhang, Yue Chen, Shijin Sun, Yikai Zhang, Lina Wang, Zhipu Luo, Meng Liu, Liang Dong, Ningzheng Dong, and Qingyu Wu. A conserved LDL-receptor motif regulates corin and CD320 membrane targeting in polarized renal epithelial cells. *eLife*, 9, 2020.

[8412] Chang Zhang, Chanjuan Hao, Guanghou Shui, and Wei Li. BLOS1 mediates kinesin switch during endosomal recycling of LDL receptor. *eLife*, 9, 2020.

[8413] Danyang Zhang, Yan Zhang, Jun Ma, Chunmei Zhu, Tongxin Niu, Wenbo Chen, Xiaoyun Pang, Yujia Zhai, and Fei Sun. Cryo-EM structures of S-OPA1 reveal its interactions with membrane and changes upon nucleotide binding. *eLife*, 9, 2020.

[8414] Fengwen Zhang and Paul D Bieniasz. HIV-1 Vpr induces cell cycle arrest and enhances viral gene expression by depleting CCDC137. *eLife*, 9, 2020.

[8415] Hao Zhang, Yang Zhang, Xinyue Zhou, Shaela Wright, Judith Hyle, Lianzhong Zhao, Jie An, Xujie Zhao, Ying Shao, Beisi Xu, Hyeong-Min Lee, Taosheng Chen, Yang Zhou, Xiang Chen, Rui Lu, and Chunliang Li. Functional interrogation of HOXA9 regulome in MLLr leukemia via reporter-based CRISPR/Cas9 screen. *eLife*, 9, 2020.

[8416] Huiliang Zhang, Nathan N Alder, Wang Wang, Hazel Szeto, David J Marcinek, and Peter S Rabinovitch. Reduction of elevated proton leak rejuvenates mitochondria in the aged cardiomyocyte. *eLife*, 9, 2020.

[8417] Jie Zhang, Lihong Song, Dennis V Pedersen, Anna Li, John D Lambris, Gregers Rom Andersen, Tom Eirik Mollnes, Ying Jie Ma, and Peter Garred. Soluble collectin-12 mediates C3-independent docking of properdin that activates the alternative pathway of complement. *eLife*, 9, 2020.

[8418] Kuan Zhang, Erica Yao, Chuwen Lin, Yu-Ting Chou, Julia Wong, Jianying Li, Paul J Wolters, and Pao-Tien Chuang. A mammalian Wnt5a–Ror2–Vangl2 axis controls the cytoskeleton and confers cellular properties required for alveologenesis. *eLife*, 9, 2020.

[8419] Li Zhang, Yohei Takahashi, Po-Kai Hsu, Hannes Kollist, Ebe Merilo, Patrick J Krysan, and Julian I Schroeder. FRET kinase sensor development reveals SnRK2/OST1 activation by ABA but not by MeJA and high CO2 during stomatal closure. *eLife*, 9, 2020.

[8420] Roger Shen Zhang, Kif Liakath-Ali, and Thomas C Südhof. Latrophilin-2 and latrophilin-3 are redundantly essential for parallel-fiber synapse function in cerebellum. *eLife*, 9, 2020.

[8421] Shi-Chao Zhang, Ming-Yu Wang, Jun-Rui Feng, Yue Chang, Shang-Rong Ji, and Yi Wu. Reversible promoter methylation determines fluctuating expression of acute phase proteins. *eLife*, 9, 2020.

[8422] Weina Zhang, Yu Chen, Jiaqing Yang, Jing Zhang, Jiayu Yu, Mengting Wang, Xiaodong Zhao, Ke Wei, Xiaoping Wan, Xiaojun Xu, Ying Jiang, Jiayu Chen, Shaorong Gao, and Zhiyong Mao. A high-throughput small molecule screen identifies farrerol as a potentiator of CRISPR/Cas9-mediated genome editing. *eLife*, 9, 2020.

[8423] Xiang Zhang, Jianguo Liu, Li Wu, and Xiaoyu Hu. MicroRNAs of the miR-17~92 family maintain adipose tissue macrophage homeostasis by sustaining il-10 expression. *eLife*, 9, 2020.

[8424] Xiaoguo Zhang, Xinxin Ding, Richard Scott Marshall, Julio Paez-Valencia, Patrick Lacey, Richard David Vierstra, and Marisa S Otegui. Reticulon proteins modulate autophagy of the endoplasmic reticulum in maize endosperm. *eLife*, 9, 2020.

[8425] Xin-Yan Zhang, Yan-Nong Dou, Lei Yuan, Qing Li, Yan-Jing Zhu, Meng Wang, and Yan-Gang Sun. Different neuronal populations mediate inflammatory pain analgesia by exogenous and endogenous opioids. *eLife*, 9, 2020.

[8426] Yanzhao Zhang, Takuya Tada, Seiya Ozono, Satoshi Kishigami, Hideaki Fujita, and Kenzo Tokunaga. MARCH8 inhibits viral infection by two different mechanisms. *eLife*, 9, 2020.

[8427] Ying Zhang, Yi Zhu, Shu-Xia Cao, Peng Sun, Jian-Ming Yang, Yan-Fang Xia, Shi-Ze Xie, Xiao-Dan Yu, Jia-Yu Fu, Chen-Jie Shen, Hai-Yang He, Hao-Qi Pan, Xiao-Juan Chen, Hao Wang, and Xiao-Ming Li. MeCP2 in cholinergic interneurons of nucleus accumbens regulates fear learning. *eLife*, 9, 2020.

[8428] Zhen-Jia Zhang, Qiang Gao, Xiao-Dong Fang, Zhi-Hang Ding, Dong-Min Gao, Wen-Ya Xu, Qing Cao, Ji-Hui Qiao, Yi-Zhou Yang, Chenggui Han, Ying Wang, Xuefeng Yuan, Dawei Li, and Xian-Bing Wang. CCR4, a RNA decay factor, is hijacked by a plant cytorhabdovirus phosphoprotein to facilitate virus replication. *eLife*, 9, 2020.

[8429] Junjun Zhao, Albert Hiu Ka Fok, Ruolin Fan, Pui-Yi Kwan, Hei-Lok Chan, Louisa Hoi-Ying Lo, Ying-Shing Chan, Wing-Ho Yung, Jiandong Huang, Cora Sau Wan Lai, and Kwok-On Lai. Specific depletion of the motor protein KIF5B leads to deficits in dendritic transport, synaptic plasticity and memory. *eLife*, 9, 2020.

[8430] Siyuan Zhao, Yevgen Yudin, and Tibor Rohacs. Disease-associated mutations in the human TRPM3 render the channel overactive via two distinct mechanisms. *eLife*, 9, 2020.

[8431] Xin Zhao, Peng Shao, Kexin Gai, Fengyin Li, Qiang Shan, and Hai-Hui Xue. $\beta$-catenin and $\gamma$-catenin are dispensable for T lymphocytes and AML leukemic stem cells. *eLife*, 9, 2020.

[8432] Zuodong Zhao, Zhuqiang Zhang, Jingjing Li, Qiang Dong, Jun Xiong, Yingfeng Li, Mengying Lan, Gang Li, and Bing Zhu. Sustained TNF-$\alpha$ stimulation leads to transcriptional memory that greatly enhances signal sensitivity and robustness. *eLife*, 9, 2020.

[8433] Leilei Zhong, Lutian Yao, Robert J Tower, Yulong Wei, Zhen Miao, Jihwan Park, Rojesh Shrestha, Luqiang Wang, Wei Yu, Nicholas Holdreith, Xiaobin Huang, Yejia Zhang, Wei Tong, Yanqing Gong, Jaimo Ahn, Katalin Susztak, Nathanial Dyment, Mingyao Li, Fanxin Long, Chider Chen, Patrick Seale, and Ling Qin. Single cell transcriptomics identifies a unique adipose lineage cell population that regulates bone marrow environment. *eLife*, 9, 2020.

[8434] Feng Zhou, Jialin Li, Weihua Zhao, Lei Xu, Xiaoxiao Zheng, Meina Fu, Shuxia Yao, Keith M Kendrick, Tor D Wager, and Benjamin Becker. Empathic pain evoked by sensory and emotional-communicative cues share common and process-specific neural representations. *eLife*, 9, 2020.

[8435] Joy Zhou, Amanda M Brown, Elizabeth P Lackey, Marife Arancillo, Tao Lin, and Roy V Sillitoe. Purkinje cell neurotransmission patterns cerebellar basket cells into zonal modules defined by distinct pinceau sizes. *eLife*, 9, 2020.

[8436] Liang Zhou, Chong-Yu Shao, Ya-Jun Xie, Na Wang, Si-Min Xu, Ben-Yan Luo, Zhi-Ying Wu, Yue Hai Ke, Mengsheng Qiu, and Ying Shen. Gab1 mediates PDGF signaling and is essential to oligodendrocyte differentiation and CNS myelination. *eLife*, 9, 2020.

[8437] Taifeng Zhou, Bo Gao, Yi Fan, Yuchen Liu, Shuhao Feng, Qian Cong, Xiaolei Zhang, Yaxing Zhou, Prem S Yadav, Jiachen Lin, Nan Wu, Liang Zhao, Dongsheng Huang, Shuanhu Zhou, Peiqiang Su, and Yingzi Yang. Piezo1/2 mediate mechanotransduction essential for bone formation through concerted activation of NFAT-YAP1-ß-catenin. *eLife*, 9, 2020.

[8438] Yuqi Zhou, Atsushi Yasumoto, Cheng Lei, Chun-Jung Huang, Hirofumi Kobayashi, Yunzhao Wu, Sheng Yan, Chia-Wei Sun, Yutaka Yatomi, and Keisuke Goda. Intelligent classification of platelet aggregates by agonist type. *eLife*, 9, 2020.

[8439] Jianxi Zhu, Gehua Zhen, Senbo An, Xiao Wang, Mei Wan, Yusheng Li, Zhiyong Chen, Yun Guan, Xinzhong Dong, Yihe Hu, and Xu Cao. Aberrant subchondral osteoblastic metabolism modifies NaV1.8 for osteoarthritis. *eLife*, 9, 2020.

[8440] Songli Zhu, Mohammadjavad Paydar, Feifei Wang, Yanqiu Li, Ling Wang, Benoit Barrette, Tadayoshi Bessho, Benjamin H Kwok, and Aimin Peng. Kinesin Kif2C in regulation of DNA double strand break dynamics and repair. *eLife*, 9, 2020.

[8441] Aleksandra Zielińska, Abigail Savietto, Anabela de Sousa Borges, Denis Martinez, Melanie Berbon, Joël R Roelofsen, Alwin M Hartman, Rinse de Boer, Ida J Van der Klei, Anna KH Hirsch, Birgit Habenstein, Marc Bramkamp, and Dirk-Jan Scheffers. Flotillin-mediated membrane fluidity controls peptidoglycan synthesis and MreB movement. *eLife*, 9, 2020.

[8442] Dario Zimmerli, Costanza Borrelli, Amaia Jauregi-Miguel, Simon Söderholm, Salome Brütsch, Nikolaos Doumpas, Jan Reichmuth, Fabienne Murphy-Seiler, MIchel Aguet, Konrad Basler, Andreas E Moor, and Claudio Cantù. TBX3 acts as tissue-specific component of the Wnt/$\beta$-catenin transcriptional complex. *eLife*, 9, 2020.

[8443] Eckart Zimmermann. Saccade suppression depends on context. *eLife*, 9, 2020.

[8444] Amanda M Zimmet, Di Cao, Amy J Bastian, and Noah J Cowan. Cerebellar patients have intact feedback control that can be leveraged to improve reaching. *eLife*, 9, 2020.

[8445] Lisa Zipper, Denise Jassmann, Sofie Burgmer, Bastian Görlich, and Tobias Reiff. Ecdysone steroid hormone remote controls intestinal stem cell fate decisions via the PPAR$\gamma$-homolog Eip75B in Drosophila. *eLife*, 9, 2020.

[8446] Javier Zorrilla de San Martin, Cristina Donato, Jérémy Peixoto, Andrea Aguirre, Vikash Choudhary, Angela Michela De Stasi, Joana LourenÇo, Marie-Claude Potier, and Alberto Bacci. Alterations of specific cortical GABAergic circuits underlie abnormal network activity in a mouse model of Down syndrome. *eLife*, 9, 2020.

[8447] Shuzhen Zuo, Lei Wang, Jung Han Shin, Yudian Cai, Boqiang Zhang, Sang Wan Lee, Kofi Appiah, Yong-di Zhou, and Sze Chai Kwok. Behavioral evidence for memory replay of video episodes in the macaque. *eLife*, 9, 2020.

[8448] Nicole Aaron, Michael J Kraakman, Qiuzhong Zhou, Qiongming Liu, Samantha Costa, Jing Yang, Longhua Liu, Lexiang Yu, Liheng Wang, Ying He, Lihong Fan, Hiroyuki Hirakawa, Lei Ding, James Lo, Weidong Wang, Baohong Zhao, Edward Guo, Lei Sun, Cliff J Rosen, and Li Qiang. Adipsin promotes bone marrow adiposity by priming mesenchymal stem cells. *eLife*, 10, 2021.

[8449] Sheryl Ann Abdukahil, Matthew D Hall, Joaquín Baruch, Gail Carson, Barbara Wanjiru Citarella, Andrew Dagens, Emmanuelle A Dankwa, Christl A Donnelly, Jake Dunning, Martina Escher, Christiana Kartsonaki, Laura Merson, Mark Pritchard, Jia Wei, Peter W Horby, Amanda Rojek, and Piero L Olliaro. Ten months of temporal variation in the clinical journey of hospitalised patients with COVID-19: An observational cohort. *eLife*, 10, 2021.

[8450] Sanjeewa Abeytunge, Francesco Gianoli, AJ Hudspeth, and Andrei S Kozlov. Rapid mechanical stimulation of inner-ear hair cells by photonic pressure. *eLife*, 10, 2021.

[8451] Emanuela Abiusi, Paola Infante, Cinzia Cagnoli, Ludovica Lospinoso Severini, Marika Pane, Giorgia Coratti, Maria Carmela Pera, Adele D'Amico, Federica Diano, Agnese Novelli, Serena Spartano, Stefania Fiori, Giovanni Baranello, Isabella Moroni, Marina Mora, Maria Barbara Pasanisi, Krizia Pocino, Loredana Le Pera, Davide D'Amico, Lorena Travaglini, Francesco Ria, Claudio Bruno, Denise Locatelli, Enrico Silvio Bertini, Lucia Ovidia Morandi, Eugenio Mercuri, Lucia Di Marcotullio, and Francesco Danilo Tiziano. SMA-miRs (miR-181a-5p, -324-5p, and -451a) are overexpressed in spinal muscular atrophy skeletal muscle and serum samples. *eLife*, 10, 2021.

[8452] Katie Abley, Pau Formosa-Jordan, Hugo Tavares, Emily YT Chan, Mana Afsharinafar, Ottoline Leyser, and James CW Locke. An ABA-GA bistable switch can account for natural variation in the variability of Arabidopsis seed germination time. *eLife*, 10, 2021.

[8453] Maria Abou Chakra, Ruth Isserlin, Thinh N Tran, and Gary D Bader. Control of tissue development and cell diversity by cell cycle-dependent transcriptional filtering. *eLife*, 10, 2021.

[8454] Joshua Abrams and Jeremy Nance. A polarity pathway for exocyst-dependent intracellular tube extension. *eLife*, 10, 2021.

[8455] Michael J Abrams, Fayth Hui Tan, Yutian Li, Ty Basinger, Martin L Heithe, Anish Sarma, Iris T Lee, Zevin J Condiotte, Misha Raffiee, John O Dabiri, David A Gold, and Lea Goentoro. A conserved strategy for inducing appendage regeneration in moon jellyfish, Drosophila, and mice. *eLife*, 10, 2021.

[8456] Shaun R Abrams and Jeremy F Reiter. Ciliary Hedgehog signaling regulates cell survival to build the facial midline. *eLife*, 10, 2021.

[8457] Mieradilijiang Abudupataer, Shichao Zhu, Shiqiang Yan, Kehua Xu, Jingjing Zhang, Shaman Luo, Wenrui Ma, Md Fazle Alam, Yuyi Tang, Hui Huang, Nan Chen, Li Wang, Guoquan Yan, Jun Li, Hao Lai, Chunsheng Wang, Kai Zhu, and Weijia Zhang. Aorta smooth muscle-on-a-chip reveals impaired mitochondrial dynamics as a therapeutic target for aortic aneurysm in bicuspid aortic valve disease. *eLife*, 10, 2021.

[8458] Alice Accorsi, Andrew C Box, Robert Peuß, Christopher Wood, Alejandro Sánchez Alvarado, and Nicolas Rohner. Image3C, a multimodal image-based and label-independent integrative method for single-cell analysis. *eLife*, 10, 2021.

[8459] Nathan P Achilly, Ling-jie He, Olivia A Kim, Shogo Ohmae, Gregory J Wojaczynski, Tao Lin, Roy V Sillitoe, Javier F Medina, and Huda Y Zoghbi. Deleting Mecp2 from the cerebellum rather than its neuronal subtypes causes a delay in motor learning in mice. *eLife*, 10, 2021.

[8460] Quadri Adewale, Ahmed F Khan, Felix Carbonell, and Yasser Iturria-Medina. Integrated transcriptomic and neuroimaging brain model decodes biological mechanisms in aging and Alzheimer's disease. *eLife*, 10, 2021.

[8461] Alison H Affinati, Paul V Sabatini, Cadence True, Abigail J Tomlinson, Melissa Kirigiti, Sarah R Lindsley, Chien Li, David P Olson, Paul Kievit, Martin G Myers, and Alan C Rupp. Cross-species analysis defines the conservation of anatomically segregated VMH neuron populations. *eLife*, 10, 2021.

[8462] Negar Afshar, Bilge Argunhan, Maierdan Palihati, Goki Taniguchi, Hideo Tsubouchi, and Hiroshi Iwasaki. A novel motif of Rad51 serves as an interaction hub for recombination auxiliary factors. *eLife*, 10, 2021.

[8463] Ipsita Agarwal and Molly Przeworski. Mutation saturation for fitness effects at human CpG sites. *eLife*, 10, 2021.

[8464] Parya Aghasafari, Pei-Chi Yang, Divya C Kernik, Kazuho Sakamoto, Yasunari Kanda, Junko Kurokawa, Igor Vorobyov, and Colleen E Clancy. A deep learning algorithm to translate and classify cardiac electrophysiology. *eLife*, 10, 2021.

[8465] Jenny I Aguilar, Mary Hongying Cheng, Josep Font, Alexandra C Schwartz, Kaitlyn Ledwitch, Amanda Duran, Samuel J Mabry, Andrea N Belovich, Yanqi Zhu, Angela M Carter, Lei Shi, Manju A Kurian, Cristina Fenollar-Ferrer, Jens Meiler, Renae Monique Ryan, Hassane S Mchaourab, Ivet Bahar, Heinrich JG Matthies, and Aurelio Galli. Psychomotor impairments and therapeutic implications revealed by a mutation associated with infantile Parkinsonism-Dystonia. *eLife*, 10, 2021.

[8466] Valeria Aguillon-Rodriguez, Dora Angelaki, Hannah Bayer, Niccolo Bonacchi, Matteo Carandini, Fanny Cazettes, Gaelle Chapuis, Anne K Churchland, Yang Dan, Eric Dewitt, Mayo Faulkner, Hamish Forrest, Laura Haetzel, Michael Häusser, Sonja B Hofer, Fei Hu, Anup Khanal, Christopher Krasniak, Ines Laranjeira, Zachary F Mainen, Guido Meijer, Nathaniel J Miska, Thomas D Mrsic-Flogel, Masayoshi Murakami, Jean-Paul Noel, Alejandro Pan-Vazquez, Cyrille Rossant, Joshua Sanders, Karolina Socha, Rebecca Terry, Anne E Urai, Hernando Vergara, Miles Wells, Christian J Wilson, Ilana B Witten, Lauren E Wool, and Anthony M Zador. Standardized and reproducible measurement of decision-making in mice. *eLife*, 10, 2021.

[8467] Regina L Agulto, Melissa M Rogers, Tracy C Tan, Amrita Ramkumar, Ashlyn M Downing, Hannah Bodin, Julia Castro, Dan W Nowakowski, and Kassandra M Ori-McKenney. Autoregulatory control of microtubule binding in doublecortin-like kinase 1. *eLife*, 10, 2021.

[8468] Jahed Ahmed, Jahed Ahmed, Oluwafemi Alaba, Gazala Ameen, Vaishali Arora, Mario A Arteaga-Vazquez, Alok Arun, Julia Bailey-Serres, Laura E Bartley, George W Bassel, Dominique C Bergmann, Edoardo Bertolini, Kaushal Kumar Bhati, Noel Blanco-Touriñán, Steven P Briggs, Javier Brumos, Benjamin Buer, Adrien Burlaocot, Sergio Alan Cervantes-Pérez, Sixue Chen, Bruno Contreras-Moreira, Francisco J CORPAS, Alfredo Cruz-Ramirez, Cesar L Cuevas-Velazquez, Josh T Cuperus, Lisa I David, Stefan de Folter, Peter H Denolf, Pingtao Ding, William P Dwyer, Matthew MS Evans, Nancy George, Pubudu P Handakumbura, Maria J Harrison, Elizabeth S Haswell, Venura Herath, Yuling Jiao, Robert E Jinkerson, Uwe John, Sanjay Joshi, Abhishek Joshi, Lydia-Marie Joubert, Ramesh Katam, Harmanpreet Kaur, Yana Kazachkova, Sunil K Kenchanmane Raju, Mather A Khan, Rajdeep Khangura, Ajay Kumar, Arun Kumar, Pankaj Kumar, Pradeep Kumar, Dhruv Lavania, Tedrick Thomas Salim Lew, Mathew G Lewsey, Chien-Yuan Lin, Dianyi Liu, Le Liu, Tie Liu, Ansul Lokdarshi, Ai My Luong, Iain C Macaulay, Sakil Mahmud, Ari Pekka Mähönen, Kamal Kumar Malukani, Alexandre P Marand, Carly A Martin, Claire D McWhite, Devang Mehta, Miguel Miñambres Martín, Jenny C Mortimer, Lachezar A Nikolov, Tatsuya Nobori, Trevor M Nolan, Aaron J Ogden, Marisa S Otegui, Mark-Christoph Ott, José M Palma, Puneet Paul, Atique U Rehman, Maida Romera-Branchat, Luis C Romero, Ronelle Roth, Saroj K Sah, Rachel Shahan, Shyam Solanki, Bao-Hua Song, Rosangela Sozzani, Gary Stacey, Anna N Stepanova, Nicolas L Taylor, Marcela K Tello-Ruiz, Tuan M Tran, Rajiv Kumar Tripathi, Batthula Vijaya Lakshmi Vadde

Vadde, Tamas Varga, Marija Vidovic, Justin W Walley, Zhiyong Wang, Renate A Weizbauer, James Whelan, Asela J Wijeratne, Tingting Xiang, Shouling Xu, Ramin Yadegari, Houlin Yu, Hai Ying Yuan, Fabio Zanini, Feng Zhao, Jie Zhu, Xiaohong Zhuang, Suryatapa Ghosh Jha, Alexander T Borowsky, Benjamin J Cole, Noah Fahlgren, Andrew Farmer, Shao-shan Carol Huang, Purva Karia, Marc Libault, Nicholas J Provart, Selena L Rice, Maite Saura-Sanchez, Pinky Agarwal, Amir H Ahkami, Christopher R Anderton, Steven P Briggs, Jennifer AN Brophy, Peter Denolf, Luigi F Di Costanzo, Moises Exposito-Alonso, Stefania Giacomello, Fabio Gomez-Cano, Kerstin Kaufmann, Dae Kwan Ko, Sagar Kumar, Andrey V Malkovskiy, Naomi Nakayama, Toshihiro Obata, Marisa S Otegui, Gergo Palfalvi, Elsa H Quezada-Rodríguez, Rajveer Singh, R Glen Uhrig, Jamie Waese, Klaas Van Wijk, R Clay Wright, David W Ehrhardt, Kenneth D Birnbaum, and Seung Y Rhee. Vision, challenges and opportunities for a Plant Cell Atlas. *eLife*, 10, 2021.

[8469] Elizabeth H Aitken, Timon Damelang, Amaya Ortega-Pajares, Agersew Alemu, Wina Hasang, Saber Dini, Holger W Unger, Maria Ome-Kaius, Morten A Nielsen, Ali Salanti, Joe Smith, Stephen Kent, P Mark Hogarth, Bruce D Wines, Julie A Simpson, Amy W Chung, and Stephen J Rogerson. Developing a multivariate prediction model of antibody features associated with protection of malaria-infected pregnant women from placental malaria. *eLife*, 10, 2021.

[8470] Kotaro Akaki, Kosuke Ogata, Yuhei Yamauchi, Noriki Iwai, Ka Man Tse, Fabian Hia, Atsushi Mochizuki, Yasushi Ishihama, Takashi Mino, and Osamu Takeuchi. IRAK1-dependent Regnase-1-14-3-3 complex formation controls Regnase-1-mediated mRNA decay. *eLife*, 10, 2021.

[8471] Katarina Akhmetova, Maxim Balasov, and Igor Chesnokov. Drosophila STING protein has a role in lipid metabolism. *eLife*, 10, 2021.

[8472] Haya Akkad, Joshua Dupont-Hadwen, Edward Kane, Carys Evans, Liam Barrett, Amba Frese, Irena Tetkovic, Sven Bestmann, and Charlotte J Stagg. Increasing human motor skill acquisition by driving theta–gamma coupling. *eLife*, 10, 2021.

[8473] Ayla Aksoy-Aksel, Andrea Gall, Anna Seewald, Francesco Ferraguti, and Ingrid Ehrlich. Midbrain dopaminergic inputs gate amygdala intercalated cell clusters by distinct and cooperative mechanisms in male mice. *eLife*, 10, 2021.

[8474] Mroj Alassaf and Mary C Halloran. Pregnancy-associated plasma protein-aa regulates endoplasmic reticulum–mitochondria associations. *eLife*, 10, 2021.

[8475] Vasyl Alba, James E Carthew, Richard W Carthew, and Madhav Mani. Global constraints within the developmental program of the Drosophila wing. *eLife*, 10, 2021.

[8476] Javier Alegre-Cortés, María Sáez, Roberto Montanari, and Ramon Reig. Medium spiny neurons activity reveals the discrete segregation of mouse dorsal striatum. *eLife*, 10, 2021.

[8477] Brian D Alford, Eduardo Tassoni-Tsuchida, Danish Khan, Jeremy J Work, Gregory Valiant, and Onn Brandman. ReporterSeq reveals genome-wide dynamic modulators of the heat shock response across diverse stressors. *eLife*, 10, 2021.

[8478] Ahmad F Alghanem, Javier Abello, Joshua M Maurer, Ashutosh Kumar, Chau My Ta, Susheel K Gunasekar, Urooj Fatima, Chen Kang, Litao Xie, Oluwaseun Adeola, Megan Riker, Macaulay Elliot-Hudson, Rachel A Minerath, Chad E Grueter, Robert F Mullins, Amber N Stratman, and Rajan Sah. The SWELL1-LRRC8 complex regulates endothelial AKT-eNOS signaling and vascular function. *eLife*, 10, 2021.

[8479] Fatima Alghoul, Schaeffer Laure, Gilbert Eriani, and Franck Martin. Translation inhibitory elements from Hoxa3 and Hoxa11 mRNAs use uORFs for translation inhibition. *eLife*, 10, 2021.

[8480] Laith Alhussein and Maurice A Smith. Motor planning under uncertainty. *eLife*, 10, 2021.

[8481] Rémi Allio, Marie-Ka Tilak, Celine Scornavacca, Nico L Avenant, Andrew C Kitchener, Erwan Corre, Benoit Nabholz, and Frédéric Delsuc. High-quality carnivoran genomes from roadkill samples enable comparative species delineation in aardwolf and bat-eared fox. *eLife*, 10, 2021.

[8482] Rawan AlSubaie, Ryan WS Wee, Anne Ritoux, Karyna Mishchanchuk, Jessica Passlack, Daniel Regester, and Andrew F MacAskill. Control of parallel hippocampal output pathways by amygdalar long-range inhibition. *eLife*, 10, 2021.

[8483] Sergio álvarez Parra, Ricardo Pérez-de la Fuente, Enrique Peñalver, Eduardo Barrón, Luis Alcalá, Jordi Pérez-Cano, Carles Martín-Closas, Khaled Trabelsi, Nieves Meléndez, Rafael López Del Valle, Rafael P Lozano, David Peris, Ana Rodrigo, Víctor Sarto i Monteys, Carlos A Bueno-Cebollada, César Menor-Salván, Marc Philippe, Alba Sánchez-García, Constanza Peña-Kairath, Antonio Arillo, Eduardo Espílez, Luis Mampel, and Xavier Delclòs. Dinosaur bonebed amber from an original swamp forest soil. *eLife*, 10, 2021.

[8484] Linda M Amarante and Mark Laubach. Coherent theta activity in the medial and orbital frontal cortices encodes reward value. *eLife*, 10, 2021.

[8485] Thomas H Ambrosi, Rahul Sinha, Holly M Steininger, Malachia Y Hoover, Matthew P Murphy, Lauren S Koepke, Yuting Wang, Wan-Jin Lu, Maurizio Morri, Norma F Neff, Irving L Weissman, Michael T Longaker, and Charles KF Chan. Distinct skeletal stem cell types orchestrate long bone skeletogenesis. *eLife*, 10, 2021.

[8486] Benjamin Amoah, Robert S McCann, Alinune N Kabaghe, Monicah Mburu, Michael G Chipeta, Paula Moraga, Steven Gowelo, Tinashe Tizifa, Henk van den Berg, Themba Mzilahowa, Willem Takken, Michele van Vugt, Kamija S Phiri, Peter J Diggle, Dianne J Terlouw, and Emanuele Giorgi. Identifying Plasmodium falciparum transmission patterns through parasite prevalence and entomological inoculation rate. *eLife*, 10, 2021.

[8487] Piero Amodio, Benjamin G Farrar, Christopher Krupenye, Ljerka Ostojić, and Nicola S Clayton. Little evidence that Eurasian jays protect their caches by responding to cues about a conspecific's desire and visual perspective. *eLife*, 10, 2021.

[8488] Anna Anchimiuk, Virginia S Lioy, Florian Patrick Bock, Anita Minnen, Frederic Boccard, and Stephan Gruber. A low Smc flux avoids collisions and facilitates chromosome organization in Bacillus subtilis. *eLife*, 10, 2021.

[8489] Eric N Anderson, Andrés A Morera, Sukhleen Kour, Jonathan D Cherry, Nandini Ramesh, Amanda Gleixner, Jacob C Schwartz, Christopher Ebmeier, William Old, Christopher J Donnelly, Jeffrey P Cheng, Anthony E Kline, Julia Kofler, Thor D Stein, and Udai Bhan Pandey. Traumatic injury compromises nucleocytoplasmic transport and leads to TDP-43 pathology. *eLife*, 10, 2021.

[8490] Jordan A Anderson, Rachel A Johnston, Amanda J Lea, Fernando A Campos, Tawni N Voyles, Mercy Y Akinyi, Susan C Alberts, Elizabeth A Archie, and Jenny Tung. High social status males experience accelerated epigenetic aging in wild baboons. *eLife*, 10, 2021.

[8491] Ariana R Andrei, Samantha Debes, Mircea Chelaru, Xiaoqin Liu, Elsa Rodarte, John L Spudich, Roger Janz, and Valentin Dragoi. Heterogeneous side effects of cortical inactivation in behaving animals. *eLife*, 10, 2021.

[8492] Rizki Tsari Andriani and Yoshihiro Kubo. Voltage-clamp fluorometry analysis of structural rearrangements of ATP-gated channel P2X2 upon hyperpolarization. *eLife*, 10, 2021.

[8493] Qi Yan Ang, Diana L Alba, Vaibhav Upadhyay, Jordan E Bisanz, Jingwei Cai, Ho Lim Lee, Eliseo Barajas, Grace Wei, Cecilia Noecker, Andrew D Patterson, Suneil K Koliwad, and Peter J Turnbaugh. The East Asian gut microbiome is distinct from colocalized White subjects and connected to metabolic health. *eLife*, 10, 2021.

[8494] Suzanne Angeli, Anna Foulger, Manish Chamoli, Tanuja Harshani Peiris, Akos Gerencser, Azar Asadi Shahmirzadi, Julie Andersen, and Gordon Lithgow. The mitochondrial permeability transition pore activates the mitochondrial unfolded protein response and promotes aging. *eLife*, 10, 2021.

[8495] Guillem Angulo, Jelena Zeleznjak, Pablo Martínez-Vicente, Joan Puñet-Ortiz, Hartmut Hengel, Martin Messerle, Annette Oxenius, Stipan Jonjic, Astrid Krmpotić, Pablo Engel, and Ana Angulo. Cytomegalovirus restricts ICOSL expression on antigen-presenting cells disabling T cell co-stimulation and contributing to immune evasion. *eLife*, 10, 2021.

[8496] Mohd Anisul, Jarrod Shilts, Jeremy Schwartzentruber, James Hayhurst, Annalisa Buniello, Elmutaz Shaikho Elhaj Mohammed, Jie Zheng, Michael Holmes, David Ochoa, Miguel Carmona, Joseph Maranville, Tom R Gaunt, Valur Emilsson, Vilmundur Gudnason, Ellen M McDonagh, Gavin J Wright, Maya Ghoussaini, and Ian Dunham. A proteome-wide genetic investigation identifies several SARS-CoV-2-exploited host targets of clinical relevance. *eLife*, 10, 2021.

[8497] Kerim Anlas and Vikas Trivedi. Studying evolution of the primary body axis in vivo and in vitro. *eLife*, 10, 2021.

[8498] Jyot D Antani, Anita X Sumali, Tanmay P Lele, and Pushkar P Lele. Asymmetric random walks reveal that the chemotaxis network modulates flagellar rotational bias in Helicobacter pylori. *eLife*, 10, 2021.

[8499] Scott M Anthony, Natalija Van Braeckel-Budimir, Steven J Moioffer, Stephanie van de Wall, Qiang Shan, Rahul Vijay, Ramakrishna Sompallae, Stacey M Hartwig, Isaac J Jensen, Steven M Varga, Noah S Butler, Hai-Hui Xue, Vladimir P Badovinac, and John T Harty. Protective function and durability of mouse lymph node-resident memory CD8+ T cells. *eLife*, 10, 2021.

[8500] Jose D Aponte, David C Katz, Daniela M Roth, Marta Vidal-García, Wei Liu, Fernando Andrade, Charles C Roseman, Steven A Murray, James Cheverud, Daniel Graf, Ralph S Marcucio, and Benedikt Hallgrímsson. Relating multivariate shapes to genescapes using phenotype-biological process associations for craniofacial shape. *eLife*, 10, 2021.

[8501] Alessandra Araujo, Alexandra Safronova, Elise Burger, Américo López-Yglesias, Shilpi Giri, Ellie T Camanzo, Andrew T Martin, Sergei Grivennikov, and Felix Yarovinsky. IFN-$\gamma$ mediates Paneth cell death via suppression of mTOR. *eLife*, 10, 2021.

[8502] Carla CM Arce, Vanitha Theepan, Bernardus CJ Schimmel, Geoffrey Jaffuel, Matthias Erb, and Ricardo AR Machado. Plant-associated CO2 mediates long-distance host location and foraging behaviour of a root herbivore. *eLife*, 10, 2021.

[8503] Sarah M Ardell and Sergey Kryazhimskiy. The population genetics of collateral resistance and sensitivity. *eLife*, 10, 2021.

[8504] J Roman Arguello, Liliane Abuin, Jan Armida, Kaan Mika, Phing Chian Chai, and Richard Benton. Targeted molecular profiling of rare olfactory sensory neurons identifies fate, wiring, and functional determinants. *eLife*, 10, 2021.

[8505] Luisa F Arias Padilla, Diana C Castañeda-Cortés, Ivana F Rosa, Omar D Moreno Acosta, Ricardo S Hattori, Rafael H Nóbrega, and Juan I Fernandino. Cystic proliferation of germline stem cells is necessary to reproductive success and normal mating behavior in medaka. *eLife*, 10, 2021.

[8506] Shalini Aricthota and Devyani Haldar. DDK/Hsk1 phosphorylates and targets fission yeast histone deacetylase Hst4 for degradation to stabilize stalled DNA replication forks. *eLife*, 10, 2021.

[8507] Muhammad Arif, Martina Klevstig, Rui Benfeitas, Stephen Doran, Hasan Turkez, Mathias Uhlén, Maryam Clausen, Johannes Wikström, Damla Etal, Cheng Zhang, Malin Levin, Adil Mardinoglu, and Jan Boren. Integrative transcriptomic analysis of tissue-specific metabolic crosstalk after myocardial infarction. *eLife*, 10, 2021.

[8508] Yoanna Ariosa-Morejon, Alberto Santos, Roman Fischer, Simon Davis, Philip Charles, Rajesh Thakker, Angus KT Wann, and Tonia L Vincent. Age-dependent changes in protein incorporation into collagen-rich tissues of mice by in vivo pulsed SILAC labelling. *eLife*, 10, 2021.

[8509] Heidi A Arjes, Lisa Willis, Haiwen Gui, Yangbo Xiao, Jason Peters, Carol Gross, and Kerwyn Casey Huang. Three-dimensional biofilm colony growth supports a mutualism involving matrix and nutrient sharing. *eLife*, 10, 2021.

[8510] Laura Arribas-Hernández, Sarah Rennie, Tino Köster, Carlotta Porcelli, Martin Lewinski, Dorothee Staiger, Robin Andersson, and Peter Brodersen. Principles of mRNA targeting via the Arabidopsis m6A-binding protein ECT2. *eLife*, 10, 2021.

[8511] Laura Arribas-Hernández, Sarah Rennie, Michael Schon, Carlotta Porcelli, Balaji Enugutti, Robin Andersson, Michael D Nodine, and Peter Brodersen. The YTHDF proteins ECT2 and ECT3 bind largely overlapping target sets and influence target mRNA abundance, not alternative polyadenylation. *eLife*, 10, 2021.

[8512] Ashley L Arthur, Amy Crawford, Anne Houdusse, and Margaret A Titus. VASP-mediated actin dynamics activate and recruit a filopodia myosin. *eLife*, 10, 2021.

[8513] Janine JG Arts, Eike K Mahlandt, Max LB Grönloh, Lilian Schimmel, Ivar Noordstra, Emma Gordon, Abraham CI van Steen, Simon Tol, Barbara Walzog, Jos van Rijssel, Martijn A Nolte, Marten Postma, Satya Khuon, John M Heddleston, Eric Wait, Teng Leong Chew, Mark Winter, Eloi Montanez, Joachim Goedhart, and Jaap D van Buul. Endothelial junctional membrane protrusions serve as hotspots for neutrophil transmigration. *eLife*, 10, 2021.

[8514] Seksiri Arttamangkul, Emily J Platt, James Carroll, and David Farrens. Functional independence of endogenous $\mu$- and $\delta$-opioid receptors co-expressed in cholinergic interneurons. *eLife*, 10, 2021.

[8515] Mária Ashaber, Yusuke Tomina, Pegah Kassraian, Eric A Bushong, William B Kristan, Mark H Ellisman, and Daniel A Wagenaar. Anatomy and activity patterns in a multifunctional motor neuron and its surrounding circuits. *eLife*, 10, 2021.

[8516] Peter Ashcroft, Sonja Lehtinen, Daniel C Angst, Nicola Low, and Sebastian Bonhoeffer. Quantifying the impact of quarantine duration on COVID-19 transmission. *eLife*, 10, 2021.

[8517] Mishal Ashraf, Sharika Mohanan, Byu Ri Sim, Anthony Tam, Kiamehr Rahemipour, Denis Brousseau, Simon Thibault, Alexander D Corbett, and Gil Bub. Random access parallel microscopy. *eLife*, 10, 2021.

[8518] Magda L Atilano, Sebastian Grönke, Teresa Niccoli, Liam Kempthorne, Oliver Hahn, Javier Morón-Oset, Oliver Hendrich, Miranda Dyson, Mirjam Lisette Adams, Alexander Hull, Marie-Therese Salcher-Konrad, Amy Monaghan, Magda Bictash, Idoia Glaria, Adrian M Isaacs, and Linda Partridge. Enhanced insulin signalling ameliorates C9orf72 hexanucleotide repeat expansion toxicity in Drosophila. *eLife*, 10, 2021.

[8519] Susann Auer, Nele A Haeltermann, Tracey L Weissgerber, Jeffrey C Erlich, Damar Susilaradeya, Magdalena Julkowska, Małgorzata Anna Gazda, Benjamin Schwessinger, Nafisa M Jadavji, Angela Abitua, Angela Abitua, Anzela Niraulu, Aparna Shah, April Clyburne-Sherinb, Benoit Guiquel, Bradly Alicea, Caroline LaManna, Diep Ganguly, Eric Perkins, Helena Jambor, Ian Man Ho Li, Jennifer Tsang, Joanne Kamens, Lenny Teytelman, Mariella Paul, Michelle Cronin, Nicolas Schmelling, Peter Crisp, Rintu Kutum, Santosh Phuyal, Sarvenaz Sarabipour, Sonali Roy, Susanna M Bachle, Tuan Tran, Tyler Ford, Vicky Steeves, Vinodh Ilangovan, Ana Baburamani, and Susanna Bachle. A community-led initiative for training in reproducible research. *eLife*, 10, 2021.

[8520] Anita E Autry, Zheng Wu, Vikrant Kapoor, Johannes Kohl, Dhananjay Bambah-Mukku, Nimrod D Rubinstein, Brenda Marin-Rodriguez, Ilaria Carta, Victoria Sedwick, Ming Tang, and Catherine Dulac. Urocortin-3 neurons in the mouse perifornical area promote infant-directed neglect and aggression. *eLife*, 10, 2021.

[8521] Adel Avetisyan, Yael Glatt, Maya Cohen, Yael Timerman, Nitay Aspis, Atalya Nachman, Naomi Halachmi, Ella Preger-Ben Noon, and Adi Salzberg. Delilah, prospero, and D-Pax2 constitute a gene regulatory network essential for the development of functional proprioceptors. *eLife*, 10, 2021.

[8522] Emir E Avilés-Pagán, Masatoshi Hara, and Terry L Orr-Weaver. The GNU subunit of PNG kinase, the developmental regulator of mRNA translation, binds BIC-C to localize to RNP granules. *eLife*, 10, 2021.

[8523] Lilach Avitan, Zac Pujic, Jan Mölter, Shuyu Zhu, Biao Sun, and Geoffrey J Goodhill. Spontaneous and evoked activity patterns diverge over development. *eLife*, 10, 2021.

[8524] Oshri Avraham, Rui Feng, Eric Edward Ewan, Justin Rustenhoven, Guoyan Zhao, and Valeria Cavalli. Profiling sensory neuron microenvironment after peripheral and central axon injury reveals key pathways for neural repair. *eLife*, 10, 2021.

[8525] Ghulam R Awab, Fahima Aaram, Natsuda Jamornthanyawat, Kanokon Suwannasin, Watcharee Pagornrat, James A Watson, Charles J Woodrow, Arjen M Dondorp, Nicholas PJ Day, Mallika Imwong, and Nicholas J White. Protective effect of Mediterranean-type glucose-6-phosphate dehydrogenase deficiency against Plasmodium vivax malaria. *eLife*, 10, 2021.

[8526] Asli Azizoglu, Roger Brent, and Fabian Rudolf. A precisely adjustable, variation-suppressed eukaryotic transcriptional controller to enable genetic discovery. *eLife*, 10, 2021.

[8527] Iris Bachmutsky, Xin Paul Wei, Adelae Durand, and Kevin Yackle. ß-arrestin 2 germline knockout does not attenuate opioid respiratory depression. *eLife*, 10, 2021.

[8528] Luka Bacic, Guillaume Gaullier, Anton Sabantsev, Laura C Lehmann, Klaus Brackmann, Despoina Dimakou, Mario Halic, Graeme Hewitt, Simon J Boulton, and Sebastian Deindl. Structure and dynamics of the chromatin remodeler ALC1 bound to a PARylated nucleosome. *eLife*, 10, 2021.

[8529] Mohamed A Badawy, Basma A Yasseen, Riem M El-Messiery, Engy A Abdel-Rahman, Aya A Elkhodiry, Azza G Kamel, Hajar El-sayed, Asmaa M Shedra, Rehab Hamdy, Mona Zidan, Diaa Al-Raawi, Mahmoud Hammad, Nahla Elsharkawy, Mohamed El Ansary, Ahmed Al-Halfawy, Alaa Elhadad, Ashraf Hatem, Sherif Abouelnaga, Laura L Dugan, and Sameh Saad Ali. Neutrophil-mediated oxidative stress and albumin structural damage predict COVID-19-associated mortality. *eLife*, 10, 2021.

[8530] Krishna N Badhiwala, Abby S Primack, Celina E Juliano, and Jacob T Robinson. Multiple neuronal networks coordinate Hydra mechanosensory behavior. *eLife*, 10, 2021.

[8531] Thomas Baeriswyl, Alexandre Dumoulin, Martina Schaettin, Georgia Tsapara, Vera Niederkofler, Denise Helbling, Evelyn Avilés, Jeannine A Frei, Nicole H Wilson, Matthias Gesemann, Beat Kunz, and Esther T Stoeckli. Endoglycan plays a role in axon guidance by modulating cell adhesion. *eLife*, 10, 2021.

[8532] Nathan A Baertsch, Nicholas E Bush, Nicholas J Burgraff, and Jan-Marino Ramirez. Dual mechanisms of opioid-induced respiratory depression in the inspiratory rhythm-generating network. *eLife*, 10, 2021.

[8533] Yang Bai, Caiyun He, Pan Chu, Junjiajia Long, Xuefei Li, and Xiongfei Fu. Spatial modulation of individual behaviors enables an ordered structure of diverse phenotypes during bacterial group migration. *eLife*, 10, 2021.

[8534] Gaurav Bajpai, Daria Amiad Pavlov, Dana Lorber, Talila Volk, and Samuel Safran. Mesoscale phase separation of chromatin in the nucleus. *eLife*, 10, 2021.

[8535] Salah A Baker, Wesley A Leigh, Guillermo Del Valle, Inigo F De Yturriaga, Sean M Ward, Caroline A Cobine, Bernard T Drumm, and Kenton M Sanders. Ca2+ signaling driving pacemaker activity in submucosal interstitial cells of Cajal in the murine colon. *eLife*, 10, 2021.

[8536] Christopher W Bakerlee, Angela M Phillips, Alex N Nguyen Ba, and Michael M Desai. Dynamics and variability in the pleiotropic effects of adaptation in laboratory budding yeast populations. *eLife*, 10, 2021.

[8537] Trygve E Bakken, Cindy TJ van Velthoven, Vilas Menon, Rebecca D Hodge, Zizhen Yao, Thuc Nghi Nguyen, Lucas T Graybuck, Gregory D Horwitz, Darren Bertagnolli, Jeff Goldy, Anna Marie Yanny, Emma Garren, Sheana Parry, Tamara Casper, Soraya I Shehata, Eliza R Barkan, Aaron Szafer, Boaz P Levi, Nick Dee, Kimberly A Smith, Susan M Sunkin, Amy Bernard, John Phillips, Michael J Hawrylycz, Christof Koch, Gabe J Murphy, Ed Lein, Hongkui Zeng, and Bosiljka Tasic. Single-cell and single-nucleus RNA-seq uncovers shared and distinct axes of variation in dorsal LGN neurons in mice, non-human primates, and humans. *eLife*, 10, 2021.

[8538] Erik Bakkeren, Joana Anuschka Herter, Jana Sanne Huisman, Yves Steiger, Ersin Gül, Joshua Patrick Mark Newson, Alexander Oliver Brachmann, Jörn Piel, Roland Regoes, Sebastian Bonhoeffer, Médéric Diard, and Wolf-Dietrich Hardt. Pathogen invasion-dependent tissue reservoirs and plasmid-encoded antibiotic degradation boost plasmid spread in the gut. *eLife*, 10, 2021.

[8539] Anahita Bakochi, Tirthankar Mohanty, Paul Theodor Pyl, Carlos Alberto Gueto-Tettay, Lars Malmström, Adam Linder, and Johan Malmström. Cerebrospinal fluid proteome maps detect pathogen-specific host response patterns in meningitis. *eLife*, 10, 2021.

[8540] Francisco de Asis Balaguer, Clara Aicart-Ramos, Gemma LM Fisher, Sara de BraganÇa, Eva M Martin-Cuevas, Cesar L Pastrana, Mark Simon Dillingham, and Fernando Moreno-Herrero. CTP promotes efficient ParB-dependent DNA condensation by facilitating one-dimensional diffusion from parS. *eLife*, 10, 2021.

[8541] Fernando R Balestra, Andrés Domínguez-Calvo, Benita Wolf, Coralie Busso, Alizée Buff, Tessa Averink, Marita Lipsanen-Nyman, Pablo Huertas, Rosa M Ríos, and Pierre Gönczy. TRIM37 prevents formation of centriolar protein assemblies by regulating Centrobin. *eLife*, 10, 2021.

[8542] Mariona Baliu-Piqué, Vera van Hoeven, Julia Drylewicz, Lotte E van der Wagen, Anke Janssen, Sigrid A Otto, Menno C van Zelm, Rob J de Boer, Jürgen Kuball, Jose AM Borghans, and Kiki Tesselaar. Cell-density independent increased lymphocyte production and loss rates post-autologous HSCT. *eLife*, 10, 2021.

[8543] Timothy S Balmer, Carolina Borges-Merjane, and Laurence O Trussell. Incomplete removal of extracellular glutamate controls synaptic transmission and integration at a cerebellar synapse. *eLife*, 10, 2021.

[8544] Tarryn Balsdon, Pascal Mamassian, and Valentin Wyart. Separable neural signatures of confidence during perceptual decisions. *eLife*, 10, 2021.

[8545] Kianoush Banaie Boroujeni, Paul Tiesinga, and Thilo Womelsdorf. Interneuron-specific gamma synchronization indexes cue uncertainty and prediction errors in lateral prefrontal and anterior cingulate cortex. *eLife*, 10, 2021.

[8546] Giulia Bandini, Sebastian Damerow, Maria Lucia Sempaio Guther, Hongjie Guo, Angela Mehlert, Jose Carlos Paredes Franco, Stephen Beverley, and Michael AJ Ferguson. An essential, kinetoplastid-specific GDP-Fuc: $\beta$-D-Gal $\alpha$-1,2-fucosyltransferase is located in the mitochondrion of Trypanosoma brucei. *eLife*, 10, 2021.

[8547] Sudeep Banjade, Yousuf H Shah, Shaogeng Tang, and Scott D Emr. Design principles of the ESCRT-III Vps24-Vps2 module. *eLife*, 10, 2021.

[8548] Beatriz Bano-Otalora, Matthew J Moye, Timothy Brown, Robert J Lucas, Casey O Diekman, and Mino DC Belle. Daily electrical activity in the master circadian clock of a diurnal mammal. *eLife*, 10, 2021.

[8549] László Bányai, Maria Trexler, Krisztina Kerekes, Orsolya Csuka, and László Patthy. Use of signals of positive and negative selection to distinguish cancer genes and passenger genes. *eLife*, 10, 2021.

[8550] Antoine Barbot, Woon Ju Park, Cherlyn J Ng, Ru-Yuan Zhang, Krystel R Huxlin, Duje Tadin, and Geunyoung Yoon. Functional reallocation of sensory processing resources caused by long-term neural adaptation to altered optics. *eLife*, 10, 2021.

[8551] Carter J Barger, Linda Chee, Mustafa Albahrani, Catalina Munoz-Trujillo, Lidia Boghean, Connor Branick, Kunle Odunsi, Ronny Drapkin, Lee Zou, and Adam R Karpf. Co-regulation and function of FOXM1/RHNO1 bidirectional genes in cancer. *eLife*, 10, 2021.

[8552] Arnab Barik, Anupama Sathyamurthy, James Thompson, Mathew Seltzer, Ariel Levine, and Alexander Chesler. A spinoparabrachial circuit defined by Tacr1 expression drives pain. *eLife*, 10, 2021.

[8553] Michael A Barnett, Geoffrey K Aguirre, and David Brainard. A quadratic model captures the human V1 response to variations in chromatic direction and contrast. *eLife*, 10, 2021.

[8554] Sierra M Barone, Alberta GA Paul, Lyndsey M Muehling, Joanne A Lannigan, William W Kwok, Ronald B Turner, Judith A Woodfolk, and Jonathan M Irish. Unsupervised machine learning reveals key immune cell subsets in COVID-19, rhinovirus infection, and cancer therapy. *eLife*, 10, 2021.

[8555] Emilie Barruet, Steven M Garcia, Jake Wu, Blanca M Morales, Stanley Tamaki, Tania Moody, Jason H Pomerantz, and Edward C Hsiao. Modeling the ACVR1R206H mutation in human skeletal muscle stem cells. *eLife*, 10, 2021.

[8556] Zachary K Barth, Maria HT Nguyen, and Kimberley D Seed. A chimeric nuclease substitutes a phage CRISPR-Cas system to provide sequence-specific immunity against subviral parasites. *eLife*, 10, 2021.

[8557] Michael I Barton, Stuart A MacGowan, Mikhail A Kutuzov, Omer Dushek, Geoffrey John Barton, and P Anton van der Merwe. Effects of common mutations in the SARS-CoV-2 Spike RBD and its ligand, the human ACE2 receptor on binding affinity and kinetics. *eLife*, 10, 2021.

[8558] Carlos AZ Bassetto, João Luis Carvalho-de Souza, and Francisco Bezanilla. Molecular basis for functional connectivity between the voltage sensor and the selectivity filter gate in Shaker K+ channels. *eLife*, 10, 2021.

[8559] André M Bastos, Jacob A Donoghue, Scott L Brincat, Meredith Mahnke, Jorge Yanar, Josefina Correa, Ayan S Waite, Mikael Lundqvist, Jefferson Roy, Emery N Brown, and Earl K Miller. Neural effects of propofol-induced unconsciousness and its reversal using thalamic stimulation. *eLife*, 10, 2021.

[8560] Aditi Batra, Roderich Roemhild, Emilie Rousseau, Sören Franzenburg, Stefan Niemann, and Hinrich Schulenburg. High potency of sequential therapy with only $\beta$-lactam antibiotics. *eLife*, 10, 2021.

[8561] Joshua Batson, Gytis Dudas, Eric Haas-Stapleton, Amy L Kistler, Lucy M Li, Phoenix Logan, Kalani Ratnasiri, and Hanna Retallack. Single mosquito metatranscriptomics identifies vectors, emerging pathogens and reservoirs in one assay. *eLife*, 10, 2021.

[8562] Rafael Bayarri-Olmos, Laust Bruun Johnsen, Manja Idorn, Line S Reinert, Anne Rosbjerg, Søren Vang, Cecilie Bo Hansen, Charlotte Helgstrand, Jais Rose Bjelke, Theresa Bak-Thomsen, Søren R Paludan, Peter Garred, and Mikkel-Ole Skjoedt. The alpha/B.1.1.7 SARS-CoV-2 variant exhibits significantly higher affinity for ACE-2 and requires lower inoculation doses to cause disease in K18-hACE2 mice. *eLife*, 10, 2021.

[8563] Amélie Beaudet, Robert C Atwood, Winfried Kockelmann, Vincent Fernandez, Thomas Connolley, Nghia Trong Vo, Ronald Clarke, and Dominic Stratford. Preliminary paleohistological observations of the StW 573 ('Little Foot') skull. *eLife*, 10, 2021.

[8564] Mikkel Malling Beck, Meaghan Elizabeth Spedden, Martin Jensen Dietz, Anke Ninija Karabanov, Mark Schram Christensen, and Jesper Lundbye-Jensen. Cortical signatures of precision grip force control in children, adolescents, and adults. *eLife*, 10, 2021.

[8565] Robert Becker, Silvia Vergarajauregui, Florian Billing, Maria Sharkova, Eleonora Lippolis, Kamel Mamchaoui, Fulvia Ferrazzi, and Felix B Engel. Myogenin controls via AKAP6 non-centrosomal microtubule-organizing center formation at the nuclear envelope. *eLife*, 10, 2021.

[8566] Joseph L Bedont, Hirofumi Toda, Mi Shi, Christine H Park, Christine Quake, Carly Stein, Anna Kolesnik, and Amita Sehgal. Short and long sleeping mutants reveal links between sleep and macroautophagy. *eLife*, 10, 2021.

[8567] Gerrit Cornelis Beekhof, Catarina Osório, Joshua J White, Scott van Zoomeren, Hannah van der Stok, Bilian Xiong, Ingo HMS Nettersheim, Willem Ashwin Mak, Marit Runge, Francesca Romana Fiocchi, Henk-Jan Boele, Freek E Hoebeek, and Martijn Schonewille. Differential spatiotemporal development of Purkinje cell populations and cerebellum-dependent sensorimotor behaviors. *eLife*, 10, 2021.

[8568] Annaliese K Beery, Sarah A Lopez, Katrina L Blandino, Nicole S Lee, and Natalie S Bourdon. Social selectivity and social motivation in voles. *eLife*, 10, 2021.

[8569] Francesco Beghini, Lauren J McIver, Aitor Blanco-Míguez, Leonard Dubois, Francesco Asnicar, Sagun Maharjan, Ana Mailyan, Paolo Manghi, Matthias Scholz, Andrew Maltez Thomas, Mireia Valles-Colomer, George Weingart, Yancong Zhang, Moreno Zolfo, Curtis Huttenhower, Eric A Franzosa, and Nicola Segata. Integrating taxonomic, functional, and strain-level profiling of diverse microbial communities with bioBakery 3. *eLife*, 10, 2021.

[8570] Hourinaz Behesti, Arif Kocabas, David E Buchholz, Thomas S Carroll, and Mary E Hatten. Altered temporal sequence of transcriptional regulators in the generation of human cerebellar granule cells. *eLife*, 10, 2021.

[8571] Inbal Ben-Ami Bartal, Jocelyn M Breton, Huanjie Sheng, Kimberly LP Long, Stella Chen, Aline Halliday, Justin W Kenney, Anne L Wheeler, Paul Frankland, Carrie Shilyansky, Karl Deisseroth, Dacher Keltner, and Daniela Kaufer. Neural correlates of ingroup bias for prosociality in rats. *eLife*, 10, 2021.

[8572] Eyal Ben-David, James Boocock, Longhua Guo, Stefan Zdraljevic, Joshua S Bloom, and Leonid Kruglyak. Whole-organism eQTL mapping at cellular resolution with single-cell sequencing. *eLife*, 10, 2021.

[8573] Hanna M Bensch, Emily A O'Connor, and Charlie Kinahan Cornwallis. Living with relatives offsets the harm caused by pathogens in natural populations. *eLife*, 10, 2021.

[8574] Noah C Benson, Eline R Kupers, Antoine Barbot, Marisa Carrasco, and Jonathan Winawer. Cortical magnification in human visual cortex parallels task performance around the visual field. *eLife*, 10, 2021.

[8575] Bidesh K Bera, Omer Tzuk, Jamie JR Bennett, and Ehud Meron. Linking spatial self-organization to community assembly and biodiversity. *eLife*, 10, 2021.

[8576] Joy Bergelson, Martin Kreitman, Dmitri A Petrov, Alvaro Sanchez, and Mikhail Tikhonov. Functional biology in its natural context: A search for emergent simplicity. *eLife*, 10, 2021.

[8577] Cathrine Bergh, Stephanie A Heusser, Rebecca Howard, and Erik Lindahl. Markov state models of proton- and pore-dependent activation in a pentameric ligand-gated ion channel. *eLife*, 10, 2021.

[8578] Emily G Berghoff, Lori Glenwinkel, Abhishek Bhattacharya, HaoSheng Sun, Erdem Varol, Nicki Mohammadi, Amelia Antone, Yi Feng, Ken Nguyen, Steven J Cook, Jordan F Wood, Neda Masoudi, Cyril C Cros, Yasmin H Ramadan, Denise M Ferkey, David H Hall, and Oliver Hobert. The Prop1-like homeobox gene unc-42 specifies the identity of synaptically connected neurons. *eLife*, 10, 2021.

[8579] Yelena Y Bernadskaya, Haicen Yue, Calina Copos, Lionel Christiaen, and Alex Mogilner. Supracellular organization confers directionality and mechanical potency to migrating pairs of cardiopharyngeal progenitor cells. *eLife*, 10, 2021.

[8580] Zachariah Bertels, Harinder Singh, Isaac Dripps, Kendra Siegersma, Alycia F Tipton, Wiktor D Witkowski, Zoie Sheets, Pal Shah, Catherine Conway, Elizaveta Mangutov, Mei Ao, Valentina Petukhova, Bhargava Karumudi, Pavel A Petukhov, Serapio M Baca, Mark M Rasenick, and Amynah A Pradhan. Neuronal complexity is attenuated in preclinical models of migraine and restored by HDAC6 inhibition. *eLife*, 10, 2021.

[8581] Tessa M Bertozzi, Nozomi Takahashi, Geula Hanin, Anastasiya Kazachenka, and Anne C Ferguson-Smith. A spontaneous genetically induced epiallele at a retrotransposon shapes host genome function. *eLife*, 10, 2021.

[8582] Anton Bespalov, René Bernard, Anja Gilis, Björn Gerlach, Javier Guillén, Vincent Castagné, Isabel A Lefevre, Fiona Ducrey, Lee Monk, Sandrine Bongiovanni, Bruce Altevogt, María Arroyo-Araujo, Lior Bikovski, Natasja de Bruin, Esmeralda Castaños-Vélez, Alexander Dityatev, Christoph H Emmerich, Raafat Fares, Chantelle Ferland-Beckham, Christelle Froger-Colléaux, Valerie Gailus-Durner, Sabine M Hölter, Martine CJ Hofmann, Patricia Kabitzke, Martien JH Kas, Claudia Kurreck, Paul Moser, Malgorzata Pietraszek, Piotr Popik, Heidrun Potschka, Ernesto Prado Montes de Oca, Leonardo Restivo, Gernot Riedel, Merel Ritskes-Hoitinga, Janko Samardzic, Michael Schunn, Claudia Stöger, Vootele Voikar, Jan Vollert, Kimberley E Wever, Kathleen Wuyts, Malcolm R MacLeod, Ulrich Dirnagl, and Thomas Steckler. Introduction to the EQIPD quality system. *eLife*, 10, 2021.

[8583] Benjamin Bessières, Emmanuel Cruz, and Cristina M Alberini. Metabolomic profiling reveals a differential role for hippocampal glutathione reductase in infantile memory formation. *eLife*, 10, 2021.

[8584] Katherine T Best, Antonion Korcari, Keshia E Mora, Anne EC Nichols, Samantha N Muscat, Emma Knapp, Mark R Buckley, and Alayna E Loiselle. Scleraxis-lineage cell depletion improves tendon healing and disrupts adult tendon homeostasis. *eLife*, 10, 2021.

[8585] Nikhil R Bhagwat, Shannon N Owens, Masaru Ito, Jay V Boinapalli, Philip Poa, Alexander Ditzel, Srujan Kopparapu, Meghan Mahalawat, Owen Richard Davies, Sean R Collins, Jeffrey R Johnson, Nevan J Krogan, and Neil Hunter. SUMO is a pervasive regulator of meiosis. *eLife*, 10, 2021.

[8586] Pradeep Bhandari, David Vandael, Diego Fernández-Fernández, Thorsten Fritzius, David Kleindienst, Cihan önal, Jacqueline Montanaro, Martin Gassmann, Peter Jonas, Akos Kulik, Bernhard Bettler, Ryuichi Shigemoto, and Peter Koppensteiner. GABAB receptor auxiliary subunits modulate Cav2.3-mediated release from medial habenula terminals. *eLife*, 10, 2021.

[8587] Shreyas Bhat, Marco Niello, Klaus Schicker, Christian Pifl, Harald H Sitte, Michael Freissmuth, and Walter Sandtner. Handling of intracellular K+ determines voltage dependence of plasmalemmal monoamine transporter function. *eLife*, 10, 2021.

[8588] Shipra Bhatia, Dirk Jan Kleinjan, Kirsty Uttley, Anita Mann, Nefeli Dellepiane, and Wendy A Bickmore. Quantitative spatial and temporal assessment of regulatory element activity in zebrafish. *eLife*, 10, 2021.

[8589] Celia Biane, Florian Rückerl, Therese Abrahamsson, Cécile Saint-Cloment, Jean Mariani, Ryuichi Shigemoto, David A DiGregorio, Rachel M Sherrard, and Laurence Cathala. Developmental emergence of two-stage nonlinear synaptic integration in cerebellar interneurons. *eLife*, 10, 2021.

[8590] Aleksandra Białas, Thorsten Langner, Adeline Harant, Mauricio P Contreras, Clare EM Stevenson, David M Lawson, Jan Sklenar, Ronny Kellner, Matthew J Moscou, Ryohei Terauchi, Mark J Banfield, and Sophien Kamoun. Two NLR immune receptors acquired high-affinity binding to a fungal effector through convergent evolution of their integrated domain. *eLife*, 10, 2021.

[8591] Cornelia Biehler, Katheryn E Rothenberg, Alexandra Jette, Helori-Mael Gaude, Rodrigo Fernandez-Gonzalez, and Patrick Laprise. Pak1 and PP2A antagonize aPKC function to support cortical tension induced by the Crumbs-Yurt complex. *eLife*, 10, 2021.

[8592] Savannah F Bifulco, Griffin D Scott, Sakher Sarairah, Zeinab Birjandian, Caroline H Roney, Steven A Niederer, Christian Mahnkopf, Peter Kuhnlein, Marcel Mitlacher, David Tirschwell, WT Longstreth, Nazem Akoum, and Patrick M Boyle. Computational modeling identifies embolic stroke of undetermined source patients with potential arrhythmic substrate. *eLife*, 10, 2021.

[8593] Patrick G Bissett, McKenzie P Hagen, Henry M Jones, and Russell A Poldrack. Design issues and solutions for stop-signal data from the Adolescent Brain Cognitive Development (ABCD) study. *eLife*, 10, 2021.

[8594] Edoardo Bistaffa, Alba Marín-Moreno, Juan Carlos Espinosa, Chiara Maria Giulia De Luca, Federico Angelo Cazzaniga, Sara Maria Portaleone, Luigi Celauro, Giuseppe Legname, Giorgio Giaccone, Juan Maria Torres, and Fabio Moda. PMCA-generated prions from the olfactory mucosa of patients with Fatal Familial Insomnia cause prion disease in mice. *eLife*, 10, 2021.

[8595] Sean R Bittner, Agostina Palmigiano, Alex T Piet, Chunyu A Duan, Carlos D Brody, Kenneth D Miller, and John Cunningham. Interrogating theoretical models of neural computation with emergent property inference. *eLife*, 10, 2021.

[8596] Stephen M Blazie, Seika Takayanagi-Kiya, Katherine A McCulloch, and Yishi Jin. Eukaryotic initiation factor EIF-3.G augments mRNA translation efficiency to regulate neuronal activity. *eLife*, 10, 2021.

[8597] Edgar E Boczek, Julius Fürsch, Marie Laura Niedermeier, Louise Jawerth, Marcus Jahnel, Martine Ruer-Gruß, Kai-Michael Kammer, Peter Heid, Laura Mediani, Jie Wang, Xiao Yan, Andrej Pozniakovski, Ina Poser, Daniel Mateju, Lars Hubatsch, Serena Carra, Simon Alberti, Anthony A Hyman, and Florian Stengel. HspB8 prevents aberrant phase transitions of FUS by chaperoning its folded RNA-binding domain. *eLife*, 10, 2021.

[8598] Juliette Boeri, Claude Meunier, Hervé Le Corronc, Pascal Branchereau, Yulia Timofeeva, FranÇois-Xavier Lejeune, Christine Mouffle, Hervé Arulkandarajah, Jean Marie Mangin, Pascal Legendre, and Antonny Czarnecki. Two opposite voltage-dependent currents control the unusual early development pattern of embryonic Renshaw cell electrical activity. *eLife*, 10, 2021.

[8599] Miriam Böhm, Kerstin Killinger, Alexander Dudziak, Pradeep Pant, Karolin Jänen, Simone Hohoff, Karl Mechtler, Mihkel örd, Mart Loog, Elsa Sanchez-Garcia, and Stefan Westermann. Cdc4 phosphodegrons allow differential regulation of Ame1CENP-U protein stability across the cell cycle. *eLife*, 10, 2021.

[8600] Sebastian Bohm, Falk Mersmann, Alessandro Santuz, Arno Schroll, and Adamantios Arampatzis. Muscle-specific economy of force generation and efficiency of work production during human running. *eLife*, 10, 2021.

[8601] James P Bohnslav, Nivanthika K Wimalasena, Kelsey J Clausing, Yu Y Dai, David A Yarmolinsky, Tomás Cruz, Adam D Kashlan, M Eugenia Chiappe, Lauren L Orefice, Clifford J Woolf, and Christopher D Harvey. DeepEthogram, a machine learning pipeline for supervised behavior classification from raw pixels. *eLife*, 10, 2021.

[8602] Tisha E Bohr, Divya A Shiroor, and Carolyn E Adler. Planarian stem cells sense the identity of the missing pharynx to launch its targeted regeneration. *eLife*, 10, 2021.

[8603] Krista Bond, Kyle Dunovan, Alexis Porter, Jonathan E Rubin, and Timothy Verstynen. Dynamic decision policy reconfiguration under outcome uncertainty. *eLife*, 10, 2021.

[8604] Matthew C Bond, Lucia Vidakovic, Praveen K Singh, Knut Drescher, and Carey D Nadell. Matrix-trapped viruses can prevent invasion of bacterial biofilms by colonizing cells. *eLife*, 10, 2021.

[8605] Giulio Bondanelli, Thomas Deneux, Brice Bathellier, and Srdjan Ostojic. Network dynamics underlying OFF responses in the auditory cortex. *eLife*, 10, 2021.

[8606] Alessandro Bonfini, Adam J Dobson, David Duneau, Jonathan Revah, Xi Liu, Philip Houtz, and Nicolas Buchon. Multiscale analysis reveals that diet-dependent midgut plasticity emerges from alterations in both stem cell niche coupling and enterocyte size. *eLife*, 10, 2021.

[8607] Braulio Bonilla, Alexander J Brown, Sarah R Hengel, Kyle S Rapchak, Debra Mitchell, Catherine A Pressimone, Adeola A Fagunloye, Thong T Luong, Reagan A Russell, Rudri K Vyas, Tony M Mertz, Hani S Zaher, Nima Mosammaparast, Ewa P Malc, Piotr A Mieczkowski, Steven A Roberts, and Kara A Bernstein. The Shu complex prevents mutagenesis and cytotoxicity of single-strand specific alkylation lesions. *eLife*, 10, 2021.

[8608] Michael Borg, Ranjith K Papareddy, Rodolphe Dombey, Elin Axelsson, Michael D Nodine, David Twell, and Frédéric Berger. Epigenetic reprogramming rewires transcription during the alternation of generations in Arabidopsis. *eLife*, 10, 2021.

[8609] V Borghesani, CL Dale, S Lukic, LBN Hinkley, M Lauricella, W Shwe, D Mizuiri, S Honma, Z Miller, B Miller, JF Houde, ML Gorno-Tempini, and SS Nagarajan. Neural dynamics of semantic categorization in semantic variant of primary progressive aphasia. *eLife*, 10, 2021.

[8610] Madeleine Bossaert, Angélique Pipier, Jean-Francois Riou, Céline Noirot, Linh-Trang Nguyên, Remy-Felix Serre, Olivier Bouchez, Eric Defrancq, Patrick Calsou, Sébastien Britton, and Dennis Gomez. Transcription-associated topoisomerase 2$\alpha$ (TOP2A) activity is a major effector of cytotoxicity induced by G-quadruplex ligands. *eLife*, 10, 2021.

[8611] Maude Bouchard, Jean-Marc Lina, Pierre-Olivier Gaudreault, Alexandre Lafrenière, Jonathan Dubé, Nadia Gosselin, and Julie Carrier. Sleeping at the switch. *eLife*, 10, 2021.

[8612] Benoît Boulan, Charlotte Ravanello, Amandine Peyrel, Christophe Bosc, Christian Delphin, Florence Appaix, Eric Denarier, Alexandra Kraut, Muriel Jacquier-Sarlin, Alyson Fournier, Annie Andrieux, Sylvie Gory-Fauré, and Jean-Christophe Deloulme. CRMP4-mediated fornix development involves Semaphorin-3E signaling pathway. *eLife*, 10, 2021.

[8613] Rozlyn CT Boutin, Charisse Petersen, Sarah E Woodward, Antonio Serapio-Palacios, Tahereh Bozorgmehr, Rachelle Loo, Alina Chalanuchpong, Mihai Cirstea, Bernard Lo, Kelsey E Huus, Weronika Barcik, Meghan B Azad, Allan B Becker, Piush J Mandhane, Theo J Moraes, Malcolm R Sears, Padmaja Subbarao, Kelly M McNagny, Stuart E Turvey, and B Brett Finlay. Bacterial–fungal interactions in the neonatal gut influence asthma outcomes later in life. *eLife*, 10, 2021.

[8614] Clara Bouyx, Marion Schiavone, Marie-Ange Teste, Etienne Dague, Nathalie Sieczkowski, Anne Julien, and Jean Marie FranÇois. The dual role of amyloid-$\beta$-sheet sequences in the cell surface properties of FLO11-encoded flocculins in Saccharomyces cerevisiae. *eLife*, 10, 2021.

[8615] Samuel Bowerman, Jeff Wereszczynski, and Karolin Luger. Archaeal chromatin 'slinkies' are inherently dynamic complexes with deflected DNA wrapping pathways. *eLife*, 10, 2021.

[8616] Corey JA Bradshaw, Christopher N Johnson, John Llewelyn, Vera Weisbecker, Giovanni Strona, and Frédérik Saltré. Relative demographic susceptibility does not explain the extinction chronology of Sahul's megafauna. *eLife*, 10, 2021.

[8617] Jan W Brascamp, Gilles de Hollander, Michael D Wertheimer, Ashley N DePew, and Tomas Knapen. Separable pupillary signatures of perception and action during perceptual multistability. *eLife*, 10, 2021.

[8618] Alejandro J Brenes, Maud Vandereyken, Olivia J James, Harriet Watt, Jens Hukelmann, Laura Spinelli, Dina Dikovskaya, Angus I Lamond, and Mahima Swamy. Tissue environment, not ontogeny, defines murine intestinal intraepithelial T lymphocytes. *eLife*, 10, 2021.

[8619] Ellen KW Brennan, Izabela Jedrasiak-Cape, Sameer Kailasa, Sharena P Rice, Shyam Kumar Sudhakar, and Omar J Ahmed. Thalamus and claustrum control parallel layer 1 circuits in retrosplenial cortex. *eLife*, 10, 2021.

[8620] Assaf Breska and Richard B Ivry. The human cerebellum is essential for modulating perceptual sensitivity based on temporal expectations. *eLife*, 10, 2021.

[8621] Andrew A Bridges and Bonnie L Bassler. Inverse regulation of Vibrio cholerae biofilm dispersal by polyamine signals. *eLife*, 10, 2021.

[8622] Axel F Brilot, Andrew S Lyon, Alex Zelter, Shruthi Viswanath, Alison Maxwell, Michael J MacCoss, Eric G Muller, Andrej Sali, Trisha N Davis, and David A Agard. CM1-driven assembly and activation of yeast $\gamma$-tubulin small complex underlies microtubule nucleation. *eLife*, 10, 2021.

[8623] Ben J Brintz, Benjamin Haaland, Joel Howard, Dennis L Chao, Joshua L Proctor, Ashraful I Khan, Sharia M Ahmed, Lindsay T Keegan, Tom Greene, Adama Mamby Keita, Karen L Kotloff, James A Platts-Mills, Eric J Nelson, Adam C Levine, Andrew T Pavia, and Daniel T Leung. A modular approach to integrating multiple data sources into real-time clinical prediction for pediatric diarrhea. *eLife*, 10, 2021.

[8624] Silvia Brochet, Andrew Quinn, Ruben AT Mars, Nicolas Neuschwander, Uwe Sauer, and Philipp Engel. Niche partitioning facilitates coexistence of closely related honey bee gut bacteria. *eLife*, 10, 2021.

[8625] Savannah G Brovero, Julia C Fortier, Hongru Hu, Pamela C Lovejoy, Nicole R Newell, Colleen M Palmateer, Ruei-Ying Tzeng, Pei-Tseng Lee, Kai Zinn, and Michelle N Arbeitman. Investigation of Drosophila fruitless neurons that express Dpr/DIP cell adhesion molecules. *eLife*, 10, 2021.

[8626] Brian Brown, Sahana Mitra, Finnegan D Roach, Deepika Vasudevan, and Hyung Don Ryoo. The transcription factor Xrp1 is required for PERK-mediated antioxidant gene induction in Drosophila. *eLife*, 10, 2021.

[8627] Elizabeth B Brown, Kreesha D Shah, Justin Palermo, Manali Dey, Anupama Dahanukar, and Alex C Keene. Ir56d-dependent fatty acid responses in Drosophila uncover taste discrimination between different classes of fatty acids. *eLife*, 10, 2021.

[8628] Alexander P Browning, Jesse A Sharp, Ryan J Murphy, Gency Gunasingh, Brodie Lawson, Kevin Burrage, Nikolas K Haass, and Matthew Simpson. Quantitative analysis of tumour spheroid structure. *eLife*, 10, 2021.

[8629] Thibaut Brunet, Marvin Albert, William Roman, Maxwell C Coyle, Danielle C Spitzer, and Nicole King. A flagellate-to-amoeboid switch in the closest living relatives of animals. *eLife*, 10, 2021.

[8630] Michael Bucher, Stephan Niebling, Yuhao Han, Dmitry Molodenskiy, Fatemeh Hassani Nia, Hans-Jürgen Kreienkamp, Dmitri Svergun, Eunjoon Kim, Alla S Kostyukova, Michael R Kreutz, and Marina Mikhaylova. Autism-associated SHANK3 missense point mutations impact conformational fluctuations and protein turnover at synapses. *eLife*, 10, 2021.

[8631] Janet C Buckner, Robert C Sanders, Brant C Faircloth, and Prosanta Chakrabarty. The critical importance of vouchers in genomics. *eLife*, 10, 2021.

[8632] Vince Buffalo. Quantifying the relationship between genetic diversity and population size suggests natural selection cannot explain Lewontin's Paradox. *eLife*, 10, 2021.

[8633] Antimo Buonocore, Xiaoguang Tian, Fatemeh Khademi, and Ziad M Hafed. Instantaneous movement-unrelated midbrain activity modifies ongoing eye movements. *eLife*, 10, 2021.

[8634] Joshua B Burt, Katrin H Preller, Murat Demirtas, Jie Lisa Ji, John H Krystal, Franz X Vollenweider, Alan Anticevic, and John D Murray. Transcriptomics-informed large-scale cortical model captures topography of pharmacological neuroimaging effects of LSD. *eLife*, 10, 2021.

[8635] Nicholas O Burton, Alexandra Willis, Kinsey Fisher, Fabian Braukmann, Jonathan Price, Lewis Stevens, L Ryan Baugh, Aaron Reinke, and Eric A Miska. Intergenerational adaptations to stress are evolutionarily conserved, stress-specific, and have deleterious trade-offs. *eLife*, 10, 2021.

[8636] Shawn D Burton and Nathaniel N Urban. Cell and circuit origins of fast network oscillations in the mammalian main olfactory bulb. *eLife*, 10, 2021.

[8637] Joseph D Butner, Geoffrey V Martin, Zhihui Wang, Bruna Corradetti, Mauro Ferrari, Nestor Esnaola, Caroline Chung, David S Hong, James W Welsh, Naomi Hasegawa, Elizabeth A Mittendorf, Steven A Curley, Shu-Hsia Chen, Ping-Ying Pan, Steven K Libutti, Shridar Ganesan, Richard L Sidman, Renata Pasqualini, Wadih Arap, Eugene J Koay, and Vittorio Cristini. Early prediction of clinical response to checkpoint inhibitor therapy in human solid tumors through mathematical modeling. *eLife*, 10, 2021.

[8638] Terkild B Buus, Alberto Herrera, Ellie Ivanova, Eleni Mimitou, Anthony Cheng, Ramin S Herati, Thales Papagiannakopoulos, Peter Smibert, Niels Odum, and Sergei B Koralov. Improving oligo-conjugated antibody signal in multimodal single-cell analysis. *eLife*, 10, 2021.

[8639] Marie Buysse, Anna Maria Floriano, Yuval Gottlieb, Tiago Nardi, Francesco Comandatore, Emanuela Olivieri, Alessia Giannetto, Ana M Palomar, Benjamin L Makepeace, Chiara Bazzocchi, Alessandra Cafiso, Davide Sassera, and Olivier Duron. A dual endosymbiosis supports nutritional adaptation to hematophagy in the invasive tick Hyalomma marginatum. *eLife*, 10, 2021.

[8640] Filomena Caccavale, Giovanni Annona, Lucie Subirana, Hector Escriva, Stephanie Bertrand, and Salvatore D'Aniello. Crosstalk between nitric oxide and retinoic acid pathways is essential for amphioxus pharynx development. *eLife*, 10, 2021.

[8641] Ana J Caetano, Val Yianni, Ana Volponi, Veronica Booth, Eleanor M D'Agostino, and Paul Sharpe. Defining human mesenchymal and epithelial heterogeneity in response to oral inflammatory disease. *eLife*, 10, 2021.

[8642] Rodrigo Caetano, Yaroslav Ispolatov, and Michael Doebeli. Evolution of diversity in metabolic strategies. *eLife*, 10, 2021.

[8643] Jinfeng Cai, Hongbo Gao, Jiacong Zhao, Shujing Hu, Xinyu Liang, Yanyan Yang, Zhuanglin Dai, Zhongsi Hong, and Kai Deng. Infection with a newly designed dual fluorescent reporter HIV-1 effectively identifies latently infected CD4+ T cells. *eLife*, 10, 2021.

[8644] Xinying Cai and Camillo Padoa-Schioppa. Neuronal activity in dorsal anterior cingulate cortex during economic choices under variable action costs. *eLife*, 10, 2021.

[8645] Christophe Caillat, Delphine Guilligay, Johana Torralba, Nikolas Friedrich, Jose L Nieva, Alexandra Trkola, Christophe J Chipot, FranÇois L Dehez, and Winfried Weissenhorn. Structure of HIV-1 gp41 with its membrane anchors targeted by neutralizing antibodies. *eLife*, 10, 2021.

[8646] Joaquín Calatayud, Magnus Neuman, Alexis Rojas, Anton Eriksson, and Martin Rosvall. Regularities in species' niches reveal the world's climate regions. *eLife*, 10, 2021.

[8647] Marcial Camacho, Bradley Quade, Thorsten Trimbuch, Junjie Xu, Levent Sari, Josep Rizo, and Christian Rosenmund. Control of neurotransmitter release by two distinct membrane-binding faces of the Munc13-1 C1C2B region. *eLife*, 10, 2021.

[8648] Ewan Cameron, Alyssa J Young, Katherine A Twohig, Emilie Pothin, Darlene Bhavnani, Amber Dismer, Jean Baptiste Merilien, Karen Hamre, Phoebe Meyer, Arnaud Le Menach, Justin M Cohen, Samson Marseille, Jean Frantz Lemoine, Marc-Aurèle Telfort, Michelle A Chang, Kimberly Won, Alaine Knipes, Eric Rogier, Punam Amratia, Daniel J Weiss, Peter W Gething, and Katherine E Battle. Mapping the endemicity and seasonality of clinical malaria for intervention targeting in Haiti using routine case data. *eLife*, 10, 2021.

[8649] Sydney Campbell, Clementina Mesaros, Luke Izzo, Hayley Affronti, Michael Noji, Bethany E Schaffer, Tiffany Tsang, Kathryn Sun, Sophie Trefely, Salisa Kruijning, John Blenis, Ian A Blair, and Kathryn E Wellen. Glutamine deprivation triggers NAGK-dependent hexosamine salvage. *eLife*, 10, 2021.

[8650] Cesar P Canales, Myka L Estes, Karol Cichewicz, Kartik Angara, John Paul Aboubechara, Scott Cameron, Kathryn Prendergast, Linda Su-Feher, Iva Zdilar, Ellie J Kreun, Emma C Connolly, Jin Myeong Seo, Jack B Goon, Kathleen Farrelly, Tyler W Stradleigh, Deborah van der List, Lori Haapanen, Judy Van de Water, Daniel Vogt, A Kimberley McAllister, and Alex S Nord. Sequential perturbations to mouse corticogenesis following in utero maternal immune activation. *eLife*, 10, 2021.

[8651] Robin Cao, Alexander Pastukhov, Stepan Aleshin, Maurizio Mattia, and Jochen Braun. Binocular rivalry reveals an out-of-equilibrium neural dynamics suited for decision-making. *eLife*, 10, 2021.

[8652] Jesusa Capera, Mireia Pérez-Verdaguer, Roberta Peruzzo, María Navarro-Pérez, Juan Martínez-Pinna, Armando Alberola-Die, Andrés Morales, Luigi Leanza, Ildiko Szabó, and Antonio Felipe. A novel mitochondrial Kv1.3–caveolin axis controls cell survival and apoptosis. *eLife*, 10, 2021.

[8653] Jean-Pascal Capp, James DeGregori, Aurora M Nedelcu, Antoine M Dujon, Justine Boutry, Pascal Pujol, Catherine Alix-Panabières, Rodrigo Hamede, Benjamin Roche, Beata Ujvari, Andriy Marusyk, Robert Gatenby, and Frédéric Thomas. Group phenotypic composition in cancer. *eLife*, 10, 2021.

[8654] Tümay Capraz, Nikolaus F Kienzl, Elisabeth Laurent, Jan W Perthold, Esther Föderl-Höbenreich, Clemens Grünwald-Gruber, Daniel Maresch, Vanessa Monteil, Janine Niederhöfer, Gerald Wirnsberger, Ali Mirazimi, Kurt Zatloukal, Lukas Mach, Josef M Penninger, Chris Oostenbrink, and Johannes Stadlmann. Structure-guided glyco-engineering of ACE2 for improved potency as soluble SARS-CoV-2 decoy receptor. *eLife*, 10, 2021.

[8655] Riccardo Caramellino, Eugenio Piasini, Andrea Buccellato, Anna Carboncino, Vijay Balasubramanian, and Davide Zoccolan. Rat sensitivity to multipoint statistics is predicted by efficient coding of natural scenes. *eLife*, 10, 2021.

[8656] Alessia Caramello, Christophe Galichet, Karine Rizzoti, and Robin Lovell-Badge. Dentate gyrus development requires a cortical hem-derived astrocytic scaffold. *eLife*, 10, 2021.

[8657] Romain Cardis, Sandro Lecci, Laura MJ Fernandez, Alejandro Osorio-Forero, Paul Chu Sin Chung, Stephany Fulda, Isabelle Decosterd, and Anita Lüthi. Cortico-autonomic local arousals and heightened somatosensory arousability during NREMS of mice in neuropathic pain. *eLife*, 10, 2021.

[8658] Emma Carley, Rachel M Stewart, Abigail Zieman, Iman Jalilian, Diane E King, Amanda Zubek, Samantha Lin, Valerie Horsley, and Megan C King. The LINC complex transmits integrin-dependent tension to the nuclear lamina and represses epidermal differentiation. *eLife*, 10, 2021.

[8659] Amado Carreras-Sureda, Laurence Abrami, Kim Ji-Hee, Wen-An Wang, Christopher Henry, Maud Frieden, Monica Didier, F Gisou van der Goot, and Nicolas Demaurex. S-acylation by ZDHHC20 targets ORAI1 channels to lipid rafts for efficient Ca2+ signaling by Jurkat T cell receptors at the immune synapse. *eLife*, 10, 2021.

[8660] Brian A Cary and Gina G Turrigiano. Stability of neocortical synapses across sleep and wake states during the critical period in rats. *eLife*, 10, 2021.

[8661] Arturo Casadevall, Quigly Dragotakes, Patrick W Johnson, Jonathon W Senefeld, Stephen A Klassen, R Scott Wright, Michael J Joyner, Nigel Paneth, and Rickey E Carter. Convalescent plasma use in the USA was inversely correlated with COVID-19 mortality. *eLife*, 10, 2021.

[8662] Melanie Castro-Mollo, Sakshi Gera, Marc Ruiz-Martinez, Maria Feola, Anisa Gumerova, Marina Planoutene, Cara Clementelli, Veena Sangkhae, Carla Casu, Se-Min Kim, Vaughn Ostland, Huiling Han, Elizabeta Nemeth, Robert Fleming, Stefano Rivella, Daria Lizneva, Tony Yuen, Mone Zaidi, and Yelena Ginzburg. The hepcidin regulator erythroferrone is a new member of the erythropoiesis-iron-bone circuitry. *eLife*, 10, 2021.

[8663] Davide Cavalieri, Alexandra Angelova, Anas Islah, Catherine Lopez, Marco Bocchio, Yannick Bollmann, Agnès Baude, and Rosa Cossart. CA1 pyramidal cell diversity is rooted in the time of neurogenesis. *eLife*, 10, 2021.

[8664] Jack F Cazet, Adrienne Cho, and Celina E Juliano. Generic injuries are sufficient to induce ectopic Wnt organizers in Hydra. *eLife*, 10, 2021.

[8665] Arantxa Cebrian-Silla, Marcos Assis Nascimento, Stephanie A Redmond, Benjamin Mansky, David Wu, Kirsten Obernier, Ricardo Romero Rodriguez, Susana Gonzalez-Granero, Jose Manuel García-Verdugo, Daniel A Lim, and Arturo álvarez Buylla. Single-cell analysis of the ventricular-subventricular zone reveals signatures of dorsal and ventral adult neurogenesis. *eLife*, 10, 2021.

[8666] Evan Cesanek, Zhaoran Zhang, James N Ingram, Daniel M Wolpert, and J Randall Flanagan. Motor memories of object dynamics are categorically organized. *eLife*, 10, 2021.

[8667] Rahul Chadda, Nathan Bernhardt, Elizabeth G Kelley, Susana CM Teixeira, Kacie Griffith, Alejandro Gil-Ley, Tuğba N öztürk, Lauren E Hughes, Ana Forsythe, Venkatramanan Krishnamani, José D Faraldo-Gómez, and Janice L Robertson. Membrane transporter dimerization driven by differential lipid solvation energetics of dissociated and associated states. *eLife*, 10, 2021.

[8668] Boris P Chagnaud, Jonathan T Perelmuter, Paul M Forlano, and Andrew H Bass. Gap junction-mediated glycinergic inhibition ensures precise temporal patterning in vocal behavior. *eLife*, 10, 2021.

[8669] Jinwei Chai, Xin Chen, Tiaofei Ye, Baishuang Zeng, Qingye Zeng, Jiena Wu, Barbora Kascakova, Larissa Almeida Martins, Tatyana Prudnikova, Ivana Kuta Smatanova, Michail Kotsyfakis, and Xueqing Xu. Characterization and functional analysis of cathelicidin-MH, a novel frog-derived peptide with anti-septicemic properties. *eLife*, 10, 2021.

[8670] Kasturi Chakraborty, Palapuravan Anees, Sunaina Surana, Simona Martin, Jihad Aburas, Sandrine Moutel, Franck Perez, Sandhya P Koushika, Paschalis Kratsios, and Yamuna Krishnan. Tissue-specific targeting of DNA nanodevices in a multicellular living organism. *eLife*, 10, 2021.

[8671] Diptiman Chanda, Mohammad Rehan, Samuel R Smith, Kevin G Dsouza, Yong Wang, Karen Bernard, Deepali Kurundkar, Vinayak Memula, Kyoko Kojima, James A Mobley, Gloria A Benavides, Victor Darley-Usmar, Young-iL Kim, Jaroslaw W Zmijewski, Jessy S Deshane, Stijn De Langhe, and Victor J Thannickal. Mesenchymal stromal cell aging impairs the self-organizing capacity of lung alveolar epithelial stem cells. *eLife*, 10, 2021.

[8672] Viswanathan Chandrasekaran, Nirupa Desai, Nicholas O Burton, Hanting Yang, Jon Price, Eric A Miska, and V Ramakrishnan. Visualizing formation of the active site in the mitochondrial ribosome. *eLife*, 10, 2021.

[8673] Hsiang-Chun Chang, Jason Solomon Shapiro, Xinghang Jiang, Grant Senyei, Teruki Sato, Justin Geier, Konrad T Sawicki, and Hossein Ardehali. Augmenter of liver regeneration regulates cellular iron homeostasis by modulating mitochondrial transport of ATP-binding cassette B8. *eLife*, 10, 2021.

[8674] Kae-Jiun Chang, Ira Agrawal, Anna Vainshtein, Wan Yun Ho, Wendy Xin, Greg Tucker-Kellogg, Keiichiro Susuki, Elior Peles, Shuo-Chien Ling, and Jonah R Chan. TDP-43 maximizes nerve conduction velocity by repressing a cryptic exon for paranodal junction assembly in Schwann cells. *eLife*, 10, 2021.

[8675] Shih-Hsin Chang, Yi-Ching Su, Mien Chang, and Jun-An Chen. MicroRNAs mediate precise control of spinal interneuron populations to exert delicate sensory-to-motor outputs. *eLife*, 10, 2021.

[8676] Dimitrios Chantzichristos, Per-Arne Svensson, Terence Garner, Camilla AM Glad, Brian R Walker, Ragnhildur Bergthorsdottir, Oskar Ragnarsson, Penelope Trimpou, Roland H Stimson, Stina W Borresen, Ulla Feldt-Rasmussen, Per-Anders Jansson, Stanko Skrtic, Adam Stevens, and Gudmundur Johannsson. Identification of human glucocorticoid response markers using integrated multi-omic analysis from a randomized crossover trial. *eLife*, 10, 2021.

[8677] Aniruddha Chattaraj, Michael L Blinov, and Leslie M Loew. The solubility product extends the buffering concept to heterotypic biomolecular condensates. *eLife*, 10, 2021.

[8678] Karina Chaudhari, Madhavi Gorla, Chao Chang, Artur Kania, and Greg J Bashaw. Robo recruitment of the Wave regulatory complex plays an essential and conserved role in midline repulsion. *eLife*, 10, 2021.

[8679] Shivesh Chaudhary, Sol Ah Lee, Yueyi Li, Dhaval S Patel, and Hang Lu. Graphical-model framework for automated annotation of cell identities in dense cellular images. *eLife*, 10, 2021.

[8680] Lakshya Chauhan, Uday Ram, Kishore Hari, and Mohit Kumar Jolly. Topological signatures in regulatory network enable phenotypic heterogeneity in small cell lung cancer. *eLife*, 10, 2021.

[8681] Albert Tian Chen, Kevin Altschuler, Shing Hei Zhan, Yujia Alina Chan, and Benjamin E Deverman. COVID-19 CG enables SARS-CoV-2 mutation and lineage tracking by locations and dates of interest. *eLife*, 10, 2021.

[8682] Bo-Ruei Chen, Yinan Wang, Anthony Tubbs, Dali Zong, Faith C Fowler, Nicholas Zolnerowich, Wei Wu, Amelia Bennett, Chun-Chin Chen, Wendy Feng, Andre Nussenzweig, Jessica K Tyler, and Barry P Sleckman. LIN37-DREAM prevents DNA end resection and homologous recombination at DNA double-strand breaks in quiescent cells. *eLife*, 10, 2021.

[8683] Cathy S Chen, Evan Knep, Autumn Han, R Becket Ebitz, and Nicola M Grissom. Sex differences in learning from exploration. *eLife*, 10, 2021.

[8684] Chang-Rui Chen, Yu-Heng Zhong, Shan Jiang, Wei Xu, Lei Xiao, Zan Wang, Wei-Min Qu, and Zhi-Li Huang. Dysfunctions of the paraventricular hypothalamic nucleus induce hypersomnia in mice. *eLife*, 10, 2021.

[8685] Chen Chen, Hongyi Chen, Shijie Huang, Taoshan Jiang, Chuanhong Wang, Zhen Tao, Chen He, Qingfeng Tang, and Peijin Li. Volatile DMNT directly protects plants against Plutella xylostella by disrupting the peritrophic matrix barrier in insect midgut. *eLife*, 10, 2021.

[8686] Gukui Chen, Jiashen Zhou, Yili Zuo, Weiping Huo, Juan Peng, Meng Li, Yani Zhang, Tietao Wang, Lin Zhang, Liang Zhang, and Haihua Liang. Structural basis for diguanylate cyclase activation by its binding partner in Pseudomonas aeruginosa. *eLife*, 10, 2021.

[8687] Jie Chen, Sihui Jin, Dandan Chen, Jie Cao, Xiaoxiao Ji, Qionglin Peng, and Yufeng Pan. fruitless tunes functional flexibility of courtship circuitry during development. *eLife*, 10, 2021.

[8688] Kepu Chen, Yuting Ye, Nikolaus F Troje, and Wen Zhou. Oxytocin modulates human chemosensory decoding of sex in a dose-dependent manner. *eLife*, 10, 2021.

[8689] Lu Chen, Jun-Jie Gu, Qiang Yang, Dong Ren, Alexander Blanke, and Olivier Béthoux. Ovipositor and mouthparts in a fossil insect support a novel ecological role for early orthopterans in 300 million years old forests. *eLife*, 10, 2021.

[8690] Min Chen, Fangfang Dong, Min Chen, Zhiming Shen, Haowei Wu, Changhuo Cen, Xiuhong Cui, Shilai Bao, and Fei Gao. PRMT5 regulates ovarian follicle development by facilitating Wt1 translation. *eLife*, 10, 2021.

[8691] Paul Z Chen, Niklas Bobrovitz, Zahra Premji, Marion Koopmans, David N Fisman, and Frank X Gu. Heterogeneity in transmissibility and shedding SARS-CoV-2 via droplets and aerosols. *eLife*, 10, 2021.

[8692] Paul Z Chen, Niklas Bobrovitz, Zahra A Premji, Marion Koopmans, David N Fisman, and Frank X Gu. SARS-CoV-2 shedding dynamics across the respiratory tract, sex, and disease severity for adult and pediatric COVID-19. *eLife*, 10, 2021.

[8693] Qian Chen, Yuyan Liu, Jiping Ren, Panpan Zhong, Manni Chen, Dongsheng Jia, Hongyan Chen, and Taiyun Wei. Exosomes mediate horizontal transmission of viral pathogens from insect vectors to plant phloem. *eLife*, 10, 2021.

[8694] Weijie Chen, Neha Mani, Hamid Karani, Hao Li, Sridhar Mani, and Jay X Tang. Confinement discerns swarmers from planktonic bacteria. *eLife*, 10, 2021.

[8695] Xiangrong Chen, Yusuf I Ali, Charlotte EL Fisher, Raquel Arribas-Bosacoma, Mohan B Rajasekaran, Gareth Williams, Sarah Walker, Jessica R Booth, Jessica JR Hudson, S Mark Roe, Laurence H Pearl, Simon E Ward, Frances MG Pearl, and Antony W Oliver. Uncovering an allosteric mode of action for a selective inhibitor of human Bloom syndrome protein. *eLife*, 10, 2021.

[8696] Xue Chen, Haidong Yao, Muhammad Kashif, Gwladys Revêchon, Maria Eriksson, Jianjiang Hu, Ting Wang, Yiran Liu, Elin Tüksammel, Staffan Strömblad, Ian M Ahearn, Mark R Philips, Clotilde Wiel, Mohamed X Ibrahim, and Martin O Bergo. A small-molecule ICMT inhibitor delays senescence of Hutchinson-Gilford progeria syndrome cells. *eLife*, 10, 2021.

[8697] Yanan Chen, Rejani B Kunjamma, Molly Weiner, Jonah R Chan, and Brian Popko. Prolonging the integrated stress response enhances CNS remyelination in an inflammatory environment. *eLife*, 10, 2021.

[8698] Yao Chen, Weiya Xu, Shuwei Yu, Kang Ni, Guangbiao She, Xiaodong Ye, Qiong Xing, Jian Zhao, and Chengdong Huang. Assembly status transition offers an avenue for activity modulation of a supramolecular enzyme. *eLife*, 10, 2021.

[8699] Zhongwen Chen, Dongmyung Oh, Kabir Hassan Biswas, Ronen Zaidel-Bar, and Jay T Groves. Probing the effect of clustering on EphA2 receptor signaling efficiency by subcellular control of ligand-receptor mobility. *eLife*, 10, 2021.

[8700] Luqi Cheng, Yuanchao Zhang, Gang Li, Jiaojian Wang, Chet Sherwood, Gaolang Gong, Lingzhong Fan, and Tianzi Jiang. Connectional asymmetry of the inferior parietal lobule shapes hemispheric specialization in humans, chimpanzees, and rhesus macaques. *eLife*, 10, 2021.

[8701] Shuhua Cheng, Wei Zhang, Giorgio Inghirami, and Wayne Tam. Mutation analysis links angioimmunoblastic T-cell lymphoma to clonal hematopoiesis and smoking. *eLife*, 10, 2021.

[8702] Xiaohang Cheng and Magdalena Bezanilla. SABRE populates ER domains essential for cell plate maturation and cell expansion influencing cell and tissue patterning. *eLife*, 10, 2021.

[8703] Jingyi Chi, Zeran Lin, William Barr, Audrey Crane, Xiphias Ge Zhu, and Paul Cohen. Early postnatal interactions between beige adipocytes and sympathetic neurites regulate innervation of subcutaneous fat. *eLife*, 10, 2021.

[8704] Chin-Hsuan Chia, Xin-Wei Tang, Yue Cao, Hua-Teng Cao, Wei Zhang, Jun-Fa Wu, Yu-Lian Zhu, Ying Chen, Yi Lin, Yi Wu, Zhe Zhang, Ti-Fei Yuan, and Rui-Ping Hu. Cortical excitability signatures for the degree of sleepiness in human. *eLife*, 10, 2021.

[8705] Joanne Chia, Shyi-Chyi Wang, Sheena Wee, David James Gill, Felicia Tay, Srinivasaraghavan Kannan, Chandra S Verma, Jayantha Gunaratne, and Frederic A Bard. Src activates retrograde membrane traffic through phosphorylation of GBF1. *eLife*, 10, 2021.

[8706] Myron Barber Child, Jack R Bateman, Amir Jahangiri, Armando Reimer, Nicholas C Lammers, Nica Sabouni, Diego Villamarin, Grace C McKenzie-Smith, Justine E Johnson, Daniel Jost, and Hernan G Garcia. Live imaging and biophysical modeling support a button-based mechanism of somatic homolog pairing in Drosophila. *eLife*, 10, 2021.

[8707] Florence WJ Chioh, Siew-Wai Fong, Barnaby E Young, Kan-Xing Wu, Anthony Siau, Shuba Krishnan, Yi-Hao Chan, Guillaume Carissimo, Louis LY Teo, Fei Gao, Ru San Tan, Liang Zhong, Angela S Koh, Seow-Yen Tan, Paul A Tambyah, Laurent Renia, Lisa FP Ng, David C Lye, and Christine Cheung. Convalescent COVID-19 patients are susceptible to endothelial dysfunction due to persistent immune activation. *eLife*, 10, 2021.

[8708] Jian-geng Chiou, Kyle D Moran, and Daniel J Lew. How cells determine the number of polarity sites. *eLife*, 10, 2021.

[8709] Peter H Chipman, Chi Chung Alan Fung, Alejandra Pazo Fernandez, Abhilash Sawant, Angelo Tedoldi, Atsushi Kawai, Sunita Ghimire Gautam, Mizuki Kurosawa, Manabu Abe, Kenji Sakimura, Tomoki Fukai, and Yukiko Goda. Astrocyte GluN2C NMDA receptors control basal synaptic strengths of hippocampal CA1 pyramidal neurons in the stratum radiatum. *eLife*, 10, 2021.

[8710] Charlotte P Choi, Rebecca J Tay, Margaret R Starostik, Suhua Feng, James J Moresco, Brooke E Montgomery, Emily Xu, Maya A Hammonds, Michael C Schatz, Taiowa A Montgomery, John R Yates, Steven E Jacobsen, and John K Kim. SNPC-1.3 is a sex-specific transcription factor that drives male piRNA expression in C. elegans. *eLife*, 10, 2021.

[8711] Jaemyung Choi, David B Lyons, and Daniel Zilberman. Histone H1 prevents non-CG methylation-mediated small RNA biogenesis in Arabidopsis heterochromatin. *eLife*, 10, 2021.

[8712] Jung-Hwa Choi, Erik R Duboue, Michelle Macurak, Jean-Michel Chanchu, and Marnie E Halpern. Specialized neurons in the right habenula mediate response to aversive olfactory cues. *eLife*, 10, 2021.

[8713] Pavan Choppakatla, Bastiaan Dekker, Erin E Cutts, Alessandro Vannini, Job Dekker, and Hironori Funabiki. Linker histone H1.8 inhibits chromatin binding of condensins and DNA topoisomerase II to tune chromosome length and individualization. *eLife*, 10, 2021.

[8714] Shu-Min Chou, Ke-Xin Li, Ming-Yueh Huang, Chao Chen, Yuan-Hung Lin King, Grant Guangnan Li, Wei Zhou, Chin Fen Teo, Yuh Nung Jan, Lily Yeh Jan, and Shi-Bing Yang. Kv1.1 channels regulate early postnatal neurogenesis in mouse hippocampus via the TrkB signaling pathway. *eLife*, 10, 2021.

[8715] Jenna R Christensen, Agnieszka A Kendrick, Joey B Truong, Adriana Aguilar-Maldonado, Vinit Adani, Monika Dzieciatkowska, and Samara L Reck-Peterson. Cytoplasmic dynein-1 cargo diversity is mediated by the combinatorial assembly of FTS–Hook–FHIP complexes. *eLife*, 10, 2021.

[8716] Katherine Christian, Carolyn Johnstone, Jo-ann Larkins, Wendy Wright, and Michael R Doran. A survey of early-career researchers in Australia. *eLife*, 10, 2021.

[8717] Chanwoo Chun, Tirthabir Biswas, and Vikas Bhandawat. Drosophila uses a tripod gait across all walking speeds, and the geometry of the tripod is important for speed control. *eLife*, 10, 2021.

[8718] Timothy W Church, Parul Tewatia, Saad Hannan, João Antunes, Olivia Eriksson, Trevor G Smart, Jeanette Hellgren Kotaleski, and Matthew G Gold. AKAP79 enables calcineurin to directly suppress protein kinase A activity. *eLife*, 10, 2021.

[8719] Elisa Ciaramelli, Flavia De Luca, Donna Kwan, Jenkin Mok, Francesca Bianconi, Violetta Knyagnytska, Carl Craver, Leonard Green, Joel Myerson, and Shayna Rosenbaum. The role of ventromedial prefrontal cortex in reward valuation and future thinking during intertemporal choice. *eLife*, 10, 2021.

[8720] Giancarlo Cicconofri, Giovanni Noselli, and Antonio DeSimone. The biomechanical role of extra-axonemal structures in shaping the flagellar beat of Euglena gracilis. *eLife*, 10, 2021.

[8721] Patricia Giselle Cipriani, Olivia Bay, John Zinno, Michelle Gutwein, Hin Hark Gan, Vinay K Mayya, George Chung, Jia-Xuan Chen, Hala Fahs, Yu Guan, Thomas F Duchaine, Matthias Selbach, Fabio Piano, and Kristin C Gunsalus. Novel LOTUS-domain proteins are organizational hubs that recruit C. elegans Vasa to germ granules. *eLife*, 10, 2021.

[8722] Andrea Cipriano, Martina Macino, Giulia Buonaiuto, Tiziana Santini, Beatrice Biferali, Giovanna Peruzzi, Alessio Colantoni, Chiara Mozzetta, and Monica Ballarino. Epigenetic regulation of Wnt7b expression by the cis-acting long noncoding RNA Lnc-Rewind in muscle stem cells. *eLife*, 10, 2021.

[8723] Maria-Daniela Cirnaru, Sicheng Song, Kizito-Tshitoko Tshilenge, Chuhyon Corwin, Justyna Mleczko, Carlos Galicia Aguirre, Houda Benlhabib, Jaroslav Bendl, Pasha Apontes, John Fullard, Jordi Creus-Muncunill, Azadeh Reyahi, Ali M Nik, Peter Carlsson, Panos Roussos, Sean D Mooney, Lisa M Ellerby, and Michelle E Ehrlich. Unbiased identification of novel transcription factors in striatal compartmentation and striosome maturation. *eLife*, 10, 2021.

[8724] Deniz Cizmeci, Giuseppe Lofano, Evan Rossignol, Anne-Sophie Dugast, Dongkyoon Kim, Guy Cavet, Ngan Nguyen, Yann Chong Tan, Michael S Seaman, Galit Alter, and Boris Julg. Distinct clonal evolution of B-cells in HIV controllers with neutralizing antibody breadth. *eLife*, 10, 2021.

[8725] Joseph T Clark, David A Christian, Jodi A Gullicksrud, Joseph A Perry, Jeongho Park, Maxime Jacquet, James C Tarrant, Enrico Radaelli, Jonathan Silver, and Christopher A Hunter. IL-33 promotes innate lymphoid cell-dependent IFN-$\gamma$ production required for innate immunity to Toxoplasma gondii. *eLife*, 10, 2021.

[8726] Federico Claudi, Adam L Tyson, Luigi Petrucco, Troy W Margrie, Ruben Portugues, and Tiago Branco. Visualizing anatomically registered data with brainrender. *eLife*, 10, 2021.

[8727] Colin M Cleary, Brenda M Milla, Fu-Shan Kuo, Shaun James, William F Flynn, Paul Robson, and Daniel K Mulkey. Somatostatin-expressing parafacial neurons are CO2/H+ sensitive and regulate baseline breathing. *eLife*, 10, 2021.

[8728] Jan Clemens, Stefan Schöneich, Konstantinos Kostarakos, R Matthias Hennig, and Berthold Hedwig. A small, computationally flexible network produces the phenotypic diversity of song recognition in crickets. *eLife*, 10, 2021.

[8729] Zachary Clemens, Sruthi Sivakumar, Abish Pius, Amrita Sahu, Sunita Shinde, Hikaru Mamiya, Nathaniel Luketich, Jian Cui, Purushottam Dixit, Joerg D Hoeck, Sebastian Kreuz, Michael Franti, Aaron Barchowsky, and Fabrisia Ambrosio. The biphasic and age-dependent impact of klotho on hallmarks of aging and skeletal muscle function. *eLife*, 10, 2021.

[8730] Justine C Cléry, Yuki Hori, David J Schaeffer, Ravi S Menon, and Stefan Everling. Neural network of social interaction observation in marmosets. *eLife*, 10, 2021.

[8731] Steven J Clipman, Shruti H Mehta, Aylur K Srikrishnan, Katie JC Zook, Priya Duggal, Shobha Mohapatra, Saravanan Shanmugam, Paneerselvam Nandagopal, Muniratnam S Kumar, Elizabeth Ogburn, Gregory M Lucas, Carl A Latkin, and Sunil S Solomon. Role of direct and indirect social and spatial ties in the diffusion of HIV and HCV among people who inject drugs: a cross-sectional community-based network analysis in New Delhi, India. *eLife*, 10, 2021.

[8732] Jonathon Alexis Coates, Elliot Brooks, Amy Louise Brittle, Emma Louise Armitage, Martin Peter Zeidler, and Iwan Robert Evans. Identification of functionally distinct macrophage subpopulations in Drosophila. *eLife*, 10, 2021.

[8733] Maria M Cobo, Caroline Hartley, Deniz Gursul, Foteini Andritsou, Marianne van der Vaart, Gabriela Schmidt Mellado, Luke Baxter, Eugene P Duff, Miranda Buckle, Ria Evans Fry, Gabrielle Green, Amy Hoskin, Richard Rogers, Eleri Adams, Fiona Moultrie, and Rebeccah Slater. Quantifying noxious-evoked baseline sensitivity in neonates to optimise analgesic trials. *eLife*, 10, 2021.

[8734] Olivier Cochet-Escartin, Mete Demircigil, Satomi Hirose, Blandine Allais, Philippe Gonzalo, Ivan Mikaelian, Kenichi Funamoto, Christophe Anjard, Vincent Calvez, and Jean-Paul Rieu. Hypoxia triggers collective aerotactic migration in Dictyostelium discoideum. *eLife*, 10, 2021.

[8735] Alexandra Colin, Gabriele Micali, Louis Faure, Marco Cosentino Lagomarsino, and Sven van Teeffelen. Two different cell-cycle processes determine the timing of cell division in Escherichia coli. *eLife*, 10, 2021.

[8736] Ewan Colman, Vittoria Colizza, Ephraim M Hanks, David P Hughes, and Shweta Bansal. Social fluidity mobilizes contagion in human and animal populations. *eLife*, 10, 2021.

[8737] Marco Colnaghi, Andrew Pomiankowski, and Nick Lane. The need for high-quality oocyte mitochondria at extreme ploidy dictates mammalian germline development. *eLife*, 10, 2021.

[8738] Alessio Vittorio Colombo, Rebecca Katie Sadler, Gemma Llovera, Vikramjeet Singh, Stefan Roth, Steffanie Heindl, Laura Sebastian Monasor, Aswin Verhoeven, Finn Peters, Samira Parhizkar, Frits Kamp, Mercedes Gomez de Aguero, Andrew J MacPherson, Edith Winkler, Jochen Herms, Corinne Benakis, Martin Dichgans, Harald Steiner, Martin Giera, Christian Haass, Sabina Tahirovic, and Arthur Liesz. Microbiota-derived short chain fatty acids modulate microglia and promote A$\beta$ plaque deposition. *eLife*, 10, 2021.

[8739] Megan M Colonnetta, Lauren R Lym, Lillian Wilkins, Gretchen Kappes, Elias A Castro, Pearl V Ryder, Paul Schedl, Dorothy A Lerit, and Girish Deshpande. Antagonism between germ cell-less and Torso receptor regulates transcriptional quiescence underlying germline/soma distinction. *eLife*, 10, 2021.

[8740] Jordi Comelles, Soumya SS, Linjie Lu, Emilie Le Maout, S Anvitha, Guillaume Salbreux, Frank Jülicher, Mandar M Inamdar, and Daniel Riveline. Epithelial colonies in vitro elongate through collective effects. *eLife*, 10, 2021.

[8741] María J Conde-Dusman, Partha N Dey, óscar Elía-Zudaire, Luis G Rabaneda, Carmen García-Lira, Teddy Grand, Victor Briz, Eric R Velasco, Raül Andero, Sergio Niñerola, Angel Barco, Pierre Paoletti, John F Wesseling, Fabrizio Gardoni, Steven J Tavalin, and Isabel Perez-Otaño. Control of protein synthesis and memory by GluN3A-NMDA receptors through inhibition of GIT1/mTORC1 assembly. *eLife*, 10, 2021.

[8742] Ian Cone and Harel Z Shouval. Learning precise spatiotemporal sequences via biophysically realistic learning rules in a modular, spiking network. *eLife*, 10, 2021.

[8743] Sonia Coni, Federica A Falconio, Marta Marzullo, Marzia Munafò, Benedetta Zuliani, Federica Mosti, Alessandro Fatica, Zaira Ianniello, Rosa Bordone, Alberto Macone, Enzo Agostinelli, Alessia Perna, Tanja Matkovic, Stephan Sigrist, Gabriella Silvestri, Gianluca Canettieri, and Laura Ciapponi. Translational control of polyamine metabolism by CNBP is required for Drosophila locomotor function. *eLife*, 10, 2021.

[8744] Federica Contò, Grace Edwards, Sarah Tyler, Danielle Parrott, Emily Grossman, and Lorella Battelli. Attention network modulation via tRNS correlates with attention gain. *eLife*, 10, 2021.

[8745] Guy Alexander Cooper, Hadleigh Frost, Ming Liu, and Stuart Andrew West. The evolution of division of labour in structured and unstructured groups. *eLife*, 10, 2021.

[8746] Brendan Cornwell, Katrina Armstrong, Nia S Walker, Marilla Lippert, Victor Nestor, Yimnang Golbuu, and Stephen R Palumbi. Widespread variation in heat tolerance and symbiont load are associated with growth tradeoffs in the coral Acropora hyacinthus in Palau. *eLife*, 10, 2021.

[8747] Aurelio Cortese, Asuka Yamamoto, Maryam Hashemzadeh, Pradyumna Sepulveda, Mitsuo Kawato, and Benedetto De Martino. Value signals guide abstraction during learning. *eLife*, 10, 2021.

[8748] Kevin R Costello, Amy Leung, Candi Trac, Michael Lee, Mudaser Basam, J Andrew Pospisilik, and Dustin E Schones. Sequence features of retrotransposons allow for epigenetic variability. *eLife*, 10, 2021.

[8749] Tyler Couch, Kyle D Berger, Dana L Kneisley, Tyler W McCullock, Paul Kammermeier, and David M Maclean. Topography and motion of acid-sensing ion channel intracellular domains. *eLife*, 10, 2021.

[8750] Robert Coukos, David Yao, Mateo I Sanchez, Eric T Strand, Meagan E Olive, Namrata D Udeshi, Jonathan S Weissman, Steven A Carr, Michael C Bassik, and Alice Y Ting. An engineered transcriptional reporter of protein localization identifies regulators of mitochondrial and ER membrane protein trafficking in high-throughput CRISPRi screens. *eLife*, 10, 2021.

[8751] Lisa I Couper, Johannah E Farner, Jamie M Caldwell, Marissa L Childs, Mallory J Harris, Devin G Kirk, Nicole Nova, Marta Shocket, Eloise B Skinner, Lawrence H Uricchio, Moises Exposito-Alonso, and Erin A Mordecai. How will mosquitoes adapt to climate warming? *eLife*, 10, 2021.

[8752] Jamie L Courtland, Tyler WA Bradshaw, Greg Waitt, Erik J Soderblom, Tricia Ho, Anna Rajab, Ricardo Vancini, Il Hwan Kim, and Scott H Soderling. Genetic disruption of WASHC4 drives endo-lysosomal dysfunction and cognitive-movement impairments in mice and humans. *eLife*, 10, 2021.

[8753] Louise Cramer. Alternative strategies for closing the award gap between white and minority ethnic students. *eLife*, 10, 2021.

[8754] Blake A Creighton, Simone Afriyie, Deepa Ajit, Cristine R Casingal, Kayleigh M Voos, Joan Reger, April M Burch, Eric Dyne, Julia Bay, Jeffrey K Huang, ES Anton, Meng-Meng Fu, and Damaris N Lorenzo. Giant ankyrin-B mediates transduction of axon guidance and collateral branch pruning factor sema 3A. *eLife*, 10, 2021.

[8755] Stephanie E Crilly, Wooree Ko, Zara Y Weinberg, and Manojkumar A Puthenveedu. Conformational specificity of opioid receptors is determined by subcellular location irrespective of agonist. *eLife*, 10, 2021.

[8756] Emanuela Cristiani, Anita Radini, Andrea Zupancich, Angelo Gismondi, Alessia D'Agostino, Claudio Ottoni, Marialetizia Carra, Snežana Vukojičić, Mihai Constantinescu, Dragana Antonović, T Douglas Price, and DuŠan Borić. Wild cereal grain consumption among Early Holocene foragers of the Balkans predates the arrival of agriculture. *eLife*, 10, 2021.

[8757] Christine E Cucinotta, Rachel H Dell, Keean CA Braceros, and Toshio Tsukiyama. RSC primes the quiescent genome for hypertranscription upon cell-cycle re-entry. *eLife*, 10, 2021.

[8758] Emanuel Cura Costa, Leo Otsuki, Aida Rodrigo Albors, Elly M Tanaka, and Osvaldo Chara. Spatiotemporal control of cell cycle acceleration during axolotl spinal cord regeneration. *eLife*, 10, 2021.

[8759] Rafael Custodio, Rhian M Ford, Cara J Ellison, Guangyu Liu, Gerda Mickute, Christoph M Tang, and Rachel M Exley. Type VI secretion system killing by commensal Neisseria is influenced by expression of type four pili. *eLife*, 10, 2021.

[8760] Fabio Da Silva, Fariba Jian Motamedi, Lahiru Chamara Weerasinghe Arachchige, Amelie Tison, Stephen T Bradford, Jonathan Lefebvre, Pascal Dolle, Norbert B Ghyselinck, Kay D Wagner, and Andreas Schedl. Retinoic acid signaling is directly activated in cardiomyocytes and protects mouse hearts from apoptosis after myocardial infarction. *eLife*, 10, 2021.

[8761] Torsten Dahlén, Mark Clements, Jingcheng Zhao, Martin L Olsson, and Gustaf Edgren. An agnostic study of associations between ABO and RhD blood group and phenome-wide disease risk. *eLife*, 10, 2021.

[8762] Jian Dai, Ying Li, Fuyuki Kametani, Xiaoran Cui, Yuichi Igarashi, Jia Huo, Hiroki Miyahara, Masayuki Mori, and Keiichi Higuchi. Curcumin promotes AApoAII amyloidosis and peroxisome proliferation in mice by activating the PPAR$\alpha$ signaling pathway. *eLife*, 10, 2021.

[8763] Martina Dal Bello, Alfonso Pérez-Escudero, Frank C Schroeder, and Jeff Gore. Inversion of pheromone preference optimizes foraging in C. elegans. *eLife*, 10, 2021.

[8764] Jasbir Singh Dalal, Kellen Diamond Winden, Catherine Lourdes Salussolia, Maria Sundberg, Achint Singh, Truc Thanh Pham, Pingzhu Zhou, William T Pu, Meghan T Miller, and Mustafa Sahin. Loss of Tsc1 in cerebellar Purkinje cells induces transcriptional and translation changes in FMRP target transcripts. *eLife*, 10, 2021.

[8765] Saeed Daneshmandi, Teresa Cassel, Richard M Higashi, Teresa W-M Fan, and Pankaj Seth. 6-Phosphogluconate dehydrogenase (6PGD), a key checkpoint in reprogramming of regulatory T cells metabolism and function. *eLife*, 10, 2021.

[8766] Zdravka Daneva, Matteo Ottolini, Yen Lin Chen, Eliska Klimentova, Maniselvan Kuppusamy, Soham A Shah, Richard D Minshall, Cheikh I Seye, Victor E Laubach, Brant E Isakson, and Swapnil K Sonkusare. Endothelial pannexin 1–TRPV4 channel signaling lowers pulmonary arterial pressure in mice. *eLife*, 10, 2021.

[8767] Zharko Daniloski, Tristan X Jordan, Juliana K Ilmain, Xinyi Guo, Gira Bhabha, Benjamin R tenOever, and Neville E Sanjana. The Spike D614G mutation increases SARS-CoV-2 infection of multiple human cell types. *eLife*, 10, 2021.

[8768] Anil K Dasanna, Sebastian Hillringhaus, Gerhard Gompper, and Dmitry A Fedosov. Effect of malaria parasite shape on its alignment at erythrocyte membrane. *eLife*, 10, 2021.

[8769] Noopur Dave, Ugur Cetiner, Daniel Arroyo, Joshua Fonbuena, Megna Tiwari, Patricia Barrera, Noelia Lander, Andriy Anishkin, Sergei Sukharev, and Veronica Jimenez. A novel mechanosensitive channel controls osmoregulation, differentiation, and infectivity in Trypanosoma cruzi. *eLife*, 10, 2021.

[8770] Lloyd Davis, Inja Radman, Angeliki Goutou, Ailish Tynan, Kieran Baxter, Zhiyan Xi, Jack M O'Shea, Jason W Chin, and Sebastian Greiss. Precise optical control of gene expression in C elegans using improved genetic code expansion and Cre recombinase. *eLife*, 10, 2021.

[8771] Anamika Dayal, Monica L Fernández-Quintero, Klaus R Liedl, and Manfred Grabner. Pore mutation N617D in the skeletal muscle DHPR blocks Ca2+ influx due to atypical high-affinity Ca2+ binding. *eLife*, 10, 2021.

[8772] Roberto de Filippo, Benjamin R Rost, Alexander Stumpf, Claire Cooper, John J Tukker, Christoph Harms, Prateep Beed, and Dietmar Schmitz. Somatostatin interneurons activated by 5-HT2A receptor suppress slow oscillations in medial entorhinal cortex. *eLife*, 10, 2021.

[8773] Lorraine De Jesús-Kim, Larry J Friedman, Marko Lõoke, Christian K Ramsoomair, Jeff Gelles, and Stephen P Bell. DDK regulates replication initiation by controlling the multiplicity of Cdc45-GINS binding to Mcm2-7. *eLife*, 10, 2021.

[8774] Rose De Kock, Weiwei Zhou, Wilsaan M Joiner, and Martin Wiener. Slowing the body slows down time perception. *eLife*, 10, 2021.

[8775] Juan Carlos De la Concepcion, Javier Vega Benjumea, Aleksandra Bialas, Ryohei Terauchi, Sophien Kamoun, and Mark J Banfield. Functional diversification gave rise to allelic specialization in a rice NLR immune receptor pair. *eLife*, 10, 2021.

[8776] Saskia MA de Man, Gooitzen Zwanenburg, Tanne van der Wal, Mark A Hink, and Renée van Amerongen. Quantitative live-cell imaging and computational modeling shed new light on endogenous WNT/CTNNB1 signaling dynamics. *eLife*, 10, 2021.

[8777] Marcos H de Moraes, FoSheng Hsu, Dean Huang, Dustin E Bosch, Jun Zeng, Matthew C Radey, Noah Simon, Hannah E Ledvina, Jacob P Frick, Paul A Wiggins, S Brook Peterson, and Joseph D Mougous. An interbacterial DNA deaminase toxin directly mutagenizes surviving target populations. *eLife*, 10, 2021.

[8778] Benedetta De Ponte Conti, Annarita Miluzio, Fabio Grassi, Sergio Abrignani, Stefano Biffo, and Sara Ricciardi. mTOR-dependent translation drives tumor infiltrating CD8+ effector and CD4+ Treg cells expansion. *eLife*, 10, 2021.

[8779] Aurelie de Rus Jacquet, Jenna L Tancredi, Andrew L Lemire, Michael C DeSantis, Wei-Ping Li, and Erin K O'Shea. The LRRK2 G2019S mutation alters astrocyte-to-neuron communication via extracellular vesicles and induces neuron atrophy in a human iPSC-derived model of Parkinson's disease. *eLife*, 10, 2021.

[8780] Dasmanthie De Silva, Lucas Ferguson, Grant H Chin, Benjamin E Smith, Ryan A Apathy, Theodore L Roth, Franziska Blaeschke, Marek Kudla, Alexander Marson, Nicholas T Ingolia, and Jamie HD Cate. Robust T cell activation requires an eIF3-driven burst in T cell receptor translation. *eLife*, 10, 2021.

[8781] Katherine A Deets, Randilea Nichols Doyle, Isabella Rauch, and Russell E Vance. Inflammasome activation leads to cDC1-independent cross-priming of CD8 T cells by epithelial cell-derived antigen. *eLife*, 10, 2021.

[8782] Florian A Dehmelt, Rebecca Meier, Julian Hinz, Takeshi Yoshimatsu, Clara A Simacek, Ruoyu Huang, Kun Wang, Tom Baden, and Aristides B Arrenberg. Spherical arena reveals optokinetic response tuning to stimulus location, size, and frequency across entire visual field of larval zebrafish. *eLife*, 10, 2021.

[8783] Stefanie Deinhardt-Emmer, Daniel Wittschieber, Juliane Sanft, Sandra Kleemann, Stefan Elschner, Karoline Frieda Haupt, Vanessa Vau, Clio Häring, Jürgen Rödel, Andreas Henke, Christina Ehrhardt, Michael Bauer, Mike Philipp, Nikolaus Gaßler, Sandor Nietzsche, Bettina Löffler, and Gita Mall. Early postmortem mapping of SARS-CoV-2 RNA in patients with COVID-19 and the correlation with tissue damage. *eLife*, 10, 2021.

[8784] Teresa del Peso Santos, Laura Alvarez, Brandon Sit, Oihane Irazoki, Jonathon Blake, Benjamin R Warner, Alyson R Warr, Anju Bala, Vladimir Benes, Matthew K Waldor, Kurt Fredrick, and Felipe Cava. BipA exerts temperature-dependent translational control of biofilm-associated colony morphology in Vibrio cholerae. *eLife*, 10, 2021.

[8785] Steven J Del Signore, Charlotte F Kelley, Emily M Messelaar, Tania Lemos, Michelle F Marchan, Biljana Ermanoska, Markus Mund, Thomas G Fai, Marko Kaksonen, and Avital Adah Rodal. An autoinhibitory clamp of actin assembly constrains and directs synaptic endocytosis. *eLife*, 10, 2021.

[8786] Gustavo Della-Flora Nunes, Emma R Wilson, Edward Hurley, Bin He, Bert W O'Malley, Yannick Poitelon, Lawrence Wrabetz, and M Laura Feltri. Activation of mTORC1 and c-Jun by Prohibitin1 loss in Schwann cells may link mitochondrial dysfunction to demyelination. *eLife*, 10, 2021.

[8787] Keith F DeLuca, Jeanne E Mick, Amy H Ide, Wanessa C Lima, Lori Sherman, Kristin L Schaller, Steven M Anderson, Ning Zhao, Timothy J Stasevich, Dileep Varma, Jakob Nilsson, and Jennifer G DeLuca. Generation and diversification of recombinant monoclonal antibodies. *eLife*, 10, 2021.

[8788] Xinna Deng, Huiqing Hou, Xiaoxi Wang, Qingxia Li, Xiuyuan Li, Zhaohua Yang, and Haijiang Wu. Development and validation of a nomogram to better predict hypertension based on a 10-year retrospective cohort study in China. *eLife*, 10, 2021.

[8789] Sven Dennerlein, Sabine Poerschke, Silke Oeljeklaus, Cong Wang, Ricarda Richter-Dennerlein, Johannes Sattmann, Diana Bauermeister, Elisa Hanitsch, Stefan Stoldt, Thomas Langer, Stefan Jakobs, Bettina Warscheid, and Peter Rehling. Defining the interactome of the human mitochondrial ribosome identifies SMIM4 and TMEM223 as respiratory chain assembly factors. *eLife*, 10, 2021.

[8790] Eric L Denovellis, Anna K Gillespie, Michael E Coulter, Marielena Sosa, Jason E Chung, Uri T Eden, and Loren M Frank. Hippocampal replay of experience at real-world speeds. *eLife*, 10, 2021.

[8791] Tanvi Deora, Siddharth S Sane, and Sanjay P Sane. Wings and halteres act as coupled dual oscillators in flies. *eLife*, 10, 2021.

[8792] Astrid Deryckere, Ruth Styfhals, Ali Murat Elagoz, Gregory E Maes, and Eve Seuntjens. Identification of neural progenitor cells and their progeny reveals long distance migration in the developing octopus brain. *eLife*, 10, 2021.

[8793] Mitul Desai, Jitendra Sharma, Adrian L Slusarczyk, Ashley A Chapin, Robert Ohlendorf, Agata Wisniowska, Mriganka Sur, and Alan Jasanoff. Hemodynamic molecular imaging of tumor-associated enzyme activity in the living brain. *eLife*, 10, 2021.

[8794] Kobe Desender, K Richard Ridderinkhof, and Peter R Murphy. Understanding neural signals of post-decisional performance monitoring: An integrative review. *eLife*, 10, 2021.

[8795] Lorenz Deserno, Rani Moran, Jochen Michely, Ying Lee, Peter Dayan, and Raymond J Dolan. Dopamine enhances model-free credit assignment through boosting of retrospective model-based inference. *eLife*, 10, 2021.

[8796] Manon Destalminil-Letourneau, Ismaël Morin-Poulard, Yushun Tian, Nathalie Vanzo, and Michele Crozatier. The vascular niche controls Drosophila hematopoiesis via fibroblast growth factor signaling. *eLife*, 10, 2021.

[8797] Andrew W DeVilbiss, Zhiyu Zhao, Misty S Martin-Sandoval, Jessalyn M Ubellacker, Alpaslan Tasdogan, Michalis Agathocleous, Thomas P Mathews, and Sean J Morrison. Metabolomic profiling of rare cell populations isolated by flow cytometry from tissues. *eLife*, 10, 2021.

[8798] Chenlu Di, Jesus Murga Moreno, Diego F Salazar-Tortosa, M Elise Lauterbur, and David Enard. Decreased recent adaptation at human mendelian disease genes as a possible consequence of interference between advantageous and deleterious variants. *eLife*, 10, 2021.

[8799] Christian A Di Buduo, Pierre-Alexandre Laurent, Carlo Zaninetti, Larissa Lordier, Paolo M Soprano, Aikaterini Ntai, Serena Barozzi, Alberto La Spada, Ida Biunno, Hana Raslova, James B Bussel, David L Kaplan, Carlo L Balduini, Alessandro Pecci, and Alessandra Balduini. Miniaturized 3D bone marrow tissue model to assess response to Thrombopoietin-receptor agonists in patients. *eLife*, 10, 2021.

[8800] Jacopo Di Russo, Jennifer L Young, Julian WR Wegner, Timmy Steins, Horst Kessler, and Joachim P Spatz. Integrin $\alpha 5 \beta 1$ nano-presentation regulates collective keratinocyte migration independent of substrate rigidity. *eLife*, 10, 2021.

[8801] E Mika Diamanti, Charu Bai Reddy, Sylvia Schröder, Tomaso Muzzu, Kenneth D Harris, Aman B Saleem, and Matteo Carandini. Spatial modulation of visual responses arises in cortex with active navigation. *eLife*, 10, 2021.

[8802] Carlos Manlio Díaz-García, Dylan J Meyer, Nidhi Nathwani, Mahia Rahman, Juan Ramón Martínez-FranÇois, and Gary Yellen. The distinct roles of calcium in rapid control of neuronal glycolysis and the tricarboxylic acid cycle. *eLife*, 10, 2021.

[8803] Paloma Diaz-Maroto, Alba Rey-Iglesia, Isabel Cartajena, Lautaro Núñez, Michael V Westbury, Valeria Varas, Mauricio Moraga, Paula F Campos, Pablo Orozco-terWengel, Juan Carlos Marin, and Anders J Hansen. Ancient DNA reveals the lost domestication history of South American camelids in Northern Chile and across the Andes. *eLife*, 10, 2021.

[8804] Francisco Díaz-Pascual, Martin Lempp, Kazuki Nosho, Hannah Jeckel, Jeanyoung K Jo, Konstantin Neuhaus, Raimo Hartmann, Eric Jelli, Mads Frederik Hansen, Alexa Price-Whelan, Lars EP Dietrich, Hannes Link, and Knut Drescher. Spatial alanine metabolism determines local growth dynamics of Escherichia coli colonies. *eLife*, 10, 2021.

[8805] Ilhame Diboun, Sachin Wani, Stuart H Ralston, and Omar ME Albagha. Epigenetic analysis of Paget's disease of bone identifies differentially methylated loci that predict disease status. *eLife*, 10, 2021.

[8806] Allison L Didychuk, Stephanie N Gates, Matthew R Gardner, Lisa M Strong, Andreas Martin, and Britt A Glaunsinger. A pentameric protein ring with novel architecture is required for herpesviral packaging. *eLife*, 10, 2021.

[8807] Peter J Diebold, Felicia N New, Michael Hovan, Michael J Satlin, and Ilana L Brito. Linking plasmid-based beta-lactamases to their bacterial hosts using single-cell fusion PCR. *eLife*, 10, 2021.

[8808] Tom Dierschke, Eduardo Flores-Sandoval, Madlen I Rast-Somssich, Felix Althoff, Sabine Zachgo, and John L Bowman. Gamete expression of TALE class HD genes activates the diploid sporophyte program in Marchantia polymorpha. *eLife*, 10, 2021.

[8809] Darcy A Diesburg, Jeremy DW Greenlee, and Jan R Wessel. Cortico-subcortical $\beta$ burst dynamics underlying movement cancellation in humans. *eLife*, 10, 2021.

[8810] Maria Eugenia Dieterle, Carles Solà-Riera, Chunyan Ye, Samuel M Goodfellow, Eva Mittler, Ezgi Kasikci, Steven B Bradfute, Jonas Klingström, Rohit K Jangra, and Kartik Chandran. Genetic depletion studies inform receptor usage by virulent hantaviruses in human endothelial cells. *eLife*, 10, 2021.

[8811] Ramile Dilshat, Valerie Fock, Colin Kenny, Ilse Gerritsen, Romain Maurice Jacques Lasseur, Jana Travnickova, Ossia M Eichhoff, Philipp Cerny, Katrin Möller, Sara Sigurbjörnsdóttir, Kritika Kirty, Berglind ósk Einarsdottir, Phil F Cheng, Mitchell Levesque, Robert A Cornell, E Elizabeth Patton, Lionel Larue, Marie de Tayrac, Erna Magnúsdóttir, Margrét Helga ögmundsdóttir, and Eirikur Steingrimsson. MITF reprograms the extracellular matrix and focal adhesion in melanoma. *eLife*, 10, 2021.

[8812] Jennifer Ding, Albert Chen, Janet Chung, Hector Acaron Ledesma, Mofei Wu, David M Berson, Stephanie E Palmer, and Wei Wei. Spatially displaced excitation contributes to the encoding of interrupted motion by a retinal direction-selective circuit. *eLife*, 10, 2021.

[8813] Yanyan Ding, Yuzhe Li, Ziqian Zhao, Qiangfeng Cliff Zhang, and Feng Liu. The chromatin-remodeling enzyme Smarca5 regulates erythrocyte aggregation via Keap1-Nrf2 signaling. *eLife*, 10, 2021.

[8814] Adam S Dingens, Payal Pratap, Keara Malone, Sarah K Hilton, Thomas Ketas, Christopher A Cottrell, Julie Overbaugh, John P Moore, PJ Klasse, Andrew B Ward, and Jesse D Bloom. High-resolution mapping of the neutralizing and binding specificities of polyclonal sera post-HIV Env trimer vaccination. *eLife*, 10, 2021.

[8815] Tra Thi Huong Dinh, Hiroyoshi Iseki, Seiya Mizuno, Saori Iijima-Mizuno, Yoko Tanimoto, Yoko Daitoku, Kanako Kato, Yuko Hamada, Ammar Shaker Hamed Hasan, Hayate Suzuki, Kazuya Murata, Masafumi Muratani, Masatsugu Ema, Jun-Dal Kim, Junji Ishida, Akiyoshi Fukamizu, Mitsuyasu Kato, Satoru Takahashi, Ken-ichi Yagami, Valerie Wilson, Ruth M Arkell, and Fumihiro Sugiyama. Disruption of entire Cables2 locus leads to embryonic lethality by diminished Rps21 gene expression and enhanced p53 pathway. *eLife*, 10, 2021.

[8816] Giada Dirupo, Sabrina Totaro, Jeanne Richard, and Corrado Corradi-Dell'Acqua. Medical education and distrust modulate the response of insular-cingulate network and ventral striatum in pain diagnosis. *eLife*, 10, 2021.

[8817] Zachary Dobson, Safa Ahad, Jackson Vanlandingham, Hila Toporik, Natalie Vaughn, Michael Vaughn, Dewight Williams, Michael Reppert, Petra Fromme, and Yuval Mazor. The structure of photosystem I from a high-light-tolerant cyanobacteria. *eLife*, 10, 2021.

[8818] Esteban Dodero-Rojas, Jose N Onuchic, and Paul Charles Whitford. Sterically confined rearrangements of SARS-CoV-2 Spike protein control cell invasion. *eLife*, 10, 2021.

[8819] Hilje M Doekes, Glenn A Mulder, and Rutger Hermsen. Repeated outbreaks drive the evolution of bacteriophage communication. *eLife*, 10, 2021.

[8820] Laura E Doepker, Sonja Danon, Elias Harkins, Duncan K Ralph, Zak Yaffe, Meghan E Garrett, Amrit Dhar, Cassia Wagner, Megan M Stumpf, Dana Arenz, James A Williams, Walter Jaoko, Kishor Mandaliya, Kelly K Lee, Frederick A Matsen, and Julie M Overbaugh. Development of antibody-dependent cell cytotoxicity function in HIV-1 antibodies. *eLife*, 10, 2021.

[8821] Monika E Dolega, Sylvain Monnier, Benjamin Brunel, Jean-FranÇois Joanny, Pierre Recho, and Giovanni Cappello. Extracellular matrix in multicellular aggregates acts as a pressure sensor controlling cell proliferation and motility. *eLife*, 10, 2021.

[8822] Soledad Domínguez, Liang Ma, Han Yu, Gabrielle Pouchelon, Christian Mayer, George D Spyropoulos, Claudia Cea, György Buzsáki, Gordon Fishell, Dion Khodagholy, and Jennifer N Gelinas. A transient postnatal quiescent period precedes emergence of mature cortical dynamics. *eLife*, 10, 2021.

[8823] Jorge Domínguez-Andrés, Yunus Kuijpers, Olivier B Bakker, Martin Jaeger, Cheng-Jian Xu, Jos WM Van der Meer, Mattias Jakobsson, Jaume Bertranpetit, Leo AB Joosten, Yang Li, and Mihai G Netea. Evolution of cytokine production capacity in ancient and modern European populations. *eLife*, 10, 2021.

[8824] Rafal Donczew and Steven Hahn. BET family members Bdf1/2 modulate global transcription initiation and elongation in Saccharomyces cerevisiae. *eLife*, 10, 2021.

[8825] Jinrui Dong, Sivakumar Viswanathan, Eleonora Adami, Sebastian Schafer, Fathima F Kuthubudeen, Anissa A Widjaja, and Stuart A Cook. The pro-regenerative effects of hyperIL6 in drug-induced liver injury are unexpectedly due to competitive inhibition of IL11 signaling. *eLife*, 10, 2021.

[8826] Drake A Donovan, Johnathan G Crandall, Vi N Truong, Abigail L Vaaler, Thomas B Bailey, Devin Dinwiddie, Orion GB Banks, Laura E McKnight, and Jeffrey N McKnight. Basis of specificity for a conserved and promiscuous chromatin remodeling protein. *eLife*, 10, 2021.

[8827] Shuaijie Dou, Guoxian Li, Ge Li, Chunmei Hou, Yang Zheng, Lili Tang, Yang Gao, Rongliang Mo, Yuxiang Li, Renxi Wang, Beifen Shen, Jun Zhang, and Gencheng Han. Ubiquitination and degradation of NF90 by Tim-3 inhibits antiviral innate immunity. *eLife*, 10, 2021.

[8828] Jessica Douthit, Ariel Hairston, Gina Lee, Carolyn A Morrison, Isabel Holguera, and Jessica E Treisman. R7 photoreceptor axon targeting depends on the relative levels of lost and found expression in R7 and its synaptic partners. *eLife*, 10, 2021.

[8829] Robert AR Drake, Kenneth A Steel, Richard Apps, Bridget M Lumb, and Anthony E Pickering. Loss of cortical control over the descending pain modulatory system determines the development of the neuropathic pain state in rats. *eLife*, 10, 2021.

[8830] Rossana Droghetti, Nicolas Agier, Gilles Fischer, Marco Gherardi, and Marco Cosentino Lagomarsino. An evolutionary model identifies the main evolutionary biases for the evolution of genome-replication profiles. *eLife*, 10, 2021.

[8831] Natascha Ingrid Drude, Lorena Martinez Gamboa, Meggie Danziger, Ulrich Dirnagl, and Ulf Toelch. Improving preclinical studies through replications. *eLife*, 10, 2021.

[8832] Jingyue Duan, Leila Rieder, Megan M Colonnetta, Annie Huang, Mary Mckenney, Scott Watters, Girish Deshpande, William Jordan, Nicolas Fawzi, and Erica Larschan. CLAMP and Zelda function together to promote Drosophila zygotic genome activation. *eLife*, 10, 2021.

[8833] Magda Dubois, Johanna Habicht, Jochen Michely, Rani Moran, Ray J Dolan, and Tobias U Hauser. Human complex exploration strategies are enriched by noradrenaline-modulated heuristics. *eLife*, 10, 2021.

[8834] Jennifer E Dumaine, Adam Sateriale, Alexis R Gibson, Amita G Reddy, Jodi A Gullicksrud, Emma N Hunter, Joseph T Clark, and Boris Striepen. The enteric pathogen Cryptosporidium parvum exports proteins into the cytosol of the infected host cell. *eLife*, 10, 2021.

[8835] Christopher Duncan-Lewis, Ella Hartenian, Valeria King, and Britt A Glaunsinger. Cytoplasmic mRNA decay represses RNA polymerase II transcription during early apoptosis. *eLife*, 10, 2021.

[8836] Valentin Dunsing, Annett Petrich, and Salvatore Chiantia. Multicolor fluorescence fluctuation spectroscopy in living cells via spectral detection. *eLife*, 10, 2021.

[8837] Hélène Duplus-Bottin, Martin Spichty, Gérard Triqueneaux, Christophe Place, Philippe Emmanuel Mangeot, Théophile Ohlmann, Franck Vittoz, and Gaël Yvert. A single-chain and fast-responding light-inducible Cre recombinase as a novel optogenetic switch. *eLife*, 10, 2021.

[8838] Bercem Dutagaci, Grzegorz Nawrocki, Joyce Goodluck, Ali Akbar Ashkarran, Charles G Hoogstraten, Lisa J Lapidus, and Michael Feig. Charge-driven condensation of RNA and proteins suggests broad role of phase separation in cytoplasmic environments. *eLife*, 10, 2021.

[8839] Abhishek Dutta, Fabien Dutreux, and Joseph Schacherer. Loss of heterozygosity results in rapid but variable genome homogenization across yeast genetic backgrounds. *eLife*, 10, 2021.

[8840] Fabien Duveau, Petra Vande Zande, Brian PH Metzger, Crisandra J Diaz, Elizabeth A Walker, Stephen Tryban, Mohammad A Siddiq, Bing Yang, and Patricia J Wittkopp. Mutational sources of trans-regulatory variation affecting gene expression in Saccharomyces cerevisiae. *eLife*, 10, 2021.

[8841] Hany KM Dweck, Gaëlle JS Talross, Wanyue Wang, and John R Carlson. Evolutionary shifts in taste coding in the fruit pest Drosophila suzukii. *eLife*, 10, 2021.

[8842] Natalie A Dye, Marko Popović, K Venkatesan Iyer, Jana F Fuhrmann, Romina Piscitello-Gómez, Suzanne Eaton, and Frank Jülicher. Self-organized patterning of cell morphology via mechanosensitive feedback. *eLife*, 10, 2021.

[8843] Vinay V Eapen, Sharan Swarup, Melissa J Hoyer, Joao A Paulo, and J Wade Harper. Quantitative proteomics reveals the selectivity of ubiquitin-binding autophagy receptors in the turnover of damaged lysosomes by lysophagy. *eLife*, 10, 2021.

[8844] Evelyn L Eastwood, Kayla A Jara, Susanne Bornelöv, Marzia Munafò, Vasileios Frantzis, Emma Kneuss, Elisar J Barbar, Benjamin Czech, and Gregory J Hannon. Dimerisation of the PICTS complex via LC8/Cut-up drives co-transcriptional transposon silencing in Drosophila. *eLife*, 10, 2021.

[8845] Emily R Ebel, Frans A Kuypers, Carrie Lin, Dmitri A Petrov, and Elizabeth S Egan. Common host variation drives malaria parasite fitness in healthy human red cells. *eLife*, 10, 2021.

[8846] Erich R Eberts, Christopher G Guglielmo, and Kenneth C Welch. Reversal of the adipostat control of torpor during migration in hummingbirds. *eLife*, 10, 2021.

[8847] Risa Ebina-Shibuya, Erin E West, Rosanne Spolski, Peng Li, Jangsuk Oh, Majid Kazemian, Daniel Gromer, Phillip Swanson, Ning Du, Dorian B McGavern, and Warren J Leonard. Thymic stromal lymphopoietin limits primary and recall CD8+ T-cell anti-viral responses. *eLife*, 10, 2021.

[8848] Evelien Eenjes, Marjon Buscop-van Kempen, Anne Boerema-de Munck, Gabriela G Edel, Floor Benthem, Lisette de Kreij-de Bruin, Marco Schnater, Dick Tibboel, Jennifer Collins, and Robbert J Rottier. SOX21 modulates SOX2-initiated differentiation of epithelial cells in the extrapulmonary airways. *eLife*, 10, 2021.

[8849] Katharina Eitzen, Priyamedha Sengupta, Samuel Kroll, Eric Kemen, and Gunther Doehlemann. A fungal member of the Arabidopsis thaliana phyllosphere antagonizes Albugo laibachii via a GH25 lysozyme. *eLife*, 10, 2021.

[8850] Johannes Elferich, Sarah Clark, Jingpeng Ge, April Goehring, Aya Matsui, and Eric Gouaux. Molecular structures and conformations of protocadherin-15 and its complexes on stereocilia elucidated by cryo-electron tomography. *eLife*, 10, 2021.

[8851] Jamie M Ellingford, Ryan George, John H McDermott, Shazaad Ahmad, Jonathan J Edgerley, David Gokhale, William G Newman, Stephen Ball, Nicholas Machin, and Graeme CM Black. Genomic and healthcare dynamics of nosocomial SARS-CoV-2 transmission. *eLife*, 10, 2021.

[8852] Amicia D Elliott, Adama Berndt, Matthew Houpert, Snehashis Roy, Robert L Scott, Carson C Chow, Hari Shroff, and Benjamin H White. Pupal behavior emerges from unstructured muscle activity in response to neuromodulation in Drosophila. *eLife*, 10, 2021.

[8853] Yasmin ElMaghloob, Begoña Sot, Michael J McIlwraith, Esther Garcia, Tamas Yelland, and Shehab Ismail. ARL3 activation requires the co-GEF BART and effector-mediated turnover. *eLife*, 10, 2021.

[8854] Rania Elsabrouty, Youngah Jo, Seonghwan Hwang, Dong-Jae Jun, and Russell A DeBose-Boyd. Type 1 polyisoprenoid diphosphate phosphatase modulates geranylgeranyl-mediated control of HMG CoA reductase and UBIAD1. *eLife*, 10, 2021.

[8855] Javier Emperador-Melero, Giovanni de Nola, and Pascal S Kaeser. Intact synapse structure and function after combined knockout of PTP$\delta$, PTP$\sigma$, and LAR. *eLife*, 10, 2021.

[8856] Taylor P Enrico, Wayne Stallaert, Elizaveta T Wick, Peter Ngoi, Xianxi Wang, Seth M Rubin, Nicholas G Brown, Jeremy E Purvis, and Michael J Emanuele. Cyclin F drives proliferation through SCF-dependent degradation of the retinoblastoma-like tumor suppressor p130/RBL2. *eLife*, 10, 2021.

[8857] Neta Erez, Lena Israitel, Eliya Bitman-Lotan, Wing H Wong, Gal Raz, Dayanne V Cornelio-Parra, Salwa Danial, Na'ama Flint Brodsly, Elena Belova, Oksana Maksimenko, Pavel Georgiev, Todd Druley, Ryan D Mohan, and Amir Orian. A Non-stop identity complex (NIC) supervises enterocyte identity and protects from premature aging. *eLife*, 10, 2021.

[8858] Maria Ermakova, Hannah Osborn, Michael Groszmann, Soumi Bala, Andrew Bowerman, Samantha McGaughey, Caitlin Byrt, Hugo Alonso-cantabrana, Steve Tyerman, Robert T Furbank, Robert E Sharwood, and Susanne von Caemmerer. Expression of a CO2-permeable aquaporin enhances mesophyll conductance in the C4 species Setaria viridis. *eLife*, 10, 2021.

[8859] Timothy M Errington, Alexandria Denis, Anne B Allison, Renee Araiza, Pedro Aza-Blanc, Lynette R Bower, Jessica Campos, Heidi Chu, Sarah Denson, Cristine Donham, Kaitlyn Harr, Babette Haven, Elizabeth Iorns, Jennie Kwok, Elysia McDonald, Steven Pelech, Nicole Perfito, Amanda Pike, Darryl Sampey, Michael Settles, David A Scott, Vidhu Sharma, Todd Tolentino, Angela Trinh, Rachel Tsui, Brandon Willis, Joshua Wood, and Lisa Young. Experiments from unfinished Registered Reports in the Reproducibility Project: Cancer Biology. *eLife*, 10, 2021.

[8860] Timothy M Errington, Alexandria Denis, Nicole Perfito, Elizabeth Iorns, and Brian A Nosek. Challenges for assessing replicability in preclinical cancer biology. *eLife*, 10, 2021.

[8861] Timothy M Errington, Maya Mathur, Courtney K Soderberg, Alexandria Denis, Nicole Perfito, Elizabeth Iorns, and Brian A Nosek. Investigating the replicability of preclinical cancer biology. *eLife*, 10, 2021.

[8862] Sarah R Erwin, Brianna N Bristow, Kaitlin E Sullivan, Rennie M Kendrick, Brian Marriott, Lihua Wang, Jody Clements, Andrew L Lemire, Jesse Jackson, and Mark S Cembrowski. Spatially patterned excitatory neuron subtypes and projections of the claustrum. *eLife*, 10, 2021.

[8863] Claire Eschbach, Akira Fushiki, Michael Winding, Bruno Afonso, Ingrid V Andrade, Benjamin T Cocanougher, Katharina Eichler, Ruben Gepner, Guangwei Si, Javier Valdes-Aleman, Richard D Fetter, Marc Gershow, Gregory SXE Jefferis, Aravinthan DT Samuel, James W Truman, Albert Cardona, and Marta Zlatic. Circuits for integrating learned and innate valences in the insect brain. *eLife*, 10, 2021.

[8864] Abdelmoneim Eshra, Hartmut Schmidt, Jens Eilers, and Stefan Hallermann. Calcium dependence of neurotransmitter release at a high fidelity synapse. *eLife*, 10, 2021.

[8865] Roland Esser, Christoph W Korn, Florian Ganzer, and Jan Haaker. L-DOPA modulates activity in the vmPFC, nucleus accumbens, and VTA during threat extinction learning in humans. *eLife*, 10, 2021.

[8866] Jordi Estefa, Paul Tafforeau, Alice M Clement, Jozef Klembara, Grzegorz Niedźwiedzki, Camille Berruyer, and Sophie Sanchez. New light shed on the early evolution of limb-bone growth plate and bone marrow. *eLife*, 10, 2021.

[8867] Chris Estell, Lee Davidson, Pieter C Steketee, Adam Monier, and Steven West. ZC3H4 restricts non-coding transcription in human cells. *eLife*, 10, 2021.

[8868] Thomas J Etheridge, Desiree Villahermosa, Eduard Campillo-Funollet, Alex David Herbert, Anja Irmisch, Adam T Watson, Hung Q Dang, Mark A Osborne, Antony W Oliver, Antony M Carr, and Johanne M Murray. Live-cell single-molecule tracking highlights requirements for stable Smc5/6 chromatin association in vivo. *eLife*, 10, 2021.

[8869] Evangelos Evangelou, Hideaki Suzuki, Wenjia Bai, Raha Pazoki, He Gao, Paul M Matthews, and Paul Elliott. Alcohol consumption in the general population is associated with structural changes in multiple organ systems. *eLife*, 10, 2021.

[8870] Nicholas J Everetts, Melanie I Worley, Riku Yasutomi, Nir Yosef, and Iswar K Hariharan. Single-cell transcriptomics of the Drosophila wing disc reveals instructive epithelium-to-myoblast interactions. *eLife*, 10, 2021.

[8871] Chelsea L Faber, Jennifer D Deem, Bao Anh Phan, Tammy P Doan, Kayoko Ogimoto, Zaman Mirzadeh, Michael W Schwartz, and Gregory J Morton. Leptin receptor neurons in the dorsomedial hypothalamus regulate diurnal patterns of feeding, locomotion, and metabolism. *eLife*, 10, 2021.

[8872] Martin H Fabry, Federica A Falconio, Fadwa Joud, Emily K Lythgoe, Benjamin Czech, and Gregory J Hannon. Maternally inherited piRNAs direct transient heterochromatin formation at active transposons during early Drosophila embryogenesis. *eLife*, 10, 2021.

[8873] Seyi Falekun, Jaime Sepulveda, Yasaman Jami-Alahmadi, Hahnbeom Park, James A Wohlschlegel, and Paul A Sigala. Divergent acyl carrier protein decouples mitochondrial Fe-S cluster biogenesis from fatty acid synthesis in malaria parasites. *eLife*, 10, 2021.

[8874] Chiara Fallerini, Sergio Daga, Stefania Mantovani, Elisa Benetti, Nicola Picchiotti, Daniela Francisci, Francesco Paciosi, Elisabetta Schiaroli, Margherita Baldassarri, Francesca Fava, Maria Palmieri, Serena Ludovisi, Francesco Castelli, Eugenia Quiros-Roldan, Massimo Vaghi, Stefano Rusconi, Matteo Siano, Maria Bandini, Ottavia Spiga, Katia Capitani, Simone Furini, Francesca Mari, Floriana Valentino, Floriana Valentino, Gabriella Doddato, Annarita Giliberti, Rossella Tita, Sara Amitrano, Mirella Bruttini, Susanna Croci, Ilaria Meloni, Maria Antonietta Mencarelli, Caterina Lo Rizzo, Anna Maria Pinto, Laura Di Sarno, Giada Beligni, Andrea Tommasi, Nicola Iuso, Francesca Montagnani, Massimiliano Fabbiani, Barbara Rossetti, Giacomo Zanelli, Elena Bargagli, Laura Bergantini, Miriana D'Alessandro, Paolo Cameli, David Bennett, Federico Anedda, Simona Marcantonio, Sabino Scolletta, Federico Franchi, Maria Antonietta Mazzei, Susanna Guerrini, Edoardo Conticini, Luca Cantarini, Bruno Frediani, Danilo Tacconi, Chiara Spertilli, Marco Feri, Alice Donati, Raffaele Scala, Luca Guidelli, Genni Spargi, Marta Corridi, Cesira Nencioni, Leonardo Croci, Gian Piero Caldarelli, Maurizio Spagnesi, Davide Romani, Paolo Piacentini, Elena Desanctis, Silvia Cappelli,

Anna Canaccini, Agnese Verzuri, Valentina Anemoli, Agostino Ognibene, Antonella D'Arminio Monforte, Federica Gaia Miraglia, Massimo Girardis, Sophie Venturelli, Stefano Busani, Andrea Cossarizza, Andrea Antinori, Alessandra Vergori, Arianna Emiliozzi, Arianna Gabrieli, Agostino Riva, Pier Giorgio Scotton, Francesca Andretta, Sandro Panese, Renzo Scaggiante, Francesca Gatti, Saverio Giuseppe Parisi, Stefano Baratti, Melania Degli Antoni, Matteo Della Monica, Carmelo Piscopo, Mario Capasso, Roberta Russo, Immacolata Andolfo, Achille Iolascon, Giuseppe Fiorentino, Massimo Carella, Marco Castori, Giuseppe Merla, Gabriella Maria Squeo, Filippo Aucella, Pamela Raggi, Carmen Marciano, Rita Perna, Matteo Bassetti, Antonio Di Biagio, Maurizio Sanguinetti, Luca Masucci, Serafina Valente, Marco Mandalà, Alessia Giorli, Lorenzo Salerni, Patrizia Zucchi, Pierpaolo Parravicini, Elisabetta Menatti, Tullio Trotta, Ferdinando Giannattasio, Gabriella Coiro, Fabio Lena, Domenico A Coviello, Cristina Mussini, Giancarlo Bosio, Enrico Martinelli, Sandro Mancarella, Luisa Tavecchia, Marco Gori, Lia Crotti, Gianfranco Parati, Chiara Gabbi, Isabella Zanella, Marco Rizzi, Franco Maggiolo, Diego Ripamonti, Tiziana Bachetti, Maria Teresa La Rovere, Simona Sarzi-Braga, Maurizio Bussotti, Mario Chiariello, Mary Ann Belli, Simona Dei, Alessandra Renieri, Mario U Mondelli, and Elisa Frullanti. Association of Toll-like receptor 7 variants with life-threatening COVID-19 disease in males: findings from a nested case-control study. *eLife*, 10, 2021.

[8875] Teresa WM Fan, Richard M Higashi, Huan Song, Saeed Daneshmandi, Angela L Mahan, Matthew S Purdom, Therese J Bocklage, Thomas A Pittman, Daheng He, Chi Wang, and Andrew N Lane. Innate immune activation by checkpoint inhibition in human patient-derived lung cancer tissues. *eLife*, 10, 2021.

[8876] Wei Fan, Shuang Tang, Xiaojuan Fan, Yi Fang, Xiaojiang Xu, Leping Li, Jian Xu, Jian-Liang Li, Zefeng Wang, and Xiaoling Li. SIRT1 regulates sphingolipid metabolism and neural differentiation of mouse embryonic stem cells through c-Myc-SMPDL3B. *eLife*, 10, 2021.

[8877] Xiaochen Fan, V Pragathi Masamsetti, Jane QJ Sun, Kasper Engholm-Keller, Pierre Osteil, Joshua Studdert, Mark E Graham, Nicolas Fossat, and Patrick PL Tam. TWIST1 and chromatin regulatory proteins interact to guide neural crest cell differentiation. *eLife*, 10, 2021.

[8878] Yana Fandakova, Elliott G Johnson, and Simona Ghetti. Distinct neural mechanisms underlie subjective and objective recollection and guide memory-based decision making. *eLife*, 10, 2021.

[8879] Huaqiang Fang, Alexei M Bygrave, Richard H Roth, Richard C Johnson, and Richard L Huganir. An optimized CRISPR/Cas9 approach for precise genome editing in neurons. *eLife*, 10, 2021.

[8880] Wentong Fang, Chengheng Liao, Rachel Shi, Jeremy M Simon, Travis S Ptacek, Giada Zurlo, Youqiong Ye, Leng Han, Cheng Fan, Lei Bao, Christopher Llynard Ortiz, Hong-Rui Lin, Ujjawal Manocha, Weibo Luo, Yan Peng, William Y Kim, Lee-Wei Yang, and Qing Zhang. ZHX2 promotes HIF1$\alpha$ oncogenic signaling in triple-negative breast cancer. *eLife*, 10, 2021.

[8881] Dayana C Farhat, Matthew W Bowler, Guillaume Communie, Dominique Pontier, Lucid Belmudes, Caroline Mas, Charlotte Corrao, Yohann Couté, Alexandre Bougdour, Thierry Lagrange, Mohamed-Ali Hakimi, and Christopher Swale. A plant-like mechanism coupling m6A reading to polyadenylation safeguards transcriptome integrity and developmental gene partitioning in Toxoplasma. *eLife*, 10, 2021.

[8882] Isabella Farhy-Tselnicker, Matthew M Boisvert, Hanqing Liu, Cari Dowling, Galina A Erikson, Elena Blanco-Suarez, Chen Farhy, Maxim N Shokhirev, Joseph R Ecker, and Nicola J Allen. Activity-dependent modulation of synapse-regulating genes in astrocytes. *eLife*, 10, 2021.

[8883] Eva M Fast, Audrey Sporrij, Margot Manning, Edroaldo Lummertz Rocha, Song Yang, Yi Zhou, Jimin Guo, Ninib Baryawno, Nikolaos Barkas, David Scadden, Fernando Camargo, and Leonard I Zon. External signals regulate continuous transcriptional states in hematopoietic stem cells. *eLife*, 10, 2021.

[8884] Agnieszka Fatalska, Emma Stepinac, Magdalena Richter, Levente Kovacs, Zbigniew Pietras, Martin Puchinger, Gang Dong, Michal Dadlez, and David M Glover. The dimeric Golgi protein Gorab binds to Sas6 as a monomer to mediate centriole duplication. *eLife*, 10, 2021.

[8885] Nikhil Faulkner, Kevin W Ng, Mary Y Wu, Ruth Harvey, Marios Margaritis, Stavroula Paraskevopoulou, Catherine Houlihan, Saira Hussain, Maria Greco, William Bolland, Scott Warchal, Judith Heaney, Hannah Rickman, Moria Spyer, Daniel Frampton, Matthew Byott, Tulio de Oliveira, Alex Sigal, Svend Kjaer, Charles Swanton, Sonia Gandhi, Rupert Beale, Steve J Gamblin, John W McCauley, Rodney Stuart Daniels, Michael Howell, David Bauer, Eleni Nastouli, and George Kassiotis. Reduced antibody cross-reactivity following infection with B.1.1.7 than with parental SARS-CoV-2 strains. *eLife*, 10, 2021.

[8886] Julie Favre, Emilie Vessieres, Anne-Laure Guihot, Coralyne Proux, Linda Grimaud, Jordan Rivron, Manuela CL Garcia, Léa Réthoré, Rana Zahreddine, Morgane Davezac, Chanaelle Fébrissy, Marine Adlanmerini, Laurent Loufrani, Vincent Procaccio, Jean-Michel Foidart, Gilles Flouriot, FranÇoise Lenfant, Coralie Fontaine, Jean-FranÇois Arnal, and Daniel Henrion. Membrane estrogen receptor alpha (ER$\alpha$) participates in flow-mediated dilation in a ligand-independent manner. *eLife*, 10, 2021.

[8887] Maurizio Fazio, Ellen van Rooijen, Michelle Dang, Glenn van de Hoek, Julien Ablain, Jeffrey K Mito, Song Yang, Andrew Thomas, Jonathan Michael, Tania Fabo, Rodsy Modhurima, Patrizia Pessina, Charles K Kaufman, Yi Zhou, Richard M White, and Leonard I Zon. SATB2 induction of a neural crest mesenchyme-like program drives melanoma invasion and drug resistance. *eLife*, 10, 2021.

[8888] Alison F Feder, Kristin N Harper, Chanson J Brumme, and Pleuni S Pennings. Understanding patterns of HIV multi-drug resistance through models of temporal and spatial drug heterogeneity. *eLife*, 10, 2021.

[8889] Ma Feilong, J Swaroop Guntupalli, and James V Haxby. The neural basis of intelligence in fine-grained cortical topographies. *eLife*, 10, 2021.

[8890] Michele Felletti, Cédric Romilly, E Gerhart H Wagner, and Kristina Jonas. A nascent polypeptide sequence modulates DnaA translation elongation in response to nutrient availability. *eLife*, 10, 2021.

[8891] Hemin Feng, Junfeng Su, Wei Fang, Xi Chen, and Jufang He. The entorhinal cortex modulates trace fear memory formation and neuroplasticity in the mouse lateral amygdala via cholecystokinin. *eLife*, 10, 2021.

[8892] Qianxi Feng, Eric Nickels, Ivo S Muskens, Adam J de Smith, W James Gauderman, Amy C Yee, Charite Ricker, Thomas Mack, Andrew D Leavitt, Lucy A Godley, and Joseph L Wiemels. Increased burden of familial-associated early-onset cancer risk among minority Americans compared to non-Latino Whites. *eLife*, 10, 2021.

[8893] Xiaoyu Feng, Zhifang Wu, Junji Xu, Yipu Xu, Bin Zhao, Baoxing Pang, Xingmin Qu, Liang Hu, Lei Hu, Zhipeng Fan, Luyuan Jin, Dengsheng Xia, Shimin Chang, Jingsong Wang, Chunmei Zhang, and Songlin Wang. Dietary nitrate supplementation prevents radiotherapy-induced xerostomia. *eLife*, 10, 2021.

[8894] Jose A Fernandez-Leon, Douglas S Engelke, Guillermo Aquino-Miranda, Alexandria Goodson, Maria N Rasheed, and Fabricio H Do Monte. Neural correlates and determinants of approach–avoidance conflict in the prelimbic prefrontal cortex. *eLife*, 10, 2021.

[8895] Monica L Fernández-Quintero, Yousra El Ghaleb, Petronel Tuluc, Marta Campiglio, Klaus R Liedl, and Bernhard E Flucher. Structural determinants of voltage-gating properties in calcium channels. *eLife*, 10, 2021.

[8896] Mark Ferris, Rebecca Ferris, Chris Workman, Eoin O'Connor, David A Enoch, Emma Goldesgeyme, Natalie Quinnell, Parth Patel, Jo Wright, Geraldine Martell, Christine Moody, Ashley Shaw, Christopher JR Illingworth, Nicholas J Matheson, and Michael P Weekes. Efficacy of FFP3 respirators for prevention of SARS-CoV-2 infection in healthcare workers. *eLife*, 10, 2021.

[8897] Rebecca Feyh, Nadine B Waeber, Simone Prinz, Pietro Ivan Giammarinaro, Gert Bange, Georg Hochberg, Roland K Hartmann, and Florian Altegoer. Structure and mechanistic features of the prokaryotic minimal RNase P. *eLife*, 10, 2021.

[8898] Adam Filipowicz, Jonathan Lalsiamthara, and Alejandro Aballay. TRPM channels mediate learned pathogen avoidance following intestinal distention. *eLife*, 10, 2021.

[8899] Maria Carmela Filomena, Daniel L Yamamoto, Pierluigi Carullo, Roman Medvedev, Andrea Ghisleni, Nicoletta Piroddi, Beatrice Scellini, Roberta Crispino, Francesca D'Autilia, Jianlin Zhang, Arianna Felicetta, Simona Nemska, Simone Serio, Chiara Tesi, Daniele Catalucci, Wolfgang A Linke, Roman Polishchuk, Corrado Poggesi, Mathias Gautel, and Marie-Louise Bang. Myopalladin knockout mice develop cardiac dilation and show a maladaptive response to mechanical pressure overload. *eLife*, 10, 2021.

[8900] Rachel C Findlay, Mohamed Osman, Kirstin A Spence, Paul M Kaye, Pegine B Walrad, and Laurence G Wilson. High-speed, three-dimensional imaging reveals chemotactic behaviour specific to human-infective Leishmania parasites. *eLife*, 10, 2021.

[8901] Anthony S Findley, Alan Monziani, Allison L Richards, Katherine Rhodes, Michelle C Ward, Cynthia A Kalita, Adnan Alazizi, Ali Pazokitoroudi, Sriram Sankararaman, Xiaoquan Wen, David E Lanfear, Roger Pique-Regi, Yoav Gilad, and Francesca Luca. Functional dynamic genetic effects on gene regulation are specific to particular cell types and environmental conditions. *eLife*, 10, 2021.

[8902] Teresa M Findley, David G Wyrick, Jennifer L Cramer, Morgan A Brown, Blake Holcomb, Robin Attey, Dorian Yeh, Eric Monasevitch, Nelly Nouboussi, Isabelle Cullen, Jeremea O Songco, Jared F King, Yashar Ahmadian, and Matthew C Smear. Sniff-synchronized, gradient-guided olfactory search by freely moving mice. *eLife*, 10, 2021.

[8903] Paul Fischer, Shatanik Mukherjee, Enrico Schiewer, Matthias Broser, Franz Bartl, and Peter Hegemann. The inner mechanics of rhodopsin guanylyl cyclase during cGMP-formation revealed by real-time FTIR spectroscopy. *eLife*, 10, 2021.

[8904] Gemma LM Fisher, Jani R Bolla, Karthik V Rajasekar, Jarno Mäkelä, Rachel Baker, Man Zhou, Josh P Prince, Mathew Stracy, Carol V Robinson, Lidia K Arciszewska, and David J Sherratt. Competitive binding of MatP and topoisomerase IV to the MukB hinge domain. *eLife*, 10, 2021.

[8905] M'Lynn E Fisher, Elisa Bovo, Rodrigo Aguayo-Ortiz, Ellen E Cho, Marsha P Pribadi, Michael P Dalton, Nishadh Rathod, M Joanne Lemieux, L Michel Espinoza-Fonseca, Seth L Robia, Aleksey V Zima, and Howard S Young. Dwarf open reading frame (DWORF) is a direct activator of the sarcoplasmic reticulum calcium pump SERCA. *eLife*, 10, 2021.

[8906] Anders Martin Fjell, Hakon Grydeland, Yunpeng Wang, Inge K Amlien, David Bartres-Faz, Andreas M Brandmaier, Sandra Düzel, Jeremy Elman, Carol E Franz, Asta K Håberg, Tim C Kietzmann, Rogier Andrew Kievit, William S Kremen, Stine K Krogsrud, Simone Kühn, Ulman Lindenberger, Didac Macía, Athanasia Monika Mowinckel, Lars Nyberg, Matthew S Panizzon, Cristina Solé-Padullés, Øystein Sørensen, Rene Westerhausen, and Kristine Beate Walhovd. The genetic organization of longitudinal subcortical volumetric change is stable throughout the lifespan. *eLife*, 10, 2021.

[8907] David Fleck, Lina Kenzler, Nadine Mundt, Martin Strauch, Naofumi Uesaka, Robert Moosmann, Felicitas Bruentgens, Annika Missel, Artur Mayerhofer, Dorit Merhof, Jennifer Spehr, and Marc Spehr. ATP activation of peritubular cells drives testicular sperm transport. *eLife*, 10, 2021.

[8908] Oliver Fleetwood, Jens Carlsson, and Lucie Delemotte. Identification of ligand-specific G protein-coupled receptor states and prediction of downstream efficacy via data-driven modeling. *eLife*, 10, 2021.

[8909] Anja Flöser, Katharina Becker, Evi Kostenis, Gabriele König, Cornelius Krasel, Peter Kolb, and Moritz Bünemann. Disentangling bias between Gq, GRK2, and arrestin3 recruitment to the M3 muscarinic acetylcholine receptor. *eLife*, 10, 2021.

[8910] Ming-fai Fong, Kevin R Duffy, Madison P Leet, Christian T Candler, and Mark F Bear. Correction of amblyopia in cats and mice after the critical period. *eLife*, 10, 2021.

[8911] Antonio HO Fonseca, Gustavo M Santana, Gabriela M Bosque Ortiz, Sérgio Bampi, and Marcelo O Dietrich. Analysis of ultrasonic vocalizations from mice using computer vision and machine learning. *eLife*, 10, 2021.

[8912] Laura Fontenas and Sarah Kucenas. Spinal cord precursors utilize neural crest cell mechanisms to generate hybrid peripheral myelinating glia. *eLife*, 10, 2021.

[8913] Angelo Forli, Matteo Pisoni, Yoav Printz, Ofer Yizhar, and Tommaso Fellin. Optogenetic strategies for high-efficiency all-optical interrogation using blue-light-sensitive opsins. *eLife*, 10, 2021.

[8914] Nadia Formicola, Marjorie Heim, Jérémy Dufourt, Anne-Sophie Lancelot, Akira Nakamura, Mounia Lagha, and Florence Besse. Tyramine induces dynamic RNP granule remodeling and translation activation in the Drosophila brain. *eLife*, 10, 2021.

[8915] Cédric Foucault and Florent Meyniel. Gated recurrence enables simple and accurate sequence prediction in stochastic, changing, and structured environments. *eLife*, 10, 2021.

[8916] Holly Fowle, Ziran Zhao, Qifang Xu, Jason S Wasserman, Xinru Wang, Mary Adeyemi, Felicity Feiser, Alison N Kurimchak, Diba Atar, Brennan C McEwan, Arminja N Kettenbach, Rebecca Page, Wolfgang Peti, Roland L Dunbrack, and Xavier Graña. PP2A/B55$\alpha$ substrate recruitment as defined by the retinoblastoma-related protein p107. *eLife*, 10, 2021.

[8917] Ilana Fox-Fisher, Sheina Piyanzin, Bracha Lea Ochana, Agnes Klochendler, Judith Magenheim, Ayelet Peretz, Netanel Loyfer, Joshua Moss, Daniel Cohen, Yaron Drori, Nehemya Friedman, Michal Mandelboim, Marc E Rothenberg, Julie M Caldwell, Mark Rochman, Arash Jamshidi, Gordon Cann, David Lavi, Tommy Kaplan, Benjamin Glaser, Ruth Shemer, and Yuval Dor. Remote immune processes revealed by immune-derived circulating cell-free DNA. *eLife*, 10, 2021.

[8918] Daniela Fraccarollo, Jonas Neuser, Julian Möller, Christian Riehle, Paolo Galuppo, and Johann Bauersachs. Expansion of CD10neg neutrophils and CD14+HLA-DRneg/low monocytes driving proinflammatory responses in patients with acute myocardial infarction. *eLife*, 10, 2021.

[8919] Carolina Franco Nitta, Ellen W Green, Elton D Jhamba, Justine M Keth, Iraís Ortiz-Caraveo, Rachel M Grattan, David J Schodt, Aubrey C Gibson, Ashwani Rajput, Keith A Lidke, Bridget S Wilson, Mara P Steinkamp, and Diane S Lidke. EGFR transactivates RON to drive oncogenic crosstalk. *eLife*, 10, 2021.

[8920] Tom P Franken, Brian J Bondy, David B Haimes, Joshua H Goldwyn, Nace L Golding, Philip H Smith, and Philip X Joris. Glycinergic axonal inhibition subserves acute spatial sensitivity to sudden increases in sound intensity. *eLife*, 10, 2021.

[8921] Tom P Franken and John H Reynolds. Columnar processing of border ownership in primate visual cortex. *eLife*, 10, 2021.

[8922] Markus Frey, Sander Tanni, Catherine Perrodin, Alice O'Leary, Matthias Nau, Jack Kelly, Andrea Banino, Daniel Bendor, Julie Lefort, Christian F Doeller, and Caswell Barry. Interpreting wide-band neural activity using convolutional neural networks. *eLife*, 10, 2021.

[8923] Ryan Z Friedman, David M Granas, Connie A Myers, Joseph C Corbo, Barak A Cohen, and Michael A White. Information content differentiates enhancers from silencers in mouse photoreceptors. *eLife*, 10, 2021.

[8924] Amit Frishberg, Inge van den Munckhof, Rob ter Horst, Kiki Schraa, Leo AB Joosten, Joost HW Rutten, Adrian C Iancu, Ioana M Dregoesc, Bogdan A Tigu, Mihai G Netea, Niels P Riksen, and Irit Gat-Viks. An integrative model of cardiometabolic traits identifies two types of metabolic syndrome. *eLife*, 10, 2021.

[8925] Romy Frömer, Matthew R Nassar, Rasmus Bruckner, Birgit Stürmer, Werner Sommer, and Nick Yeung. Response-based outcome predictions and confidence regulate feedback processing and learning. *eLife*, 10, 2021.

[8926] Frédéric Frottin, Manuela Pérez-Berlanga, F Ulrich Hartl, and Mark S Hipp. Multiple pathways of toxicity induced by C9orf72 dipeptide repeat aggregates and G4C2 RNA in a cellular model. *eLife*, 10, 2021.

[8927] Yong Fu, Kevin M Brown, Nathaniel G Jones, Silvia NJ Moreno, and L David Sibley. Toxoplasma bradyzoites exhibit physiological plasticity of calcium and energy stores controlling motility and egress. *eLife*, 10, 2021.

[8928] Robert P Fuchs, Asako Isogawa, Joao A Paulo, Kazumitsu Onizuka, Tatsuro Takahashi, Ravindra Amunugama, Julien P Duxin, and Shingo Fujii. Crosstalk between repair pathways elicits double-strand breaks in alkylated DNA and implications for the action of temozolomide. *eLife*, 10, 2021.

[8929] Saori Fukao, Kei Haniuda, Hiromasa Tamaki, and Daisuke Kitamura. Protein kinase C$\delta$ is essential for the IgG response against T-cell-independent type 2 antigens and commensal bacteria. *eLife*, 10, 2021.

[8930] Takemichi Fukasawa, Ayumi Yoshizaki, Satoshi Ebata, Asako Yoshizaki-Ogawa, Yoshihide Asano, Atsushi Enomoto, Kiyoshi Miyagawa, Yutaka Kazoe, Kazuma Mawatari, Takehiko Kitamori, and Shinichi Sato. Single-cell-level protein analysis revealing the roles of autoantigen-reactive B lymphocytes in autoimmune disease and the murine model. *eLife*, 10, 2021.

[8931] Tomoyuki Fukuda, Fajar Sofyantoro, Yen Teng Tai, Kim Hou Chia, Takato Matsuda, Takaaki Murase, Yuichi Morozumi, Hisashi Tatebe, Tomotake Kanki, and Kazuhiro Shiozaki. Tripartite suppression of fission yeast TORC1 signaling by the GATOR1-Sea3 complex, the TSC complex, and Gcn2 kinase. *eLife*, 10, 2021.

[8932] Nadia Füllbrunn, Zehao Li, Lara Jorde, Christian P Richter, Rainer Kurre, Lars Langemeyer, Changyuan Yu, Carola Meyer, Jörg Enderlein, Christian Ungermann, Jacob Piehler, and Changjiang You. Nanoscopic anatomy of dynamic multi-protein complexes at membranes resolved by graphene-induced energy transfer. *eLife*, 10, 2021.

[8933] Kara A Fulton and Kevin L Briggman. Permeabilization-free en bloc immunohistochemistry for correlative microscopy. *eLife*, 10, 2021.

[8934] Debora Fusca and Peter Kloppenburg. Task-specific roles of local interneurons for inter- and intraglomerular signaling in the insect antennal lobe. *eLife*, 10, 2021.

[8935] Wanxia Gai, Ze Zhou, Sean Agbor-Enoh, Xiaodan Fan, Sheng Lian, Peiyong Jiang, Suk Hang Cheng, John Wong, Stephen L Chan, Moon Kyoo Jang, Yanqin Yang, Raymond HS Liang, Wai Kong Chan, Edmond SK Ma, Tak Y Leung, Rossa WK Chiu, Hannah Valantine, KC Allen Chan, and YM Dennis Lo. Applications of genetic-epigenetic tissue mapping for plasma DNA in prenatal testing, transplantation and oncology. *eLife*, 10, 2021.

[8936] Swati Gaikwad, Fardin Ghobakhlou, David J Young, Jyothsna Visweswaraiah, Hongen Zhang, and Alan G Hinnebusch. Reprogramming of translation in yeast cells impaired for ribosome recycling favors short, efficiently translated mRNAs. *eLife*, 10, 2021.

[8937] Matthew D Galbraith, Kohl T Kinning, Kelly D Sullivan, Ryan Baxter, Paula Araya, Kimberly R Jordan, Seth Russell, Keith P Smith, Ross E Granrath, Jessica R Shaw, Monika Dzieciatkowska, Tusharkanti Ghosh, Andrew A Monte, Angelo D'Alessandro, Kirk C Hansen, Tellen D Benett,

Elena WY Hsieh, and Joaquín M Espinosa. Seroconversion stages COVID19 into distinct pathophysiological states. *eLife*, 10, 2021.

[8938] Ioannis Galdadas, Luca Carlino, Richard A Ward, Samantha J Hughes, Shozeb Haider, and Francesco Luigi Gervasio. Structural basis of the effect of activating mutations on the EGF receptor. *eLife*, 10, 2021.

[8939] Ioannis Galdadas, Shen Qu, Ana Sofia F Oliveira, Edgar Olehnovics, Andrew R Mack, Maria F Mojica, Pratul K Agarwal, Catherine L Tooke, Francesco Luigi Gervasio, James Spencer, Robert A Bonomo, Adrian J Mulholland, and Shozeb Haider. Allosteric communication in class A $\beta$-lactamases occurs via cooperative coupling of loop dynamics. *eLife*, 10, 2021.

[8940] Isabel Gamache, Marc-André Legault, Jean-Christophe Grenier, Rocio Sanchez, Eric Rhéaume, Samira Asgari, Amina Barhdadi, Yassamin Feroz Zada, Holly Trochet, Yang Luo, Leonid Lecca, Megan Murray, Soumya Raychaudhuri, Jean-Claude Tardif, Marie-Pierre Dubé, and Julie Hussin. A sex-specific evolutionary interaction between ADCY9 and CETP. *eLife*, 10, 2021.

[8941] Stephanie C Gantz, Maria M Ortiz, Andrew J Belilos, and Khaled Moussawi. Excitation of medium spiny neurons by 'inhibitory' ultrapotent chemogenetics via shifts in chloride reversal potential. *eLife*, 10, 2021.

[8942] Longfei Gao, Matthew Decker, Haidee Chen, and Lei Ding. Thrombopoietin from hepatocytes promotes hematopoietic stem cell regeneration after myeloablation. *eLife*, 10, 2021.

[8943] Yuanxiao Gao, Hye Jin Park, Arne Traulsen, and Yuriy Pichugin. Evolution of irreversible somatic differentiation. *eLife*, 10, 2021.

[8944] Paolo Garagnani, Julien Marquis, Massimo Delledonne, Chiara Pirazzini, Elena Marasco, Katarzyna Malgorzata Kwiatkowska, Vincenzo Iannuzzi, Maria Giulia Bacalini, Armand Valsesia, Jerome Carayol, Frederic Raymond, Alberto Ferrarini, Luciano Xumerle, Sebastiano Collino, Daniela Mari, Beatrice Arosio, Martina Casati, Evelyn Ferri, Daniela Monti, Benedetta Nacmias, Sandro Sorbi, Donata Luiselli, Davide Pettener, Gastone Castellani, Claudia Sala, Giuseppe Passarino, Francesco De Rango, Patrizia D'Aquila, Luca Bertamini, Nicola Martinelli, Domenico Girelli, Oliviero Olivieri, Cristina Giuliani, Patrick Descombes, and Claudio Franceschi. Whole-genome sequencing analysis of semi-supercentenarians. *eLife*, 10, 2021.

[8945] Angela R Garcia, Caleb Finch, Margaret Gatz, Thomas Kraft, Daniel Eid Rodriguez, Daniel Cummings, Mia Charifson, Kenneth Buetow, Bret A Beheim, Hooman Allayee, Gregory S Thomas, Jonathan Stieglitz, Michael D Gurven, Hillard Kaplan, and Benjamin C Trumble. APOE4 is associated with elevated blood lipids and lower levels of innate immune biomarkers in a tropical Amerindian subsistence population. *eLife*, 10, 2021.

[8946] David M Garcia, Edgar A Campbell, Christopher M Jakobson, Mitsuhiro Tsuchiya, Ethan A Shaw, Acadia L DiNardo, Matt Kaeberlein, and Daniel F Jarosz. A prion accelerates proliferation at the expense of lifespan. *eLife*, 10, 2021.

[8947] Alam García-Heredia, Takehiro Kado, Caralyn E Sein, Julia Puffal, Sarah H Osman, Julius Judd, Todd A Gray, Yasu S Morita, and M Sloan Siegrist. Membrane-partitioned cell wall synthesis in mycobacteria. *eLife*, 10, 2021.

[8948] Mikel Garcia-Marcos. Complementary biosensors reveal different G-protein signaling modes triggered by GPCRs and non-receptor activators. *eLife*, 10, 2021.

[8949] Yesenia Garcia-Sifuentes and Donna L Maney. Reporting and misreporting of sex differences in the biological sciences. *eLife*, 10, 2021.

[8950] Vivek Garg, Junji Suzuki, Ishan Paranjpe, Tiffany Unsulangi, Liron Boyman, Lorin S Milescu, W Jonathan Lederer, and Yuriy Kirichok. The mechanism of MICU-dependent gating of the mitochondrial Ca2+uniporter. *eLife*, 10, 2021.

[8951] Emirena Garrafa, Marika Vezzoli, Marco Ravanelli, Davide Farina, Andrea Borghesi, Stefano Calza, and Roberto Maroldi. Early prediction of in-hospital death of COVID-19 patients: a machine-learning model based on age, blood analyses, and chest x-ray score. *eLife*, 10, 2021.

[8952] Tamara Garrido-Gomez, Nerea Castillo-Marco, Mónica Clemente-Ciscar, Teresa Cordero, Irene Muñoz-Blat, Alicia Amadoz, Jorge Jimenez-Almazan, Rogelio Monfort-Ortiz, Reyes Climent, Alfredo Perales-Marin, and Carlos Simon. Disrupted PGR-B and ESR1 signaling underlies defective decidualization linked to severe preeclampsia. *eLife*, 10, 2021.

[8953] Marissa M Gaskill, Tyler J Gibson, Elizabeth D Larson, and Melissa M Harrison. GAF is essential for zygotic genome activation and chromatin accessibility in the early Drosophila embryo. *eLife*, 10, 2021.

[8954] Benjamin D Gastfriend, Hideaki Nishihara, Scott G Canfield, Koji L Foreman, Britta Engelhardt, Sean P Palecek, and Eric V Shusta. Wnt signaling mediates acquisition of blood–brain barrier properties in naïve endothelium derived from human pluripotent stem cells. *eLife*, 10, 2021.

[8955] Alberto T Gatta, Yolanda Olmos, Caroline L Stoten, Qu Chen, Peter B Rosenthal, and Jeremy G Carlton. CDK1 controls CHMP7-dependent nuclear envelope reformation. *eLife*, 10, 2021.

[8956] Jeremie Gautheron, Christophe Morisseau, Wendy K Chung, Jamila Zammouri, Martine Auclair, Genevieve Baujat, Emilie Capel, Celia Moulin, Yuxin Wang, Jun Yang, Bruce D Hammock, Barbara Cerame, Franck Phan, Bruno Fève, Corinne Vigouroux, Fabrizio Andreelli, and Isabelle Jeru. EPHX1 mutations cause a lipoatrophic diabetes syndrome due to impaired epoxide hydrolysis and increased cellular senescence. *eLife*, 10, 2021.

[8957] Natalja Gavrilov and Andreas Nieder. Distinct neural networks for the volitional control of vocal and manual actions in the monkey homologue of Broca's area. *eLife*, 10, 2021.

[8958] Katy AM Gaythorpe, Kaja Abbas, John Huber, Andromachi Karachaliou, Niket Thakkar, Kim Woodruff, Xiang Li, Susy Echeverria-Londono, Andre Arsene Bita Fouda, Andre Arsene Bita Fouda, Felicity Cutts, Emily Dansereau, Antoine Durupt, Ulla Griffiths, Jennifer Horton, L Kendall Krause, Katrina Kretsinger, Tewodaj Mengistu, Imran Mirza, Simon R Procter, Stephanie Shendale, Matthew Ferrari, Michael L Jackson, Kevin McCarthy, T Alex Perkins, Caroline Trotter, and Mark Jit. Impact of COVID-19-related disruptions to measles, meningococcal A, and yellow fever vaccination in 10 countries. *eLife*, 10, 2021.

[8959] Matthieu Genestine, Daisy Ambriz, Gregg W Crabtree, Patrick Dummer, Anna Molotkova, Michael Quintero, Angeliki Mela, Saptarshi Biswas, Huijuan Feng, Chaolin Zhang, Peter Canoll, Gunnar Hargus, Dritan Agalliu, Joseph A Gogos, and Edmund Au. Vascular-derived SPARC and SerpinE1 regulate interneuron tangential migration and accelerate functional maturation of human stem cell-derived interneurons. *eLife*, 10, 2021.

[8960] Guangfeng Geng, Jinhua Liu, Changlu Xu, Yandong Pei, Linbo Chen, Chenglong Mu, Ding Wang, Jie Gao, Yue Li, Jing Liang, Tian Zhao, Chuanmei Zhang, Jiaxi Zhou, Quan Chen, Yushan Zhu, and Lihong Shi. Receptor-mediated mitophagy regulates EPO production and protects against renal anemia. *eLife*, 10, 2021.

[8961] Alessandra Gentile, Anabela Bensimon-Brito, Rashmi Priya, Hans-Martin Maischein, Janett Piesker, Stefan Guenther, Felix Gunawan, and Didier YR Stainier. The EMT transcription factor Snai1 maintains myocardial wall integrity by repressing intermediate filament gene expression. *eLife*, 10, 2021.

[8962] Ginto George, Satoshi Ninagawa, Hirokazu Yagi, Jun-ichi Furukawa, Noritaka Hashii, Akiko Ishii-Watabe, Ying Deng, Kazutoshi Matsushita, Tokiro Ishikawa, Yugoviandi P Mamahit, Yuta Maki, Yasuhiro Kajihara, Koichi Kato, Tetsuya Okada, and Kazutoshi Mori. Purified EDEM3 or EDEM1 alone produces determinant oligosaccharide structures from M8B in mammalian glycoprotein ERAD. *eLife*, 10, 2021.

[8963] Daniel Gerber, Jorge A Pereira, Joanne Gerber, Ge Tan, Slavica Dimitrieva, Emilio Yángüez, and Ueli Suter. Transcriptional profiling of mouse peripheral nerves to the single-cell level to build a sciatic nerve ATlas (SNAT). *eLife*, 10, 2021.

[8964] Andreas Gerondopoulos, Philipp Bräuer, Tomoaki Sobajima, Zhiyi Wu, Joanne L Parker, Philip C Biggin, Francis A Barr, and Simon Newstead. A signal capture and proofreading mechanism for the KDEL-receptor explains selectivity and dynamic range in ER retrieval. *eLife*, 10, 2021.

[8965] Samuel J Gershman, Petra EM Balbi, C Randy Gallistel, and Jeremy Gunawardena. Reconsidering the evidence for learning in single cells. *eLife*, 10, 2021.

[8966] Ryan J Geusz, Allen Wang, Joshua Chiou, Joseph J Lancman, Nichole Wetton, Samy Kefalopoulou, Jinzhao Wang, Yunjiang Qiu, Jian Yan, Anthony Aylward, Bing Ren, P Duc Si Dong, Kyle J Gaulton, and Maike Sander. Pancreatic progenitor epigenome maps prioritize type 2 diabetes risk genes with roles in development. *eLife*, 10, 2021.

[8967] Brian Q Geuther, Asaf Peer, Hao He, Gautam Sabnis, Vivek M Philip, and Vivek Kumar. Action detection using a neural network elucidates the genetics of mouse grooming behavior. *eLife*, 10, 2021.

[8968] Krishna S Ghanta, Zexiang Chen, Aamir Mir, Gregoriy A Dokshin, Pranathi M Krishnamurthy, Yeonsoo Yoon, Judith Gallant, Ping Xu, Xiao-Ou Zhang, Ahmet Rasit Ozturk, Masahiro Shin, Feston Idrizi, Pengpeng Liu, Hassan Gneid, Alireza Edraki, Nathan D Lawson, Jaime A Rivera-Pérez, Erik J Sontheimer, Jonathan K Watts, and Craig C Mello. 5'-Modifications improve potency and efficacy of DNA donors for precision genome editing. *eLife*, 10, 2021.

[8969] Alireza Ghasemizadeh, Emilie Christin, Alexandre Guiraud, Nathalie Couturier, Marie Abitbol, Valerie Risson, Emmanuelle Girard, Christophe Jagla, Cedric Soler, Lilia Laddada, Colline Sanchez, Francisco-Ignacio Jaque-Fernandez, Vincent Jacquemond, Jean-Luc Thomas, Marine Lanfranchi, Julien Courchet, Julien Gondin, Laurent Schaeffer, and Vincent Gache. MACF1 controls skeletal muscle function through the microtubule-dependent localization of extra-synaptic myonuclei and mitochondria biogenesis. *eLife*, 10, 2021.

[8970] Filippo Ghezzi, Andre Marques-Smith, Paul G Anastasiades, Daniel Lyngholm, Cristiana Vagnoni, Alexandra Rowett, Gokul Parameswaran, Anna Hoerder-Suabedissen, Yasushi Nakagawa, Zoltan Molnar, and Simon JB Butt. Non-canonical role for Lpar1-EGFP subplate neurons in early postnatal mouse somatosensory cortex. *eLife*, 10, 2021.

[8971] Shima Ghoroghi, Benjamin Mary, Annabel Larnicol, Nandini Asokan, Annick Klein, Naël Osmani, Ignacio Busnelli, FranÇois Delalande, Nicodème Paul, Sébastien Halary, Frédéric Gros, Laetitia Fouillen, Anne-Marie Haeberle, Cathy Royer, Coralie Spiegelhalter, Gwennan André-Grégoire, Vincent Mittelheisser, Alexandre Detappe, Kendelle Murphy, Paul Timpson, Raphaël Carapito, Marcel Blot-Chabaud, Julie Gavard, Christine Carapito, Nicolas Vitale, Olivier Lefebvre, Jacky G Goetz, and Vincent Hyenne. Ral GTPases promote breast cancer metastasis by controlling biogenesis and organ targeting of exosomes. *eLife*, 10, 2021.

[8972] Marie Ghraichy, Valentin von Niederhäusern, Aleksandr Kovaltsuk, Jacob D Galson, Charlotte M Deane, and Johannes Trück. Different B cell subpopulations show distinct patterns in their IgH repertoire metrics. *eLife*, 10, 2021.

[8973] Matthew JS Gibson, María de Lourdes Torres, Yaniv Brandvain, and Leonie C Moyle. Introgression shapes fruit color convergence in invasive Galápagos tomato. *eLife*, 10, 2021.

[8974] Alice Gilbert, Xabier Elorza-Vidal, Armelle Rancillac, Audrey Chagnot, Mervé Yetim, Vincent Hingot, Thomas Deffieux, Anne-Cécile Boulay, Rodrigo Alvear-Perez, Salvatore Cisternino, Sabrina Martin, Sonia Taïb, Aontoinette Gelot, Virginie Mignon, Maryline Favier, Isabelle Brunet, Xavier Declèves, Mickael Tanter, Raul Estevez, Denis Vivien, Bruno Saubaméa, and Martine Cohen-Salmon. Megalencephalic leukoencephalopathy with subcortical cysts is a developmental disorder of the gliovascular unit. *eLife*, 10, 2021.

[8975] Christopher J Gill, Christian E Gunning, William B MacLeod, Lawrence Mwananyanda, Donald M Thea, Rachel C Pieciak, Geoffrey Kwenda, Zacharia Mupila, and Pejman Rohani. Asymptomatic Bordetella pertussis infections in a longitudinal cohort of young African infants and their mothers. *eLife*, 10, 2021.

[8976] Jonathan Gilley, Oscar Jackson, Menelaos Pipis, Mehrdad A Estiar, Ammar Al-Chalabi, Matt C Danzi, Kristel R van Eijk, Stephen A Goutman, Matthew B Harms, Henry Houlden, Alfredo Iacoangeli, Julia Kaye, Leandro Lima, John Ravits, Guy A Rouleau, Rebecca Schüle, Jishu Xu, Stephan Züchner, Johnathan Cooper-Knock, Ziv Gan-Or, Mary M Reilly, and Michael P Coleman. Enrichment of SARM1 alleles encoding variants with constitutively hyperactive NADase in patients with ALS and other motor nerve disorders. *eLife*, 10, 2021.

[8977] Joanna L Gillis, Josephine A Hinneh, Natalie K Ryan, Swati Irani, Max Moldovan, Lake-Ee Quek, Raj K Shrestha, Adrienne R Hanson, Jianling Xie, Andrew J Hoy, Jeff Holst, Margaret M Centenera, Ian G Mills, David J Lynn, Luke A Selth, and Lisa M Butler. A feedback loop between the androgen receptor and 6-phosphogluoconate dehydrogenase (6PGD) drives prostate cancer growth. *eLife*, 10, 2021.

[8978] Manuel Giménez-Andrés, Tadej EmerŠič, Sandra Antoine-Bally, Juan Martin D'Ambrosio, Bruno Antonny, Jure Derganc, and Alenka čopič. Exceptional stability of a perilipin on lipid droplets depends on its polar residues, suggesting multimeric assembly. *eLife*, 10, 2021.

[8979] Andrea Giometto, David R Nelson, and Andrew W Murray. Antagonism between killer yeast strains as an experimental model for biological nucleation dynamics. *eLife*, 10, 2021.

[8980] Deborah Giordano, Angelo Facchiano, Sabato D'Auria, and Francesco Loreto. A hypothesis on the capacity of plant odorant-binding proteins to bind volatile isoprenoids based on in silico evidences. *eLife*, 10, 2021.

[8981] Juliet R Girard, Lauren M Goins, Dung M Vuu, Mark S Sharpley, Carrie M Spratford, Shreya R Mantri, and Utpal Banerjee. Paths and pathways that generate cell-type heterogeneity and developmental progression in hematopoiesis. *eLife*, 10, 2021.

[8982] Cédric Girard-Buttoz, Patrick J Tkaczynski, Liran Samuni, Pawel Fedurek, Cristina Gomes, Therese Löhrich, Virgile Manin, Anna Preis, Prince F Valé, Tobias Deschner, Roman M Wittig, and Catherine Crockford. Early maternal loss leads to short- but not long-term effects on diurnal cortisol slopes in wild chimpanzees. *eLife*, 10, 2021.

[8983] Jack Gisby, Candice L Clarke, Nicholas Medjeral-Thomas, Talat H Malik, Artemis Papadaki, Paige M Mortimer, Norzawani B Buang, Shanice Lewis, Marie Pereira, Frederic Toulza, Ester Fagnano, Marie-Anne Mawhin, Emma E Dutton, Lunnathaya Tapeng, Arianne C Richard, Paul DW Kirk, Jacques Behmoaras, Eleanor Sandhu, Stephen P McAdoo, Maria F Prendecki, Matthew C Pickering, Marina Botto, Michelle Willicombe, David C Thomas, and James E Peters. Longitudinal proteomic profiling of dialysis patients with COVID-19 reveals markers of severity and predictors of death. *eLife*, 10, 2021.

[8984] Erida Gjini and Kevin B Wood. Price equation captures the role of drug interactions and collateral effects in the evolution of multidrug resistance. *eLife*, 10, 2021.

[8985] Konstantinos Gkatzis, Paolo Panza, Sofia Peruzzo, and Didier YR Stainier. Differentiation of mouse fetal lung alveolar progenitors in serum-free organotypic cultures. *eLife*, 10, 2021.

[8986] Caleb R Glassman, Leon Su, Sonia S Majri-Morrison, Hauke Winkelmann, Fei Mo, Peng Li, Magdiel Pérez-Cruz, Peggy P Ho, Ievgen Koliesnik, Nadine Nagy, Tereza Hnizdilova, Lora K Picton, Marek Kovar, Paul Bollyky, Lawrence Steinman, Everett Meyer, Jacob Piehler, Warren J Leonard, and K Christopher Garcia. Calibration of cell-intrinsic interleukin-2 response thresholds guides design of a regulatory T cell biased agonist. *eLife*, 10, 2021.

[8987] Lori Glenwinkel, Seth R Taylor, Kasper Langebeck-Jensen, Laura Pereira, Molly B Reilly, Manasa Basavaraju, Ibnul Rafi, Eviatar Yemini, Roger Pocock, Nenad Sestan, Marc Hammarlund, David M Miller, and Oliver Hobert. In silico analysis of the transcriptional regulatory logic of neuronal identity specification throughout the C. elegans nervous system. *eLife*, 10, 2021.

[8988] Chaim Glück, Kim David Ferrari, Noemi Binini, Annika Keller, Aiman S Saab, Jillian L Stobart, and Bruno Weber. Distinct signatures of calcium activity in brain mural cells. *eLife*, 10, 2021.

[8989] Elisabeth Gludovacz, Kornelia Schuetzenberger, Marlene Resch, Katharina Tillmann, Karin Petroczi, Markus Schosserer, Sigrid Vondra, Serhii Vakal, Gerald Klanert, Jürgen Pollheimer, Tiina A Salminen, Bernd Jilma, Nicole Borth, and Thomas Boehm. Heparin-binding motif mutations of human diamine oxidase allow the development of a first-in-class histamine-degrading biopharmaceutical. *eLife*, 10, 2021.

[8990] Iris E Glykofridis, Jaco C Knol, Jesper A Balk, Denise Westland, Thang V Pham, Sander R Piersma, Sinéad M Lougheed, Sepide Derakhshan, Puck Veen, Martin A Rooimans, Saskia E van Mil, Franziska Böttger, Pino J Poddighe, Irma van de Beek, Jarno Drost, Fried JT Zwartkruis, Renee X de Menezes, Hanne EJ Meijers-Heijboer, Arjan C Houweling, Connie R Jimenez, and Rob MF Wolthuis. Loss of FLCN-FNIP1/2 induces a non-canonical interferon response in human renal tubular epithelial cells. *eLife*, 10, 2021.

[8991] Benoit G Godard, Remi Dumollard, Carl-Philipp Heisenberg, and Alex McDougall. Combined effect of cell geometry and polarity domains determines the orientation of unequal division. *eLife*, 10, 2021.

[8992] Jack Goffinet, Samuel Brudner, Richard Mooney, and John Pearson. Low-dimensional learned feature spaces quantify individual and group differences in vocal repertoires. *eLife*, 10, 2021.

[8993] Hannah C Goldbach, Bradley Akitake, Caitlin E Leedy, and Mark H Histed. Performance in even a simple perceptual task depends on mouse secondary visual areas. *eLife*, 10, 2021.

[8994] Erin J Golden, Eric D Larson, Lauren A Shechtman, G Devon Trahan, Dany Gaillard, Timothy J Fellin, Jennifer K Scott, Kenneth L Jones, and Linda A Barlow. Onset of taste bud cell renewal starts at birth and coincides with a shift in SHH function. *eLife*, 10, 2021.

[8995] Alina Goldstein-Levitin, Himanshu Pandey, Kanary Allhuzaeel, Itamar Kass, and Larisa Gheber. Intracellular functions and motile properties of bi-directional kinesin-5 Cin8 are regulated by neck linker docking. *eLife*, 10, 2021.

[8996] Harold Fernando Gómez, Mathilde Sabine Dumond, Leonie Hodel, Roman Vetter, and Dagmar Iber. 3D cell neighbour dynamics in growing pseudostratified epithelia. *eLife*, 10, 2021.

[8997] André Brás GonÇalves, Sarah Kirstine Hasselbalch, Beinta Biskopstø Joensen, Sebastian Patzke, Pernille Martens, Signe Krogh Ohlsen, Mathieu Quinodoz, Konstantinos Nikopoulos, Reem Suleiman, Magnus Per Damsø Jeppesen, Catja Weiss, Søren Tvorup Christensen, Carlo Rivolta, Jens S Andersen, Pietro Farinelli, and Lotte Bang Pedersen. CEP78 functions downstream of CEP350 to control biogenesis of primary cilia by negatively regulating CP110 levels. *eLife*, 10, 2021.

[8998] Yan Gong, Julien Alassimone, Rachel Varnau, Nidhi Sharma, Lily S Cheung, and Dominique C Bergmann. Tuning self-renewal in the Arabidopsis stomatal lineage by hormone and nutrient regulation of asymmetric cell division. *eLife*, 10, 2021.

[8999] Inés González-Calvo, Keerthana Iyer, Mélanie Carquin, Anouar Khayachi, Fernando A Giuliani, Séverine M Sigoillot, Jean Vincent, Martial Séveno, Maxime Veleanu, Sylvana Tahraoui, Mélanie Albert, Oana Vigy, Célia Bosso-Lefèvre, Yann Nadjar, Andréa Dumoulin, Antoine Triller, Jean-Louis Bessereau, Laure Rondi-Reig, Philippe Isope, and Fekrije Selimi. Sushi domain-containing protein 4 controls synaptic plasticity and motor learning. *eLife*, 10, 2021.

[9000] Alvaro Gonzalez Rajal, Kamila A Marzec, Rachael A McCloy, Max Nobis, Venessa Chin, Jordan F Hastings, Kaitao Lai, Marina Kennerson, William E Hughes, Vijesh Vaghjiani, Paul Timpson, Jason E Cain, D Neil Watkins, David R Croucher, and Andrew Burgess. A non-genetic, cell cycle-dependent mechanism of platinum resistance in lung adenocarcinoma. *eLife*, 10, 2021.

[9001] Sarah E Goodale, Nafis Ahmed, Chong Zhao, Jacco A de Zwart, Pinar S özbay, Dante Picchioni, Jeff Duyn, Dario J Englot, Victoria L Morgan, and Catie Chang. fMRI-based detection of alertness predicts behavioral response variability. *eLife*, 10, 2021.

[9002] Janani Gopalan, Mitchell H Omar, Ankita Roy, Nelly M Cruz, Jerome Falcone, Kiana N Jones, Katherine A Forbush, Jonathan Himmelfarb, Benjamin S Freedman, and John D Scott. Targeting an anchored phosphatase-deacetylase unit restores renal ciliary homeostasis. *eLife*, 10, 2021.

[9003] Sayali V Gore, Eric J James, Lin-chien Huang, Jenn J Park, Andrea Berghella, Adrian C Thompson, Hollis T Cline, and Carlos D Aizenman. Role of matrix metalloproteinase-9 in neurodevelopmental deficits and experience-dependent plasticity in Xenopus laevis. *eLife*, 10, 2021.

[9004] Ilaria Gori, Roger George, Andrew G Purkiss, Stephanie Strohbuecker, Rebecca A Randall, Roksana Ogrodowicz, Virginie Carmignac, Laurence Faivre, Dhira Joshi, Svend Kjær, and Caroline S Hill. Mutations in SKI in Shprintzen–Goldberg syndrome lead to attenuated TGF-$\beta$ responses through SKI stabilization. *eLife*, 10, 2021.

[9005] Nicole R Gould, Katrina M Williams, Humberto C Joca, Olivia M Torre, James S Lyons, Jenna M Leser, Manasa P Srikanth, Marcus Hughes, Ramzi J Khairallah, Ricardo A Feldman, Christopher W Ward, and Joseph P Stains. Disparate bone anabolic cues activate bone formation by regulating the rapid lysosomal degradation of sclerostin protein. *eLife*, 10, 2021.

[9006] Anitha P Govind, Okunola Jeyifous, Theron A Russell, Zola Yi, Aubrey V Weigel, Abhijit Ramaprasad, Luke Newell, William Ramos, Fernando M Valbuena, Jason C Casler, Jing-Zhi Yan, Benjamin S Glick, Geoffrey T Swanson, Jennifer Lippincott-Schwartz, and William N Green. Activity-dependent Golgi satellite formation in dendrites reshapes the neuronal surface glycoproteome. *eLife*, 10, 2021.

[9007] Graham Gower, Pablo Iáñez Picazo, Matteo Fumagalli, and Fernando Racimo. Detecting adaptive introgression in human evolution using convolutional neural networks. *eLife*, 10, 2021.

[9008] Ashish Goyal, Daniel B Reeves, E Fabian Cardozo-Ojeda, Joshua T Schiffer, and Bryan T Mayer. Viral load and contact heterogeneity predict SARS-CoV-2 transmission and super-spreading events. *eLife*, 10, 2021.

[9009] Olivia Gozel and Wulfram Gerstner. A functional model of adult dentate gyrus neurogenesis. *eLife*, 10, 2021.

[9010] Chad Paul Grabner and Tobias Moser. The mammalian rod synaptic ribbon is essential for Cav channel facilitation and ultrafast synaptic vesicle fusion. *eLife*, 10, 2021.

[9011] Daniel Gradeci, Anna Bove, Giulia Vallardi, Alan R Lowe, Shiladitya Banerjee, and Guillaume Charras. Cell-scale biophysical determinants of cell competition in epithelia. *eLife*, 10, 2021.

[9012] Francis Grafton, Jaclyn Ho, Sara Ranjbarvaziri, Farshad Farshidfar, Anastasiia Budan, Stephanie Steltzer, Mahnaz Maddah, Kevin E Loewke, Kristina Green, Snahel Patel, Tim Hoey, and Mohammad Ali Mandegar. Deep learning detects cardiotoxicity in a high-content screen with induced pluripotent stem cell-derived cardiomyocytes. *eLife*, 10, 2021.

[9013] Roger I Grant, Elizabeth M Doncheck, Kelsey M Vollmer, Kion T Winston, Elizaveta V Romanova, Preston N Siegler, Heather Holman, Christopher W Bowen, and James M Otis. Specialized coding patterns among dorsomedial prefrontal neuronal ensembles predict conditioned reward seeking. *eLife*, 10, 2021.

[9014] Austin R Graves, Richard H Roth, Han L Tan, Qianwen Zhu, Alexei M Bygrave, Elena Lopez-Ortega, Ingie Hong, Alina C Spiegel, Richard C Johnson, Joshua T Vogelstein, Daniel J Tward, Michael I Miller, and Richard L Huganir. Visualizing synaptic plasticity in vivo by large-scale imaging of endogenous AMPA receptors. *eLife*, 10, 2021.

[9015] Gregory G Grecco, Briana E Mork, Jui-Yen Huang, Corinne E Metzger, David L Haggerty, Kaitlin C Reeves, Yong Gao, Hunter Hoffman, Simon N Katner, Andrea R Masters, Cameron W Morris, Erin A Newell, Eric A Engleman, Anthony J Baucum, Jiuen Kim, Bryan K Yamamoto, Matthew R Allen, Yu-Chien Wu, Hui-Chen Lu, Patrick L Sheets, and Brady K Atwood. Prenatal methadone exposure disrupts behavioral development and alters motor neuron intrinsic properties and local circuitry. *eLife*, 10, 2021.

[9016] Joshua D Greenlee, Maria Lopez-Cavestany, Nerymar Ortiz-Otero, Kevin Liu, Tejas Subramanian, Burt Cagir, and Michael R King. Oxaliplatin resistance in colorectal cancer enhances TRAIL sensitivity via death receptor 4 upregulation and lipid raft localization. *eLife*, 10, 2021.

[9017] Daniel Griffith and Alex S Holehouse. PARROT is a flexible recurrent neural network framework for analysis of large protein datasets. *eLife*, 10, 2021.

[9018] Iago Grobas, Marco Polin, and Munehiro Asally. Swarming bacteria undergo localized dynamic phase transition to form stress-induced biofilms. *eLife*, 10, 2021.

[9019] Marloes Grobben, Karlijn van der Straten, Philip JM Brouwer, Mitch Brinkkemper, Pauline Maisonnasse, Nathalie Dereuddre-Bosquet, Brent Appelman, AH Ayesha Lavell, Lonneke A van Vught, Judith A Burger, Meliawati Poniman, Melissa Oomen, Dirk Eggink, Tom PL Bijl, Hugo DG van Willigen, Elke Wynberg, Bas J Verkaik, Orlane JA Figaroa, Peter J de Vries, Tessel M Boertien, Diederik van de Beek, Marije K Bomers, Jonne J Sikkens, Roger Le Grand, Menno D de Jong, Maria Prins, Amy W Chung, Godelieve J de Bree, Rogier W Sanders, and Marit J van Gils. Cross-reactive antibodies after SARS-CoV-2 infection and vaccination. *eLife*, 10, 2021.

[9020] Matthew William Grol, Nele A Haelterman, Joohyun Lim, Elda M Munivez, Marilyn Archer, David M Hudson, Sara F Tufa, Douglas R Keene, Kevin Lei, Dongsu Park, Cole D Kuzawa, Catherine G Ambrose, David R Eyre, and Brendan H Lee. Tendon and motor phenotypes in the Crtap-/- mouse model of recessive osteogenesis imperfecta. *eLife*, 10, 2021.

[9021] Franziska Gruhl, Peggy Janich, Henrik Kaessmann, and David Gatfield. Circular RNA repertoires are associated with evolutionarily young transposable elements. *eLife*, 10, 2021.

[9022] Christina Grønberg, Qiaoxia Hu, Dhani Ram Mahato, Elena Longhin, Nina Salustros, Annette Duelli, Pin Lyu, Viktoria Bågenholm, Jonas Eriksson, Komal Umashankar Rao, Domhnall Iain Henderson, Gabriele Meloni, Magnus Andersson, Tristan Croll, Gabriela Godaly, Kaituo Wang, and Pontus Gourdon. Structure and ion-release mechanism of PIB-4-type ATPases. *eLife*, 10, 2021.

[9023] Hao Gu, Xi Zeng, Liusheng Peng, Chuanying Xiang, Yangyang Zhou, Xiaomin Zhang, Jixin Zhang, Ning Wang, Gang Guo, Yan Li, Kaiyun Liu, Jiang Gu, Hao Zeng, Yuan Zhuang, Haibo Li, Jinyong Zhang, Weijun Zhang, Quanming Zou, and Yun Shi. Vaccination induces rapid protection against bacterial pneumonia via training alveolar macrophage in mice. *eLife*, 10, 2021.

[9024] Gabriel A Guerrero, Maxime J Derisbourg, Felix AMC Mayr, Laura E Wester, Marco Giorda, J Eike Dinort, Matías D Hartman, Klara Schilling, María José Alonso-De Gennaro, Ryan J Lu, Bérénice A Benayoun, and Martin S Denzel. NHR-8 and P-glycoproteins uncouple xenobiotic resistance from longevity in chemosensory C. elegans mutants. *eLife*, 10, 2021.

[9025] Paul Guilhamon, Charles Chesnelong, Michelle M Kushida, Ana Nikolic, Divya Singhal, Graham MacLeod, Seyed Ali Madani Tonekaboni, Florence MG Cavalli, Christopher Arlidge, Nishani Rajakulendran, Naghmeh Rastegar, Xiaoguang Hao, Rozina Hassam, Laura J Smith, Heather Whetstone, Fiona J Coutinho, Bettina Nadorp, Katrina I Ellestad, H Artee Luchman, Jennifer Ai-wen Chan, Molly S Shoichet, Michael D Taylor, Benjamin Haibe-Kains, Samuel Weiss, Stephane Angers,

Marco Gallo, Peter B Dirks, and Mathieu Lupien. Single-cell chromatin accessibility profiling of glioblastoma identifies an invasive cancer stem cell population associated with lower survival. *eLife*, 10, 2021.

[9026] Charlene Guillot, Yannis Djeffal, Arthur Michaut, Brian Rabe, and Olivier Pourquié. Dynamics of primitive streak regression controls the fate of neuromesodermal progenitors in the chicken embryo. *eLife*, 10, 2021.

[9027] Jian-Zhong Guo, Britton A Sauerbrei, Jeremy D Cohen, Matteo Mischiati, Austin R Graves, Ferruccio Pisanello, Kristin M Branson, and Adam W Hantman. Disrupting cortico-cerebellar communication impairs dexterity. *eLife*, 10, 2021.

[9028] Lei Guo, Xin-yu Fan, Xiaomu Qiao, Craig Montell, and Jia Huang. An octopamine receptor confers selective toxicity of amitraz on honeybees and Varroa mites. *eLife*, 10, 2021.

[9029] Monica S Guo, Ryo Kawamura, Megan L Littlehale, John F Marko, and Michael T Laub. High-resolution, genome-wide mapping of positive supercoiling in chromosomes. *eLife*, 10, 2021.

[9030] Qiuyu Guo, Albert Kim, Bin Li, Andrew Ransick, Helena Bugacov, Xi Chen, Nils Lindström, Aaron Brown, Leif Oxburgh, Bing Ren, and Andrew P McMahon. A $\beta$-catenin-driven switch in TCF/LEF transcription factor binding to DNA target sites promotes commitment of mammalian nephron progenitor cells. *eLife*, 10, 2021.

[9031] Xiaohu Guo, Annika Söderholm, Sandesh Kanchugal P, Geir V Isaksen, Omar Warsi, Ulrich Eckhard, Silvia Trigüis, Adolf Gogoll, Jon Jerlström-Hultqvist, Johan Åqvist, Dan I Andersson, and Maria Selmer. Structure and mechanism of a phage-encoded SAM lyase revises catalytic function of enzyme family. *eLife*, 10, 2021.

[9032] Yang Guo, Ze-Yan Yu, Jianxin Wu, Hutao Gong, Scott Kesteven, Siiri E Iismaa, Andrea Y Chan, Sara Holman, Silvia Pinto, Andy Pironet, Charles D Cox, Robert M Graham, Rudi Vennekens, Michael P Feneley, and Boris Martinac. The Ca2+-activated cation channel TRPM4 is a positive regulator of pressure overload-induced cardiac hypertrophy. *eLife*, 10, 2021.

[9033] Rahul Gupta, Konrad J Karczewski, Daniel Howrigan, Benjamin M Neale, and Vamsi K Mootha. Human genetic analyses of organelles highlight the nucleus in age-related trait heritability. *eLife*, 10, 2021.

[9034] Ritu Gupta and Sunil Laxman. Cycles, sources, and sinks: Conceptualizing how phosphate balance modulates carbon flux using yeast metabolic networks. *eLife*, 10, 2021.

[9035] Shalini Gupta, Larry J Friedman, Jeff Gelles, and Stephen P Bell. A helicase-tethered ORC flip enables bidirectional helicase loading. *eLife*, 10, 2021.

[9036] Manickam Gurusaran and Owen Richard Davies. A molecular mechanism for LINC complex branching by structurally diverse SUN-KASH 6:6 assemblies. *eLife*, 10, 2021.

[9037] Jenny K Gustafsson, Jazmyne E Davis, Tracy Rappai, Keely G McDonald, Devesha H Kulkarni, Kathryn A Knoop, Simon P Hogan, James AJ Fitzpatrick, Wayne I Lencer, and Rodney D Newberry. Intestinal goblet cells sample and deliver lumenal antigens by regulated endocytic uptake and transcytosis. *eLife*, 10, 2021.

[9038] Axel A Guzmán-Solís, Viridiana Villa-Islas, Miriam J Bravo-López, Marcela Sandoval-Velasco, Julie K Wesp, Jorge A Gómez-Valdés, María de la Luz Moreno-Cabrera, Alejandro Meraz, Gabriela Solís-Pichardo, Peter Schaaf, Benjamin R TenOever, Daniel Blanco-Melo, and María C ávila Arcos. Ancient viral genomes reveal introduction of human pathogenic viruses into Mexico during the transatlantic slave trade. *eLife*, 10, 2021.

[9039] Thomas Hackl, Sarah Duponchel, Karina Barenhoff, Alexa Weinmann, and Matthias G Fischer. Virophages and retrotransposons colonize the genomes of a heterotrophic flagellate. *eLife*, 10, 2021.

[9040] Sara Hägg and Juulia Jylhävä. Sex differences in biological aging with a focus on human studies. *eLife*, 10, 2021.

[9041] Lukas Alexander Hahn, Dmitry Balakhonov, Erica Fongaro, Andreas Nieder, and Jonas Rose. Working memory capacity of crows and monkeys arises from similar neuronal computations. *eLife*, 10, 2021.

[9042] Baruch Haimson, Yoav Hadas, Nimrod Bernat, Artur Kania, Monica A Daley, Yuval Cinnamon, Aharon Lev-Tov, and Avihu Klar. Spinal lumbar dI2 interneurons contribute to stability of bipedal stepping. *eLife*, 10, 2021.

[9043] Bence Hajdusits, Marcin J Suskiewicz, Nikolas Hundt, Anton Meinhart, Robert Kurzbauer, Julia Leodolter, Philipp Kukura, and Tim Clausen. McsB forms a gated kinase chamber to mark aberrant bacterial proteins for degradation. *eLife*, 10, 2021.

[9044] Azadeh HajiHosseini and Cendri A Hutcherson. Alpha oscillations and event-related potentials reflect distinct dynamics of attribute construction and evidence accumulation in dietary decision making. *eLife*, 10, 2021.

[9045] Sanja M Hakala, Marie-Pierre Meurville, Michael Stumpe, and Adria C LeBoeuf. Biomarkers in a socially exchanged fluid reflect colony maturity, behavior, and distributed metabolism. *eLife*, 10, 2021.

[9046] Caryn R Hale, Kirsty Sawicka, Kevin Mora, John J Fak, Jin Joo Kang, Paula Cutrim, Katarzyna Cialowicz, Thomas S Carroll, and Robert B Darnell. FMRP regulates mRNAs encoding distinct functions in the cell body and dendrites of CA1 pyramidal neurons. *eLife*, 10, 2021.

[9047] Eric T Hall, Miriam E Dillard, Daniel P Stewart, Yan Zhang, Ben Wagner, Rachel M Levine, Shondra M Pruett-Miller, April Sykes, Jamshid Temirov, Richard E Cheney, Motomi Mori, Camenzind G Robinson, and Stacey K Ogden. Cytoneme delivery of Sonic Hedgehog from ligand-producing cells requires Myosin 10 and a Dispatched-BOC/CDON co-receptor complex. *eLife*, 10, 2021.

[9048] Pille Hallast, Laura Kibena, Margus Punab, Elena Arciero, Siiri Rootsi, Marina Grigorova, Rodrigo Flores, Mark A Jobling, Olev Poolamets, Kristjan Pomm, Paul Korrovits, Kristiina Rull, Yali Xue, Chris Tyler-Smith, and Maris Laan. A common 1.6 mb Y-chromosomal inversion predisposes to subsequent deletions and severe spermatogenic failure in humans. *eLife*, 10, 2021.

[9049] Shawn A Hallett, Yuki Matsushita, Wanida Ono, Naoko Sakagami, Koji Mizuhashi, Nicha Tokavanich, Mizuki Nagata, Annabelle Zhou, Takao Hirai, Henry M Kronenberg, and Noriaki Ono. Chondrocytes in the resting zone of the growth plate are maintained in a Wnt-inhibitory environment. *eLife*, 10, 2021.

[9050] Fletcher W Halliday, Mikko Jalo, and Anna-Liisa Laine. The effect of host community functional traits on plant disease risk varies along an elevational gradient. *eLife*, 10, 2021.

[9051] Kelsey M Hallinen, Ross Dempsey, Monika Scholz, Xinwei Yu, Ashley Linder, Francesco Randi, Anuj K Sharma, Joshua W Shaevitz, and Andrew M Leifer. Decoding locomotion from population neural activity in moving C. elegans. *eLife*, 10, 2021.

[9052] Thierry Halter, Jingyu Wang, Delase Amesefe, Emmanuelle Lastrucci, Magali Charvin, Meenu Singla Rastogi, and Lionel Navarro. The Arabidopsis active demethylase ROS1 cis-regulates defence genes by erasing DNA methylation at promoter-regulatory regions. *eLife*, 10, 2021.

[9053] Lucy Ham, Marcel Jackson, and Michael PH Stumpf. Pathway dynamics can delineate the sources of transcriptional noise in gene expression. *eLife*, 10, 2021.

[9054] Iman Hamid, Katharine L Korunes, Sandra Beleza, and Amy Goldberg. Rapid adaptation to malaria facilitated by admixture in the human population of Cabo Verde. *eLife*, 10, 2021.

[9055] Allison E Hamilos, Giulia Spedicato, Ye Hong, Fangmiao Sun, Yulong Li, and John A Assad. Slowly evolving dopaminergic activity modulates the moment-to-moment probability of reward-related self-timed movements. *eLife*, 10, 2021.

[9056] Andrew M Hamilton, Olga A Balashova, and Laura N Borodinsky. Non-canonical Hedgehog signaling regulates spinal cord and muscle regeneration in Xenopus laevis larvae. *eLife*, 10, 2021.

[9057] William L Hamilton, Gerry Tonkin-Hill, Emily R Smith, Dinesh Aggarwal, Charlotte J Houldcroft, Ben Warne, Luke W Meredith, Myra Hosmillo, Aminu S Jahun, Martin D Curran, Surendra Parmar, Laura G Caller, Sarah L Caddy, Fahad A Khokhar, Anna Yakovleva, Grant Hall, Theresa Feltwell, Malte L Pinckert, Iliana Georgana, Yasmin Chaudhry, Colin S Brown, Sonia GonÇalves, Roberto Amato, Ewan M Harrison, Nicholas M Brown, Mathew A Beale, Michael Spencer Chapman, David K Jackson, Ian Johnston, Alex Alderton, John Sillitoe, Cordelia Langford, Gordon Dougan, Sharon J Peacock, Dominic P Kwiatowski, Ian G Goodfellow, and M Estee Torok. Genomic epidemiology of COVID-19 in care homes in the east of England. *eLife*, 10, 2021.

[9058] David W Hammers, Cora C Hart, Michael K Matheny, Ernest G Heimsath, Young il Lee, John A Hammer, Richard E Cheney, and H Lee Sweeney. Filopodia powered by class x myosin promote fusion of mammalian myoblasts. *eLife*, 10, 2021.

[9059] Reza K Hammond, Matthew C Pahl, Chun Su, Diana L Cousminer, Michelle E Leonard, Sumei Lu, Claudia A Doege, Yadav Wagley, Kenyaita M Hodge, Chiara Lasconi, Matthew E Johnson, James A Pippin, Kurt D Hankenson, Rudolph L Leibel, Alessandra Chesi, Andrew D Wells, and Struan FA Grant. Biological constraints on GWAS SNPs at suggestive significance thresholds reveal additional BMI loci. *eLife*, 10, 2021.

[9060] Alvin X Han, Zandra C Felix Garza, Matthijs RA Welkers, René M Vigeveno, Nhu Duong Tran, Thi Quynh Mai Le, Thai Pham Quang, Dinh Thoang Dang, Thi Ngoc Anh Tran, Manh Tuan Ha, Thanh Hung Nguyen, Quoc Thinh Le, Thanh Hai Le, Thi Bich Ngoc Hoang, Kulkanya Chokephaibulkit, Pilaipan Puthavathana, Van Vinh Chau Nguyen, My Ngoc Nghiem, Van Kinh Nguyen, Tuyet Trinh Dao, Tinh Hien Tran, Heiman FL Wertheim, Peter W Horby, Annette Fox, H Rogier van Doorn, Dirk Eggink, Menno D de Jong, and Colin A Russell. Within-host evolutionary dynamics of seasonal and pandemic human influenza A viruses in young children. *eLife*, 10, 2021.

[9061] Bin Han, Qiaohong Wei, Fan Wu, Han Hu, Chuan Ma, Lifeng Meng, Xufeng Zhang, Mao Feng, Yu Fang, Olav Rueppell, and Jianke Li. Tachykinin signaling inhibits task-specific behavioral responsiveness in honeybee workers. *eLife*, 10, 2021.

[9062] Jeong-Kyu Han, Sun-Ho Kwon, Yong Gyu Kim, Jaeyong Choi, Jong-Il Kim, Yong-Seok Lee, Sang-Kyu Ye, and Sang Jeong Kim. Ablation of STAT3 in Purkinje cells reorganizes cerebellar synaptic plasticity in long-term fear memory network. *eLife*, 10, 2021.

[9063] Sangyoon J Han, Evgenia V Azarova, Austin J Whitewood, Alexia Bachir, Edgar Guttierrez, Alex Groisman, Alan R Horwitz, Benjamin T Goult, Kevin M Dean, and Gaudenz Danuser. Pre-complexation of talin and vinculin without tension is required for efficient nascent adhesion maturation. *eLife*, 10, 2021.

[9064] Xia Han, Jifan Feng, Tingwei Guo, Yong-Hwee Eddie Loh, Yuan Yuan, Thach-Vu Ho, Courtney Kyeong Cho, Jingyuan Li, Junjun Jing, Eva Janeckova, Jinzhi He, Fei Pei, Jing Bi, Brian Song, and Yang Chai. Runx2-Twist1 interaction coordinates cranial neural crest guidance of soft palate myogenesis. *eLife*, 10, 2021.

[9065] Simona Hankeova, Jakub Salplachta, Tomas Zikmund, Michaela Kavkova, Noémi Van Hul, Adam Brinek, Veronika Smekalova, Jakub Laznovsky, Feven Dawit, Josef Jaros, Vítězslav Bryja, Urban Lendahl, Ewa Ellis, Antal Nemeth, Björn Fischler, Edouard Hannezo, Jozef Kaiser, and Emma Rachel Andersson. DUCT reveals architectural mechanisms contributing to bile duct recovery in a mouse model for Alagille syndrome. *eLife*, 10, 2021.

[9066] Eilis Hannon, Emma L Dempster, Georgina Mansell, Joe Burrage, Nick Bass, Marc M Bohlken, Aiden Corvin, Charles J Curtis, David Dempster, Marta Di Forti, Timothy G Dinan, Gary Donohoe, Fiona Gaughran, Michael Gill, Amy Gillespie, Cerisse Gunasinghe, Hilleke E Hulshoff, Christina M Hultman, Viktoria Johansson, René S Kahn, Jaakko Kaprio, Gunter Kenis, Kaarina Kowalec, James MacCabe, Colm McDonald, Andrew McQuillin, Derek W Morris, Kieran C Murphy, Colette J Mustard, Igor Nenadic, Michael C O'Donovan, Diego Quattrone, Alexander L Richards, Bart PF Rutten, David St Clair, Sebastian Therman, Timothea Toulopoulou, Jim Van Os, John L Waddington, Patrick Sullivan, Evangelos Vassos, Gerome Breen, David Andrew Collier, Robin M Murray, Leonard S Schalkwyk, and Jonathan Mill. DNA methylation meta-analysis reveals cellular alterations in psychosis and markers of treatment-resistant schizophrenia. *eLife*, 10, 2021.

[9067] Jesse M Hansen, Avital Horowitz, Eric M Lynch, Daniel P Farrell, Joel Quispe, Frank DiMaio, and Justin M Kollman. Cryo-EM structures of CTP synthase filaments reveal mechanism of pH-sensitive assembly during budding yeast starvation. *eLife*, 10, 2021.

[9068] Hongyan Hao, Shilpi Kalra, Laura E Jameson, Leslie A Guerrero, Natalie E Cain, Jessica Bolivar, and Daniel A Starr. The Nesprin-1/-2 ortholog ANC-1 regulates organelle positioning in C. elegans independently from its KASH or actin-binding domains. *eLife*, 10, 2021.

[9069] Yaoyao Hao, Alyse Marian Thomas, and Nuo Li. Fully autonomous mouse behavioral and optogenetic experiments in home-cage. *eLife*, 10, 2021.

[9070] Paz Har-shai Yahav and Elana Zion Golumbic. Linguistic processing of task-irrelevant speech at a cocktail party. *eLife*, 10, 2021.

[9071] Akikazu Harada, Shinji Matsumoto, Yoshiaki Yasumizu, Kensaku Shojima, Toshiyuki Akama, Hidetoshi Eguchi, and Akira Kikuchi. Localization of KRAS downstream target ARL4C to invasive pseudopods accelerates pancreatic cancer cell invasion. *eLife*, 10, 2021.

[9072] Ben J Hardcastle, Jaison J Omoto, Pratyush Kandimalla, Bao-Chau M Nguyen, Mehmet F Keleş, Natalie K Boyd, Volker Hartenstein, and Mark A Frye. A visual pathway for skylight polarization processing in Drosophila. *eLife*, 10, 2021.

[9073] Diane M Harper, Madiha Tariq, Asraa Alhawli, Nadia Syed, Minal Patel, and Ken Resnicow. Cancer risk perception and physician communication behaviors on cervical cancer and colorectal cancer screening. *eLife*, 10, 2021.

[9074] Neale J Harrison, Elizabeth Connolly, Alicia Gascón Gubieda, Zidan Yang, Benjamin Altenhein, Maria Losada Perez, Marta Moreira, Jun Sun, and Alicia Hidalgo. Regenerative neurogenic response from glia requires insulin-driven neuron-glia communication. *eLife*, 10, 2021.

[9075] Samuel Frederick Mock Hart, Chi-Chun Chen, and Wenying Shou. Pleiotropic mutations can rapidly evolve to directly benefit self and cooperative partner despite unfavorable conditions. *eLife*, 10, 2021.

[9076] William S Hart, Philip K Maini, and Robin N Thompson. High infectiousness immediately before COVID-19 symptom onset highlights the importance of continued contact tracing. *eLife*, 10, 2021.

[9077] Hayden AM Hatch, Helen M Belalcazar, Owen J Marshall, and Julie Secombe. A KDM5–Prospero transcriptional axis functions during early neurodevelopment to regulate mushroom body formation. *eLife*, 10, 2021.

[9078] Ryo Hatori, Brent M Wood, Guilherme Oliveira Barbosa, and Thomas B Kornberg. Regulated delivery controls Drosophila Hedgehog, Wingless, and Decapentaplegic signaling. *eLife*, 10, 2021.

[9079] Mark E Hauber, Matthew IM Louder, and Simon C Griffith. Neurogenomic insights into the behavioral and vocal development of the zebra finch. *eLife*, 10, 2021.

[9080] Kelsey A Haugh, Mark S Ladinsky, Irfan Ullah, Helen M Stone, Ruoxi Pi, Alexandre Gilardet, Michael W Grunst, Priti Kumar, Pamela J Bjorkman, Walther Mothes, and Pradeep D Uchil. In vivo imaging of retrovirus infection reveals a role for Siglec-1/CD169 in multiple routes of transmission. *eLife*, 10, 2021.

[9081] Fiona Haward, Magdalena M Maslon, Patricia L Yeyati, Nicolas Bellora, Jan N Hansen, Stuart Aitken, Jennifer Lawson, Alex von Kriegsheim, Dagmar Wachten, Pleasantine Mill, Ian R Adams, and Javier F Caceres. Nucleo-cytoplasmic shuttling of splicing factor SRSF1 is required for development and cilia function. *eLife*, 10, 2021.

[9082] Cassandra L Hays, Asia L Sladek, Greg D Field, and Wallace B Thoreson. Properties of multivesicular release from mouse rod photoreceptors support transmission of single-photon responses. *eLife*, 10, 2021.

[9083] Kai He, Triston G Eastman, Hannah Czolacz, Shuhao Li, Akio Shinohara, Shin-ichiro Kawada, Mark S Springer, Michael Berenbrink, and Kevin L Campbell. Myoglobin primary structure reveals multiple convergent transitions to semi-aquatic life in the world's smallest mammalian divers. *eLife*, 10, 2021.

[9084] Lizhi He, Henry Pratt, Mingshi Gao, Fengxiang Wei, Zhiping Weng, and Kevin Struhl. YAP and TAZ are transcriptional co-activators of AP-1 proteins and STAT3 during breast cellular transformation. *eLife*, 10, 2021.

[9085] Xinyi Jenny He, Janki Patel, Connor E Weiss, Xiang Ma, Brenda L Bloodgood, and Matthew R Banghart. Convergent, functionally independent signaling by mu and delta opioid receptors in hippocampal parvalbumin interneurons. *eLife*, 10, 2021.

[9086] Yujiao He, Zhiguo Yang, Chen-si Zhao, Zhihui Xiao, Yu Gong, Yun-Yi Li, Yiqi Chen, Yunting Du, Dianying Feng, Amnon Altman, and Yingqiu Li. T-cell receptor (TCR) signaling promotes the assembly of RanBP2/RanGAP1-SUMO1/Ubc9 nuclear pore subcomplex via PKC-$\theta$-mediated phosphorylation of RanGAP1. *eLife*, 10, 2021.

[9087] Christoph Heier, Oskar Knittelfelder, Harald F Hofbauer, Wolfgang Mende, Ingrid Pörnbacher, Laura Schiller, Gabriele Schoiswohl, Hao Xie, Sebastian Grönke, Andrej Shevchenko, and Ronald P Kühnlein. Hormone-sensitive lipase couples intergenerational sterol metabolism to reproductive success. *eLife*, 10, 2021.

[9088] Maren Heimhalt, Alex Berndt, Jane Wagstaff, Madhanagopal Anandapadamanaban, Olga Perisic, Sarah Maslen, Stephen McLaughlin, Conny Wing-Heng Yu, Glenn R Masson, Andreas Boland, Xiaodan Ni, Keitaro Yamashita, Garib N Murshudov, Mark Skehel, Stefan M Freund, and Roger L Williams. Bipartite binding and partial inhibition links DEPTOR and mTOR in a mutually antagonistic embrace. *eLife*, 10, 2021.

[9089] Stanley Heinze, Basil el Jundi, Bente G Berg, Uwe Homberg, Randolf Menzel, Keram Pfeiffer, Ronja Hensgen, Frederick Zittrell, Marie Dacke, Eric Warrant, Gerit Pfuhl, Jürgen Rybak, and Kevin Tedore. A unified platform to manage, share, and archive morphological and functional data in insect neuroscience. *eLife*, 10, 2021.

[9090] Mathias L Heltberg, Judith Miné-Hattab, Angela Taddei, Aleksandra M Walczak, and Thierry Mora. Physical observables to determine the nature of membrane-less cellular sub-compartments. *eLife*, 10, 2021.

[9091] Karthik Amudhala Hemanthakumar, Shentong Fang, Andrey Anisimov, Mikko I Mäyränpää, Eero Mervaala, and Riikka Kivelä. Cardiovascular disease risk factors induce mesenchymal features and senescence in mouse cardiac endothelial cells. *eLife*, 10, 2021.

[9092] Audrey M Hendley, Arjun A Rao, Laura Leonhardt, Sudipta Ashe, Jennifer A Smith, Simone Giacometti, Xianlu L Peng, Honglin Jiang, David I Berrios, Mathias Pawlak, Lucia Y Li, Jonghyun

Lee, Eric A Collisson, Mark S Anderson, Gabriela K Fragiadakis, Jen Jen Yeh, Chun Jimmie Ye, Grace E Kim, Valerie M Weaver, and Matthias Hebrok. Single-cell transcriptome analysis defines heterogeneity of the murine pancreatic ductal tree. *eLife*, 10, 2021.

[9093] Siyu Heng, Wendy P O'Meara, Ryan A Simmons, and Dylan S Small. Relationship between changing malaria burden and low birth weight in sub-Saharan Africa: A difference-in-differences study via a pair-of-pairs approach. *eLife*, 10, 2021.

[9094] Josephine Henke, Raven Bunk, Dina von Werder, Stefan Häusler, Virginia L Flanagin, and Kay Thurley. Distributed coding of duration in rodent prefrontal cortex during time reproduction. *eLife*, 10, 2021.

[9095] Philipp Henneke, Katrin Kierdorf, Lindsay J Hall, Markus Sperandio, and Mathias Hornef. Perinatal development of innate immune topology. *eLife*, 10, 2021.

[9096] Camille Henry, Laurent Loiseau, Alexandra Vergnes, Didier Vertommen, Angela Mérida-Floriano, Sindhu Chitteni-Pattu, Elizabeth A Wood, Josep Casadesús, Michael M Cox, Frédéric Barras, and Benjamin Ezraty. Redox controls RecA protein activity via reversible oxidation of its methionine residues. *eLife*, 10, 2021.

[9097] Cristina Herencias, Jerónimo Rodríguez-Beltrán, Ricardo León-Sampedro, Aida Alonso-del Valle, Jana Palkovičová, Rafael Cantón, and álvaro San Millán. Collateral sensitivity associated with antibiotic resistance plasmids. *eLife*, 10, 2021.

[9098] Damián G Hernández, Catalina Rivera, Jessica Cande, Baohua Zhou, David L Stern, and Gordon J Berman. A framework for studying behavioral evolution by reconstructing ancestral repertoires. *eLife*, 10, 2021.

[9099] Liz Hernandez Borrero, David T Dicker, John Santiago, Jennifer Sanders, Xiaobing Tian, Nagib Ahsan, Avital Lev, Lanlan Zhou, and Wafik S El-Deiry. A subset of CB002 xanthine analogs bypass p53-signaling to restore a p53 transcriptome and target an S-phase cell cycle checkpoint in tumors with mutated-p53. *eLife*, 10, 2021.

[9100] Elvira Hernandez-Lagana, Gabriella Mosca, Ethel Mendocilla-Sato, Nuno Pires, Anja Frey, Alejandro Giraldo-Fonseca, Caroline Michaud, Ueli Grossniklaus, Olivier Hamant, Christophe Godin, Arezki Boudaoud, Daniel Grimanelli, Daphné Autran, and Célia Baroux. Organ geometry channels reproductive cell fate in the Arabidopsis ovule primordium. *eLife*, 10, 2021.

[9101] Víctor M Hernández-Rocamora, Natalia Baranova, Katharina Peters, Eefjan Breukink, Martin Loose, and Waldemar Vollmer. Real-time monitoring of peptidoglycan synthesis by membrane-reconstituted penicillin-binding proteins. *eLife*, 10, 2021.

[9102] Helen Hertenstein, Ellen McMullen, Astrid Weiler, Anne Volkenhoff, Holger M Becker, and Stefanie Schirmeier. Starvation-induced regulation of carbohydrate transport at the blood–brain barrier is TGF-$\beta$-signaling dependent. *eLife*, 10, 2021.

[9103] Julia Hesse, Christoph Owenier, Tobias Lautwein, Ria Zalfen, Jonas F Weber, Zhaoping Ding, Christina Alter, Alexander Lang, Maria Grandoch, Norbert Gerdes, Jens W Fischer, Gunnar W Klau, Christoph Dieterich, Karl Köhrer, and Jürgen Schrader. Single-cell transcriptomics defines heterogeneity of epicardial cells and fibroblasts within the infarcted murine heart. *eLife*, 10, 2021.

[9104] Alexander J Hetherington, Siobhán L Bridson, Anna Lee Jones, Hagen Hass, Hans Kerp, and Liam Dolan. An evidence-based 3D reconstruction of Asteroxylon mackiei, the most complex plant preserved from the Rhynie chert. *eLife*, 10, 2021.

[9105] David Hickey, Andrej Vilfan, and Ramin Golestanian. Ciliary chemosensitivity is enhanced by cilium geometry and motility. *eLife*, 10, 2021.

[9106] Ben J Hightower, Patrick WA Wijnings, Rick Scholte, Rivers Ingersoll, Diana D Chin, Jade Nguyen, Daniel Shorr, and David Lentink. How oscillating aerodynamic forces explain the timbre of the hummingbird's hum and other animals in flapping flight. *eLife*, 10, 2021.

[9107] Cameron Hill, Elisabetta Brunello, Luca Fusi, Jesús G Ovejero, and Malcolm Irving. Myosin-based regulation of twitch and tetanic contractions in mammalian skeletal muscle. *eLife*, 10, 2021.

[9108] Danika L Hill, Carly E Whyte, Silvia Innocentin, Jia Le Lee, James Dooley, Jiong Wang, Eddie A James, James C Lee, William W Kwok, Martin S Zand, Adrian Liston, Edward J Carr, and Michelle A Linterman. Impaired HA-specific T follicular helper cell and antibody responses to influenza vaccination are linked to inflammation in humans. *eLife*, 10, 2021.

[9109] Pamela Himadewi, Xue Qing David Wang, Fan Feng, Haley Gore, Yushuai Liu, Lei Yu, Ryo Kurita, Yukio Nakamura, Gerd P Pfeifer, Jie Liu, and Xiaotian Zhang. 3'HS1 CTCF binding site in human $\beta$-globin locus regulates fetal hemoglobin expression. *eLife*, 10, 2021.

[9110] Katie Hinde, Carlos Eduardo G Amorim, Alyson F Brokaw, Nicole Burt, Mary C Casillas, Albert Chen, Tara Chestnut, Patrice K Connors, Mauna Dasari, Connor Fox Ditelberg, Jeanne Dietrick, Josh Drew, Lara Durgavich, Brian Easterling, Charon Henning, Anne Hilborn, Elinor K Karlsson, Marc Kissel, Jennifer Kobylecky, Jason Krell, Danielle N Lee, Kate M Lesciotto, Kristi L Lewton, Jessica E Light, Jessica Martin, Asia Murphy, William Nickley, Alejandra Núñez-de la Mora, Olivia Pellicer, Valeria Pellicer, Anali Maughan Perry, Stephanie G Schuttler, Anne C Stone, Brian Tanis, Jesse Weber, Melissa Wilson, Emma Willcocks, and Christopher N Anderson. March Mammal Madness and the power of narrative in science outreach. *eLife*, 10, 2021.

[9111] Amelia E Hinman, Charul Jani, Stephanie C Pringle, Wei R Zhang, Neharika Jain, Amanda J Martinot, and Amy K Barczak. Mycobacterium tuberculosis canonical virulence factors interfere with a late component of the TLR2 response. *eLife*, 10, 2021.

[9112] Tjorven Hinzke, Manuel Kleiner, Mareike Meister, Rabea Schlüter, Christian Hentschker, Jan Pané-Farré, Petra Hildebrandt, Horst Felbeck, Stefan M Sievert, Florian Bonn, Uwe Völker, Dörte Becher, Thomas Schweder, and Stephanie Markert. Bacterial symbiont subpopulations have different roles in a deep-sea symbiosis. *eLife*, 10, 2021.

[9113] Ken-ichi Hirano, Hiroyuki Hosokawa, Maria Koizumi, Yusuke Endo, Takashi Yahata, Kiyoshi Ando, and Katsuto Hozumi. LMO2 is essential to maintain the ability of progenitors to differentiate into T-cell lineage in mice. *eLife*, 10, 2021.

[9114] Tetsuya Hisanaga, Shota Fujimoto, Yihui Cui, Katsutoshi Sato, Ryosuke Sano, Shohei Yamaoka, Takayuki Kohchi, Frédéric Berger, and Keiji Nakajima. Deep evolutionary origin of gamete-directed zygote activation by KNOX/BELL transcription factors in green plants. *eLife*, 10, 2021.

[9115] Jessica SY Ho, Federico Di Tullio, Megan Schwarz, Diana Low, Danny Incarnato, Florence Gay, Tommaso Tabaglio, JingXian Zhang, Heike Wollmann, Leilei Chen, Omer An, Tim Hon Man Chan, Alexander Hall Hickman, Simin Zheng, Vladimir Roudko, Sujun Chen, Alcida Karz, Musaddeque Ahmed, Housheng Hansen He, Benjamin D Greenbaum, Salvatore Oliviero, Michela Serresi, Gaetano Gargiulo, Karen M Mann, Eva Hernando, David Mulholland, Ivan Marazzi, Dave Keng Boon Wee, and Ernesto Guccione. HNRNPM controls circRNA biogenesis and splicing fidelity to sustain cancer cell fitness. *eLife*, 10, 2021.

[9116] Thao Phuong Ho-Le, Thach S Tran, Dana Bliuc, Hanh M Pham, Steven A Frost, Jacqueline R Center, John A Eisman, and Tuan V Nguyen. Epidemiological transition to mortality and refracture following an initial fracture. *eLife*, 10, 2021.

[9117] Andreas Hober, Khue Hua Tran-Minh, Dominic Foley, Thomas McDonald, Johannes PC Vissers, Rebecca Pattison, Samantha Ferries, Sigurd Hermansson, Ingvar Betner, Mathias Uhlén, Morteza Razavi, Richard Yip, Matthew E Pope, Terry W Pearson, Leigh N Andersson, Amy Bartlett, Lisa Calton, Jessica J Alm, Lars Engstrand, and Fredrik Edfors. Rapid and sensitive detection of SARS-CoV-2 infection using quantitative peptide enrichment LC-MS analysis. *eLife*, 10, 2021.

[9118] David L Hocker, Carlos D Brody, Cristina Savin, and Christine M Constantinople. Subpopulations of neurons in lOFC encode previous and current rewards at time of choice. *eLife*, 10, 2021.

[9119] Kenneth B Hoehn, Jackson S Turner, Frederick I Miller, Ruoyi Jiang, Oliver G Pybus, Ali H Ellebedy, and Steven H Kleinstein. Human B cell lineages associated with germinal centers following influenza vaccination are measurably evolving. *eLife*, 10, 2021.

[9120] Marieke MB Hoekstra, Maxime Jan, Georgia Katsioudi, Yann Emmenegger, and Paul Franken. The sleep-wake distribution contributes to the peripheral rhythms in PERIOD-2. *eLife*, 10, 2021.

[9121] Astrid Hoermann, Sofia Tapanelli, Paolo Capriotti, Giuseppe Del Corsano, Ellen KG Masters, Tibebu Habtewold, George K Christophides, and Nikolai Windbichler. Converting endogenous genes of the malaria mosquito into simple non-autonomous gene drives for population replacement. *eLife*, 10, 2021.

[9122] Patrick C Hoffmann, Stefano L Giandomenico, Iva Ganeva, Michael R Wozny, Magdalena Sutcliffe, Madeline A Lancaster, and Wanda Kukulski. Electron cryo-tomography reveals the subcellular architecture of growing axons in human brain organoids. *eLife*, 10, 2021.

[9123] Simon M Hofmann, Felix Klotzsche, Alberto Mariola, Vadim Nikulin, Arno Villringer, and Michael Gaebler. Decoding subjective emotional arousal from EEG during an immersive virtual reality experience. *eLife*, 10, 2021.

[9124] Lindsay A Hohsfield, Allison R Najafi, Yasamine Ghorbanian, Neelakshi Soni, Joshua Crapser, Dario X Figueroa Velez, Shan Jiang, Sarah E Royer, Sung Jin Kim, Caden M Henningfield, Aileen Anderson, Sunil P Gandhi, Ali Mortazavi, Matthew A Inlay, and Kim N Green. Subventricular zone/white matter microglia reconstitute the empty adult microglial niche in a dynamic wave. *eLife*, 10, 2021.

[9125] Carl D Holmgren, Paul Stahr, Damian J Wallace, Kay-Michael Voit, Emily J Matheson, Juergen Sawinski, Giacomo Bassetto, and Jason ND Kerr. Visual pursuit behavior in mice maintains the pursued prey on the retinal region with least optic flow. *eLife*, 10, 2021.

[9126] Melanie Holmgren, Michael E Ravicz, Kenneth E Hancock, Olga Strelkova, Dorina Kallogjeri, Artur A Indzhykulian, Mark E Warchol, and Lavinia Sheets. Mechanical overstimulation causes acute injury and synapse loss followed by fast recovery in lateral-line neuromasts of larval zebrafish. *eLife*, 10, 2021.

[9127] Jesse R Holt, Wei-Zheng Zeng, Elizabeth L Evans, Seung-Hyun Woo, Shang Ma, Hamid Abuwarda, Meaghan Loud, Ardem Patapoutian, and Medha M Pathak. Spatiotemporal dynamics of PIEZO1 localization controls keratinocyte migration during wound healing. *eLife*, 10, 2021.

[9128] Yukiko Hori, Koki Mimura, Yuji Nagai, Atsushi Fujimoto, Kei Oyama, Erika Kikuchi, Ken-ichi Inoue, Masahiko Takada, Tetsuya Suhara, Barry J Richmond, and Takafumi Minamimoto. Single caudate neurons encode temporally discounted value for formulating motivation for action. *eLife*, 10, 2021.

[9129] Mahmood S Hoseini, Bryan Higashikubo, Frances S Cho, Andrew H Chang, Alexandra Clemente-Perez, Irene Lew, Agnieszka Ciesielska, Michael P Stryker, and Jeanne T Paz. Gamma rhythms and visual information in mouse V1 specifically modulated by somatostatin+ neurons in reticular thalamus. *eLife*, 10, 2021.

[9130] Li Hou, Siyuan Guo, Yuanyuan Wang, Xin Nie, Pengcheng Yang, Ding Ding, Beibei Li, Le Kang, and Xianhui Wang. Neuropeptide ACP facilitates lipid oxidation and utilization during long-term flight in locusts. *eLife*, 10, 2021.

[9131] Martin Houlard, Erin E Cutts, Muhammad S Shamim, Jonathan Godwin, David Weisz, Aviva Presser Aiden, Erez Lieberman Aiden, Lothar Schermelleh, Alessandro Vannini, and Kim Nasmyth. MCPH1 inhibits Condensin II during interphase by regulating its SMC2-Kleisin interface. *eLife*, 10, 2021.

[9132] Leah Houri-Zeevi, Guy Teichman, Hila Gingold, and Oded Rechavi. Stress resets ancestral heritable small RNA responses. *eLife*, 10, 2021.

[9133] Elise Houssin, Mathieu Pinot, Karen Bellec, and Roland Le Borgne. Par3 cooperates with Sanpodo for the assembly of Notch clusters following asymmetric division of Drosophila sensory organ precursor cells. *eLife*, 10, 2021.

[9134] Aubrey GA Howard, Phillip A Baker, Rodrigo Ibarra-García-Padilla, Joshua A Moore, Lucia J Rivas, James J Tallman, Eileen W Singleton, Jessa L Westheimer, Julia A Corteguera, and Rosa A Uribe. An atlas of neural crest lineages along the posterior developing zebrafish at single-cell resolution. *eLife*, 10, 2021.

[9135] Emily Hsu, Nathan R Zemke, and Arnold J Berk. Promoter-specific changes in initiation, elongation, and homeostasis of histone H3 acetylation during CBP/p300 inhibition. *eLife*, 10, 2021.

[9136] Tsung-Yuan Hsu, Bo Zhang, Noelle D L'Etoile, and Bi-Tzen Juang. C. elegans orthologs MUT-7/CeWRN-1 of Werner syndrome protein regulate neuronal plasticity. *eLife*, 10, 2021.

[9137] Kevin Hu, Mahmoud Ghandi, and Franklin W Huang. Integrated evaluation of telomerase activation and telomere maintenance across cancer cell lines. *eLife*, 10, 2021.

[9138] Shikai Hu, Jacquelyn O Russell, Silvia Liu, Catherine Cao, Jackson McGaughey, Ravi Rai, Karis Kosar, Junyan Tao, Edward Hurley, Minakshi Poddar, Sucha Singh, Aaron Bell, Donghun Shin, Reben Raeman, Aatur D Singhi, Kari Nejak-Bowen, Sungjin Ko, and Satdarshan P Monga. $\beta$-Catenin-NF-$\kappa$B-CFTR interactions in cholangiocytes regulate inflammation and fibrosis during ductular reaction. *eLife*, 10, 2021.

[9139] Yang Hu, Chen Hu, Edmund Derrington, Brice Corgnet, Chen Qu, and Jean-Claude Dreher. Neural basis of corruption in power-holders. *eLife*, 10, 2021.

[9140] Delai Huang, Victor M Lewis, Tarah N Foster, Matthew B Toomey, Joseph C Corbo, and David M Parichy. Development and genetics of red coloration in the zebrafish relative Danio albolineatus. *eLife*, 10, 2021.

[9141] Fang Huang, Trang TT Nguyen, Ignacia Echeverria, Ramachandran Rakesh, Daniele C Cary, Hana Paculova, Andrej Sali, Arthur Weiss, Boris Matija Peterlin, and Koh Fujinaga. Reversible phosphorylation of cyclin T1 promotes assembly and stability of P-TEFb. *eLife*, 10, 2021.

[9142] Hui Huang, Youpei Lin, Wenrui Ma, Jiannan Liu, Jing Han, Xiaoyi Hu, Meilin Tang, Shiqiang Yan, Mieradilijiang Abudupataer, Chenping Zhang, Qiang Gao, and Weijia Zhang. A pre-screening strategy to assess resected tumor margins by imaging cytoplasmic viscosity and hypoxia. *eLife*, 10, 2021.

[9143] Lawrence Huang, Peter Ledochowitsch, Ulf Knoblich, Jérôme Lecoq, Gabe J Murphy, R Clay Reid, Saskia EJ de Vries, Christof Koch, Hongkui Zeng, Michael A Buice, Jack Waters, and Lu Li. Relationship between simultaneously recorded spiking activity and fluorescence signal in GCaMP6 transgenic mice. *eLife*, 10, 2021.

[9144] Qiaoli Huang, Huihui Zhang, and Huan Luo. Sequence structure organizes items in varied latent states of working memory neural network. *eLife*, 10, 2021.

[9145] Wuqing Huang, Jun Xiao, Jianguang Ji, and Liangwan Chen. Association of lipid-lowering drugs with COVID-19 outcomes from a Mendelian randomization study. *eLife*, 10, 2021.

[9146] Yongjian Huang, Jana Ognjenovic, Deepti Karandur, Kate Miller, Alan Merk, Sriram Subramaniam, and John Kuriyan. A molecular mechanism for the generation of ligand-dependent differential outputs by the epidermal growth factor receptor. *eLife*, 10, 2021.

[9147] Yongzhi Huang, Bomin Sun, Jean Debarros, Chao Zhang, Shikun Zhan, Dianyou Li, Chencheng Zhang, Tao Wang, Peng Huang, Yijie Lai, Peter Brown, Chunyan Cao, and Huiling Tan. Increased theta/alpha synchrony in the habenula-prefrontal network with negative emotional stimuli in human patients. *eLife*, 10, 2021.

[9148] Lars Hubatsch, Louise M Jawerth, Celina Love, Jonathan Bauermann, TY Dora Tang, Stefano Bo, Anthony A Hyman, and Christoph A Weber. Quantitative theory for the diffusive dynamics of liquid condensates. *eLife*, 10, 2021.

[9149] Marinus Huber, Kosmas V Kepesidis, Liudmila Voronina, Frank Fleischmann, Ernst Fill, Jacqueline Hermann, Ina Koch, Katrin Milger-Kneidinger, Thomas Kolben, Gerald B Schulz, Friedrich Jokisch, Jürgen Behr, Nadia Harbeck, Maximilian Reiser, Christian Stief, Ferenc Krausz, and Mihaela Zigman. Infrared molecular fingerprinting of blood-based liquid biopsies for the detection of cancer. *eLife*, 10, 2021.

[9150] Meret Huber, Thomas Roder, Sandra Irmisch, Alexander Riedel, Saskia Gablenz, Julia Fricke, Peter Rahfeld, Michael Reichelt, Christian Paetz, Nicole Liechti, Lingfei Hu, Zoe Bont, Ye Meng, Wei Huang, Christelle AM Robert, Jonathan Gershenzon, and Matthias Erb. A beta-glucosidase of an insect herbivore determines both toxicity and deterrence of a dandelion defense metabolite. *eLife*, 10, 2021.

[9151] Sebastian Hückesfeld, Philipp Schlegel, Anton Miroschnikow, Andreas Schoofs, Ingo Zinke, André N Haubrich, Casey M Schneider-Mizell, James W Truman, Richard D Fetter, Albert Cardona, and Michael J Pankratz. Unveiling the sensory and interneuronal pathways of the neuroendocrine connectome in Drosophila. *eLife*, 10, 2021.

[9152] Robert J Huebner, Abdul Naseer Malmi-Kakkada, Sena Sarıkaya, Shinuo Weng, D Thirumalai, and John B Wallingford. Mechanical heterogeneity along single cell-cell junctions is driven by lateral clustering of cadherins during vertebrate axis elongation. *eLife*, 10, 2021.

[9153] Alice C Hughes, Benjamin Michael Marshall, and Colin T Strine. Gaps in global wildlife trade monitoring leave amphibians vulnerable. *eLife*, 10, 2021.

[9154] Christopher L Hughes, Sharlene N Flesher, Jeffrey M Weiss, Michael Boninger, Jennifer L Collinger, and Robert A Gaunt. Perception of microstimulation frequency in human somatosensory cortex. *eLife*, 10, 2021.

[9155] Elizabeth R Hughes, Maria G Winter, Laice Alves da Silva, Matthew K Muramatsu, Angel G Jimenez, Caroline C Gillis, Luisella Spiga, Rachael B Chanin, Renato L Santos, Wenhan Zhu, and Sebastian E Winter. Reshaping of bacterial molecular hydrogen metabolism contributes to the outgrowth of commensal E. coli during gut inflammation. *eLife*, 10, 2021.

[9156] Karthik Hullahalli and Matthew K Waldor. Pathogen clonal expansion underlies multiorgan dissemination and organ-specific outcomes during murine systemic infection. *eLife*, 10, 2021.

[9157] Katina D Hulme, Anjana C Karawita, Cassandra Pegg, Myrna JM Bunte, Helle Bielefeldt-Ohmann, Conor J Bloxham, Silvie Van den Hoecke, Yin Xiang Setoh, Bram Vrancken, Monique Spronken, Lauren E Steele, Nathalie AJ Verzele, Kyle R Upton, Alexander A Khromykh, Keng Yih Chew, Maria Sukkar, Simon Phipps, and Kirsty R Short. A paucigranulocytic asthma host environment promotes the emergence of virulent influenza viral variants. *eLife*, 10, 2021.

[9158] Ann Louise Hunter, Charlotte E Pelekanou, Nichola J Barron, Rebecca C Northeast, Magdalena Grudzien, Antony D Adamson, Polly Downton, Thomas Cornfield, Peter S Cunningham, Jean-Noel Billaud, Leanne Hodson, Andrew SI Loudon, Richard D Unwin, Mudassar Iqbal, David W Ray, and David A Bechtold. Adipocyte NR1D1 dictates adipose tissue expansion during obesity. *eLife*, 10, 2021.

[9159] Kelsey E Huntington, Anna D Louie, Chun Geun Lee, Jack A Elias, Eric A Ross, and Wafik S El-Deiry. Cytokine ranking via mutual information algorithm correlates cytokine profiles with presenting disease severity in patients infected with SARS-CoV-2. *eLife*, 10, 2021.

[9160] Phil Huss, Anthony Meger, Megan Leander, Kyle Nishikawa, and Srivatsan Raman. Mapping the functional landscape of the receptor binding domain of T7 bacteriophage by deep mutational scanning. *eLife*, 10, 2021.

[9161] Sayedeh Hussaini, Vishalini Venkatesan, Valentina Biasci, José M Romero Sepúlveda, Raul A Quiñonez Uribe, Leonardo Sacconi, Gil Bub, Claudia Richter, Valentin Krinski, Ulrich Parlitz, Rupamanjari Majumder, and Stefan Luther. Drift and termination of spiral waves in optogenetically modified cardiac tissue at sub-threshold illumination. *eLife*, 10, 2021.

[9162] Kai Hwang, James M Shine, Joel Bruss, Daniel Tranel, and Aaron Boes. Neuropsychological evidence of multi-domain network hubs in the human thalamus. *eLife*, 10, 2021.

[9163] Sun-Hee Hwang, Bandarigoda N Somatilaka, Kevin White, and Saikat Mukhopadhyay. Ciliary and extraciliary Gpr161 pools repress hedgehog signaling in a tissue-specific manner. *eLife*, 10, 2021.

[9164] Theresa Hwang, Sara S Parker, Samantha M Hill, Meucci W Ilunga, Robert A Grant, Ghassan Mouneimne, and Amy E Keating. A distributed residue network permits conformational binding specificity in a conserved family of actin remodelers. *eLife*, 10, 2021.

[9165] Baher A Ibrahim, Caitlin A Murphy, Georgiy Yudintsev, Yoshitaka Shinagawa, Matthew I Banks, and Daniel A Llano. Corticothalamic gating of population auditory thalamocortical transmission in mouse. *eLife*, 10, 2021.

[9166] Christopher Icke, Freya J Hodges, Karthik Pullela, Samantha A McKeand, Jack Alfred Bryant, Adam F Cunningham, Jeff A Cole, and Ian R Henderson. Glycine acylation and trafficking of a new class of bacterial lipoprotein by a composite secretion system. *eLife*, 10, 2021.

[9167] Shintaro Ide, Yoshihiko Kobayashi, Kana Ide, Sarah A Strausser, Koki Abe, Savannah Herbek, Lori L O'Brien, Steven D Crowley, Laura Barisoni, Aleksandra Tata, Purushothama Rao Tata, and Tomokazu Souma. Ferroptotic stress promotes the accumulation of pro-inflammatory proximal tubular cells in maladaptive renal repair. *eLife*, 10, 2021.

[9168] Claudia Igler, Jens Rolff, and Roland Regoes. Multi-step vs. single-step resistance evolution under different drugs, pharmacokinetics, and treatment regimens. *eLife*, 10, 2021.

[9169] Christopher JR Illingworth, William L Hamilton, Ben Warne, Matthew Routledge, Ashley Popay, Chris Jackson, Tom Fieldman, Luke W Meredith, Charlotte J Houldcroft, Myra Hosmillo, Aminu S Jahun, Laura G Caller, Sarah L Caddy, Anna Yakovleva, Grant Hall, Fahad A Khokhar, Theresa Feltwell, Malte L Pinckert, Iliana Georgana, Yasmin Chaudhry, Martin D Curran, Surendra Parmar, Dominic Sparkes, Lucy Rivett, Nick K Jones, Sushmita Sridhar, Sally Forrest, Tom Dymond, Kayleigh Grainger, Chris Workman, Mark Ferris, Effrossyni Gkrania-Klotsas, Nicholas M Brown, Michael P Weekes, Stephen Baker, Sharon J Peacock, Ian G Goodfellow, Theodore Gouliouris, Daniela de Angelis, and M Estée Török. Superspreaders drive the largest outbreaks of hospital onset COVID-19 infections. *eLife*, 10, 2021.

[9170] Kaushik Inamdar, Feng-Ching Tsai, Rayane Dibsy, Aurore de Poret, John Manzi, Peggy Merida, Remi Muller, Pekka Lappalainen, Philippe Roingeard, Johnson Mak, Patricia Bassereau, Cyril Favard, and Delphine Muriaux. Full assembly of HIV-1 particles requires assistance of the membrane curvature factor IRSp53. *eLife*, 10, 2021.

[9171] Naokazu Inoue, Yoshihisa Hagihara, and Ikuo Wada. Evolutionarily conserved sperm factors, DCST1 and DCST2, are required for gamete fusion. *eLife*, 10, 2021.

[9172] Nobuko Inoue, Hirofumi Nishizumi, Rumi Ooyama, Kazutaka Mogi, Katsuhiko Nishimori, Takefumi Kikusui, and Hitoshi Sakano. The olfactory critical period is determined by activity-dependent Sema7A/PlxnC1 signaling within glomeruli. *eLife*, 10, 2021.

[9173] Yusuke Inoue, Ana Nikolic, Dylan Farnsworth, Rocky Shi, Fraser D Johnson, Alvin Liu, Marc Ladanyi, Romel Somwar, Marco Gallo, and William W Lockwood. Extracellular signal-regulated kinase mediates chromatin rewiring and lineage transformation in lung cancer. *eLife*, 10, 2021.

[9174] Domenica Ippolito, Saurabh Thapliyal, and Dominique A Glauser. Ca2+/CaM binding to CaMKI promotes IMA-3 importin binding and nuclear translocation in sensory neurons to control behavioral adaptation. *eLife*, 10, 2021.

[9175] Rose Irwin, Ofrah Faust, Ivana Petrovic, Sharon Grayer Wolf, Hagen Hofmann, and Rina Rosenzweig. Hsp40s play complementary roles in the prevention of tau amyloid formation. *eLife*, 10, 2021.

[9176] Mamoru Ishii, Tomoko Tateya, Michiyuki Matsuda, and Tsuyoshi Hirashima. Retrograde ERK activation waves drive base-to-apex multicellular flow in murine cochlear duct morphogenesis. *eLife*, 10, 2021.

[9177] Hannah M Isles, Catherine A Loynes, Sultan Alasmari, Fu Chuen Kon, Katherine M Henry, Anastasia Kadochnikova, Jack Hales, Clare F Muir, Maria-Cristina Keightley, Visakan Kadirkamanathan, Noémie Hamilton, Graham J Lieschke, Stephen A Renshaw, and Philip M Elks. Pioneer neutrophils release chromatin within in vivo swarms. *eLife*, 10, 2021.

[9178] Ariel Israel, Alejandro A Schäffer, Assi Cicurel, Kuoyuan Cheng, Sanju Sinha, Eyal Schiff, Ilan Feldhamer, Ameer Tal, Gil Lavie, and Eytan Ruppin. Identification of drugs associated with reduced severity of COVID-19 – a case-control study in a large population. *eLife*, 10, 2021.

[9179] Yasmine Issah, Amruta Naik, Soon Y Tang, Kaitlyn Forrest, Thomas G Brooks, Nicholas Lahens, Katherine N Theken, Mara Mermigos, Amita Sehgal, George S Worthen, Garret A FitzGerald, and Shaon Sengupta. Loss of circadian protection against influenza infection in adult mice exposed to hyperoxia as neonates. *eLife*, 10, 2021.

[9180] Jumpei Ito, Shigemiki Omiya, Mara-Camelia Rusu, Hiromichi Ueda, Tomokazu Murakawa, Yohei Tanada, Hajime Abe, Kazuki Nakahara, Michio Asahi, Manabu Taneike, Kazuhiko Nishida, Ajay M Shah, and Kinya Otsu. Iron derived from autophagy-mediated ferritin degradation induces cardiomyocyte death and heart failure in mice. *eLife*, 10, 2021.

[9181] Maria LC Iurilli, Maria LC Iurilli, Bin Zhou, James E Bennett, Rodrigo M Carrillo-Larco, Marisa K Sophiea, Andrea Rodriguez-Martinez, Honor Bixby, Bethlehem D Solomon, Cristina Taddei, Goodarz Danaei, Mariachiara Di Cesare, Gretchen A Stevens, Leanne M Riley, Stefan Savin, Melanie J Cowan, Pascal Bovet, Albertino Damasceno, Adela Chirita-Emandi, Alison J Hayes, Nayu Ikeda, Rod T Jackson, Young-Ho Khang, Avula Laxmaiah, Jing Liu, J Jaime Miranda, Olfa Saidi, Sylvain Sebert, Maroje Sorić, Gregor Starc, Edward W Gregg, Leandra Abarca-Gómez, Ziad A Abdeen, Shynar Abdrakhmanova, Suhaila Abdul Ghaffar, Hanan F Abdul Rahim, Niveen M Abu-Rmeileh, Jamila Abubakar Garba, Benjamin Acosta-Cazares, Robert J Adams, Wichai Aekplakorn, Kaosar Afsana, Shoaib Afzal, Imelda A Agdeppa, Javad Aghazadeh-Attari, Carlos A Aguilar-Salinas, Charles Agyemang, Mohamad Hasnan Ahmad, Noor Ani Ahmad, Ali Ahmadi, Naser Ahmadi, Soheir H Ahmed, Wolfgang Ahrens, Gulmira Aitmurzaeva, Kamel Ajlouni, Hazzaa M Al-Hazzaa, Badreya Al-Lahou, Rajaa Al-Raddadi, Monira Alarouj, Fadia AlBuhairan, Shahla AlDhukair, Mohamed M Ali, Abdullah Alkandari, Ala'a Alkerwi, Kristine Allin, Mar Alvarez-Pedrerol, Eman Aly, Deepak N Amarapurkar, Parisa Amiri, Norbert Amougou, Philippe Amouyel, Lars Bo Andersen, Sigmund A Anderssen, Lars ängquist, Ranjit Mohan Anjana, Alireza Ansari-Moghaddam, Hajer Aounallah-Skhiri, Joana Araújo, Inger Ariansen, Tahir Aris, Raphael E Arku, Nimmathota Arlappa, Krishna K Aryal, Thor Aspelund, Felix K Assah, Maria Cecília F AssunÇão, May Soe Aung, Juha Auvinen, Mária Avdicová, Shina Avi, Ana Azevedo, Mohsen Azimi-Nezhad, Fereidoun Azizi,

Mehrdad Azmin, Bontha V Babu, Maja Bæksgaard Jørgensen, Azli Baharudin, Suhad Bahijri, Jennifer L Baker, Nagalla Balakrishna, Mohamed Bamoshmoosh, Maciej Banach, Piotr Bandosz, José R Banegas, Joanna Baran, Carlo M Barbagallo, Alberto Barceló, Amina Barkat, Aluisio JD Barros, Mauro Virgílio Gomes Barros, Abdul Basit, Joao Luiz D Bastos, Iqbal Bata, Anwar M Batieha, Rosangela L Batista, Zhamilya Battakova, Assembekov Batyrbek, Louise A Baur, Robert Beaglehole, Silvia Bel-Serrat, Antonisamy Belavendra, Habiba Ben Romdhane, Judith Benedics, Mikhail Benet, Ingunn Holden Bergh, Salim Berkinbayev, Antonio Bernabe-Ortiz, Gailute Bernotiene, Heloísa Bettiol, Jorge Bezerra, Aroor Bhagyalaxmi, Sumit Bharadwaj, Santosh K Bhargava, Zulfiqar A Bhutta, Hongsheng Bi, Yufang Bi, Daniel Bia, Elysée Claude Bika Lele, Mukharram M Bikbov, Bihungum Bista, Dusko J Bjelica, Peter Bjerregaard, Espen Bjertness, Marius B Bjertness, Cecilia Björkelund, Katia V Bloch, Anneke Blokstra, Simona Bo, Martin Bobak, Lynne M Boddy, Bernhard O Boehm, Heiner Boeing, Jose G Boggia, Elena Bogova, Carlos P Boissonnet, Stig E Bojesen, Marialaura Bonaccio, Vanina Bongard, Alice Bonilla-Vargas, Matthias Bopp, Herman Borghs, Lien Braeckevelt, Lutgart Braeckman, Marjolijn CE Bragt, Imperia Brajkovich, Francesco Branca, Juergen Breckenkamp, João Breda, Hermann Brenner, Lizzy M Brewster, Garry R Brian, Lacramioara Brinduse, Sinead Brophy, Graziella Bruno, H Bas Bueno-de Mesquita, Anna Bugge, Marta Buoncristiano, Genc Burazeri, Con Burns, Antonio Cabrera de León, Joseph Cacciottolo, Hui Cai, Tilema Cama, Christine Cameron, José Camolas, Günay Can, Ana Paula C Cândido, Felicia Cañete, Mario V Capanzana, Nadežda Capková, Eduardo Capuano, Vincenzo Capuano, Marloes Cardol, Viviane C Cardoso, Axel C Carlsson, Esteban Carmuega, Joana Carvalho, José A Casajús, Felipe F Casanueva, Ertugrul Celikcan, Laura Censi, Marvin Cervantes-Loaiza, Juraci A Cesar, Snehalatha Chamukuttan, Angelique W Chan, Queenie Chan, Himanshu K Chaturvedi, Nish Chaturvedi, Norsyamlina Che Abdul Rahim, Miao Li Chee, Chien-Jen Chen, Fangfang Chen, Huashuai Chen, Shuohua Chen, Zhengming Chen, Ching-Yu Cheng, Bahman Cheraghian, Angela Chetrit, Ekaterina Chikova-Iscener, Arnaud Chiolero, Shu-Ti Chiou, María-Dolores Chirlaque, Belong Cho, Kaare Christensen, Diego G Christofaro, Jerzy Chudek, Renata Cifkova, Michelle Cilia, Eliza Cinteza, Frank Claessens, Janine Clarke, Els Clays, Emmanuel Cohen, Hans Concin, Susana C Confortin, Cyrus Cooper, Tara C Coppinger, Eva Corpeleijn, Simona Costanzo, Dominique Cottel, Chris Cowell, Cora L Craig, Amelia C Crampin, Ana B Crujeiras, Semánová Csilla, Alexandra M Cucu, Liufu Cui, Felipe V Cureau, Ewelina Czenczek-Lewandowska, Graziella D'Arrigo, Eleonora d'Orsi, Liliana Dacica, María ángeles Dal Re Saavedra, Jean Dallongeville, Camilla T Damsgaard, Rachel Dankner, Thomas M Dantoft, Parasmani Dasgupta, Saeed Dastgiri, Luc Dauchet, Kairat Davletov, Guy De Backer, Dirk De Bacquer, Giovanni de Gaetano, Stefaan De Henauw, Paula Duarte de Oliveira, David De Ridder, Karin De Ridder, Susanne R de Rooij, Delphine De Smedt, Mohan Deepa, Alexander D Deev, Vincent Jr DeGennaro, Abbas Dehghan, Hélène Delisle, Francis Delpeuch, Stefaan Demarest, Elaine Dennison, Katarzyna Dereń, Valérie Deschamps, Meghnath Dhimal, Augusto F Di Castelnuovo, Juvenal Soares Dias-da Costa, María Elena Díaz-Sánchez, Alejandro Diaz, Zivka Dika, Shirin Djalalinia, Visnja Djordjic, Ha TP Do, Annette J Dobson, Maria Benedetta Donati, Chiara Donfrancesco, Silvana P Donoso, Angela Döring, Maria Dorobantu, Ahmad Reza Dorosty, Kouamelan Doua, Nico Dragano, Wojciech Drygas, Jia Li Duan, Charmaine A Duante, Priscilla Duboz, Rosemary B Duda, Vesselka Duleva, Virginija Dulskiene, Samuel C Dumith, Anar Dushpanova, Vilnis Dzerve, Elzbieta Dziankowska-Zaborszczyk, Ricky Eddie, Ebrahim Eftekhar, Eruke E Egbagbe, Robert Eggertsen, Sareh Eghtesad, Gabriele Eiben, Ulf Ekelund, Mohammad El-Khateeb, Jalila El Ati, Denise Eldemire-Shearer, Marie Eliasen, Paul Elliott, Reina Engle-Stone, Macia Enguerran, Rajiv T Erasmus, Raimund Erbel, Cihangir Erem, Louise Eriksen, Johan G Eriksson, Jorge Escobedo-de la Peña, Saeid Eslami, Ali Esmaeili, Alun Evans, David Faeh, Albina A Fakhretdinova, Caroline H Fall, Elnaz Faramarzi, Mojtaba Farjam, Victoria Farrugia Sant'Angelo, Farshad Farzadfar, Mohammad Reza Fattahi, Asher Fawwad, Francisco J Felix-Redondo, Trevor S Ferguson, Romulo A Fernandes, Daniel Fernández-Bergés, Daniel Ferrante, Thomas Ferrao, Marika Ferrari, Marco M Ferrario, Catterina Ferreccio, Eldridge Ferrer, Jean Ferrieres, Thamara Hubler Figueiró, Anna Fijalkowska, Günther Fink, Krista Fischer, Leng Huat Foo, Maria Forsner, Heba M Fouad, Damian K Francis, Maria do Carmo Franco, Ruth Frikke-Schmidt, Guillermo Frontera, Flavio D Fuchs, Sandra C Fuchs, Isti I Fujiati, Yuki Fujita, Matsuda Fumihiko, Takuro Furusawa, Zbigniew Gaciong, Mihai Gafencu, Andrzej Galbarczyk, Henrike Galenkamp, Daniela Galeone, Myriam Galfo, Fabio Galvano, Jingli Gao,

Manoli Garcia-de-la Hera, Marta García-Solano, Dickman Gareta, Sarah P Garnett, Jean-Michel Gaspoz, Magda Gasull, Adroaldo Cesar Araujo Gaya, Anelise Reis Gaya, Andrea Gazzinelli, Ulrike Gehring, Harald Geiger, Johanna M Geleijnse, Ali Ghanbari, Erfan Ghasemi, Oana-Florentina Gheorghe-Fronea, Simona Giampaoli, Francesco Gianfagna, Tiffany K Gill, Jonathan Giovannelli, Glen Gironella, Aleksander Giwercman, Konstantinos Gkiouras, Justyna Godos, Sibel Gogen, Marcel Goldberg, Rebecca A Goldsmith, David Goltzman, Santiago F Gómez, Aleksandra Gomula, Bruna Goncalves Cordeiro da Silva, Helen GonÇalves, David A Gonzalez-Chica, Marcela Gonzalez-Gross, Margot González-Leon, Juan P González-Rivas, Clicerio González-Villalpando, María-Elena González-Villalpando, Angel R Gonzalez, Frederic Gottrand, Antonio Pedro GraÇa, Sidsel Graff-Iversen, DuŠan Grafnetter, Aneta Grajda, Maria G Grammatikopoulou, Ronald D Gregor, Tomasz Grodzicki, Else Karin Grøholt, Anders Grøntved, Giuseppe Grosso, Gabriella Gruden, Dongfeng Gu, Emanuela Gualdi-Russo, Pilar Guallar-Castillón, Andrea Gualtieri, Elias F Gudmundsson, Vilmundur Gudnason, Ramiro Guerrero, Idris Guessous, Andre L Guimaraes, Martin C Gulliford, Johanna Gunnlaugsdottir, Marc J Gunter, Xiu-Hua Guo, Yin Guo, Prakash C Gupta, Rajeev Gupta, Oye Gureje, Beata Gurzkowska, Enrique Gutiérrez-González, Laura Gutierrez, Felix Gutzwiller, Seongjun Ha, Farzad Hadaegh, Charalambos A Hadjigeorgiou, Rosa Haghshenas, Hamid Hakimi, Jytte Halkjær, Ian R Hambleton, Behrooz Hamzeh, Dominique Hange, Abu AM Hanif, Sari Hantunen, Jie Hao, Rachakulla Hari Kumar, Seyed Mohammad Hashemi-Shahri, Maria Hassapidou, Jun Hata, Teresa Haugsgjerd, Jiang He, Yuan He, Yuna He, Regina Heidinger-Felso, Mirjam Heinen, Tatjana Hejgaard, Marleen Elisabeth Hendriks, Rafael dos Santos Henrique, Ana Henriques, Leticia Hernandez Cadena, Sauli Herrala, Victor M Herrera, Isabelle Herter-Aeberli, Ramin Heshmat, Allan G Hill, Sai Yin Ho, Suzanne C Ho, Michael Hobbs, Michelle Holdsworth, Reza Homayounfar, Clara Homs, Wilma M Hopman, Andrea RVR Horimoto, Claudia M Hormiga, Bernardo L Horta, Leila Houti, Christina Howitt, Thein Thein Htay, Aung Soe Htet, Maung Maung Than Htike, Yonghua Hu, José María Huerta, Ilpo Tapani Huhtaniemi, Laetitia Huiart, Constanta Huidumac Petrescu, Martijn Huisman, Abdullatif Husseini, Chinh Nguyen Huu, Inge Huybrechts, Nahla Hwalla, Jolanda Hyska, Licia Iacoviello, Jesús M Ibarluzea, Mohsen M Ibrahim, Norazizah Ibrahim Wong, M Arfan Ikram, Violeta Iotova, Vilma E Irazola, Takafumi Ishida, Muhammad Islam, Sheikh Mohammed Shariful Islam, Masanori Iwasaki, Jeremy M Jacobs, Hashem Y Jaddou, Tazeen Jafar, Kenneth James, Kazi M Jamil, Konrad Jamrozik, Imre Janszky, Edward Janus, Juel Jarani, Marjo-Riitta Jarvelin, Grazyna Jasienska, Ana Jelakovic, Bojan Jelakovic, Garry Jennings, Anjani Kumar Jha, Chao Qiang Jiang, Ramon O Jimenez, Karl-Heinz Jöckel, Michel Joffres, Mattias Johansson, Jari J Jokelainen, Jost B Jonas, Jitendra Jonnagaddala, Torben Jørgensen, Pradeep Joshi, Farahnaz Joukar, Dragana P Jovic, Jacek J Jóźwiak, Anne Juolevi, Gregor Jurak, Iulia Jurca Simina, Vesna Juresa, Rudolf Kaaks, Felix O Kaducu, Anthony Kafatos, Eero O Kajantie, Zhanna Kalmatayeva, Ofra Kalter-Leibovici, Yves Kameli, Freja B Kampmann, Kodanda R Kanala, Srinivasan Kannan, Efthymios Kapantais, Argyro Karakosta, Line L Kårhus, Khem B Karki, Marzieh Katibeh, Joanne Katz, Peter T Katzmarzyk, Jussi Kauhanen, Prabhdeep Kaur, Maryam Kavousi, Gyulli M Kazakbaeva, Ulrich Keil, Lital Keinan Boker, Sirkka Keinänen-Kiukaanniemi, Roya Kelishadi, Cecily Kelleher, Han CG Kemper, Andre P Kengne, Maryam Keramati, Alina Kerimkulova, Mathilde Kersting, Timothy Key, Yousef Saleh Khader, Davood Khalili, Kay-Tee Khaw, Bahareh Kheiri, Motahareh Kheradmand, Alireza Khosravi, Ilse MSL Khouw, Ursula Kiechl-Kohlendorfer, Stefan Kiechl, Japhet Killewo, Dong Wook Kim, Hyeon Chang Kim, Jeongseon Kim, Jenny M Kindblom, Heidi Klakk, Magdalena Klimek, Jeannette Klimont, Jurate Klumbiene, Michael Knoflach, Bhawesh Koirala, Elin Kolle, Patrick Kolsteren, Jürgen König, Raija Korpelainen, Paul Korrovits, Magdalena Korzycka, Jelena Kos, Seppo Koskinen, Katsuyasu Kouda, Viktoria A Kovacs, Sudhir Kowlessur, Slawomir Koziel, Jana Kratenova, Wolfgang Kratzer, Susi Kriemler, Peter Lund Kristensen, Steinar Krokstad, Daan Kromhout, Herculina S Kruger, Ruzena Kubinova, Renata Kuciene, Urho M Kujala, Enisa Kujundzic, Zbigniew Kulaga, R Krishna Kumar, Marie KuneŠová, Pawel Kurjata, Yadlapalli S Kusuma, Kari Kuulasmaa, Catherine Kyobutungi, Quang Ngoc La, Fatima Zahra Laamiri, Tiina Laatikainen, Carl Lachat, Youcef Laid, Tai Hing Lam, Christina-Paulina Lambrinou, Edwige Landais, Vera Lanska, Georg Lappas, Bagher Larijani, Tint Swe Latt, Laura Lauria, Maria Lazo-Porras, Gwenaëlle Le Coroller, Khanh Le Nguyen Bao, Agnès Le Port, Tuyen D Le, Jeannette Lee, Jeonghee Lee, Paul H Lee, Nils Lehmann, Terho Lehtimäki, Daniel

Lemogoum, Naomi S Levitt, Yanping Li, Merike Liivak, Christa L Lilly, Wei-Yen Lim, M Fernanda Lima-Costa, Hsien-Ho Lin, Xu Lin, Yi-Ting Lin, Lars Lind, Allan Linneberg, Lauren Lissner, Mieczyslaw Litwin, Lijuan Liu, Wei-Cheng Lo, Helle-Mai Loit, Khuong Quynh Long, Luis Lopes, Oscar Lopes, Esther Lopez-Garcia, Tania Lopez, Paulo A Lotufo, José Eugenio Lozano, Janice L Lukrafka, Dalia Luksiene, Annamari Lundqvist, Robert Lundqvist, Nuno Lunet, Charles Lunogelo, Michala Lustigová, Edyta Łuszczki, Guansheng Ma, Jun Ma, Xu Ma, George LL Machado-Coelho, Aristides M Machado-Rodrigues, Luisa M Macieira, Ahmed A Madar, Stefania Maggi, Dianna J Magliano, Sara Magnacca, Emmanuella Magriplis, Gowri Mahasampath, Bernard Maire, Marjeta Majer, Marcia Makdisse, Päivi Mäki, Fatemeh Malekzadeh, Reza Malekzadeh, Rahul Malhotra, Kodavanti Mallikharjuna Rao, Sofia K Malyutina, Lynell V Maniego, Yannis Manios, Jim I Mann, Fariborz Mansour-Ghanaei, Enzo Manzato, Paula Margozzini, Anastasia Markaki, Oonagh Markey, Eliza Markidou Ioannidou, Pedro Marques-Vidal, Larissa Pruner Marques, Jaume Marrugat, Yves Martin-Prevel, Rosemarie Martin, Reynaldo Martorell, Eva Martos, Katharina Maruszczak, Stefano Marventano, Luis P Mascarenhas, Shariq R Masoodi, Ellisiv B Mathiesen, Prashant Mathur, Alicia Matijasevich, Tandi E Matsha, Christina Mavrogianni, Artur Mazur, Jean Claude N Mbanya, Shelly R McFarlane, Stephen T McGarvey, Martin McKee, Stela McLachlan, Rachael M McLean, Scott B McLean, Breige A McNulty, Sounnia Mediene Benchekor, Jurate Medzioniene, Parinaz Mehdipour, Kirsten Mehlig, Amir Houshang Mehrparvar, Aline Meirhaeghe, Jørgen Meisfjord, Christa Meisinger, Ana Maria B Menezes, Geetha R Menon, Gert BM Mensink, Maria Teresa Menzano, Alibek Mereke, Indrapal I Meshram, Andres Metspalu, Haakon E Meyer, Jie Mi, Kim F Michaelsen, Nathalie Michels, Kairit Mikkel, Karolina Milkowska, Jody C Miller, Cláudia S Minderico, GK Mini, Juan Francisco Miquel, Mohammad Reza Mirjalili, Daphne Mirkopoulou, Erkin Mirrakhimov, Marjeta MiŠigoj-Durakovic, Antonio Mistretta, Veronica Mocanu, Pietro A Modesti, Sahar Saeedi Moghaddam, Bahram Mohajer, Mostafa K Mohamed, Shukri F Mohamed, Kazem Mohammad, Zahra Mohammadi, Noushin Mohammadifard, Reza Mohammadpourhodki, Viswanathan Mohan, Salim Mohanna, Muhammad Fadhli Mohd Yusoff, Iraj Mohebbi, Farnam Mohebi, Marie Moitry, Drude Molbo, Line T Møllehave, Niels C Møller, Dénes Molnár, Amirabbas Momenan, Charles K Mondo, Michele Monroy-Valle, Eric Monterrubio-Flores, Kotsedi Daniel K Monyeki, Jin Soo Moon, Mahmood Moosazadeh, Leila B Moreira, Alain Morejon, Luis A Moreno, Karen Morgan, Suzanne N Morin, Erik Lykke Mortensen, George Moschonis, Malgorzata Mossakowska, Aya Mostafa, Anabela Mota-Pinto, Jorge Mota, Mohammad Esmaeel Motlagh, Jorge Motta, Marcos André Moura-dos Santos, Malay K Mridha, Kelias P Msyamboza, Thet Thet Mu, Magdalena Muc, Boban MugoŠa, Maria L Muiesan, Parvina Mukhtorova, Martina Müller-Nurasyid, Neil Murphy, Jaakko Mursu, Elaine M Murtagh, Kamarul Imran Musa, Sanja Music Milanovic, Vera Musil, Norlaila Mustafa, Iraj Nabipour, Shohreh Naderimagham, Gabriele Nagel, Balkish M Naidu, Farid Najafi, Harunobu Nakamura, Jana NámeŠná, Ei Ei K Nang, Vinay B Nangia, Martin Nankap, Sameer Narake, Paola Nardone, Matthias Nauck, William A Neal, Azim Nejatizadeh, Chandini Nekkantti, Keiu Nelis, Liis Nelis, Ilona Nenko, Martin Neovius, Flavio Nervi, Chung T Nguyen, Nguyen D Nguyen, Quang Ngoc Nguyen, Ramfis E Nieto-Martínez, Yury P Nikitin, Guang Ning, Toshiharu Ninomiya, Sania Nishtar, Marianna Noale, Oscar A Noboa, Helena Nogueira, Teresa Norat, Maria Nordendahl, Børge G Nordestgaard, Davide Noto, Natalia Nowak-Szczepanska, Mohannad Al Nsour, Irfan Nuhoglu, Eha Nurk, Terence W O'Neill, Dermot O'Reilly, Galina Obreja, Caleb Ochimana, Angélica M Ochoa-Avilés, Eiji Oda, Kyungwon Oh, Kumiko Ohara, Claes Ohlsson, Ryutaro Ohtsuka, örn Olafsson, Maria Teresa A Olinto, Isabel O Oliveira, Mohd Azahadi Omar, Altan Onat, Sok King Ong, Lariane M Ono, Pedro Ordunez, Rui Ornelas, Ana P Ortiz, Pedro J Ortiz, Merete Osler, Clive Osmond, Sergej M Ostojic, Afshin Ostovar, Johanna A Otero, Kim Overvad, Ellis Owusu-Dabo, Fred Michel Paccaud, Cristina Padez, Ioannis Pagkalos, Elena Pahomova, Karina Mary de Paiva, Andrzej Pajak, Domenico Palli, Alberto Palloni, Luigi Palmieri, Wen-Harn Pan, Songhomitra Panda-Jonas, Arvind Pandey, Francesco Panza, Dimitrios Papandreou, Soon-Woo Park, Suyeon Park, Winsome R Parnell, Mahboubeh Parsaeian, Ionela M Pascanu, Patrick Pasquet, Nikhil D Patel, Ivan Pecin, Mangesh S Pednekar, Nasheeta Peer, Gao Pei, Sergio Viana Peixoto, Markku Peltonen, Alexandre C Pereira, Marco A Peres, Napoleón Pérez-Farinós, Cynthia M Pérez, Valentina Peterkova, Annette Peters, Astrid Petersmann, Janina Petkeviciene, Ausra Petrauskiene, Emanuela Pettenuzzo, Niloofar Peykari, Son Thai Pham, Rafael N Pichardo, Daniela Pierannunzio,

Iris Pigeot, Hynek Pikhart, Aida Pilav, Lorenza Pilotto, Francesco Pistelli, Freda Pitakaka, Aleksandra Piwonska, Andreia N Pizarro, Pedro Plans-Rubió, Bee Koon Poh, Hermann Pohlabeln, Raluca M Pop, Stevo R Popovic, Miquel Porta, Georg Posch, Anil Poudyal, Dimitrios Poulimeneas, Hamed Pouraram, Farhad Pourfarzi, Akram Pourshams, Hossein Poustchi, Rajendra Pradeepa, Alison J Price, Jacqueline F Price, Rui Providencia, Jardena J Puder, Iveta Pudule, Soile E Puhakka, Maria Puiu, Margus Punab, Radwan F Qasrawi, Mostafa Qorbani, Tran Quoc Bao, Ivana Radic, Ricardas Radisauskas, Salar Rahimikazerooni, Mahfuzar Rahman, Mahmudur Rahman, Olli Raitakari, Manu Raj, Ellina Rakhimova, Sherali Rakhmatulloev, Ivo Rakovac, Sudha Ramachandra Rao, Ambady Ramachandran, Jacqueline Ramke, Elisabete Ramos, Rafel Ramos, Lekhraj Rampal, Sanjay Rampal, Vayia Rarra, Ramon A Rascon-Pacheco, Mette Rasmussen, Cassiano Ricardo Rech, Josep Redon, Paul Ferdinand M Reganit, Valéria Regecová, Luis Revilla, Abbas Rezaianzadeh, Lourdes Ribas-Barba, Robespierre Ribeiro, Elio Riboli, Adrian Richter, Fernando Rigo, Natascia Rinaldo, Tobias F Rinke de Wit, Ana Rito, Raphael M Ritti-Dias, Juan A Rivera, Cynthia Robitaille, Romana Roccaldo, Daniela Rodrigues, Fernando Rodríguez-Artalejo, María del Cristo Rodriguez-Perez, Laura A Rodríguez-Villamizar, Ulla Roggenbuck, Rosalba Rojas-Martinez, Nipa Rojroongwasinkul, Dora Romaguera, Elisabetta L Romeo, Rafaela V Rosario, Annika Rosengren, Ian Rouse, Joel GR Roy, Adolfo Rubinstein, Frank J Rühli, Jean-Bernard Ruidavets, Blanca Sandra Ruiz-Betancourt, Maria Ruiz-Castell, Emma Ruiz Moreno, Iuliia A Rusakova, Kenisha Russell Jonsson, Paola Russo, Petra Rust, Marcin Rutkowski, Charumathi Sabanayagam, Elena Sacchini, Harshpal S Sachdev, Alireza Sadjadi, Ali Reza Safarpour, Saeid Safiri, Nader Saki, Benoit Salanave, Eduardo Salazar Martinez, Diego Salmerón, Veikko Salomaa, Jukka T Salonen, Massimo Salvetti, Margarita Samoutian, Jose Sánchez-Abanto, Sandjaja, Susana Sans, Loreto Santa Marina, Diana A Santos, Ina S Santos, Lèlita C Santos, Maria Paula Santos, Osvaldo Santos, Rute Santos, Sara Santos Sanz, Jouko L Saramies, Luis B Sardinha, Nizal Sarrafzadegan, Thirunavukkarasu Sathish, Kai-Uwe Saum, Savvas Savva, Mathilde Savy, Norie Sawada, Mariana Sbaraini, Marcia Scazufca, Beatriz D Schaan, Angelika Schaffrath Rosario, Herman Schargrodsky, Anja Schienkiewitz, Sabine Schipf, Carsten O Schmidt, Ida Maria Schmidt, Peter Schnohr, Ben Schöttker, Sara Schramm, Stine Schramm, Helmut Schröder, Constance Schultsz, Aletta E Schutte, Aye Aye Sein, Rusidah Selamat, Vedrana Sember, Abhijit Sen, Idowu O Senbanjo, Sadaf G Sepanlou, Victor Sequera, Luis Serra-Majem, Jennifer Servais, Ludmila Ševcíková, Svetlana A Shalnova, Teresa Shamah-Levy, Morteza Shamshirgaran, Coimbatore Subramaniam Shanthirani, Maryam Sharafkhah, Sanjib K Sharma, Jonathan E Shaw, Amaneh Shayanrad, Ali Akbar Shayesteh, Lela Shengelia, Zumin Shi, Kenji Shibuya, Hana Shimizu-Furusawa, Dong Wook Shin, Majid Shirani, Rahman Shiri, Namuna Shrestha, Khairil Si-Ramlee, Alfonso Siani, Rosalynn Siantar, Abla M Sibai, Antonio M Silva, Diego Augusto Santos Silva, Mary Simon, Judith Simons, Leon A Simons, Agneta Sjöberg, Michael Sjöström, Gry Skodje, Jolanta Slowikowska-Hilczer, Przemyslaw Slusarczyk, Liam Smeeth, Hung-Kwan So, Fernanda Cunha Soares, Grzegorz Sobek, Eugène Sobngwi, Morten Sodemann, Stefan Söderberg, Moesijanti YE Soekatri, Agustinus Soemantri, Reecha Sofat, Vincenzo Solfrizzi, Mohammad Hossein Somi, Emily Sonestedt, Yi Song, Thorkild IA Sørensen, Elin P Sørgjerd, Charles Sossa Jérome, Victoria E Soto-Rojas, Aïcha Soumaré, Slavica Sovic, Bente Sparboe-Nilsen, Karen Sparrenberger, Angela Spinelli, Igor Spiroski, Jan A Staessen, Hanspeter Stamm, Maria G Stathopoulou, Kaspar Staub, Bill Stavreski, Jostein Steene-Johannessen, Peter Stehle, Aryeh D Stein, George S Stergiou, Jochanan Stessman, Ranko Stevanovic, Jutta Stieber, Doris Stöckl, Tanja Stocks, Jakub Stokwiszewski, Ekaterina Stoyanova, Gareth Stratton, Karien Stronks, Maria Wany Strufaldi, Lela Sturua, Ramón Suárez-Medina, Machi Suka, Chien-An Sun, Johan Sundström, Yn-Tz Sung, Jordi Sunyer, Paibul Suriyawongpaisal, Boyd A Swinburn, Rody G Sy, Holly E Syddall, René Charles Sylva, Moyses Szklo, Lucjan Szponar, E Shyong Tai, Mari-Liis Tammesoo, Abdonas Tamosiunas, Eng Joo Tan, Xun Tang, Maya Tanrygulyyeva, Frank Tanser, Yong Tao, Mohammed Rasoul Tarawneh, Jakob Tarp, Carolina B Tarqui-Mamani, Radka Taxová Braunerová, Anne Taylor, Julie Taylor, Félicité Tchibindat, William R Tebar, Grethe S Tell, Tania Tello, Yih Chung Tham, KR Thankappan, Holger Theobald, Xenophon Theodoridis, Lutgarde Thijs, Nihal Thomas, Betina H Thuesen, Lubica Tichá, Erik J Timmermans, Anne Tjonneland, Hanna K Tolonen, Janne S Tolstrup, Murat Topbas, Roman Topór-Madry, Liv Elin Torheim, María José Tormo, Michael J Tornaritis, Maties Torrent, Laura Torres-Collado, Stefania Toselli, Giota Touloumi, Pierre Traissac, Thi Tuyet-Hanh

Tran, Dimitrios Trichopoulos, Antonia Trichopoulou, Oanh TH Trinh, Atul Trivedi, Lechaba Tshepo, Maria Tsigga, Shoichiro Tsugane, Azaliia M Tuliakova, Marshall K Tulloch-Reid, Fikru Tullu, Tomi-Pekka Tuomainen, Jaakko Tuomilehto, Maria L Turley, Gilad Twig, Per Tynelius, Themistoklis Tzotzas, Christophe Tzourio, Peter Ueda, Eunice Ugel, Flora AM Ukoli, Hanno Ulmer, Belgin Unal, Zhamyila Usupova, Hannu MT Uusitalo, Nalan Uysal, Justina Vaitkeviciute, Gonzalo Valdivia, Susana Vale, Damaskini Valvi, Rob M van Dam, Johan Van der Heyden, Yvonne T van der Schouw, Koen Van Herck, Hoang Van Minh, Natasja M Van Schoor, Irene GM van Valkengoed, Dirk Vanderschueren, Diego Vanuzzo, Anette Varbo, Gregorio Varela-Moreiras, Patricia Varona-Pérez, Senthil K Vasan, Tomas Vega, Toomas Veidebaum, Gustavo Velasquez-Melendez, Biruta Velika, Giovanni Veronesi, WM Monique Verschuren, Cesar G Victora, Giovanni Viegi, Lucie Viet, Salvador Villalpando, Paolo Vineis, Jesus Vioque, Jyrki K Virtanen, Marjolein Visser, Sophie Visvikis-Siest, Bharathi Viswanathan, Mihaela Vladulescu, Tiina Vlasoff, Dorja Vocanec, Peter Vollenweider, Henry Völzke, Ari Voutilainen, Sari Voutilainen, Martine Vrijheid, Tanja GM Vrijkotte, Alisha N Wade, Aline Wagner, Thomas Waldhör, Janette Walton, Elvis OA Wambiya, Wan Mohamad Wan Bebakar, Wan Nazaimoon Wan Mohamud, Rildo de Souza Wanderley Júnior, Ming-Dong Wang, Ningli Wang, Qian Wang, Xiangjun Wang, Ya Xing Wang, Ying-Wei Wang, S Goya Wannamethee, Nicholas Wareham, Adelheid Weber, Niels Wedderkopp, Deepa Weerasekera, Daniel Weghuber, Wenbin Wei, Aneta Weres, Bo Werner, Peter H Whincup, Kurt Widhalm, Indah S Widyahening, Andrzej Wiecek, Rainford J Wilks, Johann Willeit, Peter Willeit, Julianne Williams, Tom Wilsgaard, Bogdan Wojtyniak, Roy A Wong-McClure, Andrew Wong, Jyh Eiin Wong, Tien Yin Wong, Jean Woo, Mark Woodward, Frederick C Wu, Jianfeng Wu, Li Juan Wu, Shouling Wu, Haiquan Xu, Liang Xu, Nor Azwany Yaacob, Uruwan Yamborisut, Weili Yan, Ling Yang, Xiaoguang Yang, Yang Yang, Nazan Yardim, Mehdi Yaseri, Tabara Yasuharu, Xingwang Ye, Panayiotis K Yiallouros, Moein Yoosefi, Akihiro Yoshihara, Qi Sheng You, San-Lin You, Novie O Younger-Coleman, Safiah Md Yusof, Ahmad Faudzi Yusoff, Luciana Zaccagni, Vassilis Zafiropulos, Ahmad A Zainuddin, Seyed Rasoul Zakavi, Farhad Zamani, Sabina Zambon, Antonis Zampelas, Hana Zamrazilová, Maria Elisa Zapata, Abdul Hamid Zargar, Ko Ko Zaw, Tomasz Zdrojewski, Kristyna Zejglicova, Tajana Zeljkovic Vrkic, Yi Zeng, Luxia Zhang, Zhen-Yu Zhang, Dong Zhao, Ming-Hui Zhao, Wenhua Zhao, Shiqi Zhen, Wei Zheng, Yingfeng Zheng, Bekbolat Zholdin, Maigeng Zhou, Dan Zhu, Marie Zins, Emanuel Zitt, Yanina Zocalo, Julio Zuñiga Cisneros, Monika Zuziak, Majid Ezzati, and Sarah Filippi. Heterogeneous contributions of change in population distribution of body mass index to change in obesity and underweight. *eLife*, 10, 2021.

[9182] Motoko Iwashita and Masato Yoshizawa. Social-like responses are inducible in asocial Mexican cavefish despite the exhibition of strong repetitive behavior. *eLife*, 10, 2021.

[9183] Maryam Izadi, Eric Seemann, Dirk Schlobinski, Lukas Schwintzer, Britta Qualmann, and Michael M Kessels. Functional interdependence of the actin nucleator Cobl and Cobl-like in dendritic arbor development. *eLife*, 10, 2021.

[9184] Christopher G Jacob, Nguyen Thuy-Nhien, Mayfong Mayxay, Richard J Maude, Huynh Hong Quang, Bouasy Hongvanthong, Viengxay Vanisaveth, Thang Ngo Duc, Huy Rekol, Rob van der Pluijm, Lorenz von Seidlein, Rick Fairhurst, FranÇois Nosten, Md Amir Hossain, Naomi Park, Scott Goodwin, Pascal Ringwald, Keobouphaphone Chindavongsa, Paul Newton, Elizabeth Ashley, Sonexay Phalivong, Rapeephan Maude, Rithea Leang, Cheah Huch, Le Thanh Dong, Kim-Tuyen Nguyen, Tran Minh Nhat, Tran Tinh Hien, Hoa Nguyen, Nicole Zdrojewski, Sara Canavati, Abdullah Abu Sayeed, Didar Uddin, Caroline Buckee, Caterina I Fanello, Marie Onyamboko, Thomas Peto, Rupam Tripura, Chanaki Amaratunga, Aung Myint Thu, Gilles Delmas, Jordi Landier, Daniel M Parker, Nguyen Hoang Chau, Dysoley Lek, Seila Suon, James Callery, Podjanee Jittamala, Borimas Hanboonkunupakarn, Sasithon Pukrittayakamee, Aung Pyae Phyo, Frank Smithuis, Khin Lin, Myo Thant, Tin Maung Hlaing, Parthasarathi Satpathi, Sanghamitra Satpathi, Prativa K Behera, Amar Tripura, Subrata Baidya, Neena Valecha, Anupkumar R Anvikar, Akhter Ul Islam, Abul Faiz, Chanon Kunasol, Eleanor Drury, Mihir Kekre, Mozam Ali, Katie Love, Shavanthi Rajatileka, Anna E Jeffreys, Kate Rowlands, Christina S Hubbart, Mehul Dhorda, Ranitha Vongpromek, Namfon Kotanan, Phrutsamon Wongnak, Jacob Almagro Garcia, Richard D Pearson, Cristina V Ariani, Thanat Chookajorn, Cinzia Malangone, T Nguyen, Jim Stalker, Ben Jeffery, Jonathan Keatley, Kimberly J Johnson, Dawn Muddyman, Xin Hui S Chan, John Sillitoe, Roberto Amato, Victoria Simpson, Sonia GonÇalves, Kirk Rockett, Nicholas P Day, Arjen M Dondorp, Dominic P Kwiatkowski, and Olivo Miotto. Genetic surveillance in the Greater Mekong subregion and South Asia to support malaria control and elimination. *eLife*, 10, 2021.

[9185] Georgin Jacob, Harish Katti, Thomas Cherian, Jhilik Das, KA Zhivago, and SP Arun. A naturalistic environment to study visual cognition in unrestrained monkeys. *eLife*, 10, 2021.

[9186] Reeba Susan Jacob, Cédric Eichmann, Alessandro Dema, Davide Mercadante, and Philipp Selenko. $\alpha$-Synuclein plasma membrane localization correlates with cellular phosphatidylinositol polyphosphate levels. *eLife*, 10, 2021.

[9187] Sem H Jacobs, Eva Dóró, Ffion R Hammond, Mai E Nguyen-Chi, Georges Lutfalla, Geert F Wiegertjes, and Maria Forlenza. Occurrence of foamy macrophages during the innate response of zebrafish to trypanosome infections. *eLife*, 10, 2021.

[9188] Basile Jacquel, Théo Aspert, Damien Laporte, Isabelle Sagot, and Gilles Charvin. Monitoring single-cell dynamics of entry into quiescence during an unperturbed life cycle. *eLife*, 10, 2021.

[9189] Adrian V Jaeggi, Aaron D Blackwell, Christopher von Rueden, Benjamin C Trumble, Jonathan Stieglitz, Angela R Garcia, Thomas S Kraft, Bret A Beheim, Paul L Hooper, Hillard Kaplan, and Michael Gurven. Do wealth and inequality associate with health in a small-scale subsistence society? *eLife*, 10, 2021.

[9190] Polona Jager, Gerald Moore, Padraic Calpin, Xhuljana Durmishi, Irene Salgarella, Lucy Menage, Yoshiaki Kita, Yan Wang, Dong Won Kim, Seth Blackshaw, Simon R Schultz, Stephen Brickley, Tomomi Shimogori, and Alessio Delogu. Dual midbrain and forebrain origins of thalamic inhibitory interneurons. *eLife*, 10, 2021.

[9191] Mihaela Jagrić, Patrik Risteski, Jelena Martinčić, Ana Milas, and Iva M Tolić. Optogenetic control of PRC1 reveals its role in chromosome alignment on the spindle by overlap length-dependent forces. *eLife*, 10, 2021.

[9192] Michael Jahn, Nick Crang, Markus Janasch, Andreas Hober, Björn Forsström, Kyle Kimler, Alexander Mattausch, Qi Chen, Johannes Asplund-Samuelsson, and Elton Paul Hudson. Protein allocation and utilization in the versatile chemolithoautotroph Cupriavidus necator. *eLife*, 10, 2021.

[9193] Adam SB Jalal, Ngat T Tran, Clare EM Stevenson, Afroze Chimthanawala, Anjana Badrinarayanan, David M Lawson, and Tung BK Le. A CTP-dependent gating mechanism enables ParB spreading on DNA. *eLife*, 10, 2021.

[9194] Jennifer E James, Sara M Willis, Paul G Nelson, Catherine Weibel, Luke J Kosinski, and Joanna Masel. Universal and taxon-specific trends in protein sequences as a function of age. *eLife*, 10, 2021.

[9195] Anthony I Jang, Ravi Sharma, and Jan Drugowitsch. Optimal policy for attention-modulated decisions explains human fixation behavior. *eLife*, 10, 2021.

[9196] Radek Jankele, Rob Jelier, and Pierre Gönczy. Physically asymmetric division of the C. elegans zygote ensures invariably successful embryogenesis. *eLife*, 10, 2021.

[9197] Danielle Janosevic, Jered Myslinski, Thomas W McCarthy, Amy Zollman, Farooq Syed, Xiaoling Xuei, Hongyu Gao, Yun-Long Liu, Kimberly S Collins, Ying-Hua Cheng, Seth Winfree, Tarek M El-Achkar, Bernhard Maier, Ricardo Melo Ferreira, Michael T Eadon, Takashi Hato, and Pierre C Dagher. The orchestrated cellular and molecular responses of the kidney to endotoxin define a precise sepsis timeline. *eLife*, 10, 2021.

[9198] Rick Jansen, Laura KM Han, Josine E Verhoeven, Karolina A Aberg, Edwin CGJ van den Oord, Yuri Milaneschi, and Brenda WJH Penninx. An integrative study of five biological clocks in somatic and mental health. *eLife*, 10, 2021.

[9199] Fanny Janssen, Anastasios Bardoutsos, Shady El Gewily, and Joop De Beer. Future life expectancy in Europe taking into account the impact of smoking, obesity, and alcohol. *eLife*, 10, 2021.

[9200] Przemyslaw Jarzebowski, Clara S Tang, Ole Paulsen, and Y Audrey Hay. Impaired spatial learning and suppression of sharp wave ripples by cholinergic activation at the goal location. *eLife*, 10, 2021.

[9201] Elizabeth Jaworski, Rose M Langsjoen, Brooke Mitchell, Barbara Judy, Patrick Newman, Jessica A Plante, Kenneth S Plante, Aaron L Miller, Yiyang Zhou, Daniele Swetnam, Stephanea Sotcheff, Victoria Morris, Nehad Saada, Rafael RG Machado, Allan McConnell, Steven G Widen, Jill Thompson, Jianli Dong, Ping Ren, Rick B Pyles, Thomas G Ksiazek, Vineet D Menachery, Scott C Weaver, and Andrew L Routh. Tiled-ClickSeq for targeted sequencing of complete coronavirus genomes with simultaneous capture of RNA recombination and minority variants. *eLife*, 10, 2021.

[9202] Philip Jean-Richard-dit Bressel, Jessica C Lee, Shi Xian Liew, Gabrielle Weidemann, Peter F Lovibond, and Gavan P McNally. Punishment insensitivity in humans is due to failures in instrumental contingency learning. *eLife*, 10, 2021.

[9203] Isaac J Jensen, Xiang Li, Patrick W McGonagill, Qiang Shan, Micaela G Fosdick, Mikaela M Tremblay, Jon CD Houtman, Hai-Hui Xue, Thomas S Griffith, Weiqun Peng, and Vladimir P Badovinac. Sepsis leads to lasting changes in phenotype and function of memory CD8 T cells. *eLife*, 10, 2021.

[9204] Thomas P Jensen, Olga Kopach, James P Reynolds, Leonid P Savtchenko, and Dmitri A Rusakov. Release probability increases towards distal dendrites boosting high-frequency signal transfer in the rodent hippocampus. *eLife*, 10, 2021.

[9205] Dongjin Jeong, Hye Sung Kim, Hye Young Kim, Min Jueng Kang, Hyeryeon Jung, Yumi Oh, Donghyun Kim, Jaemoon Koh, Sung-Yup Cho, Yoon Kyung Jeon, Eun Bong Lee, Seung Hyo Lee, Eui-Cheol Shin, Ho Min Kim, Eugene C Yi, and Doo Hyun Chung. Soluble Fas ligand drives autoantibody-induced arthritis by binding to DR5/TRAIL-R2. *eLife*, 10, 2021.

[9206] Yong Dam Jeong, Keisuke Ejima, Kwang Su Kim, Shoya Iwanami, Ana I Bento, Yasuhisa Fujita, Il Hyo Jung, Kazuyuki Aihara, Koichi Watashi, Taiga Miyazaki, Takaji Wakita, Shingo Iwami, and Marco Ajelli. Revisiting the guidelines for ending isolation for COVID-19 patients. *eLife*, 10, 2021.

[9207] Daisy X Ji, Kristen C Witt, Dmitri I Kotov, Shally R Margolis, Alexander Louie, Victoria Chevée, Katherine J Chen, Moritz M Gaidt, Harmandeep S Dhaliwal, Angus Y Lee, Stephen L Nishimura, Dario S Zamboni, Igor Kramnik, Daniel A Portnoy, K Heran Darwin, and Russell E Vance. Role of the transcriptional regulator SP140 in resistance to bacterial infections via repression of type I interferons. *eLife*, 10, 2021.

[9208] Hongfei Ji, Anthony D Fouad, Shelly Teng, Alice Liu, Pilar Alvarez-Illera, Bowen Yao, Zihao Li, and Christopher Fang-Yen. Phase response analyses support a relaxation oscillator model of locomotor rhythm generation in Caenorhabditis elegans. *eLife*, 10, 2021.

[9209] Jie Lisa Ji, Markus Helmer, Clara Fonteneau, Joshua B Burt, Zailyn Tamayo, Jure DemŠar, Brendan D Adkinson, Aleksandar Savić, Katrin H Preller, Flora Moujaes, Franz X Vollenweider, William J Martin, Grega RepovŠ, Youngsun T Cho, Christopher Pittenger, John D Murray, and Alan Anticevic. Mapping brain-behavior space relationships along the psychosis spectrum. *eLife*, 10, 2021.

[9210] Ni Ji, Gurrein K Madan, Guadalupe I Fabre, Alyssa Dayan, Casey M Baker, Talya S Kramer, Ijeoma Nwabudike, and Steven W Flavell. A neural circuit for flexible control of persistent behavioral states. *eLife*, 10, 2021.

[9211] Ni Ji, Vivek Venkatachalam, Hillary Denise Rodgers, Wesley Hung, Taizo Kawano, Christopher M Clark, Maria Lim, Mark J Alkema, Mei Zhen, and Aravinthan DT Samuel. Corollary discharge promotes a sustained motor state in a neural circuit for navigation. *eLife*, 10, 2021.

[9212] Zhejun Ji, Jacky Chuen, Marianthi Kiparaki, and Nicholas Baker. Cell competition removes segmental aneuploid cells from Drosophila imaginal disc-derived tissues based on ribosomal protein gene dose. *eLife*, 10, 2021.

[9213] Xiaoxuan Jia, Ha Hong, and James J DiCarlo. Unsupervised changes in core object recognition behavior are predicted by neural plasticity in inferior temporal cortex. *eLife*, 10, 2021.

[9214] Rundong Jiang, Ian Max Andolina, Ming Li, and Shiming Tang. Clustered functional domains for curves and corners in cortical area V4. *eLife*, 10, 2021.

[9215] Zhong-Jiao Jiang, Wenping Li, Li-Hua Yao, Badeia Saed, Yan Rao, Brian S Grewe, Andrea McGinley, Kelly Varga, Simon Alford, Ying S Hu, and Liang-Wei Gong. TRPM7 is critical for short-term synaptic depression by regulating synaptic vesicle endocytosis. *eLife*, 10, 2021.

[9216] Sungho Jin, Nal Ae Yoon, Zhong-Wu Liu, Jae Eun Song, Tamas L Horvath, Jung Dae Kim, and Sabrina Diano. Drp1 is required for AgRP neuronal activity and feeding. *eLife*, 10, 2021.

[9217] Junjun Jing, Jifan Feng, Jingyuan Li, Hu Zhao, Thach-Vu Ho, Jinzhi He, Yuan Yuan, Tingwei Guo, Jiahui Du, Mark Urata, Paul Sharpe, and Yang Chai. Reciprocal interaction between mesenchymal stem cells and transit amplifying cells regulates tissue homeostasis. *eLife*, 10, 2021.

[9218] Jacob L Johansen, Lauren E Nadler, Adam Habary, Alyssa J Bowden, and Jodie Rummer. Thermal acclimation of tropical coral reef fishes to global heat waves. *eLife*, 10, 2021.

[9219] Corbin SC Johnson, Carol A Shively, Kristofer T Michalson, Amanda J Lea, Ryne J DeBo, Timothy D Howard, Gregory A Hawkins, Susan E Appt, Yongmei Liu, Charles E McCall, David M Herrington, Edward H Ip, Thomas C Register, and Noah Snyder-Mackler. Contrasting effects of Western vs Mediterranean diets on monocyte inflammatory gene expression and social behavior in a primate model. *eLife*, 10, 2021.

[9220] Mark C Johnson, Geylani Can, Miguel Monteiro Santos, Diana Alexander, and Philip Zegerman. Checkpoint inhibition of origin firing prevents inappropriate replication outside of S-phase. *eLife*, 10, 2021.

[9221] Milo S Johnson, Shreyas Gopalakrishnan, Juhee Goyal, Megan E Dillingham, Christopher W Bakerlee, Parris T Humphrey, Tanush Jagdish, Elizabeth R Jerison, Katya Kosheleva, Katherine R Lawrence, Jiseon Min, Alief Moulana, Angela M Phillips, Julia C Piper, Ramya Purkanti, Artur Rego-Costa, Michael J McDonald, Alex N Nguyen Ba, and Michael M Desai. Phenotypic and molecular evolution across 10,000 generations in laboratory budding yeast populations. *eLife*, 10, 2021.

[9222] Rachel A Johnston, Philippe Vullioud, Jack Thorley, Henry Kirveslahti, Leyao Shen, Sayan Mukherjee, Courtney M Karner, Tim Clutton-Brock, and Jenny Tung. Morphological and genomic shifts in mole-rat 'queens' increase fecundity but reduce skeletal integrity. *eLife*, 10, 2021.

[9223] Stephen Johnston, Sarah L Parylak, Stacy Kim, Nolan Mac, Christina Lim, Iryna Gallina, Cooper Bloyd, Alexander Newberry, Christian D Saavedra, Ondrej Novak, J Tiago GonÇalves, Fred H Gage, and Matthew Shtrahman. AAV ablates neurogenesis in the adult murine hippocampus. *eLife*, 10, 2021.

[9224] Jennifer E Jones, Valerie Le Sage, Gabriella H Padovani, Michael Calderon, Erik S Wright, and Seema S Lakdawala. Parallel evolution between genomic segments of seasonal human influenza viruses reveals RNA-RNA relationships. *eLife*, 10, 2021.

[9225] Joshua M Jones, Ilana Grinberg, Avigdor Eldar, and Alan D Grossman. A mobile genetic element increases bacterial host fitness by manipulating development. *eLife*, 10, 2021.

[9226] Jakob Jordan, Maximilian Schmidt, Walter Senn, and Mihai A Petrovici. Evolving interpretable plasticity for spiking networks. *eLife*, 10, 2021.

[9227] Asha Mary Joseph, Saheli Daw, Ismath Sadhir, and Anjana Badrinarayanan. Coordination between nucleotide excision repair and specialized polymerase DnaE2 action enables DNA damage survival in non-replicating bacteria. *eLife*, 10, 2021.

[9228] Marco Jost, Amy N Jacobson, Jeffrey A Hussmann, Giana Cirolia, Michael A Fischbach, and Jonathan S Weissman. CRISPR-based functional genomics in human dendritic cells. *eLife*, 10, 2021.

[9229] Elena N Judd, Samantha M Lewis, and Abigail L Person. Diverse inhibitory projections from the cerebellar interposed nucleus. *eLife*, 10, 2021.

[9230] Linda Julian, Gregory Naylor, Grant R Wickman, Nicola Rath, Giovanni Castino, David Stevenson, Sheila Bryson, June Munro, Lynn McGarry, Margaret Mullin, Alistair Rice, Armandodel Del Río Hernández, and Michael F Olson. Defective apoptotic cell contractility provokes sterile inflammation, leading to liver damage and tumour suppression. *eLife*, 10, 2021.

[9231] Sergi Junyent, Joshua C Reeves, James LA Szczerkowski, Clare L Garcin, Tung-Jui Trieu, Matthew Wilson, Jethro Lundie-Brown, and Shukry J Habib. Wnt- and glutamate-receptors orchestrate stem cell dynamics and asymmetric cell division. *eLife*, 10, 2021.

[9232] Radhia Kacher, FranÇois-Xavier Lejeune, Sandrine Noël, Cécile Cazeneuve, Alexis Brice, Sandrine Humbert, and Alexandra Durr. Propensity for somatic expansion increases over the course of life in Huntington disease. *eLife*, 10, 2021.

[9233] Shereen R Kadir, Andrew Lilja, Nick Gunn, Campbell Strong, Rowan T Hughes, Benjamin J Bailey, James Rae, Robert G Parton, and John McGhee. Nanoscape, a data-driven 3D real-time interactive virtual cell environment. *eLife*, 10, 2021.

[9234] Morgan P Kain, Eloise B Skinner, Andrew F van den Hurk, Hamish McCallum, and Erin A Mordecai. Physiology and ecology combine to determine host and vector importance for Ross River virus. *eLife*, 10, 2021.

[9235] Hari Teja Kalidindi, Kevin P Cross, Timothy P Lillicrap, Mohsen Omrani, Egidio Falotico, Philip N Sabes, and Stephen H Scott. Rotational dynamics in motor cortex are consistent with a feedback controller. *eLife*, 10, 2021.

[9236] Varun Kamat, Brian M Robbings, Seung-Ryoung Jung, John Kelly, James B Hurley, Kenneth P Bube, and Ian R Sweet. Fluidics system for resolving concentration-dependent effects of dissolved gases on tissue metabolism. *eLife*, 10, 2021.

[9237] Tsukushi Kamiya, Nicole M Davis, Megan A Greischar, David Schneider, and Nicole Mideo. Linking functional and molecular mechanisms of host resilience to malaria infection. *eLife*, 10, 2021.

[9238] Nikolay P Kandul, Junru Liu, Jared B Bennett, John M Marshall, and Omar S Akbari. A confinable home-and-rescue gene drive for population modification. *eLife*, 10, 2021.

[9239] Patrick Bodilly Kane and Jonathan Kimmelman. Is preclinical research in cancer biology reproducible enough? *eLife*, 10, 2021.

[9240] Jin Young Kang, Eliza Llewellyn, James Chen, Paul Dominic B Olinares, Joshua Brewer, Brian T Chait, Elizabeth A Campbell, and Seth A Darst. Structural basis for transcription complex disruption by the Mfd translocase. *eLife*, 10, 2021.

[9241] Victor Kang, Robin T White, Simon Chen, and Walter Federle. Extreme suction attachment performance from specialised insects living in mountain streams (Diptera: Blephariceridae). *eLife*, 10, 2021.

[9242] Yul HR Kang, Anne Löffler, Danique Jeurissen, Ariel Zylberberg, Daniel M Wolpert, and Michael N Shadlen. Multiple decisions about one object involve parallel sensory acquisition but time-multiplexed evidence incorporation. *eLife*, 10, 2021.

[9243] Georgia Eleni Kapetaniou, Matthias A Reinhard, Patricia Christian, Andrea Jobst, Philippe N Tobler, Frank Padberg, and Alexander Soutschek. The role of oxytocin in delay of gratification and flexibility in non-social decision making. *eLife*, 10, 2021.

[9244] Mohammed Kaplan, Georges Chreifi, Lauren Ann Metskas, Janine Liedtke, Cecily R Wood, Catherine M Oikonomou, William J Nicolas, Poorna Subramanian, Lori A Zacharoff, Yuhang Wang, Yi-Wei Chang, Morgan Beeby, Megan J Dobro, Yongtao Zhu, Mark J McBride, Ariane Briegel, Carrie L Shaffer, and Grant J Jensen. In situ imaging of bacterial outer membrane projections and associated protein complexes using electron cryo-tomography. *eLife*, 10, 2021.

[9245] Prathitha Kar, Sriram Tiruvadi-Krishnan, Jaana Männik, Jaan Männik, and Ariel Amir. Distinguishing different modes of growth using single-cell data. *eLife*, 10, 2021.

[9246] Usha Pallabi Kar, Himani Dey, and Abdur Rahaman. Cardiolipin targets a dynamin-related protein to the nuclear membrane. *eLife*, 10, 2021.

[9247] Anastassios Karagiannis, Thierry Gallopin, Alexandre Lacroix, Fabrice Plaisier, Juliette Piquet, Hélène Geoffroy, Régine Hepp, Jérémie Naudé, Benjamin Le Gac, Richard Egger, Bertrand Lambolez, Dongdong Li, Jean Rossier, Jochen F Staiger, Hiromi Imamura, Susumu Seino, Jochen Roeper, and Bruno Cauli. Lactate is an energy substrate for rodent cortical neurons and enhances their firing activity. *eLife*, 10, 2021.

[9248] Farina Karim, Inbal Gazy, Sandile Cele, Yenzekile Zungu, Robert Krause, Mallory Bernstein, Khadija Khan, Yashica Ganga, Hylton Rodel, Ntombifuthi Mthabela, Matilda Mazibuko, Daniel Muema, Dirhona Ramjit, Thumbi Ndung'u, Willem Hanekom, Bernadett Gosnell, Moherndran Archary, Moherndran Archary, Kaylesh J Dullabh, Jennifer Giandhari, Philip Goulder, Guy Harling, Rohen Harrichandparsad, Kobus Herbst, Prakash Jeena, Thandeka Khoza, Nigel Klein, Rajhmun Madansein, Mohlopheni Marakalala, Mosa Moshabela, Kogie Naidoo, Zaza Ndhlovu, Kennedy Nyamande, Nesri Padayatchi, Vinod Patel, Theresa Smit, Adrie Steyn, Richard J Lessells, Emily B Wong, Tulio de Oliveira, Mahomed-Yunus S Moosa, Gil Lustig, Alasdair Leslie, Henrik Kløverpris, and Alex Sigal. HIV status alters disease severity and immune cell responses in Beta variant SARS-CoV-2 infection wave. *eLife*, 10, 2021.

[9249] Hamid Karimi-Rouzbahani, Alexandra Woolgar, and Anina N Rich. Neural signatures of vigilance decrements predict behavioural errors before they occur. *eLife*, 10, 2021.

[9250] Samuel Kariuki, Zoe A Dyson, Cecilia Mbae, Ronald Ngetich, Susan M Kavai, Celestine Wairimu, Stephen Anyona, Naomi Gitau, Robert Sanaya Onsare, Beatrice Ongandi, Sebastian Duchene, Mohamed Ali, John David Clemens, Kathryn E Holt, and Gordon Dougan. Multiple introductions of multidrug-resistant typhoid associated with acute infection and asymptomatic carriage, Kenya. *eLife*, 10, 2021.

[9251] Ariel Karlinsky and Dmitry Kobak. Tracking excess mortality across countries during the COVID-19 pandemic with the World Mortality Dataset. *eLife*, 10, 2021.

[9252] Maria Rita Karlocai, Judit Heredi, Tünde Benedek, Noemi Holderith, Andrea Lorincz, and Zoltan Nusser. Variability in the Munc13-1 content of excitatory release sites. *eLife*, 10, 2021.

[9253] Maren Kasper, Michael Heming, David Schafflick, Xiaolin Li, Tobias Lautwein, Melissa Meyer zu Horste, Dirk Bauer, Karoline Walscheid, Heinz Wiendl, Karin Loser, Arnd Heiligenhaus, and Gerd Meyer zu Hörste. Intraocular dendritic cells characterize HLA-B27-associated acute anterior uveitis. *eLife*, 10, 2021.

[9254] Eugene A Katrukha, Daphne Jurriens, Desiree M Salas Pastene, and Lukas C Kapitein. Quantitative mapping of dense microtubule arrays in mammalian neurons. *eLife*, 10, 2021.

[9255] Hiroko Katsuno-Kambe, Jessica L Teo, Robert J Ju, James Hudson, Samantha J Stehbens, and Alpha S Yap. Collagen polarization promotes epithelial elongation by stimulating locoregional cell proliferation. *eLife*, 10, 2021.

[9256] Ella Katz, Jia-Jie Li, Benjamin Jaegle, Haim Ashkenazy, Shawn R Abrahams, Clement Bagaza, Samuel Holden, Chris J Pires, Ruthie Angelovici, and Daniel J Kliebenstein. Genetic variation, environment and demography intersect to shape Arabidopsis defense metabolite variation across Europe. *eLife*, 10, 2021.

[9257] Spencer R Katz, Maksim A Yakovlev, Daniel J Vanselow, Yifu Ding, Alex Y Lin, Dilworth Y Parkinson, Yuxin Wang, Victor A Canfield, Khai C Ang, and Keith C Cheng. Whole-organism 3D quantitative characterization of zebrafish melanin by silver deposition micro-CT. *eLife*, 10, 2021.

[9258] Alaattin Kaya, Cheryl Zi Jin Phua, Mitchell Lee, Lu Wang, Alexander Tyshkovskiy, Siming Ma, Benjamin Barre, Weiqiang Liu, Benjamin R Harrison, Xiaqing Zhao, Xuming Zhou, Brian M Wasko, Theo K Bammler, Daniel EL Promislow, Matt Kaeberlein, and Vadim N Gladyshev. Evolution of natural lifespan variation and molecular strategies of extended lifespan in yeast. *eLife*, 10, 2021.

[9259] Aynur Kaya-Çopur, Fabio Marchiano, Marco Y Hein, Daniel Alpern, Julie Russeil, Nuno Miguel Luis, Matthias Mann, Bart Deplancke, Bianca H Habermann, and Frank Schnorrer. The Hippo pathway controls myofibril assembly and muscle fiber growth by regulating sarcomeric gene expression. *eLife*, 10, 2021.

[9260] Haneen Kayyal, Sailendrakumar Kolatt Chandran, Adonis Yiannakas, Nathaniel Gould, Mohammad Khamaisy, and Kobi Rosenblum. Insula to mPFC reciprocal connectivity differentially underlies novel taste neophobic response and learning in mice. *eLife*, 10, 2021.

[9261] Alison L Kearney, Dougall M Norris, Milad Ghomlaghi, Martin Kin Lok Wong, Sean J Humphrey, Luke Carroll, Guang Yang, Kristen C Cooke, Pengyi Yang, Thomas A Geddes, Sungyoung Shin, Daniel J Fazakerley, Lan K Nguyen, David E James, and James G Burchfield. Akt phosphorylates insulin receptor substrate to limit PI3K-mediated PIP3 synthesis. *eLife*, 10, 2021.

[9262] Ellen A Kearney, Paul A Agius, Victor Chaumeau, Julia C Cutts, Julie A Simpson, and Freya JI Fowkes. Anopheles salivary antigens as serological biomarkers of vector exposure and malaria transmission: A systematic review with multilevel modelling. *eLife*, 10, 2021.

[9263] Madeline M Keenen, David Brown, Lucy D Brennan, Roman Renger, Harrison Khoo, Christopher R Carlson, Bo Huang, Stephan W Grill, Geeta J Narlikar, and Sy Redding. HP1 proteins compact DNA into mechanically and positionally stable phase separated domains. *eLife*, 10, 2021.

[9264] Shai Kellner, Abeer Abbasi, Ido Carmi, Ronit Heinrich, Tali Garin-Shkolnik, Tova Hershkovitz, Moshe Giladi, Yoni Haitin, Katrine M Johannesen, Rikke Steensbjerre Møller, and Shai Berlin. Two de novo GluN2B mutations affect multiple NMDAR-functions and instigate severe pediatric encephalopathy. *eLife*, 10, 2021.

[9265] Justin W Kenney, Patrick E Steadman, Olivia Young, Meng Ting Shi, Maris Polanco, Saba Dubaishi, Kristopher Covert, Thomas Mueller, and Paul W Frankland. A 3D adult zebrafish brain atlas (AZBA) for the digital age. *eLife*, 10, 2021.

[9266] Hanna Keren, Charles Zheng, David C Jangraw, Katharine Chang, Aria Vitale, Robb B Rutledge, Francisco Pereira, Dylan M Nielson, and Argyris Stringaris. The temporal representation of experience in subjective mood. *eLife*, 10, 2021.

[9267] Nadja Kern, Rui Dong, Shawn M Douglas, Ronald D Vale, and Meghan A Morrissey. Tight nanoscale clustering of Fc$\gamma$ receptors using DNA origami promotes phagocytosis. *eLife*, 10, 2021.

[9268] S Jordan Kerns, Chaitra Belgur, Debora Petropolis, Marianne Kanellias, Riccardo Barrile, Johannes Sam, Tina Weinzierl, Tanja Fauti, Anne Freimoser-Grundschober, Jan Eckmann, Carina Hage, Martina Geiger, Patrick Ray Ng, William Tien-Street, Dimitris V Manatakis, Virginie Micallef, Regine Gerard, Michael Bscheider, Ekaterina Breous-Nystrom, Anneliese Schneider, Anna Maria Giusti, Cristina Bertinetti-Lapatki, Heather Shannon Grant, Adrian B Roth, Geraldine A Hamilton, Thomas Singer, Katia Karalis, Annie Moisan, Peter Bruenker, Christian Klein, Marina Bacac,

Nikolce Gjorevski, and Lauriane Cabon. Human immunocompetent Organ-on-Chip platforms allow safety profiling of tumor-targeted T-cell bispecific antibodies. *eLife*, 10, 2021.

[9269] Aziz Khan. A call to eradicate non-inclusive terms from the life sciences. *eLife*, 10, 2021.

[9270] Shahanshah Khan, Mahnoush S Shafiei, Christopher Longoria, John W Schoggins, Rashmin C Savani, and Hasan Zaki. SARS-CoV-2 spike protein induces inflammation via TLR2-dependent activation of the NF-$\kappa$B pathway. *eLife*, 10, 2021.

[9271] Shahzad S Khan, Yuriko Sobu, Herschel S Dhekne, Francesca Tonelli, Kerryn Berndsen, Dario R Alessi, and Suzanne R Pfeffer. Pathogenic LRRK2 control of primary cilia and Hedgehog signaling in neurons and astrocytes of mouse brain. *eLife*, 10, 2021.

[9272] Kanika Khanna, Javier Lopez-Garrido, Joseph Sugie, Kit Pogliano, and Elizabeth Villa. Asymmetric localization of the cell division machinery during Bacillus subtilis sporulation. *eLife*, 10, 2021.

[9273] Nandish Khanra, Patricia MGE Brown, Amanda M Perozzo, Derek Bowie, and Joel R Meyerson. Architecture and structural dynamics of the heteromeric GluK2/K5 kainate receptor. *eLife*, 10, 2021.

[9274] Michelina Kierzek, Parker E Deal, Evan W Miller, Shatanik Mukherjee, Dagmar Wachten, Arnd Baumann, U Benjamin Kaupp, Timo Strünker, and Christoph Brenker. Simultaneous recording of multiple cellular signaling events by frequency- and spectrally-tuned multiplexing of fluorescent probes. *eLife*, 10, 2021.

[9275] Sanne Kikkert, Dario Pfyffer, Michaela Verling, Patrick Freund, and Nicole Wenderoth. Finger somatotopy is preserved after tetraplegia but deteriorates over time. *eLife*, 10, 2021.

[9276] Angela Kim, Jakob G Knudsen, Joseph C Madara, Anna Benrick, Thomas G Hill, Lina Abdul Kadir, Joely A Kellard, Lisa Mellander, Caroline Miranda, Haopeng Lin, Timothy James, Kinga Suba, Aliya F Spigelman, Yanling Wu, Patrick E MacDonald, Ingrid Wernstedt Asterholm, Tore Magnussen, Mikkel Christensen, Tina Vilsbøll, Victoria Salem, Filip K Knop, Patrik Rorsman, Bradford B Lowell, and Linford JB Briant. Arginine-vasopressin mediates counter-regulatory glucagon release and is diminished in type 1 diabetes. *eLife*, 10, 2021.

[9277] Angela Kim, Joseph C Madara, Chen Wu, Mark L Andermann, and Bradford B Lowell. Neural basis for regulation of vasopressin secretion by anticipated disturbances in osmolality. *eLife*, 10, 2021.

[9278] Bernard Y Kim, Jeremy R Wang, Danny E Miller, Olga Barmina, Emily Delaney, Ammon Thompson, Aaron A Comeault, David Peede, Emmanuel RR D'Agostino, Julianne Pelaez, Jessica M Aguilar, Diler Haji, Teruyuki Matsunaga, Ellie E Armstrong, Molly Zych, Yoshitaka Ogawa, Marina Stamenković-Radak, Mihailo Jelić, Marija Savić Veselinović, Marija Tanasković, Pavle Erić, Jian-Jun Gao, Takehiro K Katoh, Masanori J Toda, Hideaki Watabe, Masayoshi Watada, Jeremy S Davis, Leonie C Moyle, Giulia Manoli, Enrico Bertolini, Vladimír KoŠťál, R Scott Hawley, Aya Takahashi, Corbin D Jones, Donald K Price, Noah Whiteman, Artyom Kopp, Daniel R Matute, and Dmitri A Petrov. Highly contiguous assemblies of 101 drosophilid genomes. *eLife*, 10, 2021.

[9279] Dong Hyun Kim, Hee Young Kim, Sunjung Cho, Su-Jin Yoo, Won-Ju Kim, Hye Ran Yeon, Kyungho Choi, Je-Min Choi, Seong Wook Kang, and Won-Woo Lee. Induction of the IL-1RII decoy receptor by NFAT/FOXP3 blocks IL-1$\beta$-dependent response of Th17 cells. *eLife*, 10, 2021.

[9280] Gha-Hyun J Kim, Han Mo, Harrison Liu, Zhihao Wu, Steven Chen, Jiashun Zheng, Xiang Zhao, Daryl Nucum, James Shortland, Longping Peng, Mannuel Elepano, Benjamin Tang, Steven Olson, Nick Paras, Hao Li, Adam R Renslo, Michelle R Arkin, Bo Huang, Bingwei Lu, Marina Sirota, and Su Guo. A zebrafish screen reveals Renin-angiotensin system inhibitors as neuroprotective via mitochondrial restoration in dopamine neurons. *eLife*, 10, 2021.

[9281] Heesun Kim, Yue-He Ding, Shan Lu, Mei-Qing Zuo, Wendy Tan, Darryl Conte, Meng-Qiu Dong, and Craig C Mello. PIE-1 SUMOylation promotes germline fates and piRNA-dependent silencing in C. elegans. *eLife*, 10, 2021.

[9282] Heesun Kim, Yue-He Ding, Gangming Zhang, Yong-Hong Yan, Darryl Conte, Meng-Qiu Dong, and Craig C Mello. HDAC1 SUMOylation promotes Argonaute-directed transcriptional silencing in C. elegans. *eLife*, 10, 2021.

[9283] Jee Min Kim, Pat Visanpattanasin, Vivian Jou, Sheng Liu, Xiaona Tang, Qinsi Zheng, Kai Yu Li, Jonathan Snedeker, Luke D Lavis, Timothee Lionnet, and Carl Wu. Single-molecule imaging of chromatin remodelers reveals role of ATPase in promoting fast kinetics of target search and dissociation from chromatin. *eLife*, 10, 2021.

[9284] Minseong Kim, Carmen Reinhard, and Christof Niehrs. A MET-PTPRK kinase-phosphatase rheostat controls ZNRF3 and Wnt signaling. *eLife*, 10, 2021.

[9285] Peter K Kim, Christopher J Halbrook, Samuel A Kerk, Megan Radyk, Stephanie Wisner, Daniel M Kremer, Peter Sajjakulnukit, Anthony Andren, Sean W Hou, Ayush Trivedi, Galloway Thurston, Abhinav Anand, Liang Yan, Lucia Salamanca-Cardona, Samuel D Welling, Li Zhang, Matthew R Pratt, Kayvan R Keshari, Haoqiang Ying, and Costas A Lyssiotis. Hyaluronic acid fuels pancreatic cancer cell growth. *eLife*, 10, 2021.

[9286] Soeun Kim, Guk-Yeol Park, Jong Seok Park, Jiho Park, Hyebeen Hong, and Yoontae Lee. Regulation of positive and negative selection and TCR signaling during thymic T cell development by capicua. *eLife*, 10, 2021.

[9287] Suil Kim and Douglas G McMahon. Light sets the brain's daily clock by regional quickening and slowing of the molecular clockworks at dawn and dusk. *eLife*, 10, 2021.

[9288] Maia Kinnebrew, Giovanni Luchetti, Ria Sircar, Sara Frigui, Lucrezia Vittoria Viti, Tomoki Naito, Francis Beckert, Yasunori Saheki, Christian Siebold, Arun Radhakrishnan, and Rajat Rohatgi. Patched 1 reduces the accessibility of cholesterol in the outer leaflet of membranes. *eLife*, 10, 2021.

[9289] Holly E Kinser, Matthew C Mosley, Isaac B Plutzer, and Zachary Pincus. Global, cell non-autonomous gene regulation drives individual lifespan among isogenic C. elegans. *eLife*, 10, 2021.

[9290] Michael C Kiritsy, Laurisa M Ankley, Justin Trombley, Gabrielle P Huizinga, Audrey E Lord, Pontus Orning, Roland Elling, Katherine A Fitzgerald, and Andrew J Olive. A genetic screen in macrophages identifies new regulators of IFN$\gamma$-inducible MHCII that contribute to T cell activation. *eLife*, 10, 2021.

[9291] Michael C Kiritsy, Katelyn McCann, Daniel Mott, Steven M Holland, Samuel M Behar, Christopher M Sassetti, and Andrew J Olive. Mitochondrial respiration contributes to the interferon gamma response in antigen-presenting cells. *eLife*, 10, 2021.

[9292] Nina Kirstein, Alexander Buschle, Xia Wu, Stefan Krebs, Helmut Blum, Elisabeth Kremmer, Ina M Vorberg, Wolfgang Hammerschmidt, Laurent Lacroix, Olivier Hyrien, Benjamin Audit, and Aloys Schepers. Human ORC/MCM density is low in active genes and correlates with replication time but does not delimit initiation zones. *eLife*, 10, 2021.

[9293] Kathryn E Kistler and Trevor Bedford. Evidence for adaptive evolution in the receptor-binding domain of seasonal coronaviruses OC43 and 229e. *eLife*, 10, 2021.

[9294] Katarzyna Kita, Catarina Albergaria, Ana S Machado, Megan R Carey, Martin Müller, and Igor Delvendahl. GluA4 facilitates cerebellar expansion coding and enables associative memory formation. *eLife*, 10, 2021.

[9295] Amrutha Kizhedathu, Piyush Chhajed, Lahari Yeramala, Deblina Sain Basu, Tina Mukherjee, Kutti R Vinothkumar, and Arjun Guha. Duox-generated reactive oxygen species activate ATR/Chk1 to induce G2 arrest in Drosophila tracheoblasts. *eLife*, 10, 2021.

[9296] Jan L Klee, Bryan C Souza, and Francesco P Battaglia. Learning differentially shapes prefrontal and hippocampal activity during classical conditioning. *eLife*, 10, 2021.

[9297] Amanda Klein, Hans Jürgen Solinski, Nathalie M Malewicz, Hada Fong-ha Ieong, Elizabeth I Sypek, Steven G Shimada, Timothy V Hartke, Matthew Wooten, Gang Wu, Xinzhong Dong, Mark A Hoon, Robert H LaMotte, and Matthias Ringkamp. Pruriception and neuronal coding in nociceptor subtypes in human and nonhuman primates. *eLife*, 10, 2021.

[9298] Lukas Klimmasch, Johann Schneider, Alexander Lelais, Maria Fronius, Bertram Emil Shi, and Jochen Triesch. The development of active binocular vision under normal and alternate rearing conditions. *eLife*, 10, 2021.

[9299] P Christiaan Klink, Xing Chen, Wim Vanduffel, and Pieter R Roelfsema. Population receptive fields in nonhuman primates from whole-brain fMRI and large-scale neurophysiology in visual cortex. *eLife*, 10, 2021.

[9300] Daniel S Kluger, Elio Balestrieri, Niko A Busch, and Joachim Gross. Respiration aligns perception with neural excitability. *eLife*, 10, 2021.

[9301] Barbara C Klump, James JH St Clair, and Christian Rutz. New Caledonian crows keep 'valuable' hooked tools safer than basic non-hooked tools. *eLife*, 10, 2021.

[9302] Sven Klumpe, Herman KH Fung, Sara K Goetz, Ievgeniia Zagoriy, Bernhard Hampoelz, Xiaojie Zhang, Philipp S Erdmann, Janina Baumbach, Christoph W Müller, Martin Beck, Jürgen M Plitzko, and Julia Mahamid. A modular platform for automated cryo-FIB workflows. *eLife*, 10, 2021.

[9303] Jackie Knee, Trent Sumner, Zaida Adriano, Claire Anderson, Farran Bush, Drew Capone, Veronica Casmo, David Holcomb, Pete Kolsky, Amy MacDougall, Evgeniya Molotkova, Judite Monteiro Braga, Celina Russo, Wolf Peter Schmidt, Jill Stewart, Winnie Zambrana, Valentina Zuin, Rassul Nalá, Oliver Cumming, and Joe Brown. Effects of an urban sanitation intervention on childhood enteric infection and diarrhea in Maputo, Mozambique: A controlled before-and-after trial. *eLife*, 10, 2021.

[9304] Daniel R Knight, Korakrit Imwattana, Brian Kullin, Enzo Guerrero-Araya, Daniel Paredes-Sabja, Xavier Didelot, Kate E Dingle, David W Eyre, César Rodríguez, and Thomas V Riley. Major genetic discontinuity and novel toxigenic species in Clostridioides difficile taxonomy. *eLife*, 10, 2021.

[9305] Gwenan M Knight, Rebecca E Glover, C Finn McQuaid, Ioana D Olaru, Karin Gallandat, Quentin J Leclerc, Naomi M Fuller, Sam J Willcocks, Rumina Hasan, Esther van Kleef, and Clare IR Chandler. Antimicrobial resistance and COVID-19: Intersections and implications. *eLife*, 10, 2021.

[9306] John RP Knight, Nikola Vlahov, David M Gay, Rachel A Ridgway, William James Faller, Christopher Proud, Giovanna R Mallucci, Tobias von der Haar, Christopher Mark Smales, Anne E Willis, and Owen J Sansom. Rpl24Bst mutation suppresses colorectal cancer by promoting eEF2 phosphorylation via eEF2K. *eLife*, 10, 2021.

[9307] Rachel Knoener, Edward Evans, Jordan T Becker, Mark Scalf, Bayleigh Benner, Nathan M Sherer, and Lloyd M Smith. Identification of host proteins differentially associated with HIV-1 RNA splice variants. *eLife*, 10, 2021.

[9308] Kwang Woo Ko, Laura Devault, Yo Sasaki, Jeffrey Milbrandt, and Aaron DiAntonio. Live imaging reveals the cellular events downstream of SARM1 activation. *eLife*, 10, 2021.

[9309] Allison Koenecke, Michael Powell, Ruoxuan Xiong, Zhu Shen, Nicole Fischer, Sakibul Huq, Adham M Khalafallah, Marco Trevisan, Pär Sparen, Juan J Carrero, Akihiko Nishimura, Brian Caffo, Elizabeth A Stuart, Renyuan Bai, Verena Staedtke, David L Thomas, Nickolas Papadopoulos, Ken W Kinzler, Bert Vogelstein, Shibin Zhou, Chetan Bettegowda, Maximilian F Konig, Brett D Mensh, Joshua T Vogelstein, and Susan Athey. Alpha-1 adrenergic receptor antagonists to prevent hyperinflammation and death from lower respiratory tract infection. *eLife*, 10, 2021.

[9310] Friederike E Kohrs, Ilsa-Maria Daumann, Bojana Pavlovic, Eugene Jennifer Jin, F Ridvan Kiral, Shih-Ching Lin, Filip Port, Heike Wolfenberg, Thomas F Mathejczyk, Gerit A Linneweber, Chih-Chiang Chan, Michael Boutros, and P Robin Hiesinger. Systematic functional analysis of rab GTPases reveals limits of neuronal robustness to environmental challenges in flies. *eLife*, 10, 2021.

[9311] Keiichiro Koiwai, Takashi Koyama, Soichiro Tsuda, Atsushi Toyoda, Kiyoshi Kikuchi, Hiroaki Suzuki, and Ryuji Kawano. Single-cell RNA-seq analysis reveals penaeid shrimp hemocyte subpopulations and cell differentiation process. *eLife*, 10, 2021.

[9312] Piotr Kolata and Rouslan G Efremov. Structure of Escherichia coli respiratory complex I reconstituted into lipid nanodiscs reveals an uncoupled conformation. *eLife*, 10, 2021.

[9313] Philipp Kolb, Katja Hoffmann, Annika Sievert, Henrike Reinhard, Eva Merce-Maldonado, Vu Thuy Khanh Le-Trilling, Anne Halenius, Dominique Gütle, and Hartmut Hengel. Human cytomegalovirus antagonizes activation of Fc$\gamma$ receptors by distinct and synergizing modes of IgG manipulation. *eLife*, 10, 2021.

[9314] Astrid Kollewe, Vladimir Chubanov, Fong Tsuen Tseung, Leonor Correia, Eva Schmidt, Anna Rössig, Susanna Zierler, Alexander Haupt, Catrin Swantje Müller, Wolfgang Bildl, Uwe Schulte, Annette Nicke, Bernd Fakler, and Thomas Gudermann. The molecular appearance of native TRPM7 channel complexes identified by high-resolution proteomics. *eLife*, 10, 2021.

[9315] Mi-Seon Kong, Eun Joo Kim, Sanggeon Park, Larry S Zweifel, Yeowool Huh, Jeiwon Cho, and Jeansok J Kim. 'Fearful-place' coding in the amygdala-hippocampal network. *eLife*, 10, 2021.

[9316] Georgina Kontou, Pantelis Antonoudiou, Marina Podpolny, Blanka R Szulc, I Lorena Arancibia-Carcamo, Nathalie F Higgs, Guillermo Lopez-Domenech, Patricia C Salinas, Edward O Mann, and Josef T Kittler. Miro1-dependent mitochondrial dynamics in parvalbumin interneurons. *eLife*, 10, 2021.

[9317] Renée S Koolschijn, Anna Shpektor, William T Clarke, I Betina Ip, David Dupret, Uzay E Emir, and Helen C Barron. Memory recall involves a transient break in excitatory-inhibitory balance. *eLife*, 10, 2021.

[9318] Aleksandra Kołodziej, Mikołaj Magnuski, Anastasia Ruban, and Aneta Brzezicka. No relationship between frontal alpha asymmetry and depressive disorders in a multiverse analysis of five studies. *eLife*, 10, 2021.

[9319] Ian Krajbich, Andres Mitsumasu, Rafael Polania, Christian C Ruff, and Ernst Fehr. A causal role for the right frontal eye fields in value comparison. *eLife*, 10, 2021.

[9320] Manuela Kramer, Melvin Rodriguez-Heredia, Francesco Saccon, Laura Mosebach, Manuel Twachtmann, Anja Krieger-Liszkay, Chris Duffy, Robert J Knell, Giovanni Finazzi, and Guy Thomas Hanke. Regulation of photosynthetic electron flow on dark to light transition by ferredoxin:NADP(H) oxidoreductase interactions. *eLife*, 10, 2021.

[9321] FranÇois Kroll, Gareth T Powell, Marcus Ghosh, Gaia Gestri, Paride Antinucci, Timothy J Hearn, Hande Tunbak, Sumi Lim, Harvey W Dennis, Joseph M Fernandez, David Whitmore, Elena Dreosti, Stephen W Wilson, Ellen J Hoffman, and Jason Rihel. A simple and effective F0 knockout method for rapid screening of behaviour and other complex phenotypes. *eLife*, 10, 2021.

[9322] Alena Kroupova, Fabian Ackle, Igor Asanović, Stefan Weitzer, Franziska M Boneberg, Marco Faini, Alexander Leitner, Alessia Chui, Ruedi Aebersold, Javier Martinez, and Martin Jinek. Molecular architecture of the human tRNA ligase complex. *eLife*, 10, 2021.

[9323] Lara Katharina Krüger, Matthieu Gélin, Liang Ji, Carlos Kikuti, Anne Houdusse, Manuel Théry, Laurent Blanchoin, and Phong T Tran. Kinesin-6 Klp9 orchestrates spindle elongation by regulating microtubule sliding and growth. *eLife*, 10, 2021.

[9324] Julia A Kuhn, Ilia D Vainchtein, Joao Braz, Katherine Hamel, Mollie Bernstein, Veronica Craik, Madelene W Dahlgren, Jorge Ortiz-Carpena, Ari B Molofsky, Anna V Molofsky, and Allan I Basbaum. Regulatory T-cells inhibit microglia-induced pain hypersensitivity in female mice. *eLife*, 10, 2021.

[9325] Alexandra Kühnlein, Simon A Lanzmich, and Dieter Braun. tRNA sequences can assemble into a replicator. *eLife*, 10, 2021.

[9326] Saurabh Kulkarni, Jonathan Marquez, Priya Date, Rosa Ventrella, Brian J Mitchell, and Mustafa K Khokha. Mechanical stretch scales centriole number to apical area via Piezo1 in multiciliated cells. *eLife*, 10, 2021.

[9327] Shiri Kult, Tsviya Olender, Marco Osterwalder, Svetalana Markman, Dena Leshkowitz, Sharon Krief, Ronnie Blecher-Gonen, Shani Ben-Moshe, Lydia Farack, Hadas Keren-Shaul, Tomer-Meir Salame, Terence D Capellini, Shalev Itzkovitz, Ido Amit, Axel Visel, and Elazar Zelzer. Bi-fated tendon-to-bone attachment cells are regulated by shared enhancers and KLF transcription factors. *eLife*, 10, 2021.

[9328] Amit Kumar, Edi Barkai, and Jackie Schiller. Plasticity of olfactory bulb inputs mediated by dendritic NMDA-spikes in rodent piriform cortex. *eLife*, 10, 2021.

[9329] Amit Kumar, Manisha Vaish, Saravanan S Karuppagounder, Irina Gazaryan, John W Cave, Anatoly A Starkov, Elizabeth T Anderson, Sheng Zhang, John T Pinto, Austin M Rountree, Wang Wang, Ian R Sweet, and Rajiv R Ratan. HIF1$\alpha$ stabilization in hypoxia is not oxidant-initiated. *eLife*, 10, 2021.

[9330] Charanya Kumar, Sahil Batra, Jack D Griffith, and Dirk Remus. The interplay of RNA:DNA hybrid structure and G-quadruplexes determines the outcome of R-loop-replisome collisions. *eLife*, 10, 2021.

[9331] Manoj Kumar, Sneha Gulati, Asgar H Ansari, Rhythm Phutela, Sundaram Acharya, Mohd Azhar, Jayaram Murthy, Poorti Kathpalia, Akshay Kanakan, Ranjeet Maurya, Janani Srinivasa Vasudevan, Aparna S, Rajesh Pandey, Souvik Maiti, and Debojyoti Chakraborty. FnCas9-based CRISPR diagnostic for rapid and accurate detection of major SARS-CoV-2 variants on a paper strip. *eLife*, 10, 2021.

[9332] Rajnish Kumar Singh, Yonggang Pei, Dipayan Bose, Zachary L Lamplugh, Kunfeng Sun, Yan Yuan, Paul Lieberman, Jianxin You, and Erle S Robertson. KSHV-encoded vCyclin can modulate HIF1$\alpha$ levels to promote DNA replication in hypoxia. *eLife*, 10, 2021.

[9333] Jennifer M Kunselman, Achla Gupta, Ivone Gomes, Lakshmi A Devi, and Manojkumar A Puthenveedu. Compartment-specific opioid receptor signaling is selectively modulated by different dynorphin peptides. *eLife*, 10, 2021.

[9334] Marco Künzli, Peter Reuther, Daniel D Pinschewer, and Carolyn G King. Opposing effects of T cell receptor signal strength on CD4 T cells responding to acute versus chronic viral infection. *eLife*, 10, 2021.

[9335] Mizuki Kurashina, Jane Wang, Jeffrey Lin, Kathy Kyungeun Lee, Arpun Johal, and Kota Mizumoto. Sustained expression of unc-4 homeobox gene and unc-37/Groucho in postmitotic neurons specifies the spatial organization of the cholinergic synapses in C. elegans. *eLife*, 10, 2021.

[9336] Lisa E Kursel, Henry D Cope, and Ofer Rog. Unconventional conservation reveals structure-function relationships in the synaptonemal complex. *eLife*, 10, 2021.

[9337] Hyae Yon Kweon, Mi-Ni Lee, Max Dorfel, Seungwoon Seo, Leah Gottlieb, Thomas PaPazyan, Nina McTiernan, Rasmus Ree, David Bolton, Andrew Garcia, Michael Flory, Jonathan Crain, Alison Sebold, Scott Lyons, Ahmed Ismail, Elaine Marchi, Seong-keun Sonn, Se-Jin Jeong, Sejin Jeon, Shinyeong Ju, Simon J Conway, Taesoo Kim, Hyun-Seok Kim, Cheolju Lee, Tae-Young Roh, Thomas Arnesen, Ronen Marmorstein, Goo Taeg Oh, and Gholson J Lyon. Naa12 compensates for Naa10 in mice in the amino-terminal acetylation pathway. *eLife*, 10, 2021.

[9338] Hyeogsun Kwon, Mubasher Mohammed, Oscar Franzén, Johan Ankarklev, and Ryan C Smith. Single-cell analysis of mosquito hemocytes identifies signatures of immune cell subtypes and cell differentiation. *eLife*, 10, 2021.

[9339] Jonas Hansen Kymre, XiaoLan Liu, Elena Ian, Christoffer Nerland Berge, GuiRong Wang, Bente Gunnveig Berg, XinCheng Zhao, and Xi Chu. Distinct protocerebral neuropils associated with attractive and aversive female-produced odorants in the male moth brain. *eLife*, 10, 2021.

[9340] Evdoxia Kyriazopoulou, Periklis Panagopoulos, Symeon Metallidis, George N Dalekos, Garyphallia Poulakou, Nikolaos Gatselis, Eleni Karakike, Maria Saridaki, Georgia Loli, Aggelos Stefos, Danai Prasianaki, Sarah Georgiadou, Olga Tsachouridou, Vasileios Petrakis, Konstantinos Tsiakos, Maria Kosmidou, Vassiliki Lygoura, Maria Dareioti, Haralampos Milionis, Ilias C Papanikolaou, Karolina Akinosoglou, Dimitra-Melia Myrodia, Areti Gravvani, Aliki Stamou, Theologia Gkavogianni, Konstantina Katrini, Theodoros Marantos, Ioannis P Trontzas, Konstantinos Syrigos, Loukas Chatzis, Stamatios Chatzis, Nikolaos Vechlidis, Christina Avgoustou, Stamatios Chalvatzis, Miltiades Kyprianou, Jos WM van der Meer, Jesper Eugen-Olsen, Mihai G Netea, and Evangelos J Giamarellos-Bourboulis. An open label trial of anakinra to prevent respiratory failure in COVID-19. *eLife*, 10, 2021.

[9341] Gaspare La Rocca, Bryan King, Bing Shui, Xiaoyi Li, Minsi Zhang, Kemal M Akat, Paul Ogrodowski, Chiara Mastroleo, Kevin Chen, Vincenzo Cavalieri, Yilun Ma, Viviana Anelli, Doron Betel, Joana Vidigal, Thomas Tuschl, Gunter Meister, Craig B Thompson, Tullia Lindsten, Kevin Haigis, and Andrea Ventura. Inducible and reversible inhibition of miRNA-mediated gene repression in vivo. *eLife*, 10, 2021.

[9342] Sharon R Ladyman, Kirsten M Carter, Matt L Gillett, Zin Khant Aung, and David R Grattan. A reduction in voluntary physical activity in early pregnancy in mice is mediated by prolactin. *eLife*, 10, 2021.

[9343] Fanny Lafouresse, Romain Jugele, Sabina Müller, Marine Doineau, Valérie Duplan-Eche, Eric Espinosa, Marie-Pierre Puisségur, Sébastien Gadat, and Salvatore Valitutti. Stochastic asymmetric repartition of lytic machinery in dividing CD8+ T cells generates heterogeneous killing behavior. *eLife*, 10, 2021.

[9344] Ines Lahmann, Joscha Griger, Jie-Shin Chen, Yao Zhang, Markus Schuelke, and Carmen Birchmeier. Met and Cxcr4 cooperate to protect skeletal muscle stem cells against inflammation-induced damage during regeneration. *eLife*, 10, 2021.

[9345] Jean-Benoît Lalanne and Gene-Wei Li. First-principles model of optimal translation factors stoichiometry. *eLife*, 10, 2021.

[9346] Maureen C Lamb, Chathuri P Kaluarachchi, Thiranjeewa I Lansakara, Samuel Q Mellentine, Yiling Lan, Alexei V Tivanski, and Tina L Tootle. Fascin limits Myosin activity within Drosophila border cells to control substrate stiffness and promote migration. *eLife*, 10, 2021.

[9347] Jason T Lambert, Linda Su-Feher, Karol Cichewicz, Tracy L Warren, Iva Zdilar, Yurong Wang, Kenneth J Lim, Jessica L Haigh, Sarah J Morse, Cesar P Canales, Tyler W Stradleigh, Erika Castillo Palacios, Viktoria Haghani, Spencer D Moss, Hannah Parolini, Diana Quintero, Diwash

Shrestha, Daniel Vogt, Leah C Byrne, and Alex S Nord. Parallel functional testing identifies enhancers active in early postnatal mouse brain. *eLife*, 10, 2021.

[9348] Wout S Lamers, Kevin Boyack, Vincent Larivière, Cassidy R Sugimoto, Nees Jan van Eck, Ludo Waltman, and Dakota Murray. Investigating disagreement in the scientific literature. *eLife*, 10, 2021.

[9349] Dan Landayan, Brian P Wang, Jennifer Zhou, and Fred W Wolf. Thirst interneurons that promote water seeking and limit feeding behavior in Drosophila. *eLife*, 10, 2021.

[9350] Elizabeth A Landis, Angela M Oliverio, Erin A McKenney, Lauren M Nichols, Nicole Kfoury, Megan Biango-Daniels, Leonora K Shell, Anne A Madden, Lori Shapiro, Shravya Sakunala, Kinsey Drake, Albert Robbat, Matthew Booker, Robert R Dunn, Noah Fierer, and Benjamin E Wolfe. The diversity and function of sourdough starter microbiomes. *eLife*, 10, 2021.

[9351] Veronika Lang, Sebastian Ferencik, Bharath Ananthasubramaniam, Achim Kramer, and Bert Maier. Susceptibility rhythm to bacterial endotoxin in myeloid clock-knockout mice. *eLife*, 10, 2021.

[9352] Merlin Lange, AhmetCan Solak, Shruthi Vijay Kumar, Hirofumi Kobayashi, Bin Yang, and Loïc Alain Royer. ZAF, the first open source fully automated feeder for aquatic facilities. *eLife*, 10, 2021.

[9353] Lukas M Langer, Fabien Bonneau, Yair Gat, and Elena Conti. Cryo-EM reconstructions of inhibitor-bound SMG1 kinase reveal an autoinhibitory state dependent on SMG8. *eLife*, 10, 2021.

[9354] Ina Lantzsch, Che-Hang Yu, Yu-Zen Chen, Vitaly Zimyanin, Hossein Yazdkhasti, Norbert Lindow, Erik Szentgyoergyi, Ariel M Pani, Steffen Prohaska, Martin Srayko, Sebastian Fürthauer, and Stefanie Redemann. Microtubule reorganization during female meiosis in C. elegans. *eLife*, 10, 2021.

[9355] Jacqueline A Larouche, Mahir Mohiuddin, Jeongmoon J Choi, Peter J Ulintz, Paula Fraczek, Kaitlyn Sabin, Sethuramasundaram Pitchiaya, Sarah J Kurpiers, Jesus Castor-Macias, Wenxuan Liu, Robert Louis Hastings, Lemuel A Brown, James F Markworth, Kanishka De Silva, Benjamin Levi, Sofia D Merajver, Gregorio Valdez, Joe V Chakkalakal, Young C Jang, Susan V Brooks, and Carlos A Aguilar. Murine muscle stem cell response to perturbations of the neuromuscular junction are attenuated with aging. *eLife*, 10, 2021.

[9356] Daniel B Larremore, Bailey K Fosdick, Kate M Bubar, Sam Zhang, Stephen M Kissler, C Jessica E Metcalf, Caroline O Buckee, and Yonatan H Grad. Estimating SARS-CoV-2 seroprevalence and epidemiological parameters with uncertainty from serological surveys. *eLife*, 10, 2021.

[9357] Abagael M Lasseigne, Fabio A Echeverry, Sundas Ijaz, Jennifer Carlisle Michel, E Anne Martin, Audrey J Marsh, Elisa Trujillo, Kurt C Marsden, Alberto E Pereda, and Adam C Miller. Electrical synaptic transmission requires a postsynaptic scaffolding protein. *eLife*, 10, 2021.

[9358] Eva On-Chai Lau, Devid Damiani, Georges Chehade, Nuria Ruiz-Reig, Rana Saade, Yves Jossin, Mohamed Aittaleb, Olivier Schakman, Nicolas Tajeddine, Philippe Gailly, and Fadel Tissir. DIAPH3 deficiency links microtubules to mitotic errors, defective neurogenesis, and brain dysfunction. *eLife*, 10, 2021.

[9359] Michael S Lauer, Jamie Doyle, Joy Wang, and Deepshikha Roychowdhury. Associations of topic-specific peer review outcomes and institute and center award rates with funding disparities at the National Institutes of Health. *eLife*, 10, 2021.

[9360] Michael S Lauer and Deepshikha Roychowdhury. Inequalities in the distribution of National Institutes of Health research project grant funding. *eLife*, 10, 2021.

[9361] Knut H Lauritzen, Maria Belland Olsen, Mohammed Shakil Ahmed, Kuan Yang, Johanne Egge Rinholm, Linda H Bergersen, Qin Ying Esbensen, Lars Jansen Sverkeli, Mathias Ziegler, Håvard Attramadal, Bente Halvorsen, Pål Aukrust, and Arne Yndestad. Instability in NAD+ metabolism leads to impaired cardiac mitochondrial function and communication. *eLife*, 10, 2021.

[9362] Anders Laustsen, Renée M van der Sluis, Albert Gris-Oliver, Sabina Sánchez Hernández, Ena Cemalovic, Hai Q Tang, Lars Henning Pedersen, Niels Uldbjerg, Martin R Jakobsen, and Rasmus O Bak. Ascorbic acid supports ex vivo generation of plasmacytoid dendritic cells from circulating hematopoietic stem cells. *eLife*, 10, 2021.

[9363] Thomas Laval, Laura Pedró-Cos, Wladimir Malaga, Laure Guenin-Macé, Alexandre Pawlik, Véronique Mayau, Hanane Yahia-Cherbal, Océane Delos, Wafa Frigui, Justine Bertrand-Michel, Christophe Guilhot, and Caroline Demangel. De novo synthesized polyunsaturated fatty acids operate as both host immunomodulators and nutrients for Mycobacterium tuberculosis. *eLife*, 10, 2021.

[9364] Elizabeth J Lawrence, Goker Arpag, Cayetana Arnaiz, and Marija Zanic. SSNA1 stabilizes dynamic microtubules and detects microtubule damage. *eLife*, 10, 2021.

[9365] Daniel J Laydon, Ilaria Dorigatti, Wes R Hinsley, Gemma Nedjati-Gilani, Laurent Coudeville, and Neil M Ferguson. Efficacy profile of the CYD-TDV dengue vaccine revealed by Bayesian survival analysis of individual-level phase III data. *eLife*, 10, 2021.

[9366] Aurel A Lazar, Tingkai Liu, Mehmet Kerem Turkcan, and Yiyin Zhou. Accelerating with FlyBrainLab the discovery of the functional logic of the Drosophila brain in the connectomic and synaptomic era. *eLife*, 10, 2021.

[9367] Laura Le Pelletier, Matthieu Mantecon, Jennifer Gorwood, Martine Auclair, Roberta Foresti, Roberto Motterlini, Mireille Laforge, Michael Atlan, Bruno Fève, Jacqueline Capeau, Claire Lagathu, and Veronique Bereziat. Metformin alleviates stress-induced cellular senescence of aging human adipose stromal cells and the ensuing adipocyte dysfunction. *eLife*, 10, 2021.

[9368] Maxence Le Vasseur, Jonathan Friedman, Marco Jost, Jiawei Xu, Justin Yamada, Martin Kampmann, Max A Horlbeck, Michelle R Salemi, Brett S Phinney, Jonathan S Weissman, and Jodi Nunnari. Genome-wide CRISPRi screening identifies OCIAD1 as a prohibitin client and regulatory determinant of mitochondrial Complex III assembly in human cells. *eLife*, 10, 2021.

[9369] Katherine B LeClair, Kenny L Chan, Manuella P Kaster, Lyonna F Parise, Charles Joseph Burnett, and Scott J Russo. Individual history of winning and hierarchy landscape influence stress susceptibility in mice. *eLife*, 10, 2021.

[9370] Brian R Lee, Agata Budzillo, Kristen Hadley, Jeremy A Miller, Tim Jarsky, Katherine Baker, DiJon Hill, Lisa Kim, Rusty Mann, Lindsay Ng, Aaron Oldre, Ram Rajanbabu, Jessica Trinh, Sara Vargas, Thomas Braun, Rachel A Dalley, Nathan W Gouwens, Brian E Kalmbach, Tae Kyung Kim, Kimberly A Smith, Gilberto Soler-Llavina, Staci Sorensen, Bosiljka Tasic, Jonathan T Ting, Ed Lein, Hongkui Zeng, Gabe J Murphy, and Jim Berg. Scaled, high fidelity electrophysiological, morphological, and transcriptomic cell characterization. *eLife*, 10, 2021.

[9371] Caroline S Lee, Mariam Aly, and Christopher Baldassano. Anticipation of temporally structured events in the brain. *eLife*, 10, 2021.

[9372] Douglas G Lee and Jean Daunizeau. Trading mental effort for confidence in the metacognitive control of value-based decision-making. *eLife*, 10, 2021.

[9373] Eric Kenji Lee, Hymavathy Balasubramanian, Alexandra Tsolias, Stephanie Udochukwu Anakwe, Maria Medalla, Krishna V Shenoy, and Chandramouli Chandrasekaran. Non-linear dimensionality reduction on extracellular waveforms reveals cell type diversity in premotor cortex. *eLife*, 10, 2021.

[9374] Frank K Lee, Jane C Lee, Bo Shui, Shaun Reining, Megan Jibilian, David M Small, Jason S Jones, Nathaniel H Allan-Rahill, Michael RE Lamont, Megan A Rizzo, Sendoa Tajada, Manuel F Navedo, Luis Fernando Santana, Nozomi Nishimura, and Michael I Kotlikoff. Genetically engineered mice for combinatorial cardiovascular optobiology. *eLife*, 10, 2021.

[9375] Gavin Lee, Jane W Liang, Qing Zhang, Theodore Huang, Christine Choirat, Giovanni Parmigiani, and Danielle Braun. Multi-syndrome, multi-gene risk modeling for individuals with a family history of cancer with the novel R package PanelPRO. *eLife*, 10, 2021.

[9376] John Lee, Kosuke Kawai, Jeffrey R Holt, and Gwenaëlle SG Géléoc. Sensory transduction is required for normal development and maturation of cochlear inner hair cell synapses. *eLife*, 10, 2021.

[9377] Kristen Lee and Chris Q Doe. A locomotor neural circuit persists and functions similarly in larvae and adult Drosophila. *eLife*, 10, 2021.

[9378] Yo Han Lee, Hyun-Jun Jang, Sounkou Kim, Sun Sil Choi, Keon Woo Khim, Hye-jin Eom, Jimin Hyun, Kyeong Jin Shin, Young Chan Chae, Hongtae Kim, Jiyoung Park, Neung Hwa Park, Chang-Yun Woo, Chung Hwan Hong, Eun Hee Koh, Dougu Nam, and Jang Hyun Choi. Hepatic MIR20B promotes nonalcoholic fatty liver disease by suppressing PPARA. *eLife*, 10, 2021.

[9379] Solène N Lefebvre, Antoine Taly, Anaïs Menny, Karima Medjebeur, and Pierre-Jean Corringer. Mutational analysis to explore long-range allosteric couplings involved in a pentameric channel receptor pre-activation and activation. *eLife*, 10, 2021.

[9380] Loïc Léger, Deborah Byrne, Paul Guiraud, Elsa Germain, and Etienne Maisonneuve. NirD curtails the stringent response by inhibiting RelA activity in Escherichia coli. *eLife*, 10, 2021.

[9381] Michael Jonathan Lehrke, Michael Jeremy Shapiro, Matthew J Rajcula, Madeleine M Kennedy, Shaylene A McCue, Kay L Medina, and Virginia Smith Shapiro. The mitochondrial iron transporter ABCB7 is required for B cell development, proliferation, and class switch recombination in mice. *eLife*, 10, 2021.

[9382] Jaakko I Lehtimäki, Eeva Kaisa Rajakylä, Sari Tojkander, and Pekka Lappalainen. Generation of stress fibers through myosin-driven reorganization of the actin cortex. *eLife*, 10, 2021.

[9383] Ienglam Lei, Shuo Tian, Wenbin Gao, Liu Liu, Yijing Guo, Paul Tang, Eugene Chen, and Zhong Wang. Acetyl-CoA production by specific metabolites promotes cardiac repair after myocardial infarction via histone acetylation. *eLife*, 10, 2021.

[9384] Joël Lemière, Yuan Ren, and Julien Berro. Rapid adaptation of endocytosis, exocytosis, and eisosomes after an acute increase in membrane tension in yeast cells. *eLife*, 10, 2021.

[9385] Stefan M Lemke, Dhakshin S Ramanathan, David Darevksy, Daniel Egert, Joshua D Berke, and Karunesh Ganguly. Coupling between motor cortex and striatum increases during sleep over long-term skill learning. *eLife*, 10, 2021.

[9386] Todd Lencz, Daniel Backenroth, Einat Granot-Hershkovitz, Adam Green, Kyle Gettler, Judy H Cho, Omer Weissbrod, Or Zuk, and Shai Carmi. Utility of polygenic embryo screening for disease depends on the selection strategy. *eLife*, 10, 2021.

[9387] Maximilian Lenz, Amelie Eichler, Pia Kruse, Julia Muellerleile, Thomas Deller, Peter Jedlicka, and Andreas Vlachos. All-trans retinoic acid induces synaptopodin-dependent metaplasticity in mouse dentate granule cells. *eLife*, 10, 2021.

[9388] Maximilian Lenz, Pia Kruse, Amelie Eichler, Jakob Straehle, Jürgen Beck, Thomas Deller, and Andreas Vlachos. All-trans retinoic acid induces synaptic plasticity in human cortical neurons. *eLife*, 10, 2021.

[9389] Silvia Leon, Rajae Talbi, Elizabeth A McCarthy, Kaitlin Ferrari, Chrysanthi Fergani, Lydie Naule, Ji Hae Choi, Rona S Carroll, Ursula B Kaiser, Carlos F Aylwin, Alejandro Lomniczi, and Víctor M Navarro. Sex-specific pubertal and metabolic regulation of Kiss1 neurons via Nhlh2. *eLife*, 10, 2021.

[9390] Calvin Jon A Leonen, Miho Shimada, Caroline E Weller, Tomoyoshi Nakadai, Peter L Hsu, Elizabeth L Tyson, Arpit Mishra, Patrick MM Shelton, Martin Sadilek, R David Hawkins, Ning Zheng, Robert G Roeder, and Champak Chatterjee. Sumoylation of the human histone H4 tail inhibits p300-mediated transcription by RNA polymerase II in cellular extracts. *eLife*, 10, 2021.

[9391] Eitan Lerner, Anders Barth, Jelle Hendrix, Benjamin Ambrose, Victoria Birkedal, Scott C Blanchard, Richard Börner, Hoi Sung Chung, Thorben Cordes, Timothy D Craggs, Ashok A Deniz, Jiajie Diao, Jingyi Fei, Ruben L Gonzalez, Irina V Gopich, Taekjip Ha, Christian A Hanke, Gilad Haran, Nikos S Hatzakis, Sungchul Hohng, Seok-Cheol Hong, Thorsten Hugel, Antonino Ingargiola, Chirlmin Joo, Achillefs N Kapanidis, Harold D Kim, Ted Laurence, Nam Ki Lee, Tae-Hee Lee, Edward A Lemke, Emmanuel Margeat, Jens Michaelis, Xavier Michalet, Sua Myong, Daniel Nettels, Thomas-Otavio Peulen, Evelyn Ploetz, Yair Razvag, Nicole C Robb, Benjamin Schuler, Hamid Soleimaninejad, Chun Tang, Reza Vafabakhsh, Don C Lamb, Claus AM Seidel, and Shimon Weiss. FRET-based dynamic structural biology: Challenges, perspectives and an appeal for open-science practices. *eLife*, 10, 2021.

[9392] Amanda Lesar, Javan Tahir, Jason Wolk, and Marc Gershow. Switch-like and persistent memory formation in individual Drosophila larvae. *eLife*, 10, 2021.

[9393] Mor Levi-Ferber, Rewayd Shalash, Adrien Le-Thomas, Yehuda Salzberg, Maor Shurgi, Jennifer IC Benichou, Avi Ashkenazi, and Sivan Henis-Korenblit. Neuronal regulated ire-1-dependent mRNA decay controls germline differentiation in Caenorhabditis elegans. *eLife*, 10, 2021.

[9394] Amanda H Lewis and Jörg Grandl. Piezo1 ion channels inherently function as independent mechanotransducers. *eLife*, 10, 2021.

[9395] Alexander R Leydon, Wei Wang, Hardik P Gala, Sabrina Gilmour, Samuel Juarez-Solis, Mollye L Zahler, Joseph E Zemke, Ning Zheng, and Jennifer L Nemhauser. Repression by the Arabidopsis TOPLESS corepressor requires association with the core mediator complex. *eLife*, 10, 2021.

[9396] Chun-Yang Li, Xiu-Juan Wang, Xiu-Lan Chen, Qi Sheng, Shan Zhang, Peng Wang, Mussa Quareshy, Branko Rihtman, Xuan Shao, Chao Gao, Fuchuan Li, Shengying Li, Weipeng Zhang, Xiao-Hua Zhang, Gui-Peng Yang, Jonathan D Todd, Yin Chen, and Yu-Zhong Zhang. A novel ATP dependent dimethylsulfoniopropionate lyase in bacteria that releases dimethyl sulfide and acryloyl-CoA. *eLife*, 10, 2021.

[9397] Dianrong Li, Jie Chen, Jia Guo, Lin Li, Gaihong Cai, She Chen, Jia Huang, Hui Yang, Yinhua Zhuang, Fengchao Wang, and Xiaodong Wang. A phosphorylation of RIPK3 kinase initiates an intracellular apoptotic pathway that promotes prostaglandin2$\alpha$-induced corpus luteum regression. *eLife*, 10, 2021.

[9398] Guorong Li, Chanyoung Lee, A Thomas Read, Ke Wang, Jungmin Ha, Megan Kuhn, Iris Navarro, Jenny Cui, Katherine Young, Rahul Gorijavolu, Todd Sulchek, Casey Kopczynski, Sina Farsiu, John Samples, Pratap Challa, C Ross Ethier, and W Daniel Stamer. Anti-fibrotic activity of a rho-kinase inhibitor restores outflow function and intraocular pressure homeostasis. *eLife*, 10, 2021.

[9399] Hai Li, Chia-Ying Huang, Elena G Govorunova, Oleg A Sineshchekov, Adrian Yi, Kenneth J Rothschild, Meitian Wang, Lei Zheng, and John L Spudich. The crystal structure of bromide-bound GtACR1 reveals a pre-activated state in the transmembrane anion tunnel. *eLife*, 10, 2021.

[9400] Hailun Li, Lorenzo Lones, and Aaron DiAntonio. Bidirectional regulation of glial potassium buffering – glioprotection versus neuroprotection. *eLife*, 10, 2021.

[9401] Jia Li, Xuan Qin, Jie Shi, Xiaoshuang Wang, Tong Li, Mengyao Xu, Xiaosu Chen, Yujia Zhao, Jiahao Han, Yongjun Piao, Wenwen Zhang, Pengpeng Qu, Longlong Wang, Rong Xiang, and Yi Shi. A systematic CRISPR screen reveals an IL-20/IL20RA-mediated immune crosstalk to prevent the ovarian cancer metastasis. *eLife*, 10, 2021.

[9402] Jiajianghui Li, Tianjia Guan, Qian Guo, Guannan Geng, Huiyu Wang, Fuyu Guo, Jiwei Li, and Tao Xue. Exposure to landscape fire smoke reduced birthweight in low- and middle-income countries: findings from a siblings-matched case-control study. *eLife*, 10, 2021.

[9403] Jie Li, Lei Yue, Zhihua Li, Wenting Zhang, Bing Zhang, Fangqing Zhao, and Xiuzhu Dong. aCPSF1 cooperates with terminator U-tract to dictate archaeal transcription termination efficacy. *eLife*, 10, 2021.

[9404] Jing Li, Jiabin Yan, and Timothy A Springer. Low-affinity integrin states have faster ligand-binding kinetics than the high-affinity state. *eLife*, 10, 2021.

[9405] Jingxiang Li, Shupan Guo, Fangni Chai, Qi Sun, Pan Li, Li Gao, Lunzhi Dai, Xiaoxiao Ouyang, Zhihui Zhou, Li Zhou, Wei Cheng, Shiqian Qi, Kefeng Lu, and Haiyan Ren. Genetically incorporated crosslinkers reveal NleE attenuates host autophagy dependent on PSMD10. *eLife*, 10, 2021.

[9406] Laifu Li, Li-Zi Zhang, Zhi-Xiong He, Huan Ma, Yu-Ting Zhang, Yu-Feng Xun, Wei Yuan, Wen-Juan Hou, Yi-Tong Li, Zi-Jian Lv, Rui Jia, and Fa-Dao Tai. Dorsal raphe nucleus to anterior cingulate cortex 5-HTergic neural circuit modulates consolation and sociability. *eLife*, 10, 2021.

[9407] Meijing Li, Jianfei Ma, Xueming Li, and Sen-Fang Sui. In situ cryo-ET structure of phycobilisome–photosystem II supercomplex from red alga. *eLife*, 10, 2021.

[9408] Na Li, Shangang Zhao, Zhuzhen Zhang, Yi Zhu, Christy M Gliniak, Lavanya Vishvanath, Yu A An, May-yun Wang, Yingfeng Deng, Qingzhang Zhu, Bo Shan, Amber Sherwood, Toshiharu Onodera, Orhan K Oz, Ruth Gordillo, Rana K Gupta, Ming Liu, Tamas L Horvath, Vishwa Deep Dixit, and Philipp E Scherer. Adiponectin preserves metabolic fitness during aging. *eLife*, 10, 2021.

[9409] Qiuling Li, Hyunsoo Jang, Kayla Y Lim, Alexie Lessing, and Nicholas Stavropoulos. insomniac links the development and function of a sleep-regulatory circuit. *eLife*, 10, 2021.

[9410] Siqi Li and Christopher M Counter. Signaling levels mold the RAS mutation tropism of urethane. *eLife*, 10, 2021.

[9411] Wan Hua Li, Ke Huang, Yang Cai, Qian Wen Wang, Wen Jie Zhu, Yun Nan Hou, Sujing Wang, Sheng Cao, Zhi Ying Zhao, Xu Jie Xie, Yang Du, Chi-Sing Lee, Hon Cheung Lee, Hongmin Zhang, and Yong Juan Zhao. Permeant fluorescent probes visualize the activation of SARM1 and uncover an anti-neurodegenerative drug candidate. *eLife*, 10, 2021.

[9412] Weihan Li, Kelly Crotty, Diego Garrido Ruiz, Mark Voorhies, Carlos Rivera, Anita Sil, R Dyche Mullins, Matthew P Jacobson, Jirka Peschek, and Peter Walter. Protomer alignment modulates specificity of RNA substrate recognition by Ire1. *eLife*, 10, 2021.

[9413] Wenqing Li, Virginia Tran, Iftach Shaked, Belinda Xue, Thomas Moore, Rhonda Lightle, David Kleinfeld, Issam A Awad, and Mark H Ginsberg. Abortive intussusceptive angiogenesis causes multi-cavernous vascular malformations. *eLife*, 10, 2021.

[9414] Xiaoling Li, Richard Fetter, Tina Schwabe, Christophe Jung, Liren Liu, Hermann Steller, and Ulrike Gaul. The cAMP effector PKA mediates Moody GPCR signaling in Drosophila blood–brain barrier formation and maturation. *eLife*, 10, 2021.

[9415] Yunbo Li, Erin M Ritchie, Christopher L Steinke, Cai Qi, Lizhen Chen, Binhai Zheng, and Yishi Jin. Activation of MAP3K DLK and LZK in Purkinje cells causes rapid and slow degeneration depending on signaling strength. *eLife*, 10, 2021.

[9416] Fei Liang, Bo Wang, Juan Geng, Guoling You, Jingjing Fa, Min Zhang, Hunying Sun, Huiwen Chen, Qihua Fu, Xiaoqing Zhang, and Zhen Zhang. SORBS2 is a genetic factor contributing to cardiac malformation of 4q deletion syndrome patients. *eLife*, 10, 2021.

[9417] Jormay Lim, Hsiao-Hsin Tai, Wei-Hao Liao, Ya-Cherng Chu, Chen-Ming Hao, Yueh-Chun Huang, Cheng-Han Lee, Shao-Shien Lin, Sherry Hsu, Ya-Chih Chien, Dar-Ming Lai, Wen-Shiang Chen, Chih-Cheng Chen, and Jaw-Lin Wang. ASIC1a is required for neuronal activation via low-intensity ultrasound stimulation in mouse brain. *eLife*, 10, 2021.

[9418] Soyeon Lim, You-Joung Kim, Sooyeon Park, Ji-heon Choi, Young Hoon Sung, Katsuhiko Nishimori, Zbynek Kozmik, Han-Woong Lee, and Jin Woo Kim. mTORC1-induced retinal progenitor cell overproliferation leads to accelerated mitotic aging and degeneration of descendent Müller glia. *eLife*, 10, 2021.

[9419] Tony KY Lim and Edward S Ruthazer. Microglial trogocytosis and the complement system regulate axonal pruning in vivo. *eLife*, 10, 2021.

[9420] Audrie Lin, Andrew N Mertens, Benjamin F Arnold, Sophia Tan, Jue Lin, Christine P Stewart, Alan E Hubbard, Shahjahan Ali, Jade Benjamin-Chung, Abul K Shoab, Md Ziaur Rahman, Syeda L Famida, Md Saheen Hossen, Palash Mutsuddi, Salma Akther, Mahbubur Rahman, Leanne Unicomb, Ruchira Tabassum Naved, Md Mahfuz Al Mamun, Kausar Parvin, Firdaus S Dhabhar, Patricia Kariger, Lia CH Fernald, Stephen P Luby, and John M Colford. Telomere length is associated with growth in children in rural Bangladesh. *eLife*, 10, 2021.

[9421] Jian-You Lin, Narendra Mukherjee, Max J Bernstein, and Donald B Katz. Perturbation of amygdala-cortical projections reduces ensemble coherence of palatability coding in gustatory cortex. *eLife*, 10, 2021.

[9422] Xiao-Tong Lin, Hong-Qiang Yu, Lei Fang, Ye Tan, Ze-Yu Liu, Di Wu, Jie Zhang, Hao-Jun Xiong, and Chuan-Ming Xie. Elevated FBXO45 promotes liver tumorigenesis through enhancing IGF2BP1 ubiquitination and subsequent PLK1 upregulation. *eLife*, 10, 2021.

[9423] Yu-En Lin, Chin-Hsien Lin, En-Peng Ho, Yi-Ci Ke, Stavroula Petridi, Christopher JH Elliott, Lee-Yan Sheen, and Cheng-Ting Chien. Glial Nrf2 signaling mediates the neuroprotection exerted by Gastrodia elata Blume in Lrrk2-G2019S Parkinson's disease. *eLife*, 10, 2021.

[9424] Axel Lindner, Daniel Wiesen, and Hans-Otto Karnath. Lying in a 3T MRI scanner induces neglect-like spatial attention bias. *eLife*, 10, 2021.

[9425] Ross T Lindsay, Sophie Dieckmann, Dominika Krzyzanska, Dominic Manetta-Jones, James A West, Cecilia Castro, Julian L Griffin, and Andrew J Murray. $\beta$-hydroxybutyrate accumulates in the rat heart during low-flow ischaemia with implications for functional recovery. *eLife*, 10, 2021.

[9426] Pablo J Lituma, Hyung-Bae Kwon, Karina Alviña, Rafael Luján, and Pablo E Castillo. Presynaptic NMDA receptors facilitate short-term plasticity and BDNF release at hippocampal mossy fiber synapses. *eLife*, 10, 2021.

[9427] Bingyu Liu, Qingyang Tian, and Yong Gu. Robust vestibular self-motion signals in macaque posterior cingulate region. *eLife*, 10, 2021.

[9428] Fangyu Liu, James Lee, and Jue Chen. Molecular structures of the eukaryotic retinal importer ABCA4. *eLife*, 10, 2021.

[9429] Hui Liu, Junyi Xin, Sheng Cai, and Xia Jiang. Mendelian randomization analysis provides causality of smoking on the expression of ACE2, a putative SARS-CoV-2 receptor. *eLife*, 10, 2021.

[9430] Jiaqi Liu, Xianzun Tao, Yi Zhu, Chong Li, Kai Ruan, Zoraida Diaz-Perez, Priyamvada Rai, Hongbo Wang, and R Grace Zhai. NMNAT promotes glioma growth through regulating post-translational modifications of P53 to inhibit apoptosis. *eLife*, 10, 2021.

[9431] Ka-Cheuk Liu, Alethia Villasenor, Maria Bertuzzi, Nicole Schmitner, Niki Radros, Linn Rautio, Kenny Mattonet, Ryota L Matsuoka, Sven Reischauer, Didier YR Stainier, and Olov Andersson. Insulin-producing $\beta$-cells regenerate ectopically from a mesodermal origin under the perturbation of hemato-endothelial specification. *eLife*, 10, 2021.

[9432] Lei Liu, Zhiran Fan, Xavier Rovira, Li Xue, Salomé Roux, Isabelle Brabet, Mingxia Xin, Jean-Philippe Pin, Philippe Rondard, and Jianfeng Liu. Allosteric ligands control the activation of a class C GPCR heterodimer by acting at the transmembrane interface. *eLife*, 10, 2021.

[9433] Li Liu, Christoph Sasse, Benedict Dirnberger, Oliver Valerius, Enikő Fekete-Szücs, Rebekka Harting, Daniela E Nordzieke, Stefanie Pöggeler, Petr Karlovsky, Jennifer Gerke, and Gerhard H Braus. Secondary metabolites of Hülle cells mediate protection of fungal reproductive and overwintering structures against fungivorous animals. *eLife*, 10, 2021.

[9434] Menghan Liu, Joseph C Devlin, Jiyuan Hu, Angelina Volkova, Thomas W Battaglia, Melody Ho, John R Asplin, Allyson Byrd, P'ng Loke, Huilin Li, Kelly V Ruggles, Aristotelis Tsirigos, Martin J Blaser, and Lama Nazzal. Microbial genetic and transcriptional contributions to oxalate degradation by the gut microbiota in health and disease. *eLife*, 10, 2021.

[9435] Peng Liu, Anastasia Chrysidou, Juliane Doehler, Martin N Hebart, Thomas Wolbers, and Esther Kuehn. The organizational principles of de-differentiated topographic maps in somatosensory cortex. *eLife*, 10, 2021.

[9436] Qiang Liu, Jin Wang, Xin Wei, Juan Hu, Conghui Ping, Yue Gao, Chang Xie, Peiyu Wang, Peng Cao, Zhengyu Cao, Ye Yu, Dongdong Li, and Jing Yao. Therapeutic inhibition of keratinocyte TRPV3 sensory channel by local anesthetic dyclonine. *eLife*, 10, 2021.

[9437] Qiaoling Liu, Qi Xiao, Zhen Sun, Bo Wang, Lina Wang, Na Wang, Kai Wang, Chengli Song, and Qingkai Yang. Exosome component 1 cleaves single-stranded DNA and sensitizes human kidney renal clear cell carcinoma cells to poly(ADP-ribose) polymerase inhibitor. *eLife*, 10, 2021.

[9438] Qiuying Liu, Xiaoli Chen, Mariah K Novak, Shaojie Zhang, and Wenqian Hu. Repressing Ago2 mRNA translation by Trim71 maintains pluripotency through inhibiting let-7 microRNAs. *eLife*, 10, 2021.

[9439] Qiuying Liu, Mariah K Novak, Rachel M Pepin, Taylor Eich, and Wenqian Hu. microRNA-mediated regulation of microRNA machinery controls cell fate decisions. *eLife*, 10, 2021.

[9440] Shou Liu, Wenjian Cao, Yichi Niu, Jiayi Luo, Yanhua Zhao, Zhiying Hu, and Chenghang Zong. Single-PanIN-seq unveils that ARID1A deficiency promotes pancreatic tumorigenesis by attenuating KRAS-induced senescence. *eLife*, 10, 2021.

[9441] Tengyuan Liu, Tingting Zhang, Maya Nicolas, Lydie Boussicault, Heather Rice, Alessia Soldano, Annelies Claeys, Iveta Petrova, Lee Fradkin, Bart De Strooper, Marie-Claude Potier, and Bassem A Hassan. The amyloid precursor protein is a conserved Wnt receptor. *eLife*, 10, 2021.

[9442] Xia Liu, Jingjing Yu, Longyong Xu, Katharine Umphred-Wilson, Fanglue Peng, Yao Ding, Brendan M Barton, Xiangdong Lv, Michael Y Zhao, Shengyi Sun, Yuning Hong, Ling Qi, Stanley Adoro, and Xi Chen. Notch-induced endoplasmic reticulum-associated degradation governs mouse thymocyte $\beta$-selection. *eLife*, 10, 2021.

[9443] Xiao-Man Liu, Liang Ma, and Randy Schekman. Selective sorting of microRNAs into exosomes by phase-separated YBX1 condensates. *eLife*, 10, 2021.

[9444] Xinhuai Liu, Shel-Hwa Yeo, H James McQuillan, Michel K Herde, Sabine Hessler, Isaiah Cheong, Robert Porteous, and Allan E Herbison. Highly redundant neuropeptide volume co-transmission underlying episodic activation of the GnRH neuron dendron. *eLife*, 10, 2021.

[9445] Yan-Xia Liu, Bin Xue, Wei-Yue Sun, Jenna L Wingfield, Jun Sun, Mingfu Wu, Karl F Lechtreck, Zhenlong Wu, and Zhen-Chuan Fan. Bardet–Biedl syndrome 3 protein promotes ciliary exit of the signaling protein phospholipase D via the BBSome. *eLife*, 10, 2021.

[9446] Yanyan Liu, Pu Ke, Yi-Chun Kuo, Yuxiao Wang, Xuewu Zhang, Chen Song, and Yibing Shan. A putative structural mechanism underlying the antithetic effect of homologous RND1 and RhoD GTPases in mammalian plexin regulation. *eLife*, 10, 2021.

[9447] Yi Liu, Nicolas Basty, Brandon Whitcher, Jimmy D Bell, Elena P Sorokin, Nick van Bruggen, E Louise Thomas, and Madeleine Cule. Genetic architecture of 11 organ traits derived from abdominal MRI using deep learning. *eLife*, 10, 2021.

[9448] Yunzhe Liu, Raymond J Dolan, Cameron Higgins, Hector Penagos, Mark W Woolrich, H Freyja ólafsdóttir, Caswell Barry, Zeb Kurth-Nelson, and Timothy E Behrens. Temporally delayed linear modelling (TDLM) measures replay in both animals and humans. *eLife*, 10, 2021.

[9449] Zelin Liu, Changyu Tao, Shiwei Li, Minghao Du, Yongtai Bai, Xueyan Hu, Yu Li, Jian Chen, and Ence Yang. circFL-seq reveals full-length circular RNAs with rolling circular reverse transcription and nanopore sequencing. *eLife*, 10, 2021.

[9450] Zhaoyang Liu, Amro A Hussien, Yunjia Wang, Terry Heckmann, Roberto Gonzalez, Courtney M Karner, Jess G Snedeker, and Ryan S Gray. An adhesion G protein-coupled receptor is required in cartilaginous and dense connective tissues to maintain spine alignment. *eLife*, 10, 2021.

[9451] Luca Livraghi, Joseph J Hanly, Steven M Van Bellghem, Gabriela Montejo-Kovacevich, Eva SM van der Heijden, Ling Sheng Loh, Anna Ren, Ian A Warren, James J Lewis, Carolina Concha, Laura Hebberecht, Charlotte J Wright, Jonah M Walker, Jessica Foley, Zachary H Goldberg, Henry Arenas-Castro, Camilo Salazar, Michael W Perry, Riccardo Papa, Arnaud Martin, W Owen McMillan, and Chris D Jiggins. Cortex cis-regulatory switches establish scale colour identity and pattern diversity in Heliconius. *eLife*, 10, 2021.

[9452] Line S Löken, Joao M Braz, Alexander Etlin, Mahsa Sadeghi, Mollie Bernstein, Madison Jewell, Marilyn Steyert, Julia Kuhn, Katherine Hamel, Ida J Llewellyn-Smith, and Allan Basbaum. Contribution of dorsal horn CGRP-expressing interneurons to mechanical sensitivity. *eLife*, 10, 2021.

[9453] GonÇalo Lopes, Karolina Farrell, Edward AB Horrocks, Chi-Yu Lee, Mai M Morimoto, Tomaso Muzzu, Amalia Papanikolaou, Fabio R Rodrigues, Thomas Wheatcroft, Stefano Zucca, Samuel G Solomon, and Aman B Saleem. Creating and controlling visual environments using BonVision. *eLife*, 10, 2021.

[9454] M Fernanda López-Gutiérrez, Zeus Gracia-Tabuenca, Juan J Ortiz, Francisco J Camacho, Larry J Young, Raúl G Paredes, Néstor F Díaz, Wendy Portillo, and Sarael Alcauter. Brain functional networks associated with social bonding in monogamous voles. *eLife*, 10, 2021.

[9455] José Fabricio López Hernández, Rachel M Helston, Jeffrey J Lange, R Blake Billmyre, Samantha H Schaffner, Michael T Eickbush, Scott McCroskey, and Sarah E Zanders. Diverse mating phenotypes impact the spread of wtf meiotic drivers in Schizosaccharomyces pombe. *eLife*, 10, 2021.

[9456] Nathan D Lord, Adam N Carte, Philip B Abitua, and Alexander F Schier. The pattern of nodal morphogen signaling is shaped by co-receptor expression. *eLife*, 10, 2021.

[9457] Andrea Loreto, Carlo Angeletti, Weixi Gu, Andrew Osborne, Bart Nieuwenhuis, Jonathan Gilley, Elisa Merlini, Peter Arthur-Farraj, Adolfo Amici, Zhenyao Luo, Lauren Hartley-Tassell, Thomas Ve, Laura M Desrochers, Qi Wang, Bostjan Kobe, Giuseppe Orsomando, and Michael P Coleman. Neurotoxin-mediated potent activation of the axon degeneration regulator SARM1. *eLife*, 10, 2021.

[9458] Heather S Loring, Victoria L Czech, Janneke D Icso, Lauren O'Connor, Sangram S Parelkar, Alexandra B Byrne, and Paul R Thompson. A phase transition enhances the catalytic activity of SARM1, an NAD+ glycohydrolase involved in neurodegeneration. *eLife*, 10, 2021.

[9459] Justin D Lormand, Soo-Kyoung Kim, George A Walters-Marrah, Bryce A Brownfield, J Christopher Fromme, Wade C Winkler, Jonathan R Goodson, Vincent T Lee, and Holger Sondermann. Structural characterization of NrnC identifies unifying features of dinucleases. *eLife*, 10, 2021.

[9460] Maxime Louet, Marina Casiraghi, Marjorie Damian, Mauricio GS Costa, Pedro Renault, Antoniel AS Gomes, Paulo R Batista, Céline M'Kadmi, Sophie Mary, Sonia Cantel, Severine Denoyelle, Khoubaib Ben Haj Salah, David Perahia, Paulo M Bisch, Jean-Alain Fehrentz, Laurent J Catoire, Nicolas Floquet, and Jean-Louis Banères. Concerted conformational dynamics and water movements in the ghrelin G protein-coupled receptor. *eLife*, 10, 2021.

[9461] Eduardo Loureiro-Campos, António Mateus-Pinheiro, Patrícia Patrício, Carina Soares-Cunha, Joana Silva, Vanessa Morais Sardinha, Bárbara Mendes-Pinheiro, Tiago Silveira-Rosa, Ana Verónica Domingues, Ana João Rodrigues, João Oliveira, Nuno Sousa, Nuno Dinis Alves, and Luísa Pinto. Constitutive deficiency of the neurogenic hippocampal modulator AP2$\gamma$ promotes anxiety-like behavior and cumulative memory deficits in mice from juvenile to adult periods. *eLife*, 10, 2021.

[9462] Cecilia Lövkvist, Pawel Mikulski, Svenja Reeck, Matthew Hartley, Caroline Dean, and Martin Howard. Hybrid protein assembly-histone modification mechanism for PRC2-based epigenetic switching and memory. *eLife*, 10, 2021.

[9463] Brandon R Lowe, Rajesh K Yadav, Ryan A Henry, Patrick Schreiner, Atsushi Matsuda, Alfonso G Fernandez, David Finkelstein, Margaret Campbell, Satish Kallappagoudar, Carolyn M Jablonowski, Andrew J Andrews, Yasushi Hiraoka, and Janet F Partridge. Surprising phenotypic diversity of cancer-associated mutations of Gly 34 in the histone H3 tail. *eLife*, 10, 2021.

[9464] Elijah D Lowenstein, Aleksandra Rusanova, Jonas Stelzer, Marc Hernaiz-Llorens, Adrian E Schroer, Ekaterina Epifanova, Francesca Bladt, Eser Göksu Isik, Sven Buchert, Shiqi Jia, Victor Tarabykin, and Luis R Hernandez-Miranda. Olig3 regulates early cerebellar development. *eLife*, 10, 2021.

[9465] Yu-Xuan Lu, Jennifer C Regan, Jacqueline Eßer, Lisa F Drews, Thomas Weinseis, Julia Stinn, Oliver Hahn, Richard A Miller, Sebastian Grönke, and Linda Partridge. A TORC1-histone axis regulates chromatin organisation and non-canonical induction of autophagy to ameliorate ageing. *eLife*, 10, 2021.

[9466] Lawrence Lubyayi, Harriet Mpairwe, Gyaviira Nkurunungi, Swaib A Lule, Angela Nalwoga, Emily L Webb, Jonathan Levin, and Alison M Elliott. Infection-exposure in infancy is associated with reduced allergy-related disease in later childhood in a Ugandan cohort. *eLife*, 10, 2021.

[9467] Federica Lucantonio, Eunyoung Kim, Zhixiao Su, Anna J Chang, Bilal A Bari, and Jeremiah Y Cohen. Aversive stimuli bias corticothalamic responses to motivationally significant cues. *eLife*, 10, 2021.

[9468] Bronwyn A Lucas, Benjamin A Himes, Liang Xue, Timothy Grant, Julia Mahamid, and Nikolaus Grigorieff. Locating macromolecular assemblies in cells by 2D template matching with cisTEM. *eLife*, 10, 2021.

[9469] Matthias Luft, Johanna Klepetko, Silvia Muceli, Jaime Ibáñez, Vlad Tereshenko, Christopher Festin, Gregor Laengle, Olga Politikou, Udo Maierhofer, Dario Farina, Oskar C Aszmann, and Konstantin Davide Bergmeister. Proof of concept for multiple nerve transfers to a single target muscle. *eLife*, 10, 2021.

[9470] Nikolay Lukoyanov, Hiroyuki Watanabe, Liliana S Carvalho, Olga Kononenko, Daniil Sarkisyan, Mengliang Zhang, Marlene Storm Andersen, Elena A Lukoyanova, Vladimir Galatenko, Alex Tonevitsky, Igor Bazov, Tatiana Iakovleva, Jens Schouenborg, and Georgy Bakalkin. Left-right side-specific endocrine signaling complements neural pathways to mediate acute asymmetric effects of brain injury. *eLife*, 10, 2021.

[9471] Dianne Lumaquin, Eleanor Johns, Emily Montal, Joshua M Weiss, David Ola, Abderhman Abuhashem, and Richard M White. An in vivo reporter for tracking lipid droplet dynamics in transparent zebrafish. *eLife*, 10, 2021.

[9472] Derek S Lundberg, Pratchaya Pramoj Na Ayutthaya, Annett Strauß, Gautam Shirsekar, Wen-Sui Lo, Thomas Lahaye, and Detlef Weigel. Host-associated microbe PCR (hamPCR) enables convenient measurement of both microbial load and community composition. *eLife*, 10, 2021.

[9473] Teresa Romeo Luperchio, Leandros Boukas, Li Zhang, Genay Pilarowski, Jenny Jiang, Allison Kalinousky, Kasper D Hansen, and Hans T Bjornsson. Leveraging the Mendelian disorders of the epigenetic machinery to systematically map functional epigenetic variation. *eLife*, 10, 2021.

[9474] Lydia K Lutes, Zoë Steier, Laura L McIntyre, Shraddha Pandey, James Kaminski, Ashley R Hoover, Silvia Ariotti, Aaron Streets, Nir Yosef, and Ellen A Robey. T cell self-reactivity during thymic development dictates the timing of positive selection. *eLife*, 10, 2021.

[9475] Xiao Lv, Feng Gao, Tuo Peter Li, Peng Xue, Xiao Wang, Mei Wan, Bo Hu, Hao Chen, Amit Jain, Zengwu Shao, and Xu Cao. Skeleton interoception regulates bone and fat metabolism through hypothalamic neuroendocrine NPY. *eLife*, 10, 2021.

[9476] Evan H Lyall, Daniel P Mossing, Scott R Pluta, Yun Wen Chu, Amir Dudai, and Hillel Adesnik. Synthesis of a comprehensive population code for contextual features in the awake sensory cortex. *eLife*, 10, 2021.

[9477] Yang Lyu, Kristina J Weaver, Humza A Shaukat, Marta L Plumoff, Maria Tjilos, Daniel EL Promislow, and Scott D Pletcher. Drosophila serotonin 2A receptor signaling coordinates central metabolic processes to modulate aging in response to nutrient choice. *eLife*, 10, 2021.

[9478] Chien-Hui Ma, Kamyab Javanmardi, Ilya J Finkelstein, and Makkuni Jayaram. Disintegration promotes protospacer integration by the Cas1-Cas2 complex. *eLife*, 10, 2021.

[9479] Dingbang Ma, Dariusz Przybylski, Katharine C Abruzzi, Matthias Schlichting, Qunlong Li, Xi Long, and Michael Rosbash. A transcriptomic taxonomy of Drosophila circadian neurons around the clock. *eLife*, 10, 2021.

[9480] Heng Ma, Pengcheng Li, Jiaming Hu, Xingya Cai, Qianling Song, and Haidong D Lu. Processing of motion boundary orientation in macaque V2. *eLife*, 10, 2021.

[9481] Jiajia Ma, Claire A Scott, Ying Na Ho, Harsha Mahabaleshwar, Katherine S Marsay, Changqing Zhang, Christopher KJ Teow, Ser Sue Ng, Weibin Zhang, Vinay Tergaonkar, Lynda J Partridge, Sudipto Roy, Enrique Amaya, and Tom J Carney. Matriptase activation of Gq drives epithelial disruption and inflammation via RSK and DUOX. *eLife*, 10, 2021.

[9482] Kevin C Ma, Tigist F Menkir, Stephen Kissler, Yonatan H Grad, and Marc Lipsitch. Modeling the impact of racial and ethnic disparities on COVID-19 epidemic dynamics. *eLife*, 10, 2021.

[9483] Peijun Ma, Lorrie L He, Alejandro Pironti, Hannah H Laibinis, Christoph M Ernst, Abigail L Manson, Roby P Bhattacharyya, Ashlee M Earl, Jonathan Livny, and Deborah T Hung. Genetic determinants facilitating the evolution of resistance to carbapenem antibiotics. *eLife*, 10, 2021.

[9484] Weirui Ma, Gang Zhen, Wei Xie, and Christine Mayr. In vivo reconstitution finds multivalent RNA–RNA interactions as drivers of mesh-like condensates. *eLife*, 10, 2021.

[9485] Heather E Machado, Alan O Bergland, Ryan Taylor, Susanne Tilk, Emily Behrman, Kelly Dyer, Daniel K Fabian, Thomas Flatt, Josefa González, Talia L Karasov, Bernard Kim, Iryna Kozeretska, Brian P Lazzaro, Thomas JS Merritt, John E Pool, Katherine O'Brien, Subhash Rajpurohit, Paula R Roy, Stephen W Schaeffer, Svitlana Serga, Paul Schmidt, and Dmitri A Petrov. Broad geographic sampling reveals the shared basis and environmental correlates of seasonal adaptation in Drosophila. *eLife*, 10, 2021.

[9486] Kirsty ML Mackinlay, Bailey AT Weatherbee, Viviane Souza Rosa, Charlotte E Handford, George Hudson, Tim Coorens, Lygia V Pereira, Sam Behjati, Ludovic Vallier, Marta N Shahbazi, and Magdalena Zernicka-Goetz. An in vitro stem cell model of human epiblast and yolk sac interaction. *eLife*, 10, 2021.

[9487] Owen Mackwood, Laura B Naumann, and Henning Sprekeler. Learning excitatory-inhibitory neuronal assemblies in recurrent networks. *eLife*, 10, 2021.

[9488] Alex Macpherson, Maisem Laabei, Zainab Ahdash, Melissa A Graewert, James R Birtley, Monika-Sarah ED Schulze, Susan Crennell, Sarah A Robinson, Ben Holmes, Vladas Oleinikovas, Per H Nilsson, James Snowden, Victoria Ellis, Tom Eirik Mollnes, Charlotte M Deane, Dmitri Svergun,

Alastair DG Lawson, and Jean MH van den Elsen. The allosteric modulation of complement C5 by knob domain peptides. *eLife*, 10, 2021.

[9489] Timurs Maculins, Erik Verschueren, Trent Hinkle, Meena Choi, Patrick Chang, Cecile Chalouni, Shilpa Rao, Youngsu Kwon, Junghyun Lim, Anand Kumar Katakam, Ryan C Kunz, Brian K Erickson, Ting Huang, Tsung-Heng Tsai, Olga Vitek, Mike Reichelt, Yasin Senbabaoglu, Brent Mckenzie, John R Rohde, Ivan Dikic, Donald S Kirkpatrick, and Aditya Murthy. Multiplexed proteomics of autophagy-deficient murine macrophages reveals enhanced antimicrobial immunity via the oxidative stress response. *eLife*, 10, 2021.

[9490] Fernando Y Maeda, Jurriaan JH van Haaren, David B Langley, Daniel Christ, Norma W Andrews, and Wenxia Song. Surface-associated antigen induces permeabilization of primary mouse B-cells and lysosome exocytosis facilitating antigen uptake and presentation to T-cells. *eLife*, 10, 2021.

[9491] Kaisei Maeda, Yukiko Okuda, Gen Enomoto, Satoru Watanabe, and Masahiko Ikeuchi. Biosynthesis of a sulfated exopolysaccharide, synechan, and bloom formation in the model cyanobacterium Synechocystis sp. strain PCC 6803. *eLife*, 10, 2021.

[9492] Taro Maeda, Shunichi Takahashi, Takao Yoshida, Shigeru Shimamura, Yoshihiro Takaki, Yukiko Nagai, Atsushi Toyoda, Yutaka Suzuki, Asuka Arimoto, Hisaki Ishii, Nori Satoh, Tomoaki Nishiyama, Mitsuyasu Hasebe, Tadashi Maruyama, Jun Minagawa, Junichi Obokata, and Shuji Shigenobu. Chloroplast acquisition without the gene transfer in kleptoplastic sea slugs, Plakobranchus ocellatus. *eLife*, 10, 2021.

[9493] Laura E Maglio, José A Noriega-Prieto, Irene B Maroto, Jesús Martin-Cortecero, Antonio Muñoz-Callejas, Marta Callejo-Móstoles, and David Fernández de Sevilla. IGF-1 facilitates extinction of conditioned fear. *eLife*, 10, 2021.

[9494] Joseph O Magliozzi and James B Moseley. Pak1 kinase controls cell shape through ribonucleoprotein granules. *eLife*, 10, 2021.

[9495] Jens P Magnusson, Antonio Ray Rios, Lingling Wu, and Lei S Qi. Enhanced Cas12a multi-gene regulation using a CRISPR array separator. *eLife*, 10, 2021.

[9496] ádám Magó, Noémi Kis, Balázs Lükő, and Judit K Makara. Distinct dendritic Ca2+ spike forms produce opposing input-output transformations in rat CA3 pyramidal cells. *eLife*, 10, 2021.

[9497] Lisa Mahler, Sarah P Niehs, Karin Martin, Thomas Weber, Kirstin Scherlach, Christian Hertweck, Martin Roth, and Miriam A Rosenbaum. Highly parallelized droplet cultivation and prioritization of antibiotic producers from natural microbial communities. *eLife*, 10, 2021.

[9498] G Larisa Maier, Nikita Komarov, Felix Meyenhofer, Jae Young Kwon, and Simon G Sprecher. Taste sensing and sugar detection mechanisms in Drosophila larval primary taste center. *eLife*, 10, 2021.

[9499] Roni O Maimon-Mor, Hunter R Schone, David Henderson Slater, A Aldo Faisal, and Tamar R Makin. Early life experience sets hard limits on motor learning as evidenced from artificial arm use. *eLife*, 10, 2021.

[9500] Rabina Mainali, Manal Zabalawi, David Long, Nancy Buechler, Ellen Quillen, Chia-Chi Key, Xuewei Zhu, John S Parks, Cristina Furdui, Peter W Stacpoole, Jennifer Martinez, Charles E McCall, and Matthew A Quinn. Dichloroacetate reverses sepsis-induced hepatic metabolic dysfunction. *eLife*, 10, 2021.

[9501] Giulia Malaguti and Pieter Rein ten Wolde. Theory for the optimal detection of time-varying signals in cellular sensing systems. *eLife*, 10, 2021.

[9502] Elise LV Malavasi, Aniket Ghosh, Daniel G Booth, Michele Zagnoni, Diane L Sherman, and Peter J Brophy. Dynamic early clusters of nodal proteins contribute to node of Ranvier assembly during myelination of peripheral neurons. *eLife*, 10, 2021.

[9503] Maria Maldonado, Fei Guo, and James A Letts. Atomic structures of respiratory complex III2, complex IV, and supercomplex III2-IV from vascular plants. *eLife*, 10, 2021.

[9504] Marta Maltese, Jeffrey R March, Alexander G Bashaw, and Nicolas X Tritsch. Dopamine differentially modulates the size of projection neuron ensembles in the intact and dopamine-depleted striatum. *eLife*, 10, 2021.

[9505] Eugenio Mancera, Isabel Nocedal, Stephen Hammel, Megha Gulati, Kaitlin F Mitchell, David R Andes, Clarissa J Nobile, Geraldine Butler, and Alexander D Johnson. Evolution of the complex transcription network controlling biofilm formation in Candida species. *eLife*, 10, 2021.

[9506] Christopher P Mancuso, Hyunseok Lee, Clare I Abreu, Jeff Gore, and Ahmad S Khalil. Environmental fluctuations reshape an unexpected diversity-disturbance relationship in a microbial community. *eLife*, 10, 2021.

[9507] Zachary F Mandell, Reid T Oshiro, Alexander V Yakhnin, Rishi Vishwakarma, Mikhail Kashlev, Daniel B Kearns, and Paul Babitzke. NusG is an intrinsic transcription termination factor that stimulates motility and coordinates gene expression with NusA. *eLife*, 10, 2021.

[9508] Nandini Mani, Sithara S Wijeratne, and Radhika Subramanian. Micron-scale geometrical features of microtubules as regulators of microtubule organization. *eLife*, 10, 2021.

[9509] Martial Marbouty, Agnès Thierry, Gaël A Millot, and Romain Koszul. MetaHiC phage-bacteria infection network reveals active cycling phages of the healthy human gut. *eLife*, 10, 2021.

[9510] Aurélien Marc, Marion Kerioui, FranÇois Blanquart, Julie Bertrand, Oriol Mitjà, Marc Corbacho-Monné, Michael Marks, and Jeremie Guedj. Quantifying the relationship between SARS-CoV-2 viral load and infectiousness. *eLife*, 10, 2021.

[9511] Marco Marconi, Marcal Gallemi, Eva Benkova, and Krzysztof Wabnik. A coupled mechano-biochemical model for cell polarity guided anisotropic root growth. *eLife*, 10, 2021.

[9512] Isabelle J Marié, Lara Brambilla, Doua Azzouz, Ze Chen, Gisele V Baracho, Azlann Arnett, Haiyan S Li, Weiguo Liu, Luisa Cimmino, Pratip Chattopadhyay, Gregg Silverman, Stephanie S Watowich, Bernard Khor, and David E Levy. Tonic interferon restricts pathogenic IL-17-driven inflammatory disease via balancing the microbiome. *eLife*, 10, 2021.

[9513] Mirna Marinić, Katelyn Mika, Sravanthi Chigurupati, and Vincent J Lynch. Evolutionary transcriptomics implicates HAND2 in the origins of implantation and regulation of gestation length. *eLife*, 10, 2021.

[9514] Brandon Mark, Sen-Lin Lai, Aref Arzan Zarin, Laurina Manning, Heather Q Pollington, Ashok Litwin-Kumar, Albert Cardona, James W Truman, and Chris Q Doe. A developmental framework linking neurogenesis and circuit formation in the Drosophila CNS. *eLife*, 10, 2021.

[9515] Ross D Markello, Aurina Arnatkeviciute, Jean-Baptiste Poline, Ben D Fulcher, Alex Fornito, and Bratislav Misic. Standardizing workflows in imaging transcriptomics with the abagen toolbox. *eLife*, 10, 2021.

[9516] Christopher J Markiewicz, Krzysztof J Gorgolewski, Franklin Feingold, Ross Blair, Yaroslav O Halchenko, Eric Miller, Nell Hardcastle, Joe Wexler, Oscar Esteban, Mathias Goncavles, Anita Jwa, and Russell Poldrack. The OpenNeuro resource for sharing of neuroscience data. *eLife*, 10, 2021.

[9517] Marie Marotel, Marine Villard, Annabelle Drouillard, Issam Tout, Laurie Besson, Omran Allatif, Marine Pujol, Yamila Rocca, Michelle Ainouze, Guillaume Roblot, Sébastien Viel, Melissa Gomez, Veronique Loustaud, Sophie Alain, David Durantel, Thierry Walzer, Uzma Hasan, and Antoine MarÇais. Peripheral natural killer cells in chronic hepatitis B patients display multiple molecular features of T cell exhaustion. *eLife*, 10, 2021.

[9518] Carolina Marques, Thomas Unterkircher, Paula Kroon, Barbara Oldrini, Annalisa Izzo, Yuliia Dramaretska, Roberto Ferrarese, Eva Kling, Oliver Schnell, Sven Nelander, Erwin F Wagner, Latifa Bakiri, Gaetano Gargiulo, Maria Stella Carro, and Massimo Squatrito. NF1 regulates mesenchymal glioblastoma plasticity and aggressiveness through the AP-1 transcription factor FOSL1. *eLife*, 10, 2021.

[9519] Karla Marie Márquez-Nogueras, Miryam Andrea Hortua Triana, Nathan M Chasen, Ivana Y Kuo, and Silvia NJ Moreno. Calcium signaling through a transient receptor channel is important for Toxoplasma gondii growth. *eLife*, 10, 2021.

[9520] Briana J Martiszus, Timur Tsintsadze, Wenhan Chang, and Stephen M Smith. Enhanced excitability of cortical neurons in low-divalent solutions is primarily mediated by altered voltage-dependence of voltage-gated sodium channels. *eLife*, 10, 2021.

[9521] Takahisa Maruno, Akihisa Fukuda, Norihiro Goto, Motoyuki Tsuda, Kozo Ikuta, Yukiko Hiramatsu, Satoshi Ogawa, Yuki Nakanishi, Yuichi Yamaga, Takuto Yoshioka, Kyoichi Takaori, Shinji Uemoto, Dieter Saur, Tsutomu Chiba, and Hiroshi Seno. Visualization of stem cell activity in pancreatic cancer expansion by direct lineage tracing with live imaging. *eLife*, 10, 2021.

[9522] George A Mashour, Ben JA Palanca, Mathias Basner, Duan Li, Wei Wang, Stefanie Blain-Moraes, Nan Lin, Kaitlyn Maier, Maxwell Muench, Vijay Tarnal, Giancarlo Vanini, E Andrew Ochroch, Rosemary Hogg, Marlon Schwartz, Hannah Maybrier, Randall Hardie, Ellen Janke, Goodarz Golmirzaie, Paul Picton, Andrew R McKinstry-Wu, Michael S Avidan, and Max B Kelz. Recovery of consciousness and cognition after general anesthesia in humans. *eLife*, 10, 2021.

[9523] Alan E Mast, Alisa S Wolberg, David Gailani, Michael R Garvin, Christiane Alvarez, J Izaak Miller, Bruce Aronow, and Daniel Jacobson. SARS-CoV-2 suppresses anticoagulant and fibrinolytic gene expression in the lung. *eLife*, 10, 2021.

[9524] Fred D Mast, Peter C Fridy, Natalia E Ketaren, Junjie Wang, Erica Y Jacobs, Jean Paul Olivier, Tanmoy Sanyal, Kelly R Molloy, Fabian Schmidt, Magdalena Rutkowska, Yiska Weisblum, Lucille M Rich, Elizabeth R Vanderwall, Nicholas Dambrauskas, Vladimir Vigdorovich, Sarah Keegan, Jacob B Jiler, Milana E Stein, Paul Dominic B Olinares, Louis Herlands, Theodora Hatziioannou, D Noah Sather, Jason S Debley, David Fenyö, Andrej Sali, Paul D Bieniasz, John D Aitchison, Brian T Chait, and Michael P Rout. Highly synergistic combinations of nanobodies that target SARS-CoV-2 and are resistant to escape. *eLife*, 10, 2021.

[9525] Isao Masuda, Jae-Yeon Hwang, Thomas Christian, Sunita Maharjan, Fuad Mohammad, Howard Gamper, Allen R Buskirk, and Ya-Ming Hou. Loss of N1-methylation of G37 in tRNA induces ribosome stalling and reprograms gene expression. *eLife*, 10, 2021.

[9526] Haritha Mathsyaraja, Jonathen Catchpole, Brian Freie, Emily Eastwood, Ekaterina Babaeva, Michael Geuenich, Pei Feng Cheng, Jessica Ayers, Ming Yu, Nan Wu, Sitapriya Moorthi, Kumud R Poudel, Amanda Koehne, William Grady, A McGarry Houghton, Alice H Berger, Yuzuru Shiio, David MacPherson, and Robert N Eisenman. Loss of MGA repression mediated by an atypical polycomb complex promotes tumor progression and invasiveness. *eLife*, 10, 2021.

[9527] Bridget A Matikainen-Ankney, Thomas Earnest, Mohamed Ali, Eric Casey, Justin G Wang, Amy K Sutton, Alex A Legaria, Kia M Barclay, Laura B Murdaugh, Makenzie R Norris, Yu-Hsuan Chang, Katrina P Nguyen, Eric Lin, Alex Reichenbach, Rachel E Clarke, Romana Stark, Sineadh M Conway, Filipe Carvalho, Ream Al-Hasani, Jordan G McCall, Meaghan C Creed, Victor Cazares, Matthew W Buczynski, Michael J Krashes, Zane B Andrews, and Alexxai V Kravitz. An open-source device for measuring food intake and operant behavior in rodent home-cages. *eLife*, 10, 2021.

[9528] Brya G Matthews, Sanja Novak, Francesca V Sbrana, Jessica L Funnell, Ye Cao, Emma J Buckels, Danka Grcevic, and Ivo Kalajzic. Heterogeneity of murine periosteum progenitors involved in fracture healing. *eLife*, 10, 2021.

[9529] David Mauduit, Ibrahim Ihsan Taskiran, Liesbeth Minnoye, Maxime de Waegeneer, Valerie Christiaens, Gert Hulselmans, Jonas Demeulemeester, Jasper Wouters, and Stein Aerts. Analysis of long and short enhancers in melanoma cell states. *eLife*, 10, 2021.

[9530] Olga A Maximova, Daniel E Sturdevant, John C Kash, Kishore Kanakabandi, Yongli Xiao, Mahnaz Minai, Ian N Moore, Jeff Taubenberger, Craig Martens, Jeffrey I Cohen, and Alexander G Pletnev. Virus infection of the CNS disrupts the immune-neural-synaptic axis via induction of pleiotropic gene regulation of host responses. *eLife*, 10, 2021.

[9531] Koshlan Mayer-Blackwell, Stefan Schattgen, Liel Cohen-Lavi, Jeremy C Crawford, Aisha Souquette, Jessica A Gaevert, Tomer Hertz, Paul G Thomas, Philip Bradley, and Andrew Fiore-Gartland. TCR meta-clonotypes for biomarker discovery with tcrdist3 enabled identification of public, HLA-restricted clusters of SARS-CoV-2 TCRs. *eLife*, 10, 2021.

[9532] Stephanie A Maynard, Philippe Rostaing, Natascha Schaefer, Olivier Gemin, Adrien Candat, Andréa Dumoulin, Carmen Villmann, Antoine Triller, and Christian G Specht. Identification of a stereotypic molecular arrangement of endogenous glycine receptors at spinal cord synapses. *eLife*, 10, 2021.

[9533] Abhishek Mazumder, Richard H Ebright, and Achillefs N Kapanidis. Transcription initiation at a consensus bacterial promoter proceeds via a 'bind-unwind-load-and-lock' mechanism. *eLife*, 10, 2021.

[9534] Allison W McClure and John FX Diffley. Rad53 checkpoint kinase regulation of DNA replication fork rate via Mrc1 phosphorylation. *eLife*, 10, 2021.

[9535] James W McCormick, Marielle AX Russo, Samuel Thompson, Aubrie Blevins, and Kimberly A Reynolds. Structurally distributed surface sites tune allosteric regulation. *eLife*, 10, 2021.

[9536] Shawn McGuirk, Yannick Audet-Delage, Matthew G Annis, Yibo Xue, Mathieu Vernier, Kaiqiong Zhao, Catherine St-Louis, Lucía Minarrieta, David A Patten, Geneviève Morin, Celia MT Greenwood, Vincent Giguère, Sidong Huang, Peter M Siegel, and Julie St-Pierre. Resistance to different anthracycline chemotherapeutics elicits distinct and actionable primary metabolic dependencies in breast cancer. *eLife*, 10, 2021.

[9537] Benjamin C McIlwain, Roja Gundepudi, B Ben Koff, and Randy B Stockbridge. The fluoride permeation pathway and anion recognition in Fluc family fluoride channels. *eLife*, 10, 2021.

[9538] Shane G McInally, Jane Kondev, and Bruce L Goode. Scaling of subcellular actin structures with cell length through decelerated growth. *eLife*, 10, 2021.

[9539] Colleen N McLaughlin, Maria Brbić, Qijing Xie, Tongchao Li, Felix Horns, Sai Saroja Kolluru, Justus M Kebschull, David Vacek, Anthony Xie, Jiefu Li, Robert C Jones, Jure Leskovec, Stephen R Quake, Liqun Luo, and Hongjie Li. Single-cell transcriptomes of developing and adult olfactory receptor neurons in Drosophila. *eLife*, 10, 2021.

[9540] David V McLeod and Sylvain Gandon. Understanding the evolution of multiple drug resistance in structured populations. *eLife*, 10, 2021.

[9541] Kerrie-Ann McMahon, David A Stroud, Yann Gambin, Vikas Tillu, Michele Bastiani, Emma Sierecki, Mark E Polinkovsky, Thomas E Hall, Guillermo A Gomez, Yeping Wu, Marie-Odile Parat, Nick Martel, Harriet P Lo, Kum Kum Khanna, Kirill Alexandrov, Roger Daly, Alpha Yap, Michael T Ryan, and Robert G Parton. Cavin3 released from caveolae interacts with BRCA1 to regulate the cellular stress response. *eLife*, 10, 2021.

[9542] Kirsty J McMillan, Paul J Banks, Francesca LN Hellel, Ruth E Carmichael, Thomas Clairfeuille, Ashley J Evans, Kate J Heesom, Philip Lewis, Brett M Collins, Zafar I Bashir, Jeremy M Henley, Kevin A Wilkinson, and Peter J Cullen. Sorting nexin-27 regulates AMPA receptor trafficking through the synaptic adhesion protein LRFN2. *eLife*, 10, 2021.

[9543] Andrew S McNeal, Rachel L Belote, Hanlin Zeng, Marcus Urquijo, Kendra Barker, Rodrigo Torres, Meghan Curtin, A Hunter Shain, Robert HI Andtbacka, Sheri Holmen, David H Lum, Timothy H McCalmont, Matt W VanBrocklin, Douglas Grossman, Maria L Wei, Ursula E Lang, and Robert L Judson-Torres. BRAFV600E induces reversible mitotic arrest in human melanocytes via microRNA-mediated suppression of AURKB. *eLife*, 10, 2021.

[9544] Helen R McPherson, Cedric Duval, Stephen R Baker, Matthew S Hindle, Lih T Cheah, Nathan L Asquith, Marco M Domingues, Victoria C Ridger, Simon DA Connell, Khalid M Naseem, Helen Philippou, Ramzi A Ajjan, and Robert AS Ariëns. Fibrinogen $\alpha$C-subregions critically contribute blood clot fibre growth, mechanical stability, and resistance to fibrinolysis. *eLife*, 10, 2021.

[9545] Jacob Graves McPherson and Maria F Bandres. Spontaneous neural synchrony links intrinsic spinal sensory and motor networks during unconsciousness. *eLife*, 10, 2021.

[9546] Lucile Megret, Barbara Gris, Satish Sasidharan Nair, Jasmin Cevost, Mary Wertz, Jeff Aaronson, Jim Rosinski, Thomas F Vogt, Hilary Wilkinson, Myriam Heiman, and Christian Neri. Shape deformation analysis reveals the temporal dynamics of cell-type-specific homeostatic and pathogenic responses to mutant huntingtin. *eLife*, 10, 2021.

[9547] Raphaël Méheust, Shuo Huang, Rafael Rivera-Lugo, Jillian F Banfield, and Samuel H Light. Post-translational flavinylation is associated with diverse extracytosolic redox functionalities throughout bacterial life. *eLife*, 10, 2021.

[9548] Larissa Melo do Nascimento, Franziska Egler, Katharina Arnold, Nina Papavasiliou, Christine Clayton, and Esteban Erben. Functional insights from a surface antigen mRNA-bound proteome. *eLife*, 10, 2021.

[9549] Monique S Mendes, Linh Le, Jason Atlas, Zachary Brehm, Antonio Ladron-de Guevara, Evelyn Matei, Cassandra Lamantia, Matthew N McCall, and Ania K Majewska. The role of P2Y12 in the kinetics of microglial self-renewal and maturation in the adult visual cortex in vivo. *eLife*, 10, 2021.

[9550] Oscar A Mendez, Emiliano Flores Machado, Jing Lu, and Anita A Koshy. Injection with Toxoplasma gondii protein affects neuron health and survival. *eLife*, 10, 2021.

[9551] Qing-Tao Meng, Xian-Yu Liu, Xue-Ting Liu, Juan Liu, Admire Munanairi, Devin M Barry, Benlong Liu, Hua Jin, Yu Sun, Qianyi Yang, Fang Gao, Li Wan, Jiahang Peng, Jin-Hua Jin, Kai-Feng Shen, Ray Kim, Jun Yin, Ailin Tao, and Zhou-Feng Chen. BNP facilitates NMB-encoded histaminergic itch via NPRC-NMBR crosstalk. *eLife*, 10, 2021.

[9552] Hannah R Meredith, John R Giles, Javier Perez-Saez, Théophile Mande, Andrea Rinaldo, Simon Mutembo, Elliot N Kabalo, Kabondo Makungo, Caroline O Buckee, Andrew J Tatem, C Jessica E Metcalf, and Amy Wesolowski. Characterizing human mobility patterns in rural settings of sub-Saharan Africa. *eLife*, 10, 2021.

[9553] Nicole Merritt, Keith Garcia, Dushyandi Rajendran, Zhen-Yuan Lin, Xiaomeng Zhang, Katrina A Mitchell, Nicholas Borcherding, Colleen Fullenkamp, Michael S Chimenti, Anne-Claude Gingras, Kieran F Harvey, and Munir R Tanas. TAZ-CAMTA1 and YAP-TFE3 alter the TAZ/YAP transcriptome by recruiting the ATAC histone acetyltransferase complex. *eLife*, 10, 2021.

[9554] Jasmin Mertins, Jérôme Finke, Ricarda Sies, Kerstin M Rink, Jan Hasenauer, and Thorsten Lang. The mesoscale organization of syntaxin 1A and SNAP25 is determined by SNARE–SNARE interactions. *eLife*, 10, 2021.

[9555] Joris JB Messelink, Fabian Meyer, Marc Bramkamp, and Chase P Broedersz. Single-cell growth inference of Corynebacterium glutamicum reveals asymptotically linear growth. *eLife*, 10, 2021.

[9556] Piotr Michaluk, Janosch Peter Heller, and Dmitri A Rusakov. Rapid recycling of glutamate transporters on the astroglial surface. *eLife*, 10, 2021.

[9557] Yusuke Mii, Kenichi Nakazato, Chan-Gi Pack, Takafumi Ikeda, Yasushi Sako, Atsushi Mochizuki, Masanori Taira, and Shinji Takada. Quantitative analyses reveal extracellular dynamics of Wnt ligands in Xenopus embryos. *eLife*, 10, 2021.

[9558] Katelyn Mika, Mirna Marinić, Manvendra Singh, Joanne Muter, Jan Joris Brosens, and Vincent J Lynch. Evolutionary transcriptomics implicates new genes and pathways in human pregnancy and adverse pregnancy outcomes. *eLife*, 10, 2021.

[9559] Steven Miletić, Russell J Boag, Anne C Trutti, Niek Stevenson, Birte U Forstmann, and Andrew Heathcote. A new model of decision processing in instrumental learning tasks. *eLife*, 10, 2021.

[9560] Pierre Millard, Brice Enjalbert, Sandrine Uttenweiler-Joseph, Jean-Charles Portais, and Fabien Létisse. Control and regulation of acetate overflow in Escherichia coli. *eLife*, 10, 2021.

[9561] Justin J Miller, Ishaan T Shah, Jayda Hatten, Yasaman Barekatain, Elizabeth A Mueller, Ahmed M Moustafa, Rachel L Edwards, Cynthia S Dowd, Geoffrey C Hoops, R Jeremy Johnson, Paul J Planet, Florian L Muller, Joseph M Jez, and Audrey R Odom John. Structure-guided microbial targeting of antistaphylococcal prodrugs. *eLife*, 10, 2021.

[9562] Karl N Miller, Nirmalya Dasgupta, Tianhui Liu, Peter D Adams, and Maria Grazia Vizioli. Cytoplasmic chromatin fragments—from mechanisms to therapeutic potential. *eLife*, 10, 2021.

[9563] Jason W Millington, George P Brownrigg, Charlotte Chao, Ziwei Sun, Paige J Basner-Collins, Lianna W Wat, Bruno Hudry, Irene Miguel-Aliaga, and Elizabeth J Rideout. Female-biased upregulation of insulin pathway activity mediates the sex difference in Drosophila body size plasticity. *eLife*, 10, 2021.

[9564] Kathryn Milne, Alasdair Ivens, Adam J Reid, Magda E Lotkowska, Aine O'Toole, Geetha Sankaranarayanan, Diana Munoz Sandoval, Wiebke Nahrendorf, Clement Regnault, Nick J Edwards, Sarah E Silk, Ruth O Payne, Angela M Minassian, Navin Venkatraman, Mandy J Sanders, Adrian VS Hill, Michael Barrett, Matthew Berriman, Simon J Draper, J Alexandra Rowe, and Philip J Spence. Mapping immune variation and var gene switching in naive hosts infected with Plasmodium falciparum. *eLife*, 10, 2021.

[9565] Aaron D Milstein, Yiding Li, Katie C Bittner, Christine Grienberger, Ivan Soltesz, Jeffrey C Magee, and Sandro Romani. Bidirectional synaptic plasticity rapidly modifies hippocampal representations. *eLife*, 10, 2021.

[9566] Andrea Milzi, Enrico Domenico Lemma, Rosalia Dettori, Kathrin Burgmaier, Nikolaus Marx, Sebastian Reith, and Mathias Burgmaier. Coronary plaque composition influences biomechanical stress and predicts plaque rupture in a morpho-mechanic OCT analysis. *eLife*, 10, 2021.

[9567] Soohong Min, Yangkyun Oh, Pushpa Verma, Samuel C Whitehead, Nilay Yapici, David Van Vactor, Greg SB Suh, and Stephen Liberles. Control of feeding by Piezo-mediated gut mechanosensation in Drosophila. *eLife*, 10, 2021.

[9568] Luca Minati, Claudia Firrito, Alessia Del Piano, Alberto Peretti, Simone Sidoli, Daniele Peroni, Romina Belli, Francesco Gandolfi, Alessandro Romanel, Paola Bernabo, Jacopo Zasso, Alessandro Quattrone, Graziano Guella, Fabio Lauria, Gabriella Viero, and Massimiliano Clamer. One-shot analysis of translated mammalian lncRNAs with AHARIBO. *eLife*, 10, 2021.

[9569] Judith Miné-Hattab, Mathias Heltberg, Marie Villemeur, Chloé Guedj, Thierry Mora, Aleksandra M Walczak, Maxime Dahan, and Angela Taddei. Single molecule microscopy reveals key physical features of repair foci in living cells. *eLife*, 10, 2021.

[9570] Anastasia A Minervina, Ekaterina A Komech, Aleksei Titov, Meriem Bensouda Koraichi, Elisa Rosati, Ilgar Z Mamedov, Andre Franke, Grigory A Efimov, Dmitriy M Chudakov, Thierry Mora, Aleksandra M Walczak, Yuri B Lebedev, and Mikhail V Pogorelyy. Longitudinal high-throughput TCR repertoire profiling reveals the dynamics of T-cell memory formation after mild COVID-19 infection. *eLife*, 10, 2021.

[9571] Nadiatou T Miningou Zobon, Joanna Jędrzejewska-Szmek, and Kim T Blackwell. Temporal pattern and synergy influence activity of ERK signaling pathways during L-LTP induction. *eLife*, 10, 2021.

[9572] Ila Mishra, Clemens Duerrschmid, Zhiqiang Ku, Yang He, Wei Xie, Elizabeth Sabath Silva, Jennifer Hoffman, Wei Xin, Ningyan Zhang, Yong Xu, Zhiqiang An, and Atul R Chopra. Asprosin-neutralizing antibodies as a treatment for metabolic syndrome. *eLife*, 10, 2021.

[9573] Jessica Mitchell, Carlas S Smith, Josh Titlow, Nils Otto, Pieter van Velde, Martin Booth, Ilan Davis, and Scott Waddell. Selective dendritic localization of mRNA in Drosophila mushroom body output neurons. *eLife*, 10, 2021.

[9574] Sibylle Mitschka and Christine Mayr. Endogenous p53 expression in human and mouse is not regulated by its 3'UTR. *eLife*, 10, 2021.

[9575] Divyansh Mittal and Rishikesh Narayanan. Resonating neurons stabilize heterogeneous grid-cell networks. *eLife*, 10, 2021.

[9576] Briana E Mittleman, Sebastian Pott, Shane Warland, Kenneth Barr, Claudia Cuevas, and Yoav Gilad. Divergence in alternative polyadenylation contributes to gene regulatory differences between humans and chimpanzees. *eLife*, 10, 2021.

[9577] Ryo Miyamoto, Akinori Kanai, Hiroshi Okuda, Yosuke Komata, Satoshi Takahashi, Hirotaka Matsui, Toshiya Inaba, and Akihiko Yokoyama. HOXA9 promotes MYC-mediated leukemogenesis by maintaining gene expression for multiple anti-apoptotic pathways. *eLife*, 10, 2021.

[9578] Ryoji Miyazaki, Tetsuro Watanabe, Kohei Yoshitani, and Yoshinori Akiyama. Edge-strand of BepA interacts with immature LptD on the $\beta$-barrel assembly machine to direct it to on- and off-pathways. *eLife*, 10, 2021.

[9579] Taisuke Miyazaki, Megumi Morimoto-Tomita, Coralie Berthoux, Kotaro Konno, Yoav Noam, Tokiwa Yamasaki, Matthijs Verhage, Pablo E Castillo, Masahiko Watanabe, and Susumu Tomita. Excitatory and inhibitory receptors utilize distinct post- and trans-synaptic mechanisms in vivo. *eLife*, 10, 2021.

[9580] Mohammad Mofatteh, Fabio Echegaray-Iturra, Andrew Alamban, Francesco Dalla Ricca, Anand Bakshi, and Mustafa G Aydogan. Autonomous clocks that regulate organelle biogenesis, cytoskeletal organization, and intracellular dynamics. *eLife*, 10, 2021.

[9581] Homa MohammadiPeyhani, Anush Chiappino-Pepe, Kiandokht Haddadi, Jasmin Hafner, Noushin Hadadi, and Vassily Hatzimanikatis. NICEdrug.ch, a workflow for rational drug design and systems-level analysis of drug metabolism. *eLife*, 10, 2021.

[9582] Jugal Mohapatra, Kyuto Tashiro, Ryan L Beckner, Jorge Sierra, Jessica A Kilgore, Noelle S Williams, and Glen Liszczak. Serine ADP-ribosylation marks nucleosomes for ALC1-dependent chromatin remodeling. *eLife*, 10, 2021.

[9583] Louise H Moncla, Allison Black, Chas DeBolt, Misty Lang, Nicholas R Graff, Ailyn C Pérez-Osorio, Nicola F Müller, Dirk Haselow, Scott Lindquist, and Trevor Bedford. Repeated introductions and intensive community transmission fueled a mumps virus outbreak in Washington State. *eLife*, 10, 2021.

[9584] Debasmita Mondal, Ameya G Prabhune, Sriram Ramaswamy, and Prerna Sharma. Strong confinement of active microalgae leads to inversion of vortex flow and enhanced mixing. *eLife*, 10, 2021.

[9585] Jorrit S Montijn, Koen Seignette, Marcus H Howlett, J Leonie Cazemier, Maarten Kamermans, Christiaan N Levelt, and J Alexander Heimel. A parameter-free statistical test for neuronal responsiveness. *eLife*, 10, 2021.

[9586] Benjamin M Moran, Cheyenne Payne, Quinn Langdon, Daniel L Powell, Yaniv Brandvain, and Molly Schumer. The genomic consequences of hybridization. *eLife*, 10, 2021.

[9587] Laura Morcom, Timothy J Edwards, Eric Rider, Dorothy Jones-Davis, Jonathan WC Lim, Kok-Siong Chen, Ryan J Dean, Jens Bunt, Yunan Ye, Ilan Gobius, Rodrigo Suárez, Simone Mandelstam, Elliott H Sherr, and Linda J Richards. DRAXIN regulates interhemispheric fissure remodelling to influence the extent of corpus callosum formation. *eLife*, 10, 2021.

[9588] Laura Morcom, Ilan Gobius, Ashley PL Marsh, Rodrigo Suárez, Jonathan WC Lim, Caitlin Bridges, Yunan Ye, Laura R Fenlon, Yvrick Zagar, Amelia M Douglass, Amber-Lee S Donahoo, Thomas Fothergill, Samreen Shaikh, Peter Kozulin, Timothy J Edwards, Helen M Cooper, Elliott H Sherr, Alain Chédotal, Richard J Leventer, Paul J Lockhart, and Linda J Richards. DCC regulates astroglial development essential for telencephalic morphogenesis and corpus callosum formation. *eLife*, 10, 2021.

[9589] Carlos G Moreira, Christian R Baumann, Maurizio Scandella, Sergio I Nemirovsky, Sven Leach, Reto Huber, and Daniela Noain. Closed-loop auditory stimulation method to modulate sleep slow waves and motor learning performance in rats. *eLife*, 10, 2021.

[9590] Yunuen Moreno-Lopez, Charlotte Bichara, Gilles Delbecq, Philippe Isope, and Matilde Cordero-Erausquin. The corticospinal tract primarily modulates sensory inputs in the mouse lumbar cord. *eLife*, 10, 2021.

[9591] Tania Moreno-Mármol, Mario Ledesma-Terrón, Noemi Tabanera, Maria Jesús Martin-Bermejo, Marcos J Cardozo, Florencia Cavodeassi, and Paola Bovolenta. Stretching of the retinal pigment epithelium contributes to zebrafish optic cup morphogenesis. *eLife*, 10, 2021.

[9592] Sophie M Morgani, Jie Su, Jennifer Nichols, Joan Massagué, and Anna-Katerina Hadjantonakis. The transcription factor Rreb1 regulates epithelial architecture, invasiveness, and vasculogenesis in early mouse embryos. *eLife*, 10, 2021.

[9593] Dylan H Morris, Kwe Claude Yinda, Amandine Gamble, Fernando W Rossine, Qishen Huang, Trenton Bushmaker, Robert J Fischer, M Jeremiah Matson, Neeltje Van Doremalen, Peter J Vikesland, Linsey C Marr, Vincent J Munster, and James O Lloyd-Smith. Mechanistic theory predicts the effects of temperature and humidity on inactivation of SARS-CoV-2 and other enveloped viruses. *eLife*, 10, 2021.

[9594] Amy Morris-Drake, Julie M Kern, and Andrew N Radford. Experimental evidence for delayed post-conflict management behaviour in wild dwarf mongooses. *eLife*, 10, 2021.

[9595] Robin E Morrison, Winnie Eckardt, Fernando Colchero, Veronica Vecellio, and Tara S Stoinski. Social groups buffer maternal loss in mountain gorillas. *eLife*, 10, 2021.

[9596] Martin Steen Mortensen, Morten Arendt Rasmussen, Jakob Stokholm, Asker Daniel Brejnrod, Christina Balle, Jonathan Thorsen, Karen Angeliki Krogfelt, Hans Bisgaard, and Søren Johannes Sørensen. Modeling transfer of vaginal microbiota from mother to infant in early life. *eLife*, 10, 2021.

[9597] Michal Motyka, Dominik Kusy, Matej Bocek, Renata Bilkova, and Ladislav Bocak. Phylogenomic and mitogenomic data can accelerate inventorying of tropical beetles during the current biodiversity crisis. *eLife*, 10, 2021.

[9598] Isobel C Mouat, Zachary J Morse, Iryna Shanina, Kelly L Brown, and Marc S Horwitz. Latent gammaherpesvirus exacerbates arthritis through modification of age-associated B cells. *eLife*, 10, 2021.

[9599] Rebecca A Mount, Sudiksha Sridhar, Kyle R Hansen, Ali I Mohammed, Moona Abdulkerim, Robb Kessel, Bobak Nazer, Howard J Gritton, and Xue Han. Distinct neuronal populations contribute to trace conditioning and extinction learning in the hippocampal CA1. *eLife*, 10, 2021.

[9600] Andria Mousa, Peter Winskill, Oliver John Watson, Oliver Ratmann, Mélodie Monod, Marco Ajelli, Aldiouma Diallo, Peter J Dodd, Carlos G Grijalva, Moses Chapa Kiti, Anand Krishnan, Rakesh Kumar, Supriya Kumar, Kin O Kwok, Claudio F Lanata, Olivier Le Polain de Waroux, Kathy Leung, Wiriya Mahikul, Alessia Melegaro, Carl D Morrow, Joël Mossong, Eleanor FG Neal, D James Nokes, Wirichada Pan-ngum, Gail E Potter, Fiona M Russell, Siddhartha Saha, Jonathan D Sugimoto, Wan In Wei, Robin R Wood, Joseph Wu, Juanjuan Zhang, Patrick Walker, and Charles Whittaker. Social contact patterns and implications for infectious disease transmission – a systematic review and meta-analysis of contact surveys. *eLife*, 10, 2021.

[9601] Ibrahim T Mughrabi, Jordan Hickman, Naveen Jayaprakash, Dane Thompson, Umair Ahmed, Eleni S Papadoyannis, Yao-Chuan Chang, Adam Abbas, Timir Datta-Chaudhuri, Eric H Chang, Theodoros P Zanos, Sunhee C Lee, Robert C Froemke, Kevin J Tracey, Cristin Welle, Yousef Al-Abed, and Stavros Zanos. Development and characterization of a chronic implant mouse model for vagus nerve stimulation. *eLife*, 10, 2021.

[9602] Diptendu Mukherjee, Ben Jerry Gonzales, Reut Ashwal-Fluss, Hagit Turm, Maya Groysman, and Ami Citri. Egr2 induction in spiny projection neurons of the ventrolateral striatum contributes to cocaine place preference in mice. *eLife*, 10, 2021.

[9603] Kaustav Mukherjee, Li Xue, Antanas Planutis, Merlin Nithya Gnanapragasam, Andrew Chess, and James J Bieker. EKLF/KLF1 expression defines a unique macrophage subset during mouse erythropoiesis. *eLife*, 10, 2021.

[9604] Haleigh N Mulholland, Bettina Hein, Matthias Kaschube, and Gordon B Smith. Tightly coupled inhibitory and excitatory functional networks in the developing primary visual cortex. *eLife*, 10, 2021.

[9605] Thorsten G Müller, Vojtech Zila, Kyra Peters, Sandra Schifferdecker, Mia Stanic, Bojana Lucic, Vibor Laketa, Marina Lusic, Barbara Müller, and Hans-Georg Kräusslich. HIV-1 uncoating by release of viral cDNA from capsid-like structures in the nucleus of infected cells. *eLife*, 10, 2021.

[9606] Marzia Munafò, Victoria R Lawless, Alessandro Passera, Serena MacMillan, Susanne Bornelöv, Irmgard U Haussmann, Matthias Soller, Gregory J Hannon, and Benjamin Czech. Channel nuclear pore complex subunits are required for transposon silencing in Drosophila. *eLife*, 10, 2021.

[9607] Marcia A Munoz, Emma K Fletcher, Oliver P Skinner, Julie Jurczyluk, Esther Kristianto, Mark P Hodson, Shuting Sun, Frank H Ebetino, David R Croucher, Philip M Hansbro, Jacqueline R Center, and Michael J Rogers. Bisphosphonate drugs have actions in the lung and inhibit the mevalonate pathway in alveolar macrophages. *eLife*, 10, 2021.

[9608] Leon P Munting, Marc Derieppe, Ernst Suidgeest, Lydiane Hirschler, Matthias JP van Osch, Baudouin Denis de Senneville, and Louise van der Weerd. Cerebral blood flow and cerebrovascular reactivity are preserved in a mouse model of cerebral microvascular amyloidosis. *eLife*, 10, 2021.

[9609] Dinavahi VPS Murty, Keerthana Manikandan, Wupadrasta Santosh Kumar, Ranjini Garani Ramesh, Simran Purokayastha, Bhargavi Nagendra, Abhishek ML, Aditi Balakrishnan, Mahendra Javali, Naren Prahalada Rao, and Supratim Ray. Stimulus-induced gamma rhythms are weaker in human elderly with mild cognitive impairment and Alzheimer's disease. *eLife*, 10, 2021.

[9610] Jessica H Myers, Kirsten Denman, Chris DuPont, Ahmed A Hawash, Kevin R Novak, Andrew Koesters, Manfred Grabner, Anamika Dayal, Andrew A Voss, and Mark M Rich. The mechanism underlying transient weakness in myotonia congenita. *eLife*, 10, 2021.

[9611] Anna Z Mykytyn, Tim I Breugem, Samra Riesebosch, Debby Schipper, Petra B van den Doel, Robbert J Rottier, Mart M Lamers, and Bart L Haagmans. SARS-CoV-2 entry into human airway organoids is serine protease-mediated and facilitated by the multibasic cleavage site. *eLife*, 10, 2021.

[9612] Soojung Na, Dongil Chung, Andreas Hula, Ofer Perl, Jennifer Jung, Matthew Heflin, Sylvia Blackmore, Vincenzo G Fiore, Peter Dayan, and Xiaosi Gu. Humans use forward thinking to exploit social controllability. *eLife*, 10, 2021.

[9613] Farha Naaz, Megha Agrawal, Soumyadeep Chakraborty, Mahesh S Tirumkudulu, and KV Venkatesh. Ligand sensing enhances bacterial flagellar motor output via stator recruitment. *eLife*, 10, 2021.

[9614] Asieh Naderi, Elham Soltanmaohammadi, Vimala Kaza, Shayne Barlow, Ioulia Chatzistamou, and Hippokratis Kiaris. Persistent effects of pair bonding in lung cancer cell growth in monogamous Peromyscus californicus. *eLife*, 10, 2021.

[9615] Nana Naetar, Konstantina Georgiou, Christian Knapp, Irena Bronshtein, Elisabeth Zier, Petra Fichtinger, Thomas Dechat, Yuval Garini, and Roland Foisner. LAP2alpha maintains a mobile and low assembly state of A-type lamins in the nuclear interior. *eLife*, 10, 2021.

[9616] Suman Nag and Darshan V Trivedi. To lie or not to lie: Super-relaxing with myosins. *eLife*, 10, 2021.

[9617] Shivathmihai Nagappan and Kevin M Franks. Parallel processing by distinct classes of principal neurons in the olfactory cortex. *eLife*, 10, 2021.

[9618] Jenna Nagy, Briana Ebbinghaus, Mrinalini Hoon, and Raunak Sinha. GABAA presynaptic inhibition regulates the gain and kinetics of retinal output neurons. *eLife*, 10, 2021.

[9619] Anna Nagy-Staron, Kathrin Tomasek, Caroline Caruso Carter, Elisabeth Sonnleitner, Bor Kavčič, Tiago Paixão, and Calin C Guet. Local genetic context shapes the function of a gene regulatory network. *eLife*, 10, 2021.

[9620] Wiebke Nahrendorf, Alasdair Ivens, and Philip J Spence. Inducible mechanisms of disease tolerance provide an alternative strategy of acquired immunity to malaria. *eLife*, 10, 2021.

[9621] Rohini R Nair, Dmitry Zabezhinsky, Rita Gelin-Licht, Brian J Haas, Michael CA Dyhr, Hannah S Sperber, Chad Nusbaum, and Jeffrey E Gerst. Multiplexed mRNA assembly into ribonucleoprotein particles plays an operon-like role in the control of yeast cell physiology. *eLife*, 10, 2021.

[9622] Richard K Nakamura, Lee S Mann, Mark D Lindner, Jeremy Braithwaite, Mei-Ching Chen, Adrian Vancea, Noni Byrnes, Valerie Durrant, and Bruce Reed. An experimental test of the effects of redacting grant applicant identifiers on peer review outcomes. *eLife*, 10, 2021.

[9623] Joji Nakayama, Lora Tan, Yan Li, Boon Cher Goh, Shu Wang, Hideki Makinoshima, and Zhiyuan Gong. A zebrafish embryo screen utilizing gastrulation identifies the HTR2C inhibitor pizotifen as a suppressor of EMT-mediated metastasis. *eLife*, 10, 2021.

[9624] Yuta Nakazawa, Nanako Nishiyama, Hitoshi Koizumi, Kazumasa Kanemaru, Chigusa Nakahashi-Oda, and Akira Shibuya. Tumor-derived extracellular vesicles regulate tumor-infiltrating regulatory T cells via the inhibitory immunoreceptor CD300a. *eLife*, 10, 2021.

[9625] Alice Namias, Ndey Bassin Jobe, Krijn Petrus Paaijmans, and Silvie Huijben. The need for practical insecticide-resistance guidelines to effectively inform mosquito-borne disease control programs. *eLife*, 10, 2021.

[9626] Ashwin Nandagiri, Avinash Satish Gaikwad, David L Potter, Reza Nosrati, Julio Soria, Moira K O'Bryan, Sameer Jadhav, and Ranganathan Prabhakar. Flagellar energetics from high-resolution imaging of beating patterns in tethered mouse sperm. *eLife*, 10, 2021.

[9627] Adishesh K Narahari, Alex JB Kreutzberger, Pablo S Gaete, Yu-Hsin Chiu, Susan A Leonhardt, Christopher B Medina, Xueyao Jin, Patrycja W Oleniacz, Volker Kiessling, Paula Q Barrett, Kodi S Ravichandran, Mark Yeager, Jorge E Contreras, Lukas K Tamm, and Douglas A Bayliss. ATP and large signaling metabolites flux through caspase-activated Pannexin 1 channels. *eLife*, 10, 2021.

[9628] Máté Nászai, Karen Bellec, Yachuan Yu, Alvaro Román-Fernández, Emma Sandilands, Joel Johansson, Andrew D Campbell, Jim C Norman, Owen J Sansom, David M Bryant, and Julia B Cordero. RAL GTPases mediate EGFR-driven intestinal stem cell proliferation and tumourigenesis. *eLife*, 10, 2021.

[9629] Salwa Naushin, Viren Sardana, Rajat Ujjainiya, Nitin Bhatheja, Rintu Kutum, Akash Kumar Bhaskar, Shalini Pradhan, Satyartha Prakash, Raju Khan, Birendra Singh Rawat, Karthik Bharadwaj Tallapaka, Mahesh Anumalla, Giriraj Ratan Chandak, Amit Lahiri, Susanta Kar, Shrikant Ramesh Mulay, Madhav Nilakanth Mugale, Mrigank Srivastava, Shaziya Khan, Anjali Srivastava, Bhawana Tomar, Murugan Veerapandian, Ganesh Venkatachalam, Selvamani Raja Vijayakumar, Ajay Agarwal, Dinesh Gupta, Prakash M Halami, Muthukumar Serva Peddha, Gopinath M Sundaram, Ravindra P Veeranna, Anirban Pal, Vinay Kumar Agarwal, Anil Ku Maurya, Ranvijay Kumar Singh, Ashok Kumar Raman, Suresh Kumar Anandasadagopan, Parimala Karuppanan, Subramanian Venkatesan, Harish Kumar Sardana, Anamika Kothari, Rishabh Jain, Anupama Thakur, Devendra Singh Parihar, Anas Saifi, Jasleen Kaur, Virendra Kumar, Avinash Mishra, Iranna Gogeri, Geethavani Rayasam, Praveen Singh, Rahul Chakraborty, Gaura Chaturvedi, Pinreddy Karunakar, Rohit Yadav, Sunanda Singhmar, Dayanidhi Singh, Sharmistha Sarkar, Purbasha Bhattacharya, Sundaram Acharya, Vandana Singh, Shweta Verma, Drishti Soni, Surabhi Seth, Sakshi Vashisht, Sarita Thakran, Firdaus Fatima, Akash Pratap Singh, Akanksha Sharma, Babita Sharma, Manikandan Subramanian, Yogendra S Padwad, Vipin Hallan, Vikram Patial, Damanpreet Singh, Narendra Vijay Tripude, Partha Chakrabarti, Sujay Krishna Maity, Dipyaman Ganguly, Jit Sarkar, Sistla Ramakrishna, Balthu Narender Kumar, Kiran A Kumar, Sumit G Gandhi, Piyush Singh Jamwal, Rekha Chouhan, Vijay Lakshmi Jamwal, Nitika Kapoor, Debashish Ghosh, Ghanshyam Thakkar, Umakanta Subudhi, Pradip Sen, Saumya Ray Chaudhury, Rashmi Kumar, Pawan Gupta, Amit Tuli, Deepak Sharma, Rajesh P Ringe, Amarnarayan D, Mahesh Kulkarni, Dhansekaran Shanmugam, Mahesh S Dharne, Sayed G Dastager, Rakesh Joshi, Amita P Patil, Sachin N Mahajan, Abujunaid Habib Khan, Vasudev Wagh, Rakesh Kumar Yadav, Ajinkya Khilari, Mayuri Bhadange, Arvindkumar H Chaurasiya, Shabda E Kulsange, Krishna Khairnar, Shilpa Paranjape, Jatin Kalita, Narahari G Sastry, Tridip Phukan, Prasenjit Manna, Wahengbam Romi, Pankaj Bharali, Dibyajyoti Ozah, Ravi Kumar Sahu, Elapavalooru VSSK Babu, Rajeev Sukumaran, Aiswarya R Nair, Prajeesh Kooloth Valappil, Anoop Puthiyamadam, Adarsh Velayudhanpillai, Kalpana Chodankar, Samir Damare, Yennapu Madhavi, Ved Varun Aggarwal, Sumit Dahiya, Anurag Agrawal, Debasis Dash, and Shantanu Sengupta. Insights from a Pan India Sero-Epidemiological survey (Phenome-India Cohort) for SARS-CoV2. *eLife*, 10, 2021.

[9630] Cesar Nava Gonzales, Quintyn McKaughan, Eric A Bushong, Kalyani Cauwenberghs, Renny Ng, Matthew Madany, Mark H Ellisman, and Chih-Ying Su. Systematic morphological and morphometric analysis of identified olfactory receptor neurons in Drosophila melanogaster. *eLife*, 10, 2021.

[9631] Derek Evan Nee. Integrative frontal-parietal dynamics supporting cognitive control. *eLife*, 10, 2021.

[9632] Jason Neidleman, Xiaoyu Luo, Matthew McGregor, Guorui Xie, Victoria Murray, Warner C Greene, Sulggi A Lee, and Nadia R Roan. mRNA vaccine-induced T cells respond identically to SARS-CoV-2 variants of concern but differ in longevity and homing properties depending on prior infection status. *eLife*, 10, 2021.

[9633] Margaret AM Nelson, Kelsey L McLaughlin, James T Hagen, Hannah S Coalson, Cameron Schmidt, Miki Kassai, Kimberly A Kew, Joseph M McClung, P Darrell Neufer, Patricia Brophy, Nasreen A Vohra, Darla Liles, Myles C Cabot, and Kelsey H Fisher-Wellman. Intrinsic OXPHOS limitations underlie cellular bioenergetics in leukemia. *eLife*, 10, 2021.

[9634] Peter Neofotis, Joshua Temple, Oliver L Tessmer, Jacob Bibik, Nicole Norris, Eric Pollner, Ben Lucker, Sarathi M Weraduwage, Alecia Withrow, Barbara Sears, Greg Mogos, Melinda Frame, David Hall, Joseph Weissman, and David M Kramer. The induction of pyrenoid synthesis by hyperoxia and its implications for the natural diversity of photosynthetic responses in Chlamydomonas. *eLife*, 10, 2021.

[9635] Nguyen Thi Thuy Ngan, Nhat Thanh Hoang Le, Nguyen Ngo Vi Vi, Ninh Thi Thanh Van, Nguyen Thi Hoang Mai, Duong Van Anh, Phan Hai Trieu, Nguyen Phu Huong Lan, Nguyen Hoan Phu, Nguyen Van Vinh Chau, David G Lalloo, William Hope, Justin Beardsley, Nicholas J White, Ronald Geskus, Guy E Thwaites, Damian Krysan, Luong Thi Hue Tai, Evelyne Kestelyn, Tran Quang Binh, Le Quoc Hung, Nguyen Le Nhu Tung, and Jeremy N Day. An open label randomized controlled trial of tamoxifen combined with amphotericin B and fluconazole for cryptococcal meningitis. *eLife*, 10, 2021.

[9636] Huyen Nguyen, Christian Wandell Thorball, Jacques Fellay, Jürg Böni, Sabine Yerly, Matthieu Perreau, Hans H Hirsch, Katharina Kusejko, Maria Christine Thurnheer, Manuel Battegay, Matthias Cavassini, Christian R Kahlert, Enos Bernasconi, Huldrych F Günthard, and Roger D Kouyos. Systematic screening of viral and human genetic variation identifies antiretroviral resistance and immune escape link. *eLife*, 10, 2021.

[9637] Khanh Dinh Quoc Nguyen, Michael Vigers, Eric Sefah, Susanna Seppälä, Jennifer Paige Hoover, Nicole Star Schonenbach, Blake Mertz, Michelle Ann O'Malley, and Songi Han. Homo-oligomerization of the human adenosine A2A receptor is driven by the intrinsically disordered C-terminus. *eLife*, 10, 2021.

[9638] Minh Q Nguyen, Lars J von Buchholtz, Ashlie N Reker, Nicholas JP Ryba, and Steve Davidson. Single-nucleus transcriptomic analysis of human dorsal root ganglion neurons. *eLife*, 10, 2021.

[9639] Robert J Nichols, Benjamin LaFrance, Naiya R Phillips, Devon R Radford, Luke M Oltrogge, Luis E Valentin-Alvarado, Amanda J Bischoff, Eva Nogales, and David F Savage. Discovery and characterization of a novel family of prokaryotic nanocompartments involved in sulfur metabolism. *eLife*, 10, 2021.

[9640] Alba Nicolas-Boluda, Javier Vaquero, Lene Vimeux, Thomas Guilbert, Sarah Barrin, Chahrazade Kantari-Mimoun, Matteo Ponzo, Gilles Renault, Piotr Deptula, Katarzyna Pogoda, Robert Bucki, Ilaria Cascone, José Courty, Laura Fouassier, Florence Gazeau, and Emmanuel Donnadieu. Tumor stiffening reversion through collagen crosslinking inhibition improves T cell migration and anti-PD-1 treatment. *eLife*, 10, 2021.

[9641] Felix Nicolaus, Ane Metola, Daphne Mermans, Amanda Liljenström, Ajda Krč, Salmo Mohammed Abdullahi, Matthew Zimmer, Thomas F Miller III, and Gunnar von Heijne. Residue-by-residue analysis of cotranslational membrane protein integration in vivo. *eLife*, 10, 2021.

[9642] Markus Niebuhr, Julia Belde, Anke Fähnrich, Arnauld Serge, Magali Irla, Christoph T Ellebrecht, Christoph M Hammers, Katja Bieber, Jürgen Westermann, and Kathrin Kalies. Receptor repertoires of murine follicular T helper cells reveal a high clonal overlap in separate lymph nodes in autoimmunity. *eLife*, 10, 2021.

[9643] Rene Niehus, Nuno M Oliveira, Aming Li, Alexander G Fletcher, and Kevin R Foster. The evolution of strategy in bacterial warfare via the regulation of bacteriocins and antibiotics. *eLife*, 10, 2021.

[9644] Mathias Wullum Nielsen, Christine Friis Baker, Emer Brady, Michael Bang Petersen, and Jens Peter Andersen. Weak evidence of country- and institution-related status bias in the peer review of abstracts. *eLife*, 10, 2021.

[9645] Lukasz Nierzwicki, Kyle W East, Uriel N Morzan, Pablo R Arantes, Victor S Batista, George P Lisi, and Giulia Palermo. Enhanced specificity mutations perturb allosteric signaling in CRISPR-Cas9. *eLife*, 10, 2021.

[9646] Nader Nikbakht and Mathew E Diamond. Conserved visual capacity of rats under red light. *eLife*, 10, 2021.

[9647] Thushani D Nilaweera, David A Nyenhuis, and David S Cafiso. Structural intermediates observed only in intact Escherichia coli indicate a mechanism for TonB-dependent transport. *eLife*, 10, 2021.

[9648] Takeshi Ninchoji, Dominic T Love, Ross O Smith, Marie Hedlund, Dietmar Vestweber, William C Sessa, and Lena Claesson-Welsh. eNOS-induced vascular barrier disruption in retinopathy by c-Src activation and tyrosine phosphorylation of VE-cadherin. *eLife*, 10, 2021.

[9649] Hiroshi Nishida, Morihiro Okada, Lynna Yang, Tomomi Takano, Sho Tabata, Tomoyoshi Soga, Diana M Ho, Jongkyeong Chung, Yasuhiro Minami, and Sa Kan Yoo. Methionine restriction breaks obligatory coupling of cell proliferation and death by an oncogene Src in Drosophila. *eLife*, 10, 2021.

[9650] Yiming Niu, Xiao Tao, George Vaisey, Paul Dominic B Olinares, Hanan Alwaseem, Brian T Chait, and Roderick MacKinnon. Analysis of the mechanosensor channel functionality of TACAN. *eLife*, 10, 2021.

[9651] Luke M Noble, John Yuen, Lewis Stevens, Nicolas Moya, Riaad Persaud, Marc Moscatelli, Jacqueline L Jackson, Gaotian Zhang, Rojin Chitrakar, L Ryan Baugh, Christian Braendle, Erik C Andersen, Hannah S Seidel, and Matthew V Rockman. Selfing is the safest sex for Caenorhabditis tropicalis. *eLife*, 10, 2021.

[9652] Klara Katarina Nordén, Chad M Eliason, and Mary Caswell Stoddard. Evolution of brilliant iridescent feather nanostructures. *eLife*, 10, 2021.

[9653] Aaron J Norris, Jordan R Shaker, Aaron L Cone, Imeh B Ndiokho, and Michael R Bruchas. Parabrachial opioidergic projections to preoptic hypothalamus mediate behavioral and physiological thermal defenses. *eLife*, 10, 2021.

[9654] Joseph W Nors, Shipra Gupta, and Marcel P Goldschen-Ohm. A critical residue in the $\alpha$1M2–M3 linker regulating mammalian GABAA receptor pore gating by diazepam. *eLife*, 10, 2021.

[9655] Behrad Noudoost, Kelsey Lynne Clark, and Tirin Moore. Working memory gates visual input to primate prefrontal neurons. *eLife*, 10, 2021.

[9656] Reuben W Nowell, Christopher G Wilson, Pedro Almeida, Philipp H Schiffer, Diego Fontaneto, Lutz Becks, Fernando Rodriguez, Irina R Arkhipova, and Timothy G Barraclough. Evolutionary dynamics of transposable elements in bdelloid rotifers. *eLife*, 10, 2021.

[9657] Urszula Nowicka, Piotr Chroscicki, Karen Stroobants, Maria Sladowska, Michal Turek, Barbara Uszczynska-Ratajczak, Rishika Kundra, Tomasz Goral, Michele Perni, Christopher M Dobson, Michele Vendruscolo, and Agnieszka Chacinska. Cytosolic aggregation of mitochondrial proteins disrupts cellular homeostasis by stimulating the aggregation of other proteins. *eLife*, 10, 2021.

[9658] Ole Numssen, Danilo Bzdok, and Gesa Hartwigsen. Functional specialization within the inferior parietal lobes across cognitive domains. *eLife*, 10, 2021.

[9659] Kate Nussenbaum and Catherine A Hartley. Developmental change in prefrontal cortex recruitment supports the emergence of value-guided memory. *eLife*, 10, 2021.

[9660] Kazumasa Oda, Takashi Nomura, Takanori Nakane, Keitaro Yamashita, Keiichi Inoue, Shota Ito, Johannes Vierock, Kunio Hirata, Andrés D Maturana, Kota Katayama, Tatsuya Ikuta, Itsuki Ishigami, Tamaki Izume, Rie Umeda, Ryuun Eguma, Satomi Oishi, Go Kasuya, Takafumi Kato, Tsukasa Kusakizako, Wataru Shihoya, Hiroto Shimada, Tomoyuki Takatsuji, Mizuki Takemoto, Reiya Taniguchi, Atsuhiro Tomita, Ryoki Nakamura, Masahiro Fukuda, Hirotake Miyauchi, Yongchan Lee, Eriko Nango, Rie Tanaka, Tomoyuki Tanaka, Michihiro Sugahara, Tetsunari Kimura, Tatsuro Shimamura, Takaaki Fujiwara, Yasuaki Yamanaka, Shigeki Owada, Yasumasa Joti, Kensuke Tono, Ryuichiro Ishitani, Shigehiko Hayashi, Hideki Kandori, Peter Hegemann, So Iwata, Minoru Kubo, Tomohiro Nishizawa, and Osamu Nureki. Time-resolved serial femtosecond crystallography reveals early structural changes in channelrhodopsin. *eLife*, 10, 2021.

[9661] Pascal D Odermatt, Teemu P Miettinen, Joël Lemière, Joon Ho Kang, Emrah Bostan, Scott R Manalis, Kerwyn Casey Huang, and Fred Chang. Variations of intracellular density during the cell cycle arise from tip-growth regulation in fission yeast. *eLife*, 10, 2021.

[9662] Takashi Ogino, Naoya Matsunaga, Takahiro Tanaka, Tomohito Tanihara, Hideki Terajima, Hikari Yoshitane, Yoshitaka Fukada, Akito Tsuruta, Satoru Koyanagi, and Shigehiro Ohdo. Post-transcriptional repression of circadian component CLOCK regulates cancer-stemness in murine breast cancer cells. *eLife*, 10, 2021.

[9663] Shinya Ohara, Stefan Blankvoort, Rajeevkumar Raveendran Nair, Maximiliano J Nigro, Eirik S Nilssen, Clifford Kentros, and Menno P Witter. Local projections of layer Vb-to-Va are more prominent in lateral than in medial entorhinal cortex. *eLife*, 10, 2021.

[9664] Kazuhide S Okuda, Mikaela S Keyser, David B Gurevich, Caterina Sturtzel, Elizabeth A Mason, Scott Paterson, Huijun Chen, Mark Scott, Nicholas D Condon, Paul Martin, Martin Distel, and Benjamin M Hogan. Live-imaging of endothelial Erk activity reveals dynamic and sequential signalling events during regenerative angiogenesis. *eLife*, 10, 2021.

[9665] Gabriela C Olivera, Emily C Ross, Christiane Peuckert, and Antonio Barragan. Blood-brain barrier-restricted translocation of Toxoplasma gondii from cortical capillaries. *eLife*, 10, 2021.

[9666] Julia Eve Olivieri, Roozbeh Dehghannasiri, Peter L Wang, SoRi Jang, Antoine de Morree, Serena Y Tan, Jingsi Ming, Angela Ruohao Wu, Robert C Jones, Stephen R Quake, Mark A Krasnow, and Julia Salzman. RNA splicing programs define tissue compartments and cell types at single-cell resolution. *eLife*, 10, 2021.

[9667] Deike J Omnus, Matthias J Fink, Klaudia Szwedo, and Kristina Jonas. The Lon protease temporally restricts polar cell differentiation events during the Caulobacter cell cycle. *eLife*, 10, 2021.

[9668] Shataakshi Dube O'Neil, Bence Rácz, Walter Evan Brown, Yudong Gao, Erik J Soderblom, Ryohei Yasuda, and Scott H Soderling. Action potential-coupled Rho GTPase signaling drives presynaptic plasticity. *eLife*, 10, 2021.

[9669] Alan M O'Neill, Kate A Worthing, Nikhil Kulkarni, Fengwu Li, Teruaki Nakatsuji, Dominic McGrosso, Robert H Mills, Gayathri Kalla, Joyce Y Cheng, Jacqueline M Norris, Kit Pogliano, Joe Pogliano, David J Gonzalez, and Richard L Gallo. Antimicrobials from a feline commensal bacterium inhibit skin infection by drug-resistant S. pseudintermedius. *eLife*, 10, 2021.

[9670] Koh Onimaru, Kaori Tatsumi, Chiharu Tanegashima, Mitsutaka Kadota, Osamu Nishimura, and Shigehiro Kuraku. Developmental hourglass and heterochronic shifts in fin and limb development. *eLife*, 10, 2021.

[9671] Tarik Seref Onur, Andrew Laitman, He Zhao, Ryan Keyho, Hyemin Kim, Jennifer Wang, Megan Mair, Huilan Wang, Lifang Li, Alma Perez, Maria de Haro, Ying-Wooi Wan, Genevera Allen, Boxun Lu, Ismael Al-Ramahi, Zhandong Liu, and Juan Botas. Downregulation of glial genes involved in synaptic function mitigates Huntington's disease pathogenesis. *eLife*, 10, 2021.

[9672] Chet Huan Oon and Kenneth E Prehoda. Phases of cortical actomyosin dynamics coupled to the neuroblast polarity cycle. *eLife*, 10, 2021.

[9673] James W Opzoomer, Jessica A Timms, Kevin Blighe, Thanos P Mourikis, Nicolas Chapuis, Richard Bekoe, Sedigeh Kareemaghay, Paola Nocerino, Benedetta Apollonio, Alan G Ramsay, Mahvash Tavassoli, Claire Harrison, Francesca Ciccarelli, Peter Parker, Michaela Fontenay, Paul R Barber, James N Arnold, and Shahram Kordasti. ImmunoCluster provides a computational framework for the nonspecialist to profile high-dimensional cytometry data. *eLife*, 10, 2021.

[9674] Jasmien Orije, Emilie Cardon, Julie Hamaide, Elisabeth Jonckers, Veerle M Darras, Marleen Verhoye, and Annemie Van der Linden. Uncovering a 'sensitive window' of multisensory and motor neuroplasticity in the cerebrum and cerebellum of male and female starlings. *eLife*, 10, 2021.

[9675] Pedro Ortega, José Antonio Mérida-Cerro, Ana G Rondón, Belén Gómez-González, and Andrés Aguilera. DNA-RNA hybrids at DSBs interfere with repair by homologous recombination. *eLife*, 10, 2021.

[9676] Carolina Ortiz-Cordero, Alessandro Magli, Neha R Dhoke, Taylor Kuebler, Sridhar Selvaraj, Nelio AJ Oliveira, Haowen Zhou, Yuk Y Sham, Anne G Bang, and Rita CR Perlingeiro. NAD+ enhances ribitol and ribose rescue of $\alpha$-dystroglycan functional glycosylation in human FKRP-mutant myotubes. *eLife*, 10, 2021.

[9677] Yuki Osawa, Kazuya Murata, Miho Usui, Yumeno Kuba, Hoai Thu Le, Natsuki Mikami, Toshinori Nakagawa, Yoko Daitoku, Kanako Kato, Hossam Hassan Shawki, Yoshihisa Ikeda, Akihiro Kuno, Kento Morimoto, Yoko Tanimoto, Tra Thi Huong Dinh, Ken-ichi Yagami, Masatsugu Ema, Shosei Yoshida, Satoru Takahashi, Seiya Mizuno, and Fumihiro Sugiyama. EXOC1 plays an integral role in spermatogonia pseudopod elongation and spermatocyte stable syncytium formation in mice. *eLife*, 10, 2021.

[9678] Anna Osnato, Stephanie Brown, Christel Krueger, Simon Andrews, Amanda J Collier, Shota Nakanoh, Mariana Quiroga Londoño, Brandon T Wesley, Daniele Muraro, A Sophie Brumm, Kathy K Niakan, Ludovic Vallier, Daniel Ortmann, and Peter J Rugg-Gunn. TGF$\beta$ signalling is required to maintain pluripotency of human naïve pluripotent stem cells. *eLife*, 10, 2021.

[9679] Ratan Othayoth and Chen Li. Propelling and perturbing appendages together facilitate strenuous ground self-righting. *eLife*, 10, 2021.

[9680] Kurataka Otsuka, Akihito Mamiya, Mineko Konishi, Mamoru Nozaki, Atsuko Kinoshita, Hiroaki Tamaki, Masaki Arita, Masato Saito, Kayoko Yamamoto, Takushi Hachiya, Ko Noguchi, Takashi Ueda, Yusuke Yagi, Takehito Kobayashi, Takahiro Nakamura, Yasushi Sato, Takashi Hirayama, and Munetaka Sugiyama. Temperature-dependent fasciation mutants provide a link between mitochondrial RNA processing and lateral root morphogenesis. *eLife*, 10, 2021.

[9681] Maria C Ow, Alexandra M Nichitean, and Sarah E Hall. Somatic aging pathways regulate reproductive plasticity in Caenorhabditis elegans. *eLife*, 10, 2021.

[9682] Ethan Ozment, Arianna N Tamvacakis, Jianhong Zhou, Pablo Yamild Rosiles-Loeza, Esteban Elías Escobar-Hernandez, Selene L Fernandez-Valverde, and Nagayasu Nakanishi. Cnidarian hair cell development illuminates an ancient role for the class IV POU transcription factor in defining mechanoreceptor identity. *eLife*, 10, 2021.

[9683] Attila Ozsvár, Gergely Komlósi, Gáspár Oláh, Judith Baka, Gábor Molnár, and Gábor Tamás. Predominantly linear summation of metabotropic postsynaptic potentials follows coactivation of neurogliaform interneurons. *eLife*, 10, 2021.

[9684] Bilge E öztürk, Molly E Johnson, Michael Kleyman, Serhan TurunÇ, Jing He, Sara Jabalameli, Zhouhuan Xi, Meike Visel, Valérie L Dufour, Simone Iwabe, Luis Felipe L Pompeo Marinho, Gustavo D Aguirre, José-Alain Sahel, David V Schaffer, Andreas R Pfenning, John G Flannery, William A Beltran, William R Stauffer, and Leah C Byrne. scAAVengr, a transcriptome-based pipeline for quantitative ranking of engineered AAVs with single-cell resolution. *eLife*, 10, 2021.

[9685] Alba Pacheco-Moreno, Francesca L Stefanato, Jonathan J Ford, Christine Trippel, Simon Uszkoreit, Laura Ferrafiat, Lucia Grenga, Ruth Dickens, Nathan Kelly, Alexander DH Kingdon, Liana Ambrosetti, Sergey A Nepogodiev, Kim C Findlay, Jitender Cheema, Martin Trick, Govind Chandra, Graham Tomalin, Jacob G Malone, and Andrew W Truman. Pan-genome analysis identifies intersecting roles for Pseudomonas specialized metabolites in potato pathogen inhibition. *eLife*, 10, 2021.

[9686] Fabiola Paciello, Marco Rinaudo, Valentina Longo, Sara Cocco, Giulia Conforto, Anna Pisani, Maria Vittoria Podda, Anna Rita Fetoni, Gaetano Paludetti, and Claudio Grassi. Auditory sensory deprivation induced by noise exposure exacerbates cognitive decline in a mouse model of Alzheimer's disease. *eLife*, 10, 2021.

[9687] Periklis Paganos, Danila Voronov, Jacob M Musser, Detlev Arendt, and Maria Ina Arnone. Single-cell RNA sequencing of the Strongylocentrotus purpuratus larva reveals the blueprint of major cell types and nervous system of a non-chordate deuterostome. *eLife*, 10, 2021.

[9688] Virender Kumar Pal, Ragini Agrawal, Srabanti Rakshit, Pooja Shekar, Diwakar Tumkur Narasimha Murthy, Annapurna Vyakarnam, and Amit Singh. Hydrogen sulfide blocks HIV rebound by maintaining mitochondrial bioenergetics and redox homeostasis. *eLife*, 10, 2021.

[9689] Saravanan Palani, Sayantika Ghosh, Esther Ivorra-Molla, Scott Clarke, Andrejus Suchenko, Mohan K Balasubramanian, and Darius Vasco Köster. Calponin-homology domain mediated bending of membrane-associated actin filaments. *eLife*, 10, 2021.

[9690] Lavanya H Palavalli Parsons, Sridevi Challa, Bryan A Gibson, Tulip Nandu, MiKayla S Stokes, Dan Huang, Jayanthi S Lea, and W Lee Kraus. Identification of PARP-7 substrates reveals a role for MARylation in microtubule control in ovarian cancer cells. *eLife*, 10, 2021.

[9691] Julius Palme, Jue Wang, and Michael Springer. Variation in the modality of a yeast signaling pathway is mediated by a single regulator. *eLife*, 10, 2021.

[9692] Daniel J Paluh, Karina Riddell, Catherine M Early, Maggie M Hantak, Gregory FM Jongsma, Rachel M Keeffe, Fernanda Magalhães Silva, Stuart V Nielsen, María Camila Vallejo-Pareja, Edward L Stanley, and David C Blackburn. Rampant tooth loss across 200 million years of frog evolution. *eLife*, 10, 2021.

[9693] Qiaowei Pan, Romain Feron, Elodie Jouanno, Hugo Darras, Amaury Herpin, Ben Koop, Eric Rondeau, Frederick W Goetz, Wesley A Larson, Louis Bernatchez, Mike Tringali, Stephen S Curran, Eric Saillant, Gael PJ Denys, Frank A von Hippel, Songlin Chen, J Andrés López, Hugo Verreycken, Konrad Ocalewicz, Rene Guyomard, Camille Eche, Jerome Lluch, Celine Roques, Hongxia Hu, Roger Tabor, Patrick DeHaan, Krista M Nichols, Laurent Journot, Hugues Parrinello, Christophe Klopp, Elena A Interesova, Vladimir Trifonov, Manfred Schartl, John Postlethwait, and Yann Guiguen. The rise and fall of the ancient northern pike master sex-determining gene. *eLife*, 10, 2021.

[9694] Susu Pan, Kaili Yin, Zhiwei Tang, Shuren Wang, Zhuo Chen, Yirong Wang, Hongxia Zhu, Yunyun Han, Mei Liu, Man Jiang, Ningzhi Xu, and Guo Zhang. Stimulation of hypothalamic oxytocin neurons suppresses colorectal cancer progression in mice. *eLife*, 10, 2021.

[9695] Xingyu Pan, Jingrong Zhao, Zhiying Zhou, Jijun Chen, Zhenxing Yang, Yuxuan Wu, Meizhu Bai, Yang Jiao, Yun Yang, Xuye Hu, Tianling Cheng, Qianyun Lu, Bin Wang, Chang-Lin Li, Ying-Jin Lu, Lei Diao, Yan-Qing Zhong, Jing Pan, Jianmin Zhu, Hua-Sheng Xiao, Zi-Long Qiu, Jinsong Li, Zefeng Wang, Jingyi Hui, Lan Bao, and Xu Zhang. 5′-UTR SNP of FGF13 causes translational defect and intellectual disability. *eLife*, 10, 2021.

[9696] Manish Pandey, Sakshi Bansal, Sudipta Bar, Amit Kumar Yadav, Nicholas S Sokol, Jason M Tennessen, Pankaj Kapahi, and Geetanjali Chawla. miR-125-chinmo pathway regulates dietary restriction-dependent enhancement of lifespan in Drosophila. *eLife*, 10, 2021.

[9697] Pooja Pandey, Alexandre Y Leary, Yasin Tumtas, Zachary Savage, Bayantes Dagvadorj, Cian Duggan, Enoch LH Yuen, Nattapong Sanguankiattichai, Emily Tan, Virendrasinh Khandare, Amber J Connerton, Temur Yunusov, Mathias Madalinski, Federico Gabriel Mirkin, Sebastian Schornack, Yasin Dagdas, Sophien Kamoun, and Tolga O Bozkurt. An oomycete effector subverts host vesicle trafficking to channel starvation-induced autophagy to the pathogen interface. *eLife*, 10, 2021.

[9698] Prithvi R Pandey, Bartosz Różycki, Reinhard Lipowsky, and Thomas R Weikl. Structural variability and concerted motions of the T cell receptor – CD3 complex. *eLife*, 10, 2021.

[9699] Veethika Pandey, Alicia Fleming-Martinez, Ligia Bastea, Heike R Doeppler, Jillian Eisenhauer, Tam Le, Brandy Edenfield, and Peter Storz. CXCL10/CXCR3 signaling contributes to an inflammatory microenvironment and its blockade enhances progression of murine pancreatic precancerous lesions. *eLife*, 10, 2021.

[9700] Swapnesh Panigrahi, Dorothée Murat, Antoine Le Gall, Eugénie Martineau, Kelly Goldlust, Jean-Bernard Fiche, Sara Rombouts, Marcelo Nöllmann, Leon Espinosa, and Tâm Mignot. Misic, a general deep learning-based method for the high-throughput cell segmentation of complex bacterial communities. *eLife*, 10, 2021.

[9701] Ranjith K Papareddy, Katalin Páldi, Anna D Smolka, Patrick Hüther, Claude Becker, and Michael D Nodine. Repression of CHROMOMETHYLASE 3 prevents epigenetic collateral damage in Arabidopsis. *eLife*, 10, 2021.

[9702] Casey Paquola, Jessica Royer, Lindsay B Lewis, Claude Lepage, Tristan Glatard, Konrad Wagstyl, Jordan DeKraker, Paule-J Toussaint, Sofie L Valk, Louis Collins, Ali R Khan, Katrin Amunts, Alan C Evans, Timo Dickscheid, and Boris Bernhardt. The BigBrainWarp toolbox for integration of BigBrain 3D histology with multimodal neuroimaging. *eLife*, 10, 2021.

[9703] Diogo Paramos-de Carvalho, Antonio Jacinto, and Leonor Saúde. The right time for senescence. *eLife*, 10, 2021.

[9704] Smriti Parashar, Ravi Chidambaram, Shuliang Chen, Christina R Liem, Eric Griffis, Gerard G Lambert, Nathan C Shaner, Matthew Wortham, Jesse C Hay, and Susan Ferro-Novick. Endoplasmic reticulum tubules limit the size of misfolded protein condensates. *eLife*, 10, 2021.

[9705] Bo-yong Park, Richard AI Bethlehem, Casey Paquola, Sara Larivière, Raul Rodríguez-Cruces, Reinder Vos de Wael, Edward Bullmore, Edward Bullmore, Raymond Dolan, Ian Goodyer, Peter Fonagy, Peter Jones, Michael Moutoussis, Tobias Hauser, Sharon Neufeld, Rafael Romero-Garcia, Michelle St Clair, Petra Vértes, Kirstie Whitaker, Becky Inkster, Gita Prabhu, Cinly Ooi, Umar Toseeb, Barry Widmer, Junaid Bhatti, Laura Villis, Ayesha Alrumaithi, Sarah Birt, Aislinn Bowler, Kalia Cleridou, Hina Dadabhoy, Emma Davies, Ashlyn Firkins, Sian Granville, Elizabeth Harding, Alexandra Hopkins, Daniel Isaacs, Janchai King, Danae Kokorikou, Christina Maurice, Cleo McIntosh, Jessica Memarzia, Harriet Mills, Ciara O'Donnell, Sara Pantaleone, Jenny Scott, Beatrice Kiddle, Ela Polek, Pasco Fearon, John Suckling, Anne-Laura van Harmelen, Rogier Kievit, Sam Chamberlain, Edward T Bullmore, and Boris C Bernhardt. An expanding manifold in transmodal regions characterizes adolescent reconfiguration of structural connectome organization. *eLife*, 10, 2021.

[9706] Chanhyun Park, Yuki Sakurai, Hirofumi Sato, Shinji Kanda, Yuichi Iino, and Hirofumi Kunitomo. Roles of the ClC chloride channel CLH-1 in food-associated salt chemotaxis behavior of C. elegans. *eLife*, 10, 2021.

[9707] Jaeeun Park, Misung Kim, Bora Shin, Mingyeong Kang, Jihye Yang, Tae Kwon Lee, and Woojun Park. A novel decoy strategy for polymyxin resistance in Acinetobacter baumannii. *eLife*, 10, 2021.

[9708] Ki-Sun Park, Beenish Rahat, Hyung Chul Lee, Zu-Xi Yu, Jacob Noeker, Apratim Mitra, Connor M Kean, Russell H Knutsen, Danielle Springer, Claudia M Gebert, Beth A Kozel, and Karl Pfeifer. Cardiac pathologies in mouse loss of imprinting models are due to misexpression of H19 long noncoding RNA. *eLife*, 10, 2021.

[9709] Seongjin Park, Karine Prévost, Emily M Heideman, Marie-Claude Carrier, Muhammad S Azam, Matthew A Reyer, Wei Liu, Eric Massé, and Jingyi Fei. Dynamic interactions between the RNA chaperone Hfq, small regulatory RNAs, and mRNAs in live bacterial cells. *eLife*, 10, 2021.

[9710] Matthew T Parker, Katarzyna Knop, Vasiliki Zacharaki, Anna V Sherwood, Daniel Tomé, Xuhong Yu, Pascal GP Martin, Jim Beynon, Scott D Michaels, Geoffrey J Barton, and Gordon G Simpson. Widespread premature transcription termination of Arabidopsis thaliana NLR genes by the spen protein FPA. *eLife*, 10, 2021.

[9711] Matthew W Parker, Jonchee A Kao, Alvin Huang, James M Berger, and Michael R Botchan. Molecular determinants of phase separation for Drosophila DNA replication licensing factors. *eLife*, 10, 2021.

[9712] Eloy Parra-Barrero, Kamran Diba, and Sen Cheng. Neuronal sequences during theta rely on behavior-dependent spatial maps. *eLife*, 10, 2021.

[9713] Helen Parry, Gokhan Tut, Rachel Bruton, Sian Faustini, Christine Stephens, Philip Saunders, Christopher Bentley, Katherine Hilyard, Kevin Brown, Gayatri Amirthalingam, Sue Charlton, Stephanie Leung, Emily Chiplin, Naomi S Coombes, Kevin R Bewley, Elizabeth J Penn, Cathy Rowe, Ashley Otter, Rosie Watts, Silvia D'Arcangelo, Bassam Hallis, Andrew Makin, Alex Richter, Jianmin Zuo, and Paul Moss. mRNA vaccination in people over 80 years of age induces strong humoral immune responses against SARS-CoV-2 with cross neutralization of P.1 Brazilian variant. *eLife*, 10, 2021.

[9714] Natalya Pashkova, Lokesh Gakhar, Liping Yu, Nicholas J Schnicker, Annabel Y Minard, Stanley Winistorfer, Ivan E Johnson, and Robert C Piper. ANTH domains within CALM, HIP1R, and Sla2 recognize ubiquitin internalization signals. *eLife*, 10, 2021.

[9715] Alexander O Pasternak, Jelmer Vroom, Neeltje A Kootstra, Ferdinand WNM Wit, Marijn de Bruin, Davide De Francesco, Margreet Bakker, Caroline A Sabin, Alan Winston, Jan M Prins, Peter Reiss, Ben Berkhout, P Reiss, P Reiss, FWNM Wit, J Schouten, KW Kooij, RA van Zoest, BC Elsenga, FR Janssen, M Heidenrijk, W Zikkenheiner, M van der Valk, NA Kootstra, T Booiman, AM Harskamp-Holwerda, I Maurer, MM Mangas Ruiz, AF Girigorie, J Villaudy, E Frankin, AO Pasternak, B Berkhout, T van der Kuyl, P Portegies, BA Schmand, GJ Geurtsen, JA ter Stege, M Klein Twennaar, CBLM Majoie, MWA Caan, T Su, K Weijer, PHLT Bisschop, A Kalsbeek, M Wezel, I Visser, HG Ruhé, C Franceschi, P Garagnani, C Pirazzini, M Capri, F Dall'Olio, M Chiricolo, S Salvioli, J Hoeijmakers, J Pothof, M Prins, M Martens, S Moll, J Berkel, M Totté, S Kovalev, M Gisslén, D Fuchs, H Zetterberg, A Winston, J Underwood, L McDonald, M Stott, K Legg, A Lovell, O Erlwein, N Doyle, C Kingsley, DJ Sharp, R Leech, JH Cole, S Zaheri, MMJ Hillebregt, YMC Ruijs, DP Benschop, D Burger, M de Graaff-Teulen, G Guaraldi, A Bürkle, T Sindlinger, M Moreno-Villanueva, A Keller, C Sabin, D de Francesco, C Libert, and S Dewaele. Non-nucleoside reverse transcriptase inhibitor-based combination antiretroviral therapy is associated with lower cell-associated HIV RNA and DNA levels compared to protease inhibitor-based therapy. *eLife*, 10, 2021.

[9716] Ankur V Patel, Robert D Turner, Aline Rifflet, Adelina E Acosta-Martin, Andrew Nichols, Milena M Awad, Dena Lyras, Ivo Gomperts Boneca, Marshall Bern, Mark O Collins, and Stéphane Mesnage. PGFinder, a novel analysis pipeline for the consistent, reproducible, and high-resolution structural analysis of bacterial peptidoglycans. *eLife*, 10, 2021.

[9717] Vishwa Patel and Nishad Matange. Adaptation and compensation in a bacterial gene regulatory network evolving under antibiotic selection. *eLife*, 10, 2021.

[9718] Gajanan S Patil, Priyadarshan Kinatukara, Sudipta Mondal, Sakshi Shambhavi, Ketan D Patel, Surabhi Pramanik, Noopur Dubey, Subhash Narasimhan, Murali Krishna Madduri, Biswajit Pal, Rajesh S Gokhale, and Rajan Sankaranarayanan. A universal pocket in fatty acyl-AMP ligases ensures redirection of fatty acid pool away from coenzyme A-based activation. *eLife*, 10, 2021.

[9719] James Oliver Patterson, Souradeep Basu, Paul Rees, and Paul Nurse. CDK control pathways integrate cell size and ploidy information to control cell division. *eLife*, 10, 2021.

[9720] Maarten W Paul, Arshdeep Sidhu, Yongxin Liang, Sarah E van Rossum-Fikkert, Hanny Odijk, Alex N Zelensky, Roland Kanaar, and Claire Wyman. Role of BRCA2 DNA-binding and C-terminal domain in its mobility and conformation in DNA repair. *eLife*, 10, 2021.

[9721] Dennis V Pedersen, Martin Nors Pedersen, Sofia MM Mazarakis, Yong Wang, Kresten Lindorff-Larsen, Lise Arleth, and Gregers R Andersen. Properdin oligomers adopt rigid extended conformations supporting function. *eLife*, 10, 2021.

[9722] Renata O Pereira, Alex Marti, Angela Crystal Olvera, Satya Murthy Tadinada, Sarah Hartwick Bjorkman, Eric Thomas Weatherford, Donald A Morgan, Michael Westphal, Pooja H Patel, Ana Karina Kirby, Rana Hewezi, William Bùi Trân, Luis Miguel García-Peña, Rhonda A Souvenir, Monika Mittal, Christopher M Adams, Kamal Rahmouni, Matthew J Potthoff, and E Dale Abel. OPA1 deletion in brown adipose tissue improves thermoregulation and systemic metabolism via FGF21. *eLife*, 10, 2021.

[9723] Harvey Perez, May F Abdallah, Jose I Chavira, Angelina S Norris, Martin T Egeland, Karen L Vo, Callan L Buechsenschuetz, Valentina Sanghez, Jeannie L Kim, Molly Pind, Kotoka Nakamura, Geoffrey G Hicks, Richard A Gatti, Joaquin Madrenas, Michelina Iacovino, Peter J McKinnon, and Paul J Mathews. A novel, ataxic mouse model of ataxia telangiectasia caused by a clinically relevant nonsense mutation. *eLife*, 10, 2021.

[9724] Julio D Perez, Susanne tom Dieck, Beatriz Alvarez-Castelao, Georgi Tushev, Ivy CW Chan, and Erin M Schuman. Subcellular sequencing of single neurons reveals the dendritic transcriptome of GABAergic interneurons. *eLife*, 10, 2021.

[9725] Vicente Perez-Garcia, Georgia Lea, Pablo Lopez-Jimenez, Hanneke Okkenhaug, Graham J Burton, Ashley Moffett, Margherita Y Turco, and Myriam Hemberger. BAP1/ASXL complex modulation regulates epithelial-mesenchymal transition during trophoblast differentiation and invasion. *eLife*, 10, 2021.

[9726] Jesús Pérez-Ortega, Tzitzitlini Alejandre-García, and Rafael Yuste. Long-term stability of cortical ensembles. *eLife*, 10, 2021.

[9727] Jimena Pérez-Vargas, Elin Teppa, Fouzia Amirache, Bertrand Boson, Rémi Pereira de Oliveira, Christophe Combet, Anja Böckmann, Floriane Fusil, Natalia Freitas, Alessandra Carbone, and FranÇois-Loïc Cosset. A fusion peptide in preS1 and the human protein disulfide isomerase ERp57 are involved in hepatitis B virus membrane fusion process. *eLife*, 10, 2021.

[9728] Jessamyn I Perlmutter, Jane E Meyers, and Seth R Bordenstein. A single synonymous nucleotide change impacts the male-killing phenotype of prophage WO gene wmk. *eLife*, 10, 2021.

[9729] Stefano Perni and Kurt Beam. Neuronal junctophilins recruit specific CaV and RyR isoforms to ER-PM junctions and functionally alter CaV2.1 and CaV2.2. *eLife*, 10, 2021.

[9730] George H Perry. Evolutionary medicine. *eLife*, 10, 2021.

[9731] Rita de Cassia Pessotti, Bridget L Hansen, Jewel N Reaso, Javier A Ceja-Navarro, Laila El-Hifnawi, Eoin L Brodie, and Matthew F Traxler. Multiple lineages of Streptomyces produce antimicrobials within passalid beetle galleries across eastern North America. *eLife*, 10, 2021.

[9732] Christopher J Petell, Kathyrn Randene, Michael Pappas, Diego Sandoval, Brian D Strahl, Joseph S Harrison, and Joshua P Steimel. Mechanically transduced immunosorbent assay to measure protein-protein interactions. *eLife*, 10, 2021.

[9733] Michael A Petr, Irene Alfaras, Melissa Krawcyzk, Woei-Nan Bair, Sarah J Mitchell, Christopher H Morrell, Stephanie A Studenski, Nathan L Price, Kenneth W Fishbein, Richard G Spencer, Morten Scheibye-Knudsen, Edward G Lakatta, Luigi Ferrucci, Miguel A Aon, Michel Bernier, and Rafael de Cabo. A cross-sectional study of functional and metabolic changes during aging through the lifespan in male mice. *eLife*, 10, 2021.

[9734] Natalia Petrenko and Kevin Struhl. Comparison of transcriptional initiation by RNA polymerase II across eukaryotic species. *eLife*, 10, 2021.

[9735] Johannes Pettmann, Anna Huhn, Enas Abu Shah, Mikhail A Kutuzov, Daniel B Wilson, Michael L Dustin, Simon J Davis, P Anton van der Merwe, and Omer Dushek. The discriminatory power of the T cell receptor. *eLife*, 10, 2021.

[9736] Gordon H Petty, Amanda K Kinnischtzke, Y Kate Hong, and Randy M Bruno. Effects of arousal and movement on secondary somatosensory and visual thalamus. *eLife*, 10, 2021.

[9737] Markus Pfenninger, Friederike Reuss, Angelika Kiebler, Philipp Schönnenbeck, Cosima Caliendo, Susanne Gerber, Berardino Cocchiararo, Sabrina Reuter, Nico Blüthgen, Karsten Mody, Bagdevi Mishra, Miklós Bálint, Marco Thines, and Barbara Feldmeyer. Genomic basis for drought resistance in European beech forests threatened by climate change. *eLife*, 10, 2021.

[9738] Ragini Phansalkar, Josephine Krieger, Mingming Zhao, Sai Saroja Kolluru, Robert C Jones, Stephen R Quake, Irving Weissman, Daniel Bernstein, Virginia D Winn, Gaetano D'Amato, and Kristy Red-Horse. Coronary blood vessels from distinct origins converge to equivalent states during mouse and human development. *eLife*, 10, 2021.

[9739] Chansavath Phetsouphanh, Prabhjeet Phalora, Carl-Philipp Hackstein, John Thornhill, C Mee Ling Munier, Jodi Meyerowitz, Lyle Murray, Cloete VanVuuren, Dominique Goedhals, Linnea Drexhage, Rebecca A Russell, Quentin J Sattentau, Jeffrey YW Mak, David P Fairlie, Sarah Fidler, Anthony D Kelleher, John Frater, and Paul Klenerman. Human MAIT cells respond to and suppress HIV-1. *eLife*, 10, 2021.

[9740] Angela M Phillips, Katherine R Lawrence, Alief Moulana, Thomas Dupic, Jeffrey Chang, Milo S Johnson, Ivana Cvijovic, Thierry Mora, Aleksandra M Walczak, and Michael M Desai. Binding affinity landscapes constrain the evolution of broadly neutralizing anti-influenza antibodies. *eLife*, 10, 2021.

[9741] Daniel A Phillips, Lori A Zacharoff, Cheri M Hampton, Grace W Chong, Anthony P Malanoski, Lauren Ann Metskas, Shuai Xu, Lina J Bird, Brian J Eddie, Aleksandr E Miklos, Grant J Jensen, Lawrence F Drummy, Mohamed Y El-Naggar, and Sarah M Glaven. A bacterial membrane sculpting protein with BAR domain-like activity. *eLife*, 10, 2021.

[9742] Stephen R Piccolo, Zachary E Ence, Elizabeth C Anderson, Jeffrey T Chang, and Andrea H Bild. Simplifying the development of portable, scalable, and reproducible workflows. *eLife*, 10, 2021.

[9743] Alexa Pichet Binette, Guillaume Theaud, FranÇois Rheault, Maggie Roy, D Louis Collins, Johannes Levin, Hiroshi Mori, Jae Hong Lee, Martin Rhys Farlow, Peter Schofield, Jasmeer P Chhatwal, Colin L Masters, Tammie Benzinger, John Morris, Randall Bateman, John CS Breitner, Judes Poirier, Julie Gonneaud, Maxime Descoteaux, and Sylvia Villeneuve. Bundle-specific associations between white matter microstructure and A$\beta$ and tau pathology in preclinical Alzheimer's disease. *eLife*, 10, 2021.

[9744] Domenico Pimpinella, Valentina Mastrorilli, Corinna Giorgi, Silke Coemans, Salvatore Lecca, Arnaud L Lalive, Hannah Ostermann, Elke C Fuchs, Hannah Monyer, Andrea Mele, Enrico Cherubini, and Marilena Griguoli. Septal cholinergic input to CA2 hippocampal region controls social novelty discrimination via nicotinic receptor-mediated disinhibition. *eLife*, 10, 2021.

[9745] David G Pina, Ivan Buljan, Darko Hren, and Ana MaruŠić. A retrospective analysis of the peer review of more than 75,000 Marie Curie proposals between 2007 and 2018. *eLife*, 10, 2021.

[9746] Charline Sophie Pinna, Maëlle Vilbert, Stephan Borensztajn, Willy Daney de Marcillac, Florence Piron-Prunier, Aaron Pomerantz, Nipam H Patel, Serge Berthier, Christine Andraud, Doris Gomez, and Marianne Elias. Mimicry can drive convergence in structural and light transmission features of transparent wings in Lepidoptera. *eLife*, 10, 2021.

[9747] Rosa Pipitone, Simona Eicke, Barbara Pfister, Gaetan Glauser, Denis Falconet, Clarisse Uwizeye, Thibaut Pralon, Samuel C Zeeman, Felix Kessler, and Emilie Demarsy. A multifaceted analysis reveals two distinct phases of chloroplast biogenesis during de-etiolation in Arabidopsis. *eLife*, 10, 2021.

[9748] Sashank Pisupati, Lital Chartarifsky-Lynn, Anup Khanal, and Anne K Churchland. Lapses in perceptual decisions reflect exploration. *eLife*, 10, 2021.

[9749] Laura Plassart, Ramtin Shayan, Christian Montellese, Dana Rinaldi, Natacha Larburu, Carole Pichereaux, Carine Froment, Simon Lebaron, Marie-FranÇoise O'Donohue, Ulrike Kutay, Julien Marcoux, Pierre-Emmanuel Gleizes, and Celia Plisson-Chastang. The final step of 40S ribosomal subunit maturation is controlled by a dual key lock. *eLife*, 10, 2021.

[9750] Jason D Plummer, Spike DL Postnikoff, Jessica K Tyler, and Jay E Johnson. Selenium supplementation inhibits IGF-1 signaling and confers methionine restriction-like healthspan benefits to mice. *eLife*, 10, 2021.

[9751] Maria Podinovskaia, Cristina Prescianotto-Baschong, Dominik P Buser, and Anne Spang. A novel live-cell imaging assay reveals regulation of endosome maturation. *eLife*, 10, 2021.

[9752] Ella Podvalny, Leana E King, and Biyu J He. Spectral signature and behavioral consequence of spontaneous shifts of pupil-linked arousal in human. *eLife*, 10, 2021.

[9753] Martin Pofahl, Negar Nikbakht, André N Haubrich, Theresa Nguyen, Nicola Masala, Fabian Distler, Oliver Braganza, Jakob H Macke, Laura A Ewell, Kurtulus Golcuk, and Heinz Beck. Synchronous activity patterns in the dentate gyrus during immobility. *eLife*, 10, 2021.

[9754] Matthias Pöge, Julia Mahamid, Sanae S Imanishi, Jürgen M Plitzko, Krzysztof Palczewski, and Wolfgang Baumeister. Determinants shaping the nanoscale architecture of the mouse rod outer segment. *eLife*, 10, 2021.

[9755] Theresa Pohlkamp, Xunde Xian, Connie H Wong, Murat S Durakoglugil, Gordon Chandler Werthmann, Takaomi C Saido, Bret M Evers, Charles L White, Jade Connor, Robert E Hammer, and Joachim Herz. NHE6 depletion corrects ApoE4-mediated synaptic impairments and reduces amyloid plaque load. *eLife*, 10, 2021.

[9756] Jacob T Polaski, Dylan B Udy, Luisa F Escobar-Hoyos, Gokce Askan, Steven D Leach, Andrea Ventura, Ram Kannan, and Robert K Bradley. The origins and consequences of UPF1 variants in pancreatic adenosquamous carcinoma. *eLife*, 10, 2021.

[9757] Melissa J Polonenko and Ross K Maddox. Exposing distinct subcortical components of the auditory brainstem response evoked by continuous naturalistic speech. *eLife*, 10, 2021.

[9758] Tessa M Popay, Jing Wang, Clare M Adams, Gregory Caleb Howard, Simona G Codreanu, Stacy D Sherrod, John A McLean, Lance R Thomas, Shelly L Lorey, Yuichi J Machida, April M Weissmiller, Christine M Eischen, Qi Liu, and William P Tansey. MYC regulates ribosome biogenesis and mitochondrial gene expression programs through its interaction with host cell factor–1. *eLife*, 10, 2021.

[9759] Eric C Porges, Greg Jensen, Brent Foster, Richard AE Edden, and Nicolaas AJ Puts. The trajectory of cortical GABA across the lifespan, an individual participant data meta-analysis of edited MRS studies. *eLife*, 10, 2021.

[9760] Xavier Portillo, Yu-Ting Huang, Ronald R Breaker, David P Horning, and Gerald F Joyce. Witnessing the structural evolution of an RNA enzyme. *eLife*, 10, 2021.

[9761] Daniel Porto, Yohei Matsunaga, Barbara Franke, Rhys M Williams, Hiroshi Qadota, Olga Mayans, Guy M Benian, and Hang Lu. Conformational changes in twitchin kinase in vivo revealed by FRET imaging of freely moving C. elegans. *eLife*, 10, 2021.

[9762] Sabrina Pospich, H Lee Sweeney, Anne Houdusse, and Stefan Raunser. High-resolution structures of the actomyosin-V complex in three nucleotide states provide insights into the force generation mechanism. *eLife*, 10, 2021.

[9763] Christian H Poth. Urgency forces stimulus-driven action by overcoming cognitive control. *eLife*, 10, 2021.

[9764] Daniel Powell, Sara A Haddad, Srinivas Gorur-Shandilya, and Eve Marder. Coupling between fast and slow oscillator circuits in Cancer borealis is temperature-compensated. *eLife*, 10, 2021.

[9765] Anjali M Prabhat, Catherine K Miller, Thomas Cody Prang, Jeffrey Spear, Scott A Williams, and Jeremy M DeSilva. Homoplasy in the evolution of modern human-like joint proportions in Australopithecus afarensis. *eLife*, 10, 2021.

[9766] Hanumanthu BD Prasada Rao, Takeshi Sato, Kiran Challa, Yurika Fujita, Miki Shinohara, and Akira Shinohara. Phosphorylation of luminal region of the SUN-domain protein Mps3 promotes nuclear envelope localization during meiosis. *eLife*, 10, 2021.

[9767] Cosimo Prestigio, Daniele Ferrante, Antonella Marte, Alessandra Romei, Gabriele Lignani, Franco Onofri, Pierluigi Valente, Fabio Benfenati, and Pietro Baldelli. REST/NRSF drives homeostatic plasticity of inhibitory synapses in a target-dependent fashion. *eLife*, 10, 2021.

[9768] Ian F Price, Hannah L Hertz, Benjamin Pastore, Jillian Wagner, and Wen Tang. Proximity labeling identifies LOTUS domain proteins that promote the formation of perinuclear germ granules in C. elegans. *eLife*, 10, 2021.

[9769] Michael F Priest, Elizabeth EL Lee, and Francisco Bezanilla. Tracking the movement of discrete gating charges in a voltage-gated potassium channel. *eLife*, 10, 2021.

[9770] Lucia L Prieto-Godino, Hayden R Schmidt, and Richard Benton. Molecular reconstruction of recurrent evolutionary switching in olfactory receptor specificity. *eLife*, 10, 2021.

[9771] Luigi Prisco, Stephan Hubertus Deimel, Hanna Yeliseyeva, André Fiala, and Gaia Tavosanis. The anterior paired lateral neuron normalizes odour-evoked activity in the Drosophila mushroom body calyx. *eLife*, 10, 2021.

[9772] Carl Procko, Swetha Murthy, William T Keenan, Seyed Ali Reza Mousavi, Tsegaye Dabi, Adam Coombs, Erik Procko, Lisa Baird, Ardem Patapoutian, and Joanne Chory. Stretch-activated ion channels identified in the touch-sensitive structures of carnivorous Droseraceae plants. *eLife*, 10, 2021.

[9773] Mira I Pronobis, Susan Zheng, Sumeet Pal Singh, Joseph A Goldman, and Kenneth D Poss. In vivo proximity labeling identifies cardiomyocyte protein networks during zebrafish heart regeneration. *eLife*, 10, 2021.

[9774] Eduardo Pulgar, Cornelia Schwayer, Néstor Guerrero, Loreto López, Susana Márquez, Steffen Härtel, Rodrigo Soto, Carl-Philipp Heisenberg, and Miguel L Concha. Apical contacts stemming from incomplete delamination guide progenitor cell allocation through a dragging mechanism. *eLife*, 10, 2021.

[9775] Neha Puri, Amy J Fernandez, Valerie L O'Shea Murray, Sarah McMillan, James L Keck, and James M Berger. The molecular coupling between substrate recognition and ATP turnover in a AAA+ hexameric helicase loader. *eLife*, 10, 2021.

[9776] Cheng Qian, Zhaofa Wu, Rongbo Sun, Huasheng Yu, Jianzhi Zeng, Yi Rao, and Yulong Li. Localization, proteomics, and metabolite profiling reveal a putative vesicular transporter for UDP-glucose. *eLife*, 10, 2021.

[9777] Kang-Ying Qian, Wan-Xin Zeng, Yue Hao, Xian-Ting Zeng, Haowen Liu, Lei Li, Lili Chen, Fumin Tian, Cindy Chang, Qi Hall, Chun-Xue Song, Shangbang Gao, Zhitao Hu, Joshua M Kaplan, Qian Li, and Xia-Jing Tong. Male pheromones modulate synaptic transmission at the C. elegans neuromuscular junction in a sexually dimorphic manner. *eLife*, 10, 2021.

[9778] Xinbo Qiao, Yixiao Zhang, Lisha Sun, Qingtian Ma, Jie Yang, Liping Ai, Jinqi Xue, Guanglei Chen, Hao Zhang, Ce Ji, Xi Gu, Haixin Lei, Yongliang Yang, and Caigang Liu. Association of human breast cancer CD44-/CD24- cells with delayed distant metastasis. *eLife*, 10, 2021.

[9779] Qiang Qiu, Yunming Wu, Limei Ma, Wenjing Xu, Max Hills, Vivekanandan Ramalingam, and C Ron Yu. Acquisition of innate odor preference depends on spontaneous and experiential activities during critical period. *eLife*, 10, 2021.

[9780] Markus Quante, Jasper Iske, Timm Heinbokel, Bhavna N Desai, Hector Rodriguez Cetina Biefer, Yeqi Nian, Felix Krenzien, Tomohisa Matsunaga, Hirofumi Uehara, Ryoichi Maenosono, Haruhito Azuma, Johann Pratschke, Christine S Falk, Tammy Lo, Eric Sheu, Ali Tavakkoli, Reza Abdi, David Perkins, Maria-Luisa Alegre, Alexander S Banks, Hao Zhou, Abdallah Elkhal, and Stefan G Tullius. Restored TDCA and valine levels imitate the effects of bariatric surgery. *eLife*, 10, 2021.

[9781] Tooba Quidwai, Jiaolong Wang, Emma A Hall, Narcis A Petriman, Weihua Leng, Petra Kiesel, Jonathan N Wells, Laura C Murphy, Margaret A Keighren, Joseph A Marsh, Esben Lorentzen, Gaia Pigino, and Pleasantine Mill. A WDR35-dependent coat protein complex transports ciliary membrane cargo vesicles to cilia. *eLife*, 10, 2021.

[9782] Eyad A Qunaibi, Mohamed Helmy, Iman Basheti, and Iyad Sultan. A high rate of COVID-19 vaccine hesitancy in a large-scale survey on Arabs. *eLife*, 10, 2021.

[9783] Georges Raad, Fabrizio Serra, Luc Martin, Marie-Alix Derieppe, Jérôme Gilleron, Vera L Costa, Didier F Pisani, Ez-Zoubir Amri, Michele Trabucchi, and Valerie Grandjean. Paternal multigenerational exposure to an obesogenic diet drives epigenetic predisposition to metabolic diseases in mice. *eLife*, 10, 2021.

[9784] Tomasz Radaszkiewicz, Michaela Nosková, Kristína Gömöryová, Olga Vondálová Blanářová, Katarzyna Anna Radaszkiewicz, Markéta Picková, Ráchel Víchová, TomáŠ Gybeľ, Karol Kaiser, Lucia Demková, Lucia Kučerová, TomáŠ Bárta, David PotěŠil, Zbyněk Zdráhal, Karel Souček, and Vítězslav Bryja. RNF43 inhibits WNT5A-driven signaling and suppresses melanoma invasion and resistance to the targeted therapy. *eLife*, 10, 2021.

[9785] Viktor J Radermacher, Vincent Fernandez, Emma R Schachner, Richard J Butler, Emese M Bordy, Michael Naylor Hudgins, William J de Klerk, Kimberley EJ Chapelle, and Jonah N Choiniere. A new Heterodontosaurus specimen elucidates the unique ventilatory macroevolution of ornithischian dinosaurs. *eLife*, 10, 2021.

[9786] James Rae, Charles Ferguson, Nicholas Ariotti, Richard I Webb, Han-Hao Cheng, James L Mead, James D Riches, Dominic JB Hunter, Nick Martel, Joanne Baltos, Arthur Christopoulos, Nicole S Bryce, Maria Lastra Cagigas, Sachini Fonseka, Marcel E Sayre, Edna C Hardeman, Peter W Gunning, Yann Gambin, Thomas E Hall, and Robert G Parton. A robust method for particulate detection of a genetic tag for 3D electron microscopy. *eLife*, 10, 2021.

[9787] Aditya Raghunandan, Antonio Ladron-de Guevara, Jeffrey Tithof, Humberto Mestre, Ting Du, Maiken Nedergaard, John H Thomas, and Douglas H Kelley. Bulk flow of cerebrospinal fluid observed in periarterial spaces is not an artifact of injection. *eLife*, 10, 2021.

[9788] Palur V Raghuvamsi, Nikhil K Tulsian, Firdaus Samsudin, Xinlei Qian, Kiren Purushotorman, Gu Yue, Mary M Kozma, Wong Y Hwa, Julien Lescar, Peter J Bond, Paul A MacAry, and Ganesh S Anand. SARS-CoV-2 S protein:ACE2 interaction reveals novel allosteric targets. *eLife*, 10, 2021.

[9789] Stephan Raiders, Erik Calvin Black, Andrea Bae, Stephen MacFarlane, Mason Klein, Shai Shaham, and Aakanksha Singhvi. Glia actively sculpt sensory neurons by controlled phagocytosis to tune animal behavior. *eLife*, 10, 2021.

[9790] Surender Rajasekaran, Caleb P Bupp, Mara Leimanis-Laurens, Ankit Shukla, Christopher Russell, Joseph Junewick, Emily Gleason, Elizabeth A VanSickle, Yvonne Edgerly, Bryan M Wittmann, Jeremy W Prokop, and André S Bachmann. Repurposing eflornithine to treat a patient with a rare ODC1 gain-of-function variant disease. *eLife*, 10, 2021.

[9791] Shankar Ramachandran, Navonil Banerjee, Raja Bhattacharya, Michele L Lemons, Jeremy Florman, Christopher M Lambert, Denis Touroutine, Kellianne Alexander, Liliane Schoofs, Mark J Alkema, Isabel Beets, and Michael M Francis. A conserved neuropeptide system links head and body motor circuits to enable adaptive behavior. *eLife*, 10, 2021.

[9792] Dhruva V Raman and Timothy O'Leary. Optimal plasticity for memory maintenance during ongoing synaptic change. *eLife*, 10, 2021.

[9793] Parvathy Ramesh, Nidhi Sharma Dey, Aditya Kanwal, Sudip Mandal, and Lolitika Mandal. Relish plays a dynamic role in the niche to modulate Drosophila blood progenitor homeostasis in development and infection. *eLife*, 10, 2021.

[9794] Vaidhiswaran Ramesh and J Krishnan. Symmetry breaking meets multisite modification. *eLife*, 10, 2021.

[9795] M Ines Pascoal Ramos, Linjie Tian, Emma J de Ruiter, Chang Song, Ana Paucarmayta, Akashdip Singh, Eline Elshof, Saskia V Vijver, Jahangheer Shaik, Jason Bosiacki, Zachary Cusumano, Christina Jensen, Nicholas Willumsen, Morten A Karsdal, Linda Liu, Sol Langermann, Stefan Willems, Dallas Flies, and Linde Meyaard. Cancer immunotherapy by NC410, a LAIR-2 Fc protein blocking human LAIR-collagen interaction. *eLife*, 10, 2021.

[9796] Félix Ramos-León, Matthew J Bush, Joseph W Sallmen, Govind Chandra, Jake Richardson, Kim C Findlay, Joseph R McCormick, and Susan Schlimpert. A conserved cell division protein directly regulates FtsZ dynamics in filamentous and unicellular actinobacteria. *eLife*, 10, 2021.

[9797] David Ranava, Yiying Yang, Luis Orenday-Tapia, FranÇois Rousset, Catherine Turlan, Violette Morales, Lun Cui, Cyril Moulin, Carine Froment, Gladys Munoz, Jérôme Rech, Julien Marcoux, Anne Caumont-Sarcos, Cécile Albenne, David Bikard, and Raffaele Ieva. Lipoprotein DolP supports proper folding of BamA in the bacterial outer membrane promoting fitness upon envelope stress. *eLife*, 10, 2021.

[9798] Nishtha Ranawat and Ichiro Masai. Mechanisms underlying microglial colonization of developing neural retina in zebrafish. *eLife*, 10, 2021.

[9799] Gisela Rangel-Yescas, Cecilia Cervantes, Miguel A Cervantes-Rocha, Esteban Suárez-Delgado, Anastazia T Banaszak, Ernesto Maldonado, Ian Scott Ramsey, Tamara Rosenbaum, and Leon D Islas. Discovery and characterization of Hv1-type proton channels in reef-building corals. *eLife*, 10, 2021.

[9800] Chaitra Rao, Danielle E Frodyma, Siddesh Southekal, Robert A Svoboda, Adrian R Black, Chittibabu Guda, Tomohiro Mizutani, Hans Clevers, Keith R Johnson, Kurt W Fisher, and Robert E Lewis. KSR1- and ERK-dependent translational regulation of the epithelial-to-mesenchymal transition. *eLife*, 10, 2021.

[9801] Brandon G Rasman, Patrick A Forbes, Ryan M Peters, Oscar Ortiz, Ian Franks, J Timothy Inglis, Romeo Chua, and Jean-Sébastien Blouin. Learning to stand with unexpected sensorimotor delays. *eLife*, 10, 2021.

[9802] Manoj K Rathinaswamy, Zied Gaieb, Kaelin D Fleming, Chiara Borsari, Noah J Harris, Brandon E Moeller, Matthias P Wymann, Rommie E Amaro, and John E Burke. Disease-related mutations in PI3K$\gamma$ disrupt regulatory C-terminal dynamics and reveal a path to selective inhibitors. *eLife*, 10, 2021.

[9803] Shannon Rausser, Caroline Trumpff, Marlon A McGill, Alex Junker, Wei Wang, Siu-Hong Ho, Anika Mitchell, Kalpita R Karan, Catherine Monk, Suzanne C Segerstrom, Rebecca G Reed, and Martin Picard. Mitochondrial phenotypes in purified human immune cell subtypes and cell mixtures. *eLife*, 10, 2021.

[9804] Rossana Rauti, Meishar Shahoha, Yael Leichtmann-Bardoogo, Rami Nasser, Eyal Paz, Rina Tamir, Victoria Miller, Tal Babich, Kfir Shaked, Avner Ehrlich, Konstantinos Ioannidis, Yaakov Nahmias, Roded Sharan, Uri Ashery, and Ben Meir Maoz. Effect of SARS-CoV-2 proteins on vascular permeability. *eLife*, 10, 2021.

[9805] Primoz Ravbar, Neil Zhang, and Julie H Simpson. Behavioral evidence for nested central pattern generator control of Drosophila grooming. *eLife*, 10, 2021.

[9806] Kavita Rawat, Anita Tewari, Madeline J Morrisson, Tor D Wager, and Claudia V Jakubzick. Redefining innate natural antibodies as important contributors to anti-tumor immunity. *eLife*, 10, 2021.

[9807] Thomas M Rawlings, Komal Makwana, Deborah M Taylor, Matteo A Molè, Katherine J Fishwick, Maria Tryfonos, Joshua Odendaal, Amelia Hawkes, Magdalena Zernicka-Goetz, Geraldine M Hartshorne, Jan J Brosens, and Emma S Lucas. Modelling the impact of decidual senescence on embryo implantation in human endometrial assembloids. *eLife*, 10, 2021.

[9808] Adria Razzauti and Patrick Laurent. Ectocytosis prevents accumulation of ciliary cargo in C. elegans sensory neurons. *eLife*, 10, 2021.

[9809] William R Reay, Sahar I El Shair, Michael P Geaghan, Carlos Riveros, Elizabeth G Holliday, Mark A McEvoy, Stephen Hancock, Roseanne Peel, Rodney J Scott, John R Attia, and Murray J Cairns. Genetic association and causal inference converge on hyperglycaemia as a modifiable factor to improve lung function. *eLife*, 10, 2021.

[9810] Aspen T Reese, Katia S Chadaideh, Caroline E Diggins, Laura D Schell, Mark Beckel, Peggy Callahan, Roberta Ryan, Melissa Emery Thompson, and Rachel N Carmody. Effects of domestication on the gut microbiota parallel those of human industrialization. *eLife*, 10, 2021.

[9811] Eric Torsten Reifenstein, Ikhwan Bin Khalid, and Richard Kempter. Synaptic learning rules for sequence learning. *eLife*, 10, 2021.

[9812] Amy Reilein, Helen V Kogan, Rachel Misner, Karen Sophia Park, and Daniel Kalderon. Adult stem cells and niche cells segregate gradually from common precursors that build the adult Drosophila ovary during pupal development. *eLife*, 10, 2021.

[9813] Ganna Reint, Zhuokun Li, Kornel Labun, Salla Keskitalo, Inkeri Soppa, Katariina Mamia, Eero Tolo, Monika Szymanska, Leonardo A Meza-Zepeda, Susanne Lorenz, Artur Cieslar-Pobuda, Xian Hu, Diana L Bordin, Judith Staerk, Eivind Valen, Bernhard Schmierer, Markku Varjosalo, Jussi Taipale, and Emma Haapaniemi. Rapid genome editing by CRISPR-Cas9-POLD3 fusion. *eLife*, 10, 2021.

[9814] Fernando MCV Reis, Johannes Y Lee, Sandra Maesta-Pereira, Peter J Schuette, Meghmik Chakerian, Jinhan Liu, Mimi Q La-Vu, Brooke C Tobias, Juliane M Ikebara, Alexandre Hiroaki Kihara, Newton S Canteras, Jonathan C Kao, and Avishek Adhikari. Dorsal periaqueductal gray ensembles represent approach and avoidance states. *eLife*, 10, 2021.

[9815] Juan Carlos Rengifo-Gonzalez, Krystel El Hage, Marie-Jeanne Clément, Emilie Steiner, Vandana Joshi, Pierrick Craveur, Dominique Durand, David Pastré, and Ahmed Bouhss. The cooperative binding of TDP-43 to GU-rich RNA repeats antagonizes TDP-43 aggregation. *eLife*, 10, 2021.

[9816] Rafael Renteria, Christian Cazares, Emily T Baltz, Drew C Schreiner, Ege A Yalcinbas, Thomas Steinkellner, Thomas S Hnasko, and Christina M Gremel. Mechanism for differential recruitment of orbitostriatal transmission during actions and outcomes following chronic alcohol exposure. *eLife*, 10, 2021.

[9817] Francois Rerolle, Emily Dantzer, Andrew A Lover, John M Marshall, Bouasy Hongvanthong, Hugh JW Sturrock, and Adam Bennett. Spatio-temporal associations between deforestation and malaria incidence in Lao PDR. *eLife*, 10, 2021.

[9818] Justyna A Resztak, Allison K Farrell, Henriette Mair-Meijers, Adnan Alazizi, Xiaoquan Wen, Derek E Wildman, Samuele Zilioli, Richard B Slatcher, Roger Pique-Regi, and Francesca Luca. Psychosocial experiences modulate asthma-associated genes through gene-environment interactions. *eLife*, 10, 2021.

[9819] Roger Revilla-i Domingo, Vinoth Babu Veedin Rajan, Monika Waldherr, Günther Prohaczka, Hugo Musset, Lukas Orel, Elliot Gerrard, Moritz Smolka, Alexander Stockinger, Matthias Farlik, Robert J Lucas, Florian Raible, and Kristin Tessmar-Raible. Characterization of cephalic and non-cephalic sensory cell types provides insight into joint photo- and mechanoreceptor evolution. *eLife*, 10, 2021.

[9820] Sequoyah Reynoso, Vanessa Castillo, Gajanan Dattatray Katkar, Inmaculada Lopez-Sanchez, Sahar Taheri, Celia Espinoza, Cristina Rohena, Debashis Sahoo, Pascal Gagneux, and Pradipta Ghosh. GIV/Girdin, a non-receptor modulator for G$\alpha$i/s, regulates spatiotemporal signaling during sperm capacitation and is required for male fertility. *eLife*, 10, 2021.

[9821] William F Richter, Rohan N Shah, and Alexander J Ruthenburg. Non-canonical H3K79me2-dependent pathways promote the survival of MLL-rearranged leukemia. *eLife*, 10, 2021.

[9822] Ekemini AU Riley and Randy Schekman. Open science takes on Parkinson's disease. *eLife*, 10, 2021.

[9823] Regina Rillo-Bohn, Renzo Adilardi, Therese Mitros, Barış Avşaroğlu, Lewis Stevens, Simone Köhler, Joshua Bayes, Clara Wang, Sabrina Lin, K Alienor Baskevitch, Daniel S Rokhsar, and Abby F Dernburg. Analysis of meiosis in Pristionchus pacificus reveals plasticity in homolog pairing and synapsis in the nematode lineage. *eLife*, 10, 2021.

[9824] Nicholas Rinkenberger, Michael E Abrams, Sumit K Matta, John W Schoggins, Neal M Alto, and L David Sibley. Overexpression screen of interferon-stimulated genes identifies RARRES3 as a restrictor of Toxoplasma gondii infection. *eLife*, 10, 2021.

[9825] Brian Ritchey, Qimin Hai, Juying Han, John Barnard, and Jonathan D Smith. Genetic variant in $3'$ untranslated region of the mouse pycard gene regulates inflammasome activity. *eLife*, 10, 2021.

[9826] Blaise Robert, Eyal Y Kimchi, Yurika Watanabe, Tatenda Chakoma, Miao Jing, Yulong Li, and Daniel B Polley. A functional topography within the cholinergic basal forebrain for encoding sensory cues and behavioral reinforcement outcomes. *eLife*, 10, 2021.

[9827] Vincent Robert, Ludivine Therreau, Vivien Chevaleyre, Eude Lepicard, Cécile Viollet, Julie Cognet, Arthur JY Huang, Roman Boehringer, Denis Polygalov, Thomas J McHugh, and Rebecca Ann Piskorowski. Local circuit allowing hypothalamic control of hippocampal area CA2 activity and consequences for CA1. *eLife*, 10, 2021.

[9828] DeElegant Robinson, Michael Place, James Hose, Adam Jochem, and Audrey P Gasch. Natural variation in the consequences of gene overexpression and its implications for evolutionary trajectories. *eLife*, 10, 2021.

[9829] Elektra K Robinson, Pratibha Jagannatha, Sergio Covarrubias, Matthew Cattle, Valeriya Smaliy, Rojin Safavi, Barbara Shapleigh, Robin Abu-Shumays, Miten Jain, Suzanne M Cloonan, Mark Akeson, Angela N Brooks, and Susan Carpenter. Inflammation drives alternative first exon usage to regulate immune genes including a novel iron-regulated isoform of Aim2. *eLife*, 10, 2021.

[9830] Jennifer C Robinson and Mark P Brandon. Skipping ahead: A circuit for representing the past, present, and future. *eLife*, 10, 2021.

[9831] Benoit Roch, Vincent Abramowski, Olivier Etienne, Stefania Musilli, Pierre David, Jean-Baptiste Charbonnier, Isabelle Callebaut, FranÇois D Boussin, and Jean-Pierre de Villartay. An XRCC4 mutant mouse, a model for human X4 syndrome, reveals interplays with Xlf, PAXX, and ATM in lymphoid development. *eLife*, 10, 2021.

[9832] Pierre-Luc Rochon, Catherine Theriault, Aline Giselle Rangel Olguin, and Arjun Krishnaswamy. The cell adhesion molecule Sdk1 shapes assembly of a retinal circuit that detects localized edges. *eLife*, 10, 2021.

[9833] Diana Rodrigues, Yoan Renaud, K VijayRaghavan, Lucas Waltzer, and Maneesha S Inamdar. Differential activation of JAK-STAT signaling reveals functional compartmentalization in Drosophila blood progenitors. *eLife*, 10, 2021.

[9834] Kelsie M Rodriguez, Sara C Buch-Larsen, Ilsa T Kirby, Ivan Rodriguez Siordia, David Hutin, Marit Rasmussen, Denis M Grant, Larry L David, Jason Matthews, Michael L Nielsen, and Michael S Cohen. Chemical genetics and proteome-wide site mapping reveal cysteine MARylation by PARP-7 on immune-relevant protein targets. *eLife*, 10, 2021.

[9835] Alan Rodriguez Carvajal, Irina Grishkovskaya, Carlos Gomez Diaz, Antonia Vogel, Adar Sonn-Segev, Manish S Kushwah, Katrin Schodl, Luiza Deszcz, Zsuzsanna Orban-Nemeth, Shinji Sakamoto, Karl Mechtler, Philipp Kukura, Tim Clausen, David Haselbach, and Fumiyo Ikeda. The linear ubiquitin chain assembly complex (LUBAC) generates heterotypic ubiquitin chains. *eLife*, 10, 2021.

[9836] Sean Rogers, Hanaa Hariri, N Ezgi Wood, Natalie Ortiz Speer, and W Mike Henne. Glucose restriction drives spatial reorganization of mevalonate metabolism. *eLife*, 10, 2021.

[9837] Timothy T Rogers, Christopher R Cox, Qihong Lu, Akihiro Shimotake, Takayuki Kikuchi, Takeharu Kunieda, Susumu Miyamoto, Ryosuke Takahashi, Akio Ikeda, Riki Matsumoto, and Matthew A Lambon Ralph. Evidence for a deep, distributed and dynamic code for animacy in human ventral anterior temporal cortex. *eLife*, 10, 2021.

[9838] Michèle Romanos, Guillaume Allio, Myriam Roussigné, Léa Combres, Nathalie Escalas, Cathy Soula, FranÇois Médevielle, Benjamin Steventon, Ariane Trescases, and Bertrand Bénazéraf. Cell-to-cell heterogeneity in Sox2 and Bra expression guides progenitor motility and destiny. *eLife*, 10, 2021.

[9839] Nicolas Romeo, Alasdair Hastewell, Alexander Mietke, and Jörn Dunkel. Learning developmental mode dynamics from single-cell trajectories. *eLife*, 10, 2021.

[9840] Gabriel E Romero and Laurence O Trussell. Distinct forms of synaptic plasticity during ascending vs descending control of medial olivocochlear efferent neurons. *eLife*, 10, 2021.

[9841] Tahel Ronel, Matthew Harries, Kate Wicks, Theres Oakes, Helen Singleton, Rebecca Dearman, Gavin Maxwell, and Benny Chain. The clonal structure and dynamics of the human T cell response to an organic chemical hapten. *eLife*, 10, 2021.

[9842] Yao Rong, Jinghui Jiang, Yiwei Gao, Jianli Guo, Danfeng Song, Wenhao Liu, Mingmin Zhang, Yan Zhao, Bailong Xiao, and Zhenfeng Liu. TMEM120A contains a specific coenzyme A-binding site and might not mediate poking- or stretch-induced channel activities in cells. *eLife*, 10, 2021.

[9843] Remi Ronzano, Camille Lancelin, Gardave Singh Bhumbra, Robert M Brownstone, and Marco Beato. Proximal and distal spinal neurons innervating multiple synergist and antagonist motor pools. *eLife*, 10, 2021.

[9844] Marion Rosello, Juliette Vougny, FranÇois Czarny, Marina C Mione, Jean-Paul Concordet, Shahad Albadri, and Filippo Del Bene. Precise base editing for the in vivo study of developmental signaling and human pathologies in zebrafish. *eLife*, 10, 2021.

[9845] Ayelet M Rosenberg, Shannon Rausser, Junting Ren, Eugene V Mosharov, Gabriel Sturm, R Todd Ogden, Purvi Patel, Rajesh Kumar Soni, Clay Lacefield, Desmond J Tobin, Ralf Paus, and Martin Picard. Quantitative mapping of human hair greying and reversal in relation to life stress. *eLife*, 10, 2021.

[9846] Matthew Rosenberg, Tony Zhang, Pietro Perona, and Markus Meister. Mice in a labyrinth show rapid learning, sudden insight, and efficient exploration. *eLife*, 10, 2021.

[9847] Sina C Rosenkranz, Artem A Shaposhnykov, Simone Träger, Jan Broder Engler, Maarten E Witte, Vanessa Roth, Vanessa Vieira, Nanne Paauw, Simone Bauer, Celina Schwencke-Westphal, Charlotte Schubert, Lukas Can Bal, Benjamin Schattling, Ole Pless, Jack van Horssen, Marc Freichel, and Manuel A Friese. Enhancing mitochondrial activity in neurons protects against neurodegeneration in a mouse model of multiple sclerosis. *eLife*, 10, 2021.

[9848] Nicole Rosskothen-Kuhl, Alexa N Buck, Kongyan Li, and Jan WH Schnupp. Microsecond interaural time difference discrimination restored by cochlear implants after neonatal deafness. *eLife*, 10, 2021.

[9849] Nina Rouhani and Yael Niv. Signed and unsigned reward prediction errors dynamically enhance learning and memory. *eLife*, 10, 2021.

[9850] Dorota Rousová, Vaishnavi Nivsarkar, Veronika Altmannova, Vivek B Raina, Saskia K Funk, David Liedtke, Petra Janning, Franziska Müller, Heidi Reichle, Gerben Vader, and John R Weir. Novel mechanistic insights into the role of Mer2 as the keystone of meiotic DNA break formation. *eLife*, 10, 2021.

[9851] Yann Roussel, Stephanie F Gaudreau, Emily R Kacer, Mohini Sengupta, and Tuan V Bui. Modeling spinal locomotor circuits for movements in developing zebrafish. *eLife*, 10, 2021.

[9852] Anirban Roy, Meiricris Tomaz da Silva, Raksha Bhat, Kyle R Bohnert, Takao Iwawaki, and Ashok Kumar. The IRE1/XBP1 signaling axis promotes skeletal muscle regeneration through a cell non-autonomous mechanism. *eLife*, 10, 2021.

[9853] Anthony A Ruberto, Aline Gréchez-Cassiau, Sophie Guérin, Luc Martin, Johana S Revel, Mohamed Mehiri, Malayannan Subramaniam, Franck Delaunay, and Michèle Teboul. KLF10 integrates circadian timing and sugar signaling to coordinate hepatic metabolism. *eLife*, 10, 2021.

[9854] Johannes Rudolph, Genevieve Roberts, Uma M Muthurajan, and Karolin Luger. HPF1 and nucleosomes mediate a dramatic switch in activity of PARP1 from polymerase to hydrolase. *eLife*, 10, 2021.

[9855] E Josue Ruiz, Adan Pinto-Fernandez, Andrew P Turnbull, Linxiang Lan, Thomas M Charlton, Hannah C Scott, Andreas Damianou, George Vere, Eva M Riising, Clive Da Costa, Wojciech W Krajewski, David Guerin, Jeffrey D Kearns, Stephanos Ioannidis, Marie Katz, Crystal McKinnon, Jonathan O'Connell, Natalia Moncaut, Ian Rosewell, Emma Nye, Neil Jones, Claire Heride, Malte Gersch, Min Wu, Christopher J Dinsmore, Tim R Hammonds, Sunkyu Kim, David Komander, Sylvie Urbe, Michael J Clague, Benedikt M Kessler, and Axel Behrens. USP28 deletion and small-molecule inhibition destabilizes c-MYC and elicits regression of squamous cell lung carcinoma. *eLife*, 10, 2021.

[9856] Anuththara Rupasinghe, Nikolas Francis, Ji Liu, Zac Bowen, Patrick O Kanold, and Behtash Babadi. Direct extraction of signal and noise correlations from two-photon calcium imaging of ensemble neuronal activity. *eLife*, 10, 2021.

[9857] John P Russell, Xinhong Lim, Alice Santambrogio, Val Yianni, Yasmine Kemkem, Bruce Wang, Matthew Fish, Scott Haston, Anaëlle Grabek, Shirleen Hallang, Emily J Lodge, Amanda L Patist, Andreas Schedl, Patrice Mollard, Roel Nusse, and Cynthia L Andoniadou. Pituitary stem cells produce paracrine WNT signals to control the expansion of their descendant progenitor cells. *eLife*, 10, 2021.

[9858] Emilie V Russler-Germain, Jaeu Yi, Shannon Young, Katherine Nutsch, Harikesh S Wong, Teresa L Ai, Jiani N Chai, Vivek Durai, Daniel H Kaplan, Ronald N Germain, Kenneth M Murphy, and Chyi-Song Hsieh. Gut Helicobacter presentation by multiple dendritic cell subsets enables context-specific regulatory T cell generation. *eLife*, 10, 2021.

[9859] Dylan Gerard Ryan, Ming Yang, Hiran A Prag, Giovanny Rodriguez Blanco, Efterpi Nikitopoulou, Marc Segarra-Mondejar, Christopher A Powell, Tim Young, Nils Burger, Jan Lj Miljkovic, Michal Minczuk, Michael P Murphy, Alex von Kriegsheim, and Christian Frezza. Disruption of the TCA cycle reveals an ATF4-dependent integration of redox and amino acid metabolism. *eLife*, 10, 2021.

[9860] Pia Ryt-Hansen, Jesper Schak Krog, Solvej Østergaard Breum, Charlotte Kristiane Hjulsager, Anders Gorm Pedersen, Ramona Trebbien, and Lars Erik Larsen. Co-circulation of multiple influenza A reassortants in swine harboring genes from seasonal human and swine influenza viruses. *eLife*, 10, 2021.

[9861] Seungjin Ryu, Irina Shchukina, Yun-Hee Youm, Hua Qing, Brandon Hilliard, Tamara Dlugos, Xinbo Zhang, Yuki Yasumoto, Carmen J Booth, Carlos Fernández-Hernando, Yajaira Suárez, Kamal Khanna, Tamas L Horvath, Marcelo O Dietrich, Maxim Artyomov, Andrew Wang, and Vishwa Deep Dixit. Ketogenic diet restrains aging-induced exacerbation of coronavirus infection in mice. *eLife*, 10, 2021.

[9862] Paul V Sabatini, Jine Wang, Alan C Rupp, Alison H Affinati, Jonathan N Flak, Chien Li, David P Olson, and Martin G Myers. tTARGIT AAVs mediate the sensitive and flexible manipulation of intersectional neuronal populations in mice. *eLife*, 10, 2021.

[9863] Akshay Sabnis, Katheryn LH Hagart, Anna Klöckner, Michele Becce, Lindsay E Evans, R Christopher D Furniss, Despoina AI Mavridou, Ronan Murphy, Molly M Stevens, Jane C Davies, Gérald J Larrouy-Maumus, Thomas B Clarke, and Andrew M Edwards. Colistin kills bacteria by targeting lipopolysaccharide in the cytoplasmic membrane. *eLife*, 10, 2021.

[9864] Andrea Sacchetti, Miriam Teeuwssen, Mathijs Verhagen, Rosalie Joosten, Tong Xu, Roberto Stabile, Berdine van der Steen, Martin M Watson, Alem Gusinac, Won Kyu Kim, Inge Ubink, Harmen JG Van de Werken, Arianna Fumagalli, Madelon Paauwe, Jacco Van Rheenen, Owen J Sansom, Onno Kranenburg, and Riccardo Fodde. Phenotypic plasticity underlies local invasion and distant metastasis in colon cancer. *eLife*, 10, 2021.

[9865] Daniela Saderi, Zachary P Schwartz, Charles R Heller, Jacob R Pennington, and Stephen V David. Dissociation of task engagement and arousal effects in auditory cortex and midbrain. *eLife*, 10, 2021.

[9866] Polona Safaric Tepes, Debjani Pal, Trine Lindsted, Ingrid Ibarra, Amaia Lujambio, Vilma Jimenez Sabinina, Serif Senturk, Madison Miller, Navya Korimerla, Jiahao Huang, Lawrence Glassman, Paul Lee, David Zeltsman, Kevin Hyman, Michael Esposito, Gregory J Hannon, and Raffaella Sordella. An epigenetic switch regulates the ontogeny of AXL-positive/EGFR-TKi-resistant cells by modulating miR-335 expression. *eLife*, 10, 2021.

[9867] Arne Sahm, Matthias Platzer, Philipp Koch, Yoshiyuki Henning, Martin Bens, Marco Groth, Hynek Burda, Sabine Begall, Saskia Ting, Moritz Goetz, Paul Van Daele, Magdalena Staniszewska, Jasmin Mona Klose, Pedro Fragoso Costa, Steve Hoffmann, Karol Szafranski, and Philip Dammann. Increased longevity due to sexual activity in mole-rats is associated with transcriptional changes in the HPA stress axis. *eLife*, 10, 2021.

[9868] Darjan Salaj, Anand Subramoney, Ceca Kraisnikovic, Guillaume Bellec, Robert Legenstein, and Wolfgang Maass. Spike frequency adaptation supports network computations on temporally dispersed information. *eLife*, 10, 2021.

[9869] Maria Danielle Sallee, Melissa A Pickett, and Jessica L Feldman. Apical PAR complex proteins protect against programmed epithelial assaults to create a continuous and functional intestinal lumen. *eLife*, 10, 2021.

[9870] Madhuja Samaddar, Jérôme Goudeau, Melissa Sanchez, David H Hall, K Adam Bohnert, Maria Ingaramo, and Cynthia Kenyon. A genetic screen identifies new steps in oocyte maturation that enhance proteostasis in the immortal germ lineage. *eLife*, 10, 2021.

[9871] Adrian L Sanborn, Benjamin T Yeh, Jordan T Feigerle, Cynthia V Hao, Raphael JL Townshend, Erez Lieberman Aiden, Ron O Dror, and Roger D Kornberg. Simple biochemical features underlie transcriptional activation domain diversity and dynamic, fuzzy binding to Mediator. *eLife*, 10, 2021.

[9872] Julia Sánchez-Ceinos, Rocío Guzmán-Ruiz, Oriol Alberto Rangel-Zúñiga, Jaime López-Alcalá, Elena Moreno-Caño, Mercedes Del Río-Moreno, Juan Luis Romero-Cabrera, Pablo Pérez-Martínez, Elsa Maymo-Masip, Joan Vendrell, Sonia Fernández-Veledo, José Manuel Fernández-Real, Jurga Laurencikiene, Mikael Rydén, Antonio Membrives, Raul M Luque, José López-Miranda, and María M Malagón. Impaired mRNA splicing and proteostasis in preadipocytes in obesity-related metabolic disease. *eLife*, 10, 2021.

[9873] Yara E Sánchez-Corrales, Guy B Blanchard, and Katja Röper. Correct regionalization of a tissue primordium is essential for coordinated morphogenesis. *eLife*, 10, 2021.

[9874] David W Sanders, Chanelle C Jumper, Paul J Ackerman, Dan Bracha, Anita Donlic, Hahn Kim, Devin Kenney, Ivan Castello-Serrano, Saori Suzuki, Tomokazu Tamura, Alexander H Tavares, Mohsan Saeed, Alex S Holehouse, Alexander Ploss, Ilya Levental, Florian Douam, Robert F Padera, Bruce D Levy, and Clifford P Brangwynne. SARS-CoV-2 requires cholesterol for viral entry and pathological syncytia formation. *eLife*, 10, 2021.

[9875] Richard Sando and Thomas C Südhof. Latrophilin GPCR signaling mediates synapse formation. *eLife*, 10, 2021.

[9876] Steven R Sando, Nikhil Bhatla, Eugene LQ Lee, and H Robert Horvitz. An hourglass circuit motif transforms a motor program via subcellularly localized muscle calcium signaling and contraction. *eLife*, 10, 2021.

[9877] Madeline Sandoval, Zhe Ying, and Slobodan Beronja. Interplay of opposing fate choices stalls oncogenic growth in murine skin epithelium. *eLife*, 10, 2021.

[9878] Achinta Sannigrahi, Sourav Chowdhury, Bidisha Das, Amrita Banerjee, Animesh Halder, Amaresh Kumar, Mohammed Saleem, Athi N Naganathan, Sanat Karmakar, and Krishnananda Chattopadhyay. The metal cofactor zinc and interacting membranes modulate SOD1 conformation-aggregation landscape in an in vitro ALS model. *eLife*, 10, 2021.

[9879] Andrea Santeford, Aaron Y Lee, Abdoulaye Sene, Lynn M Hassman, Alexey A Sergushichev, Ekaterina Loginicheva, Maxim N Artyomov, Philip A Ruzycki, and Rajendra S Apte. Loss of Mir146b with aging contributes to inflammation and mitochondrial dysfunction in thioglycollate-elicited peritoneal macrophages. *eLife*, 10, 2021.

[9880] Ricardo M Santos and Anton Sirota. Phasic oxygen dynamics confounds fast choline-sensitive biosensor signals in the brain of behaving rodents. *eLife*, 10, 2021.

[9881] Alfonso Santos-Lopez, Christopher W Marshall, Allison L Haas, Caroline Turner, Javier Rasero, and Vaughn S Cooper. The roles of history, chance, and natural selection in the evolution of antibiotic resistance. *eLife*, 10, 2021.

[9882] Gabriela Santos-Rodriguez, Irina Voineagu, and Robert J Weatheritt. Evolutionary dynamics of circular RNAs in primates. *eLife*, 10, 2021.

[9883] Richa Sardana, Carolyn M Highland, Beth E Straight, Christopher F Chavez, J Christopher Fromme, and Scott D Emr. Golgi membrane protein Erd1 Is essential for recycling a subset of Golgi glycosyltransferases. *eLife*, 10, 2021.

[9884] Esther Sasson, Shira Anzi, Batia Bell, Oren Yakovian, Meshi Zorsky, Urban Deutsch, Britta Engelhardt, Eilon Sherman, Gad Vatine, Ron Dzikowski, and Ayal Ben-Zvi. Nano-scale architecture of blood-brain barrier tight-junctions. *eLife*, 10, 2021.

[9885] Hikaru Sato, Juan Santos-González, and Claudia Köhler. Combinations of maternal-specific repressive epigenetic marks in the endosperm control seed dormancy. *eLife*, 10, 2021.

[9886] Dominik Saul, David G Monroe, Jennifer L Rowsey, Robyn Laura Kosinsky, Stephanie J Vos, Madison L Doolittle, Joshua N Farr, and Sundeep Khosla. Modulation of fracture healing by the transient accumulation of senescent cells. *eLife*, 10, 2021.

[9887] Daniel Sauter and Frank Kirchhoff. Evolutionary conflicts and adverse effects of antiviral factors. *eLife*, 10, 2021.

[9888] Chad W Sauvola, Yulia Akbergenova, Karen L Cunningham, Nicole A Aponte-Santiago, and J Troy Littleton. The decoy SNARE Tomosyn sets tonic versus phasic release properties and is required for homeostatic synaptic plasticity. *eLife*, 10, 2021.

[9889] Kristin M Scaplen, Mustafa Talay, John D Fisher, Raphael Cohn, Altar SorkaÇ, Yoshi Aso, Gilad Barnea, and Karla R Kaun. Transsynaptic mapping of Drosophila mushroom body output neurons. *eLife*, 10, 2021.

[9890] Anna M Scarborough, Juliana N Flaherty, Olga V Hunter, Kuanqing Liu, Ashwani Kumar, Chao Xing, Benjamin P Tu, and Nicholas K Conrad. SAM homeostasis is regulated by CFIm-mediated splicing of MAT2A. *eLife*, 10, 2021.

[9891] Mathias Scharmann, Anthony G Rebelo, and John R Pannell. High rates of evolution preceded shifts to sex-biased gene expression in Leucadendron, the most sexually dimorphic angiosperms. *eLife*, 10, 2021.

[9892] Michael Schaum, Edoardo Pinzuti, Alexandra Sebastian, Klaus Lieb, Pascal Fries, Arian Mobascher, Patrick Jung, Michael Wibral, and Oliver Tüscher. Right inferior frontal gyrus implements motor inhibitory control via beta-band oscillations in humans. *eLife*, 10, 2021.

[9893] Isabelle Schiffer, Birgit Gerisch, Kazuto Kawamura, Raymond Laboy, Jennifer Hewitt, Martin Sebastian Denzel, Marcelo A Mori, Siva Vanapalli, Yidong Shen, Orsolya Symmons, and Adam Antebi. miR-1 coordinately regulates lysosomal v-ATPase and biogenesis to impact proteotoxicity and muscle function during aging. *eLife*, 10, 2021.

[9894] Morten Schiøtt and Jacobus J Boomsma. Proteomics reveals synergy between biomass degrading enzymes and inorganic Fenton chemistry in leaf-cutting ant colonies. *eLife*, 10, 2021.

[9895] Domino K Schlegel, Srinivasagan Ramkumar, Johannes von Lintig, and Stephan CF Neuhauss. Disturbed retinoid metabolism upon loss of rlbp1a impairs cone function and leads to subretinal lipid deposits and photoreceptor degeneration in the zebrafish retina. *eLife*, 10, 2021.

[9896] Jenia Schlegel, Gregoire Denay, Rene Wink, Karine Gustavo Pinto, Yvonne Stahl, Julia Schmid, Patrick Blümke, and Rüdiger GW Simon. Control of Arabidopsis shoot stem cell homeostasis by two antagonistic CLE peptide signalling pathways. *eLife*, 10, 2021.

[9897] Markus Frederik Schliffka, Anna Francesca Tortorelli, özge özgüÇ, Ludmilla de Plater, Oliver Polzer, Diane Pelzer, and Jean-Léon Maître. Multiscale analysis of single and double maternal-zygotic Myh9 and Myh10 mutants during mouse preimplantation development. *eLife*, 10, 2021.

[9898] Franca Schmid, Giulia Conti, Patrick Jenny, and Bruno Weber. The severity of microstrokes depends on local vascular topology and baseline perfusion. *eLife*, 10, 2021.

[9899] Calvin M Schmidt and Christina D Smolke. A convolutional neural network for the prediction and forward design of ribozyme-based gene-control elements. *eLife*, 10, 2021.

[9900] Helen Schmidt, Andrea Putnam, Dominique Rasoloson, and Geraldine Seydoux. Protein-based condensation mechanisms drive the assembly of RNA-rich P granules. *eLife*, 10, 2021.

[9901] Ruth Schmidt, Stephen Curry, and Anna Hatch. Creating SPACE to evolve academic assessment. *eLife*, 10, 2021.

[9902] Oskar H Schnaack and Armita Nourmohammad. Optimal evolutionary decision-making to store immune memory. *eLife*, 10, 2021.

[9903] Casey M Schneider-Mizell, Agnes L Bodor, Forrest Collman, Derrick Brittain, Adam Bleckert, Sven Dorkenwald, Nicholas L Turner, Thomas Macrina, Kisuk Lee, Ran Lu, Jingpeng Wu, Jun Zhuang, Anirban Nandi, Brian Hu, JoAnn Buchanan, Marc M Takeno, Russel Torres, Gayathri Mahalingam, Daniel J Bumbarger, Yang Li, Thomas Chartrand, Nico Kemnitz, William M Silversmith, Dodam Ih, Jonathan Zung, Aleksandar Zlateski, Ignacio Tartavull, Sergiy Popovych, William Wong, Manuel Castro, Chris S Jordan, Emmanouil Froudarakis, Lynne Becker, Shelby Suckow, Jacob Reimer, Andreas S Tolias, Costas A Anastassiou, H Sebastian Seung, R Clay Reid, and Nuno MaÇarico da Costa. Structure and function of axo-axonic inhibition. *eLife*, 10, 2021.

[9904] Michael Schoof, Morgane Boone, Lan Wang, Rosalie Lawrence, Adam Frost, and Peter Walter. eIF2B conformation and assembly state regulate the integrated stress response. *eLife*, 10, 2021.

[9905] Maria Schörnig, Xiangchun Ju, Luise Fast, Sebastian Ebert, Anne Weigert, Sabina Kanton, Theresa Schaffer, Nael Nadif Kasri, Barbara Treutlein, Benjamin Marco Peter, Wulf Hevers, and Elena Taverna. Comparison of induced neurons reveals slower structural and functional maturation in humans than in apes. *eLife*, 10, 2021.

[9906] Ara Schorscher-Petcu, Flóra Takács, and Liam E Browne. Scanned optogenetic control of mammalian somatosensory input to map input-specific behavioral outputs. *eLife*, 10, 2021.

[9907] Karin Schrieber, Sarah Catherine Paul, Levke Valena Höche, Andrea Cecilia Salas, Rabi Didszun, Jakob Mößnang, Caroline Müller, Alexandra Erfmeier, and Elisabeth Johanna Eilers. Inbreeding in a dioecious plant has sex- and population origin-specific effects on its interactions with pollinators. *eLife*, 10, 2021.

[9908] Cornelius Schröder, Jonathan Oesterle, Philipp Berens, Takeshi Yoshimatsu, and Tom Baden. Distinct synaptic transfer functions in same-type photoreceptors. *eLife*, 10, 2021.

[9909] Courtney M Schroeder, Sarah A Tomlin, Isabel Mejia Natividad, John R Valenzuela, Janet M Young, and Harmit S Malik. An actin-related protein that is most highly expressed in Drosophila testes is critical for embryonic development. *eLife*, 10, 2021.

[9910] Simon Schug, Frederik Benzing, and Angelika Steger. Presynaptic stochasticity improves energy efficiency and helps alleviate the stability-plasticity dilemma. *eLife*, 10, 2021.

[9911] Auguste Schulz, Christoph Miehl, Michael J Berry, and Julijana Gjorgjieva. The generation of cortical novelty responses through inhibitory plasticity. *eLife*, 10, 2021.

[9912] Dominik Schumacher, Andrea Harms, Silke Bergeler, Erwin Frey, and Lotte Søgaard-Andersen. PomX, a ParA/MinD ATPase activating protein, is a triple regulator of cell division in Myxococcus xanthus. *eLife*, 10, 2021.

[9913] Marc A Schureck, Joseph E Darling, Alan Merk, Jinfeng Shao, Geervani Daggupati, Prakash Srinivasan, Paul Dominic B Olinares, Michael P Rout, Brian T Chait, Kurt Wollenberg, Sriram Subramaniam, and Sanjay A Desai. Malaria parasites use a soluble RhopH complex for erythrocyte invasion and an integral form for nutrient uptake. *eLife*, 10, 2021.

[9914] Sarah Schuster, Jaime Lisack, Ines Subota, Henriette Zimmermann, Christian Reuter, Tobias Mueller, Brooke Morriswood, and Markus Engstler. Unexpected plasticity in the life cycle of Trypanosoma brucei. *eLife*, 10, 2021.

[9915] Alex R Schuurman, Tom DY Reijnders, Anno Saris, Ivan Ramirez Moral, Michiel Schinkel, Justin de Brabander, Christine van Linge, Louis Vermeulen, Brendon P Scicluna, W Joost Wiersinga, Felipe A Vieira Braga, and Tom van der Poll. Integrated single-cell analysis unveils diverging immune features of COVID-19, influenza, and other community-acquired pneumonia. *eLife*, 10, 2021.

[9916] Philipp Schwenk, David J Sheerin, Jathish Ponnu, Anne-Marie Staudt, Klara L Lesch, Elisabeth Lichtenberg, Katalin F Medzihradszky, Ute Hoecker, Eva Klement, András Viczián, and Andreas Hiltbrunner. Uncovering a novel function of the CCR4-NOT complex in phytochrome A-mediated light signalling in plants. *eLife*, 10, 2021.

[9917] Srijan Seal, Guha Dharmarajan, and Imroze Khan. Evolution of pathogen tolerance and emerging infections: A missing experimental paradigm. *eLife*, 10, 2021.

[9918] Sofiene Seef, Julien Herrou, Paul de Boissier, Laetitia My, Gael Brasseur, Donovan Robert, Rikesh Jain, Romain Mercier, Eric Cascales, Bianca H Habermann, and Tâm Mignot. A Tad-like apparatus is required for contact-dependent prey killing in predatory social bacteria. *eLife*, 10, 2021.

[9919] Cristina Segalin, Jalani Williams, Tomomi Karigo, May Hui, Moriel Zelikowsky, Jennifer J Sun, Pietro Perona, David J Anderson, and Ann Kennedy. The Mouse Action Recognition System (MARS) software pipeline for automated analysis of social behaviors in mice. *eLife*, 10, 2021.

[9920] Mona Seifert, Subhas C Bera, Pauline van Nies, Robert N Kirchdoerfer, Ashleigh Shannon, Thi-Tuyet-Nhung Le, Xiangzhi Meng, Hongjie Xia, James M Wood, Lawrence D Harris, Flavia S Papini, Jamie J Arnold, Steven Almo, Tyler L Grove, Pei-Yong Shi, Yan Xiang, Bruno Canard, Martin Depken, Craig E Cameron, and David Dulin. Inhibition of SARS-CoV-2 polymerase by nucleotide analogs from a single-molecule perspective. *eLife*, 10, 2021.

[9921] Yaniv Sela, Marieke MB Hoekstra, and Paul Franken. Sub-minute prediction of brain temperature based on sleep–wake state in the mouse. *eLife*, 10, 2021.

[9922] Yogev Sela, Jinyang Li, Paola Kuri, Allyson J Merrell, Ning Li, Chris Lengner, Pantelis Rompolas, and Ben Z Stanger. Dissecting phenotypic transitions in metastatic disease via photoconversion-based isolation. *eLife*, 10, 2021.

[9923] Ali Seleit, Alexander Aulehla, and Alexandre Paix. Endogenous protein tagging in medaka using a simplified CRISPR/Cas9 knock-in approach. *eLife*, 10, 2021.

[9924] Titas Sengupta, Noelle L Koonce, Nabor Vázquez-Martínez, Mark W Moyle, Leighton H Duncan, Sarah E Emerson, Xiaofei Han, Lin Shao, Yicong Wu, Anthony Santella, Li Fan, Zhirong Bao, William A Mohler, Hari Shroff, and Daniel A Colón-Ramos. Differential adhesion regulates neurite placement via a retrograde zippering mechanism. *eLife*, 10, 2021.

[9925] Alexander J Sercel, Alexander N Patananan, Tianxing Man, Ting-Hsiang Wu, Amy K Yu, Garret W Guyot, Shahrooz Rabizadeh, Kayvan R Niazi, Pei-Yu Chiou, and Michael A Teitell. Stable transplantation of human mitochondrial DNA by high-throughput, pressurized isolated mitochondrial delivery. *eLife*, 10, 2021.

[9926] Juana Serrano-Lopez, Shailaja Hegde, Sachin Kumar, Josefina Serrano, Jing Fang, Ashley M Wellendorf, Paul A Roche, Yamileth Rangel, Leolene J Carrington, Hartmut Geiger, H Leighton Grimes, Sanjiv Luther, Ivan Maillard, Joaquin Sanchez-Garcia, Daniel T Starczynowski, and Jose A Cancelas. Inflammation rapidly recruits mammalian GMP and MDP from bone marrow into regional lymphatics. *eLife*, 10, 2021.

[9927] Saumil Sethna, Wadih M Zein, Sehar Riaz, Arnaud PJ Giese, Julie M Schultz, Todd Duncan, Robert B Hufnagel, Carmen C Brewer, Andrew J Griffith, T Michael Redmond, Saima Riazuddin, Thomas B Friedman, and Zubair M Ahmed. Proposed therapy, developed in a Pcdh15-deficient mouse, for progressive loss of vision in human Usher syndrome. *eLife*, 10, 2021.

[9928] Mireia Seuma, Andre J Faure, Marta Badia, Ben Lehner, and Benedetta Bolognesi. The genetic landscape for amyloid beta fibril nucleation accurately discriminates familial Alzheimer's disease mutations. *eLife*, 10, 2021.

[9929] Bikash Shakya, Saurabh D Patel, Yoshihiko Tani, and Elizabeth S Egan. Erythrocyte CD55 mediates the internalization of Plasmodium falciparum parasites. *eLife*, 10, 2021.

[9930] Viplendra PS Shakya, William A Barbeau, Tianyao Xiao, Christina S Knutson, Max H Schuler, and Adam L Hughes. A nuclear-based quality control pathway for non-imported mitochondrial proteins. *eLife*, 10, 2021.

[9931] Zhao Shan, Leike Li, Constance Lynn Atkins, Meng Wang, Yankai Wen, Jongmin Jeong, Nicolas F Moreno, Dechun Feng, Xun Gui, Ningyan Zhang, Chun Geun Lee, Jack A Elias, William M Lee, Bin Gao, Fong Wilson Lam, Zhiqiang An, and Cynthia Ju. Chitinase 3-like-1 contributes to acetaminophen-induced liver injury by promoting hepatic platelet recruitment. *eLife*, 10, 2021.

[9932] Ophir Shani, Yael Raz, Lea Monteran, Ye'ela Scharff, Oshrat Levi-Galibov, Or Megides, Hila Shacham, Noam Cohen, Dana Silverbush, Camilla Avivi, Roded Sharan, Asaf Madi, Ruth Scherz-Shouval, Iris Barshack, Ilan Tsarfaty, and Neta Erez. Evolution of fibroblasts in the lung metastatic microenvironment is driven by stage-specific transcriptional plasticity. *eLife*, 10, 2021.

[9933] Naima G Sharaf, Mona Shahgholi, Esther Kim, Jeffrey Y Lai, David G VanderVelde, Allen T Lee, and Douglas C Rees. Characterization of the ABC methionine transporter from Neisseria meningitidis reveals that lipidated MetQ is required for interaction. *eLife*, 10, 2021.

[9934] Abhinav Sharma, Diego Vidaurre, Jan Vesper, Alfons Schnitzler, and Esther Florin. Differential dopaminergic modulation of spontaneous cortico–subthalamic activity in Parkinson's disease. *eLife*, 10, 2021.

[9935] Deepika Sharma, Yilin Yu, Leyao Shen, Guo-Fang Zhang, and Courtney M Karner. SLC1A5 provides glutamine and asparagine necessary for bone development in mice. *eLife*, 10, 2021.

[9936] Kunal Sharma, Neeraj Dhar, Vivek V Thacker, Thomas M Simonet, Francois Signorino-Gelo, Graham W Knott, and John D McKinney. Dynamic persistence of UPEC intracellular bacterial communities in a human bladder-chip model of urinary tract infection. *eLife*, 10, 2021.

[9937] Medha Sharma, Tao Jiang, Zi Chen Jiang, Carlos E Moguel-Lehmer, and Tony JC Harris. Emergence of a smooth interface from growth of a dendritic network against a mechanosensitive contractile material. *eLife*, 10, 2021.

[9938] Swati Sharma and Richa Rikhy. Spatiotemporal recruitment of RhoGTPase protein GRAF inhibits actomyosin ring constriction in Drosophila cellularization. *eLife*, 10, 2021.

[9939] Simon A Sharples and Gareth B Miles. Maturation of persistent and hyperpolarization-activated inward currents shapes the differential activation of motoneuron subtypes during postnatal development. *eLife*, 10, 2021.

[9940] Louis Shekhtman, Miquel Navasa, Natasha Sansone, Gonzalo Crespo, Gitanjali Subramanya, Tje Lin Chung, E Fabian Cardozo-Ojeda, Sofía Pérez-del Pulgar, Alan S Perelson, Scott J Cotler, Xavier Forns, Susan L Uprichard, and Harel Dahari. Modeling hepatitis C virus kinetics during liver transplantation reveals the role of the liver in virus clearance. *eLife*, 10, 2021.

[9941] Jianpeng Sheng, Qi Chen, Xiaoting Wu, Yu Wen Dong, Johannes Mayer, Junlei Zhang, Lin Wang, Xueli Bai, Tingbo Liang, Yang Ho Sung, Wilson Wen Bin Goh, Franca Ronchese, and Christiane Ruedl. Fate mapping analysis reveals a novel murine dermal migratory Langerhans-like cell population. *eLife*, 10, 2021.

[9942] Wanhui Sheng, Scott W Harden, Yalun Tan, Eric G Krause, and Charles J Frazier. Dendritic osmosensors modulate activity-induced calcium influx in oxytocinergic magnocellular neurons of the mouse PVN. *eLife*, 10, 2021.

[9943] Courtney T Shepard, Amanda M Pocratsky, Brandon L Brown, Morgan A Van Rijswijck, Rachel M Zalla, Darlene A Burke, Johnny R Morehouse, Amberley S Riegler, Scott R Whittemore, and David SK Magnuson. Silencing long ascending propriospinal neurons after spinal cord injury improves hindlimb stepping in the adult rat. *eLife*, 10, 2021.

[9944] Kristin M Sherrard, Maureen Cetera, and Sally Horne-Badovinac. DAAM mediates the assembly of long-lived, treadmilling stress fibers in collectively migrating epithelial cells in Drosophila. *eLife*, 10, 2021.

[9945] Jue Shi, Jui-Hsia Weng, and Timothy J Mitchison. Immunomodulatory drug discovery from herbal medicines: Insights from organ-specific activity and xenobiotic defenses. *eLife*, 10, 2021.

[9946] Ching-Hua Shih and Justin Fay. Cis-regulatory variants affect gene expression dynamics in yeast. *eLife*, 10, 2021.

[9947] Kenichi Shimada, John A Bachman, Jeremy L Muhlich, and Timothy J Mitchison. shinyDepMap, a tool to identify targetable cancer genes and their functional connections from Cancer Dependency Map data. *eLife*, 10, 2021.

[9948] Ortal Shimon-Raz, Roy Salomon, Miki Bloch, Gabi Aisenberg Romano, Yaara Yeshurun, Adi Ulmer Yaniv, Orna Zagoory-Sharon, and Ruth Feldman. Mother brain is wired for social moments. *eLife*, 10, 2021.

[9949] Daisuke Shimura, Esther Nuebel, Rachel Baum, Steven E Valdez, Shaohua Xiao, Junco S Warren, Joseph A Palatinus, TingTing Hong, Jared Rutter, and Robin M Shaw. Protective mitochondrial fission induced by stress-responsive protein GJA1-20k. *eLife*, 10, 2021.

[9950] Eun Ju Shin, Yunsil Jang, Soyoun Kim, Hoseok Kim, Xinying Cai, Hyunjung Lee, Jung Hoon Sul, Sung-Hyun Lee, Yeonseung Chung, Daeyeol Lee, and Min Whan Jung. Robust and distributed neural representation of action values. *eLife*, 10, 2021.

[9951] Mackenzie M Shipley, Vidya Mangala Prasad, Laura E Doepker, Adam Dingens, Duncan K Ralph, Elias Harkins, Amrit Dhar, Dana Arenz, Vrasha Chohan, Haidyn Weight, Kishor Mandaliya, Jesse D Bloom, Frederick A Matsen, Kelly K Lee, and Julie M Overbaugh. Functional development of a V3/glycan-specific broadly neutralizing antibody isolated from a case of HIV superinfection. *eLife*, 10, 2021.

[9952] Yekaterina Shulgina and Sean R Eddy. A computational screen for alternative genetic codes in over 250,000 genomes. *eLife*, 10, 2021.

[9953] Jiahui Si, Jiachen Li, Canqing Yu, Yu Guo, Zheng Bian, Iona Millwood, Ling Yang, Robin Walters, Yiping Chen, Huaidong Du, Li Yin, Jianwei Chen, Junshi Chen, Zhengming Chen, Liming Li, Liming Liang, and Jun Lv. Improved lipidomic profile mediates the effects of adherence to healthy lifestyles on coronary heart disease. *eLife*, 10, 2021.

[9954] Jiahui Si, Songchun Yang, Dianjianyi Sun, Canqing Yu, Yu Guo, Yifei Lin, Iona Y Millwood, Robin G Walters, Ling Yang, Yiping Chen, Huaidong Du, Yujie Hua, Jingchao Liu, Junshi Chen, Zhengming Chen, Wei Chen, Jun Lv, Liming Liang, and Liming Li. Epigenome-wide analysis of DNA methylation and coronary heart disease: a nested case-control study. *eLife*, 10, 2021.

[9955] Ana C Sias, Ashleigh K Morse, Sherry Wang, Venuz Y Greenfield, Caitlin M Goodpaster, Tyler M Wrenn, Andrew M Wikenheiser, Sandra M Holley, Carlos Cepeda, Michael S Levine, and Kate M Wassum. A bidirectional corticoamygdala circuit for the encoding and retrieval of detailed reward memories. *eLife*, 10, 2021.

[9956] Sumit Siddharth, Sheetal Parida, Nethaji Muniraj, Shawn Hercules, David Lim, Arumugam Nagalingam, Chenguang Wang, Balazs Gyorffy, Juliet M Daniel, and Dipali Sharma. Concomitant activation of GLI1 and Notch1 contributes to racial disparity of human triple negative breast cancer progression. *eLife*, 10, 2021.

[9957] Jessica M Sidisky, Daniel Weaver, Sarrah Hussain, Meryem Okumus, Russell Caratenuto, and Daniel Babcock. Mayday sustains trans-synaptic BMP signaling required for synaptic maintenance with age. *eLife*, 10, 2021.

[9958] Joshua H Siegle, Peter Ledochowitsch, Xiaoxuan Jia, Daniel J Millman, Gabriel K Ocker, Shiella Caldejon, Linzy Casal, Andy Cho, Daniel J Denman, Séverine Durand, Peter A Groblewski, Gregg Heller, India Kato, Sara Kivikas, Jérôme Lecoq, Chelsea Nayan, Kiet Ngo, Philip R Nicovich, Kat North, Tamina K Ramirez, Jackie Swapp, Xana Waughman, Ali Williford, Shawn R Olsen, Christof Koch, Michael A Buice, and Saskia EJ de Vries. Reconciling functional differences in populations of neurons recorded with two-photon imaging and electrophysiology. *eLife*, 10, 2021.

[9959] Justin N Siemian, Miguel A Arenivar, Sarah Sarsfield, Cara B Borja, Lydia J Erbaugh, Andrew L Eagle, Alfred J Robison, Gina Leinninger, and Yeka Aponte. An excitatory lateral hypothalamic circuit orchestrating pain behaviors in mice. *eLife*, 10, 2021.

[9960] Catia AP Silva, Annik Yalnizyan-Carson, M Victoria Fernández Busch, Mike van Zwieten, Matthijs Verhage, and Christian Lohmann. Activity-dependent regulation of mitochondrial motility in developing cortical dendrites. *eLife*, 10, 2021.

[9961] João L Silva-Filho, João CK Dos-Santos, Carla Judice, Dario Beraldi, Kannan Venugopal, Diogenes Lima, Helder I Nakaya, Erich V De Paula, Stefanie CP Lopes, Marcus VG Lacerda, Matthias Marti, and Fabio TM Costa. Total parasite biomass but not peripheral parasitaemia is associated with endothelial and haematological perturbations in Plasmodium vivax patients. *eLife*, 10, 2021.

[9962] Dina Simkin, Kelly A Marshall, Carlos G Vanoye, Reshma R Desai, Bernabe I Bustos, Brandon N Piyevsky, Juan A Ortega, Marc Forrest, Gabriella L Robertson, Peter Penzes, Linda C Laux, Steven J Lubbe, John J Millichap, Alfred L George, and Evangelos Kiskinis. Dyshomeostatic modulation of Ca2+-activated K+ channels in a human neuronal model of KCNQ2 encephalopathy. *eLife*, 10, 2021.

[9963] Eléanor Simon, Carlos Jiménez-Jiménez, Irene Seijo-Barandiarán, Gustavo Aguilar, David Sánchez-Hernández, Adrián Aguirre-Tamaral, Laura González-Méndez, Pedro Ripoll, and Isabel Guerrero. Glypicans define unique roles for the Hedgehog co-receptors boi and ihog in cytoneme-mediated gradient formation. *eLife*, 10, 2021.

[9964] Amanjot Singh, Joern Hulsmeier, Arvind Reddy Kandi, Sai Shruti Pothapragada, Jens Hillebrand, Arnas Petrauskas, Khushboo Agrawal, Krishnan RT, Devasena Thiagarajan, Deepa Jayaprakashappa, K VijayRaghavan, Mani Ramaswami, and Baskar Bakthavachalu. Antagonistic roles for Ataxin-2 structured and disordered domains in RNP condensation. *eLife*, 10, 2021.

[9965] Francois Singh, Alan R Prescott, Philippa Rosewell, Graeme Ball, Alastair D Reith, and Ian G Ganley. Pharmacological rescue of impaired mitophagy in Parkinson's disease-related LRRK2 G2019S knock-in mice. *eLife*, 10, 2021.

[9966] Moirangthem Kiran Singh, Parisa Zangoui, Yuki Yamanaka, and Linda J Kenney. Genetic code expansion enables visualization of Salmonella type three secretion system components and secreted effectors. *eLife*, 10, 2021.

[9967] Raushan K Singh, McKenzie Jonely, Evan Leslie, Nick A Rejali, Rodrigo Noriega, and Brenda L Bass. Transient kinetic studies of the antiviral Drosophila Dicer-2 reveal roles of ATP in self–nonself discrimination. *eLife*, 10, 2021.

[9968] Sumnima Singh, Patricia Bastos-Amador, Jessica Ann Thompson, Mauro Truglio, Bahtiyar Yilmaz, Silvia Cardoso, Daniel Sobral, and Miguel P Soares. Glycan-based shaping of the microbiota during primate evolution. *eLife*, 10, 2021.

[9969] Nasa Sinnott-Armstrong, Sahin Naqvi, Manuel Rivas, and Jonathan K Pritchard. GWAS of three molecular traits highlights core genes and pathways alongside a highly polygenic background. *eLife*, 10, 2021.

[9970] Sahana Sitaraman, Gnaneshwar Yadav, Vandana Agarwal, Shaista Jabeen, Shivangi Verma, Meha Jadhav, and Vatsala Thirumalai. Gjd2b-mediated gap junctions promote glutamatergic synapse formation and dendritic elaboration in Purkinje neurons. *eLife*, 10, 2021.

[9971] Lise M Sjøgaard-Frich, Andreas Prestel, Emilie S Pedersen, Marc Severin, Kristian Kølby Kristensen, Johan G Olsen, Birthe B Kragelund, and Stine Falsig Pedersen. Dynamic Na+/H+ exchanger 1 (NHE1) – calmodulin complexes of varying stoichiometry and structure regulate Ca2+-dependent NHE1 activation. *eLife*, 10, 2021.

[9972] Katalin Skrapits, Miklós Sárvári, Imre Farkas, Balázs Göcz, Szabolcs Takács, éva Rumpler, Viktória Váczi, Csaba Vastagh, Gergely Rácz, András Matolcsy, Norbert Solymosi, Szilárd Póliska, Blanka Tóth, Ferenc Erdélyi, Gábor Szabó, Michael D Culler, Cecile Allet, Ludovica Cotellessa, Vincent Prévot, Paolo Giacobini, and Erik Hrabovszky. The cryptic gonadotropin-releasing hormone neuronal system of human basal ganglia. *eLife*, 10, 2021.

[9973] David Smith, Geetha Kannan, Isabelle Coppens, Fengrong Wang, Hoa Mai Nguyen, Aude Cerutti, Einar B Olafsson, Patrick A Rimple, Tracey L Schultz, Nayanna M Mercado Soto, Manlio Di Cristina, Sébastien Besteiro, and Vern B Carruthers. Toxoplasma TgATG9 is critical for autophagy and long-term persistence in tissue cysts. *eLife*, 10, 2021.

[9974] David RM Smith, Laura Temime, and Lulla Opatowski. Microbiome-pathogen interactions drive epidemiological dynamics of antibiotic resistance: A modeling study applied to nosocomial pathogen control. *eLife*, 10, 2021.

[9975] Logan K Smith, Kareem Fawaz, and Bebhinn Treanor. Galectin-9 regulates the threshold of B cell activation and autoimmunity. *eLife*, 10, 2021.

[9976] Quentin M Smith, Alessio V Inchingolo, Madalina-Daniela Mihailescu, Hongsheng Dai, and Neil M Kad. Single-molecule imaging reveals the concerted release of myosin from regulated thin filaments. *eLife*, 10, 2021.

[9977] Aaron Snoberger, Bipasha Barua, Jennifer L Atherton, Henry Shuman, Eva Forgacs, Yale E Goldman, Donald A Winkelmann, and E Michael Ostap. Myosin with hypertrophic cardiac mutation R712L has a decreased working stroke which is rescued by omecamtiv mecarbil. *eLife*, 10, 2021.

[9978] Tatiana P Soares da Costa, Cody J Hall, Santosh Panjikar, Jessica A Wyllie, Rebecca M Christoff, Saadi Bayat, Mark D Hulett, Belinda M Abbott, Anthony R Gendall, and Matthew A Perugini. Towards novel herbicide modes of action by inhibiting lysine biosynthesis in plants. *eLife*, 10, 2021.

[9979] Filip Sobczak, Patricia Pais-Roldán, Kengo Takahashi, and Xin Yu. Decoding the brain state-dependent relationship between pupil dynamics and resting state fMRI signal fluctuation. *eLife*, 10, 2021.

[9980] Sacha Sokoloski, Amir Aschner, and Ruben Coen-Cagli. Modelling the neural code in large populations of correlated neurons. *eLife*, 10, 2021.

[9981] David Sokolov, Emily R Sechrest, Yekai Wang, Connor Nevin, Jianhai Du, and Saravanan Kolandaivelu. Nuclear NAD+-biosynthetic enzyme NMNAT1 facilitates development and early survival of retinal neurons. *eLife*, 10, 2021.

[9982] Rene Solano Fonseca, Patrick Metang, Nathan Egge, Yingjian Liu, Kielen R Zuurbier, Karthigayini Sivaprakasam, Shawn Shirazi, Ashleigh Chuah, Sonja LB Arneaud, Genevieve Konopka, Dong Qian, and Peter M Douglas. Glycolytic preconditioning in astrocytes mitigates trauma-induced neurodegeneration. *eLife*, 10, 2021.

[9983] Magdalena Solyga and Tania Rinaldi Barkat. Emergence and function of cortical offset responses in sound termination detection. *eLife*, 10, 2021.

[9984] Robert A Sommer, Jerry T DeWitt, Raymond Tan, and Douglas R Kellogg. Growth-dependent signals drive an increase in early G1 cyclin concentration to link cell cycle entry with cell growth. *eLife*, 10, 2021.

[9985] Aixin Song, Zachary Hazlett, Dulith Abeykoon, Jeremy Dortch, Andrew Dillon, Justin Curtiss, Sarah Bollinger Martinez, Christopher P Hill, Clinton Yu, Lan Huang, David Fushman, Robert E Cohen, and Tingting Yao. Branched ubiquitin chain binding and deubiquitination by UCH37 facilitate proteasome clearance of stress-induced inclusions. *eLife*, 10, 2021.

[9986] Hongki Song, Thomas L Torng, Amy S Orr, Axel T Brunger, and William T Wickner. Sec17/Sec18 can support membrane fusion without help from completion of SNARE zippering. *eLife*, 10, 2021.

[9987] Hongki Song and William T Wickner. Fusion of tethered membranes can be driven by Sec18/NSF and Sec17/$\alpha$SNAP without HOPS. *eLife*, 10, 2021.

[9988] Huan Song, Fang Fang, Henrik Larsson, Nancy L Pedersen, Patrik KE Magnusson, Catarina Almqvist, and Unnur A Valdimarsdóttir. Loss of a co-twin at birth and subsequent risk of psychiatric disorders. *eLife*, 10, 2021.

[9989] Kangcheng Song, Miao Wei, Wenjun Guo, Li Quan, Yunlu Kang, Jing-Xiang Wu, and Lei Chen. Structural basis for human TRPC5 channel inhibition by two distinct inhibitors. *eLife*, 10, 2021.

[9990] Minmin Song, Guangfeng Zhao, Haixiang Sun, Simin Yao, Zhenhua Zhou, Peipei Jiang, Qianwen Wu, Hui Zhu, Huiyan Wang, Chenyan Dai, Jingmei Wang, Ruotian Li, Yun Cao, Haining Lv, Dan Liu, Jianwu Dai, Yan Zhou, and Yali Hu. circPTPN12/miR-21–5 p/$\Delta$Np63$\alpha$ pathway contributes to human endometrial fibrosis. *eLife*, 10, 2021.

[9991] Yonghyun Song and Changbong Hyeon. Cost-precision trade-off relation determines the optimal morphogen gradient for accurate biological pattern formation. *eLife*, 10, 2021.

[9992] Aylesse Sordillo and Cornelia I Bargmann. Behavioral control by depolarized and hyperpolarized states of an integrating neuron. *eLife*, 10, 2021.

[9993] Elliott D SoRelle, Joanne Dai, Emmanuela N Bonglack, Emma M Heckenberg, Jeffrey Y Zhou, Stephanie N Giamberardino, Jeffrey A Bailey, Simon G Gregory, Cliburn Chan, and Micah A Luftig. Single-cell RNA-seq reveals transcriptomic heterogeneity mediated by host–pathogen dynamics in lymphoblastoid cell lines. *eLife*, 10, 2021.

[9994] Pierpaolo Sorrentino, Caio Seguin, Rosaria Rucco, Marianna Liparoti, Emahnuel Troisi Lopez, Simona Bonavita, Mario Quarantelli, Giuseppe Sorrentino, Viktor Jirsa, and Andrew Zalesky. The structural connectome constrains fast brain dynamics. *eLife*, 10, 2021.

[9995] Célia Souque, José Antonio Escudero, and R Craig MacLean. Integron activity accelerates the evolution of antibiotic resistance. *eLife*, 10, 2021.

[9996] Alexander Soutschek, Susanna C Weber, Thorsten Kahnt, Boris B Quednow, and Philippe N Tobler. Opioid antagonism modulates wanting-related frontostriatal connectivity. *eLife*, 10, 2021.

[9997] Frank W Soveg, Johannes Schwerk, Nandan S Gokhale, Karen Cerosaletti, Julian R Smith, Erola Pairo-Castineira, Alison M Kell, Adriana Forero, Shivam A Zaver, Katharina Esser-Nobis, Justin A Roby, Tien-Ying Hsiang, Snehal Ozarkar, Jonathan M Clingan, Eileen T McAnarney, Amy EL Stone, Uma Malhotra, Cate Speake, Joseph Perez, Chiraag Balu, Eric J Allenspach, Jennifer L Hyde, Vineet D Menachery, Saumendra N Sarkar, Joshua J Woodward, Daniel B Stetson, John Kenneth Baillie, Jane H Buckner, Michael Gale, and Ram Savan. Endomembrane targeting of human OAS1 p46 augments antiviral activity. *eLife*, 10, 2021.

[9998] Davide Spalla, Isabel Maria Cornacchia, and Alessandro Treves. Continuous attractors for dynamic memories. *eLife*, 10, 2021.

[9999] Brian Spurlock, Danitra Parker, Malay Kumar Basu, Anita Hjelmeland, Sajina GC, Shanrun Liu, Gene P Siegal, Alan Gunter, Aida Moran, and Kasturi Mitra. Fine-tuned repression of Drp1-driven mitochondrial fission primes a 'stem/progenitor-like state' to support neoplastic transformation. *eLife*, 10, 2021.

[10000] Lotte Victoria Winther Stagsted, Eoghan Thomas O'Leary, Karoline Kragh Ebbesen, and Thomas Birkballe Hansen. The RNA-binding protein SFPQ preserves long-intron splicing and regulates circRNA biogenesis in mammals. *eLife*, 10, 2021.

[10001] Sara N Stahley, Lena P Basta, Rishabh Sharan, and Danelle Devenport. Celsr1 adhesive interactions mediate the asymmetric organization of planar polarity complexes. *eLife*, 10, 2021.

[10002] Fernanda I Staquicini, Amin Hajitou, Wouter HP Driessen, Bettina Proneth, Marina Cardó-Vila, Daniela I Staquicini, Christopher Markosian, Maria Hoh, Mauro Cortez, Anupama Hooda-Nehra, Mohammed Jaloudi, Israel T Silva, Jaqueline Buttura, Diana N Nunes, Emmanuel Dias-Neto, Bedrich Eckhardt, Javier Ruiz-Ramírez, Prashant Dogra, Zhihui Wang, Vittorio Cristini, Martin Trepel, Robin Anderson, Richard L Sidman, Juri G Gelovani, Massimo Cristofanilli, Gabriel N Hortobagyi, Zaver M Bhujwalla, Stephen K Burley, Wadih Arap, and Renata Pasqualini. Targeting a cell surface vitamin D receptor on tumor-associated macrophages in triple-negative breast cancer. *eLife*, 10, 2021.

[10003] Benjamin J Stauch, Alina Peter, Heike Schuler, and Pascal Fries. Stimulus-specific plasticity in human visual gamma-band activity and functional connectivity. *eLife*, 10, 2021.

[10004] Oskar Staufer, Jacqueline A De Lora, Eleonora Bailoni, Alisina Bazrafshan, Amelie S Benk, Kevin Jahnke, Zachary A Manzer, Lado Otrin, Telmo Díez Pérez, Judee Sharon, Jan Steinkühler, Katarzyna P Adamala, Bruna Jacobson, Marileen Dogterom, Kerstin Göpfrich, Darko Stefanovic, Susan R Atlas, Michael Grunze, Matthew R Lakin, Andrew P Shreve, Joachim P Spatz, and Gabriel P López. Building a community to engineer synthetic cells and organelles from the bottom-up. *eLife*, 10, 2021.

[10005] Elizabeth M Steenkiste, Jason D Berndt, Carissa Pilling, Christopher Simpkins, and Jonathan A Cooper. A Cas-BCAR3 co-regulatory circuit controls lamellipodia dynamics. *eLife*, 10, 2021.

[10006] Erich Stefan, Richard Obexer, Susanne Hofmann, Khanh Vu Huu, Yichao Huang, Nina Morgner, Hiroaki Suga, and Robert Tampé. De novo macrocyclic peptides dissect energy coupling of a heterodimeric ABC transporter by multimode allosteric inhibition. *eLife*, 10, 2021.

[10007] Elsa Steinfath, Adrian Palacios-Muñoz, Julian R Rottschäfer, Deniz Yuezak, and Jan Clemens. Fast and accurate annotation of acoustic signals with deep neural networks. *eLife*, 10, 2021.

[10008] Barbara Stekas, Steve Yeo, Alice Troitskaia, Masayoshi Honda, Sei Sho, Maria Spies, and Yann R Chemla. Switch-like control of helicase processivity by single-stranded DNA binding protein. *eLife*, 10, 2021.

[10009] Vita Stepanova, Kaja Ewa Moczulska, Guido N Vacano, Ilia Kurochkin, Xiangchun Ju, Stephan Riesenberg, Dominik Macak, Tomislav Maricic, Linda Dombrowski, Maria Schörnig, Konstantinos Anastassiadis, Oliver Baker, Ronald Naumann, Ekaterina Khrameeva, Anna Vanushkina, Elena Stekolshchikova, Alina Egorova, Anna Tkachev, Randall Mazzarino, Nathan Duval, Dmitri Zubkov, Patrick Giavalisco, Terry G Wilkinson, David Patterson, Philipp Khaitovich, and Svante Pääbo. Reduced purine biosynthesis in humans after their divergence from Neandertals. *eLife*, 10, 2021.

[10010] Tilman Stephani, Alice Hodapp, Mina Jamshidi Idaji, Arno Villringer, and Vadim V Nikulin. Neural excitability and sensory input determine intensity perception with opposing directions in initial cortical responses. *eLife*, 10, 2021.

[10011] Gabriella R Sterne, Hideo Otsuna, Barry J Dickson, and Kristin Scott. Classification and genetic targeting of cell types in the primary taste and premotor center of the adult Drosophila brain. *eLife*, 10, 2021.

[10012] Sharon R Stevens, Colleen M Longley, Yuki Ogawa, Lindsay H Teliska, Anithachristy S Arumanayagam, Supna Nair, Juan A Oses-Prieto, Alma L Burlingame, Matthew D Cykowski, Mingshan Xue, and Matthew N Rasband. Ankyrin-R regulates fast-spiking interneuron excitability through perineuronal nets and Kv3.1b K+ channels. *eLife*, 10, 2021.

[10013] Alessandro Stirpe, Nora Guidotti, Sarah J Northall, Sinan Kilic, Alexandre Hainard, Oscar Vadas, Beat Fierz, and Thomas Schalch. SUV39 SET domains mediate crosstalk of heterochromatic histone marks. *eLife*, 10, 2021.

[10014] Oliver Stirrup, Joseph Hughes, Matthew Parker, David G Partridge, James G Shepherd, James Blackstone, Francesc Coll, Alexander Keeley, Benjamin B Lindsey, Aleksandra Marek, Christine Peters, Joshua B Singer, Asif Tamuri, Thushan I de Silva, Emma C Thomson, and Judith Breuer. Rapid feedback on hospital onset SARS-CoV-2 infections combining epidemiological and sequencing data. *eLife*, 10, 2021.

[10015] Anika Stockert, Michael Schwartze, David Poeppel, Alfred Anwander, and Sonja A Kotz. Temporo-cerebellar connectivity underlies timing constraints in audition. *eLife*, 10, 2021.

[10016] Barbara H Stokes, Satish K Dhingra, Kelly Rubiano, Sachel Mok, Judith Straimer, Nina F Gnädig, Ioanna Deni, Kyra A Schindler, Jade R Bath, Kurt E Ward, Josefine Striepen, Tomas Yeo, Leila S Ross, Eric Legrand, Frédéric Ariey, Clark H Cunningham, Issa M Souleymane, Adama Gansané, Romaric Nzoumbou-Boko, Claudette Ndayikunda, Abdunoor M Kabanywanyi, Aline Uwimana, Samuel J Smith, Olimatou Kolley, Mathieu Ndounga, Marian Warsame, Rithea Leang, FranÇois Nosten, Timothy JC Anderson, Philip J Rosenthal, Didier Ménard, and David A Fidock. Plasmodium falciparum K13 mutations in Africa and Asia impact artemisinin resistance and parasite fitness. *eLife*, 10, 2021.

[10017] Julia Stokes, Arielle Freed, Rebecca Bornstein, Kevin N Su, John Snell, Amanda Pan, Grace X Sun, Kyung Yeon Park, Sangwook Jung, Hailey Worstman, Brittany M Johnson, Philip G Morgan, Margaret M Sedensky, and Simon C Johnson. Mechanisms underlying neonate-specific metabolic effects of volatile anesthetics. *eLife*, 10, 2021.

[10018] Elitsa Stoyanova, Michael Riad, Anjana Rao, and Nathaniel Heintz. 5-Hydroxymethylcytosine-mediated active demethylation is required for mammalian neuronal differentiation and function. *eLife*, 10, 2021.

[10019] Nicholas Strash, Sophia DeLuca, Geovanni L Janer Carattini, Soon Chul Heo, Ryne Gorsuch, and Nenad Bursac. Human Erbb2-induced Erk activity robustly stimulates cycling and functional remodeling of rat and human cardiomyocytes. *eLife*, 10, 2021.

[10020] Monique S Straub, Carolina Alvadia, Marta Sawicka, and Raimund Dutzler. Cryo-EM structures of the caspase-activated protein XKR9 involved in apoptotic lipid scrambling. *eLife*, 10, 2021.

[10021] John R Strikwerda and Steven M Sine. Unmasking coupling between channel gating and ion permeation in the muscle nicotinic receptor. *eLife*, 10, 2021.

[10022] Lior Strinkovsky, Evgeny Havkin, Ruby Shalom-Feuerstein, and Yonatan Savir. Spatial correlations constrain cellular lifespan and pattern formation in corneal epithelium homeostasis. *eLife*, 10, 2021.

[10023] Lauren K Strohacker, Douglas R Mackay, Madeline A Whitney, Genevieve C Couldwell, Wesley I Sundquist, and Katharine S Ullman. Identification of abscission checkpoint bodies as structures that regulate ESCRT factors to control abscission timing. *eLife*, 10, 2021.

[10024] Amy R Strom, Ronald J Biggs, Edward J Banigan, Xiaotao Wang, Katherine Chiu, Cameron Herman, Jimena Collado, Feng Yue, Joan C Ritland Politz, Leah J Tait, David Scalzo, Agnes Telling, Mark Groudine, Clifford P Brangwynne, John F Marko, and Andrew D Stephens. HP1$\alpha$ is a chromatin crosslinker that controls nuclear and mitotic chromosome mechanics. *eLife*, 10, 2021.

[10025] Lisa M Strong, Chunmei Chang, Julia F Riley, C Alexander Boecker, Thomas G Flower, Cosmo Z Buffalo, Xuefeng Ren, Andrea KH Stavoe, Erika LF Holzbaur, and James H Hurley. Structural basis for membrane recruitment of ATG16L1 by WIPI2 in autophagy. *eLife*, 10, 2021.

[10026] Andreas Strube, Michael Rose, Sepideh Fazeli, and Christian Büchel. The temporal and spectral characteristics of expectations and prediction errors in pain and thermoception. *eLife*, 10, 2021.

[10027] Jeremie Subrini and James Turner. Y chromosome functions in mammalian spermatogenesis. *eLife*, 10, 2021.

[10028] Victoria J Sugrue, Joseph Alan Zoller, Pritika Narayan, Ake T Lu, Oscar J Ortega-Recalde, Matthew J Grant, C Simon Bawden, Skye R Rudiger, Amin Haghani, Donna M Bond, Reuben R Hore, Michael Garratt, Karen E Sears, Nan Wang, Xiangdong William Yang, Russell G Snell, Timothy A Hore, and Steve Horvath. Castration delays epigenetic aging and feminizes DNA methylation at androgen-regulated loci. *eLife*, 10, 2021.

[10029] Jody A Summers and Elizabeth Martinez. Visually induced changes in cytokine production in the chick choroid. *eLife*, 10, 2021.

[10030] Michael Chas Sumner, Steven B Torrisi, Donna G Brickner, and Jason H Brickner. Random sub-diffusion and capture of genes by the nuclear pore reduces dynamics and coordinates interchromosomal movement. *eLife*, 10, 2021.

[10031] Dawei Sun, Lewis Evans, Francesca Perrone, Vanesa Sokleva, Kyungtae Lim, Saba Rezakhani, Matthias Lutolf, Matthias Zilbauer, and Emma L Rawlins. A functional genetic toolbox for human tissue-derived organoids. *eLife*, 10, 2021.

[10032] Qi Sun, Gehua Zhen, Tuo Peter Li, Qiaoyue Guo, Yusheng Li, Weiping Su, Peng Xue, Xiao Wang, Mei Wan, Yun Guan, Xinzhong Dong, Shaohua Li, Ming Cai, and Xu Cao. Parathyroid hormone attenuates osteoarthritis pain by remodeling subchondral bone in mice. *eLife*, 10, 2021.

[10033] Xuelong Sun, Shigang Yue, and Michael Mangan. How the insect central complex could coordinate multimodal navigation. *eLife*, 10, 2021.

[10034] Zhen Sun, Yingpeng Yao, Menghao You, Jingjing Liu, Wenhui Guo, Zhihong Qi, Zhao Wang, Fang Wang, Weiping Yuan, and Shuyang Yu. The kinase PDK1 is critical for promoting T follicular helper cell differentiation. *eLife*, 10, 2021.

[10035] Molly C Sutherland, Deanna L Mendez, Shalon E Babbitt, Dustin E Tillman, Olga Melnikov, Nathan L Tran, Noah T Prizant, Andrea L Collier, and Robert G Kranz. In vitro reconstitution reveals major differences between human and bacterial cytochrome c synthases. *eLife*, 10, 2021.

[10036] Mototaka Suzuki, Jaan Aru, and Matthew E Larkum. Double-$\mu$Periscope, a tool for multilayer optical recordings, optogenetic stimulations or both. *eLife*, 10, 2021.

[10037] Sho W Suzuki, Akihiko Oishi, Nadia Nikulin, Jeff R Jorgensen, Matthew G Baile, and Scott D Emr. A PX-BAR protein Mvp1/SNX8 and a dynamin-like GTPase Vps1 drive endosomal recycling. *eLife*, 10, 2021.

[10038] Sarah Lauren Svensson and Cynthia Mira Sharma. RNase III-mediated processing of a trans-acting bacterial sRNA and its cis-encoded antagonist. *eLife*, 10, 2021.

[10039] Elliott Swanson, Cara Lord, Julian Reading, Alexander T Heubeck, Palak C Genge, Zachary Thomson, Morgan DA Weiss, Xiao-jun Li, Adam K Savage, Richard R Green, Troy R Torgerson, Thomas F Bumol, Lucas T Graybuck, and Peter J Skene. Simultaneous trimodal single-cell measurement of transcripts, epitopes, and chromatin accessibility using TEA-seq. *eLife*, 10, 2021.

[10040] S Zachary Swartz, Hieu T Nguyen, Brennan C McEwan, Mark E Adamo, Iain M Cheeseman, and Arminja N Kettenbach. Selective dephosphorylation by PP2A-B55 directs the meiosis I-meiosis II transition in oocytes. *eLife*, 10, 2021.

[10041] Sarah G Swygert, Dejun Lin, Stephanie Portillo-Ledesma, Po-Yen Lin, Dakota R Hunt, Cheng-Fu Kao, Tamar Schlick, William S Noble, and Toshio Tsukiyama. Local chromatin fiber folding represses transcription and loop extrusion in quiescent cells. *eLife*, 10, 2021.

[10042] Balázs Szigeti, Laura Kartner, Allan Blemings, Fernando Rosas, Amanda Feilding, David J Nutt, Robin L Carhart-Harris, and David Erritzoe. Self-blinding citizen science to explore psychedelic microdosing. *eLife*, 10, 2021.

[10043] Agata Szuba, Fouzia Bano, Gerard Castro-Linares, Francois Iv, Manos Mavrakis, Ralf P Richter, Aurélie Bertin, and Gijsje H Koenderink. Membrane binding controls ordered self-assembly of animal septins. *eLife*, 10, 2021.

[10044] Hannah Tabakh, Adelle P McFarland, Maureen K Thomason, Alex J Pollock, Rochelle C Glover, Shivam A Zaver, and Joshua J Woodward. 4-Hydroxy-2-nonenal antimicrobial toxicity is neutralized by an intracellular pathogen. *eLife*, 10, 2021.

[10045] Hiroaki Tachiwana, Mariko Dacher, Kazumitsu Maehara, Akihito Harada, Yosuke Seto, Ryohei Katayama, Yasuyuki Ohkawa, Hiroshi Kimura, Hitoshi Kurumizaka, and Noriko Saitoh. Chromatin structure-dependent histone incorporation revealed by a genome-wide deposition assay. *eLife*, 10, 2021.

[10046] Michael A Taffe and Nicholas W Gilpin. Racial inequity in grant funding from the US National Institutes of Health. *eLife*, 10, 2021.

[10047] Luke Stephen Tain, Robert Sehlke, Ralf Leslie Meilenbrock, Thomas Leech, Jonathan Paulitz, Manopriya Chokkalingam, Nagarjuna Nagaraj, Sebastian Grönke, Jenny Fröhlich, Ilian Atanassov, Matthias Mann, Andreas Beyer, and Linda Partridge. Tissue-specific modulation of gene expression in response to lowered insulin signalling in Drosophila. *eLife*, 10, 2021.

[10048] Tamar Tak, Giulio Prevedello, Gaël Simon, Noémie Paillon, Camélia Benlabiod, Caroline Marty, Isabelle Plo, Ken R Duffy, and Leïla Perié. HSPCs display within-family homogeneity in differentiation and proliferation despite population heterogeneity. *eLife*, 10, 2021.

[10049] Yu Takagi, Laurence Tudor Hunt, Mark W Woolrich, Timothy EJ Behrens, and Miriam C Klein-Flügge. Adapting non-invasive human recordings along multiple task-axes shows unfolding of spontaneous and over-trained choice. *eLife*, 10, 2021.

[10050] Satoshi Takahashi, Akinori Kanai, Hiroshi Okuda, Ryo Miyamoto, Yosuke Komata, Takeshi Kawamura, Hirotaka Matsui, Toshiya Inaba, Akifumi Takaori-Kondo, and Akihiko Yokoyama. HBO1-MLL interaction promotes AF4/ENL/P-TEFb-mediated leukemogenesis. *eLife*, 10, 2021.

[10051] Takahiro Takano, Amanda M Wahl, Kai-Ting Huang, Takanori Narita, John Rugis, James Sneyd, and David I Yule. Highly localized intracellular Ca2+ signals promote optimal salivary gland fluid secretion. *eLife*, 10, 2021.

[10052] Sumiko Takao, Lauren Forbes, Masahiro Uni, Shuyuan Cheng, Jose Mario Bello Pineda, Yusuke Tarumoto, Paolo Cifani, Gerard Minuesa, Celine Chen, Michael G Kharas, Robert K Bradley, Christopher R Vakoc, Richard P Koche, and Alex Kentsis. Convergent organization of aberrant MYB complex controls oncogenic gene expression in acute myeloid leukemia. *eLife*, 10, 2021.

[10053] Jun Takatoh, Jae Hong Park, Jinghao Lu, Shun Li, PM Thompson, Bao-Xia Han, Shengli Zhao, David Kleinfeld, Beth Friedman, and Fan Wang. Constructing an adult orofacial premotor atlas in Allen mouse CCF. *eLife*, 10, 2021.

[10054] Hiroki Takechi, Satoko Hakeda-Suzuki, Yohei Nitta, Yuichi Ishiwata, Riku Iwanaga, Makoto Sato, Atsushi Sugie, and Takashi Suzuki. Glial insulin regulates cooperative or antagonistic Golden goal/Flamingo interactions during photoreceptor axon guidance. *eLife*, 10, 2021.

[10055] Yuka Takemon, Joel M Chick, Isabela Gerdes Gyuricza, Daniel A Skelly, Olivier Devuyst, Steven P Gygi, Gary A Churchill, and Ron Korstanje. Proteomic and transcriptomic profiling reveal different aspects of aging in the kidney. *eLife*, 10, 2021.

[10056] Deng Tan, Rui Chen, Yuejian Mo, Shu Gu, Jiao Ma, Wei Xu, Xibin Lu, Huiyu He, Fan Jiang, Weimin Fan, Yili Wang, Xi Chen, and Wei Huang. Quantitative control of noise in mammalian gene expression by dynamic histone regulation. *eLife*, 10, 2021.

[10057] Milton Tan, Anthony K Redmond, Helen Dooley, Ryo Nozu, Keiichi Sato, Shigehiro Kuraku, Sergey Koren, Adam M Phillippy, Alistair DM Dove, and Timothy Read. The whale shark genome reveals patterns of vertebrate gene family evolution. *eLife*, 10, 2021.

[10058] Yukiko Tando, Hitoshi Hiura, Asuka Takehara, Yumi Ito-Matsuoka, Takahiro Arima, and Yasuhisa Matsui. Epi-mutations for spermatogenic defects by maternal exposure to di(2-ethylhexyl) phthalate. *eLife*, 10, 2021.

[10059] Cheng-Chia Tang, Christian D Castro Andrade, Maureen J O'Meara, Sung-Hee Yoon, Tadatoshi Sato, Daniel J Brooks, Mary L Bouxsein, Janaina da Silva Martins, Jinhua Wang, Nathanael S Gray, Barbara Misof, Paul Roschger, Stephane Blouin, Klaus Klaushofer, Annegreet Velduis-Vlug, Yosta Vegting, Clifford J Rosen, Daniel O'Connell, Thomas B Sundberg, Ramnik J Xavier, Peter Ung, Avner Schlessinger, Henry M Kronenberg, Rebecca Berdeaux, Marc Foretz, and Marc N Wein. Dual targeting of salt inducible kinases and CSF1R uncouples bone formation and bone resorption. *eLife*, 10, 2021.

[10060] Wenbo Tang, Justin D Shin, and Shantanu P Jadhav. Multiple time-scales of decision-making in the hippocampus and prefrontal cortex. *eLife*, 10, 2021.

[10061] Xiaotian Tang, Yongguo Cao, Gunjan Arora, Jesse Hwang, Andaleeb Sajid, Courtney L Brown, Sameet Mehta, Alejandro Marín-López, Yu-Min Chuang, Ming-Jie Wu, Hongwei Ma, Utpal Pal, Sukanya Narasimhan, and Erol Fikrig. The Lyme disease agent co-opts adiponectin receptor-mediated signaling in its arthropod vector. *eLife*, 10, 2021.

[10062] Wucheng Tao, Joel Lee, Xiumin Chen, Javier Díaz-Alonso, Jing Zhou, Samuel Pleasure, and Roger A Nicoll. Synaptic memory requires CaMKII. *eLife*, 10, 2021.

[10063] Alicia Tapias, David Lázaro, Bo-Kun Yin, Seyed Mohammad Mahdi Rasa, Anna Krepelova, Erika Kelmer Sacramento, Paulius Grigaravicius, Philipp Koch, Joanna Kirkpatrick, Alessandro Ori, Francesco Neri, and Zhao-Qi Wang. HAT cofactor TRRAP modulates microtubule dynamics via SP1 signaling to prevent neurodegeneration. *eLife*, 10, 2021.

[10064] Alexander J Tarashansky, Jacob M Musser, Margarita Khariton, Pengyang Li, Detlev Arendt, Stephen R Quake, and Bo Wang. Mapping single-cell atlases throughout Metazoa unravels cell type evolution. *eLife*, 10, 2021.

[10065] Nick Taubert, Michael Stettler, Ramona Siebert, Silvia Spadacenta, Louisa Sting, Peter Dicke, Peter Thier, and Martin A Giese. Shape-invariant encoding of dynamic primate facial expressions in human perception. *eLife*, 10, 2021.

[10066] Helen M Tauc, Imilce A Rodriguez-Fernandez, Jason A Hackney, Michal Pawlak, Tal Ronnen Oron, Jerome Korzelius, Hagar F Moussa, Subhra Chaudhuri, Zora Modrusan, Bruce A Edgar, and Heinrich Jasper. Age-related changes in polycomb gene regulation disrupt lineage fidelity in intestinal stem cells. *eLife*, 10, 2021.

[10067] Bassam Tawfik, Joana S Martins, Sébastien Houy, Cordelia Imig, Paulo S Pinheiro, Sonja M Wojcik, Nils Brose, Benjamin H Cooper, and Jakob Balslev Sørensen. Synaptotagmin-7 places dense-core vesicles at the cell membrane to promote Munc13-2- and Ca2+-dependent priming. *eLife*, 10, 2021.

[10068] James A Taylor, Yeonee Seol, Jagat Budhathoki, Keir C Neuman, and Kiyoshi Mizuuchi. CTP and parS coordinate ParB partition complex dynamics and ParA-ATPase activation for ParABS-mediated DNA partitioning. *eLife*, 10, 2021.

[10069] Sanne ten Oever and Andrea E Martin. An oscillating computational model can track pseudo-rhythmic speech by using linguistic predictions. *eLife*, 10, 2021.

[10070] Dmitry Ter-Ovanesyan, Maia Norman, Roey Lazarovits, Wendy Trieu, Ju-Hyun Lee, George M Church, and David R Walt. Framework for rapid comparison of extracellular vesicle isolation methods. *eLife*, 10, 2021.

[10071] Lauren Tereshko, Ya Gao, Brian A Cary, Gina G Turrigiano, and Piali Sengupta. Ciliary neuropeptidergic signaling dynamically regulates excitatory synapses in postnatal neocortical pyramidal neurons. *eLife*, 10, 2021.

[10072] Federico Tessadori, Erika Tsingos, Enrico Sandro Colizzi, Fabian Kruse, Susanne C van den Brink, Malou van den Boogaard, Vincent M Christoffels, Roeland MH Merks, and Jeroen Bakkers. Twisting of the zebrafish heart tube during cardiac looping is a tbx5-dependent and tissue-intrinsic process. *eLife*, 10, 2021.

[10073] Pratish Thakore, Michael G Alvarado, Sher Ali, Amreen Mughal, Paulo W Pires, Evan Yamasaki, Harry AT Pritchard, Brant E Isakson, Cam Ha T Tran, and Scott Earley. Brain endothelial cell TRPA1 channels initiate neurovascular coupling. *eLife*, 10, 2021.

[10074] Dibesh Thapa, Joäo de Sousa Valente, Brentton Barrett, Matthew John Smith, Fulye Argunhan, Sheng Y Lee, Sofya Nikitochkina, Xenia Kodji, and Susan D Brain. Dysfunctional TRPM8 signalling in the vascular response to environmental cold in ageing. *eLife*, 10, 2021.

[10075] Jordana K Thibado, Jean-Yves Tano, Joon Lee, Leslie Salas-Estrada, Davide Provasi, Alexa Strauss, Joao Marcelo Lamim Ribeiro, Guoqing Xiang, Johannes Broichhagen, Marta Filizola, Martin J Lohse, and Joshua Levitz. Differences in interactions between transmembrane domains tune the activation of metabotropic glutamate receptors. *eLife*, 10, 2021.

[10076] Armin W Thomas, Felix Molter, and Ian Krajbich. Uncovering the computational mechanisms underlying many-alternative choice. *eLife*, 10, 2021.

[10077] Sydney P Thomas and John M Denu. Short-chain fatty acids activate acetyltransferase p300. *eLife*, 10, 2021.

[10078] Olivier Thomine, Samuel Alizon, Corentin Boennec, Marc Barthelemy, and Mircea Sofonea. Emerging dynamics from high-resolution spatial numerical epidemics. *eLife*, 10, 2021.

[10079] Jakob Thyrring and Lloyd S Peck. Global gradients in intertidal species richness and functional groups. *eLife*, 10, 2021.

[10080] Fang Tian, Bei Tong, Liang Sun, Shengchao Shi, Bin Zheng, Zibin Wang, Xianchi Dong, and Peng Zheng. N501Y mutation of spike protein in SARS-CoV-2 strengthens its binding to receptor ACE2. *eLife*, 10, 2021.

[10081] Miaomiao Tian, Fengqi Hao, Xin Jin, Xue Sun, Ying Jiang, Yang Wang, Dan Li, Tianyi Chang, Yingying Zou, Pinghui Peng, Chaoyi Xia, Jia Liu, Yuanxi Li, Ping Wang, Yunpeng Feng, and Min Wei. ACLY ubiquitination by CUL3-KLHL25 induces the reprogramming of fatty acid metabolism to facilitate iTreg differentiation. *eLife*, 10, 2021.

[10082] Ben C Tilly, Gillian E Chalkley, Jan A van der Knaap, Yuri M Moshkin, Tsung Wai Kan, Dick HW Dekkers, Jeroen AA Demmers, and C Peter Verrijzer. In vivo analysis reveals that ATP-hydrolysis couples remodeling to SWI/SNF release from chromatin. *eLife*, 10, 2021.

[10083] Courtney Tindle, MacKenzie Fuller, Ayden Fonseca, Sahar Taheri, Stella-Rita Ibeawuchi, Nathan Beutler, Gajanan Dattatray Katkar, Amanraj Claire, Vanessa Castillo, Moises Hernandez, Hana Russo, Jason Duran, Laura E Crotty Alexander, Ann Tipps, Grace Lin, Patricia A Thistlethwaite, Ranajoy Chattopadhyay, Thomas F Rogers, Debashis Sahoo, Pradipta Ghosh, and Soumita Das. Adult stem cell-derived complete lung organoid models emulate lung disease in COVID-19. *eLife*, 10, 2021.

[10084] Ana B Tinoco, Antón Barreiro-Iglesias, Luis Alfonso Yañez Guerra, Jérôme Delroisse, Ya Zhang, Elizabeth F Gunner, Cleidiane G Zampronio, Alexandra M Jones, Michaela Egertová, and Maurice R Elphick. Ancient role of sulfakinin/cholecystokinin-type signalling in inhibitory regulation of feeding processes revealed in an echinoderm. *eLife*, 10, 2021.

[10085] Luca Tirinato, Maria Grazia Marafioti, Francesca Pagliari, Jeannette Jansen, Ilenia Aversa, Rachel Hanley, Clelia Nisticò, Daniel Garcia-Calderón, Geraldine Genard, Joana Filipa Guerreiro, Francesco Saverio Costanzo, and Joao Seco. Lipid droplets and ferritin heavy chain: a devilish liaison in human cancer cell radioresistance. *eLife*, 10, 2021.

[10086] Alexei V Tkachenko, Sergei Maslov, Tong Wang, Ahmed Elbana, George N Wong, and Nigel Goldenfeld. Stochastic social behavior coupled to COVID-19 dynamics leads to waves, plateaus, and an endemic state. *eLife*, 10, 2021.

[10087] Nicolai J Toft, Trine V Axelsen, Helene L Pedersen, Marco Mele, Mark Burton, Eva Balling, Tonje Johansen, Mads Thomassen, Peer M Christiansen, and Ebbe Boedtkjer. Acid-base transporters and pH dynamics in human breast carcinomas predict proliferative activity, metastasis, and survival. *eLife*, 10, 2021.

[10088] Sherzod A Tokamov, Ting Su, Anne Ullyot, and Richard G Fehon. Negative feedback couples Hippo pathway activation with Kibra degradation independent of Yorkie-mediated transcription. *eLife*, 10, 2021.

[10089] Jakub Tomala, Petra Weberova, Barbora Tomalova, Zuzana Jiraskova Zakostelska, Ladislav Sivak, Jirina Kovarova, and Marek Kovar. IL-2/JES6-1 mAb complexes dramatically increase sensitivity to LPS through IFN-$\gamma$ production by CD25+Foxp3- T cells. *eLife*, 10, 2021.

[10090] Ai Phuong S Tong, Alex P Vaz, John H Wittig, Sara K Inati, and Kareem A Zaghloul. Ripples reflect a spectrum of synchronous spiking activity in human anterior temporal lobe. *eLife*, 10, 2021.

[10091] Gerry Tonkin-Hill, Inigo Martincorena, Roberto Amato, Andrew RJ Lawson, Moritz Gerstung, Ian Johnston, David K Jackson, Naomi Park, Stefanie V Lensing, Michael A Quail, Sónia GonÇalves, Cristina Ariani, Michael Spencer Chapman, William L Hamilton, Luke W Meredith, Grant Hall, Aminu S Jahun, Yasmin Chaudhry, Myra Hosmillo, Malte L Pinckert, Iliana Georgana, Anna Yakovleva, Laura G Caller, Sarah L Caddy, Theresa Feltwell, Fahad A Khokhar, Charlotte J Houldcroft, Martin D Curran, Surendra Parmar, Alex Alderton, Rachel Nelson, Ewan M Harrison, John Sillitoe, Stephen D Bentley, Jeffrey C Barrett, M Estee Torok, Ian G Goodfellow, Cordelia Langford, and Dominic Kwiatkowski. Patterns of within-host genetic diversity in SARS-CoV-2. *eLife*, 10, 2021.

[10092] Jaspreet Toor, Susy Echeverria-Londono, Xiang Li, Kaja Abbas, Emily D Carter, Hannah E Clapham, Andrew Clark, Margaret J de Villiers, Kirsten Eilertson, Matthew Ferrari, Ivane Gamkrelidze, Timothy B Hallett, Wes R Hinsley, Daniel Hogan, John H Huber, Michael L Jackson, Kevin Jean, Mark Jit, Andromachi Karachaliou, Petra Klepac, Alicia Kraay, Justin Lessler, Xi Li, Benjamin A Lopman, Tewodaj Mengistu, C Jessica E Metcalf, Sean M Moore, Shevanthi Nayagam, Timos Papadopoulos, T Alex Perkins, Allison Portnoy, Homie Razavi, Devin Razavi-Shearer, Stephen Resch, Colin Sanderson, Steven Sweet, Yvonne Tam, Hira Tanvir, Quan Tran Minh, Caroline L Trotter, Shaun A Truelove, Emilia Vynnycky, Neff Walker, Amy Winter, Kim Woodruff, Neil M Ferguson, and Katy AM Gaythorpe. Lives saved with vaccination for 10 pathogens across 112 countries in a pre-COVID-19 world. *eLife*, 10, 2021.

[10093] Margaret E Torrence, Michael R MacArthur, Aaron M Hosios, Alexander J Valvezan, John M Asara, James R Mitchell, and Brendan D Manning. The mTORC1-mediated activation of ATF4 promotes protein and glutathione synthesis downstream of growth signals. *eLife*, 10, 2021.

[10094] Abel Torres-Espín, Austin Chou, J Russell Huie, Nikos Kyritsis, Pavan S Upadhyayula, and Adam R Ferguson. Reproducible analysis of disease space via principal components using the novel R package syndRomics. *eLife*, 10, 2021.

[10095] Abel Torres-Espín, Jenny Haefeli, Reza Ehsanian, Dolores Torres, Carlos A Almeida, J Russell Huie, Austin Chou, Dmitriy Morozov, Nicole Sanderson, Benjamin Dirlikov, Catherine G Suen, Jessica L Nielson, Nikos Kyritsis, Debra D Hemmerle, Jason F Talbott, Geoffrey T Manley, Sanjay S Dhall, William D Whetstone, Jacqueline C Bresnahan, Michael S Beattie, Stephen L McKenna, Jonathan Z Pan, Adam R Ferguson, and MS Beattie. Topological network analysis of patient similarity for precision management of acute blood pressure in spinal cord injury. *eLife*, 10, 2021.

[10096] Stéphanie Torrino, Victor Tiroille, Bastien Dolfi, Maeva Dufies, Charlotte Hinault, Laurent Bonesso, Sonia Dagnino, Jennifer Uhler, Marie Irondelle, Anne-sophie Gay, Lucile Fleuriot, Delphine Debayle, Sandra Lacas-Gervais, Mireille Cormont, Thomas Bertero, Frederic Bost, Jerome Gilleron, and Stephan Clavel. UBTD1 regulates ceramide balance and endolysosomal positioning to coordinate EGFR signaling. *eLife*, 10, 2021.

[10097] Chiara Tortelli, Marco Turi, David Charles Burr, and Paola Binda. Objective pupillometry shows that perceptual styles covary with autistic-like personality traits. *eLife*, 10, 2021.

[10098] Chirine Toufaily, Jérôme Fortin, Carlos AI Alonso, Evelyne Lapointe, Xiang Zhou, Yorgui Santiago-Andres, Yeu-Farn Lin, Yiming Cui, Ying Wang, Dominic Devost, Ferdinand Roelfsema, Frederik Steyn, Aylin C Hanyaloglu, Terence E Hébert, Tatiana Fiordelisio, Derek Boerboom, and Daniel J Bernard. Addition of a carboxy-terminal tail to the normally tailless gonadotropin-releasing hormone receptor impairs fertility in female mice. *eLife*, 10, 2021.

[10099] Joleen JH Traets, Servaas N van der Burght, Suzanne Rademakers, Gert Jansen, and Jeroen S van Zon. Mechanism of life-long maintenance of neuron identity despite molecular fluctuations. *eLife*, 10, 2021.

[10100] Evgeniya Trofimenko, Gianvito Grasso, Mathieu Heulot, Nadja Chevalier, Marco A Deriu, Gilles Dubuis, Yoan Arribat, Marc Serulla, Sebastien Michel, Gil Vantomme, Florine Ory, Linh Chi Dam, Julien Puyal, Francesca Amati, Anita Lüthi, Andrea Danani, and Christian Widmann. Genetic, cellular, and structural characterization of the membrane potential-dependent cell-penetrating peptide translocation pore. *eLife*, 10, 2021.

[10101] Kathryn P Trogden, Justin Lee, Kai M Bracey, Kung-Hsien Ho, Hudson McKinney, Xiaodong Zhu, Goker Arpag, Thomas G Folland, Anna B Osipovich, Mark A Magnuson, Marija Zanic, Guoqiang Gu, William R Holmes, and Irina Kaverina. Microtubules regulate pancreatic $\beta$-cell heterogeneity via spatiotemporal control of insulin secretion hot spots. *eLife*, 10, 2021.

[10102] Julianne M Troiano, Federico Perozeni, Raymundo Moya, Luca Zuliani, Kwangyrul Baek, EonSeon Jin, Stefano Cazzaniga, Matteo Ballottari, and Gabriela S Schlau-Cohen. Identification of distinct pH- and zeaxanthin-dependent quenching in LHCSR3 from Chlamydomonas reinhardtii. *eLife*, 10, 2021.

[10103] Johannes Trück, Anne Eugster, Pierre Barennes, Christopher M Tipton, Eline T Luning Prak, Davide Bagnara, Cinque Soto, Jacob S Sherkow, Aimee S Payne, Marie-Paule Lefranc, Andrew Farmer, Magnolia Bostick, and Encarnita Mariotti-Ferrandiz. Biological controls for standardization and interpretation of adaptive immune receptor repertoire profiling. *eLife*, 10, 2021.

[10104] Emily N Truckenbrod, Kristina S Burrack, Todd P Knutson, Henrique Borges da Silva, Katharine E Block, Stephen D O'Flanagan, Katie R Stagliano, Arthur A Hurwitz, Ross B Fulton, Kristin R Renkema, and Stephen C Jameson. CD8+ T cell self-tolerance permits responsiveness but limits tissue damage. *eLife*, 10, 2021.

[10105] Brian V Tsu, Christopher Beierschmitt, Andrew P Ryan, Rimjhim Agarwal, Patrick S Mitchell, and Matthew D Daugherty. Diverse viral proteases activate the NLRP1 inflammasome. *eLife*, 10, 2021.

[10106] Masaki Tsujimura, Keiichi Kojima, Shiho Kawanishi, Yuki Sudo, and Hiroshi Ishikita. Proton transfer pathway in anion channelrhodopsin-1. *eLife*, 10, 2021.

[10107] Yifan Tu, Lu Wang, Yi Rong, Vincent Tam, Taijun Yin, Song Gao, Rashim Singh, and Ming Hu. Hepatoenteric recycling is a new disposition mechanism for orally administered phenolic drugs and phytochemicals in rats. *eLife*, 10, 2021.

[10108] Yuan-Po Tu, Jameel Iqbal, and Timothy O'Leary. Sensitivity of ID NOW and RT–PCR for detection of SARS-CoV-2 in an ambulatory population. *eLife*, 10, 2021.

[10109] Srinu Tumpara, Matthias Ballmaier, Sabine Wrenger, Mandy König, Matthias Lehmann, Ralf Lichtinghagen, Beatriz Martinez-Delgado, Elena Korenbaum, David DeLuca, Nils Jedicke, Tobias Welte, Malin Fromme, Pavel Strnad, Jan Stolk, and Sabina Janciauskiene. Polymerization of misfolded Z alpha-1 antitrypsin protein lowers CX3CR1 expression in human PBMCs. *eLife*, 10, 2021.

[10110] Rachael Emily Turner, Paul F Harrison, Angavai Swaminathan, Calvin A Kraupner-Taylor, Belinda J Goldie, Michael See, Amanda L Peterson, Ralf B Schittenhelm, David R Powell, Darren J Creek, Bernhard Dichtl, and Traude H Beilharz. Genetic and pharmacological evidence for kinetic competition between alternative poly(A) sites in yeast. *eLife*, 10, 2021.

[10111] Jernej TurnŠek, John K Brunson, Maria del Pilar Martinez Viedma, Thomas J Deerinck, AleŠ Horák, Miroslav Oborník, Vincent A Bielinski, and Andrew Ellis Allen. Proximity proteomics in a marine diatom reveals a putative cell surface-to-chloroplast iron trafficking pathway. *eLife*, 10, 2021.

[10112] Miguel Turrero García, Sarah K Stegmann, Tiara E Lacey, Christopher M Reid, Sinisa Hrvatin, Caleb Weinreb, Manal A Adam, M Aurel Nagy, and Corey C Harwell. Transcriptional profiling of sequentially generated septal neuron fates. *eLife*, 10, 2021.

[10113] Benjamin A Turturice, Juliana Theorell, Mary Dawn Koenig, Lisa Tussing-Humphreys, Diane R Gold, Augusto A Litonjua, Emily Oken, Sheryl L Rifas-Shiman, David L Perkins, and Patricia W Finn. Perinatal granulopoiesis and risk of pediatric asthma. *eLife*, 10, 2021.

[10114] Jürgen Tuvikene, Eli-Eelika Esvald, Annika Rähni, Kaie Uustalu, Anna Zhuravskaya, Annela Avarlaid, Eugene V Makeyev, and Tõnis Timmusk. Intronic enhancer region governs transcript-specific Bdnf expression in rodent neurons. *eLife*, 10, 2021.

[10115] Joshua M Tworig, Chandler J Coate, and Marla B Feller. Excitatory neurotransmission activates compartmentalized calcium transients in Müller glia without affecting lateral process motility. *eLife*, 10, 2021.

[10116] Abdul Malik Tyagi, Trevor M Darby, Emory Hsu, Mingcan Yu, Subhashis Pal, Hamid Dar, Jau-Yi Li, Jonathan Adams, Rheinallt M Jones, and Roberto Pacifici. The gut microbiota is a transmissible determinant of skeletal maturation. *eLife*, 10, 2021.

[10117] Shiou-Ru Tzeng, Yin-Chu Tseng, Chien-Chu Lin, Chia-Ying Hsu, Shing-Jong Huang, Yi-Ting Kuo, and Chung-I Chang. Molecular insights into substrate recognition and discrimination by the N-terminal domain of Lon AAA+ protease. *eLife*, 10, 2021.

[10118] Hironori Uehara, Xiaohui Zhang, Felipe Pereira, Siddharth Narendran, Susie Choi, Sai Bhuvanagiri, Jinlu Liu, Sangeetha Ravi Kumar, Austin Bohner, Lara Carroll, Bonnie Archer, Yue Zhang, Wei Liu, Guangping Gao, Jayakrishna Ambati, Albert S Jun, and Balamurali K Ambati. Start codon disruption with CRISPR/Cas9 prevents murine Fuchs' endothelial corneal dystrophy. *eLife*, 10, 2021.

[10119] Masashi Ugawa, Yoko Kawamura, Keisuke Toda, Kazuki Teranishi, Hikari Morita, Hiroaki Adachi, Ryo Tamoto, Hiroko Nomaru, Keiji Nakagawa, Keiki Sugimoto, Evgeniia Borisova, Yuri An, Yusuke Konishi, Seiichiro Tabata, Soji Morishita, Misa Imai, Tomoiku Takaku, Marito Araki, Norio Komatsu, Yohei Hayashi, Issei Sato, Ryoichi Horisaki, Hiroyuki Noji, and Sadao Ota. In silico-labeled ghost cytometry. *eLife*, 10, 2021.

[10120] Eva Liliane Ujeneza, Wilfred Ndifon, Shobna Sawry, Geoffrey Fatti, Julien Riou, Mary-Ann Davies, and Martin Nieuwoudt. A mechanistic model for long-term immunological outcomes in South African HIV-infected children and adults receiving ART. *eLife*, 10, 2021.

[10121] Carolin Ulbricht, Ruth Leben, Asylkhan Rakhymzhan, Frank Kirchhoff, Lars Nitschke, Helena Radbruch, Raluca A Niesner, and Anja E Hauser. Intravital quantification reveals dynamic calcium concentration changes across B cell differentiation stages. *eLife*, 10, 2021.

[10122] Maria F Ullo and Jeremy S Logue. ADF and cofilin-1 collaborate to promote cortical actin flow and the leader bleb-based migration of confined cells. *eLife*, 10, 2021.

[10123] Ki Bum Um, Suyun Hahn, So Woon Kim, Yoon Je Lee, Lutz Birnbaumer, Hyun Jin Kim, and Myoung Kyu Park. TRPC3 and NALCN channels drive pacemaking in substantia nigra dopaminergic neurons. *eLife*, 10, 2021.

[10124] Colton M Unger, Jay Devine, Benedikt Hallgrímsson, and Campbell Rolian. Selection for increased tibia length in mice alters skull shape through parallel changes in developmental mechanisms. *eLife*, 10, 2021.

[10125] Irene Unterman, Idit Bloch, Simona Cazacu, Gila Kazimirsky, Bruria Ben-Zeev, Benjamin P Berman, Chaya Brodie, and Yuval Tabach. Expanding the MECP2 network using comparative genomics reveals potential therapeutic targets for Rett syndrome. *eLife*, 10, 2021.

[10126] Taylor B Updegrove, Jailynn Harke, Vivek Anantharaman, Jin Yang, Nikhil Gopalan, Di Wu, Grzegorz Piszczek, David M Stevenson, Daniel Amador-Noguez, Jue D Wang, L Aravind, and Kumaran S Ramamurthi. Reformulation of an extant ATPase active site to mimic ancestral GTPase activity reveals a nucleotide base requirement for function. *eLife*, 10, 2021.

[10127] Rajendra Uprety, Tao Che, Saheem A Zaidi, Steven G Grinnell, Balázs R Varga, Abdelfattah Faouzi, Samuel T Slocum, Abdullah Allaoa, András Varadi, Melissa Nelson, Sarah M Bernhard, Elizaveta Kulko, Valerie Le Rouzic, Shainnel O Eans, Chloe A Simons, Amanda Hunkele, Joan Subrath, Ying Xian Pan, Jonathan A Javitch, Jay P McLaughlin, Bryan L Roth, Gavril W Pasternak, Vsevolod Katritch, and Susruta Majumdar. Controlling opioid receptor functional selectivity by targeting distinct subpockets of the orthosteric site. *eLife*, 10, 2021.

[10128] Lara Urban, Andre Holzer, J Jotautas Baronas, Michael B Hall, Philipp Braeuninger-Weimer, Michael J Scherm, Daniel J Kunz, Surangi N Perera, Daniel E Martin-Herranz, Edward T Tipper, Susannah J Salter, and Maximilian R Stammnitz. Freshwater monitoring by nanopore sequencing. *eLife*, 10, 2021.

[10129] Koichiro Uriu, Bo-Kai Liao, Andrew C Oates, and Luis G Morelli. From local resynchronization to global pattern recovery in the zebrafish segmentation clock. *eLife*, 10, 2021.

[10130] Inigo Urrutia-Irazabal, James R Ault, Frank Sobott, Nigel J Savery, and Mark S Dillingham. Analysis of the PcrA-RNA polymerase complex reveals a helicase interaction motif and a role for PcrA/UvrD helicase in the suppression of R-loops. *eLife*, 10, 2021.

[10131] Avinash R Vaidya, Henry M Jones, Johanny Castillo, and David Badre. Neural representation of abstract task structure during generalization. *eLife*, 10, 2021.

[10132] Trisha V Vaidyanathan, Max Collard, Sae Yokoyama, Michael E Reitman, and Kira E Poskanzer. Cortical astrocytes independently regulate sleep depth and duration via separate GPCR pathways. *eLife*, 10, 2021.

[10133] Gabriele Valentini, Theodore P Pavlic, Sara Imari Walker, Stephen C Pratt, Dora Biro, and Takao Sasaki. Naïve individuals promote collective exploration in homing pigeons. *eLife*, 10, 2021.

[10134] Raphael Vallat and Matthew P Walker. An open-source, high-performance tool for automated sleep staging. *eLife*, 10, 2021.

[10135] Eljo Van Battum, Celine Heitz-Marchaland, Yvrick Zagar, Stéphane Fouquet, Rohini Kuner, and Alain Chédotal. Plexin-B2 controls the timing of differentiation and the motility of cerebellar granule neurons. *eLife*, 10, 2021.

[10136] Jasmin van den Heuvel, Caroline Ashiono, Ludovic C Gillet, Kerstin Dörner, Emanuel Wyler, Ivo Zemp, and Ulrike Kutay. Processing of the ribosomal ubiquitin-like fusion protein FUBI-eS30/FAU is required for 40S maturation and depends on USP36. *eLife*, 10, 2021.

[10137] Meike E van der Heijden, Elizabeth P Lackey, Ross Perez, Fatma S Ișleyen, Amanda M Brown, Sarah G Donofrio, Tao Lin, Huda Y Zoghbi, and Roy V Sillitoe. Maturation of Purkinje cell firing properties relies on neurogenesis of excitatory neurons. *eLife*, 10, 2021.

[10138] Mariena JA van der Plas, Jun Cai, Jitka Petrlova, Karim Saleh, Sven Kjellström, and Artur Schmidtchen. Method development and characterisation of the low-molecular-weight peptidome of human wound fluids. *eLife*, 10, 2021.

[10139] Stijn van Dorp, Ruoyi Qiu, Ucheor B Choi, Minnie M Wu, Michelle Yen, Michael Kirmiz, Axel T Brunger, and Richard S Lewis. Conformational dynamics of auto-inhibition in the ER calcium sensor STIM1. *eLife*, 10, 2021.

[10140] Jonathan G Van Vranken, Jiaming Li, Dylan C Mitchell, José Navarrete-Perea, and Steven P Gygi. Assessing target engagement using proteome-wide solvent shift assays. *eLife*, 10, 2021.

[10141] Ruud van Zessen, Yue Li, Lucile Marion-Poll, Nicolas Hulo, Jérôme Flakowski, and Christian Lüscher. Dynamic dichotomy of accumbal population activity underlies cocaine sensitization. *eLife*, 10, 2021.

[10142] Charlotte Vanacker, R Anthony Defazio, Charlene M Sykes, and Suzanne M Moenter. A role for glial fibrillary acidic protein (GFAP)-expressing cells in the regulation of gonadotropin-releasing hormone (GnRH) but not arcuate kisspeptin neuron output in male mice. *eLife*, 10, 2021.

[10143] Heather Vance, Peter T Madsen, Natacha Aguilar de Soto, Danuta Maria Wisniewska, Michael Ladegaard, Sascha Hooker, and Mark Johnson. Echolocating toothed whales use ultra-fast echo-kinetic responses to track evasive prey. *eLife*, 10, 2021.

[10144] Youna Vandaele, Magalie Lenoir, Caroline Vouillac-Mendoza, Karine Guillem, and Serge H Ahmed. Probing the decision-making mechanisms underlying choice between drug and nondrug rewards in rats. *eLife*, 10, 2021.

[10145] Servaas Vandecappelle, Lucas Deckers, Neetha Das, Amir Hossein Ansari, Alexander Bertrand, and Tom Francart. EEG-based detection of the locus of auditory attention with convolutional neural networks. *eLife*, 10, 2021.

[10146] Giulia Vanoni, Giuseppe Ercolano, Simona Candiani, Mariangela Rutigliani, Mariangela Lanata, Laurent Derré, Emanuela Marcenaro, Pascal Schneider, Pedro Romero, Camilla Jandus, and Sara Trabanelli. Human primed ILCPs support endothelial activation through NF-$\kappa$B signaling. *eLife*, 10, 2021.

[10147] Roger Vargas, Luca Freschi, Maximillian Marin, L Elaine Epperson, Melissa Smith, Irina Oussenko, David Durbin, Michael Strong, Max Salfinger, and Maha Reda Farhat. In-host population dynamics of Mycobacterium tuberculosis complex during active disease. *eLife*, 10, 2021.

[10148] Harsh Vashistha, Maryam Kohram, and Hanna Salman. Non-genetic inheritance restraint of cell-to-cell variation. *eLife*, 10, 2021.

[10149] Juan M Vazquez and Vincent J Lynch. Pervasive duplication of tumor suppressors in Afrotherians during the evolution of large bodies and reduced cancer risk. *eLife*, 10, 2021.

[10150] Marisa Vedor, Nuno Queiroz, Gonzalo Mucientes, Ana Couto, Ivo da Costa, António dos Santos, Frederic Vandeperre, Jorge Fontes, Pedro Afonso, Rui Rosa, Nicolas E Humphries, and David W Sims. Climate-driven deoxygenation elevates fishing vulnerability for the ocean's widest ranging shark. *eLife*, 10, 2021.

[10151] Pavan Vedula, Satoshi Kurosaka, Brittany MacTaggart, Qin Ni, Garegin Papoian, Yi Jiang, Dawei W Dong, and Anna Kashina. Different translation dynamics of $\beta$- and $\gamma$-actin regulates cell migration. *eLife*, 10, 2021.

[10152] Lena Veit, Lucas Y Tian, Christian J Monroy Hernandez, and Michael S Brainard. Songbirds can learn flexible contextual control over syllable sequencing. *eLife*, 10, 2021.

[10153] Vidyasiri Vemulapalli, Lily A Chylek, Alison Erickson, Anamarija Pfeiffer, Khal-Hentz Gabriel, Jonathan LaRochelle, Kartik Subramanian, Ruili Cao, Kimberley Stegmaier, Morvarid Mohseni, Matthew J LaMarche, Michael G Acker, Peter K Sorger, Steven P Gygi, and Stephen C Blacklow. Time-resolved phosphoproteomics reveals scaffolding and catalysis-responsive patterns of SHP2-dependent signaling. *eLife*, 10, 2021.

[10154] Richard Venz, Tina Pekec, Iskra Katic, Rafal Ciosk, and Collin Yvès Ewald. End-of-life targeted degradation of DAF-2 insulin/IGF-1 receptor promotes longevity free from growth-related pathologies. *eLife*, 10, 2021.

[10155] Jason D Vevea, Grant F Kusick, Kevin C Courtney, Erin Chen, Shigeki Watanabe, and Edwin R Chapman. Synaptotagmin 7 is targeted to the axonal plasma membrane through $\gamma$-secretase processing to promote synaptic vesicle docking in mouse hippocampal neurons. *eLife*, 10, 2021.

[10156] Ricardo Viais, Marcos Fariña-Mosquera, Marina Villamor-Payà, Sadanori Watanabe, Lluís Palenzuela, Cristina Lacasa, and Jens Lüders. Augmin deficiency in neural stem cells causes p53-dependent apoptosis and aborts brain development. *eLife*, 10, 2021.

[10157] Francisco Victorino, Tarin M Bigley, Eugene Park, Cong-Hui Yao, Jeanne Benoit, Li-Ping Yang, Sytse J Piersma, Elvin J Lauron, Rebecca M Davidson, Gary J Patti, and Wayne M Yokoyama. HIF1$\alpha$ is required for NK cell metabolic adaptation during virus infection. *eLife*, 10, 2021.

[10158] Didac Vidal-Pineiro, Yunpeng Wang, Stine K Krogsrud, Inge K Amlien, William FC Baaré, David Bartres-Faz, Lars Bertram, Andreas M Brandmaier, Christian A Drevon, Sandra Düzel, Klaus Ebmeier, Richard N Henson, Carme Junqué, Rogier Andrew Kievit, Simone Kühn, Esten Leonardsen, Ulman Lindenberger, Kathrine S Madsen, Fredrik Magnussen, Athanasia Monika Mowinckel, Lars Nyberg, James M Roe, Barbara Segura, Stephen M Smith, Øystein Sørensen, Sana Suri, Rene Westerhausen, Andrew Zalesky, Enikő Zsoldos, Kristine Beate Walhovd, and Anders Fjell. Individual variations in 'brain age' relate to early-life factors more than to longitudinal brain change. *eLife*, 10, 2021.

[10159] Athul Vijayan, Rachele Tofanelli, Sören Strauss, Lorenzo Cerrone, Adrian Wolny, Joanna Strohmeier, Anna Kreshuk, Fred A Hamprecht, Richard S Smith, and Kay Schneitz. A digital 3D reference atlas reveals cellular growth patterns shaping the Arabidopsis ovule. *eLife*, 10, 2021.

[10160] Margaritis Voliotis, Xiao Feng Li, Ross Alexander De Burgh, Geffen Lass, Deyana Ivanova, Caitlin McIntyre, Kevin O'Byrne, and Krasimira Tsaneva-Atanasova. Modulation of pulsatile GnRH dynamics across the ovarian cycle via changes in the network excitability and basal activity of the arcuate kisspeptin network. *eLife*, 10, 2021.

[10161] Mihály Vöröslakos, Peter C Petersen, Balázs Vöröslakos, and György Buzsáki. Metal microdrive and head cap system for silicon probe recovery in freely moving rodent. *eLife*, 10, 2021.

[10162] Daan Vorselen, Sarah R Barger, Yifan Wang, Wei Cai, Julie A Theriot, Nils C Gauthier, and Mira Krendel. Phagocytic 'teeth' and myosin-II 'jaw' power target constriction during phagocytosis. *eLife*, 10, 2021.

[10163] Sabrina Vullo, Nicolas Ambrosio, Jan P Kucera, Olivier Bignucolo, and Stephan Kellenberger. Kinetic analysis of ASIC1a delineates conformational signaling from proton-sensing domains to the channel gate. *eLife*, 10, 2021.

[10164] Nguyen Lam Vuong, Phung Khanh Lam, Damien Keng Yen Ming, Huynh Thi Le Duyen, Nguyet Minh Nguyen, Dong Thi Hoai Tam, Kien Duong Thi Hue, Nguyen VV Chau, Ngoun Chanpheaktra, Lucy Chai See Lum, Ernesto Pleités, Cameron P Simmons, Kerstin D Rosenberger, Thomas Jaenisch, David Bell, Nathalie Acestor, Christine Halleux, Piero L Olliaro, Bridget A Wills, Ronald B Geskus, and Sophie Yacoub. Combination of inflammatory and vascular markers in the febrile phase of dengue is associated with more severe outcomes. *eLife*, 10, 2021.

[10165] Thanh Thi Vuong-Brender, Sean Flynn, Yvonne Vallis, Saliha E Sönmez, and Mario de Bono. Neuronal calmodulin levels are controlled by CAMTA transcription factors. *eLife*, 10, 2021.

[10166] Natalia Vydra, Patryk Janus, Paweł Kus, Tomasz Stokowy, Katarzyna Mrowiec, Agnieszka Toma-Jonik, Aleksandra Krzywon, Alexander Jorge Cortez, Bartosz Wojtas, Bartłomiej Gielniewski, Roman Jaksik, Marek Kimmel, and Wieslawa Widlak. Heat shock factor 1 (HSF1) cooperates with estrogen receptor $\alpha$ (ER$\alpha$) in the regulation of estrogen action in breast cancer cells. *eLife*, 10, 2021.

[10167] Cameron Wagg, Yann Hautier, Sarah Pellkofer, Samiran Banerjee, Bernhard Schmid, and Marcel GA van der Heijden. Diversity and asynchrony in soil microbial communities stabilizes ecosystem functioning. *eLife*, 10, 2021.

[10168] Laura J Wagstaff, Jose A Gomez-Sanchez, Shaline V Fazal, Georg W Otto, Alastair M Kilpatrick, Kirolos Michael, Liam YN Wong, Ki H Ma, Mark Turmaine, John Svaren, Tessa Gordon, Peter Arthur-Farraj, Sergio Velasco-Aviles, Hugo Cabedo, Cristina Benito, Rhona Mirsky, and Kristjan R Jessen. Failures of nerve regeneration caused by aging or chronic denervation are rescued by restoring Schwann cell c-Jun. *eLife*, 10, 2021.

[10169] Matthias Wälchli, Karolin Berneiser, Francesca Mangia, Stefan Imseng, Louise-Marie Craigie, Edward Stuttfeld, Michael N Hall, and Timm Maier. Regulation of human mTOR complexes by DEPTOR. *eLife*, 10, 2021.

[10170] A Sarah Walker, Emma Pritchard, Thomas House, Julie V Robotham, Paul J Birrell, Iain Bell, John I Bell, John N Newton, Jeremy Farrar, Ian Diamond, Ruth Studley, Jodie Hay, Karina-Doris Vihta, Timothy EA Peto, Nicole Stoesser, Philippa C Matthews, David W Eyre, and Koen B Pouwels. Ct threshold values, a proxy for viral load in community SARS-CoV-2 cases, demonstrate wide variation across populations and over time. *eLife*, 10, 2021.

[10171] Shannon M Walsh, Ryan M Sheridan, Erin D Lucas, Thu A Doan, Brian C Ware, Johnathon Schafer, Rui Fu, Matthew A Burchill, Jay R Hesselberth, and Beth Ann Jiron Tamburini. Molecular tracking devices quantify antigen distribution and archiving in the murine lymph node. *eLife*, 10, 2021.

[10172] Dawei Wang, Zuodong Ye, Wenjie Wei, Jingting Yu, Lihong Huang, Hongmin Zhang, and Jianbo Yue. Capping protein regulates endosomal trafficking by controlling F-actin density around endocytic vesicles and recruiting RAB5 effectors. *eLife*, 10, 2021.

[10173] Feng Wang, Maria Gracia Gervasi, Ana BoŠković, Fengyun Sun, Vera D Rinaldi, Jun Yu, Mary C Wallingford, Darya A Tourzani, Jesse Mager, Lihua Julie Zhu, Oliver J Rando, Pablo E Visconti, Lara Strittmatter, and Ingolf Bach. Deficient spermiogenesis in mice lacking Rlim. *eLife*, 10, 2021.

[10174] Hsiang-Tsui Wang, Hyun-Wook Lee, Mao-wen Weng, Yan Liu, William C Huang, Herbert Lepor, Xue-Ru Wu, and Moon-shong Tang. The role of TAp63$\gamma$ and P53 point mutations in regulating DNA repair, mutational susceptibility and invasion of bladder cancer cells. *eLife*, 10, 2021.

[10175] Jie Wang, Amir Rattner, and Jeremy Nathans. A transcriptome atlas of the mouse iris at single-cell resolution defines cell types and the genomic response to pupil dilation. *eLife*, 10, 2021.

[10176] Tingting Wang, Wenju Sun, Ligang Fan, Canfeng Hua, Nan Wu, Shaorong Fan, Jilin Zhang, Xin Deng, and Jian Yan. An atlas of the binding specificities of transcription factors in Pseudomonas aeruginosa directs prediction of novel regulators in virulence. *eLife*, 10, 2021.

[10177] Weisheng Wang, Peter J Schuette, Mimi Q La-Vu, Anita Torossian, Brooke C Tobias, Marta Ceko, Philip A Kragel, Fernando MCV Reis, Shiyu Ji, Megha Sehgal, Sandra Maesta-Pereira, Meghmik Chakerian, Alcino J Silva, Newton S Canteras, Tor Wager, Jonathan C Kao, and Avishek Adhikari. Dorsal premammillary projection to periaqueductal gray controls escape vigor from innate and conditioned threats. *eLife*, 10, 2021.

[10178] Xin-Qiu Wang, Jian-sheng Guo, Dan-Ting Li, Yang Yu, Jaco Hagoort, Bernard Moussian, and Chuan-Xi Zhang. Three-dimensional reconstruction of a whole insect reveals its phloem sap-sucking mechanism at nano-resolution. *eLife*, 10, 2021.

[10179] Xuechun Wang, Weilin Zeng, Xiaodan Yang, Yongsheng Zhang, Chunyu Fang, Shaoqun Zeng, Yunyun Han, and Peng Fei. Bi-channel image registration and deep-learning segmentation (BIRDS) for efficient, versatile 3D mapping of mouse brain. *eLife*, 10, 2021.

[10180] Y Claire Wang, Elizabeth Brondolo, Rachel Monane, Michaela Kiernan, Karina W Davidson, Catherine M Alfano, Catherine M Alfano, Betty Diamond, Joan Duer-Hefele, Felicia Hill-Briggs, Jerry Kim, James Peacock, Nancy D Spector, and Sunmoo Yoon. Introducing the MAVEN Leadership Training Initiative to diversify the scientific workforce. *eLife*, 10, 2021.

[10181] Yang Wang, Yifeng Guo, Guanluan Li, Chunhong Liu, Lei Wang, Aihua Zhang, Zhiqiang Yan, and Chen Song. The push-to-open mechanism of the tethered mechanosensitive ion channel NompC. *eLife*, 10, 2021.

[10182] Yawei Wang, Osamu Toyoshima, Jun Kunimatsu, Hiroshi Yamada, and Masayuki Matsumoto. Tonic firing mode of midbrain dopamine neurons continuously tracks reward values changing moment-by-moment. *eLife*, 10, 2021.

[10183] Yiquan Wang, Ruipeng Lei, Armita Nourmohammad, and Nicholas C Wu. Antigenic evolution of human influenza H3N2 neuraminidase is constrained by charge balancing. *eLife*, 10, 2021.

[10184] Zhu Wang, Mi Cheong Cheong, Jet Tsien, Heping Deng, Tian Qin, Jonathan DC Stoltzfus, Tegegn G Jaleta, Xinshe Li, James B Lok, Steven A Kliewer, and David J Mangelsdorf. Characterization of the endogenous DAF-12 ligand and its use as an anthelmintic agent in Strongyloides stercoralis. *eLife*, 10, 2021.

[10185] Khursheed A Wani, Debanjan Goswamy, Stefan Taubert, Ramesh Ratnappan, Arjumand Ghazi, and Javier E Irazoqui. NHR-49/PPAR-$\alpha$ and HLH-30/TFEB cooperate for C. elegans host defense via a flavin-containing monooxygenase. *eLife*, 10, 2021.

[10186] Michelle C Ward, Nicholas E Banovich, Abhishek Sarkar, Matthew Stephens, and Yoav Gilad. Dynamic effects of genetic variation on gene expression revealed following hypoxic stress in cardiomyocytes. *eLife*, 10, 2021.

[10187] Richard A Warren, Qianyun Zhang, Judah R Hoffman, Edward Y Li, Y Kate Hong, Randy M Bruno, and Nathaniel B Sawtell. A rapid whisker-based decision underlying skilled locomotion in mice. *eLife*, 10, 2021.

[10188] Leonhard Waschke, Thomas Donoghue, Lorenz Fiedler, Sydney Smith, Douglas D Garrett, Bradley Voytek, and Jonas Obleser. Modality-specific tracking of attention and sensory statistics in the human electrophysiological spectral exponent. *eLife*, 10, 2021.

[10189] Joshua T Washington, Katelyn R Cavender, Ashley U Amukamara, Elizabeth C McKinney, Robert J Schmitz, and Patricia J Moore. The essential role of Dnmt1 in gametogenesis in the large milkweed bug Oncopeltus fasciatus. *eLife*, 10, 2021.

[10190] Lianna W Wat, Zahid S Chowdhury, Jason W Millington, Puja Biswas, and Elizabeth J Rideout. Sex determination gene transformer regulates the male-female difference in Drosophila fat storage via the adipokinetic hormone pathway. *eLife*, 10, 2021.

[10191] Akinobu Watanabe, Amy M Balanoff, Paul M Gignac, M Eugenia L Gold, and Mark A Norell. Novel neuroanatomical integration and scaling define avian brain shape evolution and development. *eLife*, 10, 2021.

[10192] Takamitsu Watanabe. Causal roles of prefrontal cortex during spontaneous perceptual switching are determined by brain state dynamics. *eLife*, 10, 2021.

[10193] Samuel James Watson, Na Li, Yiting Ye, Feijie Wu, Qihua Ling, and R Paul Jarvis. Crosstalk between the chloroplast protein import and SUMO systems revealed through genetic and molecular investigation in Arabidopsis. *eLife*, 10, 2021.

[10194] Daniel K Weber, U Venkateswara Reddy, Songlin Wang, Erik K Larsen, Tata Gopinath, Martin B Gustavsson, Razvan L Cornea, David D Thomas, Alfonso De Simone, and Gianluigi Veglia. Structural basis for allosteric control of the SERCA-Phospholamban membrane complex by Ca2+ and phosphorylation. *eLife*, 10, 2021.

[10195] Miriam S Weber, Matthias Eibauer, Suganya Sivagurunathan, Thomas M Magin, Robert D Goldman, and Ohad Medalia. Structural heterogeneity of cellular K5/K14 filaments as revealed by cryo-electron microscopy. *eLife*, 10, 2021.

[10196] Kevin H-C Wei, Carolus Chan, and Doris Bachtrog. Establishment of H3K9me3-dependent heterochromatin during embryogenesis in Drosophila miranda. *eLife*, 10, 2021.

[10197] Xiaolu Wei, Danna G Eickbush, Iain Speece, and Amanda M Larracuente. Heterochromatin-dependent transcription of satellite DNAs in the Drosophila melanogaster female germline. *eLife*, 10, 2021.

[10198] Yulong Wei, Hao Sun, Tao Gui, Lutian Yao, Leilei Zhong, Wei Yu, Su-Jin Heo, Lin Han, Nathaniel A Dyment, Xiaowei Sherry Liu, Yejia Zhang, Eiki Koyama, Fanxin Long, Miltiadis H Zgonis, Robert L Mauck, Jaimo Ahn, and Ling Qin. The critical role of Hedgehog-responsive mesenchymal progenitors in meniscus development and injury repair. *eLife*, 10, 2021.

[10199] Ben G Weinstein, Sergio Marconi, Stephanie A Bohlman, Alina Zare, Aditya Singh, Sarah J Graves, and Ethan P White. A remote sensing derived data set of 100 million individual tree crowns for the National Ecological Observatory Network. *eLife*, 10, 2021.

[10200] Ada Weinstock, Karishma Rahman, Or Yaacov, Hitoo Nishi, Prashanthi Menon, Cyrus A Nikain, Michela L Garabedian, Stephanie Pena, Naveed Akbar, Brian E Sansbury, Sean P Heffron, Jianhua Liu, Gregory Marecki, Dawn Fernandez, Emily J Brown, Kelly V Ruggles, Stephen A Ramsey, Chiara Giannarelli, Matthew Spite, Robin P Choudhury, P'ng Loke, and Edward A Fisher. Wnt signaling enhances macrophage responses to IL-4 and promotes resolution of atherosclerosis. *eLife*, 10, 2021.

[10201] Carly V Weiss, Lana Harshman, Fumitaka Inoue, Hunter B Fraser, Dmitri A Petrov, Nadav Ahituv, and David Gokhman. The cis-regulatory effects of modern human-specific variants. *eLife*, 10, 2021.

[10202] Kai Weißenbruch, Justin Grewe, Marc Hippler, Magdalena Fladung, Moritz Tremmel, Kathrin Stricker, Ulrich Sebastian Schwarz, and Martin Bastmeyer. Distinct roles of nonmuscle myosin II isoforms for establishing tension and elasticity during cell morphodynamics. *eLife*, 10, 2021.

[10203] Samantha Wellington Miranda, Qian Cong, Amy L Schaefer, Emily Kenna MacLeod, Angelina Zimenko, David Baker, and E Peter Greenberg. A covariation analysis reveals elements of selectivity in quorum sensing systems. *eLife*, 10, 2021.

[10204] Kaylee M Wells, Kristina Kelley, Mary Baumel, Warren A Vieira, and Catherine D McCusker. Neural control of growth and size in the axolotl limb regenerate. *eLife*, 10, 2021.

[10205] Chentao Wen, Takuya Miura, Venkatakaushik Voleti, Kazushi Yamaguchi, Motosuke Tsutsumi, Kei Yamamoto, Kohei Otomo, Yukako Fujie, Takayuki Teramoto, Takeshi Ishihara, Kazuhiro Aoki, Tomomi Nemoto, Elizabeth MC Hillman, and Koutarou D Kimura. 3DeeCellTracker, a deep learning-based pipeline for segmenting and tracking cells in 3D time lapse images. *eLife*, 10, 2021.

[10206] Zachary Werkhoven, Alyssa Bravin, Kyobi Skutt-Kakaria, Pablo Reimers, Luisa F Pallares, Julien Ayroles, and Benjamin L de Bivort. The structure of behavioral variation within a genotype. *eLife*, 10, 2021.

[10207] Darryl A Wesener, Zachary W Beller, Samantha L Peters, Amir Rajabi, Gianluca Dimartino, Richard J Giannone, Robert L Hettich, and Jeffrey I Gordon. Microbiota functional activity biosensors for characterizing nutrient metabolism in vivo. *eLife*, 10, 2021.

[10208] Laura Whitworth, Jacob Coxon, Arjan van Laarhoven, Nguyen Thuy Thuong Thuong, Sofiati Dian, Bachti Alisjahbana, Ahmad Rizal Ganiem, Reinout van Crevel, Guy E Thwaites, Mark Troll, Paul H Edelstein, Roger Sewell, and Lalita Ramakrishnan. A Bayesian analysis of the association between Leukotriene A4 Hydrolase genotype and survival in tuberculous meningitis. *eLife*, 10, 2021.

[10209] J Stephan Wichers, Gerry Tonkin-Hill, Thorsten Thye, Ralf Krumkamp, Benno Kreuels, Jan Strauss, Heidrun von Thien, Judith AM Scholz, Helle Smedegaard Hansson, Rasmus Weisel Jensen, Louise Turner, Freia-Raphaella Lorenz, Anna Schöllhorn, Iris Bruchhaus, Egbert Tannich, Rolf Fendel, Thomas D Otto, Thomas Lavstsen, Tim W Gilberger, Michael F Duffy, and Anna Bachmann. Common virulence gene expression in adult first-time infected malaria patients and severe cases. *eLife*, 10, 2021.

[10210] Gregg Wildenberg, Anastasia Sorokina, Jessica Koranda, Alexis Monical, Chad Heer, Mark Sheffield, Xiaoxi Zhuang, Daniel McGehee, and Bobby Kasthuri. Partial connectomes of labeled dopaminergic circuits reveal non-synaptic communication and axonal remodeling after exposure to cocaine. *eLife*, 10, 2021.

[10211] Drew Wilfahrt, Rachael L Philips, Jyoti Lama, Monika Kizerwetter, Michael Jeremy Shapiro, Shaylene A McCue, Madeleine M Kennedy, Matthew J Rajcula, Hu Zeng, and Virginia Smith Shapiro. Histone deacetylase 3 represses cholesterol efflux during CD4+ T-cell activation. *eLife*, 10, 2021.

[10212] Scott A Williams, Thomas Cody Prang, Marc R Meyer, Thierra K Nalley, Renier Van Der Merwe, Christopher Yelverton, Daniel García-Martínez, Gabrielle A Russo, Kelly R Ostrofsky, Jeffrey Spear, Jennifer Eyre, Mark Grabowski, Shahed Nalla, Markus Bastir, Peter Schmid, Steven E Churchill, and Lee R Berger. New fossils of Australopithecus sediba reveal a nearly complete lower back. *eLife*, 10, 2021.

[10213] Gerald Willimsky, Christin Beier, Lena Immisch, George Papafotiou, Vivian Scheuplein, Andrean Goede, Hermann-Georg Holzhütter, Thomas Blankenstein, and Peter M Kloetzel. In vitro proteasome processing of neo-splicetopes does not predict their presentation in vivo. *eLife*, 10, 2021.

[10214] Patrick Morley Willoughby, Molly Allen, Jessica Yu, Roman Korytnikov, Tianhui Chen, Yupeng Liu, Isis So, Haoyu Wan, Neil Macpherson, Jennifer A Mitchell, Rodrigo Fernandez-Gonzalez, and Ashley EE Bruce. The recycling endosome protein Rab25 coordinates collective cell movements in the zebrafish surface epithelium. *eLife*, 10, 2021.

[10215] Meredith H Wilson, Stephen C Ekker, and Steven A Farber. Imaging cytoplasmic lipid droplets in vivo with fluorescent perilipin 2 and perilipin 3 knock-in zebrafish. *eLife*, 10, 2021.

[10216] Lennart Winkler, Maria Moiron, Edward H Morrow, and Tim Janicke. Stronger net selection on males across animals. *eLife*, 10, 2021.

[10217] Miles Wischnewski and Marius V Peelen. Causal neural mechanisms of context-based object recognition. *eLife*, 10, 2021.

[10218] Anirudh Wodeyar, Mark Schatza, Alik S Widge, Uri T Eden, and Mark A Kramer. A state space modeling approach to real-time phase estimation. *eLife*, 10, 2021.

[10219] Eva Wolf, Emmanuel Gaquerel, Mathias Scharmann, Levi Yant, and Marcus A Koch. Evolutionary footprints of a cold relic in a rapidly warming world. *eLife*, 10, 2021.

[10220] Michael Roland Wolff, Andrea Schmid, Philipp Korber, and Ulrich Gerland. Effective dynamics of nucleosome configurations at the yeast PHO5 promoter. *eLife*, 10, 2021.

[10221] Timothy M Wolock, Seth Flaxman, Kathryn A Risher, Tawanda Dadirai, Simon Gregson, and Jeffrey W Eaton. Evaluating distributional regression strategies for modelling self-reported sexual age-mixing. *eLife*, 10, 2021.

[10222] Jeffrey V Wong, Max Franz, Metin Can Siper, Dylan Fong, Funda Durupinar, Christian Dallago, Augustin Luna, John Giorgi, Igor Rodchenkov, özgün Babur, John A Bachman, Benjamin M Gyori, Emek Demir, Gary D Bader, and Chris Sander. Author-sourced capture of pathway knowledge in computable form using Biofactoid. *eLife*, 10, 2021.

[10223] Jeremy D Wong, Tyler Cluff, and Arthur D Kuo. The energetic basis for smooth human arm movements. *eLife*, 10, 2021.

[10224] Michael L Wood, Colin D Veal, Rita Neumann, Nicolás M Suárez, Jenna Nichols, Andrei J Parker, Diana Martin, Simon PR Romaine, Veryan Codd, Nilesh J Samani, Adriaan A Voors, Maciej Tomaszewski, Louis Flamand, Andrew J Davison, and Nicola J Royle. Variation in human herpesvirus 6B telomeric integration, excision, and transmission between tissues and individuals. *eLife*, 10, 2021.

[10225] Inge MN Wortel and Johannes Textor. Artistoo, a library to build, share, and explore simulations of cells and tissues in the web browser. *eLife*, 10, 2021.

[10226] Marina E Wosniack, Jan H Kirchner, Ling-Ya Chao, Nawal Zabouri, Christian Lohmann, and Julijana Gjorgjieva. Adaptation of spontaneous activity in the developing visual cortex. *eLife*, 10, 2021.

[10227] Arielle Woznica, Ashwani Kumar, Carolyn R Sturge, Chao Xing, Nicole King, and Julie K Pfeiffer. STING mediates immune responses in the closest living relatives of animals. *eLife*, 10, 2021.

[10228] Beibei Wu, Laura Rice, Jennifer Shrimpton, Dylan Lawless, Kieran Walker, Clive Carter, Lynn McKeown, Rashida Anwar, Gina M Doody, Sonal Srikanth, Yousang Gwack, and Sinisa Savic. Biallelic mutations in calcium release activated channel regulator 2A (CRACR2A) cause a primary immunodeficiency disorder. *eLife*, 10, 2021.

[10229] Gang Wu, Yin Yeng Lee, Evelyn M Gulla, Andrew Potter, Joseph Kitzmiller, Marc D Ruben, Nathan Salomonis, Jeffery A Whitsett, Lauren J Francey, John B Hogenesch, and David F Smith. Short-term exposure to intermittent hypoxia leads to changes in gene expression seen in chronic pulmonary disease. *eLife*, 10, 2021.

[10230] Mingzheng Wu, Samuel Minkowicz, Vasin Dumrongprechachan, Pauline Hamilton, Lei Xiao, and Yevgenia Kozorovitskiy. Attenuated dopamine signaling after aversive learning is restored by ketamine to rescue escape actions. *eLife*, 10, 2021.

[10231] Taoyu Wu and Shihui Han. Neural mechanisms of modulations of empathy and altruism by beliefs of others' pain. *eLife*, 10, 2021.

[10232] Xiaoting Wu, Takashi Saito, Takaomi C Saido, Anna M Barron, and Christiane Ruedl. Microglia and CD206+ border-associated mouse macrophages maintain their embryonic origin during Alzheimer's disease. *eLife*, 10, 2021.

[10233] Yangyu Wu, Zhuyuan Chen, Fred J Sigworth, and Cecilia M Canessa. Structure and analysis of nanobody binding to the human ASIC1a ion channel. *eLife*, 10, 2021.

[10234] Yue Kris Wu and Friedemann Zenke. Nonlinear transient amplification in recurrent neural networks with short-term plasticity. *eLife*, 10, 2021.

[10235] Zhenyong Wu, Nadiv Dharan, Zachary A McDargh, Sathish Thiyagarajan, Ben O'Shaughnessy, and Erdem Karatekin. The neuronal calcium sensor Synaptotagmin-1 and SNARE proteins cooperate to dilate fusion pores. *eLife*, 10, 2021.

[10236] Phillip Wulfridge and Kavitha Sarma. A nuclease- and bisulfite-based strategy captures strand-specific R-loops genome-wide. *eLife*, 10, 2021.

[10237] Elsio A Wunder, Haritha Adhikarla, Camila Hamond, Katharine A Owers Bonner, Li Liang, Camila B Rodrigues, Vimla Bisht, Jarlath E Nally, David P Alt, Mitermayer G Reis, Peter J Diggle, Philip L Felgner, and Albert Ko. A live attenuated-vaccine model confers cross-protective immunity against different species of the Leptospira genus. *eLife*, 10, 2021.

[10238] Shi-Li Xia, Meng Li, Bing Chen, Chao Wang, Yong-Hong Yan, Meng-Qiu Dong, and Yingchuan B Qi. The LRR-TM protein PAN-1 interacts with MYRF to promote its nuclear translocation in synaptic remodeling. *eLife*, 10, 2021.

[10239] Kehui Xiang and David P Bartel. The molecular basis of coupling between poly(A)-tail length and translational efficiency. *eLife*, 10, 2021.

[10240] Siheng Xiang and Douglas Koshland. Cohesin architecture and clustering in vivo. *eLife*, 10, 2021.

[10241] Jordan Yupeng Xiao, Antonina Hafner, and Alistair N Boettiger. How subtle changes in 3D structure can create large changes in transcription. *eLife*, 10, 2021.

[10242] Qijing Xie, Maria Brbic, Felix Horns, Sai Saroja Kolluru, Robert C Jones, Jiefu Li, Anay R Reddy, Anthony Xie, Sayeh Kohani, Zhuoran Li, Colleen N McLaughlin, Tongchao Li, Chuanyun Xu, David Vacek, David J Luginbuhl, Jure Leskovec, Stephen R Quake, Liqun Luo, and Hongjie Li. Temporal evolution of single-cell transcriptomes of Drosophila olfactory projection neurons. *eLife*, 10, 2021.

[10243] Tianfa Xie, Sarah R St Pierre, Nonthakorn Olaranont, Lauren E Brown, Min Wu, and Yubing Sun. Condensation tendency and planar isotropic actin gradient induce radial alignment in confined monolayers. *eLife*, 10, 2021.

[10244] Victoria Cochran Xie, Jinyue Pu, Brian PH Metzger, Joseph W Thornton, and Bryan C Dickinson. Contingency and chance erase necessity in the experimental evolution of ancestral proteins. *eLife*, 10, 2021.

[10245] Yi Xie, Rashmi Budhathoki, and J Todd Blankenship. Combinatorial deployment of F-actin regulators to build complex 3D actin structures in vivo. *eLife*, 10, 2021.

[10246] Yihu Xie, Bradley P Clarke, Yong Joon Kim, Austin L Ivey, Pate S Hill, Yi Shi, and Yi Ren. Cryo-EM structure of the yeast TREX complex and coordination with the SR-like protein Gbp2. *eLife*, 10, 2021.

[10247] Zhiyong Xie, Mengdi Wang, Zeyuan Liu, Congping Shang, Changjiang Zhang, Le Sun, Huating Gu, Gengxin Ran, Qing Pei, Qiang Ma, Meizhu Huang, Junjing Zhang, Rui Lin, Youtong Zhou, Jiyao Zhang, Miao Zhao, Minmin Luo, Qian Wu, Peng Cao, and Xiaoqun Wang. Transcriptomic encoding of sensorimotor transformation in the midbrain. *eLife*, 10, 2021.

[10248] Alba Xifra-Porxas, Michalis Kassinopoulos, and Georgios D Mitsis. Physiological and motion signatures in static and time-varying functional connectivity and their subject identifiability. *eLife*, 10, 2021.

[10249] Siqi Xiong, Ajay Kumar, Shenghe Tian, Eman E Taher, Enzhi Yang, Paul R Kinchington, Xiaobo Xia, and Yiqin Du. Stem cell transplantation rescued a primary open-angle glaucoma mouse model. *eLife*, 10, 2021.

[10250] Zherui Xiong, Harriet P Lo, Kerrie-Ann McMahon, Nick Martel, Alun Jones, Michelle M Hill, Robert G Parton, and Thomas E Hall. In vivo proteomic mapping through GFP-directed proximity-dependent biotin labelling in zebrafish. *eLife*, 10, 2021.

[10251] Pengfei Xu, Haoze V Yu, Kuo-Chang Tseng, Mackenzie Flath, Peter Fabian, Neil Segil, and J Gage Crump. Foxc1 establishes enhancer accessibility for craniofacial cartilage differentiation. *eLife*, 10, 2021.

[10252] Simiao Xu, Yangyang Liu, Ruixiang Hu, Min Wang, Oliver Stöhr, Yibo Xiong, Liang Chen, Hong Kang, Lingyun Zheng, Songjie Cai, Li He, Cunchuan Wang, Kyle D Copps, Morris F White, and Ji Miao. TAZ inhibits glucocorticoid receptor and coordinates hepatic glucose homeostasis in normal physiological states. *eLife*, 10, 2021.

[10253] Xiaozheng Xu, Takeya Masubuchi, Qixu Cai, Yunlong Zhao, and Enfu Hui. Molecular features underlying differential SHP1/SHP2 binding of immune checkpoint receptors. *eLife*, 10, 2021.

[10254] Xuan Xu, Jessica Kawakami, Nuwan Indika Millagaha Gedara, Jim E Riviere, Emma Meyer, Gerald J Wyckoff, and Majid Jaberi-Douraki. Data mining methodology for response to hypertension symptomology—application to COVID-19-related pharmacovigilance. *eLife*, 10, 2021.

[10255] Yongjin Xu, Richard M Bergenstal, Timothy C Dunn, and Ramzi A Ajjan. Addressing shortfalls of laboratory HbA1c using a model that incorporates red cell lifespan. *eLife*, 10, 2021.

[10256] Zhengchao Xu, Zhao Feng, Mengting Zhao, Qingtao Sun, Lei Deng, Xueyan Jia, Tao Jiang, Pan Luo, Wu Chen, Ayizuohere Tudi, Jing Yuan, Xiangning Li, Hui Gong, Qingming Luo, and Anan Li. Whole-brain connectivity atlas of glutamatergic and GABAergic neurons in the mouse dorsal and median raphe nuclei. *eLife*, 10, 2021.

[10257] Fan Xue, Zhenlei Zhao, Yanpei Gu, Jianxin Han, Keqiang Ye, and Ying Zhang. 7,8-Dihydroxyflavone modulates bone formation and resorption and ameliorates ovariectomy-induced osteoporosis. *eLife*, 10, 2021.

[10258] Jing Xue, Yan Han, Hamid Baniasadi, Weizhong Zeng, Jimin Pei, Nick V Grishin, Junmei Wang, Benjamin P Tu, and Youxing Jiang. TMEM120A is a coenzyme A-binding membrane protein with structural similarities to ELOVL fatty acid elongase. *eLife*, 10, 2021.

[10259] Yunlu Xue, Sean K Wang, Parimal Rana, Emma R West, Christin M Hong, Helian Feng, David M Wu, and Constance L Cepko. AAV-Txnip prolongs cone survival and vision in mouse models of retinitis pigmentosa. *eLife*, 10, 2021.

[10260] Ravi Yadav, Srinivas Govindan, Courtney Daczkowski, Andrew Mesecar, Srinivas Chakravarthy, and Nicholas Noinaj. Structural insight into the dual function of LbpB in mediating Neisserial pathogenesis. *eLife*, 10, 2021.

[10261] Mayu Yamada, Hirono Ohashi, Koh Hosoda, Daisuke Kurabayashi, and Shunsuke Shigaki. Multisensory-motor integration in olfactory navigation of silkmoth, Bombyx mori, using virtual reality system. *eLife*, 10, 2021.

[10262] Masahito Yamagata, Wenjun Yan, and Joshua R Sanes. A cell atlas of the chick retina based on single-cell transcriptomics. *eLife*, 10, 2021.

[10263] Nobuhiro Yamagata, Takahiro Ezaki, Takahiro Takahashi, Hongyang Wu, and Hiromu Tanimoto. Presynaptic inhibition of dopamine neurons controls optimistic bias. *eLife*, 10, 2021.

[10264] Io Yamamoto, Kexin Zhang, Jingjing Zhang, Egor Vorontsov, and Hiroki Shibuya. Telomeric double-strand DNA-binding proteins DTN-1 and DTN-2 ensure germline immortality in Caenorhabditis elegans. *eLife*, 10, 2021.

[10265] Naoki Yamawaki, Martinna G Raineri Tapies, Austin Stults, Gregory A Smith, and Gordon MG Shepherd. Circuit organization of the excitatory sensorimotor loop through hand/forelimb S1 and M1. *eLife*, 10, 2021.

[10266] Yasuo Yamazaki, Yuka Eura, and Koichi Kokame. V-ATPase V0a1 promotes Weibel–Palade body biogenesis through the regulation of membrane fission. *eLife*, 10, 2021.

[10267] Stephanie M Yan, Rachel M Sherman, Dylan J Taylor, Divya R Nair, Andrew N Bortvin, Michael C Schatz, and Rajiv C McCoy. Local adaptation and archaic introgression shape global diversity at human structural variant loci. *eLife*, 10, 2021.

[10268] Xiaohui Yan, Ke Jiang, Hui Li, Ziyi Wang, Kyle Perkins, and Fan Cao. Convergent and divergent brain structural and functional abnormalities associated with developmental dyslexia. *eLife*, 10, 2021.

[10269] Seiga Yanagisawa and Wayne D Frasch. pH-dependent 11° F1FO ATP synthase sub-steps reveal insight into the FO torque generating mechanism. *eLife*, 10, 2021.

[10270] Dafeng Yang, Stefan Haemmig, Haoyang Zhou, Daniel Pérez-Cremades, Xinghui Sun, Lei Chen, Jie Li, Jorge Haneo-Mejia, Tianlun Yang, Ivana Hollan, and Mark W Feinberg. Methotrexate attenuates vascular inflammation through an adenosine-microRNA-dependent pathway. *eLife*, 10, 2021.

[10271] Haibin Yang, Leishu Lin, Kang Sun, Ting Zhang, Wan Chen, Lianghui Li, Yuchen Xie, Chuanyue Wu, Zhiyi Wei, and Cong Yu. Complex structures of Rsu1 and PINCH1 reveal a regulatory mechanism of the ILK/PINCH/Parvin complex for F-actin dynamics. *eLife*, 10, 2021.

[10272] Hongdian Yang, Bilal A Bari, Jeremiah Y Cohen, and Daniel H O'Connor. Locus coeruleus spiking differently correlates with S1 cortex activity and pupil diameter in a tactile detection task. *eLife*, 10, 2021.

[10273] Qing Yang, Nicholas R Meyerson, Stephen K Clark, Camille L Paige, Will T Fattor, Alison R Gilchrist, Arturo Barbachano-Guerrero, Benjamin G Healy, Emma R Worden-Sapper, Sharon S Wu, Denise Muhlrad, Carolyn J Decker, Tassa K Saldi, Erika Lasda, Patrick Gonzales, Morgan R Fink, Kimngan L Tat, Cole R Hager, Jack C Davis, Christopher D Ozeroff, Gloria R Brisson, Matthew B McQueen, Leslie A Leinwand, Roy Parker, and Sara L Sawyer. Saliva TwoStep for rapid detection of asymptomatic SARS-CoV-2 carriers. *eLife*, 10, 2021.

[10274] Shu Yang, Ya Zhang, Chuxuan Yang, Xuefeng Wu, Sarah Maria El Oud, Rongfang Chen, Xudong Cai, Xufeng S Wu, Ganhui Lan, and Xiaoyan Zheng. Competitive coordination of the dual roles of the Hedgehog co-receptor in homophilic adhesion and signal reception. *eLife*, 10, 2021.

[10275] Weiwei Yang, Yu-Cheng Lin, William Johnson, Nan Dai, Romualdas Vaisvila, Peter Weigele, Yan-Jiun Lee, Ivan R Corrêa, Ira Schildkraut, and Laurence Ettwiller. A genome-phenome association study in native microbiomes identifies a mechanism for cytosine modification in DNA and RNA. *eLife*, 10, 2021.

[10276] Xingbo Yang, Gloria Ha, and Daniel J Needleman. A coarse-grained NADH redox model enables inference of subcellular metabolic fluxes from fluorescence lifetime imaging. *eLife*, 10, 2021.

[10277] Ya-Hui Yang, Pei-Shan Ho, Trong-Neng Wu, Peng-Wei Wang, Chun-Hung Richard Lin, Jui-Hsiu Tsai, Yue Leon Guo, and Hung-Yi Chuang. Risk of motor vehicle collisions after methadone use. *eLife*, 10, 2021.

[10278] Ziwei Yang, Jun Wang, Bailin He, Xiaolin Zhang, Xiaojuan Li, and Ersheng Kuang. RTN3 inhibits RIG-I-mediated antiviral responses by impairing TRIM25-mediated K63-linked polyubiquitination. *eLife*, 10, 2021.

[10279] David J Yanofsky, Justin M Di Trani, Sylwia Król, Rana Abdelaziz, Stephanie A Bueler, Peter Imming, Peter Brzezinski, and John L Rubinstein. Structure of mycobacterial CIII2CIV2 respiratory supercomplex bound to the tuberculosis drug candidate telacebec (Q203). *eLife*, 10, 2021.

[10280] Andriy S Yatsenko, Mariya M Kucherenko, Yuanbin Xie, Henning Urlaub, and Halyna R Shcherbata. Exocyst-mediated membrane trafficking of the lissencephaly-associated ECM receptor dystroglycan is required for proper brain compartmentalization. *eLife*, 10, 2021.

[10281] Gang Ye, Joseph Gallant, Jian Zheng, Christopher Massey, Ke Shi, Wanbo Tai, Abby Odle, Molly Vickers, Jian Shang, Yushun Wan, Lanying Du, Hideki Aihara, Stanley Perlman, Aaron LeBeau, and Fang Li. The development of Nanosota-1 as anti-SARS-CoV-2 nanobody drug candidates. *eLife*, 10, 2021.

[10282] Ali Ekrem Yesilkanal, Dongbo Yang, Andrea Valdespino, Payal Tiwari, Alan U Sabino, Long Chi Nguyen, Jiyoung Lee, Xiao-He Xie, Siqi Sun, Christopher Dann, Lydia Robinson-Mailman, Ethan Steinberg, Timothy Stuhlmiller, Casey Frankenberger, Elizabeth Goldsmith, Gary L Johnson, Alexandre F Ramos, and Marsha R Rosner. Limited inhibition of multiple nodes in a driver network blocks metastasis. *eLife*, 10, 2021.

[10283] Chao Yi, Tim WGM Spitters, Ezz Al-Din Ahmed Al-Far, Sen Wang, TianLong Xiong, Simian Cai, Xin Yan, Kaomei Guan, Michael Wagner, Ali El-Armouche, and Christopher L Antos. A calcineurin-mediated scaling mechanism that controls a K+-leak channel to regulate morphogen and growth factor transcription. *eLife*, 10, 2021.

[10284] Man Yi Yim, Lorenzo A Sadun, Ila R Fiete, and Thibaud Taillefumier. Place-cell capacity and volatility with grid-like inputs. *eLife*, 10, 2021.

[10285] Nai-Hao Yin, Paul Fromme, Ian McCarthy, and Helen L Birch. Individual variation in Achilles tendon morphology and geometry changes susceptibility to injury. *eLife*, 10, 2021.

[10286] Takeshi Yoneshiro, Naoya Kataoka, Jacquelyn M Walejko, Kenji Ikeda, Zachary Brown, Momoko Yoneshiro, Scott B Crown, Tsuyoshi Osawa, Juro Sakai, Robert W McGarrah, Phillip J White, Kazuhiro Nakamura, and Shingo Kajimura. Metabolic flexibility via mitochondrial BCAA carrier SLC25A44 is required for optimal fever. *eLife*, 10, 2021.

[10287] Toshihide W Yoshioka, Takahiro Doi, Mohammad Abdolrahmani, and Ichiro Fujita. Specialized contributions of mid-tier stages of dorsal and ventral pathways to stereoscopic processing in macaque. *eLife*, 10, 2021.

[10288] David M Young, Siavash Fazel Darbandi, Grace Schwartz, Zachary Bonzell, Deniz Yuruk, Mai Nojima, Laurent C Gole, John LR Rubenstein, Weimiao Yu, and Stephan J Sanders. Constructing and optimizing 3D atlases from 2D data with application to the developing mouse brain. *eLife*, 10, 2021.

[10289] Hedi Young, Beatriz Belbut, Margarida Baeta, and Leopoldo Petreanu. Laminar-specific cortico-cortical loops in mouse visual cortex. *eLife*, 10, 2021.

[10290] Rodrigo M Young, Camila J Solis, Andres Barriga-Fehrman, Carlos Abogabir, Alvaro R Thadani, Mariana Labarca, Eva Bustamante, Cecilia V Tapia, Antonia G Sarda, Francisca Sepulveda, Nadia Pozas, Leslie C Cerpa, María A Lavanderos, Nelson M Varela, Alvaro Santibañez, Ana M Sandino, Felipe Reyes-Lopez, Garth Dixon, and Luis A Quiñones. Smartphone screen testing, a novel pre-diagnostic method to identify SARS-CoV-2 infectious individuals. *eLife*, 10, 2021.

[10291] Matt Yousefzadeh, Chathurika Henpita, Rajesh Vyas, Carolina Soto-Palma, Paul Robbins, and Laura Niedernhofer. DNA damage—how and why we age? *eLife*, 10, 2021.

[10292] Allen T Yu, Carmen Berasain, Sonam Bhatia, Keith Rivera, Bodu Liu, Frank Rigo, Darryl J Pappin, and David L Spector. PHAROH lncRNA regulates Myc translation in hepatocellular carcinoma via sequestering TIAR. *eLife*, 10, 2021.

[10293] Liping Yu, Jiawei Hu, Chenlin Shi, Li Zhou, Maozhi Tian, Jiping Zhang, and Jinghong Xu. The causal role of auditory cortex in auditory working memory. *eLife*, 10, 2021.

[10294] Qian Yu, Soma Biswas, Gang Ma, Peiquan Zhao, Baojie Li, and Jing Li. Canonical NF-$\kappa$B signaling maintains corneal epithelial integrity and prevents corneal aging via retinoic acid. *eLife*, 10, 2021.

[10295] Rosemary Yu, Egor Vorontsov, Carina Sihlbom, and Jens Nielsen. Quantifying absolute gene expression profiles reveals distinct regulation of central carbon metabolism genes in yeast. *eLife*, 10, 2021.

[10296] Xinwei Yu, Matthew S Creamer, Francesco Randi, Anuj K Sharma, Scott W Linderman, and Andrew M Leifer. Fast deep neural correspondence for tracking and identifying neurons in C. elegans using semi-synthetic training. *eLife*, 10, 2021.

[10297] Peijun Yuan, Ruichen Hu, Xue Zhang, Ying Wang, and Yi Jiang. Cortical entrainment to hierarchical contextual rhythms recomposes dynamic attending in visual perception. *eLife*, 10, 2021.

[10298] Nicole Laurencia Yuwono, Kristina Warton, and Caroline Elizabeth Ford. The influence of biological and lifestyle factors on circulating cell-free DNA in blood plasma. *eLife*, 10, 2021.

[10299] William N Zagotta, Brandon S Sim, Anthony K Nhim, Marium M Raza, Eric GB Evans, Yarra Venkatesh, Chloe M Jones, Ryan A Mehl, E James Petersson, and Sharona E Gordon. An improved fluorescent noncanonical amino acid for measuring conformational distributions using time-resolved transition metal ion FRET. *eLife*, 10, 2021.

[10300] Sebastian H Zahler, David E Taylor, Joey Y Wong, Julia M Adams, and Evan H Feinberg. Superior colliculus drives stimulus-evoked directionally biased saccades and attempted head movements in head-fixed mice. *eLife*, 10, 2021.

[10301] Jacob A Zahm, Michael G Stewart, Joseph S Carrier, Stephen C Harrison, and Matthew P Miller. Structural basis of Stu2 recruitment to yeast kinetochores. *eLife*, 10, 2021.

[10302] Shenhab Zaig, Carolina da Silveira Scarpellini, and Gaspard Montandon. Respiratory depression and analgesia by opioid drugs in freely behaving larval zebrafish. *eLife*, 10, 2021.

[10303] Brooke Zanco, Christen K Mirth, Carla M Sgrò, and Matthew DW Piper. A dietary sterol trade-off determines lifespan responses to dietary restriction in Drosophila melanogaster females. *eLife*, 10, 2021.

[10304] Jingjing Zang, Matthias Gesemann, Jennifer Keim, Marijana Samardzija, Christian Grimm, and Stephan CF Neuhauss. Circadian regulation of vertebrate cone photoreceptor function. *eLife*, 10, 2021.

[10305] Maria Paula Zappia, Ana Guarner, Nadia Kellie-Smith, Alice Rogers, Robert Morris, Brandon Nicolay, Myriam Boukhali, Wilhelm Haas, Nicholas J Dyson, and Maxim V Frolov. E2F/Dp inactivation in fat body cells triggers systemic metabolic changes. *eLife*, 10, 2021.

[10306] Taraneh Zarin, Bob Strome, Gang Peng, Iva PritiŠanac, Julie D Forman-Kay, and Alan M Moses. Identifying molecular features that are associated with biological function of intrinsically disordered protein regions. *eLife*, 10, 2021.

[10307] Peter Zatka-Haas, Nicholas A Steinmetz, Matteo Carandini, and Kenneth D Harris. Sensory coding and the causal impact of mouse cortex in a visual decision. *eLife*, 10, 2021.

[10308] Leonie Zeitler, Alessandra Fiore, Claudia Meyer, Marion Russier, Gaia Zanella, Sabine Suppmann, Marco Gargaro, Sachdev S Sidhu, Somasekar Seshagiri, Caspar Ohnmacht, Thomas Köcher, Francesca Fallarino, Andreas Linkermann, and Peter J Murray. Anti-ferroptotic mechanism of IL4i1-mediated amino acid metabolism. *eLife*, 10, 2021.

[10309] Tima Zeng, Alexa Tompary, Anna C Schapiro, and Sharon L Thompson-Schill. Tracking the relation between gist and item memory over the course of long-term memory consolidation. *eLife*, 10, 2021.

[10310] Sara Zeppilli, Tobias Ackels, Robin Attey, Nell Klimpert, Kimberly D Ritola, Stefan Boeing, Anton Crombach, Andreas T Schaefer, and Alexander Fleischmann. Molecular characterization of projection neuron subtypes in the mouse olfactory bulb. *eLife*, 10, 2021.

[10311] Rediet Zewdu, Elnaz Mirzaei Mehrabad, Kelley Ingram, Pengshu Fang, Katherine L Gillis, Soledad A Camolotto, Grace Orstad, Alex Jones, Michelle C Mendoza, Benjamin T Spike, and Eric L Snyder. An NKX2-1/ERK/WNT feedback loop modulates gastric identity and response to targeted therapy in lung adenocarcinoma. *eLife*, 10, 2021.

[10312] Jinbin Zhai, Hyukmin Kim, Seung Baek Han, Meredith Manire, Rachel Yoo, Shuhuan Pang, George M Smith, and Young-Jin Son. Co-targeting myelin inhibitors and CSPGs markedly enhances regeneration of GDNF-stimulated, but not conditioning-lesioned, sensory axons into the spinal cord. *eLife*, 10, 2021.

[10313] Bruce Zhang and David Gems. Gross ways to live long: Parasitic worms as an anti-inflammaging therapy? *eLife*, 10, 2021.

[10314] Chunbo Zhang, Tianyu Zhong, Yuanyuan Li, Xianfeng Li, Xiaopeng Yuan, Linlin Liu, Weilin Wu, Jing Wu, Ye Wu, Rui Liang, Xinhua Xie, Chuanchuan Kang, Yuwen Liu, Zhonghong Lai, Jianbo Xiao, Zhixian Tang, Riqun Jin, Yan Wang, Yongwei Xiao, Jin Zhang, Jian Li, Qian Liu, Zhongsheng Sun, and Jianing Zhong. The hepatic AMPK-TET1-SIRT1 axis regulates glucose homeostasis. *eLife*, 10, 2021.

[10315] Haitao Zhang, Xiaolei Cao, Mei Tang, Guoxuan Zhong, Yuan Si, Haidong Li, Feifeng Zhu, Qinghua Liao, Liuju Li, Jianhui Zhao, Jia Feng, Shuaifeng Li, Chenliang Wang, Manuel Kaulich, Fangwei Wang, Liangyi Chen, Li Li, Zongping Xia, Tingbo Liang, Huasong Lu, Xin-Hua Feng, and Bin Zhao. A subcellular map of the human kinome. *eLife*, 10, 2021.

[10316] Hui Zhang, Haifang Wang, Xiaoyu Shen, Xinling Jia, Shuguang Yu, Xiaoying Qiu, Yufan Wang, Jiulin Du, Jun Yan, and Jie He. The landscape of regulatory genes in brain-wide neuronal phenotypes of a vertebrate brain. *eLife*, 10, 2021.

[10317] Jiedong Zhang, Yong Jiang, Yunjie Song, Peng Zhang, and Sheng He. Spatial tuning of face part representations within face-selective areas revealed by high-field fMRI. *eLife*, 10, 2021.

[10318] Liyun Zhang, Conan Chen, Jie Fu, Brendan Lilley, Cynthia Berlinicke, Baranda Hansen, Ding Ding, Guohua Wang, Tao Wang, Daniel Shou, Ying Ye, Timothy Mulligan, Kevin Emmerich, Meera T Saxena, Kelsi R Hall, Abigail V Sharrock, Carlene Brandon, Hyejin Park, Tae-In Kam, Valina L Dawson, Ted M Dawson, Joong Sup Shim, Justin Hanes, Hongkai Ji, Jun O Liu, Jiang Qian, David F Ackerley, Baerbel Rohrer, Donald J Zack, and Jeff S Mumm. Large-scale phenotypic drug screen identifies neuroprotectants in zebrafish and mouse models of retinitis pigmentosa. *eLife*, 10, 2021.

[10319] Martin Jinye Zhang, Angela Oliveira Pisco, Spyros Darmanis, and James Zou. Mouse aging cell atlas analysis reveals global and cell type-specific aging signatures. *eLife*, 10, 2021.

[10320] Wenchao Zhang, Mei Luo, Chunying Feng, Huaqing Liu, Hong Zhang, Rachel R Bennett, Andrew S Utada, Zhi Liu, and Kun Zhao. Crash landing of Vibrio cholerae by MSHA pili-assisted braking and anchoring in a viscoelastic environment. *eLife*, 10, 2021.

[10321] Xiao Zhang, Hero Robles, Kristann L Magee, Madelyn R Lorenz, Zhaohua Wang, Charles A Harris, Clarissa S Craft, and Erica L Scheller. A bone-specific adipogenesis pathway in fat-free mice defines key origins and adaptations of bone marrow adipocytes with age and disease. *eLife*, 10, 2021.

[10322] Xingyu Zhang, Li Zou, Lanxia Meng, Min Xiong, Lina Pan, Guiqin Chen, Yongfa Zheng, Jing Xiong, Zhihao Wang, Duc M Duong, Zhaohui Zhang, Xuebing Cao, Tao Wang, Li Tang, Keqiang Ye, and Zhentao Zhang. Amphiphysin I cleavage by asparagine endopeptidase leads to tau hyperphosphorylation and synaptic dysfunction. *eLife*, 10, 2021.

[10323] Xuebin Zhang, Penghu Lian, Mingming Su, Zhigang Ji, Jianhua Deng, Guoyang Zheng, Wenda Wang, Xinyu Ren, Taijiao Jiang, Peng Zhang, and Hanzhong Li. Single-cell transcriptome analysis identifies a unique tumor cell type producing multiple hormones in ectopic ACTH and CRH secreting pheochromocytoma. *eLife*, 10, 2021.

[10324] Yang Zhang, Gerd Bobe, Cristobal L Miranda, Malcolm B Lowry, Victor L Hsu, Christiane V Lohr, Carmen P Wong, Donald B Jump, Matthew M Robinson, Thomas J Sharpton, Claudia S Maier, Jan F Stevens, and Adrian F Gombart. Tetrahydroxanthohumol, a xanthohumol derivative, attenuates high-fat diet-induced hepatic steatosis by antagonizing PPAR$\gamma$. *eLife*, 10, 2021.

[10325] Yanmin Zhang, Sourav Chowdhury, João V Rodrigues, and Eugene Shakhnovich. Development of antibacterial compounds that constrain evolutionary pathways to resistance. *eLife*, 10, 2021.

[10326] Yaojun Zhang, Bin Xu, Benjamin G Weiner, Yigal Meir, and Ned S Wingreen. Decoding the physical principles of two-component biomolecular phase separation. *eLife*, 10, 2021.

[10327] Zhe Zhang, Jay R Gibson, and Kimberly M Huber. Experience-dependent weakening of callosal synaptic connections in the absence of postsynaptic FMRP. *eLife*, 10, 2021.

[10328] Zhi Zhang, Jae Whan Park, In Sook Ahn, Graciel Diamante, Nilla Sivakumar, Douglas Arneson, Xia Yang, J Edward van Veen, and Stephanie M Correa. Estrogen receptor alpha in the brain mediates tamoxifen-induced changes in physiology in mice. *eLife*, 10, 2021.

[10329] Bohan Zhao, Jiameng Sun, Qian Li, and Yi Zhong. Differential conditioning produces merged long-term memory in Drosophila. *eLife*, 10, 2021.

[10330] Fenghui Zhao, Chao Zhang, Qingtong Zhou, Kaini Hang, Xinyu Zou, Yan Chen, Fan Wu, Qidi Rao, Antao Dai, Wanchao Yin, Dan-Dan Shen, Yan Zhang, Tian Xia, Raymond C Stevens, H Eric Xu, Dehua Yang, Lihua Zhao, and Ming-Wei Wang. Structural insights into hormone recognition by the human glucose-dependent insulinotropic polypeptide receptor. *eLife*, 10, 2021.

[10331] Kangmei Zhao, Deze Kong, Benjamin Jin, Christina D Smolke, and Seung Yon Rhee. A novel bivalent chromatin associates with rapid induction of camalexin biosynthesis genes in response to a pathogen signal in Arabidopsis. *eLife*, 10, 2021.

[10332] Qingqing Zhao, Yang Kong, Alec Kittredge, Yao Li, Yin Shen, Yu Zhang, Stephen H Tsang, and Tingting Yang. Distinct expression requirements and rescue strategies for BEST1 loss- and gain-of-function mutations. *eLife*, 10, 2021.

[10333] Qingyuan Zhao, Jingshu Wang, Zhen Miao, Nancy R Zhang, Sean Hennessy, Dylan S Small, and Daniel J Rader. A Mendelian randomization study of the role of lipoprotein subfractions in coronary artery disease. *eLife*, 10, 2021.

[10334] Ting-Ting Zhao, Yi-Jing Feng, Pham Ngoc Doanh, Somphou Sayasone, Virak Khieu, Choosak Nithikathkul, Men-Bao Qian, Yuan-Tao Hao, and Ying-Si Lai. Model-based spatial-temporal mapping of opisthorchiasis in endemic countries of Southeast Asia. *eLife*, 10, 2021.

[10335] Tingting Zhao, Irina O Vvedenskaya, William KM Lai, Shrabani Basu, B Franklin Pugh, Bryce E Nickels, and Craig D Kaplan. Ssl2/TFIIH function in transcription start site scanning by RNA polymerase II in Saccharomyces cerevisiae. *eLife*, 10, 2021.

[10336] Xianye Zhao, Yilun Yu, Matthew E Clapham, Evgeny Yan, Jun Chen, Edmund A Jarzembowski, Xiangdong Zhao, and Bo Wang. Early evolution of beetles regulated by the end-Permian deforestation. *eLife*, 10, 2021.

[10337] Xiaochuan Zhao, Umar F Shahul Hameed, Vladlena Kharchenko, Chenyi Liao, Franceline Huser, Jacob M Remington, Anand K Radhakrishnan, Mariusz Jaremko, Łukasz Jaremko, Stefan T Arold, and Jianing Li. Molecular basis for the adaptive evolution of environment-sensing by H-NS proteins. *eLife*, 10, 2021.

[10338] Yili Zhao, Lei Zhang, Markus Rütgen, Ronald Sladky, and Claus Lamm. Neural dynamics between anterior insular cortex and right supramarginal gyrus dissociate genuine affect sharing from perceptual saliency of pretended pain. *eLife*, 10, 2021.

[10339] Vladimir Zhemkov, Jonathon A Ditlev, Wan-Ru Lee, Mikaela Wilson, Jen Liou, Michael K Rosen, and Ilya Bezprozvanny. The role of sigma 1 receptor in organization of endoplasmic reticulum signaling microdomains. *eLife*, 10, 2021.

[10340] Haining Zhong, Cesar C Ceballos, Crystian I Massengill, Michael A Muniak, Lei Ma, Maozhen Qin, Stefanie Kaech Petrie, and Tianyi Mao. High-fidelity, efficient, and reversible labeling of endogenous proteins using CRISPR-based designer exon insertion. *eLife*, 10, 2021.

[10341] Jing Zhou, Yong Lin, Trung Huynh, Hirofumi Noguchi, Jeffrey O Bush, and Samuel J Pleasure. NMDA receptors control development of somatosensory callosal axonal projections. *eLife*, 10, 2021.

[10342] Julie Y Zhou, Carlos A Alvarez, and Brian A Cobb. Integration of IL-2 and IL-4 signals coordinates divergent regulatory T cell responses and drives therapeutic efficacy. *eLife*, 10, 2021.

[10343] Shan Zhou, Weiwei Wang, Xiaoting Zhou, Yuying Zhang, Yuezheng Lai, Yanting Tang, Jinxu Xu, Dongmei Li, Jianping Lin, Xiaolin Yang, Ting Ran, Hongming Chen, Luke W Guddat, Quan Wang, Yan Gao, Zihe Rao, and Hongri Gong. Structure of Mycobacterium tuberculosis cytochrome bcc in complex with Q203 and TB47, two anti-TB drug candidates. *eLife*, 10, 2021.

[10344] Xiaoxue Zhou, Wenxue Li, Yansheng Liu, and Angelika Amon. Cross-compartment signal propagation in the mitotic exit network. *eLife*, 10, 2021.

[10345] Xin Zhou, Jennifer W Li, Zirong Chen, Wei Ni, Xuehui Li, Rongqiang Yang, Huangxuan Shen, Jian Liu, Francesco J DeMayo, Jianrong Lu, Frederic J Kaye, and Lizi Wu. Dependency of human and murine LKB1-inactivated lung cancer on aberrant CRTC-CREB activation. *eLife*, 10, 2021.

[10346] Xin Zhou, Seula Shin, Chenxi He, Qiang Zhang, Matthew N Rasband, Jiangong Ren, Congxin Dai, Rocío I Zorrilla-Veloz, Takashi Shingu, Liang Yuan, Yunfei Wang, Yiwen Chen, Fei Lan, and Jian Hu. Qki regulates myelinogenesis through Srebp2-dependent cholesterol biosynthesis. *eLife*, 10, 2021.

[10347] Yang Zhou, Matthew C Rosen, Sruthi K Swaminathan, Nicolas Y Masse, Ou Zhu, and David J Freedman. Distributed functions of prefrontal and parietal cortices during sequential categorical decisions. *eLife*, 10, 2021.

[10348] Yuan Zhou, Prativa Amom, Sarah H Reeder, Byung Ha Lee, Adam Helton, and Anna A Dobritsa. Members of the ELMOD protein family specify formation of distinct aperture domains on the Arabidopsis pollen surface. *eLife*, 10, 2021.

[10349] Meng Zhu, Marta Shahbazi, Angel Martin, Chuanxin Zhang, Berna Sozen, Mate Borsos, Rachel S Mandelbaum, Richard J Paulson, Matteo A Mole, Marga Esbert, Shiny Titus, Richard T Scott, Alison Campbell, Simon Fishel, Viviana Gradinaru, Han Zhao, Keliang Wu, Zi-Jiang Chen, Emre Seli, Maria J de los Santos, and Magdalena Zernicka Goetz. Human embryo polarization requires PLC signaling to mediate trophectoderm specification. *eLife*, 10, 2021.

[10350] Xinyu Zhu, Aren Boulet, Katherine M Buckley, Casey B Phillips, Micah G Gammon, Laura E Oldfather, Stanley A Moore, Scot C Leary, and Paul A Cobine. Mitochondrial copper and phosphate transporter specificity was defined early in the evolution of eukaryotes. *eLife*, 10, 2021.

[10351] Lucie Zilova, Venera Weinhardt, Tinatini Tavhelidse, Christina Schlagheck, Thomas Thumberger, and Joachim Wittbrodt. Fish primary embryonic pluripotent cells assemble into retinal tissue mirroring in vivo early eye development. *eLife*, 10, 2021.

[10352] Valeria Zoni, Rasha Khaddaj, Pablo Campomanes, Abdou Rachid Thiam, Roger Schneiter, and Stefano Vanni. Pre-existing bilayer stresses modulate triglyceride accumulation in the ER versus lipid droplets. *eLife*, 10, 2021.

[10353] Benkun Zou, Michael Goodwin, Danial Saleem, Wei Jiang, Jianguo Tang, Yiwei Chu, Robert S Munford, and Mingfang Lu. A highly conserved host lipase deacylates oxidized phospholipids and ameliorates acute lung injury in mice. *eLife*, 10, 2021.

[10354] Magdalena Żak and Nicolas Daudet. A gradient of Wnt activity positions the neurosensory domains of the inner ear. *eLife*, 10, 2021.

[10355] Mohamed Abbas, Anne Cori, Samuel Cordey, Florian Laubscher, Tomás Robalo Nunes, Ashleigh Myall, Julien Salamun, Philippe Huber, Dina Zekry, Virginie Prendki, Anne Iten, Laure Vieux, Valérie Sauvan, Christophe E Graf, and Stephan Harbarth. Reconstruction of transmission chains of SARS-CoV-2 amidst multiple outbreaks in a geriatric acute-care hospital: a combined retrospective epidemiological and genomic study. *eLife*, 11, 2022.

[10356] Moustafa Abdalla, Mohamed Abdalla, Salwa Abdalla, Mohamed Saad, David S Jones, and Scott H Podolsky. Insights from full-text analyses of the Journal of the American Medical Association and the New England Journal of Medicine. *eLife*, 11, 2022.

[10357] Majd Abdallah, Gaston E Zanitti, Valentin Iovene, and Demian Wassermann. Functional gradients in the human lateral prefrontal cortex revealed by a comprehensive coordinate-based meta-analysis. *eLife*, 11, 2022.

[10358] Reem Abdel-Haq, Johannes CM Schlachetzki, Joseph C Boktor, Thaisa M Cantu-Jungles, Taren Thron, Mengying Zhang, John W Bostick, Tahmineh Khazaei, Sujatha Chilakala, Livia H Morais, Greg Humphrey, Ali Keshavarzian, Jonathan E Katz, Matthew Thomson, Rob Knight, Viviana Gradinaru, Bruce R Hamaker, Christopher K Glass, and Sarkis K Mazmanian. A prebiotic diet modulates microglial states and motor deficits in $\alpha$-synuclein overexpressing mice. *eLife*, 11, 2022.

[10359] Shimaa A Abdellatef, Hisashi Tadakuma, Kangmin Yan, Takashi Fujiwara, Kodai Fukumoto, Yuichi Kondo, Hiroko Takazaki, Rofia Boudria, Takuo Yasunaga, Hideo Higuchi, and Keiko Hirose. Oscillatory movement of a dynein-microtubule complex crosslinked with DNA origami. *eLife*, 11, 2022.

[10360] Rany Abend, Diana Burk, Sonia G Ruiz, Andrea L Gold, Julia L Napoli, Jennifer C Britton, Kalina J Michalska, Tomer Shechner, Anderson M Winkler, Ellen Leibenluft, Daniel S Pine, and Bruno B Averbeck. Computational modeling of threat learning reveals links with anxiety and neuroanatomy in humans. *eLife*, 11, 2022.

[10361] Aliya Mari Adefuin, Sander Lindeman, Janine Kristin Reinert, and Izumi Fukunaga. State-dependent representations of mixtures by the olfactory bulb. *eLife*, 11, 2022.

[10362] Mohammad Afsar, Rohan Narayan, Md Noor Akhtar, Deepakash Das, Huma Rahil, Santhosh Kambaiah Nagaraj, Sandeep M Eswarappa, Shashank Tripathi, and Tanweer Hussain. Drug targeting Nsp1-ribosomal complex shows antiviral activity against SARS-CoV-2. *eLife*, 11, 2022.

[10363] Sobhika Agarwala, Keun-Young Kim, Sebastien Phan, Saeyeon Ju, Ye Eun Kong, Guillaume A Castillon, Eric A Bushong, Mark H Ellisman, and Owen J Tamplin. Defining the ultrastructure of the hematopoietic stem cell niche by correlative light and electron microscopy. *eLife*, 11, 2022.

[10364] Charles N Agoti, Lynette Isabella Ochola-Oyier, Simon Dellicour, Khadija Said Mohammed, Arnold W Lambisia, Zaydah R de Laurent, John M Morobe, Maureen W Mburu, Donwilliams O Omuoyo, Edidah M Ongera, Leonard Ndwiga, Eric Maitha, Benson Kitole, Thani Suleiman, Mohamed Mwakinangu, John K Nyambu, John Otieno, Barke Salim, Jennifer Musyoki, Nickson Murunga, Edward Otieno, John N Kiiru, Kadondi Kasera, Patrick Amoth, Mercy Mwangangi, Rashid Aman, Samson Kinyanjui, George Warimwe, My Phan, Ambrose Agweyu, Matthew Cotten, Edwine Barasa, Benjamin Tsofa, D James Nokes, Philip Bejon, and George Githinji. Transmission networks of SARS-CoV-2 in Coastal Kenya during the first two waves: A retrospective genomic study. *eLife*, 11, 2022.

[10365] Ritvija Agrawal, John P Gillies, Juliana L Zang, Jingjing Zhang, Sharon R Garrott, Hiroki Shibuya, Jayakrishnan Nandakumar, and Morgan E DeSantis. The KASH5 protein involved in meiotic chromosomal movements is a novel dynein activating adaptor. *eLife*, 11, 2022.

[10366] Nazia Ahmad, Sanmati Dugad, Varsha Chauhan, Shubbir Ahmed, Kunal Sharma, Sangita Kachhap, Rana Zaidi, William R Bishai, Gyanu Lamichhane, and Pankaj Kumar. Allosteric cooperation in $\beta$-lactam binding to a non-classical transpeptidase. *eLife*, 11, 2022.

[10367] Mai Ahmed, Yutaka Kojima, and Ichiro Masai. Strip1 regulates retinal ganglion cell survival by suppressing Jun-mediated apoptosis to promote retinal neural circuit formation. *eLife*, 11, 2022.

[10368] Honoka Aida, Takamasa Hashizume, Kazuha Ashino, and Bei-Wen Ying. Machine learning-assisted discovery of growth decision elements by relating bacterial population dynamics to environmental diversity. *eLife*, 11, 2022.

[10369] Yousra Ajouaou, Abdulkader Azouz, Anaëlle Taquin, Sebastien Denanglaire, Hind Hussein, Mohammad Krayem, Fabienne Andris, Muriel Moser, Stanislas Goriely, and Oberdan Leo. The oxygen sensor prolyl hydroxylase domain 2 regulates the in vivo suppressive capacity of regulatory T cells. *eLife*, 11, 2022.

[10370] Yemsratch T Akalu, Maria E Mercau, Marleen Ansems, Lindsey D Hughes, James Nevin, Emily J Alberto, Xinran N Liu, Li-Zhen He, Diego Alvarado, Tibor Keler, Yong Kong, William M Philbrick, Marcus Bosenberg, Silvia C Finnemann, Antonio Iavarone, Anna Lasorella, Carla V Rothlin, and Sourav Ghosh. Tissue-specific modifier alleles determine Mertk loss-of-function traits. *eLife*, 11, 2022.

[10371] Thomas Akam, Andy Lustig, James M Rowland, Sampath KT Kapanaiah, Joan Esteve-Agraz, Mariangela Panniello, Cristina Márquez, Michael M Kohl, Dennis Kätzel, Rui M Costa, and Mark E Walton. Open-source, Python-based, hardware and software for controlling behavioural neuroscience experiments. *eLife*, 11, 2022.

[10372] Vahid Akbari, Jean-Michel Garant, Kieran O'Neill, Pawan Pandoh, Richard Moore, Marco A Marra, Martin Hirst, and Steven JM Jones. Genome-wide detection of imprinted differentially methylated regions using nanopore sequencing. *eLife*, 11, 2022.

[10373] Chidiebere Akusobi, Bouchra S Benghomari, Junhao Zhu, Ian D Wolf, Shreya Singhvi, Charles L Dulberger, Thomas R Ioerger, and Eric J Rubin. Transposon mutagenesis in Mycobacterium abscessus identifies an essential penicillin-binding protein involved in septal peptidoglycan synthesis and antibiotic sensitivity. *eLife*, 11, 2022.

[10374] Basel Maher Al-Barghouthi, Will T Rosenow, Kang-Ping Du, Jinho Heo, Robert Maynard, Larry Mesner, Gina Calabrese, Aaron Nakasone, Bhavya Senwar, Louis Gerstenfeld, James Larner, Virginia Ferguson, Cheryl Ackert-Bicknell, Elise Morgan, David Brautigan, and Charles R Farber. Transcriptome-wide association study and eQTL colocalization identify potentially causal genes responsible for human bone mineral density GWAS associations. *eLife*, 11, 2022.

[10375] Tzitzitlini Alejandre-García, Samuel Kim, Jesús Pérez-Ortega, and Rafael Yuste. Intrinsic excitability mechanisms of neuronal ensemble formation. *eLife*, 11, 2022.

[10376] Aleksandr Alekseev, Georgii Pobegalov, Natalia Morozova, Alexey Vedyaykin, Galina Cherevatenko, Alexander Yakimov, Dmitry Baitin, and Mikhail Khodorkovskii. A new insight into RecA filament regulation by RecX from the analysis of conformation-specific interactions. *eLife*, 11, 2022.

[10377] Marie Alexandre, Romain Marlin, Mélanie Prague, Severin Coleon, Nidhal Kahlaoui, Sylvain Cardinaud, Thibaut Naninck, Benoit Delache, Mathieu Surenaud, Mathilde Galhaut, Nathalie Dereuddre-Bosquet, Mariangela Cavarelli, Pauline Maisonnasse, Mireille Centlivre, Christine Lacabaratz, Aurelie Wiedemann, Sandra Zurawski, Gerard Zurawski, Olivier Schwartz, Rogier W Sanders, Roger Le Grand, Yves Levy, and Rodolphe Thiébaut. Modelling the response to vaccine in non-human primates to define SARS-CoV-2 mechanistic correlates of protection. *eLife*, 11, 2022.

[10378] Massimiliano Alfano, Luca De Antoni, Federica Centofanti, Virginia Veronica Visconti, Simone Maestri, Chiara Degli Esposti, Roberto Massa, Maria Rosaria D'Apice, Giuseppe Novelli, Massimo Delledonne, Annalisa Botta, and Marzia Rossato. Characterization of full-length CNBP expanded alleles in myotonic dystrophy type 2 patients by Cas9-mediated enrichment and nanopore sequencing. *eLife*, 11, 2022.

[10379] Zain Alhashem, Dylan Feldner-Busztin, Christopher Revell, Macarena Alvarez-Garcillan Portillo, Karen Camargo-Sosa, Joanna Richardson, Manuel Rocha, Anton Gauert, Tatianna Corbeaux, Martina Milanetto, Francesco Argenton, Natascia Tiso, Robert N Kelsh, Victoria E Prince, Katie Bentley, and Claudia Linker. Notch controls the cell cycle to define leader versus follower identities during collective cell migration. *eLife*, 11, 2022.

[10380] Scott R Allen, Rebeccah K Stewart, Michael Rogers, Ivan Jimenez Ruiz, Erez Cohen, Alain Laederach, Christopher M Counter, Jessica K Sawyer, and Donald T Fox. Distinct responses to rare codons in select Drosophila tissues. *eLife*, 11, 2022.

[10381] William J Allen, Robin A Corey, Daniel W Watkins, A Sofia F Oliveira, Kiel Hards, Gregory M Cook, and Ian Collinson. Rate-limiting transport of positively charged arginine residues through the Sec-machinery is integral to the mechanism of protein secretion. *eLife*, 11, 2022.

[10382] Laura Amado-Rodríguez, Estefania Salgado del Riego, Juan Gomez de Ona, Inés López Alonso, Helena Gil-Pena, Cecilia López-Martínez, Paula Martín-Vicente, Antonio Lopez-Vazquez, Adrian Gonzalez Lopez, Elias Cuesta-Llavona, Raquel Rodriguez-Garcia, Jose Antonio Boga, Marta Elena alvarez Arguelles, Juan Mayordomo-Colunga, Jose Ramon Vidal-Castineira, Irene Crespo, Margarita Fernandez, Loreto Criado, Victoria Salvadores, Francisco Jose Jimeno-Demuth, Lluis Blanch, Belen Prieto, Alejandra Fernandez-Fernandez, Carlos Lopez-Larrea, Eliecer Coto, and Guillermo M Albaiceta. Effects of IFIH1 rs1990760 variants on systemic inflammation and outcome in critically ill COVID-19 patients in an observational translational study. *eLife*, 11, 2022.

[10383] Ryoji Amamoto, Grace K Wallick, and Constance L Cepko. Retinoic acid signaling mediates peripheral cone photoreceptor survival in a mouse model of retina degeneration. *eLife*, 11, 2022.

[10384] Adi Amar-Schwartz, Vered Ben Hur, Amina Jbara, Yuval Cohen, Georgina D Barnabas, Eliran Arbib, Zahava Siegfried, Bayan Mashahreh, Fouad Hassouna, Asaf Shilo, Mohammad Abu-Odeh, Michael Berger, Reuven Wiener, Rami Aqeilan, Tamar Geiger, and Rotem Karni. S6K1 phosphorylates Cdk1 and MSH6 to regulate DNA repair. *eLife*, 11, 2022.

[10385] Giulia Ambrosi, Oksana Voloshanenko, Antonia F Eckert, Dominique Kranz, G Ulrich Nienhaus, and Michael Boutros. Allele-specific endogenous tagging and quantitative analysis of $\beta$-catenin in colorectal cancer cells. *eLife*, 11, 2022.

[10386] Rana Amini, Archit Bhatnagar, Raimund Schlüßler, Stephanie Möllmert, Jochen Guck, and Caren Norden. Amoeboid-like migration ensures correct horizontal cell layer formation in the developing vertebrate retina. *eLife*, 11, 2022.

[10387] Marissa Jeme Andersen, ChunKi Fong, Alyssa Ann La Bella, Jonathan Jesus Molina, Alex Molesan, Matthew M Champion, Caitlin Howell, and Ana L Flores-Mireles. Inhibiting host-protein deposition on urinary catheters reduces associated urinary tract infections. *eLife*, 11, 2022.

[10388] Sarah Rose Anderson, Jacqueline M Roberts, Nathaniel Ghena, Emmalyn A Irvin, Joon Schwakopf, Isabelle B Cooperstein, Alejandra Bosco, and Monica L Vetter. Neuronal apoptosis drives remodeling states of microglia and shifts in survival pathway dependence. *eLife*, 11, 2022.

[10389] Tomas Andreani, Clark Rosensweig, Shiju Sisobhan, Emmanuel Ogunlana, William Kath, and Ravi Allada. Circadian programming of the ellipsoid body sleep homeostat in Drosophila. *eLife*, 11, 2022.

[10390] Massimo Andreatta, Ariel Tjitropranoto, Zachary Sherman, Michael C Kelly, Thomas Ciucci, and Santiago J Carmona. A CD4+ T cell reference map delineates subtype-specific adaptation during acute and chronic viral infections. *eLife*, 11, 2022.

[10391] Andrey Andreev, Valerie Komatsu, Paula Almiron, Kasey Rose, Alexandria Hughes, and Maurice Y Lee. Welcome to the lab. *eLife*, 11, 2022.

[10392] Evgeniya N Andreyeva, Alexander V Emelyanov, Markus Nevil, Lu Sun, Elena Vershilova, Christina A Hill, Michael-C Keogh, Robert J Duronio, Arthur I Skoultchi, and Dmitry V Fyodorov. Drosophila SUMM4 complex couples insulator function and DNA replication control. *eLife*, 11, 2022.

[10393] Botond Antal, Liam P McMahon, Syed Fahad Sultan, Andrew Lithen, Deborah J Wexler, Bradford Dickerson, Eva-Maria Ratai, and Lilianne R Mujica-Parodi. Type 2 diabetes mellitus accelerates brain aging and cognitive decline: Complementary findings from UK Biobank and meta-analyses. *eLife*, 11, 2022.

[10394] Brendan Antiochos, Daniela Trejo-Zambrano, Paride Fenaroli, Avi Rosenberg, Alan Baer, Archit Garg, Jungsan Sohn, Jessica Li, Michelle Petri, Daniel W Goldman, Christopher Mecoli, Livia Casciola-Rosen, and Antony Rosen. The DNA sensors AIM2 and IFI16 are SLE autoantigens that bind neutrophil extracellular traps. *eLife*, 11, 2022.

[10395] Priscila C Antonello, Thomas F Varley, John Beggs, Marimélia Porcionatto, Olaf Sporns, and Jean Faber. Self-organization of in vitro neuronal assemblies drives to complex network topology. *eLife*, 11, 2022.

[10396] James W Antony, America Romero, Anthony H Vierra, Rebecca S Luenser, Robert D Hawkins, and Kelly A Bennion. Semantic relatedness retroactively boosts memory and promotes memory interdependence across episodes. *eLife*, 11, 2022.

[10397] Lyndsey Aponik-Gremillion, Yvonne Y Chen, Eleonora Bartoli, Seth R Koslov, Hernan G Rey, Kevin S Weiner, Daniel Yoshor, Benjamin Y Hayden, Sameer A Sheth, and Brett L Foster. Distinct population and single-neuron selectivity for executive and episodic processing in human dorsal posterior cingulate. *eLife*, 11, 2022.

[10398] Marissa C Applegate and Dmitriy Aronov. Flexible use of memory by food-caching birds. *eLife*, 11, 2022.

[10399] Max Jameson Aragon, Aaron T Mok, Jamien Shea, Mengran Wang, Haein Kim, Nathan Barkdull, Chris Xu, and Nilay Yapici. Multiphoton imaging of neural structure and activity in Drosophila through the intact cuticle. *eLife*, 11, 2022.

[10400] Evan L Ardiel, Andrew Lauziere, Stephen Xu, Brandon J Harvey, Ryan Patrick Christensen, Stephen Nurrish, Joshua M Kaplan, and Hari Shroff. Stereotyped behavioral maturation and rhythmic quiescence in C. elegans embryos. *eLife*, 11, 2022.

[10401] Corson Areshenkoff, Daniel J Gale, Dominic Standage, Joseph Y Nashed, J Randall Flanagan, and Jason P Gallivan. Neural excursions from manifold structure explain patterns of learning during human sensorimotor adaptation. *eLife*, 11, 2022.

[10402] Jérémy Argenty, Nelly Rouquié, Cyrielle Bories, Suzanne Mélique, Valérie Duplan-Eche, Abdelhadi Saoudi, Nicolas Fazilleau, and Renaud Lesourne. A selective LIS1 requirement for mitotic spindle assembly discriminates distinct T-cell division mechanisms within the T-cell lineage. *eLife*, 11, 2022.

[10403] Thodoris Argyriou, Sam Giles, and Matt Friedman. A Permian fish reveals widespread distribution of neopterygian-like jaw suspension. *eLife*, 11, 2022.

[10404] Giacomo Ariani, J Andrew Pruszynski, and Jörn Diedrichsen. Motor planning brings human primary somatosensory cortex into action-specific preparatory states. *eLife*, 11, 2022.

[10405] Charlotte Arlt, Roberto Barroso-Luque, Shinichiro Kira, Carissa A Bruno, Ningjing Xia, Selmaan N Chettih, Sofia Soares, Noah L Pettit, and Christopher D Harvey. Cognitive experience alters cortical involvement in goal-directed navigation. *eLife*, 11, 2022.

[10406] Manoj Arra, Gaurav Swarnkar, Naga Suresh Adapala, Syeda Kanwal Naqvi, Lei Cai, Muhammad Farooq Rai, Srikanth Singamaneni, Gabriel Mbalaviele, Robert Brophy, and Yousef Abu-Amer. Glutamine metabolism modulates chondrocyte inflammatory response. *eLife*, 11, 2022.

[10407] Brogan Ashley, Claire Simner, Antigoni Manousopoulou, Carl Jenkinson, Felicity Hey, Jennifer M Frost, Faisal I Rezwan, Cory H White, Emma M Lofthouse, Emily Hyde, Laura DF Cooke, Sheila Barton, Pamela Mahon, Elizabeth M Curtis, Rebecca J Moon, Sarah R Crozier, Hazel M Inskip, Keith M Godfrey, John W Holloway, Cyrus Cooper, Kerry S Jones, Rohan M Lewis, Martin Hewison, Spiros DD Garbis, Miguel R Branco, Nicholas C Harvey, and Jane K Cleal. Placental uptake and metabolism of 25(OH)vitamin D determine its activity within the fetoplacental unit. *eLife*, 11, 2022.

[10408] Théo Aspert, Didier Hentsch, and Gilles Charvin. DetecDiv, a generalist deep-learning platform for automated cell division tracking and survival analysis. *eLife*, 11, 2022.

[10409] Antonio Astorga-Gamaza, Judith Grau-Expósito, Joaquín Burgos, Jordi Navarro, Adrià Curran, Bibiana Planas, Paula Suanzes, VicenÇ Falcó, Meritxell Genescà, and Maria J Buzon. Identification of HIV-reservoir cells with reduced susceptibility to antibody-dependent immune response. *eLife*, 11, 2022.

[10410] Lauren Y Atlas, Troy C Dildine, Esther E Palacios-Barrios, Qingbao Yu, Richard C Reynolds, Lauren A Banker, Shara S Grant, and Daniel S Pine. Instructions and experiential learning have similar impacts on pain and pain-related brain responses but produce dissociations in value-based reversal learning. *eLife*, 11, 2022.

[10411] Bahaaeddin Attaallah, Pierre Petitet, Elista Slavkova, Vicky Turner, Youssuf Saleh, Sanjay G Manohar, and Masud Husain. Hyperreactivity to uncertainty is a key feature of subjective cognitive impairment. *eLife*, 11, 2022.

[10412] Heiner Atze, Yucheng Liang, Jean-Emmanuel Hugonnet, Arnaud Gutierrez, Filippo Rusconi, and Michel Arthur. Heavy isotope labeling and mass spectrometry reveal unexpected remodeling of bacterial cell wall expansion in response to drugs. *eLife*, 11, 2022.

[10413] Benjamin Auxier, Tamás L Czárán, and Duur K Aanen. Modeling the consequences of the dikaryotic life cycle of mushroom-forming fungi on genomic conflict. *eLife*, 11, 2022.

[10414] Charlotte Avet, Arturo Mancini, Billy Breton, Christian Le Gouill, Alexander S Hauser, Claire Normand, Hiroyuki Kobayashi, Florence Gross, Mireille Hogue, Viktoriya Lukasheva, Stéphane St-Onge, Marilyn Carrier, Madeleine Héroux, Sandra Morissette, Eric B Fauman, Jean-Philippe Fortin, Stephan Schann, Xavier Leroy, David E Gloriam, and Michel Bouvier. Effector membrane translocation biosensors reveal G protein and $\beta$arrestin coupling profiles of 100 therapeutically relevant GPCRs. *eLife*, 11, 2022.

[10415] Daniela ávila González, Wendy Portillo, Carla P Barragán-álvarez, Georgina Hernandez-Montes, Eliezer Flores-Garza, Anayansi Molina-Hernández, Néstor Emmanuel Díaz-Martínez, and Néstor F Díaz. The human amniotic epithelium confers a bias to differentiate toward the neuroectoderm lineage in human embryonic stem cells. *eLife*, 11, 2022.

[10416] Guy Avraham, Jordan A Taylor, Assaf Breska, Richard B Ivry, and Samuel D McDougle. Contextual effects in sensorimotor adaptation adhere to associative learning rules. *eLife*, 11, 2022.

[10417] Florian Aymanns, Chin-Lin Chen, and Pavan Ramdya. Descending neuron population dynamics during odor-evoked and spontaneous limb-dependent behaviors. *eLife*, 11, 2022.

[10418] Vladislav Ayzenberg and Stella Lourenco. Perception of an object's global shape is best described by a model of skeletal structure in human infants. *eLife*, 11, 2022.

[10419] Hind Baba Aïssa, Romain W Sala, Elena Laura Georgescu Margarint, Jimena Laura Frontera, Andrés Pablo Varani, Fabien Menardy, Assunta Pelosi, Denis Hervé, Clément Léna, and Daniela Popa. Functional abnormalities in the cerebello-thalamic pathways in a mouse model of DYT25 dystonia. *eLife*, 11, 2022.

[10420] ákos Babiczky and Ferenc Matyas. Molecular characteristics and laminar distribution of prefrontal neurons projecting to the mesolimbic system. *eLife*, 11, 2022.

[10421] Anamaria Babosan, David Skurnik, Anaëlle Muggeo, Gerald B Pier, Zeynep Baharoglu, Thomas Jové, Marie-Cécile Ploy, Sophie Griveau, Fethi Bedioui, Sébastien Vergnolle, Sophie Moussalih, Christophe de Champs, Didier Mazel, and Thomas Guillard. A qnr-plasmid allows aminoglycosides to induce SOS in Escherichia coli. *eLife*, 11, 2022.

[10422] Swathy Babu, Yuki Takeuchi, and Ichiro Masai. Banp regulates DNA damage response and chromosome segregation during the cell cycle in zebrafish retina. *eLife*, 11, 2022.

[10423] Martin Baccino-Calace, Katharina Schmidt, and Martin Müller. The E3 ligase Thin controls homeostatic plasticity through neurotransmitter release repression. *eLife*, 11, 2022.

[10424] Harshith Bachimanchi, Benjamin Midtvedt, Daniel Midtvedt, Erik Selander, and Giovanni Volpe. Microplankton life histories revealed by holographic microscopy and deep learning. *eLife*, 11, 2022.

[10425] Mihaly Badonyi and Joseph A Marsh. Large protein complex interfaces have evolved to promote cotranslational assembly. *eLife*, 11, 2022.

[10426] Hwan Bae, Thibault Viennet, Eunyoung Park, Nam Chu, Antonieta Salguero, Michael J Eck, Haribabu Arthanari, and Philip A Cole. PH domain-mediated autoinhibition and oncogenic activation of Akt. *eLife*, 11, 2022.

[10427] Soo Ji Baek, Jin Sung Park, Jinhyun Kim, Yukio Yamamoto, and Keiko Tanaka-Yamamoto. VTA-projecting cerebellar neurons mediate stress-dependent depression-like behaviors. *eLife*, 11, 2022.

[10428] Ling Bai, Nilla Sivakumar, Shenliang Yu, Sheyda Mesgarzadeh, Tom Ding, Truong Ly, Timothy V Corpuz, James CR Grove, Brooke C Jarvie, and Zachary A Knight. Enteroendocrine cell types that drive food reward and aversion. *eLife*, 11, 2022.

[10429] Andrew D Bailey, Jason Talkish, Hongxu Ding, Haller Igel, Alejandra Duran, Shreya Mantripragada, Benedict Paten, and Manuel Ares. Concerted modification of nucleotides at functional centers of the ribosome revealed by single-molecule RNA modification profiling. *eLife*, 11, 2022.

[10430] Raisa Bailon-Zambrano, Juliana Sucharov, Abigail Mumme-Monheit, Matthew Murry, Amanda Stenzel, Anthony T Pulvino, Jennyfer M Mitchell, Kathryn L Colborn, and James T Nichols. Variable paralog expression underlies phenotype variation. *eLife*, 11, 2022.

[10431] Patrick Bakenecker, Tobias Weingarten, Daniel Hahn, and Brent Raiteri. Residual force enhancement is affected more by quadriceps muscle length than stretch amplitude. *eLife*, 11, 2022.

[10432] EmilyClare P Baker, Ryan Sayegh, Kristin M Kohler, Wyatt Borman, Claire K Goodfellow, Eden R Brush, and Matthew F Barber. Evolution of host-microbe cell adherence by receptor domain shuffling. *eLife*, 11, 2022.

[10433] Gert-Jan Bakker, Sarah Weischer, Júlia Ferrer Ortas, Judith Heidelin, Volker Andresen, Marcus Beutler, Emmanuel Beaurepaire, and Peter Friedl. Intravital deep-tumor single-beam 3-photon, 4-photon, and harmonic microscopy. *eLife*, 11, 2022.

[10434] Priya Balasubramanian, Anne E Schaar, Grace E Gustafson, Alex B Smith, Porsha R Howell, Angela Greenman, Scott Baum, Ricki J Colman, Dudley W Lamming, Gary M Diffee, and Rozalyn M Anderson. Adiponectin receptor agonist AdipoRon improves skeletal muscle function in aged mice. *eLife*, 11, 2022.

[10435] Sandhya Balasubramanian, Matteo Andreani, Júlia Goncalves Andrade, Tannishtha Saha, Devakumar Sundaravinayagam, Javier Garzón, Wenzhu Zhang, Oliver Popp, Shin-ichiro Hiraga, Ali Rahjouei, Daniel B Rosen, Philipp Mertins, Brian T Chait, Anne D Donaldson, and Michela Di Virgilio. Protection of nascent DNA at stalled replication forks is mediated by phosphorylation of RIF1 intrinsically disordered region. *eLife*, 11, 2022.

[10436] Austin T Baldwin, Juliana H Kim, Hyemin Seo, and John B Wallingford. Global analysis of cell behavior and protein dynamics reveals region-specific roles for Shroom3 and N-cadherin during neural tube closure. *eLife*, 11, 2022.

[10437] Namrata Bali, Hyung-Kook (Peter) Lee, and Kai Zinn. Sticks and Stones, a conserved cell surface ligand for the Type IIa RPTP Lar, regulates neural circuit wiring in Drosophila. *eLife*, 11, 2022.

[10438] Jee Yoon Bang, Julia Kathryn Sunstrum, Danielle Garand, Gustavo Morrone Parfitt, Melanie Woodin, Wataru Inoue, and Junchul Kim. Hippocampal-hypothalamic circuit controls context-dependent innate defensive responses. *eLife*, 11, 2022.

[10439] Keting Bao, Wenwen Liu, Zhouzhi Song, Jiali Feng, Zhifan Mao, Lingyuan Bao, Tianyue Sun, Zelan Hu, and Jian Li. Crotamiton derivative JM03 extends lifespan and improves oxidative and hypertonic stress resistance in Caenorhabditis elegans via inhibiting OSM-9. *eLife*, 11, 2022.

[10440] Mayank Baranwal, Ryan L Clark, Jaron Thompson, Zeyu Sun, Alfred O Hero, and Ophelia S Venturelli. Recurrent neural networks enable design of multifunctional synthetic human gut microbiome dynamics. *eLife*, 11, 2022.

[10441] Paul R Barber, Rami Mustapha, Fabian Flores-Borja, Giovanna Alfano, Kenrick Ng, Gregory Weitsman, Luigi Dolcetti, Ali Abdulnabi Suwaidan, Felix Wong, Jose M Vicencio, Myria Galazi, James W Opzoomer, James N Arnold, Selvam Thavaraj, Shahram Kordasti, Jana Doyle, Jon Greenberg, Magnus T Dillon, Kevin J Harrington, Martin Forster, Anthony CC Coolen, and Tony Ng. Predicting progression-free survival after systemic therapy in advanced head and neck cancer: Bayesian regression and model development. *eLife*, 11, 2022.

[10442] Carlos-Henrique D Barbosa, Fábio B Canto, Ariel Gomes, Layza M Brandao, Jéssica R Lima, Guilherme A Melo, Alessandra Granato, Eula GA Neves, Walderez O Dutra, Ana-Carolina Oliveira, Alberto Nóbrega, and Maria Bellio. Cytotoxic CD4+ T cells driven by T-cell intrinsic IL-18R/MyD88 signaling predominantly infiltrate Trypanosoma cruzi-infected hearts. *eLife*, 11, 2022.

[10443] Nicholas W Barendregt, Joshua I Gold, KreŠimir Josić, and Zachary P Kilpatrick. Normative decision rules in changing environments. *eLife*, 11, 2022.

[10444] Natasja Barki, Daniele Bolognini, Ulf Börjesson, Laura Jenkins, John Riddell, David I Hughes, Trond Ulven, Brian D Hudson, Elisabeth Rexen Ulven, Niek Dekker, Andrew B Tobin, and Graeme Milligan. Chemogenetics defines a short-chain fatty acid receptor gut–brain axis. *eLife*, 11, 2022.

[10445] Ana Beatriz F Barletta, Banhisikha Saha, Nathanie Trisnadi, Octavio AC Talyuli, Gianmarco Raddi, and Carolina Barillas-Mury. Hemocyte differentiation to the megacyte lineage enhances mosquito immunity against Plasmodium. *eLife*, 11, 2022.

[10446] Samuel J Barnes, Georg B Keller, and Tara Keck. Homeostatic regulation through strengthening of neuronal network-correlated synaptic inputs. *eLife*, 11, 2022.

[10447] Justinn Barr, Maria Elena Gentile, Sunyoung Lee, Maya E Kotas, Maria Fernanda de Mello Costa, Nicolas P Holcomb, Abigail Jaquish, Gargi Palashikar, Marcella Soewignjo, Margaret McDaniel, Ichiro Matsumoto, Robert Margolskee, Jakob Von Moltke, Noam A Cohen, Xin Sun, and Andrew E Vaughan. Injury-induced pulmonary tuft cells are heterogenous, arise independent of key Type 2 cytokines, and are dispensable for dysplastic repair. *eLife*, 11, 2022.

[10448] Kelsey Barrasso, Denise Chac, Meti D Debela, Catherine Geigel, Anjali Steenhaut, Abigail Rivera Seda, Chelsea N Dunmire, Jason B Harris, Regina C Larocque, Firas S Midani, Firdausi Qadri, Jing Yan, Ana A Weil, and Wai-Leung Ng. Impact of a human gut microbe on Vibrio cholerae host colonization through biofilm enhancement. *eLife*, 11, 2022.

[10449] Lindy E Barrett. Why both sides of the gender equation matter. *eLife*, 11, 2022.

[10450] Benjamin Barsi-Rhyne, Aashish Manglik, and Mark von Zastrow. Discrete GPCR-triggered endocytic modes enable $\beta$-arrestins to flexibly regulate cell signaling. *eLife*, 11, 2022.

[10451] Rachael E Barton, Lucia F Massari, Daniel Robertson, and Adèle L Marston. Eco1-dependent cohesin acetylation anchors chromatin loops and cohesion to define functional meiotic chromosome domains. *eLife*, 11, 2022.

[10452] Hila Barzilai-Tutsch, Valerie Morin, Gauthier Toulouse, Oleksandr Chernyavskiy, Stephen Firth, Christophe Marcelle, and Olivier Serralbo. Transgenic quails reveal dynamic TCF/$\beta$-catenin signaling during avian embryonic development. *eLife*, 11, 2022.

[10453] Angika Basant and Michael Way. The relative binding position of Nck and Grb2 adaptors impacts actin-based motility of Vaccinia virus. *eLife*, 11, 2022.

[10454] Mustafa Basaran, Y Ilker Yaman, Tevfik Can Yüce, Roman Vetter, and Askin Kocabas. Large-scale orientational order in bacterial colonies during inward growth. *eLife*, 11, 2022.

[10455] Hauke Basedau, Lisa-Marie Sturm, Jan Mehnert, Kuan-Po Peng, Marlene Schellong, and Arne May. Migraine monoclonal antibodies against CGRP change brain activity depending on ligand or receptor target – an fMRI study. *eLife*, 11, 2022.

[10456] Lisa Baumgartner, Dominik Handler, Sebastian Wolfgang Platzer, Changwei Yu, Peter Duchek, and Julius Brennecke. The Drosophila ZAD zinc finger protein Kipferl guides Rhino to piRNA clusters. *eLife*, 11, 2022.

[10457] Elizabeth A Bearce, Zoe H Irons, Johnathan R O'Hara-Smith, Colin J Kuhns, Sophie I Fisher, William E Crow, and Daniel T Grimes. Urotensin II-related peptides, Urp1 and Urp2, control zebrafish spine morphology. *eLife*, 11, 2022.

[10458] Antoine Beauchamp, Yohan Yee, Ben C Darwin, Armin Raznahan, Rogier B Mars, and Jason P Lerch. Whole-brain comparison of rodent and human brains using spatial transcriptomics. *eLife*, 11, 2022.

[10459] Yannick Becker, Nicolas Claidière, Konstantina Margiotoudi, Damien Marie, Muriel Roth, Bruno Nazarian, Jean-Luc Anton, Olivier Coulon, and Adrien Meguerditchian. Broca's cerebral asymmetry reflects gestural communication's lateralisation in monkeys (Papio anubis). *eLife*, 11, 2022.

[10460] Danielle Beckman, Adele MH Seelke, Jeffrey Bennett, Paige Dougherty, Koen KA Van Rompay, Rebekah Keesler, Patricia A Pesavento, Lark LA Coffey, John H Morrison, and Eliza Bliss-Moreau. Neuroanatomical abnormalities in a nonhuman primate model of congenital Zika virus infection. *eLife*, 11, 2022.

[10461] Verena Behringer, Andreas Berghänel, Tobias Deschner, Sean M Lee, Barbara Fruth, and Gottfried Hohmann. Transition to siblinghood causes a substantial and long-lasting increase in urinary cortisol levels in wild bonobos. *eLife*, 11, 2022.

[10462] Kevin Beier. Modified viral-genetic mapping reveals local and global connectivity relationships of ventral tegmental area dopamine cells. *eLife*, 11, 2022.

[10463] Pearis Bellamy. A guide for writing anti-racist tenure and promotion letters. *eLife*, 11, 2022.

[10464] Saul Bello-Rojas and Martha W Bagnall. Clonally related, Notch-differentiated spinal neurons integrate into distinct circuits. *eLife*, 11, 2022.

[10465] Nicola Bellotto, Jaime Agudo-Canalejo, Remy Colin, Ramin Golestanian, Gabriele Malengo, and Victor Sourjik. Dependence of diffusion in Escherichia coli cytoplasm on protein size, environmental conditions, and cell growth. *eLife*, 11, 2022.

[10466] Daniel W Belsky, Avshalom Caspi, David L Corcoran, Karen Sugden, Richie Poulton, Louise Arseneault, Andrea Baccarelli, Kartik Chamarti, Xu Gao, Eilis Hannon, Hona Lee Harrington, Renate Houts, Meeraj Kothari, Dayoon Kwon, Jonathan Mill, Joel Schwartz, Pantel Vokonas, Cuicui Wang, Benjamin S Williams, and Terrie E Moffitt. DunedinPACE, a DNA methylation biomarker of the pace of aging. *eLife*, 11, 2022.

[10467] Corinne Belville, Flora Ponelle-Chachuat, Marion Rouzaire, Christelle Gross, Bruno Pereira, Denis Gallot, Vincent Sapin, and Loïc Blanchon. Physiological TLR4 regulation in human fetal membranes as an explicative mechanism of a pathological preterm case. *eLife*, 11, 2022.

[10468] Vladislav Belyy, Iratxe Zuazo-Gaztelu, Andrew Alamban, Avi Ashkenazi, and Peter Walter. Endoplasmic reticulum stress activates human IRE1$\alpha$ through reversible assembly of inactive dimers into small oligomers. *eLife*, 11, 2022.

[10469] Aviad Ben-Shmuel, Batel Sabag, Abhishek Puthenveetil, Guy Biber, Moria Levy, Tammir Jubany, Fatima Awwad, Roshan Kumar Roy, Noah Joseph, Omri Matalon, Jessica Kivelevitz, and Mira Barda-Saad. Inhibition of SHP-1 activity by PKC-$\theta$ regulates NK cell activation threshold and cytotoxicity. *eLife*, 11, 2022.

[10470] Corinne Benakis, Alba Simats, Sophie Tritschler, Steffanie Heindl, Simon Besson-Girard, Gemma Llovera, Kelsey Pinkham, Anna Kolz, Alessio Ricci, Fabian J Theis, Stefan Bittner, özgün Gökce, Anneli Peters, and Arthur Liesz. T cells modulate the microglial response to brain ischemia. *eLife*, 11, 2022.

[10471] Gonzalo Benegas, Jonathan Fischer, and Yun S Song. Robust and annotation-free analysis of alternative splicing across diverse cell types in mice. *eLife*, 11, 2022.

[10472] Samuel A Bentley, Hannah Laeverenz-Schlogelhofer, Vasileios Anagnostidis, Jan Cammann, Marco G Mazza, Fabrice Gielen, and Kirsty Y Wan. Phenotyping single-cell motility in microfluidic confinement. *eLife*, 11, 2022.

[10473] Jiyoon Beon, Sungwook Han, Hyeokjun Yang, Seung Eun Park, Kwangbeom Hyun, Song-Yi Lee, Hyun-Woo Rhee, Jeong Kon Seo, Jaehoon Kim, Seyun Kim, and Daeyoup Lee. Inositol polyphosphate multikinase physically binds to the SWI/SNF complex and modulates BRG1 occupancy in mouse embryonic stem cells. *eLife*, 11, 2022.

[10474] Manindra Bera, Sathish Ramakrishnan, Jeff Coleman, Shyam S Krishnakumar, and James E Rothman. Molecular determinants of complexin clamping and activation function. *eLife*, 11, 2022.

[10475] Andrew C Bergen, Rachel A Kocik, James Hose, Megan N McClean, and Audrey P Gasch. Modeling single-cell phenotypes links yeast stress acclimation to transcriptional repression and pre-stress cellular states. *eLife*, 11, 2022.

[10476] Lucie A Bergeron, Søren Besenbacher, Tychele Turner, Cyril J Versoza, Richard J Wang, Alivia Lee Price, Ellie Armstrong, Meritxell Riera, Jedidiah Carlson, Hwei-yen Chen, Matthew W Hahn, Kelley Harris, April Snøfrid Kleppe, Elora H López-Nandam, Priya Moorjani, Susanne P Pfeifer, George P Tiley, Anne D Yoder, Guojie Zhang, and Mikkel H Schierup. The Mutationathon highlights the importance of reaching standardization in estimates of pedigree-based germline mutation rates. *eLife*, 11, 2022.

[10477] Jeffrey C Berry, Mingsheng Qi, Balasaheb V Sonawane, Amy Sheflin, Asaph Cousins, Jessica Prenni, Daniel P Schachtman, Peng Liu, and Rebecca S Bart. Increased signal-to-noise ratios within experimental field trials by regressing spatially distributed soil properties as principal components. *eLife*, 11, 2022.

[10478] Amol Bhandare, Joseph van de Wiel, Reno Roberts, Ingke Braren, Robert Huckstepp, and Nicholas Dale. Analyzing the brainstem circuits for respiratory chemosensitivity in freely moving mice. *eLife*, 11, 2022.

[10479] Tapomoy Bhattacharjee, Daniel B Amchin, Ricard Alert, Jenna Anne Ott, and Sujit Sankar Datta. Chemotactic smoothing of collective migration. *eLife*, 11, 2022.

[10480] Yanxia Bi, Jian Su, Shengru Zhou, Yingjie Zhao, Yan Zhang, Huihui Zhang, Mingdong Liu, Aiwu Zhou, Jianrong Xu, Meng Pan, Yiming Zhao, and Fubin Li. Distinct impact of IgG subclass on autoantibody pathogenicity in different IgG4-mediated diseases. *eLife*, 11, 2022.

[10481] Brianna Bibel, Elad Elkayam, Steve Silletti, Elizabeth A Komives, and Leemor Joshua-Tor. Target binding triggers hierarchical phosphorylation of human Argonaute-2 to promote target release. *eLife*, 11, 2022.

[10482] Annie R Bice, Qingli Xiao, Justin Kong, Ping Yan, Zachary Pollack Rosenthal, Andrew W Kraft, Karen P Smith, Tadeusz Wieloch, Jin-Moo Lee, Joseph P Culver, and Adam Q Bauer. Homotopic contralesional excitation suppresses spontaneous circuit repair and global network reconnections following ischemic stroke. *eLife*, 11, 2022.

[10483] Emmanuelle Bioud, Corentin Tasu, and Mathias Pessiglione. A computational account of why more valuable goals seem to require more effortful actions. *eLife*, 11, 2022.

[10484] Sonja Bisch-Knaden, Michelle A Rafter, Markus Knaden, and Bill S Hansson. Unique neural coding of crucial versus irrelevant plant odors in a hawkmoth. *eLife*, 11, 2022.

[10485] Mitchell Bishop, Maximilian Weinhold, Ariana Z Turk, Afuh Adeck, and Shahriar SheikhBahaei. An open-source tool for automated analysis of breathing behaviors in common marmosets and rodents. *eLife*, 11, 2022.

[10486] Sebastian H Bitzenhofer, Elena A Westeinde, Han-Xiong Bear Zhang, and Jeffry S Isaacson. Rapid odor processing by layer 2 subcircuits in lateral entorhinal cortex. *eLife*, 11, 2022.

[10487] Davia Blake, Caleb M Radens, Max B Ferretti, Matthew R Gazzara, and Kristen W Lynch. Alternative splicing of apoptosis genes promotes human T cell survival. *eLife*, 11, 2022.

[10488] Matthew R Blake, Diana C Parrish, Melanie A Staffenson, Shanice Sueda, William R Woodward, and Beth A Habecker. Chondroitin sulfate proteoglycan 4,6 sulfation regulates sympathetic nerve regeneration after myocardial infarction. *eLife*, 11, 2022.

[10489] Matthias Blanc, Giovanni Dalmasso, Frederic Udina, and Cristina Pujades. A dynamic and expandable digital 3D-atlas maker for monitoring the temporal changes in tissue growth during hindbrain morphogenesis. *eLife*, 11, 2022.

[10490] FranÇois Blanquart, Nathanaël Hozé, Benjamin John Cowling, Florence Débarre, and Simon Cauchemez. Selection for infectivity profiles in slow and fast epidemics, and the rise of SARS-CoV-2 variants. *eLife*, 11, 2022.

[10491] Florian Bleffert, Joachim Granzin, Muttalip Caliskan, Stephan N Schott-Verdugo, Meike Siebers, Björn Thiele, Laurence Rahme, Sebastian Felgner, Peter Dörmann, Holger Gohlke, Renu Batra-Safferling, Karl-Erich Jaeger, and Filip Kovacic. Structural, mechanistic, and physiological insights into phospholipase A-mediated membrane phospholipid degradation in Pseudomonas aeruginosa. *eLife*, 11, 2022.

[10492] Alexandra Blenkinsop, Mélodie Monod, Ard van Sighem, Nikos Pantazis, Daniela Bezemer, Eline Op de Coul, Thijs van de Laar, Christophe Fraser, Maria Prins, Peter Reiss, Godelieve J de Bree, and Oliver Ratmann. Estimating the potential to prevent locally acquired HIV infections in a UNAIDS Fast-Track City, Amsterdam. *eLife*, 11, 2022.

[10493] Marilou Boddé, Alex Makunin, Diego Ayala, Lemonde Bouafou, Abdoulaye Diabaté, Uwem Friday Ekpo, Mahamadi Kientega, Gilbert Le Goff, Boris K Makanga, Marc F Ngangue, Olaitan Olamide Omitola, Nil Rahola, Frederic Tripet, Richard Durbin, and Mara KN Lawniczak. High-resolution species assignment of Anopheles mosquitoes using k-mer distances on targeted sequences. *eLife*, 11, 2022.

[10494] Chandra Sekhar Boddupalli, Shiny Nair, Glenn Belinsky, Joseph Gans, Erin Teeple, Tri-Hung Nguyen, Sameet Mehta, Lilu Guo, Martin L Kramer, Jiapeng Ruan, Honggge Wang, Matthew Davison, Dinesh Kumar, DJ Vidyadhara, Bailin Zhang, Katherine Klinger, and Pramod K Mistry. Neuroinflammation in neuronopathic Gaucher disease: Role of microglia and NK cells, biomarkers, and response to substrate reduction therapy. *eLife*, 11, 2022.

[10495] Ariane C Boehm, Anja B Friedrich, Sydney Hunt, Paul Bandow, KP Siju, Jean Francois De Backer, Julia Claussen, Marie Helen Link, Thomas F Hofmann, Corinna Dawid, and Ilona C Grunwald Kadow. A dopamine-gated learning circuit underpins reproductive state-dependent odor preference in Drosophila females. *eLife*, 11, 2022.

[10496] Jan Boelts, Jan-Matthis Lueckmann, Richard Gao, and Jakob H Macke. Flexible and efficient simulation-based inference for models of decision-making. *eLife*, 11, 2022.

[10497] Jerome Bohere, Buffy L Eldridge-Thomas, and Golnar Kolahgar. Vinculin recruitment to $\alpha$-catenin halts the differentiation and maturation of enterocyte progenitors to maintain homeostasis of the Drosophila intestine. *eLife*, 11, 2022.

[10498] Arnau Bolet, Thomas L Stubbs, Jorge A Herrera-Flores, and Michael J Benton. The Jurassic rise of squamates as supported by lepidosaur disparity and evolutionary rates. *eLife*, 11, 2022.

[10499] Elena Bollati, Niclas H Lyndby, Cecilia D'Angelo, Michael Kühl, Jörg Wiedenmann, and Daniel Wangpraseurt. Green fluorescent protein-like pigments optimise the internal light environment in symbiotic reef-building corals. *eLife*, 11, 2022.

[10500] Saskia Bollmann, Hendrik Mattern, Michaël Bernier, Simon D Robinson, Daniel Park, Oliver Speck, and Jonathan R Polimeni. Imaging of the pial arterial vasculature of the human brain in vivo using high-resolution 7T time-of-flight angiography. *eLife*, 11, 2022.

[10501] Daan B Boltje, Jacob P Hoogenboom, Arjen J Jakobi, Grant J Jensen, Caspar TH Jonker, Max J Kaag, Abraham J Koster, Mart GF Last, Cecilia de Agrela Pinto, Jürgen M Plitzko, Stefan Raunser, Sebastian Tacke, Zhexin Wang, Ernest B van der Wee, Roger Wepf, and Sander den Hoedt. A cryogenic, coincident fluorescence, electron, and ion beam microscope. *eLife*, 11, 2022.

[10502] Klaus Bonazza, Roxana E Iacob, Nathan E Hudson, Jing Li, Chafen Lu, John R Engen, and Timothy A Springer. Von Willebrand factor A1 domain stability and affinity for GPIb$\alpha$ are differentially regulated by its O-glycosylated N- and C-linker. *eLife*, 11, 2022.

[10503] Jeffrey P Bonin, Paul J Sapienza, and Andrew L Lee. Dynamic allostery in substrate binding by human thymidylate synthase. *eLife*, 11, 2022.

[10504] Elsa Bonnard, Jun Liu, Nicolina Zjacic, Luis Alvarez, and Monika Scholz. Automatically tracking feeding behavior in populations of foraging C. elegans. *eLife*, 11, 2022.

[10505] Morgane Boone, Lan Wang, Rosalie E Lawrence, Adam Frost, Peter Walter, and Michael Schoof. A point mutation in the nucleotide exchange factor eIF2B constitutively activates the integrated stress response by allosteric modulation. *eLife*, 11, 2022.

[10506] James Boot, Gabriel Rosser, Dailya Kancheva, Claire Vinel, Yau Mun Lim, Nicola Pomella, Xinyu Zhang, Loredana Guglielmi, Denise Sheer, Michael Barnes, Sebastian Brandner, Sven Nelander, Kiavash Movahedi, and Silvia Marino. Global hypo-methylation in a proportion of glioblastoma enriched for an astrocytic signature is associated with increased invasion and altered immune landscape. *eLife*, 11, 2022.

[10507] Blake Bordelon and Cengiz Pehlevan. Population codes enable learning from few examples by shaping inductive bias. *eLife*, 11, 2022.

[10508] Jazlyn P Borges, Ragnhild SR Sætra, Allen Volchuk, Marit Bugge, Pascal Devant, Bjørnar Sporsheim, Bridget R Kilburn, Charles L Evavold, Jonathan C Kagan, Neil M Goldenberg, Trude Helen Flo, and Benjamin Ethan Steinberg. Glycine inhibits NINJ1 membrane clustering to suppress plasma membrane rupture in cell death. *eLife*, 11, 2022.

[10509] Parastoo Boroumand, David C Prescott, Tapas Mukherjee, Philip J Bilan, Michael Wong, Jeff Shen, Ivan Tattoli, Yuhuan Zhou, Angela Li, Tharini Sivasubramaniyam, Nancy Shi, Lucie Y Zhu, Zhi Liu, Clinton Robbins, Dana J Philpott, Stephen E Girardin, and Amira Klip. Bone marrow adipocytes drive the development of tissue invasive Ly6Chigh monocytes during obesity. *eLife*, 11, 2022.

[10510] Alberto Borsatto, Obaeda Akkad, Ioannis Galdadas, Shumeng Ma, Shymaa Damfo, Shozeb Haider, Frank Kozielski, Carolina Estarellas, and Francesco Luigi Gervasio. Revealing druggable cryptic pockets in the Nsp1 of SARS-CoV-2 and other $\beta$-coronaviruses by simulations and crystallography. *eLife*, 11, 2022.

[10511] Guillermo Bosch, Margarita Posso, Javier Louro, Marta Roman, Miquel Porta, Xavier Castells, and Francesc Macià. Impact of the COVID-19 pandemic on breast cancer screening indicators in a Spanish population-based program: a cohort study. *eLife*, 11, 2022.

[10512] Justin A Bosch, Berrak Ugur, Israel Pichardo-Casas, Jordan Rabasco, Felipe Escobedo, Zhongyuan Zuo, Ben Brown, Susan Celniker, David A Sinclair, Hugo J Bellen, and Norbert Perrimon. Two neuronal peptides encoded from a single transcript regulate mitochondrial complex III in Drosophila. *eLife*, 11, 2022.

[10513] Anna Leabourn Boss, Tanvi Damani, Tayla J Wickman, Larry W Chamley, Joanna L James, and Anna ES Brooks. Full spectrum flow cytometry reveals mesenchymal heterogeneity in first trimester placentae and phenotypic convergence in culture, providing insight into the origins of placental mesenchymal stromal cells. *eLife*, 11, 2022.

[10514] Matthew P Bostock, Anadika R Prasad, Alicia Donoghue, and Vilaiwan M Fernandes. Photoreceptors generate neuronal diversity in their target field through a Hedgehog morphogen gradient in Drosophila. *eLife*, 11, 2022.

[10515] Flora Bouchacourt, Sina Tafazoli, Marcelo G Mattar, Timothy J Buschman, and Nathaniel D Daw. Fast rule switching and slow rule updating in a perceptual categorization task. *eLife*, 11, 2022.

[10516] Jonathan Bouvette, Qinwen Huang, Amanda A Riccio, William C Copeland, Alberto Bartesaghi, and Mario J Borgnia. Automated systematic evaluation of cryo-EM specimens with SmartScope. *eLife*, 11, 2022.

[10517] Seth Boyer, Ho-Joon Lee, Nina Steele, Li Zhang, Peter Sajjakulnukit, Anthony Andren, Matthew H Ward, Rima Singh, Venkatesha Basrur, Yaqing Zhang, Alexey I Nesvizhskii, Marina Pasca di Magliano, Christopher J Halbrook, and Costas A Lyssiotis. Multiomic characterization of pancreatic cancer-associated macrophage polarization reveals deregulated metabolic programs driven by the GM-CSF–PI3K pathway. *eLife*, 11, 2022.

[10518] Mario Bräcklein, Deren Yusuf Barsakcioglu, Jaime Ibáñez, Jonathan Eden, Etienne Burdet, Carsten Mehring, and Dario Farina. The control and training of single motor units in isometric tasks are constrained by a common input signal. *eLife*, 11, 2022.

[10519] William John Bradshaw, Michael Poeschla, Aleksandra Placzek, Samuel Kean, and Dario Riccardo Valenzano. Extensive age-dependent loss of antibody diversity in naturally short-lived turquoise killifish. *eLife*, 11, 2022.

[10520] Nathaniel R Braffman, Terry B Ruskoski, Katherine M Davis, Nathaniel R Glasser, Cassidy Johnson, C Denise Okafor, Amie K Boal, and Emily P Balskus. Structural basis for an unprecedented enzymatic alkylation in cylindrocyclophane biosynthesis. *eLife*, 11, 2022.

[10521] Ines Braga Goncalves, Amy Morris-Drake, Patrick Kennedy, and Andrew N Radford. Fitness consequences of outgroup conflict. *eLife*, 11, 2022.

[10522] Ines Braga Goncalves and Andrew N Radford. Experimental evidence that chronic outgroup conflict reduces reproductive success in a cooperatively breeding fish. *eLife*, 11, 2022.

[10523] Djamal Brahim Belhaouari, Gabriel Augusto Pires De Souza, David C Lamb, Steven L Kelly, Jared V Goldstone, John J Stegeman, Philippe Colson, Bernard La Scola, and Sarah Aherfi. Metabolic arsenal of giant viruses: Host hijack or self-use? *eLife*, 11, 2022.

[10524] Ana S Brandão, Jorge Borbinha, Telmo Pereira, Patrícia H Brito, Raquel LourenÇo, Anabela Bensimon-Brito, and Antonio Jacinto. A regeneration-triggered metabolic adaptation is necessary for cell identity transitions and cell cycle re-entry to support blastema formation and bone regeneration. *eLife*, 11, 2022.

[10525] Amanda E Brandon, Lewin Small, Tuong-Vi Nguyen, Eurwin Suryana, Henry Gong, Christian Yassmin, Sarah E Hancock, Tamara Pulpitel, Sophie Stonehouse, Letisha Prescott, Melkam A Kebede, Belinda Yau, Lake-Ee Quek, Greg M Kowalski, Clinton R Bruce, Nigel Turner, and Gregory J Cooney. Insulin sensitivity is preserved in mice made obese by feeding a high starch diet. *eLife*, 11, 2022.

[10526] Christopher J Brereton, Liudi Yao, Elizabeth R Davies, Yilu Zhou, Milica Vukmirovic, Joseph A Bell, Siyuan Wang, Robert A Ridley, Lareb SN Dean, Orestis G Andriotis, Franco Conforti, Lennart Brewitz, Soran Mohammed, Timothy Wallis, Ali Tavassoli, Rob M Ewing, Aiman Alzetani, Benjamin G Marshall, Sophie V Fletcher, Philipp J Thurner, Aurelie Fabre, Naftali Kaminski, Luca Richeldi, Atul Bhaskar, Christopher J Schofield, Matthew Loxham, Donna E Davies, Yihua Wang, and Mark G Jones. Pseudohypoxic HIF pathway activation dysregulates collagen structure-function in human lung fibrosis. *eLife*, 11, 2022.

[10527] Martin W Breuss, Xiaoxu Yang, Valentina Stanley, Jennifer McEvoy-Venneri, Xin Xu, Arlene J Morales, and Joseph G Gleeson. Unbiased mosaic variant assessment in sperm: a cohort study to test predictability of transmission. *eLife*, 11, 2022.

[10528] Derek M Britain, Jason P Town, and Orion David Weiner. Progressive enhancement of kinetic proofreading in T cell antigen discrimination from receptor activation to DAG generation. *eLife*, 11, 2022.

[10529] Christian Brodbeck, Shohini Bhattasali, Aura AL Cruz Heredia, Philip Resnik, Jonathan Z Simon, and Ellen Lau. Parallel processing in speech perception with local and global representations of linguistic context. *eLife*, 11, 2022.

[10530] Molly Brothers and Jasper Rine. Distinguishing between recruitment and spread of silent chromatin structures in Saccharomyces cerevisiae. *eLife*, 11, 2022.

[10531] Owain J Bryant, Paraminder Dhillon, Colin Hughes, and Gillian M Fraser. Recognition of discrete export signals in early flagellar subunits during bacterial type III secretion. *eLife*, 11, 2022.

[10532] S Jimmy Budiardjo, Jacqueline J Stevens, Anna L Calkins, Ayotunde P Ikujuni, Virangika K Wimalasena, Emre Firlar, David A Case, Julie S Biteen, Jason T Kaelber, and Joanna SG Slusky. Colicin E1 opens its hinge to plug TolC. *eLife*, 11, 2022.

[10533] Christoph Budjan, Shichen Liu, Adrian Ranga, Senjuti Gayen, Olivier Pourquié, and Sahand Hormoz. Paraxial mesoderm organoids model development of human somites. *eLife*, 11, 2022.

[10534] Chandima Bulumulla, Andrew T Krasley, Ben Cristofori-Armstrong, William C Valinsky, Deepika Walpita, David Ackerman, David E Clapham, and Abraham G Beyene. Visualizing synaptic dopamine efflux with a 2D composite nanofilm. *eLife*, 11, 2022.

[10535] Emily A Burger, Inge MCM de Kok, James F O'Mahony, Matejka Rebolj, Erik EL Jansen, Daniel D de Bondt, James Killen, Sharon J Hanley, Alejandra Castanon, Mary Caroline Regan, Jane J Kim, Karen Canfell, and Megan A Smith. A model-based analysis of the health impacts of COVID-19 disruptions to primary cervical screening by time since last screen for current and future disruptions. *eLife*, 11, 2022.

[10536] Rayanne B Burl, Elizabeth Ann Rondini, Hongguang Wei, Roger Pique-Regi, and James G Granneman. Deconstructing cold-induced brown adipocyte neogenesis in mice. *eLife*, 11, 2022.

[10537] Andrew A Burnim, Matthew A Spence, Da Xu, Colin J Jackson, and Nozomi Ando. Comprehensive phylogenetic analysis of the ribonucleotide reductase family reveals an ancestral clade. *eLife*, 11, 2022.

[10538] Shawn D Burton, Audrey Brown, Thomas P Eiting, Isaac A Youngstrom, Thomas C Rust, Michael Schmuker, and Matt Wachowiak. Mapping odorant sensitivities reveals a sparse but structured representation of olfactory chemical space by sensory input to the mouse olfactory bulb. *eLife*, 11, 2022.

[10539] Satvika Burugupalli, Adam Alexander T Smith, Gavriel Oshlensky, Kevin Huynh, Corey Giles, Tingting Wang, Alexandra George, Sudip Paul, Anh Nguyen, Thy Duong, Natalie Mellett, Michelle Cinel, Sartaj Ahmad Mir, Li Chen, Markus R Wenk, Neerja Karnani, Fiona Collier, Richard Saffery, Peter Vuillermin, Anne-Louise Ponsonby, David Burgner, Peter Meikle, and Peter Vuillermin. Ontogeny of circulating lipid metabolism in pregnancy and early childhood – a longitudinal population study. *eLife*, 11, 2022.

[10540] Trixia M Buscagan, Jens T Kaiser, and Douglas C Rees. Selenocyanate derived Se-incorporation into the nitrogenase Fe protein cluster. *eLife*, 11, 2022.

[10541] Tobias J Buscham, Maria A Eichel-Vogel, Anna M Steyer, Olaf Jahn, Nicola Strenzke, Rakshit Dardawal, Tor R Memhave, Sophie B Siems, Christina Müller, Martin Meschkat, Ting Sun, Torben Ruhwedel, Wiebke Möbius, Eva-Maria Krämer-Albers, Susann Boretius, Klaus-Armin Nave, and Hauke B Werner. Progressive axonopathy when oligodendrocytes lack the myelin protein CMTM5. *eLife*, 11, 2022.

[10542] Brittany J Bush, Caroline Donnay, Eva-Jeneé A Andrews, Darielle Lewis-Sanders, Cloe L Gray, Zhimei Qiao, Allison J Brager, Hadiya Johnson, Hamadi CS Brewer, Sahil Sood, Talib Saafir, Morris Benveniste, Ketema N Paul, and J Christopher Ehlen. Non-rapid eye movement sleep determines resilience to social stress. *eLife*, 11, 2022.

[10543] Deny Cabezas-Bratesco, Francisco A Mcgee, Charlotte K Colenso, Kattina Zavala, Daniele Granata, Vincenzo Carnevale, Juan C Opazo, and Sebastian E Brauchi. Sequence and structural conservation reveal fingerprint residues in TRP channels. *eLife*, 11, 2022.

[10544] Marie-Jeanne Cabirol, Laura Cardoit, Gilles Courtand, Marie-Eve Mayeur, John Simmers, Olivier Pascual, and Muriel Thoby-Brisson. Microglia shape the embryonic development of mammalian respiratory networks. *eLife*, 11, 2022.

[10545] Clotilde Cadart, Larisa Venkova, Matthieu Piel, and Marco Cosentino Lagomarsino. Volume growth in animal cells is cell cycle dependent and shows additive fluctuations. *eLife*, 11, 2022.

[10546] Dawei Cai, Siqi Zhu, Mian Gong, Naifan Zhang, Jia Wen, Qiyao Liang, Weilu Sun, Xinyue Shao, Yaqi Guo, Yudong Cai, Zhuqing Zheng, Wei Zhang, Songmei Hu, Xiaoyang Wang, He Tian, Youqian Li, Wei Liu, Miaomiao Yang, Jian Yang, Duo Wu, Ludovic Orlando, and Yu Jiang. Radiocarbon and genomic evidence for the survival of Equus Sussemionus until the late Holocene. *eLife*, 11, 2022.

[10547] Wesley L Cai, Jocelyn Fang-Yi Chen, Huacui Chen, Emily Wingrove, Sarah J Kurley, Lok Hei Chan, Meiling Zhang, Anna Arnal-Estape, Minghui Zhao, Amer Balabaki, Wenxue Li, Xufen Yu, Ethan D Krop, Yali Dou, Yansheng Liu, Jian Jin, Thomas F Westbrook, Don X Nguyen, and Qin Yan.

Human WDR5 promotes breast cancer growth and metastasis via KMT2-independent translation regulation. *eLife*, 11, 2022.

[10548] Arnault H Caillet, Andrew TM Phillips, Dario Farina, and Luca Modenese. Mathematical relationships between spinal motoneuron properties. *eLife*, 11, 2022.

[10549] Nuno Calaim, Florian A Dehmelt, Pedro J GonÇalves, and Christian K Machens. The geometry of robustness in spiking neural networks. *eLife*, 11, 2022.

[10550] Lesly Calderon, Felix D Weiss, Jonathan A Beagan, Marta S Oliveira, Radina Georgieva, Yi-Fang Wang, Thomas S Carroll, Gopuraja Dharmalingam, Wanfeng Gong, Kyoko Tossell, Vincenzo de Paola, Chad Whilding, Mark A Ungless, Amanda G Fisher, Jennifer E Phillips-Cremins, and Matthias Merkenschlager. Cohesin-dependence of neuronal gene expression relates to chromatin loop length. *eLife*, 11, 2022.

[10551] Benjamin FN Campbell, Antje Dittmann, Birgit Dreier, Andreas Plückthun, and Shiva K Tyagarajan. A DARPin-based molecular toolset to probe gephyrin and inhibitory synapse biology. *eLife*, 11, 2022.

[10552] Jaclyn Camuglia, Soline Chanet, and Adam C Martin. Morphogenetic forces planar polarize LGN/Pins in the embryonic head during Drosophila gastrulation. *eLife*, 11, 2022.

[10553] Irving Cancino-Muñoz, Mariana G López, Manuela Torres-Puente, Luis M Villamayor, Rafael Borrás, María Borrás-Máñez, Montserrat Bosque, Juan J Camarena, Caroline Colijn, Ester Colomer-Roig, Javier Colomina, Isabel Escribano, Oscar Esparcia-Rodríguez, Francisco García-García, Ana Gil-Brusola, Concepción Gimeno, Adelina Gimeno-Gascón, Bárbara Gomila-Sard, Damiana Gónzales-Granda, Nieves Gonzalo-Jiménez, María Remedios Guna-Serrano, José Luis López-Hontangas, Coral Martín-González, Rosario Moreno-Muñoz, David Navarro, María Navarro, Nieves Orta, Elvira Pérez, Josep Prat, Juan Carlos Rodríguez, Ma Montserrat Ruiz-García, Hermelinda Vanaclocha, Manuel Belda-álvarez, and Iñaki Comas. Population-based sequencing of Mycobacterium tuberculosis reveals how current population dynamics are shaped by past epidemics. *eLife*, 11, 2022.

[10554] Sarah E Canetta, Emma S Holt, Laura J Benoit, Eric Teboul, Gabriella M Sahyoun, R Todd Ogden, Alexander Z Harris, and Christoph Kellendonk. Mature parvalbumin interneuron function in prefrontal cortex requires activity during a postnatal sensitive period. *eLife*, 11, 2022.

[10555] Marella D Canny and Michael P Latham. LRET-derived HADDOCK structural models describe the conformational heterogeneity required for DNA cleavage by the Mre11-Rad50 DNA damage repair complex. *eLife*, 11, 2022.

[10556] Gerard Cantero-Recasens, Josune Alonso-Marañón, Teresa Lobo-Jarne, Marta Garrido, Mar Iglesias, Lluis Espinosa, and Vivek Malhotra. Reversing chemorefraction in colorectal cancer cells by controlling mucin secretion. *eLife*, 11, 2022.

[10557] Jose Ernesto Canton-Josh, Joanna Qin, Joseph Salvo, and Yevgenia Kozorovitskiy. Dopaminergic regulation of vestibulo-cerebellar circuits through unipolar brush cells. *eLife*, 11, 2022.

[10558] Rui Cao, John H Bladon, Stephen J Charczynski, Michael E Hasselmo, and Marc W Howard. Internally generated time in the rodent hippocampus is logarithmically compressed. *eLife*, 11, 2022.

[10559] Xi Cao, Ting-Ting Shi, Chuan-Hai Zhang, Wan-Zhu Jin, Li-Ni Song, Yi-Chen Zhang, Jing-Yi Liu, Fang-Yuan Yang, Charles N Rotimi, Aimin Xu, and Jin-Kui Yang. ACE2 pathway regulates thermogenesis and energy metabolism. *eLife*, 11, 2022.

[10560] Yinan Cao and Konstantinos Tsetsos. Clarifying the role of an unavailable distractor in human multiattribute choice. *eLife*, 11, 2022.

[10561] Claudia C Carcamo, Matthew F Poyton, Anand Ranjan, Giho Park, Robert K Louder, Xinyu A Feng, Jee Min Kim, Thuc Dzu, Carl Wu, and Taekjip Ha. ATP binding facilitates target search of SWR1 chromatin remodeler by promoting one-dimensional diffusion on DNA. *eLife*, 11, 2022.

[10562] Alberto Carignano, Dai Hua Chen, Cannon Mallory, R Clay Wright, Georg Seelig, and Eric Klavins. Modular, robust, and extendible multicellular circuit design in yeast. *eLife*, 11, 2022.

[10563] Sara A Carioscia, Kathryn J Weaver, Andrew N Bortvin, Hao Pan, Daniel Ariad, Avery Davis Bell, and Rajiv C McCoy. A method for low-coverage single-gamete sequence analysis demonstrates adherence to Mendel's first law across a large sample of human sperm. *eLife*, 11, 2022.

[10564] Caterina Carraro, Lorenzo Bonaguro, Jonas Schulte-Schrepping, Arik Horne, Marie Oestreich, Stefanie Warnat-Herresthal, Tim Helbing, Michele De Franco, Kristian Haendler, Sach Mukherjee, Thomas Ulas, Valentina Gandin, Richard Goettlich, Anna C Aschenbrenner, Joachim L Schultze, and Barbara Gatto. Decoding mechanism of action and sensitivity to drug candidates from integrated transcriptome and chromatin state. *eLife*, 11, 2022.

[10565] Germán D Carrasquilla, Mario García-Ureña, Tove Fall, Thorkild IA Sørensen, and Tuomas O Kilpeläinen. Mendelian randomization suggests a bidirectional, causal relationship between physical inactivity and adiposity. *eLife*, 11, 2022.

[10566] Arnaud Carrier, Cécile Desjobert, Loic Ponger, Laurence Lamant, Matias Bustos, Jorge Torres-Ferreira, Rui Henrique, Carmen Jeronimo, Luisa Lanfrancone, Audrey Delmas, Gilles Favre, Antoine Daunay, Florence Busato, Dave SB Hoon, Jorg Tost, Chantal Etievant, Joëlle Riond, and Paola B Arimondo. DNA methylome combined with chromosome cluster-oriented analysis provides an early signature for cutaneous melanoma aggressiveness. *eLife*, 11, 2022.

[10567] Thomas A Carter, Manvendra Singh, Gabrijela Dumbović, Jason D Chobirko, John L Rinn, and Cédric Feschotte. Mosaic cis-regulatory evolution drives transcriptional partitioning of HERVH endogenous retrovirus in the human embryo. *eLife*, 11, 2022.

[10568] Lindsay B Case, Milagros De Pasquale, Lisa Henry, and Michael K Rosen. Synergistic phase separation of two pathways promotes integrin clustering and nascent adhesion formation. *eLife*, 11, 2022.

[10569] Kai R Caspar, Fabian Pallasdies, Larissa Mader, Heitor Sartorelli, and Sabine Begall. The evolution and biological correlates of hand preferences in anthropoid primates. *eLife*, 11, 2022.

[10570] Meaghan Castledine, Daniel Padfield, Pawel Sierocinski, Jesica Soria Pascual, Adam Hughes, Lotta Mäkinen, Ville-Petri Friman, Jean-Paul Pirnay, Maya Merabishvili, Daniel de Vos, and Angus Buckling. Parallel evolution of Pseudomonas aeruginosa phage resistance and virulence loss in response to phage treatment in vivo and in vitro. *eLife*, 11, 2022.

[10571] Inês Gomes Castro, Shawn P Shortill, Samantha Katarzyna Dziurdzik, Angela Cadou, Suriakarthiga Ganesan, Rosario Valenti, Yotam David, Michael Davey, Carsten Mattes, Ffion B Thomas, Reut Ester Avraham, Hadar Meyer, Amir Fadel, Emma J Fenech, Robert Ernst, Vanina Zaremberg, Tim P Levine, Christopher Stefan, Elizabeth Conibear, and Maya Schuldiner. Systematic analysis of membrane contact sites in Saccharomyces cerevisiae uncovers modulators of cellular lipid distribution. *eLife*, 11, 2022.

[10572] Catarina Catela, Yihan Chen, Yifei Weng, Kailong Wen, and Paschalis Kratsios. Control of spinal motor neuron terminal differentiation through sustained Hoxc8 gene activity. *eLife*, 11, 2022.

[10573] Lisa AE Catsburg, Manon Westra, Annemarie ML van Schaik, and Harold D MacGillavry. Dynamics and nanoscale organization of the postsynaptic endocytic zone at excitatory synapses. *eLife*, 11, 2022.

[10574] Nathanael A Caveney, Caleb R Glassman, Kevin M Jude, Naotaka Tsutsumi, and K Christopher Garcia. Structure of the IL-27 quaternary receptor signaling complex. *eLife*, 11, 2022.

[10575] Gabriel Cavin-Meza, Michelle M Kwan, and Sarah M Wignall. Multiple motors cooperate to establish and maintain acentrosomal spindle bipolarity in C. elegans oocyte meiosis. *eLife*, 11, 2022.

[10576] Işın çakır, Pauline Lining Pan, Colleen K Hadley, Abdulrahman El-Gamal, Amina Fadel, Dina Elsayegh, Omnia Mohamed, Nasser M Rizk, and Masoud Ghamari-Langroudi. Sulforaphane reduces obesity by reversing leptin resistance. *eLife*, 11, 2022.

[10577] Romane Cecchi, Fabien Vinckier, Jiri Hammer, Petr Marusic, Anca Nica, Sylvain Rheims, Agnès Trebuchon, Emmanuel J Barbeau, Marie Denuelle, Louis Maillard, Lorella Minotti, Philippe Kahane, Mathias Pessiglione, and Julien Bastin. Intracerebral mechanisms explaining the impact of incidental feedback on mood state and risky choice. *eLife*, 11, 2022.

[10578] Benjamin J Chadwick, Tuyetnhu Pham, Xiaofeng Xie, Laura C Ristow, Damian J Krysan, and Xiaorong Lin. The RAM signaling pathway links morphology, thermotolerance, and CO2 tolerance in the global fungal pathogen Cryptococcus neoformans. *eLife*, 11, 2022.

[10579] Rituparna Chakrabarti, Lina María Jaime Tobón, Loujin Slitin, Magdalena Redondo Canales, Gerhard Hoch, Marina Slashcheva, Elisabeth Fritsch, Kai Bodensiek, özge Demet özÇete, Mehmet Gültas, Susann Michanski, Felipe Opazo, Jakob Neef, Tina Pangrsic, Tobias Moser, and Carolin Wichmann. Optogenetics and electron tomography for structure-function analysis of cochlear ribbon synapses. *eLife*, 11, 2022.

[10580] Morteza Chalabi Hajkarim, Ella Karjalainen, Mikhail Osipovitch, Konstantinos Dimopoulos, Sandra L Gordon, Francesca Ambri, Kasper Dindler Rasmussen, Kirsten Grønbæk, Kristian Helin, Krister Wennerberg, and Kyoung-Jae Won. Comprehensive and unbiased multiparameter high-throughput screening by compaRe finds effective and subtle drug responses in AML models. *eLife*, 11, 2022.

[10581] Sridevi Challa, Keun W Ryu, Amy L Whitaker, Jonathan C Abshier, Cristel V Camacho, and W Lee Kraus. Development and characterization of new tools for detecting poly(ADP-ribose) in vitro and in vivo. *eLife*, 11, 2022.

[10582] Andrea M Chambers, Kyle B Lupo, Jiao Wang, Jingming Cao, Sagar Utturkar, Nadia Lanman, Victor Bernal-Crespo, Shadia Jalal, Sharon R Pine, Sandra Torregrosa-Allen, Bennett D Elzey, and Sandro Matosevic. Engineered natural killer cells impede the immunometabolic CD73-adenosine axis in solid tumors. *eLife*, 11, 2022.

[10583] Samuel E Champer, Isabel K Kim, Andrew G Clark, Philipp W Messer, and Jackson Champer. Anopheles homing suppression drive candidates exhibit unexpected performance differences in simulations with spatial structure. *eLife*, 11, 2022.

[10584] Melanie MY Chan, Omid Sadeghi-Alavijeh, Filipa M Lopes, Alina C Hilger, Horia C Stanescu, Catalin D Voinescu, Glenda M Beaman, William G Newman, Marcin Zaniew, Stefanie Weber, Yee Mang Ho, John O Connolly, Dan Wood, Carlo Maj, Alexander Stuckey, Athanasios Kousathanas, JC Ambrose, Robert Kleta, Adrian S Woolf, Detlef Bockenhauer, Adam P Levine, and Daniel P Gale. Diverse ancestry whole-genome sequencing association study identifies TBX5 and PTK7 as susceptibility genes for posterior urethral valves. *eLife*, 11, 2022.

[10585] Marie Chancel, H Henrik Ehrsson, and Wei Ji Ma. Uncertainty-based inference of a common cause for body ownership. *eLife*, 11, 2022.

[10586] Karthik Chandiran, Jenny E Suarez-Ramirez, Yinghong Hu, Evan R Jellison, Zeynep Ugur, Jun Siong Low, Bryan McDonald, Susan M Kaech, and Linda S Cauley. SMAD4 and TGF$\beta$ are architects of inverse genetic programs during fate determination of antiviral CTLs. *eLife*, 11, 2022.

[10587] Ching-Ho Chang, Lauren E Gregory, Kathleen E Gordon, Colin D Meiklejohn, and Amanda M Larracuente. Unique structure and positive selection promote the rapid divergence of Drosophila Y chromosomes. *eLife*, 11, 2022.

[10588] Shu-Jung Chang, Yu-Ting Hsu, Yun Chen, Yen-Yi Lin, Maria Lara-Tejero, and Jorge E Galan. Typhoid toxin sorting and exocytic transport from Salmonella Typhi-infected cells. *eLife*, 11, 2022.

[10589] Suhee Chang, Diana Fulmer, Stella K Hur, Joanne L Thorvaldsen, Li Li, Yemin Lan, Eric A Rhon-Calderon, Nicolae Adrian Leu, Xiaowen Chen, Jonathan A Epstein, and Marisa S Bartolomei. Dysregulated H19/Igf2 expression disrupts cardiac-placental axis during development of Silver-Russell syndrome-like mouse models. *eLife*, 11, 2022.

[10590] Callie R Chappell, Manpreet K Dhami, Mark C Bitter, Lucas Czech, Sur Herrera Paredes, Fatoumata Binta Barrie, Yadira Calderón, Katherine Eritano, Lexi-Ann Golden, Daria Hekmat-Scafe, Veronica Hsu, Clara Kieschnick, Shyamala Malladi, Nicole Rush, and Tadashi Fukami. Wide-ranging consequences of priority effects governed by an overarching factor. *eLife*, 11, 2022.

[10591] Edouard Charlebois, Carine Fillebeen, Angeliki Katsarou, Aleksandr Rabinovich, Kazimierz Wisniewski, Vivek Venkataramani, Bernhard Michalke, Anastasia Velentza, and Kostas Pantopoulos. A crosstalk between hepcidin and IRE/IRP pathways controls ferroportin expression and determines serum iron levels in mice. *eLife*, 11, 2022.

[10592] Marie JE Charpentier, Clémence Poirotte, Berta Roura-Torres, Paul Amblard-Rambert, Eric Willaume, Peter M Kappeler, FranÇois Rousset, and Julien P Renoult. Mandrill mothers associate with infants who look like their own offspring using phenotype matching. *eLife*, 11, 2022.

[10593] Debdeep Chatterjee, Reeba S Jacob, Soumik Ray, Ambuja Navalkar, Namrata Singh, Shinjinee Sengupta, Laxmikant Gadhe, Pradeep Kadu, Debalina Datta, Ajoy Paul, Sakunthala Arunima, Surabhi Mehra, Chinmai Pindi, Santosh Kumar, Praful Singru, Sanjib Senapati, and Samir K Maji. Co-aggregation and secondary nucleation in the life cycle of human prolactin/galanin functional amyloids. *eLife*, 11, 2022.

[10594] Deeptiman Chatterjee, Caique Almeida Machado Costa, Xian-Feng Wang, Allison Jevitt, Yi-Chun Huang, and Wu-Min Deng. Single-cell transcriptomics identifies Keap1-Nrf2 regulated collective invasion in a Drosophila tumor model. *eLife*, 11, 2022.

[10595] Payel Chatterjee, Agnish Dev Prusty, Umesh Mohan, and Sanjay P Sane. Integration of visual and antennal mechanosensory feedback during head stabilization in hawkmoths. *eLife*, 11, 2022.

[10596] Susheel N Chaurasia, Mohammad Ekhlak, Geeta Kushwaha, Vipin Singh, Ram L Mallick, and Debabrata Dash. Notch signaling functions in noncanonical juxtacrine manner in platelets to amplify thrombogenicity. *eLife*, 11, 2022.

[10597] Bing Chen and Monica A Perez. Altered regulation of Ia afferent input during voluntary contraction in humans with spinal cord injury. *eLife*, 11, 2022.

[10598] Dafeng Chen, Li Hou, Jianing Wei, Siyuan Guo, Weichan Cui, Pengcheng Yang, Le Kang, and Xianhui Wang. Aggregation pheromone 4-vinylanisole promotes the synchrony of sexual maturation in female locusts. *eLife*, 11, 2022.

[10599] Fangrui Chen, Jingchao Wu, Malina K Iwanski, Daphne Jurriens, Arianna Sandron, Milena Pasolli, Gianmarco Puma, Jannes Z Kromhout, Chao Yang, Wilco Nijenhuis, Lukas C Kapitein, Florian Berger, and Anna Akhmanova. Self-assembly of pericentriolar material in interphase cells lacking centrioles. *eLife*, 11, 2022.

[10600] Fei Chen, Burcu F Darst, Ravi K Madduri, Alex A Rodriguez, Xin Sheng, Christopher T Rentsch, Caroline Andrews, Wei Tang, Adam S Kibel, Anna Plym, Kelly Cho, Mohamed Jalloh, Serigne Magueye Gueye, Lamine Niang, Olufemi J Ogunbiyi, Olufemi Popoola, Akindele O Adebiyi, Oseremen I Aisuodionoe-Shadrach, Hafees O Ajibola, Mustapha A Jamda, Olabode P Oluwole, Maxwell Nwegbu, Ben Adusei, Sunny Mante, Afua Darkwa-Abrahams, James E Mensah, Andrew Anthony Adjei, Halimatou Diop, Joseph Lachance, Timothy R Rebbeck, Stefan Ambs, J Michael Gaziano, Amy C Justice, David V Conti, and Christopher A Haiman. Validation of a

multi-ancestry polygenic risk score and age-specific risks of prostate cancer: A meta-analysis within diverse populations. *eLife*, 11, 2022.

[10601] Heng-Yi Chen, Ana Almonte-Loya, Fang-Yun Lay, Michael Hsu, Eric Johnson, Edahí González-Avalos, Jieyun Yin, Richard S Bruno, Qin Ma, Hazem E Ghoneim, Daniel J Wozniak, Fiona E Harrison, and Chan-Wang Jerry Lio. Epigenetic remodeling by vitamin C potentiates plasma cell differentiation. *eLife*, 11, 2022.

[10602] Huan-Huan Chen, Tie-Ning Zhang, Fang-Yuan Zhang, and Tao Zhang. Non-coding RNAs in drug and radiation resistance of bone and soft-tissue sarcoma: a systematic review. *eLife*, 11, 2022.

[10603] Jia Chen and Daniel St Johnston. De novo apical domain formation inside the Drosophila adult midgut epithelium. *eLife*, 11, 2022.

[10604] Leying Chen, Qing You, Min Liu, Shuaihu Li, Zhaoyu Wu, Jiajun Hu, Yurui Ma, Liangyong Xia, Ying Zhou, Nan Xu, and Shiyi Zhang. Remodeling of dermal adipose tissue alleviates cutaneous toxicity induced by anti-EGFR therapy. *eLife*, 11, 2022.

[10605] Li Chen, Zhenzhen Song, Xiaowan Cao, Mingsong Fan, Yan Zhou, and Guoying Zhang. Interleukin-33 regulates the endoplasmic reticulum stress of human myometrium via an influx of calcium during initiation of labor. *eLife*, 11, 2022.

[10606] Liqiang Chen, Chetan Nagaraja, Samuel Daniels, Zoe A Fisk, Rachel Dvorak, Lindsay Meyerdirk, Jennifer A Steiner, Martha L Escobar Galvis, Michael X Henderson, Maxime WC Rousseaux, Patrik Brundin, and Hong-Yuan Chu. Synaptic location is a determinant of the detrimental effects of $\alpha$-synuclein pathology to glutamatergic transmission in the basolateral amygdala. *eLife*, 11, 2022.

[10607] Shanwen Chen, Yunfan Jin, Siqi Wang, Shaozhen Xing, Yingchao Wu, Yuhuan Tao, Yongchen Ma, Shuai Zuo, Xiaofan Liu, Yichen Hu, Hongyan Chen, Yuandeng Luo, Feng Xia, Chuanming Xie, Jianhua Yin, Xin Wang, Zhihua Liu, Ning Zhang, Zhenjiang Zech Xu, Zhi John Lu, and Pengyuan Wang. Cancer type classification using plasma cell-free RNAs derived from human and microbes. *eLife*, 11, 2022.

[10608] Spencer Chin-Yu Chen, Giacomo Benvenuti, Yuzhi Chen, Satwant Kumar, Charu Ramakrishnan, Karl Deisseroth, Wilson S Geisler, and Eyal Seidemann. Similar neural and perceptual masking effects of low-power optogenetic stimulation in primate V1. *eLife*, 11, 2022.

[10609] Weizhao Chen, Qiongping Zheng, Qiaoying Huang, Shanshan Ma, and Mingtao Li. Repressing PTBP1 fails to convert reactive astrocytes to dopaminergic neurons in a 6-hydroxydopamine mouse model of Parkinson's disease. *eLife*, 11, 2022.

[10610] Yang S Chen, Wanfu Hou, Sharon Tracy, Alex T Harvey, Vince Harjono, Fan Xu, James J Moresco, John R Yates, and Brian M Zid. Rvb1/Rvb2 proteins couple transcription and translation during glucose starvation. *eLife*, 11, 2022.

[10611] Yu Chen, Claudia Cattoglio, Gina M Dailey, Qiulin Zhu, Robert Tjian, and Xavier Darzacq. Mechanisms governing target search and binding dynamics of hypoxia-inducible factors. *eLife*, 11, 2022.

[10612] Yu-Chi Chen, Aurina Arnatkevičiūtė, Eugene McTavish, James C Pang, Sidhant Chopra, Chao Suo, Alex Fornito, and Kevin M Aquino. The individuality of shape asymmetries of the human cerebral cortex. *eLife*, 11, 2022.

[10613] Zhenghao Chen, Anil Raj, GV Prateek, Andrea Di Francesco, Justin Liu, Brice E Keyes, Ganesh Kolumam, Vladimir Jojic, and Adam Freund. Automated, high-dimensional evaluation of physiological aging and resilience in outbred mice. *eLife*, 11, 2022.

[10614] Ningyan Cheng, Guanchen Li, Mohammed Kanchwala, Bret M Evers, Chao Xing, and Hongtao Yu. STAG2 promotes the myelination transcriptional program in oligodendrocytes. *eLife*, 11, 2022.

[10615] Pan Cheng, Xin Zhao, Lizabeth Katsnelson, Elaine M Camacho-Hernandez, Angela Mermerian, Joseph C Mays, Scott M Lippman, Reyna Edith Rosales-Alvarez, Raquel Moya, Jasmine Shwetar, Dominic Grun, David Fenyo, and Teresa Davoli. Proteogenomic analysis of cancer aneuploidy and normal tissues reveals divergent modes of gene regulation across cellular pathways. *eLife*, 11, 2022.

[10616] Kaitlin Ching, Jennifer T Wang, and Tim Stearns. Long-range migration of centrioles to the apical surface of the olfactory epithelium. *eLife*, 11, 2022.

[10617] Mattia Chini, Thomas Pfeffer, and Ileana Hanganu-Opatz. An increase of inhibition drives the developmental decorrelation of neural activity. *eLife*, 11, 2022.

[10618] Manjusha Chintalapati, Nick Patterson, and Priya Moorjani. The spatiotemporal patterns of major human admixture events during the European Holocene. *eLife*, 11, 2022.

[10619] Kelvin F Cho, Shawn M Gillespie, Nicholas A Kalogriopoulos, Michael A Quezada, Martin Jacko, Michelle Monje, and Alice Y Ting. A light-gated transcriptional recorder for detecting cell-cell contacts. *eLife*, 11, 2022.

[10620] Chan Hee J Choi, William Barr, Samir Zaman, Corey Model, Annsea Park, Mascha Koenen, Zeran Lin, Sarah K Szwed, Francois Marchildon, Audrey Crane, Thomas S Carroll, Henrik Molina, and Paul Cohen. LRG1 is an adipokine that promotes insulin sensitivity and suppresses inflammation. *eLife*, 11, 2022.

[10621] Kiri Choi, Won Kyu Kim, and Changbong Hyeon. Olfactory responses of Drosophila are encoded in the organization of projection neurons. *eLife*, 11, 2022.

[10622] Michael Chong, Pedrum Mohammadi-Shemirani, Nicolas Perrot, Walter Nelson, Robert Morton, Sukrit Narula, Ricky Lali, Irfan Khan, Mohammad Khan, Conor Judge, Tafadzwa Machipisa, Nathan Cawte, Martin O'Donnell, Marie Pigeyre, Loubna Akhabir, and Guillaume Paré. GWAS and ExWAS of blood mitochondrial DNA copy number identifies 71 loci and highlights a potential causal role in dementia. *eLife*, 11, 2022.

[10623] Yoojeong Choo, Dora Matzke, Mark D Bowren, Daniel Tranel, and Jan R Wessel. Right inferior frontal gyrus damage is associated with impaired initiation of inhibitory control, but not its implementation. *eLife*, 11, 2022.

[10624] Angelo Ferreira Chora, Dora Pedroso, Eleni Kyriakou, Nadja Pejanovic, Henrique ColaÇo, Raffaella Gozzelino, André Barros, Katharina Willmann, Tiago Velho, Catarina F Moita, Isa Santos, Pedro Pereira, Silvia Carvalho, Filipa Batalha Martins, João A Ferreira, Sérgio Fernandes de Almeida, Vladimir Benes, Josef Anrather, Sebastian Weis, Miguel P Soares, Arie Geerlof, Jacques Neefjes, Michael Sattler, Ana C Messias, Ana Neves-Costa, and Luis Ferreira Moita. DNA damage independent inhibition of NF-$\kappa$B transcription by anthracyclines. *eLife*, 11, 2022.

[10625] Shao-Pei Chou, Adriana K Alexander, Edward J Rice, Lauren A Choate, and Charles G Danko. Genetic dissection of the RNA polymerase II transcription cycle. *eLife*, 11, 2022.

[10626] Lucy Chou-Zheng and Asma Hatoum-Aslan. Critical roles for 'housekeeping' nucleases in type III CRISPR-Cas immunity. *eLife*, 11, 2022.

[10627] Gaurav S Choudhary, Andrea Pellagatti, Bogos Agianian, Molly A Smith, Tushar D Bhagat, Shanisha Gordon-Mitchell, Srabani Sahu, Sanjay Pandey, Nishi Shah, Srinivas Aluri, Ritesh Aggarwal, Sarah Aminov, Leya Schwartz, Violetta Steeples, Robert N Booher, Murali Ramachandra, Maria Samson, Milagros Carbajal, Kith Pradhan, Teresa V Bowman, Manoj M Pillai, Britta Will, Amittha Wickrema, Aditi Shastri, Robert K Bradley, Robert E Martell, Ulrich G Steidl, Evripidis Gavathiotis, Jacqueline Boultwood, Daniel T Starczynowski, and Amit Verma. Activation of targetable inflammatory immune signaling is seen in myelodysplastic syndromes with SF3B1 mutations. *eLife*, 11, 2022.

[10628] Eleni Chrysostomou, Hakima Flici, Sebastian G Gornik, Miguel Salinas-Saavedra, James M Gahan, Emma T McMahon, Kerry Thompson, Shirley Hanley, Michelle Kilcoyne, Christine E Schnitzler, Paul Gonzalez, Andreas D Baxevanis, and Uri Frank. A cellular and molecular analysis of SoxB-driven neurogenesis in a cnidarian. *eLife*, 11, 2022.

[10629] Xiaojing Chu, Anna WM Janssen, Hans Koenen, Linzhung Chang, Xuehui He, Irma Joosten, Rinke Stienstra, Yunus Kuijpers, Cisca Wijmenga, Cheng-Jian Xu, Mihai G Netea, Cees J Tack, and Yang Li. A genome-wide functional genomics approach uncovers genetic determinants of immune phenotypes in type 1 diabetes. *eLife*, 11, 2022.

[10630] Daphne Chylinski, Maxime Van Egroo, Justinas Narbutas, Vincenzo Muto, Mohamed Ali Bahri, Christian Berthomier, Eric Salmon, Christine Bastin, Christophe Phillips, Fabienne Collette, Pierre Maquet, Julie Carrier, Jean-Marc Lina, and Gilles Vandewalle. Timely coupling of sleep spindles and slow waves linked to early amyloid-$\beta$ burden and predicts memory decline. *eLife*, 11, 2022.

[10631] Małgorzata Anna Cichoń, Karin Pfisterer, Judith Leitner, Lena Wagner, Clement Staud, Peter Steinberger, and Adelheid Elbe-Bürger. Interoperability of RTN1A in dendrite dynamics and immune functions in human Langerhans cells. *eLife*, 11, 2022.

[10632] Dario Cilleros-Rodriguez, Raquel Martin-Morales, Pablo Barbeito, Abhijit Deb Roy, Abdelhalim Loukil, Belen Sierra-Rodero, Gonzalo Herranz, Olatz Pampliega, Modesto Redrejo-Rodriguez, Sarah C Goetz, Manuel Izquierdo, Takanari Inoue, and Francesc R Garcia-Gonzalo. Multiple ciliary localization signals control INPP5E ciliary targeting. *eLife*, 11, 2022.

[10633] Erik Clark, Margherita Battistara, and Matthew A Benton. A timer gene network is spatially regulated by the terminal system in the Drosophila embryo. *eLife*, 11, 2022.

[10634] Ian A Clark, Siawoosh Mohammadi, Martina F Callaghan, and Eleanor A Maguire. Conduction velocity along a key white matter tract is associated with autobiographical memory recall ability. *eLife*, 11, 2022.

[10635] Pieter Clauw, Envel Kerdaffrec, Joanna Gunis, Ilka Reichardt-Gomez, Viktoria Nizhynska, Stefanie Koemeda, Jakub Jez, and Magnus Nordborg. Locally adaptive temperature response of vegetative growth in Arabidopsis thaliana. *eLife*, 11, 2022.

[10636] Joseph M Cleary, Tae Kim, Annan SI Cook, Lauren A McCormick, William O Hancock, and Luke M Rice. Measurements and simulations of microtubule growth imply strong longitudinal interactions and reveal a role for GDP on the elongating end. *eLife*, 11, 2022.

[10637] Alice M Clement, Tom J Challands, Richard Cloutier, Laurent Houle, Per E Ahlberg, Shaun P Collin, and John A Long. Morphometric analysis of lungfish endocasts elucidates early dipnoan palaeoneurological evolution. *eLife*, 11, 2022.

[10638] Chad Steven Cockrum and Susan Strome. Maternal H3K36 and H3K27 HMTs protect germline development via regulation of the transcription factor LIN-15B. *eLife*, 11, 2022.

[10639] Jennette M Codjoe, Ryan A Richardson, Fionn McLoughlin, Richard David Vierstra, and Elizabeth S Haswell. Unbiased proteomic and forward genetic screens reveal that mechanosensitive ion channel MSL10 functions at ER–plasma membrane contact sites in Arabidopsis thaliana. *eLife*, 11, 2022.

[10640] Agnese Codutti, Mohammad A Charsooghi, Elisa Cerdá-Doñate, Hubert M Taïeb, Tom Robinson, Damien Faivre, and Stefan Klumpp. Interplay of surface interaction and magnetic torque in single-cell motion of magnetotactic bacteria in microfluidic confinement. *eLife*, 11, 2022.

[10641] Irene Cogliati Dezza, Axel Cleeremans, and William H Alexander. Independent and interacting value systems for reward and information in the human brain. *eLife*, 11, 2022.

[10642] Hadar Cohen, Noam Baram, Chaya Mushka Fridman, Liat Edry-Botzer, Dor Salomon, and Motti Gerlic. Post-phagocytosis activation of NLRP3 inflammasome by two novel T6SS effectors. *eLife*, 11, 2022.

[10643] Samantha S Cohen, Nim Tottenham, and Christopher Baldassano. Developmental changes in story-evoked responses in the neocortex and hippocampus. *eLife*, 11, 2022.

[10644] Yarden Cohen, David Aaron Nicholson, Alexa Sanchioni, Emily K Mallaber, Viktoriya Skidanova, and Timothy J Gardner. Automated annotation of birdsong with a neural network that segments spectrograms. *eLife*, 11, 2022.

[10645] John Darby Cole, Jacobo Sarabia del Castillo, Gabriele Gut, Daniel Gonzalez-Bohorquez, Lucas Pelkmans, and Sebastian Jessberger. Characterization of the neurogenic niche in the aging dentate gyrus using iterative immunofluorescence imaging. *eLife*, 11, 2022.

[10646] James E Collier and Kim A Nasmyth. DNA passes through cohesin's hinge as well as its Smc3–kleisin interface. *eLife*, 11, 2022.

[10647] Mahlon A Collins, Gemechu Mekonnen, and Frank Wolfgang Albert. Variation in ubiquitin system genes creates substrate-specific effects on proteasomal protein degradation. *eLife*, 11, 2022.

[10648] Noah J Connally, Sumaiya Nazeen, Daniel Lee, Huwenbo Shi, John Stamatoyannopoulos, Sung Chun, Chris Cotsapas, Christopher A Cassa, and Shamil R Sunyaev. The missing link between genetic association and regulatory function. *eLife*, 11, 2022.

[10649] Marika Constant, Roy Salomon, and Elisa Filevich. Judgments of agency are affected by sensory noise without recruiting metacognitive processing. *eLife*, 11, 2022.

[10650] William Conway, Robert Kiewisz, Gunar Fabig, Colm P Kelleher, Hai-Yin Wu, Maya Anjur-Dietrich, Thomas Müller-Reichert, and Daniel J Needleman. Self-organization of kinetochore-fibers in human mitotic spindles. *eLife*, 11, 2022.

[10651] Eve B Cooper, Lauren JN Brent, Noah Snyder-Mackler, Mewa Singh, Asmita Sengupta, Sunil Khatiwada, Suchinda Malaivijitnond, Zhou Qi Hai, and James P Higham. The rhesus macaque as a success story of the Anthropocene. *eLife*, 11, 2022.

[10652] Ricardo A Cordova, Jagannath Misra, Parth H Amin, Anglea J Klunk, Nur P Damayanti, Kenneth R Carlson, Andrew J Elmendorf, Hyeong-Geug Kim, Emily T Mirek, Bennet D Elzey, Marcus J Miller, X Charlie Dong, Liang Cheng, Tracy G Anthony, Roberto Pili, Ronald C Wek, and Kirk A Staschke. GCN2 eIF2 kinase promotes prostate cancer by maintaining amino acid homeostasis. *eLife*, 11, 2022.

[10653] Alex Cornean, Jakob Gierten, Bettina Welz, Juan Luis Mateo, Thomas Thumberger, and Joachim Wittbrodt. Precise in vivo functional analysis of DNA variants with base editing using ACEofBASEs target prediction. *eLife*, 11, 2022.

[10654] Kimberley RG Cortenbach, Alexander HJ Staal, Teske Schoffelen, Mark AJ Gorris, Lieke L Van der Woude, Anne FM Jansen, Paul Poyck, Robert Jan Van Suylen, Peter C Wever, Chantal P Bleeker-Rovers, Mangala Srinivas, Konnie M Hebeda, Marcel van Deuren, Jos W Van der Meer, Jolanda M De Vries, and Roland RJ Van Kimmenade. Differences in local immune cell landscape between Q fever and atherosclerotic abdominal aortic aneurysms identified by multiplex immunohistochemistry. *eLife*, 11, 2022.

[10655] Vivian Vasconcelos Costa, Michelle A Sugimoto, Josy Hubner, Caio S Bonilha, Celso Martins Queiroz-Junior, Marcela Helena GonÇalves-Pereira, Jianmin Chen, Thomas Gobbetti, Gisele Olinto Libanio Rodrigues, Jordana L Bambirra, Ingredy B Passos, Carla Elizabeth Machado Lopes, Thaiane P Moreira, Kennedy Bonjour, Rossana CN Melo, Milton AP Oliveira, Marcus Vinicius M Andrade, Lirlândia Pires Sousa, Danielle Gloria Souza, Helton da Costa Santiago, Mauro Perretti, and Mauro Martins Teixeira. Targeting the Annexin A1-FPR2/ALX pathway for host-directed therapy in dengue disease. *eLife*, 11, 2022.

[10656] Luca Costantino, Stefania Ferrari, Matteo Santucci, Outi MH Salo-Ahen, Emanuele Carosati, Silvia Franchini, Angela Lauriola, Cecilia Pozzi, Matteo Trande, Gaia Gozzi, Puneet Saxena, Giuseppe

Cannazza, Lorena Losi, Daniela Cardinale, Alberto Venturelli, Antonio Quotadamo, Pasquale Linciano, Lorenzo Tagliazucchi, Maria Gaetana Moschella, Remo Guerrini, Salvatore Pacifico, Rosaria Luciani, Filippo Genovese, Stefan Henrich, Silvia Alboni, Nuno Santarem, Anabela da Silva Cordeiro, Elisa Giovannetti, Godefridus J Peters, Paolo Pinton, Alessandro Rimessi, Gabriele Cruciani, Robert M Stroud, Rebecca C Wade, Stefano Mangani, Gaetano Marverti, Domenico D'Arca, Glauco Ponterini, and Maria Paola Costi. Destabilizers of the thymidylate synthase homodimer accelerate its proteasomal degradation and inhibit cancer growth. *eLife*, 11, 2022.

[10657] Mary Couvillion, Kevin M Harlen, Kate C Lachance, Kristine L Trotta, Erin Smith, Christian Brion, Brendan M Smalec, and L Stirling Churchman. Transcription elongation is finely tuned by dozens of regulatory factors. *eLife*, 11, 2022.

[10658] Robert A Crawford, Mark P Ashe, Simon J Hubbard, and Graham D Pavitt. Cytosolic aspartate aminotransferase moonlights as a ribosome-binding modulator of Gcn2 activity during oxidative stress. *eLife*, 11, 2022.

[10659] Elise C Croteau-Chonka, Michael S Clayton, Lalanti Venkatasubramanian, Samuel N Harris, Benjamin MW Jones, Lakshmi Narayan, Michael Winding, Jean-Baptiste Masson, Marta Zlatic, and Kristina T Klein. High-throughput automated methods for classical and operant conditioning of Drosophila larvae. *eLife*, 11, 2022.

[10660] Andrew R Crowley, Harini Natarajan, Andrew P Hederman, Carly A Bobak, Joshua A Weiner, Wendy Wieland-Alter, Jiwon Lee, Evan M Bloch, Aaron AR Tobian, Andrew D Redd, Joel N Blankson, Dana Wolf, Tessa Goetghebuer, Arnaud Marchant, Ruth I Connor, Peter F Wright, and Margaret E Ackerman. Boosting of cross-reactive antibodies to endemic coronaviruses by SARS-CoV-2 infection but not vaccination with stabilized spike. *eLife*, 11, 2022.

[10661] Antonio Cuevas-Navarro, Laura Rodriguez-Muñoz, Joaquim Grego-Bessa, Alice Cheng, Katherine A Rauen, Anatoly Urisman, Frank McCormick, Gerardo Jimenez, and Pau Castel. Cross-species analysis of LZTR1 loss-of-function mutants demonstrates dependency to RIT1 orthologs. *eLife*, 11, 2022.

[10662] Can Cui, Caroline Ingre, Li Yin, Xia Li, John Andersson, Christina Seitz, Nicolas Ruffin, Yudi Pawitan, Fredrik Piehl, and Fang Fang. Correlation between leukocyte phenotypes and prognosis of amyotrophic lateral sclerosis. *eLife*, 11, 2022.

[10663] Karen L Cunningham, Chad W Sauvola, Sara Tavana, and J Troy Littleton. Regulation of presynaptic Ca2+ channel abundance at active zones through a balance of delivery and turnover. *eLife*, 11, 2022.

[10664] Camila Cupello, Tatsuya Hirasawa, Norifumi Tatsumi, Yoshitaka Yabumoto, Pierre Gueriau, Sumio Isogai, Ryoko Matsumoto, Toshiro Saruwatari, Andrew King, Masato Hoshino, Kentaro Uesugi, Masataka Okabe, and Paulo M Brito. Lung evolution in vertebrates and the water-to-land transition. *eLife*, 11, 2022.

[10665] Brian Czaya, Kylie Heitman, Isaac Campos, Christopher Yanucil, Dominik Kentrup, David Westbrook, Orlando Gutierrez, Jodie L Babitt, Grace Jung, Isidro B Salusky, Mark Hanudel, and Christian Faul. Hyperphosphatemia increases inflammation to exacerbate anemia and skeletal muscle wasting independently of FGF23-FGFR4 signaling. *eLife*, 11, 2022.

[10666] Alwyn Dady, Lindsay Davidson, Pamela A Halley, and Kate G Storey. Human spinal cord in vitro differentiation pace is initially maintained in heterologous embryonic environments. *eLife*, 11, 2022.

[10667] Hiba Dagher, Anne-Marie Chaftari, Patricia Mulanovich, Ying Jiang, Ray Hachem, Alexandre E Malek, Jovan Borjan, George M Viola, and Issam Raad. Procalcitonin for antimicrobial stewardship among cancer patients admitted with COVID-19. *eLife*, 11, 2022.

[10668] Sumedha Dahal, Humaira Siddiqua, Shivangi Sharma, Ravi K Babu, Diksha Rathore, Sheetal Sharma, and Sathees C Raghavan. Unleashing a novel function of Endonuclease G in mitochondrial genome instability. *eLife*, 11, 2022.

[10669] Jinye Dai, Kif Liakath-Ali, Samantha Rose Golf, and Thomas C Südhof. Distinct neurexin-cerebellin complexes control AMPA- and NMDA-receptor responses in a circuit-dependent manner. *eLife*, 11, 2022.

[10670] Sarah Daimer, Lorenz L Mihatsch, Sharon AS Neufeld, Graham K Murray, and Franziska Knolle. Investigating the relationship of COVID-19 related stress and media consumption with schizotypy, depression, and anxiety in cross-sectional surveys repeated throughout the pandemic in Germany and the UK. *eLife*, 11, 2022.

[10671] Lorenza Dall'Aglio, Hannah H Kim, Sander Lamballais, Jeremy Labrecque, Ryan L Muetzel, and Henning Tiemeier. Attention-deficit hyperactivity disorder symptoms and brain morphology: Examining confounding bias. *eLife*, 11, 2022.

[10672] Marshall A Dalton, Arkiev D'Souza, Jinglei Lv, and Fernando Calamante. New insights into anatomical connectivity along the anterior–posterior axis of the human hippocampus using in vivo quantitative fibre tracking. *eLife*, 11, 2022.

[10673] Kevin G Daly, Benjamin S Arbuckle, Conor Rossi, Valeria Mattiangeli, Phoebe A Lawlor, Marjan Mashkour, Eberhard Sauer, Joséphine Lesur, Levent Atici, Cevdet Merih Erek, and Daniel G Bradley. A novel lineage of the Capra genus discovered in the Taurus Mountains of Turkey using ancient genomics. *eLife*, 11, 2022.

[10674] Hugo GJ Damstra, Boaz Mohar, Mark Eddison, Anna Akhmanova, Lukas C Kapitein, and Paul W Tillberg. Visualizing cellular and tissue ultrastructure using Ten-fold Robust Expansion Microscopy (TREx). *eLife*, 11, 2022.

[10675] Hilda Björk Daníelsdóttir, Thor Aspelund, Edda Bjork Thordardottir, Katja Fall, Fang Fang, Gunnar Tómasson, Harpa Rúnarsdóttir, Qian Yang, Karmel W Choi, Beatrice Kennedy, Thorhildur Halldorsdottir, Donghao Lu, Huan Song, Jóhanna Jakobsdóttir, Arna Hauksdóttir, and Unnur Anna Valdimarsdóttir. Adverse childhood experiences and resilience among adult women: A population-based study. *eLife*, 11, 2022.

[10676] Robin F Dard, Erwan Leprince, Julien Denis, Shrisha Rao Balappa, Dmitrii Suchkov, Richard Boyce, Catherine Lopez, Marie Giorgi-Kurz, Tom Szwagier, Théo Dumont, Hervé Rouault, Marat Minlebaev, Agnès Baude, Rosa Cossart, and Michel A Picardo. The rapid developmental rise of somatic inhibition disengages hippocampal dynamics from self-motion. *eLife*, 11, 2022.

[10677] Osman Darici and Arthur D Kuo. Humans optimally anticipate and compensate for an uneven step during walking. *eLife*, 11, 2022.

[10678] Anup Das, John Myers, Raissa Mathura, Ben Shofty, Brian A Metzger, Kelly Bijanki, Chengyuan Wu, Joshua Jacobs, and Sameer A Sheth. Spontaneous neuronal oscillations in the human insula are hierarchically organized traveling waves. *eLife*, 11, 2022.

[10679] Bhaba K Das, Lei Wang, Toshifumi Fujiwara, Jian Zhou, Nukhet Aykin-Burns, Kimberly J Krager, Renny Lan, Samuel G Mackintosh, Ricky Edmondson, Michael L Jennings, Xiaofang Wang, Jian Q Feng, Tomasa Barrientos, Jyoti Gogoi, Aarthi Kannan, Ling Gao, Weirong Xing, Subburaman Mohan, and Haibo Zhao. Transferrin receptor 1-mediated iron uptake regulates bone mass in mice via osteoclast mitochondria and cytoskeleton. *eLife*, 11, 2022.

[10680] Rakesh Das, Takahiro Sakaue, GV Shivashankar, Jacques Prost, and Tetsuya Hiraiwa. How enzymatic activity is involved in chromatin organization. *eLife*, 11, 2022.

[10681] Sudeshna Das Chakraborty, Hetan Chang, Bill S Hansson, and Silke Sachse. Higher-order olfactory neurons in the lateral horn support odor valence and odor identity coding in Drosophila. *eLife*, 11, 2022.

[10682] Reetu Daswani, Carlotta Gilardi, Michael Soutschek, Prakruti Nanda, Kerstin Weiss, Silvia Bicker, Roberto Fiore, Christoph Dieterich, Pierre-Luc Germain, Jochen Winterer, and Gerhard Schratt. MicroRNA-138 controls hippocampal interneuron function and short-term memory in mice. *eLife*, 11, 2022.

[10683] Swapneeta S Date, Peng Xu, Nathaniel L Hepowit, Nicholas S Diab, Jordan Best, Boyang Xie, Jiale Du, Eric R Strieter, Lauren P Jackson, Jason A MacGurn, and Todd R Graham. Ubiquitination drives COPI priming and Golgi SNARE localization. *eLife*, 11, 2022.

[10684] Arnab Datta, Sagnik Ghosh, and Jane Kondev. How to assemble a scale-invariant gradient. *eLife*, 11, 2022.

[10685] Chintan Dave, Shirley HJ Mei, Andrea McRae, Christine Hum, Katrina J Sullivan, Josee Champagne, Tim Ramsay, and Lauralyn McIntyre. Comparison of freshly cultured versus cryopreserved mesenchymal stem cells in animal models of inflammation: A pre-clinical systematic review. *eLife*, 11, 2022.

[10686] Charlotte E Davies, Jessica E Thomas, Sophie H Malkin, Frederico M Batista, Andrew F Rowley, and Christopher J Coates. Hematodinium sp. infection does not drive collateral disease contraction in a crustacean host. *eLife*, 11, 2022.

[10687] Thomas C Day, Stephanie S Höhn, Seyed A Zamani-Dahaj, David Yanni, Anthony Burnetti, Jennifer Pentz, Aurelia R Honerkamp-Smith, Hugo Wioland, Hannah R Sleath, William C Ratcliff, Raymond E Goldstein, and Peter J Yunker. Cellular organization in lab-evolved and extant multicellular species obeys a maximum entropy law. *eLife*, 11, 2022.

[10688] Abhishek De and Gregory D Horwitz. Coding of chromatic spatial contrast by macaque V1 neurons. *eLife*, 11, 2022.

[10689] Massimo De Agrò, Chiara Matschunas, and Tomer J Czaczkes. Bundling and segregation affect pheromone deposition, but not choice, in an ant. *eLife*, 11, 2022.

[10690] Leonardo Vinicius Monteiro de Assis, Lisbeth Harder, José Thalles Lacerda, Rex Parsons, Meike Kaehler, Ingolf Cascorbi, Inga Nagel, Oliver Rawashdeh, Jens Mittag, and Henrik Oster. Rewiring of liver diurnal transcriptome rhythms by triiodothyronine (T3) supplementation. *eLife*, 11, 2022.

[10691] Helene De Bruyn, Nikky Corthout, Sebastian Munck, Wouter Everaerts, and Thomas Voets. Machine learning-assisted fluoroscopy of bladder function in awake mice. *eLife*, 11, 2022.

[10692] Mickaël De Carvalho, Guo-Song Jia, Ananya Nidamangala Srinivasa, R Blake Billmyre, Yan-Hui Xu, Jeffrey J Lange, Ibrahim M Sabbarini, Li-Lin Du, and Sarah E Zanders. The wtf meiotic driver gene family has unexpectedly persisted for over 100 million years. *eLife*, 11, 2022.

[10693] Iago de Castro Silva, Anna Bianchi, Nilesh U Deshpande, Prateek Sharma, Siddharth Mehra, Vanessa Tonin Garrido, Shannon Jacqueline Saigh, Jonathan England, Peter Joel Hosein, Deukwoo Kwon, Nipun B Merchant, and Jashodeep Datta. Neutrophil-mediated fibroblast-tumor cell il-6/stat-3 signaling underlies the association between neutrophil-to-lymphocyte ratio dynamics and chemotherapy response in localized pancreatic cancer: A hybrid clinical-preclinical study. *eLife*, 11, 2022.

[10694] Andres De La Rossa, Marine H Laporte, Simone Astori, Thomas Marissal, Sylvie Montessuit, Preethi Sheshadri, Eva Ramos-Fernández, Pablo Mendez, Abbas Khani, Charles Quairiaux, Eric B Taylor, Jared Rutter, José Manuel Nunes, Alan Carleton, Michael R Duchen, Carmen Sandi, and Jean-Claude Martinou. Paradoxical neuronal hyperexcitability in a mouse model of mitochondrial pyruvate import deficiency. *eLife*, 11, 2022.

[10695] Alejandro de la Vega, Roberta Rocca, Ross W Blair, Christopher J Markiewicz, Jeff Mentch, James D Kent, Peer Herholz, Satrajit S Ghosh, Russell A Poldrack, and Tal Yarkoni. Neuroscout, a unified platform for generalizable and reproducible fMRI research. *eLife*, 11, 2022.

[10696] Miguel Antonio Xavier de Lima, Marcus Vinicius C Baldo, Fernando A Oliveira, and Newton Sabino Canteras. The anterior cingulate cortex and its role in controlling contextual fear memory to predatory threats. *eLife*, 11, 2022.

[10697] Marc de Manuel, Felix L Wu, and Molly Przeworski. A paternal bias in germline mutation is widespread in amniotes and can arise independently of cell division numbers. *eLife*, 11, 2022.

[10698] Laura de Nies, Susheel Bhanu Busi, Benoit Josef Kunath, Patrick May, and Paul Wilmes. Mobilome-driven segregation of the resistome in biological wastewater treatment. *eLife*, 11, 2022.

[10699] Florian V De Rop, Joy N Ismail, Carmen Bravo González-Blas, Gert J Hulselmans, Christopher Campbell Flerin, Jasper Janssens, Koen Theunis, Valerie M Christiaens, Jasper Wouters, Gabriele Marcassa, Joris de Wit, Suresh Poovathingal, and Stein Aerts. Hydrop enables droplet-based single-cell ATAC-seq and single-cell RNA-seq using dissolvable hydrogel beads. *eLife*, 11, 2022.

[10700] Didier De Saint Jan. Target-specific control of olfactory bulb periglomerular cells by GABAergic and cholinergic basal forebrain inputs. *eLife*, 11, 2022.

[10701] Jelmer P de Vries, Arash Akbarinia, Alban Flachot, and Karl R Gegenfurtner. Emergent color categorization in a neural network trained for object recognition. *eLife*, 11, 2022.

[10702] Erika Alden DeBenedictis, Dieter Söll, and Kevin M Esvelt. Measuring the tolerance of the genetic code to altered codon size. *eLife*, 11, 2022.

[10703] Jordan DeKraker, Roy AM Haast, Mohamed D Yousif, Bradley Karat, Jonathan C Lau, Stefan Köhler, and Ali R Khan. Automated hippocampal unfolding for morphometry and subfield segmentation with HippUnfold. *eLife*, 11, 2022.

[10704] Diego del Alamo, Davide Sala, Hassane S Mchaourab, and Jens Meiler. Sampling alternative conformational states of transporters and receptors with AlphaFold2. *eLife*, 11, 2022.

[10705] Pascal Demange, Etienne Joly, Julien Marcoux, Patrick RA Zanon, Dymytrii Listunov, Pauline Rullière, Cécile Barthes, Céline Noirot, Jean-Baptiste Izquierdo, Alexandrine Rozié, Karen Pradines, Romain Hee, Maria Vieira de Brito, Marlène Marcellin, Remy-Felix Serre, Olivier Bouchez, Odile Burlet-Schiltz, Maria ConceiÇão Ferreira Oliveira, Stéphanie Ballereau, Vania Bernardes-Génisson, Valérie Maraval, Patrick Calsou, Stephan M Hacker, Yves Génisson, Remi Chauvin, and Sébastien Britton. SDR enzymes oxidize specific lipidic alkynylcarbinols into cytotoxic protein-reactive species. *eLife*, 11, 2022.

[10706] Beatrice Demarchi, Meaghan Mackie, Zhiheng Li, Tao Deng, Matthew J Collins, and Julia Clarke. Survival of mineral-bound peptides into the Miocene. *eLife*, 11, 2022.

[10707] Yun Deng, Haihong Wang, Xiaohui Liu, Hao Yuan, Jin Xu, Hugues de Thé, Jun Zhou, and Jun Zhu. Zbtb14 regulates monocyte and macrophage development through inhibiting pu.1 expression in zebrafish. *eLife*, 11, 2022.

[10708] Nicolas Deperrois, Mihai A Petrovici, Walter Senn, and Jakob Jordan. Learning cortical representations through perturbed and adversarial dreaming. *eLife*, 11, 2022.

[10709] Dylane Detilleux, Peggy Raynaud, Berengere Pradet-Balade, and Dominique Helmlinger. The TRRAP transcription cofactor represses interferon-stimulated genes in colorectal cancer cells. *eLife*, 11, 2022.

[10710] Tyler C Detomasi, Adriana M Rico-Ramírez, Richard I Sayler, A Pedro GonÇalves, Michael A Marletta, and N Louise Glass. A moonlighting function of a chitin polysaccharide monooxygenase, CWR-1, in Neurospora crassa allorecognition. *eLife*, 11, 2022.

[10711] Liselot Dewachter, Julien Dénéréaz, Xue Liu, Vincent de Bakker, Charlotte Costa, Mara Baldry, Jean-Claude Sirard, and Jan-Willem Veening. Amoxicillin-resistant Streptococcus pneumoniae can be resensitized by targeting the mevalonate pathway as indicated by sCRilecs-seq. *eLife*, 11, 2022.

[10712] Richard Burkett Dewell, Ying Zhu, Margaret Eisenbrandt, Richard Morse, and Fabrizio Gabbiani. Contrast polarity-specific mapping improves efficiency of neuronal computation for collision detection. *eLife*, 11, 2022.

[10713] Subash Dhakal, Qiuting Ren, Jiangqu Liu, Bradley Akitake, Izel Tekin, Craig Montell, and Youngseok Lee. Drosophila TRP$\gamma$ is required in neuroendocrine cells for post-ingestive food selection. *eLife*, 11, 2022.

[10714] Shreya H Dhume, Steven A Connor, Fergil Mills, Parisa Karimi Tari, Sarah HM Au-Yeung, Benjamin Karimi, Shinichiro Oku, Reiko T Roppongi, Hiroshi Kawabe, Shernaz X Bamji, Yu Tian Wang, Nils Brose, Michael F Jackson, Ann Marie Craig, and Tabrez J Siddiqui. Distinct but overlapping roles of LRRTM1 and LRRTM2 in developing and mature hippocampal circuits. *eLife*, 11, 2022.

[10715] Giulia Di Felice, Giovanni Visci, Federica Teglia, Marco Angelini, and Paolo Boffetta. Effect of cancer on outcome of COVID-19 patients: a systematic review and meta-analysis of studies of unvaccinated patients. *eLife*, 11, 2022.

[10716] Raquel Dias, Doug Evans, Shang-Fu Chen, Kai-Yu Chen, Salvatore Loguercio, Leslie Chan, and Ali Torkamani. Rapid, Reference-Free human genotype imputation with denoising autoencoders. *eLife*, 11, 2022.

[10717] Marco A Díaz-Salinas, Qi Li, Monir Ejemel, Leonid Yurkovetskiy, Jeremy Luban, Kuang Shen, Yang Wang, and James B Munro. Conformational dynamics and allosteric modulation of the SARS-CoV-2 spike. *eLife*, 11, 2022.

[10718] Peter Dietrich, Jens Schumacher, Nico Eisenhauer, and Christiane Roscher. Eco-evolutionary dynamics modulate plant responses to global change depending on plant diversity and species identity. *eLife*, 11, 2022.

[10719] Noah M Dietzen, Mark J Arcario, Lawrence J Chen, John T Petroff, K Trent Moreland, Kathiresan Krishnan, Grace Brannigan, Douglas F Covey, and Wayland WL Cheng. Polyunsaturated fatty acids inhibit a pentameric ligand-gated ion channel through one of two binding sites. *eLife*, 11, 2022.

[10720] Thibaud Dieudonné, Sara Abad Herrera, Michelle Juknaviciute Laursen, Maylis Lejeune, Charlott Stock, Kahina Slimani, Christine Jaxel, Joseph A Lyons, Cédric Montigny, Thomas Günther Pomorski, Poul Nissen, and Guillaume Lenoir. Autoinhibition and regulation by phosphoinositides of ATP8B1, a human lipid flippase associated with intrahepatic cholestatic disorders. *eLife*, 11, 2022.

[10721] Brian A Dillard, Albert K Chung, Alex R Gunderson, Shane C Campbell-Staton, and Andrew H Moeller. Humanization of wildlife gut microbiota in urban environments. *eLife*, 11, 2022.

[10722] Diana C Dima, Tyler M Tomita, Christopher J Honey, and Leyla Isik. Social-affective features drive human representations of observed actions. *eLife*, 11, 2022.

[10723] Vasileios Dimakopoulos, Pierre Mégevand, Lennart H Stieglitz, Lukas Imbach, and Johannes Sarnthein. Information flows from hippocampus to auditory cortex during replay of verbal working memory items. *eLife*, 11, 2022.

[10724] Michael Dimitriou. Human muscle spindles are wired to function as controllable signal-processing devices. *eLife*, 11, 2022.

[10725] Chen Ding, Youjun Wu, Hadas Dabas, and Marc Hammarlund. Activation of the CaMKII-Sarm1-ASK1-p38 MAP kinase pathway protects against axon degeneration caused by loss of mitochondria. *eLife*, 11, 2022.

[10726] Ding Ding, Jie Zhang, Baozhen Du, Xuanzhao Wang, Li Hou, Siyuan Guo, Bing Chen, and Le Kang. Non-canonical function of an Hif-1$\alpha$ splice variant contributes to the sustained flight of locusts. *eLife*, 11, 2022.

[10727] Lingjun Ding, Giuseppe Balsamo, Hongbiao Chen, Eduardo Blanco-Hernandez, Ioannis S Zouridis, Robert Naumann, Patricia Preston-Ferrer, and Andrea Burgalossi. Juxtacellular opto-tagging of hippocampal CA1 neurons in freely moving mice. *eLife*, 11, 2022.

[10728] Peng Ding, Chuan Gao, Youshui Gao, Delin Liu, Hao Li, Jun Xu, Xiaoyi Chen, Yigang Huang, Changqing Zhang, Minghao Zheng, and Junjie Gao. Osteocytes regulate senescence of bone and bone marrow. *eLife*, 11, 2022.

[10729] Yonghe Ding, Di Lang, Jianhua Yan, Haisong Bu, Hongsong Li, Kunli Jiao, Jingchun Yang, Haibo Ni, Stefano Morotti, Tai Le, Karl J Clark, Jenna Port, Stephen C Ekker, Hung Cao, Yuji Zhang, Jun Wang, Eleonora Grandi, Zhiqiang Li, Yongyong Shi, Yigang Li, Alexey V Glukhov, and Xiaolei Xu. A phenotype-based forward genetic screen identifies Dnajb6 as a sick sinus syndrome gene. *eLife*, 11, 2022.

[10730] John G Dingus, Jonathan CY Tang, Ryoji Amamoto, Grace K Wallick, and Constance L Cepko. A general approach for stabilizing nanobodies for intracellular expression. *eLife*, 11, 2022.

[10731] Mauro DiNuzzo, Silvia Mangia, Marta Moraschi, Daniele Mascali, Gisela E Hagberg, and Federico Giove. Perception is associated with the brain's metabolic response to sensory stimulation. *eLife*, 11, 2022.

[10732] Matthew A Dixon, Peter Winskill, Wendy E Harrison, Charles Whittaker, Veronika Schmidt, Astrid Carolina Flórez Sánchez, Zulma M Cucunuba, Agnes U Edia-Asuke, Martin Walker, and María-Gloria Basáñez. Global variation in force-of-infection trends for human Taenia solium taeniasis/cysticercosis. *eLife*, 11, 2022.

[10733] Lucie Dixsaut and Johannes Gräff. Brain-wide screen of prelimbic cortex inputs reveals a functional shift during early fear memory consolidation. *eLife*, 11, 2022.

[10734] Kelsie RS Doering, Xuanjin Cheng, Luke Milburn, Ramesh Ratnappan, Arjumand Ghazi, Dana L Miller, and Stefan Taubert. Nuclear hormone receptor NHR-49 acts in parallel with HIF-1 to promote hypoxia adaptation in Caenorhabditis elegans. *eLife*, 11, 2022.

[10735] Laura M Doherty, Caitlin E Mills, Sarah A Boswell, Xiaoxi Liu, Charles Tapley Hoyt, Benjamin Gyori, Sara J Buhrlage, and Peter K Sorger. Integrating multi-omics data reveals function and therapeutic potential of deubiquitinating enzymes. *eLife*, 11, 2022.

[10736] Hisham M Dokainish, Suyong Re, Takaharu Mori, Chigusa Kobayashi, Jaewoon Jung, and Yuji Sugita. The inherent flexibility of receptor binding domains in SARS-CoV-2 spike protein. *eLife*, 11, 2022.

[10737] Kimberly A Dolan, Mandira Dutta, David M Kern, Abhay Kotecha, Gregory A Voth, and Stephen G Brohawn. Structure of SARS-CoV-2 M protein in lipid nanodiscs. *eLife*, 11, 2022.

[10738] Shauni Doms, Hanna Fokt, Malte Christoph Rühlemann, Cecilia J Chung, Axel Kuenstner, Saleh M Ibrahim, Andre Franke, Leslie M Turner, and John F Baines. Key features of the genetic architecture and evolution of host-microbe interactions revealed by high-resolution genetic mapping of the mucosa-associated gut microbiome in hybrid mice. *eLife*, 11, 2022.

[10739] Kirsten A Donald, Mahmoud Maina, Nilesh Patel, Carine Nguemeni, Wael Mohammed, Amina Abubakar, Matthew Brown, Raliza Stoyanova, Andrew Welchman, Natasha Walker, Alexis Willett, Symon M Kariuki, Anthony Figaji, Dan J Stein, Amadi O Ihunwo, William Daniels, and Charles R Newton. What is next in African neuroscience? *eLife*, 11, 2022.

[10740] Zhe Dong, William Mau, Yu Feng, Zachary T Pennington, Lingxuan Chen, Yosif Zaki, Kanaka Rajan, Tristan Shuman, Daniel Aharoni, and Denise J Cai. Minian, an open-source miniscope analysis pipeline. *eLife*, 11, 2022.

[10741] Nicholas A Donnelly, Ullrich Bartsch, Hayley A Moulding, Christopher Eaton, Hugh Marston, Jessica H Hall, Jeremy Hall, Michael J Owen, Marianne BM van den Bree, and Matt W Jones. Sleep EEG in young people with 22q11.2 deletion syndrome: A cross-sectional study of slow-waves, spindles and correlations with memory and neurodevelopmental symptoms. *eLife*, 11, 2022.

[10742] Lise-Marie Donnio, Elena Cerutti, Charlene Magnani, Damien Neuillet, Pierre-Olivier Mari, and Giuseppina Giglia-Mari. XAB2 dynamics during DNA damage-dependent transcription inhibition. *eLife*, 11, 2022.

[10743] Dennis J Doorduijn, Marie V Lukassen, Marije FL van 't Wout, Vojtech Franc, Maartje Ruyken, Bart W Bardoel, Albert JR Heck, and Suzan HM Rooijakkers. Soluble MAC is primarily released from MAC-resistant bacteria that potently convert complement component C5. *eLife*, 11, 2022.

[10744] Justin YA Doritchamou, Jonathan P Renn, Bethany Jenkins, Almahamoudou Mahamar, Alassane Dicko, Michal Fried, and Patrick E Duffy. A single full-length VAR2CSA ectodomain variant purifies broadly neutralizing antibodies against placental malaria isolates. *eLife*, 11, 2022.

[10745] Sven Dorkenwald, Nicholas L Turner, Thomas Macrina, Kisuk Lee, Ran Lu, Jingpeng Wu, Agnes L Bodor, Adam A Bleckert, Derrick Brittain, Nico Kemnitz, William M Silversmith, Dodam Ih, Jonathan Zung, Aleksandar Zlateski, Ignacio Tartavull, Szi-Chieh Yu, Sergiy Popovych, William Wong, Manuel Castro, Chris S Jordan, Alyssa M Wilson, Emmanouil Froudarakis, JoAnn Buchanan, Marc M Takeno, Russel Torres, Gayathri Mahalingam, Forrest Collman, Casey M Schneider-Mizell, Daniel J Bumbarger, Yang Li, Lynne Becker, Shelby Suckow, Jacob Reimer, Andreas S Tolias, Nuno Macarico da Costa, R Clay Reid, and H Sebastian Seung. Binary and analog variation of synapses between cortical pyramidal neurons. *eLife*, 11, 2022.

[10746] Sara G Dosil, Sheila Lopez-Cobo, Ana Rodriguez-Galan, Irene Fernandez-Delgado, Marta Ramirez-Huesca, Paula Milan-Rois, Milagros Castellanos, Alvaro Somoza, Manuel José Gómez, Hugh T Reyburn, Mar Vales-Gomez, Francisco Sánchez Madrid, and Lola Fernandez-Messina. Natural killer (NK) cell-derived extracellular-vesicle shuttled microRNAs control T cell responses. *eLife*, 11, 2022.

[10747] Dobromir Dotov, Lana Delasanta, Daniel J Cameron, Edward W Large, and Laurel Trainor. Collective dynamics support group drumming, reduce variability, and stabilize tempo drift. *eLife*, 11, 2022.

[10748] Raymond Doudlah, Ting-Yu Chang, Lowell W Thompson, Byounghoon Kim, Adhira Sunkara, and Ari Rosenberg. Parallel processing, hierarchical transformations, and sensorimotor associations along the 'where' pathway. *eLife*, 11, 2022.

[10749] Haikel Dridi, Frances Forrester, Alisa Umanskaya, Wenjun Xie, Steven Reiken, Alain Lacampagne, and Andrew Marks. Role of oxidation of excitation-contraction coupling machinery in age-dependent loss of muscle function in Caenorhabditis elegans. *eLife*, 11, 2022.

[10750] Layla Drwesh, Benjamin Heim, Max Graf, Linda Kehr, Lea Hansen-Palmus, Mirita Franz-Wachtel, Boris Macek, Hubert Kalbacher, Johannes Buchner, and Doron Rapaport. A network of cytosolic (co)chaperones promotes the biogenesis of mitochondrial signal-anchored outer membrane proteins. *eLife*, 11, 2022.

[10751] Jiale Du, Sandor Babik, Yanfeng Li, Kirandeep K Deol, Stephen J Eyles, Jasna Fejzo, Marco Tonelli, and Eric Strieter. A cryptic K48 ubiquitin chain binding site on UCH37 is required for its role in proteasomal degradation. *eLife*, 11, 2022.

[10752] Mingrui Duan, Smitha Sivapragasam, Jacob S Antony, Jenna Ulibarri, John M Hinz, Gregory MK Poon, John J Wyrick, and Peng Mao. High-resolution mapping demonstrates inhibition of DNA excision repair by transcription factors. *eLife*, 11, 2022.

[10753] Mohit Dubey, Maria Pascual-Garcia, Koke Helmes, Dennis D Wever, Mustafa S Hamada, Steven A Kushner, and Maarten HP Kole. Myelination synchronizes cortical oscillations by consolidating parvalbumin-mediated phasic inhibition. *eLife*, 11, 2022.

[10754] Vasin Dumrongprechachan, Ryan B Salisbury, Lindsey Butler, Matthew L MacDonald, and Yevgenia Kozorovitskiy. Dynamic proteomic and phosphoproteomic atlas of corticostriatal axons in neurodevelopment. *eLife*, 11, 2022.

[10755] Kai Dünser, Maria Schöller, Ann-Kathrin Rößling, Christian Löfke, Nannan Xiao, Barbora Pařízková, Stanislav Melnik, Marta Rodriguez-Franco, Eva Stöger, Ondřej Novák, and Jürgen Kleine-Vehn. Endocytic trafficking promotes vacuolar enlargements for fast cell expansion rates in plants. *eLife*, 11, 2022.

[10756] Ilaria Dutto, Julian Gerhards, Antonio Herrera, Olga Souckova, Václava Škopová, Jordann A Smak, Alexandra Junza, Oscar Yanes, Cedric Boeckx, Martin D Burkhalter, Marie Zikánová, Sebastian Pons, Melanie Philipp, Jens Lüders, and Travis H Stracker. Pathway-specific effects of ADSL deficiency on neurodevelopment. *eLife*, 11, 2022.

[10757] Ali Ebrahimi, Akshit Goyal, and Otto X Cordero. Particle foraging strategies promote microbial diversity in marine environments. *eLife*, 11, 2022.

[10758] András Ecker, Bence Bagi, Eszter Vértes, Orsolya Steinbach-Németh, Mária R Karlócai, Orsolya I Papp, István Miklós, Norbert Hájos, Tamás F Freund, Attila I Gulyás, and Szabolcs Káli. Hippocampal sharp wave-ripples and the associated sequence replay emerge from structured synaptic interactions in a network model of area CA3. *eLife*, 11, 2022.

[10759] Manuela Ecker, Richard Schregle, Natasha Kapoor-Kaushik, Pascal Rossatti, Verena M Betzler, Daryan Kempe, Maté Biro, Nicholas Ariotti, Gregory MI Redpath, and Jeremie Rossy. SNX9-induced membrane tubulation regulates CD28 cluster stability and signalling. *eLife*, 11, 2022.

[10760] Maria Katharina Eckstein, Sarah L Master, Liyu Xia, Ronald E Dahl, Linda Wilbrecht, and Anne GE Collins. The interpretation of computational model parameters depends on the context. *eLife*, 11, 2022.

[10761] Rebecca CS Edgar, Ghizal Siddiqui, Katheryn Hjerrild, Tess R Malcolm, Natalie B Vinh, Chaille T Webb, Clare Holmes, Christopher A MacRaild, Hope C Chernih, Willy W Suen, Natalie A Counihan, Darren J Creek, Peter J Scammells, Sheena McGowan, and Tania F de Koning-Ward. Genetic and chemical validation of Plasmodium falciparum aminopeptidase PfA-M17 as a drug target in the hemoglobin digestion pathway. *eLife*, 11, 2022.

[10762] Yochai Edlitz and Eran Segal. Prediction of type 2 diabetes mellitus onset using logistic regression-based scorecards. *eLife*, 11, 2022.

[10763] Michaela A Edmond, Andy Hinojo-Perez, Xiaoan Wu, Marta E Perez Rodriguez, and Rene Barro-Soria. Distinctive mechanisms of epilepsy-causing mutants discovered by measuring S4 movement in KCNQ2 channels. *eLife*, 11, 2022.

[10764] Laura R Edmondson, Alejandro Jiménez Rodríguez, and Hannes P Saal. Expansion and contraction of resource allocation in sensory bottlenecks. *eLife*, 11, 2022.

[10765] Hjörleifur Einarsson, Marco Salvatore, Christian Vaagensø, Nicolas Alcaraz, Jette Bornholdt, Sarah Rennie, and Robin Andersson. Promoter sequence and architecture determine expression variability and confer robustness to genetic variants. *eLife*, 11, 2022.

[10766] Almut S Eisele, Jason Cosgrove, Aurelie Magniez, Emilie Tubeuf, Sabrina Tenreira Bento, Cecile Conrad, Fanny Cayrac, Tamar Tak, Anne-Marie Lyne, Jos Urbanus, and Leïla Perié. Erythropoietin directly remodels the clonal composition of murine hematopoietic multipotent progenitor cells. *eLife*, 11, 2022.

[10767] Fatima El Alaoui, Ignacio Casuso, David Sanchez-Fuentes, Charlotte Arpin-Andre, Raissa Rathar, Volker Baecker, Anna Castro, Thierry Lorca, Julien Viaud, Stéphane Vassilopoulos, Adrian Carretero-Genevrier, and Laura Picas. Structural organization and dynamics of FCHo2 docking on membranes. *eLife*, 11, 2022.

[10768] Assaf Elazar, Nicholas J Chandler, Ashleigh S Davey, Jonathan Y Weinstein, Julie V Nguyen, Raphael Trenker, Ryan S Cross, Misty R Jenkins, Melissa J Call, Matthew E Call, and Sarel J Fleishman. De novo-designed transmembrane domains tune engineered receptor functions. *eLife*, 11, 2022.

[10769] Michaël Elbaz, Amalia Callado Perez, Maxime Demers, Shengli Zhao, Conrad Foo, David Kleinfeld, and Martin Deschenes. A vibrissa pathway that activates the limbic system. *eLife*, 11, 2022.

[10770] Johannes Elferich, Giulia Schiroli, David T Scadden, and Nikolaus Grigorieff. Defocus Corrected Large Area Cryo-EM (DeCo-LACE) for label-free detection of molecules across entire cell sections. *eLife*, 11, 2022.

[10771] George Elias, Pieter Meysman, Esther Bartholomeus, Nicolas De Neuter, Nina Keersmaekers, Arvid Suls, Hilde Jansens, Aisha Souquette, Hans De Reu, Marie-Paule Emonds, Evelien Smits, Eva Lion, Paul G Thomas, Geert Mortier, Pierre Van Damme, Philippe Beutels, Kris Laukens, Viggo Van Tendeloo, and Benson Ogunjimi. Preexisting memory CD4 T cells in naïve individuals confer robust immunity upon hepatitis B vaccination. *eLife*, 11, 2022.

[10772] Susan Eliazer, Xuefeng Sun, Emilie Barruet, and Andrew S Brack. Heterogeneous levels of delta-like 4 within a multinucleated niche cell maintains muscle stem cell diversity. *eLife*, 11, 2022.

[10773] Marianne E Emmert, Parul Aggarwal, Kritton Shay-Winkler, Se-Jin Lee, Qingnian Goh, and Roger Cornwall. Sex-specific role of myostatin signaling in neonatal muscle growth, denervation atrophy, and neuromuscular contractures. *eLife*, 11, 2022.

[10774] Rachel J Ende, Rebecca L Murray, Samantha K D'Spain, Isabelle Coppens, and Isabelle Derré. Phosphoregulation accommodates Type III secretion and assembly of a tether of ER-Chlamydia inclusion membrane contact sites. *eLife*, 11, 2022.

[10775] Stefanie Engert, Gabriella R Sterne, Davi D Bock, and Kristin Scott. Drosophila gustatory projections are segregated by taste modality and connectivity. *eLife*, 11, 2022.

[10776] Olivia Eriksson, Upinder Singh Bhalla, Kim T Blackwell, Sharon M Crook, Daniel Keller, Andrei Kramer, Marja-Leena Linne, Ausra Saudargienė, Rebecca C Wade, and Jeanette Hellgren Kotaleski. Combining hypothesis- and data-driven neuroscience modeling in FAIR workflows. *eLife*, 11, 2022.

[10777] Cyrrus M Espino, Cheyanne M Lewis, Serena Ortiz, Miloni S Dalal, Snigdha Garlapalli, Kaylee M Wells, Darik A O'Neil, Katherine A Wilkinson, and Theanne N Griffith. NaV1.1 is essential for proprioceptive signaling and motor behaviors. *eLife*, 11, 2022.

[10778] Max T Eyre, Fábio N Souza, Ticiana SA Carvalho-Pereira, Nivison Nery, Daiana de Oliveira, Jaqueline S Cruz, Gielson A Sacramento, Hussein Khalil, Elsio A Wunder, Kathryn P Hacker, José E Hagan, James E Childs, Mitermayer G Reis, Mike Begon, Peter J Diggle, Albert I Ko, Emanuele Giorgi, and Federico Costa. Linking rattiness, geography and environmental degradation to spillover Leptospira infections in marginalised urban settings: An eco-epidemiological community-based cohort study in Brazil. *eLife*, 11, 2022.

[10779] Julia Falo-Sanjuan and Sarah Bray. Notch-dependent and -independent transcription are modulated by tissue movements at gastrulation. *eLife*, 11, 2022.

[10780] Chengcheng Fan and Douglas C Rees. Glutathione binding to the plant AtAtm3 transporter and implications for the conformational coupling of ABC transporters. *eLife*, 11, 2022.

[10781] Mengyang Fan, Wenchao Lu, Jianwei Che, Nicholas P Kwiatkowski, Yang Gao, Hyuk-Soo Seo, Scott B Ficarro, Prafulla C Gokhale, Yao Liu, Ezekiel A Geffken, Jimit Lakhani, Kijun Song, Miljan Kuljanin, Wenzhi Ji, Jie Jiang, Zhixiang He, Jason Tse, Andrew S Boghossian, Matthew G Rees, Melissa M Ronan, Jennifer A Roth, Joseph D Mancias, Jarrod A Marto, Sirano Dhe-Paganon, Tinghu Zhang, and Nathanael S Gray. Covalent disruptor of YAP-TEAD association suppresses defective Hippo signaling. *eLife*, 11, 2022.

[10782] Deborah Fanfone, Zhichong Wu, Jade Mammi, Kevin Berthenet, David Neves, Kathrin Weber, Andrea Halaburkova, FranÇois Virard, Félix Bunel, Catherine Jamard, Hector Hernandez-Vargas, Stephen WG Tait, Ana Hennino, and Gabriel Ichim. Confined migration promotes cancer metastasis through resistance to anoikis and increased invasiveness. *eLife*, 11, 2022.

[10783] Rui Fang, Jason Hon, Mengying Zhou, and Ying Lu. An empirical energy landscape reveals mechanism of proteasome in polypeptide translocation. *eLife*, 11, 2022.

[10784] Si Fang, Michael V Holmes, Tom R Gaunt, George Davey Smith, and Tom G Richardson. Constructing an atlas of associations between polygenic scores from across the human phenome and circulating metabolic biomarkers. *eLife*, 11, 2022.

[10785] Jean-Claude Farre, Krypton Carolino, Lou Devanneaux, and Suresh Subramani. OXPHOS deficiencies affect peroxisome proliferation by downregulating genes controlled by the SNF1 signaling pathway. *eLife*, 11, 2022.

[10786] Spencer Farrell, Alice E Kane, Elise Bisset, Susan E Howlett, and Andrew D Rutenberg. Measurements of damage and repair of binary health attributes in aging mice and humans reveal that robustness and resilience decrease with age, operate over broad timescales, and are affected differently by interventions. *eLife*, 11, 2022.

[10787] John S Favate, Shun Liang, Alexander L Cope, Srujana S Yadavalli, and Premal Shah. The landscape of transcriptional and translational changes over 22 years of bacterial adaptation. *eLife*, 11, 2022.

[10788] Sophie L Fayad, Guillaume Ourties, Benjamin Le Gac, Baptiste Jouffre, Sylvain Lamoine, Antoine Fruquière, Sophie Laffray, Laila Gasmi, Bruno Cauli, Christophe Mallet, Emmanuel Bourinet, Thomas Bessaih, Régis C Lambert, and Nathalie Leresche. Centrally expressed Cav3.2 T-type calcium channel is critical for the initiation and maintenance of neuropathic pain. *eLife*, 11, 2022.

[10789] Lukas P Feilen, Shu-Yu Chen, Akio Fukumori, Regina Feederle, Martin Zacharias, and Harald Steiner. Active site geometry stabilization of a presenilin homolog by the lipid bilayer promotes intramembrane proteolysis. *eLife*, 11, 2022.

[10790] Jifan Feng, Xia Han, Yuan Yuan, Courtney Kyeong Cho, Eva Janečková, Tingwei Guo, Siddhika Pareek, Md Shaifur Rahman, Banghong Zheng, Jing Bi, Junjun Jing, Mingyi Zhang, Jian Xu, Thach-Vu Ho, and Yang Chai. TGF-$\beta$ signaling and Creb5 cooperatively regulate Fgf18 to control pharyngeal muscle development. *eLife*, 11, 2022.

[10791] Yi Feng, Rafik Neme, Leslie Y Beh, Xiao Chen, Jasper Braun, Michael W Lu, and Laura F Landweber. Comparative genomics reveals insight into the evolutionary origin of massively scrambled genomes. *eLife*, 11, 2022.

[10792] Zhanying Feng, Zhana Duren, Jingxue Xin, Qiuyue Yuan, Yaoxi He, Bing Su, Wing Hung Wong, and Yong Wang. Heritability enrichment in context-specific regulatory networks improves phenotype-relevant tissue identification. *eLife*, 11, 2022.

[10793] Sara Feola, Jacopo Chiaro, Beatriz Martins, Salvatore Russo, Manlio Fusciello, Erkko Ylösmäki, Chiara Bonini, Eliana Ruggiero, Firas Hamdan, Michaela Feodoroff, Gabriella Antignani, Tapani Viitala, Sari Pesonen, Mikaela Grönholm, Rui MM Branca, Janne Lehtiö, and Vincenzo Cerullo. A novel immunopeptidomic-based pipeline for the generation of personalized oncolytic cancer vaccines. *eLife*, 11, 2022.

[10794] Kayla M Ferko, Anna Blumenthal, Chris B Martin, Daria Proklova, Alexander N Minos, Lisa M Saksida, Timothy J Bussey, Ali R Khan, and Stefan Köhler. Activity in perirhinal and entorhinal cortex predicts perceived visual similarities among category exemplars with highest precision. *eLife*, 11, 2022.

[10795] GonÇalo Fernandes, Huy Tran, Maxime Andrieu, Youssoupha Diaw, Carmina Perez Romero, Cécile Fradin, Mathieu Coppey, Aleksandra M Walczak, and Nathalie Dostatni. Synthetic reconstruction of the hunchback promoter specifies the role of Bicoid, Zelda and Hunchback in the dynamics of its transcription. *eLife*, 11, 2022.

[10796] Keith Conrad Fernandez, Laura Feeney, Ryan M Smolkin, Wei-Feng Yen, Allysia J Matthews, William Alread, John HJ Petrini, and Jayanta Chaudhuri. The structure-selective endonucleases GEN1 and MUS81 mediate complementary functions in safeguarding the genome of proliferating B lymphocytes. *eLife*, 11, 2022.

[10797] Rafael Jesus Fernandez, Zachary JG Gardner, Katherine J Slovik, Derek C Liberti, Katrina N Estep, Wenli Yang, Qijun Chen, Garrett T Santini, Javier V Perez, Sarah Root, Ranvir Bhatia, John W Tobias, Apoorva Babu, Michael P Morley, David B Frank, Edward E Morrisey, Christopher J Lengner, and F Brad Johnson. GSK3 inhibition rescues growth and telomere dysfunction in dyskeratosis congenita iPSC-derived type II alveolar epithelial cells. *eLife*, 11, 2022.

[10798] Elena Fernández álvaro, Phat Voong Vinh, Cristina de Cozar, David R Willé, Beatriz Urones, Alvaro Cortés, Alan Price, Nhu Tran Do Hoang, Tuyen Ha Thanh, Molly McCloskey, Shareef Shaheen, Denise Dayao, Amanda Martinot, Jaime de Mercado, Pablo Castañeda, Adolfo García-Perez, Benson Singa, Patricia Pavlinac, Judd Walson, Maria Santos Martínez-Martínez, Samuel LM Arnold, Saul Tzipori, Lluis Ballell Pages, and Stephen Baker. The repurposing of Tebipenem pivoxil as alternative therapy for severe gastrointestinal infections caused by extensively drug-resistant Shigella spp. *eLife*, 11, 2022.

[10799] Elena Fernández álvaro, Phat Voong Vinh, Cristina de Cozar, David R Willé, Beatriz Urones, Alvaro Cortés, Alan Price, Nhu Tran Do Hoang, Tuyen Ha Thanh, Molly McCloskey, Shareef Shaheen, Denise Dayao, Amanda Martinot, Jaime de Mercado, Pablo Castañeda, Adolfo García-Perez, Benson Singa, Patricia Pavlinac, Judd Walson, Maria Santos Martínez-Martínez, Samuel LM Arnold, Saul Tzipori, Lluis Ballell Pages, and Stephen Baker. Correction: The repurposing of Tebipenem pivoxil as alternative therapy for severe gastrointestinal infections caused by extensively drug-resistant Shigella spp. *eLife*, 11, 2022.

[10800] Caroline B Ferreira, Talita M Silva, Phelipe E Silva, Claudio L Castro, Catherine Czeisler, José J Otero, Ana C Takakura, and Thiago S Moreira. Phox2b mutation mediated by Atoh1 expression impaired respiratory rhythm and ventilatory responses to hypoxia and hypercapnia. *eLife*, 11, 2022.

[10801] Rachel D Field and Lael Parrott. Mapping the functional connectivity of ecosystem services supply across a regional landscape. *eLife*, 11, 2022.

[10802] Edward Fielder, Tengfei Wan, Ghazaleh Alimohammadiha, Abbas Ishaq, Evon Low, B Melanie Weigand, George Kelly, Craig Parker, Brigid Griffin, Diana Jurk, Viktor I Korolchuk, Thomas von Zglinicki, and Satomi Miwa. Short senolytic or senostatic interventions rescue progression of radiation-induced frailty and premature ageing in mice. *eLife*, 11, 2022.

[10803] Ceri Alan Fielding, Pragati Sabberwal, James C Williamson, Edward JD Greenwood, Thomas WM Crozier, Wioleta Zelek, Jeffrey Seow, Carl Graham, Isabella Huettner, Jonathan D Edgeworth, David A Price, Paul B Morgan, Kristin Ladell, Matthias Eberl, Ian R Humphreys, Blair Merrick,

Katie Doores, Sam J Wilson, Paul J Lehner, Eddie CY Wang, and Richard J Stanton. SARS-CoV-2 host-shutoff impacts innate NK cell functions, but antibody-dependent NK activity is strongly activated through non-spike antibodies. *eLife*, 11, 2022.

[10804] Skye RS Fishbein, John I Robinson, Tiffany Hink, Kimberly A Reske, Erin P Newcomer, Carey-Ann D Burnham, Jeffrey P Henderson, Erik R Dubberke, and Gautam Dantas. Multi-omics investigation of Clostridioides difficile-colonized patients reveals pathogen and commensal correlates of C. difficile pathogenesis. *eLife*, 11, 2022.

[10805] Ethan S FitzGerald and Amanda M Jamieson. Comment on 'SARS-CoV-2 suppresses anticoagulant and fibrinolytic gene expression in the lung'. *eLife*, 11, 2022.

[10806] Sarah M Fixsen, Kelsey R Cone, Stephen A Goldstein, Thomas A Sasani, Aaron R Quinlan, Stefan Rothenburg, and Nels C Elde. Poxviruses capture host genes by LINE-1 retrotransposition. *eLife*, 11, 2022.

[10807] Philipp Fleig, Mirna Kramar, Michael Wilczek, and Karen Alim. Emergence of behaviour in a self-organized living matter network. *eLife*, 11, 2022.

[10808] Josué Flores-Kim, Genevieve S Dobihal, Thomas G Bernhardt, and David Z Rudner. WhyD tailors surface polymers to prevent premature bacteriolysis and direct cell elongation in Streptococcus pneumoniae. *eLife*, 11, 2022.

[10809] Julia M Flynn, Neha Samant, Gily Schneider-Nachum, David T Barkan, Nese Kurt Yilmaz, Celia A Schiffer, Stephanie A Moquin, Dustin Dovala, and Daniel NA Bolon. Comprehensive fitness landscape of SARS-CoV-2 Mpro reveals insights into viral resistance mechanisms. *eLife*, 11, 2022.

[10810] Taylor Follansbee, Dan Domocos, Eileen Nguyen, Amanda Nguyen, Aristea Bountouvas, Lauren Velasquez, Mirela Iodi Carstens, Keiko Takanami, Sarah E Ross, and Earl Carstens. Inhibition of itch by neurokinin 1 receptor (Tacr1) -expressing ON cells in the rostral ventromedial medulla in mice. *eLife*, 11, 2022.

[10811] Vincent Fontanier, Matthieu Sarazin, Frederic M Stoll, Bruno Delord, and Emmanuel Procyk. Inhibitory control of frontal metastability sets the temporal signature of cognition. *eLife*, 11, 2022.

[10812] Holly C Ford, William J Allen, GonÇalo C Pereira, Xia Liu, Mark Simon Dillingham, and Ian Collinson. Towards a molecular mechanism underlying mitochondrial protein import through the TOM and TIM23 complexes. *eLife*, 11, 2022.

[10813] Hannah R Foster, Thuong Ho, Evgeniy Potapenko, Sophia M Sdao, Shih Ming Huang, Sophie L Lewandowski, Halena R VanDeusen, Shawn M Davidson, Rebecca L Cardone, Marc Prentki, Richard G Kibbey, and Matthew J Merrins. $\beta$-cell deletion of the PKm1 and PKm2 isoforms of pyruvate kinase in mice reveals their essential role as nutrient sensors for the KATP channel. *eLife*, 11, 2022.

[10814] Marjorie Fournier, Amélie Rodrigue, Larissa Milano, Jean-Yves Bleuyard, Anthony M Couturier, Jacob Wall, Jessica Ellins, Svenja Hester, Stephen J Smerdon, László Tora, Jean-Yves Masson, and Fumiko Esashi. KAT2-mediated acetylation switches the mode of PALB2 chromatin association to safeguard genome integrity. *eLife*, 11, 2022.

[10815] Faith C Fowler, Bo-Ruei Chen, Nicholas Zolnerowich, Wei Wu, Raphael Pavani, Jacob Paiano, Chelsea Peart, Zulong Chen, André Nussenzweig, Barry P Sleckman, and Jessica K Tyler. DNA-PK promotes DNA end resection at DNA double strand breaks in G0 cells. *eLife*, 11, 2022.

[10816] Joseph Wayne M Fowler, Rong Zhang, Bo Tao, Nabil E Boutagy, and William C Sessa. Inflammatory stress signaling via NF-kB alters accessible cholesterol to upregulate SREBP2 transcriptional activity in endothelial cells. *eLife*, 11, 2022.

[10817] Philip W Fowler, Carla Wright, Helen Spiers, Tingting Zhu, Elisabeth ML Baeten, Sarah W Hoosdally, Ana L Gibertoni Cruz, Aysha Roohi, Samaneh Kouchaki, Timothy M Walker, Timothy EA Peto, Grant Miller, Chris Lintott, David Clifton, Derrick W Crook, A Sarah Walker, and Ivan Barilar. A crowd of BashTheBug volunteers reproducibly and accurately measure the minimum inhibitory concentrations of 13 antitubercular drugs from photographs of 96-well broth microdilution plates. *eLife*, 11, 2022.

[10818] Damon JA Frampton, Koushik Choudhury, Johan Nikesjö, Lucie Delemotte, and Sara I Liin. Subtype-specific responses of hKv7.4 and hKv7.5 channels to polyunsaturated fatty acids reveal an unconventional modulatory site and mechanism. *eLife*, 11, 2022.

[10819] Flavia P Franco, Pingxi Xu, Brandon J Harris, Vladimir Yarov-Yarovoy, and Walter S Leal. Single amino acid residue mediates reciprocal specificity in two mosquito odorant receptors. *eLife*, 11, 2022.

[10820] Sandra Franco Iborra, Jessica Polka, and Iratxe Puebla. Promoting constructive feedback on preprints with the FAST principles. *eLife*, 11, 2022.

[10821] Marie Francois, Isabella Canal Delgado, Nikolay Shargorodsky, Cheng-Shiun Leu, and Lori Zeltser. Assessing the effects of stress on feeding behaviors in laboratory mice. *eLife*, 11, 2022.

[10822] Felipe Freitas and David Attwell. Pericyte-mediated constriction of renal capillaries evokes no-reflow and kidney injury following ischaemia. *eLife*, 11, 2022.

[10823] Jiaqi Fu, Mowei Zhou, Marina A Gritsenko, Ernesto S Nakayasu, Lei Song, and Zhao-Qing Luo. Legionella pneumophila modulates host energy metabolism by ADP-ribosylation of ADP/ATP translocases. *eLife*, 11, 2022.

[10824] Weili Fu, Sijie Chen, Runze Yang, Chen Li, Haoxiang Gao, Jian Li, and Xuegong Zhang. Cellular features of localized microenvironments in human meniscal degeneration: a single-cell transcriptomic study. *eLife*, 11, 2022.

[10825] Xiaoming Fu, Heta P Patel, Stefano Coppola, Libin Xu, Zhixing Cao, Tineke L Lenstra, and Ramon Grima. Quantifying how post-transcriptional noise and gene copy number variation bias transcriptional parameter inference from mRNA distributions. *eLife*, 11, 2022.

[10826] Xiaoqin Fu, Lu Rao, Peijun Li, Xinglei Liu, Qi Wang, Alexander I Son, Arne Gennerich, and Judy Shih-Hwa Liu. Doublecortin and JIP3 are neural-specific counteracting regulators of dynein-mediated retrograde trafficking. *eLife*, 11, 2022.

[10827] Zhongjie Fu, Anders K Nilsson, Ann Hellstrom, and Lois EH Smith. Retinopathy of prematurity: Metabolic risk factors. *eLife*, 11, 2022.

[10828] Tanja Fuchsberger, Claudia Clopath, Przemyslaw Jarzebowski, Zuzanna Brzosko, Hongbing Wang, and Ole Paulsen. Postsynaptic burst reactivation of hippocampal neurons enables associative plasticity of temporally discontiguous inputs. *eLife*, 11, 2022.

[10829] Ryo Fujisawa, Cristian Polo Rivera, and Karim PM Labib. Multiple UBX proteins reduce the ubiquitin threshold of the mammalian p97-UFD1-NPL4 unfoldase. *eLife*, 11, 2022.

[10830] Kei Fukuda, Yoshinori Makino, Satoru Kaneko, Chikako Shimura, Yuki Okada, Kenji Ichiyanagi, and Yoichi Shinkai. Potential role of KRAB-ZFP binding and transcriptional states on DNA methylation of retroelements in human male germ cells. *eLife*, 11, 2022.

[10831] Chris Furlan, Nipa Chongdar, Pooja Gupta, Wolfgang Lubitz, Hideaki Ogata, James N Blaza, and James A Birrell. Structural insight on the mechanism of an electron-bifurcating [FeFe] hydrogenase. *eLife*, 11, 2022.

[10832] R Christopher D Furniss, Nikol Kaderabkova, Declan Barker, Patricia Bernal, Evgenia Maslova, Amanda AA Antwi, Helen E McNeil, Hannah L Pugh, Laurent Dortet, Jessica MA Blair, Gerald Larrouy-Maumus, Ronan R McCarthy, Diego Gonzalez, and Despoina AI Mavridou. Breaking antimicrobial resistance by disrupting extracytoplasmic protein folding. *eLife*, 11, 2022.

[10833] Gokul G and Jogender Singh. Dithiothreitol causes toxicity in C. elegans by modulating the methionine–homocysteine cycle. *eLife*, 11, 2022.

[10834] Christopher J Gabriel, Zachary Zeidler, Benita Jin, Changliang Guo, Caitlin M Goodpaster, Adrienne Q Kashay, Anna Wu, Molly Delaney, Jovian Cheung, Lauren E DiFazio, Melissa J Sharpe, Daniel Aharoni, Scott A Wilke, and Laura A DeNardo. BehaviorDEPOT is a simple, flexible tool for automated behavioral detection based on markerless pose tracking. *eLife*, 11, 2022.

[10835] Danni A Gadd, Robert F Hillary, Daniel L McCartney, Shaza B Zaghlool, Anna J Stevenson, Yipeng Cheng, Chloe Fawns-Ritchie, Cliff Nangle, Archie Campbell, Robin Flaig, Sarah E Harris, Rosie M Walker, Liu Shi, Elliot M Tucker-Drob, Christian Gieger, Annette Peters, Melanie Waldenberger, Johannes Graumann, Allan F McRae, Ian J Deary, David J Porteous, Caroline Hayward, Peter M Visscher, Simon R Cox, Kathryn L Evans, Andrew M McIntosh, Karsten Suhre, and Riccardo E Marioni. Epigenetic scores for the circulating proteome as tools for disease prediction. *eLife*, 11, 2022.

[10836] Michael A Gaffield, Britton A Sauerbrei, and Jason M Christie. Cerebellum encodes and influences the initiation, performance, and termination of discontinuous movements in mice. *eLife*, 11, 2022.

[10837] Kevin D Gallagher, Madhav Mani, and Richard W Carthew. Emergence of a geometric pattern of cell fates from tissue-scale mechanics in the Drosophila eye. *eLife*, 11, 2022.

[10838] Cecilia Gallego-Carracedo, Matthew G Perich, Raeed H Chowdhury, Lee E Miller, and Juan álvaro Gallego. Local field potentials reflect cortical population dynamics in a region-specific and frequency-dependent manner. *eLife*, 11, 2022.

[10839] Travis Gallo, Mason Fidino, Brian Gerber, Adam A Ahlers, Julia L Angstmann, Max Amaya, Amy L Concilio, David Drake, Danielle Gay, Elizabeth W Lehrer, Maureen H Murray, Travis J Ryan, Colleen Cassady St Clair, Carmen M Salsbury, Heather A Sander, Theodore Stankowich, Jaque Williamson, J Amy Belaire, Kelly Simon, and Seth B Magle. Mammals adjust diel activity across gradients of urbanization. *eLife*, 11, 2022.

[10840] Nicola Galvanetto, Zhongjie Ye, Arin Marchesi, Simone Mortal, Sourav Maity, Alessandro Laio, and Vincent Torre. Unfolding and identification of membrane proteins in situ. *eLife*, 11, 2022.

[10841] Nicolás MS Gálvez, Gaspar A Pacheco, Bárbara M Schultz, Felipe Melo-González, Jorge A Soto, Luisa F Duarte, Liliana A González, Daniela Rivera-Pérez, Mariana Ríos, Roslye V Berrios, Yaneisi Vázquez, Daniela Moreno-Tapia, Omar P Vallejos, Catalina A Andrade, Guillermo Hoppe-Elsholz, Carolina Iturriaga, Marcela Urzua, María S Navarrete, álvaro Rojas, Rodrigo Fasce, Jorge Fernández, Judith Mora, Eugenio Ramírez, Aracelly Gaete-Argel, Mónica L Acevedo, Fernando Valiente-Echeverría, Ricardo Soto-Rifo, Daniela Weiskopf, Alba Grifoni, Alessandro Sette, Gang Zeng, Weining Meng, José V González-Aramundiz, Marina Johnson, David Goldblatt, Pablo A González, Katia Abarca, Susan M Bueno, and Alexis M Kalergis. Differences in the immune response elicited by two immunization schedules with an inactivated SARS-CoV-2 vaccine in a randomized phase 3 clinical trial. *eLife*, 11, 2022.

[10842] Alejandro Galvez-Pol, Pavandeep Virdee, Javier Villacampa, and James Kilner. Active tactile discrimination is coupled with and modulated by the cardiac cycle. *eLife*, 11, 2022.

[10843] Feng Gao, Changgong Li, Susan M Smith, Neil Peinado, Golenaz Kohbodi, Evelyn Tran, Yong-Hwee Eddie Loh, Wei Li, Zea Borok, and Parviz Minoo. Decoding the IGF1 signaling gene regulatory network behind alveologenesis from a mouse model of bronchopulmonary dysplasia. *eLife*, 11, 2022.

[10844] Ge Gao, Shuyu Guo, Quan Zhang, Hefei Zhang, Cuizhen Zhang, and Gang Peng. Kiaa1024L/Minar2 is essential for hearing by regulating cholesterol distribution in hair bundles. *eLife*, 11, 2022.

[10845] Luna Gao, Jian Zhao, Evan Ardiel, Qi Hall, Stephen Nurrish, and Joshua M Kaplan. Shank promotes action potential repolarization by recruiting BK channels to calcium microdomains. *eLife*, 11, 2022.

[10846] Mu Gao, Davi Nakajima An, and Jeffrey Skolnick. Deep learning-driven insights into super protein complexes for outer membrane protein biogenesis in bacteria. *eLife*, 11, 2022.

[10847] Zhiyao Gao, Li Zheng, Katya Krieger-Redwood, Ajay Halai, Daniel S Margulies, Jonathan Smallwood, and Elizabeth Jefferies. Flexing the principal gradient of the cerebral cortex to suit changing semantic task demands. *eLife*, 11, 2022.

[10848] Stephanie Chow Garbern, Eric J Nelson, Sabiha Nasrin, Adama Mamby Keita, Ben J Brintz, Monique Gainey, Henry Badji, Dilruba Nasrin, Joel Howard, Mami Taniuchi, James A Platts-Mills, Karen L Kotloff, Rashidul Haque, Adam C Levine, Samba O Sow, Nur Haque Alam, and Daniel T Leung. External validation of a mobile clinical decision support system for diarrhea etiology prediction in children: A multicenter study in Bangladesh and Mali. *eLife*, 11, 2022.

[10849] Fabiola García Fernández, Etienne Almayrac, Ànnia Carré Simon, Renaud Batrin, Yasmine Khalil, Michel Boissac, and Emmanuelle Fabre. Global chromatin mobility induced by a DSB is dictated by chromosomal conformation and defines the HR outcome. *eLife*, 11, 2022.

[10850] Jonay Garcia-Luis, Hélène Bordelet, Agnès Thierry, Romain Koszul, and Luis Aragon. Depletion or cleavage of cohesin during anaphase differentially affects chromatin structure and segregation. *eLife*, 11, 2022.

[10851] Mario García-Navarrete, Merisa Avdovic, Sara Pérez-Garcia, Diego Ruiz Sanchis, and Krzysztof Wabnik. Macroscopic control of cell electrophysiology through ion channel expression. *eLife*, 11, 2022.

[10852] Daniel Garcia-Souto, Alicia L Bruzos, Seila Diaz, Sara Rocha, Ana Pequeño-Valtierra, Camila F Roman-Lewis, Juana Alonso, Rosana Rodriguez, Damian Costas, Jorge Rodriguez-Castro, Antonio Villanueva, Luis Silva, Jose Maria Valencia, Giovanni Annona, Andrea Tarallo, Fernando Ricardo, Ana BratoŠ Cetinić, David Posada, Juan Jose Pasantes, and Jose MC Tubio. Mitochondrial genome sequencing of marine leukaemias reveals cancer contagion between clam species in the Seas of Southern Europe. *eLife*, 11, 2022.

[10853] Vasiliki-Ilya Gargareta, Josefine Reuschenbach, Sophie B Siems, Ting Sun, Lars Piepkorn, Carolina Mangana, Erik Späte, Sandra Goebbels, Inge Huitinga, Wiebke Möbius, Klaus-Armin Nave, Olaf Jahn, and Hauke B Werner. Conservation and divergence of myelin proteome and oligodendrocyte transcriptome profiles between humans and mice. *eLife*, 11, 2022.

[10854] Rikki M Garner and Julie A Theriot. Leading edge maintenance in migrating cells is an emergent property of branched actin network growth. *eLife*, 11, 2022.

[10855] Michael Garratt, Ilkim Erturk, Roxann Alonzo, Frank Zufall, Trese Leinders-Zufall, Scott D Pletcher, and Richard A Miller. Lifespan extension in female mice by early, transient exposure to adult female olfactory cues. *eLife*, 11, 2022.

[10856] Meghan E Garrett, Jared G Galloway, Caitlin Wolf, Jennifer K Logue, Nicholas Franko, Helen Y Chu, Frederick A Matsen, and Julie M Overbaugh. Comprehensive characterization of the antibody responses to SARS-CoV-2 Spike protein finds additional vaccine-induced epitopes beyond those for mild infection. *eLife*, 11, 2022.

[10857] Sandra Gattas, Gabriel A Elias, John Janecek, Michael A Yassa, and Norbert J Fortin. Proximal CA1 20–40 Hz power dynamics reflect trial-specific information processing supporting nonspatial sequence memory. *eLife*, 11, 2022.

[10858] Christl Gaubitz, Xingchen Liu, Joshua Pajak, Nicholas P Stone, Janelle A Hayes, Gabriel Demo, and Brian A Kelch. Cryo-EM structures reveal high-resolution mechanism of a DNA polymerase sliding clamp loader. *eLife*, 11, 2022.

[10859] Noémie Gaudin, Paula Martin Gil, Meriem Boumendjel, Dmitry Ershov, Catherine Pioche-Durieu, Manon Bouix, Quentin Delobelle, Lucia Maniscalco, Than Bich Ngan Phan, Vincent Heyer, Bernardo Reina-San-Martin, and Juliette Azimzadeh. Evolutionary conservation of centriole rotational asymmetry in the human centrosome. *eLife*, 11, 2022.

[10860] Riem Gawish, Barbara Maier, Georg Obermayer, Martin L Watzenboeck, Anna-Dorothea Gorki, Federica Quattrone, Asma Farhat, Karin Lakovits, Anastasiya Hladik, Ana Korosec, Arman Alimohammadi, Ildiko Mesteri, Felicitas Oberndorfer, Fiona Oakley, John Brain, Louis Boon, Irene Lang, Christoph J Binder, and Sylvia Knapp. A neutrophil–B-cell axis impacts tissue damage control in a mouse model of intraabdominal bacterial infection via Cxcr4. *eLife*, 11, 2022.

[10861] Riem Gawish, Philipp Starkl, Lisabeth Pimenov, Anastasiya Hladik, Karin Lakovits, Felicitas Oberndorfer, Shane JF Cronin, Anna Ohradanova-Repic, Gerald Wirnsberger, Benedikt Agerer, Lukas Endler, Tümay Capraz, Jan W Perthold, Domagoj Cikes, Rubina Koglgruber, Astrid Hagelkruys, Nuria Montserrat, Ali Mirazimi, Louis Boon, Hannes Stockinger, Andreas Bergthaler, Chris Oostenbrink, Josef M Penninger, and Sylvia Knapp. ACE2 is the critical in vivo receptor for SARS-CoV-2 in a novel COVID-19 mouse model with TNF- and IFN$\gamma$-driven immunopathology. *eLife*, 11, 2022.

[10862] Linda Geerligs, Dora Gözükara, Djamari Oetringer, Karen L Campbell, Marcel van Gerven, and Umut GüÇlü. A partially nested cortical hierarchy of neural states underlies event segmentation in the human brain. *eLife*, 11, 2022.

[10863] Joseph V Geisberg, Zarmik Moqtaderi, Nova Fong, Benjamin Erickson, David L Bentley, and Kevin Struhl. Nucleotide-level linkage of transcriptional elongation and polyadenylation. *eLife*, 11, 2022.

[10864] Steffen Geisel, Eleonora Secchi, and Jan Vermant. The role of surface adhesion on the macroscopic wrinkling of biofilms. *eLife*, 11, 2022.

[10865] Jakub Gemperle, Thomas S Harrison, Chloe Flett, Antony D Adamson, and Patrick T Caswell. On demand expression control of endogenous genes with DExCon, DExogron and LUXon reveals differential dynamics of Rab11 family members. *eLife*, 11, 2022.

[10866] Jinli Geng, Yingjun Tang, Zhen Yu, Yunming Gao, Wenxiang Li, Yitong Lu, Bo Wang, Huiming Zhou, Ping Li, Nan Liu, Ping Wang, Yubo Fan, Yaxiong Yang, Zengcai V Guo, and Xiaodong Liu. Chronic Ca2+ imaging of cortical neurons with long-term expression of GCaMP-X. *eLife*, 11, 2022.

[10867] Damiano Gentiletti, Marco de Curtis, Vadym Gnatkovsky, and Piotr Suffczynski. Focal seizures are organized by feedback between neural activity and ion concentration changes. *eLife*, 11, 2022.

[10868] Athina Georgiadou, Claire Dunican, Pablo Soro-Barrio, Hyun Jae Lee, Myrsini Kaforou, and Aubrey J Cunnington. Comparative transcriptomic analysis reveals translationally relevant processes in mouse models of malaria. *eLife*, 11, 2022.

[10869] Tamar Gera, Felix Jonas, Roye More, and Naama Barkai. Evolution of binding preferences among whole-genome duplicated transcription factors. *eLife*, 11, 2022.

[10870] Tobias Gerber, Cristina Loureiro, Nico Schramma, Siyu Chen, Akanksha Jain, Anne Weber, Anne Weigert, Malgorzata Santel, Karen Alim, Barbara Treutlein, and J Gray Camp. Spatial transcriptomic and single-nucleus analysis reveals heterogeneity in a gigantic single-celled syncytium. *eLife*, 11, 2022.

[10871] Alexandre A Germanos, Sonali Arora, Ye Zheng, Erica T Goddard, Ilsa M Coleman, Anson T Ku, Scott Wilkinson, Hanbing Song, Nicholas J Brady, Robert A Amezquita, Michael Zager, Annalysa Long, Yu Chi Yang, Jason H Bielas, Raphael Gottardo, David S Rickman, Franklin W Huang, Cyrus M Ghajar, Peter S Nelson, Adam G Sowalsky, Manu Setty, and Andrew C Hsieh. Defining cellular population dynamics at single-cell resolution during prostate cancer progression. *eLife*, 11, 2022.

[10872] Richard Gerum, Elham Mirzahossein, Mar Eroles, Jennifer Elsterer, Astrid Mainka, Andreas Bauer, Selina Sonntag, Alexander Winterl, Johannes Bartl, Lena Fischer, Shada Abuhattum, Ruchi Goswami, Salvatore Girardo, Jochen Guck, Stefan Schrüfer, Nadine Ströhlein, Mojtaba Nosratlo, Harald Herrmann, Dorothea Schultheis, Felix Rico, Sebastian Johannes Müller, Stephan Gekle, and Ben Fabry. Viscoelastic properties of suspended cells measured with shear flow deformation cytometry. *eLife*, 11, 2022.

[10873] Hussein Ghareh, Isis Alonso-Lozares, Dustin Schetters, Rae J Herman, Tim S Heistek, Yvar Van Mourik, Philip Jean-Richard-dit Bressel, Gerald Zernig, Huibert D Mansvelder, Taco J De Vries, and Nathan J Marchant. Role of anterior insula cortex in context-induced relapse of nicotine-seeking. *eLife*, 11, 2022.

[10874] James M Gibson, Heying Cui, M Yusuf Ali, Xiaoxin Zhao, Erik W Debler, Jing Zhao, Kathleen M Trybus, Sozanne R Solmaz, and Chunyu Wang. Coil-to-$\alpha$-helix transition at the Nup358-BicD2 interface activates BicD2 for dynein recruitment. *eLife*, 11, 2022.

[10875] Allison M Gicking, Tzu-Chen Ma, Qingzhou Feng, Rui Jiang, Somayesadat Badieyan, Michael A Cianfrocco, and William O Hancock. Kinesin-1, -2, and -3 motors use family-specific mechanochemical strategies to effectively compete with dynein during bidirectional transport. *eLife*, 11, 2022.

[10876] Marc Alwin Gieselmann and Alexander Thiele. Stimulus dependence of directed information exchange between cortical layers in macaque V1. *eLife*, 11, 2022.

[10877] James J Gilchrist, Silvia N Kariuki, James A Watson, Gavin Band, Sophie Uyoga, Carolyne M Ndila, Neema Mturi, Salim Mwarumba, Shebe Mohammed, Moses Mosobo, Kaur Alasoo, Kirk A Rockett, Alexander J Mentzer, Dominic P Kwiatkowski, Adrian VS Hill, Kathryn Maitland, J Anthony G Scott, and Thomas N Williams. BIRC6 modifies risk of invasive bacterial infection in Kenyan children. *eLife*, 11, 2022.

[10878] John P Gillies, Janice M Reimer, Eva P Karasmanis, Indrajit Lahiri, Zaw Min Htet, Andres E Leschziner, and Samara L Reck-Peterson. Structural basis for cytoplasmic dynein-1 regulation by Lis1. *eLife*, 11, 2022.

[10879] Robert M Gingras, Abigail M Sulpizio, Joelle Park, and Anthony Bretscher. High-resolution secretory timeline from vesicle formation at the Golgi to fusion at the plasma membrane in S. cerevisiae. *eLife*, 11, 2022.

[10880] Sai Srinivas Panapakkam Giridharan, Guangming Luo, Pilar Rivero-Rios, Noah Steinfeld, Helene Tronchere, Amika Singla, Ezra Burstein, Daniel D Billadeau, Michael A Sutton, and Lois S Weisman. Lipid kinases VPS34 and PIKfyve coordinate a phosphoinositide cascade to regulate retriever-mediated recycling on endosomes. *eLife*, 11, 2022.

[10881] Stefano G Giulieri, Romain Guérillot, Sebastian Duchene, Abderrahman Hachani, Diane Daniel, Torsten Seemann, Joshua S Davis, Steven YC Tong, Bernadette C Young, Daniel J Wilson, Timothy P Stinear, and Benjamin P Howden. Niche-specific genome degradation and convergent evolution shaping Staphylococcus aureus adaptation during severe infections. *eLife*, 11, 2022.

[10882] Aleksander Giwercman, K Barbara Sahlin, Indira Pla Parada, Krzysztof Pawlowski, Carl Fehninger, Yvonne Lundberg Giwercman, Irene Leijonhufvud, Roger Appelqvist, György Marko-Varga, Aniel Sanchez, and Johan Malm. Novel protein markers of androgen activity in humans: proteomic study of plasma from young chemically castrated men. *eLife*, 11, 2022.

[10883] Joan Gizzio, Abhishek Thakur, Allan Haldane, and Ronald M Levy. Evolutionary divergence in the conformational landscapes of tyrosine vs serine/threonine kinases. *eLife*, 11, 2022.

[10884] Evripidis Gkanias, Li Yan McCurdy, Michael N Nitabach, and Barbara Webb. An incentive circuit for memory dynamics in the mushroom body of Drosophila melanogaster. *eLife*, 11, 2022.

[10885] Juliane Glaser, Julian Iranzo, Maud Borensztein, Mattia Marinucci, Angelica Gualtieri, Colin Jouhanneau, Aurélie Teissandier, Carles Gaston-Massuet, and Deborah Bourc'his. The imprinted Zdbf2 gene finely tunes control of feeding and growth in neonates. *eLife*, 11, 2022.

[10886] Giacomo L Glotzer, Pietro Tardivo, and Elly M Tanaka. Canonical Wnt signaling and the regulation of divergent mesenchymal Fgf8 expression in axolotl limb development and regeneration. *eLife*, 11, 2022.

[10887] Amélie Luise Godeau, Marco Leoni, Jordi Comelles, Tristan Guyomar, Michele Lieb, Hélène Delanoë-Ayari, Albrecht Ott, Sebastien Harlepp, Pierre Sens, and Daniel Riveline. 3D single cell migration driven by temporal correlation between oscillating force dipoles. *eLife*, 11, 2022.

[10888] Anne-Claire Godet, Emilie Roussel, Florian David, Fransky Hantelys, Florent Morfoisse, Joffrey Alves, FranÇoise Pujol, Isabelle Ader, Edouard Bertrand, Odile Burlet-Schiltz, Carine Froment, Anthony K Henras, Patrice Vitali, Eric Lacazette, Florence Tatin, Barbara Garmy-Susini, and Anne-Catherine Prats. Long non-coding RNA Neat1 and paraspeckle components are translational regulators in hypoxia. *eLife*, 11, 2022.

[10889] Erich J Goebel, Luisina Ongaro, Emily C Kappes, Kylie Vestal, Elitza Belcheva, Roselyne Castonguay, Ravindra Kumar, Daniel J Bernard, and Thomas B Thompson. The orphan ligand, activin C, signals through activin receptor-like kinase 7. *eLife*, 11, 2022.

[10890] Jessica N Goedhoop, Bastijn JG van den Boom, Rhiannon Robke, Felice Veen, Lizz Fellinger, Wouter van Elzelingen, Tara Arbab, and Ingo Willuhn. Nucleus accumbens dopamine tracks aversive stimulus duration and prediction but not value or prediction error. *eLife*, 11, 2022.

[10891] Antigoni Gogolou, Celine Souilhol, Ilaria Granata, Filip J Wymeersch, Ichcha Manipur, Matthew Wind, Thomas JR Frith, Maria Guarini, Alessandro Bertero, Christoph Bock, Florian Halbritter, Minoru Takasato, Mario R Guarracino, and Anestis Tsakiridis. Early anteroposterior regionalisation of human neural crest is shaped by a pro-mesodermal factor. *eLife*, 11, 2022.

[10892] Jeff Goldsmith, Tomoko Kitago, Angel Garcia de la Garza, Robinson Kundert, Andreas Luft, Cathy Stinear, Winston D Byblow, Gert Kwakkel, and John W Krakauer. Arguments for the biological and predictive relevance of the proportional recovery rule. *eLife*, 11, 2022.

[10893] Allison T Goldstein, Terrence R Stanford, and Emilio Salinas. Exogenous capture accounts for fundamental differences between pro- and antisaccade performance. *eLife*, 11, 2022.

[10894] Gustavo A Gomez, Patrick Aghajanian, Sheila Pourteymoor, Destiney Larkin, and Subburaman Mohan. Differences in pathways contributing to thyroid hormone effects on postnatal cartilage calcification versus secondary ossification center development. *eLife*, 11, 2022.

[10895] Gustavo A Gomez, Charles H Rundle, Weirong Xing, Chandrasekhar Kesavan, Sheila Pourteymoor, Robert E Lewis, David R Powell, and Subburaman Mohan. Contrasting effects of Ksr2, an obesity gene, on trabecular bone volume and bone marrow adiposity. *eLife*, 11, 2022.

[10896] Bronner P GonÇalves, Matthew Hall, Waasila Jassat, Valeria Balan, Srinivas Murthy, Christiana Kartsonaki, Malcolm G Semple, Amanda Rojek, Joaquín Baruch, Luis Felipe Reyes, Abhishek Dasgupta, Jake Dunning, Barbara Wanjiru Citarella, Mark Pritchard, Alejandro Martín-Quiros, Uluhan Sili, J Kenneth Baillie, Diptesh Aryal, Yaseen Arabi, Aasiyah Rashan, Andrea Angheben, Janice Caoili, FranÇois Martin Carrier, Ewen M Harrison, Joan Gómez-Junyent, Claudia Figueiredo-Mello, James Joshua Douglas, Mohd Basri Mat Nor, Yock Ping Chow, Xin Ci Wong, Silvia Bertagnolio, Soe Soe Thwin, Anca Streinu-Cercel, Leonardo Salazar, Asgar Rishu, Rajavardhan Rangappa, David SY Ong, Madiha Hashmi, Gail Carson, Janet Diaz, Rob Fowler, Moritz UG Kraemer, Evert-Jan Wils, Peter Horby, Laura Merson, Piero L Olliaro, and Sheryl Ann Abdukahil. An international observational study to assess the impact of the Omicron variant emergence on the clinical epidemiology of COVID-19 in hospitalised patients. *eLife*, 11, 2022.

[10897] Marina GonÇalves Antunes, Matthieu Sanial, Vincent Contremoulins, Sandra Carvalho, Anne Plessis, and Isabelle Becam. High hedgehog signaling is transduced by a multikinase-dependent switch controlling the apico-basal distribution of the GPCR smoothened. *eLife*, 11, 2022.

[10898] Mikhail M Goncharov, Ekaterina A Bryushkova, Nikita I Sharaev, Valeria D Skatova, Anastasiya M Baryshnikova, George V Sharonov, Vadim Karnaukhov, Maria T Vakhitova, Igor V Samoylenko, Lev V Demidov, Sergey Lukyanov, Dmitriy M Chudakov, and Ekaterina O Serebrovskaya. Pinpointing the tumor-specific T cells via TCR clusters. *eLife*, 11, 2022.

[10899] Mónika Gönczi, Zsolt Ráduly, László Szabó, János Fodor, Andrea Telek, Nóra Dobrosi, Norbert Balogh, Péter Szentesi, Gréta Kis, Miklós Antal, György Trencsenyi, Beatrix Dienes, and László Csernoch. Septin7 is indispensable for proper skeletal muscle architecture and function. *eLife*, 11, 2022.

[10900] Louisa Gonzalez Somermeyer, Aubin Fleiss, Alexander S Mishin, Nina G Bozhanova, Anna A Igolkina, Jens Meiler, Maria-Elisenda Alaball Pujol, Ekaterina V Putintseva, Karen S Sarkisyan, and Fyodor A Kondrashov. Heterogeneity of the GFP fitness landscape and data-driven protein design. *eLife*, 11, 2022.

[10901] Kerry Marie Goodman, Phinikoula S Katsamba, Rotem Rubinstein, Göran Ahlsén, Fabiana Bahna, Seetha Mannepalli, Hanbin Dan, Rosemary V Sampogna, Lawrence Shapiro, and Barry Honig. How clustered protocadherin binding specificity is tuned for neuronal self-/nonself-recognition. *eLife*, 11, 2022.

[10902] Vishhvaan Gopalakrishnan, Dena Crozier, Kyle J Card, Lacy D Chick, Nikhil P Krishnan, Erin McClure, Julia Pelesko, Drew FK Williamson, Daniel Nichol, Soumyajit Mandal, Robert A Bonomo, and Jacob G Scott. A low-cost, open-source evolutionary bioreactor and its educational use. *eLife*, 11, 2022.

[10903] Srinivas Gorur-Shandilya, Elizabeth M Cronin, Anna C Schneider, Sara Ann Haddad, Philipp Rosenbaum, Dirk Bucher, Farzan Nadim, and Eve Marder. Mapping circuit dynamics during function and dysfunction. *eLife*, 11, 2022.

[10904] Maxime RF Gosselin, Virginie Mournetas, Malgorzata Borczyk, Suraj Verma, Annalisa Occhipinti, Justyna Róg, Lukasz Bozycki, Michal Korostynski, Samuel C Robson, Claudio Angione, Christian Pinset, and Dariusz C Gorecki. Loss of full-length dystrophin expression results in major cell-autonomous abnormalities in proliferating myoblasts. *eLife*, 11, 2022.

[10905] Hemant N Goswami, Jay Rai, Anuska Das, and Hong Li. Molecular mechanism of active Cas7-11 in processing CRISPR RNA and interfering target RNA. *eLife*, 11, 2022.

[10906] Erwan Goy, Maxime Tomezak, Caterina Facchin, Nathalie Martin, Emmanuel Bouchaert, Jerome Benoit, Clementine de Schutter, Joe Nassour, Laure Saas, Claire Drullion, Priscille M Brodin, Alexandre Vandeputte, Olivier Molendi-Coste, Laurent Pineau, Gautier Goormachtigh, Olivier Pluquet, Albin Pourtier, Fabrizio Cleri, Eric Lartigau, Nicolas Penel, and Corinne Abbadie. The out-of-field dose in radiation therapy induces delayed tumorigenesis by senescence evasion. *eLife*, 11, 2022.

[10907] Akshit Goyal, Leonora S Bittleston, Gabriel E Leventhal, Lu Lu, and Otto X Cordero. Interactions between strains govern the eco-evolutionary dynamics of microbial communities. *eLife*, 11, 2022.

[10908] Jürgen Graf, Vahid Rahmati, Myrtill Majoros, Otto W Witte, Christian Geis, Stefan J Kiebel, Knut Holthoff, and Knut Kirmse. Network instability dynamics drive a transient bursting period in the developing hippocampus in vivo. *eLife*, 11, 2022.

[10909] Thomas GW Graham, John Joseph Ferrie, Gina M Dailey, Robert Tjian, and Xavier Darzacq. Detecting molecular interactions in live-cell single-molecule imaging with proximity-assisted photoactivation (PAPA). *eLife*, 11, 2022.

[10910] Xenia Grande, Magdalena M Sauvage, Andreas Becke, Emrah Düzel, and David Berron. Transversal functional connectivity and scene-specific processing in the human entorhinal-hippocampal circuitry. *eLife*, 11, 2022.

[10911] Nicholas M Grebe, Jean Paul Hirwa, Tara S Stoinski, Linda Vigilant, and Stacy Rosenbaum. Mountain gorillas maintain strong affiliative biases for maternal siblings despite high male reproductive skew and extensive exposure to paternal kin. *eLife*, 11, 2022.

[10912] Leah J Greenspan, Margaret de Cuevas, Kathy H Le, Jennifer M Viveiros, and Erika L Matunis. Activation of the EGFR/MAPK pathway drives transdifferentiation of quiescent niche cells to stem cells in the Drosophila testis niche. *eLife*, 11, 2022.

[10913] Alexandre Grimaldi, Glenda Comai, Sebastien Mella, and Shahragim Tajbakhsh. Identification of bipotent progenitors that give rise to myogenic and connective tissues in mouse. *eLife*, 11, 2022.

[10914] David Robert Grimes. The ellipse of insignificance, a refined fragility index for ascertaining robustness of results in dichotomous outcome trials. *eLife*, 11, 2022.

[10915] Julien Gronnier, Christina M Franck, Martin Stegmann, Thomas A DeFalco, Alicia Abarca, Michelle von Arx, Kai Dünser, Wenwei Lin, Zhenbiao Yang, Jürgen Kleine-Vehn, Christoph Ringli, and Cyril Zipfel. Regulation of immune receptor kinase plasma membrane nanoscale organization by a plant peptide hormone and its receptors. *eLife*, 11, 2022.

[10916] Ruth Großeholz, Friederike Wanke, Leander Rohr, Nina Glöckner, Luiselotte Rausch, Stefan Scholl, Emanuele Scacchi, Amelie-Jette Spazierer, Lana Shabala, Sergey Shabala, Karin Schumacher, Ursula Kummer, and Klaus Harter. Computational modeling and quantitative physiology reveal central parameters for brassinosteroid-regulated early cell physiological processes linked to elongation growth of the Arabidopsis root. *eLife*, 11, 2022.

[10917] Bon-Mi Gu and Joshua D Berke. Altered basal ganglia output during self-restraint. *eLife*, 11, 2022.

[10918] Yihui Gu, Zhichao Wang, Chengjiang Wei, Yuehua Li, Wei Feng, Wei Wang, Meiqi Chang, Yu Chen, and Qingfeng Li. Photonic hyperthermia of malignant peripheral nerve sheath tumors at the third near-infrared biowindow. *eLife*, 11, 2022.

[10919] Charles Guan, Tyson Aflalo, Carey Y Zhang, Elena Amoruso, Emily R Rosario, Nader Pouratian, and Richard A Andersen. Stability of motor representations after paralysis. *eLife*, 11, 2022.

[10920] Matthias Guggenmos. Reverse engineering of metacognition. *eLife*, 11, 2022.

[10921] Tejas Guha, Elana J Fertig, and Atul Deshpande. Generating colorblind-friendly scatter plots for single-cell data. *eLife*, 11, 2022.

[10922] Anne Günther, Matthias Hose, Hanna Abberger, Fabian Schumacher, Ylva Veith, Burkhard Kleuser, Kai Matuschewski, Karl Sebastian Lang, Erich Gulbins, Jan Buer, Astrid M Westendorf, and Wiebke Hansen. The acid ceramidase/ceramide axis controls parasitemia in Plasmodium yoelii-infected mice by regulating erythropoiesis. *eLife*, 11, 2022.

[10923] Dongsheng Guo, Katelyn Daman, Jennifer JC Chen, Meng-Jiao Shi, Jing Yan, Zdenka Matijasevic, Amanda M Rickard, Monica H Bennett, Alex Kiselyov, Haowen Zhou, Anne G Bang, Kathryn R Wagner, René Maehr, Oliver D King, Lawrence J Hayward, and Charles P Emerson. iMyoblasts for ex vivo and in vivo investigations of human myogenesis and disease modeling. *eLife*, 11, 2022.

[10924] Hanmin Guo, Lin Hou, Yu Shi, Sheng Chih Jin, Xue Zeng, Boyang Li, Richard P Lifton, Martina Brueckner, Hongyu Zhao, and Qiongshi Lu. Quantifying concordant genetic effects of de novo mutations on multiple disorders. *eLife*, 11, 2022.

[10925] Heyun Guo, Ericca L Stamper, Aya Sato-Carlton, Masa A Shimazoe, Xuan Li, Liangyu Zhang, Lewis Stevens, KC Jacky Tam, Abby F Dernburg, and Peter M Carlton. Phosphoregulation of DSB-1 mediates control of meiotic double-strand break activity. *eLife*, 11, 2022.

[10926] Liang Guo, Lufeng Zhao, Junlong Ye, Zijun Ji, Jian-Jun Tang, Keyu Bai, Sijun Zheng, Liangliang Hu, and Xin Chen. Using aquatic animals as partners to increase yield and maintain soil nitrogen in the paddy ecosystems. *eLife*, 11, 2022.

[10927] Nannan Guo, Kelsey D McDermott, Yu-Tzu Shih, Haley Zanga, Debolina Ghosh, Charlotte Herber, William R Meara, James Coleman, Alexia Zagouras, Lai Ping Wong, Ruslan Sadreyev, J Tiago GonÇalves, and Amar Sahay. Transcriptional regulation of neural stem cell expansion in the adult hippocampus. *eLife*, 11, 2022.

[10928] Tingwei Guo, Xia Han, Jinzhi He, Jifan Feng, Junjun Jing, Eva Janečková, Jie Lei, Thach-Vu Ho, Jian Xu, and Yang Chai. KDM6B interacts with TFDP1 to activate P53 signaling in regulating mouse palatogenesis. *eLife*, 11, 2022.

[10929] Animesh Gupta, Luis Zaman, Hannah M Strobel, Jenna Gallie, Alita R Burmeister, Benjamin Kerr, Einat S Tamar, Roy Kishony, and Justin R Meyer. Host-parasite coevolution promotes innovation through deformations in fitness landscapes. *eLife*, 11, 2022.

[10930] Dinesh Gupta, Katie E Shalvarjian, and Dipti D Nayak. An Archaea-specific c-type cytochrome maturation machinery is crucial for methanogenesis in Methanosarcina acetivorans. *eLife*, 11, 2022.

[10931] Kuldeepkumar R Gupta, Celena M Gwin, Kathryn C Rahlwes, Kyle J Biegas, Chunyan Wang, Jin Ho Park, Jun Liu, Benjamin M Swarts, Yasu S Morita, and E Hesper Rego. An essential periplasmic protein coordinates lipid trafficking and is required for asymmetric polar growth in mycobacteria. *eLife*, 11, 2022.

[10932] Nehal Gupta, Hanbing Song, Wei Wu, Rovingaile K Ponce, Yone K Lin, Ji Won Kim, Eric J Small, Felix Y Feng, Franklin W Huang, and Ross A Okimoto. The CIC-ERF co-deletion underlies fusion-independent activation of ETS family member, ETV1, to drive prostate cancer progression. *eLife*, 11, 2022.

[10933] J Ignacio Gutierrez, Gregory P Brittingham, Yonca Karadeniz, Kathleen D Tran, Arnob Dutta, Alex S Holehouse, Craig L Peterson, and Liam J Holt. SWI/SNF senses carbon starvation with a pH-sensitive low-complexity sequence. *eLife*, 11, 2022.

[10934] Amit Gutwillig, Nadine Santana-Magal, Leen Farhat-Younis, Diana Rasoulouniriana, Asaf Madi, Chen Luxenburg, Jonathan Cohen, Krishnanand Padmanabhan, Noam Shomron, Guy Shapira, Annette Gleiberman, Roma Parikh, Carmit Levy, Meora Feinmesser, Dov Hershkovitz, Valentina Zemser-Werner, Oran Zlotnik, Sanne Kroon, Wolf-Dietrich Hardt, Reno Debets, Nathan Edward Reticker-Flynn, Peleg Rider, and Yaron Carmi. Transient cell-in-cell formation underlies tumor relapse and resistance to immunotherapy. *eLife*, 11, 2022.

[10935] Charlotte Guyomar, Clément Bousquet, Siou Ku, John M Heumann, Gabriel Guilloux, Natacha Gaillard, Claire Heichette, Laurence Duchesne, Michel O Steinmetz, Romain Gibeaux, and Denis Chrétien. Changes in seam number and location induce holes within microtubules assembled from porcine brain tubulin and in Xenopus egg cytoplasmic extracts. *eLife*, 11, 2022.

[10936] Wilmer Cristobal Guzman-Vilca, Manuel Castillo-Cara, and Rodrigo M Carrillo-Larco. Development, validation, and application of a machine learning model to estimate salt consumption in 54 countries. *eLife*, 11, 2022.

[10937] Julian Haase, Richard Chen, Wesley M Parker, Mary Kate Bonner, Lisa M Jenkins, and Alexander E Kelly. The TFIIH complex is required to establish and maintain mitotic chromosome structure. *eLife*, 11, 2022.

[10938] Noushin Hadadi, Martina Spiljar, Karin Steinbach, Melis Çolakoğlu, Claire Chevalier, Gabriela Salinas, Doron Merkler, and Mirko Trajkovski. Comparative multi-tissue profiling reveals extensive tissue-specificity in transcriptome reprogramming during thermal adaptation. *eLife*, 11, 2022.

[10939] Travis A Hage, Alice Bosma-Moody, Christopher A Baker, Megan B Kratz, Luke Campagnola, Tim Jarsky, Hongkui Zeng, and Gabe J Murphy. Synaptic connectivity to L2/3 of primary visual cortex measured by two-photon optogenetic stimulation. *eLife*, 11, 2022.

[10940] David L Haggerty, Braulio Munoz, Taylor Pennington, Gonzalo Viana Di Prisco, Gregory G Grecco, and Brady K Atwood. The role of anterior insular cortex inputs to dorsolateral striatum in binge alcohol drinking. *eLife*, 11, 2022.

[10941] Gerald Hahn, Arvind Kumar, Helmut Schmidt, Thomas R Knösche, and Gustavo Deco. Rate and oscillatory switching dynamics of a multilayer visual microcircuit model. *eLife*, 11, 2022.

[10942] Michael A Hahn, Kathrin Bothe, Dominik Heib, Manuel Schabus, Randolph F Helfrich, and Kerstin Hoedlmoser. Slow oscillation–spindle coupling strength predicts real-life gross-motor learning in adolescents and adults. *eLife*, 11, 2022.

[10943] Shojiro Haji, Taiki Ito, Carla Guenther, Miyako Nakano, Takashi Shimizu, Daiki Mori, Yasunori Chiba, Masato Tanaka, Sushil K Mishra, Janet A Willment, Gordon D Brown, Masamichi Nagae, and Sho Yamasaki. Human Dectin-1 is O-glycosylated and serves as a ligand for C-type lectin receptor CLEC-2. *eLife*, 11, 2022.

[10944] David B Halling, Ashley E Philpo, and Richard W Aldrich. Calcium dependence of both lobes of calmodulin is involved in binding to a cytoplasmic domain of SK channels. *eLife*, 11, 2022.

[10945] Hunter E Halverson, Jinsook Kim, Andrei Khilkevich, Michael D Mauk, and George J Augustine. Feedback inhibition underlies new computational functions of cerebellar interneurons. *eLife*, 11, 2022.

[10946] Tuomas Hämälä, Weixuan Ning, Helmi Kuittinen, Nader Aryamanesh, and Outi Savolainen. Environmental response in gene expression and DNA methylation reveals factors influencing the adaptive potential of Arabidopsis lyrata. *eLife*, 11, 2022.

[10947] George L Hamilton, Nabanita Saikia, Sujit Basak, Franceine S Welcome, Fang Wu, Jakub Kubiak, Changcheng Zhang, Yan Hao, Claus AM Seidel, Feng Ding, Hugo Sanabria, and Mark E Bowen. Fuzzy supertertiary interactions within PSD-95 enable ligand binding. *eLife*, 11, 2022.

[10948] Bin Han, Qiaohong Wei, Esmaeil Amiri, Han Hu, Lifeng Meng, Micheline K Strand, David R Tarpy, Shufa Xu, Jianke Li, and Olav Rueppell. The molecular basis of socially induced egg-size plasticity in honey bees. *eLife*, 11, 2022.

[10949] Bing Han, Xing Li, Ruo-Song Ai, Si-Ying Deng, Ze-Qing Ye, Xin Deng, Wen Ma, Shun Xiao, Jing-Zhi Wang, Li-Mei Wang, Chong Xie, Yan Zhang, Yan Xu, and Yuan Zhang. Atmospheric particulate matter aggravates CNS demyelination through involvement of TLR-4/NF-kB signaling and microglial activation. *eLife*, 11, 2022.

[10950] Shuo Han, Sophia Peng, Joshua Vance, Kimberly Tran, Nhu Do, Nauy Bui, Zhenhua Gui, and Shizhen Wang. Structural dynamics determine voltage and pH gating in human voltage-gated proton channel. *eLife*, 11, 2022.

[10951] Yifu Han, Chun Chien, Pragya Goel, Kaikai He, Cristian Pinales, Christopher Buser, and Dion Dickman. Botulinum neurotoxin accurately separates tonic vs. phasic transmission and reveals heterosynaptic plasticity rules in Drosophila. *eLife*, 11, 2022.

[10952] Yongchao Han, Lei Peng, and Tao Wang. Tadr is an axonal histidine transporter required for visual neurotransmission in Drosophila. *eLife*, 11, 2022.

[10953] Jeffrey L Hansen, Kaiser J Loell, and Barak A Cohen. A test of the pioneer factor hypothesis using ectopic liver gene activation. *eLife*, 11, 2022.

[10954] Sarah R Hansen, David S White, Mark Scalf, Ivan R Corrêa, Lloyd M Smith, and Aaron A Hoskins. Multi-step recognition of potential 5′ splice sites by the Saccharomyces cerevisiae U1 snRNP. *eLife*, 11, 2022.

[10955] Scott D Hansen, Albert A Lee, Benjamin R Duewell, and Jay T Groves. Membrane-mediated dimerization potentiates PIP5K lipid kinase activity. *eLife*, 11, 2022.

[10956] Lana Hantzsch, Benjamin Parrell, and Caroline A Niziolek. A single exposure to altered auditory feedback causes observable sensorimotor adaptation in speech. *eLife*, 11, 2022.

[10957] Kawssar Harb, Melanie Richter, Nagammal Neelagandan, Elia Magrinelli, Hend Harfoush, Katrin Kuechler, Melad Henis, Irm Hermanns-Borgmeyer, Froylan Calderon de Anda, and Kent Duncan. Pum2 and TDP-43 refine area-specific cytoarchitecture post-mitotically and modulate translation of Sox5, Bcl11b, and Rorb mRNAs in developing mouse neocortex. *eLife*, 11, 2022.

[10958] Richard Hardstone, Matthew W Flounders, Michael Zhu, and Biyu J He. Frequency-specific neural signatures of perceptual content and perceptual stability. *eLife*, 11, 2022.

[10959] Kishore Hari, Varun Ullanat, Archana Balasubramanian, Aditi Gopalan, and Mohit Kumar Jolly. Landscape of epithelial–mesenchymal plasticity as an emergent property of coordinated teams in regulatory networks. *eLife*, 11, 2022.

[10960] Diane M Harper, Melissa Plegue, Masahito Jimbo, Sherri Sheinfeld Gorin, and Ananda Sen. US women screen at low rates for both cervical and colorectal cancers than a single cancer: a cross-sectional population-based observational study. *eLife*, 11, 2022.

[10961] William S Hart, Sam Abbott, Akira Endo, Joel Hellewell, Elizabeth Miller, Nick Andrews, Philip K Maini, Sebastian Funk, and Robin N Thompson. Inference of the SARS-CoV-2 generation time using UK household data. *eLife*, 11, 2022.

[10962] Ali Hashmi, Sham Tlili, Pierre Perrin, Molly Lowndes, Hanna Peradziryi, Joshua M Brickman, Alfonso Martínez Arias, and Pierre-FranÇois Lenne. Cell-state transitions and collective cell movement generate an endoderm-like region in gastruloids. *eLife*, 11, 2022.

[10963] Joshua Hatterschide, Paola Castagnino, Hee Won Kim, Steven M Sperry, Kathleen T Montone, Devraj Basu, and Elizabeth A White. YAP1 activation by human papillomavirus E7 promotes basal cell identity in squamous epithelia. *eLife*, 11, 2022.

[10964] Alexander S Hauser, Charlotte Avet, Claire Normand, Arturo Mancini, Asuka Inoue, Michel Bouvier, and David E Gloriam. Common coupling map advances GPCR-G protein selectivity. *eLife*, 11, 2022.

[10965] David W Hawman, Kimberly Meade-White, Jacob Archer, Shanna S Leventhal, Drew Wilson, Carl Shaia, Samantha Randall, Amit P Khandhar, Kyle Krieger, Tien-Ying Hsiang, Michael Gale, Peter Berglund, Deborah Heydenburg Fuller, Heinz Feldmann, and Jesse H Erasmus. SARS-CoV2 variant-specific replicating RNA vaccines protect from disease following challenge with heterologous variants of concern. *eLife*, 11, 2022.

[10966] Iain M Hay, Katie E Mulholland, Tiffany Lai, Stephen C Graham, Hayley J Sharpe, and Janet E Deane. Molecular mechanism of Afadin substrate recruitment to the receptor phosphatase PTPRK via its pseudophosphatase domain. *eLife*, 11, 2022.

[10967] James A Hay, Stephen M Kissler, Joseph R Fauver, Christina Mack, Caroline G Tai, Radhika M Samant, Sarah Connolly, Deverick J Anderson, Gaurav Khullar, Matthew MacKay, Miral Patel, Shannan Kelly, April Manhertz, Isaac Eiter, Daisy Salgado, Tim Baker, Ben Howard, Joel T Dudley, Christopher E Mason, Manoj Nair, Yaoxing Huang, John DiFiori, David D Ho, Nathan D Grubaugh, and Yonatan H Grad. Quantifying the impact of immune history and variant on SARS-CoV-2 viral kinetics and infection rebound: A retrospective cohort study. *eLife*, 11, 2022.

[10968] Masaaki Hayashi, Kohei Okuyama, Nobuaki Mizuguchi, Ryotaro Hirose, Taisuke Okamoto, Michiyuki Kawakami, and Junichi Ushiba. Spatially bivariate EEG-neurofeedback can manipulate interhemispheric inhibition. *eLife*, 11, 2022.

[10969] Elizabeth M Haynes, Korri H Burnett, Jiaye He, Marcel W Jean-Pierre, Martin Jarzyna, Kevin W Eliceiri, Jan Huisken, and Mary C Halloran. KLC4 shapes axon arbors during development and mediates adult behavior. *eLife*, 11, 2022.

[10970] Laura Katharine Hayward and Guy Sella. Polygenic adaptation after a sudden change in environment. *eLife*, 11, 2022.

[10971] Feng He, Chan Wu, and Allan Jacobson. Dcp2 C-terminal cis-binding elements control selective targeting of the decapping enzyme by forming distinct decapping complexes. *eLife*, 11, 2022.

[10972] Zan He, Zijuan Xin, Qiong Yang, Chen Wang, Meng Li, Wei Rao, Zhimin Du, Jia Bai, Zixuan Guo, Xiuyan Ruan, Zhaojun Zhang, Xiangdong Fang, and Hua Zhao. Mapping the single-cell landscape of acral melanoma and analysis of the molecular regulatory network of the tumor microenvironments. *eLife*, 11, 2022.

[10973] Anne Hebert, Annet Simons, Janneke HM Schuurs-Hoeijmakers, Hans JPM Koenen, Evelien Zonneveld-Huijssoon, Stefanie SV Henriet, Ellen JH Schatorjé, Esther PAH Hoppenreijs, Erika KSM Leenders, Etienne JM Janssen, Gijs WE Santen, Sonja A de Munnik, Simon V van Reijmersdal, Esther van Rijssen, Simone Kersten, Mihai G Netea, Ruben L Smeets, Frank L van de Veerdonk, Alexander Hoischen, and Caspar I van der Made. Trio-based whole exome sequencing in patients with suspected sporadic inborn errors of immunity: A retrospective cohort study. *eLife*, 11, 2022.

[10974] Alec Heckert, Liza Dahal, Robert Tjian, and Xavier Darzacq. Recovering mixtures of fast-diffusing states from short single-particle trajectories. *eLife*, 11, 2022.

[10975] Emily L Heckman and Chris Q Doe. Presynaptic contact and activity opposingly regulate postsynaptic dendrite outgrowth. *eLife*, 11, 2022.

[10976] Ann Kathrin Heilig, Ryohei Nakamura, Atsuko Shimada, Yuka Hashimoto, Yuta Nakamura, Joachim Wittbrodt, Hiroyuki Takeda, and Toru Kawanishi. Wnt11 acts on dermomyotome cells to guide epaxial myotome morphogenesis. *eLife*, 11, 2022.

[10977] James P Held, Gaomin Feng, Benjamin R Saunders, Claudia V Pereira, Kristopher Burkewitz, and Maulik R Patel. A tRNA processing enzyme is a key regulator of the mitochondrial unfolded protein response. *eLife*, 11, 2022.

[10978] Robert N Helsley, Tatsunori Miyata, Anagha Kadam, Venkateshwari Varadharajan, Naseer Sangwan, Emily C Huang, Rakhee Banerjee, Amanda L Brown, Kevin K Fung, William J Massey, Chase Neumann, Danny Orabi, Lucas J Osborn, Rebecca C Schugar, Megan R McMullen, Annette Bellar, Kyle L Poulsen, Adam Kim, Vai Pathak, Marko Mrdjen, James T Anderson, Belinda Willard, Craig J McClain, Mack Mitchell, Arthur J McCullough, Svetlana Radaeva, Bruce Barton, Gyongyi Szabo, Srinivasan Dasarathy, Jose Carlos Garcia-Garcia, Daniel M Rotroff, Daniela S Allende, Zeneng Wang, Stanley L Hazen, Laura E Nagy, and Jonathan Mark Brown. Gut microbial trimethylamine is elevated in alcohol-associated hepatitis and contributes to ethanol-induced liver injury in mice. *eLife*, 11, 2022.

[10979] Margaret M Henderson, Rosanne L Rademaker, and John T Serences. Flexible utilization of spatial- and motor-based codes for the storage of visuo-spatial information. *eLife*, 11, 2022.

[10980] Ardalan Hendi, Long-Gang Niu, Andrew William Snow, Richard Ikegami, Zhao-Wen Wang, and Kota Mizumoto. Channel-independent function of UNC-9/Innexin in spatial arrangement of GABAergic synapses in C. elegans. *eLife*, 11, 2022.

[10981] Marlene Henríquez-Urrutia, Rebecca Spanner, Consuelo Olivares-Yánez, Aldo Seguel-Avello, Rodrigo Pérez-Lara, Hector Guillén-Alonso, Robert Winkler, Alfredo Herrera-Estrella, Paulo Canessa, and Luis F Larrondo. Circadian oscillations in Trichoderma atroviride and the role of core clock components in secondary metabolism, development, and mycoparasitism against the phytopathogen Botrytis cinerea. *eLife*, 11, 2022.

[10982] Dongeun Heo, Jonathan P Ling, Gian C Molina-Castro, Abraham J Langseth, Ari Waisman, Klaus-Armin Nave, Wiebke Möbius, Phil C Wong, and Dwight E Bergles. Stage-specific control of oligodendrocyte survival and morphogenesis by TDP-43. *eLife*, 11, 2022.

[10983] Yunseok Heo, Eojin Yoon, Ye-Eun Jeon, Ji-Hye Yun, Naito Ishimoto, Hyeonuk Woo, Sam-Yong Park, Ji-Joon Song, and Weontae Lee. Cryo-EM structure of the human somatostatin receptor 2 complex with its agonist somatostatin delineates the ligand-binding specificity. *eLife*, 11, 2022.

[10984] Alice L Herneisen, Zhu-Hong Li, Alex W Chan, Silvia NJ Moreno, and Sebastian Lourido. Temporal and thermal profiling of the Toxoplasma proteome implicates parasite Protein Phosphatase 1 in the regulation of Ca2+-responsive pathways. *eLife*, 11, 2022.

[10985] Julia Takuno Hespanhol, Daniel Enrique Sanchez-Limache, Gianlucca GonÇalves Nicastro, Liam Mead, Edgar Enrique Llontop, Gustavo Chagas-Santos, Chuck Shaker Farah, Robson Francisco de Souza, Rodrigo da Silva Galhardo, Andrew L Lovering, and Ethel Bayer-Santos. Antibacterial T6SS effectors with a VRR-Nuc domain are structure-specific nucleases. *eLife*, 11, 2022.

[10986] Stephanie A Heusser, Christian B Borg, Janne M Colding, and Stephan A Pless. Conformational decoupling in acid-sensing ion channels uncovers mechanism and stoichiometry of PcTx1-mediated inhibition. *eLife*, 11, 2022.

[10987] Graham JM Hickey, Candice L Wike, Xichen Nie, Yixuan Guo, Mengyao Tan, Patrick J Murphy, and Bradley R Cairns. Establishment of developmental gene silencing by ordered polycomb complex recruitment in early zebrafish embryos. *eLife*, 11, 2022.

[10988] Frank Hidalgo, Laura M Nocka, Neel H Shah, Kent Gorday, Naomi R Latorraca, Pradeep Bandaru, Sage Templeton, David Lee, Deepti Karandur, Jeffrey G Pelton, Susan Marqusee, David Wemmer, and John Kuriyan. A saturation-mutagenesis analysis of the interplay between stability and activation in Ras. *eLife*, 11, 2022.

[10989] Michèle MG Hillege, Andi Shi, Ricardo A Galli, Gang Wu, Philippe Bertolino, Willem MH Hoogaars, and Richard T Jaspers. Lack of Tgfbr1 and Acvr1b synergistically stimulates myofibre hypertrophy and accelerates muscle regeneration. *eLife*, 11, 2022.

[10990] Bretta Hixson, Xiao-Li Bing, Xiaowei Yang, Alessandro Bonfini, Peter Nagy, and Nicolas Buchon. A transcriptomic atlas of Aedes aegypti reveals detailed functional organization of major body parts and gut regional specializations in sugar-fed and blood-fed adult females. *eLife*, 11, 2022.

[10991] Po-Yi Ho, Benjamin H Good, and Kerwyn Casey Huang. Competition for fluctuating resources reproduces statistics of species abundance over time across wide-ranging microbiotas. *eLife*, 11, 2022.

[10992] Christopher M Hoel, Lin Zhang, and Stephen G Brohawn. Structure of the GOLD-domain seven-transmembrane helix protein family member TMEM87A. *eLife*, 11, 2022.

[10993] Katelyn J Hoff, Jayne E Aiken, Mark A Gutierrez, Santos J Franco, and Jeffrey K Moore. TUBA1A tubulinopathy mutants disrupt neuron morphogenesis and override XMAP215/Stu2 regulation of microtubule dynamics. *eLife*, 11, 2022.

[10994] Henriette Hoffmann-Veltung, Nsoh Godwin Anabire, Michael Fokuo Ofori, Peter Janhmatz, Niklas Ahlborg, Lars Hviid, and Maria del Pilar Quintana. Analysis of allelic cross-reactivity of monoclonal IgG antibodies by a multiplexed reverse FluoroSpot assay. *eLife*, 11, 2022.

[10995] Danielle Holz, Aaron R Hall, Eiji Usukura, Sawako Yamashiro, Naoki Watanabe, and Dimitrios Vavylonis. A mechanism with severing near barbed ends and annealing explains structure and dynamics of dendritic actin networks. *eLife*, 11, 2022.

[10996] Matthias Hose, Anne Günther, Eyad Naser, Fabian Schumacher, Tina Schönberger, Julia Falkenstein, Athanasios Papadamakis, Burkhard Kleuser, Katrin Anne Becker, Erich Gulbins, Adriana Haimovitz-Friedman, Jan Buer, Astrid M Westendorf, and Wiebke Hansen. Cell-intrinsic ceramides determine T cell function during melanoma progression. *eLife*, 11, 2022.

[10997] David J Hosfield, Sandra Weber, Nan-Sheng Li, Madline Sauvage, Carstyn F Joiner, Govinda R Hancock, Emily A Sullivan, Estelle Ndukwe, Ross Han, Sydney Cush, Muriel Lainé, Sylvie C Mader, Geoffrey L Greene, and Sean W Fanning. Stereospecific lasofoxifene derivatives reveal the interplay between estrogen receptor alpha stability and antagonistic activity in ESR1 mutant breast cancer cells. *eLife*, 11, 2022.

[10998] Shoko Hososhima, Ritsu Mizutori, Rei Abe-Yoshizumi, Andrey Rozenberg, Shunta Shigemura, Alina Pushkarev, Masae Konno, Kota Katayama, Keiichi Inoue, Satoshi P Tsunoda, Oded Béjà, and Hideki Kandori. Proton-transporting heliorhodopsins from marine giant viruses. *eLife*, 11, 2022.

[10999] Morten Hostrup, Anders Krogh Lemminger, Ben Stocks, Alba Gonzalez-Franquesa, Jeppe Kjærgaard Larsen, Julia Prats Quesada, Martin Thomassen, Brian Tate Weinert, Jens Bangsbo, and Atul Shahaji Deshmukh. High-intensity interval training remodels the proteome and acetylome of human skeletal muscle. *eLife*, 11, 2022.

[11000] Thomas M Houslay, Ryan L Earley, Stephen J White, Wiebke Lammers, Andrew J Grimmer, Laura M Travers, Elizabeth L Johnson, Andrew J Young, and Alastair Wilson. Genetic integration of behavioural and endocrine components of the stress response. *eLife*, 11, 2022.

[11001] Sébastien Houy, Joana S Martins, Noa Lipstein, and Jakob Balslev Sørensen. Phorbolester-activated Munc13-1 and ubMunc13-2 exert opposing effects on dense-core vesicle secretion. *eLife*, 11, 2022.

[11002] Laurence J Howe, Ben Brumpton, Humaira Rasheed, Bjørn Olav Åsvold, George Davey Smith, and Neil M Davies. Taller height and risk of coronary heart disease and cancer: A within-sibship Mendelian randomization study. *eLife*, 11, 2022.

[11003] Mariah L Hoye, Lorenzo Calviello, Abigail J Poff, Nna-Emeka Ejimogu, Carly R Newman, Maya D Montgomery, Jianhong Ou, Stephen N Floor, and Debra L Silver. Aberrant cortical development is driven by impaired cell cycle and translational control in a DDX3X syndrome model. *eLife*, 11, 2022.

[11004] Chih-Wei Hsu, Juan Cerda, Jason M Kirk, Williamson D Turner, Tara L Rasmussen, Carlos P Flores Suarez, Mary E Dickinson, and Joshua D Wythe. EZ Clear for simple, rapid, and robust mouse whole organ clearing. *eLife*, 11, 2022.

[11005] Han Hu, Yan Wang, Paul G McDonald, Stephen Wroe, Jingmai K O'Connor, Alexander Bjarnason, Joseph J Bevitt, Xuwei Yin, Xiaoting Zheng, Zhonghe Zhou, and Roger BJ Benson. Earliest evidence for fruit consumption and potential seed dispersal by birds. *eLife*, 11, 2022.

[11006] Huan-Huan Hu, Guang-Ming Lu, Chia-Chun Chang, Yilan Li, Jiale Zhong, Chen-Jun Guo, Xian Zhou, Boqi Yin, Tianyi Zhang, and Ji-Long Liu. Filamentation modulates allosteric regulation of PRPS. *eLife*, 11, 2022.

[11007] Jinfeng Hu, Catherine A Forster, Xing Xu, Qi Zhao, Yiming He, and Fenglu Han. Computed tomographic analysis of the dental system of three Jurassic ceratopsians and implications for the evolution of tooth replacement pattern and diet in early-diverging ceratopsians. *eLife*, 11, 2022.

[11008] Lu Hu, Yang Sun, Shun Liu, Hannah Erb, Alka Singh, Junhao Mao, Xuelian Luo, and Xu Wu. Discovery of a new class of reversible TEA domain transcription factor inhibitors with a novel binding mode. *eLife*, 11, 2022.

[11009] Tuanjun Hu, Lorna Taylor, Adrian Sherman, Christian Keambou Tiambo, Steven J Kemp, Bruce Whitelaw, Rachel J Hawken, Appolinaire Djikeng, and Michael J McGrew. A low-tech, cost-effective and efficient method for safeguarding genetic diversity by direct cryopreservation of poultry embryonic reproductive cells. *eLife*, 11, 2022.

[11010] Zicheng Hu, Kattria van der Ploeg, Saborni Chakraborty, Prabhu S Arunachalam, Diego AM Mori, Karen B Jacobson, Hector Bonilla, Julie Parsonnet, Jason R Andrews, Marisa Holubar, Aruna Subramanian, Chaitan Khosla, Yvonne Maldonado, Haley Hedlin, Lauren de la Parte, Kathleen Press, Maureen Ty, Gene S Tan, Catherine Blish, Saki Takahashi, Isabel Rodriguez-Barraquer, Bryan Greenhouse, Atul J Butte, Upinder Singh, Bali Pulendran, Taia T Wang, and Prasanna Jagannathan. Early immune markers of clinical, virological, and immunological outcomes in patients with COVID-19: a multi-omics study. *eLife*, 11, 2022.

[11011] Huachao Huang, Yinshan Fang, Ming Jiang, Yihan Zhang, Jana Biermann, Johannes C Melms, Jennifer A Danielsson, Ying Yang, Li Qiang, Jia Liu, Yiwu Zhou, Manli Wang, Zhihong Hu, Timothy C Wang, Anjali Saqi, Jie Sun, Ichiro Matsumoto, Wellington V Cardoso, Charles W Emala, Jian Zhu, Benjamin Izar, Hongmei Mou, and Jianwen Que. Contribution of Trp63CreERT2-labeled cells to alveolar regeneration is independent of tuft cells. *eLife*, 11, 2022.

[11012] Meng Huang, Minjie Hong, Xinhao Hou, Chengming Zhu, Di Chen, Xiangyang Chen, Shouhong Guang, and Xuezhu Feng. H3K9me1/2 methylation limits the lifespan of daf-2 mutants in C. elegans. *eLife*, 11, 2022.

[11013] Pinxiu Huang, Wenbo Deng, Haili Bao, Zhong Lin, Mengying Liu, Jinxiang Wu, Xiaobo Zhou, Manting Qiao, Yihua Yang, Han Cai, Faiza Rao, Jingsi Chen, Dunjin Chen, Jinhua Lu, Haibin Wang, Aiping Qin, and Shuangbo Kong. SOX4 facilitates PGR protein stability and FOXO1 expression conducive for human endometrial decidualization. *eLife*, 11, 2022.

[11014] Shuya Kate Huang, Omar Almurad, Reizel J Pejana, Zachary A Morrison, Aditya Pandey, Louis-Philippe Picard, Mark Nitz, Adnan Sljoka, and R Scott Prosser. Allosteric modulation of the adenosine A2A receptor by cholesterol. *eLife*, 11, 2022.

[11015] Wei Huang, Joel Vega-Rodriguez, Chritopher Kizito, Sung-Jae Cha, and Marcelo Jacobs-Lorena. Combining transgenesis with paratransgenesis to fight malaria. *eLife*, 11, 2022.

[11016] Xiaoning Huang, Yongping Xin, and Ting Lu. A systematic, complexity-reduction approach to dissect the kombucha tea microbiome. *eLife*, 11, 2022.

[11017] Yuheng Huang, Harsh Shukla, and Yuh Chwen G Lee. Species-specific chromatin landscape determines how transposable elements shape genome evolution. *eLife*, 11, 2022.

[11018] Stefan T Huber, Edin Sarajlic, Roeland Huijink, Felix Weis, Wiel H Evers, and Arjen J Jakobi. Nanofluidic chips for cryo-EM structure determination from picoliter sample volumes. *eLife*, 11, 2022.

[11019] Amanda M Hughes, Eleanor Sanderson, Tim Morris, Ziada Ayorech, Martin Tesli, Helga Ask, Ted Reichborn-Kjennerud, Ole A Andreassen, Per Magnus, Øyvind Helgeland, Stefan Johansson, Pål Njølstad, George Davey Smith, Alexandra Havdahl, Laura D Howe, and Neil M Davies. Body mass index and childhood symptoms of depression, anxiety, and attention-deficit hyperactivity disorder: A within-family Mendelian randomization study. *eLife*, 11, 2022.

[11020] Fern A Hughes, Alexis R Barr, and Philipp Thomas. Patterns of interdivision time correlations reveal hidden cell cycle factors. *eLife*, 11, 2022.

[11021] Nadia M Huisjes, Thomas M Retzer, Matthias J Scherr, Rohit Agarwal, Lional Rajappa, Barbara Safaric, Anita Minnen, and Karl E Duderstadt. Mars, a molecule archive suite for reproducible analysis and reporting of single-molecule properties from bioimages. *eLife*, 11, 2022.

[11022] Brooke D Huisman, Zheng Dai, David K Gifford, and Michael E Birnbaum. A high-throughput yeast display approach to profile pathogen proteomes for MHC-II binding. *eLife*, 11, 2022.

[11023] Jana S Huisman, Jérémie Scire, Daniel C Angst, Jinzhou Li, Richard A Neher, Marloes H Maathuis, Sebastian Bonhoeffer, and Tanja Stadler. Estimation and worldwide monitoring of the effective reproductive number of SARS-CoV-2. *eLife*, 11, 2022.

[11024] Wouter Huiting, Suzanne L Dekker, Joris CJ van der Lienden, Rafaella Mergener, Maiara K Musskopf, Gabriel V Furtado, Emma Gerrits, David Coit, Mehrnoosh Oghbaie, Luciano H Di Stefano, Hein Schepers, Maria AWH van Waarde-Verhagen, Suzanne Couzijn, Lara Barazzuol, John LaCava, Harm H Kampinga, and Steven Bergink. Targeting DNA topoisomerases or checkpoint kinases results in an overload of chaperone systems, triggering aggregation of a metastable subproteome. *eLife*, 11, 2022.

[11025] Brian Hurwitz, Nicola Guzzi, Anita Gola, Vincent F Fiore, Ataman Sendoel, Maria Nikolova, Douglas Barrows, Thomas S Carroll, H Amalia Pasolli, and Elaine Fuchs. The integrated stress response remodels the microtubule-organizing center to clear unfolded proteins following proteotoxic stress. *eLife*, 11, 2022.

[11026] Cendri A Hutcherson and Anita Tusche. Evidence accumulation, not 'self-control', explains dorsolateral prefrontal activation during normative choice. *eLife*, 11, 2022.

[11027] Erica J Hutchins, Shashank Gandhi, Jose Chacon, Michael Piacentino, and Marianne E Bronner. RNA-binding protein Elavl1/HuR is required for maintenance of cranial neural crest specification. *eLife*, 11, 2022.

[11028] Nora Hutchinson, Hannah Moyer, Deborah A Zarin, and Jonathan Kimmelman. The proportion of randomized controlled trials that inform clinical practice. *eLife*, 11, 2022.

[11029] Minh Sang Huynh, Yogesh Hooda, Yuzi Raina Li, Maciej Jagielnicki, Christine Chieh-Lin Lai, and Trevor F Moraes. Reconstitution of surface lipoprotein translocation through the Slam translocon. *eLife*, 11, 2022.

[11030] Justin H Hwang, Rand Arafeh, Ji-Heui Seo, Sylvan C Baca, Megan Ludwig, Taylor E Arnoff, Lydia Sawyer, Camden Richter, Sydney Tape, Hannah E Bergom, Sean McSweeney, Jonathan P Rennhack, Sarah A Klingenberg, Alexander TM Cheung, Jason Kwon, Jonathan So, Steven Kregel, Eliezer M Van Allen, Justin M Drake, Matthew L Freedman, and William C Hahn. CREB5 reprograms FOXA1 nuclear interactions to promote resistance to androgen receptor-targeting therapies. *eLife*, 11, 2022.

[11031] Kai Hwang, James M Shine, Michael W Cole, and Evan Sorenson. Thalamocortical contributions to cognitive task activity. *eLife*, 11, 2022.

[11032] Liang-Dar Hwang, Gunn-Helen Moen, and David M Evans. Using adopted individuals to partition indirect maternal genetic effects into prenatal and postnatal effects on offspring phenotypes. *eLife*, 11, 2022.

[11033] Woong Y Hwang, Valentyna Kostiuk, Delfina P González, C Patrick Lusk, and Mustafa K Khokha. Kap-$\beta$2/Transportin mediates $\beta$-catenin nuclear transport in Wnt signaling. *eLife*, 11, 2022.

[11034] Olga Ibañez-Solé, Alex M Ascensión, Marcos J Araúzo-Bravo, and Ander Izeta. Lack of evidence for increased transcriptional noise in aged tissues. *eLife*, 11, 2022.

[11035] Hiroshi Ichise, Shoko Tsukamoto, Tsuyoshi Hirashima, Yoshinobu Konishi, Choji Oki, Shinya Tsukiji, Satoshi Iwano, Atsushi Miyawaki, Kenta Sumiyama, Kenta Terai, and Michiyuki Matsuda. Functional visualization of NK cell-mediated killing of metastatic single tumor cells. *eLife*, 11, 2022.

[11036] Aoi Ichiyama, Samuel Mestern, Gabriel B Benigno, Kaela E Scott, Brian L Allman, Lyle Muller, and Wataru Inoue. State-dependent activity dynamics of hypothalamic stress effector neurons. *eLife*, 11, 2022.

[11037] Metehan Ilter, Ramazan Kasmer, Farzaneh Jalalypour, Canan Atilgan, Ozan Topcu, Nihal Karakas, and Ozge Sensoy. Inhibition of mutant RAS-RAF interaction by mimicking structural and dynamic properties of phosphorylated RAS. *eLife*, 11, 2022.

[11038] Takeshi Imai, Ryuta Tobe, Koji Honda, Mai Tanaka, Jun Kawamoto, and Hisaaki Mihara. Group II truncated haemoglobin YjbI prevents reactive oxygen species-induced protein aggregation in Bacillus subtilis. *eLife*, 11, 2022.

[11039] Tsuyoshi Imasaki, Satoshi Kikkawa, Shinsuke Niwa, Yumiko Saijo-Hamano, Hideki Shigematsu, Kazuhiro Aoyama, Kaoru Mitsuoka, Takahiro Shimizu, Mari Aoki, Ayako Sakamoto, Yuri Tomabechi, Naoki Sakai, Mikako Shirouzu, Shinya Taguchi, Yosuke Yamagishi, Tomiyoshi Setsu, Yoshiaki Sakihama, Eriko Nitta, Masatoshi Takeichi, and Ryo Nitta. CAMSAP2 organizes a $\gamma$-tubulin-independent microtubule nucleation centre through phase separation. *eLife*, 11, 2022.

[11040] Kengo Inada, Kazoku Tsujimoto, Masahide Yoshida, Katsuhiko Nishimori, and Kazunari Miyamichi. Oxytocin signaling in the posterior hypothalamus prevents hyperphagic obesity in mice. *eLife*, 11, 2022.

[11041] Henrike Indrischek, Juliane Hammer, Anja Machate, Nikolai Hecker, Bogdan Kirilenko, Juliana Roscito, Stefan Hans, Caren Norden, Michael Brand, and Michael Hiller. Vision-related convergent gene losses reveal SERPINE3′s unknown role in the eye. *eLife*, 11, 2022.

[11042] Shawn Irgen-Gioro, Shawn Yoshida, Victoria Walling, and Shasha Chong. Fixation can change the appearance of phase separation in living cells. *eLife*, 11, 2022.

[11043] Munenori Ishibashi, Joyce Keung, Catherine W Morgans, Sue A Aicher, James R Carroll, Joshua H Singer, Li Jia, Wei Li, Iris Fahrenfort, Christophe P Ribelayga, and Stephen C Massey. Analysis of rod/cone gap junctions from the reconstruction of mouse photoreceptor terminals. *eLife*, 11, 2022.

[11044] Yuzuru Itoh, Vivek Singh, Anas Khawaja, Andreas Naschberger, Minh Duc Nguyen, Joanna Rorbach, and Alexey Amunts. Structure of the mitoribosomal small subunit with streptomycin reveals Fe-S clusters and physiological molecules. *eLife*, 11, 2022.

[11045] Aleksandar Z Ivanov, Andrew J King, Ben DB Willmore, Kerry MM Walker, and Nicol S Harper. Cortical adaptation to sound reverberation. *eLife*, 11, 2022.

[11046] Amrita A Iyer, Ishwar Hosamani, John D Nguyen, Tiantian Cai, Sunita Singh, Melissa M McGovern, Lisa Beyer, Hongyuan Zhang, Hsin-I Jen, Rizwan Yousaf, Onur Birol, Jenny J Sun, Russell S Ray, Yehoash Raphael, Neil Segil, and Andrew K Groves. Cellular reprogramming with ATOH1, GFI1, and POU4F3 implicate epigenetic changes and cell-cell signaling as obstacles to hair cell regeneration in mature mammals. *eLife*, 11, 2022.

[11047] Hamit Izgi, Dingding Han, Ulas Isildak, Shuyun Huang, Ece Kocabiyik, Philipp Khaitovich, Mehmet Somel, and Handan Melike Dönertaş. Inter-tissue convergence of gene expression during ageing suggests age-related loss of tissue and cellular identity. *eLife*, 11, 2022.

[11048] John Jackson, Christie Le Coeur, and Owen Jones. Life history predicts global population responses to the weather in terrestrial mammals. *eLife*, 11, 2022.

[11049] David S Jacobs, Madeleine C Allen, Junchol Park, and Bita Moghaddam. Learning of probabilistic punishment as a model of anxiety produces changes in action but not punisher encoding in the dmPFC and VTA. *eLife*, 11, 2022.

[11050] Guillaume Jacquemin, Annabelle Wurmser, Mathilde Huyghe, Wenjie Sun, Zeinab Homayed, Candice Merle, Meghan Perkins, Fairouz Qasrawi, Sophie Richon, Florent Dingli, Guillaume Arras, Damarys Loew, Danijela Vignjevic, Julie Pannequin, and Silvia Fre. Paracrine signalling between intestinal epithelial and tumour cells induces a regenerative programme. *eLife*, 11, 2022.

[11051] Corentin Jacques, Jacques Jonas, Sophie Colnat-Coulbois, Louis Maillard, and Bruno Rossion. Low and high frequency intracranial neural signals match in the human associative cortex. *eLife*, 11, 2022.

[11052] Erik Jacques, Yinni Kuang, Allison P Kann, Fabien Le Grand, Robert S Krauss, and Penney M Gilbert. The mini-IDLE 3D biomimetic culture assay enables interrogation of mechanisms governing muscle stem cell quiescence and niche repopulation. *eLife*, 11, 2022.

[11053] Luisa Maria Jaimes-Nino, Jürgen Heinze, and Jan Oettler. Late-life fitness gains and reproductive death in Cardiocondyla obscurior ants. *eLife*, 11, 2022.

[11054] Amit Jairaman, Amanda McQuade, Alberto Granzotto, You Jung Kang, Jean Paul Chadarevian, Sunil Gandhi, Ian Parker, Ian Smith, Hansang Cho, Stefano L Sensi, Shivashankar Othy, Mathew Blurton-Jones, and Michael D Cahalan. TREM2 regulates purinergic receptor-mediated calcium signaling and motility in human iPSC-derived microglia. *eLife*, 11, 2022.

[11055] Maximilian AH Jakobs, Assaf Zemel, and Kristian Franze. Unrestrained growth of correctly oriented microtubules instructs axonal microtubule orientation. *eLife*, 11, 2022.

[11056] Elham Jalalvand, Jonatan Alvelid, Giovanna Coceano, Steven Edwards, Brita Robertson, Sten Grillner, and Ilaria Testa. ExSTED microscopy reveals contrasting functions of dopamine and somatostatin CSF-c neurons along the lamprey central canal. *eLife*, 11, 2022.

[11057] Jaime James, Yifei Chen, Clara M Hernandez, Florian Forster, Markus Dagnell, Qing Cheng, Amir A Saei, Hassan Gharibi, Gonzalo Fernandez Lahore, Annika Åstrand, Rajneesh Malhotra, Bernard Malissen, Roman A Zubarev, Elias SJ Arnér, and Rikard Holmdahl. Redox regulation of PTPN22 affects the severity of T-cell-dependent autoimmune inflammation. *eLife*, 11, 2022.

[11058] Mohammed Janahi, Leon Aksman, Jonathan M Schott, Younes Mokrab, and Andre Altmann. Nomograms of human hippocampal volume shifted by polygenic scores. *eLife*, 11, 2022.

[11059] Dong Gil Jang, Keun Yeong Kwon, Yeong Cheon Kweon, Byung-gyu Kim, Kyungjae Myung, Hyun-Shik Lee, Chan Young Park, Taejoon Kwon, and Tae Joo Park. GJA1 depletion causes ciliary defects by affecting Rab11 trafficking to the ciliary base. *eLife*, 11, 2022.

[11060] Pramod K Jangir, Qiue Yang, Liam P Shaw, Julio Diaz Caballero, Lois Ogunlana, Rachel Wheatley, Timothy Walsh, and R Craig MacLean. Pre-existing chromosomal polymorphisms in pathogenic E. coli potentiate the evolution of resistance to a last-resort antibiotic. *eLife*, 11, 2022.

[11061] Kayla A Jara, Nikolaus M Loening, Patrick N Reardon, Zhen Yu, Prajna Woonnimani, Coban Brooks, Cat H Vesely, and Elisar J Barbar. Multivalency, autoinhibition, and protein disorder in the regulation of interactions of dynein intermediate chain with dynactin and the nuclear distribution protein. *eLife*, 11, 2022.

[11062] Sanja Jasek, Csaba Verasztó, Emelie Brodrick, Réza Shahidi, Tom Kazimiers, Alexandra Kerbl, and Gáspár Jékely. Desmosomal connectomics of all somatic muscles in an annelid larva. *eLife*, 11, 2022.

[11063] Jacob S Jaszczak, Laura DeVault, Lily Yeh Jan, and Yuh Nung Jan. Steroid hormone signaling activates thermal nociception during Drosophila peripheral nervous system development. *eLife*, 11, 2022.

[11064] Saskia-Larissa Jauch-Speer, Marisol Herrera-Rivero, Nadine Ludwig, Bruna Caroline Véras De Carvalho, Leonie Martens, Jonas Wolf, Achmet Imam Chasan, Anika Witten, Birgit Markus, Bernhard Schieffer, Thomas Vogl, Jan Rossaint, Monika Stoll, Johannes Roth, and Olesja Fehler. C/EBP$\delta$-induced epigenetic changes control the dynamic gene transcription of S100a8 and S100a9. *eLife*, 11, 2022.

[11065] Katarzyna Jaworska, Yuening Yan, Nicola J van Rijsbergen, Robin AA Ince, and Philippe G Schyns. Different computations over the same inputs produce selective behavior in algorithmic brain networks. *eLife*, 11, 2022.

[11066] Viraaj Jayaram, Nirag Kadakia, and Thierry Emonet. Sensing complementary temporal features of odor signals enhances navigation of diverse turbulent plumes. *eLife*, 11, 2022.

[11067] Jayson Jeganathan, Megan Campbell, Matthew Hyett, Gordon Parker, and Michael Breakspear. Quantifying dynamic facial expressions under naturalistic conditions. *eLife*, 11, 2022.

[11068] Brian B Jeon, Thomas Fuchs, Steven M Chase, and Sandra J Kuhlman. Visual experience has opposing influences on the quality of stimulus representation in adult primary visual cortex. *eLife*, 11, 2022.

[11069] Hyeonsoo Jeong, Nicole M Baran, Dan Sun, Paramita Chatterjee, Thomas S Layman, Christopher N Balakrishnan, Donna L Maney, and Soojin V Yi. Dynamic molecular evolution of a supergene with suppressed recombination in white-throated sparrows. *eLife*, 11, 2022.

[11070] Hyun-Woo Jeong, Rodrigo Diéguez-Hurtado, Hendrik Arf, Jian Song, Hongryeol Park, Kai Kruse, Lydia Sorokin, and Ralf H Adams. Single-cell transcriptomics reveals functionally specialized vascular endothelium in brain. *eLife*, 11, 2022.

[11071] Joshua Jeong, Joyce H Lee, Claudia C Carcamo, Matthew W Parker, and James M Berger. DNA-Stimulated Liquid-Liquid phase separation by eukaryotic topoisomerase ii modulates catalytic function. *eLife*, 11, 2022.

[11072] Marieke Jepma, Mathieu Roy, Kiran Ramlakhan, Monique van Velzen, and Albert Dahan. Different brain systems support learning from received and avoided pain during human pain-avoidance learning. *eLife*, 11, 2022.

[11073] Dillon Jevon, Kylie Deng, Nicole Hallahan, Krish Kumar, Jason Tong, Wan Jun Gan, Clara Tran, Marcela Bilek, and Peter Thorn. Local activation of focal adhesion kinase orchestrates the positioning of presynaptic scaffold proteins and Ca2+ signalling to control glucose-dependent insulin secretion. *eLife*, 11, 2022.

[11074] Huiru Jia, Yongqiang Liu, Xaiokang Li, Hui Li, Yunfei Pan, Chaoxing Hu, Xainyong Zhou, Kris AG Wyckhuys, and Kongming Wu. Windborne migration amplifies insect-mediated pollination services. *eLife*, 11, 2022.

[11075] Jia Jia, Guangzhao Li, and Ke-Qin Gao. Palatal morphology predicts the paleobiology of early salamanders. *eLife*, 11, 2022.

[11076] Yinjun Jia, Shuaishuai Li, Xuan Guo, Bo Lei, Junqiang Hu, Xiao-Hong Xu, and Wei Zhang. Selfee, self-supervised features extraction of animal behaviors. *eLife*, 11, 2022.

[11077] Yutian Jia, Yanming Zhang, Wenhao Wang, Jianlin Lei, Zhengxin Ying, and Guanghui Yang. Structural and functional insights of the human peroxisomal ABC transporter ALDP. *eLife*, 11, 2022.

[11078] Fu-Qing Jiang, Kun Liu, Jia-Xuan Chen, Yan Cao, Wu-Yun Chen, Wan-Ling Zhao, Guo-Hua Song, Chi-Qian Liang, Yi-Min Zhou, Huan-Lei Huang, Rui-Jin Huang, Hui Zhao, Kyu-Sang Park, Zhenyu Ju, Dongqing Cai, and Xu-Feng Qi. Mettl3-mediated m6A modification of Fgf16 restricts cardiomyocyte proliferation during heart regeneration. *eLife*, 11, 2022.

[11079] Yuwei Jiang, Misako Komatsu, Yuyan Chen, Ruoying Xie, Kaiwei Zhang, Ying Xia, Peng Gui, Zhifeng Liang, and Liping Wang. Constructing the hierarchy of predictive auditory sequences in the marmoset brain. *eLife*, 11, 2022.

[11080] Wei Jiao, Gard Spreemann, Evelyne Ruchti, Soumya Banerjee, Samuel Vernon, Ying Shi, R Steven Stowers, Kathryn Hess, and Brian D McCabe. Intact Drosophila central nervous system cellular quantitation reveals sexual dimorphism. *eLife*, 11, 2022.

[11081] Judit Jimenez-Sainz, Joshua Mathew, Gemma Moore, Sudipta Lahiri, Jennifer Garbarino, Joseph P Eder, Eli Rothenberg, and Ryan B Jensen. BRCA2 BRC missense variants disrupt RAD51-dependent DNA repair. *eLife*, 11, 2022.

[11082] Eric N Jimenez-Vazquez, Michael Arad, álvaro Macías, Maria L Vera-Pedrosa, Francisco Miguel Cruz, Lilian K Gutierrez, Ashley J Cuttitta, André Monteiro da Rocha, Todd J Herron, Daniela Ponce-Balbuena, Guadalupe Guerrero-Serna, Ofer Binah, Daniel E Michele, and José Jalife. SNTA1 gene rescues ion channel function and is antiarrhythmic in cardiomyocytes derived from induced pluripotent stem cells from muscular dystrophy patients. *eLife*, 11, 2022.

[11083] Nicolai Tidemand Johansen, Marta Bonaccorsi, Tone Bengtsen, Andreas Haahr Larsen, Frederik Grønbæk Tidemand, Martin Cramer Pedersen, Pie Huda, Jens Berndtsson, Tamim Darwish, Nageshewar Rao Yepuri, Anne Martel, Thomas Günther Pomorski, Andrea Bertarello, Mark Sansom, Mikaela Rapp, Ramon Crehuet, Tobias Schubeis, Kresten Lindorff-Larsen, Guido Pintacuda, and Lise Arleth. Mg2+-dependent conformational equilibria in CorA and an integrated view on transport regulation. *eLife*, 11, 2022.

[11084] Juliane John, Oskar Aurelius, Vivek Srinivas, Patricia Saura, In-Sik Kim, Asmit Bhowmick, Philipp S Simon, Medhanjali Dasgupta, Cindy Pham, Sheraz Gul, Kyle D Sutherlin, Pierre Aller, Agata Butryn, Allen M Orville, Mun Hon Cheah, Shigeki Owada, Kensuke Tono, Franklin D Fuller, Alexander Batyuk, Aaron S Brewster, Nicholas K Sauter, Vittal K Yachandra, Junko Yano, Ville RI Kaila, Jan Kern, Hugo Lebrette, and Martin Högbom. Redox-controlled reorganization and flavin strain within the ribonucleotide reductase R2b–NrdI complex monitored by serial femtosecond crystallography. *eLife*, 11, 2022.

[11085] JP Johnson, Thilo Focken, Kuldip Khakh, Parisa Karimi Tari, Celine Dube, Samuel J Goodchild, Jean-Christophe Andrez, Girish Bankar, David Bogucki, Kristen Burford, Elaine Chang, Sultan Chowdhury, Richard Dean, Gina de Boer, Shannon Decker, Christoph Dehnhardt, Mandy Feng, Wei Gong, Michael Grimwood, Abid Hasan, Angela Hussainkhel, Qi Jia, Stephanie Lee, Jenny Li, Sophia Lin, Andrea Lindgren, Verner Lofstrand, Janette Mezeyova, Rostam Namdari, Karen Nelkenbrecher, Noah Gregory Shuart, Luis Sojo, Shaoyi Sun, Matthew Taron, Matthew Waldbrook, Diana Weeratunge, Steven Wesolowski, Aaron Williams, Michael Wilson, Zhiwei Xie, Rhena Yoo, Clint Young, Alla Zenova, Wei Zhang, Alison J Cutts, Robin P Sherrington, Simon N Pimstone, Raymond Winquist, Charles J Cohen, and James R Empfield. NBI-921352, a first-in-class, NaV1.6 selective, sodium channel inhibitor that prevents seizures in Scn8a gain-of-function mice, and wild-type mice and rats. *eLife*, 11, 2022.

[11086] Milo S Johnson and Michael M Desai. Mutational robustness changes during long-term adaptation in laboratory budding yeast populations. *eLife*, 11, 2022.

[11087] Nathan R Johnson, Luis F Larrondo, José M álvarez, and Elena A Vidal. Comprehensive re-analysis of hairpin small RNAs in fungi reveals loci with conserved links. *eLife*, 11, 2022.

[11088] Peter S Johnstone, Maite Ogueta, Olga Akay, Inan Top, Sheyum Syed, Ralf Stanewsky, and Deniz Top. Real time, in vivo measurement of neuronal and peripheral clocks in Drosophila melanogaster. *eLife*, 11, 2022.

[11089] Jolle Wolter Jolles, Matthew MG Sosna, Geoffrey PF Mazué, Colin R Twomey, Joseph Bak-Coleman, Daniel I Rubenstein, and Iain D Couzin. Both prey and predator features predict the individual predation risk and survival of schooling prey. *eLife*, 11, 2022.

[11090] Laura Jones, Madeleine Verriotis, Robert J Cooper, Maria Pureza Laudiano-Dray, Mohammed Rupawala, Judith Meek, Lorenzo Fabrizi, and Maria Fitzgerald. Widespread nociceptive maps in the human neonatal somatosensory cortex. *eLife*, 11, 2022.

[11091] David J Jörg, Doris H Fuertinger, Alhaji Cherif, David A Bushinsky, Ariella Mermelstein, Jochen G Raimann, and Peter Kotanko. Modeling osteoporosis to design and optimize pharmacological therapies comprising multiple drug types. *eLife*, 11, 2022.

[11092] Siddhartha Joshi and Joshua I Gold. Context-dependent relationships between locus coeruleus firing patterns and coordinated neural activity in the anterior cingulate cortex. *eLife*, 11, 2022.

[11093] Manuel F Juette, Jordan D Carelli, Emily J Rundlet, Alan Brown, Sichen Shao, Angelica Ferguson, Michael R Wasserman, Mikael Holm, Jack Taunton, and Scott C Blanchard. Didemnin B and ternatin-4 differentially inhibit conformational changes in eEF1A required for aminoacyl-tRNA accommodation into mammalian ribosomes. *eLife*, 11, 2022.

[11094] Damien Jullié, Camila Benitez, Tracy A Knight, Milos S Simic, and Mark von Zastrow. Endocytic trafficking determines cellular tolerance of presynaptic opioid signaling. *eLife*, 11, 2022.

[11095] Na Young Jun, Douglas A Ruff, Lily E Kramer, Brittany Bowes, Surya T Tokdar, Marlene R Cohen, and Jennifer M Groh. Coordinated multiplexing of information about separate objects in visual cortex. *eLife*, 11, 2022.

[11096] Paula Kaanders, Pradyumna Sepulveda, Tomas Folke, Pietro Ortoleva, and Benedetto De Martino. Humans actively sample evidence to support prior beliefs. *eLife*, 11, 2022.

[11097] Koichiro Kajikawa, Brad K Hulse, Athanassios G Siapas, and Evgueniy V Lubenov. UP-DOWN states and ripples differentially modulate membrane potential dynamics across DG, CA3, and CA1 in awake mice. *eLife*, 11, 2022.

[11098] Muntasir Kamal, Levon Tokmakjian, Jessica Knox, Peter Mastrangelo, Jingxiu Ji, Hao Cai, Jakub W Wojciechowski, Michael P Hughes, Kristóf Takács, Xiaoquan Chu, Jianfeng Pei, Vince Grolmusz, Malgorzata Kotulska, Julie Deborah Forman-Kay, and Peter J Roy. A spatiotemporal reconstruction of the C. elegans pharyngeal cuticle reveals a structure rich in phase-separating proteins. *eLife*, 11, 2022.

[11099] Suchitra Kamle, Bing Ma, Chang Min Lee, Gail Schor, Yang Zhou, Chun Geun Lee, and Jack A Elias. Host chitinase 3-like-1 is a universal therapeutic target for SARS-CoV-2 viral variants in COVID-19. *eLife*, 11, 2022.

[11100] Oguz Kanca, Jonathan Zirin, Yanhui Hu, Burak Tepe, Debdeep Dutta, Wen-Wen Lin, Liwen Ma, Ming Ge, Zhongyuan Zuo, Lu-Ping Liu, Robert W Levis, Norbert Perrimon, and Hugo J Bellen. An expanded toolkit for Drosophila gene tagging using synthesized homology donor constructs for CRISPR-mediated homologous recombination. *eLife*, 11, 2022.

[11101] Genki N Kanda, Taku Tsuzuki, Motoki Terada, Noriko Sakai, Naohiro Motozawa, Tomohiro Masuda, Mitsuhiro Nishida, Chihaya T Watanabe, Tatsuki Higashi, Shuhei A Horiguchi, Taku Kudo, Motohisa Kamei, Genshiro A Sunagawa, Kenji Matsukuma, Takeshi Sakurada, Yosuke Ozawa, Masayo Takahashi, Koichi Takahashi, and Tohru Natsume. Robotic search for optimal cell culture in regenerative medicine. *eLife*, 11, 2022.

[11102] Harry Kane, Nelson M LaMarche, áine Ní Scannail, Amanda E Garza, Hui-Fern Koay, Adiba I Azad, Britta Kunkemoeller, Brenneth Stevens, Michael B Brenner, and Lydia Lynch. Longitudinal analysis of invariant natural killer T cell activation reveals a cMAF-associated transcriptional state of NKT10 cells. *eLife*, 11, 2022.

[11103] Miki Kaneshige, Kei Obara, Michiaki Suzuki, Toshiki Tazoe, and Yukio Nishimura. Tuning of motor outputs produced by spinal stimulation during voluntary control of torque directions in monkeys. *eLife*, 11, 2022.

[11104] Minsu Kang, Hee Young Na, Soomin Ahn, Ji-Won Kim, Sejoon Lee, Soyeon Ahn, Ju Hyun Lee, Jeonghwan Youk, Haesook T Kim, Kui-Jin Kim, Koung Jin Suh, Jun Suh Lee, Se Hyun Kim, Jin Won Kim, Yu Jung Kim, Keun-Wook Lee, Yoo-Seok Yoon, Jee Hyun Kim, Jin-Haeng Chung, Ho-Seong Han, and Jong Seok Lee. Gallbladder adenocarcinomas undergo subclonal diversification and selection from precancerous lesions to metastatic tumors. *eLife*, 11, 2022.

[11105] Anna Kankaanpää, Asko Tolvanen, Aino Heikkinen, Jaakko Kaprio, Miina Ollikainen, and Elina Sillanpää. The role of adolescent lifestyle habits in biological aging: A prospective twin study. *eLife*, 11, 2022.

[11106] Fumihiro Kano, Yuri Kawaguchi, and Yeow Hanling. Experimental evidence that uniformly white sclera enhances the visibility of eye-gaze direction in humans and chimpanzees. *eLife*, 11, 2022.

[11107] Tadayoshi Karasawa, Takanori Komada, Naoya Yamada, Emi Aizawa, Yoshiko Mizushina, Sachiko Watanabe, Chintogtokh Baatarjav, Takayoshi Matsumura, and Masafumi Takahashi. Cryo-sensitive aggregation triggers NLRP3 inflammasome assembly in cryopyrin-associated periodic syndrome. *eLife*, 11, 2022.

[11108] Yael Kashash, Grace Smarsh, Noga Zilkha, Yossi Yovel, and Tali Kimchi. Alone, in the dark: The extraordinary neuroethology of the solitary blind mole rat. *eLife*, 11, 2022.

[11109] Edward R Kastenhuber, Marisa Mercadante, Benjamin Nilsson-Payant, Jared L Johnson, Javier A Jaimes, Frauke Muecksch, Yiska Weisblum, Yaron Bram, Vasuretha Chandar, Gary R Whittaker, Benjamin R tenOever, Robert E Schwartz, and Lewis Cantley. Coagulation factors directly cleave SARS-CoV-2 spike and enhance viral entry. *eLife*, 11, 2022.

[11110] Go Kasuya and Koichi Nakajo. Optimized tight binding between the S1 segment and KCNE3 is required for the constitutively open nature of the KCNQ1-KCNE3 channel complex. *eLife*, 11, 2022.

[11111] Koji Kato, Tasuku Hamaguchi, Ryo Nagao, Keisuke Kawakami, Yoshifumi Ueno, Takehiro Suzuki, Hiroko Uchida, Akio Murakami, Yoshiki Nakajima, Makio Yokono, Seiji Akimoto, Naoshi Dohmae, Koji Yonekura, and Jian-Ren Shen. Structural basis for the absence of low-energy chlorophylls in a photosystem I trimer from Gloeobacter violaceus. *eLife*, 11, 2022.

[11112] Dhruva Katrekar, Yichen Xiang, Nathan Palmer, Anushka Saha, Dario Meluzzi, and Prashant Mali. Comprehensive interrogation of the ADAR2 deaminase domain for engineering enhanced RNA editing activity and specificity. *eLife*, 11, 2022.

[11113] Eva Kaulich, Trae Carroll, Brian D Ackley, Yi-Quan Tang, Iris Hardege, Keith Nehrke, William R Schafer, and Denise S Walker. Distinct roles for two Caenorhabditis elegans acid-sensing ion channels in an ultradian clock. *eLife*, 11, 2022.

[11114] Hidehiko Kawai, Ren Iwata, Shungo Ebi, Ryusei Sugihara, Shogo Masuda, Chiho Fujiwara, Shingo Kimura, and Hiroyuki Kamiya. Development of a versatile high-throughput mutagenesis assay with multiplexed short-read NGS using DNA-barcoded supF shuttle vector library amplified in E. coli. *eLife*, 11, 2022.

[11115] Lachlan Kay, Rebecca Keogh, Thomas Andrillon, and Joel Pearson. The pupillary light response as a physiological index of aphantasia, sensory and phenomenological imagery strength. *eLife*, 11, 2022.

[11116] Sinan Kaya-Zeeb, Lorenz Engelmayer, Mara Straßburger, Jasmin Bayer, Heike Bähre, Roland Seifert, Oliver Scherf-Clavel, and Markus Thamm. Octopamine drives honeybee thermogenesis. *eLife*, 11, 2022.

[11117] Kanwal Kayastha, Alexander Katsyv, Christina Himmrich, Sonja Welsch, Jan M Schuller, Ulrich Ermler, and Volker Müller. Structure-based electron-confurcation mechanism of the Ldh-EtfAB complex. *eLife*, 11, 2022.

[11118] Christian Keine, Mohammed Al-Yaari, Tamara Radulovic, Connon I Thomas, Paula Valino Ramos, Debbie Guerrero-Given, Mrinalini Ranjan, Holger Taschenberger, Naomi Kamasawa, and Samuel M Young. Presynaptic Rac1 controls synaptic strength through the regulation of synaptic vesicle priming. *eLife*, 11, 2022.

[11119] Kevin P Kelly, Mroj Alassaf, Camille E Sullivan, Ava E Brent, Zachary H Goldberg, Michelle E Poling, Julien Dubrulle, and Akhila Rajan. Fat body phospholipid state dictates hunger-driven feeding behavior. *eLife*, 11, 2022.

[11120] Lauritz Kennedy, Emilie R Glesaaen, Vuk Palibrk, Marco Pannone, Wei Wang, Ali Al-Jabri, Rajikala Suganthan, Niklas Meyer, Marie Landa Austbø, Xiaolin Lin, Linda H Bergersen, Magnar Bjørås, and Johanne E Rinholm. Lactate receptor HCAR1 regulates neurogenesis and microglia activation after neonatal hypoxia-ischemia. *eLife*, 11, 2022.

[11121] Anna Keppner, Miguel Correia, Sara Santambrogio, Teng Wei Koay, Darko Maric, Carina Osterhof, Denise V Winter, Angèle Clerc, Michael Stumpe, Frédéric Chalmel, Sylvia Dewilde, Alex Odermatt, Dieter Kressler, Thomas Hankeln, Roland H Wenger, and David Hoogewijs. Androglobin, a chimeric mammalian globin, is required for male fertility. *eLife*, 11, 2022.

[11122] Samuel A Kerk, Lin Lin, Amy L Myers, Damien J Sutton, Anthony Andren, Peter Sajjakulnukit, Li Zhang, Yaqing Zhang, Jennifer A Jiménez, Barbara S Nelson, Brandon Chen, Anthony Robinson, Galloway Thurston, Samantha B Kemp, Nina G Steele, Megan T Hoffman, Hui-Ju Wen, Daniel Long, Sarah E Ackenhusen, Johanna Ramos, Xiaohua Gao, Zeribe C Nwosu, Stefanie Galban, Christopher J Halbrook, David B Lombard, David R Piwnica-Worms, Haoqiang Ying, Marina Pasca di Magliano, Howard C Crawford, Yatrik M Shah, and Costas A Lyssiotis. Metabolic requirement for GOT2 in pancreatic cancer depends on environmental context. *eLife*, 11, 2022.

[11123] Ali A Kermani, Olive E Burata, B Ben Koff, Akiko Koide, Shohei Koide, and Randy B Stockbridge. Crystal structures of bacterial small multidrug resistance transporter EmrE in complex with structurally diverse substrates. *eLife*, 11, 2022.

[11124] Pius Kern, Micha Heilbron, Floris P de Lange, and Eelke Spaak. Cortical activity during naturalistic music listening reflects short-range predictions based on long-term experience. *eLife*, 11, 2022.

[11125] Casper Kerrén, Sander van Bree, Benjamin J Griffiths, and Maria Wimber. Phase separation of competing memories along the human hippocampal theta rhythm. *eLife*, 11, 2022.

[11126] Lauren Kershberg, Aditi Banerjee, and Pascal S Kaeser. Protein composition of axonal dopamine release sites in the striatum. *eLife*, 11, 2022.

[11127] Madhura D Ketkar, Burak Gür, Sebastian Molina-Obando, Maria Ioannidou, Carlotta Martelli, and Marion Silies. First-order visual interneurons distribute distinct contrast and luminance information across ON and OFF pathways to achieve stable behavior. *eLife*, 11, 2022.

[11128] Hina Khan, Partha Paul, Ritesh Rajesh Sevalkar, Sangita Kachhap, Balvinder Singh, and Dibyendu Sarkar. Convergence of two global regulators to coordinate expression of essential virulence determinants of Mycobacterium tuberculosis. *eLife*, 11, 2022.

[11129] Nasar Khan, Hüsnü Aslan, Henning Büttner, Holger Rohde, Thaddeus Wayne Golbek, Steven Joop Roeters, Sander Woutersen, Tobias Weidner, and Rikke Louise Meyer. The giant staphylococcal protein Embp facilitates colonization of surfaces through Velcro-like attachment to fibrillated fibronectin. *eLife*, 11, 2022.

[11130] Sabab Hasan Khan, Sean M Braet, Stephen John Koehler, Elizabeth Elacqua, Ganesh Srinivasan Anand, and C Denise Okafor. Ligand-induced shifts in conformational ensembles that describe transcriptional activation. *eLife*, 11, 2022.

[11131] Divya Khattar, Sharlene Fernandes, John Snowball, Minzhe Guo, Matthew C Gillen, Suchi Singh Jain, Debora Sinner, William Zacharias, and Daniel T Swarr. PI3K signaling specifies proximal-distal fate by driving a developmental gene regulatory network in SOX9+ mouse lung progenitors. *eLife*, 11, 2022.

[11132] Jian Ming Khor and Charles A Ettensohn. Architecture and evolution of the cis-regulatory system of the echinoderm kirrelL gene. *eLife*, 11, 2022.

[11133] Robert Kiewisz, Gunar Fabig, William Conway, Daniel Baum, Daniel Needleman, and Thomas Müller-Reichert. Three-dimensional structure of kinetochore-fibers in human mitotic spindles. *eLife*, 11, 2022.

[11134] Atsushi Kikumoto, Ulrich Mayr, and David Badre. The role of conjunctive representations in prioritizing and selecting planned actions. *eLife*, 11, 2022.

[11135] Murat Kilinc, Vineet Arora, Thomas K Creson, Camilo Rojas, Aliza A Le, Julie Lauterborn, Brent Wilkinson, Nicolas Hartel, Nicholas Graham, Adrian Reich, Gemma Gou, Yoichi Araki, Àlex Bayés, Marcelo Coba, Gary Lynch, Courtney A Miller, and Gavin Rumbaugh. Endogenous Syngap1 alpha splice forms promote cognitive function and seizure protection. *eLife*, 11, 2022.

[11136] Elisabeth A Kilroy, Amanda C Ignacz, Kaylee L Brann, Claire E Schaffer, Devon Varney, Sarah S Alrowaished, Kodey J Silknitter, Jordan N Miner, Ahmed Almaghasilah, Tashawna L Spellen, Alexandra D Lewis, Karissa Tilbury, Benjamin L King, Joshua B Kelley, and Clarissa A Henry. Beneficial impacts of neuromuscular electrical stimulation on muscle structure and function in the zebrafish model of Duchenne muscular dystrophy. *eLife*, 11, 2022.

[11137] Hyojin Kim, Eric B Gao, Adam Draper, Noah C Berens, Hanna Vihma, Xinyuan Zhang, Alexandra Higashi-Howard, Kimberly D Ritola, Jeremy M Simon, Andrew J Kennedy, and Benjamin D Philpot. Rescue of behavioral and electrophysiological phenotypes in a Pitt-Hopkins syndrome mouse model by genetic restoration of Tcf4 expression. *eLife*, 11, 2022.

[11138] HyungGoo R Kim, Dora E Angelaki, and Gregory C DeAngelis. A neural mechanism for detecting object motion during self-motion. *eLife*, 11, 2022.

[11139] Jessica H Kim, Grace H Kromm, Olivia K Barnhill, Jacob Sperber, Lauren B Heuer, Sierra Loomis, Matthew C Newman, Kenneth Han, Faris F Gulamali, Theresa B Legan, Katharine E Jensen, Samuel C Funderburk, Michael J Krashes, and Matthew E Carter. A discrete parasubthalamic nucleus subpopulation plays a critical role in appetite suppression. *eLife*, 11, 2022.

[11140] Jun Kim, David S Jimenez, Bhavana Ragipani, Bo Zhang, Lena A Street, Maxwell Kramer, Sarah E Albritton, Lara H Winterkorn, Ana K Morao, and Sevinc Ercan. Condensin DC loads and spreads from recruitment sites to create loop-anchored TADs in C. elegans. *eLife*, 11, 2022.

[11141] Jung-Min Kim, Yeon-Suk Yang, Jaehyoung Hong, Sachin Chaugule, Hyonho Chun, Marjolein CH van der Meulen, Ren Xu, Matthew B Greenblatt, and Jae-hyuck Shim. Biphasic regulation of osteoblast development via the ERK MAPK–mTOR pathway. *eLife*, 11, 2022.

[11142] Seaho Kim, CheukMan C Au, Mohd Azrin Bin Jamalruddin, Naira Essam Abou-Ghali, Eiman Mukhtar, Luigi Portella, Adeline Berger, Daniel Worroll, Prerna Vatsa, David S Rickman, David M Nanus, and Paraskevi Giannakakou. AR-V7 exhibits non-canonical mechanisms of nuclear import and chromatin engagement in castrate-resistant prostate cancer. *eLife*, 11, 2022.

[11143] Siyoung Kim, Jeeyun Chung, Henning Arlt, Alexander J Pak, Robert V Farese, Tobias C Walther, and Gregory A Voth. Seipin transmembrane segments critically function in triglyceride nucleation and lipid droplet budding from the membrane. *eLife*, 11, 2022.

[11144] Soo Jeong Kim, Youngsik Woo, Hyun Jin Kim, Bon Seong Goo, Truong Thi My Nhung, Seol-Ae Lee, Bo Kyoung Suh, Dong Jin Mun, Joung-Hun Kim, and Sang Ki Park. Retinoic acid-induced protein 14 controls dendritic spine dynamics associated with depressive-like behaviors. *eLife*, 11, 2022.

[11145] Woohyun Kim, Jae Jin Shin, Yu Jin Jeong, Kyungdeok Kim, Jung Won Bae, Young Woo Noh, Seungjoon Lee, Woochul Choi, Se-Bum Paik, Min Whan Jung, Eunee Lee, and Eunjoon Kim. Suppressed prefrontal neuronal firing variability and impaired social representation in IRSp53-mutant mice. *eLife*, 11, 2022.

[11146] Yang Joon Kim, Kaitlin Rhee, Jonathan Liu, Selene Jeammet, Meghan A Turner, Stephen J Small, and Hernan G Garcia. Predictive modeling reveals that higher-order cooperativity drives transcriptional repression in a synthetic developmental enhancer. *eLife*, 11, 2022.

[11147] Hirokazu Kimura, Raymond M Paranal, Neha Nanda, Laura D Wood, James R Eshleman, Ralph H Hruban, Michael G Goggins, Alison P Klein, Randall Brand, and Nicholas J Roberts. Functional CDKN2A assay identifies frequent deleterious alleles misclassified as variants of uncertain significance. *eLife*, 11, 2022.

[11148] Sylvia P Kinosian and Paul G Wolf. The biology of C. richardii as a tool to understand plant evolution. *eLife*, 11, 2022.

[11149] Marianthi Kiparaki, Chaitali Khan, Virginia Folgado-Marco, Jacky Chuen, Panagiotis Moulos, and Nicholas E Baker. The transcription factor Xrp1 orchestrates both reduced translation and cell competition upon defective ribosome assembly or function. *eLife*, 11, 2022.

[11150] Lisa L Kirkemo, Susanna K Elledge, Jiuling Yang, James R Byrnes, Jeff E Glasgow, Robert Blelloch, and James A Wells. Cell-surface tethered promiscuous biotinylators enable comparative small-scale surface proteomic analysis of human extracellular vesicles and cells. *eLife*, 11, 2022.

[11151] Phoebe M Kirkwood, Douglas A Gibson, Isaac Shaw, Ross Dobie, Olympia Kelepouri, Neil C Henderson, and Philippa TK Saunders. Single-cell RNA sequencing and lineage tracing confirm mesenchyme to epithelial transformation (MET) contributes to repair of the endometrium at menstruation. *eLife*, 11, 2022.

[11152] Djem U Kissiov, Alexander Ethell, Sean Chen, Natalie K Wolf, Chenyu Zhang, Susanna M Dang, Yeara Jo, Katrine N Madsen, Ishan Paranjpe, Angus Y Lee, Bryan Chim, Stefan A Muljo, and David H Raulet. Binary outcomes of enhancer activity underlie stable random monoallelic expression. *eLife*, 11, 2022.

[11153] Gordon L Klein. Is calcium a link between inflammatory bone resorption and heart disease? *eLife*, 11, 2022.

[11154] Riku Klén, Disha Purohit, Ricardo Gómez-Huelgas, José Manuel Casas-Rojo, Juan Miguel Antón-Santos, Jesús Millán Núñez-Cortés, Carlos Lumbreras, José Manuel Ramos-Rincón, Noelia García Barrio, Miguel Pedrera-Jiménez, Antonio Lalueza Blanco, María Dolores Martin-Escalante, Francisco Rivas-Ruiz, Maria ángeles Onieva-García, Pablo Young, Juan Ignacio Ramirez, Estela Edith Titto Omonte, Rosmery Gross Artega, Magdy Teresa Canales Beltrán, Pascual Ruben Valdez, Florencia Pugliese, Rosa Castagna, Ivan A Huespe, Bruno Boietti, Javier A Pollan, Nico Funke, Benjamin Leiding, and David Gómez-Varela. Development and evaluation of a machine learning-based in-hospital COVID-19 disease outcome predictor (CODOP): A multicontinental retrospective study. *eLife*, 11, 2022.

[11155] Maren Klingelhöfer-Jens, Mana R Ehlers, Manuel Kuhn, Vincent Keyaniyan, and Tina B Lonsdorf. Robust group- but limited individual-level (longitudinal) reliability and insights into cross-phases response prediction of conditioned fear. *eLife*, 11, 2022.

[11156] Janus RL Kobbersmed, Manon MM Berns, Susanne Ditlevsen, Jakob B Sørensen, and Alexander M Walter. Allosteric stabilization of calcium and phosphoinositide dual binding engages several synaptotagmins in fast exocytosis. *eLife*, 11, 2022.

[11157] Alex A Koch, James S Bagnall, Nicola J Smyllie, Nicola Begley, Antony D Adamson, Jennifer L Fribourgh, David G Spiller, Qing-Jun Meng, Carrie L Partch, Korbinian Strimmer, Thomas A House, Michael H Hastings, and Andrew SI Loudon. Quantification of protein abundance and interaction defines a mechanism for operation of the circadian clock. *eLife*, 11, 2022.

[11158] Yuta Koganezawa, Miki Umetani, Moritoshi Sato, and Yuichi Wakamoto. History-dependent physiological adaptation to lethal genetic modification under antibiotic exposure. *eLife*, 11, 2022.

[11159] Robin Köhler, Eugen Kaganovitch, and Seán M Murray. High-throughput imaging and quantitative analysis uncovers the nature of plasmid positioning by ParABS. *eLife*, 11, 2022.

[11160] Andreas Kolbeck, Peter Marhavý, Damien De Bellis, Baohai Li, Takehiro Kamiya, Toru Fujiwara, Lothar Kalmbach, and Niko Geldner. CASP microdomain formation requires cross cell wall stabilization of domains and non-cell autonomous action of LOTR1. *eLife*, 11, 2022.

[11161] Piotr Kolesinski, Kuei-Chen Wang, Yujiro Hirose, Victor Nizet, and Partho Ghosh. An M protein coiled coil unfurls and exposes its hydrophobic core to capture LL-37. *eLife*, 11, 2022.

[11162] Polina Kosillo, Kamran M Ahmed, Erin E Aisenberg, Vasiliki Karalis, Bradley M Roberts, Stephanie J Cragg, and Helen S Bateup. Dopamine neuron morphology and output are differentially controlled by mTORC1 and mTORC2. *eLife*, 11, 2022.

[11163] Michael A Kovacs, Maureen N Cowan, Isaac W Babcock, Lydia A Sibley, Katherine Still, Samantha J Batista, Sydney A Labuzan, Ish Sethi, and Tajie H Harris. Meningeal lymphatic drainage promotes T cell responses against Toxoplasma gondii but is dispensable for parasite control in the brain. *eLife*, 11, 2022.

[11164] Amanda Kowalczyk, Maria Chikina, and Nathan Clark. Complementary evolution of coding and noncoding sequence underlies mammalian hairlessness. *eLife*, 11, 2022.

[11165] Eric Kowarz, Lea Krutzke, Marius Külp, Patrick Streb, Patrizia Larghero, Jennifer Reis, Silvia Bracharz, Tatjana Engler, Stefan Kochanek, and Rolf Marschalek. Vaccine-induced COVID-19 mimicry syndrome. *eLife*, 11, 2022.

[11166] Mateusz Kozak and Marko Kaksonen. Condensation of Ede1 promotes the initiation of endocytosis. *eLife*, 11, 2022.

[11167] Nataliia Kozhemiako, Jun Wang, Chenguang Jiang, Lei A Wang, Guanchen Gai, Kai Zou, Zhe Wang, Xiaoman Yu, Lin Zhou, Shen Li, Zhenglin Guo, Robert Law, James Coleman, Dimitrios Mylonas, Lu Shen, Guoqiang Wang, Shuping Tan, Shengying Qin, Hailiang Huang, Michael Murphy, Robert Stickgold, Dara Manoach, Zhenhe Zhou, Wei Zhu, Mei-Hua Hal, Shaun M Purcell, and Jen Q Pan. Non-rapid eye movement sleep and wake neurophysiology in schizophrenia. *eLife*, 11, 2022.

[11168] Linda K Krasniewski, Papiya Chakraborty, Chang-Yi Cui, Krystyna Mazan-Mamczarz, Christopher Dunn, Yulan Piao, Jinshui Fan, Changyou Shi, Tonya Wallace, Cuong Nguyen, Isabelle A Rathbun, Rachel Munk, Dimitrios Tsitsipatis, Supriyo De, Payel Sen, Luigi Ferrucci, and Myriam Gorospe. Single-cell analysis of skeletal muscle macrophages reveals age-associated functional subpopulations. *eLife*, 11, 2022.

[11169] Robert Krause, Jumari Snyman, Hwa Shi-Hsia, Daniel Muema, Farina Karim, Yashica Ganga, Abigail Ngoepe, Yenzekile Zungu, Inbal Gazy, Mallory Bernstein, Khadija Khan, Matilda Mazibuko, Ntombifuthi Mthabela, Dirhona Ramjit, Moherndran Archary, Oliver Limbo, Joseph Jardine, Devin Sok, Ian A Wilson, Willem Hanekom, Alex Sigal, Henrik Kløverpris, Thumbi Ndung'u, and Alasdair Leslie. HIV skews the SARS-CoV-2 B cell response towards an extrafollicular maturation pathway. *eLife*, 11, 2022.

[11170] Henrik Krehenwinkel, Sven Weber, Rieke Broekmann, Anja Melcher, Julian Hans, Rüdiger Wolf, Axel Hochkirch, Susan Rachel Kennedy, Jan Koschorreck, Sven Künzel, Christoph Müller, Rebecca Retzlaff, Diana Teubner, Sonja Schanzer, Roland Klein, Martin Paulus, Thomas Udelhoven, and Michael Veith. Environmental DNA from archived leaves reveals widespread temporal turnover and biotic homogenization in forest arthropod communities. *eLife*, 11, 2022.

[11171] Philip HC Kremer, Bart Ferwerda, Hester J Bootsma, Nienke Y Rots, Alienke J Wijmenga-Monsuur, Elisabeth AM Sanders, Krzysztof Trzciński, Anne L Wyllie, Paul Turner, Arie van der Ende, Matthijs C Brouwer, Stephen D Bentley, Diederik van de Beek, and John A Lees. Pneumococcal genetic variability in age-dependent bacterial carriage. *eLife*, 11, 2022.

[11172] Lenno Krenning, Stijn Sonneveld, and Marvin E Tanenbaum. Time-resolved single-cell sequencing identifies multiple waves of mRNA decay during the mitosis-to-G1 phase transition. *eLife*, 11, 2022.

[11173] Elena Kreutzer, Walter Senn, and Mihai A Petrovici. Natural-gradient learning for spiking neurons. *eLife*, 11, 2022.

[11174] Saloni Krishnan, Gabriel J Cler, Harriet J Smith, Hanna E Willis, Salomi S Asaridou, Máiréad P Healy, Daniel Papp, and Kate E Watkins. Quantitative MRI reveals differences in striatal myelin in children with DLD. *eLife*, 11, 2022.

[11175] Vivek Krishnan, Sher Ali, Albert L Gonzales, Pratish Thakore, Caoimhin S Griffin, Evan Yamasaki, Michael G Alvarado, Martin T Johnson, Mohamed Trebak, and Scott Earley. STIM1-dependent peripheral coupling governs the contractility of vascular smooth muscle cells. *eLife*, 11, 2022.

[11176] Emil Laust Kristoffersen, Matthew Burman, Agnes Noy, and Philipp Holliger. Rolling circle RNA synthesis catalyzed by RNA. *eLife*, 11, 2022.

[11177] Virginia Kroef, Sabine Ruegenberg, Moritz Horn, Kira Allmeroth, Lena Ebert, Seyma Bozkus, Stephan Miethe, Ulrich Elling, Bernhard Schermer, Ulrich Baumann, and Martin Sebastian Denzel. GFPT2/GFAT2 and AMDHD2 act in tandem to control the hexosamine pathway. *eLife*, 11, 2022.

[11178] Marius Kröger, Jörg Scheffel, Evgeny A Shirshin, Johannes Schleusener, Martina C Meinke, Jürgen Lademann, Marcus Maurer, and Maxim E Darvin. Label-free imaging of M1 and M2 macrophage phenotypes in the human dermis in vivo using two-photon excited FLIM. *eLife*, 11, 2022.

[11179] Logesvaran Krshnan, Wingyan Skyla Siu, Michael Van de Weijer, Daniel Hayward, Elena Navarro Guerrero, Ulrike Gruneberg, and Pedro Carvalho. Regulated degradation of the inner nuclear membrane protein SUN2 maintains nuclear envelope architecture and function. *eLife*, 11, 2022.

[11180] Stefan Kruse and Ulrike Herzschuh. Regional opportunities for tundra conservation in the next 1000 years. *eLife*, 11, 2022.

[11181] Lea Krutzke, Reinhild Rösler, Ellen Allmendinger, Tatjana Engler, Sebastian Wiese, and Stefan Kochanek. Process- and product-related impurities in the ChAdOx1 nCov-19 vaccine. *eLife*, 11, 2022.

[11182] Iga Kucharska, Lamia Hossain, Danton Ivanochko, Qiren Yang, John L Rubinstein, Régis Pomès, and Jean-Philippe Julien. Structural basis of Plasmodium vivax inhibition by antibodies binding to the circumsporozoite protein repeats. *eLife*, 11, 2022.

[11183] Kevin S Kuchinski, Kara D Loos, Danae M Suchan, Jennifer N Russell, Ashton N Sies, Charles Kumakamba, Francisca Muyembe, Placide Mbala Kingebeni, Ipos Ngay Lukusa, Frida N'Kawa, Joseph Atibu Losoma, Maria Makuwa, Amethyst Gillis, Matthew LeBreton, James A Ayukekbong, Nicole A Lerminiaux, Corina Monagin, Damien O Joly, Karen Saylors, Nathan D Wolfe, Edward M Rubin, Jean J Muyembe Tamfum, Natalie A Prystajecky, David J McIver, Christian E Lange, and Andrew DS Cameron. Targeted genomic sequencing with probe capture for discovery and surveillance of coronaviruses in bats. *eLife*, 11, 2022.

[11184] Cansu Küey, Méghane Sittewelle, Gabrielle Larocque, Miguel Hernández-González, and Stephen J Royle. Recruitment of clathrin to intracellular membranes is sufficient for vesicle formation. *eLife*, 11, 2022.

[11185] Nahoko Kuga, Reimi Abe, Kotomi Takano, Yuji Ikegaya, and Takuya Sasaki. Prefrontal-amygdalar oscillations related to social behavior in mice. *eLife*, 11, 2022.

[11186] Jayashree Kumar, Lela Lackey, Justin M Waldern, Abhishek Dey, Anthony M Mustoe, Kevin M Weeks, David H Mathews, and Alain Laederach. Quantitative prediction of variant effects on alternative splicing in MAPT using endogenous pre-messenger RNA structure probing. *eLife*, 11, 2022.

[11187] Vineet Kumar, Rajesh Kumar Mishra, Debarghya Ghose, Arunima Kalita, Pulkit Dhiman, Anand Prakash, Nirja Thakur, Gopa Mitra, Vinod D Chaudhari, Amit Arora, and Dipak Dutta. Free spermidine evokes superoxide radicals that manifest toxicity. *eLife*, 11, 2022.

[11188] Vibha Kumra Ahnlide, Johannes Kumra Ahnlide, Sebastian Wrighton, Jason P Beech, and Pontus Nordenfelt. Nanoscale binding site localization by molecular distance estimation on native cell surfaces using topological image averaging. *eLife*, 11, 2022.

[11189] Charlotte Kunze, Pepijn Luijckx, Andrew L Jackson, and Ian Donohue. Alternate patterns of temperature variation bring about very different disease outcomes at different mean temperatures. *eLife*, 11, 2022.

[11190] Andrew Kuo, Antonio Checa, Colin Niaudet, Bongnam Jung, Zhongjie Fu, Craig E Wheelock, Sasha A Singh, Masanori Aikawa, Lois E Smith, Richard L Proia, and Timothy Hla. Murine endothelial serine palmitoyltransferase 1 (SPTLC1) is required for vascular development and systemic sphingolipid homeostasis. *eLife*, 11, 2022.

[11191] Jan W Kurzawski, Claudia Lunghi, Laura Biagi, Michela Tosetti, Maria Concetta Morrone, and Paola Binda. Short-term plasticity in the human visual thalamus. *eLife*, 11, 2022.

[11192] Jill T Kuwabara, Akitoshi Hara, Sumit Bhutada, Greg S Gojanovich, Jasmine Chen, Kanani Hokutan, Vikram Shettigar, Anson Y Lee, Lydia P DeAngelo, Jack R Heckl, Julia R Jahansooz, Dillon K Tacdol, Mark T Ziolo, Suneel S Apte, and Michelle D Tallquist. Consequences of PDGFR$\alpha$+ fibroblast reduction in adult murine hearts. *eLife*, 11, 2022.

[11193] Alejandro La Greca, Nicolás Bellora, FranÇois Le Dily, Rodrigo Jara, Ana Silvina Nacht, Javier Quilez Oliete, José Luis Villanueva, Enrique Vidal, Gabriela Merino, Cristóbal Fresno, Inti Tarifa Reischle, Griselda Vallejo, Guillermo Vicent, Elmer Fernández, Miguel Beato, and Patricia Saragüeta. Chromatin topology defines estradiol-primed progesterone receptor and PAX2 binding in endometrial cancer cells. *eLife*, 11, 2022.

[11194] Danilo La Terra, Ann-Sofie Bjerre, Marius Rosier, Rei Masuda, Tomás J Ryan, and Lucy M Palmer. The role of higher-order thalamus during learning and correct performance in goal-directed behavior. *eLife*, 11, 2022.

[11195] Mimi Q La-Vu, Ekayana Sethi, Sandra Maesta-Pereira, Peter J Schuette, Brooke C Tobias, Fernando MCV Reis, Weisheng Wang, Anita Torossian, Amy Bishop, Saskia J Leonard, Lilly Lin, Catherine M Cahill, and Avishek Adhikari. Sparse genetically defined neurons refine the canonical role of periaqueductal gray columnar organization. *eLife*, 11, 2022.

[11196] Jérôme Lacoste, Hédi Soula, Angélique Burg, Agnès Audibert, Pénélope Darnat, Michel Gho, and Sophie Louvet-Vallée. A neural progenitor mitotic wave is required for asynchronous axon outgrowth and morphology. *eLife*, 11, 2022.

[11197] Rebecca LaCroix, Benjamin Lin, Tae-Yun Kang, and Andre Levchenko. Complex effects of kinase localization revealed by compartment-specific regulation of protein kinase A activity. *eLife*, 11, 2022.

[11198] Mato Lagator, Srdjan Sarikas, Magdalena Steinrueck, David Toledo-Aparicio, Jonathan P Bollback, Calin C Guet, and GaŠper Tkačik. Predicting bacterial promoter function and evolution from random sequences. *eLife*, 11, 2022.

[11199] Jason KH Lai, Pearlyn JY Toh, Hamizah A Cognart, Geetika Chouhan, and Timothy E Saunders. DNA-damage induced cell death in yap1;wwtr1 mutant epidermal basal cells. *eLife*, 11, 2022.

[11200] Alexander Lalejini, Emily Dolson, Anya E Vostinar, and Luis Zaman. Artificial selection methods from evolutionary computing show promise for directed evolution of microbes. *eLife*, 11, 2022.

[11201] Paul Lambey, Omolade Otun, Xiaojing Cong, FranÇois Hoh, Luc Brunel, Pascal Verdié, Claire M Grison, Fanny Peysson, Sylvain Jeannot, Thierry Durroux, Cherine Bechara, Sébastien Granier, and Cédric Leyrat. Structural insights into recognition of chemokine receptors by Staphylococcus aureus leukotoxins. *eLife*, 11, 2022.

[11202] Colin LaMont, Jakub Otwinowski, Kanika Vanshylla, Henning Gruell, Florian Klein, and Armita Nourmohammad. Design of an optimal combination therapy with broadly neutralizing antibodies to suppress HIV-1. *eLife*, 11, 2022.

[11203] Amel Lamri, Jayneel Limbachia, Karleen M Schulze, Dipika Desai, Brian Kelly, Russell J de Souza, Guillaume Paré, Deborah A Lawlor, John Wright, and Sonia S Anand. The genetic risk of gestational diabetes in South Asian women. *eLife*, 11, 2022.

[11204] Andrew T Landau, Pojeong Park, J David Wong-Campos, He Tian, Adam E Cohen, and Bernardo L Sabatini. Dendritic branch structure compartmentalizes voltage-dependent calcium influx in cortical layer 2/3 pyramidal cells. *eLife*, 11, 2022.

[11205] Anna M Langmüller, Jackson Champer, Sandra Lapinska, Lin Xie, Matthew Metzloff, Samuel E Champer, Jingxian Liu, Yineng Xu, Jie Du, Andrew G Clark, and Philipp W Messer. Fitness effects of CRISPR endonucleases in Drosophila melanogaster populations. *eLife*, 11, 2022.

[11206] Emma Laporte, Florian Hermans, Silke De Vriendt, Annelies Vennekens, Diether Lambrechts, Charlotte Nys, Benoit Cox, and Hugo Vankelecom. Decoding the activated stem cell phenotype of the neonatally maturing pituitary. *eLife*, 11, 2022.

[11207] Valérie Laroute, Catherine Beaufrand, Pedro Gomes, Sébastien Nouaille, Valérie Tondereau, Marie-Line Daveran-Mingot, Vassilia Theodorou, Hélène Eutamene, Muriel Mercier-Bonin, and Muriel Cocaign-Bousquet. Lactococcus lactis NCDO2118 exerts visceral antinociceptive properties in rat via GABA production in the gastro-intestinal tract. *eLife*, 11, 2022.

[11208] Therese LaRue, Heike Lindner, Ankit Srinivas, Moises Exposito-Alonso, Guillaume Lobet, and José R Dinneny. Uncovering natural variation in root system architecture and growth dynamics using a robotics-assisted phenomics platform. *eLife*, 11, 2022.

[11209] Elise Laruelle, Katia Belcram, Alain Trubuil, Jean-Christophe Palauqui, and Philippe Andrey. Large-scale analysis and computer modeling reveal hidden regularities behind variability of cell division patterns in Arabidopsis thaliana embryogenesis. *eLife*, 11, 2022.

[11210] Davis Laundon, Nathan Chrismas, Kimberley Bird, Seth Thomas, Thomas Mock, and Michael Cunliffe. A cellular and molecular atlas reveals the basis of chytrid development. *eLife*, 11, 2022.

[11211] Timothée Laurent, Pravin Kumar, Susanne Liese, Farnaz Zare, Mattias Jonasson, Andreas Carlson, and Lars-Anders Carlson. Architecture of the chikungunya virus replication organelle. *eLife*, 11, 2022.

[11212] Matthew D Lauver, Ge Jin, Katelyn N Ayers, Sarah N Carey, Charles S Specht, Catherine S Abendroth, and Aron E Lukacher. T cell deficiency precipitates antibody evasion and emergence of neurovirulent polyomavirus. *eLife*, 11, 2022.

[11213] Alyssa J Lawler, Easwaran Ramamurthy, Ashley R Brown, Naomi Shin, Yeonju Kim, Noelle Toong, Irene M Kaplow, Morgan Wirthlin, Xiaoyu Zhang, BaDoi N Phan, Grant A Fox, Kirsten Wade, Jing He, Bilge Esin Ozturk, Leah C Byrne, William R Stauffer, Kenneth N Fish, and Andreas R Pfenning. Machine learning sequence prioritization for cell type-specific enhancer design. *eLife*, 11, 2022.

[11214] Charlotte Lawrenson, Elena Paci, Jasmine Pickford, Robert AR Drake, Bridget M Lumb, and Richard Apps. Cerebellar modulation of memory encoding in the periaqueductal grey and fear behaviour. *eLife*, 11, 2022.

[11215] Campbell D Lawson, Samantha Peel, Asier Jayo, Adam Corrigan, Preeti Iyer, Mabel Baxter Dalrymple, Richard J Marsh, Susan Cox, Isabel Van Audenhove, Jan Gettemans, and Maddy Parsons. Nuclear fascin regulates cancer cell survival. *eLife*, 11, 2022.

[11216] özgün Le Roux, Nicole LK Pershing, Erin Kaltenbrun, Nicole J Newman, Jeffrey I Everitt, Elisa Baldelli, Mariaelena Pierobon, Emanuel F Petricoin, and Christopher M Counter. Genetically manipulating endogenous Kras levels and oncogenic mutations in vivo influences tissue patterning of murine tumorigenesis. *eLife*, 11, 2022.

[11217] Megan Leander, Zhuang Liu, Qiang Cui, and Srivatsan Raman. Deep mutational scanning and machine learning reveal structural and molecular rules governing allosteric hotspots in homologous proteins. *eLife*, 11, 2022.

[11218] Alexis Lebecq, Mehdi Doumane, Aurelie Fangain, Vincent Bayle, Jia Xuan Leong, Frédérique Rozier, Maria del Marques-Bueno, Laia Armengot, Romain Boisseau, Mathilde Laetitia Simon, Mirita Franz-Wachtel, Boris Macek, Suayib üstün, Yvon Jaillais, and Marie-Cécile Caillaud. The Arabidopsis SAC9 enzyme is enriched in a cortical population of early endosomes and restricts PI(4,5)P2 at the plasma membrane. *eLife*, 11, 2022.

[11219] Karl F Lechtreck, Yi Liu, Jin Dai, Rama A Alkhofash, Jack Butler, Lea Alford, and Pinfen Yang. Chlamydomonas ARMC2/PF27 is an obligate cargo adapter for intraflagellar transport of radial spokes. *eLife*, 11, 2022.

[11220] Byron Lee, Nima Jaberi-Lashkari, and Eliezer Calo. A unified view of low complexity regions (LCRs) across species. *eLife*, 11, 2022.

[11221] Candice Lee, Emerson F Harkin, Xuming Yin, Richard Naud, and Simon Chen. Cell-type-specific responses to associative learning in the primary motor cortex. *eLife*, 11, 2022.

[11222] Dongjin R Lee, Christopher Rhodes, Apratim Mitra, Yajun Zhang, Dragan Maric, Ryan K Dale, and Timothy J Petros. Transcriptional heterogeneity of ventricular zone cells in the ganglionic eminences of the mouse forebrain. *eLife*, 11, 2022.

[11223] Hongmi Lee and Janice Chen. A generalized cortical activity pattern at internally generated mental context boundaries during unguided narrative recall. *eLife*, 11, 2022.

[11224] Jae-Ik Lee, Richard Seist, Stephen McInturff, Daniel J Lee, M Christian Brown, Konstantina M Stankovic, and Shelley Fried. Magnetic stimulation allows focal activation of the mouse cochlea. *eLife*, 11, 2022.

[11225] Jae-Joong Lee, Sungwoo Lee, Dong Hee Lee, and Choong-Wan Woo. Functional brain reconfiguration during sustained pain. *eLife*, 11, 2022.

[11226] Jeffrey Y Lee, Peter AC Wing, Dalia S Gala, Marko Noerenberg, Aino I Järvelin, Joshua Titlow, Xiaodong Zhuang, Natasha Palmalux, Louisa Iselin, Mary Kay Thompson, Richard M Parton, Maria Prange-Barczynska, Alan Wainman, Francisco J Salguero, Tammie Bishop, Daniel Agranoff, William James, Alfredo Castello, Jane A McKeating, and Ilan Davis. Absolute quantitation of individual SARS-CoV-2 RNA molecules provides a new paradigm for infection dynamics and variant differences. *eLife*, 11, 2022.

[11227] Mark S Lee, Peter J Tuohy, Caleb Y Kim, Katrina Lichauco, Heather L Parrish, Koenraad Van Doorslaer, and Michael S Kuhns. Enhancing and inhibitory motifs regulate CD4 activity. *eLife*, 11, 2022.

[11228] Seok Hee Lee, Xiaowei Liu, David Jimenez-Morales, and Paolo F Rinaudo. Murine blastocysts generated by in vitro fertilization show increased Warburg metabolism and altered lactate production. *eLife*, 11, 2022.

[11229] Joël Lemière, Paula Real-Calderon, Liam J Holt, Thomas G Fai, and Fred Chang. Control of nuclear size by osmotic forces in Schizosaccharomyces pombe. *eLife*, 11, 2022.

[11230] Susanna Lemmelä, Eleanor M Wigmore, Christian Benner, Aki S Havulinna, Rachel MY Ong, Tibor Kempf, Kai C Wollert, Stefan Blankenberg, Tanja Zeller, James E Peters, Veikko Salomaa, Maria Fritsch, Ruth March, Aarno Palotie, Mark Daly, Adam S Butterworth, Mervi Kinnunen, Dirk S Paul, and Athena Matakidou. Integrated analyses of growth differentiation factor-15 concentration and cardiometabolic diseases in humans. *eLife*, 11, 2022.

[11231] Sarah Lensch, Michael H Herschl, Connor H Ludwig, Joydeb Sinha, Michaela M Hinks, Adi Mukund, Taihei Fujimori, and Lacramioara Bintu. Dynamic spreading of chromatin-mediated gene silencing and reactivation between neighboring genes in single cells. *eLife*, 11, 2022.

[11232] Carrie E Leonard, Jolie Quiros, Frances Lefcort, and Lisa A Taneyhill. Loss of Elp1 disrupts trigeminal ganglion neurodevelopment in a model of familial dysautonomia. *eLife*, 11, 2022.

[11233] Manuel Lera-Ramirez, FranÇois J Nédélec, and Phong T Tran. Microtubule rescue at midzone edges promotes overlap stability and prevents spindle collapse during anaphase B. *eLife*, 11, 2022.

[11234] Xavier Leray, Jacob K Hilton, Kamsi Nwangwu, Alissa Becerril, Vedrana Mikusevic, Gabriel Fitzgerald, Anowarul Amin, Mary R Weston, and Joseph A Mindell. Tonic inhibition of the chloride/proton antiporter ClC-7 by PI(3,5)P2 is crucial for lysosomal pH maintenance. *eLife*, 11, 2022.

[11235] Clarisse Leseigneur, Laurent Boucontet, Magalie Duchateau, Javier Pizarro-Cerda, Mariette Matondo, Emma Colucci-Guyon, and Olivier Dussurget. NAD kinase promotes Staphylococcus aureus pathogenesis by supporting production of virulence factors and protective enzymes. *eLife*, 11, 2022.

[11236] Alexandria MH Lesicko, Christopher F Angeloni, Jennifer M Blackwell, Mariella De Biasi, and Maria N Geffen. Corticofugal regulation of predictive coding. *eLife*, 11, 2022.

[11237] Nicholas P Lesner, Xun Wang, Zhenkang Chen, Anderson Frank, Cameron J Menezes, Sara House, Spencer D Shelton, Andrew Lemoff, David G McFadden, Janaka Wansapura, Ralph J DeBerardinis, and Prashant Mishra. Differential requirements for mitochondrial electron transport chain components in the adult murine liver. *eLife*, 11, 2022.

[11238] Jake Leyhr, Laura Waldmann, Beata Filipek-Górniok, Hanqing Zhang, Amin Allalou, and Tatjana Haitina. A novel cis-regulatory element drives early expression of Nkx3.2 in the gnathostome primary jaw joint. *eLife*, 11, 2022.

[11239] Bochuan Li, Ting Zhang, Mengxia Liu, Zhen Cui, Yanhong Zhang, Mingming Liu, Yanan Liu, Yongqiao Sun, Mengqi Li, Yikui Tian, Ying Yang, Hongfeng Jiang, and Degang Liang. RNA N6-methyladenosine modulates endothelial atherogenic responses to disturbed flow in mice. *eLife*, 11, 2022.

[11240] Chong Li, Marie R Culhane, Declan C Schroeder, Maxim C-J Cheeran, Lucina Galina Pantoja, Micah L Jansen, and Montserrat Torremorell. Vaccination decreases the risk of influenza A virus reassortment but not genetic variation in pigs. *eLife*, 11, 2022.

[11241] Deng-Fei Li, Wen-Long Han, Susanne S Renner, and Shuang-Quan Huang. Touch-sensitive stamens enhance pollen dispersal by scaring away visitors. *eLife*, 11, 2022.

[11242] Enjie Li, Mingyue Xia, Yu Du, Kaili Long, Feng Ji, Feiyan Pan, Lingfeng He, Zhigang Hu, and Zhigang Guo. METTL3 promotes homologous recombination repair and modulates chemotherapeutic response in breast cancer by regulating the EGF/RAD51 axis. *eLife*, 11, 2022.

[11243] Hanqin Li, Oriol Busquets, Yogendra Verma, Khaja Mohieddin Syed, Nitzan Kutnowski, Gabriella R Pangilinan, Luke A Gilbert, Helen S Bateup, Donald C Rio, Dirk Hockemeyer, and Frank Soldner. Highly efficient generation of isogenic pluripotent stem cell models using prime editing. *eLife*, 11, 2022.

[11244] Jiaqian Li, Di Huang, Bingxi Lei, Jingying Huang, Linbing Yang, Man Nie, Shicheng Su, Qiyi Zhao, and Ying Wang. VLA-4 suppression by senescence signals regulates meningeal immunity and leptomeningeal metastasis. *eLife*, 11, 2022.

[11245] Jie Li, Jiayi Wu, Catherine Hall, Xiao-chen Bai, and Eunhee Choi. Molecular basis for the role of disulfide-linked $\alpha$CTs in the activation of insulin-like growth factor 1 receptor and insulin receptor. *eLife*, 11, 2022.

[11246] Junhao Li, Antonio Pinto-Duarte, Mark Zander, Michael S Cuoco, Chi-Yu Lai, Julia Osteen, Linjing Fang, Chongyuan Luo, Jacinta D Lucero, Rosa Gomez-Castanon, Joseph R Nery, Isai Silva-Garcia, Yan Pang, Terrence J Sejnowski, Susan B Powell, Joseph R Ecker, Eran A Mukamel, and M Margarita Behrens. Dnmt3a knockout in excitatory neurons impairs postnatal synapse maturation and increases the repressive histone modification H3K27me3. *eLife*, 11, 2022.

[11247] Miaomiao Li, Chih Yean Ong, Christophe J Langouët-Astrié, Lisi Tan, Ashwni Verma, Yimu Yang, Xiaoxiao Zhang, Dhaval K Shah, Eric P Schmidt, and Ding Xu. Heparan sulfate-dependent RAGE oligomerization is indispensable for pathophysiological functions of RAGE. *eLife*, 11, 2022.

[11248] Mingyang Li, Tingting Liu, Xinyi Xu, Qingqing Wen, Zhiyong Zhao, Xixi Dang, Yi Zhang, and Dan Wu. Development of visual cortex in human neonates is selectively modified by postnatal experience. *eLife*, 11, 2022.

[11249] Shasha Li, Michael D Bern, Benpeng Miao, Changxu Fan, Xiaoyun Xing, Takeshi Inoue, Sytse J Piersma, Ting Wang, Marco Colonna, Tomohiro Kurosaki, and Wayne M Yokoyama. The transcription factor Bach2 negatively regulates murine natural killer cell maturation and function. *eLife*, 11, 2022.

[11250] Tai-De Li, Peter Bieling, Julian Weichsel, R Dyche Mullins, and Daniel A Fletcher. The molecular mechanism of load adaptation by branched actin networks. *eLife*, 11, 2022.

[11251] Tianzhi Li, Qiqi Cheng, Shen Wang, and Cong Ma. Rabphilin 3A binds the N-peptide of SNAP-25 to promote SNARE complex assembly in exocytosis. *eLife*, 11, 2022.

[11252] Wenyang Li, Jennifer Y Chen, Cheng Sun, Robert P Sparks, Lorena Pantano, Raza-Ur Rahman, Sean P Moran, Joshua V Pondick, Rory Kirchner, David Wrobel, Michael Bieler, Achim Sauer, Shannan J Ho Sui, Julia F Doerner, Jörg F Rippmann, and Alan C Mullen. Nanchangmycin regulates FYN, PTK2, and MAPK1/3 to control the fibrotic activity of human hepatic stellate cells. *eLife*, 11, 2022.

[11253] Xiangyi Li, Lin Jiang, Chao Xue, Mulin Jun Li, and Miaoxin Li. A conditional gene-based association framework integrating isoform-level eQTL data reveals new susceptibility genes for schizophrenia. *eLife*, 11, 2022.

[11254] Xiaodong Li, Patrick J Gordon, John A Gaynes, Alexandra W Fuller, Randy Ringuette, Clayton P Santiago, Valerie Wallace, Seth Blackshaw, Pulin Li, and Edward M Levine. Lhx2 is a progenitor-intrinsic modulator of Sonic Hedgehog signaling during early retinal neurogenesis. *eLife*, 11, 2022.

[11255] Xin Li, Noor Singh, Camille Miller, India Washington, Bintou Sosseh, and Kacy Lynn Gordon. The C. elegans gonadal sheath Sh1 cells extend asymmetrically over a differentiating germ cell population in the proliferative zone. *eLife*, 11, 2022.

[11256] Xinfeng Li, Fang Chen, Xiaoyu Liu, Jinfeng Xiao, Binda T Andongma, Qing Tang, Xiaojian Cao, Shan-Ho Chou, Michael Y Galperin, and Jin He. Clp protease and antisense RNA jointly regulate the global regulator CarD to mediate mycobacterial starvation response. *eLife*, 11, 2022.

[11257] Xinxuan Li, Yajing Zhou, Shuai Yuan, Xuan Zhou, Lijuan Wang, Jing Sun, Lili Yu, Jinghan Zhu, Han Zhang, Nan Yang, Shuhui Dai, Peige Song, Susanna C Larsson, Evropi Theodoratou, Yimin Zhu, and Xue Li. Genetically predicted high IGF-1 levels showed protective effects on COVID-19 susceptibility and hospitalization: a Mendelian randomisation study with data from 60 studies across 25 countries. *eLife*, 11, 2022.

[11258] Yang Li, Yan Wu, Elena A Federzoni, Xiaodan Wang, Andre Dharmawan, Xiaoyi Hu, Hui Wang, Robert J Hawley, Sean Stevens, Megan Sykes, and Yong-Guang Yang. CD47 cross-dressing by extracellular vesicles expressing CD47 inhibits phagocytosis without transmitting cell death signals. *eLife*, 11, 2022.

[11259] Ye Li, Shiqi Liu, Yingdan Zhang, Zi Jing Seng, Haoran Xu, Liang Yang, and Yilin Wu. Self-organized canals enable long-range directed material transport in bacterial communities. *eLife*, 11, 2022.

[11260] Yitong Li, Vijaya Kumar Balakrishnan, Michael Rowse, Cheng-Guo Wu, Anastasia Phoebe Bravos, Vikash K Yadav, Ylva Ivarsson, Stefan Strack, Irina V Novikova, and Yongna Xing. Coupling to short linear motifs creates versatile PME-1 activities in PP2A holoenzyme demethylation and inhibition. *eLife*, 11, 2022.

[11261] Ziru Li, Emily Bowers, Junxiong Zhu, Hui Yu, Julie Hardij, Devika P Bagchi, Hiroyuki Mori, Kenneth T Lewis, Katrina Granger, Rebecca L Schill, Steven M Romanelli, Simin Abrishami, Kurt D Hankenson, Kanakadurga Singer, Clifford J Rosen, and Ormond A MacDougald. Lipolysis of bone marrow adipocytes is required to fuel bone and the marrow niche during energy deficits. *eLife*, 11, 2022.

[11262] Jie Lian, Jiajun Deng, Edward S Hui, Mohamad Koohi-Moghadam, Yunlang She, Chang Chen, and Varut Vardhanabhuti. Early stage NSCLS patients' prognostic prediction with multi-information using transformer and graph neural network model. *eLife*, 11, 2022.

[11263] Zifei Liang, Choong H Lee, Tanzil M Arefin, Zijun Dong, Piotr Walczak, Song-Hai Shi, Florian Knoll, Yulin Ge, Leslie Ying, and Jiangyang Zhang. Virtual mouse brain histology from multi-contrast MRI via deep learning. *eLife*, 11, 2022.

[11264] Brandon Wey-Hung Liauw, Arash Foroutan, Michael R Schamber, Weifeng Lu, Hamid Samareh Afsari, and Reza Vafabakhsh. Conformational fingerprinting of allosteric modulators in metabotropic glutamate receptor 2. *eLife*, 11, 2022.

[11265] Linda H Lidborg, Catharine Penelope Cross, and Lynda G Boothroyd. A meta-analysis of the association between male dimorphism and fitness outcomes in humans. *eLife*, 11, 2022.

[11266] Joshua L Lillvis, Hideo Otsuna, Xiaoyu Ding, Igor Pisarev, Takashi Kawase, Jennifer Colonell, Konrad Rokicki, Cristian Goina, Ruixuan Gao, Amy Hu, Kaiyu Wang, John Bogovic, Daniel E Milkie, Linus Meienberg, Brett D Mensh, Edward S Boyden, Stephan Saalfeld, Paul W Tillberg, and Barry J Dickson. Rapid reconstruction of neural circuits using tissue expansion and light sheet microscopy. *eLife*, 11, 2022.

[11267] Jennifer M Lin, Tyler A Mitchell, Megan Rothstein, Alison Pehl, Ed Zandro M Taroc, Raghu R Katreddi, Katherine E Parra, Damian G Zuloaga, Marcos Simoes-Costa, and Paolo Emanuele Forni. Sociosexual behavior requires both activating and repressive roles of Tfap2e/AP-2$\epsilon$ in vomeronasal sensory neurons. *eLife*, 11, 2022.

[11268] Yu-Ling Lin, Zhu-Sen Yang, Wai-Yi Wong, Shih-Che Lin, Shuu-Jiun Wang, Shih-Pin Chen, Jen-Kun Cheng, Hui Lu, and Cheng-Chang Lien. Cellular mechanisms underlying central sensitization in a mouse model of chronic muscle pain. *eLife*, 11, 2022.

[11269] Jaime Lisack, Brooke Morriswood, and Markus Engstler. Response to comment on 'Unexpected plasticity in the life cycle of Trypanosoma Brucei'. *eLife*, 11, 2022.

[11270] Fang Liu, Sekhar Kambakam, Maira P Almeida, Zhitao Ming, Jordan M Welker, Wesley A Wierson, Laura E Schultz-Rogers, Stephen C Ekker, Karl J Clark, Jeffrey J Essner, and Maura McGrail. Cre/lox regulated conditional rescue and inactivation with zebrafish UFlip alleles generated by CRISPR-Cas9 targeted integration. *eLife*, 11, 2022.

[11271] Li Liu, Steve Dodd, Ryan D Hunt, Nikorn Pothayee, Tatjana Atanasijevic, Nadia Bouraoud, Dragan Maric, E Ashley Moseman, Selamawit Gossa, Dorian B McGavern, and Alan P Koretsky. Early detection of cerebrovascular pathology and protective antiviral immunity by MRI. *eLife*, 11, 2022.

[11272] Ming Liu, Stuart Andrew West, and Geoff Wild. The evolution of manipulative cheating. *eLife*, 11, 2022.

[11273] Qiangqiang Liu, Qian Luo, Jianyu Feng, Yanping Zhao, Biao Ma, Hongcheng Cheng, Tian Zhao, Hong Lei, Chenglong Mu, Linbo Chen, Yuanyuan Meng, Jiaojiao Zhang, Yijia Long, Jingyi Su, Guo Chen, Yanjun Li, Gang Hu, Xudong Liao, Quan Chen, and Yushan Zhu. Hypoxia-induced proteasomal degradation of DBC1 by SIAH2 in breast cancer progression. *eLife*, 11, 2022.

[11274] Qingqing Liu, Catherine E Chang, Alexandra C Wooldredge, Benjamin Fong, Brian K Kennedy, and Chuankai Zhou. Tom70-based transcriptional regulation of mitochondrial biogenesis and aging. *eLife*, 11, 2022.

[11275] Rui Liu, Kangcheng Song, Jing-Xiang Wu, Xiao-Peng Geng, Liming Zheng, Xiaoyin Gao, Hailin Peng, and Lei Chen. Structure of human phagocyte NADPH oxidase in the resting state. *eLife*, 11, 2022.

[11276] Wenbo Liu, Xukun Lu, Zheng-Hui Zhao, Ruibao SU, Qian-Nan Li Li, Yue Xue, Zheng Gao, Si-Min Sun Sun, Wen-Long Lei, Lei Li, Geng An, Hanyan Liu, Zhiming Han, Ying-Chun Ouyang, Yi Hou, Zhen-Bo Wang, Qing-Yuan Sun, and Jianqiao Liu. SRSF10 is essential for progenitor spermatogonia expansion by regulating alternative splicing. *eLife*, 11, 2022.

[11277] Xiaojie Liu, Casey R Vickstrom, Hao Yu, Shuai Liu, Shana Terai Snarrenberg, Vladislav Friedman, Lianwei Mu, Bixuan Chen, Thomas J Kelly, David A Baker, and Qing-song Liu. Epac2 in midbrain dopamine neurons contributes to cocaine reinforcement via enhancement of dopamine release. *eLife*, 11, 2022.

[11278] Xingchen Liu, Christl Gaubitz, Joshua Pajak, and Brian A Kelch. A second DNA binding site on RFC facilitates clamp loading at gapped or nicked DNA. *eLife*, 11, 2022.

[11279] Yanan Liu, Juanjuan Feng, Kun Yuan, Zhengzhen Wu, Longmiao Hu, Yue Lu, Kun Li, Jiawei Guo, Jing Chen, Chengbin Ma, and Xiufeng Pang. The oncoprotein BCL6 enables solid tumor cells to evade genotoxic stress. *eLife*, 11, 2022.

[11280] Yupeng Liu, Siyu Le, Yi Liu, Huinan Jiang, Binye Ruan, Yufeng Huang, Xuemei Ao, Xudong Shi, Xiaoyi Fu, and Shuran Wang. The effect of calcium supplementation in people under 35 years old: A systematic review and meta-analysis of randomized controlled trials. *eLife*, 11, 2022.

[11281] Zaoqu Liu, Siyuan Weng, Qin Dang, Hui Xu, Yuqing Ren, Chunguang Guo, Zhe Xing, Zhenqiang Sun, and Xinwei Han. Gene interaction perturbation network deciphers a high-resolution taxonomy in colorectal cancer. *eLife*, 11, 2022.

[11282] Zhen Liu, Meng-Hua Wu, Qi-Xuan Wang, Shao-Zhen Lin, Xi-Qiao Feng, Bo Li, and Xin Liang. Drosophila mechanical nociceptors preferentially sense localized poking. *eLife*, 11, 2022.

[11283] Zhi Liu, Tian Chen, Sicheng Zhang, Tianfang Yang, Yun Gong, Hong-Wen Deng, Ding Bai, Weidong Tian, and YiPing Chen. Discovery and functional assessment of a novel adipocyte population driven by intracellular Wnt/$\beta$-catenin signaling in mammals. *eLife*, 11, 2022.

[11284] Zhihui Liu, Man Jiang, Kif Liakath-Ali, Alessandra Sclip, Jaewon Ko, Roger Shen Zhang, and Thomas C Südhof. Deletion of Calsyntenin-3, an atypical cadherin, suppresses inhibitory synapses but increases excitatory parallel-fiber synapses in cerebellum. *eLife*, 11, 2022.

[11285] Zhuang Liu, Letian Zhang, Maria A Toma, Dongqing Li, Xiaowei Bian, Irena Pastar, Marjana Tomic-Canic, Pehr Sommar, and Ning Xu Landén. Integrative small and long RNA omics analysis of human healing and nonhealing wounds discovers cooperating microRNAs as therapeutic targets. *eLife*, 11, 2022.

[11286] Arnau Llobet Rosell, Maria Paglione, Jonathan Gilley, Magdalena Kocia, Giulia Perillo, Massimiliano Gasparrini, Lucia Cialabrini, Nadia Raffaelli, Carlo Angeletti, Giuseppe Orsomando, Pei-Hsuan Wu, Michael P Coleman, Andrea Loreto, and Lukas Jakob Neukomm. The NAD+ precursor NMN activates dSarm to trigger axon degeneration in Drosophila. *eLife*, 11, 2022.

[11287] Andrew Loffer, Jasleen Singh, Akihito Fukudome, Vibhor Mishra, Feng Wang, and Craig S Pikaard. A DCL3 dicing code within Pol IV-RDR2 transcripts diversifies the siRNA pool guiding RNA-directed DNA methylation. *eLife*, 11, 2022.

[11288] J Carter Loftus, Roi Harel, Chase L Núñez, and Margaret C Crofoot. Ecological and social pressures interfere with homeostatic sleep regulation in the wild. *eLife*, 11, 2022.

[11289] Fidel-Nicolás Lolo, Dácil María Pavón, Araceli Grande-García, Alberto Elosegui-Artola, Valeria Inés Segatori, Sara Sánchez, Xavier Trepat, Pere Roca-Cusachs, and Miguel A del Pozo. Caveolae couple mechanical stress to integrin recycling and activation. *eLife*, 11, 2022.

[11290] Gina M LoMastro, Chelsea G Drown, Aubrey L Maryniak, Cayla E Jewett, Margaret A Strong, and Andrew Jon Holland. PLK4 drives centriole amplification and apical surface area expansion in multiciliated cells. *eLife*, 11, 2022.

[11291] Jason T Long, Abigail Leinroth, Yihan Liao, Yinshi Ren, Anthony J Mirando, Tuyet Nguyen, Wendi Guo, Deepika Sharma, Douglas Rouse, Colleen Wu, Kathryn Song Eng Cheah, Courtney M Karner, and Matthew J Hilton. Hypertrophic chondrocytes serve as a reservoir for marrow-associated skeletal stem and progenitor cells, osteoblasts, and adipocytes during skeletal development. *eLife*, 11, 2022.

[11292] Katrina M Longhini and Michael Glotzer. Aurora A and cortical flows promote polarization and cytokinesis by inducing asymmetric ECT-2 accumulation. *eLife*, 11, 2022.

[11293] Niklas Lonnemann, Shirin Hosseini, Melanie Ohm, Robert Geffers, Karsten Hiller, Charles A Dinarello, and Martin Korte. IL-37 expression reduces acute and chronic neuroinflammation and rescues cognitive impairment in an Alzheimer's disease mouse model. *eLife*, 11, 2022.

[11294] Noella Lopes, Nicolas Boucherit, Jérémy C Santamaria, Nathan Provin, Jonathan Charaix, Pierre Ferrier, Matthieu Giraud, and Magali Irla. Thymocytes trigger self-antigen-controlling pathways in immature medullary thymic epithelial stages. *eLife*, 11, 2022.

[11295] Jaime G Lopez and Ned S Wingreen. Noisy metabolism can promote microbial cross-feeding. *eLife*, 11, 2022.

[11296] Tania López-Hernández, Koh-ichiro Takenaka, Yasunori Mori, Pornparn Kongpracha, Shushi Nagamori, Volker Haucke, and Shigeo Takamori. Clathrin-independent endocytic retrieval of SV proteins mediated by the clathrin adaptor AP-2 at mammalian central synapses. *eLife*, 11, 2022.

[11297] Sophia S Loschky, Giovanna Maria Spano, William Marshall, Andrea Schroeder, Kelsey Marie Nemec, Shannon Sandra Schiereck, Luisa de Vivo, Michele Bellesi, Sebastian Weyn Banningh, Giulio Tononi, and Chiara Cirelli. Ultrastructural effects of sleep and wake on the parallel fiber synapses of the cerebellum. *eLife*, 11, 2022.

[11298] John T Lovell, Avinash Sreedasyam, M Eric Schranz, Melissa Wilson, Joseph W Carlson, Alex Harkess, David Emms, David M Goodstein, and Jeremy Schmutz. GENESPACE tracks regions of interest and gene copy number variation across multiple genomes. *eLife*, 11, 2022.

[11299] Sofia Lövestam, Fujiet Adrian Koh, Bart van Knippenberg, Abhay Kotecha, Alexey G Murzin, Michel Goedert, and Sjors HW Scheres. Assembly of recombinant tau into filaments identical to those of Alzheimer's disease and chronic traumatic encephalopathy. *eLife*, 11, 2022.

[11300] Qihong Lu, Uri Hasson, and Kenneth A Norman. A neural network model of when to retrieve and encode episodic memories. *eLife*, 11, 2022.

[11301] Wen Lu, Margot Lakonishok, Anna S Serpinskaya, and Vladimir I Gelfand. A novel mechanism of bulk cytoplasmic transport by cortical dynein in Drosophila ovary. *eLife*, 11, 2022.

[11302] Asael Lubotzky, Ilana Pelov, Ronen Teplitz, Daniel Neiman, Adama Smadja, Hai Zemmour, Sheina Piyanzin, Bracha-Lea Ochana, Kirsty L Spalding, Benjamin Glaser, Ruth Shemer, Yuval Dor, and Yoav Kohn. Elevated brain-derived cell-free DNA among patients with first psychotic episode – a proof-of-concept study. *eLife*, 11, 2022.

[11303] Bronwyn A Lucas, Kexin Zhang, Sarah Loerch, and Nikolaus Grigorieff. In situ single particle classification reveals distinct 60S maturation intermediates in cells. *eLife*, 11, 2022.

[11304] Daria Lukasz, Alisha Beirl, and Katie Kindt. Chronic neurotransmission increases the susceptibility of lateral-line hair cells to ototoxic insults. *eLife*, 11, 2022.

[11305] Andrea J Lund, Kristen J Wade, Zachary L Nikolakis, Kathleen N Ivey, Blair W Perry, Hamish NC Pike, Sara H Paull, Yang Liu, Todd A Castoe, David D Pollock, and Elizabeth J Carlton. Integrating genomic and epidemiologic data to accelerate progress toward schistosomiasis elimination. *eLife*, 11, 2022.

[11306] Mimmi LE Lundahl, Morgane Mitermite, Dylan Gerard Ryan, Sarah Case, Niamh C Williams, Ming Yang, Roisin I Lynch, Eimear Lagan, Filipa M Lebre, Aoife L Gorman, Bojan Stojkovic, Adrian P Bracken, Christian Frezza, Frederick J Sheedy, Eoin M Scanlan, Luke AJ O'Neill, Stephen V Gordon, and Ed C Lavelle. Macrophage innate training induced by IL-4 and IL-13 activation enhances OXPHOS driven anti-mycobacterial responses. *eLife*, 11, 2022.

[11307] Elias T Lunsford, Alexandra Paz, Alex C Keene, and James C Liao. Evolutionary convergence of a neural mechanism in the cavefish lateral line system. *eLife*, 11, 2022.

[11308] Dan Luo, Ni Fan, Xiuying Zhang, Fung Yin Ngo, Jia Zhao, Wei Zhao, Ming Huang, Ding Li, Yu Wang, and Jianhui Rong. Covalent inhibition of endoplasmic reticulum chaperone GRP78 disconnects the transduction of ER stress signals to inflammation and lipid accumulation in diet-induced obese mice. *eLife*, 11, 2022.

[11309] Wei-Jia Luo, Sung-Liang Yu, Chia-Ching Chang, Min-Hui Chien, Ya-Ling Chang, Keng-Mao Liao, Pei-Chun Lin, Kuei-Pin Chung, Ya-Hui Chuang, Jeremy JW Chen, Pan-Chyr Yang, and Kang-Yi Su. HLJ1 amplifies endotoxin-induced sepsis severity by promoting IL-12 heterodimerization in macrophages. *eLife*, 11, 2022.

[11310] Sarah M Lurie, James E Kragel, Stephan U Schuele, and Joel L Voss. Human hippocampal responses to network intracranial stimulation vary with theta phase. *eLife*, 11, 2022.

[11311] Andrew R Lynch, Nicholas L Arp, Amber S Zhou, Beth A Weaver, and Mark E Burkard. Quantifying chromosomal instability from intratumoral karyotype diversity using agent-based modeling and Bayesian inference. *eLife*, 11, 2022.

[11312] Ekaterini Maria Lyras, Karin Zimmermann, Lisa Katharina Wagner, Dorothea Dörr, Christoph SN Klose, Cornelius Fischer, Steffen Jung, Simon Yona, Avi-Hai Hovav, Werner Stenzel, Steffen Dommerich, Thomas Conrad, Achim Leutz, and Alexander Mildner. Tongue immune compartment analysis reveals spatial macrophage heterogeneity. *eLife*, 11, 2022.

[11313] Dandan Ma, Fulin Wang, Rongyi Wang, Yingchun Hu, Zhiquan Chen, Ning Huang, Yonglu Tian, Yuqing Xia, Junlin Teng, and Jianguo Chen. $\alpha$-/$\gamma$-Taxilin are required for centriolar subdistal appendage assembly and microtubule organization. *eLife*, 11, 2022.

[11314] Justin Ma, Lian Bi, James Spurlin, and Peter Lwigale. Nephronectin-integrin $\alpha 8$ signaling is required for proper migration of periocular neural crest cells during chick corneal development. *eLife*, 11, 2022.

[11315] Qing Ma, Liuyi Yang, Karen Tolentino, Guiping Wang, Yang Zhao, Ulrike M Litzenburger, Quanming Shi, Lin Zhu, Chen Yang, Huiyuan Jiao, Feng Zhang, Rui Li, Miao-Chih Tsai, Jun-An Chen, Ian Lai, Hong Zeng, Lingjie Li, and Howard Y Chang. Inducible lncRNA transgenic mice reveal continual role of HOTAIR in promoting breast cancer metastasis. *eLife*, 11, 2022.

[11316] Tongcui Ma, Matthew McGregor, Leila Giron, Guorui Xie, Ashley F George, Mohamed Abdel-Mohsen, and Nadia R Roan. Single-cell glycomics analysis by CyTOF-Lec reveals glycan features defining cells differentially susceptible to HIV. *eLife*, 11, 2022.

[11317] Lewis Macdonald, Gillian C Taylor, Jennifer Margaret Brisbane, Ersi Christodoulou, Lucy Scott, Alex von Kriegsheim, Janet Rossant, Bin Gu, and Andrew J Wood. Rapid and specific degradation of endogenous proteins in mouse models using auxin-inducible degrons. *eLife*, 11, 2022.

[11318] Juan F Macias-Velasco, Celine L St Pierre, Jessica P Wayhart, Li Yin, Larry Spears, Mario A Miranda, Caryn Carson, Katsuhiko Funai, James M Cheverud, Clay F Semenkovich, and Heather A Lawson. Parent-of-origin effects propagate through networks to shape metabolic traits. *eLife*, 11, 2022.

[11319] Charles E MacKay, Miranda Floen, M Dennis Leo, Raquibul Hasan, Tessa AC Garrud, Carlos Fernández-Peña, Purnima Singh, Kafait U Malik, and Jonathan H Jaggar. A plasma membrane-localized polycystin-1/polycystin-2 complex in endothelial cells elicits vasodilation. *eLife*, 11, 2022.

[11320] Emily RR Mackie, Andrew S Barrow, Rebecca M Christoff, Belinda M Abbott, Anthony R Gendall, and Tatiana P Soares da Costa. A dual-target herbicidal inhibitor of lysine biosynthesis. *eLife*, 11, 2022.

[11321] Emil Bargmann Madsen, Mathias Wullum Nielsen, Josefine Bjørnholm, Reshma Jagsi, and Jens Peter Andersen. Author-level data confirm the widening gender gap in publishing rates during COVID-19. *eLife*, 11, 2022.

[11322] Lerato E Magosi, Yinfeng Zhang, Tanya Golubchik, Victor DeGruttola, Eric Tchetgen Tchetgen, Vladimir Novitsky, Janet Moore, Pam Bachanas, Tebogo Segolodi, Refeletswe Lebelonyane, Molly Pretorius Holme, Sikhulile Moyo, Joseph Makhema, Shahin Lockman, Christophe Fraser, Myron Max Essex, and Marc Lipsitch. Deep-sequence phylogenetics to quantify patterns of HIV transmission in the context of a universal testing and treatment trial – BCPP/Ya Tsie trial. *eLife*, 11, 2022.

[11323] Gayathri Mahalingam, Russel Torres, Daniel Kapner, Eric T Trautman, Tim Fliss, Shamishtaa Seshamani, Eric Perlman, Rob Young, Samuel Kinn, JoAnn Buchanan, Marc M Takeno, Wenjing Yin, Daniel J Bumbarger, Ryder P Gwinn, Julie Nyhus, Ed Lein, Steven J Smith, R Clay Reid, Khaled A Khairy, Stephan Saalfeld, Forrest Collman, and Nuno Macarico da Costa. A scalable and modular automated pipeline for stitching of large electron microscopy datasets. *eLife*, 11, 2022.

[11324] Alexis DJ Makin, John Tyson-Carr, Giulia Rampone, Yiovanna Derpsch, Damien Wright, and Marco Bertamini. Lessons from a catalogue of 6674 brain recordings. *eLife*, 11, 2022.

[11325] Alexander N Malyavko, Olga A Petrova, Maria I Zvereva, Vladimir I Polshakov, and Olga A Dontsova. Telomere length regulation by Rif1 protein from Hansenula polymorpha. *eLife*, 11, 2022.

[11326] Ana MG Manea, Anna Zilverstand, Kamil Ugurbil, Sarah R Heilbronner, and Jan Zimmermann. Intrinsic timescales as an organizational principle of neural processing across the whole rhesus macaque brain. *eLife*, 11, 2022.

[11327] Pierre Mangeol, Dominique Massey-Harroche, Fabrice Richard, Jean-Paul Concordet, Pierre-FranÇois Lenne, and André Le Bivic. Super-resolution imaging uncovers the nanoscopic segregation of polarity proteins in epithelia. *eLife*, 11, 2022.

[11328] Kasra Manoocheri and Adam G Carter. Rostral and caudal basolateral amygdala engage distinct circuits in the prelimbic and infralimbic prefrontal cortex. *eLife*, 11, 2022.

[11329] Toby Mansell, Richard Saffery, Satvika Burugupalli, Anne-Louise Ponsonby, Mimi LK Tang, Martin O'Hely, Siroon Bekkering, Adam Alexander T Smith, Rebecca Rowland, Sarath Ranganathan, Peter D Sly, Peter Vuillermin, Fiona Collier, Peter Meikle, David Burgner, and John Carlin. Early life infection and proinflammatory, atherogenic metabolomic and lipidomic profiles in infancy: a population-based cohort study. *eLife*, 11, 2022.

[11330] Fatma Mansour, Christian Hinze, Narasimha Swamy Telugu, Jelena Kresoja, Iman B Shaheed, Christian Mosimann, Sebastian Diecke, and Kai M Schmidt-Ott. The centrosomal protein 83 (CEP83) regulates human pluripotent stem cell differentiation toward the kidney lineage. *eLife*, 11, 2022.

[11331] Siavash Mansouri, Daniel Heylmann, Thorsten Stiewe, Michael Kracht, and Rajkumar Savai. Cancer genome and tumor microenvironment: Reciprocal crosstalk shapes lung cancer plasticity. *eLife*, 11, 2022.

[11332] Aurelia A Mapps, Erica Boehm, Corinne Beier, William T Keenan, Jennifer Langel, Michael Liu, Michael B Thomsen, Samer Hattar, Haiqing Zhao, Emmanouil Tampakakis, and Rejji Kuruvilla. Satellite glia modulate sympathetic neuron survival, activity, and autonomic function. *eLife*, 11, 2022.

[11333] Marco Marchetti, Chenge Zhang, and Bruce A Edgar. An improved organ explant culture method reveals stem cell lineage dynamics in the adult Drosophila intestine. *eLife*, 11, 2022.

[11334] Biliana Marcheva, Benjamin J Weidemann, Akihiko Taguchi, Mark Perelis, Kathryn Moynihan Ramsey, Marsha V Newman, Yumiko Kobayashi, Chiaki Omura, Jocelyn E Manning Fox, Haopeng Lin, Patrick E Macdonald, and Joseph Bass. P2Y1 purinergic receptor identified as a diabetes target in a small-molecule screen to reverse circadian $\beta$-cell failure. *eLife*, 11, 2022.

[11335] Maxim I Maron, Alyssa D Casill, Varun Gupta, Jacob S Roth, Simone Sidoli, Charles C Query, Matthew J Gamble, and David Shechter. Type I and II PRMTs inversely regulate post-transcriptional intron detention through Sm and CHTOP methylation. *eLife*, 11, 2022.

[11336] Bryan Marsh, Yan Zhou, Mirhan Kapidzic, Susan Fisher, and Robert Blelloch. Regionally distinct trophoblast regulate barrier function and invasion in the human placenta. *eLife*, 11, 2022.

[11337] Felipe Martelli, Natalia H Hernandes, Zhongyuan Zuo, Julia Wang, Ching-On Wong, Nicholas E Karagas, Ute Roessner, Thusita Rupasinghe, Charles Robin, Kartik Venkatachalam, Trent Perry, Philip Batterham, and Hugo J Bellen. Low doses of the organic insecticide spinosad trigger lysosomal defects, elevated ROS, lipid dysregulation, and neurodegeneration in flies. *eLife*, 11, 2022.

[11338] Susan Martin, Jessica Tyrrell, E Louise Thomas, Matthew J Bown, Andrew R Wood, Robin N Beaumont, Lam C Tsoi, Philip E Stuart, James T Elder, Philip Law, Richard Houlston, Christopher Kabrhel, Nikos Papadimitriou, Marc J Gunter, Caroline J Bull, Joshua A Bell, Emma E Vincent, Naveed Sattar, Malcolm G Dunlop, Ian PM Tomlinson, Sara Lindström, Jimmy D Bell, Timothy M Frayling, and Hanieh Yaghootkar. Disease consequences of higher adiposity uncoupled from its adverse metabolic effects using Mendelian randomisation. *eLife*, 11, 2022.

[11339] Fernando Martín-Fernández, Ana Bermejo-Santos, Lorena Bragg-Gonzalo, Carlos G Briz, Esther Serrano-Saiz, and Marta Nieto. Role of Nrp1 in controlling cortical inter-hemispheric circuits. *eLife*, 11, 2022.

[11340] Guillaume Martinez, Charles Coutton, Corinne Loeuillet, Caroline Cazin, Jana Muroňová, Magalie Boguenet, Emeline Lambert, Magali Dhellemmes, Geneviève Chevalier, Jean-Pascal Hograindleur, Charline Vilpreux, Yasmine Neirijnck, Zine-Eddine Kherraf, Jessica Escoffier, Serge Nef, Pierre F Ray, and Christophe Arnoult. Oligogenic heterozygous inheritance of sperm abnormalities in mouse. *eLife*, 11, 2022.

[11341] Maria Cecilia Martinez, Camila Lidia Zold, Marcos Antonio Coletti, Mario Gustavo Murer, and Mariano Andrés Belluscio. Dorsal striatum coding for the timely execution of action sequences. *eLife*, 11, 2022.

[11342] Juan Martinez-Cervantes, Prachi Shah, Anna Phan, and Isaac Cervantes-Sandoval. Higher-order unimodal olfactory sensory preconditioning in Drosophila. *eLife*, 11, 2022.

[11343] Nuné Martiros, Vikrant Kapoor, Spencer E Kim, and Venkatesh N Murthy. Distinct representation of cue-outcome association by D1 and D2 neurons in the ventral striatum's olfactory tubercle. *eLife*, 11, 2022.

[11344] Kelly M Martyniuk, Arturo Torres-Herraez, Daniel C Lowes, Marcelo Rubinstein, Marie A Labouesse, and Christoph Kellendonk. Dopamine D2Rs coordinate cue-evoked changes in striatal acetylcholine levels. *eLife*, 11, 2022.

[11345] Kazuaki Maruyama, Sachiko Miyagawa-Tomita, Yuka Haneda, Mayuko Kida, Fumio Matsuzaki, Kyoko Imanaka-Yoshida, and Hiroki Kurihara. The cardiopharyngeal mesoderm contributes to lymphatic vessel development in mouse. *eLife*, 11, 2022.

[11346] Omid Mashinchian, Xiaotong Hong, Joris Michaud, Eugenia Migliavacca, Gregory Lefebvre, Christophe Boss, Filippo De Franceschi, Emmeran Le Moal, Jasmin Collerette-Tremblay, Joan Isern, Sylviane Metairon, Frederic Raymond, Patrick Descombes, Nicolas Bouche, Pura Muñoz-Cánoves, Jerome N Feige, and C Florian Bentzinger. In vivo transcriptomic profiling using cell encapsulation identifies effector pathways of systemic aging. *eLife*, 11, 2022.

[11347] Rolando Masís-Obando, Kenneth A Norman, and Christopher Baldassano. Schema representations in distinct brain networks support narrative memory during encoding and retrieval. *eLife*, 11, 2022.

[11348] Thiery Masserey, Tamsin Lee, Monica Golumbeanu, Andrew J Shattock, Sherrie L Kelly, Ian M Hastings, and Melissa A Penny. The influence of biological, epidemiological, and treatment factors on the establishment and spread of drug-resistant Plasmodium falciparum. *eLife*, 11, 2022.

[11349] Anand O Masson, Bryce Besler, W Brent Edwards, and Roman J Krawetz. High spatial resolution analysis using automated indentation mapping differentiates biomechanical properties of normal vs. degenerated articular cartilage in mice. *eLife*, 11, 2022.

[11350] Alan E Mast, Alisa S Wolberg, David Gailani, Michael R Garvin, Christiane Alvarez, J Izaak Miller, Piet Jones, Bruce Aronow, and Daniel Jacobson. Response to comment on 'SARS-CoV-2 suppresses anticoagulant and fibrinolytic gene expression in the lung'. *eLife*, 11, 2022.

[11351] Lior Matityahu, Jeffrey M Malgady, Meital Schirelman, Yvonne Johansson, Jennifer A Wilking, Gilad Silberberg, Joshua A Goldberg, and Joshua L Plotkin. A tonic nicotinic brake controls spike timing in striatal spiny projection neurons. *eLife*, 11, 2022.

[11352] Gabriel Machado Matos, Michael D Lewis, Carlos Talavera-López, Matthew Yeo, Edmundo C Grisard, Louisa A Messenger, Michael A Miles, and Björn Andersson. Microevolution of Trypanosoma cruzi reveals hybridization and clonal mechanisms driving rapid genome diversification. *eLife*, 11, 2022.

[11353] Mitsunori Matsuo, Jia Yuan, Yeon Sun Kim, Amanda Dewar, Hidetoshi Fujita, Sudhansu K Dey, and Xiaofei Sun. Targeted depletion of uterine glandular Foxa2 induces embryonic diapause in mice. *eLife*, 11, 2022.

[11354] Eriko Matsuura-Suzuki, Tadahiro Shimazu, Mari Takahashi, Kaoru Kotoshiba, Takehiro Suzuki, Kazuhiro Kashiwagi, Yoshihiro Sohtome, Mai Akakabe, Mikiko Sodeoka, Naoshi Dohmae, Takuhiro Ito, Yoichi Shinkai, and Shintaro Iwasaki. METTL18-mediated histidine methylation of RPL3 modulates translation elongation for proteostasis maintenance. *eLife*, 11, 2022.

[11355] Keith R Matthews and Stephen Larcombe. Comment on 'Unexpected plasticity in the life cycle of Trypanosoma brucei'. *eLife*, 11, 2022.

[11356] Stefania Mattioni, Mohamed Rezk, Ceren Battal, Jyothirmayi Vadlamudi, and Olivier Collignon. Impact of blindness onset on the representation of sound categories in occipital and temporal cortices. *eLife*, 11, 2022.

[11357] Joanna Mattis, Ala Somarowthu, Kevin M Goff, Evan Jiang, Jina Yom, Nathaniel Sotuyo, Laura M Mcgarry, Huijie Feng, Keisuke Kaneko, and Ethan M Goldberg. Corticohippocampal circuit dysfunction in a mouse model of Dravet syndrome. *eLife*, 11, 2022.

[11358] Austin K Mattox, Christopher Douville, Natalie Silliman, Janine Ptak, Lisa Dobbyn, Joy Schaefer, Maria Popoli, Cherie Blair, Kathy Judge, Kai Pollard, Christine Pratilas, Jaishri Blakeley, Fausto Rodriguez, Nickolas Papadopoulos, Allan Belzberg, and Chetan Bettegowda. Detection of malignant peripheral nerve sheath tumors in patients with neurofibromatosis using aneuploidy and mutation identification in plasma. *eLife*, 11, 2022.

[11359] Marta Matuszewska, Gemma GR Murray, Xiaoliang Ba, Rhiannon Wood, Mark A Holmes, and Lucy A Weinert. Stable antibiotic resistance and rapid human adaptation in livestock-associated MRSA. *eLife*, 11, 2022.

[11360] Alena Maul, Antje Kathrin Huebner, Nicola Strenzke, Tobias Moser, Rudolf Rübsamen, SaŠa Jovanovic, and Christian A Hübner. The Cl–channel TMEM16A is involved in the generation of cochlear Ca2+ waves and promotes the refinement of auditory brainstem networks in mice. *eLife*, 11, 2022.

[11361] Michael Sean Mauro, Gunta Celma, Vitaly Zimyanin, Magdalena M Magaj, Kimberley H Gibson, Stefanie Redemann, and Shirin Bahmanyar. Ndc1 drives nuclear pore complex assembly independent of membrane biogenesis to promote nuclear formation and growth. *eLife*, 11, 2022.

[11362] Hisham Mazal, Franz-Ferdinand Wieser, and Vahid Sandoghdar. Deciphering a hexameric protein complex with Angstrom optical resolution. *eLife*, 11, 2022.

[11363] Fulvia Mazzaferri, Massimo Mirandola, Alessia Savoldi, Pasquale De Nardo, Matteo Morra, Maela Tebon, Maddalena Armellini, Giulia De Luca, Lucrezia Calandrino, Lolita Sasset, Denise D'Elia, Emanuela Sozio, Elisa Danese, Davide Gibellini, Isabella Monne, Giovanna Scroccaro, Nicola Magrini, Annamaria Cattelan, Carlo Tascini, Francesca Simbeni, and Evelina Tacconelli. Exploratory data on the clinical efficacy of monoclonal antibodies against SARS-CoV-2 Omicron variant of concern. *eLife*, 11, 2022.

[11364] Luca Mazzucato. Neural mechanisms underlying the temporal organization of naturalistic animal behavior. *eLife*, 11, 2022.

[11365] Thomas S McAlear and Susanne Bechstedt. The mitotic spindle protein CKAP2 potently increases formation and stability of microtubules. *eLife*, 11, 2022.

[11366] Jeffrey R McArthur, Jierong Wen, Andrew Hung, Rocio K Finol-Urdaneta, and David J Adams. $\mu$-Theraphotoxin Pn3a inhibition of CaV3.3 channels reveals a novel isoform-selective drug binding site. *eLife*, 11, 2022.

[11367] Caitlyn L McCafferty, Ophelia Papoulas, Mareike A Jordan, Gabriel Hoogerbrugge, Candice Nichols, Gaia Pigino, David W Taylor, John B Wallingford, and Edward M Marcotte. Integrative modeling reveals the molecular architecture of the intraflagellar transport A (IFT-A) complex. *eLife*, 11, 2022.

[11368] Colin D McClure, Amira Hassan, Gabriel N Aughey, Khushbakht Butt, Alicia Estacio-Gómez, Aneisha Duggal, Chee Ying Sia, Annika F Barber, and Tony D Southall. An auxin-inducible, GAL4-compatible, gene expression system for Drosophila. *eLife*, 11, 2022.

[11369] Sean E McGeary, Namita Bisaria, Thy M Pham, Peter Y Wang, and David P Bartel. MicroRNA $3'-$compensatory pairing occurs through two binding modes, with affinity shaped by nucleotide identity and position. *eLife*, 11, 2022.

[11370] Matthew McGill, Ariel E Hight, Yurika L Watanabe, Aravindakshan Parthasarathy, Dongqin Cai, Kameron Clayton, Kenneth E Hancock, Anne Takesian, Sharon G Kujawa, and Daniel B Polley. Neural signatures of auditory hypersensitivity following acoustic trauma. *eLife*, 11, 2022.

[11371] James N McGregor, Abigail L Grassler, Paul I Jaffe, Amanda Louise Jacob, Michael S Brainard, and Samuel J Sober. Shared mechanisms of auditory and non-auditory vocal learning in the songbird brain. *eLife*, 11, 2022.

[11372] Andrew McKay, Emma K Costa, Jingxun Chen, Chi-Kuo Hu, Xiaoshan Chen, Claire N Bedbrook, Rishad C Khondker, Mike Thielvoldt, Param Priya Singh, Tony Wyss-Coray, and Anne Brunet. An automated feeding system for the African killifish reveals the impact of diet on lifespan and allows scalable assessment of associative learning. *eLife*, 11, 2022.

[11373] Daniel W McKay, Heather E McFarlane, Yue Qu, Apriadi Situmorang, Matthew Gilliham, and Stefanie Wege. Plant Trans-Golgi Network/Early Endosome pH regulation requires Cation Chloride Cotransporter (CCC1). *eLife*, 11, 2022.

[11374] Jennifer McKey, Dilara N Anbarci, Corey Bunce, Alejandra E Ontiveros, Richard R Behringer, and Blanche Capel. Integration of mouse ovary morphogenesis with developmental dynamics of the oviduct, ovarian ligaments, and rete ovarii. *eLife*, 11, 2022.

[11375] Shannon J McKie, Parth Rakesh Desai, Yeonee Seol, Adam MB Allen, Anthony Maxwell, and Keir C Neuman. Topoisomerase VI is a chirally-selective, preferential DNA decatenase. *eLife*, 11, 2022.

[11376] Amelia C McKitterick and Thomas G Bernhardt. Phage resistance profiling identifies new genes required for biogenesis and modification of the corynebacterial cell envelope. *eLife*, 11, 2022.

[11377] Ian G McLachlan, Talya S Kramer, Malvika Dua, Elizabeth M DiLoreto, Matthew A Gomes, Ugur Dag, Jagan Srinivasan, and Steven W Flavell. Diverse states and stimuli tune olfactory receptor expression levels to modulate food-seeking behavior. *eLife*, 11, 2022.

[11378] Angela McLaughlin, Vincent Montoya, Rachel L Miller, Gideon J Mordecai, Michael Worobey, Art FY Poon, and Jeffrey B Joy. Genomic epidemiology of the first two waves of SARS-CoV-2 in Canada. *eLife*, 11, 2022.

[11379] Timothy J Mead, Daniel R Martin, Lauren W Wang, Stuart A Cain, Cagri Gulec, Elisabeth Cahill, Joseph Mauch, Dieter Reinhardt, Cecilia Lo, Clair Baldock, and Suneel S Apte. Proteolysis of fibrillin-2 microfibrils is essential for normal skeletal development. *eLife*, 11, 2022.

[11380] Tomas Mediavilla, özgün özalay, Héctor M Estévez-Silva, Bárbara Frias, Greger Orädd, Fahad R Sultan, Claudio Brozzoli, Benjamín Garzón, Martin Lövdén, and Daniel J Marcellino. Learning-related contraction of gray matter in rodent sensorimotor cortex is associated with adaptive myelination. *eLife*, 11, 2022.

[11381] Laura Medina-Ruiz, Robin Bartolini, Gillian J Wilson, Douglas P Dyer, Francesca Vidler, Catherine E Hughes, Fabian Schuette, Samantha Love, Marieke Pingen, Alan James Hayes, Jun Fu, Adrian Francis Stewart, and Gerard J Graham. Analysis of combinatorial chemokine receptor expression dynamics using multi-receptor reporter mice. *eLife*, 11, 2022.

[11382] J Kyle Medley, Jenna Persons, Tathagata Biswas, Luke Olsen, Robert Peuß, Jaya Krishnan, Shaolei Xiong, and Nicolas Rohner. The metabolome of Mexican cavefish shows a convergent signature highlighting sugar, antioxidant, and Ageing-Related metabolites. *eLife*, 11, 2022.

[11383] Kimberly Meechan, Wei Guan, Alfons Riedinger, Vera Stankova, Azumi Yoshimura, Rosa Pipitone, Arthur Milberger, Helmuth Schaar, Inés Romero-Brey, Rachel Templin, Christopher J Peddie, Nicole L Schieber, Martin L Jones, Lucy Collinson, and Yannick Schwab. Crosshair, semi-automated targeting for electron microscopy with a motorised ultramicrotome. *eLife*, 11, 2022.

[11384] Marine Megemont, Jim McBurney-Lin, and Hongdian Yang. Pupil diameter is not an accurate real-time readout of locus coeruleus activity. *eLife*, 11, 2022.

[11385] Catherine Meh and Prabhat Jha. Trends in female-selective abortion among Asian diasporas in the United States, United Kingdom, Canada and Australia. *eLife*, 11, 2022.

[11386] Ved Mehta, Basavraj Khanppnavar, Dina Schuster, Ilayda Kantarci, Irene Vercellino, Angela Kosturanova, Tarun Iype, Sasa Stefanic, Paola Picotti, and Volodymyr M Korkhov. Structure of Mycobacterium tuberculosis Cya, an evolutionary ancestor of the mammalian membrane adenylyl cyclases. *eLife*, 11, 2022.

[11387] Shaoyi Mei, Yi Wu, Yan Wang, Yubo Cui, Miao Zhang, Tong Zhang, Xiaosheng Huang, Sejie Yu, Tao Yu, and Jun Zhao. Disruption of PIKFYVE causes congenital cataract in human and zebrafish. *eLife*, 11, 2022.

[11388] Anne C Meinema, Anna Marzelliusardottir, Mihailo Mirkovic, Théo Aspert, Sung Sik Lee, Gilles Charvin, and Yves Barral. DNA circles promote yeast ageing in part through stimulating the reorganization of nuclear pore complexes. *eLife*, 11, 2022.

[11389] Steven Lee Meisler and John DE Gabrieli. Fiber-specific structural properties relate to reading skills in children and adolescents. *eLife*, 11, 2022.

[11390] Jorge F Mejías and Xiao-Jing Wang. Mechanisms of distributed working memory in a large-scale network of macaque neocortex. *eLife*, 11, 2022.

[11391] Julian Melgar, Mads F Schou, Maud Bonato, Zanell Brand, Anel Engelbrecht, Schalk WP Cloete, and Charlie K Cornwallis. Experimental evidence that group size generates divergent benefits of cooperative breeding for male and female ostriches. *eLife*, 11, 2022.

[11392] Fabrizio Menardo. Understanding drivers of phylogenetic clustering and terminal branch lengths distribution in epidemics of Mycobacterium tuberculosis. *eLife*, 11, 2022.

[11393] Evdokia Menelaou, Sandeep Kishore, and David L McLean. Mixed synapses reconcile violations of the size principle in zebrafish spinal cord. *eLife*, 11, 2022.

[11394] Danilo Menicucci, Claudia Lunghi, Andrea Zaccaro, Maria Concetta Morrone, and Angelo Gemignani. Mutual interaction between visual homeostatic plasticity and sleep in adult humans. *eLife*, 11, 2022.

[11395] Natalie D Mercuri and Brian J Cox. The need for more research into reproductive health and disease. *eLife*, 11, 2022.

[11396] Zsolt Merényi, Máté Virágh, Emile Gluck-Thaler, Jason C Slot, Brigitta Kiss, Torda Varga, András Geösel, Botond Hegedüs, Balázs Bálint, and László G Nagy. Gene age shapes the transcriptional landscape of sexual morphogenesis in mushroom-forming fungi (Agaricomycetes). *eLife*, 11, 2022.

[11397] Timon Merk, Victoria Peterson, Witold J Lipski, Benjamin Blankertz, Robert S Turner, Ningfei Li, Andreas Horn, Robert Mark Richardson, and Wolf-Julian Neumann. Electrocorticography is superior to subthalamic local field potentials for movement decoding in Parkinson's disease. *eLife*, 11, 2022.

[11398] Emeric Merour, Hatem Hmidan, Corentine Marie, Pierre-Henri Helou, Haiyang Lu, Antoine Potel, Jean-Baptiste Hure, Adrien Clavairoly, Yi Ping Shih, Salman Goudarzi, Sebastien Dussaud, Philippe Ravassard, Sassan Hafizi, Su Hao Lo, Bassem A Hassan, and Carlos Parras. Transient regulation of focal adhesion via Tensin3 is required for nascent oligodendrocyte differentiation. *eLife*, 11, 2022.

[11399] Christina M Merrick, Tanner C Dixon, Assaf Breska, Jack Lin, Edward F Chang, David King-Stephens, Kenneth D Laxer, Peter B Weber, Jose Carmena, Robert Thomas Knight, and Richard B Ivry. Left hemisphere dominance for bilateral kinematic encoding in the human brain. *eLife*, 11, 2022.

[11400] Andrea Merseburg, Jacquelin Kasemir, Eric W Buss, Felix Leroy, Tobias Bock, Alessandro Porro, Anastasia Barnett, Simon E Tröder, Birgit Engeland, Malte Stockebrand, Anna Moroni, Steven A Siegelbaum, Dirk Isbrandt, and Bina Santoro. Seizures, behavioral deficits, and adverse drug responses in two new genetic mouse models of HCN1 epileptic encephalopathy. *eLife*, 11, 2022.

[11401] Andrea S Meseguer, Alice Michel, Pierre-Henri Fabre, Oscar A Pérez Escobar, Guillaume Chomicki, Ricarda Riina, Alexandre Antonelli, Pierre-Olivier Antoine, Frédéric Delsuc, and Fabien L Condamine. Diversification dynamics in the Neotropics through time, clades, and biogeographic regions. *eLife*, 11, 2022.

[11402] Anna Metzger and Matteo Toscani. Unsupervised learning of haptic material properties. *eLife*, 11, 2022.

[11403] Ann-Kristin Meyer and Roland G Benoit. Suppression weakens unwanted memories via a sustained reduction of neural reactivation. *eLife*, 11, 2022.

[11404] Benjamin H Meyer, Panagiotis S Adam, Ben A Wagstaff, George E Kolyfetis, Alexander J Probst, Sonja V Albers, and Helge C Dorfmueller. Agl24 is an ancient archaeal homolog of the eukaryotic N-glycan chitobiose synthesis enzymes. *eLife*, 11, 2022.

[11405] Dylan J Meyer, Carlos Manlio Díaz-García, Nidhi Nathwani, Mahia Rahman, and Gary Yellen. The Na+/K+ pump dominates control of glycolysis in hippocampal dentate granule cells. *eLife*, 11, 2022.

[11406] Gretchen A Meyer, Stavros Thomopoulos, Yousef Abu-Amer, and Karen C Shen. Tenotomy-induced muscle atrophy is sex-specific and independent of NF$\kappa$B. *eLife*, 11, 2022.

[11407] Keith J Mickolajczyk, Paul Dominic B Olinares, Brian T Chait, Shixin Liu, and Tarun M Kapoor. The MIDAS domain of AAA mechanoenzyme Mdn1 forms catch bonds with two different substrates. *eLife*, 11, 2022.

[11408] Teemu P Miettinen, Kevin S Ly, Alice Lam, and Scott R Manalis. Single-cell monitoring of dry mass and dry mass density reveals exocytosis of cellular dry contents in mitosis. *eLife*, 11, 2022.

[11409] Katelyn Mika, Camilla M Whittington, Bronwyn M McAllan, and Vincent J Lynch. Gene expression phylogenies and ancestral transcriptome reconstruction resolves major transitions in the origins of pregnancy. *eLife*, 11, 2022.

[11410] Artem Mikelov, Evgeniia I Alekseeva, Ekaterina A Komech, Dmitry B Staroverov, Maria A Turchaninova, Mikhail Shugay, Dmitriy M Chudakov, Georgii A Bazykin, and Ivan V Zvyagin. Memory persistence and differentiation into antibody-secreting cells accompanied by positive selection in longitudinal BCR repertoires. *eLife*, 11, 2022.

[11411] Alexander R Mikesell, Olena Isaeva, Francie Moehring, Katelyn E Sadler, Anthony D Menzel, and Cheryl L Stucky. Keratinocyte PIEZO1 modulates cutaneous mechanosensation. *eLife*, 11, 2022.

[11412] Nace Mikus, Sebastian Korb, Claudia Massaccesi, Christian Gausterer, Irene Graf, Matthäus Willeit, Christoph Eisenegger, Claus Lamm, Giorgia Silani, and Christoph Mathys. Effects of dopamine D2/3 and opioid receptor antagonism on the trade-off between model-based and model-free behaviour in healthy volunteers. *eLife*, 11, 2022.

[11413] Kevin J Miller, Matthew M Botvinick, and Carlos D Brody. Value representations in the rodent orbitofrontal cortex drive learning, not choice. *eLife*, 11, 2022.

[11414] Blayze Millward, Steve Maddock, and Michael Mangan. CompoundRay, an open-source tool for high-speed and high-fidelity rendering of compound eyes. *eLife*, 11, 2022.

[11415] Veronika Miskolci, Kelsey E Tweed, Michael R Lasarev, Emily C Britt, Alex J Walsh, Landon J Zimmerman, Courtney E McDougal, Mark R Cronan, Jing Fan, John-Demian Sauer, Melissa C Skala, and Anna Huttenlocher. In vivo fluorescence lifetime imaging of macrophage intracellular metabolism during wound responses in zebrafish. *eLife*, 11, 2022.

[11416] Noah P Mitchell, Dillon J Cislo, Suraj Shankar, Yuzheng Lin, Boris I Shraiman, and Sebastian J Streichan. Visceral organ morphogenesis via calcium-patterned muscle constrictions. *eLife*, 11, 2022.

[11417] Wayne Mitchell, Jeffrey D Tamucci, Emery L Ng, Shaoyi Liu, Alexander V Birk, Hazel H Szeto, Eric R May, Andrei T Alexandrescu, and Nathan N Alder. Structure-activity relationships of mitochondria-targeted tetrapeptide pharmacological compounds. *eLife*, 11, 2022.

[11418] Catalin Mitelut, Yongxu Zhang, Yuki Sekino, Jamie D Boyd, Federico Bollanos, Nicholas V Swindale, Greg Silasi, Shreya Saxena, and Timothy H Murphy. Mesoscale cortex-wide neural dynamics predict self-initiated actions in mice several seconds prior to movement. *eLife*, 11, 2022.

[11419] Noriyo Mitome, Shintaroh Kubo, Sumie Ohta, Hikaru Takashima, Yuto Shigefuji, Toru Niina, and Shoji Takada. Cooperation among c-subunits of FoF1-ATP synthase in rotation-coupled proton translocation. *eLife*, 11, 2022.

[11420] Ramkrishna Mitra, Clare M Adams, and Christine M Eischen. Systematic lncRNA mapping to genome-wide co-essential modules uncovers cancer dependency on uncharacterized lncRNAs. *eLife*, 11, 2022.

[11421] Masatoshi Miyakoshi, Teppei Morita, Asaki Kobayashi, Anna Berger, Hiroki Takahashi, Yasuhiro Gotoh, Tetsuya Hayashi, and Kan Tanaka. Glutamine synthetase mRNA releases sRNA from its 3'UTR to regulate carbon/nitrogen metabolic balance in Enterobacteriaceae. *eLife*, 11, 2022.

[11422] Takahisa Miyao, Maki Miyauchi, S Thomas Kelly, Tommy W Terooatea, Tatsuya Ishikawa, Eugene Oh, Sotaro Hirai, Kenta Horie, Yuki Takakura, Houko Ohki, Mio Hayama, Yuya Maruyama, Takao Seki, Hiroto Ishii, Haruka Yabukami, Masaki Yoshida, Azusa Inoue, Asako Sakaue-Sawano, Atsushi Miyawaki, Masafumi Muratani, Aki Minoda, Nobuko Akiyama, and Taishin Akiyama. Integrative analysis of scRNA-seq and scATAC-seq revealed transit-amplifying thymic epithelial cells expressing autoimmune regulator. *eLife*, 11, 2022.

[11423] Yuu Miyazaki, Atsuhiko Ichimura, Ryo Kitayama, Naoki Okamoto, Tomoki Yasue, Feng Liu, Takaaki Kawabe, Hiroki Nagatomo, Yohei Ueda, Ichiro Yamauchi, Takuro Hakata, Kazumasa Nakao, Sho Kakizawa, Miyuki Nishi, Yasuo Mori, Haruhiko Akiyama, Kazuwa Nakao, and Hiroshi Takeshima. C-type natriuretic peptide facilitates autonomic $Ca2+$ entry in growth plate chondrocytes for stimulating bone growth. *eLife*, 11, 2022.

[11424] Hidenobu Miyazawa, Marteinn T Snaebjornsson, Nicole Prior, Eleni Kafkia, Henrik M Hammarén, Nobuko Tsuchida-Straeten, Kiran R Patil, Martin Beck, and Alexander Aulehla. Glycolytic flux-signaling controls mouse embryo mesoderm development. *eLife*, 11, 2022.

[11425] Kouhei Mizuno, Mais Maree, Toshihiko Nagamura, Akihiro Koga, Satoru Hirayama, Soichi Furukawa, Kenji Tanaka, and Kazuya Morikawa. Novel multicellular prokaryote discovered next to an underground stream. *eLife*, 11, 2022.

[11426] Xiaoyi Mo, Peiyuan Pang, Yulin Wang, Dexiang Jiang, Mengyu Zhang, Yang Li, Peiyu Wang, Qizhi Geng, Chang Xie, Hai-Ning Du, Bo Zhong, Dongdong Li, and Jing Yao. Tyrosine phosphorylation tunes chemical and thermal sensitivity of TRPV2 ion channel. *eLife*, 11, 2022.

[11427] Maryam H Mofrad, Greydon Gilmore, Dominik Koller, Seyed M Mirsattari, Jorge G Burneo, David A Steven, Ali R Khan, Ana Suller Marti, and Lyle Muller. Waveform detection by deep learning reveals multi-area spindles that are selectively modulated by memory load. *eLife*, 11, 2022.

[11428] Jeffrey Molendijk, Ronnie Blazev, Richard J Mills, Yaan-Kit Ng, Kevin I Watt, Daryn Chau, Paul Gregorevic, Peter J Crouch, James BW Hilton, Leszek Lisowski, Peixiang Zhang, Karen Reue, Aldons J Lusis, James E Hudson, David E James, Marcus M Seldin, and Benjamin L Parker. Proteome-wide systems genetics identifies UFMylation as a regulator of skeletal muscle function. *eLife*, 11, 2022.

[11429] Nardus Mollentze, Deborah Keen, Uuriintuya Munkhbayar, Roman Biek, and Daniel G Streicker. Variation in the ACE2 receptor has limited utility for SARS-CoV-2 host prediction. *eLife*, 11, 2022.

[11430] Katrin Möller, Max Brambach, Ambra Villani, Elisa Gallo, Darren Gilmour, and Francesca Peri. A role for the centrosome in regulating the rate of neuronal efferocytosis by microglia in vivo. *eLife*, 11, 2022.

[11431] Fanny Momboisse, Giacomo Nardi, Philippe Colin, Melanie Hery, Nelia Cordeiro, Simon Blachier, Olivier Schwartz, Fernando Arenzana-Seisdedos, Nathalie Sauvonnet, Jean-Christophe Olivo-Marin, Bernard Lagane, Thibault Lagache, and Anne Brelot. Tracking receptor motions at the plasma membrane reveals distinct effects of ligands on CCR5 dynamics depending on its dimerization status. *eLife*, 11, 2022.

[11432] Gemma Moncunill, Jason Carnes, William Chad Young, Lindsay Carpp, Stephen De Rosa, Joseph J Campo, Augusto Nhabomba, Maxmillian Mpina, Chenjerai Jairoce, Greg Finak, Paige Haas, Carl Muriel, Phu Van, Héctor Sanz, Sheetij Dutta, Benjamin Mordmüller, Selidji T Agnandji, Núria Díez-Padrisa, Nana Aba Williams, John J Aponte, Clarissa Valim, Daniel E Neafsey, Claudia Daubenberger, M Juliana McElrath, Carlota Dobaño, Ken Stuart, and Raphael Gottardo. Transcriptional correlates of malaria in RTS,S/AS01-vaccinated African children: a matched case–control study. *eLife*, 11, 2022.

[11433] Sudipta Mondal, Priyadarshan Kinatukara, Shubham Singh, Sakshi Shambhavi, Gajanan S Patil, Noopur Dubey, Salam Herojeet Singh, Biswajit Pal, P Chandra Shekar, Siddhesh S Kamat, and Rajan Sankaranarayanan. DIP2 is a unique regulator of diacylglycerol lipid homeostasis in eukaryotes. *eLife*, 11, 2022.

[11434] Nicolás Mongiardino Koch, Jeffrey R Thompson, Avery S Hiley, Marina F McCowin, A Frances Armstrong, Simon E Coppard, Felipe Aguilera, Omri Bronstein, Andreas Kroh, Rich Mooi, and Greg W Rouse. Phylogenomic analyses of echinoid diversification prompt a re-evaluation of their fossil record. *eLife*, 11, 2022.

[11435] Arnau Montagud, Jonas Béal, Luis Tobalina, Pauline Traynard, Vigneshwari Subramanian, Bence Szalai, Róbert Alföldi, László Puskás, Alfonso Valencia, Emmanuel Barillot, Julio Saez-Rodriguez, and Laurence Calzone. Patient-specific Boolean models of signalling networks guide personalised treatments. *eLife*, 11, 2022.

[11436] Stefania Monterisi, Johanna Michl, Alzbeta Hulikova, Jana Koth, Esther M Bridges, Amaryllis E Hill, Gulnar Abdullayeva, Walter F Bodmer, and Pawel Swietach. Solute exchange through gap junctions lessens the adverse effects of inactivating mutations in metabolite-handling genes. *eLife*, 11, 2022.

[11437] Sean Akira Montgomery, Tetsuya Hisanaga, Nan Wang, Elin Axelsson, Svetlana Akimcheva, Milos Sramek, Chang Liu, and Frédéric Berger. Polycomb-mediated repression of paternal chromosomes maintains haploid dosage in diploid embryos of Marchantia. *eLife*, 11, 2022.

[11438] Edmund RR Moody, Tara A Mahendrarajah, Nina Dombrowski, James W Clark, Celine Petitjean, Pierre Offre, Gergely J Szöllősi, Anja Spang, and Tom A Williams. An estimate of the deepest branches of the tree of life from ancient vertically evolving genes. *eLife*, 11, 2022.

[11439] Jamie Moore-Fried, Mahinder Paul, Zhixin Jing, David Fooksman, and Gregoire Lauvau. CD169+ macrophages orchestrate plasmacytoid dendritic cell arrest and retention for optimal priming in the bone marrow of malaria-infected mice. *eLife*, 11, 2022.

[11440] Jenna A Moore-Ott, Selena Chiu, Daniel B Amchin, Tapomoy Bhattacharjee, and Sujit S Datta. A biophysical threshold for biofilm formation. *eLife*, 11, 2022.

[11441] Sofia N Moraes, Jordan T Becker, Seyed Arad Moghadasi, Nadine M Shaban, Ashley A Auerbach, Adam Z Cheng, and Reuben S Harris. Evidence linking APOBEC3B genesis and evolution of innate immune antagonism by gamma-herpesvirus ribonucleotide reductases. *eLife*, 11, 2022.

[11442] Mychel RPT Morais, Pinyuan Tian, Craig Lawless, Syed Murtuza-Baker, Louise Hopkinson, Steven Woods, Aleksandr Mironov, David A Long, Daniel P Gale, Telma MT Zorn, Susan J Kimber, Roy Zent, and Rachel Lennon. Kidney organoids recapitulate human basement membrane assembly in health and disease. *eLife*, 11, 2022.

[11443] Fernanda Morales Berstein, Daniel L McCartney, Ake T Lu, Konstantinos K Tsilidis, Emmanouil Bouras, Philip Haycock, Kimberley Burrows, Amanda I Phipps, Daniel D Buchanan, Iona Cheng, Rosalind A Eeles, Richard M Martin, George Davey Smith, Caroline L Relton, Steve Horvath, Riccardo E Marioni, Tom G Richardson, and Rebecca C Richmond. Assessing the causal role of epigenetic clocks in the development of multiple cancers: a Mendelian randomization study. *eLife*, 11, 2022.

[11444] Daniel E Morales-Mantilla, Bailee Kain, Duy Le, Anthony R Flores, Silke Paust, and Katherine Y King. Hematopoietic stem and progenitor cells improve survival from sepsis by boosting immunomodulatory cells. *eLife*, 11, 2022.

[11445] Makenna M Morck, Debanjan Bhowmik, Divya Pathak, Aminah Dawood, James Spudich, and Kathleen M Ruppel. Hypertrophic cardiomyopathy mutations in the pliant and light chain-binding regions of the lever arm of human $\beta$-cardiac myosin have divergent effects on myosin function. *eLife*, 11, 2022.

[11446] Brittany L Morgan, Mariana C Stern, Eliseo J Pérez-Stable, Monica Webb Hooper, and Laura Fejerman. Adding a One Health approach to a research framework for minority health and health disparities. *eLife*, 11, 2022.

[11447] Andréanne Morin, Britney A Helling, Seetha Krishnan, Laurie E Risner, Nykia D Walker, and Nancy B Schwartz. Surveying the experience of postdocs in the United States before and during the COVID-19 pandemic. *eLife*, 11, 2022.

[11448] Tsuyoshi Morita and Ken'ichiro Hayashi. Actin-related protein 5 functions as a novel modulator of MyoD and MyoG in skeletal muscle and in rhabdomyosarcoma. *eLife*, 11, 2022.

[11449] Ekaterina Morozova, Peter Newstein, and Eve Marder. Reciprocally inhibitory circuits operating with distinct mechanisms are differently robust to perturbation and modulation. *eLife*, 11, 2022.

[11450] Ryan J Morrill, James Bigelow, Jefferson DeKloe, and Andrea R Hasenstaub. Audiovisual task switching rapidly modulates sound encoding in mouse auditory cortex. *eLife*, 11, 2022.

[11451] Stefan Möstl, Fabian Hoffmann, Jan-Niklas Hönemann, Jose Ramon Alvero-Cruz, Jörn Rittweger, Jens Tank, and Jens Jordan. Utility of estimated pulse wave velocity for assessing vascular stiffness: comparison of methods. *eLife*, 11, 2022.

[11452] Susan M Motch Perrine, M Kathleen Pitirri, Emily L Durham, Mizuho Kawasaki, Hao Zheng, Danny Z Chen, Kazuhiko Kawasaki, and Joan T Richtsmeier. A dysmorphic mouse model reveals developmental interactions of chondrocranium and dermatocranium. *eLife*, 11, 2022.

[11453] Erick VS Motta, Alejandra Gage, Thomas E Smith, Kristin J Blake, Waldan K Kwong, Ian M Riddington, and Nancy Moran. Host-microbiome metabolism of a plant toxin in bees. *eLife*, 11, 2022.

[11454] Dallas L Mould, Mirjana Stevanovic, Alix Ashare, Daniel Schultz, and Deborah A Hogan. Metabolic basis for the evolution of a common pathogenic Pseudomonas aeruginosa variant. *eLife*, 11, 2022.

[11455] Evangelos Mourkas, Koji Yahara, Sion C Bayliss, Jessica K Calland, Håkan Johansson, Leonardos Mageiros, Zilia Y Muñoz-Ramirez, Grant Futcher, Guillaume Méric, Matthew D Hitchings, Santiago Sandoval-Motta, Javier Torres, Keith A Jolley, Martin CJ Maiden, Patrik Ellström, Jonas Waldenström, Ben Pascoe, and Samuel K Sheppard. Host ecology regulates interspecies recombination in bacteria of the genus Campylobacter. *eLife*, 11, 2022.

[11456] Khyobeni Mozhui, Ake T Lu, Caesar Z Li, Amin Haghani, Jose Vladimir Sandoval-Sierra, Yibo Wu, Robert W Williams, and Steve Horvath. Genetic loci and metabolic states associated with murine epigenetic aging. *eLife*, 11, 2022.

[11457] Lena Maria Muckenfuss, Anabel Carmen Migenda Herranz, Franziska Maria Boneberg, Marcello Clerici, and Martin Jinek. Fip1 is a multivalent interaction scaffold for processing factors in human mRNA 3' end biogenesis. *eLife*, 11, 2022.

[11458] Rasa A Muktupavela, Martin Petr, Laure Ségurel, Thorfinn Korneliussen, John Novembre, and Fernando Racimo. Modeling the spatiotemporal spread of beneficial alleles using ancient genomes. *eLife*, 11, 2022.

[11459] Christine Müller, Laura M Zidek, Sabrina Eichwald, Gertrud Kortman, Mirjam H Koster, and Cornelis F Calkhoven. Enhanced C/EBP$\beta$ function promotes hyperplastic versus hypertrophic fat tissue growth and prevents steatosis in response to high-fat diet feeding. *eLife*, 11, 2022.

[11460] Jan L Münch, Fabian Paul, Ralf Schmauder, and Klaus Benndorf. Bayesian inference of kinetic schemes for ion channels by Kalman filtering. *eLife*, 11, 2022.

[11461] Lucas E Murray, Haein Kim, Luke M Rice, and Charles L Asbury. Working strokes produced by curling protofilaments at disassembling microtubule tips can be biochemically tuned and vary with species. *eLife*, 11, 2022.

[11462] Stefano Musardo, Alessandro Contestabile, Marit Knoop, Olivier Baud, and Camilla Bellone. Oxytocin neurons mediate the effect of social isolation via the VTA circuits. *eLife*, 11, 2022.

[11463] Aftab Nadeem, Alexandra Berg, Hudson Pace, Athar Alam, Eric Toh, Jörgen Ådén, Nikola Zlatkov, Si Lhyam Myint, Karina Persson, Gerhard Gröbner, Anders Sjöstedt, Marta Bally, Jonas Barandun, Bernt Eric Uhlin, and Sun Nyunt Wai. Protein-lipid interaction at low pH induces oligomerization of the MakA cytotoxin from Vibrio cholerae. *eLife*, 11, 2022.

[11464] Kenichi Nagano, Kei Yamana, Hiroaki Saito, Riku Kiviranta, Ana Clara Pedroni, Dhairya Raval, Christof Niehrs, Francesca Gori, and Roland Baron. R-spondin 3 deletion induces Erk phosphorylation to enhance Wnt signaling and promote bone formation in the appendicular skeleton. *eLife*, 11, 2022.

[11465] Irène Nagle, Florence Delort, Sylvie Hénon, Claire Wilhelm, Sabrina Batonnet-Pichon, and Myriam Reffay. The importance of intermediate filaments in the shape maintenance of myoblast model tissues. *eLife*, 11, 2022.

[11466] Daisuke Nakane, Gen Enomoto, Heike Bähre, Yuu Hirose, Annegret Wilde, and Takayuki Nishizaka. Thermosynechococcus switches the direction of phototaxis by a c-di-GMP-dependent process with high spatial resolution. *eLife*, 11, 2022.

[11467] Akhilesh Nandan, Abhishek Das, Robert Lott, and Aneta Koseska. Cells use molecular working memory to navigate in changing chemoattractant fields. *eLife*, 11, 2022.

[11468] Nilay Nandi, Zuhair Zaidi, Charles Tracy, and Helmut Krämer. A phosphoswitch at acinus-serine437 controls autophagic responses to cadmium exposure and neurodegenerative stress. *eLife*, 11, 2022.

[11469] Jana Narasimhan, Suzanne Letinski, Stephen P Jung, Aleksey Gerasyuto, Jiashi Wang, Michael Arnold, Guangming Chen, Jean Hedrick, Melissa Dumble, Kanchana Ravichandran, Talya Levitz, Chang Cui, Catherine L Drennan, JoAnne Stubbe, Gary Karp, and Arthur Branstrom. Ribonucleotide reductase, a novel drug target for gonorrhea. *eLife*, 11, 2022.

[11470] Darshana Z Narayanan, Daniel Y Takahashi, Lauren M Kelly, Sabina I Hlavaty, Junzhou Huang, and Asif A Ghazanfar. Prenatal development of neonatal vocalizations. *eLife*, 11, 2022.

[11471] Naser Nasrollahzadeh, Peyman Karami, Jian Wang, Lida Bagheri, Yanheng Guo, Philippe Abdel-Sayed, Lee Laurent-Applegate, and Dominique P Pioletti. Temperature evolution following joint loading promotes chondrogenesis by synergistic cues via calcium signaling. *eLife*, 11, 2022.

[11472] Joris P Nassal, Fiona H Murphy, Ruud F Toonen, and Matthijs Verhage. Differential axonal trafficking of Neuropeptide Y-, LAMP1-, and RAB7-tagged organelles in vivo. *eLife*, 11, 2022.

[11473] Laura B Naumann, Joram Keijser, and Henning Sprekeler. Invariant neural subspaces maintained by feedback modulation. *eLife*, 11, 2022.

[11474] Beatriz Navarro-Dominguez, Ching-Ho Chang, Cara L Brand, Christina A Muirhead, Daven C Presgraves, and Amanda M Larracuente. Epistatic selection on a selfish Segregation Distorter supergene – drive, recombination, and genetic load. *eLife*, 11, 2022.

[11475] Alba Navarro-Romero, Lorena Galera-López, Paula Ortiz-Romero, Alberto Llorente-Ovejero, Lucía de los Reyes-Ramírez, Iker Bengoetxea de Tena, Anna Garcia-Elias, Aleksandra Mas-Stachurska, Marina Reixachs-Solé, Antoni Pastor, Rafael de la Torre, Rafael Maldonado, Begoña Benito, Eduardo Eyras, Rafael Rodríguez-Puertas, Victoria Campuzano, and Andres Ozaita. Cannabinoid signaling modulation through JZL184 restores key phenotypes of a mouse model for Williams–Beuren syndrome. *eLife*, 11, 2022.

[11476] Andrea Navas-Olive, Rodrigo Amaducci, Maria-Teresa Jurado-Parras, Enrique R Sebastian, and Liset M de la Prida. Deep learning-based feature extraction for prediction and interpretation of sharp-wave ripples in the rodent hippocampus. *eLife*, 11, 2022.

[11477] Ashok R Nayak and Montserrat Samsó. Ca2+ inactivation of the mammalian ryanodine receptor type 1 in a lipidic environment revealed by cryo-EM. *eLife*, 11, 2022.

[11478] Jakke Neiro, Divya Sridhar, Anish Dattani, and Aziz Aboobaker. Identification of putative enhancer-like elements predicts regulatory networks active in planarian adult stem cells. *eLife*, 11, 2022.

[11479] Nuno GB Neto, Sinead A O'Rourke, Mimi Zhang, Hannah K Fitzgerald, Aisling Dunne, and Michael G Monaghan. Non-invasive classification of macrophage polarisation by 2P-FLIM and machine learning. *eLife*, 11, 2022.

[11480] Kok Pin Ng, Xing Qian, Kwun Kei Ng, Fang Ji, Pedro Rosa-Neto, Serge Gauthier, Nagaendran Kandiah, and Juan Helen Zhou. Stage-dependent differential influence of metabolic and structural networks on memory across Alzheimer's disease continuum. *eLife*, 11, 2022.

[11481] Wai Hoe Ng, Elizabeth K Johnston, Jun Jie Tan, Jacqueline M Bliley, Adam W Feinberg, Donna B Stolz, Ming Sun, Piyumi Wijesekara, Finn Hawkins, Darrell N Kotton, and Xi Ren. Recapitulating human cardio-pulmonary co-development using simultaneous multilineage differentiation of pluripotent stem cells. *eLife*, 11, 2022.

[11482] Nhat TD Nguyen, Ashutosh K Pathak, and Isabella M Cattadori. Gastrointestinal helminths increase Bordetella bronchiseptica shedding and host variation in supershedding. *eLife*, 11, 2022.

[11483] Tricia T Nguyen and Gia K Voeltz. An ER phospholipid hydrolase drives ER-associated mitochondrial constriction for fission and fusion. *eLife*, 11, 2022.

[11484] Alex N Nguyen Ba, Katherine R Lawrence, Artur Rego-Costa, Shreyas Gopalakrishnan, Daniel Temko, Franziska Michor, and Michael M Desai. Barcoded bulk QTL mapping reveals highly polygenic and epistatic architecture of complex traits in yeast. *eLife*, 11, 2022.

[11485] Amy M Ni, Chengcheng Huang, Brent Doiron, and Marlene R Cohen. A general decoding strategy explains the relationship between behavior and correlated variability. *eLife*, 11, 2022.

[11486] Qin Ni, Kaustubh Wagh, Aashli Pathni, Haoran Ni, Vishavdeep Vashisht, Arpita Upadhyaya, and Garegin A Papoian. A tug of war between filament treadmilling and myosin induced contractility generates actin rings. *eLife*, 11, 2022.

[11487] Raffaele Nicastro, Hélène Gaillard, Laura Zarzuela, Marie-Pierre Péli-Gulli, Elisabet Fernández-García, Mercedes Tomé, Néstor García-Rodríguez, Raúl V Durán, Claudio De Virgilio, and Ralf Erik Wellinger. Manganese is a physiologically relevant TORC1 activator in yeast and mammals. *eLife*, 11, 2022.

[11488] Jonathan Nicholas, Nathaniel D Daw, and Daphna Shohamy. Uncertainty alters the balance between incremental learning and episodic memory. *eLife*, 11, 2022.

[11489] Aaron L Nichols, Zack Blumenfeld, Chengcheng Fan, Laura Luebbert, Annet EM Blom, Bruce N Cohen, Jonathan S Marvin, Philip M Borden, Charlene H Kim, Anand K Muthusamy, Amol V Shivange, Hailey J Knox, Hugo Rego Campello, Jonathan H Wang, Dennis A Dougherty, Loren L Looger, Timothy Gallagher, Douglas C Rees, and Henry A Lester. Fluorescence activation mechanism and imaging of drug permeation with new sensors for smoking-cessation ligands. *eLife*, 11, 2022.

[11490] Alec R Nickolls, Gabrielle S O'Brien, Sarah Shnayder, Yunxiao Zhang, Maximilian Nagel, Ardem Patapoutian, and Alexander T Chesler. Reevaluation of Piezo1 as a gut RNA sensor. *eLife*, 11, 2022.

[11491] Judith Nicolas, Bradley R King, David Levesque, Latifa Lazzouni, Emily Coffey, Stephan Swinnen, Julien Doyon, Julie Carrier, and Genevieve Albouy. Sigma oscillations protect or reinstate motor memory depending on their temporal coordination with slow waves. *eLife*, 11, 2022.

[11492] Ole Haagen Nielsen, Fernando Fernandez-Banares, Toshiro Sato, and Darrell S Pardi. Microscopic colitis: Etiopathology, diagnosis, and rational management. *eLife*, 11, 2022.

[11493] Fugui Niu, Peng Han, Jian Zhang, Yuanchu She, Lixin Yang, Jun Yu, Mengru Zhuang, Kezhen Tang, Yuwei Shi, Baisheng Yang, Chunqiao Liu, Bo Peng, and Sheng-Jian Ji. The m6A reader YTHDF2 is a negative regulator for dendrite development and maintenance of retinal ganglion cells. *eLife*, 11, 2022.

[11494] Thandeka Nkosi, Caroline Chasara, Andrea O Papadopoulos, Tiza L Nguni, Farina Karim, Mahomed-Yunus S Moosa, Inbal Gazy, Kondwani Jambo, Willem Hanekom, Alex Sigal, and Zaza M Ndhlovu. Unsuppressed HIV infection impairs T cell responses to SARS-CoV-2 infection and abrogates T cell cross-recognition. *eLife*, 11, 2022.

[11495] Isabel Nocedal and Michael T Laub. Ancestral reconstruction of duplicated signaling proteins reveals the evolution of signaling specificity. *eLife*, 11, 2022.

[11496] Jean-Paul Noel, Edoardo Balzani, Eric Avila, Kaushik J Lakshminarasimhan, Stefania Bruni, Panos Alefantis, Cristina Savin, and Dora E Angelaki. Coding of latent variables in sensory, parietal, and frontal cortices during closed-loop virtual navigation. *eLife*, 11, 2022.

[11497] Jean-Paul Noel, Sabyasachi Shivkumar, Kalpana Dokka, Ralf M Haefner, and Dora E Angelaki. Aberrant causal inference and presence of a compensatory mechanism in autism spectrum disorder. *eLife*, 11, 2022.

[11498] André Nogueira Alves, Marisa Mateus Oliveira, Takashi Koyama, Alexander Shingleton, and Christen Kerry Mirth. Ecdysone coordinates plastic growth with robust pattern in the developing wing. *eLife*, 11, 2022.

[11499] Nighat Noureen, Zhenqing Ye, Yidong Chen, Xiaojing Wang, and Siyuan Zheng. Signature-scoring methods developed for bulk samples are not adequate for cancer single-cell RNA sequencing data. *eLife*, 11, 2022.

[11500] Christopher J Nunn and Sidhartha Goyal. Contingency and selection in mitochondrial genome dynamics. *eLife*, 11, 2022.

[11501] Toshiyuki Oda, Haruaki Yanagisawa, Hideyuki Shinmori, Youichi Ogawa, and Tatsuyoshi Kawamura. Cryo-electron tomography of Birbeck granules reveals the molecular mechanism of langerin lattice formation. *eLife*, 11, 2022.

[11502] Ariel Ogran, Tal Havkin-Solomon, Shirly Becker-Herman, Keren David, Idit Shachar, and Rivka Dikstein. Polysome-CAGE of TCL1-driven chronic lymphocytic leukemia revealed multiple N-terminally altered epigenetic regulators and a translation stress signature. *eLife*, 11, 2022.

[11503] SeCheol Oh, Fabrizio Marinelli, Wenchang Zhou, Jooyeon Lee, Ho Jeong Choi, Min Kim, José D Faraldo-Gómez, and Richard K Hite. Differential ion dehydration energetics explains selectivity in the non-canonical lysosomal K+ channel TMEM175. *eLife*, 11, 2022.

[11504] Ryan P O'Hara, Edem Binka, Adityo Prakosa, Stefan L Zimmerman, Mark J Cartoski, M Roselle Abraham, Dai-Yin Lu, Patrick M Boyle, and Natalia A Trayanova. Personalized computational heart models with T1-mapped fibrotic remodeling predict sudden death risk in patients with hypertrophic cardiomyopathy. *eLife*, 11, 2022.

[11505] Philip J O'Herron, David A Hartmann, Kun Xie, Prakash Kara, and Andy Y Shih. 3D optogenetic control of arteriole diameter in vivo. *eLife*, 11, 2022.

[11506] Michael Ohh, Cassandra C Taber, Fraser G Ferens, and Daniel Tarade. Hypoxia-inducible factor underlies von Hippel-Lindau disease stigmata. *eLife*, 11, 2022.

[11507] Megan Okada, Krithika Rajaram, Russell P Swift, Amanda Mixon, John Alan Maschek, Sean T Prigge, and Paul A Sigala. Critical role for isoprenoids in apicoplast biogenesis by malaria parasites. *eLife*, 11, 2022.

[11508] Justin T Okano, Joan Ponce, Matthias Krönke, and Sally Blower. Lack of ownership of mobile phones could hinder the rollout of mHealth interventions in Africa. *eLife*, 11, 2022.

[11509] Hiroko Okawa, Takeru Kondo, Akishige Hokugo, Philip Cherian, Jesus J Campagna, Nicholas A Lentini, Eric C Sung, Samantha Chiang, Yi-Ling Lin, Frank H Ebetino, Varghese John, Shuting Sun, Charles E McKenna, and Ichiro Nishimura. Mechanism of bisphosphonate-related osteonecrosis of the jaw (BRONJ) revealed by targeted removal of legacy bisphosphonate from jawbone using competing inert hydroxymethylene diphosphonate. *eLife*, 11, 2022.

[11510] Viktor J Olah, Annie M Goettemoeller, Sruti Rayaprolu, Eric B Dammer, Nicholas T Seyfried, Srikant Rangaraju, Jordane Dimidschstein, and Matthew JM Rowan. Biophysical Kv3 channel alterations dampen excitability of cortical PV interneurons and contribute to network hyperexcitability in early Alzheimer's. *eLife*, 11, 2022.

[11511] Viktor J Oláh, Nigel P Pedersen, and Matthew JM Rowan. Ultrafast simulation of large-scale neocortical microcircuitry with biophysically realistic neurons. *eLife*, 11, 2022.

[11512] Marianne Oldehinkel, Alberto Llera, Myrthe Faber, Ismael Huertas, Jan K Buitelaar, Bastiaan R Bloem, Andre F Marquand, Rick C Helmich, Koen V Haak, and Christian F Beckmann. Mapping dopaminergic projections in the human brain with resting-state fMRI. *eLife*, 11, 2022.

[11513] Sandra Olenic, Lim Heo, Michael Feig, and Lee Kroos. Inhibitory proteins block substrate access by occupying the active site cleft of Bacillus subtilis intramembrane protease SpoIVFB. *eLife*, 11, 2022.

[11514] Valeria Oliva, Ron Hartley-Davies, Rosalyn Moran, Anthony E Pickering, and Jonathan CW Brooks. Simultaneous brain, brainstem, and spinal cord pharmacological-fMRI reveals involvement of an endogenous opioid network in attentional analgesia. *eLife*, 11, 2022.

[11515] Giorgio Oliviero, Sergey Kovalchuk, Adelina Rogowska-Wrzesinska, Veit Schwämmle, and Ole N Jensen. Distinct and diverse chromatin proteomes of ageing mouse organs reveal protein signatures that correlate with physiological functions. *eLife*, 11, 2022.

[11516] Yaara Oppenheimer-Shaanan, Gilad Jakoby, Maya L Starr, Romiel Karliner, Gal Eilon, Maxim Itkin, Sergey Malitsky, and Tamir Klein. A dynamic rhizosphere interplay between tree roots and soil bacteria under drought stress. *eLife*, 11, 2022.

[11517] Richard F Oppong, Antonio Terracciano, Martin Picard, Yong Qian, Thomas J Butler, Toshiko Tanaka, Ann Zenobia Moore, Eleanor M Simonsick, Krista Opsahl-Ong, Christopher Coletta, Angelina R Sutin, Myriam Gorospe, Susan M Resnick, Francesco Cucca, Sonja W Scholz, Bryan J Traynor, David Schlessinger, Luigi Ferrucci, and Jun Ding. Personality traits are consistently associated with blood mitochondrial DNA copy number estimated from genome sequences in two genetic cohort studies. *eLife*, 11, 2022.

[11518] Yerdos A Ordabayev, Larry J Friedman, Jeff Gelles, and Douglas L Theobald. Bayesian machine learning analysis of single-molecule fluorescence colocalization images. *eLife*, 11, 2022.

[11519] James Osei-Owusu, Zheng Ruan, Ljubica Mihaljević, Daniel S Matasic, Kevin Hong Chen, Wei Lü, and Zhaozhu Qiu. Molecular mechanism underlying desensitization of the proton-activated chloride channel PAC. *eLife*, 11, 2022.

[11520] Amy Osterman and Alfonso Mondragón. Structures of topoisomerase V in complex with DNA reveal unusual DNA-binding mode and novel relaxation mechanism. *eLife*, 11, 2022.

[11521] Siti N Othman, Spartak N Litvinchuk, Irina Maslova, Hollis Dahn, Kevin R Messenger, Desiree Andersen, Michael J Jowers, Yosuke Kojima, Dmitry V Skorinov, Kiyomi Yasumiba, Ming-Feng Chuang, Yi-Huey Chen, Yoonhyuk Bae, Jennifer Hoti, Yikweon Jang, and Amael Borzee. From Gondwana to the Yellow Sea, evolutionary diversifications of true toads Bufo sp. in the Eastern Palearctic and a revisit of species boundaries for Asian lineages. *eLife*, 11, 2022.

[11522] Simón Oxenford, Jan Roediger, Clemens Neudorfer, Luka Milosevic, Christopher Güttler, Philipp Spindler, Peter Vajkoczy, Wolf-Julian Neumann, Andrea Kühn, and Andreas Horn. Lead-OR: A multimodal platform for deep brain stimulation surgery. *eLife*, 11, 2022.

[11523] Osnat Oz, Lior Matityahu, Aviv Mizrahi-Kliger, Alexander Kaplan, Noa Berkowitz, Lior Tiroshi, Hagai Bergman, and Joshua A Goldberg. Non-uniform distribution of dendritic nonlinearities differentially engages thalamostriatal and corticostriatal inputs onto cholinergic interneurons. *eLife*, 11, 2022.

[11524] Lang Pan, Lu Chen, Jun Lv, Yuanjie Pang, Yu Guo, Pei Pei, Huaidong Du, Ling Yang, Iona Y Millwood, Robin G Walters, Yiping Chen, Weiwei Gong, Junshi Chen, Canqing Yu, Zhengming Chen, and Liming Li. Association of egg consumption, metabolic markers, and risk of cardiovascular diseases: A nested case-control study. *eLife*, 11, 2022.

[11525] Mingyao Pan and Bo Li. T cell receptor convergence is an indicator of antigen-specific T cell response in cancer immunotherapies. *eLife*, 11, 2022.

[11526] James C Pang, James K Rilling, James A Roberts, Martijn P van den Heuvel, and Luca Cocchi. Evolutionary shaping of human brain dynamics. *eLife*, 11, 2022.

[11527] Lucas C Pantaleão, Isabella Inzani, Samuel Furse, Elena Loche, Antonia Hufnagel, Thomas Ashmore, Heather L Blackmore, Benjamin Jenkins, Asha A M Carpenter, Ania Wilczynska, Martin Bushell, Albert Koulman, Denise S Fernandez-Twinn, and Susan E Ozanne. Maternal diet-induced obesity during pregnancy alters lipid supply to mouse E18.5 fetuses and changes the cardiac tissue lipidome in a sex-dependent manner. *eLife*, 11, 2022.

[11528] Alonso Javier Pardal and Andrew James Bowman. A specific role for importin-5 and NASP in the import and nuclear hand-off of monomeric H3. *eLife*, 11, 2022.

[11529] Cheon-Gyu Park, Wookyung Yu, and Byung-Chang Suh. Molecular basis of the PIP2-dependent regulation of CaV2.2 channel and its modulation by CaV $\beta$ subunits. *eLife*, 11, 2022.

[11530] Hae J Park, Mark A Gregory, Vadym Zaberezhnyy, Andrew Goodspeed, Craig T Jordan, Jeffrey S Kieft, and James DeGregori. Therapeutic resistance in acute myeloid leukemia cells is mediated by a novel ATM/mTOR pathway regulating oxidative phosphorylation. *eLife*, 11, 2022.

[11531] Jong S Park, Austin M Gabel, Polina Kassir, Lois Kang, Prableen K Chowdhary, Afia Osei-Ntansah, Neil D Tran, Soujanya Viswanathan, Bryanna Canales, Pengfei Ding, Young-Sam Lee, and Rachel Brewster. N-myc downstream regulated gene 1 (ndrg1) functions as a molecular switch for cellular adaptation to hypoxia. *eLife*, 11, 2022.

[11532] Joonkoo Park and David E Huber. A visual sense of number emerges from divisive normalization in a simple center-surround convolutional network. *eLife*, 11, 2022.

[11533] Sohyeon Park, Hyunjoong Kim, Yi Wang, Dae Seok Eom, and Jun Allard. Zebrafish airinemes optimize their shape between ballistic and diffusive search. *eLife*, 11, 2022.

[11534] Su Mi Park, Cooper E Roache, Philip H Iffland, Hans J Moldenhauer, Katia K Matychak, Amber E Plante, Abby G Lieberman, Peter B Crino, and Andrea Meredith. BK channel properties correlate with neurobehavioral severity in three KCNMA1-linked channelopathy mouse models. *eLife*, 11, 2022.

[11535] Matthew T Parker, Beth K Soanes, Jelena Kusakina, Antoine Larrieu, Katarzyna Knop, Nisha Joy, Friedrich Breidenbach, Anna V Sherwood, Geoffrey J Barton, Sebastian M Fica, Brendan H Davies, and Gordon G Simpson. m6A modification of U6 snRNA modulates usage of two major classes of pre-mRNA 5′ splice site. *eLife*, 11, 2022.

[11536] Philip RL Parker, Elliott TT Abe, Natalie T Beatie, Emmalyn SP Leonard, Dylan M Martins, Shelby L Sharp, David G Wyrick, Luca Mazzucato, and Cristopher M Niell. Distance estimation from monocular cues in an ethological visuomotor task. *eLife*, 11, 2022.

[11537] Pau Pascual-Garcia, Shawn C Little, and Maya Capelson. Nup98-dependent transcriptional memory is established independently of transcription. *eLife*, 11, 2022.

[11538] Andrius PaŠukonis, Shirley Jennifer Serrano-Rojas, Marie-Therese Fischer, Matthias-Claudio Loretto, Daniel A Shaykevich, Bibiana Rojas, Max Ringler, Alexandre B Roland, Alejandro Marcillo-Lara, Eva Ringler, Camilo Rodríguez, Luis A Coloma, and Lauren A O'Connell. Contrasting parental roles shape sex differences in poison frog space use but not navigational performance. *eLife*, 11, 2022.

[11539] Ruhi Patel, Himani Galagali, John K Kim, and Alison R Frand. Feedback between a retinoid-related nuclear receptor and the let-7 microRNAs controls the pace and number of molting cycles in C. elegans. *eLife*, 11, 2022.

[11540] Neil Paterson and Tim Lämmermann. Macrophage network dynamics depend on haptokinesis for optimal local surveillance. *eLife*, 11, 2022.

[11541] Dilan SR Patiranage, Elodie Rey, Nazgol Emrani, Gordon Wellman, Karl Schmid, Sandra M Schmöckel, Mark Tester, and Christian Jung. Genome-wide association study in quinoa reveals selection pattern typical for crops with a short breeding history. *eLife*, 11, 2022.

[11542] Jennifer Patritti Cram, Jianqiang Wu, Robert A Coover, Tilat A Rizvi, Katherine E Chaney, Ramya Ravindran, Jose A Cancelas, Robert J Spinner, and Nancy Ratner. P2RY14 cAMP signaling regulates Schwann cell precursor self-renewal, proliferation, and nerve tumor initiation in a mouse model of neurofibromatosis. *eLife*, 11, 2022.

[11543] Austin H Patton, Emilie J Richards, Katelyn J Gould, Logan K Buie, and Christopher H Martin. Hybridization alters the shape of the genotypic fitness landscape, increasing access to novel fitness peaks during adaptive radiation. *eLife*, 11, 2022.

[11544] Jordan L Pauli, Jane Y Chen, Marcus L Basiri, Sekun Park, Matthew E Carter, Elisenda Sanz, G Stanley McKnight, Garret D Stuber, and Richard D Palmiter. Molecular and anatomical characterization of parabrachial neurons and their axonal projections. *eLife*, 11, 2022.

[11545] Eric Paulissen, Nicholas J Palmisano, Joshua S Waxman, and Benjamin L Martin. Somite morphogenesis is required for axial blood vessel formation during zebrafish embryogenesis. *eLife*, 11, 2022.

[11546] Julie Paxman, Zhen Zhou, Richard O'Laughlin, Yuting Liu, Yang Li, Wanying Tian, Hetian Su, Yanfei Jiang, Shayna E Holness, Elizabeth Stasiowski, Lev S Tsimring, Lorraine Pillus, Jeff Hasty, and Nan Hao. Age-dependent aggregation of ribosomal RNA-binding proteins links deterioration in chromatin stability with challenges to proteostasis. *eLife*, 11, 2022.

[11547] Maximilian Peer, Hua Yuan, Yubo Zhang, Katharina Korbula, Peter Novick, and Gang Dong. Double NPY motifs at the N-terminus of the yeast t-SNARE Sso2 synergistically bind Sec3 to promote membrane fusion. *eLife*, 11, 2022.

[11548] Di Peng, Koji Ando, Melina Hußmann, Marleen Gloger, Renae Skoczylas, Naoki Mochizuki, Christer Betsholtz, Shigetomo Fukuhara, Stefan Schulte-Merker, Nathan D Lawson, and Katarzyna Koltowska. Proper migration of lymphatic endothelial cells requires survival and guidance cues from arterial mural cells. *eLife*, 11, 2022.

[11549] Catalina Pereira, Gerardo A Arroyo-Martinez, Matthew Z Guo, Michael S Downey, Emma R Kelly, Kathryn J Grive, Shantha K Mahadevaiah, Jennie R Sims, Vitor M Faca, Charlton Tsai, Carl J Schiltz, Niek Wit, Heinz Jacobs, Nathan L Clark, Raimundo Freire, James Turner, Amy M Lyndaker, Miguel A Brieno-Enriquez, Paula E Cohen, Marcus B Smolka, and Robert S Weiss. Multiple 9-1-1 complexes promote homolog synapsis, DSB repair, and ATR signaling during mammalian meiosis. *eLife*, 11, 2022.

[11550] Marta Perera, Silas Boye Nissen, Martin Proks, Sara Pozzi, Rita S Monteiro, Ala Trusina, and Joshua M Brickman. Transcriptional heterogeneity and cell cycle regulation as central determinants of Primitive Endoderm priming. *eLife*, 11, 2022.

[11551] Maria C Perez-Flores, Eric Verschooten, Jeong Han Lee, Hyo Jeong Kim, Philip X Joris, and Ebenezer N Yamoah. Intrinsic mechanical sensitivity of mammalian auditory neurons as a contributor to sound-driven neural activity. *eLife*, 11, 2022.

[11552] Albert Pérez-Martí, Suresh Ramakrishnan, Jiayi Li, Aurelien Dugourd, Martijn R Molenaar, Luigi R De La Motte, Kelli Grand, Anis Mansouri, Mélanie Parisot, Soeren S Lienkamp, Julio Saez-Rodriguez, and Matias Simons. Reducing lipid bilayer stress by monounsaturated fatty acids protects renal proximal tubules in diabetes. *eLife*, 11, 2022.

[11553] Nicholas D Peterson, Janneke D Icso, J Elizabeth Salisbury, Tomás Rodríguez, Paul R Thompson, and Read Pukkila-Worley. Pathogen infection and cholesterol deficiency activate the C. elegans p38 immune pathway through a TIR-1/SARM1 phase transition. *eLife*, 11, 2022.

[11554] Radmila Petric, Matina Kalcounis-Rueppell, and Catherine A Marler. Testosterone pulses paired with a location induce a place preference to the nest of a monogamous mouse under field conditions. *eLife*, 11, 2022.

[11555] Iurii Petrov and Andrey Alexeyenko. Individualized discovery of rare cancer drivers in global network context. *eLife*, 11, 2022.

[11556] Stéphane Peyrégne, Janet Kelso, Benjamin M Peter, and Svante Pääbo. The evolutionary history of human spindle genes includes back-and-forth gene flow with Neandertals. *eLife*, 11, 2022.

[11557] Omar Peña-Ramos, Lucia Chiao, Xianghua Liu, Xiaomeng Yu, Tianyou Yao, Henry He, and Zheng Zhou. Autophagosomes fuse to phagosomes and facilitate the degradation of apoptotic cells in Caenorhabditis elegans. *eLife*, 11, 2022.

[11558] Alejandro Peñin-Franch, José Antonio García-Vidal, Carlos Manuel Martínez, Pilar Escolar-Reina, Rosa M Martínez-Ojeda, Ana I Gómez, Juan M Bueno, Francisco Minaya-Muñoz, Fermín Valera-Garrido, Francesc Medina-Mirapeix, and Pablo Pelegrín. Galvanic current activates the NLRP3 inflammasome to promote Type I collagen production in tendon. *eLife*, 11, 2022.

[11559] Thomas Pfeffer, Christian Keitel, Daniel S Kluger, Anne Keitel, Alena Russmann, Gregor Thut, Tobias H Donner, and Joachim Gross. Coupling of pupil- and neuronal population dynamics reveals diverse influences of arousal on cortical processing. *eLife*, 11, 2022.

[11560] Jasmin Philip, Mihkel örd, Andriele Silva, Shaneen Singh, John FX Diffley, Dirk Remus, Mart Loog, and Amy E Ikui. Cdc6 is sequentially regulated by PP2A-Cdc55, Cdc14, and Sic1 for origin licensing in S. cerevisiae. *eLife*, 11, 2022.

[11561] Ellen Phillips, Naseer Ahmad, Li Sun, James Iben, Christopher J Walkey, Aleksandra Rusin, Tony Yuen, Clifford J Rosen, Ian M Willis, Mone Zaidi, and Deborah L Johnson. MAF1, a repressor of RNA polymerase III-dependent transcription, regulates bone mass. *eLife*, 11, 2022.

[11562] Jonathan E Phillips, Maribel Santos, Mohammed Konchwala, Chao Xing, and Duojia Pan. Genome editing in the unicellular holozoan Capsaspora owczarzaki suggests a premetazoan role for the Hippo pathway in multicellular morphogenesis. *eLife*, 11, 2022.

[11563] Ryan S Phillips, Hidehiko Koizumi, Yaroslav I Molkov, Jonathan E Rubin, and Jeffrey C Smith. Predictions and experimental tests of a new biophysical model of the mammalian respiratory oscillator. *eLife*, 11, 2022.

[11564] Ryan S Phillips and Jonathan E Rubin. Putting the theory into 'burstlet theory' with a biophysical model of burstlets and bursts in the respiratory preBötzinger complex. *eLife*, 11, 2022.

[11565] Chiara Piantoni, Manuel Paina, David Molla, Sheng Liu, Giorgia Bertoli, Hongmei Jiang, Yanyan Wang, Yi Wang, Yi Wang, Dario DiFrancesco, Andrea Barbuti, Annalisa Bucchi, and Mirko Baruscotti. Chinese natural compound decreases pacemaking of rabbit cardiac sinoatrial cells by targeting second messenger regulation of f-channels. *eLife*, 11, 2022.

[11566] Francesco M Piccolo, Nathaniel R Kastan, Tomomi Haremaki, Qingyun Tian, Tiago L Laundos, Riccardo De Santis, Andrew J Beaudoin, Thomas S Carroll, Ji-Dung Luo, Ksenia Gnedeva, Fred Etoc, AJ Hudspeth, and Ali H Brivanlou. Role of YAP in early ectodermal specification and a Huntington's Disease model of human neurulation. *eLife*, 11, 2022.

[11567] Benjamin Pillet, Alfonso Méndez-Godoy, Guillaume Murat, Sébastien Favre, Michael Stumpe, Laurent Falquet, and Dieter Kressler. Dedicated chaperones coordinate co-translational regulation of ribosomal protein production with ribosome assembly to preserve proteostasis. *eLife*, 11, 2022.

[11568] Morgan L Pimm, Xinbei Liu, Farzana Tuli, Jennifer Heritz, Ashley Lojko, and Jessica L Henty-Ridilla. Visualizing molecules of functional human profilin. *eLife*, 11, 2022.

[11569] Judith Pineau, Léa Pinon, Olivier Mesdjian, Jacques Fattaccioli, Ana-Maria Lennon Duménil, and Paolo Pierobon. Microtubules restrict F-actin polymerization to the immune synapse via GEF-H1 to maintain polarity in lymphocytes. *eLife*, 11, 2022.

[11570] Justine M Pinskey, Tyler M Hoard, Xiao-Feng Zhao, Nicole E Franks, Zoë C Frank, Alexandra N McMellen, Roman J Giger, and Benjamin L Allen. Plexins promote Hedgehog signaling through their cytoplasmic GAP activity. *eLife*, 11, 2022.

[11571] Justine M Pinskey, Adhya Lagisetty, Long Gui, Nhan Phan, Evan Reetz, Amirrasoul Tavakoli, Gang Fu, and Daniela Nicastro. Three-dimensional flagella structures from animals' closest unicellular relatives, the Choanoflagellates. *eLife*, 11, 2022.

[11572] Lucas Pinto, David W Tank, and Carlos D Brody. Multiple timescales of sensory-evidence accumulation across the dorsal cortex. *eLife*, 11, 2022.

[11573] Lukasz Piszczek, Andreea Constantinescu, Dominic Kargl, Jelena Lazovic, Anton Pekcec, Janet R Nicholson, and Wulf Haubensak. Dissociation of impulsive traits by subthalamic metabotropic glutamate receptor 4. *eLife*, 11, 2022.

[11574] Christiane Pleuger, Dingding Ai, Minea L Hoppe, Laura T Winter, Daniel Bohnert, Dominik Karl, Stefan Guenther, Slava Epelman, Crystal Kantores, Monika Fijak, Sarina Ravens, Ralf Middendorff, Johannes U Mayer, Kate L Loveland, Mark Hedger, Sudhanshu Bhushan, and Andreas Meinhardt. The regional distribution of resident immune cells shapes distinct immunological environments along the murine epididymis. *eLife*, 11, 2022.

[11575] Ignacio Polti, Matthias Nau, Raphael Kaplan, Virginie van Wassenhove, and Christian F Doeller. Rapid encoding of task regularities in the human hippocampus guides sensorimotor timing. *eLife*, 11, 2022.

[11576] Maurizio Porfiri, Peng Zhang, and Sean D Peterson. Hydrodynamic model of fish orientation in a channel flow. *eLife*, 11, 2022.

[11577] Marion Portes, Thomas Mangeat, Natacha Escallier, Ophélie Dufrancais, Brigitte Raynaud-Messina, Christophe Thibault, Isabelle Maridonneau-Parini, Christel Vérollet, and Renaud Poincloux. Nanoscale architecture and coordination of actin cores within the sealing zone of human osteoclasts. *eLife*, 11, 2022.

[11578] Dmitry Postnov, Donald J Marsh, Will A Cupples, Niels-Henrik Holstein-Rathlou, and Olga Sosnovtseva. Synchronization in renal microcirculation unveiled with high-resolution blood flow imaging. *eLife*, 11, 2022.

[11579] Cilia R Pothast, Romy C Dijkland, Melissa Thaler, Renate S Hagedoorn, Michel GD Kester, Anne K Wouters, Pieter S Hiemstra, Martijn J van Hemert, Stephanie Gras, JH Frederik Falkenburg, and Mirjam HM Heemskerk. SARS-CoV-2-specific CD4+ and CD8+ T cell responses can originate from cross-reactive CMV-specific T cells. *eLife*, 11, 2022.

[11580] Davide Potrich, Mirko Zanon, and Giorgio Vallortigara. Archerfish number discrimination. *eLife*, 11, 2022.

[11581] Emily Nicole Powers, Charlene Chan, Ella Doron-Mandel, Lidia Llacsahuanga Allcca, Jenny Kim Kim, Marko Jovanovic, and Gloria Ann Brar. Bidirectional promoter activity from expression cassettes can drive off-target repression of neighboring gene translation. *eLife*, 11, 2022.

[11582] RT Pramod, Michael A Cohen, Joshua B Tenenbaum, and Nancy Kanwisher. Invariant representation of physical stability in the human brain. *eLife*, 11, 2022.

[11583] Anadika R Prasad, Inês Lago-Baldaia, Matthew P Bostock, Zaynab Housseini, and Vilaiwan M Fernandes. Differentiation signals from glia are fine-tuned to set neuronal numbers during development. *eLife*, 11, 2022.

[11584] S Ganga Prasath, Souvik Mandal, Fabio Giardina, Jordan Kennedy, Venkatesh N Murthy, and L Mahadevan. Dynamics of cooperative excavation in ant and robot collectives. *eLife*, 11, 2022.

[11585] Hannes Preiß, Anna C Kögler, David Mörsdorf, Daniel čapek, Gary H Soh, Katherine W Rogers, Hernán Morales-Navarrete, María Almuedo-Castillo, and Patrick Müller. Regulation of Nodal signaling propagation by receptor interactions and positive feedback. *eLife*, 11, 2022.

[11586] Asher Preska Steinberg, Mingzhi Lin, and Edo Kussell. Core genes can have higher recombination rates than accessory genes within global microbial populations. *eLife*, 11, 2022.

[11587] Carlos A Prete, Lewis F Buss, Charles Whittaker, Tassila Salomon, Marcio K Oikawa, Rafael HM Pereira, Isabel CG Moura, Lucas Delerino, Manoel Barral-Netto, Natalia M Tavares, Rafael FO Franca, Viviane S Boaventura, Fabio Miyajima, Alfredo Mendrone-Junior, Cesar de Almeida-Neto, Nanci A Salles, Suzete C Ferreira, Karine A Fladzinski, Luana M de Souza, Luciane K Schier, Patricia M Inoue, Lilyane A Xabregas, Myuki AE Crispim, Nelson Fraiji, Fernando LV Araujo, Luciana MB Carlos, Veridiana Pessoa, Maisa A Ribeiro, Rosenvaldo E de Souza, Sônia MN da Silva, Anna F Cavalcante, Maria IB ValenÇa, Maria V da Silva, Esther Lopes, Luiz A Filho, Sheila OG

Mateos, Gabrielle T Nunes, Alexander L Silva-Junior, Michael P Busch, Marcia C Castro, Christopher Dye, Oliver Ratmann, Nuno R Faria, Vítor H Nascimento, and Ester C Sabino. SARS-CoV-2 antibody dynamics in blood donors and COVID-19 epidemiology in eight Brazilian state capitals: A serial cross-sectional study. *eLife*, 11, 2022.

[11588] Carlotta Pribbenow, Yi-chun Chen, M-Marcel Heim, Desiree Laber, Silas Reubold, Eric Reynolds, Isabella Balles, Tania Fernández-d V Alquicira, Raquel Suárez-Grimalt, Lisa Scheunemann, Carolin Rauch, Tanja Matkovic, Jörg Rösner, Gregor Lichtner, Sridhar R Jagannathan, and David Owald. Postsynaptic plasticity of cholinergic synapses underlies the induction and expression of appetitive and familiarity memories in Drosophila. *eLife*, 11, 2022.

[11589] Ariel Pribluda, Anneleen Daemen, Anthony Nelson Lima, Xi Wang, Marc Hafner, Chungkee Poon, Zora Modrusan, Anand Kumar Katakam, Oded Foreman, Jefferey Eastham, Jefferey Hung, Benjamin Haley, Julia T Garcia, Erica L Jackson, and Melissa R Junttila. EHMT2 methyltransferase governs cell identity in the lung and is required for KRAS G12D tumor development and propagation. *eLife*, 11, 2022.

[11590] Jacob S Prince, Ian Charest, Jan W Kurzawski, John A Pyles, Michael J Tarr, and Kendrick N Kay. Improving the accuracy of single-trial fMRI response estimates using GLMsingle. *eLife*, 11, 2022.

[11591] Felix Proulx-Giraldeau, Jan M Skotheim, and Paul FranÇois. Evolution of cell size control is canalized towards adders or sizers by cell cycle structure and selective pressures. *eLife*, 11, 2022.

[11592] Pauline Provini, Alexandre Brunet, Andréa Filippo, and Sam Van Wassenbergh. In vivo intraoral waterflow quantification reveals hidden mechanisms of suction feeding in fish. *eLife*, 11, 2022.

[11593] Christian A Pulver, Emine Celiker, Charlie Woodrow, Inga Geipel, Carl D Soulsbury, Darron A Cullen, Stephen M Rogers, Daniel Veitch, and Fernando Montealegre-Z. Ear pinnae in a neotropical katydid (Orthoptera: Tettigoniidae) function as ultrasound guides for bat detection. *eLife*, 11, 2022.

[11594] Natasha M Puri, Giovanna R Romano, Ting-Yu Lin, Quynh N Mai, and Roshanak Irannejad. The organic cation transporter 2 regulates dopamine D1 receptor signaling at the Golgi apparatus. *eLife*, 11, 2022.

[11595] Karin Purshouse, Elias T Friman, Shelagh Boyle, Pooran Singh Dewari, Vivien Grant, Alhafidz Hamdan, Gillian M Morrison, Paul M Brennan, Sjoerd V Beentjes, Steven M Pollard, and Wendy A Bickmore. Oncogene expression from extrachromosomal DNA is driven by copy number amplification and does not require spatial clustering in glioblastoma stem cells. *eLife*, 11, 2022.

[11596] Guangyao Qi, Wen Fang, Shenghao Li, Junru Li, and Liping Wang. Neural dynamics of causal inference in the macaque frontoparietal circuit. *eLife*, 11, 2022.

[11597] Jialing Qi, Annegret Rittershaus, Rashmi Priya, Shivani Mansingh, Didier YR Stainier, and Christian SM Helker. Apelin signaling dependent endocardial protrusions promote cardiac trabeculation in zebrafish. *eLife*, 11, 2022.

[11598] Shan Qi, Javier Mota, Siu-Hong Chan, Johanna Villarreal, Nan Dai, Shailee Arya, Robert A Hromas, Manjeet K Rao, Ivan R Corrêa Jr, and Yogesh K Gupta. RNA binding to human METTL3-METTL14 restricts N6-deoxyadenosine methylation of DNA in vitro. *eLife*, 11, 2022.

[11599] Pengge Qian, Xu Wang, Chuan-Qi Zhong, Jiaxu Wang, Mengya Cai, Wang Nguitragool, Jian Li, Huiting Cui, and Jing Yuan. Inner membrane complex proteomics reveals a palmitoylation regulation critical for intraerythrocytic development of malaria parasite. *eLife*, 11, 2022.

[11600] Lingxia Qiao, Zhi-Bo Zhang, Wei Zhao, Ping Wei, and Lei Zhang. Network design principle for robust oscillatory behaviors with respect to biological noise. *eLife*, 11, 2022.

[11601] Juan Qin, Jingfeng Zhang, Lianyun Lin, Omid Haji-Ghassemi, Zhi Lin, Kenneth J Woycechowsky, Filip Van Petegem, Yan Zhang, and Zhiguang Yuchi. Structures of PKA–phospholamban complexes reveal a mechanism of familial dilated cardiomyopathy. *eLife*, 11, 2022.

[11602] Fernando Racimo, Elia Valentini, Gaston Rijo De León, Teresa L Santos, Anna Norberg, Lane M Atmore, Myranda Murray, Sanja M Hakala, Frederik Appel Olsen, Charlie J Gardner, and Julia B Halder. The biospheric emergency calls for scientists to change tactics. *eLife*, 11, 2022.

[11603] Atanas Radkov, Anne L Sapiro, Sebastian Flores, Corey Henderson, Hayden Saunders, Rachel Kim, Steven Massa, Samuel Thompson, Chase Mateusiak, Jacob Biboy, Ziyi Zhao, Lea M Starita, William L Hatleberg, Waldemar Vollmer, Alistair B Russell, Jean-Pierre Simorre, Spencer Anthony-Cahill, Peter Brzovic, Beth Hayes, and Seemay Chou. Antibacterial potency of type VI amidase effector toxins is dependent on substrate topology and cellular context. *eLife*, 11, 2022.

[11604] Daniel Radtke, Natalie Thuma, Christine Schülein, Philipp Kirchner, Arif B Ekici, Kilian Schober, and David Voehringer. Th2 single-cell heterogeneity and clonal distribution at distant sites in helminth-infected mice. *eLife*, 11, 2022.

[11605] Atefeh Rafiei, Sofía Cruz Tetlalmatzi, Claire H Edrington, Linda Lee, D Alex Crowder, Daniel J Saltzberg, Andrej Sali, Gary Brouhard, and David C Schriemer. Doublecortin engages the microtubule lattice through a cooperative binding mode involving its C-terminal domain. *eLife*, 11, 2022.

[11606] Sarah M Rajtmajer, Timothy M Errington, and Frank G Hillary. How failure to falsify in high-volume science contributes to the replication crisis. *eLife*, 11, 2022.

[11607] Michael J Rale, Brianna Romer, Brian P Mahon, Sophie M Travis, and Sabine Petry. The conserved centrosomin motif, $\gamma$TuNA, forms a dimer that directly activates microtubule nucleation by the $\gamma$-tubulin ring complex ($\gamma$TuRC). *eLife*, 11, 2022.

[11608] Janani Ramachandran, Weiqiang Zhou, Anna E Bardenhagen, Talia Nasr, Ellen R Yates, Aaron M Zorn, Hongkai Ji, and Steven A Vokes. Hedgehog regulation of epithelial cell state and morphogenesis in the larynx. *eLife*, 11, 2022.

[11609] Karthik Ramanadane, Monique S Straub, Raimund Dutzler, and Cristina Manatschal. Structural and functional properties of a magnesium transporter of the SLC11/NRAMP family. *eLife*, 11, 2022.

[11610] Komal Ramani, Nirmala Mavila, Aushinie Abeynayake, Maria Lauda Tomasi, Jiaohong Wang, Michitaka Matsuda, and Eki Seki. Targeting A-kinase anchoring protein 12 phosphorylation in hepatic stellate cells regulates liver injury and fibrosis in mouse models. *eLife*, 11, 2022.

[11611] Abhinay Ramaprasad, Paul-Christian Burda, Enrica Calvani, Aaron J Sait, Susana Alejandra Palma-Duran, Chrislaine Withers-Martinez, Fiona Hackett, James Macrae, Lucy Collinson, Tim Wolf Gilberger, and Michael J Blackman. A choline-releasing glycerophosphodiesterase essential for phosphatidylcholine biosynthesis and blood stage development in the malaria parasite. *eLife*, 11, 2022.

[11612] Miguel Ramirez, Yuliya Badayeva, Joanna Yeung, Joshua Wu, Ayasha Abdalla-Wyse, Erin Yang, Brett Trost, Stephen W Scherer, and Daniel Goldowitz. Temporal analysis of enhancers during mouse cerebellar development reveals dynamic and novel regulatory functions. *eLife*, 11, 2022.

[11613] Erika K Ramos, Chia-Feng Tsai, Yuzhi Jia, Yue Cao, Megan Manu, Rokana Taftaf, Andrew D Hoffmann, Lamiaa El-Shennawy, Marina A Gritsenko, Valery Adorno-Cruz, Emma J Schuster, David Scholten, Dhwani Patel, Xia Liu, Priyam Patel, Brian Wray, Youbin Zhang, Shanshan Zhang, Ronald J Moore, Jeremy V Mathews, Matthew J Schipma, Tao Liu, Valerie L Tokars, Massimo Cristofanilli, Tujin Shi, Yang Shen, Nurmaa K Dashzeveg, and Huiping Liu. Machine learning-assisted elucidation of CD81–CD44 interactions in promoting cancer stemness and extracellular vesicle integrity. *eLife*, 11, 2022.

[11614] Yajamana Ramu, Jayden Yamakaze, Yufeng Zhou, Toshinori Hoshi, and Zhe Lu. Blocking Kir6.2 channels with SpTx1 potentiates glucose-stimulated insulin secretion from murine pancreatic $\beta$ cells and lowers blood glucose in diabetic mice. *eLife*, 11, 2022.

[11615] Kamalini G Ranasinghe, Parul Verma, Chang Cai, Xihe Xie, Kiwamu Kudo, Xiao Gao, Hannah Lerner, Danielle Mizuiri, Amelia Strom, Leonardo Iaccarino, Renaud La Joie, Bruce L Miller, Maria Luisa Gorno-Tempini, Katherine P Rankin, William J Jagust, Keith Vossel, Gil D Rabinovici, Ashish Raj, and Srikantan S Nagarajan. Altered excitatory and inhibitory neuronal subpopulation parameters are distinctly associated with tau and amyloid in Alzheimer's disease. *eLife*, 11, 2022.

[11616] Ann Rancourt, Sachiko Sato, and Masahiko S Satoh. Empirical single-cell tracking and cell-fate simulation reveal dual roles of p53 in tumor suppression. *eLife*, 11, 2022.

[11617] Lyra O Randzavola, Paige M Mortimer, Emma Garside, Elizabeth R Dufficy, Andrea Schejtman, Georgia Roumelioti, Lu Yu, Mercedes Pardo, Kerstin Spirohn, Charlotte Tolley, Cordelia Brandt, Katherine Harcourt, Esme Nichols, Mike Nahorski, Geoff Woods, James C Williamson, Shreehari Suresh, John M Sowerby, Misaki Matsumoto, Celio XC Santos, Cher Shen Kiar, Subhankar Mukhopadhyay, William M Rae, Gordon J Dougan, John Grainger, Paul J Lehner, Michael A Calderwood, Jyoti Choudhary, Simon Clare, Anneliese Speak, Giorgia Santilli, Alex Bateman, Kenneth GC Smith, Francesca Magnani, and David C Thomas. EROS is a selective chaperone regulating the phagocyte NADPH oxidase and purinergic signalling. *eLife*, 11, 2022.

[11618] Sanket Rane, Thea Hogan, Edward Lee, Benedict Seddon, and Andrew J Yates. Towards a unified model of naive T cell dynamics across the lifespan. *eLife*, 11, 2022.

[11619] Joseph L Ransdell, Jonathan D Moreno, Druv Bhagavan, Jonathan R Silva, and Jeanne M Nerbonne. Intrinsic mechanisms in the gating of resurgent Na+ currents. *eLife*, 11, 2022.

[11620] Bitya Raphael-Mizrahi, Malka Attar-Namdar, Mukesh Chourasia, Maria G Cascio, Avital Shurki, Joseph Tam, Moshe Neuman, Neta Rimmerman, Zvi Vogel, Arie Shteyer, Roger G Pertwee, Andreas Zimmer, Natalya M Kogan, Itai Bab, and Yankel Gabet. Osteogenic growth peptide is a potent anti-inflammatory and bone preserving hormone via cannabinoid receptor type 2. *eLife*, 11, 2022.

[11621] David V Rasicci, Prince Tiwari, Skylar ML Bodt, Rohini Desetty, Fredrik R Sadler, Sivaraj Sivaramakrishnan, Roger Craig, and Christopher M Yengo. Dilated cardiomyopathy mutation E525K in human beta-cardiac myosin stabilizes the interacting-heads motif and super-relaxed state of myosin. *eLife*, 11, 2022.

[11622] Liisa Raud, Christina Thunberg, and René J Huster. Partial response electromyography as a marker of action stopping. *eLife*, 11, 2022.

[11623] Parth K Raval, Sriram G Garg, and Sven B Gould. Endosymbiotic selective pressure at the origin of eukaryotic cell biology. *eLife*, 11, 2022.

[11624] Guillaume Ravel, Michel Bergmann, Alain Trubuil, Julien Deschamps, Romain Briandet, and Simon Labarthe. Inferring characteristics of bacterial swimming in biofilm matrix from time-lapse confocal laser scanning microscopy. *eLife*, 11, 2022.

[11625] Nithin Sam Ravi, Beeke Wienert, Stacia K Wyman, Henry William Bell, Anila George, Gokulnath Mahalingam, Jonathan T Vu, Kirti Prasad, Bhanu Prasad Bandlamudi, Nivedhitha Devaraju, Vignesh Rajendiran, Nazar Syedbasha, Aswin Anand Pai, Yukio Nakamura, Ryo Kurita, Muthuraman Narayanasamy, Poonkuzhali Balasubramanian, Saravanabhavan Thangavel, Srujan Marepally, Shaji R Velayudhan, Alok Srivastava, Mark A DeWitt, Merlin Crossley, Jacob E Corn, and Kumarasamypet M Mohankumar. Identification of novel HPFH-like mutations by CRISPR base editing that elevate the expression of fetal hemoglobin. *eLife*, 11, 2022.

[11626] Ethiraj Ravindran, Nobuto Arashiki, Lena-Luise Becker, Kohtaro Takizawa, Jonathan Lévy, Thomas Rambaud, Konstantin L Makridis, Yoshio Goshima, Na Li, Maaike Vreeburg, Bénédicte Demeer, Achim Dickmanns, Alexander PA Stegmann, Hao Hu, Fumio Nakamura, and Angela M Kaindl. Monoallelic CRMP1 gene variants cause neurodevelopmental disorder. *eLife*, 11, 2022.

[11627] Daniel J Rawle, Thuy T Le, Troy Dumenil, Cameron Bishop, Kexin Yan, Eri Nakayama, Phillip I Bird, and Andreas Suhrbier. Widespread discrepancy in Nnt genotypes and genetic backgrounds complicates granzyme A and other knockout mouse studies. *eLife*, 11, 2022.

[11628] Alokananda Ray and Xin Li. A Notch-dependent transcriptional mechanism controls expression of temporal patterning factors in Drosophila medulla. *eLife*, 11, 2022.

[11629] Suhita Ray, Linda Chee, Yichao Zhou, Meg A Schaefer, Michael J Naldrett, Sophie Alvarez, Nicholas T Woods, and Kyle J Hewitt. Functional requirements for a Samd14-capping protein complex in stress erythropoiesis. *eLife*, 11, 2022.

[11630] Steffen M Recktenwald, Greta Simionato, Marcelle GM Lopes, Fabia Gamboni, Monika Dzieciatkowska, Patrick Meybohm, Kai Zacharowski, Andreas von Knethen, Christian Wagner, Lars Kaestner, Angelo D'Alessandro, and Stephan Quint. Cross-talk between red blood cells and plasma influences blood flow and omics phenotypes in severe COVID-19. *eLife*, 11, 2022.

[11631] William T Redman, Nora S Wolcott, Luca Montelisciani, Gabriel Luna, Tyler D Marks, Kevin K Sit, Che-Hang Yu, Spencer Smith, and Michael J Goard. Long-term transverse imaging of the hippocampus with glass microperiscopes. *eLife*, 11, 2022.

[11632] Alex Reichenbach, Rachel E Clarke, Romana Stark, Sarah Haas Lockie, Mathieu Mequinion, Harry Dempsey, Sasha Rawlinson, Felicia Reed, Tara Sepehrizadeh, Michael DeVeer, Astrid C Munder, Juan Nunez-Iglesias, David C Spanswick, Randall Mynatt, Alexxai V Kravitz, Christopher V Dayas, Robyn Brown, and Zane B Andrews. Metabolic sensing in AgRP neurons integrates homeostatic state with dopamine signalling in the striatum. *eLife*, 11, 2022.

[11633] Felicia Reinitz, Elizabeth Y Chen, Benedetta Nicolis di Robilant, Bayarsaikhan Chuluun, Jane Antony, Robert C Jones, Neha Gubbi, Karen Lee, William Hai Dang Ho, Sai Saroja Kolluru, Dalong Qian, Maddalena Adorno, Katja Piltti, Aileen Anderson, Michelle Monje, H Craig Heller, Stephen R Quake, and Michael F Clarke. Inhibiting USP16 rescues stem cell aging and memory in an Alzheimer's model. *eLife*, 11, 2022.

[11634] Dina Rekler and Chaya Kalcheim. Completion of neural crest cell production and emigration is regulated by retinoic-acid-dependent inhibition of BMP signaling. *eLife*, 11, 2022.

[11635] Lea Rems, Xinru Tang, Fangwei Zhao, Sergio Pérez-Conesa, Ilaria Testa, and Lucie Delemotte. Identification of electroporation sites in the complex lipid organization of the plasma membrane. *eLife*, 11, 2022.

[11636] Boyang Ren, Huan Xia, Yijun Liao, Hang Zhou, Zhongnan Wang, Yaoyao Shi, and Mingzhao Zhu. Endothelial SIRP$\alpha$ signaling controls VE-cadherin endocytosis for thymic homing of progenitor cells. *eLife*, 11, 2022.

[11637] Gabriel Renaud, Maibritt Nørgaard, Johan Lindberg, Henrik Grönberg, Bram De Laere, Jørgen Bjerggaard Jensen, Michael Borre, Claus Lindbjerg Andersen, Karina Dalsgaard Sørensen, Lasse Maretty, and Søren Besenbacher. Unsupervised detection of fragment length signatures of circulating tumor DNA using non-negative matrix factorization. *eLife*, 11, 2022.

[11638] Joseph M Replogle, Jessica L Bonnar, Angela N Pogson, Christina R Liem, Nolan K Maier, Yufang Ding, Baylee J Russell, Xingren Wang, Kun Leng, Alina Guna, Thomas M Norman, Ryan A Pak, Daniel M Ramos, Michael E Ward, Luke A Gilbert, Martin Kampmann, Jonathan S Weissman, and Marco Jost. Maximizing CRISPRi efficacy and accessibility with dual-sgRNA libraries and optimal effectors. *eLife*, 11, 2022.

[11639] Vanessa Ress, Arne Traulsen, and Yuriy Pichugin. Eco-evolutionary dynamics of clonal multicellular life cycles. *eLife*, 11, 2022.

[11640] Jack Rhodes, Andra-Octavia Roman, Marta Bjornson, Benjamin Brandt, Paul Derbyshire, Michele Wyler, Marc W Schmid, Frank LH Menke, Julia Santiago, and Cyril Zipfel. Perception of a conserved family of plant signalling peptides by the receptor kinase HSL3. *eLife*, 11, 2022.

[11641] Katherine Rhodes, Kenneth A Barr, Joshua M Popp, Benjamin J Strober, Alexis Battle, and Yoav Gilad. Human embryoid bodies as a novel system for genomic studies of functionally diverse cell types. *eLife*, 11, 2022.

[11642] Maria Ribeiro and Miguel Castelo-Branco. Slow fluctuations in ongoing brain activity decrease in amplitude with ageing yet their impact on task-related evoked responses is dissociable from behavior. *eLife*, 11, 2022.

[11643] Mark Richards, Emmanuel Nwadozi, Sagnik Pal, Pernilla Martinsson, Mika Kaakinen, Marleen Gloger, Elin Sjöberg, Katarzyna Koltowska, Christer Betsholtz, Lauri Eklund, Sofia Nordling, and Lena Claesson-Welsh. Claudin5 protects the peripheral endothelial barrier in an organ and vessel-type-specific manner. *eLife*, 11, 2022.

[11644] Amy Richardson, Victoria Ciampani, Mihai Stancu, Kseniia Bondarenko, Sherylanne Newton, Joern R Steinert, Nadia Pilati, Bruce P Graham, Conny Kopp-Scheinpflug, and Ian D Forsythe. Kv3.3 subunits control presynaptic action potential waveform and neurotransmitter release at a central excitatory synapse. *eLife*, 11, 2022.

[11645] Tom G Richardson, Grace M Power, and George Davey Smith. Adiposity may confound the association between vitamin D and disease risk – a lifecourse Mendelian randomization study. *eLife*, 11, 2022.

[11646] Daniel J Richter, Romain Watteaux, Thomas Vannier, Jade Leconte, Paul Frémont, Gabriel Reygondeau, Nicolas Maillet, Nicolas Henry, Gaëtan Benoit, Ophélie Da Silva, Tom O Delmont, Antonio Fernàndez-Guerra, Samir Suweis, Romain Narci, Cédric Berney, Damien Eveillard, Frederick Gavory, Lionel Guidi, Karine Labadie, Eric Mahieu, Julie Poulain, Sarah Romac, Simon Roux, Céline Dimier, Stefanie Kandels, Marc Picheral, Sarah Searson, Silvia G Acinas, Stéphane Pesant, Jean-Marc Aury, Jennifer R Brum, Claire Lemaitre, Eric Pelletier, Peer Bork, Shinichi Sunagawa, Fabien Lombard, Lee Karp-Boss, Chris Bowler, Matthew B Sullivan, Eric Karsenti, Mahendra Mariadassou, Ian Probert, Pierre Peterlongo, Patrick Wincker, Colomban de Vargas, Maurizio Ribera d'Alcalà, Daniele Iudicone, and Olivier Jaillon. Genomic evidence for global ocean plankton biogeography shaped by large-scale current systems. *eLife*, 11, 2022.

[11647] Joyce Rigal, Ane Martin Anduaga, Elena Bitman, Emma Rivellese, Sebastian Kadener, and Michael T Marr. Artificially stimulating retrotransposon activity increases mortality and accelerates a subset of aging phenotypes in Drosophila. *eLife*, 11, 2022.

[11648] Nicola Rigolli, Nicodemo Magnoli, Lorenzo Rosasco, and Agnese Seminara. Learning to predict target location with turbulent odor plumes. *eLife*, 11, 2022.

[11649] Nicola Rigolli, Gautam Reddy, Agnese Seminara, and Massimo Vergassola. Alternation emerges as a multi-modal strategy for turbulent odor navigation. *eLife*, 11, 2022.

[11650] Lance A Riley, Xiping Zhang, Collin M Douglas, Joseph M Mijares, David W Hammers, Christopher A Wolff, Neil B Wood, Hailey R Olafson, Ping Du, Siegfried Labeit, Michael J Previs, Eric T Wang, and Karyn A Esser. The skeletal muscle circadian clock regulates titin splicing through RBM20. *eLife*, 11, 2022.

[11651] Camilo Riquelme-Guzmán, Stephanie L Tsai, Karen Carreon Paz, Congtin Nguyen, David Oriola, Maritta Schuez, Jan Brugués, Joshua D Currie, and Tatiana Sandoval-Guzmán. Osteoclast-mediated resorption primes the skeleton for successful integration during axolotl limb regeneration. *eLife*, 11, 2022.

[11652] Rafael Rivera-Lugo, David Deng, Andrea Anaya-Sanchez, Sara Tejedor-Sanz, Eugene Tang, Valeria M Reyes Ruiz, Hans B Smith, Denis V Titov, John-Demian Sauer, Eric P Skaar, Caroline M Ajo-Franklin, Daniel A Portnoy, and Samuel H Light. Listeria monocytogenes requires cellular respiration for NAD+ regeneration and pathogenesis. *eLife*, 11, 2022.

[11653] Zaigham Abbas Rizvi, Rajdeep Dalal, Srikanth Sadhu, Akshay Binayke, Jyotsna Dandotiya, Yashwant Kumar, Tripti Shrivastava, Sonu Kumar Gupta, Suruchi Aggarwal, Manas Ranjan Tripathy, Deepak Kumar Rathore, Amit Kumar Yadav, Guruprasad R Medigeshi, Amit Kumar Pandey, Sweety Samal, Shailendra Asthana, and Amit Awasthi. Golden Syrian hamster as a model to study cardiovascular complications associated with SARS-CoV-2 infection. *eLife*, 11, 2022.

[11654] Amy L Roberts, Alessandro Morea, Ariella Amar, Antonino Zito, Julia S El-Sayed Moustafa, Max Tomlinson, Ruth CE Bowyer, Xinyuan Zhang, Colette Christiansen, Ricardo Costeira, Claire J Steves, Massimo Mangino, Jordana T Bell, Chloe CY Wong, Timothy J Vyse, and Kerrin S Small. Age acquired skewed X chromosome inactivation is associated with adverse health outcomes in humans. *eLife*, 11, 2022.

[11655] Alejandro Rodriguez Gama, Tayla Miller, Jeffrey J Lange, Jay R Unruh, and Randal Halfmann. A nucleation barrier spring-loads the CBM signalosome for binary activation. *eLife*, 11, 2022.

[11656] Maria Rodriguez-Lopez, Shajahan Anver, Cristina Cotobal, Stephan Kamrad, Michal Malecki, Clara Correia-Melo, Mimoza Hoti, StJohn Townsend, Samuel Marguerat, Sheng Kai Pong, Mary Y Wu, Luis Montemayor, Michael Howell, Markus Ralser, and Jürg Bähler. Functional profiling of long intergenic non-coding RNAs in fission yeast. *eLife*, 11, 2022.

[11657] Sabela Rodríguez-Lorenzo, Lynn van Olst, Carla Rodriguez-Mogeda, Alwin Kamermans, Susanne MA van der Pol, Ernesto Rodríguez, Gijs Kooij, and Helga E de Vries. Single-cell profiling reveals periventricular CD56bright NK cell accumulation in multiple sclerosis. *eLife*, 11, 2022.

[11658] Jesse D Rogers and William J Richardson. Fibroblast mechanotransduction network predicts targets for mechano-adaptive infarct therapies. *eLife*, 11, 2022.

[11659] Shanna Romand, Hela Abdelkefi, Cécile Lecampion, Mohamed Belaroussi, Melanie Dussenne, Brigitte Ksas, Sylvie Citerne, Jose Caius, Stefano D'Alessandro, Hatem Fakhfakh, Stefano Caffarri, Michel Havaux, and Ben Field. A guanosine tetraphosphate (ppGpp) mediated brake on photosynthesis is required for acclimation to nitrogen limitation in Arabidopsis. *eLife*, 11, 2022.

[11660] Maria C Romero, Lara Merken, Peter Janssen, and Marco Davare. Neural effects of continuous theta-burst stimulation in macaque parietal neurons. *eLife*, 11, 2022.

[11661] Rafael Romero-Becerra, Alfonso Mora, Elisa Manieri, Ivana Nikolic, Ayelén Melina Santamans, Valle Montalvo-Romeral, Francisco Miguel Cruz, Elena Rodríguez, Marta León, Luis Leiva-Vega, Laura Sanz, Víctor Bondía, David Filgueiras-Rama, Luis Jesús Jiménez-Borreguero, José Jalife, Barbara Gonzalez-Teran, and Guadalupe Sabio. MKK6 deficiency promotes cardiac dysfunction through MKK3-p38$\gamma/\delta$-mTOR hyperactivation. *eLife*, 11, 2022.

[11662] Remi Ronzano, Sophie Skarlatou, Bianca K Barriga, B Anne Bannatyne, Gardave Singh Bhumbra, Joshua D Foster, Jeffrey D Moore, Camille Lancelin, Amanda M Pocratsky, Mustafa Görkem özyurt, Calvin Chad Smith, Andrew J Todd, David J Maxwell, Andrew J Murray, Samuel L Pfaff, Robert M Brownstone, Niccolò Zampieri, and Marco Beato. Spinal premotor interneurons controlling antagonistic muscles are spatially intermingled. *eLife*, 11, 2022.

[11663] Victoria Root, Dollyane Muret, Maite Arribas, Elena Amoruso, John Thornton, Aurelie Tarall-Jozwiak, Irene Tracey, and Tamar R Makin. Complex pattern of facial remapping in somatosensory cortex following congenital but not acquired hand loss. *eLife*, 11, 2022.

[11664] Fernando Rodrigo Rosas Bringas, Sonia Stinus, Pien de Zoeten, Marita Cohn, and Michael Chang. Rif2 protects Rap1-depleted telomeres from MRX-mediated degradation in Saccharomyces cerevisiae. *eLife*, 11, 2022.

[11665] Gail M Rosenbaum, Hannah L Grassie, and Catherine A Hartley. Valence biases in reinforcement learning shift across adolescence and modulate subsequent memory. *eLife*, 11, 2022.

[11666] Marion Rouault, Aurélien Weiss, Junseok K Lee, Jan Drugowitsch, Valerian Chambon, and Valentin Wyart. Controllability boosts neural and cognitive signatures of changes-of-mind in uncertain environments. *eLife*, 11, 2022.

[11667] Daniel Routledge, Sally Rogers, Yosuke Ono, Lucy Brunt, Valerie Meniel, Giusy Tornillo, Hassan Ashktorab, Toby J Phesse, and Steffen Scholpp. The scaffolding protein flot2 promotes cytoneme-based transport of wnt3 in gastric cancer. *eLife*, 11, 2022.

[11668] Frédéric Roux, George Parish, Ramesh Chelvarajah, David T Rollings, Vijay Sawlani, Hajo Hamer, Stephanie Gollwitzer, Gernot Kreiselmeyer, Marije J ter Wal, Luca Kolibius, Bernhard P Staresina, Maria Wimber, Matthew W Self, and Simon Hanslmayr. Oscillations support short latency co-firing of neurons during human episodic memory formation. *eLife*, 11, 2022.

[11669] Ruchi Roy, Janet Zayas, Sunil K Singh, Kaylee Delgado, Stephen J Wood, Mohamed F Mohamed, Dulce M Frausto, Yasmeen A Albalawi, Thea P Price, Ricardo Estupinian, Eileena F Giurini, Timothy M Kuzel, Andrew Zloza, Jochen Reiser, and Sasha H Shafikhani. Overriding impaired FPR chemotaxis signaling in diabetic neutrophil stimulates infection control in murine diabetic wound. *eLife*, 11, 2022.

[11670] Udita Roy, Shir Yaish, Ziva Weissman, Mariel Pinsky, Sunanda Dey, Guy Horev, and Daniel Kornitzer. Ferric reductase-related proteins mediate fungal heme acquisition. *eLife*, 11, 2022.

[11671] Hector M Rubiato, Mengqi Liu, Richard J O'Connell, and Mads E Nielsen. Plant SYP12 syntaxins mediate an evolutionarily conserved general immunity to filamentous pathogens. *eLife*, 11, 2022.

[11672] Alexis Ruiz, Sofia Benucci, Urs Duthaler, Christoph Bachmann, Martina Franchini, Faiza Noreen, Laura Pietrangelo, Feliciano Protasi, Susan Treves, and Francesco Zorzato. Improvement of muscle strength in a mouse model for congenital myopathy treated with HDAC and DNA methyltransferase inhibitors. *eLife*, 11, 2022.

[11673] Carlos Ruiz-Arenas, Carles Hernandez-Ferrer, Marta Vives-Usano, Sergi Marí, Ines Quintela, Dan Mason, Solène Cadiou, Maribel Casas, Sandra Andrusaityte, Kristine Bjerve Gutzkow, Marina Vafeiadi, John Wright, Johanna Lepeule, Regina Grazuleviciene, Leda Chatzi, ángel Carracedo, Xavier Estivill, Eulàlia Marti, Geòrgia Escaramís, Martine Vrijheid, Juan R González, and Mariona Bustamante. Identification of autosomal cis expression quantitative trait methylation (cis eQTMs) in children's blood. *eLife*, 11, 2022.

[11674] Charles M Russell, Katherine G Schaefer, Andrew Dixson, Amber LH Gray, Robert J Pyron, Daiane S Alves, Nicholas Moore, Elizabeth A Conley, Ryan J Schuck, Tommi A White, Thanh D Do, Gavin M King, and Francisco N Barrera. The Candida albicans virulence factor candidalysin polymerizes in solution to form membrane pores and damage epithelial cells. *eLife*, 11, 2022.

[11675] Magdalena L Russell, Aisha Souquette, David M Levine, Stefan A Schattgen, E Kaitlynn Allen, Guillermina Kuan, Noah Simon, Angel Balmaseda, Aubree Gordon, Paul G Thomas, Frederick A Matsen, and Philip Bradley. Combining genotypes and T cell receptor distributions to infer genetic loci determining V(D)J recombination probabilities. *eLife*, 11, 2022.

[11676] Marie C Russell, Catherine M Herzog, Zachary Gajewski, Chloe Ramsay, Fadoua El Moustaid, Michelle V Evans, Trishna Desai, Nicole L Gottdenker, Sara L Hermann, Alison G Power, and Andrew C McCall. Both consumptive and non-consumptive effects of predators impact mosquito populations and have implications for disease transmission. *eLife*, 11, 2022.

[11677] Saige Rutherford, Charlotte Fraza, Richard Dinga, Seyed Mostafa Kia, Thomas Wolfers, Mariam Zabihi, Pierre Berthet, Amanda Worker, Serena Verdi, Derek Andrews, Laura KM Han, Johanna MM Bayer, Paola Dazzan, Phillip McGuire, Roel T Mocking, Aart Schene, Chandra Sripada,

Ivy F Tso, Elizabeth R Duval, Soo-Eun Chang, Brenda WJH Penninx, Mary M Heitzeg, S Alexandra Burt, Luke W Hyde, David Amaral, Christine Wu Nordahl, Ole A Andreasssen, Lars T Westlye, Roland Zahn, Henricus G Ruhe, Christian Beckmann, and Andre F Marquand. Charting brain growth and aging at high spatial precision. *eLife*, 11, 2022.

[11678] Adriano Rutz, Maria Sorokina, Jakub Galgonek, Daniel Mietchen, Egon Willighagen, Arnaud Gaudry, James G Graham, Ralf Stephan, Roderic Page, Jiří VondráŠek, Christoph Steinbeck, Guido F Pauli, Jean-Luc Wolfender, Jonathan Bisson, and Pierre-Marie Allard. The LOTUS initiative for open knowledge management in natural products research. *eLife*, 11, 2022.

[11679] Alicia J Rybicki, Sophie L Sowden, Bianca Schuster, and Jennifer L Cook. Dopaminergic challenge dissociates learning from primary versus secondary sources of information. *eLife*, 11, 2022.

[11680] Vitaly Ryu, Anisa Gumerova, Funda Korkmaz, Seong Su Kang, Pavel Katsel, Sari Miyashita, Hasni Kannangara, Liam Cullen, Pokman Chan, TanChun Kuo, Ashley Padilla, Farhath Sultana, Soleil A Wizman, Natan Kramskiy, Samir Zaidi, Se-Min Kim, Maria I New, Clifford J Rosen, Ki A Goosens, Tal Frolinger, Vahram Haroutunian, Keqiang Ye, Daria Lizneva, Terry F Davies, Tony Yuen, and Mone Zaidi. Brain atlas for glycoprotein hormone receptors at single-transcript level. *eLife*, 11, 2022.

[11681] Ankit Sabharwal, Mark D Wishman, Roberto Lopez Cervera, MaKayla R Serres, Jennifer L Anderson, Shannon R Holmberg, Bibekananda Kar, Anthony J Treichel, Noriko Ichino, Weibin Liu, Jingchun Yang, Yonghe Ding, Yun Deng, Jean M Lacey, William J Laxen, Perry R Loken, Devin Oglesbee, Steven A Farber, Karl J Clark, Xiaolei Xu, and Stephen C Ekker. Genetic therapy in a mitochondrial disease model suggests a critical role for liver dysfunction in mortality. *eLife*, 11, 2022.

[11682] João C Sabino, Madalena R de Almeida, Patrícia L Abreu, Ana M Ferreira, Paulo Caldas, Marco M Domingues, Nuno C Santos, Claus M Azzalin, Ana Rita Grosso, and Sérgio Fernandes de Almeida. Epigenetic reprogramming by TET enzymes impacts co-transcriptional R-loops. *eLife*, 11, 2022.

[11683] Sadra Sadeh and Claudia Clopath. Contribution of behavioural variability to representational drift. *eLife*, 11, 2022.

[11684] Souvik Sadhukhan and Saroj Kumar Nandi. On the origin of universal cell shape variability in confluent epithelial monolayers. *eLife*, 11, 2022.

[11685] Maliheh Safari, Bhargavi Jayaraman, Henni Zommer, Shumin Yang, Cynthia Smith, Jason D Fernandes, and Alan D Frankel. Functional and structural segregation of overlapping helices in HIV-1. *eLife*, 11, 2022.

[11686] Saswati Saha, Lionel Spinelli, Jaime A Castro Mondragon, Anaïs Kervadec, Michaela Lynott, Laurent Kremmer, Laurence Roder, Sallouha Krifa, Magali Torres, Christine Brun, Georg Vogler, Rolf Bodmer, Alexandre R Colas, Karen Ocorr, and Laurent Perrin. Genetic architecture of natural variation of cardiac performance from flies to humans. *eLife*, 11, 2022.

[11687] Korin Sahinyan, Darren M Blackburn, Marie-Michelle Simon, Felicia Lazure, Tony Kwan, Guillaume Bourque, and Vahab D Soleimani. Application of ATAC-Seq for genome-wide analysis of the chromatin state at single myofiber resolution. *eLife*, 11, 2022.

[11688] Vinciane Saint-Criq, Anita Guequén, Amber R Philp, Sandra Villanueva, Tábata Apablaza, Ignacio Fernández-Moncada, Agustín Mansilla, Livia Delpiano, Iván Ruminot, Cristian Carrasco, Michael A Gray, and Carlos A Flores. Inhibition of the sodium-dependent HCO3- transporter SLC4A4, produces a cystic fibrosis-like airway disease phenotype. *eLife*, 11, 2022.

[11689] Shunsuke Saito, Tokiro Ishikawa, Satoshi Ninagawa, Tetsuya Okada, and Kazutoshi Mori. A motor neuron disease-associated mutation produces non-glycosylated Seipin that induces ER stress and apoptosis by inactivating SERCA2b. *eLife*, 11, 2022.

[11690] Yuichiro Saito, Venny Santosa, Kei-ichiro Ishiguro, and Masato T Kanemaki. MCMBP promotes the assembly of the MCM2–7 hetero-hexamer to ensure robust DNA replication in human cells. *eLife*, 11, 2022.

[11691] Masahide Sakabe, Michael Thompson, Nong Chen, Mark Verba, Aishlin Hassan, Richard Lu, and Mei Xin. Inhibition of $\beta$1-AR/G$\alpha$s signaling promotes cardiomyocyte proliferation in juvenile mice through activation of RhoA-YAP axis. *eLife*, 11, 2022.

[11692] Kazumi Sakai, Yoshinori Shichida, Yasushi Imamoto, and Takahiro Yamashita. Creation of photocyclic vertebrate rhodopsin by single amino acid substitution. *eLife*, 11, 2022.

[11693] Mohammad Ali Salehinejad, Elham Ghanavati, Jörg Reinders, Jan G Hengstler, Min-Fang Kuo, and Michael A Nitsche. Sleep-dependent upscaled excitability, saturated neuroplasticity, and modulated cognition in the human brain. *eLife*, 11, 2022.

[11694] Ana CG Salina, Douglas dos Santos, Tamara S Rodrigues, Marlon Fortes-Rocha, Edismauro G Freitas-Filho, Daniel L Alzamora-Terrel, Icaro MS Castro, Thais FC Fraga da Silva, Mikhael HF de Lima, Daniele C Nascimento, Camila M Silva, Juliana E Toller-Kawahisa, Amanda Becerra, Samuel Oliveira, Diego B Caetité, Leticia Almeida, Adriene Y Ishimoto, Thais M Lima, Ronaldo B Martins, Flavio Veras, Natália B do Amaral, Marcela C Giannini, Letícia P Bonjorno, Maria IF Lopes, Maira N Benatti, Sabrina S Batah, Rodrigo C Santana, Fernando C Vilar, Maria A Martins, Rodrigo L Assad, Sergio CL de Almeida, Fabiola R de Oliveira, Eurico Arruda Neto, Thiago M Cunha, José C Alves-Filho, Vania LD Bonato, Fernando Q Cunha, Alexandre T Fabro, Helder I Nakaya, Dario S Zamboni, Paulo Louzada-Junior, Rene DR Oliveira, and Larissa D Cunha. Efferocytosis of SARS-CoV-2-infected dying cells impairs macrophage anti-inflammatory functions and clearance of apoptotic cells. *eLife*, 11, 2022.

[11695] Kathryn A Salvati, George MPR Souza, Adam C Lu, Matthew L Ritger, Patrice Guyenet, Stephen B Abbott, and Mark P Beenhakker. Respiratory alkalosis provokes spike-wave discharges in seizure-prone rats. *eLife*, 11, 2022.

[11696] Rebeca San Martin, Priyojit Das, Jacob T Sanders, Ashtyn M Hill, and Rachel Patton McCord. Transcriptional profiling of Hutchinson-Gilford Progeria syndrome fibroblasts reveals deficits in mesenchymal stem cell commitment to differentiation related to early events in endochondral ossification. *eLife*, 11, 2022.

[11697] Héctor Sánchez-Iranzo, Aliaksandr Halavatyi, and Alba Diz-Muñoz. Strength of interactions in the Notch gene regulatory network determines patterning and fate in the notochord. *eLife*, 11, 2022.

[11698] Theo Sanderson. Taxonium, a web-based tool for exploring large phylogenetic trees. *eLife*, 11, 2022.

[11699] Guillem Santamaria, Chen Liao, Chloe Lindberg, Yanyan Chen, Zhe Wang, Kyu Rhee, Francisco Rodrigues Pinto, Jinyuan Yan, and Joao B Xavier. Evolution and regulation of microbial secondary metabolism. *eLife*, 11, 2022.

[11700] Anthony Santella, Irina Kolotuev, Caroline Kizilyaprak, and Zhirong Bao. Cross-modality synthesis of EM time series and live fluorescence imaging. *eLife*, 11, 2022.

[11701] Vishnu Muraleedharan Saraswathy, Akshai Janardhana Kurup, Priyanka Sharma, Sophie Polès, Morgane Poulain, and Maximilian Fürthauer. The E3 ubiquitin ligase mindbomb1 controls planar cell polarity-dependent convergent extension movements during zebrafish gastrulation. *eLife*, 11, 2022.

[11702] Haruka Sato, Jun Hatakeyama, Takuji Iwasato, Kimi Araki, Nobuhiko Yamamoto, and Kenji Shimamura. Thalamocortical axons control the cytoarchitecture of neocortical layers by area-specific supply of VGF. *eLife*, 11, 2022.

[11703] Tatsuya Sato, Jason Solomon Shapiro, Hsiang-Chun Chang, Richard A Miller, and Hossein Ardehali. Aging is associated with increased brain iron through cortex-derived hepcidin expression. *eLife*, 11, 2022.

[11704] Chie Satou, Rachael L Neve, Hassana K Oyibo, Pawel Zmarz, Kuo-Hua Huang, Estelle Arn Bouldoires, Takuma Mori, Shin-ichi Higashijima, Georg B Keller, and Rainer W Friedrich. A viral toolbox for conditional and transneuronal gene expression in zebrafish. *eLife*, 11, 2022.

[11705] Jonas-Frederic Sauer and Marlene Bartos. Disrupted-in-schizophrenia-1 is required for normal pyramidal cell–interneuron communication and assembly dynamics in the prefrontal cortex. *eLife*, 11, 2022.

[11706] Josh Saul, Takashi Hirose, and H Robert Horvitz. The transcriptional corepressor CTBP-1 acts with the SOX family transcription factor EGL-13 to maintain AIA interneuron cell identity in Caenorhabditis elegans. *eLife*, 11, 2022.

[11707] Rubens Sautchuk, Brianna H Kalicharan, Katherine Escalera-Rivera, Jennifer H Jonason, George A Porter, Hani A Awad, and Roman A Eliseev. Transcriptional regulation of cyclophilin D by BMP/Smad signaling and its role in osteogenic differentiation. *eLife*, 11, 2022.

[11708] Melissa Savard, Tharick A Pascoal, Stijn Servaes, Thijs Dhollander, Yasser Iturria-Medina, Min Su Kang, Paolo Vitali, Joseph Therriault, Sulantha Mathotaarachchi, Andrea Lessa Benedet, Serge Gauthier, and Pedro Rosa-Neto. Impact of long- and short-range fibre depletion on the cognitive deficits of fronto-temporal dementia. *eLife*, 11, 2022.

[11709] Ayana Sawai, Sarah Pfennig, Milica Bulajić, Alexander Miller, Alireza Khodadadi-Jamayran, Esteban O Mazzoni, and Jeremy S Dasen. PRC1 sustains the integrity of neural fate in the absence of PRC2 function. *eLife*, 11, 2022.

[11710] Shreya Saxena, Abigail A Russo, John Cunningham, and Mark M Churchland. Motor cortex activity across movement speeds is predicted by network-level strategies for generating muscle activity. *eLife*, 11, 2022.

[11711] Wasim A Sayyad and Thomas D Pollard. The number of cytokinesis nodes in mitotic fission yeast scales with cell size. *eLife*, 11, 2022.

[11712] Miranda L Scalabrino, Mishek Thapa, Lindsey A Chew, Esther Zhang, Jason Xu, Alapakkam P Sampath, Jeannie Chen, and Greg D Field. Robust cone-mediated signaling persists late into rod photoreceptor degeneration. *eLife*, 11, 2022.

[11713] Allison Schad, Rebekah L Layton, Debra Ragland, and Jeanette Gowen Cook. Mental health in medical and biomedical doctoral students during the 2020 COVID-19 pandemic and racial protests. *eLife*, 11, 2022.

[11714] Marco Schade, Nils Knötschke, Marie K Hörnig, Carina Paetzel, and Sebastian Stumpf. Neurovascular anatomy of dwarf dinosaur implies precociality in sauropods. *eLife*, 11, 2022.

[11715] Stefan L Schaefer and Gerhard Hummer. Sublytic gasdermin-D pores captured in atomistic molecular simulations. *eLife*, 11, 2022.

[11716] Kyra Schapiro, KreŠimir Josić, Zachary P Kilpatrick, and Joshua I Gold. Strategy-dependent effects of working-memory limitations on human perceptual decision-making. *eLife*, 11, 2022.

[11717] Désirée Schatton, Giada Di Pietro, Karolina Szczepanowska, Matteo Veronese, Marie-Charlotte Marx, Kristina Braunöhler, Esther Barth, Stefan Müller, Patrick Giavalisco, Thomas Langer, Aleksandra Trifunovic, and Elena I Rugarli. CLUH controls astrin-1 expression to couple mitochondrial metabolism to cell cycle progression. *eLife*, 11, 2022.

[11718] Alexej Schatz and York Winter. LabNet hardware control software for the Raspberry Pi. *eLife*, 11, 2022.

[11719] Paul E Schavemaker and Michael Lynch. Flagellar energy costs across the tree of life. *eLife*, 11, 2022.

[11720] Myriam Scherer, Clarissa Read, Gregor Neusser, Christine Kranz, Anna K Kuderna, Regina Müller, Florian Full, Sonja Wörz, Anna Reichel, Eva-Maria Schilling, Paul Walther, and Thomas Stamminger. Dual signaling via interferon and DNA damage response elicits entrapment by giant PML nuclear bodies. *eLife*, 11, 2022.

[11721] Philipp E Scherer, John P Kirwan, and Clifford J Rosen. Post-acute sequelae of COVID-19: A metabolic perspective. *eLife*, 11, 2022.

[11722] Kevin Schilling, Yuankun Zhai, Zhuang Zhou, Bin Zhou, Edward Brown, and Xinping Zhang. High-resolution imaging of the osteogenic and angiogenic interface at the site of murine cranial bone defect repair via multiphoton microscopy. *eLife*, 11, 2022.

[11723] Raimund Schlüßler, Kyoohyun Kim, Martin Nötzel, Anna Taubenberger, Shada Abuhattum, Timon Beck, Paul Müller, Shovamaye Maharana, Gheorghe Cojoc, Salvatore Girardo, Andreas Hermann, Simon Alberti, and Jochen Guck. Correlative all-optical quantification of mass density and mechanics of subcellular compartments with fluorescence specificity. *eLife*, 11, 2022.

[11724] Jonathan S Schor, Isabelle Gonzalez Montalvo, Perry WE Spratt, Rea J Brakaj, Jasmine A Stansil, Emily L Twedell, Kevin J Bender, and Alexandra B Nelson. Therapeutic deep brain stimulation disrupts movement-related subthalamic nucleus activity in parkinsonian mice. *eLife*, 11, 2022.

[11725] Marina Schrecker, Juan C Castaneda, Sujan Devbhandari, Charanya Kumar, Dirk Remus, and Richard K Hite. Multistep loading of a DNA sliding clamp onto DNA by replication factor C. *eLife*, 11, 2022.

[11726] Olga T Schubert, Joshua S Bloom, Meru J Sadhu, and Leonid Kruglyak. Genome-wide base editor screen identifies regulators of protein abundance in yeast. *eLife*, 11, 2022.

[11727] Rebecca C Schugar, Christy M Gliniak, Lucas J Osborn, William Massey, Naseer Sangwan, Anthony Horak, Rakhee Banerjee, Danny Orabi, Robert N Helsley, Amanda L Brown, Amy Burrows, Chelsea Finney, Kevin K Fung, Frederick M Allen, Daniel Ferguson, Anthony D Gromovsky, Chase Neumann, Kendall Cook, Amy McMillan, Jennifer A Buffa, James T Anderson, Margarete Mehrabian, Maryam Goudarzi, Belinda Willard, Tytus D Mak, Andrew R Armstrong, Garth Swanson, Ali Keshavarzian, Jose Carlos Garcia-Garcia, Zeneng Wang, Aldons J Lusis, Stanley L Hazen, and Jonathan Mark Brown. Gut microbe-targeted choline trimethylamine lyase inhibition improves obesity via rewiring of host circadian rhythms. *eLife*, 11, 2022.

[11728] Helena Schulz-Mirbach, Alexandra Müller, Tong Wu, Pascal Pfister, SelÇuk Aslan, Lennart Schada von Borzyskowski, Tobias J Erb, Arren Bar-Even, and Steffen N Lindner. On the flexibility of the cellular amination network in E coli. *eLife*, 11, 2022.

[11729] Erika L Schumacher and Bruce A Carlson. Convergent mosaic brain evolution is associated with the evolution of novel electrosensory systems in teleost fishes. *eLife*, 11, 2022.

[11730] Samantha C Schwager, Katherine M Young, Lauren A Hapach, Caroline M Carlson, Jenna A Mosier, Tanner J McArdle, Wenjun Wang, Curtis Schunk, Anissa L Jayathilake, Madison E Bates, Francois Bordeleau, Marc A Antonyak, Richard A Cerione, and Cynthia A Reinhart-King. Weakly migratory metastatic breast cancer cells activate fibroblasts via microvesicle-Tg2 to facilitate dissemination and metastasis. *eLife*, 11, 2022.

[11731] Ryan W Schwark, Matthew J Fuxjager, and Marc F Schmidt. Proposing a neural framework for the evolution of elaborate courtship displays. *eLife*, 11, 2022.

[11732] Aaron ZA Schwartz, Nikita Tsyba, Yusuff Abdu, Maulik R Patel, and Jeremy Nance. Independent regulation of mitochondrial DNA quantity and quality in Caenorhabditis elegans primordial germ cells. *eLife*, 11, 2022.

[11733] Madeleine Seale, Oleksandr Zhdanov, Merel B Soons, Cathal Cummins, Erika Kroll, Michael R Blatt, Hossein Zare-Behtash, Angela Busse, Enrico Mastropaolo, James M Bullock, Ignazio M Viola, and Naomi Nakayama. Environmental morphing enables informed dispersal of the dandelion diaspore. *eLife*, 11, 2022.

[11734] Elizabeth M Sefton, Mirialys Gallardo, Claire E Tobin, Brittany C Collins, Mary P Colasanto, Allyson J Merrell, and Gabrielle Kardon. Fibroblast-derived Hgf controls recruitment and expansion of muscle during morphogenesis of the mammalian diaphragm. *eLife*, 11, 2022.

[11735] Ivonne Sehring, Hossein Falah Mohammadi, Melanie Haffner-Luntzer, Anita Ignatius, Markus Huber-Lang, and Gilbert Weidinger. Zebrafish fin regeneration involves generic and regeneration-specific osteoblast injury responses. *eLife*, 11, 2022.

[11736] Ron Sender, Yinon Bar-On, Sang Woo Park, Elad Noor, Jonathan Dushoff, and Ron Milo. The unmitigated profile of COVID-19 infectiousness. *eLife*, 11, 2022.

[11737] GökÇe Senger, Stefano Santaguida, and Martin H Schaefer. Regulation of protein complex partners as a compensatory mechanism in aneuploid tumors. *eLife*, 11, 2022.

[11738] Shreoshi Sengupta, Mainak Mondal, Kaval Reddy Prasasvi, Arani Mukherjee, Prerna Magod, Serge Urbach, Dinorah Friedmann-Morvinski, Philippe Marin, and Kumaravel Somasundaram. Differentiated glioma cell-derived fibromodulin activates integrin-dependent Notch signaling in endothelial cells to promote tumor angiogenesis and growth. *eLife*, 11, 2022.

[11739] Irene Senna, Sophia Piller, Itay Ben-Zion, and Marc O Ernst. Recalibrating vision-for-action requires years after sight restoration from congenital cataracts. *eLife*, 11, 2022.

[11740] Eugene Serebryany, Sourav Chowdhury, Christopher N Woods, David C Thorn, Nicki E Watson, Arthur A McClelland, Rachel E Klevit, and Eugene I Shakhnovich. A native chemical chaperone in the human eye lens. *eLife*, 11, 2022.

[11741] Manutea C Serrero, Virginie Girault, Sebastian Weigang, Todd M Greco, Ana Ramos-Nascimento, Fenja Anderson, Antonio Piras, Ana Hickford Martinez, Jonny Hertzog, Anne Binz, Anja Pohlmann, Ute Prank, Jan Rehwinkel, Rudolf Bauerfeind, Ileana M Cristea, Andreas Pichlmair, Georg Kochs, and Beate Sodeik. The interferon-inducible GTPase MxB promotes capsid disassembly and genome release of herpesviruses. *eLife*, 11, 2022.

[11742] Francesco A Servello, Rute Fernandes, Matthias Eder, Nathan Harris, Olivier MF Martin, Natasha Oswal, Anders Lindberg, Nohelly Derosiers, Piali Sengupta, Nicholas Stroustrup, and Javier Apfeld. Neuronal temperature perception induces specific defenses that enable C. elegans to cope with the enhanced reactivity of hydrogen peroxide at high temperature. *eLife*, 11, 2022.

[11743] Amy L Seufert, James W Hickman, Ste K Traxler, Rachael M Peterson, Trent A Waugh, Sydney J Lashley, Natalia Shulzhenko, Ruth J Napier, and Brooke A Napier. Enriched dietary saturated fatty acids induce trained immunity via ceramide production that enhances severity of endotoxemia and clearance of infection. *eLife*, 11, 2022.

[11744] Osman Shabir, Ben Pendry, Llywelyn Lee, Beth Eyre, Paul S Sharp, Monica A Rebollar, David Drew, Clare Howarth, Paul R Heath, Stephen B Wharton, Sheila E Francis, and Jason Berwick. Assessment of neurovascular coupling and cortical spreading depression in mixed mouse models of atherosclerosis and Alzheimer's disease. *eLife*, 11, 2022.

[11745] Orie T Shafer, Gabrielle J Gutierrez, Kimberly Li, Amber Mildenhall, Daphna Spira, Jonathan Marty, Aurel A Lazar, and Maria de la Paz Fernandez. Connectomic analysis of the Drosophila lateral neuron clock cells reveals the synaptic basis of functional pacemaker classes. *eLife*, 11, 2022.

[11746] Sahil H Shah, Lucio M Schiapparelli, Yuanhui Ma, Satoshi Yokota, Melissa Atkins, Xin Xia, Evan G Cameron, Thanh Huang, Sarah Saturday, Catalina B Sun, Cara Knasel, Seth Blackshaw, John R Yates, Hollis T Cline, and Jeffrey L Goldberg. Quantitative transportomics identifies Kif5a as a major regulator of neurodegeneration. *eLife*, 11, 2022.

[11747] Montaser F Shaheen, Julie Y Tse, Ethan S Sokol, Margaret Masterson, Pranshu Bansal, Ian Rabinowitz, Christy A Tarleton, Andrey S Dobroff, Tracey L Smith, Thèrése J Bocklage, Brian K Mannakee, Ryan N Gutenkunst, Joyce Bischoff, Scott A Ness, Gregory M Riedlinger, Roman Groisberg, Renata Pasqualini, Shridar Ganesan, and Wadih Arap. Genomic landscape of lymphatic malformations: a case series and response to the PI3K$\alpha$ inhibitor alpelisib in an N-of-1 clinical trial. *eLife*, 11, 2022.

[11748] Rajesh Sharma, Jeong Joo Kim, Liying Qin, Philipp Henning, Madoka Akimoto, Bryan Van-Schouwen, Gundeep Kaur, Banumathi Sankaran, Kevin R MacKenzie, Giuseppe Melacini, Darren E Casteel, Friedrich W Herberg, and Choel Kim. An auto-inhibited state of protein kinase G and implications for selective activation. *eLife*, 11, 2022.

[11749] Paul B Sharp, Evan M Russek, Quentin JM Huys, Raymond J Dolan, and Eran Eldar. Humans perseverate on punishment avoidance goals in multigoal reinforcement learning. *eLife*, 11, 2022.

[11750] Karthik Shekhar, Irene E Whitney, Salwan Butrus, Yi-Rong Peng, and Joshua R Sanes. Diversification of multipotential postmitotic mouse retinal ganglion cell precursors into discrete types. *eLife*, 11, 2022.

[11751] Leyao Shen, Yilin Yu, Yunji Zhou, Shondra M Pruett-Miller, Guo-Fang Zhang, and Courtney M Karner. SLC38A2 provides proline to fulfill unique synthetic demands arising during osteoblast differentiation and bone formation. *eLife*, 11, 2022.

[11752] Qing Shen, Huan Song, Thor Aspelund, Jingru Yu, Donghao Lu, Jóhanna Jakobsdóttir, Jacob Bergstedt, Lu Yi, Patrick Sullivan, Arvid Sjölander, Weimin Ye, Katja Fall, Fang Fang, and Unnur Valdimarsdóttir. Cardiovascular disease and subsequent risk of psychiatric disorders: a nationwide sibling-controlled study. *eLife*, 11, 2022.

[11753] Xin Shen, Jonas van den Brink, Anna Bergan-Dahl, Terje R Kolstad, Einar S Norden, Yufeng Hou, Martin Laasmaa, Yuriana Aguilar-Sanchez, Ann P Quick, Emil KS Espe, Ivar Sjaastad, Xander HT Wehrens, Andrew G Edwards, Christian Soeller, and William E Louch. Prolonged $\beta$-adrenergic stimulation disperses ryanodine receptor clusters in cardiomyocytes and has implications for heart failure. *eLife*, 11, 2022.

[11754] Zeyang Shen, Rick Z Li, Thomas A Prohaska, Marten A Hoeksema, Nathan J Spann, Jenhan Tao, Gregory J Fonseca, Thomas Le, Lindsey K Stolze, Mashito Sakai, Casey E Romanoski, and Christopher K Glass. Systematic analysis of naturally occurring insertions and deletions that alter transcription factor spacing identifies tolerant and sensitive transcription factor pairs. *eLife*, 11, 2022.

[11755] Sushila A Shenoy, Sushuang Zheng, Wencheng Liu, Yuanyi Dai, Yuanxiu Liu, Zhipeng Hou, Susumu Mori, Yi Tang, Jerry Cheng, Wenzhen Duan, and Chenjian Li. A novel and accurate full-length HTT mouse model for Huntington's disease. *eLife*, 11, 2022.

[11756] Arefeh Sherafati, Noel Dwyer, Aahana Bajracharya, Mahlega Samira Hassanpour, Adam T Eggebrecht, Jill B Firszt, Joseph P Culver, and Jonathan E Peelle. Prefrontal cortex supports speech perception in listeners with cochlear implants. *eLife*, 11, 2022.

[11757] Dawafuti Sherpa, Judith Mueller, özge Karayel, Peng Xu, Yu Yao, Jakub Chrustowicz, Karthik V Gottemukkala, Christine Baumann, Annette Gross, Oliver Czarnecki, Wei Zhang, Jun Gu, Johan Nilvebrant, Sachdev S Sidhu, Peter J Murray, Matthias Mann, Mitchell J Weiss, Brenda A Schulman, and Arno F Alpi. Modular UBE2H-CTLH E2-E3 complexes regulate erythroid maturation. *eLife*, 11, 2022.

[11758] Yi-Jun Sheu, Risa Karakida Kawaguchi, Jesse Gillis, and Bruce Stillman. Prevalent and dynamic binding of the cell cycle checkpoint kinase Rad53 to gene promoters. *eLife*, 11, 2022.

[11759] Meng Shi, Hieng Chiong Tie, Mahajan Divyanshu, Xiuping Sun, Yan Zhou, Boon Kim Boh, Leah A Vardy, and Lei Lu. Arl15 upregulates the TGF$\beta$ family signaling by promoting the assembly of the Smad-complex. *eLife*, 11, 2022.

[11760] Weikang Shi, Sebastien Ballesta, and Camillo Padoa-Schioppa. Neuronal origins of reduced accuracy and biases in economic choices under sequential offers. *eLife*, 11, 2022.

[11761] Yoichiro Shibuya, Akishige Hokugo, Hiroko Okawa, Takeru Kondo, Daniel Khalil, Lixin Wang, Yvonne Roca, Adam Clements, Hodaka Sasaki, Ella Berry, Ichiro Nishimura, and Reza Jarrahy. Therapeutic downregulation of neuronal PAS domain 2 (Npas2) promotes surgical skin wound healing. *eLife*, 11, 2022.

[11762] Philip K Shiu, Gabriella R Sterne, Stefanie Engert, Barry J Dickson, and Kristin Scott. Taste quality and hunger interactions in a feeding sensorimotor circuit. *eLife*, 11, 2022.

[11763] Tikva Shore, Tgst Levi, Rachel Kalifa, Amatzia Dreifuss, Dina Rekler, Ariella Weinberg-Shukron, Yuval Nevo, Tzofia Bialistoky, Victoria Moyal, Merav Yaffa Gold, Shira Leebhoff, David Zangen, Girish Deshpande, and Offer Gerlitz. Nucleoporin107 mediates female sexual differentiation via Dsx. *eLife*, 11, 2022.

[11764] David P Shorten, Viola Priesemann, Michael Wibral, and Joseph T Lizier. Early lock-in of structured and specialised information flows during neural development. *eLife*, 11, 2022.

[11765] Shawn P Shortill, Mia S Frier, Ponthakorn Wongsangaroonsri, Michael Davey, and Elizabeth Conibear. The VINE complex is an endosomal VPS9-domain GEF and SNX-BAR coat. *eLife*, 11, 2022.

[11766] Dmitry Shvarev, Jannis Schoppe, Caroline König, Angela Perz, Nadia Füllbrunn, Stephan Kiontke, Lars Langemeyer, Dovile Januliene, Kilian Schnelle, Daniel Kümmel, Florian Fröhlich, Arne Moeller, and Christian Ungermann. Structure of the HOPS tethering complex, a lysosomal membrane fusion machinery. *eLife*, 11, 2022.

[11767] Sophie R Sichel, Benjamin P Bratton, and Nina R Salama. Distinct regions of H. pylori's bactofilin CcmA regulate protein–protein interactions to control helical cell shape. *eLife*, 11, 2022.

[11768] Dirk H Siepe, Lukas T Henneberg, Steven C Wilson, Gaelen T Hess, Michael C Bassik, Kai Zinn, and K Christopher Garcia. Identification of orphan ligand-receptor relationships using a cell-based CRISPRa enrichment screening platform. *eLife*, 11, 2022.

[11769] Jakob M Silberberg, Charlott Stock, Lisa Hielkema, Robin A Corey, Jan Rheinberger, Dorith Wunnicke, Victor RA Dubach, Phillip J Stansfeld, Inga Hänelt, and Cristina Paulino. Inhibited KdpFABC transitions into an E1 off-cycle state. *eLife*, 11, 2022.

[11770] Anita Siller, Nadja T Hofer, Giulia Tomagra, Nicole Burkert, Simon Hess, Julia Benkert, Aisylu Gaifullina, Desiree Spaich, Johanna Duda, Christina Poetschke, Kristina Vilusic, Eva Maria Fritz, Toni Schneider, Peter Kloppenburg, Birgit Liss, Valentina Carabelli, Emilio Carbone, Nadine Jasmin Ortner, and Jörg Striessnig. $\beta$2-subunit alternative splicing stabilizes Cav2.3 Ca2+ channel activity during continuous midbrain dopamine neuron-like activity. *eLife*, 11, 2022.

[11771] Sara Silva Pereira, Mariana De Niz, Karine Serre, Marie Ouarné, Joana E Coelho, Cláudio A Franco, and Luisa M Figueiredo. Immunopathology and Trypanosoma congolense parasite sequestration cause acute cerebral trypanosomiasis. *eLife*, 11, 2022.

[11772] Matt J Silver, Ayden Saffari, Noah J Kessler, Gririraj R Chandak, Caroline HD Fall, Prachand Issarapu, Akshay Dedaniya, Modupeh Betts, Sophie E Moore, Michael N Routledge, Zdenko Herceg, Cyrille Cuenin, Maria Derakhshan, Philip T James, David Monk, and Andrew M Prentice. Environmentally sensitive hotspots in the methylome of the early human embryo. *eLife*, 11, 2022.

[11773] Noah J Silverstein, Yetao Wang, Zachary Manickas-Hill, Claudia Carbone, Ann Dauphin, Brittany P Boribong, Maggie Loiselle, Jameson Davis, Maureen M Leonard, Leticia Kuri-Cervantes, Kendall Lavin-Parsons, Nuala J Meyer, Michael R Betts, Jonathan Z Li, Bruce D Walker, Xu G Yu, Lael M Yonker, and Jeremy Luban. Innate lymphoid cells and COVID-19 severity in SARS-CoV-2 infection. *eLife*, 11, 2022.

[11774] Malcolm J W Sim, Zachary Stotz, Jinghua Lu, Paul Brennan, Eric O Long, and Peter D Sun. T cells discriminate between groups C1 and C2 HLA-C. *eLife*, 11, 2022.

[11775] Dorottya Anna Simon, András Tálas, Péter István Kulcsár, Zsuzsanna Biczók, Sarah Laura Krausz, György Várady, and Ervin Welker. PEAR, a flexible fluorescent reporter for the identification and enrichment of successfully prime edited cells. *eLife*, 11, 2022.

[11776] Jennie R Sims, Vitor M FaÇa, Catalina Pereira, Carolline AscenÇão, William Comstock, Jumana Badar, Gerardo A Arroyo-Martinez, Raimundo Freire, Paula E Cohen, Robert S Weiss, and Marcus B Smolka. Phosphoproteomics of ATR signaling in mouse testes. *eLife*, 11, 2022.

[11777] Sudhuman Singh, Torri D Wilson, Spring Valdivia, Barbara Benowitz, Sarah Chaudhry, Jun Ma, Anisha P Adke, Omar Soler-Cedeño, Daniela Velasquez, Mario A Penzo, and Yarimar Carrasquillo. An inhibitory circuit from central amygdala to zona incerta drives pain-related behaviors in mice. *eLife*, 11, 2022.

[11778] Laura M Sipe, Mehdi Chaib, Emily B Korba, Heejoon Jo, Mary Camille Lovely, Brittany R Counts, Ubaid Tanveer, Jeremiah R Holt, Jared C Clements, Neena A John, Deidre Daria, Tony N Marion, Margaret S Bohm, Radhika Sekhri, Ajeeth K Pingili, Bin Teng, James A Carson, D Neil Hayes, Matthew J Davis, Katherine L Cook, Joseph F Pierre, and Liza Makowski. Response to immune checkpoint blockade improved in pre-clinical model of breast cancer after bariatric surgery. *eLife*, 11, 2022.

[11779] Edhriz Siraliev-Perez, Jordan TB Stariha, Reece M Hoffmann, Brenda RS Temple, Qisheng Zhang, Nicole Hajicek, Meredith L Jenkins, John E Burke, and John Sondek. Dynamics of allosteric regulation of the phospholipase C-$\gamma$ isozymes upon recruitment to membranes. *eLife*, 11, 2022.

[11780] Nikoloz Sirmpilatze, Judith Mylius, Michael Ortiz-Rios, Jürgen Baudewig, Jaakko Paasonen, Daniel Golkowski, Andreas Ranft, Rüdiger Ilg, Olli Gröhn, and Susann Boretius. Spatial signatures of anesthesia-induced burst-suppression differ between primates and rodents. *eLife*, 11, 2022.

[11781] Christoph J Slavetinsky, Janna N Hauser, Cordula Gekeler, Jessica Slavetinsky, André Geyer, Alexandra Kraus, Doris Heilingbrunner, Samuel Wagner, Michael Tesar, Bernhard Krismer, Sebastian Kuhn, Christoph M Ernst, and Andreas Peschel. Sensitizing Staphylococcus aureus to antibacterial agents by decoding and blocking the lipid flippase MprF. *eLife*, 11, 2022.

[11782] Carol Smith, Jill G Canestrari, Archer J Wang, Matthew M Champion, Keith M Derbyshire, Todd A Gray, and Joseph T Wade. Pervasive translation in Mycobacterium tuberculosis. *eLife*, 11, 2022.

[11783] Clare M Smith, Richard E Baker, Megan K Proulx, Bibhuti B Mishra, Jarukit E Long, Sae Woong Park, Ha-Na Lee, Michael C Kiritsy, Michelle M Bellerose, Andrew J Olive, Kenan C Murphy, Kadamba Papavinasasundaram, Frederick J Boehm, Charlotte J Reames, Rachel K Meade, Brea K Hampton, Colton L Linnertz, Ginger D Shaw, Pablo Hock, Timothy A Bell, Sabine Ehrt, Dirk Schnappinger, Fernando Pardo-Manuel de Villena, Martin T Ferris, Thomas R Ioerger, and Christopher M Sassetti. Host-pathogen genetic interactions underlie tuberculosis susceptibility in genetically diverse mice. *eLife*, 11, 2022.

[11784] Courtney J Smith, Nasa Sinnott-Armstrong, Anna Cichońska, Heli Julkunen, Eric B Fauman, Peter Würtz, and Jonathan K Pritchard. Integrative analysis of metabolite GWAS illuminates the molecular basis of pleiotropy and genetic correlation. *eLife*, 11, 2022.

[11785] Elliot H Smith, Jyun-you Liou, Edward M Merricks, Tyler Davis, Kyle Thomson, Bradley Greger, Paul House, Ronald G Emerson, Robert Goodman, Guy M McKhann, Sameer Sheth, Catherine Schevon, and John D Rolston. Human interictal epileptiform discharges are bidirectional traveling waves echoing ictal discharges. *eLife*, 11, 2022.

[11786] Richard D Smith, Ilya Kolb, Shinsuke Tanaka, Albert K Lee, Timothy D Harris, and Mladen Barbic. Robotic multi-probe single-actuator inchworm neural microdrive. *eLife*, 11, 2022.

[11787] Spenser S Smith, Daniel Chu, Tiange Qu, Jessye A Aggleton, and Richard A Schneider. Species-specific sensitivity to TGF$\beta$ signaling and changes to the Mmp13 promoter underlie avian jaw development and evolution. *eLife*, 11, 2022.

[11788] Simon Snoeck, Bradley W Abramson, Anthony GK Garcia, Ashley N Egan, Todd P Michael, and Adam D Steinbrenner. Evolutionary gain and loss of a plant pattern-recognition receptor for HAMP recognition. *eLife*, 11, 2022.

[11789] Tolulope Sokoya, Jan Parolek, Mads Møller Foged, Dmytro I Danylchuk, Manuel Bozan, Bingshati Sarkar, Angelika Hilderink, Michael Philippi, Lorenzo D Botto, Paulien A Terhal, Outi Mäkitie, Jacob Piehler, Yeongho Kim, Christopher G Burd, Andrey S Klymchenko, Kenji Maeda, and Joost CM Holthuis. Pathogenic variants of sphingomyelin synthase SMS2 disrupt lipid landscapes in the secretory pathway. *eLife*, 11, 2022.

[11790] Javier Solivan-Rivera, Zinger Yang Loureiro, Tiffany DeSouza, Anand Desai, Sabine Pallat, Qin Yang, Raziel Rojas-Rodriguez, Rachel Ziegler, Pantos Skritakis, Shannon Joyce, Denise Zhong, Tammy Nguyen, and Silvia Corvera. A neurogenic signature involving monoamine Oxidase-A controls human thermogenic adipose tissue development. *eLife*, 11, 2022.

[11791] Maja Solman, Sasja Blokzijl-Franke, Florian Piques, Chuan Yan, Qiqi Yang, Marion Strullu, Sarah M Kamel, Pakize Ak, Jeroen Bakkers, David M Langenau, Hélène Cavé, and Jeroen den Hertog. Inflammatory response in hematopoietic stem and progenitor cells triggered by activating SHP2 mutations evokes blood defects. *eLife*, 11, 2022.

[11792] Cwyn Solvi, Yonghe Zhou, Yunxiao Feng, Yuyi Lu, Mark Roper, Li Sun, Rebecca J Reid, Lars Chittka, Andrew B Barron, and Fei Peng. Bumblebees retrieve only the ordinal ranking of foraging options when comparing memories obtained in distinct settings. *eLife*, 11, 2022.

[11793] Rachana Deven Somaiya, Katelyn Stebbins, Ellen C Gingrich, Hehuang Xie, John N Campbell, A Denise R Garcia, and Michael A Fox. Sonic hedgehog-dependent recruitment of GABAergic interneurons into the developing visual thalamus. *eLife*, 11, 2022.

[11794] Markus J Sommer, Sooyoung Cha, Ales Varabyou, Natalia Rincon, Sukhwan Park, Ilia Minkin, Mihaela Pertea, Martin Steinegger, and Steven L Salzberg. Structure-guided isoform identification for the human transcriptome. *eLife*, 11, 2022.

[11795] Thomas Sonnweber, Piotr Tymoszuk, Sabina Sahanic, Anna Boehm, Alex Pizzini, Anna Luger, Christoph Schwabl, Manfred Nairz, Philipp Grubwieser, Katharina Kurz, Sabine Koppelstätter, Magdalena Aichner, Bernhard Puchner, Alexander Egger, Gregor Hoermann, Ewald Wöll, Günter Weiss, Gerlig Widmann, Ivan Tancevski, and Judith Löffler-Ragg. Investigating phenotypes of pulmonary COVID-19 recovery: A longitudinal observational prospective multicenter trial. *eLife*, 11, 2022.

[11796] Pranidhi Sood, Athena Lin, Connie Yan, Rebecca McGillivary, Ulises Diaz, Tatyana Makushok, Ambika V Nadkarni, Sindy KY Tang, and Wallace F Marshall. Modular, cascade-like transcriptional program of regeneration in Stentor. *eLife*, 11, 2022.

[11797] Trevor R Sorrells, Anjali Pandey, Adriana Rosas-Villegas, and Leslie B Vosshall. A persistent behavioral state enables sustained predation of humans by mosquitoes. *eLife*, 11, 2022.

[11798] Sol Sotillos, Isabel von der Decken, Ivan Domenech Mercadé, Sriraksha Srinivasan, Dmytro Sirokha, Ludmila Livshits, Stefano Vanni, Serge Nef, Anna Biason-Lauber, Daniel Rodríguez Gutiérrez, and James Castelli-Gair Hombría. A conserved function of Human DLC3 and Drosophila Cv-c in testis development. *eLife*, 11, 2022.

[11799] Efe Soyman, Rune Bruls, Kalliopi Ioumpa, Laura Müller-Pinzler, Selene Gallo, Chaoyi Qin, Elisabeth CW van Straaten, Matthew W Self, Judith C Peters, Jessy K Possel, Yoshiyuki Onuki, Johannes C Baayen, Sander Idema, Christian Keysers, and Valeria Gazzola. Intracranial human recordings reveal association between neural activity and perceived intensity for the pain of others in the insula. *eLife*, 11, 2022.

[11800] Martin A Spacek, Davide Crombie, Yannik Bauer, Gregory Born, Xinyu Liu, Steffen Katzner, and Laura Busse. Robust effects of corticothalamic feedback and behavioral state on movie responses in mouse dLGN. *eLife*, 11, 2022.

[11801] Carola Sparn, Eleni Dimou, Annalena Meyer, Roberto Saleppico, Sabine Wegehingel, Matthias Gerstner, Severina Klaus, Helge Ewers, and Walter Nickel. Glypican-1 drives unconventional secretion of fibroblast growth factor 2. *eLife*, 11, 2022.

[11802] Manuel Spitschan, Nayantara Santhi, Amrita Ahluwalia, Dorothee Fischer, Lilian Hunt, Natasha A Karp, Francis Lévi, Inés Pineda-Torra, Parisa Vidafar, and Rhiannon White. Sex differences and sex bias in human circadian and sleep physiology research. *eLife*, 11, 2022.

[11803] Allan C Spradling, Wanbao Niu, Qi Yin, Madhulika Pathak, and Bhawana Maurya. Conservation of oocyte development in germline cysts from Drosophila to mouse. *eLife*, 11, 2022.

[11804] Sriram Srikant, Chantal K Guegler, and Michael T Laub. The evolution of a counter-defense mechanism in a virus constrains its host range. *eLife*, 11, 2022.

[11805] Prerna Srivastava, Geoff de Rosenroll, Akihiro Matsumoto, Tracy Michaels, Zachary Turple, Varsha Jain, Santhosh Sethuramanujam, Benjamin L Murphy-Baum, Keisuke Yonehara, and Gautam Bhagwan Awatramani. Spatiotemporal properties of glutamate input support direction selectivity in the dendrites of retinal starburst amacrine cells. *eLife*, 11, 2022.

[11806] Rachel V Stadler, Shane R Nelson, David M Warshaw, and Gary E Ward. A circular zone of attachment to the extracellular matrix provides directionality to the motility of Toxoplasma gondii in 3D. *eLife*, 11, 2022.

[11807] Aaron Stahl, Nathaniel C Noyes, Tamara Boto, Valentina Botero, Connor N Broyles, Miao Jing, Jianzhi Zeng, Lanikea B King, Yulong Li, Ronald L Davis, and Seth M Tomchik. Associative learning drives longitudinally graded presynaptic plasticity of neurotransmitter release along axonal compartments. *eLife*, 11, 2022.

[11808] Desislava P Staneva, Stefan Bresson, Tatsiana Auchynnikava, Christos Spanos, Juri Rappsilber, A Arockia Jeyaprakash, David Tollervey, Keith R Matthews, and Robin C Allshire. The SPARC complex defines RNAPII promoters in Trypanosoma brucei. *eLife*, 11, 2022.

[11809] Catherine Stark, Teanna Bautista-Leung, Joanna Siegfried, and Daniel Herschlag. Systematic investigation of the link between enzyme catalysis and cold adaptation. *eLife*, 11, 2022.

[11810] Benjamin J Stauch, Alina Peter, Isabelle Ehrlich, Zora Nolte, and Pascal Fries. Human visual gamma for color stimuli. *eLife*, 11, 2022.

[11811] Akis Stavropoulos, Kaushik J Lakshminarasimhan, Jean Laurens, Xaq Pitkow, and Dora E Angelaki. Influence of sensory modality and control dynamics on human path integration. *eLife*, 11, 2022.

[11812] Laura Stefan, Nadine Engbersen, and Christian Schöb. Rapid transgenerational adaptation in response to intercropping reduces competition. *eLife*, 11, 2022.

[11813] Janina I Steinert, Henrike Sternberg, Giuseppe A Veltri, and Tim Büthe. How should COVID-19 vaccines be distributed between the Global North and South: a discrete choice experiment in six European countries. *eLife*, 11, 2022.

[11814] Simon Stenberg, Jing Li, Arne B Gjuvsland, Karl Persson, Erik Demitz-Helin, Carles González Peña, Jia-Xing Yue, Ciaran Gilchrist, Timmy ärengård, Payam Ghiaci, Lisa Larsson-Berglund, Martin Zackrisson, Silvana Smits, Johan Hallin, Johanna L Höög, Mikael Molin, Gianni Liti, Stig W Omholt, and Jonas Warringer. Genetically controlled mtDNA deletions prevent ROS damage by arresting oxidative phosphorylation. *eLife*, 11, 2022.

[11815] Donté Alexander Stevens, Christopher Beierschmitt, Swetha Mahesula, Miles R Corley, John Salogiannis, Brian V Tsu, Bryant Cao, Andrew P Ryan, Hiroyuki Hakozawki, Samara L Reck-Peterson, and Matthew D Daugherty. Antiviral function and viral antagonism of the rapidly evolving dynein activating adaptor NINL. *eLife*, 11, 2022.

[11816] Valentina Štimac, Isabella Koprivec, Martina Manenica, Juraj Simunić, and Iva M Tolić. Augmin prevents merotelic attachments by promoting proper arrangement of bridging and kinetochore fibers. *eLife*, 11, 2022.

[11817] Oliver Stirrup, James Blackstone, Fiona Mapp, Alyson MacNeil, Monica Panca, Alison Holmes, Nicholas Machin, Gee Yen Shin, Tabitha Mahungu, Kordo Saeed, Tranprit Saluja, Yusri Taha, Nikunj Mahida, Cassie Pope, Anu Chawla, Maria-Teresa Cutino-Moguel, Asif Tamuri, Rachel Williams, Alistair Darby, David L Robertson, Flavia Flaviani, Eleni Nastouli, Samuel Robson, Darren Smith, Matthew Loose, Kenneth Laing, Irene Monahan, Beatrix Kele, Sam Haldenby, Ryan George, Matthew Bashton, Adam A Witney, Matthew Byott, Francesc Coll, Michael Chapman, Sharon J Peacock, Joseph Hughes, Gaia Nebbia, David G Partridge, Matthew Parker, James Richard Price, Christine Peters, Sunando Roy, Luke B Snell, Thushan I de Silva, Emma Thomson, Paul Flowers, Andrew Copas, and Judith Breuer. Effectiveness of rapid SARS-CoV-2 genome sequencing in supporting infection control for hospital-onset COVID-19 infection: Multicentre, prospective study. *eLife*, 11, 2022.

[11818] Ohad Stoler, Alexandra Stavsky, Yana Khrapunsky, Israel Melamed, Grace Stutzmann, Daniel Gitler, Israel Sekler, and Ilya Fleidervish. Frequency- and spike-timing-dependent mitochondrial Ca2+ signaling regulates the metabolic rate and synaptic efficacy in cortical neurons. *eLife*, 11, 2022.

[11819] Anastasia V Stolyarova, Tatiana V Neretina, Elena A Zvyagina, Anna V Fedotova, Alexey S Kondrashov, and Georgii A Bazykin. Complex fitness landscape shapes variation in a hyperpolymorphic species. *eLife*, 11, 2022.

[11820] Nikos Stratakis, Alexandros P Siskos, Eleni Papadopoulou, Anh N Nguyen, Yinqi Zhao, Katerina Margetaki, Chung-Ho E Lau, Muireann Coen, Lea Maitre, Silvia Fernández-Barrés, Lydiane Agier, Sandra Andrusaityte, Xavier Basagaña, Anne Lise Brantsaeter, Maribel Casas, Serena Fossati, Regina Grazuleviciene, Barbara Heude, Rosemary RC McEachan, Helle Margrete Meltzer, Christopher Millett, Fernanda Rauber, Oliver Robinson, Theano Roumeliotaki, Eva Borras, Eduard Sabidó, Jose Urquiza, Marina Vafeiadi, Paolo Vineis, Trudy Voortman, John Wright, David V Conti, Martine Vrijheid, Hector C Keun, and Leda Chatzi. Urinary metabolic biomarkers of diet quality in European children are associated with metabolic health. *eLife*, 11, 2022.

[11821] Dominik Straub and Constantin A Rothkopf. Putting perception into action with inverse optimal control for continuous psychophysics. *eLife*, 11, 2022.

[11822] Sören Strauss, Adam Runions, Brendan Lane, Dennis Eschweiler, Namrata Bajpai, Nicola Trozzi, Anne-Lise Routier-Kierzkowska, Saiko Yoshida, Sylvia Rodrigues da Silveira, Athul Vijayan, Rachele Tofanelli, Mateusz Majda, Emillie Echevin, Constance Le Gloanec, Hana Bertrand-Rakusova, Milad Adibi, Kay Schneitz, George W Bassel, Daniel Kierzkowski, Johannes Stegmaier, Miltos Tsiantis, and Richard S Smith. Using positional information to provide context for biological image analysis with MorphoGraphX 2.0. *eLife*, 11, 2022.

[11823] Sebastian Ströh, Eric W Hammerschmith, David W Tank, H Sebastian Seung, and Adrian Andreas Wanner. In situ X-ray-assisted electron microscopy staining for large biological samples. *eLife*, 11, 2022.

[11824] Weiping Su, Guanqiao Liu, Bahram Mohajer, Jiekang Wang, Alena Shen, Weixin Zhang, Bin Liu, Ali Guermazi, Peisong Gao, Xu Cao, Shadpour Demehri, and Mei Wan. Senescent preosteoclast secretome promotes metabolic syndrome associated osteoarthritis through cyclooxygenase 2. *eLife*, 11, 2022.

[11825] Laura E Suarez, Yossi Yovel, Martijn P van den Heuvel, Olaf Sporns, Yaniv Assaf, Guillaume Lajoie, and Bratislav Misic. A connectomics-based taxonomy of mammals. *eLife*, 11, 2022.

[11826] Nikkil Sudharsanan, Caterina Favaretti, Violetta Hachaturyan, Till Bärnighausen, and Alain Vandormael. Effects of side-effect risk framing strategies on COVID-19 vaccine intentions: a randomized controlled trial. *eLife*, 11, 2022.

[11827] Ko Sugawara, Çağrı Çevrim, and Michalis Averof. Tracking cell lineages in 3D by incremental deep learning. *eLife*, 11, 2022.

[11828] Chie Sugimoto, Yukie Murakami, Eisuke Ishii, Hiroyoshi Fujita, and Hiroshi Wakao. Reprogramming and redifferentiation of mucosal-associated invariant T cells reveal tumor inhibitory activity. *eLife*, 11, 2022.

[11829] Alyson Sujkowski, Kristin Richardson, Matthew V Prifti, Robert J Wessells, and Sokol V Todi. Endurance exercise ameliorates phenotypes in Drosophila models of spinocerebellar ataxias. *eLife*, 11, 2022.

[11830] Vaishnavi Sukumar, Roland S Johansson, and J Andrew Pruszynski. Precise and stable edge orientation signaling by human first-order tactile neurons. *eLife*, 11, 2022.

[11831] Bethany Sump, Donna G Brickner, Agustina D'Urso, Seo Hyun Kim, and Jason H Brickner. Mitotically heritable, RNA polymerase II-independent H3K4 dimethylation stimulates INO1 transcriptional memory. *eLife*, 11, 2022.

[11832] Anton Sumser, Maximilian Joesch, Peter Jonas, and Yoav Ben-Simon. Fast, high-throughput production of improved rabies viral vectors for specific, efficient and versatile transsynaptic retrograde labeling. *eLife*, 11, 2022.

[11833] Kai Sun, Chun Wang, Jianqiu Xiao, Michael D Brodt, Luorongxin Yuan, Tong Yang, Yael Alippe, Huimin Hu, Dingjun Hao, Yousef Abu-Amer, Matthew J Silva, Jie Shen, and Gabriel Mbalaviele. Fracture healing is delayed in the absence of gasdermin-interleukin-1 signaling. *eLife*, 11, 2022.

[11834] Xin-Yao Sun, Xiang-Chun Ju, Yang Li, Peng-Ming Zeng, Jian Wu, Ying-Ying Zhou, Li-Bing Shen, Jian Dong, Yue-Jun Chen, and Zhen-Ge Luo. Generation of vascularized brain organoids to study neurovascular interactions. *eLife*, 11, 2022.

[11835] Derek C Sung, Xiaowen Chen, Mei Chen, Jisheng Yang, Susan Schultz, Apoorva Babu, Yitian Xu, Siqi Gao, TC Stevenson Keller, Patricia Mericko-Ishizuka, Michelle Lee, Ying Yang, Joshua P Scallan, and Mark L Kahn. VE-cadherin enables trophoblast endovascular invasion and spiral artery remodeling during placental development. *eLife*, 11, 2022.

[11836] Pooja Suresh, Vahe Galstyan, Rob Phillips, and Sophie Dumont. Modeling and mechanical perturbations reveal how spatially regulated anchorage gives rise to spatially distinct mechanics across the mammalian spindle. *eLife*, 11, 2022.

[11837] Norimitsu Suzuki, Malinda LS Tantirigama, K Phyu Aung, Helena HY Huang, and John M Bekkers. Fast and slow feedforward inhibitory circuits for cortical odor processing. *eLife*, 11, 2022.

[11838] Clara Suñer, Annarita Sibilio, Judit Martín, Chiara Lara Castellazzi, Oscar Reina, Ivan Dotu, Adrià Caballé, Elisa Rivas, Vittorio Calderone, Juana Díez, Angel R Nebreda, and Raúl Méndez. Macrophage inflammation resolution requires CPEB4-directed offsetting of mRNA degradation. *eLife*, 11, 2022.

[11839] Claire A Symanski, John H Bladon, Emi T Kullberg, Paul Miller, and Shantanu P Jadhav. Rhythmic coordination and ensemble dynamics in the hippocampal-prefrontal network during odor-place associative memory and decision making. *eLife*, 11, 2022.

[11840] Eszter Szabó, Cinzia Chiandetti, Ernő Téglás, Elisabetta Versace, Gergely Csibra, ágnes Melinda Kovács, and Giorgio Vallortigara. Young domestic chicks spontaneously represent the absence of objects. *eLife*, 11, 2022.

[11841] Zoltán Szadai, Hyun-Jae Pi, Quentin Chevy, Katalin ócsai, Dinu F Albeanu, Balázs Chiovini, Gergely Szalay, Gergely Katona, Adam Kepecs, and Balázs Rózsa. Cortex-wide response mode of VIP-expressing inhibitory neurons by reward and punishment. *eLife*, 11, 2022.

[11842] Eva M Szego, Laura Malz, Nadine Bernhardt, Angela Rösen-Wolff, Björn H Falkenburger, and Hella Luksch. Constitutively active STING causes neuroinflammation and degeneration of dopaminergic neurons in mice. *eLife*, 11, 2022.

[11843] Katarzyna Szymanska, Karsten Boldt, Clare V Logan, Matthew Adams, Philip A Robinson, Marius Ueffing, Elton Zeqiraj, Gabrielle Wheway, and Colin A Johnson. Regulation of canonical Wnt signalling by the ciliopathy protein MKS1 and the E2 ubiquitin-conjugating enzyme UBE2E1. *eLife*, 11, 2022.

[11844] Shinya Taguchi, Juri Nakano, Tsuyoshi Imasaki, Tomoki Kita, Yumiko Saijo-Hamano, Naoki Sakai, Hideki Shigematsu, Hiromichi Okuma, Takahiro Shimizu, Eriko Nitta, Satoshi Kikkawa, Satoshi Mizobuchi, Shinsuke Niwa, and Ryo Nitta. Structural model of microtubule dynamics inhibition by kinesin-4 from the crystal structure of KLP-12 –tubulin complex. *eLife*, 11, 2022.

[11845] Masak Takaine, Hiromi Imamura, and Satoshi Yoshida. High and stable ATP levels prevent aberrant intracellular protein aggregation in yeast. *eLife*, 11, 2022.

[11846] Yasuyuki Tamai, Akiko Eguchi, Ryuta Shigefuku, Hiroshi Kitamura, Mina Tempaku, Ryosuke Sugimoto, Yoshinao Kobayashi, Motoh Iwasa, Yoshiyuki Takei, and Hayato Nakagawa. Association of lithocholic acid with skeletal muscle hypertrophy through TGR5-IGF-1 and skeletal muscle mass in cultured mouse myotubes, chronic liver disease rats and humans. *eLife*, 11, 2022.

[11847] Stella Tamana, Maria Xenophontos, Anna Minaidou, Coralea Stephanou, Cornelis L Harteveld, Celeste Bento, Joanne Traeger-Synodinos, Irene Fylaktou, Norafiza Mohd Yasin, Faidatul Syazlin Abdul Hamid, Ezalia Esa, Hashim Halim-Fikri, Bin Alwi Zilfalil, Andrea C Kakouri, Marina Kleanthous, and Petros Kountouris. Evaluation of in silico predictors on short nucleotide variants in HBA1, HBA2, and HBB associated with haemoglobinopathies. *eLife*, 11, 2022.

[11848] Chao Tan, Giovanni de Nola, Claire Qiao, Cordelia Imig, Richard T Born, Nils Brose, and Pascal S Kaeser. Munc13 supports fusogenicity of non-docked vesicles at synapses with disrupted active zones. *eLife*, 11, 2022.

[11849] Hua Leonhard Tan, Stefanie Bungert-Plümke, Daniel Kortzak, Christoph Fahlke, and Gabriel Stölting. Determination of oligomeric states of proteins via dual-color colocalization with single molecule localization microscopy. *eLife*, 11, 2022.

[11850] Shoji Tane, Keishi Shintomi, Kazuhisa Kinoshita, Yuko Tsubota, Makoto M Yoshida, Tomoko Nishiyama, and Tatsuya Hirano. Cell cycle-specific loading of condensin I is regulated by the N-terminal tail of its kleisin subunit. *eLife*, 11, 2022.

[11851] Ting Tang, Naili Zhang, Franca J Bongers, Michael Staab, Andreas Schuldt, Felix Fornoff, Hong Lin, Jeannine Cavender-Bares, Andrew L Hipp, Shan Li, Yu Liang, Baocai Han, Alexandra-Maria Klein, Helge Bruelheide, Walter Durka, Bernhard Schmid, Keping Ma, and Xiaojuan Liu. Tree species and genetic diversity increase productivity via functional diversity and trophic feedbacks. *eLife*, 11, 2022.

[11852] Vi T Tang, Joseph McCormick, Bolin Xu, Yawei Wang, Huan Fang, Xiao Wang, David Siemieniak, Rami Khoriaty, Brian T Emmer, Xiao-Wei Chen, and David Ginsburg. Hepatic inactivation of murine Surf4 results in marked reduction in plasma cholesterol. *eLife*, 11, 2022.

[11853] Xin-Zi Tang, Lieselotte S M Kreuk, Cynthia Cho, Ross J Metzger, and Christopher D C Allen. Bronchus-associated macrophages efficiently capture and present soluble inhaled antigens and are capable of local Th2 cell activation. *eLife*, 11, 2022.

[11854] Yan Tang, Haihong Zong, Hyokjoon Kwon, Yunping Qiu, Jacob B Pessin, Licheng Wu, Katherine A Buddo, Ilya Boykov, Cameron A Schmidt, Chien-Te Lin, P Darrell Neufer, Gary J Schwartz, Irwin J Kurland, and Jeffrey E Pessin. TIGAR deficiency enhances skeletal muscle thermogenesis by increasing neuromuscular junction cholinergic signaling. *eLife*, 11, 2022.

[11855] Kirill V Tarasov, Khalid Chakir, Daniel R Riordon, Alexey E Lyashkov, Ismayil Ahmet, Maria Grazia Perino, Allwin Jennifa Silvester, Jing Zhang, Mingyi Wang, Yevgeniya O Lukyanenko, Jia-Hua Qu, Miguel Calvo-Rubio Barrera, Magdalena Juhaszova, Yelena S Tarasova, Bruce Ziman, Richard Telljohann, Vikas Kumar, Mark Ranek, John Lammons, Rostislav Bychkov, Rafael de Cabo, Seungho Jun, Gizem Keceli, Ashish Gupta, Dongmei Yang, Miguel A Aon, Luigi Adamo, Christopher H Morrell, Walter Otu, Cameron Carroll, Shane Chambers, Nazareno Paolocci, Thanh Huynh, Karel Pacak, Robert Weiss, Loren Field, Steven J Sollott, and Edward G Lakatta. A remarkable adaptive paradigm of heart performance and protection emerges in response to marked cardiac-specific overexpression of ADCY8. *eLife*, 11, 2022.

[11856] Darya Task, Chun-Chieh Lin, Alina Vulpe, Ali Afify, Sydney Ballou, Maria Brbic, Philipp Schlegel, Joshua Raji, Gregory SXE Jefferis, Hongjie Li, Karen Menuz, and Christopher J Potter. Chemoreceptor co-expression in Drosophila melanogaster olfactory neurons. *eLife*, 11, 2022.

[11857] Meltem Tatli, Sarah Moraïs, Omar E Tovar-Herrera, Yannick J Bomble, Edward A Bayer, Ohad Medalia, and Itzhak Mizrahi. Nanoscale resolution of microbial fiber degradation in action. *eLife*, 11, 2022.

[11858] Sonia Taïb, Noël Lamandé, Sabrina Martin, Fanny Coulpier, Piotr Topilko, and Isabelle Brunet. Myelinating Schwann cells and Netrin-1 control intra-nervous vascularization of the developing mouse sciatic nerve. *eLife*, 11, 2022.

[11859] Sara Tejedor-Sanz, Eric T Stevens, Siliang Li, Peter Finnegan, James Nelson, Andre Knoesen, Samuel H Light, Caroline M Ajo-Franklin, and Maria L Marco. Extracellular electron transfer increases fermentation in lactic acid bacteria via a hybrid metabolism. *eLife*, 11, 2022.

[11860] Sanne ten Oever, Sara Carta, Greta Kaufeld, and Andrea E Martin. Neural tracking of phrases in spoken language comprehension is automatic and task-dependent. *eLife*, 11, 2022.

[11861] Benjamin C Tendler, Taylor Hanayik, Olaf Ansorge, Sarah Bangerter-Christensen, Gregory S Berns, Mads F Bertelsen, Katherine L Bryant, Sean Foxley, Martijn P van den Heuvel, Amy FD Howard, Istvan N Huszar, Alexandre A Khrapitchev, Anna Leonte, Paul R Manger, Ricarda AL Menke, Jeroen Mollink, Duncan Mortimer, Menuka Pallebage-Gamarallage, Lea Roumazeilles, Jerome Sallet, Lianne H Scholtens, Connor Scott, Adele Smart, Martin R Turner, Chaoyue Wang, Saad Jbabdi, Rogier B Mars, and Karla L Miller. The Digital Brain Bank, an open access platform for post-mortem imaging datasets. *eLife*, 11, 2022.

[11862] Bochuan Teng, Joshua P Kaplan, Ziyu Liang, Zachary Krieger, Yu-Hsiang Tu, Batuujin Burendei, Andrew B Ward, and Emily R Liman. Structural motifs for subtype-specific pH-sensitive gating of vertebrate otopetrin proton channels. *eLife*, 11, 2022.

[11863] Nicole A Terpolilli, Reinhard Dolp, Kai Waehner, Susanne M Schwarzmaier, Elisabeth Rumbler, Boyan Todorov, Michel D Ferrari, Arn MJM van den Maagdenberg, and Nikolaus Plesnila. CaV2.1 channel mutations causing familial hemiplegic migraine type 1 increase the susceptibility for cortical spreading depolarizations and seizures and worsen outcome after experimental traumatic brain injury. *eLife*, 11, 2022.

[11864] Christian JG Tessier, Raymond M Sturgeon, Johnathon R Emlaw, Gregory D McCluskey, F Javier Pérez-Areales, and Corrie JB daCosta. Ancestral acetylcholine receptor $\beta$-subunit forms homopentamers that prime before opening spontaneously. *eLife*, 11, 2022.

[11865] Yuta Tezuka, Kenta M Hagihara, Kenichi Ohki, Tomoo Hirano, and Yoshiaki Tagawa. Developmental stage-specific spontaneous activity contributes to callosal axon projections. *eLife*, 11, 2022.

[11866] Shruti Thapliyal, Kristin L Arendt, Anthony G Lau, and Lu Chen. Retinoic acid-gated BDNF synthesis in neuronal dendrites drives presynaptic homeostatic plasticity. *eLife*, 11, 2022.

[11867] Alexandre P Thiery, Ariane SI Standing, Rory L Cooper, and Gareth J Fraser. An epithelial signalling centre in sharks supports homology of tooth morphogenesis in vertebrates. *eLife*, 11, 2022.

[11868] H Nayanga Thirimanne, Feinan Wu, Derek H Janssens, Jherek Swanger, Ahmed Diab, Heather M Feldman, Robert A Amezquita, Raphael Gottardo, Patrick J Paddison, Steven Henikoff, and Bruce E Clurman. Global and context-specific transcriptional consequences of oncogenic Fbw7 mutations. *eLife*, 11, 2022.

[11869] Mathi Thiruppathy, Peter Fabian, J Andrew Gillis, and J Gage Crump. Gill developmental program in the teleost mandibular arch. *eLife*, 11, 2022.

[11870] Christopher S Thom, Madison B Wilken, Stella T Chou, and Benjamin F Voight. Body mass index and adipose distribution have opposing genetic impacts on human blood traits. *eLife*, 11, 2022.

[11871] Alex F Thompson, Patrick R Blackburn, Noah S Arons, Sarah N Stevens, Dusica Babovic-Vuksanovic, Jane B Lian, Eric W Klee, and Jason Stumpff. Pathogenic mutations in the chromokinesin KIF22 disrupt anaphase chromosome segregation. *eLife*, 11, 2022.

[11872] Eric E Thomson, Mark Harfouche, Kanghyun Kim, Pavan C Konda, Catherine W Seitz, Colin Cooke, Shiqi Xu, Whitney S Jacobs, Robin Blazing, Yang Chen, Sunanda Sharma, Timothy W Dunn, Jaehee Park, Roarke W Horstmeyer, and Eva A Naumann. Gigapixel imaging with a novel multi-camera array microscope. *eLife*, 11, 2022.

[11873] Thomas Thumberger, Tinatini Tavhelidse-Suck, Jose Arturo Gutierrez-Triana, Alex Cornean, Rebekka Medert, Bettina Welz, Marc Freichel, and Joachim Wittbrodt. Boosting targeted genome editing using the hei-tag. *eLife*, 11, 2022.

[11874] Henning Tiedemann, Yaniv Morgenstern, Filipp Schmidt, and Roland W Fleming. One-shot generalization in humans revealed through a drawing task. *eLife*, 11, 2022.

[11875] Susanne Tilk, Svyatoslav Tkachenko, Christina Curtis, Dmitri A Petrov, and Christopher D McFarland. Most cancers carry a substantial deleterious load due to Hill-Robertson interference. *eLife*, 11, 2022.

[11876] Dorothea Ogmore Tilley, Ulrike Abuabed, Ursula Zimny Arndt, Monika Schmid, Stefan Florian, Peter R Jungblut, Volker Brinkmann, Alf Herzig, and Arturo Zychlinsky. Histone H3 clipping is a novel signature of human neutrophil extracellular traps. *eLife*, 11, 2022.

[11877] James A Timmons, Andrew Anighoro, Robert J Brogan, Jack Stahl, Claes Wahlestedt, David Gordon Farquhar, Jake Taylor-King, Claude-Henry Volmar, William E Kraus, and Stuart M Phillips. A human-based multi-gene signature enables quantitative drug repurposing for metabolic disease. *eLife*, 11, 2022.

[11878] Alba Timón-Gómez, Alexandra L Scharr, Nicholas Y Wong, Erwin Ni, Arijit Roy, Min Liu, Julisia Chau, Jack L Lampert, Homza Hireed, Noah S Kim, Masood Jan, Alexander R Gupta, Ryan W Day, James M Gardner, Richard JA Wilson, Antoni Barrientos, and Andy J Chang. Tissue-specific mitochondrial HIGD1C promotes oxygen sensitivity in carotid body chemoreceptors. *eLife*, 11, 2022.

[11879] See-Yeun Ting, Kaitlyn D LaCourse, Hannah E Ledvina, Rutan Zhang, Matthew C Radey, Hemantha D Kulasekara, Rahul Somavanshi, Savannah K Bertolli, Larry A Gallagher, Jennifer Kim, Kelsi M Penewit, Stephen J Salipante, Libin Xu, S Brook Peterson, and Joseph D Mougous. Discovery of coordinately regulated pathways that provide innate protection against interbacterial antagonism. *eLife*, 11, 2022.

[11880] Alexandra Tinnermann, Christian Sprenger, and Christian Büchel. Opioid analgesia alters corticospinal coupling along the descending pain system in healthy participants. *eLife*, 11, 2022.

[11881] Johnny M Tkach, Reuben Philip, Amit Sharma, Jonathan Strecker, Daniel Durocher, and Laurence Pelletier. Global cellular response to chemical perturbation of PLK4 activity and abnormal centrosome number. *eLife*, 11, 2022.

[11882] Marco Todesco, Natalia Bercovich, Amy Kim, Ivana Imerovski, Gregory L Owens, óscar Dorado Ruiz, Srinidhi V Holalu, Lufiani L Madilao, Mojtaba Jahani, Jean-Sébastien Légaré, Benjamin K Blackman, and Loren H Rieseberg. Genetic basis and dual adaptive role of floral pigmentation in sunflowers. *eLife*, 11, 2022.

[11883] Andrea Toledo, Mathieu Letellier, Giorgia Bimbi, Béatrice Tessier, Sophie Daburon, Alexandre Favereaux, Ingrid Chamma, Kristel Vennekens, Jeroen Vanderlinden, Matthieu Sainlos, Joris de Wit, Daniel Choquet, and Olivier Thoumine. MDGAs are fast-diffusing molecules that delay excitatory synapse development by altering neuroligin behavior. *eLife*, 11, 2022.

[11884] Theadora Tolkin, Ariz Mohammad, Todd A Starich, Ken CQ Nguyen, David H Hall, Tim Schedl, E Jane Albert Hubbard, and David Greenstein. Innexin function dictates the spatial relationship between distal somatic cells in the Caenorhabditis elegans gonad without impacting the germline stem cell pool. *eLife*, 11, 2022.

[11885] Isabella Tomanek and Călin C Guet. Adaptation dynamics between copy-number and point mutations. *eLife*, 11, 2022.

[11886] Kathrin Tomasek, Alexander Leithner, Ivana Glatzova, Michael S Lukesch, Calin C Guet, and Michael Sixt. Type 1 piliated uropathogenic Escherichia coli hijack the host immune response by binding to CD14. *eLife*, 11, 2022.

[11887] Rute A Tomaz, Ekaterini D Zacharis, Fabian Bachinger, Annabelle Wurmser, Daniel Yamamoto, Sandra Petrus-Reurer, Carola M Morell, Dominika Dziedzicka, Brandon T Wesley, Imbisaat Geti, Charis-Patricia Segeritz, Miguel C de Brito, Mariya Chhatriwala, Daniel Ortmann, Kourosh Saeb-Parsy, and Ludovic Vallier. Generation of functional hepatocytes by forward programming with nuclear receptors. *eLife*, 11, 2022.

[11888] Hideaki Tomita, Kelly M Hines, Josi M Herron, Amy Li, David W Baggett, and Libin Xu. 7-Dehydrocholesterol-derived oxysterols cause neurogenic defects in Smith-Lemli-Opitz syndrome. *eLife*, 11, 2022.

[11889] Andrea Torneri, Lander Willem, Vittoria Colizza, Cécile Kremer, Christelle Meuris, Gilles Darcis, Niel Hens, and Pieter JK Libin. Controlling SARS-CoV-2 in schools using repetitive testing strategies. *eLife*, 11, 2022.

[11890] Julia R Torvi, Jonathan Wong, Daniel Serwas, Amir Moayed, David G Drubin, and Georjana Barnes. Reconstitution of kinetochore motility and microtubule dynamics reveals a role for a kinesin-8 in establishing end-on attachments. *eLife*, 11, 2022.

[11891] Ayala Tovy, Jaime M Reyes, Linda Zhang, Yung-Hsin Huang, Carina Rosas, Alexes C Daquinag, Anna Guzman, Raghav Ramabadran, Chun-Wei Chen, Tianpeng Gu, Sinjini Gupta, Laura Ortinau, Dongsu Park, Aaron R Cox, Rachel E Rau, Sean M Hartig, Mikhail G Kolonin, and Margaret A Goodell. Constitutive loss of DNMT3A causes morbid obesity through misregulation of adipogenesis. *eLife*, 11, 2022.

[11892] Robert J Tower, Emily Busse, Josue Jaramillo, Michelle Lacey, Kevin Hoffseth, Anyonya R Guntur, Jennifer Simkin, and Mimi C Sammarco. Spatial transcriptomics reveals metabolic changes underly age-dependent declines in digit regeneration. *eLife*, 11, 2022.

[11893] Wataru Toyokawa and Wolfgang Gaissmaier. Conformist social learning leads to self-organised prevention against adverse bias in risky decision making. *eLife*, 11, 2022.

[11894] Lucas R Trambaiolli, Xiaolong Peng, Julia F Lehman, Gary Linn, Brian E Russ, Charles E Schroeder, Hesheng Liu, and Suzanne N Haber. Anatomical and functional connectivity support the existence of a salience network node within the caudal ventrolateral prefrontal cortex. *eLife*, 11, 2022.

[11895] Ngoc Lan Tran, Lorena Maria Ferreira, Blanca Alvarez-Moya, Valentina Buttiglione, Barbara Ferrini, Paola Zordan, Andrea Monestiroli, Claudio Fagioli, Eugenia Bezzecchi, Giulia Maria Scotti, Antonio Esposito, Riccardo Leone, Chiara Gnasso, Andrea Brendolan, Luca G Guidotti, and Giovanni Sitia. Continuous sensing of IFN$\alpha$ by hepatic endothelial cells shapes a vascular antimetastatic barrier. *eLife*, 11, 2022.

[11896] Quang D Tran, Eric Galiana, Philippe Thomen, Céline Cohen, FranÇois Orange, Fernando Peruani, and Xavier Noblin. Coordination of two opposite flagella allows high-speed swimming and active turning of individual zoospores. *eLife*, 11, 2022.

[11897] Sydney Trask, Jeffrey S Mogil, Fred J Helmstetter, Cheryl L Stucky, and Katelyn E Sadler. Contextual control of conditioned pain tolerance and endogenous analgesic systems. *eLife*, 11, 2022.

[11898] Laura Christin Trautenberg, Marko Brankatschk, Andrej Shevchenko, Stuart Wigby, and Klaus Reinhardt. Ecological lipidology. *eLife*, 11, 2022.

[11899] Ethan B Trepka, Shude Zhu, Ruobing Xia, Xiaomo Chen, and Tirin Moore. Functional interactions among neurons within single columns of macaque V1. *eLife*, 11, 2022.

[11900] Abhinandan M Tripathi, Yael Admoni, Arie Fridrich, Magda Lewandowska, Joachim M Surm, Reuven Aharoni, and Yehu Moran. Functional characterization of a 'plant-like' HYL1 homolog in the cnidarian Nematostella vectensis indicates a conserved involvement in microRNA biogenesis. *eLife*, 11, 2022.

[11901] Claudio Tripodo, Barbara Bassani, Elena Jachetti, Valeria Cancila, Claudia Chiodoni, Paola Portararo, Laura Botti, Cesare Valenti, Milena Perrone, Maurilio Ponzoni, Patrizia Comoli, Mara Lecchi, Paolo Verderio, Antonio Curti, Mario P Colombo, and Sabina Sangaletti. Neutrophil extracellular traps arm DC vaccination against NPM-mutant myeloproliferation. *eLife*, 11, 2022.

[11902] Julie Trolle, Ross M McBee, Andrew Kaufman, Sudarshan Pinglay, Henri Berger, Sergei German, Liyuan Liu, Michael J Shen, Xinyi Guo, J Andrew Martin, Michael E Pacold, Drew R Jones, Jef D Boeke, and Harris H Wang. Resurrecting essential amino acid biosynthesis in mammalian cells. *eLife*, 11, 2022.

[11903] Marius Tröndle, Tzvetan Popov, Sabine Dziemian, and Nicolas Langer. Decomposing the role of alpha oscillations during brain maturation. *eLife*, 11, 2022.

[11904] Jeremy M Trott, Ann N Hoffman, Irina Zhuravka, and Michael S Fanselow. Conditional and unconditional components of aversively motivated freezing, flight and darting in mice. *eLife*, 11, 2022.

[11905] Shaun Truelove, Claire P Smith, Michelle Qin, Luke C Mullany, Rebecca K Borchering, Justin Lessler, Katriona Shea, Emily Howerton, Lucie Contamin, John Levander, Jessica Kerr, Harry Hochheiser, Matt Kinsey, Kate Tallaksen, Shelby Wilson, Lauren Shin, Kaitlin Rainwater-Lovett, Joseph C Lemairtre, Juan Dent, Joshua Kaminsky, Elizabeth C Lee, Javier Perez-Saez, Alison Hill, Dean Karlen, Matteo Chinazzi, Jessica T Davis, Kunpeng Mu, Xinyue Xiong, Ana Pastore y Piontti, Alessandro Vespignani, Ajitesh Srivastava, Przemyslaw Porebski, Srinivasan Venkatramanan, Aniruddha Adiga, Bryan Lewis, Brian Klahn, Joseph Outten, Mark Orr, Galen Harrison, Benjamin Hurt, Jiangzhuo Chen, Anil Vullikanti, Madhav Marathe, Stefan Hoops, Parantapa Bhattacharya, Dustin Machi, Shi Chen, Rajib Paul, Daniel Janies, Jean-Claude Thill, Marta Galanti, Teresa K Yamana, Sen Pei, Jeffrey L Shaman, Jessica M Healy, Rachel B Slayton, Matthew Biggerstaff, Michael A Johansson, Michael C Runge, and Cecile Viboud. Projected resurgence of COVID-19 in the United States in July—December 2021 resulting from the increased transmissibility of the Delta variant and faltering vaccination. *eLife*, 11, 2022.

[11906] Jonathan S Tsay, Hyosub Kim, Adrian M Haith, and Richard B Ivry. Understanding implicit sensorimotor adaptation as a process of proprioceptive re-alignment. *eLife*, 11, 2022.

[11907] Sau Yee Tsoi, Merve öncül, Ella Svahn, Mark Robertson, Zuzanna Bogdanowicz, Christina McClure, and Gülşen Sürmeli. Telencephalic outputs from the medial entorhinal cortex are copied directly to the hippocampus. *eLife*, 11, 2022.

[11908] Wenyu Tu and Nanyin Zhang. Neural underpinning of a respiration-associated resting-state fMRI network. *eLife*, 11, 2022.

[11909] Tayfun Tumkaya, Safwan Burhanudin, Asghar Khalilnezhad, James Stewart, Hyungwon Choi, and Adam Claridge-Chang. Most primary olfactory neurons have individually neutral effects on behavior. *eLife*, 11, 2022.

[11910] Gabriela C Tunes, Eliezyer Fermino de Oliveira, Estevão UP Vieira, Marcelo S Caetano, André M Cravo, and Marcelo Bussotti Reyes. Time encoding migrates from prefrontal cortex to dorsal striatum during learning of a self-timed response duration task. *eLife*, 11, 2022.

[11911] Paul Tupper, Shraddha Pai, Urooj Khan, and Caroline Colijn. COVID-19 cluster size and transmission rates in schools from crowdsourced case reports. *eLife*, 11, 2022.

[11912] David J Turner, Alexander Saveliev, Fiamma Salerno, Louise S Matheson, Michael Screen, Hannah Lawson, David Wotherspoon, Kamil R Kranc, and Martin Turner. A functional screen of RNA binding proteins identifies genes that promote or limit the accumulation of CD138+ plasma cells. *eLife*, 11, 2022.

[11913] Hannah Twarkowski, Victor Steininger, Min Jae Kim, and Amar Sahay. A dentate gyrus-CA3 inhibitory circuit promotes evolution of hippocampal-cortical ensembles during memory consolidation. *eLife*, 11, 2022.

[11914] Ceereena Ubaida-Mohien, Sally Spendiff, Alexey Lyashkov, Ruin Moaddel, Norah J MacMillan, Marie-Eve Filion, Jose A Morais, Tanja Taivassalo, Luigi Ferrucci, and Russell T Hepple. Unbiased proteomics, histochemistry, and mitochondrial DNA copy number reveal better mitochondrial health in muscle of high-functioning octogenarians. *eLife*, 11, 2022.

[11915] Balazs B Ujfalussy and Gergő Orbán. Sampling motion trajectories during hippocampal theta sequences. *eLife*, 11, 2022.

[11916] Sinan U Umu, Hilde Langseth, Verena Zuber, Åslaug Helland, Robert Lyle, and Trine B Rounge. Serum RNAs can predict lung cancer up to 10 years prior to diagnosis. *eLife*, 11, 2022.

[11917] Ambuj Upadhyay, Nathan R Feltman, Adam Sychla, Anna Janzen, Siba R Das, Maciej Maselko, and Michael Smanski. Genetically engineered insects with sex-selection and genetic incompatibility enable population suppression. *eLife*, 11, 2022.

[11918] Lara Urban, Mariana De Niz, Florencia Fernández-Chiappe, Hedyeh Ebrahimi, Laura KM Han, Devang Mehta, Regina Mencia, Divyansh Mittal, Elizabeth Ochola, Carolina Paz Quezada, Facundo Romani, Lana Sinapayen, Andy Tay, Aalok Varma, and Lamis Yahia Mohamed Elkheir. eLife's new model and its impact on science communication. *eLife*, 11, 2022.

[11919] Sebastian GW Urchs, Angela Tam, Pierre Orban, Clara Moreau, Yassine Benhajali, Hien Duy Nguyen, Alan C Evans, and Pierre Bellec. Functional connectivity subtypes associate robustly with ASD diagnosis. *eLife*, 11, 2022.

[11920] Pantelis Vafidis, David Owald, Tiziano D'Albis, and Richard Kempter. Learning accurate path integration in ring attractor models of the head direction system. *eLife*, 11, 2022.

[11921] Alicia N Vagnozzi, Matthew T Moore, Minshan Lin, Elyse M Brozost, Ritesh KC, Aambar Agarwal, Lindsay A Schwarz, Xin Duan, Niccolò Zampieri, Lynn T Landmesser, and Polyxeni Philippidou. Coordinated cadherin functions sculpt respiratory motor circuit connectivity. *eLife*, 11, 2022.

[11922] Kotryna Vaidžiulytė, Anne-Sophie Macé, Aude Battistella, William Beng, Kristine Schauer, and Mathieu Coppey. Persistent cell migration emerges from a coupling between protrusion dynamics and polarized trafficking. *eLife*, 11, 2022.

[11923] George Vaisey, Priyam Banerjee, Alison J North, Christoph A Haselwandter, and Roderick MacKinnon. Piezo1 as a force-through-membrane sensor in red blood cells. *eLife*, 11, 2022.

[11924] Giulio Valperga and Mario de Bono. Impairing one sensory modality enhances another by reconfiguring peptidergic signalling in Caenorhabditis elegans. *eLife*, 11, 2022.

[11925] Corine M van der Weele and William R Jeffery. Cavefish cope with environmental hypoxia by developing more erythrocytes and overexpression of hypoxia-inducible genes. *eLife*, 11, 2022.

[11926] Alexandra Van Hall-Beauvais, Jesse R Poganik, Kuan-Ting Huang, Saba Parvez, Yi Zhao, Hong-Yu Lin, Xuyu Liu, Marcus John Curtis Long, and Yimon Aye. Z-REX uncovers a bifurcation in function of Keap1 paralogs. *eLife*, 11, 2022.

[11927] Eric Van Otterloo, Isaac Milanda, Hamish Pike, Jamie A Thompson, Hong Li, Kenneth L Jones, and Trevor Williams. AP-2$\alpha$ and AP-2$\beta$ cooperatively function in the craniofacial surface ectoderm to regulate chromatin and gene expression dynamics during facial development. *eLife*, 11, 2022.

[11928] Jacob A Van Winkle, Stefan T Peterson, Elizabeth A Kennedy, Michael J Wheadon, Harshad Ingle, Chandni Desai, Rachel Rodgers, David A Constant, Austin P Wright, Lena Li, Maxim N Artyomov, Sanghyun Lee, Megan T Baldridge, and Timothy J Nice. Homeostatic interferon-lambda response to bacterial microbiota stimulates preemptive antiviral defense within discrete pockets of intestinal epithelium. *eLife*, 11, 2022.

[11929] Brianna Vandrey, Jack Armstrong, Christina M Brown, Derek LF Garden, and Matthew F Nolan. Fan cells in lateral entorhinal cortex directly influence medial entorhinal cortex through synaptic connections in layer 1. *eLife*, 11, 2022.

[11930] Chiara Vanni, Matthew S Schechter, Silvia G Acinas, Albert Barberán, Pier Luigi Buttigieg, Emilio O Casamayor, Tom O Delmont, Carlos M Duarte, A Murat Eren, Robert D Finn, Renzo Kottmann, Alex Mitchell, Pablo Sánchez, Kimmo Siren, Martin Steinegger, Frank Oliver Gloeckner, and Antonio Fernàndez-Guerra. Unifying the known and unknown microbial coding sequence space. *eLife*, 11, 2022.

[11931] Sreejith Jayasree Varma, Enrica Calvani, Nana-Maria Grüning, Christoph B Messner, Nicholas Grayson, Floriana Capuano, Michael Mülleder, and Markus Ralser. Global analysis of cytosine and adenine DNA modifications across the tree of life. *eLife*, 11, 2022.

[11932] Elena Vasileva, Mikako Warren, Timothy J Triche, and James F Amatruda. Dysregulated heparan sulfate proteoglycan metabolism promotes Ewing sarcoma tumor growth. *eLife*, 11, 2022.

[11933] Juan Manuel Vazquez, Maria T Pena, Baaqeyah Muhammad, Morgan Kraft, Linda B Adams, and Vincent J Lynch. Parallel evolution of reduced cancer risk and tumor suppressor duplications in Xenarthra. *eLife*, 11, 2022.

[11934] Sara E Vazquez, Sabrina A Mann, Aaron Bodansky, Andrew F Kung, Zoe Quandt, Elise MN Ferré, Nils Landegren, Daniel Eriksson, Paul Bastard, Shen-Ying Zhang, Jamin Liu, Anthea Mitchell, Irina Proekt, David Yu, Caleigh Mandel-Brehm, Chung-Yu Wang, Brenda Miao, Gavin Sowa, Kelsey Zorn, Alice Y Chan, Veronica M Tagi, Chisato Shimizu, Adriana Tremoulet, Kara Lynch, Michael R Wilson, Olle Kämpe, Kerry Dobbs, Ottavia M Delmonte, Rosa Bacchetta, Luigi D Notarangelo, Jane C Burns, Jean-Laurent Casanova, Michail S Lionakis, Troy R Torgerson, Mark S Anderson, and Joseph L DeRisi. Autoantibody discovery across monogenic, acquired, and COVID-19-associated autoimmunity with scalable PhIP-seq. *eLife*, 11, 2022.

[11935] Sergio Velasco-Aviles, Nikiben Patel, Angeles Casillas-Bajo, Laura Frutos-Rincón, Enrique Velasco, Juana Gallar, Peter Arthur-Farraj, Jose A Gomez-Sanchez, and Hugo Cabedo. A genetic compensatory mechanism regulated by Jun and Mef2d modulates the expression of distinct class IIa Hdacs to ensure peripheral nerve myelination and repair. *eLife*, 11, 2022.

[11936] Leandro M Velez, Cassandra Van, Timothy Moore, Zhenqi Zhou, Casey Johnson, Andrea L Hevener, and Marcus M Seldin. Genetic variation of putative myokine signaling is dominated by biological sex and sex hormones. *eLife*, 11, 2022.

[11937] Griselda Velez-Aguilera, Batool Ossareh-Nazari, Lucie Van Hove, Nicolas Joly, and Lionel Pintard. Cortical microtubule pulling forces contribute to the union of the parental genomes in the Caenorhabditis elegans zygote. *eLife*, 11, 2022.

[11938] Larisa Venkova, Amit Singh Vishen, Sergio Lembo, Nishit Srivastava, Baptiste Duchamp, Artur Ruppel, Alice Williart, Stéphane Vassilopoulos, Alexandre Deslys, Juan Manuel Garcia Arcos, Alba Diz-Muñoz, Martial Balland, Jean-FranÇois Joanny, Damien Cuvelier, Pierre Sens, and Matthieu Piel. A mechano-osmotic feedback couples cell volume to the rate of cell deformation. *eLife*, 11, 2022.

[11939] Claudia A Vera-Arias, Aurel Holzschuh, Colins O Oduma, Kingsley Badu, Mutala Abdul-Hakim, Joshua Yukich, Manuel W Hetzel, Bakar S Fakih, Abdullah Ali, Marcelo U Ferreira, Simone Ladeia-Andrade, Fabián E Sáenz, Yaw Afrane, Endalew Zemene, Delenasaw Yewhalaw, James W Kazura, Guiyun Yan, and Cristian Koepfli. High-throughput Plasmodium falciparum hrp2 and hrp3 gene deletion typing by digital PCR to monitor malaria rapid diagnostic test efficacy. *eLife*, 11, 2022.

[11940] Sergio Oscar Verduzco-Flores and Erik De Schutter. Self-configuring feedback loops for sensorimotor control. *eLife*, 11, 2022.

[11941] Niels JM Verstegen, Ruth R Hagen, Jet van den Dijssel, Lisan H Kuijper, Christine Kreher, Thomas Ashhurst, Laura YL Kummer, Maurice Steenhuis, Mariel Duurland, Rivka de Jongh, Nina de Jong, C Ellen van der Schoot, Amélie V Bos, Erik Mul, Katherine Kedzierska, Koos PJ van Dam, Eileen W Stalman, Laura Boekel, Gertjan Wolbink, Sander W Tas, Joep Killestein, Zoé LE van Kempen, Luuk Wieske, Taco W Kuijpers, Filip Eftimov, Theo Rispens, S Marieke van Ham, Anja ten Brinke, and Carolien E van de Sandt. Immune dynamics in SARS-CoV-2 experienced immunosuppressed rheumatoid arthritis or multiple sclerosis patients vaccinated with mRNA-1273. *eLife*, 11, 2022.

[11942] Alice Vidal, Salvador Soto-Faraco, and Rubén Moreno-Bote. Balance between breadth and depth in human many-alternative decisions. *eLife*, 11, 2022.

[11943] Berta Vidal, Burcu Gulez, Wen Xi Cao, Eduardo Leyva-Díaz, Molly B Reilly, Tessa Tekieli, and Oliver Hobert. The enteric nervous system of the C. elegans pharynx is specified by the Sine oculis-like homeobox gene ceh-34. *eLife*, 11, 2022.

[11944] Edmundo G Vides, Ayan Adhikari, Claire Y Chiang, Pawel Lis, Elena Purlyte, Charles Limouse, Justin L Shumate, Elena Spínola-Lasso, Herschel S Dhekne, Dario R Alessi, and Suzanne R Pfeffer. A feed-forward pathway drives LRRK2 kinase membrane recruitment and activation. *eLife*, 11, 2022.

[11945] Joana B Vieira and Andreas Olsson. Neural defensive circuits underlie helping under threat in humans. *eLife*, 11, 2022.

[11946] Ryan C Vignogna, Mariateresa Allocca, Maria Monticelli, Joy W Norris, Richard Steet, Ethan O Perlstein, Giuseppina Andreotti, and Gregory I Lang. Evolutionary rescue of phosphomannomutase deficiency in yeast models of human disease. *eLife*, 11, 2022.

[11947] Nuria Vilaplana-Lopera, Vincent Cuminetti, Ruba Almaghrabi, Grigorios Papatzikas, Ashok Kumar Rout, Mark Jeeves, Elena González, Yara Alyahyawi, Alan Cunningham, Ayşegül Erdem, Frank Schnütgen, Manoj Raghavan, Sandeep Potluri, Jean-Baptiste Cazier, Jan Jacob Schuringa, Michelle AC Reed, Lorena Arranz, Ulrich L Günther, and Paloma Garcia. Crosstalk between AML and stromal cells triggers acetate secretion through the metabolic rewiring of stromal cells. *eLife*, 11, 2022.

[11948] Osvaldo Villa, Nicole L Stuhr, Chia-an Yen, Eileen M Crimmins, Thalida Em Arpawong, and Sean P Curran. Genetic variation in ALDH4A1 is associated with muscle health over the lifespan and across species. *eLife*, 11, 2022.

[11949] Alejandro Villalta, Alain Schmitt, Leandro F Estrozi, Emmanuelle RJ Quemin, Jean-Marie Alempic, Audrey Lartigue, Vojtěch Pražák, Lucid Belmudes, Daven Vasishtan, Agathe MG Colmant, Flora A Honoré, Yohann Couté, Kay Grünewald, and Chantal Abergel. The giant mimivirus 1.2 Mb genome is elegantly organized into a 30-nm diameter helical protein shield. *eLife*, 11, 2022.

[11950] Kevin Vinberg, Jörgen Rosén, Granit Kastrati, and Fredrik Ahs. Whole brain correlates of individual differences in skin conductance responses during discriminative fear conditioning to social cues. *eLife*, 11, 2022.

[11951] Stefania Viola, William Roseby, Stefano Santabarbara, Dennis Nürnberg, Ricardo AssunÇão, Holger Dau, Julien Sellés, Alain Boussac, Andrea Fantuzzi, and A William Rutherford. Impact of energy limitations on function and resilience in long-wavelength Photosystem II. *eLife*, 11, 2022.

[11952] MatouŠ Vobořil, Jiří Březina, TomáŠ Brabec, Jan DobeŠ, Ondřej Ballek, Martina DobeŠová, Jasper Manning, Richard S Blumberg, and Dominik Filipp. A model of preferential pairing between epithelial and dendritic cells in thymic antigen transfer. *eLife*, 11, 2022.

[11953] Marina Voinson, Charles L Nunn, and Amy Goldberg. Primate malarias as a model for cross-species parasite transmission. *eLife*, 11, 2022.

[11954] Andrea Volante, Juan Carlos Alonso, and Kiyoshi Mizuuchi. Distinct architectural requirements for the parS centromeric sequence of the pSM19035 plasmid partition machinery. *eLife*, 11, 2022.

[11955] Nadine Vollmuth, Lisa Schlicker, Yongxia Guo, Pargev Hovhannisyan, Sudha Janaki-Raman, Naziia Kurmasheva, Werner Schmitz, Almut Schulze, Kathrin Stelzner, Karthika Rajeeve, and Thomas Rudel. c-Myc plays a key role in IFN-$\gamma$-induced persistence of Chlamydia trachomatis. *eLife*, 11, 2022.

[11956] Allison R Wagner, Chi G Weindel, Kelsi O West, Haley M Scott, Robert O Watson, and Kristin L Patrick. SRSF6 balances mitochondrial-driven innate immune outcomes through alternative splicing of BAX. *eLife*, 11, 2022.

[11957] Julian M Wagner, C Jaco Klok, Meghan E Duell, John J Socha, Guohua Cao, Hao Gong, and Jon F Harrison. Isometric spiracular scaling in scarab beetles—implications for diffusive and advective oxygen transport. *eLife*, 11, 2022.

[11958] Petra Wahle, Eric Sobierajski, Ina Gasterstädt, Nadja Lehmann, Susanna Weber, Joachim HR Lübke, Maren Engelhardt, Claudia Distler, and Gundela Meyer. Neocortical pyramidal neurons with axons emerging from dendrites are frequent in non-primates, but rare in monkey and human. *eLife*, 11, 2022.

[11959] Sigrid Wahlen, Filip Matthijssens, Wouter Van Loocke, Sylvie Taveirne, Laura Kiekens, Eva Persyn, Els Van Ammel, Zenzi De Vos, Stijn De Munter, Patrick Matthys, Filip Van Nieuwerburgh, Tom Taghon, Bart Vandekerckhove, Pieter Van Vlierberghe, and Georges Leclercq. The transcription factor RUNX2 drives the generation of human NK cells and promotes tissue residency. *eLife*, 11, 2022.

[11960] Tom WN Walker, Konstantin Gavazov, Thomas Guillaume, Thibault Lambert, Pierre Mariotte, Devin Routh, Constant Signarbieux, Sebastián Block, Tamara Münkemüller, Hanna Nomoto, Thomas W Crowther, Andreas Richter, Alexandre Buttler, and Jake M Alexander. Lowland plant arrival in alpine ecosystems facilitates a decrease in soil carbon content under experimental climate warming. *eLife*, 11, 2022.

[11961] Ryan L Wallace, Eric Lu, Xiangxia Luo, and Andrew P Capaldi. Ait1 regulates TORC1 signaling and localization in budding yeast. *eLife*, 11, 2022.

[11962] Jeffrey Wammes, Kenneth A Norman, and Nicholas Turk-Browne. Increasing stimulus similarity drives nonmonotonic representational change in hippocampus. *eLife*, 11, 2022.

[11963] Bin Wan, şeyma Bayrak, Ting Xu, H Lina Schaare, Richard AI Bethlehem, Boris C Bernhardt, and Sofie L Valk. Heritability and cross-species comparisons of human cortical functional organization asymmetry. *eLife*, 11, 2022.

[11964] Amy Wang, Alexander R Dunn, and William I Weis. Mechanism of the cadherin–catenin F-actin catch bond interaction. *eLife*, 11, 2022.

[11965] Anmin Wang, Wanyiin Tao, Jiyu Tong, Juanzi Gao, Jinghao Wang, Gaopeng Hou, Chen Qian, Guorong Zhang, Runzhi Li, Decai Wang, Xingxing Ren, Kaiguang Zhang, Siyuan Ding, Richard A Flavell, Huabing Li, Wen Pan, and Shu Zhu. m6A modifications regulate intestinal immunity and rotavirus infection. *eLife*, 11, 2022.

[11966] Camille S Wang, Natali L Chanaday, Lisa M Monteggia, and Ege T Kavalali. Probing the segregation of evoked and spontaneous neurotransmission via photobleaching and recovery of a fluorescent glutamate sensor. *eLife*, 11, 2022.

[11967] Fengbin Wang, Chi Ho Chan, Victor Suciu, Khawla Mustafa, Madeline Ammend, Dong Si, Allon I Hochbaum, Edward H Egelman, and Daniel R Bond. Structure of Geobacter OmcZ filaments suggests extracellular cytochrome polymers evolved independently multiple times. *eLife*, 11, 2022.

[11968] Jia C Wang, Yang-In Yim, Xufeng Wu, Valentin Jaumouille, Andrew Cameron, Clare M Waterman, John H Kehrl, and John A Hammer. A B-cell actomyosin arc network couples integrin co-stimulation to mechanical force-dependent immune synapse formation. *eLife*, 11, 2022.

[11969] Jin Wang, Jiayi Xie, Daosong Wang, Xue Han, Minqi Chen, Guojun Shi, Linjia Jiang, and Meng Zhao. CXCR4high megakaryocytes regulate host-defense immunity against bacterial pathogens. *eLife*, 11, 2022.

[11970] Jingqiang Wang, Chunye Liu, Lingli He, Zhiyao Xie, Lanyue Bai, Wentao Yu, Zuoyun Wang, Yi Lu, Dong Gao, Junfen Fu, Lei Zhang, and Yi Arial Zeng. Selective YAP activation in Procr cells is essential for ovarian stem/progenitor expansion and epithelium repair. *eLife*, 11, 2022.

[11971] Lan Wang, Hannah Toutkoushian, Vladislav Belyy, Claire Y Kokontis, and Peter Walter. Conserved structural elements specialize ATAD1 as a membrane protein extraction machine. *eLife*, 11, 2022.

[11972] Libo Wang, Zaoqu Liu, Ruopeng Liang, Weijie Wang, Rongtao Zhu, Jian Li, Zhe Xing, Siyuan Weng, Xinwei Han, and Yu-ling Sun. Comprehensive machine-learning survival framework develops a consensus model in large-scale multicenter cohorts for pancreatic cancer. *eLife*, 11, 2022.

[11973] Mengyao Wang, Yu Qi, Yutao Cao, Xiaosheng Zhang, Yongsheng Wang, Qingyou Liu, Jinlong Zhang, Guangbin Zhou, Yue Ai, Shao Wei, Linli Wang, Guoshi Liu, Zhengxing Lian, and Hongbing Han. Domain fusion TLR2-4 enhances the autophagy-dependent clearance of Staphylococcus aureus in the genetic engineering goat. *eLife*, 11, 2022.

[11974] Min Wang, Thomas A Stidham, Jingmai K O'Connor, and Zhonghe Zhou. Insight into the evolutionary assemblage of cranial kinesis from a Cretaceous bird. *eLife*, 11, 2022.

[11975] Qi-Shan Wang, Kai-Jian Fan, Hui Teng, Sijia Chen, Bing-Xin Xu, Di Chen, and Ting-Yu Wang. Mir204 and Mir211 suppress synovial inflammation and proliferation in rheumatoid arthritis by targeting Ssrp1. *eLife*, 11, 2022.

[11976] Shiying Wang, Yubo Ma, Qian Wu, Min Wang, Dongyu Hu, Corwin Sullivan, and Xing Xu. Digital restoration of the pectoral girdles of two Early Cretaceous birds and implications for early-flight evolution. *eLife*, 11, 2022.

[11977] Tao Wang, Biyang Jing, Bowen Deng, Kai Shi, Jing Li, Baoxu Ma, Fengming Wu, and Chuan Zhou. Drosulfakinin signaling modulates female sexual receptivity in Drosophila. *eLife*, 11, 2022.

[11978] Wanyue Wang, Hany KM Dweck, Gaëlle JS Talross, Ali Zaidi, Joshua M Gendron, and John R Carlson. Sugar sensation and mechanosensation in the egg-laying preference shift of Drosophila suzukii. *eLife*, 11, 2022.

[11979] Weikang Wang, Dante Poe, Yaxuan Yang, Thomas Hyatt, and Jianhua Xing. Epithelial-to-mesenchymal transition proceeds through directional destabilization of multidimensional attractor. *eLife*, 11, 2022.

[11980] Xin Wang, Anjun Jiao, Lina Sun, Wenhua Li, Biao Yang, Yanhong Su, Renyi Ding, Cangang Zhang, Haiyan Liu, Xiaofeng Yang, Chenming Sun, and Baojun Zhang. Zinc finger protein Zfp335 controls early T-cell development and survival through $\beta$-selection-dependent and -independent mechanisms. *eLife*, 11, 2022.

[11981] Xing Wang, Jean Vannier, Xiaoguang Yang, Lucas Leclère, Qiang Ou, Xikun Song, Tsuyoshi Komiya, and Jian Han. Muscle systems and motility of early animals highlighted by cnidarians from the basal Cambrian. *eLife*, 11, 2022.

[11982] Xue-Ping Wang, Deidra M Balchak, Clayton Gentilcore, Nathan L Clark, and Ossama B Kashlan. Activation by cleavage of the epithelial Na+ channel $\alpha$ and $\gamma$ subunits independently coevolved with the vertebrate terrestrial migration. *eLife*, 11, 2022.

[11983] Xueyong Wang, Murad Nawaz, Chris DuPont, Jessica H Myers, Steve RA Burke, Roger A Bannister, Brent D Foy, Andrew A Voss, and Mark M Rich. The role of action potential changes in depolarization-induced failure of excitation contraction coupling in mouse skeletal muscle. *eLife*, 11, 2022.

[11984] Yajing Wang and Min Zhu. Squamation and scale morphology at the root of jawed vertebrates. *eLife*, 11, 2022.

[11985] Yan Wang, Xiaohui Liang, Yuqian Jiang, Danjiang Dong, Cong Zhang, Tianqiang Song, Ming Chen, Yong You, Han Liu, Min Ge, Haibin Dai, Fengchan Xi, Wanqing Zhou, Jian-Qun Chen, Qiang Wang, Qihan Chen, and Wenkui Yu. Novel fast pathogen diagnosis method for severe pneumonia patients in the intensive care unit: randomized clinical trial. *eLife*, 11, 2022.

[11986] Yi-wen Wang, Chris C Wreden, Maayan Levy, Julia L Meng, Zarion D Marshall, Jason MacLean, and Ellie Heckscher. Sequential addition of neuronal stem cell temporal cohorts generates a feed-forward circuit in the Drosophila larval nerve cord. *eLife*, 11, 2022.

[11987] Yifan Wang, C Richard Boland, Ajay Goel, Dominik Wodarz, and Natalia L Komarova. Aspirin's effect on kinetic parameters of cells contributes to its role in reducing incidence of advanced colorectal adenomas, shown by a multiscale computational study. *eLife*, 11, 2022.

[11988] Yingying Wang, Xi Chen, Katharina Spengler, Karoline Terberger, Marko Boehm, Jens Appel, Thomas Barske, Stefan Timm, Natalia Battchikova, Martin Hagemann, and Kirstin Gutekunst. Pyruvate:ferredoxin oxidoreductase and low abundant ferredoxins support aerobic photomixotrophic growth in cyanobacteria. *eLife*, 11, 2022.

[11989] Yonghui Wang, Shaopeng Wang, Liqing Zhao, Cunzhu Liang, Bailing Miao, Qing Zhang, Xiaxia Niu, Wenhong Ma, and Bernhard Schmid. Stability and asynchrony of local communities but less so diversity increase regional stability of Inner Mongolian grassland. *eLife*, 11, 2022.

[11990] Yuexin Wang, Hui Yang, Anna Jia, Yufei Wang, Qiuli Yang, Yingjie Dong, Yueru Hou, Yejin Cao, Lin Dong, Yujing Bi, and Guangwei Liu. Dendritic cell Piezo1 directs the differentiation of TH1 and Treg cells in cancer. *eLife*, 11, 2022.

[11991] Zhaowei Wang, Xiaoling Xia, Jiaqi Li, and Tatsushi Igaki. Tumor elimination by clustered microRNAs miR-306 and miR-79 via noncanonical activation of JNK signaling. *eLife*, 11, 2022.

[11992] Zhenzhen Wang, Ziyu Zhai, Chunyu Chen, Xuejiao Tian, Zhen Xing, Panfei Xing, Yushun Yang, Junfeng Zhang, Chunming Wang, and Lei Dong. Air pollution particles hijack peroxidasin to disrupt immunosurveillance and promote lung cancer. *eLife*, 11, 2022.

[11993] Zhongxiao Wang, Felix Yemanyi, Alexandra K Blomfield, Kiran Bora, Shuo Huang, Chi-Hsiu Liu, William R Britton, Steve S Cho, Yohei Tomita, Zhongjie Fu, Jian-xing Ma, Wen-hong Li, and Jing Chen. Amino acid transporter SLC38A5 regulates developmental and pathological retinal angiogenesis. *eLife*, 11, 2022.

[11994] Zimei Wang, Adam Romanski, Vatsal Mehra, Yunfang Wang, Matthew Brannigan, Benjamin C Campbell, Gregory A Petsko, Pantelis Tsoulfas, and Murray G Blackmore. Brain-wide analysis of the supraspinal connectome reveals anatomical correlates to functional recovery after spinal injury. *eLife*, 11, 2022.

[11995] Zinan Wang, Joseph P Receveur, Jian Pu, Haosu Cong, Cole Richards, Muxuan Liang, and Henry Chung. Desiccation resistance differences in Drosophila species can be largely explained by variations in cuticular hydrocarbons. *eLife*, 11, 2022.

[11996] Ziyue Wang, Michael Mülleder, Ihor Batruch, Anjali Chelur, Kathrin Textoris-Taube, Torsten Schwecke, Johannes Hartl, Jason Causon, Jose Castro-Perez, Vadim Demichev, Stephen Tate, and Markus Ralser. High-throughput proteomics of nanogram-scale samples with Zeno SWATH MS. *eLife*, 11, 2022.

[11997] Stephanie A Wankowicz, Saulo H de Oliveira, Daniel W Hogan, Henry van den Bedem, and James S Fraser. Ligand binding remodels protein side-chain conformational heterogeneity. *eLife*, 11, 2022.

[11998] Brandt Warecki, Simon William Abraham Titen, Mohammad Shahriyar Alam, Giovanni Vega, Nassim Lemseffer, Karen Hug, Jonathan S Minden, and William Sullivan. Wolbachia action in the sperm produces developmentally deferred chromosome segregation defects during the Drosophila mid-blastula transition. *eLife*, 11, 2022.

[11999] Robert Warneford-Thomson, Parisha P Shah, Patrick Lundgren, Jonathan Lerner, Jason Morgan, Antonio Davila, Benjamin S Abella, Kenneth Zaret, Jonathan Schug, Rajan Jain, Christoph A Thaiss, and Roberto Bonasio. A LAMP sequencing approach for high-throughput co-detection of SARS-CoV-2 and influenza virus in human saliva. *eLife*, 11, 2022.

[12000] Sam Wass, Emily Phillips, Celia Smith, Elizabeth OOB Fatimehin, and Louise Goupil. Vocal communication is tied to interpersonal arousal coupling in caregiver-infant dyads. *eLife*, 11, 2022.

[12001] Paul M Wassarman and Eveline S Litscher. Female fertility and the zona pellucida. *eLife*, 11, 2022.

[12002] Kate M Wassum. Amygdala-cortical collaboration in reward learning and decision making. *eLife*, 11, 2022.

[12003] James A Watson, Robert J Commons, Joel Tarning, Julie A Simpson, Alejandro Llanos Cuentas, Marcus VG Lacerda, Justin A Green, Gavin CKW Koh, Cindy S Chu, FranÇois H Nosten, Richard N Price, Nicholas PJ Day, and Nicholas J White. The clinical pharmacology of tafenoquine in the radical cure of Plasmodium vivax malaria: An individual patient data meta-analysis. *eLife*, 11, 2022.

[12004] Anna Isabell Weaver, Laura Alvarez, Kelly M Rosch, Asraa Ahmed, Garrett Sean Wang, Michael S van Nieuwenhze, Felipe Cava, and Tobias Dörr. Lytic transglycosylases mitigate periplasmic crowding by degrading soluble cell wall turnover products. *eLife*, 11, 2022.

[12005] Lilian Aline Weber, Sara Tomiello, Dario Schöbi, Katharina V Wellstein, Daniel Mueller, Sandra Iglesias, and Klaas Enno Stephan. Auditory mismatch responses are differentially sensitive to changes in muscarinic acetylcholine versus dopamine receptor function. *eLife*, 11, 2022.

[12006] Amy K Webster, Rojin Chitrakar, Maya Powell, Jingxian Chen, Kinsey Fisher, Robyn E Tanny, Lewis Stevens, Kathryn Evans, Angela Wei, Igor Antoshechkin, Erik C Andersen, and L Ryan Baugh. Using population selection and sequencing to characterize natural variation of starvation resistance in Caenorhabditis elegans. *eLife*, 11, 2022.

[12007] Marcus M Weera, Abigail E Agoglia, Eliza Douglass, Zhiying Jiang, Shivakumar Rajamanickam, Rosetta S Shackett, Melissa A Herman, Nicholas J Justice, and Nicholas W Gilpin. Generation of a CRF1-Cre transgenic rat and the role of central amygdala CRF1 cells in nociception and anxiety-like behavior. *eLife*, 11, 2022.

[12008] Waja Wegner, Heinz Steffens, Carola Gregor, Fred Wolf, and Katrin I Willig. Environmental enrichment enhances patterning and remodeling of synaptic nanoarchitecture as revealed by STED nanoscopy. *eLife*, 11, 2022.

[12009] Lilija Wehling, Liam Keegan, Paula Fernández-Palanca, Reham Hassan, Ahmed Ghallab, Jennifer Schmitt, Yingyue Tang, Maxime Le Marois, Stephanie Roessler, Peter Schirmacher, Ursula Kummer, Jan G Hengstler, Sven Sahle, and Kai Breuhahn. Spatial modeling reveals nuclear phosphorylation and subcellular shuttling of YAP upon drug-induced liver injury. *eLife*, 11, 2022.

[12010] Jingjing Wei, Linghui Hou, Jingtong Liu, Ziwen Wang, Siqi Gao, Tao Qi, Song Gao, Shuna Sun, and Yongming Wang. Closely related type II-C Cas9 orthologs recognize diverse PAMs. *eLife*, 11, 2022.

[12011] Kristin Weineck, Olivia Xin Wen, and Molly J Henry. Neural synchronization is strongest to the spectral flux of slow music and depends on familiarity and beat salience. *eLife*, 11, 2022.

[12012] Alison R Weiss, William A Liguore, Kristin Brandon, Xiaojie Wang, Zheng Liu, Jacqueline S Domire, Dana Button, Sathya Srinivasan, Christopher D Kroenke, and Jodi L McBride. A novel rhesus macaque model of Huntington's disease recapitulates key neuropathological changes along with motor and cognitive decline. *eLife*, 11, 2022.

[12013] Joshua M Weiss, Dianne Lumaquin-Yin, Emily Montal, Shruthy Suresh, Carl S Leonhardt, and Richard M White. Shifting the focus of zebrafish toward a model of the tumor microenvironment. *eLife*, 11, 2022.

[12014] Georg Welzel and Stefan Schuster. Connexins evolved after early chordates lost innexin diversity. *eLife*, 11, 2022.

[12015] Bingqiang Wen, Guolun Wang, Enhong Li, Olena A Kolesnichenko, Zhaowei Tu, Senad Divanovic, Tanya V Kalin, and Vladimir V Kalinichenko. In vivo generation of bone marrow from embryonic stem cells in interspecies chimeras. *eLife*, 11, 2022.

[12016] Dawn M Wenzel, Douglas R Mackay, Jack J Skalicky, Elliott L Paine, Matthew S Miller, Katharine S Ullman, and Wesley I Sundquist. Comprehensive analysis of the human ESCRT-III-MIT domain interactome reveals new cofactors for cytokinetic abscission. *eLife*, 11, 2022.

[12017] Jacob A Westerberg, Michelle S Schall, Alexander Maier, Geoffrey F Woodman, and Jeffrey D Schall. Laminar microcircuitry of visual cortex producing attention-associated electric fields. *eLife*, 11, 2022.

[12018] Sarah E Westrick, Mara Laslo, and Eva K Fischer. The big potential of the small frog Eleutherodactylus coqui. *eLife*, 11, 2022.

[12019] Joshua R Wheeler, Oscar N Whitney, Thomas O Vogler, Eric D Nguyen, Bradley Pawlikowski, Evan Lester, Alicia Cutler, Tiffany Elston, Nicole Dalla Betta, Kevin R Parker, Kathryn E Yost, Hannes Vogel, Thomas A Rando, Howard Y Chang, Aaron M Johnson, Roy Parker, and Bradley B Olwin. RNA-binding proteins direct myogenic cell fate decisions. *eLife*, 11, 2022.

[12020] Richard J White, Eirinn Mackay, Stephen W Wilson, and Elisabeth M Busch-Nentwich. Allele-specific gene expression can underlie altered transcript abundance in zebrafish mutants. *eLife*, 11, 2022.

[12021] Jane E Whitney, In-Hee Lee, Ji-Won Lee, and Sek Won Kong. Evolution of multiple omics approaches to define pathophysiology of pediatric acute respiratory distress syndrome. *eLife*, 11, 2022.

[12022] Felix C Widmer, Sean M O'Toole, and Georg B Keller. NMDA receptors in visual cortex are necessary for normal visuomotor integration and skill learning. *eLife*, 11, 2022.

[12023] Alexandra Wiesinger, Jiuru Li, Lianne Fokkert, Priscilla Bakker, Arie O Verkerk, Vincent M Christoffels, Gerard JJ Boink, and Harsha D Devalla. A single cell transcriptional roadmap of human pacemaker cell differentiation. *eLife*, 11, 2022.

[12024] Angela R Wild, Peter W Hogg, Stephane Flibotte, Glory G Nasseri, Rocio B Hollman, Danya Abazari, Kurt Haas, and Shernaz X Bamji. Exploring the expression patterns of palmitoylating and de-palmitoylating enzymes in the mouse brain using the curated RNA-seq database BrainPalmSeq. *eLife*, 11, 2022.

[12025] Sophia A Wild, Ian G Cannell, Ashley Nicholls, Katarzyna Kania, Dario Bressan, Gregory J Hannon, Gregory J Hannon, and Kirsty Sawicka. Clonal transcriptomics identifies mechanisms of chemoresistance and empowers rational design of combination therapies. *eLife*, 11, 2022.

[12026] Elizabeth T Wiles, Colleen C Mumford, Kevin J McNaught, Hideki Tanizawa, and Eric U Selker. The ACF chromatin-remodeling complex is essential for Polycomb repression. *eLife*, 11, 2022.

[12027] Martin Wilkinson, Oliver J Wilkinson, Connie Feyerherm, Emma E Fletcher, Dale B Wigley, and Mark S Dillingham. Structures of RecBCD in complex with phage-encoded inhibitor proteins reveal distinctive strategies for evasion of a bacterial immunity hub. *eLife*, 11, 2022.

[12028] Audrey Miller Williams, Seth Donoughe, Edwin Munro, and Sally Horne-Badovinac. Fat2 polarizes the WAVE complex in trans to align cell protrusions for collective migration. *eLife*, 11, 2022.

[12029] Christopher S Williams, W Kimryn Rathmell, John M Carethers, Diane M Harper, YM Dennis Lo, Peter J Ratcliffe, and Mone Zaidi. A global view of the aspiring physician-scientist. *eLife*, 11, 2022.

[12030] Darian Williams, Marwa Mahmoud, Renfa Liu, Aitor Andueza, Sandeep Kumar, Dong-Won Kang, Jiahui Zhang, Ian Tamargo, Nicolas Villa-Roel, Kyung-In Baek, Hwakyoung Lee, Yongjin An, Leran Zhang, Edward W Tate, Pritha Bagchi, Jan Pohl, Laurent O Mosnier, Eleftherios P Diamandis, Koichiro Mihara, Morley D Hollenberg, Zhifei Dai, and Hanjoong Jo. Stable flow-induced expression of KLK10 inhibits endothelial inflammation and atherosclerosis. *eLife*, 11, 2022.

[12031] Luc Edward Wilson, Jason da Silva Castanheira, and Sylvain Baillet. Time-resolved parameterization of aperiodic and periodic brain activity. *eLife*, 11, 2022.

[12032] Ashtyn T Wiltbank, Emma R Steinson, Stacey J Criswell, Melanie Piller, and Sarah Kucenas. Cd59 and inflammation regulate Schwann cell development. *eLife*, 11, 2022.

[12033] Reinhard Windoffer, Nicole Schwarz, Sungjun Yoon, Teodora Piskova, Michael Scholkemper, Johannes Stegmaier, Andrea Bönsch, Jacopo Di Russo, and Rudolf E Leube. Quantitative mapping of keratin networks in 3D. *eLife*, 11, 2022.

[12034] Jacob M Winter, Heidi L Fresenius, Corey N Cunningham, Peng Wei, Heather R Keys, Jordan Berg, Alex Bott, Tarun Yadav, Jeremy Ryan, Deepika Sirohi, Sheryl R Tripp, Paige Barta, Neeraj Agarwal, Anthony Letai, David M Sabatini, Matthew L Wohlever, and Jared Rutter. Collateral deletion of the mitochondrial AAA+ ATPase ATAD1 sensitizes cancer cells to proteasome dysfunction. *eLife*, 11, 2022.

[12035] Gregory S Wirak, Jeremy Florman, Mark J Alkema, Christopher W Connor, and Christopher V Gabel. Age-associated changes to neuronal dynamics involve a disruption of excitatory/inhibitory balance in C. elegans. *eLife*, 11, 2022.

[12036] Aemiro Bezabih Woldeyohannes, Sessen Daniel Iohannes, Mara Miculan, Leonardo Caproni, Jemal Seid Ahmed, Kauê de Sousa, Ermias Abate Desta, Carlo Fadda, Mario Enrico Pè, and Matteo Dell'Acqua. Data-driven, participatory characterization of farmer varieties discloses teff breeding potential under current and future climates. *eLife*, 11, 2022.

[12037] Parker S Woods, Lucas M Kimmig, Kaitlyn A Sun, Angelo Y Meliton, Obada R Shamaa, Yufeng Tian, Rengül Cetin-Atalay, Willard W Sharp, Robert B Hamanaka, and Gökhan M Mutlu. HIF-1$\alpha$ induces glycolytic reprograming in tissue-resident alveolar macrophages to promote cell survival during acute lung injury. *eLife*, 11, 2022.

[12038] Marina E Wosniack, Dylan Festa, Nan Hu, Julijana Gjorgjieva, and Jimena Berni. Adaptation of Drosophila larva foraging in response to changes in food resources. *eLife*, 11, 2022.

[12039] Kevin M Wright, Andrew G Deighan, Andrea Di Francesco, Adam Freund, Vladimir Jojic, Gary A Churchill, and Anil Raj. Age and diet shape the genetic architecture of body weight in diversity outbred mice. *eLife*, 11, 2022.

[12040] Chi-Hong Wu, Vedakumar Tatavarty, Pierre M Jean Beltran, Andrea A Guerrero, Hasmik Keshishian, Karsten Krug, Melanie A MacMullan, Li Li, Steven A Carr, Jeffrey R Cottrell, and Gina G Turrigiano. A bidirectional switch in the Shank3 phosphorylation state biases synapses toward up- or downscaling. *eLife*, 11, 2022.

[12041] Juan Wu, Dongna Chen, Hui Huang, Ning Luo, Huishuang Chen, Junjie Zhao, Yanyan Wang, Tian Zhao, Siyuan Huang, Yang Ren, Teng Zhai, Weibin Sun, Houxuan Li, and Wei Li. A novel gene ZNF862 causes hereditary gingival fibromatosis. *eLife*, 11, 2022.

[12042] Meiling Wu, Anda Zhao, Xingchen Yan, Hongyang Gao, Chunwang Zhang, Xiaomin Liu, Qiwen Luo, Feizhou Xie, Shanlin Liu, and Dongyun Shi. Hepatic AMPK signaling dynamic activation in response to REDOX balance are sentinel biomarkers of exercise and antioxidant intervention to improve blood glucose control. *eLife*, 11, 2022.

[12043] Yushi Wu, Arun Devotta, Diana S José-Edwards, Jamie E Kugler, Lenny J Negrón-Piñeiro, Karina Braslavskaya, Jermyn Addy, Jean-Pierre Saint-Jeannet, and Anna Di Gregorio. Xbp1 and Brachyury establish an evolutionarily conserved subcircuit of the notochord gene regulatory network. *eLife*, 11, 2022.

[12044] Feng Xian, Julia Regina Sondermann, David Gomez Varela, and Manuela Schmidt. Deep proteome profiling reveals signatures of age and sex differences in paw skin and sciatic nerve of naïve mice. *eLife*, 11, 2022.

[12045] Han Xiao, Tao Zhang, Changjun Li, Yong Cao, Linfeng Wang, Huabin Chen, Shengcan Li, Changbiao Guan, Jianzhong Hu, Di Chen, Can Chen, and Hongbin Lu. Mechanical stimulation promotes enthesis injury repair by mobilizing Prrx1+ cells via ciliary TGF-$\beta$ signaling. *eLife*, 11, 2022.

[12046] Na Xiao, Jie Wang, Ting Wang, Xingliang Xiong, Junyi Zhou, Xian Su, Jing Peng, Chao Yang, Xiaofeng Li, Ge Lin, Guangxiu Lu, Fei Gong, and Lamei Cheng. Metformin abrogates pathological TNF-$\alpha$-producing B cells through mTOR-dependent metabolic reprogramming in polycystic ovary syndrome. *eLife*, 11, 2022.

[12047] Yucheng Xiao, Jonathan W Theile, Agnes Zybura, Yanling Pan, Zhixin Lin, and Theodore R Cummins. A-type FHFs mediate resurgent currents through TTX-resistant voltage-gated sodium channels. *eLife*, 11, 2022.

[12048] Rong Xie, Dan Yi, Daofu Zeng, Qiang Jie, Qinglin Kang, Zeng Zhang, Zhenlin Zhang, Guozhi Xiao, Lin Chen, Liping Tong, and Di Chen. Specific deletion of Axin1 leads to activation of $\beta$-catenin/BMP signaling resulting in fibular hemimelia phenotype in mice. *eLife*, 11, 2022.

[12049] Yanling Xin, Qinghai He, Huilin Liang, Ke Zhang, Jingyi Guo, Qi Zhong, Dan Chen, Jinyan Li, Yizhi Liu, and Shuyi Chen. m6A epitranscriptomic modification regulates neural progenitor-to-glial cell transition in the retina. *eLife*, 11, 2022.

[12050] Le Xiong, Erik A Tolen, Jinmi Choi, Sergiy Velychko, Livia Caizzi, Taras Velychko, Kenjiro Adachi, Caitlin M MacCarthy, Michael Lidschreiber, Patrick Cramer, and Hans R Schöler. Oct4 differentially regulates chromatin opening and enhancer transcription in pluripotent stem cells. *eLife*, 11, 2022.

[12051] Cong Xu, Van Vuong Dinh, Kai Kruse, Hyun-Woo Jeong, Emma C Watson, Susanne Adams, Frank Berkenfeld, Martin Stehling, Seyed Javad Rasouli, Rui Fan, Rui Chen, Ivan Bedzhov, Qi Chen, Katsuhiro Kato, Mara E Pitulescu, and Ralf H Adams. Induction of osteogenesis by bone-targeted Notch activation. *eLife*, 11, 2022.

[12052] Jun Xu, Ah-Ram Kim, Ross W Cheloha, Fabian A Fischer, Joshua Shing Shun Li, Yuan Feng, Emily Stoneburner, Richard Binari, Stephanie E Mohr, Jonathan Zirin, Hidde L Ploegh, and Norbert Perrimon. Protein visualization and manipulation in Drosophila through the use of epitope tags recognized by nanobodies. *eLife*, 11, 2022.

[12053] Tong Xu, Mathijs Verhagen, Rosalie Joosten, Wenjie Sun, Andrea Sacchetti, Leonel Munoz Sagredo, Véronique Orian-Rousseau, and Riccardo Fodde. Alternative splicing downstream of EMT enhances phenotypic plasticity and malignant behavior in colon cancer. *eLife*, 11, 2022.

[12054] Xiaowei Xu, Xu Hua, Kyle Brown, Xiaojun Ren, and Zhiguo Zhang. Mcm2 promotes stem cell differentiation via its ability to bind H3-H4. *eLife*, 11, 2022.

[12055] Yi Xue, Laura Waller, Hillel Adesnik, and Nicolas Pégard. Three-dimensional multi-site random access photostimulation (3D-MAP). *eLife*, 11, 2022.

[12056] Alkesh Yadav, Quentin Vagne, Pierre Sens, Garud Iyengar, and Madan Rao. Glycan processing in the Golgi as optimal information coding that constrains cisternal number and enzyme specificity. *eLife*, 11, 2022.

[12057] Takayoshi Yamamoto, Yuta Kambayashi, Yuta Otsuka, Boni A Afouda, Claudiu Giuraniuc, Tatsuo Michiue, and Stefan Hoppler. Positive feedback regulation of frizzled-7 expression robustly shapes a steep Wnt gradient in Xenopus heart development, together with sFRP1 and heparan sulfate. *eLife*, 11, 2022.

[12058] Shunpei Yamauchi, Takashi Nozoe, Reiko Okura, Edo Kussell, and Yuichi Wakamoto. A unified framework for measuring selection on cellular lineages and traits. *eLife*, 11, 2022.

[12059] Mingchao Yan, Wenwen Yu, Qian Lv, Qiming Lv, Tingting Bo, Xiaoyu Chen, Yilin Liu, Yafeng Zhan, Shengyao Yan, Xiangyu Shen, Baofeng Yang, Qiming Hu, Jiangli Yu, Zilong Qiu, Yuanjing Feng, Xiao-Yong Zhang, He Wang, Fuqiang Xu, and Zheng Wang. Mapping brain-wide excitatory projectome of primate prefrontal cortex at submicron resolution and comparison with diffusion tractography. *eLife*, 11, 2022.

[12060] Hagai Yanai, Christopher Dunn, Bongsoo Park, Christopher Coletta, Ross A McDevitt, Taylor McNeely, Michael Leone, Robert P Wersto, Kathy A Perdue, and Isabel Beerman. Male rat leukocyte population dynamics predict a window for intervention in aging. *eLife*, 11, 2022.

[12061] Bingyi Yang, Bernardo García-Carreras, Justin Lessler, Jonathan M Read, Huachen Zhu, C Jessica E Metcalf, James A Hay, Kin O Kwok, Ruiyun Shen, Chao Q Jiang, Yi Guan, Steven Riley, and Derek A Cummings. Long term intrinsic cycling in human life course antibody responses to influenza A(H3N2): an observational and modeling study. *eLife*, 11, 2022.

[12062] Chen Yang, Hailin Zhang, Mengnuo Chen, Siying Wang, Ruolan Qian, Linmeng Zhang, Xiaowen Huang, Jun Wang, Zhicheng Liu, Wenxin Qin, Cun Wang, Hualian Hang, and Hui Wang. A survey of optimal strategy for signature-based drug repositioning and an application to liver cancer. *eLife*, 11, 2022.

[12063] Haomin Yang, Nirmala Bhoo-Pathy, Judith S Brand, Elham Hedayati, Felix Grassmann, Erwei Zeng, Jonas Bergh, Weiwei Bian, Jonas F Ludvigsson, Per Hall, and Kamila Czene. Risk of heart disease following treatment for breast cancer – results from a population-based cohort study. *eLife*, 11, 2022.

[12064] Jane Yang, Husain Shakil, Stéphanie Ratté, and Steven A Prescott. Minimal requirements for a neuron to coregulate many properties and the implications for ion channel correlations and robustness. *eLife*, 11, 2022.

[12065] Lei Yang, Xichen Dong, Zheng Liu, Jinjing Tan, Xiaoxi Huang, Tao Wen, Hao Qu, and Zhenjun Wang. VPS9D1-AS1 overexpression amplifies intratumoral TGF-$\beta$ signaling and promotes tumor cell escape from CD8+ T cell killing in colorectal cancer. *eLife*, 11, 2022.

[12066] Liheng Yang, Eleanor C Semmes, Cristian Ovies, Christina Megli, Sallie Permar, Jennifer B Gilner, and Carolyn B Coyne. Innate immune signaling in trophoblast and decidua organoids defines differential antiviral defenses at the maternal-fetal interface. *eLife*, 11, 2022.

[12067] Marty G Yang, Emi Ling, Christopher J Cowley, Michael E Greenberg, and Thomas Vierbuchen. Characterization of sequence determinants of enhancer function using natural genetic variation. *eLife*, 11, 2022.

[12068] Ning Yang, Xuebo Lu, Yanan Jiang, Lili Zhao, Donghao Wang, Yaxing Wei, Yin Yu, Myoung Ok Kim, Kyle Vaughn Laster, Xin Li, Baoyin Yuan, Zigang Dong, and Kangdong Liu. Arbidol inhibits human esophageal squamous cell carcinoma growth in vitro and in vivo through suppressing ataxia telangiectasia and Rad3-related protein kinase. *eLife*, 11, 2022.

[12069] Peng Yang, Yaohui Chen, Zhiqi Huang, Huidan Xia, Ling Cheng, Hao Wu, Yueling Zhang, and Fan Wang. Single-cell RNA sequencing analysis of shrimp immune cells identifies macrophage-like phagocytes. *eLife*, 11, 2022.

[12070] Qianli Yang, Zhongqiao Lin, Wenyi Zhang, Jianshu Li, Xiyuan Chen, Jiaqi Zhang, and Tianming Yang. Monkey plays Pac-Man with compositional strategies and hierarchical decision-making. *eLife*, 11, 2022.

[12071] Qixin Yang, Yuchuan Miao, Leonard J Campanello, Matt J Hourwitz, Bedri Abubaker-Sharif, Abby L Bull, Peter N Devreotes, John T Fourkas, and Wolfgang Losert. Cortical waves mediate the cellular response to electric fields. *eLife*, 11, 2022.

[12072] Shu Yang, Xuefeng Wu, Euphrosyne I Daoutidou, Ya Zhang, MaryJane Shimell, Kun-Han Chuang, Aidan J Peterson, Michael B O'Connor, and Xiaoyan Zheng. The NDNF-like factor Nord is a Hedgehog-induced extracellular BMP modulator that regulates Drosophila wing patterning and growth. *eLife*, 11, 2022.

[12073] Wan Yang and Jeffrey L Shaman. COVID-19 pandemic dynamics in South Africa and epidemiological characteristics of three variants of concern (Beta, Delta, and Omicron). *eLife*, 11, 2022.

[12074] Yang Yang, Jing Wu, Qiao Li, Jing Wang, Lixian Mu, Li Hui, Min Li, Wei Xu, Hailong Yang, and Lin Wei. A non-bactericidal cathelicidin provides prophylactic efficacy against bacterial infection by driving phagocyte influx. *eLife*, 11, 2022.

[12075] Yu Hsuan Carol Yang, Linford JB Briant, Christopher A Raab, Sri Teja Mullapudi, Hans-Martin Maischein, Koichi Kawakami, and Didier YR Stainier. Innervation modulates the functional connectivity between pancreatic endocrine cells. *eLife*, 11, 2022.

[12076] Murat Yildirim, Chloe Delepine, Danielle Feldman, Vincent A Pham, Stephanie Chou, Jacque Ip, Alexi Nott, Li-Huei Tsai, Guo-Li Ming, Peter TC So, and Mriganka Sur. Label-free three-photon imaging of intact human cerebral organoids for tracking early events in brain development and deficits in Rett syndrome. *eLife*, 11, 2022.

[12077] Vural Yilmaz, Panayiota Louca, Louiza Potamiti, Mihalis Panayiotidis, and Katerina Strati. A novel lineage-tracing mouse model for studying early MmuPV1 infections. *eLife*, 11, 2022.

[12078] Willa Wen-You Yim, Hayashi Yamamoto, and Noboru Mizushima. A pulse-chasable reporter processing assay for mammalian autophagic flux with HaloTag. *eLife*, 11, 2022.

[12079] DongAhn Yoo, Junhee Park, Chul Lee, Injun Song, Young Ho Lee, Tery Yun, Hyemin Lee, Adriana Heguy, Jae Yong Han, Jeremy S Dasen, Heebal Kim, and Myungin Baek. Little skate genome provides insights into genetic programs essential for limb-based locomotion. *eLife*, 11, 2022.

[12080] Makoto M Yoshida, Kazuhisa Kinoshita, Yuuki Aizawa, Shoji Tane, Daisuke Yamashita, Keishi Shintomi, and Tatsuya Hirano. Molecular dissection of condensin II-mediated chromosome assembly using in vitro assays. *eLife*, 11, 2022.

[12081] Nozomu Yoshioka, Masayuki Kurose, Masato Yano, Dang Minh Tran, Shujiro Okuda, Yukiko Mori-Ochiai, Masao Horie, Toshihiro Nagai, Ichizo Nishino, Shinsuke Shibata, and Hirohide Takebayashi. Isoform-specific mutation in Dystonin-b gene causes late-onset protein aggregate myopathy and cardiomyopathy. *eLife*, 11, 2022.

[12082] Remy A Yovanno, Tsung Han Chou, Sarah J Brantley, Hiro Furukawa, and Albert Y Lau. Excitatory and inhibitory D-serine binding to the NMDA receptor. *eLife*, 11, 2022.

[12083] Gongchen Yu, James P Herman, Leor N Katz, and Richard J Krauzlis. Microsaccades as a marker not a cause for attention-related modulation. *eLife*, 11, 2022.

[12084] Houqing Yu, Roarke A Kamber, and Vladimir Denic. The peroxisomal exportomer directly inhibits phosphoactivation of the pexophagy receptor Atg36 to suppress pexophagy in yeast. *eLife*, 11, 2022.

[12085] Huasheng Yu, Jingwei Xiong, Adam Yongxin Ye, Suna Li Cranfill, Tariq Cannonier, Mayank Gautam, Marina Zhang, Rayan Bilal, Jong-Eun Park, Yuji Xue, Vidhur Polam, Zora Vujovic, Daniel Dai, William Ong, Jasper Ip, Amanda Hsieh, Nour Mimouni, Alejandra Lozada, Medhini Sosale, Alex Ahn, Minghong Ma, Long Ding, Javier Arsuaga, and Wenqin Luo. Scratch-AID, a deep learning-based system for automatic detection of mouse scratching behavior with high accuracy. *eLife*, 11, 2022.

[12086] Wentao Yu, Lei Kang, Victor TC Tsang, Yan Zhang, Ivy HM Wong, and Terence TW Wong. Translational rapid ultraviolet-excited sectioning tomography for whole-organ multicolor imaging with real-time molecular staining. *eLife*, 11, 2022.

[12087] Zhou Yu, Maxwell H Turner, Jacob Baudin, and Fred Rieke. Adaptation in cone photoreceptors contributes to an unexpected insensitivity of primate On parasol retinal ganglion cells to spatial structure in natural images. *eLife*, 11, 2022.

[12088] Alex Eric Yuan and Wenying Shou. Data-driven causal analysis of observational biological time series. *eLife*, 11, 2022.

[12089] Lei Yuan, Peiyao Li, Huiru Jing, Qian Zheng, and Hui Xiao. trim-21 promotes proteasomal degradation of CED-1 for apoptotic cell clearance in C. elegans. *eLife*, 11, 2022.

[12090] Wendy Wing Sze Yue, Lin Yuan, Joao M Braz, Allan I Basbaum, and David Julius. TRPV1 drugs alter core body temperature via central projections of primary afferent sensory neurons. *eLife*, 11, 2022.

[12091] Asieh Zadbood, Samuel Nastase, Janice Chen, Kenneth A Norman, and Uri Hasson. Neural representations of naturalistic events are updated as our understanding of the past changes. *eLife*, 11, 2022.

[12092] Michela Zaffagni, Jenna M Harris, Ines L Patop, Nagarjuna Reddy Pamudurti, Sinead Nguyen, and Sebastian Kadener. SARS-CoV-2 Nsp14 mediates the effects of viral infection on the host cell transcriptome. *eLife*, 11, 2022.

[12093] Ioannis Zalachoras, Eva Ramos-Fernández, Fiona Hollis, Laura Trovo, João Rodrigues, Alina Strasser, Olivia Zanoletti, Pascal Steiner, Nicolas Preitner, Lijing Xin, Simone Astori, and Carmen Sandi. Glutathione in the nucleus accumbens regulates motivation to exert reward-incentivized effort. *eLife*, 11, 2022.

[12094] Ryan Zander, Achia Khatun, Moujtaba Y Kasmani, Yao Chen, and Weiguo Cui. Delineating the transcriptional landscape and clonal diversity of virus-specific CD4+ T cells during chronic viral infection. *eLife*, 11, 2022.

[12095] Juan P Zanin and Wilma J Friedman. p75NTR prevents the onset of cerebellar granule cell migration via RhoA activation. *eLife*, 11, 2022.

[12096] Mark A Zaydman, Alexander S Little, Fidel Haro, Valeryia Aksianiuk, William J Buchser, Aaron DiAntonio, Jeffrey I Gordon, Jeffrey Milbrandt, and Arjun S Raman. Defining hierarchical protein interaction networks from spectral analysis of bacterial proteomes. *eLife*, 11, 2022.

[12097] Yanis Zekri, Romain Guyot, Inés Garteizgogeascoa Suñer, Laurence Canaple, Amandine Gautier Stein, Justine Vily Petit, Denise Aubert, Sabine Richard, Frédéric Flamant, and Karine Gauthier. Brown adipocytes local response to thyroid hormone is required for adaptive thermogenesis in adult male mice. *eLife*, 11, 2022.

[12098] Gang Zhai, Tingting Shu, Guangqing Yu, Haipei Tang, Chuang Shi, Jingyi Jia, Qiyong Lou, Xiangyan Dai, Xia Jin, Jiangyan He, Wuhan Xiao, Xiaochun Liu, and Zhan Yin. Augmentation of progestin signaling rescues testis organization and spermatogenesis in zebrafish with the depletion of androgen signaling. *eLife*, 11, 2022.

[12099] Biao Zhang, Kai Xue, Shutong Zhou, Kui Wang, Wenjing Liu, Cong Xu, Lizhen Cui, Linfeng Li, Qinwei Ran, Zongsong Wang, Ronghai Hu, Yanbin Hao, Xiaoyong Cui, and Yanfen Wang. Environmental selection overturns the decay relationship of soil prokaryotic community over geographic distance across grassland biotas. *eLife*, 11, 2022.

[12100] Chenlu Zhang, Zehan Hu, Abdul G Lone, Methinee Artami, Marshall Edwards, Christos C Zouboulis, Maggie Stein, and Tamia A Harris-Tryon. Small proline-rich proteins (SPRRs) are epidermally produced antimicrobial proteins that defend the cutaneous barrier by direct bacterial membrane disruption. *eLife*, 11, 2022.

[12101] Chi Zhang, Ming-Wen Hu, Xue-Wei Wang, Xiang Cui, Jing Liu, Qian Huang, Xu Cao, Feng-Quan Zhou, Jiang Qian, Shao-Qiu He, and Yun Guan. scRNA-sequencing reveals subtype-specific transcriptomic perturbations in DRG neurons of PirtEGFPf mice in neuropathic pain condition. *eLife*, 11, 2022.

[12102] Dan Zhang, Roxane P Spiegelhalder, Emily B Abrash, Tiago DG Nunes, Inés Hidalgo, M Ximena Anleu Gil, Barbara Jesenofsky, Heike Lindner, Dominique C Bergmann, and Michael T Raissig. Opposite polarity programs regulate asymmetric subsidiary cell divisions in grasses. *eLife*, 11, 2022.

[12103] Feifei Zhang, Margo Chase-Topping, Chuan-Guo Guo, and Mark EJ Woolhouse. Predictors of human-infective RNA virus discovery in the United States, China, and Africa, an ecological study. *eLife*, 11, 2022.

[12104] Haiwei Zhang, Hongchen Li, Mingshun Lu, Shengnan Wang, Xueya Ma, Fei Wang, Jiaxi Liu, Xinyu Li, Haichao Yang, Fan Zhang, Haitao Shen, Noel J Buckley, Nikita Gamper, Ebenezer N Yamoah, and Ping Lv. Repressor element 1-silencing transcription factor deficiency yields profound hearing loss through Kv7.4 channel upsurge in auditory neurons and hair cells. *eLife*, 11, 2022.

[12105] Han Zhang, Natalia A Shevtsova, Dylan Deska-Gauthier, Colin Mackay, Kimberly J Dougherty, Simon M Danner, Ying Zhang, and Ilya A Rybak. The role of V3 neurons in speed-dependent interlimb coordination during locomotion in mice. *eLife*, 11, 2022.

[12106] Han-Xiong Bear Zhang, Laurel Heckman, Zachary Niday, Sooyeon Jo, Akie Fujita, Jaehoon Shim, Roshan Pandey, Hoor Al Jandal, Selwyn Jayakar, Lee B Barrett, Jennifer Smith, Clifford J Woolf, and Bruce P Bean. Cannabidiol activates neuronal Kv7 channels. *eLife*, 11, 2022.

[12107] Jianhua Zhang, Zachery R Gregorich, Ran Tao, Gina C Kim, Pratik A Lalit, Juliana L Carvalho, Yogananda Markandeya, Deane F Mosher, Sean P Palecek, and Timothy J Kamp. Cardiac differentiation of human pluripotent stem cells using defined extracellular matrix proteins reveals essential role of fibronectin. *eLife*, 11, 2022.

[12108] Jing Zhang, Zi Bo Han, Yu Liang, Xue Feng Zhang, Yu Qin Jin, Li Fang Du, Shuai Shao, Hui Wang, Jun Wei Hou, Ke Xu, Wenwen Lei, Ze Hua Lei, Zhao Ming Liu, Jin Zhang, Ya Nan Hou, Ning Liu, Fu Jie Shen, Jin Juan Wu, Xiang Zheng, Xin Yu Li, Xin Li, Wei Jin Huang, Gui Zhen Wu, Ji Guo Su, and Qi Ming Li. A mosaic-type trimeric RBD-based COVID-19 vaccine candidate induces potent neutralization against Omicron and other SARS-CoV-2 variants. *eLife*, 11, 2022.

[12109] Jingchao Zhang, Huijun Wu, Di Wang, Lanxin Wang, Yifan Cui, Chenxi Zhang, Kun Zhao, and Luyan Ma. Intracellular glycosyl hydrolase PslG shapes bacterial cell fate, signaling, and the biofilm development of Pseudomonas aeruginosa. *eLife*, 11, 2022.

[12110] Jingruo Zhang, Manuel A Riquelme, Rui Hua, Francisca M Acosta, Sumin Gu, and Jean X Jiang. Connexin 43 hemichannels regulate mitochondrial ATP generation, mobilization, and mitochondrial homeostasis against oxidative stress. *eLife*, 11, 2022.

[12111] Jingsong Zhang, Jessica Cunningham, Joel Brown, and Robert Gatenby. Evolution-based mathematical models significantly prolong response to abiraterone in metastatic castrate-resistant prostate cancer and identify strategies to further improve outcomes. *eLife*, 11, 2022.

[12112] Kuan Zhang, Erica Yao, Biao Chen, Ethan Chuang, Julia Wong, Robert I Seed, Stephen L Nishimura, Paul J Wolters, and Pao-Tien Chuang. Acquisition of cellular properties during alveolar formation requires differential activity and distribution of mitochondria. *eLife*, 11, 2022.

[12113] Ling-Qi Zhang, Nicolas P Cottaris, and David H Brainard. An image reconstruction framework for characterizing initial visual encoding. *eLife*, 11, 2022.

[12114] Meichao Zhang, Boris C Bernhardt, Xiuyi Wang, Dominika Varga, Katya Krieger-Redwood, Jessica Royer, Raúl Rodríguez-Cruces, Reinder Vos de Wael, Daniel S Margulies, Jonathan Smallwood, and Elizabeth Jefferies. Perceptual coupling and decoupling of the default mode network during mind-wandering and reading. *eLife*, 11, 2022.

[12115] Shengnan Zhang, Yi Zhu, Jinxia Lu, Zhenying Liu, Amanda G Lobato, Wen Zeng, Jiaqi Liu, Jiali Qiang, Shuyi Zeng, Yaoyang Zhang, Cong Liu, Jun Liu, Zhuohao He, R Grace Zhai, and Dan Li. Specific binding of Hsp27 and phosphorylated Tau mitigates abnormal Tau aggregation-induced pathology. *eLife*, 11, 2022.

[12116] Wujie Zhang, Maimon C Rose, and Michael M Yartsev. A unifying mechanism governing inter-brain neural relationship during social interactions. *eLife*, 11, 2022.

[12117] Xi Zhang, Xiaolin Zhang, Fangnian Tong, Yi Cai, Yujie Zhang, Haixu Song, Xiaoxiang Tian, Chenghui Yan, and Yaling Han. Gut microbiota induces high platelet response in patients with ST segment elevation myocardial infarction after ticagrelor treatment. *eLife*, 11, 2022.

[12118] Xinrui L Zhang, William C Spencer, Nobuko Tabuchi, Meagan M Kitt, and Evan S Deneris. Reorganization of postmitotic neuronal chromatin accessibility for maturation of serotonergic identity. *eLife*, 11, 2022.

[12119] Yanchun Zhang, Xuexue Xiong, Qi Zhu, Jiali Zhang, Shengmiao Chen, Yuetong Wang, Jian Cao, Li Chen, Linjun Hou, Xi Zhao, Piliang Hao, Jian Chen, Min Zhuang, Dake Li, and Gaofeng Fan. FER-mediated phosphorylation and PIK3R2 recruitment on IRS4 promotes AKT activation and tumorigenesis in ovarian cancer cells. *eLife*, 11, 2022.

[12120] Yanfeng Zhang, Kenneth Day, and Devin M Absher. STAT3-mediated allelic imbalance of novel genetic variant Rs1047643 and B-cell-specific super-enhancer in association with systemic lupus erythematosus. *eLife*, 11, 2022.

[12121] Yang Zhang, Pengfei Liang, Liheng Yang, Ke Zoe Shan, Liping Feng, Yong Chen, Wolfgang Liedtke, Carolyn B Coyne, and Huanghe Yang. Functional coupling between TRPV4 channel and TMEM16F modulates human trophoblast fusion. *eLife*, 11, 2022.

[12122] Yikui Zhang, Mengyun Li, Bo Yu, Shengjian Lu, Lujie Zhang, Senmiao Zhu, Zhonghao Yu, Tian Xia, Haoliang Huang, WenHao Jiang, Si Zhang, Lanfang Sun, Qian Ye, Jiaying Sun, Hui Zhu, Pingping Huang, Huifeng Hong, Shuaishuai Yu, Wenjie Li, Danni Ai, Jingfan Fan, Wentao Li, Hong Song, Lei Xu, Xiwen Chen, Tongke Chen, Meng Zhou, Jingxing Ou, Jian Yang, Wei Li, Yang Hu, and Wencan Wu. Cold protection allows local cryotherapy in a clinical-relevant model of traumatic optic neuropathy. *eLife*, 11, 2022.

[12123] Yixiao Zhang, Lisha Sun, Haonan Li, Liping Ai, Qingtian Ma, Xinbo Qiao, Jie Yang, Hao Zhang, Xunyan Ou, Yining Wang, Guanglei Chen, Jinqi Xue, Xudong Zhu, Yu Zhao, Yongliang Yang, and Caigang Liu. Binding blockade between TLN1 and integrin $\beta 1$ represses triple-negative breast cancer. *eLife*, 11, 2022.

[12124] Dezhi Zhao, Manuel A Riquelme, Teja Guda, Chao Tu, Huiyun Xu, Sumin Gu, and Jean X Jiang. Connexin hemichannels with prostaglandin release in anabolic function of bone to mechanical loading. *eLife*, 11, 2022.

[12125] Hongchang Zhao, Stella R Hartono, Kirtney Mae Flores de Vera, Zheyuan Yu, Krishni Satchi, Tracy Zhao, Roger Sciammas, Lionel Sanz, Frédéric Chédin, and Jacqueline Barlow. Senataxin and RNase H2 act redundantly to suppress genome instability during class switch recombination. *eLife*, 11, 2022.

[12126] Jing Zhao, Anahid A Birjandi, Mohi Ahmed, Yushi Redhead, Jose Villagomez Olea, and Paul Sharpe. Telocytes regulate macrophages in periodontal disease. *eLife*, 11, 2022.

[12127] Longhui Zhao, Jichao Wang, Haodi Zhang, Tongliang Wang, Yue Yang, Yezhong Tang, Wouter Halfwerk, and Jianguo Cui. Parasite defensive limb movements enhance acoustic signal attraction in male little torrent frogs. *eLife*, 11, 2022.

[12128] Meng Zhao, Niels Banhos Danneskiold-Samsøe, Livia Ulicna, Quennie Nguyen, Laetitia Voilquin, David E Lee, James P White, Zewen Jiang, Nickeisha Cuthbert, Shrika Paramasivam, Ewa Bielczyk-Maczynska, Capucine Van Rechem, and Katrin J Svensson. Phosphoproteomic mapping reveals distinct signaling actions and activation of muscle protein synthesis by Isthmin-1. *eLife*, 11, 2022.

[12129] Rong Zhao, Stacy D Grunke, Caleb A Wood, Gabriella A Perez, Melissa Comstock, Ming-Hua Li, Anand K Singh, Kyung-Won Park, and Joanna L Jankowsky. Activity disruption causes degeneration of entorhinal neurons in a mouse model of Alzheimer's circuit dysfunction. *eLife*, 11, 2022.

[12130] Yiheng Zhao, Linchan Yu, Shuchen Zhang, Xiangyu Su, and Xiang Zhou. Extrachromosomal circular DNA: Current status and future prospects. *eLife*, 11, 2022.

[12131] Zikai Zhao, Qi Li, Usama Ashraf, Mengjie Yang, Wenjing Zhu, Jun Gu, Zheng Chen, Changqin Gu, Youhui Si, Shengbo Cao, and Jing Ye. Zika virus causes placental pyroptosis and associated adverse fetal outcomes by activating GSDME. *eLife*, 11, 2022.

[12132] Eric B Zheng and Li Zhao. Protein evidence of unannotated ORFs in Drosophila reveals diversity in the evolution and properties of young proteins. *eLife*, 11, 2022.

[12133] Fengwei Zheng, Roxana Georgescu, Nina Y Yao, Huilin Li, and Michael E O'Donnell. Cryo-EM structures reveal that RFC recognizes both the 3'- and 5'-DNA ends to load PCNA onto gaps for DNA repair. *eLife*, 11, 2022.

[12134] Minyan Zheng, Olga Zueva, and Veronica F Hinman. Regeneration of the larval sea star nervous system by wounding induced respecification to the Sox2 lineage. *eLife*, 11, 2022.

[12135] Rui Zheng, Yonglan Du, Xintai Wang, Tailin Liao, Zhe Zhang, Na Wang, Xiumao Li, Ying Shen, Lei Shi, Jianhong Luo, Jun Xia, Ziyi Wang, and Junyu Xu. KIF2C regulates synaptic plasticity and cognition in mice through dynamic microtubule depolymerization. *eLife*, 11, 2022.

[12136] Xiang Zheng, Haimiao Yuan, and Chaoqun Ni. How parenthood contributes to gender gaps in academia. *eLife*, 11, 2022.

[12137] Xiaowei Zheng, Sampath Narayanan, Cheng Xu, Sofie Eliasson Angelstig, Jacob Grünler, Allan Zhao, Alessandro Di Toro, Luciano Bernardi, Massimiliano Mazzone, Peter Carmeliet, Marianna Del Sole, Giancarlo Solaini, Elisabete A Forsberg, Ao Zhang, Kerstin Brismar, Tomas A Schiffer, Neda Rajamand Ekberg, Ileana Ruxandra Botusan, Fredrik Palm, and Sergiu-Bogdan Catrina. Repression of hypoxia-inducible factor-1 contributes to increased mitochondrial reactive oxygen species production in diabetes. *eLife*, 11, 2022.

[12138] Jiale Zhong, Chen-Jun Guo, Xian Zhou, Chia-Chun Chang, Boqi Yin, Tianyi Zhang, Huan-Huan Hu, Guang-Ming Lu, and Ji-Long Liu. Structural basis of dynamic P5CS filaments. *eLife*, 11, 2022.

[12139] Ling Zhong, Peng Hao, Qian Zhang, Tao Jiang, Huan Li, Jialing Xiao, Chenglong Li, Lan Luo, Chunbao Xie, Jiang Hu, Liang Wang, Yuping Liu, Yi Shi, Wei Zhang, and Bo Gong. Revised International Staging System (R-ISS) stage-dependent analysis uncovers oncogenes and potential immunotherapeutic targets in multiple myeloma (MM). *eLife*, 11, 2022.

[12140] Baohua Zhou, Zifan Li, Sunnie Kim, John Lafferty, and Damon A Clark. Shallow neural networks trained to detect collisions recover features of visual loom-selective neurons. *eLife*, 11, 2022.

[12141] Shouhao Zhou, Xinyi Liu, Xinying Fang, Vernon M Chinchilli, Michael Wang, Hong-Gang Wang, Nikolay V Dokholyan, Chan Shen, and J Jack Lee. Robust and Efficient Assessment of Potency (REAP) as a quantitative tool for dose-response curve estimation. *eLife*, 11, 2022.

[12142] Tyng-An Zhou, Hsuan-Po Hsu, Yueh-Hua Tu, Hui-Kuei Cheng, Chih-Yu Lin, Nien-Jung Chen, Jin-Wu Tsai, Ellen A Robey, Hsuan-Cheng Huang, Chia-Lin Hsu, and Ivan L Dzhagalov. Thymic macrophages consist of two populations with distinct localization and origin. *eLife*, 11, 2022.

[12143] Yizhuo Zhou, Ying Yang, Lihao Guo, Jun Qian, Jian Ge, Debora Sinner, Hongxu Ding, Andrea Califano, and Wellington V Cardoso. Airway basal cells show regionally distinct potential to undergo metaplastic differentiation. *eLife*, 11, 2022.

[12144] Zheng Zhou, Valentyna Krashevska, Rahayu Widyastuti, Stefan Scheu, and Anton Potapov. Tropical land use alters functional diversity of soil food webs and leads to monopolization of the detrital energy channel. *eLife*, 11, 2022.

[12145] Haoran Zhu, Keefe T Chan, Xinran Huang, Carmelo Cerra, Shaun Blake, Anna S Trigos, Dovile Anderson, Darren J Creek, David P De Souza, Xi Wang, Caiyun Fu, Metta Jana, Elaine Sanij, Richard B Pearson, and Jian Kang. Cystathionine-$\beta$-synthase is essential for AKT-induced senescence and suppresses the development of gastric cancers with PI3K/AKT activation. *eLife*, 11, 2022.

[12146] Lu Zhu, Qing Zhang, Ciro D Cordeiro, Sudeep Banjade, Richa Sardana, Yuxin Mao, and Scott D Emr. Adaptor linked K63 di-ubiquitin activates Nedd4/Rsp5 E3 ligase. *eLife*, 11, 2022.

[12147] Seren Zhu, Kaushik J Lakshminarasimhan, Nastaran Arfaei, and Dora E Angelaki. Eye movements reveal spatiotemporal dynamics of visually-informed planning in navigation. *eLife*, 11, 2022.

[12148] Ya-Bing Zhu, Yan Wang, Xiao-Xiao Hua, Ling Xu, Ming-Zhe Liu, Rui Zhang, Peng-Fei Liu, Jin-Bao Li, Ling Zhang, and Di Mu. PBN-PVT projections modulate negative affective states in mice. *eLife*, 11, 2022.

[12149] Yannan Zhu, Yimeng Zeng, Jingyuan Ren, Lingke Zhang, Changming Chen, Guillen Fernandez, and Shaozheng Qin. Emotional learning retroactively promotes memory integration through rapid neural reactivation and reorganization. *eLife*, 11, 2022.

[12150] Zhanwu Zhu, Jingjing Liu, Huan Feng, Yanning Zhang, Ruiqi Huang, Qiaochu Pan, Jing Nan, Ruidong Miao, and Bo Cheng. Overcoming the cytoplasmic retention of GDOWN1 modulates global transcription and facilitates stress adaptation. *eLife*, 11, 2022.

[12151] Jasenko Zivanov, Joaquín Otón, Zunlong Ke, Andriko von Kügelgen, Euan Pyle, Kun Qu, Dustin Morado, Daniel Castaño Díez, Giulia Zanetti, Tanmay AM Bharat, John AG Briggs, and Sjors HW Scheres. A Bayesian approach to single-particle electron cryo-tomography in RELION-4.0. *eLife*, 11, 2022.

[12152] Omid A Zobeiri and Kathleen E Cullen. Distinct representations of body and head motion are dynamically encoded by Purkinje cell populations in the macaque cerebellum. *eLife*, 11, 2022.

[12153] Yan Zou, Miaomiao Zhang, Qiongfeng Wu, Ning Zhao, Minwei Chen, Cui Yang, Yimei Du, and Bing Han. Activation of transient receptor potential vanilloid 4 is involved in pressure overload-induced cardiac hypertrophy. *eLife*, 11, 2022.

[12154] Philipp K Zuber, Tina Daviter, Ramona Heißmann, Ulrike Persau, Kristian Schweimer, and Stefan H Knauer. Structural and thermodynamic analyses of the $\beta$-to-$\alpha$ transformation in RfaH reveal principles of fold-switching proteins. *eLife*, 11, 2022.

[12155] Stefan A Zukin, Matthew R Marunde, Irina K Popova, Katarzyna M Soczek, Eva Nogales, and Avinash B Patel. Structure and flexibility of the yeast NuA4 histone acetyltransferase complex. *eLife*, 11, 2022.

[12156] Knut Sindre Åbjørsbråten, Gry HE Syverstad Skaaraas, Céline Cunen, Daniel M Bjørnstad, Kristin M Gullestad Binder, Laura Bojarskaite, Vidar Jensen, Lars NG Nilsson, Shreyas B Rao, Wannan Tang, Gudmund Horn Hermansen, Erlend A Nagelhus, Ole Petter Ottersen, Reidun Torp, and Rune Enger. Impaired astrocytic Ca2+ signaling in awake-behaving Alzheimer's disease transgenic mice. *eLife*, 11, 2022.

[12157] Saman Abbaspoor, Ahmed T Hussin, and Kari L Hoffman. Theta- and gamma-band oscillatory uncoupling in the macaque hippocampus. *eLife*, 12, 2023.

[12158] Tooba Abbassi-Daloii, Salma el Abdellaoui, Lenard M Voortman, Thom TJ Veeger, Davy Cats, Hailiang Mei, Duncan E Meuffels, Ewoud van Arkel, Peter AC 't Hoen, Hermien E Kan, and Vered Raz. A transcriptome atlas of leg muscles from healthy human volunteers reveals molecular and cellular signatures associated with muscle location. *eLife*, 12, 2023.

[12159] Moustafa Abdalla, Salwa Abdalla, and Mohamed Abdalla. Tracing the path of 37,050 studies into practice across 18 specialties of the 2.4 million published between 2011 and 2020. *eLife*, 12, 2023.

[12160] Mahmoud EA Abdellahi, Anne CM Koopman, Matthias S Treder, and Penelope A Lewis. Targeted memory reactivation in human REM sleep elicits detectable reactivation. *eLife*, 12, 2023.

[12161] Abdirahman I Abdi, Fiona Achcar, Lauriane Sollelis, João Luiz Silva-Filho, Kioko Mwikali, Michelle Muthui, Shaban Mwangi, Hannah W Kimingi, Benedict Orindi, Cheryl Andisi Kivisi, Manon Alkema, Amrita Chandrasekar, Peter C Bull, Philip Bejon, Katarzyna Modrzynska, Teun Bousema, and Matthias Marti. Plasmodium falciparum adapts its investment into replication versus transmission according to the host environment. *eLife*, 12, 2023.

[12162] Taylor J Abele, Zachary P Billman, Lupeng Li, Carissa K Harvest, Alexia K Bryan, Gabrielle R Magalski, Joseph P Lopez, Heather N Larson, Xiao-Ming Yin, and Edward A Miao. Apoptotic signaling clears engineered Salmonella in an organ-specific manner. *eLife*, 12, 2023.

[12163] Yonathan Tamrat Aberra, Lijiang Ma, Johan LM Björkegren, and Mete Civelek. Predicting mechanisms of action at genetic loci associated with discordant effects on type 2 diabetes and abdominal fat accumulation. *eLife*, 12, 2023.

[12164] Shai Abramson, Benjamin J Kraus, John A White, Michael E Hasselmo, Dori Derdikman, and Genela Morris. Flexible coding of time or distance in hippocampal cells. *eLife*, 12, 2023.

[12165] Ellen M Acosta, Katherine A Little, Benjamin P Bratton, Jaime G Lopez, Xuming Mao, Aimee S Payne, Mohamed Donia, Danelle Devenport, and Zemer Gitai. Bacterial DNA on the skin surface overrepresents the viable skin microbiome. *eLife*, 12, 2023.

[12166] Jan Andreas Adelmann, Roman Vetter, and Dagmar Iber. Patterning precision under non-linear morphogen decay and molecular noise. *eLife*, 12, 2023.

[12167] Adedeji M Aderounmu, P Joseph Aruscavage, Bryan Kolaczkowski, and Brenda L Bass. Ancestral protein reconstruction reveals evolutionary events governing variation in Dicer helicase function. *eLife*, 12, 2023.

[12168] Yalda Afshar, Feyiang Ma, Austin Quach, Anhyo Jeong, Hannah L Sunshine, Vanessa Freitas, Yasaman Jami-Alahmadi, Raphael Helaers, Xinmin Li, Matteo Pellegrini, James A Wohlschlegel, Casey E Romanoski, Miikka Vikkula, and M Luisa Iruela-Arispe. Transcriptional drifts associated with environmental changes in endothelial cells. *eLife*, 12, 2023.

[12169] Ipsita Agarwal, Zachary L Fuller, Simon R Myers, and Molly Przeworski. Relating pathogenic loss-of-function mutations in humans to their evolutionary fitness costs. *eLife*, 12, 2023.

[12170] Dina Aggad, Nicolas Brouilly, Shizue Omi, Clara Luise Essmann, Benoit Dehapiot, Cathy Savage-Dunn, Fabrice Richard, Chantal Cazevieille, Kristin A Politi, David H Hall, Remy Pujol, and Nathalie Pujol. Meisosomes, folded membrane microdomains between the apical extracellular matrix and epidermis. *eLife*, 12, 2023.

[12171] Ahmed-Noor A Agip, Injae Chung, Alvaro Sanchez-Martinez, Alexander J Whitworth, and Judy Hirst. Cryo-EM structures of mitochondrial respiratory complex I from Drosophila melanogaster. *eLife*, 12, 2023.

[12172] Leire Aguinagalde Salazar, Maurits A den Boer, Suzanne M Castenmiller, Seline A Zwarthoff, Carla de Haas, Piet C Aerts, Frank J Beurskens, Janine Schuurman, Albert JR Heck, Kok van Kessel, and Suzan HM Rooijakkers. Promoting Fc-Fc interactions between anti-capsular antibodies provides strong immune protection against Streptococcus pneumoniae. *eLife*, 12, 2023.

[12173] Tim Aguirre, Gillian L Dornan, Sarah Hostachy, Martin Neuenschwander, Carola Seyffarth, Volker Haucke, Anja Schüetz, Jens Peter von Kries, and Dorothea Fiedler. An unconventional gatekeeper mutation sensitizes inositol hexakisphosphate kinases to an allosteric inhibitor. *eLife*, 12, 2023.

[12174] Sara Ahadi, Kenneth A Wilson, Boris Babenko, Cory Y McLean, Drew Bryant, Orion Pritchard, Ajay Kumar, Enrique M Carrera, Ricardo Lamy, Jay M Stewart, Avinash Varadarajan, Marc Berndl, Pankaj Kapahi, and Ali Bashir. Longitudinal fundus imaging and its genome-wide association analysis provide evidence for a human retinal aging clock. *eLife*, 12, 2023.

[12175] Ali Reza Ahmadi, Guang Song, Tianshun Gao, Jing Ma, Xiaomei Han, Ming-Wen Hu, Andrew M Cameron, Russell N Wesson, Benjamin Philosophe, Shane Ottmann, Elizabeth King, Ahmet Gurakar, Le Qi, Brandon Peiffer, James Burdick, Robert Anders, Zhanxiang Zhou, Hongkun Lu, Dechun Feng, Chien-Sheng Chen, Jiang Qian, Bin Gao, Heng Zhu, and Zhaoli Sun. Discovery and characterization of cross-reactive intrahepatic antibodies in severe alcoholic hepatitis. *eLife*, 12, 2023.

[12176] Ibrahim Ahmed, Shen-Hsi Yang, Samuel Ogden, Wei Zhang, Yaoyong Li, and Andrew D Sharrocks. eRNA profiling uncovers the enhancer landscape of oesophageal adenocarcinoma and reveals new deregulated pathways. *eLife*, 12, 2023.

[12177] Sharia M Ahmed, Ben J Brintz, Patricia B Pavlinac, Lubaba Shahrin, Sayeeda Huq, Adam C Levine, Eric J Nelson, James A Platts-Mills, Karen L Kotloff, and Daniel T Leung. Derivation and external validation of clinical prediction rules identifying children at risk of linear growth faltering. *eLife*, 12, 2023.

[12178] Sana Ahmed-Seghir, Manisha Jalan, Helen E Grimsley, Aman Sharma, Shyam Twayana, Settapong T Kosiyatrakul, Christopher Thompson, Carl L Schildkraut, and Simon N Powell. A local ATR-dependent checkpoint pathway is activated by a site-specific replication fork block in human cells. *eLife*, 12, 2023.

[12179] Ji-Eun Ahn and Hubert Amrein. Opposing chemosensory functions of closely related gustatory receptors. *eLife*, 12, 2023.

[12180] Sophie Aimon, Karen Y Cheng, Julijana Gjorgjieva, and Ilona C Grunwald Kadow. Global change in brain state during spontaneous and forced walk in Drosophila is composed of combined activity patterns of different neuron classes. *eLife*, 12, 2023.

[12181] Marina Ainciburu, Teresa Ezponda, Nerea Berastegui, Ana Alfonso-Pierola, Amaia Vilas-Zornoza, Patxi San Martin-Uriz, Diego Alignani, Jose Lamo-Espinosa, Mikel San-Julian, Tamara Jiménez-Solas, Felix Lopez, Sandra Muntion, Fermin Sanchez-Guijo, Antonieta Molero, Julia Montoro, Guillermo Serrano, Aintzane Diaz-Mazkiaran, Miren Lasaga, David Gomez-Cabrero, Maria Diez-Campelo, David Valcarcel, Mikel Hernaez, Juan P Romero, and Felipe Prosper. Uncovering perturbations in human hematopoiesis associated with healthy aging and myeloid malignancies at single-cell resolution. *eLife*, 12, 2023.

[12182] Kyle Aitken and Stefan Mihalas. Neural population dynamics of computing with synaptic modulations. *eLife*, 12, 2023.

[12183] Ngozi D Akingbesote, Brooks P Leitner, Daniel G Jovin, Reina Desrouleaux, Dennis Owusu, Wanling Zhu, Zongyu Li, Michael N Pollak, and Rachel J Perry. Gene and protein expression and metabolic flux analysis reveals metabolic scaling in liver ex vivo and in vivo. *eLife*, 12, 2023.

[12184] Hiroki Akizawa, Emily M Lopes, and Rafael A Fissore. Zn2+ is essential for Ca2+ oscillations in mouse eggs. *eLife*, 12, 2023.

[12185] Mastura Akter, Mahadi Hasan, Aruna Surendran Ramkrishnan, Zafar Iqbal, Xianlin Zheng, Zhongqi Fu, Zhuogui Lei, Anwarul Karim, and Ying Li. Astrocyte and L-lactate in the anterior cingulate cortex modulate schema memory and neuronal mitochondrial biogenesis. *eLife*, 12, 2023.

[12186] Andrea Alamia, Lucie Terral, Malo Renaud D'ambra, and Rufin VanRullen. Distinct roles of forward and backward alpha-band waves in spatial visual attention. *eLife*, 12, 2023.

[12187] Mark Borris D Aldonza, Junghwa Cha, Insung Yong, Jayoung Ku, Pavel Sinitcyn, Dabin Lee, Ryeong-Eun Cho, Roben D Delos Reyes, Dongwook Kim, Soyeon Kim, Minjeong Kang, Yongsuk Ku, Geonho Park, Hye-Jin Sung, Han Suk Ryu, Sukki Cho, Tae Min Kim, Pilnam Kim, Je-Yoel Cho, and Yoosik Kim. Multi-targeted therapy resistance via drug-induced secretome fucosylation. *eLife*, 12, 2023.

[12188] Reemy Ali Nasser, Yuval Harel, and Shay Stern. Early-life experience reorganizes neuromodulatory regulation of stage-specific behavioral responses and individuality dimensions during development. *eLife*, 12, 2023.

[12189] David Allard, Isabelle Cousineau, Eric H Ma, Bertrand Allard, Yacine Bareche, Hubert Fleury, and John Stagg. The CD73 immune checkpoint promotes tumor cell metabolic fitness. *eLife*, 12, 2023.

[12190] Hooman Allayee, Charles R Farber, Marcus M Seldin, Evan Graehl Williams, David E James, and Aldons J Lusis. Systems genetics approaches for understanding complex traits with relevance for human disease. *eLife*, 12, 2023.

[12191] Adam R Almeida and Wendy B Macklin. Early myelination involves the dynamic and repetitive ensheathment of axons which resolves through a low and consistent stabilization rate. *eLife*, 12, 2023.

[12192] Aurora Alvarez-Buylla, Marie-Therese Fischer, Maria Dolores Moya Garzon, Alexandra E Rangel, Elicio E Tapia, Julia T Tanzo, H Tom Soh, Luis A Coloma, Jonathan Z Long, and Lauren A O'Connell. Binding and sequestration of poison frog alkaloids by a plasma globulin. *eLife*, 12, 2023.

[12193] Andrew J Aman, Lauren M Saunders, August A Carr, Sanjay Srivatasan, Colten Eberhard, Blake Carrington, Dawn Watkins-Chow, William J Pavan, Cole Trapnell, and David M Parichy. Transcriptomic profiling of tissue environments critical for post-embryonic patterning and morphogenesis of zebrafish skin. *eLife*, 12, 2023.

[12194] Mathieu Amand, Philipp Adams, Rafaela Schober, Gilles Iserentant, Jean-Yves Servais, Michel Moutschen, and Carole Seguin-Devaux. The anti-caspase 1 inhibitor VX-765 reduces immune activation, CD4+ T cell depletion, viral load, and total HIV-1 DNA in HIV-1 infected humanized mice. *eLife*, 12, 2023.

[12195] Wenbin An, Maria Feola, Maayan Levy, Srinivas Aluri, Marc Ruiz-Martinez, Ashwin Sridharan, Eitan Fibach, Xiaofan Zhu, Amit Verma, and Yelena Ginzburg. Iron chelation improves ineffective erythropoiesis and iron overload in myelodysplastic syndrome mice. *eLife*, 12, 2023.

[12196] James T Anderson, Steven Henikoff, and Kami Ahmad. Chromosome-specific maturation of the epigenome in the Drosophila male germline. *eLife*, 12, 2023.

[12197] Nigel K Anderson, Doris Preininger, and Matthew J Fuxjager. Comment on 'Parasite defensive limb movements enhance acoustic signal attraction in male little torrent frogs'. *eLife*, 12, 2023.

[12198] Warren Anderson, Fariba Barahmand-pour Whitman, Peter S Linsley, Karen Cerosaletti, Jane H Buckner, and David J Rawlings. PTPN22 R620W gene editing in T cells enhances low-avidity TCR responses. *eLife*, 12, 2023.

[12199] Francesco Andreata, Marc Clément, Robert A Benson, Juliette Hadchouel, Emanuele Procopio, Guillaume Even, Julie Vorbe, Samira Benadda, Véronique Ollivier, Benoit Ho-Tin-Noe, Marie Le Borgne, Pasquale Maffia, Antonino Nicoletti, and Giuseppina Caligiuri. CD31 signaling promotes the detachment at the uropod of extravasating neutrophils allowing their migration to sites of inflammation. *eLife*, 12, 2023.

[12200] Lilya Andrianova, Steliana Yanakieva, Gabriella Margetts-Smith, Shivali Kohli, Erica S Brady, John P Aggleton, and Michael T Craig. No evidence from complementary data sources of a direct glutamatergic projection from the mouse anterior cingulate area to the hippocampal formation. *eLife*, 12, 2023.

[12201] Khai C Ang, Victor A Canfield, Tiffany C Foster, Thaddeus D Harbaugh, Kathryn A Early, Rachel L Harter, Katherine P Reid, Shou Ling Leong, Yuka Kawasawa, Dajiang Liu, John W Hawley, and Keith C Cheng. Native American genetic ancestry and pigmentation allele contributions to skin color in a Caribbean population. *eLife*, 12, 2023.

[12202] Juan M Angueyra, Vincent P Kunze, Laura K Patak, Hailey Kim, Katie Kindt, and Wei Li. Transcription factors underlying photoreceptor diversity. *eLife*, 12, 2023.

[12203] Asgar H Ansari, Manoj Kumar, Sajal Sarkar, Souvik Maiti, and Debojyoti Chakraborty. CriSNPr, a single interface for the curated and de novo design of gRNAs for CRISPR diagnostics using diverse Cas systems. *eLife*, 12, 2023.

[12204] Niki Anthoney, Lucy Tainton-Heap, Hang Luong, Eleni Notaras, Amber B Kewin, Qiongyi Zhao, Trent Perry, Philip Batterham, Paul J Shaw, and Bruno van Swinderen. Experimentally induced active and quiet sleep engage non-overlapping transcriptional programs in Drosophila. *eLife*, 12, 2023.

[12205] Rea L Antoniou-Kourounioti, Anis Meschichi, Svenja Reeck, Scott Berry, Govind Menon, Yusheng Zhao, John Fozard, Terri Holmes, Lihua Zhao, Huamei Wang, Matthew Hartley, Caroline Dean, Stefanie Rosa, and Martin Howard. Integrating analog and digital modes of gene expression at Arabidopsis FLC. *eLife*, 12, 2023.

[12206] Juan J Apiz Saab, Lindsey N Dzierozynski, Patrick B Jonker, Roya AminiTabrizi, Hardik Shah, Rosa Elena Menjivar, Andrew J Scott, Zeribe C Nwosu, Zhou Zhu, Riona N Chen, Moses Oh, Colin Sheehan, Daniel R Wahl, Marina Pasca di Magliano, Costas A Lyssiotis, Kay F Macleod, Christopher R Weber, and Alexander Muir. Pancreatic tumors exhibit myeloid-driven amino acid stress and upregulate arginine biosynthesis. *eLife*, 12, 2023.

[12207] Nicole Aponte Santiago, Stephanie Konecki, Lauren Porter, and Harvinder Virk. Tales of the unexpected. *eLife*, 12, 2023.

[12208] Alber Aqil, Leo Speidel, Pavlos Pavlidis, and Omer Gokcumen. Balancing selection on genomic deletion polymorphisms in humans. *eLife*, 12, 2023.

[12209] Hirotaka Araki, Shinjiro Hino, Kotaro Anan, Kanji Kuribayashi, Kan Etoh, Daiki Seko, Ryuta Takase, Kensaku Kohrogi, Yuko Hino, Yusuke Ono, Eiichi Araki, and Mitsuyoshi Nakao. LSD1 defines the fiber type-selective responsiveness to environmental stress in skeletal muscle. *eLife*, 12, 2023.

[12210] Raul Araya-Secchi, Katrine Bugge, Pernille Seiffert, Amalie Petry, Gitte W Haxholm, Kresten Lindorff-Larsen, Stine Falsig Pedersen, Lise Arleth, and Birthe B Kragelund. The prolactin receptor scaffolds Janus kinase 2 via co-structure formation with phosphoinositide-4,5-bisphosphate. *eLife*, 12, 2023.

[12211] Edwin Ardiansyah, Julian Avila-Pacheco, Le Thanh Hoang Nhat, Sofiati Dian, Dao Nguyen Vinh, Hoang Thanh Hai, Kevin Bullock, Bachti Alisjahbana, Mihai G Netea, Riwanti Estiasari, Trinh Thi Bich Tram, Joseph Donovan, Dorothee Heemskerk, Tran Thi Hong Chau, Nguyen Duc Bang, Ahmad Rizal Ganiem, Rovina Ruslami, Valerie ACM Koeken, Raph L Hamers, Darma Imran, Kartika Maharani, Vinod Kumar, Clary B Clish, Reinout van Crevel, Guy Thwaites, Arjan van Laarhoven, and Nguyen Thuy Thuong Thuong. Tryptophan metabolism determines outcome in tuberculous meningitis: a targeted metabolomic analysis. *eLife*, 12, 2023.

[12212] Sahab Arinrad, Constanze Depp, Sophie B Siems, Andrew Octavian Sasmita, Maria A Eichel, Anja Ronnenberg, Kurt Hammerschmidt, Katja A Lüders, Hauke B Werner, Hannelore Ehrenreich, and Klaus-Armin Nave. Isolated catatonia-like executive dysfunction in mice with forebrain-specific loss of myelin integrity. *eLife*, 12, 2023.

[12213] Daniel A Armendariz, Sean C Goetsch, Anjana Sundarrajan, Sushama Sivakumar, Yihan Wang, Shiqi Xie, Nikhil V Munshi, and Gary C Hon. CHD-associated enhancers shape human cardiomyocyte lineage commitment. *eLife*, 12, 2023.

[12214] Daniel A Armendariz, Anjana Sundarrajan, and Gary C Hon. Breaking enhancers to gain insights into developmental defects. *eLife*, 12, 2023.

[12215] Amandeep S Arora, Hsiang-Ling Huang, Ramanpreet Singh, Yoshie Narui, Andrejus Suchenko, Tomoyuki Hatano, Sarah M Heissler, Mohan K Balasubramanian, and Krishna Chinthalapudi. Structural insights into actin isoforms. *eLife*, 12, 2023.

[12216] Samantha DM Arras, Nellie Sibaeva, Ryan J Catchpole, Nobuyuki Horinouchi, Dayong Si, Alannah M Rickerby, Kengo Deguchi, Makoto Hibi, Koichi Tanaka, Michiki Takeuchi, Jun Ogawa, and Anthony M Poole. Characterisation of an Escherichia coli line that completely lacks ribonucleotide reduction yields insights into the evolution of parasitism and endosymbiosis. *eLife*, 12, 2023.

[12217] Diego M Arribas, Antonia Marin-Burgin, and Luis G Morelli. Adult-born granule cells improve stimulus encoding and discrimination in the dentate gyrus. *eLife*, 12, 2023.

[12218] Thiruma V Arumugam, Asfa Alli-Shaik, Elisa A Liehn, Sharmelee Selvaraji, Luting Poh, Vismitha Rajeev, Yoonsuk Cho, Yongeun Cho, Jongho Kim, Joonki Kim, Hannah LF Swa, David Tan Zhi Hao, Chutima Rattanasopa, David Yang-Wei Fann, David Castano Mayan, Gavin Yong-Quan Ng, Sang-Ha Baik, Karthik Mallilankaraman, Mathias Gelderblom, Grant R Drummond, Christopher G Sobey, Brian K Kennedy, Roshni R Singaraja, Mark P Mattson, Dong-Gyu Jo, and Jayantha Gunaratne. Multiomics analyses reveal dynamic bioenergetic pathways and functional remodeling of the heart during intermittent fasting. *eLife*, 12, 2023.

[12219] Takuma Asahi, Shinya Abe, Guangwei Cui, Akihiro Shimba, Tsukasa Nabekura, Hitoshi Miyachi, Satsuki Kitano, Keizo Ohira, Johannes M Dijkstra, Masaki Miyazaki, Akira Shibuya, Hiroshi Ohno, and Koichi Ikuta. Liver type 1 innate lymphoid cells lacking IL-7 receptor are a native killer cell subset fostered by parenchymal niches. *eLife*, 12, 2023.

[12220] Angelo Ascani, Sara Torstensson, Sanjiv Risal, Haojiang Lu, Gustaw Eriksson, Congru Li, Sabrina Teschl, Joana Menezes, Katalin Sandor, Claes Ohlsson, Camilla I Svensson, Mikael CI Karlsson, Martin Helmut Stradner, Barbara Obermayer-Pietsch, and Elisabet Stener-Victorin. The role of B cells in immune cell activation in polycystic ovary syndrome. *eLife*, 12, 2023.

[12221] Yoshinori Aso, Daichi Yamada, Daniel Bushey, Karen L Hibbard, Megan Sammons, Hideo Otsuna, Yichun Shuai, and Toshihide Hige. Neural circuit mechanisms for transforming learned olfactory valences into wind-oriented movement. *eLife*, 12, 2023.

[12222] Nanako Atsumi, Keiko Yasumatsu, Yuriko Takashina, Chiaki Ito, Norihisa Yasui, Robert F Margolskee, and Atsuko Yamashita. Chloride ions evoke taste sensations by binding to the extracellular ligand-binding domain of sweet/umami taste receptors. *eLife*, 12, 2023.

[12223] Amélie Aussel, Ian C Fiebelkorn, Sabine Kastner, Nancy J Kopell, and Benjamin Rafael Pittman-Polletta. Interacting rhythms enhance sensitivity of target detection in a fronto-parietal computational model of visual attention. *eLife*, 12, 2023.

[12224] Chiara Auwerx, Marie C Sadler, Tristan Woh, Alexandre Reymond, Zoltán Kutalik, and Eleonora Porcu. Exploiting the mediating role of the metabolome to unravel transcript-to-phenotype associations. *eLife*, 12, 2023.

[12225] Riham Ayoubi, Joel Ryan, Michael S Biddle, Walaa Alshafie, Maryam Fotouhi, Sara Gonzalez Bolivar, Vera Ruiz Moleon, Peter Eckmann, Donovan Worrall, Ian McDowell, Kathleen Southern, Wolfgang Reintsch, Thomas M Durcan, Claire Brown, Anita Bandrowski, Harvinder Virk, Aled M Edwards, Peter McPherson, and Carl Laflamme. Scaling of an antibody validation procedure enables quantification of antibody performance in major research applications. *eLife*, 12, 2023.

[12226] Arianne M Babina, Serhiy Surkov, Weihua Ye, Jon Jerlström-Hultqvist, Mårten Larsson, Erik Holmqvist, Per Jemth, Dan I Andersson, and Michael Knopp. Rescue of Escherichia coli auxotrophy by de novo small proteins. *eLife*, 12, 2023.

[12227] Ludovica Bachschmid-Romano, Nicholas G Hatsopoulos, and Nicolas Brunel. Interplay between external inputs and recurrent dynamics during movement preparation and execution in a network model of motor cortex. *eLife*, 12, 2023.

[12228] David J Bacsik, Bernadeta Dadonaite, Andrew Butler, Allison J Greaney, Nicholas S Heaton, and Jesse D Bloom. Influenza virus transcription and progeny production are poorly correlated in single cells. *eLife*, 12, 2023.

[12229] Brett C Baggett, Kevin R Murphy, Elif Sengun, Eric Mi, Yueming Cao, Nilufer N Turan, Yichun Lu, Lorraine Schofield, Tae Yun Kim, Anatoli Y Kabakov, Peter Bronk, Zhilin Qu, Patrizia Camelliti, Patrycja Dubielecka, Dmitry Terentyev, Federica del Monte, Bum-Rak Choi, John Sedivy, and Gideon Koren. Myofibroblast senescence promotes arrhythmogenic remodeling in the aged infarcted rabbit heart. *eLife*, 12, 2023.

[12230] Elmehdi Bahafid, Imke Bradtmöller, Ann M Thies, Thi TON Nguyen, Crisanto Gutierrez, Bénédicte Desvoyes, Yvonne Stahl, Ikram Blilou, and Rüdiger GW Simon. The Arabidopsis SHORTROOT network coordinates shoot apical meristem development with auxin-dependent lateral organ initiation. *eLife*, 12, 2023.

[12231] Amichai Baichman-Kass, Tingting Song, and Jonathan Friedman. Competitive interactions between culturable bacteria are highly non-additive. *eLife*, 12, 2023.

[12232] Sydney M Bailes, Daniel EP Gomez, Beverly Setzer, and Laura D Lewis. Resting-state fMRI signals contain spectral signatures of local hemodynamic response timing. *eLife*, 12, 2023.

[12233] Sameer S Bajikar, Ashley G Anderson, Jian Zhou, Mark A Durham, Alexander J Trostle, Ying-Wooi Wan, Zhandong Liu, and Huda Y Zoghbi. MeCP2 regulates Gdf11, a dosage-sensitive gene critical for neurological function. *eLife*, 12, 2023.

[12234] Zachary Baker, Molly Przeworski, and Guy Sella. Down the Penrose stairs, or how selection for fewer recombination hotspots maintains their existence. *eLife*, 12, 2023.

[12235] Katarína Balážová, Hans Clevers, and Antonella FM Dost. The role of macrophages in non-small cell lung cancer and advancements in 3D co-cultures. *eLife*, 12, 2023.

[12236] Valentina Baldazzi, Delphine Ropers, Jean-Luc Gouzé, Tomas Gedeon, and Hidde de Jong. Resource allocation accounts for the large variability of rate-yield phenotypes across bacterial strains. *eLife*, 12, 2023.

[12237] Amy Louise Ball, Carol E Jolly, Mark G Lennon, Jonathan J Lyon, Ana Alfirevic, and Amy E Chadwick. The generation of HepG2 transmitochondrial cybrids to reveal the role of mitochondrial genotype in idiosyncratic drug-induced liver injury. *eLife*, 12, 2023.

[12238] Georgia Balsevich, Gavin N Petrie, Daniel E Heinz, Arashdeep Singh, Robert J Aukema, Avery C Hunker, Haley A Vecchiarelli, Hiulan Yau, Martin Sticht, Roger J Thompson, Francis S Lee, Larry S Zweifel, Prasanth K Chelikani, Nils C Gassen, and Matthew N Hill. A genetic variant of fatty acid amide hydrolase (FAAH) exacerbates hormone-mediated orexigenic feeding in mice. *eLife*, 12, 2023.

[12239] Lior Baltiansky, Guy Frankel, and Ofer Feinerman. Emergent regulation of ant foraging frequency through a computationally inexpensive forager movement rule. *eLife*, 12, 2023.

[12240] Rachel A Banks, Vahe Galstyan, Heun Jin Lee, Soichi Hirokawa, Athena Ierokomos, Tyler D Ross, Zev Bryant, Matt Thomson, and Rob Phillips. Motor processivity and speed determine structure and dynamics of microtubule-motor assemblies. *eLife*, 12, 2023.

[12241] David Bann, Liam Wright, Neil M Davies, and Vanessa Moulton. Weakening of the cognition and height association from 1957 to 2018: Findings from four British birth cohort studies. *eLife*, 12, 2023.

[12242] Mengjing Bao, Ruth E Dörig, Paula Maria Vazquez-Pianzola, Dirk Beuchle, and Beat Suter. Differential modification of the C-terminal tails of different $\alpha$-tubulins and their importance for microtubule function in vivo. *eLife*, 12, 2023.

[12243] Noa Barak-Gavish, Bareket Dassa, Constanze Kuhlisch, Inbal Nussbaum, Alexander Brandis, Gili Rosenberg, Roi Avraham, and Assaf Vardi. Bacterial lifestyle switch in response to algal metabolites. *eLife*, 12, 2023.

[12244] Natasja Barki, Laura Jenkins, Sara Marsango, Domonkos Dedeo, Daniele Bolognini, Louis Dwomoh, Aisha M Abdelmalik, Margaret Nilsen, Manon Stoffels, Falko Nagel, Stefan Schulz, Andrew B Tobin, and Graeme Milligan. Phosphorylation bar-coding of free fatty acid receptor 2 is generated in a tissue-specific manner. *eLife*, 12, 2023.

[12245] Ida L Barlow, Eirinn Mackay, Emily Wheater, Aimee Goel, Sumi Lim, Steve Zimmerman, Ian Woods, David A Prober, and Jason Rihel. The zebrafish mutant dreammist implicates sodium homeostasis in sleep regulation. *eLife*, 12, 2023.

[12246] Jose Gabriel Nino Barreat and Aris Katzourakis. A billion years arms-race between viruses, virophages, and eukaryotes. *eLife*, 12, 2023.

[12247] Joseph L Basalla, Claudia A Mak, Jordan A Byrne, Maria Ghalmi, Y Hoang, and Anthony G Vecchiarelli. Dissecting the phase separation and oligomerization activities of the carboxysome positioning protein McdB. *eLife*, 12, 2023.

[12248] Elizaveta V Bashkirova, Nell Klimpert, Kevin Monahan, Christine E Campbell, Jason Osinski, Longzhi Tan, Ira Schieren, Ariel Pourmorady, Beka Stecky, Gilad Barnea, Xiaoliang Sunney Xie, Ishmail Abdus-Saboor, Benjamin M Shykind, Bianca J Marlin, Richard M Gronostajski, Alexander Fleischmann, and Stavros Lomvardas. Opposing, spatially-determined epigenetic forces impose restrictions on stochastic olfactory receptor choice. *eLife*, 12, 2023.

[12249] Mohsen Basiri, Mohammad E Ghaffari, Jiapeng Ruan, Vagishwari Murugesan, Nathaniel Kleytman, Glenn Belinsky, Amir Akhavan, Andrew Lischuk, Lilu Guo, Katherine Klinger, and Pramod K Mistry. Osteonecrosis in Gaucher disease in the era of multiple therapies: Biomarker set for risk stratification from a tertiary referral center. *eLife*, 12, 2023.

[12250] Jordan T Bateman and Erica S Levitt. Opioid suppression of an excitatory pontomedullary respiratory circuit by convergent mechanisms. *eLife*, 12, 2023.

[12251] Sophie Bavard and Stefano Palminteri. The functional form of value normalization in human reinforcement learning. *eLife*, 12, 2023.

[12252] Sion C Bayliss, Rebecca K Locke, Claire Jenkins, Marie Anne Chattaway, Timothy J Dallman, and Lauren A Cowley. Rapid geographical source attribution of Salmonella enterica serovar Enteritidis genomes using hierarchical machine learning. *eLife*, 12, 2023.

[12253] Amélie Beaudet and Edwin de Jager. Broca's area, variation and taxic diversity in early Homo from Koobi Fora (Kenya). *eLife*, 12, 2023.

[12254] Kristina B Beck, Ben C Sheldon, and Josh A Firth. Social learning mechanisms shape transmission pathways through replicate local social networks of wild birds. *eLife*, 12, 2023.

[12255] Florian Becker, Simon Fuchs, Lukas Refisch, Friedel Drepper, Wolfgang Bildl, Uwe Schulte, Shuo Liang, Jonas Immanuel Heinicke, Sierra C Hansen, Clemens Kreutz, Bettina Warscheid, Bernd Fakler, Evgeny V Mymrikov, and Carola Hunte. Conformational regulation and target-myristoyl switch of calcineurin B homologous protein 3. *eLife*, 12, 2023.

[12256] Patrick Becker, Fiona Naughton, Deborah Brotherton, Raul Pacheco-Gomez, Oliver Beckstein, and Alexander D Cameron. Mechanism of substrate binding and transport in BASS transporters. *eLife*, 12, 2023.

[12257] Claire N Bedbrook, Ravi D Nath, Rahul Nagvekar, Karl Deisseroth, and Anne Brunet. Rapid and precise genome engineering in a naturally short-lived vertebrate. *eLife*, 12, 2023.

[12258] Celine Bellegarda, Guillaume Zavard, Lionel Moisan, FranÇoise Brochard-Wyart, Jean-FranÇois Joanny, Ryan S Gray, Yasmine Cantaut-Belarif, and Claire Wyart. The Reissner fiber under tension in vivo shows dynamic interaction with ciliated cells contacting the cerebrospinal fluid. *eLife*, 12, 2023.

[12259] Tal Ben Yaakov, Tanya Wasserman, Eliel Aknin, and Yonatan Savir. Single-cell analysis of the aged ovarian immune system reveals a shift towards adaptive immunity and attenuated cell function. *eLife*, 12, 2023.

[12260] Lucas Benjamin, Ana Fló, Fosca Al Roumi, and Ghislaine Dehaene-Lambertz. Humans parsimoniously represent auditory sequences by pruning and completing the underlying network structure. *eLife*, 12, 2023.

[12261] Richard Benton and Nathaniel J Himmel. Structural screens identify candidate human homologs of insect chemoreceptors and cryptic Drosophila gustatory receptor-like proteins. *eLife*, 12, 2023.

[12262] Cory A Berger and Ann M Tarrant. Sensory conflict disrupts circadian rhythms in the sea anemone Nematostella vectensis. *eLife*, 12, 2023.

[12263] Andreas Berghaenel, Jeroen MG Stevens, Gottfried Hohmann, Tobias Deschner, and Verena Behringer. Evidence for adolescent length growth spurts in bonobos and other primates highlights the importance of scaling laws. *eLife*, 12, 2023.

[12264] Alexander Berne, Tom Zhang, Joseph Shomar, Anggie J Ferrer, Aaron Valdes, Tomoko Ohyama, and Mason Klein. Mechanical vibration patterns elicit behavioral transitions and habituation in crawling Drosophila larvae. *eLife*, 12, 2023.

[12265] Cecile Berne, Sébastien Zappa, and Yves V Brun. eDNA-stimulated cell dispersion from Caulobacter crescentus biofilms upon oxygen limitation is dependent on a toxin–antitoxin system. *eLife*, 12, 2023.

[12266] Martin H Berryer, Gizem Rizki, Anna Nathanson, Jenny A Klein, Darina Trendafilova, Sara G Susco, Daisy Lam, Angelica Messana, Kristina M Holton, Kyle W Karhohs, Beth A Cimini, Kathleen Pfaff, Anne E Carpenter, Lee L Rubin, and Lindy E Barrett. High-content synaptic phenotyping in human cellular models reveals a role for BET proteins in synapse assembly. *eLife*, 12, 2023.

[12267] Carsten Beta, Leah Edelstein-Keshet, Nir Gov, and Arik Yochelis. From actin waves to mechanism and back: How theory aids biological understanding. *eLife*, 12, 2023.

[12268] Anshuman Bhanja, Margaret K Seeley-Fallen, Michelle Lazzaro, Arpita Upadhyaya, and Wenxia Song. N-WASP-dependent branched actin polymerization attenuates B-cell receptor signaling by increasing the molecular density of receptor clusters. *eLife*, 12, 2023.

[12269] Shreyas Bhat, Ali El-Kasaby, Ameya Kasture, Danila Boytsov, Julian B Reichelt, Thomas Hummel, Sonja Sucic, Christian Pifl, Michael Freissmuth, and Walter Sandtner. A mechanism of uncompetitive inhibition of the serotonin transporter. *eLife*, 12, 2023.

[12270] Rahul Bhattacharjee, Aaron R Hall, MariaSanta C Mangione, Maya G Igarashi, Rachel H Roberts-Galbraith, Jun-Song Chen, Dimitrios Vavylonis, and Kathleen L Gould. Multiple polarity kinases inhibit phase separation of F-BAR protein Cdc15 and antagonize cytokinetic ring assembly in fission yeast. *eLife*, 12, 2023.

[12271] Vikrant K Bhosle, Chunxiang Sun, Sajedabanu Patel, Tse Wing Winnie Ho, Johannes Westman, Dustin A Ammendolia, Fatemeh Mirshafiei Langari, Noah Fine, Nicole Toepfner, Zhubing Li, Manraj Sharma, Judah Glogauer, Mariana I Capurro, Nicola L Jones, Jason T Maynes, Warren L Lee, Michael Glogauer, Sergio Grinstein, and Lisa A Robinson. The chemorepellent, SLIT2, bolsters innate immunity against Staphylococcus aureus. *eLife*, 12, 2023.

[12272] Xiling Bian, Jiemin Zhu, Xiaobo Jia, Wenjun Liang, Sihan Yu, Zhiqiang Li, Wenxia Zhang, and Yi Rao. Suggestion of creatine as a new neurotransmitter by approaches ranging from chemical analysis and biochemistry to electrophysiology. *eLife*, 12, 2023.

[12273] Flavie Bidel, Yaron Meirovitch, Richard Lee Schalek, Xiaotang Lu, Elisa Catherine Pavarino, Fuming Yang, Adi Peleg, Yuelong Wu, Tal Shomrat, Daniel Raimund Berger, Adi Shaked, Jeff William Lichtman, and Binyamin Hochner. Connectomics of the Octopus vulgaris vertical lobe provides insight into conserved and novel principles of a memory acquisition network. *eLife*, 12, 2023.

[12274] Merve Bilgic, Quan Wu, Taeko Suetsugu, Atsunori Shitamukai, Yuji Tsunekawa, Tomomi Shimogori, Mitsutaka Kadota, Osamu Nishimura, Shigehiro Kuraku, Hiroshi Kiyonari, and Fumio Matsuzaki. Truncated radial glia as a common precursor in the late corticogenesis of gyrencephalic mammals. *eLife*, 12, 2023.

[12275] Katja Birker, Shuchao Ge, Natalie J Kirkland, Jeanne L Theis, James Marchant, Zachary C Fogarty, Maria A Missinato, Sreehari Kalvakuri, Paul Grossfeld, Adam J Engler, Karen Ocorr, Timothy J Nelson, Alexandre R Colas, Timothy M Olson, Georg Vogler, and Rolf Bodmer. Mitochondrial MICOS complex genes, implicated in hypoplastic left heart syndrome, maintain cardiac contractility and actomyosin integrity. *eLife*, 12, 2023.

[12276] Lasse M Blaabjerg, Maher M Kassem, Lydia L Good, Nicolas Jonsson, Matteo Cagiada, Kristoffer E Johansson, Wouter Boomsma, Amelie Stein, and Kresten Lindorff-Larsen. Rapid protein stability prediction using deep learning representations. *eLife*, 12, 2023.

[12277] Holly H Black, Jessica L Hanson, Julia E Roberts, Shannon N Leslie, Will Campodonico, Christopher C Ebmeier, G Aaron Holling, Jian Wei Tay, Autumn M Matthews, Elizabeth Ung, Cristina I Lau, and Alexandra M Whiteley. UBQLN2 restrains the domesticated retrotransposon PEG10 to maintain neuronal health in ALS. *eLife*, 12, 2023.

[12278] Garrett J Blair, Changliang Guo, Shiyun Wang, Michael S Fanselow, Peyman Golshani, Daniel Aharoni, and Hugh T Blair. Hippocampal place cell remapping occurs with memory storage of aversive experiences. *eLife*, 12, 2023.

[12279] Oscar Blanch-Lombarte, Dan Ouchi, Esther Jimenez-Moyano, Julieta Carabelli, Miguel Angel Marin, Ruth Peña, Adam Pelletier, Aarthi Talla, Ashish Sharma, Judith Dalmau, José Ramón Santos, Rafick-Pierre Sékaly, Bonaventura Clotet, and Julia G Prado. Selective loss of CD107a TIGIT+ memory HIV-1-specific CD8+ T cells in PLWH over a decade of ART. *eLife*, 12, 2023.

[12280] Jadna Bogado Lopes, Anna N Senko, Klaas Bahnsen, Daniel Geisler, Eugene Kim, Michel Bernanos, Diana Cash, Stefan Ehrlich, Anthony C Vernon, and Gerd Kempermann. Individual behavioral trajectories shape whole-brain connectivity in mice. *eLife*, 12, 2023.

[12281] Katarzyna Bogucka-Janczi, Gregory Harms, Marie-May Coissieux, Mohamed Bentires-Alj, Bernd Thiede, and Krishnaraj Rajalingam. ERK3/MAPK6 dictates CDC42/RAC1 activity and ARP2/3-dependent actin polymerization. *eLife*, 12, 2023.

[12282] Briana M Bohannon, Jessica J Jowais, Leif Nyberg, Vanessa Olivier-Meo, Valentina Corradi, D Peter Tieleman, Sara I Liin, and H Peter Larsson. Mechanistic insights into robust cardiac IKs potassium channel activation by aromatic polyunsaturated fatty acid analogues. *eLife*, 12, 2023.

[12283] Christopher H Bohrer and Daniel R Larson. Synthetic analysis of chromatin tracing and live-cell imaging indicates pervasive spatial coupling between genes. *eLife*, 12, 2023.

[12284] Tom Boissonnet, Matteo Tripodi, and Hiroki Asari. Awake responses suggest inefficient dense coding in the mouse retina. *eLife*, 12, 2023.

[12285] Ashley C Bolte, Daniel A Shapiro, Arun B Dutta, Wei Feng Ma, Katherine R Bruch, Michael A Kovacs, Ana Royo Marco, Hannah E Ennerfelt, and John R Lukens. The meningeal transcriptional response to traumatic brain injury and aging. *eLife*, 12, 2023.

[12286] David Bolumar, Javier Moncayo-Arlandi, Javier Gonzalez-Fernandez, Ana Ochando, Inmaculada Moreno, Ana Monteagudo-Sanchez, Carlos Marin, Antonio Diez, Paula Fabra, Miguel Angel Checa, Juan Jose Espinos, David K Gardner, Carlos Simon, and Felipe Vilella. Vertical transmission of maternal DNA through extracellular vesicles associates with altered embryo bioenergetics during the periconception period. *eLife*, 12, 2023.

[12287] Tyler Bonnen and Mark AG Eldridge. Inconsistencies between human and macaque lesion data can be resolved with a stimulus-computable model of the ventral visual stream. *eLife*, 12, 2023.

[12288] Caroline Bonnet, Justine Charpentier, Natacha Retailleau, Daniel Choquet, and FranÇoise Coussen. Regulation of different phases of AMPA receptor intracellular transport by 4.1N and SAP97. *eLife*, 12, 2023.

[12289] Jacopo Bono, Sara Zannone, Victor Pedrosa, and Claudia Clopath. Learning predictive cognitive maps with spiking neurons during behavior and replays. *eLife*, 12, 2023.

[12290] Mandy Y Boontanrart, Elia Mächler, Simone Ponta, Jan C Nelis, Viviana G Preiano, and Jacob E Corn. Engineering of the endogenous HBD promoter increases HbA2. *eLife*, 12, 2023.

[12291] Bavat Bornstein, Lia Heinemann-Yerushalmi, Sharon Krief, Ruth Adler, Bareket Dassa, Dena Leshkowitz, Minchul Kim, Guy Bewick, Robert W Banks, and Elazar Zelzer. Molecular characterization of the intact mouse muscle spindle using a multi-omics approach. *eLife*, 12, 2023.

[12292] Fernanda M Bosada, Karel van Duijvenboden, Alexandra E Giovou, Mathilde R Rivaud, Jae-Sun Uhm, Arie O Verkerk, Bastiaan J Boukens, and Vincent M Christoffels. An atrial fibrillation-associated regulatory region modulates cardiac Tbx5 levels and arrhythmia susceptibility. *eLife*, 12, 2023.

[12293] Christopher T Boughter and Martin Meier-Schellersheim. Conserved biophysical compatibility among the highly variable germline-encoded regions shapes TCR-MHC interactions. *eLife*, 12, 2023.

[12294] Manon Boulet, Guerric Gilbert, Yoan Renaud, Martina Schmidt-Dengler, Emilie Plantié, Romane Bertrand, Xinsheng Nan, Tomasz Jurkowski, Mark Helm, Laurence Vandel, and Lucas Waltzer. Adenine methylation is very scarce in the Drosophila genome and not erased by the ten-eleven translocation dioxygenase. *eLife*, 12, 2023.

[12295] Kieran A Boyle, Erika Polgar, Maria Gutierrez-Mecinas, Allen C Dickie, Andrew H Cooper, Andrew M Bell, Evelline Jumolea, Adrian Casas-Benito, Masahiko Watanabe, David I Hughes, Gregory A Weir, John S Riddell, and Andrew J Todd. Neuropeptide Y-expressing dorsal horn inhibitory interneurons gate spinal pain and itch signalling. *eLife*, 12, 2023.

[12296] Philip Bradley. Structure-based prediction of T cell receptor:peptide-MHC interactions. *eLife*, 12, 2023.

[12297] Sean M Braet, Theresa SC Buckley, Varun Venkatakrishnan, Kim-Marie A Dam, Pamela J Bjorkman, and Ganesh S Anand. Timeline of changes in spike conformational dynamics in emergent SARS-CoV-2 variants reveal progressive stabilization of trimer stalk with altered NTD dynamics. *eLife*, 12, 2023.

[12298] Anke Braun and Tobias H Donner. Adaptive biasing of action-selective cortical build-up activity by stimulus history. *eLife*, 12, 2023.

[12299] Barbara Bravi, Andrea Di Gioacchino, Jorge Fernandez-de Cossio-Diaz, Aleksandra M Walczak, Thierry Mora, Simona Cocco, and Rémi Monasson. A transfer-learning approach to predict antigen immunogenicity and T-cell receptor specificity. *eLife*, 12, 2023.

[12300] Ignacio Bravo-Plaza, Victor G Tagua, Herbert N Arst, Ana Alonso, Mario Pinar, Begoña Monterroso, Antonio Galindo, and Miguel A Peñalva. The Uso1 globular head interacts with SNAREs to maintain viability even in the absence of the coiled-coil domain. *eLife*, 12, 2023.

[12301] Melanie N Brien, Anna Orteu, Eugenie C Yen, Juan A Galarza, Jimi Kirvesoja, Hannu Pakkanen, Kazumasa Wakamatsu, Chris D Jiggins, and Johanna Mappes. Colour polymorphism associated with a gene duplication in male wood tiger moths. *eLife*, 12, 2023.

[12302] Emma M Briggs, Catarina A Marques, Guy R Oldrieve, Jihua Hu, Thomas D Otto, and Keith R Matthews. Profiling the bloodstream form and procyclic form Trypanosoma brucei cell cycle using single-cell transcriptomics. *eLife*, 12, 2023.

[12303] Jennifer K Briggs, Anne Gresch, Isabella Marinelli, JaeAnn M Dwulet, David J Albers, Vira Kravets, and Richard KP Benninger. $\beta$-cell intrinsic dynamics rather than gap junction structure dictates subpopulations in the islet functional network. *eLife*, 12, 2023.

[12304] Michele Brischigliaro, Alfredo Cabrera-Orefice, Susanne Arnold, Carlo Viscomi, Massimo Zeviani, and Erika Fernández-Vizarra. Structural rather than catalytic role for mitochondrial respiratory chain supercomplexes. *eLife*, 12, 2023.

[12305] DuŠica BriŠevac, Carolina M Peralta, and Tobias S Kaiser. An oligogenic architecture underlying ecological and reproductive divergence in sympatric populations. *eLife*, 12, 2023.

[12306] Léa Broca-Brisson, Rania Harati, Clémence Disdier, Orsolya Mozner, Romane Gaston-Breton, Auriane Maïza, Narciso Costa, Anne-Cécile Guyot, Balazs Sarkadi, Agota Apati, Matthew R Skelton, Lucie Madrange, Frank Yates, Jean Armengaud, Rifat Hamoudi, and Aloïse Mabondzo. Deciphering neuronal deficit and protein profile changes in human brain organoids from patients with creatine transporter deficiency. *eLife*, 12, 2023.

[12307] Rebeca Brocal-Ruiz, Ainara Esteve-Serrano, Carlos Mora-Martínez, Maria Luisa Franco-Rivadeneira, Peter Swoboda, Juan J Tena, MarÇal Vilar, and Nuria Flames. Forkhead transcription factor FKH-8 cooperates with RFX in the direct regulation of sensory cilia in Caenorhabditis elegans. *eLife*, 12, 2023.

[12308] Christian Brodbeck, Proloy Das, Marlies Gillis, Joshua P Kulasingham, Shohini Bhattasali, Phoebe Gaston, Philip Resnik, and Jonathan Z Simon. Eelbrain, a Python toolkit for time-continuous analysis with temporal response functions. *eLife*, 12, 2023.

[12309] Brigitte M Browe, Ying-Jie Peng, Jayasri Nanduri, Nanduri R Prabhakar, and Alfredo J Garcia. Gasotransmitter modulation of hypoglossal motoneuron activity. *eLife*, 12, 2023.

[12310] Nora C Brown, Benjamin Gordon, Caitlin E McDonough-Goldstein, Snigdha Misra, Geoffrey D Findlay, Andrew G Clark, and Mariana Federica Wolfner. The seminal odorant binding protein Obp56g is required for mating plug formation and male fertility in Drosophila melanogaster. *eLife*, 12, 2023.

[12311] Tanya L Brown, Emma C Horton, Evan W Craig, Camille EA Goo, Erik C Black, Madeleine N Hewitt, Nathaniel G Yee, Everett T Fan, David W Raible, and Jeffrey P Rasmussen. Dermal appendage-dependent patterning of zebrafish atoh1a+ Merkel cells. *eLife*, 12, 2023.

[12312] Natalia S Brunetti, Gustavo G Davanzo, Diogo de Moraes, Allan JR Ferrari, Gabriela F Souza, Stéfanie Primon Muraro, Thiago L Knittel, Vinicius O Boldrini, Lauar B Monteiro, João Victor Virgílio-da Silva, Gerson S Profeta, Natália S Wassano, Luana Nunes Santos, Victor C Carregari, Artur HS Dias, Flavio P Veras, Lucas A Tavares, Julia Forato, Icaro MS Castro, Lícia C Silva-Costa, André C Palma, Eli Mansour, Raisa G Ulaf, Ana F Bernardes, Thyago A Nunes, Luciana C Ribeiro, Marcus V

Agrela, Maria Luiza Moretti, Lucas I Buscaratti, Fernanda Crunfli, Raissa G Ludwig, Jaqueline A Gerhardt, Natália Munhoz-Alves, Ana Maria Marques, Renata Sesti-Costa, Mariene R Amorim, Daniel A Toledo-Teixeira, Pierina Lorencini Parise, Matheus Cavalheiro Martini, Karina Bispos-dos Santos, Camila L Simeoni, Fabiana Granja, Virgínia C Silvestrini, Eduardo B de Oliveira, Vitor M Faca, Murilo Carvalho, Bianca G Castelucci, Alexandre B Pereira, Laís D Coimbra, Marieli MG Dias, Patricia B Rodrigues, Arilson Bernardo SP Gomes, Fabricio B Pereira, Leonilda MB Santos, Louis-Marie Bloyet, Spencer Stumpf, Marjorie C Pontelli, Sean Whelan, Andrei C Sposito, Robson F Carvalho, André S Vieira, Marco AR Vinolo, André Damasio, Licio Velloso, Ana Carolina M Figueira, Luis LP da Silva, Thiago Mattar Cunha, Helder I Nakaya, Henrique Marques-Souza, Rafael E Marques, Daniel Martins-de Souza, Munir S Skaf, Jose Luiz Proenca-Modena, Pedro MM Moraes-Vieira, Marcelo A Mori, and Alessandro S Farias. SARS-CoV-2 uses CD4 to infect T helper lymphocytes. *eLife*, 12, 2023.

[12313] Clément Brunner, Gabriel Montaldo, and Alan Urban. Functional ultrasound imaging of stroke in awake rats. *eLife*, 12, 2023.

[12314] Peter Bruun-Rasmussen, Morten Hanefeld Dziegiel, Karina Banasik, Pär Ingemar Johansson, and Søren Brunak. Associations of ABO and Rhesus D blood groups with phenome-wide disease incidence: A 41-year retrospective cohort study of 482,914 patients. *eLife*, 12, 2023.

[12315] Jiawen Bu, Yixiao Zhang, Nan Niu, Kewei Bi, Lisha Sun, Xinbo Qiao, Yimin Wang, Yinan Zhang, Xiaofan Jiang, Dan Wang, Qingtian Ma, Huajun Li, and Caigang Liu. Dalpiciclib partially abrogates ER signaling activation induced by pyrotinib in HER2+HR+ breast cancer. *eLife*, 12, 2023.

[12316] Christian Büchel. The role of expectations, control and reward in the development of pain persistence based on a unified model. *eLife*, 12, 2023.

[12317] Eva Budinská, Martina Hrivňáková, Tina Catela Ivkovic, Marie Madrzyk, Rudolf Nenutil, Beatrix Bencsiková, Dagmar Al Tukmachi, Michaela Ručková, Lenka Zdražilová Dubská, Ondřej Slabý, Josef Feit, Mihnea-Paul Dragomir, Petra Borilova Linhartova, Sabine Tejpar, and Vlad Popovici. Molecular portraits of colorectal cancer morphological regions. *eLife*, 12, 2023.

[12318] Danielle B Buglak, Pauline Bougaran, Molly R Kulikauskas, Ziqing Liu, Elizabeth Monaghan-Benson, Ariel L Gold, Allison P Marvin, Andrew Burciu, Natalie T Tanke, Morgan Oatley, Shea N Ricketts, Karina Kinghorn, Bryan N Johnson, Celia E Shiau, Stephen Rogers, Christophe Guilluy, and Victoria L Bautch. Nuclear SUN1 stabilizes endothelial cell junctions via microtubules to regulate blood vessel formation. *eLife*, 12, 2023.

[12319] Caroline Bull, Emma Hazelwood, Joshua A Bell, Vanessa Tan, Andrei-Emil Constantinescu, Carolina Borges, Danny Legge, Kimberley Burrows, Jeroen R Huyghe, Hermann Brenner, Sergi Castellvi-Bel, Andrew T Chan, Sun-Seog Kweon, Loic Le Marchand, Li Li, Iona Cheng, Rish K Pai, Jane C Figueiredo, Neil Murphy, Marc J Gunter, Nicholas J Timpson, and Emma E Vincent. Identifying metabolic features of colorectal cancer liability using Mendelian randomization. *eLife*, 12, 2023.

[12320] Maya L Bulos, Edyta M Grzelak, Chloris Li-Ma, Emily Chen, Mitchell Hull, Kristen A Johnson, and Michael J Bollong. Pharmacological inhibition of CLK2 activates YAP by promoting alternative splicing of AMOTL2. *eLife*, 12, 2023.

[12321] Lydie Burglen, Evelien Van Hoeymissen, Leila Qebibo, Magalie Barth, Newell Belnap, Felix Boschann, Christel Depienne, Katrien De Clercq, Andrew GL Douglas, Mark P Fitzgerald, Nicola Foulds, Catherine Garel, Ingo Helbig, Katharina Held, Denise Horn, Annelies Janssen, Angela M Kaindl, Vinodh Narayanan, Christina Prager, Mailys Rupin-Mas, Alexandra Afenjar, Siyuan Zhao, Vincent Th Ramaekers, Sarah M Ruggiero, Simon Thomas, Stéphanie Valence, Lionel Van Maldergem, Tibor Rohacs, Diana Rodriguez, David Dyment, Thomas Voets, and Joris Vriens. Gain-of-function variants in the ion channel gene TRPM3 underlie a spectrum of neurodevelopmental disorders. *eLife*, 12, 2023.

[12322] Annamarie E Bustion, Renuka R Nayak, Ayushi Agrawal, Peter J Turnbaugh, and Katherine S Pollard. SIMMER employs similarity algorithms to accurately identify human gut microbiome species and enzymes capable of known chemical transformations. *eLife*, 12, 2023.

[12323] Roberta Cacioppo, Hesna Begum Akman, Taner Tuncer, Ayse Elif Erson-Bensan, and Catherine Lindon. Differential translation of mRNA isoforms underlies oncogenic activation of cell cycle kinase Aurora A. *eLife*, 12, 2023.

[12324] Ana J Caetano, Yushi Redhead, Farah Karim, Pawan Dhami, Shichina Kannambath, Rosamond Nuamah, Ana A Volponi, Luigi Nibali, Veronica Booth, Eleanor M D'Agostino, and Paul T Sharpe. Spatially resolved transcriptomics reveals pro-inflammatory fibroblast involved in lymphocyte recruitment through CXCL8 and CXCL10. *eLife*, 12, 2023.

[12325] Cori K Cahoon, Colette M Richter, Amelia E Dayton, and Diana E Libuda. Sexual dimorphic regulation of recombination by the synaptonemal complex in C. elegans. *eLife*, 12, 2023.

[12326] Marco Caligaris, Raffaele Nicastro, Zehan Hu, Farida Tripodi, Johannes Erwin Hummel, Benjamin Pillet, Marie-Anne Deprez, Joris Winderickx, Sabine Rospert, Paola Coccetti, Jörn Dengjel, and Claudio De Virgilio. Snf1/AMPK fine-tunes TORC1 signaling in response to glucose starvation. *eLife*, 12, 2023.

[12327] Francesco Camaglia, Arie Ryvkin, Erez Greenstein, Shlomit Reich-Zeliger, Benny Chain, Thierry Mora, Aleksandra M Walczak, and Nir Friedman. Quantifying changes in the T cell receptor repertoire during thymic development. *eLife*, 12, 2023.

[12328] Diego Candia-Rivera, Federico Raimondo, Pauline Pérez, Lionel Naccache, Catherine Tallon-Baudry, and Jacobo D Sitt. Conscious processing of global and local auditory irregularities causes differentiated heartbeat-evoked responses. *eLife*, 12, 2023.

[12329] Luyang Cao, Lixiang Ma, Juan Zhao, Xiangyu Wang, Xinzou Fang, Wei Li, Yawen Qi, Yingkui Tang, Jieya Liu, Shengxian Peng, Li Yang, Liangxue Zhou, Li Li, Xiaobo Hu, Yuan Ji, Yingyong Hou, Yi Zhao, Xianming Zhang, You-yang Zhao, Yong Zhao, Yuquan Wei, Asrar B Malik, Hexige Saiyin, and Jingsong Xu. An unexpected role of neutrophils in clearing apoptotic hepatocytes in vivo. *eLife*, 12, 2023.

[12330] Megan E Carey, Zoe A Dyson, Danielle J Ingle, Afreenish Amir, Mabel K Aworh, Marie Anne Chattaway, Ka Lip Chew, John A Crump, Nicholas A Feasey, Benjamin P Howden, Karen H Keddy, Mailis Maes, Christopher M Parry, Sandra Van Puyvelde, Hattie E Webb, Ayorinde Oluwatobiloba Afolayan, Anna P Alexander, Shalini Anandan, Jason R Andrews, Philip M Ashton, Buddha Basnyat, Ashish Bavdekar, Isaac I Bogoch, John D Clemens, Kesia Esther da Silva, Anuradha De, Joep de Ligt, Paula Lucia Diaz Guevara, Christiane Dolecek, Shanta Dutta, Marthie M Ehlers, Louise Francois Watkins, Denise O Garrett, Gauri Godbole, Melita A Gordon, Andrew R Greenhill, Chelsey Griffin, Madhu Gupta, Rene S Hendriksen, Robert S Heyderman, Yogesh Hooda, Juan Carlos Hormazabal, Odion O Ikhimiukor, Junaid Iqbal, Jobin John Jacob, Claire Jenkins, Dasaratha Ramaiah Jinka, Jacob John, Gagandeep Kang, Abdoulie Kanteh, Arti Kapil, Abhilasha Karkey, Samuel Kariuki, Robert A Kingsley, Roshine Mary Koshy, AC Lauer, Myron M Levine, Ravikumar Kadahalli Lingegowda, Stephen P Luby, Grant Austin Mackenzie, Tapfumanei Mashe, Chisomo Msefula, Ankur Mutreja, Geetha Nagaraj, Savitha Nagaraj, Satheesh Nair, Take K Naseri, Susana Nimarota-Brown, Elisabeth Njamkepo, Iruka N Okeke, Sulochana Putli Bai Perumal, Andrew J Pollard, Agila Kumari Pragasam, Firdausi Qadri, Farah N Qamar, Sadia Isfat Ara Rahman, Savitra Devi Rambocus, David A Rasko, Pallab Ray, Roy Robins-Browne, Temsunaro Rongsen-Chandola, Jean Pierre Rutanga, Samir K Saha, Senjuti Saha, Karnika Saigal, Mohammad Saiful Islam Sajib, Jessica C Seidman, Jivan Shakya, Varun Shamanna, Jayanthi Shastri, Rajeev Shrestha, Sonia Sia, Michael J Sikorski, Ashita Singh, Anthony M Smith, Kaitlin A Tagg, Dipesh Tamrakar, Arif Mohammed Tanmoy, Maria Thomas, Mathew S Thomas, Robert Thomsen, Nicholas R Thomson, Siaosi Tupua, Krista Vaidya, Mary Valcanis, Balaji Veeraraghavan, FranÇois-Xavier Weill, Jackie Wright, Gordon Dougan, Silvia Argimón, Jacqueline A Keane, David M Aanensen, Stephen Baker, Kathryn E Holt,

and Peter Aaby. Global diversity and antimicrobial resistance of typhoid fever pathogens: Insights from a meta-analysis of 13,000 Salmonella Typhi genomes. *eLife*, 12, 2023.

[12331] Rebecca Elizabeth Carlisle and Arthur D Kuo. Optimization of energy and time predicts dynamic speeds for human walking. *eLife*, 12, 2023.

[12332] Pablo M Casillas-Espinosa, Alison Anderson, Anna Harutyunyan, Crystal Li, Jiyoon Lee, Emma L Braine, Rhys D Brady, Mujun Sun, Cheng Huang, Christopher K Barlow, Anup D Shah, Ralf B Schittenhelm, Richelle Mychasiuk, Nigel C Jones, Sandy R Shultz, and Terence J O'Brien. Disease-modifying effects of sodium selenate in a model of drug-resistant, temporal lobe epilepsy. *eLife*, 12, 2023.

[12333] Ido Caspy, Maria Fadeeva, Yuval Mazor, and Nathan Nelson. Structure of Dunaliella photosystem II reveals conformational flexibility of stacked and unstacked supercomplexes. *eLife*, 12, 2023.

[12334] Raúl Castanera, Noemia Morales-Díaz, Sonal Gupta, Michael Purugganan, and Josep M Casacuberta. Transposons are important contributors to gene expression variability under selection in rice populations. *eLife*, 12, 2023.

[12335] Anna Castano, Margaux Silvestre, Carrow I Wells, Jennifer L Sanderson, Carla A Ferrer, Han Wee Ong, Yi Lang, William Richardson, Josie A Silvaroli, Frances M Bashore, Jeffery L Smith, Isabelle M Genereux, Kelvin Dempster, David H Drewry, Navlot S Pabla, Alex N Bullock, Tim A Benke, Sila K Ultanir, and Alison D Axtman. Discovery and characterization of a specific inhibitor of serine-threonine kinase cyclin-dependent kinase-like 5 (CDKL5) demonstrates role in hippocampal CA1 physiology. *eLife*, 12, 2023.

[12336] Hugo G Castelán-Sánchez, Luis Delaye, Rhys PD Inward, Simon Dellicour, Bernardo Gutierrez, Natalia Martinez de la Vina, Celia Boukadida, Oliver G Pybus, Guillermo de Anda Jáuregui, Plinio Guzmán, Marisol Flores-Garrido, óscar Fontanelli, Maribel Hernández Rosales, Amilcar Meneses, Gabriela Olmedo-Alvarez, Alfredo Heriberto Herrera-Estrella, Alejandro Sánchez-Flores, José Esteban Muñoz-Medina, Andreu Comas-García, Bruno Gómez-Gil, Selene Zárate, Blanca Taboada, Susana López, Carlos F Arias, Moritz UG Kraemer, Antonio Lazcano, and Marina Escalera Zamudio. Comparing the evolutionary dynamics of predominant SARS-CoV-2 virus lineages co-circulating in Mexico. *eLife*, 12, 2023.

[12337] Maikel Castellano-Pozo, Georgios Sioutas, Consuelo Barroso, Josh P Prince, Pablo Lopez-Jimenez, Joseph Davy, Angel-Luis Jaso-Tamame, Oliver Crawley, Nan Shao, Jesus Page, and Enrique Martinez-Perez. The kleisin subunit controls the function of C. elegans meiotic cohesins by determining the mode of DNA binding and differential regulation by SCC-2 and WAPL-1. *eLife*, 12, 2023.

[12338] Jesus A Castor-Macias, Jacqueline A Larouche, Emily C Wallace, Bonnie D Spence, Alec Eames, Pamela Duran, Benjamin A Yang, Paula M Fraczek, Carol A Davis, Susan V Brooks, Krishna Rao Maddipati, James F Markworth, and Carlos A Aguilar. Maresin 1 repletion improves muscle regeneration after volumetric muscle loss. *eLife*, 12, 2023.

[12339] Carlos Castrillon, Lea Simoni, Theo van den Broek, Cees van der Poel, Elliot H Akama-Garren, Minghe Ma, and Michael C Carroll. Complex subsets but redundant clonality after B cells egress from spontaneous germinal centers. *eLife*, 12, 2023.

[12340] Nathanael A Caveney, Naotaka Tsutsumi, and K Christopher Garcia. Structural insight into guanylyl cyclase receptor hijacking of the kinase–Hsp90 regulatory mechanism. *eLife*, 12, 2023.

[12341] Erika Cecon, Atsuro Oishi, Marine Luka, Delphine Ndiaye-Lobry, Arnaud FranÇois, Mathias Lescuyer, Fany Panayi, Julie Dam, Patricia Machado, and Ralf Jockers. Novel repertoire of tau biosensors to monitor pathological tau transformation and seeding activity in living cells. *eLife*, 12, 2023.

[12342] Lucydalila Cedillo, Fasih M Ahsan, Sainan Li, Nicole L Stuhr, Yifei Zhou, Yuyao Zhang, Adebanjo Adedoja, Luke M Murphy, Armen Yerevanian, Sinclair Emans, Khoi Dao, Zhaozhi Li, Nicholas D Peterson, Jeramie Watrous, Mohit Jain, Sudeshna Das, Read Pukkila-Worley, Sean P Curran, and Alexander A Soukas. Ether lipid biosynthesis promotes lifespan extension and enables diverse pro-longevity paradigms in Caenorhabditis elegans. *eLife*, 12, 2023.

[12343] Laura Celotto, Fabian Rost, Anja Machate, Juliane Bläsche, Andreas Dahl, Anke Weber, Stefan Hans, and Michael Brand. Single-cell RNA sequencing unravels the transcriptional network underlying zebrafish retina regeneration. *eLife*, 12, 2023.

[12344] Juan Carlos Cerpa, Alessandro Piccin, Margot Dehove, Marina Lavigne, Eric J Kremer, Mathieu Wolff, Shauna L Parkes, and Etienne Coutureau. Inhibition of noradrenergic signalling in rodent orbitofrontal cortex impairs the updating of goal-directed actions. *eLife*, 12, 2023.

[12345] Valentin Chabert, Geun-Don Kim, Danye Qiu, Guizhen Liu, Lydie Michaillat Mayer, Muhammed Jamsheer K, Henning J Jessen, and Andreas Mayer. Inositol pyrophosphate dynamics reveals control of the yeast phosphate starvation program through 1,5-IP8 and the SPX domain of Pho81. *eLife*, 12, 2023.

[12346] Sílvia Chafino, Panagiotis Giannios, Jordi Casanova, David Martín, and Xavier Franch-Marro. Antagonistic role of the BTB-zinc finger transcription factors Chinmo and Broad-Complex in the juvenile/pupal transition and in growth control. *eLife*, 12, 2023.

[12347] Pragnya Chakraborty, Bipan Kumar Deb, Vikas Arige, Thasneem Musthafa, Sundeep Malik, David I Yule, Colin W Taylor, and Gaiti Hasan. Regulation of store-operated Ca2+ entry by IP3 receptors independent of their ability to release Ca2+. *eLife*, 12, 2023.

[12348] Alex W Chan, Malgorzata Broncel, Eden Yifrach, Nicole R Haseley, Sundeep Chakladar, Elena Andree, Alice L Herneisen, Emily Shortt, Moritz Treeck, and Sebastian Lourido. Analysis of CDPK1 targets identifies a trafficking adaptor complex that regulates microneme exocytosis in Toxoplasma. *eLife*, 12, 2023.

[12349] Gary KL Chan, Samantha Maisel, Yeonjoo C Hwang, Bryan C Pascual, Rebecca RB Wolber, Phuong Vu, Krushna C Patra, Mehdi Bouhaddou, Heidi L Kenerson, Huat C Lim, Donald Long, Raymond S Yeung, Praveen Sethupathy, Danielle L Swaney, Nevan J Krogan, Rigney E Turnham, Kimberly J Riehle, John D Scott, Nabeel Bardeesy, and John D Gordan. Oncogenic PKA signaling increases c-MYC protein expression through multiple targetable mechanisms. *eLife*, 12, 2023.

[12350] Magnus Chan, Harutyun Sahakyan, Jodene Eldstrom, Daniel Sastre, Yundi Wang, Ying Dou, Marc Pourrier, Vitya Vardanyan, and David Fedida. A generic binding pocket for small molecule IKs activators at the extracellular inter-subunit interface of KCNQ1 and KCNE1 channel complexes. *eLife*, 12, 2023.

[12351] Abel Chandra, Laura Tünnermann, Tommy Löfstedt, and Regina Gratz. Transformer-based deep learning for predicting protein properties in the life sciences. *eLife*, 12, 2023.

[12352] Ching-Ho Chang, Isabel Mejia Natividad, and Harmit S Malik. Expansion and loss of sperm nuclear basic protein genes in Drosophila correspond with genetic conflicts between sex chromosomes. *eLife*, 12, 2023.

[12353] Nathan J Cheetham, Milla Kibble, Andrew Wong, Richard J Silverwood, Anika Knuppel, Dylan M Williams, Olivia KL Hamilton, Paul H Lee, Charis Bridger Staatz, Giorgio Di Gessa, Jingmin Zhu, Srinivasa Vittal Katikireddi, George B Ploubidis, Ellen J Thompson, Ruth CE Bowyer, Xinyuan Zhang, Golboo Abbasian, Maria Paz Garcia, Deborah Hart, Jeffrey Seow, Carl Graham, Neophytos Kouphou, Sam Acors, Michael H Malim, Ruth E Mitchell, Kate Northstone, Daniel Major-Smith, Sarah Matthews, Thomas Breeze, Michael Crawford, Lynn Molloy, Alex SF Kwong, Katie Doores, Nishi Chaturvedi, Emma L Duncan, Nicholas J Timpson, and Claire J Steves. Antibody levels following vaccination against SARS-CoV-2: associations with post-vaccination infection and risk factors in two UK longitudinal studies. *eLife*, 12, 2023.

[12354] Nicholas G Chehade and Omar A Gharbawie. Motor actions are spatially organized in motor and dorsal premotor cortex. *eLife*, 12, 2023.

[12355] Mohamad Ibrahim Cheikh, Joel Tchoufag, Miriam Osterfield, Kevin Dean, Swayamdipta Bhaduri, Chuzhong Zhang, Kranthi Kiran Mandadapu, and Konstantin Doubrovinski. A comprehensive model of Drosophila epithelium reveals the role of embryo geometry and cell topology in mechanical responses. *eLife*, 12, 2023.

[12356] Karthickeyan Chella Krishnan, Elie-Julien El Hachem, Mark P Keller, Sanjeet G Patel, Luke Carroll, Alexis Diaz Vegas, Isabela Gerdes Gyuricza, Christine Light, Yang Cao, Calvin Pan, Karolina Elżbieta Kaczor-Urbanowicz, Varun Shravah, Diana Anum, Matteo Pellegrini, Chi Fung Lee, Marcus M Seldin, Nadia A Rosenthal, Gary A Churchill, Alan D Attie, Benjamin Parker, David E James, and Aldons J Lusis. Genetic architecture of heart mitochondrial proteome influencing cardiac hypertrophy. *eLife*, 12, 2023.

[12357] Benjamin Chen, Boris Julg, Sindhu Mohandas, Steven B Bradfute, and Zaki A Sherif. Viral persistence, reactivation, and mechanisms of long COVID. *eLife*, 12, 2023.

[12358] Guanglei Chen, Xi Gu, Jinqi Xue, Xu Zhang, Xiaopeng Yu, Yu Zhang, Ailin Li, Yi Zhao, Guijin He, Meiyue Tang, Fei Xing, Jianqiao Yin, Xiaobo Bian, Ye Han, Shuo Cao, Chao Liu, Xiaofan Jiang, Keliang Zhang, Yan Xia, Huajun Li, Nan Niu, and Caigang Liu. Effects of neoadjuvant stereotactic body radiotherapy plus adebrelimab and chemotherapy for triple-negative breast cancer: A pilot study. *eLife*, 12, 2023.

[12359] Holly Y Chen, Manju Swaroop, Samantha Papal, Anupam K Mondal, Hyun Beom Song, Laura Campello, Gregory J Tawa, Florian Regent, Hiroko Shimada, Kunio Nagashima, Natalia de Val, Samuel G Jacobson, Wei Zheng, and Anand Swaroop. Reserpine maintains photoreceptor survival in retinal ciliopathy by resolving proteostasis imbalance and ciliogenesis defects. *eLife*, 12, 2023.

[12360] Hongwen Chen, Yu Qin, Marissa Chou, Jason G Cyster, and Xiaochun Li. Transmembrane protein CD69 acts as an S1PR1 agonist. *eLife*, 12, 2023.

[12361] Hsiao-Fan Chen, Wei-Jan Wang, Chung-Yu Chen, Wei-Chao Chang, Po-Ren Hsueh, Shin-Lei Peng, Chen-Shiou Wu, Yeh Chen, Hsin-Yu Huang, Wan-Jou Shen, Shao-Chun Wang, and Mien-Chie Hung. The natural tannins oligomeric proanthocyanidins and punicalagin are potent inhibitors of infection by SARS-CoV-2. *eLife*, 12, 2023.

[12362] Jiangfei Chen, Kunal Baxi, Amanda E Lipsitt, Nicole Rae Hensch, Long Wang, Prethish Sreenivas, Paulomi Modi, Xiang Ru Zhao, Antoine Baudin, Daniel G Robledo, Abhik Bandyopadhyay, Aaron Sugalski, Anil K Challa, Dias Kurmashev, Andrea R Gilbert, Gail E Tomlinson, Peter Houghton, Yidong Chen, Madeline N Hayes, Eleanor Y Chen, David S Libich, and Myron S Ignatius. Defining function of wild-type and three patient-specific TP53 mutations in a zebrafish model of embryonal rhabdomyosarcoma. *eLife*, 12, 2023.

[12363] Kehan Chen, Lie Wang, Jiemin Shen, Ah-Lim Tsai, Ming Zhou, and Gang Wu. Mechanism of stepwise electron transfer in six-transmembrane epithelial antigen of the prostate (STEAP) 1 and 2. *eLife*, 12, 2023.

[12364] Kevin S Chen, Rui Wu, Marc H Gershow, and Andrew M Leifer. Continuous odor profile monitoring to study olfactory navigation in small animals. *eLife*, 12, 2023.

[12365] Po Jui Chen, Anna B McMullin, Bryan J Visser, Qian Mei, Susan M Rosenberg, and David Bates. Interdependent progression of bidirectional sister replisomes in E. coli. *eLife*, 12, 2023.

[12366] Si-Ting Chen, Wen-Wen Shi, Yu-Qian Lin, Zhen-Shan Yang, Ying Wang, Meng-Yuan Li, Yue Li, Ai-Xia Liu, Yali Hu, and Zeng-Ming Yang. Embryo-derive TNF promotes decidualization via fibroblast activation. *eLife*, 12, 2023.

[12367] Vivian Chen, Milo S Johnson, Lucas Hérissant, Parris T Humphrey, David C Yuan, Yuping Li, Atish Agarwala, Samuel B Hoelscher, Dmitri A Petrov, Michael M Desai, and Gavin Sherlock. Evolution of haploid and diploid populations reveals common, strong, and variable pleiotropic effects in non-home environments. *eLife*, 12, 2023.

[12368] Xudong Chen, Bowen Jia, Shihan Zhu, and Mingjie Zhang. Phase separation-mediated actin bundling by the postsynaptic density condensates. *eLife*, 12, 2023.

[12369] Yiwang Chen, Qi Jiang, Mijiti Peierdun, Howard E Takiff, and Qian Gao. The mutational signatures of poor treatment outcomes on the drug-susceptible Mycobacterium tuberculosis genome. *eLife*, 12, 2023.

[12370] Yiya Chen, Yige Gao, Zhifen He, Zhouyuan Sun, Yu Mao, Robert F Hess, Peng Zhang, and Jiawei Zhou. Internal neural states influence the short-term effect of monocular deprivation in human adults. *eLife*, 12, 2023.

[12371] Yuanshu Chen, Haochen Zou, Xin Hou, Chuimei Lan, Jing Wang, Yanan Qing, Wangjun Chen, Shuxia Yao, and Keith M Kendrick. Oxytocin administration enhances pleasantness and neural responses to gentle stroking but not moderate pressure social touch by increasing peripheral concentrations. *eLife*, 12, 2023.

[12372] Zhe Chen, Garrett J Blair, Changliang Guo, Jim Zhou, Juan-Luis Romero-Sosa, Alicia Izquierdo, Peyman Golshani, Jason Cong, Daniel Aharoni, and Hugh T Blair. A hardware system for real-time decoding of in vivo calcium imaging data. *eLife*, 12, 2023.

[12373] Danni Cheng, Ke Qiu, Yufang Rao, Minzi Mao, Li Li, Yan Wang, Yao Song, Junren Chen, Xiaowei Yi, Xiuli Shao, Shao Hui Huang, Yi Zhang, Xuemei Chen, Sisi Wu, Shuaishuai Yu, Jun Liu, Haiyang Wang, Xingchen Peng, Daibo Li, Lin Yang, Li Chen, Zhiye Ying, Yongbo Zheng, Meijun Zheng, Binwu Ying, Xiaoxi Zeng, Wei Zhang, Wei Xu, Geoffrey Liu, Fei Chen, Haopeng Yu, Yu Zhao, and Jianjun Ren. Proliferative exhausted CD8+ T cells exacerbate long-lasting anti-tumor effects in human papillomavirus-positive head and neck squamous cell carcinoma. *eLife*, 12, 2023.

[12374] Amy Cheung, Kotaro Konno, Yuka Imamura, Aya Matsui, Manabu Abe, Kenji Sakimura, Toshikuni Sasaoka, Takeshi Uemura, Masahiko Watanabe, and Kensuke Futai. Neurexins in serotonergic neurons regulate neuronal survival, serotonin transmission, and complex mouse behaviors. *eLife*, 12, 2023.

[12375] Foo Cheung, Richard Apps, Lesia Dropulic, Yuri Kotliarov, Jinguo Chen, Tristan Jordan, Marc Langweiler, Julian Candia, Angelique Biancotto, Kyu Lee Han, Nicholas Rachmaninoff, Harlan Pietz, Kening Wang, John S Tsang, and Jeffrey I Cohen. Sex and prior exposure jointly shape innate immune responses to a live herpesvirus vaccine. *eLife*, 12, 2023.

[12376] Yajing Chi, Mu Su, Dongdong Zhou, Fangchao Zheng, Baoxuan Zhang, Ling Qiang, Guohua Ren, Lihua Song, Bing Bu, Shu Fang, Bo Yu, Jinxing Zhou, Jinming Yu, and Huihui Li. Dynamic analysis of circulating tumor DNA to predict the prognosis and monitor the treatment response of patients with metastatic triple-negative breast cancer: A prospective study. *eLife*, 12, 2023.

[12377] Rupesh K Chillale, Shihab Shamma, Srdjan Ostojic, and Yves Boubenec. Dynamics and maintenance of categorical responses in primary auditory cortex during task engagement. *eLife*, 12, 2023.

[12378] Kavitha Chinnaiya, Sarah Burbridge, Aragorn Jones, Dong Won Kim, Elsie Place, Elizabeth Manning, Ian Groves, Changyu Sun, Matthew Towers, Seth Blackshaw, and Marysia Placzek. A neuroepithelial wave of BMP signalling drives anteroposterior specification of the tuberal hypothalamus. *eLife*, 12, 2023.

[12379] Chandramohan Chitraju, Alexander W Fischer, Yohannes A Ambaw, Kun Wang, Bo Yuan, Sheng Hui, Tobias C Walther, and Robert V Farese. Mice lacking triglyceride synthesis enzymes in adipose tissue are resistant to diet-induced obesity. *eLife*, 12, 2023.

[12380] Eunjin Cho, Xiangguo Che, Mary Jasmin Ang, Seongmin Cheon, Jinkyung Lee, Kwang Soo Kim, Chang Hoon Lee, Sang-Yeop Lee, Hee-Young Yang, Changjong Moon, Chungoo Park, Je-Yong Choi, and Tae-Hoon Lee. Peroxiredoxin 5 regulates osteogenic differentiation through interaction with hnRNPK during bone regeneration. *eLife*, 12, 2023.

[12381] Ukrae H Cho and Martin W Hetzer. Caspase-mediated nuclear pore complex trimming in cell differentiation and endoplasmic reticulum stress. *eLife*, 12, 2023.

[12382] Seung Choi, Mark E Hauber, Lucas J Legendre, Noe-Heon Kim, Yuong-Nam Lee, and David J Varricchio. Microstructural and crystallographic evolution of palaeognath (Aves) eggshells. *eLife*, 12, 2023.

[12383] Budhaditya Chowdhury, Lakshman Abhilash, Antonio Ortega, Sha Liu, and Orie Shafer. Homeostatic control of deep sleep and molecular correlates of sleep pressure in Drosophila. *eLife*, 12, 2023.

[12384] Nahian Shahmat Chowdhury, Alan KI Chiang, Samantha K Millard, Patrick Skippen, Wei-Ju Chang, David A Seminowicz, and Siobhan M Schabrun. Combined transcranial magnetic stimulation and electroencephalography reveals alterations in cortical excitability during pain. *eLife*, 12, 2023.

[12385] Tsz Long Chu, Peikai Chen, Anna Xiaodan Yu, Mingpeng Kong, Zhijia Tan, Kwok Yeung Tsang, Zhongjun Zhou, and Kathryn Song Eng Cheah. MMP14 cleaves PTH1R in the chondrocyte-derived osteoblast lineage, curbing signaling intensity for proper bone anabolism. *eLife*, 12, 2023.

[12386] Bryce Chung, Muneeb Zia, Kyle A Thomas, Jonathan A Michaels, Amanda Jacob, Andrea Pack, Matthew J Williams, Kailash Nagapudi, Lay Heng Teng, Eduardo Arrambide, Logan Ouellette, Nicole Oey, Rhuna Gibbs, Philip Anschutz, Jiaao Lu, Yu Wu, Mehrdad Kashefi, Tomomichi Oya, Rhonda Kersten, Alice C Mosberger, Sean O'Connell, Runming Wang, Hugo Marques, Ana Rita Mendes, Constanze Lenschow, Gayathri Kondakath, Jeong Jun Kim, William Olson, Kiara N Quinn, Pierce Perkins, Graziana Gatto, Ayesha Thanawalla, Susan Coltman, Taegyo Kim, Trevor Smith, Ben Binder-Markey, Martin Zaback, Christopher K Thompson, Simon Giszter, Abigail Person, Martyn Goulding, Eiman Azim, Nitish Thakor, Daniel O'Connor, Barry Trimmer, Susana Q Lima, Megan R Carey, Chethan Pandarinath, Rui M Costa, J Andrew Pruszynski, Muhannad Bakir, and Samuel J Sober. Myomatrix arrays for high-definition muscle recording. *eLife*, 12, 2023.

[12387] Griffin Chure and Jonas Cremer. An optimal regulation of fluxes dictates microbial growth in and out of steady state. *eLife*, 12, 2023.

[12388] Ernesto Ciabatti, Ana González-Rueda, Daniel de Malmazet, Hassal Lee, Fabio Morgese, and Marco Tripodi. Genomic stability of self-inactivating rabies. *eLife*, 12, 2023.

[12389] Erin J Ciampa, Padraich Flahardy, Harini Srinivasan, Christopher Jacobs, Linus Tsai, S Ananth Karumanchi, and Samir M Parikh. Hypoxia-inducible factor 1 signaling drives placental aging and can provoke preterm labor. *eLife*, 12, 2023.

[12390] Leslie D Claar, Irene Rembado, Jacqulyn R Kuyat, Simone Russo, Lydia C Marks, Shawn R Olsen, and Christof Koch. Cortico-thalamo-cortical interactions modulate electrically evoked EEG responses in mice. *eLife*, 12, 2023.

[12391] Jenny Clarkson, Siew Hoong Yip, Robert Porteous, Alexia Kauff, Alison K Heather, and Allan E Herbison. CRISPR-Cas9 knockdown of ESR1 in preoptic GABA-kisspeptin neurons suppresses the preovulatory surge and estrous cycles in female mice. *eLife*, 12, 2023.

[12392] Khalyd J Clay, Yongzhi Yang, Christina Clark, and Michael Petrascheck. Proteostasis is differentially modulated by inhibition of translation initiation or elongation. *eLife*, 12, 2023.

[12393] Mathew Clement, Kristin Ladell, Kelly L Miners, Morgan Marsden, Lucy Chapman, Anna Cardus Figueras, Jake Scott, Robert Andrews, Simon Clare, Valeriia V Kriukova, Ksenia R Lupyr, Olga V Britanova, David R Withers, Simon A Jones, Dmitriy M Chudakov, David A Price, and

Ian R Humphreys. Inhibitory IL-10-producing CD4+ T cells are T-bet-dependent and facilitate cytomegalovirus persistence via coexpression of arginase-1. *eLife*, 12, 2023.

[12394] Marcella M Cline, Barbara Juarez, Avery Hunker, Ernesto G Regiarto, Bryan Hariadi, Marta E Soden, and Larry S Zweifel. Netrin-1 regulates the balance of synaptic glutamate signaling in the adult ventral tegmental area. *eLife*, 12, 2023.

[12395] Hudson W Coates, Isabelle M Capell-Hattam, Ellen M Olzomer, Ximing Du, Rhonda Farrell, Hongyuan Yang, Frances L Byrne, and Andrew J Brown. Hypoxia truncates and constitutively activates the key cholesterol synthesis enzyme squalene monooxygenase. *eLife*, 12, 2023.

[12396] Kelly D Cobey, Christophe A Fehlmann, Marina Christ Franco, Ana Patricia Ayala, Lindsey Sikora, Danielle B Rice, Chenchen Xu, John PA Ioannidis, Manoj M Lalu, Alixe Ménard, Andrew Neitzel, Bea Nguyen, Nino Tsertsvadze, and David Moher. Epidemiological characteristics and prevalence rates of research reproducibility across disciplines: A scoping review of articles published in 2018-2019. *eLife*, 12, 2023.

[12397] Olivier Codol, Mehrdad Kashefi, Christopher J Forgaard, Joseph M Galea, J Andrew Pruszynski, and Paul L Gribble. Sensorimotor feedback loops are selectively sensitive to reward. *eLife*, 12, 2023.

[12398] Nikki Coleman-Gosser, Yanhui Hu, Shiva Raghuvanshi, Shane Stitzinger, Weihang Chen, Arthur Luhur, Daniel Mariyappa, Molly Josifov, Andrew Zelhof, Stephanie E Mohr, Norbert Perrimon, and Amanda Simcox. Continuous muscle, glial, epithelial, neuronal, and hemocyte cell lines for Drosophila research. *eLife*, 12, 2023.

[12399] Léonard Colin, Celine Reyes, Julien Berthezene, Laetitia Maestroni, Laurent Modolo, Esther Toselli, Nicolas Chanard, Stephane Schaak, Olivier Cuvier, Yannick Gachet, Stephane Coulon, Pascal Bernard, and Sylvie Tournier. Condensin positioning at telomeres by shelterin proteins drives sister-telomere disjunction in anaphase. *eLife*, 12, 2023.

[12400] Lindsay Collins, John Francis, Brett Emanuel, and David A McCormick. Cholinergic and noradrenergic axonal activity contains a behavioral-state signal that is coordinated across the dorsal cortex. *eLife*, 12, 2023.

[12401] Megan M Colonnetta, Paul Schedl, and Girish Deshpande. Germline/soma distinction in Drosophila embryos requires regulators of zygotic genome activation. *eLife*, 12, 2023.

[12402] Bradley M Colquitt, Kelly Li, Foad Green, Robert Veline, and Michael S Brainard. Neural circuit-wide analysis of changes to gene expression during deafening-induced birdsong destabilization. *eLife*, 12, 2023.

[12403] Crescent L Combe, Carol M Upchurch, Carmen C Canavier, and Sonia Gasparini. Cholinergic modulation shifts the response of CA1 pyramidal cells to depolarizing ramps via TRPM4 channels with potential implications for place field firing. *eLife*, 12, 2023.

[12404] Ely Contreras, Jacob D Bhoi, Takuma Sonoda, Lutz Birnbaumer, and Tiffany M Schmidt. Melanopsin activates divergent phototransduction pathways in intrinsically photosensitive retinal ganglion cell subtypes. *eLife*, 12, 2023.

[12405] Anna A Cook, Tsz Chui Sophia Leung, Max Rice, Maya Nachman, élyse Zadigue-Dube, and Alanna Jean Watt. Endosomal dysfunction contributes to cerebellar deficits in spinocerebellar ataxia type 6. *eLife*, 12, 2023.

[12406] Daniel C Cook and Timothy A Ryan. GABABR silencing of nerve terminals. *eLife*, 12, 2023.

[12407] Ian Coombs, Cécile Bats, Craig A Sexton, Dorota Studniarczyk, Stuart G Cull-Candy, and Mark Farrant. Enhanced functional detection of synaptic calcium-permeable AMPA receptors using intracellular NASPM. *eLife*, 12, 2023.

[12408] Courtney A Copeland, Benjamin A Olenchock, David Ziehr, Sarah McGarrity, Kevin Leahy, Jamey D Young, Joseph Loscalzo, and William M Oldham. MYC overrides HIF-1$\alpha$ to regulate proliferating primary cell metabolism in hypoxia. *eLife*, 12, 2023.

[12409] Romain Coppée, FranÇois Blanquart, Aude Jary, Valentin Leducq, Valentine Marie Ferré, Anna Maria Franco Yusti, Léna Daniel, Charlotte Charpentier, Samuel Lebourgeois, Karen Zafilaza, Vincent Calvez, Diane Descamps, Anne-Geneviève Marcelin, Benoit Visseaux, and Antoine Bridier-Nahmias. Phylodynamics of SARS-CoV-2 in France, Europe, and the world in 2020. *eLife*, 12, 2023.

[12410] Elaine A Corbett, L Alexandra Martinez-Rodriguez, Cian Judd, Redmond G O'Connell, and Simon P Kelly. Multiphasic value biases in fast-paced decisions. *eLife*, 12, 2023.

[12411] Olivier Corneille, Jo Havemann, Emma L Henderson, Hans IJzerman, Ian Hussey, Jean-Jacques Orban de Xivry, Lee Jussim, Nicholas P Holmes, Artur Pilacinski, Brice Beffara, Harriet Carroll, Nicholas Otieno Outa, Peter Lush, and Leon D Lotter. Beware 'persuasive communication devices' when writing and reading scientific articles. *eLife*, 12, 2023.

[12412] Ricardo Cortez Cardoso Penha, Karl Smith-Byrne, Joshua R Atkins, Philip C Haycock, Siddhartha Kar, Veryan Codd, Nilesh J Samani, Christopher Nelson, Maja Milojevic, Aurélie AG Gabriel, Christopher Amos, Paul Brennan, Rayjean J Hung, Linda Kachuri, and James D Mckay. Common genetic variations in telomere length genes and lung cancer: a Mendelian randomisation study and its novel application in lung tumour transcriptome. *eLife*, 12, 2023.

[12413] Jamie D Costabile, Kaarthik A Balakrishnan, Sina Schwinn, and Martin Haesemeyer. Model discovery to link neural activity to behavioral tasks. *eLife*, 12, 2023.

[12414] Kara K Cover, Abby G Lieberman, Morgan M Heckman, and Brian N Mathur. The rostral intralaminar nuclear complex of the thalamus supports striatally mediated action reinforcement. *eLife*, 12, 2023.

[12415] Matthew V Cowley, Johanna Pruller, Massimo Ganassi, Peter S Zammit, and Christopher RS Banerji. An in silico FSHD muscle fiber for modeling DUX4 dynamics and predicting the impact of therapy. *eLife*, 12, 2023.

[12416] Liam P Coyne, Xiaowen Wang, Jiyao Song, Ebbing de Jong, Karin Schneider, Paul T Massa, Frank A Middleton, Thomas Becker, and Xin Jie Chen. Mitochondrial protein import clogging as a mechanism of disease. *eLife*, 12, 2023.

[12417] Lachlan Cribb, Ramon Sha, Stephanie Yiallourou, Natalie A Grima, Marina Cavuoto, Andree-Ann Baril, and Matthew P Pase. Sleep regularity and mortality: a prospective analysis in the UK Biobank. *eLife*, 12, 2023.

[12418] Wayne Croft, Hayden Pearce, Sandra Margielewska-Davies, Lindsay Lim, Samantha M Nicol, Fouzia Zayou, Daniel Blakeway, Francesca Marcon, Sarah Powell-Brett, Brinder Mahon, Reena Merard, Jianmin Zuo, Gary Middleton, Keith Roberts, Rachel M Brown, and Paul Moss. Spatial determination and prognostic impact of the fibroblast transcriptome in pancreatic ductal adenocarcinoma. *eLife*, 12, 2023.

[12419] Adeline Cros, Alba De Juan, Renaud Leclère, Julio L Sampaio, Mabel San Roman, Mathieu Maurin, Sandrine Heurtebise-Chrétien, and Elodie Segura. Homeostatic activation of aryl hydrocarbon receptor by dietary ligands dampens cutaneous allergic responses by controlling Langerhans cells migration. *eLife*, 12, 2023.

[12420] Megan M Cullinan, Robert C Klipp, Abigail Camenisch, and John R Bankston. Dynamic landscape of the intracellular termini of acid-sensing ion channel 1a. *eLife*, 12, 2023.

[12421] Irene Cumplido-Mayoral, Marina García-Prat, Grégory Operto, Carles Falcon, Mahnaz Shekari, Raffaele Cacciaglia, Marta Milà-Alomà, Luigi Lorenzini, Silvia Ingala, Alle Meije Wink, Henk JMM Mutsaerts, Carolina Minguillón, Karine Fauria, José Luis Molinuevo, Sven Haller, Gael Chetelat, Adam Waldman, Adam J Schwarz, Frederik Barkhof, Ivonne Suridjan, Gwendlyn Kollmorgen, Anna Bayfield, Henrik Zetterberg, Kaj Blennow, Marc Suárez-Calvet, Verónica Vilaplana, and Juan Domingo Gispert. Biological brain age prediction using machine learning on structural neuroimaging data: Multi-cohort validation against biomarkers of Alzheimer's disease and neurodegeneration stratified by sex. *eLife*, 12, 2023.

[12422] Pedro P Cunha, Eleanor Minogue, Lena CM Krause, Rita M Hess, David Bargiela, Brennan J Wadsworth, Laura Barbieri, Carolin Brombach, Iosifina P Foskolou, Ivan Bogeski, Pedro Velica, and Randall S Johnson. Oxygen levels at the time of activation determine T cell persistence and immunotherapeutic efficacy. *eLife*, 12, 2023.

[12423] Ashton J Curtis, Jian Zhu, Christopher J Penny, and Matthew G Gold. Molecular basis of interactions between CaMKII and $\alpha$-actinin-2 that underlie dendritic spine enlargement. *eLife*, 12, 2023.

[12424] Alayne Cuzick, James Seager, Valerie Wood, Martin Urban, Kim Rutherford, and Kim E Hammond-Kosack. A framework for community curation of interspecies interactions literature. *eLife*, 12, 2023.

[12425] Victoria L Czech, Lauren C O'Connor, Brendan Philippon, Emily Norman, and Alexandra B Byrne. TIR-1/SARM1 inhibits axon regeneration and promotes axon degeneration. *eLife*, 12, 2023.

[12426] Denis Dacheux, Guillaume Martinez, Christine E Broster Reix, Julie Beurois, Patrick Lores, Magamba Tounkara, Jean-William Dupuy, Derrick Roy Robinson, Corinne Loeuillet, Emeline Lambert, Zeina Wehbe, Jessica Escoffier, Amir Amiri-Yekta, Abbas Daneshipour, Seyedeh-Hanieh Hosseini, Raoudha Zouari, Selima Fourati Ben Mustapha, Lazhar Halouani, Xiaohui Jiang, Ying Shen, Chunyu Liu, Nicolas Thierry-Mieg, Amandine Septier, Marie Bidart, Véronique Satre, Caroline Cazin, Zine Eddine Kherraf, Christophe Arnoult, Pierre F Ray, Aminata Toure, Mélanie Bonhivers, and Charles Coutton. Novel axonemal protein ZMYND12 interacts with TTC29 and DNAH1, and is required for male fertility and flagellum function. *eLife*, 12, 2023.

[12427] Hiba Dagher, Anne-Marie Chaftari, Ishwaria M Subbiah, Alexandre E Malek, Ying Jiang, Peter Lamie, Bruno Granwehr, Teny John, Eduardo Yepez, Jovan Borjan, Cielito Reyes-Gibby, Mary Flores, Fareed Khawaja, Mala Pande, Noman Ali, Raniv Rojo, Daniel D Karp, Patrick Chaftari, Ray Hachem, and Issam I Raad. Long COVID in cancer patients: preponderance of symptoms in majority of patients over long time period. *eLife*, 12, 2023.

[12428] Carole Daly, Akim Abdul Guseinov, Hyunggu Hahn, Adam Wright, Irina G Tikhonova, Alex Rojas Bie Thomsen, and Bianca Plouffe. $\beta$-Arrestin-dependent and -independent endosomal G protein activation by the vasopressin type 2 receptor. *eLife*, 12, 2023.

[12429] Christelle Damon-Soubeyrand, Antonino Bongiovanni, Areski Chorfa, Chantal Goubely, Nelly Pirot, Luc Pardanaud, Laurence Piboin-Fragner, Caroline Vachias, Stephanie Bravard, Rachel Guiton, Jean-Leon Thomas, Fabrice Saez, Ayhan Kocer, Meryem Tardivel, Joël R Drevet, and Joelle Henry-Berger. Three-dimensional imaging of vascular development in the mouse epididymis. *eLife*, 12, 2023.

[12430] Salvatore D'Aniello, Stephanie Bertrand, and Hector Escriva. Amphioxus as a model to study the evolution of development in chordates. *eLife*, 12, 2023.

[12431] Michael JD Daniels, Lucas Lefevre, Stefan Szymkowiak, Alice Drake, Laura McCulloch, Makis Tzioras, Jack Barrington, Owen R Dando, Xin He, Mehreen Mohammad, Hiroki Sasaguri, Takashi Saito, Takaomi C Saido, Tara L Spires-Jones, and Barry W McColl. Cystatin F (Cst7) drives sex-dependent changes in microglia in an amyloid-driven model of Alzheimer's disease. *eLife*, 12, 2023.

[12432] Sandipan Dasgupta, Daniella Y Dayagi, Gal Haimovich, Emanuel Wyler, Tsviya Olender, Robert H Singer, Markus Landthaler, and Jeffrey E Gerst. Global analysis of contact-dependent human-to-mouse intercellular mRNA and lncRNA transfer in cell culture. *eLife*, 12, 2023.

[12433] Heledd Davies, Hugo Belda, Malgorzata Broncel, Jill Dalimot, and Moritz Treeck. PerTurboID, a targeted in situ method reveals the impact of kinase deletion on its local protein environment in the cytoadhesion complex of malaria-causing parasites. *eLife*, 12, 2023.

[12434] Kia L Davis, Nicole Ackermann, Lisa M Klesges, Nora Leahy, Callie Walsh-Bailey, Sarah Humble, Bettina Drake, and Vetta L Sanders Thompson. Understanding disruptions in cancer care to reduce increased cancer burden. *eLife*, 12, 2023.

[12435] Zachary W Davis, Nicholas M Dotson, Tom P Franken, Lyle Muller, and John H Reynolds. Spike-phase coupling patterns reveal laminar identity in primate cortex. *eLife*, 12, 2023.

[12436] Svenja de Buhr and Frauke Gräter. Myristoyl's dual role in allosterically regulating and localizing Abl kinase. *eLife*, 12, 2023.

[12437] Roberto De Filippo and Dietmar Schmitz. Differential ripple propagation along the hippocampal longitudinal axis. *eLife*, 12, 2023.

[12438] Steven De Gieter, Casey I Gallagher, Eveline Wijckmans, Diletta Pasini, Chris Ulens, and Rouslan G Efremov. Sterol derivative binding to the orthosteric site causes conformational changes in an invertebrate Cys-loop receptor. *eLife*, 12, 2023.

[12439] Saskia EJ de Vries, Joshua H Siegle, and Christof Koch. Sharing neurophysiology data from the Allen Brain Observatory. *eLife*, 12, 2023.

[12440] Julia U Deere, Arvin A Sarkissian, Meifeng Yang, Hannah A Uttley, Nicole Martinez Santana, Lam Nguyen, Kaushiki Ravi, and Anita V Devineni. Selective integration of diverse taste inputs within a single taste modality. *eLife*, 12, 2023.

[12441] Jean-Philippe Defour, Emilie Leroy, Sharmila Dass, Thomas Balligand, Gabriel Levy, Ian C Brett, Nicolas Papadopoulos, Céline Mouton, Lidvine Genet, Christian Pecquet, Judith Staerk, Steven O Smith, and Stefan N Constantinescu. Constitutive activation and oncogenicity are mediated by loss of helical structure at the cytosolic boundary of thrombopoietin receptor mutant dimers. *eLife*, 12, 2023.

[12442] Mark M Dekker, Luc E Coffeng, Frank P Pijpers, Debabrata Panja, and Sake J de Vlas. Reducing societal impacts of SARS-CoV-2 interventions through subnational implementation. *eLife*, 12, 2023.

[12443] Jordan DeKraker, Nicola Palomero-Gallagher, Olga Kedo, Neda Ladbon-Bernasconi, Sascha EA Muenzing, Markus Axer, Katrin Amunts, Ali R Khan, Boris C Bernhardt, and Alan C Evans. Evaluation of surface-based hippocampal registration using ground-truth subfield definitions. *eLife*, 12, 2023.

[12444] Claudia Dell'Amico, Marilyn M Angulo Salavarria, Yutaka Takeo, Ichiko Saotome, Maria Teresa Dell'Anno, Maura Galimberti, Enrica Pellegrino, Elena Cattaneo, Angeliki Louvi, and Marco Onorati. Microcephaly-associated protein WDR62 shuttles from the Golgi apparatus to the spindle poles in human neural progenitors. *eLife*, 12, 2023.

[12445] Alessandro Dema, Rabab Charafeddine, Shima Rahgozar, Jeffrey van Haren, and Torsten Wittmann. Growth cone advance requires EB1 as revealed by genomic replacement with a light-sensitive variant. *eLife*, 12, 2023.

[12446] Nisarg Desai, Praneet Bala, Rebecca Richardson, Jessica Raper, Jan Zimmermann, and Benjamin Hayden. OpenApePose, a database of annotated ape photographs for pose estimation. *eLife*, 12, 2023.

[12447] Simon Desch, Petra Schweinhardt, Ben Seymour, Herta Flor, and Susanne Becker. Evidence for dopaminergic involvement in endogenous modulation of pain relief. *eLife*, 12, 2023.

[12448] Carlos P Descovich, Kendall J Lough, Akankshya Jena, Jessica J Wu, Jina Yom, Danielle C Spitzer, Manuela Uppalapati, Katarzyna M Kedziora, and Scott E Williams. AGS3 antagonizes LGN to balance oriented cell divisions and cell fate choices in mammalian epidermis. *eLife*, 12, 2023.

[12449] Damien Detraux, Marino Caruso, Louise Feller, Maude Fransolet, Sébastien Meurant, Julie Mathieu, Thierry Arnould, and Patricia Renard. A critical role for heme synthesis and succinate in the regulation of pluripotent states transitions. *eLife*, 12, 2023.

[12450] Arun Devotta, Hugo Juraver-Geslin, Casey Griffin, and Jean-Pierre Saint-Jeannet. Npr3 regulates neural crest and cranial placode progenitors formation through its dual function as clearance and signaling receptor. *eLife*, 12, 2023.

[12451] Nihav Dhawale and Madhusudhan Venkadesan. How human runners regulate footsteps on uneven terrain. *eLife*, 12, 2023.

[12452] Sandeep S Dhawan, Carl Pinter, Andy CH Lee, and Rutsuko Ito. Male rodent perirhinal cortex, but not ventral hippocampus, inhibition induces approach bias under object-based approach-avoidance conflict. *eLife*, 12, 2023.

[12453] Herschel S Dhekne, Francesca Tonelli, Wondwossen M Yeshaw, Claire Y Chiang, Charles Limouse, Ebsy Jaimon, Elena Purlyte, Dario R Alessi, and Suzanne R Pfeffer. Genome-wide screen reveals Rab12 GTPase as a critical activator of Parkinson's disease-linked LRRK2 kinase. *eLife*, 12, 2023.

[12454] Alexis Diaz-Vegas, Søren Madsen, Kristen C Cooke, Luke Carroll, Jasmine XY Khor, Nigel Turner, Xin Y Lim, Miro A Astore, Jonathan C Morris, Anthony S Don, Amanda Garfield, Simona Zarini, Karin A Zemski Berry, Andrew P Ryan, Bryan C Bergman, Joseph T Brozinick, David E James, and James G Burchfield. Mitochondrial electron transport chain, ceramide, and coenzyme Q are linked in a pathway that drives insulin resistance in skeletal muscle. *eLife*, 12, 2023.

[12455] Amanda R Dicks, Grigory I Maksaev, Zainab Harissa, Alireza Savadipour, Ruhang Tang, Nancy Steward, Wolfgang Liedtke, Colin G Nichols, Chia-Lung Wu, and Farshid Guilak. Skeletal dysplasia-causing TRPV4 mutations suppress the hypertrophic differentiation of human iPSC-derived chondrocytes. *eLife*, 12, 2023.

[12456] Geoffrey W Diehl and A David Redish. Differential processing of decision information in subregions of rodent medial prefrontal cortex. *eLife*, 12, 2023.

[12457] Nicolas Diekmann and Sen Cheng. A model of hippocampal replay driven by experience and environmental structure facilitates spatial learning. *eLife*, 12, 2023.

[12458] Sydney Dimmock, Cian O'Donnell, and Conor Houghton. Bayesian analysis of phase data in EEG and MEG. *eLife*, 12, 2023.

[12459] Jason P Dinh and SN Patek. Tradeoffs explain scaling, sex differences, and seasonal oscillations in the remarkable weapons of snapping shrimp (Alpheus spp.). *eLife*, 12, 2023.

[12460] Claudia Dominici, Oscar D Villarreal, Junio Dort, Emilie Heckel, Yu Chang Wang, Ioannis Ragoussis, Jean-Sebastien Joyal, Nicolas Dumont, and Stéphane Richard. Inhibition of type I PRMTs reforms muscle stem cell identity enhancing their therapeutic capacity. *eLife*, 12, 2023.

[12461] Maxime Donadieu, Nathanael J Lee, María I Gaitán, Seung-Kwon Ha, Nicholas J Luciano, Snehashis Roy, Benjamin Ineichen, Emily C Leibovitch, Cecil C Yen, Dzung L Pham, Afonso C Silva, Mac Johnson, Steve Jacobson, Pascal Sati, and Daniel S Reich. In vivo MRI is sensitive to remyelination in a nonhuman primate model of multiple sclerosis. *eLife*, 12, 2023.

[12462] Yang Dong, Jiali Wang, Rachel-Ann Garibsingh, Keino Hutchinson, Yueyue Shi, Gilad Eisenberg, Xiaozhen Yu, Avner Schlessinger, and Christof Grewer. Conserved allosteric inhibition mechanism in SLC1 transporters. *eLife*, 12, 2023.

[12463] Narges Doostani, Gholam-Ali Hossein-Zadeh, and Maryam Vaziri-Pashkam. The normalization model predicts responses in the human visual cortex during object-based attention. *eLife*, 12, 2023.

[12464] Edward JA Douglas, Nathanael Palk, Tarcisio Brignoli, Dina Altwiley, Marcia Boura, Maisem Laabei, Mario Recker, Gordon CY Cheung, Ryan Liu, Roger C Hseih, Michael Otto, Eoin O'Brien, Rachel M McLoughlin, and Ruth C Massey. Extensive remodelling of the cell wall during the development of Staphylococcus aureus bacteraemia. *eLife*, 12, 2023.

[12465] Leonard Drees, Susi Schneider, Dietmar Riedel, Reinhard Schuh, and Matthias Behr. The proteolysis of ZP proteins is essential to control cell membrane structure and integrity of developing tracheal tubes in Drosophila. *eLife*, 12, 2023.

[12466] Chunxiu Du, Hu Xu, Cong Cao, Jiahui Cao, Yufei Zhang, Cong Zhang, Rongfang Qiao, Wenhua Ming, Yaqing Li, Huiwen Ren, Xiaohui Cui, Zhilin Luan, Youfei Guan, and Xiaoyan Zhang. Neutral amino acid transporter SLC38A2 protects renal medulla from hyperosmolarity-induced ferroptosis. *eLife*, 12, 2023.

[12467] Charles Ducrot, Gregory de Carvalho, Benoît Delignat-Lavaud, Constantin VL Delmas, Priyabrata Halder, Nicolas Giguère, Consiglia Pacelli, Sriparna Mukherjee, Marie-Josée Bourque, Martin Parent, Lulu Y Chen, and Louis-Eric Trudeau. Conditional deletion of neurexins dysregulates neurotransmission from dopamine neurons. *eLife*, 12, 2023.

[12468] Loïc Duffet, Elyse T Williams, Andrea Gresch, Simin Chen, Musadiq A Bhat, Dietmar Benke, Nina Hartrampf, and Tommaso Patriarchi. Optical tools for visualizing and controlling human GLP-1 receptor activation with high spatiotemporal resolution. *eLife*, 12, 2023.

[12469] Emilie Dujardin, André Marjolaine, Aurelie Dewaele, Béatrice Mandon-Pépin, Francis Poulat, Anne Frambourg, Dominique Thépot, Luc Jouneau, Geneviève Jolivet, Eric Pailhoux, and Maelle Pannetier. DMRT1 is a testis-determining gene in rabbits and is also essential for female fertility. *eLife*, 12, 2023.

[12470] Vanessa Dumeaux, Samira Massahi, Van Bettauer, Austin Mottola, Anna Dukovny, Sanny Singh Khurdia, Anna Carolina Borges Pereira Costa, Raha Parvizi Omran, Shawn Simpson, Jinglin Lucy Xie, Malcolm Whiteway, Judith Berman, and Michael T Hallett. Candida albicans exhibits heterogeneous and adaptive cytoprotective responses to antifungal compounds. *eLife*, 12, 2023.

[12471] Maud Dumoux, Thomas Glen, Jake LR Smith, Elaine ML Ho, Luis MA Perdigão, Avery Pennington, Sven Klumpe, Neville BY Yee, David Andrew Farmer, Pui YA Lai, William Bowles, Ron Kelley, Jürgen M Plitzko, Liang Wu, Mark Basham, Daniel K Clare, C Alistair Siebert, Michele C Darrow, James H Naismith, and Michael Grange. Cryo-plasma FIB/SEM volume imaging of biological specimens. *eLife*, 12, 2023.

[12472] Pierre Dupuy, Shreya Ghosh, Allison Fay, Oyindamola Adefisayo, Richa Gupta, Stewart Shuman, and Michael S Glickman. Roles for mycobacterial DinB2 in frameshift and substitution mutagenesis. *eLife*, 12, 2023.

[12473] Audrey Dureux, Alessandro Zanini, Janahan Selvanayagam, Ravi S Menon, and Stefan Everling. Gaze patterns and brain activations in humans and marmosets in the Frith-Happé theory-of-mind animation task. *eLife*, 12, 2023.

[12474] Anna Durner, Ellis Durner, and Annette Nicke. Improved ANAP incorporation and VCF analysis reveal details of P2X7 current facilitation and a limited conformational interplay between ATP binding and the intracellular ballast domain. *eLife*, 12, 2023.

[12475] Jules Duruz, Marta Sprecher, Jenifer C Kaldun, Al-Sayed Al-Soudy, Heidi EL Lischer, Geert van Geest, Pamela Nicholson, Rémy Bruggmann, and Simon G Sprecher. Molecular characterization of cell types in the squid Loligo vulgaris. *eLife*, 12, 2023.

[12476] éléonore Duvelle, Roddy M Grieves, and Matthijs AA van der Meer. Temporal context and latent state inference in the hippocampal splitter signal. *eLife*, 12, 2023.

[12477] Niamh Eastwood, Jiarui Zhou, Romain Derelle, Mohamed Abou-Elwafa Abdallah, William A Stubbings, Yunlu Jia, Sarah E Crawford, Thomas A Davidson, John K Colbourne, Simon Creer, Holly Bik, Henner Hollert, and Luisa Orsini. 100 years of anthropogenic impact causes changes in freshwater functional biodiversity. *eLife*, 12, 2023.

[12478] Jan Eckhardt, Alexis Ruiz, Stéphane Koenig, Maud Frieden, Hervé Meier, Alexander Schmidt, Susan Treves, and Francesco Zorzato. Quantitative proteomic analysis of skeletal muscles from wild-type and transgenic mice carrying recessive Ryr1 mutations linked to congenital myopathies. *eLife*, 12, 2023.

[12479] Carol A Edwards, William MD Watkinson, Stephanie B Telerman, Lisa C Hulsmann, Russell S Hamilton, and Anne C Ferguson-Smith. Reassessment of weak parent-of-origin expression bias shows it rarely exists outside of known imprinted regions. *eLife*, 12, 2023.

[12480] Avner Ehrlich, Konstantinos Ioannidis, Makram Nasar, Ismaeel Abu Alkian, Yuval Daskal, Nofar Atari, Limor Kliker, Nir Rainy, Matan Hofree, Sigal Shafran Tikva, Inbal Houri, Arrigo Cicero, Chiara Pavanello, Cesare R Sirtori, Jordana B Cohen, Julio A Chirinos, Lisa Deutsch, Merav Cohen, Amichai Gottlieb, Adina Bar-Chaim, Oren Shibolet, Michal Mandelboim, Shlomo L Maayan, and Yaakov Nahmias. Efficacy and safety of metabolic interventions for the treatment of severe COVID-19: in vitro, observational, and non-randomized open-label interventional study. *eLife*, 12, 2023.

[12481] Matthias Ekman, Sarah Kusch, and Floris P de Lange. Successor-like representation guides the prediction of future events in human visual cortex and hippocampus. *eLife*, 12, 2023.

[12482] Krystel El Hage, Nicolas Babault, Olek Maciejak, Bénédicte Desforges, Pierrick Craveur, Emilie Steiner, Juan Carlos Rengifo-Gonzalez, Hélène Henrie, Marie-Jeanne Clement, Vandana Joshi, Ahmed Bouhss, Liya Wang, Cyril Bauvais, and David Pastré. Targeting RNA:protein interactions with an integrative approach leads to the identification of potent YBX1 inhibitors. *eLife*, 12, 2023.

[12483] Elma El Khouri, Farah Diab, Camille Louvrier, Eman Assrawi, Aphrodite Daskalopoulou, Alexandre Nguyen, William Piterboth, Samuel Deshayes, Alexandra Desdoits, Bruno Copin, Florence Dastot Le Moal, Sonia Athina Karabina, Serge Amselem, Achille Aouba, and Irina Giurgea. A critical region of A20 unveiled by missense TNFAIP3 variations that lead to autoinflammation. *eLife*, 12, 2023.

[12484] Mariam El-Zein, Rami Ali, Eliya Farah, Sarah Botting-Provost, Eduardo L Franco, and Aisha Lofters. Pan-Canadian survey on the impact of the COVID-19 pandemic on cervical cancer screening and management: cross-sectional survey of healthcare professionals. *eLife*, 12, 2023.

[12485] Ahmad Elbahnsi, John Cowgill, Verena Burtscher, Linda Wedemann, Luise Zeckey, Baron Chanda, and Lucie Delemotte. Interplay between VSD, pore, and membrane lipids in electromechanical coupling in HCN channels. *eLife*, 12, 2023.

[12486] Sarah Elbaz-Hayoun, Batya Rinsky, Shira Hagbi-Levi, Michelle Grunin, and Itay Chowers. CCR1 mediates Müller cell activation and photoreceptor cell death in macular and retinal degeneration. *eLife*, 12, 2023.

[12487] Miriam Elfström, Penelope Grace Gray, and Joakim Dillner. Cervical cancer screening improvements with self-sampling during the COVID-19 pandemic. *eLife*, 12, 2023.

[12488] Vlad Elgart and Joseph Loscalzo. Local generation and efficient evaluation of numerous drug combinations in a single sample. *eLife*, 12, 2023.

[12489] Chad M Eliason, Jenna M McCullough, Shannon J Hackett, and Michael J Andersen. Complex plumages spur rapid color diversification in kingfishers (Aves: Alcedinidae). *eLife*, 12, 2023.

[12490] David Elkind, Hannah Hochgerner, Etay Aloni, Noam Shental, and Amit Zeisel. Sex, strain, and lateral differences in brain cytoarchitecture across a large mouse population. *eLife*, 12, 2023.

[12491] Nicole A Ellis, Kevin S Myers, Jessica Tung, Anne Davidson Ward, Kathryn Johnston, Katherine E Bonnington, Timothy J Donohue, and Matthias P Machner. A randomized multiplex CRISPRi-Seq approach for the identification of critical combinations of genes. *eLife*, 12, 2023.

[12492] Carolyn Elya, Danylo Lavrentovich, Emily Lee, Cassandra Pasadyn, Jasper Duval, Maya Basak, Valerie Saykina, and Benjamin de Bivort. Neural mechanisms of parasite-induced summiting behavior in 'zombie' Drosophila. *eLife*, 12, 2023.

[12493] Jon C Emery, Peter J Dodd, Sayera Banu, Beatrice Frascella, Frances L Garden, Katherine C Horton, Shahed Hossain, Irwin Law, Frank van Leth, Guy B Marks, Hoa Binh Nguyen, Hai Viet Nguyen, Ikushi Onozaki, Maria Imelda D Quelapio, Alexandra S Richards, Nabila Shaikh, Edine W Tiemersma, Richard G White, Khalequ Zaman, Frank Cobelens, and Rein MGJ Houben. Estimating the contribution of subclinical tuberculosis disease to transmission: An individual patient data analysis from prevalence surveys. *eLife*, 12, 2023.

[12494] Christopher H Emfinger, Lauren E Clark, Brian Yandell, Kathryn L Schueler, Shane P Simonett, Donnie S Stapleton, Kelly A Mitok, Matthew J Merrins, Mark P Keller, and Alan D Attie. Novel regulators of islet function identified from genetic variation in mouse islet Ca2+ oscillations. *eLife*, 12, 2023.

[12495] Alexandriya MX Emonds, Ramanujan Srinath, Kristina J Nielsen, and Charles E Connor. Object representation in a gravitational reference frame. *eLife*, 12, 2023.

[12496] Scott M Emrich, Ryan E Yoast, Xuexin Zhang, Adam J Fike, Yin-Hu Wang, Kristen N Bricker, Anthony Y Tao, Ping Xin, Vonn Walter, Martin T Johnson, Trayambak Pathak, Adam C Straub, Stefan Feske, Ziaur SM Rahman, and Mohamed Trebak. Orai3 and Orai1 mediate CRAC channel function and metabolic reprogramming in B cells. *eLife*, 12, 2023.

[12497] Markus Engstler and Tom Beneke. Gene editing and scalable functional genomic screening in Leishmania species using the CRISPR/Cas9 cytosine base editor toolbox LeishBASEedit. *eLife*, 12, 2023.

[12498] Imke Ensinck, Alexander Maman, Waleed S Albihlal, Michelangelo Lassandro, Giulia Salzano, Theodora Sideri, Steven A Howell, Enrica Calvani, Harshil Patel, Guy Bushkin, Markus Ralser, Ambrosius P Snijders, Mark Skehel, Ana Casañal, Schraga Schwartz, and Folkert J van Werven. The yeast RNA methylation complex consists of conserved yet reconfigured components with m6A-dependent and independent roles. *eLife*, 12, 2023.

[12499] Berra Erkosar, Cindy Dupuis, Fanny Cavigliasso, Loriane Savary, Laurent Kremmer, Hector Gallart-Ayala, Julijana Ivanisevic, and Tadeusz J Kawecki. Evolutionary adaptation to juvenile malnutrition impacts adult metabolism and impairs adult fitness in Drosophila. *eLife*, 12, 2023.

[12500] Ayca Ersoy, Bengi Altintel, Nurit Livnat Levanon, Nir Ben-Tal, Turkan Haliloglu, and Oded Lewinson. Computational analysis of long-range allosteric communications in CFTR. *eLife*, 12, 2023.

[12501] Alejandra Escós, Ester Diaz-Mora, Michael Pattison, Pilar Fajardo, Diego González-Romero, Ana Risco, José Martín-Gómez, éric Bonneil, Nahum Sonenberg, Seyed Mehdi Jafarnejad, Juan José Sanz-Ezquerro, Steven C Ley, and Ana Cuenda. p38$\gamma$ and p38$\delta$ modulate innate immune response by regulating MEF2D activation. *eLife*, 12, 2023.

[12502] Hiroaki Eshima, Justin L Shahtout, Piyarat Siripoksup, MacKenzie J Pearson, Ziad S Mahmassani, Patrick J Ferrara, Alexis W Lyons, John Alan Maschek, Alek D Peterlin, Anthony RP Verkerke, Jordan M Johnson, Anahy Salcedo, Jonathan J Petrocelli, Edwin R Miranda, Ethan J Anderson, Sihem Boudina, Qitao Ran, James E Cox, Micah J Drummond, and Katsuhiko Funai. Lipid hydroperoxides promote sarcopenia through carbonyl stress. *eLife*, 12, 2023.

[12503] Jamal Esmaily, Sajjad Zabbah, Reza Ebrahimpour, and Bahador Bahrami. Interpersonal alignment of neural evidence accumulation to social exchange of confidence. *eLife*, 12, 2023.

[12504] Chandler B Est and Regina M Murphy. An in vitro model for vitamin A transport across the human blood–brain barrier. *eLife*, 12, 2023.

[12505] Edmée Eyraud, Elise Maurat, Jean-Marc Sac-Epée, Pauline Henrot, Maeva Zysman, Pauline Esteves, Thomas Trian, Jean-William Dupuy, Alexander Leipold, Antoine-Emmanuel Saliba, Hugues Begueret, Pierre-Olivier Girodet, Matthieu Thumerel, Romain Hustache-Castaing, Roger Marthan, Florian Levet, Pierre Vallois, Cécile Contin-Bordes, Patrick Berger, and Isabelle Dupin. Short-range interactions between fibrocytes and CD8+ T cells in COPD bronchial inflammatory response. *eLife*, 12, 2023.

[12506] Yi Fan, Ping Lyu, Ruiye Bi, Chen Cui, Ruoshi Xu, Clifford J Rosen, Quan Yuan, and Chenchen Zhou. Creating an atlas of the bone microenvironment during oral inflammatory-related bone disease using single-cell profiling. *eLife*, 12, 2023.

[12507] Ching Fang, Dmitriy Aronov, LF Abbott, and Emily L Mackevicius. Neural learning rules for generating flexible predictions and computing the successor representation. *eLife*, 12, 2023.

[12508] Huan Fang, Wenlong Ren, Qiuxia Cui, Huichun Liang, Chuanyu Yang, Wenjing Liu, Xinye Wang, Xue Liu, Yujie Shi, Jing Feng, and Ceshi Chen. Integrin $\beta 4$ promotes DNA damage-related drug resistance in triple-negative breast cancer via TNFAIP2/IQGAP1/RAC1. *eLife*, 12, 2023.

[12509] Payam E Farahani, Xiaoyu Yang, Emily V Mesev, Kaylan A Fomby, Ellen H Brumbaugh-Reed, Caleb J Bashor, Celeste M Nelson, and Jared E Toettcher. pYtags enable spatiotemporal measurements of receptor tyrosine kinase signaling in living cells. *eLife*, 12, 2023.

[12510] Talia Fargason, Naiduwadura Ivon Upekala De Silva, Erin Powell, Zihan Zhang, Trenton Paul, Jamal Shariq, Steve Zaharias, and Jun Zhang. Peptides that Mimic RS repeats modulate phase separation of SRSF1, revealing a reliance on combined stacking and electrostatic interactions. *eLife*, 12, 2023.

[12511] Elena Farmaki, Aritro Nath, Rena Emond, Kimya L Karimi, Vince K Grolmusz, Patrick A Cosgrove, and Andrea H Bild. ONC201/TIC10 enhances durability of mTOR inhibitor everolimus in metastatic ER+ breast cancer. *eLife*, 12, 2023.

[12512] Mariah Farrell, Heather Fairfield, Michelle Karam, Anastasia D'Amico, Connor S Murphy, Carolyne Falank, Romanos Sklavenitis Pistofidi, Amanda Cao, Catherine R Marinac, Julie A Dragon, Lauren McGuinness, Carlos G Gartner, Reagan Di Iorio, Edward Jachimowicz, Victoria DeMambro, Calvin Vary, and Michaela R Reagan. Targeting the fatty acid binding proteins disrupts multiple myeloma cell cycle progression and MYC signaling. *eLife*, 12, 2023.

[12513] John S Favate, Kyle S Skalenko, Eric Chiles, Xiaoyang Su, Srujana Samhita Yadavalli, and Premal Shah. Linking genotypic and phenotypic changes in the E. coli long-term evolution experiment using metabolomics. *eLife*, 12, 2023.

[12514] Xing Feng, Ruifeng Sun, Moonyoung Lee, Xinyue Chen, Shangqin Guo, Huimin Geng, Marcus Müschen, Jungmin Choi, and Joao Pedro Pereira. Cell circuits between leukemic cells and mesenchymal stem cells block lymphopoiesis by activating lymphotoxin beta receptor signaling. *eLife*, 12, 2023.

[12515] Brielle Ferguson, Cameron Glick, and John R Huguenard. Prefrontal PV interneurons facilitate attention and are linked to attentional dysfunction in a mouse model of absence epilepsy. *eLife*, 12, 2023.

[12516] Corey Fernandez, Jiefeng Jiang, Shao-Fang Wang, Hannah Lee Choi, and Anthony D Wagner. Representational integration and differentiation in the human hippocampus following goal-directed navigation. *eLife*, 12, 2023.

[12517] Justin M Fine, Archana S Mysore, Maria E Fini, William J Tyler, and Marco Santello. Transcranial focused ultrasound to human rIFG improves response inhibition through modulation of the P300 onset latency. *eLife*, 12, 2023.

[12518] Louis Fisher, Helen J Curtis, Richard Croker, Milan Wiedemann, Victoria Speed, Christopher Wood, Andrew Brown, Lisa EM Hopcroft, Rose Higgins, Jon Massey, Peter Inglesby, Caroline E Morton, Alex J Walker, Jessica Morley, Amir Mehrkar, Seb Bacon, George Hickman, Orla Macdonald, Tom Lewis, Marion Wood, Martin Myers, Miriam Samuel, Robin Conibere, Wasim Baqir, Harpreet Sood, Charles Drury, Kiren Collison, Chris Bates, David Evans, Iain Dillingham, Tom Ward, Simon Davy, Rebecca M Smith, William Hulme, Amelia Green, John Parry, Frank Hester, Sam Harper, Jonathan Cockburn, Shaun O'Hanlon, Alex Eavis, Richard Jarvis, Dima Avramov, Paul Griffiths, Aaron Fowles, Nasreen Parkes, Brian MacKenna, and Ben Goldacre. Eleven key measures for monitoring general practice clinical activity during COVID-19: A retrospective cohort study using 48 million adults' primary care records in England through OpenSAFELY. *eLife*, 12, 2023.

[12519] Swann Floc'hlay, Ramya Balaji, Dimitrije Stanković, Valerie M Christiaens, Carmen Bravo González-Blas, Seppe De Winter, Gert J Hulselmans, Maxime De Waegeneer, Xiaojiang Quan, Duygu Koldere, Mardelle Atkins, Georg Halder, Mirka Uhlirova, Anne-Kathrin Classen, and Stein Aerts. Shared enhancer gene regulatory networks between wound and oncogenic programs. *eLife*, 12, 2023.

[12520] Barnaby Flower, Le Manh Hung, Leanne Mccabe, M Azim Ansari, Chau Le Ngoc, Thu Vo Thi, Hang Vu Thi Kim, Phuong Nguyen Thi Ngoc, Le Thanh Phuong, Vo Minh Quang, Thuan Dang Trong, Thao Le Thi, Tran Nguyen Bao, Cherry Kingsley, David Smith, Richard M Hoglund, Joel Tarning, Evelyne Kestelyn, Sarah L Pett, Rogier van Doorn, Jennifer Ilo Van Nuil, Hugo Turner, Guy E Thwaites, Eleanor Barnes, Motiur Rahman, Ann Sarah Walker, Jeremy N Day, Nguyen VV Chau, and Graham S Cooke. Efficacy of ultra-short, response-guided sofosbuvir and daclatasvir therapy for hepatitis C in a single-arm mechanistic pilot study. *eLife*, 12, 2023.

[12521] Sagen Flowers, Rushali Kothari, Yamila N Torres Cleuren, Melissa R Alcorn, Chee Kiang Ewe, Geneva Alok, Samantha L Fiallo, Pradeep M Joshi, and Joel H Rothman. Regulation of defective mitochondrial DNA accumulation and transmission in C. elegans by the programmed cell death and aging pathways. *eLife*, 12, 2023.

[12522] Virginia Folgado-Marco, Kristina Ames, Jacky Chuen, Kira Gritsman, and Nicholas E Baker. Haploinsufficiency of the essential gene Rps12 causes defects in erythropoiesis and hematopoietic stem cell maintenance. *eLife*, 12, 2023.

[12523] Mary F Fontana, Erica Ollmann Saphire, and Marion Pepper. Plasmodium infection disrupts the T follicular helper cell response to heterologous immunization. *eLife*, 12, 2023.

[12524] Hugh Z Ford, Angelika Manhart, and Jonathan R Chubb. Controlling periodic long-range signalling to drive a morphogenetic transition. *eLife*, 12, 2023.

[12525] Haleh Fotowat and Florian Engert. Neural circuits underlying habituation of visually evoked escape behaviors in larval zebrafish. *eLife*, 12, 2023.

[12526] Celine Ann Fox, Chi Tak Lee, Anna Kathleen Hanlon, Tricia XF Seow, Kevin Lynch, Siobhán Harty, Derek Richards, Jorge Palacios, Veronica O'Keane, Klaas Enno Stephan, and Claire M Gillan. An observational treatment study of metacognition in anxious-depression. *eLife*, 12, 2023.

[12527] Kai J Fox, Daniel Birman, and Justin L Gardner. Gain, not concomitant changes in spatial receptive field properties, improves task performance in a neural network attention model. *eLife*, 12, 2023.

[12528] John A Fozard, Chris Morgan, and Martin Howard. Coarsening dynamics can explain meiotic crossover patterning in both the presence and absence of the synaptonemal complex. *eLife*, 12, 2023.

[12529] Antonietta Franco, Jiajia Li, Daniel P Kelly, Ray E Hershberger, Ali J Marian, Renate M Lewis, Moshi Song, Xiawei Dang, Alina D Schmidt, Mary E Mathyer, John R Edwards, Cristina de Guzman Strong, and Gerald W Dorn. A human mitofusin 2 mutation can cause mitophagic cardiomyopathy. *eLife*, 12, 2023.

[12530] Alexandre Francou, Kathryn V Anderson, and Anna-Katerina Hadjantonakis. A ratchet-like apical constriction drives cell ingression during the mouse gastrulation EMT. *eLife*, 12, 2023.

[12531] Daniel Frank, Maria Bergamasco, Michael J Mlodzianoski, Andrew Kueh, Ellen Tsui, Cathrine Hall, Georgios Kastrappis, Anne Kathrin Voss, Catriona McLean, Maree Faux, Kelly L Rogers, Bang Tran, Elizabeth Vincan, David Komander, Grant Dewson, and Hoanh Tran. Trabid patient mutations impede the axonal trafficking of adenomatous polyposis coli to disrupt neurite growth. *eLife*, 12, 2023.

[12532] Michelle M Frank, Austen A Sitko, Kirupa Suthakar, Lester Torres Cadenas, Mackenzie Hunt, Mary Caroline Yuk, Catherine JC Weisz, and Lisa V Goodrich. Experience-dependent flexibility in a molecularly diverse central-to-peripheral auditory feedback system. *eLife*, 12, 2023.

[12533] William Tyler Frantz, Sharanya Iyengar, James Neiswender, Alyssa Cousineau, René Maehr, and Craig J Ceol. Pigment cell progenitor heterogeneity and reiteration of developmental signaling underlie melanocyte regeneration in zebrafish. *eLife*, 12, 2023.

[12534] Giovanni Frighetto and Mark A Frye. Columnar neurons support saccadic bar tracking in Drosophila. *eLife*, 12, 2023.

[12535] Felix F Fuchsberger, Dongyoon Kim, Natalia Baranova, Hanka Vrban, Marten Kagelmacher, Robert Wawrzinek, and Christoph Rademacher. Information transfer in mammalian glycan-based communication. *eLife*, 12, 2023.

[12536] Jingyi Guo Fuglstad, Pearl Saldanha, Jacopo Paglia, and Jonathan R Whitlock. Histological E-data Registration in rodent Brain Spaces. *eLife*, 12, 2023.

[12537] Nico Fuhrmann, Celine Prakash, and Tobias S Kaiser. Polygenic adaptation from standing genetic variation allows rapid ecotype formation. *eLife*, 12, 2023.

[12538] Yuzo Fujino, Morio Ueyama, Taro Ishiguro, Daisaku Ozawa, Hayato Ito, Toshihiko Sugiki, Asako Murata, Akira Ishiguro, Tania Gendron, Kohji Mori, Eiichi Tokuda, Tomoya Taminato, Takuya Konno, Akihide Koyama, Yuya Kawabe, Toshihide Takeuchi, Yoshiaki Furukawa, Toshimichi Fujiwara, Manabu Ikeda, Toshiki Mizuno, Hideki Mochizuki, Hidehiro Mizusawa, Keiji Wada, Kinya Ishikawa, Osamu Onodera, Kazuhiko Nakatani, Leonard Petrucelli, Hideki Taguchi, and Yoshitaka Nagai. FUS regulates RAN translation through modulating the G-quadruplex structure of GGGGCC repeat RNA in C9orf72-linked ALS/FTD. *eLife*, 12, 2023.

[12539] Daniel Fuks, Yoel Melamed, Dafna Langgut, Tali Erickson-Gini, Yotam Tepper, Guy Bar-Oz, and Ehud Weiss. Unprecedented yet gradual nature of first millennium CE intercontinental crop plant dispersal revealed in ancient Negev desert refuse. *eLife*, 12, 2023.

[12540] Hironori Funabiki, Isabel E Wassing, Qingyuan Jia, Ji-Dung Luo, and Thomas Carroll. Coevolution of the CDCA7-HELLS ICF-related nucleosome remodeling complex and DNA methyltransferases. *eLife*, 12, 2023.

[12541] Lindsay Fuzzell, Naomi C Brownstein, Holly B Fontenot, Paige W Lake, Alexandra Michel, Ashley Whitmer, Sarah L Rossi, McKenzie McIntyre, Susan T Vadaparampil, and Rebecca B Perkins. Examining the association of clinician characteristics with perceived changes in cervical cancer screening and colposcopy practice during the COVID-19 pandemic: a mixed methods assessment. *eLife*, 12, 2023.

[12542] Lindsay Fuzzell, Paige Lake, Naomi C Brownstein, Holly B Fontenot, Ashley Whitmer, Alexandra Michel, McKenzie McIntyre, Sarah L Rossi, Sidika Kajtezovic, Susan T Vadaparampil, and Rebecca Perkins. Examining the perceived impact of the COVID-19 pandemic on cervical cancer screening practices among clinicians practicing in Federally Qualified Health Centers: A mixed methods study. *eLife*, 12, 2023.

[12543] Yafit Gabay, Avi Karni, and Lori L Holt. Memory for incidentally learned categories evolves in the post-learning interval. *eLife*, 12, 2023.

[12544] Francisco X Galdos, Carissa Lee, Soah Lee, Sharon Paige, William Goodyer, Sidra Xu, Tahmina Samad, Gabriela V Escobar, Adrija Darsha, Aimee Beck, Rasmus O Bak, Matthew H Porteus, and Sean M Wu. Combined lineage tracing and scRNA-seq reveals unexpected first heart field predominance of human iPSC differentiation. *eLife*, 12, 2023.

[12545] Louis Gall, Carrie Duckworth, Ferran Jardi, Lieve Lammens, Aimee Parker, Ambra Bianco, Holly Kimko, David Mark Pritchard, and Carmen Pin. Homeostasis, injury, and recovery dynamics at multiple scales in a self-organizing mouse intestinal crypt. *eLife*, 12, 2023.

[12546] Robert Philip Ganley, Marilia Magalhaes de Sousa, Kira Werder, Tugce öztürk, Raquel Mendes, Matteo Ranucci, Hendrik Wildner, and Hanns Ulrich Zeilhofer. Targeted anatomical and functional identification of antinociceptive and pronociceptive serotonergic neurons that project to the spinal dorsal horn. *eLife*, 12, 2023.

[12547] Claire Gao, Chiraag A Gohel, Yan Leng, Jun Ma, David Goldman, Ariel J Levine, and Mario A Penzo. Molecular and spatial profiling of the paraventricular nucleus of the thalamus. *eLife*, 12, 2023.

[12548] Huanqing Gao, Yiming Zhong, Liang Zhou, Sixiong Lin, Xiaoting Hou, Zhen Ding, Yan Li, Qing Yao, Huiling Cao, Xuenong Zou, Di Chen, Xiaochun Bai, and Guozhi Xiao. Kindlin-2 inhibits TNF/NF-$\kappa$B-Caspase 8 pathway in hepatocytes to maintain liver development and function. *eLife*, 12, 2023.

[12549] Xin Gao, Kaiming Luo, Diya Wang, Yunbo Wei, Yin Yao, Jun Deng, Yang Yang, Qunxiong Zeng, Xiaoru Dong, Le Xiong, Dongcheng Gong, Lin Lin, Kai Pohl, Shaoling Liu, Yu Liu, Lu Liu, Thi HO Nguyen, Lilith F Allen, Katherine Kedzierska, Yanliang Jin, Mei-Rong Du, Wanping Chen, Liangjing Lu, Nan Shen, Zheng Liu, Ian A Cockburn, Wenjing Luo, and Di Yu. T follicular helper 17 (Tfh17) cells are superior for immunological memory maintenance. *eLife*, 12, 2023.

[12550] Amanda K Garcia, Derek F Harris, Alex J Rivier, Brooke M Carruthers, Azul Pinochet-Barros, Lance C Seefeldt, and Betül KaÇar. Nitrogenase resurrection and the evolution of a singular enzymatic mechanism. *eLife*, 12, 2023.

[12551] Gilberto Garcia, Hanlin Zhang, Sophia Moreno, C Kimberly Tsui, Brant Michael Webster, Ryo Higuchi-Sanabria, and Andrew Dillin. Lipid homeostasis is essential for a maximal ER stress response. *eLife*, 12, 2023.

[12552] José Manuel García-Martínez, Ana Chocarro-Calvo, Javier Martínez-Useros, María Jesús Fernández-Aceñero, M Carmen Fiuza, José Cáceres-Rentero, Antonio De la Vieja, Antonio Barbáchano, Alberto Muñoz, María Jesús Larriba, and Custodia García Jiménez. Vitamin D induces SIRT1 activation through K610 deacetylation in colon cancer. *eLife*, 12, 2023.

[12553] Rania Garde, Abhyudai Singh, Asif Ali, and David Pincus. Transcriptional regulation of Sis1 promotes fitness but not feedback in the heat shock response. *eLife*, 12, 2023.

[12554] Maire Gavagan, Noel Jameson, and Jesse G Zalatan. The Axin scaffold protects the kinase GSK3$\beta$ from cross-pathway inhibition. *eLife*, 12, 2023.

[12555] John A Gaynes, Samuel A Budoff, Michael J Grybko, and Alon Poleg-Polsky. Heterogeneous presynaptic receptive fields contribute to directional tuning in starburst amacrine cells. *eLife*, 12, 2023.

[12556] Gaoran Ge, Qianping Guo, Ying Zhou, Wenming Li, Wei Zhang, Jiaxiang Bai, Qing Wang, Huaqiang Tao, Wei Wang, Zhen Wang, Minfeng Gan, Yaozeng Xu, Huilin Yang, Bin Li, and Dechun Geng. GLI1 facilitates collagen-induced arthritis in mice by collaborative regulation of DNA methyltransferases. *eLife*, 12, 2023.

[12557] Mengyuan Ge, Judith Molina, Jin-Ju Kim, Shamroop K Mallela, Anis Ahmad, Javier Varona Santos, Hassan Al-Ali, Alla Mitrofanova, Kumar Sharma, Flavia Fontanesi, Sandra Merscher, and Alessia Fornoni. Empagliflozin reduces podocyte lipotoxicity in experimental Alport syndrome. *eLife*, 12, 2023.

[12558] Carolin Gehr, Jeremie Sibille, and Jens Kremkow. Retinal input integration in excitatory and inhibitory neurons in the mouse superior colliculus in vivo. *eLife*, 12, 2023.

[12559] Florian Geisler, Sanne Remmelzwaal, Vera Jankowski, Ruben Schmidt, Mike Boxem, and Rudolf E Leube. Intermediate filament network perturbation in the C. elegans intestine causes systemic dysfunctions. *eLife*, 12, 2023.

[12560] Qibin Geng, Yushun Wan, Fu-Chun Hsueh, Jian Shang, Gang Ye, Fan Bu, Morgan Herbst, Rowan Wilkens, Bin Liu, and Fang Li. Lys417 acts as a molecular switch that regulates the conformation of SARS-CoV-2 spike protein. *eLife*, 12, 2023.

[12561] Jason T George and Herbert Levine. Optimal cancer evasion in a dynamic immune microenvironment generates diverse post-escape tumor antigenicity profiles. *eLife*, 12, 2023.

[12562] Gaspard Gerschenfeld, Fanny Coulpier, Aurélie Gresset, Pernelle Pulh, Bastien Job, Thomas Topilko, Julie Siegenthaler, Maria Eleni Kastriti, Isabelle Brunet, Patrick Charnay, and Piotr Topilko. Neural tube-associated boundary caps are a major source of mural cells in the skin. *eLife*, 12, 2023.

[12563] Sophia Gessner, Zela Alexandria-Mae Martin, Michael A Reiche, Joana A Santos, Ryan Dinkele, Atondaho Ramudzuli, Neeraj Dhar, Timothy J de Wet, Saber Anoosheh, Dirk M Lang, Jesse Aaron, Teng-Leong Chew, Jennifer Herrmann, Rolf Müller, John D McKinney, Roger Woodgate, Valerie Mizrahi, česlovas Venclovas, Meindert H Lamers, and Digby F Warner. Investigating the composition and recruitment of the mycobacterial ImuA'-ImuB-DnaE2 mutasome. *eLife*, 12, 2023.

[12564] Anna C Geuzebroek, Hannah Craddock, Redmond G O'Connell, and Simon P Kelly. Balancing true and false detection of intermittent sensory targets by adjusting the inputs to the evidence accumulation process. *eLife*, 12, 2023.

[12565] Valentin Gfeller, Jan Waelchli, Stephanie Pfister, Gabriel Deslandes-Hérold, Fabio Mascher, Gaetan Glauser, Yvo Aeby, Adrien Mestrot, Christelle AM Robert, Klaus Schlaeppi, and Matthias Erb. Plant secondary metabolite-dependent plant-soil feedbacks can improve crop yield in the field. *eLife*, 12, 2023.

[12566] Sue Ghosh, Jackie Fantes, Karin Leschly, Julio Mazul, and Rebecca B Perkins. Electronic data review, client reminders, and expanded clinic hours for improving cervical cancer screening rates after the COVID-19 pandemic shutdowns: A multicomponent quality improvement program. *eLife*, 12, 2023.

[12567] Yelena Ginzburg, Xiuli An, Stefano Rivella, and Adam Goldfarb. Normal and dysregulated crosstalk between iron metabolism and erythropoiesis. *eLife*, 12, 2023.

[12568] Paolo Giorgi Rossi, Giuliano Carrozzi, Patrizia Falini, Letizia Sampaolo, Giuseppe Gorini, Manuel Zorzi, Paola Armaroli, Carlo Senore, Priscilla Sassoli de Bianchi, Maria Masocco, Marco Zappa, Francesca Battisti, and Paola Mantellini. The impact of the COVID-19 pandemic on Italian population-based cancer screening activities and test coverage: Results from national cross-sectional repeated surveys in 2020. *eLife*, 12, 2023.

[12569] Laura K Globig, Nora Holtz, and Tali Sharot. Changing the incentive structure of social media platforms to halt the spread of misinformation. *eLife*, 12, 2023.

[12570] Aaron Gochman, Xiao-Feng Tan, Chanhyung Bae, Helen Chen, Kenton J Swartz, and Andres Jara-Oseguera. Cannabidiol sensitizes TRPV2 channels to activation by 2-APB. *eLife*, 12, 2023.

[12571] Godber M Godbersen, Sebastian Klug, Wolfgang Wadsak, Verena Pichler, Julia Raitanen, Anna Rieckmann, Lars Stiernman, Luca Cocchi, Michael Breakspear, Marcus Hacker, Rupert Lanzenberger, and Andreas Hahn. Task-evoked metabolic demands of the posteromedial default mode network are shaped by dorsal attention and frontoparietal control networks. *eLife*, 12, 2023.

[12572] Adwait A Godbole, Sneha Gopalan, Thien-Kim Nguyen, Alexander L Munden, Dominique S Lui, Matthew J Fanelli, Paula Vo, Caroline A Lewis, Jessica B Spinelli, Thomas G Fazzio, and Amy K Walker. S-adenosylmethionine synthases specify distinct H3K4me3 populations and gene expression patterns during heat stress. *eLife*, 12, 2023.

[12573] Shivani Sujay Godbole and Nikolay V Dokholyan. Allosteric regulation of kinase activity in living cells. *eLife*, 12, 2023.

[12574] Baira Godneeva, Maria Ninova, Katalin Fejes-Toth, and Alexei Aravin. SUMOylation of Bonus, the Drosophila homolog of Transcription Intermediary Factor 1, safeguards germline identity by recruiting repressive chromatin complexes to silence tissue-specific genes. *eLife*, 12, 2023.

[12575] Rafael Soares Godoy, Nicholas D Cober, David P Cook, Emma McCourt, Yupu Deng, Liyuan Wang, Kenny Schlosser, Katelynn Rowe, and Duncan J Stewart. Single-cell transcriptomic atlas of lung microvascular regeneration after targeted endothelial cell ablation. *eLife*, 12, 2023.

[12576] Lisa Goebel, Tonia Kirschner, Sandra Koska, Amrita Rai, Petra Janning, Stefano Maffini, Helge Vatheuer, Paul Czodrowski, Roger S Goody, Matthias P Müller, and Daniel Rauh. Targeting oncogenic KRasG13C with nucleotide-based covalent inhibitors. *eLife*, 12, 2023.

[12577] Raeann Goering, Ankita Arora, Megan C Pockalny, and J Matthew Taliaferro. RNA localization mechanisms transcend cell morphology. *eLife*, 12, 2023.

[12578] Lex J Gómez, James C Dooley, and Mark S Blumberg. Activity in developing prefrontal cortex is shaped by sleep and sensory experience. *eLife*, 12, 2023.

[12579] Sofia Gomez, Lionel Bureau, Karin John, Elise-Noëlle Chêne, Delphine Débarre, and Sigolene Lecuyer. Substrate stiffness impacts early biofilm formation by modulating Pseudomonas aeruginosa twitching motility. *eLife*, 12, 2023.

[12580] Ryszard Stefan Gomolka, Lauren M Hablitz, Humberto Mestre, Michael Giannetto, Ting Du, Natalie Linea Hauglund, Lulu Xie, Weiguo Peng, Paula Melero Martinez, Maiken Nedergaard, and Yuki Mori. Loss of aquaporin-4 results in glymphatic system dysfunction via brain-wide interstitial fluid stagnation. *eLife*, 12, 2023.

[12581] Joaquin Gonzalez, Pablo Torterolo, and Adriano BL Tort. Mechanisms and functions of respiration-driven gamma oscillations in the primary olfactory cortex. *eLife*, 12, 2023.

[12582] L Sofia Gonzalez, Austen A Fisher, Shane P D'Souza, Evelin M Cotella, Richard A Lang, and J Elliott Robinson. Ventral striatum dopamine release encodes unique properties of visual stimuli in mice. *eLife*, 12, 2023.

[12583] Amanda J González Segarra, Gina Pontes, Nicholas Jourjine, Alexander Del Toro, and Kristin Scott. Hunger- and thirst-sensing neurons modulate a neuroendocrine network to coordinate sugar and water ingestion. *eLife*, 12, 2023.

[12584] Adam Gordon-Fennell, Joumana M Barbakh, MacKenzie T Utley, Shreya Singh, Paula Bazzino, Raajaram Gowrishankar, Michael R Bruchas, Mitchell F Roitman, and Garret D Stuber. An open-source platform for head-fixed operant and consummatory behavior. *eLife*, 12, 2023.

[12585] Mark Gormley, Tom Dudding, Steven J Thomas, Jessica Tyrrell, Andrew R Ness, Miranda Pring, Danny Legge, George Davey Smith, Rebecca C Richmond, Emma E Vincent, and Caroline Bull. Evaluating the effect of metabolic traits on oral and oropharyngeal cancer risk using Mendelian randomization. *eLife*, 12, 2023.

[12586] Teppei Goto, Mitsue Hagihara, and Kazunari Miyamichi. Dynamics of pulsatile activities of arcuate kisspeptin neurons in aging female mice. *eLife*, 12, 2023.

[12587] Flavia Venetucci Gouveia, Jurgen Germann, Gavin JB Elias, Alexandre Boutet, Aaron Loh, Adriana Lucia Lopez Rios, Cristina Torres Diaz, William Omar Contreras Lopez, Raquel Chacon Ruiz Martinez, Erich Talamoni Fonoff, Juan Carlos Benedetti-Isaac, Peter Giacobbe, Pablo M Arango Pava, Han Yan, George M Ibrahim, Nir Lipsman, Andres Lozano, and Clement Hamani. Multi-centre analysis of networks and genes modulated by hypothalamic stimulation in patients with aggressive behaviours. *eLife*, 12, 2023.

[12588] Shiri Graff van Creveld, Sacha N Coesel, Stephen Blaskowski, Ryan D Groussman, Megan J Schatz, and E Virginia Armbrust. Divergent functions of two clades of flavodoxin in diatoms mitigate oxidative stress and iron limitation. *eLife*, 12, 2023.

[12589] Alejandro Granados, Alejandro A Granados, Simon Bucher, Hanbing Song, Aditi Agrawal, Ann T Chen, Tien Peng, Norma Neff, Angela Oliveira Pisco, Franklin Huang, and Bruce Wang. Single-nuclei characterization of pervasive transcriptional signatures across organs in response to COVID-19. *eLife*, 12, 2023.

[12590] Raquel Granell, John A Curtin, Sadia Haider, Negusse Tadesse Kitaba, Sara A Mathie, Lisa G Gregory, Laura L Yates, Mauro Tutino, Jenny Hankinson, Mauro Perretti, Judith M Vonk, Hasan S Arshad, Paul Cullinan, Sara Fontanella, Graham C Roberts, Gerard H Koppelman, Angela Simpson, Steve W Turner, Clare S Murray, Clare M Lloyd, John W Holloway, and Adnan Custovic. A meta-analysis of genome-wide association studies of childhood wheezing phenotypes identifies ANXA1 as a susceptibility locus for persistent wheezing. *eLife*, 12, 2023.

[12591] Franziska Grathwol, Christian Roos, Dietmar Zinner, Benjamin Hume, Stéphanie M Porcier, Didier Berthet, Jacques Cuisin, Stefan Merker, Claudio Ottoni, Wim Van Neer, Nathaniel J Dominy, and Gisela H Kopp. Adulis and the transshipment of baboons during classical antiquity. *eLife*, 12, 2023.

[12592] Emily I Green, Etienne Jaouen, Dennis Klug, Roenick Proveti Olmo, Amandine Gautier, Stéphanie Blandin, and Eric Marois. A population modification gene drive targeting both Saglin and Lipophorin impairs Plasmodium transmission in Anopheles mosquitoes. *eLife*, 12, 2023.

[12593] Maura Greiser, Mariusz Karbowski, Aaron David Kaplan, Andrew Kyle Coleman, Nicolas Verhoeven, Carmen A Mannella, W Jonathan Lederer, and Liron Boyman. Calcium and bicarbonate signaling pathways have pivotal, resonating roles in matching ATP production to demand. *eLife*, 12, 2023.

[12594] Théodore Grenier, Jessika Consuegra, Mariana G Ferrarini, Houssam Akherraz, Longwei Bai, Yves Dusabyinema, Isabelle Rahioui, Pedro Da Silva, Benjamin Gillet, Sandrine Hughes, Cathy I Ramos, Renata C Matos, and FranÇois Leulier. Intestinal GCN2 controls Drosophila systemic growth in response to Lactiplantibacillus plantarum symbiotic cues encoded by r/tRNA operons. *eLife*, 12, 2023.

[12595] Elizabeth Griggs, Kyle Trageser, Sean Naughton, Eun-Jeong Yang, Brian Mathew, Grace Van Hyfte, Linh Hellmers, Nathalie Jette, Molly Estill, Li Shen, Tracy Fischer, and Giulio Maria Pasinetti. Recapitulation of pathophysiological features of AD in SARS-CoV-2-infected subjects. *eLife*, 12, 2023.

[12596] Xiaochun Gu, Zhong Zhao, Xueli Chen, Lifeng Zhang, Huaqiang Fang, Ting Zhao, Shenghong Ju, Weizheng Gao, Xiaoyu Qian, Xianhua Wang, Jue Zhang, and Heping Cheng. Imaging microglia surveillance during sleep-wake cycles in freely behaving mice. *eLife*, 12, 2023.

[12597] Jian Guan, J David Peske, Michael Manoharan Valerio, Chansu Park, Ellen A Robey, and Scheherazade Sadegh-Nasseri. Commensal bacteria maintain a Qa-1b-restricted unconventional CD8+ T population in gut epithelium. *eLife*, 12, 2023.

[12598] Nina Gubensäk, Theo Sagmeister, Christoph Buhlheller, Bruno Di Geronimo, Gabriel E Wagner, Lukas Petrowitsch, Melissa A Gräwert, Markus Rotzinger, Tamara M Ismael Berger, Jan Schäfer, Isabel Usón, Joachim Reidl, Pedro A Sánchez-Murcia, Klaus Zangger, and Tea Pavkov-Keller. Vibrio cholerae's ToxRS bile sensing system. *eLife*, 12, 2023.

[12599] Zhannetta V Gugel, Elizabeth G Maurais, and Elizabeth J Hong. Chronic exposure to odors at naturally occurring concentrations triggers limited plasticity in early stages of Drosophila olfactory processing. *eLife*, 12, 2023.

[12600] Shiyu Gui, Boaz Yuval, Tobias Engl, Yongyue Lu, and Daifeng Cheng. Protein feeding mediates sex pheromone biosynthesis in an insect. *eLife*, 12, 2023.

[12601] Rafaela M Guimarães, ConceiÇão E Aníbal-Silva, Marcela Davoli-Ferreira, Francisco Isaac F Gomes, Atlante Mendes, Maria CM Cavallini, Miriam M Fonseca, Samara Damasceno, Larissa P Andrade, Marco Colonna, Cyril Rivat, Fernando Q Cunha, José C Alves-Filho, and Thiago M Cunha. Neuron-associated macrophage proliferation in the sensory ganglia is associated with peripheral nerve injury-induced neuropathic pain involving CX3CR1 signaling. *eLife*, 12, 2023.

[12602] Benjamin Guinet, David Lepetit, Sylvain Charlat, Peter N Buhl, David G Notton, Astrid Cruaud, Jean-Yves Rasplus, Julia Stigenberg, Damien M de Vienne, Bastien Boussau, and Julien Varaldi. Endoparasitoid lifestyle promotes endogenization and domestication of dsDNA viruses. *eLife*, 12, 2023.

[12603] Yuqi Guo, Jianlu Wei, Chuanju Liu, Xin Li, and Wenbo Yan. Metformin regulates bone marrow stromal cells to accelerate bone healing in diabetic mice. *eLife*, 12, 2023.

[12604] Pratik Rajan Gupte, Gregory F Albery, Jakob Gismann, Amy Sweeny, and Franz J Weissing. Novel pathogen introduction triggers rapid evolution in animal social movement strategies. *eLife*, 12, 2023.

[12605] Ellen J Guss, Yulia Akbergenova, Karen L Cunningham, and J Troy Littleton. Loss of the extracellular matrix protein Perlecan disrupts axonal and synaptic stability during Drosophila development. *eLife*, 12, 2023.

[12606] Enric Gutiérrez-Martínez, Susana Benet Garrabé, Nicolas Mateos, Itziar Erkizia, Jon Ander Nieto-Garai, Maier Lorizate, Kyra JE Borgman, Carlo Manzo, Felix Campelo, Nuria Izquierdo-Useros, Javier Martinez-Picado, and Maria F Garcia-Parajo. Actin-regulated Siglec-1 nanoclustering influences HIV-1 capture and virus-containing compartment formation in dendritic cells. *eLife*, 12, 2023.

[12607] Utsav Gyawali, David A Martin, Fangmiao Sun, Yulong Li, and Donna Calu. Dopamine in the dorsal bed nucleus of stria terminalis signals Pavlovian sign-tracking and reward violations. *eLife*, 12, 2023.

[12608] Zsuzsanna Gyöngy, Gábor Mocsár, éva Hegedűs, Thomas Stockner, Zsuzsanna Ritter, László Homolya, Anita Schamberger, Tamás I Orbán, Judit Remenyik, Gergely Szakacs, and Katalin Goda. Nucleotide binding is the critical regulator of ABCG2 conformational transitions. *eLife*, 12, 2023.

[12609] Abderrahman Hachani, Stefano G Giulieri, Romain Guérillot, Calum J Walsh, Marion Herisse, Ye Mon Soe, Sarah L Baines, David R Thomas, Shane Doris Cheung, Ashleigh S Hayes, Ellie Cho, Hayley J Newton, Sacha Pidot, Ruth C Massey, Benjamin P Howden, and Timothy P Stinear. A high-throughput cytotoxicity screening platform reveals agr-independent mutations in bacteraemia-associated Staphylococcus aureus that promote intracellular persistence. *eLife*, 12, 2023.

[12610] Farah Haddad, Amro M Soliman, Michael E Wong, Emilie H Albers, Shawna L Semple, Débora Torrealba, Ryan D Heimroth, Asif Nashiry, Keith B Tierney, and Daniel R Barreda. Fever integrates antimicrobial defences, inflammation control, and tissue repair in a cold-blooded vertebrate. *eLife*, 12, 2023.

[12611] Daniel Haenelt, Robert Trampel, Shahin Nasr, Jonathan R Polimeni, Roger BH Tootell, Martin I Sereno, Kerrin J Pine, Luke J Edwards, Saskia Helbling, and Nikolaus Weiskopf. High-resolution quantitative and functional MRI indicate lower myelination of thin and thick stripes in human secondary visual cortex. *eLife*, 12, 2023.

[12612] Omar A Hafez, Benjamin Escribano, Rouven L Ziegler, Jan J Hirtz, Ernst Niebur, and Jan Pielage. The cellular architecture of memory modules in Drosophila supports stochastic input integration. *eLife*, 12, 2023.

[12613] Paul Hage, In Kyu Jang, Vivian Looi, Mohammad Amin Fakharian, Simon P Orozco, Jay S Pi, Ehsan Sedaghat-Nejad, and Reza Shadmehr. Effort cost of harvest affects decisions and movement vigor of marmosets during foraging. *eLife*, 12, 2023.

[12614] Hanako Hagio, Wataru Koyama, Shiori Hosaka, Aysenur Deniz Song, Janchiv Narantsatsral, Koji Matsuda, Takashi Shimizu, Shoko Hososhima, Satoshi P Tsunoda, Hideki Kandori, and Masahiko Hibi. Optogenetic manipulation of neuronal and cardiomyocyte functions in zebrafish using microbial rhodopsins and adenylyl cyclases. *eLife*, 12, 2023.

[12615] Hanako Hagio, Wataru Koyama, Shiori Hosaka, Aysenur Deniz Song, Janchiv Narantsatsral, Koji Matsuda, Tomohiro Sugihara, Takashi Shimizu, Mitsumasa Koyanagi, Akihisa Terakita, and Masahiko Hibi. Optogenetic manipulation of Gq- and Gi/o-coupled receptor signaling in neurons and heart muscle cells. *eLife*, 12, 2023.

[12616] Emma A Hall, Dhivya Kumar, Suzanna L Prosser, Patricia L Yeyati, Vicente Herranz-Pérez, Jose Manuel García-Verdugo, Lorraine Rose, Lisa McKie, Daniel O Dodd, Peter A Tennant, Roly Megaw, Laura C Murphy, Marisa F Ferreira, Graeme Grimes, Lucy Williams, Tooba Quidwai, Laurence Pelletier, Jeremy F Reiter, and Pleasantine Mill. Centriolar satellites expedite mother centriole remodeling to promote ciliogenesis. *eLife*, 12, 2023.

[12617] Nicole Xiao Han and Miguel Patricio Eckstein. Inferential eye movement control while following dynamic gaze. *eLife*, 12, 2023.

[12618] Yong Han, Yang He, Lauren Harris, Yong Xu, and Qi Wu. Identification of a GABAergic neural circuit governing leptin signaling deficiency-induced obesity. *eLife*, 12, 2023.

[12619] Linda Handlin, Giovanni Novembre, Helene Lindholm, Robin Kämpe, Elisabeth Paul, and India Morrison. Human endogenous oxytocin and its neural correlates show adaptive responses to social touch based on recent social context. *eLife*, 12, 2023.

[12620] Benjamin Hänisch, Justine Y Hansen, Boris C Bernhardt, Simon B Eickhoff, Juergen Dukart, Bratislav Misic, and Sofie Louise Valk. Cerebral chemoarchitecture shares organizational traits with brain structure and function. *eLife*, 12, 2023.

[12621] Thomas Hansen, Jocelyn Lee, Naama Reicher, Gil Ovadia, Shuaiqi Guo, Wangbiao Guo, Jun Liu, Ido Braslavsky, Yinon Rudich, and Peter L Davies. Ice nucleation proteins self-assemble into large fibres to trigger freezing at near 0 °C. *eLife*, 12, 2023.

[12622] Yuchen Hao, Estelle Toulmé, Benjamin König, Christian Rosenmund, and Andrew JR Plested. Targeted sensors for glutamatergic neurotransmission. *eLife*, 12, 2023.

[12623] Yuichiro Hara and Shigehiro Kuraku. The impact of local genomic properties on the evolutionary fate of genes. *eLife*, 12, 2023.

[12624] Yoichiro Harada, Yu Mizote, Takehiro Suzuki, Akiyoshi Hirayama, Satsuki Ikeda, Mikako Nishida, Toru Hiratsuka, Ayaka Ueda, Yusuke Imagawa, Kento Maeda, Yuki Ohkawa, Junko Murai, Hudson H Freeze, Eiji Miyoshi, Shigeki Higashiyama, Heiichiro Udono, Naoshi Dohmae, Hideaki Tahara, and Naoyuki Taniguchi. Metabolic clogging of mannose triggers dNTP loss and genomic instability in human cancer cells. *eLife*, 12, 2023.

[12625] Emerson F Harkin, Michael B Lynn, Alexandre Payeur, Jean-FranÇois Boucher, Léa Caya-Bissonnette, Dominic Cyr, Chloe Stewart, André Longtin, Richard Naud, and Jean-Claude Béïque. Temporal derivative computation in the dorsal raphe network revealed by an experimentally driven augmented integrate-and-fire modeling framework. *eLife*, 12, 2023.

[12626] David J Harning, Samuel Sacco, Kesara Anamthawat-Jónsson, Nicolò Ardenghi, Thor Thordarson, Jonathan H Raberg, Julio Sepúlveda, áslaug Geirsdóttir, Beth Shapiro, and Gifford H Miller. Delayed postglacial colonization of Betula in Iceland and the circum North Atlantic. *eLife*, 12, 2023.

[12627] Em P Harrington, Riley B Catenacci, Matthew D Smith, Dongeun Heo, Cecilia E Miller, Keya R Meyers, Jenna Glatzer, Dwight E Bergles, and Peter A Calabresi. MHC class I and MHC class II reporter mice enable analysis of immune oligodendroglia in mouse models of multiple sclerosis. *eLife*, 12, 2023.

[12628] Noah J Harris, Meredith L Jenkins, Sung-Eun Nam, Manoj K Rathinaswamy, Matthew AH Parson, Harish Ranga-Prasad, Udit Dalwadi, Brandon E Moeller, Eleanor Sheeky, Scott D Hansen, Calvin K Yip, and John E Burke. Allosteric activation or inhibition of PI3K$\gamma$ mediated through conformational changes in the p110$\gamma$ helical domain. *eLife*, 12, 2023.

[12629] Scott C Harris and Felice A Dunn. Asymmetric retinal direction tuning predicts optokinetic eye movements across stimulus conditions. *eLife*, 12, 2023.

[12630] Madeleine L Hart, Evan Quon, Anna-Lena BG Vigil, Ian A Engstrom, Oliver J Newsom, Kristian Davidsen, Pia Hoellerbauer, Samantha M Carlisle, and Lucas B Sullivan. Mitochondrial redox adaptations enable alternative aspartate synthesis in SDH-deficient cells. *eLife*, 12, 2023.

[12631] Eva Hasel de Carvalho, Shivani S Dharmadhikari, Kateryna Shkarina, Jingwei Rachel Xiong, Bruno Reversade, Petr Broz, and Maria Leptin. The Opto-inflammasome in zebrafish as a tool to study cell and tissue responses to speck formation and cell death. *eLife*, 12, 2023.

[12632] Josue Haubrich and Karim Nader. Network-level changes in the brain underlie fear memory strength. *eLife*, 12, 2023.

[12633] Tamarinde L Haven, Susan Abunijela, and Nicole Hildebrand. Biomedical supervisors' role modeling of open science practices. *eLife*, 12, 2023.

[12634] Marito Hayashi, Judith A Kaye, Ella R Douglas, Narendra R Joshi, Fiona M Gribble, Frank Reimann, and Stephen D Liberles. Enteroendocrine cell lineages that differentially control feeding and gut motility. *eLife*, 12, 2023.

[12635] Rebecca Frank Hayward, F Phil Brooks, Shang Yang, Shiqiang Gao, and Adam E Cohen. Diminishing neuronal acidification by channelrhodopsins with low proton conduction. *eLife*, 12, 2023.

[12636] Lily S He, Yujia Qi, Corey AH Allard, Wendy A Valencia-Montoya, Stephanie P Krueger, Keiko Weir, Agnese Seminara, and Nicholas W Bellono. Molecular tuning of sea anemone stinging. *eLife*, 12, 2023.

[12637] Yang He, Taohui Liu, Quansheng He, Wei Ke, Xiaoyu Li, Jinjin Du, Suixin Deng, Zhenfeng Shu, Jialin Wu, Baozhi Yang, Yuqing Wang, Ying Mao, Yanxia Rao, Yousheng Shu, and Bo Peng. Microglia facilitate and stabilize the response to general anesthesia via modulating the neuronal network in a brain region-specific manner. *eLife*, 12, 2023.

[12638] Yi He, Yanhua H Huang, Oliver M Schlüter, and Yan Dong. Cue- versus reward-encoding basolateral amygdala projections to nucleus accumbens. *eLife*, 12, 2023.

[12639] Martin N Hebart, Oliver Contier, Lina Teichmann, Adam H Rockter, Charles Y Zheng, Alexis Kidder, Anna Corriveau, Maryam Vaziri-Pashkam, and Chris I Baker. THINGS-data, a multimodal collection of large-scale datasets for investigating object representations in human brain and behavior. *eLife*, 12, 2023.

[12640] David B Heisler, Kristen A Johnson, Duo H Ma, Maikke B Ohlson, Lishu Zhang, Michelle Tran, Chase D Corley, Michael E Abrams, Jeffrey G McDonald, John W Schoggins, Neal M Alto, and Arun Radhakrishnan. A concerted mechanism involving ACAT and SREBPs by which oxysterols deplete accessible cholesterol to restrict microbial infection. *eLife*, 12, 2023.

[12641] Stewart Heitmann, Jamie I Vandenberg, and Adam P Hill. Assessing drug safety by identifying the axis of arrhythmia in cardiomyocyte electrophysiology. *eLife*, 12, 2023.

[12642] Clair R Henthorn, Paul M Airs, Emma K Neumann, and Mostafa Zamanian. Resolving the origins of secretory products and anthelmintic responses in a human parasitic nematode at single-cell resolution. *eLife*, 12, 2023.

[12643] Kyra Hermanns, Marco Marklewitz, Florian Zirkel, Anne Kopp, Stephanie Kramer-Schadt, and Sandra Junglen. Mosquito community composition shapes virus prevalence patterns along anthropogenic disturbance gradients. *eLife*, 12, 2023.

[12644] Ines Hernandez-Perez, Javier Rubio, Adrian Baumann, Henrique Girao, Miriam Ferrando, Elena Rebollo, Anna M Aragay, and María Isabel Geli. Kazrin promotes dynein/dynactin-dependent traffic from early to recycling endosomes. *eLife*, 12, 2023.

[12645] R Samuel Herron, Alexander K Kunisky, Jessica R Madden, Vivian I Anyaeche, May Z Maung, and Hun-Way Hwang. A twin UGUA motif directs the balance between gene isoforms through CFIm and the mTORC1 signaling pathway. *eLife*, 12, 2023.

[12646] Ann-Kathrin Herzfeldt, Marta Puig Gamez, Eva Martin, Lukasz Miloslaw Boryn, Praveen Baskaran, Heinrich J Huber, Michael Schuler, John E Park, and Lee Kim Swee. Complementary CRISPR screen highlights the contrasting role of membrane-bound and soluble ICAM-1 in regulating antigen-specific tumor cell killing by cytotoxic T cells. *eLife*, 12, 2023.

[12647] Katja Heuer, Nicolas Traut, Alexandra Allison de Sousa, Sofie Louise Valk, Julien Clavel, and Roberto Toro. Diversity and evolution of cerebellar folding in mammals. *eLife*, 12, 2023.

[12648] Joshua R Heyza, Mariia Mikhova, Aastha Bahl, David G Broadbent, and Jens C Schmidt. Systematic analysis of the molecular and biophysical properties of key DNA damage response factors. *eLife*, 12, 2023.

[12649] Sanne Hiddingh, Aridaman Pandit, Fleurieke Verhagen, Rianne Rijken, Nila Hendrika Servaas, Rina CGK Wichers, Ninette H ten Dam-van Loon, Saskia M Imhof, Timothy RDJ Radstake, Joke H de Boer, and Jonas JW Kuiper. Transcriptome network analysis implicates CX3CR1-positive type 3 dendritic cells in non-infectious uveitis. *eLife*, 12, 2023.

[12650] Satoshi Higurashi, Sachio Tsukada, Binta Maria Aleogho, Joo Hyun Park, Yana Al-Hebri, Masaru Tanaka, Shunji Nakano, Ikue Mori, and Kentaro Noma. Bacterial diet affects the age-dependent decline of associative learning in Caenorhabditis elegans. *eLife*, 12, 2023.

[12651] Nathaniel J Himmel, Akira Sakurai, Atit A Patel, Shatabdi Bhattacharjee, Jamin M Letcher, Maggie N Benson, Thomas R Gray, Gennady S Cymbalyuk, and Daniel N Cox. Chloride-dependent mechanisms of multimodal sensory discrimination and nociceptive sensitization in Drosophila. *eLife*, 12, 2023.

[12652] Elodie Hinnekens, Marianne Barbu-Roth, Manh-Cuong Do, Bastien Berret, and Caroline Teulier. Generating variability from motor primitives during infant locomotor development. *eLife*, 12, 2023.

[12653] Minato Hirano, Gaddiel Galarza-Muñoz, Chloe Nagasawa, Geraldine Schott, Liuyang Wang, Alejandro L Antonia, Vaibhav Jain, Xiaoying Yu, Steven G Widen, Farren BS Briggs, Simon G Gregory, Dennis C Ko, William S Fagg, Shelton Bradrick, and Mariano A Garcia-Blanco. The RNA helicase DDX39B activates FOXP3 RNA splicing to control T regulatory cell fate. *eLife*, 12, 2023.

[12654] Jessica Xin Hjaltelin, Sif Ingibergsdóttir Novitski, Isabella Friis Jørgensen, Troels Siggaard, Siri Amalie Vulpius, David Westergaard, Julia Sidenius Johansen, Inna M Chen, Lars Juhl Jensen, and Søren Brunak. Pancreatic cancer symptom trajectories from Danish registry data and free text in electronic health records. *eLife*, 12, 2023.

[12655] Brandon Ho, Ethan J Sanford, Raphael Loll-Krippleber, Nikko P Torres, Marcus B Smolka, and Grant W Brown. Mec1-independent activation of the Rad53 checkpoint kinase revealed by quantitative analysis of protein localization dynamics. *eLife*, 12, 2023.

[12656] Kevin Yueh Lin Ho, Rosalyn Leigh Carr, Alexandra Dmitria Dvoskin, and Guy Tanentzapf. Kinetics of blood cell differentiation during hematopoiesis revealed by quantitative long-term live imaging. *eLife*, 12, 2023.

[12657] Peh Joo Ho, Iain BeeHuat Tan, Dawn Qingqing Chong, Chiea Chuen Khor, Jian-Min Yuan, Woon-Puay Koh, Rajkumar Dorajoo, and Jingmei Li. Polygenic risk scores for the prediction of common cancers in East Asians: A population-based prospective cohort study. *eLife*, 12, 2023.

[12658] Huu Hoang, Shinichiro Tsutsumi, Masanori Matsuzaki, Masanobu Kano, Mitsuo Kawato, Kazuo Kitamura, and Keisuke Toyama. Dynamic organization of cerebellar climbing fiber response and synchrony in multiple functional components reduces dimensions for reinforcement learning. *eLife*, 12, 2023.

[12659] Daniel Hoces, Giorgia Greter, Markus Arnoldini, Melanie L Stäubli, Claudia Moresi, Anna Sintsova, Sara Berent, Isabel Kolinko, Florence Bansept, Aurore Woller, Janine Häfliger, Eric Martens, Wolf-Dietrich Hardt, Shinichi Sunagawa, Claude Loverdo, and Emma Slack. Fitness advantage of Bacteroides thetaiotaomicron capsular polysaccharide in the mouse gut depends on the resident microbiota. *eLife*, 12, 2023.

[12660] Flavia Hodel, Zhi Ming Xu, Christian Wandall Thorball, Roxane de La Harpe, Prunelle Letang-Mathieu, Nicole Brenner, Julia Butt, Noemi Bender, Tim Waterboer, Pedro Manuel Marques-Vidal, Peter Vollenweider, Julien Vaucher, and Jacques Fellay. Associations of genetic and infectious risk factors with coronary heart disease. *eLife*, 12, 2023.

[12661] Thomas Hoffmann, Alexandra Hörmann, Maja Corcokovic, Jakub Zmajkovic, Matthias Hinterndorfer, Jasko Salkanovic, Fiona Spreitzer, Anna Köferle, Katrin Gitschtaler, Alexandra Popa, Sarah Oberndorfer, Florian Andersch, Markus Schaefer, Michaela Fellner, Nicole Budano, Jan G Ruppert, Paolo Chetta, Melanie Wurm, Johannes Zuber, and Ralph A Neumüller. Precision RNAi using synthetic shRNAmir target sites. *eLife*, 12, 2023.

[12662] Jonas Maurice Holzinger, Martina Toelge, Maren Werner, Katharina Ursula Ederer, Heiko Ingo Siegmund, David Peterhoff, Stefan Helmut Blaas, Nicolas Gisch, Christoph Brochhausen, André Gessner, and Sigrid Bülow. Scorpionfish BPI is highly active against multiple drug-resistant Pseudomonas aeruginosa isolates from people with cystic fibrosis. *eLife*, 12, 2023.

[12663] Sebastian Hormigo, Ji Zhou, Dorian Chabbert, Sarmad Sajid, Natan Busel, and Manuel Castro-Alamancos. Zona incerta distributes a broad movement signal that modulates behavior. *eLife*, 12, 2023.

[12664] Isabella Horton, Conor J Kelly, Adam Dziulko, David M Simpson, and Edward B Chuong. Mouse B2 SINE elements function as IFN-inducible enhancers. *eLife*, 12, 2023.

[12665] Changkun Hu, Taylor Bugbee, Rachel Palinski, Ibukun A Akinyemi, Michael T McIntosh, Thomas MacCarthy, Sumita Bhaduri-McIntosh, and Nicholas Wallace. Beta human papillomavirus 8E6 promotes alternative end joining. *eLife*, 12, 2023.

[12666] Hang Hu, Rachel E Hostetler, and Ariel Agmon. Ultrafast (400 Hz) network oscillations induced in mouse barrel cortex by optogenetic activation of thalamocortical axons. *eLife*, 12, 2023.

[12667] Jie Hu, Arkady Konovalov, and Christian C Ruff. A unified neural account of contextual and individual differences in altruism. *eLife*, 12, 2023.

[12668] Jing Hu, Bingna Zhou, Xiaoyun Lin, Qian Zhang, Feifei Guan, Lei Sun, Jiayi Liu, Ou Wang, Yan Jiang, Wei-bo Xia, Xiaoping Xing, and Mei Li. Impaired bone strength and bone microstructure in a novel early-onset osteoporotic rat model with a clinically relevant PLS3 mutation. *eLife*, 12, 2023.

[12669] Jun Hu, Zhe Zhang, Sai Xiao, Yalei Cao, Yinghong Chen, Jiaming Weng, Hui Jiang, Wei Li, Jia-Yu Chen, and Chao Liu. Microhomology-mediated circular DNA formation from oligonucleosomal fragments during spermatogenesis. *eLife*, 12, 2023.

[12670] Yucen Hu, Jingxiu Xu, Erqing Gao, Xueyuan Fan, Jieli Wei, Bingcheng Ye, Suhong Xu, and Weirui Ma. Enhanced single RNA imaging reveals dynamic gene expression in live animals. *eLife*, 12, 2023.

[12671] Haoyan Huang, Meng Qian, Yue Liu, Shang Chen, Huifang Li, Zhibo Han, Zhong-Chao Han, Xiang-Mei Chen, Qiang Zhao, and Zongjin Li. Genetically engineered mesenchymal stem cells as a nitric oxide reservoir for acute kidney injury therapy. *eLife*, 12, 2023.

[12672] Jiawen Huang, Isabel Velarde, Wei Ji Ma, and Christopher Baldassano. Schema-based predictive eye movements support sequential memory encoding. *eLife*, 12, 2023.

[12673] Kaixin Huang, Laura K Milton, Harry Dempsey, Stephen J Power, Kyna-Anne Conn, Zane B Andrews, and Claire J Foldi. Rapid, automated, and experimenter-free touchscreen testing reveals reciprocal interactions between cognitive flexibility and activity-based anorexia in female rats. *eLife*, 12, 2023.

[12674] Lingli Huang, Wenqing Li, Xingxing Dai, Shuai Zhao, Bo Xu, Fengsong Wang, Ren-Tao Jin, Lihua Luo, Limin Wu, Xue Jiang, Yu Cheng, Jiaqi Zou, Caoling Xu, Xianhong Tong, Heng-Yu Fan, Han Zhao, and Jianqiang Bao. Biallelic variants in MAD2L1BP (p31comet) cause female infertility characterized by oocyte maturation arrest. *eLife*, 12, 2023.

[12675] Long Shuang Huang, Mohammad Anas, Jingsong Xu, Bisheng Zhou, Peter T Toth, Yamuna Krishnan, Anke Di, and Asrar B Malik. Endosomal trafficking of two-pore K+ efflux channel TWIK2 to plasmalemma mediates NLRP3 inflammasome activation and inflammatory injury. *eLife*, 12, 2023.

[12676] Mi Huang, Daniel Coral, Hamidreza Ardalani, Peter Spegel, Alham Saadat, Melina Claussnitzer, Hindrik Mulder, Paul W Franks, and Sebastian Kalamajski. Identification of a weight loss-associated causal eQTL in MTIF3 and the effects of MTIF3 deficiency on human adipocyte function. *eLife*, 12, 2023.

[12677] Rong Huang, Jiahui Shao, Zhihui Xu, Yuqi Chen, Yunpeng Liu, Dandan Wang, Haichao Feng, Weibing Xun, Qirong Shen, Nan Zhang, and Ruifu Zhang. A toxin-mediated policing system in Bacillus optimizes division of labor via penalizing cheater-like nonproducers. *eLife*, 12, 2023.

[12678] Xin Huang, Tian-Neng Zhu, Ying-Chao Liu, Guo-An Qi, Jian-Nan Zhang, and Guo-Bo Chen. Efficient estimation for large-scale linkage disequilibrium patterns of the human genome. *eLife*, 12, 2023.

[12679] Alyssa Huff, Marlusa Karlen-Amarante, Luiz M Oliveira, and Jan-Marino Ramirez. Role of the postinspiratory complex in regulating swallow–breathing coordination and other laryngeal behaviors. *eLife*, 12, 2023.

[12680] Brandon W Hughes, Benjamin M Siemsen, Evgeny Tsvetkov, Stefano Berto, Jaswinder Kumar, Rebecca G Cornbrooks, Rose Marie Akiki, Jennifer Y Cho, Jordan S Carter, Kirsten K Snyder, Ahlem Assali, Michael D Scofield, Christopher W Cowan, and Makoto Taniguchi. NPAS4 in the medial prefrontal cortex mediates chronic social defeat stress-induced anhedonia-like behavior and reductions in excitatory synapses. *eLife*, 12, 2023.

[12681] Victoria T Hunniford, Agnes Grudniewicz, Dean A Fergusson, Joshua Montroy, Emma Grigor, Casey Lansdell, and Manoj M Lalu. A systematic assessment of preclinical multilaboratory studies and a comparison to single laboratory studies. *eLife*, 12, 2023.

[12682] Adel Hussein, Shujie Fan, Maria Lopez-Redondo, Ian Kenney, Xihui Zhang, Oliver Beckstein, and David L Stokes. Energy coupling and stoichiometry of Zn2+/H+ antiport by the prokaryotic cation diffusion facilitator YiiP. *eLife*, 12, 2023.

[12683] Melina Hußmann, Dörte Schulte, Sarah Weischer, Claudia Carlantoni, Hiroyuki Nakajima, Naoki Mochizuki, Didier YR Stainier, Thomas Zobel, Manuel Koch, and Stefan Schulte-Merker. Svep1 is a binding ligand of Tie1 and affects specific aspects of facial lymphatic development in a Vegfc-independent manner. *eLife*, 12, 2023.

[12684] Jae Yeon Hwang, Pengxin Chai, Shoaib Nawaz, Jungmin Choi, Francesc Lopez-Giraldez, Shabir Hussain, Kaya Bilguvar, Shrikant Mane, Richard P Lifton, Wasim Ahmad, Kai Zhang, and Jean-Ju Chung. LRRC23 truncation impairs radial spoke 3 head assembly and sperm motility underlying male infertility. *eLife*, 12, 2023.

[12685] Alexandre Hyafil, Jaime de la Rocha, Cristina Pericas, Leor N Katz, Alexander C Huk, and Jonathan W Pillow. Temporal integration is a robust feature of perceptual decisions. *eLife*, 12, 2023.

[12686] Consuelo Ibar, Krishna Chinthalapudi, Sarah M Heissler, and Kenneth D Irvine. Competition between myosin II and $\beta$H-spectrin regulates cytoskeletal tension. *eLife*, 12, 2023.

[12687] Airat Ibragimov, Xin Yang Bing, Yulii V Shidlovskii, Michael Levine, Pavel Georgiev, and Paul Schedl. lncRNA read-through regulates the BX-C insulator Fub-1. *eLife*, 12, 2023.

[12688] Leena Ali Ibrahim, Brie Wamsley, Norah Alghamdi, Nusrath Yusuf, Elaine Sevier, Ariel Hairston, Mia Sherer, Xavier Hubert Jaglin, Qing Xu, Lihua Guo, Alireza Khodadadi-Jamayran, Emilia Favuzzi, Yuan Yuan, Jordane Dimidschstein, Robert B Darnell, and Gordon Fishell. Nova proteins direct synaptic integration of somatostatin interneurons through activity-dependent alternative splicing. *eLife*, 12, 2023.

[12689] Sotaro Ichinose, Yoshihiro Susuki, Nobutake Hosoi, Ryosuke Kaneko, Mizuho Ebihara, Hirokazu Hirai, and Hirohide Iwasaki. Interaction between Teneurin-2 and microtubules via EB proteins provides a platform for GABAA receptor exocytosis. *eLife*, 12, 2023.

[12690] Dinesh C Indurthi and Anthony Auerbach. Agonist efficiency links binding and gating in a nicotinic receptor. *eLife*, 12, 2023.

[12691] Kazuki Inoue, Yongli Qin, Yuhan Xia, Jie Han, Ruoxi Yuan, Jun Sun, Ren Xu, Jean X Jiang, Matthew B Greenblatt, and Baohong Zhao. Bone marrow Adipoq-lineage progenitors are a major cellular source of M-CSF that dominates bone marrow macrophage development, osteoclastogenesis, and bone mass. *eLife*, 12, 2023.

[12692] Domenica Ippolito and Dominique A Glauser. Multiple antagonist calcium-dependent mechanisms control CaM kinase-1 subcellular localization in a C. elegans thermal nociceptor. *eLife*, 12, 2023.

[12693] Asako Ishii, Jianyu Shan, Xin Sheng, Eunchul Kim, Akimasa Watanabe, Makio Yokono, Chiyo Noda, Chihong Song, Kazuyoshi Murata, Zhenfeng Liu, and Jun Minagawa. The photosystem I supercomplex from a primordial green alga Ostreococcus tauri harbors three light-harvesting complex trimers. *eLife*, 12, 2023.

[12694] Yuu Ishii, Hironori Ishii, Takeshi Kuroha, Ryusuke Yokoyama, Ryusaku Deguchi, Kazuhiko Nishitani, Jun Minagawa, Masakado Kawata, Shunichi Takahashi, and Shinichiro Maruyama. Environmental pH signals the release of monosaccharides from cell wall in coral symbiotic alga. *eLife*, 12, 2023.

[12695] Hiroaki Ishikawa, Jeremy Moore, Dennis R Diener, Markus Delling, and Wallace F Marshall. Testing the ion-current model for flagellar length sensing and IFT regulation. *eLife*, 12, 2023.

[12696] Igor Y Iskusnykh, Nikolai Fattakhov, Yiran Li, Laure Bihannic, Matthew K Kirchner, Ekaterina Y Steshina, Paul A Northcott, and Victor V Chizhikov. Lmx1a is a master regulator of the cortical hem. *eLife*, 12, 2023.

[12697] Yasuko Isoe, Ryohei Nakamura, Shigenori Nonaka, Yasuhiro Kamei, Teruhiro Okuyama, Naoyuki Yamamoto, Hideaki Takeuchi, and Hiroyuki Takeda. Epigenetically distinct synaptic architecture in clonal compartments in the teleostean dorsal pallium. *eLife*, 12, 2023.

[12698] Yurika Ito, Mari Yamagata, Takuya Yamamoto, Katsuya Hirasaka, Takeshi Nikawa, and Takahiko Sato. The reciprocal regulation between mitochondrial-associated membranes and Notch signaling in skeletal muscle atrophy. *eLife*, 12, 2023.

[12699] Krishnan S Iyer, Chaitra Prabhakara, Satyajit Mayor, and Madan Rao. Cellular compartmentalisation and receptor promiscuity as a strategy for accurate and robust inference of position during morphogenesis. *eLife*, 12, 2023.

[12700] William D Jackson, Chiara Giacomassi, Sophie Ward, Amber Owen, Tiago C Luis, Sarah Spear, Kevin J Woollard, Cecilia Johansson, Jessica Strid, and Marina Botto. TLR7 activation at epithelial barriers promotes emergency myelopoiesis and lung antiviral immunity. *eLife*, 12, 2023.

[12701] Jessica Jacobs-Li, Weiyi Tang, Can Li, and Marianne E Bronner. Single-cell profiling coupled with lineage analysis reveals vagal and sacral neural crest contributions to the developing enteric nervous system. *eLife*, 12, 2023.

[12702] Evelyn Jagoda, Davide Marnetto, Gayani Senevirathne, Victoria Gonzalez, Kaushal Baid, Francesco Montinaro, Daniel Richard, Darryl Falzarano, Emmanuelle V LeBlanc, Che C Colpitts, Arinjay Banerjee, Luca Pagani, and Terence D Capellini. Regulatory dissection of the severe COVID-19 risk locus introgressed by Neanderthals. *eLife*, 12, 2023.

[12703] Kanishk Jain, Matthew R Marunde, Jonathan M Burg, Susan L Gloor, Faith M Joseph, Karl F Poncha, Zachary B Gillespie, Keli L Rodriguez, Irina K Popova, Nathan W Hall, Anup Vaidya, Sarah A Howard, Hailey F Taylor, Laylo Mukhsinova, Ugochi C Onuoha, Emily F Patteson, Spencer W Cooke, Bethany C Taylor, Ellen N Weinzapfel, Marcus A Cheek, Matthew J Meiners, Geoffrey C Fox, Kevin EW Namitz, Martis W Cowles, Krzysztof Krajewski, Zu-Wen Sun, Michael S Cosgrove, Nicolas L Young, Michael-Christopher Keogh, and Brian D Strahl. An acetylation-mediated chromatin switch governs H3K4 methylation read-write capability. *eLife*, 12, 2023.

[12704] Anthony MV Jakob and Samuel J Gershman. Rate-distortion theory of neural coding and its implications for working memory. *eLife*, 12, 2023.

[12705] Binta Jalloh, Carly L Lancaster, J Christopher Rounds, Brianna E Brown, Sara W Leung, Ayan Banerjee, Derrick J Morton, Rick S Bienkowski, Milo B Fasken, Isaac J Kremsky, Matthew Tegowski, Kate Meyer, Anita Corbett, and Ken Moberg. The Drosophila Nab2 RNA binding protein inhibits m6A methylation and male-specific splicing of Sex lethal transcript in female neuronal tissue. *eLife*, 12, 2023.

[12706] Bhagyshree Jamge, Zdravko J Lorković, Elin Axelsson, Akihisa Osakabe, Vikas Shukla, Ramesh Yelagandula, Svetlana Akimcheva, Annika Luisa Kuehn, and Frédéric Berger. Histone variants shape chromatin states in Arabidopsis. *eLife*, 12, 2023.

[12707] Abhishek Jamwal, Cristina F Constantin, Stephan Hirschi, Sebastian Henrich, Wolfgang Bildl, Bernd Fakler, Simon J Draper, Uwe Schulte, and Matthew K Higgins. Erythrocyte invasion-neutralising antibodies prevent Plasmodium falciparum RH5 from binding to basigin-containing membrane protein complexes. *eLife*, 12, 2023.

[12708] Pramod K Jangir, Lois Ogunlana, Petra Szili, Marton Czikkely, Liam P Shaw, Emily J Stevens, Yang Yu, Qiue Yang, Yang Wang, Csaba Pál, Timothy R Walsh, and Craig R MacLean. The evolution of colistin resistance increases bacterial resistance to host antimicrobial peptides and virulence. *eLife*, 12, 2023.

[12709] Alana Jaskir and Michael J Frank. On the normative advantages of dopamine and striatal opponency for learning and choice. *eLife*, 12, 2023.

[12710] Alagie Jassey, James Logue, Stuart Weston, Michael A Wagner, Ganna Galitska, Katelyn Miller, Matthew Frieman, and William T Jackson. SIRT-1 is required for release of enveloped enteroviruses. *eLife*, 12, 2023.

[12711] Suvi Jauhiainen, Henna Ilmonen, Pia Vuola, Heta Rasinkangas, Heidi H Pulkkinen, Sara Keränen, Miika Kiema, Jade J Liikkanen, Nihay Laham-Karam, Svetlana Laidinen, Mustafa Beter, Einari Aavik, Kimmo Lappalainen, Jouko Lohi, Johanna Aronniemi, Tiit örd, Minna U Kaikkonen, Päivi Salminen, Erkki Tukiainen, Seppo Ylä-Herttuala, and Johanna P Laakkonen. ErbB signaling is a potential therapeutic target for vascular lesions with fibrous component. *eLife*, 12, 2023.

[12712] Sehrish Javed, Ya-Ting Chang, Yoobin Cho, Yu-Ju Lee, Hao-Cheng Chang, Minza Haque, Yu Cheng Lin, and Wei-Hsiang Huang. Smith–Magenis syndrome protein RAI1 regulates body weight homeostasis through hypothalamic BDNF-producing neurons and neurotrophin downstream signalling. *eLife*, 12, 2023.

[12713] Fabrice Jean-Pierre, Thomas H Hampton, Daniel Schultz, Deborah A Hogan, Marie-Christine Groleau, Eric Déziel, and George A O'Toole. Community composition shapes microbial-specific phenotypes in a cystic fibrosis polymicrobial model system. *eLife*, 12, 2023.

[12714] Meghan Jelen, Pierre-Yves Musso, Pierre Junca, and Michael D Gordon. Optogenetic induction of appetitive and aversive taste memories in Drosophila. *eLife*, 12, 2023.

[12715] Jayashree Jena, Luis Miguel García-Peña, Eric T Weatherford, Alex Marti, Sarah H Bjorkman, Kevin Kato, Jivan Koneru, Jason H Chen, Randy J Seeley, E Dale Abel, and Renata O Pereira. GDF15 is required for cold-induced thermogenesis and contributes to improved systemic metabolic health following loss of OPA1 in brown adipocytes. *eLife*, 12, 2023.

[12716] Myeongjune Jeon, Goowon Jeong, Yupeng Yang, Xiao Luo, Daesong Jeong, Jinseul Kyung, Youbong Hyun, Yuehui He, and Ilha Lee. Vernalization-triggered expression of the antisense transcript COOLAIR is mediated by CBF genes. *eLife*, 12, 2023.

[12717] Cayla E Jewett, Bailey L McCurdy, Eileen T O'Toole, Alexander J Stemm-Wolf, Katherine S Given, Carrie H Lin, Valerie Olsen, Whitney Martin, Laura Reinholdt, Joaquín M Espinosa, Kelly D Sullivan, Wendy B Macklin, Rytis Prekeris, and Chad G Pearson. Trisomy 21 induces pericentrosomal crowding delaying primary ciliogenesis and mouse cerebellar development. *eLife*, 12, 2023.

[12718] Shihai Jia, Evan M Ratzan, Ellison J Goodrich, Raisa Abrar, Luke Heiland, Basile Tarchini, and Michael R Deans. The dark kinase STK32A regulates hair cell planar polarity opposite of EMX2 in the developing mouse inner ear. *eLife*, 12, 2023.

[12719] Xiaobo Jia, Jiemin Zhu, Xiling Bian, Sulin Liu, Sihan Yu, Wenjun Liang, Lifen Jiang, Renbo Mao, Wenxia Zhang, and Yi Rao. Importance of glutamine in synaptic vesicles revealed by functional studies of SLC6A17 and its mutations pathogenic for intellectual disability. *eLife*, 12, 2023.

[12720] Guo Jiahui, Ma Feilong, Samuel A Nastase, James V Haxby, and M Ida Gobbini. Cross-movie prediction of individualized functional topography. *eLife*, 12, 2023.

[12721] Lan Jiang, Ge Liu, Annette Oeser, Andrea Ihegword, Alyson L Dickson, Laura L Daniel, Adriana M Hung, Nancy J Cox, Cecilia P Chung, Wei-Qi Wei, C Michael Stein, and Qiping Feng. Association between APOL1 risk variants and the occurrence of sepsis in Black patients hospitalized with infections: a retrospective cohort study. *eLife*, 12, 2023.

[12722] Yufei Jiang, Yan He, Songting Liu, Gaizhen Li, Dunjin Chen, Wenbo Deng, Ping Li, Ying Zhang, Jinxiang Wu, Jianing Li, Longmei Wang, Jiajing Lin, Haibin Wang, Shuangbo Kong, and Guixiu Shi. G$\alpha$q-PKD/PKC$\mu$ signal regulating the nuclear export of HDAC5 to induce the I$\kappa$B expression and limit the NF-$\kappa$B-mediated inflammatory response essential for early pregnancy. *eLife*, 12, 2023.

[12723] Michael Jigo, Daniel Tavdy, Marc M Himmelberg, and Marisa Carrasco. Cortical magnification eliminates differences in contrast sensitivity across but not around the visual field. *eLife*, 12, 2023.

[12724] Blanca Jimeno and Simon Verhulst. Meta-analysis reveals glucocorticoid levels reflect variation in metabolic rate, not 'stress'. *eLife*, 12, 2023.

[12725] Hui-Juan Jin, Tiechao Ruan, Siyu Dai, Xin-Yan Geng, Yihong Yang, Ying Shen, and Su-Ren Chen. Identification of CFAP52 as a novel diagnostic target of male infertility with defects of sperm head-tail connection and flagella development. *eLife*, 12, 2023.

[12726] Xavier Job and Konstantina Kilteni. Action does not enhance but attenuates predicted touch. *eLife*, 12, 2023.

[12727] Ann-Kathrin Joechner, Michael A Hahn, Georg Gruber, Kerstin Hoedlmoser, and Markus Werkle-Bergner. Sleep spindle maturity promotes slow oscillation-spindle coupling across child and adolescent development. *eLife*, 12, 2023.

[12728] Philippa Anne Johnson, Tessel Blom, Simon van Gaal, Daniel Feuerriegel, Stefan Bode, and Hinze Hogendoorn. Position representations of moving objects align with real-time position in the early visual response. *eLife*, 12, 2023.

[12729] Hannah Jones, Jenny A Willis, Lucy C Firth, Carlo NG Giachello, and Giorgio F Gilestro. A reductionist paradigm for high-throughput behavioural fingerprinting in Drosophila melanogaster. *eLife*, 12, 2023.

[12730] Sabrina A Jones, Jacob H Barfield, V Kindler Norman, and Woodrow L Shew. Scale-free behavioral dynamics directly linked with scale-free cortical dynamics. *eLife*, 12, 2023.

[12731] Elisabeth Jongsma, Anita Goyala, José Maria Mateos, and Collin Yvès Ewald. Removal of extracellular human amyloid beta aggregates by extracellular proteases in C. elegans. *eLife*, 12, 2023.

[12732] Rebecca Jordan and Georg B Keller. The locus coeruleus broadcasts prediction errors across the cortex to promote sensorimotor plasticity. *eLife*, 12, 2023.

[12733] Pooja Joshi, Shanea Estes, Deeptankar DeMazumder, Bjorn C Knollmann, and Swati Dey. Ryanodine receptor 2 inhibition reduces dispersion of cardiac repolarization, improves contractile function, and prevents sudden arrhythmic death in failing hearts. *eLife*, 12, 2023.

[12734] Thomas H Julian, Johnathan Cooper-Knock, Stuart MacGregor, Hui Guo, Tariq Aslam, Eleanor Sanderson, Graeme CM Black, and Panagiotis I Sergouniotis. Phenome-wide Mendelian randomisation analysis identifies causal factors for age-related macular degeneration. *eLife*, 12, 2023.

[12735] Aaron W Jurrjens, Marcus M Seldin, Corey Giles, Peter J Meikle, Brian G Drew, and Anna C Calkin. The potential of integrating human and mouse discovery platforms to advance our understanding of cardiometabolic diseases. *eLife*, 12, 2023.

[12736] Olivia Justynski, Kate Bridges, Will Krause, Maria Fernanda Forni, Quan M Phan, Teresa Sandoval-Schaefer, Kristyn Carter, Diane E King, Henry C Hsia, Michael I Gazes, Steven D Vyce, Ryan R Driskell, Kathryn Miller-Jensen, and Valerie Horsley. Apoptosis recognition receptors regulate skin tissue repair in mice. *eLife*, 12, 2023.

[12737] Takehiro Kado, Zarina Akbary, Daisuke Motooka, Ian L Sparks, Emily S Melzer, Shota Nakamura, Enrique R Rojas, Yasu S Morita, and M Sloan Siegrist. A cell wall synthase accelerates plasma membrane partitioning in mycobacteria. *eLife*, 12, 2023.

[12738] Sevim Kahraman, Kimitaka Shibue, Dario F De Jesus, Hyunki Kim, Jiang Hu, Debasish Manna, Bridget Wagner, Amit Choudhary, and Rohit N Kulkarni. Fluorescein-based sensors to purify human $\alpha$-cells for functional and transcriptomic analyses. *eLife*, 12, 2023.

[12739] Alicia Kairouani, Dominique Pontier, Claire Picart, Fabien Mounet, Yves Martinez, Lucie Le-Bot, Mathieu Fanuel, Philippe Hammann, Lucid Belmudes, Remy Merret, Jacinthe Azevedo, Marie-Christine Carpentier, Dominique Gagliardi, Yohann Couté, Richard Sibout, Natacha Bies-Etheve, and Thierry Lagrange. Cell-type-specific control of secondary cell wall formation by Musashi-type translational regulators in Arabidopsis. *eLife*, 12, 2023.

[12740] Yasutetsu Kanaoka, Koun Onodera, Kaori Watanabe, Yusaku Hayashi, Tadao Usui, Tadashi Uemura, and Yukako Hattori. Inter-organ Wingless/Ror/Akt signaling regulates nutrient-dependent hyperarborization of somatosensory neurons. *eLife*, 12, 2023.

[12741] Matheswaran Kandasamy, Uzi Gileadi, Pramila Rijal, Tiong Kit Tan, Lian N Lee, Jili Chen, Gennaro Prota, Paul Klenerman, Alain Townsend, and Vincenzo Cerundolo. Recombinant single-cycle influenza virus with exchangeable pseudotypes allows repeated immunization to augment anti-tumour immunity with immune checkpoint inhibitors. *eLife*, 12, 2023.

[12742] Tej Kandola, Shriram Venkatesan, Jiahui Zhang, Brooklyn T Lerbakken, Alex Von Schulze, Jillian F Blanck, Jianzheng Wu, Jay R Unruh, Paula Berry, Jeffrey J Lange, Andrew C Box, Malcolm Cook, Celeste Sagui, and Randal Halfmann. Pathologic polyglutamine aggregation begins with a self-poisoning polymer crystal. *eLife*, 12, 2023.

[12743] Kota Kaneko, Yan Liang, Qing Liu, Shuo Zhang, Alexander Scheiter, Dan Song, and Gen-Sheng Feng. Identification of CD133+ intercellsomes in intercellular communication to offset intracellular signal deficit. *eLife*, 12, 2023.

[12744] Ling Kang, Jonas Ranft, and Vincent Hakim. Beta oscillations and waves in motor cortex can be accounted for by the interplay of spatially structured connectivity and fluctuating inputs. *eLife*, 12, 2023.

[12745] Piotr Karaś, Klaudia Kochanowicz, Marcin Pitek, Przemyslaw Domanski, Igor Obuchowski, Barlomiej Tomiczek, and Krzysztof Liberek. Evolution towards simplicity in bacterial small heat shock protein system. *eLife*, 12, 2023.

[12746] Masha Karelina, Joseph J Noh, and Ron O Dror. How accurately can one predict drug binding modes using AlphaFold models? *eLife*, 12, 2023.

[12747] Javad Karimi Abadchi, Zahra Rezaei, Thomas Knöpfel, Bruce L McNaughton, and Majid H Mohajerani. Inhibition is a prevalent mode of activity in the neocortex around awake hippocampal ripples in mice. *eLife*, 12, 2023.

[12748] Silvia N Kariuki, Alexander W Macharia, Johnstone Makale, Wilfred Nyamu, Stephen L Hoffman, Melissa C Kapulu, Philip Bejon, Julian C Rayner, and Thomas N Williams. The Dantu blood group prevents parasite growth in vivo: Evidence from a controlled human malaria infection study. *eLife*, 12, 2023.

[12749] Clement Karsenty, Celine Guilbeau-Frugier, Gaël Genet, Marie-Helene Seguelas, Philippe Alzieu, Olivier Cazorla, Alexandra Montagner, Yuna Blum, Caroline Dubroca, Julile Maupoint, Blandine Tramunt, Marie Cauquil, Thierry Sulpice, Sylvain Richard, Silvia Arcucci, Remy Flores-Flores, Nicolas Pataluch, Romain Montoriol, Pierre Sicard, Antoine Deney, Thierry Couffinhal, Jean-Michel Senard, and Celine Galés. Ephrin-B1 regulates the adult diastolic function through a late postnatal maturation of cardiomyocyte surface crests. *eLife*, 12, 2023.

[12750] Nadja S Katheder, Kristen C Browder, Diana Chang, Ann De Maziere, Pekka Kujala, Suzanne van Dijk, Judith Klumperman, Tzu-Chiao Lu, Hongjie Li, Zijuan Lai, Dewakar Sangaraju, and Heinrich Jasper. Nicotinic acetylcholine receptor signaling maintains epithelial barrier integrity. *eLife*, 12, 2023.

[12751] Takuya Kato, Robert P Jenkins, Stefanie Derzsi, Melda Tozluoglu, Antonio Rullan, Steven Hooper, Raphaël AG Chaleil, Holly Joyce, Xiao Fu, Selvam Thavaraj, Paul A Bates, and Erik Sahai. Interplay of adherens junctions and matrix proteolysis determines the invasive pattern and growth of squamous cell carcinoma. *eLife*, 12, 2023.

[12752] Yusuke Kato, Hiroshi Kuroda, Shin-Ichiro Ozawa, Keisuke Saito, Vivek Dogra, Martin Scholz, Guoxian Zhang, Catherine de Vitry, Hiroshi Ishikita, Chanhong Kim, Michael Hippler, Yuichiro Takahashi, and Wataru Sakamoto. Characterization of tryptophan oxidation affecting D1 degradation by FtsH in the photosystem II quality control of chloroplasts. *eLife*, 12, 2023.

[12753] Yuuki Kawabata, Hideyuki Akada, Ken-ichiro Shimatani, Gregory Naoki Nishihara, Hibiki Kimura, Nozomi Nishiumi, and Paolo Domenici. Multiple preferred escape trajectories are explained by a geometric model incorporating prey's turn and predator attack endpoint. *eLife*, 12, 2023.

[12754] Fang Ke, Zachary L Benet, Mitra P Maz, Jianhua Liu, Alexander L Dent, Joanne Michelle Kahlenberg, and Irina L Grigorova. Germinal center B cells that acquire nuclear proteins are specifically suppressed by follicular regulatory T cells. *eLife*, 12, 2023.

[12755] Megan E Kelley, Adi Y Berman, David R Stirling, Beth A Cimini, Yu Han, Shantanu Singh, Anne E Carpenter, Tarun M Kapoor, and Gregory P Way. High-content microscopy reveals a morphological signature of bortezomib resistance. *eLife*, 12, 2023.

[12756] CJ Kelly, Reid K Couch, Vivian T Ha, Camille M Bodart, Judy Wu, Sydney Huff, Nicole T Herrel, Hyunsung D Kim, Azaad O Zimmermann, Jessica Shattuck, Yu-Chen Pan, Matt Kaeberlein, and Anthony S Grillo. Iron status influences mitochondrial disease progression in Complex I-deficient mice. *eLife*, 12, 2023.

[12757] Jimmy J Kelly, Hua Wen, and Paul Brehm. Single-cell RNAseq analysis of spinal locomotor circuitry in larval zebrafish. *eLife*, 12, 2023.

[12758] Christopher Kesten, Valentin Leitner, Susanne Dora, James W Sims, Julian Dindas, Cyril Zipfel, Consuelo M De Moraes, and Clara Sanchez-Rodriguez. Soil-borne fungi alter the apoplastic purinergic signaling in plants by deregulating the homeostasis of extracellular ATP and its metabolite adenosine. *eLife*, 12, 2023.

[12759] Valentina Khalil, Islam Faress, Noëmie Mermet-Joret, Peter Kerwin, Keisuke Yonehara, and Sadegh Nabavi. Subcortico-amygdala pathway processes innate and learned threats. *eLife*, 12, 2023.

[12760] Nazir M Khan, Martha Elena Diaz-Hernandez, Samir Chihab, Priyanka Priyadarshani, Pallavi Bhattaram, Luke J Mortensen, Rosa M Guzzo, and Hicham Drissi. Differential chondrogenic differentiation between iPSC derived from healthy and OA cartilage is associated with changes in epigenetic regulation and metabolic transcriptomic signatures. *eLife*, 12, 2023.

[12761] Diana Khoromskaia and Guillaume Salbreux. Active morphogenesis of patterned epithelial shells. *eLife*, 12, 2023.

[12762] Chelsea U Kidwell, Joseph R Casalini, Soorya Pradeep, Sandra D Scherer, Daniel Greiner, Defne Bayik, Dionysios C Watson, Gregory S Olson, Justin D Lathia, Jarrod S Johnson, Jared Rutter, Alana L Welm, Thomas A Zangle, and Minna Roh-Johnson. Transferred mitochondria accumulate reactive oxygen species, promoting proliferation. *eLife*, 12, 2023.

[12763] Brian Kim, Seth Haney, Ana P Milan, Shruti Joshi, Zane Aldworth, Nikolai Rulkov, Alexander T Kim, Maxim Bazhenov, and Mark A Stopfer. Olfactory receptor neurons generate multiple response motifs, increasing coding space dimensionality. *eLife*, 12, 2023.

[12764] C Kim, Benjamin Chen, Sindhu Mohandas, Jalees Rehman, Zaki A Sherif, K Coombs, and Boris Julg. The importance of patient-partnered research in addressing long COVID: Takeaways for biomedical research study design from the RECOVER Initiative's Mechanistic Pathways taskforce. *eLife*, 12, 2023.

[12765] Jung-Hoon Kim, Josepheen De Asis-Cruz, Dhineshvikram Krishnamurthy, and Catherine Limperopoulos. Toward a more informative representation of the fetal–neonatal brain connectome using variational autoencoder. *eLife*, 12, 2023.

[12766] Mean-Hwan Kim, Cristina Radaelli, Elliot R Thomsen, Deja Monet, Thomas Chartrand, Nikolas L Jorstad, Joseph T Mahoney, Michael J Taormina, Brian Long, Katherine Baker, Trygve E Bakken, Luke Campagnola, Tamara Casper, Michael Clark, Nick Dee, Florence D'Orazi, Clare Gamlin, Brian E Kalmbach, Sara Kebede, Brian R Lee, Lindsay Ng, Jessica Trinh, Charles Cobbs, Ryder P Gwinn, C Dirk Keene, Andrew L Ko, Jeffrey G Ojemann, Daniel L Silbergeld, Staci A Sorensen, Jim Berg, Kimberly A Smith, Philip R Nicovich, Tim Jarsky, Hongkui Zeng, Jonathan T Ting, Boaz P Levi, and Ed Lein. Target cell-specific synaptic dynamics of excitatory to inhibitory neuron connections in supragranular layers of human neocortex. *eLife*, 12, 2023.

[12767] Sang Hwa Kim, Kye D Nichols, Eric N Anderson, Yining Liu, Nandini Ramesh, Weiyan Jia, Connor J Kuerbis, Mark Scalf, Lloyd M Smith, Udai Bhan Pandey, and Randal S Tibbetts. Axon guidance genes modulate neurotoxicity of ALS-associated UBQLN2. *eLife*, 12, 2023.

[12768] Seoyoon Kim, Daehyo Lee, WC Bhashini Wijesinghe, and Duyoung Min. Robust membrane protein tweezers reveal the folding speed limit of helical membrane proteins. *eLife*, 12, 2023.

[12769] Yong-Il Kim, Rebecca O'Rourke, and Charles G Sagerström. scMultiome analysis identifies embryonic hindbrain progenitors with mixed rhombomere identities. *eLife*, 12, 2023.

[12770] Michael T Kimble, Matthew J Johnson, Mattie R Nester, and Lorraine S Symington. Long-range DNA end resection supports homologous recombination by checkpoint activation rather than extensive homology generation. *eLife*, 12, 2023.

[12771] Maedbh King, Ladan Shahshahani, Richard B Ivry, and Jörn Diedrichsen. A task-general connectivity model reveals variation in convergence of cortical inputs to functional regions of the cerebellum. *eLife*, 12, 2023.

[12772] Michael Kintscher, Olexiy Kochubey, and Ralf Schneggenburger. A striatal circuit balances learned fear in the presence and absence of sensory cues. *eLife*, 12, 2023.

[12773] Sina B Kirchhofer, Victor Jun Yu Lim, Sebastian Ernst, Noemi Karsai, Julia G Ruland, Meritxell Canals, Peter Kolb, and Moritz Bünemann. Differential interaction patterns of opioid analgesics with $\mu$ opioid receptors correlate with ligand-specific voltage sensitivity. *eLife*, 12, 2023.

[12774] Manfred G Kitzbichler, Daniel Martins, Richard AI Bethlehem, Richard Dear, Rafael Romero-Garcia, Varun Warrier, Jakob Seidlitz, Ottavia Dipasquale, Federico Turkheimer, Mara Cercignani, Edward T Bullmore, and Neil A Harrison. Two human brain systems micro-structurally associated with obesity. *eLife*, 12, 2023.

[12775] Katharina MC Klee, Michael W Hess, Michael Lohmüller, Sebastian Herzog, Kristian Pfaller, Thomas Müller, Georg F Vogel, and Lukas A Huber. A CRISPR screen in intestinal epithelial cells identifies novel factors for polarity and apical transport. *eLife*, 12, 2023.

[12776] Constance Kleijwegt, Florent Bressac, Coline Seurre, Wilhelm Bouchereau, Camille Cohen, Pascale Texier, Thomas Simonet, Laurent Schaeffer, Patrick Lomonte, and Armelle Corpet. Interplay between PML NBs and HIRA for H3.3 dynamics following type I interferon stimulus. *eLife*, 12, 2023.

[12777] Riku Klén, Ivan A Huespe, Felipe Aníbal Gregalio, Antonio Lalueza Lalueza Blanco, Miguel Pedrera Jimenez, Noelia Garcia Barrio, Pascual Ruben Valdez, Matias A Mirofsky, Bruno Boietti, Ricardo Gómez-Huelgas, José Manuel Casas-Rojo, Juan Miguel Antón-Santos, Javier Alberto Pollan, and David Gómez-Varela. Development and validation of COEWS (COVID-19 Early Warning Score) for hospitalized COVID-19 with laboratory features: A multicontinental retrospective study. *eLife*, 12, 2023.

[12778] Jackie Kleynhans, Lorenzo Dall'Amico, Laetitia Gauvin, Michele Tizzoni, Lucia Maloma, Sibongile Walaza, Neil A Martinson, Anne von Gottberg, Nicole Wolter, Mvuyo Makhasi, Cheryl Cohen, Ciro Cattuto, Stefano Tempia, and Amelia Buys. Association of close-range contact patterns with SARS-CoV-2: a household transmission study. *eLife*, 12, 2023.

[12779] Zane Kliesmete, Lucas Esteban Wange, Beate Vieth, Miriam Esgleas, Jessica Radmer, Matthias Hülsmann, Johanna Geuder, Daniel Richter, Mari Ohnuki, Magdelena Götz, Ines Hellmann, and Wolfgang Enard. Regulatory and coding sequences of TRNP1 co-evolve with brain size and cortical folding in mammals. *eLife*, 12, 2023.

[12780] Jonas R Knudsen, Kaspar W Persson, Carlos Henriquez-Olguin, Zhencheng Li, Nicolas Di Leo, Sofie A Hesselager, Steffen H Raun, Janne R Hingst, Raphaël Trouillon, Martin Wohlwend, Jørgen FP Wojtaszewski, Martin AM Gijs, and Thomas Elbenhardt Jensen. Microtubule-mediated GLUT4 trafficking is disrupted in insulin-resistant skeletal muscle. *eLife*, 12, 2023.

[12781] Cassandra Koh, Lionel Frangeul, Hervé Blanc, Carine Ngoagouni, Sébastien Boyer, Philippe Dussart, Nina Grau, Romain Girod, Jean-Bernard Duchemin, and Maria-Carla Saleh. Ribosomal RNA (rRNA) sequences from 33 globally distributed mosquito species for improved metagenomics and species identification. *eLife*, 12, 2023.

[12782] Jaemoon Koh, Yeon Duk Woo, Hyun Jung Yoo, Jun-Pyo Choi, Sae Hoon Kim, Yoon-Seok Chang, Kyeong Cheon Jung, Ji Hyung Kim, Yoon Kyung Jeon, Hye Young Kim, and Doo Hyun Chung. De novo fatty-acid synthesis protects invariant NKT cells from cell death, thereby promoting their homeostasis and pathogenic roles in airway hyperresponsiveness. *eLife*, 12, 2023.

[12783] Carmen Kohl, Michelle XM Wong, Jing Jun Wong, Matthew FS Rushworth, and Bolton KH Chau. Intraparietal stimulation disrupts negative distractor effects in human multi-alternative decision-making. *eLife*, 12, 2023.

[12784] Friederike E Kohrs, Susann Auer, Alexandra Bannach-Brown, Susann Fiedler, Tamarinde Laura Haven, Verena Heise, Constance Holman, Flavio Azevedo, René Bernard, Arnim Bleier, Nicole Bössel, Brian Patrick Cahill, Leyla Jael Castro, Adrian Ehrenhofer, Kristina Eichel, Maximillian Frank, Claudia Frick, Malte Friese, Anne Gärtner, Kerstin Gierend, David Joachim Grüning, Lena Hahn, Maren Hülsemann, Malika Ihle, Sabrina Illius, Laura König, Matthias König, Louisa Kulke, Anton Kutlin, Fritjof Lammers, David MA Mehler, Christoph Miehl, Anett Müller-Alcazar, Claudia Neuendorf, Helen Niemeyer, Florian Pargent, Aaron Peikert, Christina U Pfeuffer, Robert Reinecke, Jan Philipp Röer, Jessica L Rohmann, Alfredo Sánchez-Tójar, Stefan Scherbaum, Elena Sixtus, Lisa Spitzer, Vera Maren Straßburger, Marcel Weber, Clarissa J Whitmire, Josephine Zerna, Dilara Zorbek, Philipp Zumstein, and Tracey L Weissgerber. Eleven strategies for making reproducible research and open science training the norm at research institutions. *eLife*, 12, 2023.

[12785] Yoshinao Koike, Masahiko Takahata, Masahiro Nakajima, Nao Otomo, Hiroyuki Suetsugu, Xiaoxi Liu, Tsutomu Endo, Shiro Imagama, Kazuyoshi Kobayashi, Takashi Kaito, Satoshi Kato, Yoshiharu Kawaguchi, Masahiro Kanayama, Hiroaki Sakai, Takashi Tsuji, Takeshi Miyamoto, Hiroyuki Inose, Toshitaka Yoshii, Masafumi Kashii, Hiroaki Nakashima, Kei Ando, Yuki Taniguchi, Kazuhiro Takeuchi, Shuji Ito, Kohei Tomizuka, Keiko Hikino, Yusuke Iwasaki, Yoichiro Kamatani, Shingo Maeda, Hideaki Nakajima, Kanji Mori, Atsushi Seichi, Shunsuke Fujibayashi, Tsukasa Kanchiku, Kei Watanabe, Toshihiro Tanaka, Kazunobu Kida, Sho Kobayashi, Masahito Takahashi, Kei Yamada, Hiroshi Takuwa, Hsing-Fang Lu, Shumpei Niida, Kouichi Ozaki, Yukihide Momozawa, Tsuji Takashi, Masashi Yamazaki, Atsushi Okawa, Morio Matsumoto, Norimasa Iwasaki, Chikashi Terao,

and Shiro Ikegawa. Genetic insights into ossification of the posterior longitudinal ligament of the spine. *eLife*, 12, 2023.

[12786] Takeru Kondo, Annie Gleason, Hiroko Okawa, Akishige Hokugo, and Ichiro Nishimura. Mouse gingival single-cell transcriptomic atlas identified a novel fibroblast subpopulation activated to guide oral barrier immunity in periodontitis. *eLife*, 12, 2023.

[12787] Xiangmudong Kong, Wensheng Yan, Wenqiang Sun, Yanhong Zhang, Hee Jung Yang, Mingyi Chen, Hongwu Chen, Ralph W de Vere White, Jin Zhang, and Xinbin Chen. Isoform-specific disruption of the TP73 gene reveals a critical role for TAp73$\gamma$ in tumorigenesis via leptin. *eLife*, 12, 2023.

[12788] Antonion Korcari, Anne EC Nichols, Mark R Buckley, and Alayna E Loiselle. Scleraxis-lineage cells are required for tendon homeostasis and their depletion induces an accelerated extracellular matrix aging phenotype. *eLife*, 12, 2023.

[12789] Maria Körner, Susanne R Meyer, Gabriella Marincola, Maximilian J Kern, Clemens Grimm, Christina Schuelein-Voelk, Utz Fischer, Kay Hofmann, and Alexander Buchberger. The FAM104 proteins VCF1/2 promote the nuclear localization of p97/VCP. *eLife*, 12, 2023.

[12790] Eleftheria Koropouli, Qiang Wang, Rebeca Mejías, Randal Hand, Tao Wang, David D Ginty, and Alex L Kolodkin. Palmitoylation regulates neuropilin-2 localization and function in cortical neurons and conveys specificity to semaphorin signaling via palmitoyl acyltransferases. *eLife*, 12, 2023.

[12791] Oron Kotler, Yana Khrapunsky, Arik Shvartsman, Hui Dai, Leigh D Plant, Steven AN Goldstein, and Ilya Fleidervish. SUMOylation of NaV1.2 channels regulates the velocity of backpropagating action potentials in cortical pyramidal neurons. *eLife*, 12, 2023.

[12792] Lenka Koudelková, Markéta Pelantová, Zuzana Brůhová, Martin Sztacho, Vojtěch Pavlík, Dalibor Pánek, Jakub Gemperle, Pavel Talacko, Jan Brábek, and Daniel Rösel. Phosphorylation of tyrosine 90 in SH3 domain is a new regulatory switch controlling Src kinase. *eLife*, 12, 2023.

[12793] Robert A Kozol, Andrew J Conith, Anders Yuiska, Alexia Cree-Newman, Bernadeth Tolentino, Kasey Benesh, Alexandra Paz, Evan Lloyd, Johanna E Kowalko, Alex C Keene, Craig Albertson, and Erik R Duboue. A brain-wide analysis maps structural evolution to distinct anatomical module. *eLife*, 12, 2023.

[12794] Daniel A Kramer, Heidy Y Narvaez-Ortiz, Urval Patel, Rebecca Shi, Kang Shen, Brad J Nolen, Julien Roche, and Baoyu Chen. The intrinsically disordered cytoplasmic tail of a dendrite branching receptor uses two distinct mechanisms to regulate the actin cytoskeleton. *eLife*, 12, 2023.

[12795] Friedrich Krohn, Manuele Novello, Ruben S van der Giessen, Chris I De Zeeuw, Johan JM Pel, and Laurens WJ Bosman. The integrated brain network that controls respiration. *eLife*, 12, 2023.

[12796] Carly M Krull, Haiyi Li, and Amit Pathak. Nuclear export inhibition jumbles epithelial–mesenchymal states and gives rise to migratory disorder in healthy epithelia. *eLife*, 12, 2023.

[12797] Marc Kschonsak, Christine C Jao, Christopher P Arthur, Alexis L Rohou, Philippe Bergeron, Daniel F Ortwine, Steven J McKerrall, David H Hackos, Lunbin Deng, Jun Chen, Tianbo Li, Peter S Dragovich, Matthew Volgraf, Matthew R Wright, Jian Payandeh, Claudio Ciferri, and John C Tellis. Cryo-EM reveals an unprecedented binding site for NaV1.7 inhibitors enabling rational design of potent hybrid inhibitors. *eLife*, 12, 2023.

[12798] Subhash Kulkarni, Monalee Saha, Jared Slosberg, Alpana Singh, Sushma Nagaraj, Laren Becker, Chengxiu Zhang, Alicia Bukowski, Zhuolun Wang, Guosheng Liu, Jenna M Leser, Mithra Kumar, Shriya Bakhshi, Matthew J Anderson, Mark Lewandoski, Elizabeth Vincent, Loyal A Goff, and Pankaj Jay Pasricha. Age-associated changes in lineage composition of the enteric nervous system regulate gut health and disease. *eLife*, 12, 2023.

[12799] Jessie Kulsuptrakul, Elizabeth A Turcotte, Michael Emerman, and Patrick S Mitchell. A human-specific motif facilitates CARD8 inflammasome activation after HIV-1 infection. *eLife*, 12, 2023.

[12800] Amit Kumar, Oscar Vadas, Nicolas Dos Santos Pacheco, Xu Zhang, Kin Chao, Nicolas Darvill, Helena Ø Rasmussen, Yingqi Xu, Gloria Meng-Hsuan Lin, Fisentzos A Stylianou, Jan Skov Pedersen, Sarah L Rouse, Marc L Morgan, Dominique Soldati-Favre, and Stephen Matthews. Structural and regulatory insights into the glideosome-associated connector from Toxoplasma gondii. *eLife*, 12, 2023.

[12801] Saurav Kumar, Amanda Stainer, Julien Dubrulle, Christopher Simpkins, and Jonathan A Cooper. Cas phosphorylation regulates focal adhesion assembly. *eLife*, 12, 2023.

[12802] Zeyneb Kurt, Jenny Cheng, Rio Barrere-Cain, Caden N McQuillen, Zara Saleem, Neil Hsu, Nuoya Jiang, Calvin Pan, Oscar Franzén, Simon Koplev, Susanna Wang, Johan Björkegren, Aldons J Lusis, Montgomery Blencowe, and Xia Yang. Shared and distinct pathways and networks genetically linked to coronary artery disease between human and mouse. *eLife*, 12, 2023.

[12803] James ZJ Kwan, Thomas F Nguyen, Anuli C Uzozie, Marek A Budzynski, Jieying Cui, Joseph MC Lee, Filip Van Petegem, Philipp F Lange, and Sheila S Teves. RNA Polymerase II transcription independent of TBP in murine embryonic stem cells. *eLife*, 12, 2023.

[12804] Sheldon JJ Kwok, Daniel T Montoro, Adam L Haber, Seok-Hyun Yun, and Vladimir Vinarsky. Single-cell transcriptomics of a dynamic cell behavior in murine airways. *eLife*, 12, 2023.

[12805] Daniel R Labuz, Giavonni Lewis, Irma D Fleming, Callie M Thompson, Yan Zhai, Matthew A Firpo, and Daniel T Leung. Targeted multi-omic analysis of human skin tissue identifies alterations of conventional and unconventional T cells associated with burn injury. *eLife*, 12, 2023.

[12806] Beatrice Lace, Chao Su, Daniel Invernot Perez, Marta Rodriguez-Franco, Tatiana Vernié, Morgane Batzenschlager, Sabrina Egli, Cheng-Wu Liu, and Thomas Ott. RPG acts as a central determinant for infectosome formation and cellular polarization during intracellular rhizobial infections. *eLife*, 12, 2023.

[12807] Michael J Lafferty, Nil Aygün, Niyanta K Patel, Oleh Krupa, Dan Liang, Justin M Wolter, Daniel H Geschwind, Luis de la Torre-Ubieta, and Jason L Stein. MicroRNA-eQTLs in the developing human neocortex link miR-4707-3p expression to brain size. *eLife*, 12, 2023.

[12808] Amrita Lamba, Matthew R Nassar, and Oriel FeldmanHall. Prefrontal cortex state representations shape human credit assignment. *eLife*, 12, 2023.

[12809] Laurie Anne Lamiré, Martin Haesemeyer, Florian Engert, Michael Granato, and Owen Randlett. Functional and pharmacological analyses of visual habituation learning in larval zebrafish. *eLife*, 12, 2023.

[12810] Laurenz Lammer, Frauke Beyer, Melanie Luppa, Christian Sanders, Ronny Baber, Christoph Engel, Kerstin Wirkner, Markus Loffler, Steffi G Riedel-Heller, Arno Villringer, and A Veronica Witte. Impact of social isolation on grey matter structure and cognitive functions: A population-based longitudinal neuroimaging study. *eLife*, 12, 2023.

[12811] Alexandro Landshammer, Adriano Bolondi, Helene Kretzmer, Christian Much, René Buschow, Alina Rose, Hua-Jun Wu, Sebastian D Mackowiak, Bjoern Braendl, Pay Giesselmann, Rosaria Tornisiello, Krishna Mohan Parsi, Jack Huey, Thorsten Mielke, David Meierhofer, René Maehr, Denes Hnisz, Franziska Michor, John L Rinn, and Alexander Meissner. T-REX17 is a transiently expressed non-coding RNA essential for human endoderm formation. *eLife*, 12, 2023.

[12812] Joseph S Latshaw, Reece E Mazade, Mary Petersen, Julie A Mustard, Irina Sinakevitch, Lothar Wissler, Xiaojiao Guo, Chelsea Cook, Hong Lei, Jürgen Gadau, and Brian Smith. Tyramine and its Amtyr1 receptor modulate attention in honey bees (Apis mellifera). *eLife*, 12, 2023.

[12813] Meghan Laturney, Gabriella R Sterne, and Kristin Scott. Mating activates neuroendocrine pathways signaling hunger in Drosophila females. *eLife*, 12, 2023.

[12814] Michael S Lauer, Joy Wang, and Deepshikha Roychowdhury. National Institutes of Health research project grant inflation 1998 to 2021. *eLife*, 12, 2023.

[12815] Richard C Lauer, Marc Barry, Tracey L Smith, Andrew Maltez Thomas, Jin Wu, Ruofei Du, Ji-Hyun Lee, Arpit Rao, Andrey S Dobroff, Marco A Arap, Diana N Nunes, Israel T Silva, Emmanuel Dias-Neto, Isan Chen, Dennis J McCance, Webster K Cavenee, Renata Pasqualini, and Wadih Arap. Dysregulation of the PRUNE2/PCA3 genetic axis in human prostate cancer: from experimental discovery to validation in two independent patient cohorts. *eLife*, 12, 2023.

[12816] Rimma Laufer Britva, Aviad Keren, Marta Bertolini, Yehuda Ullmann, Ralf Paus, and Amos Gilhar. Involvement of ILC1-like innate lymphocytes in human autoimmunity, lessons from alopecia areata. *eLife*, 12, 2023.

[12817] Romain Laurent, Zachary A Szpiech, Sergio S da Costa, Valentin Thouzeau, Cesar A Fortes-Lima, FranÇoise Dessarps-Freichey, Laure Lémée, José Utgé, Noah A Rosenberg, Marlyse Baptista, and Paul Verdu. A genetic and linguistic analysis of the admixture histories of the islands of Cabo Verde. *eLife*, 12, 2023.

[12818] Peter M Lauro, Shane Lee, Daniel E Amaya, David D Liu, Umer Akbar, and Wael F Asaad. Concurrent decoding of distinct neurophysiological fingerprints of tremor and bradykinesia in Parkinson's disease. *eLife*, 12, 2023.

[12819] M Elise Lauterbur, Maria Izabel A Cavassim, Ariella L Gladstein, Graham Gower, Nathaniel S Pope, Georgia Tsambos, Jeffrey Adrion, Saurabh Belsare, Arjun Biddanda, Victoria Caudill, Jean Cury, Ignacio Echevarria, Benjamin C Haller, Ahmed R Hasan, Xin Huang, Leonardo Nicola Martin Iasi, Ekaterina Noskova, Jana Obsteter, Vitor Antonio Correa Pavinato, Alice Pearson, David Peede, Manolo F Perez, Murillo F Rodrigues, Chris CR Smith, Jeffrey P Spence, Anastasia Teterina, Silas Tittes, Per Unneberg, Juan Manuel Vazquez, Ryan K Waples, Anthony Wilder Wohns, Yan Wong, Franz Baumdicker, Reed A Cartwright, Gregor Gorjanc, Ryan N Gutenkunst, Jerome Kelleher, Andrew D Kern, Aaron P Ragsdale, Peter L Ralph, Daniel R Schrider, and Ilan Gronau. Expanding the stdpopsim species catalog, and lessons learned for realistic genome simulations. *eLife*, 12, 2023.

[12820] Maria I Lazaro-Pena, Adam B Cornwell, Carlos A Diaz-Balzac, Ritika Das, Zachary C Ward, Nicholas Macoretta, Juilee Thakar, and Andrew V Samuelson. Homeodomain-interacting protein kinase maintains neuronal homeostasis during normal Caenorhabditis elegans aging and systemically regulates longevity from serotonergic and GABAergic neurons. *eLife*, 12, 2023.

[12821] Oscar Marcelo Lazo and Giampietro Schiavo. Rab10 regulates the sorting of internalised TrkB for retrograde axonal transport. *eLife*, 12, 2023.

[12822] Thao DV Le, Dianxin Liu, Gai-Linn K Besing, Ritika Raghavan, Blair J Ellis, Ryan P Ceddia, Sheila Collins, and Julio E Ayala. Glucagon-like peptide-1 receptor activation stimulates PKA-mediated phosphorylation of Raptor and this contributes to the weight loss effect of liraglutide. *eLife*, 12, 2023.

[12823] Kristian Kyle Le Vay, Elia Salibi, Basusree Ghosh, TY Dora Tang, and Hannes Mutschler. Ribozyme activity modulates the physical properties of RNA–peptide coacervates. *eLife*, 12, 2023.

[12824] Romane Leboutet, Céline Largeau, Leonie Müller, Magali Prigent, Grégoire Quinet, Manuel S Rodriguez, Marie-Hélène Cuif, Thorsten Hoppe, Emmanuel Culetto, Christophe Lefebvre, and Renaud Legouis. LGG-1/GABARAP lipidation is not required for autophagy and development in Caenorhabditis elegans. *eLife*, 12, 2023.

[12825] Ivan Lebovka, Bruno Hay Mele, Xiaomin Liu, Alexandra Zakieva, Theresa Schlamp, Nial Rau Gursanscky, Roeland MH Merks, Ruth Großeholz, and Thomas Greb. Computational modeling of

cambium activity provides a regulatory framework for simulating radial plant growth. *eLife*, 12, 2023.

[12826] Owen Leddy, Forest M White, and Bryan D Bryson. Immunopeptidomics reveals determinants of Mycobacterium tuberculosis antigen presentation on MHC class I. *eLife*, 12, 2023.

[12827] Dongwon Lee, Wu Chen, Heet Naresh Kaku, Xinming Zhuo, Eugene S Chao, Armand Soriano, Allen Kuncheria, Stephanie Flores, Joo Hyun Kim, Armando Rivera, Frank Rigo, Paymaan Jafar-nejad, Arthur L Beaudet, Matthew S Caudill, and Mingshan Xue. Antisense oligonucleotide therapy rescues disturbed brain rhythms and sleep in juvenile and adult mouse models of Angelman syndrome. *eLife*, 12, 2023.

[12828] Haksoo Lee, Hyunkoo Kang, Changjong Moon, and BuHyun Youn. PAK3 downregulation induces cognitive impairment following cranial irradiation. *eLife*, 12, 2023.

[12829] Han Nim Lee, Jenu Varghese Chacko, Ariadna Gonzalez Solís, Kuo-En Chen, Jessica AS Barros, Santiago Signorelli, A Harvey Millar, Richard David Vierstra, Kevin W Eliceiri, and Marisa S Otegui. The autophagy receptor NBR1 directs the clearance of photodamaged chloroplasts. *eLife*, 12, 2023.

[12830] Kiran GL Lee, Adele Mennerat, Dieter Lukas, Hannah L Dugdale, and Antica Culina. The effect of the COVID-19 pandemic on the gender gap in research productivity within academia. *eLife*, 12, 2023.

[12831] Peter H Lee, Michael Anaya, Mark S Ladinsky, Justin M Reitsma, and Kai Zinn. An extracellular vesicle targeting ligand that binds to Arc proteins and facilitates Arc transport in vivo. *eLife*, 12, 2023.

[12832] Reagan Lee, Wei Xu, Marshall Dozier, Ruth McQuillan, Evropi Theodoratou, and Jonine Figueroa. A rapid review of COVID-19's global impact on breast cancer screening participation rates and volumes from January to December 2020. *eLife*, 12, 2023.

[12833] Wha-Eum Lee, Sang Woo Park, Daniel M Weinberger, Donald Olson, Lone Simonsen, Bryan T Grenfell, and Cécile Viboud. Direct and indirect mortality impacts of the COVID-19 pandemic in the United States, March 1, 2020 to January 1, 2022. *eLife*, 12, 2023.

[12834] Yunjeong Lee, In Seo Yeo, Namhee Kim, Dong-Keun Lee, Kyung-Tai Kim, Jiyoung Yoon, Jawoon Yi, Young Bin Hong, Byung-Ok Choi, Yoichi Kosodo, Daesoo Kim, Jihwan Park, and Mi-Ryoung Song. Transcriptional control of motor pool formation and motor circuit connectivity by the LIM-HD protein Isl2. *eLife*, 12, 2023.

[12835] Bonnibelle K Leeds, Katelyn F Kostello, Yuna Y Liu, Christian R Nelson, Sue Biggins, and Charles L Asbury. Mechanical coupling coordinates microtubule growth. *eLife*, 12, 2023.

[12836] Matthew F Lefebvre, Nikolas H Claussen, Noah P Mitchell, Hannah J Gustafson, and Sebastian J Streichan. Geometric control of myosin II orientation during axis elongation. *eLife*, 12, 2023.

[12837] Elena Farah Lehmann, Márton Liziczai, Katarzyna Drożdżyk, Patrick Altermatt, Cassiano Langini, Vania Manolova, Hanna Sundstrom, Franz Dürrenberger, Raimund Dutzler, and Cristina Manatschal. Structures of ferroportin in complex with its specific inhibitor vamifeport. *eLife*, 12, 2023.

[12838] Xiao Lei, Daniel GonÇalves-Carneiro, Trinity M Zang, and Paul D Bieniasz. Initiation of HIV-1 Gag lattice assembly is required for recognition of the viral genome packaging signal. *eLife*, 12, 2023.

[12839] Marco Leibinger, Charlotte Zeitler, Miriam Paulat, Philipp Gobrecht, Alexander Hilla, Anastasia Andreadaki, Rainer Guthoff, and Dietmar Fischer. Inhibition of microtubule detyrosination by parthenolide facilitates functional CNS axon regeneration. *eLife*, 12, 2023.

[12840] Agnieszka Lemanska, Colm Andrews, Louis Fisher, Seb Bacon, Adam E Frampton, Amir Mehrkar, Peter Inglesby, Simon Davy, Keith Roberts, Praveetha Patalay, Ben Goldacre, Brian MacKenna, Alex J Walker, and Alex J Walker. Healthcare in England was affected by the COVID-19 pandemic across the pancreatic cancer pathway: A cohort study using OpenSAFELY-TPP. *eLife*, 12, 2023.

[12841] René Lemcke, Christine Egebjerg, Nicolai T Berendtsen, Kristoffer L Egerod, Allan R Thomsen, Tune H Pers, Jan P Christensen, and Birgitte R Kornum. Molecular consequences of peripheral Influenza A infection on cell populations in the murine hypothalamus. *eLife*, 12, 2023.

[12842] Eline Lemerle, Jeanne Lainé, Marion Benoist, Gilles Moulay, Anne Bigot, Clémence Labasse, Angéline Madelaine, Alexis Canette, Perrine Aubin, Jean-Michel Vallat, Norma B Romero, Marc Bitoun, Vincent Mouly, Isabelle Marty, Bruno Cadot, Laura Picas, and Stéphane Vassilopoulos. Caveolae and Bin1 form ring-shaped platforms for T-tubule initiation. *eLife*, 12, 2023.

[12843] Lukas Leopold Lengersdorff, Isabella C Wagner, Gloria Mittmann, David Sastre-Yagüe, Andre Lüttig, Andreas Olsson, Pedrag Petrovic, and Claus Lamm. Neuroimaging and behavioral evidence that violent video games exert no negative effect on human empathy for pain and emotional reactivity to violence. *eLife*, 12, 2023.

[12844] Movitz Lenninger, Mikael Skoglund, Pawel Andrzej Herman, and Arvind Kumar. Are single-peaked tuning curves tuned for speed rather than accuracy? *eLife*, 12, 2023.

[12845] Joaquín Letelier, Lorena Buono, María Almuedo-Castillo, Jingjing Zang, Constanza Mounieres, Sergio González-Díaz, Rocío Polvillo, Estefanía Sanabria-Reinoso, Jorge Corbacho, Ana Sousa-Ortega, Ruth Diez del Corral, Stephan CF Neuhauss, and Juan R Martínez-Morales. Mutation of vsx genes in zebrafish highlights the robustness of the retinal specification network. *eLife*, 12, 2023.

[12846] Gidon Levakov, Alon Kaplan, Anat Yaskolka Meir, Ehud Rinott, Gal Tsaban, Hila Zelicha, Matthias Blüher, Uta Ceglarek, Michael Stumvoll, Ilan Shelef, Galia Avidan, and Iris Shai. The effect of weight loss following 18 months of lifestyle intervention on brain age assessed with resting-state functional connectivity. *eLife*, 12, 2023.

[12847] Tylor R Lewis, Mustafa S Makia, Carson M Castillo, Ying Hao, Muayyad R Al-Ubaidi, Nikolai P Skiba, Shannon M Conley, Vadim Y Arshavsky, and Muna I Naash. ROM1 is redundant to PRPH2 as a molecular building block of photoreceptor disc rims. *eLife*, 12, 2023.

[12848] Tylor R Lewis, Sebastien Phan, Carson M Castillo, Keun-Young Kim, Kelsey Coppenrath, William Thomas, Ying Hao, Nikolai P Skiba, Marko E Horb, Mark H Ellisman, and Vadim Y Arshavsky. Photoreceptor disc incisures form as an adaptive mechanism ensuring the completion of disc enclosure. *eLife*, 12, 2023.

[12849] Allyson Li, Rashmi Voleti, Minhee Lee, Dejan Gagoski, and Neel H Shah. High-throughput profiling of sequence recognition by tyrosine kinases and SH2 domains using bacterial peptide display. *eLife*, 12, 2023.

[12850] Fu Li, Gregory Artiushin, and Amita Sehgal. Modulation of sleep by trafficking of lipids through the Drosophila blood-brain barrier. *eLife*, 12, 2023.

[12851] Gen Li, Binshi Bo, Puxin Wang, Peixing Qian, Mingzhe Li, Yuyan Li, Chuanjun Tong, Kaiwei Zhang, Baogui Zhang, Tianzi Jiang, Zhifeng Liang, and Xiaojie Duan. Instantaneous antidepressant effect of lateral habenula deep brain stimulation in rats studied with functional MRI. *eLife*, 12, 2023.

[12852] Hao Li and Xin Jin. Multiple dynamic interactions from basal ganglia direct and indirect pathways mediate action selection. *eLife*, 12, 2023.

[12853] Hongzhe Li, Sandro Bräunig, Parashar Dhapolar, Göran Karlsson, Stefan Lang, and Stefan Scheding. Identification of phenotypically, functionally, and anatomically distinct stromal niche populations in human bone marrow based on single-cell RNA sequencing. *eLife*, 12, 2023.

[12854] Jiang Li, Rebecca E Ruggiero-Ruff, Yuxin He, Xinru Qiu, Nancy Lainez, Pedro Villa, Adam Godzik, Djurdjica Coss, and Meera G Nair. Sexual dimorphism in obesity is governed by RELM$\alpha$ regulation of adipose macrophages and eosinophils. *eLife*, 12, 2023.

[12855] Jingxuan Li, Chunlan Yang, Alexandre Jousset, Keming Yang, Xiaofang Wang, Zhihui Xu, Tianjie Yang, Xinlan Mei, Zengtao Zhong, Yangchun Xu, Qirong Shen, Ville-Petri Friman, and Zhong Wei. Engineering multifunctional rhizosphere probiotics using consortia of Bacillus amyloliquefaciens transposon insertion mutants. *eLife*, 12, 2023.

[12856] Kai Li, Yuma Tsukasa, Misato Kurio, Kaho Maeta, Akimitsu Tsumadori, Shumpei Baba, Risa Nishimura, Akira Murakami, Koun Onodera, Takako Morimoto, Tadashi Uemura, and Tadao Usui. Belly roll, a GPI-anchored Ly6 protein, regulates Drosophila melanogaster escape behaviors by modulating the excitability of nociceptive peptidergic interneurons. *eLife*, 12, 2023.

[12857] Kexin Li, Qingji Huo, Nathan H Dimmitt, Guofan Qu, Junjie Bao, Pankita H Pandya, M Reza Saadatzadeh, Khadijeh Bijangi-Vishehsaraei, Melissa A Kacena, Karen E Pollok, Chien-Chi Lin, Bai-Yan Li, and Hiroki Yokota. Osteosarcoma-enriched transcripts paradoxically generate osteosarcoma-suppressing extracellular proteins. *eLife*, 12, 2023.

[12858] Shuxian Li, Lingbing Li, Changqing Zhang, Huaxuan Fu, Shuping Yu, Meijuan Zhou, Junjun Guo, Zhenya Fang, Anna Li, Man Zhao, Meihua Zhang, and Xietong Wang. PM2.5 leads to adverse pregnancy outcomes by inducing trophoblast oxidative stress and mitochondrial apoptosis via KLF9/CYP1A1 transcriptional axis. *eLife*, 12, 2023.

[12859] Wei Li, Xinqi Lou, Yingjie Zha, Yinyin Qin, Jun Zha, Lei Hong, Zhanli Xie, Shudi Yang, Chen Wang, Jianzhong An, Zhenhao Zhang, and Shigang Qiao. Single-cell RNA-seq of heart reveals intercellular communication drivers of myocardial fibrosis in diabetic cardiomyopathy. *eLife*, 12, 2023.

[12860] Xiaoxu Li, Jean-David Morel, Giorgia Benegiamo, Johanne Poisson, Alexis Bachmann, Alexis Rapin, Jonathan Sulc, Evan Williams, Alessia Perino, Kristina Schoonjans, Maroun Bou Sleiman, and Johan Auwerx. Genetic and dietary modulators of the inflammatory response in the gastrointestinal tract of the BXD mouse genetic reference population. *eLife*, 12, 2023.

[12861] Ya-tang Li and Markus Meister. Functional cell types in the mouse superior colliculus. *eLife*, 12, 2023.

[12862] Yanqiu Li, Feifei Wang, Xin Li, Ling Wang, Zheng Yang, Zhongsheng You, and Aimin Peng. The ATM-E6AP-MASTL axis mediates DNA damage checkpoint recovery. *eLife*, 12, 2023.

[12863] Yongjun Li, Paula Haynes, Shirley L Zhang, Zhifeng Yue, and Amita Sehgal. Ecdysone acts through cortex glia to regulate sleep in Drosophila. *eLife*, 12, 2023.

[12864] Yu Li, Pablo Guaman Tipan, Hilary J Selden, Jayashree Srinivasan, Laura P Hale, and Lauren IR Ehrlich. CCR4 and CCR7 differentially regulate thymocyte localization with distinct outcomes for central tolerance. *eLife*, 12, 2023.

[12865] Yuanyuan Li, Wenyan Xu, Svetlana Makova, Martina Brueckner, and Zhaoxia Sun. Inactivation of Invs/Nphp2 in renal epithelial cells drives infantile nephronophthisis like phenotypes in mouse. *eLife*, 12, 2023.

[12866] Yutian Li, Anish A Sarma, Iris T Lee, Fayth Hui Tan, Michael J Abrams, Zevin J Condiotte, Martin Heithe, Misha Raffiee, John O Dabiri, David A Gold, and Lea Goentoro. Response to comment on 'A conserved strategy for inducing appendage regeneration in moon jellyfish, Drosophila, and mice'. *eLife*, 12, 2023.

[12867] Jiurong Liang, Guanling Huang, Xue Liu, Ningshan Liu, Forough Taghavifar, Kristy Dai, Changfu Yao, Nan Deng, Yizhou Wang, Peter Chen, Cory Hogaboam, Barry R Stripp, William C Parks, Paul W Noble, and Dianhua Jiang. Reciprocal interactions between alveolar progenitor dysfunction and aging promote lung fibrosis. *eLife*, 12, 2023.

[12868] Wenbo Liao, Ying Jiang, Long Jin, and Stefan Lüpold. How hibernation in frogs drives brain and reproductive evolution in opposite directions. *eLife*, 12, 2023.

[12869] Guang Lin, Burak Tepe, Geoff McGrane, Regine C Tipon, Gist Croft, Leena Panwala, Amanda Hope, Agnes JH Liang, Zhongyuan Zuo, Seul Kee Byeon, Lily Wang, Akhilesh Pandey, and Hugo J Bellen. Exploring therapeutic strategies for infantile neuronal axonal dystrophy (INAD/PARK14). *eLife*, 12, 2023.

[12870] Jiamin Lin, Pingping Zhang, Wei Liu, Guorong Liu, Juan Zhang, Min Yan, Yuyou Duan, and Na Yang. A positive feedback loop between ZEB2 and ACSL4 regulates lipid metabolism to promote breast cancer metastasis. *eLife*, 12, 2023.

[12871] Rixing Lin, Aron Kos, Juan Pablo Lopez, Julien Dine, Laura M Fiori, Jennie Yang, Yair Ben-Efraim, Zahia Aouabed, Pascal Ibrahim, Haruka Mitsuhashi, Tak Pan Wong, El Cherif Ibrahim, Catherine Belzung, Pierre Blier, Faranak Farzan, Benicio N Frey, Raymond W Lam, Roumen Milev, Daniel J Muller, Sagar V Parikh, Claudio Soares, Rudolf Uher, Corina Nagy, Naguib Mechawar, Jane A Foster, Sidney H Kennedy, Alon Chen, and Gustavo Turecki. SNORD90 induces glutamatergic signaling following treatment with monoaminergic antidepressants. *eLife*, 12, 2023.

[12872] Xiaoxuan Lin, Patrick R Haller, Navid Bavi, Nabil Faruk, Eduardo Perozo, and Tobin R Sosnick. Folding of prestin's anion-binding site and the mechanism of outer hair cell electromotility. *eLife*, 12, 2023.

[12873] Yunfeng Lin, Jia Li, Haichao Zhao, Anne McMahon, Kelly McGhee, and Shan Yan. APE1 recruits ATRIP to ssDNA in an RPA-dependent and -independent manner to promote the ATR DNA damage response. *eLife*, 12, 2023.

[12874] Zong-Shin Lin, Chiao-Chen Chung, Yu-Chia Liu, Chu-Han Chang, Hui-Chia Liu, Yung-Yi Liang, Teng-Le Huang, Tsung-Ming Chen, Che-Hsin Lee, Chih-Hsin Tang, Mien-Chie Hung, and Ya-Huey Chen. EZH2/hSULF1 axis mediates receptor tyrosine kinase signaling to shape cartilage tumor progression. *eLife*, 12, 2023.

[12875] Max Lindmark, Malin Karlsson, and Anna Gårdmark. Larger but younger fish when growth outpaces mortality in heated ecosystem. *eLife*, 12, 2023.

[12876] Caigang Liu, Hong Yu, Guanglei Chen, Qichao Yang, Zichu Wang, Nan Niu, Ling Han, Dongyu Zhao, Manji Wang, Yuanyuan Liu, and Yongliang Yang. An herbal drug combination identified by knowledge graph alleviates the clinical symptoms of plasma cell mastitis patients: A nonrandomized controlled trial. *eLife*, 12, 2023.

[12877] Can Liu, Jiawei Zhou, Stephan Kudlacek, Timothy Qi, Tyler Dunlap, and Yanguang Cao. Population dynamics of immunological synapse formation induced by bispecific T cell engagers predict clinical pharmacodynamics and treatment resistance. *eLife*, 12, 2023.

[12878] Cong Liu, Milena Schönke, Borah Spoorenberg, Joost M Lambooij, Hendrik JP van der Zande, Enchen Zhou, Maarten E Tushuizen, Anne-Christine Andreasson, Andrew Park, Stephanie Oldham, Martin Uhrbom, Ingela Ahlstedt, Yasuhiro Ikeda, Kristina Wallenius, Xiao-Rong Peng, Bruno Guigas, Mariëtte R Boon, Yanan Wang, and Patrick CN Rensen. FGF21 protects against hepatic lipotoxicity and macrophage activation to attenuate fibrogenesis in nonalcoholic steatohepatitis. *eLife*, 12, 2023.

[12879] Hang Liu, Zhuoran Zhang, Yifan Gu, Changsheng Dai, Guanqiao Shan, Haocong Song, Daniel Li, Wenyuan Chen, Ge Lin, and Yu Sun. Development and evaluation of a live birth prediction model for evaluating human blastocysts from a retrospective study. *eLife*, 12, 2023.

[12880] Hanghang Liu, Akane Wada, Isabella Le, Phuong T Le, Andrew WF Lee, Jun Zhou, Francesca Gori, Roland Baron, and Clifford J Rosen. PTH regulates osteogenesis and suppresses adipogenesis through Zfp467 in a feed-forward, PTH1R-cyclic AMP-dependent manner. *eLife*, 12, 2023.

[12881] He Liu, Meisuo Liu, Xibao Tian, Haina Wang, Jiujiao Gao, Hanrui Li, Zhehuan Zhao, Yu Liu, Caigang Liu, Xuan Chen, and Yongliang Yang. Discovery and biological evaluation of a potent

small molecule CRM1 inhibitor for its selective ablation of extranodal NK/T cell lymphoma. *eLife*, 12, 2023.

[12882] Jin Liu, Hyesang Chang, Daniel A Abrams, Julia Boram Kang, Lang Chen, Miriam Rosenberg-Lee, and Vinod Menon. Atypical cognitive training-induced learning and brain plasticity and their relation to insistence on sameness in children with autism. *eLife*, 12, 2023.

[12883] Jingnan Liu, Yuanbing Zhang, Qiao-Qi Wang, Youfang Zhou, and Ji-Long Liu. Fat body-specific reduction of CTPS alleviates HFD-induced obesity. *eLife*, 12, 2023.

[12884] Linjie Liu, Xiaojie Chen, and Attila Szolnoki. Coevolutionary dynamics via adaptive feedback in collective-risk social dilemma game. *eLife*, 12, 2023.

[12885] Wei Liu, Rupendra Shrestha, Albert Lowe, Xusheng Zhang, and Ludovic Spaeth. Self-formation of concentric zones of telencephalic and ocular tissues and directional retinal ganglion cell axons. *eLife*, 12, 2023.

[12886] Xiao-Lan Liu, Yulin Yang, Yue Hu, Jingjing Wu, Chuqiao Han, Qiaojia Lu, Xihui Gan, Shaohua Qi, Jinhu Guo, Qun He, Yi Liu, and Xiao Liu. The nutrient-sensing GCN2 signaling pathway is essential for circadian clock function by regulating histone acetylation under amino acid starvation. *eLife*, 12, 2023.

[12887] Xiaoliang Liu, Ting Yu, Xiaoqin Tan, Daqing Jin, Wenmin Yang, Jiangping Zhang, Lu Dai, Zhongwei He, Dongliang Li, Yunfeng Zhang, Shuyi Liao, Jinghong Zhao, Tao P Zhong, and Chi Liu. Renal interstitial cells promote nephron regeneration by secreting prostaglandin E2. *eLife*, 12, 2023.

[12888] Xiaozhen Liu, Jing Qiao, Ruixuan Jia, Fan Zhang, Xiang Meng, Yang Li, and Liping Yang. Allele-specific gene-editing approach for vision loss restoration in RHO-associated retinitis pigmentosa. *eLife*, 12, 2023.

[12889] Yingtao Liu, Eri Hasegawa, Akinao Nose, Maarten F Zwart, and Hiroshi Kohsaka. Synchronous multi-segmental activity between metachronal waves controls locomotion speed in Drosophila larvae. *eLife*, 12, 2023.

[12890] Beth Lloyd, Lycia D de Voogd, Verónica Mäki-Marttunen, and Sander Nieuwenhuis. Pupil size reflects activation of subcortical ascending arousal system nuclei during rest. *eLife*, 12, 2023.

[12891] Alexander Lobanov, Samantha Dyckman, Helen Kurkjian, and Babak Momeni. Spatial structure favors microbial coexistence except when slower mediator diffusion weakens interactions. *eLife*, 12, 2023.

[12892] Anne Löffler, Ariel Zylberberg, Michael N Shadlen, and Daniel M Wolpert. Judging the difficulty of perceptual decisions. *eLife*, 12, 2023.

[12893] Claudia Londoño-Nieto, Roberto García-Roa, Clara Garcia-Co, Paula González, and Pau Carazo. Thermal phenotypic plasticity of pre- and post-copulatory male harm buffers sexual conflict in wild Drosophila melanogaster. *eLife*, 12, 2023.

[12894] Lijiang Long, Wen Xu, Francisco Valencia, Annalise B Paaby, and Patrick T McGrath. A toxin-antidote selfish element increases fitness of its host. *eLife*, 12, 2023.

[12895] Vincent Loreau, Renate Rees, Eunice HoYee Chan, Waltraud Taxer, Kathrin Gregor, Bianka Mußil, Christophe Pitaval, Nuno Miguel Luis, Pierre Mangeol, Frank Schnorrer, and Dirk Görlich. A nanobody toolbox to investigate localisation and dynamics of Drosophila titins and other key sarcomeric proteins. *eLife*, 12, 2023.

[12896] Elena Lorenzi, Stefano Tambalo, Giorgio Vallortigara, and Angelo Bifone. Manganese-enhanced magnetic resonance imaging reveals light-induced brain asymmetry in embryo. *eLife*, 12, 2023.

[12897] Isabel IC Low, Lisa M Giocomo, and Alex H Williams. Remapping in a recurrent neural network model of navigation and context inference. *eLife*, 12, 2023.

[12898] Sebastián Loyola, Tycho M Hoogland, Hugo Hoedemaker, Vincenzo Romano, Mario Negrello, and Chris I De Zeeuw. How inhibitory and excitatory inputs gate output of the inferior olive. *eLife*, 12, 2023.

[12899] Anthony Lozano, Francois-Régis Souche, Carine Chavey, Valérie Dardalhon, Christel Ramirez, Serena Vegna, Guillaume Desandre, Anaïs Riviere, Amal Zine El Aabidine, Philippe Fort, Leila Akkari, Urszula Hibner, and Damien Grégoire. Ras/MAPK signalling intensity defines subclonal fitness in a mouse model of hepatocellular carcinoma. *eLife*, 12, 2023.

[12900] Jinping Lu, Ingo Dreyer, Miles Sasha Dickinson, Sabine Panzer, Dawid Jaślan, Carlos Navarro-Retamal, Dietmar Geiger, Ulrich Terpitz, Dirk Becker, Robert M Stroud, Irene Marten, and Rainer Hedrich. Vicia faba SV channel VfTPC1 is a hyperexcitable variant of plant vacuole Two Pore Channels. *eLife*, 12, 2023.

[12901] Junyan Lu, Britta Velten, Bernd Klaus, Mauricio Ramm, Wolfgang Huber, and Rachel Coulthard-Graf. The changing career paths of PhDs and postdocs trained at EMBL. *eLife*, 12, 2023.

[12902] Ke Lu, Ming Zhang, Guizheng Wei, Guozhi Xiao, Liping Tong, and Di Chen. Multiple cullin-associated E3 ligases regulate cyclin D1 protein stability. *eLife*, 12, 2023.

[12903] Haitao Luan, Timothy A Bielecki, Bhopal C Mohapatra, Namista Islam, Insha Mushtaq, Aaqib M Bhat, Sameer Mirza, Sukanya Chakraborty, Mohsin Raza, Matthew D Storck, Michael S Toss, Jane L Meza, Wallace B Thoreson, Donald W Coulter, Emad A Rakha, Vimla Band, and Hamid Band. EHD2 overexpression promotes tumorigenesis and metastasis in triple-negative breast cancer by regulating store-operated calcium entry. *eLife*, 12, 2023.

[12904] Bronwyn A Lucas, Benjamin A Himes, and Nikolaus Grigorieff. Baited reconstruction with 2D template matching for high-resolution structure determination in vitro and in vivo without template bias. *eLife*, 12, 2023.

[12905] Eric Lucas, Raul Murillo, Silvina Arrossi, Martin Bárcena, Youssef Chami, Ashrafun Nessa, Suraj Perera, Padmaka Silva, Suleeporn Sangrajrang, Richard Muwonge, and Partha Basu. Quantification of impact of COVID-19 pandemic on cancer screening programmes – a case study from Argentina, Bangladesh, Colombia, Morocco, Sri Lanka, and Thailand. *eLife*, 12, 2023.

[12906] Rebecca Luckett, Sarah Feldman, Yin Ling Woo, Anna-Barbara Moscicki, Anna R Giuliano, Silvia de Sanjosé, Andreas M Kaufmann, Shuk On Annie Leung, Francisco Garcia, Karen Chan, Neerja Bhatla, Margaret Stanley, Julia Brotherton, Joel Palefsky, and Suzanne Garland. COVID-19 as a catalyst for reimagining cervical cancer prevention. *eLife*, 12, 2023.

[12907] Alison M Luckey, Lauren S McLeod, Yuefeng Huang, Anusha Mohan, and Sven Vanneste. Making memories last using the peripheral effect of direct current stimulation. *eLife*, 12, 2023.

[12908] Agustin Leonardo Lujan, Ombretta Foresti, Conor Sugden, Nathalie Brouwers, Alex Mateo Farre, Alessio Vignoli, Mahshid Azamian, Alicia Turner, Jose Wojnacki, and Vivek Malhotra. Defects in lipid homeostasis reflect the function of TANGO2 in phospholipid and neutral lipid metabolism. *eLife*, 12, 2023.

[12909] Cihang Luo, George O Poinar, Chunpeng Xu, De Zhuo, Edmund A Jarzembowski, and Bo Wang. Widespread mermithid nematode parasitism of Cretaceous insects. *eLife*, 12, 2023.

[12910] Zhao-Bo Luo, Shengzhong Han, Xi-Jun Yin, Hongye Liu, Junxia Wang, Meifu Xuan, Chunyun Hao, Danqi Wang, Yize Liu, Shuangyan Chang, Dongxu Li, Kai Gao, Huiling Li, Biaohu Quan, Lin-Hu Quan, and Jin-Dan Kang. Fecal transplant from myostatin deletion pigs positively impacts the gut-muscle axis. *eLife*, 12, 2023.

[12911] Francisco J Luongo, Lu Liu, Chun Lum Andy Ho, Janis K Hesse, Joseph B Wekselblatt, Frank F Lanfranchi, Daniel Huber, and Doris Y Tsao. Mice and primates use distinct strategies for visual segmentation. *eLife*, 12, 2023.

[12912] Shira M Lupkin and Vincent B McGinty. Monkeys exhibit human-like gaze biases in economic decisions. *eLife*, 12, 2023.

[12913] Kennedy Lushasi, Kirstyn Brunker, Malavika Rajeev, Elaine A Ferguson, Gurdeep Jaswant, Laurie Louise Baker, Roman Biek, Joel Changalucha, Sarah Cleaveland, Anna Czupryna, Anthony R Fooks, Nicodemus J Govella, Daniel T Haydon, Paul CD Johnson, Rudovick Kazwala, Tiziana Lembo, Denise Marston, Msanif Masoud, Matthew Maziku, Eberhard Mbunda, Geofrey Mchau, Ally Z Mohamed, Emmanuel Mpolya, Chanasa Ngeleja, Kija Ng'habi, Hezron Nonga, Kassim Omar, Kristyna Rysava, Maganga Sambo, Lwitiko Sikana, Rachel Steenson, and Katie Hampson. Integrating contact tracing and whole-genome sequencing to track the elimination of dog-mediated rabies: An observational and genomic study. *eLife*, 12, 2023.

[12914] Kevin Luxem, Jennifer J Sun, Sean P Bradley, Keerthi Krishnan, Eric Yttri, Jan Zimmermann, Talmo D Pereira, and Mark Laubach. Open-source tools for behavioral video analysis: Setup, methods, and best practices. *eLife*, 12, 2023.

[12915] Huiyun Lyu, Guohua Yuan, Xinyi Liu, Xiaobo Wang, Shuang Geng, Tie Xia, Xuyu Zhou, Yinqing Li, Xiaoyu Hu, and Yan Shi. Sustained store-operated calcium entry utilizing activated chromatin state leads to instability in iTregs. *eLife*, 12, 2023.

[12916] Liang Ma, Jasleen Singh, and Randy Schekman. Two RNA-binding proteins mediate the sorting of miR223 from mitochondria into exosomes. *eLife*, 12, 2023.

[12917] Xuan Ma, Fabio Rizzoglio, Kevin L Bodkin, Eric Perreault, Lee E Miller, and Ann Kennedy. Using adversarial networks to extend brain computer interface decoding accuracy over time. *eLife*, 12, 2023.

[12918] HaDi MaBouDi, James AR Marshall, Neville Dearden, and Andrew B Barron. How honey bees make fast and accurate decisions. *eLife*, 12, 2023.

[12919] Nicolas Macaisne, Laura Bellutti, Kimberley Laband, Frances Edwards, Laras Pitayu-Nugroho, Alison Gervais, Thadshagine Ganeswaran, Hélène Geoffroy, Gilliane Maton, Julie C Canman, Benjamin Lacroix, and Julien Dumont. Synergistic stabilization of microtubules by BUB-1, HCP-1, and CLS-2 controls microtubule pausing and meiotic spindle assembly. *eLife*, 12, 2023.

[12920] Robert Machold, Shlomo Dellal, Manuel Valero, Hector Zurita, Ilya Kruglikov, John Hongyu Meng, Jessica L Hanson, Yoshiko Hashikawa, Benjamin Schuman, György Buzsáki, and Bernardo Rudy. Id2 GABAergic interneurons comprise a neglected fourth major group of cortical inhibitory cells. *eLife*, 12, 2023.

[12921] Emily L Mackevicius, Shijie Gu, Natalia I Denisenko, and Michale S Fee. Self-organization of songbird neural sequences during social isolation. *eLife*, 12, 2023.

[12922] Maria-Bernadette Madel, Julia Halper, Lidia Ibáñez, Lozano Claire, Matthieu Rouleau, Antoine Boutin, Adrien Mahler, Rodolphe Pontier-Bres, Thomas Ciucci, Majlinda Topi, Christophe Hue, Jerome Amiaud, Salvador Iborra, David Sancho, Dominique Heymann, Henri-Jean Garchon, Dorota Czerucka, Florence Apparailly, Isabelle Duroux-Richard, Abdelilah Wakkach, and Claudine Blin-Wakkach. Specific targeting of inflammatory osteoclastogenesis by the probiotic yeast S. boulardii CNCM I-745 reduces bone loss in osteoporosis. *eLife*, 12, 2023.

[12923] Bhoomi Madhu, Mohammed Farhan Lakdawala, and Tina L Gumienny. The DBL-1/TGF-$\beta$ signaling pathway tailors behavioral and molecular host responses to a variety of bacteria in Caenorhabditis elegans. *eLife*, 12, 2023.

[12924] Naïma Madi, Daisy Chen, Richard Wolff, B Jesse Shapiro, and Nandita R Garud. Community diversity is associated with intra-species genetic diversity and gene loss in the human gut microbiome. *eLife*, 12, 2023.

[12925] Luba Mahbub, Guennadi Kozlov, Pengyu Zong, Emma L Lee, Sandra Tetteh, Thushara Nethramangalath, Caroline Knorn, Jianning Jiang, Ashkan Shahsavan, Lixia Yue, Loren Runnels, and Kalle Gehring. Structural insights into regulation of CNNM-TRPM7 divalent cation uptake by the small GTPase ARL15. *eLife*, 12, 2023.

[12926] Eike K Mahlandt, Sebastián Palacios Martínez, Janine JG Arts, Simon Tol, Jaap D van Buul, and Joachim Goedhart. Opto-RhoGEFs, an optimized optogenetic toolbox to reversibly control Rho GTPase activity on a global to subcellular scale, enabling precise control over vascular endothelial barrier strength. *eLife*, 12, 2023.

[12927] Josephine HR Maidment, Motoki Shimizu, Adam R Bentham, Sham Vera, Marina Franceschetti, Apinya Longya, Clare EM Stevenson, Juan Carlos De la Concepcion, Aleksandra Białas, Sophien Kamoun, Ryohei Terauchi, and Mark J Banfield. Effector target-guided engineering of an integrated domain expands the disease resistance profile of a rice NLR immune receptor. *eLife*, 12, 2023.

[12928] Robert Maier, Pavel Flegontov, Olga Flegontova, Ulaş Işıldak, Piya Changmai, and David Reich. On the limits of fitting complex models of population history to f-statistics. *eLife*, 12, 2023.

[12929] Samarpan Maiti, Kaushik Bhattacharya, Diana Wider, Dina Hany, Olesya Panasenko, Lilia Bernasconi, Nicolas Hulo, and Didier Picard. Hsf1 and the molecular chaperone Hsp90 support a 'rewiring stress response' leading to an adaptive cell size increase in chronic stress. *eLife*, 12, 2023.

[12930] Gaurav Majumdar, Timothy A Liddle, Calum Stewart, Christopher J Marshall, Maureen Bain, and Tyler Stevenson. FSH$\beta$ links photoperiodic signaling to seasonal reproduction in Japanese quail. *eLife*, 12, 2023.

[12931] Victoria P Mak, Kami White, Lynne R Wilkens, Iona Cheng, Christopher A Haiman, and Loic Le Marchand. The impact of COVID-19 on cancer screening and treatment in older adults: The Multiethnic Cohort Study. *eLife*, 12, 2023.

[12932] Tamar R Makin and John W Krakauer. Against cortical reorganisation. *eLife*, 12, 2023.

[12933] Gerardo Malagon, Jongyun Myeong, and Vitaly A Klyachko. Two forms of asynchronous release with distinctive spatiotemporal dynamics in central synapses. *eLife*, 12, 2023.

[12934] Hugo Malagon-Vina, Stéphane Ciocchi, and Thomas Klausberger. Firing patterns of ventral hippocampal neurons predict the exploration of anxiogenic locations. *eLife*, 12, 2023.

[12935] FranÇois Mallard, Bruno Afonso, and Henrique Teotónio. Selection and the direction of phenotypic evolution. *eLife*, 12, 2023.

[12936] Stefano Mammola, Martino Adamo, Dragan Antić, Jacopo Calevo, Tommaso Cancellario, Pedro Cardoso, Dan Chamberlain, Matteo Chialva, Furkan Durucan, Diego Fontaneto, Duarte Goncalves, Alejandro Martínez, Luca Santini, Iñigo Rubio-Lopez, Ronaldo Sousa, David Villegas-Rios, Aida Verdes, and Ricardo A Correia. Drivers of species knowledge across the tree of life. *eLife*, 12, 2023.

[12937] Irene Man, Damien Georges, Maxime Bonjour, and Iacopo Baussano. Approximating missing epidemiological data for cervical cancer through Footprinting: A case study in India. *eLife*, 12, 2023.

[12938] Irene Man, Damien Georges, Rengaswamy Sankaranarayanan, Partha Basu, and Iacopo Baussano. Building resilient cervical cancer prevention through gender-neutral HPV vaccination. *eLife*, 12, 2023.

[12939] Irina AD Mancini, Riccardo Levato, Marlena M Ksiezarczyk, Miguel Dias Castilho, Michael Chen, Mattie HP van Rijen, Lonneke L IJsseldijk, Marja Kik, P René van Weeren, and Jos Malda. Microstructural differences in the osteochondral unit of terrestrial and aquatic mammals. *eLife*, 12, 2023.

[12940] Rebecca EK Mandt, Madeline R Luth, Mark A Tye, Ralph Mazitschek, Sabine Ottilie, Elizabeth A Winzeler, Maria Jose Lafuente-Monasterio, Francisco Javier Gamo, Dyann F Wirth, and Amanda K Lukens. Diverse evolutionary pathways challenge the use of collateral sensitivity as a strategy to suppress resistance. *eLife*, 12, 2023.

[12941] Jake P Mann, Xiaowen Duan, Satish Patel, Luis Carlos Tábara, Fabio Scurria, Anna Alvarez-Guaita, Afreen Haider, Ineke Luijten, Matthew Page, Margherita Protasoni, Koini Lim, Sam Virtue, Stephen O'Rahilly, Martin Armstrong, Julien Prudent, Robert K Semple, and David B Savage. A mouse model of human mitofusin-2-related lipodystrophy exhibits adipose-specific mitochondrial stress and reduced leptin secretion. *eLife*, 12, 2023.

[12942] Omer Mano, Minseung Choi, Ryosuke Tanaka, Matthew S Creamer, Natalia CB Matos, Joseph W Shomar, Bara A Badwan, Thomas R Clandinin, and Damon A Clark. Long-timescale anti-directional rotation in Drosophila optomotor behavior. *eLife*, 12, 2023.

[12943] Arian Mansur, Remi Joseph, Euri S Kim, Pierre M Jean-Beltran, Namrata D Udeshi, Cadence Pearce, Hanjie Jiang, Reina Iwase, Miroslav P Milev, Hashem A Almousa, Elyshia McNamara, Jeffrey Widrick, Claudio Perez, Gianina Ravenscroft, Michael Sacher, Philip A Cole, Steven A Carr, and Vandana A Gupta. Dynamic regulation of inter-organelle communication by ubiquitylation controls skeletal muscle development and disease onset. *eLife*, 12, 2023.

[12944] Wei Mao, Lars D Renner, Charlène Cornilleau, Ines Li de la Sierra-Gallay, Sana Afensiss, Sarah Benlamara, Yoan Ah-Seng, Herman Van Tilbeurgh, Sylvie Nessler, Aurélie Bertin, Arnaud Chastanet, and Rut Carballido-Lopez. On the role of nucleotides and lipids in the polymerization of the actin homolog MreB from a Gram-positive bacterium. *eLife*, 12, 2023.

[12945] Yanbo Mao, Thomas Nakel, Isil Erbasol Serbes, Saurabh Joshi, Dawit G Tekleyohans, Thomas Baum, and Rita Groß-Hardt. ECS1 and ECS2 suppress polyspermy and the formation of haploid plants by promoting double fertilization. *eLife*, 12, 2023.

[12946] Sophie Marbach, Noah Ziethen, Leonie Bastin, Felix K Bäuerle, and Karen Alim. Vein fate determined by flow-based but time-delayed integration of network architecture. *eLife*, 12, 2023.

[12947] Emmeline Marchal-Duval, Méline Homps-Legrand, Antoine Froidure, Madeleine Jaillet, Mada Ghanem, Deneuville Lou, Aurélien Justet, Arnaud Maurac, Aurelie Vadel, Emilie Fortas, Aurelie Cazes, Audrey Joannes, Laura Giersh, Herve Mal, Pierre Mordant, Tristan Piolot, Marin Truchin, Carine M Mounier, Ksenija Schirduan, Martina Korfei, Andreas Gunther, Bernard Mari, Frank Jaschinski, Bruno Crestani, and Arnaud A Mailleux. Identification of Paired-related Homeobox Protein 1 as a key mesenchymal transcription factor in pulmonary fibrosis. *eLife*, 12, 2023.

[12948] Miguel A Maria-Solano and Sun Choi. Dynamic allosteric networks drive adenosine A1 receptor activation and G-protein coupling. *eLife*, 12, 2023.

[12949] Marija Markicevic, Oliver Sturman, Johannes Bohacek, Markus Rudin, Valerio Zerbi, Ben D Fulcher, and Nicole Wenderoth. Neuromodulation of striatal D1 cells shapes BOLD fluctuations in anatomically connected thalamic and cortical regions. *eLife*, 12, 2023.

[12950] Odeya Marmor, Yael Pollak, Chen Doron, Fritjof Helmchen, and Ariel Gilad. History information emerges in the cortex during learning. *eLife*, 12, 2023.

[12951] Carine M Marshall, Veronica L Thompson, Nicky M Creux, and Stacey L Harmer. The circadian clock controls temporal and spatial patterns of floral development in sunflower. *eLife*, 12, 2023.

[12952] Anatolie Marta, TomáŠ Tichopád, Oldřich BartoŠ, Jiří Klíma, Mujahid Ali Shah, Vendula Šlechtová Bohlen, Joerg Bohlen, Karel Halačka, LukáŠ Choleva, Matthias Stöck, Dmitrij Dedukh, and Karel Janko. Genetic and karyotype divergence between parents affect clonality and sterility in hybrids. *eLife*, 12, 2023.

[12953] Josep Martí-Solans, Aina Børve, Paul Bump, Andreas Hejnol, and Timothy Lynagh. Peripheral and central employment of acid-sensing ion channels during early bilaterian evolution. *eLife*, 12, 2023.

[12954] Kendall E Martin, Padmapriyadarshini Ravisankar, Manu Beerens, Calum A MacRae, and Joshua S Waxman. Nr2f1a maintains atrial nkx2.5 expression to repress pacemaker identity within venous atrial cardiomyocytes of zebrafish. *eLife*, 12, 2023.

[12955] Carolina A Martinez-Gutierrez, Josef C Uyeda, and Frank O Aylward. A timeline of bacterial and archaeal diversification in the ocean. *eLife*, 12, 2023.

[12956] Julieta Martino, Sebastián Omar Siri, Nicolás Luis Calzetta, Natalia Soledad Paviolo, Cintia Garro, Maria F Pansa, Sofía Carbajosa, Aaron C Brown, José Luis Bocco, Israel Gloger, Gerard Drewes, Kevin P Madauss, Gastón Soria, and Vanesa Gottifredi. Inhibitors of Rho kinases (ROCK) induce multiple mitotic defects and synthetic lethality in BRCA2-deficient cells. *eLife*, 12, 2023.

[12957] William Duncan Martinson, Rebecca McLennan, Jessica M Teddy, Mary C McKinney, Lance A Davidson, Ruth E Baker, Helen M Byrne, Paul M Kulesa, and Philip K Maini. Dynamic fibronectin assembly and remodeling by leader neural crest cells prevents jamming in collective cell migration. *eLife*, 12, 2023.

[12958] Emily Martyn, Sarah Eisen, Nicky Longley, Philippa Harris, Julian Surey, James Norman, Michael Brown, Binta Sultan, Tongai G Maponga, Collins Iwuji, Stuart Flanagan, Indrajit Ghosh, Alistair Story, and Philippa C Matthews. The forgotten people: Hepatitis B virus (HBV) infection as a priority for the inclusion health agenda. *eLife*, 12, 2023.

[12959] Alex Mas-Sandoval, Sara Mathieson, and Matteo Fumagalli. The genomic footprint of social stratification in admixing American populations. *eLife*, 12, 2023.

[12960] Javier Masís, Travis Chapman, Juliana Y Rhee, David D Cox, and Andrew M Saxe. Strategically managing learning during perceptual decision making. *eLife*, 12, 2023.

[12961] Peter M Masschelin, Pradip Saha, Scott A Ochsner, Aaron R Cox, Kang Ho Kim, Jessica B Felix, Robert Sharp, Xin Li, Lin Tan, Jun Hyoung Park, Liping Wang, Vasanta Putluri, Philip L Lorenzi, Alli M Nuotio-Antar, Zheng Sun, Benny Abraham Kaipparettu, Nagireddy Putluri, David D Moore, Scott A Summers, Neil J McKenna, and Sean M Hartig. Vitamin B2 enables regulation of fasting glucose availability. *eLife*, 12, 2023.

[12962] Stewart WC Masson, Søren Madsen, Kristen C Cooke, Meg Potter, Alexis Diaz Vegas, Luke Carroll, Senthil Thillainadesan, Harry B Cutler, Ken R Walder, Gregory J Cooney, Grant Morahan, Jacqueline Stöckli, and David E James. Leveraging genetic diversity to identify small molecules that reverse mouse skeletal muscle insulin resistance. *eLife*, 12, 2023.

[12963] Francesca Mastrogiuseppe, Naoki Hiratani, and Peter Latham. Evolution of neural activity in circuits bridging sensory and abstract knowledge. *eLife*, 12, 2023.

[12964] Giuseppina Mastrototaro, Pierluigi Carullo, Jianlin Zhang, Beatrice Scellini, Nicoletta Piroddi, Simona Nemska, Maria Carmela Filomena, Simone Serio, Carol A Otey, Chiara Tesi, Fabian Emrich, Wolfgang A Linke, Corrado Poggesi, Simona Boncompagni, and Marie-Louise Bang. Ablation of palladin in adult heart causes dilated cardiomyopathy associated with intercalated disc abnormalities. *eLife*, 12, 2023.

[12965] Surjeet Mastwal, Xinjian Li, Rianne Stowell, Matthew Manion, Wenyu Zhang, Nam-Shik Kim, Ki-Jun Yoon, Hongjun Song, Guo-Li Ming, and Kuan Hong Wang. Adolescent neurostimulation of dopamine circuit reverses genetic deficits in frontal cortex function. *eLife*, 12, 2023.

[12966] Ivona Mateska, Anke Witt, Eman Hagag, Anupam Sinha, Canelif Yilmaz, Evangelia Thanou, Na Sun, Ourania Kolliniati, Maria Patschin, Heba Abdelmegeed, Holger Henneicke, Waldemar Kanczkowski, Ben Wielockx, Christos Tsatsanis, Andreas Dahl, Axel Karl Walch, Ka Wan Li, Mirko Peitzsch, Triantafyllos Chavakis, and Vasileia Ismini Alexaki. Succinate mediates inflammation-induced adrenocortical dysfunction. *eLife*, 12, 2023.

[12967] William Matlock, Samuel Lipworth, Kevin K Chau, Manal AbuOun, Leanne Barker, James Kavanagh, Monique Andersson, Sarah Oakley, Marcus Morgan, Derrick W Crook, Daniel S Read, Muna Anjum, Liam P Shaw, Nicole Stoesser, and Manal AbuOun. Enterobacterales plasmid sharing amongst human bloodstream infections, livestock, wastewater, and waterway niches in Oxfordshire, UK. *eLife*, 12, 2023.

[12968] Christine Mau, Heike Rudolf, Frederic Strobl, Benjamin Schmid, Timo Regensburger, Ralf Palmisano, Ernst HK Stelzer, Leila Taher, and Ezzat El-Sherif. How enhancers regulate wavelike gene expression patterns. *eLife*, 12, 2023.

[12969] Lampros Mavrommatis, Hyun-Woo Jeong, Urs Kindler, Gemma Gomez-Giro, Marie-Cecile Kienitz, Martin Stehling, Olympia E Psathaki, Dagmar Zeuschner, M Gabriele Bixel, Dong Han, Gabriela Morosan-Puopolo, Daniela Gerovska, Ji Hun Yang, Jeong Beom Kim, Marcos J Arauzo-Bravo, Jens C Schwamborn, Stephan A Hahn, Ralf H Adams, Hans R Schöler, Matthias Vorgerd, Beate Brand-Saberi, and Holm Zaehres. Human skeletal muscle organoids model fetal myogenesis and sustain uncommitted PAX7 myogenic progenitors. *eLife*, 12, 2023.

[12970] Dennis May, Sangwon Yun, David G Gonzalez, Sangbum Park, Yanbo Chen, Elizabeth Lathrop, Biao Cai, Tianchi Xin, Hongyu Zhao, Siyuan Wang, Lauren E Gonzalez, Katie Cockburn, and Valentina Greco. Live imaging reveals chromatin compaction transitions and dynamic transcriptional bursting during stem cell differentiation in vivo. *eLife*, 12, 2023.

[12971] Gemma E May, Christina Akirtava, Matthew Agar-Johnson, Jelena Micic, John Woolford, and Joel McManus. Unraveling the influences of sequence and position on yeast uORF activity using massively parallel reporter systems and machine learning. *eLife*, 12, 2023.

[12972] Marianne Mazevet, Anissa Belhadef, Maxance Ribeiro, Delphine Dayde, Anna Llach, Marion Laudette, Tiphaine Belleville, Philippe Mateo, Mélanie Gressette, Florence Lefebvre, Ju Chen, Christilla Bachelot-Loza, Catherine Rucker-Martin, Frank Lezoualch, Bertrand Crozatier, Jean-Pierre Benitah, Marie-Catherine Vozenin, Rodolphe Fischmeister, Ana-Maria Gomez, Christophe Lemaire, and Eric Morel. EPAC1 inhibition protects the heart from doxorubicin-induced toxicity. *eLife*, 12, 2023.

[12973] Sarah E McComas, Tom Reichenbach, Darko Mitrovic, Claudia Alleva, Marta Bonaccorsi, Lucie Delemotte, and David Drew. Determinants of sugar-induced influx in the mammalian fructose transporter GLUT5. *eLife*, 12, 2023.

[12974] Sharon A McGrath-Morrow, Jarrett Venezia, Roland Ndeh, Nigel Michki, Javier Perez, Benjamin David Singer, Raffaello Cimbro, Mark Soloski, and Alan L Scott. Cellular and molecular dynamics in the lungs of neonatal and juvenile mice in response to E. coli. *eLife*, 12, 2023.

[12975] Andrew McKinney, Ming Hu, Amber Hoskins, Arian Mohammadyar, Nabeeha Naeem, Junzhan Jing, Saumil S Patel, Bhavin R Sheth, and Xiaolong Jiang. Cellular composition and circuit organization of the locus coeruleus of adult mice. *eLife*, 12, 2023.

[12976] Andrea McQuate, Sharmon Knecht, and David W Raible. Activity regulates a cell type-specific mitochondrial phenotype in zebrafish lateral line hair cells. *eLife*, 12, 2023.

[12977] Taylor N Medwig-Kinney, Brian A Kinney, Michael AQ Martinez, Callista Yee, Sydney S Sirota, Angelina A Mullarkey, Neha Somineni, Justin Hippler, Wan Zhang, Kang Shen, Christopher Hammell, Ariel M Pani, and David Q Matus. Dynamic compartmentalization of the pro-invasive transcription factor NHR-67 reveals a role for Groucho in regulating a proliferative-invasive cellular switch in C. elegans. *eLife*, 12, 2023.

[12978] Geoffrey W Meissner, Aljoscha Nern, Zachary Dorman, Gina M DePasquale, Kaitlyn Forster, Theresa Gibney, Joanna H Hausenfluck, Yisheng He, Nirmala A Iyer, Jennifer Jeter, Lauren Johnson, Rebecca M Johnston, Kelley Lee, Brian Melton, Brianna Yarbrough, Christopher T Zugates, Jody Clements, Cristian Goina, Hideo Otsuna, Konrad Rokicki, Robert R Svirskas, Yoshinori Aso,

Gwyneth M Card, Barry J Dickson, Erica Ehrhardt, Jens Goldammer, Masayoshi Ito, Dagmar Kainmueller, Wyatt Korff, Lisa Mais, Ryo Minegishi, Shigehiro Namiki, Gerald M Rubin, Gabriella R Sterne, Tanya Wolff, and Oz Malkesman. A searchable image resource of Drosophila GAL4 driver expression patterns with single neuron resolution. *eLife*, 12, 2023.

[12979] Sahar Melamed, Aixia Zhang, Michal Jarnik, Joshua Mills, Aviezer Silverman, Hongen Zhang, and Gisela Storz. $\sigma$28-dependent small RNA regulation of flagella biosynthesis. *eLife*, 12, 2023.

[12980] Artur Meller, Jeffrey M Lotthammer, Louis G Smith, Borna Novak, Lindsey A Lee, Catherine C Kuhn, Lina Greenberg, Leslie A Leinwand, Michael J Greenberg, and Gregory R Bowman. Drug specificity and affinity are encoded in the probability of cryptic pocket opening in myosin motor domains. *eLife*, 12, 2023.

[12981] Mariana R Melo, Alexander D Wykes, Angela A Connelly, Jaspreet K Bassi, Shane D Cheung, Stuart J McDougall, Clément Menuet, Ross AD Bathgate, and Andrew M Allen. Selective transduction and photoinhibition of pre-Bötzinger complex neurons that project to the facial nucleus in rats affects nasofacial activity. *eLife*, 12, 2023.

[12982] Daniël P Melters, Keir C Neuman, Reda S Bentahar, Tatini Rakshit, and Yamini Dalal. Single molecule analysis of CENP-A chromatin by high-speed atomic force microscopy. *eLife*, 12, 2023.

[12983] Matthew J Mender, Samuel R Nason-Tomaszewski, Hisham Temmar, Joseph T Costello, Dylan M Wallace, Matthew S Willsey, Nishant Ganesh Kumar, Theodore A Kung, Parag Patil, and Cynthia A Chestek. The impact of task context on predicting finger movements in a brain-machine interface. *eLife*, 12, 2023.

[12984] Gaurav Mendiratta and Edward Stites. Theoretical analysis reveals a role for RAF conformational autoinhibition in paradoxical activation. *eLife*, 12, 2023.

[12985] Rosa E Menjivar, Zeribe C Nwosu, Wenting Du, Katelyn L Donahue, Hanna S Hong, Carlos Espinoza, Kristee Brown, Ashley Velez-Delgado, Wei Yan, Fatima Lima, Allison Bischoff, Padma Kadiyala, Daniel Salas-Escabillas, Howard C Crawford, Filip Bednar, Eileen Carpenter, Yaqing Zhang, Christopher J Halbrook, Costas A Lyssiotis, and Marina Pasca di Magliano. Arginase 1 is a key driver of immune suppression in pancreatic cancer. *eLife*, 12, 2023.

[12986] Miriam Menzel, David Gräßel, Ivan Rajkovic, Michael M Zeineh, and Marios Georgiadis. Using light and X-ray scattering to untangle complex neuronal orientations and validate diffusion MRI. *eLife*, 12, 2023.

[12987] Christian Merkel, Luise Burgmann, Mandy Viktoria Bartsch, Mircea Ariel Schoenfeld, and Jens-Max Hopf. Serial attentional resource allocation during parallel feature value tracking. *eLife*, 12, 2023.

[12988] Ljubica Mihaljević, Zheng Ruan, James Osei-Owusu, Wei Lü, and Zhaozhu Qiu. Inhibition of the proton-activated chloride channel PAC by PIP2. *eLife*, 12, 2023.

[12989] Gerdien Mijnheer, Nila Hendrika Servaas, Jing Yao Leong, Arjan Boltjes, Eric Spierings, Phyllis Chen, Liyun Lai, Alessandra Petrelli, Sebastiaan Vastert, Rob J de Boer, Salvatore Albani, Aridaman Pandit, and Femke van Wijk. Compartmentalization and persistence of dominant (regulatory) T cell clones indicates antigen skewing in juvenile idiopathic arthritis. *eLife*, 12, 2023.

[12990] Flora Mikaeloff, Marco Gelpi, Rui Benfeitas, Andreas D Knudsen, Beate Vestad, Julie Høgh, Johannes R Hov, Thomas Benfield, Daniel Murray, Christian G Giske, Adil Mardinoglu, Marius Trøseid, Susanne D Nielsen, and Ujjwal Neogi. Network-based multi-omics integration reveals metabolic at-risk profile within treated HIV-infection. *eLife*, 12, 2023.

[12991] Peter R Millar, Brian A Gordon, Patrick H Luckett, Tammie LS Benzinger, Carlos Cruchaga, Anne M Fagan, Jason J Hassenstab, Richard J Perrin, Suzanne E Schindler, Ricardo F Allegri, Gregory S Day, Martin R Farlow, Hiroshi Mori, Georg Nübling, Adam Sarah, Randall J Bateman, John C Morris, and Beau M Ances. Multimodal brain age estimates relate to Alzheimer disease biomarkers and cognition in early stages: a cross-sectional observational study. *eLife*, 12, 2023.

[12992] Alexandria N Miller, Patrick R Houlihan, Ella Matamala, Deny Cabezas-Bratesco, Gi Young Lee, Ben Cristofori-Armstrong, Tanya L Dilan, Silvia Sanchez-Martinez, Doreen Matthies, Rui Yan, Zhiheng Yu, Dejian Ren, Sebastian E Brauchi, and David E Clapham. The SARS-CoV-2 accessory protein Orf3a is not an ion channel, but does interact with trafficking proteins. *eLife*, 12, 2023.

[12993] Shashwat Mishra, Mohamed Dabaja, Asra Akhlaq, Bianca Pereira, Kelsey Marbach, Mediha Rovcanin, Rashmi Chandra, Antonio Caballero, Diana Fernandes de Abreu, QueeLim Ch'ng, and Joy Alcedo. Specific sensory neurons and insulin-like peptides modulate food type-dependent oogenesis and fertilization in Caenorhabditis elegans. *eLife*, 12, 2023.

[12994] Darko Mitrovic, Sarah E McComas, Claudia Alleva, Marta Bonaccorsi, David Drew, and Lucie Delemotte. Reconstructing the transport cycle in the sugar porter superfamily using coevolution-powered machine learning. *eLife*, 12, 2023.

[12995] Valentin Mitterer, Matthias Thoms, Robert Buschauer, Otto Berninghausen, Ed Hurt, and Roland Beckmann. Concurrent remodelling of nucleolar 60S subunit precursors by the Rea1 ATPase and Spb4 RNA helicase. *eLife*, 12, 2023.

[12996] Akihiro Miura, Hemanta Sarmah, Junichi Tanaka, Youngmin Hwang, Anri Sawada, Yuko Shimamura, Takehiro Otoshi, Yuri Kondo, Yinshan Fang, Dai Shimizu, Zurab Ninish, Jake Le Suer, Nicole C Dubois, Jennifer Davis, Shinichi Toyooka, Jun Wu, Jianwen Que, Finn J Hawkins, Chyuan-Sheng Lin, and Munemasa Mori. Conditional blastocyst complementation of a defective Foxa2 lineage efficiently promotes the generation of the whole lung. *eLife*, 12, 2023.

[12997] Ryota Miyashita, Atsuya Nishiyama, Weihua Qin, Yoshie Chiba, Satomi Kori, Norie Kato, Chieko Konishi, Soichiro Kumamoto, Hiroko Kozuka-Hata, Masaaki Oyama, Yoshitaka Kawasoe, Toshiki Tsurimoto, Tatsuro S Takahashi, Heinrich Leonhardt, Kyohei Arita, and Makoto Nakanishi. The termination of UHRF1-dependent PAF15 ubiquitin signaling is regulated by USP7 and ATAD5. *eLife*, 12, 2023.

[12998] Kaoru Mizukami, Hiroki Higashiyama, Yuichiro Arima, Koji Ando, Norihiro Okada, Katsumi Kose, Shigehito Yamada, Jun K Takeuchi, Kazuko Koshiba-Takeuchi, Shigetomo Fukuhara, Sachiko Miyagawa-Tomita, and Hiroki Kurihara. Coronary artery established through amniote evolution. *eLife*, 12, 2023.

[12999] Brijesh Modi, Matteo Guardamagna, Federico Stella, Marilena Griguoli, Enrico Cherubini, and Francesco P Battaglia. State-dependent coupling of hippocampal oscillations. *eLife*, 12, 2023.

[13000] Mehrab N Modi, Adithya E Rajagopalan, Hervé Rouault, Yoshinori Aso, and Glenn C Turner. Flexible specificity of memory in Drosophila depends on a comparison between choices. *eLife*, 12, 2023.

[13001] Sebastian Moeller, Anton M Unakafov, Julia Fischer, Alexander Gail, Stefan Treue, and Igor Kagan. Human and macaque pairs employ different coordination strategies in a transparent decision game. *eLife*, 12, 2023.

[13002] Fatma F Mohamed, Chunxi Ge, Shawn A Hallett, Alec C Bancroft, Randy T Cowling, Noriaki Ono, Abdul-Aziz Binrayes, Barry Greenberg, Benjamin Levi, Vesa M Kaartinen, and Renny T Franceschi. Control of craniofacial development by the collagen receptor, discoidin domain receptor 2. *eLife*, 12, 2023.

[13003] Sindhu Mohandas, Prasanna Jagannathan, Timothy J Henrich, Zaki A Sherif, Christian Bime, Erin Quinlan, Michael A Portman, Marila Gennaro, Jalees Rehman, and Steven B Bradfute. Immune mechanisms underlying COVID-19 pathology and post-acute sequelae of SARS-CoV-2 infection (PASC). *eLife*, 12, 2023.

[13004] Ali Mohebi, Val L Collins, and Joshua D Berke. Accumbens cholinergic interneurons dynamically promote dopamine release and enable motivation. *eLife*, 12, 2023.

[13005] Sarah Mondoloni, Claire Nguyen, Eléonore Vicq, Maria Ciscato, Joachim Jehl, Romain Durand-de Cuttoli, Nicolas Torquet, Stefania Tolu, Stéphanie Pons, Uwe Maskos, Fabio Marti, Philippe Faure, and Alexandre Mourot. Prolonged nicotine exposure reduces aversion to the drug in mice by altering nicotinic transmission in the interpeduncular nucleus. *eLife*, 12, 2023.

[13006] Siavash Monfared, Guruswami Ravichandran, José Andrade, and Amin Doostmohammadi. Mechanical basis and topological routes to cell elimination. *eLife*, 12, 2023.

[13007] Mahdi Moradi Marjaneh, Edwin P Kirk, Ralph Patrick, Dimuthu Alankarage, David T Humphreys, Gonzalo Del Monte-Nieto, Paola Cornejo-Paramo, Vaibhao Janbandhu, Tram B Doan, Sally L Dunwoodie, Emily S Wong, Chris Moran, Ian CA Martin, Peter C Thomson, and Richard P Harvey. Quantitative trait and transcriptome analysis of genetic complexity underpinning cardiac interatrial septation in mice using an advanced intercross line. *eLife*, 12, 2023.

[13008] Edoardo Moretto, Federico Miozzo, Anna Longatti, Caroline Bonnet, Francoise Coussen, Fanny Jaudon, Lorenzo A Cingolani, and Maria Passafaro. The tetraspanin TSPAN5 regulates AMPAR exocytosis by interacting with the AP4 complex. *eLife*, 12, 2023.

[13009] Kimberly J Morgan, Karen Doggett, Fansuo Geng, Stephen Mieruszynski, Lachlan Whitehead, Kelly A Smith, Benjamin M Hogan, Cas Simons, Gregory J Baillie, Ramyar Molania, Anthony T Papenfuss, Thomas E Hall, Elke A Ober, Didier YR Stainier, Zhiyuan Gong, and Joan K Heath. ahctf1 and kras mutations combine to amplify oncogenic stress and restrict liver overgrowth in a zebrafish model of hepatocellular carcinoma. *eLife*, 12, 2023.

[13010] Eva Morgun, Jennifer Zhu, Sultan Almunif, Sharan Bobbala, Melissa S Aguilar, Junzhong Wang, Kathleen Conner, Yongyong Cui, Liang Cao, Chetan Seshadri, Evan A Scott, and Chyung-Ru Wang. Vaccination with mycobacterial lipid loaded nanoparticle leads to lipid antigen persistence and memory differentiation of antigen-specific T cells. *eLife*, 12, 2023.

[13011] Eitan Moses, Roman Franek, and Itamar Harel. A scalable and tunable platform for functional interrogation of peptide hormones in fish. *eLife*, 12, 2023.

[13012] Ali Motahharynia, Ahmad Pourmohammadi, Armin Adibi, Vahid Shaygannejad, Fereshteh Ashtari, Iman Adibi, and Mehdi Sanayei. A mechanistic insight into sources of error of visual working memory in multiple sclerosis. *eLife*, 12, 2023.

[13013] Alief Moulana, Thomas Dupic, Angela M Phillips, Jeffrey Chang, Anne A Roffler, Allison J Greaney, Tyler N Starr, Jesse D Bloom, and Michael M Desai. The landscape of antibody binding affinity in SARS-CoV-2 Omicron BA.1 evolution. *eLife*, 12, 2023.

[13014] Laura Moutard, Caroline Goudin, Catherine Jaeger, Céline Duparc, Estelle Louiset, Tony Pereira, FranÇois Fraissinet, Marion Delessard, Justine Saulnier, Aurélie Rives-Feraille, Christelle Delalande, Hervé Lefebvre, Nathalie Rives, Ludovic Dumont, and Christine Rondanino. Steroidogenesis and androgen/estrogen signaling pathways are altered in in vitro matured testicular tissues of prepubertal mice. *eLife*, 12, 2023.

[13015] Guillermo Moya-Alvarado, Reynaldo Tiburcio-Felix, María Raquel Ibáñez, Alejandro A Aguirre-Soto, Miguel V Guerra, Chengbiao Wu, William C Mobley, Eran Perlson, and Francisca C Bronfman. BDNF/TrkB signaling endosomes in axons coordinate CREB/mTOR activation and protein synthesis in the cell body to induce dendritic growth in cortical neurons. *eLife*, 12, 2023.

[13016] Brian D Mueller, Sean A Merrill, Shigeki Watanabe, Ping Liu, Longgang Niu, Anish Singh, Pablo Maldonado-Catala, Alex Cherry, Matthew S Rich, Malan Silva, Andres Villu Maricq, Zhao-Wen Wang, and Erik M Jorgensen. CaV1 and CaV2 calcium channels mediate the release of distinct pools of synaptic vesicles. *eLife*, 12, 2023.

[13017] Taulant Muka, Joshua JX Li, Sahar J Farahani, and John PA Ioannidis. An umbrella review of systematic reviews on the impact of the COVID-19 pandemic on cancer prevention and management, and patient needs. *eLife*, 12, 2023.

[13018] Karl S Muller, Jonathan Matthis, Kathryn Bonnen, Lawrence K Cormack, Alex C Huk, and Mary Hayhoe. Retinal motion statistics during natural locomotion. *eLife*, 12, 2023.

[13019] Ayumi Mure, Yuki Sugiura, Rae Maeda, Kohei Honda, Nozomu Sakurai, Yuuki Takahashi, Masayoshi Watada, Toshihiko Katoh, Aina Gotoh, Yasuhiro Gotoh, Itsuki Taniguchi, Keiji Nakamura, Tetsuya Hayashi, Takane Katayama, Tadashi Uemura, and Yukako Hattori. Identification of key yeast species and microbe–microbe interactions impacting larval growth of Drosophila in the wild. *eLife*, 12, 2023.

[13020] Alan E Murphy, Nurun Fancy, and Nathan Skene. Avoiding false discoveries in single-cell RNA-seq by revisiting the first Alzheimer's disease dataset. *eLife*, 12, 2023.

[13021] David A Murphy, Eyal Elyashiv, Guy Amster, and Guy Sella. Broad-scale variation in human genetic diversity levels is predicted by purifying selection on coding and non-coding elements. *eLife*, 12, 2023.

[13022] Sae-Il Murtada, Nicole Mikush, Mo Wang, Pengwei Ren, Yuki Kawamura, Abhay B Ramachandra, David S Li, Demetrios T Braddock, George Tellides, Leslie B Gordon, and Jay D Humphrey. Lonafarnib improves cardiovascular function and survival in a mouse model of Hutchinson-Gilford progeria syndrome. *eLife*, 12, 2023.

[13023] Muniesh Muthaiyan Shanmugam, Jyotiska Chaudhuri, Durai Sellegounder, Amit Kumar Sahu, Sanjib Guha, Manish Chamoli, Brian Hodge, Neelanjan Bose, Charis Amber, Dominique O Farrera, Gordon Lithgow, Richmond Sarpong, James J Galligan, and Pankaj Kapahi. Methylglyoxal-derived hydroimidazolone, MG-H1, increases food intake by altering tyramine signaling via the GATA transcription factor ELT-3 in Caenorhabditis elegans. *eLife*, 12, 2023.

[13024] Musleh M Muthana, Xuexiang Du, Mingyue Liu, Xu Wang, Wei Wu, Chunxia Ai, Lishan Su, Pan Zheng, and Yang Liu. CTLA-4 antibody-drug conjugate reveals autologous destruction of B-lymphocytes associated with regulatory T cell impairment. *eLife*, 12, 2023.

[13025] Daniel Muñoz-Reyes, Levi J McClelland, Sandra Arroyo-Urea, Sonia Sánchez-Yepes, Juan Sabín, Sara Pérez-Suárez, Margarita Menendez, Alicia Mansilla, Javier García-Nafría, Stephen Sprang, and Maria Jose Sanchez-Barrena. The neuronal calcium sensor NCS-1 regulates the phosphorylation state and activity of the G$\alpha$ chaperone and GEF Ric-8A. *eLife*, 12, 2023.

[13026] Sana Naderi, Peter E Chen, Carmen Lia Murall, Raphael Poujol, Susanne Kraemer, Bradley S Pickering, Selena M Sagan, and B Jesse Shapiro. Zooanthroponotic transmission of SARS-CoV-2 and host-specific viral mutations revealed by genome-wide phylogenetic analysis. *eLife*, 12, 2023.

[13027] Fares Z Najar, Evan Linde, Chelsea L Murphy, Veniamin A Borin, Huan Wang, Shozeb Haider, and Pratul K Agarwal. Future COVID19 surges prediction based on SARS-CoV-2 mutations surveillance. *eLife*, 12, 2023.

[13028] Yaeko Nakajima-Takagi, Motohiko Oshima, Junichiro Takano, Shuhei Koide, Naoki Itokawa, Shun Uemura, Masayuki Yamashita, Shohei Andoh, Kazumasa Aoyama, Yusuke Isshiki, Daisuke Shinoda, Atsunori Saraya, Fumio Arai, Kiyoshi Yamaguchi, Yoichi Furukawa, Haruhiko Koseki, Tomokatsu Ikawa, and Atsushi Iwama. Polycomb repressive complex 1.1 coordinates homeostatic and emergency myelopoiesis. *eLife*, 12, 2023.

[13029] Yuka Nakamura, Miyuki Kurabe, Mami Matsumoto, Tokiharu Sato, Satoshi Miyashita, Kana Hoshina, Yoshinori Kamiya, Kazuki Tainaka, Hitoshi Matsuzawa, Nobuhiko Ohno, and Masaki Ueno. Cerebrospinal fluid-contacting neuron tracing reveals structural and functional connectivity for locomotion in the mouse spinal cord. *eLife*, 12, 2023.

[13030] Pedro Nascimento de Lima, Rosita van den Puttelaar, Anne I Hahn, Matthias Harlass, Nicholson Collier, Jonathan Ozik, Ann G Zauber, Iris Lansdorp-Vogelaar, and Carolyn M Rutter. Projected long-term effects of colorectal cancer screening disruptions following the COVID-19 pandemic. *eLife*, 12, 2023.

[13031] Nurul Jannah Mohamed Nasir, Hans Heemskerk, Julia Jenkins, Nur Hidayah Hamadee, Ralph Bunte, and Lisa Tucker-Kellogg. Myoglobin-derived iron causes wound enlargement and impaired regeneration in pressure injuries of muscle. *eLife*, 12, 2023.

[13032] Kim A Nasmyth, Byung-Gil Lee, Maurici Brunet Roig, and Jan Löwe. What AlphaFold tells us about cohesin's retention on and release from chromosomes. *eLife*, 12, 2023.

[13033] Irene Navarro Lobato, Adrian Aleman-Zapata, Anumita Samanta, Milan Bogers, Shekhar Narayanan, Abdelrahman Rayan, Alejandra Alonso, Jacqueline van der Meij, Mehdi Khamassi, Zafar U Khan, and Lisa Genzel. Increased cortical plasticity leads to memory interference and enhanced hippocampal-cortical interactions. *eLife*, 12, 2023.

[13034] Janna C Nawroth, Christoph Giez, Alexander Klimovich, Eva Kanso, and Thomas CG Bosch. Spontaneous body wall contractions stabilize the fluid microenvironment that shapes host–microbe associations. *eLife*, 12, 2023.

[13035] Stephan Nebe, Mario Reutter, Daniel H Baker, Jens Bölte, Gregor Domes, Matthias Gamer, Anne Gärtner, Carsten Gießing, Caroline Gurr, Kirsten Hilger, Philippe Jawinski, Louisa Kulke, Alexander Lischke, Sebastian Markett, Maria Meier, Christian J Merz, Tzvetan Popov, Lara MC Puhlmann, Daniel S Quintana, Tim Schäfer, Anna-Lena Schubert, Matthias FJ Sperl, Antonia Vehlen, Tina B Lonsdorf, and Gordon B Feld. Enhancing precision in human neuroscience. *eLife*, 12, 2023.

[13036] Felix Christopher Nebeling, Stefanie Poll, Lena Christine Justus, Julia Steffen, Kevin Keppler, Manuel Mittag, and Martin Fuhrmann. Microglial motility is modulated by neuronal activity and correlates with dendritic spine plasticity in the hippocampus of awake mice. *eLife*, 12, 2023.

[13037] Julie C Necarsulmer, Jeremy M Simon, Baggio A Evangelista, Youjun Chen, Xu Tian, Sara Nafees, Ariana B Marquez, Huijun Jiang, Ping Wang, Deepa Ajit, Viktoriya D Nikolova, Kathryn M Harper, J Ashley Ezzell, Feng-Chang Lin, Adriana S Beltran, Sheryl S Moy, and Todd J Cohen. RNA-binding deficient TDP-43 drives cognitive decline in a mouse model of TDP-43 proteinopathy. *eLife*, 12, 2023.

[13038] Abigail C Neininger-Castro, James B Hayes, Zachary C Sanchez, Nilay Taneja, Aidan M Fenix, Satish Moparthi, Stéphane Vassilopoulos, and Dylan Tyler Burnette. Independent regulation of Z-lines and M-lines during sarcomere assembly in cardiac myocytes revealed by the automatic image analysis software sarcApp. *eLife*, 12, 2023.

[13039] Alexey Dmitrievich Neverov, Gennady Fedonin, Anfisa Popova, Daria Bykova, and Georgii Bazykin. Coordinated evolution at amino acid sites of SARS-CoV-2 spike. *eLife*, 12, 2023.

[13040] Kahlan E Newman, Sarah N Tindall, Sophie L Mader, Syma Khalid, Gavin H Thomas, and Marjan W Van Der Woude. A novel fold for acyltransferase-3 (AT3) proteins provides a framework for transmembrane acyl-group transfer. *eLife*, 12, 2023.

[13041] Kim Bich Nguyen, Malte Roerden, Christopher J Copeland, Coralie M Backlund, Nory G Klop-Packel, Tanaka Remba, Byungji Kim, Nishant K Singh, Michael E Birnbaum, Darrell J Irvine, and Stefani Spranger. Decoupled neoantigen cross-presentation by dendritic cells limits anti-tumor immunity against tumors with heterogeneous neoantigen expression. *eLife*, 12, 2023.

[13042] Van Thien Chi Nguyen, Trong Hieu Nguyen, Nhu Nhat Tan Doan, Thi Mong Quynh Pham, Giang Thi Huong Nguyen, Thanh Dat Nguyen, Thuy Thi Thu Tran, Duy Long Vo, Thanh Hai Phan, Thanh Xuan Jasmine, Van Chu Nguyen, Huu Thinh Nguyen, Trieu Vu Nguyen, Thi Hue Hanh Nguyen, Le Anh Khoa Huynh, Trung Hieu Tran, Quang Thong Dang, Thuy Nguyen Doan, Anh Minh Tran, Viet Hai Nguyen, Vu Tuan Anh Nguyen, Le Minh Quoc Ho, Quang Dat Tran, Thi Thu Thuy Pham, Tan Dat Ho, Bao Toan Nguyen, Thanh Nhan Vo Nguyen, Thanh Dang Nguyen, Dung Thai Bieu Phu, Boi Hoan Huu Phan, Thi Loan Vo, Thi Huong Thoang Nai, Thuy Trang Tran, My Hoang Truong, Ngan Chau Tran, Trung Kien Le, Thanh Huong Thi Tran, Minh Long Duong, Hoai Phuong Thi Bach, Van Vu Kim, The Anh Pham, Duc Huy Tran, Trinh Ngoc An Le, Truong Vinh Ngoc Pham, Minh Triet Le, Dac Ho Vo, Thi Minh Thu Tran, Minh Nguyen Nguyen, Thi

Tuong Vi Van, Anh Nhu Nguyen, Thi Trang Tran, Vu Uyen Tran, Minh Phong Le, Thi Thanh Do, Thi Van Phan, Hong-Dang Luu Nguyen, Duy Sinh Nguyen, Van Thinh Cao, Thanh-Thuy Thi Do, Dinh Kiet Truong, Hung Sang Tang, Hoa Giang, Hoai-Nghia Nguyen, Minh-Duy Phan, and Le Son Tran. Multimodal analysis of methylomics and fragmentomics in plasma cell-free DNA for multi-cancer early detection and localization. *eLife*, 12, 2023.

[13043] Carolyn Nickson, Megan A Smith, Eleonora Feletto, Louiza S Velentzis, Kate Broun, Sabine Deij, Paul Grogan, Michaela Hall, Emily He, D James St John, Jie-Bin Lew, Pietro Procopio, Kate T Simms, Joachim Worthington, G Bruce Mann, and Karen Canfell. A modelled evaluation of the impact of COVID-19 on breast, bowel, and cervical cancer screening programmes in Australia. *eLife*, 12, 2023.

[13044] Terren K Niethamer, Lillian I Levin, Michael P Morley, Apoorva Babu, Su Zhou, and Edward E Morrisey. Atf3 defines a population of pulmonary endothelial cells essential for lung regeneration. *eLife*, 12, 2023.

[13045] Nikos Nikolopoulos, Renata C Matos, Stephanie Ravaud, Pascal Courtin, Houssam Akherraz, Simon Palussiere, Virginie Gueguen-Chaignon, Marie Salomon-Mallet, Alain Guillot, Yann Guerardel, Marie-Pierre Chapot-Chartier, Christophe Grangeasse, and FranÇois Leulier. Structure–function analysis of Lactiplantibacillus plantarum DltE reveals D-alanylated lipoteichoic acids as direct cues supporting Drosophila juvenile growth. *eLife*, 12, 2023.

[13046] Gai Nishikawa, Yu Sugo, Keisuke Saito, and Hiroshi Ishikita. Absence of electron-transfer-associated changes in the time-dependent X-ray free-electron laser structures of the photosynthetic reaction center. *eLife*, 12, 2023.

[13047] Nan Niu, Yinan Zhang, Yang Bai, Xin Wang, Shunchao Yan, Dong Song, Hong Xu, Tong Liu, Bin Hua, Yingchao Zhang, Jinchi Liu, Xinbo Qiao, Jiaxiang Liu, Xinyu Zheng, Hongyi Cao, and Caigang Liu. Efficacy and safety of endocrine therapy after mastectomy in patients with hormone receptor positive breast ductal carcinoma in situ: Retrospective cohort study. *eLife*, 12, 2023.

[13048] Laura M Nocka, Timothy J Eisen, Anthony T Iavarone, Jay T Groves, and John Kuriyan. Stimulation of the catalytic activity of the tyrosine kinase Btk by the adaptor protein Grb2. *eLife*, 12, 2023.

[13049] Gregory M Noetscher, Peter J Serano, Marc Horner, Alexander Prokop, Jonathan Hanson, Kyoko Fujimoto, James Brown, Ara Nazarian, Jerome Ackerman, and Sergey N Makaroff. An in silico testbed for fast and accurate MR labeling of orthopedic implants. *eLife*, 12, 2023.

[13050] Hirofumi Noguchi, Jessica Chelsea Arela, Thomas Ngo, Laura Cocas, and Samuel Pleasure. Shh from mossy cells contributes to preventing NSC pool depletion after seizure-induced neurogenesis and in aging. *eLife*, 12, 2023.

[13051] Mette Hartmann Nonboe, George Napolitano, Jeppe Bennekou Schroll, Ilse Vejborg, Marianne Waldstrøm, and Elsebeth Lynge. Impact of COVID-19 pandemic on breast and cervical cancer screening in Denmark: A register-based study. *eLife*, 12, 2023.

[13052] Stijn A Nuiten, Jan Willem de Gee, Jasper B Zantvoord, Johannes J Fahrenfort, and Simon van Gaal. Catecholaminergic neuromodulation and selective attention jointly shape perceptual decision-making. *eLife*, 12, 2023.

[13053] Kate Nussenbaum, Rebecca E Martin, Sean Maulhardt, Yi (Jen) Yang, Greer Bizzell-Hatcher, Naiti S Bhatt, Maximilian Koenig, Gail M Rosenbaum, John P O'Doherty, Jeffrey Cockburn, and Catherine A Hartley. Novelty and uncertainty differentially drive exploration across development. *eLife*, 12, 2023.

[13054] Eider Nuñez, Frederick Jones, Arantza Muguruza-Montero, Janire Urrutia, Alejandra Aguado, Covadonga Malo, Ganeko Bernardo-Seisdedos, Carmen Domene, Oscar Millet, Nikita Gamper, and

Alvaro Villarroel. Redox regulation of KV7 channels through EF3 hand of calmodulin. *eLife*, 12, 2023.

[13055] Izumi Oikawa, Shu Kondo, Kao Hashimoto, Akiho Yoshida, Megumi Hamajima, Hiromu Tanimoto, Katsuo Furukubo-Tokunaga, and Ken Honjo. A descending inhibitory mechanism of nociception mediated by an evolutionarily conserved neuropeptide system in Drosophila. *eLife*, 12, 2023.

[13056] Hidehiko Okuma, Jeffrey M Hord, Ishita Chandel, David Venzke, Mary E Anderson, Ameya S Walimbe, Soumya Joseph, Zeita Gastel, Yuji Hara, Fumiaki Saito, Kiichiro Matsumura, and Kevin P Campbell. N-terminal domain on dystroglycan enables LARGE1 to extend matriglycan on $\alpha$-dystroglycan and prevents muscular dystrophy. *eLife*, 12, 2023.

[13057] Laure Olazcuaga, Raymonde Baltenweck, Nicolas Leménager, Alessandra Maia-Grondard, Patricia Claudel, Philippe Hugueney, and Julien Foucaud. Metabolic consequences of various fruit-based diets in a generalist insect species. *eLife*, 12, 2023.

[13058] Tina Bech Olesen, Henry Jensen, Henrik Møller, Jens Winther Jensen, Berit Andersen, and Morten Rasmussen. Nationwide participation in FIT-based colorectal cancer screening in Denmark during the COVID-19 pandemic: An observational study. *eLife*, 12, 2023.

[13059] Tina Bech Olesen, Henry Jensen, Henrik Møller, Jens Winther Jensen, Berit Andersen, Ilse Vejborg, and Sisse H Njor. Nationwide mammography screening participation in Denmark during the COVID-19 pandemic: An observational study. *eLife*, 12, 2023.

[13060] Tina Bech Olesen, Henry Jensen, Henrik Møller, Jens Winther Jensen, Marianne Waldstrøm, and Berit Andersen. Participation in the nationwide cervical cancer screening programme in Denmark during the COVID-19 pandemic: An observational study. *eLife*, 12, 2023.

[13061] Eli Olson, Theadora Ceccarelli, and Malini Raghavan. Endo-lysosomal assembly variations among human leukocyte antigen class I (HLA class I) allotypes. *eLife*, 12, 2023.

[13062] Pei-Wen Ong, Ya-Ping Lin, Hung-Wei Chen, Cheng-Yu Lo, Marina Burlyaeva, Thomas Noble, Ramakrishnan Madhavan Nair, Roland Schafleitner, Margarita Vishnyakova, Eric Bishop-von Wettberg, Maria Samsonova, Sergey Nuzhdin, Chau-Ti Ting, and Cheng-Ruei Lee. Environment as a limiting factor of the historical global spread of mungbean. *eLife*, 12, 2023.

[13063] James DB O'Sullivan, Thomas S Blacker, Claire Scott, Weise Chang, Mohi Ahmed, Val Yianni, and Zoe F Mann. Gradients of glucose metabolism regulate morphogen signalling required for specifying tonotopic organisation in the chicken cochlea. *eLife*, 12, 2023.

[13064] David J Ottenheimer, Madelyn M Hjort, Anna J Bowen, Nicholas A Steinmetz, and Garret D Stuber. A stable, distributed code for cue value in mouse cortex during reward learning. *eLife*, 12, 2023.

[13065] Ian R Outhwaite, Sukrit Singh, Benedict-Tilman Berger, Stefan Knapp, John D Chodera, and Markus A Seeliger. Death by a thousand cuts through kinase inhibitor combinations that maximize selectivity and enable rational multitargeting. *eLife*, 12, 2023.

[13066] John F Ouyang, Kunal Mishra, Yi Xie, Harry Park, Kevin Y Huang, Enrico Petretto, and Jacques Behmoaras. Systems level identification of a matrisome-associated macrophage polarisation state in multi-organ fibrosis. *eLife*, 12, 2023.

[13067] Dai Owaki, Volker Dürr, and Josef Schmitz. A hierarchical model for external electrical control of an insect, accounting for inter-individual variation of muscle force properties. *eLife*, 12, 2023.

[13068] Danielle J Owen, Elisa Aguilar-Martinez, Zongling Ji, Yaoyong Li, and Andrew D Sharrocks. ZMYM2 controls human transposable element transcription through distinct co-regulatory complexes. *eLife*, 12, 2023.

[13069] Maruthi Kumar Pabba, Christian Ritter, Vadim O Chagin, Janis Meyer, Kerem Celikay, Jeffrey H Stear, Dinah Loerke, Ksenia Kolobynina, Paulina Prorok, Alice Kristin Schmid, Heinrich Leonhardt, Karl Rohr, and M Cristina Cardoso. Replisome loading reduces chromatin motion independent of DNA synthesis. *eLife*, 12, 2023.

[13070] Abhilash Padavannil, Anjaneyulu Murari, Shauna-Kay Rhooms, Edward Owusu-Ansah, and James A Letts. Resting mitochondrial complex I from Drosophila melanogaster adopts a helix-locked state. *eLife*, 12, 2023.

[13071] Elliott L Paine, Jack J Skalicky, Frank G Whitby, Douglas R Mackay, Katharine S Ullman, Christopher P Hill, and Wesley I Sundquist. The Calpain-7 protease functions together with the ESCRT-III protein IST1 within the midbody to regulate the timing and completion of abscission. *eLife*, 12, 2023.

[13072] Sinisa Pajevic, Dietmar Plenz, Peter J Basser, and R Douglas Fields. Oligodendrocyte-mediated myelin plasticity and its role in neural synchronization. *eLife*, 12, 2023.

[13073] Camryn Pajon, Marla C Fortoul, Gabriela Diaz-Tang, Estefania Marin Meneses, Ariane R Kalifa, Elinor Sevy, Taniya Mariah, Brandon Toscan, Maili Marcelin, Daniella M Hernandez, Melissa M Marzouk, Allison J Lopatkin, Omar Tonsi Eldakar, and Robert P Smith. Interactions between metabolism and growth can determine the co-existence of Staphylococcus aureus and Pseudomonas aeruginosa. *eLife*, 12, 2023.

[13074] Lin Pan, Amelia Trimarco, Alice J Zhang, Ko Fujimori, Yoshihiro Urade, Lu O Sun, Carla Taveggia, and Ye Zhang. Oligodendrocyte-lineage cell exocytosis and L-type prostaglandin D synthase promote oligodendrocyte development and myelination. *eLife*, 12, 2023.

[13075] Wenfei Pan, Vladimir A Meshcheryakov, Tianjie Li, Yi Wang, Gourisankar Ghosh, and Vivien Ya-Fan Wang. Structures of NF-$\kappa$B p52 homodimer-DNA complexes rationalize binding mechanisms and transcription activation. *eLife*, 12, 2023.

[13076] Xueyang Pan, Albert N Alvarez, Mengqi Ma, Shenzhao Lu, Michael W Crawford, Lauren C Briere, Oguz Kanca, Shinya Yamamoto, David A Sweetser, Jenny L Wilson, Ruth J Napier, Jonathan N Pruneda, and Hugo J Bellen. Allelic strengths of encephalopathy-associated UBA5 variants correlate between in vivo and in vitro assays. *eLife*, 12, 2023.

[13077] Liang-Yu Pang, Steven DeLuca, Haolong Zhu, John M Urban, and Allan C Spradling. Chromatin and gene expression changes during female Drosophila germline stem cell development illuminate the biology of highly potent stem cells. *eLife*, 12, 2023.

[13078] Janani Parameswaran, Nancy Zhang, Elke Braems, Kedamawit Tilahun, Devesh C Pant, Keena Yin, Seneshaw Asress, Kara Heeren, Anwesha Banerjee, Emma Davis, Samantha L Schwartz, Graeme L Conn, Gary J Bassell, Ludo Van Den Bosch, and Jie Jiang. Antisense, but not sense, repeat expanded RNAs activate PKR/eIF2$\alpha$-dependent ISR in C9ORF72 FTD/ALS. *eLife*, 12, 2023.

[13079] Joshua K Park, Shantanu Bafna, Iain S Forrest, áine Duffy, Carla Marquez-Luna, Ben O Petrazzini, Ha My Vy, Daniel M Jordan, Marie Verbanck, Jagat Narula, Robert S Rosenson, Ghislain Rocheleau, and Ron Do. Phenome-wide Mendelian randomization study of plasma triglyceride levels and 2600 disease traits. *eLife*, 12, 2023.

[13080] Kieop Park, Ranjay Jayadev, Sara G Payne, Isabel W Kenny-Ganzert, Qiuyi Chi, Daniel S Costa, William Ramos-Lewis, Siddharthan B Thendral, and David R Sherwood. Reciprocal discoidin domain receptor signaling strengthens integrin adhesion to connect adjacent tissues. *eLife*, 12, 2023.

[13081] Yein Christina Park, Bharat Reddy, Navid Bavi, Eduardo Perozo, and José D Faraldo-Gómez. State-specific morphological deformations of the lipid bilayer explain mechanosensitive gating of MscS ion channels. *eLife*, 12, 2023.

[13082] Matthew T Parker, Sebastian M Fica, Geoffrey J Barton, and Gordon G Simpson. Inter-species association mapping links splice site evolution to METTL16 and SNRNP27K. *eLife*, 12, 2023.

[13083] Susan Parker, Ashish A Deshmukh, Baojiang Chen, David R Lairson, Maria Daheri, Sally W Vernon, and Jane R Montealegre. Perceived barriers to cervical cancer screening and motivators for at-home human papillomavirus self-sampling during the COVID-19 pandemic: Results from a telephone survey. *eLife*, 12, 2023.

[13084] Rachel H Parkinson, Jennifer Scott, Anna L Dorling, Hannah Jones, Martha Haslam, Alex E McDermott-Roberts, and Geraldine A Wright. Mouthparts of the bumblebee (Bombus terrestris) exhibit poor acuity for the detection of pesticides in nectar. *eLife*, 12, 2023.

[13085] Benjamin Pasquereau and Robert S Turner. Neural dynamics underlying self-control in the primate subthalamic nucleus. *eLife*, 12, 2023.

[13086] Maribel Patiño, Willian N Lagos, Neelakshi S Patne, Paula A Miyazaki, Sai Krishna Bhamidipati, Forrest Collman, and Edward M Callaway. Postsynaptic cell type and synaptic distance do not determine efficiency of monosynaptic rabies virus spread measured at synaptic resolution. *eLife*, 12, 2023.

[13087] Daniel T Pederick, Nicole A Perry-Hauser, Huyan Meng, Zhigang He, Jonathan A Javitch, and Liqun Luo. Context-dependent requirement of G protein coupling for Latrophilin-2 in target selection of hippocampal axons. *eLife*, 12, 2023.

[13088] Benjamin D Pedigo, Mike Powell, Eric W Bridgeford, Michael Winding, Carey E Priebe, and Joshua T Vogelstein. Generative network modeling reveals quantitative definitions of bilateral symmetry exhibited by a whole insect brain connectome. *eLife*, 12, 2023.

[13089] James M Pemberton, Dang Nguyen, Elizabeth J Osterlund, Wiebke Schormann, Justin P Pogmore, Nehad Hirmiz, Brian Leber, and David W Andrews. The carboxyl-terminal sequence of PUMA binds to both anti-apoptotic proteins and membranes. *eLife*, 12, 2023.

[13090] Mihir Pendse, Haley De Selle, Nguyen Vo, Gabriella Quinn, Chaitanya Dende, Yun Li, Cristine N Salinas, Tarun Srinivasan, Daniel C Propheter, Alexander A Crofts, Eugene Koo, Brian Hassell, Kelly A Ruhn, Prithvi Raj, Yuuki Obata, and Lora V Hooper. Macrophages regulate gastrointestinal motility through complement component 1q. *eLife*, 12, 2023.

[13091] Jennifer T Pentz, Kathryn MacGillivray, James G DuBose, Peter L Conlin, Emma Reinhardt, Eric Libby, and William C Ratcliff. Evolutionary consequences of nascent multicellular life cycles. *eLife*, 12, 2023.

[13092] Valerie Ann Perez, David W Sanders, Ayde Mendoza-Oliva, Barbara Elena Stopschinski, Vishruth Mullapudi, Charles L White, Lukasz A Joachimiak, and Marc I Diamond. DnaJC7 specifically regulates tau seeding. *eLife*, 12, 2023.

[13093] Catherine Perrodin, Colombine Verzat, and Daniel Bendor. Courtship behaviour reveals temporal regularity is a critical social cue in mouse communication. *eLife*, 12, 2023.

[13094] Konstantinos-Evangelos Petousakis, Jiyoung Park, Athanasia Papoutsi, Stelios Smirnakis, and Panayiota Poirazi. Modeling apical and basal tree contribution to orientation selectivity in a mouse primary visual cortex layer 2/3 pyramidal cell. *eLife*, 12, 2023.

[13095] Maurice A Petroccione, Lianna Y D'Brant, Nurat Affinnih, Patrick H Wehrle, Gabrielle C Todd, Shergil Zahid, Haley E Chesbro, Ian L Tschang, and Annalisa Scimemi. Neuronal glutamate transporters control reciprocal inhibition and gain modulation in D1 medium spiny neurons. *eLife*, 12, 2023.

[13096] Elijah A Petter, Isabella P Fallon, Ryan N Hughes, Glenn DR Watson, Warren H Meck, Francesco Paolo Ulloa Severino, and Henry H Yin. Elucidating a locus coeruleus-dentate gyrus dopamine pathway for operant reinforcement. *eLife*, 12, 2023.

[13097] Thomas-O Peulen, Carola S Hengstenberg, Ralf Biehl, Mykola Dimura, Charlotte Lorenz, Alessandro Valeri, Julian Folz, Christian A Hanke, Semra Ince, Tobias Vöpel, Bela Farago, Holger Gohlke, Johann P Klare, Andreas M Stadler, Claus AM Seidel, and Christian Herrmann. Integrative dynamic structural biology unveils conformers essential for the oligomerization of a large GTPase. *eLife*, 12, 2023.

[13098] Wesley A Phelps, Matthew D Hurton, Taylor N Ayers, Anne E Carlson, Joel C Rosenbaum, and Miler T Lee. Hybridization led to a rewired pluripotency network in the allotetraploid Xenopus laevis. *eLife*, 12, 2023.

[13099] Angela M Phillips, Daniel P Maurer, Caelan Brooks, Thomas Dupic, Aaron G Schmidt, and Michael M Desai. Hierarchical sequence-affinity landscapes shape the evolution of breadth in an anti-influenza receptor binding site antibody. *eLife*, 12, 2023.

[13100] Robert A Phillips, Ethan Wan, Jennifer J Tuscher, David Reid, Olivia R Drake, Lara Ianov, and Jeremy J Day. Temporally specific gene expression and chromatin remodeling programs regulate a conserved Pdyn enhancer. *eLife*, 12, 2023.

[13101] Merrick D Pierson Smela, Christian C Kramme, Patrick RJ Fortuna, Jessica L Adams, Rui Su, Edward Dong, Mutsumi Kobayashi, Garyk Brixi, Venkata Srikar Kavirayuni, Emma Tysinger, Richie E Kohman, Toshi Shioda, Pranam Chatterjee, and George M Church. Directed differentiation of human iPSCs to functional ovarian granulosa-like cells via transcription factor overexpression. *eLife*, 12, 2023.

[13102] Julian MJ Pietsch, Alán F Muñoz, Diane-Yayra A Adjavon, Iseabail Farquhar, Ivan BN Clark, and Peter S Swain. Determining growth rates from bright-field images of budding cells through identifying overlaps. *eLife*, 12, 2023.

[13103] Delphine Pinatel, Edouard Pearlstein, Giulia Bonetto, Laurence Goutebroze, Domna Karagogeos, Valérie Crepel, and Catherine Faivre-Sarrailh. A class-specific effect of dysmyelination on the excitability of hippocampal interneurons. *eLife*, 12, 2023.

[13104] Ana Sofia Pinheiro, Vasilios Tsarouhas, Kirsten André Senti, Badrul Arefin, and Christos Samakovlis. Scavenger receptor endocytosis controls apical membrane morphogenesis in the Drosophila airways. *eLife*, 12, 2023.

[13105] Romain Pintore, Raphaël Cornette, Alexandra Houssaye, and Ronan Allain. Femora from an exceptionally large population of coeval ornithomimosaurs yield evidence of sexual dimorphism in extinct theropod dinosaurs. *eLife*, 12, 2023.

[13106] Romina Piscitello-Gómez, Franz S Gruber, Abhijeet Krishna, Charlie Duclut, Carl D Modes, Marko Popović, Frank Jülicher, Natalie A Dye, and Suzanne Eaton. Core PCP mutations affect short-time mechanical properties but not tissue morphogenesis in the Drosophila pupal wing. *eLife*, 12, 2023.

[13107] Alexandra Polyák, Leila Topal, Noémi Zombori-Tóth, Noémi Tóth, János Prorok, Zsófia Kohajda, Szilvia Déri, Vivien Demeter-Haludka, Péter Hegyi, Viktória Venglovecz, Gergely ágoston, Zoltán Husti, Péter Gazdag, Jozefina Szlovák, Tamás árpádffy Lovas, Muhammad Naveed, Annamária Sarusi, Norbert Jost, László Virág, Norbert Nagy, István Baczkó, Attila S Farkas, and András Varró. Cardiac electrophysiological remodeling associated with enhanced arrhythmia susceptibility in a canine model of elite exercise. *eLife*, 12, 2023.

[13108] Anton A Polyansky, Laura D Gallego, Roman G Efremov, Alwin Köhler, and Bojan Zagrovic. Protein compactness and interaction valency define the architecture of a biomolecular condensate across scales. *eLife*, 12, 2023.

[13109] Jörn K Pomper, Mohammad Shams, Shengjun Wen, Friedemann Bunjes, and Peter Thier. Non-shared coding of observed and executed actions prevails in macaque ventral premotor mirror neurons. *eLife*, 12, 2023.

[13110] Margarita Pons-Salort, Ben Lambert, Everlyn Kamau, Richard Pebody, Heli Harvala, Peter Simmonds, and Nicholas C Grassly. Changes in transmission of Enterovirus D68 (EV-D68) in England inferred from seroprevalence data. *eLife*, 12, 2023.

[13111] Galina Popova, Hanna Retallack, Chang N Kim, Albert Wang, David Shin, Joseph L DeRisi, and Tomasz Nowakowski. Rubella virus tropism and single-cell responses in human primary tissue and microglia-containing organoids. *eLife*, 12, 2023.

[13112] Rémi Porte, Rita Silva-Gomes, Charlotte Theroude, Raffaella Parente, Fatemeh Asgari, Marina Sironi, Fabio Pasqualini, Sonia Valentino, Rosanna Asselta, Camilla Recordati, Marta Noemi Monari, Andrea Doni, Antonio Inforzato, Carlos Rodriguez-Gallego, Ignacio Obando, Elena Colino, Barbara Bottazzi, and Alberto Mantovani. Regulation of inflammation and protection against invasive pneumococcal infection by the long pentraxin PTX3. *eLife*, 12, 2023.

[13113] Hayley Porter, Yang Li, Maria Victoria Neguembor, Manuel Beltran, Wazeer Varsally, Laura Martin, Manuel Tavares Cornejo, Dubravka Pezić, Amandeep Bhamra, Silvia Surinova, Richard G Jenner, Maria Pia Cosma, and Suzana Hadjur. Cohesin-independent STAG proteins interact with RNA and R-loops and promote complex loading. *eLife*, 12, 2023.

[13114] Sol Pose-Méndez, Paul Schramm, Barbara Winter, Jochen C Meier, Konstantinos Ampatzis, and Reinhard W Köster. Lifelong regeneration of cerebellar Purkinje cells after induced cell ablation in zebrafish. *eLife*, 12, 2023.

[13115] Tamara A Potapova, Jay R Unruh, Juliana Conkright-Fincham, Charles AS Banks, Laurence Florens, David Alan Schneider, and Jennifer L Gerton. Distinct states of nucleolar stress induced by anticancer drugs. *eLife*, 12, 2023.

[13116] Helle Rus Povlsen, Amalie Kai Bentzen, Mohammad Kadivar, Leon Eyrich Jessen, Sine Reker Hadrup, and Morten Nielsen. Improved T cell receptor antigen pairing through data-driven filtering of sequencing information from single cells. *eLife*, 12, 2023.

[13117] Léa Pradier and Stéphanie Bedhomme. Ecology, more than antibiotics consumption, is the major predictor for the global distribution of aminoglycoside-modifying enzymes. *eLife*, 12, 2023.

[13118] Amy Pribadi, Michael A Rieger, Kaila Rosales, Kirthi C Reddy, and Sreekanth H Chalasani. Dopamine signaling regulates predator-driven changes in Caenorhabditis elegans' egg laying behavior. *eLife*, 12, 2023.

[13119] Francisco Prieto-Ruiz, Elisa Gómez-Gil, Rebeca Martín-García, Armando Jesús Pérez-Díaz, Jero Vicente-Soler, Alejandro Franco, Teresa Soto, Pilar Pérez, Marisa Madrid, and José Cansado. Myosin II regulatory light chain phosphorylation and formin availability modulate cytokinesis upon changes in carbohydrate metabolism. *eLife*, 12, 2023.

[13120] Christian Pritz, Eyal Itskovits, Eduard Bokman, Rotem Ruach, Vladimir Gritsenko, Tal Nelken, Mai Menasherof, Aharon Azulay, and Alon Zaslaver. Principles for coding associative memories in a compact neural network. *eLife*, 12, 2023.

[13121] Mikhail Proskurin, Maxim Manakov, and Alla Karpova. ACC neural ensemble dynamics are structured by strategy prevalence. *eLife*, 12, 2023.

[13122] Andrea E Prota, Daniel Lucena-Agell, Yuntao Ma, Juan Estevez-Gallego, Shuo Li, Katja Bargsten, Fernando Josa-Prado, Karl-Heinz Altmann, Natacha Gaillard, Shinji Kamimura, Tobias Mühlethaler, Federico Gago, Maria A Oliva, Michel O Steinmetz, Wei-Shuo Fang, and J Fernando Díaz. Structural insight into the stabilization of microtubules by taxanes. *eLife*, 12, 2023.

[13123] Chinmay Purandare and Mayank Mehta. Mega-scale movie-fields in the mouse visuo-hippocampal network. *eLife*, 12, 2023.

[13124] Chao Qi, Silvia Acosta Gutierrez, Pia Lavriha, Alaa Othman, Diego Lopez-Pigozzi, Erva Bayraktar, Dina Schuster, Paola Picotti, Nicola Zamboni, Mario Bortolozzi, Francesco Luigi Gervasio, and Volodymyr M Korkhov. Structure of the connexin-43 gap junction channel in a putative closed state. *eLife*, 12, 2023.

[13125] Wesley W Qian, Jennifer N Wei, Benjamin Sanchez-Lengeling, Brian K Lee, Yunan Luo, Marnix Vlot, Koen Dechering, Jian Peng, Richard C Gerkin, and Alexander B Wiltschko. Metabolic activity organizes olfactory representations. *eLife*, 12, 2023.

[13126] Yulong Qiao, Qiang Sun, Xiaona Chen, Liangqiang He, Di Wang, Ruibao Su, Yuanchao Xue, Hao Sun, and Huating Wang. Nuclear m6A reader YTHDC1 promotes muscle stem cell activation/proliferation by regulating mRNA splicing and nuclear export. *eLife*, 12, 2023.

[13127] Yi Qin, Mehran Ahmadlou, Samuel Suhai, Paul Neering, Leander de Kraker, J Alexander Heimel, and Christiaan N Levelt. Thalamic regulation of ocular dominance plasticity in adult visual cortex. *eLife*, 12, 2023.

[13128] Maofeng Qing, Dan Yang, Qianhui Shang, Jiakuan Peng, Jiaxin Deng, Jiang Lu, Jing Li, HongXia Dan, Yu Zhou, Hao Xu, and Qianming Chen. CD8+ tissue-resident memory T cells induce oral lichen planus erosion via cytokine network. *eLife*, 12, 2023.

[13129] Weiwei Qiu, Chelsea R Hutch, Yi Wang, Jennifer Wloszek, Rachel A Rucker, Martin G Myers, and Darleen Sandoval. Multiple NTS neuron populations cumulatively suppress food intake. *eLife*, 12, 2023.

[13130] Valentin Quiniou, Pierre Barennes, Vanessa Mhanna, Paul Stys, Helene Vantomme, Zhicheng Zhou, Federica Martina, Nicolas Coatnoan, Michele Barbie, Hang-Phuong Pham, Béatrice Clémenceau, Henri Vie, Mikhail Shugay, Adrien Six, Barbara Brandao, Roberto Mallone, Encarnita Mariotti-Ferrandiz, and David Klatzmann. Human thymopoiesis produces polyspecific CD8+ $\alpha/\beta$ T cells responding to multiple viral antigens. *eLife*, 12, 2023.

[13131] Xarxa Quiroga, Nikhil Walani, Andrea Disanza, Albert Chavero, Alexandra Mittens, Francesc Tebar, Xavier Trepat, Robert G Parton, María Isabel Geli, Giorgio Scita, Marino Arroyo, Anabel-Lise Le Roux, and Pere Roca-Cusachs. A mechanosensing mechanism controls plasma membrane shape homeostasis at the nanoscale. *eLife*, 12, 2023.

[13132] Issam I Raad, Ray Hachem, Nigo Masayuki, Tarcila Datoguia, Hiba Dagher, Ying Jiang, Vivek Subbiah, Bilal Siddiqui, Arnaud Bayle, Robert Somer, Ana Fernández Cruz, Edward Gorak, Arvinder Bhinder, Nobuyoshi Mori, Nelson Hamerschlak, Samuel Shelanski, Tomislav Dragovich, Yee Elise Vong Kiat, Suha Fakhreddine, Abi Hanna Pierre, Roy F Chemaly, Victor Mulanovich, Javier Adachi, Jovan Borjan, Fareed Khawaja, Bruno Granwehr, Teny John, Eduardo Yepez Yepez, Harrys A Torres, Natraj Reddy Ammakkanavar, Marcel Yibirin, Cielito C Reyes-Gibby, Mala Pande, Noman Ali, Raniv Dawey Rojo, Shahnoor M Ali, Rita E Deeba, Patrick Chaftari, Takahiro Matsuo, Kazuhiro Ishikawa, Ryo Hasegawa, Ramón Aguado-Noya, Alvaro Garcia García, Cristina Traseira Puchol, Dong Gun Lee, Monica Slavin, Benjamin Teh, Cesar A Arias, Dimitrios P Kontoyiannis, Alexandre E Malek, and Anne-Marie Chaftari. International multicenter study comparing COVID-19 in patients with cancer to patients without cancer: Impact of risk factors and treatment modalities on survivorship. *eLife*, 12, 2023.

[13133] Enrico Radaelli, Charles-Antoine Assenmacher, Jillian Verrelle, Esha Banerjee, Florence Manero, Salim Khiati, Anais Girona, Guillermo Lopez-Lluch, Placido Navas, and Marco Spinazzi. Mitochondrial defects caused by PARL deficiency lead to arrested spermatogenesis and ferroptosis. *eLife*, 12, 2023.

[13134] Milos Radivojevic and Anna Rostedt Punga. Functional imaging of conduction dynamics in cortical and spinal axons. *eLife*, 12, 2023.

[13135] Bahar Rahsepar, Jacob F Norman, Jad Noueihed, Benjamin Lahner, Melanie H Quick, Kevin Ghaemi, Aashna Pandya, Fernando R Fernandez, Steve Ramirez, and John A White. Theta-phase-specific modulation of dentate gyrus memory neurons. *eLife*, 12, 2023.

[13136] Arjun Rajan, Lucas Anhezini, Noemi Rives-Quinto, Jay Y Chhabra, Megan C Neville, Elizabeth D Larson, Stephen F Goodwin, Melissa M Harrison, and Cheng-Yu Lee. Low-level repressive histone marks fine-tune gene transcription in neural stem cells. *eLife*, 12, 2023.

[13137] Karthik Ramanadane, Márton Liziczai, Dragana Markovic, Monique S Straub, Gian T Rosalen, Anto Udovcic, Raimund Dutzler, and Cristina Manatschal. Structural and functional properties of a plant NRAMP-related aluminum transporter. *eLife*, 12, 2023.

[13138] Vaidhiswaran Ramesh and J Krishnan. A unified approach to dissecting biphasic responses in cell signaling. *eLife*, 12, 2023.

[13139] Vijyendra Ramesh, Fang Liu, Melyssa S Minto, Urann Chan, and Anne E West. Bidirectional regulation of postmitotic H3K27me3 distributions underlie cerebellar granule neuron maturation dynamics. *eLife*, 12, 2023.

[13140] Ricardo Omar Ramirez Flores, Jan David Lanzer, Daniel Dimitrov, Britta Velten, and Julio Saez-Rodriguez. Multicellular factor analysis of single-cell data for a tissue-centric understanding of disease. *eLife*, 12, 2023.

[13141] Andy Rampersaud, Jeannette Connerney, and David J Waxman. Plasma growth hormone pulses induce male-biased pulsatile chromatin opening and epigenetic regulation in adult mouse liver. *eLife*, 12, 2023.

[13142] Dingyu Rao, Hua Lu, Xiongwei Wang, Zhonghong Lai, Jiali Zhang, and Zhixian Tang. Tissue-derived exosome proteomics identifies promising diagnostic biomarkers for esophageal cancer. *eLife*, 12, 2023.

[13143] Longfeng Rao, Nils Horn, Nadja Meister, Stefan Preiss, William R Taylor, Alessandro Santuz, and Pascal Schütz. Comparable in vivo joint kinematics between self-reported stable and unstable knees after TKA can be explained by muscular adaptation strategies: A retrospective observational study. *eLife*, 12, 2023.

[13144] Lucija Rapan, Sean Froudist-Walsh, Meiqi Niu, Ting Xu, Ling Zhao, Thomas Funck, Xiao-Jing Wang, Katrin Amunts, and Nicola Palomero-Gallagher. Cytoarchitectonic, receptor distribution and functional connectivity analyses of the macaque frontal lobe. *eLife*, 12, 2023.

[13145] Parth K Raval, Wing Yui Ngan, Jenna Gallie, and Deepa Agashe. The layered costs and benefits of translational redundancy. *eLife*, 12, 2023.

[13146] Mukulika Ray, Ashley Mae Conard, Jennifer Urban, Pranav Mahableshwarkar, Joseph Aguilera, Annie Huang, Smriti Vaidyanathan, and Erica Larschan. Sex-specific splicing occurs genome-wide during early Drosophila embryogenesis. *eLife*, 12, 2023.

[13147] Shamayeeta Ray, Samuel P Berry, Eric A Wilson, Casey H Zhang, Mrinal Shekhar, Abhishek Singharoy, and Rachelle Gaudet. High-resolution structures with bound Mn2+ and Cd2+ map the metal import pathway in an Nramp transporter. *eLife*, 12, 2023.

[13148] Lyudmil Raykov, Manon Mottet, Jahn Nitschke, and Thierry Soldati. A TRAF-like E3 ubiquitin ligase TrafE coordinates ESCRT and autophagy in endolysosomal damage response and cell-autonomous immunity to Mycobacterium marinum. *eLife*, 12, 2023.

[13149] Davide Reato, Raphael Steinfeld, André Tacão Monteiro, and Alfonso Renart. Response outcome gates the effect of spontaneous cortical state fluctuations on perceptual decisions. *eLife*, 12, 2023.

[13150] Jenaid M Rees, Victoria A Sleight, Stephen J Clark, Tetsuya Nakamura, and J Andrew Gillis. Ectodermal Wnt signaling, cell fate determination, and polarity of the skate gill arch skeleton. *eLife*, 12, 2023.

[13151] Stella Reichling, Peter F Doubleday, Tomas Germade, Ariane Bergmann, Robbie Loewith, Uwe Sauer, and Duncan Holbrook-Smith. Dynamic metabolome profiling uncovers potential TOR signaling genes. *eLife*, 12, 2023.

[13152] Janice M Reimer, Morgan E DeSantis, Samara L Reck-Peterson, and Andres E Leschziner. Structures of human dynein in complex with the lissencephaly 1 protein, LIS1. *eLife*, 12, 2023.

[13153] Ronja Reinhardt and Thomas A Leonard. A critical evaluation of protein kinase regulation by activation loop autophosphorylation. *eLife*, 12, 2023.

[13154] Zeinab Rekad, Michaël Ruff, Agata Radwanska, Dominique Grall, Delphine Ciais, and Ellen Van Obberghen-Schilling. Coalescent RNA-localizing and transcriptional activities of SAM68 modulate adhesion and subendothelial basement membrane assembly. *eLife*, 12, 2023.

[13155] Pascal Dominic Rem, Vita Sereikaite, Diego Fernández-Fernández, Sebastian Reinartz, Daniel Ulrich, Thorsten Fritzius, Luca Trovo, Salomé Roux, Ziyang Chen, Philippe Rondard, Jean-Philippe Pin, Jochen Schwenk, Bernd Fakler, Martin Gassmann, Tania Rinaldi Barkat, Kristian Strømgaard, and Bernhard Bettler. Soluble amyloid-$\beta$ precursor peptide does not regulate GABAB receptor activity. *eLife*, 12, 2023.

[13156] Olaya Rendueles, Jorge AM de Sousa, and Eduardo PC Rocha. Competition between lysogenic and sensitive bacteria is determined by the fitness costs of the different emerging phage-resistance strategies. *eLife*, 12, 2023.

[13157] Simone Rey, Henrike Ohm, Frederieke Moschref, Dagmar Zeuschner, Marit Praetz, and Christian Klämbt. Glial-dependent clustering of voltage-gated ion channels in Drosophila precedes myelin formation. *eLife*, 12, 2023.

[13158] Fernanda Lenita Ribeiro, Ashley York, Elizabeth Zavitz, Steffen Bollmann, Marcello GP Rosa, and Alexander Puckett. Variability of visual field maps in human early extrastriate cortex challenges the canonical model of organization of V2 and V3. *eLife*, 12, 2023.

[13159] Janet W Rich-Edwards and Donna L Maney. Best practices to promote rigor and reproducibility in the era of sex-inclusive research. *eLife*, 12, 2023.

[13160] Daniel Richard, Steven Pregizer, Divya Venkatasubramanian, Rosanne M Raftery, Pushpanathan Muthuirulan, Zun Liu, Terence D Capellini, and April M Craft. Lineage-specific differences and regulatory networks governing human chondrocyte development. *eLife*, 12, 2023.

[13161] Eve Richardson, Špela Binter, Miha Kosmac, Marie Ghraichy, Valentin von Niederhäusern, Aleksandr Kovaltsuk, Jacob D Galson, Johannes Trück, Dominic F Kelly, Charlotte M Deane, Paul Kellam, and Simon J Watson. Characterisation of the immune repertoire of a humanised transgenic mouse through immunophenotyping and high-throughput sequencing. *eLife*, 12, 2023.

[13162] Tom G Richardson, Genevieve M Leyden, and George Davey Smith. Time-varying and tissue-dependent effects of adiposity on leptin levels: A Mendelian randomization study. *eLife*, 12, 2023.

[13163] Manuela Richter, Lila Neahring, Jinghui Tao, Renaldo Sutanto, Nathan H Cho, and Sophie Dumont. Kinetochore-fiber lengths are maintained locally but coordinated globally by poles in the mammalian spindle. *eLife*, 12, 2023.

[13164] Jana Riegger and Rolf E Brenner. Increase of cell surface vimentin is associated with vimentin network disruption and subsequent stress-induced premature senescence in human chondrocytes. *eLife*, 12, 2023.

[13165] Atle E Rimehaug, Alexander J Stasik, Espen Hagen, Yazan N Billeh, Josh H Siegle, Kael Dai, Shawn R Olsen, Christof Koch, Gaute T Einevoll, and Anton Arkhipov. Uncovering circuit mechanisms of current sinks and sources with biophysical simulations of primary visual cortex. *eLife*, 12, 2023.

[13166] Vera Rinaldi, Kathleen Messemer, Kathleen Desevin, Fengyun Sun, Bethany C Berry, Shweta Kukreja, Andrew R Tapper, Amy J Wagers, and Oliver J Rando. Evidence for RNA or protein transport from somatic tissues to the male reproductive tract in mouse. *eLife*, 12, 2023.

[13167] Alan V Rincon, Bridget M Waller, Julie Duboscq, Alexander Mielke, Claire Pérez, Peter R Clark, and Jérôme Micheletta. Higher social tolerance is associated with more complex facial behavior in macaques. *eLife*, 12, 2023.

[13168] Juan Luis Riquelme, Mike Hemberger, Gilles Laurent, and Julijana Gjorgjieva. Single spikes drive sequential propagation and routing of activity in a cortical network. *eLife*, 12, 2023.

[13169] Milena Rmus, Mingjian He, Beth Baribault, Edward G Walsh, Elena K Festa, Anne GE Collins, and Matthew R Nassar. Age-related differences in prefrontal glutamate are associated with increased working memory decay that gives the appearance of learning deficits. *eLife*, 12, 2023.

[13170] Jessica M Roberts, James D Beck, Tanner B Pollock, Devin P Bendixsen, and Eric J Hayden. RNA sequence to structure analysis from comprehensive pairwise mutagenesis of multiple self-cleaving ribozymes. *eLife*, 12, 2023.

[13171] Matthew Roberts, Julia Ogden, AS Mukarram Hossain, Anshuman Chaturvedi, Alastair RW Kerr, Caroline Dive, Jennifer Ellen Beane, and Carlos Lopez-Garcia. Interrogating the precancerous evolution of pathway dysfunction in lung squamous cell carcinoma using XTABLE. *eLife*, 12, 2023.

[13172] Oliver Robinson, ChungHo E Lau, Sungyeon Joo, Sandra Andrusaityte, Eva Borras, Paula de Prado-Bert, Lida Chatzi, Hector C Keun, Regina Grazuleviciene, Kristine B Gutzkow, Lea Maitre, Dries S Martens, Eduard Sabido, Valérie Siroux, Jose Urquiza, Marina Vafeiadi, John Wright, Tim S Nawrot, Mariona Bustamante, and Martine Vrijheid. Associations of four biological age markers with child development: A multi-omic analysis in the European HELIX cohort. *eLife*, 12, 2023.

[13173] Kimberly E Roche, Johannes R Bjork, Mauna R Dasari, Laura Grieneisen, David Jansen, Trevor J Gould, Laurence R Gesquiere, Luis B Barreiro, Susan C Alberts, Ran Blekhman, Jack A Gilbert, Jenny Tung, Sayan Mukherjee, and Elizabeth A Archie. Universal gut microbial relationships in the gut microbiome of wild baboons. *eLife*, 12, 2023.

[13174] Ellen M Rodberg, Carolina R den Hartog, Emma S Dauster, and Elena M Vazey. Sex-dependent noradrenergic modulation of premotor cortex during decision-making. *eLife*, 12, 2023.

[13175] Yuri Elias Rodrigues, Cezar M Tigaret, Hélène Marie, Cian O'Donnell, and Romain Veltz. A stochastic model of hippocampal synaptic plasticity with geometrical readout of enzyme dynamics. *eLife*, 12, 2023.

[13176] Jonathan Rodriguez, Abdon Iniguez, Nilamani Jena, Prasanthi Tata, Zhong-Ying Liu, Arthur D Lander, John Lowengrub, and Richard A Van Etten. Predictive nonlinear modeling of malignant myelopoiesis and tyrosine kinase inhibitor therapy. *eLife*, 12, 2023.

[13177] María Rodríguez-López, Nicola Bordin, Jon Lees, Harry Scholes, Shaimaa Hassan, Quentin Saintain, Stephan Kamrad, Christine Orengo, and Jürg Bähler. Broad functional profiling of fission yeast proteins using phenomics and machine learning. *eLife*, 12, 2023.

[13178] James M Roe, Didac Vidal-Pineiro, Inge K Amlien, Mengyu Pan, Markus H Sneve, Michel Thiebaut de Schotten, Patrick Friedrich, Zhiqiang Sha, Clyde Francks, Espen M Eilertsen, Yunpeng Wang, Kristine B Walhovd, Anders M Fjell, and René Westerhausen. Tracing the development and lifespan change of population-level structural asymmetry in the cerebral cortex. *eLife*, 12, 2023.

[13179] Satish Rojekar, Anusha R Pallapati, Judit Gimenez-Roig, Funda Korkmaz, Farhath Sultana, Damini Sant, Clement M Haeck, Anne Macdonald, Se-Min Kim, Clifford J Rosen, Orly Barak, Marcia Meseck, John Caminis, Daria Lizneva, Tony Yuen, and Mone Zaidi. Development and biophysical characterization of a humanized FSH–blocking monoclonal antibody therapeutic formulated at an ultra-high concentration. *eLife*, 12, 2023.

[13180] Romain Rollin, Jean-FranÇois Joanny, and Pierre Sens. Physical basis of the cell size scaling laws. *eLife*, 12, 2023.

[13181] Justin L Roncaioli, Janet Peace Babirye, Roberto A Chavez, Fitty L Liu, Elizabeth A Turcotte, Angus Y Lee, Cammie F Lesser, and Russell E Vance. A hierarchy of cell death pathways confers layered resistance to shigellosis in mice. *eLife*, 12, 2023.

[13182] Natalia C Rosas, Jonathan Wilksch, Jake Barber, Jiahui Li, Yanan Wang, Zhewei Sun, Andrea Rocker, Chaille T Webb, Laura Perlaza-Jiménez, Christopher J Stubenrauch, Vijaykrishna Dhanasekaran, Jiangning Song, George Taiaroa, Mark Davies, Richard A Strugnell, Qiyu Bao, Tieli Zhou, Michael J McDonald, and Trevor Lithgow. The evolutionary mechanism of non-carbapenemase carbapenem-resistant phenotypes in Klebsiella spp. *eLife*, 12, 2023.

[13183] Ruben Rosas, Rhiannon R Aguilar, Nina Arslanovic, Anna Seck, Duncan J Smith, Jessica K Tyler, and Mair EA Churchill. A novel single alpha-helix DNA-binding domain in CAF-1 promotes gene silencing and DNA damage survival through tetrasome-length DNA selectivity and spacer function. *eLife*, 12, 2023.

[13184] Noah H Rose, Athanase Badolo, Massamba Sylla, Jewelna Akorli, Sampson Otoo, Andrea Gloria-Soria, Jeffrey R Powell, Bradley J White, Jacob E Crawford, and Carolyn S McBride. Dating the origin and spread of specialization on human hosts in Aedes aegypti mosquitoes. *eLife*, 12, 2023.

[13185] Greta Rossi, Gabriele Ordazzo, Niccolò N Vanni, Valerio Castoldi, Angelo Iannielli, Dario Di Silvestre, Edoardo Bellini, Letizia Bernardo, Serena G Giannelli, Mirko Luoni, Sharon Muggeo, Letizia Leocani, PierLuigi Mauri, and Vania Broccoli. MCT1-dependent energetic failure and neuroinflammation underlie optic nerve degeneration in Wolfram syndrome mice. *eLife*, 12, 2023.

[13186] Martina Rossi, Carlos Anerillas, Maria Laura Idda, Rachel Munk, Chang Hoon Shin, Stefano Donega, Dimitrios Tsitsipatis, Allison B Herman, Jennifer L Martindale, Xiaoling Yang, Yulan Piao, Krystyna Mazan-Mamczarz, Jinshui Fan, Luigi Ferrucci, Peter F Johnson, Supriyo De, Kotb Abdelmohsen, and Myriam Gorospe. Pleiotropic effects of BAFF on the senescence-associated secretome and growth arrest. *eLife*, 12, 2023.

[13187] Sofia Rossini, Marco Gargaro, Giulia Scalisi, Elisa Bianconi, Sara Ambrosino, Eleonora Panfili, Claudia Volpi, Ciriana Orabona, Antonio Macchiarulo, Francesca Fallarino, and Giada Mondanelli. A back-door insight into the modulation of Src kinase activity by the polyamine spermidine. *eLife*, 12, 2023.

[13188] Priscila S Rothier, Anne-Claire Fabre, Julien Clavel, Roger BJ Benson, and Anthony Herrel. Mammalian forelimb evolution is driven by uneven proximal-to-distal morphological diversity. *eLife*, 12, 2023.

[13189] Jean-Philippe Rousseau, Andreea Furdui, Carolina da Silveira Scarpellini, Richard L Horner, and Gaspard Montandon. Medullary tachykinin precursor 1 neurons promote rhythmic breathing. *eLife*, 12, 2023.

[13190] Roshni Roy, Pei-Lun Kuo, Julián Candia, Dimitra Sarantopoulou, Ceereena Ubaida-Mohien, Dena Hernandez, Mary Kaileh, Sampath Arepalli, Amit Singh, Arsun Bektas, Jaekwan Kim, Ann Z Moore, Toshiko Tanaka, Julia McKelvey, Linda Zukley, Cuong Nguyen, Tonya Wallace, Christopher Dunn, William Wood, Yulan Piao, Christopher Coletta, Supriyo De, Jyoti Sen, Nan-ping Weng, Ranjan Sen, and Luigi Ferrucci. Epigenetic signature of human immune aging in the GESTALT study. *eLife*, 12, 2023.

[13191] Lars N Royall, Diana Machado, Sebastian Jessberger, and Annina Denoth-Lippuner. Asymmetric inheritance of centrosomes maintains stem cell properties in human neural progenitor cells. *eLife*, 12, 2023.

[13192] Xiaohao Ruan, Da Huang, Yongle Zhan, Jingyi Huang, Jinlun Huang, Ada Tsui-Lin Ng, James Hok-Leung Tsu, and Rong Na. Risk of second primary cancers after a diagnosis of first primary cancer: A pan-cancer analysis and Mendelian randomization study. *eLife*, 12, 2023.

[13193] Mika Rubinov. Circular and unified analysis in network neuroscience. *eLife*, 12, 2023.

[13194] Maria Ruesseler, Lilian Aline Weber, Tom Rhys Marshall, Jill O'Reilly, and Laurence Tudor Hunt. Quantifying decision-making in dynamic, continuously evolving environments. *eLife*, 12, 2023.

[13195] Artur Ruppel, Dennis Wörthmüller, Vladimir Misiak, Manasi Kelkar, Irène Wang, Philippe Moreau, Adrien Méry, Jean Révilloud, Guillaume Charras, Giovanni Cappello, Thomas Boudou, Ulrich S Schwarz, and Martial Balland. Force propagation between epithelial cells depends on active coupling and mechano-structural polarization. *eLife*, 12, 2023.

[13196] Magdalena L Russell, Noah Simon, Philip Bradley, and Frederick A Matsen. Statistical inference reveals the role of length, GC content, and local sequence in V(D)J nucleotide trimming. *eLife*, 12, 2023.

[13197] Mark A Rutherford, Atri Bhattacharyya, Maolei Xiao, Hou-Ming Cai, Indra Pal, and Maria Eulalia Rubio. GluA3 subunits are required for appropriate assembly of AMPAR GluA2 and GluA4 subunits on cochlear afferent synapses and for presynaptic ribbon modiolar–pillar morphology. *eLife*, 12, 2023.

[13198] Saige Rutherford, Pieter Barkema, Ivy F Tso, Chandra Sripada, Christian F Beckmann, Henricus G Ruhe, and Andre F Marquand. Evidence for embracing normative modeling. *eLife*, 12, 2023.

[13199] Charumathi Sabanayagam, Feng He, Simon Nusinovici, Jialiang Li, Cynthia Lim, Gavin Tan, and Ching Yu Cheng. Prediction of diabetic kidney disease risk using machine learning models: A population-based cohort study of Asian adults. *eLife*, 12, 2023.

[13200] Shievanie Sabesan, Andreas Fragner, Ciaran Bench, Fotios Drakopoulos, and Nicholas A Lesica. Large-scale electrophysiology and deep learning reveal distorted neural signal dynamics after hearing loss. *eLife*, 12, 2023.

[13201] Lopamudra Sadhu, Nikolaos Tsopoulidis, Md Hasanuzzaman, Vibor Laketa, Michael Way, and Oliver T Fackler. ARPC5 isoforms and their regulation by calcium-calmodulin-N-WASP drive distinct Arp2/3-dependent actin remodeling events in CD4 T cells. *eLife*, 12, 2023.

[13202] Julie M Sadino, Xander G Bradeen, Conor J Kelly, Liza E Brusman, Deena M Walker, and Zoe R Donaldson. Prolonged partner separation erodes nucleus accumbens transcriptional signatures of pair bonding in male prairie voles. *eLife*, 12, 2023.

[13203] Lina Sagert, Christian Winter, Ina Ruppert, Maximilian Zehetmaier, Christoph Thomas, and Robert Tampé. The ER folding sensor UGGT1 acts on TAPBPR-chaperoned peptide-free MHC I. *eLife*, 12, 2023.

[13204] Pankaj Sahai-Hernandez, Claire Pouget, Shai Eyal, Ondrej Svoboda, Jose Chacon, Lin Grimm, Tor Gjøen, and David Traver. Dermomyotome-derived endothelial cells migrate to the dorsal aorta to support hematopoietic stem cell emergence. *eLife*, 12, 2023.

[13205] Jiro Sakai, Jiyeon Yang, Chao-Kai Chou, Wells W Wu, and Mustafa Akkoyunlu. B cell receptor-induced IL-10 production from neonatal mouse CD19+CD43- cells depends on STAT5-mediated IL-6 secretion. *eLife*, 12, 2023.

[13206] Nehuén Salas, Manuela Blasco Pedreros, Tuanne dos Santos Melo, Vanina G Maguire, Jihui Sha, James A Wohlschlegel, Antonio Pereira-Neves, and Natalia de Miguel. Role of cytoneme structures and extracellular vesicles in Trichomonas vaginalis parasite-parasite communication. *eLife*, 12, 2023.

[13207] Federico Salas-Lucia, Csaba Fekete, Richárd Sinkó, Péter Egri, Kristóf Rada, Yvette Ruska, Balázs Gereben, and Antonio C Bianco. Axonal T3 uptake and transport can trigger thyroid hormone signaling in the brain. *eLife*, 12, 2023.

[13208] Pedro Sampaio, Sara Pestana, Catarina Bota, Adán Guerrero, Ivo A Telley, David Smith, and Susana Santos Lopes. Fluid extraction from the left-right organizer uncovers mechanical properties needed for symmetry breaking. *eLife*, 12, 2023.

[13209] Kai J Sandbrink, Pranav Mamidanna, Claudio Michaelis, Matthias Bethge, Mackenzie Weygandt Mathis, and Alexander Mathis. Contrasting action and posture coding with hierarchical deep neural network models of proprioception. *eLife*, 12, 2023.

[13210] Theo Sanderson, Maxwell L Bileschi, David Belanger, and Lucy J Colwell. ProteInfer, deep neural networks for protein functional inference. *eLife*, 12, 2023.

[13211] Dasha A Sandra, Jay A Olson, Ellen J Langer, and Mathieu Roy. Presenting a sham treatment as personalised increases the placebo effect in a randomised controlled trial. *eLife*, 12, 2023.

[13212] Sreedhara Sangadala, Chi Heon Kim, Lorenzo M Fernandes, Pooja Makkar, George R Beck, Scott D Boden, Hicham Drissi, and Steven M Presciutti. Sclerostin small-molecule inhibitors promote osteogenesis by activating canonical Wnt and BMP pathways. *eLife*, 12, 2023.

[13213] Fumiya K Sano, Hiroaki Akasaka, Wataru Shihoya, and Osamu Nureki. Cryo-EM structure of the endothelin-1-ETB-Gi complex. *eLife*, 12, 2023.

[13214] Marianna Santopaolo, Michaela Gregorova, Fergus Hamilton, David Arnold, Anna Long, Aurora Lacey, Elizabeth Oliver, Alice Halliday, Holly Baum, Kristy Hamilton, Rachel Milligan, Olivia Pearce, Lea Knezevic, Begonia Morales Aza, Alice Milne, Emily Milodowski, Eben Jones, Rajeka Lazarus, Anu Goenka, Adam Finn, Nicholas Maskell, Andrew D Davidson, Kathleen Gillespie, Linda Wooldridge, and Laura Rivino. Prolonged T-cell activation and long COVID symptoms independently associate with severe COVID-19 at 3 months. *eLife*, 12, 2023.

[13215] Yonatan Sanz Perl, Sol Fittipaldi, Cecilia Gonzalez Campo, Sebastián Moguilner, Josephine Cruzat, Matias E Fraile-Vazquez, Rubén Herzog, Morten L Kringelbach, Gustavo Deco, Pavel Prado, Agustin Ibanez, and Enzo Tagliazucchi. Model-based whole-brain perturbational landscape of neurodegenerative diseases. *eLife*, 12, 2023.

[13216] Anne L Sapiro, Beth M Hayes, Regan F Volk, Jenny Y Zhang, Diane M Brooks, Calla Martyn, Atanas Radkov, Ziyi Zhao, Margie Kinnersley, Patrick R Secor, Balyn W Zaro, and Seemay Chou. Longitudinal map of transcriptome changes in the Lyme pathogen Borrelia burgdorferi during tickborne transmission. *eLife*, 12, 2023.

[13217] Raphael Sarfati, Kunaal Joshi, Owen Martin, Julie C Hayes, Srividya Iyer-Biswas, and Orit Peleg. Emergent periodicity in the collective synchronous flashing of fireflies. *eLife*, 12, 2023.

[13218] Abby Sarkar, Yinhua Jin, Brian C DeFelice, Catriona Y Logan, Yan Yang, Teni Anbarchian, Peng Wu, Maurizio Morri, Norma F Neff, Huy Nguyen, Eric Rulifson, Matthew Fish, Avi Gurion Kaye, Azalia M Martínez Jaimes, and Roel Nusse. Intermittent fasting induces rapid hepatocyte proliferation to restore the hepatostat in the mouse liver. *eLife*, 12, 2023.

[13219] Akshat Sarkari, Sophie Korenfeld, Karina Deniz, Katherine Ladner, Phillip Wong, Sanyukta Padmanabhan, Rachel I Vogel, Laura A Sherer, Naomi Courtemanche, Clifford Steer, Kerem Wainer-Katsir, and Emil Lou. Treatment with tumor-treating fields (TTFields) suppresses intercellular tunneling nanotube formation in vitro and upregulates immuno-oncologic biomarkers in vivo in malignant mesothelioma. *eLife*, 12, 2023.

[13220] Dylan C Sarver, Cheng Xu, Susana Rodriguez, Susan Aja, Andrew E Jaffe, Feng J Gao, Michael Delannoy, Muthu Periasamy, Yasuhiro Kazuki, Mitsuo Oshimura, Roger H Reeves, and G William Wong. Hypermetabolism in mice carrying a near-complete human chromosome 21. *eLife*, 12, 2023.

[13221] Takashi Sasaki, Yoshinori Nishimoto, Takumi Hirata, Yukiko Abe, Nobuyoshi Hirose, Michiyo Takayama, Toru Takebayashi, Hideyuki Okano, and Yasumichi Arai. Status and physiological significance of circulating adiponectin in the very old and centenarians: an observational study. *eLife*, 12, 2023.

[13222] Serkan Sayin, Brittany Rosener, Carmen G Li, Bao Ho, Olga Ponomarova, Doyle V Ward, Albertha JM Walhout, and Amir Mitchell. Evolved bacterial resistance to the chemotherapy gemcitabine modulates its efficacy in co-cultured cancer cells. *eLife*, 12, 2023.

[13223] Valeria Scagliotti, Maria Lillina Vignola, Thea Willis, Mark Howard, Eugenia Marinelli, Carles Gaston-Massuet, Cynthia Andoniadou, and Marika Charalambous. Imprinted Dlk1 dosage as a size determinant of the mammalian pituitary gland. *eLife*, 12, 2023.

[13224] Elias Scheer and Cornelia I Bargmann. Sensory neurons couple arousal and foraging decisions in Caenorhabditis elegans. *eLife*, 12, 2023.

[13225] Louise Schenberg, Aïda Palou, FranÇois Simon, Tess Bonnard, Charles-Elliot Barton, Desdemona Fricker, Michele Tagliabue, Jordi Llorens, and Mathieu Beraneck. Multisensory gaze stabilization in response to subchronic alteration of vestibular type I hair cells. *eLife*, 12, 2023.

[13226] Van Schepler-Luu, Coline Sciallano, Melissa Stiebner, Chonghui Ji, Gabriel Boulard, Amadou Diallo, Florence Auguy, Si Nian Char, Yugander Arra, Kyrylo Schenstnyi, Marcel Buchholzer, Eliza PI Loo, Atugonza L Bilaro, David Lihepanyama, Mohammed Mkuya, Rosemary Murori, Ricardo Oliva, Sebastien Cunnac, Bing Yang, Boris Szurek, and Wolf B Frommer. Genome editing of an African elite rice variety confers resistance against endemic and emerging Xanthomonas oryzae pv. oryzae strains. *eLife*, 12, 2023.

[13227] Andria C Schibler, Predrag Jevtic, Gianluca Pegoraro, Daniel L Levy, and Tom Misteli. Identification of epigenetic modulators as determinants of nuclear size and shape. *eLife*, 12, 2023.

[13228] Judith Schick, Meltem Altunay, Matthew Lacorcia, Nathalie Marschner, Stefanie Westermann, Julia Schluckebier, Christoph Schubart, Barbara Bodendorfer, Dennis Christensen, Christian Alexander, Stefan Wirtz, David Voehringer, Clarissa Prazeres da Costa, and Roland Lang. IL-4 and helminth infection downregulate MINCLE-dependent macrophage response to mycobacteria and Th17 adjuvanticity. *eLife*, 12, 2023.

[13229] Erik S Schild, Shivam Gupta, Clément Dubois, Euclides E Fernandes Póvoa, Marie-Anne Félix, Andrew Mugler, and Hendrik C Korswagen. Precise temporal control of neuroblast migration through combined regulation and feedback of a Wnt receptor. *eLife*, 12, 2023.

[13230] William HK Schilling, Podjanee Jittamala, James A Watson, Maneerat Ekkapongpisit, Tanaya Siripoon, Thundon Ngamprasertchai, Viravarn Luvira, Sasithorn Pongwilai, Cintia Cruz, James J Callery, Simon Boyd, Varaporn Kruabkontho, Thatsanun Ngernseng, Jaruwan Tubprasert, Mohammad Yazid Abdad, Nattaporn Piaraksa, Kanokon Suwannasin, Pongtorn Hanboonkunupakarn, Borimas Hanboonkunupakarn, Sakol Sookprome, Kittiyod Poovorawan, Janjira Thaipadungpanit, Stuart Blacksell, Mallika Imwong, Joel Tarning, Walter RJ Taylor, Vasin Chotivanich, Chunlanee Sangketchon, Wiroj Ruksakul, Kesinee Chotivanich, Mauro Martins Teixeira, Sasithon Pukrittayakamee, Arjen M Dondorp, Nicholas PJ Day, Watcharapong Piyaphanee, Weerapong Phumratanaprapin, and Nicholas J White. Pharmacometrics of high-dose ivermectin in early COVID-19 from an open label, randomized, controlled adaptive platform trial (PLATCOV). *eLife*, 12, 2023.

[13231] Philip Schmassmann, Julien Roux, Steffen Dettling, Sabrina Hogan, Tala Shekarian, Tomás A Martins, Marie-FranÇoise Ritz, Sylvia Herter, Marina Bacac, and Gregor Hutter. Single-cell characterization of human GBM reveals regional differences in tumor-infiltrating leukocyte activation. *eLife*, 12, 2023.

[13232] Konstantin Schmidt, GonÇalo Santos-Matos, Stefan Leopold-Messer, Yassine El Chazli, Olivier Emery, Théodora Steiner, Joern Piel, and Philipp Engel. Integration host factor regulates colonization factors in the bee gut symbiont Frischella perrara. *eLife*, 12, 2023.

[13233] Ekaterina Schneider, Dora Hopf, Corina Aguilar-Raab, Dirk Scheele, Andreas B Neubauer, Uta Sailer, René Hurlemann, Monika Eckstein, and Beate Ditzen. Affectionate touch and diurnal oxytocin levels: An ecological momentary assessment study. *eLife*, 12, 2023.

[13234] Simon Schneider, Andjela Kovacevic, Michelle Mayer, Ann-Kristin Dicke, Lena Arévalo, Sophie A Koser, Jan N Hansen, Samuel Young, Christoph Brenker, Sabine Kliesch, Dagmar Wachten, Gregor Kirfel, Timo Struenker, Frank Tüttelmann, and Hubert Schorle. Cylicins are a structural component of the sperm calyx being indispensable for male fertility in mice and human. *eLife*, 12, 2023.

[13235] Anna Elisabeth Schnell, Maarten Leemans, Kasper Vinken, and Hans Op de Beeck. A computationally informed comparison between the strategies of rodents and humans in visual object recognition. *eLife*, 12, 2023.

[13236] Florian Schueder, Pierre Mangeol, Eunice HoYee Chan, Renate Rees, Jürgen Schünemann, Ralf Jungmann, Dirk Görlich, and Frank Schnorrer. Nanobodies combined with DNA-PAINT super-resolution reveal a staggered titin nanoarchitecture in flight muscles. *eLife*, 12, 2023.

[13237] Heiko H Schütt, Alexander D Kipnis, Jörn Diedrichsen, and Nikolaus Kriegeskorte. Statistical inference on representational geometries. *eLife*, 12, 2023.

[13238] Irene Schwartz, Milica Vunjak, Valentina Budroni, Adriana Cantoran García, Marialaura Mastrovito, Adrian Soderholm, Matthias Hinterndorfer, Melanie de Almeida, Kathrin Hacker, Jingkui Wang, Kimon Froussios, Julian Jude, Thomas Decker, Johannes Zuber, and Gijs A Versteeg. SPOP targets the immune transcription factor IRF1 for proteasomal degradation. *eLife*, 12, 2023.

[13239] Sara Scinicariello, Adrian Soderholm, Markus Schäfer, Alexandra Shulkina, Irene Schwartz, Kathrin Hacker, Rebeca Gogova, Robert Kalis, Kimon Froussios, Valentina Budroni, Annika Bestehorn, Tim Clausen, Pavel Kovarik, Johannes Zuber, and Gijs A Versteeg. HUWE1 controls tristetraprolin proteasomal degradation by regulating its phosphorylation. *eLife*, 12, 2023.

[13240] Kwi Shan Seah and Vinodkumar Saranathan. Hierarchical morphogenesis of swallowtail butterfly wing scale nanostructures. *eLife*, 12, 2023.

[13241] Federico G Segala, Aurelio Bruno, Joel T Martin, Myat T Aung, Alex R Wade, and Daniel H Baker. Different rules for binocular combination of luminance flicker in cortical and subcortical pathways. *eLife*, 12, 2023.

[13242] Effie Segas, Sébastien Mick, Vincent Leconte, Océane Dubois, Rémi Klotz, Daniel Cattaert, and Aymar de Rugy. Intuitive movement-based prosthesis control enables arm amputees to reach naturally in virtual reality. *eLife*, 12, 2023.

[13243] Dario Segura-Peña, Oda Hovet, Hemanga Gogoi, Jennine Dawicki-McKenna, Stine Malene Hansen Wøien, Manuel Carrer, Ben E Black, Michele Cascella, and Nikolina Sekulic. The structural basis of the multi-step allosteric activation of Aurora B kinase. *eLife*, 12, 2023.

[13244] Yasunari Seita, Keren Cheng, John R McCarrey, Nomesh Yadu, Ian H Cheeseman, Alec Bagwell, Corinna N Ross, Isamar Santana Toro, Li-hua Yen, Sean Vargas, Christopher S Navara, Brian P Hermann, and Kotaro Sasaki. Efficient generation of marmoset primordial germ cell-like cells using induced pluripotent stem cells. *eLife*, 12, 2023.

[13245] Uri Seroussi, Andrew Lugowski, Lina Wadi, Robert X Lao, Alexandra R Willis, Winnie Zhao, Adam E Sundby, Amanda G Charlesworth, Aaron W Reinke, and Julie M Claycomb. A comprehensive survey of C. elegans argonaute proteins reveals organism-wide gene regulatory networks and functions. *eLife*, 12, 2023.

[13246] Nelson BC Serre, DaŠa Wernerová, Pruthvi Vittal, Shiv Mani Dubey, Eva Medvecká, Adriana Jelínková, Jan PetráŠek, Guido Grossmann, and MatyáŠ Fendrych. The AUX1-AFB1-CNGC14 module establishes a longitudinal root surface pH profile. *eLife*, 12, 2023.

[13247] Damiano Sgarbossa, Umberto Lupo, and Anne-Florence Bitbol. Generative power of a protein language model trained on multiple sequence alignments. *eLife*, 12, 2023.

[13248] Keanu Shadron and José Luis Peña. Development of frequency tuning shaped by spatial cue reliability in the barn owl's auditory midbrain. *eLife*, 12, 2023.

[13249] Viral S Shah, Jue Hou, Vladimir Vinarsky, Jiajie Xu, Manalee V Surve, Charles P Lin, and Jayaraj Rajagopal. Autofluorescence imaging permits label-free cell type assignment and reveals the dynamic formation of airway secretory cell associated antigen passages (SAPs). *eLife*, 12, 2023.

[13250] Hossein Shahabi, Dileep R Nair, and Richard M Leahy. Multilayer brain networks can identify the epileptogenic zone and seizure dynamics. *eLife*, 12, 2023.

[13251] Naeem Yusuf Shaikh and Kartik Sunagar. The deep-rooted origin of disulfide-rich spider venom toxins. *eLife*, 12, 2023.

[13252] Deepthi Shankaran, Anjali Singh, Stanzin Dawa, Prabhakar Arumugam, Sheetal Gandotra, and Vivek Rao. The antidepressant sertraline provides a novel host directed therapy module for augmenting TB therapy. *eLife*, 12, 2023.

[13253] Sachin Sharma, Russell Littman, John D Tompkins, Douglas Arneson, Jaime Contreras, Al-Hassan Dajani, Kaitlyn Ang, Amit Tsanhani, Xin Sun, Patrick Y Jay, Herbert Herzog, Xia Yang, and Olujimi A Ajijola. Tiered sympathetic control of cardiac function revealed by viral tracing and single cell transcriptome profiling. *eLife*, 12, 2023.

[13254] Chun Shen, Edmund T Rolls, Shitong Xiang, Christelle Langley, Barbara J Sahakian, Wei Cheng, and Jianfeng Feng. Brain and molecular mechanisms underlying the nonlinear association between close friendships, mental health, and cognition in children. *eLife*, 12, 2023.

[13255] Jiemin Shen, Azaan Saalim Wilbon, Ming Zhou, and Yaping Pan. Mechanism of Ca2+ transport by ferroportin. *eLife*, 12, 2023.

[13256] Zeyu Shen, Bowen Jia, Yang Xu, Jonas Wessén, Tanmoy Pal, Hue Sun Chan, Shengwang Du, and Mingjie Zhang. Biological condensates form percolated networks with molecular motion properties distinctly different from dilute solutions. *eLife*, 12, 2023.

[13257] Courtney T Shepard, Brandon L Brown, Morgan A Van Rijswijck, Rachel M Zalla, Darlene A Burke, Johnny R Morehouse, Amberly S Riegler, Scott R Whittemore, and David SK Magnuson. Silencing long-descending inter-enlargement propriospinal neurons improves hindlimb stepping after contusive spinal cord injuries. *eLife*, 12, 2023.

[13258] Katharine Sherratt, Hugo Gruson, Rok Grah, Helen Johnson, Rene Niehus, Bastian Prasse, Frank Sandmann, Jannik Deuschel, Daniel Wolffram, Sam Abbott, Alexander Ullrich, Graham Gibson, Evan L Ray, Nicholas G Reich, Daniel Sheldon, Yijin Wang, Nutcha Wattanachit, Lijing Wang, Jan Trnka, Guillaume Obozinski, Tao Sun, Dorina Thanou, Loic Pottier, Ekaterina Krymova, Jan H Meinke, Maria Vittoria Barbarossa, Neele Leithauser, Jan Mohring, Johanna Schneider, Jaroslaw Wlazlo, Jan Fuhrmann, Berit Lange, Isti Rodiah, Prasith Baccam, Heidi Gurung, Steven Stage, Bradley Suchoski, Jozef Budzinski, Robert Walraven, Inmaculada Villanueva, Vit Tucek, Martin Smid, Milan Zajicek, Cesar Perez Alvarez, Borja Reina, Nikos I Bosse, Sophie R Meakin, Lauren Castro, Geoffrey Fairchild, Isaac Michaud, Dave Osthus, Pierfrancesco Alaimo Di Loro, Antonello Maruotti, Veronika Eclerova, Andrea Kraus, David Kraus, Lenka Pribylova, Bertsimas Dimitris, Michael Lingzhi Li, Soni Saksham, Jonas Dehning, Sebastian Mohr, Viola Priesemann, Grzegorz Redlarski, Benjamin Bejar, Giovanni Ardenghi, Nicola Parolini, Giovanni Ziarelli, Wolfgang Bock, Stefan Heyder, Thomas Hotz, David E Singh, Miguel Guzman-Merino, Jose L Aznarte, David Morina, Sergio Alonso, Enric Alvarez, Daniel Lopez, Clara Prats, Jan Pablo Burgard, Arne Rodloff, Tom Zimmermann, Alexander Kuhlmann, Janez Zibert, Fulvia Pennoni, Fabio Divino, Marti Catala, Gianfranco Lovison, Paolo Giudici, Barbara Tarantino, Francesco Bartolucci, Giovanna Jona Lasinio, Marco Mingione, Alessio Farcomeni, Ajitesh Srivastava, Pablo Montero-Manso, Aniruddha Adiga,

Benjamin Hurt, Bryan Lewis, Madhav Marathe, Przemyslaw Porebski, Srinivasan Venkatramanan, Rafal P Bartczuk, Filip Dreger, Anna Gambin, Krzysztof Gogolewski, Magdalena Gruziel-Slomka, Bartosz Krupa, Antoni Moszyński, Karol Niedzielewski, Jedrzej Nowosielski, Maciej Radwan, Franciszek Rakowski, Marcin Semeniuk, Ewa Szczurek, Jakub Zielinski, Jan Kisielewski, Barbara Pabjan, Kirsten Holger, Yuri Kheifetz, Markus Scholz, Biecek Przemyslaw, Marcin Bodych, Maciej Filinski, Radoslaw Idzikowski, Tyll Krueger, Tomasz Ozanski, Johannes Bracher, and Sebastian Funk. Predictive performance of multi-model ensemble forecasts of COVID-19 across European nations. *eLife*, 12, 2023.

[13259] Dasvit Shetty and Linda J Kenney. A pH-sensitive switch activates virulence in Salmonella. *eLife*, 12, 2023.

[13260] Meng Shi, Jiaqi Zhao, Simin Zhang, Wei Huang, Mengfei Li, Xue Bai, Wenxue Zhang, Kai Zhang, Xuefeng Chen, and Song Xiang. Structural basis for the Rad6 activation by the Bre1 N-terminal domain. *eLife*, 12, 2023.

[13261] Tuo Shi, Marielle O Beaulieu, Lauren M Saunders, Peter Fabian, Cole Trapnell, Neil Segil, J Gage Crump, and David W Raible. Single-cell transcriptomic profiling of the zebrafish inner ear reveals molecularly distinct hair cell and supporting cell subtypes. *eLife*, 12, 2023.

[13262] Mitsugu Shimobayashi, Amandine Thomas, Sunil Shetty, Irina C Frei, Bettina K Wölnerhanssen, Diana Weissenberger, Anke Vandekeere, Mélanie Planque, Nikolaus Dietz, Danilo Ritz, Anne Christin Meyer-Gerspach, Timm Maier, Nissim Hay, Ralph Peterli, Sarah-Maria Fendt, Nicolas Rohner, and Michael N Hall. Diet-induced loss of adipose hexokinase 2 correlates with hyperglycemia. *eLife*, 12, 2023.

[13263] Heon Shin, Amy Leung, Kevin R Costello, Parijat Senapati, Hiroyuki Kato, Roger E Moore, Michael Lee, Dimitri Lin, Xiaofang Tang, Patrick Pirrotte, Zhen Bouman Chen, and Dustin E Schones. Inhibition of DNMT1 methyltransferase activity via glucose-regulated O-GlcNAcylation alters the epigenome. *eLife*, 12, 2023.

[13264] Paul Shin, Qi Pian, Hidehiro Ishikawa, Gen Hamanaka, Emiri T Mandeville, Shuzhen Guo, Buyin Fu, Mohammed Alfadhel, Srinivasa Rao Allu, Ikbal şencan Eğilmez, Baoqiang Li, Chongzhao Ran, Sergei A Vinogradov, Cenk Ayata, Eng Lo, Ken Arai, Anna Devor, and Sava Sakadžić. Aerobic exercise reverses aging-induced depth-dependent decline in cerebral microcirculation. *eLife*, 12, 2023.

[13265] Rabina Shrestha, Tess McCann, Harini Saravanan, Jaret Lieberth, Prashanna Koirala, and Joshua Bloomekatz. The myocardium utilizes a platelet-derived growth factor receptor alpha (Pdgfra)–phosphoinositide 3-kinase (PI3K) signaling cascade to steer toward the midline during zebrafish heart tube formation. *eLife*, 12, 2023.

[13266] Ashelyn E Sidders, Katarzyna M Kedziora, Melina Arts, Jan-Martin Daniel, Stefania de Benedetti, Jenna E Beam, Duyen T Bui, Joshua B Parsons, Tanja Schneider, Sarah E Rowe, and Brian P Conlon. Antibiotic-induced accumulation of lipid II synergizes with antimicrobial fatty acids to eradicate bacterial populations. *eLife*, 12, 2023.

[13267] Maheen Siddiqui, Paola Pinti, Sabrina Brigadoi, Sarah Lloyd-Fox, Clare E Elwell, Mark H Johnson, Ilias Tachtsidis, and Emily JH Jones. Using multi-modal neuroimaging to characterise social brain specialisation in infants. *eLife*, 12, 2023.

[13268] Jaydeep Sidhaye, Philipp Trepte, Natalie Sepke, Maria Novatchkova, Michael Schutzbier, Gerhard Dürnberger, Karl Mechtler, and Jürgen A Knoblich. Integrated transcriptome and proteome analysis reveals posttranscriptional regulation of ribosomal genes in human brain organoids. *eLife*, 12, 2023.

[13269] Anthony V Signore, Phillip R Morrison, Colin J Brauner, Angela Fago, Roy E Weber, and Kevin L Campbell. Evolution of an extreme hemoglobin phenotype contributed to the sub-Arctic specialization of extinct Steller's sea cows. *eLife*, 12, 2023.

[13270] Srinjoy Sil, Sarah Keegan, Farida Ettefa, Lance T Denes, Jef D Boeke, and Liam J Holt. Condensation of LINE-1 is critical for retrotransposition. *eLife*, 12, 2023.

[13271] Brian Silver, Kevin Gerrish, and Erik Tokar. Cell-free DNA as a potential biomarker of differentiation and toxicity in cardiac organoids. *eLife*, 12, 2023.

[13272] Patrik Simmler, Eleonora I Ioannidi, Tamara Mengis, Kim Fabiano Marquart, Simran Asawa, Kjong Van-Lehmann, Andre Kahles, Tinu Thomas, Cornelia Schwerdel, Nicola Aceto, Gunnar Rätsch, Markus Stoffel, and Gerald Schwank. Mutant SF3B1 promotes malignancy in PDAC. *eLife*, 12, 2023.

[13273] Márton A Simon, Iordan Iordanov, Andras Szollosi, and László Csanády. Estimating the true stability of the prehydrolytic outward-facing state in an ABC protein. *eLife*, 12, 2023.

[13274] Yosef Singer, Luke Taylor, Ben DB Willmore, Andrew J King, and Nicol S Harper. Hierarchical temporal prediction captures motion processing along the visual pathway. *eLife*, 12, 2023.

[13275] Pragyan Singh, Inbal Gazy, and Martin Kupiec. Control of telomere length in yeast by SUMOylated PCNA and the Elg1 PCNA unloader. *eLife*, 12, 2023.

[13276] Tamar Skaist Mehlman, Justin T Biel, Syeda Maryam Azeem, Elliot R Nelson, Sakib Hossain, Louise Dunnett, Neil G Paterson, Alice Douangamath, Romain Talon, Danny Axford, Helen Orins, Frank von Delft, and Daniel A Keedy. Room-temperature crystallography reveals altered binding of small-molecule fragments to PTP1B. *eLife*, 12, 2023.

[13277] Rastko Sknepnek, Ilyas Djafer-Cherif, Manli Chuai, Cornelis Weijer, and Silke Henkes. Generating active T1 transitions through mechanochemical feedback. *eLife*, 12, 2023.

[13278] Patryk Slusarczyk, Pratik Kumar Mandal, Gabriela Zurawska, Marta Niklewicz, Komal Chouhan, Raghunandan Mahadeva, Aneta Jończy, Matylda Macias, Aleksandra Szybinska, Magdalena Cybulska-Lubak, Olga Krawczyk, Sylwia Herman, Michal Mikula, Remigiusz Serwa, Małgorzata Lenartowicz, Wojciech Pokrzywa, and Katarzyna Mleczko-Sanecka. Impaired iron recycling from erythrocytes is an early hallmark of aging. *eLife*, 12, 2023.

[13279] Anastasia O Smirnova, Anna M Miroshnichenkova, Yulia V Olshanskaya, Michael A Maschan, Yuri B Lebedev, Dmitriy M Chudakov, Ilgar Z Mamedov, and Alexander Komkov. The use of non-functional clonotypes as a natural calibrator for quantitative bias correction in adaptive immune receptor repertoire profiling. *eLife*, 12, 2023.

[13280] Samuel P Smithers, Yulong Shao, James Altham, and Peter J Bex. Large depth differences between target and flankers can increase crowding: Evidence from a multi-depth plane display. *eLife*, 12, 2023.

[13281] Mateo Sokač, Asbjørn Kjær, Lars Dyrskjøt, Benjamin Haibe-Kains, Hugo JWL Aerts, and Nicolai J Birkbak. Spatial transformation of multi-omics data unlocks novel insights into cancer biology. *eLife*, 12, 2023.

[13282] Fangfang Song, Won Dong Lee, Tyler Marmo, Xing Ji, Chao Song, Xueyang Liao, Rebecca Seeley, Lutian Yao, Haoran Liu, and Fanxin Long. Osteoblast-intrinsic defect in glucose metabolism impairs bone formation in type II diabetic male mice. *eLife*, 12, 2023.

[13283] Hayoung Song, Won Mok Shim, and Monica D Rosenberg. Large-scale neural dynamics in a shared low-dimensional state space reflect cognitive and attentional dynamics. *eLife*, 12, 2023.

[13284] Janet Song, Justin T Okano, Joan Ponce, Lesego Busang, Khumo Seipone, Eugenio Valdano, and Sally Blower. The role of migration networks in the development of Botswana's generalized HIV epidemic. *eLife*, 12, 2023.

[13285] Jinyeop Song, Bo-Seong Jeong, Seong-Woo Kim, Seong-Bin Im, Seonghoon Kim, Chih-Jen Lai, Wonki Cho, Jae U Jung, Myung-Ju Ahn, and Byung-Ha Oh. Noncovalent antibody catenation on a target surface greatly increases the antigen-binding avidity. *eLife*, 12, 2023.

[13286] Yoshifumi Sonobe, Soojin Lee, Gopinath Krishnan, Yuanzheng Gu, Deborah Y Kwon, Fen-Biao Gao, Raymond P Roos, and Paschalis Kratsios. Translation of dipeptide repeat proteins in C9ORF72 ALS/FTD through unique and redundant AUG initiation codons. *eLife*, 12, 2023.

[13287] Altar SorkaÇ, Rareș A Moșneanu, Anthony M Crown, Doruk Savaş, Angel M Okoro, Ezgi Memiş, Mustafa Talay, and Gilad Barnea. retro-Tango enables versatile retrograde circuit tracing in Drosophila. *eLife*, 12, 2023.

[13288] Shaghayegh Soudi, Mojtaba Jahani, Marco Todesco, Gregory L Owens, Natalia Bercovich, Loren H Rieseberg, and Sam Yeaman. Repeatability of adaptation in sunflowers reveals that genomic regions harbouring inversions also drive adaptation in species lacking an inversion. *eLife*, 12, 2023.

[13289] Alexander Soutschek and Philippe N Tobler. A process model account of the role of dopamine in intertemporal choice. *eLife*, 12, 2023.

[13290] Breanne Sparta, Nont Kosaisawe, Michael Pargett, Madhura Patankar, Nicholaus DeCuzzi, and John G Albeck. Continuous sensing of nutrients and growth factors by the mTORC1-TFEB axis. *eLife*, 12, 2023.

[13291] John P Spencer, Samuel H Forbes, Sophie Naylor, Vinay P Singh, Kiara Jackson, Sean Deoni, Madhuri Tiwari, and Aarti Kumar. Poor air quality is associated with impaired visual cognition in the first two years of life: A longitudinal investigation. *eLife*, 12, 2023.

[13292] Amy E Spens, Nicholas A Sutliff, Sean R Bennett, Amy E Campbell, and Stephen J Tapscott. Human DUX4 and mouse Dux interact with STAT1 and broadly inhibit interferon-stimulated gene induction. *eLife*, 12, 2023.

[13293] Ean Spielvogel, Sook-Kyung Lee, Shuntai Zhou, Gordon J Lockbaum, Mina Henes, Amy Sondgeroth, Klajdi Kosovrasti, Ellen A Nalivaika, Akbar Ali, Nese Kurt Yilmaz, Celia A Schiffer, and Ronald Swanstrom. Selection of HIV-1 for resistance to fifth-generation protease inhibitors reveals two independent pathways to high-level resistance. *eLife*, 12, 2023.

[13294] George A Spirou, Matthew Kersting, Sean Carr, Bayan Razzaq, Carolyna Yamamoto Alves Pinto, Mariah Dawson, Mark H Ellisman, and Paul B Manis. High-resolution volumetric imaging constrains compartmental models to explore synaptic integration and temporal processing by cochlear nucleus globular bushy cells. *eLife*, 12, 2023.

[13295] Gabriel J Starrett, Kelly Yu, Yelena Golubeva, Petra Lenz, Mary L Piaskowski, David Petersen, Michael Dean, Ajay Israni, Brenda Y Hernandez, Thomas C Tucker, Iona Cheng, Lou Gonsalves, Cyllene R Morris, Shehnaz K Hussain, Charles F Lynch, Reuben S Harris, Ludmila Prokunina-Olsson, Paul S Meltzer, Christopher B Buck, and Eric A Engels. Evidence for virus-mediated oncogenesis in bladder cancers arising in solid organ transplant recipients. *eLife*, 12, 2023.

[13296] Michael R Stehnach, Richard J Henshaw, Sheri A Floge, and Jeffrey S Guasto. Multiplexed microfluidic screening of bacterial chemotaxis. *eLife*, 12, 2023.

[13297] Max L Sterling, Ruben Teunisse, and Bernhard Englitz. Rodent ultrasonic vocal interaction resolved with millimeter precision using hybrid beamforming. *eLife*, 12, 2023.

[13298] Merav Stern, Nicolae Istrate, and Luca Mazzucato. A reservoir of timescales emerges in recurrent circuits with heterogeneous neural assemblies. *eLife*, 12, 2023.

[13299] Zachary C Stevenson, Megan J Moerdyk-Schauwecker, Stephen A Banse, Dhaval S Patel, Hang Lu, and Patrick C Phillips. High-throughput library transgenesis in Caenorhabditis elegans via Transgenic Arrays Resulting in Diversity of Integrated Sequences (TARDIS). *eLife*, 12, 2023.

[13300] Laura Stidsholt, Antoniya Hubancheva, Stefan Greif, Holger R Goerlitz, Mark Johnson, Yossi Yovel, and Peter T Madsen. Echolocating bats prefer a high risk-high gain foraging strategy to increase prey profitability. *eLife*, 12, 2023.

[13301] Sebastian Strauss, Julia Acker, Guido Papa, Daniel Desirò, Florian Schueder, Alexander Borodavka, and Ralf Jungmann. Principles of RNA recruitment to viral ribonucleoprotein condensates in a segmented dsRNA virus. *eLife*, 12, 2023.

[13302] Peter R Strege, Luke M Cowan, Constanza Alcaino, Amelia Mazzone, Christopher A Ahern, Lorin S Milescu, Gianrico Farrugia, and Arthur Beyder. Mechanosensitive pore opening of a prokaryotic voltage-gated sodium channel. *eLife*, 12, 2023.

[13303] Stefan Strütt, Thibaut Sellinger, Sylvain Glémin, Aurélien Tellier, and Stefan Laurent. Joint inference of evolutionary transitions to self-fertilization and demographic history using whole-genome sequences. *eLife*, 12, 2023.

[13304] Alina Studenova, Carina Forster, Denis Alexander Engemann, Tilman Hensch, Christian Sanders, Nicole Mauche, Ulrich Hegerl, Markus Loffler, Arno Villringer, and Vadim Nikulin. Event-related modulation of alpha rhythm explains the auditory P300-evoked response in EEG. *eLife*, 12, 2023.

[13305] Esteban Suárez-Delgado, Maru Orozco-Contreras, Gisela E Rangel-Yescas, and Leon D Islas. Activation-pathway transitions in human voltage-gated proton channels revealed by a non-canonical fluorescent amino acid. *eLife*, 12, 2023.

[13306] Aygul Subkhangulova, Miguel A Gonzalez-Lozano, Alexander JA Groffen, Jan RT van Weering, August B Smit, Ruud F Toonen, and Matthijs Verhage. Tomosyn affects dense core vesicle composition but not exocytosis in mammalian neurons. *eLife*, 12, 2023.

[13307] Karla J Suchacki, Benjamin J Thomas, Yoshiko M Ikushima, Kuan-Chan Chen, Claire Fyfe, Adriana AS Tavares, Richard J Sulston, Andrea Lovdel, Holly J Woodward, Xuan Han, Domenico Mattiucci, Eleanor J Brain, Carlos J Alcaide-Corral, Hiroshi Kobayashi, Gillian A Gray, Phillip D Whitfield, Roland H Stimson, Nicholas M Morton, Alexandra M Johnstone, and William P Cawthorn. The effects of caloric restriction on adipose tissue and metabolic health are sex- and age-dependent. *eLife*, 12, 2023.

[13308] Hoi Ching Suen, Shitao Rao, Alfred Chun Shui Luk, Ruoyu Zhang, Lele Yang, Huayu Qi, Hon Cheong So, Robin M Hobbs, Tin-lap Lee, and Jinyue Liao. The single-cell chromatin accessibility landscape in mouse perinatal testis development. *eLife*, 12, 2023.

[13309] Jini Sugatha, Amulya Priya, Prateek Raj, Ebsy Jaimon, Uma Swaminathan, Anju Jose, Thomas John Pucadyil, and Sunando Datta. Insights into cargo sorting by SNX32 and its role in neurite outgrowth. *eLife*, 12, 2023.

[13310] Wei-Sheng Sun, Lena Lassinantti, Michael Järvå, Andreas Schmitt, Josy ter Beek, and Ronnie P-A Berntsson. Structural foundation for the role of enterococcal PrgB in conjugation, biofilm formation, and virulence. *eLife*, 12, 2023.

[13311] Weiwei Sun, Yelizaveta Rassadkina, Ce Gao, Sarah Isabel Collens, Xiaodong Lian, Isaac H Solomon, Shibani S Mukerji, Xu G Yu, and Mathias Lichterfeld. Persistence of intact HIV-1 proviruses in the brain during antiretroviral therapy. *eLife*, 12, 2023.

[13312] Weizhao Sun, Michelle Ina Lange, Jürgen Gadau, and Jan Buellesbach. Decoding the genetic and chemical basis of sexual attractiveness in parasitic wasps. *eLife*, 12, 2023.

[13313] Xiaomei Sun, Yuxin Ye, Naofumi Sakurai, Hang Wang, Koji Kato, Jian Yu, Keizo Yuasa, Akihiko Tsuji, and Min Yao. Structural basis of EHEP-mediated offense against phlorotannin-induced defense from brown algae to protect akuBGL activity. *eLife*, 12, 2023.

[13314] Xun Sun, Jing Yu, Cheng Zhu, Xinreng Mo, Qiangqiang Sun, Dandan Yang, Chang Su, and Yang Lu. Recognition of galactose by a scaffold protein recruits a transcriptional activator for the GAL regulon induction in Candida albicans. *eLife*, 12, 2023.

[13315] Hayeon Sung, Anoumid Vaziri, Daniel Wilinski, Riley KR Woerner, Lydia Freddolino, and Monica Dus. Nutrigenomic regulation of sensory plasticity. *eLife*, 12, 2023.

[13316] Suhas Sureshchandra, Brianna M Doratt, Norma Mendza, Oleg Varlamov, Monica Rincon, Nicole E Marshall, and Ilhem Messaoudi. Maternal obesity blunts antimicrobial responses in fetal monocytes. *eLife*, 12, 2023.

[13317] Ericka D Surrett, Kirsten R Guckes, Shyan Cousins, Terry B Ruskoski, Andrew G Cecere, Denise A Ludvik, C Denise Okafor, Mark J Mandel, and Tim I Miyashiro. Two enhancer binding proteins activate $\sigma$54-dependent transcription of a quorum regulatory RNA in a bacterial symbiont. *eLife*, 12, 2023.

[13318] Gautam Susarla, Priyanka Kataria, Amrita Kundu, and Patrick D'Silva. Saccharomyces cerevisiae DJ-1 paralogs maintain genome integrity through glycation repair of nucleic acids and proteins. *eLife*, 12, 2023.

[13319] Zachary Susswein, Eva C Rest, and Shweta Bansal. Disentangling the rhythms of human activity in the built environment for airborne transmission risk: An analysis of large-scale mobility data. *eLife*, 12, 2023.

[13320] Anne Sustar and John C Tuthill. Comment on 'A conserved strategy for inducing appendage regeneration in moon jellyfish, Drosophila, and mice'. *eLife*, 12, 2023.

[13321] Jelena Sučević and Anna C Schapiro. A neural network model of hippocampal contributions to category learning. *eLife*, 12, 2023.

[13322] Russell P Swift, Rubayet Elahi, Krithika Rajaram, Hans B Liu, and Sean T Prigge. The Plasmodium falciparum apicoplast cysteine desulfurase provides sulfur for both iron-sulfur cluster assembly and tRNA modification. *eLife*, 12, 2023.

[13323] Sahla Syed, Yifei Duan, and Bomyi Lim. Modulation of protein-DNA binding reveals mechanisms of spatiotemporal gene control in early Drosophila embryos. *eLife*, 12, 2023.

[13324] Anita Szőke, Orsolya Sárkány, Géza Schermann, Orsolya Kapuy, Axel CR Diernfellner, Michael Brunner, Norbert Gyöngyösi, and Krisztina Káldi. Adaptation to glucose starvation is associated with molecular reorganization of the circadian clock in Neurospora crassa. *eLife*, 12, 2023.

[13325] Simon Grund Sørensen, Amruta Shrikhande, Gustav Alexander Poulsgaard, Mikkel Hovden Christensen, Johanna Bertl, Britt Elmedal Laursen, Eva R Hoffmann, and Jakob Skou Pedersen. Pancancer association of DNA repair deficiencies with whole-genome mutational patterns. *eLife*, 12, 2023.

[13326] Hirohide Takahashi, Toshiki Yamada, Jerod S Denton, Kevin Strange, and Erkan Karakas. Cryo-EM structures of an LRRC8 chimera with native functional properties reveal heptameric assembly. *eLife*, 12, 2023.

[13327] Valeria Taliani, Giulia Buonaiuto, Fabio Desideri, Adriano Setti, Tiziana Santini, Silvia Galfrè, Leonardo Schirone, Davide Mariani, Giacomo Frati, Valentina Valenti, Sebastiano Sciarretta, Emerald Perlas, Carmine Nicoletti, Antonio Musarò, and Monica Ballarino. The long noncoding RNA Charme supervises cardiomyocyte maturation by controlling cell differentiation programs in the developing heart. *eLife*, 12, 2023.

[13328] Stephanie W Tam, Kate Huffer, Mufeng Li, and Kenton J Swartz. Ion permeation pathway within the internal pore of P2X receptor channels. *eLife*, 12, 2023.

[13329] Eshwar R Tammineni, Lourdes Figueroa, Carlo Manno, Disha Varma, Natalia Kraeva, Carlos A Ibarra, Amira Klip, Sheila Riazi, and Eduardo Rios. Muscle calcium stress cleaves junctophilin1, unleashing a gene regulatory program predicted to correct glucose dysregulation. *eLife*, 12, 2023.

[13330] Hui Zhen Tan, Justin JFJ Jansen, Gary A Allport, Kritika M Garg, Balaji Chattopadhyay, Martin Irestedt, Sean EH Pang, Glen Chilton, Chyi Yin Gwee, and Frank E Rheindt. Megafaunal extinctions, not climate change, may explain Holocene genetic diversity declines in Numenius shorebirds. *eLife*, 12, 2023.

[13331] Zhenyu Tan, Yang Yue, Felipe Leprevost, Sarah Haynes, Venkatesha Basrur, Alexey I Nesvizhskii, Kristen J Verhey, and Michael A Cianfrocco. Autoinhibited kinesin-1 adopts a hierarchical folding pattern. *eLife*, 12, 2023.

[13332] Zhi Yang Tan, Shujun Cai, Alex J Noble, Jon K Chen, Jian Shi, and Lu Gan. Heterogeneous non-canonical nucleosomes predominate in yeast cells in situ. *eLife*, 12, 2023.

[13333] Annick FN Tanguay, Daniela J Palombo, Brittany Love, Rafael Glikstein, Patrick SR Davidson, and Louis Renoult. The shared and unique neural correlates of personal semantic, general semantic, and episodic memory. *eLife*, 12, 2023.

[13334] Helen Taylor, Arash Zaghi, and Sara Rankin. Marginalising dyslexic researchers is bad for science. *eLife*, 12, 2023.

[13335] Samuel JP Taylor, Laura Bel Borja, Flavie Soubigou, Jack Houston, Dhanya K Cheerambathur, and Federico Pelisch. BUB-1 and CENP-C recruit PLK-1 to control chromosome alignment and segregation during meiosis I in C. elegans oocytes. *eLife*, 12, 2023.

[13336] Seth R Taylor, Mariko Kobayashi, Antonietta Vilella, Durgesh Tiwari, Norjin Zolboot, Jessica X Du, Kathryn R Spencer, Andrea Hartzell, Carol Girgiss, Yusuf T Abaci, Yufeng Shao, Claudia De Sanctis, Gian Carlo Bellenchi, Robert B Darnell, Christina Gross, Michele Zoli, Darwin K Berg, and Giordano Lippi. MicroRNA-218 instructs proper assembly of hippocampal networks. *eLife*, 12, 2023.

[13337] Nydia Tejeda-Munoz, Yagmur Azbazdar, Julia Monka, Grace Binder, Alex Dayrit, Raul Ayala, Neil O'Brien, and Edward M De Robertis. The PMA phorbol ester tumor promoter increases canonical Wnt signaling via macropinocytosis. *eLife*, 12, 2023.

[13338] Godfrey S Temba, Nadira Vadaq, Vesla Kullaya, Tal Pecht, Paolo Lionetti, Duccio Cavalieri, Joachim L Schultze, Reginald Kavishe, Leo AB Joosten, Andre J van der Ven, Blandina T Mmbaga, Mihai G Netea, and Quirijn de Mast. Differences in the inflammatory proteome of East African and Western European adults and associations with environmental and dietary factors. *eLife*, 12, 2023.

[13339] Benjamin C Tendler, Maddie Welland, Karla L Miller, and Melanie Alexis-Butler. Why every lab needs a handbook. *eLife*, 12, 2023.

[13340] Bochuan Teng, Joshua P Kaplan, Ziyu Liang, Kevin Saejin Chyung, Marcel P Goldschen-Ohm, and Emily R Liman. Zinc activation of OTOP proton channels identifies structural elements of the gating apparatus. *eLife*, 12, 2023.

[13341] Dmitry Ter-Ovanesyan, Tal Gilboa, Bogdan Budnik, Adele Nikitina, Sara Whiteman, Roey Lazarovits, Wendy Trieu, David Kalish, George M Church, and David R Walt. Improved isolation of extracellular vesicles by removal of both free proteins and lipoproteins. *eLife*, 12, 2023.

[13342] Mügen Terzioglu, Kristo Veeroja, Toni Montonen, Teemu O Ihalainen, Tiina S Salminen, Paule Bénit, Pierre Rustin, Young-Tae Chang, Takeharu Nagai, and Howard T Jacobs. Mitochondrial temperature homeostasis resists external metabolic stresses. *eLife*, 12, 2023.

[13343] Filiz Tezcan, Hugo Weissbart, and Andrea E Martin. A tradeoff between acoustic and linguistic feature encoding in spoken language comprehension. *eLife*, 12, 2023.

[13344] Astha Thakkar, Kith Pradhan, Benjamin Duva, Juan Manuel Carreno, Srabani Sahu, Victor Thiruthuvanathan, Sean Campbell, Sonia Gallego, Tushar D Bhagat, Johanna Rivera, Gaurav Choudhary, Raul Olea, Maite Sabalza, Lauren C Shapiro, Matthew Lee, Ryann Quinn, Ioannis Mantzaris, Edward Chu, Britta Will, Liise-anne Pirofski, Florian Krammer, Amit Verma, and Balazs Halmos. Study of efficacy and longevity of immune response to third and fourth doses of COVID-19 vaccines in patients with cancer: A single arm clinical trial. *eLife*, 12, 2023.

[13345] Friederike Thams and Stefanie Brassen. The need to change: Is there a critical role of midlife adaptation in mental health later in life? *eLife*, 12, 2023.

[13346] Robin Thandiackal and George Lauder. In-line swimming dynamics revealed by fish interacting with a robotic mechanism. *eLife*, 12, 2023.

[13347] Keshav Thapa, James J Kadiri, Karla Saukkonen, Iida Pennanen, Bishwa Ghimire, Minying Cai, Eriika Savontaus, and Petteri Rinne. Melanocortin 1 receptor regulates cholesterol and bile acid metabolism in the liver. *eLife*, 12, 2023.

[13348] Yuttapong Thawornwattana, Fernando Seixas, Ziheng Yang, and James Mallet. Major patterns in the introgression history of Heliconius butterflies. *eLife*, 12, 2023.

[13349] Alexandre P Thiery, Ailin Leticia Buzzi, Eva Hamrud, Chris Cheshire, Nicholas M Luscombe, James Briscoe, and Andrea Streit. scRNA-sequencing in chick suggests a probabilistic model for cell fate allocation at the neural plate border. *eLife*, 12, 2023.

[13350] Connon I Thomas, Melissa A Ryan, Naomi Kamasawa, and Benjamin Scholl. Postsynaptic mitochondria are positioned to support functional diversity of dendritic spines. *eLife*, 12, 2023.

[13351] F Emil Thomasen, Matthew J Cuneo, Tanja Mittag, and Kresten Lindorff-Larsen. Conformational and oligomeric states of SPOP from small-angle X-ray scattering and molecular dynamics simulations. *eLife*, 12, 2023.

[13352] Basil T Thommen, Jerzy M Dziekan, Fiona Achcar, Seth Tjia, Armin Passecker, Katarzyna Buczak, Christin Gumpp, Alexander Schmidt, Matthias Rottmann, Christof Grüring, Matthias Marti, Zbynek Bozdech, and Nicolas MB Brancucci. Genetic validation of PfFKBP35 as an antimalarial drug target. *eLife*, 12, 2023.

[13353] Jeffrey Thompson, Yidi Wang, Tobias Dreischulte, Olga Barreiro, Rodrigo J Gonzalez, Pavel Hanč, Colette Matysiak, Harold R Neely, Marietta Rottenkolber, Thomas Haskell, Stefan Endres, and Ulrich H von Andrian. Association between bisphosphonate use and COVID-19 related outcomes. *eLife*, 12, 2023.

[13354] Jakob Thyrring, Colin D Macleod, Katie E Marshall, Jessica Kennedy, Réjean Tremblay, and Christopher DG Harley. Ocean acidification increases susceptibility to sub-zero air temperatures in ecosystem engineers and limits poleward range shifts. *eLife*, 12, 2023.

[13355] Lucas Y Tian, Timothy L Warren, William H Mehaffey, and Michael S Brainard. Dynamic top-down biasing implements rapid adaptive changes to individual movements. *eLife*, 12, 2023.

[13356] Qiaomu Tian, Peng Zhang, Yihan Wang, Youhui Si, Dengping Yin, Christopher R Weber, Melissa L Fishel, Karen E Pollok, Bo Qiu, Fei Xiao, and Anita S Chong. A novel triptolide analog downregulates NF-$\kappa$B and induces mitochondrial apoptosis pathways in human pancreatic cancer. *eLife*, 12, 2023.

[13357] Ruben A Tikidji-Hamburyan, Gubbi Govindaiah, William Guido, and Matthew T Colonnese. Synaptic and circuit mechanisms prevent detrimentally precise correlation in the developing mammalian visual system. *eLife*, 12, 2023.

[13358] David N Tippett, Colum Breen, Stephen J Butler, Marta Sawicka, and Raimund Dutzler. Structural and functional properties of the transporter SLC26A6 reveal mechanism of coupled anion exchange. *eLife*, 12, 2023.

[13359] Wilma Tixi, Maricela Maldonado, Ya-Ting Chang, Amy Chiu, Wilson Yeung, Nazia Parveen, Michael S Nelson, Ryan Hart, Shihao Wang, Wu Jih Hsu, Patrick Fueger, Janel L Kopp, Mark O Huising, Sangeeta Dhawan, and Hung Ping Shih. Coordination between ECM and cell-cell adhesion regulates the development of islet aggregation, architecture, and functional maturation. *eLife*, 12, 2023.

[13360] Elena Tomas Bort, Megan D Joseph, Qiaoying Wang, Edward P Carter, Nicolas J Roth, Jessica Gibson, Ariana Samadi, Hemant M Kocher, Sabrina Simoncelli, Peter J McCormick, and Richard P Grose. Purinergic GPCR-integrin interactions drive pancreatic cancer cell invasion. *eLife*, 12, 2023.

[13361] Francesca G Tomasi, Satoshi Kimura, Eric J Rubin, and Matthew K Waldor. A tRNA modification in Mycobacterium tuberculosis facilitates optimal intracellular growth. *eLife*, 12, 2023.

[13362] Parker Tope, Eliya Farah, Rami Ali, Mariam El-Zein, Wilson H Miller, and Eduardo L Franco. The impact of lag time to cancer diagnosis and treatment on clinical outcomes prior to the COVID-19 pandemic: A scoping review of systematic reviews and meta-analyses. *eLife*, 12, 2023.

[13363] Guillem Torcal Garcia, Elisabeth Kowenz-Leutz, Tian V Tian, Antonis Klonizakis, Jonathan Lerner, Luisa De Andres-Aguayo, Valeriia Sapozhnikova, Clara Berenguer, Marcos Plana Carmona, Maria Vila Casadesus, Romain Bulteau, Mirko Francesconi, Sandra Peiro, Philipp Mertins, Kenneth Zaret, Achim Leutz, and Thomas Graf. Carm1-arginine methylation of the transcription factor C/EBP$\alpha$ regulates transdifferentiation velocity. *eLife*, 12, 2023.

[13364] Valerie A Tornini, Liyun Miao, Ho-Joon Lee, Timothy Gerson, Sarah E Dube, Valeria Schmidt, FranÇois Kroll, Yin Tang, Katherine Du, Manik Kuchroo, Charles E Vejnar, Ariel Alejandro Bazzini, Smita Krishnaswamy, Jason Rihel, and Antonio J Giraldez. linc-mipep and linc-wrb encode micropeptides that regulate chromatin accessibility in vertebrate-specific neural cells. *eLife*, 12, 2023.

[13365] David J Torres, Paulus Mrass, Janie Byrum, Arrick Gonzales, Dominick N Martinez, Evelyn Juarez, Emily Thompson, Vaiva Vezys, Melanie E Moses, and Judy L Cannon. Quantitative analyses of T cell motion in tissue reveals factors driving T cell search in tissues. *eLife*, 12, 2023.

[13366] Arturo Torres Ortiz, Michelle Kendall, Nathaniel Storey, James Hatcher, Helen Dunn, Sunando Roy, Rachel Williams, Charlotte Williams, Richard A Goldstein, Xavier Didelot, Kathryn Harris, Judith Breuer, and Louis Grandjean. Within-host diversity improves phylogenetic and transmission reconstruction of SARS-CoV-2 outbreaks. *eLife*, 12, 2023.

[13367] Abdoulie O Touray, Rishi Rajesh, Tony Isebe, Tamara Sternlieb, Mira Loock, Oksana Kutova, and Igor Cestari. PI(3,4,5)P3 allosteric regulation of repressor activator protein 1 controls antigenic variation in trypanosomes. *eLife*, 12, 2023.

[13368] Thach Tran, Thao Ho-Le, Dana Bliuc, Bo Abrahamsen, Louise Hansen, Peter Vestergaard, Jacqueline R Center, and Tuan V Nguyen. 'Skeletal Age' for mapping the impact of fracture on mortality. *eLife*, 12, 2023.

[13369] Janine Traut, Jose Prius Mengual, Elise J Meijer, Laura E McKillop, Hannah Alfonsa, Anna Hoerder-Suabedissen, Seo Ho Song, Kristoffer D Fehér, Dieter Riemann, Zoltan Molnar, Colin J Akerman, Vladyslav V Vyazovskiy, and Lukas B Krone. Effects of clozapine-N-oxide and compound 21 on sleep in laboratory mice. *eLife*, 12, 2023.

[13370] Michaela E Trautman, Leah N Braucher, Christian Elliehausen, Wenyuan G Zhu, Esther Zelenovskiy, Madelyn Green, Michelle M Sonsalla, Chung-Yang Yeh, Troy A Hornberger, Adam R Konopka, and Dudley W Lamming. Resistance exercise protects mice from protein-induced fat accretion. *eLife*, 12, 2023.

[13371] Amanda M Travis, Samiya Manocha, Jason R Willer, Timothy S Wessler, Nikolai P Skiba, and Jillian N Pearring. Disrupting the ciliary gradient of active Arl3 affects rod photoreceptor nuclear migration. *eLife*, 12, 2023.

[13372] Tarah Elizabeth Trebino, Borna Markusic, Haihan Nan, Shrhea Banerjee, and Zhihong Wang. Unveiling the domain-specific and RAS isoform-specific details of BRAF kinase regulation. *eLife*, 12, 2023.

[13373] Katherine E Trevers, Hui-Chun Lu, Youwen Yang, Alexandre P Thiery, Anna C Strobl, Claire Anderson, Božena PálinkáŠová, Nidia MM de Oliveira, Irene M de Almeida, Mohsin AF Khan, Natalia Moncaut, Nicholas M Luscombe, Leslie Dale, Andrea Streit, and Claudio D Stern. A gene regulatory network for neural induction. *eLife*, 12, 2023.

[13374] Federico Trigo and Shin-ya Kawaguchi. Analogue signaling of somatodendritic synaptic activity to axon enhances GABA release in young cerebellar molecular layer interneurons. *eLife*, 12, 2023.

[13375] James W Truman, Jacquelyn Price, Rosa L Miyares, and Tzumin Lee. Metamorphosis of memory circuits in Drosophila reveals a strategy for evolving a larval brain. *eLife*, 12, 2023.

[13376] Tim K Tsang, Xiaotong Huang, Can Wang, Sijie Chen, Bingyi Yang, Simon Cauchemez, and Benjamin John Cowling. The effect of variation of individual infectiousness on SARS-CoV-2 transmission in households. *eLife*, 12, 2023.

[13377] Oksana Tsyklauri, Tereza Chadimova, Veronika Niederlova, Jirina Kovarova, Juraj Michalik, Iva Malatova, Sarka Janusova, Olha Ivashchenko, Helene Rossez, Ales Drobek, Hana Vecerova, Virginie Galati, Marek Kovar, and Ondrej Stepanek. Regulatory T cells suppress the formation of potent KLRK1 and IL-7R expressing effector CD8 T cells by limiting IL-2. *eLife*, 12, 2023.

[13378] Leo Uesaka, Yusuke Goto, Masaru Naruoka, Henri Weimerskirch, Katsufumi Sato, and Kentaro Q Sakamoto. Wandering albatrosses exert high take-off effort only when both wind and waves are gentle. *eLife*, 12, 2023.

[13379] Rupali Ugrankar-Banerjee, Son Tran, Jade Bowerman, Anastasiia Kovalenko, Blessy Paul, and W Mike Henne. The fat body cortical actin network regulates Drosophila inter-organ nutrient trafficking, signaling, and adipose cell size. *eLife*, 12, 2023.

[13380] Lara Urban, Allison K Miller, Daryl Eason, Deidre Vercoe, Megan Shaffer, Shaun P Wilkinson, Gert-Jan Jeunen, Neil J Gemmell, and Andrew Digby. Non-invasive real-time genomic monitoring of the critically endangered kākāpō. *eLife*, 12, 2023.

[13381] Masayuki Ushio, Hiroki Saito, Motoaki Tojo, and Atsushi J Nagano. An ecological network approach for detecting and validating influential organisms for rice growth. *eLife*, 12, 2023.

[13382] Vera Valakh, Derek Wise, Xiaoyue Aelita Zhu, Mingqi Sha, Jaidyn Fok, Stephen D Van Hooser, Robin Schectman, Isabel Cepeda, Ryan Kirk, Sean M O'Toole, and Sacha B Nelson. A transcriptional constraint mechanism limits the homeostatic response to activity deprivation in mammalian neocortex. *eLife*, 12, 2023.

[13383] Sofie Louise Valk, Philipp Kanske, Bo-yong Park, Seok-Jun Hong, Anne Böckler, Fynn-Mathis Trautwein, Boris C Bernhardt, and Tania Singer. Functional and microstructural plasticity following social and interoceptive mental training. *eLife*, 12, 2023.

[13384] Miguel Vallebueno-Estrada, Guillermo G Hernández-Robles, Eduardo González-Orozco, Ivan Lopez-Valdivia, Teresa Rosales Tham, Víctor Vásquez Sánchez, Kelly Swarts, Tom D Dillehay, Jean-Philippe Vielle-Calzada, and Rafael Montiel. Domestication and lowland adaptation of coastal preceramic maize from Paredones, Peru. *eLife*, 12, 2023.

[13385] Daan F van den Berg, Baltus A van der Steen, Ana Rita Costa, and Stan JJ Brouns. Phage tRNAs evade tRNA-targeting host defenses through anticodon loop mutations. *eLife*, 12, 2023.

[13386] Ruben van den Bosch, Frank H Hezemans, Jessica I Määttä, Lieke Hofmans, Danae Papadopetraki, Robbert-Jan Verkes, Andre F Marquand, Jan Booij, and Roshan Cools. Evidence for absence of links between striatal dopamine synthesis capacity and working memory capacity, spontaneous eye-blink rate, and trait impulsivity. *eLife*, 12, 2023.

[13387] Thijs L van der Plas, Jérôme Tubiana, Guillaume Le Goc, Geoffrey Migault, Michael Kunst, Herwig Baier, Volker Bormuth, Bernhard Englitz, and Georges Debrégeas. Neural assemblies uncovered by generative modeling explain whole-brain activity statistics and reflect structural connectivity. *eLife*, 12, 2023.

[13388] Jenny van Dongen, Gonneke Willemsen, Bastiaan T Heijmans, Eco JC de Geus, Dorret I Boomsma, and Michael C Neale. Effects of smoking on genome-wide DNA methylation profiles: A study of discordant and concordant monozygotic twin pairs. *eLife*, 12, 2023.

[13389] Laura C Van Eyndhoven, Vincent PG Verberne, Carlijn VC Bouten, Abhyudai Singh, and Jurjen Tel. Transiently heritable fates and quorum sensing drive early IFN-I response dynamics. *eLife*, 12, 2023.

[13390] Jasper van Oort, Alberto Llera, Nils Kohn, Ting Mei, Rose M Collard, Fleur A Duyser, Janna N Vrijsen, Christian F Beckmann, Aart H Schene, Guillén Fernández, Indira Tendolkar, and Philip FP van Eijndhoven. Brain structure and function link to variation in biobehavioral dimensions across the psychopathological continuum. *eLife*, 12, 2023.

[13391] Koen Vandelannoote, Andrew H Buultjens, Jessica L Porter, Anita Velink, John R Wallace, Kim R Blasdell, Michael Dunn, Victoria Boyd, Janet AM Fyfe, Ee Laine Tay, Paul DR Johnson, Saras M Windecker, Nick Golding, and Timothy P Stinear. Statistical modeling based on structured surveys of Australian native possum excreta harboring Mycobacterium ulcerans predicts Buruli ulcer occurrence in humans. *eLife*, 12, 2023.

[13392] Sindhu Vangeti, Sara Falck-Jones, Meng Yu, Björn österberg, Sang Liu, Muhammad Asghar, Klara Sondén, Clare Paterson, Penn Whitley, Jan Albert, Niclas Johansson, Anna Färnert, and Anna Smed-Sörensen. Human influenza virus infection elicits distinct patterns of monocyte and dendritic cell mobilization in blood and the nasopharynx. *eLife*, 12, 2023.

[13393] Louiza S Velentzis, Megan A Smith, James Killen, Julia ML Brotherton, Rebecca Guy, and Karen Canfell. A modelled analysis of the impact of COVID-19-related disruptions to HPV vaccination. *eLife*, 12, 2023.

[13394] Aarya Venkat, Grace Watterson, Dominic P Byrne, Brady O'Boyle, Safal Shrestha, Nathan Gravel, Emma E Fairweather, Leonard A Daly, Claire Bunn, Wayland Yeung, Ishan Aggarwal, Samiksha Katiyar, Claire E Eyers, Patrick A Eyers, and Natarajan Kannan. Mechanistic and evolutionary insights into isoform-specific 'supercharging' in DCLK family kinases. *eLife*, 12, 2023.

[13395] Krithika Venkataraman, Nadav Shai, Priyanka Lakhiani, Sarah Zylka, Jieqing Zhao, Margaret Herre, Joshua Zeng, Lauren A Neal, Henrik Molina, Li Zhao, and Leslie B Vosshall. Two novel, tightly linked, and rapidly evolving genes underlie Aedes aegypti mosquito reproductive resilience during drought. *eLife*, 12, 2023.

[13396] Roxane Verdikt, Abigail A Armstrong, Jenny Cheng, Young Sun Hwang, Amander T Clark, Xia Yang, and Patrick Allard. Metabolic memory of $\delta$9-tetrahydrocannabinol exposure in pluripotent stem cells and primordial germ cells-like cells. *eLife*, 12, 2023.

[13397] Dániel Veréb, Mite Mijalkov, Anna Canal-Garcia, Yu-Wei Chang, Emiliano Gomez-Ruiz, Blanca Zufiria Gerboles, Miia Kivipelto, Per Svenningsson, Henrik Zetterberg, Giovanni Volpe, Matthew Betts, Heidi IL Jacobs, and Joana B Pereira. Age-related differences in the functional topography of the locus coeruleus and their implications for cognitive and affective functions. *eLife*, 12, 2023.

[13398] Jeroen PH Verharen. ChatGPT identifies gender disparities in scientific peer review. *eLife*, 12, 2023.

[13399] Niels JM Verstegen, Sabrina Pollastro, Peter-Paul A Unger, Casper Marsman, George Elias, Tineke Jorritsma, Marij Streutker, Kevin Bassler, Kristian Haendler, Theo Rispens, Joachim L Schultze, Anja ten Brinke, Marc Beyer, and S Marieke van Ham. Single-cell analysis reveals dynamics of human B cell differentiation and identifies novel B and antibody-secreting cell intermediates. *eLife*, 12, 2023.

[13400] Ben Vezina, Stephen C Watts, Jane Hawkey, Helena B Cooper, Louise M Judd, Adam WJ Jenney, Jonathan M Monk, Kathryn E Holt, and Kelly L Wyres. Bactabolize is a tool for high-throughput generation of bacterial strain-specific metabolic models. *eLife*, 12, 2023.

[13401] Maria Vias, Lena Morrill Gavarró, Carolin M Sauer, Deborah A Sanders, Anna M Piskorz, Dominique-Laurent Couturier, Stéphane Ballereau, Bárbara Hernando, Michael P Schneider, James Hall, Filipe Correia-Martins, Florian Markowetz, Geoff Macintyre, and James D Brenton. High-grade serous ovarian carcinoma organoids as models of chromosomal instability. *eLife*, 12, 2023.

[13402] Guillaume Viejo, Daniel Levenstein, Sofia Skromne Carrasco, Dhruv Mehrotra, Sara Mahallati, Gilberto R Vite, Henry Denny, Lucas Sjulson, Francesco P Battaglia, and Adrien Peyrache. Pynapple, a toolbox for data analysis in neuroscience. *eLife*, 12, 2023.

[13403] Ilja Vietor, Domagoj Cikes, Kati Piironen, Theodora Vasakou, David Heimdörfer, Ronald Gstir, Matthias David Erlacher, Ivan Tancevski, Philipp Eller, Egon Demetz, Michael W Hess, Volker Kuhn, Gerald Degenhart, Jan Rozman, Martin Klingenspor, Martin Hrabe de Angelis, Taras Valovka, and Lukas A Huber. The negative adipogenesis regulator Dlk1 is transcriptionally regulated by Ifrd1 (TIS7) and translationally by its orthologue Ifrd2 (SKMc15). *eLife*, 12, 2023.

[13404] Stefanny Villalobos-Cantor, Ruth M Barrett, Alec F Condon, Alicia Arreola-Bustos, Kelsie M Rodriguez, Michael S Cohen, and Ian Martin. Rapid cell type-specific nascent proteome labeling in Drosophila. *eLife*, 12, 2023.

[13405] Hélène Vitet, Julie Bruyère, Hao Xu, Claire Séris, Jacques Brocard, Yah-Sé Abada, Benoît Delatour, Chiara Scaramuzzino, Laurent Venance, and Frédéric Saudou. Huntingtin recruits KIF1A to transport synaptic vesicle precursors along the mouse axon to support synaptic transmission and motor skill learning. *eLife*, 12, 2023.

[13406] Christiane Voufo, Andy Quaen Chen, Benjamin E Smith, Rongshan Yan, Marla B Feller, and Alexandre Tiriac. Circuit mechanisms underlying embryonic retinal waves. *eLife*, 12, 2023.

[13407] Emily K Vraga, Sonya S Brady, Chloe Gansen, Euna Mehnaz Khan, Sarah L Bennis, Madalyn Nones, Rongwei Tang, Jaideep Srivastava, and Shalini Kulasingam. A review of HPV and HBV vaccine hesitancy, intention, and uptake in the era of social media and COVID-19. *eLife*, 12, 2023.

[13408] Ziva Vuckovic, Jinan Wang, Vi Pham, Jesse I Mobbs, Matthew J Belousoff, Apurba Bhattarai, Wessel AC Burger, Geoff Thompson, Mahmuda Yeasmin, Vindhya Nawaratne, Katie Leach, Emma T van der Westhuizen, Elham Khajehali, Yi-Lynn Liang, Alisa Glukhova, Denise Wootten, Craig W Lindsley, Andrew Tobin, Patrick Sexton, Radostin Danev, Celine Valant, Yinglong Miao, Arthur Christopoulos, and David M Thal. Pharmacological hallmarks of allostery at the M4 muscarinic receptor elucidated through structure and dynamics. *eLife*, 12, 2023.

[13409] Elizabeth L Wagner, Jun-Sub Im, Stefano Sala, Maura I Nakahata, Terence E Imbery, Sihan Li, Daniel Chen, Katherine Nimchuk, Yael Noy, David W Archer, Wenhao Xu, George Hashisaki, Karen B Avraham, Patrick W Oakes, and Jung-Bum Shin. Repair of noise-induced damage to stereocilia F-actin cores is facilitated by XIRP2 and its novel mechanosensor domain. *eLife*, 12, 2023.

[13410] Rohan S Wakade, Laura C Ristow, Melanie Wellington, and Damian J Krysan. Intravital imaging-based genetic screen reveals the transcriptional network governing Candida albicans filamentation during mammalian infection. *eLife*, 12, 2023.

[13411] Lucas M Walker, James RJ Haycocks, Julia C Van Kessel, Triana N Dalia, Ankur B Dalia, and David C Grainger. A simple mechanism for integration of quorum sensing and cAMP signalling in Vibrio cholerae. *eLife*, 12, 2023.

[13412] Klara M Wanelik, Mike Begon, Janette E Bradley, Ida M Friberg, Joseph A Jackson, Christopher H Taylor, and Steve Paterson. Effects of an IgE receptor polymorphism acting on immunity, susceptibility to infection, and reproduction in a wild rodent. *eLife*, 12, 2023.

[13413] Bo Wang, Zheyong Liang, Tan Tan, Miao Zhang, Yina Jiang, Yangyang Shang, Xiaoqian Gao, Shaoran Song, Ruiqi Wang, He Chen, Jie Liu, Juan Li, Yu Ren, and Peijun Liu. CRB3 navigates Rab11 trafficking vesicles to promote $\gamma$TuRC assembly during ciliogenesis. *eLife*, 12, 2023.

[13414] Chaoming Wang, Tianqiu Zhang, Xiaoyu Chen, Sichao He, Shangyang Li, and Si Wu. BrainPy, a flexible, integrative, efficient, and extensible framework for general-purpose brain dynamics programming. *eLife*, 12, 2023.

[13415] Dong Wang, Tiemei Zheng, Songlin Zhou, Mingwen Liu, Yaobo Liu, Xiaosong Gu, Susu Mao, and Bin Yu. Promoting axon regeneration by inhibiting RNA N6-methyladenosine demethylase ALKBH5. *eLife*, 12, 2023.

[13416] Dongyang Wang, Xiaojing Zheng, Lihong Chai, Junli Zhao, Jiuling Zhu, Yanqing Li, Peiyan Yang, Qinwen Mao, and Haibin Xia. FAM76B regulates NF-$\kappa$B-mediated inflammatory pathway by influencing the translocation of hnRNPA2B1. *eLife*, 12, 2023.

[13417] Fan Wang, Yuge Chen, Yuxin Lin, Xuze Wang, Kaiyuan Li, Yong Han, Jintao Wu, Xingyi Shi, Zhenggang Zhu, Chaoying Long, Xiaojun Hu, Shumin Duan, and Zhihua Gao. A parabrachial to hypothalamic pathway mediates defensive behavior. *eLife*, 12, 2023.

[13418] Fan Wang, Han Zhang, Zhengwang Wu, Dan Hu, Zhen Zhou, Jessica B Girault, Li Wang, Weili Lin, and Gang Li. Fine-grained functional parcellation maps of the infant cerebral cortex. *eLife*, 12, 2023.

[13419] Haisheng Wang, Xiaofeng Wang, Liuyan Huang, Chenglin Wang, Fanyuan Yu, and Ling Ye. Overburdened ferroptotic stress impairs tooth morphogenesis. *eLife*, 12, 2023.

[13420] Huan Wang, Syed RA Rizvi, Danni Dong, Jiaqi Lou, Qian Wang, Watanyoo Sopipong, Yufeng Su, Fares Najar, Pratul K Agarwal, Frank Kozielski, and Shozeb Haider. Emerging variants of SARS-CoV-2 NSP10 highlight strong functional conservation of its binding to two non-structural proteins, NSP14 and NSP16. *eLife*, 12, 2023.

[13421] Jie Wang, Amir Rattner, and Jeremy Nathans. Bacterial meningitis in the early postnatal mouse studied at single-cell resolution. *eLife*, 12, 2023.

[13422] Jing Wang, Hamid Azimi, Yilei Zhao, Melanie Kaeser, Pilar Vaca Sánchez, Abraham Vazquez-Guardado, John A Rogers, Michael Harvey, and Gregor Rainer. Optogenetic activation of visual thalamus generates artificial visual percepts. *eLife*, 12, 2023.

[13423] Jinyang Wang, Yaxin Wen, Wentao Zhao, Yan Zhang, Furong Lin, Cong Ouyang, Huihui Wang, Lizheng Yao, Huanhuan Ma, Yue Zhuo, Huiying Huang, Xiulin Shi, Liubin Feng, Donghai Lin, Bin Jiang, and Qinxi Li. Hepatic conversion of acetyl-CoA to acetate plays crucial roles in energy stress. *eLife*, 12, 2023.

[13424] Mingkun Wang, Belle Yanyu Lin, Shuofei Sun, Charles Dai, Feifei Long, and Jonathan T Butcher. Shear and hydrostatic stress regulate fetal heart valve remodeling through YAP-mediated mechanotransduction. *eLife*, 12, 2023.

[13425] Nan Wang, Jing He, Xiaoyu Feng, Shengyou Liao, Yi Zhao, Fuchou Tang, and Kehkooi Kee. Single-cell profiling of lncRNAs in human germ cells and molecular analysis reveals transcriptional regulation of LNC1845 on LHX8. *eLife*, 12, 2023.

[13426] Shumin Wang, Eri O Maruyama, John Martinez, Justin Lopes, Trunee Hsu, Wencheng Wu, Wei Hsu, and Takamitsu Maruyama. miRNA-27a is essential for bone remodeling by modulating p62-mediated osteoclast signaling. *eLife*, 12, 2023.

[13427] Xiang Wang, Vitaliy V Bondar, Oliver B Davis, Michael T Maloney, Maayan Agam, Marcus Y Chin, Audrey Cheuk-Nga Ho, Rajarshi Ghosh, Dara E Leto, David Joy, Meredith EK Calvert, Joseph W Lewcock, Gilbert Di Paolo, Robert G Thorne, Zachary K Sweeney, and Anastasia G Henry. Rab12 is a regulator of LRRK2 and its activation by damaged lysosomes. *eLife*, 12, 2023.

[13428] Xiaosha Wang, Bijun Wang, and Yanchao Bi. Early language exposure affects neural mechanisms of semantic representations. *eLife*, 12, 2023.

[13429] Xin-Tai Wang, Lin Zhou, Bin-Bin Dong, Fang-Xiao Xu, De-Juan Wang, En-Wei Shen, Xin-Yu Cai, Yin Wang, Na Wang, Sheng-Jian Ji, Wei Chen, Martijn Schonewille, J Julius Zhu, Chris I De Zeeuw, and Ying Shen. cAMP-EPAC-PKC$\epsilon$-RIM1$\alpha$ signaling regulates presynaptic long-term potentiation and motor learning. *eLife*, 12, 2023.

[13430] Yaoxuan Wang, Mengda Jiang, Yuting Zhu, Lu Xue, Wenying Shu, Xiang Li, Hongsai Chen, Yun Li, Ying Chen, Yongchuan Chai, Yu Zhang, Yinghua Chu, Yang Song, Xiaofeng Tao, Zhaoyan Wang, and Hao Wu. Impact of inner ear malformation and cochlear nerve deficiency on the development of auditory-language network in children with profound sensorineural hearing loss. *eLife*, 12, 2023.

[13431] Yu Wang, Meghan Lee Arnold, Anna Joelle Smart, Guoqiang Wang, Rebecca J Androwski, Andres Morera, Ken CQ Nguyen, Peter J Schweinsberg, Ge Bai, Jason Cooper, David H Hall, Monica Driscoll, and Barth D Grant. Large vesicle extrusions from C. elegans neurons are consumed and stimulated by glial-like phagocytosis activity of the neighboring cell. *eLife*, 12, 2023.

[13432] Yuanyuan Wang, Yinhe Mao, Xiaoqing Chen, Xinhuang Huang, Zhongyi Jiang, Kaiyan Yang, Lixing Tian, Tong Jiang, Yun Zou, Xiaoyuan Ma, Chaoyue Xu, Zili Zhou, Xianwei Wu, Lei Pan, Huaping Liang, Lin Zhong, and Changbin Chen. Homeostatic control of an iron repressor in a GI tract resident. *eLife*, 12, 2023.

[13433] Yuanyuan Wang, Tiffany Wu, Ming-Chi Tsai, Mitchell G Rezzonico, Alyaa M Abdel-Haleem, Luke Xie, Vineela D Gandham, Hai Ngu, Kimberly Stark, Caspar Glock, Daqi Xu, Oded Foreman, Brad A Friedman, Morgan Sheng, and Jesse E Hanson. TPL2 kinase activity regulates microglial inflammatory responses and promotes neurodegeneration in tauopathy mice. *eLife*, 12, 2023.

[13434] Yueyang Wang, Lee D Troughton, Fan Xu, Aritra Chatterjee, Chang Ding, Han Zhao, Laura Pulido Cifuentes, Ryan B Wagner, Tianqi Wang, Shelly Tan, Jingjuan Chen, Linlin Li, David Umulis, Shihuan Kuang, Daniel M Suter, Chongli Yuan, Deva Chan, Fang Huang, Patrick W Oakes, and Qing Deng. Atypical peripheral actin band formation via overactivation of RhoA and nonmuscle myosin II in mitofusin 2-deficient cells. *eLife*, 12, 2023.

[13435] Zhipeng Wang, Cheng Jin, Pengyu Li, Yiran Li, Jielin Tang, Zhixin Yu, Tao Jiao, Jinhuan Ou, Han Wang, Dingfeng Zou, Mengzhen Li, Xinyu Mang, Jun Liu, Yan Lu, Kai Li, Ning Zhang, Jia Yu, Shiying Miao, Linfang Wang, and Wei Song. Identification of quiescent FOXC2+ spermatogonial stem cells in adult mammals. *eLife*, 12, 2023.

[13436] Rachel Warren, Handeng Lyu, Kylie Klinkhammer, and Stijn P De Langhe. Hippo signaling impairs alveolar epithelial regeneration in pulmonary fibrosis. *eLife*, 12, 2023.

[13437] Emma T Watson, Michaela M Pauers, Michael J Seibert, Jason D Vevea, and Edwin R Chapman. Synaptic vesicle proteins are selectively delivered to axons in mammalian neurons. *eLife*, 12, 2023.

[13438] Kristina J Weaver, Sonakshi Raju, Rachel A Rucker, Tuhin Chakraborty, Robert A Holt, and Scott D Pletcher. Behavioral dissection of hunger states in Drosophila. *eLife*, 12, 2023.

[13439] Bryn D Webb, Sara M Nowinski, Ashley Solmonson, Jaya Ganesh, Richard J Rodenburg, Joao Leandro, Anthony Evans, Hieu S Vu, Thomas P Naidich, Bruce D Gelb, Ralph J DeBerardinis, Jared Rutter, and Sander M Houten. Recessive pathogenic variants in MCAT cause combined oxidative phosphorylation deficiency. *eLife*, 12, 2023.

[13440] Emily R Webb, Georgia L Dodd, Michaela Noskova, Esme Bullock, Morwenna Muir, Margaret C Frame, Alan Serrels, and Valerie G Brunton. Kindlin-1 regulates IL-6 secretion and modulates the immune environment in breast cancer models. *eLife*, 12, 2023.

[13441] Ke-Hsuan Wei, I-Ting Lin, Kaushik Chowdhury, Khai Lone Lim, Kuan-Ting Liu, Tai-Ming Ko, Yao-Ming Chang, Kai-Chien Yang, and Shih-Lei (Ben) Lai. Comparative single-cell profiling reveals distinct cardiac resident macrophages essential for zebrafish heart regeneration. *eLife*, 12, 2023.

[13442] Xinzhu Wei, Christopher R Robles, Ali Pazokitoroudi, Andrea Ganna, Alexander Gusev, Arun Durvasula, Steven Gazal, Po-Ru Loh, David Reich, and Sriram Sankararaman. The lingering effects of Neanderthal introgression on human complex traits. *eLife*, 12, 2023.

[13443] Yujian Wen, Jielong Huang, Shuhui Guo, Yehezqel Elyahu, Alon Monsonego, Hai Zhang, Yanqing Ding, and Hao Zhu. Applying causal discovery to single-cell analyses using CausalCell. *eLife*, 12, 2023.

[13444] Zhuangfeng Weng, Jiefu Zheng, Yiyi Zhou, Zuer Lu, Yixi Wu, Dongyi Xu, Huanhuan Li, Huanhuan Liang, and Yingfang Liu. Structural and mechanistic insights into the MCM8/9 helicase complex. *eLife*, 12, 2023.

[13445] Valentin Wernet, Marius Kriegler, Vojtech Kumpost, Ralf Mikut, Lennart Hilbert, and Reinhard Fischer. Synchronization of oscillatory growth prepares fungal hyphae for fusion. *eLife*, 12, 2023.

[13446] Robert West, Hadrien Delattre, Elad Noor, Elisenda Feliu, and Orkun S Soyer. Dynamics of co-substrate pools can constrain and regulate metabolic fluxes. *eLife*, 12, 2023.

[13447] Benjamin D Wheeler, John D Gagnon, Wandi S Zhu, Priscila Muñoz-Sandoval, Simon K Wong, Dimitre S Simeonov, Zhongmei Li, Rachel DeBarge, Matthew H Spitzer, Alexander Marson, and K Mark Ansel. The lncRNA Malat1 inhibits miR-15/16 to enhance cytotoxic T cell activation and memory cell formation. *eLife*, 12, 2023.

[13448] Szczepan Wiecha, Przemysław Seweryn Kasiak, Piotr Szwed, Tomasz Kowalski, Igor Cieśliński, Marek Postuła, and Andrzej Klusiewicz. VO2max prediction based on submaximal cardiorespiratory relationships and body composition in male runners and cyclists: a population study. *eLife*, 12, 2023.

[13449] Christoph Wiest, Flavie Torrecillos, Alek Pogosyan, Manuel Bange, Muthuraman Muthuraman, Sergiu Groppa, Natasha Hulse, Harutomo Hasegawa, Keyoumars Ashkan, Fahd Baig, Francesca Morgante, Erlick A Pereira, Nicolas Mallet, Peter J Magill, Peter Brown, Andrew Sharott, and Huiling Tan. The aperiodic exponent of subthalamic field potentials reflects excitation/inhibition balance in Parkinsonism. *eLife*, 12, 2023.

[13450] Hannah J Williams, Vivek H Sridhar, Edward Hurme, Gabriella EC Gall, Natalia Borrego, Genevieve E Finerty, Iain D Couzin, C Giovanni Galizia, Nathaniel J Dominy, Hannah M Rowland, Mark E Hauber, James P Higham, Ariana Strandburg-Peshkin, and Amanda D Melin. Sensory collectives in natural systems. *eLife*, 12, 2023.

[13451] Justin Krish Williams, Jordan Matthew Ngo, Isabelle Madeline Lehman, and Randy Schekman. Annexin A6 mediates calcium-dependent exosome secretion during plasma membrane repair. *eLife*, 12, 2023.

[13452] Siân Wilson, Maximilian Pietsch, Lucilio Cordero-Grande, Daan Christiaens, Alena Uus, Vyacheslav R Karolis, Vanessa Kyriakopoulou, Kathleen Colford, Anthony N Price, Jana Hutter, Mary A Rutherford, Emer J Hughes, Serena J Counsell, Jacques-Donald Tournier, Joseph V Hajnal, A David Edwards, Jonathan O'Muircheartaigh, and Tomoki Arichi. Spatiotemporal tissue maturation of thalamocortical pathways in the human fetal brain. *eLife*, 12, 2023.

[13453] Sebastien Wolf, Guillaume Le Goc, Georges Debrégeas, Simona Cocco, and Rémi Monasson. Emergence of time persistence in a data-driven neural network model. *eLife*, 12, 2023.

[13454] Andrzej S Wolniewicz, Yuefeng Shen, Qiang Li, Yuanyuan Sun, Yu Qiao, Yajie Chen, Yi-Wei Hu, and Jun Liu. An armoured marine reptile from the Early Triassic of South China and its phylogenetic and evolutionary implications. *eLife*, 12, 2023.

[13455] Pieter J Wolters, Doret Wouters, Yury M Tikunov, Shimlal Ayilalath, Linda P Kodde, Miriam F Strijker, Lotte Caarls, Richard GF Visser, and Vivianne GAA Vleeshouwers. Tetraose steroidal glycoalkaloids from potato provide resistance against Alternaria solani and Colorado potato beetle. *eLife*, 12, 2023.

[13456] Andrew I Wong, Tiago Beites, Kyle A Planck, Rachael A Fieweger, Kathryn A Eckartt, Shuqi Li, Nicholas C Poulton, Brian C VanderVen, Kyu Y Rhee, Dirk Schnappinger, Sabine Ehrt, and Jeremy Rock. Cyclic AMP is a critical mediator of intrinsic drug resistance and fatty acid metabolism in M. tuberculosis. *eLife*, 12, 2023.

[13457] Kenneth Kin Lam Wong, Tongchao Li, Tian-Ming Fu, Gaoxiang Liu, Cheng Lyu, Sayeh Kohani, Qijing Xie, David J Luginbuhl, Srigokul Upadhyayula, Eric Betzig, and Liqun Luo. Origin of wiring specificity in an olfactory map revealed by neuron type–specific, time-lapse imaging of dendrite targeting. *eLife*, 12, 2023.

[13458] Shannon E Wright and Peter K Todd. Native functions of short tandem repeats. *eLife*, 12, 2023.

[13459] Meng Wu, Rongyu Zhang, Zixiong Zhang, Ning Zhang, Chenfan Li, Yongli Xie, Haoran Xia, Fangjiao Huang, Ruoying Zhang, Ming Liu, Xiaoyu Li, Shan Cen, and Jinming Zhou. Selective androgen receptor degrader (SARD) to overcome antiandrogen resistance in castration-resistant prostate cancer. *eLife*, 12, 2023.

[13460] Mengyao Wu, Tak-Ho Lo, Liping Li, Jia Sun, Chujun Deng, Ka-Ying Chan, Xiang Li, Steve Ting-Yuan Yeh, Jimmy Tsz Hang Lee, Pauline Po Yee Lui, Aimin Xu, and Chi-Ming Wong. Amelioration of non-alcoholic fatty liver disease by targeting adhesion G protein-coupled receptor F1 (Adgrf1). *eLife*, 12, 2023.

[13461] Shenjie Wu, Nancy C Hernandez Villegas, Daniel W Sirkis, Iona Thomas-Wright, Richard Wade-Martins, and Randy Schekman. Unconventional secretion of $\alpha$-synuclein mediated by palmitoylated DNAJC5 oligomers. *eLife*, 12, 2023.

[13462] Timothy Wu, Jennifer M Deger, Hui Ye, Caiwei Guo, Justin Dhindsa, Brandon T Pekarek, Rami Al-Ouran, Zhandong Liu, Ismael Al-Ramahi, Juan Botas, and Joshua M Shulman. Tau polarizes an aging transcriptional signature to excitatory neurons and glia. *eLife*, 12, 2023.

[13463] Xiangkun Wu, Hong Yan, Mingxing Qiu, Xiaoping Qu, Jing Wang, Shaowan Xu, Yiran Zheng, Minghui Ge, Linlin Yan, and Li Liang. Comprehensive characterization of tumor microenvironment in colorectal cancer via molecular analysis. *eLife*, 12, 2023.

[13464] Yeyao Wu, Wangmi Fu, Ning Zang, and Chun Zhou. Structural characterization of human RPA70N association with DNA damage response proteins. *eLife*, 12, 2023.

[13465] Yuan-hao Wu, Ella Podvalny, and Biyu J He. Spatiotemporal neural dynamics of object recognition under uncertainty in humans. *eLife*, 12, 2023.

[13466] Meghan E Wynne, Oluwaseun Ogunbona, Alicia R Lane, Avanti Gokhale, Stephanie A Zlatic, Chongchong Xu, Zhexing Wen, Duc M Duong, Sruti Rayaprolu, Anna Ivanova, Eric A Ortlund, Eric B Dammer, Nicholas T Seyfried, Blaine R Roberts, Amanda Crocker, Vinit Shanbhag, Michael Petris, Nanami Senoo, Selvaraju Kandasamy, Steven Michael Claypool, Antoni Barrientos, Aliza Wingo, Thomas S Wingo, Srikant Rangaraju, Allan I Levey, Erica Werner, and Victor Faundez. APOE expression and secretion are modulated by mitochondrial dysfunction. *eLife*, 12, 2023.

[13467] Baolong Xia, Raghuvir Viswanatha, Yanhui Hu, Stephanie E Mohr, and Norbert Perrimon. Pooled genome-wide CRISPR activation screening for rapamycin resistance genes in Drosophila cells. *eLife*, 12, 2023.

[13468] Dongsheng Xiao, Yuhao Yan, and Timothy H Murphy. Mesotrode chronic simultaneous mesoscale cortical imaging and subcortical or peripheral nerve spiking activity recording in mice. *eLife*, 12, 2023.

[13469] Michael Xiao, Chia-Hua Wu, Graham Meek, Brian Kelly, Dara Buendia Castillo, Lyndsay EA Young, Sara Martire, Sajina Dhungel, Elizabeth McCauley, Purbita Saha, Altair L Dube, Matthew S Gentry, Laura A Banaszynski, Ramon C Sun, and Chintan K Kikani. PASK links cellular energy metabolism with a mitotic self-renewal network to establish differentiation competence. *eLife*, 12, 2023.

[13470] Shuyun Xiao, Valerie Michael, and Richard Mooney. Nested circuits mediate the decision to vocalize. *eLife*, 12, 2023.

[13471] Tong Xiao, Kaijie Wu, Peiliang Wang, Yali Ding, Xiao Yang, Chao Chang, and Yan Yang. Sensory input-dependent gain modulation of the optokinetic nystagmus by mid-infrared stimulation in pigeons. *eLife*, 12, 2023.

[13472] Juan Xie, Weilin Zhang, Xiaolei Zhu, Minghua Deng, and Luhua Lai. Coevolution-based prediction of key allosteric residues for protein function regulation. *eLife*, 12, 2023.

[13473] Marjorie Xie, Samuel P Muscinelli, Kameron Decker Harris, and Ashok Litwin-Kumar. Task-dependent optimal representations for cerebellar learning. *eLife*, 12, 2023.

[13474] Weizhen Xie, Marcus Cappiello, Michael A Yassa, Edward Ester, Kareem A Zaghloul, and Weiwei Zhang. The entorhinal-DG/CA3 pathway in the medial temporal lobe retains visual working memory of a simple surface feature. *eLife*, 12, 2023.

[13475] Jiyu Xin, Yang Shi, Xin Zhang, Xinyi Yuan, Yueyong Xin, Huimin He, Jiejie Shen, Robert E Blankenship, and Xiaoling Xu. Carotenoid assembly regulates quinone diffusion and the Roseiflexus castenholzii reaction center-light harvesting complex architecture. *eLife*, 12, 2023.

[13476] Adele Francis Xu, Rut Molinuevo, Elisa Fazzari, Harrison Tom, Zijian Zhang, Julien Menendez, Kerriann M Casey, Davide Ruggero, Lindsay Hinck, Jonathan K Pritchard, and Maria Barna. Subfunctionalized expression drives evolutionary retention of ribosomal protein paralogs Rps27 and Rps27l in vertebrates. *eLife*, 12, 2023.

[13477] Han Xu, Lian Xia, John R Spence, Mingxian Lin, Chunyang Lu, Yanpeng Li, Jie Chen, Tushou Luo, Yide Li, and Suqin Fang. Flying squirrels use a mortise-tenon structure to fix nuts on understory twigs. *eLife*, 12, 2023.

[13478] Huaming Xu, Zhijian Li, Chao-Chung Kuo, Katrin Götz, Thomas Look, Marcelo AS de Toledo, Kristin Seré, Ivan G Costa, and Martin Zenke. A lncRNA identifies Irf8 enhancer element in negative feedback control of dendritic cell differentiation. *eLife*, 12, 2023.

[13479] Ke Xu, Xianwei Su, Kailun Fang, Yue Lv, Tao Huang, Mengjing Li, Ziqi Wang, Yingying Yin, Tahir Muhammad, Shangming Liu, Xiangfeng Chen, Jing Jiang, Jinsong Li, Wai-Yee Chan, Jinlong Ma, Gang Lu, Zi-Jiang Chen, and Hongbin Liu. The Slingshot phosphatase 2 is required for acrosome biogenesis during spermatogenesis in mice. *eLife*, 12, 2023.

[13480] Lijuan Xu, Xinyao Zhang, Yin Zhao, Xiaorui Gang, Tao Zhou, Jialing Han, Yang Cao, Binyan Qi, Shuning Song, Xiaojie Wang, and Yuanbo Liang. Metformin protects trabecular meshwork against oxidative injury via activating integrin/ROCK signals. *eLife*, 12, 2023.

[13481] Tianze Xu, Jing Cai, Lei Wang, Li Xu, Hongting Zhao, Fudi Wang, Esther G Meyron-Holtz, Fanis Missirlis, Tong Qiao, and Kuanyu Li. Hormone replacement therapy for postmenopausal atherosclerosis is offset by late age iron deposition. *eLife*, 12, 2023.

[13482] Wenjie Xu, Chenglong Zhu, Xueli Gao, Baosheng Wu, Han Xu, Mingliang Hu, Honghui Zeng, Xiaoni Gan, Chenguang Feng, Jiangmin Zheng, Jing Bo, Li-Sheng He, Qiang Qiu, Wen Wang, Shunping He, and Kun Wang. Chromosome-level genome assembly of hadal snailfish reveals mechanisms of deep-sea adaptation in vertebrates. *eLife*, 12, 2023.

[13483] Yang Xu, Tian-liang Cui, Jia-yi Li, Bingchen Chen, and Jin-Hui Wang. Associative memory neurons of encoding multi-modal signals are recruited by neuroligin-3-mediated new synapse formation. *eLife*, 12, 2023.

[13484] Surabhi Yadav, Olivia Delau, Adam J Bonner, Daniela Markovic, William Patterson, Sasha Ottey, Richard P Buyalos, and Ricardo Azziz. Direct economic burden of mental health disorders associated with polycystic ovary syndrome: Systematic review and meta-analysis. *eLife*, 12, 2023.

[13485] Daichi Yamada, Daniel Bushey, Feng Li, Karen L Hibbard, Megan Sammons, Jan Funke, Ashok Litwin-Kumar, Toshihide Hige, and Yoshinori Aso. Hierarchical architecture of dopaminergic circuits enables second-order conditioning in Drosophila. *eLife*, 12, 2023.

[13486] Tetsushi Yamagata, Ikuo Ogiwara, Tetsuya Tatsukawa, Toshimitsu Suzuki, Yuka Otsuka, Nao Imaeda, Emi Mazaki, Ikuyo Inoue, Natsuko Tokonami, Yurina Hibi, Shigeyoshi Itohara, and Kazuhiro Yamakawa. Scn1a-GFP transgenic mouse revealed Nav1.1 expression in neocortical pyramidal tract projection neurons. *eLife*, 12, 2023.

[13487] Atomu Yamaguchi, Noriaki Maeshige, Hikari Noguchi, Jiawei Yan, Xiaoqi Ma, Mikiko Uemura, Dongming Su, Hiroyo Kondo, Kristopher Sarosiek, and Hidemi Fujino. Pulsed ultrasound promotes secretion of anti-inflammatory extracellular vesicles from skeletal myotubes via elevation of intracellular calcium level. *eLife*, 12, 2023.

[13488] Ayako Yamaguchi and Manon Peltier. Two conserved vocal central pattern generators broadly tuned for fast and slow rates generate species-specific vocalizations in Xenopus clawed frogs. *eLife*, 12, 2023.

[13489] Hiroshi Yamaguchi, Motohiro Morikawa, and Masahide Kikkawa. Calaxin stabilizes the docking of outer arm dyneins onto ciliary doublet microtubule in vertebrates. *eLife*, 12, 2023.

[13490] Yumeya Yamamori, Oliver J Robinson, and Jonathan P Roiser. Approach-avoidance reinforcement learning as a translational and computational model of anxiety-related avoidance. *eLife*, 12, 2023.

[13491] Nao Yamamoto, Keisuke Ejima, Roger S Zoh, and Andrew W Brown. Bias in nutrition-health associations is not eliminated by excluding extreme reporters in empirical or simulation studies. *eLife*, 12, 2023.

[13492] Chaojun Yan, Jingjing Xiong, Zirui Zhou, Qifang Li, Chuan Gao, Mengyao Zhang, Liya Yu, Jinpeng Li, Ming-Ming Hu, Chen-Song Zhang, Cheguo Cai, Haojian Zhang, and Jing Zhang. A cleaved METTL3 potentiates the METTL3–WTAP interaction and breast cancer progression. *eLife*, 12, 2023.

[13493] Ryuji Yanase, Flávia Moreira-Leite, Edward Rea, Lauren Wilburn, Jovana Sádlová, Barbora Vojtkova, Katerina Pružinová, Atsushi Taniguchi, Shigenori Nonaka, Petr Volf, and Jack D Sunter. Formation and three-dimensional architecture of Leishmania adhesion in the sand fly vector. *eLife*, 12, 2023.

[13494] Jie Yang, Ruijun Tang, Shiye Chen, Yinan Chen, Kai Yuan, Rui Huang, and Liming Wang. Exposure to high-sugar diet induces transgenerational changes in sweet sensitivity and feeding behavior via H3K27me3 reprogramming. *eLife*, 12, 2023.

[13495] Qiming Yang, Te-Wen Lo, KatjuŠa Brejc, Caitlin Schartner, Edward J Ralston, Denise M Lapidus, and Barbara J Meyer. X-chromosome target specificity diverged between dosage compensation mechanisms of two closely related Caenorhabditis species. *eLife*, 12, 2023.

[13496] Rui-Qiu Yang, Yong-Hong Chen, Qin-yi Wu, Jie Tang, Shan-Zhuang Niu, Qiu Zhao, Yi-Cheng Ma, and Cheng-Gang Zou. Indole produced during dysbiosis mediates host–microorganism chemical communication. *eLife*, 12, 2023.

[13497] Runze Yang, Tianhao Xu, Lei Zhang, Minghao Ge, Liwei Yan, Jian Li, and Weili Fu. A single-cell atlas depicting the cellular and molecular features in human anterior cruciate ligamental degeneration: A single cell combined spatial transcriptomics study. *eLife*, 12, 2023.

[13498] Xiaoyan Yang, Hsiang-Chun Chang, Yuki Tatekoshi, Amir Mahmoodzadeh, Maryam Balibegloo, Zeinab Najafi, Rongxue Wu, Chunlei Chen, Tatsuya Sato, Jason Shapiro, and Hossein Ardehali. SIRT2 inhibition protects against cardiac hypertrophy and ischemic injury. *eLife*, 12, 2023.

[13499] Xiong Yang, Rong Wan, Zhiwen Liu, Su Feng, Jiaxin Yang, Naihe Jing, and Ke Tang. The differentiation and integration of the hippocampal dorsoventral axis are controlled by two nuclear receptor genes. *eLife*, 12, 2023.

[13500] Lutian Yao, Jiawei Lu, Leilei Zhong, Yulong Wei, Tao Gui, Luqiang Wang, Jaimo Ahn, Joel D Boerckel, Danielle Rux, Christina Mundy, Ling Qin, and Maurizio Pacifici. Activin A marks a novel progenitor cell population during fracture healing and reveals a therapeutic strategy. *eLife*, 12, 2023.

[13501] Yiqi Yao and Alejandro Ochoa. Limitations of principal components in quantitative genetic association models for human studies. *eLife*, 12, 2023.

[13502] Zihao Yao, Weizhong Qi, Hongbo Zhang, Zhicheng Zhang, Liangliang Liu, Yan Shao, Hua Zeng, Jianbin Yin, Haoyan Pan, Xiongtian Guo, Anling Liu, Daozhang Cai, Xiaochun Bai, and Haiyan Zhang. Down-regulated GAS6 impairs synovial macrophage efferocytosis and promotes obesity-associated osteoarthritis. *eLife*, 12, 2023.

[13503] Yi Tian Yap, Wei Li, Qian Huang, Qi Zhou, David Zhang, Yi Sheng, Ljljiana Mladenovic-Lucas, Siu-Pok Yee, Kyle E Orwig, James G Granneman, David C Williams, Rex A Hess, Aminata Toure, and Zhibing Zhang. DNALI1 interacts with the MEIG1/PACRG complex within the manchette and is required for proper sperm flagellum assembly in mice. *eLife*, 12, 2023.

[13504] Yujie Ye, Shumpei Morita, Justin J Chang, Patrick M Buckley, Kiera B Wilhelm, Daniel DiMaio, Jay T Groves, and Francisco N Barrera. Allosteric inhibition of the T cell receptor by a designed membrane ligand. *eLife*, 12, 2023.

[13505] Priscilla S-W Yeung, Megumi Yamashita, and Murali Prakriya. A pathogenic human Orai1 mutation unmasks STIM1-independent rapid inactivation of Orai1 channels. *eLife*, 12, 2023.

[13506] Ece Yildiz, Gaby El Alam, Alessia Perino, Antoine Jalil, Pierre-Damien Denechaud, Katharina Huber, Lluis Fajas, Johan Auwerx, Giovanni Sorrentino, and Kristina Schoonjans. Hepatic lipid overload triggers biliary epithelial cell activation via E2Fs. *eLife*, 12, 2023.

[13507] Yuk-Hoi Yiu and Christian Leibold. A theory of hippocampal theta correlations accounting for extrinsic and intrinsic sequences. *eLife*, 12, 2023.

[13508] Gehad Youssef, Luke Gammon, Leah Ambler, Sophia Lunetto, Alice Scemama, Hannah Cottom, Kim Piper, Ian C Mackenzie, Michael P Philpott, and Adrian Biddle. Disseminating cells in human oral tumours possess an EMT cancer stem cell marker profile that is predictive of metastasis in image-based machine learning. *eLife*, 12, 2023.

[13509] James Yu, Stephanie Dancausse, Maria Paz, Tolu Faderin, Melissa Gaviria, Joseph W Shomar, Dave Zucker, Vivek Venkatachalam, and Mason Klein. Continuous, long-term crawling behavior characterized by a robotic transport system. *eLife*, 12, 2023.

[13510] Qinlin Yu, Minhui Ouyang, John Detre, Huiying Kang, Di Hu, Bo Hong, Fang Fang, Yun Peng, and Hao Huang. Infant brain regional cerebral blood flow increases supporting emergence of the default-mode network. *eLife*, 12, 2023.

[13511] Xiaoyu Yu, Jinsai Shang, and Douglas J Kojetin. Molecular basis of ligand-dependent Nurr1-RXR$\alpha$ activation. *eLife*, 12, 2023.

[13512] Erliang Yuan, Huijuan Guo, Weiyao Chen, Bingru Du, Yingjie Mi, Zhaorui Qi, Yiyang Yuan, Keyan Zhu-Salzman, Feng Ge, and Yucheng Sun. A novel gene REPTOR2 activates the autophagic degradation of wing disc in pea aphid. *eLife*, 12, 2023.

[13513] Shuai Yuan, Jie Chen, Xixian Ruan, Yuhao Sun, Ke Zhang, Xiaoyan Wang, Xue Li, Dipender Gill, Stephen Burgess, Edward Giovannucci, and Susanna C Larsson. Smoking, alcohol consumption, and 24 gastrointestinal diseases: Mendelian randomization analysis. *eLife*, 12, 2023.

[13514] John Se Kit Yuen Jr, Michael K Saad, Ning Xiang, Brigid M Barrick, Hailey DiCindio, Chunmei Li, Sabrina W Zhang, Miriam Rittenberg, Emily T Lew, Kevin Lin Zhang, Glenn Leung, Jaymie A Pietropinto, and David L Kaplan. Aggregating in vitro-grown adipocytes to produce macroscale cell-cultured fat tissue with tunable lipid compositions for food applications. *eLife*, 12, 2023.

[13515] George Zacharopoulos, Francesco Sella, Uzay Emir, and Roi Cohen Kadosh. Dissecting the chain of information processing and its interplay with neurochemicals and fluid intelligence across development. *eLife*, 12, 2023.

[13516] Mone Zaidi, Se-Min Kim, Mehr Mathew, Funda Korkmaz, Farhath Sultana, Sari Miyashita, Anisa Azatovna Gumerova, Tal Frolinger, Ofer Moldavski, Orly Barak, Anusha Pallapati, Satish Rojekar, John Caminis, Yelena Ginzburg, Vitaly Ryu, Terry F Davies, Daria Lizneva, Clifford J Rosen, and Tony Yuen. Bone circuitry and interorgan skeletal crosstalk. *eLife*, 12, 2023.

[13517] Mohammad Saeed Zarzor, Ingmar Blumcke, and Silvia Budday. Exploring the role of the outer subventricular zone during cortical folding through a physics-based model. *eLife*, 12, 2023.

[13518] Anastasiya Zaytseva, Evelina Bouckova, McKennon J Wiles, Madison H Wustrau, Isabella G Schmidt, Hadassah Mendez-Vazquez, Latika Khatri, and Seonil Kim. Ketamine's rapid antidepressant effects are mediated by Ca2+-permeable AMPA receptors. *eLife*, 12, 2023.

[13519] Benjamin M Zemel, Alexander A Nevue, Leonardo ES Tavares, Andre Dagostin, Peter V Lovell, Dezhe Z Jin, Claudio V Mello, and Henrique von Gersdorff. Motor cortex analogue neurons in songbirds utilize Kv3 channels to generate ultranarrow spikes. *eLife*, 12, 2023.

[13520] Fu Zeng, Adam Zaidel, and Aihua Chen. Contrary neuronal recalibration in different multisensory cortical areas. *eLife*, 12, 2023.

[13521] Yu Zeng, David C Plachetzki, Kristen Nieders, Hannah Campbell, Marissa Cartee, M Sabrina Pankey, Kennedy Guillen, and Douglas Fudge. Epidermal threads reveal the origin of hagfish slime. *eLife*, 12, 2023.

[13522] Chen Zhang, Zhi-Fu Guo, Wennan Liu, Kyosuke Kazama, Louis Hu, Xiaobo Sun, Lu Wang, Hyoungjoo Lee, Lin Lu, Xiao-Feng Yang, Ross Summer, and Jianxin Sun. PIMT is a novel and potent suppressor of endothelial activation. *eLife*, 12, 2023.

[13523] Cheng Zhang, Shasha Zhao, Huan Deng, Shihua Zhang, Juan Wang, Xiaoye Song, Deen Yu, Yue Zhang, and Wensheng Deng. STAT3 promotes RNA polymerase III-directed transcription by controlling the miR-106a-5p/TP73 axis. *eLife*, 12, 2023.

[13524] Chi Zhang, Xianfu Yi, Mengfan Hou, Qingyang Li, Xueying Li, Lu Lu, Enlin Qi, Mingxin Wu, Lin Qi, Huan Jian, Zhangyang Qi, Yigang Lv, Xiaohong Kong, Mingjun Bi, Shiqing Feng, and Hengxing Zhou. The landscape of m1A modification and its posttranscriptional regulatory functions in primary neurons. *eLife*, 12, 2023.

[13525] Congyan Zhang and Randy Schekman. Syncytin-mediated open-ended membrane tubular connections facilitate the intercellular transfer of cargos including Cas9 protein. *eLife*, 12, 2023.

[13526] Fan Zhang, Yuanyuan Wang, Ying Zhao, Manqi Wang, Bin Zhou, Bin Zhou, and Xianpeng Ge. NFATc1 marks articular cartilage progenitors and negatively determines articular chondrocyte differentiation. *eLife*, 12, 2023.

[13527] Hangjie Zhang, Qianhui Hua, Nani Nani Xu, Xinpei Zhang, Bo Chen, Xijun Ma, Jie Hu, Zhongbing Chen, Pengfei Yu, Huijun Lei, Shenyu Wang, Linling Ding, Jian Fu, Yuting Liao, Juan Yang, Jianmin Jiang, and Huakun Lv. Evaluation of antibody kinetics and durability in healthy individuals vaccinated with inactivated COVID-19 vaccine (CoronaVac): A cross-sectional and cohort study in Zhejiang, China. *eLife*, 12, 2023.

[13528] Hui Zhang, Chan Li, Qingyang Liu, Jingmei Li, Hao Wu, Rui Xu, Yidan Sun, Ming Cheng, Xiaoe Zhao, Menghao Pan, Qiang Wei, and Baohua Ma. C-type natriuretic peptide improves maternally aged oocytes quality by inhibiting excessive PINK1/Parkin-mediated mitophagy. *eLife*, 12, 2023.

[13529] Liangyu Zhang, Weston T Stauffer, John S Wang, Fan Wu, Zhouliang Yu, Chenshu Liu, Hyung Jun Kim, and Abby F Dernburg. Recruitment of Polo-like kinase couples synapsis to meiotic progression via inactivation of CHK-2. *eLife*, 12, 2023.

[13530] Qinrong Zhang, Yuhan Yang, Kevin J Cao, Wei Chen, Santosh Paidi, Chun-hong Xia, Richard H Kramer, Xiaohua Gong, and Na Ji. Retinal microvascular and neuronal pathologies probed in vivo by adaptive optical two-photon fluorescence microscopy. *eLife*, 12, 2023.

[13531] Rui Zhang, Qingxi Liu, Sa Zhou, Hongpeng He, Mingfeng Zhao, and Wenjian Ma. Mesenchymal stem cell suppresses the efficacy of CAR-T toward killing lymphoma cells by modulating the microenvironment through stanniocalcin-1. *eLife*, 12, 2023.

[13532] Ruolin Zhang, Weibing Mao, Lumeng Niu, Wendai Bao, Yiqi Wang, Ying Wang, Yasha Zhu, Zhihao Yang, Jincao Chen, Jiawen Dong, Meng Cai, Zilong Yuan, Haikun Song, Guangqiang Li, Min Zhang, Nanxiang Xiong, Jun Wei, and Zhiqiang Dong. NSC-derived exosomes enhance therapeutic effects of NSC transplantation on cerebral ischemia in mice. *eLife*, 12, 2023.

[13533] Songdou Zhang, Jianying Li, Dongyue Zhang, Zhixian Zhang, Shili Meng, Zhen Li, and Xiaoxia Liu. miR-252 targeting temperature receptor CcTRPM to mediate the transition from summer-form to winter-form of Cacopsylla chinensis. *eLife*, 12, 2023.

[13534] Tao Zhang, Liyang Wan, Han Xiao, Linfeng Wang, Jianzhong Hu, and Hongbin Lu. Single-cell RNA sequencing reveals cellular and molecular heterogeneity in fibrocartilaginous enthesis formation. *eLife*, 12, 2023.

[13535] Weixin Zhang, Kathleen Noller, Janet Crane, Mei Wan, Xiaojun Wu, Patrick Cahan, and Xu Cao. RANK+TLR2+ myeloid subpopulation converts autoimmune to joint destruction in rheumatoid arthritis. *eLife*, 12, 2023.

[13536] Wenxin Zhang, Taki Nishimura, Deepanshi Gahlot, Chieko Saito, Colin Davis, Harold BJ Jefferies, Anne Schreiber, Lipi Thukral, and Sharon A Tooze. Autophagosome membrane expansion is mediated by the N-terminus and cis-membrane association of human ATG8s. *eLife*, 12, 2023.

[13537] Yijun Zhang, Tao Wu, Zhimin He, Wenlong Lai, Xiangyi Shen, Jiaoyan Lv, Yuanhao Wang, and Li Wu. Regulation of pDC fate determination by histone deacetylase 3. *eLife*, 12, 2023.

[13538] Ying Zhang, Kenneth E Schriver, Jia Ming Hu, and Anna Wang Roe. Spatial frequency representation in V2 and V4 of macaque monkey. *eLife*, 12, 2023.

[13539] Yingnan Zhang, Kelly Zhang, Adityo Prakosa, Cynthia James, Stefan L Zimmerman, Richard Carrick, Eric Sung, Alessio Gasperetti, Crystal Tichnell, Brittney Murray, Hugh Calkins, and Natalia A Trayanova. Predicting ventricular tachycardia circuits in patients with arrhythmogenic right ventricular cardiomyopathy using genotype-specific heart digital twins. *eLife*, 12, 2023.

[13540] Yunpei Zhang, Lingling Neng, Kushal Sharma, Zhiqiang Hou, Anatasiya Johnson, Junha Song, Alain Dabdoub, and Xiaorui Shi. Pericytes control vascular stability and auditory spiral ganglion neuron survival. *eLife*, 12, 2023.

[13541] Zengdi Zhang, Zan Huang, Mohamed Awad, Mohammed Elsalanty, James Cray, Lauren E Ball, Jason C Maynard, Alma L Burlingame, Hu Zeng, Kim C Mansky, and Hai-Bin Ruan. O-GlcNAc glycosylation orchestrates fate decision and niche function of bone marrow stromal progenitors. *eLife*, 12, 2023.

[13542] Zhiwang Zhang, Qichao Liao, Tingli Pan, Lin Yu, Zupeng Luo, Songtao Su, Shi Liu, Menglong Hou, Yixing Li, Turtushikh Damba, Yunxiao Liang, and Lei Zhou. BATF relieves hepatic steatosis by inhibiting PD1 and promoting energy metabolism. *eLife*, 12, 2023.

[13543] Jianjian Zhao, Xuchen Zhang, Bohan Zhao, Wantong Hu, Tongxin Diao, Liyuan Wang, Yi Zhong, and Qian Li. Genetic dissection of mutual interference between two consecutive learning tasks in Drosophila. *eLife*, 12, 2023.

[13544] Jie Zhao, Bohan Fan, Jian Huang, Benjamin John Cowling, Shiu Lun Ryan Au Yeung, Andrea Baccarelli, Gabriel M Leung, and C Mary Schooling. Environment- and epigenome-wide association study of obesity in 'Children of 1997' birth cohort. *eLife*, 12, 2023.

[13545] Kunli Zhao, Kohichi Matsunaga, Kouichi Mizuno, Hao Wang, Katsuhide Okunishi, and Tetsuro Izumi. Functional hierarchy among different Rab27 effectors involved in secretory granule exocytosis. *eLife*, 12, 2023.

[13546] Lifen Zhao, Fangyu Wei, Xinheng He, Antao Dai, Dehua Yang, Hualiang Jiang, Liuqing Wen, and Xi Cheng. Identification of a carbohydrate recognition motif of purinergic receptors. *eLife*, 12, 2023.

[13547] Longhui Zhao, Wouter Halfwerk, and Jianguo Cui. Response to comment on 'Parasite defensive limb movements enhance acoustic signal attraction in male little torrent frogs'. *eLife*, 12, 2023.

[13548] Zhuoran Zhao, Xiayu Shen, Shuang Chen, Jing Gu, Haun Wang, Maria F Mojica, Moumita Samanta, Debsindhu Bhowmik, Alejandro J Vila, Robert A Bonomo, and Shozeb Haider. Gating interactions steer loop conformational changes in the active site of the L1 metallo-$\beta$-lactamase. *eLife*, 12, 2023.

[13549] Hong Zheng, Qianjin Li, Shanhu Li, Zhiguo Li, Marco Brotto, Daiana Weiss, Domenick Prosdocimo, Chunhui Xu, Ashruth Reddy, Michelle Puchowicz, Xinyang Zhao, M Neale Weitzmann, Mukesh K Jain, and Cheng-Kui Qu. Loss of Ptpmt1 limits mitochondrial utilization of carbohydrates and leads to muscle atrophy and heart failure in tissue-specific knockout mice. *eLife*, 12, 2023.

[13550] Yiqiao Zheng, Chi Sun, Xiaodong Zhang, Philip A Ruzycki, and Shiming Chen. Missense mutations in CRX homeodomain cause dominant retinopathies through two distinct mechanisms. *eLife*, 12, 2023.

[13551] Zhichao Zheng, Lihong Wu, Zhicong Li, Ruoshu Tang, Hongtao Li, Yinyin Huang, Tianqi Wang, Shaofen Xu, Haoyu Cheng, Zhitong Ye, Dong Xiao, Xiaolin Lin, Gang Wu, Richard T Jaspers, and Janak L Pathak. Mir155 regulates osteogenesis and bone mass phenotype via targeting S1pr1 gene. *eLife*, 12, 2023.

[13552] Leilei Zhong, Jiawei Lu, Jiankang Fang, Lutian Yao, Wei Yu, Tao Gui, Michael Duffy, Nicholas Holdreith, Catherine A Bautista, Xiaobin Huang, Shovik Bandyopadhyay, Kai Tan, Chider Chen, Yongwon Choi, Jean X Jiang, Shuying Yang, Wei Tong, Nathanial Dyment, and Ling Qin. Csf1 from marrow adipogenic precursors is required for osteoclast formation and hematopoiesis in bone. *eLife*, 12, 2023.

[13553] Coral Y Zhou, Bastiaan Dekker, Ziyuan Liu, Hilda Cabrera, Joel Ryan, Job Dekker, and Rebecca Heald. Mitotic chromosomes scale to nuclear-cytoplasmic ratio and cell size in Xenopus. *eLife*, 12, 2023.

[13554] Jeff Jiajing Zhou, Jin Sun Cho, Han Han, Ira L Blitz, Wenqi Wang, and Ken WY Cho. Histone deacetylase 1 maintains lineage integrity through histone acetylome refinement during early embryogenesis. *eLife*, 12, 2023.

[13555] Ling-Yun Zhou, Chen-Xi Jin, Wen-Xiao Wang, Lei Song, Jung-Bum Shin, Ting-Ting Du, and Hao Wu. Differential regulation of hair cell actin cytoskeleton mediated by SRF and MRTFB. *eLife*, 12, 2023.

[13556] Mengmeng Zhou, Yuhong Han, and Jin Jiang. Ulk4 promotes Shh signaling by regulating Stk36 ciliary localization and Gli2 phosphorylation. *eLife*, 12, 2023.

[13557] Yufeng Zhou, John H Lewis, and Zhe Lu. Tracking multiple conformations occurring on angstrom-and-millisecond scales in single amino-acid-transporter molecules. *eLife*, 12, 2023.

[13558] Changyu Zhu, Yadira M Soto-Feliciano, John P Morris, Chun-Hao Huang, Richard P Koche, Yu-jui Ho, Ana Banito, Chun-Wei Chen, Aditya Shroff, Sha Tian, Geulah Livshits, Chi-Chao Chen, Myles Fennell, Scott A Armstrong, C David Allis, Darjus F Tschaharganeh, and Scott W Lowe. MLL3 regulates the CDKN2A tumor suppressor locus in liver cancer. *eLife*, 12, 2023.

[13559] Tianyu Zhu, Yuxin Zhang, Xunlun Sheng, Xiangzheng Zhang, Yu Chen, Hongjing Zhu, Yueshuai Guo, Yaling Qi, Yichen Zhao, Qi Zhou, Xue Chen, Xuejiang Guo, and Chen Zhao. Absence of CEP78 causes photoreceptor and sperm flagella impairments in mice and a human individual. *eLife*, 12, 2023.

[13560] Xiaoquan Zhu, Chao Chen, Dong Wei, Yong Xu, Siying Liang, Wenlong Jia, Jian Li, Yanchun Qu, Jianpo Zhai, Yaoguang Zhang, Pengjie Wu, Qiang Hao, Linlin Zhang, Wei Zhang, Xinyu Yang, Lin Pan, Ruomei Qi, Yao Li, Feiliang Wang, Rui Yi, Ze Yang, Jianye Wang, and Yanyang Zhao. FOXP2 confers oncogenic effects in prostate cancer. *eLife*, 12, 2023.

[13561] Aojia Zhuang, Aobo Zhuang, Yijiao Chen, Zhaoyu Qin, Dexiang Zhu, Li Ren, Ye Wei, Pengyang Zhou, Xuetong Yue, Fuchu He, Jianmin Xu, and Chen Ding. Proteomic characteristics reveal the signatures and the risks of T1 colorectal cancer metastasis to lymph nodes. *eLife*, 12, 2023.

[13562] Catharina Zich, Andrew J Quinn, James J Bonaiuto, George O'Neill, Lydia C Mardell, Nick S Ward, and Sven Bestmann. Spatiotemporal organisation of human sensorimotor beta burst activity. *eLife*, 12, 2023.

[13563] Valentyna Zinchenko, Johannes Hugger, Virginie Uhlmann, Detlev Arendt, and Anna Kreshuk. MorphoFeatures for unsupervised exploration of cell types, tissues, and organs in volume electron microscopy. *eLife*, 12, 2023.

[13564] Luke H Ziolkowski, Elena O Gracheva, and Sviatoslav N Bagriantsev. Mechanotransduction events at the physiological site of touch detection. *eLife*, 12, 2023.

[13565] Erika Zodda, Olga Tura-Ceide, Nicholas L Mills, Josep Tarragó-Celada, Marina Carini, Timothy M Thomson, and Marta Cascante. Autonomous metabolic reprogramming and oxidative stress characterize endothelial dysfunction in acute myocardial infarction. *eLife*, 12, 2023.

[13566] Dan-Dan Zou, Ya-Zhou Sun, Xin-Jie Li, Wen-Juan Wu, Dan Xu, Yu-Tong He, Jue Qi, Ying Tu, Yang Tang, Yun-Hua Tu, Xiao-Li Wang, Xing Li, Feng-Yan Lu, Ling Huang, Heng Long, Li He, and Xin Li. Single-cell sequencing highlights heterogeneity and malignant progression in actinic keratosis and cutaneous squamous cell carcinoma. *eLife*, 12, 2023.

[13567] Jiajie Zou, Yuran Zhang, Jialu Li, Xing Tian, and Nai Ding. Human attention during goal-directed reading comprehension relies on task optimization. *eLife*, 12, 2023.

[13568] Qin Zou, Rong Yuan, Yu Zhang, Yifei Wang, Ting Zheng, Rui Shi, Mei Zhang, Yujing Li, Kaixin Fei, Ran Feng, Binyun Pan, Xinyue Zhang, Zhengyin Gong, Li Zhu, Guoqing Tang, Mingzhou Li, Xuewei Li, and Yanzhi Jiang. A single-cell transcriptome atlas of pig skin characterizes anatomical positional heterogeneity. *eLife*, 12, 2023.

[13569] Xinle Zou, Sri Karthika Shanmugam, Scott A Kanner, Kevin J Sampson, Robert S Kass, and Henry M Colecraft. Divergent regulation of KCNQ1/E1 by targeted recruitment of protein kinase A to distinct sites on the channel complex. *eLife*, 12, 2023.

[13570] Dalen Zuidema, Alexis Jones, Won-Hee Song, Michal Zigo, and Peter Sutovsky. Identification of candidate mitochondrial inheritance determinants using the mammalian cell-free system. *eLife*, 12, 2023.

[13571] Lorena Zuzic, Jan K Marzinek, Ganesh S Anand, Jim Warwicker, and Peter J Bond. A pH-dependent cluster of charges in a conserved cryptic pocket on flaviviral envelopes. *eLife*, 12, 2023.

[13572] Theodore J Zwang, Rachel E Bennett, Maria Lysandrou, Benjamin Woost, Anqi Zhang, Charles M Lieber, Douglas S Richardson, and Bradley T Hyman. Tissue libraries enable rapid determination of conditions that preserve antibody labeling in cleared mouse and human tissue. *eLife*, 12, 2023.

[13573] Asaph Zylbertal and Isaac H Bianco. Recurrent network interactions explain tectal response variability and experience-dependent behavior. *eLife*, 12, 2023.

[13574] Keene L Abbott, Ahmed Ali, Bradley I Reinfeld, Amy Deik, Sonu Subudhi, Madelyn D Landis, Rachel A Hongo, Kirsten L Young, Tenzin Kunchok, Christopher S Nabel, Kayla D Crowder, Johnathan R Kent, Maria Lucia L Madariaga, Rakesh K Jain, Kathryn E Beckermann, Caroline A Lewis, Clary B Clish, Alexander Muir, W Kimryn Rathmell, Jeffrey Rathmell, and Matthew G Vander Heiden. Metabolite profiling of human renal cell carcinoma reveals tissue-origin dominance in nutrient availability. *eLife*, 13, 2024.

[13575] Reham Abdelaziz, Adam P Tomczak, Andreas Neef, and Luis A Pardo. Revealing a hidden conducting state by manipulating the intracellular domains in KV10.1 exposes the coupling between two gating mechanisms. *eLife*, 12, 2024.

[13576] Kenta Abe, Yuki Kambe, Kei Majima, Zijing Hu, Makoto Ohtake, Ali Momennezhad, Hideki Izumi, Takuma Tanaka, Ashley Matunis, Emma Stacy, Takahide Itokazu, Takashi R Sato, and Tatsuo Sato. Functional diversity of dopamine axons in prefrontal cortex during classical conditioning. *eLife*, 12, 2024.

[13577] Mitsuhiro Abe, Masataka Yanagawa, Michio Hiroshima, Toshihide Kobayashi, and Yasushi Sako. Bilateral regulation of EGFR activity and local PI(4,5)P2 dynamics in mammalian cells observed with superresolution microscopy. *eLife*, 13, 2024.

[13578] Syeda Nayab Fatima Abidi, Sara Chan, Kerstin Seidel, Daniel Lafkas, Louis Vermeulen, Frank Peale, and Christian W Siebel. The Jag2/Notch1 signaling axis promotes sebaceous gland differentiation and controls progenitor proliferation. *eLife*, 13, 2024.

[13579] Adlin Abramian, Rein I Hoogstraaten, Fiona H Murphy, Kathryn F McDaniel, Ruud F Toonen, and Matthijs Verhage. Rabphilin-3A negatively regulates neuropeptide release, through its SNAP25 interaction. *eLife*, 13, 2024.

[13580] Emily M Adamic, Adam R Teed, Jason Avery, Feliberto de la Cruz, and Sahib Khalsa. Hemispheric divergence of interoceptive processing across psychiatric disorders. *eLife*, 13, 2024.

[13581] Maciek Adamowski, Ivana Matijević, and Jiří Friml. Developmental patterning function of GNOM ARF-GEF mediated from the cell periphery. *eLife*, 13, 2024.

[13582] Maria L Adelus, Jiacheng Ding, Binh T Tran, Austin C Conklin, Anna K Golebiewski, Lindsey K Stolze, Michael B Whalen, Darren A Cusanovich, and Casey E Romanoski. Single-cell 'omic profiles of human aortic endothelial cells in vitro and human atherosclerotic lesions ex vivo reveal heterogeneity of endothelial subtype and response to activating perturbations. *eLife*, 12, 2024.

[13583] Bikash Adhikari, Katharina Schneider, Mathias Diebold, Christoph Sotriffer, and Elmar Wolf. Identification of suitable target/E3 ligase pairs for PROTAC development using a rapamycin-induced proximity assay (RiPA). *eLife*, 13, 2024.

[13584] Moneeza A Agha, Sandeep Kishore, and David L McLean. Cell-type-specific origins of locomotor rhythmicity at different speeds in larval zebrafish. *eLife*, 13, 2024.

[13585] Raphael Aguillon, Mieka Rinsky, Noa Simon-Blecher, Tirza Doniger, Lior Appelbaum, and Oren Levy. CLOCK evolved in cnidaria to synchronize internal rhythms with diel environmental cues. *eLife*, 12, 2024.

[13586] Saher Ahmed, David J Adams, Muzlifah Haniffa, Aidan Maartens, and Maria Augusta Arruda. Targeted action to increase inclusion at the Wellcome Sanger Institute. *eLife*, 13, 2024.

[13587] Munir Akkaya, Jafar Al Souz, Daniel Williams, Rahul Kamdar, Olena Kamenyeva, Juraj Kabat, Ethan Shevach, and Billur Akkaya. Illuminating T cell-dendritic cell interactions in vivo by FlAsHing antigens. *eLife*, 12, 2024.

[13588] Daniel C Akuma, Kimberly A Wodzanowski, Ronit Schwartz Wertman, Patrick M Exconde, Víctor R Vázquez Marrero, Chukwuma E Odunze, Daniel Grubaugh, Sunny Shin, Cornelius Taabazuing, and Igor E Brodsky. Catalytic activity and autoprocessing of murine caspase-11 mediate noncanonical inflammasome assembly in response to cytosolic LPS. *eLife*, 13, 2024.

[13589] David Alais, Jacob Coorey, Randolph Blake, and Matthew J Davidson. A new 'CFS tracking' paradigm reveals uniform suppression depth regardless of target complexity or salience. *eLife*, 12, 2024.

[13590] Danielle Algranati, Roni Oren, Bareket Dassa, Liat Fellus-Alyagor, Alexander Plotnikov, Haim Barr, Alon Harmelin, Nir London, Guy Ron, Noa Furth, and Efrat Shema. Dual targeting of histone deacetylases and MYC as potential treatment strategy for H3-K27M pediatric gliomas. *eLife*, 13, 2024.

[13591] Megan E Amason, Cole J Beatty, Carissa K Harvest, Daniel R Saban, and Edward A Miao. Chemokine expression profile of an innate granuloma. *eLife*, 13, 2024.

[13592] Tommaso Amico, Samuel Toluwanimi Dada, Andrea Lazzari, Michaela Brezinova, Antonio Trovato, Michele Vendruscolo, Monika Fuxreiter, and Amos Maritan. A scale-invariant log-normal droplet size distribution below the critical concentration for protein phase separation. *eLife*, 13, 2024.

[13593] Saira Amir, Olatunbosun Arowolo, Ekaterina Mironova, Joseph McGaunn, Oladele Oluwayiose, Oleg Sergeyev, J Richard Pilsner, and Alexander Suvorov. Mechanistic target of rapamycin (mTOR) pathway in Sertoli cells regulates age-dependent changes in sperm DNA methylation. *eLife*, 13, 2024.

[13594] Jake W Anderson, David Vaisar, David N Jones, Laurel M Pegram, Guy P Vigers, Huifen Chen, John G Moffat, and Natalie G Ahn. Conformation selection by ATP-competitive inhibitors and allosteric communication in ERK2. *eLife*, 12, 2024.

[13595] Chiara Andolina, Wouter Graumans, Moussa Guelbeogo, Geert-Jan van Gemert, Jordache Ramijth, Soré Harouna, Zongo Soumanaba, Rianne Stoter, Marga Vegte-Bolmer, Martina Pangos, Photini Sinnis, Katharine Collins, Sarah G Staedke, Alfred B Tiono, Chris Drakeley, Kjerstin Lanke, and Teun Bousema. Quantification of sporozoite expelling by Anopheles mosquitoes infected with laboratory and naturally circulating P. falciparum gametocytes. *eLife*, 12, 2024.

[13596] Clare Andradi-Brown, Jan Stephan Wichers-Misterek, Heidrun von Thien, Yannick D Höppner, Judith AM Scholz, Helle Hansson, Emma Filtenborg Hocke, Tim Wolf Gilberger, Michael F Duffy, Thomas Lavstsen, Jake Baum, Thomas D Otto, Aubrey J Cunnington, and Anna Bachmann. A novel computational pipeline for var gene expression augments the discovery of changes in the Plasmodium falciparum transcriptome during transition from in vivo to short-term in vitro culture. *eLife*, 12, 2024.

[13597] Arman Angaji, Michel Owusu, Christoph Velling, Nicola Dick, Donate Weghorn, and Johannes Berg. High-density sampling reveals volume growth in human tumours. *eLife*, 13, 2024.

[13598] Giovanni Anobile, Irene Petrizzo, Daisy Paiardini, David Burr, and Guido Marco Cicchini. Sensorimotor mechanisms selective to numerosity derived from individual differences. *eLife*, 12, 2024.

[13599] Marwan Anoud, Emmanuelle Delagoutte, Quentin Helleu, Alice Brion, Evelyne Duvernois-Berthet, Marie As, Xavier Marques, Khadija Lamribet, Catherine Senamaud-Beaufort, Laurent Jourdren, Annie Adrait, Sophie Heinrich, Geraldine Toutirais, Sahima Hamlaoui, Giacomo Gropplero, Ilaria Giovannini, Loic Ponger, Marc Geze, Corinne Blugeon, Yohann Couté, Roberto Guidetti, Lorena Rebecchi, Carine Giovannangeli, Anne De Cian, and Jean-Paul Concordet. Comparative transcriptomics reveal a novel tardigrade-specific DNA-binding protein induced in response to ionizing radiation. *eLife*, 13, 2024.

[13600] Lina Antenucci, Salla Virtanen, Chandan Thapa, Minne Jartti, Ilona Pitkänen, Helena Tossavainen, and Perttu Permi. Reassessing the substrate specificities of the major Staphylococcus aureus peptidoglycan hydrolases lysostaphin and LytM. *eLife*, 13, 2024.

[13601] Margaret L Antonio, Clemens L Weiß, Ziyue Gao, Susanna Sawyer, Victoria Oberreiter, Hannah M Moots, Jeffrey P Spence, Olivia Cheronet, Brina Zagorc, Elisa Praxmarer, Kadir Toykan özdoğan, Lea Demetz, Pere Gelabert, Daniel Fernandes, Michaela Lucci, Timka Alihodžić, Selma Amrani, Pavel Avetisyan, Christèle Baillif-Ducros, željka Bedić, Audrey Bertrand, Maja Bilić, Luca Bondioli, Paulina Borówka, Emmanuel Botte, Josip Burmaz, Domagoj Bužanić, Francesca Candilio, Mirna Cvetko, Daniela De Angelis, Ivan Drnić, Kristián Elschek, Mounir Fantar, Andrej Gaspari, Gabriella Gasperetti, Francesco Genchi, Snežana Golubović, Zuzana Hukeľová, Rimantas Jankauskas, Kristina Jelinčić Vučković, Gordana Jeremić, Iva Kaić, Kevin Kazek, Hamazasp Khachatryan, Anahit Khudaverdyan, Sylvia Kirchengast, Miomir Korać, Valérie Kozlowski, Mária KroŠláková, Dora KuŠan Špalj, Francesco La Pastina, Marie Laguardia, Sandra Legrand, Tino Leleković, Tamara Leskovar, Wiesław Lorkiewicz, Dženi Los, Ana Maria Silva, Rene Masaryk, Vinka Matijević, Yahia Mehdi Seddik Cherifi, Nicolas Meyer, Ilija Mikić, NataŠa Miladinović-Radmilović, Branka MiloŠević Zakić, Lina Nacouzi, Magdalena Natuniewicz-Sekuła, Alessia Nava, Christine Neugebauer-Maresch, Jan Nováček, Anna Osterholtz, Julianne Paige, Lujana Paraman, Dominique Pieri, Karol Pieta, Stefan Pop-Lazić, Matej Ruttkay, Mirjana Sanader, Arkadiusz Sołtysiak, Alessandra Sperduti, Tijana Stankovic Pesterac, Maria Teschler-Nicola, Iwona Teul, Domagoj Tončinić, Julien Trapp, Dragana Vulović, Tomasz Waliszewski, Diethard Walter, MiloŠ živanović, Mohamed el Mostefa Filah, Morana čauŠević Bully, Mario Šlaus, DuŠan Borić, Mario Novak, Alfredo Coppa, Ron Pinhasi, and Jonathan K Pritchard. Stable population structure in Europe since the Iron Age, despite high mobility. *eLife*, 13, 2024.

[13602] Alessandra Machado Araujo, Joseph D Dekker, Kendra Garrison, Zhe Su, Catherine Rhee, Zicheng Hu, Bum-Kyu Lee, Daniel Osorio, Jiwon Lee, Vishwanath R Iyer, Lauren IR Ehrlich, George Georgiou, Gregory Ippolito, Stephen Yi, and Haley O Tucker. Lymphoid origin of intrinsically activated plasmacytoid dendritic cells in mice. *eLife*, 13, 2024.

[13603] Josep-Maria Armengol-Collado, Livio Nicola Carenza, and Luca Giomi. Hydrodynamics and multiscale order in confluent epithelia. *eLife*, 13, 2024.

[13604] Erick E Arroyo-Pérez, John C Hook, Alejandra Alvarado, Stephan Wimmi, Timo Glatter, Kai Thormann, and Simon Ringgaard. A conserved cell-pole determinant organizes proper polar flagellum formation. *eLife*, 13, 2024.

[13605] Arvind Arul Nambi Rajan, Ryuta Asada, and Ben Montpetit. Gle1 is required for tRNA to stimulate Dbp5 ATPase activity in vitro and promote Dbp5-mediated tRNA export in vivo in Saccharomyces cerevisiae. *eLife*, 12, 2024.

[13606] Rieko Asai, Vivek N Prakash, Shubham Sinha, Manu Prakash, and Takashi Mikawa. Coupling and uncoupling of midline morphogenesis and cell flow in amniote gastrulation. *eLife*, 12, 2024.

[13607] Carolline AscenÇão, Jennie R Sims, Alexis Dziubek, William Comstock, Elizabeth A Fogarty, Jumana Badar, Raimundo Freire, Andrew Grimson, Robert S Weiss, Paula E Cohen, and Marcus B Smolka. A TOPBP1 allele causing male infertility uncouples XY silencing dynamics from sex body formation. *eLife*, 12, 2024.

[13608] Olgerta Asko, Alejandro Omar Blenkmann, Sabine Liliana Leske, Maja Dyhre Foldal, Anais LLorens, Ingrid Funderud, Torstein R Meling, Robert T Knight, Tor Endestad, and Anne-Kristin Solbakk. Altered hierarchical auditory predictive processing after lesions to the orbitofrontal cortex. *eLife*, 13, 2024.

[13609] Hasiba Asma, Ellen Tieke, Kevin D Deem, Jabale Rahmat, Tiffany Dong, Xinbo Huang, Yoshinori Tomoyasu, and Marc S Halfon. Regulatory genome annotation of 33 insect species. *eLife*, 13, 2024.

[13610] Keisuke Atsumi, Yuusuke Nishida, Masayuki Ushio, Hirotaka Nishi, Takanori Genroku, and Shogoro Fujiki. Boosting biodiversity monitoring using smartphone-driven, rapidly accumulating community-sourced data. *eLife*, 13, 2024.

[13611] Teresa Attenborough, Kate A Rawlinson, Carmen L Diaz Soria, Kirsty Ambridge, Geetha Sankaranarayanan, Jennie Graham, James A Cotton, Stephen R Doyle, Gabriel Rinaldi, and Matthew Berriman. A single-cell atlas of the miracidium larva of Schistosoma mansoni reveals cell types, developmental pathways, and tissue architecture. *eLife*, 13, 2024.

[13612] Nicolas Aubert, Madeleine Purcarea, Julien Novarino, Julien Schopp, Alexis Audibert, Wangtianrui Li, Marie Fornier, Léonie Cagnet, Marie Naturel, Armanda Casrouge, Marie-Caroline Dieu-Nosjean, Nicolas Blanchard, Gilles Dietrich, Cedric Peirs, and Gilles Marodon. Enkephalin-mediated modulation of basal somatic sensitivity by regulatory T cells in mice. *eLife*, 13, 2024.

[13613] Matias I Autio, Efthymios Motakis, Arnaud Perrin, Talal Bin Amin, Zenia Tiang, Dang Vinh Do, Jiaxu Wang, Joanna Tan, Shirley Suet Lee Ding, Wei Xuan Tan, Chang Jie Mick Lee, Adrian Kee Keong Teo, and Roger SY Foo. Computationally defined and in vitro validated putative genomic safe harbour loci for transgene expression in human cells. *eLife*, 13, 2024.

[13614] Simon Avrillon, FranÇois Hug, Roger M Enoka, Arnault HD Caillet, and Dario Farina. The identification of extensive samples of motor units in human muscles reveals diverse effects of neuromodulatory inputs on the rate coding. *eLife*, 13, 2024.

[13615] Alexandra a Aybar-Torres, Lennon A Saldarriaga, Ann T Pham, Amir M Emtiazjoo, Ashish K Sharma, Andrew j Bryant, and Lei Jin. The common Sting1 HAQ, AQ alleles rescue CD4 T cellpenia, restore T-regs, and prevent SAVI (N153S) inflammatory disease in mice. *eLife*, 13, 2024.

[13616] Valentin Babosha, Natalia Klimenko, Anastasia Revel-Muroz, Evgeniya Tikhonova, Pavel Georgiev, and Oksana Maksimenko. N-terminus of Drosophila melanogaster MSL1 is critical for dosage compensation. *eLife*, 13, 2024.

[13617] Sugith Badugu, Kshitiza Mohan Dhyani, Manoj Thakur, and Kalappa Muniyappa. Saccharomyces cerevisiae Rev7 promotes non-homologous end-joining by blocking Mre11 nuclease and Rad50's ATPase activities and homologous recombination. *eLife*, 13, 2024.

[13618] Seo-Gyeong Bae, Guo Nan Yin, Jiyeon Ock, Jun-Kyu Suh, Ji-Kan Ryu, and Jihwan Park. Single-cell transcriptome analysis of cavernous tissues reveals the key roles of pericytes in diabetic erectile dysfunction. *eLife*, 12, 2024.

[13619] Alexander S Baier, Nathan Gioacchini, Priit Eek, Erik M Leith, Song Tan, and Craig L Peterson. Dual engagement of the nucleosomal acidic patches is essential for deposition of histone H2A.Z by SWR1C. *eLife*, 13, 2024.

[13620] Ioannis Bakoyiannis, Eva Gunnel Ducourneau, Mateo N'diaye, Alice Fermigier, Celine Ducroix-Crepy, Clementine Bosch-Bouju, Etienne Coutureau, Pierre Trifilieff, and Guillaume Ferreira. Obesogenic diet induces circuit-specific memory deficits in mice. *eLife*, 13, 2024.

[13621] Revathi Balasubramanian, Krishnakumar Kizhatil, Taibo Li, Nicholas Tolman, Aakriti Bhandari, Graham Clark, Violet Bupp-Chickering, Ruth A Kelly, Sally Zhou, John Peregrin, Marina Simón, Christa Montgomery, W Daniel Stamer, Jiang Qian, and Simon WM John. Transcriptomic profiling of Schlemm's canal cells reveals a lymphatic-biased identity and three major cell states. *eLife*, 13, 2024.

[13622] Daniel Ballmer and Bungo Akiyoshi. Dynamic localization of the chromosomal passenger complex in trypanosomes is controlled by the orphan kinesins KIN-A and KIN-B. *eLife*, 13, 2024.

[13623] Gbolahan Bamgbose, Guillaume Bordet, Niraj Lodhi, and Alexei Tulin. Mono-methylated histones control PARP-1 in chromatin and transcription. *eLife*, 13, 2024.

[13624] Yi Ban, Yue Zou, Yingzhuo Liu, Sharrel Lee, Robert B Bednarczyk, Jianting Sheng, Yuliang Cao, Stephen TC Wong, and Dingcheng Gao. Targeting ribosome biogenesis as a novel therapeutic approach to overcome EMT-related chemoresistance in breast cancer. *eLife*, 12, 2024.

[13625] Paula Banca, Maria Herrojo Ruiz, Miguel Fernando Gonzalez-Zalba, Marjan Biria, Aleya A Marzuki, Thomas Piercy, Akeem Sule, Naomi A Fineberg, and Trevor W Robbins. Action sequence learning, habits, and automaticity in obsessive-compulsive disorder. *eLife*, 12, 2024.

[13626] Mischa Vance Bandet and Ian Robert Winship. Aberrant cortical activity, functional connectivity, and neural assembly architecture after photothrombotic stroke in mice. *eLife*, 12, 2024.

[13627] Rajdeep Banerjee, Thomas J Meyer, Margaret C Cam, Sukhbir Kaur, and David D Roberts. Differential regulation by CD47 and thrombospondin-1 of extramedullary erythropoiesis in mouse spleen. *eLife*, 12, 2024.

[13628] Jaskamaljot Kaur Banwait, Liana Islam, and Aaron L Lucius. Single turnover transient state kinetics reveals processive protein unfolding catalyzed by Escherichia coli ClpB. *eLife*, 13, 2024.

[13629] Mahta Barekatain, Yameng Liu, Ashley Archambeau, Vadim Cherezov, Scott Fraser, Kate L White, and Mark A Hayes. Insulator-based dielectrophoresis-assisted separation of insulin secretory vesicles. *eLife*, 13, 2024.

[13630] Jeffrey Barr, Austin Walz, Anthony C Restaino, Moran Amit, Sarah M Barclay, Elisabeth G Vichaya, William C Spanos, Robert Dantzer, Sebastien Talbot, and Paola D Vermeer. Tumor-infiltrating nerves functionally alter brain circuits and modulate behavior in a mouse model of head-and-neck cancer. *eLife*, 13, 2024.

[13631] Pierre Barrat-Charlaix and Richard A Neher. Eco-evolutionary dynamics of adapting pathogens and host immunity. *eLife*, 13, 2024.

[13632] Bethany M Bartlett, Yatendra Kumar, Shelagh Boyle, Tamoghna Chowdhury, Andrea Quintanilla, Charlene Boumendil, Juan Carlos Acosta, and Wendy A Bickmore. TPR is required for cytoplasmic chromatin fragment formation during senescence. *eLife*, 13, 2024.

[13633] Aleksandar Bartolome, Julia C Heiby, Domenico Di Fraia, Ivonne Heinze, Hannah Knaudt, Ellen Spaeth, Omid Omrani, Alberto Minetti, Maleen Hofmann, Joanna M Kirkpatrick, Therese Dau, and Alessandro Ori. Quantitative mapping of proteasome interactomes and substrates using ProteasomeID. *eLife*, 13, 2024.

[13634] Robert J Bastidas, Mateusz Kędzior, Robert K Davidson, Stephen C Walsh, Lee Dolat, Barbara S Sixt, Jonathan N Pruneda, Jorn Coers, and Raphael H Valdivia. The acetylase activity of Cdu1 regulates bacterial exit from infected cells by protecting Chlamydia effectors from degradation. *eLife*, 12, 2024.

[13635] Lisa Baumgartner, Jonathan J Ipsaro, Ulrich Hohmann, Dominik Handler, Alexander Schleiffer, Peter Duchek, and Julius Brennecke. Evolutionary adaptation of an HP1-protein chromodomain integrates chromatin and DNA sequence signals. *eLife*, 13, 2024.

[13636] Elizabeth A Beath, Cynthia Bailey, Meghana Mahantesh Magadam, Shuyan Qiu, Karen L McNally, and Francis J McNally. Katanin, kinesin-13, and ataxin-2 inhibit premature interaction between maternal and paternal genomes in C. elegans zygotes. *eLife*, 13, 2024.

[13637] Timothy E Behrens, Olujimi A Ajijola, and Mayank Chugh. How and why eLife selects papers for peer review. *eLife*, 13, 2024.

[13638] Clara Bekirian, Isabel Valsecchi, Sophie Bachellier-Bassi, Cyril Scandola, J Inaki Guijarro, Murielle Chauvel, Thierry Mourer, Neil AR Gow, Vishu Kumar Aimanianda, Christophe d'Enfert, and Thierry Fontaine. $\beta$-1,6-Glucan plays a central role in the structure and remodeling of the bilaminate fungal cell wall. *eLife*, 13, 2024.

[13639] Ryan T Bell, Harutyun Sahakyan, Kira S Makarova, Yuri I Wolf, and Eugene V Koonin. CoCoNuTs are a diverse subclass of Type IV restriction systems predicted to target RNA. *eLife*, 13, 2024.

[13640] Tristan A Bell, Bridget E Luce, Pusparanee Hakim, Virly Y Ananda, Hiba Dardari, Tran H Nguyen, Arezu Monshizadeh, and Luke H Chao. Prominin 1 and Tweety Homology 1 both induce extracellular vesicle formation. *eLife*, 13, 2024.

[13641] Julia Belyaeva and Matthias Elgeti. Exploring protein structural ensembles: Integration of sparse experimental data from electron paramagnetic resonance spectroscopy with molecular modeling methods. *eLife*, 13, 2024.

[13642] Roee Ben Nissan, Eliya Milshtein, Vanessa Pahl, Benoit de Pins, Ghil Jona, Dikla Levi, Hadas Yung, Noga Nir, Dolev Ezra, Shmuel Gleizer, Hannes Link, Elad Noor, and Ron Milo. Autotrophic growth of Escherichia coli is achieved by a small number of genetic changes. *eLife*, 12, 2024.

[13643] Sabrina Benas, Ximena Fernandez, and Emilio Kropff. Modeled grid cells aligned by a flexible attractor. *eLife*, 12, 2024.

[13644] Florian Bénitière, Anamaria Necsulea, and Laurent Duret. Random genetic drift sets an upper limit on mRNA splicing accuracy in metazoans. *eLife*, 13, 2024.

[13645] Leif Benner, Savannah Muron, Jillian G Gomez, and Brian Oliver. OVO positively regulates essential maternal pathways by binding near the transcriptional start sites in the Drosophila female germline. *eLife*, 13, 2024.

[13646] Kelly A Bennion, Jade Phong, Mytien Le, Kunhua Cheng, Christopher N Wahlheim, and James W Antony. Semantic relatedness proactively benefits learning, memory, and interdependence across episodes. *eLife*, 13, 2024.

[13647] Marco Bergamini, Alberto Dalla Volta, Carlotta Palumbo, Stefania Zamboni, Luca Triggiani, Manuel Zamparini, Marta Laganà, Luca Rinaudo, Nunzia Di Meo, Irene Caramella, Roberto Bresciani, Francesca Valcamonico, Paolo Borghetti, Andrea Guerini, Davide Farina, Alessandro Antonelli, Claudio Simeone, Gherardo Mazziotti, and Alfredo Berruti. Relationship between circulating FSH levels and body composition and bone health in patients with prostate cancer who undergo androgen deprivation therapy: The BLADE study. *eLife*, 13, 2024.

[13648] Michael Berger, Michèle Fraatz, Katrin Auweiler, Katharina Dorn, Tanna El Khadrawe, and Henrike Scholz. Octopamine integrates the status of internal energy supply into the formation of food-related memories. *eLife*, 12, 2024.

[13649] Cathrine Bergh, UrŠka RovŠnik, Rebecca Howard, and Erik Lindahl. Discovery of lipid binding sites in a ligand-gated ion channel by integrating simulations and cryo-EM. *eLife*, 12, 2024.

[13650] Corentin Bernou, Marc-André Mouthon, Mathieu Daynac, Thierry Kortulewski, Benjamin Demaille, Vilma Barroca, Sebastien Couillard-Despres, Nathalie Dechamps, Véronique Ménard, Léa Bellenger, Christophe Antoniewski, Alexandra Déborah Chicheportiche, and FranÇois Dominique Boussin. Switching of RNA splicing regulators in immature neuroblasts during adult neurogenesis. *eLife*, 12, 2024.

[13651] Augusto Berrocal, Nicholas C Lammers, Hernan G Garcia, and Michael B Eisen. Unified bursting strategies in ectopic and endogenous even-skipped expression patterns. *eLife*, 12, 2024.

[13652] Michele Bertacchi, Gwendoline Maharaux, Agnès Loubat, Matthieu Jung, and Michèle Studer. FGF8-mediated gene regulation affects regional identity in human cerebral organoids. *eLife*, 13, 2024.

[13653] Elazar Besser, Anat Gelfand, Shiri Procaccia, Paula Berman, and David Meiri. Cannabinoid combination targets NOTCH1-mutated T-cell acute lymphoblastic leukemia through the integrated stress response pathway. *eLife*, 12, 2024.

[13654] Yelyzaveta Bessonova and Baranidharan Raman. Serotonergic amplification of odor-evoked neural responses maps onto flexible behavioral outcomes. *eLife*, 12, 2024.

[13655] Mitchell Bestry, Alexander N Larcombe, Nina Kresoje, Emily K Chivers, Chloe Bakker, James P Fitzpatrick, Elizabeth J Elliott, Jeffrey M Craig, Evelyne Muggli, Jane Halliday, Delyse Hutchinson, Sam Buckberry, Ryan Lister, Martyn Symons, and David Martino. Early moderate prenatal alcohol exposure and maternal diet impact offspring DNA methylation across species. *eLife*, 12, 2024.

[13656] Luis Alberto Bezares Calderón, Réza Shahidi, and Gáspár Jékely. Mechanism of barotaxis in marine zooplankton. *eLife*, 13, 2024.

[13657] Lijun Bian, Zhimin Ma, Xiangjin Fu, Chen Ji, Tianpei Wang, Caiwang Yan, Juncheng Dai, Hongxia Ma, Zhibin Hu, Hongbing Shen, Lu Wang, Meng Zhu, and Guangfu Jin. Associations of combined phenotypic aging and genetic risk with incident cancer: A prospective cohort study. *eLife*, 13, 2024.

[13658] Rob Bierman, Jui M Dave, Daniel M Greif, and Julia Salzman. Statistical analysis supports pervasive RNA subcellular localization and alternative 3′ UTR regulation. *eLife*, 12, 2024.

[13659] Zachary Paul Billman, Stephen Bela Kovacs, Bo Wei, Kidong Kang, Ousmane H Cissé, and Edward A Miao. Caspase-1 activates gasdermin A in non-mammals. *eLife*, 12, 2024.

[13660] Ornella Bimai, Ipsita Banerjee, Inna Rozman Grinberg, Ping Huang, Lucas Hultgren, Simon Ekström, Daniel Lundin, Britt-Marie Sjöberg, and Derek T Logan. Nucleotide binding to the ATP-cone in anaerobic ribonucleotide reductases allosterically regulates activity by modulating substrate binding. *eLife*, 12, 2024.

[13661] Ikhwan Bin Khalid, Eric T Reifenstein, Naomi Auer, Lukas Kunz, and Richard Kempter. Quantitative modeling of the emergence of macroscopic grid-like representations. *eLife*, 13, 2024.

[13662] Xinyang Bing, Wenfan Ke, Miki Fujioka, Amina Kurbidaeva, Sarah Levitt, Mike Levine, Paul Schedl, and James B Jaynes. Chromosome structure in Drosophila is determined by boundary pairing not loop extrusion. *eLife*, 13, 2024.

[13663] Helmut Bischof, Selina Maier, Piotr Koprowski, Bogusz Kulawiak, Sandra Burgstaller, Joanna Jasińska, Kristian Serafimov, Monika Zochowska, Dominic Gross, Werner Schroth, Lucas Matt, David Arturo Juarez Lopez, Ying Zhang, Irina Bonzheim, Florian A Büttner, Falko Fend, Matthias Schwab, Andreas L Birkenfeld, Roland Malli, Michael Lämmerhofer, Piotr Bednarczyk, Adam Szewczyk, and Robert Lukowski. mitoBKCa is functionally expressed in murine and human breast cancer cells and potentially contributes to metabolic reprogramming. *eLife*, 12, 2024.

[13664] Parijat Biswas, Priyanka Roy, Subhamoy Jana, Dipanjan Ray, Jibitesh Das, Bipasa Chaudhuri, Ridita Ray Basunia, Bidisha Sinha, and Deepak Kumar Sinha. Exploring the role of macromolecular crowding and TNFR1 in cell volume control. *eLife*, 13, 2024.

[13665] Sara Bitar, Timo Baumann, Christopher Weber, Majd Abusaada, Liliana Rojas-Charry, Patrick Ziegler, Thomas Schettgen, Isabella Eva Randerath, Vivek Venkataramani, Bernhard Michalke, Eva-Maria Hanschmann, Giuseppe Arena, Rejko Krueger, Li Zhang, and Axel Methner. Iron-sulfur cluster loss in mitochondrial CISD1 mediates PINK1 loss-of-function phenotypes. *eLife*, 13, 2024.

[13666] Amanda Mixon Blackwell, Yasaman Jami-Alahmadi, Armiyaw S Nasamu, Shota Kudo, Akinobu Senoo, Celine Slam, Kouhei Tsumoto, James A Wohlschlegel, Jose Manuel Martinez Caaveiro, Daniel E Goldberg, and Paul A Sigala. Malaria parasites require a divergent heme oxygenase for apicoplast gene expression and biogenesis. *eLife*, 13, 2024.

[13667] Andrew S Blaeser, Jun Zhao, Arthur U Sugden, Simone Carneiro-Nascimento, Mark L Andermann, and Dan Levy. Sensitization of meningeal afferents to locomotion-related meningeal deformations in a migraine model. *eLife*, 12, 2024.

[13668] Maryline Blin, Louis Valay, Manon Kuratko, Marie Pavie, and Sylvie Rétaux. The evolution of olfactory sensitivity, preferences, and behavioral responses in Mexican cavefish is influenced by fish personality. *eLife*, 12, 2024.

[13669] David B Blumenthal, Marta Lucchetta, Linda Kleist, Sándor P Fekete, Markus List, and Martin H Schaefer. Emergence of power law distributions in protein-protein interaction networks through study bias. *eLife*, 13, 2024.

[13670] Desiree Böck, Maria Wilhelm, Jonas Mumenthaler, Daniel Fabio Carpanese, Peter I Kulcsár, Simon d'Aquin, Alessio Cremonesi, Anahita Rassi, Johannes Häberle, Tommaso Patriarchi, and Gerald Schwank. Base editing of Ptbp1 in neurons alleviates symptoms in a mouse model of Parkinson's disease. *eLife*, 13, 2024.

[13671] Bruno Bockorny, Lakshmi Muthuswamy, Ling Huang, Marco Hadisurya, Christine Maria Lim, Leo L Tsai, Ritu R Gill, Jesse L Wei, Andrea J Bullock, Joseph E Grossman, Robert J Besaw, Supraja Narasimhan, Weiguo Andy Tao, Sofia Perea, Mandeep S Sawhney, Steven D Freedman, Manuel Hildago, Anton Iliuk, and Senthil K Muthuswamy. A large-scale proteomics resource of circulating extracellular vesicles for biomarker discovery in pancreatic cancer. *eLife*, 12, 2024.

[13672] Juan Carlos Boffi, Brice Bathellier, Hiroki Asari, and Robert Prevedel. Noisy neuronal populations effectively encode sound localization in the dorsal inferior colliculus of awake mice. *eLife*, 13, 2024.

[13673] Nicholas M Boffi, Yipei Guo, Chris H Rycroft, and Ariel Amir. How microscopic epistasis and clonal interference shape the fitness trajectory in a spin glass model of microbial long-term evolution. *eLife*, 12, 2024.

[13674] Valentin Bohl, Nele Merret Hollmann, Tobias Melzer, Panagiotis Katikaridis, Lena Meins, Bernd Simon, Dirk Flemming, Irmgard Sinning, Janosch Hennig, and Axel Mogk. The Listeria monocytogenes persistence factor ClpL is a potent stand-alone disaggregase. *eLife*, 12, 2024.

[13675] Laura Boi, Yvonne Johansson, Raffaella Tonini, Rosario Moratalla, Gilberto Fisone, and Gilad Silberberg. Serotonergic and dopaminergic neurons in the dorsal raphe are differentially altered in a mouse model for parkinsonism. *eLife*, 12, 2024.

[13676] Simona Bolamperti, Hiroaki Saito, Sarah Heerdmann, Eric Hesse, and Hanna Taipaleenmäki. Tgif1-deficiency impairs cytoskeletal architecture in osteoblasts by activating PAK3 signaling. *eLife*, 13, 2024.

[13677] Sara Bolívar, Elisenda Sanz, David Ovelleiro, Douglas W Zochodne, and Esther Udina. Neuron-specific RNA-sequencing reveals different responses in peripheral neurons after nerve injury. *eLife*, 12, 2024.

[13678] Ewa K Bomba-Warczak, Karen M Velez, Luhan T Zhou, Christelle Guillermier, Seby Edassery, Matthew L Steinhauser, Jeffrey N Savas, and Francesca E Duncan. Exceptional longevity of mammalian ovarian and oocyte macromolecules throughout the reproductive lifespan. *eLife*, 13, 2024.

[13679] Artem N Bonchuk, Konstantin I Balagurov, Rozbeh Baradaran, Konstantin M Boyko, Nikolai N Sluchanko, Anastasia M Khrustaleva, Anna D Burtseva, Olga V Arkova, Karina K Khalisova, Vladimir O Popov, Andreas Naschberger, and Pavel G Georgiev. The Arthropoda-specific Tramtrack group BTB protein domains use previously unknown interface to form hexamers. *eLife*, 13, 2024.

[13680] Sonia A Boor, Joshua D Meisel, and Dennis H Kim. Neuroendocrine gene expression coupling of interoceptive bacterial food cues to foraging behavior of C. elegans. *eLife*, 12, 2024.

[13681] Swagata Bose, Satya Ranjan Sahu, Abinash Dutta, and Narottam Acharya. A chemically induced attenuated strain of Candida albicans generates robust protective immune responses and prevents systemic candidiasis development. *eLife*, 13, 2024.

[13682] Sebastien Bouret, Emmanuel Paradis, Sandrine Prat, Laurie Castro, Pauline Perez, Emmanuel Gilissen, and Cecile Garcia. Linking the evolution of two prefrontal brain regions to social and foraging challenges in primates. *eLife*, 12, 2024.

[13683] Caroline M Boyd, Sundharraman Subramanian, Drew T Dunham, Kristin N Parent, and Kimberley D Seed. A Vibrio cholerae viral satellite maximizes its spread and inhibits phage by remodeling hijacked phage coat proteins into small capsids. *eLife*, 12, 2024.

[13684] Carly Boye, Cynthia A Kalita, Anthony S Findley, Adnan Alazizi, Julong Wei, Xiaoquan Wen, Roger Pique-Regi, and Francesca Luca. Characterization of caffeine response regulatory variants in vascular endothelial cells. *eLife*, 13, 2024.

[13685] Kathryn Branam, Joshua I Gold, and Long Ding. The subthalamic nucleus contributes causally to perceptual decision-making in monkeys. *eLife*, 13, 2024.

[13686] Rion Brattig-Correia, Joana M Almeida, Margot Julia Wyrwoll, Irene Julca, Daniel Sobral, Chandra Shekhar Misra, Sara Di Persio, Leonardo Gastón Guilgur, Hans-Christian Schuppe, Neide Silva, Pedro Prudêncio, Ana Nóvoa, Ana S Leocádio, Joana Bom, Sandra Laurentino, Moises Mallo, Sabine Kliesch, Marek Mutwil, Luis M Rocha, Frank Tüttelmann, Jörg D Becker, and Paulo Navarro-Costa. The conserved genetic program of male germ cells uncovers ancient regulators of human spermatogenesis. *eLife*, 13, 2024.

[13687] Iliana E Bray, Stephen E Clarke, Kerriann M Casey, and Paul Nuyujukian. Neuroelectrophysiology-compatible electrolytic lesioning. *eLife*, 12, 2024.

[13688] Alexis J Breen and Dominik Deffner. Risk-sensitive learning is a winning strategy for leading an urban invasion. *eLife*, 12, 2024.

[13689] Marie Breeur, George Stepaniants, Pekka Keski-Rahkonen, Philippe Rigollet, and Vivian Viallon. Optimal transport for automatic alignment of untargeted metabolomic data. *eLife*, 12, 2024.

[13690] Jordan Breffle, Hannah Germaine, Justin D Shin, Shantanu P Jadhav, and Paul Miller. Intrinsic dynamics of randomly clustered networks generate place fields and preplay of novel environments. *eLife*, 13, 2024.

[13691] Alejandro J Brenes, Eva Griesser, Linda V Sinclair, Lindsay Davidson, Alan R Prescott, Francois Singh, Elizabeth KJ Hogg, Carmen Espejo-Serrano, Hao Jiang, Harunori Yoshikawa, Melpomeni Platani, Jason R Swedlow, Greg M Findlay, Doreen A Cantrell, and Angus I Lamond. Proteomic and functional comparison between human induced and embryonic stem cells. *eLife*, 13, 2024.

[13692] Marian Brenner, Christoph Zink, Linda Witzinger, Angelika Keller, Kerstin Hadamek, Sebastian Bothe, Martin Neuenschwander, Carmen Villmann, Jens Peter von Kries, Hermann Schindelin, Elisabeth Jeanclos, and Antje Gohla. 7,8-Dihydroxyflavone is a direct inhibitor of human and murine pyridoxal phosphatase. *eLife*, 13, 2024.

[13693] Deborah H Brotherton, Sarbjit Nijjar, Christos G Savva, Nicholas Dale, and Alexander David Cameron. Structures of wild-type and a constitutively closed mutant of connexin26 shed light on channel regulation by CO2. *eLife*, 13, 2024.

[13694] Emma Brown, Gemma Swinscoe, Daniella A Lefteri, Ravi Singh, Amy Moran, Rebecca F Thompson, Daniel Maskell, Hannah Beaumont, Matthew J Bentham, Claire Donald, Alain Kohl, Andrew Macdonald, Neil Ranson, Richard Foster, Clive S McKimmie, Antreas C Kalli, and Stephen Griffin. Inhibitors of the small membrane (M) protein viroporin prevent Zika virus infection. *eLife*, 13, 2024.

[13695] Patrick E Brown, Sze Hang Fu, Leslie Newcombe, Xuyang Tang, Nico Nagelkerke, H Chaim Birnboim, Aiyush Bansal, Karen Colwill, Geneviève Mailhot, Melanie Delgado-Brand, Tulunay Tursun, Freda Qi, Anne-Claude Gingras, Arthur S Slutsky, Maria D Pasic, Jeffrey Companion, Isaac I Bogoch, Ed Morawski, Teresa Lam, Angus Reid, and Prabhat Jha. Hybrid immunity from severe acute respiratory syndrome coronavirus 2 infection and vaccination in Canadian adults: A cohort study. *eLife*, 13, 2024.

[13696] Nicolas G Brukman, Clari Valansi, and Benjamin Podbilewicz. Sperm induction of somatic cell-cell fusion as a novel functional test. *eLife*, 13, 2024.

[13697] James D Brunner and Nicholas Chia. Metabolic model-based ecological modeling for probiotic design. *eLife*, 13, 2024.

[13698] Nikaela W Bryan, Aamir Ali, Ewa Niedzialkowska, Leland Mayne, P Todd Stukenberg, and Ben E Black. Structural basis for the phase separation of the chromosome passenger complex. *eLife*, 13, 2024.

[13699] J Barney Bryson, Alexandra Kourgiantaki, Dai Jiang, Andreas Demosthenous, and Linda Greensmith. An optogenetic cell therapy to restore control of target muscles in an aggressive mouse model of amyotrophic lateral sclerosis. *eLife*, 12, 2024.

[13700] Yuanyuan Bu, Xingye Dong, Rongrong Zhang, Xianglian Shen, Yan Liu, Shu Wang, Tetsuo Takano, and Shenkui Liu. Unraveling the role of urea hydrolysis in salt stress response during seed germination and seedling growth in Arabidopsis thaliana. *eLife*, 13, 2024.

[13701] Lara Busby, Guillermo Serrano Nájera, and Benjamin John Steventon. Intrinsic and extrinsic cues time somite progenitor contribution to the vertebrate primary body axis. *eLife*, 13, 2024.

[13702] Francesca Butera, Julia E Sero, Lucas G Dent, and Chris Bakal. Actin networks modulate heterogeneous NF-$\kappa$B dynamics in response to TNF$\alpha$. *eLife*, 13, 2024.

[13703] Anamarija Butkovic, Thomas James Ellis, Ruben Gonzalez, Benjamin Jaegle, Magnus Nordborg, and Santiago F Elena. Genetic basis of Arabidopsis thaliana responses to infection by naïve and adapted isolates of turnip mosaic virus. *eLife*, 12, 2024.

[13704] Yuranny Cabral-Calderin, Daniela van Hinsberg, Axel Thielscher, and Molly J Henry. Behavioral entrainment to rhythmic auditory stimulation can be modulated by tACS depending on the electrical stimulation field properties. *eLife*, 12, 2024.

[13705] Birol Cabukusta, Shalom Borst Pauwels, Jimmy JLL Akkermans, Niek Blomberg, Aat A Mulder, Roman I Koning, Martin Giera, and Jacques Neefjes. The ORP9-ORP11 dimer promotes sphingomyelin synthesis. *eLife*, 12, 2024.

[13706] Cristina Cacho-Navas, Carmen López-Pujante, Natalia Reglero-Real, Natalia Colás-Algora, Ana Cuervo, Jose Javier Conesa, Susana Barroso, Gema de Rivas, Sergio Ciordia, Alberto Paradela, Gianluca D'Agostino, Carlo Manzo, Jorge Feito, Germán Andrés, Francisca Molina-Jiménez, Pedro Majano, Isabel Correas, José-Maria Carazo, Sussan Nourshargh, Meritxell Huch, and Jaime Millán. ICAM-1 nanoclusters regulate hepatic epithelial cell polarity by leukocyte adhesion-independent control of apical actomyosin. *eLife*, 12, 2024.

[13707] Yuqi Cai, Zhirong Zhao, Mingyue Shi, Mingfang Zheng, Ling Gong, and Miao He. Embryonic origins of forebrain oligodendrocytes revisited by combinatorial genetic fate mapping. *eLife*, 13, 2024.

[13708] Islay Campbell, Roya Sharifpour, Jose Fermin Balda Aizpurua, Elise Beckers, Ilenia Paparella, Alexandre Berger, Ekaterina Koshmanova, Nasrin Mortazavi, John Read, Mikhail Zubkov, Puneet Talwar, Fabienne Collette, Siya Sherif, Christophe Phillips, Laurent Lamalle, and Gilles Vandewalle. Regional response to light illuminance across the human hypothalamus. *eLife*, 13, 2024.

[13709] Maria Cristina Cannarsa, Filippo Liguori, Nicola Pellicciotta, Giacomo Frangipane, and Roberto Di Leonardo. Light-driven synchronization of optogenetic clocks. *eLife*, 13, 2024.

[13710] Liyu Cao. A spatial-attentional mechanism underlies action-related distortions of time judgment. *eLife*, 12, 2024.

[13711] Silvia Cardani, Tara A Janes, William Betzner, and Silvia Pagliardini. Knockdown of PHOX2B in the retrotrapezoid nucleus reduces the central CO2 chemoreflex in rats. *eLife*, 13, 2024.

[13712] Sandra P Cárdenas-García, Sundas Ijaz, and Alberto E Pereda. The components of an electrical synapse as revealed by expansion microscopy of a single synaptic contact. *eLife*, 13, 2024.

[13713] Marcelo A Carignano, Martin Kroeger, Luay M Almassalha, Vasundhara Agrawal, Wing Shun Li, Emily M Pujadas-Liwag, Rikkert J Nap, Vadim Backman, and Igal Szleifer. Local volume concentration, packing domains, and scaling properties of chromatin. *eLife*, 13, 2024.

[13714] Shannon H Carroll, Sogand Schafer, Kenta Kawasaki, Casey Tsimbal, Amelie M Jule, Shawn A Hallett, Edward Li, and Eric C Liao. Genetic requirement of dact1/2 to regulate noncanonical Wnt signaling and calpain 8 during embryonic convergent extension and craniofacial morphogenesis. *eLife*, 13, 2024.

[13715] Oceane Cassan, Lea-Lou Pimpare, Timothy Mozzanino, Cecile Fizames, Sebastien Devidal, Fabrice Roux, Alexandru Milcu, Sophie Lebre, Alain Gojon, and Antoine Martin. Natural genetic variation underlying the negative effect of elevated CO2 on ionome composition in Arabidopsis thaliana. *eLife*, 12, 2024.

[13716] Ivan Castello-Serrano, Frederick A Heberle, Barbara Diaz-Rohrer, Rossana Ippolito, Carolyn R Shurer, Pablo Lujan, Felix Campelo, Kandice R Levental, and Ilya Levental. Partitioning to ordered membrane domains regulates the kinetics of secretory traffic. *eLife*, 12, 2024.

[13717] João Paulo Portela Catani, Anouk Smet, Tine Ysenbaert, Marnik Vuylsteke, Guy Bottu, Janick Mathys, Alexander Botzki, Guadalupe Cortes-Garcia, Tod Strugnell, Raul Gomila, John Hamberger, John Catalan, Irina V Ustyugova, Timothy Farrell, Svetlana Stegalkina, Satyajit Ray, Lauren LaRue, Xavier Saelens, and Thorsten U Vogel. The antigenic landscape of human influenza N2 neuraminidases from 2009 until 2017. *eLife*, 12, 2024.

[13718] Anna Cattani, Don B Arnold, Michelle McCarthy, and Nancy Kopell. Basolateral amygdala oscillations enable fear learning in a biophysical model. *eLife*, 12, 2024.

[13719] J Leonie Cazemier, Robin Haak, TK Loan Tran, Ann TY Hsu, Medina Husic, Brandon D Peri, Lisa Kirchberger, Matthew W Self, Pieter Roelfsema, and J Alexander Heimel. Involvement of superior colliculus in complex figure detection of mice. *eLife*, 13, 2024.

[13720] Cenk Celik, Stella Tue Ting Lee, Frederick Reinhart Tanoto, Mark Veleba, Kimberly Kline, and Guillaume Thibault. Decoding the complexity of delayed wound healing following Enterococcus faecalis infection. *eLife*, 13, 2024.

[13721] Antonio Cerdán Cerdá, Nicola Toschi, Constantina A Treaba, Valeria Barletta, Elena Herranz, Ambica Mehndiratta, Jose A Gomez-Sanchez, Caterina Mainero, and Silvia De Santis. A translational MRI approach to validate acute axonal damage detection as an early event in multiple sclerosis. *eLife*, 13, 2024.

[13722] Zeinab M Chahine, Mohit Gupta, Todd Lenz, Thomas Hollin, Steven Abel, Charles Banks, Anita Saraf, Jacques Prudhomme, Suhani Bhanvadia, Laurence A Florens, and Karine G Le Roch. PfMORC protein regulates chromatin accessibility and transcriptional repression in the human malaria parasite, Plasmodium falciparum. *eLife*, 12, 2024.

[13723] Arijit Chakraborty, Arunava Bandyopadhaya, Vijay K Singh, Filip Kovacic, Sujin Cha, William M Oldham, A Aria Tzika, and Laurence G Rahme. The bacterial quorum sensing signal 2'-aminoacetophenone rewires immune cell bioenergetics through the Ppargc1a/Esrra axis to mediate tolerance to infection. *eLife*, 13, 2024.

[13724] Michael James Chambers, Sophia B Scobell, and Meru J Sadhu. Systematic genetic characterization of the human PKR kinase domain highlights its functional malleability to escape a poxvirus substrate mimic. *eLife*, 13, 2024.

[13725] Laura M Chamness, Charles P Kuntz, Andrew G McKee, Wesley D Penn, Christopher M Hemmerich, Douglas B Rusch, Hope Woods, Dyotima, Jens Meiler, and Jonathan P Schlebach. Divergent folding-mediated epistasis among unstable membrane protein variants. *eLife*, 12, 2024.

[13726] Marina Chan, Yuqi Kang, Shannon Osborne, Michael Zager, and Taranjit S Gujral. A kinase to cytokine explorer to identify molecular regulators and potential therapeutic opportunities. *eLife*, 12, 2024.

[13727] Caleb Chang, Grace Zhou, and Yang Gao. Observing one-divalent-metal-ion-dependent and histidine-promoted His-Me family I-PpoI nuclease catalysis in crystallo. *eLife*, 13, 2024.

[13728] Chao Chang, Sara L Banerjee, Sung Soon Park, Xiao Lei Zhang, David Cotnoir-White, Karla J Opperman, Muriel Desbois, Brock Grill, and Artur Kania. Ubiquitin ligase and signalling hub MYCBP2 is required for efficient EPHB2 tyrosine kinase receptor function. *eLife*, 12, 2024.

[13729] Ray Chang, Ari Davydov, Pattana Jaroenlak, Breane Budaitis, Damian C Ekiert, Gira Bhabha, and Manu Prakash. Energetics of the mokicrosporidian polar tube invasion machinery. *eLife*, 12, 2024.

[13730] Yi-Shin Chang, Kai Huang, Jessica M Lee, Christen L Vagts, Christian Ascoli, Md-Ruhul Amin, Mahmood Ghassemi, Claudia M Lora, Russell Edafetanure-Ibeh, Yue Huang, Ruth A Cherian, Nandini Sarup, Samantha R Warpecha, Sunghyun Hwang, Rhea Goel, Benjamin A Turturice, Cody Schott, Montserrat Hernandez, Yang Chen, Julianne Jorgensen, Wangfei Wang, Mladen Rasic, Richard M Novak, Patricia W Finn, and David L Perkins. Altered transcriptomic immune responses of maintenance hemodialysis patients to the COVID-19 mRNA vaccine. *eLife*, 13, 2024.

[13731] Yi-Ting Chang, Eric A Finkel, Duo Xu, and Daniel H O'Connor. Rule-based modulation of a sensorimotor transformation across cortical areas. *eLife*, 12, 2024.

[13732] Yuan Chang, Savannah Lusk, Andersen Chang, Christopher S Ward, and Russell S Ray. Vglut2-based glutamatergic signaling in central noradrenergic neurons is dispensable for normal breathing and chemosensory reflexes. *eLife*, 12, 2024.

[13733] Ana C Chang-Gonzalez, Robert J Mallis, Matthew J Lang, Ellis L Reinherz, and Wonmuk Hwang. Asymmetric framework motion of TCR$\alpha\beta$ controls load-dependent peptide discrimination. *eLife*, 13, 2024.

[13734] Charlotte F Chao, Yanina-Yasmin Pesch, Huaxu Yu, Chenjingyi Wang, Maria J Aristizabal, Tao Huan, Guy Tanentzapf, and Elizabeth Rideout. An important role for triglyceride in regulating spermatogenesis. *eLife*, 12, 2024.

[13735] Lin Chao, Chun Kuen Chan, Chao Shi, and Ulla Camilla Rang. Spatial and temporal distribution of ribosomes in single cells reveals aging differences between old and new daughters of Escherichia coli. *eLife*, 12, 2024.

[13736] Shuo Chao, Jun Lu, Li-Jun Li, Hong-Yan Guo, Kuipeng Xu, Ning Wang, Shu-Xian Zhao, Xiao-Wen Jin, Shao-Ge Wang, Shen Yin, Wei Shen, Ming-Hui Zhao, Gui-An Huang, Qing-Yuan Sun, and Zhao-Jia Ge. Maternal obesity may disrupt offspring metabolism by inducing oocyte genome hyper-methylation via increased DNMTs. *eLife*, 13, 2024.

[13737] Nyasha Charura, Ernesto Llamas, Concetta De Quattro, David Vilchez, Moritz K Nowack, and Alga Zuccaro. Root cap cell corpse clearance limits microbial colonization in Arabidopsis thaliana. *eLife*, 13, 2024.

[13738] Sara Chelaghma, Huiling Ke, Konstantin Barylyuk, Thomas Krueger, Ludek Koreny, and Ross F Waller. Apical annuli are specialised sites of post-invasion secretion of dense granules in Toxoplasma. *eLife*, 13, 2024.

[13739] Claire C Chen, Joseph Han, Carlene A Chinn, Jacob S Rounds, Xiang Li, Mehran Nikan, Marie Myszka, Liqi Tong, Luiz FM Passalacqua, Timothy Bredy, Marcelo A Wood, and Andrej Luptak. Inhibition of Cpeb3 ribozyme elevates CPEB3 protein expression and polyadenylation of its target mRNAs and enhances object location memory. *eLife*, 13, 2024.

[13740] Hedi Chen, Xiaoyu Fan, Shuqian Zhu, Yuchan Pei, Xiaochun Zhang, Xiaonan Zhang, Lihang Liu, Feng Qian, and Boxue Tian. Accurate prediction of CDR-H3 loop structures of antibodies with deep learning. *eLife*, 12, 2024.

[13741] Hong Chen, Lijun Sun, Lu Feng, Xue Han, Yunhua Zhang, Wenbo Zhai, Zehe Zhang, Michael Mulholland, Weizhen Zhang, and Yue Yin. Intermittent fasting promotes type 3 innate lymphoid cells secreting IL-22 contributing to the beigeing of white adipose tissue. *eLife*, 12, 2024.

[13742] Hui Chen, Sung-Jin Lee, Ryan Li, Asmiti Sura, Nicholas Suen, Archana Dilip, Yan Pomogov, Meghah Vuppalapaty, Timothy T Suen, Chenggang Lu, Yorick Post, and Yang Li. BRAIDing receptors for cell-specific targeting. *eLife*, 12, 2024.

[13743] Jiali Chen, Qinghe Zeng, Xu Wang, Rui Xu, Weidong Wang, Yuliang Huang, Qi Sun, Wenhua Yuan, Pinger Wang, Di Chen, Peijian Tong, and Hongting Jin. Aberrant methylation and expression of TNXB promote chondrocyte apoptosis and extracullar matrix degradation in hemophilic arthropathy via AKT signaling. *eLife*, 13, 2024.

[13744] Jianwen Chen, Bao Zhao, Hong Dong, Tianliang Li, Xiang Cheng, Wang Gong, Jing Wang, Junran Zhang, Gang Xin, Yanbao Yu, Yu L Lei, Jennifer D Black, Zihai Li, and Haitao Wen. Inhibition of O-GlcNAc transferase activates type I interferon-dependent antitumor immunity by bridging cGAS-STING pathway. *eLife*, 13, 2024.

[13745] Jinjing Chen, Ruoyu Wang, Feng Xiong, Hao Sun, Byron Kemper, Wenbo Li, and Jongsook Kemper. Hammerhead-type FXR agonists induce an enhancer RNA Fincor that ameliorates nonalcoholic steatohepatitis in mice. *eLife*, 13, 2024.

[13746] Jiyun Chen, Hong Chen, Shanshan Li, Xiaofeng Lin, Rong Hu, Kaiming Zhang, and Liang Liu. Structural and mechanistic insights into ribosomal ITS2 RNA processing by nuclease-kinase machinery. *eLife*, 12, 2024.

[13747] Ming-Ming Chen, Yue Zhao, Kun Yu, Xue-Ling Xu, Xiao-Sheng Zhang, Jin-Long Zhang, Su-Jun Wu, Zhi-Mei Liu, Yi-Ming Yuan, Xiao-Fei Guo, Shi-Yu Qi, Guang Yi, Shu-Qi Wang, Huang-Xiang Li, Ao-Wu Wu, Guo-Shi Liu, Shou-Long Deng, Hong-Bing Han, Feng-Hua Lv, Di Lian, and Zheng-Xing Lian. A MSTNDel73C mutation with FGF5 knockout sheep by CRISPR/Cas9 promotes skeletal muscle myofiber hyperplasia. *eLife*, 12, 2024.

[13748] Peigen Chen, Haicheng Chen, Ziyu Liu, Xinyi Pan, Qianru Liu, and Xing Yang. Fungal–bacteria interactions provide shelter for bacteria in Caesarean section scar diverticulum. *eLife*, 12, 2024.

[13749] Qiang Chen, Norianne T Ingram, Jacob Baudin, Juan M Angueyra, Raunak Sinha, and Fred Rieke. Predictably manipulating photoreceptor light responses to reveal their role in downstream visual responses. *eLife*, 13, 2024.

[13750] Spencer Chin-Yu Chen, Yuzhi Chen, Wilson S Geisler, and Eyal Seidemann. Neural correlates of perceptual similarity masking in primate V1. *eLife*, 12, 2024.

[13751] Ting-Wen Chen, Hsiao-Wei Liao, Michelle Noble, Jing-Yi Siao, Yu-Hsuan Cheng, Wei-Chung Chiang, Yi-Tzu Lo, and Chung-Te Chang. Human DCP1 is crucial for mRNA decapping and possesses paralog-specific gene regulating functions. *eLife*, 13, 2024.

[13752] Wei Chen, Yun Lu, Yan Zhang, Jinjin Wu, Abigail McVicar, Yilin Chen, Siyu Zhu, Guochun Zhu, You Lu, Jiayang Zhang, Matthew McConnell, and Yi-Ping Li. Cbf$\beta$ regulates Wnt/$\beta$-catenin, Hippo/Yap, and Tgf$\beta$ signaling pathways in articular cartilage homeostasis and protects from ACLT surgery-induced osteoarthritis. *eLife*, 13, 2024.

[13753] Xianwei Chen, Yanfeng Li, Jin Xu, Yong Cui, Qian Wu, Haidi Yin, Yuying Li, Chuan Gao, Liwen Jiang, Huating Wang, Zilong Wen, Zhongping Yao, and Zhenguo Wu. Styxl2 regulates de novo sarcomere assembly by binding to non-muscle myosin IIs and promoting their degradation. *eLife*, 12, 2024.

[13754] Yan Chen, Ruizhi Yang, Bin Qi, and Zhao Shan. Peptidoglycan-Chi3l1 interaction shapes gut microbiota in intestinal mucus layer. *eLife*, 13, 2024.

[13755] Yao Chi Chen, Karen Sargsyan, Jon D Wright, Yu-Hsien Chen, Yi-Shuian Huang, and Carmay Lim. PPI-hotspotID for detecting protein–protein interaction hot spots from the free protein structure. *eLife*, 13, 2024.

[13756] Yi-Ju Chen, Shun-Cheng Tseng, Peng-Tzu Chen, and Eric Hwang. The non-mitotic role of HMMR in regulating the localization of TPX2 and the dynamics of microtubules in neurons. *eLife*, 13, 2024.

[13757] Yiyi Chen, Laimdota Zizmare, Victor Calbiague, Lan Wang, Shirley Yu, Fritz W Herberg, Oliver Schmachtenberg, Francois Paquet-Durand, and Christoph Trautwein. Retinal metabolism displays evidence for uncoupling of glycolysis and oxidative phosphorylation via Cori-, Cahill-, and mini-Krebs-cycle. *eLife*, 12, 2024.

[13758] Zhenhui Chen, Lu Yu, Zhihao Zheng, Xusheng Wang, Qiqing Guo, Yuchuan Chen, Yaowei Zhang, Yuqin Zhang, Jianbiao Xiao, Keli Chen, Hongying Fan, and Yi Ding. CPT1A mediates radiation sensitivity in colorectal cancer. *eLife*, 13, 2024.

[13759] Zulong Chen, Xin Wang, Xinlei Gao, Nina Arslanovic, Kaifu Chen, and Jessica K Tyler. Transcriptional inhibition after irradiation occurs preferentially at highly expressed genes in a manner dependent on cell cycle progression. *eLife*, 13, 2024.

[13760] Bing Liang Alvin Chew, AN Qi Ngoh, Wint Wint Phoo, Kitti Wing Ki Chan, Zheng Ser, Nikhil K Tulsian, Shiao See Lim, Mei Jie Grace Weng, Satoru Watanabe, Milly M Choy, Jenny Low, Eng Eong Ooi, Christiane Ruedl, Radoslaw M Sobota, Subhash G Vasudevan, and Dahai Luo. Secreted dengue virus NS1 from infection is predominantly dimeric and in complex with high-density lipoprotein. *eLife*, 12, 2024.

[13761] Meera Chikermane, Liz Weerdmeester, Nanditha Rajamani, Richard M Köhler, Timon Merk, Jojo Vanhoecke, Andreas Horn, and Wolf Julian Neumann. Cortical beta oscillations map to shared brain networks modulated by dopamine. *eLife*, 13, 2024.

[13762] Shek Man Chim, Kristen Howell, John Dronzek, Weizhen Wu, Cristopher Van Hout, Manuel AR Ferreira, Bin Ye, Alexander Li, Susannah Brydges, Vinayagam Arunachalam, Anthony Marcketta, Adam E Locke, Jonas Bovijn, Niek Verweij, Tanima De, Luca Lotta, Lyndon Mitnaul, Michelle LeBlanc, Regeneron Genetics Center, David J Carey, Olle Melander, Alan Shuldiner, Katia Karalis, Aris N Economides, and Harikiran Nistala. Genetic inactivation of zinc transporter SLC39A5 improves liver function and hyperglycemia in obesogenic settings. *eLife*, 12, 2024.

[13763] Po-Lin Chiu, Juan D Orjuela, Bert L de Groot, Camilo Aponte Santamaría, and Thomas Walz. Structure and dynamics of cholesterol-mediated aquaporin-0 arrays and implications for lipid rafts. *eLife*, 12, 2024.

[13764] Hohyun Cho, Markus Adamek, Jon T Willie, and Peter Brunner. Novel cyclic homogeneous oscillation detection method for high accuracy and specific characterization of neural dynamics. *eLife*, 12, 2024.

[13765] Doo Eun Choi, Jun Wan Shin, Sophia Zeng, Eun Pyo Hong, Jae-Hyun Jang, Jacob M Loupe, Vanessa C Wheeler, Hannah E Stutzman, Ben Kleinstiver, and Jong-Min Lee. Base editing strategies to convert CAG to CAA diminish the disease-causing mutation in Huntington's disease. *eLife*, 12, 2024.

[13766] Jessica Y Chotiner, N Adrian Leu, Fang Yang, Isabella G Cossu, Yongjuan Guan, Huijuan Lin, and P Jeremy Wang. TRIP13 localizes to synapsed chromosomes and functions as a dosage-sensitive regulator of meiosis. *eLife*, 12, 2024.

[13767] Rupam Choudhury, Anuroop Venkateswaran Venkatasubramani, Jie Hua, Marco Borsò, Celeste Franconi, Sarah Kinkley, Ignasi Forné, and Axel Imhof. The role of RNA in the maintenance of chromatin domains as revealed by antibody-mediated proximity labelling coupled to mass spectrometry. *eLife*, 13, 2024.

[13768] Priya M Christensen, Jonathan Martin, Aparna Uppuluri, Luke R Joyce, Yahan Wei, Ziqiang Guan, Faruck Morcos, and Kelli L Palmer. Lipid discovery enabled by sequence statistics and machine learning. *eLife*, 13, 2024.

[13769] Amanda Chu, Nicholas T Gordon, Aleah M DuBois, Christa B Michel, Katherine E Hanrahan, David C Williams, Stefano Anzellotti, and Michael A McDannald. A fear conditioned cue orchestrates a suite of behaviors in rats. *eLife*, 13, 2024.

[13770] Tianhao Chu, Zilong Ji, Junfeng Zuo, Yuanyuan Mi, Wen-hao Zhang, Tiejun Huang, Daniel Bush, Neil Burgess, and Si Wu. Firing rate adaptation affords place cell theta sweeps, phase precession, and procession. *eLife*, 12, 2024.

[13771] Xiao Chu, Lidong Wang, Yiheng Zhu, Zhengshan Feng, Qingtian Guan, Lei Song, and Zhaoqing Luo. A unique cell division protein critical for the assembly of the bacterial divisome. *eLife*, 12, 2024.

[13772] Chi-Ning Chuang, Hou-Cheng Liu, Tai-Ting Woo, Ju-Lan Chao, Chiung-Ya Chen, Hisao-Tang Hu, Yi-Ping Hsueh, and Ting-Fang Wang. Noncanonical usage of stop codons in ciliates expands proteins with structurally flexible Q-rich motifs. *eLife*, 12, 2024.

[13773] Taegon Chung, Iksoo Chang, and Sangyeol Kim. Development of equation of motion deciphering locomotion including omega turns of Caenorhabditis elegans. *eLife*, 12, 2024.

[13774] Steven A Cincotta, Nainoa Richardson, Mariko H Foecke, and Diana J Laird. Differential susceptibility of male and female germ cells to glucocorticoid-mediated signaling. *eLife*, 12, 2024.

[13775] Harry Clark and Matthew F Nolan. Task-anchored grid cell firing is selectively associated with successful path integration-dependent behaviour. *eLife*, 12, 2024.

[13776] Bradley P Clarke, Alexia E Angelos, Menghan Mei, Pate S Hill, Yihu Xie, and Yi Ren. Cryo-EM structure of the CBC-ALYREF complex. *eLife*, 12, 2024.

[13777] Tom Clegg and Samraat Pawar. Variation in thermal physiology can drive the temperature-dependence of microbial community richness. *eLife*, 13, 2024.

[13778] Fanny-Mei Cloarec-Ung, Jamie Beaulieu, Arunan Suthananthan, Bernhard Lehnertz, Guy Sauvageau, Hilary M Sheppard, and David JHF Knapp. Near-perfect precise on-target editing of human hematopoietic stem and progenitor cells. *eLife*, 12, 2024.

[13779] Olivier Codol, Jonathan A Michaels, Mehrdad Kashefi, J Andrew Pruszynski, and Paul L Gribble. MotorNet, a Python toolbox for controlling differentiable biomechanical effectors with artificial neural networks. *eLife*, 12, 2024.

[13780] Reto B Cola, Salome N Niethammer, Preethi Rajamannar, Andrea Gresch, Musadiq A Bhat, Kevin Assoumou, Elyse T Williams, Patrick Hauck, Nina Hartrampf, Dietmar Benke, Miriam Stoeber, Gil Levkowitz, Sarah Melzer, and Tommaso Patriarchi. Probing PAC1 receptor activation across species with an engineered sensor. *eLife*, 13, 2024.

[13781] Reto B Cola, Diana M Roccaro-Waldmeyer, Samara Naim, Alexandre Babalian, Petra Seebeck, Gonzalo Alvarez-Bolado, and Marco R Celio. Chemo- and optogenetic activation of hypothalamic Foxb1-expressing neurons and their terminal endings in the rostral-dorsolateral PAG leads to tachypnea, bradycardia, and immobility. *eLife*, 12, 2024.

[13782] Alison G Cole, Tamar Hashimshony, Zhuo Du, and Itai Yanai. Gene regulatory patterning codes in early cell fate specification of the C. elegans embryo. *eLife*, 12, 2024.

[13783] Florent Colomb, Abhishek Jamwal, Adefunke Ogunkanbi, Tania Frangova, Alice R Savage, Sarah Kelly, Gavin J Wright, Matthew K Higgins, and Henry J McSorley. Heparan sulphate binding controls in vivo half-life of the HpARI protein family. *eLife*, 13, 2024.

[13784] Etienne Combrisson, Ruggero Basanisi, Maelle CM Gueguen, Sylvain Rheims, Philippe Kahane, Julien Bastin, and Andrea Brovelli. Neural interactions in the human frontal cortex dissociate reward and punishment learning. *eLife*, 12, 2024.

[13785] Typhaine Comyn, Thomas Preat, Alice Pavlowsky, and Pierre-Yves PlaÇais. PKC$\delta$ is an activator of neuronal mitochondrial metabolism that mediates the spacing effect on memory consolidation. *eLife*, 13, 2024.

[13786] Wei Cong, Jia Li, Charlotte Hacker, Ye Li, Yu Zhang, Lixiao Jin, Yi Zhang, Diqiang Li, Yadong Xue, and Yuguang Zhang. Different coexistence patterns between apex carnivores and mesocarnivores based on temporal, spatial, and dietary niche partitioning analysis in Qilian Mountain National Park, China. *eLife*, 13, 2024.

[13787] Sean V Connelly, Nicholas F Brazeau, Mwinyi Msellem, Billy E Ngasala, Ozkan Aydemir, Varun Goel, Karamoko Niaré, David J Giesbrecht, Zachary R Popkin-Hall, Chris Hennelly, Zackary Park, Ann M Moormann, John M Ong'echa, Robert Verity, Safia Mohammed, Shija J Shija, Lwidiko E Mhamilawa, Ulrika Morris, Andreas Mårtensson, Jessica T Lin, Anders Björkman, Jonathan J Juliano, and Jeffrey A Bailey. Strong isolation by distance and evidence of population microstructure reflect ongoing Plasmodium falciparum transmission in Zanzibar. *eLife*, 12, 2024.

[13788] Claudia D Consalvo, Adedeji M Aderounmu, Helen M Donelick, P Joseph Aruscavage, Debra M Eckert, Peter S Shen, and Brenda L Bass. Caenorhabditis elegans Dicer acts with the RIG-I-like helicase DRH-1 and RDE-4 to cleave dsRNA. *eLife*, 13, 2024.

[13789] Théo Constant, F Stephen Dobson, Caroline Habold, and Sylvain Giroud. Evolutionary trade-offs in dormancy phenology. *eLife*, 12, 2024.

[13790] Rossella Conti and Céline Auger. Associative plasticity of granule cell inputs to cerebellar Purkinje cells. *eLife*, 13, 2024.

[13791] Alexander D Cook, Mark Carrington, and Matthew K Higgins. Molecular mechanism of complement inhibition by the trypanosome receptor ISG65. *eLife*, 12, 2024.

[13792] Ivan Corbeski, Pablo Andrés Vargas-Rosales, Rajiv Kumar Bedi, Jiahua Deng, Dylan Coelho, Emmanuelle Braud, Laura Iannazzo, Yaozong Li, Danzhi Huang, Mélanie Ethève-Quelquejeu, Qiang Cui, and Amedeo Caflisch. The catalytic mechanism of the RNA methyltransferase METTL3. *eLife*, 12, 2024.

[13793] Allison Coté, Aoife O'Farrell, Ian Dardani, Margaret Dunagin, Chris Coté, Yihan Wan, Sareh Bayatpour, Heather L Drexler, Katherine A Alexander, Fei Chen, Asmamaw T Wassie, Rohan Patel, Kenneth Pham, Edward S Boyden, Shelly Berger, Jennifer Phillips-Cremins, L Stirling Churchman, and Arjun Raj. Post-transcriptional splicing can occur in a slow-moving zone around the gene. *eLife*, 12, 2024.

[13794] Donal J Cox, Sarah A Connolly, Cilian ó Maoldomhnaigh, Aenea AI Brugman, Olivia Sandby Thomas, Emily Duffin, Karl M Gogan, Oisin ó Gallchobhair, Dearbhla M Murphy, Sinead A O'Rourke, Finbarr O'Connell, Parthiban Nadarajan, James J Phelan, Laura E Gleeson, Sharee A Basdeo, and Joseph Keane. Human airway macrophages are metabolically reprogrammed by IFN-$\gamma$ resulting in glycolysis-dependent functional plasticity. *eLife*, 13, 2024.

[13795] Lucie Crhak Khaitova, Pavlina Mikulkova, Jana Pecinkova, Manikandan Kalidass, Stefan Heckmann, Inna Lermontova, and Karel Riha. Heat stress impairs centromere structure and segregation of meiotic chromosomes in Arabidopsis. *eLife*, 12, 2024.

[13796] David A Crowe, Andrew Willow, Rachael K Blackman, Adele L DeNicola, Matthew V Chafee, and Bagrat Amirikian. A prefrontal network model operating near steady and oscillatory states links spike desynchronization and synaptic deficits in schizophrenia. *eLife*, 13, 2024.

[13797] Lucia Csepregi, Kenneth Hoehn, Daniel Neumeier, Joseph M Taft, Simon Friedensohn, Cédric R Weber, Arkadij Kummer, Fabian Sesterhenn, Bruno E Correia, and Sai T Reddy. The physiological landscape and specificity of antibody repertoires are consolidated by multiple immunizations. *eLife*, 13, 2024.

[13798] Mengqiao Cui, Xiaoyuan Pan, Zhijie Fan, Shulin Wu, Ran Ji, Xianlei Wang, Xiangxi Kong, Zhou Wu, Lingzhen Song, Weiyi Song, Jun-Xia Yang, Hongjie Zhang, Hongxing Zhang, Hai-Lei Ding, and Jun-Li Cao. Dysfunctional S1P/S1PR1 signaling in the dentate gyrus drives vulnerability of chronic pain-related memory impairment. *eLife*, 13, 2024.

[13799] Yaning Cui, Hongping Qian, Jinhuan Yin, Changwen Xu, Pengyun Luo, Xi Zhang, Meng Yu, Bodan Su, Xiaojuan Li, and Jinxing Lin. Single-molecule analysis reveals the phosphorylation of FLS2 governs its spatiotemporal dynamics and immunity. *eLife*, 12, 2024.

[13800] Erin R Cullen, Mona Safari, Isabelle Mittelstadt, and Matthew C Weston. Hyperactivity of mTORC1- and mTORC2-dependent signaling mediates epilepsy downstream of somatic PTEN loss. *eLife*, 12, 2024.

[13801] Bridget M Curran, Kelsey R Nickerson, Andrea R Yung, Lisa V Goodrich, Alexander Jaworski, Marc Tessier-Lavigne, and Le Ma. Multiple guidance mechanisms control axon growth to generate precise T-shaped bifurcation during dorsal funiculus development in the spinal cord. *eLife*, 13, 2024.

[13802] Michael J Currie, James S Davies, Mariafrancesca Scalise, Ashutosh Gulati, Joshua D Wright, Michael C Newton-Vesty, Gayan S Abeysekera, Ramaswamy Subramanian, Weixiao Y Wahlgren, Rosmarie Friemann, Jane R Allison, Peter D Mace, Michael DW Griffin, Borries Demeler, Soichi Wakatsuki, David Drew, Cesare Indiveri, Renwick CJ Dobson, and Rachel A North. Structural and biophysical analysis of a Haemophilus influenzae tripartite ATP-independent periplasmic (TRAP) transporter. *eLife*, 12, 2024.

[13803] Asher D Cutter. Beyond Haldane's rule: Sex-biased hybrid dysfunction for all modes of sex determination. *eLife*, 13, 2024.

[13804] Ignacy Czajewski, Bijayalaxmi Swain, Jiawei Xu, Laurin McDowall, Andrew T Ferenbach, and Daan MF van Aalten. Rescuable sleep and synaptogenesis phenotypes in a Drosophila model of O-GlcNAc transferase intellectual disability. *eLife*, 13, 2024.

[13805] Anna Dai and Martine Ben Amar. Cyclic muscle contractions reinforce the actomyosin motors and mediate the full elongation of C. elegans embryo. *eLife*, 12, 2024.

[13806] Marina Dajka, Tobias Rath, Nina Morgner, and Benesh Joseph. Dynamic basis of lipopolysaccharide export by LptB2FGC. *eLife*, 13, 2024.

[13807] Michael P Dalton, Mary Hongying Cheng, Ivet Bahar, and Jonathan A Coleman. Structural mechanisms for VMAT2 inhibition by tetrabenazine. *eLife*, 12, 2024.

[13808] Stephen D'Amico, Varvara Kirillov, Oleksi Petrenko, and Nancy C Reich. STAT3 is a genetic modifier of TGF-beta induced EMT in KRAS mutant pancreatic cancer. *eLife*, 13, 2024.

[13809] Camille Dantzer, Justine Vaché, Aude Brunel, Isabelle Mahouche, Anne-Aurélie Raymond, Jean-William Dupuy, Melina Petrel, Paulette Bioulac-Sage, David Perrais, Nathalie Dugot-Senant,

Mireille Verdier, Barbara Bessette, Clotilde Billottet, and Violaine Moreau. Emerging role of oncogenic ß-catenin in exosome biogenesis as a driver of immune escape in hepatocellular carcinoma. *eLife*, 13, 2024.

[13810] Showkat Ahmad Dar, Sulochan Malla, Vlastimil Martinek, Matthew John Payea, Christopher Tai-Yi Lee, Jessica Martin, Aditya Jignesh Khandeshi, Jennifer L Martindale, Cedric Belair, and Manolis Maragkakis. Full-length direct RNA sequencing uncovers stress granule-dependent RNA decay upon cellular stress. *eLife*, 13, 2024.

[13811] Anup Das and Vinod Menon. Electrophysiological dynamics of salience, default mode, and frontoparietal networks during episodic memory formation and recall revealed through multi-experiment iEEG replication. *eLife*, 13, 2024.

[13812] Priyanka Das, Alejandro Aballay, and Jogender Singh. Calcineurin inhibition enhances Caenorhabditis elegans lifespan by defecation defects-mediated calorie restriction and nuclear hormone signaling. *eLife*, 12, 2024.

[13813] Proloy Das, Mingjian He, and Patrick L Purdon. A dynamic generative model can extract interpretable oscillatory components from multichannel neurophysiological recordings. *eLife*, 13, 2024.

[13814] Rajsekhar Das, Sumit Sinha, Xin Li, TR Kirkpatrick, and D Thirumalai. Free volume theory explains the unusual behavior of viscosity in a non-confluent tissue during morphogenesis. *eLife*, 12, 2024.

[13815] Subhradip Das, Sushmitha Hegde, Neel Wagh, Jyothish Sudhakaran, Adheena Elsa Roy, Girish Deshpande, and Girish S Ratnaparkhi. Caspar specifies primordial germ cell count and identity in Drosophila melanogaster. *eLife*, 13, 2024.

[13816] Agnik Dasgupta, Caleb C Reagor, Sang Peter Paik, Lauren M Snow, Adrian Jacobo, and AJ Hudspeth. Semaphorin7A patterns neural circuitry in the lateral line of the zebrafish. *eLife*, 12, 2024.

[13817] Yuki Date, Yukiko Sasazawa, Mitsuhiro Kitagawa, Kentaro Gejima, Ayami Suzuki, Hideyuki Saya, Yasuyuki Kida, Masaya Imoto, Eisuke Itakura, Nobutaka Hattori, and Shinji Saiki. Novel autophagy inducers by accelerating lysosomal clustering against Parkinson's disease. *eLife*, 13, 2024.

[13818] Csaba Dávid, Kristóf Giber, Katalin Kerti-Szigeti, Mihály Köllő, Zoltan Nusser, and Laszlo Acsady. A novel image segmentation method based on spatial autocorrelation identifies A-type potassium channel clusters in the thalamus. *eLife*, 12, 2024.

[13819] Kristian Davidsen, Jonathan S Marvin, Abhi Aggarwal, Timothy A Brown, and Lucas B Sullivan. An engineered biosensor enables dynamic aspartate measurements in living cells. *eLife*, 12, 2024.

[13820] Kristian Davidsen and Lucas B Sullivan. A robust method for measuring aminoacylation through tRNA-Seq. *eLife*, 12, 2024.

[13821] Natalie R Davidson and Casey S Greene. Analysis of science journalism reveals gender and regional disparities in coverage. *eLife*, 12, 2024.

[13822] John H Day, Catherine M Della Santina, Pema Maretich, Alexander L Auld, Kirsten K Schnieder, Tay Shin, Edward S Boyden, and Laurie A Boyer. High-throughput expansion microscopy enables scalable super-resolution imaging. *eLife*, 13, 2024.

[13823] Jorge de Carvalho, Sham Tlili, Timothy E Saunders, and Ivo A Telley. The positioning mechanics of microtubule asters in Drosophila embryo explants. *eLife*, 12, 2024.

[13824] Toni de Dios Martinez, Claudia Fontsere, Pere Renom, Josefin Stiller, Laia Llovera, Marcela Uliano-Silva, Alejandro Sánchez-Gracia, Charlotte Wright, Esther Lizano, Berta Caballero, Arcadi Navarro, Sergi Civit, Robert K Robbins, Mark Blaxter, Tomàs Marquès, Roger Vila, and Carles Lalueza-Fox. Whole genomes from the extinct Xerces Blue butterfly can help identify declining insect species. *eLife*, 12, 2024.

[13825] Rita De Gasperi, Laszlo Csernoch, Beatrix Dienes, Monika Gonczi, Jayanta K Chakrabarty, Shahar Goeta, Abdurrahman Aslan, Carlos A Toro, David Karasik, Lewis M Brown, Marco Brotto, and Christopher P Cardozo. Septin 7 interacts with Numb to preserve sarcomere structural organization and muscle contractile function. *eLife*, 12, 2024.

[13826] Giordano de Guglielmo, Lieselot Carrette, Marsida Kallupi, Molly Brennan, Brent Boomhower, Lisa Maturin, Dana Conlisk, Sharona Sedighim, Lani Tieu, McKenzie J Fannon, Angelica R Martinez, Nathan Velarde, Dyar Othman, Benjamin Sichel, Jarryd Ramborger, Justin Lau, Jenni Kononoff, Adam Kimbrough, Sierra Simpson, Lauren C Smith, Kokila Shankar, Selene Bonnet-Zahedi, Elizabeth A Sneddon, Alicia Avelar, Sonja Lorean Plasil, Joseph Mosquera, Caitlin Crook, Lucas Chun, Ashley Vang, Kristel K Milan, Paul Schweitzer, Bonnie Lin, Beverly Peng, Apurva S Chitre, Oksana Polesskaya, Leah C Solberg Woods, Abraham A Palmer, and Olivier George. Large-scale characterization of cocaine addiction-like behaviors reveals that escalation of intake, aversion-resistant responding, and breaking-points are highly correlated measures of the same construct. *eLife*, 12, 2024.

[13827] Lizbeth de La Cruz, Derek Bui, Claudia M Moreno, and Oscar Vivas. Sympathetic motor neuron dysfunction is a missing link in age-associated sympathetic overactivity. *eLife*, 12, 2024.

[13828] Carla de la Fuente, Alexandre Grondin, Bassirou Sine, Marilyne Debieu, Christophe Belin, Amir Hajjarpoor, Jonathan A Atkinson, Sixtine Passot, Marine Salson, Julie Orjuela, Christine Tranchant-Dubreuil, Jean-Rémy Brossier, Maxime Steffen, Charlotte Morgado, Hang Ngan Dinh, Bipin K Pandey, Julie Darmau, Antony Champion, Anne-Sophie Petitot, Celia Barrachina, Marine Pratlong, Thibault Mounier, Princia Nakombo-Gbassault, Pascal Gantet, Prakash Gangashetty, Yann Guedon, Vincent Vadez, Jean-Philippe Reichheld, Malcolm J Bennett, Ndjido Ardo Kane, Soazig Guyomarc'h, Darren M Wells, Yves Vigouroux, and Laurent Laplaze. Glutaredoxin regulation of primary root growth is associated with early drought stress tolerance in pearl millet. *eLife*, 12, 2024.

[13829] Silvia de Sanjosé, Rebecca B Perkins, Nicole Campos, Federica Inturrisi, Didem Egemen, Brian Befano, Ana Cecilia Rodriguez, Jose Jerónimo, Li C Cheung, Kanan Desai, Paul Han, Akiva P Novetsky, Abigail Ukwuani, Jenna Marcus, Syed Rakin Ahmed, Nicolas Wentzensen, Jayashree Kalpathy-Cramer, and Mark Schiffman. Design of the HPV-automated visual evaluation (PAVE) study: Validating a novel cervical screening strategy. *eLife*, 12, 2024.

[13830] Gaetan De Waele, Willem Waegeman, and Gerben Menschaert. An antimicrobial drug recommender system using MALDI-TOF MS and dual-branch neural networks. *eLife*, 13, 2024.

[13831] Emily L Dearlove, Chatrin Chatrin, Lori Buetow, Syed F Ahmed, Tobias Schmidt, Martin Bushell, Brian O Smith, and Danny T Huang. DTX3L ubiquitin ligase ubiquitinates single-stranded nucleic acids. *eLife*, 13, 2024.

[13832] Nincy Debeuf, Sahine Lameire, Manon Vanheerswynghels, Julie Deckers, Caroline De Wolf, Wendy Toussaint, Rein Verbeke, Kevin Verstaen, Hamida Hammad, Stijn Vanhee, and Bart N Lambrecht. TCR transgenic clone selection guided by immune receptor analysis and single-cell RNA expression of polyclonal responders. *eLife*, 13, 2024.

[13833] Lies Deceuninck and Fabian Kloosterman. Disruption of awake sharp-wave ripples does not affect memorization of locations in repeated-acquisition spatial memory tasks. *eLife*, 13, 2024.

[13834] Marc E Deetjen, Diana D Chin, Ashley M Heers, Bret W Tobalske, and David Lentink. Small deviations in kinematics and body form dictate muscle performances in the finely tuned avian downstroke. *eLife*, 12, 2024.

[13835] F Javier DeHaro-Arbona, Charalambos Roussos, Sarah Baloul, Jonathan Townson, María J Gómez Lamarca, and Sarah Bray. Dynamic modes of Notch transcription hubs conferring memory and stochastic activation revealed by live imaging the co-activator Mastermind. *eLife*, 12, 2024.

[13836] Sebastian Deichsel, Lisa Frankenreiter, Johannes Fechner, Bernd M Gahr, Mirjam Zimmermann, Helena Mastel, Irina Preis, Anette Preiss, and Anja C Nagel. Inhibition of the Notch signal transducer CSL by Pkc53E-mediated phosphorylation to fend off parasitic immune challenge in Drosophila. *eLife*, 12, 2024.

[13837] Mathilde Delacoux and Fumihiro Kano. Fine-scale tracking reveals visual field use for predator detection and escape in collective foraging of pigeon flocks. *eLife*, 13, 2024.

[13838] Geoffroy Delamare, Yosif Zaki, Denise J Cai, and Claudia Clopath. Drift of neural ensembles driven by slow fluctuations of intrinsic excitability. *eLife*, 12, 2024.

[13839] Pablo J Delclos, Kiran Adhikari, Alexander B Mai, Oluwatomi Hassan, Alexander A Oderhowho, Vyshnika Sriskantharajah, Tammie Trinh, and Richard Meisel. Trans regulation of an odorant binding protein by a proto-Y chromosome affects male courtship in house fly. *eLife*, 13, 2024.

[13840] Pan-Yue Deng, Ajeet Kumar, Valeria Cavalli, and Vitaly A Klyachko. Circuit-based intervention corrects excessive dentate gyrus output in the fragile X mouse model. *eLife*, 12, 2024.

[13841] Wei Q Deng, Nathan Cawte, Natalie Campbell, Sandi M Azab, Russell J de Souza, Amel Lamri, Katherine M Morrison, Stephanie A Atkinson, Padmaja Subbarao, Stuart E Turvey, Theo J Moraes, Koon K Teo, Piush J Mandhane, Meghan B Azad, Elinor Simons, Guillaume Paré, and Sonia S Anand. Maternal smoking DNA methylation risk score associated with health outcomes in offspring of European and South Asian ancestry. *eLife*, 13, 2024.

[13842] Brian DePasquale, Carlos D Brody, and Jonathan W Pillow. Neural population dynamics underlying evidence accumulation in multiple rat brain regions. *eLife*, 13, 2024.

[13843] Simon Desiderio, Frederick Schwaller, Kevin Tartour, Kiran Padmanabhan, Gary R Lewin, Patrick Carroll, and Frederic Marmigere. Touch receptor end-organ innervation and function require sensory neuron expression of the transcription factor Meis2. *eLife*, 12, 2024.

[13844] GVS Devakinandan, Mark Terasaki, and Adish Dani. Single-cell transcriptomics of vomeronasal neuroepithelium reveals a differential endoplasmic reticulum environment amongst neuronal subtypes. *eLife*, 13, 2024.

[13845] Debabrata Dey, Shir Marciano, Anna Poryval, Ondřej Groborz, Lucie Wohlrabova, Tomás Slanina, and Gideon Schreiber. Reversing protonation of weakly basic drugs greatly enhances intracellular diffusion and decreases lysosomal sequestration. *eLife*, 13, 2024.

[13846] Surbhi Dhingra, Prachi M Chopade, Rajesh Vinnakota, and Janesh Kumar. Functional implications of the exon 9 splice insert in GluK1 kainate receptors. *eLife*, 12, 2024.

[13847] Roberto Efraín Díaz, Andrew K Ecker, Galen J Correy, Pooja Asthana, Iris D Young, Bryan Faust, Michael C Thompson, Ian B Seiple, Steven Van Dyken, Richard M Locksley, and James S Fraser. Structural characterization of ligand binding and pH-specific enzymatic activity of mouse Acidic Mammalian Chitinase. *eLife*, 12, 2024.

[13848] Gretchen Diffendall, Aurelie Claes, Anna Barcons-Simon, Prince Nyarko, Florent Dingli, Miguel M Santos, Damarys Loew, Antoine Claessens, and Artur Scherf. RNA polymerase III is involved in regulating Plasmodium falciparum virulence. *eLife*, 13, 2024.

[13849] Xingyu Ding, Sean Froudist-Walsh, Jorge Jaramillo, Junjie Jiang, and Xiao-Jing Wang. Cell type-specific connectome predicts distributed working memory activity in the mouse brain. *eLife*, 13, 2024.

[13850] Yufang Ding, Karin Yanagi, Fang Yang, Evelyn Callaway, Clint Cheng, Martha E Hensel, Rani Menon, Robert C Alaniz, Kyongbum Lee, and Arul Jayaraman. Oral supplementation of gut microbial metabolite indole-3-acetate alleviates diet-induced steatosis and inflammation in mice. *eLife*, 12, 2024.

[13851] Silvia Diz-de Almeida, Raquel Cruz, Andre D Luchessi, José M Lorenzo-Salazar, Miguel López de Heredia, Inés Quintela, Rafaela González-Montelongo, Vivian Nogueira Silbiger, Marta Sevilla Porras, Jair Antonio Tenorio Castaño, Julian Nevado, Jose María Aguado, Carlos Aguilar, Sergio Aguilera-Albesa, Virginia Almadana, Berta Almoguera, Nuria Alvarez, álvaro Andreu-Bernabeu, Eunate Arana-Arri, Celso Arango, María J Arranz, Maria-Jesus Artiga, Raúl C Baptista-Rosas, María Barreda-Sánchez, Moncef Belhassen-Garcia, Joao F Bezerra, Marcos AC Bezerra, Lucía Boix-Palop, María Brion, Ramón Brugada, Matilde Bustos, Enrique J Calderón, Cristina Carbonell, Luis Castano, Jose E Castelao, Rosa Conde-Vicente, M Lourdes Cordero-Lorenzana, Jose L Cortes-Sanchez, Marta Corton, M Teresa Darnaude, Alba De Martino-Rodríguez, Victor del Campo-Pérez, Aranzazu Diaz de Bustamante, Elena Domínguez-Garrido, Rocío Eirós, María Carmen Fariñas, María J Fernandez-Nestosa, Uxía Fernández-Robelo, Amanda Fernández-Rodríguez, Tania Fernández-Villa, Manuela Gago-Dominguez, Belén Gil-Fournier, Javier Gómez-Arrue, Beatriz González álvarez, Fernan Gonzalez Bernaldo de Quirós, Anna González-Neira, Javier González-Peñas, Juan F Gutiérrez-Bautista, María José Herrero, Antonio Herrero-Gonzalez, María A Jimenez-Sousa, María Claudia Lattig, Anabel Liger Borja, Rosario Lopez-Rodriguez, Esther Mancebo, Caridad Martín-López, Vicente Martín, Oscar Martinez-Nieto, Iciar Martinez-Lopez, Michel F Martinez-Resendez, Angel Martinez-Perez, Juliana F Mazzeu, Eleuterio Merayo Macías, Pablo Minguez, Victor Moreno Cuerda, Silviene F Oliveira, Eva Ortega-Paino, Mara Parellada, Estela Paz-Artal, Ney PC Santos, Patricia Pérez-Matute, Patricia Perez, M Elena Pérez-Tomás, Teresa Perucho, Mellina Pinsach-Abuin, Guillermo Pita, Ericka N Pompa-Mera, Gloria L Porras-Hurtado, Aurora Pujol, Soraya Ramiro León, Salvador Resino, Marianne R Fernandes, Emilio Rodríguez-Ruiz, Fernando Rodriguez-Artalejo, José A Rodriguez-Garcia, Francisco Ruiz-Cabello, Javier Ruiz-Hornillos, Pablo Ryan, José Manuel Soria, Juan Carlos Souto, Eduardo Tamayo, Alvaro Tamayo-Velasco, Juan Carlos Taracido-Fernandez, Alejandro Teper, Lilian Torres-Tobar, Miguel Urioste, Juan Valencia-Ramos, Zuleima Yáñez, Ruth Zarate, Itziar de Rojas, Agustín Ruiz, Pascual Sánchez, Luis Miguel Real, Encarna Guillen-Navarro, Carmen Ayuso, Esteban Parra, José A Riancho, Augusto Rojas-Martinez, Carlos Flores, Pablo Lapunzina, and ángel Carracedo. Novel risk loci for COVID-19 hospitalization among admixed American populations. *eLife*, 13, 2024.

[13852] Morgane Djebar, Isabelle Anselme, Guillaume Pezeron, Pierre-Luc Bardet, Yasmine Cantaut-Belarif, Alexis Eschstruth, Diego López-Santos, Hélène Le Ribeuz, Arnim Jenett, Hanane Khoury, Joelle Veziers, Caroline Parmentier, Aurélie Hirschler, Christine Carapito, Ruxandra Bachmann-Gagescu, Sylvie Schneider-Maunoury, and Christine Vesque. Astrogliosis and neuroinflammation underlie scoliosis upon cilia dysfunction. *eLife*, 13, 2024.

[13853] Olivera Djuric, Elisabetta Larosa, Mariateresa Cassinadri, Silvia Cilloni, Eufemia Bisaccia, Davide Pepe, Laura Bonvicini, Massimo Vicentini, Francesco Venturelli, Paolo Giorgi Rossi, Patrizio Pezzotti, Alberto Mateo Urdiales, and Emanuela Bedeschi. Effect of an enhanced public health contact tracing intervention on the secondary transmission of SARS-CoV-2 in educational settings: The four-way decomposition analysis. *eLife*, 13, 2024.

[13854] Greg J Dodge, Alyssa J Anderson, Yi He, Weijing Liu, Rosa Viner, and Barbara Imperiali. Mapping the architecture of the initiating phosphoglycosyl transferase from S. enterica O-antigen biosynthesis in a liponanoparticle. *eLife*, 12, 2024.

[13855] Roswitha Dolcemascolo, María Heras-Hernández, Lucas Goiriz, Roser Montagud-Martínez, Alejandro Requena-Menéndez, Raúl Ruiz, Anna Pérez-Ràfols, R Anahí Higuera-Rodríguez, Guillermo Pérez-Ropero, Wim F Vranken, Tommaso Martelli, Wolfgang Kaiser, Jos Buijs, and Guillermo Rodrigo. Repurposing the mammalian RNA-binding protein Musashi-1 as an allosteric translation repressor in bacteria. *eLife*, 12, 2024.

[13856] Natalia Dolgova, Eva-Maria E Uhlemann, Michal T Boniecki, Frederick S Vizeacoumar, Anjuman Ara, Paria Nouri, Martina Ralle, Marco Tonelli, Syed A Abbas, Jaala Patry, Hussain Elhasasna, Andrew Freywald, Franco J Vizeacoumar, and Oleg Y Dmitriev. MEMO1 binds iron and modulates iron homeostasis in cancer cells. *eLife*, 13, 2024.

[13857] Hui Dong, Jiong Yang, Kai He, Wen-Bin Zheng, De-Hua Lai, Jing Liu, Hui-Yong Ding, Rui-Bin Wu, Kevin M Brown, Geoff Hide, Zhao-Rong Lun, Xing-Quan Zhu, and Shaojun Long. The Toxoplasma monocarboxylate transporters are involved in the metabolism within the apicoplast and are linked to parasite survival. *eLife*, 12, 2024.

[13858] Pooja Dongre, Gaëlle Lanté, Mathieu Cantat, Charlotte Canteloup, and Erica van de Waal. Role of immigrant males and muzzle contacts in the uptake of a novel food by wild vervet monkeys. *eLife*, 13, 2024.

[13859] Narges Doostani, Gholam-Ali Hossein-Zadeh, Radoslaw M Cichy, and Maryam Vaziri-Pashkam. Attention modulates human visual responses to objects by tuning sharpening. *eLife*, 12, 2024.

[13860] Rachel L Doser, Kaz M Knight, Ennis W Deihl, and Frederic J Hoerndli. Activity-dependent mitochondrial ROS signaling regulates recruitment of glutamate receptors to synapses. *eLife*, 13, 2024.

[13861] Anne Drougard, Eric H Ma, Vanessa Wegert, Ryan Sheldon, Ilaria Panzeri, Naman Vatsa, Stefanos Apostle, Luca Fagnocchi, Judith Schaf, Klaus Gossens, Josephine Völker, Shengru Pang, Anna Bremser, Erez Dror, Francesca Giacona, Sagar Sagar, Michael X Henderson, Marco Prinz, Russell G Jones, and John Andrew Pospisilik. An acute microglial metabolic response controls metabolism and improves memory. *eLife*, 12, 2024.

[13862] Katarzyna Drożdżyk, Martina Peter, and Raimund Dutzler. Structural features of heteromeric channels composed of CALHM2 and CALHM4 paralogs. *eLife*, 13, 2024.

[13863] Changlin Du, Chuanting Xu, Pengcheng Jia, Na Cai, Zhenming Zhang, Wenna Meng, Lu Chen, Zhongnan Zhou, Qi Wang, Rui Feng, Jun Li, Xiaoming Meng, Cheng Huang, and Taotao Ma. PST-PIP2 ameliorates aristolochic acid nephropathy by suppressing interleukin-19-mediated neutrophil extracellular trap formation. *eLife*, 13, 2024.

[13864] Kun-sheng Du, Jin Guo, Sarah R Losso, Stephen Pates, Ming Li, and Ai-lin Chen. Multiple origins of dorsal ecdysial sutures in trilobites and their relatives. *eLife*, 12, 2024.

[13865] Zhanwei Du, Lin Wang, Yuan Bai, Yunhu Liu, Eric HY Lau, Alison P Galvani, Robert M Krug, Benjamin John Cowling, and Lauren A Meyers. A retrospective cohort study of Paxlovid efficacy depending on treatment time in hospitalized COVID-19 patients. *eLife*, 13, 2024.

[13866] Haojing Duan, Runye Shi, Jujiao Kang, Tobias Banaschewski, Arun LW Bokde, Christian Büchel, Sylvane Desrivières, Herta Flor, Antoine Grigis, Hugh Garavan, Penny A Gowland, Andreas Heinz, Rüdiger Brühl, Jean-Luc Martinot, Marie-Laure Paillère Martinot, Eric Artiges, Frauke Nees, Dimitri Papadopoulos Orfanos, Luise Poustka, Sarah Hohmann, Nathalie Nathalie Holz, Juliane Fröhner, Michael N Smolka, Nilakshi Vaidya, Henrik Walter, Robert Whelan, Gunter Schumann, Xiaolei Lin, and Jianfeng Feng. Population clustering of structural brain aging and its association with brain development. *eLife*, 13, 2024.

[13867] Tianlin Duan, Adrien Sicard, Sylvain Glémin, and Martin Lascoux. Separating phases of allopolyploid evolution with resynthesized and natural Capsella bursa-pastoris. *eLife*, 12, 2024.

[13868] Ziyi Duan and Clayton E Curtis. Visual working memories are abstractions of percepts. *eLife*, 13, 2024.

[13869] Zofia Dubicka, Jarosław Tyszka, Agnieszka Pałczyńska, Michelle Höhne, Jelle Bijma, Max Jense, Nienke Klerks, and Ulf Bickmeyer. Biocalcification in porcelaneous foraminifera. *eLife*, 13, 2024.

[13870] Benjamin R Duewell, Naomi E Wilson, Gabriela M Bailey, Sarah E Peabody, and Scott D Hansen. Molecular dissection of PI3K$\beta$ synergistic activation by receptor tyrosine kinases, G$\beta$G$\gamma$, and Rho-family GTPases. *eLife*, 12, 2024.

[13871] Fabien Duveau, Céline Cordier, Lionel Chiron, Matthias Le Bec, Sylvain Pouzet, Julie Séguin, Artémis Llamosi, Benoit Sorre, Jean-Marc Di Meglio, and Pascal Hersen. Yeast cell responses and survival during periodic osmotic stress are controlled by glucose availability. *eLife*, 12, 2024.

[13872] Joshua D Eaton, Jessica Board, Lee Davidson, Chris Estell, and Steven West. Human promoter directionality is determined by transcriptional initiation and the opposing activities of INTS11 and CDK9. *eLife*, 13, 2024.

[13873] Pierce Eggan, Sharona E Gordon, and William N Zagotta. Ligand-coupled conformational changes in a cyclic nucleotide-gated ion channel revealed by time-resolved transition metal ion FRET. *eLife*, 13, 2024.

[13874] Kristina Ehring, Sophia Friederike Ehlers, Jurij Froese, Fabian Gude, Janna Puschmann, and Kay Grobe. Two-way Dispatched function in Sonic hedgehog shedding and transfer to high-density lipoproteins. *eLife*, 12, 2024.

[13875] Katharina Eichler, Stefanie Hampel, Adrián Alejandro-García, Steven A Calle-Schuler, Alexis Santana-Cruz, Lucia Kmecova, Jonathan M Blagburn, Eric D Hoopfer, and Andrew M Seeds. Somatotopic organization among parallel sensory pathways that promote a grooming sequence in Drosophila. *eLife*, 12, 2024.

[13876] Mohamed El Amri, Abhay Pandit, and Gerhard Schlosser. Marcks and Marcks-like 1 proteins promote spinal cord development and regeneration in Xenopus. *eLife*, 13, 2024.

[13877] Johannes Elferich, Lingli Kong, Ximena Zottig, and Nikolaus Grigorieff. CTFFIND5 provides improved insight into quality, tilt, and thickness of TEM samples. *eLife*, 13, 2024.

[13878] Christina Elliott and Patrick Brundell. How we stepped up to support others. *eLife*, 13, 2024.

[13879] Arne Elofsson, Ling Han, Enrica Bianchi, Gavin J Wright, and Luca Jovine. Deep learning insights into the architecture of the mammalian egg-sperm fusion synapse. *eLife*, 13, 2024.

[13880] Natsuko Emura, Florence DM Wavreil, Annaliese Fries, and Mamiko Yajima. The evolutionary modifications of a GoLoco motif in the AGS protein facilitate micromere formation in the sea urchin embryo. *eLife*, 13, 2024.

[13881] Anthony English, Fleur Uittenbogaard, Alexa Torrens, Dennis Sarroza, Anna Veronica Elizabeth Slaven, Daniele Piomelli, Michael R Bruchas, Nephi Stella, and Benjamin Bruce Land. A preclinical model of THC edibles that produces high-dose cannabimimetic responses. *eLife*, 12, 2024.

[13882] Bernhard Englitz, Sahar Akram, Mounya Elhilali, and Shihab Shamma. Decoding contextual influences on auditory perception from primary auditory cortex. *eLife*, 13, 2024.

[13883] Christoph Erbacher, Sebastian Britz, Philine Dinkel, Thomas Klein, Markus Sauer, Christian Stigloher, and Nurcan üÇeyler. Interaction of human keratinocytes and nerve fiber terminals at the neuro-cutaneous unit. *eLife*, 13, 2024.

[13884] Anna L Erdei, Aneth B David, Eleni C Savvidou, Vaida Džemedžionaitė, Advaith Chakravarthy, Béla P Molnár, and Teun Dekker. The push–pull intercrop Desmodium does not repel, but intercepts and kills pests. *eLife*, 13, 2024.

[13885] Phillip A Erice, Xinyan Huang, Matthew J Seasock, Matthew J Robertson, Hui-Ying Tung, Melissa A Perez-Negron, Shivani L Lotlikar, David B Corry, Farrah Kheradmand, and Antony Rodriguez. Downregulation of Mirlet7 miRNA family promotes Tc17 differentiation and emphysema via derepression of ROR$\gamma$t. *eLife*, 13, 2024.

[13886] Abraham Escobedo, Salli-Ann Holloway, Megan Votoupal, Aaron L Cone, Hannah Skelton, Alex A Legaria, Imeh Ndiokho, Tasheia Floyd, Alexxai V Kravitz, Michael R Bruchas, and Aaron J Norris. Glutamatergic supramammillary nucleus neurons respond to threatening stressors and promote active coping. *eLife*, 12, 2024.

[13887] Gabriella O Estevam, Edmond M Linossi, Christian B Macdonald, Carla A Espinoza, Jennifer M Michaud, Willow Coyote-Maestas, Eric A Collisson, Natalia Jura, and James S Fraser. Conserved regulatory motifs in the juxtamembrane domain and kinase N-lobe revealed through deep mutational scanning of the MET receptor tyrosine kinase domain. *eLife*, 12, 2024.

[13888] Giulio Facchini, Alann Rathery, Stéphane Douady, David Sillam-Dussès, and Andrea Perna. Substrate evaporation drives collective construction in termites. *eLife*, 12, 2024.

[13889] Anne Fajac, Iva Simeonova, Julia Leemput, Marc Gabriel, Aurélie Morin, Vincent Lejour, Annaïg Hamon, Jeanne Rakotopare, Wilhelm Vaysse-Zinkhöfer, Eliana Eldawra, Marina Pinskaya, Antonin Morillon, Jean-Christophe Bourdon, Boris Bardot, and Franck Toledo. Mutant mice lacking alternatively spliced p53 isoforms unveil Ackr4 as a male-specific prognostic factor in Myc-driven B-cell lymphomas. *eLife*, 13, 2024.

[13890] Johannes Falck, Lei Zhang, Laurel Raffington, Johannes Julius Mohn, Jochen Triesch, Christine Heim, and Yee Lee Shing. Hippocampus and striatum show distinct contributions to longitudinal changes in value-based learning in middle childhood. *eLife*, 12, 2024.

[13891] Rongxin Fang, Aaron Halpern, Mohammed Mostafizur Rahman, Zhengkai Huang, Zhiyun Lei, Sebastian J Hell, Catherine Dulac, and Xiaowei Zhuang. Three-dimensional single-cell transcriptome imaging of thick tissues. *eLife*, 12, 2024.

[13892] Zeming Fang, Meihua Zhao, Ting Xu, Yuhang Li, Hanbo Xie, Peng Quan, Haiyang Geng, and Ru-Yuan Zhang. Individuals with anxiety and depression use atypical decision strategies in an uncertain world. *eLife*, 13, 2024.

[13893] Zepeng Fang, Yuanyuan Dang, Zhipei Ling, Yongzheng Han, Hulin Zhao, Xin Xu, and Mingsha Zhang. The involvement of the human prefrontal cortex in the emergence of visual awareness. *eLife*, 12, 2024.

[13894] Tumininu S Faniyan, Xinyi Zhang, Donald A Morgan, Jorge Robles, Siresha Bathina, Paul S Brookes, Kamal Rahmouni, Rachel J Perry, and Kavaljit H Chhabra. A kidney-hypothalamus axis promotes compensatory glucose production in response to glycosuria. *eLife*, 12, 2024.

[13895] Islam Faress, Valentina Khalil, Wen-Hsien Hou, Andrea Moreno, Niels Andersen, Rosalina Fonseca, Joaquin Piriz, Marco Capogna, and Sadegh Nabavi. Non-Hebbian plasticity transforms transient experiences into lasting memories. *eLife*, 12, 2024.

[13896] Leen Farhat-Younis, Manho Na, Amichai Zarfin, Aseel Khateeb, Nadine Santana-Magal, Alon Richter, Amit Gutwillig, Diana Rasoulouniriana, Annette Gleiberman, Lir Beck, Tamar Giger, Avraham Ashkenazi, Adi Barzel, Peleg Rider, and Yaron Carmi. Expression of modified Fc$\gamma$RI enables myeloid cells to elicit robust tumor-specific cytotoxicity. *eLife*, 12, 2024.

[13897] Megan E Farquhar, Qianqian Yang, and Viktor Vegh. Robust, fast and accurate mapping of diffusional mean kurtosis. *eLife*, 12, 2024.

[13898] KC Farrell, Jennifer T Wang, and Tim Stearns. Spindle assembly checkpoint-dependent mitotic delay is required for cell division in absence of centrosomes. *eLife*, 12, 2024.

[13899] Amirhossein Farzmahdi, Wilbert Zarco, Winrich A Freiwald, Nikolaus Kriegeskorte, and Tal Golan. Emergence of brain-like mirror-symmetric viewpoint tuning in convolutional neural networks. *eLife*, 13, 2024.

[13900] Luisa Fassi, Shachar Hochman, Zafiris J Daskalakis, Daniel M Blumberger, and Roi Cohen Kadosh. The importance of individual beliefs in assessing treatment efficacy. *eLife*, 12, 2024.

[13901] Sami Fawaz, Severine Marti, Melody Dufossee, Yann Pucheu, Astrid Gaufroy, Jean Broitman, Audrey Bidet, Aicha Soumare, Gaëlle Munsch, Christophe Tzourio, Stephanie Debette, David-Alexandre Trégouët, Chloe James, Olivier Mansier, and Thierry Couffinhal. Evaluation of clonal

hematopoiesis and mosaic loss of Y chromosome in cardiovascular risk: An analysis in prospective studies. *eLife*, 13, 2024.

[13902] KM Rifat Faysal, James C Walsh, Nadine Renner, Chantal L Márquez, Vaibhav B Shah, Andrew J Tuckwell, Michelle P Christie, Michael W Parker, Stuart G Turville, Greg J Towers, Leo C James, David A Jacques, and Till Böcking. Pharmacologic hyperstabilisation of the HIV-1 capsid lattice induces capsid failure. *eLife*, 13, 2024.

[13903] Frederick Federer, Justin Balsor, Alexander Ingold, David P Babcock, Jordane Dimidschstein, and Alessandra Angelucci. Laminar specificity and coverage of viral-mediated gene expression restricted to GABAergic interneurons and their parvalbumin subclass in marmoset primary visual cortex. *eLife*, 13, 2024.

[13904] Guozheng Feng, Yiwen Wang, Weijie Huang, Haojie Chen, Jian Cheng, and Ni Shu. Spatial and temporal pattern of structure–function coupling of human brain connectome with development. *eLife*, 13, 2024.

[13905] Tong Feng, Qi Zhang, Si-Hui Li, Yan-ling Ping, Mu-qiu Tian, Shuan-hu Zhou, Xin Wang, Jun-Meng Wang, Fan-Rang Liang, Shu-Guang Yu, and Qiao-Feng Wu. Inhibition of miR-199b-5p reduces pathological alterations in osteoarthritis by potentially targeting Fzd6 and Gcnt2. *eLife*, 12, 2024.

[13906] Xinran Feng, Shan Xu, Yuannan Li, and Jia Liu. Body size as a metric for the affordable world. *eLife*, 12, 2024.

[13907] Giulia Ferraretti, Paolo Abondio, Marta Alberti, Agnese Dezi, Phurba T Sherpa, Paolo Cocco, Massimiliano Tiriticco, Marco Di Marcello, Guido Alberto Gnecchi-Ruscone, Luca Natali, Angela Corcelli, Giorgio Marinelli, Davide Peluzzi, Stefania Sarno, and Marco Sazzini. Archaic introgression contributed to shape the adaptive modulation of angiogenesis and cardiovascular traits in human high-altitude populations from the Himalayas. *eLife*, 12, 2024.

[13908] Alexandra M Fister, Adam Horn, Michael R Lasarev, and Anna Huttenlocher. Damage-induced basal epithelial cell migration modulates the spatial organization of redox signaling and sensory neuron regeneration. *eLife*, 13, 2024.

[13909] Stephen E Flaherty, Olivier Bezy, Brianna LaCarubba Paulhus, LouJin Song, Mary Piper, Jincheng Pang, Yoson Park, Shoh Asano, Yu-Chin Lien, John D Griffin, Andrew Robertson, Alan Opsahl, Dinesh Hirenallur Shanthappa, Youngwook Ahn, Evanthia Pashos, Rebecca A Simmons, Morris J Birnbaum, and Zhidan Wu. SPAG7 deletion causes intrauterine growth restriction, resulting in adulthood obesity and metabolic dysfunction. *eLife*, 12, 2024.

[13910] Sophie A Fleck, Puja Biswas, Emily D DeWitt, Rebecca L Knuteson, Robert C Eisman, Travis Nemkov, Angelo D'Alessandro, Jason M Tennessen, Elizabeth Rideout, and Lesley N Weaver. Auxin exposure disrupts feeding behavior and fatty acid metabolism in adult Drosophila. *eLife*, 12, 2024.

[13911] Paul AG Forbes, Gökhan Aydogan, Julia Braunstein, Boryana Todorova, Isabella C Wagner, Patricia L Lockwood, Matthew AJ Apps, Christian C Ruff, and Claus Lamm. Acute stress reduces effortful prosocial behaviour. *eLife*, 12, 2024.

[13912] Pascal Forcella, Niklas Ifflander, Chiara Rolando, Elli-Anna Balta, Aikaterini Lampada, Claudio Giachino, Tanzila Mukhtar, Thomas Bock, and Verdon Taylor. SAFB regulates hippocampal stem cell fate by targeting Drosha to destabilize Nfib mRNA. *eLife*, 13, 2024.

[13913] Eric J Foss, Carmina Lichauco, Tonibelle Gatbonton-Schwager, Sara J Gonske, Brandon Lofts, Uyen Lao, and Antonio Bedalov. Identification of 1600 replication origins in S. cerevisiae. *eLife*, 12, 2024.

[13914] Katrin Franke, Chenchen Cai, Kayla Ponder, Jiakun Fu, Sacha Sokoloski, Philipp Berens, and Andreas Savas Tolias. Asymmetric distribution of color-opponent response types across mouse visual cortex supports superior color vision in the sky. *eLife*, 12, 2024.

[13915] Cody A Freas, Ajay Narenda, Trevor Murray, and Ken Cheng. Polarised moonlight guides nocturnal bull ants home. *eLife*, 13, 2024.

[13916] Lukas Frey, Dhiman Ghosh, Bilal M Qureshi, David Rhyner, Ricardo Guerrero-Ferreira, Aditya Pokharna, Witek Kwiatkowski, Tetiana Serdiuk, Paola Picotti, Roland Riek, and Jason Greenwald. On the pH-dependence of $\alpha$-synuclein amyloid polymorphism and the role of secondary nucleation in seed-based amyloid propagation. *eLife*, 12, 2024.

[13917] Yang Fu, Xiao-Dan Luo, Jin-Ze Li, Qian-Yuan Mo, Xue Wang, Yue Zhao, You-Ming Zhang, Hao-Tong Luo, Dai-Yang Xia, Wei-Qing Ma, Jian-Ying Chen, Li-Hau Wang, Qiu-Yi Deng, Lukuyu Ben, Muhammad Kashif Saleemi, Xian-Zhi Jiang, Juan Chen, Kai Miao, Zhen-Ping Lin, Peng Zhang, Hui Ye, Qing-Yun Cao, Yong-Wen Zhu, Lin Yang, Qiang Tu, and Wence Wang. Host-derived Lactobacillus plantarum alleviates hyperuricemia by improving gut microbial community and hydrolase-mediated degradation of purine nucleosides. *eLife*, 13, 2024.

[13918] Yumin Fu, Xinyu Guo, Linmao Sun, Tianming Cui, Chenghui Wu, Jiabei Wang, Yao Liu, and Lianxin Liu. Exploring the role of the immune microenvironment in hepatocellular carcinoma: Implications for immunotherapy and drug resistance. *eLife*, 13, 2024.

[13919] Dietmar Funck, Malte Sinn, Giuseppe Forlani, and Jörg S Hartig. Guanidine production by plant homoarginine-6-hydroxylases. *eLife*, 12, 2024.

[13920] Timothy Fuqua, Yiqiao Sun, and Andreas Wagner. The emergence and evolution of gene expression in genome regions replete with regulatory motifs. *eLife*, 13, 2024.

[13921] Aurélie Anne-Gaëlle Gabriel, Julien Racle, Maryline Falquet, Camilla Jandus, and David Gfeller. Robust estimation of cancer and immune cell-type proportions from bulk tumor ATAC-Seq data. *eLife*, 13, 2024.

[13922] Sachin P Gadani, Saumitra Singh, Sophia Kim, Jingwen Hu, Matthew D Smith, Peter A Calabresi, and Pavan Bhargava. Spatial transcriptomics of meningeal inflammation reveals inflammatory gene signatures in adjacent brain parenchyma. *eLife*, 12, 2024.

[13923] Swati Gaikwad, Fardin Ghobakhlou, Hongen Zhang, and Alan G Hinnebusch. Yeast eIF2A has a minimal role in translation initiation and uORF-mediated translational control in vivo. *eLife*, 12, 2024.

[13924] Dario Galanti, Jun Hee Jung, Caroline Müller, and Oliver Bossdorf. Discarded sequencing reads uncover natural variation in pest resistance in Thlaspi arvense. *eLife*, 13, 2024.

[13925] Christian Galicia, Giambattista Guaitoli, Marcus Fislage, Christian Johannes Gloeckner, and Wim Versées. Structural insights into the GTP-driven monomerization and activation of a bacterial LRRK2 homolog using allosteric nanobodies. *eLife*, 13, 2024.

[13926] Lavinia Gambelli, Mathew McLaren, Rebecca Conners, Kelly Sanders, Matthew C Gaines, Lewis Clark, Vicki AM Gold, Daniel Kattnig, Mateusz Sikora, Cyril Hanus, Michail N Isupov, and Bertram Daum. Structure of the two-component S-layer of the archaeon Sulfolobus acidocaldarius. *eLife*, 13, 2024.

[13927] Ninghai Gan, Yan Han, Weizhong Zeng, and Youxing Jiang. TRPML1 gating modulation by allosteric mutations and lipids. *eLife*, 13, 2024.

[13928] Chenxi Gao, Huaibin Ge, Shih-Fan Kuan, Chunhui Cai, Xinghua Lu, Farzad Esni, Robert E Schoen, Jing H Wang, Edward Chu, and Jing Hu. FAK loss reduces BRAFV600E-induced ERK phosphorylation to promote intestinal stemness and cecal tumor formation. *eLife*, 13, 2024.

[13929] Lingzhi Gao, Dian Chen, and Yu Liu. Ligand response of guanidine-IV riboswitch at single-molecule level. *eLife*, 13, 2024.

[13930] Yuan Gao, Yong-Chun Cai, Dong-Yu Liu, Juan Yu, Jue Wang, Ming Li, Bin Xu, Tengfei Wang, Gang Chen, Georg Northoff, Ruiliang Bai, and Xue Mei Song. GABAergic inhibition in human hMT+ predicts visuo-spatial intelligence mediated through the frontal cortex. *eLife*, 13, 2024.

[13931] Eva M García-Cuesta, Pablo Martínez, Karthik Selvaraju, Gabriel Ulltjärn, Adrián Miguel Gómez Pozo, Gianluca D'Agostino, Sofia Gardeta, Adriana Quijada-Freire, Patricia Blanco Gabella, Carlos Roca, Daniel del Hoyo, Rodrigo Jiménez-Saiz, Alfonso García-Rubia, Blanca Soler Palacios, Pilar Lucas, Rosa Ayala-Bueno, Noelia Santander Acerete, Yolanda Carrasco, Carlos Oscar Sorzano, Ana Martinez, Nuria E Campillo, Lasse D Jensen, Jose Miguel Rodriguez Frade, César Santiago, and Mario Mellado. Allosteric modulation of the CXCR4:CXCL12 axis by targeting receptor nanoclustering via the TMV-TMVI domain. *eLife*, 13, 2024.

[13932] Pamela Garcia-Saldivar, Cynthia de León, Felipe A Mendez Salcido, Luis Concha, and Hugo Merchant. White matter structural bases for phase accuracy during tapping synchronization. *eLife*, 13, 2024.

[13933] Vincent Gardeux, Roel PJ Bevers, Fabrice PA David, Emily Rosschaert, Romain Rochepeau, and Bart Deplancke. DGRPool, a web tool leveraging harmonized Drosophila Genetic Reference Panel phenotyping data for the study of complex traits. *eLife*, 12, 2024.

[13934] Josep Garnica, Patricia Sole, Jun Yamanouchi, Joel Moro, Debajyoti Mondal, Cesar Fandos, Pau Serra, and Pere Santamaria. T-follicular helper cells are epigenetically poised to transdifferentiate into T-regulatory type 1 cells. *eLife*, 13, 2024.

[13935] Preksha Gaur, Yesheswini Rajendran, Bhagyashree Srivastava, Manasvini Markandey, Vered Fishbain-Yoskovitz, Gayatree Mohapatra, Aamir Suhail, Shikha Chaudhary, Shaifali Tyagi, Subhash Chandra Yadav, Amit Kumar Pandey, Yifat Merbl, Avinash Bajaj, Vineet Ahuja, and Chittur Srikanth. Rab7-dependent regulation of goblet cell protein CLCA1 modulates gastrointestinal homeostasis. *eLife*, 12, 2024.

[13936] David Geller-McGrath, Kishori M Konwar, Virginia P Edgcomb, Maria Pachiadaki, Jack W Roddy, Travis J Wheeler, and Jason E McDermott. Predicting metabolic modules in incomplete bacterial genomes with MetaPathPredict. *eLife*, 13, 2024.

[13937] Shang Geng, Xing Lv, Weiwei Zheng, and Tianjun Xu. An arms race between 5′ppp-RNA virus and its alternative recognition receptor MDA5 in RIG-I-lost teleost fish. *eLife*, 13, 2024.

[13938] Yoav Ger, Moni Shahar, and Nitzan Shahar. Using recurrent neural network to estimate irreducible stochasticity in human choice behavior. *eLife*, 13, 2024.

[13939] Carolin Gerke, Liane Bauersfeld, Ivo Schirmeister, Chiara Noemi-Marie Mireisz, Valerie Oberhardt, Lea Mery, Di Wu, Christopher Sebastian Jürges, Robbert M Spaapen, Claudio Mussolino, Vu Thuy Khanh Le-Trilling, Mirko Trilling, Lars Dölken, Wolfgang Paster, Florian Erhard, Maike Hofmann, Andreas Schlosser, Hartmut Hengel, Frank Momburg, and Anne Halenius. Multimodal HLA-I genotype regulation by human cytomegalovirus US10 and resulting surface patterning. *eLife*, 13, 2024.

[13940] Sila Gerlevik, Nogayhan Seymen, Shan Hama, Warisha Mumtaz, I Richard Thompson, Seyed R Jalili, Deniz E Kaya, Alfredo Iacoangeli, Andrea Pellagatti, Jacqueline Boultwood, Giorgio Napolitani, Ghulam J Mufti, and Mohammad M Karimi. Identification of novel myelodysplastic syndromes prognostic subgroups by integration of inflammation, cell-type composition, and immune signatures in the bone marrow. *eLife*, 13, 2024.

[13941] Edward M Germany, Nakajohn Thewasano, Kenichiro Imai, Yuki Maruno, Rebecca S Bamert, Christopher J Stubenrauch, Rhys A Dunstan, Yue Ding, Yukari Nakajima, XiangFeng Lai, Chaille T Webb, Kentaro Hidaka, Kher Shing Tan, Hsinhui Shen, Trevor Lithgow, and Takuya Shiota. Dual recognition of multiple signals in bacterial outer membrane proteins enhances assembly and maintains membrane integrity. *eLife*, 12, 2024.

[13942] Thomas RM Germe, Natassja G Bush, Victoria M Baskerville, Dominik Saman, Justin LP Benesch, and Anthony Maxwell. Rapid, DNA-induced interface swapping by DNA gyrase. *eLife*, 12, 2024.

[13943] Theresa Gewering, Deepali Waghray, Kristian Parey, Hendrik Jung, Nghi NB Tran, Joel Zapata, Pengyi Zhao, Hao Chen, Dovile Januliene, Gerhard Hummer, Ina Urbatsch, Arne Moeller, and Qinghai Zhang. Tracing the substrate translocation mechanism in P-glycoprotein. *eLife*, 12, 2024.

[13944] Tara Ghafari, Cecilia Mazzetti, Kelly Garner, Tjerk Gutteling, and Ole Jensen. Modulation of alpha oscillations by attention is predicted by hemispheric asymmetry of subcortical regions. *eLife*, 12, 2024.

[13945] Zahra Ghasemahmad, Aaron Mrvelj, Rishitha Panditi, Bhavya Sharma, Karthic Drishna Perumal, and Jeffrey J Wenstrup. Emotional vocalizations alter behaviors and neurochemical release into the amygdala. *eLife*, 12, 2024.

[13946] Phaedra C Ghazi, Kayla T O'Toole, Sanjana Srinivas Boggaram, Michael T Scherzer, Mark R Silvis, Yun Zhang, Madhumita Bogdan, Bryan D Smith, Guillermina Lozano, Daniel L Flynn, Eric L Snyder, Conan G Kinsey, and Martin McMahon. Inhibition of ULK1/2 and KRASG12C controls tumor growth in preclinical models of lung cancer. *eLife*, 13, 2024.

[13947] Carlo Giannangelo, Matthew P Challis, Ghizal Siddiqui, Rebecca Edgar, Tess R Malcolm, Chaille T Webb, Nyssa Drinkwater, Natalie Vinh, Christopher Macraild, Natalie Counihan, Sandra Duffy, Sergio Wittlin, Shane M Devine, Vicky M Avery, Tania De Koning-Ward, Peter Scammells, Sheena McGowan, and Darren J Creek. Chemoproteomics validates selective targeting of Plasmodium M1 alanyl aminopeptidase as an antimalarial strategy. *eLife*, 13, 2024.

[13948] Tristan Gicquel, Fabio Marchiano, Gabriela Reyes-Castellanos, Stephane Audebert, Luc Camoin, Bianca H Habermann, Benoit Giannesini, and Alice Carrier. Integrative study of skeletal muscle mitochondrial dysfunction in a murine pancreatic cancer-induced cachexia model. *eLife*, 13, 2024.

[13949] Thierry Gilbert, Camille Gorlt, Merlin Barbier, Benjamin Duployer, Marianna Plozza, Ophélie Dufrancais, Laure-Elene Martet, Elisa Dalbard, Loelia Segot, Christophe Tenailleau, Laurence Haren, Christel Vérollet, Christiane Bierkamp, and Andreas Merdes. Loss of ninein interferes with osteoclast formation and causes premature ossification. *eLife*, 13, 2024.

[13950] Anna K Gillespie, Daniela Astudillo Maya, Eric L Denovellis, Sachi Desse, and Loren M Frank. Neurofeedback training can modulate task-relevant memory replay rate in rats. *eLife*, 12, 2024.

[13951] Maxwell Gillett and Nicolas Brunel. Dynamic control of sequential retrieval speed in networks with heterogeneous learning rules. *eLife*, 12, 2024.

[13952] Bernd K Gilsbach, Franz Y Ho, Benjamin Riebenbauer, Xiaojuan Zhang, Giambattista Guaitoli, Arjan Kortholt, and Christian Johannes Gloeckner. Intramolecular feedback regulation of the LRRK2 Roc G domain by a LRRK2 kinase-dependent mechanism. *eLife*, 12, 2024.

[13953] Alon Avraham Ginatt, Maria Berihu, Einam Castel, Shlomit Medina, Gon Carmi, Adi Faigenboim-Doron, Itai Sharon, Ofir Tal, Samir Droby, Tracey Somera, Mark Mazzola, Hanan Eizenberg, and Shiri Freilich. A metabolic modeling-based framework for predicting trophic dependencies in native rhizobiomes of crop plants. *eLife*, 13, 2024.

[13954] Sebastián Giunti, María Gabriela Blanco, María José De Rosa, and Diego Rayes. The ketone body $\beta$-hydroxybutyrate ameliorates neurodevelopmental deficits in the GABAergic system of daf-18/PTEN Caenorhabditis elegans mutants. *eLife*, 13, 2024.

[13955] Siena J Glenn, Zealon Gentry-Lear, Michael Shavlik, Michael J Harms, Thomas J Asaki, and Arden Baylink. Bacterial vampirism mediated through taxis to serum. *eLife*, 12, 2024.

[13956] Vasileios Glykos and Shigeyoshi Fujisawa. Memory-specific encoding activities of the ventral tegmental area dopamine and GABA neurons. *eLife*, 12, 2024.

[13957] Simei Go, Constantinos Demetriou, Giampiero Valenzano, Sophie Hughes, Simone Lanfredini, Helen Ferry, Edward Arbe-Barnes, Shivan Sivakumar, Rachel Bashford-Rogers, Mark R Middleton, Somnath Mukherjee, Jennifer Morton, Keaton Jones, and Eric O Neill. Tissue-resident natural killer cells support survival in pancreatic cancer through promotion of cDC1-CD8 T activity. *eLife*, 13, 2024.

[13958] Andrew Goetz, Hoda Akl, and Purushottam Dixit. The ability to sense the environment is heterogeneously distributed in cell populations. *eLife*, 12, 2024.

[13959] Chetan Gohil, Rukuang Huang, Evan Roberts, Mats WJ van Es, Andrew J Quinn, Diego Vidaurre, and Mark W Woolrich. osl-dynamics, a toolbox for modeling fast dynamic brain activity. *eLife*, 12, 2024.

[13960] Dena Goldblatt, Basak Rosti, Kyla Rose Hamling, Paige Leary, Harsh Panchal, Marlyn Li, Hannah Gelnaw, Stephanie Huang, Cheryl Quainoo, and David Schoppik. Motor neurons are dispensable for the assembly of a sensorimotor circuit for gaze stabilization. *eLife*, 13, 2024.

[13961] Allison T Goldstein, Terrence R Stanford, and Emilio Salinas. Coupling of saccade plans to endogenous attention during urgent choices. *eLife*, 13, 2024.

[13962] Alessia Golluscio, Jodene Eldstrom, Jessica J Jowais, Marta Elena Perez, Kevin Peter Cunningham, Alicia De La Cruz, Xiaoan Wu, Valentina Corradi, D Peter Tieleman, David Fedida, and H Peter Larsson. PUFA stabilizes a conductive state of the selectivity filter in IKs channels. *eLife*, 13, 2024.

[13963] Arkadiy K Golov, Alexey A Gavrilov, Noam Kaplan, and Sergey V Razin. A genome-wide nucleosome-resolution map of promoter-centered interactions in human cells corroborates the enhancer-promoter looping model. *eLife*, 12, 2024.

[13964] Juan Manuel Gomez, Hendrik Nolte, Elisabeth Vogelsang, Bipasha Dey, Michiko Takeda, Girolamo Giudice, Miriam Faxel, Theresa Haunold, Alina Cepraga, Robert P Zinzen, Marcus Krüger, Evangelia Petsalaki, Yu-Chiun Wang, and Maria Leptin. Differential regulation of the proteome and phosphoproteome along the dorso-ventral axis of the early Drosophila embryo. *eLife*, 13, 2024.

[13965] Alexander Gonzalez and Lisa M Giocomo. Parahippocampal neurons encode task-relevant information for goal-directed navigation. *eLife*, 12, 2024.

[13966] Samuel J Gonzalez, Julia M Heckel, Rebecca R Goldblum, Taylor A Reid, Mark McClellan, and Melissa K Gardner. Rapid binding to protofilament edge sites facilitates tip tracking of EB1 at growing microtubule plus-ends. *eLife*, 13, 2024.

[13967] Stephen Gonzalez, Joseph Swift, Adi Yaaran, Jiaying Xu, Charlotte Miller, Natanella Illouz-Eliaz, Joseph R Nery, Wolfgang Busch, Yotam Zait, and Joseph R Ecker. Arabidopsis transcriptome responses to low water potential using high-throughput plate assays. *eLife*, 12, 2024.

[13968] Carlos Gonzalez-Islas, Zahraa Sabra, Ming-fai Fong, Pernille Yilmam, Nicholas Au Yong, Kathrin Engisch, and Peter Wenner. GABAergic synaptic scaling is triggered by changes in spiking activity rather than AMPA receptor activation. *eLife*, 12, 2024.

[13969] Rachel S Gormal, Ramon Martinez-Marmol, Andrew J Brooks, and Frédéric A Meunier. Location, location, location: Protein kinase nanoclustering for optimised signalling output. *eLife*, 13, 2024.

[13970] Sofia Granados-Aparici, Qin Yang, and Hugh J Clarke. SMAD4 promotes somatic-germline contact during murine oocyte growth. *eLife*, 13, 2024.

[13971] Thomas Grandits, Christoph M Augustin, Gundolf Haase, Norbert Jost, Gary R Mirams, Steven A Niederer, Gernot Plank, András Varró, László Virág, and Alexander Jung. Neural network emulation of the human ventricular cardiomyocyte action potential for more efficient computations in pharmacological studies. *eLife*, 12, 2024.

[13972] Alberto Granzotto, Bryce Vissel, and Stefano L Sensi. Lost in translation: Inconvenient truths on the utility of mouse models in Alzheimer's disease research. *eLife*, 13, 2024.

[13973] Jessica Gray, Von Vergel L Torres, Emily Goodall, Samantha A McKeand, Danielle Scales, Christy Collins, Laura Wetherall, Zheng Jie Lian, Jack A Bryant, Matthew T Milner, Karl A Dunne, Christopher Icke, Jessica L Rooke, Thamarai Schneiders, Peter A Lund, Adam F Cunningham, Jeff A Cole, and Ian R Henderson. Transposon mutagenesis screen in Klebsiella pneumoniae identifies genetic determinants required for growth in human urine and serum. *eLife*, 12, 2024.

[13974] Ari Grele, Tara J Massad, Kathryn A Uckele, Lee A Dyer, Yasmine Antonini, Laura Braga, Matthew L Forister, Lidia Sulca, Massuo Kato, Humberto G Lopez, André R Nascimento, Thomas Parchman, Wilmer R Simbaña, Angela M Smilanich, John O Stireman, Eric J Tepe, Thomas Walla, and Lora A Richards. Intra- and interspecific diversity in a tropical plant clade alter herbivory and ecosystem resilience. *eLife*, 12, 2024.

[13975] Johannes N Greve, Anja Marquardt, Robin Heiringhoff, Theresia Reindl, Claudia Thiel, Nataliya Di Donato, Manuel H Taft, and Dietmar J Manstein. The non-muscle actinopathy-associated mutation E334Q in cytoskeletal $\gamma$-actin perturbs interaction of actin filaments with myosin and ADF/cofilin family proteins. *eLife*, 12, 2024.

[13976] Samuel C Griffiths, Jia Tan, Armin Wagner, Levi L Blazer, Jarrett J Adams, Srisathya Srinivasan, Shayan Moghisaei, Sachdev S Sidhu, Christian Siebold, and Hsin-Yi Henry Ho. Structure and function of the ROR2 cysteine-rich domain in vertebrate noncanonical WNT5A signaling. *eLife*, 13, 2024.

[13977] David Robert Grimes. Region of Attainable Redaction, an extension of Ellipse of Insignificance analysis for gauging impacts of data redaction in dichotomous outcome trials. *eLife*, 13, 2024.

[13978] Anna-Maria Grob, Hendrik Heinbockel, Branka Milivojevic, Christian F Doeller, and Lars Schwabe. Causal role of the angular gyrus in insight-driven memory reconfiguration. *eLife*, 12, 2024.

[13979] John P Grogan, Matthias Raemaekers, Maaike MH van Swieten, Alexander L Green, Martin J Gillies, and Sanjay G Manohar. Muscarinic receptors mediate motivation via preparatory neural activity in humans. *eLife*, 13, 2024.

[13980] Matthew Grove, Hyukmin Kim, Shuhuan Pang, Jose Paz Amaya, Guoqing Hu, Jiliang Zhou, Michel Lemay, and Young-Jin Son. TEAD1 is crucial for developmental myelination, Remak bundles, and functional regeneration of peripheral nerves. *eLife*, 13, 2024.

[13981] Marta Grzonka and Hisham Bazzi. Mouse SAS-6 is required for centriole formation in embryos and integrity in embryonic stem cells. *eLife*, 13, 2024.

[13982] LeYuan Gu, WeiHui Shao, Lu Liu, Qing Xu, YuLing Wang, JiaXuan Gu, Yue Yang, ZhuoYue Zhang, YaXuan Wu, Yue Shen, Qian Yu, XiTing Lian, HaiXiang Ma, YuanLi Zhang, and HongHai Zhang. NE contribution to rebooting unconsciousness caused by midazolam. *eLife*, 13, 2024.

[13983] Shaohua Gu, Yuanzhe Shao, Karoline Rehm, Laurent Bigler, Di Zhang, Ruolin He, Ruichen Xu, Jiqi Shao, Alexandre Jousset, Ville-Petri Friman, Xiaoying Bian, Zhong Wei, Rolf Kümmerli, and Zhiyuan Li. Feature sequence-based genome mining uncovers the hidden diversity of bacterial siderophore pathways. *eLife*, 13, 2024.

[13984] Wenyue Guan, Ziyan Nie, Anne LaurenÇon, Mathilde Bouchet, Christophe Godin, Chérif Kabir, Aurelien Darnas, and Jonathan Enriquez. The role of Imp and Syp RNA-binding proteins in precise neuronal elimination by apoptosis through the regulation of transcription factors. *eLife*, 12, 2024.

[13985] Célia Guérin and David Tulasne. Recording and classifying MET receptor mutations in cancers. *eLife*, 13, 2024.

[13986] Rafael F Guerrero, Tandin Dorji, Ra'Mal M Harris, Matthew D Shoulders, and C Brandon Ogbunugafor. Evolutionary druggability for low-dimensional fitness landscapes toward new metrics for antimicrobial applications. *eLife*, 12, 2024.

[13987] Stephanie Guillet, Tomi Lazarov, Natasha Jordan, Bertrand Boisson, Maria Tello, Barbara Craddock, Ting Zhou, Chihiro Nishi, Rohan Bareja, Hairu Yang, Frederic Rieux-Laucat, Rosa Irene Fregel Lorenzo, Sabrina D Dyall, David Isenberg, David D'Cruz, Nico Lachmann, Olivier Elemento, Agnes Viale, Nicholas D Socci, Laurent Abel, Shigekazu Nagata, Morgan Huse, W Todd Miller, Jean-Laurent Casanova, and Frédéric Geissmann. ACK1 and BRK non-receptor tyrosine kinase deficiencies are associated with familial systemic lupus and involved in efferocytosis. *eLife*, 13, 2024.

[13988] Elisa Guma, Antoine Beauchamp, Siyuan Liu, Elizabeth Levitis, Jacob Ellegood, Linh Pham, Rogier B Mars, Armin Raznahan, and Jason P Lerch. Comparative neuroimaging of sex differences in human and mouse brain anatomy. *eLife*, 13, 2024.

[13989] Ankita Gumaste, Keeley L Baker, Michelle Izydorczak, Aaron C True, Ganesh Vasan, John P Crimaldi, and Justus Verhagen. Behavioral discrimination and olfactory bulb encoding of odor plume intermittency. *eLife*, 13, 2024.

[13990] Pengfei Guo, Rebecca C Lim, Keshari Rajawasam, Tiffany Trinh, Hong Sun, and Hui Zhang. A methylation-phosphorylation switch controls EZH2 stability and hematopoiesis. *eLife*, 13, 2024.

[13991] Youjia Guo, Tomohiro Kitano, Kimiko Inoue, Kensaku Murano, Michiko Hirose, Ten D Li, Akihiko Sakashita, Hirotsugu Ishizu, Narumi Ogonuki, Shogo Matoba, Masayuki Sato, Atsuo Ogura, and Haruhiko Siomi. Obox4 promotes zygotic genome activation upon loss of Dux. *eLife*, 13, 2024.

[13992] Morgan L Gustison, Rodrigo Muñoz-Castañeda, Pavel Osten, and Steven M Phelps. Sexual coordination in a whole-brain map of prairie vole pair bonding. *eLife*, 12, 2024.

[13993] Nicolas Gutierrez-Castellanos, Dario Sarra, Beatriz S Godinho, and Zachary F Mainen. Maturation of cortical input to dorsal raphe nucleus increases behavioral persistence in mice. *eLife*, 13, 2024.

[13994] Megan Haase, Tien Comlekoglu, Alexa Petrucciani, Shayn M Peirce, and Silvia S Blemker. Agent-based model demonstrates the impact of nonlinear, complex interactions between cytokinces on muscle regeneration. *eLife*, 13, 2024.

[13995] Sachiko Haga-Yamanaka, Rogelio Nunez-Flores, Christi A Scott, Sarah Perry, Stephanie Turner Chen, Crystal Pontrello, Meera G Nair, and Anandasankar Ray. Plasticity of gene expression in the nervous system by exposure to environmental odorants that inhibit HDACs. *eLife*, 12, 2024.

[13996] David L Haggerty and Brady K Atwood. Sex-dependent, lateralized engagement of anterior insular cortex inputs to the dorsolateral striatum in binge alcohol drinking. *eLife*, 13, 2024.

[13997] Hideo Hagihara, Hirotaka Shoji, Satoko Hattori, Giovanni Sala, Yoshihiro Takamiya, Mika Tanaka, Masafumi Ihara, Mihiro Shibutani, Izuho Hatada, Kei Hori, Mikio Hoshino, Akito Nakao, Yasuo Mori, Shigeo Okabe, Masayuki Matsushita, Anja Urbach, Yuta Katayama, Akinobu Matsumoto, Keiichi I Nakayama, Shota Katori, Takuya Sato, Takuji Iwasato, Haruko Nakamura, Yoshio Goshima, Matthieu Raveau, Tetsuya Tatsukawa, Kazuhiro Yamakawa, Noriko Takahashi, Haruo Kasai, Johji Inazawa, Ikuo Nobuhisa, Tetsushi Kagawa, Tetsuya Taga, Mohamed Darwish, Hirofumi Nishizono, Keizo Takao, Kiran Sapkota, Kazutoshi Nakazawa, Tsuyoshi Takagi, Haruki Fujisawa, Yoshihisa Sugimura, Kyosuke Yamanishi, Lakshmi Rajagopal, Nanette Deneen Hannah, Herbert Y Meltzer, Tohru Yamamoto, Shuji Wakatsuki, Toshiyuki Araki, Katsuhiko Tabuchi, Tadahiro Numakawa, Hiroshi Kunugi, Freesia L Huang, Atsuko Hayata-Takano, Hitoshi Hashimoto, Kota Tamada, Toru Takumi, Takaoki Kasahara, Tadafumi Kato, Isabella A Graef, Gerald R Crabtree, Nozomi Asaoka, Hikari Hatakama, Shuji Kaneko, Takao Kohno, Mitsuharu Hattori, Yoshio Hoshiba, Ryuhei Miyake, Kisho Obi-Nagata, Akiko Hayashi-Takagi, Léa J Becker, Ipek Yalcin, Yoko Hagino, Hiroko Kotajima-Murakami, Yuki Moriya, Kazutaka Ikeda, Hyopil Kim, Bong-Kiun Kaang, Hikari Otabi, Yuta Yoshida, Atsushi Toyoda, Noboru H Komiyama, Seth GN Grant, Michiru Ida-Eto, Masaaki Narita,

Ken-ichi Matsumoto, Emiko Okuda-Ashitaka, Iori Ohmori, Tadayuki Shimada, Kanato Yamagata, Hiroshi Ageta, Kunihiro Tsuchida, Kaoru Inokuchi, Takayuki Sassa, Akio Kihara, Motoaki Fukasawa, Nobuteru Usuda, Tayo Katano, Teruyuki Tanaka, Yoshihiro Yoshihara, Michihiro Igarashi, Takashi Hayashi, Kaori Ishikawa, Satoshi Yamamoto, Naoya Nishimura, Kazuto Nakada, Shinji Hirotsune, Kiyoshi Egawa, Kazuma Higashisaka, Yasuo Tsutsumi, Shoko Nishihara, Noriyuki Sugo, Takeshi Yagi, Naoto Ueno, Tomomi Yamamoto, Yoshihiro Kubo, Rie Ohashi, Nobuyuki Shiina, Kimiko Shimizu, Sayaka Higo-Yamamoto, Katsutaka Oishi, Hisashi Mori, Tamio Furuse, Masaru Tamura, Hisashi Shirakawa, Daiki X Sato, Yukiko U Inoue, Takayoshi Inoue, Yuriko Komine, Tetsuo Yamamori, Kenji Sakimura, and Tsuyoshi Miyakawa. Large-scale animal model study uncovers altered brain pH and lactate levels as a transdiagnostic endophenotype of neuropsychiatric disorders involving cognitive impairment. *eLife*, 12, 2024.

[13998] Lisa Hahn, Simon B Eickhoff, Karsten Mueller, Leonhard Schilbach, Henryk Barthel, Klaus Fassbender, Klaus Fliessbach, Johannes Kornhuber, Johannes Prudlo, Matthis Synofzik, Jens Wiltfang, Janine Diehl-Schmid, Markus Otto, Juergen Dukart, and Matthias L Schroeter. Resting-state alterations in behavioral variant frontotemporal dementia are related to the distribution of monoamine and GABA neurotransmitter systems. *eLife*, 13, 2024.

[13999] Hoang Thanh Hai, Le Thanh Hoang Nhat, Trinh Thi Bich Tram, Do Dinh Vinh, Artika P Nath, Joseph Donovan, Nguyen Thi Anh Thu, Dang Van Thanh, Nguyen Duc Bang, Dang Thi Minh Ha, Nguyen Hoan Phu, Ho Dang Trung Nghia, Le Hong Van, Michael Inouye, Guy E Thwaites, and Nguyen Thuy Thuong Thuong. Whole blood transcriptional profiles and the pathogenesis of tuberculous meningitis. *eLife*, 13, 2024.

[14000] Dipasree Hajra, Raju S Rajmani, Ayushi Devendrasingh Chaudhary, Shashi Kumar Gupta, and Dipshikha Chakravortty. Salmonella-induced SIRT1 and SIRT3 are crucial for maintaining the metabolic switch in bacteria and host for successful pathogenesis. *eLife*, 13, 2024.

[14001] Michael B Hall, Ryan R Wick, Louise M Judd, An N Nguyen, Eike J Steinig, Ouli Xie, Mark Davies, Torsten Seemann, Timothy P Stinear, and Lachlan Coin. Benchmarking reveals superiority of deep learning variant callers on bacterial nanopore sequence data. *eLife*, 13, 2024.

[14002] Ffion R Hammond, Amy Lewis, Gabriele Pollara, Gillian S Tomlinson, Mahdad Noursadeghi, Endre Kiss-Toth, and Philip M Elks. Tribbles1 is host protective during in vivo mycobacterial infection. *eLife*, 13, 2024.

[14003] Hui Han, Rong-Hua Luo, Xin-Yan Long, Li-Qiong Wang, Qian Zhu, Xin-Yue Tang, Rui Zhu, Yi-Cheng Ma, Yong-Tang Zheng, and Cheng-Gang Zou. Transcriptional regulation of SARS-CoV-2 receptor ACE2 by SP1. *eLife*, 13, 2024.

[14004] MeiQing Han, Xia Wang, Lin Su, Shiqi Pan, Ningning Liu, Duan Li, Liang Liu, JunWei Cui, Huajie Zhao, and Fan Yang. Intestinal microbiome dysbiosis increases Mycobacteria pulmonary colonization in mice by regulating the Nos2-associated pathways. *eLife*, 13, 2024.

[14005] Kazuki Hanaoka, Kensuke Nishikawa, Atsuko Ikeda, Philipp Schlarmann, Saku Sasaki, Sotaro Fujii, Sayumi Yamashita, Aya Nakaji, and Kouichi Funato. Membrane contact sites regulate vacuolar fission via sphingolipid metabolism. *eLife*, 12, 2024.

[14006] Sang-Hyeon Hann, Seon-Yong Kim, Ye Lynne Kim, Young-Woo Jo, Jong-Seol Kang, Hyerim Park, Se-Young Choi, and Young-Yun Kong. Depletion of SMN protein in mesenchymal progenitors impairs the development of bone and neuromuscular junction in spinal muscular atrophy. *eLife*, 12, 2024.

[14007] Colleen E Hannon and Michael B Eisen. Intrinsic protein disorder is insufficient to drive subnuclear clustering in embryonic transcription factors. *eLife*, 12, 2024.

[14008] Chelsea L Hansen, Cécile Viboud, and Lone Simonsen. Disentangling the relationship between cancer mortality and COVID-19 in the US. *eLife*, 13, 2024.

[14009] Qinyu Hao, Minxue Liu, Swapna Vidhur Daulatabad, Saba Gaffari, You Jin Song, Rajneesh Srivastava, Shivang Bhaskar, Anurupa Moitra, Hazel Mangan, Elizabeth Tseng, Rachel B Gilmore, Susan M Frier, Xin Chen, Chengliang Wang, Sui Huang, Stormy Chamberlain, Hong Jin, Jonas Korlach, Brian McStay, Saurabh Sinha, Sarath Chandra Janga, Supriya G Prasanth, and Kannanganattu V Prasanth. Monoallelically expressed noncoding RNAs form nucleolar territories on NOR-containing chromosomes and regulate rRNA expression. *eLife*, 13, 2024.

[14010] Masaya Harada, Laia Serratosa Capdevila, Maria Wilhelm, Denis Burdakov, and Tommaso Patriarchi. Stimulation of VTA dopamine inputs to LH upregulates orexin neuronal activity in a DRD2-dependent manner. *eLife*, 12, 2024.

[14011] Samyogita Hardikar, Bronte Mckeown, H Lina Schaare, Raven Star Wallace, Ting Xu, Mark Edgar Lauckener, Sofie Louise Valk, Daniel S Margulies, Adam Turnbull, Boris C Bernhardt, Reinder Vos de Wael, Arno Villringer, and Jonathan Smallwood. Macro-scale patterns in functional connectivity associated with ongoing thought patterns and dispositional traits. *eLife*, 13, 2024.

[14012] Harsh N Hariani, A Brynn Algstam, Christian T Candler, Isabelle F Witteveen, Jasmeen K Sidhu, and Timothy S Balmer. A system of feed-forward cerebellar circuits that extend and diversify sensory signaling. *eLife*, 12, 2024.

[14013] Parameswaran Hariharan, Yuqi Shi, Satoshi Katsube, Katleen Willibal, Nathan D Burrows, Patrick Mitchell, Amirhossein Bakhtiiari, Samantha Stanfield, Els Pardon, H Ronald Kaback, Ruibin Liang, Jan Steyaert, Rosa Viner, and Lan Guan. Mobile barrier mechanisms for Na+-coupled symport in an MFS sugar transporter. *eLife*, 12, 2024.

[14014] C Jake Harris, Zhenhui Zhong, Lucia Ichino, Suhua Feng, and Steven E Jacobsen. H1 restricts euchromatin-associated methylation pathways from heterochromatic encroachment. *eLife*, 12, 2024.

[14015] Nathan D Harry and Christina Zakas. The role of heterochronic gene expression and regulatory architecture in early developmental divergence. *eLife*, 13, 2024.

[14016] Nicholas J Hathaway, Isaac E Kim, Neeva WernsmanYoung, Sin Ting Hui, Rebecca Crudale, Emily Y Liang, Christian P Nixon, David Giesbrecht, Jonathan J Juliano, Jonathan B Parr, and Jeffrey A Bailey. Interchromosomal segmental duplication drives translocation and loss of P. falciparum histidine-rich protein 3. *eLife*, 13, 2024.

[14017] Marleen Haupt, Monika Graumann, Santani Teng, Carina Kaltenbach, and Radoslaw Cichy. The transformation of sensory to perceptual braille letter representations in the visually deprived brain. *eLife*, 13, 2024.

[14018] Kelsey A Hauser, Christina N Garvey, Ryley S Crow, Muhammad RH Hossainey, Dustin T Howard, Netra Ranganathan, Lindsey K Gentry, Amulya Yaparla, Namarta Kalia, Mira Zelle, Elizabeth J Jones, Anju N Duttargi, Louise A Rollins-Smith, Carly R Muletz-Wolz, and Leon Grayfer. Amphibian mast cells serve as barriers to chytrid fungus infections. *eLife*, 12, 2024.

[14019] Kourosh Hayatigolkhatmi, Chiara Soriani, Emanuel Soda, Elena Ceccacci, Oualid El Menna, Sebastiano Peri, Ivan Negrelli, Giacomo Bertolini, Gian Martino Franchi, Roberta Carbone, Saverio Minucci, and Simona Rodighiero. Automated workflow for the cell cycle analysis of (non-)adherent cells using a machine learning approach. *eLife*, 13, 2024.

[14020] Brandon H Hayes, Mai Wang, Hui Zhu, Steven H Phan, Lawrence J Dooling, Jason C Andrechak, Alexander H Chang, Michael P Tobin, Nicholas M Ontko, Tristan Marchena, and Dennis E Discher. Chromosomal instability induced in cancer can enhance macrophage-initiated immune responses that include anti-tumor IgG. *eLife*, 12, 2024.

[14021] Jitian He, Le Gao, Peiying Wang, Wing Keung Chan, Yiran Zheng, Yumo Zhang, Jiaman Sun, Xue Li, Jiming Wang, Xiao-Hong Li, Huaiyong Chen, Zhouxin Yang, and Youwei Wang. Prdm1 positively regulates liver Group 1 ILCs cancer immune surveillance and preserves functional heterogeneity. *eLife*, 13, 2024.

[14022] Qiong He, Miao-Miao Zhao, Ming-Jia Li, Xiao-Ya Li, Jian-Min Jin, Ying-Mei Feng, Li Zhang, Wei Jin Huang, Fangyuan Yang, and Jin-Kui Yang. Hyperglycemia induced cathepsin L maturation linked to diabetic comorbidities and COVID-19 mortality. *eLife*, 13, 2024.

[14023] Qixin He, John K Chaillet, and Frédéric Labbé. Antigenic strain diversity predicts different biogeographic patterns of maintenance and decline of antimalarial drug resistance. *eLife*, 12, 2024.

[14024] Chad Heer and Mark Sheffield. Distinct catecholaminergic pathways projecting to hippocampal CA1 transmit contrasting signals during navigation in familiar and novel environments. *eLife*, 13, 2024.

[14025] Matthias Heindorf and Georg B Keller. Antipsychotic drugs selectively decorrelate long-range interactions in deep cortical layers. *eLife*, 12, 2024.

[14026] Idan Hekselman, Assaf Vital, Maya Ziv-Agam, Lior Kerber, Ido Yairi, and Esti Yeger-Lotem. Affected cell types for hundreds of Mendelian diseases revealed by analysis of human and mouse single-cell data. *eLife*, 13, 2024.

[14027] Charles R Heller, Gregory R Hamersky, and Stephen V David. Task-specific invariant representation in auditory cortex. *eLife*, 12, 2024.

[14028] Mathieu Hénault, Souhir Marsit, Guillaume Charron, and Christian R Landry. The genomic landscape of transposable elements in yeast hybrids is shaped by structural variation and genotype-specific modulation of transposition rate. *eLife*, 12, 2024.

[14029] Gonzalo Hernandez-Hernandez, Samantha C O'Dwyer, Pei-Chi Yang, Collin Matsumoto, Mindy Tieu, Zhihui Fong, Timothy J Lewis, L Fernando Santana, and Colleen E Clancy. A computational model predicts sex-specific responses to calcium channel blockers in mammalian mesenteric vascular smooth muscle. *eLife*, 12, 2024.

[14030] Alice L Herneisen, Michelle L Peters, Tyler A Smith, Emily Shortt, and Sebastian Lourido. SPARK regulates AGC kinases central to the Toxoplasma gondii asexual cycle. *eLife*, 13, 2024.

[14031] Anna Herrmannová, Jan Jelínek, Klára PospíŠilová, Farkas Kerényi, TomáŠ Vomastek, Kathleen Watt, Jan Brábek, Mahabub Pasha Mohammad, Susan Wagner, Ivan Topisirovic, and LeoŠ Shivaya ValáŠek. Perturbations in eIF3 subunit stoichiometry alter expression of ribosomal proteins and key components of the MAPK signaling pathways. *eLife*, 13, 2024.

[14032] Fabian Hersperger, Tim Meyring, Pia Weber, Chintan Chhatbar, Gianni Monaco, Marc S Dionne, Katrin Paeschke, Marco Prinz, Olaf Groß, Anne-Kathrin Classen, and Katrin Kierdorf. DNA damage signaling in Drosophila macrophages modulates systemic cytokine levels in response to oxidative stress. *eLife*, 12, 2024.

[14033] Nadine Herzog, Hendrik Hartmann, Lieneke Katharina Janssen, Arsene Kanyamibwa, Maria Waltmann, Peter Kovacs, Lorenz Deserno, Sean Fallon, Arno Villringer, and Annette Horstmann. Working memory gating in obesity is moderated by striatal dopaminergic gene variants. *eLife*, 13, 2024.

[14034] Sina Heydari, Haotian Hang, and Eva Kanso. Mapping spatial patterns to energetic benefits in groups of flow-coupled swimmers. *eLife*, 13, 2024.

[14035] Ines Heyndrickx, Kim Deswarte, Kenneth Verstraete, Koen HG Verschueren, Ursula Smole, Helena Aegerter, Ann Dansercoer, Hamida Hammad, Savvas N Savvides, and Bart N Lambrecht. Ym1 protein crystals promote type 2 immunity. *eLife*, 12, 2024.

[14036] Eman Hijaze, Tsvia Gildor, Ronald Seidel, Majed Layous, Mark Winter, Luca Bertinetti, Yael Politi, and Smadar Ben-Tabou de Leon. ROCK and the actomyosin network control biomineral growth and morphology during sea urchin skeletogenesis. *eLife*, 12, 2024.

[14037] Max Henry Hills, Limei Ma, Ai Fang, Thelma Chiremba, Seth Malloy, Allison R Scott, Anoja G Perera, and C Ron Yu. Molecular, cellular, and developmental organization of the mouse vomeronasal organ at single cell resolution. *eLife*, 13, 2024.

[14038] Himawari Hirai, Kohtarou Konno, Miwako Yamasaki, Masahiko Watanabe, Takeshi Sakaba, and Yuki Hashimotodani. Distinct release properties of glutamate/GABA co-transmission serve as a frequency-dependent filtering of supramammillary inputs. *eLife*, 13, 2024.

[14039] Kenya Hitomi, Yoichiro Ishii, and Bei-Wen Ying. Experimental evolution for the recovery of growth loss due to genome reduction. *eLife*, 13, 2024.

[14040] David V Ho, Duncan Tormey, Aaron Odell, Aracely A Newton, Robert R Schnittker, Diana P Baumann, William B Neaves, Morgan R Schroeder, Rutendo F Sigauke, Anthony J Barley, and Peter Baumann. Post-meiotic mechanism of facultative parthenogenesis in gonochoristic whiptail lizard species. *eLife*, 13, 2024.

[14041] Larissa Höfling, Klaudia P Szatko, Christian Behrens, Yuyao Deng, Yongrong Qiu, David Alexander Klindt, Zachary Jessen, Gregory W Schwartz, Matthias Bethge, Philipp Berens, Katrin Franke, Alexander S Ecker, and Thomas Euler. A chromatic feature detector in the retina signals visual context changes. *eLife*, 13, 2024.

[14042] Lian Hollander-Cohen, Omer Cohen, Miriam Shulman, Tomer Aiznkot, Pierre Fontanaud, Omer Revah, Patrice Mollard, Matan Golan, and Berta Levavi-Sivan. The satiety hormone cholecystokinin gates reproduction in fish by controlling gonadotropin secretion. *eLife*, 13, 2024.

[14043] Corey D Holman, Alexander P Sakers, Ryan P Calhoun, Lan Cheng, Ethan C Fein, Christopher Jacobs, Linus Tsai, Evan D Rosen, and Patrick Seale. Aging impairs cold-induced beige adipogenesis and adipocyte metabolic reprogramming. *eLife*, 12, 2024.

[14044] Kentaro Homan, Tomohiro Onodera, Hisatoshi Hanamatsu, Jun-ichi Furukawa, Daisuke Momma, Masatake Matsuoka, and Norimasa Iwasaki. Articular cartilage corefucosylation regulates tissue resilience in osteoarthritis. *eLife*, 12, 2024.

[14045] Chien-Tai Hong, Chen-Chih Chung, Ruan-Ching Yu, and Lung Chan. Plasma extracellular vesicle synaptic proteins as biomarkers of clinical progression in patients with Parkinson's disease. *eLife*, 12, 2024.

[14046] Karolina Honzejkova, Dalibor Kosek, Veronika Obsilova, and Tomas Obsil. The cryo-EM structure of ASK1 reveals an asymmetric architecture allosterically modulated by TRX1. *eLife*, 13, 2024.

[14047] Daniel Hoops, Robert Kyne, Samer Salameh, Del MacGowan, Radu Gabriel Avramescu, Elise Ewing, Alina Tao He, Taylor Orsini, Anais Durand, Christina Popescu, Janet Mengyi Zhao, Kelcie Shatz, LiPing Li, Quinn Carroll, Guofa Liu, Matthew J Paul, and Cecilia Flores. The scheduling of adolescence with Netrin-1 and UNC5C. *eLife*, 12, 2024.

[14048] Michael J Hopkins, Brittni N Moore, Jasmin L Jeffery, and Andrea S Young. Demystifying the 'hidden curriculum' for minoritized graduate students. *eLife*, 13, 2024.

[14049] Paul Horn, Jenny Norlin, Kasper Almholt, Birgitte M Viuff, Elisabeth D Galsgaard, Andreas Hald, Franziska Zosel, Helle Demuth, Svend Poulsen, Peder L Norby, Morten G Rasch, Mogens Vyberg, Jan Fleckner, Mikkel Parsberg Werge, Lise Lotte Gluud, Marco R Rink, Emma Shepherd, Ellie Northall, Patricia F Lalor, Chris J Weston, Morten Fog-Tonnesen, and Philip N Newsome. Evaluation of Gremlin-1 as a therapeutic target in metabolic dysfunction-associated steatohepatitis. *eLife*, 13, 2024.

[14050] Robert Horvath, Nikolaos Minadakis, Yann Bourgeois, and Anne C Roulin. The evolution of transposable elements in Brachypodium distachyon is governed by purifying selection, while neutral and adaptive processes play a minor role. *eLife*, 12, 2024.

[14051] Steven S Hou, Yuya Ikegawa, Yeseo Kwon, Natalia Wieckiewicz, Mei CQ Houser, Brianna Lundin, Brian J Bacskai, Oksana Berezovska, and Masato Maesako. Recording $\gamma$-secretase activity in living mouse brains. *eLife*, 13, 2024.

[14052] Xinlin Hou, Peng Zhang, Licheng Mo, Cheng Peng, and Dandan Zhang. Sensitivity to vocal emotions emerges in newborns at 37 weeks gestational age. *eLife*, 13, 2024.

[14053] Julian JA Hoving, Elizabeth Harford-Wright, Patrick Wingfield-Digby, Anne-Laure Cattin, Mariana Campana, Alex Power, Toby Morgan, Erica Torchiaro, Victor Quereda, and Alison C Lloyd. N-cadherin directs the collective Schwann cell migration required for nerve regeneration through Slit2/3-mediated contact inhibition of locomotion. *eLife*, 13, 2024.

[14054] Gregory Caleb Howard, Jing Wang, Kristie L Rose, Camden Jones, Purvi Patel, Tina Tsui, Andrea C Florian, Logan Vlach, Shelly L Lorey, Brian C Grieb, Brianna N Smith, Macey J Slota, Elizabeth M Reynolds, Soumita Goswami, Michael R Savona, Frank M Mason, Taekyu Lee, Stephen Fesik, Qi Liu, and William P Tansey. Ribosome subunit attrition and activation of the p53–MDM4 axis dominate the response of MLL-rearranged cancer cells to WDR5 WIN site inhibition. *eLife*, 12, 2024.

[14055] Pyae Hein Htet, Edward Avezov, and Eric Lauga. Fluid mechanics of luminal transport in actively contracting endoplasmic reticulum. *eLife*, 13, 2024.

[14056] Can Hu, Xue-Ting Zhu, Ming-Hong He, Yangyang Shao, Zhongjun Qin, Zhi-Jing Wu, and Jin-Qiu Zhou. Elimination of subtelomeric repeat sequences exerts little effect on telomere essential functions in Saccharomyces cerevisiae. *eLife*, 12, 2024.

[14057] Chongbin Hu, Nan Zhang, Yun Hong, Ruxiu Tie, Dongdong Fan, Aifu Lin, Ye Chen, Li-xin Xiang, and Jian-zhong Shao. Single-cell RNA sequencing unveils the hidden powers of zebrafish kidney for generating both hematopoiesis and adaptive antiviral immunity. *eLife*, 13, 2024.

[14058] Yue Hu, Wenjie Du, Jiangtao Qi, Huoqing Luo, Zhao Zhang, Mengqiang Luo, and Yingwei Wang. Comparative brain-wide mapping of ketamine- and isoflurane-activated nuclei and functional networks in the mouse brain. *eLife*, 12, 2024.

[14059] Xiaojiao Hua, Chen Zhao, Jianbo Tian, Junbao Wang, Xiaoping Miao, Gen Zheng, Min Wu, Mei Ye, Ying Liu, and Yan Zhou. A Ctnnb1 enhancer transcriptionally regulates Wnt signaling dosage to balance homeostasis and tumorigenesis of intestinal epithelia. *eLife*, 13, 2024.

[14060] Fang Huang, Zhenwei Dai, Jinmiao Yu, Kainan Wang, Chaoqun Chen, Dan Chen, Jinrui Zhang, Jinyao Zhao, Mei Li, Wenjing Zhang, Xiaojie Li, Yangfan Qi, and Yang Wang. RBM7 deficiency promotes breast cancer metastasis by coordinating MFGE8 splicing switch and NF-kB pathway. *eLife*, 13, 2024.

[14061] Kai-Ting Huang, Larry E Wagner, Takahiro Takano, Xiao-Xuan Lin, Harini Bagavant, Umesh Deshmukh, and David I Yule. Dysregulated Ca2+ signaling, fluid secretion, and mitochondrial function in a mouse model of early Sjögren's disease. *eLife*, 13, 2024.

[14062] Li-Wen Huang, Derek LF Garden, Christina McClure, and Matthew F Nolan. Synaptic interactions between stellate cells and parvalbumin interneurons in layer 2 of the medial entorhinal cortex are organized at the scale of grid cell clusters. *eLife*, 12, 2024.

[14063] Qiaoli Huang and Huan Luo. Shared structure facilitates working memory of multiple sequences. *eLife*, 12, 2024.

[14064] Sin-Wei Huang, See-Khai Lim, Yao-An Yu, Yi-Chung Pan, Wan-Ju Lien, Chung-Yuan Mou, Che-Ming Jack Hu, and Kurt Yun Mou. Overcoming the nutritional immunity by engineering iron-scavenging bacteria for cancer therapy. *eLife*, 12, 2024.

[14065] Taicheng Huang and Jia Liu. A stochastic world model on gravity for stability inference. *eLife*, 12, 2024.

[14066] Tengteng Huang, Xiaoling Chen, Jun He, Ping Zheng, Yuheng Luo, Aimin Wu, Hui Yan, Bing Yu, Daiwen Chen, and Zhiqing Huang. Eugenol mimics exercise to promote skeletal muscle fiber remodeling and myokine IL-15 expression by activating TRPV1 channel. *eLife*, 12, 2024.

[14067] Yanling Huang, Haocong Mo, Jie Yang, Luyang Gao, Tian Tao, Qing Shu, Wenying Guo, Yawen Zhao, Jingya Lyu, Qimeng Wang, Jinghui Guo, Hening Zhai, Linyan Zhu, Hui Chen, and Geyang Xu. Mechano-regulation of GLP-1 production by Piezo1 in intestinal L cells. *eLife*, 13, 2024.

[14068] Yiwei Huang, Gujie Wu, Guoshu Bi, Lin Cheng, Jiaqi Liang, Ming Li, Huan Zhang, Guangyao Shan, Zhengyang Hu, Zhencong Chen, Zongwu Lin, Wei Jiang, Qun Wang, Junjie Xi, Shanye Yin, and Cheng Zhan. Unveiling chemotherapy-induced immune landscape remodeling and metabolic reprogramming in lung adenocarcinoma by scRNA-sequencing. *eLife*, 13, 2024.

[14069] Paul Huchede, Swann Meyer, Clement Berthelot, Maud Hamadou, Adrien Bertrand-Chapel, Andria Rakotomalala, Line Manceau, Julia Tomine, Nicolas Lespinasse, Paul Lewandowski, Martine Cordier-Bussat, Laura Broutier, Aurelie Dutour, Isabelle Rochet, Jean-Yves Blay, Cyril Degletagne, Valery Attignon, Angel Montero-Carcaboso, Marion Le Grand, Eddy Pasquier, Alexandre Vasiljevic, Pascale Gilardi-Hebenstreit, Samuel Meignan, Pierre Leblond, Vanessa Ribes, Erika Cosset, and Marie Castets. BMP2 and BMP7 cooperate with H3.3K27M to promote quiescence and invasiveness in pediatric diffuse midline gliomas. *eLife*, 12, 2024.

[14070] Sanjarbek Hudaiberdiev and Ivan Ovcharenko. Functional characteristics and computational model of abundant hyperactive loci in the human genome. *eLife*, 13, 2024.

[14071] Roger Huerlimann, Natacha Roux, Ken Maeda, Polina Pilieva, Saori Miura, Hsiao-chian Chen, Michael Izumiyama, Vincent Laudet, and Timothy Ravasi. The transcriptional landscape underlying larval development and metamorphosis in the Malabar grouper (Epinephelus malabaricus). *eLife*, 13, 2024.

[14072] Alyssa D Huff, Marlusa Karlen-Amarante, Luiz M Oliveira, and Jan-Marino Ramirez. Chronic intermittent hypoxia reveals role of the Postinspiratory Complex in the mediation of normal swallow production. *eLife*, 12, 2024.

[14073] Kate Huffer, Matthew CS Denley, Elisabeth V Oskoui, and Kenton J Swartz. Conservation of the cooling agent binding pocket within the TRPM subfamily. *eLife*, 13, 2024.

[14074] Anne E Hultgren, Nicole MF Patras, and Jenna Hicks. Blinding reduces institutional prestige bias during initial review of applications for a young investigator award. *eLife*, 13, 2024.

[14075] Iain Hunter, Bramwell Coulson, Tom Pettini, Jacob J Davies, Jill Parkin, Matthias Landgraf, and Richard A Baines. Balance of activity during a critical period tunes a developing network. *eLife*, 12, 2024.

[14076] Sung Won Hur, Karen Safaryan, Long Yang, Hugh T Blair, Sotiris C Masmanidis, Paul J Mathews, Daniel Aharoni, and Peyman Golshani. Correlated signatures of social behavior in cerebellum and anterior cingulate cortex. *eLife*, 12, 2024.

[14077] Mathieu C Husser, Nhat P Pham, Chris Law, Flavia RB Araujo, Vincent JJ Martin, and Alisa Piekny. Endogenous tagging using split mNeonGreen in human iPSCs for live imaging studies. *eLife*, 12, 2024.

[14078] Judith Hüttemeister, Franziska Rudolph, Michael H Radke, Claudia Fink, Dhana Friedrich, Stephan Preibisch, Martin Falcke, Eva Wagner, Stephan E Lehnart, and Michael Gotthardt. Visualizing sarcomere and cellular dynamics in skeletal muscle to improve cell therapies. *eLife*, 13, 2024.

[14079] Hyeri Hwang, Seung-Woo Jin, and Inah Lee. Differential functions of the dorsal and intermediate regions of the hippocampus for optimal goal-directed navigation in VR space. *eLife*, 13, 2024.

[14080] Ran-Der Hwang, YuNing Lu, Qing Tang, Goran Periz, Giho Park, Xiangning Li, Qiwang Xiang, Yang Liu, Tao Zhang, and Jiou Wang. DBT is a metabolic switch for maintenance of proteostasis under proteasomal impairment. *eLife*, 12, 2024.

[14081] Soo-Yeon Hwang, Kyung-Hwa Jeon, Hwa-Jong Lee, Inhye Moon, Sehyun Jung, Seul-Ah Kim, Hyunji Jo, Seojeong Park, Misun Ahn, Soo-Yeon Kwak, Younghwa Na, and Youngjoo Kwon. Synthesis and biological assessment of chalcone and pyrazoline derivatives as novel inhibitor for ELF3-MED23 interaction. *eLife*, 13, 2024.

[14082] Melody C Iacino, Taylor A Stowe, Elizabeth G Pitts, Lacey L Sexton, Shannon L Macauley, and Mark J Ferris. A unique multi-synaptic mechanism involving acetylcholine and GABA regulates dopamine release in the nucleus accumbens through early adolescence in male rats. *eLife*, 13, 2024.

[14083] Sara Ibañez, Nilapratim Sengupta, Jennifer I Luebke, Klaus Wimmer, and Christina M Weaver. Myelin dystrophy impairs signal transmission and working memory in a multiscale model of the aging prefrontal cortex. *eLife*, 12, 2024.

[14084] Ibtisam Ibtisam and Alexei F Kisselev. Early recovery of proteasome activity in cells pulse-treated with proteasome inhibitors is independent of DDI2. *eLife*, 12, 2024.

[14085] Toshiharu Ichinose, Shu Kondo, Mai Kanno, Yuichi Shichino, Mari Mito, Shintaro Iwasaki, and Hiromu Tanimoto. Translational regulation enhances distinction of cell types in the nervous system. *eLife*, 12, 2024.

[14086] Jasper Iske, Rachid El Fatimy, Yeqi Nian, Amina Ghouzlani, Siawosh K Eskandari, Hector Rodriguez Cetina Biefer, Anju Vasudevan, and Abdallah Elkhal. NAD+ prevents septic shock-induced death by non-canonical inflammasome blockade and IL-10 cytokine production in macrophages. *eLife*, 12, 2024.

[14087] Atikul Islam, Yu-Chun Chang, Xiao-Chi Chen, Chia-Wei Weng, Chien-Yu Chen, Che-Wei Wang, Mu-Kuan Chen, Alexander S Tikhomirov, Andrey E Shchekotikhin, and Pin Ju Chueh. Water-soluble 4-(dimethylaminomethyl)heliomycin exerts greater antitumor effects than parental heliomycin by targeting the tNOX-SIRT1 axis and apoptosis in oral cancer cells. *eLife*, 12, 2024.

[14088] Callum M Ives, Linh Nguyen, Carl A Fogarty, Aoife M Harbison, Yves Durocher, John Klassen, and Elisa Fadda. Role of N343 glycosylation on the SARS-CoV-2 S RBD structure and co-receptor binding across variants of concern. *eLife*, 13, 2024.

[14089] Sudhanvan Iyer, Kathryn Maxson Jones, Jill O Robinson, Nicole R Provenza, Dominique Duncan, Gabriel Lázaro-Muñoz, Amy L McGuire, Sameer A Sheth, and Mary A Majumder. The BRAIN Initiative data-sharing ecosystem: Characteristics, challenges, benefits, and opportunities. *eLife*, 13, 2024.

[14090] Masanori Izumi, Sakuya Nakamura, Kohei Otomo, Hiroyuki Ishida, Jun Hidema, Tomomi Nemoto, and Shinya Hagihara. Autophagosome development and chloroplast segmentation occur synchronously for piecemeal degradation of chloroplasts. *eLife*, 12, 2024.

[14091] Carolyn M Jablonowski, Waise Quarni, Shivendra Singh, Haiyan Tan, Dhanushka Hewa Bostanthirige, Hongjian Jin, Jie Fang, Ti-Cheng Chang, David Finkelstein, Ji-Hoon Cho, Dongli Hu, Vishwajeeth Pagala, Sadie Miki Sakurada, Shondra M Pruett-Miller, Ruoning Wang, Andrew Murphy, Kevin Freeman, Junmin Peng, Andrew M Davidoff, Gang Wu, and Jun Yang. Metabolic reprogramming of cancer cells by JMJD6-mediated pre-mRNA splicing associated with therapeutic response to splicing inhibitor. *eLife*, 12, 2024.

[14092] Martina Jabloñski, Guillermina M Luque, Matias Gomez Elias, Claudia Sanchez Cardenas, Xinran Xu, Jose L de La Vega Beltran, Gabriel Corkidi, Alejandro Linares, Victor Abonza, Aquetzalli Arenas-Hernandez, María DP Ramos-Godinez, Alejandro López-Saavedra, Dario Krapf, Diego Krapf, Alberto Darszon, Adán Guerrero, and Mariano G Buffone. Reorganization of the flagellum scaffolding induces a sperm standstill during fertilization. *eLife*, 13, 2024.

[14093] Hamidreza Jafarinia, Erik van der Giessen, and Patrick R Onck. C9orf72 polyPR directly binds to various nuclear transport components. *eLife*, 12, 2024.

[14094] N Suhas Jagannathan, Javier Yu Peng Koh, Younghwan Lee, Radoslaw Mikolaj Sobota, Aaron T Irving, Lin-fa Wang, Yoko Itahana, Koji Itahana, and Lisa Tucker-Kellogg. Multi-omic analysis of bat versus human fibroblasts reveals altered central metabolism. *eLife*, 13, 2024.

[14095] Natalia Jagielska, Thomas G Kaye, Michael B Habib, Tatsuya Hirasawa, and Michael Pittman. New soft tissue data of pterosaur tail vane reveals sophisticated, dynamic tensioning usage and expands its evolutionary origins. *eLife*, 13, 2024.

[14096] Lina María Jaime Tobón and Tobias Moser. Bridging the gap between presynaptic hair cell function and neural sound encoding. *eLife*, 12, 2024.

[14097] Nina K Jain, Pierce J Ogden, and George M Church. Comprehensive mutagenesis maps the effect of all single-codon mutations in the AAV2 rep gene on AAV production. *eLife*, 12, 2024.

[14098] Swati Jain, John J LaFrancois, Kasey Gerencer, Justin J Botterill, Meghan Kennedy, Chiara Criscuolo, and Helen E Scharfman. Increasing adult-born neurons protects mice from epilepsy. *eLife*, 12, 2024.

[14099] Swati Jain, Gherman Uritskiy, Marthandan Mahalingam, Himanshu Batra, Subhash Chand, Hung V Trinh, Charles Beck, Woong-Hee Shin, Wadad Alsalmi, Gustavo Kijak, Leigh A Eller, Jerome Kim, Daisuke Kihara, Sodsai Tovanabutra, Guido Ferrari, Merlin L Robb, Mangala Rao, and Venigalla B Rao. A remarkable genetic shift in a transmitted/founder virus broadens antibody responses against HIV-1. *eLife*, 13, 2024.

[14100] Maëliss Jallais and Marco Palombo. Introducing $\mu$GUIDE for quantitative imaging via generalized uncertainty-driven inference using deep learning. *eLife*, 13, 2024.

[14101] Sara Jamali, Sophie Bagur, Enora Bremont, Timo Van Kerkoerle, Stanislas Dehaene, and Brice Bathellier. Parallel mechanisms signal a hierarchy of sequence structure violations in the auditory cortex. *eLife*, 13, 2024.

[14102] Alex James, Franca Buelow, Liam Gibson, and Ann Brower. Female-dominated disciplines have lower evaluated research quality and funding success rates, for men and women. *eLife*, 13, 2024.

[14103] Samarjit Jana, Mainak Mondal, Sagar Mahale, Bhavana Gupta, Kaval Reddy Prasasvi, Lekha Kandasami, Neha Jha, Abhishek Chowdhury, Vani Santosh, Chandrasekhar Kanduri, and Kumaravel Somasundaram. PITAR, a DNA damage-inducible cancer/testis long noncoding RNA, inactivates p53 by binding and stabilizing TRIM28 mRNA. *eLife*, 12, 2024.

[14104] Wonjo Jang, Kanishka Senarath, Gavin Feinberg, Sumin Lu, and Nevin A Lambert. Visualization of endogenous G proteins on endosomes and other organelles. *eLife*, 13, 2024.

[14105] Serena Janho dit Hreich, Thierry Juhel, Sylvie Leroy, Alina Ghinet, Frederic Brau, Veronique Hofman, Paul Hofman, and Valerie Vouret-Craviari. Activation of the P2RX7/IL-18 pathway in immune cells attenuates lung fibrosis. *eLife*, 12, 2024.

[14106] Luka Jarc, Manuj Bandral, Elisa Zanfrini, Mathias Lesche, Vida Kufrin, Raquel Sendra, Daniela Pezzolla, Ioannis Giannios, Shahryar Khattak, Katrin Neumann, Barbara Ludwig, and Anthony Gavalas. Regulation of multiple signaling pathways promotes the consistent expansion of human pancreatic progenitors in defined conditions. *eLife*, 12, 2024.

[14107] Amandine Jarysta, Abigail LD Tadenev, Matthew Day, Barry Krawchuk, Benjamin E Low, Michael V Wiles, and Basile Tarchini. Inhibitory G proteins play multiple roles to polarize sensory hair cell morphogenesis. *eLife*, 12, 2024.

[14108] Alexandra L Jellinger, Rebecca L Suthard, Bingbing Yuan, Michelle Surets, Evan A Ruesch, Albit J Caban, Shawn Liu, Monika Shpokayte, and Steve Ramirez. Chronic activation of a negative engram induces behavioral and cellular abnormalities. *eLife*, 13, 2024.

[14109] Mathias Fynbo Jensen and Morten Nielsen. Enhancing TCR specificity predictions by combined pan- and peptide-specific training, loss-scaling, and sequence similarity integration. *eLife*, 12, 2024.

[14110] Davin Jeong, Guang Shi, Xin Li, and D Thirumalai. Structural basis for the preservation of a subset of topologically associating domains in interphase chromosomes upon cohesin depletion. *eLife*, 12, 2024.

[14111] Jiwon Jeong, Kujin Kwon, Terezia Klaudia Geisseova, Jongbin Lee, Taejoon Kwon, and Chunghun Lim. Drosulfakinin signaling encodes early-life memory for adaptive social plasticity. *eLife*, 13, 2024.

[14112] Qi Jia, Drew Young, Qixin Zhang, and Derek Sieburth. Endogenous hydrogen peroxide positively regulates secretion of a gut-derived peptide in neuroendocrine potentiation of the oxidative stress response in Caenorhabditis elegans. *eLife*, 13, 2024.

[14113] He-Hai Jiang, Ruoxuan Xu, Xiupeng Nie, Zhenghui Su, Xiaoshan Xu, Ruiqi Pang, Yi Zhou, and Fujun Luo. Neurexins control the strength and precise timing of glycinergic inhibition in the auditory brainstem. *eLife*, 13, 2024.

[14114] Shan Jiang, Lu Chen, Wei-Min Qu, Zhi-Li Huang, and Chang-Rui Chen. Hypothalamic corticotropin-releasing hormone neurons modulate sevoflurane anesthesia and the post-anesthesia stress responses. *eLife*, 12, 2024.

[14115] Dongyue Jiao, Huiru Sun, Xiaying Zhao, Yingji Chen, Zeheng Lv, Qing Shi, Yao Li, Chenji Wang, and Kun Gao. mTORC1/S6K1 signaling promotes sustained oncogenic translation through modulating CRL3IBTK-mediated ubiquitination of eIF4A1 in cancer cells. *eLife*, 12, 2024.

[14116] Hui-Juan Jin, Yong Fan, Xiaoyu Yang, Yue Dong, Xiao-Zhen Zhang, Xin-Yan Geng, Zheng Yan, Ling Wu, Meng Ma, Bin Li, Qifeng Lyu, Yun Pan, Mingxi Liu, Yanping Kuang, and Su-Ren Chen. Disruption in CYLC1 leads to acrosome detachment, sperm head deformity, and male in/subfertility in humans and mice. *eLife*, 13, 2024.

[14117] Yong Jin, Jiayu Xing, Chenyu Dai, Lei Jin, Wanying Zhang, Qianqian Tao, Mei Hou, Ziyi Li, Wen Yang, Qiyu Feng, Hongyang Wang, and Qingsheng Yu. NK cell exhaustion in Wilson's disease revealed by single-cell RNA sequencing predicts the prognosis of cholecystitis. *eLife*, 13, 2024.

[14118] Zhixin Jing, Phillip Galbo, Luis Ovando, Megan Demouth, Skylar Welte, Rosa Park, Kartik Chandran, Yinghao Wu, Thomas MacCarthy, Deyou Zheng, and David Fooksman. Fine-tuning spatial-temporal dynamics and surface receptor expression support plasma cell-intrinsic longevity. *eLife*, 12, 2024.

[14119] Emilia Johnson, Reuben Sunil Kumar Sharma, Pablo Ruiz Cuenca, Isabel Byrne, Milena Salgado-Lynn, Zarith Suraya Shahar, Lee Col Lin, Norhadila Zulkifli, Nor Dilaila Mohd Saidi, Chris Drakeley, Jason Matthiopoulos, Luca Nelli, and Kimberly Fornace. Landscape drives zoonotic malaria prevalence in non-human primates. *eLife*, 12, 2024.

[14120] Sean R Johnson, Meghana Peshwa, and Zhiyi Sun. Sensitive remote homology search by local alignment of small positional embeddings from protein language models. *eLife*, 12, 2024.

[14121] Rachel A Johnston, Katherine A Aracena, Luis B Barreiro, Amanda J Lea, and Jenny Tung. DNA methylation-environment interactions in the human genome. *eLife*, 12, 2024.

[14122] Sebastian Jojoa-Cruz, Adrienne E Dubin, Wen-Hsin Lee, and Andrew B Ward. Structure-guided mutagenesis of OSCAs reveals differential activation to mechanical stimuli. *eLife*, 12, 2024.

[14123] Kent W Jorgenson, Jamie E Hibbert, Ramy KA Sayed, Anthony N Lange, Joshua S Godwin, Paulo HC Mesquita, Bradley A Ruple, Mason C McIntosh, Andreas N Kavazis, Michael D Roberts, and Troy A Hornberger. A novel imaging method (FIM-ID) reveals that myofibrillogenesis plays a major role in the mechanically induced growth of skeletal muscle. *eLife*, 12, 2024.

[14124] Antony M Jose. Heritable epigenetic changes are constrained by the dynamics of regulatory architectures. *eLife*, 12, 2024.

[14125] Raji E Joseph, Thomas E Wales, Sandrine Jayne, Robert G Britton, D Bruce Fulton, John R Engen, Martin JS Dyer, and Amy H Andreotti. Impact of the clinically approved BTK inhibitors on the conformation of full-length BTK and analysis of the development of BTK resistance mutations in chronic lymphocytic leukemia. *eLife*, 13, 2024.

[14126] Jyotsna, Binayak Sarkar, Mohit Yadav, Alvina Deka, Manasvini Markandey, Priyadarshini Sanyal, Perumal Nagarajan, Nilesh Gaikward, Vineet Ahuja, Debasisa Mohanty, Soumen Basak, and Rajesh S Gokhale. A hepatocyte-specific transcriptional program driven by Rela and Stat3 exacerbates experimental colitis in mice by modulating bile synthesis. *eLife*, 12, 2024.

[14127] Anna Kádková, Jacqeline Murach, Maiken Østergaard, Andrea Malsam, Jörg Malsam, Fabio Lolicato, Walter Nickel, Thomas H Söllner, and Jakob Balslev Sørensen. SNAP25 disease mutations change the energy landscape for synaptic exocytosis due to aberrant SNARE interactions. *eLife*, 12, 2024.

[14128] Richard A Kahn, Harvinder Virk, Carl Laflamme, Douglas W Houston, Nicole K Polinski, Rob Meijers, Allan I Levey, Clifford B Saper, Timothy M Errington, Rachel E Turn, Anita Bandrowski, James S Trimmer, Meghan Rego, Leonard P Freedman, Fortunato Ferrara, Andrew RM Bradbury, Hannah Cable, and Skye Longworth. Antibody characterization is critical to enhance reproducibility in biomedical research. *eLife*, 13, 2024.

[14129] Yaroslav Kainov, Fursham Hamid, and Eugene V Makeyev. Recurrent disruption of tumour suppressor genes in cancer by somatic mutations in cleavage and polyadenylation signals. *eLife*, 13, 2024.

[14130] Hari Teja Kalidindi and Frederic Crevecoeur. Task-dependent coarticulation of movement sequences. *eLife*, 13, 2024.

[14131] Archana Kamalakar, Brendan Tobin, Sundus Kaimari, M Hope Robinson, Afra I Toma, Timothy Cha, Samir Chihab, Irica Moriarity, Surabhi Gautam, Pallavi Bhattaram, Shelly Abramowicz, Hicham Drissi, Andres Garcia, Levi Wood, and Steven L Goudy. Delivery of a Jagged1-PEG-MAL hydrogel with pediatric human bone cells regenerates critically sized craniofacial bone defects. *eLife*, 13, 2024.

[14132] Sangeetha Kandoi, Cassandra Martinez, Kevin Xu Chen, Miika Mehine, L Vinod K Reddy, Brian C Mansfield, Jacque L Duncan, and Deepak A Lamba. Disease modeling and pharmacological rescue of autosomal dominant retinitis pigmentosa associated with RHO copy number variation. *eLife*, 12, 2024.

[14133] Yuki Kaneda, Haruhiko Miyata, Zoulan Xu, Keisuke Shimada, Maki Kamoshita, Tatsuya Nakagawa, Chihiro Emori, and Masahito Ikawa. FBXO24 deletion causes abnormal accumulation of membraneless electron-dense granules in sperm flagella and male infertility. *eLife*, 13, 2024.

[14134] Chenlu Kang, Pengcheng Chen, Xin Yi, Dong Li, Yiping Hu, Yihong Yang, Huaqing Cai, Bo Li, and Congying Wu. Amoeboid cells undergo durotaxis with soft end polarized NMIIA. *eLife*, 13, 2024.

[14135] Hyereen Kang, Seong Woo Choi, Joo Young Kim, Soo-Jin Oh, Sung Joon Kim, and Myung-Shik Lee. ER-to-lysosome Ca2+ refilling followed by K+ efflux-coupled store-operated Ca2+ entry in inflammasome activation and metabolic inflammation. *eLife*, 12, 2024.

[14136] Jongkyun Kang, Guodong Huang, Long Ma, Youren Tong, Anu Shahapal, Phoenix Chen, and Jie Shen. Cell-autonomous role of leucine-rich repeat kinase in the protection of dopaminergic neuron survival. *eLife*, 12, 2024.

[14137] Ju Kang, Shijie Zhang, Yiyuan Niu, Fan Zhong, and Xin Wang. Intraspecific predator interference promotes biodiversity in ecosystems. *eLife*, 13, 2024.

[14138] Tae-Yun Kang, Federico Bocci, Qing Nie, José N Onuchic, and Andre Levchenko. Spatial–temporal order–disorder transition in angiogenic NOTCH signaling controls cell fate specification. *eLife*, 12, 2024.

[14139] Foteini Karapanagioti, Úlfur águst Atlason, Dirk J Slotboom, Bert Poolman, and Sebastian Obermaier. Fitness landscape of substrate-adaptive mutations in evolved amino acid-polyamine-organocation transporters. *eLife*, 13, 2024.

[14140] Ryo Kariyazono and Takashi Osanai. CyAbrB2 is a nucleoid-associated protein in Synechocystis controlling hydrogenase expression during fermentation. *eLife*, 13, 2024.

[14141] Kelly Karl, Nuala Del Piccolo, Taylor Light, Tanaya Roy, Pooja Deduja, Vlad-Constantin Ursachi, Bohumil Fafilek, Pavel Krejci, and Kalina Hristova. Ligand bias underlies differential signaling of multiple FGFs via FGFR1. *eLife*, 12, 2024.

[14142] Joakim W Karlsson, Vasu R Sah, Roger Olofsson Bagge, Irina Kuznetsova, Munir Iqba, Samuel Alsen, Sofia Stenqvist, Alka Saxena, Lars Ny, Lisa M Nilsson, and Jonas A Nilsson. Patient-derived xenografts and single-cell sequencing identifies three subtypes of tumor-reactive lymphocytes in uveal melanoma metastases. *eLife*, 12, 2024.

[14143] Mehrdad Kashefi, Sasha Reschechtko, Giacomo Ariani, Mahdiyar Shahbazi, Alice Tan, Jörn Diedrichsen, and J Andrew Pruszynski. Future movement plans interact in sequential arm movements. *eLife*, 13, 2024.

[14144] Makiko Kashio, Sandra Derouiche, Reiko U Yoshimoto, Kenji Sano, Jing Lei, Mizuho A Kido, and Makoto Tominaga. Involvement of TRPV4 in temperature-dependent perspiration in mice. *eLife*, 13, 2024.

[14145] Koji Kato, Yoshiki Nakajima, Jian Xing, Minoru Kumazawa, Haruya Ogawa, Jian-Ren Shen, Kentaro Ifuku, and Ryo Nagao. Structural basis for molecular assembly of fucoxanthin chlorophyll a/c-binding proteins in a diatom photosystem I supercomplex. *eLife*, 13, 2024.

[14146] Maiko Kawasaki, Katsushige Kawasaki, Finsa Tisna Sari, Takehisa Kudo, Jun Nihara, Madoka Kitamura, Takahiro Nagai, Vanessa Utama, Yoko Ishida, Fumiya Meguro, Alex Kesuma, Akira Fujita, Takayuki Nishimura, Yuan Kogure, Satoshi Maruyama, Jun-ichi Tanuma, Yoshito Kakihara, Takeyasu Maeda, Sarah Ghafoor, Roman H Khonsari, Pierre Corre, Paul T Sharpe, Martyn Cobourne, Brunella Franco, and Atsushi Ohazama. Cell-cell interaction determines cell fate of mesoderm-derived cell in tongue development through Hh signaling. *eLife*, 13, 2024.

[14147] Ece Kaya, Sonja A Kotz, and Molly J Henry. A novel method for estimating properties of attentional oscillators reveals an age-related decline in flexibility. *eLife*, 12, 2024.

[14148] Amir Hossein Kayvanjoo, Iva Splichalova, David Alejandro Bejarano, Hao Huang, Katharina Mauel, Nikola Makdissi, David Heider, Hui Ming Tew, Nora Reka Balzer, Eric Greto, Collins Osei-Sarpong, Kevin Baßler, Joachim L Schultze, Stefan Uderhardt, Eva Kiermaier, Marc Beyer, Andreas Schlitzer, and Elvira Mass. Fetal liver macrophages contribute to the hematopoietic stem cell niche by controlling granulopoiesis. *eLife*, 13, 2024.

[14149] Skander Kazdaghli, Iordanis Kerenidis, Jens Kieckbusch, and Philip Teare. Improved clinical data imputation via classical and quantum determinantal point processes. *eLife*, 12, 2024.

[14150] Shraddha KC, Kenny H Nguyen, Vincent Nicholson, Annie Walgren, Tony Trent, Edith Gollub, Paulette Sofia Romero-Perez, Alex S Holehouse, Shahar Sukenik, and Thomas C Boothby. Disordered proteins interact with the chemical environment to tune their protective function during drying. *eLife*, 13, 2024.

[14151] Wenfan Ke, Miki Fujioka, Paul Schedl, and James B Jaynes. Stem-loop and circle-loop TADs generated by directional pairing of boundary elements have distinct physical and regulatory properties. *eLife*, 13, 2024.

[14152] Moran Kelbert, Antonio Jordán-Pla, Lola de Miguel-Jiménez, José García-Martínez, Michael Selitrennik, Adi Guterman, Noa Henig, Sander Granneman, José E Pérez-Ortín, Sebastián Chávez, and Mordechai Choder. The zinc-finger transcription factor Sfp1 imprints specific classes of mRNAs and links their synthesis to cytoplasmic decay. *eLife*, 12, 2024.

[14153] Bryan Kennedy, Sarala N Malladi, Roger BH Tootell, and Shahin Nasr. A previously undescribed scene-selective site is the key to encoding ego-motion in naturalistic environments. *eLife*, 13, 2024.

[14154] Nicholas GW Kennedy, Jessica C Lee, Simon Killcross, R Fred Westbrook, and Nathan M Holmes. Prediction error determines how memories are organized in the brain. *eLife*, 13, 2024.

[14155] Athina Keramidioti, Sandra Schneid, Christina Busse, Christoph Cramer von Laue, Bianca Bertulat, Willi Salvenmoser, Martin Hess, Olga Alexandrova, Kristine M Glauber, Robert E Steele, Bert Hobmayer, Thomas W Holstein, and Charles N David. A new look at the architecture and dynamics of the Hydra nerve net. *eLife*, 12, 2024.

[14156] Simon Kern, Juliane Nagel, Martin F Gerchen, Çağatay Gürsoy, Andreas Meyer-Lindenberg, Peter Kirsch, Raymond J Dolan, Steffen Gais, and Gordon B Feld. Reactivation strength during cued recall is modulated by graph distance within cognitive maps. *eLife*, 12, 2024.

[14157] Natalia E Ketaren, Fred D Mast, Peter C Fridy, Jean Paul Olivier, Tanmoy Sanyal, Andrej Sali, Brian T Chait, Michael P Rout, and John D Aitchison. Nanobody repertoire generated against the spike protein of ancestral SARS-CoV-2 remains efficacious against the rapidly evolving virus. *eLife*, 12, 2024.

[14158] Viktor Nikolaus Kewenig, Gabriella Vigliocco, and Jeremy I Skipper. When abstract becomes concrete, naturalistic encoding of concepts in the brain. *eLife*, 13, 2024.

[14159] Hina Khan, Partha Paul, Harsh Goar, Bhanwar Bamnia, Navin Baid, and Dibyendu Sarkar. Correction: Mycobacterium tuberculosis PhoP integrates stress response to intracellular survival by regulating cAMP level. *eLife*, 13, 2024.

[14160] Hina Khan, Partha Paul, Harsh Goar, Bhanwar Bamniya, Navin Baid, and Dibyendu Sarkar. Mycobacterium tuberculosis PhoP integrates stress response to intracellular survival by regulating cAMP level. *eLife*, 13, 2024.

[14161] Imran S Khan, Christopher Molina, Xin Ren, Vincent C Auyeung, Max Cohen, Tatsuya Tsukui, Amha Atakilit, and Dean Sheppard. Impaired myofibroblast proliferation is a central feature of pathologic post-natal alveolar simplification. *eLife*, 13, 2024.

[14162] Ayse Kilic, Arda Halu, Margherita De Marzio, Enrico Maiorino, Melody G Duvall, Thayse Brueggemann, Joselyn J Rojas Quintero, Robert Chase, Hooman Mirzakhani, Ayse özge Sungur, Janine Koepke, Taiji Nakano, Hong Yong Peh, Nandini Krishnamoorthy, Raja-Elie Abdulnour, Katia Georgopoulos, Augusto A Litonjua, Marie Demay, Harald Renz, Bruce D Levy, and Scott T Weiss. Vitamin D constrains inflammation by modulating the expression of key genes on Chr17q12-21.1. *eLife*, 12, 2024.

[14163] Eun Joo Kim, Mi-Seon Kong, Sanggeon Park, Jeiwon Cho, and Jeansok John Kim. Periaqueductal gray activates antipredatory neural responses in the amygdala of foraging rats. *eLife*, 12, 2024.

[14164] Hee Young Kim, Yeon Jun Kang, Dong Hyun Kim, Jiyeon Jang, Su Jeong Lee, Gwanghun Kim, Hee Byung Koh, Ye Eun Ko, Hyun Mu Shin, Hajeong Lee, Tae-Hyun Yoo, and Won-Woo Lee. Uremic toxin indoxyl sulfate induces trained immunity via the AhR-dependent arachidonic acid pathway in end-stage renal disease (ESRD). *eLife*, 12, 2024.

[14165] Jaebin Kim, Edwin Bustamante, Peter Sotonyi, Nicholas Maxwell, Pooja Parameswaran, Julie K Kent, William C Wetsel, Erik J Soderblom, Bence Rácz, and Scott H Soderling. Presynaptic Rac1 in the hippocampus selectively regulates working memory. *eLife*, 13, 2024.

[14166] Jee Min Kim, Claudia C Carcamo, Sina Jazani, Zepei Xie, Xinyu A Feng, Maryam Yamadi, Matthew Poyton, Katie L Holland, Jonathan B Grimm, Luke D Lavis, Taekjip Ha, and Carl Wu. Dynamic 1D search and processive nucleosome translocations by RSC and ISW2 chromatin remodelers. *eLife*, 12, 2024.

[14167] Jeongsik Kim, Dahyun Kim, Dong-Kyun Kim, Sang-Hee Lee, Wonyul Jang, and Dae-Sik Lim. Formation of a giant unilocular vacuole via macropinocytosis-like process confers anoikis resistance. *eLife*, 13, 2024.

[14168] Namsuk Kim, Yan Li, Ri Yu, Hyo-Shin Kwon, Anji Song, Mi-Hee Jun, Jin-Young Jeong, Ji Hyun Lee, Hyun-Ho Lim, Mi-Jin Kim, Jung-Woong Kim, and Won-Jong Oh. Repulsive Sema3E-Plexin-D1 signaling coordinates both axonal extension and steering via activating an autoregulatory factor, Mtss1. *eLife*, 13, 2024.

[14169] Nayoung Kim, Sehhoon Park, Areum Jo, Hye Hyeon Eum, Hong Kwan Kim, Kyungjong Lee, Jong Ho Cho, Bo Mi Ku, Hyun Ae Jung, Jong-Mu Sun, Se-Hoon Lee, Jin Seok Ahn, Jung-Il Lee, Jung Won Choi, Dasom Jeong, Minsu Na, Huiram Kang, Jeong Yeon Kim, Jung Kyoon Choi, Hae-Ock Lee, and Myung-Ju Ahn. Unveiling the influence of tumor and immune signatures on immune checkpoint therapy in advanced lung cancer. *eLife*, 13, 2024.

[14170] Seung Mi Kim, Yaw Quagraine, Monika Singh, and Jung Hwan Kim. Rab11 suppresses neuronal stress signaling by localizing dual leucine zipper kinase to axon terminals for protein turnover. *eLife*, 13, 2024.

[14171] Eyal Y Kimchi, Anthony Burgos-Robles, Gillian A Matthews, Tatenda Chakoma, Makenzie Patarino, Javier C Weddington, Cody Siciliano, Wannan Yang, Shaun Foutch, Renee Simons, Ming-fai Fong, Miao Jing, Yulong Li, Daniel B Polley, and Kay M Tye. Reward contingency gates selective cholinergic suppression of amygdala neurons. *eLife*, 12, 2024.

[14172] Georg Kislinger, Gunar Fabig, Antonia Wehn, Lucia Rodriguez, Hanyi Jiang, Cornelia Niemann, Andrey S Klymchenko, Nikolaus Plesnila, Thomas Misgeld, Thomas Müller-Reichert, Igor Khalin, and Martina Schifferer. Combining array tomography with electron tomography provides insights into leakiness of the blood-brain barrier in mouse cortex. *eLife*, 12, 2024.

[14173] Tomoki Kita, Kyoko Chiba, Jiye Wang, Atsushi Nakagawa, and Shinsuke Niwa. Comparative analysis of two Caenorhabditis elegans kinesins KLP-6 and UNC-104 reveals a common and distinct activation mechanism in kinesin-3. *eLife*, 12, 2024.

[14174] Arndt-Lukas Klaassen and Björn Rasch. Difficulty in artificial word learning impacts targeted memory reactivation and its underlying neural signatures. *eLife*, 12, 2024.

[14175] Gordon L Klein. Phosphate as an adjunct to calcium in promoting coronary vascular calcification in chronic inflammatory states. *eLife*, 13, 2024.

[14176] Maksim Kleverov, Daria Zenkova, Vladislav Kamenev, Margarita Sablina, Maxim N Artyomov, and Alexey A Sergushichev. Phantasus, a web application for visual and interactive gene expression analysis. *eLife*, 13, 2024.

[14177] Nils Klughammer, Anders Barth, Maurice Dekker, Alessio Fragasso, Patrick R Onck, and Cees Dekker. Diameter dependence of transport through nuclear pore complex mimics studied using optical nanopores. *eLife*, 12, 2024.

[14178] Steven Klupt, Kyong Tkhe Fam, Xing Zhang, Pavan Kumar Chodisetti, Abeera Mehmood, Tumara Boyd, Danielle Grotjahn, Donghyun Park, and Howard C Hang. Secreted antigen A peptidoglycan hydrolase is essential for Enterococcus faecium cell separation and priming of immune checkpoint inhibitor therapy. *eLife*, 13, 2024.

[14179] Paul Knabl, Alexandra Schauer, Autumn P Pomreinke, Bob Zimmermann, Katherine W Rogers, Daniel čapek, Patrick Müller, and Grigory Genikhovich. Analysis of SMAD1/5 target genes in a sea anemone reveals ZSWIM4-6 as a novel BMP signaling modulator. *eLife*, 13, 2024.

[14180] Filip Knop, Apolena Zounarová, Vojtěch Šabata, Teije Corneel Middelkoop, and Marie Macůrková. Caenorhabditis elegans SEL-5/AAK1 regulates cell migration and cell outgrowth independently of its kinase activity. *eLife*, 13, 2024.

[14181] Toshiki Kobayashi and Daichi Nozaki. Implicit motor adaptation patterns in a redundant motor task manipulating a stick with both hands. *eLife*, 13, 2024.

[14182] Duk-Su Koh, Anastasiia Stratiievska, Subhashis Jana, Shauna C Otto, Teresa M Swanson, Anthony Nhim, Sara Carlson, Marium Raza, Ligia Araujo Naves, Eric N Senning, Ryan A Mehl, and Sharona E Gordon. Genetic code expansion, click chemistry, and light-activated PI3K reveal details of membrane protein trafficking downstream of receptor tyrosine kinases. *eLife*, 12, 2024.

[14183] Seiichi Koike, Masashi Tachikawa, Motosuke Tsutsumi, Takuya Okada, Tomomi Nemoto, Kazuko Keino-Masu, and Masayuki Masu. Actin dynamics switches two distinct modes of endosomal fusion in yolk sac visceral endoderm cells. *eLife*, 13, 2024.

[14184] Nada Kojovic, Sezen Cekic, Santiago Herce Castañón, Martina Franchini, Holger Franz Sperdin, Corrado Sandini, Reem Kais Jan, Daniela Zöller, Lylia Ben Hadid, Daphné Bavelier, and Marie Schaer. Unraveling the developmental dynamic of visual exploration of social interactions in autism. *eLife*, 13, 2024.

[14185] Anastasia D Komarova, Snezhana D Sinyushkina, Ilia D Shchechkin, Irina N Druzhkova, Sofia A Smirnova, Vitaliy M Terekhov, Artem M Mozherov, Nadezhda I Ignatova, Elena E Nikonova, Evgeny A Shirshin, Liubov E Shimolina, Sergey V Gamayunov, Vladislav I Shcheslavskiy, and Marina V Shirmanova. Insights into metabolic heterogeneity of colorectal cancer gained from fluorescence lifetime imaging. *eLife*, 13, 2024.

[14186] Benjamin R Kop, Yazan Shamli Oghli, Talyta C Grippe, Tulika Nandi, Judith Lefkes, Sjoerd W Meijer, Soha Farboud, Marwan Engels, Michelle Hamani, Melissa Null, Angela Radetz, Umair Hassan, Ghazaleh Darmani, Andrey Chetverikov, Hanneke EM den Ouden, Til Ole Bergmann, Robert Chen, and Lennart Verhagen. Auditory confounds can drive online effects of transcranial ultrasonic stimulation in humans. *eLife*, 12, 2024.

[14187] Avani Koparkar, Timothy L Warren, Jonathan D Charlesworth, Sooyoon Shin, Michael S Brainard, and Lena Veit. Lesions in a songbird vocal circuit increase variability in song syntax. *eLife*, 13, 2024.

[14188] Kevin P Koster and S Murray Sherman. Convergence of inputs from the basal ganglia with layer 5 of motor cortex and cerebellum in mouse motor thalamus. *eLife*, 13, 2024.

[14189] Tiantian Kou, Lan Kang, Bin Zhang, Jiaqi Li, Baohong Zhao, Wenwen Zeng, and Xiaoyu Hu. RBP-J regulates homeostasis and function of circulating Ly6Clo monocytes. *eLife*, 12, 2024.

[14190] Artemis Koumoundourou, Märt Rannap, Elodie De Bruyckere, Sigrun Nestel, Carsten Reissner, Alexei V Egorov, Pengtao Liu, Markus Missler, Bernd Heimrich, Andreas Draguhn, and Stefan Britsch. Regulation of hippocampal mossy fiber-CA3 synapse function by a Bcl11b/C1ql2/Nrxn3(25b+) pathway. *eLife*, 12, 2024.

[14191] Aleksandra Kovacevic, David RM Smith, Eve Rahbé, Sophie Novelli, Paul Henriot, Emmanuelle Varon, Robert Cohen, Corinne Levy, Laura Temime, and Lulla Opatowski. Exploring factors shaping antibiotic resistance patterns in Streptococcus pneumoniae during the 2020 COVID-19 pandemic. *eLife*, 13, 2024.

[14192] Caroline Krag, Maria Saur Svane, Sten Madsbad, Susanne Boel Graversen, Jesper Frank Christensen, Thorkild IA Sørensen, Louise Lang Lehrskov, and Tinne Laurberg. The impact of surgery and oncological treatment on risk of type 2 diabetes onset in patients with colorectal cancer: nationwide cohort study in Denmark. *eLife*, 12, 2024.

[14193] Christin Krause, Jan H Britsemmer, Miriam Bernecker, Anna Molenaar, Natalie Taege, Nuria Lopez-Alcantara, Cathleen Geißler, Meike Kaehler, Katharina Iben, Anna Judycka, Jonas Wagner, Stefan Wolter, Oliver Mann, Paul Pfluger, Ingolf Cascorbi, Hendrik Lehnert, Kerstin Stemmer, Sonja C Schriever, and Henriette Kirchner. Liver microRNA transcriptome reveals miR-182 as link between type 2 diabetes and fatty liver disease in obesity. *eLife*, 12, 2024.

[14194] Paul G Kremer, Elizabeth A Lampros, Allison M Blocker, and Adam W Barb. One N-glycan regulates natural killer cell antibody-dependent cell-mediated cytotoxicity and modulates Fc $\gamma$ receptor IIIa/CD16a structure. *eLife*, 13, 2024.

[14195] Tomoko Kubori, Kohei Arasaki, Hiromu Oide, Tomoe Kitao, and Hiroki Nagai. Multi-tiered actions of Legionella effectors to modulate host Rab10 dynamics. *eLife*, 12, 2024.

[14196] Kiwamu Kudo, Kamalini G Ranasinghe, Hirofumi Morise, Faatimah Syed, Kensuke Sekihara, Katherine P Rankin, Bruce L Miller, Joel H Kramer, Gil D Rabinovici, Keith Vossel, Heidi E Kirsch, and Srikantan S Nagarajan. Neurophysiological trajectories in Alzheimer's disease progression. *eLife*, 12, 2024.

[14197] Lauren Kuffler, Daniel A Skelly, Anne Czechanski, Haley J Fortin, Steven C Munger, Christopher L Baker, Laura G Reinholdt, and Gregory W Carter. Imputation of 3D genome structure by genetic–epigenetic interaction modeling in mice. *eLife*, 12, 2024.

[14198] Valentina Kugler, Selina Schwaighofer, Andreas Feichtner, Florian Enzler, Jakob Fleischmann, Sophie Strich, Sarah Schwarz, Rebecca Wilson, Philipp Tschaikner, Jakob Troppmair, Veronika Sexl, Pascal Meier, Teresa Kaserer, and Eduard Stefan. Impact of protein and small molecule interactions on kinase conformations. *eLife*, 13, 2024.

[14199] Ivan Kulich, Julia Schmid, Anastasia Teplova, Linlin Qi, and Jiří Friml. Rapid translocation of NGR proteins driving polarization of PIN-activating D6 protein kinase during root gravitropism. *eLife*, 12, 2024.

[14200] Pradeep Kumar, Ankit Roy, Shivapura Jagadeesha Mukul, Avinash Kumar Singh, Dipesh Kumar Singh, Aswan Nalli, Pujaita Banerjee, Kandhalu Sagadevan Dinesh Babu, Bakthisaran Raman, Shobha P Kruparani, Imran Siddiqi, and Rajan Sankaranarayanan. A translation proofreader of archaeal origin imparts multi-aldehyde stress tolerance to land plants. *eLife*, 12, 2024.

[14201] Miki Kume, Hanako Koguchi-Yoshioka, Shuichi Nakai, Yutaka Matsumura, Atsushi Tanemura, Kazunori Yokoi, Shoichi Matsuda, Yuumi Nakamura, Naoya Otani, Mifue Taminato, Koichi Tomita, Tateki Kubo, Mari Wataya-Kaneda, Atsushi Kumanogoh, Manabu Fujimoto, and Rei Watanabe. Downregulation of semaphorin 4A in keratinocytes reflects the features of non-lesional psoriasis. *eLife*, 13, 2024.

[14202] Suman Kundu, Advait Shetty, and Maria Gomes-Solecki. Exposure to live saprophytic Leptospira before challenge with a pathogenic serovar prevents severe leptospirosis and promotes kidney homeostasis. *eLife*, 13, 2024.

[14203] Maximilian Kurjahn, Antaran Deka, Antoine Girot, Leila Abbaspour, Stefan Klumpp, Maike Lorenz, Oliver Bäumchen, and Stefan Karpitschka. Quantifying gliding forces of filamentous cyanobacteria by self-buckling. *eLife*, 12, 2024.

[14204] Hyo Jun Kwon, Devi Santhosh, and Zhen Huang. A novel monomeric amyloid $\beta$-activated signaling pathway regulates brain development via inhibition of microglia. *eLife*, 13, 2024.

[14205] Nicholas LaBerge, Kenneth Hunter Wapman, Aaron Clauset, and Daniel B Larremore. Gendered hiring and attrition on the path to parity for academic faculty. *eLife*, 13, 2024.

[14206] Blake J Laham, Isha R Gore, Casey J Brown, and Elizabeth Gould. Adult-born granule cells modulate CA2 network activity during retrieval of developmental memories of the mother. *eLife*, 12, 2024.

[14207] Vilde Olsson Lalun, Maike Breiden, Sergio Galindo-Trigo, Elwira Smakowska-Luzan, Rüdiger GW Simon, and Melinka A Butenko. A dual function of the IDA peptide in regulating cell separation and modulating plant immunity at the molecular level. *eLife*, 12, 2024.

[14208] Adriano R Lameira, Madeleine E Hardus, Andrea Ravignani, Teresa Raimondi, and Marco Gamba. Recursive self-embedded vocal motifs in wild orangutans. *eLife*, 12, 2024.

[14209] Qiang Lan, Ewelina Trela, Riitta Lindström, Jyoti Prabha Satta, Beata Kaczyńska, Mona M Christensen, Martin Holzenberger, Jukka Jernvall, and Marja L Mikkola. Mesenchyme instructs growth while epithelium directs branching in the mouse mammary gland. *eLife*, 13, 2024.

[14210] Marius Landau, Sherif Elsabbagh, Harald Gross, Adrian CD Fuchs, Anita CF Schultz, and Joachim E Schultz. The membrane domains of mammalian adenylyl cyclases are lipid receptors. *eLife*, 13, 2024.

[14211] Nicolas Langer, Maurice Weber, Bruno Hebling Vieira, Dawid Strzelczyk, Lukas Wolf, Andreas Pedroni, Jonathan Heitz, Stephan Müller, Christoph Schultheiss, Marius Troendle, Juan Carlos Arango Lasprilla, Diego Rivera, Federica Scarpina, Qianhua Zhao, Rico Leuthold, Flavia Wehrle, Oskar Jenni, Peter Brugger, Tino Zaehle, Romy Lorenz, and Ce Zhang. A deep learning approach for automated scoring of the Rey–Osterrieth complex figure. *eLife*, 13, 2024.

[14212] Felix Lankester, Tito J Kibona, Kathryn J Allan, William de Glanville, Joram J Buza, Frank Katzer, Jo E Halliday, Blandina T Mmbaga, Nick Wheelhouse, Elisabeth A Innes, Kate M Thomas, Obed M Nyasebwa, Emanuel Swai, John R Claxton, and Sarah Cleaveland. Livestock abortion surveillance in Tanzania reveals disease priorities and importance of timely collection of vaginal swab samples for attribution. *eLife*, 13, 2024.

[14213] Zhuohan Lao, Kartik D Kamat, Zhongling Jiang, and Bin Zhang. OpenNucleome for high-resolution nuclear structural and dynamical modeling. *eLife*, 13, 2024.

[14214] Damien Laporte, Aurelie Massoni-Laporte, Charles Lefranc, Jim Dompierre, David Mauboules, Emmanuel T Nsamba, Anne Royou, Lihi Gal, Maya Schuldiner, Mohan L Gupta, and Isabelle Sagot. A stable microtubule bundle formed through an orchestrated multistep process controls quiescence exit. *eLife*, 12, 2024.

[14215] Megan Larmore, Orhi Esarte Palomero, Neha Kamat, and Paul G DeCaen. A synthetic method to assay polycystin channel biophysics. *eLife*, 13, 2024.

[14216] Jean-David Larouche, Céline M Laumont, Assya Trofimov, Krystel Vincent, Leslie Hesnard, Sylvie Brochu, Caroline Côté, Juliette F Humeau, éric Bonneil, Joel Lanoix, Chantal Durette, Patrick Gendron, Jean-Philippe Laverdure, Ellen R Richie, Sébastien Lemieux, Pierre Thibault, and Claude Perreault. Transposable elements regulate thymus development and function. *eLife*, 12, 2024.

[14217] Mart GF Last, Leoni Abendstein, Lenard M Voortman, and Thomas H Sharp. Streamlining segmentation of cryo-electron tomography datasets with Ais. *eLife*, 13, 2024.

[14218] Sara Latini, Veronica Venafra, Giorgia Massacci, Valeria Bica, Simone Graziosi, Giusj Monia Pugliese, Marta Iannuccelli, Filippo Frioni, Gessica Minnella, John Donald Marra, Patrizia Chiusolo, Gerardo Pepe, Manuela Helmer Citterich, Dimitros Mougiakakos, Martin Böttcher, Thomas Fischer, Livia Perfetto, and Francesca Sacco. Unveiling the signaling network of FLT3-ITD AML improves drug sensitivity prediction. *eLife*, 12, 2024.

[14219] Olujolagbe Layinka, Luca D Hargitai, Punit Shah, Lucy H Waldren, and Florence YN Leung. Five interdisciplinary tensions and opportunities in neurodiversity research. *eLife*, 13, 2024.

[14220] Audrey Le Veve, Mathieu Genete, Christelle Lepers-Blassiau, Chloé Ponitzki, Céline Poux, Xavier Vekemans, Eleonore Durand, and Vincent Castric. The genetic architecture of the load linked to dominant and recessive self-incompatibility alleles in Arabidopsis halleri and Arabidopsis lyrata. *eLife*, 13, 2024.

[14221] Giulia Leanza, Francesca Cannata, Malak Faraj, Claudio Pedone, Viola Viola, Flavia Tramontana, Niccolò Pellegrini, Gianluca Vadalà, Alessandra Piccoli, Rocky Strollo, Francesca Zalfa, Alec T Beeve, Erica L Scheller, Simon Y Tang, Roberto Civitelli, Mauro Maccarrone, Rocco Papalia, and Nicola Napoli. Bone canonical Wnt signaling is downregulated in type 2 diabetes and associates with higher advanced glycation end-products (AGEs) content and reduced bone strength. *eLife*, 12, 2024.

[14222] Sophie Leclercq, Hany Ahmed, Camille Amadieu, Géraldine Petit, Ville Koistinen, Quentin Leyrolle, Marie Poncin, Peter Stärkel, Eloise Kok, Pekka J Karhunen, Philippe de Timary, Sophie Laye, Audrey M Neyrinck, Olli K Kärkkäinen, Kati Hanhineva, and Nathalie Delzenne. Blood metabolomic profiling reveals new targets in the management of psychological symptoms associated with severe alcohol use disorder. *eLife*, 13, 2024.

[14223] Donghyung Lee, Nathan Lau, Lillian Liu, and Cory M Root. Transformation of valence signaling in a mouse striatopallidal circuit. *eLife*, 12, 2024.

[14224] Hyun-Chae Lee, Sun-Hee Park, Hye Min Jeong, Goeun Shin, Sung In Lim, Jeongtae Kim, Jaewon Shim, Yeong-Min Park, and Kyoung Seob Song. LPS-induced systemic inflammation is suppressed by the PDZ motif peptide of ZO-1 via regulation of macrophage M1/M2 polarization. *eLife*, 13, 2024.

[14225] James Lee, Leman Simpson, Yi Li, Samuel Becker, Fan Zou, Xin Zhang, and Lu Bai. Transcription factor condensates, 3D clustering, and gene expression enhancement of the MET regulon. *eLife*, 13, 2024.

[14226] Ju-Young Lee, Dahee Jung, and Sebastien Royer. Stochastic characterization of navigation strategies in an automated variant of the Barnes maze. *eLife*, 12, 2024.

[14227] Kyung-Tae Lee, Inez KA Pranoto, Soon-Young Kim, Hee-Joo Choi, Ngoc Bao To, Hansong Chae, Jeong-Yeon Lee, Jung-Eun Kim, Young V Kwon, and Jin-Wu Nam. Comparative interactome analysis of $\alpha$-arrestin families in human and Drosophila. *eLife*, 12, 2024.

[14228] Mark S Lee, Peter J Tuohy, Caleb Y Kim, Philip P Yost, Katrina Lichauco, Heather L Parrish, Koenraad Van Doorslaer, and Michael S Kuhns. The CD4 transmembrane GGXXG and juxtamembrane (C/F)CV+C motifs mediate pMHCII-specific signaling independently of CD4-LCK interactions. *eLife*, 12, 2024.

[14229] MinHyuk Lee, Se Hoon Park, Kyeung Min Joo, Jae Young Kwon, Kyung-Hoon Lee, and KyeongJin Kang. Drosophila HCN mediates gustatory homeostasis by preserving sensillar transepithelial potential in sweet environments. *eLife*, 13, 2024.

[14230] Selene Seoyun Lee, Livia Civitelli, and Laura Parkkinen. Brain-derived and in vitro-seeded alpha-synuclein fibrils exhibit distinct biophysical profiles. *eLife*, 13, 2024.

[14231] Tai-Ying Lee, Yves Weissenberger, Andrew J King, and Johannes C Dahmen. Midbrain encodes sound detection behavior without auditory cortex. *eLife*, 12, 2024.

[14232] Sindre Lee-Ødegård, Marit Hjorth, Thomas Olsen, Gunn-Helen Moen, Emily Daubney, David M Evans, Andrea L Hevener, Aldons J Lusis, Mingqi Zhou, Marcus M Seldin, Hooman Allayee, James Hilser, Jonas Krag Viken, Hanne Gulseth, Frode Norheim, Christian A Drevon, and Kåre Inge Birkeland. Serum proteomic profiling of physical activity reveals CD300LG as a novel exerkine with a potential causal link to glucose homeostasis. *eLife*, 13, 2024.

[14233] Jake D Lehle, Yu-Huey Lin, Amanda Gomez, Laura Chavez, and John R McCarrey. An in vitro approach reveals molecular mechanisms underlying endocrine disruptor-induced epimutagenesis. *eLife*, 13, 2024.

[14234] Jade Leiba, Tamara Sipka, Christina Begon-Pescia, Matteo Bernardello, Sofiane Tairi, Lionello Bossi, Anne-Alicia Gonzalez, Xavier Mialhe, Emilio J Gualda, Pablo Loza-Alvarez, Anne Blanc-Potard, Georges Lutfalla, and Mai E Nguyen-Chi. Dynamics of macrophage polarization support Salmonella persistence in a whole living organism. *eLife*, 13, 2024.

[14235] Alexandra H Leighton, Juliette E Cheyne, and Christian Lohmann. Clustered synapses develop in distinct dendritic domains in visual cortex before eye opening. *eLife*, 12, 2024.

[14236] Evgenia Leikina, Jarred M Whitlock, Kamran Melikov, Wendy Zhang, Michael P Bachmann, and Leonid Chernomordik. Formation of multinucleated osteoclasts depends on an oxidized species of cell surface-associated La protein. *eLife*, 13, 2024.

[14237] Domingos Leite de Castro, Miguel Aroso, A Pedro Aguiar, David B Grayden, and Paulo Aguiar. Disrupting abnormal neuronal oscillations with adaptive delayed feedback control. *eLife*, 13, 2024.

[14238] Dipti Ranjan Lenka, Shakti Virendra Dahe, Odetta Antico, Pritiranjan Sahoo, Alan R Prescott, Miratul MK Muqit, and Atul Kumar. Additional feedforward mechanism of Parkin activation via binding of phospho-UBL and RING0 in trans. *eLife*, 13, 2024.

[14239] Marco Leonti, Joanna Baker, Peter Staub, Laura Casu, and Julie Hawkins. Taste shaped the use of botanical drugs. *eLife*, 12, 2024.

[14240] Christopher TA Lewis, Elise G Melhedegaard, Marija M Ognjanovic, Mathilde S Olsen, Jenni Laitila, Robert AE Seaborne, Magnus Gronset, Changxin Zhang, Hiroyuki Iwamoto, Anthony L Hessel, Michel N Kuehn, Carla Merino, Nuria Amigo, Ole Frobert, Sylvain Giroud, James F Staples, Anna V Goropashnaya, Vadim B Fedorov, Brian Barnes, Oivind Toien, Kelly Drew, Ryan J Sprenger, and Julien Ochala. Remodeling of skeletal muscle myosin metabolic states in hibernating mammals. *eLife*, 13, 2024.

[14241] Aedan Yue Li, Natalia Ladyka-Wojcik, Heba Qazilbash, Ali Golestani, Dirk B Walther, Chris B Martin, and Morgan D Barense. Experience transforms crossmodal object representations in the anterior temporal lobes. *eLife*, 13, 2024.

[14242] Ang Li, Jianxun Yi, Xuejun Li, Li Dong, Lyle W Ostrow, Jianjie Ma, and Jingsong Zhou. Distinct transcriptomic profile of satellite cells contributes to preservation of neuromuscular junctions in extraocular muscles of ALS mice. *eLife*, 12, 2024.

[14243] Bin Li, Jin Zhang, Taojun He, Hanmei Yuan, Hui Wu, Peng Wang, and Chao Wu. PRR adjuvants restrain high stability peptides presentation on APCs. *eLife*, 13, 2024.

[14244] Bing Li, Prasanna Suresh, Jack Brelstaff, Shekhar Kedia, Clare E Bryant, and David Klenerman. The delayed kinetics of Myddosome formation explains why amyloid-beta aggregates trigger Toll-like receptor 4 less efficiently than lipopolysaccharide. *eLife*, 13, 2024.

[14245] Bingrui Li, Fernanda G Kugeratski, and Raghu Kalluri. A novel machine learning algorithm selects proteome signature to specifically identify cancer exosomes. *eLife*, 12, 2024.

[14246] Chunxiao Li, Tao Deng, Yang Wang, Fajun Sun, Burt Wolff, Qigao Jiangzuo, Jiao Ma, Luda Xing, Jiao Fu, Ji Zhang, and Shiqi Wang. The trunk replaces the longer mandible as the main feeding organ in elephant evolution. *eLife*, 12, 2024.

[14247] Hao Li, Jingyu Feng, Mengying Chen, Min Xin, Xi Chen, Wenhao Liu, Liping Wang, Kuan Hong Wang, and Jufang He. Cholecystokinin facilitates motor skill learning by modulating neuroplasticity in the motor cortex. *eLife*, 13, 2024.

[14248] Jialin Li, Feihong Yang, Yu Tian, Ziwu Wang, Dashi Qi, Zhengang Yang, Jiangang Song, Jing Ding, Xin Wang, and Zhuangzhi Zhang. Lateral/caudal ganglionic eminence makes limited contribution to cortical oligodendrocytes. *eLife*, 13, 2024.

[14249] Jiang Li, Yuefeng Yu, Ying Sun, Yanqi Fu, Wenqi Shen, Lingli Cai, Xiao Tan, Yan Cai, Ningjian Wang, Yingli Lu, and Bin Wang. Nuclear magnetic resonance-based metabolomics with machine learning for predicting progression from prediabetes to diabetes. *eLife*, 13, 2024.

[14250] Jing Li, Chao Ning, Yaohua Liu, Bowen Deng, Bingcai Wang, Kai Shi, Rencong Wang, Ruixin Fang, and Chuan Zhou. The function of juvenile–adult transition axis in female sexual receptivity of Drosophila melanogaster. *eLife*, 12, 2024.

[14251] Lu Li, Yin Li, Caihong Huang, Wenjuan Hou, Zijian Lv, Lizi Zhang, Yishan Qu, Yahan Sun, Kaizhe Huang, Xiao Han, Zhixiong He, and Fadao Tai. PVN-mPFC OT projections modulate pup-directed pup care or attacking in virgin mandarin voles. *eLife*, 13, 2024.

[14252] Ming Li, Nikolay P Kandul, Ruichen Sun, Ting Yang, Elena D Benetta, Daniel J Brogan, Igor Antoshechkin, Héctor M Sánchez C, Yinpeng Zhan, Nicolas A DeBeaubien, YuMin M Loh, Matthew P Su, Craig Montell, John M Marshall, and Omar S Akbari. Targeting sex determination to suppress mosquito populations. *eLife*, 12, 2024.

[14253] Shengjie Li, Jun Ren, Zhendong Jiang, Yichao Qiu, Mingxi Shao, Yingzhu Li, Jianing Wu, Yunxiao Song, Xinghuai Sun, Shunxiang Gao, and Wenjun Cao. Metabolomics identifies and validates serum androstenedione as novel biomarker for diagnosing primary angle closure glaucoma and predicting the visual field progression. *eLife*, 12, 2024.

[14254] Thomas Morgan Li, Victoria Zyulina, Ethan S Seltzer, Marija Dacic, Yurii Chinenov, Andrea R Daamen, Keila R Veiga, Noa Schwartz, David J Oliver, Pamela Cabahug-Zuckerman, Jose Lora, Yong Liu, William D Shipman, William G Ambler, Sarah F Taber, Karen B Onel, Jonathan H Zippin, Mehdi Rashighi, James G Krueger, Niroshana Anandasabapathy, Inez Rogatsky, Ali Jabbari, Carl P Blobel, Peter E Lipsky, and Theresa T Lu. The interferon-rich skin environment regulates Langerhans cell ADAM17 to promote photosensitivity in lupus. *eLife*, 13, 2024.

[14255] Tingting Li, Wenwen Shi, Margaret S Ho, and Yong Q Zhang. A Pvr–AP-1–Mmp1 signaling pathway is activated in astrocytes upon traumatic brain injury. *eLife*, 12, 2024.

[14256] Xuankun Li, Adriana E Marvaldi, Rolf G Oberprieler, Dave Clarke, Brian D Farrell, Andrea Sequeira, M Silvia Ferrer, Charles O'Brien, Shayla Salzman, Seunggwan Shin, William Tang, and Duane D McKenna. The evolutionary history of the ancient weevil family Belidae (Coleoptera: Curculionoidea) reveals the marks of Gondwana breakup and major floristic turnovers, including the rise of angiosperms. *eLife*, 13, 2024.

[14257] Xueyou Li, William V Bleisch, Wenqiang Hu, Quan Li, Hongjiao Wang, Zhongzheng Chen, Ru Bai, and Xue-Long Jiang. Human disturbance increases spatiotemporal associations among mountain forest terrestrial mammal species. *eLife*, 12, 2024.

[14258] Yangang Li, Xinyun Zhu, Yu Qi, and Yueming Wang. Revealing unexpected complex encoding but simple decoding mechanisms in motor cortex via separating behaviorally relevant neural signals. *eLife*, 12, 2024.

[14259] Yi-Hong Li, Yi-Sen Yang, Yan-Bo Xue, Hao Lei, Sai-Sai Zhang, Junbin Qian, Yushi Yao, Ruhong Zhou, and Liquan Huang. G protein subunit G$\gamma$13-mediated signaling pathway is critical to the inflammation resolution and functional recovery of severely injured lungs. *eLife*, 12, 2024.

[14260] Yiting Li, Wenqu Yin, Xin Wang, Jiawen Li, Shanglin Zhou, Chaolin Ma, Peng Yuan, and Baoming Li. Stable sequential dynamics in prefrontal cortex represents subjective estimation of time. *eLife*, 13, 2024.

[14261] Yongjun Li, Nitin S Chouhan, Shirley L Zhang, Rebecca S Moore, Sara B Noya, Joy Shon, Zhifeng Yue, and Amita Sehgal. Modulation of RNA processing genes during sleep-dependent memory. *eLife*, 12, 2024.

[14262] Yue Li, Qinyao Sun, Shunli Zhu, Congying Chu, and Jiaojian Wang. Cross-species alignment along the chronological axis reveals evolutionary effect on structural development of the human brain. *eLife*, 13, 2024.

[14263] Zhi Li, Yuedi Wang, Qiqian Qin, Lanchun Chen, Xiaoqun Dang, Zhengang Ma, and Zeyang Zhou. Imidacloprid disrupts larval molting regulation and nutrient energy metabolism, causing developmental delay in honey bee Apis mellifera. *eLife*, 12, 2024.

[14264] Zhiming Li, Xingping Liu, Yan Zhang, Yuanyuan Li, Liquan Zhou, and Shuiqiao Yuan. FBXO24 modulates mRNA alternative splicing and MIWI degradation and is required for normal sperm formation and male fertility. *eLife*, 12, 2024.

[14265] Jie Liang, Yang Pan, Wenya Zhang, Darui Gao, Yongqian Wang, Wuxiang Xie, and Fanfan Zheng. Associations of age at diagnosis of breast cancer with incident myocardial infarction and heart failure: A prospective cohort study. *eLife*, 13, 2024.

[14266] Yucheng Liang, Jean-Emmanuel Hugonnet, Filippo Rusconi, and Michel Arthur. Peptidoglycan-tethered and free forms of the Braun lipoprotein are in dynamic equilibrium in Escherichia coli. *eLife*, 12, 2024.

[14267] Zilu Liang, Simeng Wu, Jie Wu, Wen-Xu Wang, Shaozheng Qin, and Chao Liu. Distance and grid-like codes support the navigation of abstract social space in the human brain. *eLife*, 12, 2024.

[14268] Hebin Liao, Xiaodan Yan, Chenyi Wang, Chun Huang, Wei Zhang, Leyi Xiao, Jun Jiang, Yongjia Bao, Tao Huang, Hanbo Zhang, Chunming Guo, Yufeng Zhang, and Yingying Pu. Cyclic di-GMP as an antitoxin regulates bacterial genome stability and antibiotic persistence in biofilms. *eLife*, 13, 2024.

[14269] Zian Liao, Suni Tang, Kaori Nozawa, Keisuke Shimada, Masahito Ikawa, Diana Monsivais, and Martin Matzuk. Affinity-tagged SMAD1 and SMAD5 mouse lines reveal transcriptional reprogramming mechanisms during early pregnancy. *eLife*, 12, 2024.

[14270] Simon M Lichtinger, Joanne L Parker, Simon Newstead, and Philip C Biggin. The mechanism of mammalian proton-coupled peptide transporters. *eLife*, 13, 2024.

[14271] Wanyoung Lim, Inwoo Hwang, Jiande Zhang, Zhenzhong Chen, Jeonghun Han, Jaehyung Jeon, Bon-Kyoung Koo, Sangmin Kim, Jeong Eon Lee, Youngkwan Kim, Kenneth J Pienta, Sarah R Amend, Robert H Austin, Jee-Yin Ahn, and Sungsu Park. Exploration of drug resistance mechanisms in triple negative breast cancer cells using a microfluidic device and patient tissues. *eLife*, 12, 2024.

[14272] David Yin-wei Lin, Lauren E Kueffer, Puneet Juneja, Thomas E Wales, John R Engen, and Amy H Andreotti. Conformational heterogeneity of the BTK PHTH domain drives multiple regulatory states. *eLife*, 12, 2024.

[14273] Miin S Lin, Se-Young Jo, Jens Luebeck, Howard Y Chang, Sihan Wu, Paul S Mischel, and Vineet Bafna. Transcriptional immune suppression and up-regulation of double-stranded DNA damage and repair repertoires in ecDNA-containing tumors. *eLife*, 12, 2024.

[14274] Shijia Lin, Yiwen Zhang, Shupei Zhang, Yijie Wei, Mengxue Han, Yamei Deng, Jiayi Guo, Biying Zhu, Tianyuan Yang, Enhua Xia, Xiaochun Wan, William J Lucas, and Zhaoliang Zhang. Root-specific theanine metabolism and regulation at the single-cell level in tea plants (Camellia sinensis). *eLife*, 13, 2024.

[14275] Xingcheng Lin and Bin Zhang. Explicit ion modeling predicts physicochemical interactions for chromatin organization. *eLife*, 12, 2024.

[14276] Ya-Hui Lin, Li-Wen Wang, Yen-Hui Chen, Yi-Chieh Chan, Shang-Hsiu Hu, Sheng-Yan Wu, Chi-Shiun Chiang, Guan-Jie Huang, Shang-Da Yang, Shi-Wei Chu, Kuo-Chuan Wang, Chin-Hsien Lin, Pei-Hsin Huang, Hwai-Jong Cheng, Bi-Chang Chen, and Li-An Chu. Revealing intact neuronal circuitry in centimeter-sized formalin-fixed paraffin-embedded brain. *eLife*, 13, 2024.

[14277] Yiechang Lin, Elaine Tao, James P Champion, and Ben Corry. A binding site for phosphoinositides described by multiscale simulations explains their modulation of voltage-gated sodium channels. *eLife*, 12, 2024.

[14278] James Lincoff, Cole VM Helsell, Frank V Marcoline, Andrew M Natale, and Michael Grabe. Membrane curvature sensing and symmetry breaking of the M2 proton channel from Influenza A. *eLife*, 13, 2024.

[14279] Jack W Lindsey and Elias B Issa. Factorized visual representations in the primate visual system and deep neural networks. *eLife*, 13, 2024.

[14280] Jack W Lindsey and Ashok Litwin-Kumar. Selective consolidation of learning and memory via recall-gated plasticity. *eLife*, 12, 2024.

[14281] Justin Lines, Andres Baraibar, Carmen Nanclares, Eduardo D Martin, Juan Aguilar, Paulo Kofuji, Marta Navarrete, and Alfonso Araque. A spatial threshold for astrocyte calcium surge. *eLife*, 12, 2024.

[14282] Harshavardhan Lingegowda, Katherine B Zutautas, Yuhong Wei, Priyanka Yolmo, Danielle J Sisnett, Alison McCallion, Madhuri Koti, and Chandrakant Tayade. Endocannabinoids and their receptors modulate endometriosis pathogenesis and immune response. *eLife*, 13, 2024.

[14283] Fabian Link, Alyssa Borges, Oliver Karo, Marvin Jungblut, Thomas Müller, Elisabeth Meyer-Natus, Timothy Krüger, Stefan Sachs, Nicola G Jones, Mary Morphew, Markus Sauer, Christian Stigloher, J Richard McIntosh, and Markus Engstler. Continuous endosomes form functional subdomains and orchestrate rapid membrane trafficking in trypanosomes. *eLife*, 12, 2024.

[14284] Fabian Link, Sisco Jung, Xenia Malzer, Felix Zierhut, Antonia Konle, Alyssa Borges, Christopher Batters, Monika Weiland, Mara Poellmann, An Binh Nguyen, Johannes Kullmann, Claudia Veigel, Markus Engstler, and Brooke Morriswood. The actomyosin system is essential for the integrity of the endosomal system in bloodstream form Trypanosoma brucei. *eLife*, 13, 2024.

[14285] John P Liska, Declan P Rowley, Trevor Thai Kim Nguyen, Jens-Oliver Muthmann, Daniel A Butts, Jacob Yates, and Alexander C Huk. Running modulates primate and rodent visual cortex differently. *eLife*, 12, 2024.

[14286] Jordan Little, Maria Chikina, and Nathan L Clark. Evolutionary rate covariation is a reliable predictor of co-functional interactions but not necessarily physical interactions. *eLife*, 12, 2024.

[14287] Andrew Liu, Jessica O'Connell, Farley Wall, and Richard W Carthew. Scaling between cell cycle duration and wing growth is regulated by Fat-Dachsous signaling in Drosophila. *eLife*, 12, 2024.

[14288] Baiwei Liu, Zampeta-Sofia Alexopoulou, and Freek van Ede. Jointly looking to the past and the future in visual working memory. *eLife*, 12, 2024.

[14289] Chinky Shiu Chen Liu, Tithi Mandal, Parijat Biswas, Md Asmaul Hoque, Purbita Bandopadhyay, Bishnu Prasad Sinha, Jafar Sarif, Ranit D'Rozario, Deepak Kumar Sinha, Bidisha Sinha, and Dipyaman Ganguly. Piezo1 mechanosensing regulates integrin-dependent chemotactic migration in human T cells. *eLife*, 12, 2024.

[14290] Gang Liu, Yunxuan Hou, Xin Jin, Yixue Zhang, Chaoyue Sun, Chengquan Huang, Yujie Ren, Jianmin Gao, Xiuli Wang, and Xiumei Jiang. PI3K/HSCB axis facilitates FOG1 nuclear translocation to promote erythropoiesis and megakaryopoiesis. *eLife*, 13, 2024.

[14291] Jian Liu, YuYing Jiang, ZiYue Diao, DanDan Chen, RuiYuan Xia, BingWei Wang, Shuo Yang, and ZhiQiang Yin. Gasdermin D-mediated neutrophil pyroptosis drives inflammation in psoriasis. *eLife*, 13, 2024.

[14292] Jingjing Liu, Yuedong Wang, Xian Liu, Junhai Han, and Yao Tian. Spatiotemporal changes in Netrin/Dscam1 signaling dictate axonal projection direction in Drosophila small ventral lateral clock neurons. *eLife*, 13, 2024.

[14293] Jinxin Liu, Haoyue He, Yanglingxi Wang, Jun Du, Kaixin Liang, Jun Xue, Yidan Liang, Peng Chen, Shanshan Tian, and Yongbing Deng. Predictive models for secondary epilepsy in patients with acute ischemic stroke within one year. *eLife*, 13, 2024.

[14294] Juan Liu, Morgan W Tingley, Qiang Wu, Peng Ren, Tinghao Jin, Ping Ding, and Xingfeng Si. Habitat fragmentation mediates the mechanisms underlying long-term climate-driven thermophilization in birds. *eLife*, 13, 2024.

[14295] Pengfei Liu, Xinyi Liu, and Bin Qi. UPRER–immunity axis acts as physiological food evaluation system that promotes aversion behavior in sensing low-quality food. *eLife*, 13, 2024.

[14296] Qiuling Liu, Hua Yang, Jianxian Luo, Cheng Peng, Ke Wang, Guowei Zhang, Hongsheng Lin, and Zhisheng Ji. 14-3-3 protein augments the protein stability of phosphorylated spastin and promotes the recovery of spinal cord injury through its agonist intervention. *eLife*, 12, 2024.

[14297] Silvia Liu, Yan-Ping Yu, Bao-Guo Ren, Tuval Ben-Yehezkel, Caroline Obert, Mat Smith, Wenjia Wang, Alina Ostrowska, Alejandro Soto-Gutierrez, and Jian-Hua Luo. Long-read single-cell sequencing reveals expressions of hypermutation clusters of isoforms in human liver cancer cells. *eLife*, 12, 2024.

[14298] Suying Liu, Sarah E Adams, Haotian Zheng, Juliana Ehnot, Seul K Jung, Greer Jeffrey, Theresa Menna, Louise Purton, Hongzhe Lee, and Peter Kurre. Dynamic tracking of native precursors in adult mice. *eLife*, 13, 2024.

[14299] Ting Liu, Xing Shen, Yijia Ren, Hongyu Lu, Yu Liu, Chong Chen, Lin Yu, and Zhihong Xue. Genome-wide mapping of native co-localized G4s and R-loops in living cells. *eLife*, 13, 2024.

[14300] Wenjing Liu, Shujin Li, Mu Yang, Jie Ma, Lu Liu, Ping Fei, Qianchun Xiang, Lulin Huang, Peiquan Zhao, Zhenglin Yang, and Xianjun Zhu. Dysfunction of Calcyphosine-Like gene impairs retinal angiogenesis through the MYC axis and is associated with familial exudative vitreoretinopathy. *eLife*, 13, 2024.

[14301] Yang Liu, Yi Jiang, Peng Qiu, Tie Ma, Yang Bai, Jiawen Bu, Yueting Hu, Ming Jin, Tong Zhu, and Xi Gu. RGS10 deficiency facilitates distant metastasis by inducing epithelial–mesenchymal transition in breast cancer. *eLife*, 13, 2024.

[14302] Yanqi Liu, Pol Bech, Keita Tamura, Lucas T Délez, Sylvain Crochet, and Carl CH Petersen. Cell class-specific long-range axonal projections of neurons in mouse whisker-related somatosensory cortices. *eLife*, 13, 2024.

[14303] Youxi Liu, Meimei Yin, Xiaoting Mao, Shuai Wu, Shuangping Wei, Shujun Heng, Yichun Yang, Jinwen Huang, Zhuolin Guo, Chuan Li, Chao Ji, Liu Hu, Wenjie Liu, and Ling-juan Zhang. Defining cell type-specific immune responses in a mouse model of allergic contact dermatitis by single-cell transcriptomics. *eLife*, 13, 2024.

[14304] Zheng Liu. A new potential strategy for cutaneous squamous cell carcinoma treatment by generating serum-based antibodies from tumor-exposed mice. *eLife*, 13, 2024.

[14305] Zhuang Liu, Thomas G Gillis, Srivatsan Raman, and Qiang Cui. A parameterized two-domain thermodynamic model explains diverse mutational effects on protein allostery. *eLife*, 12, 2024.

[14306] Sergi Llambrich, Birger Tielemans, Ellen Saliën, Marta Atzori, Kaat Wouters, Vicky Van Bulck, Mark Platt, Laure Vanherp, Nuria Gallego Fernandez, Laura Grau de la Fuente, Harish Poptani, Lieve Verlinden, Uwe Himmelreich, Anca Croitor, Catia Attanasio, Zsuzsanna Callaerts-Vegh, Willy

Gsell, Neus Martínez-Abadías, and Greetje Vande Velde. Pleiotropic effects of trisomy and pharmacologic modulation on structural, functional, molecular, and genetic systems in a Down syndrome mouse model. *eLife*, 12, 2024.

[14307] Fabio Lolicato, Julia P Steringer, Roberto Saleppico, Daniel Beyer, Jaime Fernandez-Sobaberas, Sebastian Unger, Steffen Klein, Petra Riegerová, Sabine Wegehingel, Hans-Michael Müller, Xiao J Schmitt, Shreyas Kaptan, Christian Freund, Martin Hof, Radek Šachl, Petr Chlanda, Ilpo Vattulainen, and Walter Nickel. Disulfide bridge-dependent dimerization triggers FGF2 membrane translocation into the extracellular space. *eLife*, 12, 2024.

[14308] Andrew T Lombardo, Cameron AR Mitchell, Riasat Zaman, David J McDermitt, and Anthony Bretscher. ARHGAP18-ezrin functions as an autoregulatory module for RhoA in the assembly of distinct actin-based structures. *eLife*, 13, 2024.

[14309] Filipa M Lopes, Celine Grenier, Benjamin W Jarvis, Sara Al Mahdy, Adrian Lène-McKay, Alison M Gurney, William G Newman, Simon N Waddington, Adrian S Woolf, and Neil A Roberts. Human HPSE2 gene transfer ameliorates bladder pathophysiology in a mutant mouse model of urofacial syndrome. *eLife*, 13, 2024.

[14310] Jon R Lorsch, Lawrence A Tabak, and Monica M Bertagnolli. Applied research won't flourish without basic science. *eLife*, 13, 2024.

[14311] Jörn Lötsch, Khayal Gasimli, Sebastian Malkusch, Lisa Hahnefeld, Carlo Angioni, Yannick Schreiber, Sandra Trautmann, Saskia Wedel, Dominique Thomas, Nerea Ferreiros Bouzas, Christian H Brandts, Benjamin Schnappauf, Christine Solbach, Gerd Geisslinger, and Marco Sisignano. Machine learning and biological validation identify sphingolipids as potential mediators of paclitaxel-induced neuropathy in cancer patients. *eLife*, 13, 2024.

[14312] Augustus J Lowry, Pengfei Liang, Mo Song, Yuichun Wan, Zhen-Ming Pei, Huanghe Yang, and Yang Zhang. TMEM16 and OSCA/TMEM63 proteins share a conserved potential to permeate ions and phospholipids. *eLife*, 13, 2024.

[14313] Nicolas Loyer, Elizabeth KJ Hogg, Hayley G Shaw, Anna Pasztor, David H Murray, Greg M Findlay, and Jens Januschke. A CDK1 phosphorylation site on Drosophila PAR-3 regulates neuroblast polarisation and sensory organ formation. *eLife*, 13, 2024.

[14314] Jiansen Lu, Jiahuan Zhang, Huaji Jiang, Zhiqiang Hu, Yufen Zhang, Lian He, Jianwu Yang, Yingchao Xie, Dan Wu, Hongyu Li, Ke Zeng, Peng Tan, Qingyue Xiao, Zijing Song, Chenglong Pan, Xiaochun Bai, and Xiao Yu. Vangl2 suppresses NF-$\kappa$B signaling and ameliorates sepsis by targeting p65 for NDP52-mediated autophagic degradation. *eLife*, 12, 2024.

[14315] Xiuyuan Lu, Hiroki Hayashi, Eri Ishikawa, Yukiko Takeuchi, Julian Vincent Tabora Dychiao, Hironori Nakagami, and Sho Yamasaki. Early acquisition of S-specific Tfh clonotypes after SARS-CoV-2 vaccination is associated with the longevity of anti-S antibodies. *eLife*, 12, 2024.

[14316] Yu-Ming Lu, Shan Yan, Shih-Chieh Ti, and Chaogu Zheng. Editing of endogenous tubulins reveals varying effects of tubulin posttranslational modifications on axonal growth and regeneration. *eLife*, 13, 2024.

[14317] Simon Lui, Ashleigh K Brink, and Laura H Corbit. Optogenetic stimulation of the locus coeruleus enhances appetitive extinction in rats. *eLife*, 12, 2024.

[14318] Pablo Lujan, Carla Garcia-Cabau, Yuichi Wakana, Javier Vera Lillo, Carmen Rodilla-Ramírez, Hideaki Sugiura, Vivek Malhotra, Xavier Salvatella, Maria F Garcia-Parajo, and Felix Campelo. Sorting of secretory proteins at the trans-Golgi network by human TGN46. *eLife*, 12, 2024.

[14319] Jie Luo and Jeff Ranish. Isobaric crosslinking mass spectrometry technology for studying conformational and structural changes in proteins and complexes. *eLife*, 13, 2024.

[14320] Yin Luo, Huanmeng Hao, Zhangjie Wang, Chih Yean Ong, Robert Dutcher, Yongmei Xu, Jian Liu, Lars C Pedersen, and Ding Xu. Heparan sulfate promotes TRAIL-induced tumor cell apoptosis. *eLife*, 12, 2024.

[14321] Yongfeng Luo, Ke Cao, Joanne Chiu, Hui Chen, Hong-Jun Wang, Matthew E Thornton, Brendan H Grubbs, Martin Kolb, Michael S Parmacek, Yuji Mishina, and Wei Shi. Defective mesenchymal Bmpr1a-mediated BMP signaling causes congenital pulmonary cysts. *eLife*, 12, 2024.

[14322] Andrea I Luppi, Pedro AM Mediano, Fernando E Rosas, Judith Allanson, John Pickard, Robin L Carhart-Harris, Guy B Williams, Michael M Craig, Paola Finoia, Adrian M Owen, Lorina Naci, David K Menon, Daniel Bor, and Emmanuel A Stamatakis. A synergistic workspace for human consciousness revealed by Integrated Information Decomposition. *eLife*, 12, 2024.

[14323] Bingjiang Lyu, William D Marslen-Wilson, Yuxing Fang, and Lorraine K Tyler. Finding structure during incremental speech comprehension. *eLife*, 12, 2024.

[14324] Bo Lyu and Qisheng Song. The intricate relationship of G-Quadruplexes and bacterial pathogenicity islands. *eLife*, 12, 2024.

[14325] Jixing Lyu, Tianqi Zhang, Michael T Marty, David Clemmer, David H Russell, and Arthur Laganowsky. Double and triple thermodynamic mutant cycles reveal the basis for specific MsbA-lipid interactions. *eLife*, 12, 2024.

[14326] Tongtong Ma, Ruimin Ren, Jianqi Lv, Ruipeng Yang, Xinyi Zheng, Yang Hu, Guiyu Zhu, and Heng Wang. Transdifferentiation of fibroblasts into muscle cells to constitute cultured meat with tunable intramuscular fat deposition. *eLife*, 13, 2024.

[14327] Wenxin Ma, Lian Zhao, Biying Xu, Robert N Fariss, T Michael Redmond, Jizhong Zou, Wai T Wong, and Wei Li. Human-induced pluripotent stem cell-derived microglia integrate into mouse retina and recapitulate features of endogenous microglia. *eLife*, 12, 2024.

[14328] Zhixin Ma, Wenshu Wang, Xiaojing Yang, Menglong Rui, and Su Wang. Glial ferritin maintains neural stem cells via transporting iron required for self-renewal in Drosophila. *eLife*, 13, 2024.

[14329] Thi Thom Mac, Teddy Fauquier, Nicolas Jullien, Pauline Romanet, Heather Etchevers, Anne Barlier, Frederic Castinetti, and Thierry Brue. Modeling corticotroph deficiency with pituitary organoids supports the functional role of NFKB2 in human pituitary differentiation. *eLife*, 12, 2024.

[14330] Matt MacAinsh, Souvik Dey, and Huan-Xiang Zhou. Direct and indirect salt effects on homotypic phase separation. *eLife*, 13, 2024.

[14331] Alastair MacDonald, Alina Hebling, Xin Paul Wei, and Kevin Yackle. The breath shape controls intonation of mouse vocalizations. *eLife*, 13, 2024.

[14332] J Wesley Maddox, Gregory J Ordemann, Juan AM de la Rosa Vázquez, Angie Huang, Christof Gault, Serena R Wisner, Kate Randall, Daiki Futagi, Nihal A Salem, Dayne Mayfield, Boris V Zemelman, Steven DeVries, Mrinalini Hoon, and Amy Lee. A non-conducting role of the Cav1.4 Ca2+ channel drives homeostatic plasticity at the cone photoreceptor synapse. *eLife*, 13, 2024.

[14333] Renan Maestri, Benoît Perez-Lamarque, Anna Zhukova, and Hélène Morlon. Recent evolutionary origin and localized diversity hotspots of mammalian coronaviruses. *eLife*, 13, 2024.

[14334] Silvia Maggi, Rebecca M Hock, Martin O'Neill, Mark Buckley, Paula M Moran, Tobias Bast, Musa Sami, and Mark D Humphries. Tracking subjects' strategies in behavioural choice experiments at trial resolution. *eLife*, 13, 2024.

[14335] Satyajit Mahapatra and Tomoyuki Takahashi. Physiological roles of endocytosis and presynaptic scaffold in vesicle replenishment at fast and slow central synapses. *eLife*, 12, 2024.

[14336] Rabina Mainali, Nancy Buechler, Cristian Otero, Laken Edwards, Chia-Chi Key, Cristina Furdui, and Matthew A Quinn. Itaconate stabilizes CPT1a to enhance lipid utilization during inflammation. *eLife*, 12, 2024.

[14337] Mariano Maio, Joaquina Barros, Marine Joly, Zoi Vahlas, José Luis Marín Franco, Melanie Genoula, Sarah C Monard, María Belén Vecchione, Federico Fuentes, Virginia Gonzalez Polo, María Florencia Quiroga, Mónica Vermeulen, Thien-Phong Vu Manh, Rafael J Argüello, Sandra Inwentarz, Rosa Musella, Lorena Ciallella, Pablo González Montaner, Domingo Palmero, Geanncarlo Lugo Villarino, María del Carmen Sasiain, Olivier Neyrolles, Christel Vérollet, and Luciana Balboa. Elevated glycolytic metabolism of monocytes limits the generation of HIF1A-driven migratory dendritic cells in tuberculosis. *eLife*, 12, 2024.

[14338] Ritwik Maity, Xuepei Zhang, Francesca Romana Liberati, Chiara Scribani Rossi, Francesca Cutruzzolá, Serena Rinaldo, Massimiliano Gaetani, José Antonio Aínsa, and Javier Sancho. Merging multiomics with proteome integral solubility alteration unveils antibiotic mode of action. *eLife*, 13, 2024.

[14339] Maryam Majeed, Haejun Han, Keren Zhang, Wen Xi Cao, Chien-Po Liao, Oliver Hobert, and Hang Lu. Toolkits for detailed and high-throughput interrogation of synapses in C. elegans. *eLife*, 12, 2024.

[14340] Jarno Mäkelä, Alexandros Papagiannakis, Wei-Hsiang Lin, Michael Charles Lanz, Skye Glenn, Matthew Swaffer, Georgi K Marinov, Jan M Skotheim, and Christine Jacobs-Wagner. Genome concentration limits cell growth and modulates proteome composition in Escherichia coli. *eLife*, 13, 2024.

[14341] James Malkin, Cian O'Donnell, Conor J Houghton, and Laurence Aitchison. Signatures of Bayesian inference emerge from energy-efficient synapses. *eLife*, 12, 2024.

[14342] Eugenio Manassero, Giulia Concina, Maria Clarissa Chantal Caraig, Pietro Sarasso, Adriana Salatino, Raffaella Ricci, and Benedetto Sacchetti. Medial anterior prefrontal cortex stimulation downregulates implicit reactions to threats and prevents the return of fear. *eLife*, 13, 2024.

[14343] Chenchen Mao, Yanyu Chen, Dong Xing, Teming Zhang, Yangxuan Lin, Cong Long, Jiaye Yu, Yunhui Luo, Tao Ming, Wangkai Xie, Zheng Han, Dianfeng Mei, Dan Xiang, Mingdong Lu, Xian Shen, and Xiangyang Xue. Resting natural killer cells promote the progress of colon cancer liver metastasis by elevating tumor-derived stem cell factor. *eLife*, 13, 2024.

[14344] Renbo Mao, Jianjun Yu, Bowen Deng, Xihuimin Dai, Yuyao Du, Sujie Du, Wenxia Zhang, and Yi Rao. Conditional chemoconnectomics (cCCTomics) as a strategy for efficient and conditional targeting of chemical transmission. *eLife*, 12, 2024.

[14345] Eleanor Martin, Rossana Girardello, Gunnar Dittmar, and Alexander Ludwig. Time-resolved proximity proteomics uncovers a membrane tension-sensitive caveolin-1 interactome at the rear of migrating cells. *eLife*, 13, 2024.

[14346] Hannah R Martin, Anna Lysakowski, and Ruth Anne Eatock. The potassium channel subunit KV1.8 (Kcna10) is essential for the distinctive outwardly rectifying conductances of type I and II vestibular hair cells. *eLife*, 13, 2024.

[14347] Nuria Martin Flores, Marina Podpolny, Faye McLeod, Isaac Workman, Karen Crawford, Dobril Ivanov, Ganna Leonenko, Valentina Escott-Price, and Patricia C Salinas. Downregulation of Dickkopf-3, a Wnt antagonist elevated in Alzheimer's disease, restores synapse integrity and memory in a disease mouse model. *eLife*, 12, 2024.

[14348] Miguel Martinez-Ara, Federico Comoglio, and Bas van Steensel. Large-scale analysis of the integration of enhancer-enhancer signals by promoters. *eLife*, 12, 2024.

[14349] Irene Martínez-Gallego, Heriberto Coatl-Cuaya, and Antonio Rodriguez-Moreno. Astrocytes mediate two forms of spike timing-dependent depression at entorhinal cortex-hippocampal synapses. *eLife*, 13, 2024.

[14350] Matthew R Marunde, Harrison A Fuchs, Jonathan M Burg, Irina K Popova, Anup Vaidya, Nathan W Hall, Ellen N Weinzapfel, Matthew J Meiners, Rachel Watson, Zachary B Gillespie, Hailey F Taylor, Laylo Mukhsinova, Ugochi C Onuoha, Sarah A Howard, Katherine Novitzky, Eileen T McAnarney, Krzysztof Krajewski, Martis W Cowles, Marcus A Cheek, Zu-Wen Sun, Bryan J Venters, Michael-C Keogh, and Catherine A Musselman. Nucleosome conformation dictates the histone code. *eLife*, 13, 2024.

[14351] Nicola Masala, Manuel Mittag, Eleonora Ambrad Giovannetti, Darik A O'Neil, Fabian J Distler, Peter Rupprecht, Fritjof Helmchen, Rafael Yuste, Martin Fuhrmann, Heinz Beck, Michael Wenzel, and Tony Kelly. Aberrant hippocampal Ca2+ microwaves following synapsin-dependent adeno-associated viral expression of Ca2+ indicators. *eLife*, 13, 2024.

[14352] Elena Massai, Marco Bonizzato, Isley De Jesus, Roxanne Drainville, and Marina Martinez. Cortical neuroprosthesis-mediated functional ipsilateral control of locomotion in rats with spinal cord hemisection. *eLife*, 12, 2024.

[14353] Laura Massoz, David Bergemann, Arnaud Lavergne, Célia Reynders, Caroline Désiront, Chiara Goossens, Lydie Flasse, Bernard Peers, Marianne M Voz, and Isabelle Manfroid. Negative cell cycle regulation by calcineurin is necessary for proper beta cell regeneration in zebrafish. *eLife*, 12, 2024.

[14354] Aya Matsui, Cathy Spangler, Johannes Elferich, Momoko Shiozaki, Nikki Jean, Xiaowei Zhao, Maozhen Qin, Haining Zhong, Zhiheng Yu, and Eric Gouaux. Cryo-electron tomographic investigation of native hippocampal glutamatergic synapses. *eLife*, 13, 2024.

[14355] Mami Matsuo-Takasaki, Sho Kambayashi, Yasuko Hemmi, Tamami Wakabayashi, Tomoya Shimizu, Yuri An, Hidenori Ito, Kazuhiro Takeuchi, Masato Ibuki, Terasu Kawashima, Rio Masayasu, Manami Suzuki, Yoshikazu Kawai, Masafumi Umekage, Tomoaki M Kato, Michiya Noguchi, Koji Nakade, Yukio Nakamura, Tomoyuki Nakaishi, Naoki Nishishita, Masayoshi Tsukahara, and Yohei Hayashi. Complete suspension culture of human induced pluripotent stem cells supplemented with suppressors of spontaneous differentiation. *eLife*, 12, 2024.

[14356] John J Maurer, Alexandra Lin, Xi Jin, Jiso Hong, Nicholas Sathi, Romain Cardis, Alejandro Osorio-Forero, Anita Lüthi, Franz Weber, and Shinjae Chung. Homeostatic regulation of rapid eye movement sleep by the preoptic area of the hypothalamus. *eLife*, 12, 2024.

[14357] Jonathan Mayzel and Elad Schneidman. Homeostatic synaptic normalization optimizes learning in network models of neural population codes. *eLife*, 13, 2024.

[14358] Clément Mazeaud, Stefan Pfister, Jonathan E Owen, Higor Sette Pereira, Flavie Charbonneau, Zachary E Robinson, Anaïs Anton, Cheyanne L Bemis, Aïssatou Aïcha Sow, Trushar R Patel, Christopher J Neufeldt, Pietro Scaturro, and Laurent Chatel-Chaix. Zika virus remodels and hijacks IGF2BP2 ribonucleoprotein complex to promote viral replication organelle biogenesis. *eLife*, 13, 2024.

[14359] Audrey T Medeiros, Scott J Gratz, Ambar Delgado, Jason T Ritt, and Kate M O'Connor-Giles. Ca2+ channel and active zone protein abundance intersects with input-specific synapse organization to shape functional synaptic diversity. *eLife*, 12, 2024.

[14360] Matias A Medina, Francisco Fuentes-Villalobos, Claudio Quevedo, Felipe Aguilera, Raul Riquelme, Maria Luisa Rioseco, Sebastian Barria, Yazmin Pinos, Mario Calvo, Ian Burbulis, Camila Kossack, Raymond A Alvarez, Jose Luis Garrido, and Maria Ines Barria. Longitudinal transcriptional changes reveal genes from the natural killer cell-mediated cytotoxicity pathway as critical players underlying COVID-19 progression. *eLife*, 13, 2024.

[14361] Olivia C Meisner, Weikang Shi, Nicholas A Fagan, Joel Greenwood, Monika P Jadi, Anirvan S Nandy, and Steve WC Chang. Development of a Marmoset Apparatus for Automated Pulling to study cooperative behaviors. *eLife*, 13, 2024.

[14362] Natalia Mendes, Ariane Zanesco, Cristhiane Aguiar, Gabriela F Rodrigues-Luiz, Dayana Silva, Jonathan Campos, Niels Olsen Saraiva Camara, Pedro Moraes-Vieira, Eliana Araujo, and Licio A Velloso. CXCR3-expressing myeloid cells recruited to the hypothalamus protect against diet-induced body mass gain and metabolic dysfunction. *eLife*, 13, 2024.

[14363] Yang Meng, Yue Ni, Zhuoran Li, Tianhang Jiang, Tianshu Sun, Yanjian Li, Xindi Gao, Hailong Li, Chenhao Suo, Chao Li, Sheng Yang, Tian Lan, Guojian Liao, Tongbao Liu, Ping Wang, and Chen Ding. Interplay between acetylation and ubiquitination of imitation switch chromatin remodeler Isw1 confers multidrug resistance in Cryptococcus neoformans. *eLife*, 13, 2024.

[14364] Debashish U Menon, Prabuddha Chakraborty, Noel Murcia, and Terry Magnuson. ARID1A governs the silencing of sex-linked transcription during male meiosis in the mouse. *eLife*, 12, 2024.

[14365] Sneha Menon, Subinoy Adhikari, and Jagannath Mondal. An integrated machine learning approach delineates an entropic expansion mechanism for the binding of a small molecule to $\alpha$-synuclein. *eLife*, 13, 2024.

[14366] Jonathan AC Menzies, André Maia Chagas, Tom Baden, and Claudio R Alonso. A microRNA that controls the emergence of embryonic movement. *eLife*, 13, 2024.

[14367] Salima Messaoudi, Ada Allam, Julie Stoufflet, Theo Paillard, Anaïs Le Ven, Coralie Fouquet, Mohamed Doulazmi, Alain Trembleau, and Isabelle Caille. FMRP regulates postnatal neuronal migration via MAP1B. *eLife*, 12, 2024.

[14368] Brian PH Metzger, Yeonwoo Park, Tyler N Starr, and Joseph W Thornton. Epistasis facilitates functional evolution in an ancient transcription factor. *eLife*, 12, 2024.

[14369] Nathaniel Paul Meyer, Tania Singh, Matthew L Kutys, Todd G Nystul, and Diane L Barber. Arp2/3 complex activity enables nuclear YAP for naïve pluripotency of human embryonic stem cells. *eLife*, 13, 2024.

[14370] Yue Miao, Yongtao Du, Baolei Wang, Jingjing Liang, Yu Liang, Song Dang, Jiahao Liu, Dong Li, Kangmin He, and Mei Ding. Spatiotemporal recruitment of the ubiquitin-specific protease USP8 directs endosome maturation. *eLife*, 13, 2024.

[14371] Felix Michaud, Ruggiero Francavilla, Dimitry Topolnik, Parisa Iloun, Suhel Tamboli, Frederic Calon, and Lisa Topolnik. Altered firing output of VIP interneurons and early dysfunctions in CA1 hippocampal circuits in the 3xTg mouse model of Alzheimer's disease. *eLife*, 13, 2024.

[14372] Matthew Millard, David W Franklin, and Walter Herzog. A three filament mechanistic model of musculotendon force and impedance. *eLife*, 12, 2024.

[14373] Case Vincent Miller, Jen A Bright, Xiaoli Wang, Xiaoting Zheng, and Michael Pittman. Synthetic analysis of trophic diversity and evolution in Enantiornithes with new insights from Bohaiornithidae. *eLife*, 12, 2024.

[14374] Ana Milovic, Jonathan V Duong, and Alan G Barbour. The infection-tolerant white-footed deermouse tempers interferon responses to endotoxin in comparison to the mouse and rat. *eLife*, 12, 2024.

[14375] Yoshihisa Mimura, Tomoya Yasujima, Katsuhisa Inoue, Shogo Akino, Chitaka Namba, Hiroyuki Kusuhara, Yutaro Sekiguchi, Kinya Ohta, Takahiro Yamashiro, and Hiroaki Yuasa. SLC35G1 is a highly chloride-sensitive transporter responsible for the basolateral membrane transport in intestinal citrate absorption. *eLife*, 13, 2024.

[14376] Kyounghee Min, Batuhan Yenilmez, Mark Kelly, Dimas Echeverria, Michael Elleby, Lawrence M Lifshitz, Naideline Raymond, Emmanouela Tsagkaraki, Shauna M Harney, Chloe DiMarzio, Hui Wang, Nicholas McHugh, Brianna Bramato, Brett Morrison, Jeffery D Rothstein, Anastasia Khvorova, and Michael P Czech. Lactate transporter MCT1 in hepatic stellate cells promotes fibrotic collagen expression in nonalcoholic steatohepatitis. *eLife*, 12, 2024.

[14377] Li Min, Fanqin Bu, Jingxin Meng, Xiang Liu, Qingdong Guo, Libo Zhao, Zhi Li, Xiangji Li, Shengtao Zhu, and Shutian Zhang. Circulating small extracellular vesicle RNA profiling for the detection of T1a stage colorectal cancer and precancerous advanced adenoma. *eLife*, 12, 2024.

[14378] Mina MiŠić, Noah Lee, Francesca Zidda, Kyungjin Sohn, Katrin Usai, Martin Löffler, Md Nasir Uddin, Arsalan Farooqi, Giovanni Schifitto, Zhengwu Zhang, Frauke Nees, Paul Geha, and Herta Flor. A multisite validation of brain white matter pathways of resilience to chronic back pain. *eLife*, 13, 2024.

[14379] Somen K Mistri, Brianna M Hilton, Katherine J Horrigan, Emma S Andretta, Remi Savard, Oliver Dienz, Kenneth J Hampel, Diana L Gerrard, Joshua T Rose, Nikoletta Sidiropoulos, Dev Majumdar, and Jonathan E Boyson. SLAM/SAP signaling regulates discrete $\gamma\delta$ T cell developmental checkpoints and shapes the innate-like $\gamma\delta$ TCR repertoire. *eLife*, 13, 2024.

[14380] Wayne Mitchell, Ludger JE Goeminne, Alexander Tyshkovskiy, Sirui Zhang, Julie Y Chen, Joao A Paulo, Kerry A Pierce, Angelina H Choy, Clary B Clish, Steven P Gygi, and Vadim N Gladyshev. Multi-omics characterization of partial chemical reprogramming reveals evidence of cell rejuvenation. *eLife*, 12, 2024.

[14381] Rishav Mitra, Shlesha Richhariya, and Gaiti Hasan. Orai-mediated calcium entry determines activity of central dopaminergic neurons by regulation of gene expression. *eLife*, 12, 2024.

[14382] Rinako Miyano, Hirokazu Sakamoto, Kenzo Hirose, and Takeshi Sakaba. RIM-BP2 regulates Ca2+ channel abundance and neurotransmitter release at hippocampal mossy fiber terminals. *eLife*, 12, 2024.

[14383] Ayumi Mizuno, Malgorzata Lagisz, Pietro Pollo, Yefeng Yang, Masayo Soma, and Shinichi Nakagawa. A systematic review and meta-analysis of eyespot anti-predator mechanisms. *eLife*, 13, 2024.

[14384] Marius E Moeller, Nathaniel V Mon Père, Benjamin Werner, and Weini Huang. Measures of genetic diversification in somatic tissues at bulk and single-cell resolution. *eLife*, 12, 2024.

[14385] Jaaved Mohammed, Neha Arora, Harold S Matthews, Karissa Hansen, Maram Bader, Susan Walsh, John R Shaffer, Seth M Weinberg, Tomek Swigut, Peter Claes, Licia Selleri, and Joanna Wysocka. A common cis-regulatory variant impacts normal-range and disease-associated human facial shape through regulation of PKDCC during chondrogenesis. *eLife*, 13, 2024.

[14386] Shakur Mohibi, Yanhong Zhang, Vivian Perng, Mingyi Chen, Jin Zhang, and Xinbin Chen. Ferredoxin 1 is essential for embryonic development and lipid homeostasis. *eLife*, 13, 2024.

[14387] Robyn D Moir, Emilio Merheb, Violeta Chitu, E Richard Stanley, and Ian M Willis. Molecular basis of neurodegeneration in a mouse model of Polr3-related disease. *eLife*, 13, 2024.

[14388] Dániel Molnár, éva Viola Surányi, Tamás Trombitás, Dóra Füzesi, Rita Hirmondó, and Judit Toth. Genetic stability of Mycobacterium smegmatis under the stress of first-line antitubercular agents. *eLife*, 13, 2024.

[14389] Shanka Subhra Mondal, Steven Frankland, Taylor W Webb, and Jonathan D Cohen. Determinantal point process attention over grid cell code supports out of distribution generalization. *eLife*, 12, 2024.

[14390] Armando Montoya-Garcia, Idaira M Guerrero-Fonseca, Sandra D Chanez-Paredes, Karina B Hernandez-Almaraz, Iliana I Leon-Vega, Angelica Silva-Olivares, Abigail Betanzos, Monica Mondragon-Castelan, Ricardo Mondragon-Flores, Citlaltepetl Salinas-Lara, Hilda Vargas-Robles, and Michael Schnoor. Arpin deficiency increases actomyosin contractility and vascular permeability. *eLife*, 12, 2024.

[14391] Merlin Monzel, Pitshaporn Leelaarporn, Teresa Lutz, Johannes Schultz, Sascha Brunheim, Martin Reuter, and Cornelia McCormick. Hippocampal-occipital connectivity reflects autobiographical memory deficits in aphantasia. *eLife*, 13, 2024.

[14392] Camille Morel, Eline Lemerle, Feng-Ching Tsai, Thomas Obadia, Nishit Srivastava, Maud Marechal, Audrey Salles, Marvin Albert, Caroline Stefani, Yvonne Benito, FranÇois Vandenesch, Christophe Lamaze, Stéphane Vassilopoulos, Matthieu Piel, Patricia Bassereau, David Gonzalez-Rodriguez, Cecile Leduc, and Emmanuel Lemichez. Caveolin-1 protects endothelial cells from extensive expansion of transcellular tunnel by stiffening the plasma membrane. *eLife*, 12, 2024.

[14393] Marcos Moreno-Aguilera, Alba M Neher, Mónica B Mendoza, Martin Dodel, Faraz K Mardakheh, Raúl Ortiz, and Carme Gallego. KIS counteracts PTBP2 and regulates alternative exon usage in neurons. *eLife*, 13, 2024.

[14394] Mia C Morrell, Ilya Nemenman, and Audrey Sederberg. Neural criticality from effective latent variables. *eLife*, 12, 2024.

[14395] Katie Morris, Edita Bulovaite, Takeshi Kaizuka, Sebastian Schnorrenberg, Candace T Adams, Noboru Komiyama, Lorena Mendive-Tapia, Seth GN Grant, and Mathew H Horrocks. Sequential replacement of PSD95 subunits in postsynaptic supercomplexes is slowest in the cortex. *eLife*, 13, 2024.

[14396] Mitchell P Morton, Sachira Denagamage, Isabel J Blume, John H Reynolds, Monika P Jadi, and Anirvan S Nandy. Brain state and cortical layer-specific mechanisms underlying perception at threshold. *eLife*, 12, 2024.

[14397] Benedikt M Mortzfeld, Shakti K Bhattarai, and Vanni Bucci. Novel class IIb microcins show activity against Gram-negative ESKAPE and plant pathogens. *eLife*, 13, 2024.

[14398] Piermatteo Morucci, Sanjeev Nara, Mikel Lizarazu, Clara Martin, and Nicola Molinaro. Language experience shapes predictive coding of rhythmic sound sequences. *eLife*, 12, 2024.

[14399] Flora Moujaes, Jie Lisa Ji, Masih Rahmati, Joshua B Burt, Charles Schleifer, Brendan D Adkinson, Aleksandar Savic, Nicole Santamauro, Zailyn Tamayo, Caroline Diehl, Antonija Kolobaric, Morgan Flynn, Nathalie Rieser, Clara Fonteneau, Terry Camarro, Junqian Xu, Youngsun Cho, Grega Repovs, Sarah K Fineberg, Peter T Morgan, Erich Seifritz, Franz X Vollenweider, John H Krystal, John D Murray, Katrin H Preller, and Alan Anticevic. Ketamine induces multiple individually distinct whole-brain functional connectivity signatures. *eLife*, 13, 2024.

[14400] Ayumu Mubuchi, Mina Takechi, Shunsuke Nishio, Tsukasa Matsuda, Yoshifumi Itoh, Chihiro Sato, Ken Kitajima, Hiroshi Kitagawa, and Shinji Miyata. Assembly of neuron- and radial glial-cell-derived extracellular matrix molecules promotes radial migration of developing cortical neurons. *eLife*, 12, 2024.

[14401] Henning Mühlenbeck, Yuko Tsutsui, Mark A Lemmon, Kyle W Bender, and Cyril Zipfel. Allosteric activation of the co-receptor BAK1 by the EFR receptor kinase initiates immune signaling. *eLife*, 12, 2024.

[14402] Ananda Kishore Mukherjee, Subhajit Dutta, Ankita Singh, Shalu Sharma, Shuvra Shekhar Roy, Antara Sengupta, Megha Chatterjee, Soujanya Vinayagamurthy, Sulochana Bagri, Divya Khanna, Meenakshi Verma, Dristhi Soni, Anshul Budharaja, Sagar Kailasrao Bhisade, Vivek Anand, Ahmad Perwez, Nija George, Mohammed Faruq, Ishaan Gupta, Radhakrishnan Sabarinathan, and Shantanu

Chowdhury. Telomere length sensitive regulation of interleukin receptor 1 type 1 (IL1R1) by the shelterin protein TRF2 modulates immune signalling in the tumour microenvironment. *eLife*, 13, 2024.

[14403] Nicholas J Mullen, Surendra K Shukla, Ravi Thakur, Sai Sundeep Kollala, Dezhen Wang, Nina Chaika, Juan F Santana, William R Miklavcic, Drew A LaBreck, Jayapal Reddy Mallareddy, David H Price, Amarnath Natarajan, Kamiya Mehla, David B Sykes, Michael A Hollingsworth, and Pankaj K Singh. DHODH inhibition enhances the efficacy of immune checkpoint blockade by increasing cancer cell antigen presentation. *eLife*, 12, 2024.

[14404] Karl S Muller, Kathryn Bonnen, Stephanie M Shields, Daniel P Panfili, Jonathan Matthis, and Mary M Hayhoe. Analysis of foothold selection during locomotion using terrain reconstruction. *eLife*, 12, 2024.

[14405] Yuho Murata, Tsubasa Nishi, Izumi Kaneko, Shiroh Iwanaga, and Masao Yuda. Coordinated regulation of gene expression in Plasmodium female gametocytes by two transcription factors. *eLife*, 12, 2024.

[14406] Jana Muroňová, Zine Eddine Kherraf, Elsa Giordani, Emeline Lambert, Simon Eckert, Caroline Cazin, Amir Amiri-Yekta, Magali Court, Geneviève Chevalier, Guillaume Martinez, Yasmine Neirijnck, Francoise Kühne, Lydia Wehrli, Nikolai Klena, Virginie Hamel, Lisa De Macedo, Jessica Escoffier, Paul Guichard, Charles Coutton, Selima Fourati Ben Mustapha, Mahmoud Kharouf, Anne-Pacale Bouin, Raoudha Zouari, Nicolas Thierry-Mieg, Serge Nef, Stefan Geimer, Corinne Loeuillet, Pierre F Ray, and Christophe Arnoult. Lack of CCDC146, a ubiquitous centriole and microtubule-associated protein, leads to non-syndromic male infertility in human and mouse. *eLife*, 12, 2024.

[14407] Pinanong Na-Phatthalung, Shumin Sun, Enjun Xie, Jia Wang, Junxia Min, and Fudi Wang. The zinc transporter Slc30a1 (ZnT1) in macrophages plays a protective role against attenuated Salmonella. *eLife*, 13, 2024.

[14408] Maximilian Nagel, Marco Niestroj, Rohini Bansal, David Fleck, Angelika Lampert, Romana Stopkova, Pavel Stopka, Yoram Ben-Shaul, and Marc Spehr. Deciphering the chemical language of inbred and wild mouse conspecific scents. *eLife*, 12, 2024.

[14409] Tamas L Nagy, Evelyn Strickland, and Orion D Weiner. Neutrophils actively swell to potentiate rapid migration. *eLife*, 12, 2024.

[14410] Sudershana Nair and Nicholas E Baker. Extramacrochaetae regulates Notch signaling in the Drosophila eye through non-apoptotic caspase activity. *eLife*, 12, 2024.

[14411] Yoko Nakai-Futatsugi, Jianshi Jin, Taisaku Ogawa, Noriko Sakai, Akiko Maeda, Ken-ichi Hironaka, Masakazu Fukuda, Hiroki Danno, Yuji Tanaka, Seiji Hori, Katsuyuki Shiroguchi, and Masayo Takahashi. Pigmentation level of human iPSC-derived RPE does not indicate a specific gene expression profile. *eLife*, 12, 2024.

[14412] Arya Y Nakhe, Prasanna K Dadi, Jinsun Kim, Matthew T Dickerson, Soma Behera, Jordyn R Dobson, Shristi Shrestha, Jean-Philippe Cartailler, Leesa Sampson, Mark A Magnuson, and David A Jacobson. The MODY-associated KCNK16 L114P mutation increases islet glucagon secretion and limits insulin secretion resulting in transient neonatal diabetes and glucose dyshomeostasis in adults. *eLife*, 12, 2024.

[14413] Tatyana O Nakonechnaya, Bruno Moltedo, Ekaterina V Putintseva, Sofya Leyn, Dmitry A Bolotin, Olga V Britanova, Mikhail Shugay, and Dmitriy M Chudakov. Convergence, plasticity, and tissue residence of regulatory T cell response via TCR repertoire prism. *eLife*, 12, 2024.

[14414] Matteo Napoli, Roland Immler, Ina Rohwedder, Valerio Lupperger, Johannes Pfabe, Mariano Gonzalez Pisfil, Anna Yevtushenko, Thomas Vogl, Johannes Roth, Melanie Salvermoser, Steffen Dietzel, Marjan Slak Rupnik, Carsten Marr, Barbara Walzog, Markus Sperandio, and Monika Pruenster. Cytosolic S100A8/A9 promotes Ca2+ supply at LFA-1 adhesion clusters during neutrophil recruitment. *eLife*, 13, 2024.

[14415] Ardalan Naseri, Degui Zhi, and Shaojie Zhang. Discovery of runs-of-homozygosity diplotype clusters and their associations with diseases in UK Biobank. *eLife*, 13, 2024.

[14416] Vikas Navratna, Arvind Kumar, Jaimin K Rana, and Shyamal Mosalaganti. Structure of the human heparan-$\alpha$-glucosaminide N-acetyltransferase (HGSNAT). *eLife*, 13, 2024.

[14417] Hitendra Negi, Aravind Ravichandran, Pritha Dasgupta, Shridivya Reddy, and Ranabir Das. Plasticity of the proteasome-targeting signal Fat10 enhances substrate degradation. *eLife*, 13, 2024.

[14418] Kien Xuan Ngo, Huong T Vu, Kenichi Umeda, Minh-Nhat Trinh, Noriyuki Kodera, and Taro Uyeda. Deciphering the actin structure-dependent preferential cooperative binding of cofilin. *eLife*, 13, 2024.

[14419] Thuy TM Ngo, Bailey Liu, Feng Wang, Aakash Basu, Carl Wu, and Taekjip Ha. Dependence of nucleosome mechanical stability on DNA mismatches. *eLife*, 13, 2024.

[14420] Tenzin Ngodup, Tomohiko Irie, Seán P Elkins, and Laurence O Trussell. The Na+ leak channel NALCN controls spontaneous activity and mediates synaptic modulation by $\alpha$2-adrenergic receptors in auditory neurons. *eLife*, 12, 2024.

[14421] Ai Nguyen, Huaying Zhao, Dulguun Myagmarsuren, Sanjana Srinivasan, Di Wu, Jiji Chen, Grzegorz Piszczek, and Peter Schuck. Modulation of biophysical properties of nucleocapsid protein in the mutant spectrum of SARS-CoV-2. *eLife*, 13, 2024.

[14422] Lena H Nguyen, Youfen Xu, Maanasi Nair, and Angelique Bordey. The mTOR pathway genes MTOR, Rheb, Depdc5, Pten, and Tsc1 have convergent and divergent impacts on cortical neuron development and function. *eLife*, 12, 2024.

[14423] Phuong-Khanh Nguyen and Louise Y Cheng. Drosophila medulla neuroblast termination via apoptosis, differentiation, and gliogenic switch is scheduled by the depletion of the neuroepithelial stem cell pool. *eLife*, 13, 2024.

[14424] Quynh Anh Thi Nguyen, Andrea Rocha, Ricky Chhor, Yuna Yamashita, Christian Stadler, Crystal Pontrello, Hongdian Yang, and Sachiko Haga-Yamanaka. Hypothalamic representation of the imminence of predator threat detected by the vomeronasal organ in mice. *eLife*, 12, 2024.

[14425] Nguyen Thi Khanh Nhu, Minh-Duy Phan, Steven J Hancock, Kate M Peters, Laura Alvarez-Fraga, Brian M Forde, Stacey B Andersen, Thyl Miliya, Patrick NA Harris, Scott A Beatson, Sanmarie Schlebusch, Haakon Bergh, Paul Turner, Annelie Brauner, Benita Westerlund-Wikström, Adam D Irwin, and Mark A Schembri. High-risk Escherichia coli clones that cause neonatal meningitis and association with recrudescent infection. *eLife*, 12, 2024.

[14426] Qasem Nick, Daniel J Gale, Corson Areshenkoff, Anouk De Brouwer, Joseph Nashed, Jeffrey Wammes, Tianyao Zhu, Randy Flanagan, Jonny Smallwood, and Jason Gallivan. Reconfigurations of cortical manifold structure during reward-based motor learning. *eLife*, 12, 2024.

[14427] Litong Nie, Chao Wang, Min Huang, Xiaoguang Liu, Xu Feng, Mengfan Tang, Siting Li, Qinglei Hang, Hongqi Teng, Xi Shen, Li Ma, Boyi Gan, and Junjie Chen. DePARylation is critical for S phase progression and cell survival. *eLife*, 12, 2024.

[14428] Savvas Nikolaou, Amelie Juin, Jamie A Whitelaw, Nikki R Paul, Loic Fort, Colin Nixon, Heather J Spence, Sheila Bryson, and Laura M Machesky. CYRI-B-mediated macropinocytosis drives metastasis via lysophosphatidic acid receptor uptake. *eLife*, 13, 2024.

[14429] Satoshi Ninagawa, Masaki Matsuo, Deng Ying, Shuichiro Oshita, Shinya Aso, Kazutoshi Matsushita, Mai Taniguchi, Akane Fueki, Moe Yamashiro, Kaoru Sugasawa, Shunsuke Saito, Koshi Imami, Yasuhiko Kizuka, Tetsushi Sakuma, Takashi Yamamoto, Hirokazu Yagi, Koichi Kato, and Kazutoshi Mori. UGGT1-mediated reglucosylation of N-glycan competes with ER-associated degradation of unstable and misfolded glycoproteins. *eLife*, 12, 2024.

[14430] Yohei Nitta, Jiro Osaka, Ryuto Maki, Satoko Hakeda-Suzuki, Emiko Suzuki, Satoshi Ueki, Takashi Suzuki, and Atsushi Sugie. Drosophila model to clarify the pathological significance of OPA1 in autosomal dominant optic atrophy. *eLife*, 12, 2024.

[14431] Haley R Noonan, Alexandra M Thornock, Julia Barbano, Michael E Xifaras, Chloe S Baron, Song Yang, Katherine Koczirka, Alicia M McConnell, and Leonard I Zon. A chronic signaling TGFb zebrafish reporter identifies immune response in melanoma. *eLife*, 13, 2024.

[14432] Roberto Notario Manzano, Thibault Chaze, Eric Rubinstein, Esthel Penard, Mariette Matondo, Chiara Zurzolo, and Christel Brou. Proteomic landscape of tunneling nanotubes reveals CD9 and CD81 tetraspanins as key regulators. *eLife*, 13, 2024.

[14433] Serena Notartomaso, Nico Antenucci, Mariacristina Mazzitelli, Xavier Rovira, Serena Boccella, Flavia Ricciardi, Francesca Liberatore, Xavier Gomez-Santacana, Tiziana Imbriglio, Milena Cannella, Charleine Zussy, Livio Luongo, Sabatino Maione, Cyril Goudet, Giuseppe Battaglia, Amadeu Llebaria, Ferdinando Nicoletti, and Volker Neugebauer. A 'double-edged' role for type-5 metabotropic glutamate receptors in pain disclosed by light-sensitive drugs. *eLife*, 13, 2024.

[14434] Elham Nourani, Louise Faure, Hester Brønnvik, Martina Scacco, Enrico Bassi, Wolfgang Fiedler, Martin U Grüebler, Julia S Hatzl, David Jenny, Andrea Roverselli, Petra Sumasgutner, Matthias Tschumi, Martin Wikelski, and Kamran Safi. Developmental stage shapes the realized energy landscape for a flight specialist. *eLife*, 13, 2024.

[14435] Senem Ntourmas, Martin Sachs, Petra Paclíková, Martina Brückner, Vítězslav Bryja, Jürgen Behrens, and Dominic B Bernkopf. Endogenous oligomer formation underlies DVL2 condensates and promotes Wnt/$\beta$-catenin signaling. *eLife*, 13, 2024.

[14436] Veronica Teresa Ober, George Boniface Githure, Yuri Volpato Santos, Sidney Becker, Gabriel Moya Munoz, Jérôme Basquin, Frank Schwede, Esben Lorentzen, and Michael Boshart. Purine nucleosides replace cAMP in allosteric regulation of PKA in trypanosomatid pathogens. *eLife*, 12, 2024.

[14437] Ruby Oberin, Sigrid Petautschnig, Ellen G Jarred, Zhipeng Qu, Tesha Tsai, Neil A Youngson, Gabrielle Pulsoni, Thi T Truong, Dilini Fernando, Heidi Bildsoe, Rheannon O Blücher, Maarten van den Buuse, David K Gardner, Natalie A Sims, David L Adelson, and Patrick S Western. Fetal growth delay caused by loss of non-canonical imprinting is resolved late in pregnancy and culminates in offspring overgrowth. *eLife*, 13, 2024.

[14438] Johanna Odenwald, Bernardo Gabiatti, Silke Braune, Siqi Shen, Martin Zoltner, and Susanne Kramer. Detection of TurboID fusion proteins by fluorescent streptavidin outcompetes antibody signals and visualises targets not accessible to antibodies. *eLife*, 13, 2024.

[14439] Mariann Oemisch and Hyojung Seo. Therapeutic doses of ketamine acutely attenuate the aversive effect of losses during decision-making. *eLife*, 12, 2024.

[14440] David Oestreicher, Shashank Chepurwar, Kathrin Kusch, Vladan Rankovic, Sangyong Jung, Nicola Strenzke, and Tina Pangrsic. CaBP1 and 2 enable sustained CaV1.3 calcium currents and synaptic transmission in inner hair cells. *eLife*, 13, 2024.

[14441] Kristine Oevel, Svea Hohensee, Atul Kumar, Irving Rosas-Brugada, Francesca Bartolini, Tolga Soykan, and Volker Haucke. Rho GTPase signaling and mDia facilitate endocytosis via presynaptic actin. *eLife*, 12, 2024.

[14442] Izumi Ohigashi, Andrea J White, Mei-Ting Yang, Sayumi Fujimori, Yu Tanaka, Alison Jacques, Hiroshi Kiyonari, Yosuke Matsushita, Sevilay Turan, Michael C Kelly, Graham Anderson, and Yousuke Takahama. Developmental conversion of thymocyte-attracting cells into self-antigen-displaying cells in embryonic thymus medulla epithelium. *eLife*, 12, 2024.

[14443] Trine Line Hauge Okholm, Andreas Bjerregaard Kamstrup, Morten Muhlig Nielsen, Anne Kruse Hollensen, Mette Laugesen Graversgaard, Matilde Helbo Sørensen, Lasse Sommer Kristensen, Søren Vang, Samuel S Park, Eugene Yeo, Lars Dyrskjøt, Jørgen Kjems, Jakob Skou Pedersen, and Christian Kroun Damgaard. circHIPK3 nucleates IGF2BP2 and functions as a competing endogenous RNA. *eLife*, 13, 2024.

[14444] James D O'Leary, Rasmus Bruckner, Livia Autore, and Tomás J Ryan. Natural forgetting reversibly modulates engram expression. *eLife*, 12, 2024.

[14445] Maxim Oleynikov and Samie R Jaffrey. RNA tertiary structure and conformational dynamics revealed by BASH MaP. *eLife*, 13, 2024.

[14446] Cristina Olivieri, Yingjie Wang, Caitlin Walker, Manu Veliparambil Subrahmanian, Kim N Ha, David Bernlohr, Jiali Gao, Carlo Camilloni, Michele Vendruscolo, Susan S Taylor, and Gianluigi Veglia. The $\alpha$C-$\beta$4 loop controls the allosteric cooperativity between nucleotide and substrate in the catalytic subunit of protein kinase A. *eLife*, 12, 2024.

[14447] Krzysztof Hubert Olszyński, Rafał Polowy, Agnieszka Diana Wardak, Izabela Anna Łaska, Aneta Wiktoria Grymanowska, Wojciech Puławski, Olga Gawryś, Michał Koliński, and Robert Kuba Filipkowski. Male rats emit aversive 44-kHz ultrasonic vocalizations during prolonged Pavlovian fear conditioning. *eLife*, 12, 2024.

[14448] Kazuya Ono, Amandine Jarysta, Natasha C Hughes, Alma Jukic, Hui Ho Vanessa Chang, Michael R Deans, Ruth Anne Eatock, Kathleen E Cullen, Katie S Kindt, and Basile Tarchini. Contributions of mirror-image hair cell orientation to mouse otolith organ and zebrafish neuromast function. *eLife*, 13, 2024.

[14449] Jakub Onysk, Nicholas Gregory, Mia Whitefield, Maeghal Jain, Georgia Turner, Ben Seymour, and Flavia Mancini. Statistical learning shapes pain perception and prediction independently of external cues. *eLife*, 12, 2024.

[14450] Johannes Oppermann, Andrey Rozenberg, Thomaz Fabrin, Cristian González-Cabrera, Rafael Parker, Oded Béjà, Matthias Prigge, and Peter Hegemann. Robust optogenetic inhibition with red-light-sensitive anion-conducting channelrhodopsins. *eLife*, 12, 2024.

[14451] David O'Reilly and Ioannis Delis. Dissecting muscle synergies in the task space. *eLife*, 12, 2024.

[14452] Yuval Orsher, Ariel Rom, Rotem Perel, Yoav Lahini, Pablo Blinder, and Mark Shein-Idelson. Sequentially activated discrete modules appear as traveling waves in neuronal measurements with limited spatiotemporal sampling. *eLife*, 12, 2024.

[14453] Nicholas James Ose, Paul Campitelli, Tushar Modi, I Can Kazan, Sudhir Kumar, and Sefika Banu Ozkan. Some mechanistic underpinnings of molecular adaptations of SARS-COV-2 spike protein by integrating candidate adaptive polymorphisms with protein dynamics. *eLife*, 12, 2024.

[14454] Matthew Osmond and Graham Coop. Estimating dispersal rates and locating genetic ancestors with genome-wide genealogies. *eLife*, 13, 2024.

[14455] Benson Otarigho, Anna Frances Butts, and Alejandro Aballay. Neuronal NPR-15 modulates molecular and behavioral immune responses via the amphid sensory neuron-intestinal axis in C. elegans. *eLife*, 12, 2024.

[14456] Santiago Otero-Coronel, Thomas Preuss, and Violeta Medan. Multisensory integration enhances audiovisual responses in the Mauthner cell. *eLife*, 13, 2024.

[14457] Javier I Ottaviani, Virag Sagi-Kiss, Hagen Schroeter, and Gunter GC Kuhnle. Reliance on self-reports and estimated food composition data in nutrition research introduces significant bias that can only be addressed with biomarkers. *eLife*, 13, 2024.

[14458] Fouad Ouasti, Maxime Audin, Karine Fréon, Jean-Pierre Quivy, Mehdi Tachekort, Elizabeth Cesard, Aurélien Thureau, Virginie Ropars, Paloma Fernández Varela, Gwenaelle Moal, Ibrahim Soumana-Amadou, Aleksandra Uryga, Pierre Legrand, Jessica Andreani, Raphaël Guerois, Geneviève Almouzni, Sarah Lambert, and Francoise Ochsenbein. Disordered regions and folded modules in CAF-1 promote histone deposition in Schizosaccharomyces pombe. *eLife*, 12, 2024.

[14459] Yeyun Ouyang, Mi-Young Jeong, Corey N Cunningham, Jordan A Berg, Ashish G Toshniwal, Casey E Hughes, Kristina Seiler, Jonathan G Van Vranken, Ahmad A Cluntun, Geanette Lam, Jacob M Winter, Emel Akdogan, Katja K Dove, Sara M Nowinski, Matthew West, Greg Odorizzi, Steven P Gygi, Cory D Dunn, Dennis R Winge, and Jared Rutter. Phosphate starvation signaling increases mitochondrial membrane potential through respiration-independent mechanisms. *eLife*, 13, 2024.

[14460] Muge Ozker, Leyao Yu, Patricia Dugan, Werner Doyle, Daniel Friedman, Orrin Devinsky, and Adeen Flinker. Speech-induced suppression and vocal feedback sensitivity in human cortex. *eLife*, 13, 2024.

[14461] Kamil Pabis, Diogo Barardo, Olga Sirbu, Kumar Selvarajoo, Jan Gruber, and Brian K Kennedy. A concerted increase in readthrough and intron retention drives transposon expression during aging and senescence. *eLife*, 12, 2024.

[14462] Mátyás Paczkó, Eörs Szathmáry, and András Szilágyi. Stochastic parabolic growth promotes coexistence and a relaxed error threshold in RNA-like replicator populations. *eLife*, 13, 2024.

[14463] Zahid Padamsey, Danai Katsanevaki, Patricia Maeso, Manuela Rizzi, Emily E Osterweil, and Nathalie L Rochefort. Sex-specific resilience of neocortex to food restriction. *eLife*, 12, 2024.

[14464] Matthew C Pahl, Prabhat Sharma, Rajan M Thomas, Zachary Thompson, Zachary Mount, James A Pippin, Peter A Morawski, Peng Sun, Chun Su, Daniel Campbell, Struan FA Grant, and Andrew D Wells. Dynamic chromatin architecture identifies new autoimmune-associated enhancers for IL2 and novel genes regulating CD4+ T cell activation. *eLife*, 13, 2024.

[14465] Veronika Pak, Quadri Adewale, Danilo Bzdok, Mahsa Dadar, Yashar Zeighami, and Yasser Iturria-Medina. Distinctive whole-brain cell types predict tissue damage patterns in thirteen neurodegenerative conditions. *eLife*, 12, 2024.

[14466] Johannes Paladini, Annalena Maier, Judith Maria Habazettl, Ines Hertel, Rajesh Sonti, and Stephan Grzesiek. The molecular basis of Abelson kinase regulation by its $\alpha$I-helix. *eLife*, 12, 2024.

[14467] Dayu Pan, Kheiria Gamal Benkato, Xuequan Han, Jinjian Zheng, Vijay Kumar, Mei Wan, Junying Zheng, and Xu Cao. Senescence of endplate osteoclasts induces sensory innervation and spinal pain. *eLife*, 12, 2024.

[14468] Jiabin Pan, Rui Zhou, Lin-Lin Yao, Jie Zhang, Ning Zhang, Qing-Juan Cao, Shaopeng Sun, and Xiang-dong Li. Identification of a third myosin-5a-melanophilin interaction that mediates the association of myosin-5a with melanosomes. *eLife*, 13, 2024.

[14469] Meng-Hao Pan, Kun-Huan Zhang, Si-Le Wu, Zhen-Nan Pan, Ming-Hong Sun, Xiao-Han Li, Jia-Qian Ju, Shi-Ming Luo, Xiang-Hong Ou, and Shao-Chen Sun. FMNL2 regulates actin for endoplasmic reticulum and mitochondria distribution in oocyte meiosis. *eLife*, 12, 2024.

[14470] Yali Pan, Steven Frisson, Kara D Federmeier, and Ole Jensen. Early parafoveal semantic integration in natural reading. *eLife*, 12, 2024.

[14471] Lan Pang, Gangqi Fang, Zhiguo Liu, Zhi Dong, Jiani Chen, Ting Feng, Qichao Zhang, Yifeng Sheng, Yueqi Lu, Ying Wang, Yixiang Zhang, Guiyun Li, Xuexin Chen, Shuai Zhan, and Jianhua Huang. Coordinated molecular and ecological adaptations underlie a highly successful parasitoid. *eLife*, 13, 2024.

[14472] Marco Paoli, Antoine Wystrach, Brice Ronsin, and Martin Giurfa. Analysis of fast calcium dynamics of honey bee olfactory coding. *eLife*, 13, 2024.

[14473] Irini Papazian, Maria Kourouvani, Anastasia Dagkonaki, Vasileios Gouzouasis, Lila Dimitrakopoulou, Nikolaos Markoglou, Fotis Badounas, Theodore Tselios, Maria Anagnostouli, and Lesley Probert. Spontaneous human CD8 T cell and autoimmune encephalomyelitis-induced CD4/CD8 T cell lesions in the brain and spinal cord of HLA-DRB1*15-positive multiple sclerosis humanized immune system mice. *eLife*, 12, 2024.

[14474] Chan-Su Park, Jian Guan, Peter Rhee, Federico Gonzalez, Hee-sung Lee, Ji-hyun Park, Laurent Coscoy, Ellen A Robey, Nilabh Shastri, and Scheherazade Sadegh-Nasseri. Fam49b dampens TCR signal strength to regulate survival of positively selected thymocytes and peripheral T cells. *eLife*, 13, 2024.

[14475] Chung Park, Il-Young Hwang, Serena Li-Sue Yan, Sinmanus Vimonpatranon, Danlan Wei, Don Van Ryk, Alexandre Girard, Claudia Cicala, James Arthos, and John H Kehrl. Murine alveolar macrophages rapidly accumulate intranasally administered SARS-CoV-2 Spike protein leading to neutrophil recruitment and damage. *eLife*, 12, 2024.

[14476] Jongkeun Park, WonJong Choi, Do Young Seong, Seungpil Jeong, Ju Young Lee, Hyo Jeong Park, Dae Sun Chung, Kijong Yi, Uijin Kim, Ga-Yeon Yoon, Hyeran Kim, Taehoon Kim, Sooyeon Ko, Eun Jeong Min, Hyun-Soo Cho, Nam-Hyeok Cho, and Dongwan Hong. Accurate predictions of SARS-CoV-2 infectivity from comprehensive analysis. *eLife*, 13, 2024.

[14477] Kiwon Park, Dohoon Lee, Jiseok Jeong, Sungwon Lee, Sun Kim, and Kwangseog Ahn. Human immunodeficiency virus-1 induces host genomic R-loops and preferentially integrates its genome near the R-loop regions. *eLife*, 13, 2024.

[14478] Sanggeon Park, Yeowool Huh, Jeansok J Kim, and Jeiwon Cho. Bidirectional fear modulation by discrete anterior insular circuits in male mice. *eLife*, 13, 2024.

[14479] Won Yeong Park, Jieun Yun, Jinho Shin, Byung Ho Oh, Gilsuk Yoon, Seung-Mo Hong, and Ki Hean Kim. Open-top Bessel beam two-photon light sheet microscopy for three-dimensional pathology. *eLife*, 12, 2024.

[14480] Gina Partipilo, Yang Gao, Alexis J Holwerda, Yassir Lekbach, Ismar E Miniel Mahfoud, Trevor R Simmons, Bailey M Tibbett, Rebecca E Wilen, Marcus S Benyamin, and Benjamin K Keitz. Teaching troubleshooting skills to graduate students. *eLife*, 13, 2024.

[14481] Firdousi Parvez, Devika Sangpal, Harshad Paithankar, Zainab Amin, and Jeetender Chugh. Differential conformational dynamics in two type-A RNA-binding domains drive the double-stranded RNA recognition and binding. *eLife*, 13, 2024.

[14482] Stefan Passlick, Ghanim Ullah, and Christian Henneberger. Bidirectional dysregulation of synaptic glutamate signaling after transient metabolic failure. *eLife*, 13, 2024.

[14483] Krutik Patel, Matt Barter, Jamie Soul, Peter Clark, Carole Proctor, Ian Clark, David Young, and Daryl P Shanley. Systems analysis of miR-199a/b-5p and multiple miR-199a/b-5p targets during chondrogenesis. *eLife*, 12, 2024.

[14484] Samuel Pattillo Smith, Gregory Darnell, Dana Udwin, Julian Stamp, Arbel Harpak, Sohini Ramachandran, and Lorin Crawford. Discovering non-additive heritability using additive GWAS summary statistics. *eLife*, 13, 2024.

[14485] Samuel Pawel, Rachel Heyard, Charlotte Micheloud, and Leonhard Held. Replication of null results: Absence of evidence or evidence of absence? *eLife*, 12, 2024.

[14486] Shristi Pawnikar, Brenda S Magenheimer, Keya Joshi, Ericka Nevarez-Munoz, Allan Haldane, Robin L Maser, and Yinglong Miao. Activation of polycystin-1 signaling by binding of stalk-derived peptide agonists. *eLife*, 13, 2024.

[14487] Hannah L Payne, Jennifer L Raymond, and Mark S Goldman. Interactions between circuit architecture and plasticity in a closed-loop cerebellar system. *eLife*, 13, 2024.

[14488] Eyal Paz, Sahil Jain, Irit Gottfried, Orna Staretz-Chacham, Muhammad Mahajnah, Pritha Bagchi, Nicholas T Seyfried, Uri Ashery, and Abdussalam Azem. Biochemical and neurophysiological effects of deficiency of the mitochondrial import protein TIMM50. *eLife*, 13, 2024.

[14489] Wenyu Peng, Pan Wang, Chaoyang Tan, Han Zhao, Kun Chen, Huaxing Si, Yuchen Tian, Anxin Lou, Zhi Zhu, Yifang Yuan, Kaijie Wu, Chao Chang, Yuanming Wu, and Tao Chen. High-frequency terahertz stimulation alleviates neuropathic pain by inhibiting the pyramidal neuron activity in the anterior cingulate cortex of mice. *eLife*, 13, 2024.

[14490] Yunhui Peng, Wei Song, Vladimir B Teif, Ivan Ovcharenko, David Landsman, and Anna R Panchenko. Detection of new pioneer transcription factors as cell-type-specific nucleosome binders. *eLife*, 12, 2024.

[14491] Anne-Sophie Pepin, Patrycja A Jazwiec, Vanessa Dumeaux, Deborah M Sloboda, and Sarah Kimmins. Determining the effects of paternal obesity on sperm chromatin at histone H3 lysine 4 tri-methylation in relation to the placental transcriptome and cellular composition. *eLife*, 13, 2024.

[14492] Edgar M Pera, Josefine Nilsson-De Moura, Yuriy Pomeshchik, Laurent Roybon, and Ivana Milas. Inhibition of the serine protease HtrA1 by SerpinE2 suggests an extracellular proteolytic pathway in the control of neural crest migration. *eLife*, 12, 2024.

[14493] Pirunthan Perampalam, James I MacDonald, Komila Zakirova, Daniel T Passos, Sumaiyah Wasif, Yudith Ramos-Valdes, Maeva Hervieu, Patrick Mehlen, Rob Rottapel, Benjamin Gibert, Rohann JM Correa, Trevor G Shepherd, and Frederick A Dick. Netrin signaling mediates survival of dormant epithelial ovarian cancer cells. *eLife*, 12, 2024.

[14494] David Pérez-González, Ana Belén Lao-Rodríguez, Cristian Aedo-Sánchez, and Manuel S Malmierca. Acetylcholine modulates the precision of prediction error in the auditory cortex. *eLife*, 12, 2024.

[14495] Amanda C Perofsky, John Huddleston, Chelsea L Hansen, John R Barnes, Thomas Rowe, Xiyan Xu, Rebecca Kondor, David E Wentworth, Nicola Lewis, Lynne Whittaker, Burcu Ermetal, Ruth Harvey, Monica Galiano, Rodney Stuart Daniels, John W McCauley, Seiichiro Fujisaki, Kazuya Nakamura, Noriko Kishida, Shinji Watanabe, Hideki Hasegawa, Sheena G Sullivan, Ian G Barr, Kanta Subbarao, Florian Krammer, Trevor Bedford, and Cécile Viboud. Antigenic drift and subtype interference shape A(H3N2) epidemic dynamics in the United States. *eLife*, 13, 2024.

[14496] Simon Perrin, Maria Ethel, Vincent Bretegnier, Cassandre Goachet, Cécile-Aurore Wotawa, Marine Luka, Fanny Coulpier, Cécile Masson, Mickael Ménager, and Céline Colnot. Single-nucleus transcriptomics reveal the differentiation trajectories of periosteal skeletal/stem progenitor cells in bone regeneration. *eLife*, 13, 2024.

[14497] Jacques Pesnot Lerousseau and Christopher Summerfield. Space as a scaffold for rotational generalisation of abstract concepts. *eLife*, 13, 2024.

[14498] Paolo Petazzi, Telma Ventura, Francesca Paola Luongo, Heather McClafferty, Alisha May, Helen Alice Taylor, Michael J Shipston, Nicola Romanò, Lesley M Forrester, Pablo Menendez, and Antonella Fidanza. A novel human pluripotent stem cell gene activation system identifies IGFBP2 as a mediator in the production of haematopoietic progenitors in vitro. *eLife*, 13, 2024.

[14499] Colin H Peters, Rohit K Singh, Avery A Langley, William G Nichols, Hannah R Ferris, Danielle A Jeffrey, Catherine Proenza, and John R Bankston. LRMP inhibits cAMP potentiation of HCN4 channels by disrupting intramolecular signal transduction. *eLife*, 12, 2024.

[14500] E Nicholas Petersen, Mahmud Arif Pavel, Samuel S Hansen, Manasa Gudheti, Hao Wang, Zixuan Yuan, Keith R Murphy, William Ja, Heather A Ferris, Erik Jorgensen, and Scott B Hansen. Mechanical activation of TWIK-related potassium channel by nanoscopic movement and rapid second messenger signaling. *eLife*, 12, 2024.

[14501] Marvin Petersen, Felix Hoffstaedter, Felix L Nägele, Carola Mayer, Maximilian Schell, D Leander Rimmele, Birgit-Christiane Zyriax, Tanja Zeller, Simone Kühn, Jürgen Gallinat, Jens Fiehler, Raphael Twerenbold, Amir Omidvarnia, Kaustubh R Patil, Simon B Eickhoff, Goetz Thomalla, and Bastian Cheng. A latent clinical-anatomical dimension relating metabolic syndrome to brain structure and cognition. *eLife*, 12, 2024.

[14502] Ralph E Peterson, Aman Choudhri, Catalin Mitelut, Aramis Tanelus, Athena Capo-Battaglia, Alex H Williams, David M Schneider, and Dan H Sanes. Unsupervised discovery of family specific vocal usage in the Mongolian gerbil. *eLife*, 12, 2024.

[14503] Sophie Peterson, Amanda Maheras, Brenda Wu, Jose Chavira, and Ronald Keiflin. Sex differences in discrimination behavior and orbitofrontal engagement during context-gated reward prediction. *eLife*, 12, 2024.

[14504] Isabelle Petit-Hartlein, Annelise Vermot, Michel Thepaut, Anne-Sophie Humm, Florine Dupeux, Jerome Dupuy, Vincent Chaptal, Jose Antonio Marquez, Susan ME Smith, and Franck Fieschi. X-ray structure and enzymatic study of a bacterial NADPH oxidase highlight the activation mechanism of eukaryotic NOX. *eLife*, 13, 2024.

[14505] Céline Petitgas, Laurent Seugnet, Amina Dulac, Giorgio Matassi, Ali Mteyrek, Rebecca Fima, Marion Strehaiano, Joana Dagorret, Baya Chérif-Zahar, Sandrine Marie, Irène Ceballos-Picot, and Serge Birman. Metabolic and neurobehavioral disturbances induced by purine recycling deficiency in Drosophila. *eLife*, 12, 2024.

[14506] Gordon H Petty and Randy M Bruno. Attentional modulation of secondary somatosensory and visual thalamus of mice. *eLife*, 13, 2024.

[14507] Laurie Peverini, Sophie Shi, Karima Medjebeur, and Pierre-Jean Corringer. Mapping the molecular motions of 5-HT3 serotonin-gated channel by voltage-clamp fluorometry. *eLife*, 12, 2024.

[14508] Cuong Pham, Yuji Komaki, Anna Deàs-Just, Benjamin Le Gac, Christine Mouffle, Clara Franco, Agnès Chaperon, Vincent Vialou, Tomokazu Tsurugizawa, Bruno Cauli, and Dongdong Li. Astrocyte aquaporin mediates a tonic water efflux maintaining brain homeostasis. *eLife*, 13, 2024.

[14509] Thi Mong Quynh Pham, Thanh Nhan Nguyen, Bui Que Tran Nguyen, Thi Phuong Diem Tran, Nguyen My Diem Pham, Hoang Thien Phuc Nguyen, Thi Kim Cuong Ho, Dinh Viet Linh Nguyen, Huu Thinh Nguyen, Duc Huy Tran, Thanh Sang Tran, Truong Vinh Ngoc Pham, Minh Triet Le, Thi Tuong Vy Nguyen, Minh-Duy Phan, Hoa Giang, Hoai-Nghia Nguyen, and Le Son Tran. The T cell receptor $\beta$ chain repertoire of tumor infiltrating lymphocytes improves neoantigen prediction and prioritization. *eLife*, 13, 2024.

[14510] Tien M Phan, Young C Kim, Galia T Debelouchina, and Jeetain Mittal. Interplay between charge distribution and DNA in shaping HP1 paralog phase separation and localization. *eLife*, 12, 2024.

[14511] Jonathan E Phillips and Duojia Pan. The Hippo kinase cascade regulates a contractile cell behavior and cell density in a close unicellular relative of animals. *eLife*, 12, 2024.

[14512] Amber R Philp, Carolina R Reyes, Josselyne Mansilla, Amar Sharma, Chao Zhao, Carlos Valenzuela-Krugmann, Khalil S Rawji, Ginez A Gonzalez Martinez, Penelope Dimas, Bryan Hinrichsen, César Ulloa-Leal, Amie K Waller, Diana M Bessa de Sousa, Maite A Castro, Ludwig Aigner, Pamela Ehrenfeld, Maria Elena Silva, Ilias Kazanis, Cedric Ghevaert, Robin JM Franklin, and Francisco J Rivera. Circulating platelets modulate oligodendrocyte progenitor cell differentiation during remyelination. *eLife*, 12, 2024.

[14513] Marie-Eve Picard, Miriam Kunz, Jen-I Chen, Michel-Pierre Coll, Etienne Vachon-Presseau, Tor D Wager, and Pierre Rainville. A distributed brain response predicting the facial expression of acute nociceptive pain. *eLife*, 12, 2024.

[14514] Jose Mario Bello Pineda and Robert K Bradley. DUX4 is a common driver of immune evasion and immunotherapy failure in metastatic cancers. *eLife*, 12, 2024.

[14515] Ivan Pinos, Johana Coronel, Asma'a Albakri, Amparo Blanco, Patrick McQueen, Donald Molina, JaeYoung Sim, Edward A Fisher, and Jaume Amengual. $\beta$-Carotene accelerates the resolution of atherosclerosis in mice. *eLife*, 12, 2024.

[14516] Lenore Pipes and Rasmus Nielsen. A rapid phylogeny-based method for accurate community profiling of large-scale metabarcoding datasets. *eLife*, 13, 2024.

[14517] Annette Pisanski, Mitchell Prostebby, Clayton T Dickson, and Silvia Pagliardini. Mapping responses to focal injections of bicuculline in the lateral parafacial region identifies core regions for maximal generation of active expiration. *eLife*, 13, 2024.

[14518] Magdalena Podkowik, Andrew I Perault, Gregory Putzel, Andrew Pountain, Jisun Kim, Ashley L DuMont, Erin E Zwack, Robert J Ulrich, Theodora K Karagounis, Chunyi Zhou, Andreas F Haag, Julia Shenderovich, Gregory A Wasserman, Junbeom Kwon, John Chen, Anthony R Richardson, Jeffrey N Weiser, Carla R Nowosad, Desmond S Lun, Dane Parker, Alejandro Pironti, Xilin Zhao, Karl Drlica, Itai Yanai, Victor J Torres, and Bo Shopsin. Quorum-sensing agr system of Staphylococcus aureus primes gene expression for protection from lethal oxidative stress. *eLife*, 12, 2024.

[14519] Amy R Poe, Lucy Zhu, Si Hao Tang, Ella Valencia, and Matthew S Kayser. Energetic demands regulate sleep-wake rhythm circuit development. *eLife*, 13, 2024.

[14520] Sebastian Pöhl, Manuel Osorio-Valeriano, Emöke Cserti, Jannik Harberding, Rogelio Hernandez-Tamayo, Jacob Biboy, Patrick Sobetzko, Waldemar Vollmer, Peter L Graumann, and Martin Thanbichler. A dynamic bactofilin cytoskeleton cooperates with an M23 endopeptidase to control bacterial morphogenesis. *eLife*, 12, 2024.

[14521] Ivan Pokrovsky, Teja Curk, Andreas Dietz, Ivan Fufachev, Olga Kulikova, Sebastian Rößler, and Martin Wikelski. Foxtrot migration and dynamic over-wintering range of an Arctic raptor. *eLife*, 12, 2024.

[14522] Denise M Poltavski, Alexander T Cunha, Jaime Tan, Henry M Sucov, and Takako Makita. Lineage-specific intersection of endothelin and GDNF signaling in enteric nervous system development. *eLife*, 13, 2024.

[14523] Antonella Pomè and Eckart Zimmermann. Visuo-motor updating in individuals with heightened autistic traits. *eLife*, 13, 2024.

[14524] Josse Poppinga, Nolan J Barrett, L Niels Cornelisse, Matthijs Verhage, and Jan RT van Weering. Endosomal sorting protein SNX4 limits synaptic vesicle docking and release. *eLife*, 13, 2024.

[14525] Giuseppina Porciello, Alessandro Monti, Maria Serena Panasiti, and Salvatore Maria Aglioti. Ingestible pills reveal gastric correlates of emotions. *eLife*, 13, 2024.

[14526] Julia R Port, Dylan H Morris, Jade C Riopelle, Claude Kwe Yinda, Victoria A Avanzato, Myndi G Holbrook, Trenton Bushmaker, Jonathan E Schulz, Taylor A Saturday, Kent Barbian, Colin A Russell, Rose Perry-Gottschalk, Carl Shaia, Craig Martens, James O Lloyd-Smith, Robert J Fischer, and Vincent J Munster. Host and viral determinants of airborne transmission of SARS-CoV-2 in the Syrian hamster. *eLife*, 12, 2024.

[14527] Joanna C Porter, Jamie Inshaw, Vincente Joel Solis, Emma Denneny, Rebecca Evans, Mia I Temkin, Nathalia De Vasconcelos, Iker Valle Aramburu, Dennis Hoving, Donna Basire, Tracey Crissell, Jesusa Guinto, Alison Webb, Hanif Esmail, Victoria Johnston, Anna Last, Thomas Rampling, Lena Lippert, Elisa Theresa Helbig, Florian Kurth, Bryan Williams, Aiden Flynn, Pauline T Lukey, Veronique Birault, and Venizelos Papayannopoulos. Anti-inflammatory therapy with nebulized dornase alfa for severe COVID-19 pneumonia: a randomized unblinded trial. *eLife*, 12, 2024.

[14528] Stephanie Portillo-Ledesma, Suckwoo Chung, Jill Hoffman, and Tamar Schlick. Regulation of chromatin architecture by transcription factor binding. *eLife*, 12, 2024.

[14529] Kevin Portmann, Aline Linder, and Klaus Eyer. Stimulation-induced cytokine polyfunctionality as a dynamic concept. *eLife*, 12, 2024.

[14530] Arthur Prat-Carrabin, Florent Meyniel, and Rava Azeredo da Silveira. Resource-rational account of sequential effects in human prediction. *eLife*, 13, 2024.

[14531] Smrithi Prem, Bharati Dev, Cynthia Peng, Monal Mehta, Rohan Alibutud, Robert J Connacher, Madeline St Thomas, Xiaofeng Zhou, Paul Matteson, Jinchuan Xing, James H Millonig, and Emanuel DiCicco-Bloom. Dysregulation of mTOR signaling mediates common neurite and migration defects in both idiopathic and 16p11.2 deletion autism neural precursor cells. *eLife*, 13, 2024.

[14532] Alejandro Prieto, Luïsa Miró, Yago Margolles, Manuel Bernabeu, David Salguero, Susana Merino, Joan Tomas, Juan Alberto Corbera, Anna Perez-Bosque, Mario Huttener, Luis ángel Fernández, and Antonio Juarez. Targeting plasmid-encoded proteins that contain immunoglobulin-like domains to combat antimicrobial resistance. *eLife*, 13, 2024.

[14533] Mattia Privitera, Lukas M von Ziegler, Amalia Floriou-Servou, Sian N Duss, Runzhong Zhang, Rebecca Waag, Sebastian Leimbacher, Oliver Sturman, Fabienne K Roessler, Annelies Heylen, Yannick Vermeiren, Debby Van Dam, Peter P De Deyn, Pierre-Luc Germain, and Johannes Bohacek. Noradrenaline release from the locus coeruleus shapes stress-induced hippocampal gene expression. *eLife*, 12, 2024.

[14534] Marcel Proske, Robert Janowski, Sabrina Bacher, Hyun-Seo Kang, Thomas Monecke, Tony Koehler, Saskia Hutten, Jana Tretter, Anna Crois, Lena Molitor, Alejandro Varela-Rial, Roberto Fino, Elisa Donati, Gianni De Fabritiis, Dorothee Dormann, Michael Sattler, and Dierk Niessing. PURA syndrome-causing mutations impair PUR-domain integrity and affect P-body association. *eLife*, 13, 2024.

[14535] Sasithon Pukrittayakamee, Podjanee Jittamala, James A Watson, Borimas Hanboonkunupakarn, Pawanrat Leungsinsiri, Kittiyod Poovorawan, Kesinee Chotivanich, Germana Bancone, Cindy S Chu, Mallika Imwong, Nicholas PJ Day, Walter RJ Taylor, and Nicholas J White. Primaquine in glucose-6-phosphate dehydrogenase deficiency: an adaptive pharmacometric assessment of ascending dose regimens in healthy volunteers. *eLife*, 12, 2024.

[14536] Alain Pulfer, Diego Ulisse Pizzagalli, Paolo Armando Gagliardi, Lucien Hinderling, Paul Lopez, Romaniya Zayats, Pau Carrillo-Barberà, Paola Antonello, Miguel Palomino-Segura, Benjamin Grädel, Mariaclaudia Nicolai, Alessandro Giusti, Marcus Thelen, Luca Maria Gambardella, Thomas T Murooka, Olivier Pertz, Rolf Krause, and Santiago Fernandez Gonzalez. Transformer-based spatial–temporal detection of apoptotic cell death in live-cell imaging. *eLife*, 12, 2024.

[14537] Swarang Sachin Pundlik, Alok Barik, Ashwin Venkateshvaran, Snehasudha Subhadarshini Sahoo, Mahapatra Anshuman Jaysingh, Raviswamy GH Math, Heera Lal, Maroof Athar Hashmi, and Arvind Ramanathan. Senescent cells inhibit mouse myoblast differentiation via the SASP-lipid 15d-PGJ2 mediated modification and control of HRas. *eLife*, 13, 2024.

[14538] Eva Pyrihová, Martin S King, Alannah C King, M Rey Toleco, Mark van der Giezen, and Edmund RS Kunji. A mitochondrial carrier transports glycolytic intermediates to link cytosolic and mitochondrial glycolysis in the human gut parasite Blastocystis. *eLife*, 13, 2024.

[14539] Mu Qiao. Deciphering the genetic code of neuronal type connectivity through bilinear modeling. *eLife*, 12, 2024.

[14540] Jierui Qin, Tingting Yang, Kexin Li, Ting Liu, and Wei Zhang. Pharyngeal mechanosensory neurons control food swallow in Drosophila melanogaster. *eLife*, 12, 2024.

[14541] Lianhua Qin, Junfang Xu, Jianxia Chen, Sen Wang, Ruijuan Zheng, Zhenling Cui, Zhonghua Liu, Xiangyang Wu, Jie Wang, Xiaochen Huang, Zhaohui Wang, Mingqiao Wang, Rong Pan, Stefan HE Kaufmann, Xun Meng, Lu Zhang, Wei Sha, and Haipeng Liu. Cell-autonomous targeting of arabinogalactan by host immune factors inhibits mycobacterial growth. *eLife*, 13, 2024.

[14542] Peipei Qin, Qiuhui Bi, Zeya Guo, Liyuan Yang, Haokun Li, Peng Li, Xinyu Liang, Junhao Luo, Xiangyu Kong, Yirong Xiong, Bo Sun, Sebastian Ocklenburg, and Gaolang Gong. Microstructural asymmetries of the planum temporale predict functional lateralization of auditory-language processing. *eLife*, 13, 2024.

[14543] Sen Qin, Yawei Xu, Shimiao Yu, Wencong Han, Shiheng Fan, Wenxiang Ai, Kenan Zhang, Yizhou Wang, Xuehong Zhou, Qi Shen, Kan Gong, Luyang Sun, and Zheng Zhang. Molecular classification and tumor microenvironment characteristics in pheochromocytomas. *eLife*, 12, 2024.

[14544] Yongli Qin, Jumpei Shirakawa, Cheng Xu, Ruge Chen, Xu Yang, Courtney Ng, Shinichi Nakano, Mahmoud Elguindy, Zhonghao Deng, Kannanganattu V Prasanth, Moritz F Eissmann, Shinichi Nakagawa, William M Ricci, and Baohong Zhao. Long non-coding RNA Malat1 fine-tunes bone homeostasis and repair by orchestrating cellular crosstalk and $\beta$-catenin-OPG/Jagged1 pathway. *eLife*, 13, 2024.

[14545] Jian Qiu, Margaritis Voliotis, Martha A Bosch, Xiao Feng Li, Larry S Zweifel, Krasimira Tsaneva-Atanasova, Kevin T O'Byrne, Oline K Rønnekleiv, and Martin J Kelly. Estradiol elicits distinct firing patterns in arcuate nucleus kisspeptin neurons of females through altering ion channel conductances. *eLife*, 13, 2024.

[14546] Jia-Hua Qu, Khalid Chakir, Kirill V Tarasov, Daniel R Riordon, Maria Grazia Perino, Allwin Jennifa Silvester, and Edward G Lakatta. Reprogramming of cardiac phosphoproteome, proteome, and transcriptome confers resilience to chronic adenylyl cyclase-driven stress. *eLife*, 12, 2024.

[14547] Sebastian Quiroz Monnens, Casper Peters, Luuk Willem Hesselink, Kasper Smeets, and Bernhard Englitz. The recurrent temporal restricted Boltzmann machine captures neural assembly dynamics in whole-brain activity. *eLife*, 13, 2024.

[14548] Angela L Rachubinski, Elizabeth Wallace, Emily Gurnee, Belinda A Enriquez-Estrada, Kayleigh R Worek, Keith P Smith, Paula Araya, Katherine A Waugh, Ross E Granrath, Eleanor Britton, Hannah R Lyford, Micah G Donovan, Neetha Paul Eduthan, Amanda A Hill, Barry Martin, Kelly D Sullivan, Lina Patel, Deborah J Fidler, Matthew D Galbraith, Cory A Dunnick, David A Norris, and Joaquín M Espinosa. JAK inhibition decreases the autoimmune burden in Down syndrome. *eLife*, 13, 2024.

[14549] Maryam Rahimi-Balaei, Shayan Amiri, Thomas Lamonerie, Sih-Rong Wu, Huda Y Zoghbi, G Giacomo Consalez, Daniel Goldowitz, and Hassan Marzban. Identification of an early subset of cerebellar nuclei neurons in mice. *eLife*, 13, 2024.

[14550] Maruti Nandan Rai, Qing Lan, Chirag Parsania, Rikky Rai, Niranjan Shirgaonkar, Ruiwen Chen, Li Shen, Kaeling Tan, and Koon Ho Wong. Temporal transcriptional response of Candida glabrata during macrophage infection reveals a multifaceted transcriptional regulator CgXbp1 important for macrophage response and fluconazole resistance. *eLife*, 13, 2024.

[14551] Achuthan Raja Venkatesh, Kathy H Le, David M Weld, and Onn Brandman. Diffusive lensing as a mechanism of intracellular transport and compartmentalization. *eLife*, 12, 2024.

[14552] Prithviraj Rajebhosale, Mala R Ananth, Ronald Kim, Richard Crouse, Li Jiang, Gretchen López-Hernández, Chongbo Zhong, Christian Arty, Shaohua Wang, Alice Jone, Niraj S Desai, Yulong Li, Marina R Picciotto, Lorna W Role, and David A Talmage. Functionally refined encoding of threat memory by distinct populations of basal forebrain cholinergic projection neurons. *eLife*, 13, 2024.

[14553] Carlos A Ramirez Carbo, Olalekan G Faromiki, and Beiyan Nan. A lytic transglycosylase connects bacterial focal adhesion complexes to the peptidoglycan cell wall. *eLife*, 13, 2024.

[14554] Sabhya Rana, Firoj Alom, Robert C Martinez, David D Fuller, and Aaron D Mickle. Acute ampakines increase voiding function and coordination in a rat model of SCI. *eLife*, 12, 2024.

[14555] Damien M Rasmussen, Manny M Semonis, Joseph T Greene, Joseph M Muretta, Andrew R Thompson, Silvia Toledo Ramos, David D Thomas, William CK Pomerantz, Tanya S Freedman, and Nicholas M Levinson. Allosteric coupling asymmetry mediates paradoxical activation of BRAF by type II inhibitors. *eLife*, 13, 2024.

[14556] Aviv Ratzon, Dori Derdikman, and Omri Barak. Representational drift as a result of implicit regularization. *eLife*, 12, 2024.

[14557] Ricardo Raudales, Gukhan Kim, Sean M Kelly, Joshua Hatfield, Wuqiang Guan, Shengli Zhao, Anirban Paul, Yongjun Qian, Bo Li, and Z Josh Huang. Specific and comprehensive genetic targeting reveals brain-wide distribution and synaptic input patterns of GABAergic axo-axonic interneurons. *eLife*, 13, 2024.

[14558] Sandeep K Ravala, Sendi Rafael Adame-Garcia, Sheng Li, Chun-Liang Chen, Michael A Cianfrocco, J Silvio Gutkind, Jennifer N Cash, and John JG Tesmer. Structural and dynamic changes in P-Rex1 upon activation by PIP3 and inhibition by IP4. *eLife*, 12, 2024.

[14559] Joshua Reeves, Pierre Tournier, Pierre Becquart, Robert Carton, Yin Tang, Alessandra Vigilante, Dong Fang, and Shukry J Habib. Rejuvenating aged osteoprogenitors for bone repair. *eLife*, 13, 2024.

[14560] Josue M Regalado, Ariadna Corredera Asensio, Theresa Haunold, Andrew C Toader, Yan Ran Li, Lauren A Neal, and Priyamvada Rajasethupathy. Neural activity ramps in frontal cortex signal extended motivation during learning. *eLife*, 13, 2024.

[14561] Marius Regin, Yingnan Lei, Edouard Couvreu De Deckersberg, Charlotte Janssens, Anfien Huyghebaert, Yves Guns, Pieter Verdyck, Greta Verheyen, Hilde Van de Velde, Karen Sermon, and Claudia Spits. Complex aneuploidy triggers autophagy and p53-mediated apoptosis and impairs the second lineage segregation in human preimplantation embryos. *eLife*, 12, 2024.

[14562] Lisa Reisinger, Gianpaolo Demarchi, Jonas Obleser, William Sedley, Marta Partyka, Juliane Schubert, Quirin Gehmacher, Sebastian Roesch, Nina Suess, Eugen Trinka, Winfried Schlee, and Nathan Weisz. Aberrant auditory prediction patterns robustly characterize tinnitus. *eLife*, 13, 2024.

[14563] Yasmina Reisser, Franziska Hornung, Antje Häder, Thurid Lauf, Sandor Nietzsche, Bettina Löffler, and Stefanie Deinhardt-Emmer. Telomerase RNA component knockout exacerbates Staphylococcus aureus pneumonia by extensive inflammation and dysfunction of T cells. *eLife*, 13, 2024.

[14564] Rosalío Reyes, Arthur D Lander, and Marcos Nahmad. Dynamic readout of the Hh gradient in the Drosophila wing disc reveals pattern-specific tradeoffs between robustness and precision. *eLife*, 13, 2024.

[14565] Luke M Rice, Lauren A McCormick, Joseph M Cleary, and William O Hancock. Interface-acting nucleotide controls polymerization dynamics at microtubule plus- and minus-ends. *eLife*, 12, 2024.

[14566] Reese Richardson, Heliodoro Tejedor Navarro, Luis A Nunes Amaral, and Thomas Stoeger. Meta-Research: Understudied genes are lost in a leaky pipeline between genome-wide assays and reporting of results. *eLife*, 12, 2024.

[14567] Katherine Rickelton, Trisha M Zintel, Jason Pizzollo, Emily Miller, John J Ely, Mary Ann Raghanti, William D Hopkins, Patrick R Hof, Chet C Sherwood, Amy L Bauernfeind, and Courtney C Babbitt. Tempo and mode of gene expression evolution in the brain across primates. *eLife*, 13, 2024.

[14568] Lukas Rier, Natalie Rhodes, Daisie O Pakenham, Elena Boto, Niall Holmes, Ryan M Hill, Gonzalo Reina Rivero, Vishal Shah, Cody Doyle, James Osborne, Richard W Bowtell, Margot Taylor, and Matthew J Brookes. Tracking the neurodevelopmental trajectory of beta band oscillations with optically pumped magnetometer-based magnetoencephalography. *eLife*, 13, 2024.

[14569] Charlotte Rimbault, Christelle Breillat, Benjamin Compans, Estelle Toulmé, Filipe Nunes Vicente, Monica Fernandez-Monreal, Patrice Mascalchi, Camille Genuer, Virginia Puente-Muñoz, Isabel Gauthereau, Eric Hosy, Stéphane Claverol, Gregory Giannone, Ingrid Chamma, Cameron D Mackereth, Christel Poujol, Daniel Choquet, and Matthieu Sainlos. Engineering paralog-specific PSD-95 recombinant binders as minimally interfering multimodal probes for advanced imaging techniques. *eLife*, 13, 2024.

[14570] Victoria JH Ritvo, Alex Nguyen, Nicholas B Turk-Browne, and Kenneth A Norman. A neural network model of differentiation and integration of competing memories. *eLife*, 12, 2024.

[14571] Juan Jose Rodriguez Gotor, Kashif Mahfooz, Isabel Perez-Otano, and John F Wesseling. Parallel processing of quickly and slowly mobilized reserve vesicles in hippocampal synapses. *eLife*, 12, 2024.

[14572] Tristan Roget, Claire Macmurray, Pierre Jolivet, Sylvie Meleard, and Michael Rera. A scenario for an evolutionary selection of ageing. *eLife*, 13, 2024.

[14573] Laurel A Rohde, Arianne Bercowsky-Rama, Guillaume Valentin, Sundar Ram Naganathan, Ravi A Desai, Petr Strnad, Daniele Soroldoni, and Andrew C Oates. Cell-autonomous timing drives the vertebrate segmentation clock's wave pattern. *eLife*, 13, 2024.

[14574] Miriam Romagosa, Sharon Nieukirk, Irma Cascão, Tiago A Marques, Robert Dziak, Jean-Yves Royer, Joanne O'Brien, David K Mellinger, Andreia Pereira, Arantza Ugalde, Elena Papale, Sofia Aniceto, Giuseppa Buscaino, Marianne Rasmussen, Luis Matias, Rui Prieto, and Mónica A Silva. Fin whale song evolution in the North Atlantic. *eLife*, 13, 2024.

[14575] Julien Rossato, FranÇois Hug, Kylie Tucker, Ciara Gibbs, Lilian Lacourpaille, Dario Farina, and Simon Avrillon. I-Spin live, an open-source software based on blind-source separation for real-time decoding of motor unit activity in humans. *eLife*, 12, 2024.

[14576] Fani Roumelioti, Christos Tzaferis, Dimitris Konstantopoulos, Dimitra Papadopoulou, Alejandro Prados, Maria Sakkou, Anastasios Liakos, Panagiotis Chouvardas, Theodore Meletakos, Yiannis Pandis, Niki Karagianni, Maria C Denis, Maria Fousteri, Maria Armaka, and George Kollias. Mir221/222 drive synovial hyperplasia and arthritis by targeting cell cycle inhibitors and chromatin remodeling components. *eLife*, 13, 2024.

[14577] Jasmine Rowell, Ching-In Lau, Susan Ross, Diana C Yanez, Oscar A Peña, Benny Chain, and Tessa Crompton. Distinct T-cell receptor (TCR) gene segment usage and MHC-restriction between foetal and adult thymus. *eLife*, 13, 2024.

[14578] Shamayita Roy, Hemanta Adhikary, Sarah Isler, and Damien D'Amours. The Smc5/6 complex counteracts R-loop formation at highly transcribed genes in cooperation with RNase H2. *eLife*, 13, 2024.

[14579] Shuvra Shekhar Roy, Sulochana Bagri, Soujanya Vinayagamurthy, Avik Sengupta, Claudia Regina Then, Rahul Kumar, Sriram Sridharan, and Shantanu Chowdhury. Artificially inserted strong promoter containing multiple G-quadruplexes induces long-range chromatin modification. *eLife*, 13, 2024.

[14580] Jessica Royer, Valeria Kebets, Camille Piguet, Jianzhong Chen, Leon Qi Rong Ooi, Matthias Kirschner, Vanessa Siffredi, Bratislav Misic, BT Thomas Yeo, and Boris C Bernhardt. Multimodal neural correlates of childhood psychopathology. *eLife*, 13, 2024.

[14581] Ann-Kathrin Rößling, Kai Dünser, Chenlu Liu, Susan Lauw, Marta Rodriguez-Franco, Lothar Kalmbach, Elke Barbez, and Jürgen Kleine-Vehn. Pectin methylesterase activity is required for RALF1 peptide signalling output. *eLife*, 13, 2024.

[14582] Tiechao Ruan, Ruixi Zhou, Yihong Yang, Junchen Guo, Chuan Jiang, Xiang Wang, Gan Shen, Siyu Dai, Suren Chen, and Ying Shen. Deficiency of IQCH causes male infertility in humans and mice. *eLife*, 12, 2024.

[14583] Yang Ruan, Ning Ling, Shengjing Jiang, Xin Jing, Jin-Sheng He, Qirong Shen, and Zhibiao Nan. Warming and altered precipitation independently and interactively suppress alpine soil microbial growth in a decadal-long experiment. *eLife*, 12, 2024.

[14584] Zheng Ruan, Junuk Lee, Yangyang Li, Juan Du, and Wei Lü. Human pannexin 1 channel is not phosphorylated by Src tyrosine kinase at Tyr199 and Tyr309. *eLife*, 13, 2024.

[14585] Gerald M Rubin and Yoshinori Aso. New genetic tools for mushroom body output neurons in Drosophila. *eLife*, 12, 2024.

[14586] Sarah Rubin, Ankit Agrawal, Anne Seewald, Meng-Jia Lian, Olivia Gottdenker, Paul Villoutreix, Adrian Baule, Tomer Stern, and Elazar Zelzer. Limited column formation in the embryonic growth plate implies divergent growth mechanisms during pre- and postnatal bone development. *eLife*, 13, 2024.

[14587] Linda S Rubio, Suman Mohajan, and David S Gross. Heat Shock Factor 1 forms nuclear condensates and restructures the yeast genome before activating target genes. *eLife*, 12, 2024.

[14588] Julia Ruge, Mana R Ehlers, Alexandros Kastrinogiannis, Maren Klingelhöfer-Jens, Alina Koppold, Rany Abend, and Tina B Lonsdorf. How adverse childhood experiences get under the skin: A systematic review, integration and methodological discussion on threat and reward learning mechanisms. *eLife*, 13, 2024.

[14589] Rafael Naime Ruggiero, Danilo Benette Marques, Matheus Teixeira Rossignoli, Jana Batista De Ross, Tamiris Prizon, Ikaro Jesus Silva Beraldo, Lezio Soares Bueno-Junior, Ludmyla Kandratavicius, Jose Eduardo Peixoto-Santos, Cleiton Lopes-Aguiar, and Joao Pereira Leite. Dysfunctional hippocampal-prefrontal network underlies a multidimensional neuropsychiatric phenotype following early-life seizure. *eLife*, 12, 2024.

[14590] Javier Ruiz-Navarro, Sara Fernández-Hermira, Irene Sanz-Fernández, Pablo Barbeito, Alfonso Navarro-Zapata, Antonio Pérez-Martínez, Francesc R Garcia-Gonzalo, Víctor Calvo, and Manuel Izquierdo Pastor. Formin-like 1$\beta$ phosphorylation at S1086 is necessary for secretory polarized traffic of exosomes at the immune synapse in Jurkat T lymphocytes. *eLife*, 13, 2024.

[14591] Ilya A Rybak, Natalia A Shevtsova, Sergey N Markin, Boris I Prilutsky, and Alain Frigon. Operation regimes of spinal circuits controlling locomotion and the role of supraspinal drives and sensory feedback. *eLife*, 13, 2024.

[14592] Heungjin Ryu, Kibum Nam, Byeong Eun Lee, Yundon Jeong, Seunghun Lee, Jeongmo Kim, Young-Min Hyun, Jae-Ick Kim, and Jung-Hoon Park. The sperm hook as a functional adaptation for migration and self-organized behavior. *eLife*, 13, 2024.

[14593] Ji Eun Ryu, Kyu-Won Shim, Hyun Woong Roh, Minsung Park, Jae-Hyung Lee, and Eun Young Kim. Circadian regulation of endoplasmic reticulum calcium response in cultured mouse astrocytes. *eLife*, 13, 2024.

[14594] Vitaly Ryu, Anisa Azatovna Gumerova, Ronit Witztum, Funda Korkmaz, Liam Cullen, Hasni Kannangara, Ofer Moldavski, Orly Barak, Daria Lizneva, Ki A Goosens, Sarah Stanley, Se-Min Kim, Tony Yuen, and Mone Zaidi. An atlas of brain-bone sympathetic neural circuits in mice. *eLife*, 13, 2024.

[14595] Michael D Sacco, Lauren R Hammond, Radwan E Noor, Dipanwita Bhattacharya, Lily J McKnight, Jesper J Madsen, Xiujun Zhang, Shane G Butler, M Trent Kemp, Aiden C Jaskolka-Brown, Sebastian J Khan, Ioannis Gelis, Prahathees Eswara, and Yu Chen. Staphylococcus aureus FtsZ and PBP4 bind to the conformationally dynamic N-terminal domain of GpsB. *eLife*, 13, 2024.

[14596] Jayanarayanan Sadanandan, Sithara Thomas, Iny Elizabeth Mathew, Zhen Huang, Spiros L Blackburn, Nitin Tandon, Hrishikesh Lokhande, Pierre D McCrea, Emery H Bresnick, Pramod K Dash,

Devin W McBride, Arif Harmanci, Lalit K Ahirwar, Dania Jose, Ari C Dienel, Hussein A Zeineddine, Sungha Hong, and Peeyush Kumar T. Key epigenetic and signaling factors in the formation and maintenance of the blood-brain barrier. *eLife*, 12, 2024.

[14597] Mohsen Sadeghi, Reza Sharif Razavian, Salah Bazzi, Raeed H Chowdhury, Aaron P Batista, Patrick J Loughlin, and Dagmar Sternad. Inferring control objectives in a virtual balancing task in humans and monkeys. *eLife*, 12, 2024.

[14598] Cristina Sáenz de Miera, Nicole Bellefontaine, Susan J Allen, Martin G Myers, and Carol F Elias. Glutamate neurotransmission from leptin receptor cells is required for typical puberty and reproductive function in female mice. *eLife*, 13, 2024.

[14599] Georgia Maria Sagia, Xenia Georgiou, Georgios Chamilos, George Diallinas, and Sofia Dimou. Distinct trafficking routes of polarized and non-polarized membrane cargoes in Aspergillus nidulans. *eLife*, 13, 2024.

[14600] Andrea Salazar Lázaro, Thorsten Trimbuch, GülÇin Vardar, and Christian Rosenmund. The stability of the primed pool of synaptic vesicles and the clamping of spontaneous neurotransmitter release rely on the integrity of the C-terminal half of the SNARE domain of syntaxin-1A. *eLife*, 12, 2024.

[14601] Monica Salinas-Pena, Elena Rebollo, and Albert Jordan. Imaging analysis of six human histone H1 variants reveals universal enrichment of H1.2, H1.3, and H1.5 at the nuclear periphery and nucleolar H1X presence. *eLife*, 12, 2024.

[14602] Zeina Salloum, Kristin Dauner, Yun-feng Li, Neha Verma, David Valdivieso-González, Víctor Almendro-Vedia, John D Zhang, Kiran Nakka, Mei Xi Chen, Jeffrey McDonald, Chase D Corley, Alexander Sorisky, Bao-Liang Song, Iván López-Montero, Jie Luo, Jeffrey F Dilworth, and Xiaohui Zha. Statin-mediated reduction in mitochondrial cholesterol primes an anti-inflammatory response in macrophages by upregulating Jmjd3. *eLife*, 13, 2024.

[14603] Parthasarathy Sampathkumar, Heekyung Jung, Hui Chen, Zhengjian Zhang, Nicholas Suen, Yiran Yang, Zhong Huang, Tom Lopez, Robert Benisch, Sung-Jin Lee, Jay Ye, Wen-Chen Yeh, and Yang Li. Targeted protein degradation systems to enhance Wnt signaling. *eLife*, 13, 2024.

[14604] Pablo Sanchez Bosch, Bomsoo Cho, and Jeffrey D Axelrod. Flamingo participates in multiple models of cell competition. *eLife*, 13, 2024.

[14605] Luis Sánchez-Guardado, Peyman Callejas Razavi, Bo Wang, Antuca Callejas-Marín, and Carlos Lois. Projection neurons are necessary for the maintenance of the mouse olfactory circuit. *eLife*, 13, 2024.

[14606] Jiun Sang, Subash Dhakal, Bhanu Shrestha, Dharmendra Kumar Nath, Yunjung Kim, Anindya Ganguly, Craig Montell, and Youngseok Lee. A single pair of pharyngeal neurons functions as a commander to reject high salt in Drosophila melanogaster. *eLife*, 12, 2024.

[14607] Chananchida Sang-aram, Robin Browaeys, Ruth Seurinck, and Yvan Saeys. Spotless, a reproducible pipeline for benchmarking cell type deconvolution in spatial transcriptomics. *eLife*, 12, 2024.

[14608] Daljit Sangar, Elizabeth Hill, Kezia Jack, Mark Batchelor, Beenaben Mistry, Juan M Ribes, Graham S Jackson, Simon Mead, and Jan Bieschke. Syntaxin-6 delays prion protein fibril formation and prolongs the presence of toxic aggregation intermediates. *eLife*, 13, 2024.

[14609] Chloe Santos, Abigail R Marshall, Ailish Murray, Kate Metcalfe, Priyanka Narayan, Sandra CP de Castro, Eirini Maniou, Nicholas DE Greene, Gabriel L Galea, and Andrew J Copp. Spinal neural tube formation and tail development in human embryos. *eLife*, 12, 2024.

[14610] Susmita Sarkar, Saurabh Gupta, Chiranjit Mahato, Dibyendu Das, and Jagannath Mondal. Elucidating ATP's role as solubilizer of biomolecular aggregate. *eLife*, 13, 2024.

[14611] Dylan C Sarver, Jean Garcia-Diaz, Muzna Saqib, Ryan C Riddle, and G William Wong. Tmem263 deletion disrupts the GH/IGF-1 axis and causes dwarfism and impairs skeletal acquisition. *eLife*, 12, 2024.

[14612] Dylan C Sarver, Muzna Saqib, Fangluo Chen, and G William Wong. Mitochondrial respiration atlas reveals differential changes in mitochondrial function across sex and age. *eLife*, 13, 2024.

[14613] Daichi Sasaki, Ken Imai, Yoko Ikoma, and Ko Matsui. Plastic vasomotion entrainment. *eLife*, 13, 2024.

[14614] Thomas A Sasani, Aaron R Quinlan, and Kelley Harris. Epistasis between mutator alleles contributes to germline mutation spectrum variability in laboratory mice. *eLife*, 12, 2024.

[14615] Rosalyn W Sayaman, Masaru Miyano, Eric G Carlson, Parijat Senapati, Arrianna Zirbes, Sundus F Shalabi, Michael E Todhunter, Victoria E Seewaldt, Susan L Neuhausen, Martha R Stampfer, Dustin E Schones, and Mark A LaBarge. Luminal epithelial cells integrate variable responses to aging into stereotypical changes that underlie breast cancer susceptibility. *eLife*, 13, 2024.

[14616] Yannick Schäfer, Katja Palitzsch, Maria Leptin, Andrew R Whiteley, Thomas Wiehe, and Jaanus Suurväli. Copy number variation and population-specific immune genes in the model vertebrate zebrafish. *eLife*, 13, 2024.

[14617] Anna-Maria Schaffer, Gina Jasmin Fiala, Miriam Hils, Eriberto Natali, Lmar Babrak, Laurenz Alexander Herr, Mari Carmen Romero-Mulero, Nina Cabezas-Wallscheid, Marta Rizzi, Enkelejda Miho, Wolfgang WA Schamel, and Susana Minguet. Kidins220 regulates the development of B cells bearing the $\lambda$ light chain. *eLife*, 13, 2024.

[14618] Kim Schalcher, Estelle Milliet, Robin Séchaud, Roman Bühler, Bettina Almasi, Simon Potier, Paolo Becciu, Alexandre Roulin, and Emily LC Shepard. Landing force reveals new form of motion-induced sound camouflage in a wild predator. *eLife*, 12, 2024.

[14619] Kyra Schapiro, JD Rittenberg, Max Kenngott, and Eve Marder. Ih block reveals separation of timescales in pyloric rhythm response to temperature changes in Cancer borealis. *eLife*, 13, 2024.

[14620] Philipp H Schiffer, Paschalis Natsidis, Daniel J Leite, Helen E Robertson, FranÇois Lapraz, Ferdinand Marlétaz, Bastian Fromm, Liam Baudry, Fraser Simpson, Eirik Høye, Anne C Zakrzewski, Paschalia Kapli, Katharina J Hoff, Steven Müller, Martial Marbouty, Heather Marlow, Richard R Copley, Romain Koszul, Peter Sarkies, and Maximilian J Telford. Insights into early animal evolution from the genome of the xenacoelomorph worm Xenoturbella bocki. *eLife*, 13, 2024.

[14621] Marine Schimel, Ta-Chu Kao, and Guillaume Hennequin. When and why does motor preparation arise in recurrent neural network models of motor control? *eLife*, 12, 2024.

[14622] Meredith N Schmehl, Valeria C Caruso, Yunran Chen, Na Young Jun, Shawn M Willett, Jeff T Mohl, Douglas A Ruff, Marlene Cohen, Akinori F Ebihara, Winrich A Freiwald, Surya T Tokdar, and Jennifer M Groh. Multiple objects evoke fluctuating responses in several regions of the visual pathway. *eLife*, 13, 2024.

[14623] Flavio J Schmidig, Simon Ruch, and Katharina Henke. Episodic long-term memory formation during slow-wave sleep. *eLife*, 12, 2024.

[14624] Kara Schmidlin, Sam Apodaca, Daphne Newell, Alexander Sastokas, Grant Kinsler, and Kerry Geiler-Samerotte. Distinguishing mutants that resist drugs via different mechanisms by examining fitness tradeoffs. *eLife*, 13, 2024.

[14625] Solveig G Schmidt, Andreas Nygaard, Joseph A Mindell, and Claus J Loland. Exploring the K+ binding site and its coupling to transport in the neurotransmitter:sodium symporter LeuT. *eLife*, 12, 2024.

[14626] Nora Schmit, Hillary M Topazian, Matteo Pianella, Giovanni D Charles, Peter Winskill, Michael T White, Katharina Hauck, and Azra C Ghani. Modeling resource allocation strategies for insecticide-treated bed nets to achieve malaria eradication. *eLife*, 12, 2024.

[14627] Arthur L Schneider, Rita Martins-Silva, Alexandre Kaizeler, Nuno Saraiva-Agostinho, and Nuno L Barbosa-Morais. voyAGEr, a free web interface for the analysis of age-related gene expression alterations in human tissues. *eLife*, 12, 2024.

[14628] Daniel R Schonhaut, Aditya M Rao, Ashwin G Ramayya, Ethan A Solomon, Nora A Herweg, Itzhak Fried, and Michael J Kahana. MTL neurons phase-lock to human hippocampal theta. *eLife*, 13, 2024.

[14629] Friedrich Schuessler, Francesca Mastrogiuseppe, Srdjan Ostojic, and Omri Barak. Aligned and oblique dynamics in recurrent neural networks. *eLife*, 13, 2024.

[14630] Haiko Schurz, Vivek Naranbhai, Tom A Yates, James J Gilchrist, Tom Parks, Peter J Dodd, Marlo Möller, Eileen G Hoal, Andrew P Morris, Adrian VS Hill, and Haiko Schurz. Multi-ancestry meta-analysis of host genetic susceptibility to tuberculosis identifies shared genetic architecture. *eLife*, 13, 2024.

[14631] Jens Schuster, Xi Lu, Yonglong Dang, Joakim Klar, Amelie Wenz, Niklas Dahl, and Xingqi Chen. Epigenetic insights into GABAergic development in Dravet Syndrome iPSC and therapeutic implications. *eLife*, 12, 2024.

[14632] Jessica E Schwarz, Antonijo Mrčela, Nicholas F Lahens, Yongjun Li, Cynthia Hsu, Gregory R Grant, Carsten Skarke, Shirley L Zhang, and Amita Sehgal. Evidence for a role of human blood-borne factors in mediating age-associated changes in molecular circadian rhythms. *eLife*, 12, 2024.

[14633] Mohammed Seba, Frederic Boccard, and Stéphane Duigou. Activity of MukBEF for chromosome management in E. coli and its inhibition by MatP. *eLife*, 12, 2024.

[14634] Peter Andreas Seeber, Laura Batke, Yury Dvornikov, Alexandra Schmidt, Yi Wang, Kathleen Stoof-Leichsenring, Katie Moon, Samuel H Vohr, Beth Shapiro, and Laura S Epp. Mitochondrial genomes of Pleistocene megafauna retrieved from recent sediment layers of two Siberian lakes. *eLife*, 12, 2024.

[14635] Mina O Seedhom, Devin Dersh, Jaroslav Holly, Mariana Pavon-Eternod, Jiajie Wei, Matthew Angel, Lucas Shores, Alexandre David, Jefferson Santos, Heather Hickman, and Jonathan W Yewdell. Paradoxical imbalance between activated lymphocyte protein synthesis capacity and rapid division rate. *eLife*, 12, 2024.

[14636] Benjamin Seelbinder, Susan Wagner, Manavi Jain, Elena Erben, Sergei Klykov, Iliya Dimitrov Stoev, Venkat Raghavan Krishnaswamy, and Moritz Kreysing. Probe-free optical chromatin deformation and measurement of differential mechanical properties in the nucleus. *eLife*, 13, 2024.

[14637] Magali Seguret, Patricia Davidson, Stijn Robben, Charlène Jouve, Celine Pereira, Quitterie Lelong, Lucille Deshayes, Cyril Cerveau, Maël Le Berre, Rita S Rodrigues Ribeiro, and Jean-Sébastien Hulot. A versatile high-throughput assay based on 3D ring-shaped cardiac tissues generated from human induced pluripotent stem cell-derived cardiomyocytes. *eLife*, 12, 2024.

[14638] Koen Seignette, Nora Jamann, Paolo Papale, Huub Terra, Ralph O Porneso, Leander de Kraker, Chris van der Togt, Maaike van der Aa, Paul Neering, Emma Ruimschotel, Pieter R Roelfsema, Jorrit S Montijn, Matthew W Self, Maarten HP Kole, and Christiaan N Levelt. Experience shapes chandelier cell function and structure in the visual cortex. *eLife*, 12, 2024.

[14639] Richard Sejour, Janet Leatherwood, Alisa Yurovsky, and Bruce Futcher. Enrichment of rare codons at $5'$ ends of genes is a spandrel caused by evolutionary sequence turnover and does not improve translation. *eLife*, 12, 2024.

[14640] Nikola Sekulovski, Jenna C Wettstein, Amber E Carleton, Lauren N Juga, Linnea E Taniguchi, Xiaolong Ma, Sridhar Rao, Jenna K Schmidt, Thaddeus G Golos, Chien-Wei Lin, and Kenichiro Taniguchi. Temporally resolved early bone morphogenetic protein-driven transcriptional cascade during human amnion specification. *eLife*, 12, 2024.

[14641] Thibaut Sellinger, Frank Johannes, and Aurélien Tellier. Improved inference of population histories by integrating genomic and epigenomic data. *eLife*, 12, 2024.

[14642] Lesia Semenova, Yingfan Wang, Shane Falcinelli, Nancie Archin, Alicia D Cooper-Volkheimer, David M Margolis, Nilu Goonetilleke, David M Murdoch, Cynthia D Rudin, and Edward P Browne. Machine learning approaches identify immunologic signatures of total and intact HIV DNA during long-term antiretroviral therapy. *eLife*, 13, 2024.

[14643] Ron Sender, Elad Noor, Ron Milo, and Yuval Dor. What fraction of cellular DNA turnover becomes cfDNA? *eLife*, 12, 2024.

[14644] Katherine A Senn, Karli A Lipinski, Natalie J Zeps, Amory F Griffin, Max E Wilkinson, and Aaron A Hoskins. Control of 3' splice site selection by the yeast splicing factor Fyv6. *eLife*, 13, 2024.

[14645] Walter Senn, Dominik Dold, Akos F Kungl, Benjamin Ellenberger, Jakob Jordan, Yoshua Bengio, João Sacramento, and Mihai A Petrovici. A neuronal least-action principle for real-time learning in cortical circuits. *eLife*, 12, 2024.

[14646] Rula Sha, Ruochen Guo, Huimin Duan, Qian Peng, Ningyang Yuan, Zhenzhen Wang, Zhigang Li, Zhiqin Xie, Xue You, and Ying Feng. SRSF2 is a key player in orchestrating the directional migration and differentiation of MyoD progenitors during skeletal muscle development. *eLife*, 13, 2024.

[14647] Zachary Shaffer, Roberto Romero, Adi L Tarca, Jose Galaz, Marcia Arenas-Hernandez, Dereje W Gudicha, Tinnakorn Chaiworapongsa, Eunjung Jung, Manaphat Suksai, Kevin R Theis, and Nardhy Gomez-Lopez. The vaginal immunoproteome for the prediction of spontaneous preterm birth: A retrospective longitudinal study. *eLife*, 13, 2024.

[14648] Nishal Pradeepbhai Shah, AJ Phillips, Sasidhar Madugula, Amrith Lotlikar, Alex R Gogliettino, Madeline Rose Hays, Lauren Grosberg, Jeff Brown, Aditya Dusi, Pulkit Tandon, Pawel Hottowy, Wladyslaw Dabrowski, Alexander Sher, Alan M Litke, Subhasish Mitra, and EJ Chichilnisky. Precise control of neural activity using dynamically optimized electrical stimulation. *eLife*, 13, 2024.

[14649] Ladan Shahshahani, Maedbh King, Caroline Nettekoven, Richard B Ivry, and Jörn Diedrichsen. Selective recruitment of the cerebellum evidenced by task-dependent gating of inputs. *eLife*, 13, 2024.

[14650] Amin MD Shakhawat, Jacqueline G Foltz, Adam B Nance, Jaydev Bhateja, and Jennifer L Raymond. Systemic pharmacological suppression of neural activity reverses learning impairment in a mouse model of Fragile X syndrome. *eLife*, 12, 2024.

[14651] Moagi Tube Shaku, Peter K Um, Karl L Ocius, Alexis J Apostolos, Marcos M Pires, William R Bishai, and Bavesh D Kana. A modified BCG with depletion of enzymes associated with peptidoglycan amidation induces enhanced protection against tuberculosis in mice. *eLife*, 13, 2024.

[14652] Jinsai Shang and Douglas J Kojetin. Unanticipated mechanisms of covalent inhibitor and synthetic ligand cobinding to PPAR$\gamma$. *eLife*, 13, 2024.

[14653] Julia Shangguan and Ronald S Rock. Hundreds of myosin 10s are pushed to the tips of filopodia and could cause traffic jams on actin. *eLife*, 12, 2024.

[14654] Xingfeng Shao, Qinyang Shou, Kimberly Felix, Brandon Ojogho, Xuejuan Jiang, Brian T Gold, Megan M Herting, Eric L Goldwaser, Peter Kochunov, Elliot Hong, Ioannis Pappas, Meredith Braskie, Hosung Kim, Steven Cen, Kay Jann, and Danny JJ Wang. Age-related decline in blood-brain barrier function is more pronounced in males than females in parietal and temporal regions. *eLife*, 13, 2024.

[14655] Raman Sharma, Chao Chen, Lionel Tan, Katie Rolfe, Ioana-Gabriela Fiţa, Siôn Jones, Anup Pingle, Rachel A Gibson, Navin Goyal, Hema Sharma, and Panayota Bird. Comment on 'The clinical pharmacology of tafenoquine in the radical cure of Plasmodium vivax malaria: An individual patient data meta-analysis'. *eLife*, 13, 2024.

[14656] Vartika Sharma, Nalani Sachan, Bappi Sarkar, Mousumi Mutsuddi, and Ashim Mukherjee. E3 ubiquitin ligase Deltex facilitates the expansion of Wingless gradient and antagonizes Wingless signaling through a conserved mechanism of transcriptional effector Armadillo/$\beta$-catenin degradation. *eLife*, 12, 2024.

[14657] Rachel Sharninghausen, Jiwon Hwang, Devon D Dennison, and Ryan D Baldridge. Identification of ERAD-dependent degrons for the endoplasmic reticulum lumen. *eLife*, 12, 2024.

[14658] Huishang She, Yan Hao, Gang Song, Xu Luo, Fumin Lei, Weiwei Zhai, and Yanhua Qu. Gene expression plasticity followed by genetic change during colonization in a high-elevation environment. *eLife*, 12, 2024.

[14659] Salman Shehzada, Tomoko Noto, Julie Saksouk, and Kazufumi Mochizuki. A SUMO E3 ligase promotes long non-coding RNA transcription to regulate small RNA-directed DNA elimination. *eLife*, 13, 2024.

[14660] Xia Shen, Xiang Peng, YueGui Guo, Zhujiang Dai, Long Cui, Wei Yu, Yun Liu, and Chen-Ying Liu. YAP/TAZ enhances P-body formation to promote tumorigenesis. *eLife*, 12, 2024.

[14661] Danqi Sheng, Chen-Xi Yue, Fei Jin, Yao Wang, Muneyoshi Ichikawa, Ye Yu, Chang-Run Guo, and Motoyuki Hattori. Structural insights into the orthosteric inhibition of P2X receptors by non-ATP analog antagonists. *eLife*, 12, 2024.

[14662] Chenggang Shi, Shuang Chen, Huimin Liu, Rongrong Pan, Shiqi Li, Yanhui Wang, Xiaotong Wu, Jingjing Li, Xuewen Li, Chaofan Xing, Xian Liu, Yiquan Wang, Qingming Qu, and Guang Li. Evolution of the gene regulatory network of body axis by enhancer hijacking in amphioxus. *eLife*, 13, 2024.

[14663] Ahmed A Shibl, Michael A Ochsenkühn, Amin R Mohamed, Ashley Isaac, Lisa SY Coe, Yejie Yun, Grzegorz Skrzypek, Jean-Baptiste Raina, Justin R Seymour, Ahmed J Afzal, and Shady A Amin. Molecular mechanisms of microbiome modulation by the eukaryotic secondary metabolite azelaic acid. *eLife*, 12, 2024.

[14664] Anika Shimonty, Fabrizio Pin, Matthew Prideaux, Gang Peng, Joshua Huot, Hyeonwoo Kim, Clifford J Rosen, Bruce M Spiegelman, and Lynda F Bonewald. Deletion of FNDC5/irisin modifies murine osteocyte function in a sex-specific manner. *eLife*, 12, 2024.

[14665] Jae Hun Shin, Jooyoung Park, Jaechul Lim, Jaekwang Jeong, Ravi K Dinesh, Stephen E Maher, Jeonghyun Kim, Soyeon Park, Jun Young Hong, John Wysolmerski, Jungmin Choi, and Alfred LM Bothwell. Metastasis of colon cancer requires Dickkopf-2 to generate cancer cells with Paneth cell properties. *eLife*, 13, 2024.

[14666] Jeong-Oh Shin, Jong-Bin Lee, Sihoon Lee, and Jin-Woo Kim. Enhancing bone regeneration and osseointegration using rhPTH(1-34) and dimeric R25CPTH(1-34) in an osteoporotic beagle model. *eLife*, 13, 2024.

[14667] Saori Shinoda, Yuji Sakai, Takahide Matsui, Masaaki Uematsu, Ikuko Koyama-Honda, Jun-ichi Sakamaki, Hayashi Yamamoto, and Noboru Mizushima. Syntaxin 17 recruitment to mature autophagosomes is temporally regulated by PI4P accumulation. *eLife*, 12, 2024.

[14668] Rauf Shiraliyev and Mehmet A Orman. Metabolic disruption impairs ribosomal protein levels, resulting in enhanced aminoglycoside tolerance. *eLife*, 13, 2024.

[14669] William R Shoemaker and Jacopo Grilli. Investigating macroecological patterns in coarse-grained microbial communities using the stochastic logistic model of growth. *eLife*, 12, 2024.

[14670] Amy N Shore, Keyong Li, Mona Safari, Alshaima'a M Qunies, Brittany D Spitznagel, C David Weaver, Kyle Emmitte, Wayne Frankel, and Matthew C Weston. Heterozygous expression of a Kcnt1 gain-of-function variant has differential effects on somatostatin- and parvalbumin-expressing cortical GABAergic neurons. *eLife*, 13, 2024.

[14671] Bhanu Shrestha, Jiun Sang, Suman Rimal, and Youngseok Lee. Pharyngeal neuronal mechanisms governing sour taste perception in Drosophila melanogaster. *eLife*, 13, 2024.

[14672] Yunda Si and Chengfei Yan. Protein language model-embedded geometric graphs power inter-protein contact prediction. *eLife*, 12, 2024.

[14673] Berit Siedentop, Viacheslav N Kachalov, Christopher Witzany, Matthias Egger, Roger D Kouyos, and Sebastian Bonhoeffer. The effect of combining antibiotics on resistance: A systematic review and meta-analysis. *eLife*, 13, 2024.

[14674] Hugo Siegfried, Georges Farkouh, Rémi Le Borgne, Catherine Pioche-Durieu, Thaïs De Azevedo Laplace, Agathe Verraes, Lucien Daunas, Jean-Marc Verbavatz, and Mélina L Heuzé. The ER tether VAPA is required for proper cell motility and anchors ER-PM contact sites to focal adhesions. *eLife*, 13, 2024.

[14675] Laura-Marie Silbermann, Benjamin Vermeer, Sonja Schmid, and Katarzyna Tych. The known unknowns of the Hsp90 chaperone. *eLife*, 13, 2024.

[14676] Melissa Silva, Van Tran, and Alain Marty. A maximum of two readily releasable vesicles per docking site at a cerebellar single active zone synapse. *eLife*, 12, 2024.

[14677] Hyungtai Sim, Hyun Jung Park, Geun-Ho Park, Yeon Jeong Kim, Woong-Yang Park, Se-Hoon Lee, and Murim Choi. Increased inflammatory signature in myeloid cells of non-small cell lung cancer patients with high clonal hematopoiesis burden. *eLife*, 13, 2024.

[14678] Niklas Simon, Abu Safyan, George Pyrowolakis, and Shinya Matsuda. Dally is not essential for Dpp spreading or internalization but for Dpp stability by antagonizing Tkv-mediated Dpp internalization. *eLife*, 12, 2024.

[14679] Akashdip Singh, Alberto Miranda Bedate, Helen J von Richthofen, Saskia V Vijver, Michiel van der Vlist, Raphael Kuhn, Alexander Yermanos, Jürgen J Kuball, Can Kesmir, M Ines Pascoal Ramos, and Linde Meyaard. A novel bioinformatics pipeline for the identification of immune inhibitory receptors as potential therapeutic targets. *eLife*, 13, 2024.

[14680] Maneesh Kumar Singh, Victoria Ann Bonnell, Israel Tojal Da Silva, Verônica Feijoli Santiago, Miriam Santos Moraes, Jack Adderley, Christian Doerig, Giuseppe Palmisano, Manuel Llinas, and Celia RS Garcia. A Plasmodium falciparum MORC protein complex modulates epigenetic control of gene expression through interaction with heterochromatin. *eLife*, 12, 2024.

[14681] Mrityunjay Singh, Dinesh C Indurthi, Lovika Mittal, Anthony Auerbach, and Shailendra Asthana. Conformational dynamics of a nicotinic receptor neurotransmitter site. *eLife*, 13, 2024.

[14682] Surya Bansi Singh, Shatruhan Singh Rajput, Aditya Sharma, Sujal Kataria, Priyanka Dutta, Vaishnavi Ananthanarayanan, Amitabha Nandi, Shivprasad Patil, Amitabha Majumdar, and Deepa Subramanyam. Pathogenic Huntingtin aggregates alter actin organization and cellular stiffness resulting in stalled clathrin-mediated endocytosis. *eLife*, 13, 2024.

[14683] Nasa Sinnott-Armstrong, Stanley Fields, Frederick Roth, Lea M Starita, Cole Trapnell, Judit Villen, Douglas M Fowler, and Christine Queitsch. Understanding genetic variants in context. *eLife*, 13, 2024.

[14684] Margaux Sivori, Bowen Dempsey, Zoubida Chettouh, Franck Boismoreau, Maïlys Ayerdi, Annaliese Eymael, Sylvain Baulande, Sonia Lameiras, Fanny Coulpier, Olivier Delattre, Hermann Rohrer, Olivier Mirabeau, and Jean-FranÇois Brunet. The pelvic organs receive no parasympathetic innervation. *eLife*, 12, 2024.

[14685] Ekaterina Smirnova, Emmanuelle Bignon, Patrick Schultz, Gabor Papai, and Adam Ben Shem. Binding to nucleosome poises human SIRT6 for histone H3 deacetylation. *eLife*, 12, 2024.

[14686] Tanya M Smith, Manish Arora, Christine Austin, Janaína Nunes ávila, Mathieu Duval, Tze Tshen Lim, Philip J Piper, Petra Vaiglova, John de Vos, Ian S Williams, Jian-xin Zhao, and Daniel R Green. Oxygen isotopes in orangutan teeth reveal recent and ancient climate variation. *eLife*, 12, 2024.

[14687] Shuhei So, Masayo Asakawa, and Hitoshi Sawa. Distinct functions of three Wnt proteins control mirror-symmetric organogenesis in the C. elegans gonad. *eLife*, 13, 2024.

[14688] Fangxing Song, Xue Dong, Jiaxu Zhao, Jue Wang, Xiaohui Sang, Xin He, and Min Bao. Causal role of the frontal eye field in attention-induced ocular dominance plasticity. *eLife*, 12, 2024.

[14689] Weichen Song, Yongyong Shi, and Guan ning Lin. Haplotype function score improves biological interpretation and cross-ancestry polygenic prediction of human complex traits. *eLife*, 12, 2024.

[14690] Chhavi Sood, Md Ausrafuggaman Nahid, Kendall R Branham, Matt Pahl, Susan E Doyle, and Sarah E Siegrist. Delta-dependent Notch activation closes the early neuroblast temporal program to promote lineage progression and neurogenesis termination in Drosophila. *eLife*, 12, 2024.

[14691] Aisha Souquette and Paul G Thomas. Variation in the basal immune state and implications for disease. *eLife*, 13, 2024.

[14692] Thomas P Spargo, Lachlan Gilchrist, Guy P Hunt, Richard JB Dobson, Petroula Proitsi, Ammar Al-Chalabi, Oliver Pain, and Alfredo Iacoangeli. Statistical examination of shared loci in neuropsychiatric diseases using genome-wide association study summary statistics. *eLife*, 12, 2024.

[14693] Daniel Spari, Annina Schmid, Daniel Sanchez-Taltavull, Shaira Murugan, Keely Keller, Nadia Ennaciri, Lilian Salm, Deborah Stroka, and Guido Beldi. Released bacterial ATP shapes local and systemic inflammation during abdominal sepsis. *eLife*, 13, 2024.

[14694] Megan M Sperry, Berenice Charrez, Haleh Fotowat, Erica Gardner, Kanoelani Pilobello, Zohreh Izadifar, Tiffany Lin, Abigail Kuelker, Sahith Kaki, Michael Lewandowski, Shanda Lightbown, Ramses Martinez, Susan Marquez, Joel Moore, Maria Plaza-Oliver, Adama M Sesay, Kostyantyn Shcherbina, Katherine Sheehan, Takako Takeda, Daniela Del Campo, Kristina Andrijauskaite, Exal Cisneros, Riley Lopez, Isabella Cano, Zachary Maxwell, Israel Jessop, Rafa Veraza, Leon Bunegin, Thomas J Percival, Jaclyn Yracheta, Jorge J Pena, Diandra M Wood, Zachary T Homas, Cody J Hinshaw, Jennifer Cox-Hinshaw, Olivia G Parry, Justin J Sleeter, Erik K Weitzel, Michael Levin, Michael Super, Richard Novak, and Donald E Ingber. Identification of pharmacological inducers of a reversible hypometabolic state for whole organ preservation. *eLife*, 13, 2024.

[14695] Emma D Spikol, Ji Cheng, Michelle Macurak, Abhignya Subedi, and Marnie E Halpern. Genetically defined nucleus incertus neurons differ in connectivity and function. *eLife*, 12, 2024.

[14696] Natanael Spisak, Gabriel Athènes, Thomas Dupic, Thierry Mora, and Aleksandra M Walczak. Combining mutation and recombination statistics to infer clonal families in antibody repertoires. *eLife*, 13, 2024.

[14697] Kiran Sriram, Zhijie Qi, Dongqiang Yuan, Naseeb Kaur Malhi, Xuejing Liu, Riccardo Calandrelli, Yingjun Luo, Alonso Tapia, Shengyan Jin, Ji Shi, Martha Salas, Runrui Dang, Brian Armstrong, Saul J Priceman, Ping H Wang, Jiayu Liao, Rama Natarajan, Sheng Zhong, and Zhen Bouman Chen. Regulation of nuclear transcription by mitochondrial RNA in endothelial cells. *eLife*, 13, 2024.

[14698] Alexandra Stavsky, Leonardo A Parra-Rivas, Shani Tal, Jen Riba, Kayalvizhi Madhivanan, Subhojit Roy, and Daniel Gitler. Synapsin E-domain is essential for $\alpha$-synuclein function. *eLife*, 12, 2024.

[14699] Noah J Steinberg, Zvi N Roth, J Anthony Movshon, and Elisha Merriam. Brain representations of motion and position in the double-drift illusion. *eLife*, 13, 2024.

[14700] Elisabet Stener-Victorin, Gustaw Eriksson, Man Mohan Shrestha, Valentina Rodriguez Paris, Haojiang Lu, Jasmine Banks, Manisha Samad, Charlène Perian, Baptiste Jude, Viktor Engman, Roberto Boi, Emma Nilsson, Charlotte Ling, Jenny Nyström, Ingrid Wernstedt Asterholm, Nigel Turner, Johanna Lanner, and Anna Benrick. Proteomic analysis shows decreased type I fibers and ectopic fat accumulation in skeletal muscle from women with PCOS. *eLife*, 12, 2024.

[14701] Petteri Stenroos, Isabelle Guillemain, Federico Tesler, Olivier Montigon, Nora Collomb, Vasile Stupar, Alain Destexhe, Veronique Coizet, Olivier David, and Emmanuel L Barbier. EEG-fMRI in awake rat and whole-brain simulations show decreased brain responsiveness to sensory stimulations during absence seizures. *eLife*, 12, 2024.

[14702] Harlan P Stevens, Carly V Winegar, Arwen F Oakley, and Stephen R Piccolo. Identifying images in the biology literature that are problematic for people with a color-vision deficiency. *eLife*, 13, 2024.

[14703] Valentina Stolz, Rafael de Freitas e Silva, Ramona Rica, Ci Zhu, Teresa Preglej, Patricia Hamminger, Daniela Hainberger, Marlis Alteneder, Lena Müller, Monika Waldherr, Darina Waltenberger, Anastasiya Hladik, Benedikt Agerer, Michael Schuster, Tobias Frey, Thomas Krausgruber, Sylvia Knapp, Clarissa Campbell, Klaus Schmetterer, Michael Trauner, Andreas Bergthaler, Christoph Bock, Nicole Boucheron, and Wilfried Ellmeier. Nuclear receptor corepressor 1 controls regulatory T cell subset differentiation and effector function. *eLife*, 13, 2024.

[14704] John J Stout, Allison E George, Suhyeong Kim, Henry L Hallock, and Amy L Griffin. Using synchronized brain rhythms to bias memory-guided decisions. *eLife*, 12, 2024.

[14705] Sima Stroganov, Talia Harris, Liat Fellus-Alyagor, Lital Ben Moyal, Romina Plitman Mayo, Ofra Golani, Dana Hirsch, Shifra Ben-Dor, Alexander Brandis, Tevie Mehlman, Michal Kovo, Tal Biron-Shental, Nava Dekel, and Michal Neeman. The differential regulation of placenta trophoblast bisphosphoglycerate mutase in fetal growth restriction: preclinical study in mice and observational histological study of human placenta. *eLife*, 13, 2024.

[14706] Astrid Katharina Maria Stubbusch, Johannes M Keegstra, Julia Schwartzman, Sammy Pontrelli, Estelle E Clerc, Samuel Charlton, Roman Stocker, Cara Magnabosco, Olga T Schubert, Martin Ackermann, and Glen G D'Souza. Polysaccharide breakdown products drive degradation-dispersal cycles of foraging bacteria through changes in metabolism and motility. *eLife*, 13, 2024.

[14707] Sofía Suárez Freire, Sebastián Perez-Pandolfo, Sabrina Micaela Fresco, Julián Valinoti, Eleonora Sorianello, Pablo Wappner, and Mariana Melani. The exocyst complex controls multiple events in the pathway of regulated exocytosis. *eLife*, 12, 2024.

[14708] Sanjay Kumar Sukumar, Vimala Antonydhason, Linnea Molander, Jawdat Sandakly, Malak Kleit, Ganesh Umapathy, Patricia Mendoza-Garcia, Tafheem Masudi, Andreas Schlosser, Dick R Nässel, Christian Wegener, Margret Shirinian, and Ruth H Palmer. The Alk receptor tyrosine kinase regulates Sparkly, a novel activity regulating neuropeptide precursor in the Drosophila central nervous system. *eLife*, 12, 2024.

[14709] Rashmi Sukumaran, Achuthsankar S Nair, and Moinak Banerjee. Ethnic and region-specific genetic risk variants of stroke and its comorbid conditions can define the variations in the burden of stroke and its phenotypic traits. *eLife*, 13, 2024.

[14710] Srdjan Sumarac, Kiah A Spencer, Leon A Steiner, Conor Fearon, Emily A Haniff, Andrea A Kühn, Mojgan Hodaie, Suneil K Kalia, Andres Lozano, Alfonso Fasano, William Duncan Hutchison, and Luka Milosevic. Interrogating basal ganglia circuit function in people with Parkinson's disease and dystonia. *eLife*, 12, 2024.

[14711] Congnan Sun, Yoel Hassin, Arjan Boonman, Assaf Shwartz, and Yossi Yovel. Species and habitat specific changes in bird activity in an urban environment during Covid 19 lockdown. *eLife*, 12, 2024.

[14712] Fan Sun, Pengrong Yan, Yadong Xiao, Hongqiao Zhang, Steven D Shapiro, Gutian Xiao, and Zhaoxia Qu. Improving PD-1 blockade plus chemotherapy for complete remission of lung cancer by nanoPDLIM2. *eLife*, 12, 2024.

[14713] Jing Sun, Desmond Choy, Nicolas Sompairac, Shirin Jamshidi, Michele Mishto, and Shahram Kordasti. ImmCellTyper facilitates systematic mass cytometry data analysis for deep immune profiling. *eLife*, 13, 2024.

[14714] Jun Sun, Francisca Rojo-Cortes, Suzana Ulian-Benitez, Manuel G Forero, Guiyi Li, Deepanshu ND Singh, Xiaocui Wang, Sebastian Cachero, Marta Moreira, Dean Kavanagh, Gregory SXE Jefferis, Vincent Croset, and Alicia Hidalgo. A neurotrophin functioning with a Toll regulates structural plasticity in a dopaminergic circuit. *eLife*, 13, 2024.

[14715] Li Sun, Xuejin Chen, Chunlin Song, Wenjing Shi, Libo Liu, Shuang Bai, Xi Wang, Jiali Chen, Chengyu Jiang, Shuang-min Wang, Zhou-qing Luo, Ruiwen Wang, Yamei Wang, and Quan-wen Jin. Negative regulation of APC/C activation by MAPK-mediated attenuation of Cdc20Slp1 under stress. *eLife*, 13, 2024.

[14716] Li Sun, Libo Liu, Chunlin Song, Yamei Wang, and Quan-wen Jin. Heat stress-induced activation of MAPK pathway attenuates Atf1-dependent epigenetic inheritance of heterochromatin in fission yeast. *eLife*, 13, 2024.

[14717] Longjie Sun, Zheng Lv, Xuexue Chen, Rong Ye, Shuang Tian, Chaofan Wang, Xiaomei Xie, Lu Yan, Xiaohong Yao, Yujing Shao, Sheng Cui, Juan Chen, and Jiali Liu. Splicing factor SRSF1 is essential for homing of precursor spermatogonial stem cells in mice. *eLife*, 12, 2024.

[14718] Qi Sun, Kimberley Huynh, Dzenita Muratovic, Nicholas J Gunn, Anja R Zelmer, Lucian Bogdan Solomon, Gerald J Atkins, and Dongqing Yang. Rapid bacterial evaluation beyond the colony forming unit in osteomyelitis. *eLife*, 13, 2024.

[14719] Wenjian Sun, Haohao Wu, Yujie Peng, Xuejiao Zheng, Jing Li, Dingxuan Zeng, Peng Tang, Ming Zhao, Hemin Feng, Hao Li, Ye Liang, Junfeng Su, Xi Chen, Tomas Hökfelt, and Jufang He. Heterosynaptic plasticity of the visuo-auditory projection requires cholecystokinin released from entorhinal cortex afferents. *eLife*, 13, 2024.

[14720] Yi Sun, Zhe Chen, Minjun Jin, Haibo Xie, and Chengtian Zhao. Ciliary length regulation by intraflagellar transport in zebrafish. *eLife*, 13, 2024.

[14721] Yingjie Sun, Changheng Li, Xiaofei Deng, Wenjie Li, Xiaoyi Deng, Weiqi Ge, Miaoyuan Shi, Ying Guo, Yanxun V Yu, Hai-bing Zhou, and Youngnam N Jin. Target protein identification in live cells and organisms with a non-diffusive proximity tagging system. *eLife*, 13, 2024.

[14722] Evyatar Swissa, Uri Monsonego, Lynn T Yang, Lior Schori, Lyna Kamintsky, Sheida Mirloo, Itamar Burger, Sarit Uzzan, Rishi Patel, Peter H Sudmant, Ofer Prager, Daniela Kaufer, and Alon Friedman. Cortical plasticity is associated with blood–brain barrier modulation. *eLife*, 12, 2024.

[14723] Wiktoria Sztangierska, Hubert Wyszkowski, Maria Pokornowska, Klaudia Kochanowicz, Michal Rychłowski, Krzysztof Liberek, and Agnieszka Kłosowska. Early steps of protein disaggregation by Hsp70 chaperone and class B J-domain proteins are shaped by Hsp110. *eLife*, 13, 2024.

[14724] Iustin V Tabarean. Opposing actions of co-released GABA and neurotensin on the activity of preoptic neurons and on body temperature. *eLife*, 13, 2024.

[14725] Nathan Tague, Cristian Coriano-Ortiz, Michael B Sheets, and Mary J Dunlop. Light-inducible protein degradation in E. coli with the LOVdeg tag. *eLife*, 12, 2024.

[14726] Masahiro Takigawa, Marta Huelin Gorriz, Margot Tirole, and Daniel Bendor. Evaluating hippocampal replay without a ground truth. *eLife*, 13, 2024.

[14727] Stav Talal, Jon F Harrison, Ruth Farington, Jacob P Youngblood, Hector E Medina, Rick Overson, and Arianne J Cease. Body mass and growth rates predict protein intake across animals. *eLife*, 13, 2024.

[14728] Deepali Taneja, Poornima Viswanathan, and Sahana V Rajan. Navigating neurodiversity in higher education in India. *eLife*, 13, 2024.

[14729] Junhui Tang, Rui Yang, Hui Li, Xiaodong Wei, Zhen Yang, Wenbin Cai, Yao Jiang, Ga Zhuo, Li Meng, and Yali Xu. Derivation and internal validation of prediction models for pulmonary hypertension risk assessment in a cohort inhabiting Tibet, China. *eLife*, 13, 2024.

[14730] Ruijing Tang, Luobin Guo, Tingyu Wei, Tingting Chen, Huan Yang, Honghao Ye, Fangzhou Lin, Yongyi Zeng, Haijun Yu, Zhixiong Cai, and Xiaolong Liu. Engineering PEG10assembled endogenous virus-like particles with genetically encoded neoantigen peptides for cancer vaccination. *eLife*, 13, 2024.

[14731] James Taniguchi, Riccardo Melani, Lynne Chantranupong, Michelle J Wen, Ali Mohebi, Joshua D Berke, Bernardo L Sabatini, and Nicolas X Tritsch. Comment on 'Accumbens cholinergic interneurons dynamically promote dopamine release and enable motivation'. *eLife*, 13, 2024.

[14732] Daniyal Tariq, Nicole Maurici, Bradley M Bartholomai, Siddarth Chandrasekaran, Jay C Dunlap, Alaji Bah, and Brian R Crane. Phosphorylation, disorder, and phase separation govern the behavior of Frequency in the fungal circadian clock. *eLife*, 12, 2024.

[14733] Rebecca D Tarvin, Jeffrey L Coleman, David A Donoso, Mileidy Betancourth-Cundar, Karem López-Hervas, Kimberly S Gleason, J Ryan Sanders, Jacqueline M Smith, Santiago R Ron, Juan C Santos, Brian E Sedio, David C Cannatella, and Richard W Fitch. Passive accumulation of alkaloids in inconspicuously colored frogs refines the evolutionary paradigm of acquired chemical defenses. *eLife*, 13, 2024.

[14734] Yuki Tatekoshi, Chunlei Chen, Jason Solomon Shapiro, Hsiang-Chun Chang, Malorie Blancard, Davi M Lyra-Leite, Paul W Burridge, Matthew Feinstein, Richard D'Aquila, Priscilla Hsue, and Hossein Ardehali. Human induced pluripotent stem cell-derived cardiomyocytes to study inflammation-induced aberrant calcium transient. *eLife*, 13, 2024.

[14735] Noémie te Rietmolen, Manuel R Mercier, Agnès Trébuchon, Benjamin Morillon, and Daniele Schön. Speech and music recruit frequency-specific distributed and overlapping cortical networks. *eLife*, 13, 2024.

[14736] Liliana R Teixeira, Radha Akella, John M Humphreys, Haixia He, and Elizabeth J Goldsmith. Water and chloride as allosteric inhibitors in WNK kinase osmosensing. *eLife*, 12, 2024.

[14737] Amruta Tendolkar, Anyi Mazo-Vargas, Luca Livraghi, Joseph J Hanly, Kelsey C Van Horne, Lawrence E Gilbert, and Arnaud Martin. Cis-regulatory modes of Ultrabithorax inactivation in butterfly forewings. *eLife*, 12, 2024.

[14738] Alina Tetereva and Narun Pat. Brain age has limited utility as a biomarker for capturing fluid cognition in older individuals. *eLife*, 12, 2024.

[14739] Sabrina Tetzlaff, Arne Hillebrand, Nikiforos Drakoulis, Zala Gluhic, Sascha Maschmann, Peter Lyko, Susann Wicke, and Christian Schmitz-Linneweber. Small RNAs from mitochondrial genome recombination sites are incorporated into T. gondii mitoribosomes. *eLife*, 13, 2024.

[14740] Mathew Thayer, Michael B Heskett, Leslie G Smith, Paul T Spellman, and Phillip A Yates. ASAR lncRNAs control DNA replication timing through interactions with multiple hnRNP/RNA binding proteins. *eLife*, 13, 2024.

[14741] Daniel Thiel, Luis Alfonso Yañez Guerra, Amanda Kieswetter, Alison G Cole, Liesbet Temmerman, Ulrich Technau, and Gáspár Jékely. Large-scale deorphanization of Nematostella vectensis neuropeptide G protein-coupled receptors supports the independent expansion of bilaterian and cnidarian peptidergic systems. *eLife*, 12, 2024.

[14742] Ryan Thiermann, Michael Sandler, Gursharan Ahir, John T Sauls, Jeremy Schroeder, Steven Brown, Guillaume Le Treut, Fangwei Si, Dongyang Li, Jue D Wang, and Suckjoon Jun. Tools and methods for high-throughput single-cell imaging with the mother machine. *eLife*, 12, 2024.

[14743] Rajan M Thomas, Matthew C Pahl, Liqing Wang, Struan FA Grant, Wayne W Hancock, and Andrew D Wells. Foxp3 depends on Ikaros for control of regulatory T cell gene expression and function. *eLife*, 12, 2024.

[14744] Surya Chandra Rao Thumu, Monika Jain, Sumitha Soman, Soumen Das, Vijaya Verma, Arnab Nandi, David H Gutmann, Balaji Jayaprakash, Deepak Nair, James P Clement, Swananda Marathe, and Narendrakumar Ramanan. SRF-deficient astrocytes provide neuroprotection in mouse models of excitotoxicity and neurodegeneration. *eLife*, 13, 2024.

[14745] Junbin Tian, Fang Yang, Ying Wang, Li Wang, Ning Wang, Yi Jiang, and Li Yang. Atypical local and global biological motion perception in children with attention deficit hyperactivity disorder. *eLife*, 12, 2024.

[14746] Maojin Tian, Le Yang, Ziqian Zhao, Jigang Li, Lianqing Wang, Qingqing Yin, Wei Hu, Yunwei Lou, Jianxin Du, and Peiqing Zhao. TIPE drives a cancer stem-like phenotype by promoting glycolysis via PKM2/HIF-1$\alpha$ axis in melanoma. *eLife*, 13, 2024.

[14747] Mengxue Tian, Zhenjia Wang, Zhangli Su, Etsuko Shibata, Yoshiyuki Shibata, Anindya Dutta, and Chongzhi Zang. Integrative analysis of DNA replication origins and ORC-/MCM-binding sites in human cells reveals a lack of overlap. *eLife*, 12, 2024.

[14748] Johanna KS Tiemann, Magdalena Szczuka, Lisa Bouarroudj, Mohamed Oussaren, Steven Garcia, Rebecca J Howard, Lucie Delemotte, Erik Lindahl, Marc Baaden, Kresten Lindorff-Larsen, Matthieu Chavent, and Pierre Poulain. MDverse, shedding light on the dark matter of molecular dynamics simulations. *eLife*, 12, 2024.

[14749] Pénélope Tignard, Karen Pottin, Audrey Geeverding, Mohamed Doulazmi, Mélody Cabrera, Coralie Fouquet, Mathilde Liffran, Jonathan Fouchard, Marion Rosello, Shahad Albadri, Filippo Del Bene, Alain Trembleau, and Marie Anne Breau. Basement membranes are crucial for proper olfactory placode shape, position and boundary with the brain, and for olfactory axon development. *eLife*, 12, 2024.

[14750] Susanne Tilk, Judith Frydman, Christina Curtis, and Dmitri A Petrov. Cancers adapt to their mutational load by buffering protein misfolding stress. *eLife*, 12, 2024.

[14751] Alexei V Tkachenko and Sergei Maslov. Emergence of catalytic function in prebiotic information-coding polymers. *eLife*, 12, 2024.

[14752] Fatima Tleiss, Martina Montanari, Romane Milleville, Olivier Pierre, Julien Royet, Dani Osman, Armel Gallet, and C Leopold Kurz. Spatial and temporal coordination of Duox/TrpA1/Dh31 and IMD pathways is required for the efficient elimination of pathogenic bacteria in the intestine of Drosophila larvae. *eLife*, 13, 2024.

[14753] Takuro Tojima, Yasuyuki Suda, Natsuko Jin, Kazuo Kurokawa, and Akihiko Nakano. Spatiotemporal dissection of the Golgi apparatus and the ER-Golgi intermediate compartment in budding yeast. *eLife*, 13, 2024.

[14754] Daniel Toker, Eli Müller, Hiroyuki Miyamoto, Maurizio S Riga, Laia Lladó-Pelfort, Kazuhiro Yamakawa, Francesc Artigas, James M Shine, Andrew E Hudson, Nader Pouratian, and Martin M Monti. Criticality supports cross-frequency cortical-thalamic information transfer during conscious states. *eLife*, 13, 2024.

[14755] Alexa Tompary and Lila Davachi. Integration of overlapping sequences emerges with consolidation through medial prefrontal cortex neural ensembles and hippocampal–cortical connectivity. *eLife*, 13, 2024.

[14756] Erik Toraason, Alina Salagean, David E Almanzar, Jordan E Brown, Colette M Richter, Nicole A Kurhanewicz, Ofer Rog, and Diana E Libuda. BRCA1/BRC-1 and SMC-5/6 regulate DNA repair pathway engagement during Caenorhabditis elegans meiosis. *eLife*, 13, 2024.

[14757] Léa Torcq, Sara Majello, Catherine Vivier, and Anne A Schmidt. Tuning apicobasal polarity and junctional recycling in the hemogenic endothelium orchestrates the morphodynamic complexity of emerging pre-hematopoietic stem cells. *eLife*, 12, 2024.

[14758] Francesca Torelli, Diogo M da Fonseca, Simon W Butterworth, Joanna C Young, and Moritz Treeck. Paracrine rescue of MYR1-deficient Toxoplasma gondii mutants reveals limitations of pooled in vivo CRISPR screens. *eLife*, 13, 2024.

[14759] Paul A Torrillo and Tami D Lieberman. Reversions mask the contribution of adaptive evolution in microbiomes. *eLife*, 13, 2024.

[14760] Viraj R Torsekar, Nevo Sagi, J Alfred Daniel, Yael Hawlena, Efrat Gavish-Regev, and Dror Hawlena. Contrasting responses to aridity by different-sized decomposers cause similar decomposition rates across a precipitation gradient. *eLife*, 13, 2024.

[14761] Werner Treptow, Yichen Liu, Carlos AZ Bassetto, Bernardo I Pinto, Joao Antonio Alves Nunes, Ramon Mendoza Uriarte, Christophe J Chipot, Francisco Bezanilla, and Benoit Roux. Isoleucine gate blocks K+ conduction in C-type inactivation. *eLife*, 13, 2024.

[14762] Amy Tresenrider, Marcus Hooper, Levi Todd, Faith Kierney, Nicolai A Blasdel, Cole Trapnell, and Thomas A Reh. A multiplexed, single-cell sequencing screen identifies compounds that increase neurogenic reprogramming of murine Muller glia. *eLife*, 12, 2024.

[14763] Yi-Ting Tsai, Nogayhan Seymen, I Richard Thompson, Xinchen Zou, Warisha Mumtaz, Sila Gerlevik, Ghulam J Mufti, and Mohammad M Karimi. Expression of most retrotransposons in human blood correlates with biological aging. *eLife*, 13, 2024.

[14764] Jonathan S Tsay, Hyosub E Kim, Samuel D McDougle, Jordan A Taylor, Adrian Haith, Guy Avraham, John W Krakauer, Anne GE Collins, and Richard B Ivry. Fundamental processes in sensorimotor learning: Reasoning, refinement, and retrieval. *eLife*, 13, 2024.

[14765] Kazushi Tsutsui, Ryoya Tanaka, Kazuya Takeda, and Keisuke Fujii. Collaborative hunting in artificial agents with deep reinforcement learning. *eLife*, 13, 2024.

[14766] Wenyu Tu, Samuel R Cramer, and Nanyin Zhang. Disparity in temporal and spatial relationships between resting-state electrophysiological and fMRI signals. *eLife*, 13, 2024.

[14767] Jennifer Tullet, Jennifer Leigh, Brandon Coke, David Fisher, Johanna Haszczyn, Steven Houghton, Johanna Fish, Laura Freeman, Isabella Garcia, Stefan Penman, and Emma Hargreaves. Using reflective practice to support PhD students in the biosciences. *eLife*, 13, 2024.

[14768] Sarah Tune and Jonas Obleser. Neural attentional filters and behavioural outcome follow independent individual trajectories over the adult lifespan. *eLife*, 12, 2024.

[14769] Joanne Tung, Lei Huang, Ginto George, Heather P Harding, David Ron, and Adriana Ordonez. A genome-wide CRISPR/Cas9 screen identifies calreticulin as a selective repressor of ATF6$\alpha$. *eLife*, 13, 2024.

[14770] Jake Turley, Isaac V Chenchiah, Paul Martin, Tanniemola B Liverpool, and Helen Weavers. Deep learning for rapid analysis of cell divisions in vivo during epithelial morphogenesis and repair. *eLife*, 12, 2024.

[14771] Hanna Tutaj, Katarzyna Tomala, Adrian Pirog, Marzena Marszałek, and Ryszard Korona. Extreme positive epistasis for fitness in monosomic yeast strains. *eLife*, 12, 2024.

[14772] Yujiro Umezaki, Sergio Hidalgo, Erika Nguyen, Tiffany Nguyen, Jay Suh, Sheena S Uchino, Joanna Chiu, and Fumika Hamada. Taste triggers a homeostatic temperature control in hungry flies. *eLife*, 13, 2024.

[14773] ElÇin ünal, Amanda J Su, and Siri C Yendluri. Control of meiotic entry by dual inhibition of a key mitotic transcription factor. *eLife*, 12, 2024.

[14774] Mark W Urban, Brittany A Charsar, Nicolette M Heinsinger, Shashirekha S Markandaiah, Lindsay Sprimont, Wei Zhou, Eric V Brown, Nathan T Henderson, Samantha J Thomas, Biswarup Ghosh, Rachel E Cain, Davide Trotti, Piera Pasinelli, Megan C Wright, Matthew B Dalva, and Angelo C Lepore. EphrinB2 knockdown in cervical spinal cord preserves diaphragm innervation in a mutant SOD1 mouse model of ALS. *eLife*, 12, 2024.

[14775] Alexandra Urbancokova, Terezie Hornofova, Josef Novak, Sarka Andrs Salajkova, Sona Stemberkova Hubackova, Alena Uvizl, Tereza Buchtova, Martin Mistrik, Brian McStay, Zdenek Hodny, Jiri Bartek, and Pavla Vasicova. Topological stress triggers persistent DNA lesions in ribosomal DNA with ensuing formation of PML-nucleolar compartment. *eLife*, 12, 2024.

[14776] Pasquale Valenzisi, Veronica Marabitti, Pietro Pichierri, and Annapaola Franchitto. WRNIP1 prevents transcription-associated genomic instability. *eLife*, 12, 2024.

[14777] Sofie Louise Valk, Veronika Engert, Lara Puhlmann, Roman Linz, Benoit Caldairou, Andrea Bernasconi, Neda Bernasconi, Boris C Bernhardt, and Tania Singer. Differential increase of hippocampal subfield volume after socio-affective mental training relates to reductions in diurnal cortisol. *eLife*, 12, 2024.

[14778] Marco van den Noort, Panagiotis Drougkas, Cristina Paulino, and Bert Poolman. The substrate-binding domains of the osmoregulatory ABC importer OpuA transiently interact. *eLife*, 12, 2024.

[14779] Marie-Sophie H van der Goes, Jakob Voigts, Jonathan P Newman, Enrique HS Toloza, Norma J Brown, Pranav Murugan, and Mark T Harnett. Coordinated head direction representations in mouse anterodorsal thalamic nucleus and retrosplenial cortex. *eLife*, 13, 2024.

[14780] Maureen van der Grinten, Jaap de Ruyter van Steveninck, Antonio Lozano, Laura Pijnacker, Bodo Rueckauer, Pieter Roelfsema, Marcel van Gerven, Richard van Wezel, Umut GüÇlü, and Yağmur GüÇlütürk. Towards biologically plausible phosphene simulation for the differentiable optimization of visual cortical prostheses. *eLife*, 13, 2024.

[14781] Meike E van der Heijden, Amanda M Brown, Dominic J Kizek, and Roy V Sillitoe. Cerebellar nuclei cells produce distinct pathogenic spike signatures in mouse models of ataxia, dystonia, and tremor. *eLife*, 12, 2024.

[14782] Julian van Gerwen, Stewart WC Masson, Harry B Cutler, Alexis Diaz Vegas, Meg Potter, Jacqueline Stöckli, Søren Madsen, Marin E Nelson, Sean J Humphrey, and David E James. The genetic and dietary landscape of the muscle insulin signalling network. *eLife*, 12, 2024.

[14783] Chris van Hoorn and Andrew P Carter. A cryo-electron tomography study of ciliary rootlet organization. *eLife*, 12, 2024.

[14784] Jonathan G Van Vranken, Jiaming Li, Julian Mintseris, Ting-Yu Wei, Catherine M Sniezek, Meagan Gadzuk-Shea, Steven P Gygi, and Devin K Schweppe. Large-scale characterization of drug mechanism of action using proteome-wide thermal shift assays. *eLife*, 13, 2024.

[14785] Jake VanBelzen, Bennet Sakelaris, Donna G Brickner, Nikita Marcou, Hermann Riecke, Niall M Mangan, and Jason H Brickner. Chromatin endogenous cleavage provides a global view of yeast RNA polymerase II transcription kinetics. *eLife*, 13, 2024.

[14786] Venkateshwari Varadharajan, Iyappan Ramachandiran, William J Massey, Raghav Jain, Rakhee Banerjee, Anthony J Horak, Megan R McMullen, Emily Huang, Annette Bellar, Shuhui W Lorkowski, Kailash Gulshan, Robert N Helsley, Isabella James, Vai Pathak, Jaividhya Dasarathy, Nicole Welch, Srinivasan Dasarathy, David Streem, Ofer Reizes, Daniela S Allende, Jonathan D Smith, Judith Simcox, Laura E Nagy, and J Mark Brown. Membrane-bound O-acyltransferase 7 (MBOAT7) shapes lysosomal lipid homeostasis and function to control alcohol-associated liver injury. *eLife*, 12, 2024.

[14787] Nikolaos Vardalakis, Amélie Aussel, Nicolas P Rougier, and Fabien B Wagner. A dynamical computational model of theta generation in hippocampal circuits to study theta-gamma oscillations during neurostimulation. *eLife*, 12, 2024.

[14788] Aivaras VaŠkevičius, Denis Baronas, Janis Leitans, Agnė Kvietkauskaitė, Audronė RukŠėnaitė, Elena Manakova, Zigmantas Toleikis, Algirdas Kaupinis, Andris Kazaks, Marius Gedgaudas, Aurelija Mickevičiūtė, Vaida Juozapaitienė, Helgi B Schiöth, Kristaps Jaudzems, Mindaugas Valius, Kaspars Tars, Saulius Gražulis, Franz-Josef Meyer-Almes, Jurgita Matulienė, Asta Zubrienė, Virginija Dudutienė, and Daumantas Matulis. Targeted anticancer pre-vinylsulfone covalent inhibitors of carbonic anhydrase IX. *eLife*, 13, 2024.

[14789] Brian G Vassallo, Noemie Scheidel, Sylvia E J Fischer, and Dennis H Kim. Bacteria are a major determinant of Orsay virus transmission and infection in Caenorhabditis elegans. *eLife*, 12, 2024.

[14790] Ester Vazquez-Fernandez, Jing Yang, Ziguo Zhang, Antonina E Andreeva, Paul Emsley, and David Barford. A comparative study of the cryo-EM structures of Saccharomyces cerevisiae and human anaphase-promoting complex/cyclosome (APC/C). *eLife*, 13, 2024.

[14791] John P Veillette, Fan Gao, and Howard C Nusbaum. Cardiac afferent signals can facilitate visual dominance in binocular rivalry. *eLife*, 13, 2024.

[14792] Vineeth Vengayil, Shreyas Niphadkar, Swagata Adhikary, Sriram Varahan, and Sunil Laxman. The deubiquitinase Ubp3/Usp10 constrains glucose-mediated mitochondrial repression via phosphate budgeting. *eLife*, 12, 2024.

[14793] Anil Verma, Chase E Hawes, Sonny R Elizaldi, Justin C Smith, Dhivyaa Rajasundaram, Gabriel Kristian Pedersen, Xiaoying Shen, LaTonya D Williams, Georgia D Tomaras, Pamela A Kozlowski, Rama R Amara, and Smita S Iyer. Tailoring Tfh profiles enhances antibody persistence to a clade C HIV-1 vaccine in rhesus macaques. *eLife*, 12, 2024.

[14794] Mayank Verma, Yoko Asakura, Xuerui Wang, Kasey Zhou, Mahmut ünverdi, Allison P Kann, Robert S Krauss, and Atsushi Asakura. Endothelial cell signature in muscle stem cells validated by VEGFA-FLT1-AKT1 axis promoting survival of muscle stem cell. *eLife*, 13, 2024.

[14795] Raphael Vidal, Eoin Leen, Steffi Herold, Mareike Müller, Daniel Fleischhauer, Christina Schülein-Völk, Dimitrios Papadopoulos, Isabelle Röschert, Leonie Uhl, Carsten P Ade, Peter Gallant, Richard Bayliss, Martin Eilers, and Gabriele Büchel. Association with TFIIIC limits MYCN localisation in hubs of active promoters and chromatin accumulation of non-phosphorylated RNA polymerase II. *eLife*, 13, 2024.

[14796] Diego Vidaurre. A generative model of electrophysiological brain responses to stimulation. *eLife*, 12, 2024.

[14797] Srinivasan Vijay, Nguyen Le Hoai Bao, Dao Nguyen Vinh, Le Thanh Hoang Nhat, Do Dang Anh Thu, Nguyen Le Quang, Le Pham Tien Trieu, Hoang Ngoc Nhung, Vu Thi Ngoc Ha, Phan Vuong Khac Thai, Dang Thi Minh Ha, Nguyen Huu Lan, Maxine Caws, Guy E Thwaites, Babak Javid, and Nguyen Thuy Thuong. Rifampicin tolerance and growth fitness among isoniazid-resistant clinical Mycobacterium tuberculosis isolates from a longitudinal study. *eLife*, 13, 2024.

[14798] Pau Vilimelis Aceituno, Dominic Dall'Osto, and Ioannis Pisokas. Theoretical principles explain the structure of the insect head direction circuit. *eLife*, 13, 2024.

[14799] Esteban Villar-Rodríguez, Cristina Cano-Melle, Lidón Marin-Marin, Maria Antònia Parcet, and César Avila. What happens to the inhibitory control functions of the right inferior frontal cortex when this area is dominant for language? *eLife*, 12, 2024.

[14800] Vera Vollenweider, Karoline Rehm, Clara Chepkirui, Manuela Pérez-Berlanga, Magdalini Polymenidou, Jörn Piel, Laurent Bigler, and Rolf Kümmerli. Antimicrobial activity of iron-depriving pyoverdines against human opportunistic pathogens. *eLife*, 13, 2024.

[14801] Svetlana Volotsky, Opher Donchin, and Ronen Segev. The archerfish uses motor adaptation in shooting to correct for changing physical conditions. *eLife*, 12, 2024.

[14802] Nguyen Lam Vuong, Nguyen Than Ha Quyen, Nguyen Thi Hanh Tien, Kien Duong Thi Hue, Huynh Thi Le Duyen, Phung Khanh Lam, Dong Thi Hoai Tam, Tran Van Ngoc, Thomas Jaenisch, Cameron P Simmons, Sophie Yacoub, Bridget A Wills, and Ronald Geskus. Dengue viremia kinetics and effects on platelet count and clinical outcomes: An analysis of 2340 patients from Vietnam. *eLife*, 13, 2024.

[14803] Brennan J Wadsworth, Marina Leiwe, Eleanor A Minogue, Pedro P Cunha, Viktor Engman, Carolin Brombach, Christos Asvestis, Shiv K Sah-Teli, Emilia Marklund, Peppi Karppinen, Jorge L Ruas, Helene Rundqvist, Johanna T Lanner, and Randall S Johnson. A 2-hydroxybutyrate-mediated feedback loop regulates muscular fatigue. *eLife*, 12, 2024.

[14804] Konrad Wagstyl, Sophie Adler, Jakob Seidlitz, Simon Vandekar, Travis T Mallard, Richard Dear, Alex R DeCasien, Theodore D Satterthwaite, Siyuan Liu, Petra E Vértes, Russell T Shinohara, Aaron Alexander-Bloch, Daniel H Geschwind, and Armin Raznahan. Transcriptional cartography integrates multiscale biology of the human cortex. *eLife*, 12, 2024.

[14805] Kristine B Walhovd, Stine K Krogsrud, Inge K Amlien, Øystein Sørensen, Yunpeng Wang, Anne Cecilie S Bråthen, Knut Overbye, Jonas Kransberg, Athanasia M Mowinckel, Fredrik Magnussen, Martine Herud, Asta K Håberg, Anders Martin Fjell, and Didac Vidal-Pineiro. Fetal influence on the human brain through the lifespan. *eLife*, 12, 2024.

[14806] Timothy J Walker, Eduardo Reyes-Alvarez, Brandy D Hyndman, Michael G Sugiyama, Larissa CB Oliveira, Aisha N Rekab, Mathieu JF Crupi, Rebecca Cabral-Dias, Qianjin Guo, Patricia LM Dahia, Douglas S Richardson, Costin N Antonescu, and Lois M Mulligan. Loss of tumor suppressor TMEM127 drives RET-mediated transformation through disrupted membrane dynamics. *eLife*, 12, 2024.

[14807] Kevin Walsh, David P McGovern, Jessica Dully, Simon P Kelly, and Redmond G O'Connell. Prior probability cues bias sensory encoding with increasing task exposure. *eLife*, 12, 2024.

[14808] Ban Wang, Alexander L Starr, and Hunter B Fraser. Cell-type-specific cis-regulatory divergence in gene expression and chromatin accessibility revealed by human-chimpanzee hybrid cells. *eLife*, 12, 2024.

[14809] Chong Wang, Rikuan Zheng, Tianhang Zhang, and Chaomin Sun. Polysaccharides induce deep-sea Lentisphaerae strains to release chronic bacteriophages. *eLife*, 13, 2024.

[14810] Chuchu Wang, Chunyu Zhao, Hu Xiao, Jiali Qiang, Zhenying Liu, Jinge Gu, Shengnan Zhang, Dan Li, Yaoyang Zhang, Jacqueline Burré, Jiajia Diao, and Cong Liu. N-acetylation of $\alpha$-synuclein enhances synaptic vesicle clustering mediated by $\alpha$-synuclein and lysophosphatidylcholine. *eLife*, 13, 2024.

[14811] Chun Wang, Khushpreet Kaur, Canxin Xu, Yousef Abu-Amer, and Gabriel Mbalaviele. Chemotherapy activates inflammasomes to cause inflammation-associated bone loss. *eLife*, 13, 2024.

[14812] Deming Wang, Jiangnan Yang, Le Liu, Yi Zhou, Peng Xu, Min Qin, and Pu Huang. Alasemenia, the earliest ovule with three wings and without cupule. *eLife*, 13, 2024.

[14813] Deng Wang, Yaqin Qiang, Junfeng Guo, Jean Vannier, Zuchen Song, Jiaxin Peng, Boyao Zhang, Jie Sun, Yilun Yu, Yiheng Zhang, Tao Zhang, Xiaoguang Yang, and Jian Han. Early evolution of the ecdysozoan body plan. *eLife*, 13, 2024.

[14814] Fangyan Wang, Xiujie Liu, Furong Huang, Yan Zhou, Xinyu Wang, Zhengyang Song, Sisi Wang, Xiaoting Wang, Dibang Shi, Gaoyi Ruan, Xiawei Ji, Eryao Zhang, Zenglin Tan, Yuqing Ye, Chuang Wang, Jesse Zhu, and Wantie Wang. Gut microbiota-derived gamma-aminobutyric acid from metformin treatment reduces hepatic ischemia/reperfusion injury through inhibiting ferroptosis. *eLife*, 12, 2024.

[14815] Guangqin Wang, Yunpeng Gu, and Zhiyong Liu. Deciphering the genetic interactions between Pou4f3, Gfi1, and Rbm24 in maintaining mouse cochlear hair cell survival. *eLife*, 12, 2024.

[14816] Guoqiang Wang, Ryan J Guasp, Sangeena Salam, Edward Chuang, Andrés Morera, Anna J Smart, David Jimenez, Sahana Shekhar, Emily Friedman, Ilija Melentijevic, Ken C Nguyen, David H Hall, Barth D Grant, and Monica Driscoll. Mechanical force of uterine occupation enables large vesicle extrusion from proteostressed maternal neurons. *eLife*, 13, 2024.

[14817] Hao Wang, Kai He, Huan Zhang, Quanyong Zhang, Lei Cao, Jing Li, Zhaoshan Zhong, Hao Chen, Li Zhou, Chao Lian, Minxiao Wang, Kai Chen, Pei-Yuan Qian, and Chaolun Li. Deciphering deep-sea chemosynthetic symbiosis by single-nucleus RNA-sequencing. *eLife*, 12, 2024.

[14818] Hao Wang, Chen Ye, Qi Lu, Zhijie Jiang, Chao Jiang, Chun Zhou, Na Li, Caiqiao Zhang, Guoping Zhao, Min Yue, and Yan Li. Bacterial exonuclease III expands its enzymatic activities on single-stranded DNA. *eLife*, 13, 2024.

[14819] Hao Wang, Biying Zhu, Siming Qiao, Chunxia Dong, Xiaochun Wan, Weimin Gong, and Zhaoliang Zhang. Structure and evolution of alanine/serine decarboxylases and the engineering of theanine production. *eLife*, 12, 2024.

[14820] Huishan Wang, Xingyan Liu, Yamin Liu, Chencheng Yang, Yaxin Ye, Xiaomei Yu, Nengyin Sheng, Shihua Zhang, Bingyu Mao, and Pengcheng Ma. The E3 ubiquitin ligase RNF220 maintains hindbrain Hox expression patterns through regulation of WDR5 stability. *eLife*, 13, 2024.

[14821] Jing Wang, Min Su, Dongmei Zhang, Ludi Zhang, Chenxu Niu, Chaoyi Li, Shuangzhu You, Yuqi Sang, Yongxue Zhang, Xiaona Du, and Hailin Zhang. The cation channel mechanisms of subthreshold inward depolarizing currents in the mice VTA dopaminergic neurons and their roles in the chronic-stress-induced depression-like behavior. *eLife*, 12, 2024.

[14822] Jing Wang, Yingying Yin, Lei Yang, Junchao Qin, Zixiang Wang, Chunhong Qiu, Yuan Gao, Gang Lu, Fei Gao, Zi-Jiang Chen, Xiyu Zhang, Hongbin Liu, and Zhaojian Liu. TMC7 deficiency causes acrosome biogenesis defects and male infertility in mice. *eLife*, 13, 2024.

[14823] Jing-Ping Wang, Chun-Hao Hung, Yae-Huei Liou, Ching-Chen Liu, Kun-Hai Yeh, Keh-Yang Wang, Zheng-Sheng Lai, Biswanath Chatterjee, Tzu-Chi Hsu, Tung-Liang Lee, Yu-Chiau Shyu, Pei-Wen Hsiao, Liuh-Yow Chen, Trees-Juen Chuang, Chen-Hsin Albert Yu, Nan-Shih Liao, and C-K James Shen. Long-term hematopoietic transfer of the anti-cancer and lifespan-extending capabilities of a genetically engineered blood system by transplantation of bone marrow mononuclear cells. *eLife*, 12, 2024.

[14824] Linkang Wang, Haiyan Wang, Xinxin Li, Mengyuan Zhu, Dongyang Gao, Dayue Hu, Zhixuan Xiong, Xiangmin Li, and Ping Qian. Bacillus velezensis HBXN2020 alleviates Salmonella Typhimurium infection in mice by improving intestinal barrier integrity and reducing inflammation. *eLife*, 13, 2024.

[14825] Pinhua Wang, Sunayana Sarkar, Menghuan Zhang, Tingting Xiao, Fenhua Kong, Zhe Zhang, Deepa Balasubramanian, Nandan Jayaram, Sayantan Datta, Ruyu He, Ping Wu, Peng Chao, Ying Zhang, Michael Washburn, Laurence A Florens, Sonal Nagarkar-Jaiswal, Manish Jaiswal, and Man Mohan.

DYRK1A interacts with the tuberous sclerosis complex and promotes mTORC1 activity. *eLife*, 12, 2024.

[14826] Wen Wang, Xiao Zhang, Ning Zhao, Ze-Hua Xu, Kangxin Jin, and Zi-Bing Jin. RNA fusion in human retinal development. *eLife*, 13, 2024.

[14827] Xiang Wang, Gan Shen, Yihong Yang, Chuan Jiang, Tiechao Ruan, Xue Yang, Liangchai Zhuo, Yingteng Zhang, Yangdi Ou, Xinya Zhao, Shunhua Long, Xiangrong Tang, Tingting Lin, and Ying Shen. DNAH3 deficiency causes flagellar inner dynein arm loss and male infertility in humans and mice. *eLife*, 13, 2024.

[14828] Xiaoning Wang, Jinxiang Zhao, Jiehuan Xu, Bowen Li, Xia Liu, Gangcai Xie, Xuchu Duan, and Dong Liu. Noncaloric monosaccharides induce excessive sprouting angiogenesis in zebrafish via foxo1a-marcksl1a signal. *eLife*, 13, 2024.

[14829] Ya-Juan Wang, Xiao-Jing Di, Pei-Pei Zhang, Xi Chen, Marnie P Williams, Dong-Yun Han, Raad Nashmi, Brandon J Henderson, Fraser J Moss, and Ting-Wei Mu. Hsp47 promotes biogenesis of multi-subunit neuroreceptors in the endoplasmic reticulum. *eLife*, 13, 2024.

[14830] Yuanyuan Wang, Fan Xu, Guangyi Li, Chen Cheng, Bowen Yu, Ze Zhang, Dandan Kong, Fabao Chen, Yali Liu, Zhen Fang, Longxing Cao, Yang Yu, Yijun Gu, and Yongning He. Structure of scavenger receptor SCARF1 and its interaction with lipoproteins. *eLife*, 13, 2024.

[14831] Yuedan Wang, Ying Li, Jiaqing Feng, Chuansen Wang, Yuwei Wan, Bingyang Lv, Yinming Li, Hao Xie, Ting Chen, Faxi Wang, Ziyue Li, Anhuai Yang, and Xuan Xiao. Transcriptional responses in a mouse model of silicone wire embolization induced acute retinal artery ischemia and reperfusion. *eLife*, 13, 2024.

[14832] Yuhao Wang, Linhao Ruan, Jin Zhu, Xi Zhang, Alexander Chih-Chieh Chang, Alexis Tomaszewski, and Rong Li. Metabolic regulation of misfolded protein import into mitochondria. *eLife*, 12, 2024.

[14833] Yujiang Wang, Karoline Leiberg, Nathan Kindred, Christopher R Madan, Colline Poirier, Christopher I Petkov, Peter Neal Taylor, and Bruno Mota. Neuro-evolutionary evidence for a universal fractal primate brain shape. *eLife*, 12, 2024.

[14834] Zhuqing Wang, Yue Wang, Tong Zhou, Sheng Chen, Dayton Morris, Rubens Daniel Miserani Magalhães, Musheng Li, Shawn Wang, Hetan Wang, Yeming Xie, Hayden McSwiggin, Daniel Oliver, Shuiqiao Yuan, Huili Zheng, Jaaved Mohammed, Eric C Lai, John R McCarrey, and Wei Yan. The rapidly evolving X-linked MIR-506 family fine-tunes spermatogenesis to enhance sperm competition. *eLife*, 13, 2024.

[14835] Ziguang Wang, Shuai Li, Sheng Zhang, Tianyu Zhang, Yujie Wu, Anqi Liu, Kui Wang, Xiaowen Ji, Haiqun Cao, Yinglao Zhang, Eng King Tan, Yongcheng Wang, Yirong Wang, and Wei Liu. Hosts manipulate lifestyle switch and pathogenicity heterogeneity of opportunistic pathogens in the single-cell resolution. *eLife*, 13, 2024.

[14836] Zijing Wang, Bihan Xia, Shaochong Qi, Xian Zhang, Xiaoshuang Zhang, Yan Li, Huimin Wang, Miao Zhang, Ziyi Zhao, David Kerr, Li Yang, Shijie Cai, and Jilin Yang. Bestrophin-4 relays HES4 and interacts with TWIST1 to suppress epithelial-to-mesenchymal transition in colorectal cancer cells. *eLife*, 12, 2024.

[14837] Stephanie A Wankowicz, Ashraya Ravikumar, Shivani Sharma, Blake Riley, Akshay Raju, Daniel W Hogan, Jessica Flowers, Henry van den Bedem, Daniel A Keedy, and James S Fraser. Automated multiconformer model building for X-ray crystallography and cryo-EM. *eLife*, 12, 2024.

[14838] Rebecca Warfvinge, Linda Geironson Ulfsson, Parashar Dhapola, Fatemeh Safi, Mikael Sommarin, Shamit Soneji, Henrik Hjorth-Hansen, Satu Mustjoki, Johan Richter, Ram Krishna Thakur, and Göran Karlsson. Single-cell multiomics analysis of chronic myeloid leukemia links cellular heterogeneity to therapy response. *eLife*, 12, 2024.

[14839] Kurt Warnhoff, Sushila Bhattacharya, Jennifer Snoozy, Peter C Breen, and Gary Ruvkun. Hypoxia-inducible factor induces cysteine dioxygenase and promotes cysteine homeostasis in Caenorhabditis elegans. *eLife*, 12, 2024.

[14840] Abdul Wasim, Sneha Menon, and Jagannath Mondal. Modulation of $\alpha$-synuclein aggregation amid diverse environmental perturbation. *eLife*, 13, 2024.

[14841] Tetsuya Watabe, Shinya Yamahira, Kanako Takakura, Dean Thumkeo, Shuh Narumiya, Michiyuki Matsuda, and Kenta Terai. Calcium transients trigger switch-like discharge of prostaglandin E2 in an extracellular signal-regulated kinase-dependent manner. *eLife*, 12, 2024.

[14842] Kiichi Watanabe, Hui Chiu, and David J Anderson. Whole-brain in situ mapping of neuronal activation in Drosophila during social behaviors and optogenetic stimulation. *eLife*, 12, 2024.

[14843] Takashi Watanabe, Hikaru Hata, Yoshiki Mochizuki, Fumie Yokoyama, Tomoko Hasegawa, Naveen Kumar, Tomohiro Kurosaki, Osamu Ohara, and Hidehiro Fukuyama. Development of a new genotype–phenotype linked antibody screening system. *eLife*, 13, 2024.

[14844] Shintaro Watanuki, Hiroshi Kobayashi, Yuki Sugiura, Masamichi Yamamoto, Daiki Karigane, Kohei Shiroshita, Yuriko Sorimachi, Shinya Fujita, Takayuki Morikawa, Shuhei Koide, Motohiko Oshima, Akira Nishiyama, Koichi Murakami, Miho Haraguchi, Shinpei Tamaki, Takehiro Yamamoto, Tomohiro Yabushita, Yosuke Tanaka, Go Nagamatsu, Hiroaki Honda, Shinichiro Okamoto, Nobuhito Goda, Tomohiko Tamura, Ayako Nakamura-Ishizu, Makoto Suematsu, Atsushi Iwama, Toshio Suda, and Keiyo Takubo. Context-dependent modification of PFKFB3 in hematopoietic stem cells promotes anaerobic glycolysis and ensures stress hematopoiesis. *eLife*, 12, 2024.

[14845] Jamie Mitchel Waterman, Tristan Michael Cofer, Lei Wang, Gaetan Glauser, and Matthias Erb. High-resolution kinetics of herbivore-induced plant volatile transfer reveal clocked response patterns in neighboring plants. *eLife*, 12, 2024.

[14846] James A Watson, Robert J Commons, Joel Tarning, Julie A Simpson, Alejandro Llanos Cuentas, Marcus VG Lacerda, Justin A Green, Gavin CKW Koh, Cindy S Chu, FranÇois H Nosten, Richard N Price, Nicholas PJ Day, and Nicholas J White. Response to comment on 'The clinical pharmacology of tafenoquine in the radical cure of Plasmodium vivax malaria: An individual patient data meta-analysis'. *eLife*, 13, 2024.

[14847] Lukas M Weber, Heena R Divecha, Matthew N Tran, Sang Ho Kwon, Abby Spangler, Kelsey D Montgomery, Madhavi Tippani, Rahul Bharadwaj, Joel E Kleinman, Stephanie C Page, Thomas M Hyde, Leonardo Collado-Torres, Kristen R Maynard, Keri Martinowich, and Stephanie C Hicks. The gene expression landscape of the human locus coeruleus revealed by single-nucleus and spatially-resolved transcriptomics. *eLife*, 12, 2024.

[14848] Ramona Weber and Chung-Te Chang. Human DDX6 regulates translation and decay of inefficiently translated mRNAs. *eLife*, 13, 2024.

[14849] Xujin Wei, Jie Liu, Jia Cheng, Wangyu Cai, Wen Xie, Kang Wang, Lingyun Lin, Jingjing Hou, Jianchun Cai, and Huiqin Zhuo. Super-enhancer-driven ZFP36L1 promotes PD-L1 expression in infiltrative gastric cancer. *eLife*, 13, 2024.

[14850] Catherine A Weibel, Andrew L Wheeler, Jennifer E James, Sara M Willis, Hanon McShea, and Joanna Masel. The protein domains of vertebrate species in which selection is more effective have greater intrinsic structural disorder. *eLife*, 12, 2024.

[14851] Tobias Weinberger, Messerer Denise, Markus Joppich, Maximilian Fischer, Clarisabel Garcia Rodriguez, Konda Kumaraswami, Vanessa Wimmler, Sonja Ablinger, Saskia Räuber, Jiahui Fang, Lulu Liu, Wing Han Liu, Julia Winterhalter, Johannes Lichti, Lukas Thomas, Dena Esfandyari, Guelce Percin, Sandra Matin, Andrés Hidalgo, Claudia Waskow, Stefan Engelhardt, Andrei Todica, Ralf Zimmer, Clare Pridans, Elisa Gomez Perdiguero, and Christian Schulz. Resident and recruited macrophages differentially contribute to cardiac healing after myocardial ischemia. *eLife*, 12, 2024.

[14852] Alex Weiss, Cassandra D'Amata, Bret J Pearson, and Madeline N Hayes. A syngeneic spontaneous zebrafish model of tp53-deficient, EGFRvIII, and PI3KCAH1047R-driven glioblastoma reveals inhibitory roles for inflammation during tumor initiation and relapse in vivo. *eLife*, 13, 2024.

[14853] Chao Wen, Yuyi Lu, Cwyn Solvi, Shunping Dong, Cai Wang, Xiujun Wen, Haijun Xiao, Shikui Dong, Junbao Wen, Fei Peng, and Lars Chittka. Does bumblebee preference of continuous over interrupted strings in string-pulling tasks indicate means-end comprehension? *eLife*, 13, 2024.

[14854] Shijie Wen, Hiroshi Arakawa, Shigeru Yokoyama, Yoshiyuki Shirasaka, Haruhiro Higashida, and Ikumi Tamai. Functional identification of soluble uric acid as an endogenous inhibitor of CD38. *eLife*, 13, 2024.

[14855] Yifei Weng, Shiyi Zhou, Katherine Morillo, Rachel Kaletsky, Sarah Lin, and Coleen T Murphy. The neuron-specific IIS/FOXO transcriptome in aged animals reveals regulatory mechanisms of cognitive aging. *eLife*, 13, 2024.

[14856] Yihe Weng, Johann Kruschwitz, Laura M Rueda-Delgado, Kathy L Ruddy, Rory Boyle, Luisa Franzen, Emin Serin, Tochukwu Nweze, Jamie Hanson, Alannah Smyth, Tom Farnan, Tobias Banaschewski, Arun LW Bokde, Sylvane Desrivières, Herta Flor, Antoine Grigis, Hugh Garavan, Penny A Gowland, Andreas Heinz, Rüdiger Brühl, Jean-Luc Martinot, Marie-Laure Paillère Martinot, Eric Artiges, Jane McGrath, Frauke Nees, Dimitri Papadopoulos Orfanos, Tomas Paus, Luise Poustka, Nathalie Holz, Juliane Fröhner, Michael N Smolka, Nilakshi Vaidya, Gunter Schumann, Henrik Walter, and Robert Whelan. A robust brain network for sustained attention from adolescence to adulthood that predicts later substance use. *eLife*, 13, 2024.

[14857] Diek W Wheeler, Jeffrey D Kopsick, Nate Sutton, Carolina Tecuatl, Alexander O Komendantov, Kasturi Nadella, and Giorgio A Ascoli. Hippocampome.org 2.0 is a knowledge base enabling data-driven spiking neural network simulations of rodent hippocampal circuits. *eLife*, 12, 2024.

[14858] Kantapon Pum Wiboonsaksakul, Olivia ME Leavitt Brown, and Kathleen E Cullen. Restoring vestibular function during natural self-motion: Progress and challenges. *eLife*, 13, 2024.

[14859] Daniel M Williams and Andrew A Peden. S-acylation of NLRP3 provides a nigericin sensitive gating mechanism that controls access to the Golgi. *eLife*, 13, 2024.

[14860] Zachary H Williams, Alvaro Dafonte Imedio, Lea Gaucherand, Derek C Lee, Salwa Mohd Mostafa, James P Phelan, John M Coffin, and Welkin E Johnson. Recombinant origin and interspecies transmission of a HERV-K(HML-2)-related primate retrovirus with a novel RNA transport element. *eLife*, 13, 2024.

[14861] Jacob H Wilmot, Cassiano RAF Diniz, Ana P Crestani, Kyle R Puhger, Jacob Roshgadol, Lin Tian, and Brian Joseph Wiltgen. Phasic locus coeruleus activity enhances trace fear conditioning by increasing dopamine release in the hippocampus. *eLife*, 12, 2024.

[14862] Volker Winstel, Evan R Abt, Thuc M Le, and Caius G Radu. Targeting host deoxycytidine kinase mitigates Staphylococcus aureus abscess formation. *eLife*, 12, 2024.

[14863] Matthew F Wipperman, Allen Z Lin, Kaitlyn M Gayvert, Benjamin Lahner, Selin Somersan-Karakaya, Xuefang Wu, Joseph Im, Minji Lee, Bharatkumar Koyani, Ian Setliff, Malika Thakur, Daoyu Duan, Aurora Breazna, Fang Wang, Wei Keat Lim, Gabor Halasz, Jacek Urbanek, Yamini Patel, Gurinder S Atwal, Jennifer D Hamilton, Samuel Stuart, Oren Levy, Andreja Avbersek, Rinol Alaj, Sara C Hamon, and Olivier Harari. Digital wearable insole-based identification of knee arthropathies and gait signatures using machine learning. *eLife*, 13, 2024.

[14864] Pattama Wiriyasermkul, Satomi Moriyama, Masataka Suzuki, Pornparn Kongpracha, Nodoka Nakamae, Saki Takeshita, Yoko Tanaka, Akina Matsuda, Masaki Miyasaka, Kenji Hamase, Tomonori Kimura, Masashi Mita, Jumpei Sasabe, and Shushi Nagamori. A multi-hierarchical approach reveals d-serine as a hidden substrate of sodium-coupled monocarboxylate transporters. *eLife*, 12, 2024.

[14865] Jenny L Witten, Veronika Lukyanova, and Wolf M Harmening. Sub-cone visual resolution by active, adaptive sampling in the human foveola. *eLife*, 13, 2024.

[14866] Christoph Arne Wittkamp, Maren-Isabel Wolf, and Michael Rose. The neural dynamics of positive and negative expectations of pain. *eLife*, 13, 2024.

[14867] Martin Wohlwend, Pirkka-Pekka Laurila, Ludger JE Goeminne, Tanes Lima, Ioanna Daskalaki, Xiaoxu Li, Giacomo von Alvensleben, Barbara Crisol, Renata Mangione, Hector Gallart-Ayala, Olivier Burri, Stephen Butler, Jonathan Morris, Nigel Turner, Julijana Ivanisevic, and Johan Auwerx. Inhibition of CERS1 in skeletal muscle exacerbates age-related muscle dysfunction. *eLife*, 12, 2024.

[14868] Nicole C Woitowich, Sarah R Hengel, Christopher Solis, Tauras P Vilgalys, Joel Babdor, and Daniel J Tyrrell. Analysis of NIH K99/R00 awards and the career progression of awardees. *eLife*, 12, 2024.

[14869] Jing Jun Wong, Alessandro Bongioanni, Matthew FS Rushworth, and Bolton KH Chau. Distractor effects in decision making are related to the individual's style of integrating choice attributes. *eLife*, 12, 2024.

[14870] Victor C Wong, Patrick R Houlihan, Hui Liu, Deepika Walpita, Michael C DeSantis, Zhe Liu, and Erin K O'Shea. Plasticity-induced actin polymerization in the dendritic shaft regulates intracellular AMPA receptor trafficking. *eLife*, 13, 2024.

[14871] Andrew E Worthy, Joanna T Anderson, Alicia R Lane, Laura J Gomez-Perez, Anthony A Wang, Ronald W Griffith, Andre F Rivard, Jay B Bikoff, and Francisco J Alvarez. Spinal V1 inhibitory interneuron clades differ in birthdate, projections to motoneurons, and heterogeneity. *eLife*, 13, 2024.

[14872] Bingbing Wu, Chenghong Long, Yuzhuo Yang, Zhe Zhang, Shuang Ma, Yanjie Ma, Huafang Wei, Jinghe Li, Hui Jiang, Wei Li, and Chao Liu. CCDC113 stabilizes sperm axoneme and head-tail coupling apparatus to ensure male fertility. *eLife*, 13, 2024.

[14873] Haoyang Wu, Peiyuan Lv, Jinyu Wang, Brian Bennett, Jiajia Wang, Pishun Li, Yi Peng, Guang Hu, and Jiaji Lin. Genetic screen identified PRMT5 as a neuroprotection target against cerebral ischemia. *eLife*, 12, 2024.

[14874] Jian Wu, Nisha A Jonniya, Sophia P Hirakis, Cristina Olivieri, Gianluigi Veglia, Alexandr P Kornev, and Susan S Taylor. Role of the $\alpha$C-$\beta$4 loop in protein kinase structure and dynamics. *eLife*, 12, 2024.

[14875] Shuai Wu, Archit Garg, Zachary Mazanek, Gretchen Belotte, Jeffery J Zhou, Christina M Stallings, Jacob Lueck, Aubrey Roland, Michael A Chattergoon, and Jungsan Sohn. Design principles for inflammasome inhibition by pyrin-only-proteins. *eLife*, 13, 2024.

[14876] Shuting Wu, Asem Wardak, Mehak M Khan, Christopher H Chen, and Wade G Regehr. Implications of variable synaptic weights for rate and temporal coding of cerebellar outputs. *eLife*, 13, 2024.

[14877] Songyuan Wu, Xiaoling Tong, Chenxing Peng, Jiangwen Luo, Chenghao Zhang, Kunpeng Lu, Chunlin Li, Xin Ding, Xiaohui Duan, Yaru Lu, Hai Hu, Duan Tan, and Fangyin Dai. The BTB-ZF gene Bm-mamo regulates pigmentation in silkworm caterpillars. *eLife*, 12, 2024.

[14878] Yifan Wu, Kunhua Qin, Yi Xu, Shreya Rajhans, Truong Vo, Kevin M Lopez, Jun Liu, Michael H Nipper, Janice Deng, Xue Yin, Logan R Ramjit, Zhenqing Ye, Yu Luan, H Efsun Arda, and Pei Wang. Hippo pathway-mediated YAP1/TAZ inhibition is essential for proper pancreatic endocrine specification and differentiation. *eLife*, 13, 2024.

[14879] Zhenyong Wu, Grant F Kusick, Manon MM Berns, Sumana Raychaudhuri, Kie Itoh, Alexander M Walter, Edwin R Chapman, and Shigeki Watanabe. Synaptotagmin 7 docks synaptic vesicles to support facilitation and Doc2$\alpha$-triggered asynchronous release. *eLife*, 12, 2024.

[14880] Chenjie Xia, Huihui Xu, Liang Fang, Jiali Chen, Wenhua Yuan, Danqing Fu, Xucheng Wang, Bangjian He, Luwei Xiao, Chengliang Wu, Peijian Tong, Di Chen, Pinger Wang, and Hongting Jin. $\beta$-catenin inhibition disrupts the homeostasis of osteogenic/adipogenic differentiation leading to the development of glucocorticoid-induced osteonecrosis of the femoral head. *eLife*, 12, 2024.

[14881] Jing Xie, Yafei Xu, Xuhua Liu, Li Long, Ji Chen, Chunyan Huang, Yan Shao, Zhiqing Cai, Zhimin Zhang, Ruixin Zhou, Jiarong Leng, Xiaochun Bai, and Qiancheng Song. Mechanically stimulated osteocytes maintain tumor dormancy in bone metastasis of non-small cell lung cancer by releasing small extracellular vesicles. *eLife*, 12, 2024.

[14882] Xianghong Xie, Mingyue Gao, Wei Zhao, Chunmei Li, Weihong Zhang, Jiahui Yang, Yinliang Zhang, Enhui Chen, Yanfang Guo, Zeyu Guo, Minglong Zhang, Ebenezeri Erasto Ngowi, Heping Wang, Xiaoman Wang, Yinghan Zhu, Yiting Wang, Xiaolu Li, Hong Yao, Li Yan, Fude Fang, Meixia Li, Aijun Qiao, and Xiaojun Liu. LncRNA Snhg3 aggravates hepatic steatosis via PPAR$\gamma$ signaling. *eLife*, 13, 2024.

[14883] Xufeng Xie, Xi Chen, Shilei Zhang, Jiuxi Liu, Wenlong Zhang, and Yongguo Cao. Neutralizing gut-derived lipopolysaccharide as a novel therapeutic strategy for severe leptospirosis. *eLife*, 13, 2024.

[14884] Yeming Xie, Fengying Ruan, Yaning Li, Meng Luo, Chen Zhang, Zhichao Chen, Zhe Xie, Zhe Weng, Weitian Chen, Wenfang Chen, Yitong Fang, Yuxin Sun, Mei Guo, Juan Wang, Shouping Xu, Hongqi Wang, and Chong Tang. Spatial chromatin accessibility sequencing resolves high-order spatial interactions of epigenomic markers. *eLife*, 12, 2024.

[14885] Yu-Feng Xie, Jane Yang, Stéphanie Ratté, and Steven A Prescott. Similar excitability through different sodium channels and implications for the analgesic efficacy of selective drugs. *eLife*, 12, 2024.

[14886] Zhongyun Xie, Yongping Chai, Zhiwen Zhu, Zijie Shen, Zhengyang Guo, Zhiguang Zhao, Long Xiao, Zhuo Du, Guangshuo Ou, and Wei Li. Vacuolar H+-ATPase determines daughter cell fates through asymmetric segregation of the nucleosome remodeling and deacetylase complex. *eLife*, 12, 2024.

[14887] Luyang Xiong, Irina Zhevlakova, Xiaoxia Z West, Detao Gao, Rakhilya Murtazina, Anthony Horak, J Mark Brown, Iuliia Molokotina, Eugene A Podrez, and Tatiana V Byzova. TLR2 regulates hair follicle cycle and regeneration via BMP signaling. *eLife*, 12, 2024.

[14888] Weihong Xiong, Maozhen Qin, and Haining Zhong. PKA regulation of neuronal function requires the dissociation of catalytic subunits from regulatory subunits. *eLife*, 13, 2024.

[14889] Chundi Xu, Tyler B Ramos, Owen J Marshall, and Chris Q Doe. Notch signaling and Bsh homeodomain activity are integrated to diversify Drosophila lamina neuron types. *eLife*, 12, 2024.

[14890] Chundi Xu, Tyler B Ramos, Edward M Rogers, Michael B Reiser, and Chris Q Doe. Homeodomain proteins hierarchically specify neuronal diversity and synaptic connectivity. *eLife*, 12, 2024.

[14891] Hang Xu, Zihao Yuan, Kunpeng Qin, Shuai Jiang, and Li Sun. The molecular mechanism and evolutionary divergence of caspase 3/7-regulated gasdermin E activation. *eLife*, 12, 2024.

[14892] Jiayun Xu, Mauricio Girardi-Schappo, Jean-Claude Beique, André Longtin, and Leonard Maler. Shortcutting from self-motion signals reveals a cognitive map in mice. *eLife*, 13, 2024.

[14893] Longjiang Xu, Zan Yuan, Jiafeng Zhou, Yuan Zhao, Wei Liu, Shuaiyao Lu, Zhanlong He, Boqin Qiang, Pengcheng Shu, Yang Chen, and Xiaozhong Peng. Temporal transcriptomic dynamics in developing macaque neocortex. *eLife*, 12, 2024.

[14894] Mengyuan Xu, Torben Neelands, Alexander S Powers, Yan Liu, Steven D Miller, Grigore D Pintilie, J Du Bois, Ron O Dror, Wah Chiu, and Merritt Maduke. CryoEM structures of the human CLC-2 voltage-gated chloride channel reveal a ball-and-chain gating mechanism. *eLife*, 12, 2024.

[14895] Peibo Xu, Jian Peng, Tingli Yuan, Zhaoqin Chen, Hui He, Ziyan Wu, Ting Li, Xiaodong Li, Luyue Wang, Le Gao, Jun Yan, Wu Wei, Chengyu T Li, Zhen-Ge Luo, and Yuejun Chen. High-throughput mapping of single-neuron projection and molecular features by retrograde barcoded labeling. *eLife*, 13, 2024.

[14896] Zhimin Xu, Zhao Wang, Lifang Wang, and Yingchuan B Qi. Essential function of transmembrane transcription factor MYRF in promoting transcription of miRNA lin-4 during C. elegans development. *eLife*, 12, 2024.

[14897] Gang Xue, Xiaoyi Zhang, Wanqi Li, Lu Zhang, Zongxu Zhang, Xiaolin Zhou, Di Zhang, Lei Zhang, and Zhiyuan Li. A logic-incorporated gene regulatory network deciphers principles in cell fate decisions. *eLife*, 12, 2024.

[14898] Yunlu Xue, Yimin Zhou, and Constance L Cepko. Txnip deletions and missense alleles prolong the survival of cones in a retinitis pigmentosa mouse model. *eLife*, 12, 2024.

[14899] Dilip Kumar Yadav, Andrew C Chang, Noa WF Grooms, Samuel H Chung, and Christopher V Gabel. O-GlcNAc signaling increases neuron regeneration through one-carbon metabolism in Caenorhabditis elegans. *eLife*, 13, 2024.

[14900] Behzad Yaghmaeian Salmani, Laura Lahti, Linda Gillberg, Jesper Kjaer Jacobsen, Ioannis Mantas, Per Svenningsson, and Thomas Perlmann. Transcriptomic atlas of midbrain dopamine neurons uncovers differential vulnerability in a Parkinsonism lesion model. *eLife*, 12, 2024.

[14901] Donghui Yan, Bowen Hu, Burcu F Darst, Shubhabrata Mukherjee, Brian W Kunkle, Yuetiva Deming, Logan Dumitrescu, Yunling Wang, Adam Naj, Amanda Kuzma, Yi Zhao, Hyunseung Kang, Sterling C Johnson, Cruchaga Carlos, Timothy J Hohman, Paul K Crane, Corinne D Engelman, and Qiongshi Lu. Biobank-wide association scan identifies risk factors for late-onset Alzheimer's disease and endophenotypes. *eLife*, 12, 2024.

[14902] Guanxiong Yan, Yang Ma, Yanfang Wang, Jing Zhang, Haoming Cheng, Fanjie Tan, Su Wang, Delin Zhang, Jie Xiong, Ping Yin, and Wei Miao. A seven-sex species recognizes self and non-self mating-type via a novel protein complex. *eLife*, 13, 2024.

[14903] Hui Yan, Yao Liu, Xipeng Li, Bing Yu, Jun He, Xiangbing Mao, Jie Yu, Zhiqing Huang, Yuheng Luo, Junqiu Luo, Aimin Wu, and Daiwen Chen. Leucine alleviates cytokine storm syndrome by regulating macrophage polarization via the mTORC1/LXR$\alpha$ signaling pathway. *eLife*, 12, 2024.

[14904] Xiaodan Yan, Hebin Liao, Chenyi Wang, Chun Huang, Wei Zhang, Chunming Guo, and Yingying Pu. An improved bacterial single-cell RNA-seq reveals biofilm heterogeneity. *eLife*, 13, 2024.

[14905] Xiaoqian Yan, Sarah Shi Tung, Bella Fascendini, Yulan Diana Chen, Anthony M Norcia, and Kalanit Grill-Spector. The emergence of visual category representations in infants' brains. *eLife*, 13, 2024.

[14906] Yongzhi Yan, Scott Jarvie, and Qing Zhang. Habitat loss weakens the positive relationship between grassland plant richness and above-ground biomass. *eLife*, 12, 2024.

[14907] Toyoshi Yanagihara, Kentaro Hata, Keisuke Matsubara, Kazufumi Kunimura, Kunihiro Suzuki, Kazuya Tsubouchi, Satoshi Ikegame, Yoshihiro Baba, Yoshinori Fukui, and Isamu Okamoto. Exploratory mass cytometry analysis reveals immunophenotypes of cancer treatment-related pneumonitis. *eLife*, 12, 2024.

[14908] Fan Nils Yang, Dante Picchioni, Jacco A de Zwart, Yicun Wang, Peter van Gelderen, and Jeff H Duyn. Reproducible, data-driven characterization of sleep based on brain dynamics and transitions from whole-night fMRI. *eLife*, 13, 2024.

[14909] Jun Yang, Hanqi Zhang, and Sukbin Lim. Sensory-memory interactions via modular structure explain errors in visual working memory. *eLife*, 13, 2024.

[14910] Jun Yang, Shitian Zou, Zeyou Qiu, Mingqiang Lai, Qing Long, Huan Chen, Ping lin Lai, Sheng Zhang, Zhi Rao, Xiaoling Xie, Yan Gong, Anling Liu, Mangmang Li, and Xiaochun Bai. Mecp2 fine-tunes quiescence exit by targeting nuclear receptors. *eLife*, 12, 2024.

[14911] Lei Yang, Jingtao Zhang, Sen Liu, Yanning Zhang, Li Wang, Xiaotong Wang, Shanshan Wang, Ke Li, Mengping Wei, and Chen Zhang. Establishment of transgenic fluorescent mice for labeling synapses and screening synaptogenic adhesion molecules. *eLife*, 13, 2024.

[14912] Long Yang, Deepak Singla, Alexander K Wu, Katy A Cross, and Sotiris C Masmanidis. Dopamine lesions alter the striatal encoding of single-limb gait. *eLife*, 12, 2024.

[14913] Lu Yang, Zijin Lin, Ruijing Mu, Wenhan Wu, Hao Zhi, Xiaodong Liu, Hanyu Yang, and Li Liu. Neurons enhance blood–brain barrier function via upregulating claudin-5 and VE-cadherin expression due to glial cell line-derived neurotrophic factor secretion. *eLife*, 13, 2024.

[14914] Luting Yang, Chunqing Hu, Xiaowen Chen, Jie Zhang, Zhe Feng, Yanxin Xiao, Weitai He, Tingting Cui, Xin Zhang, Yang Yang, Yaling Zhang, and Yaping Yan. Upregulated expression of ubiquitin ligase TRIM21 promotes PKM2 nuclear translocation and astrocyte activation in experimental autoimmune encephalomyelitis. *eLife*, 13, 2024.

[14915] Ruichen Yang, Hongshang Chu, Hua Yue, Yuji Mishina, Zhenlin Zhang, Huijuan Liu, and Baojie Li. BMP signaling maintains auricular chondrocyte identity and prevents microtia development by inhibiting protein kinase A. *eLife*, 12, 2024.

[14916] Shun Kai Yang, Shintaroh Kubo, Corbin Steven Black, Katya Peri, Daniel Dai, Thibault Legal, Melissa Valente-Paterno, Jacek Gaertig, and Khanh Huy Bui. Effect of $\alpha$-tubulin acetylation on the doublet microtubule structure. *eLife*, 12, 2024.

[14917] Tingting Yang, Marko S Chavez, Christina M Niman, Shuai Xu, and Mohamed Y El-Naggar. Long-distance electron transport in multicellular freshwater cable bacteria. *eLife*, 12, 2024.

[14918] Wan-ping Yang, Mei-qi Li, Jie Ding, Jia-yan Li, Gang Wu, Bao Liu, Yu-qi Gao, Guo-hua Wang, and Qian-qian Luo. High-altitude hypoxia exposure inhibits erythrophagocytosis by inducing macrophage ferroptosis in the spleen. *eLife*, 12, 2024.

[14919] Xue Yang, Chuyi Han, Changhao Yu, Bin Zhou, Ling Ye, Feifei Li, and Fanyuan Yu. Inclusive, exclusive and hierarchical atlas of NFATc1+/PDGFR-$\alpha$+ cells in dental and periodontal mesenchyme. *eLife*, 13, 2024.

[14920] Yue Yang, Bin Huang, Yimeng Qin, Danwei Wang, Yinuo Jin, Linmin Su, Qingxin Wang, Yi Pan, Yanfeng Zhang, Yumeng Shen, Wenjun Hu, Zhengyu Cao, Liang Jin, and Fangfang Zhang. Adipocyte microRNA-802 promotes adipose tissue inflammation and insulin resistance by modulating macrophages in obesity. *eLife*, 13, 2024.

[14921] Zongju Yang, Tianqi Bai, Zhiguo E, Baixiao Niu, and Chen Chen. OsNF-YB7 inactivates OsGLK1 to inhibit chlorophyll biosynthesis in rice embryo. *eLife*, 13, 2024.

[14922] Junjun Yao, Shaoxing Dai, Ran Zhu, Ju Tan, Qiancheng Zhao, Yu Yin, Jiansen Sun, Xuewei Du, Longjiao Ge, Jianhua Xu, Chunli Hou, Nan Li, Jun Li, Weizhi Ji, Chuhong Zhu, Runrui Zhang, and Tianqing Li. Deciphering molecular heterogeneity and dynamics of human hippocampal neural stem cells at different ages and injury states. *eLife*, 12, 2024.

[14923] Changtian Ye, Ryan Ho, Kenneth H Moberg, and James Q Zheng. Adverse impact of female reproductive signaling on age-dependent neurodegeneration after mild head trauma in Drosophila. *eLife*, 13, 2024.

[14924] Xinjian Ye, Yijing Bai, Mengjun Li, Yuhang Ye, Yitong Chen, Bin Liu, Yuwei Dai, Shan Wang, Weiyi Pan, Zhiyong Wang, Yingying Mao, and Qianming Chen. Genetic associations between circulating immune cells and periodontitis highlight the prospect of systemic immunoregulation in periodontal care. *eLife*, 12, 2024.

[14925] Zhenbang Ye, Ning Huang, Yongliang Fu, Rongle Tian, Liming Wang, and Wenting Huang. Tumor purity-related genes for predicting the prognosis and drug sensitivity of DLBCL patients. *eLife*, 13, 2024.

[14926] Mighten C Yip, Mercedes M Gonzalez, Colby F Lewallen, Corey R Landry, Ilya Kolb, Bo Yang, William M Stoy, Ming-fai Fong, Matthew JM Rowan, Edward S Boyden, and Craig R Forest. Patch-walking, a coordinated multi-pipette patch clamp for efficiently finding synaptic connections. *eLife*, 13, 2024.

[14927] Baba Yogesh and Georg B Keller. Cholinergic input to mouse visual cortex signals a movement state and acutely enhances layer 5 responsiveness. *eLife*, 12, 2024.

[14928] John Yong, Jacqueline E Villalta, Ngoc Vu, Matthew A Kukurugya, Niclas Olsson, Magdalena Preciado López, Julia R Lazzari-Dean, Kayley Hake, Fiona E McAllister, Bryson D Bennett, and Calvin H Jan. Impairment of lipid homeostasis causes lysosomal accumulation of endogenous protein aggregates through ESCRT disruption. *eLife*, 12, 2024.

[14929] Kyusang Yoo, Young-Woo Jo, Takwon Yoo, Sang-Hyeon Hann, Inkuk Park, Yea-Eun Kim, Ye Lynne Kim, Joonwoo Rhee, In-Wook Song, Ji-Hoon Kim, Daehyun Baek, and Young-Yun Kong. Muscle-resident mesenchymal progenitors sense and repair peripheral nerve injury via the GDNF-BDNF axis. *eLife*, 13, 2024.

[14930] Matthew Yorek, Xingshan Jiang, Shanshan Liu, Jiaqing Hao, Jianyu Yu, Anthony Avellino, Zhanxu Liu, Melissa Curry, Henry Keen, Jianqiang Shao, Anand Kanagasabapathy, Maying Kong, Yiqin Xiong, Edward R Sauter, Sonia L Sugg, and Bing Li. FABP4-mediated lipid accumulation and lipolysis in tumor-associated macrophages promote breast cancer metastasis. *eLife*, 13, 2024.

[14931] Teruhiko Yoshida, Khun Zaw Latt, Avi Z Rosenberg, Briana A Santo, Komuraiah Myakala, Yu Ishimoto, Yongmei Zhao, Shashi Shrivastav, Bryce A Jones, Xiaoping Yang, Xiaoxin X Wang, Vincent M Tutino, Pinaki Sarder, Moshe Levi, Koji Okamoto, Cheryl A Winkler, and Jeffrey B Kopp. PKR activation-induced mitochondrial dysfunction in HIV-transgenic mice with nephropathy. *eLife*, 12, 2024.

[14932] Bo Yu, Congzhou Liu, Sean C Proll, Enna Manhardt, Shuying Liang, Sujatha Srinivasan, Elizabeth Swisher, and David N Fredricks. Identification of fallopian tube microbiota and its association with ovarian cancer. *eLife*, 12, 2024.

[14933] Daniel S Yu, Megan A Outram, Ashley Smith, Carl L McCombe, Pravin B Khambalkar, Sharmin A Rima, Xizhe Sun, Lisong Ma, Daniel J Ericsson, David A Jones, and Simon J Williams. The structural repertoire of Fusarium oxysporum f. sp. lycopersici effectors revealed by experimental and computational studies. *eLife*, 12, 2024.

[14934] Haibin Yu, Dandan Liu, Yaowen Zhang, Ruijun Tang, Xunan Fan, Song Mao, Lu Lv, Fang Chen, Hongtao Qin, Zhuohua Zhang, Daan MF van Aalten, Bing Yang, and Kai Yuan. Tissue-specific O-GlcNAcylation profiling identifies substrates in translational machinery in Drosophila mushroom body contributing to olfactory learning. *eLife*, 13, 2024.

[14935] Hao Yu, Anas M Khanshour, Aki Ushiki, Nao Otomo, Yoshinao Koike, Elisabet Einarsdottir, Yanhui Fan, Lilian Antunes, Yared H Kidane, Reuel Cornelia, Rory R Sheng, Yichi Zhang, Jimin Pei, Nick V Grishin, Bret M Evers, Jason Pui Yin Cheung, John A Herring, Chikashi Terao, You-qiang Song, Christina A Gurnett, Paul Gerdhem, Shiro Ikegawa, Jonathan J Rios, Nadav Ahituv, and Carol A Wise. Association of genetic variation in COL11A1 with adolescent idiopathic scoliosis. *eLife*, 12, 2024.

[14936] Hong Yu, Hiroshi Nishio, Joseph Barbi, Marisa Mitchell-Flack, Paolo DA Vignali, Ying Zheng, Andriana Lebid, Kwang-Yu Chang, Juan Fu, Makenzie Higgins, Ching-Tai Huang, Xuehong Zhang, Zhiguang Li, Lee Blosser, Ada Tam, Charles Drake, and Drew Pardoll. Neurotrophic factor Neuritin modulates T cell electrical and metabolic state for the balance of tolerance and immunity. *eLife*, 13, 2024.

[14937] Xiaolin Yu, Guangfei Duan, Pengfei Pei, Long Chen, Renji Gu, Wenrui Hu, Hongli Zhang, Yan-Dong Wang, Lili Gong, Lihong Liu, Ting-Ting Chu, Jin-Ping Li, and Shi-Zhong Luo. Heparan sulfate-dependent phase separation of CCL5 and its chemotactic activity. *eLife*, 13, 2024.

[14938] Xiaoxin Yu, Roger S Zoh, David A Fluharty, Luis M Mestre, Danny Valdez, Carmen D Tekwe, Colby J Vorland, Yasaman Jamshidi-Naeini, Sy Han Chiou, Stella T Lartey, and David B Allison. Misstatements, misperceptions, and mistakes in controlling for covariates in observational research. *eLife*, 13, 2024.

[14939] Xiaozhuo Yu, Wen Zhou, Xiaodong Chen, Shunyu He, Mengting Qin, Meng Yuan, Yang Wang, Woodvine Otieno Odhiambo, Yinsha Miao, and Yanhong Ji. RAG1 and RAG2 non-core regions are implicated in leukemogenesis and off-target V(D)J recombination in BCR-ABL1-driven B-cell lineage lymphoblastic leukemia. *eLife*, 12, 2024.

[14940] Yong Yu, Shihong M Gao, Youchen Guan, Pei-Wen Hu, Qinghao Zhang, Jiaming Liu, Bentian Jing, Qian Zhao, David M Sabatini, Monther Abu-Remaileh, Sung Yun Jung, and Meng C Wang. Organelle proteomic profiling reveals lysosomal heterogeneity in association with longevity. *eLife*, 13, 2024.

[14941] Augustine Xiaoran Yuan, Jennifer Colonell, Anna Lebedeva, Michael Okun, Adam S Charles, and Timothy D Harris. Multi-day neuron tracking in high-density electrophysiology recordings using earth mover's distance. *eLife*, 12, 2024.

[14942] Qianmu Yuan, Chong Tian, and Yuedong Yang. Genome-scale annotation of protein binding sites via language model and geometric deep learning. *eLife*, 13, 2024.

[14943] Tian Yuan, Li Wang, and Yi Jiang. Multi-level processing of emotions in life motion signals revealed through pupil responses. *eLife*, 12, 2024.

[14944] Tian Yuan, Yifan Wang, Yuchen Jin, Hui Yang, Shuai Xu, Heng Zhang, Qian Chen, Na Li, Xinyue Ma, Huifang Song, Chao Peng, Ze Geng, Jie Dong, Guifang Duan, Qi Sun, Yang Yang, Fan Yang, and Zhuo Huang. Coupling of Slack and NaV1.6 sensitizes Slack to quinidine blockade and guides anti-seizure strategy development. *eLife*, 12, 2024.

[14945] Tian Yue, Yongbo Guo, Xuebin Qi, Wangshan Zheng, Hui Zhang, Bin Wang, Kai Liu, Bin Zhou, Xuerui Zeng, Ouzhuluobu, Yaoxi He, and Bing Su. Sex-biased regulatory changes in the placenta of native highlanders contribute to adaptive fetal development. *eLife*, 12, 2024.

[14946] Ernie Yulyaningsih, Jung H Suh, Melania Fanok, Roni Chau, Hilda Solanoy, Ryan Takahashi, Anna I Bakardjiev, Isabel Becerra, N Butch Benitez, Chi-Lu Chiu, Sonnet S Davis, William E Dowdle, Timothy Earr, Anthony A Estrada, Audrey Gill, Connie Ha, Patrick CG Haddick, Kirk R Henne, Martin Larhammar, Amy W-S Leung, Romeo Maciuca, Bahram Memarzadeh, Hoang N Nguyen, Alicia A Nugent, Maksim Osipov, Yingqing Ran, Kevin Rebadulla, Elysia Roche, Thomas Sandman, Jing Wang, Joseph W Lewcock, Kimberly Scearce-Levie, Lesley A Kane, and Pascal E Sanchez. DNL343 is an investigational CNS penetrant eukaryotic initiation factor 2B activator that prevents and reverses the effects of neurodegeneration caused by the integrated stress response. *eLife*, 12, 2024.

[14947] Minsik Yun, Do-Hyoung Kim, Tal Soo Ha, Kang-Min Lee, Eungyu Park, Markus Knaden, Bill S Hansson, and Young-Joon Kim. Male cuticular pheromones stimulate removal of the mating plug and promote re-mating through pC1 neurons in Drosophila females. *eLife*, 13, 2024.

[14948] Anja T Zai, Anna E Stepien, Nicolas Giret, and Richard HR Hahnloser. Goal-directed vocal planning in a songbird. *eLife*, 12, 2024.

[14949] Tariq Zaman, Daniel Vogt, Jeremy Prokop, Qusai Abdulkhaliq Alsabia, Gabriel Simms, April Stafford, Bryan W Luikart, and Michael R Williams. Kit Ligand and Kit receptor tyrosine kinase sustain synaptic inhibition of Purkinje cells. *eLife*, 12, 2024.

[14950] Boglarka Zambo, Evelina Edelweiss, Bastien Morlet, Luc Negroni, Matyas Pajkos, Zsuzsanna Dosztanyi, Soren Ostergaard, Gilles Trave, Jocelyn Laporte, and Gergo Gogl. Uncovering the BIN1-SH3 interactome underpinning centronuclear myopathy. *eLife*, 13, 2024.

[14951] Mirela Zaneva, Tao Coll-Martín, Yseult Héjja-Brichard, Tamara Kalandadze, Andrea Kis, Alicja Koperska, Marie Adrienne Robles Manalili, Adrien Mathy, Christopher J Graham, Anna Hollis, Robert M Ross, Siu Kit Yeung, Veronica Allen, Flavio Azevedo, Emily Friedel, Stephanie Fuller, Vaitsa Giannouli, Biljana Gjoneska, Helena Hartmann, Max Korbmacher, Mahmoud M Elsherif, and Alyssa Hillary Zisk. An annotated introductory reading list for neurodiversity. *eLife*, 13, 2024.

[14952] Jie Zang, Shenquan Liu, Pascal Helson, and Arvind Kumar. Structural constraints on the emergence of oscillations in multi-population neural networks. *eLife*, 12, 2024.

[14953] Ando Christian Zehrer, Ana Martin-Villalba, Benedict Diederich, and Helge Ewers. An open-source, high-resolution, automated fluorescence microscope. *eLife*, 12, 2024.

[14954] Ruijie Zeng, Yuying Ma, Lijun Zhang, Dongling Luo, Rui Jiang, Huihuan Wu, Zewei Zhuo, Qi Yang, Jingwei Li, Felix W Leung, Chongyang Duan, Weihong Sha, and Hao Chen. Associations of proton pump inhibitors with susceptibility to influenza, pneumonia, and COVID-19: Evidence from a large population-based cohort study. *eLife*, 13, 2024.

[14955] Zhao-Ying Zeng, Jun-Rong Huang, Zi-Qing Liu, Ai-Ling Yang, Yu-Xuan Li, Yong-Lan Wang, and Han-Bo Zhang. Distinct effects of phyllosphere and rhizosphere microbes on invader Ageratina adenophora during its early life stages. *eLife*, 13, 2024.

[14956] Liting Zhai, Amber C Bonds, Clyde A Smith, Hannah Oo, Jonathan Chiu-Chun Chou, Paula V Welander, and Laura MK Dassama. Novel sterol binding domains in bacteria. *eLife*, 12, 2024.

[14957] Chi Zhang, Qian Huang, Neil C Ford, Nathachit Limjunyawong, Qing Lin, Fei Yang, Xiang Cui, Ankit Uniyal, Jing Liu, Megha Mahabole, Hua He, Xuewei Wang, Irina Duff, Yiru Wang, Jieru Wan, Guangwu Zhu, Srinivasa N Raja, Hongpeng Jia, Dazhi Yang, Xinzhong Dong, Xu Cao, Scheffer C Tseng, Shaoqiu He, and Yun Guan. Human birth tissue products as a non-opioid medicine to inhibit post-surgical pain. *eLife*, 13, 2024.

[14958] Chi Zhang, Rongjing Zhang, and Junhua Yuan. Potassium-mediated bacterial chemotactic response. *eLife*, 12, 2024.

[14959] Donghang Zhang, Yali Chen, Yiyong Wei, Hongjun Chen, Yujie Wu, Lin Wu, Jin Li, Qiyang Ren, Changhong Miao, Tao Zhu, Jin Liu, Bowen Ke, and Cheng Zhou. Spatial transcriptomics and single-nucleus RNA sequencing reveal a transcriptomic atlas of adult human spinal cord. *eLife*, 12, 2024.

[14960] Fan Zhang, Annie Lee, Anna V Freitas, Jake T Herb, Zong-Heng Wang, Snigdha Gupta, Zhe Chen, and Hong Xu. A transcription network underlies the dual genomic coordination of mitochondrial biogenesis. *eLife*, 13, 2024.

[14961] Honglei Zhang, Chao Liu, Shuting Wang, Qing Wang, Xu Feng, Huawei Jiang, Li Xiao, Chao Luo, Lu Zhang, Fei Hou, Minjun Zhou, Zhiyong Deng, Heng Li, Yong Zhang, Xiaosan Su, and Gaofeng Li. Proteogenomic analysis of air-pollution-associated lung cancer reveals prevention and therapeutic opportunities. *eLife*, 13, 2024.

[14962] Houyu Zhang, Yan Li, Carlos F Ibáñez, and Meng Xie. Perirenal adipose tissue contains a subpopulation of cold-inducible adipocytes derived from brown-to-white conversion. *eLife*, 13, 2024.

[14963] Kejia Zhang, Patrick Eldin, Jessica H Ciesla, Laurence Briant, Jenna M Lentini, Jillian Ramos, Justin Cobb, Joshua Munger, and Dragony Fu. Proteolytic cleavage and inactivation of the TRMT1 tRNA modification enzyme by SARS-CoV-2 main protease. *eLife*, 12, 2024.

[14964] Li Zhang, Fen Hu, Zirong Zhao, Xinfeng Li, Mingyue Zhong, Jiajun He, Fangfang Yao, Xiaomei Zhang, Yuxuan Mao, Hongping Wei, Jin He, and Hang Yang. Dimer-monomer transition defines a hyper-thermostable peptidoglycan hydrolase mined from bacterial proteome by lysin-derived antimicrobial peptide-primed screening. *eLife*, 13, 2024.

[14965] Linda Zhang, Joanne I Hsu, Etienne D Braekeleer, Chun-Wei Chen, Tajhal D Patel, Alejandra G Martell, Anna G Guzman, Katharina Wohlan, Sarah M Waldvogel, Hidetaka Uryu, Ayala Tovy, Elsa Callen, Rebecca L Murdaugh, Rosemary Richard, Sandra Jansen, Lisenka Vissers, Bert BA de Vries, Andre Nussenzweig, Shixia Huang, Cristian Coarfa, Jamie Anastas, Koichi Takahashi, George Vassiliou, and Margaret A Goodell. SOD1 is a synthetic-lethal target in PPM1D-mutant leukemia cells. *eLife*, 12, 2024.

[14966] Lingjie Zhang, Tong Deng, Zhongqi Liufu, Xiangnyu Chen, Shijie Wu, Xueyu Liu, Changhao Shi, Bingjie Chen, Zheng Hu, Qichun Cai, Chenli Liu, Mengfeng Li, Miles E Tracy, Xuemei Lu, Chung-I Wu, and Hai-Jun Wen. Characterization of cancer-driving nucleotides (CDNs) across genes, cancer types, and patients. *eLife*, 13, 2024.

[14967] Ming Zhang, Guangyi Du, Lianghua Xie, Yang Xu, and Wei Chen. Circular RNA HMGCS1 sponges MIR4521 to aggravate type 2 diabetes-induced vascular endothelial dysfunction. *eLife*, 13, 2024.

[14968] Qinyu Zhang, Rasmus Olofzon, Anna Konturek-Ciesla, Ouyang Yuan, and David Bryder. Ex vivo expansion potential of murine hematopoietic stem cells is a rare property only partially predicted by phenotype. *eLife*, 12, 2024.

[14969] Sheng-Hui Zhang, Xing-Nan Zhao, Dan-Qing Jiang, Shi-Ming Tang, and Cong Yu. Ocular dominance-dependent binocular combination of monocular neuronal responses in macaque V1. *eLife*, 13, 2024.

[14970] Shu-Ting Zhang, Shi-Kai Deng, Tao Li, Megan E Maloney, De-Feng Li, Jim C Spain, and Ning-Yi Zhou. Discovery of the 1-naphthylamine biodegradation pathway reveals a broad-substrate-spectrum enzyme catalyzing 1-naphthylamine glutamylation. *eLife*, 13, 2024.

[14971] Shuai-Shuai Zhang, Pei-Chao Wang, Chao Ning, Ke Yang, Guo-Cheng Li, Lin-Lin Cao, Ling-Qiao Huang, and Chen-Zhu Wang. The larva and adult of Helicoverpa armigera use differential gustatory receptors to sense sucrose. *eLife*, 12, 2024.

[14972] Songyao Zhang, Tuo Zhang, Guannan Cao, Jingchao Zhou, Zhibin He, Xiao Li, Yudan Ren, Tao Liu, Xi Jiang, Lei Guo, Junwei Han, and Tianming Liu. Species -shared and -unique gyral peaks on human and macaque brains. *eLife*, 12, 2024.

[14973] Ting Zhang, Alisa Ambrodji, Huixing Huang, Kelly J Bouchonville, Amy S Etheridge, Remington E Schmidt, Brianna M Bembenek, Zoey B Temesgen, Zhiquan Wang, Federico Innocenti, Deborah Stroka, Robert B Diasio, Carlo R Largiadèr, and Steven M Offer. Germline cis variant determines epigenetic regulation of the anti-cancer drug metabolism gene dihydropyrimidine dehydrogenase (DPYD). *eLife*, 13, 2024.

[14974] Tony Zhang, Matthew Rosenberg, Zeyu Jing, Pietro Perona, and Markus Meister. Endotaxis: A neuromorphic algorithm for mapping, goal-learning, navigation, and patrolling. *eLife*, 12, 2024.

[14975] Weiting Zhang, Karl Petri, Junyan Ma, Hyunho Lee, Chia-Lun Tsai, J Keith Joung, and Jing-Ruey Joanna Yeh. Enhancing CRISPR prime editing by reducing misfolded pegRNA interactions. *eLife*, 12, 2024.

[14976] Yangfan Zhang and George V Lauder. Energy conservation by collective movement in schooling fish. *eLife*, 12, 2024.

[14977] Yaojun Zhang, Andrew GT Pyo, Ross Kliegman, Yoyo Jiang, Clifford P Brangwynne, Howard A Stone, and Ned S Wingreen. The exchange dynamics of biomolecular condensates. *eLife*, 12, 2024.

[14978] Yuchen Zhang, Yitang Sun, Qi Yu, Suhang Song, J Thomas Brenna, Ye Shen, and Kaixiong Ye. Higher ratio of plasma omega-6/omega-3 fatty acids is associated with greater risk of all-cause, cancer, and cardiovascular mortality: A population-based cohort study in UK Biobank. *eLife*, 12, 2024.

[14979] Yuting Zhang, Min Zhang, Haiyan Wu, Xinwei Wang, Hang Zheng, Junjuan Feng, Jing Wang, Longlong Luo, He Xiao, Chunxia Qiao, Xinying Li, Yuanqiang Zheng, Weijin Huang, Youchun Wang, Yi Wang, Yanchun Shi, Jiannan Feng, and Guojiang Chen. A novel MARV glycoprotein-specific antibody with potentials of broad-spectrum neutralization to filovirus. *eLife*, 12, 2024.

[14980] Zhaoran Zhang, Huijun Wang, Tianyang Zhang, Zixuan Nie, and Kunlin Wei. Perceptual error based on Bayesian cue combination drives implicit motor adaptation. *eLife*, 13, 2024.

[14981] Zhe Zhang, Xu Hong, Peng Xiong, Junfeng Wang, Yaoqi Zhou, and Jian Zhan. Minimal twister sister-like self-cleaving ribozymes in the human genome revealed by deep mutational scanning. *eLife*, 12, 2024.

[14982] Zhiliang Zhang, Zhifei Zhang, Lars Holmer, Timothy P Topper, Bing Pan, and Guoxiang Li. Evolution and diversity of biomineralized columnar architecture in early Cambrian phosphatic-shelled brachiopods. *eLife*, 12, 2024.

[14983] Zhixian Zhang, Jianying Li, Yilin Wang, Zhen Li, Xiaoxia Liu, and Songdou Zhang. Neuropeptide Bursicon and its receptor-mediated the transition from summer-form to winter-form of Cacopsylla chinensis. *eLife*, 13, 2024.

[14984] Dianyuan Zhao, Ziwei Huang, Xiaoyu Li, Huan Wang, Qingwei Hou, Yuyao Wang, Fang Yan, Wenting Yang, Di Liu, Shaoqiong Yi, Chunguang Han, Yanan Hao, and Li Tang. GDF2 and BMP10 coordinate liver cellular crosstalk to maintain liver health. *eLife*, 13, 2024.

[14985] Lei Zhao, Wei Zhou, Jun He, De-Zhu Li, and Hong-Tao Li. Positive selection and relaxed purifying selection contribute to rapid evolution of male-biased genes in a dioecious flowering plant. *eLife*, 12, 2024.

[14986] Mengya Zhao, Beiying Dai, Xiaodong Li, Yixin Zhang, Chun Qiao, Yaru Qin, Zhao Li, Qingmei Li, Shuzhen Wang, Yong Yang, and Yijun Chen. RAPSYN-mediated neddylation of BCR-ABL alternatively determines the fate of Philadelphia chromosome-positive leukemia. *eLife*, 12, 2024.

[14987] Pianpian Zhao, Zhifeng Sheng, Lin Xu, Peng Li, Wenjin Xiao, Chengda Yuan, Zhanwei Xu, Mengyuan Yang, Yu Qian, Jiadong Zhong, Jiaxuan Gu, David Karasik, and Hou-Feng Zheng. Deciphering the complex relationship between type 2 diabetes and fracture risk with both genetic and observational evidence. *eLife*, 12, 2024.

[14988] Qing Zhao, Yanjing Zhu, Yilong Ren, Lijuan Zhao, Jingwei Zhao, Shuai Yin, Haofei Ni, Rongrong Zhu, Liming Cheng, and Ning Xie. Targeting resident astrocytes attenuates neuropathic pain after spinal cord injury. *eLife*, 13, 2024.

[14989] Ruonan Zhao, Emma L Moore, Madelaine M Gogol, Jay R Unruh, Zulin Yu, Allison R Scott, Yan Wang, Naresh K Rajendran, and Paul A Trainor. Identification and characterization of intermediate states in mammalian neural crest cell epithelial to mesenchymal transition and delamination. *eLife*, 13, 2024.

[14990] Tianyu Zhao, Hui Li, Meishuang Zhang, Yang Xu, Ming Zhang, and Li Chen. Systematic evaluation of multifactorial causal associations for Alzheimer's disease and an interactive platform MRAD developed based on Mendelian randomization analysis. *eLife*, 13, 2024.

[14991] Tiesuo Zhao, Pengkun Wei, Congli Zhang, Shijie Zhou, Lirui Liang, Shuoshuo Guo, Zhinan Yin, Sichang Cheng, Zerui Gan, Yuanling Xia, Yongxi Zhang, Sheng Guo, Jiateng Zhong, Zishan Yang, Fei Tu, Qianqing Wang, Jin Bai, Feng Ren, Zhiwei Feng, and Huijie Jia. Nifuroxazide suppresses PD-L1 expression and enhances the efficacy of radiotherapy in hepatocellular carcinoma. *eLife*, 12, 2024.

[14992] Xingsen Zhao, Qihang Sun, Yikai Shou, Weijun Chen, Mengxuan Wang, Wenzheng Qu, Xiaoli Huang, Ying Li, Chao Wang, Yan Gu, Chai Ji, Qiang Shu, and Xuekun Li. A human forebrain organoid model reveals the essential function of GTF2IRD1-TTR-ERK axis for the neurodevelopmental deficits of Williams syndrome. *eLife*, 13, 2024.

[14993] Yangzi Zhao, Lijun Ren, Tingting Zhao, Hanli You, Yongjie Miao, Huixin Liu, Lei Cao, Bingxin Wang, Yi Shen, Yafei Li, Ding Tang, and Zhukuan Cheng. SCC3 is an axial element essential for homologous chromosome pairing and synapsis. *eLife*, 13, 2024.

[14994] Zewei Zhao, Longyun Hu, Bigui Song, Tao Jiang, Qian Wu, Jiejing Lin, Xiaoxiao Li, Yi Cai, Jin Li, Bingxiu Qian, Siqi Liu, Jilu Lang, and Zhonghan Yang. Constitutively active receptor ADGRA3 signaling induces adipose thermogenesis. *eLife*, 13, 2024.

[14995] Lirong Zheng, Bingxin Zhou, Banghao Wu, Yang Tan, Juan Huang, Madhusudan Tyagi, Victoria García Sakai, Takeshi Yamada, Hugh O'Neill, Qiu Zhang, and Liang Hong. Decoupling of the onset of anharmonicity between a protein and its surface water around 200 K. *eLife*, 13, 2024.

[14996] Rikuan Zheng, Chong Wang, Rui Liu, Ruining Cai, and Chaomin Sun. Physiological and metabolic insights into the first cultured anaerobic representative of deep-sea Planctomycetes bacteria. *eLife*, 12, 2024.

[14997] Xueao Zheng, Hanchen Chen, Zhiping Deng, Yujing Wu, Linlin Zhong, Chong Wu, Xiaodan Yu, Qiansi Chen, and Shunping Yan. The tRNA thiolation-mediated translational control is essential for plant immunity. *eLife*, 13, 2024.

[14998] Felix Y Zhou, David P Waterman, Marissa Ashton, Suhaily Caban-Penix, Gonen Memisoglu, Vinay V Eapen, and James E Haber. Prolonged cell cycle arrest in response to DNA damage in yeast requires the maintenance of DNA damage signaling and the spindle assembly checkpoint. *eLife*, 13, 2024.

[14999] Fujun Zhou, Julie M Bocetti, Meizhen Hou, Daoming Qin, Alan G Hinnebusch, and Jon R Lorsch. Transcriptome-wide analysis of the function of Ded1 in translation preinitiation complex assembly in a reconstituted in vitro system. *eLife*, 13, 2024.

[15000] Ling Zhou, Chengyang Ni, Ruixue Liao, Xiaoqin Tang, Taian Yi, Mei Ran, Miao Huang, Rui Liao, Xiaogang Zhou, Dalian Qin, Long Wang, Feihong Huang, Xiang Xie, Ying Wan, Jiesi Luo, Yiwei Wang, and Jianming Wu. Activating SRC/MAPK signaling via 5-HT1A receptor contributes to the effect of vilazodone on improving thrombocytopenia. *eLife*, 13, 2024.

[15001] Liu Zhou, Wei Wei, Teng Leng Ooi, and Zijiang J He. An allocentric human odometer for perceiving distances on the ground plane. *eLife*, 12, 2024.

[15002] Mingqi Zhou, Ian Tamburini, Cassandra Van, Jeffrey Molendijk, Christy M Nguyen, Ivan Yao-Yi Chang, Casey Johnson, Leandro M Velez, Youngseo Cheon, Reichelle Yeo, Hosung Bae, Johnny Le, Natalie Larson, Ron Pulido, Carlos HV Nascimento-Filho, Cholsoon Jang, Ivan Marazzi, Jamie Justice, Nicholas Pannunzio, Andrea L Hevener, Lauren Sparks, Erin Kershaw, Dequina Nicholas, Benjamin L Parker, Selma Masri, and Marcus M Seldin. Leveraging inter-individual transcriptional correlation structure to infer discrete signaling mechanisms across metabolic tissues. *eLife*, 12, 2024.

[15003] Wei Zhou, Kezhang He, Chiyin Wang, Pengqi Wang, Dan Wang, Bowen Wang, Han Geng, Hong Lian, Tianhua Ma, Yu Nie, and Sheng Ding. Pharmacologically inducing regenerative cardiac cells by small molecule drugs. *eLife*, 13, 2024.

[15004] Xin Zhou, Zhinuo Jenny Wang, Julia Camps, Jakub Tomek, Alfonso Santiago, Adria Quintanas, Mariano Vazquez, Marmar Vaseghi, and Blanca Rodriguez. Clinical phenotypes in acute and chronic infarction explained through human ventricular electromechanical modelling and simulations. *eLife*, 13, 2024.

[15005] Bohan Zhu, Richard I Ainsworth, Zengmiao Wang, Zhengzhi Liu, Salvador Sierra, Chengyu Deng, Luis F Callado, J Javier Meana, Wei Wang, Chang Lu, and Javier González-Maeso. Antipsychotic-induced epigenomic reorganization in frontal cortex of individuals with schizophrenia. *eLife*, 12, 2024.

[15006] Jiayi Zhu, Jean-Christophe Boivin, Alastair Garner, Jing Ning, Yi Q Zhao, and Tomoko Ohyama. Feedback inhibition by a descending GABAergic neuron regulates timing of escape behavior in Drosophila larvae. *eLife*, 13, 2024.

[15007] Lin Zhu, Jing Jin, Tingting Wang, Yong Hu, Hainan Liu, Ting Gao, Qincai Dong, Yanwen Jin, Ping Li, Zijing Liu, Yi Huang, Xuan Liu, and Cheng Cao. Ebola virus sequesters IRF3 in viral inclusion bodies to evade host antiviral immunity. *eLife*, 12, 2024.

[15008] Yi-jun Zhu, Cai-yun Deng, Liu Fan, Ya-Qian Wang, Hui Zhou, and Hua-tai Xu. Combinatorial expression of $\gamma$-protocadherins regulates synaptic connectivity in the mouse neocortex. *eLife*, 12, 2024.

[15009] Jonathan Zirin, Barbara Jusiak, Raphael Lopes, Benjamin Ewen-Campen, Justin A Bosch, Alexandria Risbeck, Corey Forman, Christians Villalta, Yanhui Hu, and Norbert Perrimon. Expanding the Drosophila toolkit for dual control of gene expression. *eLife*, 12, 2024.

[15010] Malik Zohaib Ali, Taru S Dutt, Amy MacNeill, Amanda Walz, Camron Pearce, Ha Lam, Jamie S Philp, Johnathan Patterson, Marcela Henao-Tamayo, Richard Lee, Jiuyu Liu, Gregory T Robertson, Anthony J Hickey, Bernd Meibohm, and Mercedes Gonzalez Juarrero. A modified BPaL regimen for tuberculosis treatment replaces linezolid with inhaled spectinamides. *eLife*, 13, 2024.

[15011] Katarzyna Marta Zoltowska, Utpal Das, Sam Lismont, Thomas Enzlein, Masato Maesako, Mei CQ Houser, Maria Luisa Franco, Burcu özcan, Diana Gomes Moreira, Dmitry Karachentsev, Ann Becker, Carsten Hopf, MarÇal Vilar, Oksana Berezovska, William Mobley, and Lucía Chávez-Gutiérrez. Alzheimer's disease linked A$\beta$42 exerts product feedback inhibition on $\gamma$-secretase impairing downstream cell signaling. *eLife*, 12, 2024.

[15012] Ariel Zylberberg, Akram Bakkour, Daphna Shohamy, and Michael N Shadlen. Value construction through sequential sampling explains serial dependencies in decision making. *eLife*, 13, 2024.

[15013] Muad Y Abd El Hay, Gretel B Kamm, Alejandro Tlaie Boria, and Jan Siemens. Diverging roles of TRPV1 and TRPM2 in warm-temperature detection. *eLife*, 13, 2025.

[15014] Luis Abdala-Roberts, Adriana Puentes, Deborah L Finke, Robert J Marquis, Marta Montserrat, Erik H Poelman, Sergio Rasmann, Arnaud Sentis, Celia C Symons, Nicole M van Dam, Gina Wimp, Christer Björkman, and Kailen A Mooney. Connecting the dots: Managing species interaction networks to mitigate the impacts of global change. *eLife*, 14, 2025.

[15015] Mia L Abramsson, Robin A Corey, Jan L Skerle, Louise J Persson, Olivia Anden, Abraham O Oluwole, Rebecca J Howard, Erik Lindahl, Carol V Robinson, Kvido Strisovsky, Erik G Marklund, David Drew, Phillip J Stansfeld, and Michael Landreh. Engineering cardiolipin binding to an artificial membrane protein reveals determinants for lipid-mediated stabilization. *eLife*, 14, 2025.

[15016] Timothy J Abreo, Emma C Thompson, Anuraag Madabushi, Kristen L Park, Heun Soh, Nissi Varghese, Carlos G Vanoye, Kristen Springer, Jim Johnson, Scotty Sims, Zhigang Ji, Ana G Chavez, Miranda J Jankovic, Bereket Habte, Aamir R Zuberi, Cathleen M Lutz, Zhao Wang, Vaishnav Krishnan, Lisa Dudler, Stephanie Einsele-Scholz, Jeffrey L Noebels, Alfred L George, Atul Maheshwari, Anastasios Tzingounis, and Edward C Cooper. Plural molecular and cellular mechanisms of pore domain KCNQ2 encephalopathy. *eLife*, 13, 2025.

[15017] Cyril Achard, Timokleia Kousi, Markus Frey, Maxime Vidal, Yves Paychere, Colin Hofmann, Asim Iqbal, Sebastien B Hausmann, Stéphane Pagès, and Mackenzie Weygandt Mathis. CellSeg3D, Self-supervised 3D cell segmentation for fluorescence microscopy. *eLife*, 13, 2025.

[15018] Joseph Ackermann, Chiara Bernard, Philemon Sirven, Helene Salmon, Massimiliano Fraldi, and Martine D Ben Amar. Mechanistic insight for T-cell exclusion by cancer-associated fibroblasts in human lung cancer. *eLife*, 13, 2025.

[15019] Manuela C Aguirre-Botero, Olga Pacios, Susanna Celli, Eduardo Aliprandini, Anisha Gladston, Jean-Michel Thiberge, Pauline Formaglio, and Rogerio Amino. Late killing of Plasmodium berghei sporozoites in the liver by an anti-circumsporozoite protein antibody. *eLife*, 14, 2025.

[15020] Christine Ahrends, Mark W Woolrich, and Diego Vidaurre. Predicting individual traits from models of brain dynamics accurately and reliably using the Fisher kernel. *eLife*, 13, 2025.

[15021] Hailin Ai, Weiru Lin, Chengwen Liu, Nihong Chen, and Peng Zhang. Mesoscale functional organization and connectivity of color, disparity, and naturalistic texture in human second visual area. *eLife*, 13, 2025.

[15022] Emmanuel Akabuogu, Victor Carneiro da Cunha Martorelli, Rok KraŠovec, Ian S Roberts, and Thomas A Waigh. Emergence of ion-channel-mediated electrical oscillations in Escherichia coli biofilms. *eLife*, 13, 2025.

[15023] Aalimah Akinosho, Joseph Alexander, Kyle Floyd, and Andres Gabriel Vidal-Gadea. Independent validation of transgenerational inheritance of learned pathogen avoidance in Caenorhabditis elegans. *eLife*, 14, 2025.

[15024] Adriana K Alexander, Karina F Rodriguez, Yu-Ying Chen, Ciro Amato, Martin A Estermann, Barbara Nicol, Xin Xu, and Humphrey HC Yao. Single-nucleus multiomics reveals the gene regulatory networks underlying sex determination of murine primordial germ cells. *eLife*, 13, 2025.

[15025] Lena Alexopoulou and Magali Irla. Toll-like receptors (TLRs) in the trained immunity era. *eLife*, 14, 2025.

[15026] Gabriel Magno de Freitas Almeida, Iker Arriaga, Bruna Luiza de Azevedo, Miika Leppänen, Jonatas S Abrahão, Julien Andreani, Davide Zabeo, Janne J Ravantti, Nicola GA Abrescia, and Lotta-Riina Sundberg. Genomic and structural insights into Jyvaskylavirus, the first giant virus isolated from Finland. *eLife*, 13, 2025.

[15027] Ecem Altan, Catherine A Morgan, Steven C Dakin, and D Samuel Schwarzkopf. Spatial frequency adaptation modulates population receptive field sizes. *eLife*, 13, 2025.

[15028] Lenia Amaral, Xiaosha Wang, Yanchao Bi, and Ella Striem-Amit. Unraveling the impact of congenital deafness on individual brain organization. *eLife*, 13, 2025.

[15029] Miguel Amaral, Felix Frey, Xiuyun Jiang, Buzz Baum, and Anđela Šarić. Balancing stability and flexibility when reshaping archaeal membranes. *eLife*, 14, 2025.

[15030] Axelle Amen, Randy Yoo, Amanda Fabra-García, Judith Bolscher, William JR Stone, Isabelle Bally, Sebastián Dergan-Dylon, Iga Kucharska, Roos M de Jong, Marloes de Bruijni, Teun Bousema, C Richter King, Randall S MacGill, Robert W Sauerwein, Jean-Philippe Julien, Pascal Poignard, and Matthijs M Jore. Target-agnostic identification of human antibodies to Plasmodium falciparum sexual forms reveals cross-stage recognition of glutamate-rich repeats. *eLife*, 13, 2025.

[15031] Kenza Amroune, Lorenzo Fontolan, Agnès Baude, David Robbe, and Ingrid Bureau. Sparse innervation and local heterogeneity in the vibrissal corticostriatal projection. *eLife*, 14, 2025.

[15032] Dilara N Anbarci, Jennifer McKey, Daniel S Levic, Michel Bagnat, and Blanche Capel. Rediscovering the rete ovarii, a secreting auxiliary structure to the ovary. *eLife*, 13, 2025.

[15033] Parnian Arafi, Sujan Devkota, Emily Williams, Masato Maesako, and Michael S Wolfe. Alzheimer-mutant $\gamma$-secretase complexes stall amyloid $\beta$-peptide production. *eLife*, 13, 2025.

[15034] Hiroshi Arai, Susumu Katsuma, Noriko Matsuda-Imai, Shiou-Ruei Lin, Maki N Inoue, and Daisuke Kageyama. Prophage-encoded Hm-oscar gene recapitulates Wolbachia-induced male-killing in the tea tortrix moth Homona magnanima. *eLife*, 13, 2025.

[15035] Maria Ardaya, Marie-Catherine Tiveron, Harold Cremer, Esther Rubio-López, Abraham Martín, Benjamin Dehay, Fernando Pérez-Cerdá, Carlos Matute, Federico N Soria, and Fabio Cavaliere. Gliogenesis from the subventricular zone modulates the extracellular matrix at the glial scar after brain ischemia. *eLife*, 13, 2025.

[15036] Yasuhiro Arimura, Hide A Konishi, and Hironori Funabiki. MagIC-Cryo-EM, structural determination on magnetic beads for scarce macromolecules in heterogeneous samples. *eLife*, 13, 2025.

[15037] Toshitake Asabuki and Claudia Clopath. Embedding stochastic dynamics of the environment in spontaneous activity by prediction-based plasticity. *eLife*, 13, 2025.

[15038] Toshitake Asabuki and Tomoki Fukai. Predictive learning rules generate a cortical-like replay of probabilistic sensory experiences. *eLife*, 13, 2025.

[15039] Oghenevwogaga Joseph Atake, Fidji Berio, Melanie Debiais Thibaud, and B Frank Eames. The holocephalan ratfish endoskeleton shares trabecular and areolar mineralization patterns, but not tesserae, with elasmobranchs little skate and catshark. *eLife*, 13, 2025.

[15040] Franziska Auer, Katherine Nardone, Koji Matsuda, Masahiko Hibi, and David Schoppik. Cerebellar Purkinje cells control posture in larval zebrafish (Danio rerio). *eLife*, 13, 2025.

[15041] Hans Auer, Donna Gift Cabalo, Raúl Rodríguez-Cruces, Oualid Benkarim, Casey Paquola, Jordan DeKraker, Yezhou Wang, Sofie Louise Valk, Boris C Bernhardt, and Jessica Royer. From histology to macroscale function in the human amygdala. *eLife*, 13, 2025.

[15042] Joonas A Autio, Ikko Kimura, Takayuki Ose, Yuki Matsumoto, Masahiro Ohno, Yuta Urushibata, Takuro Ikeda, Matthew F Glasser, David C van Essen, and Takuya Hayashi. Mapping vascular network architecture in primate brain using ferumoxytol-weighted laminar MRI. *eLife*, 13, 2025.

[15043] Jessel Ayra Plasencia, Sara Medina-Suárez, Esperanza Hernández-Carralero, Jonay García-Luis, Lorraine S Symington, and Félix Machín. Cohesin reconstitution and homologous recombination repair of DNA double-strand breaks in late mitosis. *eLife*, 13, 2025.

[15044] Benura Azeroglu, Wei Wu, Raphael Pavani, Ranjodh Singh Sandhu, Tadahiko Matsumoto, André Nussenzweig, and Eros Lazzerini-Denchi. Conserved and unique features of terminal telomeric sequences in ALT-positive cancer cells. *eLife*, 14, 2025.

[15045] Azali Azlan, Li Zhu, and Ryuya Fukunaga. Female-germline specific protein Sakura interacts with Otu and is crucial for germline stem cell renewal and differentiation and oogenesis. *eLife*, 13, 2025.

[15046] Farah Bader, Clayton Bingham, Karen K David, Hermon Gebrehiwet, Crystal L Lantz, Grace CY Peng, Mauricio Rangel-Gomez, and James Gnadt. The NIH BRAIN Initiative's impacts in systems and computational neuroscience and team-scale research 2014–2023. *eLife*, 14, 2025.

[15047] Akanksha Bafna, Gareth Banks, Vadim Vasilyev, Robert Dallmann, Michael H Hastings, and Patrick M Nolan. Zinc finger homeobox-3 (ZFHX3) orchestrates genome-wide daily gene expression in the suprachiasmatic nucleus. *eLife*, 14, 2025.

[15048] M Reza Bahranifard, Jessica Chan, A Thomas Read, Guorong Li, Lin Cheng, Babak N Safa, Seyed Mohammad Siadat, Anamik Jhunjhunwala, Hans E Grossniklaus, Stanislav Y Emelianov, W Daniel Stamer, Markus H Kuehn, and C Ross Ethier. Magnetically steered cell therapy for reduction of intraocular pressure as a treatment strategy for open-angle glaucoma. *eLife*, 13, 2025.

[15049] Thomas Balligand, Claire Carpenet, Sergi Olivé Palau, Tom Jaspers, Pavana Suresh, Xin Liu, Himadri Medhi, Yoon Ho Lee, Mohammad Rashidian, Bart De Strooper, Hidde L Ploegh, and Maarten Dewilde. A pair of congenic mice for imaging of transplants by positron emission tomography using anti-transferrin receptor nanobodies. *eLife*, 14, 2025.

[15050] Andrew J Balmer, Nina FD White, Eyyüb S ünlü, Chiyun Lee, Richard D Pearson, Jacob Almagro-Garcia, and Cristina Ariani. Understanding the global rise of artemisinin resistance: Insights from over 100,000 Plasmodium falciparum samples. *eLife*, 14, 2025.

[15051] Deb Sankar Banerjee and Shiladitya Banerjee. Catalytic growth in a shared enzyme pool ensures robust control of centrosome size. *eLife*, 12, 2025.

[15052] Soumita Banerjee, Soniya Sanjay Shende, Laxmi Kata, Relestina Simon Lopes, Swathika Praveen, Ruchi Joshi, Naveen Kumar Khare, Gorantla V Raghuram, Snehal Shabrish, and Indraneel Mittra. Horizontally transferred cell-free chromatin particles function as autonomous 'satellite genomes' and vehicles for transposable elements within host cells. *eLife*, 13, 2025.

[15053] Kush Banga, Julius Benson, Jai Bhagat, Dan Biderman, Daniel Birman, Niccolò Bonacchi, Sebastian A Bruijns, Kelly Buchanan, Robert AA Campbell, Matteo Carandini, Gaelle A Chapuis, Anne K Churchland, M Felicia Davatolhagh, Hyun Dong Lee, Mayo Faulkner, Berk GerÇek, Fei Hu, Julia Huntenburg, Cole Lincoln Hurwitz, Anup Khanal, Christopher Krasniak, Petrina Lau, Christopher Langfield, Nancy Mackenzie, Guido T Meijer, Nathaniel J Miska, Zeinab Mohammadi, Jean-Paul Noel, Liam Paninski, Alejandro Pan-Vazquez, Cyrille Rossant, Noam Roth, Michael Schartner, Karolina Z Socha, Nicholas A Steinmetz, Karel Svoboda, Marsa Taheri, Anne E Urai, Shuqi Wang, Miles Wells, Steven J West, Matthew R Whiteway, Olivier Winter, Ilana B Witten, and Yizi Zhang. Reproducibility of in vivo electrophysiological measurements in mice. *eLife*, 13, 2025.

[15054] Gabriele Baniulyte and Joseph T Wade. A bacterial regulatory uORF senses multiple classes of ribosome-targeting antibiotics. *eLife*, 13, 2025.

[15055] Adriana Bankston. Campaigning for science and scientists. *eLife*, 14, 2025.

[15056] Vibhavari Aysha Bansal, Jia Min Tan, Hui Rong Soon, Norliyana Zainolabidin, Takaomi Saido, and Toh Hean Ch'ng. A$\beta$-driven nuclear pore complex dysfunction alters activation of necroptosis proteins in a mouse model of Alzheimer's disease. *eLife*, 13, 2025.

[15057] Hanbin Bao, Yanan Wang, Haoxing Li, Qiang Wang, Yutao Lei, Ying Ye, Syed F Wadood, Hui Zhu, Christian Staehelin, Gary Stacey, Shutong Xu, and Yangrong Cao. The rhizobial effector NopT targets Nod factor receptors to regulate symbiosis in Lotus japonicus. *eLife*, 13, 2025.

[15058] Rishika Baral, Kristin Ho, Ramasamy P Kumar, Jesse B Hopkins, Maxwell B Watkins, Salvatore LaRussa, Suhaily Caban-Penix, Logan A Calderone, and Niels Bradshaw. A general mechanism for initiating the bacterial general stress response. *eLife*, 13, 2025.

[15059] Alison G Barber, Cynthia M Quintero, Michael Hamilton, Nirakar Rajbhandari, Roman Sasik, Yan Zhang, Carla Kim, Hatim Husain, Xin Sun, and Tannishtha Reya. Regulation of lung cancer initiation and progression by the stem cell determinant Musashi. *eLife*, 13, 2025.

[15060] Aida Bareghamyan, Changfeng Deng, Sarah Daoudi, Shubhash C Yadav, Xiaocen Lu, Wei Zhang, Robert E Campbell, Richard H Kramer, David M Chenoweth, and Don B Arnold. A toolbox for ablating excitatory and inhibitory synapses. *eLife*, 13, 2025.

[15061] Graham L Barlow, Christian M Schürch, Salil S Bhate, Darci J Phillips, Arabella Young, Shen Dong, Hunter A Martinez, Gernot Kaber, Nadine Nagy, Sasvath Ramachandran, Janet Meng, Eva Korpos, Jeffrey A Bluestone, Garry P Nolan, and Paul L Bollyky. The extra-islet pancreas supports autoimmunity in human type 1 diabetes. *eLife*, 13, 2025.

[15062] Joseph M Barnby, Jen Nguyen, Julia Griem, Magdalena Wloszek, Henry Burgess, Linda J Richards, Jessica Kingston, Gavin Cooper, P Read Montague, Peter Dayan, Tobias Nolte, and Peter Fonagy. Self-other generalisation shapes social interaction and is disrupted in borderline personality disorder. *eLife*, 14, 2025.

[15063] Kelsey R Baron, Samantha Oviedo, Sophia Krasny, Mashiat Zaman, Rama Aldakhlallah, Prerona Bora, Prakhyat Mathur, Gerald Pfeffer, Michael J Bollong, Timothy E Shutt, Danielle A Grotjahn, and R Luke Wiseman. Pharmacologic activation of integrated stress response kinases inhibits pathologic mitochondrial fragmentation. *eLife*, 13, 2025.

[15064] Claudia Barth, Liisa AM Galea, Emily G Jacobs, Bonnie H Lee, Lars T Westlye, and Ann-Marie G de Lange. Menopausal hormone therapy and the female brain: Leveraging neuroimaging and prescription registry data from the UK Biobank cohort. *eLife*, 13, 2025.

[15065] Pál Barzó, Ildikó Szöts, Martin Tóth, éva Adrienn Csajbók, Gábor Molnár, and Gábor Tamás. Electrophysiology and morphology of human cortical supragranular pyramidal cells in a wide age range. *eLife*, 13, 2025.

[15066] Lisa M Bas, Ian D Roberts, Cendri A Hutcherson, and Anita Tusche. A neurocomputational account of the link between social perception and social action. *eLife*, 12, 2025.

[15067] Diellor Basha, Amirmohammad Azarmehri, Elian Proulx, Sylvain Chauvette, Maryam Ghorbani, and Igor Timofeev. The reuniens nucleus of the thalamus facilitates hippocampo-cortical dialogue during sleep. *eLife*, 12, 2025.

[15068] Gianpaolo Antonio Basile, Augusto Ielo, Lilla Bonanno, Antonio Cerasa, Giuseppe Santoro, Demetrio Milardi, Giuseppe Pio Anastasi, Ambra Torre, Sergio Baldari, Riccardo Laudicella, Michele Gaeta, Marina Quartu, Maria Pina Serra, Marcello Trucas, Angelo Quartarone, Manojkumar Saranathan, and Alberto Cacciola. Shared functional organization between pulvinar-cortical and cortico-cortical connectivity and its structural and molecular imaging correlates. *eLife*, 13, 2025.

[15069] Morgane Batzenschlager, Beatrice Lace, Ning Zhang, Chao Su, Anna Boiger, Sabrina Egli, Pascal Krohn, Jule Salfeld, Franck Anicet Ditengou, Thomas Laux, and Thomas Ott. Competence for transcellular infection in the root cortex involves a post-replicative, cell-cycle exit decision in Medicago truncatula. *eLife*, 12, 2025.

[15070] Helen B Belato, Alexa L Knight, Alexandra M D'Ordine, Chinmai Pindi, Zhiqiang Fan, Jinping Luo, Giulia Palermo, Gerwald Jogl, and George P Lisi. Structural and dynamic impacts of single-atom disruptions to guide RNA interactions within the recognition lobe of Geobacillus stearothermophilus Cas9. *eLife*, 13, 2025.

[15071] Christopher Bell, Lukas Kilo, Daniel Gottschalk, Jashar Arian, Lea Deneke, Hanna Kern, Christof Rickert, Oliver Kobler, Julia Strauß, Martin Heine, Carsten Duch, and Stefanie Ryglewski. Specific presynaptic functions require distinct Drosophila Cav2 splice isoforms. *eLife*, 13, 2025.

[15072] George N Bendzunas, Dominic P Byrne, Safal Shrestha, Leonard A Daly, Sally O Oswald, Samiksha Katiyar, Aarya Venkat, Wayland Yeung, Claire E Eyers, Patrick A Eyers, and Natarajan Kannan. Redox regulation and dynamic control of brain-selective kinases BRSK1/2 in the AMPK family through cysteine-based mechanisms. *eLife*, 13, 2025.

[15073] Ethan S Benevides, Prajwal P Thakre, Sabhya Rana, Michael D Sunshine, Victoria N Jensen, Karim Oweiss, and David D Fuller. Chemogenetic stimulation of phrenic motor output and diaphragm activity. *eLife*, 13, 2025.

[15074] Lee R Berger, John Hawks, Agustín Fuentes, Dirk Van Rooyen, Mathabela Tsikoane, Maropeng Mpete, Samuel Nkwe, and Keneiloe Molopyane. An initial report of circa 241,000- to 335,000-year-old rock engravings and their relation to Homo naledi in the Rising Star cave system, South Africa. *eLife*, 12, 2025.

[15075] Bruno Gideon Bergheim, Alison G Cole, Mandy Rettel, Frank Stein, Stefan Redl, Michael W Hess, Aissam Ikmi, and Suat özbek. Molecular dynamics of the matrisome across sea anemone life history. *eLife*, 14, 2025.

[15076] Sreetama Bhadra, Gabriella Damasceno, Daniela Hoss, and Alexandra Weyrich. Improving gender equity in academia depends on the workplace environment. *eLife*, 14, 2025.

[15077] Arkajyoti Bhattacharya, Thijs S Stutvoet, Mirela Perla, Stefan Loipfinger, Mathilde Jalving, Anna KL Reyners, Paola D Vermeer, Ronny Drapkin, Marco de Bruyn, Elisabeth GE de Vries, Steven de Jong, and Rudolf SN Fehrmann. Transcriptional pattern enriched for synaptic signaling is associated with shorter survival of patients with high-grade serous ovarian cancer. *eLife*, 13, 2025.

[15078] Namrata Bhattacharya, Anja Rockstroh, Sanket Suhas Deshpande, Sam Koshy Thomas, Anunay Yadav, Chitrita Goswami, Smriti Chawla, Pierre Solomon, Cynthia Fourgeux, Gaurav Ahuja, Brett Hollier, Himanshu Kumar, Antoine Roquilly, Jeremie Poschmann, Melanie Lehman, Colleen C Nelson, and Debarka Sengupta. Artificial intelligence approaches for tumor phenotype stratification from single-cell transcriptomic data. *eLife*, 13, 2025.

[15079] Ronja Bigge, Rebecca Grittner, and Anna Lisa Stöckl. Integration of parallel pathways for flight control in a hawkmoth reflects prevalence and relevance of natural visual cues. *eLife*, 14, 2025.

[15080] Célian Bimbard, Flóra Takács, Joana A Catarino, Julie MJ Fabre, Sukriti Gupta, Stephen C Lenzi, Maxwell D Melin, Nathanael O'Neill, Ivana Orsolic, Magdalena Robacha, James S Street, José M Gomes Teixeira, Simon Townsend, Enny H van Beest, Arthur M Zhang, Anne K Churchland, Chunyu A Duan, Kenneth D Harris, Dimitri Michael Kullmann, Gabriele Lignani, Zachary F Mainen, Troy W Margrie, Nathalie L Rochefort, Andrew Wikenheiser, Matteo Carandini, and Philip Coen. An adaptable, reusable, and light implant for chronic Neuropixels probes. *eLife*, 13, 2025.

[15081] Rituja S Bisen, Fathima Mukthar Iqbal, Federico Cascino-Milani, Till Bockemühl, and Jan M Ache. Nutritional state-dependent modulation of insulin-producing cells in Drosophila. *eLife*, 13, 2025.

[15082] Joseph A Bisson, Miriam Gordillo, Ritu Kumar, Neranjan de Silva, Ellen Yang, Kelly M Banks, Zhong-Dong Shi, Kihyun Lee, Dapeng Yang, Wendy K Chung, Danwei Huangfu, and Todd Evans. GATA6 regulates WNT and BMP programs to pattern precardiac mesoderm during the earliest stages of human cardiogenesis. *eLife*, 13, 2025.

[15083] Jorge Blanco Mendana, Margaret Donovan, Lindsey Gengelbach O'Brien, Benjamin Auch, John Garbe, and Daryl M Gohl. Deterministic genetic barcoding for multiplexed behavioral and single-cell transcriptomic studies. *eLife*, 12, 2025.

[15084] Itai Bloch, David Troupin, Sivan Toledo, Ran Nathan, and Nir Sapir. Combining radio-telemetry and radar measurements to test optimal foraging in an aerial insectivore bird. *eLife*, 13, 2025.

[15085] Ilona M Bloem, Leah Bakst, Joseph T McGuire, and Sam Ling. Dynamic estimation of the attentional field from visual cortical activity. *eLife*, 14, 2025.

[15086] Attila Boda, Villő Balázs, Anikó Nagy, Dávid Hargitai, Mónika Lippai, Zsófia Simon-Vecsei, Márton Molnár, Fanni Fürstenhoffer, Gábor Juhász, and Péter Lőrincz. The Rab7-Epg5 and Rab39-ema modules cooperatively position autophagosomes for efficient lysosomal fusions. *eLife*, 13, 2025.

[15087] Ida Marie Boisen, Nadia Krarup Knudsen, John E Nielsen, Ireen Kooij, Mathilde Louise Bagger, Jovanna Kaludjerovic, Peter O'Shaughnessy, Peter W Andrews, Noriko Ide, Birgitte G Toft, Anders Juul, Arnela Mehmedbasic, Anne Jørgensen, Lee B Smith, Richard Norman, Ewa Rajpert-De Meyts, Beate Lanske, and Martin Blomberg Jensen. Changes in local mineral homeostasis facilitate the formation of benign and malignant testicular microcalcifications. *eLife*, 13, 2025.

[15088] Annika Boldt, Celine Ann Fox, Claire M Gillan, and Sam Gilbert. Transdiagnostic compulsivity is associated with reduced reminder setting, only partially attributable to overconfidence. *eLife*, 13, 2025.

[15089] James Boocock, Noah Alexander, Leslie Alamo Tapia, Laura Walter-McNeill, Shivani Prashant Patel, Chetan Munugala, Joshua S Bloom, and Leonid Kruglyak. Single-cell eQTL mapping in yeast reveals a tradeoff between growth and reproduction. *eLife*, 13, 2025.

[15090] Prateeka Borar, Tapan Biswas, Ankur Chaudhuri, Pallavi T Rao, Swasti Raychaudhuri, Tom Huxford, Saikat Chakrabarti, Gourisankar Ghosh, and Smarajit Polley. Dual-specific autophosphorylation of kinase IKK2 enables phosphorylation of substrate I$\kappa$B$\alpha$ through a phosphoenzyme intermediate. *eLife*, 13, 2025.

[15091] Ilaria Borghi, Lucia Mencarelli, Michele Maiella, Elias Paolo Casula, Matteo Ferraresi, Francesca Candeo, Elena Savastano, Martina Assogna, Sonia Bonnì, and Giacomo Koch. Dual transcranial electromagnetic stimulation of the precuneus boosts human long-term memory. *eLife*, 14, 2025.

[15092] Hannah Bos, Christoph Miehl, Anne-Marie Michelle Oswald, and Brent Doiron. Untangling stability and gain modulation in cortical circuits with multiple interneuron classes. *eLife*, 13, 2025.

[15093] Thomas A Bos, Elizaveta Polyakova, Janine Maria van Gils, Antoine AF de Vries, Marie-José Goumans, Christian Freund, Marco C DeRuiter, and Monique RM Jongbloed. A systematic review and embryological perspective of pluripotent stem cell-derived autonomic postganglionic neuron differentiation for human disease modeling. *eLife*, 14, 2025.

[15094] Mahima Bose, Ishita Talwar, Varun Suresh, Urvi Mishra, Shiona Biswas, Anuradha Yadav, Shital T Suryavanshi, Simon Hippenmeyer, and Shubha Tole. Dual role of FOXG1 in regulating gliogenesis in the developing neocortex via the FGF signalling pathway. *eLife*, 13, 2025.

[15095] Justine Boutry, Océane Rieu, Lena Guimard, Jordan Meliani, Aurora M Nedelcu, Sophie Tissot, Nikita Stepanskyy, Beata Ujvari, Rodrigo Hamede, Antoine M Dujon, Jácint Tökölyi, and Fréderic Thomas. First evidence for the evolution of host manipulation by tumors during the long-term vertical transmission of tumor cells in Hydra oligactis. *eLife*, 13, 2025.

[15096] John C Bowler, George Zakka, Hyun Choong Yong, Wenke Li, Bovey Rao, Zhenrui Liao, James B Priestley, and Attila Losonczy. An Intranet of Things approach for adaptable control of behavioral and navigation-based experiments. *eLife*, 13, 2025.

[15097] Kayleigh Bozon, Hartmut Cuny, Delicia Z Sheng, Ella MMA Martin, Alena Sipka, Paul Young, David T Humphreys, and Sally L Dunwoodie. Impaired yolk sac NAD metabolism disrupts murine embryogenesis with relevance to human birth defects. *eLife*, 13, 2025.

[15098] Kai M Bracey, Margret A Fye, Alisa Cario, Kung-Hsien Ho, Pi'illani Noguchi, Guoqiang Gu, and Irina Kaverina. Glucose-stimulated KIF5B-driven microtubule sliding organizes microtubule networks in mouse pancreatic $\beta$ cells. *eLife*, 12, 2025.

[15099] Patrick Brandt, Dawayne Whittington, Kimberley D Wood, Christopher Holmquist, Ana T Nogueira, Christiann H Gaines, Patrick Brennwald, and Rebekah L Layton. Development and assessment of a sustainable PhD internship program supporting diverse biomedical career outcomes. *eLife*, 12, 2025.

[15100] Anamari Brdar, Christian Osterburg, Philipp Münick, Anne Christin Machel, Rajeshwari Rathore, Susanne Osterburg, Büşra Yüksel, Birgit Schäfer, Kristina Desch, Julian D Langer, Ivan Dikic, and Volker Dötsch. p53 isoforms have a high aggregation propensity, interact with chaperones and lack binding to p53 interaction partners. *eLife*, 13, 2025.

[15101] Andrea Brenna, Micaela Borsa, Gabriella Saro, Jürgen A Ripperger, Dominique A Glauser, Zhihong Yang, Antoine Adamantidis, and Urs Albrecht. Cyclin-dependent kinase 5 (Cdk5) activity is modulated by light and gates rapid phase shifts of the circadian clock. *eLife*, 13, 2025.

[15102] Kira Breunig, Xuifen Lei, Mauro Montalbano, Gabriela DA Guardia, Shiva Ostadrahimi, Victoria Alers, Adam Kosti, Jennifer Chiou, Nicole Klein, Corina Vinarov, Lily Wang, Mujia Li, Weidan Song, W Lee Kraus, David S Libich, Stefano Tiziani, Susan T Weintraub, Pedro AF Galante, and Luiz O Penalva. SERBP1 interacts with PARP1 and is present in PARylation-dependent protein complexes regulating splicing, cell division, and ribosome biogenesis. *eLife*, 13, 2025.

[15103] Martin Breyton, Jan Fousek, Giovanni Rabuffo, Pierpaolo Sorrentino, Lionel Kusch, Marcello Massimini, Spase Petkoski, and Viktor Jirsa. Spatiotemporal brain complexity quantifies consciousness outside of perturbation paradigms. *eLife*, 13, 2025.

[15104] Eric R Brooks, Andrew R Moorman, Bhaswati Bhattacharya, Ian S Prudhomme, Max Land, Heather L Alcorn, Roshan Sharma, Dana Pe'er, and Jennifer A Zallen. A single-cell atlas of spatial and temporal gene expression in the mouse cranial neural plate. *eLife*, 13, 2025.

[15105] Merissa Brousseau, Da Teng, Nathan E Thomas, Gregory A Voth, and Katherine A Henzler-Wildman. The C-terminus of the multi-drug efflux pump EmrE prevents proton leak by gating transport. *eLife*, 14, 2025.

[15106] Robert A Bruce, Matthew Weber, Alexandra Bova, Rachael Volkman, Casey Jacobs, Kartik Sivakumar, Hannah Stutt, Youngcho Kim, Rodica Curtu, and Kumar Narayanan. Complementary cognitive roles for D2-MSNs and D1-MSNs during interval timing. *eLife*, 13, 2025.

[15107] Jean-FranÇois Brunet. Gaskell, Langley, and the "para-sympathetic" idea. *eLife*, 14, 2025.

[15108] Alexandra V Bruter, Ekaterina A Varlamova, Nina I Stavskaya, Zoia G Antysheva, Vasily N Manskikh, Anna V Tvorogova, Diana S Korshunova, Alvina I Khamidullina, Marina V Utkina, Viktor P Bogdanov, Iuliia P Baikova, Alyona I Nikiforova, Eugene A Albert, Denis O Maksimov, Jing Li, Mengqian Chen, Gary P Schools, Alexey V Feoktistov, Alexander A Shtil, Igor B Roninson, Vladislav A Mogila, Yulia Y Silaeva, and Victor V Tatarskiy. Knockout of cyclin-dependent kinases 8 and 19 leads to depletion of cyclin C and suppresses spermatogenesis and male fertility in mice. *eLife*, 13, 2025.

[15109] Xiangning Bu, Nathanael Ashby, Teresa Vitali, Sulgi Lee, Ananya Gottumukkala, Kangsun Yun, Sana Tabbara, Patricia Latham, Christine Teal, and Inhee Chung. Cell crowding activates pro-invasive mechanotransduction pathway in high-grade DCIS via TRPV4 inhibition and cell volume reduction. *eLife*, 13, 2025.

[15110] John A Buglino, Yaprak Ozakman, Chad E Hatch, Anna Benjamin, Derek S Tan, and Michael S Glickman. Chalkophore-mediated respiratory oxidase flexibility controls M. tuberculosis virulence. *eLife*, 14, 2025.

[15111] Justin Burdge, Anissa Jhumka, Ashar Khan, Simon Ogundare, Nicholas Baer, Sasha Fulton, Alexander Kaplan, Brittany Bistis, William Foster, Joshua Thackray, Andre Toussaint, Miao Li, Yosuke M Morizawa, Jake Nazarian, Leah Yadessa, Arlene J George, Abednego Delinois, Wadzanayi Mayiseni, Noah Loran, Guang Yang, David J Margolis, Victoria E Abraira, and Ishmail Abdus-Saboor. Remote automated delivery of mechanical stimuli coupled to brain recordings in behaving mice. *eLife*, 13, 2025.

[15112] Ludwig Burger and Ulrich Gerland. Toward stable replication of genomic information in pools of RNA molecules. *eLife*, 14, 2025.

[15113] Ozanna Burnicka-Turek, Katy A Trampel, Brigitte Laforest, Michael T Broman, Xinan H Yang, Zoheb Khan, Eric Rytkin, Binjie Li, Ella Schaffer, Margaret Gadek, Kaitlyn M Shen, Igor R Efimov, and Ivan P Moskowitz. Coordinated Tbx3/Tbx5 transcriptional control of the adult ventricular conduction system. *eLife*, 13, 2025.

[15114] Silas E Busch and Christian Hansel. Non-allometric expansion and enhanced compartmentalization of Purkinje cell dendrites in the human cerebellum. *eLife*, 14, 2025.

[15115] Ole Bæk, Tik Muk, Ziyuan Wu, Yongxin Ye, Bekzod Khakimov, Alessandra Maria Casano, Bagirath Gangadharan, Ivan Bilic, Anders Brunse, Per Torp Sangild, and Duc Ninh Nguyen. Altered hepatic metabolism mediates sepsis preventive effects of reduced glucose supply in infected preterm newborns. *eLife*, 13, 2025.

[15116] Lin Cai, Taiyu Fan, Xuzhong Qu, Ying Zhang, Xianyu Gou, Quanwei Ding, Weihua Feng, Tingting Cao, Xiaohua Lv, Xiuli Liu, Qing Huang, Tingwei Quan, and Shaoqun Zeng. Automatic and accurate reconstruction of long-range axonal projections of single-neuron in mouse brain. *eLife*, 13, 2025.

[15117] Sihui Cai, Jie Zhou, Xiaotong Luo, Chenqiu Zhang, Shouheng Jin, Jian Ren, and Jun Cui. Phase transition of WTAP regulates m6A modification of interferon-stimulated genes. *eLife*, 13, 2025.

[15118] Alexander S Campbell, Martin Minařík, Roman Franěk, Michaela Vazačová, MiloŠ Havelka, David Gela, Martin PŠenička, and Clare VH Baker. Opposing roles for Bmp signalling during the development of electrosensory lateral line organs. *eLife*, 14, 2025.

[15119] Justin M Campbell, Rhiannon L Cowan, Krista L Wahlstrom, Martina K Hollearn, Dylan Jensen, Tyler Davis, Shervin Rahimpour, Ben Shofty, Amir Arain, John D Rolston, Stephan Hamann, Shuo Wang, Lawrence N Eisenman, James Swift, Tao Xie, Peter Brunner, Joseph Manns, Cory Inman, Elliot H Smith, and Jon Timothy Willie. Human single-neuron activity is modulated by intracranial theta burst stimulation of the basolateral amygdala. *eLife*, 14, 2025.

[15120] Feilin Cao, Zhaosheng Ma, Zenggui Wu, Weizhu Wu, Ouchen Wang, Binbin Cui, Xiaotao Zhu, Jing Hao, Xiaochun Ji, Zhanwen Li, Deyou Tao, Qingjing Feng, Wei Lin, Dongbo Shi, Jingde Shu, Jichun Zhou, and Shifen Huang. Pyrotinib after trastuzumab-based adjuvant therapy in patients with HER2-positive breast cancer (PERSIST): A multicenter phase II trial. *eLife*, 13, 2025.

[15121] Li-Jun Cao, Jin-Cui Chen, Joshua A Thia, Thomas L Schmidt, Richard Ffrench-Constant, Lin-Xi Zhang, Yu Yang, Meng-Chu Yuan, Jia-Yue Zhang, Xiao-Yang Zhang, Qiong Yang, Ya-Jun Gong, Hu Li, Xuexin Chen, Ary A Hoffmann, and Shu-Jun Wei. Recurrent mutations drive the rapid evolution of pesticide resistance in the two-spotted spider mite Tetranychus urticae. *eLife*, 14, 2025.

[15122] Zhiping Cao, Wing-Ho Yung, and Ya Ke. Repeated activation of preoptic area recipient neurons in posterior paraventricular nucleus mediates chronic heat-induced negative emotional valence and hyperarousal states. *eLife*, 13, 2025.

[15123] Aurelie Carabalona, Henna Kallo, Maryanne Gonzalez, Liliia Andriichuk, Ellinoora Elomaa, Florence Molinari, Christiana Fragkou, Pekka Lappalainen, Marja W Wessels, Juha Saarikangas, and Claudio Rivera. Identification of novel human microcephaly-linked protein Mtss2 that mediates cortical progenitor cell division and corticogenesis through Nedd9-RhoA. *eLife*, 13, 2025.

[15124] Eugenio Antonio Carrera Silva, Juliana Puyssegur, and Andrea Emilse Errasti. Coevolutionary interplay: Helminths-trained immunity and its impact on the rise of inflammatory diseases. *eLife*, 14, 2025.

[15125] Sergio Casas-Tinto, Nuria Garcia-Guillen, and María Losada-Perez. Adult neurogenesis through glial transdifferentiation in a CNS injury paradigm. *eLife*, 13, 2025.

[15126] Bastien Casu, Joseph W Sallmen, Peter E Haas, Govind Chandra, Pavel Afanasyev, Jingwei Xu, Martin Pilhofer, and Susan Schlimpert. Function and firing of the Streptomyces coelicolor contractile injection system requires the membrane protein CisA. *eLife*, 14, 2025.

[15127] Çağrı Çevrim, Béryl Laplace-Builhé, Ko Sugawara, Maria Lorenza Rusciano, Nicolas Labert, Jacques Brocard, Alba Almazán, and Michalis Averof. Long-term live imaging, cell identification and cell tracking in regenerating crustacean legs. *eLife*, 14, 2025.

[15128] Jacob H Cecil, Cristina M Padilla, Austin A Lipinski, Paul Langlais, Xiangxia Luo, and Andrew P Capaldi. The molecular logic of Gtr1/2- and Pib2-dependent TORC1 regulation in budding yeast. *eLife*, 13, 2025.

[15129] Chrispin Chaguza, Daan W Arends, Stephanie W Lo, Indri Hapsari Putri, Anna York, John A Lees, Anne L Wyllie, Daniel M Weinberger, Stephen D Bentley, Marien I de Jonge, and Amelieke JH Cremers. The capsule and genetic background, rather than specific individual loci, strongly influence in vitro pneumococcal growth kinetics. *eLife*, 14, 2025.

[15130] Jodie Chandler, M Elise Bullock, Arpit C Swain, Cayman Williams, Christiaan H van Dorp, Benedict Seddon, and Andrew J Yates. Tissue-resident memory CD4+ T cells are sustained by site-specific levels of self-renewal and continuous replacement. *eLife*, 14, 2025.

[15131] Iksoo Chang, Taegon Chung, and Sangyeol Kim. Wavenumber-dependent transmission of subthreshold waves on electrical synapses network model of Caenorhabditis elegans. *eLife*, 13, 2025.

[15132] Song Chang, Beilin Zheng, Les Keniston, Jinghong Xu, and Liping Yu. Auditory cortex learns to discriminate audiovisual cues through selective multisensory enhancement. *eLife*, 13, 2025.

[15133] Ana Cristina Chang-Gonzalez, Aoi Akitsu, Robert J Mallis, Matthew J Lang, Ellis L Reinherz, and Wonmuk Hwang. Load-based divergence in the dynamic allostery of two TCRs recognizing the same pMHC. *eLife*, 13, 2025.

[15134] Kuan-Hao Chao, Alan Mao, Anqi Liu, Steven L Salzberg, and Mihaela Pertea. OpenSpliceAI provides an efficient modular implementation of SpliceAI enabling easy retraining across nonhuman species. *eLife*, 14, 2025.

[15135] Martik Chatterjee, Xin Yi Yu, Noah K Brady, Connor Amendola, Gabriel C Hatto, and Robert D Reed. mirror determines the far posterior domain in butterfly wings. *eLife*, 13, 2025.

[15136] Hiam Chemaitelly, Houssein H Ayoub, Peter Coyle, Patrick Tang, Mohammad R Hasan, Hadi M Yassine, Asmaa A Al Thani, Zaina Al-Kanaani, Einas Al-Kuwari, Andrew Jeremijenko, Anvar Hassan Kaleeckal, Ali Nizar Latif, Riyazuddin Mohammad Shaik, Hanan F Abdul-Rahim, Gheyath K Nasrallah, Mohamed Ghaith Al-Kuwari, Hamad Eid Al-Romaihi, Mohamed H Al-Thani, Abdullatif Al-Khal, Roberto Bertollini, Adeel A Butt, and Laith J Abu-Raddad. Assessing healthy vaccinee effect in COVID-19 vaccine effectiveness studies: a national cohort study in Qatar. *eLife*, 14, 2025.

[15137] Chong Chen, Qian Liu, Cheng Tang, Yu Rong, Xinyi Zhao, Dandan Li, Fan Lu, Jia Qu, and Xinting Liu. ZC3H11A mutations cause high myopia by triggering PI3K-AKT and NF-$\kappa$B-mediated signaling pathway in humans and mice. *eLife*, 12, 2025.

[15138] Christopher H Chen, Zhiyi Yao, Shuting Wu, and Wade G Regehr. Characterization of direct Purkinje cell outputs to the brainstem. *eLife*, 13, 2025.

[15139] Cong Chen, Tong-Yao Gao, Hua-Wei Yi, Yi Zhang, Tong Wang, Zhi-Ling Lou, Tao-Feng Wei, Yun-Bi Lu, Tingting Li, Chun Tang, and Wei-Ping Zhang. Elevated ubiquitin phosphorylation by PINK1 contributes to proteasomal impairment and promotes neurodegeneration. *eLife*, 14, 2025.

[15140] En-Li Chen, Tsai-Wen Chen, Eric R Schreiter, and Bei-Jung Lin. Synchronous ensembles of hippocampal CA1 pyramidal neurons during novel exploration. *eLife*, 13, 2025.

[15141] Fangluo Chen, Dylan C Sarver, Muzna Saqib, Leandro M Velez, Susan Aja, Marcus M Seldin, and G William Wong. Loss of CTRP10 results in female obesity with preserved metabolic health. *eLife*, 13, 2025.

[15142] Guanghan Chen, Zhengyang Guo, Zhiwen Zhu, Shanshan Xie, Tianhua Zhou, and Guangshuo Ou. Kinesin-2 autoinhibition requires elbow phosphorylation. *eLife*, 13, 2025.

[15143] Hongqian Chen, Hui-Qing Fang, Jin-Tao Liu, Shi-Yu Chang, Li-Ben Cheng, Ming-Xin Sun, Jian-Rui Feng, Ze-Min Liu, Yong-Hong Zhang, Clifford J Rosen, and Peng Liu. Atlas of Fshr expression from novel reporter mice. *eLife*, 13, 2025.

[15144] Huanhuan Joyce Chen, Eric E Gardner, Yajas Shah, Kui Zhang, Abhimanyu Thakur, Chen Zhang, Olivier Elemento, and Harold Varmus. Metastatic small cell lung cancer arises from TP53/RB1-deficient and MYC overproduction hESC-derived PNECs. *eLife*, 13, 2025.

[15145] Jenny Chen, Phoebe R Richardson, Christopher Kirby, Sean R Eddy, and Hopi E Hoekstra. Cellular evolution of the hypothalamic preoptic area of behaviorally divergent deer mice. *eLife*, 13, 2025.

[15146] Pengbo Chen, Bo Li, Zeyu Lu, Qingyin Xu, Huoliang Zheng, Shengdan Jiang, Leisheng Jiang, and Xinfeng Zheng. PCBP2 as an intrinsic agi ng factor regulates the senescence of hBMSCs through the ROS-FGF2 signaling axis. *eLife*, 13, 2025.

[15147] Shihui Chen and Carolyn Marie Phillips. Nuclear Argonaute protein NRDE-3 switches small RNA partners during embryogenesis to mediate temporal-specific gene regulatory activity. *eLife*, 13, 2025.

[15148] Shuo Chen, Jinxiu Liang, Jie Yin, Weijia Zhang, Peijun Jiang, Wenyuan Wang, Xiaoying Chen, Yuanhong Zhou, Peng Xia, Fan Yang, Ying Gu, Ruilin Zhang, and Peidong Han. Contraction-induced endocardial id2b plays a dual role in regulating myocardial contractility and valve formation. *eLife*, 13, 2025.

[15149] Siyu Chen, Tamar Basiashvili, Joshua Hutchings, Marta Sanz Murillo, Amalia Villagran Suarez, Erica Xiong, Jaime Alegrio Louro, Andres E Leschziner, and Elizabeth Villa. Cryo-electron tomography reveals the microtubule-bound form of inactive LRRK2. *eLife*, 13, 2025.

[15150] Wanxia Chen, Jiaqi Zhao, Zhanxia Tao, Shan Zhang, Xiujuan Bei, Wen Lu, and Xiaoting Qi. Short heat shock factor A2 regulates heat resistance and growth balance in Arabidopsis. *eLife*, 13, 2025.

[15151] Xiaowen Chen, Maciej Winiarksi, Alicja Puścian, Ewelina Knapska, Thierry Mora, and Aleksandra M Walczak. Modeling collective behavior in groups of mice housed under semi-naturalistic conditions. *eLife*, 13, 2025.

[15152] Xue Chen, Deng Pan, Jia-Jia Liu, and Yanrui Yang. Endophilin A1 facilitates organization of the GABAergic postsynaptic machinery to maintain excitation-inhibition balance. *eLife*, 13, 2025.

[15153] Yao-Chung Chen, Arnaud Maupas, and Katja Nowick. Regulatory networks of KRAB zinc finger genes and transposable elements changed during human brain evolution and disease. *eLife*, 14, 2025.

[15154] Yilin Chen, Taosheng Liu, Ke Jia, Jan Theeuwes, and Mengyuan Gong. Dual-format attentional template during preparation in human visual cortex. *eLife*, 13, 2025.

[15155] Yu-Chia Chen, Mamoru Takada, Aerica Nagornyuk, Muhan Yu, Hideyuki Yamada, Takeshi Nagashima, Masayuki Ohtsuka, Jennifer G DeLuca, Steven M Markus, Motoki Takaku, and Aussie Suzuki. Inhibition of p38-MK2 pathway enhances the efficacy of microtubule inhibitors in breast cancer cells. *eLife*, 13, 2025.

[15156] Ziqi Chen, Magdalena Żak, Shuting Xu, Javier de Andrés, and Nicolas Daudet. A tissue boundary orchestrates the segregation of inner ear sensory organs. *eLife*, 14, 2025.

[15157] Huiyong Cheng, Dawson Miller, Nneka Southwell, Paola Porcari, Joshua L Fischer, Isobel Taylor, J Michael Salbaum, Claudia Kappen, Fenghua Hu, Cha Yang, Kayvan R Keshari, Steven S Gross, Marilena D'Aurelio, and Qiuying Chen. Untargeted pixel-by-pixel metabolite ratio imaging as a novel tool for biomedical discovery in mass spectrometry imaging. *eLife*, 13, 2025.

[15158] Ling Cheng, Ian Meliala, Yidi Kong, Jingyuan Chen, Christopher G Proud, and Mikael Björklund. PEBP1 amplifies mitochondrial dysfunction-induced integrated stress response. *eLife*, 13, 2025.

[15159] Susan F Cheng, Wan Lin Yue, Kwun Kei Ng, Xing Qian, Siwei Liu, Trevor WK Tan, Kim-Ngan Nguyen, Ruth LF Leong, Saima Hilal, Ching-Yu Cheng, Ai Peng Tan, Evelyn C Law, Peter D Gluckman, Christopher Li-Hsian Chen, Yap Seng Chong, Michael J Meaney, Michael WL Chee, BT Thomas Yeo, and Juan Helen Zhou. Rate of brain aging associates with future executive function in Asian children and older adults. *eLife*, 13, 2025.

[15160] Laura Childers, Jieun Park, Siyao Wang, Richard Liu, Robert Barry, Stephen A Watts, John F Rawls, and Michel Bagnat. Protein absorption in the zebrafish gut is regulated by interactions between lysosome rich enterocytes and the microbiome. *eLife*, 13, 2025.

[15161] Hui Chiu, Alice A Robie, Kristin Branson, Tanvi Vippa, Samantha Epstein, Gerald M Rubin, David J Anderson, and Catherine E Schretter. Cell type-specific contributions to a persistent aggressive internal state in female Drosophila. *eLife*, 12, 2025.

[15162] Yuma Cho, Akari Taniguchi, Akiharu Kubo, and Junichi Ikenouchi. Rho-ROCK liberates sequestered claudin for rapid de novo tight junction formation. *eLife*, 13, 2025.

[15163] Junhong Choi, Wei Chen, Hanna Liao, Xiaoyi Li, and Jay Shendure. A molecular proximity sensor based on an engineered, dual-component guide RNA. *eLife*, 13, 2025.

[15164] Jonathan Choy, Shadi Charara, Kalyani Cauwenberghs, Quintyn McKaughan, Keun-Young Kim, Mark H Ellisman, and Chih-Ying Su. Population-level morphological analysis of paired CO2- and odor-sensing olfactory neurons in D. melanogaster via volume electron microscopy. *eLife*, 14, 2025.

[15165] Janelle JY Chuah, Madalena R Daugherty, and David M Smith. Occupancy of the HbYX hydrophobic pocket is sufficient to induce gate opening in the archaeal 20S proteasomes. *eLife*, 14, 2025.

[15166] Julie N Chuong, Nadav Ben Nun, Ina Suresh, Julia Cano Matthews, Titir De, Grace Avecilla, Farah Abdul-Rahman, Nathan Brandt, Yoav Ram, and David Gresham. Template switching during DNA replication is a prevalent source of adaptive gene amplification. *eLife*, 13, 2025.

[15167] Matthew A Churgin, Danylo O Lavrentovich, Matthew A-Y Smith, Ruixuan Gao, Edward S Boyden, and Benjamin L de Bivort. A neural correlate of individual odor preference in Drosophila. *eLife*, 12, 2025.

[15168] Joseph Clayton, Aarion Romany, Evangelia Matenoglou, Evripidis Gavathiotis, Poulikos I Poulikakos, and Jana Shen. Mechanism of dimer selectivity and binding cooperativity of BRAF inhibitors. *eLife*, 13, 2025.

[15169] Agnese Codutti, Mohammad A Charsooghi, Konrad Marx, Elisa Cerdá-Doñate, Omar Muñoz, Paul Zaslansky, Vitali Telezki, Tom Robinson, Damien Faivre, and Stefan Klumpp. Physiological magnetic field strengths help magnetotactic bacteria navigate in simulated sediments. *eLife*, 13, 2025.

[15170] Oksana Cohen and Yoram Ben-Shaul. Representation of male features in the female mouse accessory olfactory bulb, and their stability during the estrus cycle. *eLife*, 14, 2025.

[15171] Brooke A Conti, Leo Novikov, Deyan Tong, Qing Xiang, Savon Vigil, Thomas J McLellan, Chuong Nguyen, Nancy De La Cruz, Reshma T Veettil, Prashant Pradhan, Parag Sahasrabudhe, Jason D Arroyo, Lei Shang, Benjamin R Sabari, David J Shields, and Mariano Oppikofer. N6-methyladenosine in DNA promotes genome stability. *eLife*, 13, 2025.

[15172] Ashley L Cook, Surojit Sur, Laura Dobbyn, Evangeline Watson, Joshua D Cohen, Blair Ptak, Bum Seok Lee, Suman Paul, Emily Hsiue, Maria Popoli, Bert Vogelstein, Nickolas Papadopoulos, Chetan Bettegowda, Kathy Gabrielson, Shibin Zhou, Kenneth W Kinzler, and Nicolas Wyhs. Identification of nonsense-mediated decay inhibitors that alter the tumor immune landscape. *eLife*, 13, 2025.

[15173] Laura E Cook, Charles Y Feigin, John D Hills, Davide M Vespasiani, Andrew J Pask, and Irene Gallego Romero. Gene regulatory dynamics during craniofacial development in a carnivorous marsupial. *eLife*, 14, 2025.

[15174] Claire Cooper, Daniel Parthier, Jeremie Sibille, John J Tukker, Nicolas Tritsch, and Dietmar Schmitz. Ultraslow serotonin oscillations in the hippocampus delineate substates across NREM and waking. *eLife*, 13, 2025.

[15175] Flavie Coquel, Sing-Zong Ho, Keng-Chang Tsai, Chun-Yen Yang, Antoine Aze, Julie Devin, Ting-Hsiang Chang, Marie Kong-Hap, Audrey Bioteau, Jerome Moreaux, Domenico Maiorano, Philippe Pourquier, Wen-Chin Yang, Yea-Lih Lin, and Philippe Pasero. Synergistic effect of inhibiting CHK2 and DNA replication on cancer cell growth. *eLife*, 13, 2025.

[15176] Katherine R Cording, Emilie M Tu, Hongli Wang, Alexander HCW Agopyan-Miu, and Helen S Bateup. Cntnap2 loss drives striatal neuron hyperexcitability and behavioral inflexibility. *eLife*, 13, 2025.

[15177] Ximena Corso Diaz, Xulong Liang, Kiam Preston, Bilguun Tegshee, Milton A English, Jacob Nellissery, Sharda Prasad Yadav, Claire Marchal, and Anand Swaroop. Maf-family bZIP transcription factor NRL interacts with RNA-binding proteins and R-loops in retinal photoreceptors. *eLife*, 13, 2025.

[15178] Ana Catarina Costa, Blanca R Murillo, Rita Bessa, Ricardo Ribeiro, Tiago Ferreira da Silva, Patrícia Porfírio-Rodrigues, Gabriel G Martins, Pedro Brites, Matthias Kneussel, Thomas Misgeld, Monika S Brill, and Monica M Sousa. Axon-specific microtubule regulation drives asymmetric regeneration of sensory neuron axons. *eLife*, 13, 2025.

[15179] Kira A Cozzolino, Lynn Sanford, Samuel Hunter, Kayla Molison, Benjamin Erickson, Meaghan CS Courvan, Taylor Jones, Deepa Ajit, Matthew D Galbraith, Joaquín M Espinosa, David Bentley, Mary Ann Allen, Robin D Dowell, and Dylan J Taatjes. Mediator kinase inhibition suppresses hyperactive interferon signaling in Down syndrome. *eLife*, 13, 2025.

[15180] Slaven Crnkovic, Helene Thekkekara Puthenparampil, Shirin Mulch, Valentina Biasin, Nemanja Radic, Jochen Wilhelm, Marek Bartkuhn, Ehsan Bonyadi Rad, Alicja Wawrzen, Ingrid Matzer, Ankita Mitra, Ryan D Leib, Bence Miklos Nagy, Anita Sahu-Osen, Francesco Valzano, Natalie Bordag, Matthias Evermann, Konrad Hoetzenecker, Andrea Olschewski, Senka Ljubojevic-Holzer, Malgorzata Wygrecka, Kurt Stenmark, Leigh M Marsh, Vinicio de Jesus Perez, and Grazyna Kwapiszewska. Adventitial fibroblasts direct smooth muscle cell-state transition in pulmonary vascular disease. *eLife*, 13, 2025.

[15181] Luuk Croijmans, Fogelina Cuperus, Dirk F van Apeldoorn, Felix JJA Bianchi, Walter AH Rossing, and Erik H Poelman. Strip cropping shows promising increases in ground beetle community diversity compared to monocultures. *eLife*, 14, 2025.

[15182] Dayna Adrienne Croock, Yolandi Swart, Haiko Schurz, Desiree C Petersen, Marlo Möller, and Caitlin Uren. Confirmation of HLA-II associations with TB susceptibility in admixed African samples. *eLife*, 13, 2025.

[15183] Bruno Cuevas Zuviría, Franka Detemple, Kaustubh Amritkar, Amanda K Garcia, Lance Seefeldt, Oliver Einsle, and Betül KaÇar. Structural evolution of nitrogenase over 3 billion years. *eLife*, 14, 2025.

[15184] Jian Cui, Hua Li, Dien Ye, Guoying Zhang, Yan Zhang, Ling Yang, Martha MS Sim, Jeremy P Wood, Yinan Wei, Zhenyu Li, and Congqing Wu. Inhibiting NINJ1-dependent plasma membrane rupture protects against inflammasome-induced blood coagulation and inflammation. *eLife*, 12, 2025.

[15185] Yan Cui, Evgeny Bondarenko, Carolina Thörn Perez, Delia N Chiu, and Jack L Feldman. Sigh generation in preBötzinger complex. *eLife*, 13, 2025.

[15186] Salam Dabsan, Gali Zur, Naim Abu-Freha, Shahar Sofer, Iris Grossman-Haham, Ayelet Gilad, and Aeid Igbaria. Cytosolic and endoplasmic reticulum chaperones inhibit wt-p53 to increase cancer cells' survival by refluxing ER-proteins to the cytosol. *eLife*, 14, 2025.

[15187] Liza Dahal, Thomas GW Graham, Gina M Dailey, Alec Heckert, Robert Tjian, and Xavier Darzacq. Surprising features of nuclear receptor interaction networks revealed by live-cell single-molecule imaging. *eLife*, 12, 2025.

[15188] Christoph D Dahl and Yaling Cheng. Individual recognition in a jumping spider (Phidippus regius). *eLife*, 13, 2025.

[15189] Vikram Dalal, Brandon K Tan, Hanrui Xu, and Wayland WL Cheng. Cryo-EM structures of a pentameric ligand-gated ion channel in liposomes. *eLife*, 14, 2025.

[15190] Leonardo Daniel, Eli Vakil, and William Saban. High cognitive violation of expectations is compromised in cerebellar ataxia. *eLife*, 14, 2025.

[15191] Priyojit Das, Rebeca San Martin, Tian Hong, and Rachel Patton McCord. Rearrangement of 3D genome organization in breast cancer epithelial to mesenchymal transition and metastasis organotropism. *eLife*, 13, 2025.

[15192] Mauna R Dasari, Kimberly E Roche, David Jansen, Jordan Anderson, Susan C Alberts, Jenny Tung, Jack A Gilbert, Ran Blekhman, Sayan Mukherjee, and Elizabeth A Archie. Social and environmental predictors of gut microbiome age in wild baboons. *eLife*, 13, 2025.

[15193] Debadatta Dash, Fumiaki Iwane, William Hayward, Roberto F Salamanca-Giron, Marlene Bönstrup, Ethan R Buch, and Leonardo G Cohen. Sequence action representations contextualize during early skill learning. *eLife*, 13, 2025.

[15194] Matthew Davenport, Ha Na Choe, Hiroaki Matsunami, and Erich Jarvis. Sex chromosome gene expression associated with vocal learning following hormonal manipulation in female zebra finches. *eLife*, 12, 2025.

[15195] Monique Marylin Alves de Almeida, Yves De Repentigny, Sabrina Gagnon, Emma R Sutton, and Rashmi Kothary. Impact of liver-specific survival motor neuron (SMN) depletion on central nervous system and peripheral tissue pathology. *eLife*, 13, 2025.

[15196] Sarah De Beuckeleer, Tim Van De Looverbosch, Johanna Van Den Daele, Peter Ponsaerts, and Winnok H De Vos. Unbiased identification of cell identity in dense mixed neural cultures. *eLife*, 13, 2025.

[15197] Sebastien De La Forest Divonne, Juliette Pouzadoux, Oceane Romatif, Caroline Montagnani, Guillaume Mitta, Delphine Destoumieux-Garzón, Benjamin Gourbal, Guillaume M Charriere, and Emmanuel Vignal. Diversity and functional specialization of oyster immune cells uncovered by integrative single-cell level investigations. *eLife*, 13, 2025.

[15198] Daniel de la Fuente, Maria Maroto, Yulia N Cajas, Karina Canon-Beltran, Raul Fernandez-Gonzalez, Ana Munoz-Maceda, Juana M Sanchez-Puig, Rafael Blasco, Paula Cots-Rodríguez, Manuel Avilés, Dimitrios Rizos, and Alfonso Gutierrez-Adan. Oviductin sets the species-specificity of the mammalian zona pellucida. *eLife*, 13, 2025.

[15199] Rebeca de la Fuente, Wladimiro Díaz-Villanueva, Vicente Arnau, and Andres Moya. Alternative splicing across the tree of life. *eLife*, 13, 2025.

[15200] Anja de Lange, Hayley Tomes, Joshua S Selfe, Ulrich Fabien Prodjinotho, Matthijs B Verhoog, Siddhartha Mahanty, Katherine Ann Smith, William Horsnell, Chummy Sikasunge, Clarissa Prazeres da Costa, and Joseph V Raimondo. Cestode larvae excite host neuronal circuits via glutamatergic signalling. *eLife*, 12, 2025.

[15201] Jean de Seze, Maud Bongaerts, Benoit Boulevard, and Mathieu Coppey. Optogenetic control of a GEF of RhoA uncovers a signaling switch from retraction to protrusion. *eLife*, 12, 2025.

[15202] Sophie Debaenst, Tamara Jarayseh, Hanna De Saffel, Jan Willem Bek, Matthieu Boone, Ivan Josipovic, Pierre Kibleur, Ronald Y Kwon, Paul J Coucke, and Andy Willaert. Crispant analysis in zebrafish as a tool for rapid functional screening of disease-causing genes for bone fragility. *eLife*, 13, 2025.

[15203] Humberto Debat, Izuchukwu Azuka Okafor, Mercury Shitindo, Olavo B Amaral, Nurul Izzati, Chandana Basu Mallick, Awika O Henry, Roseline Dzekem Dine, and Carolina Santacruz-Perez. How the Global South is reshaping scholarly communication. *eLife*, 14, 2025.

[15204] Giuseppe Deganutti, Ludovico Pipito, Roxana Maria Rujan, Tal Weizmann, Peter Griffin, Antonella Ciancetta, Stefano Moro, and Christopher Arthur Reynolds. Hidden GPCR structural transitions addressed by multiple walker supervised molecular dynamics (mwSuMD). *eLife*, 13, 2025.

[15205] Jonas Karolis Degutis, Simon Weber, Joram Soch, and John-Dylan Haynes. Neural dynamics of visual working memory representation during sensory distraction. *eLife*, 13, 2025.

[15206] Emily Delgouffe, Samuel Madureira Silva, Frédéric Chalmel, Wilfried Cools, Camille Raets, Kelly Tilleman, Guy T'Sjoen, Yoni Baert, and Ellen Goossens. Partial rejuvenation of the spermatogonial stem cell niche after gender-affirming hormone therapy in trans women. *eLife*, 13, 2025.

[15207] Cora Demler, John C Lawlor, Ronit Yelin, Dhana Llivichuzcha-Loja, Lihi Shaulov, David Kim, Megan Stewart, Frank K Lee, Natalia Shylo, Paul A Trainor, Thomas M Schultheiss, and Natasza A Kurpios. An atypical basement membrane forms a midline barrier during left-right asymmetric gut development in the chicken embryo. *eLife*, 12, 2025.

[15208] Yiting Deng, Yuanhang He, Juan Xu, Haoting He, Manling Zhang, and Guang Li. Cardiac fibroblasts regulate myocardium and coronary vasculature development in the murine heart via the collagen signaling pathway. *eLife*, 13, 2025.

[15209] Souleymane Diallo, Kateřina KaŠparová, Josef Šulc, Jibin Johny, Jan Křivánek, Jana Nebesářová, David Sillam-Dussès, Pavlína Kyjaková, Jiří VondráŠek, AleŠ Machara, Ondřej LukŠan, Ewald Grosse-Wilde, and Robert Hanus. Identification of the trail-following pheromone receptor in termites. *eLife*, 13, 2025.

[15210] David Dierks, Ran Shachar, Ronit Nir, Miguel Angel Garcia-Campos, Anna Uzonyi, David Wiener, Ursula Toth, Walter Rossmanith, Lior Lasman, Boris Slobodin, Jacob H Hanna, Yaron Antebi, Ruth Scherz-Shouval, and Schraga Schwartz. Passive shaping of intra- and intercellular m6A dynamics via mRNA metabolism. *eLife*, 13, 2025.

[15211] George Dimitriadis, Ella Svahn, Andrew F MacAskill, and Athena Akrami. Heron, a Knowledge Graph editor for intuitive implementation of Python-based experimental pipelines. *eLife*, 13, 2025.

[15212] Shihang Ding, Pei-En Chou, Shinji Deguchi, and Taeyoon Kim. Impacts of structural properties of myosin II filaments on force generation. *eLife*, 14, 2025.

[15213] Lucian DiPeso, Sriram Pendyala, Heather Z Huang, Douglas M Fowler, and Emily M Hatch. Image-based identification and isolation of micronucleated cells to dissect cellular consequences. *eLife*, 13, 2025.

[15214] Jérome Dockès, Kendra M Oudyk, Mohammad Torabi, Alejandro I de la Vega, and Jean-Baptiste Poline. Mining the neuroimaging literature. *eLife*, 13, 2025.

[15215] Joel Dokmegang, Emmanuel Faure, Patrick Lemaire, Edwin Munro, and Madhav Mani. Spectral decomposition unlocks ascidian morphogenesis. *eLife*, 13, 2025.

[15216] Angel D'Oliviera, Xuhang Dai, Saba Mottaghinia, Sophie Olson, Evan P Geissler, Lucie Etienne, Yingkai Zhang, and Jeffrey S Mugridge. Recognition and cleavage of human tRNA methyltransferase TRMT1 by the SARS-CoV-2 main protease. *eLife*, 12, 2025.

[15217] Jian Dong, Mian Chen, Jan RT van Weering, Ka Wan Li, August B Smit, Ruud F Toonen, and Matthijs Verhage. Rab10 regulates neuropeptide release by maintaining Ca2+ homeostasis and protein synthesis. *eLife*, 13, 2025.

[15218] Pengzhen Dong, Yue Li, Yuying Wang, Qiang Zhao, Tianfeng Lu, Jian Chen, Tianyu Guo, Jun Ma, Bing Yang, Honggang Wu, and Hai Huang. Fat body-derived cytokine Upd2 controls disciplined migration of tracheal stem cells in Drosophila. *eLife*, 13, 2025.

[15219] Ruihan Dong, Rongrong Liu, Ziyu Liu, Yangang Liu, Gaomei Zhao, Honglei Li, Shiyuan Hou, Xiaohan Ma, Huarui Kang, Jing Liu, Fei Guo, Ping Zhao, Junping Wang, Cheng Wang, Xingan Wu, Sheng Ye, and Cheng Zhu. Exploring the repository of de novo-designed bifunctional antimicrobial peptides through deep learning. *eLife*, 13, 2025.

[15220] Kaili Du, Hongyu Chen, Zhaonan Pan, Mengli Zhao, Shixue Cheng, Yu Luo, Wenhe Zhang, and Dan Li. Small-molecule activation of TFEB alleviates Niemann–Pick disease type C via promoting lysosomal exocytosis and biogenesis. *eLife*, 13, 2025.

[15221] Shoukang Du, Yuhan Wang, Bowen Chen, Shuangshuang Xie, Kuan Yoow Chan, David C Hay, and Ting Gang Chew. Clearance of protein aggregates during cell division. *eLife*, 13, 2025.

[15222] Xinyu Du, Haoxuan Xu, Peirun Song, Yuying Zhai, Hangting Ye, Xuehui Bao, Qianyue Huang, Hisashi Tanigawa, Zhiyi Tu, Pei Chen, Xuan Zhao, Josef P Rauschecker, and Xiongjie Yu. The multifaceted role of the inferior colliculus in sensory prediction, reward processing, and decision-making. *eLife*, 13, 2025.

[15223] Jingze Duan and Chen Song. Twist is the key to the gating of mechanosensitive ion channel NOMPC. *eLife*, 13, 2025.

[15224] David Duneau, Pierre DM Lafont, Christine Lauzeral, Nathalie Parthuisot, Christian Faucher, Xuerong Jin, Nicolas Buchon, and Jean-Baptiste Ferdy. A within-host infection model to explore tolerance and resistance. *eLife*, 14, 2025.

[15225] Soumajit Dutta and Diwakar Shukla. Characterization of binding kinetics and intracellular signaling of new psychoactive substances targeting cannabinoid receptor using transition-based reweighting method. *eLife*, 13, 2025.

[15226] Rohan Eapen, Cynthia Okoye, Christopher Stubbs, Marianne Schimpl, Thomas Tischer, Eileen J Fisher, Maria Zacharopoulou, Fernando Ferrer, David Barford, David R Spring, Catherine Lindon, Christopher Phillips, and Laura S Itzhaki. Development of D-box peptides to inhibit the anaphase-promoting complex/cyclosome. *eLife*, 14, 2025.

[15227] Sreesankar Easwaran and Denise J Montell. A genome-wide association study implicates the olfactory system in Drosophila melanogaster diapause-associated lifespan extension and fecundity. *eLife*, 13, 2025.

[15228] András Ecker, Daniela Egas Santander, Marwan Abdellah, Jorge Blanco Alonso, Sirio Bolaños-Puchet, Giuseppe Chindemi, Dhuruva Priyan Gowri Mariyappan, James B Isbister, James King, Pramod Kumbhar, Ioannis Magkanaris, Eilif B Muller, and Michael W Reimann. Assemblies, synapse clustering, and network topology interact with plasticity to explain structure-function relationships of the cortical connectome. *eLife*, 13, 2025.

[15229] Marisa S Egan, Emily A O'Rourke, Shrawan Kumar Mageswaran, Biao Zuo, Inna Martynyuk, Tabitha Demissie, Emma N Hunter, Antonia R Bass, Yi-Wei Chang, Igor E Brodsky, and Sunny Shin. Inflammasomes primarily restrict cytosolic Salmonella replication within human macrophages. *eLife*, 12, 2025.

[15230] Pierce Eggan, Sharona E Gordon, and William N Zagotta. Domain coupling in allosteric regulation of SthK measured using time-resolved transition metal ion FRET. *eLife*, 14, 2025.

[15231] Fanny Eggeler, Jonathan Boulanger-Weill, Flavia De Santis, Laura Belleri, Karine Duroure, Thomas O Auer, Shahad Albadri, and Filippo Del Bene. Meteorins regulate the formation of the left-right organizer and the establishment of vertebrate body asymmetry. *eLife*, 14, 2025.

[15232] Cameron T Ellis, Tristan S Yates, Michael J Arcaro, and Nicholas Turk-Browne. Movies reveal the fine-grained organization of infant visual cortex. *eLife*, 12, 2025.

[15233] Roger Eloiflin, Elena Pérez-Antón, Aïssata Camara, Annick Dujeancourt-Henry, Salimatou Boiro, Martial N Djetchi, Mélika Barkissa Traoré, Mathurin Koffi, Dramane Kaba, Yann Le Pennec, Bakary Doukouré, Abdoulaye Dansy Camara, Moïse Kagbadouno, Pascal Campagne, Mamadou Camara, Vincent Jamonneau, Sophie Thévenon, Jean-Mathieu Bart, Lucy Glover, and Brice Rotureau. A SHERLOCK toolbox for eco-epidemiological surveillance of African trypanosomes in domestic pigs from Western Africa. *eLife*, 14, 2025.

[15234] Assmaa Elsheikh, Camden M Driggers, Ha H Truong, Zhongying Yang, John Allen, Niel M Henriksen, Katarzyna Walczewska-Szewc, and Show-Ling Shyng. AI-based discovery and cryoEM structural elucidation of a KATP channel pharmacochaperone. *eLife*, 13, 2025.

[15235] Alaknanda Emery, Orest W Blaschuk, Doan T Dinh, Tim McPhee, Rouven Becker, Andrew D Abell, Krzysztof M Mrozik, Andrew CW Zannettino, Rebecca L Robker, and Darryl L Russell. N-cadherin mechanosensing in ovarian follicles controls oocyte maturation and ovulation. *eLife*, 13, 2025.

[15236] Akinori Endo, Chikage Takahashi, Naoko Ishibashi, Yasumasa Nishito, Koji Yamano, Keiji Tanaka, and Yukiko Yoshida. Ubiquitination-activated TAB–TAK1–IKK–NF-$\kappa$B axis modulates gene expression for cell survival in the lysosomal damage response. *eLife*, 14, 2025.

[15237] Melanie Lianne Engelfriet, Yanwu Guo, Andreas Arnold, Eivind Valen, and Rafal Ciosk. Reprograming gene expression in 'hibernating' C. elegans involves the IRE-1/XBP-1 pathway. *eLife*, 13, 2025.

[15238] Sinead English, M Emília Santos, Clare Buckley, Chrissy L Hammond, Sarah Lloyd-Fox, and Nina F Ockendon-Powell. To be, or not to be, part-time in academia. *eLife*, 14, 2025.

[15239] Derek G Epiney, Gonzalo Morales Chaya, Noah R Dillon, Sen-Lin Lai, and Chris Q Doe. Single nuclei RNA-sequencing of adult brain neurons derived from type 2 neuroblasts reveals transcriptional complexity in the insect central complex. *eLife*, 14, 2025.

[15240] Mayalen Etcheverry, Clément Moulin-Frier, Pierre-Yves Oudeyer, and Michael Levin. AI-driven automated discovery tools reveal diverse behavioral competencies of biological networks. *eLife*, 13, 2025.

[15241] Cécile Fabio and Christoph Kayser. Mixed evidence for the rhythmicity of auditory perceptual judgements in humans. *eLife*, 14, 2025.

[15242] Johannes Jacobus Fahrenfort, Philippa A Johnson, Niels Kloosterman, Timo Stein, and Simon van Gaal. Response to comment on 'Criterion placement threatens the construct validity of neural measures of consciousness'. *eLife*, 14, 2025.

[15243] Johannes Jacobus Fahrenfort, Philippa A Johnson, Niels A Kloosterman, Timo Stein, and Simon van Gaal. Criterion placement threatens the construct validity of neural measures of consciousness. *eLife*, 13, 2025.

[15244] Kayson Fakhar, Fatemeh Hadaeghi, Caio Seguin, Shrey Dixit, Arnaud Messé, Gorka Zamora-López, Bratislav Misic, and Claus C Hilgetag. A general framework for characterizing optimal communication in brain networks. *eLife*, 13, 2025.

[15245] Michel Fallah, Kenea C Udobi, Aleksandra E Swiatek, Chelsea B Scott, and Rebekah C Evans. Inhibitory basal ganglia nuclei differentially innervate pedunculopontine nucleus subpopulations and evoke differential motor and valence behaviors. *eLife*, 13, 2025.

[15246] Jiayi Fan, Nalini Dhingra, Tammy Yang, Vicki Yang, and Xiaolan Zhao. Srs2 binding to proliferating cell nuclear antigen (PCNA) and its sumoylation contribute to replication protein A (RPA) antagonism during the DNA damage response. *eLife*, 13, 2025.

[15247] Huan Fang, Huichun Liang, Chuanyu Yang, Dewei Jiang, Qianmei Luo, Wen-Ming Cao, Huifeng Zhang, and Ceshi Chen. STAMBPL1 activates the GRHL3/HIF1A/VEGFA axis through interaction with FOXO1 to promote angiogenesis in triple-negative breast cancer. *eLife*, 13, 2025.

[15248] Jie Fang, Shivendra Singh, Brennan Wells, Qiong Wu, Hongjian Jin, Laura J Janke, Shibiao Wan, Jacob A Steele, Jon P Connelly, Andrew J Murphy, Ruoning Wang, Andrew Davidoff, Margaret Ashcroft, Shondra M Pruett-Miller, and Jun Yang. The context-dependent epigenetic and organogenesis programs determine 3D vs. 2D cellular fitness of MYC-driven murine liver cancer cells. *eLife*, 14, 2025.

[15249] Laura Fargeot, Camille Poesy, Maxim Lefort, Jerome G Prunier, Madoka Krick, Rik Verdonck, Charlotte Veyssiere, Murielle Richard, Delphine Legrand, Geraldine Loot, and Blanchet Simon. Genetic diversity affects ecosystem functions across trophic levels as much as species diversity, but in an opposite direction. *eLife*, 13, 2025.

[15250] Elizabeth J Fay, Kolya Isterabadi, Charles M Rezanka, Jessica Le, and Matthew D Daugherty. Evolutionary and functional analyses reveal a role for the RHIM in tuning RIPK3 activity across vertebrates. *eLife*, 13, 2025.

[15251] Roman O Fedoryshchak, Karim El-Bouri, Dhira Joshi, Stephane Mouilleron, and Richard Treisman. PDZ-directed substrate recruitment is the primary determinant of specific 4E-BP1 dephosphorylation by PP1-Neurabin. *eLife*, 13, 2025.

[15252] Haiyi Fei, Xiaowen Lu, Zhan Shi, Xiu Liu, Cuiyu Yang, Xiaohong Zhu, Yuhan Lin, Ziqun Jiang, Jianmin Wang, Dong Huang, Liu Liu, Songying Zhang, and Lingling Jiang. Deciphering the preeclampsia-specific immune microenvironment and the role of pro-inflammatory macrophages at the maternal–fetal interface. *eLife*, 13, 2025.

[15253] Rui Feng, Sarah F Rosen, Irshad Ansari, Sebastian John, Michael B Thomsen, Oshri Avraham, Cedric G Geoffroy, and Valeria Cavalli. Endothelin B receptor inhibition rescues aging-dependent neuronal regenerative decline. *eLife*, 13, 2025.

[15254] Lotfi Ferhat, Rabia Soussi, Maxime Masse, Grigorios Kyriatzis, Stéphane Girard, Fanny Gassiot, Nicolas Gaudin, Mathieu Laurencin, Anne Bernard, Angélique Bôle, Géraldine Ferracci, Maria Smirnova, FranÇois Roman, Vincent Dive, Salvatore Cisternino, Jamal Temsamani, Marion David, Pascaline Lécorché, Guillaume Jacquot, and Michel Khrestchatisky. A peptide-neurotensin conjugate that crosses the blood-brain barrier induces pharmacological hypothermia associated with anticonvulsant, neuroprotective, and anti-inflammatory properties following status epilepticus in mice. *eLife*, 13, 2025.

[15255] Ahana Aurora Fernandez, Nora Serve, Sarah-Cecil Fabian, and Mirjam Knörnschild. Maternal behavior influences vocal practice and learning processes in the greater sac-winged bat. *eLife*, 13, 2025.

[15256] David Ferreiro, Luis Daniel González-Vázquez, Ana Prado-Comesaña, and Miguel Arenas. Forecasting protein evolution by integrating birth-death population models with structurally constrained substitution models. *eLife*, 14, 2025.

[15257] Ryan M Finnerty, Daniel J Carulli, Akshata Hedge, Yanli Wang, Frimpong Boadu, Sarayut Winuthayanon, Jianlin Jack Cheng, and Wipawee Winuthayanon. Multi-omics analyses and machine learning prediction of oviductal responses in the presence of gametes and embryos. *eLife*, 13, 2025.

[15258] Cora Fischer, Jochen Kaiser, and Christoph Bledowski. A direct neural signature of serial dependence in working memory. *eLife*, 13, 2025.

[15259] Marie-Therese Fischer, Katherine S Xue, Elizabeth K Costello, Mai Dvorak, Gaelle Raboisson, Anna Robaczewska, Stephanie N Caty, David A Relman, and Lauren A O'Connell. Effects of parental care on skin microbial community composition in poison frogs. *eLife*, 14, 2025.

[15260] Josep Fita-Torró, José Luis Garrido-Huarte, Lucía López-Gil, Agnès H Michel, Benoit Kornmann, Amparo Pascual-Ahuir, and Markus Proft. Inhibition of mitochondrial protein import and proteostasis by a pro-apoptotic lipid. *eLife*, 13, 2025.

[15261] Nicolas Flaugnatti, Loriane Bader, Mary Croisier-Coeytaux, and Melanie Blokesch. Capsular polysaccharide restrains type VI secretion in Acinetobacter baumannii. *eLife*, 14, 2025.

[15262] Nico A Flierman, Sue Ann Koay, Willem S van Hoogstraten, Tom JH Ruigrok, Pieter Roelfsema, Aleksandra Badura, and Chris I De Zeeuw. Encoding of cerebellar dentate neuron activity during visual attention in rhesus macaques. *eLife*, 13, 2025.

[15263] Ana Fló, Lucas Benjamin, Marie Palu, and Ghislaine Dehaene-Lambertz. Statistical learning beyond words in human neonates. *eLife*, 13, 2025.

[15264] Joseph R Floeder, Huijeong Jeong, Ali Mohebi, and Vijay Mohan K Namboodiri. Mesolimbic dopamine ramps reflect environmental timescales. *eLife*, 13, 2025.

[15265] Jessica Flowers, Nathaniel Echols, Galen J Correy, Priyadarshini Jaishankar, Takaya Togo, Adam R Renslo, Henry van den Bedem, James S Fraser, and Stephanie A Wankowicz. Expanding automated multiconformer ligand modeling to macrocycles and fragments. *eLife*, 14, 2025.

[15266] Catherine Suzanne Forconi, Christina Nixon, Hannah W Wu, Boaz Odwar, Sunthorn Pond-Tor, John M Ong'echa, Jonathan D Kurtis, and Ann M Moormann. T-follicular helper cell profiles differ by malaria antigen and for children compared to adults. *eLife*, 13, 2025.

[15267] Alyssa Lyn Fortier and Jonathan K Pritchard. Ancient trans-species polymorphism at the Major Histocompatibility Complex in primates. *eLife*, 14, 2025.

[15268] Zachary Fournier, Leandro M Alonso, and Eve Marder. Rhythmic circuit function is more robust to changes in synaptic than intrinsic conductances. *eLife*, 13, 2025.

[15269] Ruggiero Francavilla, Bidisha Chattopadhyaya, Jorelle Linda Damo Kamda, Vidya Jadhav, Said Kourrich, Jacques L Michaud, and Graziella Di Cristo. Syngap1 regulates the synaptic drive and membrane excitability of Parvalbumin-positive interneurons in mouse auditory cortex. *eLife*, 13, 2025.

[15270] Lawrence R Frank, Vitaly L Galinsky, Olave Krigolson, Susan Tapert, Stephan Bickel, and Antigona Martinez. Imaging of brain electric field networks with spatially resolved EEG. *eLife*, 13, 2025.

[15271] Tamar Frankovits, Prakash Varkey Cherian, Yarden Yesharim, Simon Dobler, and Omri Wurtzel. Stochastic cell-intrinsic stem cell decisions control colony growth in planarians. *eLife*, 13, 2025.

[15272] Tisa Frelih, Andraž Matkovič, TjaŠa Mlinarič, Jurij Bon, and Grega RepovŠ. Modulation of aperiodic EEG activity provides sensitive index of cognitive state changes during working memory task. *eLife*, 13, 2025.

[15273] Louna Fruchard, Anamaria Babosan, Andre Carvalho, Manon Lang, Blaise Li, Magalie Duchateau, Quentin Giai Gianetto, Mariette Matondo, Frederic Bonhomme, Isabelle Hatin, Hugo Arbes, Céline Fabret, Enora Corler, Guillaume Sanchez, Virginie Marchand, Yuri Motorin, Olivier Namy, Valérie de Crécy-Lagard, Didier Mazel, and Zeynep Baharoglu. Aminoglycoside tolerance in Vibrio cholerae engages translational reprogramming associated with queuosine tRNA modification. *eLife*, 13, 2025.

[15274] Liangliang Fu, Chenghu Yin, Qin Zhao, Shuling Guo, Wenjun Shao, Ting Xia, Quan Sun, Liangwen Chen, Jinghan Li, Min Wang, and Haibin Xia. CD81+ senescent-like fibroblasts exaggerate inflammation and activate neutrophils via C3/C3aR1 axis in periodontitis. *eLife*, 13, 2025.

[15275] Adrian CD Fuchs. Enzymatic protein fusions with 100% product yield. *eLife*, 13, 2025.

[15276] Tanja Fuchsberger, Imogen Stockwell, Matty Woods, Zuzanna Brzosko, Ingo H Greger, and Ole Paulsen. Dopamine increases protein synthesis in hippocampal neurons enabling dopamine-dependent LTP. *eLife*, 13, 2025.

[15277] Agustín Fuentes, Marc Kissel, Penny Spikins, Keneiloe Molopyane, John Hawks, and Lee R Berger. Meaning-making behavior in a small-brained hominin, Homo naledi, from the late Pleistocene: contexts and evolutionary implications. *eLife*, 12, 2025.

[15278] Lucas Fuentes Valenzuela, Paul Francois, and Jan M Skotheim. The Product neutrality function defining genetic interactions emerges from mechanistic models of cell growth. *eLife*, 14, 2025.

[15279] Yuri Fujita, Shotaro Namba, Yoshiaki Kamada, and Hisao Moriya. Impact of maximal overexpression of a non-toxic protein on yeast cell physiology. *eLife*, 13, 2025.

[15280] Zachary Gaertner, Cameron Oram, Amanda Schneeweis, Elan Schonfeld, Cyril Bolduc, Chuyu Chen, Daniel Dombeck, Loukia Parisiadou, Jean-Francois Poulin, and Rajeshwar Awatramani. Molecular and spatial transcriptomic classification of midbrain dopamine neurons and their alterations in a LRRK2G2019S model of Parkinson's disease. *eLife*, 13, 2025.

[15281] Daniel Patrick Gainey, Andrey V Shubin, and Craig P Hunter. Reported transgenerational responses to Pseudomonas aeruginosa in Caenorhabditis elegans are not robust. *eLife*, 13, 2025.

[15282] Elisabetta Gamen, Eleanor L Price, Daniela Pezzolla, Carla De Villiers, Mala Gunadasa-Rohling, Adam B Lokman, Maria-Alexa Cosma, Judith Sayers, Carolina Roque Silva, Rafik Salama, David Robert Mole, Tammie Bishop, Chris W Pugh, Robin P Choudhury, Carolyn A Carr, Joaquim Miguel Vieira, and Paul R Riley. Stabilisation of HIF signalling in the mouse epicardium extends embryonic potential and neonatal heart regeneration. *eLife*, 14, 2025.

[15283] Johanna Ganssauge, Sophie Hawkins, Seema Chandramohan Namboori, Szi Kay Leung, Jonathan Mill, and Akshay Bhinge. Rapid and inducible mislocalization of endogenous TDP43 in a novel human model of amyotrophic lateral sclerosis. *eLife*, 13, 2025.

[15284] Can Gao, Mingze Ma, Jie Chen, Xiaoxiao Ji, Qionglin Peng, and Yufeng Pan. Social experience shapes fighting strategies in Drosophila. *eLife*, 13, 2025.

[15285] Jiayin Gao, Jiawei Wang, Shiyu Liu, Jinzhu Song, Chuanxin Zhang, Boyang Liu, and Keliang Wu. The asymmetric expression of HSPA2 in blastomeres governs the first embryonic cell-fate decision. *eLife*, 13, 2025.

[15286] Yang Gao, Shao-Cong Su, Ji-Yang Xing, Zhao-Yu Liu, Dick R Nässel, Chris Bass, Congfen Gao, and Shun-Fan Wu. Pesticide-induced resurgence in brown planthoppers is mediated by action on a suite of genes that promote juvenile hormone biosynthesis and female fecundity. *eLife*, 12, 2025.

[15287] Marco Garbelli, Stephanie Niklaus, and Stephan CF Neuhauss. Characterization of postsynaptic glutamate transporter functionality in the zebrafish retinal first synapse across different wavelengths. *eLife*, 13, 2025.

[15288] Dominik Garber and József Fiser. Structure transfer and consolidation in visual implicit learning. *eLife*, 13, 2025.

[15289] Joshua D Garcia, Chenyu Wang, Ryan PD Alexander, Emmie Banks, Timothy Fenton, Jean-Marc DeKeyser, Tatiana V Abramova, Alfred L George, Roy Ben-Shalom, David H Hackos, and Kevin J Bender. Differential roles of NaV1.2 and NaV1.6 in neocortical pyramidal cell excitability. *eLife*, 14, 2025.

[15290] Nelson García-Vázquez, Tania J González-Robles, Ethan Lane, Daria Spasskaya, Qingyue Zhang, Marc A Kerzhnerman, YeonTae Jeong, Marta Collu, Daniele Simoneschi, Kelly V Ruggles, Gergely Róna, Sharon Kaisari, and Michele Pagano. Stabilization of GTSE1 by cyclin D1–CDK4/6-mediated phosphorylation promotes cell proliferation with implications for cancer prognosis. *eLife*, 13, 2025.

[15291] Suma Garibova, Eva Stickler, Fatima AlAli, Maha A Abdulla, Abbirami Sathappan, Sahar I Da'as, Lillian Ghanem, Mohamed Nadhir Djekidel, Rick Portman, and Matteo Avella. Cross-species insemination reveals mouse sperm ability to enter and cross the fish micropyle. *eLife*, 14, 2025.

[15292] Haruna Gekko, Ruri Nomura, Daiki Kuzuhara, Masato Kaneyasu, Genpei Koseki, Deepak Adhikari, Yasuyuki Mio, John Carroll, Tomohiro Kono, Hiroaki Funahashi, and Takuya Wakai. Redistribution of fragmented mitochondria ensures symmetric organelle partitioning and faithful chromosome segregation in mitotic mouse zygotes. *eLife*, 13, 2025.

[15293] Eléonore Genzoni, Tanja Schwander, and Laurent Keller. Trophic eggs affect caste determination in the ant Pogonomyrmex rugosus. *eLife*, 12, 2025.

[15294] Jayati Gera, Marishia Agard, Hannah Nave, Austin B Baldridge, Farwa Sajadi, Leena Thorat, Theresa H McKim, Shu Kondo, Dick R Nässel, Mitchell H Omar, Jean-Paul Paluzzi, and Meet Zandawala. Anti-diuretic hormone ITP signals via a guanylate cyclase receptor to modulate systemic homeostasis in Drosophila. *eLife*, 13, 2025.

[15295] Dhaval Ghone, Edward L Evans, Madison Bandini, Kaelyn G Stephenson, Nathan M Sherer, and Aussie Suzuki. HIV-1 Vif disrupts phosphatase feedback regulation at the kinetochore, leading to a pronounced pseudo-metaphase arrest. *eLife*, 13, 2025.

[15296] Giuliana Giamundo, Daniela Intartaglia, Eugenio Del Prete, Elena Polishchuk, Fabrizio Andreone, Marzia Ognibene, Sara Buonocore, Bruno Hay Mele, Francesco Giuseppe Salierno, Jlenia Monfregola, Dario Antonini, Paolo Grumati, Alessandra Eva, Rossella De Cegli, and Ivan Conte. Ezrin defines TSC complex activation at endosomal compartments through EGFR–AKT signaling. *eLife*, 13, 2025.

[15297] Arnaud PJ Giese, Wei-Hsiang Weng, Katie S Kindt, Hui Ho Vanessa Chang, Jonathan S Montgomery, Evan M Ratzan, Alisha J Beirl, Roberto Aponte Rivera, Jeffrey M Lotthammer, Sanket Walujkar, Mark P Foster, Omid A Zobeiri, Jeffrey R Holt, Saima Riazuddin, Kathleen E Cullen, Marcos Sotomayor, and Zubair M Ahmed. Complexes of vertebrate TMC1/2 and CIB2/3 proteins form hair-cell mechanotransduction cation channels. *eLife*, 12, 2025.

[15298] Cristina Gil Avila, Elisabeth S May, Felix S Bott, Laura Tiemann, Vanessa Hohn, Henrik Heitmann, Paul Theo Zebhauser, Joachim Gross, and Markus Ploner. Assessing the balance between excitation and inhibition in chronic pain through the aperiodic component of EEG. *eLife*, 13, 2025.

[15299] Karen Giménez-Orenga, Eva Martín-Martínez, Lubov Nathanson, and Elisa Oltra. HERV activation segregates ME/CFS from fibromyalgia while defining a novel nosologic entity. *eLife*, 14, 2025.

[15300] Eloise Giraud, Michael Lynn, Philippe Vincent-Lamarre, Jean-Claude Beique, and Jean-Philippe Thivierge. Unsupervised pipeline for the identification of cortical excitatory and inhibitory neurons in high-density multielectrode arrays with ground-truth validation. *eLife*, 14, 2025.

[15301] Adam Glaser, Jayaram Chandrashekar, Sonya Vasquez, Cameron Arshadi, Rajvi Javeri, Naveen Ouellette, Xiaoyun Jiang, Judith Baka, Gabor Kovacs, Micah Woodard, Shamishtaa Seshamani, Kevin Cao, Nathan Clack, Andrew Recknagel, Anna Grim, Pooja Balaram, Emily Turschak, Marcus Hooper, Alan Liddell, John Rohde, Ayana Hellevik, Kevin Takasaki, Lindsey Erion Barner, Molly Logsdon, Chris Chronopoulos, Saskia EJ de Vries, Jonathan T Ting, Steven Perlmutter, Brian E Kalmbach, Nikolai Dembrow, Bosiljka Tasic, R Clay Reid, David Feng, and Karel Svoboda. Expansion-assisted selective plane illumination microscopy for nanoscale imaging of centimeter-scale tissues. *eLife*, 12, 2025.

[15302] Agnieszka Glica, Katarzyna Wasilewska, Julia Jurkowska, Jarosław Żygierewicz, Bartosz Kossowski, and Katarzyna Jednoróg. Reevaluating the neural noise in dyslexia using biomarkers from electroencephalography and high-resolution magnetic resonance spectroscopy. *eLife*, 13, 2025.

[15303] Wesley H Godfrey, Judy J Lee, Shruthi Shanmukha, Kaho Cho, Xiaojing Deng, Chandra Shekar R Ambati, Vasanta Putluri, Abu Hena Mostafa Kamal, Paul M Kim, Nagireddy Putluri, and Michael D Kornberg. Phosphoglycerate mutase regulates Treg differentiation through control of serine synthesis and one-carbon metabolism. *eLife*, 14, 2025.

[15304] Julieta Gomez-Frittelli, Gabrielle Frederique Devienne, Lee Travis, Melinda A Kyloh, Xin Duan, Tim J Hibberd, Nick J Spencer, John R Huguenard, and Julia A Kaltschmidt. Synaptic cell adhesion molecule Cdh6 identifies a class of sensory neurons with novel functions in colonic motility. *eLife*, 13, 2025.

[15305] Xuantong Gong, Lishuang Gu, Di Yang, Yu He, Qian Li, Hao Qin, and Yong Wang. Ferroptosis-related genes mediate tumor microenvironment and prognosis in triple-negative breast cancer via integrated RNA-seq analysis. *eLife*, 13, 2025.

[15306] Dominic Gonschorek, Matías A Goldin, Jonathan Oesterle, Tom Schwerd-Kleine, Ryan Arlinghaus, Zhijian Zhao, Timm Schubert, Olivier Marre, and Thomas Euler. Nitric oxide modulates contrast suppression in a subset of mouse retinal ganglion cells. *eLife*, 13, 2025.

[15307] Tirso RJ Gonzalez Alam, Katya Krieger-Redwood, Dominika Varga, Zhiyao Gao, Aidan J Horner, Tom Hartley, Michel Thiebaut de Schotten, Magdalena Sliwinska, David Pitcher, Daniel S Margulies, Jonathan Smallwood, and Elizabeth Jefferies. A double dissociation between semantic and spatial cognition in visual to default network pathways. *eLife*, 13, 2025.

[15308] Moloy T Goswami, Eric Weh, Shubha Subramanya, Katherine M Weh, Hima Bindu Durumutla, Heather Hager, Nicholas Miller, Sraboni Chaudhury, Anthony Andren, Peter Sajjakulnukit, Li Zhang, Cagri Besirli, Costas A Lyssiotis, and Thomas J Wubben. Glutamine catabolism supports amino acid biosynthesis and suppresses the integrated stress response to promote photoreceptor survival. *eLife*, 13, 2025.

[15309] Victoire Gouirand, Sean Clancy, Courtney Macon, Jose Valle, Mariela Pauli, Hong-An Truong, Jarish Cohen, Maxime Kinet, Margaret M Lowe, Samuel J Lord, Kristen Skruber, Hobart Harris, Esther Kim, Isaac Neuhaus, Karin Reif, Ali A Zarrin, R Dyche Mullins, and Michael D Rosenblum. Layilin regulates Treg motility and suppressive capacity in skin. *eLife*, 14, 2025.

[15310] Ashwin Govindan and Nicholas K Conrad. SFSWAP is a negative regulator of OGT intron detention and global pre-mRNA splicing. *eLife*, 13, 2025.

[15311] Elena G Govorunova, Oleg A Sineshchekov, Hai Li, Yueyang Gou, Hongmei Chen, Shuyuan Yang, Yumei Wang, Stephen Mitchell, Alyssa Palmateer, Leonid S Brown, FranÇois St-Pierre, Mingshan G Xue, and John L Spudich. Blue-shifted ancyromonad channelrhodopsins for multiplex optogenetics. *eLife*, 14, 2025.

[15312] Parveen Goyal, KanagaVijayan Dhanabalan, Mariafrancesca Scalise, Rosmarie Friemann, Cesare Indiveri, Renwick CJ Dobson, Kutti R Vinothkumar, and Subramanian Ramaswamy. Molecular determinants of Neu5Ac binding to a tripartite ATP independent periplasmic (TRAP) transporter. *eLife*, 13, 2025.

[15313] Saloni Goyal, Divya Rajendran, Anup Kumar Mani, and Athi N Naganathan. Polyphosphate discriminates protein conformational ensembles more efficiently than DNA promoting diverse assembly and maturation behaviors. *eLife*, 14, 2025.

[15314] Ana Patrícia Graça, Vadim Nikitushkin, Mark Ellerhorst, Cláudia Vilhena, Tilman E Klassert, Andreas Starick, Malte Siemers, Walid K Al-Jammal, Ivan Vilotijevic, Hortense Slevogt, Kai Papenfort, and Gerald Lackner. MftG is crucial for ethanol metabolism of mycobacteria by linking mycofactocin oxidation to respiration. *eLife*, 13, 2025.

[15315] Carley N Gray, Manickam Ashokkumar, Derek H Janssens, Jennifer L Kirchherr, Brigitte Allard, Emily Hsieh, Terry L Hafer, Nancie M Archin, Edward P Browne, and Michael Emerman. Integrator complex subunit 12 knockout overcomes a transcriptional block to HIV latency reversal. *eLife*, 13, 2025.

[15316] Vitka Gres, Merve Göcer, Julia Kolter, and Philipp Henneke. Trained immunity in skin infections: Macrophages and beyond. *eLife*, 14, 2025.

[15317] Simonas Griesius, Amy Richardson, and Dimitri Michael Kullmann. Supralinear dendritic integration in murine dendrite-targeting interneurons. *eLife*, 13, 2025.

[15318] Sophie V Griswold and Stephen D Van Hooser. Premature vision drives aberrant development of response properties in primary visual cortex. *eLife*, 14, 2025.

[15319] Miaoqing Gu, Shanshan Liang, Jiahui Zhu, Ruijie Li, Ke Liu, Xuanyue Wang, Frank W Ohl, Yun Zhang, Xiang Liao, Chunqing Zhang, Hongbo Jia, Yi Zhou, Jianxiong Zhang, and Xiaowei Chen. Tonotopy is not preserved in a descending stage of auditory cortex. *eLife*, 13, 2025.

[15320] Mathias Guayasamin, Lewis R Depaauw-Holt, Ifeoluwa I Adedipe, Ossama Ghenissa, Juliette Vaugeois, Manon Duquenne, Benjamin Rogers, Jade Latraverse-Arquilla, Sarah Peyrard, Anthony Bosson, and Ciaran Murphy-Royal. Early-life stress induces persistent astrocyte dysfunction associated with fear generalisation. *eLife*, 13, 2025.

[15321] Marc-Antoine Guery, Sukai Ceesay, Sainabou Drammeh, Fatou K Jaiteh, Umberto D'Alessandro, Teun Bousema, David J Conway, and Antoine Claessens. Household clustering and seasonal genetic variation of Plasmodium falciparum at the community-level in The Gambia. *eLife*, 13, 2025.

[15322] Kate Gunther, Teagan Fisher, Dongli Liu, Jason Abbott, and Caroline Elizabeth Ford. Endometriosis is not the endometrium: Reviewing the over-representation of eutopic endometrium in endometriosis research. *eLife*, 14, 2025.

[15323] Torsten Gunther, Jacob Chisausky, ángeles M Galindo-Pellicena, Eneko Iriarte, Oscar Cortes Gardyn, Paulina G Eusebi, Rebeca García-González, Irene Ureña, Marta Moreno-García, Alfonso Alday, Manuel Rojo, Amalia Pérez, Cristina Tejedor Rodríguez, Iñigo García Martínez de Lagrán, Juan Luis Arsuaga, José-Miguel Carretero, Anders Götherström, Colin Smith, and Cristina Valdiosera. The genomic legacy of aurochs hybridisation in ancient and modern Iberian cattle. *eLife*, 13, 2025.

[15324] Bing Guo, Shannon Takala-Harrison, and Timothy D O'Connor. Benchmarking and optimization of methods for the detection of identity-by-descent in high-recombining Plasmodium falciparum genomes. *eLife*, 14, 2025.

[15325] Susu Guo, Xiaodi Hu, Jennifer L Cotton, Lifang Ma, Qi Li, Jiangtao Cui, Yongjie Wang, Ritesh P Thakare, Zhipeng Tao, Y Tony Ip, Xu Wu, Jiayi Wang, and Junhao Mao. VGLL2 and TEAD1 fusion proteins identified in human sarcoma drive YAP/TAZ-independent tumorigenesis by engaging EP300. *eLife*, 13, 2025.

[15326] Yan Guo, Yong Zhang, Guizhen Wang, Hongtao Liu, Jianfeng Wang, Xuming Deng, Liuqin He, and Jiazhang Qiu. Caffeic acid phenethyl ester protects Clostridioides difficile infection by toxin inhibition and microbiota modulation. *eLife*, 13, 2025.

[15327] Hans Tobias Gustafsson, Lucas Ferguson, Carolina Galan, Tianxiong Yu, Heather Upton, Ebru Kaymak, Zhiping Weng, Kathleen Collins, and Oliver J Rando. Deep sequencing of yeast and mouse tRNAs and tRNA fragments using OTTR. *eLife*, 14, 2025.

[15328] Bjorn Gylemo, Maike Bensberg, and Colm E Nestor. A whole-organism landscape of X-inactivation in humans. *eLife*, 14, 2025.

[15329] Eleni Hackwell, Sharon R Ladyman, Jenny Clarkson, H James McQullian, Ulrich Boehm, Allan Edward Herbison, Rosemary Brown, and David R Grattan. Prolactin-mediates a lactation-induced suppression of arcuate kisspeptin neuronal activity necessary for lactational infertility in mice. *eLife*, 13, 2025.

[15330] Hyunggu Hahn, Carole Daly, John Little, Nicole A Perry-Hauser, Emmanuel Flores-Espinoza, Asuka Inoue, Bianca Plouffe, and Alex RB Thomsen. Endosomal chemokine receptor signalosomes regulate central mechanisms underlying cell migration. *eLife*, 13, 2025.

[15331] Swagata Halder, Deepak Velgapuni Raya, and Devarajan Sridharan. Distinct neural bases of subcomponents of the attentional blink. *eLife*, 13, 2025.

[15332] Malika Hale, Kennidy K Takehara, Christopher D Thouvenel, Dina A Moustafa, Andrea Repele, Mary F Fontana, Jason Netland, Sharon McNamara, Ronald L Gibson, Joanna B Goldberg, David J Rawlings, and Marion Pepper. Monoclonal antibodies derived from B cells in subjects with cystic fibrosis reduce Pseudomonas aeruginosa burden in mice. *eLife*, 13, 2025.

[15333] Jessica A Haley, Tianyi Chen, Mikio Aoi, and Sreekanth H Chalasani. Accept–reject decision-making revealed via a quantitative and ethological study of C. elegans foraging. *eLife*, 13, 2025.

[15334] Yutaro Hama, Yuko Fujioka, Hayashi Yamamoto, Noboru Mizushima, and Nobuo Noda. The triad interaction of ULK1, ATG13, and FIP200 is required for ULK complex formation and autophagy. *eLife*, 13, 2025.

[15335] James E Hammond, Ruth E Baker, and Berta Verd. Modularity of the segmentation clock and morphogenesis. *eLife*, 14, 2025.

[15336] Han V Han, Richard Efem, Barbara Rosati, Kevin Lu, Sara Maimouni, Ya-Ping Jiang, Valeria Montoya, Ando Van Der Velden, Wei-Xing Zong, and Richard Z Lin. Propionyl-CoA carboxylase subunit B regulates anti-tumor T cells in a pancreatic cancer mouse model. *eLife*, 13, 2025.

[15337] Su Young Han, Shel-Hwa Yeo, Jae-Chang Kim, Ziyue Zhou, and Allan E Herbison. Multi-dimensional oscillatory activity of mouse GnRH neurons in vivo. *eLife*, 13, 2025.

[15338] Joseph Hanna, Yacine Touahri, Alissa Pak, Lauren Belfiore, Edwin van Oosten, Luke Ajay David, Sisu Han, Yaroslav Ilnytskyy, Igor Kovalchuk, Deborah Kurrasch, Satoshi Okawa, Antonio del Sol, Robert A Screaton, Isabelle Aubert, and Carol Schuurmans. Glycolytic flux controls retinal progenitor cell differentiation via regulating Wnt signaling. *eLife*, 13, 2025.

[15339] Ameya Harmalkar, Sergey Lyskov, and Jeffrey J Gray. Reliable protein–protein docking with AlphaFold, Rosetta, and replica exchange. *eLife*, 13, 2025.

[15340] McKenna Harpring, Junghoon Lee, Guangming Zhong, Scot P Ouellette, and John V Cox. FtsK is critical for the assembly of the unique divisome complex of the FtsZ-less Chlamydia trachomatis. *eLife*, 13, 2025.

[15341] Miriam Hauptman and Marina Bedny. Animacy semantic network supports causal inferences about illness. *eLife*, 13, 2025.

[15342] Sarah Hawes, Bo Liang, Braden Oldham, Breanna T Sullivan, Lupeng Wang, Bin Song, Lisa Chang, Da-Ting Lin, and Huaibin Cai. Patchy striatonigral neurons modulate locomotor vigor in response to environmental valence. *eLife*, 14, 2025.

[15343] Li He, Jiayu Wen, and Qi Dai. PRDM16 functions as a co-repressor in the BMP pathway to suppress neural stem cell proliferation. *eLife*, 14, 2025.

[15344] Meina He, Yongxin Tao, Kexin Mu, Haoqi Feng, Ying Fan, Tong Liu, Qiaoyun Huang, Yujie Xiao, and Wenli Chen. Coordinated regulation of chemotaxis and resistance to copper by CsoR in Pseudomonas putida. *eLife*, 13, 2025.

[15345] Shuo He, Lei Huang, Zhuozheng Chen, Ze Yuan, Yue Zhao, Lingfang Zeng, Yi Zhu, and Jinlong He. Disruption of the novel nested gene Aff3ir mediates disturbed flow-induced atherosclerosis in mice. *eLife*, 13, 2025.

[15346] Tao He, Xizi Gong, Qian Wang, Xinyi Zhu, Yunzhe Liu, and Fang Fang. Non-feature-specific elevated responses and feature-specific backward replay in human brain induced by visual sequence exposure. *eLife*, 13, 2025.

[15347] Hope M Healey, Hayden B Penn, Clayton M Small, Susan Bassham, Vithika Goyal, Micah A Woods, and William A Cresko. Single-cell sequencing provides clues about the developmental genetic basis of evolutionary adaptations in syngnathid fishes. *eLife*, 13, 2025.

[15348] Gabriel O Heckerman, Eileen Tzng, Arely Campos-Melendez, Chisomaga Ekwueme, and Adrienne Mueller. Transparency of research practices in cardiovascular literature. *eLife*, 14, 2025.

[15349] Hendrik Heinbockel, Gregor Leicht, Anthony D Wagner, and Lars Schwabe. Post-retrieval noradrenergic activation impairs subsequent memory depending on cortico-hippocampal reactivation. *eLife*, 13, 2025.

[15350] Zach Hensel. Secondary structure of the SARS-CoV-2 genome is predictive of nucleotide substitution frequency. *eLife*, 13, 2025.

[15351] Erik Henze, Russell N Burkhardt, Bennett William Fox, Tyler J Schwertfeger, Eric Gelsleichter, Kevin Michalski, Lydia Kramer, Margret Lenfest, Jordyn M Boesch, Hening Lin, Frank C Schroeder, and Toshimitsu Kawate. ATP-release pannexin channels are gated by lysophospholipids. *eLife*, 14, 2025.

[15352] Eva Herdering, Tristan Reif-Trauttmansdorff, Anuj Kumar, Tim Habenicht, Georg Hochberg, Stefan Bohn, Jan Schuller, and Ruth Anne Schmitz. 2-oxoglutarate triggers assembly of active dodecameric Methanosarcina mazei glutamine synthetase. *eLife*, 13, 2025.

[15353] Björn Herrmann. Enhanced neural speech tracking through noise indicates stochastic resonance in humans. *eLife*, 13, 2025.

[15354] Nicole Herrmann May, Anh Cao, Annika Schmid, Fabian Link, Jorge Arias-del Angel, Elisabeth Meiser, and Tom Beneke. Improved base editing and functional screening in Leishmania via co-expression of the AsCas12a ultra variant, a T7 RNA polymerase, and a cytosine base editor. *eLife*, 13, 2025.

[15355] Cecilia Hes, Abigail J Tomlinson, Lieke Michielsen, Hunter J Murdoch, Fatemeh Soltani, Maia V Kokoeva, and Paul V Sabatini. A unified rodent atlas reveals the cellular complexity and evolutionary divergence of the dorsal vagal complex. *eLife*, 14, 2025.

[15356] David Hike, Xiaochen Liu, Zeping Xie, Bei Zhang, Sangcheon Choi, Xiaoqing Alice Zhou, Andy Liu, Alyssa Murstein, Yuanyuan Jiang, Anna Devor, and Xin Yu. High-resolution awake mouse fMRI at 14 tesla. *eLife*, 13, 2025.

[15357] Connor Hill, Alexandra Indeglia, Francis Picone, Maureen E Murphy, Cara Cipriano, Robert G Maki, and Alessandro Gardini. NAB2-STAT6 drives an EGR1-dependent neuroendocrine program in solitary fibrous tumors. *eLife*, 13, 2025.

[15358] Nayo M Hill, Haley M Tripp, Daniel M Wolpert, Laura A Malone, and Amy J Bastian. Age-dependent predictors of effective reinforcement motor learning across childhood. *eLife*, 13, 2025.

[15359] Yusuke Himeoka and Chikara Furusawa. Perturbation-response analysis of in silico metabolic dynamics revealed hard-coded responsiveness in the cofactors and network sparsity. *eLife*, 13, 2025.

[15360] Masakatsu Hironaka and Tomonari Sumi. A neural network model that generates salt concentration memory-dependent chemotaxis in Caenorhabditis elegans. *eLife*, 14, 2025.

[15361] Ruoya Ho, Purushotham Pallinti, Louis FL Wilson, Yueping Wan, and Jochen Zimmer. Structure, function and assembly of soybean primary cell wall cellulose synthases. *eLife*, 13, 2025.

[15362] Yi-Yun Ho, Qiuwei Yang, Priyanka Boddu, David A Bulkin, and Melissa R Warden. Infralimbic parvalbumin neural activity facilitates cued threat avoidance. *eLife*, 12, 2025.

[15363] William Hockeimer, Ruo-Yah Lai, Maanasa Natrajan, William Snider, and James J Knierim. Leveraging place field repetition to understand positional versus nonpositional inputs to hippocampal field CA1. *eLife*, 14, 2025.

[15364] Guylaine Hoffner, Pablo Castro, Lynn Uhrig, Camilo Miguel Signorelli, Morgan Dupont, Jordy Tasserie, Alain Destexhe, Rodrigo Cofre, Jacobo Sitt, and Béchir Jarraya. Transcranial direct current stimulation modulates primate brain dynamics across states of consciousness. *eLife*, 13, 2025.

[15365] Edwin A Homan, Ankit Gilani, Alfonso Rubio-Navarro, Maya A Johnson, Odin M Schaepkens, Eric Cortada, Renan Pereira de Lima, Lisa Stoll, and James C Lo. Complement 3a receptor 1 on macrophages and Kupffer cells is not required for the pathogenesis of metabolic dysfunction-associated steatotic liver disease. *eLife*, 13, 2025.

[15366] Juken Hong, Wenzhi Xue, and Teng Wang. Emergence of alternative stable states in microbial communities undergoing horizontal gene transfer. *eLife*, 13, 2025.

[15367] Victoria B Hosack and Fritzie Arce-McShane. 3D directional tuning in the orofacial sensorimotor cortex during natural feeding and drinking. *eLife*, 13, 2025.

[15368] Kawsar Hossain, Madeline Smith, Karlin E Rufenacht, Rebecca O'Rourke, and Stephen W Santoro. In mice, discrete odors can selectively promote the neurogenesis of sensory neuron subtypes that they stimulate. *eLife*, 13, 2025.

[15369] Kazi A Hossain, Lukasz Nierzwicki, Modesto Orozco, Jacek Czub, and Giulia Palermo. Flexibility in PAM recognition expands DNA targeting in xCas9. *eLife*, 13, 2025.

[15370] Julia A Hotinger, Ian W Campbell, Karthik Hullahalli, Akina Osaki, and Matthew K Waldor. Quantification of Salmonella enterica serovar Typhimurium population dynamics in murine infection using a highly diverse barcoded library. *eLife*, 13, 2025.

[15371] Yanlin Hou, Zhengwen Nie, Qi Jiang, Sergiy Velychko, Sandra Heising, Ivan Bedzhov, Guangming Wu, Kenjiro Adachi, and Hans R Scholer. Emerging cooperativity between Oct4 and Sox2 governs the pluripotency network in early mouse embryos. *eLife*, 13, 2025.

[15372] Conor J Howard, Nathan S Abell, Beatriz A Osuna, Eric M Jones, Leon Y Chan, Henry Chan, Dean R Artis, Jonathan B Asfaha, Joshua S Bloom, Aaron R Cooper, Andrew Liao, Eden Mahdavi, Nabil Mohammed, Alan L Su, Giselle A Uribe, Sriram Kosuri, Diane E Dickel, and Nathan B Lubock. High-resolution deep mutational scanning of the melanocortin-4 receptor enables target characterization for drug discovery. *eLife*, 13, 2025.

[15373] Jesse Howe, Douglas Walker, Kyle Tengler, Maya Sonpatki, Patrick N Reardon, Justin WC Leung, and Elisar J Barbar. LC8 enhances 53BP1 foci through heterogeneous bridging of 53BP1 oligomers. *eLife*, 14, 2025.

[15374] Akiko Hozumi, Nozomu M Totsuka, Arata Onodera, Yanbin Wang, Mayuko Hamada, Akira Shiraishi, Honoo Satake, Takeo Horie, Kohji Hotta, and Yasunori Sasakura. Stimulatory and inhibitory G-protein signaling relays drive cAMP accumulation for timely metamorphosis in the chordate Ciona. *eLife*, 13, 2025.

[15375] Louise Tzung-Harn Hsieh, Belinda S Hall, Jane Newcombe, Tom A Mendum, Sonia Santana Varela, Yagnesh Umrania, Michael J Deery, Wei Q Shi, Josué Diaz-Delgado, Francisco J Salguero, and Rachel E Simmonds. The Mycobacterium ulcerans toxin mycolactone causes destructive Sec61-dependent loss of the endothelial glycocalyx and vessel basement membrane to drive skin necrosis. *eLife*, 12, 2025.

[15376] Chen-wei Hsu, Hao Ho, Ching-Hsin Yang, Yan-wei Wang, Ker-Chau Li, and Bon-chu Chung. Single-cell transcriptomes of zebrafish germline reveal progenitor types and feminization by Foxl2l. *eLife*, 14, 2025.

[15377] Yijia Hua, Ningjun Duan, Chunxiao Sun, Fan Yang, Min Tian, Yanting Sun, Shuhan Zhao, Jue Gong, Qian Liu, Xiang Huang, Yan Liang, Ziyi Fu, Wei Li, and Yongmei Yin. Targeting SLC7A11-mediated cysteine metabolism for the treatment of trastuzumab-resistant HER2-positive breast cancer. *eLife*, 14, 2025.

[15378] Changrun Huang, Dirk van Moorselaar, Joshua Foster, Mieke Donk, and Jan Theeuwes. Neural mechanisms of learned suppression uncovered by probing the hidden attentional priority map. *eLife*, 13, 2025.

[15379] Hong Huang, Ying Yang, Qiuju Zhang, Yongfeng Yang, Zhenqi Xiong, Shengqiang Mao, Tingting Song, Yilong Wang, Zhiqiang Liu, Hong Bu, Li Zhang, and Le Zhang. S100a4+ alveolar macrophages accelerate the progression of precancerous atypical adenomatous hyperplasia by promoting the angiogenic function regulated by fatty acid metabolism. *eLife*, 13, 2025.

[15380] Jiadai Huang, Fang Chen, Beifang Lu, Yue Sun, Youyue Li, Canfeng Hua, and Xin Deng. DNA methylome regulates virulence and metabolism in Pseudomonas syringae. *eLife*, 13, 2025.

[15381] Junjun Huang, Rong Jia, and Jihua Guo. RUNX2 isoform II protects cancer cells from ferroptosis and apoptosis by promoting PRDX2 expression in oral squamous cell carcinoma. *eLife*, 13, 2025.

[15382] Mei-Lun Huang, Yusuke Suzuki, Hiroki Sasaguri, Takashi Saito, Takaomi C Saido, and Itaru Imayoshi. Misclassification in memory modification in AppNL-G-F knock-in mouse model of Alzheimer's disease. *eLife*, 14, 2025.

[15383] Shih-Wen Huang, Yein-Gei Lai, Hao-Ting Liao, Chin-Ling Chang, Ruo-Yu Ma, Yung-Hsiang Chen, Yae-Huei Liou, Zhen-Qi Wu, Yu-Chen Wu, Ko-Jiunn Liu, Yen-Tsung Huang, Jen-Lung Yang, Ming-Shen Dai, and Nan-Shih Liao. Syngeneic natural killer cell therapy activates dendritic and T cells in metastatic lungs and effectively treats low-burden metastases. *eLife*, 13, 2025.

[15384] Yan Huang, Nina Bucevic, Carmen Coves, Natalia Felipe-Medina, Marina Marcet-Ortega, Nikoleta Nikou, Cristina Madrid-Sandín, Maria López-Panadés, Carolina Buza, Neus Ferrer Miralles, Antoni Iborra, Anna Pujol, Alberto M Pendás, and Ignasi Roig. The full-length BEND2 protein is dispensable for spermatogenesis but required for setting the ovarian reserve in mice. *eLife*, 13, 2025.

[15385] David E Huber. A memory model of rodent spatial navigation in which place cells are memories arranged in a grid and grid cells are non-spatial. *eLife*, 13, 2025.

[15386] Daniel Hui, Scott Dudek, Krzysztof Kiryluk, Theresa L Walunas, Iftikhar J Kullo, Wei-Qi Wei, Hemant Tiwari, Josh F Peterson, Wendy K Chung, Brittney H Davis, Atlas Khan, Leah C Kottyan, Nita A Limdi, Qiping Feng, Megan J Puckelwartz, Chunhua Weng, Johanna L Smith, Elizabeth W Karlson, Gail P Jarvik, and Marylyn D Ritchie. Risk factors affecting polygenic score performance across diverse cohorts. *eLife*, 12, 2025.

[15387] Alicia Hurtado, Victor López-Soriano, Miguel Lao, M Angeles Celis-Barroso, Pilar Lazúen, Alejandro Chacón-de Castro, Yolanda Ramírez-Casas, Miguel Alaminos, John Martin Collinson, Miguel Burgos, Rafael Jiménez, F David Carmona, and Francisco Javier Barrionuevo. Sox9 prevents retinal degeneration and is required for limbal stem cell differentiation in the adult mouse eye. *eLife*, 13, 2025.

[15388] Vincent Huson and Wade G Regehr. Realistic mossy fiber input patterns to unipolar brush cells evoke a continuum of temporal responses comprised of components mediated by different glutamate receptors. *eLife*, 13, 2025.

[15389] Saman Hussain, Katherine Pinter, Mara Uhl, Hiu-Tung Wong, and Katie S Kindt. Microtubule networks in zebrafish hair cells facilitate presynapse transport and fusion during development. *eLife*, 13, 2025.

[15390] Baher A Ibrahim, Yoshitaka Shinagawa, Austin Douglas, Gang Xiao, Alexander R Asilador, and Daniel A Llano. Microprism-based two-photon imaging of the mouse inferior colliculus reveals novel organizational principles of the auditory midbrain. *eLife*, 12, 2025.

[15391] Taro Ichimura, Taishi Kakizuka, Yoshitsugu Taniguchi, Satoshi Ejima, Yuki Sato, Keiko Itano, Kaoru Seiriki, Hitoshi Hashimoto, Ko Sugawara, Hiroya Itoga, Shuichi Onami, and Takeharu Nagai. Volumetric trans-scale imaging of massive quantity of heterogeneous cell populations in centimeter-wide tissue and embryo. *eLife*, 13, 2025.

[15392] Jittoku Ihara, Yibin Huang, Yoichi Takami, Yoichi Nozato, Toshimasa Takahashi, Akemi Kakino, Cheng Wang, Ziwei Wang, Yu Guo, Weidong Liu, Nanxiang Yin, Ryoichi Ohara, Taku Fujimoto, Shino Yoshida, Kazuhiro Hongyo, Hiroshi Koriyama, Hiroshi Akasaka, Hikari Takeshita, Shinsuke Sakai, Kazunori Inoue, Yoshitaka Isaka, Hiromi Rakugi, Tatsuya Sawamura, and Koichi Yamamoto. Oxidized low-density lipoprotein potentiates angiotensin II-induced Gq activation through the AT1-LOX1 receptor complex. *eLife*, 13, 2025.

[15393] Sharon Inberg, Yael Iosilevskii, Alba Calatayud-Sanchez, Hagar Setty, Meital Oren-Suissa, Michael Krieg, and Benjamin Podbilewicz. Sensory experience controls dendritic structure and behavior by distinct pathways involving degenerins. *eLife*, 14, 2025.

[15394] Lucia Inchauste, Elif Nurtop, Lissete Bautista Machicado, Yanine Leigue Roth, Shirley Lenz Gonzales, Maria Luisa Herrera, Katty Mina Villafan, Pedro Mamani Mamani, Marcelo Ramos Espinoza, Juan Carlos Pavel Suarez, Juan Cansio Garcia Copa, Yitzhak Leigue Zabala, Etzel Arancibia Cardozo, Pierre Gallian, Xavier de Lamballerie, and Stéphane Priet. Improving SARS-CoV-2 variants monitoring in the absence of genomic surveillance capabilities: a serological study in Bolivian blood donors in October 2021 and June 2022. *eLife*, 13, 2025.

[15395] Tudor M Ionescu, Mario Amend, Rakibul Hafiz, Andreas Maurer, Bharat Biswal, Hans F Wehrl, and Kristina Herfert. Mapping serotonergic dynamics using drug-modulated molecular connectivity in rats. *eLife*, 13, 2025.

[15396] Yevheniia Ishchenko, Amanda T Jeng, Shufang Feng, Timothy Nottoli, Cindy Manriquez-Rodriguez, Khanh K Nguyen, Melissa G Carrizales, Matthew J Vitarelli, Ellen E Corcoran, Charles A Greer, Samuel A Myers, and Anthony J Koleske. Heterozygosity for neurodevelopmental disorder-associated TRIO variants yields distinct deficits in behavior, neuronal development, and synaptic transmission in mice. *eLife*, 13, 2025.

[15397] Scott Isherwood, Sarah A Kemp, Steven Miletić, Niek Stevenson, Pierre-Louis Bazin, and Birte Forstmann. Multi-study fMRI outlooks on subcortical BOLD responses in the stop-signal paradigm. *eLife*, 12, 2025.

[15398] Kentaro K Ishii, Koichi Hashikawa, Jane Chea, Shihan Yin, Rebecca Erin Fox, Suyang Kan, Meha Shah, Zhe Charles Zhou, Jovana Navarrete, Alexandria D Murry, Eric R Szelenyi, Sam A Golden, and Garret D Stuber. Post-ejaculatory inhibition of female sexual drive via heterogeneous neuronal ensembles in the medial preoptic area. *eLife*, 12, 2025.

[15399] Ryosuke Isotani, Masaki Igarashi, Masaomi Miura, Kyoko Naruse, Satoshi Kuranami, Manami Katoh, Seitaro Nomura, and Toshimasa Yamauchi. Nicotine enhances the stemness and tumorigenicity in intestinal stem cells via Hippo-YAP/TAZ and Notch signal pathway. *eLife*, 13, 2025.

[15400] Ayaka Itani, Haruto Motomura, Ken Oda, Hideyuki Yamashita, Kanae Sakai, Ken-ichi Kusumoto, Shinsuke Shigeto, Takehiko Ichikawa, Hosain Mohammad Mubarak, Takeshi Fukuma, Takuya Katayama, Jun-ichi Maruyama, Shunsuke Masuo, Naoki Takaya, and Norio Takeshita. The increase in cell volume and nuclear number of the koji-fungus Aspergillus oryzae contributes to its high enzyme productivity. *eLife*, 14, 2025.

[15401] Yuna Izumi-Mishima, Rie Tsutsumi, Tetsuya Shiuchi, Saori Fujimoto, Momoka Taniguchi, Mizuki Sugiuchi, Manaka Tsutsumi, Yuko Okamatsu-Ogura, Takeshi Yoneshiro, Masashi Kuroda, Kazuhiro Nomura, and Hiroshi Sakaue. Brown adipose tissue and skeletal muscle coordinately contribute to thermogenesis in mice. *eLife*, 13, 2025.

[15402] Sara Jaber, Eliana Eldawra, Jeanne Rakotopare, Iva Simeonova, Vincent Lejour, Marc Gabriel, Tatiana Cañeque, Vitalina Volochtchouk, Monika Licaj, Anne Fajac, Raphaël Rodriguez, Antonin Morillon, Boris Bardot, and Franck Toledo. Oncogenic and teratogenic effects of Trp53Y217C, an inflammation-prone mouse model of the human hotspot mutant TP53Y220C. *eLife*, 13, 2025.

[15403] Georgin Jacob, RT Pramod, and SP Arun. Visual homogeneity computations in the brain enable solving property-based visual tasks. *eLife*, 13, 2025.

[15404] Paul I Jaffe, Gustavo X Santiago-Reyes, Robert J Schafer, Patrick G Bissett, and Russell A Poldrack. An image-computable model of speeded decision-making. *eLife*, 13, 2025.

[15405] Rupinder Singh Jandu, Ashim Bhattacharya, Frank Antony, Mohammed Al-Seragi, Hiroyuki Aoki, Mohan Babu, and Franck Duong van Hoa. Membrane-mimetic thermal proteome profiling (MM-TPP) toward mapping membrane protein–ligand dynamic interactions. *eLife*, 14, 2025.

[15406] Jasper Janssens, Pierre Mangeol, Nikolai Hecker, Gabriele Partel, Katina I Spanier, Joy N Ismail, Gert J Hulselmans, Stein Aerts, and Frank Schnorrer. Spatial transcriptomics in the adult Drosophila brain and body. *eLife*, 13, 2025.

[15407] Gabriel E Jara, Francesco Pontiggia, Renee Otten, Roman V Agafonov, Marcelo A Martí, and Dorothee Kern. Wide transition-state ensemble as key component for enzyme catalysis. *eLife*, 12, 2025.

[15408] Chinmaya Jena, Saillesh Chinnaraj, Soham Deolankar, and Nishad Matange. Proteostasis modulates gene dosage evolution in antibiotic-resistant bacteria. *eLife*, 13, 2025.

[15409] Hey-Kyeong Jeong, Estibaliz Gonzalez-Fernandez, Ilan Crawley, Julia M Coakley, Jinha Hwang, Dale DO Martin, Shernaz X Bamji, Jong-Il Kim, Shin H Kang, and Gareth M Thomas. Micro-scale control of oligodendrocyte morphology and myelination by the intellectual disability-linked protein acyltransferase ZDHHC9. *eLife*, 13, 2025.

[15410] Wencheng Ji, Ori Hachmo, Naama Barkai, and Ariel Amir. Design principles of transcription factors with intrinsically disordered regions. *eLife*, 14, 2025.

[15411] Zhiguang Jia and Jianhan Chen. Atomistic simulation of voltage activation of a truncated BK channel. *eLife*, 14, 2025.

[15412] Yalan Jiang, Pingping He, Ke Sheng, Yongmiao Peng, Huilan Wu, Songwei Qian, Weiping Ji, Xiaoling Guo, and Xiaoou Shan. The protective roles of eugenol on type 1 diabetes mellitus through NRF2-mediated oxidative stress pathway. *eLife*, 13, 2025.

[15413] Yixuan Jiang, Ying Liang, Fujie Zhao, Zhenguo Lu, Siyu Wang, Yao Meng, Zhanxiang Liu, Jing Zhang, and Youbao Zhao. Rtf1 HMD domain facilitates global histone H2B monoubiquitination and regulates morphogenesis and virulence in the meningitis-causing pathogen Cryptococcus neoformans. *eLife*, 13, 2025.

[15414] Yong Jiang, Stephen J Mayor, Xiuli Chu, Xiaoqi Ye, Rongzhou Man, Jing Tao, and Qing-Lai Dang. Assessing plant phenological changes based on drivers of spring phenology. *eLife*, 14, 2025.

[15415] Dongyue Jiao, Kun Chang, Jiamin Jin, Yingji Chen, Mo Ren, Yucong Zhang, Kun Gao, Yaoting Xu, Lixin Wang, and Chenji Wang. Targeting the WSB2–NOXA axis in cancer cells for enhanced sensitivity to BCL-2 family protein inhibitors. *eLife*, 13, 2025.

[15416] Zhu Liduzi Jiesisibieke, Io Ieong Chan, Jack Chun Man Ng, and C Mary Schooling. A sex-specific Mendelian randomization-phenome-wide association study of body mass index. *eLife*, 13, 2025.

[15417] Erli Jin, Jennifer K Briggs, Richard KP Benninger, and Matthew J Merrins. Glucokinase activity controls peripherally located subpopulations of $\beta$-cells that lead islet Ca2+ oscillations. *eLife*, 13, 2025.

[15418] Michelle Jin, Simon O Ogundare, Marcos Lanio, Sophia Sorid, Alicia Ruth Whye, Sofia Leal Santos, Alessandra Franceschini, and Christine Ann Denny. A SMARTTR workflow for multi-ensemble atlas mapping and brain-wide network analysis. *eLife*, 13, 2025.

[15419] Marketa Jirouskova, Karel Harant, Pavel Cejnar, Srikant Ojha, Katerina Korelova, Lenka Sarnova, Eva Sticova, Christoph H Mayr, Herbert B Schiller, and Martin Gregor. Dynamics of compartment-specific proteomic landscapes of hepatotoxic and cholestatic models of liver fibrosis. *eLife*, 13, 2025.

[15420] Marie-Dominique Jolivet, Anne Flore Deroubaix, Marie Boudsocq, Nikolaj B Abel, Marion Rocher, Terezinha Robbe, Valérie Wattelet-Boyer, Jennifer Huard, Dorian Lefebvre, Yi-Ju Lu, Brad Day, Grégoire Saias, Jahed Ahmed, Valérie Cotelle, Nathalie Giovinazzo, Jean-Luc Gallois, Yasuyuki Yamaji, Sylvie German-Retana, Julien Gronnier, Thomas Ott, Sébastien Mongrand, and Véronique Germain. Interdependence of plasma membrane nanoscale dynamics of a kinase and its cognate substrate underlies Arabidopsis response to viral infection. *eLife*, 12, 2025.

[15421] Divyesh Joshi, Preeti Jindal, Ronak K Shetty, and Maneesha S Inamdar. Rudhira-mediated microtubule stability controls TGF$\beta$ signaling during mouse vascular development. *eLife*, 13, 2025.

[15422] Luay Joudeh, Robert E Appleby, Joseph D Maman, and Luca Pellegrini. Structural mechanism of strand exchange by the RAD51 filament. *eLife*, 14, 2025.

[15423] Qianqian Ju, Wenjing Sheng, Meichen Zhang, Jing Chen, Liucheng Wu, Xiaoyu Liu, Wentao Fang, Hui Shi, and Cheng Sun. TAK1-mediated phosphorylation of PLCE1 represses PIP2 hydrolysis to impede esophageal squamous cancer metastasis. *eLife*, 13, 2025.

[15424] Nicholas Judd and Rogier Kievit. No effect of additional education on long-term brain structure, a preregistered natural experiment in thousands of individuals. *eLife*, 13, 2025.

[15425] JeYoung Jung, Steve Williams, and Matthew A Lambon Ralph. The role of GABA in semantic memory and its neuroplasticity. *eLife*, 12, 2025.

[15426] Yoshihisa Kaizuka and Rika Machida. Cell membrane glycan contents are biochemical factors that constitute a kinetic barrier to viral particle uptake in a protein-nonspecific manner. *eLife*, 14, 2025.

[15427] Márton Kajtor, Viktor András Billes, Bálint Király, Patricia Karkusova, Tibor Kovács, Hannah Stabb, Katalin Sviatkó, Andor Vizi, Eszter Ujvári, Diána Balázsfi, Sophie E Seidenbecher, Duda Kvitsiani, Tibor Vellai, and Balázs Hangya. Altered reactivity to threatening stimuli in Drosophila models of Parkinson's disease, revealed by a trial-based assay. *eLife*, 13, 2025.

[15428] Jenifer C Kaldun, Emanuele Calia, Ganesh Chinmai Bangalore Mukunda, Cornelia Fritsch, Nikita Komarov, and Simon G Sprecher. A temporally restricted function of the dopamine receptor Dop1R2 during memory formation. *eLife*, 13, 2025.

[15429] Rachel Kaletsky, Rebecca S Moore, Titas Sengupta, Renee Seto, Borja Ceballos-Llera, and Coleen T Murphy. Molecular requirements for C. elegans transgenerational epigenetic inheritance of pathogen avoidance. *eLife*, 14, 2025.

[15430] Irina Kalita, Ira Alexandra Iosub, Lorna McLaren, Louise Goossens, Sander Granneman, and Meriem El Karoui. An Hfq-dependent post-transcriptional mechanism fine tunes RecB expression in Escherichia coli. *eLife*, 13, 2025.

[15431] Lena Kallweit, Eric Daniel Hamlett, Hannah Saternos, Anah Gilmore, Ann-Charlotte Granholm, and Scott Horowitz. Chronic RNA G-quadruplex accumulation in aging and Alzheimer's disease. *eLife*, 14, 2025.

[15432] Jingbo Kan, Adrian Morales-Amador, Yozen Hernandez, Melinda A Ternei, Christophe Lemetre, Logan W Maclntyre, Nicolas Biais, and Sean F Brady. Oxydifficidin, a potent Neisseria gonorrhoeae antibiotic due to DedA-assisted uptake and ribosomal protein RplL sensitivity. *eLife*, 13, 2025.

[15433] Liqi Kang, Banghao Wu, Bingxin Zhou, Pan Tan, Yun (Kenneth) Kang, Yongzhen Yan, Yi Zong, Shuang Li, Zhuo Liu, and Liang Hong. AI-enabled alkaline-resistant evolution of protein to apply in mass production. *eLife*, 13, 2025.

[15434] Tomoharu Kanie, Beibei Liu, Julia F Love, Saxton D Fisher, Anna-Karin Gustavsson, and Peter K Jackson. A hierarchical pathway for assembly of the distal appendages that organize primary cilia. *eLife*, 14, 2025.

[15435] Tomoharu Kanie, Roy Ng, Keene L Abbott, Niaj Mohammad Tanvir, Esben Lorentzen, Olaf Pongs, and Peter K Jackson. Myristoylated Neuronal Calcium Sensor-1 captures the preciliary vesicle at distal appendages. *eLife*, 14, 2025.

[15436] Delaney Kaper, UroŠ Radović, Per-Olof Bergh, August Qvist, Marcus Henricsson, Jan Borén, and Marc Pilon. Forward genetics in C. elegans reveals genetic adaptations to polyunsaturated fatty acid deficiency. *eLife*, 13, 2025.

[15437] Mehmet Mahsum Kaplan, Erika Hudacova, Miroslav Matejcek, Haneen Tuaima, Jan Křivánek, and Ondrej Machon. Mesenchymal Meis2 controls whisker development independently from trigeminal sensory innervation. *eLife*, 13, 2025.

[15438] Yogita Kapoor, Himani Khurana, Debatri Dutta, Arnab Chakraborty, Anshu Priya, Archana Singh, Siddhesh S Kamat, Neeraj Dhar, Thomas John Pucadyil, and Vinay Kumar Nandicoori. Wag31, a membrane tether, is crucial for lipid homeostasis in mycobacteria. *eLife*, 14, 2025.

[15439] Alexander N Kapustin, Sofia Serena Tsakali, Meredith Whitehead, George Chennell, Meng-Ying Wu, Chris Molenaar, Anton Kutikhin, Yimeng Chen, Sadia Ahmad, Leo Bogdanov, Maxim Sinitsky, Kseniya Rubina, Aled Clayton, Frederik J Verweij, Dirk Michiel Pegtel, Simona Zingaro, Arseniy Lobov, Bozhana Zainullina, Dylan Owen, Maddy Parsons, Richard E Cheney, Derek T Warren, Martin James Humphries, Thomas Iskratsch, Mark Holt, and Catherine M Shanahan. Matrix-associated extracellular vesicles modulate human smooth muscle cell adhesion and directionality by presenting collagen VI. *eLife*, 12, 2025.

[15440] Nare Karagulyan, Maja überegger, Yumeng Qi, Norbert Babai, Florian Hofer, Lejo Johnson Chacko, Fangfang Wang, Maria Luque, Rudolf Glueckert, Anneliese Schrott-Fischer, Yunfeng Hua, Tobias Moser, and Christine Bandtlow. Probing the role of synaptic adhesion molecule RTN4RL2 in setting up cochlear connectivity. *eLife*, 14, 2025.

[15441] Lili Karashchuk, Jing Shuang Li, Grant M Chou, Sarah Walling-Bell, Steven L Brunton, John C Tuthill, and Bingni W Brunton. Sensorimotor delays constrain robust locomotion in a 3D kinematic model of fly walking. *eLife*, 13, 2025.

[15442] Rezwana Karim, Wendi Teng, Cameron D Behram, and Hening Lin. SIRT2-mediated ACSS2 K271 deacetylation suppresses lipogenesis under nutrient stress. *eLife*, 13, 2025.

[15443] Diana M Karosas, Leslie Gonzales, Yingxuan Wang, Christopher Bergevin, Laurel H Carney, and Kenneth S Henry. Otoacoustic emissions but not behavioral measurements predict cochlear nerve frequency tuning in an avian vocal communication specialist. *eLife*, 13, 2025.

[15444] Gülnihal Kavaklioglu, Alexandra Podhornik, Terezia Vcelkova, Jelena Marjanovic, Mirjam A Beck, Trinh Phan-Canh, Theresia Mair, Claudia Miccolo, Aleksej Drino, Mirko Doni, Gerda Egger, Susanna Chiocca, Miha Modic, and Christian Seiser. The domesticated transposon protein L1TD1 associates with its ancestor L1 ORF1p to promote LINE-1 retrotransposition. *eLife*, 13, 2025.

[15445] Shinichi Kawaguchi, Xin Xu, Takashi Soga, Kenta Yamaguchi, Ryuuya Kawasaki, Ryota Shimouchi, Susumu Date, and Toshie Kai. In silico screening by AlphaFold2 program revealed the potential binding partners of nuage-localizing proteins and piRNA-related proteins. *eLife*, 13, 2025.

[15446] Toshihiro Kawasaki, Toshiya Nishimura, Naoki Tani, Carina Ramos, Emil Karaulanov, Minori Shinya, Kenji Saito, Emily Taylor, René F Ketting, Kei-ichiro Ishiguro, Minoru Tanaka, Kellee R Siegfried, and Noriyoshi Sakai. Meioc-Piwil1 complexes regulate rRNA transcription for differentiation of spermatogonial stem cells. *eLife*, 14, 2025.

[15447] Ekin Kaya, Evan Wegienka, Alexandra Akhtarzandi-Das, Hanh Do, Ada Eban-Rothschild, and Gideon Rothschild. Food intake enhances hippocampal sharp wave-ripples. *eLife*, 14, 2025.

[15448] Madeline M Keenen, Liheng Yang, Huan Liang, Veronica J Farmer, Rizban E Worota, Rohit Singh, Amy S Gladfelter, and Carolyn B Coyne. Comparative analysis of the syncytiotrophoblast in placenta tissue and trophoblast organoids using snRNA sequencing. *eLife*, 13, 2025.

[15449] Olga Kepinska, Josue Dalboni da Rocha, Carola Tuerk, Alexis Hervais-Adelman, Florence Bouhali, David W Green, Cathy J Price, and Narly Golestani. Auditory cortex anatomy reflects multilingual phonological experience. *eLife*, 12, 2025.

[15450] David A Kessler and Nadav M Shnerb. The dominant–egalitarian transition in species-rich communities. *eLife*, 14, 2025.

[15451] Iraj Khalid, Orphee Morlaas, Hugo Bottemanne, Lisa Thonon, Thomas Da Costa, Philippe Fossati, and Liane Schmidt. Effects of experiencing the COVID-19 pandemic on optimistically biased belief updating. *eLife*, 13, 2025.

[15452] Vladimir Khayenko, Cihan Makbul, Clemens Schulte, Naomi Hemmelmann, Sonja Kachler, Bettina Böttcher, and Hans Michael Maric. Induction of hepatitis B core protein aggregation targeting an unconventional binding site. *eLife*, 13, 2025.

[15453] Mohsen Khosravi-Maharlooei, Andrea Vecchione, Nichole Danzl, Hao Wei Li, Grace Nauman, Rachel Madley, Elizabeth Waffarn, Robert Winchester, Amanda Ruiz, Xiaolan Ding, Georgia Fousteri, and Megan Sykes. Follicular helper- and peripheral helper-like T cells drive autoimmune disease in human immune system mice. *eLife*, 13, 2025.

[15454] Sami Kilpinen, Lassi Virtanen, Silvana Bodington Celma, Amos Bonsdorff, Heidi Heliölä, Kaia Achim, and Juha Partanen. Gene regulatory mechanisms guiding bifurcation of inhibitory and excitatory neuron lineages in the mouse anterior brainstem. *eLife*, 14, 2025.

[15455] Ji Hyun Kim, Muhammad Irfan, Sreelekshmi Sreekumar, Atsawasuwan Phimon, Stephanie Kim, and Seung Chung. CRISPR-edited DPSCs constitutively expressing BDNF enhance dentin regeneration in injured teeth. *eLife*, 14, 2025.

[15456] Linda H Kim, Adam Lognon, Sandeep Sharma, Michelle A Tran, Cecilia Badenhorst, Taylor Chomiak, Stephanie Tam, Claire McPherson, Todd E Stang, Shane EA Eaton, Zelma HT Kiss, and Patrick J Whelan. Restoration of locomotor function following stimulation of the A13 region in Parkinson's mouse models. *eLife*, 12, 2025.

[15457] Tackhoon Kim, Byung-Sun Park, Soobeen Heo, Heeju Jeon, Jaeyeal Kim, Donghwa Kim, Sang Kook Lee, So-Youn Jung, Sun-Young Kong, and Timothy Lu. Combinatorial CRISPR screen reveals FYN and KDM4 as targets for synergistic drug combination for treating triple negative breast cancer. *eLife*, 13, 2025.

[15458] Hirokazu Kimura, Kamel Lahouel, Cristian Tomasetti, and Nicholas Jason Roberts. Functional characterization of all CDKN2A missense variants and comparison to in silico models of pathogenicity. *eLife*, 13, 2025.

[15459] Jan H Kirchner, Lucas Euler, Ingo Fritz, André Ferreira Castro, and Julijana Gjorgjieva. Dendritic growth and synaptic organization from activity-independent cues and local activity-dependent plasticity. *eLife*, 12, 2025.

[15460] Hans Kirschner, Hanna M Molla, Matthew R Nassar, Harriet de Wit, and Markus Ullsperger. Methamphetamine-induced adaptation of learning rate dynamics depend on baseline performance. *eLife*, 13, 2025.

[15461] Matthew R Kleinman and David J Foster. Spatial localization of hippocampal replay requires dopamine signaling. *eLife*, 13, 2025.

[15462] Jacob W Klemm, Chloe Van Hazel, and Robin E Harris. Regeneration following tissue necrosis is mediated by non-apoptotic caspase activity. *eLife*, 13, 2025.

[15463] Maren Klingelhöfer-Jens, Katharina Hutterer, Miriam A Schiele, Elisabeth J Leehr, Dirk Schümann, Karoline Rosenkranz, Joscha Böhnlein, Jonathan Repple, Jürgen Deckert, Katharina Domschke, Udo Dannlowski, Ulrike Lueken, Andreas Reif, Marcel Romanos, Peter Zwanzger, Paul Pauli, Matthias Gamer, and Tina B Lonsdorf. Reduced discrimination between signals of danger and safety but not overgeneralization is linked to exposure to childhood adversity in healthy adults. *eLife*, 12, 2025.

[15464] Jason R Klug, Xunyi Yan, Hilary Hoffman, Max D Engelhardt, Fumitaka Osakada, Edward M Callaway, and Xin Jin. Asymmetric cortical projections to striatal direct and indirect pathways distinctly control actions. *eLife*, 12, 2025.

[15465] Abigail Knecht, Denise Sirias, Daniel R Utter, and Karine A Gibbs. Non-cognate immunity proteins provide broader defenses against interbacterial effectors in microbial communities. *eLife*, 12, 2025.

[15466] Hannah Riley Knight, Marie Kim, Nisha Kannan, Hannah Taylor, Hailey Main, Emily Azcue, and Aaron Esser-Kahn. Bioengineering approaches to trained immunity: Physiologic targets and therapeutic strategies. *eLife*, 14, 2025.

[15467] Alexander J Knights, Easton C Farrell, Olivia M Ellis, Michelle J Song, C Thomas Appleton, and Tristan Maerz. Synovial macrophage diversity and activation of M-CSF signaling in post-traumatic osteoarthritis. *eLife*, 12, 2025.

[15468] Ethan Knights, Richard N Henson, Alexa Morcom, Daniel J Mitchell, and Kamen A Tsvetanov. Neural evidence of functional compensation for fluid intelligence in healthy ageing. *eLife*, 13, 2025.

[15469] Hisato Kobayashi, Tatsushi Igaki, Soichiro Kumamoto, Keisuke Tanaka, Tomoya Takashima, So I Nagaoka, Shunsuke Suzuki, Masaaki Hayashi, Marilyn B Renfree, Manabu Kawahara, Shun Saito, Toshihiro Kobayashi, Hiroshi Nagashima, Hitomi Matsunari, Kazuaki Nakano, Ayuko Uchikura, Hiroshi Kiyonari, Mari Kaneko, Hiroo Imai, Kazuhiko Nakabayashi, Matthew Lorincz, and Kazuki Kurimoto. Post-fertilization transcription initiation in an ancestral LTR retrotransposon drives lineage-specific genomic imprinting of ZDBF2. *eLife*, 13, 2025.

[15470] Andrew G Koesters, Mark M Rich, and Kathrin Engisch. Homeostatic synaptic plasticity of miniature excitatory postsynaptic currents in mouse cortical cultures requires neuronal Rab3a. *eLife*, 12, 2025.

[15471] Damian Koevoet, Laura Van Zantwijk, Marnix Naber, Sebastiaan Mathôt, Stefan van der Stigchel, and Christoph Strauch. Effort drives saccade selection. *eLife*, 13, 2025.

[15472] Keita Kohno, Ryoji Shirasaka, Keita Hirose, Takahiro Masuda, and Makoto Tsuda. Interleukin-4 induces CD11c+ microglia leading to amelioration of neuropathic pain in mice. *eLife*, 14, 2025.

[15473] Yoshinori Kohwi, Xianrong Wong, Mari Grange, Thomas Sexton, Hunter W Richards, Yohko Kitagawa, Shimon Sakaguchi, Ya-Chen Liang, Cheng-Ming Chuong, Vladimir A Botchkarev, Ichiro Taniuchi, Karen L Reddy, and Terumi Kohwi-Shigematsu. Genome organization by SATB1 binding to base-unpairing regions (BURs) provides a scaffold for SATB1-regulated gene expression. *eLife*, 14, 2025.

[15474] Igor Kondrychyn, Liqun He, Haymar Wint, Christer Betsholtz, and Li-Kun Phng. Combined forces of hydrostatic pressure and actin polymerization drive endothelial tip cell migration and sprouting angiogenesis. *eLife*, 13, 2025.

[15475] Ling-Wei Kong, Wenjia Shi, Xiao-Jun Tian, and Ying-Cheng Lai. Dynamical mechanisms of growth-feedback effects on adaptive gene circuits. *eLife*, 12, 2025.

[15476] Mi-Seon Kong, Ethan Ancell, Daniela M Witten, and Larry S Zweifel. Valence and salience encoding in the central amygdala. *eLife*, 13, 2025.

[15477] Weiguang Kong, Xinyou Wang, Guangyi Ding, Peng Yang, Yong Shi, Chang Cai, Xinyi Yang, Gaofeng Cheng, Fumio Takizawa, and Zhen Xu. Newly discovered and conserved role of IgM against viral infection in an early vertebrate. *eLife*, 14, 2025.

[15478] Veronika Koren, Simone Blanco Malerba, Tilo Schwalger, and Stefano Panzeri. Efficient coding in biophysically realistic excitatory-inhibitory spiking networks. *eLife*, 13, 2025.

[15479] Joanna Kosinska, Julian C Assmann, Julica Inderhees, Helge Müller-Fielitz, Kristian Händler, Sven Geisler, Axel Künstner, Hauke Busch, Anna Worthmann, Joerg Heeren, Christian D Sadik, Matthias Gunzer, Vincent Prévot, Ruben Nogueiras, Misa Hirose, Malte Spielmann, Stefan Offermanns, Nina Wettschureck, and Markus Schwaninger. Diet modulates the therapeutic effects of dimethyl fumarate mediated by the immunometabolic neutrophil receptor HCAR2. *eLife*, 14, 2025.

[15480] Sofia V Krasik, Ekaterina A Bryushkova, George V Sharonov, Daria S Myalik, Elizaveta V Shurganova, Dmitry V Komarov, Irina A Shagina, Polina S Shpudeiko, Maria A Turchaninova, Maria T Vakhitova, Igor V Samoylenko, Dimitr T Marinov, Lev V Demidov, Vladimir E Zagaynov, Dmitriy M Chudakov, and Ekaterina O Serebrovskaya. Systematic evaluation of intratumoral and peripheral BCR repertoires in three cancers. *eLife*, 13, 2025.

[15481] Martin Krause, Wolfram Schulze, and Stefan Schuster. Learning and cognition in highspeed decision making. *eLife*, 13, 2025.

[15482] Lauren J Kreeger, Suraj Honnuraiah, Sydney Maeker, Siobhan Shea, Gordon Fishell, and Lisa Goodrich. An anatomical and physiological basis for flexible coincidence detection in the auditory system. *eLife*, 13, 2025.

[15483] Gabriela Krejčová and Adam Bajgar. Current insights into insect immune memory. *eLife*, 14, 2025.

[15484] Laura Sophie Kremer, Guanbin Gao, Giovanni Rigoni, Roberta Filograna, Mara Mennuni, Rolf Wibom, ákos Végvári, Camilla Koolmeister, and Nils-Göran Larsson. Tissue-specific responses to TFAM and mtDNA copy number manipulation in prematurely ageing mice. *eLife*, 14, 2025.

[15485] Seetha Krishnan, Can Dong, Heather Ratigan, Denisse Morales-Rodriguez, Chery Cherian, and Mark Sheffield. A contextual fear conditioning paradigm in head-fixed mice exploring virtual reality. *eLife*, 14, 2025.

[15486] Lisa M Kroell and Martin Rolfs. The magnitude and time course of pre-saccadic foveal prediction depend on the conspicuity of the saccade target. *eLife*, 12, 2025.

[15487] FranÇois Kroll, Joshua Donnelly, Güliz Gürel özcan, Eirinn Mackay, and Jason Rihel. Behavioural pharmacology predicts disrupted signalling pathways and candidate therapeutics from zebrafish mutants of Alzheimer's disease risk genes. *eLife*, 13, 2025.

[15488] Rachael Kuintzle, Leah A Santat, and Michael B Elowitz. Diversity in Notch ligand-receptor signaling interactions. *eLife*, 12, 2025.

[15489] Jessie Kulsuptrakul, Michael Emerman, and Patrick S Mitchell. CARD8 inflammasome activation during HIV-1 cell-to-cell transmission. *eLife*, 13, 2025.

[15490] Abhishek Kumar, Keiichiro Tanaka, and Martin A Schwartz. Focal adhesion-derived liquid-liquid phase separations regulate mRNA translation. *eLife*, 13, 2025.

[15491] Angelika Kunkel, Katharina Schmidt, Helena Hartmann, Torben Strietzel, Jens-Lennart Sperzel, Katja Wiech, and Ulrike Bingel. Nocebo effects are stronger and more persistent than placebo effects in healthy individuals. *eLife*, 14, 2025.

[15492] Raymond Laboy, Marjana Ndoci, Shamsh Tabrez Syed, Maximilian Vonolfen, Eugen Ballhysa, Tim Droth, Klara Schilling, Anna Loehrke, Ilian Atanassov, and Adam Antebi. Hexokinase regulates Mondo-mediated longevity via the PPP and organellar dynamics. *eLife*, 12, 2025.

[15493] Ludovica Labruna, Christina Merrick, Angel V Peterchev, Ben Inglis, Richard B Ivry, and Daniel Sheltraw. Kilohertz transcranial magnetic perturbation (kTMP) as a new non-invasive method to modulate cortical excitability. *eLife*, 13, 2025.

[15494] Kip D Lacy, Jina Lee, Kathryn Rozen-Gagnon, Wei Wang, Thomas S Carroll, and Daniel JC Kronauer. Heterozygosity at a conserved candidate sex determination locus is associated with female development in the clonal raider ant (Ooceraea biroi). *eLife*, 14, 2025.

[15495] Wei-Yun Lai, Sheng-Kai Hsu, Andreas Futschik, and Christian Schlötterer. Pleiotropy increases parallel selection signatures during adaptation from standing genetic variation. *eLife*, 13, 2025.

[15496] Poortata Lalwani, Thad Polk, and Douglas D Garrett. Modulation of brain signal variability in visual cortex reflects aging, GABA, and behavior. *eLife*, 14, 2025.

[15497] Agnès Landemard, Célian Bimbard, and Yves Boubenec. Hierarchical encoding of natural sound mixtures in ferret auditory cortex. *eLife*, 14, 2025.

[15498] Charles-francois V Latchoumane, Joon-Hyuk Lee, Seong-Wook Kim, Jinhyun Kim, and Hee-Sup Shin. Mediodorsal thalamic nucleus mediates resistance to ethanol through Cav3.1 T-type Ca2+ regulation of neural activity. *eLife*, 13, 2025.

[15499] Andrew P Latham, Longchen Zhu, Dina A Sharon, Songtao Ye, Adam P Willard, Xin Zhang, and Bin Zhang. Microphase separation produces interfacial environment within diblock biomolecular condensates. *eLife*, 12, 2025.

[15500] Marie-Alphée Laurent, Corentin Jacques, Xiaoqian Yan, Pauline Jurczynski, Sophie Colnat-Coulbois, Louis Maillard, Steven Le Cam, Radu Ranta, Benoit R Cottereau, Laurent Koessler, Jacques Jonas, and Bruno Rossion. A tight relationship between BOLD fMRI activation/deactivation and increase/decrease in single neuron responses in human association cortex. *eLife*, 14, 2025.

[15501] Irina Lazar-Contes, Rodrigo G Arzate-Mejia, Deepak K Tanwar, Leonard C Steg, Kerem Uzel, Olivier Ulrich Feudjio, Marion Crespo, Pierre-Luc Germain, and Isabelle M Mansuy. Dynamics of transcriptional programs and chromatin accessibility in mouse spermatogonial cells from early postnatal to adult life. *eLife*, 12, 2025.

[15502] Nguyet Le, Sherine Awad, Isabella Palazzo, Thanh Hoang, and Seth Blackshaw. Viral-mediated Pou5f1 (Oct4) overexpression and inhibition of Notch signaling synergistically induce neurogenic competence in mammalian Müller glia. *eLife*, 14, 2025.

[15503] Benjamin Le Gac, Marine Tournissac, Esther Belzic, Sandrine Picaud, Isabelle Dusart, Hédi Soula, Dongdong Li, Serge Charpak, and Bruno Cauli. Elevated pyramidal cell firing orchestrates arteriolar vasoconstriction through COX-2-derived prostaglandin E2 signaling. *eLife*, 13, 2025.

[15504] Théo Le Moigne, Martina Santoni, Lucile Jomat, Stéphane D Lemaire, Mirko Zaffagnini, Nicolas Chéron, and Julien Henri. Structure of the Calvin-Benson-Bassham sedoheptulose-1,7-bisphosphatase from the model microalga Chlamydomonas reinhardtii. *eLife*, 12, 2025.

[15505] Camille Le Roy, Nina Tervelde, Thomas Engels, and Florian T Muijres. Adaptations in wing morphology rather than wingbeat kinematics enable flight in small hoverfly species. *eLife*, 13, 2025.

[15506] Joséphine Ledamoisel, Bruno Buatois, Rémi Mauxion, Christine Andraud, Melanie McClure, Vincent Debat, and Violaine Llaurens. Convergent iridescence and divergent chemical signals in sympatric sister-species of Amazonian butterflies. *eLife*, 14, 2025.

[15507] Choongheon Lee, Mohammad Shokrian, Kenneth S Henry, Laurel H Carney, J Christopher Holt, and Jong-Hoon Nam. Outer hair cells stir cochlear fluids. *eLife*, 13, 2025.

[15508] Christopher R Lee, Gillian A Matthews, Mackenzie E Lemieux, Elizabeth M Wasserlein, Matilde Borio, Raymundo L Miranda, Laurel R Keyes, Gates P Schneider, Caroline Jia, Andrea Tran, Faith Aloboudi, May G Chan, Enzo Peroni, Grace Pereira, Alba López-Moraga, Anna Pallé, Eyal Y Kimchi, Nancy Padilla-Coreano, Romy Wichmann, and Kay M Tye. Separable dorsal raphe dopamine projections mimic the facets of a loneliness-like state. *eLife*, 14, 2025.

[15509] Dayeon Lee, Sin Young Choi, So-I Shin, Hyunsu An, Byeong-Sun Choi, and Jihwan Park. Multi-omics single-cell analysis reveals key regulators of HIV-1 persistence and aberrant host immune responses in early infection. *eLife*, 14, 2025.

[15510] Hanee Lee, Junsu Kang, Sang-Hee Lee, Dowoon Lee, Christine H Chung, and Junho Lee. Neuroprotective role of Hippo signaling by microtubule stability control in Caenorhabditis elegans. *eLife*, 13, 2025.

[15511] Hyun Jee Lee, Jingting Liang, Shivesh Chaudhary, Sihoon Moon, Zikai Yu, Taihong Wu, He Liu, Myung-Kyu Choi, Yun Zhang, and Hang Lu. Automated cell annotation in multi-cell images using an improved CRF_ID algorithm. *eLife*, 12, 2025.

[15512] Juhee Lee, Wenying Shou, and Hye Jin Park. The success of artificial selection for collective composition hinges on initial and target values. *eLife*, 13, 2025.

[15513] Junghoon Lee and Scot P Ouellette. The second messenger signaling molecule cyclic di-AMP drives developmental cycle progression in Chlamydia trachomatis. *eLife*, 14, 2025.

[15514] Ka Kiu Lee, Urszula Łapińska, Giulia Tolle, Maureen Micaletto, Bing Zhang, Wanida Phetsang, Anthony D Verderosa, Brandon M Invergo, Joseph Westley, Attila Bebes, Raif Yuecel, Paul A O'Neill, Audrey Farbos, Aaron R Jeffries, Stineke van Houte, Pierluigi Caboni, Mark AT Blaskovich, Benjamin E Housden, Krasimira Tsaneva-Atanasova, and Stefano Pagliara. Heterogeneous efflux pump expression underpins phenotypic resistance to antimicrobial peptides. *eLife*, 13, 2025.

[15515] Kacie Lee and Reuben Rideaux. The influence of temporal context on vision over multiple time scales. *eLife*, 14, 2025.

[15516] Raz Leib and David Franklin. Error prediction determines the coordinate system used for the representation of novel dynamics. *eLife*, 14, 2025.

[15517] Hans C Leier, Alexander J Foden, Darren A Jindal, Abigail J Wilkov, Paola Van der Linden Costello, Pamela J Vanderzalm, Jaeda Coutinho-Budd, Masashi Tabuchi, and Heather T Broihier. Glia control experience-dependent plasticity in an olfactory critical period. *eLife*, 13, 2025.

[15518] Alannah Lejeune, Chunyi Zhou, Defne Ercelen, Gregory Putzel, Xiaomin Yao, Alyson R Guy, Miranda Pawline, Magdalena Podkowik, Alejandro Pironti, Victor J Torres, Bo Shopsin, and Ken Cadwell. Sex-dependent gastrointestinal colonization resistance to MRSA is microbiota and Th17 dependent. *eLife*, 13, 2025.

[15519] Orel Levy, Shirley Libman Hackmon, Yair Zvilichovsky, Adi Korisky, Aurelie Bidet-Caulet, Julie B Schweitzer, and Elana Zion Golumbic. Selective attention and sensitivity to auditory disturbances in a virtually real classroom. *eLife*, 13, 2025.

[15520] Hang Li, Shengwei Ji, Nanang R Ariefta, Eloiza May S Galon, Shimaa AES El-Sayed, Thom Do, Lijun Jia, Miako Sakaguchi, Masahito Asada, Yoshifumi Nishikawa, Xin Qin, Mingming Liu, and Xuenan Xuan. Efficacy and mechanism of action of cipargamin as an antibabesial drug candidate. *eLife*, 13, 2025.

[15521] Hao Li, Izumi Ishizaki, Koji Kato, Xiaomei Sun, Sachiko Muramatsu, Hiroshi Itou, Toyoyuki Ose, Hiroyuki Araki, and Min Yao. Sld3CBD–Cdc45 structural insights into Cdc45 recruitment for CMG complex formation during DNA replication. *eLife*, 13, 2025.

[15522] Hong-Fei Li, Bao Dong, Yuan-Yuan Peng, Hao-Yue Luo, Xiao-Lan Ou, Zheng-Lin Ren, Yoonseong Park, Jin-Jun Wang, and Hong-Bo Jiang. The neuropeptide sulfakinin is a peripheral regulator of insect behavioral switch between mating and foraging. *eLife*, 13, 2025.

[15523] Hongchen Li, Xinchao Zhang, Xiaoben Liang, Shuyan Li, Ziyi Cui, Xinyu Zhao, Kai Wang, Bingbing Zha, Haijie Ma, Ming Xu, Lei Lv, and Yanping Xu. HNF4$\alpha$-TET2-FBP1 axis contributes to gluconeogenesis and type 2 diabetes. *eLife*, 13, 2025.

[15524] Hui Li, Jun Yang, Su-fang Kuang, Huan-zhe Fu, Hui-yin Lin, and Bo Peng. Magnesium modulates phospholipid metabolism to promote bacterial phenotypic resistance to antibiotics. *eLife*, 13, 2025.

[15525] Jingjing Li, Xinyue Wang, Laia Jordana, éric Bonneil, Victoria Ginestet, Momina Ahmed, Mohammed Bourouh, Cristina Mirela Pascariu, T Martin Schmeing, Pierre Thibault, and Vincent Archambault. Mechanisms of PP2A-Ankle2 dependent nuclear reassembly after mitosis. *eLife*, 13, 2025.

[15526] Kailai Li, Junyi Liang, Nan Li, Jianbo Fang, Xinyi Zhou, Jian Zhang, Anqi Lin, Peng Luo, and Hui Meng. A multi-gene predictive model for the radiation sensitivity of nasopharyngeal carcinoma based on machine learning. *eLife*, 13, 2025.

[15527] Keva Li, Nicholas Tolman, Ayellet V Segrè, Kelsey V Stuart, Oana A Zeleznik, Neeru A Vallabh, Kuang Hu, Nazlee Zebardast, Akiko Hanyuda, Yoshihiko Raita, Christa Montgomery, Chi Zhang, Pirro G Hysi, Ron Do, Anthony P Khawaja, Janey L Wiggs, Jae H Kang, Simon WM John, and Louis R Pasquale. Pyruvate and related energetic metabolites modulate resilience against high genetic risk for glaucoma. *eLife*, 14, 2025.

[15528] Lei Li, Guanghao Wu, Xiaolei Xu, Junling Yang, Lirong Yi, Ziqing Yang, Zheng Mo, Li Xing, Ying Shan, Zhuo Yu, and Yinchuan Li. The effects of 17$\alpha$-estradiol treatment on endocrine system revealed by single-nucleus transcriptomic sequencing of hypothalamus. *eLife*, 13, 2025.

[15529] Luhe Li, Fangfang Hong, Stephanie Badde, and Michael S Landy. Precision-based causal inference modulates audiovisual temporal recalibration. *eLife*, 13, 2025.

[15530] Mengjia Li, Hengchao Zhang, Xiuyun Wu, Mengqi Yu, Qianqian Yang, Lei Sun, Wei Li, Zhongxing Jiang, Fumin Xue, Ting Wang, Xuili An, and Lixiang Chen. IDH1 regulates human erythropoiesis by eliciting chromatin state reprogramming. *eLife*, 13, 2025.

[15531] Mengjie Li, Aiguo Tian, and Jin Jiang. Numb provides a fail-safe mechanism for intestinal stem cell self-renewal in adult Drosophila midgut. *eLife*, 14, 2025.

[15532] Mengyun Li, Huan Jin, Yongxiang Liu, Zining Wang, Lin Li, Tiantian Wang, Xiaojuan Wang, Hongxia Zhang, Bitao Huo, Tiantian Yu, Shoujie Wang, Wei Zhao, Jinyun Liu, Peng Huang, Jun Cui, and Xiaojun Xia. Aurora kinase A promotes trained immunity via regulation of endogenous S-adenosylmethionine metabolism. *eLife*, 14, 2025.

[15533] Peng Li, Sree Pulugulla, Sonali Das, Jangsuk Oh, Rosanne Spolski, Jian-Xin Lin, and Warren J Leonard. A new pipeline SPICE identifies novel JUN-IKZF1 composite elements. *eLife*, 12, 2025.

[15534] Shu Li, Yi Zou, Austin McMasters, Fuxiang Chen, and Jun Yan. Trained immunity: A new player in cancer immunotherapy. *eLife*, 14, 2025.

[15535] Tianhan Li, Liaoxun Lu, Juanjuan Qiu, Xin Dong, Le Yang, Kexin He, Yanrong Gu, Binhui Zhou, Tingting Jia, Toby Lawrence, Marie Malissen, Guixue Wang, Rong Huang, Hui Wang, Bernard Malissen, Yinming Liang, and Lichen Zhang. Xcr1+ type 1 conventional dendritic cells are essential mediators for atherosclerosis progression. *eLife*, 14, 2025.

[15536] Tingbo Li, Ling Yang, Jiaqi Tu, Yufan Hao, Zhu Zhu, Yingjie Xiong, Qingzhu Gao, Lili Zhou, Guanglei Xie, Dongdong Zhang, Xuzhao Li, Yuxiao Jin, Yiyi Zhang, Bingrui Zhao, Nan Li, Xi Wang, and Jie-Min Jia. The myeloid cell-driven transdifferentiation of endothelial cells into pericytes promotes the restoration of BBB function and brain self-repair after stroke. *eLife*, 14, 2025.

[15537] Wenqing Li, Sara McCurdy, Miguel A Lopez-Ramirez, Ho-Sup Lee, and Mark H Ginsberg. Genetic inactivation of the $\beta$1 adrenergic receptor prevents cerebral cavernous malformations in zebrafish. *eLife*, 13, 2025.

[15538] Yafen Li, Yixuan Lin, Qing Li, Yongqiang Chen, Zhifang Li, and Antao Chen. Temporal dynamics analysis reveals that concurrent working memory load eliminates the Stroop effect through disrupting stimulus-response mapping. *eLife*, 13, 2025.

[15539] Yan-Xue Li, Xin-Le Kang, Yan-Li Li, Xiao-Pei Wang, Qiao Yan, Jin-Xing Wang, and Xiao-Fan Zhao. Receptor tyrosine kinases CAD96CA and FGFR1 function as the cell membrane receptors of insect juvenile hormone. *eLife*, 13, 2025.

[15540] Yao Li, Hui Xin, Zhexun Lian, and Wei Zhang. Exploration of the metabolomic mechanisms of postmenopausal hypertension induced by low estrogen state. *eLife*, 13, 2025.

[15541] Yi Li, Long Gong, Jing Wu, Wesley Hung, Mei Zhen, and Shangbang Gao. UBR-1 deficiency leads to ivermectin resistance in Caenorhabditis elegans. *eLife*, 13, 2025.

[15542] Zhiheng Li, Jinsheng Hu, Thomas A Stidham, Mao Ye, Min Wang, Yanhong Pan, Tao Zhao, Jingshu Li, Zhonghe Zhou, and Julia A Clarke. Iridescent structural coloration in a crested Cretaceous enantiornithine bird from the Jehol Biota. *eLife*, 14, 2025.

[15543] Zhu-Hong Li, Beejan Asady, Le Chang, Myriam Andrea Hortua Triana, Catherine Li, Isabelle Coppens, and Silvia NJ Moreno. Calcium transfer from the ER to other organelles for optimal signaling in Toxoplasma gondii. *eLife*, 13, 2025.

[15544] Zongze Li, Lucia Fernandez Cardo, Michal Rokicki, Jimena Monzón-Sandoval, Viola Volpato, Frank Wessely, Caleb Webber, and Meng Li. Changes in neural progenitor lineage composition during astrocytic differentiation of human iPSCs. *eLife*, 13, 2025.

[15545] Sergiy Libert, Alex Chekholko, and Cynthia Kenyon. A mathematical model that predicts human biological age from physiological traits identifies environmental and genetic factors that influence aging. *eLife*, 13, 2025.

[15546] Carmina Lichauco, Eric J Foss, Tonibelle Gatbonton-Schwager, Nelson F Athow, Brandon Lofts, Robin Acob, Erin Taylor, James J Marquez, Uyen Lao, Shawna Miles, and Antonio Bedalov. Sir2 and Fun30 regulate ribosomal DNA replication timing via MCM helicase positioning and nucleosome occupancy. *eLife*, 13, 2025.

[15547] Clémence Ligneul, Lily Qiu, William T Clarke, Saad Jbabdi, Marco Palombo, and Jason P Lerch. Diffusion MRS tracks distinct trajectories of neuronal development in the cerebellum and thalamus of rat neonates. *eLife*, 13, 2025.

[15548] Tony KY Lim, Anne Ritoux, Luke W Paine, Larissa Ferguson, Tawab Abdul, Laura J Grundy, and Ewan St John Smith. Cap-independent co-expression of dsRNA-sensing and NF-$\kappa$B pathway inhibitors enables controllable self-amplifying RNA expression with reduced immunotoxicity. *eLife*, 14, 2025.

[15549] Susan Lin, Aravind R Gade, Hong-Gang Wang, James E Niemeyer, Allison Galante, Isabella DiStefano, Patrick Towers, Jorge Nunez, Maiko Matsui, Theodore H Schwartz, Anjali Rajadhyaksha, and Geoffrey S Pitt. Interneuron FGF13 regulates seizure susceptibility via a sodium channel-independent mechanism. *eLife*, 13, 2025.

[15550] Yi-Hsuan Lin, Tae Hun Kim, Suman Das, Tanmoy Pal, Jonas Wessén, Atul Kaushik Rangadurai, Lewis E Kay, Julie D Forman-Kay, and Hue Sun Chan. Electrostatics of salt-dependent reentrant phase behaviors highlights diverse roles of ATP in biomolecular condensates. *eLife*, 13, 2025.

[15551] Jack W Lindsey, Jeffrey Markowitz, Winthrop F Gillis, Sandeep R Datta, and Ashok Litwin-Kumar. Dynamics of striatal action selection and reinforcement learning. *eLife*, 13, 2025.

[15552] Doris Ling, Lijun Zhang, Debajit Saha, Alex Bo-Yuan Chen, and Baranidharan Raman. Adaptation invariant concentration discrimination in an insect olfactory system. *eLife*, 12, 2025.

[15553] Haibin Liu, Lulu Yu, Vladimir Majerciak, Thomas J Meyer, Ming Yi, Peter F Johnson, Maggie Cam, Douglas R Lowy, and Zhi-Ming Zheng. The long noncoding RNA lnc-FANCI-2 intrinsically restricts RAS signaling in human papillomavirus type 16-infected cervical cancer cells. *eLife*, 13, 2025.

[15554] Haowen Liu, Lei Li, Jiafan Wang, Jiayi Hu, Jingyao Xia, Xiaochun Yu, Jing Tang, Huisheng Liu, Xiaofei Yang, Cong Ma, Lijun Kang, and Zhitao Hu. Mechanisms that regulate the C1-C2B mutual inhibition control functional switch of UNC-13. *eLife*, 14, 2025.

[15555] Jianheng Fox Liu, Ben R Hawley, Luke S Nicholson, and Samie R Jaffrey. Decoding m6Am by simultaneous transcription-start mapping and methylation quantification. *eLife*, 13, 2025.

[15556] Jianing Liu, Anokhi Shah, Xinyu Liu, Joshua L Wort, Yue Ma, Katie Hardman, Niklas G Johansson, Orquidea Ribeiro, Adam Brookfield, Alice Bowen, Jari Yli-Kauhaluoma, Henri Xhaard, Lars JC Jeuken, Adrian Goldman, Christos Pliotas, and Keni Vidilaseris. Conformational dynamics and asymmetry in multimodal inhibition of membrane-bound pyrophosphatases. *eLife*, 13, 2025.

[15557] Lanfang Liu, Jiahao Jiang, Hehui Li, and Guosheng Ding. Tripartite organization of brain state dynamics underlying spoken narrative comprehension. *eLife*, 13, 2025.

[15558] Lilong Liu, Zhenghao Liu, Lei Fan, Zhipeng Yao, Junyi Hu, Yaxin Hou, Yang Li, Yuhong Ding, Yingchun Kuang, Ke Chen, Yi Hao, and Zheng Liu. An evaluation of the tumor microenvironment through CALR, IL1R1, IFNB1, and IFNG to assess prognosis and immunotherapy response in bladder cancer patients. *eLife*, 13, 2025.

[15559] Meiqiu Liu, Yue Chen, and Rongqing Chen. Social rank-order stability of mice revealed by a novel food competition paradigm in combination with available space competition paradigms. *eLife*, 14, 2025.

[15560] Ning Liu, Wenan Qiang, Philip W Jordan, John F Marko, and Huanyu Qiao. Cell cycle and age-related modulations of mouse chromosome stiffness. *eLife*, 13, 2025.

[15561] Sainan Liu, Jiepin Huang, Suhao Chen, Michael L Platt, and Yan Yang. Multi-dimensional social relationships shape social attention in monkeys. *eLife*, 14, 2025.

[15562] Shixuan Liu, Ceryl Tan, Chloe Melo-Gavin, Miriam B Ginzberg, Ron Blutrich, Nish Patel, Michael Rape, Kevin G Mark, and Ran Kafri. Oversized cells activate global proteasome-mediated protein degradation to maintain cell size homeostasis. *eLife*, 14, 2025.

[15563] Xu Liu, Yuchen Xie, Shengzhi Yang, Cong Jiang, Ke Shang, Jinxia Luo, Lin Zhang, Gang Hu, Qinghua Liu, Bisong Yue, Zhenxin Fan, Zhanlong He, and Jing Li. Multi-omics investigation of spontaneous T2DM macaque emphasizes gut microbiota could up-regulate the absorption of excess palmitic acid in the T2DM progression. *eLife*, 14, 2025.

[15564] Yating Liu, Guojing Tian, Ziyi Wang, Junkang Zheng, Huimin Liu, Sucheng Zhu, Zhao Shan, and Bin Qi. Neuronal detection triggers systemic digestive shutdown in response to adverse food sources in Caenorhabditis elegans. *eLife*, 14, 2025.

[15565] Ying Liu, Rajani Karmakar, Maria Billini, Wieland Steinchen, Saumyak Mukherjee, Rogelio Hernandez-Tamayo, Thomas Heimerl, Gert Bange, Lars V Schäfer, and Martin Thanbichler. Membrane binding properties of the cytoskeletal protein bactofilin. *eLife*, 13, 2025.

[15566] Yisi Liu, Pu Wang, Junjie Zou, and Hongwei Zhou. A novel method (RIM-Deep) for enhancing imaging depth and resolution stability of deep cleared tissue in inverted confocal microscopy. *eLife*, 13, 2025.

[15567] Julianna Locantore, Yijun Liu, Jesse White, Janet Berrios Wallace, Celia Beron, Emily Kraft, Bernardo Sabatini, and Michael Wallace. Mixed representations of choice direction and outcome by GABA/glutamate cotransmitting neurons in the entopeduncular nucleus. *eLife*, 13, 2025.

[15568] Gabriel Loewinger, Erjia Cui, David Lovinger, and Francisco Pereira. A statistical framework for analysis of trial-level temporal dynamics in fiber photometry experiments. *eLife*, 13, 2025.

[15569] Maxine K Loh, Samantha J Hurh, Paula Bazzino, Rachel M Donka, Alexandra T Keinath, Jamie D Roitman, and Mitchell F Roitman. Dopamine activity encodes the changing valence of the same stimulus in conditioned taste aversion paradigms. *eLife*, 13, 2025.

[15570] Juan Long, Meng Ma, Yuting Chen, Bo Gong, Yi Zheng, Hao Li, and Jing Yang. Deletion of sulfate transporter SUL1 extends yeast replicative lifespan via reduced PKA signaling instead of decreased sulfate uptake. *eLife*, 13, 2025.

[15571] Adam D Longhurst, Kyle Wang, Harsha Garadi Suresh, Mythili Ketavarapu, Henry N Ward, Ian R Jones, Vivek Narayan, Frances V Hundley, Arshia Zernab Hassan, Charles Boone, Chad L Myers, Yin Shen, Vijay Ramani, Brenda J Andrews, and David P Toczyski. The PRC2.1 subcomplex opposes G1 progression through regulation of CCND1 and CCND2. *eLife*, 13, 2025.

[15572] Ana Teresa López-Jiménez, Dominik Brokatzky, Kamla Pillay, Tyrese Williams, Gizem özbaykal Güler, and Serge Mostowy. High-content high-resolution microscopy and deep learning-assisted analysis reveals host and bacterial heterogeneity during Shigella infection. *eLife*, 13, 2025.

[15573] Anastasiia Lozovska, Ana Casaca, Ana Novoa, Ying-Yi Kuo, Arnon D Jurberg, Gabriel G Martins, Anna-Katerina Hadjantonakis, and Moises Mallo. Tgfbr1 regulates lateral plate mesoderm and endoderm reorganization during the trunk to tail transition. *eLife*, 13, 2025.

[15574] Han Lu, Sandra Diaz-Pier, Maximilian Lenz, and Andreas Vlachos. The interplay between homeostatic synaptic scaling and homeostatic structural plasticity maintains the robust firing rate of neural networks. *eLife*, 12, 2025.

[15575] Jing Lu, Hao Xu, Dongyue Wang, Yanlu Chen, Takeshi Inoue, Liang Gao, and Kai Lei. 3D reconstruction of neuronal allometry and neuromuscular projections in asexual planarians using expansion tiling light sheet microscopy. *eLife*, 13, 2025.

[15576] Hella Luksch, Felix Schulze, David Geißler-Lösch, David Sprott, Lennart Höfs, Eva M Szegö, Wulf Tonnus, Stefan Winkler, Claudia Günther, Andreas Linkermann, Rayk Behrendt, Lino L Teichmann, Björn H Falkenburger, and Angela Rösen-Wolff. Tissue inflammation induced by constitutively active STING is mediated by enhanced TNF signaling. *eLife*, 14, 2025.

[15577] Dong Luo, Jing Zheng, Shuning Lv, Ren Sheng, Maorong Chen, Xi He, and Xinjun Zhang. Wnt induces FZD5/8 endocytosis and degradation and the involvement of RSPO-ZNRF3/RNF43 and DVL. *eLife*, 14, 2025.

[15578] Menguxn Luo, Jia Zhou, Cailu Sun, Wanjia Chen, Chaoying Fu, Chenfang Si, Yaoyang Zhang, Yang Geng, and Yelin Chen. APP $\beta$-CTF triggers cell-autonomous synaptic toxicity independent of A$\beta$. *eLife*, 13, 2025.

[15579] Wencong Lyu, Haochen Wang, Zhehao Du, Ran Wei, Jianuo He, Fanju Meng, Jinlong Bi, Lijun Zhang, Chao Zhang, Yiting Guan, and Wei Tao. Enalapril mitigates senescence and aging-related phenotypes in human cells and mice via pSmad1/5/9-driven antioxidative genes. *eLife*, 14, 2025.

[15580] Congcong Ma, Zhaoxing Jia, Tianxiang Jiang, Qian Cai, Jinding Yang, Lin Gan, Kecheng Qian, Zixin Pan, Qinyu Ye, Mengyuan Dai, and Xianming Lin. Single-cell transcriptome sequencing for opening the blood-brain barrier through specific mode electroacupuncture stimulation. *eLife*, 14, 2025.

[15581] Mengqi Ma, Yiming Zheng, Mingxi Deng, Shenzhao Lu, Xueyang Pan, Xi Luo, Michelle Etoundi, David Li-Kroeger, Kim C Worley, Lindsay C Burrage, Lauren S Blieden, Aimee Allworth, Wei-Liang Chen, Giuseppe Merla, Barbara Mandriani, Catherine E Otten, Pierre Blanc, Jill A Rosenfeld, Debdeep Dutta, Shinya Yamamoto, Michael F Wangler, Ian A Glass, Jingheng Chen, Elizabeth Blue, Paolo Prontera, Jeremie Rosain, Sandrine Marlin, Seema R Lalani, and Hugo J Bellen. Heterozygous variants in PLCG1 affect hearing, vision, cardiac, and immune function. *eLife*, 13, 2025.

[15582] Pingchuan Ma, Peter Chen, Scott Sternson, and Yao Chen. The promise and peril of comparing fluorescence lifetime in biology revealed by simulations. *eLife*, 13, 2025.

[15583] Shuai Ma, Bin Yang, Yuyang Sun, Xinyue Wang, Houliang Guo, Ruiying Liu, Ting Ye, Chenbo Kang, Jingnan Chen, and Lingyan Jiang. Salmonella exploits host- and bacterial-derived $\beta$-alanine for replication inside host macrophages. *eLife*, 13, 2025.

[15584] Xin Ma, Meng Li, Yuanyuan Zhang, Tingting Xu, Xinchen Zhou, Mengqi Qian, Zhiren Yang, and Xinyan Han. Akkermansia muciniphila identified as key strain to alleviate gut barrier injury through Wnt signaling pathway. *eLife*, 12, 2025.

[15585] Fabienne Maassen, Vu Thuy Khanh Le-Trilling, Luisa Betke, Thilo Bracht, Corinna Siegmund, Malte Bayer, Benjamin Katschinski, Antonia Belter, Tanja Becker, Denise Mennerich, Sebastian Voigt, Lori Frappier, Barbara Sitek, Katharina Fleischhauer, and Mirko Trilling. The human cytomegalovirus-encoded pUS28 antagonizes CD4+ T cell recognition by targeting CIITA. *eLife*, 14, 2025.

[15586] HaDi MaBouDi, Jasmin Richter, Marie-Geneviève Guiraud, Mark Roper, James AR Marshall, and Lars Chittka. Active vision of bees in a simple pattern discrimination task. *eLife*, 14, 2025.

[15587] HaDi MaBouDi, Mark Roper, Marie-Geneviève Guiraud, Mikko Juusola, Lars Chittka, and James AR Marshall. A neuromorphic model of active vision shows how spatiotemporal encoding in lobula neurons can aid pattern recognition in bees. *eLife*, 14, 2025.

[15588] Matt MacAinsh, Sanbo Qin, and Huan-Xiang Zhou. DIRseq as a method for predicting drug-interacting residues of intrinsically disordered proteins from sequences. *eLife*, 14, 2025.

[15589] Donald Iain MacDonald, Monessha Jayabalan, Jonathan T Seaman, Rakshita Balaji, Alec R Nickolls, and Alexander Theodore Chesler. Pain persists in mice lacking both Substance P and CGRP$\alpha$ signaling. *eLife*, 13, 2025.

[15590] Pragyansree Machhua, Vignesh Gopalakrishnan Unnithan, Yu Liu, Yiping Jiang, Lingfeng Zhang, and Zhihong Guo. Daptomycin forms a stable complex with phosphatidylglycerol for selective uptake to bacterial membrane. *eLife*, 13, 2025.

[15591] Nagatoshi Machii, Ryo Hatashima, Tatsuya Niwa, Hideki Taguchi, Ismael A Kimirei, Hillary DJ Mrosso, Mitsuto Aibara, Tatsuki Nagasawa, and Masato Nikaido. Pronounced expression of extracellular matrix proteoglycans regulated by Wnt pathway underlies the parallel evolution of lip hypertrophy in East African cichlids. *eLife*, 13, 2025.

[15592] Ricardo J Mack, Natasha M Flores, Geoffrey C Fox, Hanyang Dong, Metehan Cebeci, Simone Hausmann, Tourkian Chasan, Jill M Dowen, Brian D Strahl, Pawel K Mazur, and Or Gozani. SETD2 suppresses tumorigenesis in a KRASG12C-driven lung cancer model, and its catalytic activity is regulated by histone acetylation. *eLife*, 14, 2025.

[15593] Marisa Mackie, Vivian Vy Le, Heather R Carstensen, Nicole R Kushnir, Dylan L Castro, Ivan M Dimov, Kathleen T Quach, Steven J Cook, Oliver Hobert, Sreekanth H Chalasani, and Ray L Hong. Evolution of lateralized gustation in nematodes. *eLife*, 14, 2025.

[15594] Pranav Mahajan, Shuangyi Tong, Sang Wan Lee, and Ben Seymour. Balancing safety and efficiency in human decision-making. *eLife*, 13, 2025.

[15595] Colleen A Maillie, Kiana Golden, Ian A Wilson, Andrew B Ward, and Marco Mravic. Ab initio prediction of specific phospholipid complexes and membrane association of HIV-1 MPER antibodies by multi-scale simulations. *eLife*, 12, 2025.

[15596] Anastasia A Makarova, Nicholas J Chua, Anna V Diakova, Inna A Desyatirkina, Pat Gunn, Song Pang, C Shan Xu, Harald F Hess, Dmitri B Chklovskii, and Alexey A Polilov. The first complete 3D reconstruction and morphofunctional mapping of an insect eye. *eLife*, 14, 2025.

[15597] Alina Malita, Anne H Skakkebaek, Olga Kubrak, Xiaokang Chen, Takashi Koyama, Elizabeth C Connolly, Nadja Ahrentloev, Ditte S Andersen, Michael J Texada, Kenneth Halberg, and Kim Rewitz. Glia-mediated gut–brain cytokine signaling couples sleep to intestinal inflammatory responses induced by oxidative stress. *eLife*, 13, 2025.

[15598] Jennifer March and Sebastian Gluth. Hunger shifts attention and attribute weighting in dietary choice. *eLife*, 13, 2025.

[15599] Tony Marchand, Kemi E Akinnola, Shoichiro Takeishi, Maria Maryanovich, Sandra Pinho, Julien Saint-Vanne, Alexander Birbrair, Thierry Lamy, Karin Tarte, Paul Frenette, and Kira Gritsman. Periosteal skeletal stem cells can migrate into the bone marrow and support hematopoiesis after injury. *eLife*, 13, 2025.

[15600] Julia M Marchingo, Laura Spinelli, Shalini Pathak, and Doreen A Cantrell. PIM kinase control of CD8 T cell protein synthesis and cell trafficking. *eLife*, 13, 2025.

[15601] Andrew Margolis and Andrew Gordus. A stochastic explanation for observed local-to-global foraging states in Caenorhabditis elegans. *eLife*, 14, 2025.

[15602] Alba Marino, Gautier Debaecker, Anna-Sophie Fiston-Lavier, Annabelle Haudry, and Benoit Nabholz. Effective population size does not explain long-term variation in genome size and transposable element content in animals. *eLife*, 13, 2025.

[15603] M Julia Maristany, Anne Aguirre Gonzalez, Jorge R Espinosa, Jan Huertas, Rosana Collepardo-Guevara, and Jerelle A Joseph. Decoding phase separation of prion-like domains through data-driven scaling laws. *eLife*, 13, 2025.

[15604] Francesco Marocco, Sabrina Garbo, Claudia Montaldo, Alessio Colantoni, Luca Quattrocchi, Gioele Gaboardi, Giovanna Sabarese, Carla Cicchini, Mario Lecce, Alessia Carnevale, Rossella Paolini, Gian Gaetano Tartaglia, Cecilia Battistelli, and Marco Tripodi. Negative regulation of miRNA sorting into EVs is mediated by the capacity of RBP PCBP2 to impair the SYNCRIP-dependent miRNA loading. *eLife*, 14, 2025.

[15605] Benita Martin-Castaño, Patricia Diez-Echave, Jorge García-García, Laura Hidalgo-García, Antonio Jesús Ruiz-Malagon, José Alberto Molina-Tijeras, María Jesús Rodríguez-Sojo, Anaïs Redruello-Romero, Margarita Martínez-Zaldívar, Emilio Mota, Fernando Cobo, Xando Díaz-Villamarin, Marta Alvarez-Estevez, Federico García, Concepción Morales-García, Silvia Merlos, Paula Garcia-Flores, Manuel Colmenero-Ruiz, José Hernández-Quero, Maria Nuñez, Maria Elena Rodriguez-Cabezas, Angel Carazo, Javier Martin, Rocio Moron, Alba Rodríguez Nogales, and Julio Galvez. The relationship between gut and nasopharyngeal microbiome composition can predict the severity of COVID-19. *eLife*, 13, 2025.

[15606] Daisuke Mashiko, Shingo Tonai, Haruhiko Miyata, Martin M Matzuk, and Masahito Ikawa. SLC35G3 is a UDP-N-acetylglucosamine transporter for sperm glycoprotein formation and underpins male fertility in mice. *eLife*, 14, 2025.

[15607] Claire Masson, Cyril Scandola, Jean-Yves Rinckel, Fabienne Proamer, Emily Janus-Bell, Fareeha Batool, Naël Osmani, Jacky G Goetz, Léa Mallo, Nathalie Brouard, Catherine Leon, Alicia Bornert, Renaud Poincloux, Olivier Destaing, Alma Mansson, Hong Qian, Maxime Lehmann, and Anita Eckly. Megakaryocytes assemble a three-dimensional cage of extracellular matrix that controls their maturation and anchoring to the vascular niche. *eLife*, 14, 2025.

[15608] Christopher S McAllester and John E Pool. The potential of inversions to accumulate balanced sexual antagonism is supported by simulations and Drosophila experiments. *eLife*, 12, 2025.

[15609] Sean McCracken, Liam McCoy, Ziyi Hu, Julie A Hodges, Katia Valkova, Philip R Williams, and Josh L Morgan. Mistargeted retinal axons induce a synaptically independent subcircuit in the visual thalamus of albino mice. *eLife*, 13, 2025.

[15610] Matthew B McDougal, Ian N Boys, Anthony M De Maria, Emi Nakahara, and John W Schoggins. IFIT1 is rapidly evolving and exhibits disparate antiviral activities across 11 mammalian orders. *eLife*, 13, 2025.

[15611] David Trombley McSwiggen, Helen Liu, Ruensern Tan, Sebastia Agramunt Puig, Lakshmi B Akella, Russell Berman, Mason Bretan, Hanzhe Chen, Xavier Darzacq, Kelsey Ford, Ruth Godbey, Eric Gonzalez, Adi Hanuka, Alec Heckert, Jaclyn J Ho, Stephanie L Johnson, Reed Kelso, Aaron Klammer, Ruchira Krishnamurthy, Jifu Li, Kevin Lin, Brian Margolin, Patrick McNamara, Laurence

Meyer, Sarah E Pierce, Akshay Sule, Connor Stashko, Yangzhong Tang, Daniel J Anderson, and Hilary P Beck. A high-throughput platform for single-molecule tracking identifies drug interaction and cellular mechanisms. *eLife*, 12, 2025.

[15612] Duncan S Mearns, Sydney A Hunt, Martin W Schneider, Ash V Parker, Manuel Stemmer, and Herwig Baier. Diverse prey capture strategies in teleost larvae. *eLife*, 13, 2025.

[15613] Joshua G Medina-Feliciano, Griselle Valentín-Tirado, Kiara Luna-Martínez, Alejandra Beltran-Rivera, Yamil Miranda-Negrón, and José E Garcia-Arraras. Single-cell RNA sequencing of the holothurian regenerating intestine reveals the pluripotency of the coelomic epithelium. *eLife*, 13, 2025.

[15614] Sara Z Mehrhof and Camilla L Nord. A common alteration in effort-based decision-making in apathy, anhedonia, and late circadian rhythm. *eLife*, 13, 2025.

[15615] Iti Mehta, Jacob B Hogins, Sydney R Hall, Gabrielle Vragel, Sankalya Ambagaspitiye, Philippe E Zimmern, and Larry Reitzer. Control of pili synthesis and putrescine homeostasis in Escherichia coli. *eLife*, 13, 2025.

[15616] Stefanie SM Meier, Michael Hörzing, Cornelia Böhm, Emma LR Düthorn, Heikki Takala, René Uebe, and Andreas Möglich. Engineering NIR-sighted bacteria. *eLife*, 14, 2025.

[15617] Geoffrey W Meissner, Allison Vannan, Jennifer Jeter, Kari Close, Gina M DePasquale, Zachary Dorman, Kaitlyn Forster, Jaye Anne Beringer, Theresa Gibney, Joanna H Hausenfluck, Yisheng He, Kristin Henderson, Lauren Johnson, Rebecca M Johnston, Gudrun Ihrke, Nirmala A Iyer, Rachel Lazarus, Kelley Lee, Hsing-Hsi Li, Hua-Peng Liaw, Brian Melton, Scott Miller, Reeham Motaher, Alexandra Novak, Omotara Ogundeyi, Alyson Petruncio, Jacquelyn Price, Sophia Protopapas, Susana Tae, Jennifer Taylor, Rebecca Vorimo, Brianna Yarbrough, Kevin Xiankun Zeng, Christopher T Zugates, Heather Dionne, Claire Angstadt, Kelly Ashley, Amanda Cavallaro, Tam Dang, Guillermo A Gonzalez, Karen L Hibbard, Cuizhen Huang, Jui-Chun Kao, Todd Laverty, Monti Mercer, Brenda Perez, Scarlett Rose Pitts, Danielle Ruiz, Viruthika Vallanadu, Grace Zhiyu Zheng, Cristian Goina, Hideo Otsuna, Konrad Rokicki, Robert R Svirskas, Han SJ Cheong, Michael-John Dolan, Erica Ehrhardt, Kai Feng, Basel EI Galfi, Jens Goldammer, Stephen J Huston, Nan Hu, Masayoshi Ito, Claire McKellar, Ryo Minegishi, Shigehiro Namiki, Aljoscha Nern, Catherine E Schretter, Gabriella R Sterne, Lalanti Venkatasubramanian, Kaiyu Wang, Tanya Wolff, Ming Wu, Reed George, Oz Malkesman, Yoshinori Aso, Gwyneth M Card, Barry J Dickson, Wyatt Korff, Kei Ito, James W Truman, Marta Zlatic, and Gerald M Rubin. A split-GAL4 driver line resource for Drosophila neuron types. *eLife*, 13, 2025.

[15618] Claire Meissner-Bernard, Friedemann Zenke, and Rainer W Friedrich. Geometry and dynamics of representations in a precisely balanced memory network related to olfactory cortex. *eLife*, 13, 2025.

[15619] Jessica Mella, Regan F Volk, Balyn W Zaro, and Abigail Buchwalter. C-terminal tagging, transmembrane domain hydrophobicity, and an ER retention motif influence the secretory trafficking of the inner nuclear membrane protein emerin. *eLife*, 14, 2025.

[15620] Almudena Mendez-Perez, Andres M Acosta-Moreno, Carlos Wert-Carvajal, Pilar Ballesteros-Cuartero, Ruben Sánchez-García, Jose R Macias, Rebeca Sanz-Pamplona, Ramon Alemany, Carlos Oscar Sorzano, Arrate Munoz-Barrutia, and Esteban Veiga. Unraveling the power of NAP-CNB's machine learning-enhanced tumor neoantigen prediction. *eLife*, 13, 2025.

[15621] Perenkita J Mendiola, Philip O'Herron, Kun Xie, Michael W Brands, Weston Bush, Rachel E Patterson, Valeria Di Stefano, and Jessica A Filosa. Blood pressure variability compromises vascular function in middle-aged mice. *eLife*, 14, 2025.

[15622] Haley N Mendoza-Romero, Jessica E Biddinger, Michelle N Bedenbaugh, and Richard Simerly. Microglia are required for developmental specification of AgRP innervation in the hypothalamus of offspring exposed to maternal high-fat diet during lactation. *eLife*, 13, 2025.

[15623] Fansen Meng, Jong Hwan Kim, Chang-Ru Tsai, Jeffrey D Steimle, Jun Wang, Yufeng Shi, Rich G Li, Bing Xie, Vaibhav Deshmukh, Shijie Liu, Xiao Li, and James F Martin. SARS-CoV-2 NSP13 interacts with TEAD to suppress Hippo-YAP signaling. *eLife*, 13, 2025.

[15624] Xiaoyu Meng, Yezhang Zhu, Kuai Liu, Yuxi Wang, Xiaoqian Liu, Chenxin Liu, Yan Zeng, Shuai Wang, Xianzhi Gao, Xin Shen, Jing Chen, Sijue Tao, Qianying Xu, Linjia Dong, Li Shen, and Lie Wang. CXXC-finger protein 1 associates with FOXP3 to stabilize homeostasis and suppressive functions of regulatory T cells. *eLife*, 13, 2025.

[15625] Ismail M Meraz, Mourad Majidi, Renduo Song, Feng Meng, Lihui Gao, Qi Wang, Jing Wang, Elizabeth J Shpall, and Jack A Roth. NPRL2 gene therapy induces effective antitumor immunity in KRAS/STK11 mutant anti-PD1 resistant metastatic non-small cell lung cancer (NSCLC) in a humanized mouse model. *eLife*, 13, 2025.

[15626] Adrián Merino-Salomón, Jonathan Scheneider, Leon Babl, Jan-Hagen Krohn, Marta Sobrinos-Sanguino, Tillman Schaefer, Juan Ramon Luque-Ortega, Carlos Alfonso, Mercedes Jiménez, Marion Jasnin, Petra Schwille, and German Rivas. Crosslinking by ZapD drives the assembly of short FtsZ filaments into toroidal structures in solution. *eLife*, 13, 2025.

[15627] Noëmie Mermet-Joret, Andrea Moreno, Agnieszka Zbela, Milad Nazari, Bárður Eyjólfsson Ellendersen, Raquel Comaposada Baro, Nathalie Krauth, Anne von Philipsborn, Andreas Toft Sørensen, Joaquin Piriz, John Yu-luen Lin, and Sadegh Nabavi. Dual-color optical activation and suppression of neurons with high temporal precision. *eLife*, 12, 2025.

[15628] Santi Mestre-Fos, Lucas Ferguson, Marena I Trinidad, Nicholas T Ingolia, and Jamie HD Cate. eIF3 engages with 3′-UTR termini of highly translated mRNAs. *eLife*, 13, 2025.

[15629] Dominique Meunier, Ricardo Corona-Torres, Kay Boulton, Zhiguang Wu, Maeve Ballantyne, Laura Glendinning, Anum Ali Ahmad, Dominika Borowska, Lorna Taylor, Lonneke Vervelde, Jorge del Pozo, Marili Vasilogianni, José Jaramillo-Ortiz, Gonzalo Sanchez-Arsuaga, Androniki Psifidi, Fiona Tomley, Kellie A Watson, Michael J McGrew, Mark P Stevens, Damer P Blake, and David A Hume. Interleukin 10 controls the balance between tolerance, pathogen elimination, and immunopathology in birds. *eLife*, 14, 2025.

[15630] Elise Miedlar, Grace E Hamilton, Samuel R Witus, Sara J Gonske, Michael Riffle, Alex Zelter, Rachel E Klevit, Charles L Asbury, Yoana N Dimitrova, and Trisha N Davis. Force transmission through the inner kinetochore is enhanced by centromeric DNA sequences. *eLife*, 14, 2025.

[15631] Ljiljana Mihajlovic, Bharat Ravi Iyengar, Florian Baier, IÇvara Barbier, Justyna Iwaszkiewicz, Vincent Zoete, Andreas Wagner, and Yolanda Schaerli. A direct experimental test of Ohno's hypothesis. *eLife*, 13, 2025.

[15632] Marwa O Mikati, Petra Erdmann-Gilmore, Rose Connors, Sineadh M Conway, Jim Malone, Justin Woods, Robert W Sprung, Reid R Townsend, and Ream Al-Hasani. Highly sensitive in vivo detection of dynamic changes in enkephalins following acute stress in mice. *eLife*, 12, 2025.

[15633] Hyeonyoung Min, Yale Y Yang, and Yunlei Yang. Cold induces brain region-selective cell activity-dependent lipid metabolism. *eLife*, 13, 2025.

[15634] Kyeongseon Min, Sungkwon Chung, Seung-Kyun Lee, Jongho Lee, Phan Tan Toi, Daehong Kim, Jung Seung Lee, and Jang-Yeon Park. Responses to membrane potential-modulating ionic solutions measured by magnetic resonance imaging of cultured cells and in vivo rat cortex. *eLife*, 13, 2025.

[15635] Martin Minařík, Alexander S Campbell, Roman Franěk, Michaela Vazačová, MiloŠ Havelka, David Gela, Martin PŠenička, and Clare VH Baker. Atoh1 is required for the formation of lateral line electroreceptors and hair cells, whereas FoxG1 represses an electrosensory fate. *eLife*, 14, 2025.

[15636] Mihailo Mirkovic, Jordan McCarthy, Anne Cornelis Meinema, Julie Parenteau, Sung Sik Lee, Sherif Abou Elela, and Yves Barral. Dissociation of the nuclear basket triggers chromosome loss in aging yeast. *eLife*, 14, 2025.

[15637] Michi Miura, Naho Kiuchi, Siu-Ying Lau, Bobo Wing-Yee Mok, Hiroshi Ushirogawa, Tadasuke Naito, Honglin Chen, and Mineki Saito. A statistical framework for quantifying the nuclear export rate of influenza viral mRNAs. *eLife*, 12, 2025.

[15638] Daigo Miyazaki, Mitsuto Sato, Naoko Shiba, Takahiro Yoshizawa, and Akinori Nakamura. Becker muscular dystrophy mice showed site-specific decay of type IIa fibers with capillary change in skeletal muscle. *eLife*, 13, 2025.

[15639] Yaroslav Molkov, Anke Borgmann, Hidehiko Koizumi, Noriyuki Hama, Ruli Zhang, and Jeffrey Smith. Inference technique for the synaptic conductances in rhythmically active networks and application to respiratory central pattern generation circuits. *eLife*, 13, 2025.

[15640] Rubén Mollá-Albaladejo, Manuel Jiménez-Caballero, and Juan Antonio Sanchez-Alcaniz. Molecular characterization of gustatory second-order neurons reveals integrative mechanisms of gustatory and metabolic information. *eLife*, 13, 2025.

[15641] Yugarshi Mondal, Ronald L Calabrese, and Eve Marder. Relative timescale of channel voltage dependence and channel density regulation impacts assembly and recovery of activity. *eLife*, 14, 2025.

[15642] Fernanda Morales Berstein, Jasmine Khouja, Mark Gormley, Elmira Ebrahimi, Shama Virani, James D McKay, Paul Brennan, Tom G Richardson, Caroline L Relton, George Davey Smith, M Carolina Borges, Tom Dudding, and Rebecca C Richmond. Reassessing the link between adiposity and head and neck cancer: a Mendelian randomization study. *eLife*, 14, 2025.

[15643] Laura Morano, Nadia Vezzio-Vié, Adam Aissanou, Tom Egger, Antoine Aze, Solène Fiachetti, Benoit Bordignon, Cedric Hassen-khodja, Hervé Seitz, Louis-Antoine Milazzo, Véronique Garambois, Laurent Chaloin, Nathalie Bonnefoy, Céline Gongora, Angelos Constantinou, and Jihane Basbous. TopBP1 biomolecular condensates as a new therapeutic target in advanced-stage colorectal cancer. *eLife*, 14, 2025.

[15644] Adrián Moreno Borrallo, Sarahi Jaramillo Ortiz, Christine Schaeffer-Reiss, Benoît Quintard, Benjamin Rey, Pierre Bize, Vincent A Viblanc, Thierry Boulinier, Olivier Chastel, Jorge S Gutiérrez, José A Masero, Fabrice Bertile, and Francois Criscuolo. Variation in albumin glycation rates in birds suggests resistance to relative hyperglycaemia rather than conformity to the pace of life syndrome hypothesis. *eLife*, 13, 2025.

[15645] Parham Mostame, Jonathan Wirsich, Thomas Alderson, Ben Ridley, Anne-Lise Giraud, David W Carmichael, Serge Vulliemoz, Maxime Guye, Louis Lemieux, and Sepideh Sadaghiani. A multiplex of connectome trajectories enables several connectivity patterns in parallel. *eLife*, 13, 2025.

[15646] Catia Mota, Kiseung Kim, Ye Ji Son, Eun Jung Thak, Su-Bin Lee, Ju-El Kim, Jeong-Kee Yoon, Min-Ho Kang, Heeyoun Hwang, Yong-Sun Bahn, J Andrew Alspaugh, and Hyun Ah Kang. Evolutionary unique N-glycan-dependent protein quality control system plays pivotal roles in cellular fitness and extracellular vesicle transport in Cryptococcus neoformans. *eLife*, 13, 2025.

[15647] Xiang Mou and Daoyun Ji. Observational activation of anterior cingulate cortical neurons coordinates hippocampal replay in social learning. *eLife*, 13, 2025.

[15648] Shahriar Mowla, Linda Farahani, Tharu Tharakan, Rhianna Davies, Goncalo DS Correia, Yun S Lee, Samit Kundu, Shirin Khanjani, Emad Sindi, Raj Rai, Lesley Regan, Dalia Khalifa, Ralf Henkel, Suks Minhas, Waljit S Dhillo, Jara Ben Nagi, Phillip Bennett, David A MacIntyre, and Channa N Jayasena. Characterisation and comparison of semen microbiota and bacterial load in men with infertility, recurrent miscarriage, or proven fertility. *eLife*, 13, 2025.

[15649] Yunfei Mu, Shijia Hu, Xiangyang Liu, Xin Tang, Jiayi Lin, and Hongjun Shi. Mechanical forces pattern endocardial Notch activation via mTORC2-PKC pathway. *eLife*, 13, 2025.

[15650] Sromona D Mukherjee, Carlos Batagello, Ava Adler, Jose Agudelo, Anna Zampini, Mangesh Suryavanshi, Andrew Nguyen, Terry Orr, Denise Dearing, Manoj Monga, and Aaron W Miller. Complex system modeling reveals oxalate homeostasis is driven by diverse oxalate-degrading bacteria. *eLife*, 14, 2025.

[15651] Soham Mukhopadhyay, Muhammad Asim Javed, Jiaxu Wu, and Edel Perez-Lopez. Structure-guided secretome analysis of gall-forming microbes offers insights into effector diversity and evolution. *eLife*, 14, 2025.

[15652] Sarah R Mullinax, Andrea M Darby, Anjali Gupta, Patrick Chan, Brittny R Smith, and Robert L Unckless. A suite of selective pressures supports the maintenance of alleles of a Drosophila immune peptide. *eLife*, 12, 2025.

[15653] James D Munday, Alicia Rosello, John Edmunds, and Sebastian Funk. Forecasting the spatial spread of an Ebola epidemic in real time: Comparing predictions of mathematical models and experts. *eLife*, 13, 2025.

[15654] Masaya Muramoto, Nozomi Hanawa, Misako Okumura, Takahiro Chihara, Masayuki Miura, and Natsuki Shinoda. Executioner caspase is proximal to Fasciclin 3 which facilitates non-lethal activation in Drosophila olfactory receptor neurons. *eLife*, 13, 2025.

[15655] Peter J Murphy, Juliette E McGregor, Zhengyang Xu, Qiang Yang, William Merigan, and David R Williams. Optogenetic stimulation of single ganglion cells in the living primate fovea. *eLife*, 12, 2025.

[15656] Nancy Nader, Lama Assaf, Lubna Zarif, Anna Halama, Sharan Yadav, Maya Dib, Nabeel Attarwala, Qiuying Chen, Karsten Suhre, Steven Gross, and Khaled Machaca. Progesterone induces meiosis through two obligate co-receptors with PLA2 activity. *eLife*, 13, 2025.

[15657] Yuji Nagai, Yukiko Hori, Ken-ichi Inoue, Toshiyuki Hirabayashi, Koki Mimura, Kei Oyama, Naohisa Miyakawa, Yuki Hori, Haruhiko Iwaoki, Katsushi Kumata, Ming-Rong Zhang, Masahiko Takada, Makoto Higuchi, and Takafumi Minamimoto. Longitudinal assessment of DREADD expression and efficacy in the monkey brain. *eLife*, 14, 2025.

[15658] Gurueswar Nagarajan, Denis Matrov, Anna C Pearson, Cecil C Yen, Sean P Bradley, and Yogita Chudasama. Cingulate cortex shapes early postnatal development of social vocalizations. *eLife*, 13, 2025.

[15659] Faez Amokrane Nait Mohamed and Daniel Lingwood. Innate immunity and training to subvert original antigenic sin by the humoral immune response. *eLife*, 14, 2025.

[15660] Shinichi Nakagawa, William K Cornwell, and Corey T Callaghan. Decoupling both local and global abundance from global range size, challenging the abundance-occupancy relationship in birds. *eLife*, 13, 2025.

[15661] Makaela Nartker, Chaz Firestone, Howard Egeth, and Ian Phillips. Sensitivity to visual features in inattentional blindness. *eLife*, 13, 2025.

[15662] Ojas Natarajan, Susanne L Gibboney, Morgan N Young, Shen Jean Lim, Felicia Nguyen, Natalia Pluta, Celine GF Atkinson, Assunta Liberti, Eric D Kees, Brittany A Leigh, Mya Breitbart, Jeffrey A Gralnick, and Larry J Dishaw. Prophage regulation of Shewanella fidelis 3313 motility and biofilm formation with implications for gut colonization dynamics in Ciona robusta. *eLife*, 14, 2025.

[15663] Dharmendra Kumar Nath, Subash Dhakal, and Youngseok Lee. TRP$\gamma$ regulates lipid metabolism through Dh44 neuroendocrine cells. *eLife*, 13, 2025.

[15664] Pavan K Nayak, Arul Subramanian, and Thomas F Schilling. Transcriptome profiling of tendon fibroblasts at the onset of embryonic muscle contraction reveals novel force-responsive genes. *eLife*, 14, 2025.

[15665] Pavla Nedbalová, Nikola Kaislerova, Lenka Chodakova, Martin Moos, and TomáŠ Doležal. SAM transmethylation pathway and adenosine recycling to ATP are essential for systemic regulation and immune response. *eLife*, 13, 2025.

[15666] Kara A Nelson, Kari F Lenhart, Lauren Anllo, and Stephen DiNardo. The Drosophila hematopoietic niche assembles through collective cell migration controlled by neighbor tissues and Slit-Robo signaling. *eLife*, 13, 2025.

[15667] Maximilian Nentwich, Marcin Leszczynski, Charles E Schroeder, Stephan Bickel, and Lucas C Parra. Intrinsic dynamic shapes responses to external stimulation in the human brain. *eLife*, 14, 2025.

[15668] Thea Ng, Eunsol Noh, and Rebecca MC Spencer. Bayesian meta-analysis reveals the mechanistic role of slow oscillation-spindle coupling in sleep-dependent memory consolidation. *eLife*, 13, 2025.

[15669] Han G Ngo, Sayed Golam Mohiuddin, Aina Ananda, and Mehmet Orman. Unraveling CRP/cAMP-mediated metabolic regulation in Escherichia coli persister cells. *eLife*, 13, 2025.

[15670] Khoa Ngo, Pei-Chi Yang, Vladimir Yarov-Yarovoy, Colleen E Clancy, and Igor Vorobyov. Harnessing AlphaFold to reveal hERG channel conformational state secrets. *eLife*, 13, 2025.

[15671] Arnaud N'Guessan, Wen Yuan Tong, Hamed Heydari, and Alex N Nguyen Ba. Refining the resolution of the yeast genotype–phenotype map using single-cell RNA-sequencing. *eLife*, 13, 2025.

[15672] Mai Nguyen, Elda Bauda, Célia Boyat, Cédric Laguri, Céline Freton, Anne Chouquet, Benoit Gallet, Morgane Baudoin, Yung-Sing Wong, Christophe Grangeasse, Christine Moriscot, Claire Durmort, André Zapun, and Cecile Morlot. Teichoic acids in the periplasm and cell envelope of Streptococcus pneumoniae. *eLife*, 14, 2025.

[15673] Roshani Nhuchhen Pradhan, Craig Montell, and Youngseok Lee. Cholesterol taste avoidance in Drosophila melanogaster. *eLife*, 14, 2025.

[15674] Ev L Nichols, Joo Lee, and Kang Shen. UNC-6/Netrin promotes both adhesion and directed growth within a single axon. *eLife*, 13, 2025.

[15675] Giorgio Nicoletti, Matteo Bruzzone, Samir Suweis, Marco dal Maschio, and Daniel Maria Busiello. Optimal information gain at the onset of habituation to repeated stimuli. *eLife*, 13, 2025.

[15676] Yamato Niitani, Kohei Matsuzaki, Erik Jonsson, Ronald D Vale, and Michio Tomishige. Kinetic regulation of kinesin's two motor domains coordinates its stepping along microtubules. *eLife*, 14, 2025.

[15677] Guozhu Ning, Yu Lin, Haixia Ma, Jiaqi Zhang, Liping Yang, Zhengyu Liu, Lei Li, Xinyu He, and Qiang Wang. BCAS2 promotes primitive hematopoiesis by sequestering $\beta$-catenin within the nucleus. *eLife*, 13, 2025.

[15678] Ke Ning, Yuanyuan Xie, Wen Sun, Lingke Feng, Can Fang, Rong Pan, Yan Li, and Ling Yu. Non-destructive in situ monitoring of structural changes of 3D tumor spheroids during the formation, migration, and fusion process. *eLife*, 13, 2025.

[15679] Nikolaos Nirakis, Sofia Dimothyra, Eleftheria Karadima, and Vasileia Ismini Alexaki. Metabolic regulation of immune memory and function of microglia. *eLife*, 14, 2025.

[15680] Katsuyuki Nishi, Taro Sakamaki, Akiomi Nagasaka, Kevin Shuolong Kao, Kay Sadaoka, Masahide Asano, Nobuyuki Yamamoto, Akifumi Takaori-Kondo, and Masanori Miyanishi. Alteration of long- and short-term hematopoietic stem cell ratio causes myeloid-biased hematopoiesis. *eLife*, 13, 2025.

[15681] Minsoo Noh, Xiangguo Che, Xian Jin, Dong-Kyo Lee, Hyun-Ju Kim, Doo Ri Park, Soo Young Lee, Hunsang Lee, Thomas J Gardella, Je-Yong Choi, and Sihoon Lee. Dimeric R25CPTH(1–34) activates the parathyroid hormone-1 receptor in vitro and stimulates bone formation in osteoporotic female mice. *eLife*, 13, 2025.

[15682] Janne Ina Nold, Tahmine Fadai, and Christian Büchel. Acute aerobic exercise intensity does not modulate pain potentially due to differences in fitness levels and sex effects: results from a pharmacological fMRI study. *eLife*, 14, 2025.

[15683] Sara A Nolin, Mary E Faulkner, Paul Stewart, Leland L Fleming, Stacy Merritt, Roxanne F Rezaei, Pradyumna K Bharadwaj, Mary Kate Franchetti, David A Raichlen, Cortney J Jessup, Lloyd Edwards, G Alex Hishaw, Emily J Van Etten, Theodore P Trouard, David Geldmacher, Virginia G Wadley, Noam Alperin, Eric S Porges, Adam J Woods, Ron A Cohen, Bonnie E Levin, Tatjana Rundek, Gene E Alexander, and Kristina M Visscher. Network segregation is associated with processing speed in the cognitively healthy oldest-old. *eLife*, 14, 2025.

[15684] Samuel Noorman, Timo Stein, Johannes Jacobus Fahrenfort, and Simon van Gaal. Perceptual and attentional impairments of conscious access involve distinct neural mechanisms despite equal task performance. *eLife*, 13, 2025.

[15685] Samuel Noorman, Timo Stein, Jasper Zantvoord, Johannes Fahrenfort, and Simon van Gaal. A causal role of the NMDA receptor in recurrent processing during perceptual integration. *eLife*, 13, 2025.

[15686] Kristin Nordin, Robin Pedersen, Farshad Falahati, Jarkko Johansson, Filip Grill, Micael Andersson, Saana M Korkki, Lars Bäckman, Andrew Zalesky, Anna Rieckmann, Lars Nyberg, and Alireza Salami. Two long-axis dimensions of hippocampal-cortical integration support memory function across the adult lifespan. *eLife*, 13, 2025.

[15687] Nobuyuki Nosaka, Vanessa Borges, Daisy Martinon, Timothy R Crother, Moshe Arditi, and Kenichi Shimada. Hypothermia protects against ventilator-induced lung injury by limiting IL-1$\beta$ release and NETs formation. *eLife*, 14, 2025.

[15688] Livio Oboti, Federico Pedraja, Marie Ritter, Marlena Lohse, Lennart Klette, and Rüdiger Krahe. Why the brown ghost chirps at night. *eLife*, 12, 2025.

[15689] J Daniel Obray, Erik T Wilkes, Mike Scofield, and L Judson Chandler. Adolescent alcohol exposure promotes mechanical allodynia and alters synaptic function at inputs from the basolateral amygdala to the prelimbic cortex. *eLife*, 13, 2025.

[15690] Thomas J O'Brien, Ida L Barlow, Luigi Feriani, and André EX Brown. High-throughput tracking enables systematic phenotyping and drug repurposing in C. elegans disease models. *eLife*, 12, 2025.

[15691] Daria M Odermatt, Frank Chidawanyika, Daniel M Mutyambai, Bernhard Schmid, Luiz A Domeignoz Horta, Collins O Onjura, Amanuel Tamiru, and Meredith C Schuman. Desmodium volatiles in 'push-pull' cropping systems and protection against the fall armyworm, Spodoptera frugiperda. *eLife*, 13, 2025.

[15692] Takashi Ogino, Akari Saito, Masato Sawada, Shoko Takemura, Yuzuki Hara, Kanami Yoshimura, Jiro Nagase, Honomi Kawase, Takamasa Sato, Hiroyuki Inada, Vicente Herranz-Pérez, Yoh-suke Mukouyama, Masatsugu Ema, José Manuel García-Verdugo, Junichi Nabekura, and Kazunobu Sawamoto. Neuronal migration depends on blood flow in the adult mammalian brain. *eLife*, 13, 2025.

[15693] Sven Ohl and Martin Rolfs. Visual routines for detecting causal interactions are tuned to motion direction. *eLife*, 13, 2025.

[15694] Gáspár Oláh, Rajmund Lákovics, Sapir Shapira, Yonatan Leibner, Attila Szücs, éva Adrienn Csajbók, Pál Barzó, Gábor Molnár, Idan Segev, and Gábor Tamás. Accelerated signal propagation speed in human neocortical dendrites. *eLife*, 13, 2025.

[15695] A Sofia F Oliveira, Fiona L Kearns, Mia A Rosenfeld, Lorenzo Casalino, Lorenzo Tulli, Imre Berger, Christiane Schaffitzel, Andrew D Davidson, Rommie E Amaro, and Adrian J Mulholland. Allosteric modulation by the fatty acid site in the glycosylated SARS-CoV-2 spike. *eLife*, 13, 2025.

[15696] Philipp S O'Neill, Martín Baccino-Calace, Peter Rupprecht, Sungmoo Lee, Yukun A Hao, Michael Z Lin, Rainer W Friedrich, Martin Mueller, and Igor Delvendahl. A deep learning framework for automated and generalized synaptic event analysis. *eLife*, 13, 2025.

[15697] Masayuki Ono, Masaya Yamaguchi, Daisuke Motooka, Yujiro Hirose, Kotaro Higashi, Tomoko Sumitomo, Tohru Miyoshi-Akiyama, Rumi Okuno, Takahiro Yamaguchi, Ryuji Kawahara, Hitoshi Otsuka, Noriko Nakanishi, Yu Kazawa, Chikara Nakagawa, Ryo Yamaguchi, Hiroo Sakai, Yuko Matsumoto, Tadayoshi Ikebe, and Shigetada Kawabata. Identifying genetic variations in emm89 Streptococcus pyogenes associated with severe invasive infections. *eLife*, 14, 2025.

[15698] Emily E Oor, Emilio Salinas, and Terrence R Stanford. Location- and feature-based selection histories make independent, qualitatively distinct contributions to urgent visuomotor performance. *eLife*, 13, 2025.

[15699] Affiong Ika Oqua, Kin Chao, Liliane El Eid, Lisa Casteller, Billy P Baxter, Alba Miguéns-Gómez, Sebastian Barg, Ben Jones, Jorge Bernardino de la Serna, Sarah L Rouse, and Alejandra Tomas. Molecular mapping and functional validation of GLP-1R cholesterol binding sites in pancreatic beta cells. *eLife*, 13, 2025.

[15700] Lucie Oriol, Melody Chao, Grace J Kollman, Dina S Dowlat, Sarthak M Singhal, Thomas Steinkellner, and Thomas S Hnasko. Ventral tegmental area interneurons revisited: GABA and glutamate projection neurons make local synapses. *eLife*, 13, 2025.

[15701] Zigmunds Orlovskis, Archana Singh, Adi Kliot, Weijie Huang, and Saskia A Hogenhout. The phytoplasma SAP54 effector acts as a molecular matchmaker for leafhopper vectors by targeting plant MADS-box factor SVP. *eLife*, 13, 2025.

[15702] FranÇois Osiurak, Giovanni Federico, Arnaud Fournel, Vivien Gaujoux, Franck Lamberton, Danièle Ibarrola, Yves Rossetti, and Mathieu Lesourd. Shaping the physical world to our ends through the left PF technical-cognition area. *eLife*, 13, 2025.

[15703] Zuzana Outla, Gizem Oyman-Eyrilmez, Katerina Korelova, Magdalena Prechova, Lukas Frick, Lenka Sarnova, Piyush Bisht, Petra Novotna, Jan Kosla, Patricia Bortel, Yasmin Borutzki, Andrea Bileck, Christopher Gerner, Mohammad Rahbari, Nuh Rahbari, Emrullah Birgin, Bibiana Kvasnicova, Andrea Galisova, Katerina Sulkova, Andreas Bauer, Njainday Jobe, Ondrej Tolde, Eva Sticova, Daniel Rösel, Tracy O'Connor, Martin Otahal, Daniel Jirak, Mathias Heikenwälder, Gerhard Wiche, Samuel M Meier-Menches, and Martin Gregor. Plectin-mediated cytoskeletal crosstalk as a target for inhibition of hepatocellular carcinoma growth and metastasis. *eLife*, 13, 2025.

[15704] Masroor Ahmad Paddar, Fulong Wang, Einar S Trosdal, Emily Hendrix, Yi He, Michelle R Salemi, Michal Mudd, Jingyue Jia, Thabata Duque, Ruheena Javed, Brett S Phinney, and Vojo Deretic. Noncanonical roles of ATG5 and membrane atg8ylation in retromer assembly and function. *eLife*, 13, 2025.

[15705] Marina Padilha, Victor Nahuel Keller, Paula Normando, Raquel M Schincaglia, Nathalia C Freitas-Costa, Samary SR Freire, Felipe M Delpino, Inês RR de Castro, Elisa MA Lacerda, Dayana R Farias, Zachary Kroezen, Meera Shanmuganathan, Philip Britz-Mckibbin, and Gilberto Kac. Serum metabolome indicators of early childhood development in the Brazilian National Survey on Child Nutrition (ENANI-2019). *eLife*, 14, 2025.

[15706] Cristina Paissoni, Sarita Puri, Luca Broggini, Manoj K Sriramoju, Martina Maritan, Rosaria Russo, Valentina Speranzini, Federico Ballabio, Mario Nuvolone, Giampaolo Merlini, Giovanni Palladini, Shang-Te Danny Hsu, Stefano Ricagno, and Carlo Camilloni. A conformational fingerprint for amyloidogenic light chains. *eLife*, 13, 2025.

[15707] Anna Pakula, Salsabiel El Nagar, N Sumru Bayin, Jens Bager Christensen, Daniel Stephen, Adam James Reid, Richard P Koche, and Alexandra L Joyner. An increase in reactive oxygen species underlies neonatal cerebellum repair. *eLife*, 14, 2025.

[15708] Luis G Palma, Daniel Alvarez-Villanueva, Maria Maqueda, Mercedes Barrero, Arnau Iglesias, Joan Bertran, Damiana Alvarez, Carlos A Garcia-Prieto, Cecilia Ballare, Virginia Rodriguez-Cortez, Clara Bueno, August Vidal, Alberto Villanueva, Pablo Menendez, Gregoire Stik, Luciano Di Croce, Bernhard Payer, Manel Esteller, Lluis Espinosa, and Anna Bigas. Chromatin activity of I$\kappa$B$\alpha$ mediates the exit from naïve pluripotency. *eLife*, 14, 2025.

[15709] Chengfang Pan, Ying Liu, Liangya Wang, Wen-Yong Fan, Yunzhi Ni, Xuefeng Zhang, Di Wu, Chenyang Li, Jin Li, Zhaoyang Li, Rui Liu, and Changlong Hu. The Kv2.2 channel mediates the inhibition of prostaglandin E2 on glucose-stimulated insulin secretion in pancreatic $\beta$-cells. *eLife*, 13, 2025.

[15710] Deepak Pant, Parik Kakani, Rushikesh Joshi, Abin Sabu, Shruti Agrawal, Atul Samaiya, and Sanjeev Shukla. Interplay of YEATS2 and GCDH regulates histone crotonylation and drives EMT in head and neck cancer. *eLife*, 14, 2025.

[15711] Alexandros Papagiannakis, Qiwei Yu, Sander K Govers, Wei-Hsiang Lin, Ned S Wingreen, and Christine Jacobs-Wagner. Nonequilibrium polysome dynamics promote chromosome segregation and its coupling to cell growth in Escherichia coli. *eLife*, 14, 2025.

[15712] Adrián Páramo, Pedro Mocho, Fernando Escaso, and Francisco Ortega. Evolution of hind limb morphology of Titanosauriformes (Dinosauria, Sauropoda) analyzed via 3D geometric morphometrics reveals wide-gauge posture as an exaptation for gigantism. *eLife*, 13, 2025.

[15713] Cesare V Parise. Correlation detection as a stimulus computable account for audiovisual perception, causal inference, and saliency maps in mammals. *eLife*, 14, 2025.

[15714] Cesare V Parise and Marc O Ernst. Multisensory integration operates on correlated input from unimodal transient channels. *eLife*, 12, 2025.

[15715] Eun Hye Park, Kally C O'Reilly Sparks, Griffin Grubbs, David Taborga, Kyndall Nicholas, Armaan S Ahmed, Natalie Ruiz-Péreza, Natalie Kim, Simon Segura-Carrillo, and André A Fenton. Cognitive control of behavior and hippocampal information processing without medial prefrontal cortex. *eLife*, 13, 2025.

[15716] Jiho Park, Grayson O Sipe, Xin Tang, Prachi Ojha, Giselle Fernandes, Yi Ning Leow, Caroline Zhang, Yuma Osako, Arundhati Natesan, Gabrielle T Drummond, Rudolf Jaenisch, and Mriganka Sur. Astrocytic modulation of population encoding in mouse visual cortex via GABA transporter 3 revealed by multiplexed CRISPR/Cas9 gene editing. *eLife*, 14, 2025.

[15717] Atit A Patel, Albert Cardona, and Daniel N Cox. Neural substrates of cold nociception in Drosophila larva. *eLife*, 12, 2025.

[15718] Bhumil Patel, Maryke Grobler, Alberto Herrera, Elias Logari, Valery Ortiz, and Needhi Bhalla. The conserved ATPase PCH-2 controls the number and distribution of crossovers by antagonizing their formation in Caenorhabditis elegans. *eLife*, 13, 2025.

[15719] Ulrike Pech, Jasper Janssens, Nils Schoovaerts, Sabine Kuenen, Carles Calatayud Aristoy, Sandra F Gallego, Samira Makhzami, Gert J Hulselmans, Suresh Poovathingal, Kristofer Davie, Adekunle T Bademosi, Jef Swerts, Sven Vilain, Stein Aerts, and Patrik Verstreken. Synaptic deregulation of cholinergic projection neurons causes olfactory dysfunction across five fly Parkinsonism models. *eLife*, 13, 2025.

[15720] María Pedraza, Valentina Grampa, Sophie Scotto-Lomassese, Julien Puech, Aude Muzerelle, Azka Mohammad, Sophie Lebon, Nicolas Renier, Christine Metin, and Justine Masson. The ciliary kinesin KIF7 controls the development of the cerebral cortex by acting differentially on SHH signaling in dorsal and ventral forebrain. *eLife*, 13, 2025.

[15721] Oana Pelea, Sarah Mayes, Quentin RV Ferry, Tudor A Fulga, and Tatjana Sauka-Spengler. Specific modulation of CRISPR transcriptional activators through RNA-sensing guide RNAs in mammalian cells and zebrafish embryos. *eLife*, 12, 2025.

[15722] David Peles, Shai Netser, Natalie Ray, Taghreed Suliman, and Shlomo Wagner. DeePosit, an AI-based tool for detecting mouse urine and fecal depositions from thermal video clips of behavioral experiments. *eLife*, 13, 2025.

[15723] Yang Peng, Jing Yang, Jixing Ao, Yilin Li, Jia Shen, Xiang He, Dihong Tang, Chaonan Chu, Congrong Liu, and Liang Weng. Single-cell profiling reveals the intratumor heterogeneity and immunosuppressive microenvironment in cervical adenocarcinoma. *eLife*, 13, 2025.

[15724] Aitana Perea-Gomez, Natividad Bellido Carreras, Magali Dhellemmes, Furong Tang, Coralie Le Gallo, and Marie-Christine Chaboissier. NR2F2 is required in the embryonic testis for fetal Leydig cell development. *eLife*, 14, 2025.

[15725] Michael Pereira, Nathan Faivre, Fosco Bernasconi, Nicholas Brandmeir, Jacob E Suffridge, Kaylee Tran, Shuo Wang, Victor Finomore, Peter Konrad, Ali Rezai, and Olaf Blanke. Subcortical correlates of consciousness with human single neuron recordings. *eLife*, 13, 2025.

[15726] Sarah Perlee, Yilun Ma, Miranda V Hunter, Jacob B Swanson, Nelly M Cruz, Zhitao Ming, Julia Xia, Timothee Lionnet, Maura McGrail, and Richard M White. Identifying in vivo genetic dependencies of melanocyte and melanoma development. *eLife*, 13, 2025.

[15727] Linsey Peters, Leonida Rakateli, Rosanna Huchzermeier, Andrea Bonnin-Marquez, Sanne L Maas, Cheng Lin, Alexander Jans, Yana Geng, Alan Gorter, Marion Gijbels, Sander Rensen, Peter Olinga, Tim Hendrikx, Marcin Krawczyk, Malvina Brisbois, Joachim Jankowski, Kiril Bidzhekov, Christian Weber, Erik AL Biessen, Ronit Shiri-Sverdlov, Tom Houben, Yvonne Doering, Matthias Bartneck, and Emiel van der Vorst. MicroRNA-26b protects against MASH development in mice and can be efficiently targeted with lipid nanoparticles. *eLife*, 13, 2025.

[15728] Alexander P Petroff, Julia Hernandez, Vladislav Kelin, and Nina Radchenko-Hannafin. Magnetotactic bacteria optimally navigate natural pore networks. *eLife*, 14, 2025.

[15729] Markus Borud Pettersen, Vemund Schøyen, Anders Malthe-Sørenssen, and Mikkel E Lepperød. Learning place cells and remapping by decoding the cognitive map. *eLife*, 13, 2025.

[15730] Julia Peukes, Charlie Lovatt, Conny Leistner, Jerome Boulanger, Dustin R Morado, Martin JG Fuller, Wanda Kukulski, Fei Zhu, Noboru H Komiyama, John AG Briggs, Seth GN Grant, and René AW Frank. The molecular infrastructure of glutamatergic synapses in the mammalian forebrain. *eLife*, 13, 2025.

[15731] Patricia E Phelps, Sung Min Ha, Rana R Khankan, Mahlet A Mekonnen, Giovanni Juarez, Kaitlin L Ingraham Dixie, Yen-Wei Chen, and Xia Yang. Olfactory ensheathing cells from adult female rats are hybrid glia that promote neural repair. *eLife*, 13, 2025.

[15732] Emily AM Phillips, Louise Goupil, Megan Whitehorn, Emma Bruce-Gardyne, Florian A Csolsim, Navsheen Kaur, Emily Greenwood, Ira Marriott Haresign, and Sam V Wass. Endogenous oscillatory rhythms and interactive contingencies jointly influence infant attention during early infant-caregiver interaction. *eLife*, 12, 2025.

[15733] Alessandro Piccin, Anne-Emilie Allain, Jérôme M Baufreton, Sandrine S Bertrand, and Angelo Contarino. Disruption of the CRF1 receptor eliminates morphine-induced sociability deficits and firing of oxytocinergic neurons in male mice. *eLife*, 13, 2025.

[15734] Sytse J Piersma, Shasha Li, Pamela Wong, Michael D Bern, Jennifer Poursine-Laurent, Liping Yang, Diana L Beckman, Bijal A Parikh, and Wayne M Yokoyama. Expression of a single inhibitory member of the Ly49 receptor family is sufficient to license NK cells for effector functions. *eLife*, 13, 2025.

[15735] Bárbara Pinto-Correia, Patrícia Caldeira-Bernardo, and Miguel Remondes. Optogenetic silencing of hippocampal inputs to the retrosplenial cortex causes a prolonged disruption of spatial working memory. *eLife*, 13, 2025.

[15736] Joar Esteban Pinto Torres, Mathieu Claes, Rik Hendrickx, Meng Yuan, Natalia Smiejkowska, Pieter Van Wielendaele, Aysima Hacisuleyman, Hans De Winter, Serge Muyldermans, Paul AM Michels, Malcolm D Walkinshaw, Wim Versées, Guy Caljon, Stefan Magez, and Yann G-J Sterckx. Allosteric inhibition of trypanosomatid pyruvate kinases by a camelid single-domain antibody. *eLife*, 13, 2025.

[15737] Daniela Piña Novo, Mang Gao, Rita Fischer, Louis Richevaux, Jianing Yu, John M Barrett, and Gordon MG Shepherd. Cortical dynamics in hand/forelimb S1 and M1 evoked by brief photostimulation of the mouse's hand. *eLife*, 14, 2025.

[15738] Sergio Plaza-Alonso, Nicolas Cano-Astorga, Javier DeFelipe, and Lidia Alonso-Nanclares. Volume electron microscopy reveals unique laminar synaptic characteristics in the human entorhinal cortex. *eLife*, 14, 2025.

[15739] Pooja Popli, Arin K Oestreich, Vineet K Maurya, Marina N Rowen, Yong Zhang, Michael J Holtzman, Ramya Masand, John P Lydon, Shizuo Akira, Kelle Moley, and Ramakrishna Kommagani. The autophagy protein ATG14 safeguards against unscheduled pyroptosis activation to enable embryo transport during early pregnancy. *eLife*, 13, 2025.

[15740] Duilio M Potenza, Xin Cheng, Guillaume Ajalbert, Andrea Brenna, Marie-Noelle Giraud, Aurelien Frobert, Stephane Cook, Kirsten D Mertz, Zhihong Yang, and Xiu-Fen Ming. Cell-autonomous and non-cell-autonomous effects of Arginase 2 on cardiac aging. *eLife*, 13, 2025.

[15741] Saugat Poudel, Jason Hyun, Ying Hefner, Jon Monk, Victor Nizet, and Bernhard O Palsson. Interpreting roles of mutations associated with the emergence of S. aureus USA300 strains using transcriptional regulatory network reconstruction. *eLife*, 12, 2025.

[15742] Derek Power, Justin Elstrott, and Jesse Schallek. Photoreceptor loss does not recruit neutrophils despite strong microglial activation. *eLife*, 13, 2025.

[15743] Supratim Pradhan, Dhruba Dhar, Debolina Manna, Shubhangi Chakraborty, Arkapriya Bhattacharyya, Khushi Chauhan, Rimi Mukherjee, Abhik Sen, Krishna Pandey, Soumen Das, and Budhaditya Mukherjee. Scrutinized lipid utilization disrupts Amphotericin-B responsiveness in clinical isolates of Leishmania donovani. *eLife*, 14, 2025.

[15744] Alexandra Prado-Mantilla, Wenxiu Ning, and Terry Lechler. Molecular and mechanical signatures contributing to mouse epidermal differentiation and barrier formation. *eLife*, 13, 2025.

[15745] Benedikt Praegel, Feng Chen, Adria Dym, Amichai Lavi-Rudel, Shaul Druckmann, and Adi Mizrahi. Age and learning shapes sound representations in auditory cortex during adolescence. *eLife*, 14, 2025.

[15746] Renaud Prevel, Erwan Pernet, Kim A Tran, Abderrahmane Sadek, Mina Sadeghi, Elizabeth Lapshina, Leonardo F Jurado, Arnold S Kristof, Mohieddine Moumni, Jeremie Poschmann, and Maziar Divangahi. $\beta$-Glucan reprograms alveolar macrophages via neutrophil/IFN$\gamma$ axis in a murine model of lung injury. *eLife*, 13, 2025.

[15747] Sujeethkumar Prithiviraj, Alejandro Garcia Garcia, Karin Linderfalk, Bai Yiguang, Sonia Ferveur, Ludvig Nilsén Falck, Agatheeswaran Subramaniam, Sofie Mohlin, David Hidalgo Gil, Steven J Dupard, Dimitra Zacharaki, Deepak Bushan Raina, and Paul E Bourgine. Compositional editing of extracellular matrices by CRISPR/Cas9 engineering of human mesenchymal stem cell lines. *eLife*, 13, 2025.

[15748] Tatiana Proboste, Abigail Turnlund, Andrew Bengsen, Matthew Gentle, Cameron Wilson, Lana Harriott, Richard A Fuller, Darren Marshall, and Ricardo J Soares-Magalhaes. Quantifying feral pig interactions to inform disease transmission networks. *eLife*, 13, 2025.

[15749] Aleksandra Prochera, Anoohya N Muppirala, Gavin A Kuziel, Salima Soualhi, Amy Shepherd, Liang Sun, Biju Issac, Harry J Rosenberg, Farah Karim, Kristina Perez, Kyle H Smith, Tonora H Archibald, Seth Rakoff-Nahoum, Susan J Hagen, and Meenakshi Rao. Enteric glia regulate Paneth cell secretion and intestinal microbial ecology. *eLife*, 13, 2025.

[15750] Róża K Przanowska, Yuechuan Chen, Takayuki-Okano Uchida, Etsuko Shibata, Xiaoxiao Hao, Isaac Segura Rueda, Kate Jensen, Piotr Przanowski, Anthony Trimboli, Yoshiyuki Shibata, Gustavo Leone, and Anindya Dutta. DNA replication in primary hepatocytes without the six-subunit ORC. *eLife*, 13, 2025.

[15751] Rachel Pudlowski, Lingyi Xu, Ljiljana Milenkovic, Chandan Kumar, Katherine Hemsworth, Zayd Aqrabawi, Tim Stearns, and Jennifer T Wang. A delta-tubulin/epsilon-tubulin/Ted protein complex is required for centriole architecture. *eLife*, 13, 2025.

[15752] Marcia Puig-Lluch, Mary Williams, Eric Wada, Imeña Valdes, Carrie Tribble, Andrew Read, Kanwardeep S Rawale, Chelsea L Newbold, Bathabile Mthombeni, Michael Moody, Laura Minero, Annarita Marrano, Melanie Link-Perez, Roger W Innes, Cody Coyotee Howard, Adriana Hernandez, Corri Hamilton, Denita Hadziabdic, Morgan Gostel, Joanna Friesner, John E Fowler, Mindy Findlater, Sakina Elshibli, Steven J Burgess, Hank W Bass, Burcu Alptekin, R Shawn Abrahams, and Patricia Baldrich. Making conferences in the plant sciences more inclusive through community recommendations. *eLife*, 14, 2025.

[15753] Erdem Pulcu and Michael Browning. Humans adapt rationally to approximate estimates of uncertainty. *eLife*, 14, 2025.

[15754] Gyeong Hee Pyeon, Hyewon Cho, Byung Min Chung, June-Seek Choi, and Yong Sang Jo. Parabrachial CGRP neurons modulate active defensive behavior under a naturalistic threat. *eLife*, 14, 2025.

[15755] Cheng Qi, Cheng Qian, Emma Steijvers, Robert A Colvin, and Daewoo Lee. A pair of dopaminergic neurons DAN-c1 mediate Drosophila larval aversive olfactory learning through D2-like receptors. *eLife*, 13, 2025.

[15756] Weiwei Qi, Yanlan Long, Ziming Li, Zhen Zhao, Jinhui Shi, Wanting Xie, Laijian Wang, Yandan Tan, Ti Zhou, Minting Liang, Ping Jiang, Bin Jiang, Xia Yang, and Guoquan Gao. The novel role of Kallistatin in linking metabolic syndromes and cognitive memory deterioration by inducing amyloid-$\beta$ plaques accumulation and tau protein hyperphosphorylation. *eLife*, 13, 2025.

[15757] Yan Qian, Qiannv Liu, Xiangyun Cheng, Chunlei Wang, Chun Kong, Mengqian Li, Chao Ren, Dong Jiang, Shuo Wang, and Pengyan Xia. A VgrG2b fragment cleaved by caspase-11/4 promotes Pseudomonas aeruginosa infection through suppressing the NLRP3 inflammasome. *eLife*, 13, 2025.

[15758] Lulu Qin, Chongbin Hu, Qiong Zhao, Yong Wang, Dongdong Fan, Aifu Lin, Lixin Xiang, Ye Chen, and Jianzhong Shao. Unraveling the role of Ctla-4 in intestinal immune homeostasis through a novel Zebrafish model of inflammatory bowel disease. *eLife*, 13, 2025.

[15759] Kathleen T Quach, Gillian A Hughes, and Sreekanth H Chalasani. Interdependence between SEB-3 receptor and NLP-49 peptides shifts across predator-induced defensive behavioral modes in Caenorhabditis elegans. *eLife*, 13, 2025.

[15760] Adi Rachum, Lee M Harten, Reut Assa, Aya Goldshtein, Xing Chen, Nesim Gonceer, and Yossi Yovel. Early experience affects foraging behavior of wild fruit bats more than their original behavioral predispositions. *eLife*, 14, 2025.

[15761] Katerina Rademacher, Zak Doric, Dominik Haddad, Aphroditi Mamaligas, Szu-Chi Liao, Rose Creed, Kohei Kano, Zac Chatterton, Yuhong Fu, Joseph H Garcia, Victoria M Vance, Yoshitaka J Sei, Anatol Kreitzer, Glenda Halliday, Alexandra B Nelson, Elyssa Margolis, and Ken Nakamura. Chronic hyperactivation of midbrain dopamine neurons causes preferential dopamine neuron degeneration. *eLife*, 13, 2025.

[15762] Denise Ragusa, Chun Wai Suen, Gabriel Torregrosa Cortes, Fabio Pastorino, Ayona Johns, Ylenia Cicirò, Liza Dijkhuis, Susanne van den Brink, Michele Cilli, Connor Byrne, Giulia-Andreea Ionescu, Joana Cerveira, Kamil R Kranc, Victor Hernandez-Hernandez, Mirco Ponzoni, Anna Bigas, Jordi Garcia-Ojalvo, Alfonso Martínez Arias, and Cristina Pina. Dissecting infant leukemia developmental origins with a hemogenic gastruloid model. *eLife*, 14, 2025.

[15763] Maryam Rahmati Ishka, Hayley Sussman, Yunfei Hu, Mashael Daghash Alqahtani, Eric Craft, Ronell Sicat, Minmin Wang, Li'ang Yu, Rachid Ait-Haddou, Bo Li, Georgia Drakakaki, Andrew DL Nelson, Miguel Pineros, Arthur Korte, Łukasz Jaremko, Christa Testerink, Mark Tester, and Magdalena M Julkowska. Natural variation in salt-induced changes in root:shoot ratio reveals SR3G as a negative regulator of root suberization and salt resilience in Arabidopsis. *eLife*, 13, 2025.

[15764] Marco Rando, Martin James, Alessandro Verri, Lorenzo Rosasco, and Agnese Seminara. Q-learning with temporal memory to navigate turbulence. *eLife*, 13, 2025.

[15765] Rajneesh Rao, Alejandro Aballay, and Jogender Singh. Inhibition of the UFD-1-NPL-4 complex triggers an aberrant immune response in Caenorhabditis elegans. *eLife*, 13, 2025.

[15766] Maha Rashid, Sarah Thomas, Jennifer Isaac, Sonia Corbett Karkare, Hannah Klein, and Malavika Murugan. A ventral hippocampal-lateral septum pathway regulates social novelty preference. *eLife*, 13, 2025.

[15767] Marietta M Ravesloot-Chavez, Erik Van Dis, Douglas Fox, Andrea Anaya-Sanchez, Scott Espich, Xammy Huu Nguyenla, Sagar Rawal, Helia Samani, Mallory Ballinger, Henry F Thomas, Dmitri I Kotov, Russell E Vance, Michael W Nachman, and Sarah A Stanley. Tuberculosis susceptibility in genetically diverse mice reveals functional diversity of neutrophils. *eLife*, 13, 2025.

[15768] Aleksandr Rayshubskiy, Stephen L Holtz, Alexander S Bates, Quinn X Vanderbeck, Laia Serratosa Capdevila, Victoria Rockwell, and Rachel Wilson. Neural circuit mechanisms for steering control in walking Drosophila. *eLife*, 13, 2025.

[15769] Kavidha Reddy, Guinevere Q Lee, Nicole Reddy, Tatenda JB Chikowore, Kathy Baisley, Krista L Dong, Bruce D Walker, Xu G Yu, Mathias Lichterfeld, and Thumbi Ndung'u. Differences in HIV-1 reservoir size, landscape characteristics, and decay dynamics in acute and chronic treated HIV-1 Clade C infection. *eLife*, 13, 2025.

[15770] William T Redman, Santiago Acosta-Mendoza, Xue-Xin Wei, and Michael J Goard. Robust variability of grid cell properties within individual grid modules enhances encoding of local space. *eLife*, 13, 2025.

[15771] Barbora Rehak Buckova, Charlotte Fraza, Rastislav Rehák, Marián Kolenič, Christian F Beckmann, Filip Španiel, Andre F Marquand, and Jaroslav Hlinka. Using normative models pre-trained on cross-sectional data to evaluate intra-individual longitudinal changes in neuroimaging data. *eLife*, 13, 2025.

[15772] Jun Ren, Ying Zhou, Yudi Hu, Jing Yang, Hongkun Fang, Xuejing Lyu, Jintao Guo, Xiaodong Shi, and Qiyuan Li. A model-based factorization method for scRNA data unveils bifurcating transcriptional modules underlying cell fate determination. *eLife*, 13, 2025.

[15773] Simon Rethemeier, Sonja Fritzsche, Dominik Mühlen, Gregor Bucher, and Vera S Hunnekuhl. Differences in size and number of embryonic type II neuroblast lineages correlate with divergent timing of central complex development between beetle and fly. *eLife*, 13, 2025.

[15774] Rebecca RG, Giorgio A Ascoli, Nate M Sutton, and Holger Dannenberg. Spatial periodicity in grid cell firing is explained by a neural sequence code of 2-D trajectories. *eLife*, 13, 2025.

[15775] David Richter, Dirk van Moorselaar, and Jan Theeuwes. Proactive distractor suppression in early visual cortex. *eLife*, 13, 2025.

[15776] Sean A Ridout, Priyathama Vellanki, and Ilya Nemenman. A mathematical model for ketosis-prone diabetes suggests the existence of multiple pancreatic $\beta$-cell inactivation mechanisms. *eLife*, 13, 2025.

[15777] Nicolas N Rieser, Milena Ronchetti, Adriana Lea Lea Hotz, and Stephan CF Neuhauss. Multifaceted role of galanin in brain excitability. *eLife*, 13, 2025.

[15778] Krishna Rijal and Pankaj Mehta. A differentiable Gillespie algorithm for simulating chemical kinetics, parameter estimation, and designing synthetic biological circuits. *eLife*, 14, 2025.

[15779] Jordina Rincon-Torroella, Marco Dal Molin, Brian Mog, Gyuri Han, Evangeline Watson, Nicolas Wyhs, Shun Ishiyama, Taha Ahmedna, Il Minn, Nilofer Azad, Chetan Bettegowda, Nickolas Papadopoulos, Kenneth W Kinzler, Shibin Zhou, Bert Vogelstein, Kathleen Gabrielson, and Surojit Sur. ME3BP-7 is a targeted cytotoxic agent that rapidly kills pancreatic cancer cells expressing high levels of monocarboxylate transporter MCT1. *eLife*, 13, 2025.

[15780] Israel Ríos, Cristina Herrero, Mónica Torres-Torresano, Baltasar López-Navarro, María Teresa Schiaffino, Francisco Díaz Crespo, Alicia Nieto-Valle, Rafael Samaniego, Yolanda Sierra-Palomares, Eduardo Oliver, Fernando Revuelta-Salgado, Ricardo García-Luján, Paloma Sánchez-Mateos, Rafael Delgado, Amaya Puig-Kröger, and Angel L Corbí. Reprogramming of GM-CSF-dependent alveolar macrophages through GSK3 activity modulation. *eLife*, 14, 2025.

[15781] Emilio Rios-Jimenez, Anoek Zomer, Raphael Collot, Mario Barrera Román, Sandra F Archidona, Hendrikus Ariese, Ravian van Ineveld, Michiel Kleinnijenhuis, Nils Bessler, Hannah Johnson, Caleb A Dawson, Anne Rios, and Maria Alieva. BEHAV3D Tumor Profiler to map heterogeneous cancer cell behavior in the tumor microenvironment. *eLife*, 13, 2025.

[15782] Erin M Ritchie, Dilan Acar, Siming Zhong, Qianyi Pu, Yunbo Li, Binhai Zheng, and Yishi Jin. Translatome analysis reveals cellular network in DLK-dependent hippocampal glutamatergic neuron degeneration. *eLife*, 13, 2025.

[15783] Mykola Roiuk, Marilena Neff, and Aurelio A Teleman. Human eIF2A has a minimal role in translation initiation and in uORF-mediated translational control in HeLa cells. *eLife*, 14, 2025.

[15784] Astrid Rollenhagen, Akram Sadeghi, Bernd Walkenfort, Claus C Hilgetag, Kurt Sätzler, and Joachim HR Lübke. Ultrastructural sublaminar-specific diversity of excitatory synaptic boutons in layer 1 of the adult human temporal lobe neocortex. *eLife*, 13, 2025.

[15785] Alejandro Rosell, Agata Adelajda Krygowska, Marta Alcón Pérez, Cristina Cuesta, Mathieu-Benoit Voisin, Juan de Paz, Héctor Sanz-Fraile, Vinothini Rajeeve, Ana Carreras-González, Alberto Berral-González, Ottilie Swinyard, Enrique Gabandé-Rodríguez, Julian Downward, Jordi Alcaraz, Juan Anguita, Carmen García-Macías, Javier De Las Rivas, Pedro R Cutillas, and Esther Castellano Sanchez. RAS–p110$\alpha$ signalling in macrophages is required for effective inflammatory response and resolution of inflammation. *eLife*, 13, 2025.

[15786] Shelby A Roseman, Allison P Siegenfeld, Ceejay Lee, Nicholas Z Lue, Amanda L Waterbury, and Brian B Liau. DNA methylation insulates genic regions from CTCF loops near nuclear speckles. *eLife*, 13, 2025.

[15787] Matthew Rosenblatt, Saloni Mehta, Hannah Peterson, Javid Dadashkarimi, Raimundo Rodriguez, Maya L Foster, Brendan D Adkinson, Qinghao Liang, Violet M Kimble, Jean Ye, Marie C McCusker, Michael C Farruggia, Max J Rolison, Margaret L Westwater, Rongtao Jiang, Stephanie Noble, and Dustin Scheinost. Trends in self-citation rates in high-impact neurology, neuroscience, and psychiatry journals. *eLife*, 12, 2025.

[15788] Yevgenia Rosenblum, Mahdad Jafarzadeh Esfahani, Nico Adelhöfer, Paul Zerr, Melanie Furrer, Reto Huber, Famke F Roest, Axel Steiger, Marcel Zeising, Csenge G Horváth, Bence Schneider, Róbert Bódizs, and Martin Dresler. Fractal cycles of sleep, a new aperiodic activity-based definition of sleep cycles. *eLife*, 13, 2025.

[15789] Jessica E Rosien, Luke D Fannin, Justin D Yeakel, Suchinda Malaivijitnond, Nathaniel J Dominy, and Amanda Tan. Food-washing monkeys recognize the law of diminishing returns. *eLife*, 13, 2025.

[15790] Jerko Rosko, Rebecca N Poon, Kelsey Cremin, Emanuele Locatelli, Mary Coates, Sarah JN Duxbury, Kieran Randall, Katie Croft, Chantal Valeriani, Marco Polin, and Orkun S Soyer. Cellular coordination underpins rapid reversals in gliding filamentous cyanobacteria and its loss results in plectonemes. *eLife*, 13, 2025.

[15791] Gitanjali Roy, Rameesha Syed, Olivia Lazaro, Sylvia Robertson, Sean D McCabe, Daniela Rodriguez, Alex M Mawla, Travis S Johnson, and Michael A Kalwat. Identification of type 2 diabetes- and obesity-associated human $\beta$-cells using deep transfer learning. *eLife*, 13, 2025.

[15792] Kevin Alen Rucci, Gabriel Barco, Andrea Onorato, Mauricio Beranek, Mariana Pueta, and Adrián Díaz. Effects of blood meal source and seasonality on reproductive traits of Culex quinquefasciatus (Diptera: Culicidae). *eLife*, 12, 2025.

[15793] Martina Rudgalvyte, Zehan Hu, Dieter Kressler, Jörn Dengjel, and Dominique A Glauser. Antagonist actions of CMK-1/CaMKI and TAX-6/calcineurin along the C. elegans thermal avoidance circuit orchestrate adaptation of nociceptive response to repeated stimuli. *eLife*, 14, 2025.

[15794] Christopher Nass Rudzitis, Monika Lakk, Ayushi Singh, Sarah N Redmon, Denisa Kirdajová, Yun-Ting Tseng, Michael L De Ieso, W Daniel Stamer, Samuel Herberg, and David Krizaj. TRPV4 activation by TGF$\beta$2 enhances cellular contractility and drives ocular hypertension. *eLife*, 14, 2025.

[15795] Rosa Rugani, Matteo Macchinizzi, Yujia Zhang, and Lucia Regolin. Prenatal light exposure affects number sense and the mental number line in young domestic chicks. *eLife*, 14, 2025.

[15796] Carmen Ruger-Herreros, Lucia Svoboda, Gurranna Male, Aseem Shrivastava, Markus Höpfner, Katharina Jetzinger, Jiří Koubek, Günter Kramer, Fabian den Brave, Axel Mogk, David S Gross, and Bernd Bukau. Nuclear and cytosolic J-domain proteins provide synergistic control of Hsf1 at distinct phases of the heat shock response. *eLife*, 14, 2025.

[15797] Brian C Ruyle, Sarah Masud, Rohith Kesaraju, Mubariz Tahirkheli, Juhi Modh, Caroline G Roth, Sofia Angulo-Lopera, Tania Lintz, Jessica A Higginbotham, Nicolas Massaly, and Jose A Morón. Peripheral opioid receptor antagonism alleviates fentanyl-induced cardiorespiratory depression and is devoid of aversive behavior. *eLife*, 13, 2025.

[15798] Veronica H Ryan, Sydney Lawton, Joel F Reyes, James Hawrot, Ashley M Frankenfield, Sahba Seddighi, Daniel M Ramos, Jacob Epstein, Faraz Faghri, Nicholas L Johnson, Jizhong Zou, Martin Kampmann, John Replogle, Yue Andy Qi, Hebao Yuan, Kory Johnson, Dragan Maric, Ling Hao, Mike A Nalls, and Michael Emmerson Ward. Maintenance of neuronal TDP-43 expression requires axonal lysosome transport. *eLife*, 14, 2025.

[15799] Ilya A Rybak, Natalia A Shevtsova, Johannie Audet, Sirine Yassine, Sergey N Markin, Boris I Prilutsky, and Alain Frigon. Operation of spinal sensorimotor circuits controlling phase durations during tied-belt and split-belt locomotion after a lateral thoracic hemisection. *eLife*, 13, 2025.

[15800] Faustine Ryckebusch, Yao Tian, Mylene Rapin, Fanny Schüpfer, Mark Austin Hanson, and Bruno Lemaitre. Layers of immunity: Deconstructing the Drosophila effector response. *eLife*, 14, 2025.

[15801] Hannes P Saal, Ingvars Birznieks, and Roland S Johansson. Fingertip viscoelasticity enables human tactile neurons to encode loading history alongside current force. *eLife*, 12, 2025.

[15802] David Saeb, Emma E Lietzke, Daisy I Fuchs, Emma C Aldrich, Kimberley D Bruce, and Kayla G Sprenger. The flexible stalk domain of sTREM2 modulates its interactions with brain-based phospholipids. *eLife*, 13, 2025.

[15803] Akiko Saiki-Ishikawa, Mark Agrios, Sajishnu Savya, Adam Forrest, Hannah Sroussi, Sarah Hsu, Diya Basrai, Feihong Xu, and Andrew Miri. Hierarchy between forelimb premotor and primary motor cortices and its manifestation in their firing patterns. *eLife*, 13, 2025.

[15804] Yusuke Sakai, Saumik Sen, Tomohiro Sugihara, Yukiya Kakeyama, Makoto Iwasaki, Gebhard FX Schertler, Xavier Deupi, Mitsumasa Koyanagi, and Akihisa Terakita. Coral anthozoan-specific opsins employ a novel chloride counterion for spectral tuning. *eLife*, 14, 2025.

[15805] Bennet Sakelaris and Hermann Riecke. Adult neurogenesis reconciles flexibility and stability of olfactory perceptual memory. *eLife*, 14, 2025.

[15806] Rosanna P Sammons, Stefano Masserini, Laura Moreno Velasquez, Verjinia D Metodieva, Gaspar Cano, Andrea Sannio, Marta Orlando, Nikolaus Maier, Richard Kempter, and Dietmar Schmitz. Subtype specific connectivity between CA3 pyramidal neurons may underlie their sequential activation during sharp waves. *eLife*, 13, 2025.

[15807] CJ San Felipe, Jyoti Batra, Monita Muralidharan, Shivali Malpotra, Durga Anand, Rachel Bauer, Kliment A Verba, Danielle L Swaney, Nevan J Krogan, Michael Grabe, and James S Fraser. Coupled equilibria of dimerization and lipid binding modulate SARS Cov 2 Orf9b interactions and interferon response. *eLife*, 14, 2025.

[15808] Carlos A Sánchez-León, Guillermo Sánchez-Garrido Campos, Marta Fernández, álvaro Sánchez-López, Javier F Medina, and Javier Márquez-Ruiz. Somatodendritic orientation determines tDCS-induced neuromodulation of Purkinje cell activity in awake mice. *eLife*, 13, 2025.

[15809] Estefania Sanchez-Vasquez, Marianne E Bronner, and Magdalena Zernicka-Goetz. HIF1A-mediated pathways promote euploid cell survival in chromosomally mosaic embryos. *eLife*, 13, 2025.

[15810] Kristian Sandberg and Morten Overgaard. Comment on 'Criterion placement threatens the construct validity of neural measures of consciousness'. *eLife*, 14, 2025.

[15811] Raphael Sarfati, Kunaal Joshi, Owen Martin, Julie C Hayes, Srividya Iyer-Biswas, and Orit Peleg. Correction: Emergent periodicity in the collective synchronous flashing of fireflies. *eLife*, 14, 2025.

[15812] Yuhkoh Satouh, Takaki Tatebe, Isei Tanida, Junji Yamaguchi, Yasuo Uchiyama, and Ken Sato. Endosomal-lysosomal organellar assembly (ELYSA) structures coordinate lysosomal degradation systems through mammalian oocyte-to-embryo transition. *eLife*, 13, 2025.

[15813] Andrea Sattin, Chiara Nardin, Simon Daste, Monica Moroni, Innem Reddy, Carlo Liberale, Stefano Panzeri, Alexander Fleischmann, and Tommaso Fellin. Aberration correction in long GRIN lens-based microendoscopes for extended field-of-view two-photon imaging in deep brain regions. *eLife*, 13, 2025.

[15814] Khund Sayeed, Sreeja Parameswaran, Matthew J Beucler, Lee E Edsall, Andrew VonHandorf, Audrey Crowther, Omer A Donmez, Matthew R Hass, Scott Richards, Carmy R Forney, Hayley K Hesse, Sydney H Jones, Katelyn A Dunn, Jay Wright, Merrin Man Long Leong, Laura A Murray-Nerger, Vijay Yechoor, Ben E Gewurz, Kenneth M Kaufman, John B Harley, Bo Zhao, William E Miller, Leah C Kottyan, and Matthew T Weirauch. Human cytomegalovirus infection coopts chromatin organization to diminish TEAD1 transcription factor activity. *eLife*, 13, 2025.

[15815] Christopher T Schafer, Raymond F Pauszek, Martin Gustavsson, Tracy M Handel, and David P Millar. Distinct activation mechanisms of CXCR4 and ACKR3 revealed by single-molecule analysis of their conformational landscapes. *eLife*, 13, 2025.

[15816] Moritz Schloetter, Georg U Maret, and Christoph J Kleineidam. Annihilation of action potentials induces electrical coupling between neurons. *eLife*, 12, 2025.

[15817] Marc W Schmid, Klara KropivŠek, Samuel E Wuest, Bernhard Schmid, and Ueli Grossniklaus. Weak evidence for heritable changes in response to selection by aphids in Arabidopsis accessions. *eLife*, 14, 2025.

[15818] Fabian Schmidt, Sarah K Danböck, Eugen Trinka, Dominic P Klein, Gianpaolo Demarchi, and Nathan Weisz. Age-related changes in 'cortical' 1/f dynamics are linked to cardiac activity. *eLife*, 13, 2025.

[15819] Douwe Schulte, Marta Šiborová, Lukas Käll, and Joost Snijder. Simultaneous polyclonal antibody sequencing and epitope mapping by cryo electron microscopy and mass spectrometry. *eLife*, 14, 2025.

[15820] Isaïh Schwab-Mohamed, Manuel R Mercier, Agnès Trebuchon, Benjamin Morillon, Leonardo Lancia, and Daniele Schön. Bridging verbal coordination and neural dynamics. *eLife*, 13, 2025.

[15821] Philipp Schwintek, Emre Eren, Christof Bernhard Mast, and Dieter Braun. Prebiotic gas flow environment enables isothermal nucleic acid replication. *eLife*, 13, 2025.

[15822] Ruben Sebastian-Perez, Shoma Nakagawa, Xiaochuan Tu, Sergi Aranda, Martina Pesaresi, Pablo Aurelio Gomez-Garcia, Marc Alcoverro-Bertran, Jose Luis Gomez-Vazquez, Davide Carnevali, Eva Borràs, Eduard Sabidó, Laura Martin, Malka Nissim-Rafinia, Eran Meshorer, Maria Victoria Neguembor, Luciano Di Croce, and Maria Pia Cosma. SMARCAD1 and TOPBP1 contribute to heterochromatin maintenance at the transition from the 2C-like to the pluripotent state. *eLife*, 12, 2025.

[15823] Marius Seidenthal, Jasmina Redzovic, Jana F Liewald, Dennis Rentsch, Stepan Shapiguzov, Noah Schuh, Nils Rosenkranz, Stefan Eimer, and Alexander Gottschalk. Flower/FLWR-1 regulates neuronal activity via the plasma membrane Ca2+ ATPase to promote recycling of synaptic vesicles. *eLife*, 13, 2025.

[15824] Antara Sengupta, Soujanya Vinayagamurthy, Dristhi Soni, Rajlekha Deb, Ananda Kishore Mukherjee, Subhajit Dutta, Jushta Jaiswal, Mukta Yadav, Shalu Sharma, Sulochana Bagri, Shuvra Shekhar Roy, Priya Poonia, Ankita Singh, Divya Khanna, Amit Kumar Kumar Bhatt, Akshay Sharma, Suman Saurav, Rajender K Motiani, and Shantanu Chowdhury. Telomeres control human telomerase (TERT) expression through non-telomeric TRF2. *eLife*, 14, 2025.

[15825] Jewoo Seo, Seung Ha Kim, Jaegeon Lee, Min Seok Kim, Yong-Seok Lee, and Sang Jeong Kim. Acquisition phase-specific contribution of climbing fiber transmission to cerebellum-dependent motor memory in mice. *eLife*, 13, 2025.

[15826] HeeYoung Seon and Dongil Chung. Policy shaping based on the learned preferences of others accounts for risky decision-making under social observation. *eLife*, 13, 2025.

[15827] Susumu Setogawa, Takashi Okauchi, Di Hu, Yasuhiro Wada, Keigo Hikishima, Hirotaka Onoe, Kayo Nishizawa, Nobuyuki Sakayori, Hiroyuki Miyawaki, Takuma Kitanishi, Kenji Mizuseki, Yilong Cui, and Kazuto Kobayashi. Acquisition of auditory discrimination mediated by different processes through two distinct circuits linked to the lateral striatum. *eLife*, 13, 2025.

[15828] Devanshi Piyush Shah, Pallavi Raj Sharma, Rachit Agarwal, and Arnab Barik. A septo-hypothalamic-medullary circuit directs stress-induced analgesia. *eLife*, 13, 2025.

[15829] Sakshi Shambhavi, Sudipta Mondal, Arnab Chakraborty, Nikita Shukla, Bapin Kumar Panda, Santhosh Kumar, Priyadarshan Kinatukara, Biswajit Pal, Siddhesh S Kamat, and Rajan Sankaranarayanan. Emergence of Dip2-mediated specific DAG-based PKC signalling axis in eukaryotes. *eLife*, 14, 2025.

[15830] Yuanyue Shan, Xinyi Guo, Mengmeng Zhang, Meiyu Chen, Ying Li, Mingfeng Zhang, and Duanqing Pei. Structure of human PIEZO1 and its slow-inactivating channelopathy mutants. *eLife*, 13, 2025.

[15831] Xiexiang Shao, Xingzuan Lin, Hao Zhou, Minghui Wang, Lili Han, Xin Fu, Sheng Li, Siyuan Zhu, Shenao Zhou, Wenjun Yang, Jianhua Wang, Zhanghua Li, and Ping Hu. Human CD29+/CD56+ myogenic progenitors display tenogenic differentiation potential and facilitate tendon regeneration. *eLife*, 13, 2025.

[15832] Zhujun Shao, Mengya Zhang, and Qing Yu. Stimulus representation in human frontal cortex supports flexible control in working memory. *eLife*, 13, 2025.

[15833] Ayni Sharif, Matthew S Jeffers, Dean A Fergusson, Raj Bapuji, Stuart G Nicholls, John Humphrey, Warren Johnston, Ed Mitchell, Mary-Ann Speirs, Laura Stronghill, Michele Vuckovic, Susan Wulf, Risa Shorr, Dar Dowlatshahi, Dale Corbett, and Manoj M Lalu. Preclinical systematic review of CCR5 antagonists as cerebroprotective and stroke recovery enhancing agents. *eLife*, 14, 2025.

[15834] Dominic WH Shayler, Kevin Stachelek, Linda Cambier, Sunhye Lee, Jinlun Bai, Bhavana Bhat, Mark W Reid, Daniel J Weisenberger, Jennifer G Aparicio, Yeha Kim, Mitali Singh, Maxwell Bay, Matthew E Thornton, Eamon K Doyle, Zachary Fouladian, Stephan G Erberich, Brendan H Grubbs, Michael A Bonaguidi, Cheryl Mae Craft, Hardeep P Singh, and David Cobrinik. Identification and characterization of early human photoreceptor states and cell-state-specific retinoblastoma-related features. *eLife*, 13, 2025.

[15835] Andrew M Shelton, David K Oliver, Ivan P Lazarte, Joachim S Grimstvedt, Ishaan Kapoor, Jake A Swann, Caitlin A Ashcroft, Simon N Williams, Niall Conway, Selma Tir, Amy Robinson, Stuart Peirson, Thomas Akam, Clifford G Kentros, Menno P Witter, Simon JB Butt, and Adam Max Packer. Single neurons and networks in the mouse claustrum integrate input from widespread cortical sources. *eLife*, 13, 2025.

[15836] Li Shen, Shuo Li, Yuhao Tian, Ying Wang, and Yi Jiang. Cortical tracking of hierarchical rhythms orchestrates the multisensory processing of biological motion. *eLife*, 13, 2025.

[15837] Darryl Shibata. Human brain ancestral barcodes. *eLife*, 13, 2025.

[15838] Kai S Shimagaki, Rebecca M Lynch, and John P Barton. Parallel HIV-1 fitness landscapes shape viral dynamics in humans and macaques that develop broadly neutralizing antibodies. *eLife*, 14, 2025.

[15839] Yichun Shuai, Megan Sammons, Gabriella R Sterne, Karen L Hibbard, He Yang, Ching-Po Yang, Claire Managan, Igor Siwanowicz, Tzumin Lee, Gerald M Rubin, Glenn C Turner, and Yoshinori Aso. Driver lines for studying associative learning in Drosophila. *eLife*, 13, 2025.

[15840] Nathan M Shugarts Devanapally, Aishwarya Sathya, Andrew L Yi, Winnie M Chan, Julia A Marre, and Antony M Jose. Intergenerational transport of double-stranded RNA in C. elegans can limit heritable epigenetic changes. *eLife*, 13, 2025.

[15841] Cheryll M Sia, Rebecca L Ambrose, Mary Valcanis, Patiyan Andersson, Susan A Ballard, Benjamin P Howden, Deborah A Williamson, Jaclyn S Pearson, and Danielle J Ingle. Distinct adaptation and epidemiological success of different genotypes within Salmonella enterica serovar Dublin. *eLife*, 13, 2025.

[15842] Rita Sianga-Mete, Penelope Hartnady, Wimbai Caroline Mandikumba, Kayleigh Rutherford, Christopher Brian Currin, Florence Phelanyane, Sabina Stefan, Steven Weaver, Sergei L Kosakovsky Pond, and Darren P Martin. Viral genome sequence datasets display pervasive evidence of strand-specific substitution biases that are best described using non-reversible nucleotide substitution models. *eLife*, 12, 2025.

[15843] Julia D Sigwart, Yunlong Li, Zeyuan Chen, Katarzyna Vončina, and Jin Sun. Still waters run deep in large-scale genome rearrangements of morphologically conservative Polyplacophora. *eLife*, 13, 2025.

[15844] Rui Vasco Simoes, Rafael Neto Henriques, Jonas L Olesen, Beatriz M Cardoso, Francisca F Fernandes, Mariana AV Monteiro, Sune N Jespersen, Tânia Carvalho, and Noam Shemesh. Deuterium metabolic imaging phenotypes mouse glioblastoma heterogeneity through glucose turnover kinetics. *eLife*, 13, 2025.

[15845] Franck Simon, Maria Colomba Comes, Tiziana Tocci, Louise Dupuis, Vincent Cabeli, Nikita Lagrange, Arianna Mencattini, Maria Carla Parrini, Eugenio Martinelli, and Herve Isambert. CausalXtract, a flexible pipeline to extract causal effects from live-cell time-lapse imaging data. *eLife*, 13, 2025.

[15846] Nelson V Simwela, Eleni Jaecklein, Christopher M Sassetti, and David G Russell. Impaired fatty acid import or catabolism in macrophages restricts intracellular growth of Mycobacterium tuberculosis. *eLife*, 13, 2025.

[15847] Davinder Singh, Yajun Liu, Yi-Hua Zhu, Sha Zhang, Shelby M Naegele, and Jian-Qiu Wu. Septins function in exocytosis via physical interactions with the exocyst complex in fission yeast cytokinesis. *eLife*, 13, 2025.

[15848] Vandana Singh and Scot P Ouellette. Altering the redox status of Chlamydia trachomatis directly impacts its developmental cycle progression. *eLife*, 13, 2025.

[15849] Ankur Sinha, Padraig Gleeson, Bóris Marin, Salvador Dura-Bernal, Sotirios Panagiotou, Sharon Crook, Matteo Cantarelli, Robert C Cannon, Andrew P Davison, Harsha Gurnani, and Robin Angus Silver. The NeuroML ecosystem for standardized multi-scale modeling in neuroscience. *eLife*, 13, 2025.

[15850] Nirvik Sinha, Sharon Israely, Ora Ben Harosh, Ran Harel, Julius PA Dewald, and Yifat Prut. Disentangling acute motor deficits and adaptive responses evoked by the loss of cerebellar output. *eLife*, 14, 2025.

[15851] Vendula Bohlen Šlechtová, TomáŠ Dvořák, Jörg Freyhof, Maurice Kottelat, Boris Levin, Alexander Golubtsov, Vlastimil Šlechta, and Joerg Bohlen. Reconstructing the phylogeny and evolutionary history of freshwater fishes (Nemacheilidae) across Eurasia since early Eocene. *eLife*, 13, 2025.

[15852] Nikolaos Smit and Martha M Robbins. Risk-taking incentives predict aggression heuristics in female gorillas. *eLife*, 14, 2025.

[15853] Laura J Smithson, Juliana L Zang, Lucas Junginger, Thomas J Waller, Lauren Reilly-Jankowiak, Sophia A Khan, Ye Li, Dawen Cai, and Catherine A Collins. Axonal injury signaling is restrained by a spared synaptic branch. *eLife*, 13, 2025.

[15854] Jisun So, Olivia Strobel, Jamie Wann, Kyungchan Kim, Avishek Paul, Dominic J Acri, Luke C Dabin, Jungsu Kim, Gang Peng, and Hyun Cheol Roh. Robust single-nucleus RNA sequencing reveals depot-specific cell population dynamics in adipose tissue remodeling during obesity. *eLife*, 13, 2025.

[15855] Octavia Soegyono, Elise Pepin, Beatrice K Leung, Billy Chieng, Bernard W Balleine, and Vincent Laurent. The influence of nucleus accumbens shell D1 and D2 neurons on outcome-specific Pavlovian instrumental transfer. *eLife*, 14, 2025.

[15856] Jelly HM Soffers, Erin Beck, Daniel J Sytkowski, Marianne E Maughan, Devasri Devarakonda, Yi Zhu, Beth A Wilson, Yu-Chieh David Chen, Ted Erclik, James W Truman, James B Skeath, and Haluk Lacin. A library of lineage-specific driver lines connects developing neuronal circuits to behavior in the Drosophila ventral nerve cord. *eLife*, 14, 2025.

[15857] Upneet K Sokhi, Ruoxi Yuan, Bikash Mishra, Yurii Chinenov, Anvita Singaraju, Karmela K Chan, Anne Bass, Richard D Bell, Laura Donlin, and Lionel B Ivashkiv. Opposing regulation of TNF responses by IFN-$\gamma$ and a PGE2-cAMP axis that is apparent in rheumatoid and immune checkpoint inhibitor-induced arthritis human IL-1$\beta$+ macrophages. *eLife*, 14, 2025.

[15858] Aryeh Solomon, Noa Bossel Ben-Moshe, Dotan Hoffman, Sébastien Trzebanski, Dror Yehezkel, Leia Vainman, Mihai G Netea, and Roi Avraham. Early and delayed STAT1-dependent responses drive local trained immunity of macrophages in the spleen. *eLife*, 13, 2025.

[15859] Magdalena Solyga and Georg B Keller. Multimodal mismatch responses in mouse auditory cortex. *eLife*, 13, 2025.

[15860] Bhanu Priya Somashekar and Upinder Singh Bhalla. Discriminating neural ensemble patterns through dendritic computations in randomly connected feedforward networks. *eLife*, 13, 2025.

[15861] Einav Somech, Debdatta Halder, Avishay Spitzer, Chaya Barbolin, Michael Tyler, Reut Halperin, Moshe Biton, Amit Tirosh, and Itay Tirosh. Subtypes and proliferation patterns of small intestine neuroendocrine tumors revealed by single-cell RNA sequencing. *eLife*, 13, 2025.

[15862] Hyun Beom Song, Laura Campello, Anupam Mondal, Holly Y Chen, Milton A English, Michael Glen, Phillip Vanlandingham, Rafal Farjo, and Anand Swaroop. Sex-specific attenuation of photoreceptor degeneration by reserpine in a rhodopsin P23H rat model of autosomal dominant retinitis pigmentosa. *eLife*, 14, 2025.

[15863] Ning Song, Hang Gao, Jianhao Li, Yi Liu, Mingze Wang, Zhiming Ma, Naisheng Zhang, and Wenlong Zhang. Microbiota from young mice counteracts susceptibility to age-related gout through modulating butyric acid levels in aged mice. *eLife*, 13, 2025.

[15864] Yan Song, Linda J Fothergill, Kari S Lee, Brandon Y Liu, Ada Koo, Mark Perelis, Shanti Diwakarla, Brid Callaghan, Jie Huang, Jill Wykosky, John B Furness, and Gene W Yeo. Stratification of enterochromaffin cells by single-cell expression analysis. *eLife*, 12, 2025.

[15865] Aneri Soni and Michael J Frank. Adaptive chunking improves effective working memory capacity in a prefrontal cortex and basal ganglia circuit. *eLife*, 13, 2025.

[15866] Tade Souaiaia, Hei Man Wu, Clive Hoggart, and Paul F O'Reilly. Sibling similarity can reveal key insights into genetic architecture. *eLife*, 12, 2025.

[15867] Riccardo Spott, Mathias W Pletz, Carolin Fleischmann-Struzek, Aurelia Kimmig, Christiane Hadlich, Matthias Hauert, Mara Lohde, Mateusz Jundzill, Mike Marquet, Petra Dickmann, Ruben Schüchner, Martin Hölzer, Denise Kühnert, and Christian Brandt. Leveraging mobility data to analyze persistent SARS-CoV-2 mutations and inform targeted genomic surveillance. *eLife*, 13, 2025.

[15868] Galen T Squiers, Chun Wan, James Gorder, Harrison Puscher, and Jingshi Shen. A Commander-independent function of COMMD3 in endosomal trafficking. *eLife*, 14, 2025.

[15869] Tino Stauber, Greta Moschini, Amro A Hussien, Patrick Klaus Jaeger, Katrien De Bock, and Jess G Snedeker. Il-6 signaling exacerbates hallmarks of chronic tendon disease by stimulating reparative fibroblasts. *eLife*, 12, 2025.

[15870] Philip Stavrides, Chris N Goulbourne, James Peddy, Chunfeng Huo, Mala Rao, Vinod Khetarpal, Deanna M Marchionini, Ralph A Nixon, and Dun-Sheng Yang. mTOR inhibition in Q175 Huntington's disease model mice facilitates neuronal autophagy and mutant huntingtin clearance. *eLife*, 14, 2025.

[15871] Christina Alexandra Stephens, Niek van Hilten, Lisa Zheng, and Michael Grabe. Simulation-based survey of TMEM16 family reveals that robust lipid scrambling requires an open groove. *eLife*, 14, 2025.

[15872] Robert G Stewart, Matthew James Marquis, Sooyeon Jo, Brandon J Harris, Aman S Aberra, Verity Cook, Zachary Whiddon, Vladimir Yarov-Yarovoy, Michael Ferns, and Jon T Sack. A Kv2 inhibitor combination reveals native neuronal conductances consistent with Kv2/KvS heteromers. *eLife*, 13, 2025.

[15873] William Stribling, Lindsey R Hall, Aubrey Powell, Casey Harless, Melissa J Martin, Brendan W Corey, Erik Snesrud, Ana Ong, Rosslyn Maybank, Jason Stam, Katelyn V Bartlett, Brendan T Jones, Lan N Preston, Katherine F Lane, Bernadette Thompson, Lynn M Young, Yoon I Kwak, Alice E Barsoumian, Ana Elizabeth Markelz, John L Kiley, Robert J Cybulski, Jason W Bennett, Patrick T Mc Gann, and Francois Lebreton. Detecting, mapping, and suppressing the spread of a decade-long Pseudomonas aeruginosa nosocomial outbreak with genomics. *eLife*, 13, 2025.

[15874] Eric V Strobl and Eric Gamazon. Discovering root causal genes with high-throughput perturbations. *eLife*, 13, 2025.

[15875] Jake Patrick Stroud, Michal Wojcik, Kristopher Torp Jensen, Makoto Kusunoki, Mikiko Kadohisa, Mark J Buckley, John Duncan, Mark G Stokes, and Mate Lengyel. Effects of noise and metabolic cost on cortical task representations. *eLife*, 13, 2025.

[15876] Hairui Su, Yong Sun, Han Guo, Chiao-Wang Sun, Qiuying Chen, Szumam Liu, Anlun Li, Min Gao, Rui Zhao, Glen Raffel, Jian Jin, Cheng-Kui Qu, Michael Yu, Christopher A Klug, George Y Zheng, Scott Ballinger, Matthew Kutny, Long X Zheng, Zechen Chong, Chamara Senevirathne, Steven Gross, Yabing Chen, Minkui Luo, and Xinyang Zhao. PRMT1-mediated metabolic reprogramming promotes leukemogenesis. *eLife*, 14, 2025.

[15877] Tsz Kin Suen, Burcu Al, Alice Scarpa, Anca Dorhoi, Mihai G Netea, and Katarzyna Placek. A dual nature of $\gamma\delta$ T cell immune memory responses. *eLife*, 14, 2025.

[15878] Mitsuru Sugimoto, Tadayuki Takagi, Tomohiro Suzuki, Hiroshi Shimizu, Goro Shibukawa, Yuki Nakajima, Yutaro Takeda, Yuki Noguchi, Reiko Kobayashi, Hidemichi Imamura, Hiroyuki Asama, Naoki Konno, Yuichi Waragai, Hidenobu Akatsuka, Rei Suzuki, Takuto Hikichi, and Hiromasa Ohira. A new preprocedural predictive risk model for post-endoscopic retrograde cholangiopancreatography pancreatitis: The SuPER model. *eLife*, 13, 2025.

[15879] Alisa Sukhina, Clemence Queriault, Saptarshi Roy, Elise Hall, Kelly Rome, Muskaan Aggarwal, Elizabeth Nunn, Ashley Weiss, Janet Nguyen, F Chris Bennett, and Will Bailis. Malnutrition drives infection susceptibility and dysregulated myelopoiesis that persists after refeeding intervention. *eLife*, 13, 2025.

[15880] Kelly G Sullivan and Greg J Bashaw. Commissureless acts as a substrate adapter in a conserved Nedd4 E3 ubiquitin ligase pathway to promote axon growth across the midline. *eLife*, 13, 2025.

[15881] Valeria Sulzyk, Ludmila Curci, Lucas N González, Abril Rebagliati Cid, Mariana Weigel Muñoz, and Patricia S Cuasnicu. The epididymis contributes to sperm DNA integrity and early embryo development through Cysteine-Rich Secretory Proteins. *eLife*, 13, 2025.

[15882] Mate Sumegi, Gaspar Olah, Istvan Paul Lukacs, Martin Blazsek, Judit K Makara, and Zoltan Nusser. Diverse calcium dynamics underlie place field formation in hippocampal CA1 pyramidal cells. *eLife*, 13, 2025.

[15883] Yuanqiang Sun, Yuange Duan, Peixiang Gao, Chenlu Liu, Kaichun Jin, Shengqian Dou, Wenxiong Tang, Hong Zhang, and Jian Lu. Upstream open reading frames buffer translational variability during Drosophila evolution and development. *eLife*, 14, 2025.

[15884] Yue Sun, Jingwei Li, Jiadai Huang, Shumin Li, Youyue Li, Beifang Lu, and Xin Deng. Architecture of genome-wide transcriptional regulatory network reveals dynamic functions and evolutionary trajectories in Pseudomonas syringae. *eLife*, 13, 2025.

[15885] Kevin Sung, Mackenzie M Johnson, Will Dumm, Noah Simon, Hugh Haddox, Julia Fukuyama, and Frederick A Matsen. Thrifty wide-context models of B cell receptor somatic hypermutation. *eLife*, 14, 2025.

[15886] Swastika Sur, Maggie Kerwin, Silvia Pineda, Poonam Sansanwal, Tara K Sigdel, Marina Sirota, and Minnie M Sarwal. Novel mechanism for tubular injury in nephropathic cystinosis. *eLife*, 13, 2025.

[15887] Elissa Sutlief, Charlie Walters, Tanya Marton, and Marshall G Hussain Shuler. The value of initiating a pursuit in temporal decision-making. *eLife*, 13, 2025.

[15888] Cassandra C Taber, Wenguang He, Geneviève MC Gasmi-Seabrook, Mia Hubert, Fraser G Ferens, Mitsuhiko Ikura, Jeffrey E Lee, and Michael Ohh. Erythrocytosis-inducing PHD2 mutations implicate biological role for N-terminal prolyl-hydroxylation in HIF1$\alpha$ oxygen-dependent degradation domain. *eLife*, 14, 2025.

[15889] Yurika Tagui, Shingo Takeda, Hiroyo Waida, Shoichi Kitahara, Tomoki Kimura, Shinji Kanda, Yoshitaka Oka, Yu Hayashi, and Chie Umatani. AgRP1 modulates breeding season-dependent feeding behavior in female medaka. *eLife*, 13, 2025.

[15890] Hiroshi Takagi, Shogo Ito, Jae Sung Shim, Akane Kubota, Andrew K Hempton, Nayoung Lee, Takamasa Suzuki, Jared S Wong, Chansie Yang, Christine T Nolan, Kerry L Bubb, Cristina M Alexandre, Daisuke Kurihara, Yoshikatsu Sato, Yasuomi Tada, Takatoshi Kiba, Jose L Pruneda-Paz, Christine Quietsch, Josh T Cuperus, and Takato Imaizumi. Companion cells with high florigen production express other small proteins and reveal a nitrogen-sensitive FT repressor. *eLife*, 14, 2025.

[15891] Suguru Takagi, Shiina Takano, Tomohiro Kubo, Yusaku Hashimoto, Shu Morise, Xiangsunze Zeng, and Akinao Nose. Segment-specific axon guidance by Wnt/Fz signaling diversifies motor commands in Drosophila larvae. *eLife*, 13, 2025.

[15892] Rajae Talbi, Todd L Stincic, Kaitlin Ferrari, Choi Ji Hae, Karol Walec, Elizabeth Medve, Achi Gerutshang, Silvia Leon, Elizabeth A McCarthy, Oline K Rønnekleiv, Martin J Kelly, and Victor M Navarro. POMC neurons control fertility through differential signaling of MC4R in kisspeptin neurons. *eLife*, 13, 2025.

[15893] Dandan Tan, Yidan Liu, Huaxia Luo, Qiang Shen, Xingbo Long, Luzheng Xu, Jieyu Liu, Nanbert A Zhong, Hong Zhang, and Hui Xiong. A novel mouse model for LAMA2-related muscular dystrophy with analysis of molecular pathogenesis and clinical phenotype. *eLife*, 13, 2025.

[15894] Gansheng Tan, Anna L Huguenard, Kara M Donovan, Phillip Demarest, Xiaoxuan Liu, Ziwei Li, Markus Adamek, Kory Lavine, Ananthv K Vellimana, Terrance T Kummer, Joshua W Osbun, Gregory J Zipfel, Peter Brunner, and Eric C Leuthardt. The effect of transcutaneous auricular vagus nerve stimulation on cardiovascular function in subarachnoid hemorrhage patients: A randomized trial. *eLife*, 13, 2025.

[15895] Yang Tan, Bingxin Zhou, Lirong Zheng, Guisheng Fan, and Liang Hong. Semantical and geometrical protein encoding toward enhanced bioactivity and thermostability. *eLife*, 13, 2025.

[15896] Toru Tanaka, Naoto Sasaki, Aga Krisnanda, Hilman Zulkifli Amin, Ken Ito, Sayo Horibe, Kazuhiko Matsuo, Ken-ichi Hirata, Takashi Nakayama, and Yoshiyuki Rikitake. C-C chemokine receptor 4 deficiency exacerbates early atherosclerosis in mice. *eLife*, 13, 2025.

[15897] Guoling Tang, Yaning Han, Xing Sun, Ruonan Zhang, Ming-Hu Han, Quanying Liu, and Pengfei Wei. Anti-drift pose tracker (ADPT), a transformer-based network for robust animal pose estimation cross-species. *eLife*, 13, 2025.

[15898] Tao Tang, Weiming Zhong, Puyu Tang, Rongsi Dai, Jiajing Guo, and Zhipeng Gao. Linalool combats Saprolegnia parasitica infections through direct killing of microbes and modulation of host immune system. *eLife*, 13, 2025.

[15899] Jing Tao, Charles A Nock, Eric B Searle, Shongming Huang, Rongzhou Man, Hua Yang, Grégoire T Freschet, Cyrille Violle, and Ji Zheng. Partitioning changes in ecosystem productivity by effects of species interactions in biodiversity experiments. *eLife*, 13, 2025.

[15900] Hiroki Taoka, Tadayoshi Murakawa, Kohei Kawaguchi, Michiko Koizumi, Tatsuya Kaminishi, Yuriko Sakamaki, Kaori Tanaka, Akihito Harada, Keiichi Inoue, Tomotake Kanki, Yasuyuki Ohkawa, and Naonobu Fujita. Transcriptional dynamics uncover the role of BNIP3 in mitophagy during muscle remodeling in Drosophila. *eLife*, 14, 2025.

[15901] Nathan Tardiff, Jiwon Kang, and Joshua I Gold. Normative evidence weighing and accumulation in correlated environments. *eLife*, 13, 2025.

[15902] Gillian AM Tarr, Linda Chui, Kim Stanford, Emmanuel W Bumunang, Rahat Zaheer, Vincent Li, Stephen B Freedman, Chad R Laing, and Tim A McAllister. Persistent cross-species transmission systems dominate Shiga toxin-producing Escherichia coli O157:H7 epidemiology in a high incidence region: A genomic epidemiology study. *eLife*, 13, 2025.

[15903] Olivia B Taylor, Nicholas DeGroff, Heithem M El-Hodiri, Chengyu Gao, and Andy J Fischer. Sphingosine-1-phosphate signaling regulates the ability of Müller glia to become neurogenic, proliferating progenitor-like cells. *eLife*, 13, 2025.

[15904] Xiangbin Teng and Ru-Yuan Zhang. Sequential temporal anticipation characterized by neural power modulation and in recurrent neural networks. *eLife*, 13, 2025.

[15905] Alexander L Tesmer, Paulius Viskaitis, Dane Donegan, Eva F Bracey, Nikola Grujic, Tommaso Patriarchi, Daria Peleg-Raibstein, and Denis Burdakov. Orexin population activity precisely reflects net body movement across behavioral and metabolic states. *eLife*, 14, 2025.

[15906] William R Thomas, Troy Richter, Erin T O'Neil, Cecilia Baldoni, Angelique Corthals, Dominik von Elverfeldt, John D Nieland, Dina Dechmann, Richard Hunter, and Liliana M Davalos. Seasonal and comparative evidence of adaptive gene expression in mammalian brain size plasticity. *eLife*, 13, 2025.

[15907] Aurore Thomazeau, Sabine Rannio, Jennifer A Brock, Hovy Ho-Wai Wong, and Per Jesper Sjöström. Neocortical layer-5 tLTD relies on non-ionotropic presynaptic NMDA receptor signaling. *eLife*, 14, 2025.

[15908] Christian Thome, Jan Maximilian Janssen, Seda Karabulut, Claudio Acuna, Elisa D'Este, Stella J Soyka, Konrad Baum, Michael Bock, Nadja Lehmann, Johannes Roos, Nikolas A Stevens, Masashi Hasegawa, Dan A Ganea, Chloé M Benoit, Jan Gründemann, Lia Y Min, Kalynn M Bird, Christian Schultz, Vann Bennett, Paul Jenkins, and Maren Engelhardt. Live imaging of excitable axonal microdomains in ankyrin-G-GFP mice. *eLife*, 12, 2025.

[15909] Li Tian, Zongzheng Zhao, Wenying Gao, Zirui Liu, Xiao Li, Wenyan Zhang, and Zhaolong Li. SARS-CoV-2 nsp16 is regulated by host E3 ubiquitin ligases, UBR5 and MARCHF7. *eLife*, 13, 2025.

[15910] Zhengming Tian, Yuning Li, Feiyang Jin, Zirui Xu, Yakun Gu, Mengyuan Guo, Qianqian Shao, Yingxia Liu, Hanjiang Luo, Yue Wang, Suyu Zhang, Chenlu Yang, Xin Liu, Xunming Ji, and Jia Liu. Brain-derived exosomal hemoglobin transfer contributes to neuronal mitochondrial homeostasis under hypoxia. *eLife*, 13, 2025.

[15911] Hieng Chiong Tie, Haiyun Wang, Divyanshu Mahajan, Hilbert Yuen In Lam, Xiuping Sun, Bing Chen, Yuguang Mu, and Lei Lu. Quantitative intra-Golgi transport and organization data suggest the stable compartment nature of the Golgi. *eLife*, 13, 2025.

[15912] Kathryn E Tiedje, Qi Zhan, Shazia Ruybal-Pésantez, Gerry Tonkin-Hill, Qixin He, Mun Hua Tan, Dionne C Argyropoulos, Samantha Deed, Anita Ghansah, Oscar Bangre, Abraham R Oduro,

Kwadwo A Koram, Mercedes Pascual, and Karen P Day. Measuring changes in Plasmodium falciparum census population size in response to sequential malaria control interventions. *eLife*, 12, 2025.

[15913] Silas Tittes, Anne Lorant, Sean P McGinty, James B Holland, Jose de Jesus Sánchez-González, Arun Seetharam, Maud Tenaillon, and Jeffrey Ross-Ibarra. The population genetics of convergent adaptation in maize and teosinte is not locally restricted. *eLife*, 12, 2025.

[15914] Gemechu Bekele Tolossa, Aidan M Schneider, Eva Dyer, and Keith B Hengen. Neurons throughout the brain embed robust signatures of their anatomical location into spike trains. *eLife*, 13, 2025.

[15915] Patsy R Tomlinson, Rachel G Knox, Olga Perisic, Helen Su, Gemma V Brierley, Roger L Williams, and Robert K Semple. Paradoxical dominant negative activity of an immunodeficiency-associated activating PIK3R1 variant. *eLife*, 13, 2025.

[15916] Ramin Toosi, Behnam Karami, Roxana Koushki, Farideh Shakerian, Jalaledin Noroozi, Ehsan Rezayat, Abdol-Hossein Vahabie, Mohammad Ali Akhaee, and Mohammad-Reza A Dehaqani. The spatial frequency representation predicts category coding in the inferior temporal cortex. *eLife*, 13, 2025.

[15917] Balázs István Tóth, Bahar Bazeli, Annelies Janssens, Erika Lisztes, Márk Racskó, Balázs Kelemen, Mihály Herczeg, Tamás Milán Nagy, Katalin E Kövér, Argha Mitra, Attila Borics, Tamás Bíró, and Thomas Voets. Direct modulation of TRPM8 ion channels by rapamycin and analog macrolide immunosuppressants. *eLife*, 13, 2025.

[15918] Hiroki Toyoda, Doyun Kim, Byeong Geon Koh, Tomomi Sano, Takashi Kanematsu, Seog Bae Oh, and Youngnam Kang. Chronic stress impairs autoinhibition in neurons of the locus coeruleus to increase asparagine endopeptidase activity. *eLife*, 14, 2025.

[15919] Quang Thinh Trac, Yue Huang, Tom Erkers, Päivi östling, Anna Bohlin, Albin Osterroos, Mattias Vesterlund, Rozbeh Jafari, Ioannis Siavelis, Helena Backvall, Santeri Kiviluoto, Lukas Orre, Mattias Rantalainen, Janne Lehtiö, Soren Lehmann, Olli Kallioniemi, Yudi Pawitan, and Trung Nghia Vu. Pathway activation model for personalized prediction of drug synergy. *eLife*, 13, 2025.

[15920] Brandon T Tran, Vidthiya Jeyanathan, Ruoqiong Cao, Eva Kaufmann, and Katherine Y King. Hematopoietic stem and progenitor cells as a reservoir for trained immunity. *eLife*, 14, 2025.

[15921] My Tran, Angel J Hernandez Viera, Patricia Q Tran, Erick D Nilsen, Lily Tran, and Charlie Y Mo. Bacteriophage infection drives loss of $\beta$-lactam resistance in methicillin-resistant Staphylococcus aureus. *eLife*, 13, 2025.

[15922] Khanh B Trang, Matthew C Pahl, James A Pippin, Chun Su, Sheridan H Littleton, Prabhat Sharma, Nikhil N Kulkarni, Louis R Ghanem, Natalie A Terry, Joan M O'Brien, Yadav Wagley, Kurt D Hankenson, Ashley Jermusyk, Jason Hoskins, Laufey T Amundadottir, Mai Xu, Kevin Brown, Stewart Anderson, Wenli Yang, Paul Titchenell, Patrick Seale, Klaus H Kaestner, Laura Cook, Megan Levings, Babette S Zemel, Alessandra Chesi, Andrew D Wells, and Struan FA Grant. 3D genomic features across >50 diverse cell types reveal insights into the genomic architecture of childhood obesity. *eLife*, 13, 2025.

[15923] Laura Troyer, Yu-Huan Wang, Shobhna Shobhna, Seunghyeon Kim, Brooke Ramsey, Jeechul Woo, Emad Tajkhorshid, and Sangjin Kim. Roles of the membrane-binding motif and the C-terminal domain of RNase E in localization and diffusion in E. coli. *eLife*, 14, 2025.

[15924] Massimo Trusel, Ziran Zhao, Danyal H Alam, Ethan S Marks, Maaya Z Ikeda, and Todd F Roberts. Synaptic connectivity of sensorimotor circuits for vocal imitation in the songbird. *eLife*, 14, 2025.

[15925] Jean Marc Tsagmo Ngoune, Parul Sharma, Aline Crouzols, Nathalie Petiot, and Brice Rotureau. Stumpy forms are the predominant transmissible forms of Trypanosoma brucei. *eLife*, 12, 2025.

[15926] Erika Tsingos, Advaita M Dick, and Baubak Bajoghli. Altered thymic niche synergistically drives the massive proliferation of malignant thymocytes. *eLife*, 13, 2025.

[15927] Kaima Tsukada, Rikiya Imamura, Tomoko Miyake, Kotaro Saikawa, Mizuki Saito, Naoya Kase, Lingyan Fu, Masamichi Ishiai, Yoshihisa Matsumoto, and Mikio Shimada. CDK-mediated phosphorylation of PNKP is required for end-processing of single-strand DNA gaps on Okazaki fragments and genome stability. *eLife*, 14, 2025.

[15928] Takayuki Tsurumi, Ayaka Kato, Arvind Kumar, and Kenji Morita. Online reinforcement learning of state representation in recurrent network supported by the power of random feedback and biological constraints. *eLife*, 14, 2025.

[15929] Gaqi Tu, Peiying Wen, Adel Halawa, and Kaori Takehara-Nishiuchi. Acetylcholine modulates prefrontal outcome coding during threat learning under uncertainty. *eLife*, 13, 2025.

[15930] Cecilia Tubert, Rodrigo Manuel Paz, Agostina Mónica Stahl, Kianny Miroslava Sanchez Armijos, Lorena Rela, and Mario Gustavo Murer. Striatal cholinergic interneuron pause response requires Kv1 channels, is absent in dyskinetic mice, and is restored by dopamine D5 receptor inverse agonism. *eLife*, 13, 2025.

[15931] Angela M Tuckowski, Safa Beydoun, Elizabeth S Kitto, Ajay Bhat, Marshall B Howington, Aditya Sridhar, Mira Bhandari, Kelly Chambers, and Scott F Leiser. fmo-4 promotes longevity and stress resistance via ER to mitochondria calcium regulation in C. elegans. *eLife*, 13, 2025.

[15932] Markus R Tünte, Stefanie Hoehl, Moritz Wunderwald, Johannes Bullinger, Asena Boyadziheva, Lara Maister, Birgit Elsner, Manos Tsakiris, and Ezgi Kayhan. Respiratory and cardiac interoceptive sensitivity in the first two years of life. *eLife*, 12, 2025.

[15933] Gergely F Turi, Sasa Teng, Xinyue Chen, Emily CY Lim, Carla Dias, Ruining Hu, Ruizhi Wang, Fenghua Zhen, and Yueqing Peng. Serotonin modulates infraslow oscillation in the dentate gyrus during non-REM sleep. *eLife*, 13, 2025.

[15934] Kevin Turner, Dakota Brockway, Md Shakhawat Hossain, Keith Griffith, Denver Greenawalt, Qingguang Zhang, Kyle Gheres, Nicole Crowley, and Patrick J Drew. Type-I nNOS neurons orchestrate cortical neural activity and vasomotion. *eLife*, 14, 2025.

[15935] Katarzyna Tutak, Izabela Broniarek, Andrzej Zielezinski, Daria Niewiadomska, Tomasz Skrzypczak, Anna Baud, and Krzysztof Sobczak. Insufficiency of 40S ribosomal proteins, RPS26 and RPS25, negatively affects biosynthesis of polyglycine-containing proteins in fragile-X associated conditions. *eLife*, 13, 2025.

[15936] Masaaki Uematsu and Jeremy M Baskin. Barcode-free multiplex plasmid sequencing using Bayesian analysis and nanopore sequencing. *eLife*, 12, 2025.

[15937] Tomohiro Umeda, Ayumi Sakai, Rumi Uekado, Keiko Shigemori, Ryota Nakajima, Kei Yamana, and Takami Tomiyama. Simply crushed zizyphi spinosi semen prevents neurodegenerative diseases and reverses age-related cognitive decline in mice. *eLife*, 13, 2025.

[15938] Nobuhisa Umeki, Yoshiyuki Kabashima, and Yasushi Sako. Evaluation of information flows in the RAS-MAPK system using transfer entropy measurements. *eLife*, 14, 2025.

[15939] Miki Umetani, Miho Fujisawa, Reiko Okura, Takashi Nozoe, Shoichi Suenaga, Hidenori Nakaoka, Edo Kussell, and Yuichi Wakamoto. Observation of persister cell histories reveals diverse modes of survival in antibiotic persistence. *eLife*, 14, 2025.

[15940] Marta Urbanska, Yan Ge, Maria Winzi, Shada Abuhattum, Syed Shafat Ali, Maik Herbig, Martin Kräter, Nicole Toepfner, Joanne Durgan, Oliver Florey, Martina Dori, Federico Calegari, Fidel-Nicolás Lolo, Miguel ángel del Pozo, Anna Taubenberger, Carlo Vittorio Cannistraci, and Jochen Guck. De novo identification of universal cell mechanics gene signatures. *eLife*, 12, 2025.

[15941] Saima Usman, William Andrew Yeudall, Muy-Teck Teh, Fatemah Ghloum, Hemanth Tummala, and Ahmad Waseem. A single cysteine residue in vimentin regulates long non-coding RNA XIST to suppress epithelial–mesenchymal transition and stemness in breast cancer. *eLife*, 14, 2025.

[15942] Julie Uveira, Antoine Donati, Marvin Léria, Marion Lechable, FranÇois Lahaye, Christine Vesque, Evelyn Houliston, and Tsuyoshi Momose. Planar cell polarity coordination in a cnidarian embryo provides clues to animal body axis evolution. *eLife*, 14, 2025.

[15943] Dylan A Valencia, Angela N Koeberlein, Haruko Nakano, Akos Rudas, Aanand A Patel, Airi Harui, Cassandra Spencer, Atsushi Nakano, and Margot E Quinlan. Human formin FHOD3-mediated actin elongation is required for sarcomere integrity in cardiomyocytes. *eLife*, 13, 2025.

[15944] José Antonio Valer, Alexandre Deber, Marius Wits, Carolina Pimenta-Lope, Marie-José Goumans, Jose Luis Rosa, Gonzalo Sánchez-Duffhues, and Francesc Ventura. PI3K$\alpha$ inhibition blocks osteochondroprogenitor specification and the hyper-inflammatory response to prevent heterotopic ossification. *eLife*, 12, 2025.

[15945] Timo van Kerkoerle, Louise Pape, Milad Ekramnia, Xiaoxia Feng, Jordy Tasserie, Morgan Dupont, Xiaolian Li, Béchir Jarraya, Wim Vanduffel, Stanislas Dehaene, and Ghislaine Dehaene-Lambertz. Brain areas for reversible symbolic reference, a potential singularity of the human brain. *eLife*, 12, 2025.

[15946] Matthijs van Veelen. The general version of Hamilton's rule. *eLife*, 14, 2025.

[15947] Artem K Velichko, Nadezhda V Petrova, Dmitry A Deriglazov, Anastasia P Kovina, Artem V Luzhin, Eugene P Kazakov, Igor I Kireev, Sergey Razin, and Omar L Kantidze. Treacle's ability to form liquid-like phase condensates is essential for nucleolar fibrillar center assembly, efficient rRNA transcription and processing, and rRNA gene repair. *eLife*, 13, 2025.

[15948] Aaron M Ver Heul, Madison Mack, Lydia Zamidar, Masato Tamari, Ting-Lin Yang, Anna M Trier, Do-Hyun Kim, Hannah Janzen-Meza, Steven J Van Dyken, Chyi-Song Hsieh, Jenny M Karo, Joseph C Sun, and Brian S Kim. RAG suppresses group 2 innate lymphoid cells. *eLife*, 13, 2025.

[15949] Csaba Verasztó, Sanja Jasek, Martin Gühmann, Luis Alberto Bezares-Calderón, Elizabeth A Williams, Réza Shahidi, and Gáspár Jékely. Whole-body connectome of a segmented annelid larva. *eLife*, 13, 2025.

[15950] Anil K Verma Rodriguez, Josue O Ramírez-Jarquin, Román Rossi-Pool, and Fatuel Tecuapetla. Basal ganglia output (entopeduncular nucleus) coding of contextual kinematics and reward in the freely moving mouse. *eLife*, 13, 2025.

[15951] Iva Veseli, Yiqun T Chen, Matthew S Schechter, Chiara Vanni, Emily C Fogarty, Andrea R Watson, Bana Jabri, Ran Blekhman, Amy D Willis, Michael K Yu, Antonio Fernàndez-Guerra, Jessika Füssel, and A Murat Eren. Microbes with higher metabolic independence are enriched in human gut microbiomes under stress. *eLife*, 12, 2025.

[15952] Rocio Vicario, Stamatina Fragkogianni, Leslie Weber, Tomi Lazarov, Yang Hu, Samantha Y Hayashi, Barbara Craddock, Nicholas D Socci, Araitz Alberdi, Ann Baako, Oyku Ay, Masato Ogishi, Estibaliz Lopez-Rodrigo, Rajya Kappagantula, Agnes Viale, Christine A Iacobuzio-Donahue, Ting Zhou, Richard M Ransohoff, Richard Chesworth, Omar Abdel-Wahab, Bertrand Boisson, Olivier Elemento, Jean-Laurent Casanova, W Todd Miller, and Frédéric Geissmann. A microglia clonal inflammatory disorder in Alzheimer's disease. *eLife*, 13, 2025.

[15953] Federico A Vignale, Andrea Hernandez Garcia, Carlos P Modenutti, Ezequiel J Sosa, Lucas A Defelipe, Renato Oliveira, Gisele L Nunes, Raúl M Acevedo, German F Burguener, Sebastian M Rossi, Pedro D Zapata, Dardo A Marti, Pedro Sansberro, Guilherme Oliveira, Emily M Catania, Madeline N Smith, Nicole M Dubs, Satish Nair, Todd J Barkman, and Adrian G Turjanski. Yerba mate (Ilex paraguariensis) genome provides new insights into convergent evolution of caffeine biosynthesis. *eLife*, 14, 2025.

[15954] Mykhailo Vladymyrov, Luca Marchetti, Sidar Aydin, Sasha GN Soldati, Adrien Mossu, Arindam Pal, Laurent Gueissaz, Akitaka Ariga, and Britta Engelhardt. UFMTrack, an Under-Flow Migration Tracker enabling analysis of the entire multi-step immune cell extravasation cascade across the blood-brain barrier in microfluidic devices. *eLife*, 13, 2025.

[15955] Kaspar E Vogt, Ashwinikumar Kulkarni, Richa Pandi, Mantre Dehnad, Genevieve Konopka, and Robert W Greene. Sleep need driven oscillation of glutamate synaptic phenotype. *eLife*, 13, 2025.

[15956] Mehul Vora, Jonathan Dietz, Zachary Wing, Karen George, Jun Kelly Liu, Christopher Rongo, and Cathy Savage-Dunn. Genome-wide analysis of Smad and Schnurri transcription factors in C. elegans demonstrates widespread interaction and a function in collagen secretion. *eLife*, 13, 2025.

[15957] Mihály Vöröslakos, Yunchang Zhang, Kathryn McClain, Roman Huszár, Aryeh Rothstein, and György Buzsáki. ThermoMaze behavioral paradigm for assessing immobility-related brain events in rodents. *eLife*, 12, 2025.

[15958] Gabriel Wainstein, Christopher J Whyte, Kaylena A Ehgoetz Martens, Eli J Müller, Vicente Medel, Britt Anderson, Elisabeth Stöttinger, James Danckert, Brandon R Munn, and James M Shine. Evidence from pupillometry, fMRI, and RNN modelling shows that gain neuromodulation mediates task-relevant perceptual switches. *eLife*, 13, 2025.

[15959] Melanie Rose Walker, Preston Leung, Elizabeth Keoshkerian, Mehdi R Pirozyan, Andrew Lloyd, Fabio Luciani, and Rowena A Bull. Temporal dynamics of viral fitness and the adaptive immune response in HCV infection. *eLife*, 13, 2025.

[15960] Raven Star Wallace, Bronte Mckeown, Ian Goodall-Halliwell, Louis Chitiz, Philippe Forest, Theodoros Karapanagiotidis, Bridget Mulholland, Adam Turnbull, Tamara Vanderwal, Samyogita Hardikar, Tirso RJ Gonzalez Alam, Boris C Bernhardt, Hao-Ting Wang, Will Strawson, Michael Milham, Ting Xu, Daniel S Margulies, Giulia L Poerio, Elizabeth Jefferies, Jeremy I Skipper, Jeffrey D Wammes, Robert Leech, and Jonathan Smallwood. Mapping patterns of thought onto brain activity during movie-watching. *eLife*, 13, 2025.

[15961] Brenna Walton, Serena Abbodante, Michaela Ellen Marshall, Justyna M Dobruchowska, Amani Alvi, Larry A Gallagher, Nikhil Vallikat, Zhemin Zhang, Daniel J Wozniak, Edward W Yu, Geert-Jan Boons, Eric Pearlman, and Arne Rietsch. A biofilm-tropic Pseudomonas aeruginosa bacteriophage uses the exopolysaccharide Psl as receptor. *eLife*, 13, 2025.

[15962] Baikui Wang, Xianqi Peng, Xiao Zhou, Xiuyan Jin, Abubakar Siddique, Jiayun Yao, Haiqi Zhang, Weifen Li, Yan Li, and Min Yue. Lacticaseibacillus rhamnosus P118 enhances host tolerance to Salmonella infection by promoting microbe-derived indole metabolites. *eLife*, 13, 2025.

[15963] Gaohang Wang, Wei Qi, Rui Huang, and Liming Wang. Sexual failure decreases sweet taste perception in male Drosophila via dopaminergic signaling. *eLife*, 14, 2025.

[15964] Huazhen Wang, Ludivine Bertonnier-Brouty, Isabella Artner, Jiayu Wen, and Qi Dai. Drosophila hamlet mediates epithelial tissue assembly of the reproductive system. *eLife*, 13, 2025.

[15965] Huei-Bin Wang, Natalie E Smale, Sarah H Brown, Sophia AMB Villanueva, David Zhou, Aly Mulji, Deap S Bhandal, Kyle Nguyen-Ngo, John R Harvey, Cristina A Ghiani, and Christopher S Colwell. Scheduled feeding improves behavioral outcomes and reduces inflammation in a mouse model of fragile X syndrome. *eLife*, 14, 2025.

[15966] Jie Wang, Jun Nishiyama, Paula Parra-Bueno, Elwy Okaz, Goksu Oz, Xiaodan Liu, Tetsuya Watabe, Irena Suponitsky-Kroyter, Timothy E McGraw, Erzsebet M Szatmari, and Ryohei Yasuda. Rab10 inactivation promotes AMPAR trafficking and spine enlargement during long-term potentiation. *eLife*, 13, 2025.

[15967] Jiongke Wang, Yujie Shi, Ying Wang, Yingqiang Shen, Huan Liu, Silu Sun, Yimei Wang, Xikun Zhou, Yu Zhou, Xin Zeng, Jing Li, and Qianming Chen. PA28$\gamma$ promotes the malignant progression of tumor by elevating mitochondrial function via C1QBP. *eLife*, 13, 2025.

[15968] Junxiu Wang, Matthana Klakong, Qiuyu Zhu, Jinting Pan, Yudie Duan, Lirong Wang, Yong Li, Jiangbo Dang, Danlong Jing, and Hong Zhou. Insight into the bioactivity and action mode of betulin, a candidate aphicide from plant metabolite, against aphids. *eLife*, 14, 2025.

[15969] Lianlei Wang, Xinyu Yang, Sen Zhao, Pengfei Zheng, Wen Wen, Kexin Xu, Xi Cheng, Qing Li, Anas M Khanshour, Yoshinao Koike, Junjun Liu, Xin Fan, Nao Otomo, Zefu Chen, Yaqi Li, Lulu Li, Haibo Xie, Panpan Zhu, Xiaoxin Li, Yuchen Niu, Shengru Wang, Sen Liu, Suomao Yuan, Chikashi Terao, Ziquan Li, Shaoke Chen, Xiuli Zhao, Pengfei Liu, Jennifer E Posey, Zhihong Wu, Guixing Qiu, Shiro Ikegawa, James R Lupski, Jonathan J Rios, Carol A Wise, Jianguo T Zhang, Chengtian Zhao, and Nan Wu. EPHA4 signaling dysregulation links abnormal locomotion and the development of idiopathic scoliosis. *eLife*, 13, 2025.

[15970] Liyi Wang, Shiqi Liu, Shu Zhang, Yizhen Wang, Yanbing Zhou, and Tizhong Shan. Single-nucleus transcriptomics reveal the cytological mechanism of conjugated linoleic acids in regulating intramuscular fat deposition. *eLife*, 13, 2025.

[15971] Long Wang, Christo Morison, and Weini Huang. Cancer–immune coevolution dictated by antigenic mutation accumulation. *eLife*, 14, 2025.

[15972] Ming-Qiang Wang, Shi-Kun Guo, Peng-Fei Guo, Juan-Juan Yang, Guo-Ai Chen, Douglas Chesters, Michael C Orr, Ze-Qing Niu, Michael Staab, Jing-Ting Chen, Yi Li, Qing-Song Zhou, Felix Fornoff, Xiaoyu Shi, Shan Li, Massimo Martini, Alexandra-Maria Klein, Andreas Schuldt, Xiaojuan Liu, Keping Ma, Helge Bruelheide, Arong Luo, and Chao-Dong Zhu. Multidimensionality of tree communities structure host-parasitoid networks and their phylogenetic composition. *eLife*, 13, 2025.

[15973] Ning Wang, Yimeng Wang, Mingkun Guo, Ling Wang, Xueling Wang, Nan Zhu, Jiajia Yang, Lei Wang, Chenguang Zheng, and Dong Ming. Dynamic gamma modulation of hippocampal place cells predominates development of theta sequences. *eLife*, 13, 2025.

[15974] Qian Wang, Hongge Li, Yingyu Mao, Ankur Garg, Eun Sil Park, Yihua Wu, Alyssa Chow, John Peregrin, and Xin Zhang. Shc1 cooperates with Frs2 and Shp2 to recruit Grb2 in FGF-induced lens development. *eLife*, 13, 2025.

[15975] Shimin Wang, Hongxian Wang, Bicheng Jin, Hongli Yan, Qingliang Zheng, and Dong Zhao. scRNA-seq and scATAC-seq reveal that Sertoli cell mediates spermatogenesis disorders through stage-specific communications in non-obstructive azoospermia. *eLife*, 13, 2025.

[15976] Siqi Wang, Danyue Dong, Xin Li, and Zefeng Wang. Pan-tissue transcriptome analysis reveals sex-dimorphic human aging. *eLife*, 13, 2025.

[15977] Weihua Wang, Junqiao Xing, Xiqi Zhang, Hongni Liu, Xingyu Liu, Haochen Jiang, Cheng Xu, Xue Zhao, and Zhangfeng Hu. Control of ciliary transcriptional programs during spermatogenesis by antagonistic transcription factors. *eLife*, 13, 2025.

[15978] Xin Wang. Overflow metabolism originates from growth optimization and cell heterogeneity. *eLife*, 13, 2025.

[15979] Yi Wang, Thomas Nguyen, Qingan He, Oliver Has, Kiarash Forouzesh, and Dae Seok Eom. Cytoneme-mediated intercellular signaling in keratinocytes is essential for epidermal remodeling in zebrafish. *eLife*, 13, 2025.

[15980] Yike Wang, Matthew R Lowerison, Zhe Huang, Qi You, Bing-Ze Lin, Daniel A Llano, and Pengfei Song. Longitudinal awake imaging of mouse deep brain microvasculature with super-resolution ultrasound localization microscopy. *eLife*, 13, 2025.

[15981] Yuqian Wang, Guibin Wang, Lishan Zhang, Qilan Cai, Meizhen Lin, Dongping Huang, Yuyue Xie, Wenxiong Lin, and Xiangmin Lin. Aeromonas hydrophila CobQ is a new type of NAD+- and Zn2+-independent protein lysine deacetylase. *eLife*, 13, 2025.

[15982] Zeng-Hang Wang, Ying An, Ting Zhao, Tong-Tong Pei, Dora Yuping Wang, Xiaoye Liang, Wenming Qin, and Tao Dong. Amidase and lysozyme dual functions in TseP reveal a new family of chimeric effectors in the type VI secretion system. *eLife*, 13, 2025.

[15983] Zezhen Wang, Weihao Mai, Yuming Chai, Kexin Qi, Hongtai Ren, Chen Shen, Shiwu Zhang, Guodong Tan, Yu Hu, and Quan Wen. The geometry and dimensionality of brain-wide activity. *eLife*, 14, 2025.

[15984] Zhen Wang, Jie Zhu, Mengda Xu, Xuyuan Ma, Maozheng Shen, Jingyu Yan, Guosheng Gan, and Xiang Zhou. Transplantation of exogenous mitochondria mitigates myocardial dysfunction after cardiac arrest. *eLife*, 13, 2025.

[15985] Zhiyan Wang, Nore Ojogun, Yiling Liu, Lu Gan, Zeling Xiao, Jintao Feng, Wei Jiang, Yeying Chen, Benkun Zou, ChengYun Yu, Changshun Li, Asha Ashuo, Xiaobo Li, Mingsheng Fu, Jian Wu, Yiwei Chu, Robert S Munford, and Mingfang Lu. A host enzyme reduces metabolic dysfunction-associated steatotic liver disease (MASLD) by inactivating intestinal lipopolysaccharide. *eLife*, 13, 2025.

[15986] Cody J Warren, Arturo Barbachano-Guerrero, Vanessa L Bauer, Alex Stabell, Obaiah Dirasantha, Qing Yang, and Sara L Sawyer. Adaptation of CD4 in gorillas and chimpanzees conveyed resistance to simian immunodeficiency viruses. *eLife*, 13, 2025.

[15987] Denise Weber, Katrin Vogt, Anton Miroschnikow, Michael J Pankratz, and Andreas S Thum. Four individually identified paired dopamine neurons signal taste punishment in larval Drosophila. *eLife*, 12, 2025.

[15988] Amy K Webster, John H Willis, Erik Johnson, Peter Sarkies, and Patrick C Phillips. Gene expression variation across genetically identical individuals predicts reproductive traits. *eLife*, 14, 2025.

[15989] Hongping Wei, Thomas Ka Chung Lam, and Hokto Kazama. Odors drive feeding through gustatory receptor neurons in Drosophila. *eLife*, 13, 2025.

[15990] Lin Wei, Hai-Yan Cao, Ruyi Zou, Min Du, Qingdong Zhang, Danrong Lu, Xiangyu Xu, Yingying Xu, Wenshuang Wang, Xiu-Lan Chen, Yu-Zhong Zhang, and Fuchuan Li. Crystal structure and catalytic mechanism of PL35 family glycosaminoglycan lyases with an ultrabroad substrate spectrum. *eLife*, 13, 2025.

[15991] Simon Weiler, Manuel Teichert, and Troy W Margrie. Layer 6 corticocortical neurons are a major route for intra- and interhemispheric feedback. *eLife*, 13, 2025.

[15992] Eric Weine, Samuel Pattillo Smith, Rebecca Kathryn Knowlton, and Arbel Harpak. Trade-offs in modeling context dependency in complex trait genetics. *eLife*, 13, 2025.

[15993] Ziyu Wen, Pingchao Li, Yue Yuan, Congcong Wang, Minchao Li, Haohang Wang, Minjuan Shi, Yizi He, Mingting Cui, Ling Chen, and Caijun Sun. Purging viral latency by a bifunctional HSV-vectored therapeutic vaccine in chronically SIV-infected macaques. *eLife*, 13, 2025.

[15994] Brooke E White, Carolyn L Hodo, Sarah Hamer, Ashley B Saunders, Susana A Laucella, Daniel B Hall, and Rick L Tarleton. Serial 'deep-sampling' PCR of fragmented DNA reveals the wide range of Trypanosoma cruzi burden among chronically infected human, macaque, and canine hosts, and allows accurate monitoring of parasite load following treatment. *eLife*, 14, 2025.

[15995] Wojciech Wietrzynski, Lorenz Lamm, William HJ Wood, Matina-Jasemi Loukeri, Lorna Malone, Tingying Peng, Matthew P Johnson, and Benjamin D Engel. Molecular architecture of thylakoid membranes within intact spinach chloroplasts. *eLife*, 14, 2025.

[15996] Anne L Willems, Lukas Van Oudenhove, and Bram Vervliet. Omissions of threat trigger subjective relief and prediction error-like signaling in the human reward and salience systems. *eLife*, 12, 2025.

[15997] David C Williams, Amanda Chu, Nicholas T Gordon, Aleah M DuBois, Suhui Qian, Genevieve Valvo, Selena Shen, Jacob B Boyce, Anaise C Fitzpatrick, Mahsa Moaddab, Emma L Russell, Liliuokalani H Counsman, and Michael A McDannald. Ethograms predict visual fear conditioning status in rats. *eLife*, 14, 2025.

[15998] Katharina Anna Wilmes, Mihai A Petrovici, Shankar Sachidhanandam, and Walter Senn. Uncertainty-modulated prediction errors in cortical microcircuits. *eLife*, 13, 2025.

[15999] Lucy A Winder, Mirre JP Simons, and Terry Burke. No evidence for a trade-off between reproduction and survival in a meta-analysis across birds. *eLife*, 12, 2025.

[16000] Kevin Winker and Kira Delmore. Seasonally migratory songbirds have different historic population size characteristics than resident relatives. *eLife*, 12, 2025.

[16001] Lucie Winkler, Markus Butz, Abhinav Sharma, Jan Vesper, Alfons Schnitzler, Petra Fischer, and Jan Hirschmann. Context-dependent modulations of subthalamo-cortical synchronization during rapid reversals of movement direction in Parkinson's disease. *eLife*, 13, 2025.

[16002] Tanya Wolff, Mark Eddison, Nan Chen, Aljoscha Nern, Preeti Sundaramurthi, Divya Sitaraman, and Gerald M Rubin. Cell type-specific driver lines targeting the Drosophila central complex and their use to investigate neuropeptide expression and sleep regulation. *eLife*, 14, 2025.

[16003] David W Wolfson, Joshua A Hull, Yongwu Li, Trevor J Gonzalez, Mourya D Jayaram, Garth W Devlin, Valentina Cigliola, Kelsey A Oonk, Alan Rosales, Nenad Bursac, Aravind Asokan, and Kenneth D Poss. Spatial and longitudinal tracking of enhancer-AAV vectors that target transgene expression to injured mouse myocardium. *eLife*, 14, 2025.

[16004] Francesca S Wong, Alina B Thomas, Simon Killcross, Vincent Laurent, R Fred Westbrook, and Nathan M Holmes. Integration of sensory and fear memories in the rat medial temporal lobe. *eLife*, 13, 2025.

[16005] Wei Jie Wong, Yi Wen Zhu, Hai Ting Wang, Jia Wen Qian, Ziyi Li, Song Li, Zhao Yuan Liu, Wei Guo, Shuang Yan Zhang, Bing Su, Fang Ping He, Kang Wang, and Florent Ginhoux. Modeling hereditary diffuse leukoencephalopathy with axonal spheroids using microglia-sufficient brain organoids. *eLife*, 13, 2025.

[16006] J Luke Woodward, Jeffrey Matthew, Rutuparna Joshi, Vishakha Vishwakarma, Ying Xiao, and SeYeon Chung. Sulfation affects apical extracellular matrix organization during development of the Drosophila embryonic salivary gland tube. *eLife*, 14, 2025.

[16007] Dan Wu, Venkateswararao Eeda, Zahra Maria, Komal Rawal, Audrey Wang, Oana Herlea-Pana, Ram Babu Undi, Hui-Ying Lim, and Weidong Wang. Targeting IRE1$\alpha$ improves insulin sensitivity and thermogenesis and suppresses metabolically active adipose tissue macrophages in male obese mice. *eLife*, 13, 2025.

[16008] Kingsley Y Wu, Ta I Hung, and Chia-en A Chang. PROTAC-induced protein structural dynamics in targeted protein degradation. *eLife*, 13, 2025.

[16009] San Pin Steve Wu, Elvis Quiroz, Tianyuan Wang, Skylar G Montague Redecke, Xin Xu, Lin Lin, Matthew L Anderson, and Francesco J DeMayo. Assessment of the histone mark-based epigenomic landscape in human myometrium at term pregnancy. *eLife*, 13, 2025.

[16010] Wei Wu, Qianzhuo Mao, Zhuang-Xin Ye, Zhenfeng Liao, Hong-Wei Shan, Jun-Min Li, Chuan-Xi Zhang, and Jian-Ping Chen. Brochosomes as an antireflective camouflage coating for leafhoppers. *eLife*, 13, 2025.

[16011] Weijie Wu, Miao Yu, Qianchen Li, Yiqian Zhao, Lei Zhang, Yi Sun, Zhenbin Wang, Yuqing Gong, Wenjing Wang, Chenying Liu, Jing Zhang, Yan Tang, Xiaojie Xu, Xiaojing Guo, Jun Zhan, and Hongquan Zhang. Loss function of tumor suppressor FRMD8 confers resistance to tamoxifen therapy via a dual mechanism. *eLife*, 13, 2025.

[16012] Xuanyu Wu and Yang Zhou. Nonlinear feedback modulation contributes to the optimization of flexible decision-making. *eLife*, 13, 2025.

[16013] Yangyu Wu, Yangyang Yan, Youshan Yang, Shumin Bian, Alberto Rivetta, Ken Allen, and Fred J Sigworth. CryoEM structures of Kv1.2 potassium channels, conducting and non-conducting. *eLife*, 12, 2025.

[16014] Yi Wu, Weilin Guo, Haoyue Kuang, Xiaohai Wu, Thi Huong Trinh, Yuexin Wang, Shizheng Zhao, Zilong Wen, and Tao Yu. Pu.1/Spi1 dosage controls the turnover and maintenance of microglia in zebrafish and mammals. *eLife*, 14, 2025.

[16015] Zhengyu Wu, Maojin Tian, Sanyuan Fu, Min Chen, Rongjing Zhang, and Junhua Yuan. Spatial integration of sensory input and motor output in Pseudomonas aeruginosa chemotaxis through colocalized distribution. *eLife*, 13, 2025.

[16016] Zhifei Wu, Baoshan Liao, Julia Ying, Jan Keung, Zongli Zheng, Virpi Ahola, and Wenjun Xiong. Simultaneous cyclin D1 overexpression and p27kip1 knockdown enable robust Müller glia cell cycle reactivation in uninjured mouse retina. *eLife*, 13, 2025.

[16017] Zhiyuan Wu, Lindi Liu, Chenchen He, Lin Xiao, Duo Yun, Junliang Chen, Zhihao Liu, Wenjun Li, Qingjie Lv, and Xiaodong Tan. CD131 contributes to ulcerative colitis pathogenesis by promoting macrophage infiltration. *eLife*, 13, 2025.

[16018] Moritz F Wurm and Doruk Yiğit ErigüÇ. Decoding the physics of observed actions in the human brain. *eLife*, 13, 2025.

[16019] Baolong Xia, Ah-Ram Kim, Feimei Liu, Myeonghoon Han, Emily Stoneburner, Stephanie Makdissi, Francesca Di Cara, Stephanie E Mohr, Aaron Ring, and Norbert Perrimon. Phage-displayed synthetic library and screening platform for nanobody discovery. *eLife*, 14, 2025.

[16020] Yifan Xiang, Vineeta Tanwar, Parminder Singh, Lizellen La Follette, Vikram Pratap Narayan, and Pankaj Kapahi. Early menarche and childbirth accelerate aging-related outcomes and age-related diseases: Evidence for antagonistic pleiotropy in humans. *eLife*, 13, 2025.

[16021] Mengmeng Xiao, Xiangji Li, Fanqin Bu, Shixiang Ma, Xiaohan Yang, Jun Chen, Yu Zhao, Ferdinando Cananzi, Chenghua Luo, and Li Min. Molecular feature-based classification of retroperitoneal liposarcoma: a prospective cohort study. *eLife*, 14, 2025.

[16022] Xing Xiao, Gagik Yeghiazaryan, Fynn Eggersmann, Anna Lena Cremer, Heiko Backes, Peter Kloppenburg, and Anne Christine Hausen. Deficiency of orexin receptor type 1 in dopaminergic neurons increases novelty-induced locomotion and exploration. *eLife*, 12, 2025.

[16023] Chen Xie, Sven Künzel, and Diethard Tautz. Fast evolutionary turnover and overlapping variances of sex-biased gene expression patterns defy a simple binary sex classification of somatic tissues. *eLife*, 13, 2025.

[16024] Enhui Xie, Shuyi Zha, Yiyang Xu, and Xianchun Li. Group identification drives brain integration for collective performance. *eLife*, 13, 2025.

[16025] Changhao Xiong, Nathan M Petro, Ke Bo, Lihan Cui, Andreas Keil, and Mingzhou Ding. Rhythmic sampling and competition of target and distractor in a motion detection task. *eLife*, 14, 2025.

[16026] Jianying Xu, Wenjie Shi, Yi Zhu, Chao Zhang, Julia Nagelschmitz, Maximilian Doelling, Sara Al-Madhi, Ujjwal Mukund Mahajan, Maciej Pech, Georg Rose, Roland Siegfried Croner, Guoliang Zheng, Christoph Kahlert, and Ulf Dietrich Kahlert. Human multiethnic radiogenomics reveals low-abundancy microRNA signature in plasma-derived extracellular vesicles for early diagnosis and molecular subtyping of pancreatic cancer. *eLife*, 14, 2025.

[16027] Manming Xu, Sarath Chandra Dantu, James A Garnett, Robert A Bonomo, Alessandro Pandini, and Shozeb Haider. Functionally important residues from graph analysis of coevolved dynamic couplings. *eLife*, 14, 2025.

[16028] Peipei Xu, Rui Zhang, Zhengrong Zhou, Honglin Xu, Yuejia Li, Mengge Yang, Ruifan Lin, Yingchun Wang, Xiahe Huang, Qi Xie, and Wenxiang Meng. MARK2 regulates Golgi apparatus reorientation by phosphorylation of CAMSAP2 in directional cell migratio. *eLife*, 14, 2025.

[16029] Yuanyuan Xu, Qi Peng, Xiaoping Lu, Long Ma, Jun Li, and Xinsheng Yao. scRNA+TCR-seq reveals the proportion and characteristics of dual TCR Treg cells in mouse lymphoid and non-lymphoid tissues. *eLife*, 14, 2025.

[16030] Genlong Xue, Jiming Yang, Yang Zhang, Ying Yang, Ruixin Zhang, Desheng Li, Tao Tian, Jialiang Li, Xiaofang Zhang, Changzhu Li, Xingda Li, Jiqin Yang, Kewei Shen, Yang Guo, Xuening Liu, Guohui Yang, Lina Xuan, Hongli Shan, Yanjie Lu, Yang Baofeng, and Zhenwei Pan. Binding of LncDACH1 to dystrophin impairs the membrane trafficking of Nav1.5 protein and increases ventricular arrhythmia susceptibility. *eLife*, 12, 2025.

[16031] Jing Xue, Weizhong Zeng, Scott John, Nicole Attiq, Michela Ottolia, and Youxing Jiang. Structural mechanisms of PIP2 activation and SEA0400 inhibition in human cardiac sodium-calcium exchanger NCX1. *eLife*, 14, 2025.

[16032] Shivraj M Yabaji, Vadim Zhernovkov, Prasanna Babu Araveti, Suruchi Lata, Oleksii S Rukhlenko, Salam Al Abdullatif, Arthur Vanvalkenburg, Yuriy O Alekseyev, Qicheng Ma, Gargi Dayama, Nelson C Lau, W Evan Johnson, William R Bishai, Nicholas A Crossland, Joshua D Campbell, Boris N Kholodenko, Alexander A Gimelbrant, Lester Kobzik, and Igor Kramnik. Lipid peroxidation and type I interferon coupling fuels pathogenic macrophage activation causing tuberculosis susceptibility. *eLife*, 14, 2025.

[16033] Odessa R Yabut, Jessica Arela, Hector G Gomez, Jesse Garcia Castillo, Thomas Ngo, and Samuel J Pleasure. Aberrant FGF signaling promotes granule neuron precursor expansion in SHH subgroup infantile medulloblastoma. *eLife*, 13, 2025.

[16034] Rei Yagasaki, Ryo Nakamura, Yuuki Shikaya, Ryosuke Tadokoro, Ruolin Hao, Zhe Wang, Mototsugu Eiraku, Masafumi Inaba, and Yoshiko Takahashi. The gut contractile organoid for studying the gut motility regulated by coordinating signals between interstitial cells of Cajal and smooth muscles. *eLife*, 13, 2025.

[16035] Risa Yamada, Giovanni B Brandani, and Shoji Takada. Multiphase separation in postsynaptic density regulated by membrane geometry via interaction valency and volume. *eLife*, 14, 2025.

[16036] Takayuki Yamaguchi, Kei Okatsu, Masato Kubota, Ayuka Mitsumori, Atsushi Yamagata, Yuko Fukata, Masaki Fukata, Mikihiro Shibata, and Shuya Fukai. Structural insights into heterohexameric assembly of epilepsy-related ligand–receptor complex LGI1–ADAM22. *eLife*, 14, 2025.

[16037] Takaki Yamamoto, Tomoki Ishibashi, Yuko Mimori-Kiyosue, Sylvain Hiver, Naoko Tokushige, Mitsusuke Tarama, Masatoshi Takeichi, and Tatsuo Shibata. Epithelial cell chirality emerges through the dynamic concentric pattern of actomyosin cytoskeleton. *eLife*, 14, 2025.

[16038] Takashi Yamamoto, Kayoko Ueji, Haruno Mizuta, Chizuko Inui-Yamamoto, Natsuko Kumamoto, Yasuhiro Shibata, and Shinya Ugawa. GPRC6A as a novel kokumi receptor responsible for enhanced taste preferences by ornithine. *eLife*, 13, 2025.

[16039] Satoshi Yamashita, Shuji Ishihara, and FranÇois Graner. Apical constriction requires patterned apical surface remodeling to synchronize cellular deformation. *eLife*, 13, 2025.

[16040] Naoki Yamawaki, Hande Login, Solbjørg Østergaard Feld-Jakobsen, Bernadett Mercedesz Molnar, Mads Zippor Kirkegaard, Maria Moltesen, Aleksandra Okrasa, Jelena Radulovic, and Asami Tanimura. Endopiriform neurons projecting to ventral CA1 are a critical node for recognition memory. *eLife*, 13, 2025.

[16041] Hong Yang, Cheng Zhang, Woonghee Kim, Mengnan Shi, Metin Kiliclioglu, Cemil Bayram, Ismail Bolar, özlem özdemir Tozlu, Cem Baba, Nursena Yuksel, Serkan Yildirim, Shazia Iqbal, Jihad Sebhaoui, Ahmet Hacımuftuoglu, Matthias Uhlen, Jan Boren, Hasan Turkez, and Adil Mardinoglu. Multi-tissue network analysis reveals the effect of JNK inhibition on dietary sucrose-induced metabolic dysfunction in rats. *eLife*, 13, 2025.

[16042] Menglei Yang, Hafiz Muhammad Jafar Hussain, Manan Khan, Zubair Muhammad, Jianteng Zhou, Ao Ma, Xiongheng Huang, Jingwei Ye, Min Chen, Aoran Zhi, Tao Liu, Ranjha Khan, Ali Asim, Wasim Shah, Aurang Zeb, Nisar Ahmad, Huan Zhang, Bo Xu, Hui Ma, Qinghua Shi, and Baolu Shi. Deficiency in DNAH12 causes male infertility by impairing DNAH1 and DNALI1 recruitment in humans and mice. *eLife*, 13, 2025.

[16043] Shujing Yang, Huanyao Liu, Jieshi Hu, Binjun Chen, Wanlu An, Xuwen Song, Yi Yang, and Fang He. ACVR2A facilitates trophoblast cell invasion through TCF7/c-JUN pathway in pre-eclampsia progression. *eLife*, 14, 2025.

[16044] Lan Yao, Wanwan Zhang, Xiaogang Yang, Meisheng Yi, and Kuntong Jia. Myosin light chain 3 serves as a receptor for nervous necrosis virus entry into host cells via the macropinocytosis pathway. *eLife*, 13, 2025.

[16045] Shengze Yao, Yesu Jeon, Barry Kesner, and Jeannie T Lee. Xist RNA binds select autosomal genes and depends on Repeat B to regulate their expression. *eLife*, 13, 2025.

[16046] Yiyang Ye, Qirun Wang, and Jie Lin. Physical constraints and biological regulations underlie universal osmoresponses. *eLife*, 13, 2025.

[16047] Qi Yin and Allan C Spradling. Distinct waves of ovarian follicles contribute to mouse oocyte production. *eLife*, 14, 2025.

[16048] Zhuomin Yin, Tao Feng, Qing Xu, Wumin Dai, Maowei Ni, Juan Ni, and Hanmei Lou. Monitoring circulating cell-free HPV DNA in metastatic or recurrent cervical cancer: clinical significance and treatment implications. *eLife*, 13, 2025.

[16049] Baba Yogesh, Matthias Heindorf, Rebecca Jordan, and Georg B Keller. Quantification of the effect of hemodynamic occlusion in two-photon imaging of mouse cortex. *eLife*, 14, 2025.

[16050] Alex J Yonk, Ivan Linares-García, Logan Pasternak, Sofia E Juliani, Mark A Gradwell, Arlene J George, and David J Margolis. Role of posterior medial thalamus in the modulation of striatal circuitry and choice behavior. *eLife*, 13, 2025.

[16051] Jung Hoon Yoon, Karthi Sellamuthu, Louise Prakash, and Satya Prakash. WRN and WRNIP1 ATPases impose high fidelity on translesion synthesis by Y-family DNA polymerases. *eLife*, 14, 2025.

[16052] Ryusei Yoshida, Kazuma Korogi, Qinfei Wu, Shogo Ozaki, and Tsutomu Katayama. Primosomal protein PriC rescues replication initiation stress by bypassing the DnaA-DnaB interaction step for DnaB helicase loading at oriC. *eLife*, 13, 2025.

[16053] Jiro Yoshino, Sonali S Mali, Claire R Williams, Takeshi Morita, Chloe E Emerson, Christopher J Arp, Sophie E Miller, Chang Yin, Lydia Thé, Chikayo Hemmi, Mana Motoyoshi, Kenichi Ishii, Kazuo Emoto, Diana M Bautista, and Jay Z Parrish. Drosophila epidermal cells are intrinsically mechanosensitive and modulate nociceptive behavioral outputs. *eLife*, 13, 2025.

[16054] Bowen Yu, Penghai Li, Haoze Xu, Yueming Wang, Kedi Xu, and Yaoyao Hao. Novel and optimized mouse behavior enabled by fully autonomous HABITS: Home-cage assisted behavioral innovation and testing system. *eLife*, 14, 2025.

[16055] Zhou Yu, Andrian Gutu, Namsoo Kim, and Erin K O'Shea. Activity-dependent synapse elimination requires caspase-3 activation. *eLife*, 13, 2025.

[16056] Kenichi Yuasa, Iris IA Groen, Giovanni Piantoni, Stephanie Montenegro, Adeen Flinker, Sasha Devore, Orrin Devinsky, Werner Doyle, Patricia Dugan, Daniel Friedman, Nick F Ramsey, Natalia Petridou, and Jonathan Winawer. Precise spatial tuning of visually driven alpha oscillations in human visual cortex. *eLife*, 12, 2025.

[16057] Paul Richard J Yulo, Nicolas Desprat, Monica L Gerth, Barbara Ritzl-Rinkenberger, Andrew D Farr, Yunhao Liu, Xue-Xian Zhang, Michael Miller, Felipe Cava, Paul B Rainey, and Heather L Hendrickson. Evolutionary rescue of spherical mreB deletion mutants of the rod-shape bacterium Pseudomonas fluorescens SBW25. *eLife*, 13, 2025.

[16058] Yordanka Zafirova and Rufin Vogels. Integration of head and body orientations in the macaque superior temporal sulcus is stronger for upright bodies. *eLife*, 14, 2025.

[16059] Scott D Zawieja, Grace A Pea, Sarah E Broyhill, Advaya Patro, Karen H Bromert, Charles E Norton, Hae Jin Kim, Sathesh Kumar Sivasankaran, Min Li, Jorge A Castorena-Gonzalez, Bernard T Drumm, and Michael J Davis. Cellular characterization of the mouse collecting lymphatic vessels reveals that lymphatic muscle cells are the innate pacemaker cells. *eLife*, 12, 2025.

[16060] Mohammad Zeeshan, Igor Blatov, Ryuji Yanase, David JP Ferguson, Sarah L Pashley, Zeinab Chahine, Yoshiki Yamaryo-Botté, Akancha Mishra, Baptiste Marche, Suhani Bhanvadia, Molly Hair, Sagar Batra, Robert Markus, Declan Brady, Andrew R Bottrill, Sue Vaughan, Cyrille Y Botté, Karine G Le Roch, Anthony A Holder, Eelco Tromer, and Rita Tewari. A novel SUN1-ALLAN complex coordinates segregation of the bipartite MTOC across the nuclear envelope during rapid closed mitosis in Plasmodium berghei. *eLife*, 14, 2025.

[16061] Qi Zeng, Chen Yao, Shimeng Zhang, Yizhi Mao, Jing Wang, Ziyang Wang, Chunjie Sheng, and Shuai Chen. ORMDL3 restrains type I interferon signaling and anti-tumor immunity by promoting RIG-I degradation. *eLife*, 13, 2025.

[16062] Yajun Zhai, Peiyi Liu, Xueqin Hu, Changjian Fan, Xiaodie Cui, Qibiao He, Dandan He, Xiaoyuan Ma, and Gongzheng Hu. Artesunate, EDTA, and colistin work synergistically against MCR-negative and -positive colistin-resistant Salmonella. *eLife*, 13, 2025.

[16063] Baihong Zhang, Shuqin Huang, Shuyu Guo, Yixuan Meng, Yuzhen Tian, Yue Zhou, Hang Chen, Xue Li, Jun Zhou, and Wenli Chen. ATG6 interacting with NPR1 increases Arabidopsis thaliana resistance to Pst DC3000/avrRps4 by increasing its nuclear accumulation and stability. *eLife*, 13, 2025.

[16064] Changli Zhang, Martha Elena Diaz-Hernandez, Takanori Fukunaga, Sreekala Shenoy, Sangwook Tim Yoon, Lisbet Haglund, and Hicham Drissi. Therapeutic effects of PDGF-AB/BB against cellular senescence in human intervertebral disc. *eLife*, 13, 2025.

[16065] Chenghang Zhang, Tarlan Vatan, and Colenso M Speer. Eye-specific differences in active zone addition during synaptic competition in the developing visual system. *eLife*, 12, 2025.

[16066] Chong Zhang, Zuoyan Zhang, Yueyi Wu, Yuchen Wu, Jing Cheng, Kaizhi Luo, Zhidi Li, Manman Zhang, Jian Wang, Xuesen Zhang, and Yangling Li. Phosphodiesterase 1A physically interacts with YTHDF2 and reinforces the progression of non-small cell lung cancer. *eLife*, 13, 2025.

[16067] Fan Zhang, Hao-Ze Chen, Bo Zheng, Liang Huang, Ye Xiang, Jing-Ren Zhang, and Jia-Feng Liu. Heat stress induces phage tolerance in Enterobacteriaceae. *eLife*, 14, 2025.

[16068] Fan Zhang, Yuan Wang, Luyang Zhang, Chunjie Wang, Deping Chen, Haibo Liu, Ren Xu, Cole M Haynes, Jae-Hyuck Shim, and Xianpeng Ge. The ESCRT protein CHMP5 restricts bone formation by controlling endolysosome-mitochondrion-mediated cell senescence. *eLife*, 13, 2025.

[16069] Hong Zhang, Yinmei Ni, and Jian Li. Oxytocin restores context-specific hyperaltruistic preference. *eLife*, 13, 2025.

[16070] Jiayao Zhang, Juan Li, Jiehong Huang, Xuerui Xiang, Ruoyu Li, Yun Zhai, Shuxian Lin, and Weicai Liu. Psychological stress disturbs bone metabolism via miR-335-3p/Fos signaling in osteoclast. *eLife*, 13, 2025.

[16071] Jie Zhang, Hui Tang, Haoming Wu, Xuewen Pang, Rong Jin, and Yu Zhang. Thymic dendritic cell-derived IL-27p28 promotes the establishment of functional bias against IFN-$\gamma$ production in newly generated CD4+ T cells through STAT1-related epigenetic mechanisms. *eLife*, 13, 2025.

[16072] Le Zhang, Yunpeng Guan, YuenYee Cheng, Nural N Cokcetin, Amy L Bottomley, Andrew Robinson, Elizabeth J Harry, Antoine M van Oijen, Qian Peter Su, and Dayong Jin. Fast evolution of SOS-independent multi-drug resistance in bacteria. *eLife*, 13, 2025.

[16073] Lizi Zhang, Yishan Qu, Larry J Young, Wenjuan Hou, Limin Liu, Jing Liu, Yuqian Wang, Lu Li, Xing Guo, Yin Li, Caihong Huang, Zijian Lv, Yi-Tong Li, Rui Jia, Ting Lian, Hao Feng, Hui Qiao, Zhixiong He, and Fa-Dao Tai. Different roles of D1/D2 medium spiny neurons in the nucleus accumbens in pair bond formation of male mandarin voles. *eLife*, 13, 2025.

[16074] Shiyu Zhang, Min Zhou, Chi Shao, Yu Zhao, Mingzhe Liu, Lei Ni, Zhiyao Bao, Qiurui Zhang, Ting Zhang, Qun Luo, Jieming Qu, Zuojun Xu, and Wei Zuo. Autologous P63+ lung progenitor cell transplantation in idiopathic pulmonary fibrosis: a phase 1 clinical trial. *eLife*, 13, 2025.

[16075] Shuo Zhang, Yan Tian, Quanying Liu, and Haiyan Wu. The neural correlates of novelty and variability in human decision-making under an active inference framework. *eLife*, 13, 2025.

[16076] Weigao Zhang, Hu Liu, Danyang Zhang, Yuguo Yi, Liang Tao, Yunfeng Zhu, Shuxian Huang, Xunan Zhao, Qianchao Shao, Peiqi Li, Yiwen Weng, Wei Lu, Jianfa Zhang, Haibing Zhang, Yuxin Chen, and Dan Weng. Role of hepatocyte RIPK1 in maintaining liver homeostasis during metabolic challenges. *eLife*, 13, 2025.

[16077] Wenyuan Zhang, Zhongru Gu, Yangkang Chen, Ran Zhang, and Xiangjiang Zhan. Birds migrate longitudinally in response to the resultant Asian monsoons of the Qinghai-Tibet Plateau uplift. *eLife*, 14, 2025.

[16078] Xin Zhang, Guangyu Zhang, Shuli Sang, Yang Fei, Xiaopeng Cao, Wenge Song, Feide Liu, Jinze Che, Haoxia Tao, Hongwei Wang, Lihua Zhang, Yiyan Guan, Shipeng Rong, Lijuan Pei, Sheng Yao, Yanchun Wang, Min Zhang, and Chunjie Liu. Single-cell dissection of prognostic architecture and immunotherap response in Helicobacter pylori infection-associated gastric cancer. *eLife*, 13, 2025.

[16079] Yafan Zhang, Irene Silvernail, Zhuyang Lin, and Xingcheng Lin. Interpretable protein-DNA interactions captured by structure-sequence optimization. *eLife*, 14, 2025.

[16080] Yan Zhang, Jingwan Han, Xie Dejian, Wenlong Shen, Ping Li, Jian You Lau, Jingyun Li, Lin Li, Grzegorz Kudla, and Zhihu Zhao. Mapping HIV-1 RNA structure, homodimers, long-range interactions and persistent domains by HiCapR. *eLife*, 13, 2025.

[16081] Yiheng Zhang, Xing Wang, Jian Han, Juyue Xiao, Yuanyuan Yong, Chiyang Yu, Ning Yue, Jie Sun, Kaiyue He, Wenjing Hao, Tao Zhang, Bin Wang, Deng Wang, and Xiaoguang Yang. Dynamic simulations of feeding and respiration of the early Cambrian periderm-bearing cnidarian polyps. *eLife*, 12, 2025.

[16082] Yiran Zhang, Hanyang Guan, Shijin Liu, Haoquan Li, Zili Bian, Jiashuai He, Zhan Zhao, Shenghui Qiu, Tianmu Mo, Xiangwei Zhang, Zuyang Chen, Hui Ding, Xiaoxu Zhao, Liang Wang, Yunlong Pan, and Jinghua Pan. Gamma Knife stereotactic radiotherapy combined with tislelizumab as later-line therapy in pMMR/MSS/MSI-L metastatic colorectal cancer: a phase II trial analysis. *eLife*, 14, 2025.

[16083] Yiteng Zhang, Jianfeng Feng, and Bin Min. Elucidating the selection mechanisms in context-dependent computation through low-rank neural network modeling. *eLife*, 13, 2025.

[16084] Yu Zhang, Shuo Cao, Nan Niu, Huilian Shan, Jinqi Xue, Guanglei Chen, Yongqing Xu, Jianqiao Yin, Chao Liu, Lisha Sun, Xiaofan Jiang, Meiyue Tang, Qianshi Xu, Mingxuan Jia, Xu Zhang, Zhenyong Zhang, Qingfu Zhang, Jianfei Wang, Ailin Li, Yongliang Yang, and Caigang Liu. Efficacy and safety of neoadjuvant stereotactic body radiation therapy plus dalpiciclib and exemestane for hormone receptor-positive, HER2-negative breast cancer: A prospective pilot study. *eLife*, 14, 2025.

[16085] Yuan Zhang, Dan Tang, Ning Zhang, Yi Xiang, Yifan Hu, Wen Qian, Yangji Baima, Xianbin Ding, Ziyun Wang, Jianzhong Yin, Xiong Xiao, and Xing Zhao. Lifestyles and their relative contribution to biological aging across multiple-organ systems: Change analysis from the China Multi-Ethnic Cohort study. *eLife*, 13, 2025.

[16086] Yueliang Zhang, Sara B Noya, Yongjun Li, Jichao Fang, and Amita Sehgal. The microbiome interacts with the circadian clock and dietary composition to regulate metabolite cycling in the Drosophila gut. *eLife*, 13, 2025.

[16087] Yumeng Zhang, Jared Zheng, and Bin Zhang. Protein language model identifies disordered, conserved motifs implicated in phase separation. *eLife*, 14, 2025.

[16088] Zongwang Zhang, Lixia Zheng, Yang Chen, Yuanyuan Chen, Junjie Hou, Chenglu Xiao, Xiaojun Zhu, Shi-Min Zhao, and Jing-Wei Xiong. AARS2 ameliorates myocardial ischemia via fine-tuning PKM2-mediated metabolism. *eLife*, 13, 2025.

[16089] Liuqun Zhao, Tanel Punga, and Suparna Sanyal. $\delta$133p53$\alpha$ and $\delta$160p53$\alpha$ isoforms of the tumor suppressor protein p53 exert dominant-negative effect primarily by co-aggregation. *eLife*, 14, 2025.

[16090] Shengsheng Zhao, Ji Lin, Bingzi Zhu, Yin Jin, Qiantong Dong, Xiaojiao Ruan, Dan Jin, Yongdong Yi, Binglong Bai, Hongzheng Li, Danna Liang, Jianhua Lu, Letian Meng, Xiang Wang, Yuekai Cui, Yuyang Gu, Xian Shen, Xufeng Lu, Shangrui Rao, and Weijian Sun. NOLC1 suppresses immunochemotherapy by inhibiting p53-mediated ferroptosis in gastric cancer. *eLife*, 13, 2025.

[16091] Xin Zhao, Yurim Chae, Destiny Smith, Valerie Chen, Dylan DeFelipe, Joshua W Sokol, Archana Sadangi, and Katherine Tschida. Short-term social isolation acts on hypothalamic neurons to promote social behavior in a sex- and context-dependent manner. *eLife*, 13, 2025.

[16092] Yan Zhao, Hanshuo Zhu, Jinqian Li, Hang Xu, and Li Sun. T3SS translocon induces pyroptosis by direct interaction with NLRC4/NAIP inflammasome. *eLife*, 13, 2025.

[16093] Zhonghao Zhao and Marc H Schieber. Progressively shifting patterns of co-modulation among premotor cortex neurons carry dynamically similar signals during action execution and observation. *eLife*, 13, 2025.

[16094] Bo Zheng, Bronner P GonÇalves, Pengfei Deng, Weibing Wang, Jie Tian, Xueyao Liang, Ye Yao, and Caoyi Xue. Protection afforded by post-infection SARS-CoV-2 vaccine doses: A cohort study in Shanghai. *eLife*, 13, 2025.

[16095] Xu Zheng, Shi Yu, Yanqiu Zhou, Kuai Yu, Yuhui Gao, Mengdan Chen, Dong Duan, Yunyi Li, Xiaoxian Cui, Jiabin Mou, Yuying Yang, Xun Wang, Min Chen, Yaming Jiu, Jincun Zhao, and Guangxun Meng. Interleukin-1 prevents SARS-CoV-2-induced membrane fusion to restrict viral transmission via induction of actin bundles. *eLife*, 13, 2025.

[16096] Shilin Zhong, Ruiyu Wang, Xinwei Gao, Qingchun Guo, Rui Lin, and Minmin Luo. Modular DNA barcoding of nanobodies enables multiplexed in situ protein imaging and high-throughput biomolecule detection. *eLife*, 14, 2025.

[16097] Han Zhou, Tonglian Zhou, Wenli Yu, Liping Liu, Yeonjin Ko, Kristen A Johnson, Ian A Wilson, Peter G Schultz, and Michael J Bollong. Targeted protein degradation by KLHDC2 ligands identified by high-throughput screening. *eLife*, 14, 2025.

[16098] Jiale Zhou, Ding Zhao, Jinze Li, Deqiang Kong, Xiangrui Li, Renquan Zhang, Yuru Liang, Xun Gao, Yuqiang Qian, Di Wang, Jiahui Chen, Liangxue Lai, Yang Han, and Zhanjun Li. Transcriptome-wide identification of 5-methylcytosine by deaminase and reader protein-assisted sequencing. *eLife*, 13, 2025.

[16099] Yuwei Zhou, Wenwen Liu, Ziru Huang, Yushu Gou, Siqi Liu, Lixu Jiang, Yue Yang, and Jian Huang. A comprehensive antigen-antibody complex database unlocking insights into interaction interface. *eLife*, 14, 2025.

[16100] Zhiheng Zhou and Joy Geng. Preparatory attentional templates in prefrontal and sensory cortex encode target-associated information. *eLife*, 14, 2025.

[16101] Ziyue Zhou, Su Young Han, Maria Pardo-Navarro, Ellen G Wall, Reena Desai, Szilvia Vas, David J Handelsman, and Allan E Herbison. GnRH pulse generator activity in mouse models of polycystic ovary syndrome. *eLife*, 13, 2025.

[16102] Bin Zhu, Rui Wei, Wenjuan Hua, Lu Li, Wenlin Zhang, and Pei Liang. A-to-I RNA editing of CYP18A1 mediates transgenerational wing dimorphism in aphids. *eLife*, 13, 2025.

[16103] Yuqing Zhu, Kevin Cho, Haluk Lacin, Yi Zhu, Jose T DiPaola, Beth A Wilson, Gary Patti, and James B Skeath. Loss of dihydroceramide desaturase drives neurodegeneration by disrupting endoplasmic reticulum and lipid droplet homeostasis in glial cells. *eLife*, 13, 2025.

[16104] Samantha J Ziegler, Brandon C Knott, Josephine N Gruber, Neal N Hengge, Qi Xu, Daniel G Olson, Eduardo E Romero, Lydia-Marie Joubert, and Yannick J Bomble. Structural characterization and dynamics of AdhE ultrastructures from Clostridium thermocellum show a containment strategy for toxic intermediates. *eLife*, 13, 2025.

[16105] Maggie E Zink, Leslie Zhen, Jacie R McHaney, Jennifer Klara, Kimberly Yurasits, Victoria E Cancel, Olivia Flemm, Claire Mitchell, Jyotishka Datta, Bharath Chandresekaran, and Aravindakshan Parthasarathy. Increased listening effort and cochlear neural degeneration underlie speech-in-noise deficits in normal-hearing middle-aged adults. *eLife*, 13, 2025.

[16106] Magdalena Ziółkowska, Narges Sotoudeh, Anna Cały, Monika Puchalska, Roberto Pagano, Malgorzata Alicja Śliwińska, Ahmad Salamian, and Kasia Radwanska. Projections from thalamic nucleus reuniens to hippocampal CA1 area participate in context fear extinction by affecting extinction-induced molecular remodeling of excitatory synapses. *eLife*, 13, 2025.
